# Upscaling biodiversity monitoring: Metabarcoding estimates 31,846 insect species from Malaise traps across Germany

**DOI:** 10.1101/2023.05.04.539402

**Authors:** Dominik Buchner, James S. Sinclair, Manfred Ayasse, Arne Beermann, Jörn Buse, Frank Dziock, Julian Enss, Mark Frenzel, Thomas Hörren, Yuanheng Li, Michael T. Monaghan, Carsten Morkel, Jörg Müller, Steffen U. Pauls, Ronny Richter, Tobias Scharnweber, Martin Sorg, Stefan Stoll, Sönke Twietmeyer, Wolfgang W. Weisser, Benedikt Wiggering, Martin Wilmking, Gerhard Zotz, Mark O. Gessner, Peter Haase, Florian Leese

**Affiliations:** Aquatic Ecosystem Research, University of Duisburg Essen, Universitätsstraße 5, 45141 Essen, Germany; Senckenberg Research Institute and Natural History Museum Frankfurt, Clamecystraße 12, 63571 Gelnhausen, Germany; Institute of Evolutionary Ecology and Conservation Genomics, University of Ulm, Albert-Einstein Allee 11, 89069 Ulm, Germany; Centre for Water and Environmental Research, Universitätsstraße 2, 45141 Essen, Germany; Black Forest National Park, Kniebisstraße 67, 72250 Freudenstadt, Germany; University of Applied Sciences HTW Dresden, Pillnitzer Platz 2, 01326 Dresden; Entomological Society Krefeld, Magdeburger Straße 38-40, 47800 Krefeld, Germany; Faculty of Biology, University of Duisburg Essen, Universitätsstraße 5, 45141 Essen, Germany; Helmholtz Centre for Environmental Research – UFZ, Department of Community Ecology, Theodor-Lieser-Straße 4, 06120 Halle, Germany; Department of Plankton and Microbial Ecology, Leibniz Institute of Freshwater Ecology & Inland Fisheries (IGB), Zur alten Fischerhütte 2, 16775 Stechlin, Germany; Department of Ecology, Berlin Institute of Technology (TU Berlin), Ernst-Reuter-Platz 1, 10587 Berlin, Germany; Department of Evolutionary and Integrative Ecology, Leibniz Institute of Freshwater Ecology and Inland Fisheries (IGB), Müggelseedamm 301, 12587 Berlin, Germany; Institute of Biology, Freie Universität Berlin, Königin-Luise-Straße 2/4, Gartenhaus 14195 Berlin, Germany; Kellerwald-Edersee National Park, Laustraße 8, 34537 Bad Wildungen, Germany; Field Station Fabrikschleichach, Department of Animal Ecology and Tropical Biology, Julius-Maximilians-Universität Würzburg, 97070 Würzburg, Germany; Bavarian Forest National Park, 94481 Grafenau, Germany; Senckenberg Research Institute and Natural History Museum Frankfurt, Senckenberganlage 25, 60325 Frankfurt am Main, Germany; LOEWE Centre for Translational Biodiversity Genomics, Senckenberganlage 25, 60325 Frankfurt am Main, Germany; Institute for Insect Biotechnology, Justus-Liebig-University Gießen, Ludwigstraße 23, 35390 Gießen, Germany; German Centre for Integrative Biodiversity Research (iDiv) Halle-Jena-Leipzig, Puschstraße 4, 04103 Leipzig, Germany; Systematic Botany and Functional Biodiversity, Institute for Biology, Leipzig University, Johannisallee 21, 04103, Leipzig, Germany; Institute for Botany and Landscape Ecology, Greifswald University, Soldmannstraße 15, 17487 Greifswald, Germany; University of Applied Sciences Trier, Umwelt-Campus Birkenfeld, Campusallee, 55768 Hoppstädten-Weiersbach; Eifel National Park, 53937 Schleiden-Gemünd, Germany; Chair for Terrestrial Ecology, Department of Life Science Systems, School of Life Sciences, Technische Universität München, Hans-Carl-von-Carlowitz-Platz 2, 85354 Freising-Weihenstephan, Germany; Lower Saxon Wadden Sea National Park Authority, Virchowstraße 1, 26382 Wilhelmshaven, Germany; Institute of Biology and Environmental Sciences, Carl von Ossietzky Universität Oldenburg, Ammerländer Heerstraße 114-118, 26129 Oldenburg, Germany

## Abstract

Mitigating ongoing losses of insects and their key functions (e.g., pollination) requires accurately tracking large-scale and long-term community changes. However, doing so has been notoriously hindered by uniquely high insect species diversity that requires prohibitively high investments of time, funding, and taxonomic expertise. Here, we show that these concerns can be addressed through a comprehensive, scalable and cost-efficient DNA metabarcoding workflow. We use 1,815 samples from 75 Malaise traps across Germany from 2019 and 2020 to demonstrate how metabarcoding can be incorporated into large-scale insect monitoring networks for less than 50 € per sample, including supplies, labor and maintenance. With on average 1.4M sequence reads per sample we uncovered 10,803 validated insect species, of which 83.9% were represented by a single OTU. We estimated another 21,403 plausible species, which likely either lack a reference barcode or are undescribed. The total of 31,846 species is similar to the number of insect species known for Germany (∼35,500). Because Malaise traps capture only a subset of insects, our approach identified many species likely unknown from Germany or new to science. Our reproducible workflow (∼80% OTU-similarity among years) provides a blueprint for large-scale biodiversity monitoring of insects and other biodiversity components in near real time.

## Introduction

Insects are the most diverse taxonomic group on Earth and contribute to essential ecosystem processes and services, such as pollination, nutrient cycling and organic matter decomposition (Cardoso et al. 2020). However, insect populations are declining (Wagner 2020) and our ability to mitigate these declines is hindered by poor understanding of spatial distributions, habitat requirements, biotic interactions, dynamics, and even the overall number of extant species. About 1 million insect species have been described to date, but recent estimates of total species numbers stand at about 5.5 million (Stork 2018) or even more (IPBES 2019). Surprisingly, many new species are being reported even for extremely well-studied and generally species-poor areas with a long history of entomological research. For example, based on 4,000 species caught with Malaise traps in Sweden, Karlsson et al. (2020) reported almost 700 insect species new to science and ∼1,300 new to Sweden. Similarly, in a DNA barcode analysis of about 62,000 specimens collected with Malaise traps, Chimeno et al. (2022) estimated over 2,000 dipteran species new to Germany, raising the total number of 33,341 known insect species (Klausnitzer 2005) to approximately 35,500. Thus, while many new insect species are being continuously reported, the vast majority still remain unknown.

A key constraint to closing the current information gap on insects is the vast number of species and samples that must be processed. Terrestrial and aquatic monitoring methods, like Malaise, canopy, or light traps, as well as sediment (benthos) samples from freshwater, can collect thousands of specimens (e.g., Resh and Jackson 1993, Karlsson et al. 2020, Habel et al. 2023), many of which are small and difficult to identify even for experts, resulting in taxonomic neglect (Srivathsan et al. 2023) and extensive time needed for identification that by far outweighs the time required for collection. National and international monitoring networks, such as the Global Malaise Trap program (Geiger et al. 2016), BioScan (Hobern 2021), LifePlan (www.helsinki.fi/en/projects/lifeplan), or the Swedish Malaise Trap program (Karlsson et al. 2020), have started coordinated large-scale insect sampling initiatives based on standardized traps and procedures (e.g., Hallmann et al. 2017). However, classical taxonomic analyses of these samples cannot keep pace with the rate of collection, particularly given the low and globally declining number of taxonomic experts (European Commission 2022). While the importance of taxonomic expertise remains undisputed, there is a clear need for alternative methods to assess insect species diversity both efficiently and reliably (van Klink et al. 2020, Chua et al. 2023).

Following more than a decade of research and trial applications, DNA metabarcoding has now reached a high technology readiness level, presenting a promising solution for examining a fuller range of insect biodiversity in large-scale monitoring programs. A range of suitable protocols for specimen collection, processing and data analysis are now available (Buchner et al. 2021c, Montgomery et al. 2021), as are suitable primer pairs (Braukmann et al. 2019, Elbrecht et al. 2019) and an accessible platform for insect barcode reference data (Ratnasingham and Hebert 2007). The key strength of DNA metabarcoding is that it rapidly delivers taxonomically highly resolved taxa lists, whereas quantitative information on species abundance or biomass remains a challenge (Sickel et al. 2023).

Despite its potential, DNA metabarcoding has not yet been implemented in large-scale, long-term insect biodiversity monitoring programs. There are several likely reasons. In particular sample throughput is still constrained by the need for significant manual labor input, expensive DNA kits and reagents, and difficulties in accessing information on lab and analysis procedures (McGee et al. 2019), which all present roadblocks to large-scale implementation. Furthermore, incomplete and partly inconsistent reference databases still limit the accuracy and quantity of species-level assignments and thus the completeness and validity of the resulting species lists. This problem remains particularly pervasive for highly diverse groups like Diptera and Hymenoptera where species that are often difficult to distinguish based on morphological criteria and thus tend to be underrepresented in reference libraries. The unknown (i.e., undescribed) species in these poorly explored groups are often referred to as “dark taxa” (Hartop 2021). However, in the absence of reference barcodes for species, or even when formal species descriptions are lacking, DNA metabarcoding can still overcome these limitations using genetic distance thresholds to approximate entities that roughly reflect species, such as molecular Operational Taxonomic Units (OTUs) or Barcode Index Numbers (Ratnasingham and Hebert 2013) (BINs). Even in the absence of species in databases, these distance-based entities can approximate species numbers and have been applied in ecological and ecotoxicological research for many years (Sturmbauer et al. 1999, Hoppeler et al. 2016, Beermann et al. 2018).

DNA metabarcoding analyses to date have been limited to either a specific region within a country (e.g., Geiger et al. 2016, Uhler et al. 2021, Habel et al. 2023), short time spans (e.g., Huang et al. 2022, Li et al. 2023) or specific taxonomic groups (e.g., Chimeno et al. 2022, Huang et al. 2022). None has quantified the full extent of insect diversity – including dark taxa – at the whole-country scale and through time. Thus, the present study fills this gap by presenting a robust DNA metabarcoding workflow for application to large-scale insect monitoring programs, combined with a new multi-level procedure to assess the validity of species records. The underlying dataset consists of 1,815 Malaise trap samples collected in 2019 and 2020 from 75 individual traps across Germany (Figure 1). Our aims were to: (1) present the workflow and provide detailed and publicly available laboratory procedures and programs; (2) quantify the time and cost investment required per sample; and (3) evaluate the reliability of the approach in terms of number of known species detected, number of dark taxa and their likely validity, and variation in species detected between years. The results obtained with the new workflow show that DNA metabarcoding is feasible for large-scale and long-term insect monitoring, and providing insight into insect diversity at scales that have been challenging to study so far.

**Figure 1:**
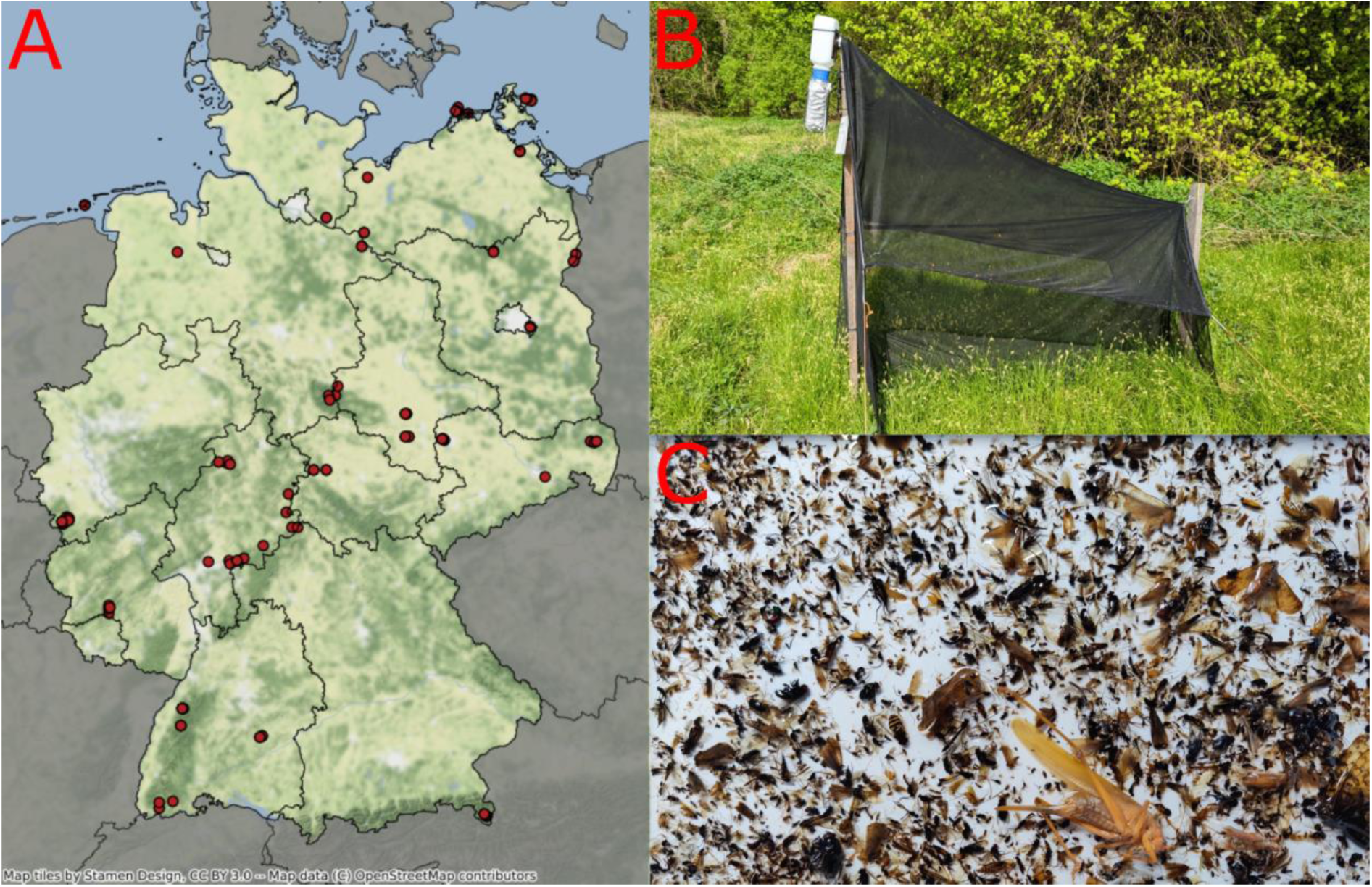
**A:** Location of the in total 75 Malaise traps across Germany**, B: Lateral view** of a Malaise trap with collection bottle protected from sunlight at the upper left end, and **C: top view** of a preserved Malaise trap sample spread in a white tray.

## Materials and Methods

### Sampling

Sampling was conducted as part of the nationwide German Malaise trap monitoring program (Welti et al. 2022) comprising forests, grassland, agricultural and urban areas (https://www.ufz.de/lter-d/index.php?de=46285). Most of the 31 sites in which the Malaise traps were placed belong either to the German LTER-D network (Haase et al. 2016, Mirtl et al. 2018) (Long-Term Ecological Research) or to the network of national natural landscapes (https://nationale-naturlandschaften.de). At each site, one to six Malaise traps have been operated to monitor biomass and species composition of flying insects in different habitats. In the present study, we used a total of 1,815 Malaise trap samples from 56 locations across Germany during 2019, with 19 locations added in 2020 (Figure 1). Traps were emptied every two weeks from the beginning of April until the end of October in both years (approx. 15 samples per year, depending on site-specific climatic conditions). Insects were caught in 80% ethanol and their wet biomass was measured following Welti et al.(2022). Samples were kept in 96% ethanol and protected from light until later genetic analysis.

### Sample processing

All samples were divided into two size classes (small ≤ 4 mm; large > 4 mm) to increase taxon recovery rates of small taxa (Elbrecht et al. 2021). Samples spread on a perforated plate sieve (4 mm hole diameter) were stirred using a magnetic stirrer (750 rpm) in ethanol (Figure S1), so that small individuals passed through the holes, whereas the large ones were retained on top of the sieve. The size fractions were homogenized following the protocol described in Buchner et al. (Buchner et al. 2021a), except that the homogenization time was reduced to 30 s. The two size fractions were subsequently pooled at a ratio of 1:4 (large 200 µL: small 800 µL) as recommended by Elbrecht et al. (2021).

### DNA extraction

Generally, the laboratory steps followed the workflow described in Buchner et al. (Buchner et al. 2021c). All procedures are available as step-by-step protocols in the protocols.io repository (Buchner 2022b). All subsequent steps after size-sorting and homogenization were completed on a Biomek FX^P^ liquid handling workstation (Beckman Coulter, Brea, CA, USA). After sample lysis (Buchner 2022e), samples were processed in duplicate during the entire library preparation to control for possible cross-contamination. Additionally, in each 96-well plate twelve negative controls were included. DNA was extracted using a magnetic bead protocol (Buchner 2022a). Extraction success was verified on a 1% agarose gel. For all samples that did not amplify, the extraction was repeated with silica spin columns (Buchner 2022a), which were always successful.

### Sequencing library preparation

The PCR for the metabarcoding library followed a two-step PCR protocol (Zizka et al. 2019) targeting a 205 bp fragment (Vamos et al. 2017) of the cytochrome oxidase c subunit I (COI) gene. DNA was amplified in a first PCR using the Qiagen Multiplex Plus Kit (Qiagen, Hilden, Germany) with a final concentration of 1x Multiplex Mastermix, 200 nM of each primer (fwh2F, fwhR2n (Vamos et al. 2017)) and 1 µL of DNA, filled up with PCR-grade water to a final volume of 10 µL. The amplification protocol was 5 min of initial denaturation at 95 °C, 20 cycles of 30 s denaturation at 95 °C, 90 s of annealing at 58 °C and 30 s of extension at 72 °C, followed by a final elongation step of 10 min at 68 °C. Each of the PCR plates used in the first step was tagged with a unique combination of inline tags. Additionally, the primers contained a universal binding site for the primer used in the second PCR step to anneal (Table S1). The PCR products were purified using a bead-based protocol and a ratio of 0.8x and an elution volume of 40 µL to remove remaining primers and potential primer dimers (Buchner 2022c).

In the second PCR, DNA was amplified at a final concentration of 1x Multiplex Mastermix, 100 nM of each primer (Table S1), 1x Corralload Loading Dye, 2 µL of the cleaned-up product of the first PCR in a final volume of 10 µL. The amplification protocol was 5 min of initial denaturation at 95°C, 25 cycles of 30 s denaturation at 95°C, 90 s of annealing at 61°C and 30 s of extension at 72°C, followed by a final elongation step of 10 min at 68 °C. PCR success was visualized on a 1% agarose gel.

To achieve a similar sequencing depth, the PCR products were normalized to equal concentrations. Normalization was achieved with a bead-based protocol and a ratio of 0.7x (Buchner 2022d) and an elution volume of 40 µL. The whole volume of the normalized samples was then pooled into the final libraries. Libraries were then concentrated using a silica spin-column protocol (Buchner 2022a). Library concentrations were quantified on a Fragment Analyzer (High Sensitivity NGS Fragment Analysis Kit; Advanced Analytical, Ankeny, IA, USA). The libraries were sequenced at Macrogen Europe using the HiSeq X platform with a paired-end (2x150 bp, 15 lanes) kit or at CeGaT (Tübingen, Germany) using the MiSeq V2 platform (2x150 bp, 1 lane).

### Bioinformatics

Raw data of the sequencing runs were delivered demultiplexed by index reads. Since no differences were detected between sequencing runs, they were all pooled before subsequent analyses. Additional demultiplexing of the inline tags was achieved with the Python package “demultiplexer” (v1.1.0, https://github.com/DominikBuchner/demultiplexer). Reads were further processed with the APSCALE pipeline (Buchner et al. 2022) (v1.4.0, https://github.com/DominikBuchner/apscale using default settings. Briefly, paired-end reads were first merged using vsearch (Rognes et al. 2016) (v2.21.1) before the primer sequences were trimmed using cutadapt (Martin 2011) (v3.5). Only reads with a length of 205 bp (±10) and with a maximum expected error of 1 were retained. Identical reads less abundant than 4 were discarded and all others dereplicated before OTUs were clustered based on a similarity threshold of 97% and mapped to OTUs including singletons. The resulting OTU table was filtered for erroneous OTUs with the LULU algorithm (Frøslev et al. 2017) as implemented in APSCALE. Taxonomic assignment was performed using BOLDigger (Buchner and Leese 2020) (v1.5.4, https://github.com/DominikBuchner/BOLDigger). The best hit was determined with the BOLDigger method and the API verification method. This resulted in a raw OTU table (Table S2) that was used in subsequent analysis.

### Data filtering

To control for possible contamination during the laboratory workflow, the technical replicates of each sample, as well as the negative controls, were merged by summing up the reads, provided that the reads were present in both replicates. Subsequently, the maximum number of reads per OTU present in all of the negative controls was subtracted from the respective OTU (Table S3). All OTUs were analyzed for stop-codons, and any OTUs containing stop-codons were removed. We analyzed two datasets: 1) OTUs assigned at the species level and 2) OTUs assigned to insects. To further clean dataset 1, all OTUs sharing species assignments were merged by summing up their reads. Retrieved species names containing numbers or punctuation marks were also removed (e.g., incomplete database records). The resulting final species list (Table S4) for all samples was used for the validation procedure.

### Validation of taxonomic assignment

To validate the resulting species-level list, three different approaches were used. First, the occurrences of all species were checked by taxonomic experts at the Entomological Society Krefeld, Germany. As a basis, the experts used the digitally available insect species catalogue from the Entomofauna Germanica (Klausnitzer 2005), which is best resembled through the list from the German Barcode of Life portal (https://gbol.bolgermany.de). All records of species found were then validated against known changes in synonymies as well as new records of insect species from Germany reported in scientific primarily literature available to them. Second, all detected species were checked for GBIF records within a 200 km radius around the given trap. This value was selected to be sufficiently high to accommodate also for the often patchy and rare species records in GBIF, the large size of the study area and the fact that many flying insect species are highly mobile. To do so, a polygon was drawn around all trap locations with occurrences of the respective species, and the border of this polygon was then extended by 200 km (Figure S2). Records were extracted from the GBIF database using the “rgbif” package (Chamberlain et al. 2022). Lastly, all species were checked against the German Barcode of Life database, which was downloaded with a custom Python script (Script S1). A record was accepted as valid when two of the three validation criteria were met, which also helps to control for false positives (Table S5).

### Statistical analysis

To assess if sequencing depth was sufficient across all samples, we conducted a read-based rarefaction using a custom python script. This approach involved randomly sampling reads without replacement from each sample in increments of 0.1% (with 50 iterations per step) of the total read count. Subsequently, we fitted a Michaelis-Menten-type equation to the resulting rarefaction curve. Using this function, we computed OTU richness when doubling the sequencing depth. If OTU richness increased by less than 5% when doubling the sequencing depth, the sequencing depth was considered sufficient (Figure S3, Table S6).

In addition to examining sequencing rarefaction curves, we also examined rarefaction curves for both validated and plausible species using the iChao2 (Chiu et al. 2014) estimator. We did so to determine the potential influence of collecting more samples from each site, such as by sampling for a longer time period or more frequently, on the number of detected insect species. This was done by randomly drawing subsets of samples (50 iterations each) from the dataset without replacement in increments of 5 from 5 to 1815.

#### Plausible species and dark taxa estimation

To address the potential over-estimation of species diversity based on OTU numbers we computed the mean number of OTUs per validated species for each of the 20 insect orders within the dataset. This mean was then used to normalize the OTU count for each order. Specifically, we divided the number of OTUs per order by the calculated mean, providing an estimate of additional species present in the dataset that have not yet been assigned a species name. We refer to these additional species as plausible species. Furthermore, we obtained data from the Barcode of Life datasystems (accession date: 14^th^ of April, 2023), including the number of species barcoded with a voucher specimen collected in Germany, along with the total number insect species known from Germany (Klausnitzer 2005). This additional information allowed us to distinguish plausible species lacking a reference sequence from potential dark data within the dataset. The described correction was performed on order as well as on family level, but we simplicity we focused the subsequent analysis on the order level.

### Time and cost estimations

To estimate the time and costs needed for our nationwide insect diversity assessment via metabarcoding, we used vendor list prices for materials, as well as runtimes on the liquid handler plus estimates for laboratory set-up before and after each step of the workflow. These costs include all labware and chemicals needed to complete the respective step of the protocol. Incubation times are not included in the time estimates, because the time can be used to process other samples. The calculated price per sample is based on 1,815 samples including replicates and negative controls, resulting in a total number of 4,149 individual reactions spread across 44 96-well PCR plates. A total of 10 blenders were used for homogenization, which are included in the cost estimate. For sequencing, cost estimates are based on the current costs of 1,100 € for 110 Gb output at a commercial service provider (Macrogen Europe) using the NovaSeq 6000 S4, which has replaced the HiSeq platform, and a sequencing depth of at least 1.5 million reads per sample (∼ 817 Gb). Labor costs were estimated at 60 € per hour, corresponding to an experienced scientist in Germany. The costs for the use of the liquid handler were estimated based on a linear depreciation over a total expected lifetime of 10 years. Annual gross maintenance costs of 40,000 € were assumed for instruments, resulting in total depreciation and maintenance costs of approximately 4,000 € for the present study. No rental and auxiliary costs needed to be applied in this study.

## Results

### Sequencing results and species validation

We performed high-throughput DNA metabarcoding including replicates and negative controls on 1,815 Malaise trap insect samples (775 for 2019 and 1,040 for 2020; Table 1) using a 205-bp fragment of the mitochondrial cytochrome *c* oxidase I (COI) gene as a marker. All samples and sequencing runs combined yielded 3,999,082,169 demultiplexed read pairs. The average read number per sample in the final read table was 1,401,469 (±SD of 618,631). Sequencing depth was sufficient for all samples, with a mean increase in richness of 0.17% by doubling the sequencing depth (0% – 1.46%; Table S6).

**Table 1:** Summary statistics of OTUs, assigned insect species and OTUs assigned to validated insect species according to the two-out-of-three validation criterion. Plausible species calculated from the average number of OTUs per validated species and family. n = number of Malaise trap samples.

|  | **2019 (n=775)** | **2020 (n=1,040)** | **$\Sigma$ (n=1,815)** |
| --- | --- | --- | --- |
| **Raw OTUs** | 41,189 | 50,166 | 52,981 |
| **Insect OTUs** | 39,367 | 47,652 | 50,087 |
| **Insect species** | 9,728 | 11,183 | 11,776 |
| **Validated species according to:** |  |  |  |
| **Expert validation** | 9,132 | 10,475 | 11,030 |
| **GBIF validation** | 7,969 | 8,867 | 9,254 |
| **GBOL validation** | 8,777 | 10,056 | 10,574 |
| **Validated species** | **8,983** | **10,276** | **10,803** |
| **Plausible species** |  |  | **21,403** |
| **Total species** |  |  | **31,846** |

Sequencing yielded a total of 52,981 raw OTUs, 50,087 of which were assigned to insects. We were able to assign 11,776 species names to a total of 15,042 OTUs. Two thirds of these species (10,803) were validated via three different criteria involving i) expert judgment from entomologists with particular knowledge of long-term Malaise trap community data from German Malaise traps, ii) a comparison with an online database that includes the known German species (Klausnitzer 2005) (GBOL), and iii) a GBIF record check within a 200 km radius. Species were regarded as ‘validated’ when two of the three validation criteria supported them. The three different validation criteria showed a pairwise agreement exceeding 80%. Taxonomic experts validated an additional 355 species not yet not in the list of species reported from Germany (Table 1), leading to an overlap of 97% with the GBOL database. GBIF and taxonomic expert validation had an overlap of 84%, and GBIF and GBOL of 82% (Table S7). Consequently, a total of 35,045 insect OTUs remained unassigned, either because species-level reference data are lacking or the species are truly unknown (i.e., dark taxa; Table 1), resulting in a large discrepancy between the number of named species and recorded insect OTUs. Although sometimes several OTUs were assigned to the same species, 9,061 (83.9%) of our validated species were represented by a single OTU (Table S8).

### Species richness estimation

Our sampling effort captured the majority of OTUs. Based on rarefaction curves, a greatly increased sampling effort in each site, for instance by sampling more frequently or for a longer period at all sites, may have only resulted in detecting an additional 4,725 OTUs (+9.4%) and 934 validated species (+8.6%; Figure 2). Additionally, most of the OTUs were found in both sampling years (36,932 = 73.7%) with more insect OTUs occurring exclusively in 2020 (10,720) than in 2019 (2,435). The same is true for the validated insect species. Most were found in both years (8,456 = 78.3%) but more than three times as many occurred exclusively in 2020 (1,820) compared to 2019 (527).

**Figure 2:**
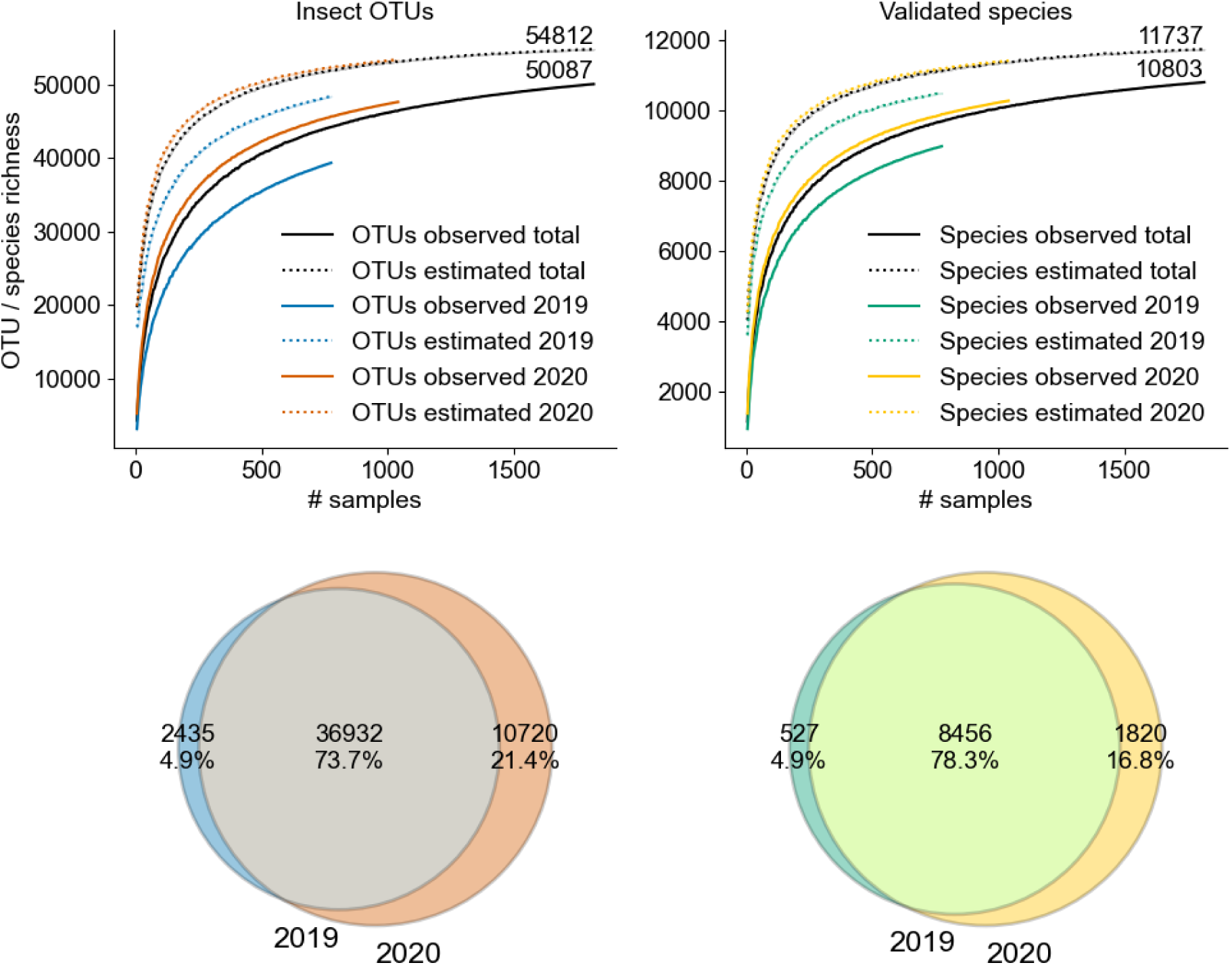
Top row: Rarefaction curves showing. richness of insect OTUs and validated species richness that were either observed (solid lines) or estimated (dotted lines). **Bottom row:** Shared and unique absolute numbers and proportions of the insect OTUs (left circles) and validated insect species (right circles) collected in 2019 and 2020.

### OTU distribution across insect groups and unknown taxa

The four most diverse insect orders in Germany (Diptera, Hymenoptera, Lepidoptera, and Coleoptera) were well represented in the dataset (Figure 3, top row). Diptera and Hymenoptera were the most common orders both at the OTU (22,732 = 45.4% and 12,823 = 25.6%) and species level (3,851 = 35.6% and 2,370 = 21.9%). The 10,803 validated insect species represent 33.5% of the 35,500 insect species known in Germany (Klausnitzer 2005) and 82.6% of the 13,076 barcoded insect species recorded in the country (Table S9).

**Figure 3:**
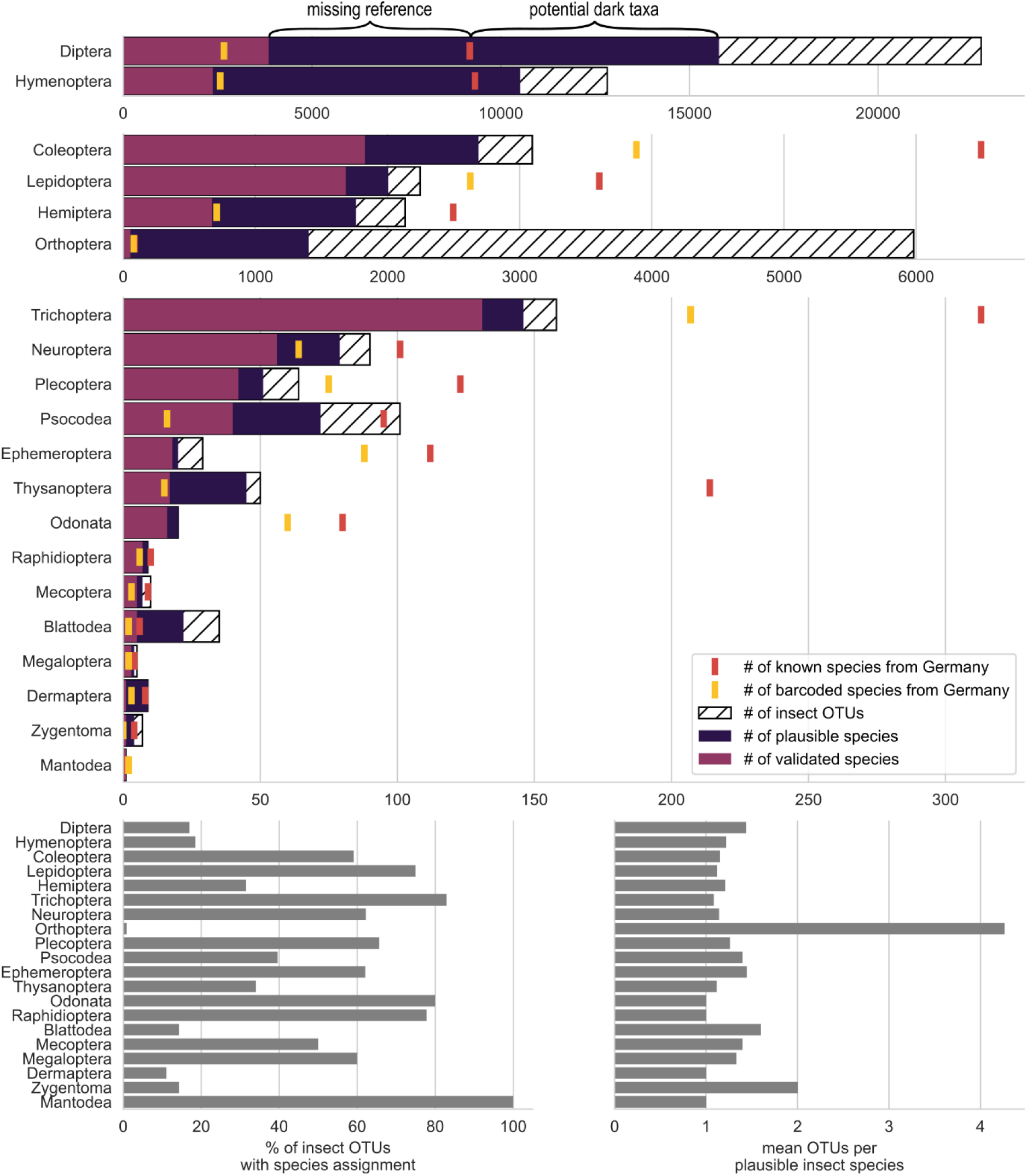
Top panels: Number of total insect OTUs, plausible and validated insect species per order identified in the present study compared to the total number of species reported from Germany or sampled in Germany and included in the BOLD database (data accessed on 14^th^ of April, 2023). **Bottom left panel**: Percentage of OTUs assigned to species. **Bottom right panel:** Mean number of distinct OTUs assigned to a given plausible species.

The percentage of OTUs assigned to species differed considerably among insect orders, being highest for Lepidoptera (75%), followed by Coleoptera (59%). Less than a fifth of the OTUs identified as Diptera and Hymenoptera could be assigned at the species-level (17% and 18%, respectively). The lowest percentage of OTUs assigned to the species level was for Orthoptera (<1%) (Figure 3, bottom left panel). The mean number of OTUs per validated insect species varied slightly among insect orders and families, typically between 1 and 1.5, but Orthoptera (specifically the Acrididae) were represented by >4 and Zygentoma (specifically Lepismatidae) by ∼2 OTUs per species (Figure 3, bottom right panel).

Based on the mean number of OTUs per validated insect species calculated at either the order or family level, we estimated that our data set respectively included either an additional 22,496 or 21,403 plausible insect species (Table S10). All OTUs belonging to Orthoptera were removed for these estimates to avoid artificially inflating the total number of species (Information S1). The lower of the two numbers is a more conservative estimate, but since we identified 435 different families (see Table S11 for further family level information), our analysis focuses, for simplicity reasons, on the order level. The additional plausible species could be those that either a) could not be identified to species level due to a lack of a reference in the present reference database, b) had not previously been recorded from Germany, or c) represent new species to science. Examples of species not reported from Germany or new to science (‘potential dark taxa’ in Figure 3, Table S10) were particularly evident in the Diptera and Hymenoptera, for which we respectively found ∼6,600 and ∼1,200 species while the missing reference data aspect affected the assignment in all orders except the Mantodea (with 1 species only).

### Time and cost estimation

The total costs of the laboratory workflow for duplicate sample processing and negative controls -including all needed labware, chemicals, sequencing and salaries - were estimated at 88,000 €, equivalent to about 46 € per sample (Tables 2, S12). Costs for laboratory materials accounted for 12 € (26%) of the total costs per sample, and salaries for 34 € (74%). Sequencing was the most expensive step (contributing 4.85 € per sample), followed by enzymatic steps such as PCR (1.65 €) and sample lysis (1.54 €). The total processing time for all 1,815 samples was 1,030 working hours or 27.5 weeks, equivalent to 141 samples within 2 weeks for one person working full time and supported by one liquid handling robot. Most of the processing time was needed for sample size-sorting and homogenization. All subsequent steps were completed within 6 weeks.

**Table 2:**
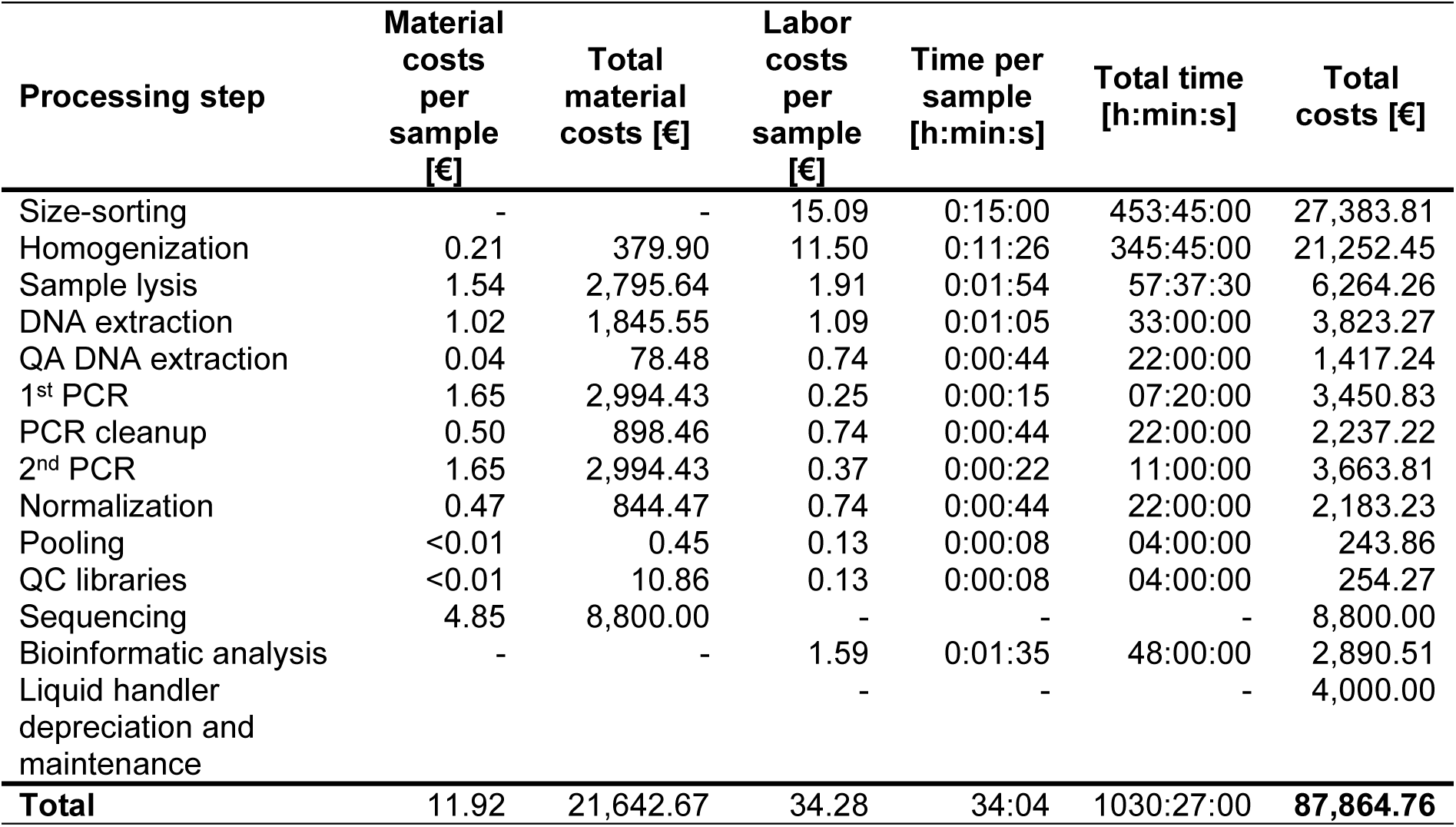
Cost estimates for the proposed workflow: Cost estimates are based on vendor list prices in 2022 where available. Time estimates are either hands-on time in the laboratory or mean runtimes of the robotic protocols. Costs and time required per sample are based on 1,815 samples. Total time and total costs are based on the respective number of replicates processed in each step.

## Discussion

### A metabarcoding workflow for large-scale biodiversity monitoring

We here present a comprehensive, scalable, and cost-efficient DNA metabarcoding workflow which allows analyzing thousands of specimen-and species-rich samples within several weeks to months depending on the available workforce. The proposed DNA metabarcoding workflow differs from others (e.g. Braukmann et al. 2019, Hardulak et al. 2020, Hausmann et al. 2020) in several important aspects that are key to high sample throughput at reduced costs and processing time while ensuring high-quality data. Key differences include: i) homogenization of samples in preservative liquid to avoid a time-consuming drying step (Buchner et al. 2021a); ii) processing of all samples in duplicate before DNA extraction as an essential quality assurance measure to reduce the probability of false positive signals; iii) completion of all laboratory procedures by an automated liquid handling robot to minimize processing time (Buchner et al. 2021c) (except for the gel electrophoresis) and maximize consistency; iv) a mean sequencing depth increased to ∼1.4 M reads per sample to boost species detectability; and v) publication of all laboratory procedures as open-source protocols (Buchner 2022b) or programs (Buchner and Leese 2020, Buchner et al. 2022) to ensure full transparency and reproducibility following the FAIR principles (Wilkinson et al. 2016).

Labor and material costs of metabarcoding protocols are often considered prohibitively high for large-scale monitoring programs (Borrell et al. 2017, Montgomery et al. 2021), frequently exceeding 200 € per sample and up to 400 € (Ji et al. 2013, Elbrecht et al. 2017, Aylagas et al. 2018). The workflow presented here considerably reduces these costs, down to <50 €. This is primarily achieved by automating crucial laboratory steps and by preparing all required solutions instead of purchasing expensive commercial kits. Costs could be further cut in future large-scale programs by ordering chemicals and consumables in bulk and by further reducing reaction volumes for all laboratory steps involving enzymes, wherever possible (Buchner et al. 2021b). An important consideration to lower labor costs is to reduce the processing time per sample, notably for size-sorting and homogenization, ideally by automating these steps as well. Furthermore, expenses related to sampling in the field are not included in our analysis, although they can easily match laboratory costs. For example, in our monitoring program, costs for Malaise trap installation, maintenance and biweekly sampling from April to October (15 samples per site and year in total) amounted to about 12 working hours or 240 € for a student aid. Adding travel time (highly variable but assumed here to average 15 hours or 300 €) and material (approx. 450 €) would result in total costs for field sampling of 66 € per sample, unless much of the work is accomplished by volunteers. This sum ignores expenses for rent and additional costs (heating, electricity etc.), which need to be added when commercial providers are solicited.

Another crucial but often neglected step in metabarcoding workflows is species validation. Validating species records poses a significant challenge for many taxa due to the dearth of reference sequences, few checklists and experts, and diverse algorithmic approaches (Hleap et al. 2021). However, these challenges could be partly addressed by using publicly available species databases, as suggested by the remarkably close agreement we found among our three validation procedures of insect experts, GBIF, and GBOL matching. The taxonomic experts added further species to the list of known species from Germany based on recent scientific evidence. This agreement suggests that relying solely on public databases, like GBIF (Telenius 2011) and the GBOL database, which most closely resembles the reports of the Entomofauna Germanica (Klausnitzer 2005), is sufficiently accurate for large-scale monitoring where expert validation tends to be prohibitive. This is not to advocate disregarding expert knowledge, which is critical to minimize the well-known errors inherent in automated validation methods (e.g. Meiklejohn et al. 2019). Instead, we would like to raise awareness that acceptable alternatives exist to overcome time constraints, as they typically arise in large biodiversity monitoring programs. FAIR and curated reference databases with up-to-date taxonomic names and integration of synonyms is essential for harmonized biodiversity monitoring and establishing such systems a key challenge (Keck and Altermatt 2023).

### A reliable account of biodiversity

Rarefaction indicates that the 10,803 insect species identified (without considering unassigned OTUs) with our protocol would increase by only 934 species (8.6%) if sampling efforts were ramped up at the established study sites. However, we acknowledge that higher species numbers would be obtained if more sites were sampled, in particular including new habitat types. This expectation is consistent with the larger number of validated insect species in 2020 (+14.4%) when 19 more sites were sampled compared to 2019. The 10,803 detected species represent one third of all insect species reported from Germany and ∼83.1% of all barcoded insects from this country. This includes nearly 100% of the up-to-date barcoded Hymenoptera and Hemiptera, indicating that with just 75 sampling sites our approach captures a large portion of the known insect biodiversity of Germany. Other metabarcoding studies from regions within Germany have identified ∼5,900 species (Uhler et al. 2021) and ∼ 11,984 insect OTUs (Habel et al. 2023). Similarly, a DNA barcoding study investigating 1% of the samples from the Swedish Malaise trap program found >4,000 species (Karlsson et al. 2020), and another recent DNA barcoding study based on Malaise traps across eight countries from four continents found >25,000 species (Srivathsan et al. 2023). While we acknowledge that comparisons of insect species and OTUs numbers among studies are difficult due to differences in protocols, these examples indicate, along with the high proportion of known or barcoded insects from Germany, that rapid and cost effective metabarcoding can provide a robust inventory of biodiversity.

At the scale of our nationwide sampling network, we consistently found the same species in two consecutive years (∼75% species and ∼80% OTU-similarity among years), highlighting the spatiotemporal reliability of taxa lists derived from metabarcoding when used in large-scale monitoring. One of the principal concerns in large-scale sampling is that high community variability in space and time, and sampling idiosyncrasies, make it difficult to detect the same species even in spatially or temporally proximate samples. This issue is particularly relevant when using methods, like metabarcoding, that can identify thousands of species, most of which will be rare and so may only occur sporadically in a single site (Jeliazkov et al. 2022). For example, traps just tens of meters apart (e.g., Steinke et al. 2021) or in successive sampling periods (e.g., Sinclair et al., in prep, Information S1) can capture very different insect communities. However, the between-year consistency of our data suggests this issue may diminish as the spatial scale of sampling for metabarcoding increases, such as in larger-scale biodiversity monitoring conducted across whole countries.

### Assessing dark taxa biodiversity

A key finding of our study is the discovery of approximately 5 times more insect OTUs than validated insect species, demonstrating the potential value of cost-effective metabarcoding for uncovering as yet unknown biodiversity in large-scale monitoring. One explanation for the higher number of OTUs we found is that many unassigned OTUs represent described species that cannot be assigned to species level because reference sequences are lacking. Alternatively, many of these OTUs could represent dark taxa, i.e. species new to science, as highlighted by Karlsson et al. (2020) and Srivathsan et al. (2023). While it is difficult to estimate the number of OTUs attributed to either known species without a reference barcode or to new species, we can roughly quantify this by comparing OTU-to-species ratios. Our mean number of OTUs per validated insect species of 1.0-1.5 suggests that our chosen cutoff of 3% sequence identity generally delineates different species accurately. The only exception is the Orthoptera, which are known to exhibit many pseudogenes (see Information S1) and that would inflate the OUT-to-species ratio. Specific correction factors for different taxonomic groups can also be estimated to infer the portion of undescribed species in the OTU datasets, which our results suggest is a large proportion. This is particularly true for the Hymenoptera and Diptera, where, after correcting the OTU numbers, we estimate about 6,600 and 1,200 potential new species to science for the two groups. This result agrees with reports from the Swedish Malaise Trap Program where most of the observed 700 species new to science (Karlsson et al. 2020) belonged to Diptera and Hymenoptera (Ronquist et al. 2020). However, our estimate of putatively undiscovered species in Germany is obviously a significant underrepresentation for two main reasons: First, while Malaise traps capture many different insect species, they likely miss many non-flying insects, insects flying high above ground, e.g. in tree canopies, or avoiding or escaping Malaise traps (e.g. Habel et al. 2023). Second, while our 75 Malaise traps reflect almost entirely the north-south and east-west gradients of Germany, they do not cover all regions in the country. Consequently, there is likely a trove of undiscovered species, even in countries like Germany and Sweden with a relatively species-poor fauna and a long taxonomic tradition. DNA metabarcoding can help discover the extent of this unknown biodiversity.

### Technical challenges and ways forward

Metabarcoding comes with some drawbacks and technical challenges that must be considered. Firstly, our approach is a compromise between improving feasibility for large-scale monitoring and the chances of detecting all taxa. Additional sample processing, such as complementary mild lysis (Marquina et al. 2022, Iwaszkiewicz-Eggebrecht et al. 2023), greater replication (Zizka et al. 2022b) and using multiple gene markers or primers (Elbrecht et al. 2019, Hajibabaei et al. 2019), as well as increasing sequencing depth (this study), would undoubtedly recover more species. However, the additional effort required must be weighed against the extra information gained, which may not be substantial (Buchner et al. 2021a) Secondly, the incompleteness of regional and interregional reference sequence databases, and the generation of reference sequences of unknown species, remains a persistent challenge (e.g. Karlsson et al. 2020, Chua et al. 2023). Poorly studied groups present the largest hurdle, such as Diptera, Hymenoptera and various beetle families present the largest impediment. Improved collaboration between taxonomists, molecular ecologists, and experts from other fields, such as computer vision and deep learning (Høye et al. 2021, Wührl et al. 2022), will be needed to fill these gaps and to improve biodiversity assessments in general. Specifically, sequence data need to be linked with valid taxonomic names and undescribed species need to be described. This could be done by re-sampling those sites, where an unknown or undescribed species has been identified by metabarcoding. Alternatively, techniques like mild lysis need to be further developed, to allow a similar high coverage of species while also allowing morphological identification. Vouchers would also help obtain the ecological trait information needed link the biodiversity changes with ecosystem function changes. Lastly, a key advantage of molecular methods is that tissue and DNA samples can easily be stored in miniaturized formats and be reanalyzed in the future (Zizka et al. 2022a). Emerging methods such as metagenomics can thus be applied in the future to stored samples to gain additional insights through reanalysis. This also makes direct intercalibration of different methods possible enabling harmonized biodiversity monitoring despite methodological advancements.

### Implications beyond insect monitoring

We used insects collected with Malaise traps to demonstrate the value of a new DNA metabarcoding workflow that is reliable, scalable, fast, cost-effective, and particularly well-suited for large-scale monitoring of highly diverse taxonomic groups. The approach is, however, by no means limited to insects from Malaise traps, given that many of the key advances we highlight (e.g., automated workflow, robust species validation) are applicable to a variety of sampling methods, other invertebrate and vertebrate taxa, and even environmental DNA from various sources including soil and sediment (Pawlowski et al. 2022), by aligning all post-sampling processing steps with the requirements for robotic high sample throughput (Buchner et al. 2021c). The presented workflow thus moves us closer to realizing the overall vision for metabarcoding, i.e., to generate and link high-throughput biodiversity analyses with large-scale monitoring (Bush et al. 2017). Such integration would greatly enhance assessments of the massive ongoing changes in global biodiversity experienced at the present (e.g., Sinclair et al., in prep, Information S2) and biodiversity protection (e.g., the Kunming-Montreal Global Biodiversity Framework of the CBD), including Red List and invasive species assessments as part of policy frameworks on biodiversity conservation (e.g., Wetzel et al. 2015). As demonstrated here, integrating metabarcoding into large-scale monitoring networks is a powerful means to improving our understanding of biodiversity change and supporting conservation actions.

## Acknowledgements

This project received funding from the Landes-Offensive zur Entwicklung Wissenschaftlich-ökonomischer Exzellenz of the German federal State of Hesse (Center for Translational Biodiversity Genomics; LOEWE-TBG), the Hessische Landesamt für Naturschutz, Umwelt und Geologie (HLNUG) and the EU Horizon 2020 project eLTER PLUS (grant agreement no. 871128). We are particular grateful to the participating LTER-D and NNL sites. We furthermore want to thank Michael Sachtleben, Monika Degebrodt and Katrin Preuß (IGB), Thomas Hahn and Petra Möhl (Ranger Nature Park Stechling-Ruppiner Land), Dr. Mario Schrumpf (Nature Park Stechlin-Ruppiner Land), Sebastian Flinkerbusch, Michael Hinz, Enno Klipp and Roland Wollgarten (Eifel National Park) and Günter Hoenselaar (Kellerwald-Edersee National Park) and all members involved in the operation of traps and the sample collection for their assistance and support of the project.

## Data accessibility statement

Demultiplexed raw read data for this study has been uploaded to the European Nucleotide Archive under the accession number PRJEB71324.

## Author contributions

DB, FL and PH conceived the study. DB performed the analyses and wrote the manuscript with contributions from all authors. DB performed the laboratory processing and the sequence analysis with help from FL and YL. TH and MS checked the taxalist for plausibility. MOG, JM, SUP, SSt, JB, JE, TH, CM, RR, TS, ST, WW, BW, MW and GZ set up Malaise traps in their respective regions, maintained traps and collected samples.

## Competing interest

The authors declare no competing interests.

## Notes

### Competing Interest Statement

The authors have declared no competing interest.

### Summary of Updates

Updated the full manuscript as well as key figures to the current submission process.

