## Supplementary material for "Upscaling biodiversity monitoring: Metabarcoding estimates 31,846 insect species from Malaise traps across Germany": Figure S3.pdf

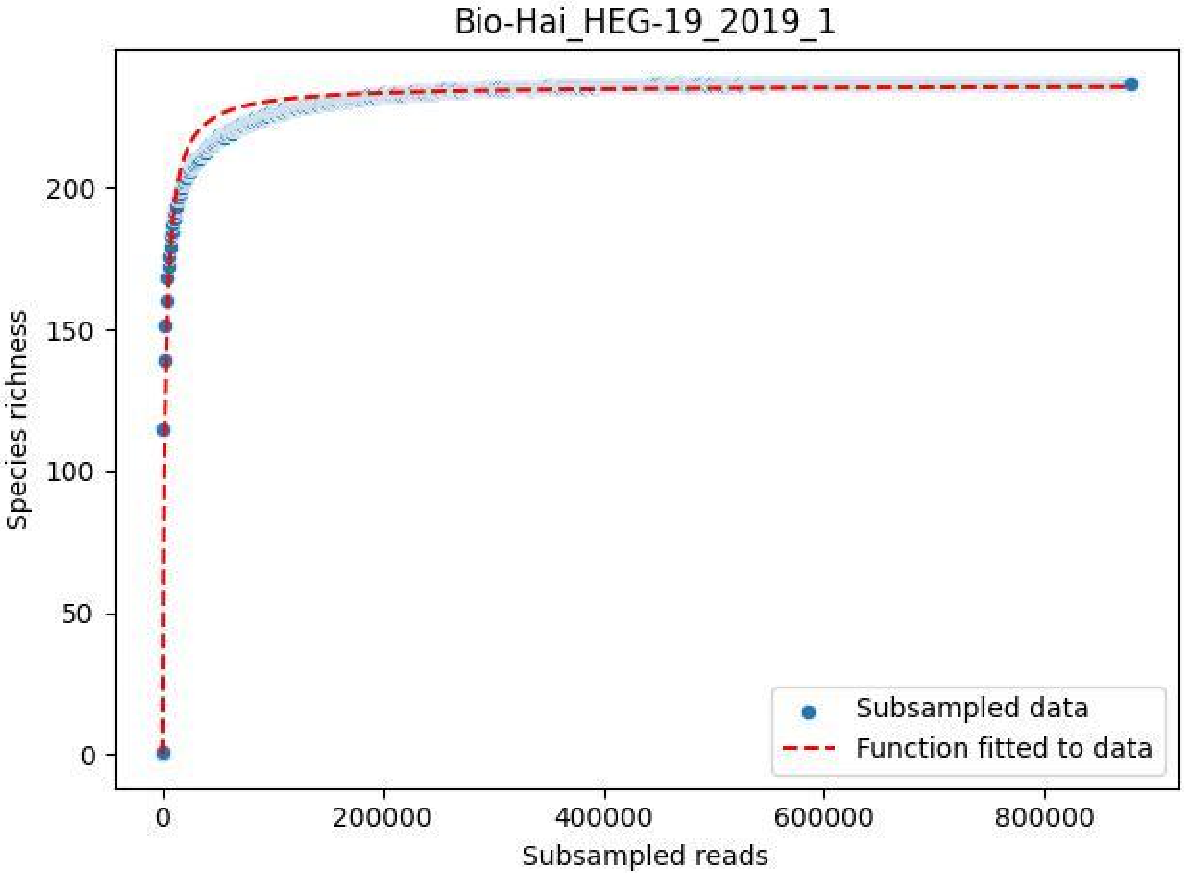

Bio-Hai_HEG-19_2019_1
Species richness
0
50
100
150
200
Subsampled reads
0
200000
400000
600000
800000
Subsampled data
Function fitted to data

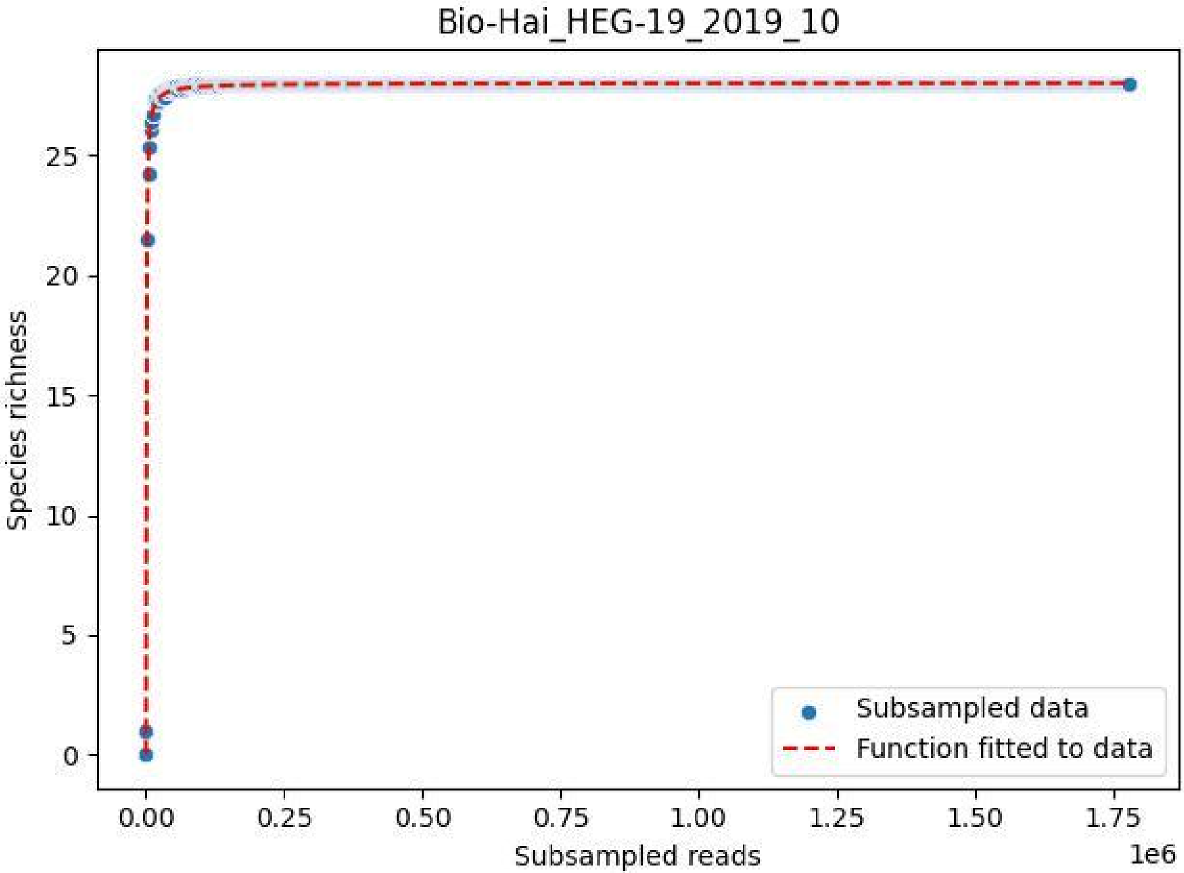
Bio-Hai_HEG-19_2019_10
Species richness
0
5
10
15
20
25
0.00
0.25
0.50
0.75
1.00
1.25
1.50
1.75
1e6
Subsampled reads
Subsampled data
Function fitted to data

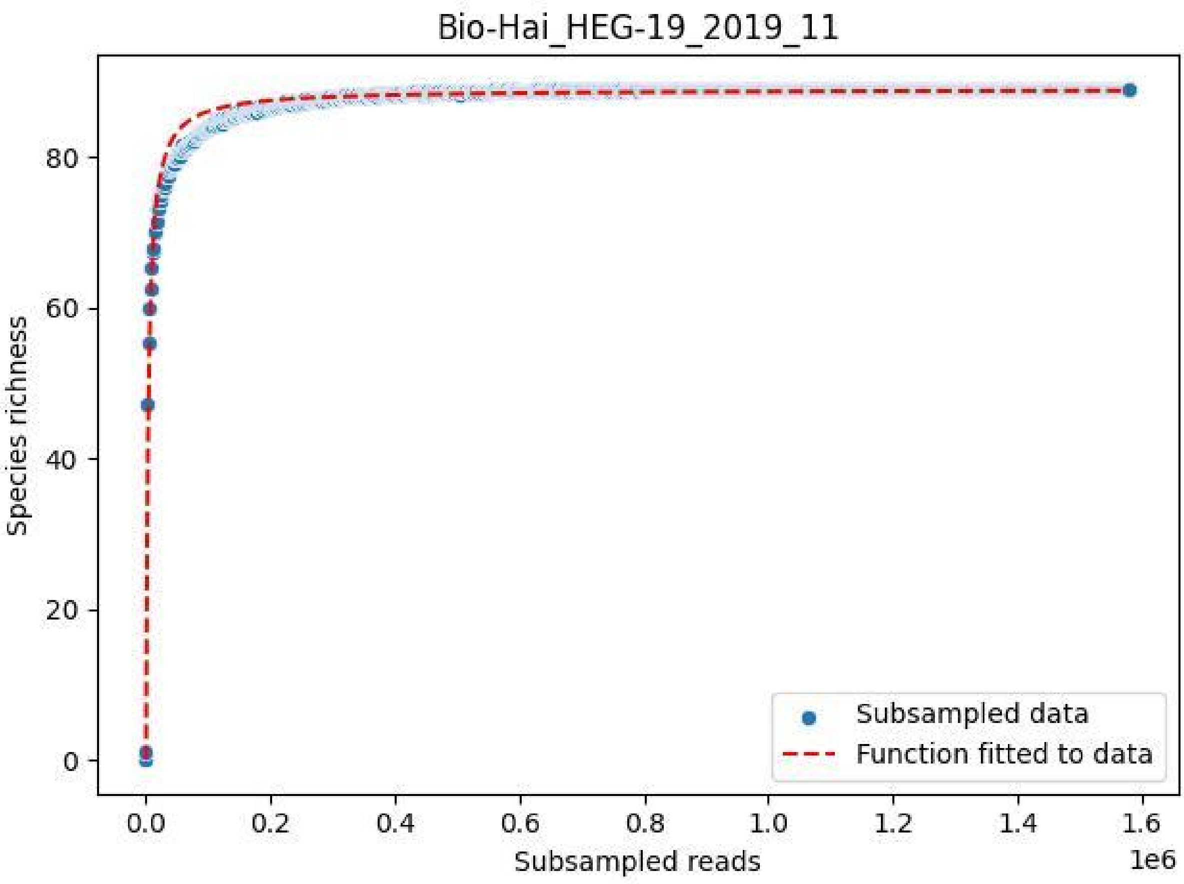

Bio-Hai_HEG-19_2019_11
Species richness
Subsampled reads
1e6
0
20
40
60
80
0.0
0.2
0.4
0.6
0.8
1.0
1.2
1.4
1.6
Subsampled data
Function fitted to data

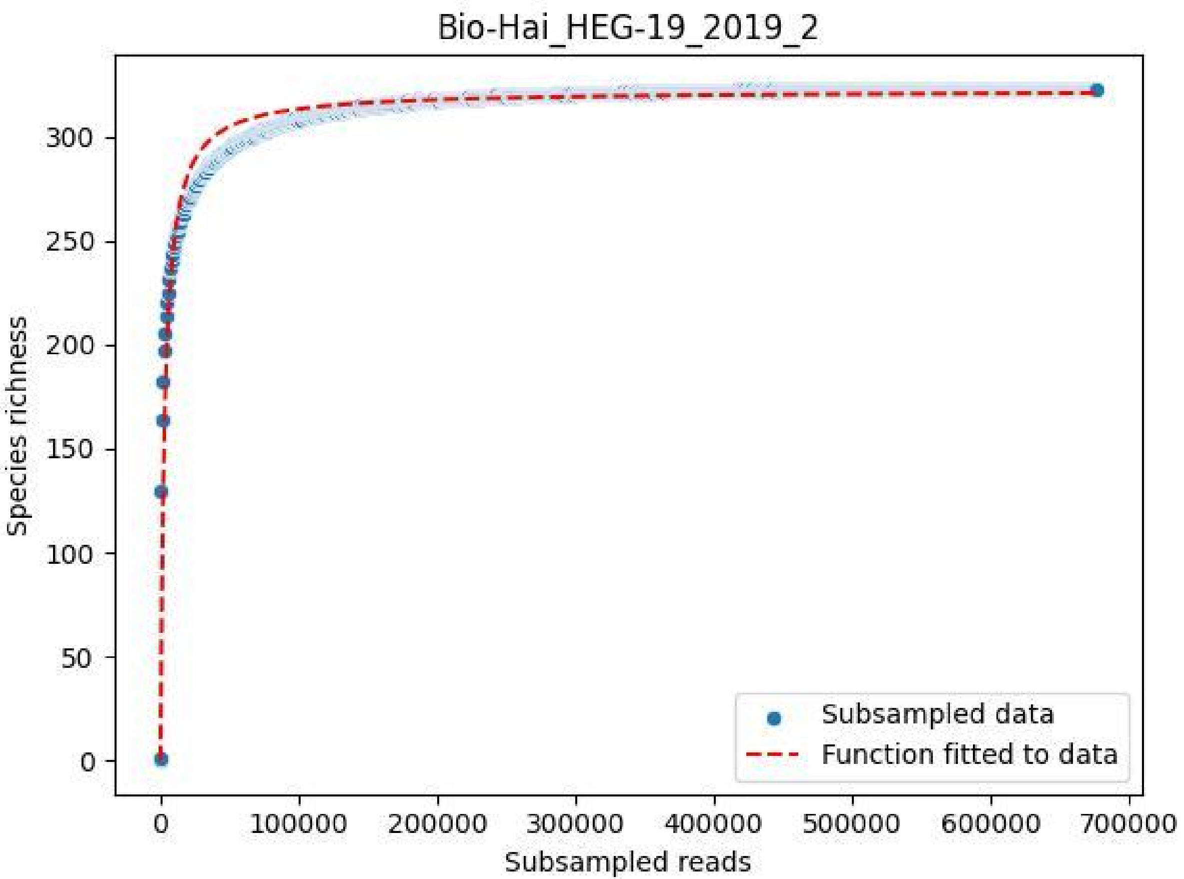
Bio-Hai_HEG-19_2019_2
Species richness
Subsampled reads
0
50
100
150
200
250
300
0
100000
200000
300000
400000
500000
600000
700000
Subsampled data
Function fitted to data

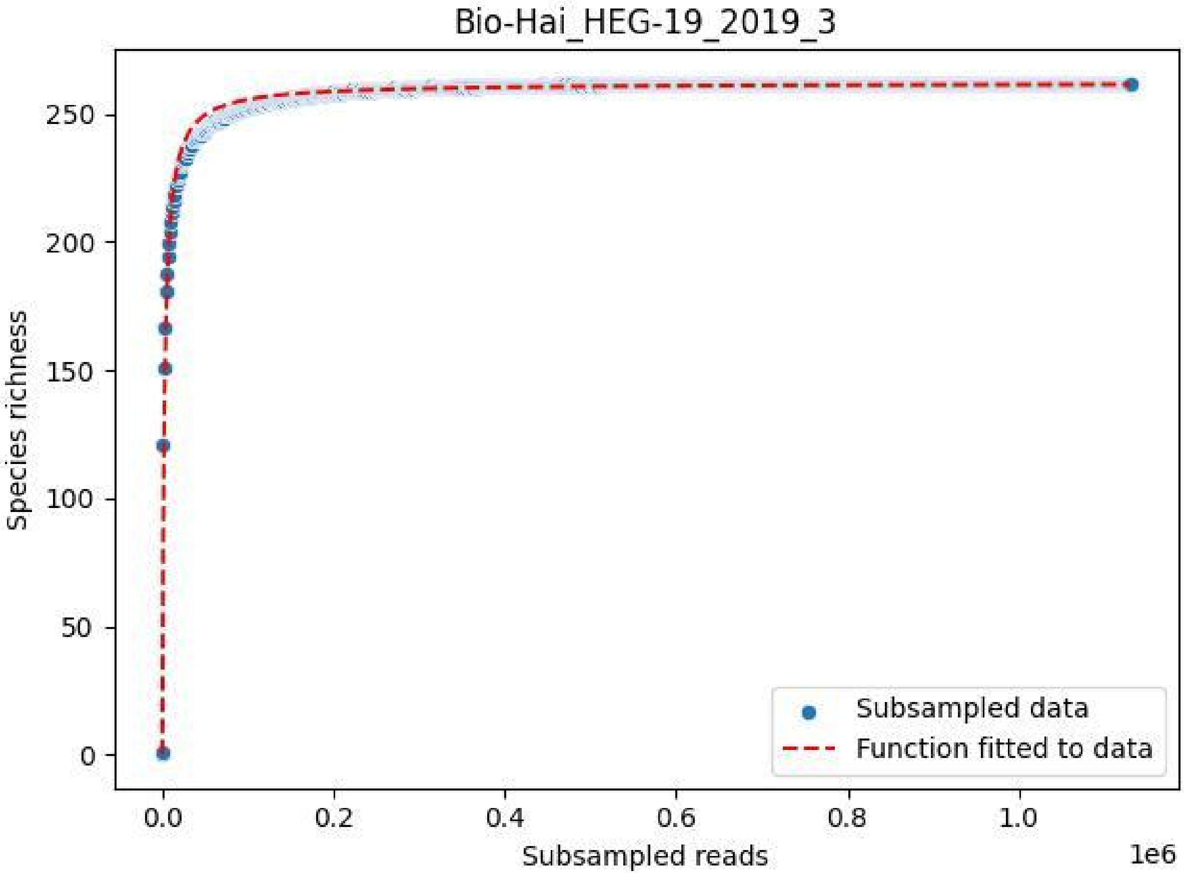

Bio-Hai_HEG-19_2019_3
Species richness
250
200
150
100
50
0
0.0
0.2
0.4
0.6
0.8
1.0
Subsampled reads
1e6
Subsampled data
Function fitted to data

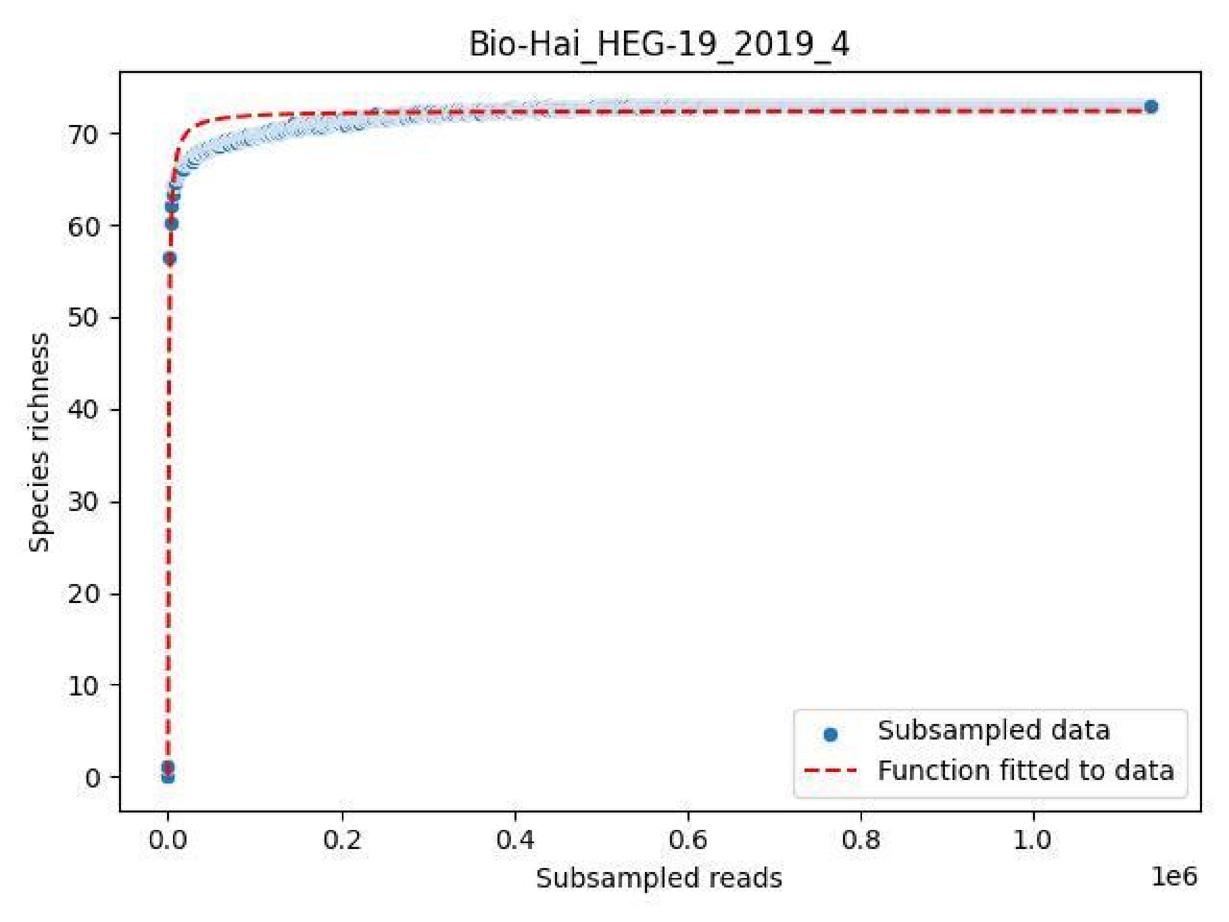

Bio-Hai_HEG-19_2019_4
Species richness
0
10
20
30
40
50
60
70
0.0
0.2
0.4
0.6
0.8
1.0
1e6
Subsampled reads
Subsampled data
Function fitted to data

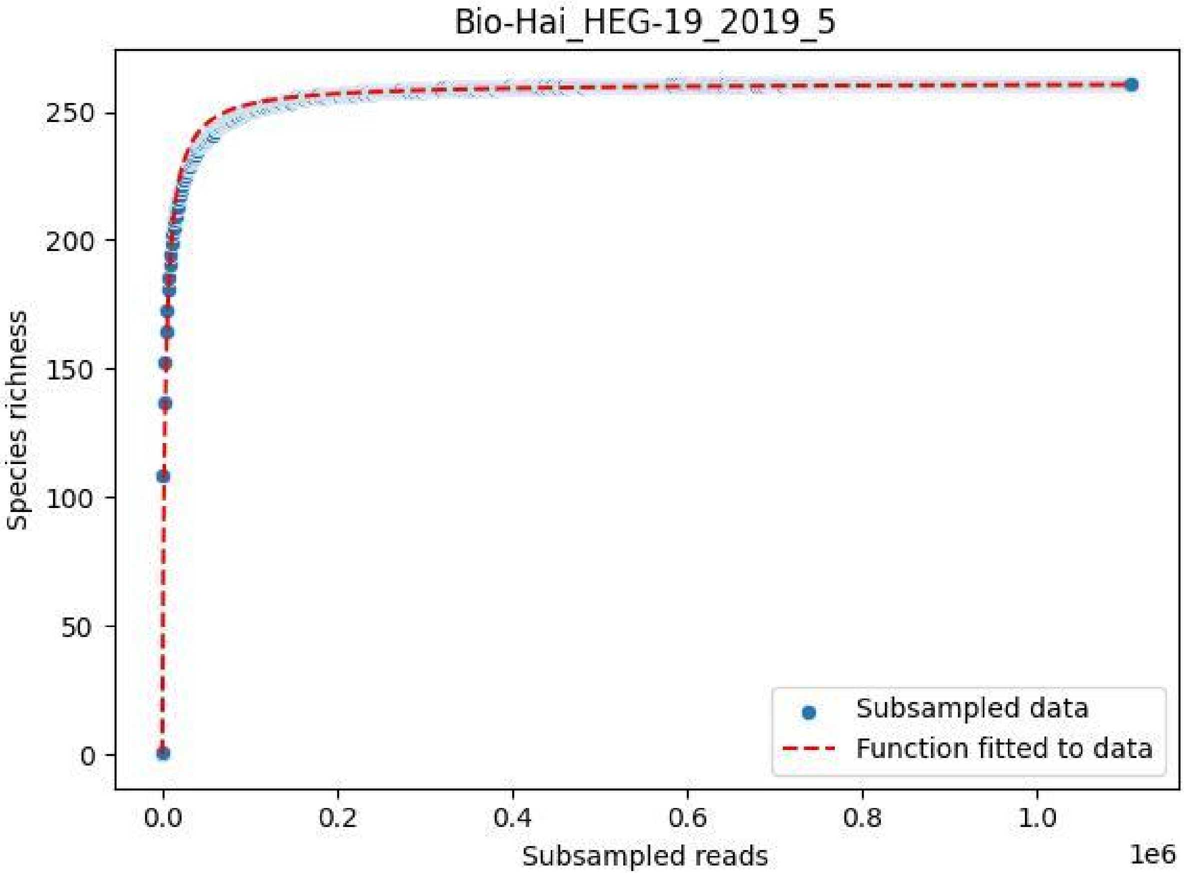
Bio-Hai_HEG-19_2019_5
Species richness
250
200
150
100
50
0
0.0
0.2
0.4
0.6
0.8
1.0
1e6
Subsampled reads
Subsampled data
Function fitted to data

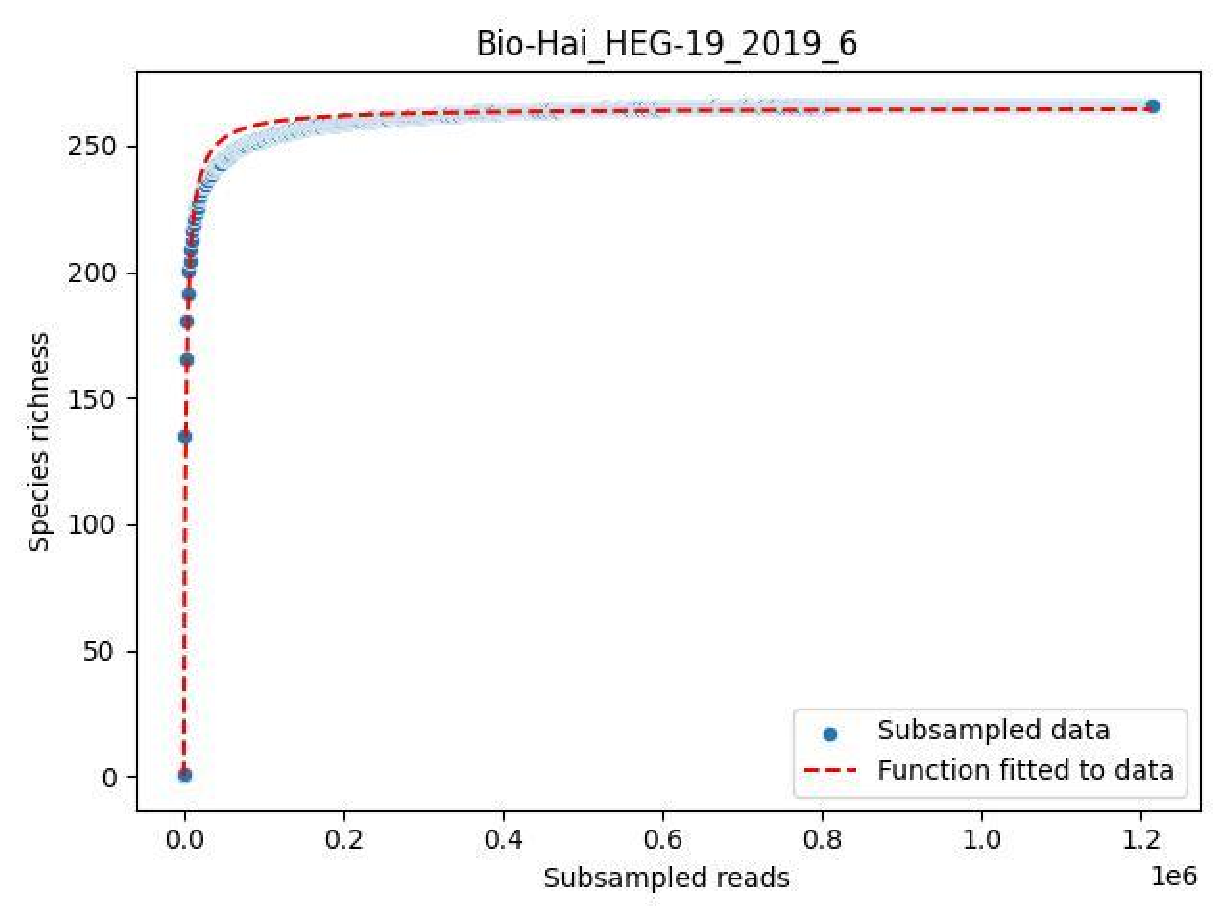

Bio-Hai_HEG-19_2019_6
Species richness
Subsampled reads
1e6
0
50
100
150
200
250
0.0
0.2
0.4
0.6
0.8
1.0
1.2
Subsampled data
Function fitted to data

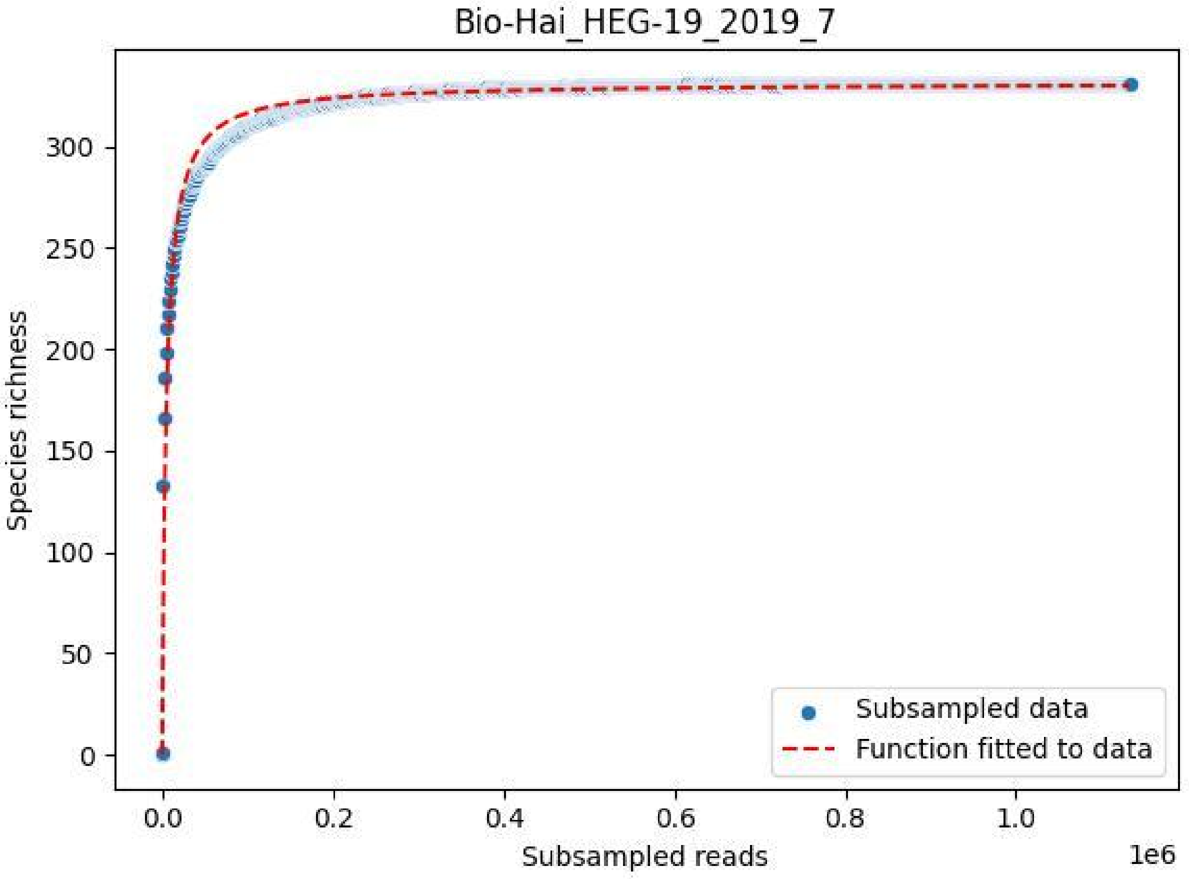
Bio-Hai_HEG-19_2019_7
Species richness
Subsampled reads
1e6
Subsampled data
Function fitted to data
0
50
100
150
200
250
300
0.0
0.2
0.4
0.6
0.8
1.0

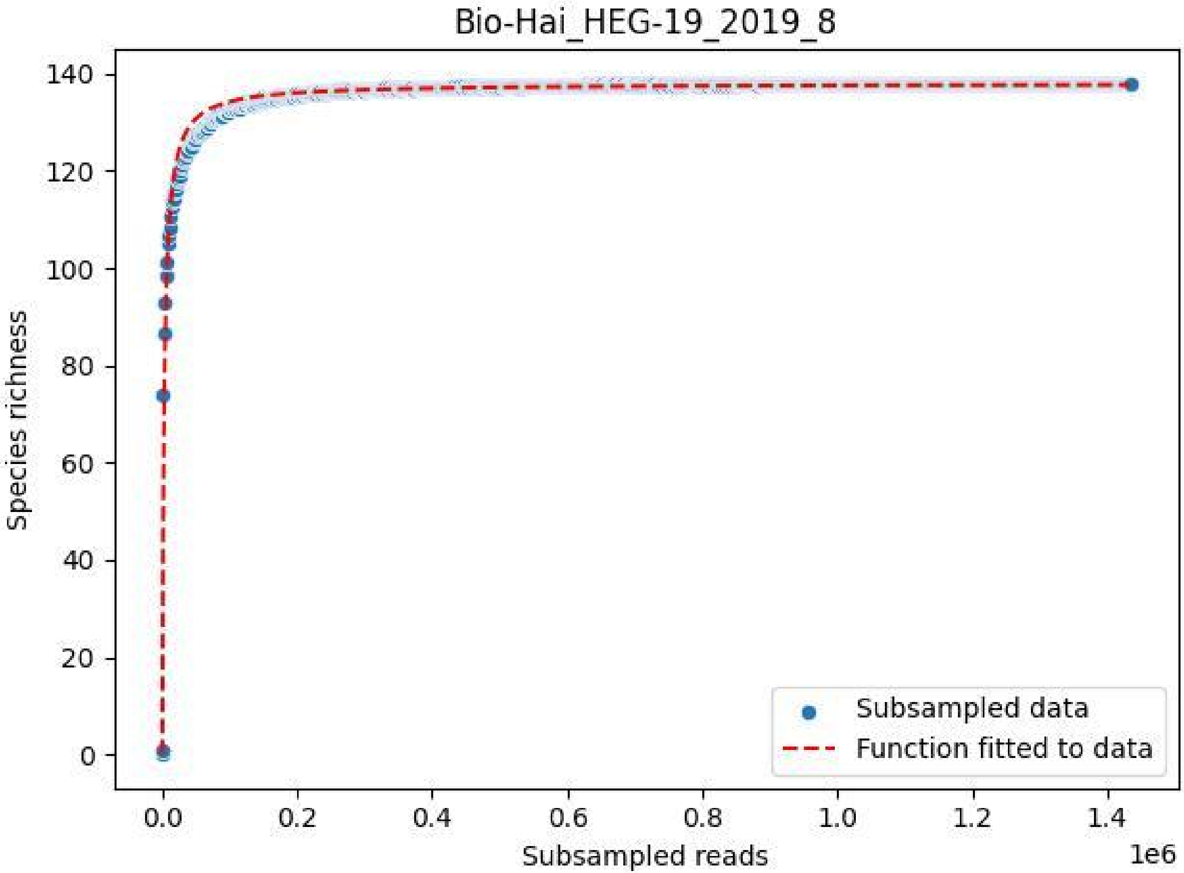
Bio-Hai_HEG-19_2019_8
Species richness
Subsampled reads
1e6
0
20
40
60
80
100
120
140
0.0
0.2
0.4
0.6
0.8
1.0
1.2
1.4
Subsampled data
Function fitted to data

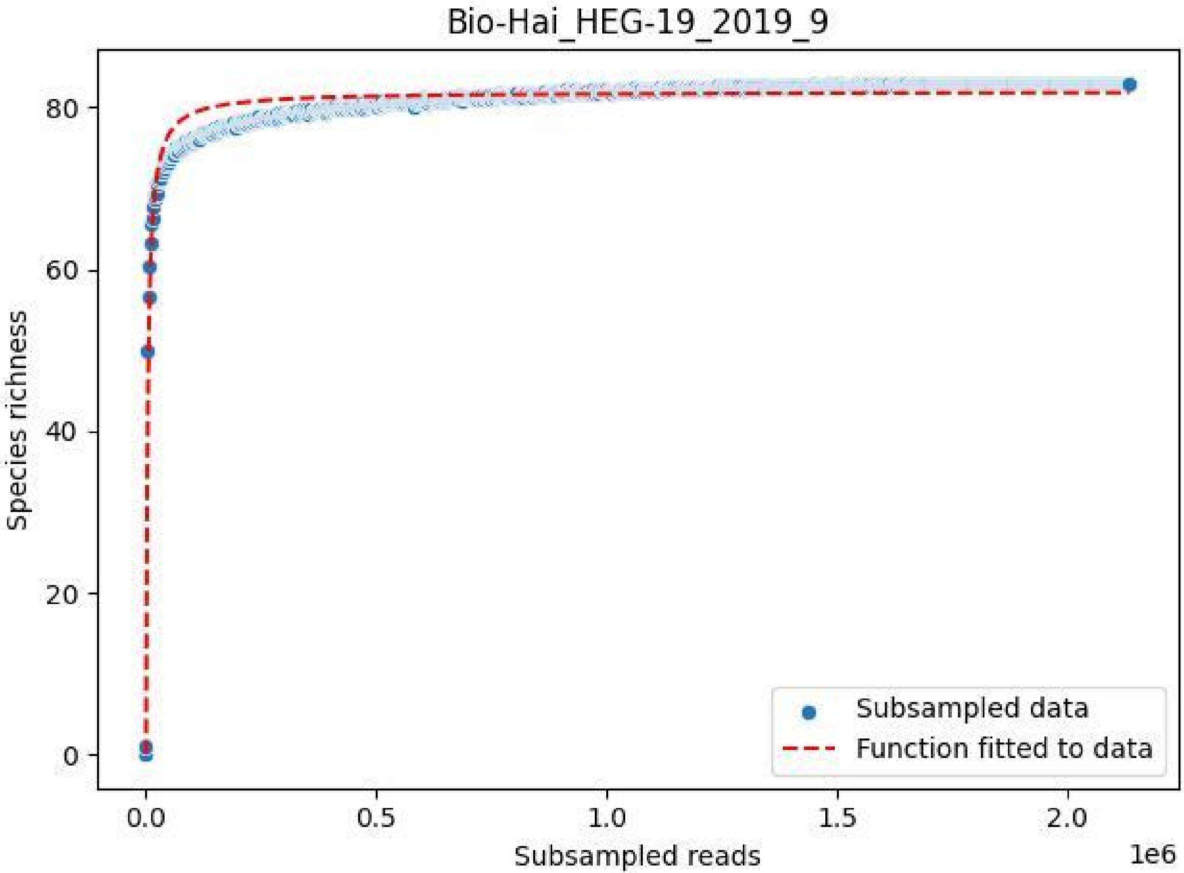
Bio-Hai_HEG-19_2019_9
Species richness
80
60
40
20
0
0.0
0.5
1.0
1.5
2.0
1e6
Subsampled reads
Subsampled data
Function fitted to data

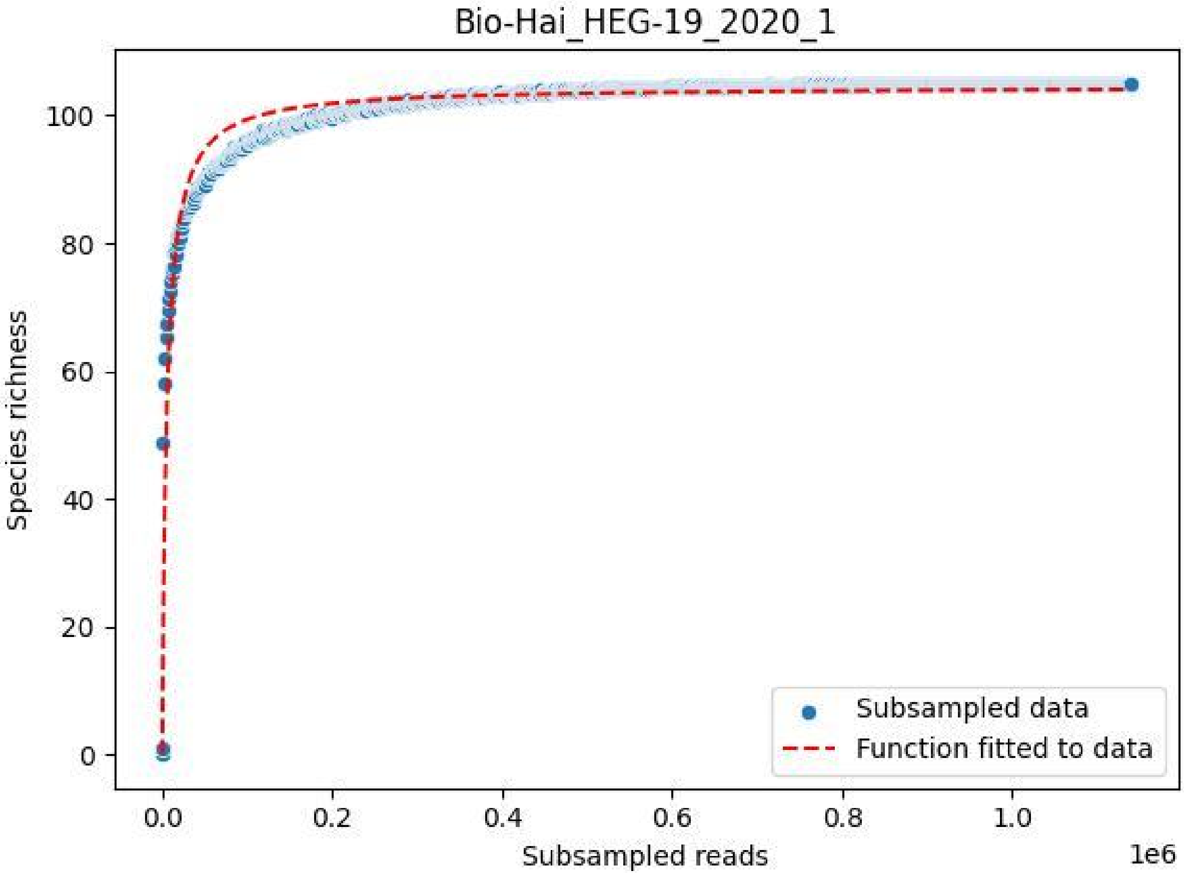
Bio-Hai_HEG-19_2020_1
Species richness
Subsampled reads
1e6
0
20
40
60
80
100
0.0
0.2
0.4
0.6
0.8
1.0
Subsampled data
Function fitted to data

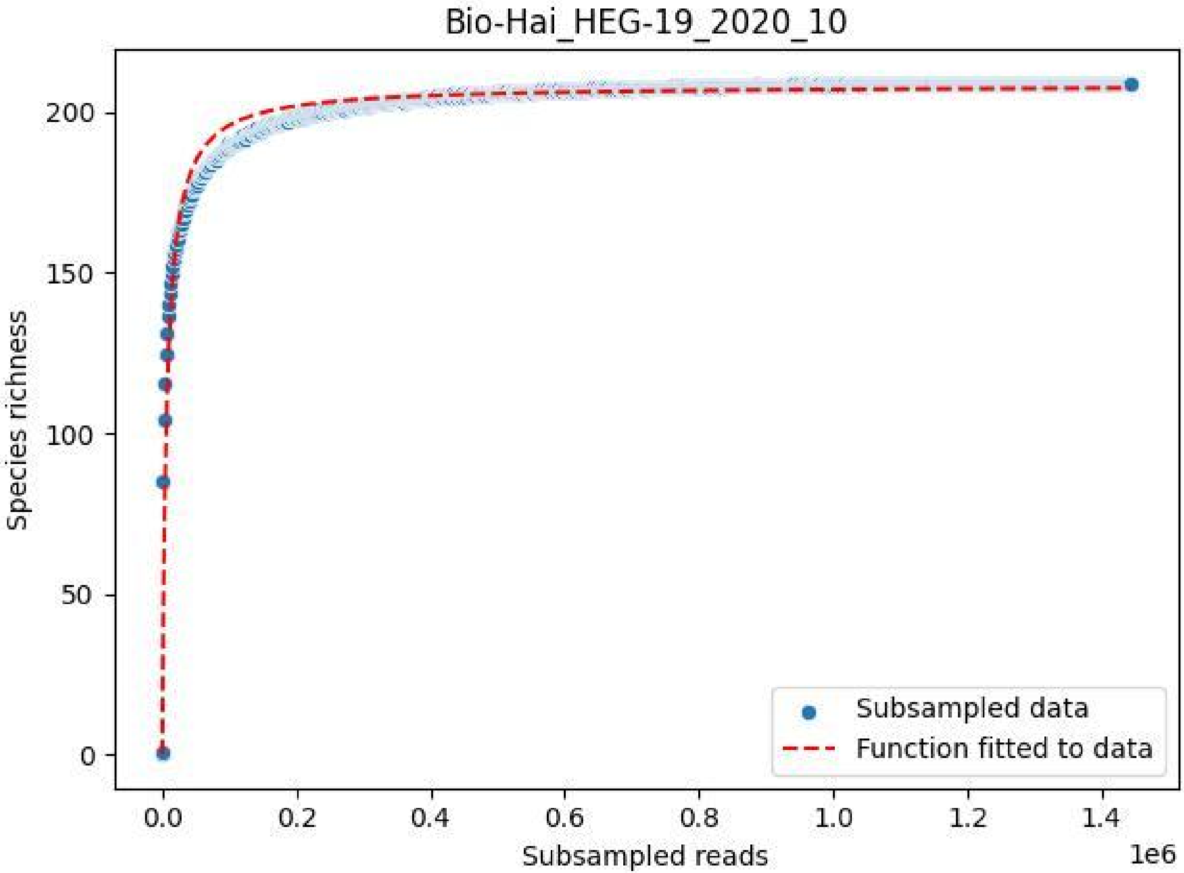

Bio-Hai_HEG-19_2020_10
Species richness
Subsampled reads
1e6
0
50
100
150
200
0.0
0.2
0.4
0.6
0.8
1.0
1.2
1.4
Subsampled data
Function fitted to data

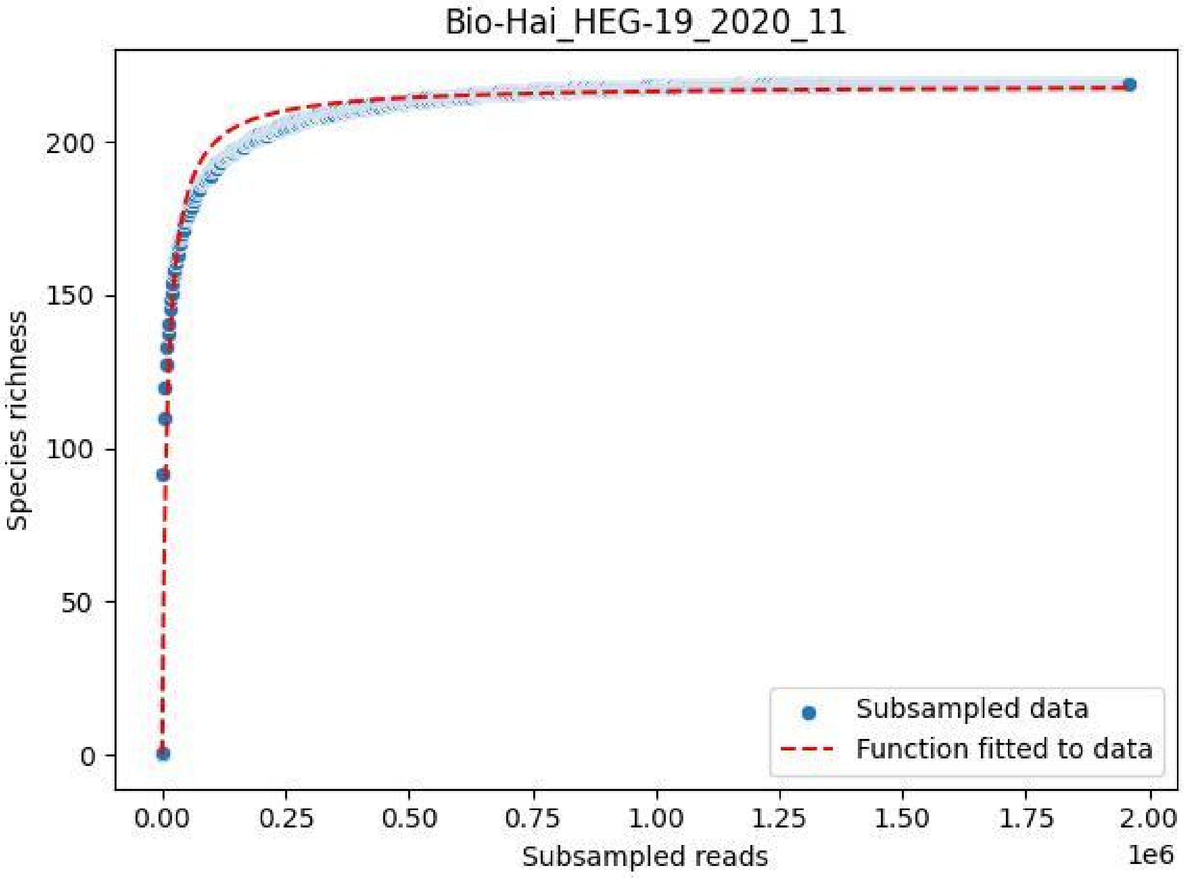
Bio-Hai_HEG-19_2020_11
Species richness
Subsampled reads
1e6
0
50
100
150
200
0.00
0.25
0.50
0.75
1.00
1.25
1.50
1.75
2.00
Subsampled data
Function fitted to data

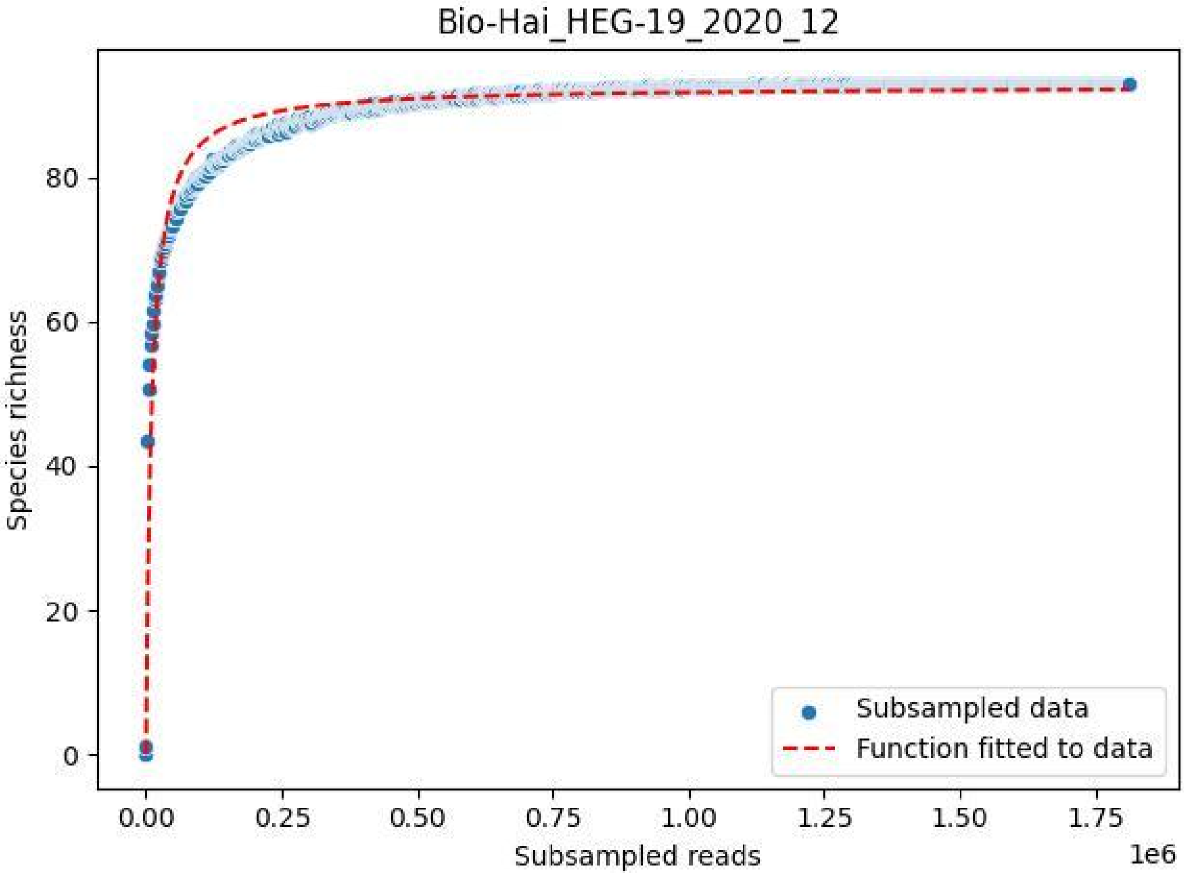
Bio-Hai_HEG-19_2020_12
Species richness
0
20
40
60
80
0.00
0.25
0.50
0.75
1.00
1.25
1.50
1.75
1e6
Subsampled reads
Subsampled data
Function fitted to data

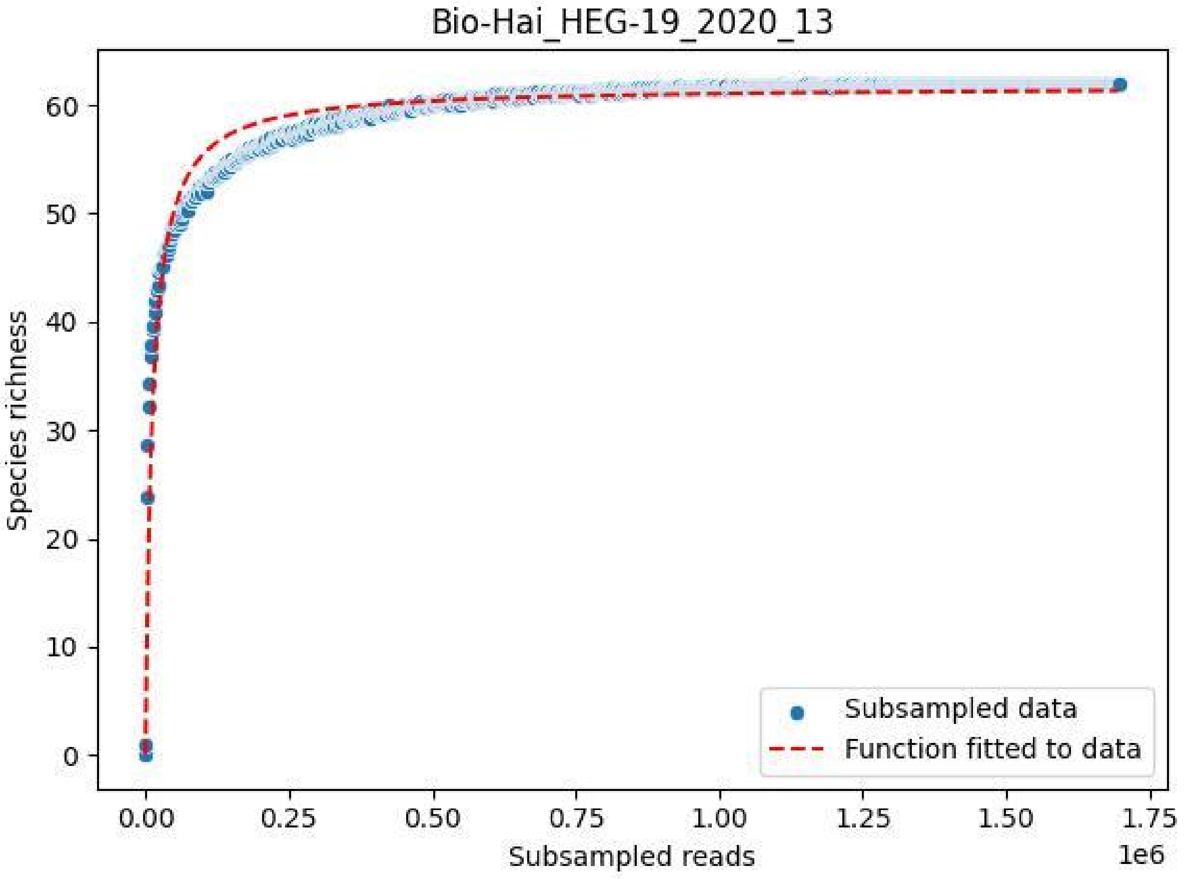

Bio-Hai_HEG-19_2020_13
Species richness
Subsampled reads
1e6
0
10
20
30
40
50
60
0.00
0.25
0.50
0.75
1.00
1.25
1.50
1.75
Subsampled data
Function fitted to data

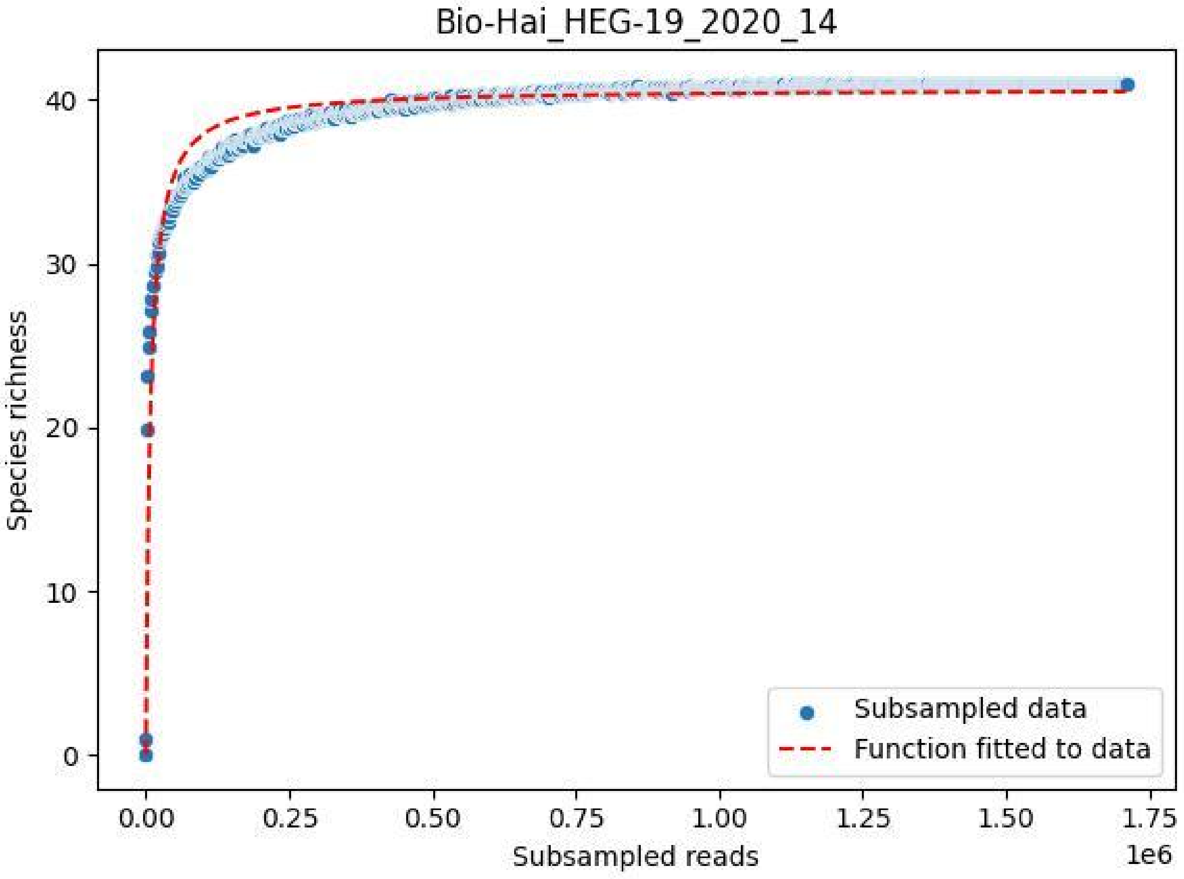

Bio-Hai_HEG-19_2020_14
Species richness
Subsampled reads
1e6
0
10
20
30
40
0.00
0.25
0.50
0.75
1.00
1.25
1.50
1.75
Subsampled data
Function fitted to data

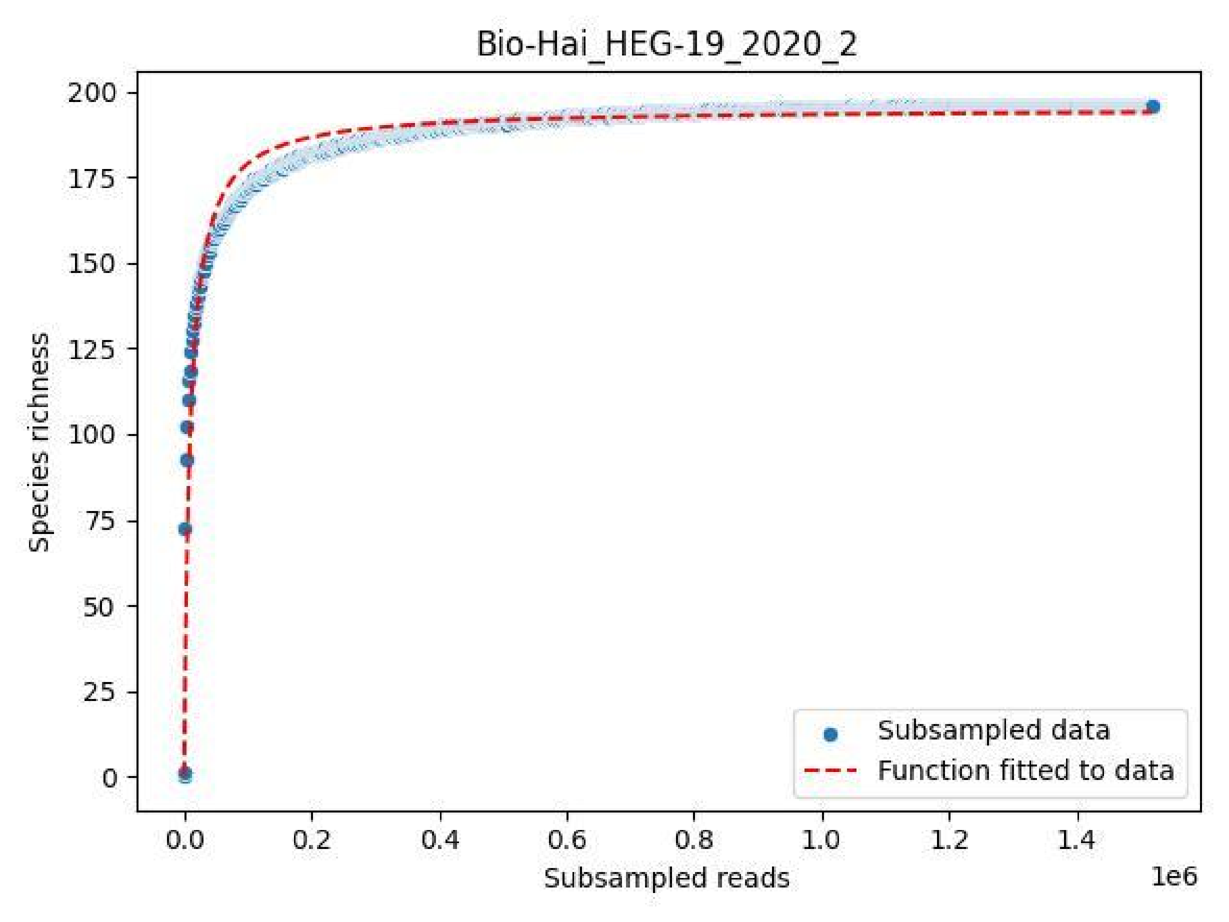

Bio-Hai_HEG-19_2020_2
Species richness
200
175
150
125
100
75
50
25
0
0.0
0.2
0.4
0.6
0.8
1.0
1.2
1.4
1e6
Subsampled reads
Subsampled data
Function fitted to data

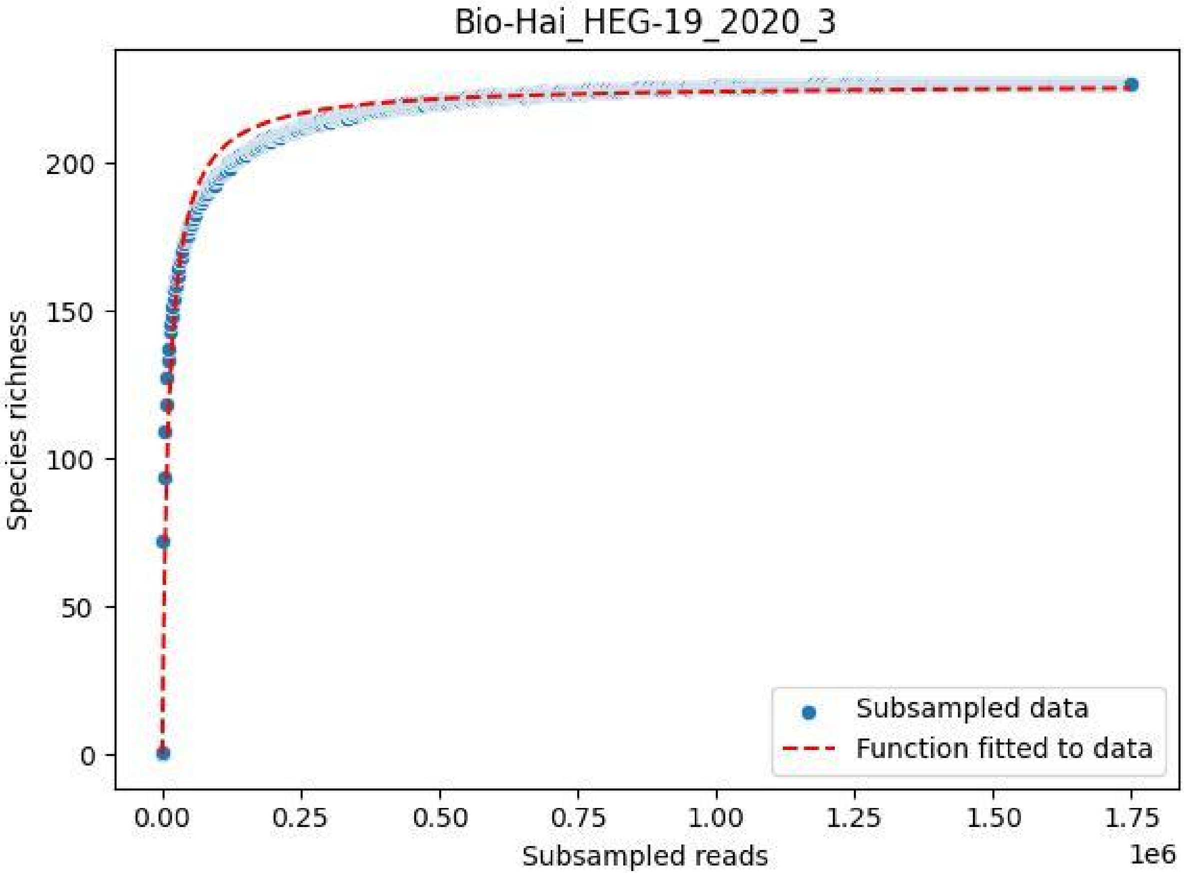

Bio-Hai_HEG-19_2020_3
Species richness
Subsampled reads
1e6
0
50
100
150
200
0.00
0.25
0.50
0.75
1.00
1.25
1.50
1.75
Subsampled data
Function fitted to data

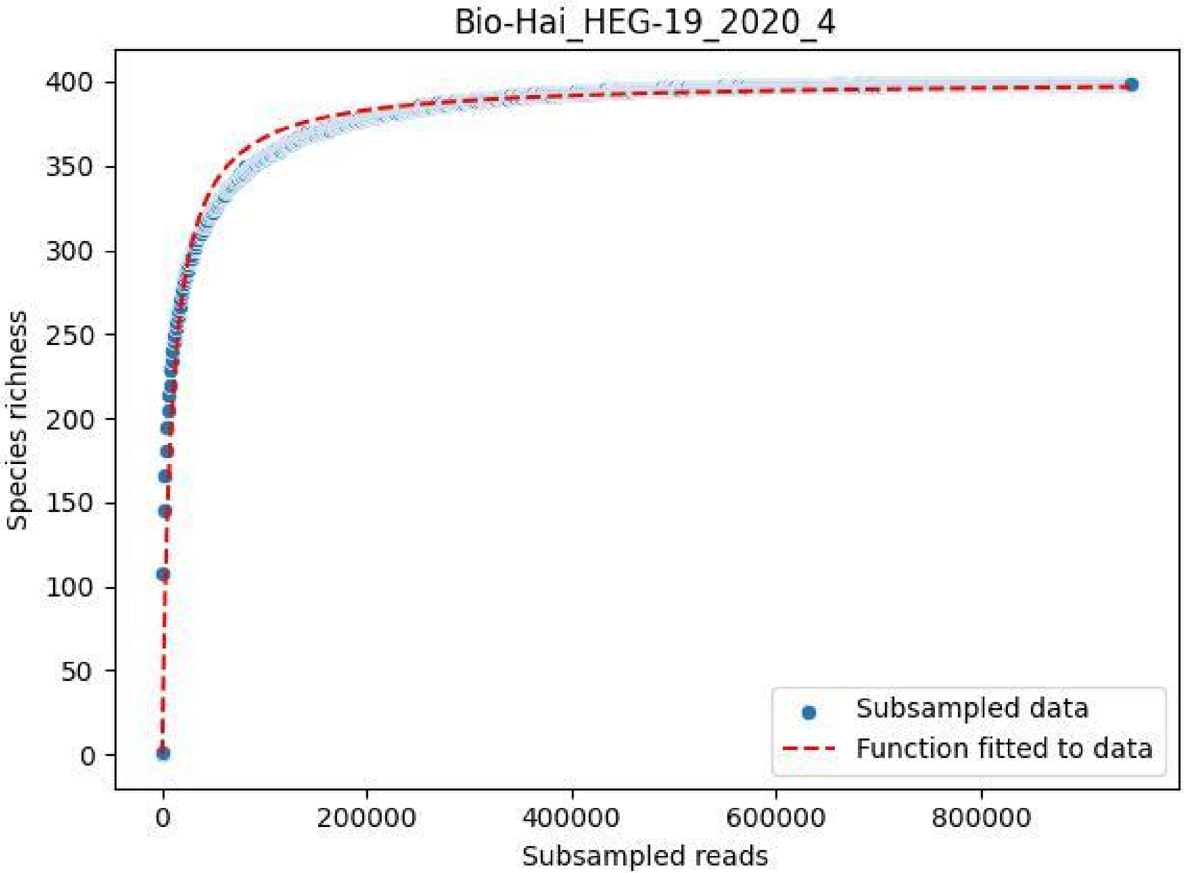
Bio-Hai_HEG-19_2020_4
Species richness
Subsampled reads
0
50
100
150
200
250
300
350
400
0
200000
400000
600000
800000
Subsampled data
Function fitted to data

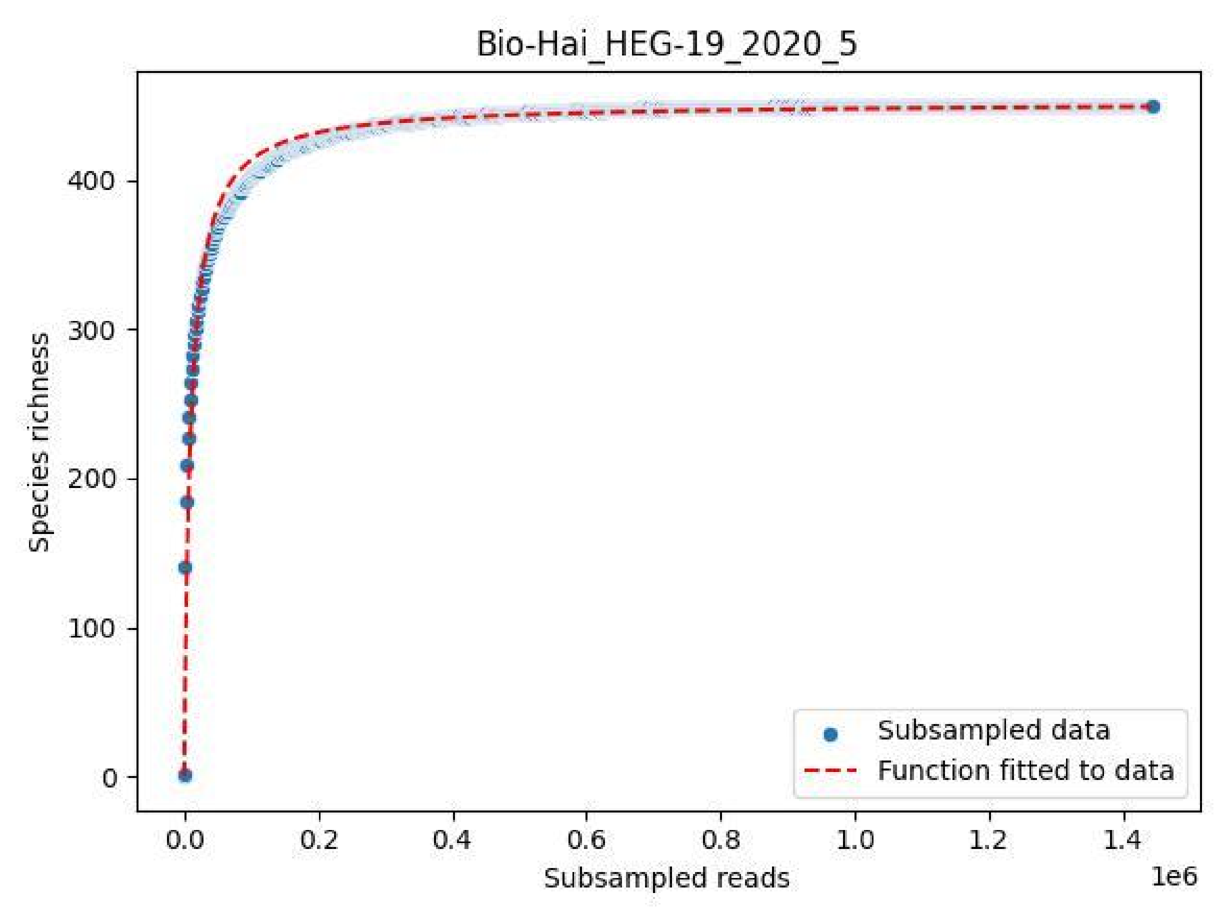

Bio-Hai_HEG-19_2020_5
Species richness
0
100
200
300
400
0.0
0.2
0.4
0.6
0.8
1.0
1.2
1.4
1e6
Subsampled reads
Subsampled data
Function fitted to data

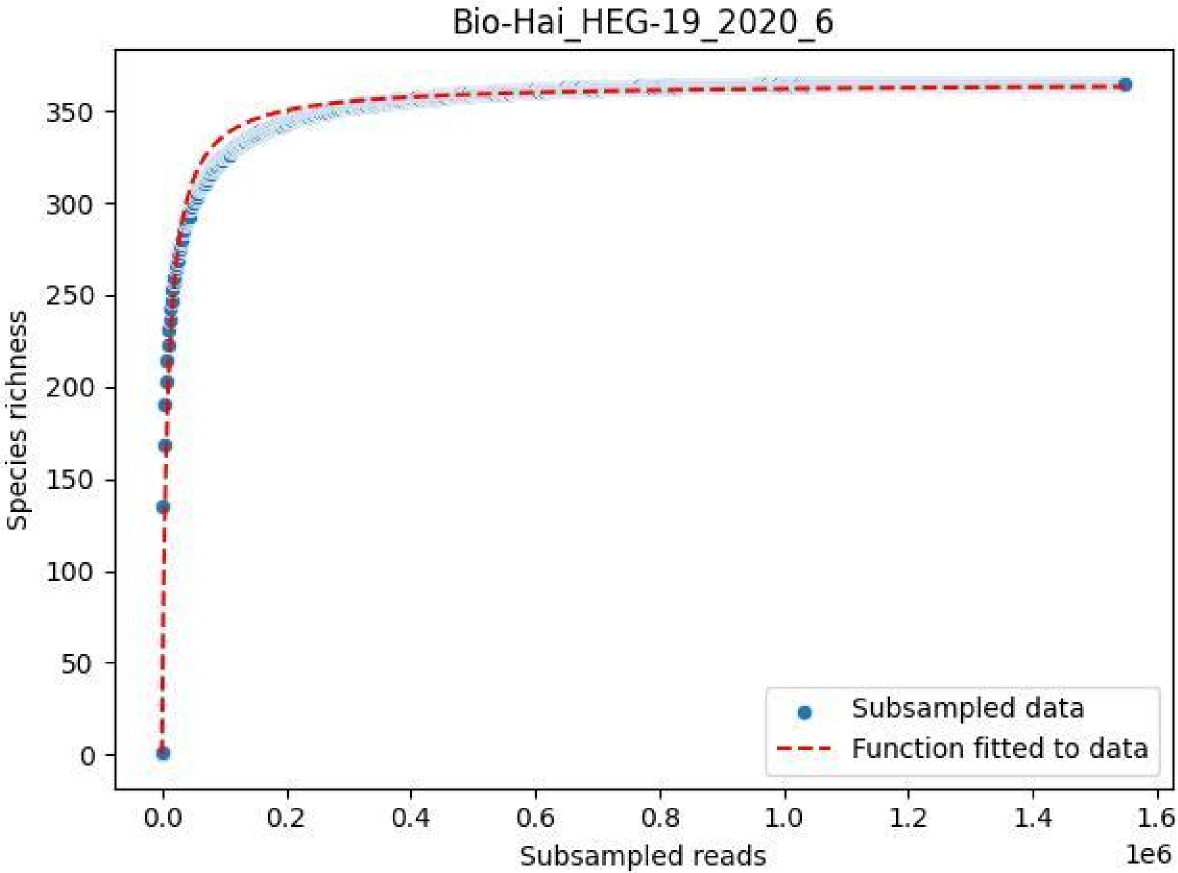
Bio-Hai_HEG-19_2020_6
Species richness
Subsampled reads
1e6
0
50
100
150
200
250
300
350
0.0
0.2
0.4
0.6
0.8
1.0
1.2
1.4
1.6
Subsampled data
Function fitted to data

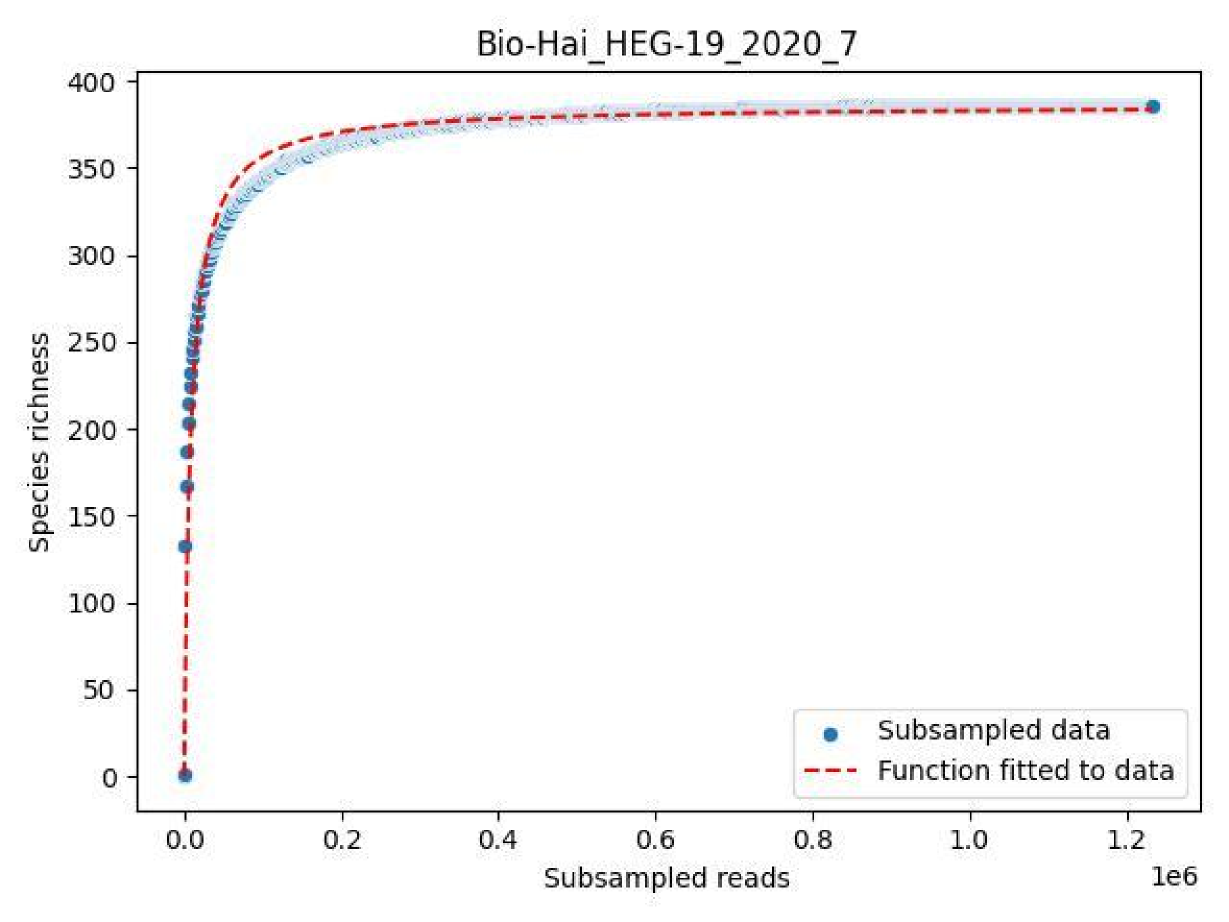
Bio-Hai_HEG-19_2020_7
Species richness
Subsampled reads
1e6
0
50
100
150
200
250
300
350
400
0.0
0.2
0.4
0.6
0.8
1.0
1.2
Subsampled data
Function fitted to data

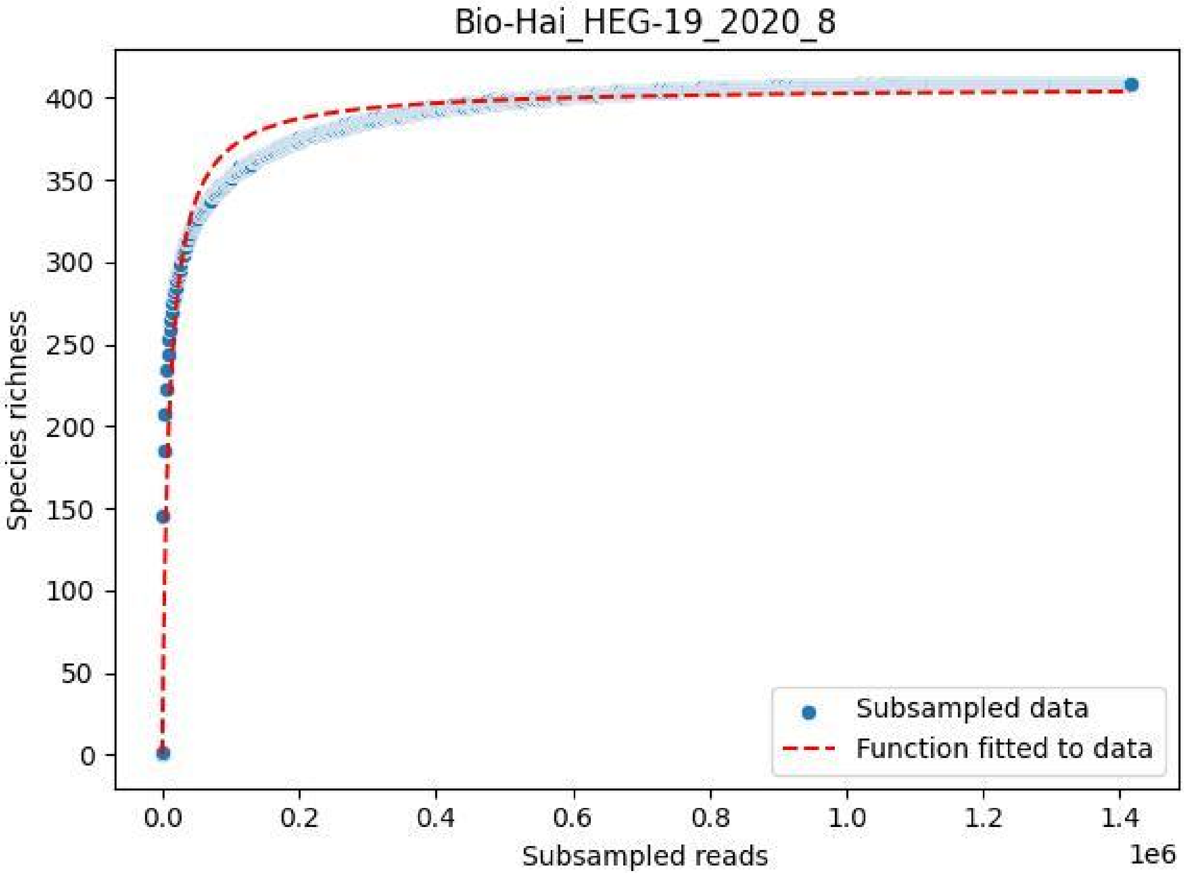

Bio-Hai_HEG-19_2020_8
Species richness
0
50
100
150
200
250
300
350
400
Subsampled reads
0.0
0.2
0.4
0.6
0.8
1.0
1.2
1.4
1e6
Subsampled data
Function fitted to data

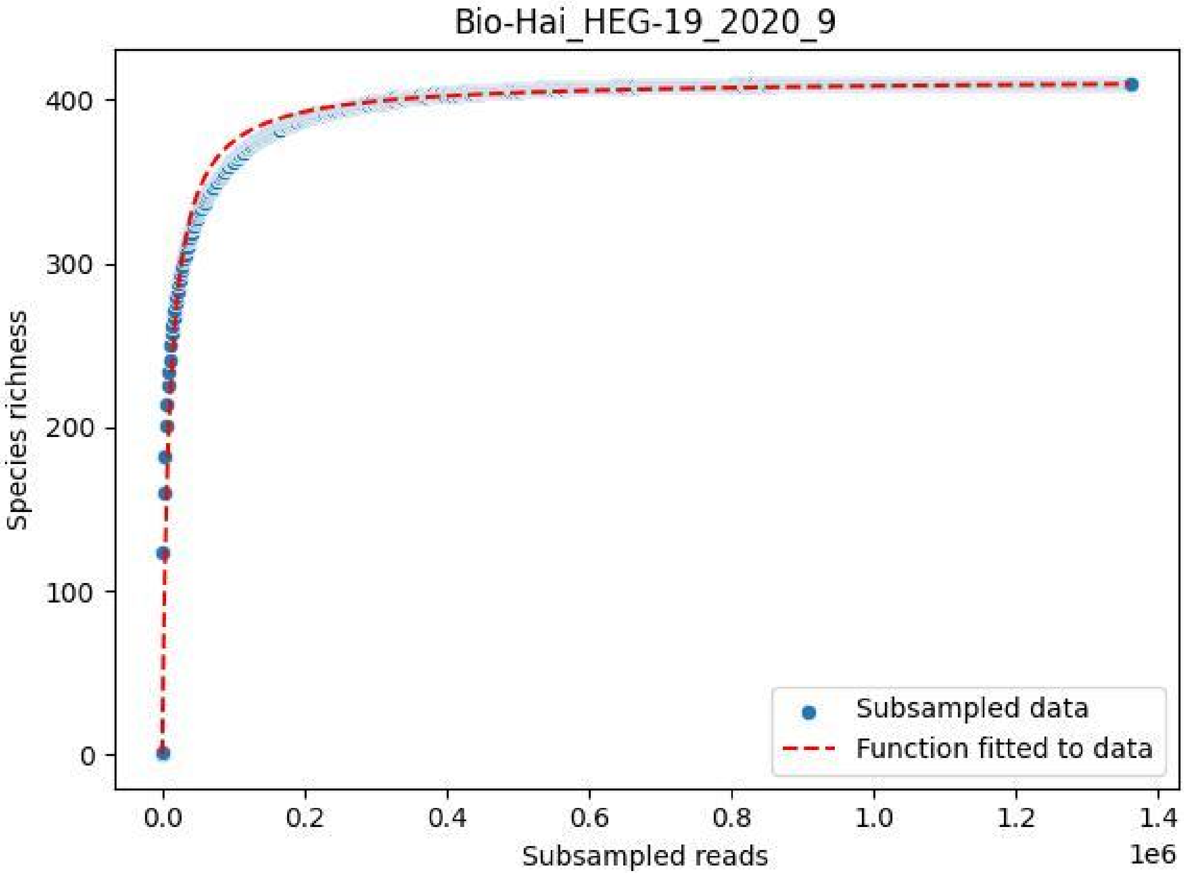

Bio-Hai_HEG-19_2020_9
Species richness
400
300
200
100
0
0.0
0.2
0.4
0.6
0.8
1.0
1.2
1.4
1e6
Subsampled reads
Subsampled data
Function fitted to data

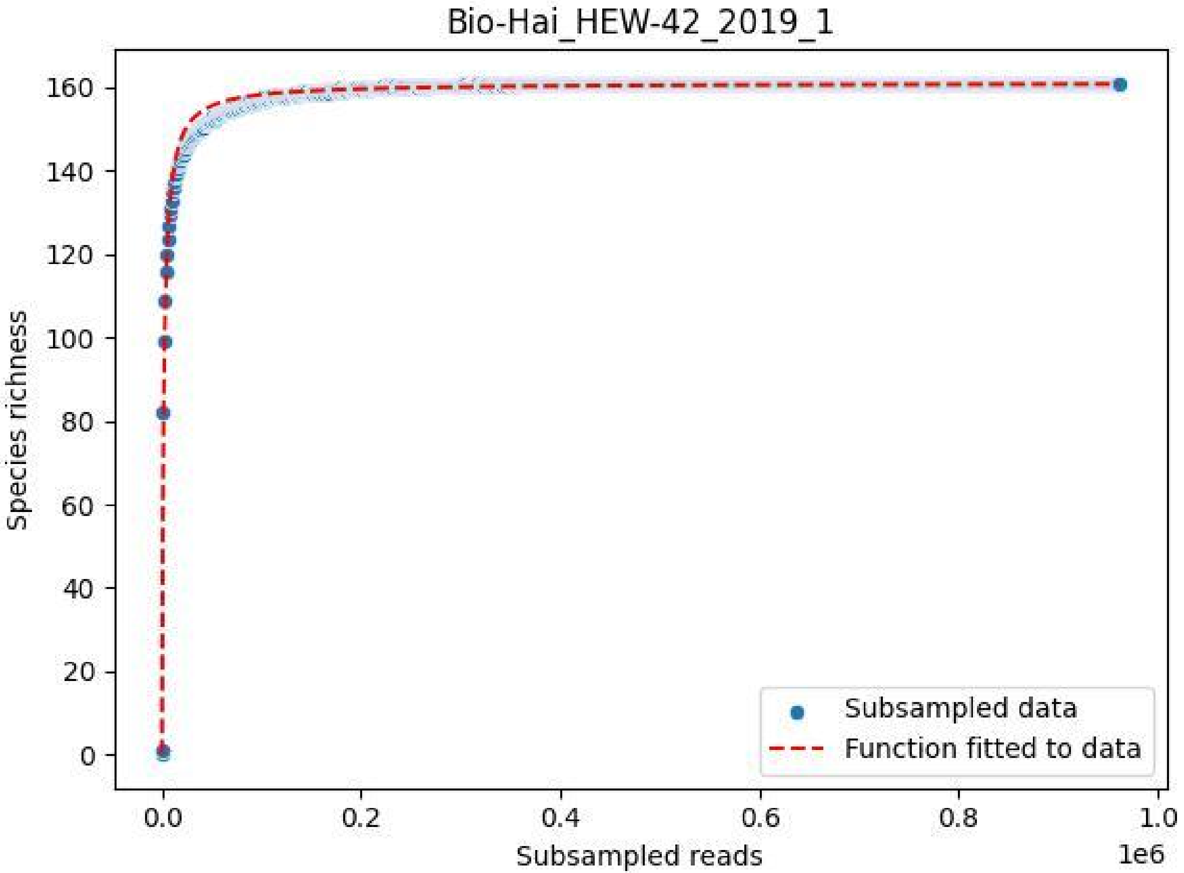
Bio-Hai_HEW-42_2019_1
Species richness
Subsampled reads
1e6
Subsampled data
Function fitted to data
0
20
40
60
80
100
120
140
160
0.0
0.2
0.4
0.6
0.8
1.0

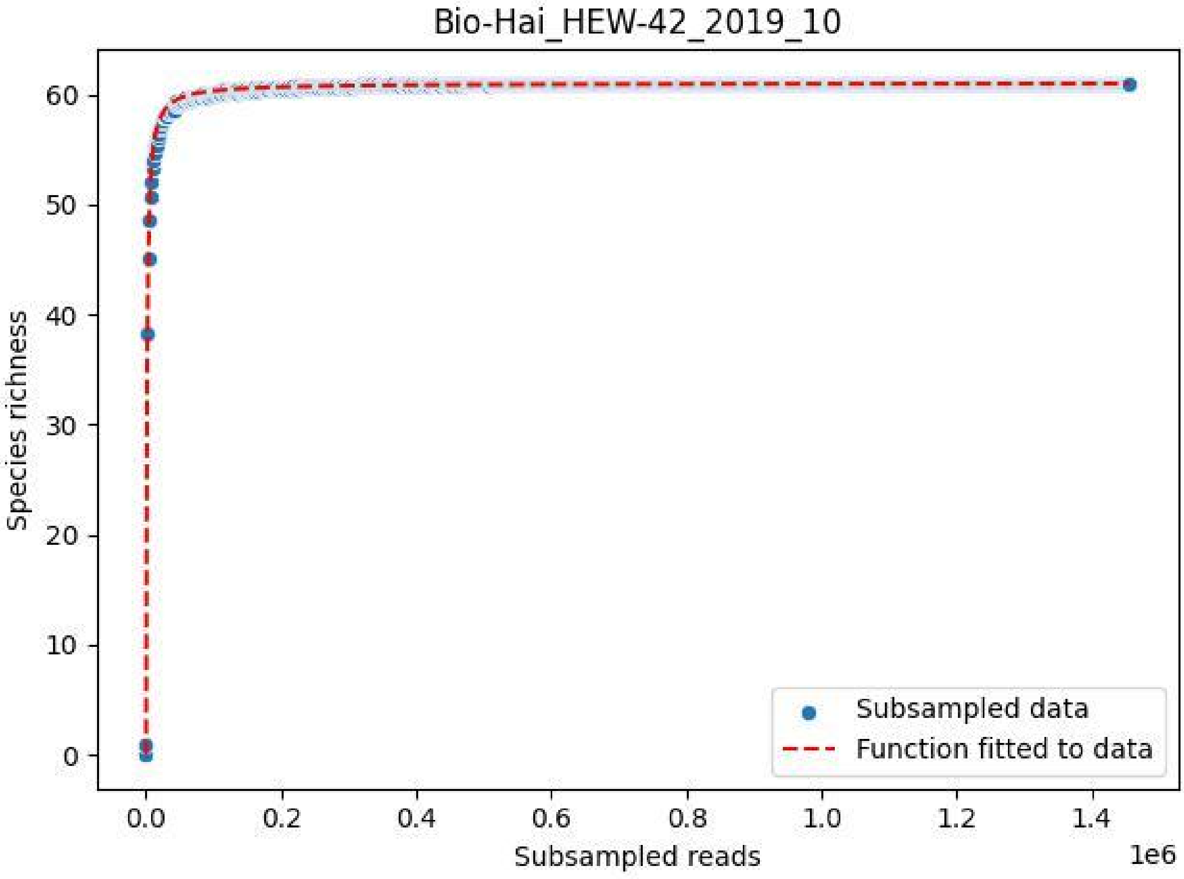

Bio-Hai_HEW-42_2019_10
Species richness
0
10
20
30
40
50
60
0.0
0.2
0.4
0.6
0.8
1.0
1.2
1.4
1e6
Subsampled reads
Subsampled data
Function fitted to data

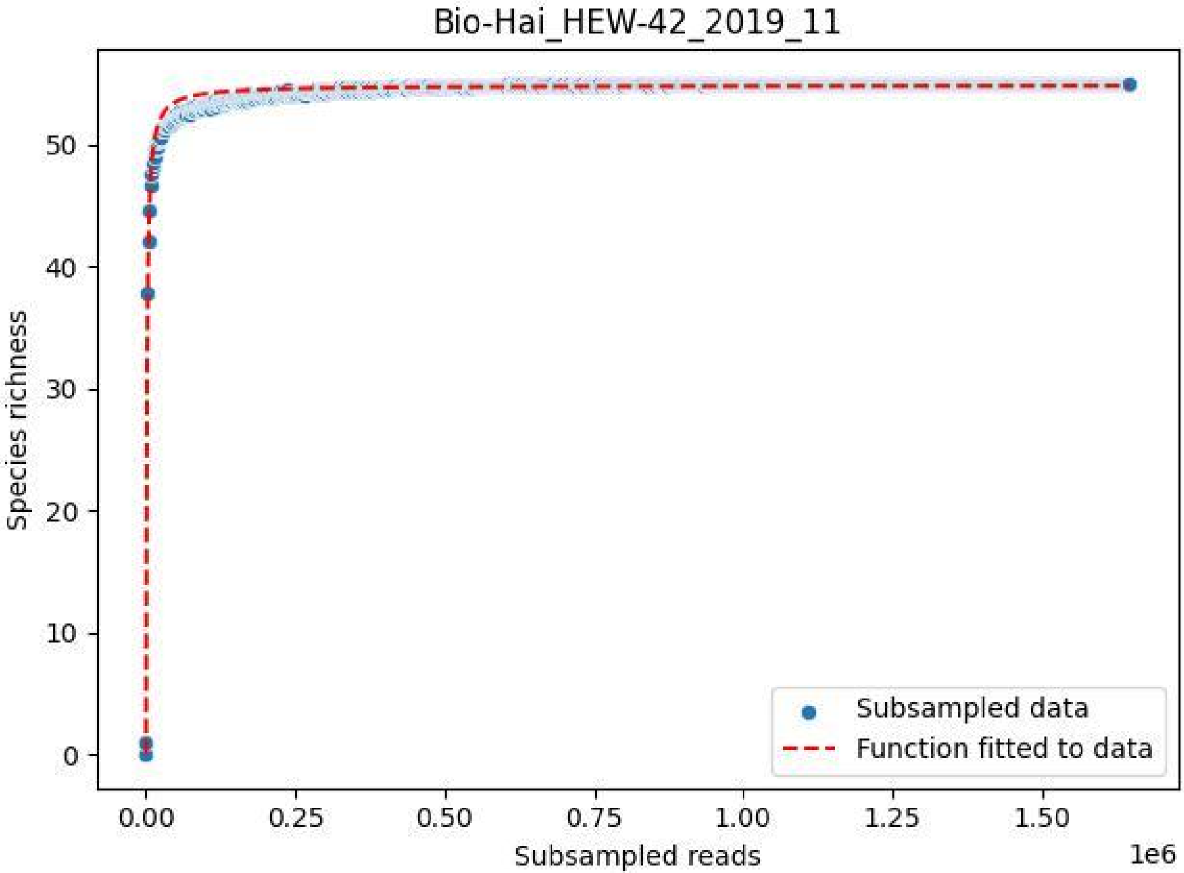
Bio-Hai_HEW-42_2019_11
Species richness
Subsampled reads
1e6
Subsampled data
Function fitted to data
0
10
20
30
40
50
0.00
0.25
0.50
0.75
1.00
1.25
1.50

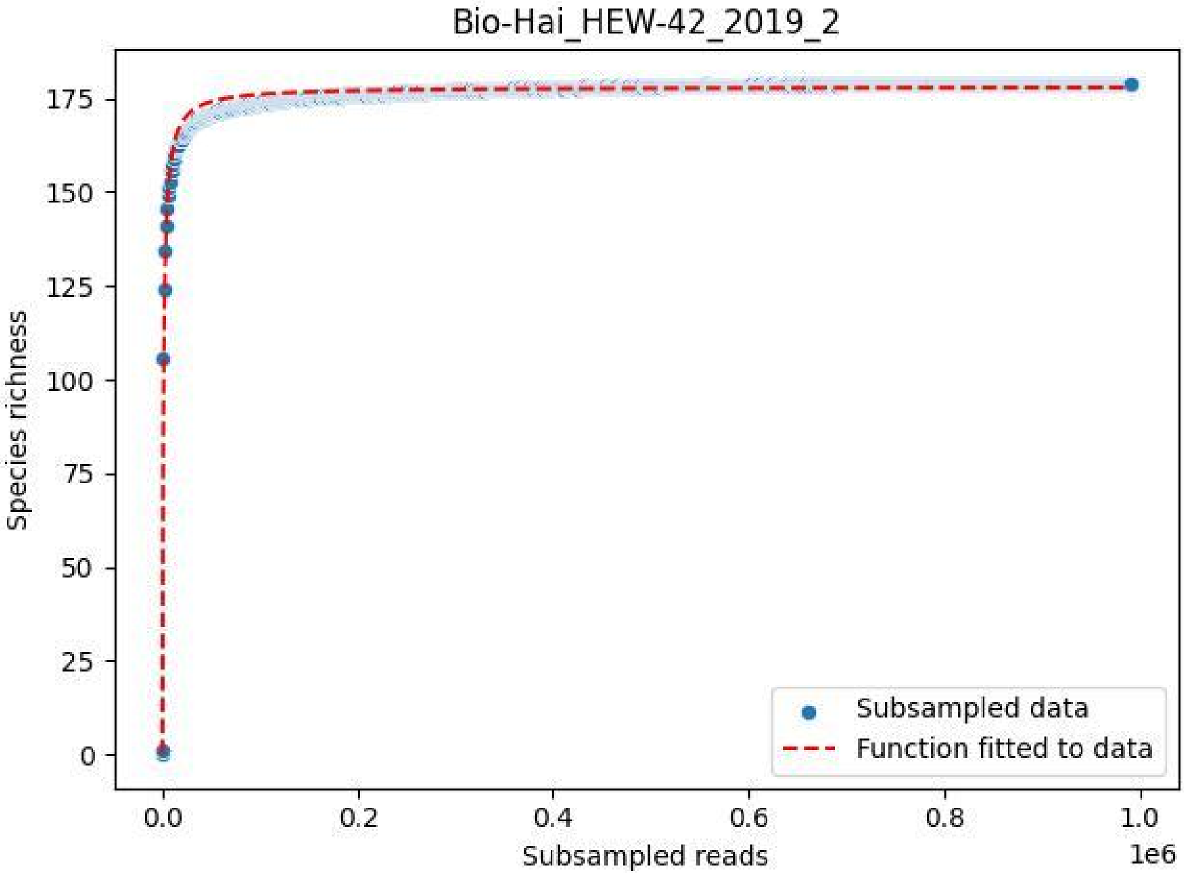
Bio-Hai_HEW-42_2019_2
Species richness
Subsampled reads
1e6
0.0
0.2
0.4
0.6
0.8
1.0
0
25
50
75
100
125
150
175
Subsampled data
Function fitted to data

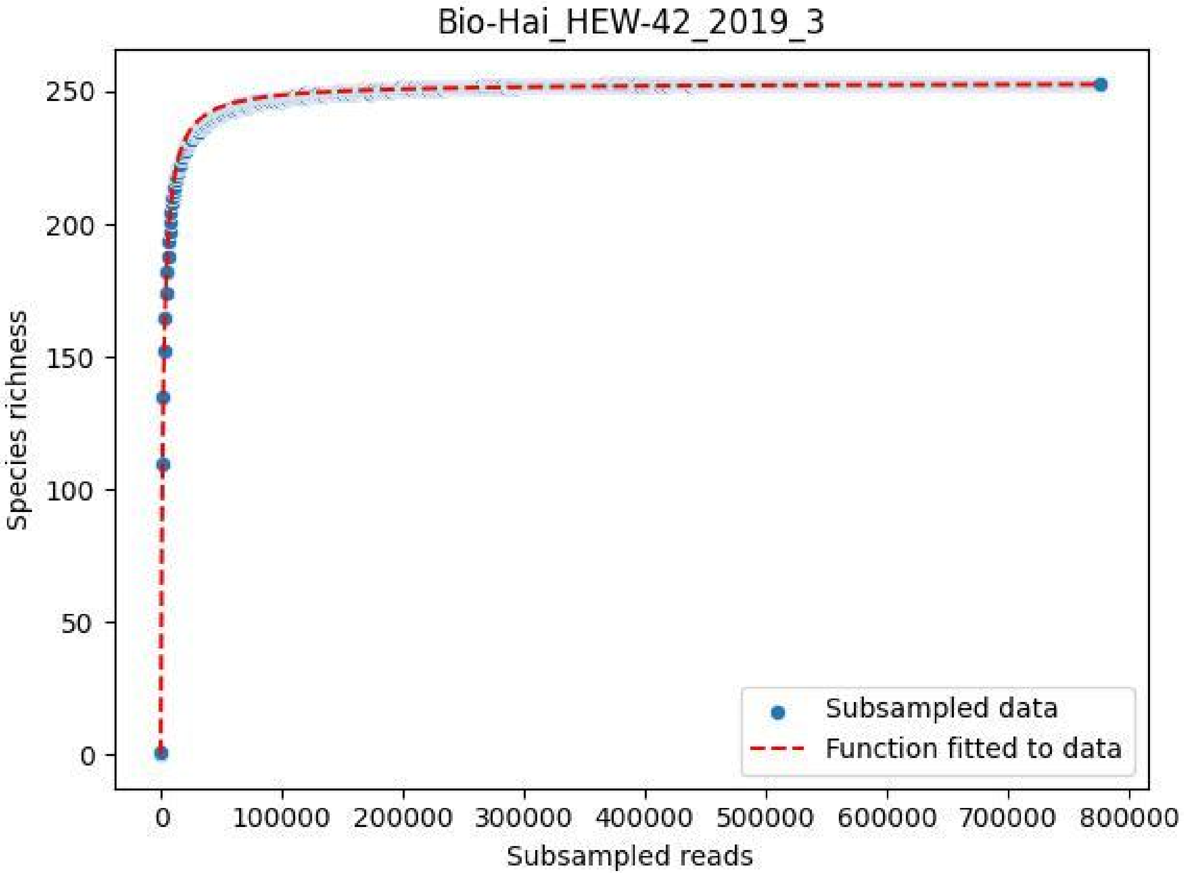
Bio-Hai_HEW-42_2019_3
Species richness
0
50
100
150
200
250
Subsampled reads
0
100000
200000
300000
400000
500000
600000
700000
800000
Subsampled data
Function fitted to data

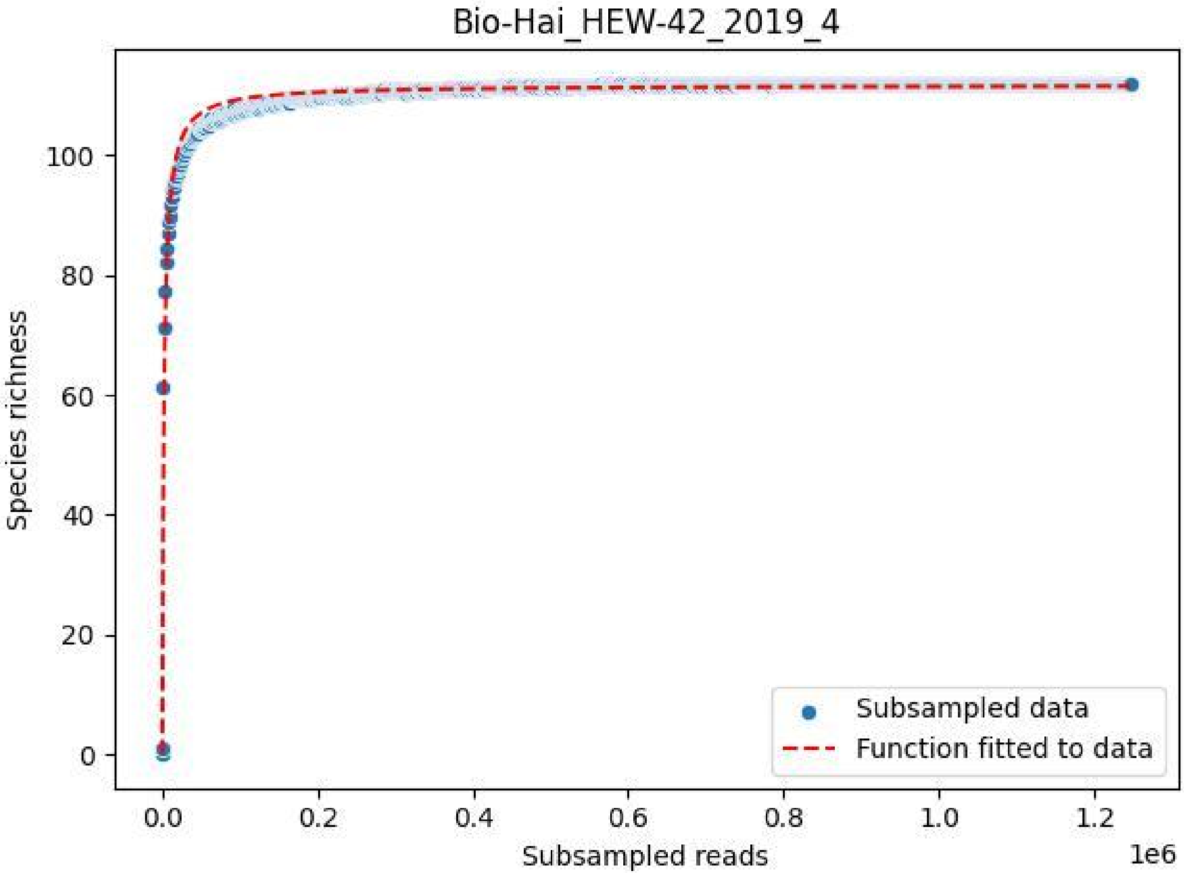
Bio-Hai_HEW-42_2019_4
Species richness
Subsampled reads
1e6
0
20
40
60
80
100
0.0
0.2
0.4
0.6
0.8
1.0
1.2
Subsampled data
Function fitted to data

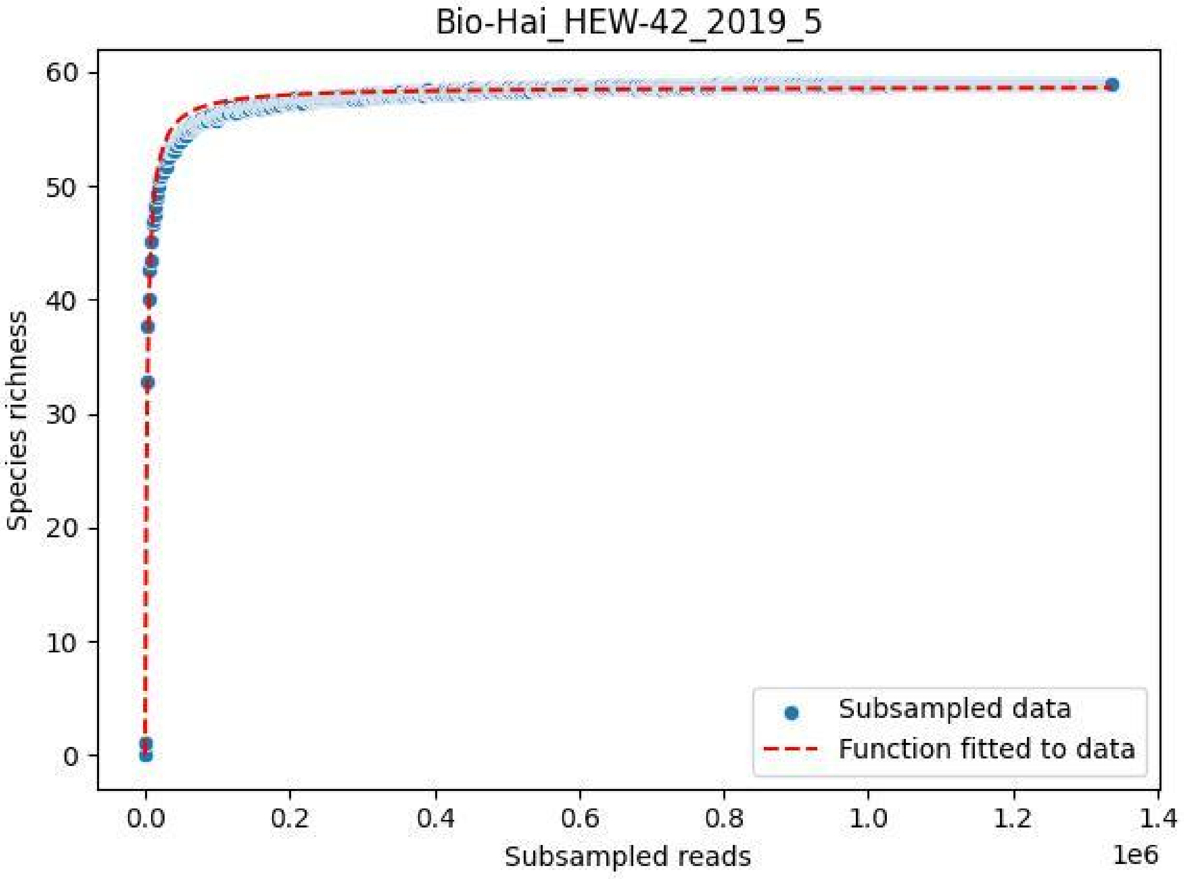
Bio-Hai_HEW-42_2019_5
Species richness
Subsampled reads
1e6
0
10
20
30
40
50
60
0.0
0.2
0.4
0.6
0.8
1.0
1.2
1.4
Subsampled data
Function fitted to data

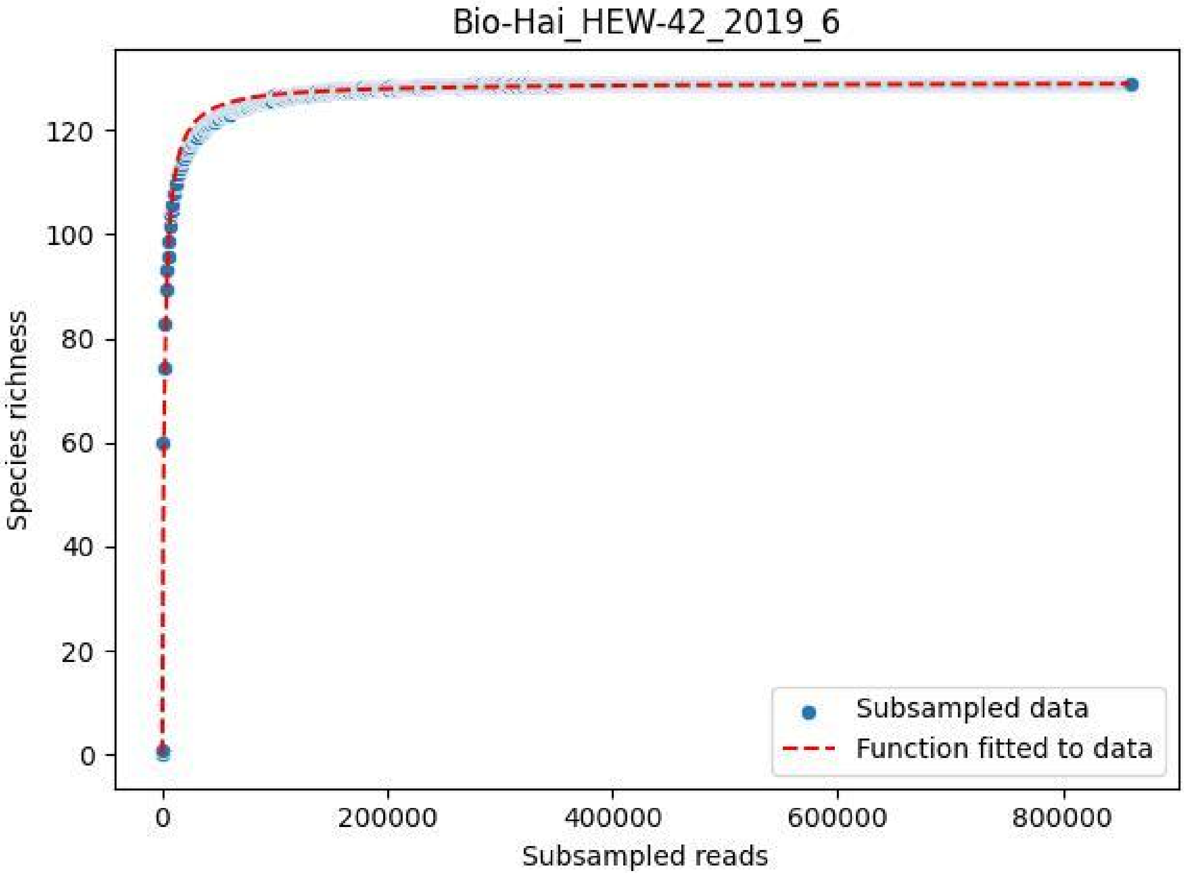
Bio-Hai_HEW-42_2019_6
Species richness
Subsampled reads
0
20
40
60
80
100
120
0
200000
400000
600000
800000
Subsampled data
Function fitted to data

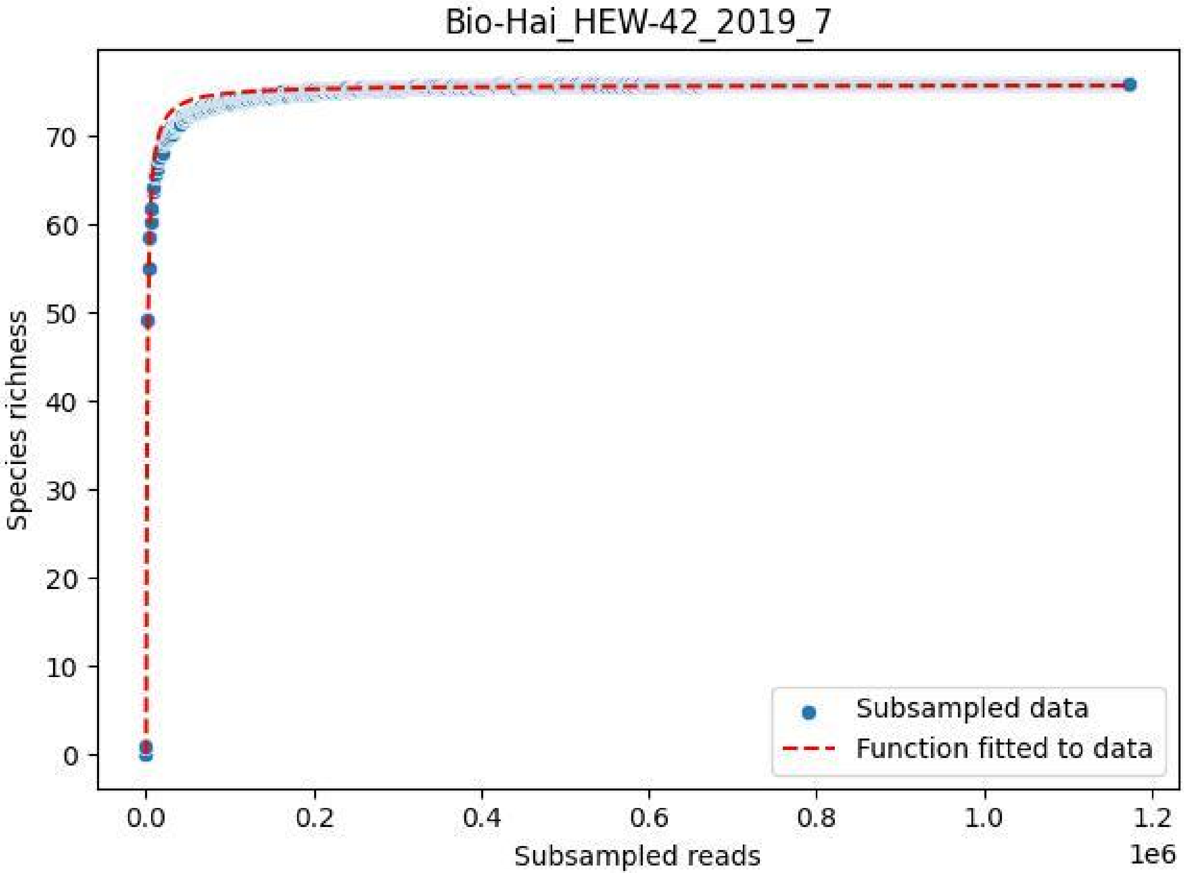
Bio-Hai_HEW-42_2019_7
Species richness
Subsampled reads
1e6
Subsampled data
Function fitted to data
0
10
20
30
40
50
60
70
0.0
0.2
0.4
0.6
0.8
1.0
1.2

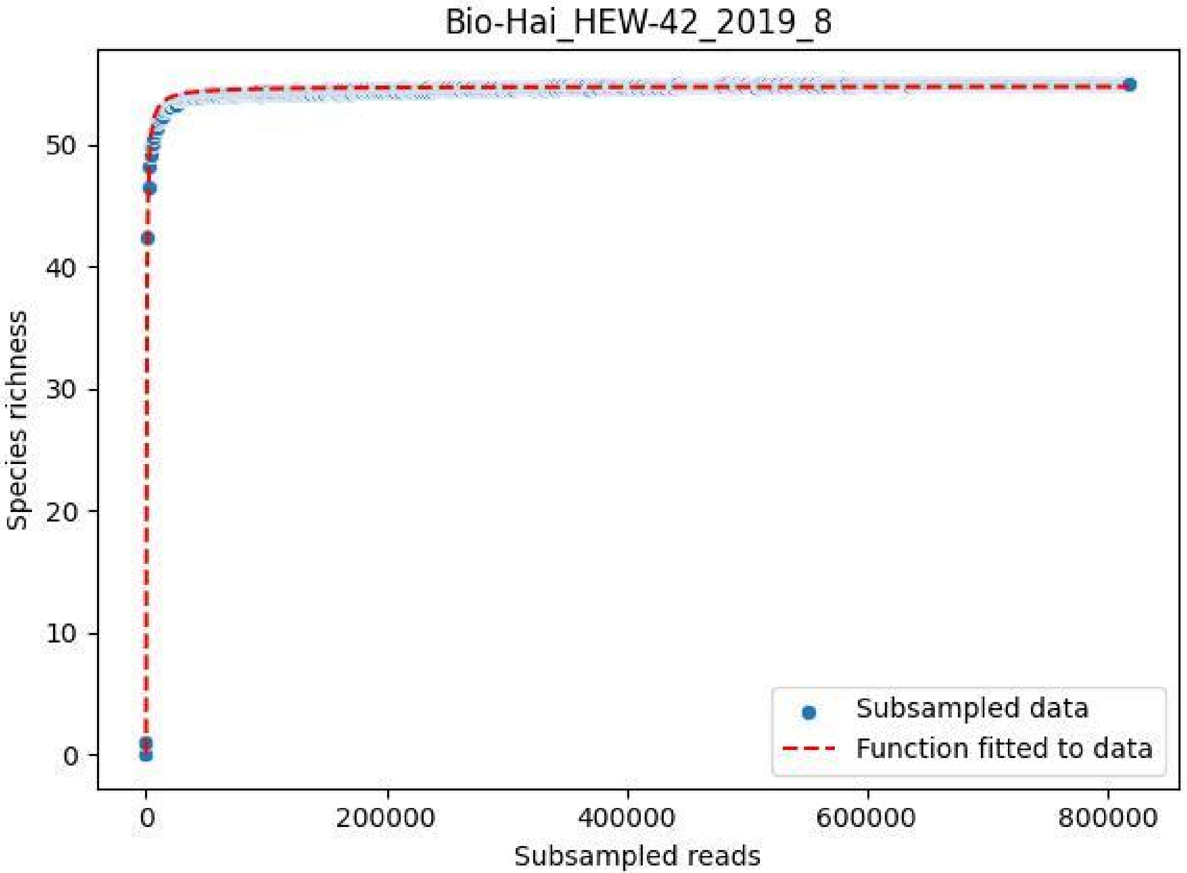
Bio-Hai_HEW-42_2019_8
Species richness
Subsampled reads
0
10
20
30
40
50
0
200000
400000
600000
800000
Subsampled data
Function fitted to data

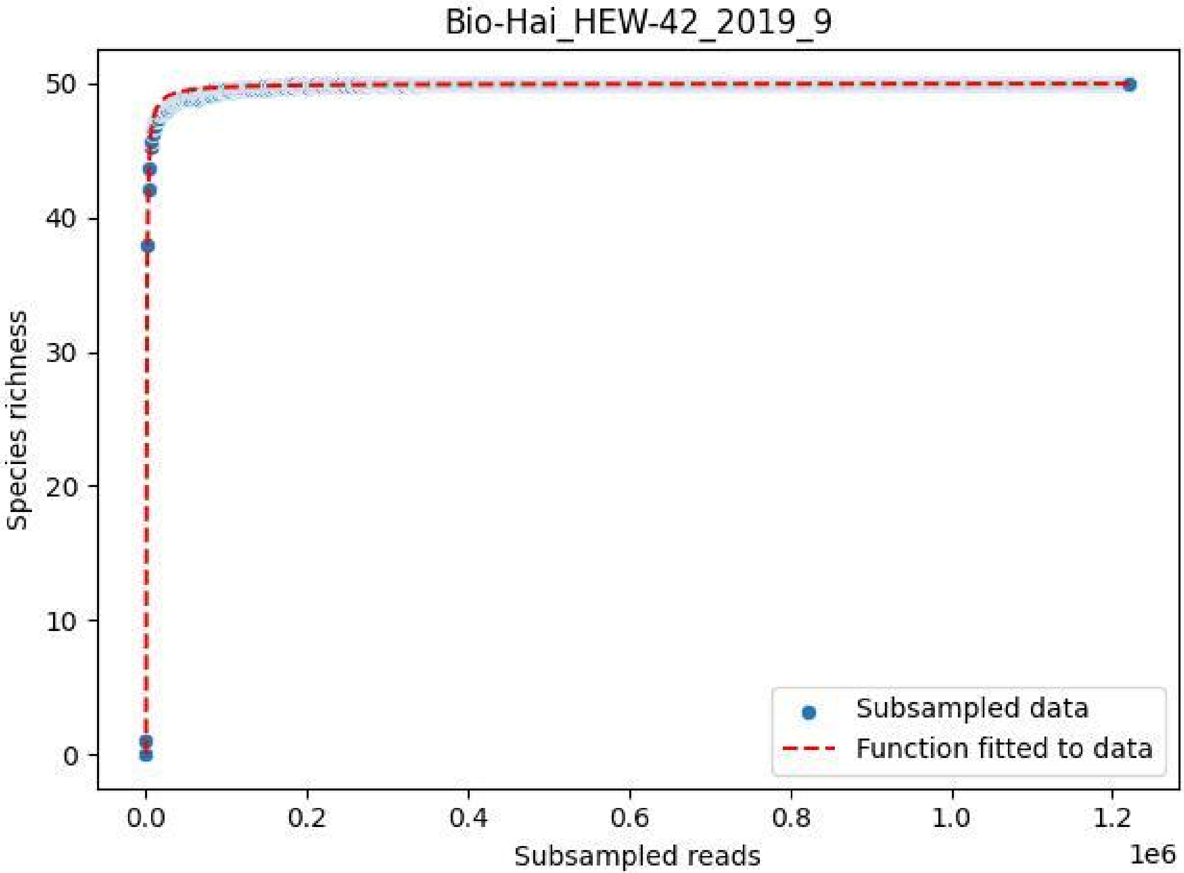
Bio-Hai_HEW-42_2019_9
Species richness
Subsampled reads
1e6
Subsampled data
Function fitted to data
0
10
20
30
40
50
0.0
0.2
0.4
0.6
0.8
1.0
1.2

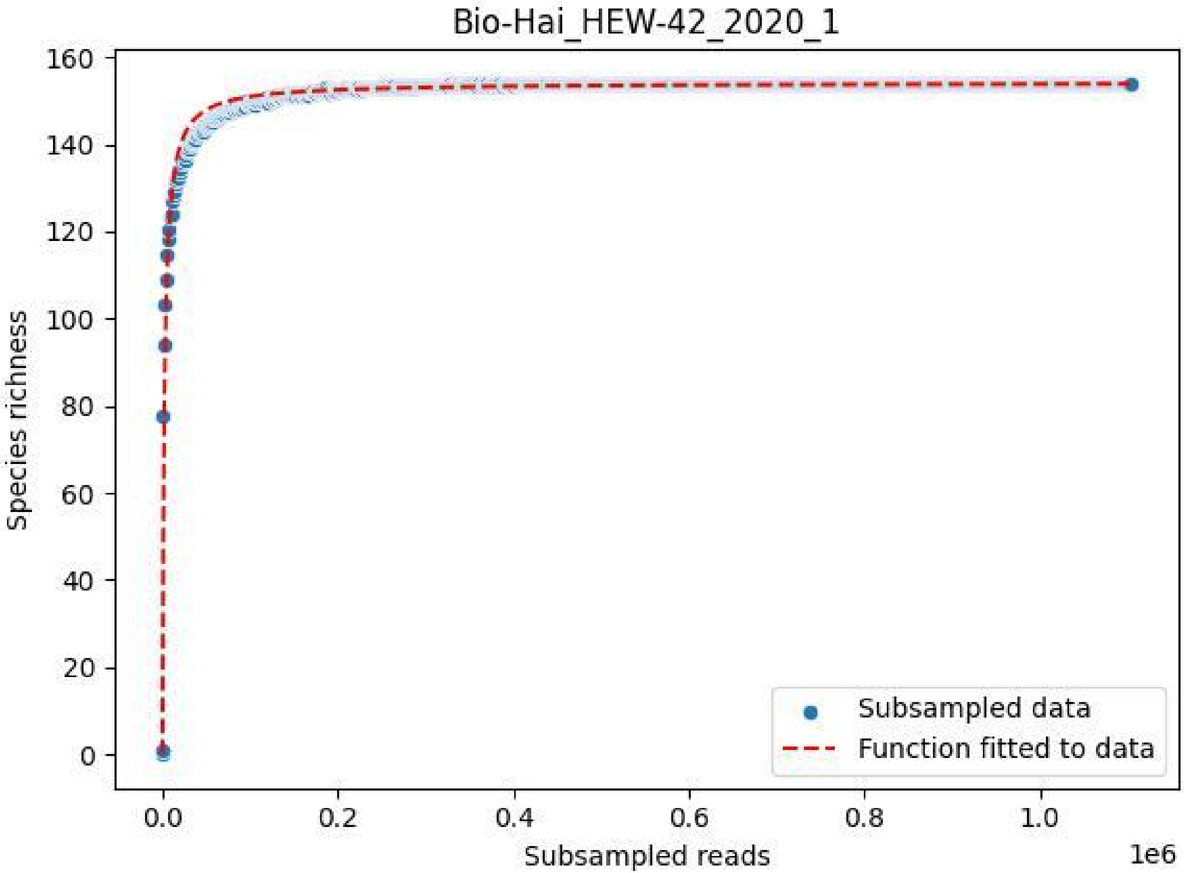
Bio-Hai_HEW-42_2020_1
Species richness
Subsampled reads
1e6
0
20
40
60
80
100
120
140
160
0.0
0.2
0.4
0.6
0.8
1.0
Subsampled data
Function fitted to data

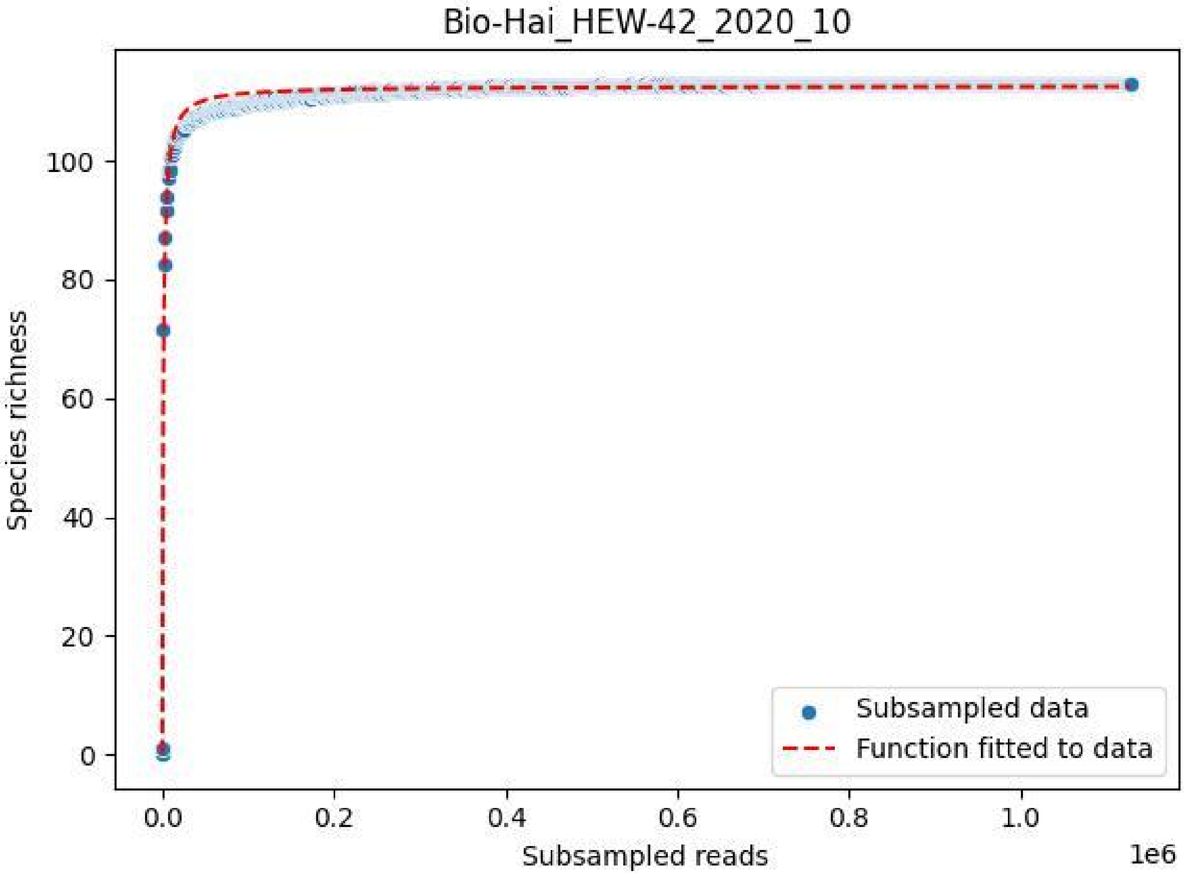
Bio-Hai_HEW-42_2020_10
Species richness
Subsampled reads
1e6
0
20
40
60
80
100
0.0
0.2
0.4
0.6
0.8
1.0
Subsampled data
Function fitted to data

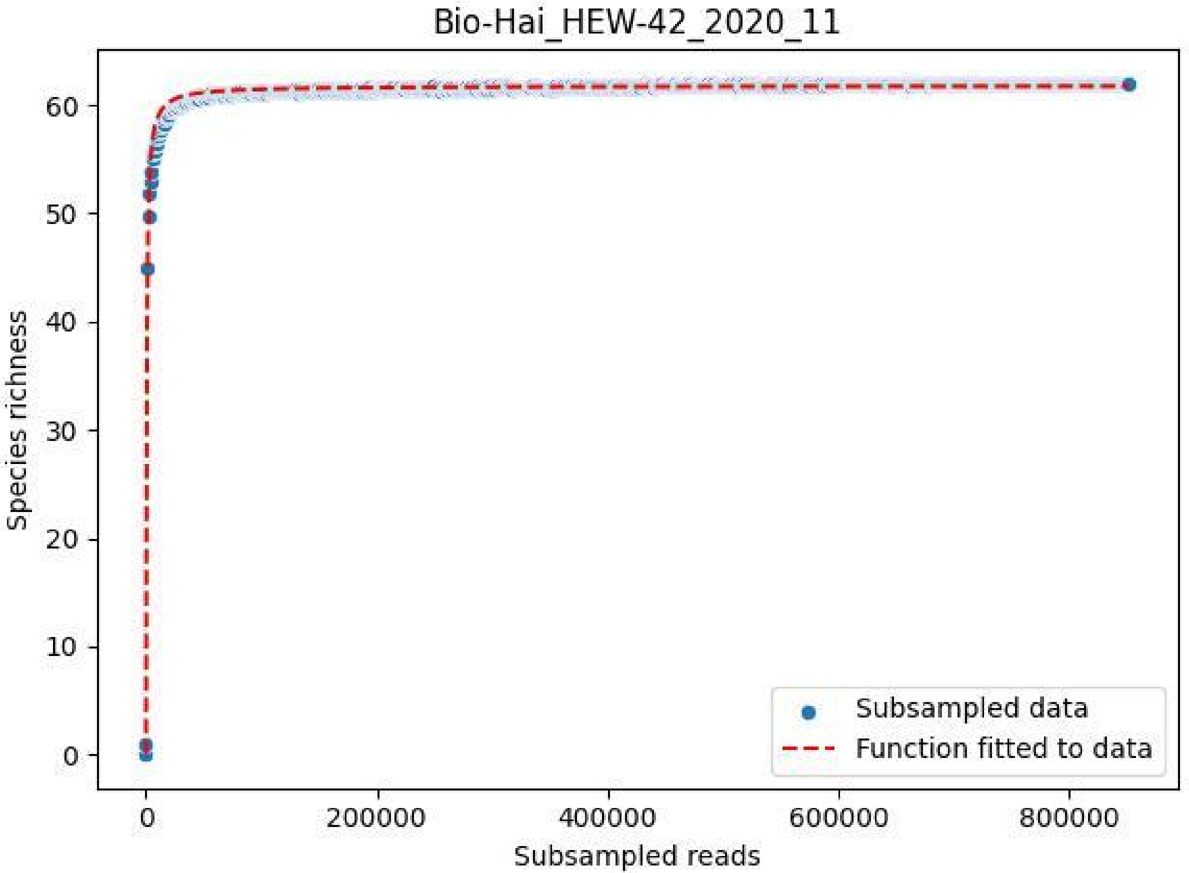
Bio-Hai_HEW-42_2020_11
Species richness
Subsampled reads
Subsampled data
Function fitted to data
0
10
20
30
40
50
60
0
200000
400000
600000
800000

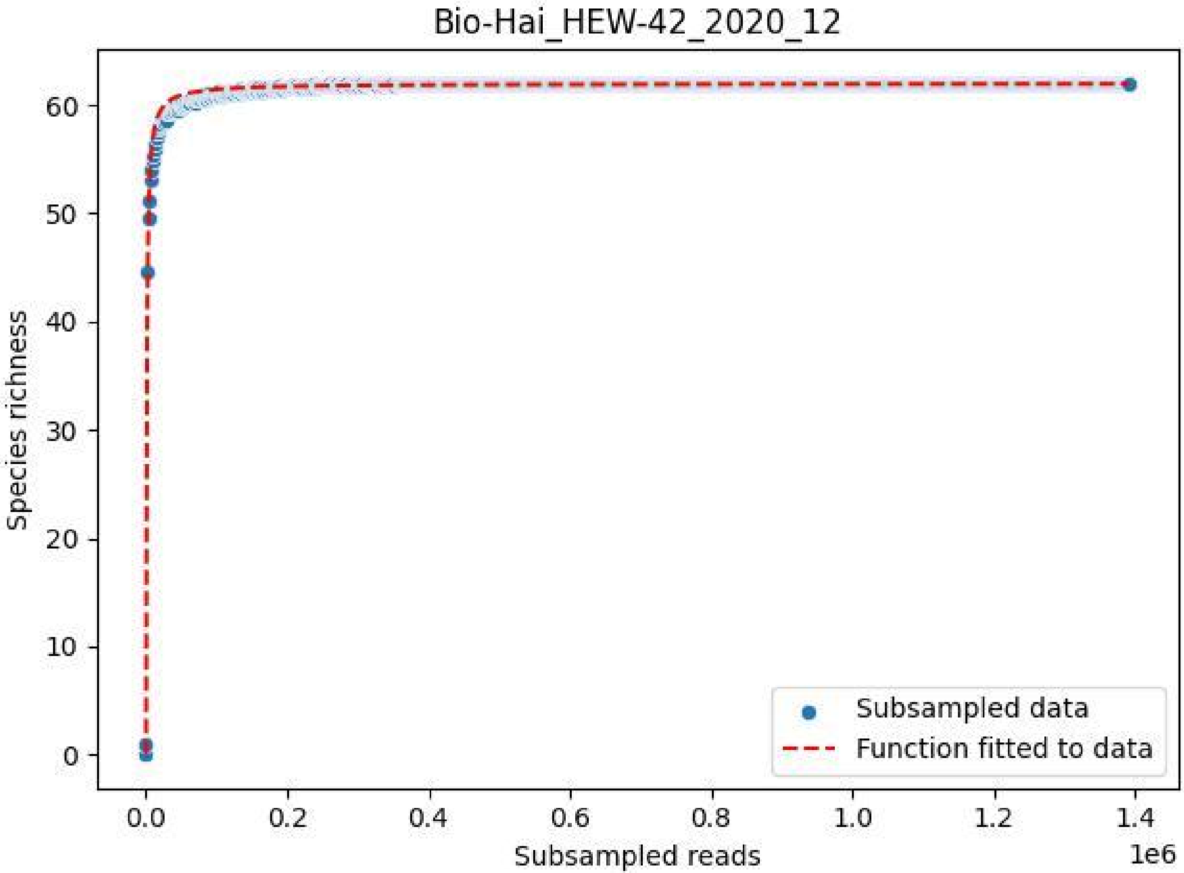
Bio-Hai_HEW-42_2020_12
Species richness
0
10
20
30
40
50
60
0.0
0.2
0.4
0.6
0.8
1.0
1.2
1.4
1e6
Subsampled reads
Subsampled data
Function fitted to data

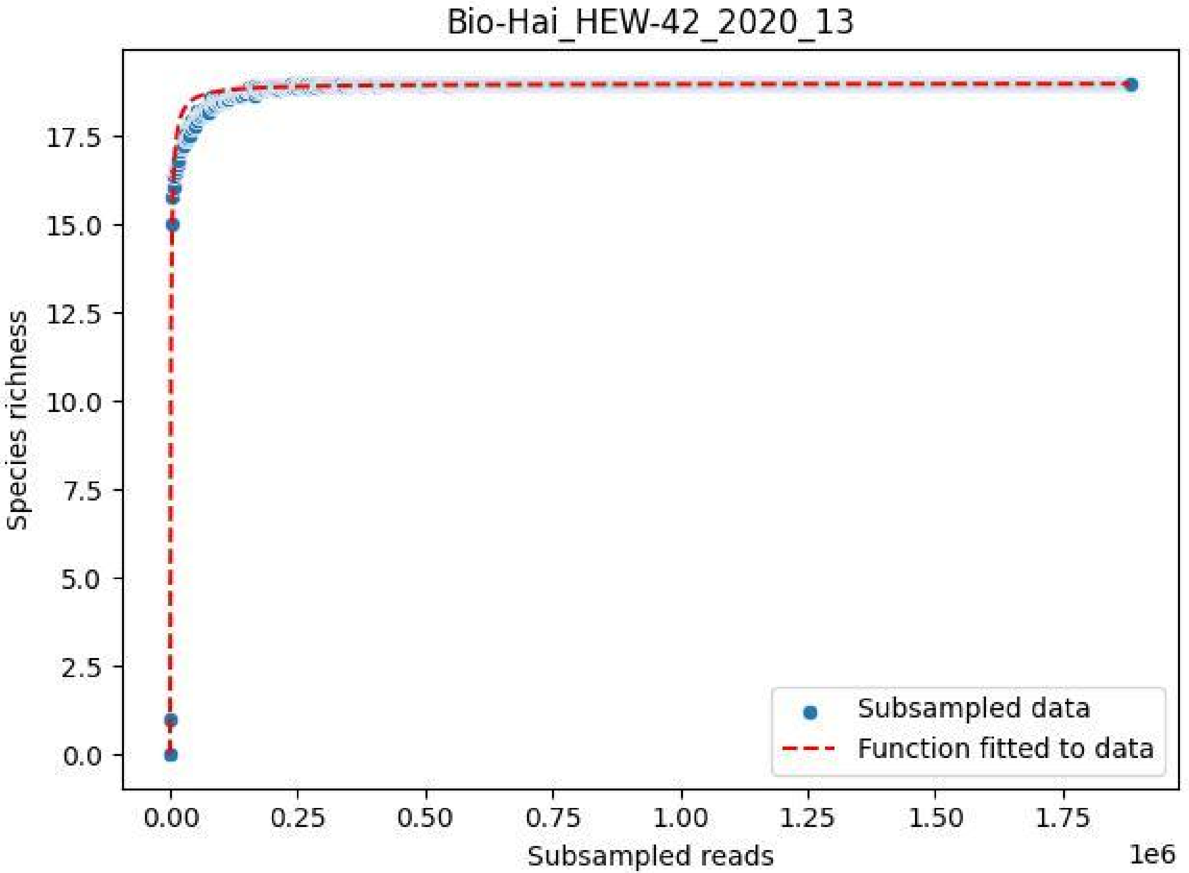

Bio-Hai_HEW-42_2020_13
Species richness
Subsampled reads
1e6
0.0
2.5
5.0
7.5
10.0
12.5
15.0
17.5
0.00
0.25
0.50
0.75
1.00
1.25
1.50
1.75
Subsampled data
Function fitted to data

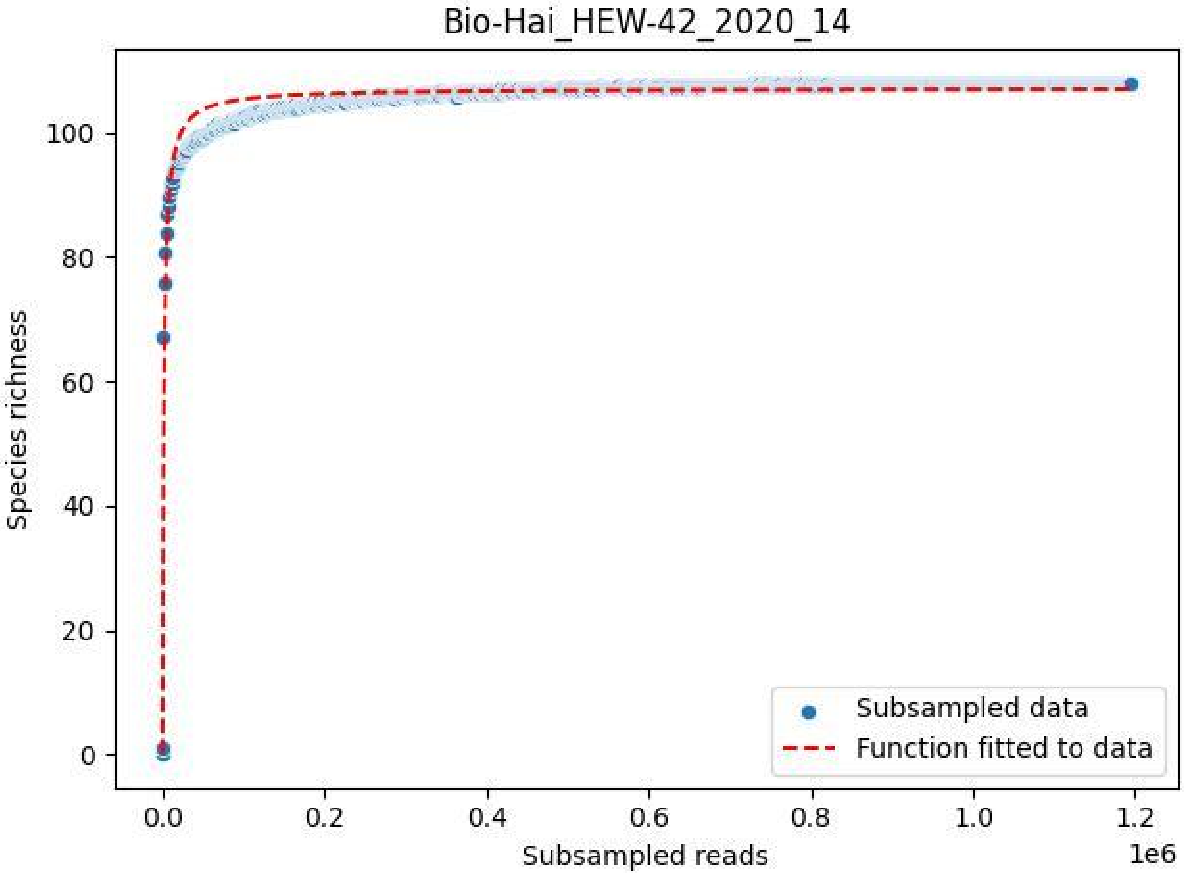
Bio-Hai_HEW-42_2020_14
Species richness
0
20
40
60
80
100
0.0
0.2
0.4
0.6
0.8
1.0
1.2
1e6
Subsampled reads
Subsampled data
Function fitted to data

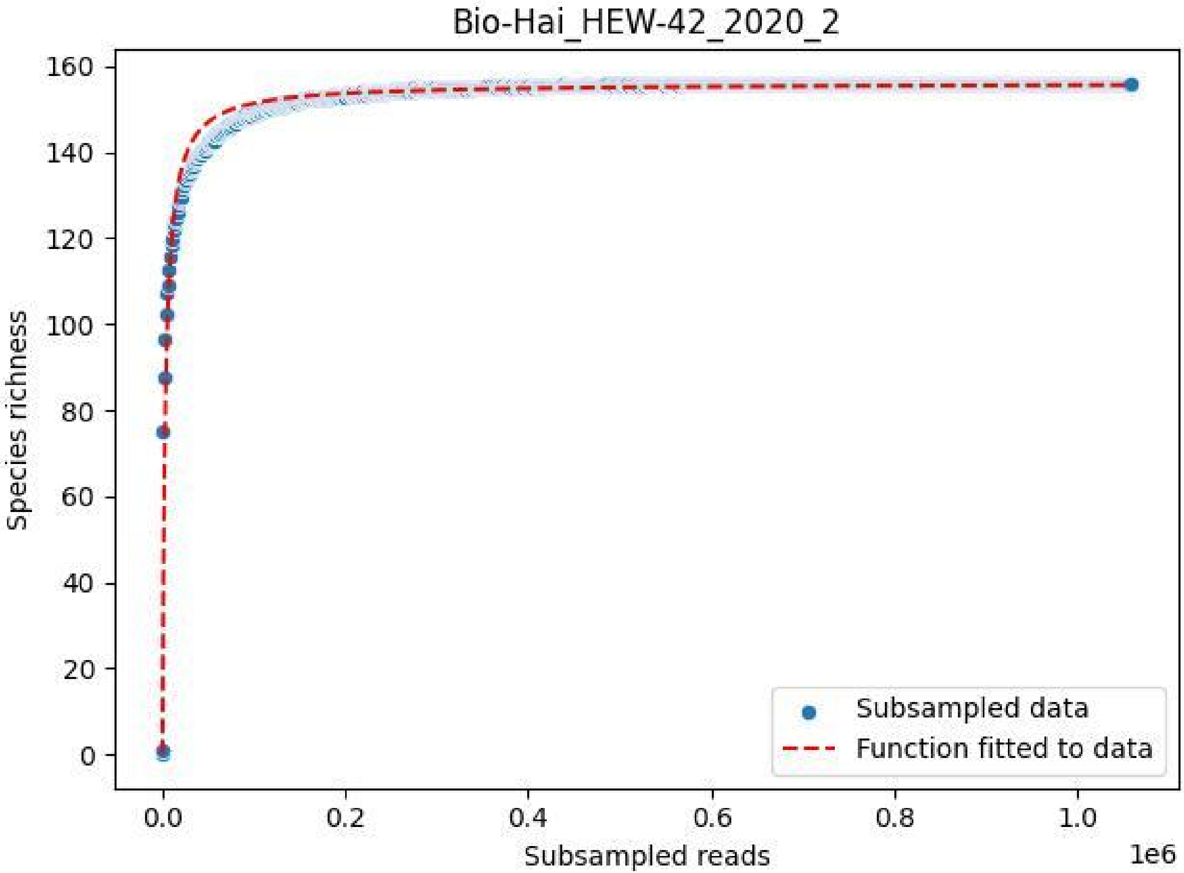
Bio-Hai_HEW-42_2020_2
Species richness
Subsampled reads
1e6
0
20
40
60
80
100
120
140
160
0.0
0.2
0.4
0.6
0.8
1.0
Subsampled data
Function fitted to data

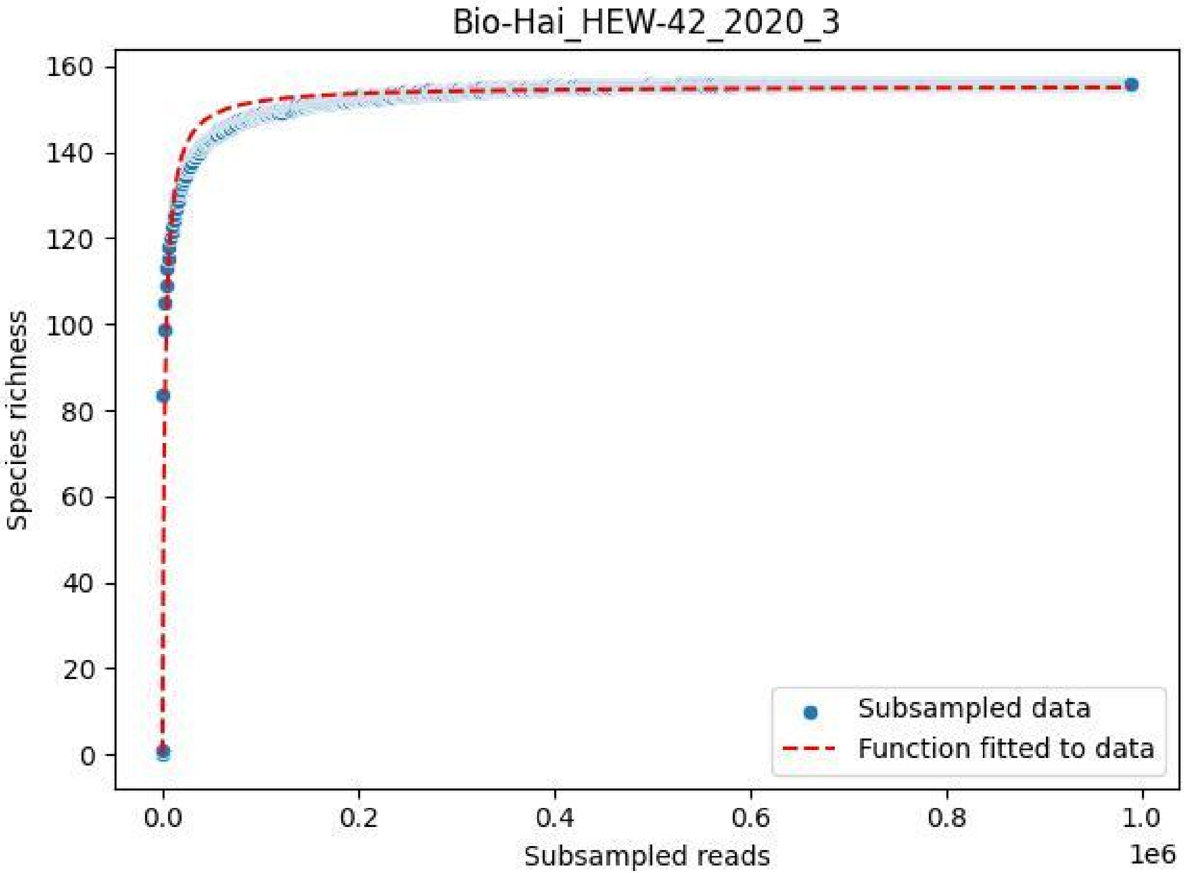
Bio-Hai_HEW-42_2020_3
Species richness
160
140
120
100
80
60
40
20
0
0.0
0.2
0.4
0.6
0.8
1.0
1e6
Subsampled reads
Subsampled data
Function fitted to data

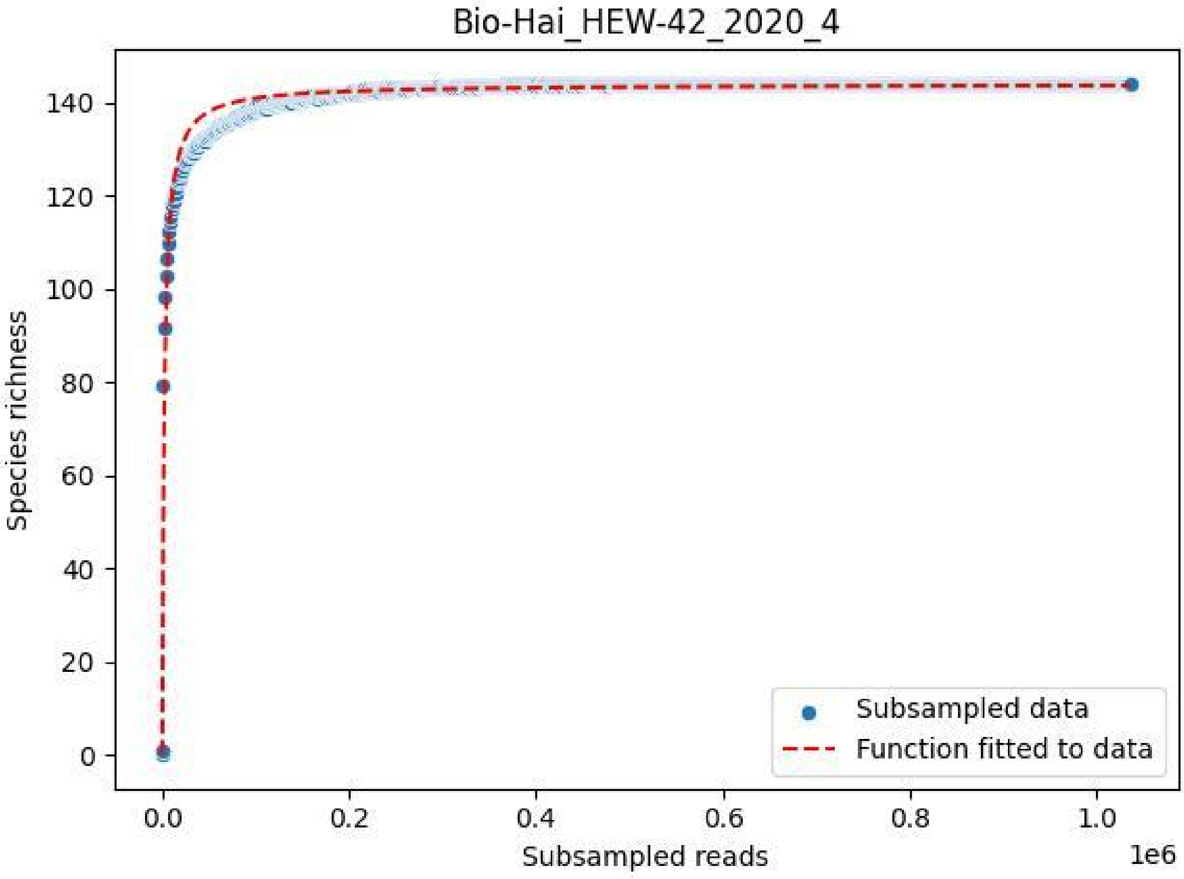

Bio-Hai_HEW-42_2020_4
Species richness
Subsampled reads
1e6
0
20
40
60
80
100
120
140
0.0
0.2
0.4
0.6
0.8
1.0
Subsampled data
Function fitted to data

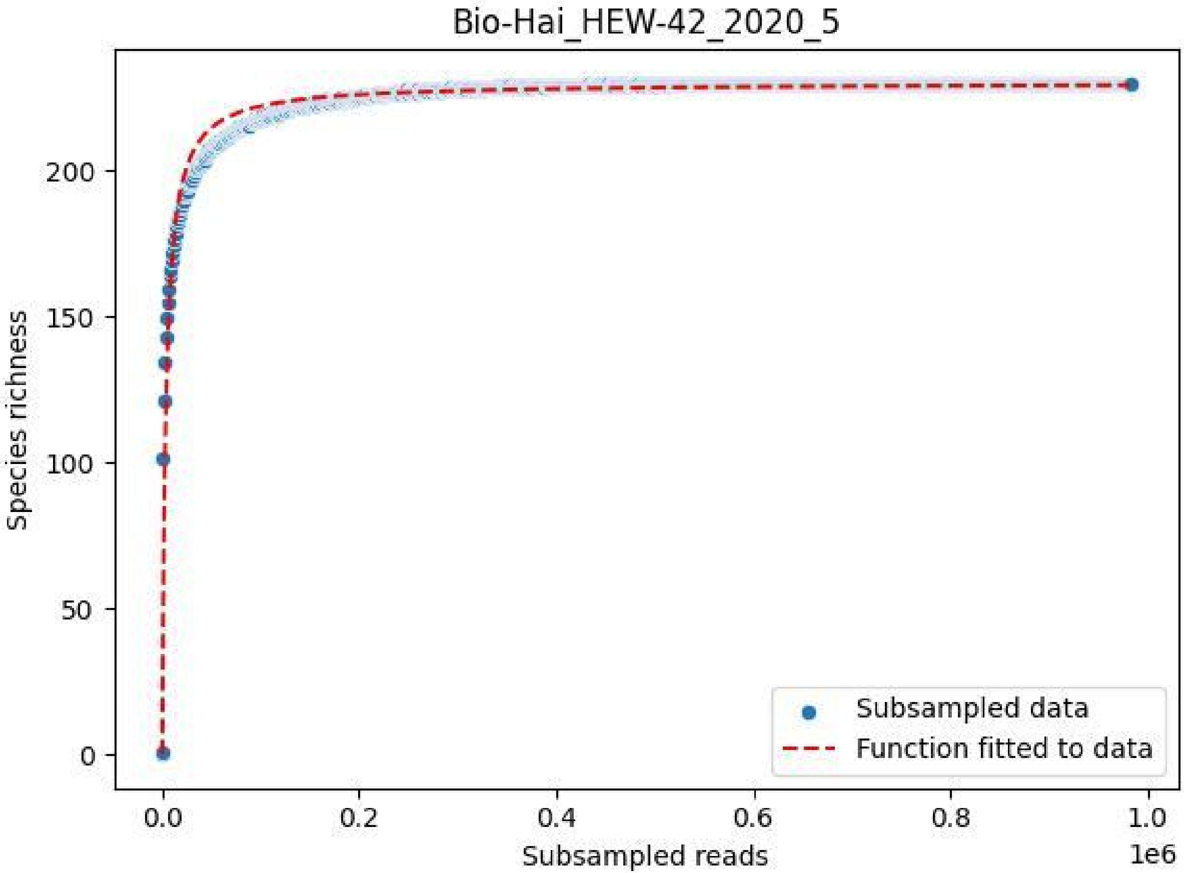

Bio-Hai_HEW-42_2020_5
Species richness
Subsampled reads
1e6
0
50
100
150
200
0.0
0.2
0.4
0.6
0.8
1.0
Subsampled data
Function fitted to data

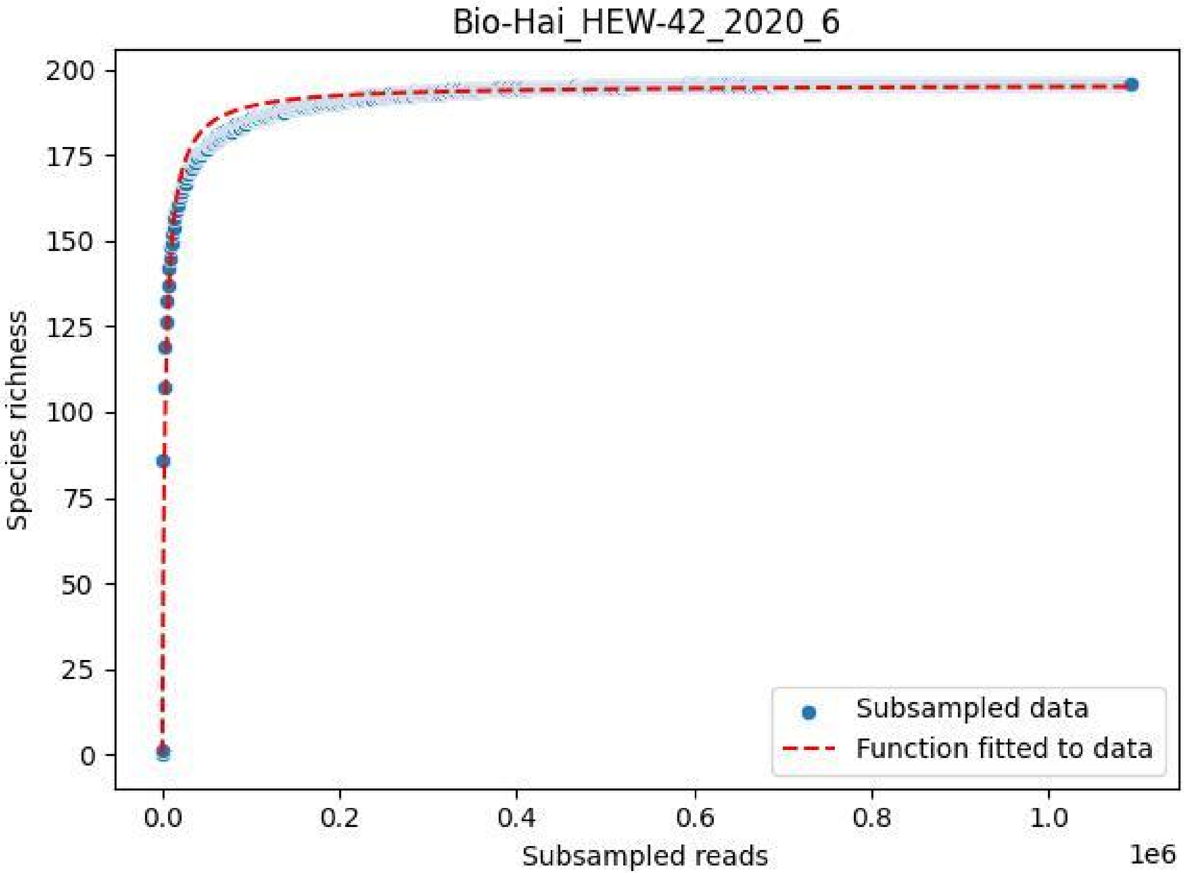

Bio-Hai_HEW-42_2020_6
Species richness
Subsampled reads
1e6
0
25
50
75
100
125
150
175
200
0.0
0.2
0.4
0.6
0.8
1.0
Subsampled data
Function fitted to data

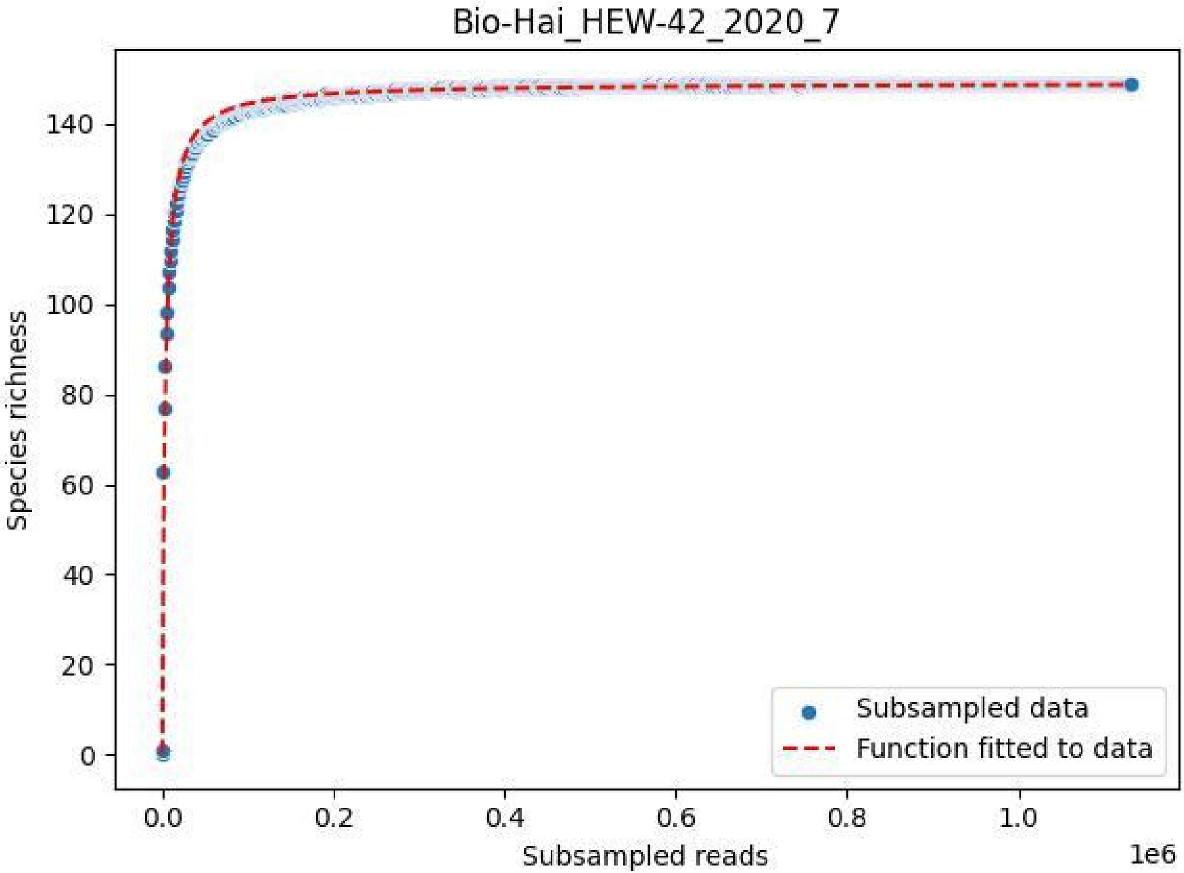
Bio-Hai_HEW-42_2020_7
Species richness
0
20
40
60
80
100
120
140
0.0
0.2
0.4
0.6
0.8
1.0
1e6
Subsampled reads
Subsampled data
Function fitted to data

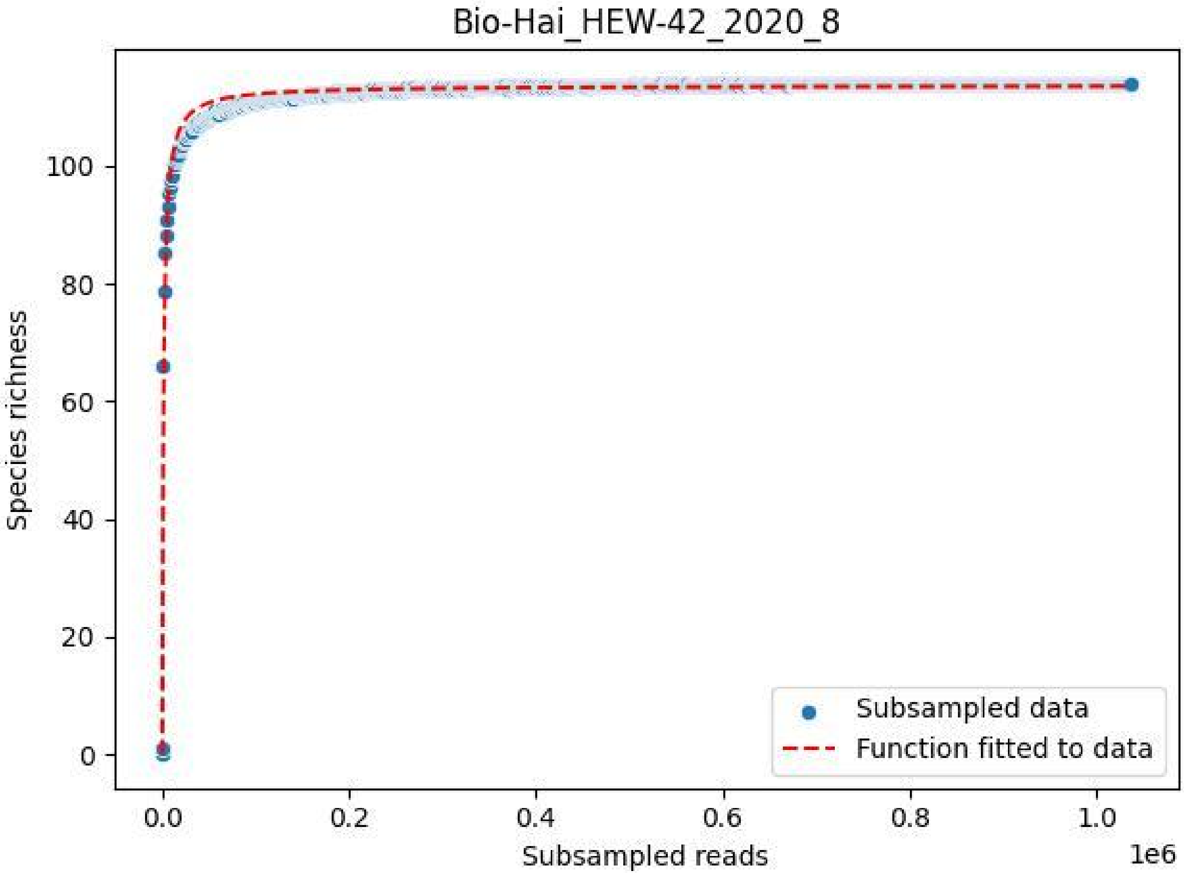
Bio-Hai_HEW-42_2020_8
Species richness
0
20
40
60
80
100
0.0
0.2
0.4
0.6
0.8
1.0
1e6
Subsampled reads
Subsampled data
Function fitted to data

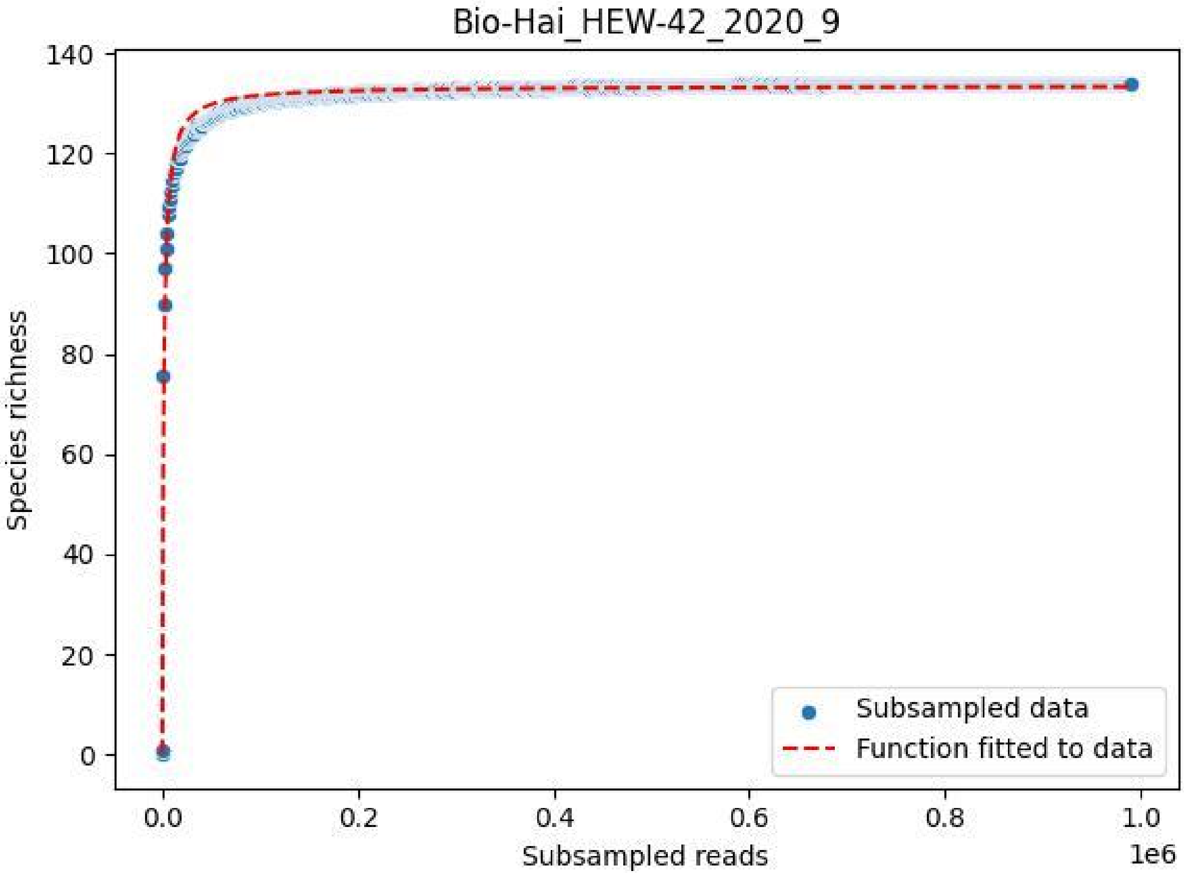
Bio-Hai_HEW-42_2020_9
Species richness
Subsampled reads
1e6
0
20
40
60
80
100
120
140
0.0
0.2
0.4
0.6
0.8
1.0
Subsampled data
Function fitted to data

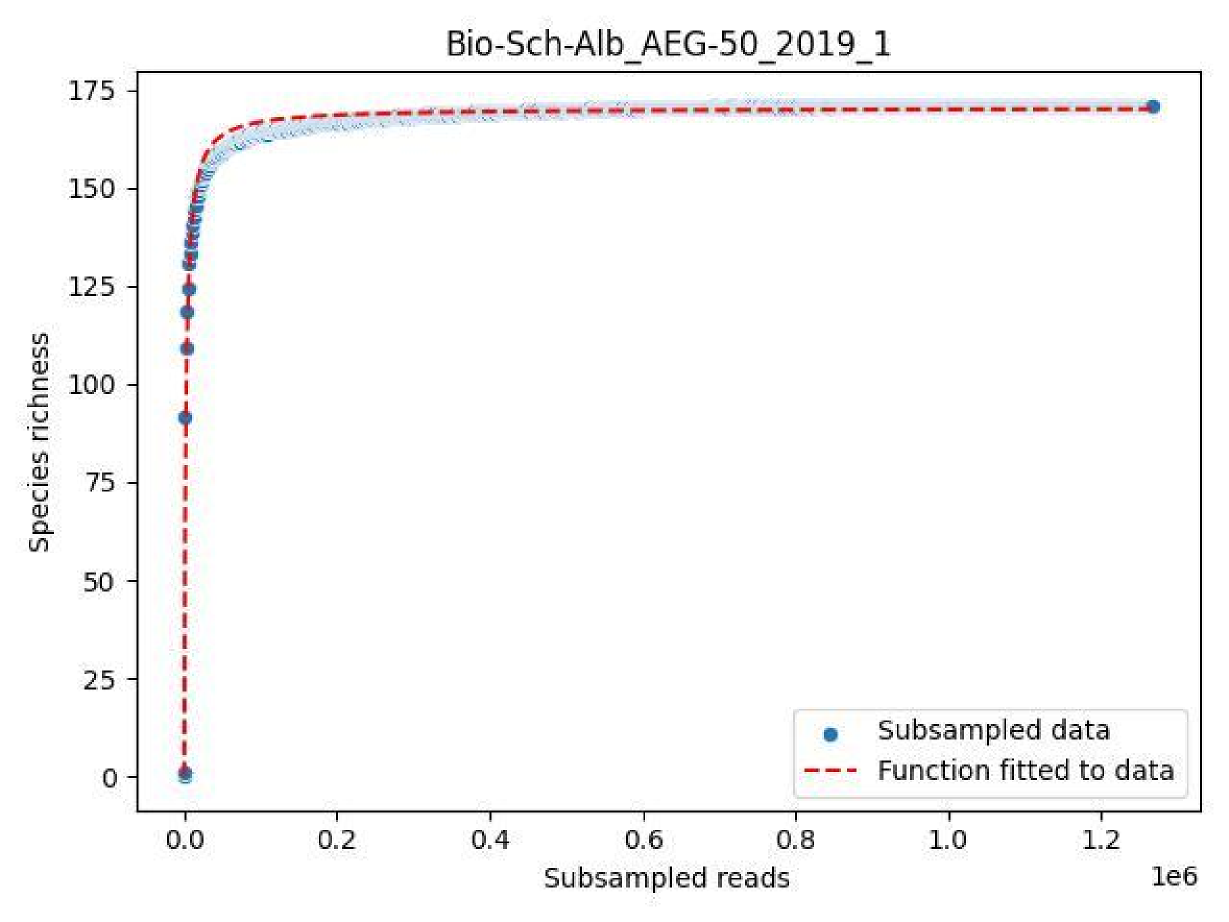

Bio-Sch-Alb_AEG-50_2019_1
Species richness
175
150
125
100
75
50
25
0
0.0
0.2
0.4
0.6
0.8
1.0
1.2
Subsampled reads
1e6
Subsampled data
Function fitted to data

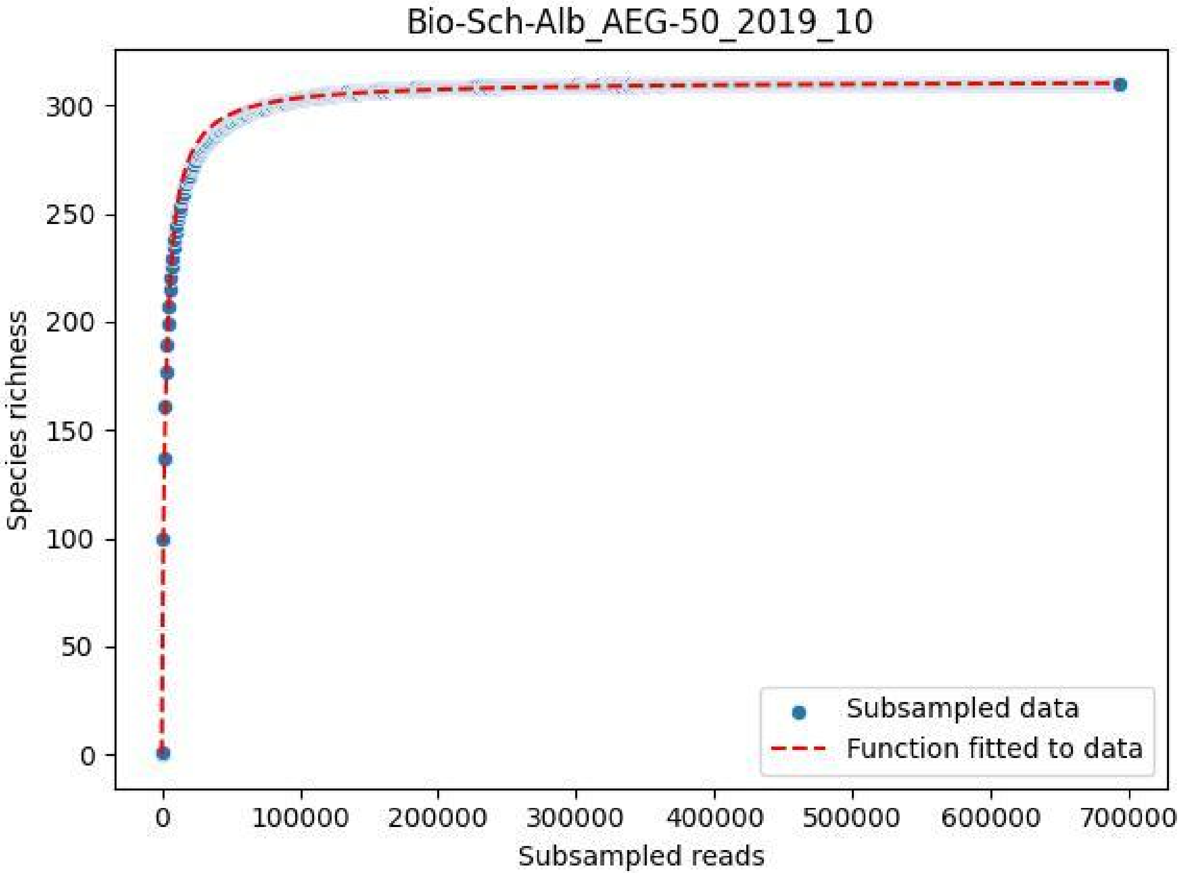

Bio-Sch-Alb_AEG-50_2019_10
Species richness
Subsampled reads
0
50
100
150
200
250
300
100000
200000
300000
400000
500000
600000
700000
Subsampled data
Function fitted to data

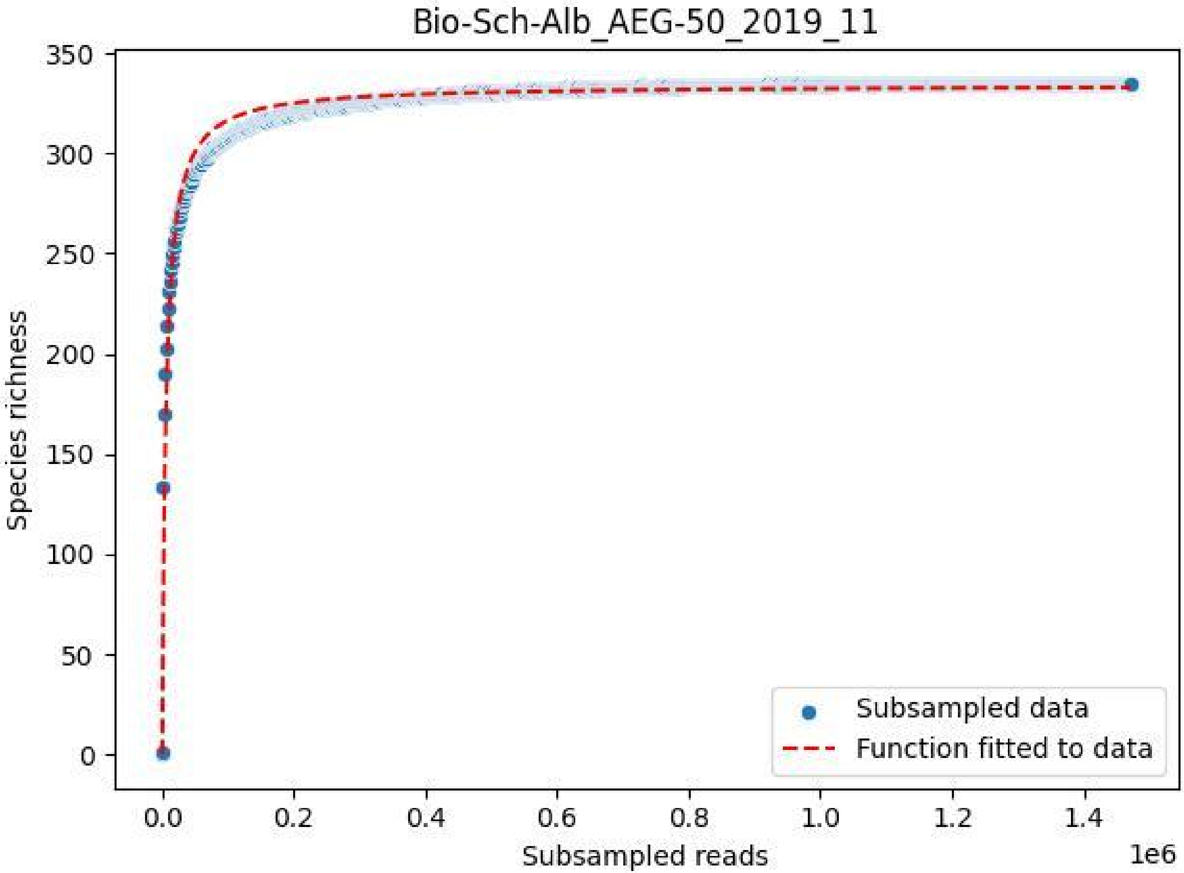
Bio-Sch-Alb_AEG-50_2019_11
Species richness
Subsampled reads
1e6
0
50
100
150
200
250
300
350
0.0
0.2
0.4
0.6
0.8
1.0
1.2
1.4
Subsampled data
Function fitted to data

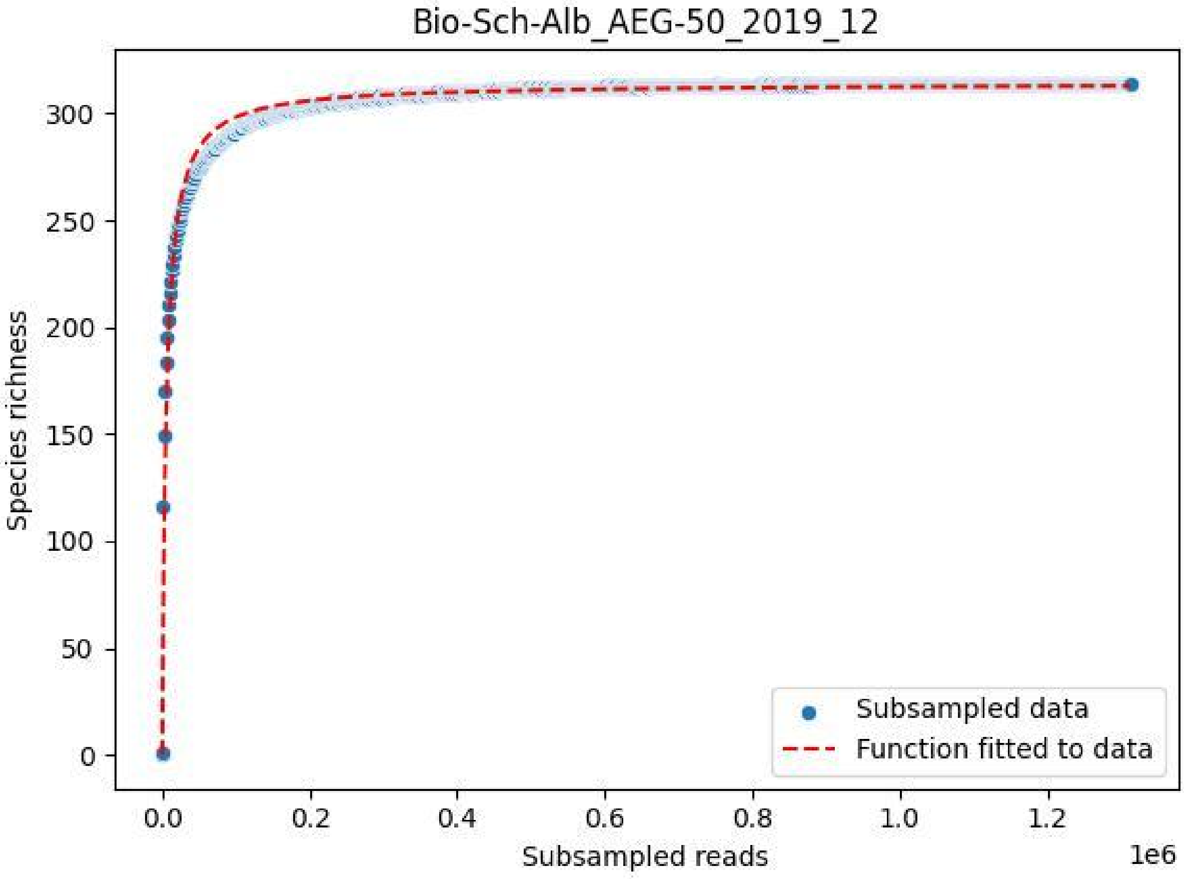

Bio-Sch-Alb_AEG-50_2019_12
Species richness
Subsampled reads
1e6
0
50
100
150
200
250
300
0.0
0.2
0.4
0.6
0.8
1.0
1.2
Subsampled data
Function fitted to data

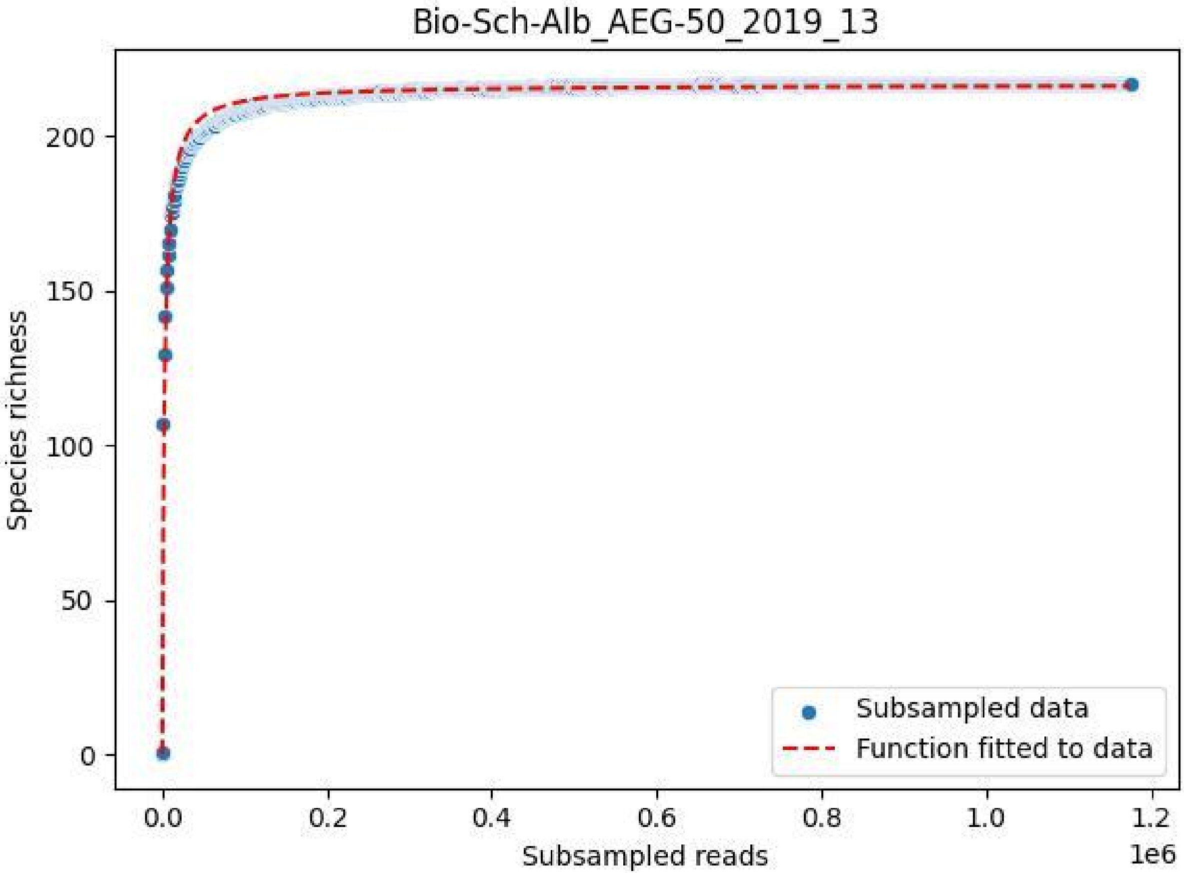
Bio-Sch-Alb_AEG-50_2019_13
Species richness
Subsampled reads
1e6
0
50
100
150
200
0.0
0.2
0.4
0.6
0.8
1.0
1.2
Subsampled data
Function fitted to data

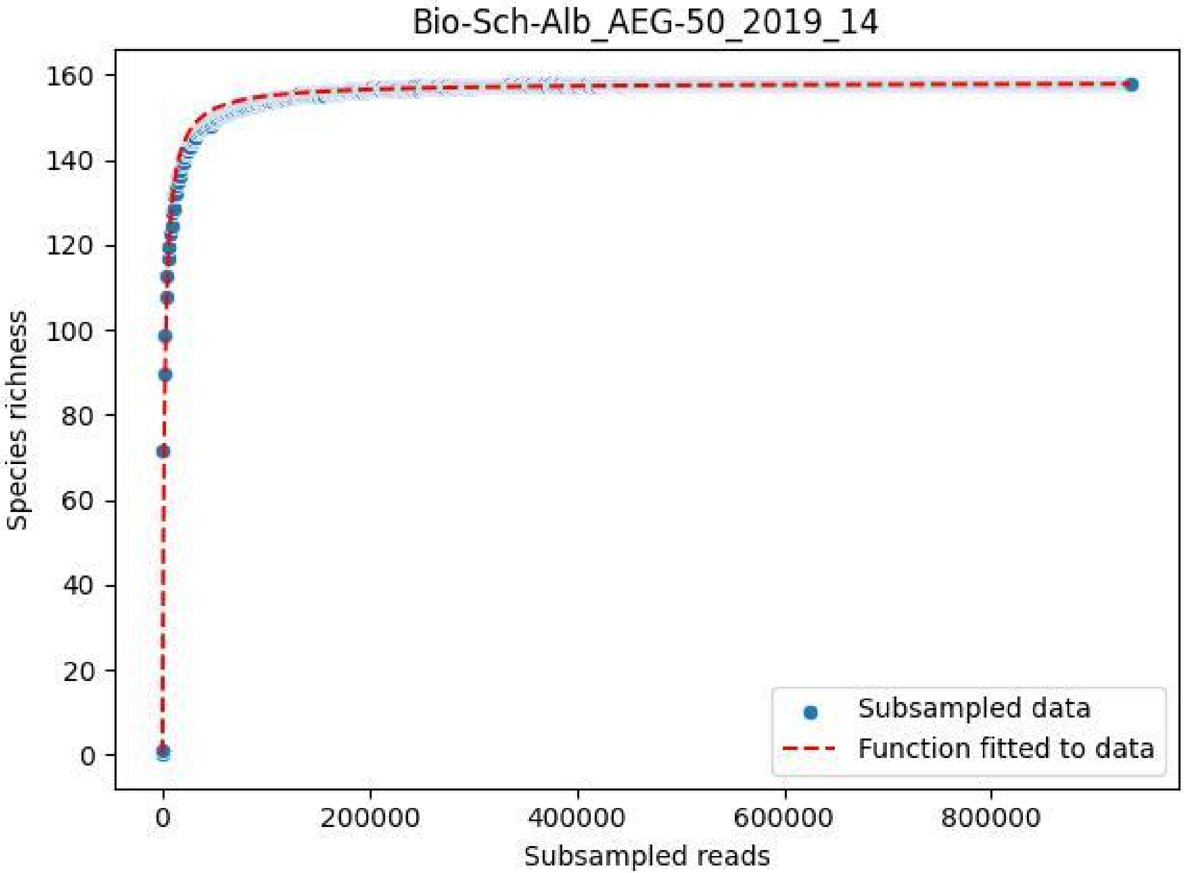
Bio-Sch-Alb_AEG-50_2019_14
Species richness
Subsampled reads
0
20
40
60
80
100
120
140
160
0
200000
400000
600000
800000
Subsampled data
Function fitted to data

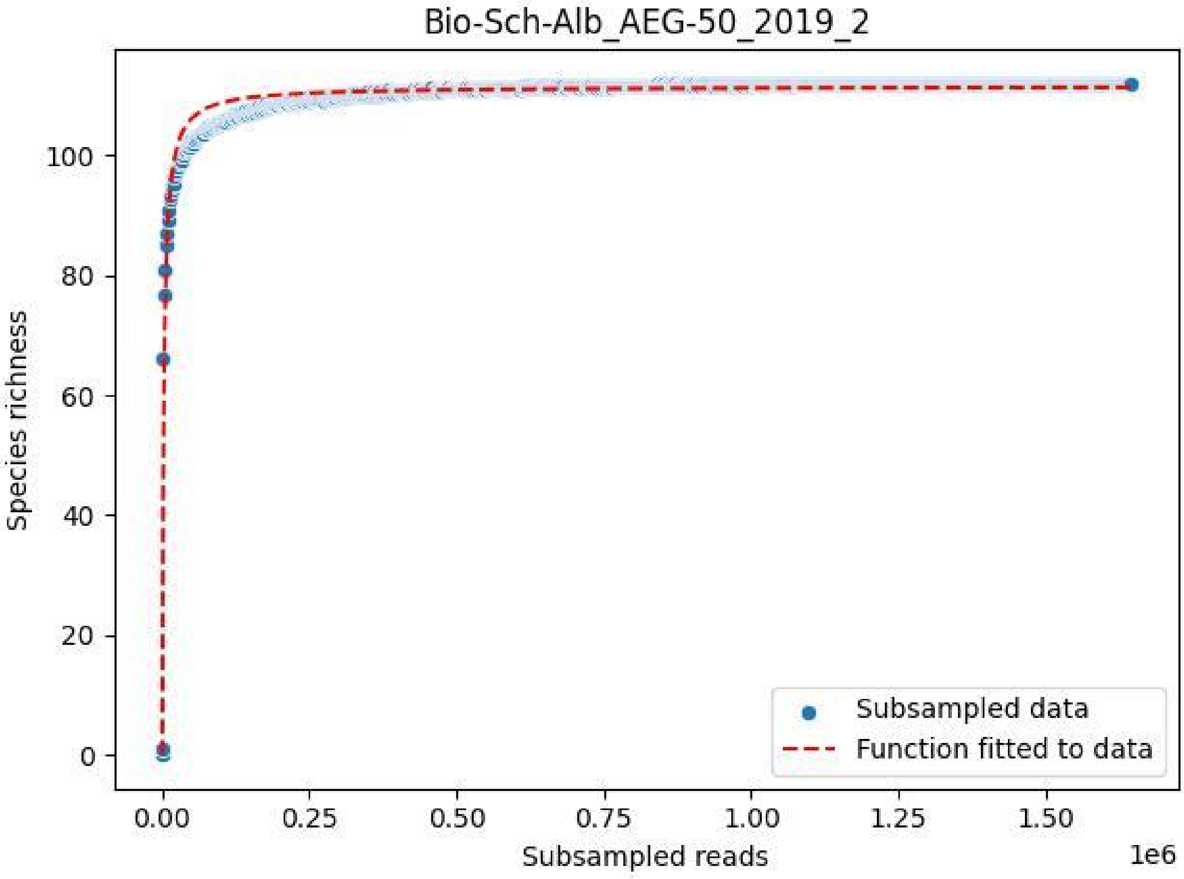

Bio-Sch-Alb_AEG-50_2019_2
Species richness
Subsampled reads
1e6
0
20
40
60
80
100
0.00
0.25
0.50
0.75
1.00
1.25
1.50
Subsampled data
Function fitted to data

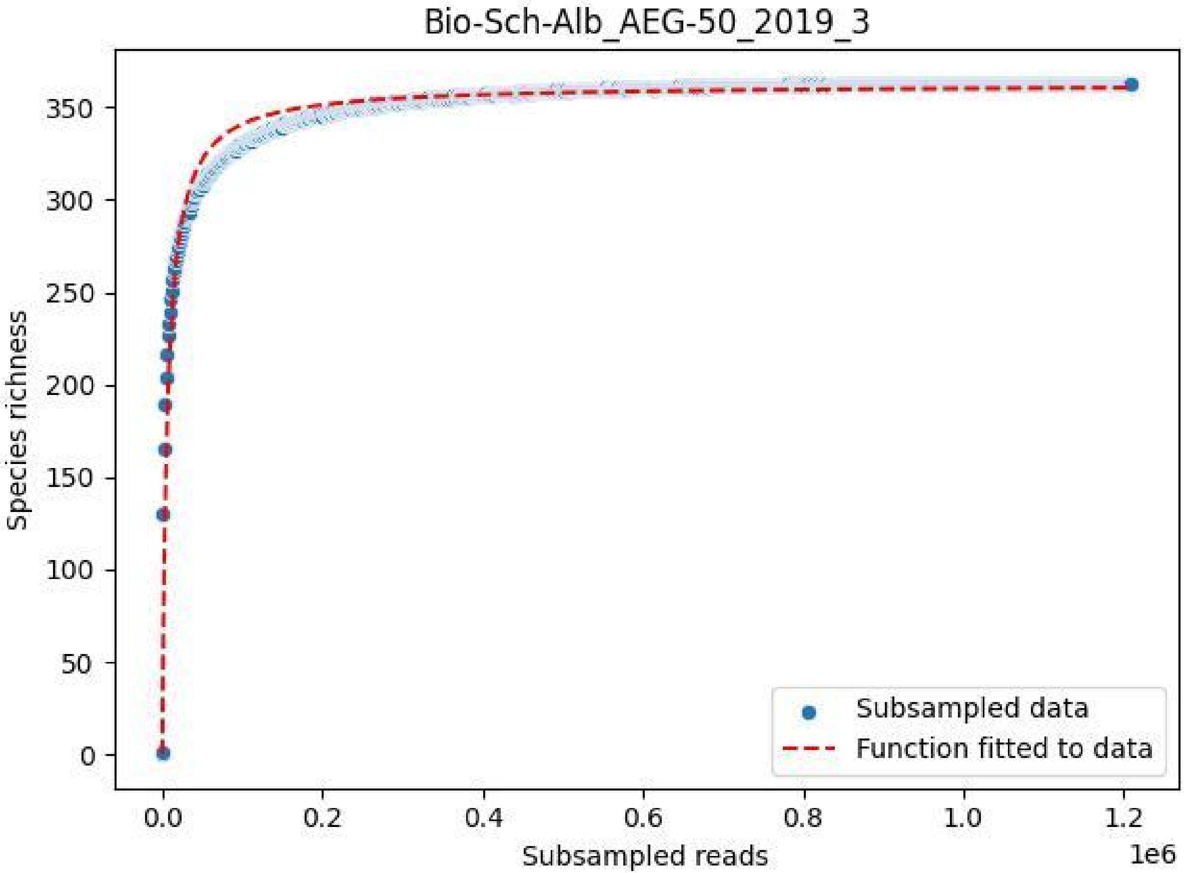
Bio-Sch-Alb_AEG-50_2019_3
Species richness
Subsampled reads
1e6
0
50
100
150
200
250
300
350
0.0
0.2
0.4
0.6
0.8
1.0
1.2
Subsampled data
Function fitted to data

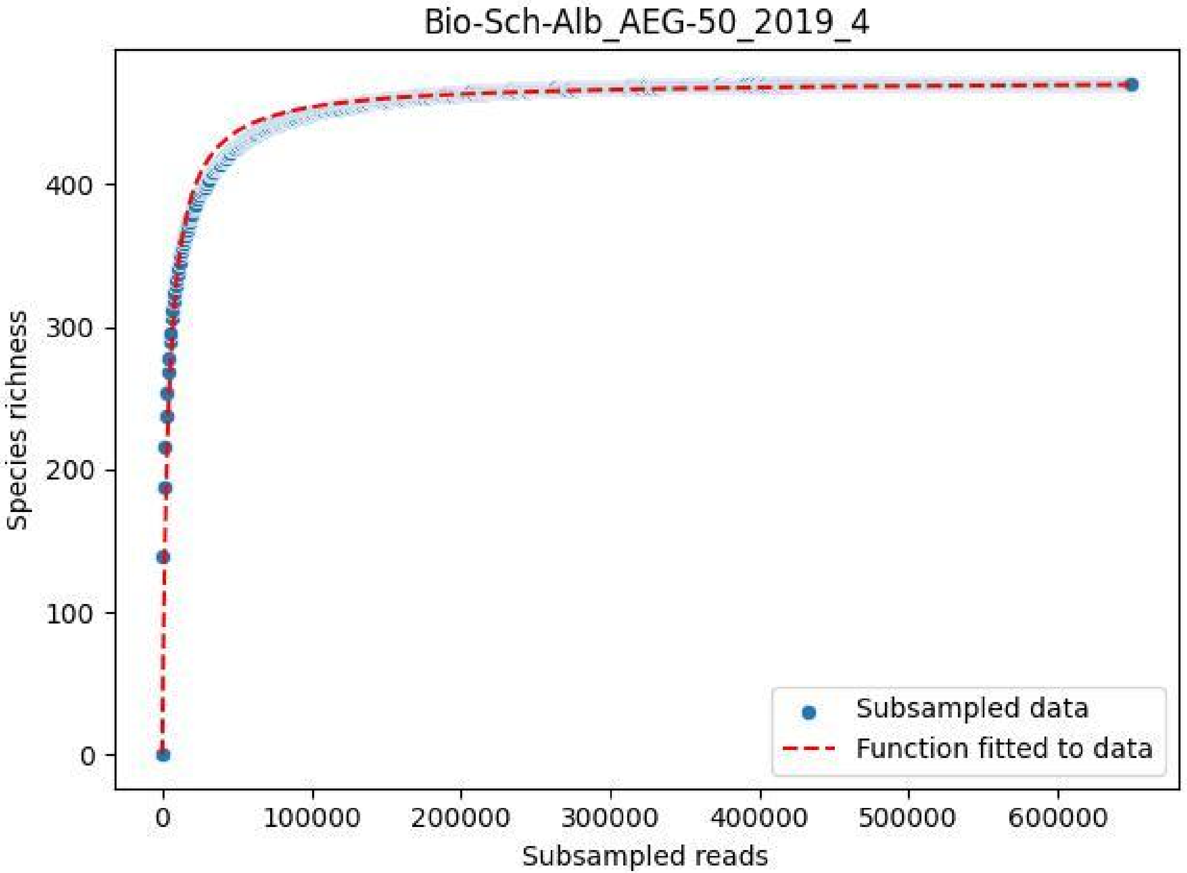
Bio-Sch-Alb_AEG-50_2019_4
Species richness
Subsampled reads
0
100
200
300
400
100000
200000
300000
400000
500000
600000
Subsampled data
Function fitted to data

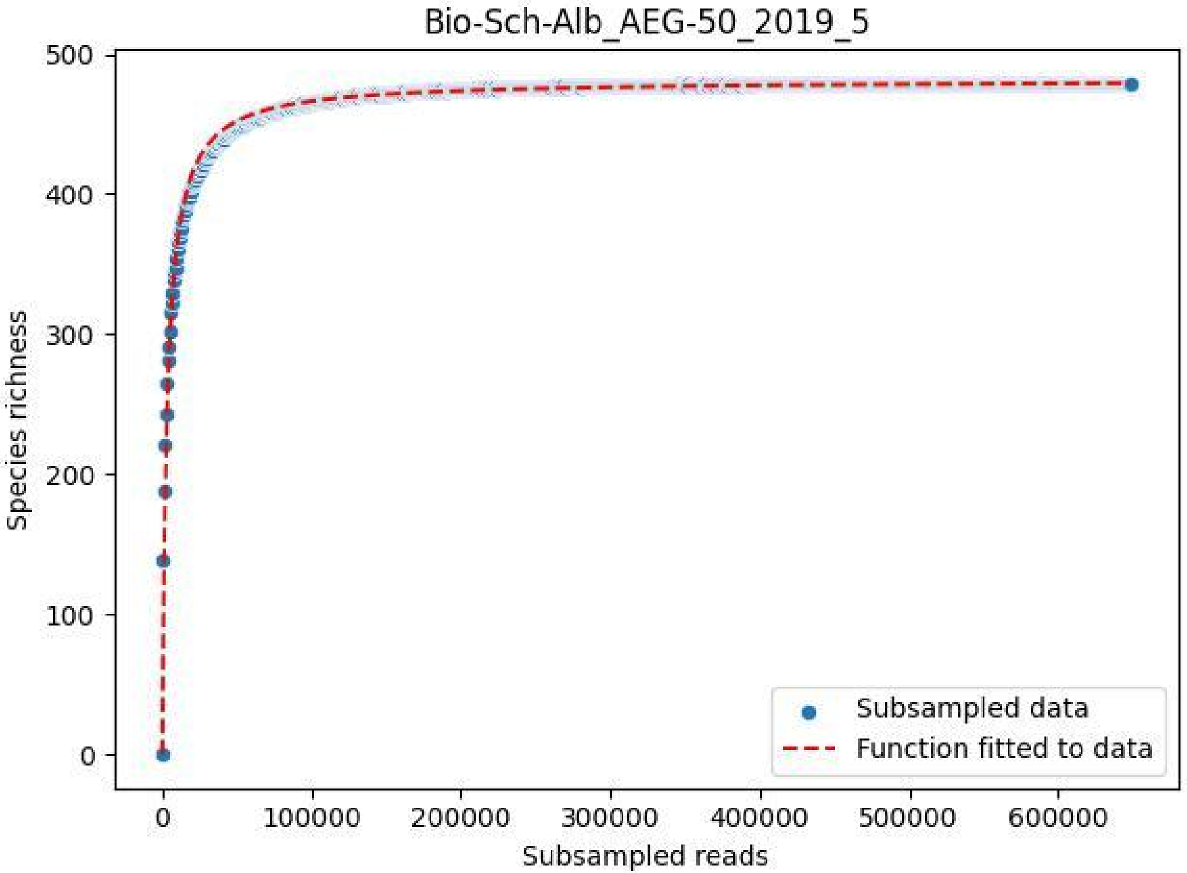
Bio-Sch-Alb_AEG-50_2019_5
Species richness
0
100
200
300
400
500
Subsampled reads
0
100000
200000
300000
400000
500000
600000
Subsampled data
Function fitted to data

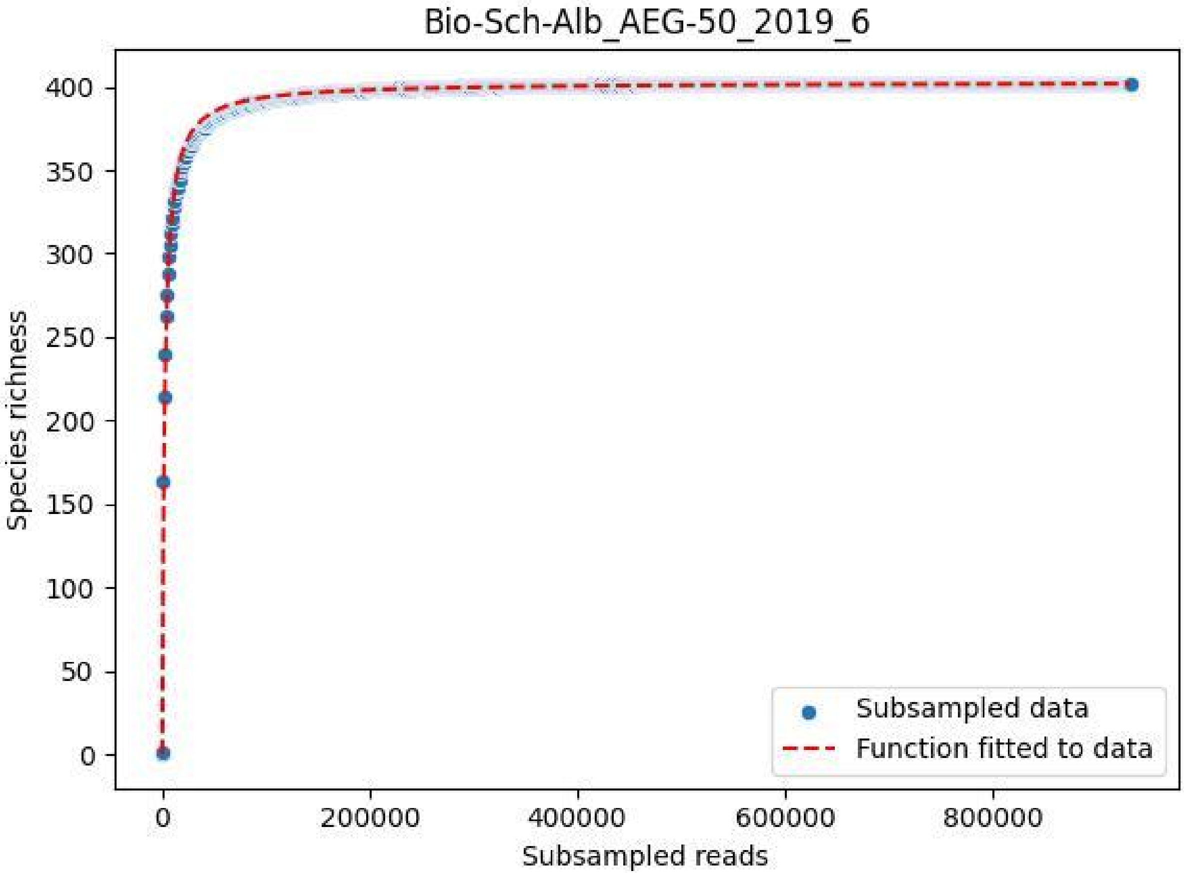
Bio-Sch-Alb_AEG-50_2019_6
Species richness
0
50
100
150
200
250
300
350
400
Subsampled reads
0
200000
400000
600000
800000
Subsampled data
Function fitted to data

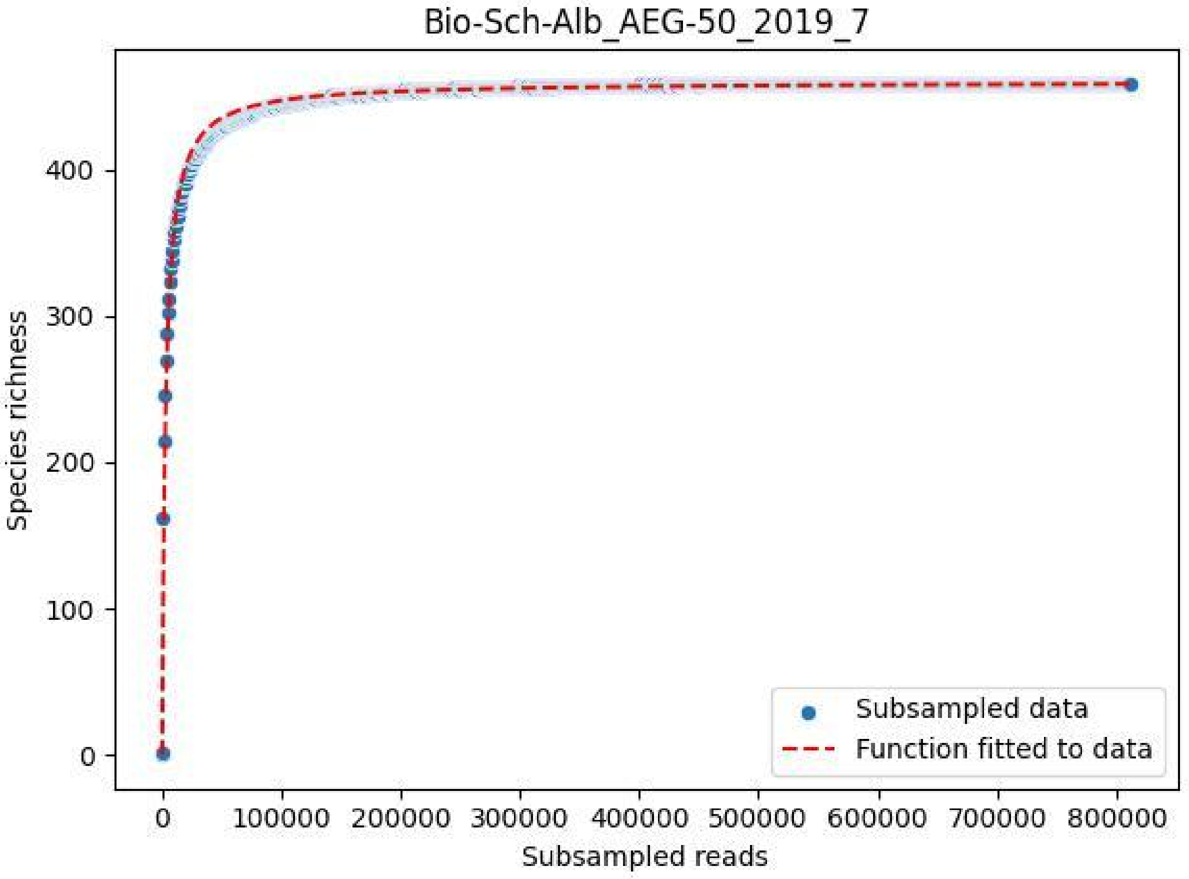

Bio-Sch-Alb_AEG-50_2019_7
Species richness
Subsampled reads
0
100
200
300
400
0
100000
200000
300000
400000
500000
600000
700000
800000
Subsampled data
Function fitted to data

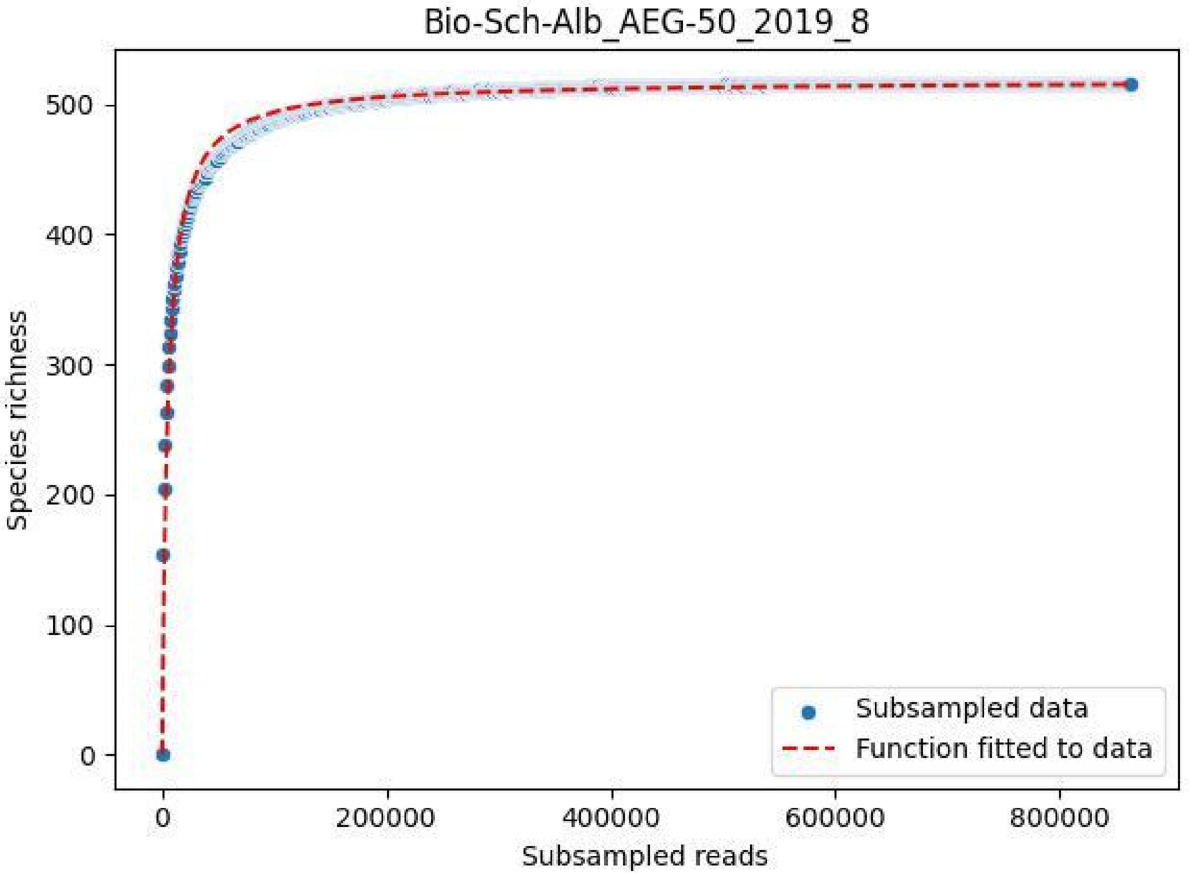
Bio-Sch-Alb_AEG-50_2019_8
Species richness
Subsampled reads
0
100
200
300
400
500
0
200000
400000
600000
800000
Subsampled data
Function fitted to data

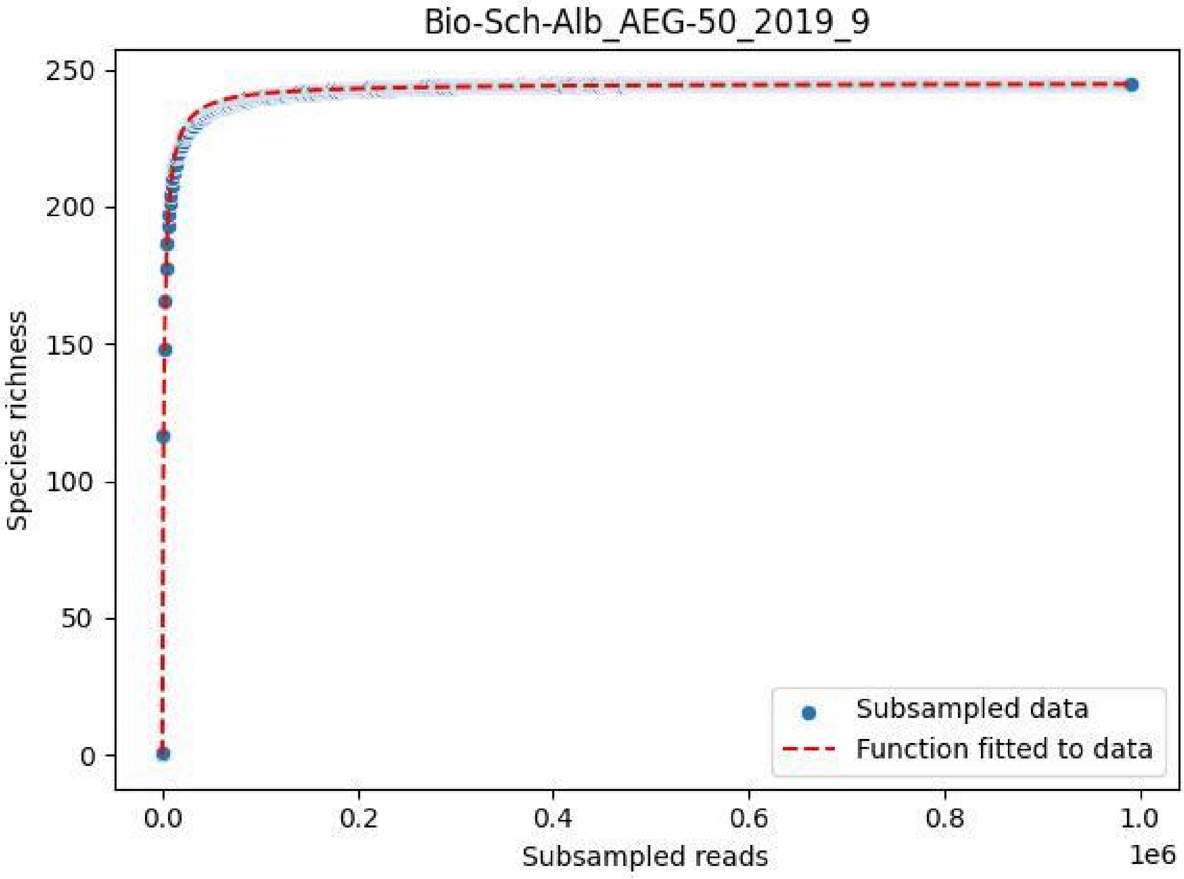

Bio-Sch-Alb_AEG-50_2019_9
250
200
150
100
50
0
Species richness
0.0
0.2
0.4
0.6
0.8
1.0
1e6
Subsampled reads
Subsampled data
Function fitted to data

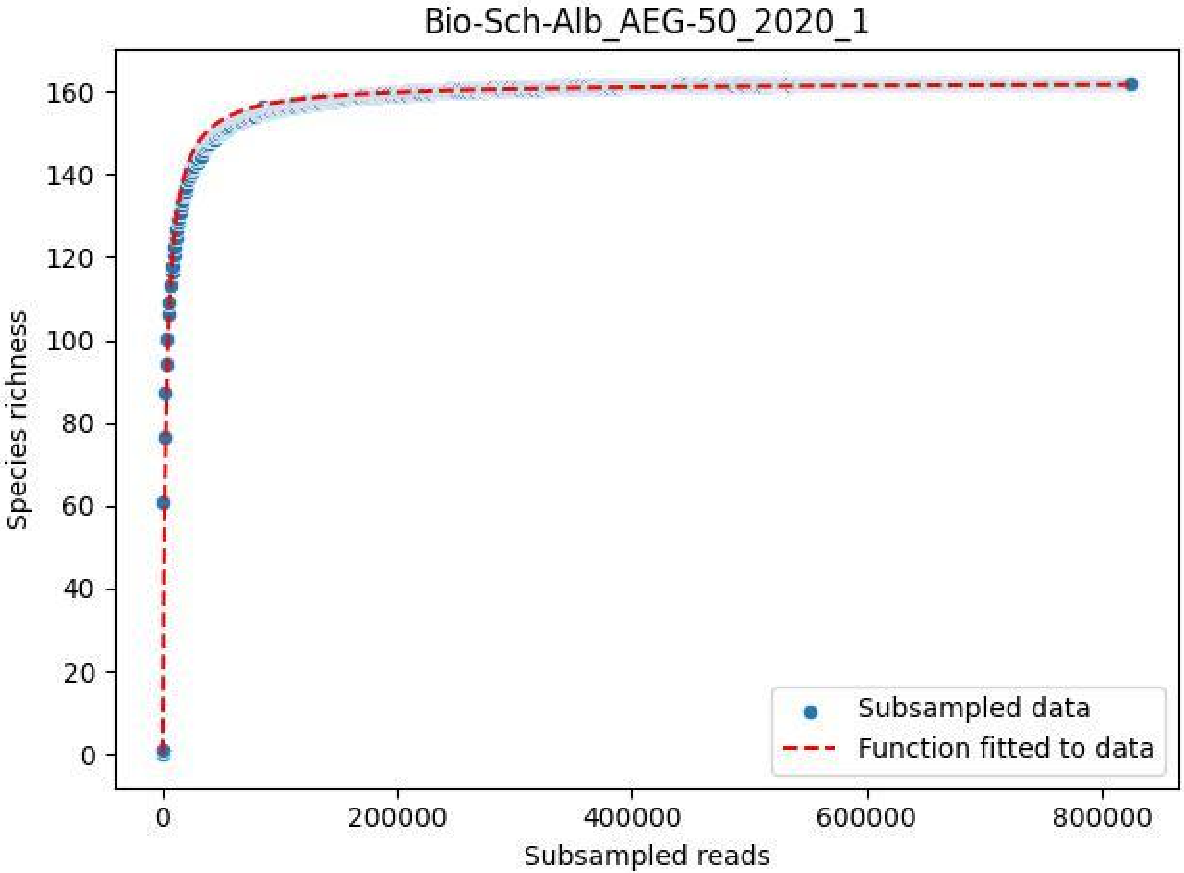

Bio-Sch-Alb_AEG-50_2020_1
Species richness
Subsampled reads
0
20
40
60
80
100
120
140
160
200000
400000
600000
800000
Subsampled data
Function fitted to data

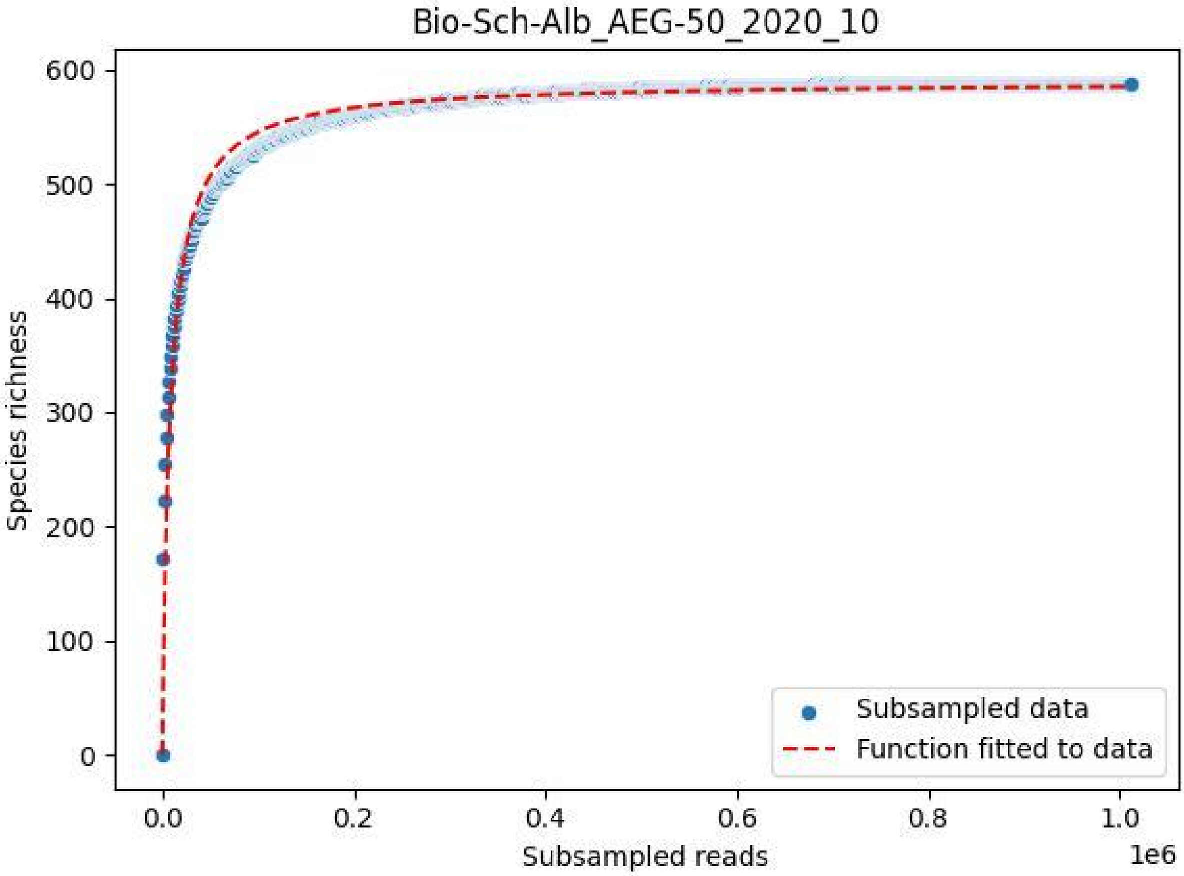
Bio-Sch-Alb_AEG-50_2020_10
Species richness
600
500
400
300
200
100
0
0.0
0.2
0.4
0.6
0.8
1.0
Subsampled reads
1e6
Subsampled data
Function fitted to data

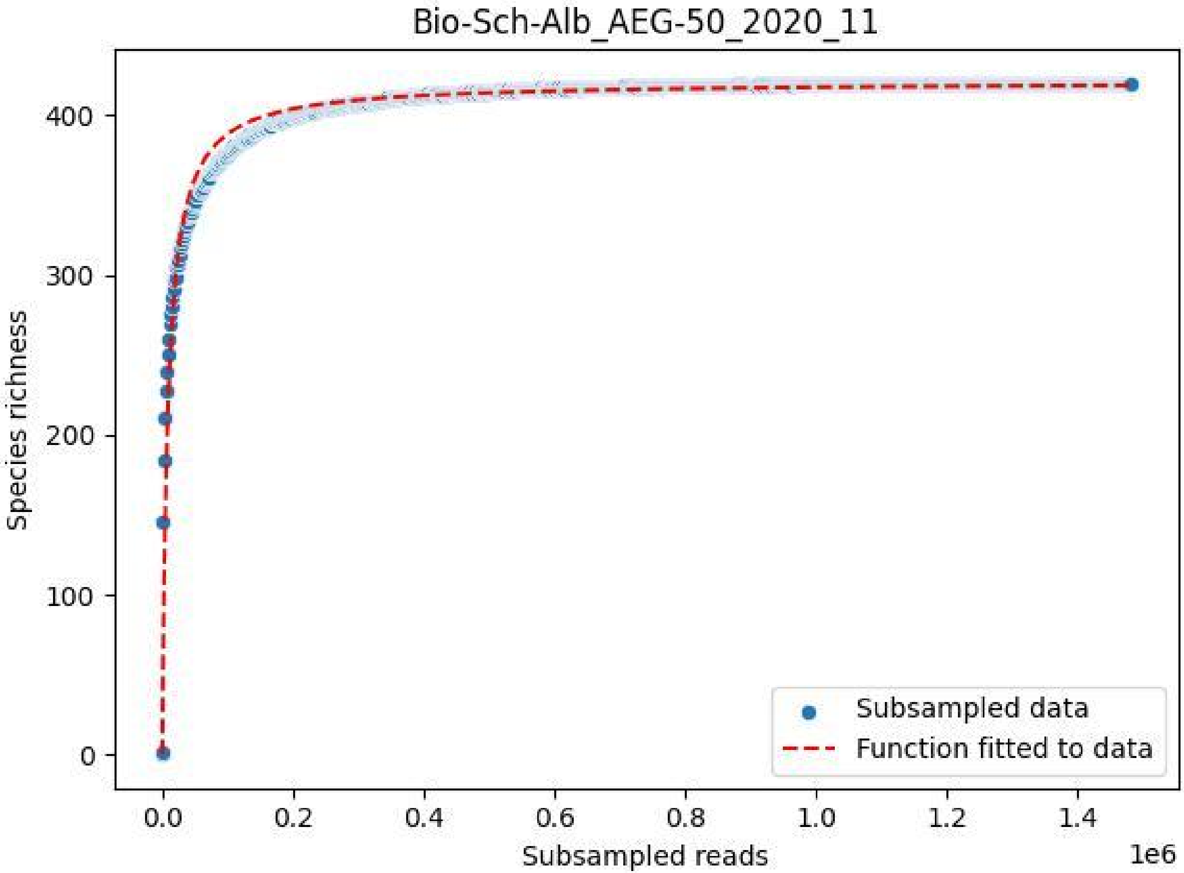
Bio-Sch-Alb_AEG-50_2020_11
Species richness
Subsampled reads
1e6
0
100
200
300
400
0.0
0.2
0.4
0.6
0.8
1.0
1.2
1.4
Subsampled data
Function fitted to data

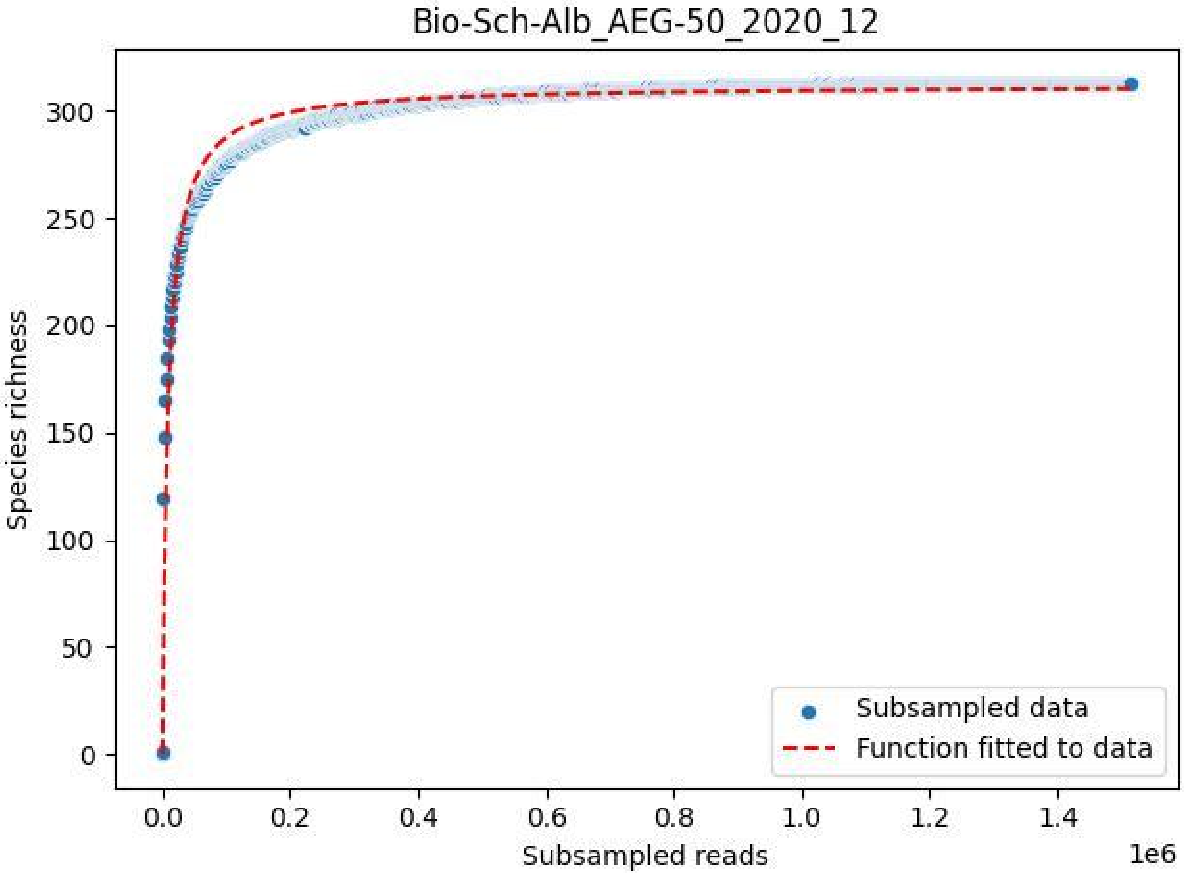

Bio-Sch-Alb_AEG-50_2020_12
Species richness
Subsampled reads
1e6
0
50
100
150
200
250
300
0.0
0.2
0.4
0.6
0.8
1.0
1.2
1.4
Subsampled data
Function fitted to data

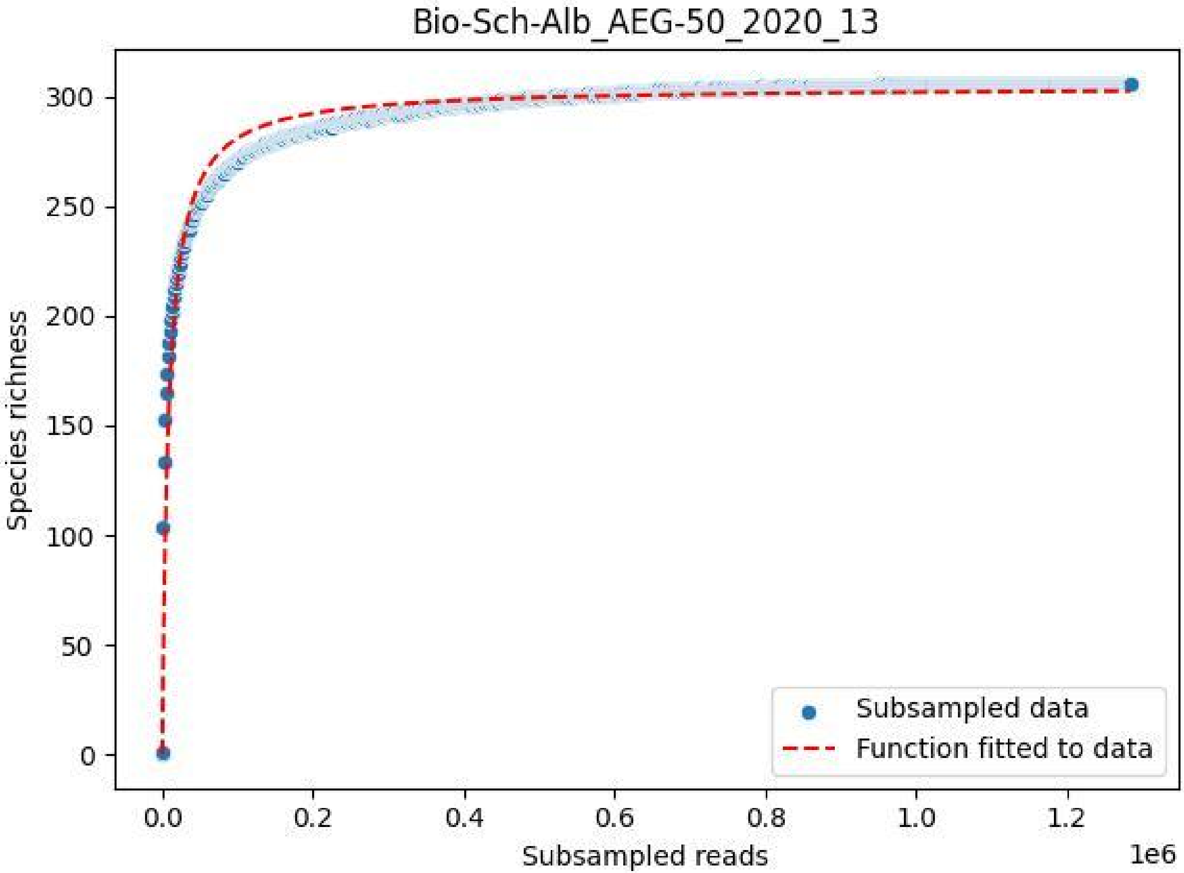

Bio-Sch-Alb_AEG-50_2020_13
Species richness
Subsampled reads
1e6
0
50
100
150
200
250
300
0.0
0.2
0.4
0.6
0.8
1.0
1.2
Subsampled data
Function fitted to data

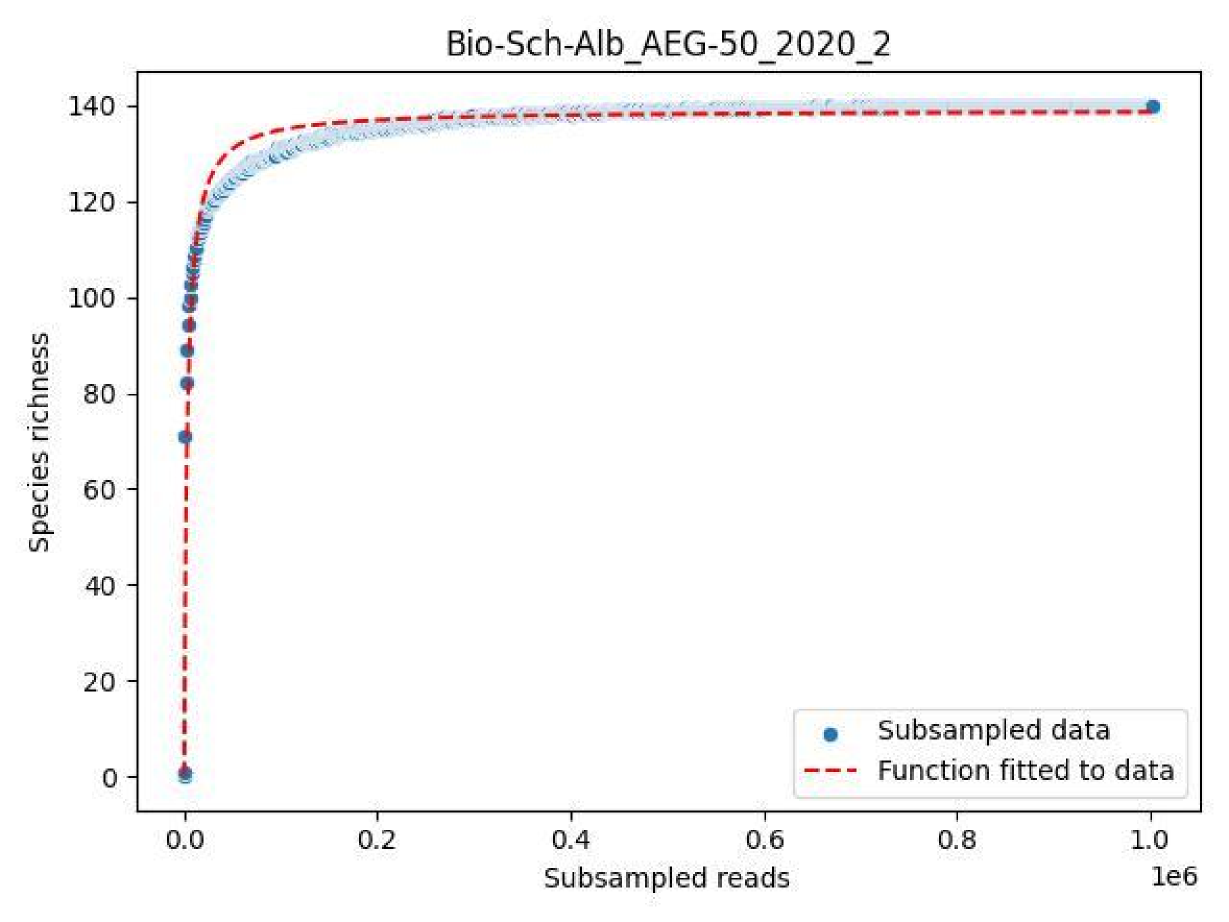
Bio-Sch-Alb_AEG-50_2020_2
Species richness
Subsampled reads
1e6
Subsampled data
Function fitted to data
0
20
40
60
80
100
120
140
0.0
0.2
0.4
0.6
0.8
1.0

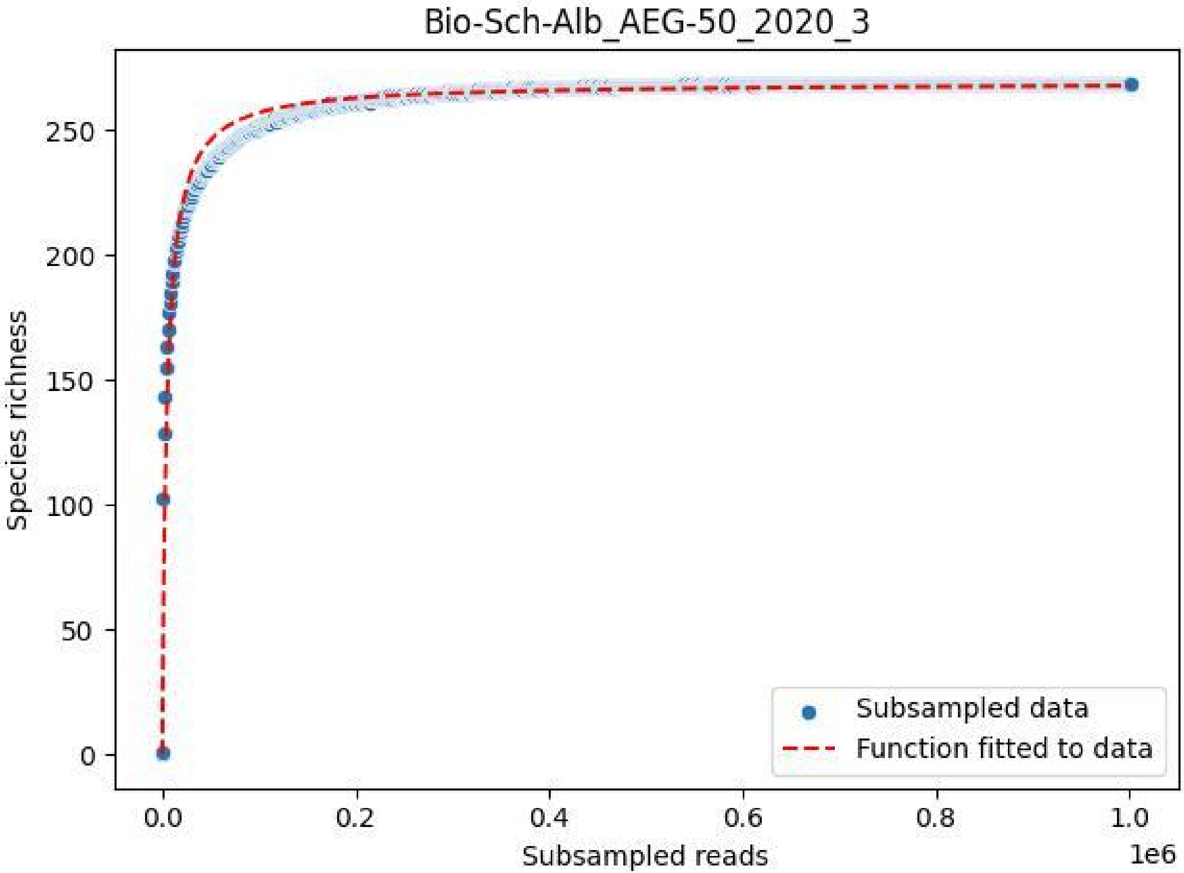
Bio-Sch-Alb_AEG-50_2020_3
Species richness
Subsampled reads
1e6
0
50
100
150
200
250
0.0
0.2
0.4
0.6
0.8
1.0
Subsampled data
Function fitted to data

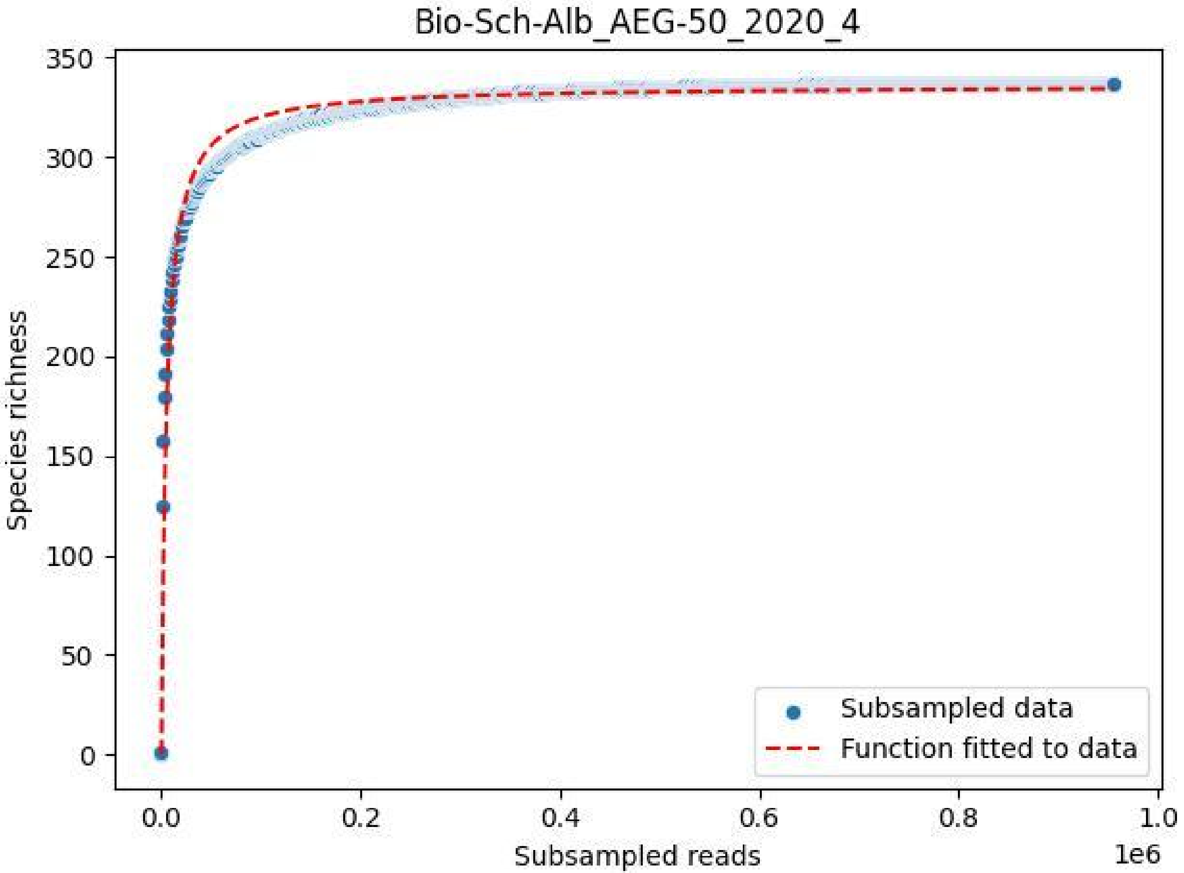
Bio-Sch-Alb_AEG-50_2020_4
Species richness
Subsampled reads
1e6
0.0
0.2
0.4
0.6
0.8
1.0
0
50
100
150
200
250
300
350
Subsampled data
Function fitted to data

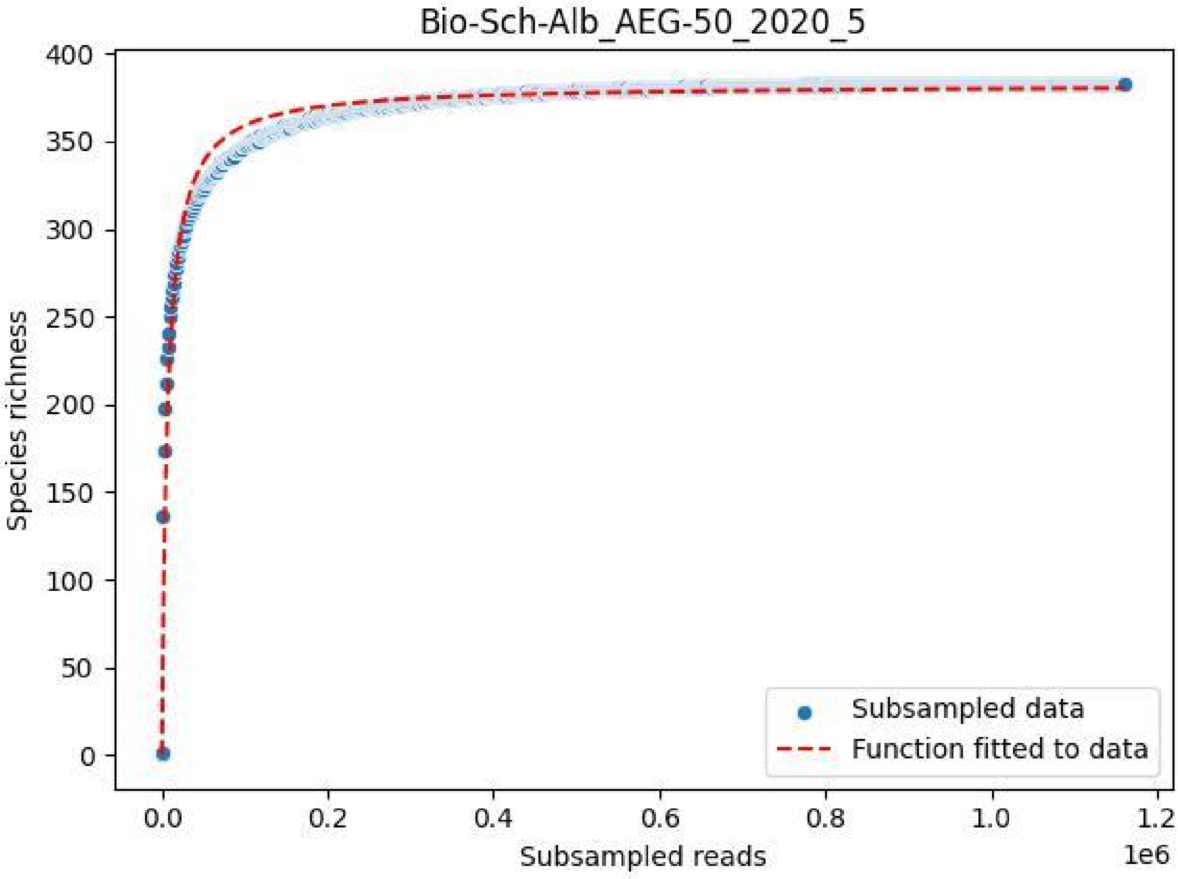
Bio-Sch-Alb_AEG-50_2020_5
Species richness
Subsampled reads
1e6
Subsampled data
Function fitted to data
400
350
300
250
200
150
100
50
0
0.0
0.2
0.4
0.6
0.8
1.0
1.2

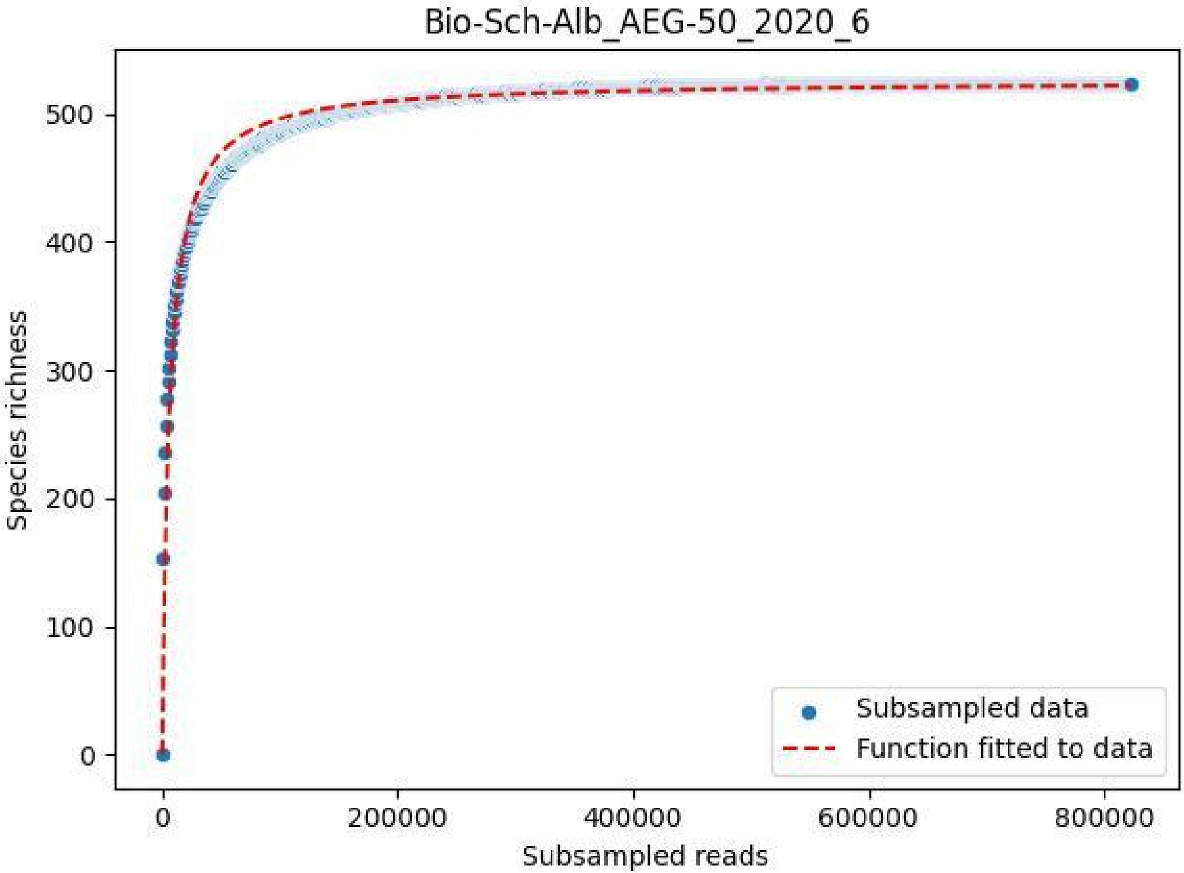

Bio-Sch-Alb_AEG-50_2020_6
Species richness
Subsampled reads
0
100
200
300
400
500
0
200000
400000
600000
800000
Subsampled data
Function fitted to data

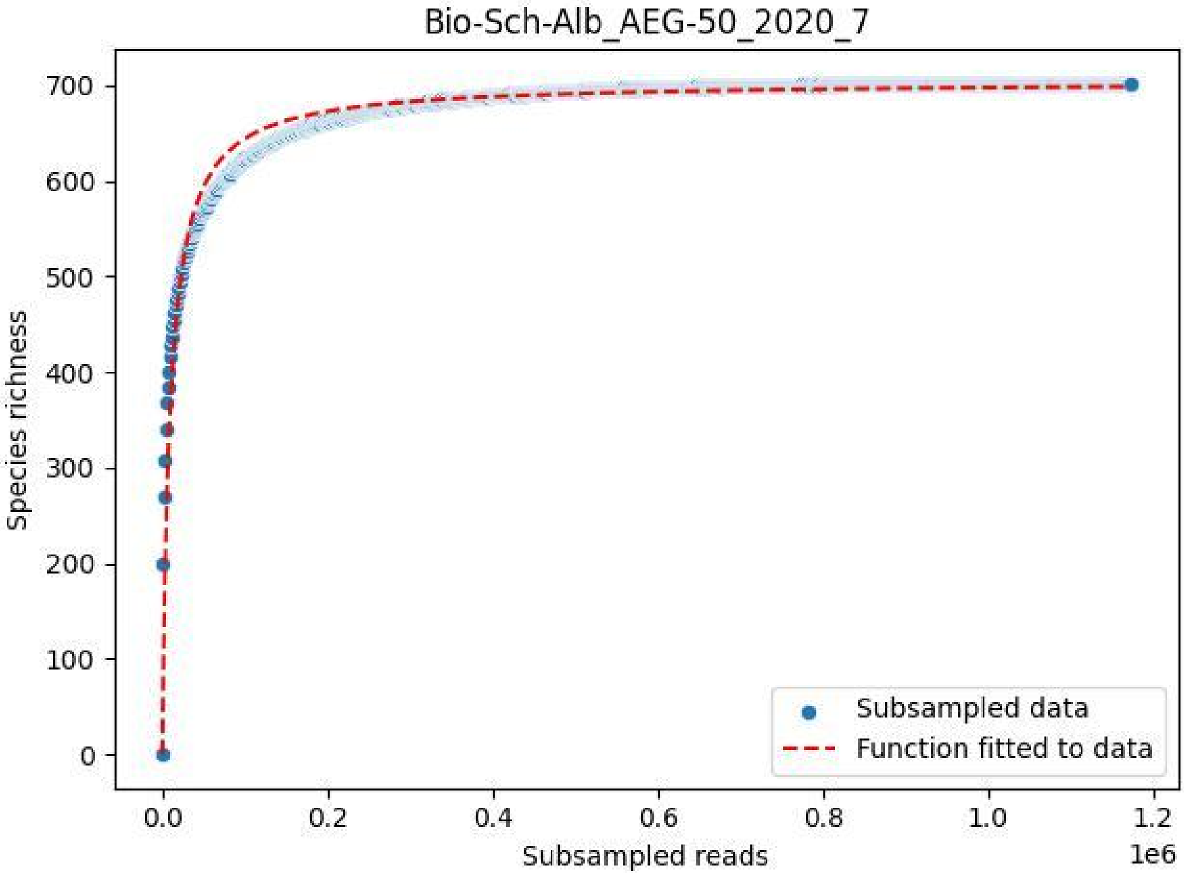

Bio-Sch-Alb_AEG-50_2020_7
Species richness
0
100
200
300
400
500
600
700
Subsampled reads
0.0
0.2
0.4
0.6
0.8
1.0
1.2
1e6
Subsampled data
Function fitted to data

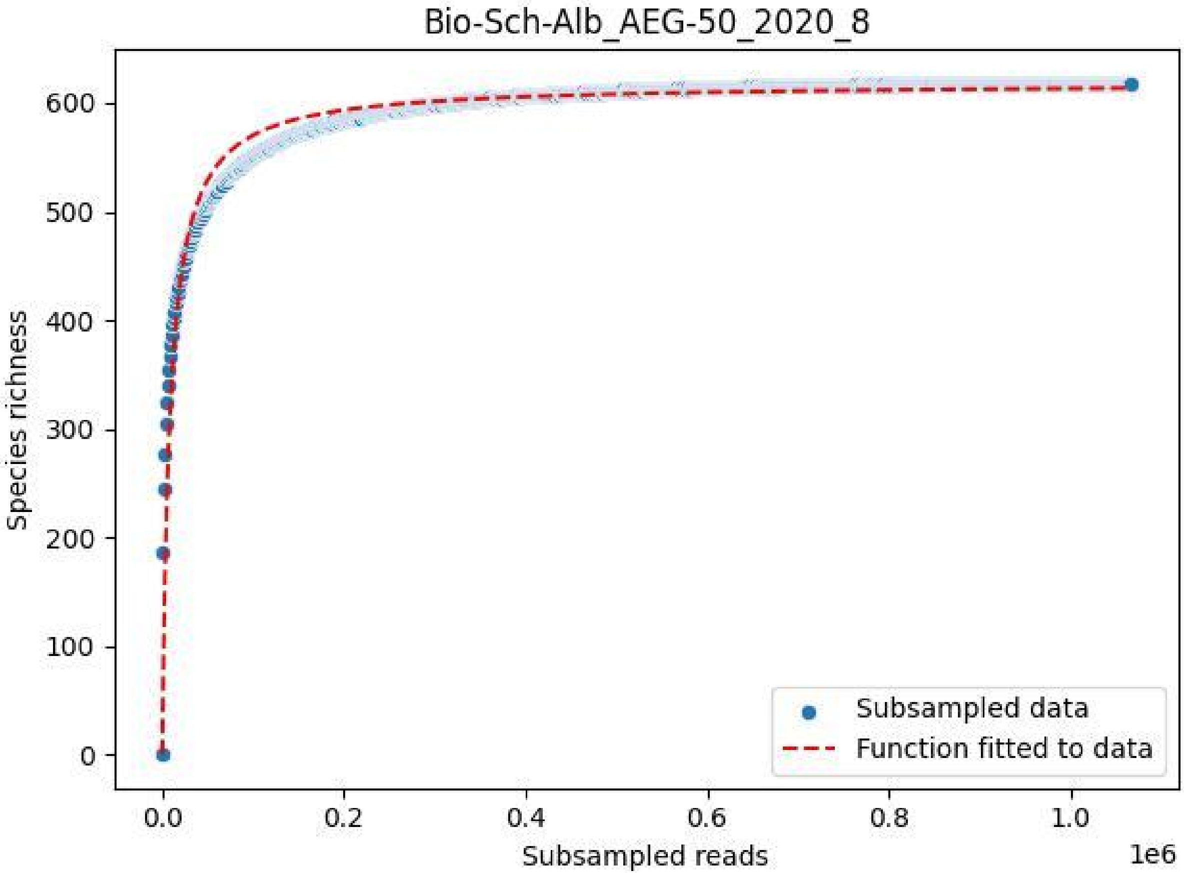
Bio-Sch-Alb_AEG-50_2020_8
Species richness
Subsampled reads
1e6
0
100
200
300
400
500
600
0.0
0.2
0.4
0.6
0.8
1.0
Subsampled data
Function fitted to data

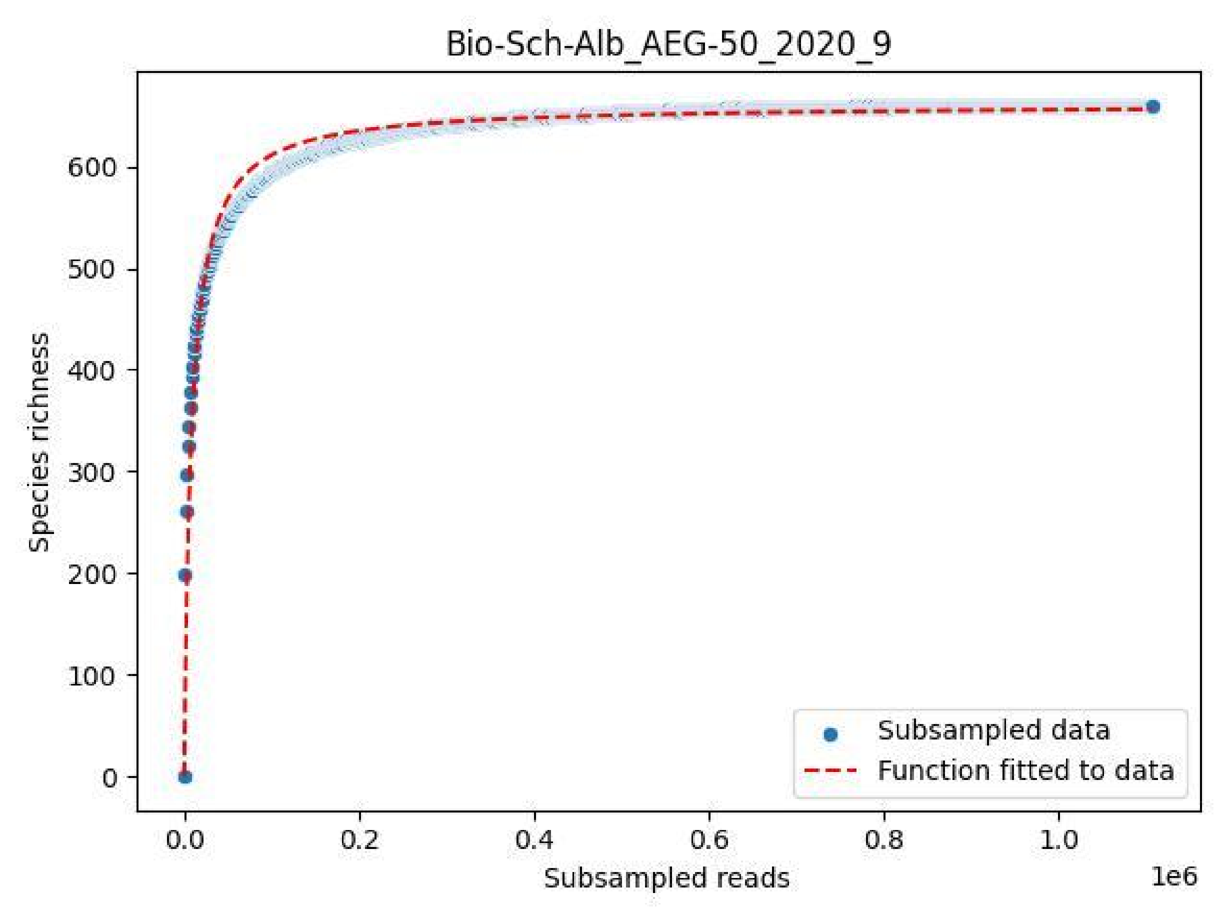

Bio-Sch-Alb_AEG-50_2020_9
Species richness
Subsampled reads
1e6
0
100
200
300
400
500
600
0.0
0.2
0.4
0.6
0.8
1.0
Subsampled data
Function fitted to data

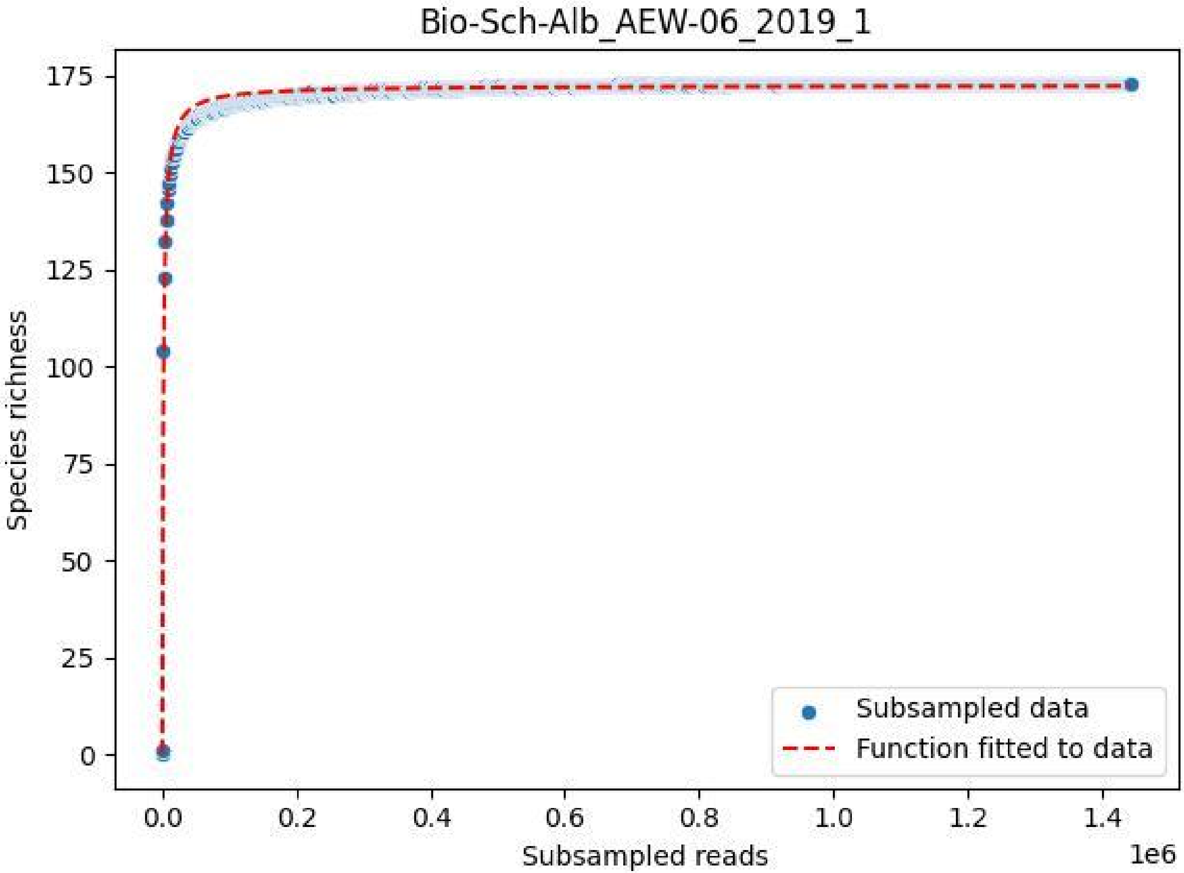
Bio-Sch-Alb_AEW-06_2019_1
Species richness
Subsampled reads
1e6
175
150
125
100
75
50
25
0
0.0
0.2
0.4
0.6
0.8
1.0
1.2
1.4
Subsampled data
Function fitted to data

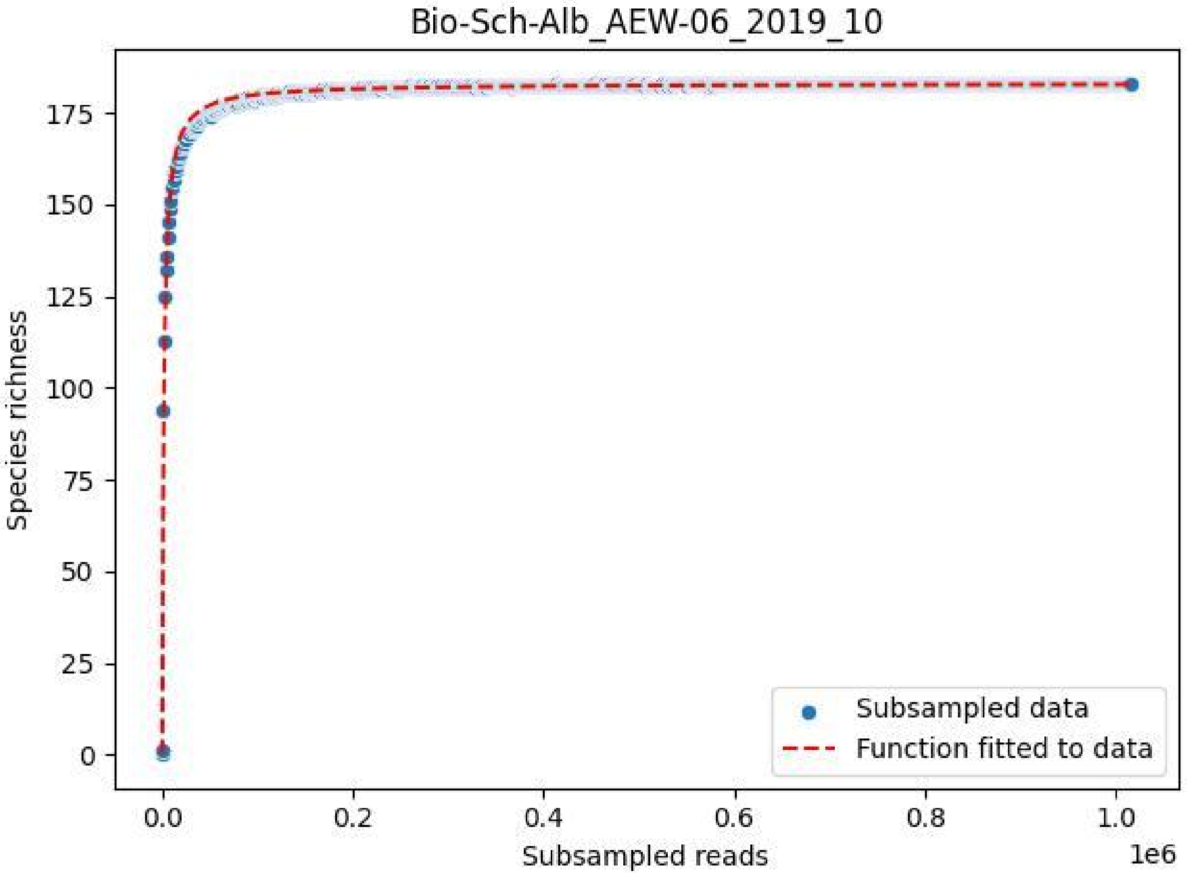
Bio-Sch-Alb_AEW-06_2019_10
Species richness
Subsampled reads
1e6
0
25
50
75
100
125
150
175
0.0
0.2
0.4
0.6
0.8
1.0
Subsampled data
Function fitted to data

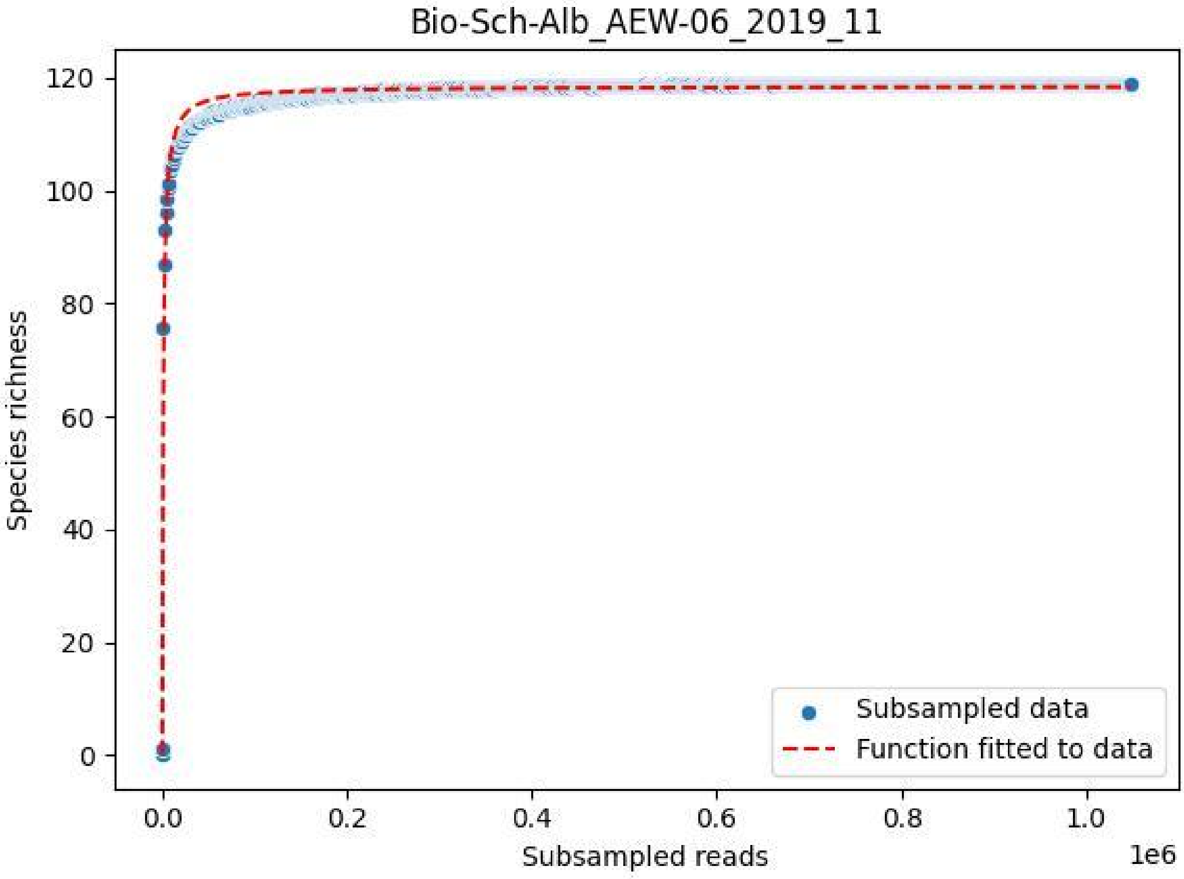

Bio-Sch-Alb_AEW-06_2019_11
Species richness
Subsampled reads
1e6
0
20
40
60
80
100
120
0.0
0.2
0.4
0.6
0.8
1.0
Subsampled data
Function fitted to data

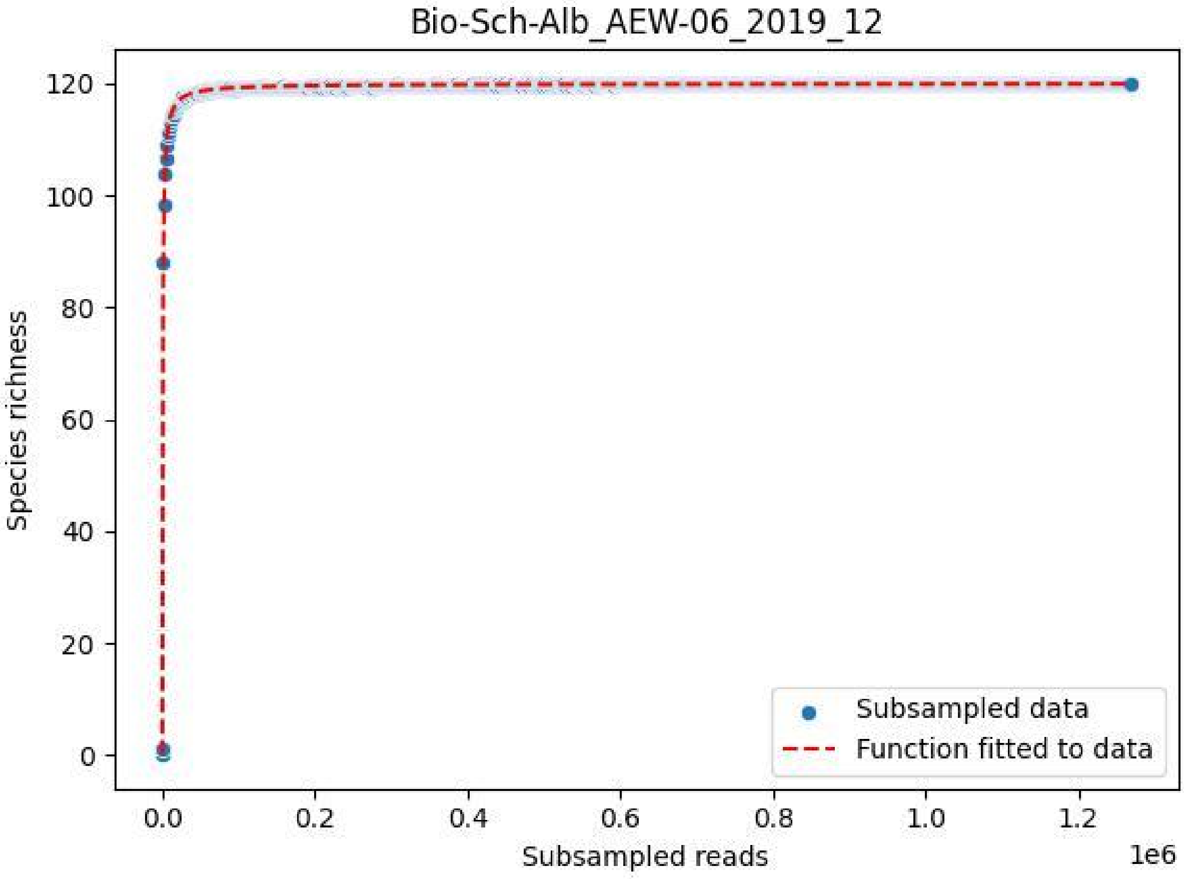
Bio-Sch-Alb_AEW-06_2019_12
Species richness
0
20
40
60
80
100
120
0.0
0.2
0.4
0.6
0.8
1.0
1.2
1e6
Subsampled reads
Subsampled data
Function fitted to data

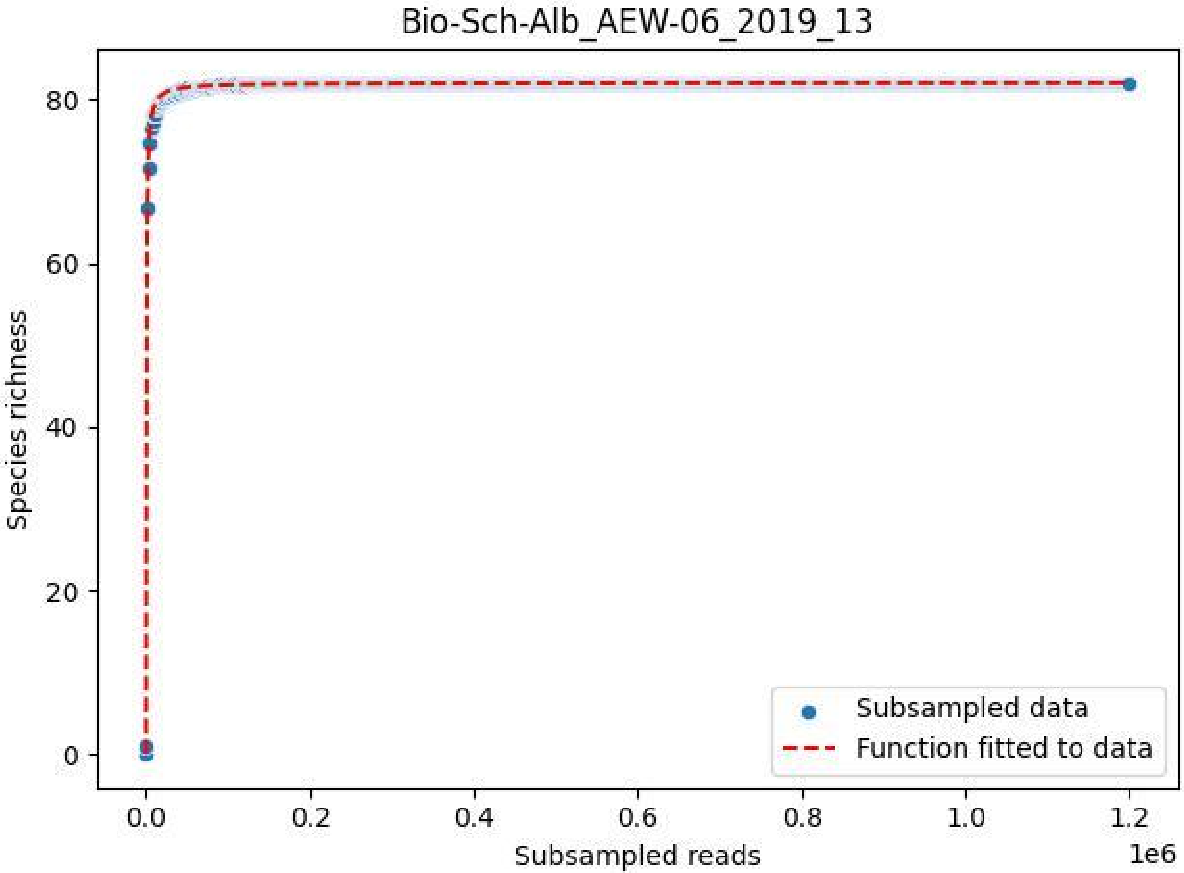

Bio-Sch-Alb_AEW-06_2019_13
Species richness
Subsampled reads
1e6
0
20
40
60
80
0.0
0.2
0.4
0.6
0.8
1.0
1.2
Subsampled data
Function fitted to data

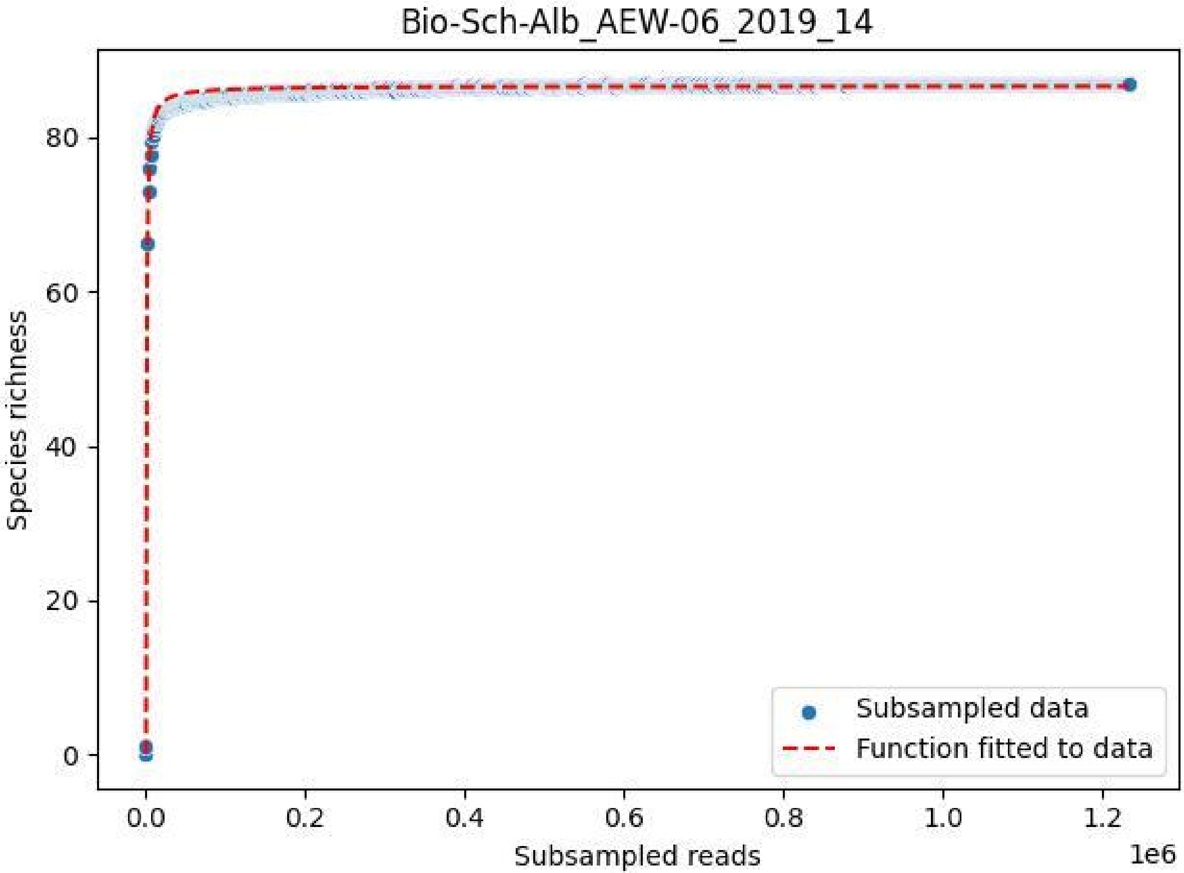
Bio-Sch-Alb_AEW-06_2019_14
Species richness
Subsampled reads
1e6
0
20
40
60
80
0.0
0.2
0.4
0.6
0.8
1.0
1.2
Subsampled data
Function fitted to data

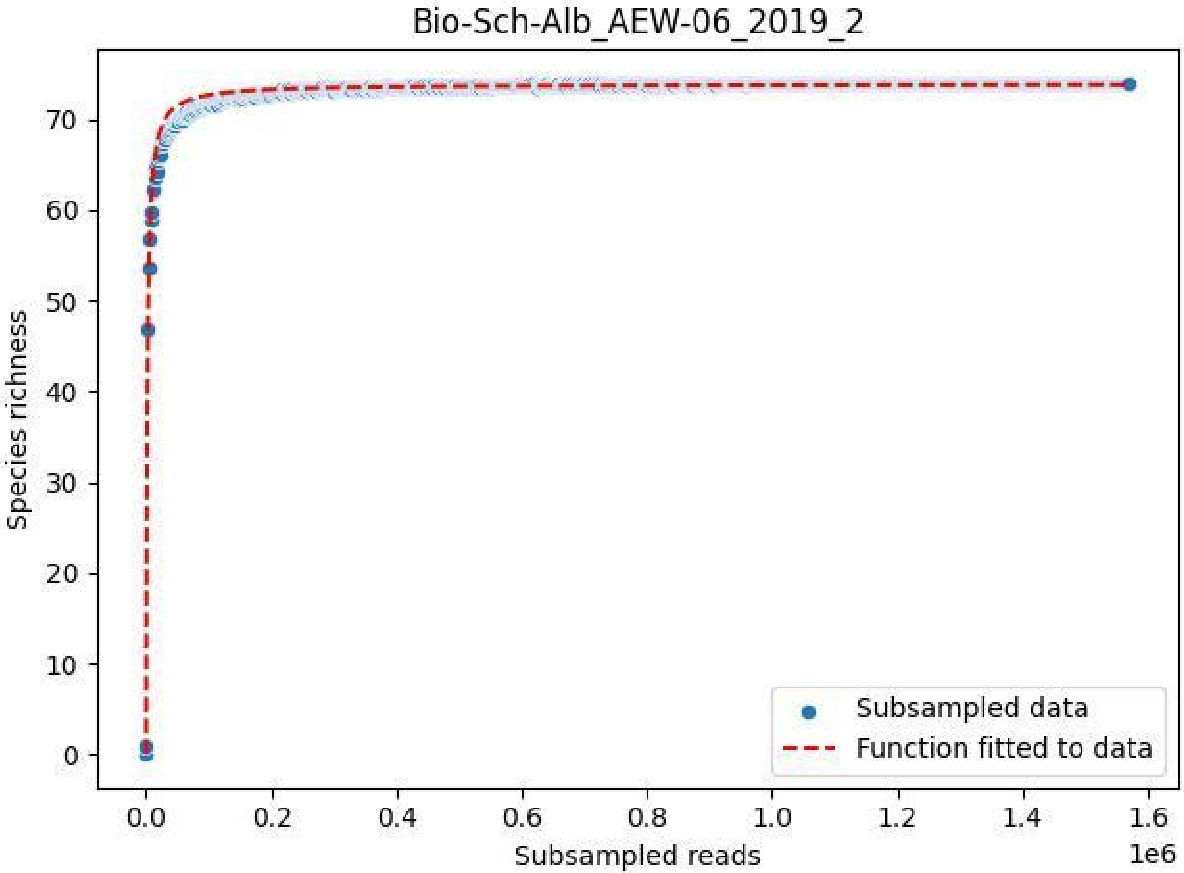

Bio-Sch-Alb_AEW-06_2019_2
Species richness
Subsampled reads
1e6
0
10
20
30
40
50
60
70
0.0
0.2
0.4
0.6
0.8
1.0
1.2
1.4
1.6
Subsampled data
Function fitted to data

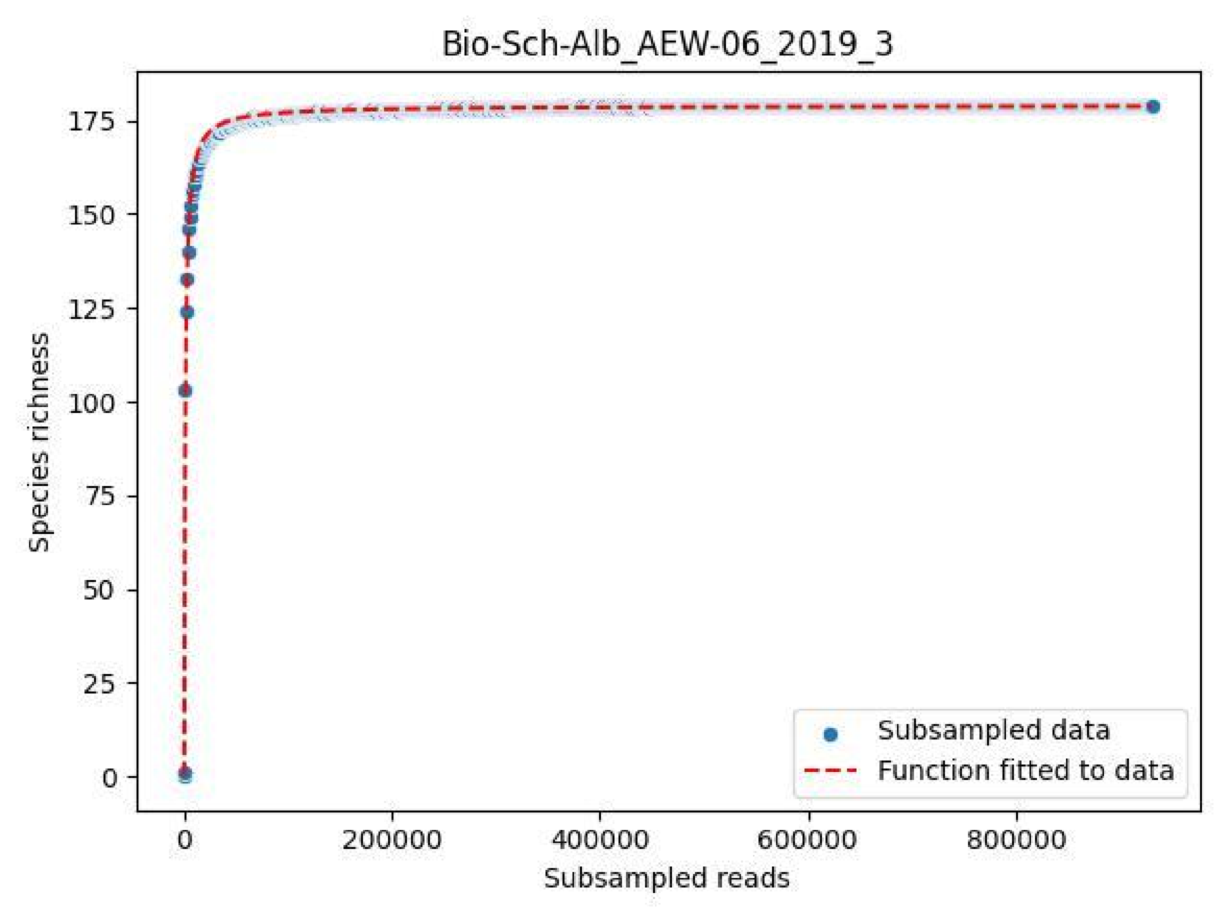
Bio-Sch-Alb_AEW-06_2019_3
Species richness
0
25
50
75
100
125
150
175
Subsampled reads
0
200000
400000
600000
800000
Subsampled data
Function fitted to data

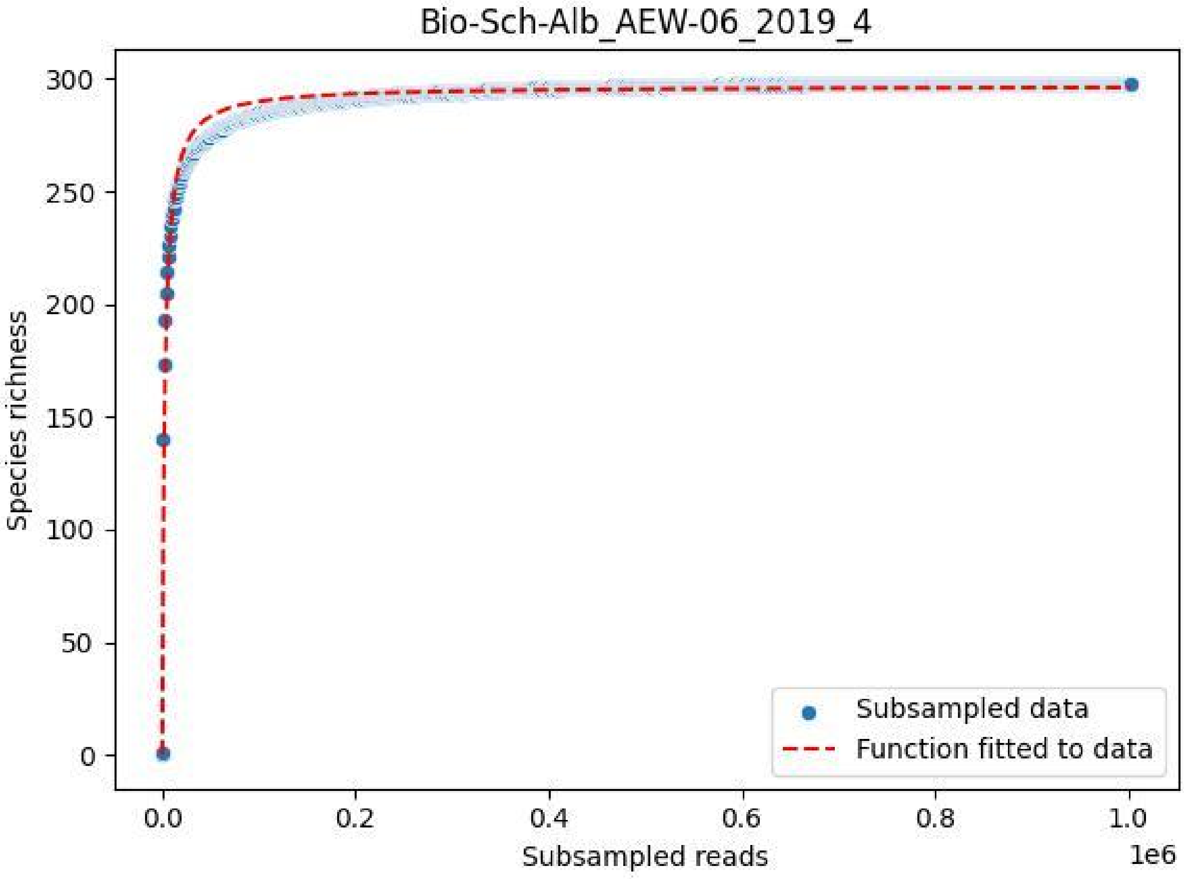

Bio-Sch-Alb_AEW-06_2019_4
300
250
200
150
100
50
0
Species richness
0.0
0.2
0.4
0.6
0.8
1.0
1e6
Subsampled reads
Subsampled data
Function fitted to data

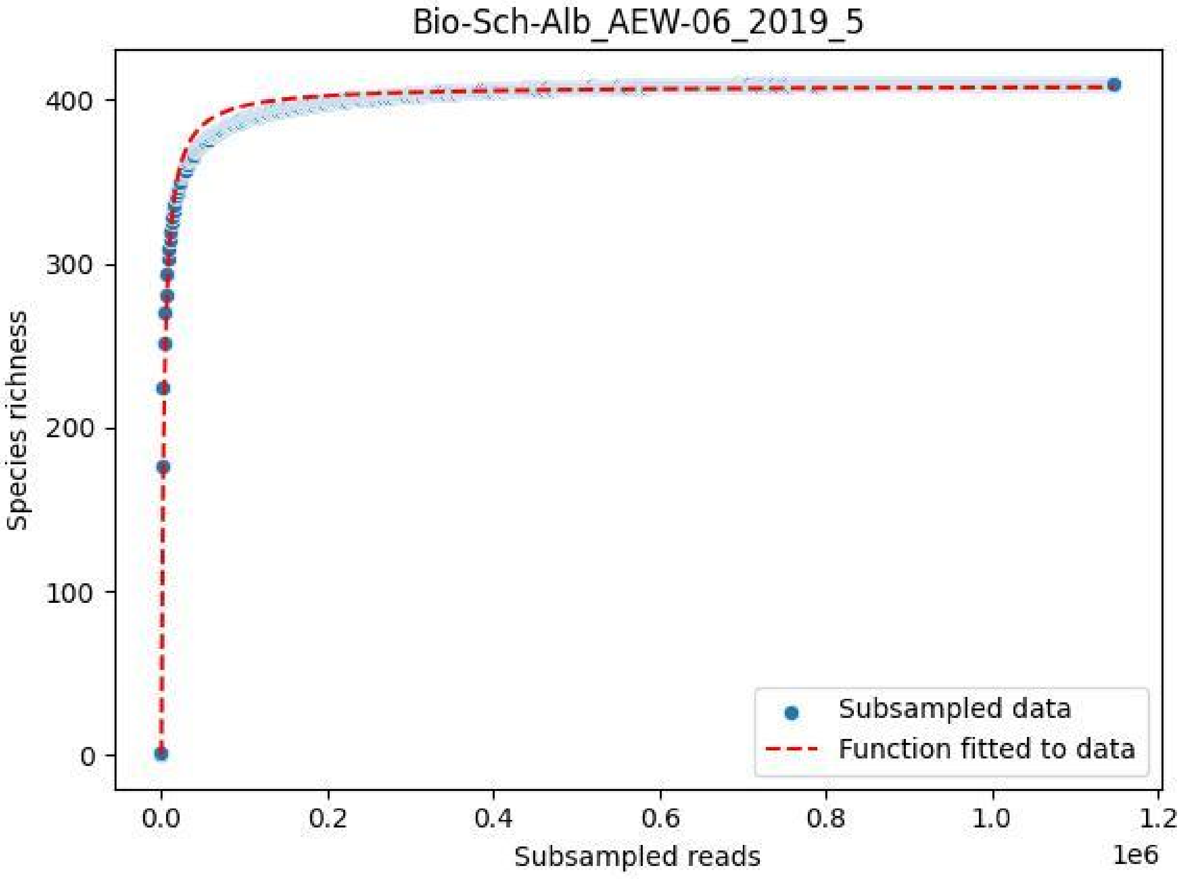
Bio-Sch-Alb_AEW-06_2019_5
Species richness
0
100
200
300
400
Subsampled reads
0.0
0.2
0.4
0.6
0.8
1.0
1.2
1e6
Subsampled data
Function fitted to data

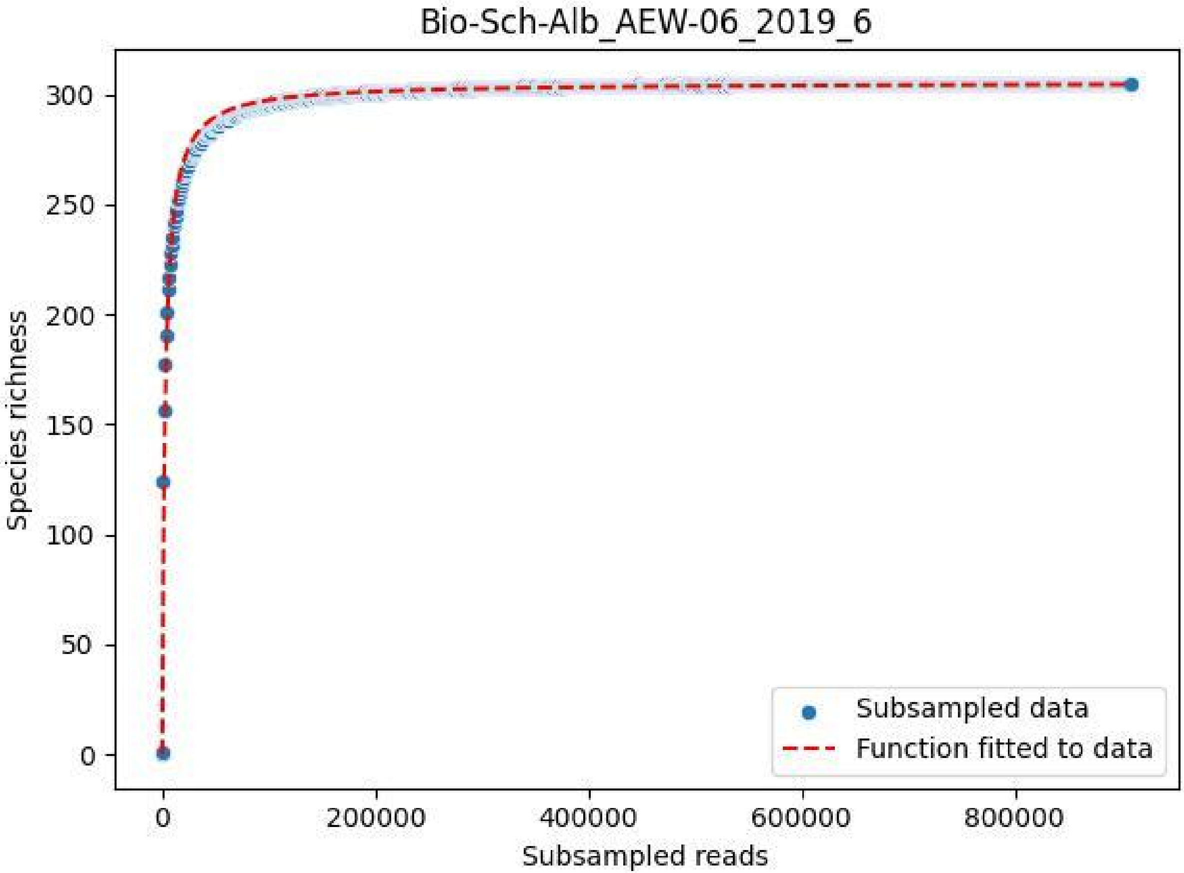

Bio-Sch-Alb_AEW-06_2019_6
Species richness
Subsampled reads
0
50
100
150
200
250
300
0
200000
400000
600000
800000
Subsampled data
Function fitted to data

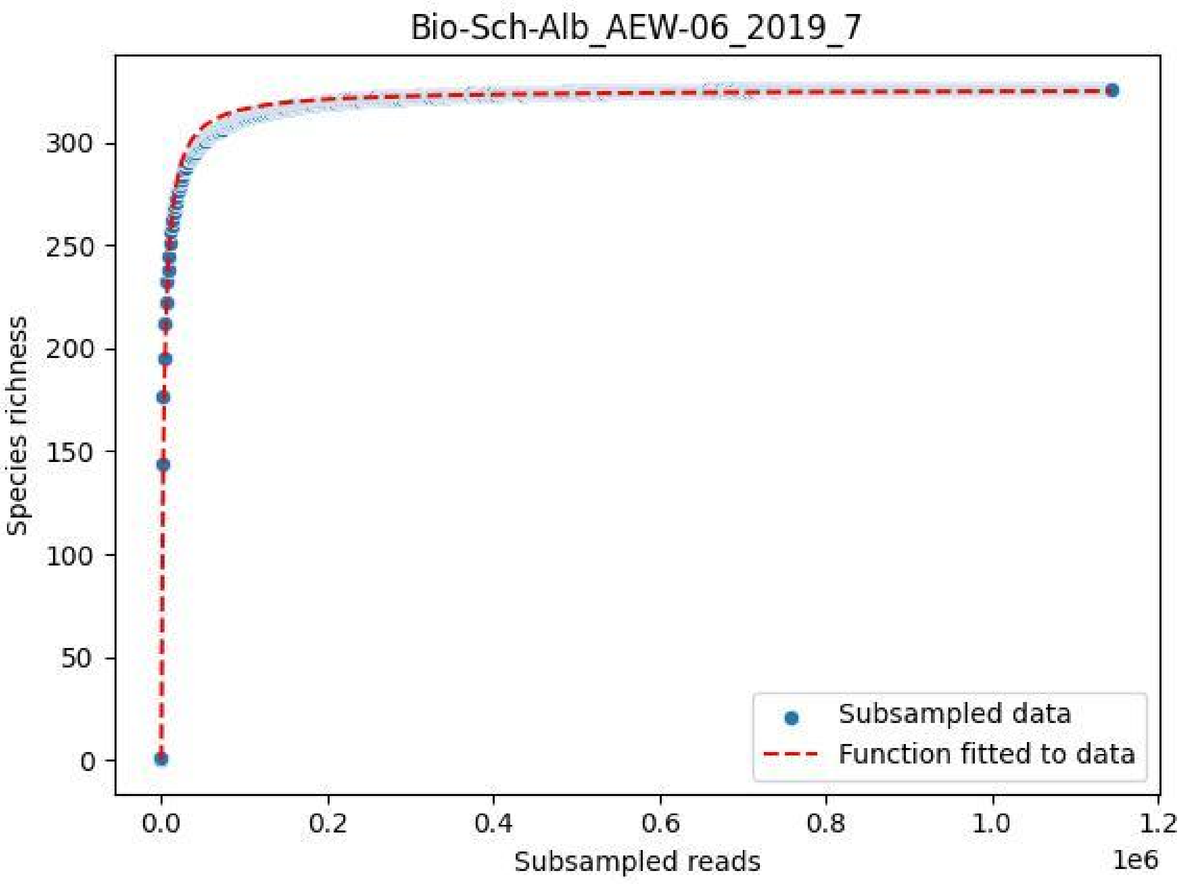
Bio-Sch-Alb_AEW-06_2019_7
Species richness
Subsampled reads
1e6
0
50
100
150
200
250
300
0.0
0.2
0.4
0.6
0.8
1.0
1.2
Subsampled data
Function fitted to data

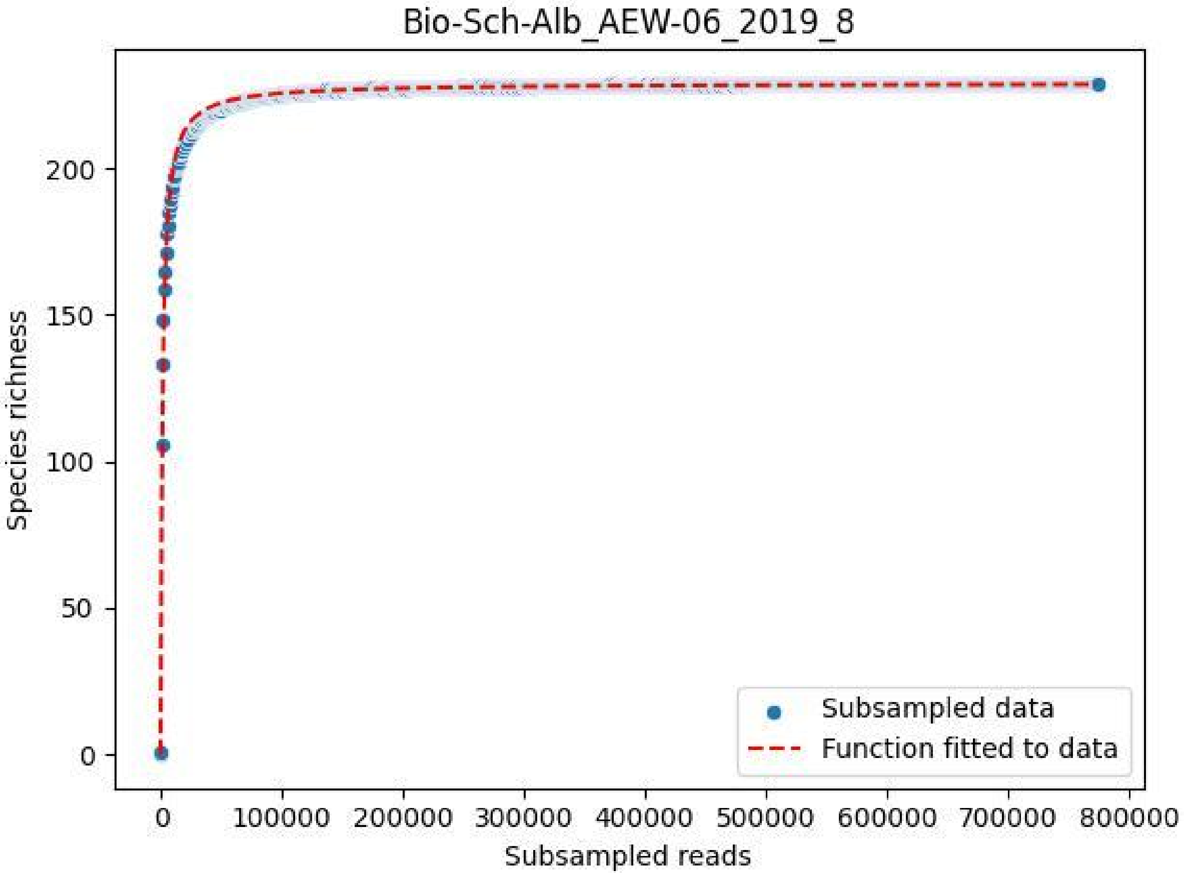
Bio-Sch-Alb_AEW-06_2019_8
Species richness
Subsampled reads
0
50
100
150
200
0
100000
200000
300000
400000
500000
600000
700000
800000
Subsampled data
Function fitted to data

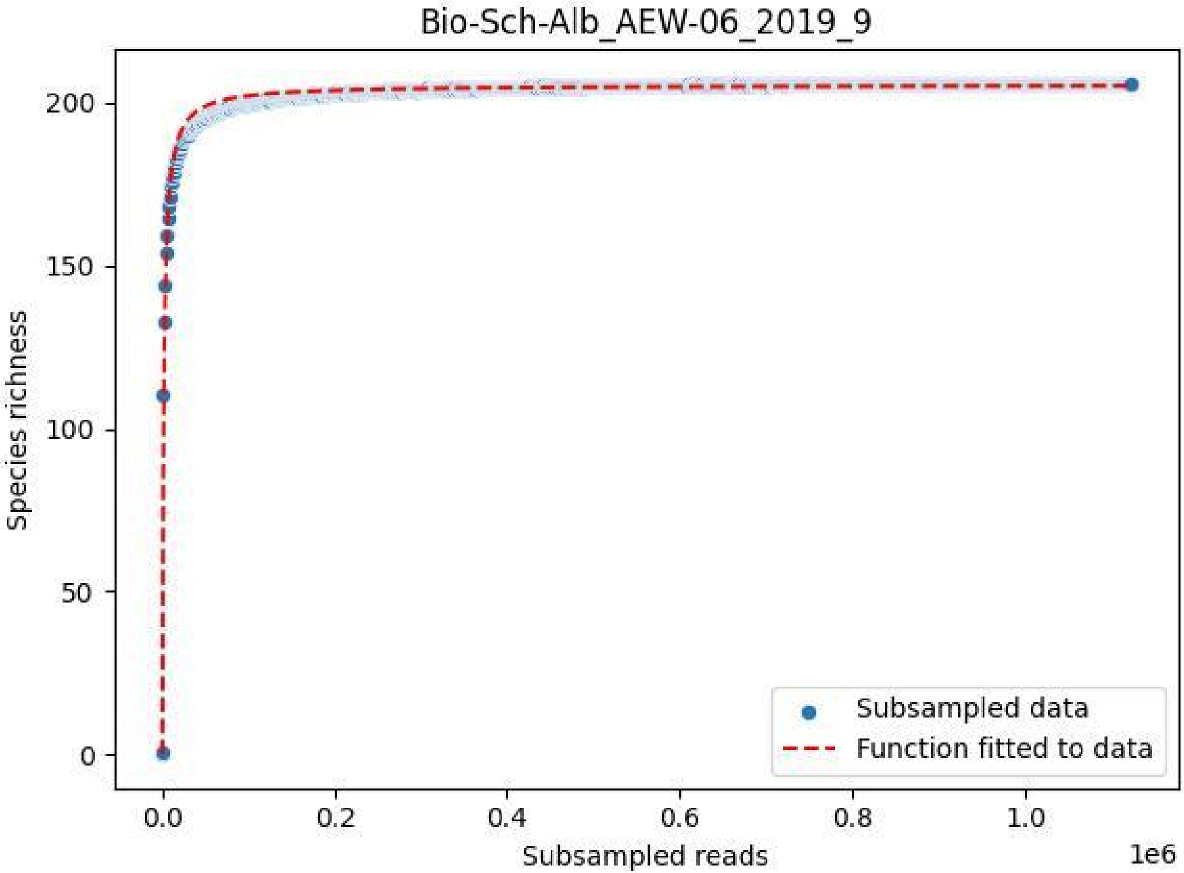
Bio-Sch-Alb_AEW-06_2019_9
Species richness
Subsampled reads
1e6
0
50
100
150
200
0.0
0.2
0.4
0.6
0.8
1.0
Subsampled data
Function fitted to data

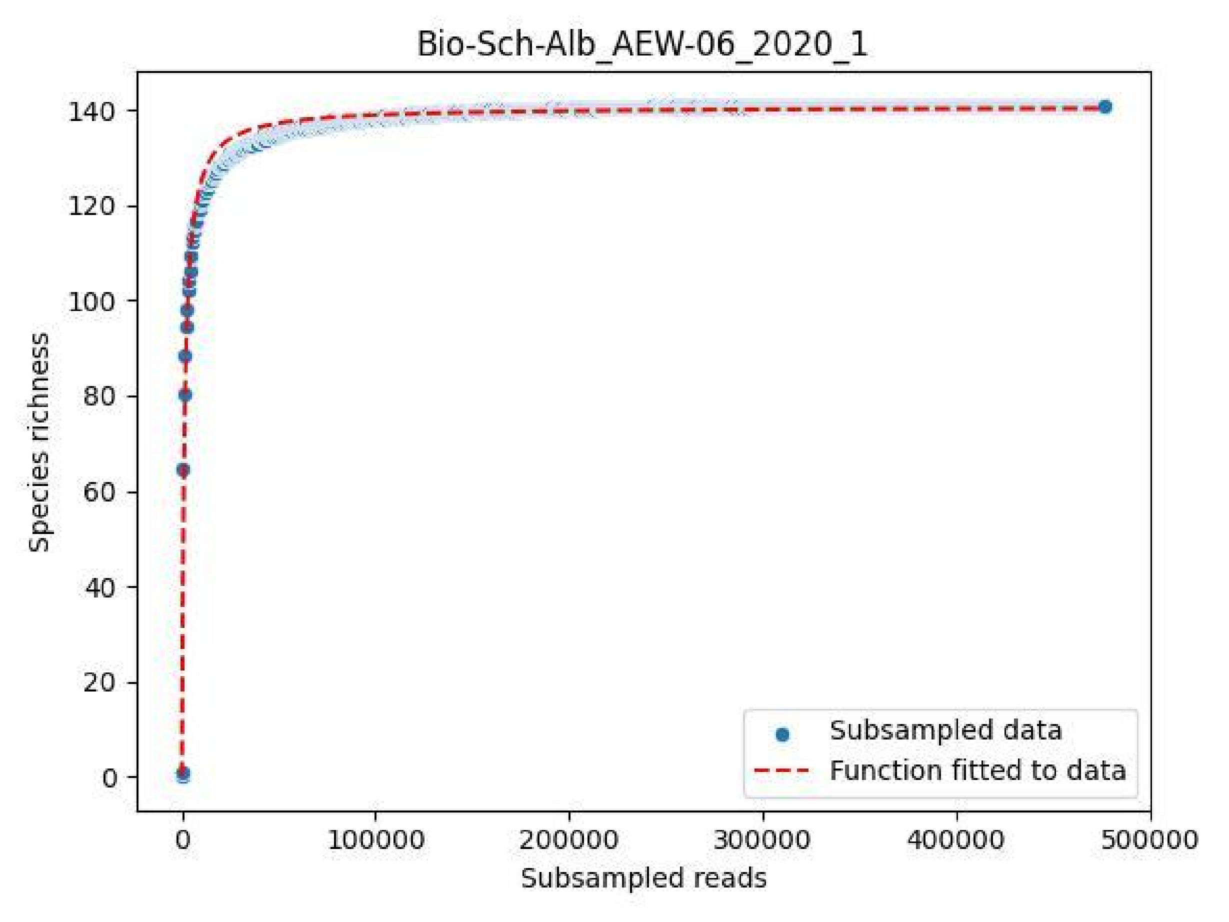
Bio-Sch-Alb_AEW-06_2020_1
Species richness
Subsampled reads
0
20
40
60
80
100
120
140
0
100000
200000
300000
400000
500000
Subsampled data
Function fitted to data

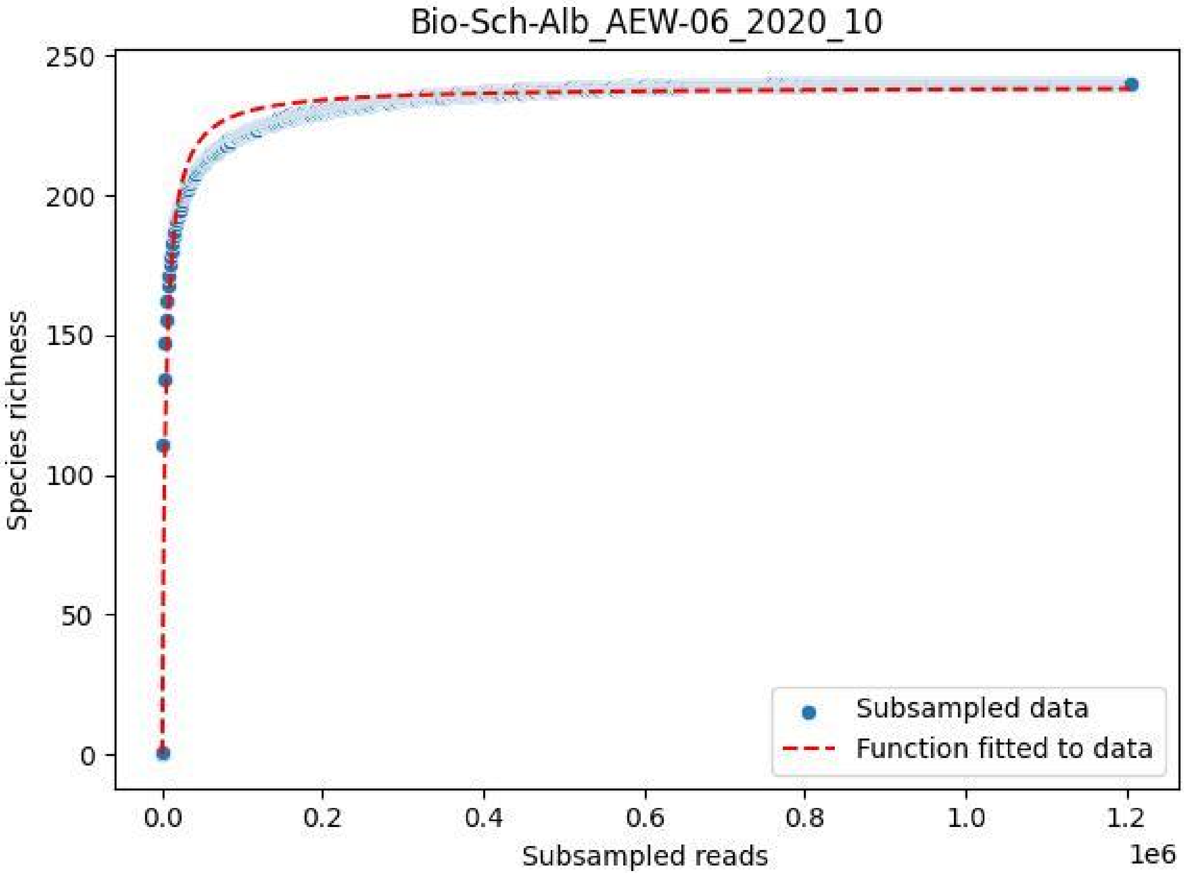
Bio-Sch-Alb_AEW-06_2020_10
Species richness
Subsampled reads
1e6
Subsampled data
Function fitted to data
0
50
100
150
200
250
0.0
0.2
0.4
0.6
0.8
1.0
1.2

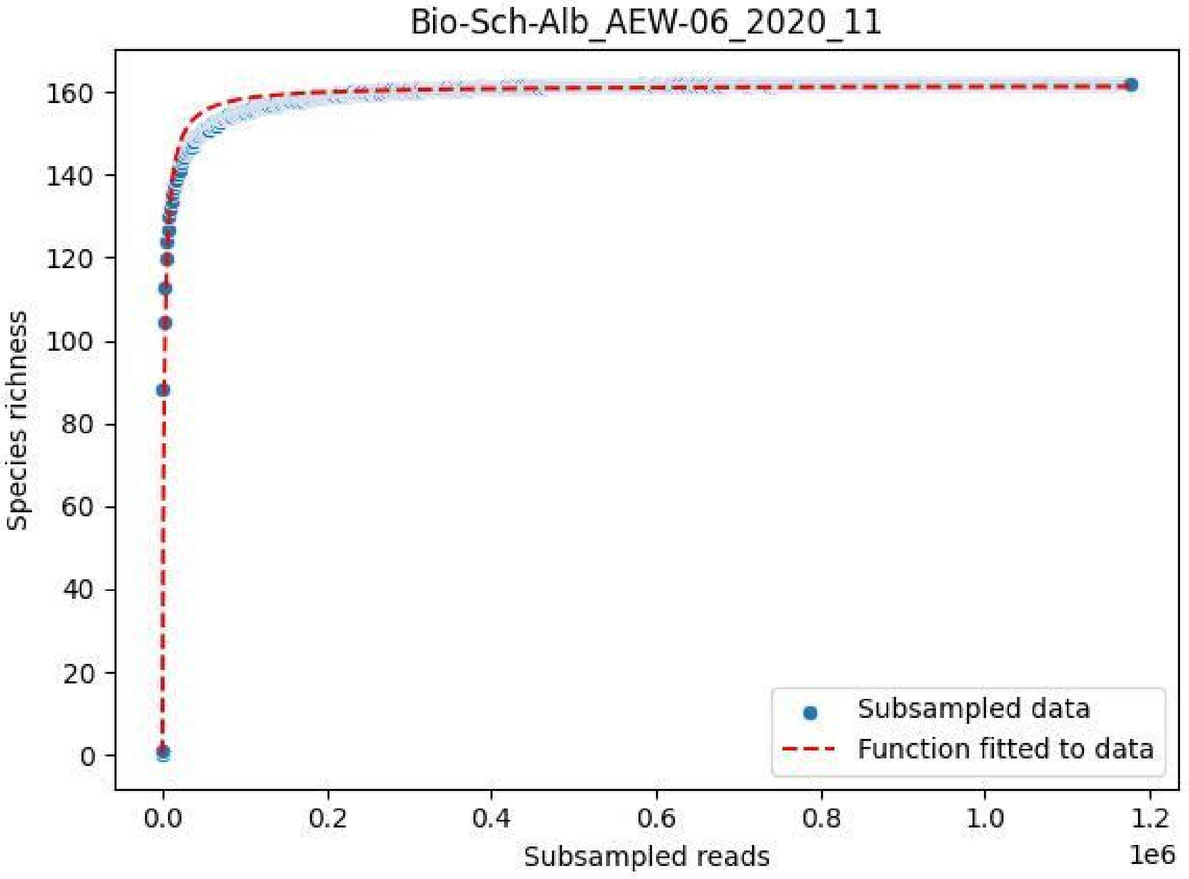

Bio-Sch-Alb_AEW-06_2020_11
Species richness
Subsampled reads
1e6
0
20
40
60
80
100
120
140
160
0.0
0.2
0.4
0.6
0.8
1.0
1.2
Subsampled data
Function fitted to data

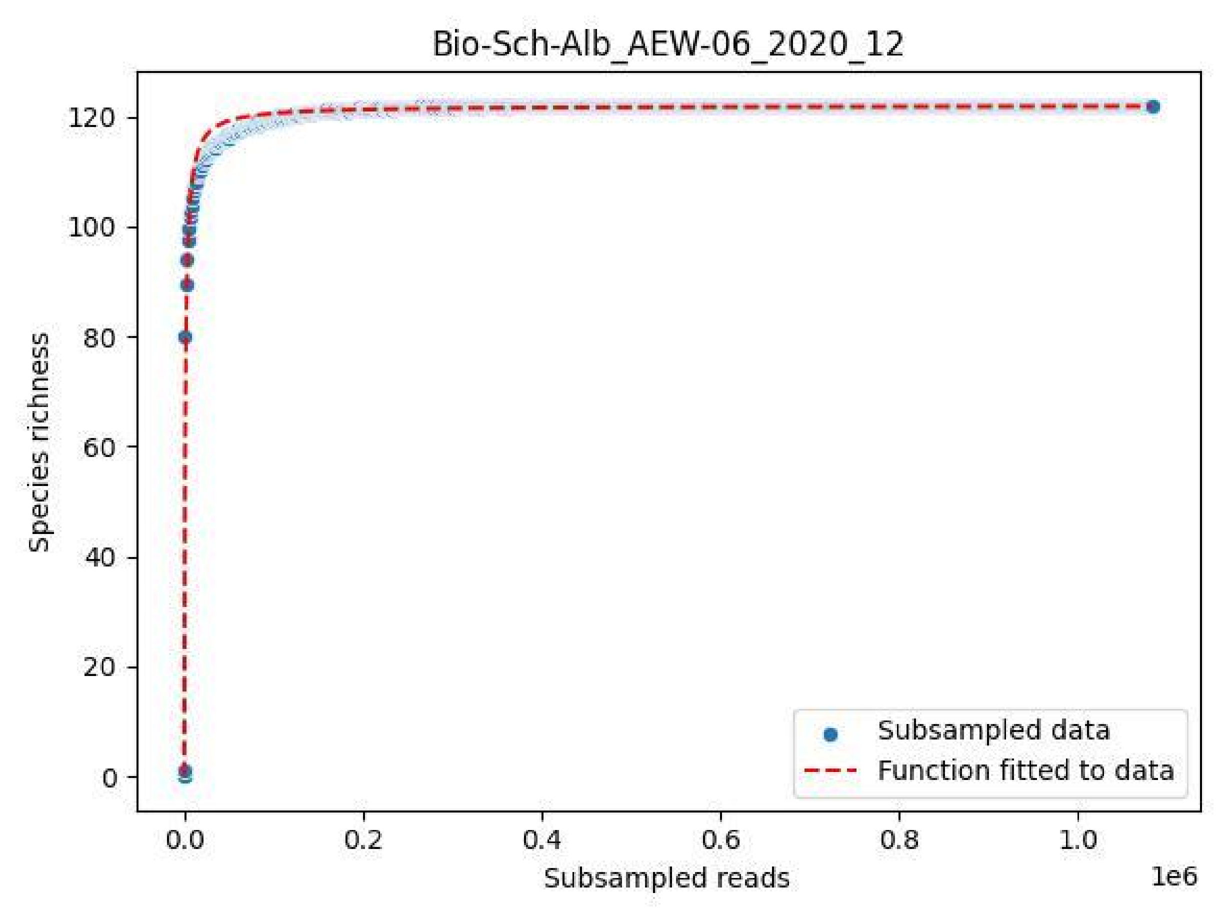

Bio-Sch-Alb_AEW-06_2020_12
Species richness
0
20
40
60
80
100
120
0.0
0.2
0.4
0.6
0.8
1.0
1e6
Subsampled reads
Subsampled data
Function fitted to data

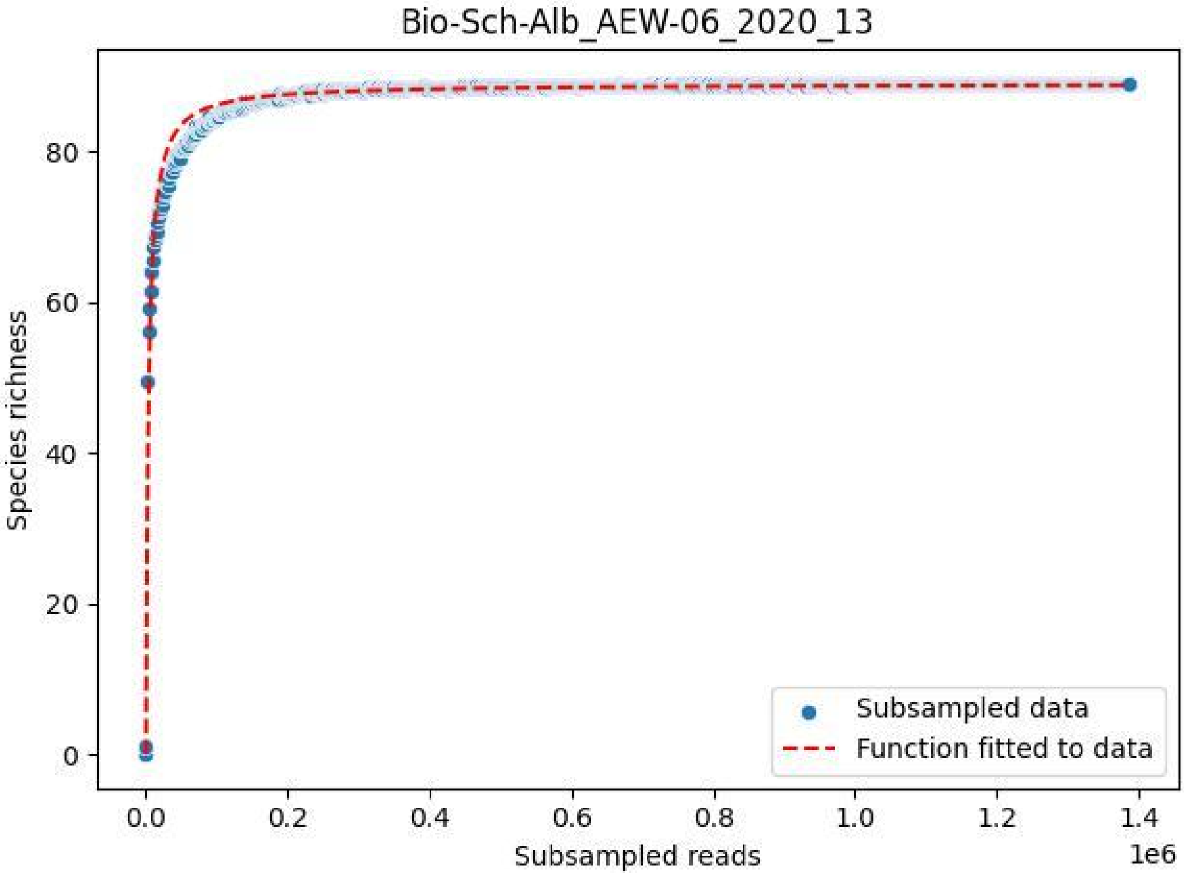
Bio-Sch-Alb_AEW-06_2020_13
Species richness
0
20
40
60
80
Subsampled reads
0.0
0.2
0.4
0.6
0.8
1.0
1.2
1.4
1e6
Subsampled data
Function fitted to data

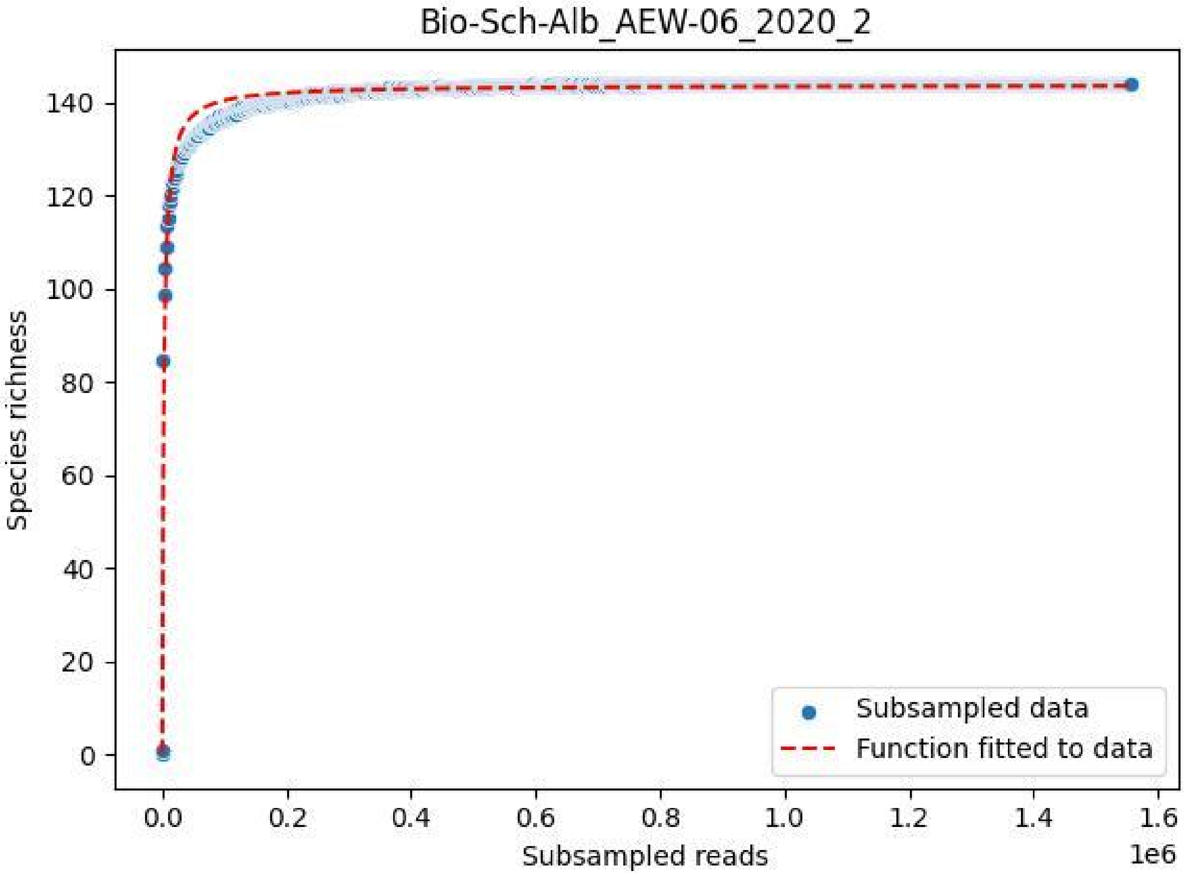
Bio-Sch-Alb_AEW-06_2020_2
Species richness
0
20
40
60
80
100
120
140
Subsampled reads
0.0
0.2
0.4
0.6
0.8
1.0
1.2
1.4
1.6
1e6
Subsampled data
Function fitted to data

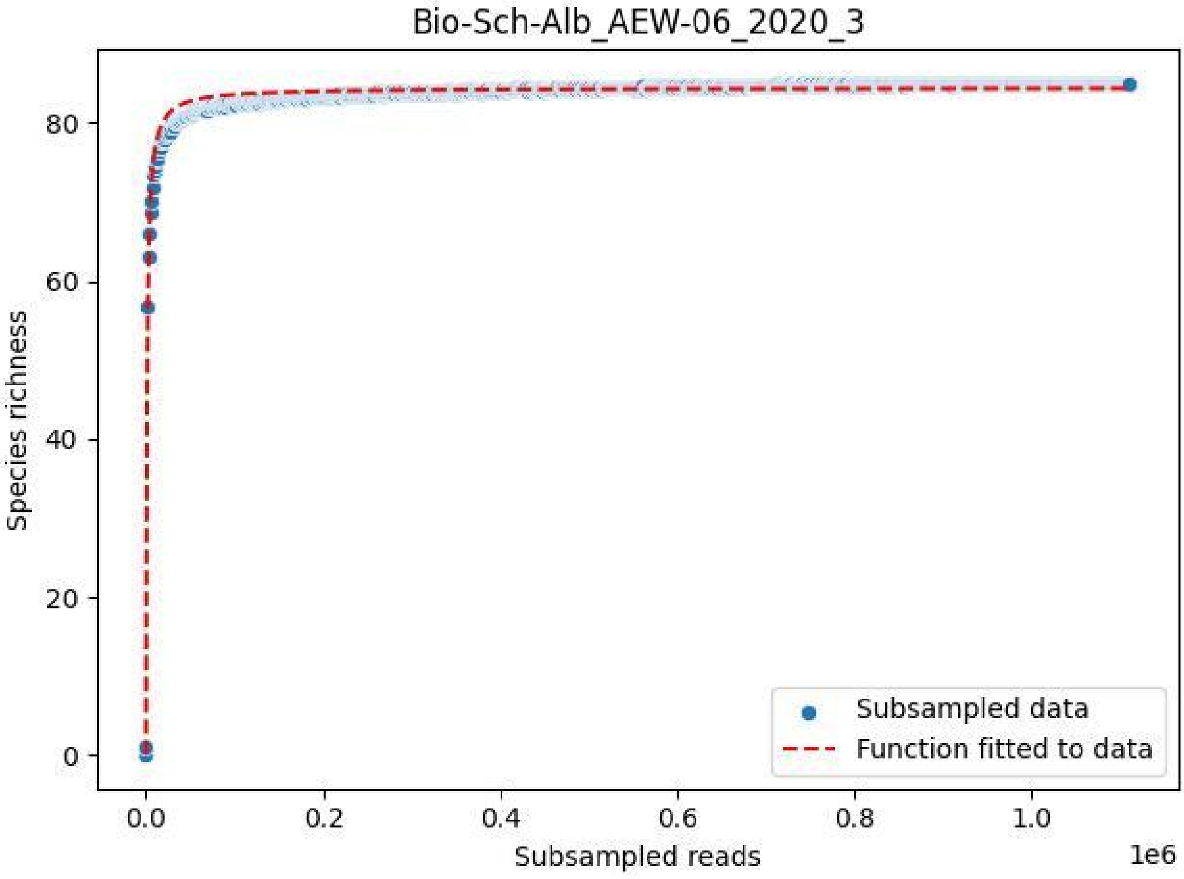

Bio-Sch-Alb_AEW-06_2020_3
Species richness
80
60
40
20
0
0.0
0.2
0.4
0.6
0.8
1.0
Subsampled reads
1e6
Subsampled data
Function fitted to data

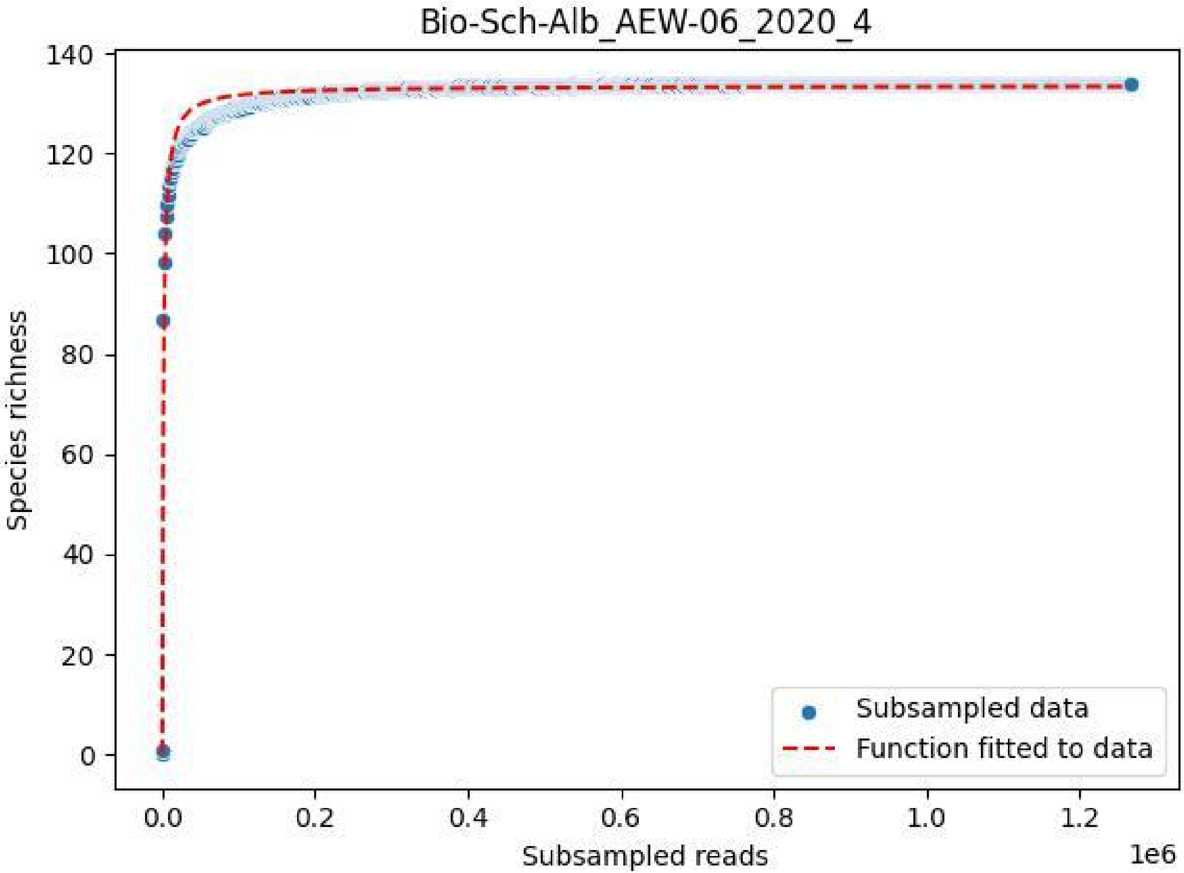
Bio-Sch-Alb_AEW-06_2020_4
Species richness
Subsampled reads
1e6
0
20
40
60
80
100
120
140
0.0
0.2
0.4
0.6
0.8
1.0
1.2
Subsampled data
Function fitted to data

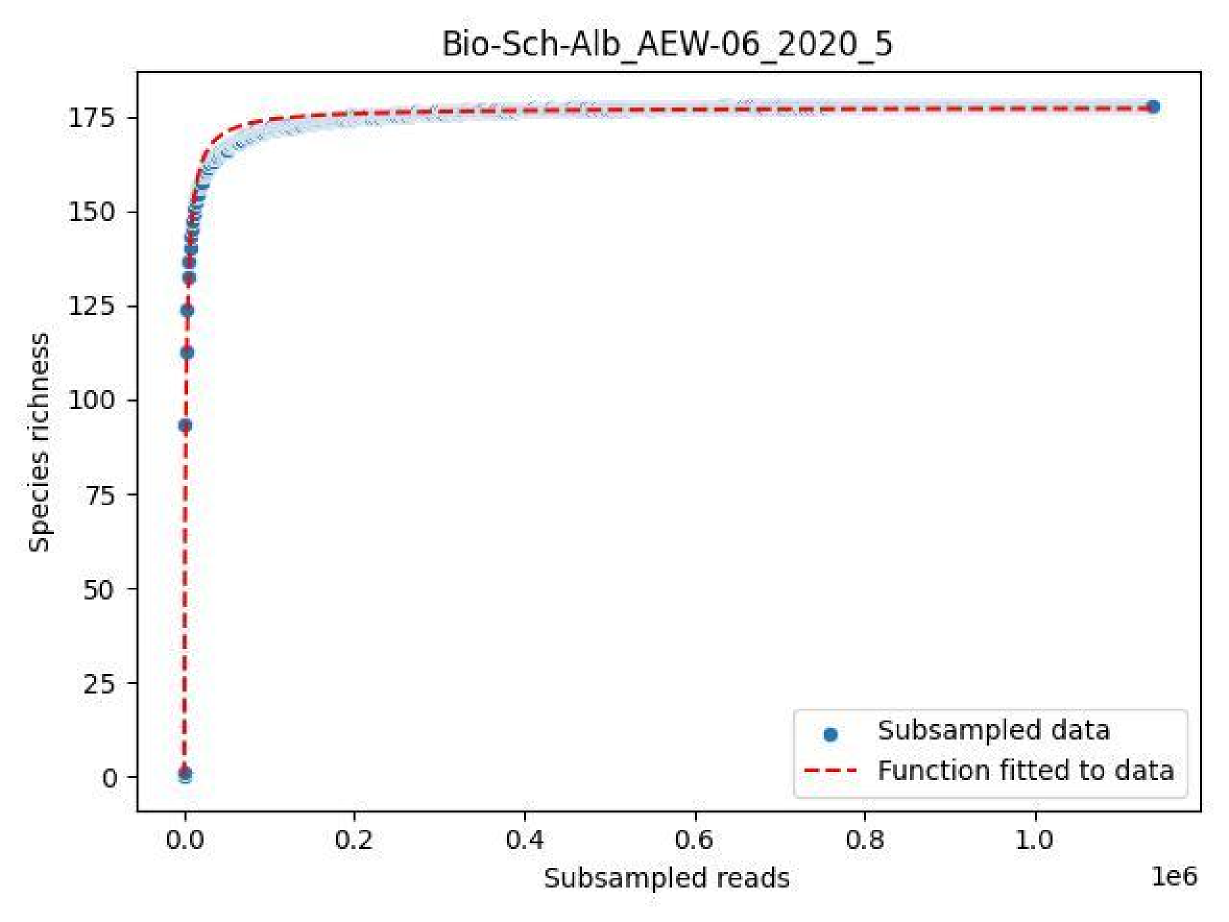
Bio-Sch-Alb_AEW-06_2020_5
Species richness
0
25
50
75
100
125
150
175
Subsampled reads
0.0
0.2
0.4
0.6
0.8
1.0
1e6
Subsampled data
Function fitted to data

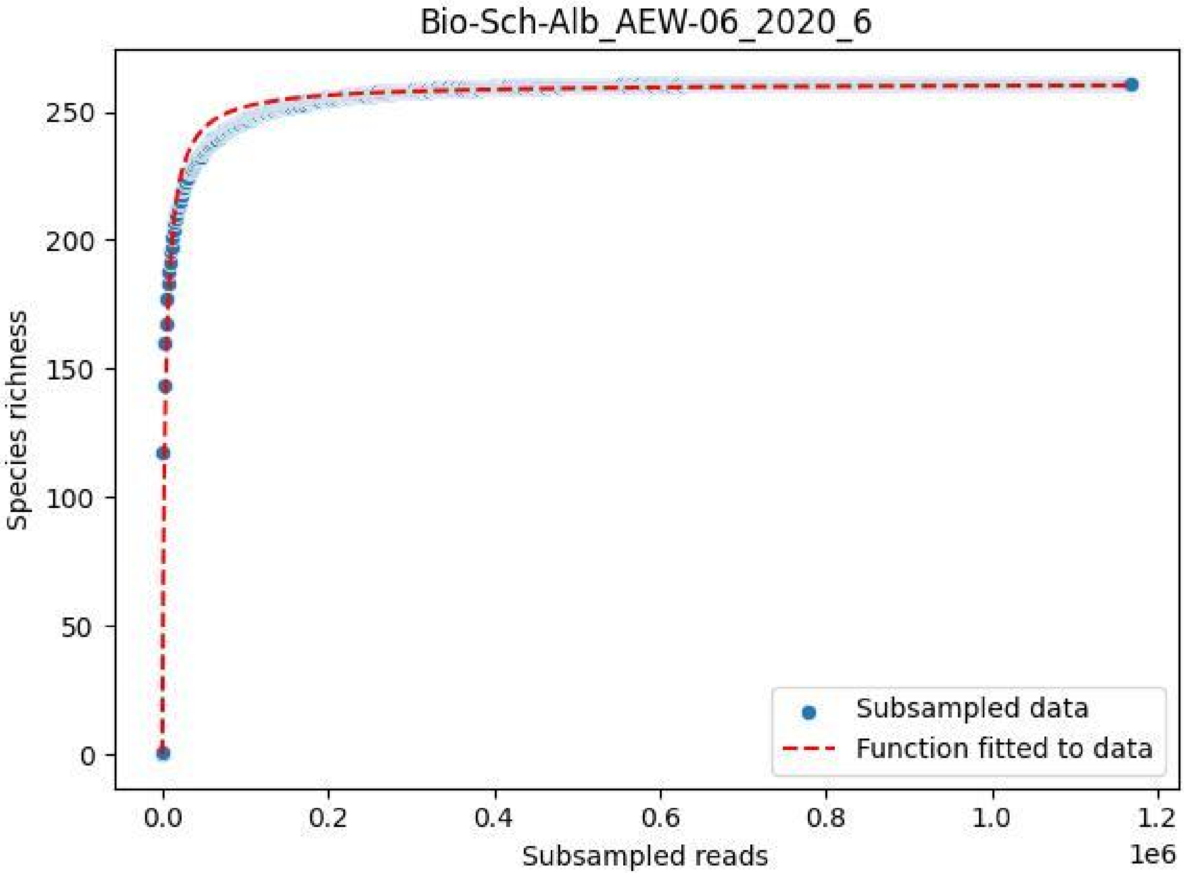
Bio-Sch-Alb_AEW-06_2020_6
Species richness
Subsampled reads
1e6
0
50
100
150
200
250
0.0
0.2
0.4
0.6
0.8
1.0
1.2
Subsampled data
Function fitted to data

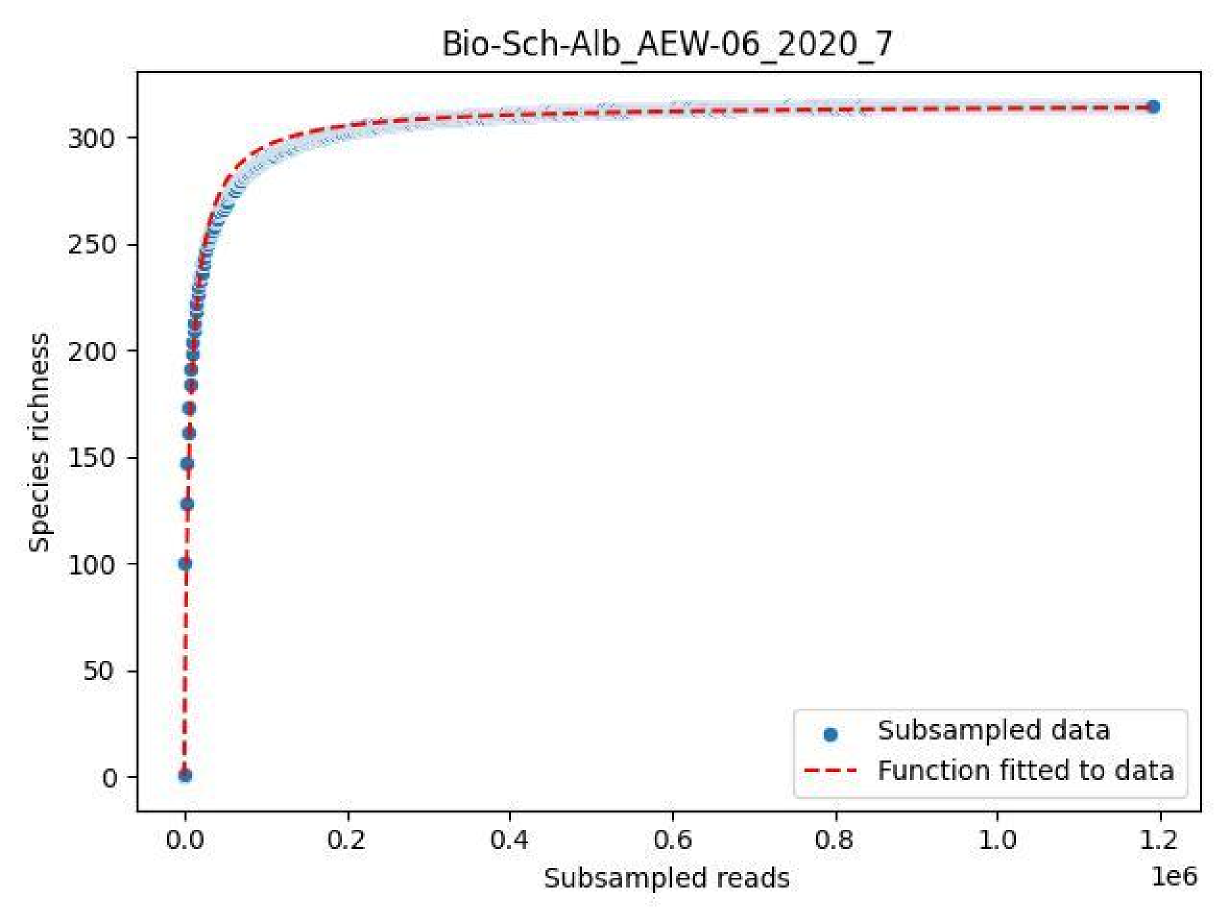
Bio-Sch-Alb_AEW-06_2020_7
Species richness
Subsampled reads
1e6
0
50
100
150
200
250
300
0.0
0.2
0.4
0.6
0.8
1.0
1.2
Subsampled data
Function fitted to data

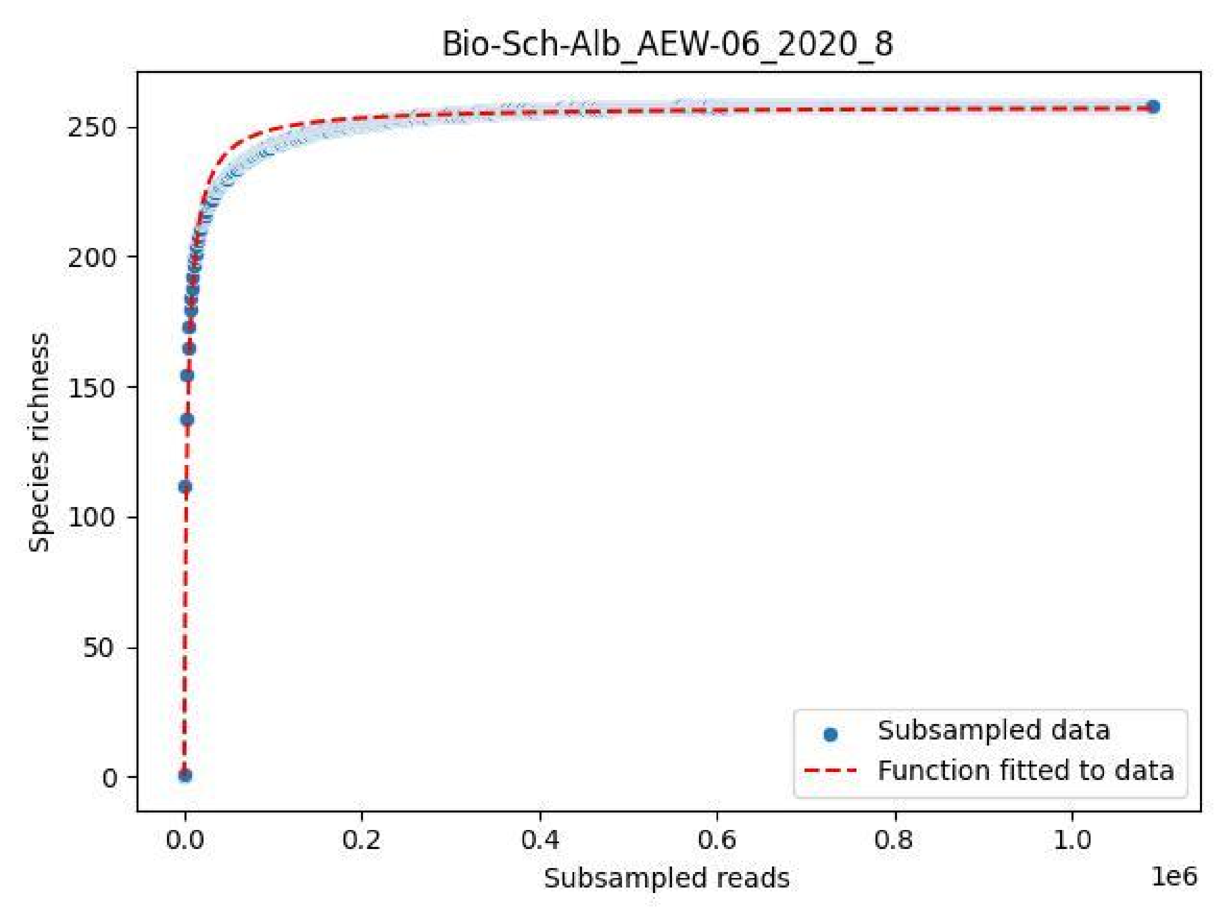

Bio-Sch-Alb_AEW-06_2020_8
Species richness
0
50
100
150
200
250
0.0
0.2
0.4
0.6
0.8
1.0
1e6
Subsampled reads
Subsampled data
Function fitted to data

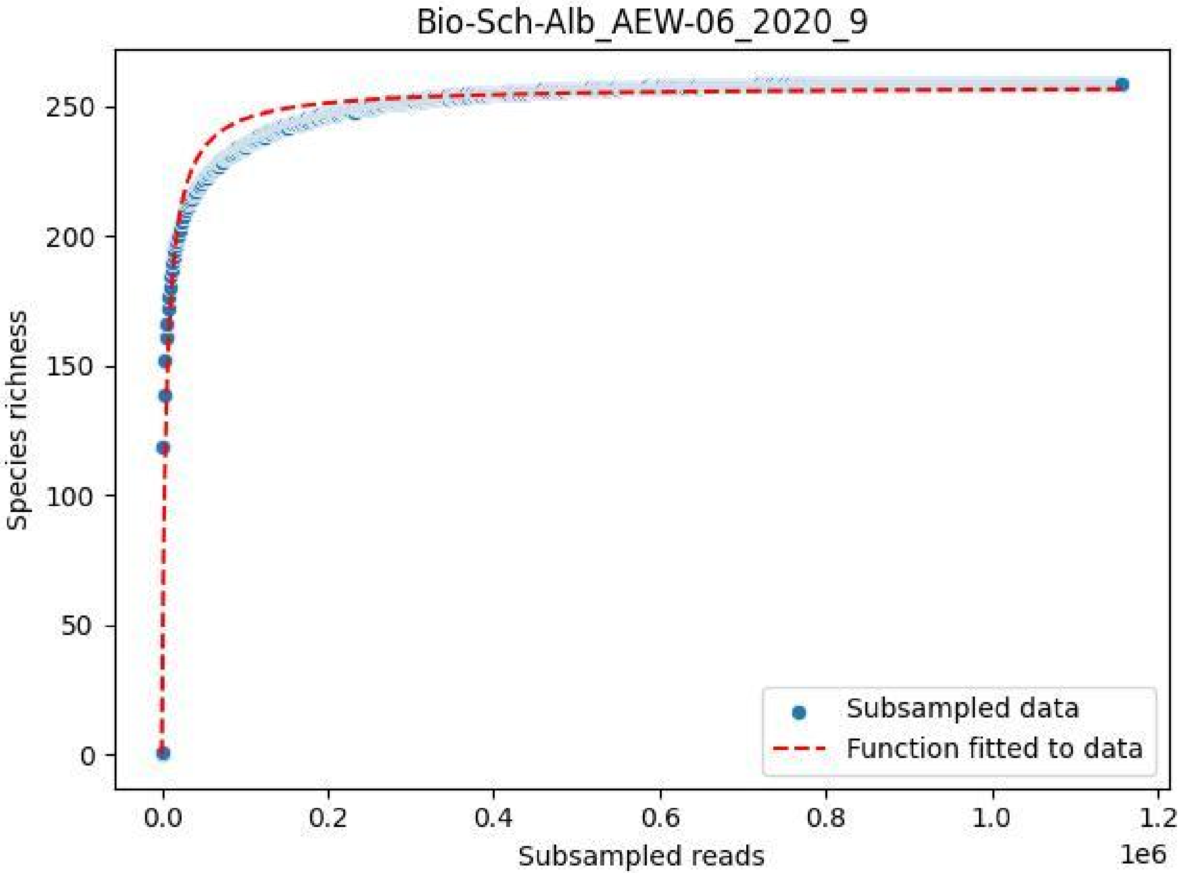
Bio-Sch-Alb_AEW-06_2020_9
Species richness
Subsampled reads
1e6
Subsampled data
Function fitted to data
0
50
100
150
200
250
0.0
0.2
0.4
0.6
0.8
1.0
1.2

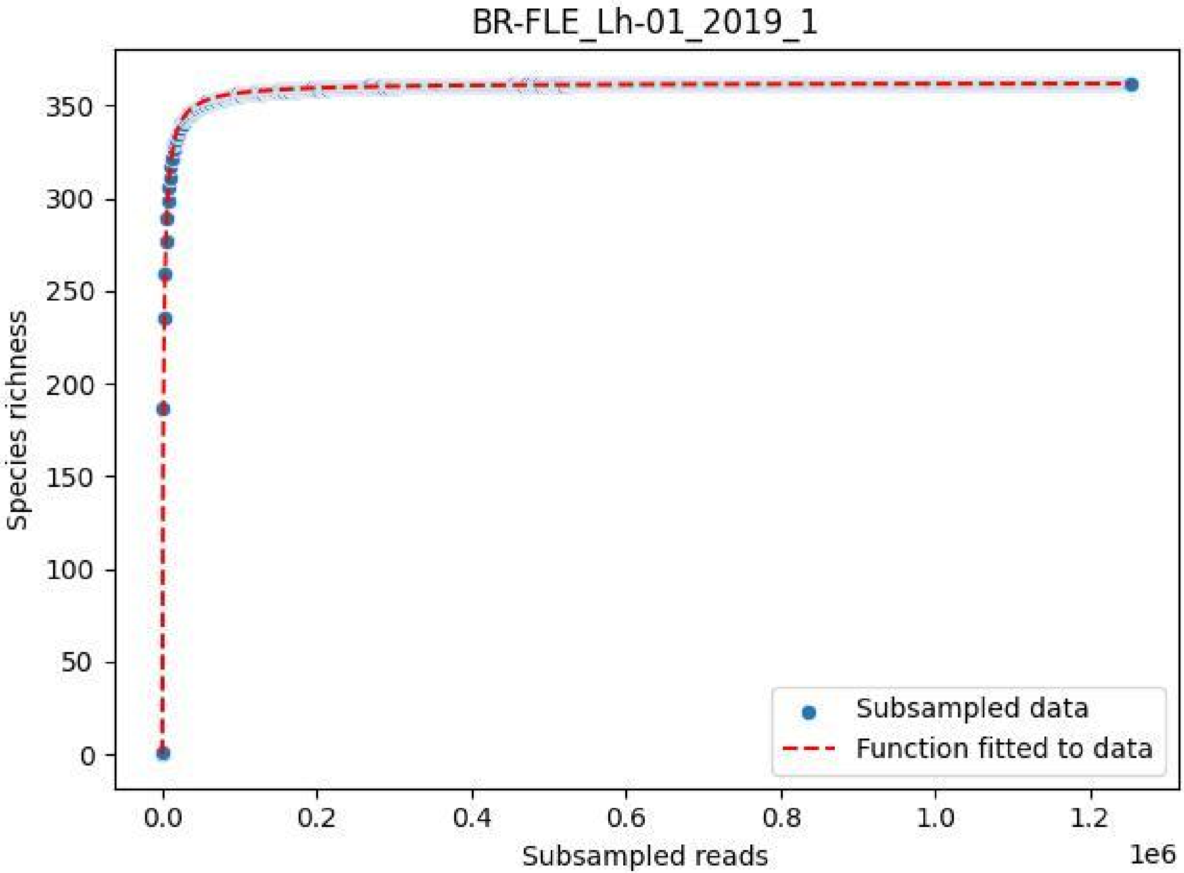
BR-FLE_Lh-01_2019_1
Species richness
Subsampled reads
1e6
0
50
100
150
200
250
300
350
0.0
0.2
0.4
0.6
0.8
1.0
1.2
Subsampled data
Function fitted to data

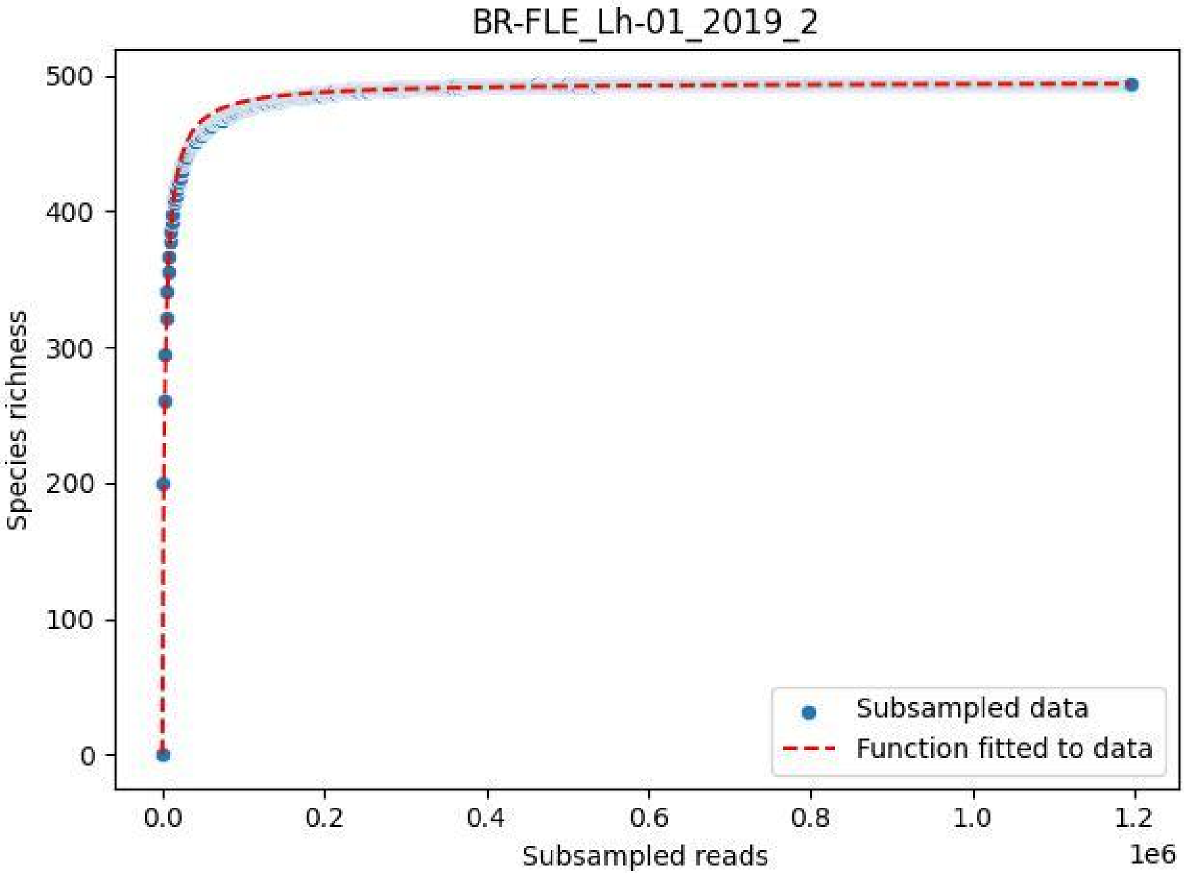
BR-FLE_Lh-01_2019_2
Species richness
Subsampled reads
1e6
0
100
200
300
400
500
0.0
0.2
0.4
0.6
0.8
1.0
1.2
Subsampled data
Function fitted to data

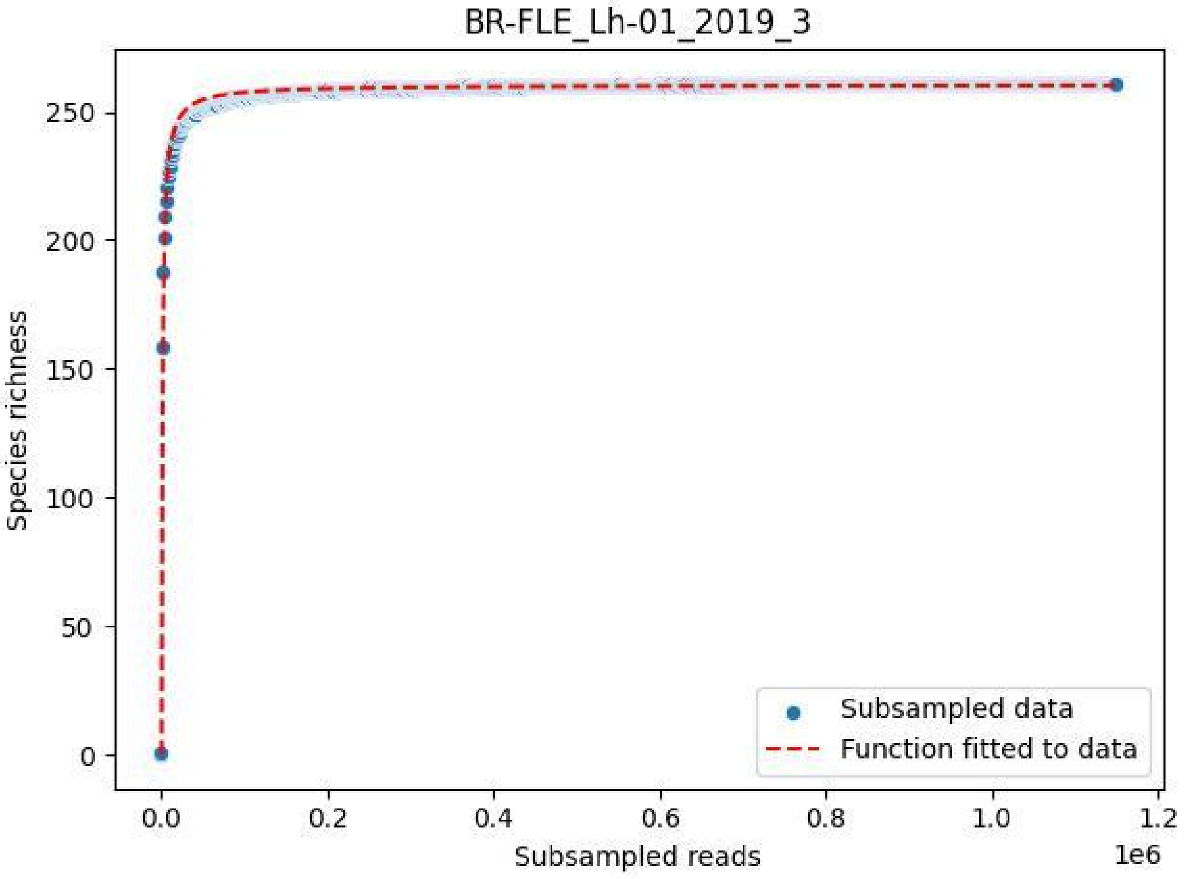
BR-FLE_Lh-01_2019_3
Species richness
Subsampled reads
1e6
0
50
100
150
200
250
0.0
0.2
0.4
0.6
0.8
1.0
1.2
Subsampled data
Function fitted to data

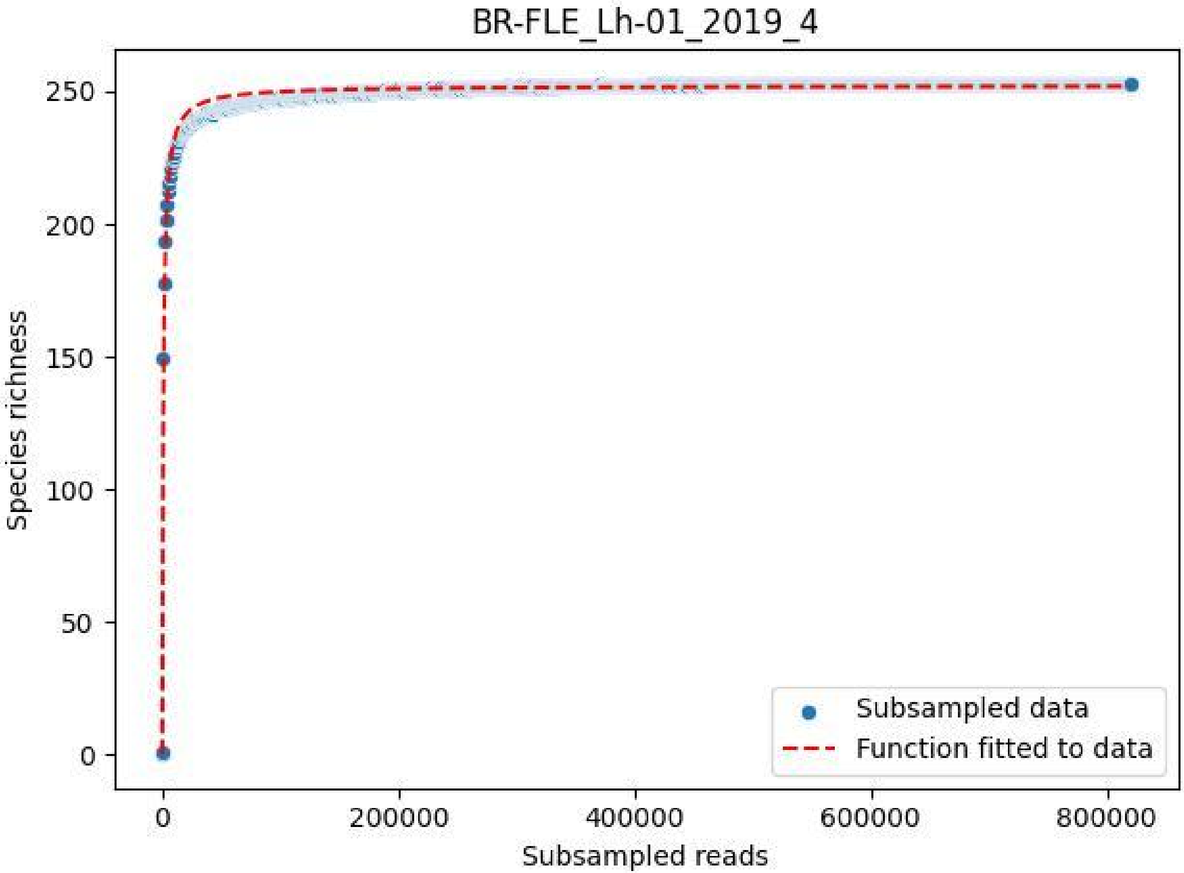
BR-FLE_Lh-01_2019_4
Species richness
Subsampled reads
0
50
100
150
200
250
0
200000
400000
600000
800000
Subsampled data
Function fitted to data

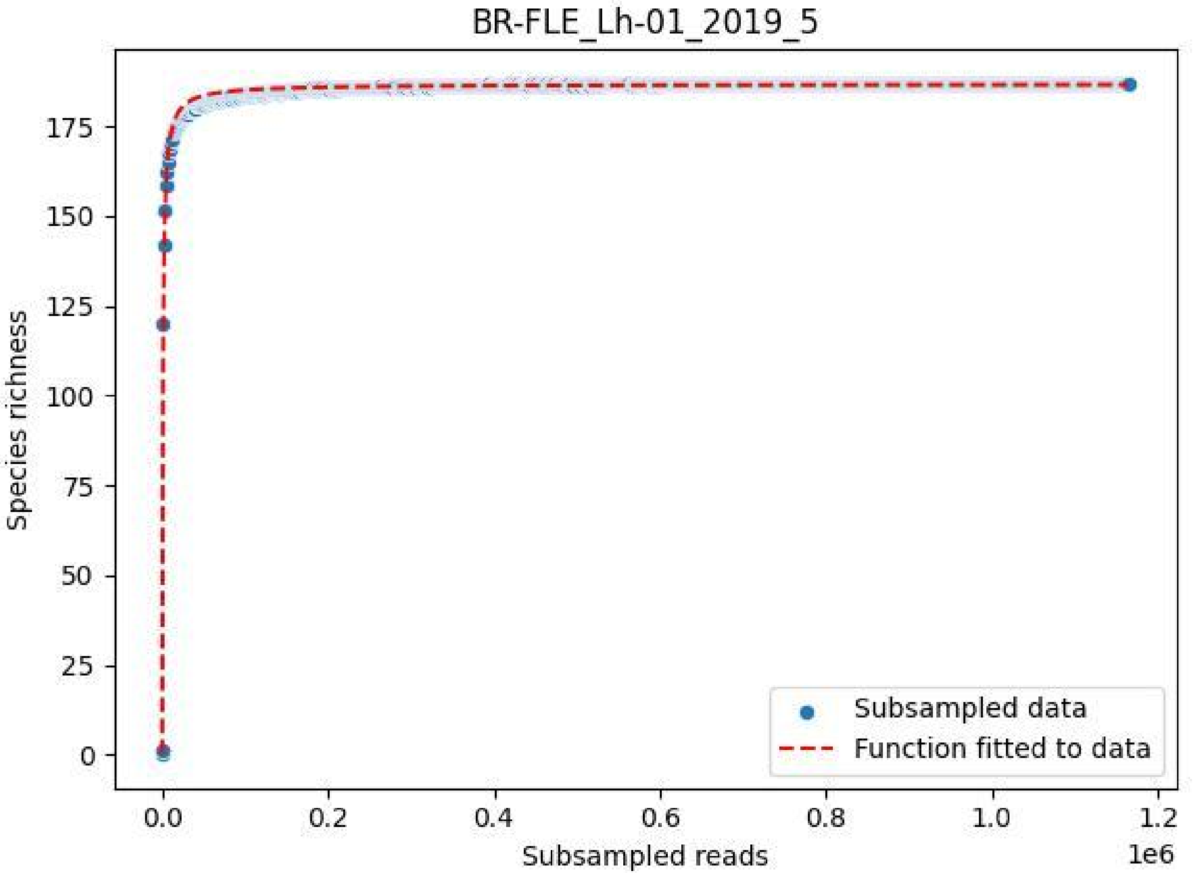
BR-FLE_Lh-01_2019_5
Species richness
Subsampled reads
1e6
0
25
50
75
100
125
150
175
0.0
0.2
0.4
0.6
0.8
1.0
1.2
Subsampled data
Function fitted to data

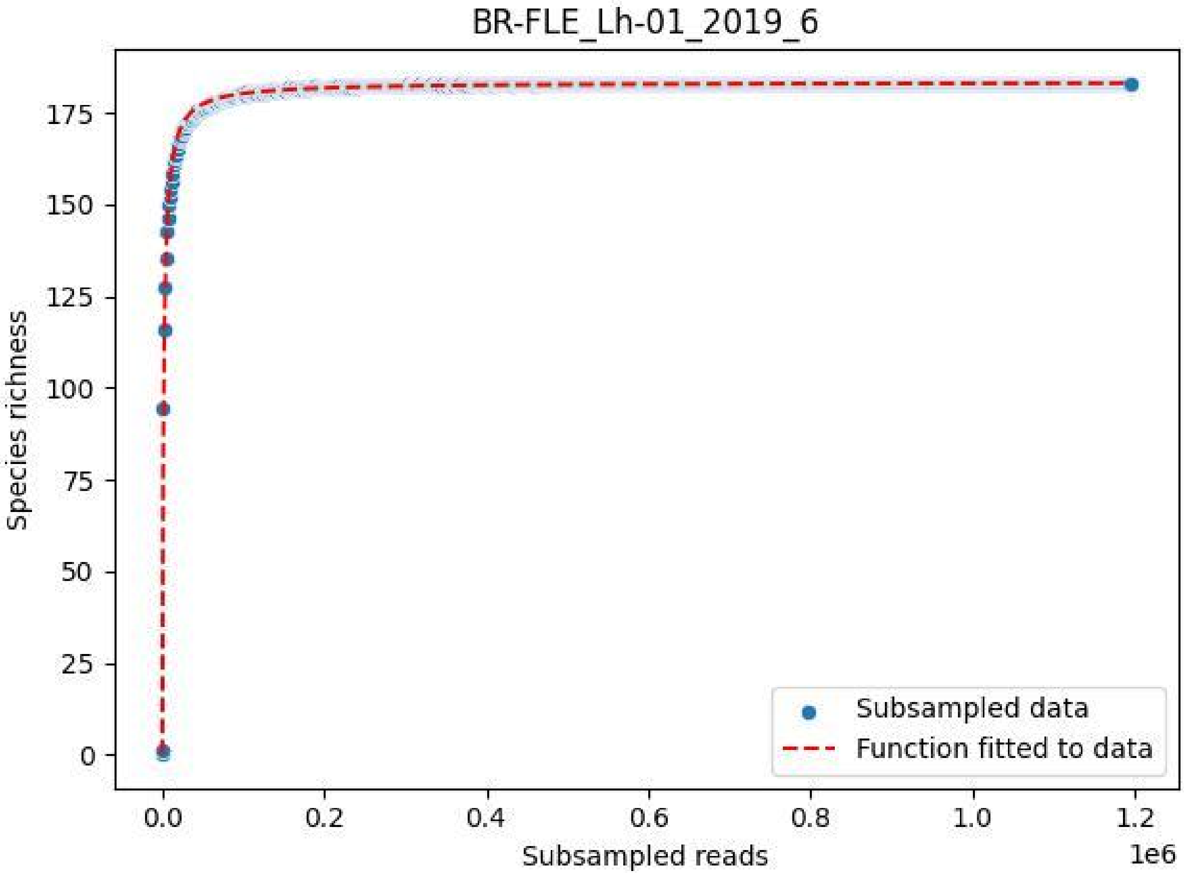

BR-FLE_Lh-01_2019_6
Species richness
Subsampled reads
1e6
0
25
50
75
100
125
150
175
0.0
0.2
0.4
0.6
0.8
1.0
1.2
Subsampled data
Function fitted to data

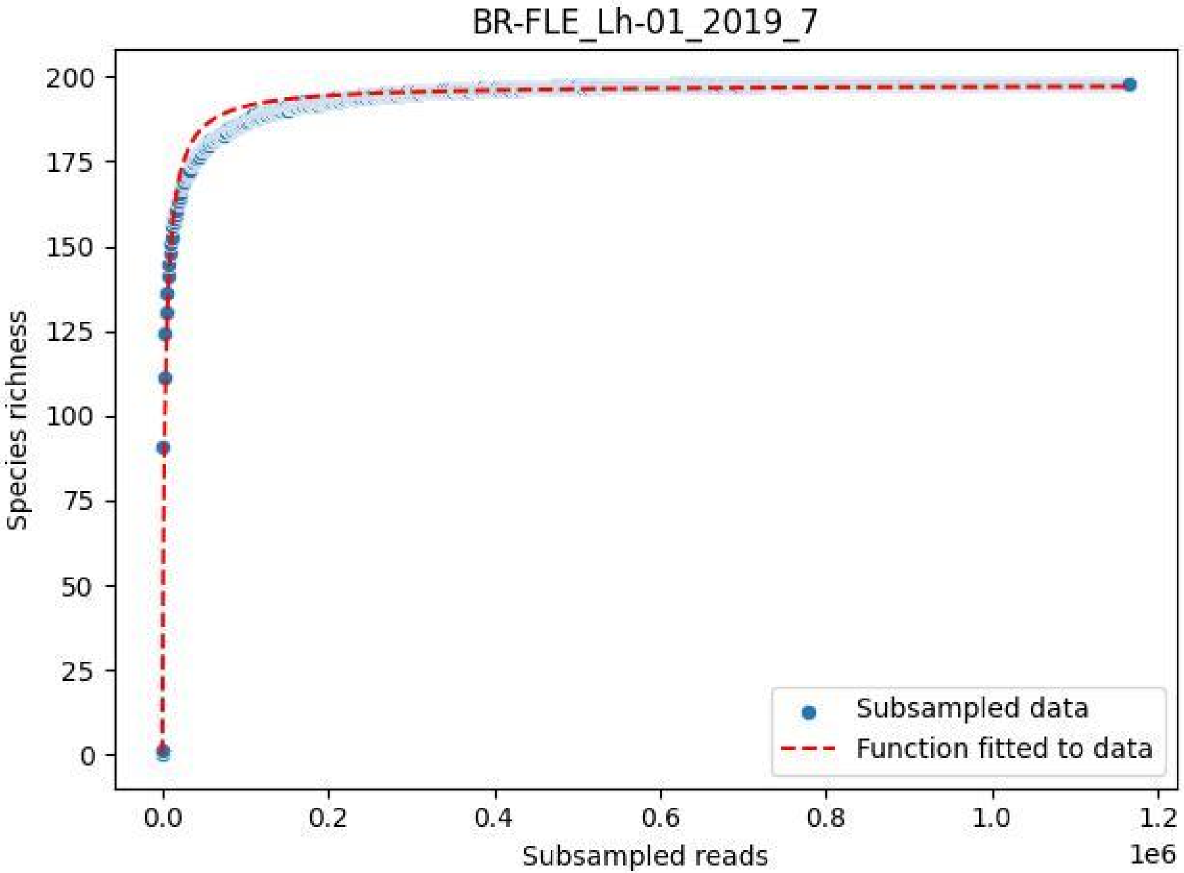
BR-FLE_Lh-01_2019_7
Species richness
Subsampled reads
1e6
0
25
50
75
100
125
150
175
200
0.0
0.2
0.4
0.6
0.8
1.0
1.2
Subsampled data
Function fitted to data

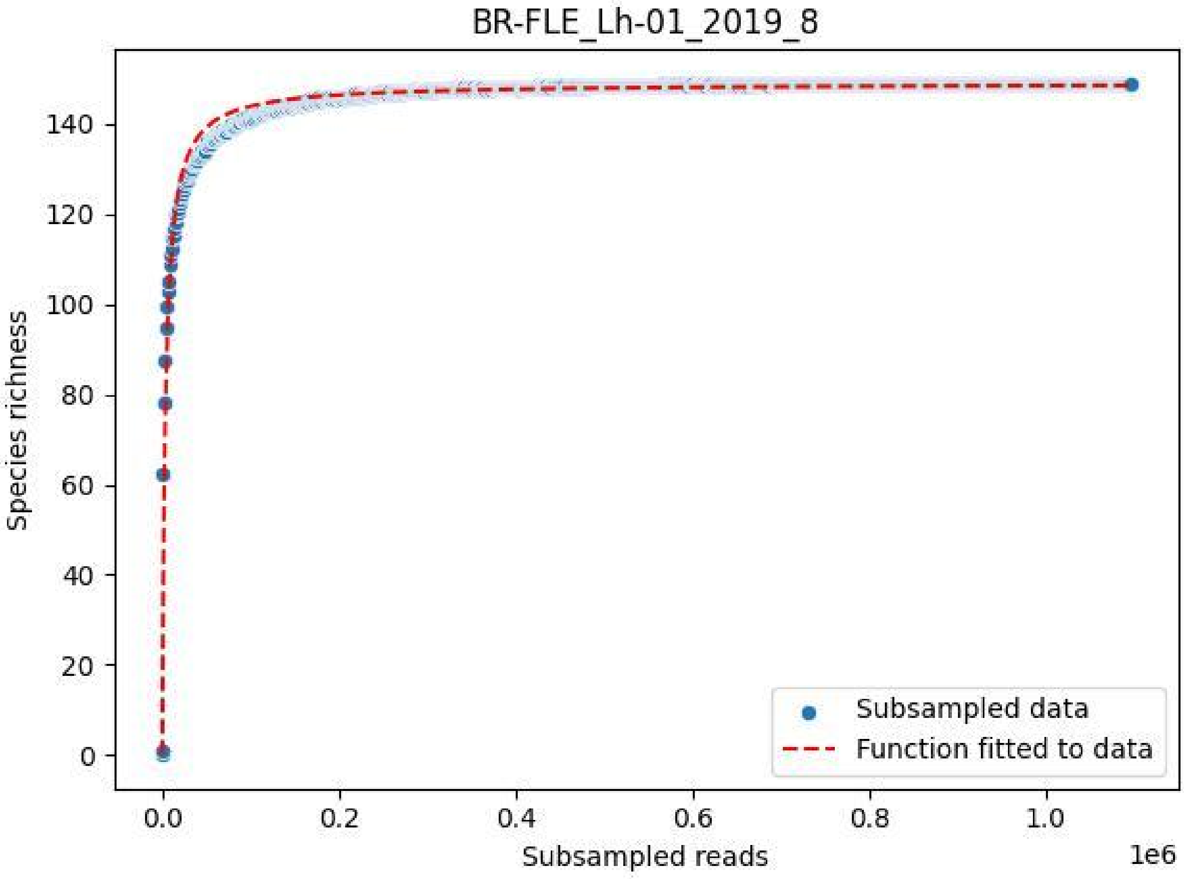
BR-FLE_Lh-01_2019_8
Species richness
Subsampled reads
1e6
Subsampled data
Function fitted to data
0
20
40
60
80
100
120
140
0.0
0.2
0.4
0.6
0.8
1.0

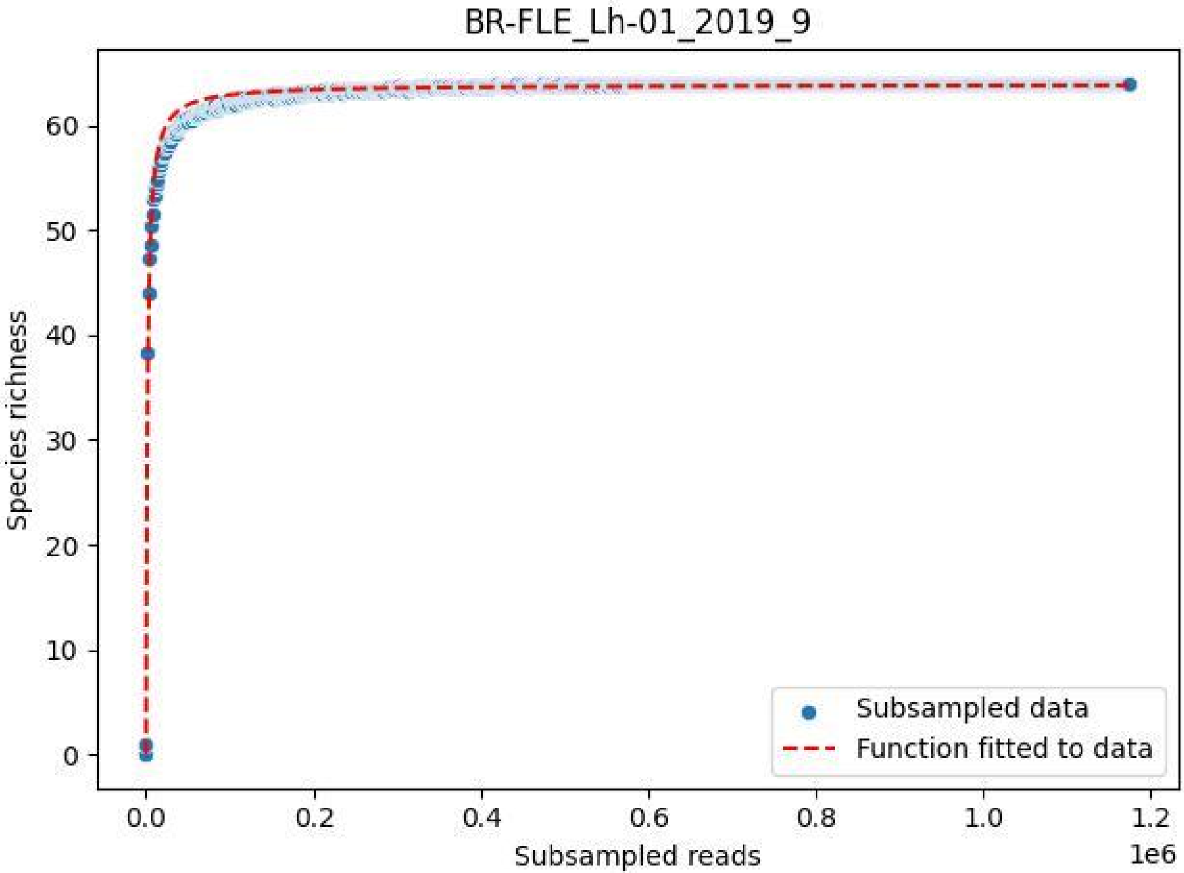
BR-FLE_Lh-01_2019_9
Species richness
Subsampled reads
1e6
0
10
20
30
40
50
60
0.0
0.2
0.4
0.6
0.8
1.0
1.2
Subsampled data
Function fitted to data

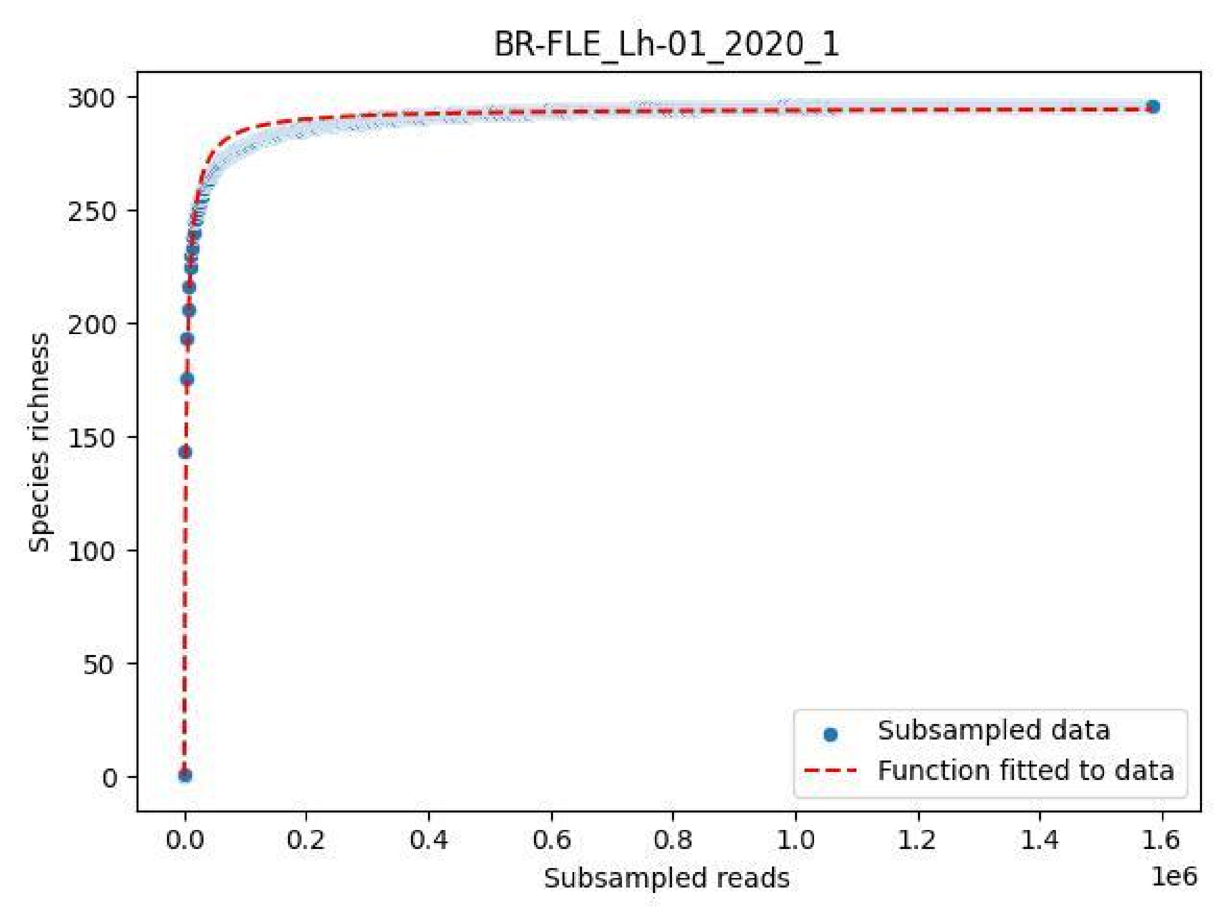
BR-FLE_Lh-01_2020_1
Species richness
Subsampled reads
1e6
0
50
100
150
200
250
300
0.0
0.2
0.4
0.6
0.8
1.0
1.2
1.4
1.6
Subsampled data
Function fitted to data

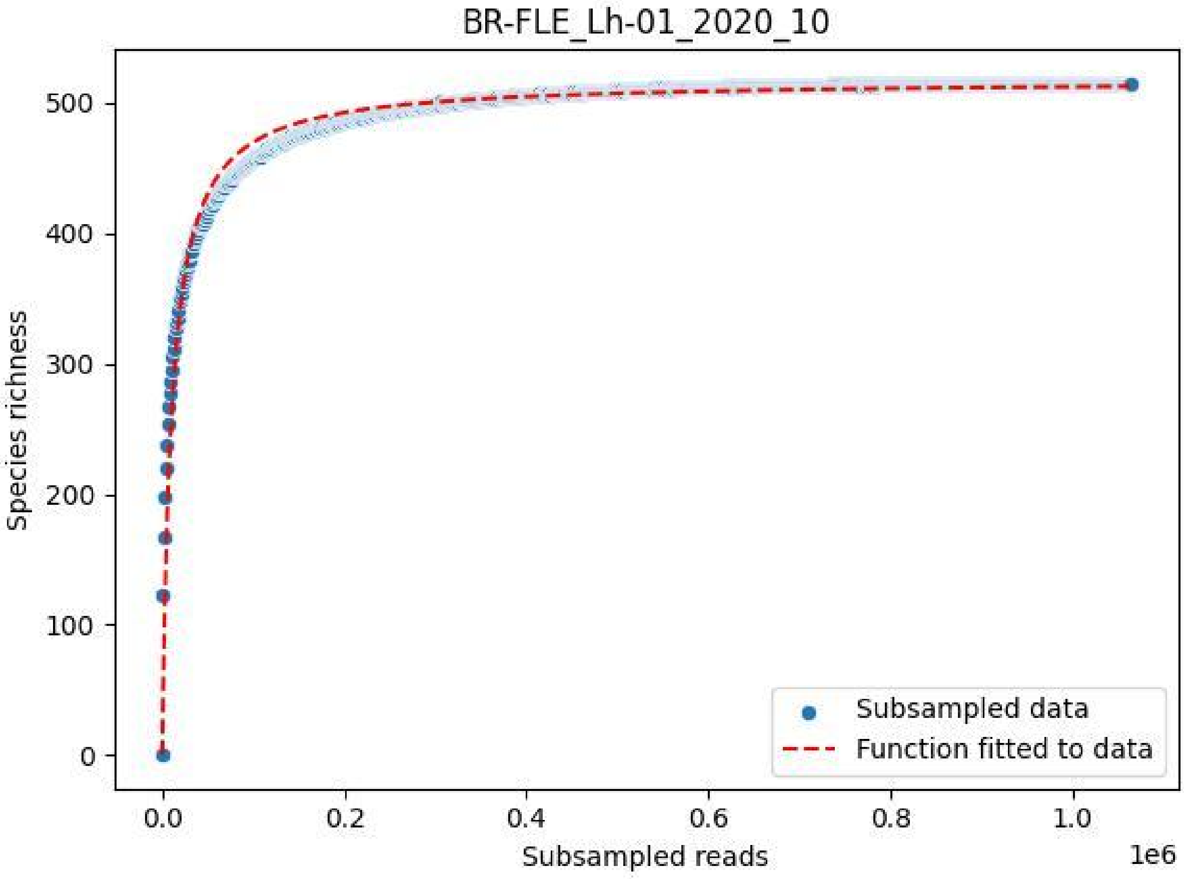
BR-FLE_Lh-01_2020_10
Species richness
Subsampled reads
1e6
0
100
200
300
400
500
0.0
0.2
0.4
0.6
0.8
1.0
Subsampled data
Function fitted to data

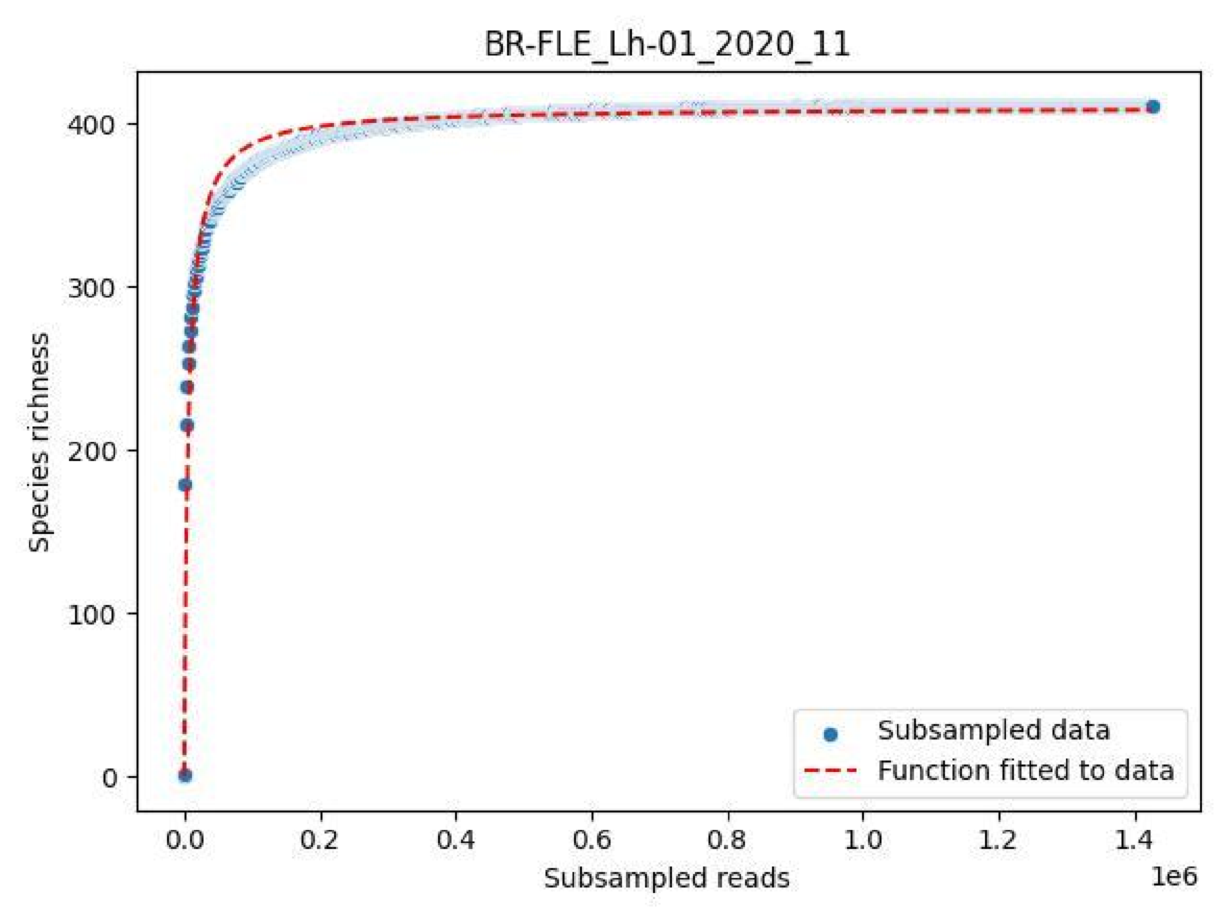

BR-FLE_Lh-01_2020_11
Species richness
Subsampled reads
1e6
0
100
200
300
400
0.0
0.2
0.4
0.6
0.8
1.0
1.2
1.4
Subsampled data
Function fitted to data

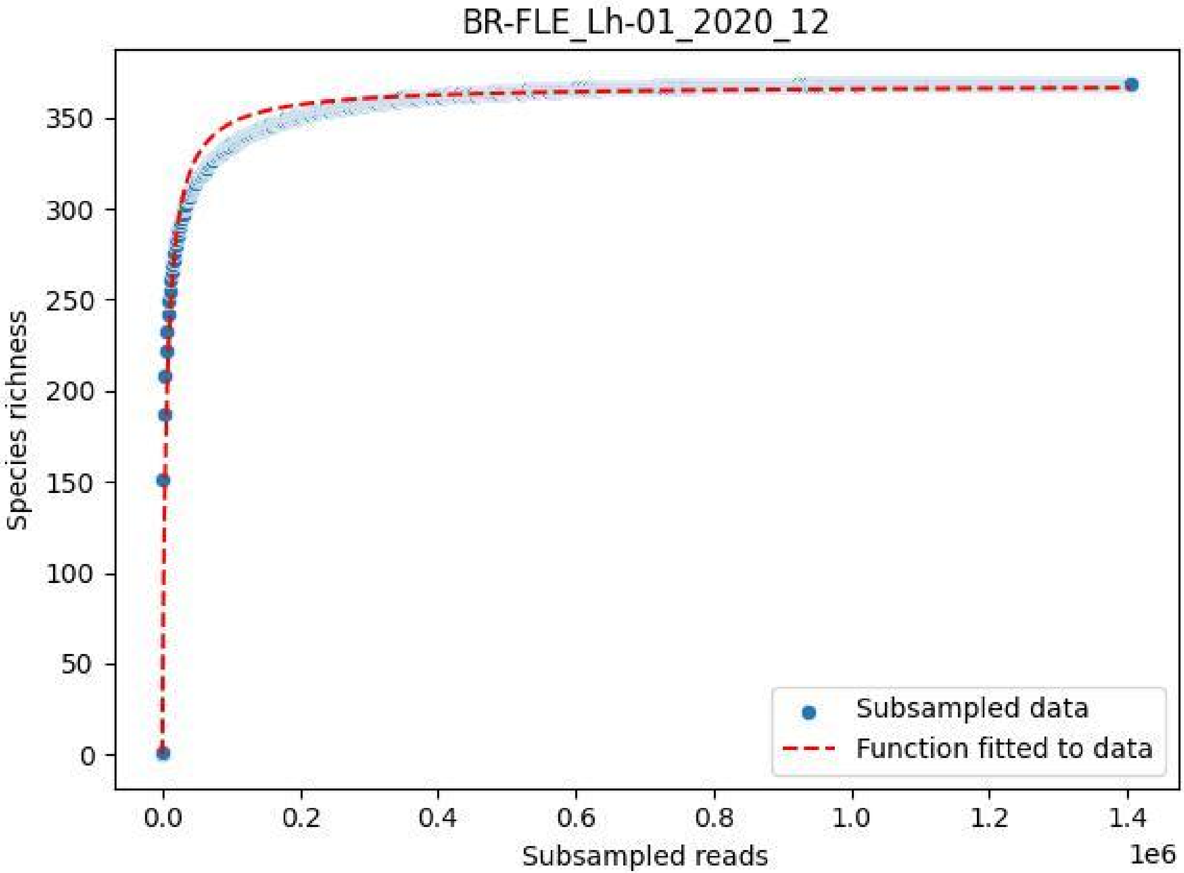
BR-FLE_Lh-01_2020_12
Species richness
Subsampled reads
1e6
0
50
100
150
200
250
300
350
0.0
0.2
0.4
0.6
0.8
1.0
1.2
1.4
Subsampled data
Function fitted to data

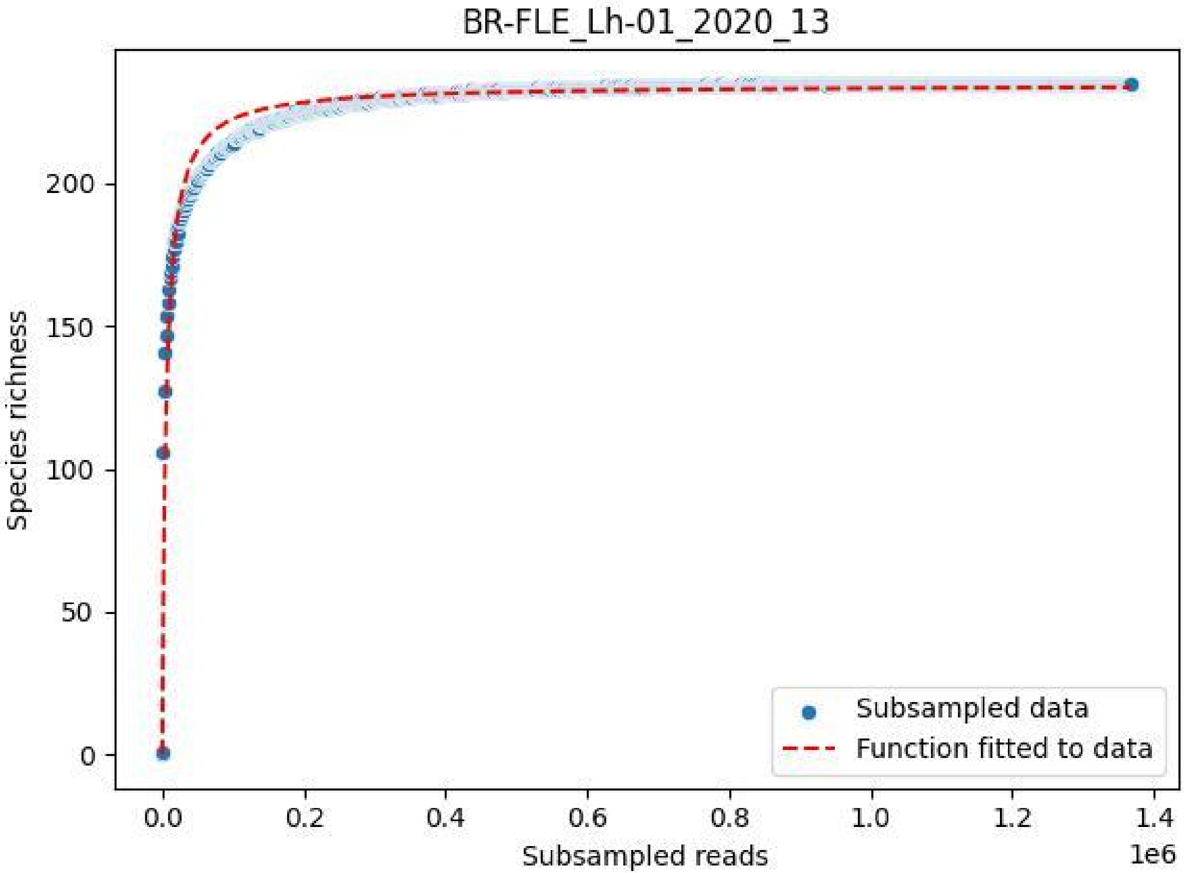
BR-FLE_Lh-01_2020_13
Species richness
Subsampled reads
1e6
0
50
100
150
200
0.0
0.2
0.4
0.6
0.8
1.0
1.2
1.4
Subsampled data
Function fitted to data

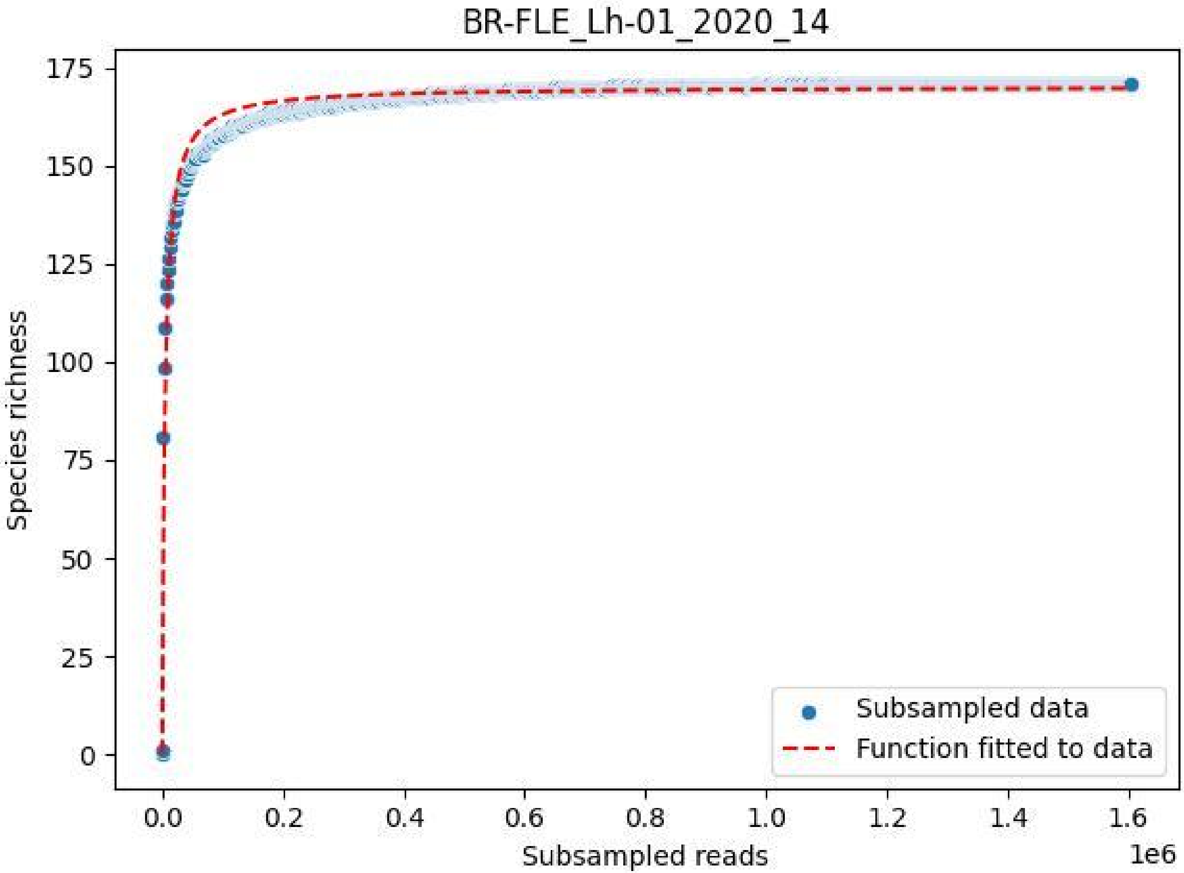

BR-FLE_Lh-01_2020_14
Species richness
Subsampled reads
1e6
Subsampled data
Function fitted to data
175
150
125
100
75
50
25
0
0.0
0.2
0.4
0.6
0.8
1.0
1.2
1.4
1.6

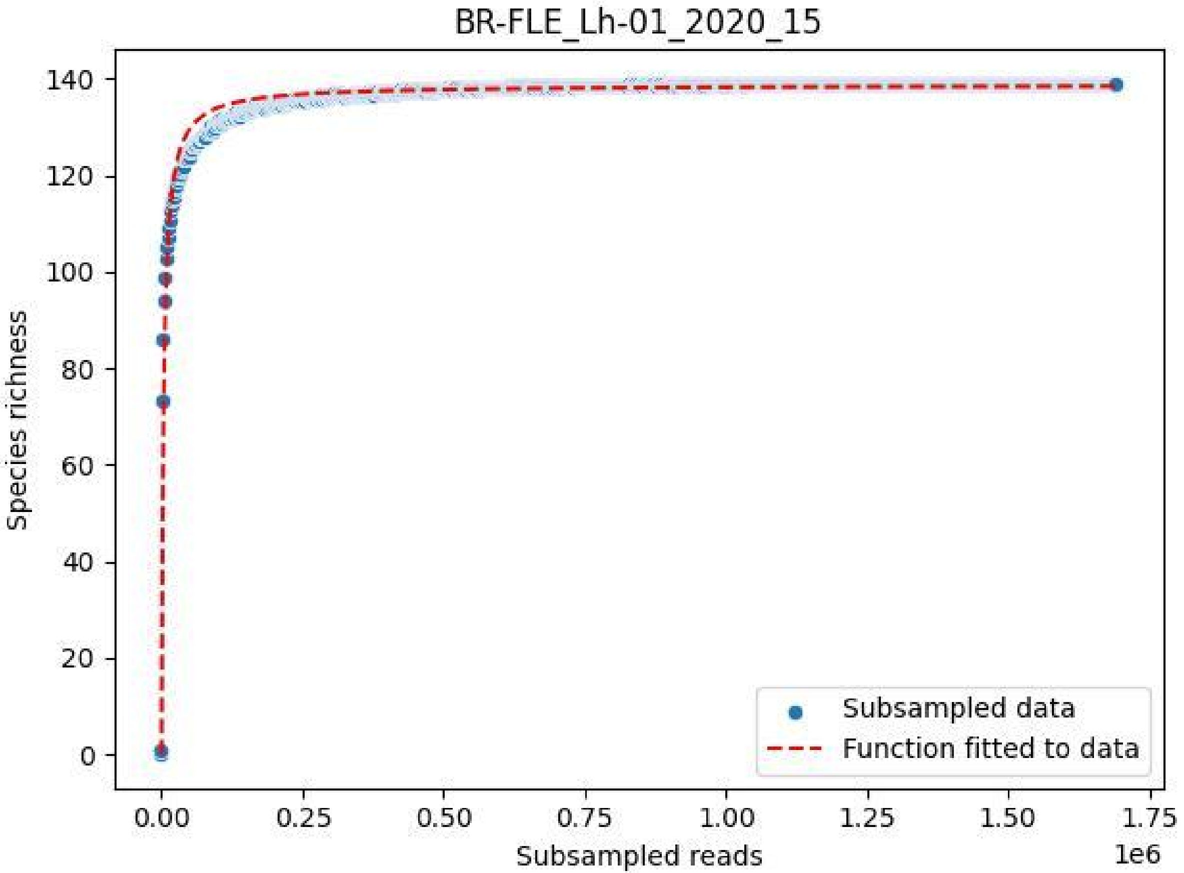
BR-FLE_Lh-01_2020_15
Species richness
Subsampled reads
1e6
0
20
40
60
80
100
120
140
0.00
0.25
0.50
0.75
1.00
1.25
1.50
1.75
Subsampled data
Function fitted to data

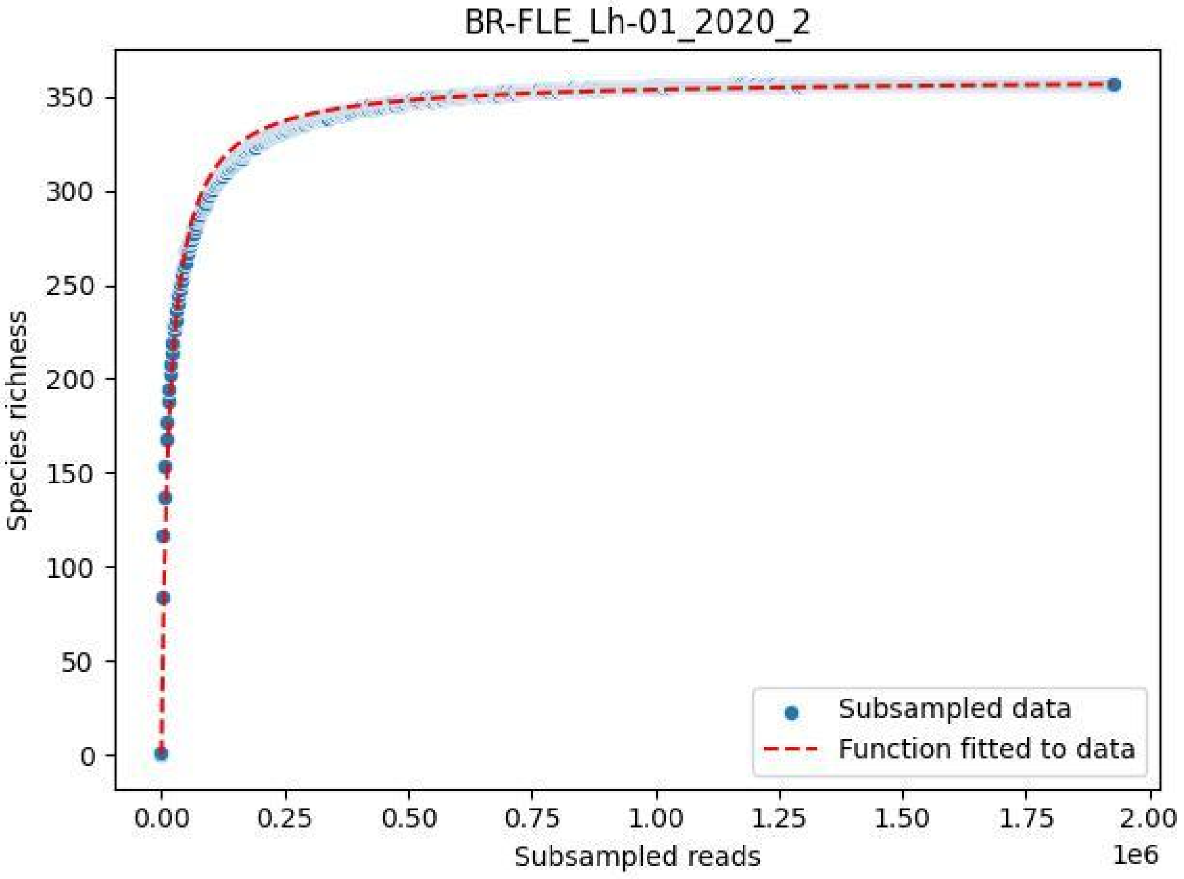

BR-FLE_Lh-01_2020_2
Species richness
Subsampled reads
1e6
0
50
100
150
200
250
300
350
0.00
0.25
0.50
0.75
1.00
1.25
1.50
1.75
2.00
Subsampled data
Function fitted to data

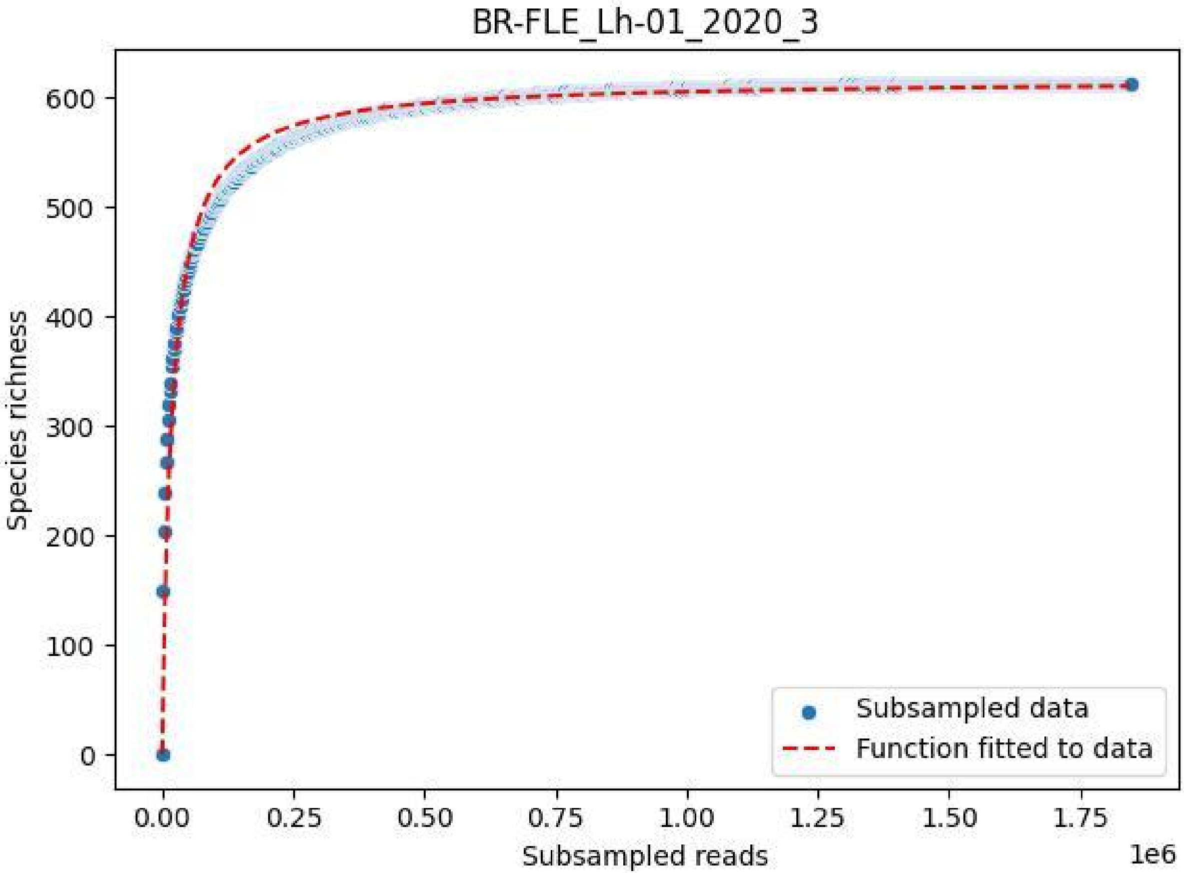

BR-FLE_Lh-01_2020_3
Species richness
Subsampled reads
1e6
0
100
200
300
400
500
600
0.00
0.25
0.50
0.75
1.00
1.25
1.50
1.75
Subsampled data
Function fitted to data

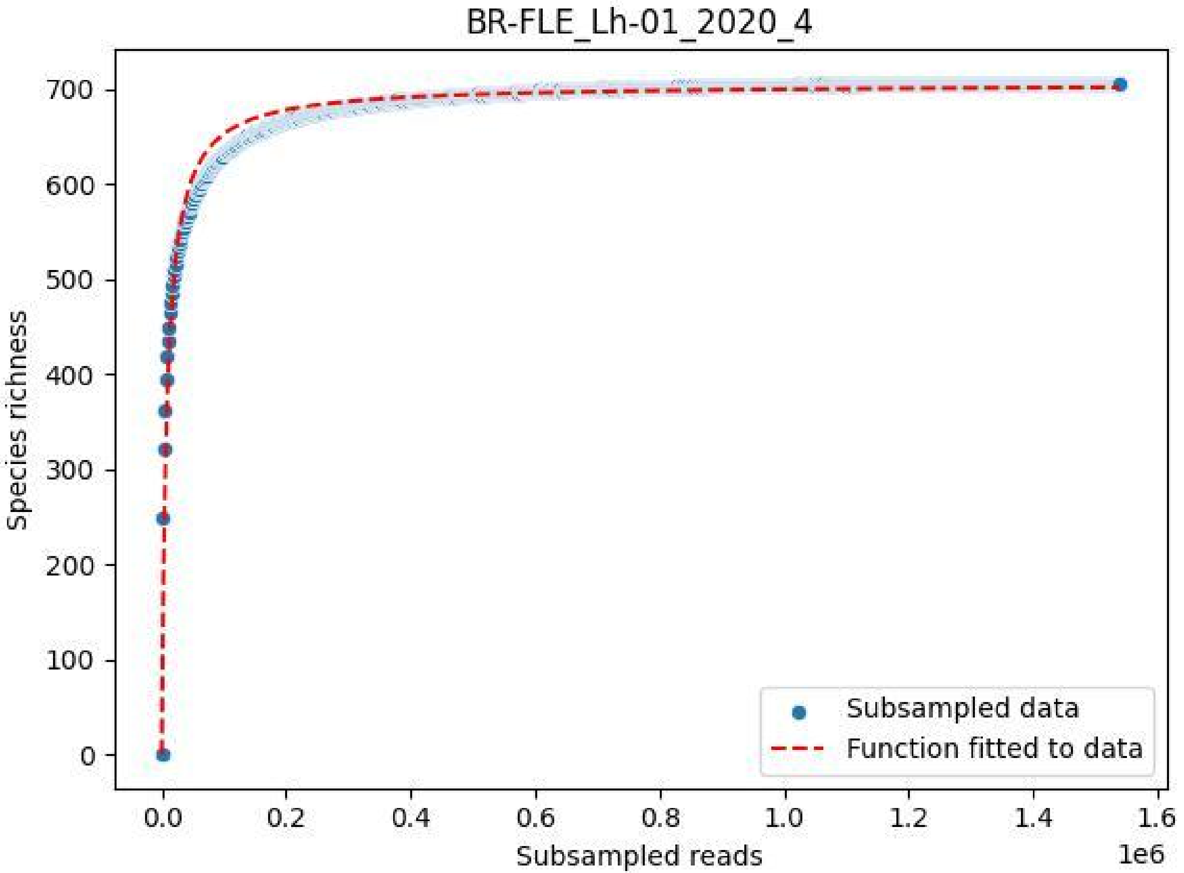
BR-FLE_Lh-01_2020_4
Species richness
Subsampled reads
1e6
0
100
200
300
400
500
600
700
0.0
0.2
0.4
0.6
0.8
1.0
1.2
1.4
1.6
Subsampled data
Function fitted to data

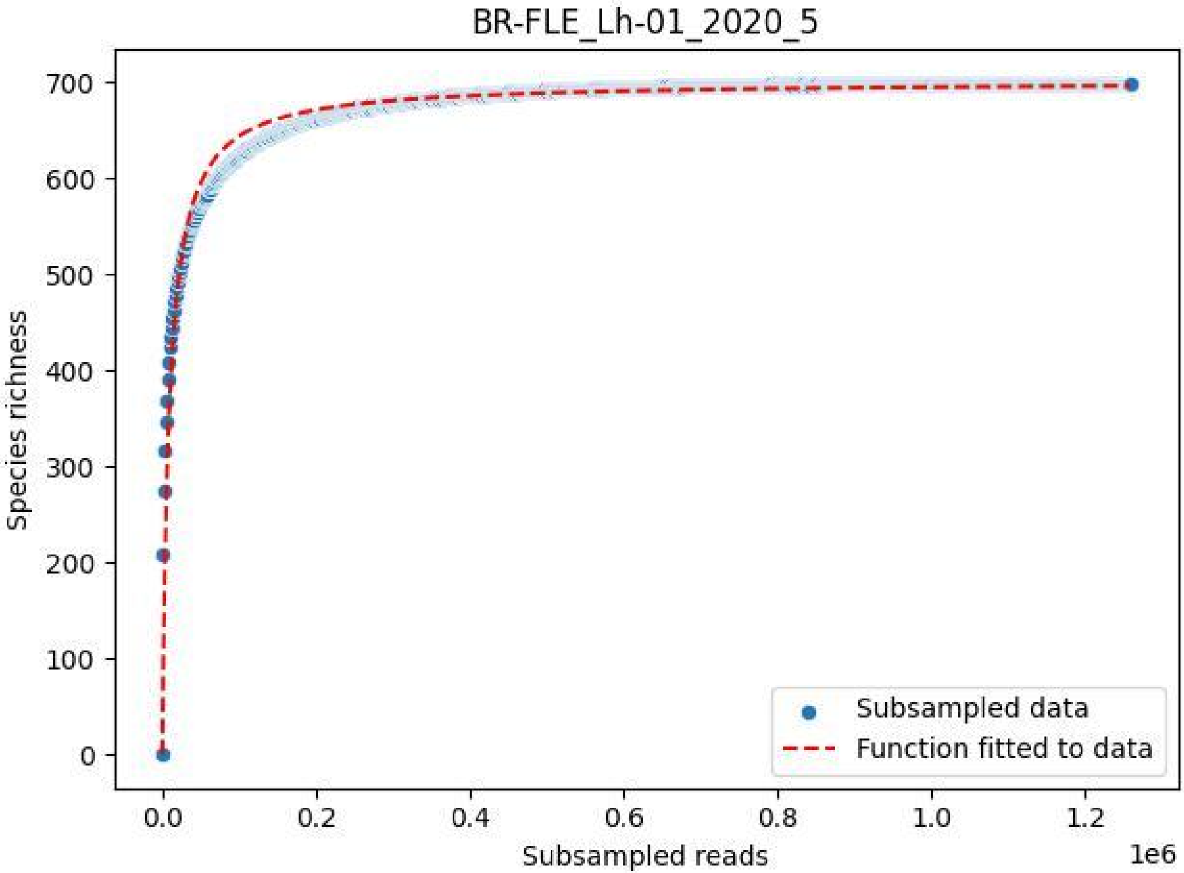
BR-FLE_Lh-01_2020_5
Species richness
Subsampled reads
1e6
0
100
200
300
400
500
600
700
0.0
0.2
0.4
0.6
0.8
1.0
1.2
Subsampled data
Function fitted to data

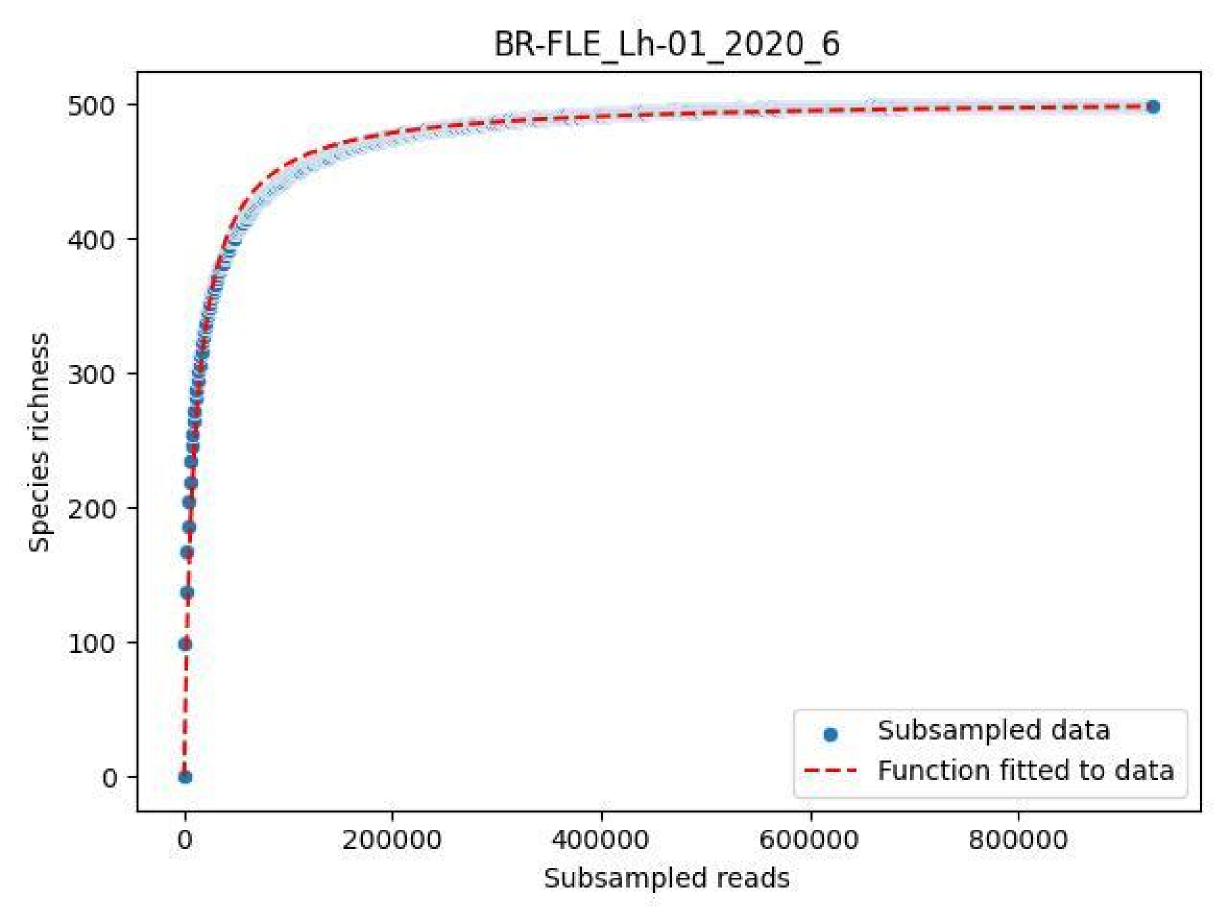
BR-FLE_Lh-01_2020_6
Species richness
Subsampled reads
0
100
200
300
400
500
0
200000
400000
600000
800000
Subsampled data
Function fitted to data

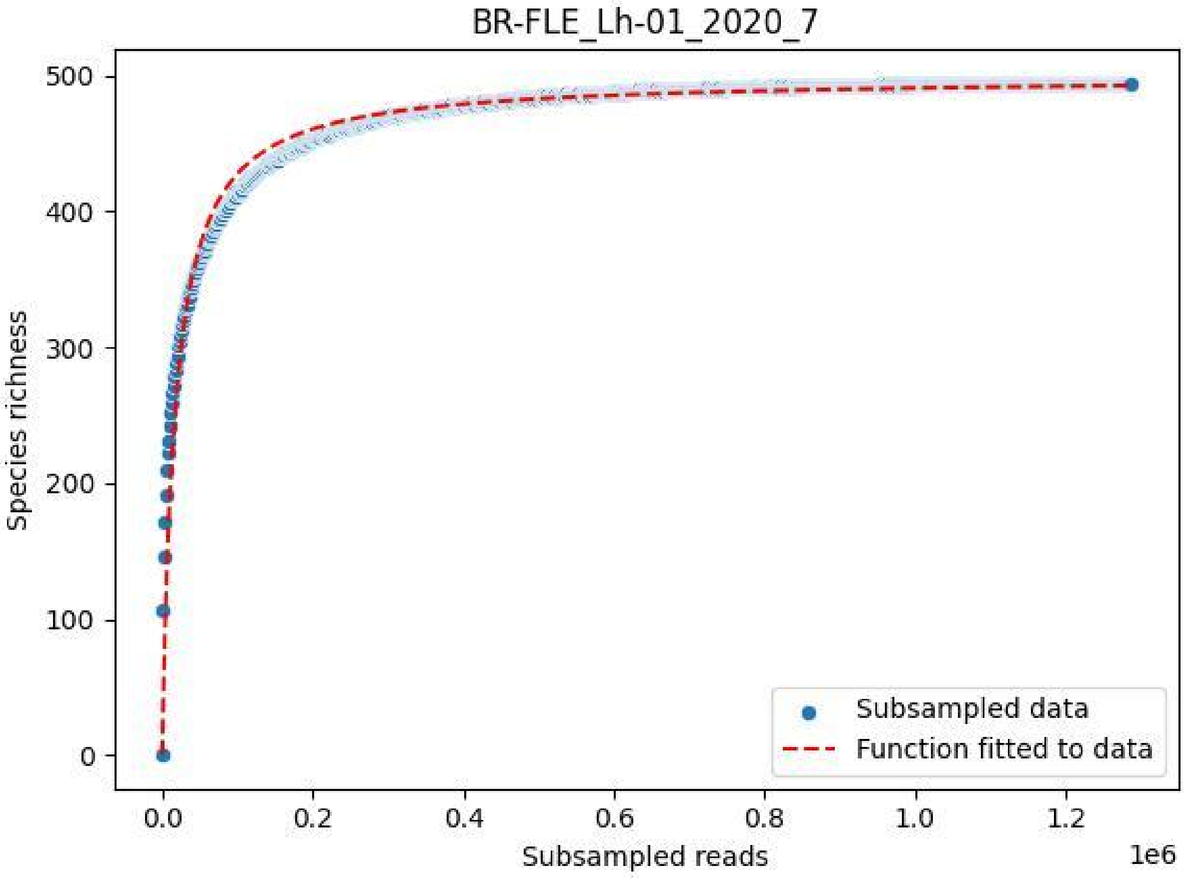

BR-FLE_Lh-01_2020_7
Species richness
500
400
300
200
100
0
0.0
0.2
0.4
0.6
0.8
1.0
1.2
1e6
Subsampled reads
Subsampled data
Function fitted to data

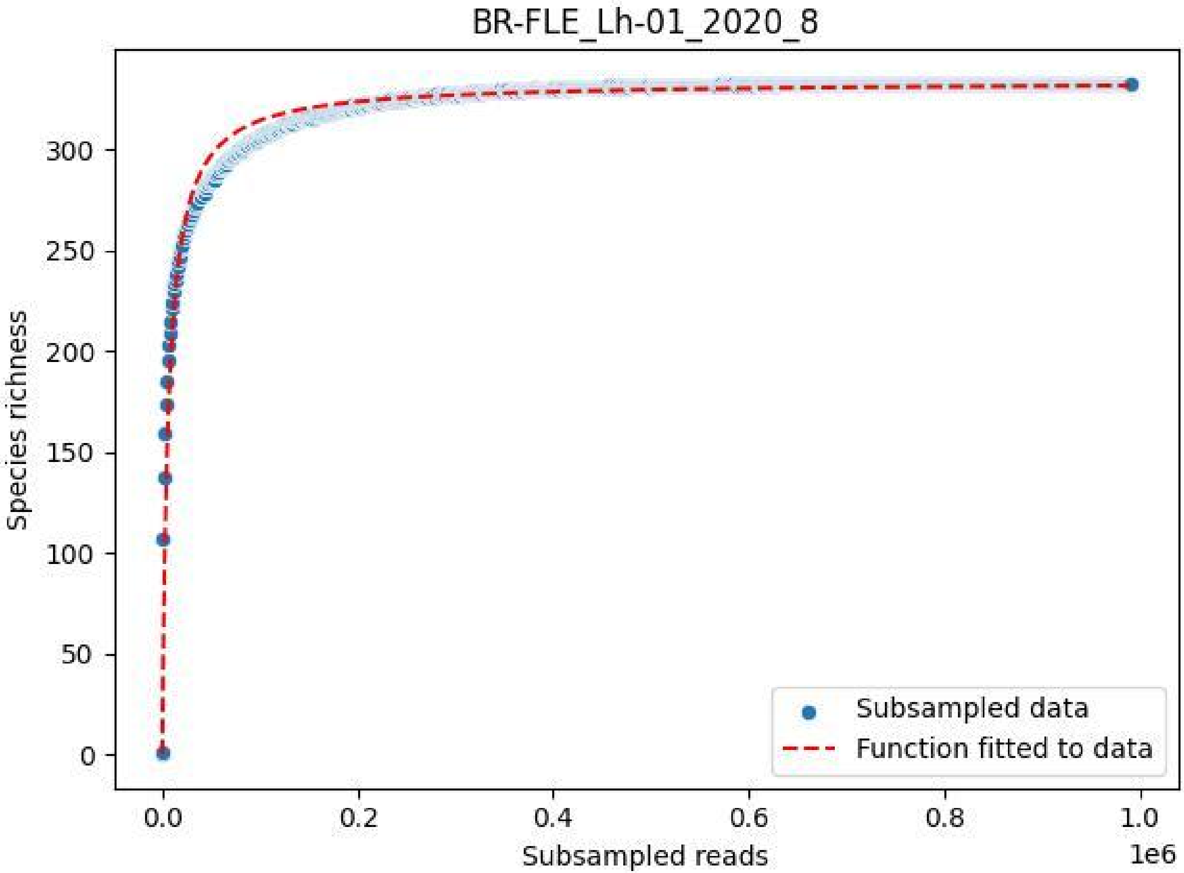
BR-FLE_Lh-01_2020_8
Species richness
Subsampled reads
1e6
0
50
100
150
200
250
300
0.0
0.2
0.4
0.6
0.8
1.0
Subsampled data
Function fitted to data

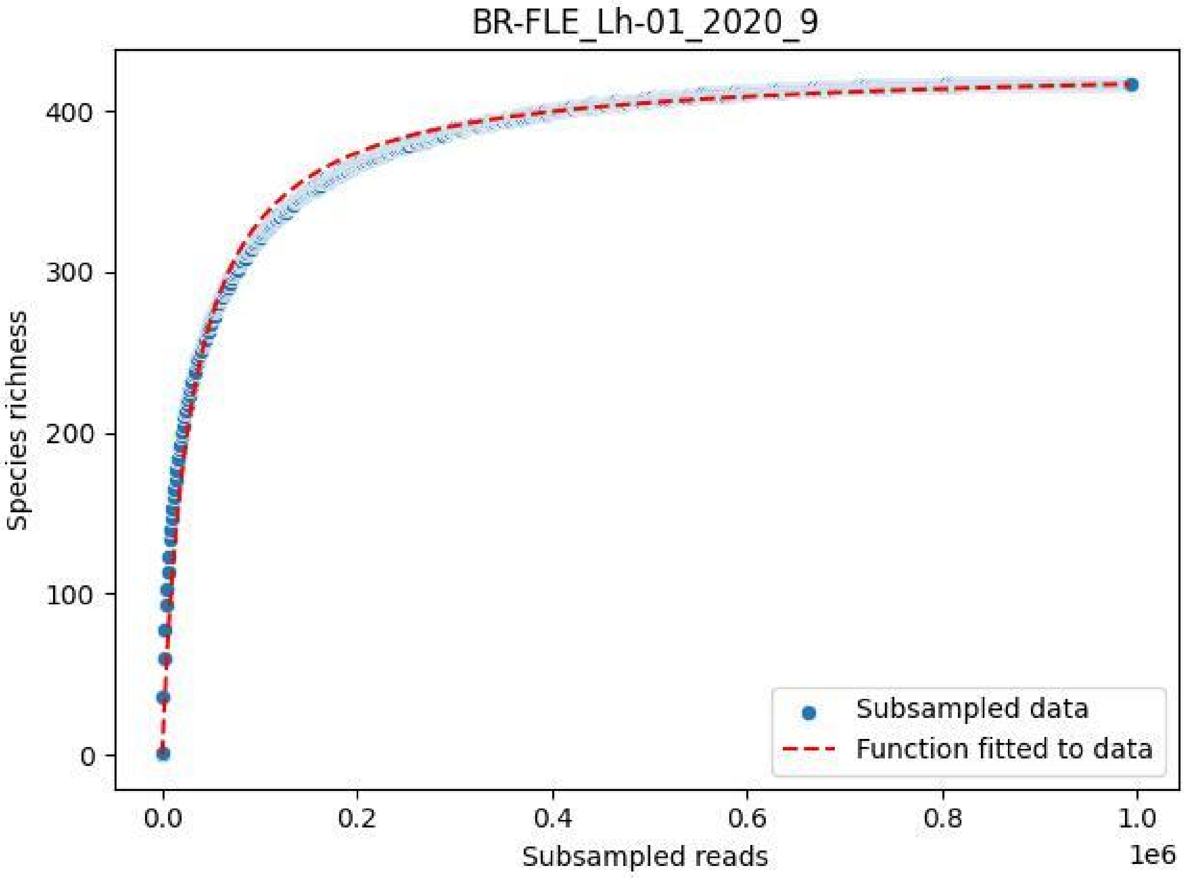
BR-FLE_Lh-01_2020_9
Species richness
Subsampled reads
1e6
0
100
200
300
400
0.0
0.2
0.4
0.6
0.8
1.0
Subsampled data
Function fitted to data

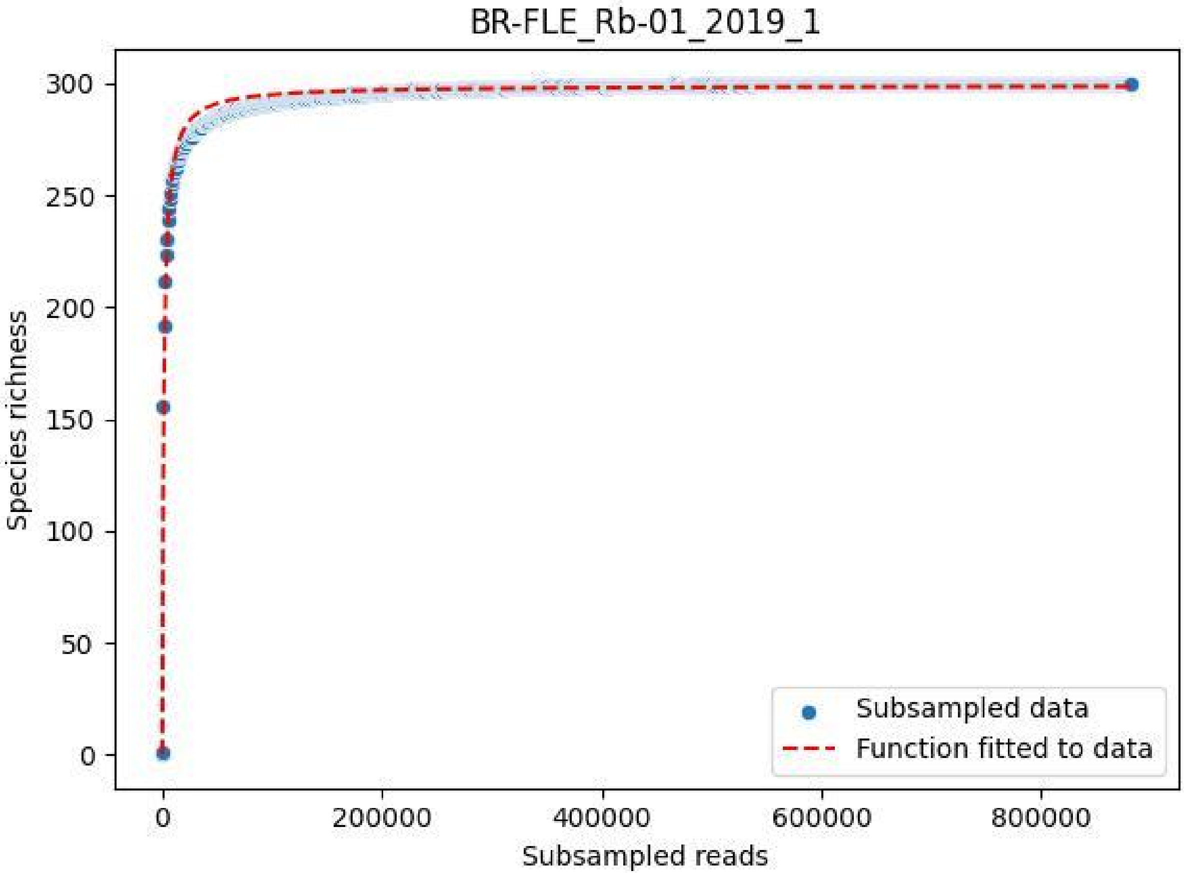
BR-FLE_Rb-01_2019_1
Species richness
Subsampled reads
0
50
100
150
200
250
300
0
200000
400000
600000
800000
Subsampled data
Function fitted to data

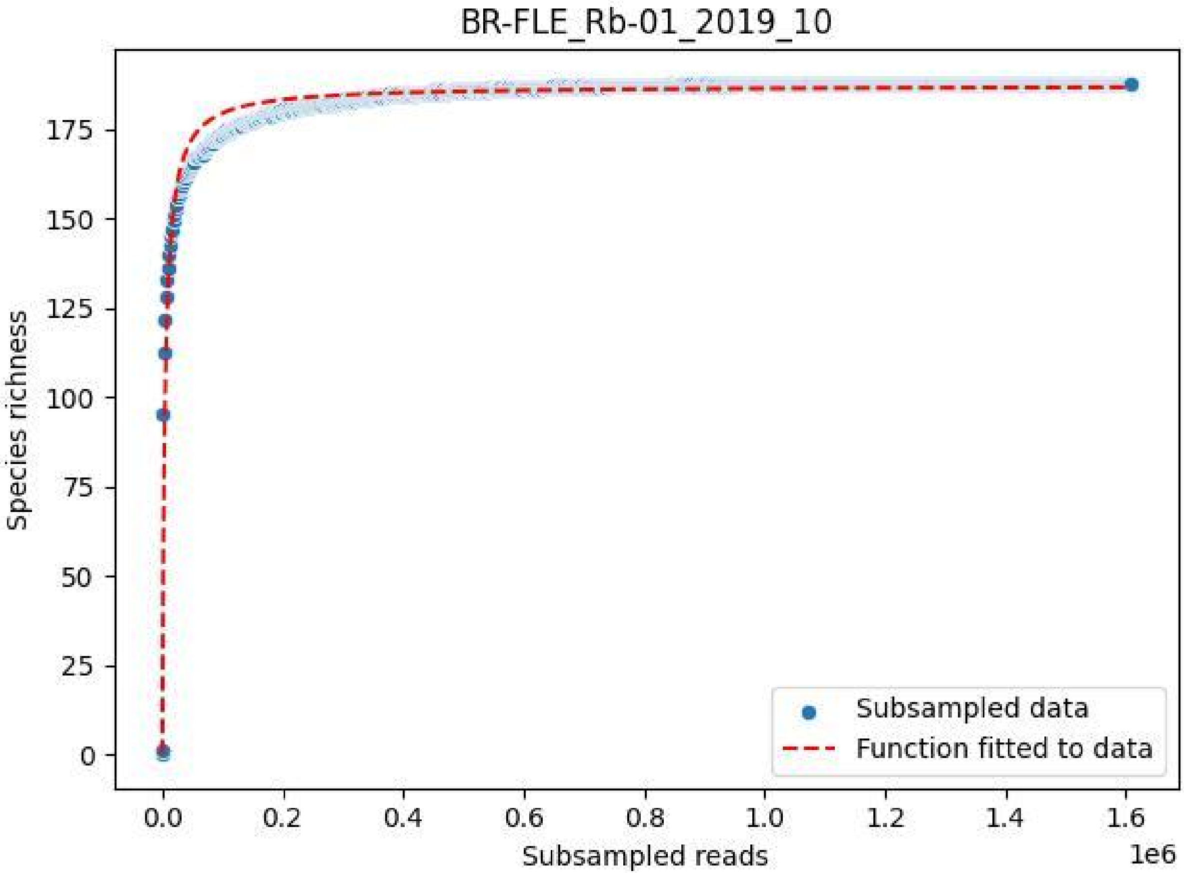
BR-FLE_Rb-01_2019_10
Species richness
Subsampled reads
1e6
0
25
50
75
100
125
150
175
0.0
0.2
0.4
0.6
0.8
1.0
1.2
1.4
1.6
Subsampled data
Function fitted to data

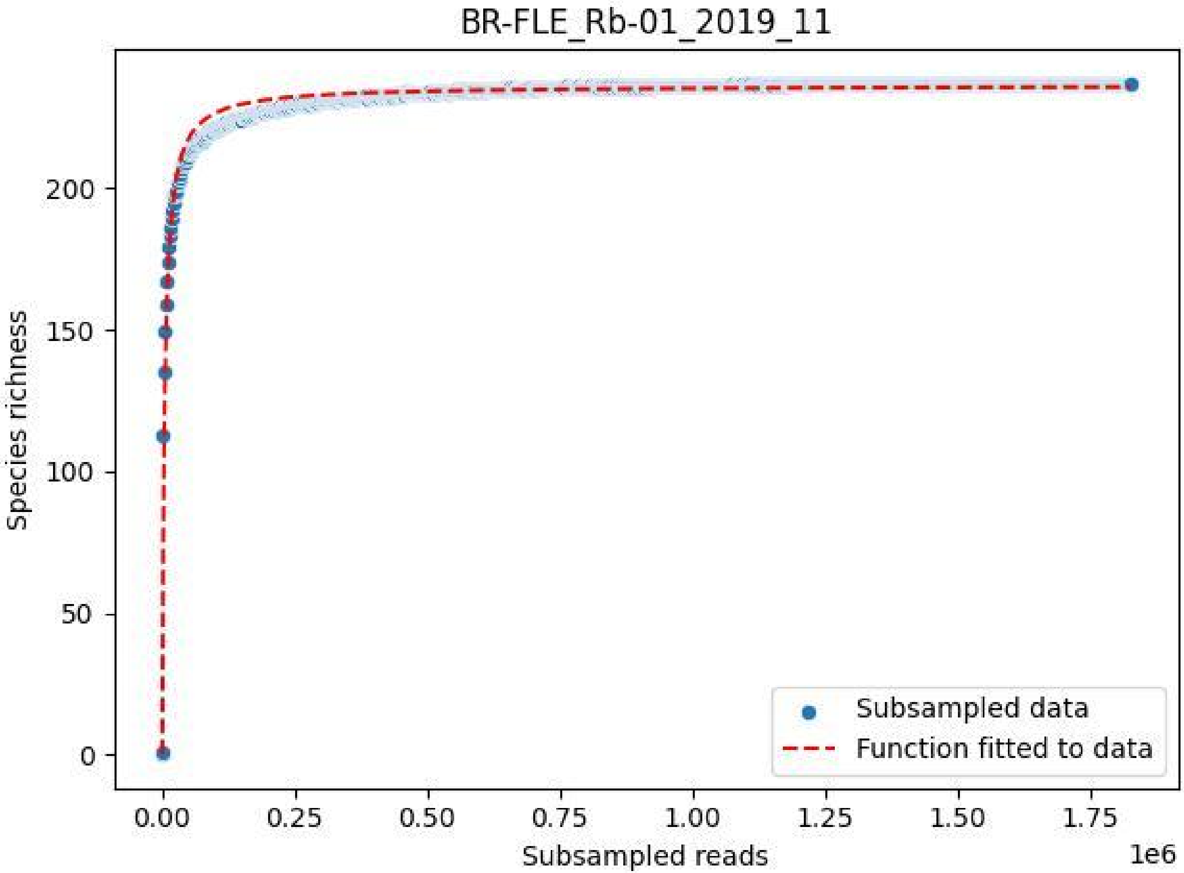
BR-FLE_Rb-01_2019_11
Species richness
Subsampled reads
1e6
0
50
100
150
200
0.00
0.25
0.50
0.75
1.00
1.25
1.50
1.75
Subsampled data
Function fitted to data

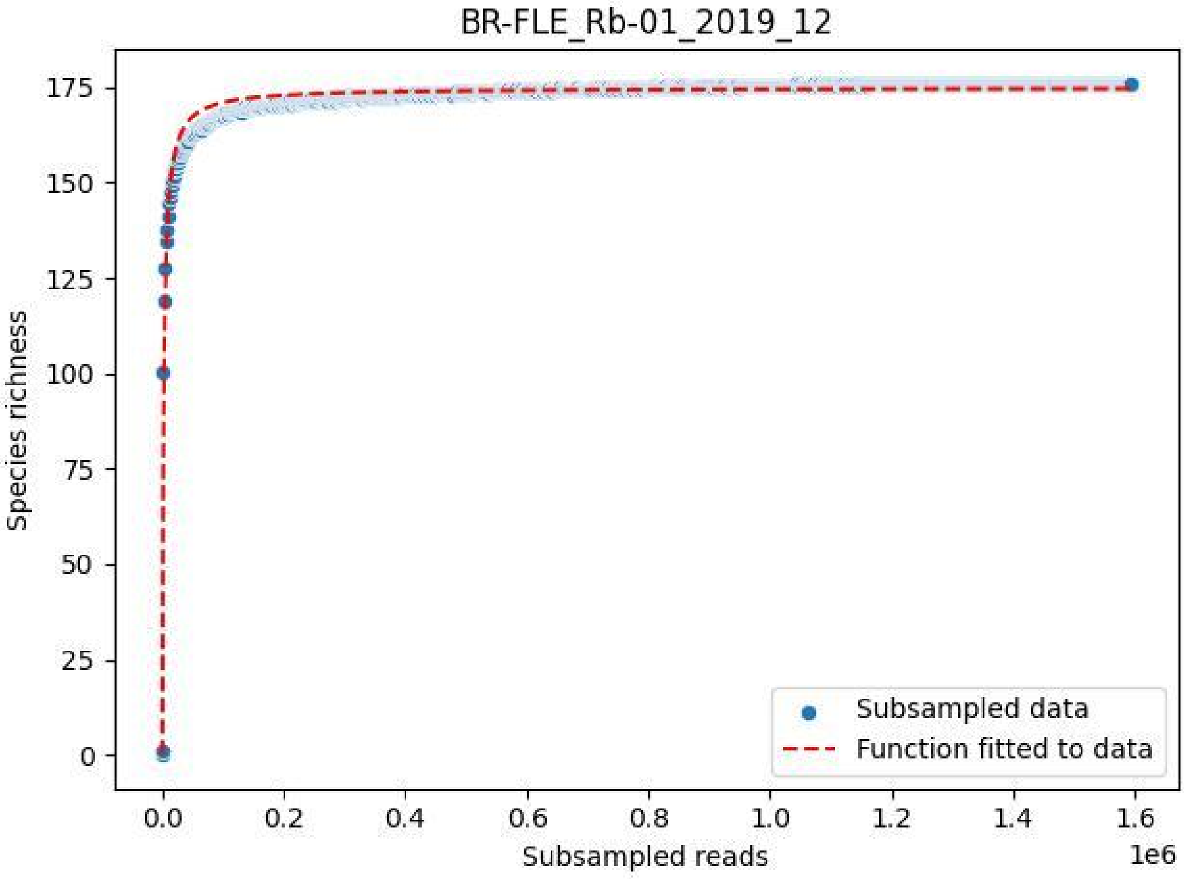

BR-FLE_Rb-01_2019_12
Species richness
Subsampled reads
1e6
0
25
50
75
100
125
150
175
0.0
0.2
0.4
0.6
0.8
1.0
1.2
1.4
1.6
Subsampled data
Function fitted to data

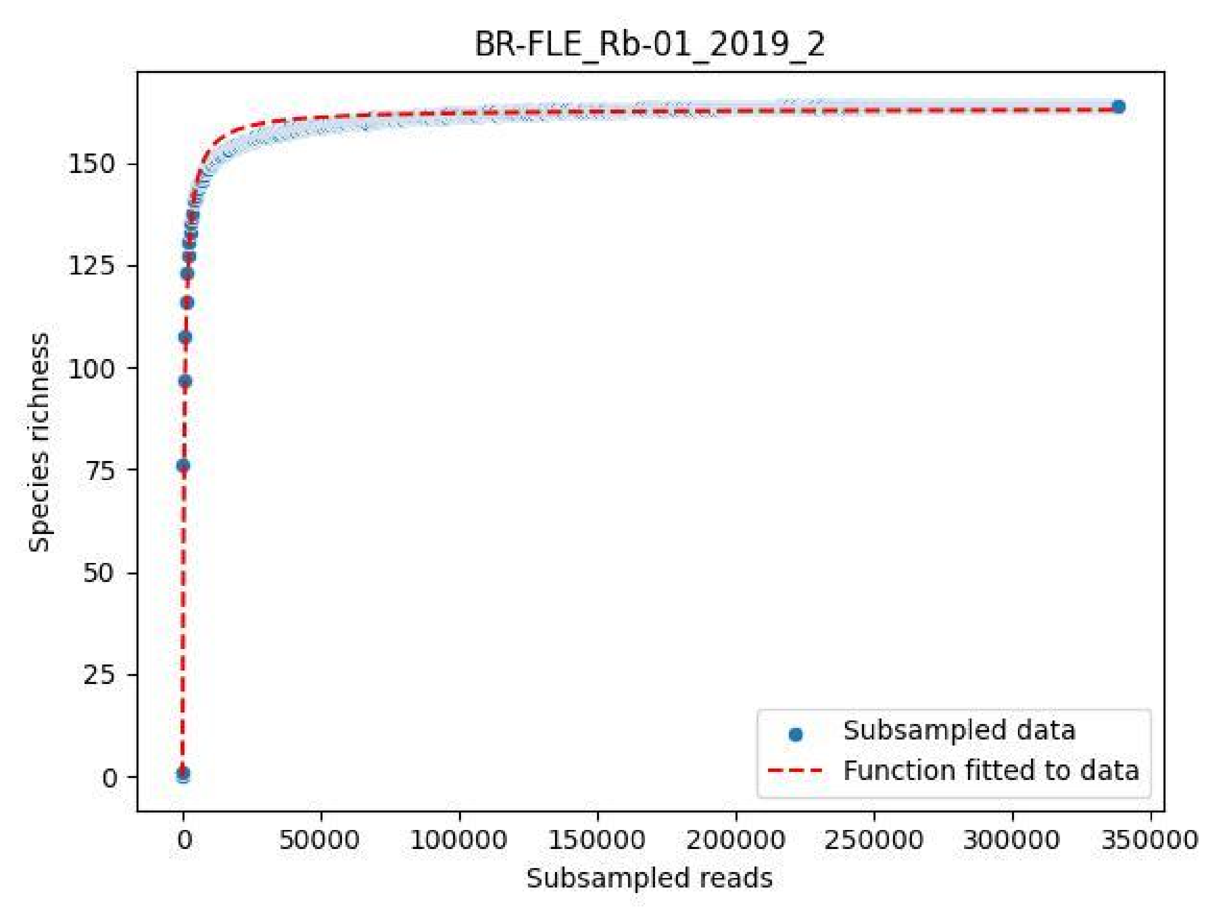
BR-FLE_Rb-01_2019_2
Species richness
Subsampled reads
0
25
50
75
100
125
150
0
50000
100000
150000
200000
250000
300000
350000
Subsampled data
Function fitted to data

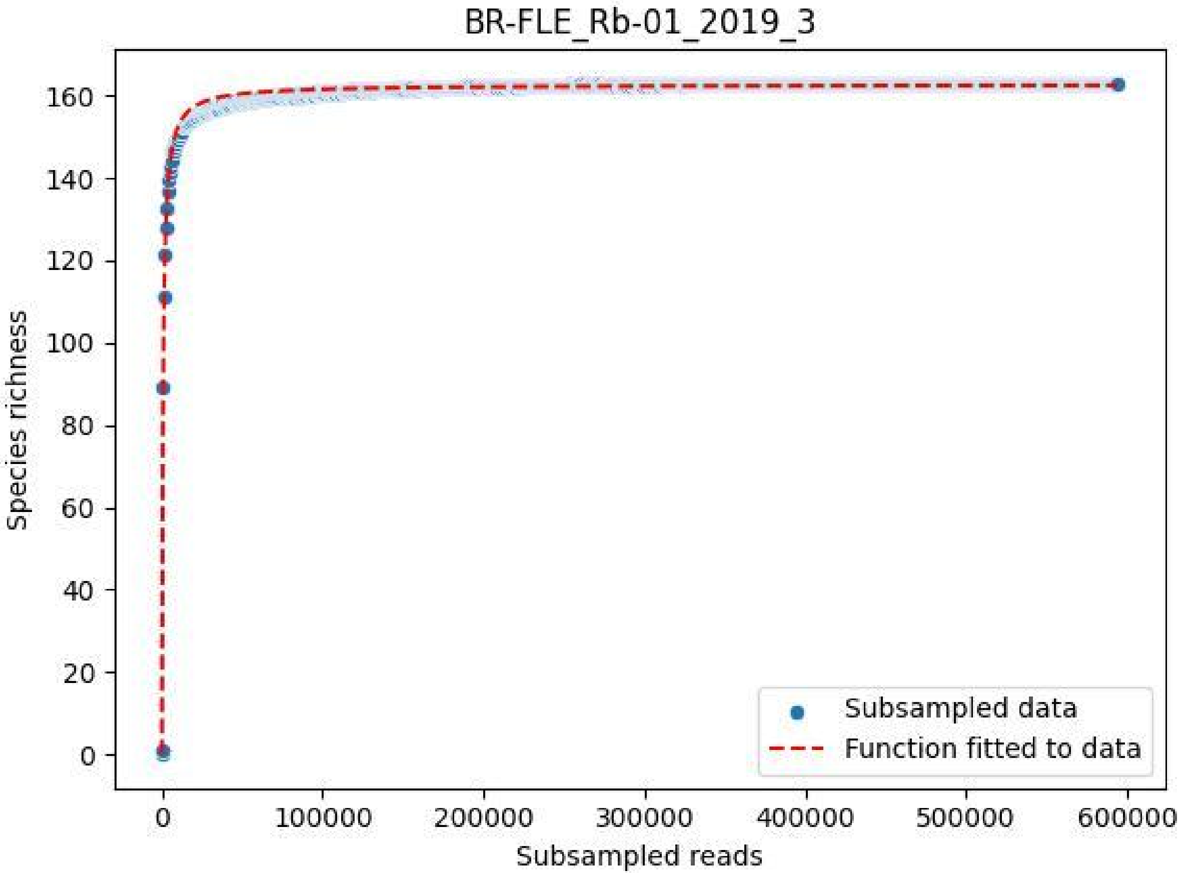
BR-FLE_Rb-01_2019_3
Species richness
Subsampled reads
0
20
40
60
80
100
120
140
160
0
100000
200000
300000
400000
500000
600000
Subsampled data
Function fitted to data

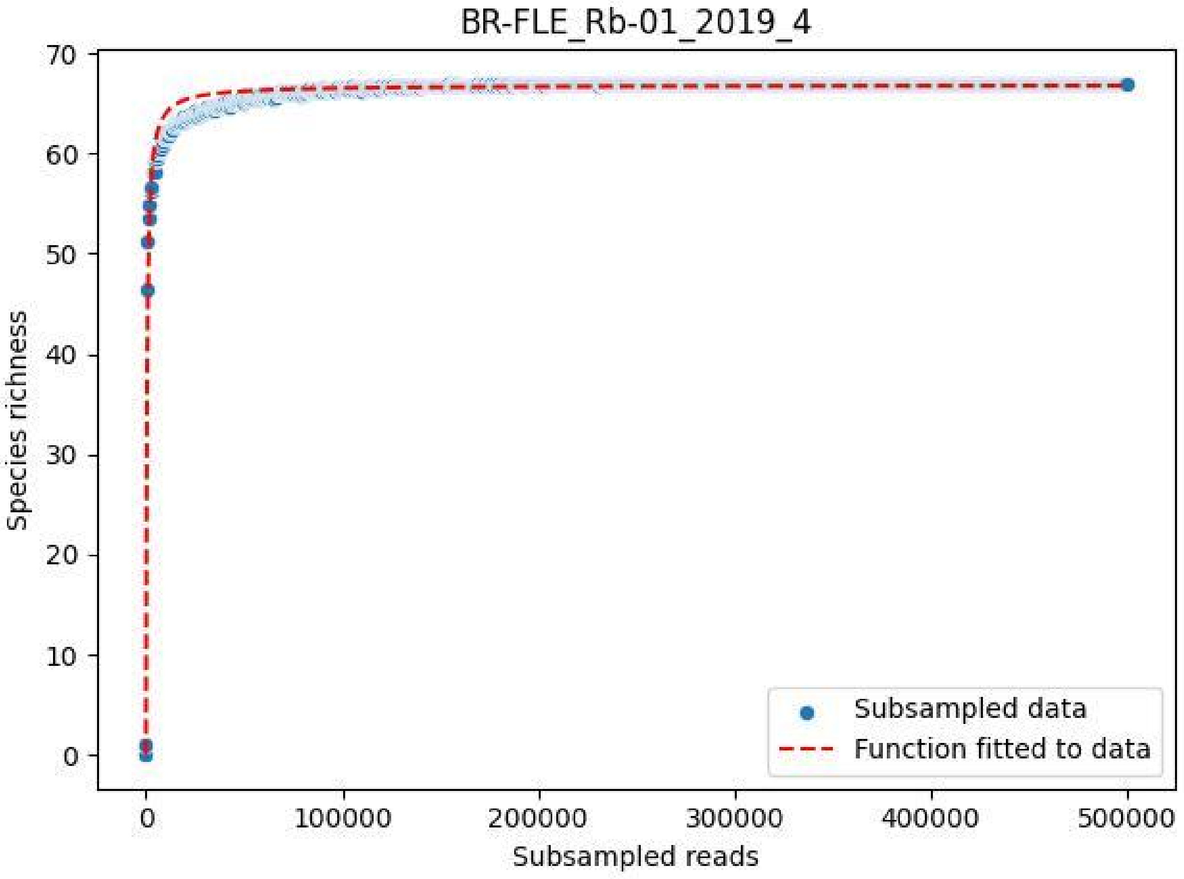
BR-FLE_Rb-01_2019_4
Species richness
Subsampled reads
0
10
20
30
40
50
60
70
0
100000
200000
300000
400000
500000
Subsampled data
Function fitted to data

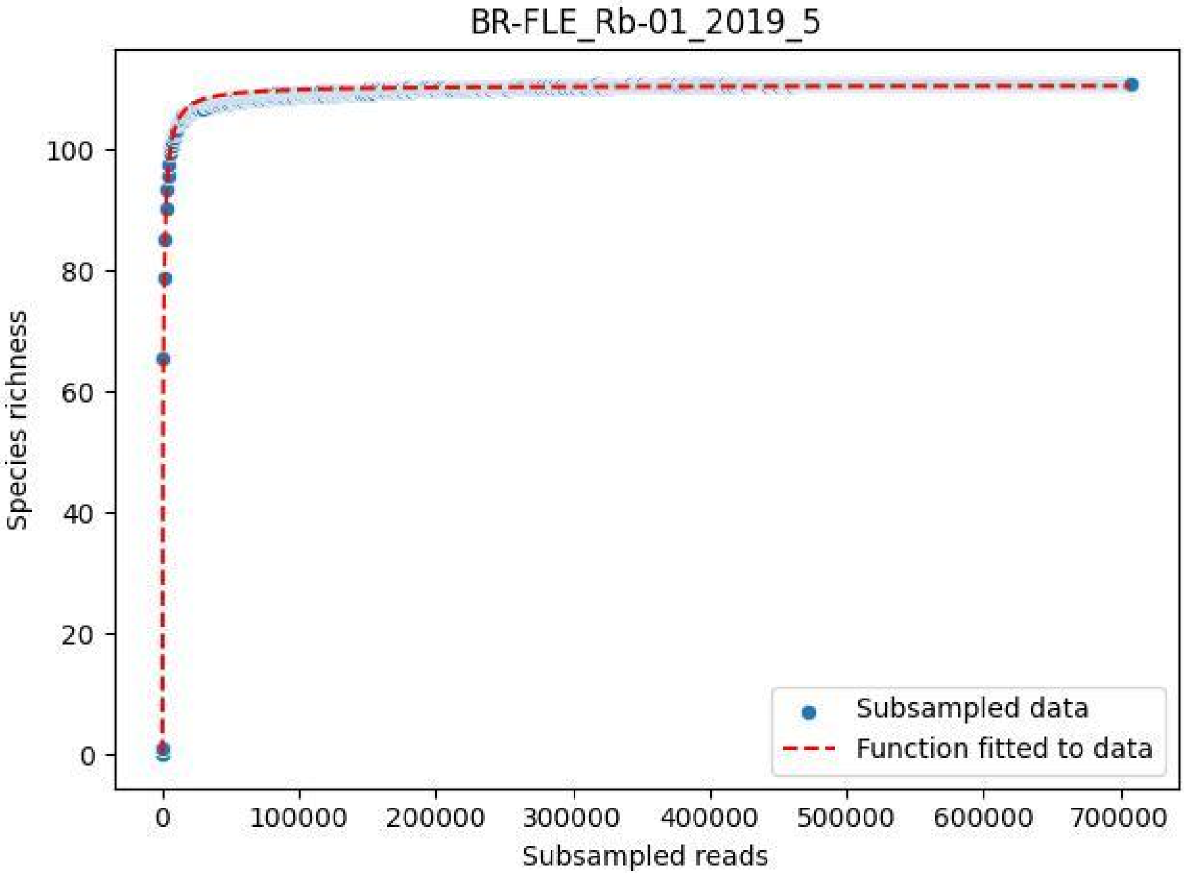
BR-FLE_Rb-01_2019_5
Species richness
0
20
40
60
80
100
0
100000
200000
300000
400000
500000
600000
700000
Subsampled reads
Subsampled data
Function fitted to data

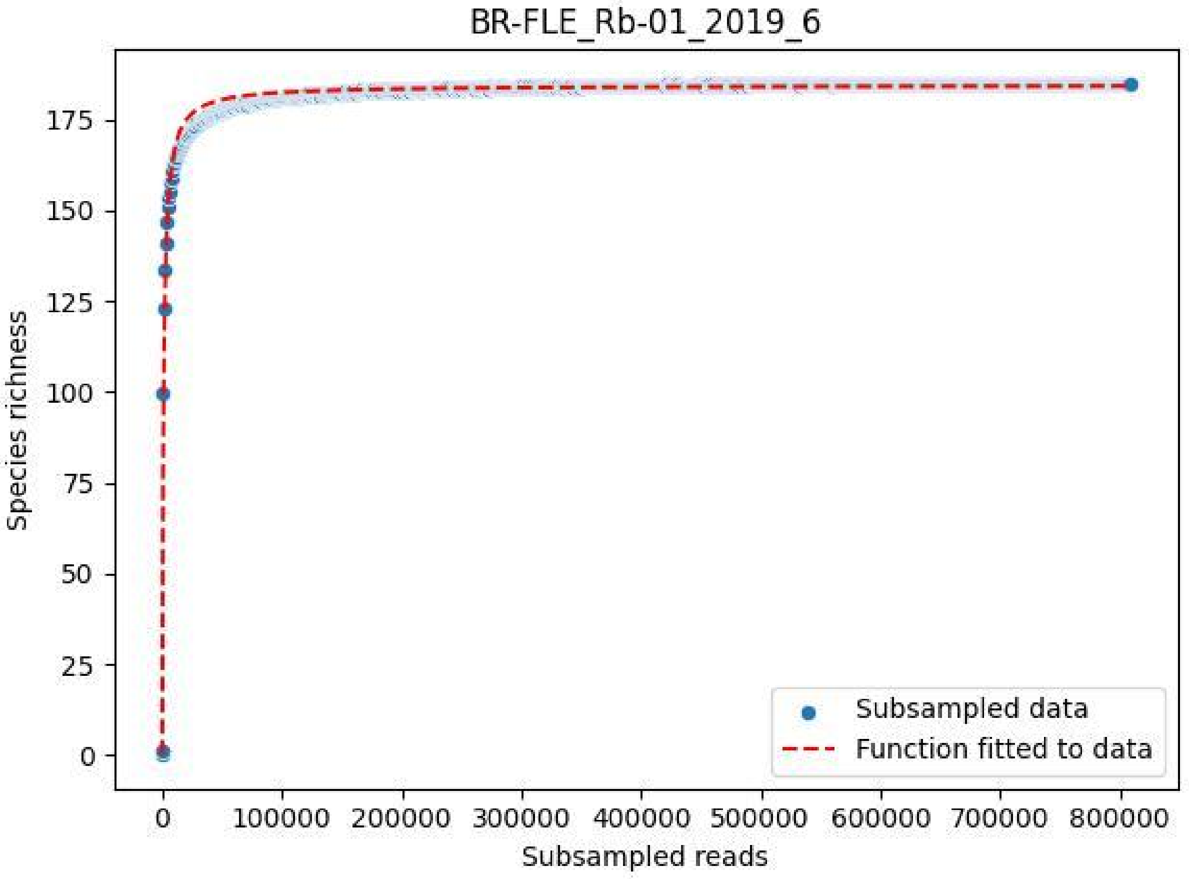
BR-FLE_Rb-01_2019_6
Species richness
Subsampled reads
Subsampled data
Function fitted to data
0
25
50
75
100
125
150
175
0
100000
200000
300000
400000
500000
600000
700000
800000

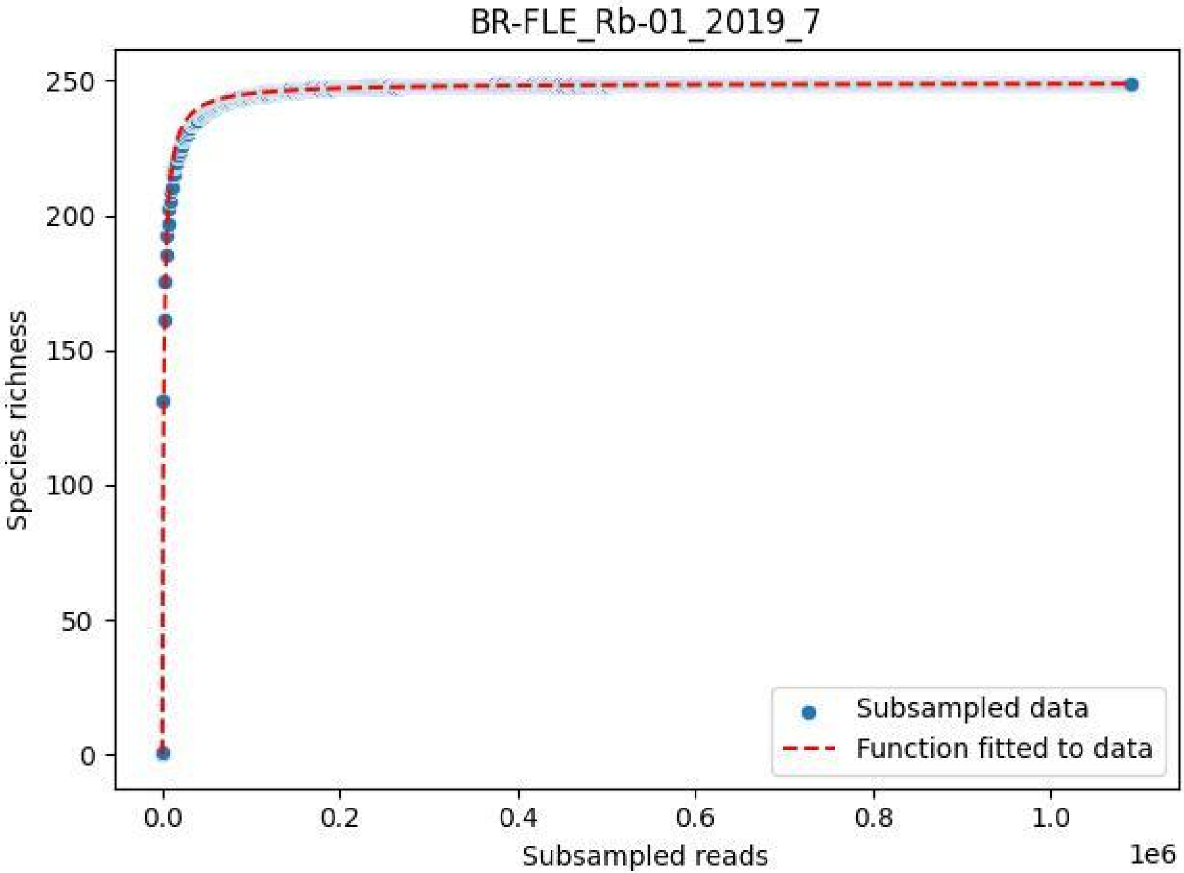
BR-FLE_Rb-01_2019_7
Species richness
250
200
150
100
50
0
0.0
0.2
0.4
0.6
0.8
1.0
1e6
Subsampled reads
Subsampled data
Function fitted to data

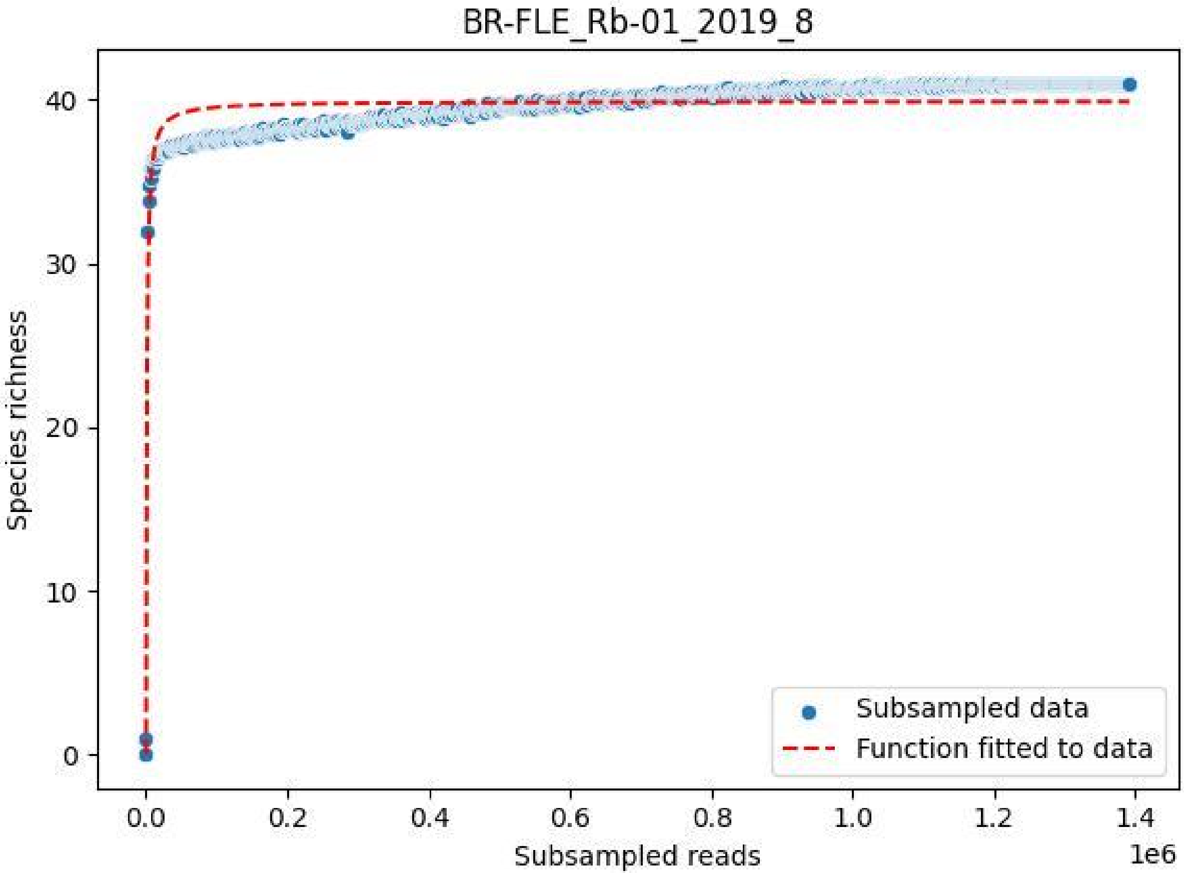
BR-FLE_Rb-01_2019_8
Species richness
Subsampled reads
1e6
0
10
20
30
40
0.0
0.2
0.4
0.6
0.8
1.0
1.2
1.4
Subsampled data
Function fitted to data

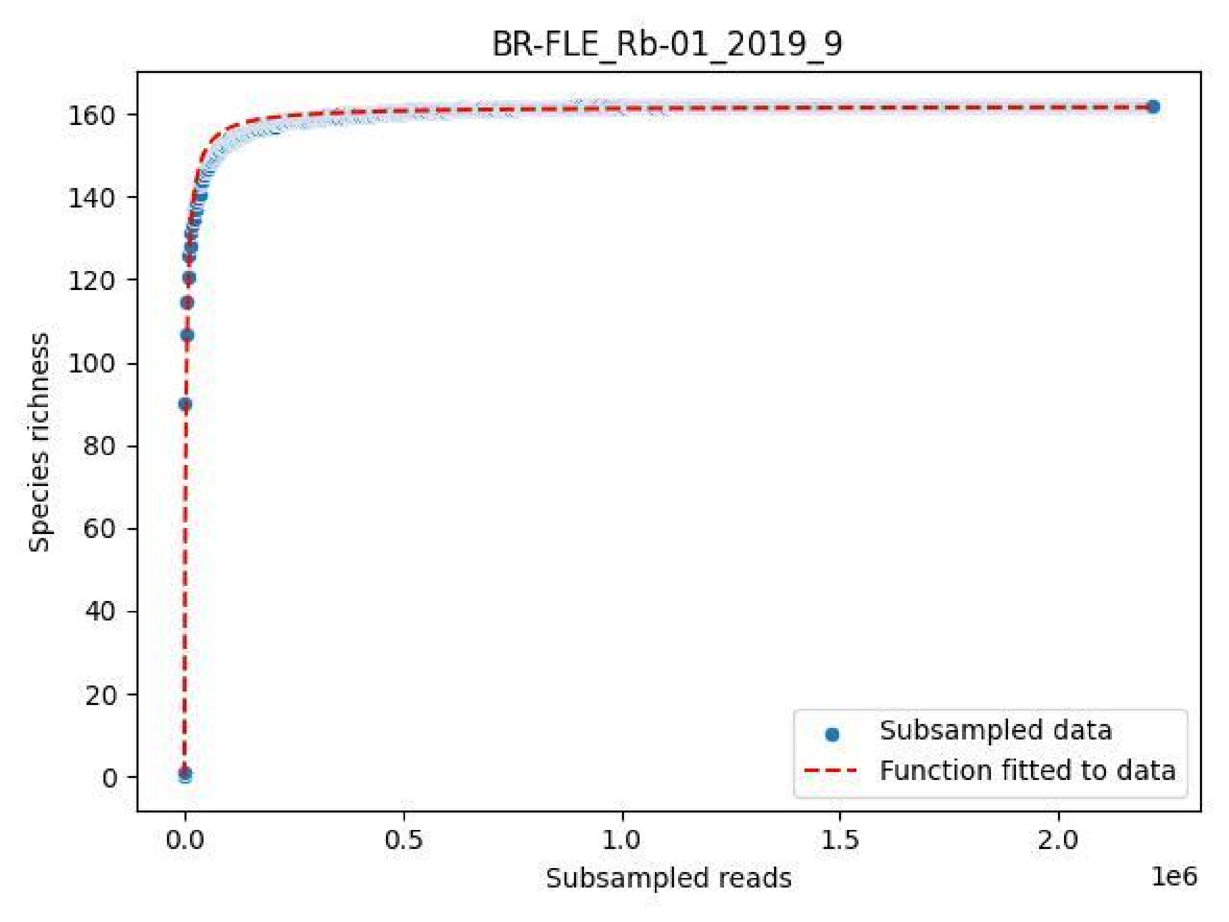
BR-FLE_Rb-01_2019_9
Species richness
160
140
120
100
80
60
40
20
0
0.0
0.5
1.0
1.5
2.0
1e6
Subsampled reads
Subsampled data
Function fitted to data

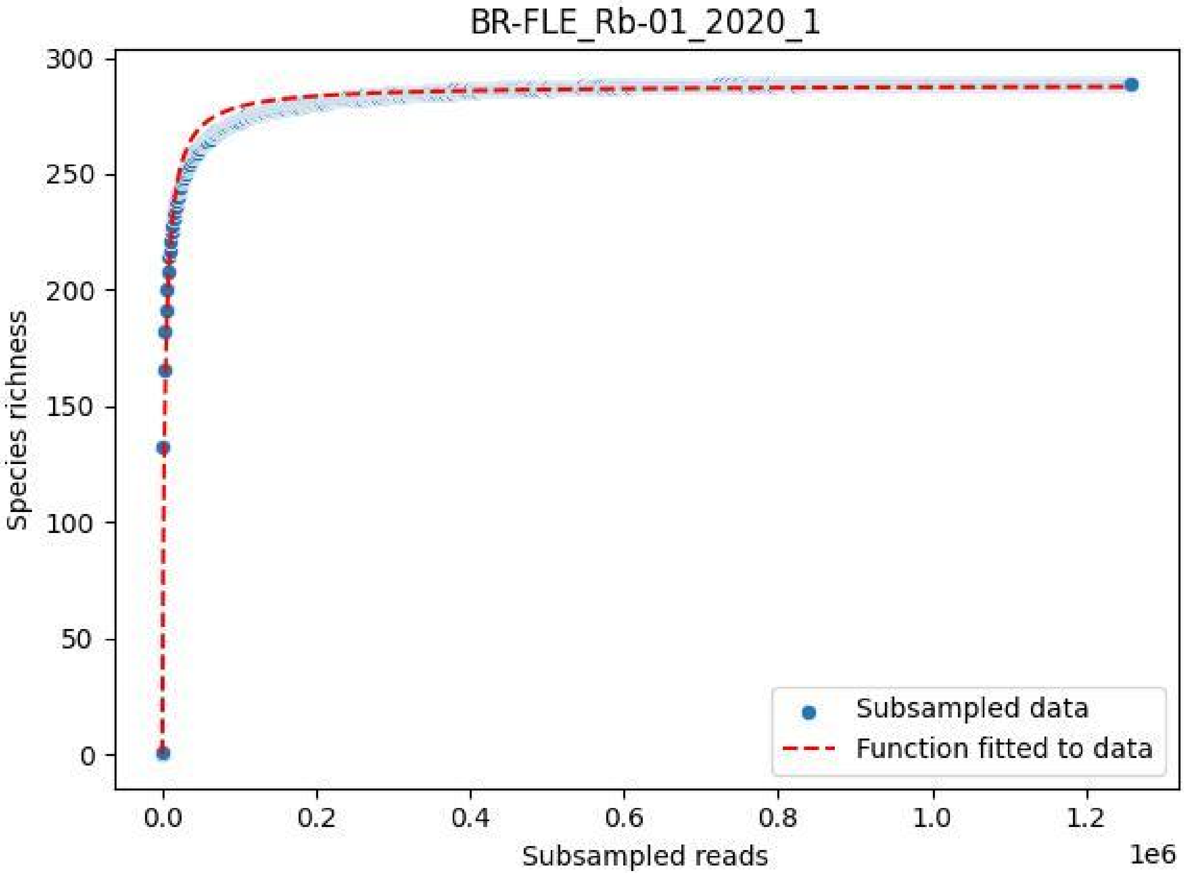
BR-FLE_Rb-01_2020_1
Species richness
Subsampled reads
1e6
Subsampled data
Function fitted to data
300
250
200
150
100
50
0
0.0
0.2
0.4
0.6
0.8
1.0
1.2

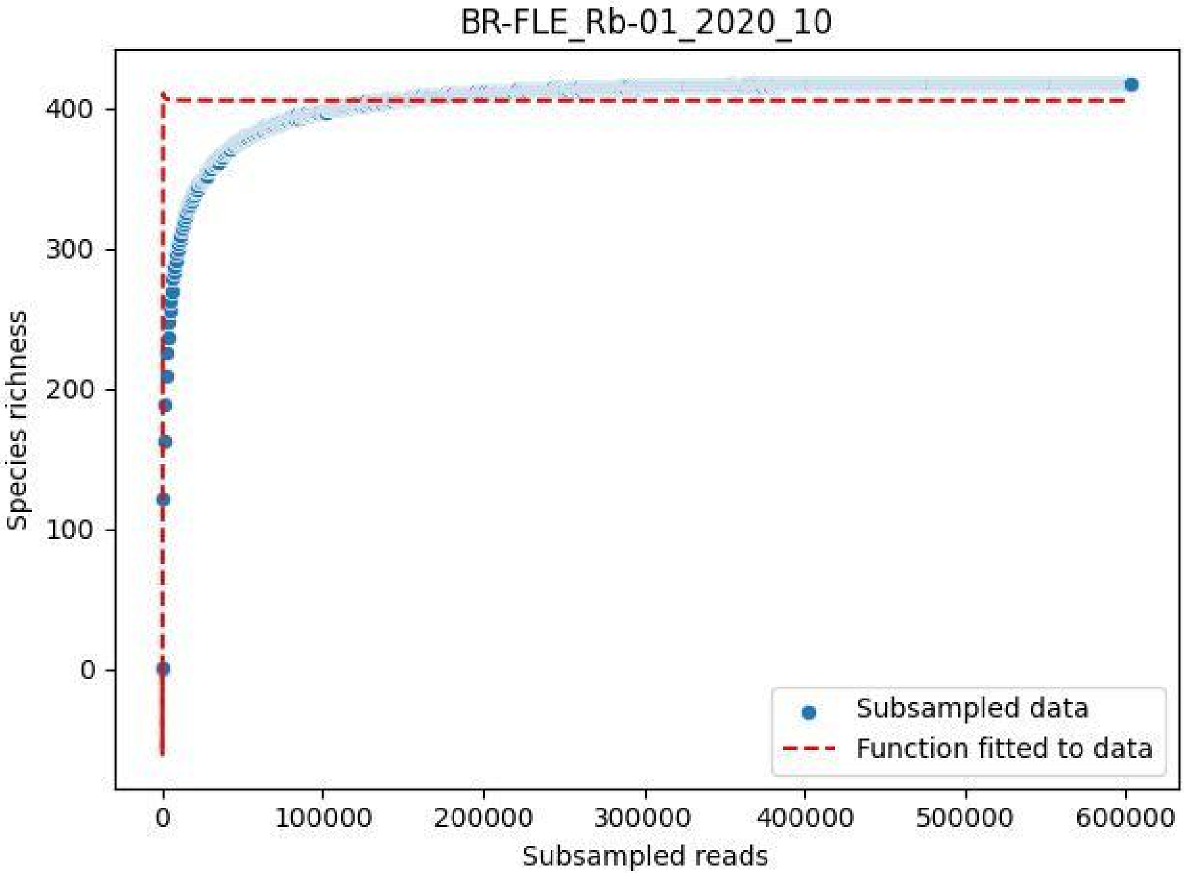
BR-FLE_Rb-01_2020_10
Species richness
Subsampled reads
0
100
200
300
400
0
100000
200000
300000
400000
500000
600000
Subsampled data
Function fitted to data

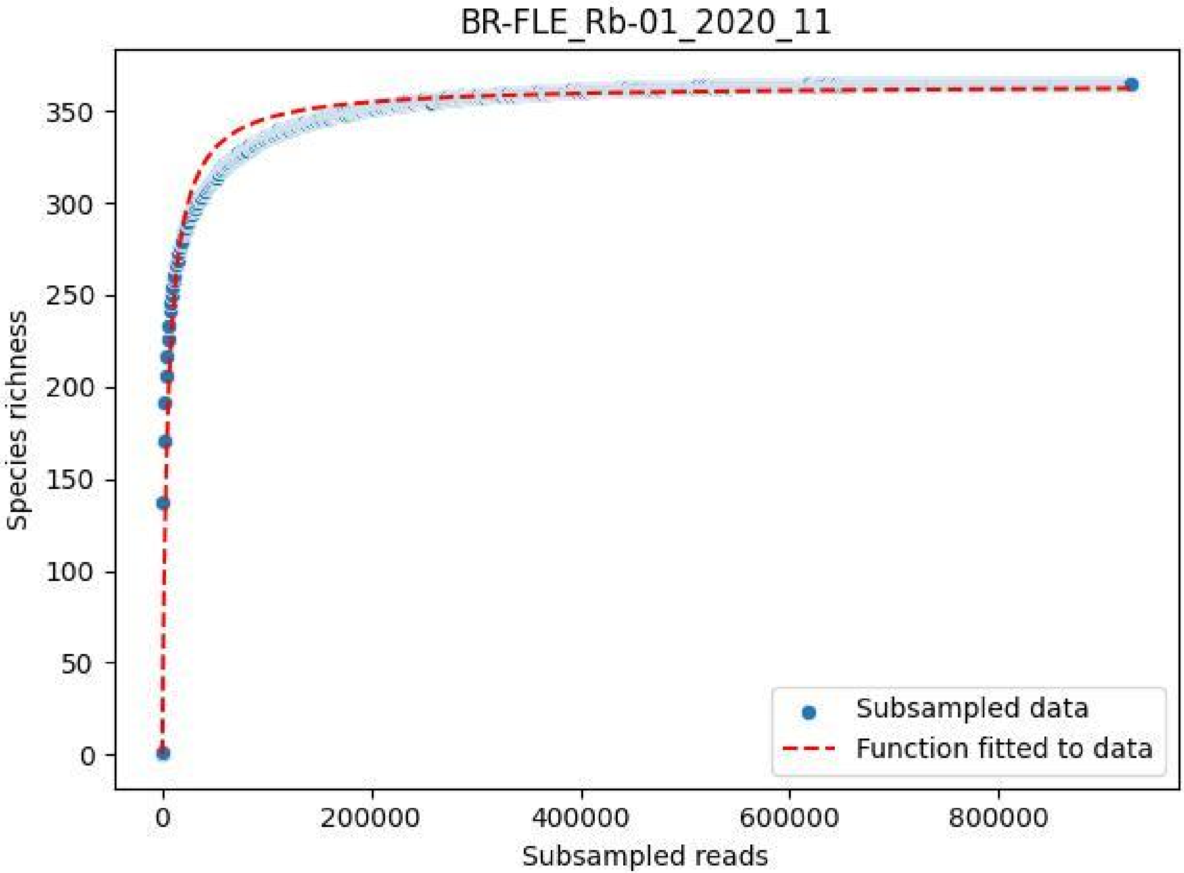
BR-FLE_Rb-01_2020_11
Species richness
0
50
100
150
200
250
300
350
Subsampled reads
0
200000
400000
600000
800000
Subsampled data
Function fitted to data

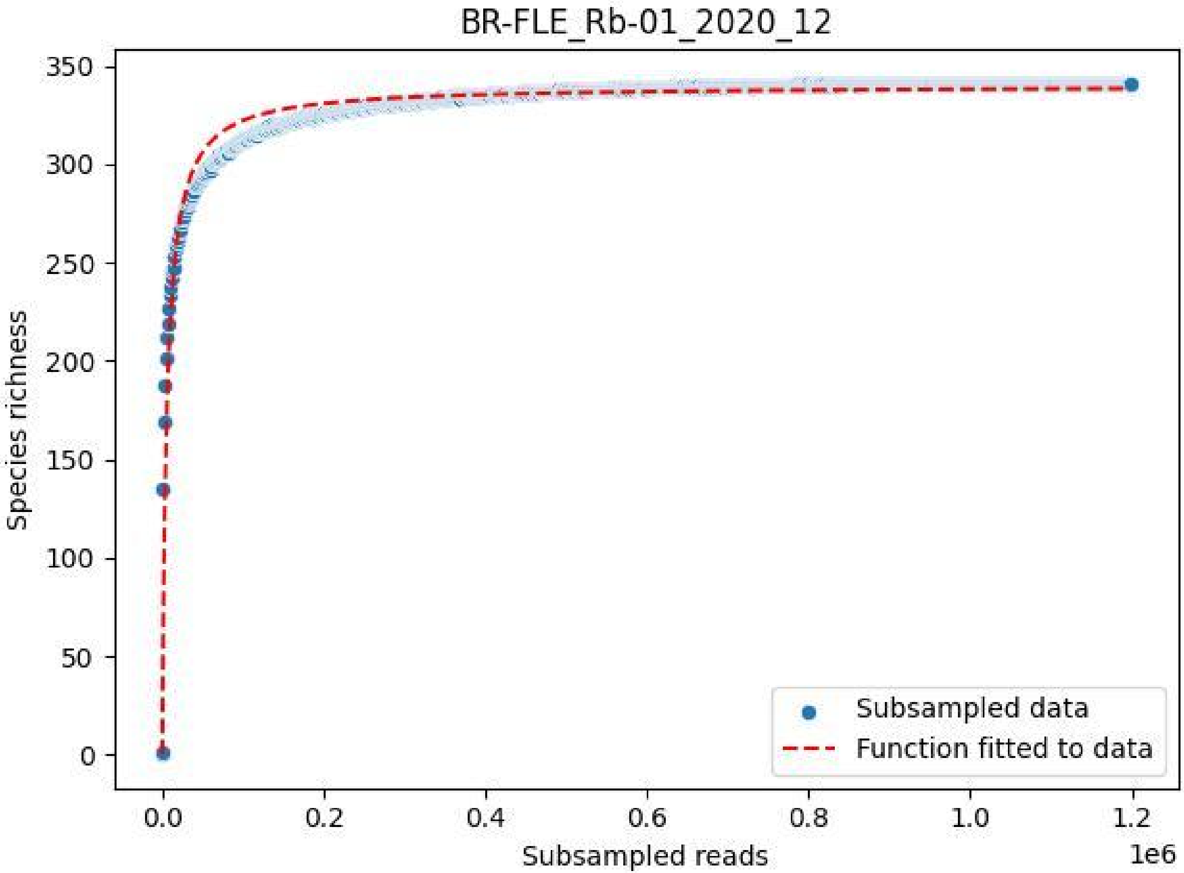
BR-FLE_Rb-01_2020_12
Species richness
350
300
250
200
150
100
50
0
0.0
0.2
0.4
0.6
0.8
1.0
1.2
Subsampled reads
1e6
Subsampled data
Function fitted to data

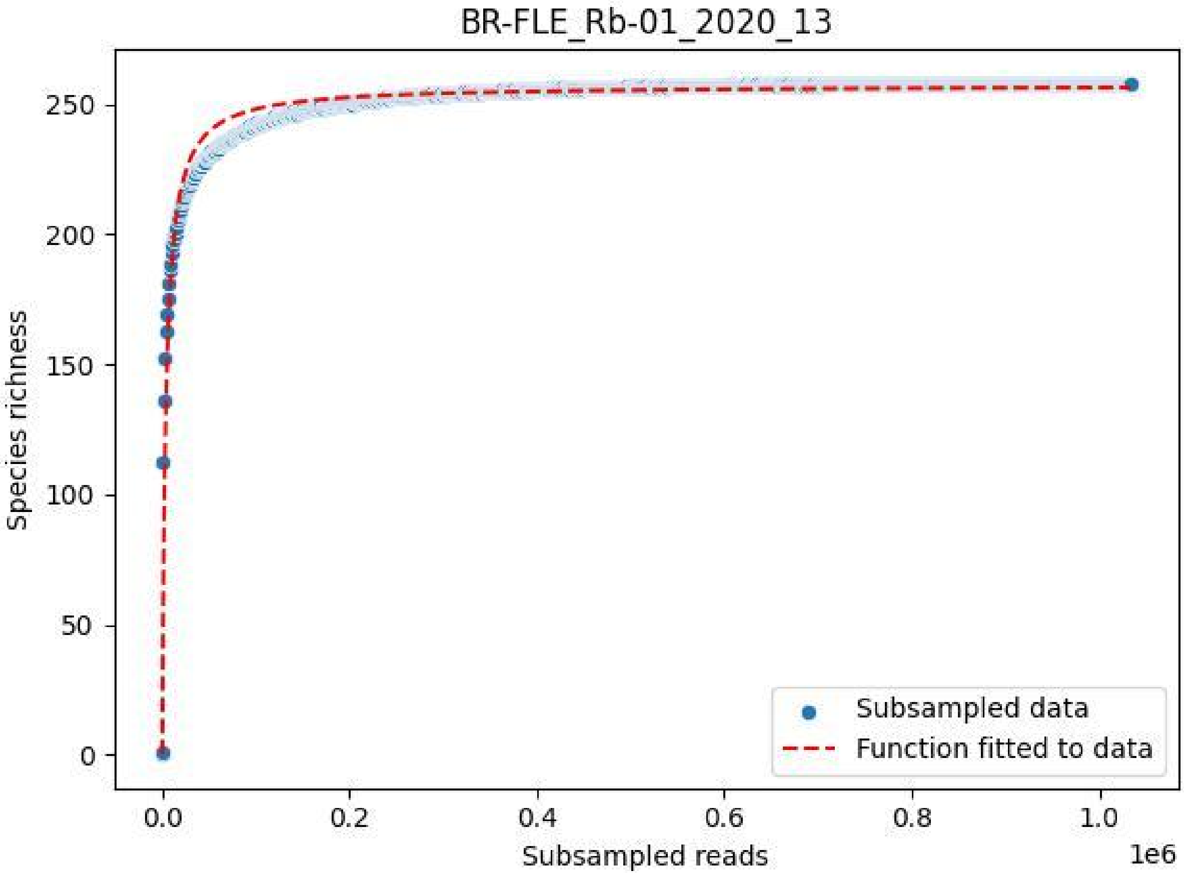

BR-FLE_Rb-01_2020_13
Species richness
Subsampled reads
1e6
0
50
100
150
200
250
0.0
0.2
0.4
0.6
0.8
1.0
Subsampled data
Function fitted to data

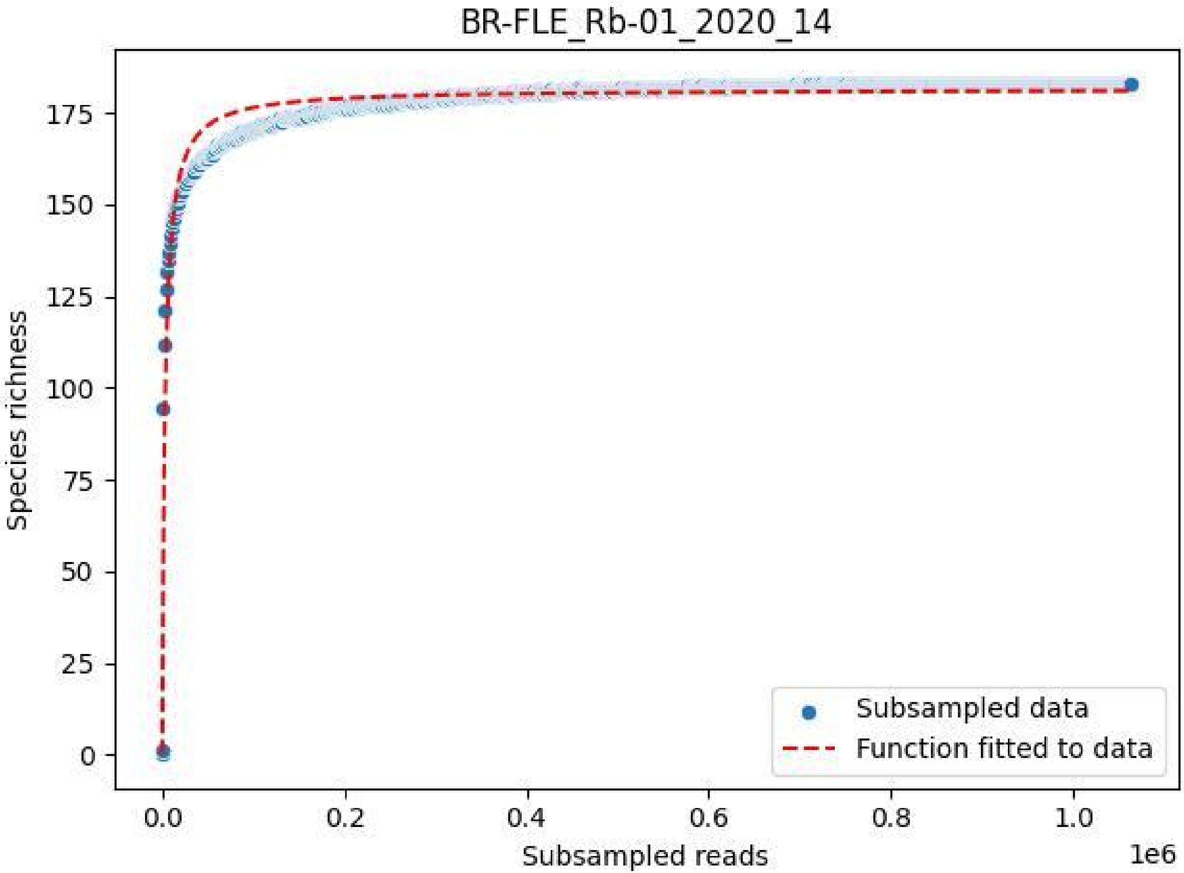
BR-FLE_Rb-01_2020_14
Species richness
Subsampled reads
1e6
0
25
50
75
100
125
150
175
0.0
0.2
0.4
0.6
0.8
1.0
Subsampled data
Function fitted to data

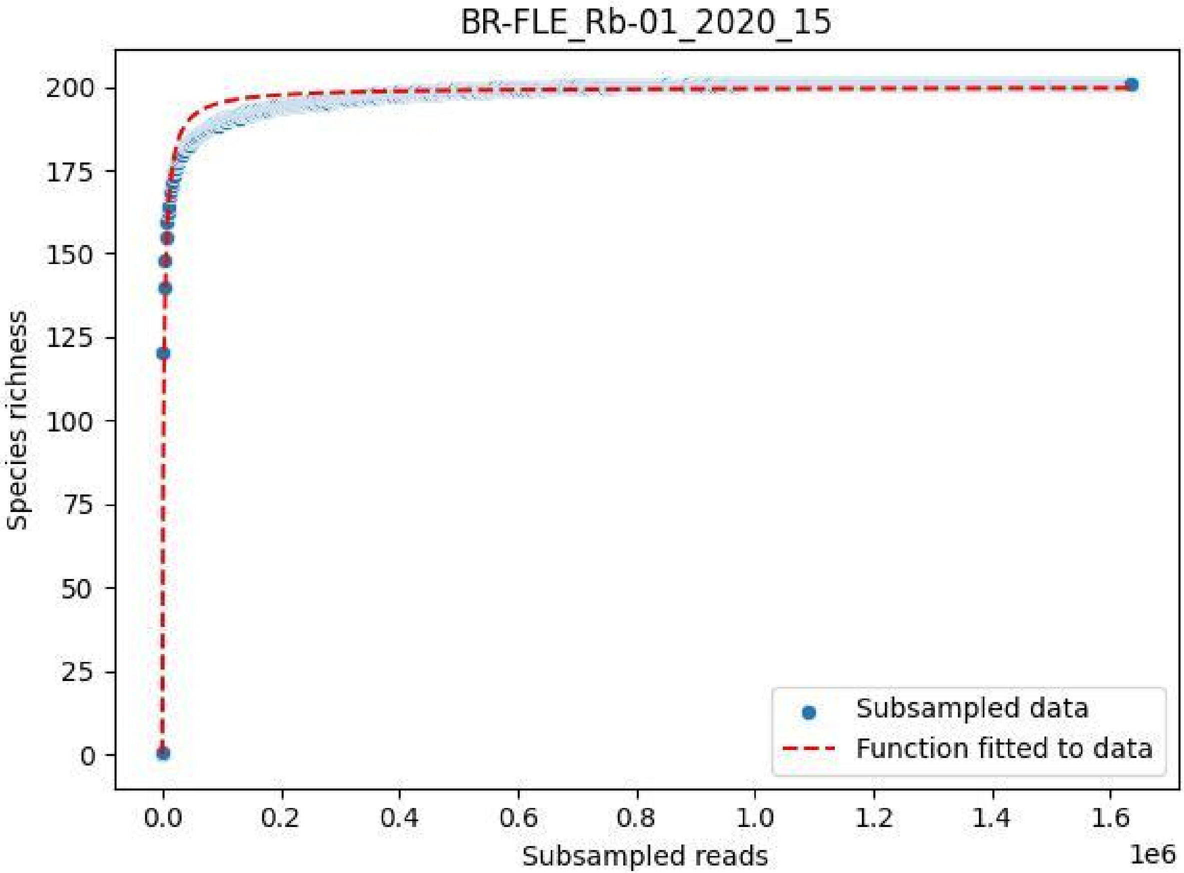
BR-FLE_Rb-01_2020_15
Species richness
Subsampled reads
1e6
0
25
50
75
100
125
150
175
200
0.0
0.2
0.4
0.6
0.8
1.0
1.2
1.4
1.6
Subsampled data
Function fitted to data

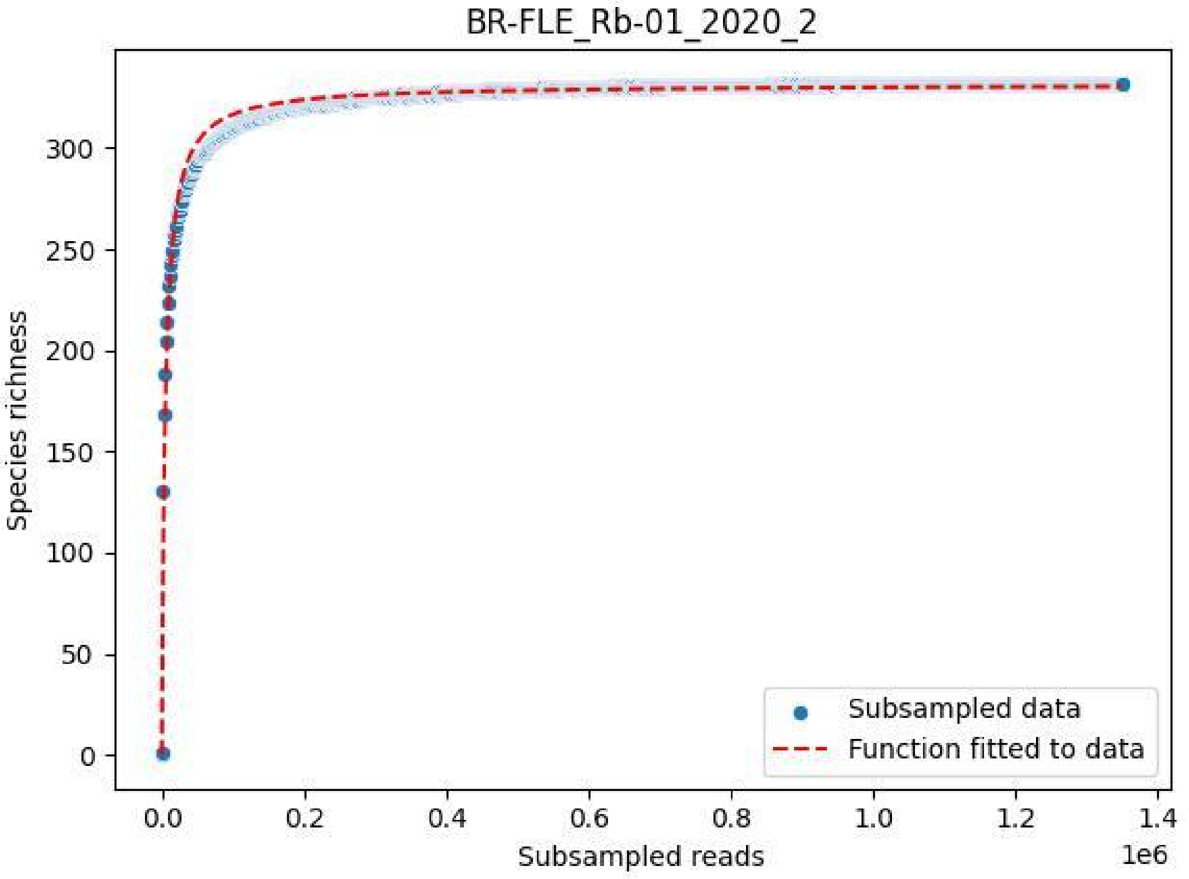
BR-FLE_Rb-01_2020_2
Species richness
Subsampled reads
0
50
100
150
200
250
300
0.0
0.2
0.4
0.6
0.8
1.0
1.2
1.4
1e6
Subsampled data
Function fitted to data

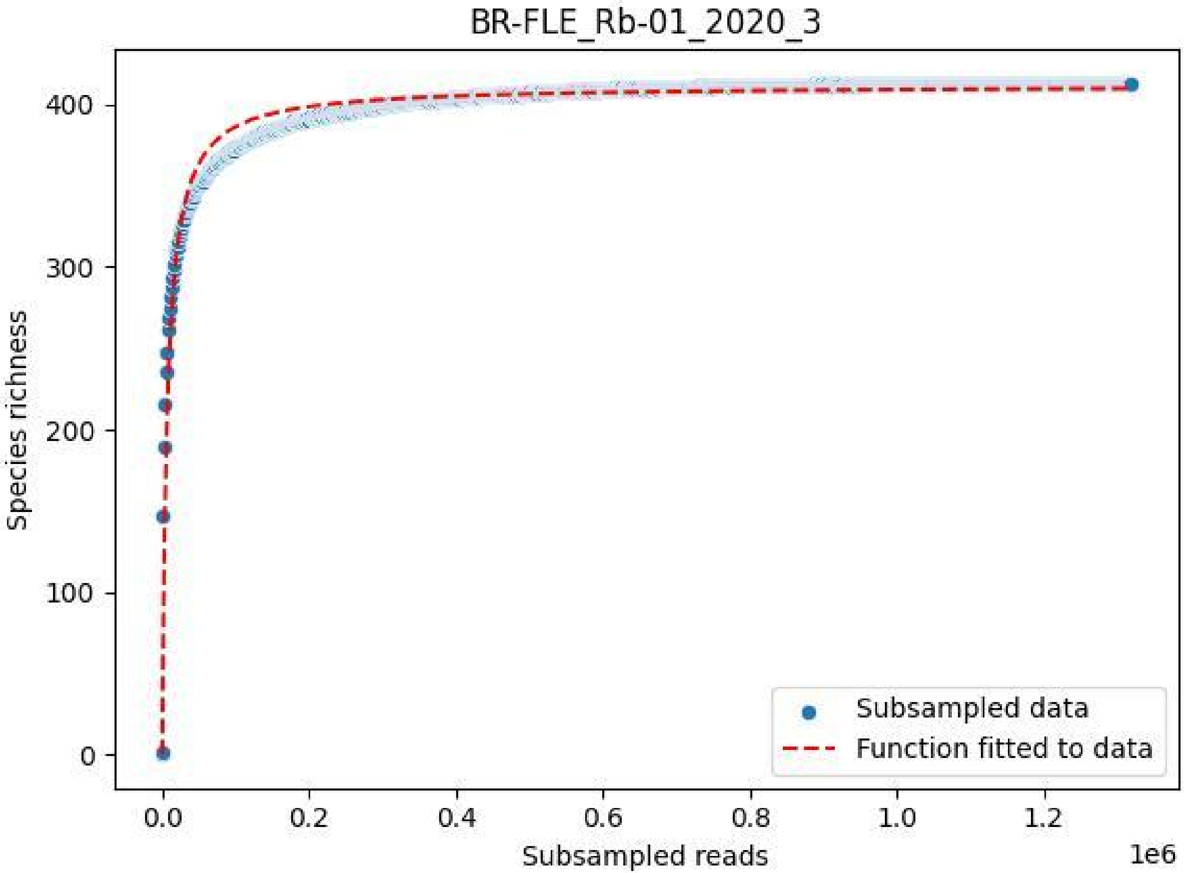

BR-FLE_Rb-01_2020_3
Species richness
Subsampled reads
1e6
0
100
200
300
400
0.0
0.2
0.4
0.6
0.8
1.0
1.2
Subsampled data
Function fitted to data

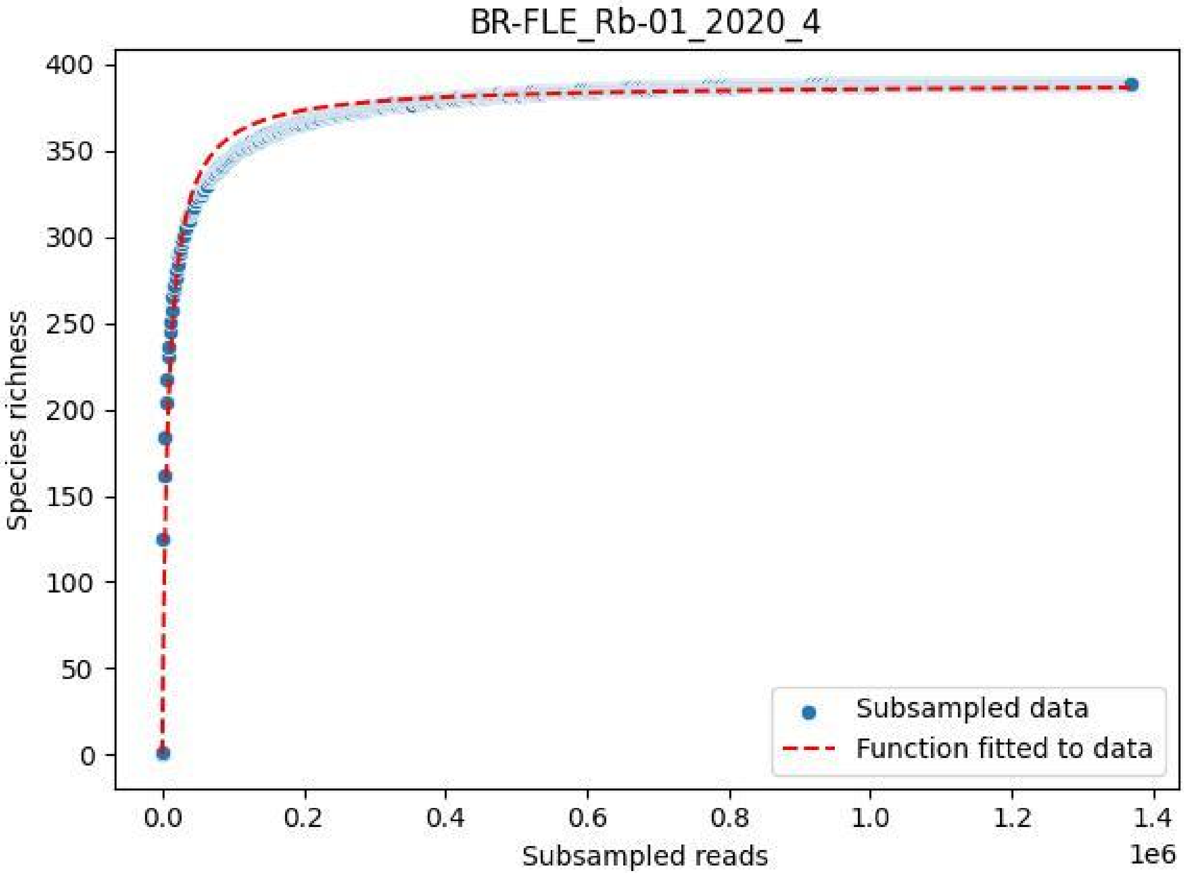
BR-FLE_Rb-01_2020_4
Species richness
400
350
300
250
200
150
100
50
0
0.0
0.2
0.4
0.6
0.8
1.0
1.2
1.4
1e6
Subsampled reads
Subsampled data
Function fitted to data

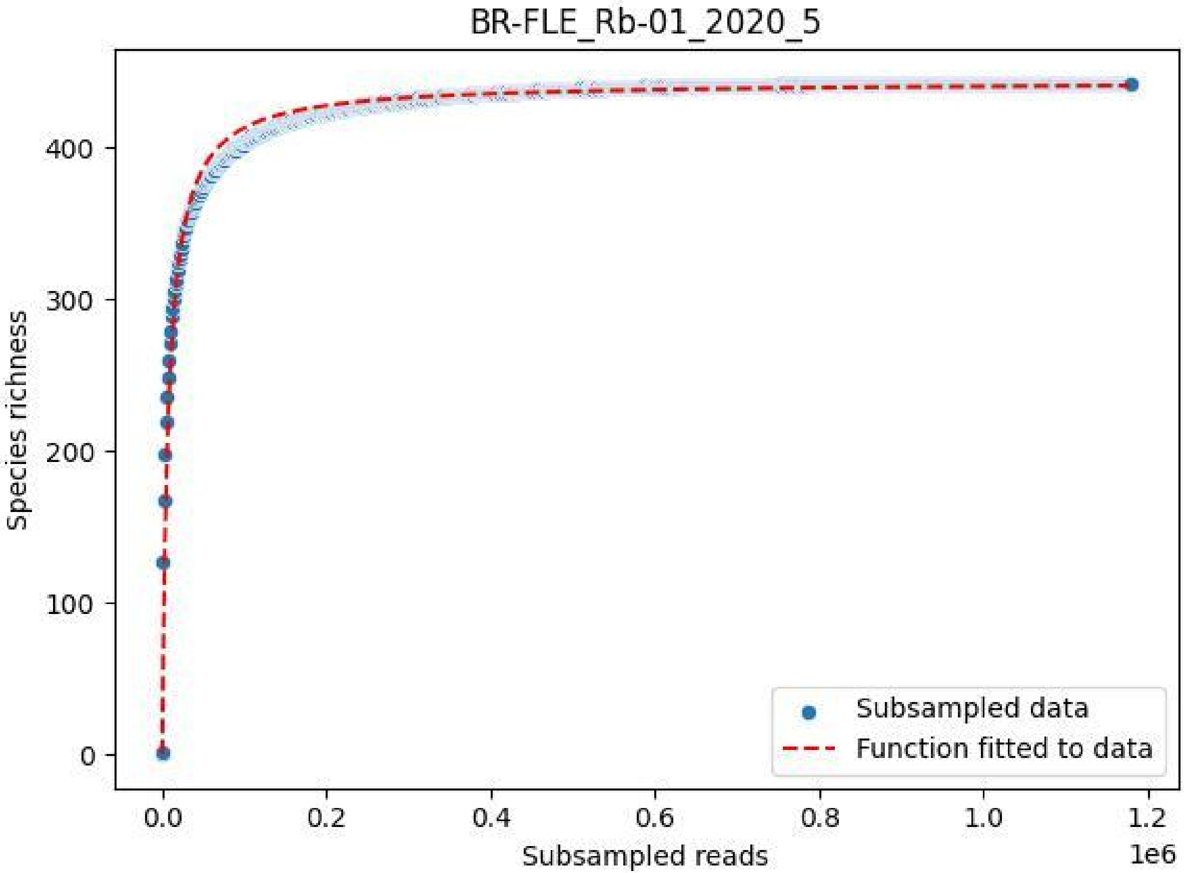
BR-FLE_Rb-01_2020_5
Species richness
Subsampled reads
1e6
0
100
200
300
400
0.0
0.2
0.4
0.6
0.8
1.0
1.2
Subsampled data
Function fitted to data

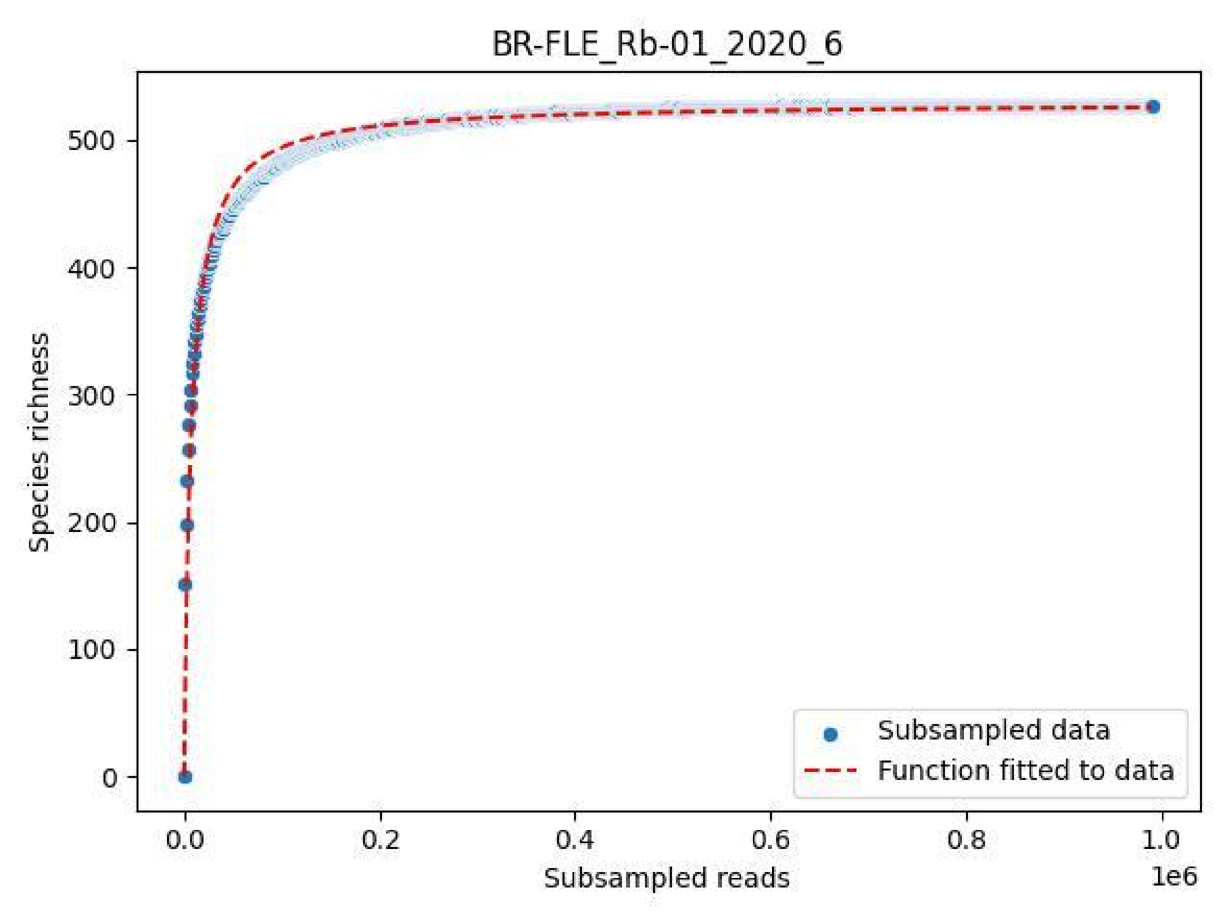

BR-FLE_Rb-01_2020_6
Species richness
Subsampled reads
1e6
0
100
200
300
400
500
0.0
0.2
0.4
0.6
0.8
1.0
Subsampled data
Function fitted to data

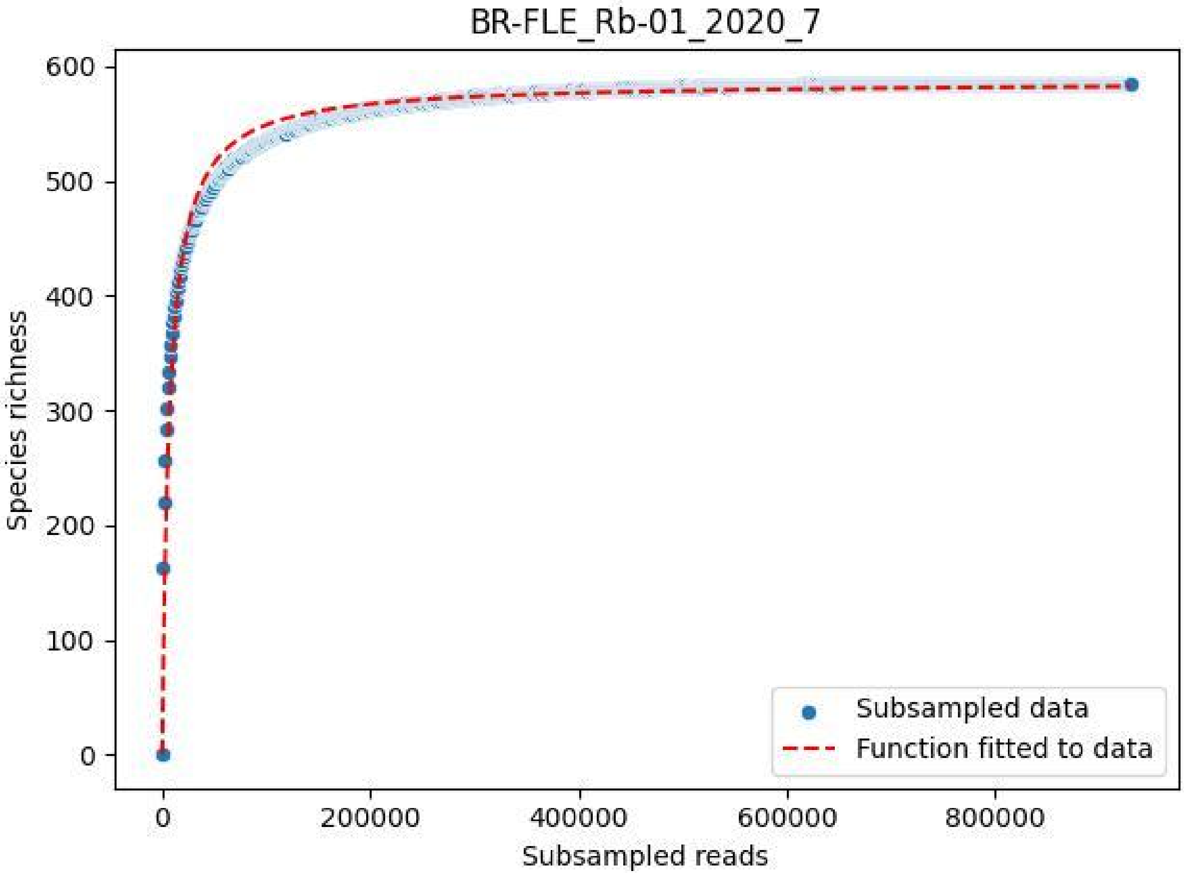

BR-FLE_Rb-01_2020_7
Species richness
Subsampled reads
0
100
200
300
400
500
600
0
200000
400000
600000
800000
Subsampled data
Function fitted to data

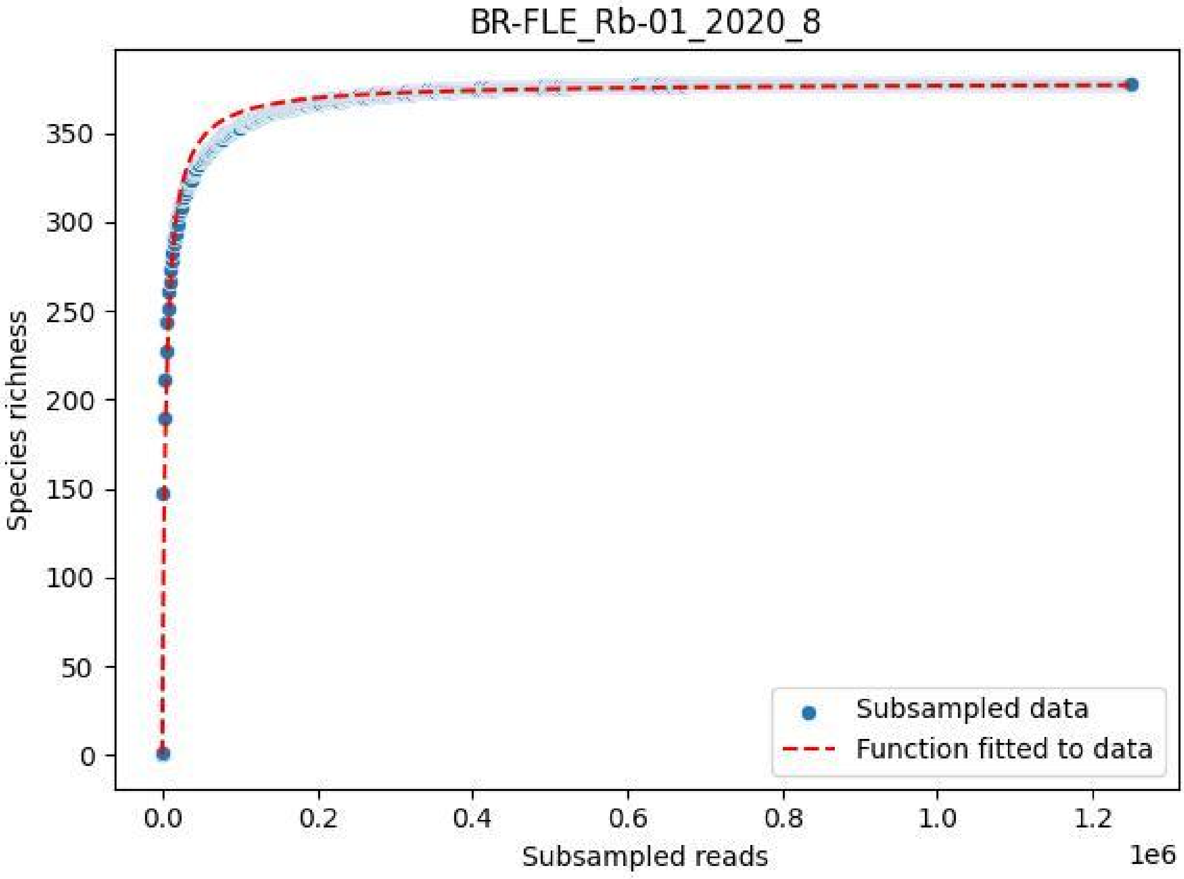
BR-FLE_Rb-01_2020_8
Species richness
Subsampled reads
1e6
0
50
100
150
200
250
300
350
0.0
0.2
0.4
0.6
0.8
1.0
1.2
Subsampled data
Function fitted to data

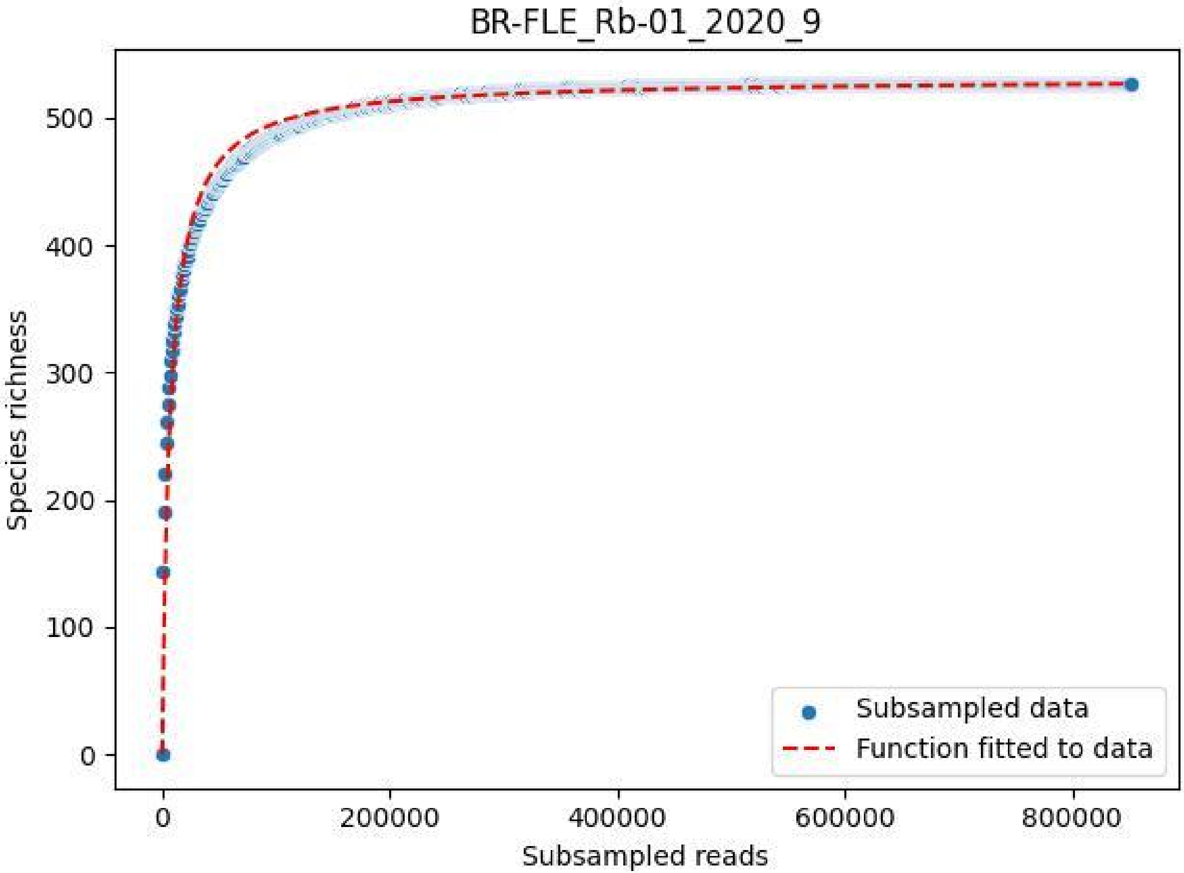
BR-FLE_Rb-01_2020_9
Species richness
Subsampled reads
0
100
200
300
400
500
0
200000
400000
600000
800000
Subsampled data
Function fitted to data

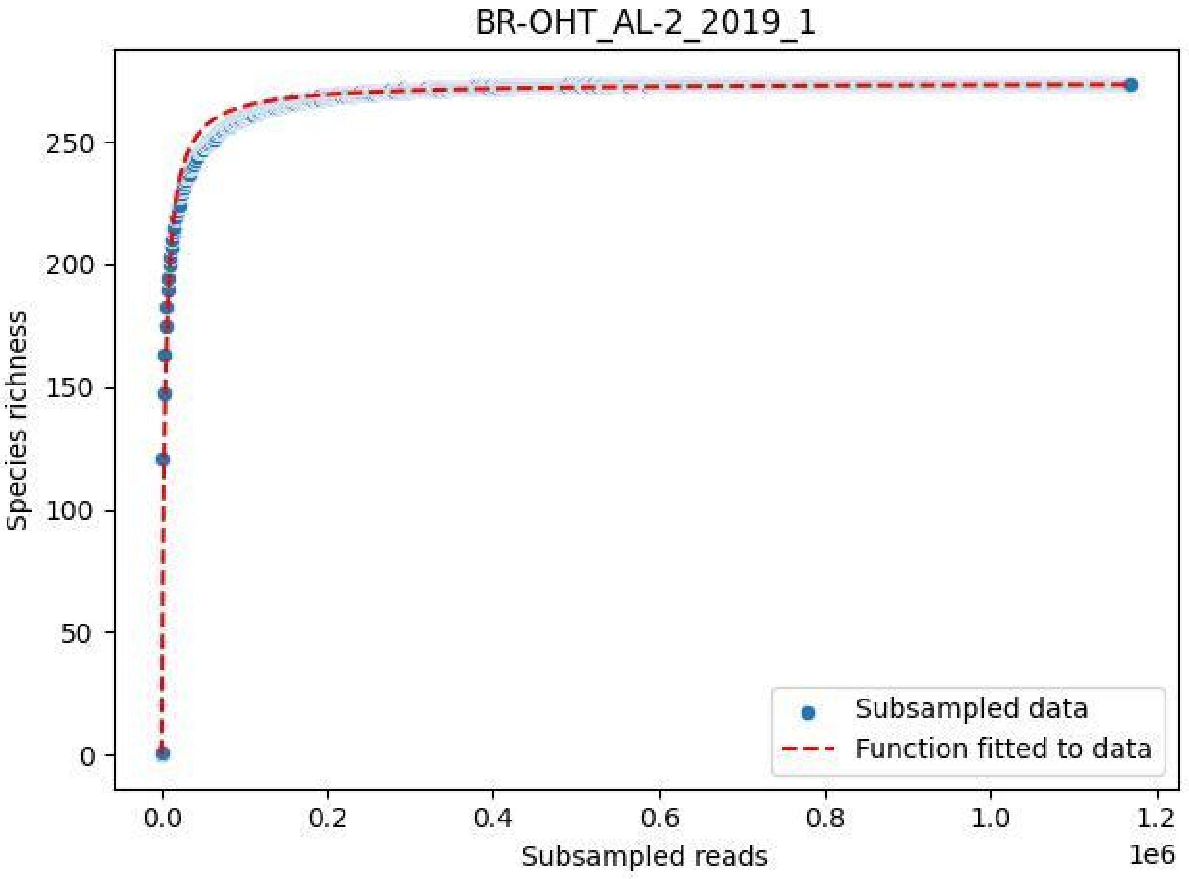
BR-OHT_AL-2_2019_1
Species richness
Subsampled reads
1e6
0
50
100
150
200
250
0.0
0.2
0.4
0.6
0.8
1.0
1.2
Subsampled data
Function fitted to data

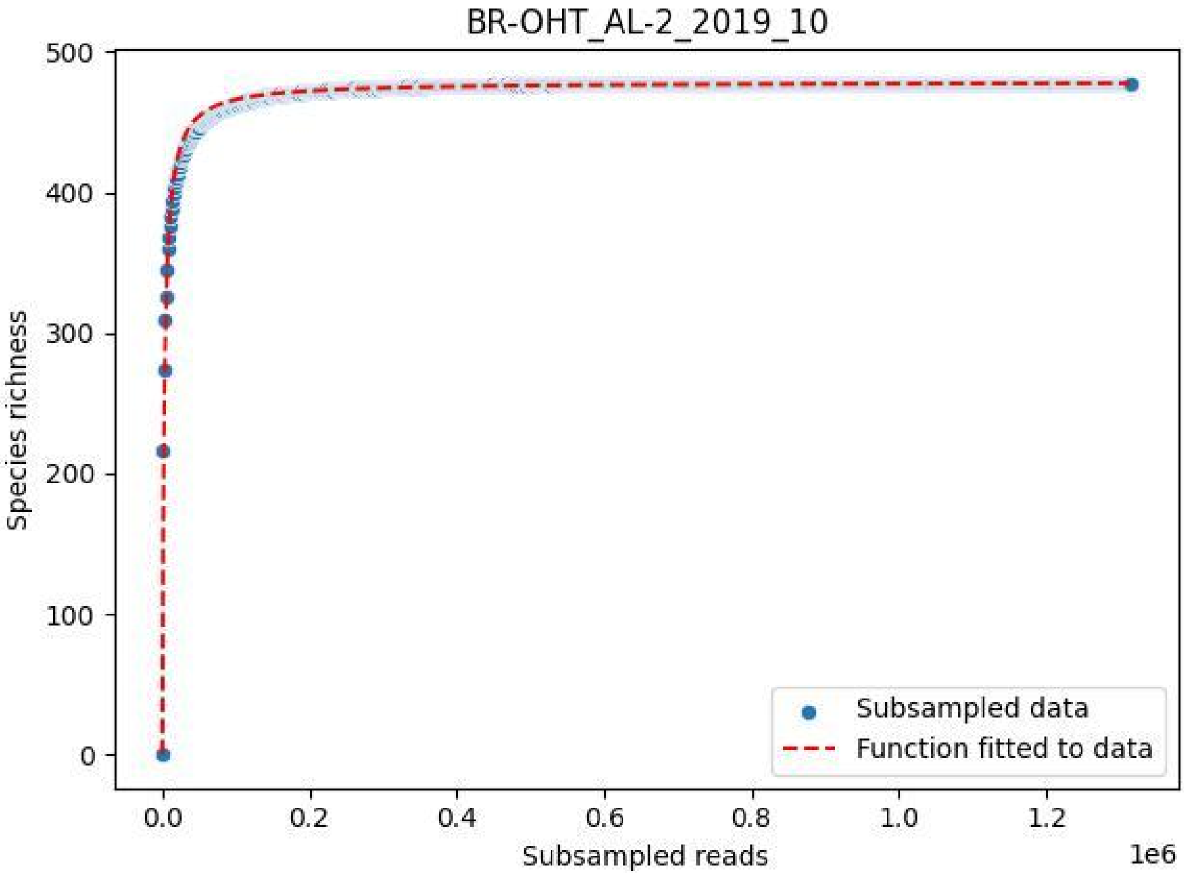

BR-OHT_AL-2_2019_10
Species richness
Subsampled reads
1e6
0
100
200
300
400
500
0.0
0.2
0.4
0.6
0.8
1.0
1.2
Subsampled data
Function fitted to data

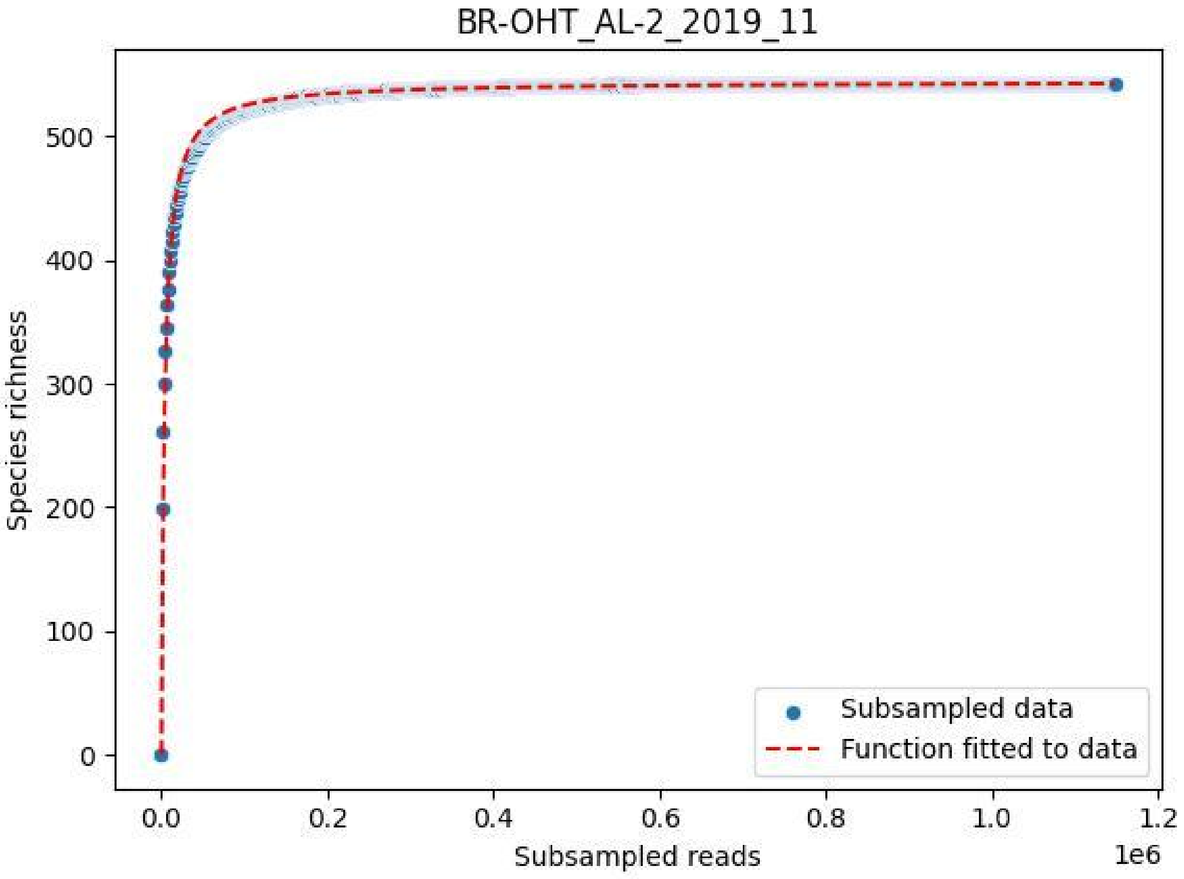
BR-OHT_AL-2_2019_11
Species richness
Subsampled reads
1e6
0
100
200
300
400
500
0.0
0.2
0.4
0.6
0.8
1.0
1.2
Subsampled data
Function fitted to data

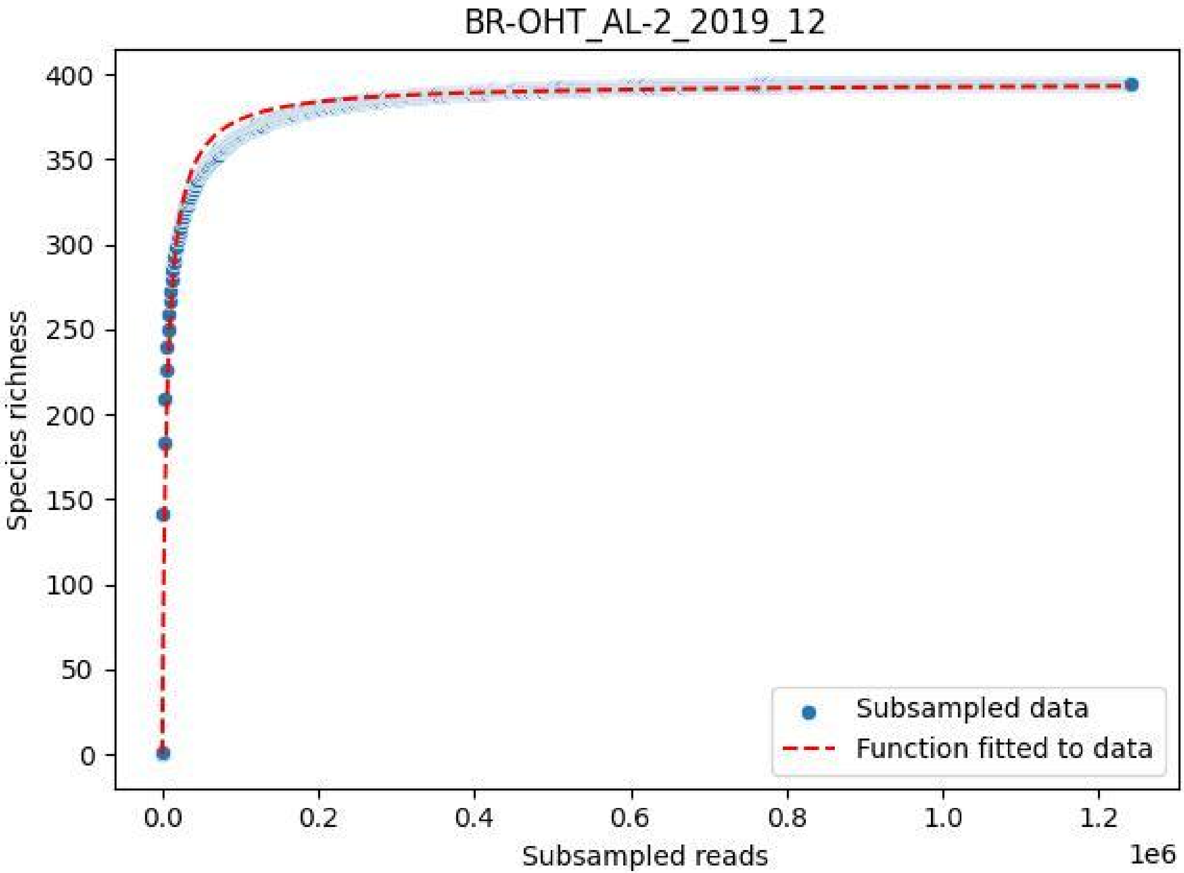

BR-OHT_AL-2_2019_12
Species richness
Subsampled reads
1e6
0
50
100
150
200
250
300
350
400
0.0
0.2
0.4
0.6
0.8
1.0
1.2
Subsampled data
Function fitted to data

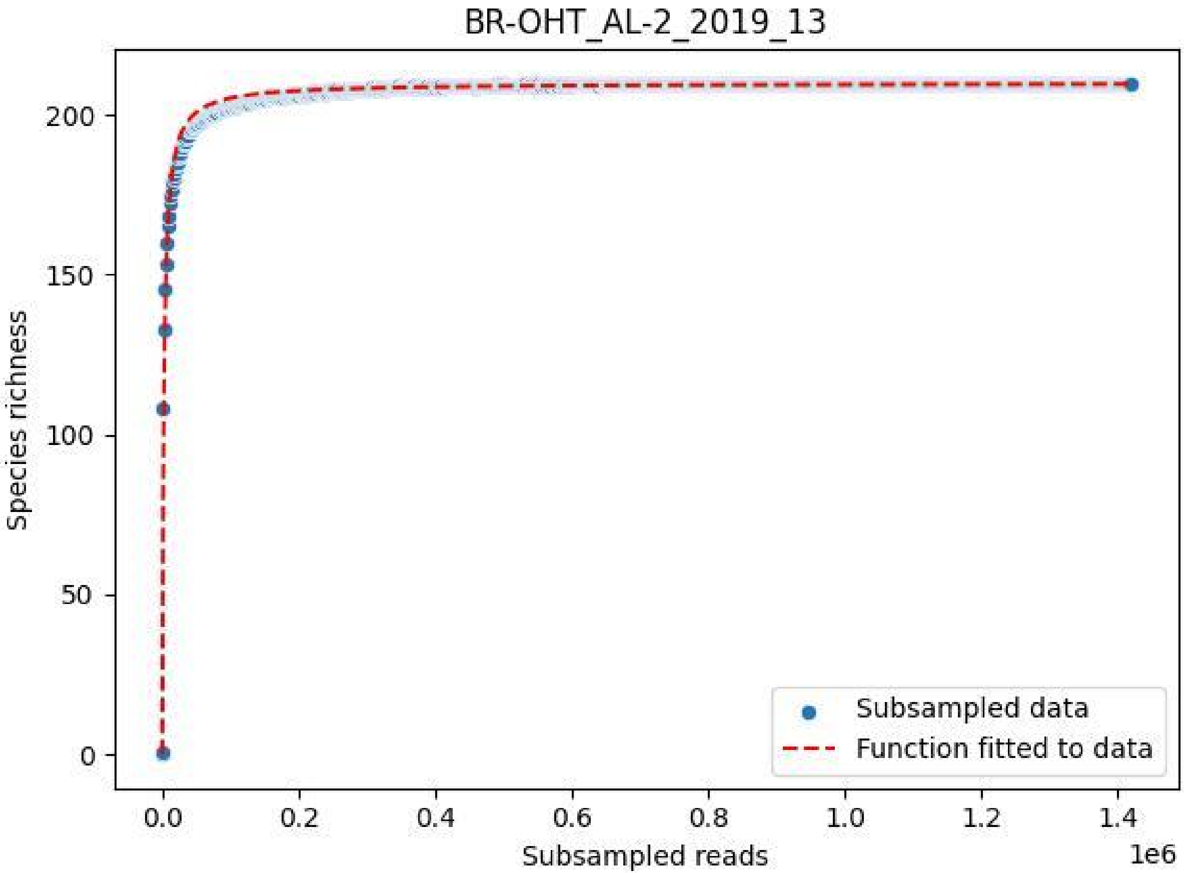

BR-OHT_AL-2_2019_13
Species richness
Subsampled reads
1e6
0
50
100
150
200
0.0
0.2
0.4
0.6
0.8
1.0
1.2
1.4
Subsampled data
Function fitted to data

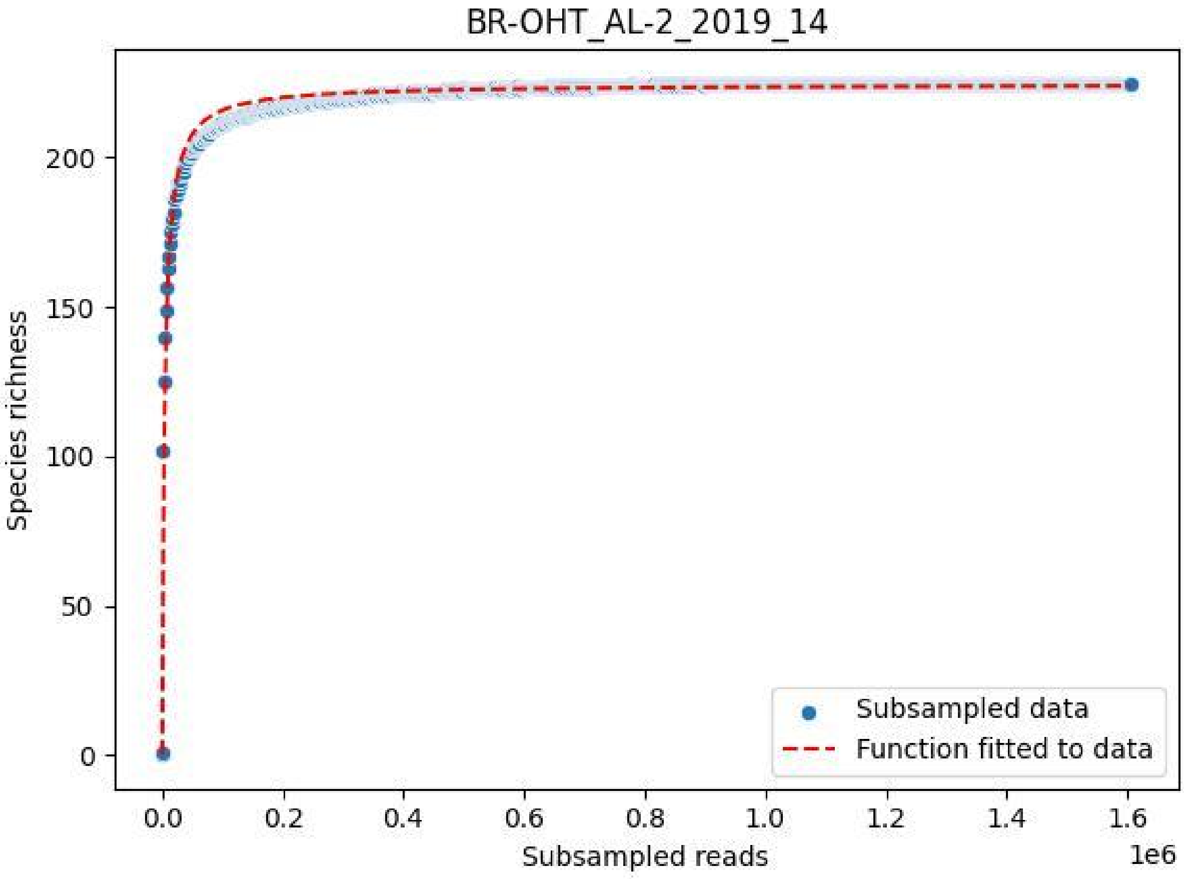
BR-OHT_AL-2_2019_14
Species richness
Subsampled reads
1e6
0
50
100
150
200
0.0
0.2
0.4
0.6
0.8
1.0
1.2
1.4
1.6
Subsampled data
Function fitted to data

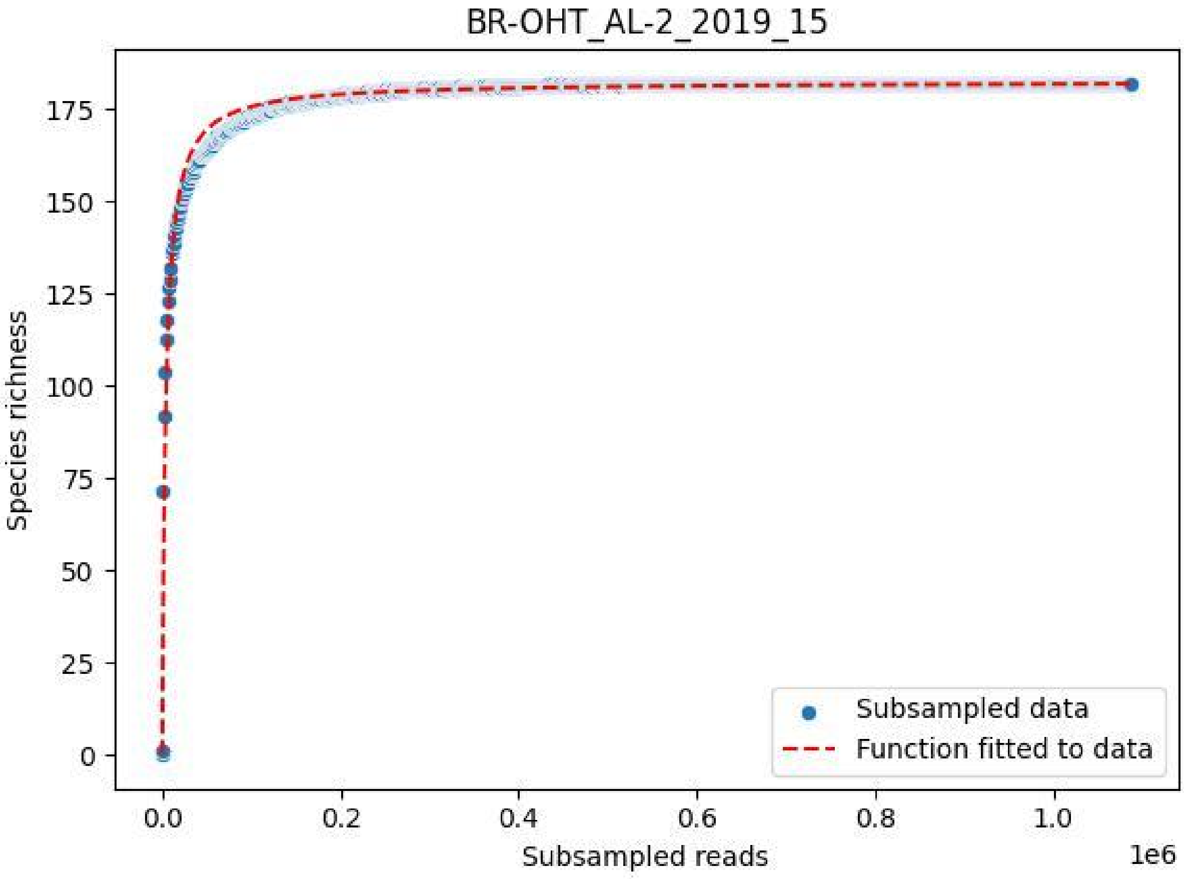
BR-OHT_AL-2_2019_15
Species richness
Subsampled reads
1e6
0
25
50
75
100
125
150
175
0.0
0.2
0.4
0.6
0.8
1.0
Subsampled data
Function fitted to data

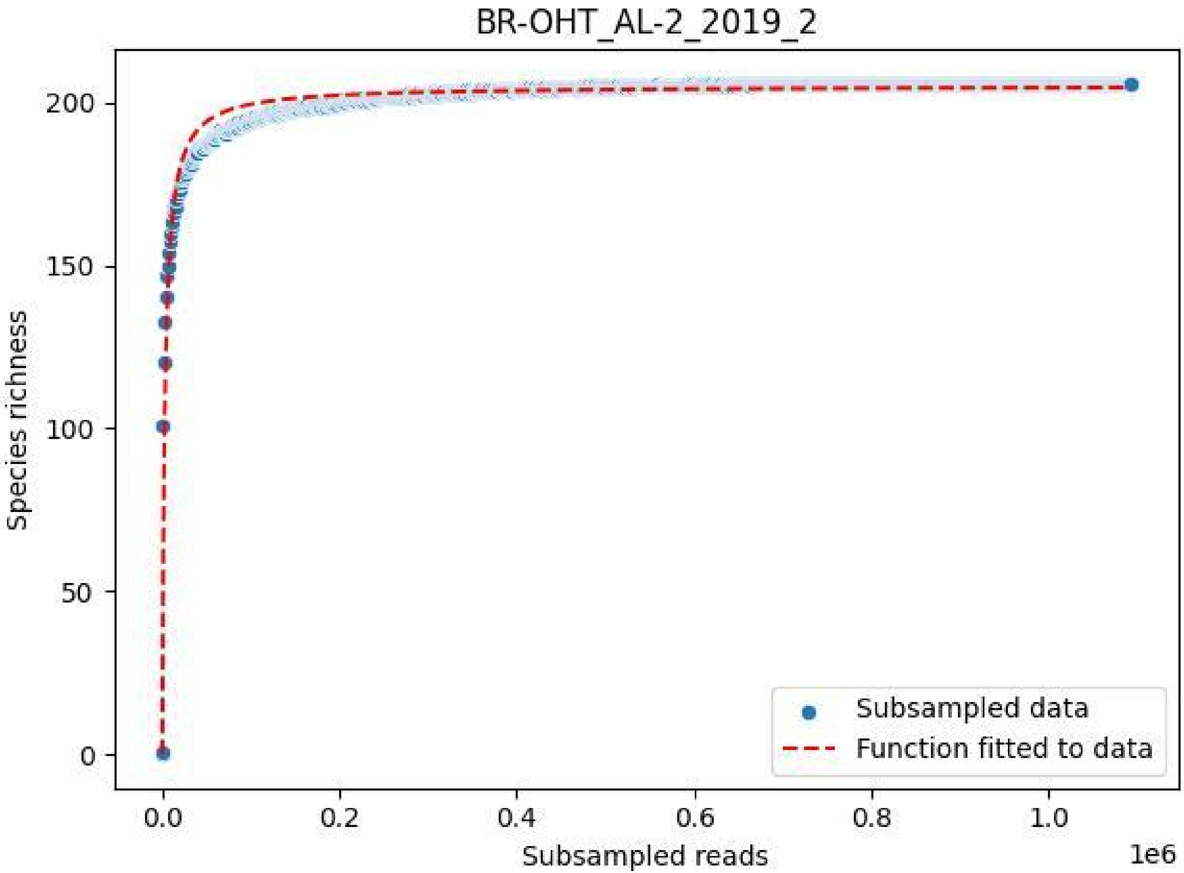

BR-OHT_AL-2_2019_2
Species richness
Subsampled reads
1e6
0
50
100
150
200
0.0
0.2
0.4
0.6
0.8
1.0
Subsampled data
Function fitted to data

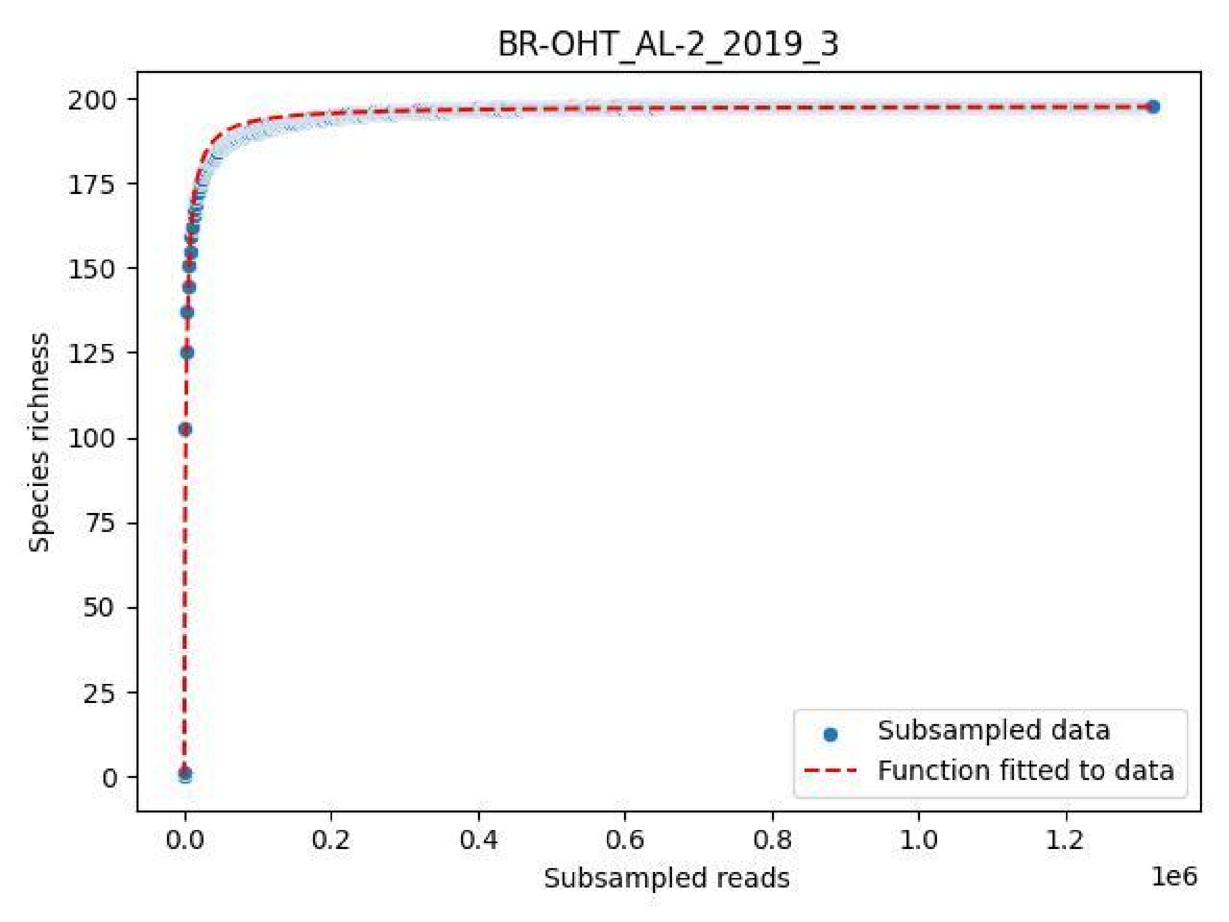
BR-OHT_AL-2_2019_3
Species richness
Subsampled reads
1e6
200
175
150
125
100
75
50
25
0
0.0
0.2
0.4
0.6
0.8
1.0
1.2
Subsampled data
Function fitted to data

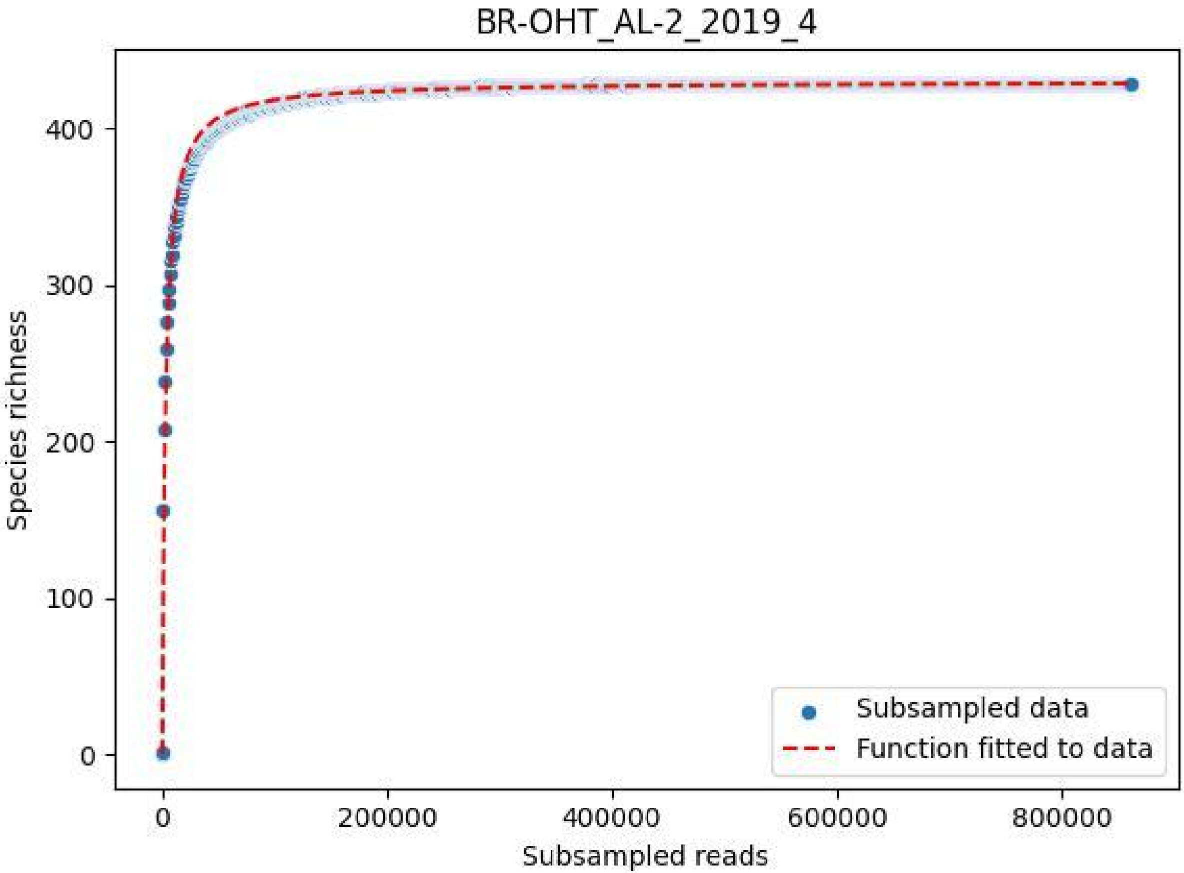
BR-OHT_AL-2_2019_4
Species richness
0
100
200
300
400
0
200000
400000
600000
800000
Subsampled reads
Subsampled data
Function fitted to data

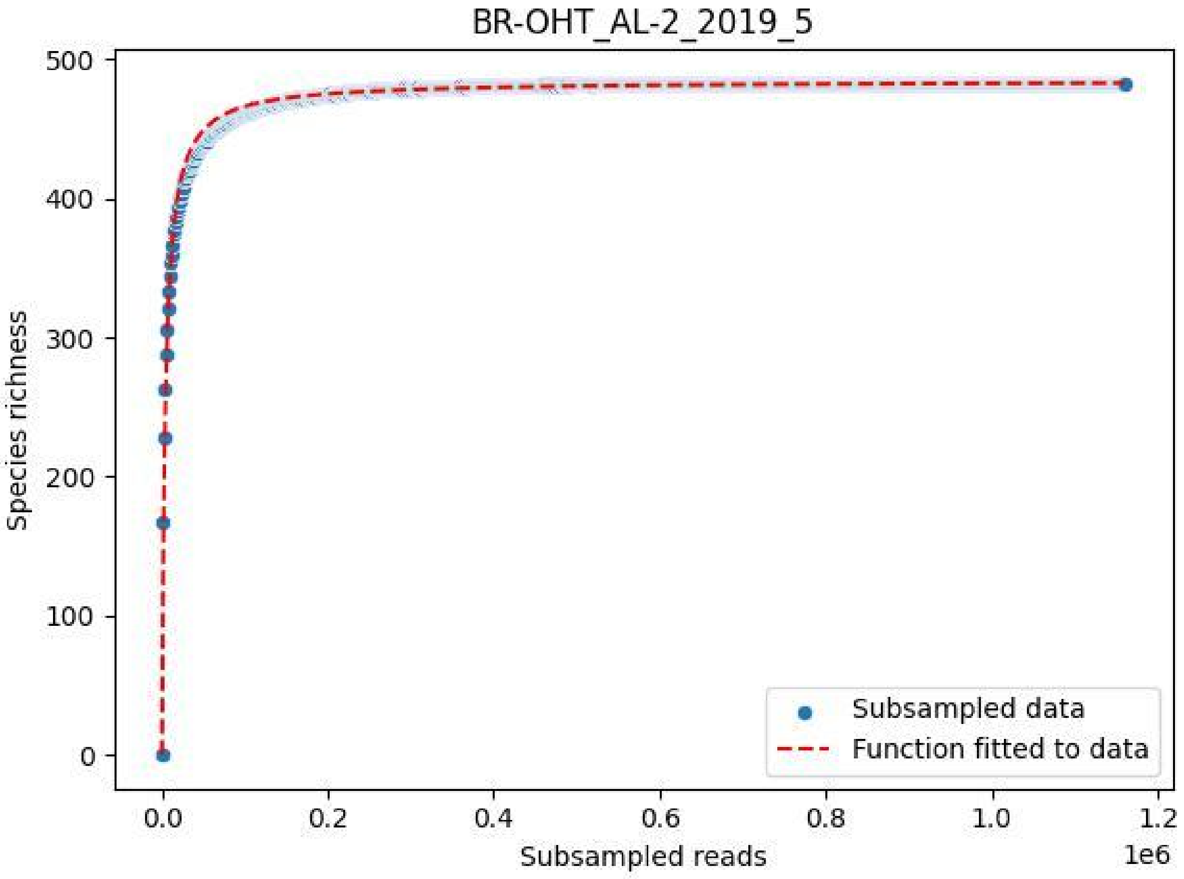
BR-OHT_AL-2_2019_5
Species richness
Subsampled reads
1e6
0
100
200
300
400
500
0.0
0.2
0.4
0.6
0.8
1.0
1.2
Subsampled data
Function fitted to data

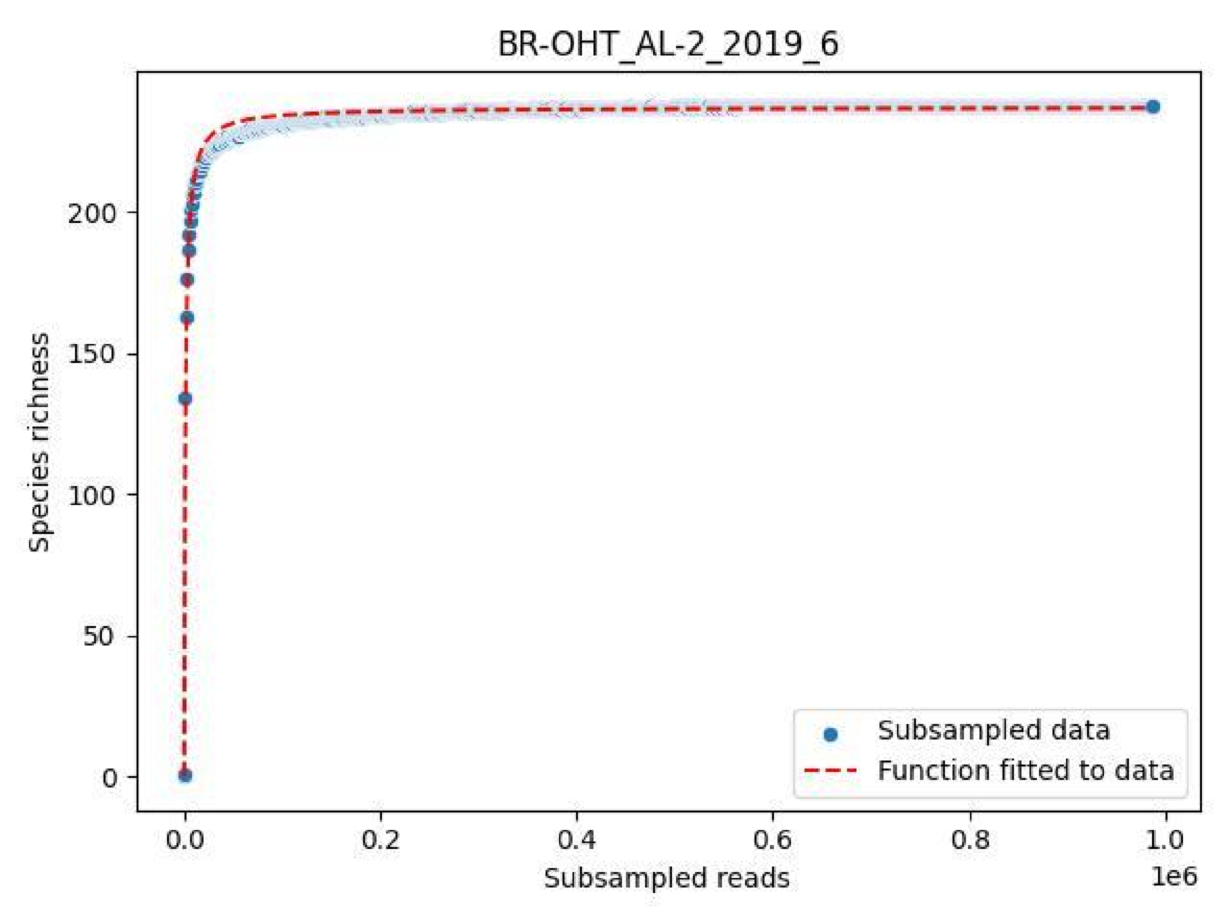
BR-OHT_AL-2_2019_6
Species richness
0
50
100
150
200
0.0
0.2
0.4
0.6
0.8
1.0
1e6
Subsampled reads
Subsampled data
Function fitted to data

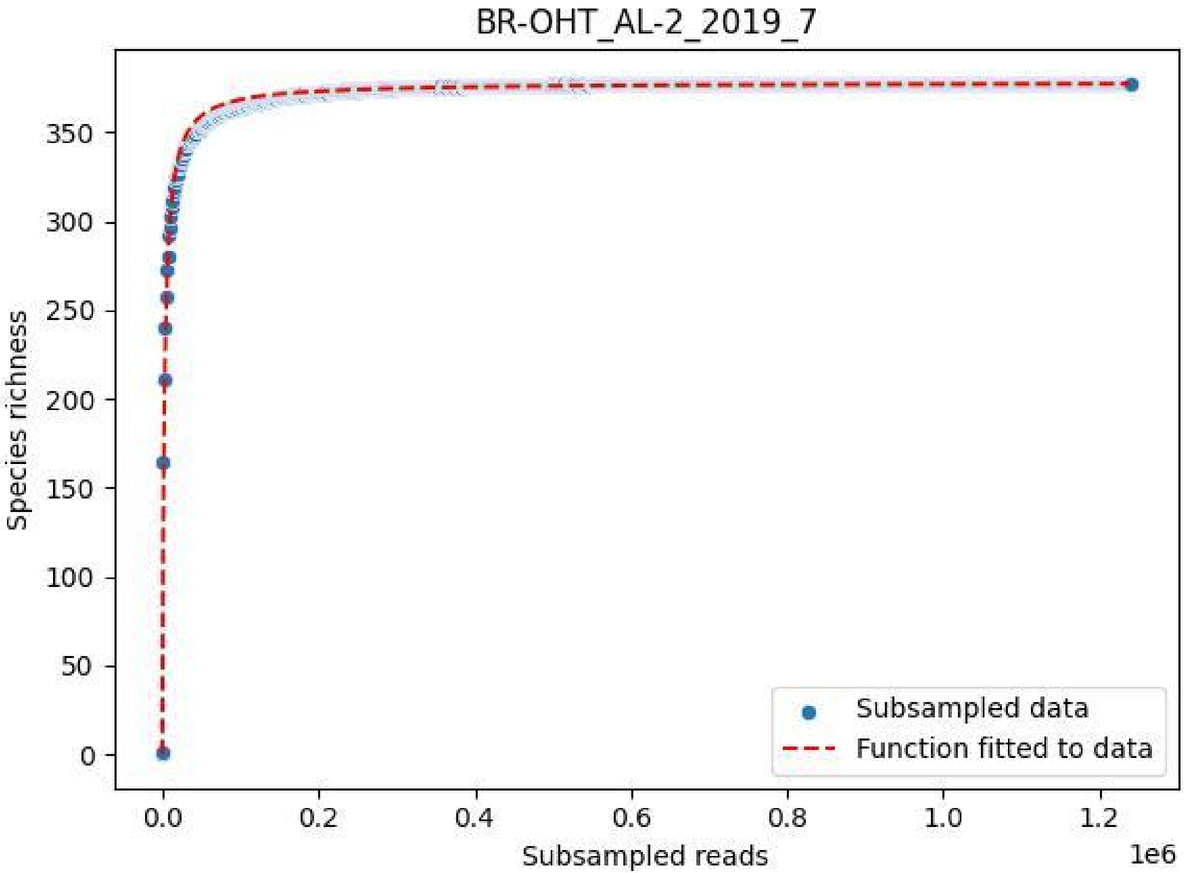

BR-OHT_AL-2_2019_7
Species richness
Subsampled reads
1e6
0
50
100
150
200
250
300
350
0.0
0.2
0.4
0.6
0.8
1.0
1.2
Subsampled data
Function fitted to data

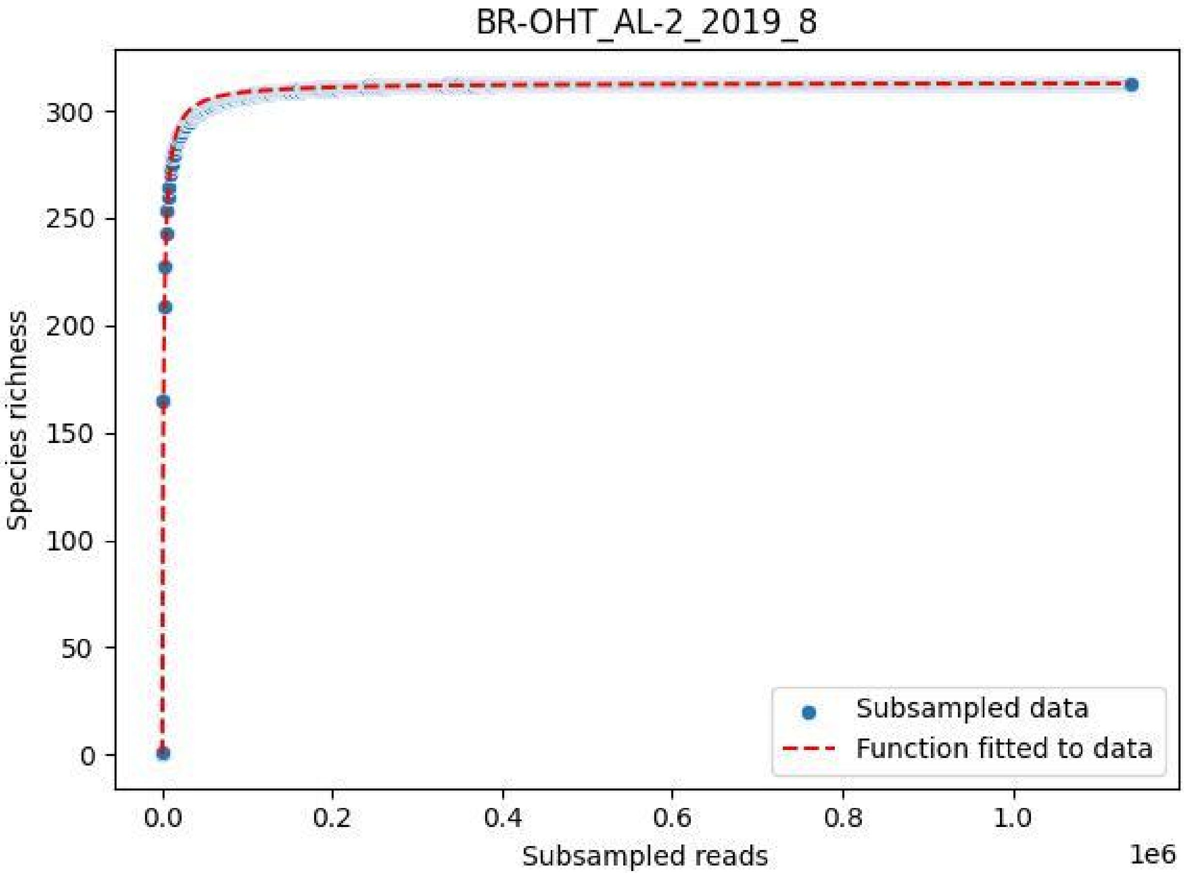
BR-OHT_AL-2_2019_8
Species richness
Subsampled reads
1e6
0.0
0.2
0.4
0.6
0.8
1.0
0
50
100
150
200
250
300
Subsampled data
Function fitted to data

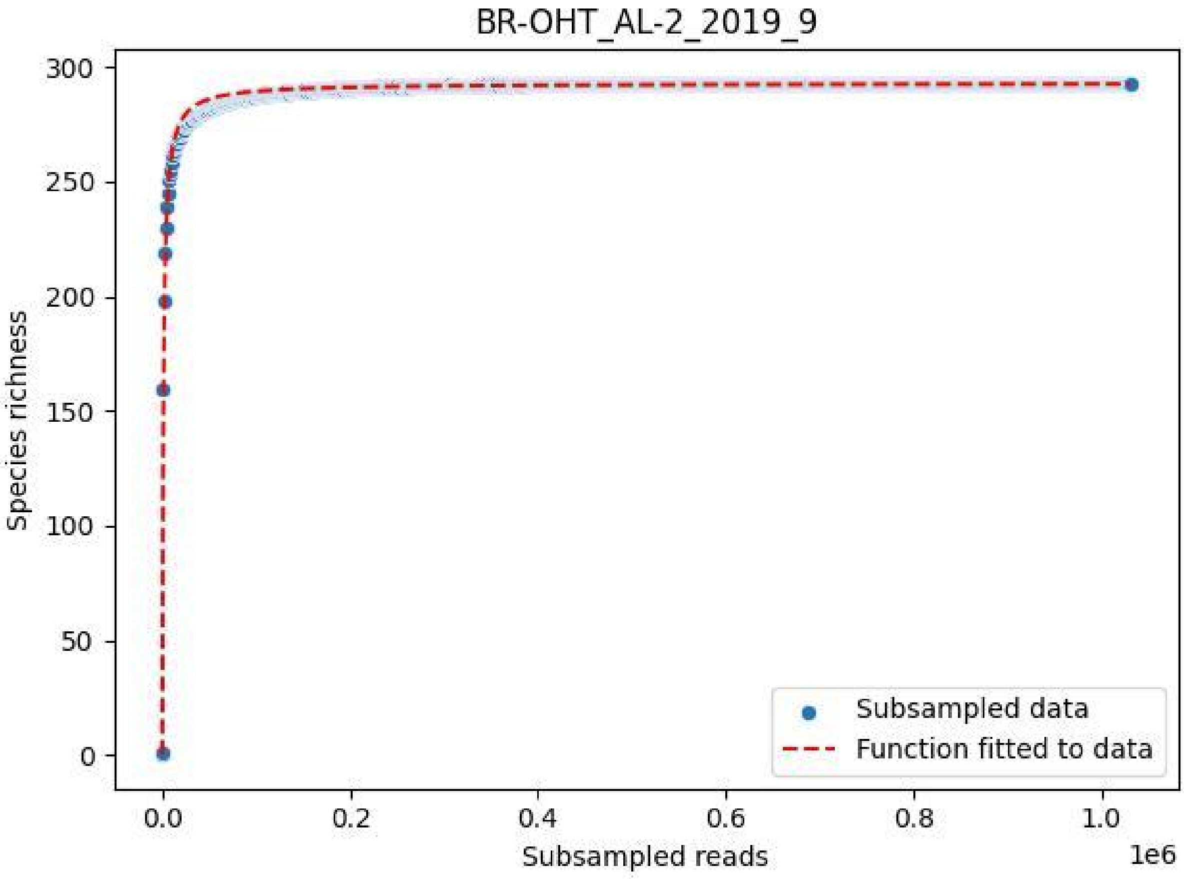
BR-OHT_AL-2_2019_9
Species richness
300
250
200
150
100
50
0
0.0
0.2
0.4
0.6
0.8
1.0
1e6
Subsampled reads
Subsampled data
Function fitted to data

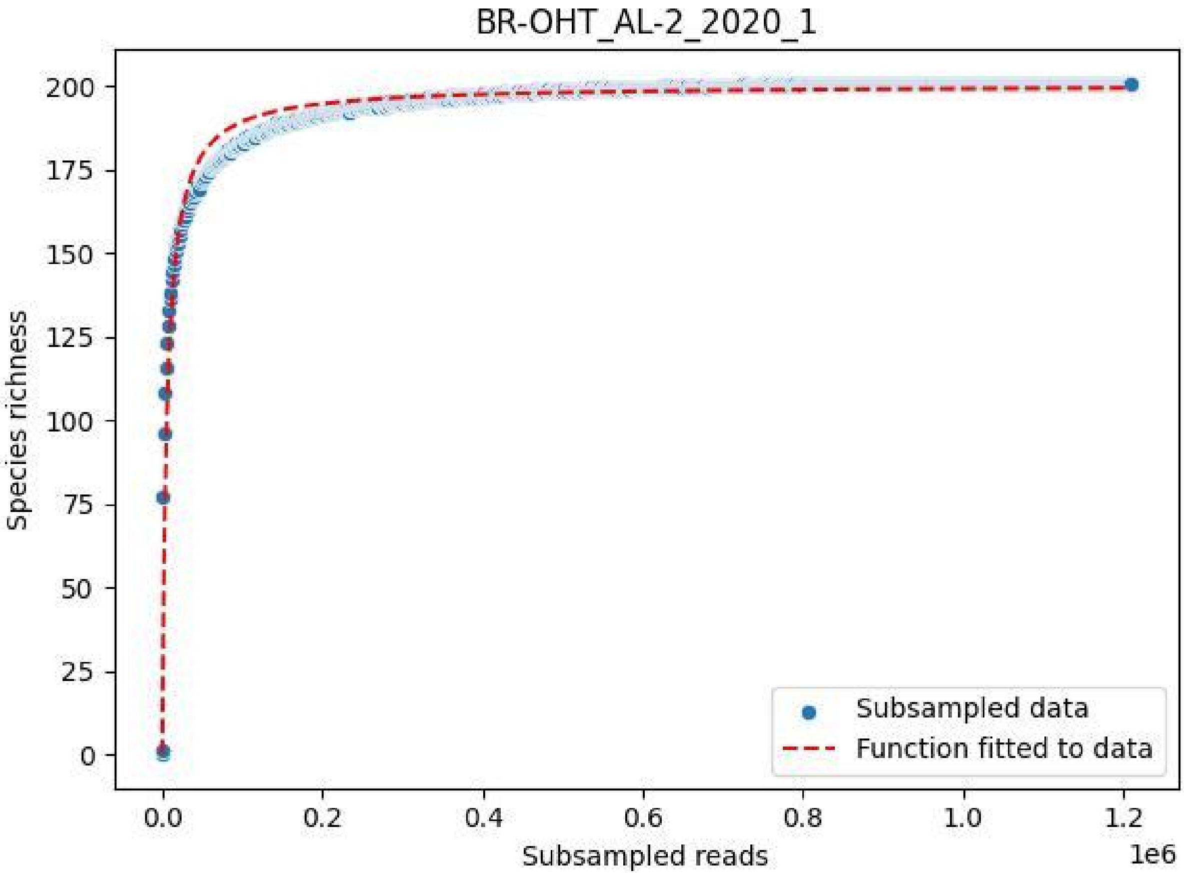

BR-OHT_AL-2_2020_1
Species richness
Subsampled reads
1e6
0
25
50
75
100
125
150
175
200
0.0
0.2
0.4
0.6
0.8
1.0
1.2
Subsampled data
Function fitted to data

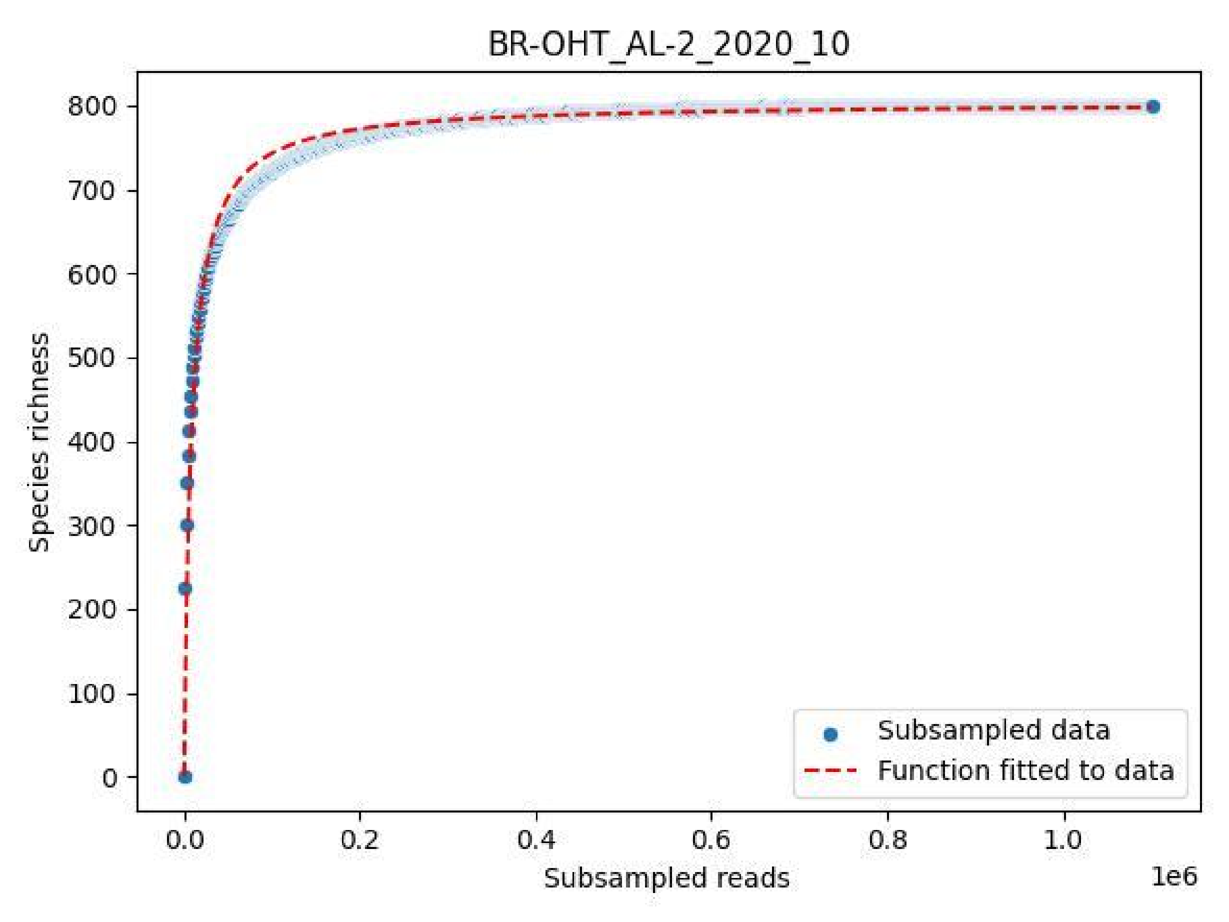

BR-OHT_AL-2_2020_10
Species richness
0
100
200
300
400
500
600
700
800
Subsampled reads
0.0
0.2
0.4
0.6
0.8
1.0
1e6
Subsampled data
Function fitted to data

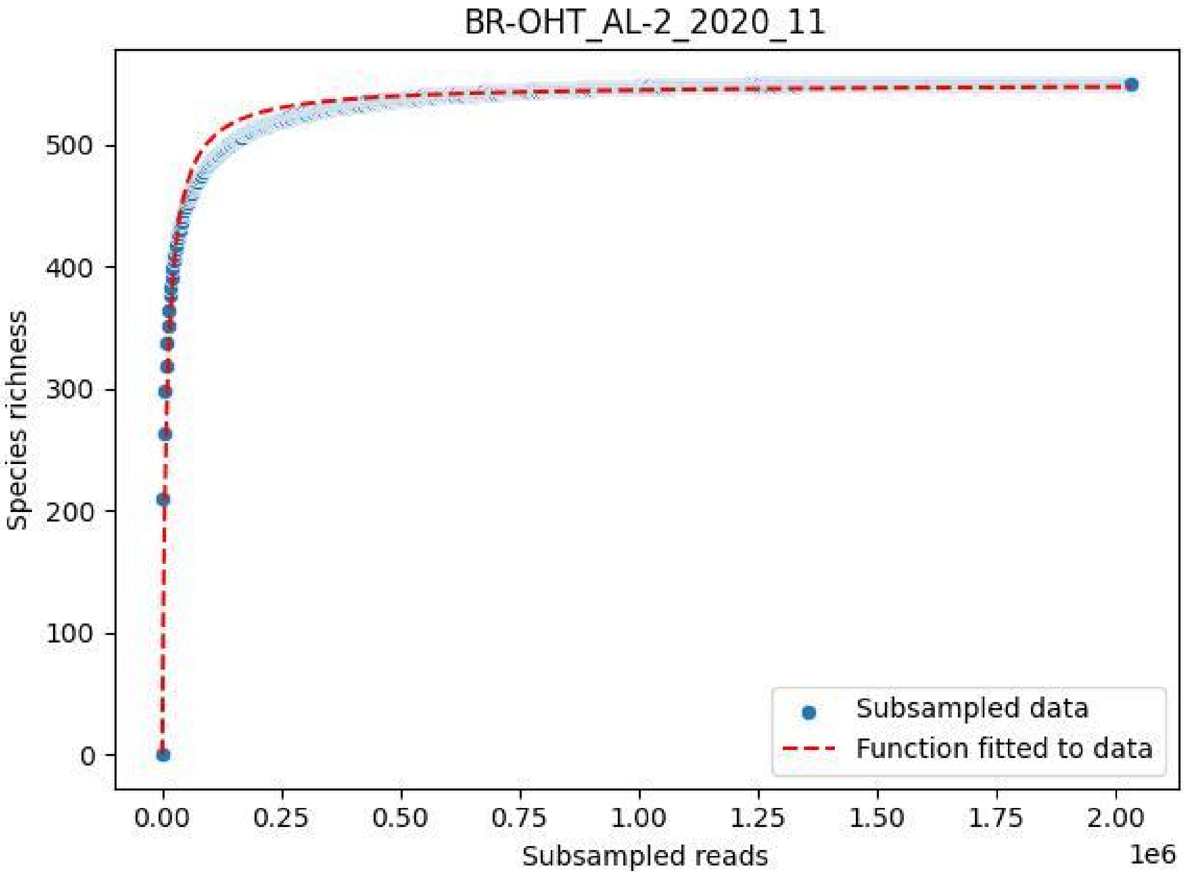
BR-OHT_AL-2_2020_11
Species richness
0
100
200
300
400
500
0.00
0.25
0.50
0.75
1.00
1.25
1.50
1.75
2.00
1e6
Subsampled reads
Subsampled data
Function fitted to data

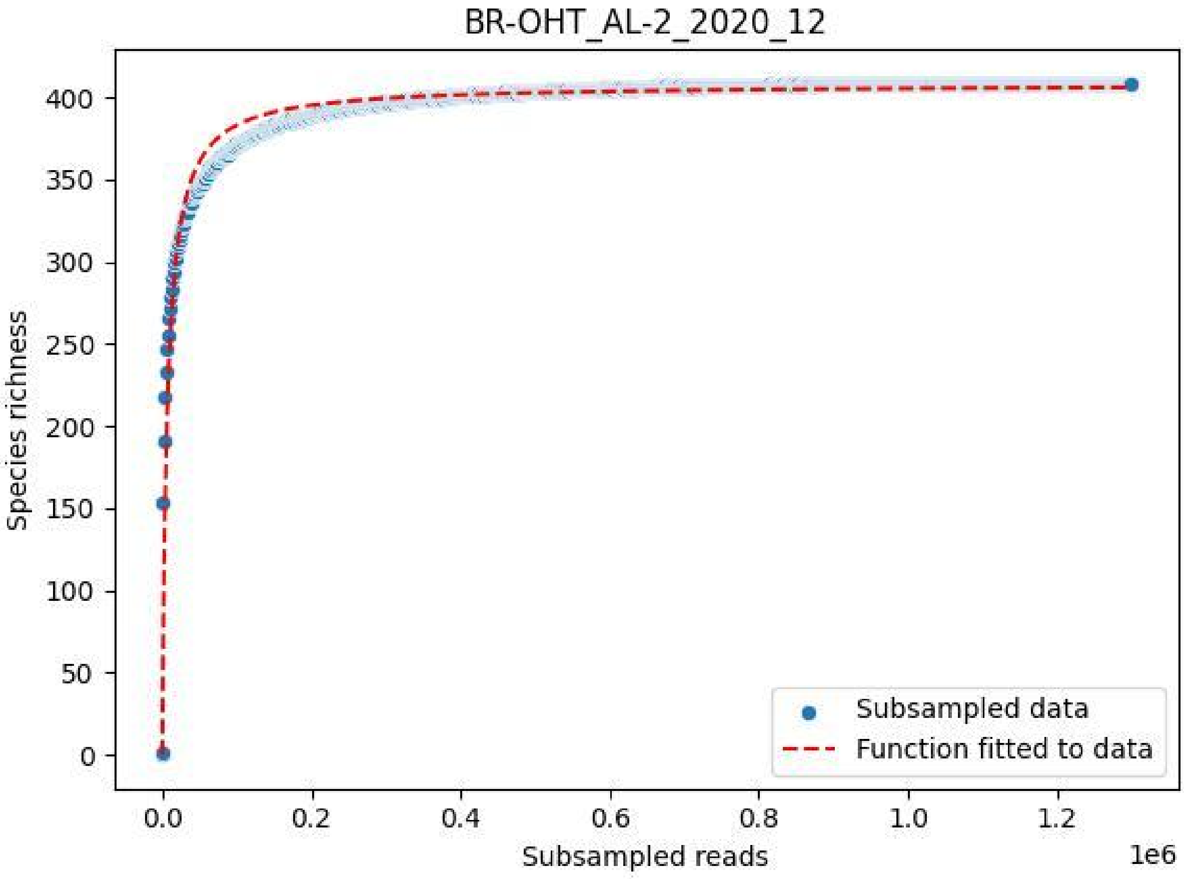

BR-OHT_AL-2_2020_12
Species richness
Subsampled reads
1e6
0
50
100
150
200
250
300
350
400
0.0
0.2
0.4
0.6
0.8
1.0
1.2
Subsampled data
Function fitted to data

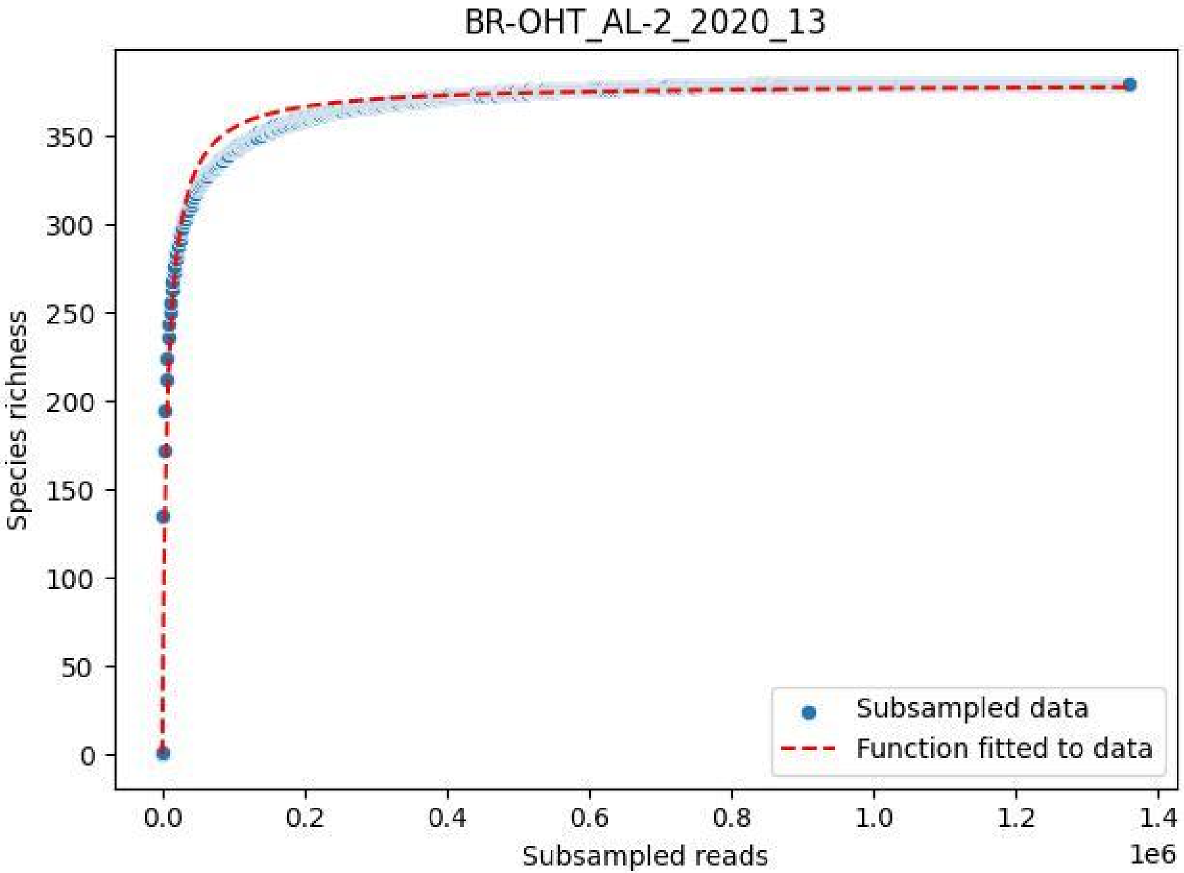

BR-OHT_AL-2_2020_13
Species richness
Subsampled reads
1e6
0
50
100
150
200
250
300
350
0.0
0.2
0.4
0.6
0.8
1.0
1.2
1.4
Subsampled data
Function fitted to data

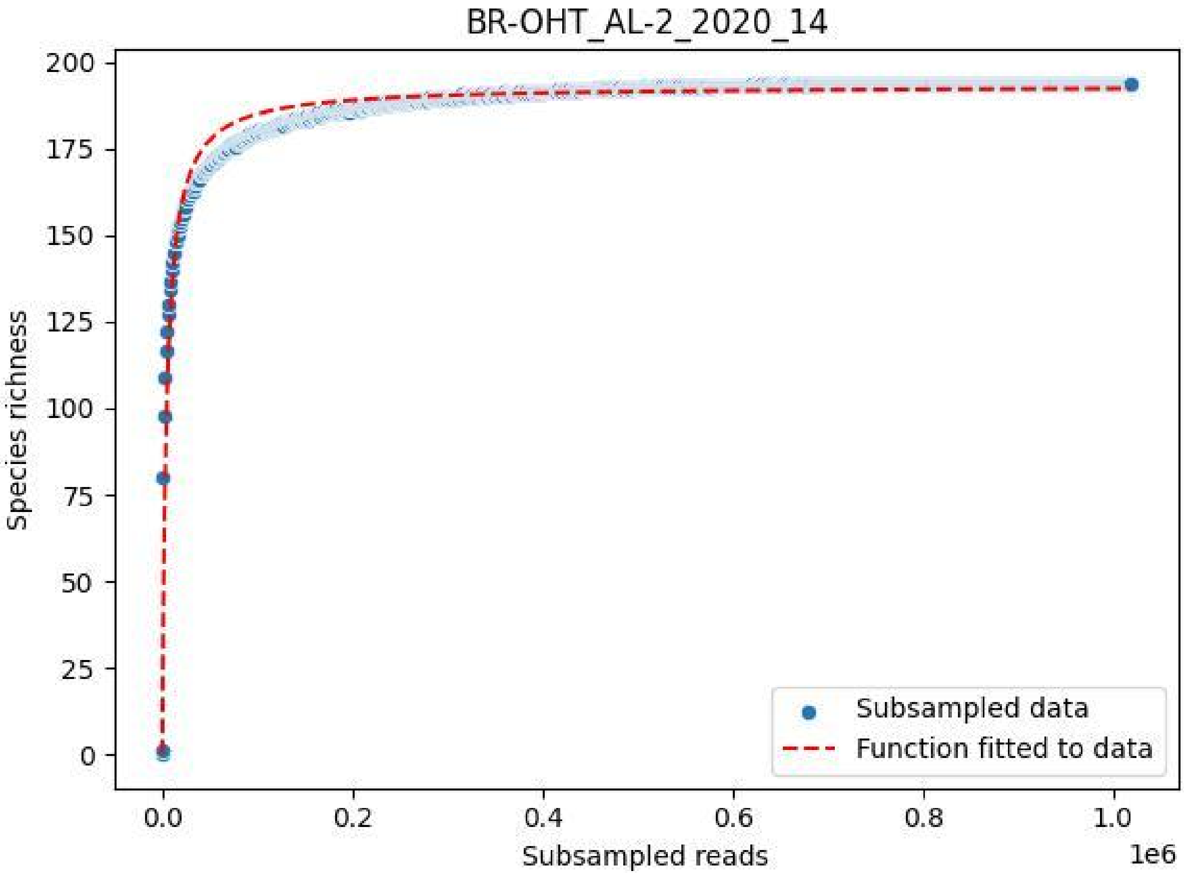
BR-OHT_AL-2_2020_14
Species richness
Subsampled reads
1e6
0.0
0.2
0.4
0.6
0.8
1.0
0
25
50
75
100
125
150
175
200
Subsampled data
Function fitted to data

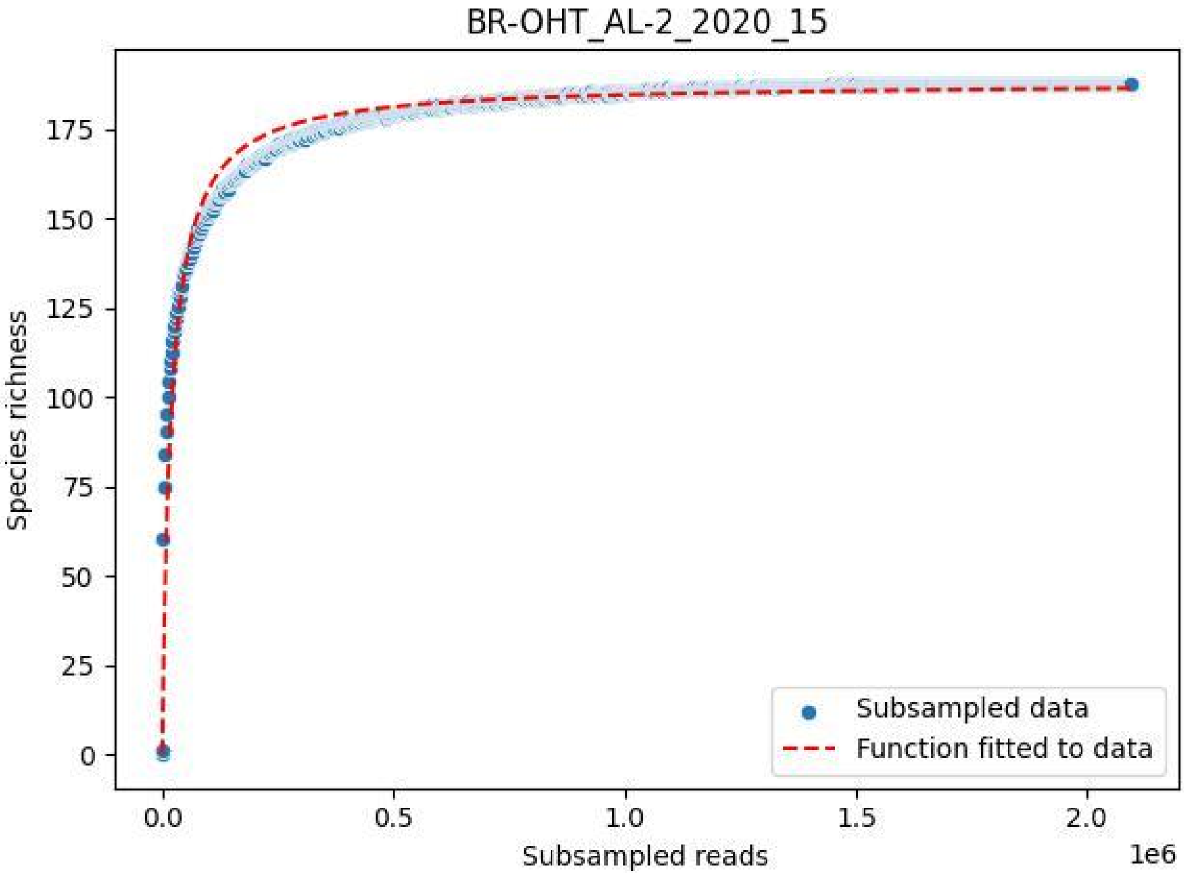
BR-OHT_AL-2_2020_15
Species richness
Subsampled reads
1e6
Subsampled data
Function fitted to data
0
25
50
75
100
125
150
175
0.0
0.5
1.0
1.5
2.0

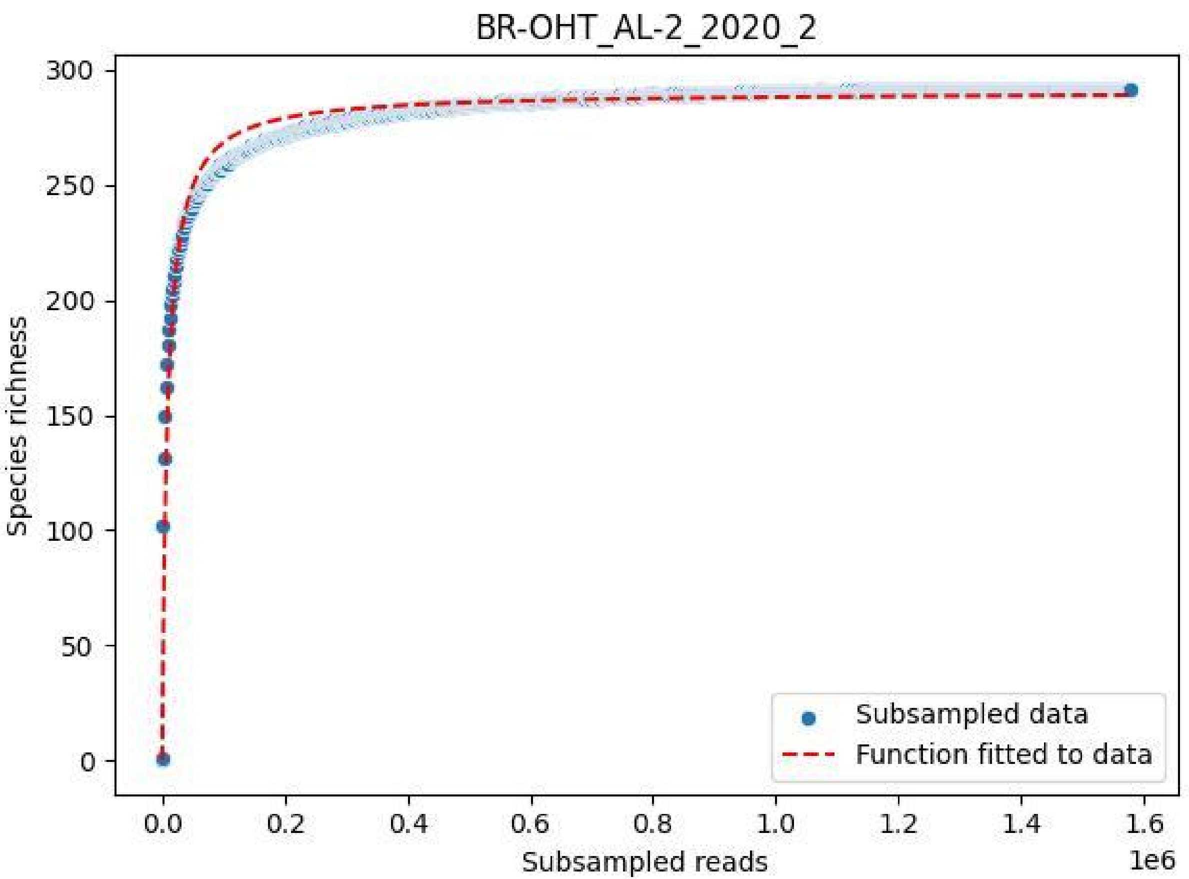
BR-OHT_AL-2_2020_2
Species richness
Subsampled reads
1e6
0
50
100
150
200
250
300
0.0
0.2
0.4
0.6
0.8
1.0
1.2
1.4
1.6
Subsampled data
Function fitted to data

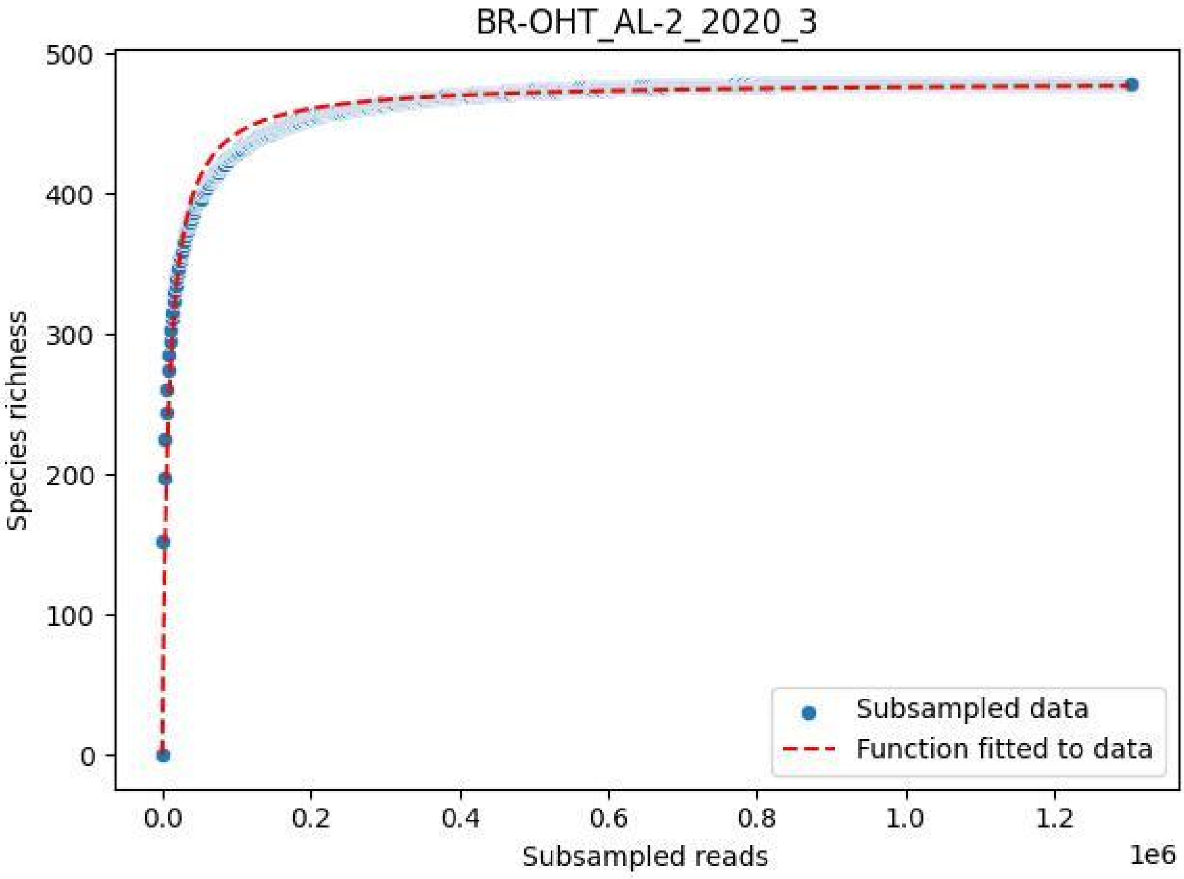
BR-OHT_AL-2_2020_3
Species richness
Subsampled reads
1e6
0
100
200
300
400
500
0.0
0.2
0.4
0.6
0.8
1.0
1.2
Subsampled data
Function fitted to data

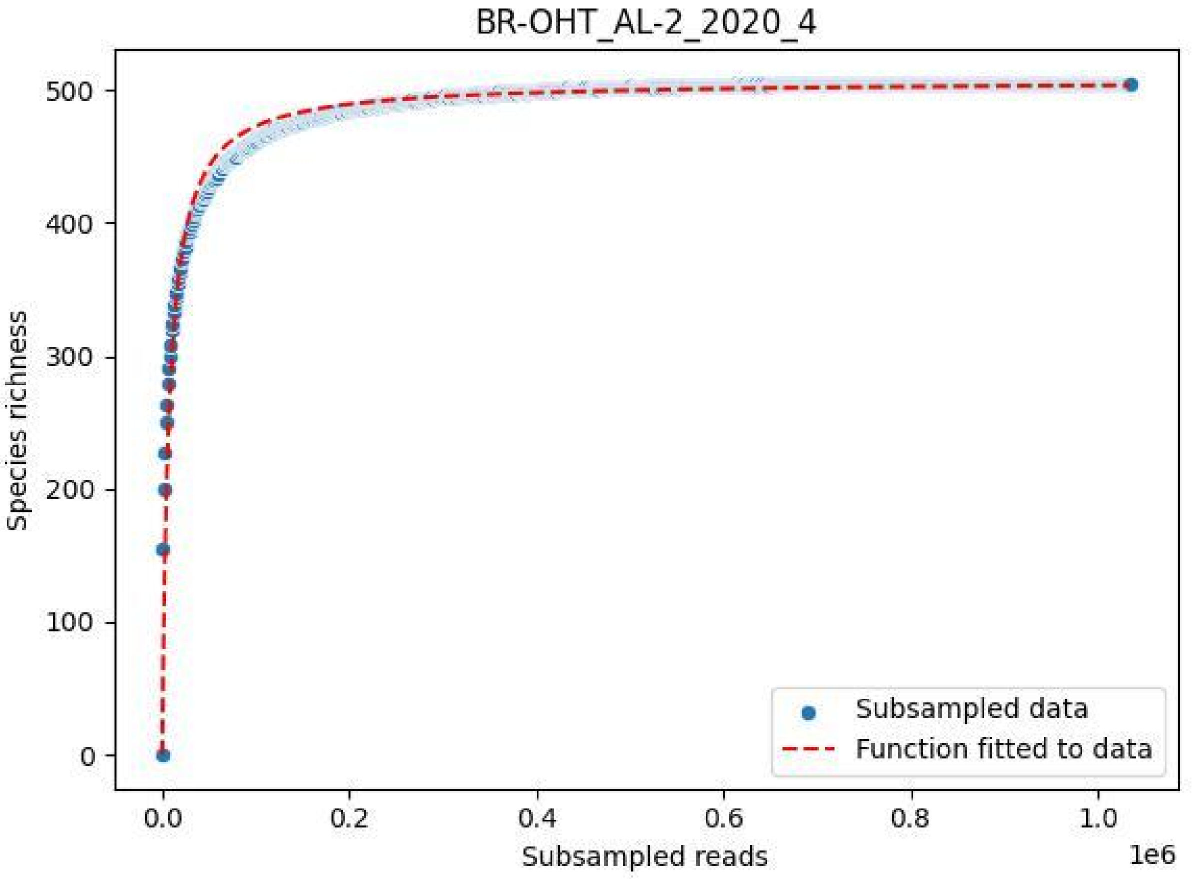

BR-OHT_AL-2_2020_4
Species richness
Subsampled reads
1e6
0
100
200
300
400
500
0.0
0.2
0.4
0.6
0.8
1.0
Subsampled data
Function fitted to data

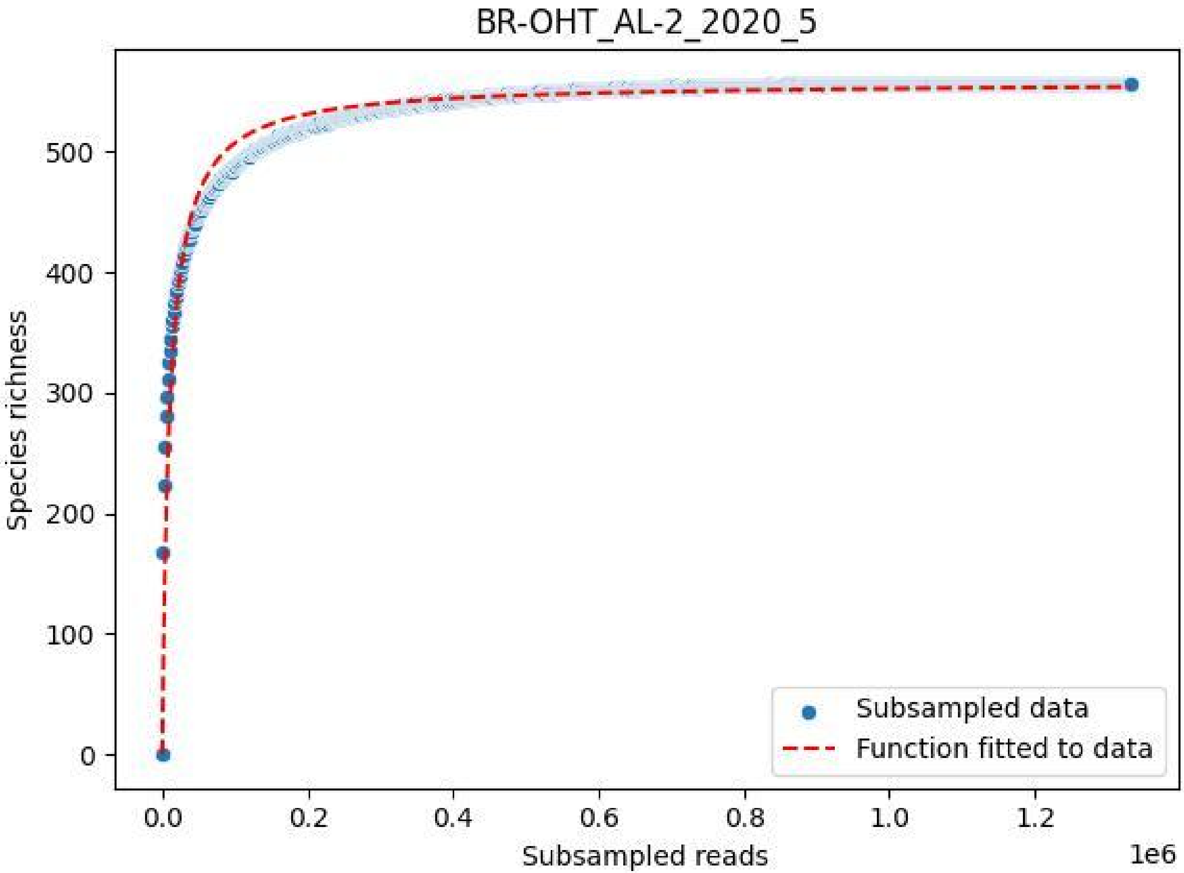
BR-OHT_AL-2_2020_5
Species richness
Subsampled reads
1e6
0
100
200
300
400
500
0.0
0.2
0.4
0.6
0.8
1.0
1.2
Subsampled data
Function fitted to data

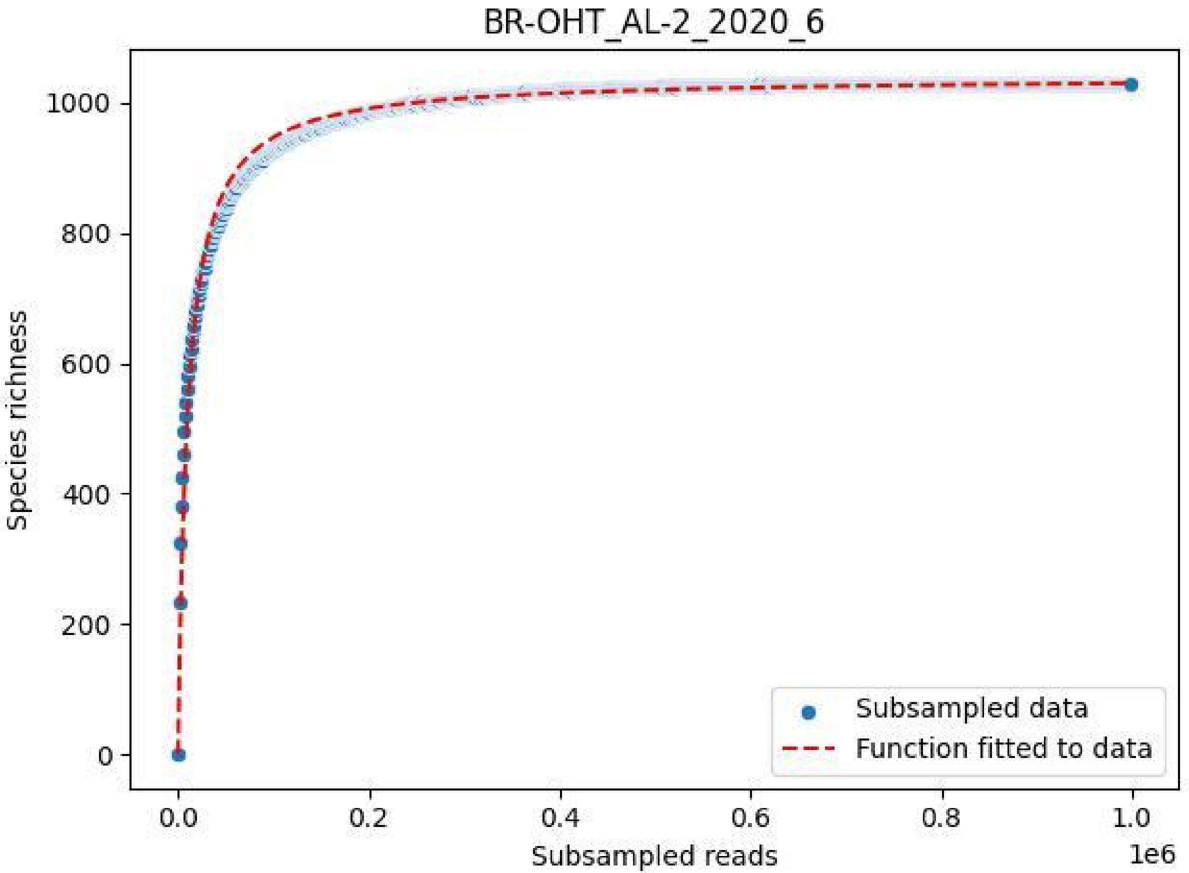
BR-OHT_AL-2_2020_6
Species richness
Subsampled reads
1e6
0
200
400
600
800
1000
0.0
0.2
0.4
0.6
0.8
1.0
Subsampled data
Function fitted to data

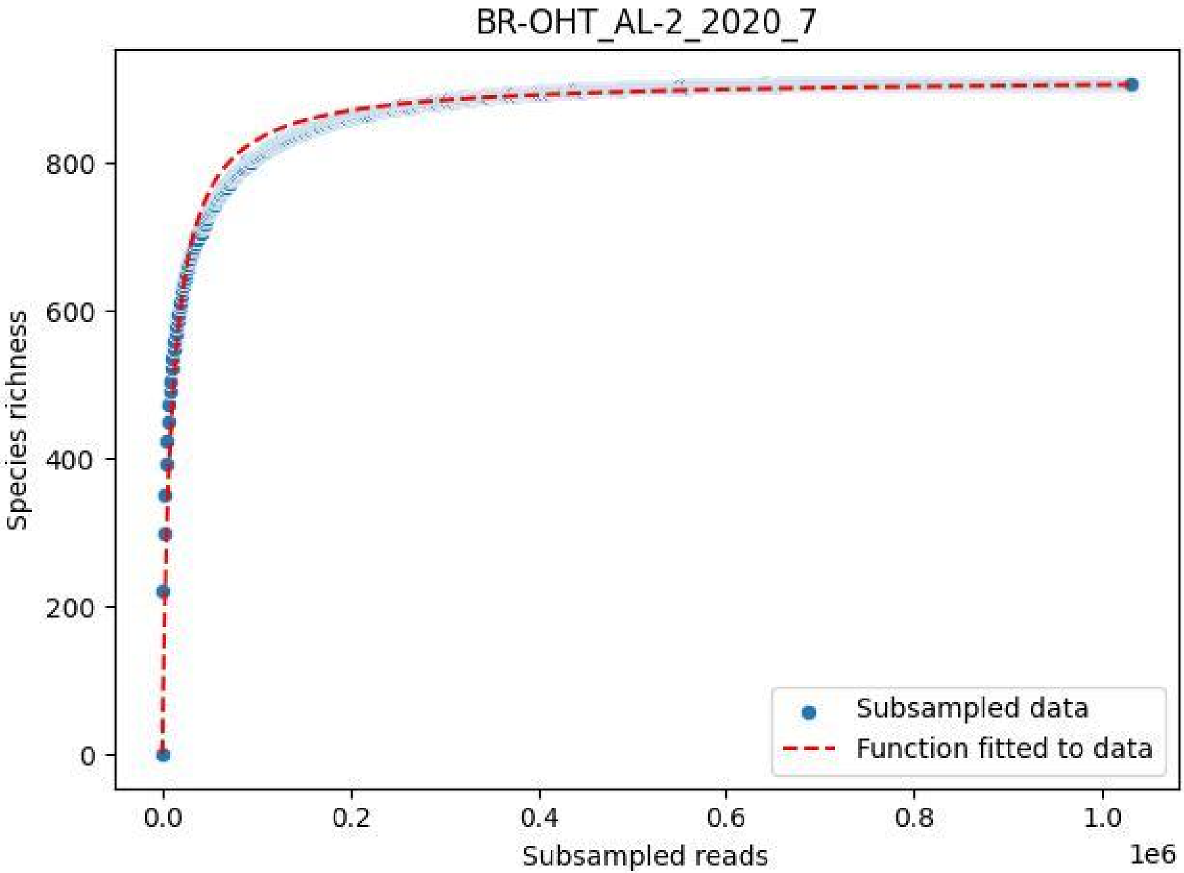

BR-OHT_AL-2_2020_7
Species richness
0
200
400
600
800
0.0
0.2
0.4
0.6
0.8
1.0
1e6
Subsampled reads
Subsampled data
Function fitted to data

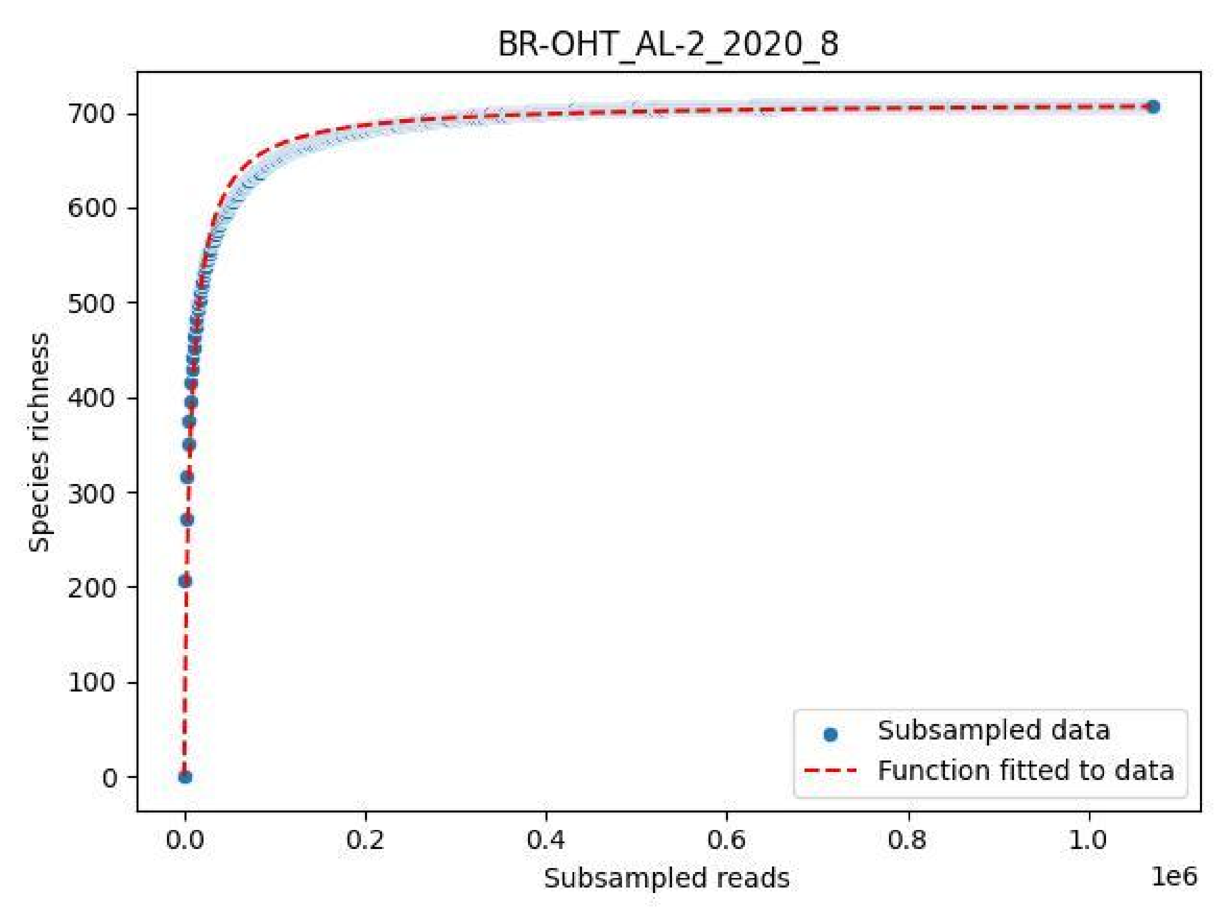

BR-OHT_AL-2_2020_8
Species richness
Subsampled reads
1e6
0
100
200
300
400
500
600
700
0.0
0.2
0.4
0.6
0.8
1.0
Subsampled data
Function fitted to data

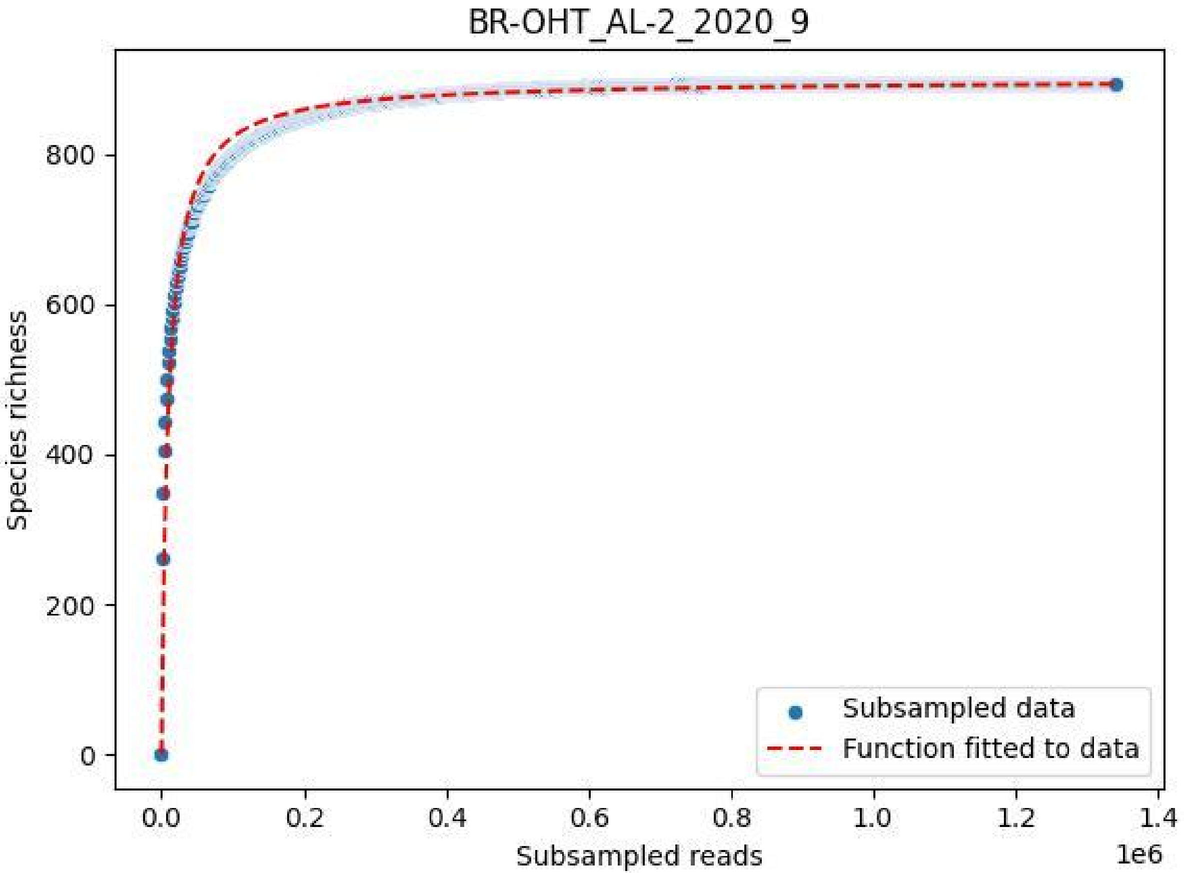
BR-OHT_AL-2_2020_9
Species richness
Subsampled reads
1e6
0
200
400
600
800
0.0
0.2
0.4
0.6
0.8
1.0
1.2
1.4
Subsampled data
Function fitted to data

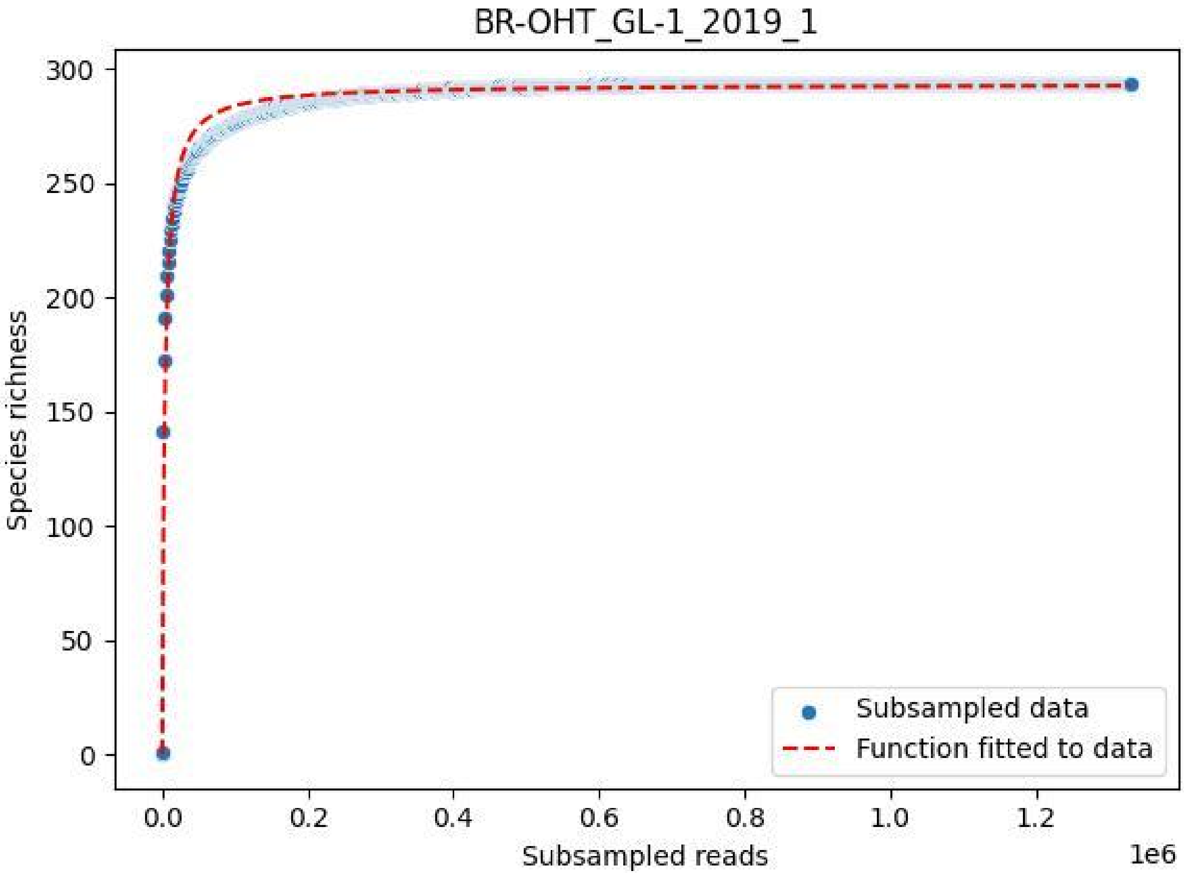
BR-OHT_GL-1_2019_1
300
250
200
150
100
50
0
Species richness
0.0
0.2
0.4
0.6
0.8
1.0
1.2
1e6
Subsampled reads
Subsampled data
Function fitted to data

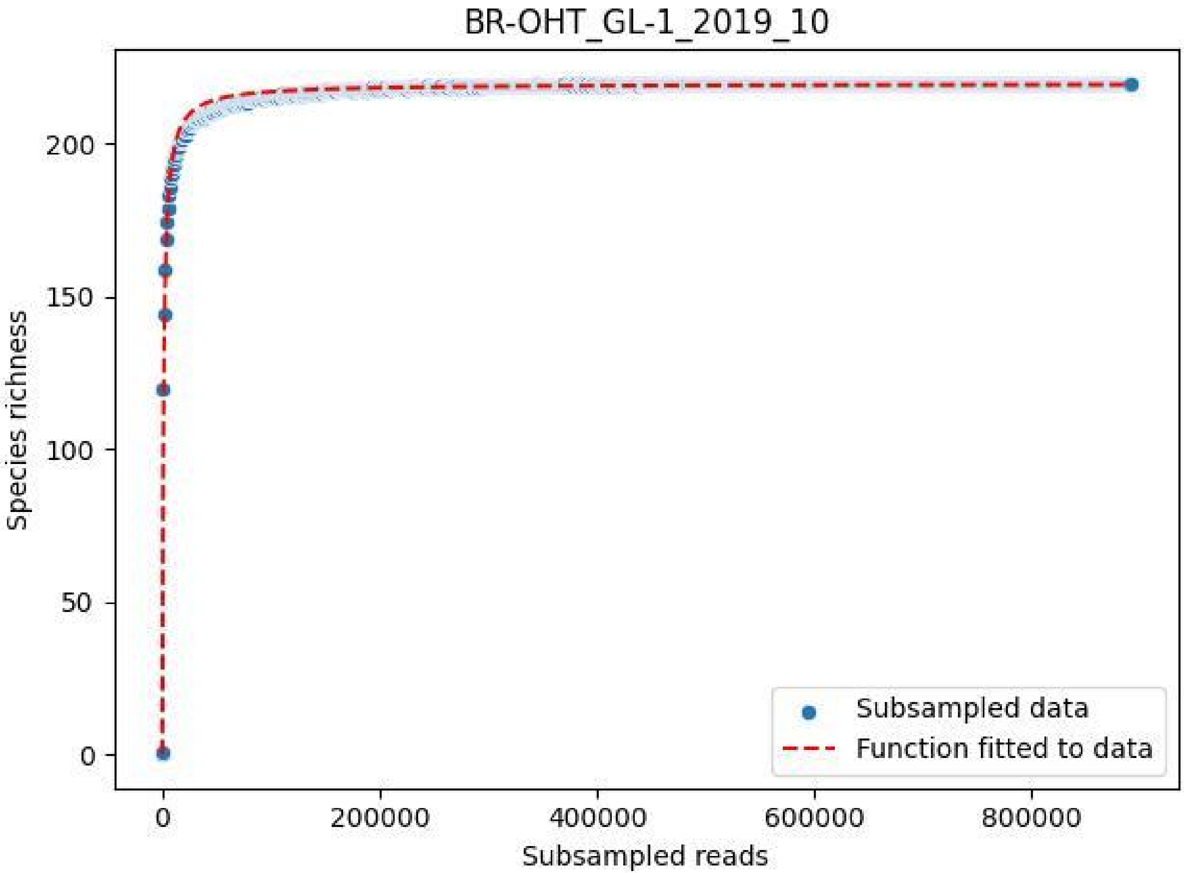
BR-OHT_GL-1_2019_10
Species richness
Subsampled reads
0
50
100
150
200
0
200000
400000
600000
800000
Subsampled data
Function fitted to data

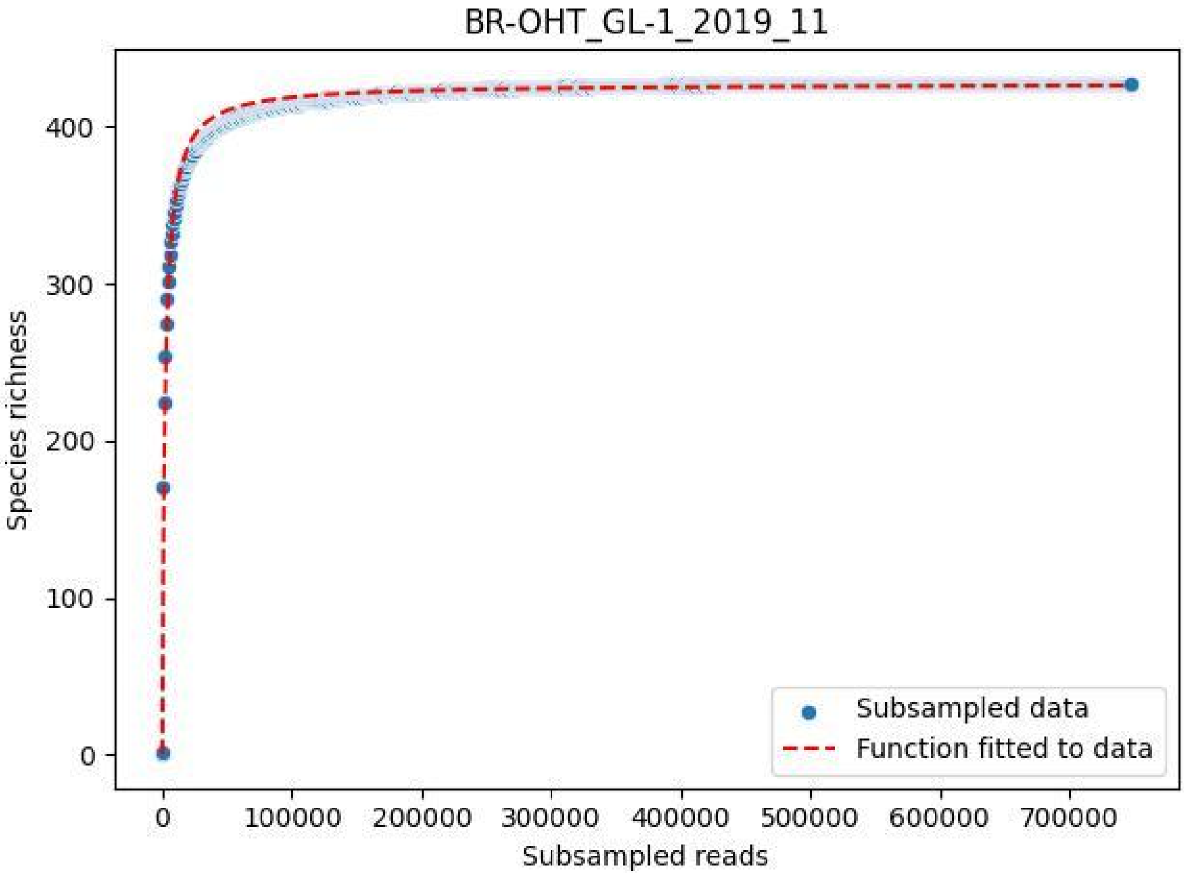
BR-OHT_GL-1_2019_11
Species richness
0
100
200
300
400
Subsampled reads
0
100000
200000
300000
400000
500000
600000
700000
Subsampled data
Function fitted to data

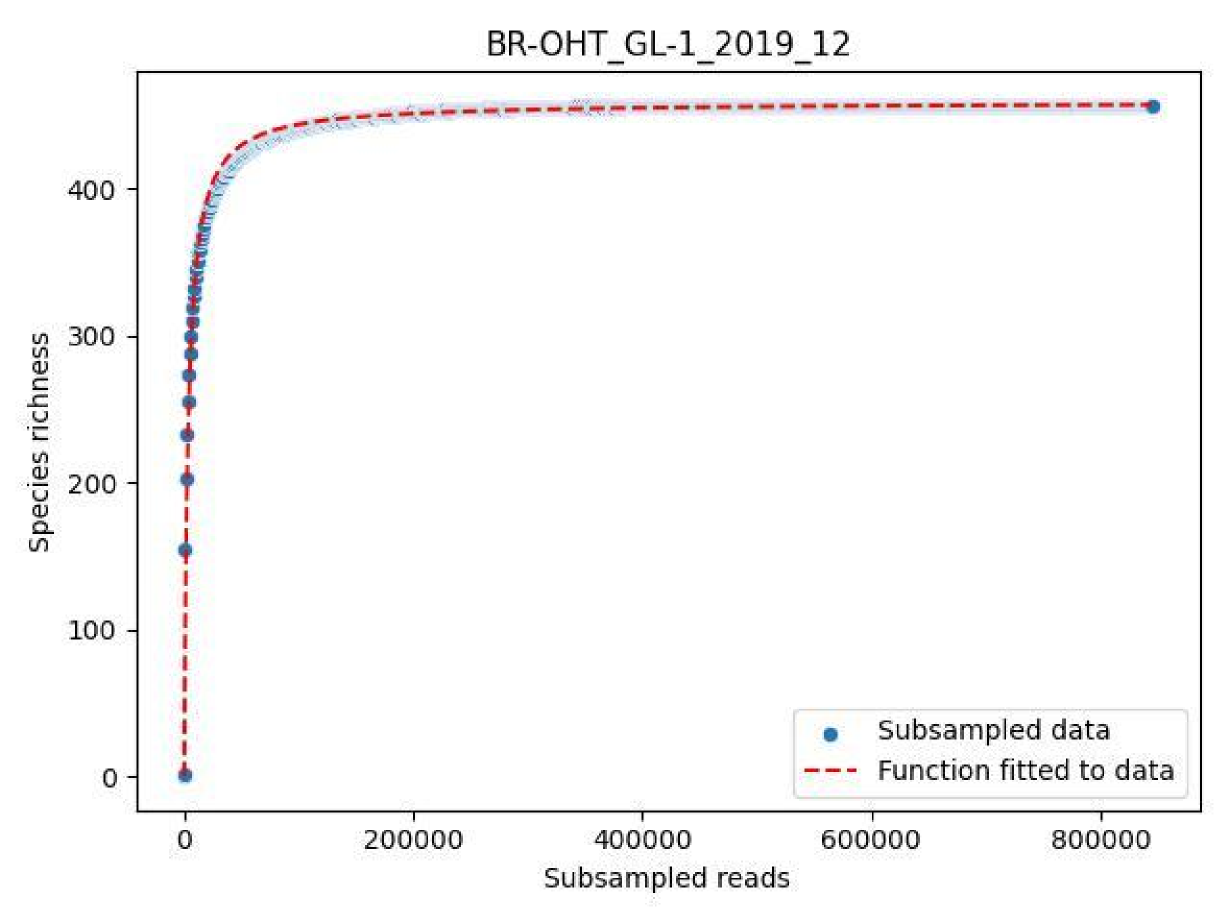

BR-OHT_GL-1_2019_12
Species richness
Subsampled reads
0
100
200
300
400
0
200000
400000
600000
800000
Subsampled data
Function fitted to data

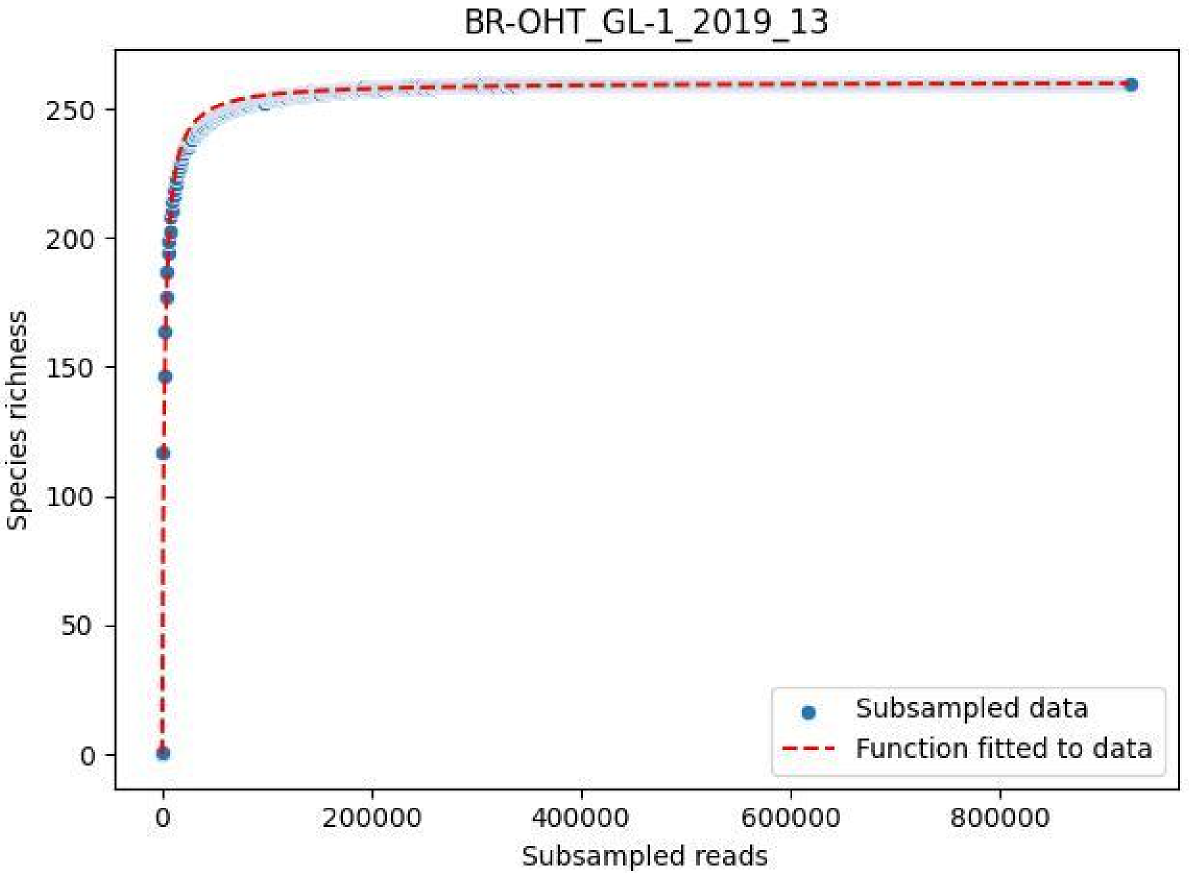
BR-OHT_GL-1_2019_13
Species richness
Subsampled reads
0
50
100
150
200
250
0
200000
400000
600000
800000
Subsampled data
Function fitted to data

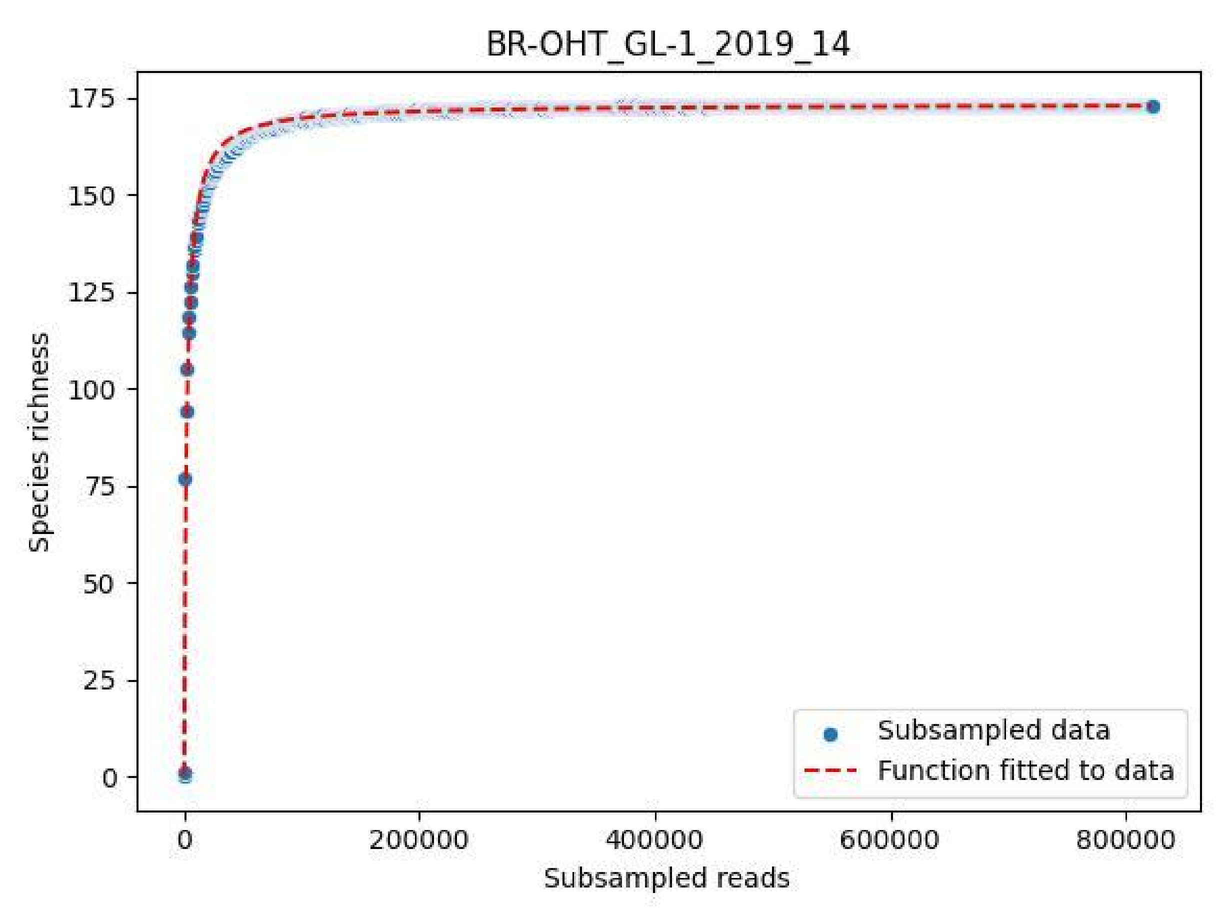

BR-OHT_GL-1_2019_14
Species richness
Subsampled reads
Subsampled data
Function fitted to data
0
25
50
75
100
125
150
175
0
200000
400000
600000
800000

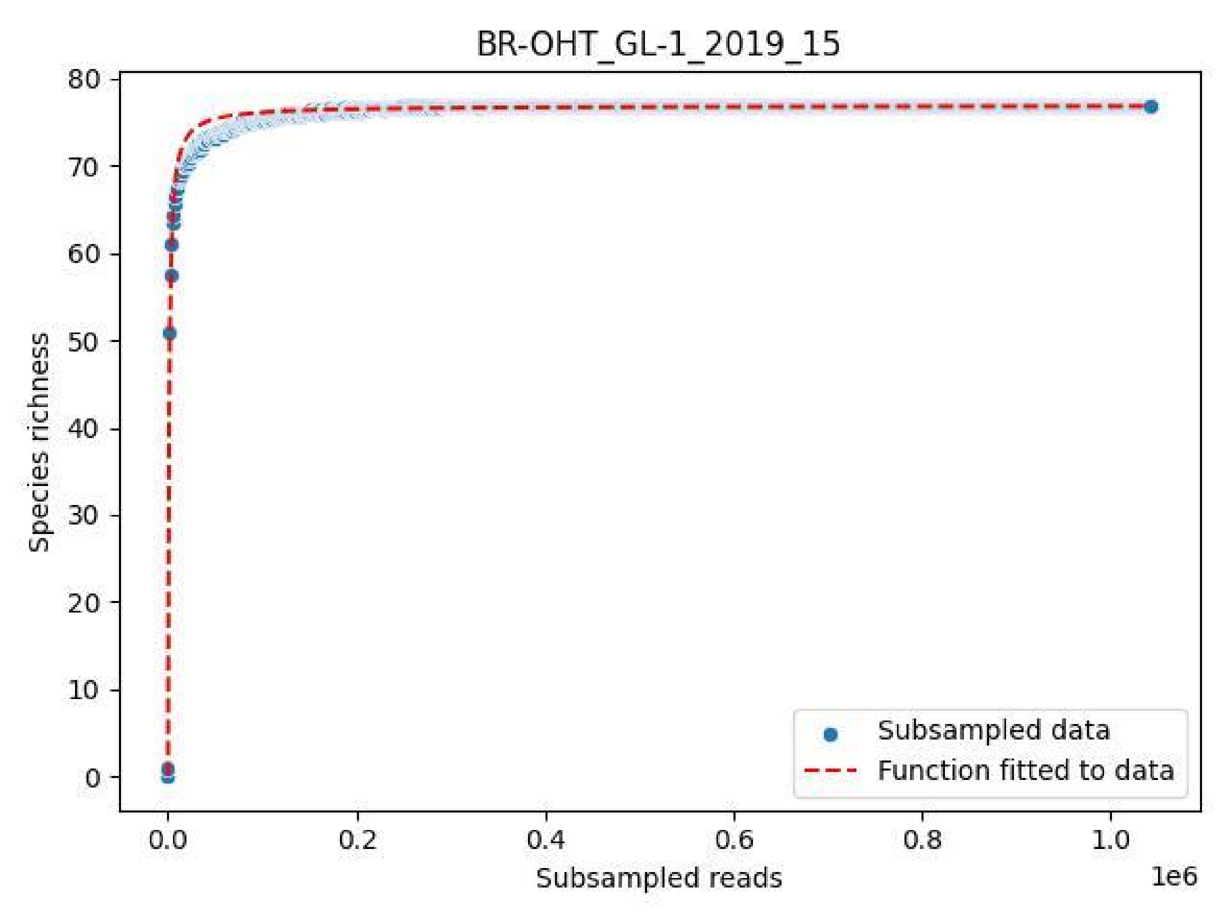
BR-OHT_GL-1_2019_15
Species richness
Subsampled reads
1e6
0
10
20
30
40
50
60
70
80
0.0
0.2
0.4
0.6
0.8
1.0
Subsampled data
Function fitted to data

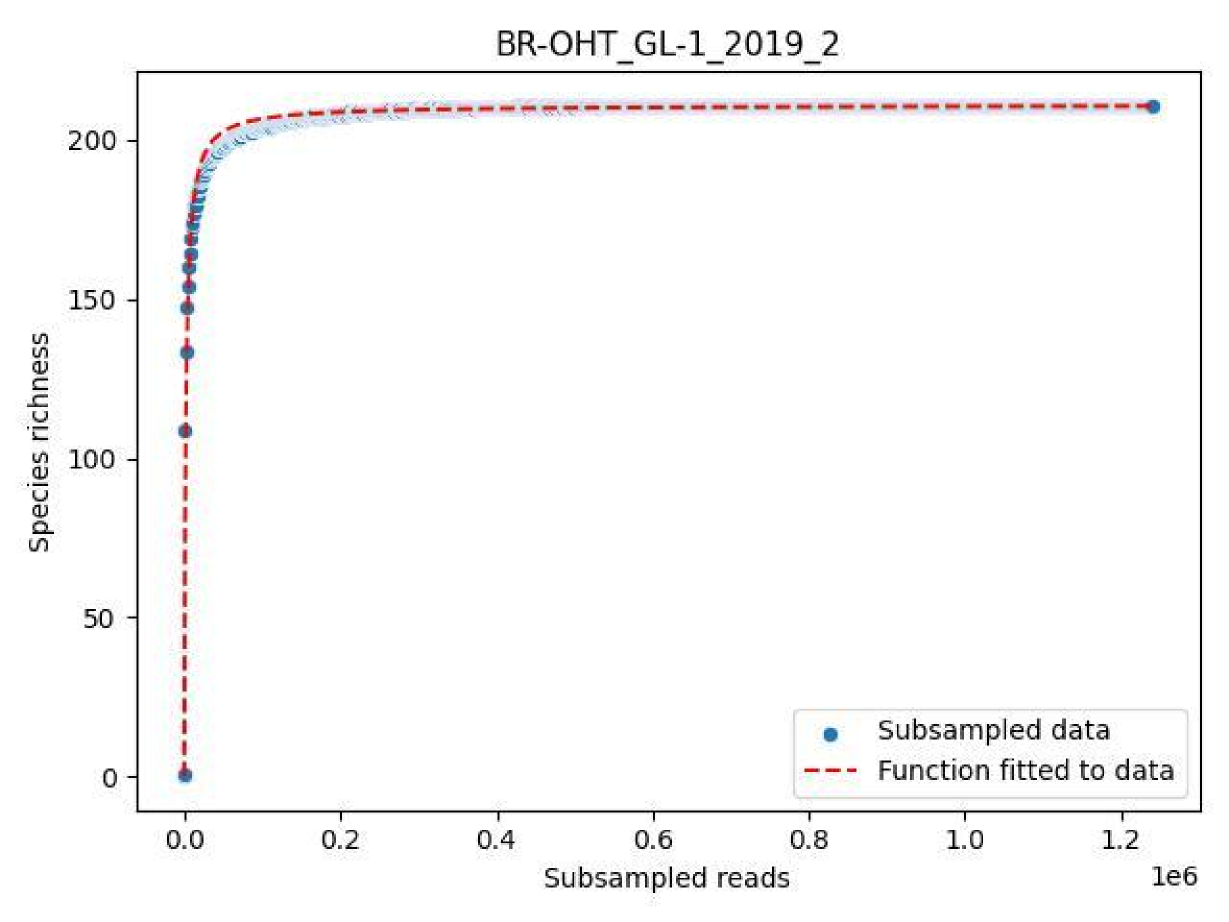

BR-OHT_GL-1_2019_2
Species richness
Subsampled reads
1e6
0
50
100
150
200
0.0
0.2
0.4
0.6
0.8
1.0
1.2
Subsampled data
Function fitted to data

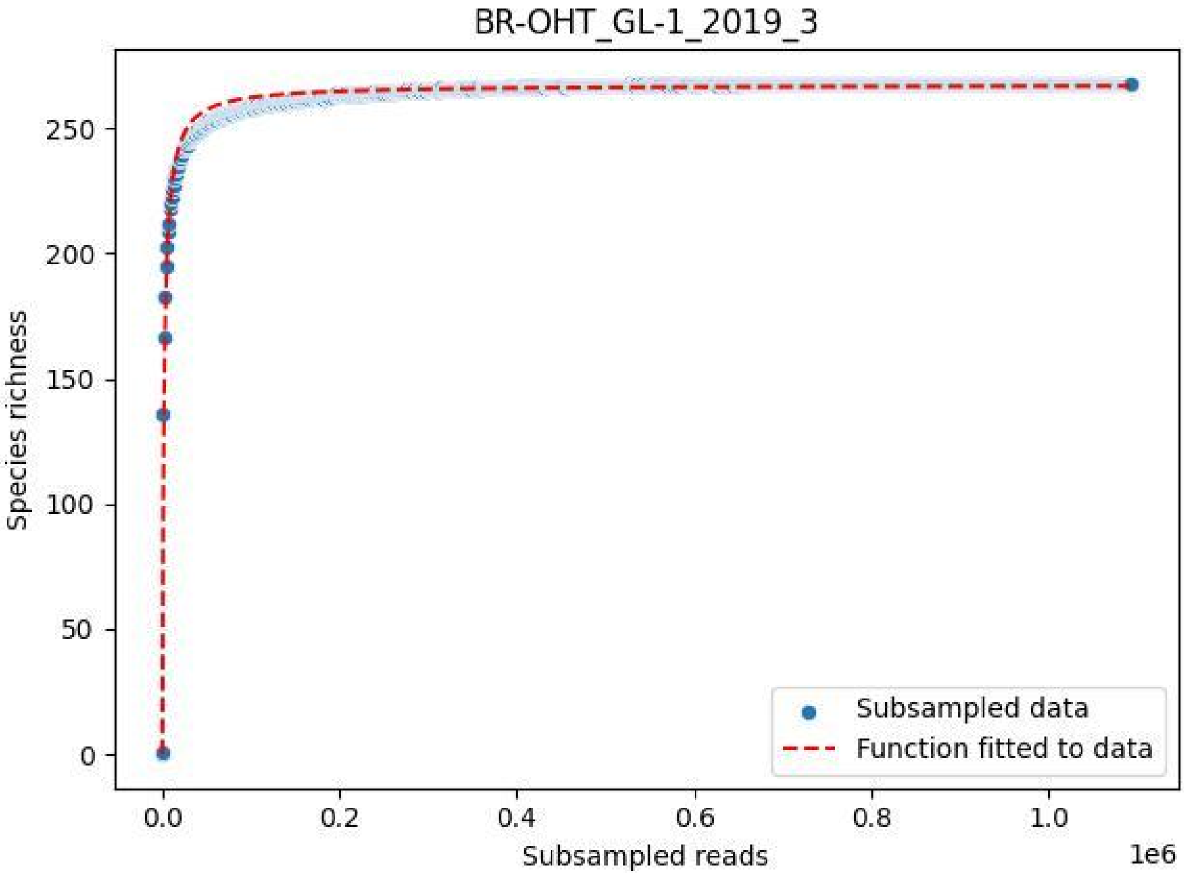
BR-OHT_GL-1_2019_3
Species richness
Subsampled reads
1e6
Subsampled data
Function fitted to data
0
50
100
150
200
250
0.0
0.2
0.4
0.6
0.8
1.0

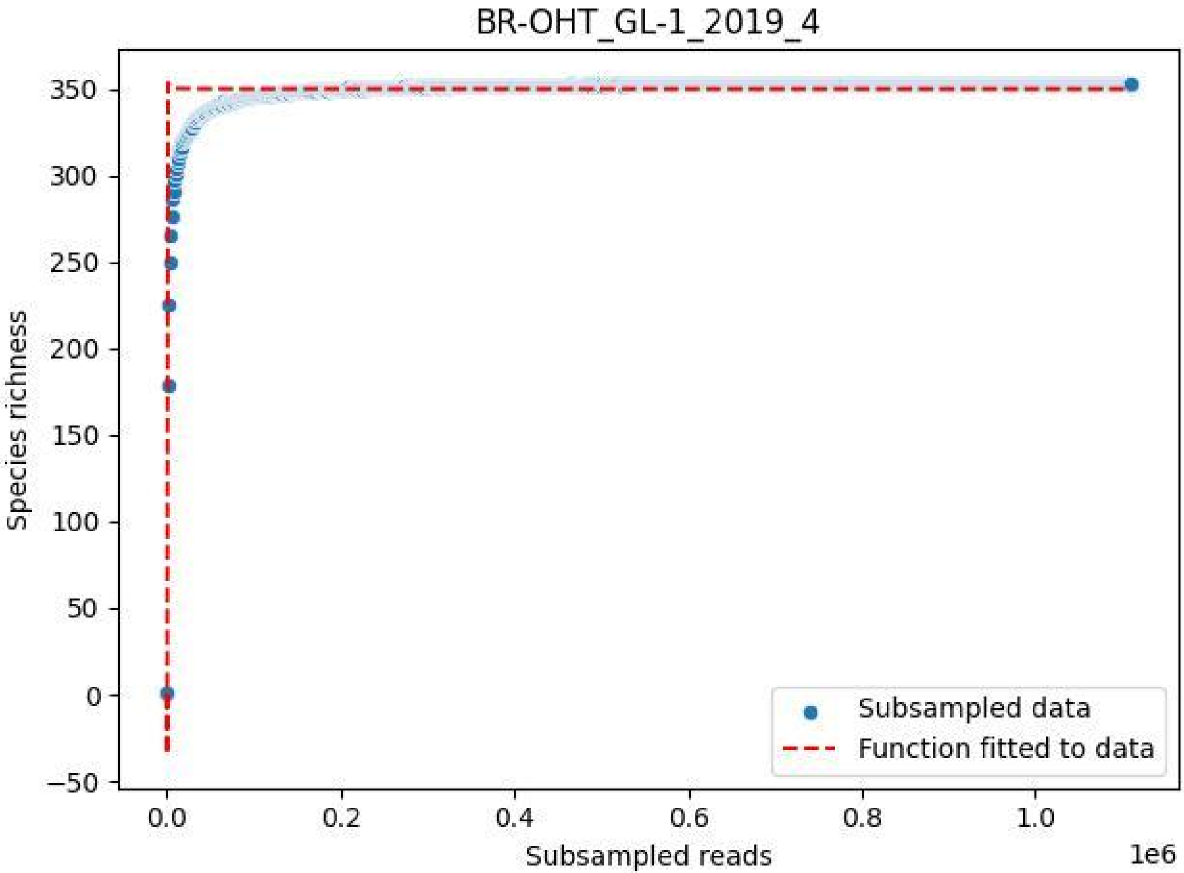
BR-OHT_GL-1_2019_4
Species richness
Subsampled reads
1e6
350
300
250
200
150
100
50
0
−50
0.0
0.2
0.4
0.6
0.8
1.0
Subsampled data
Function fitted to data

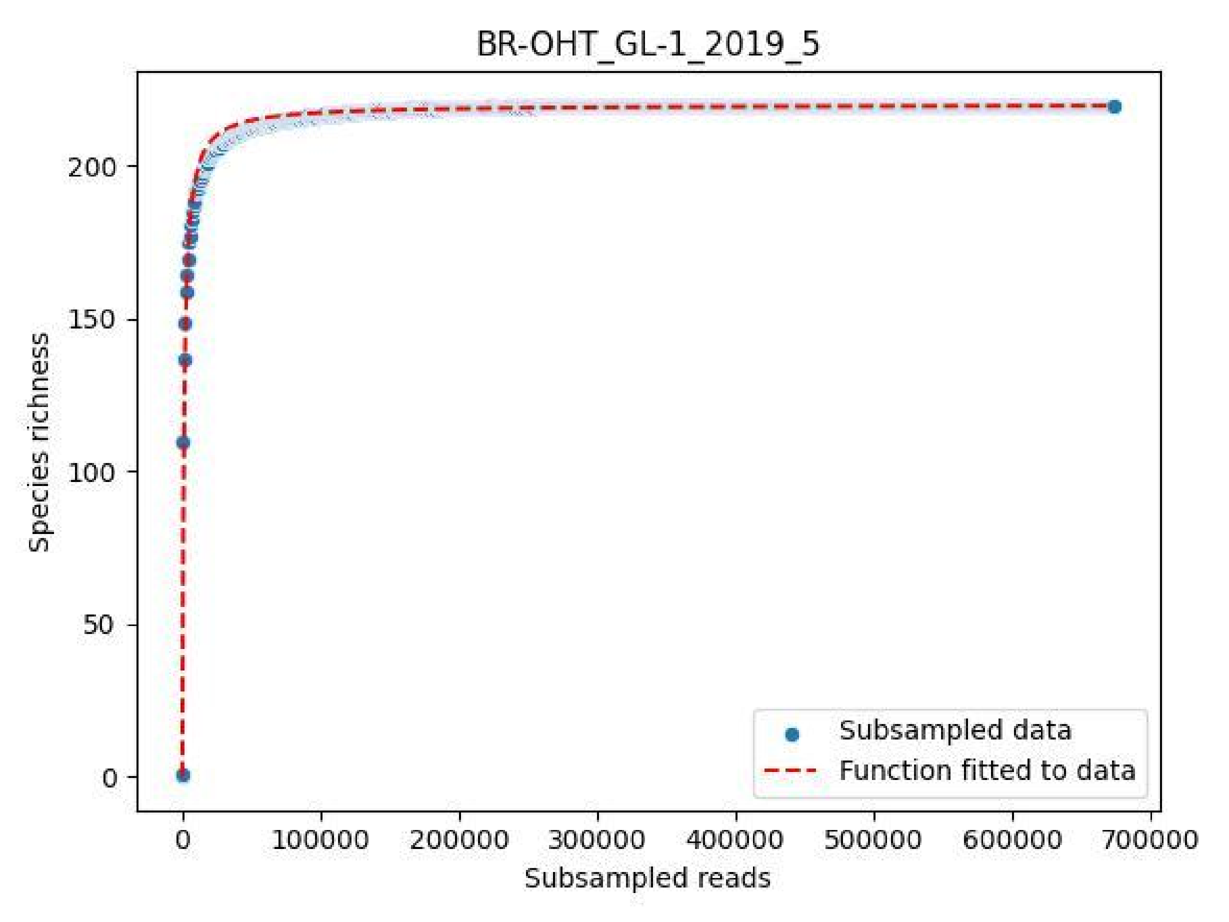

BR-OHT_GL-1_2019_5
Species richness
Subsampled reads
0
50
100
150
200
0
100000
200000
300000
400000
500000
600000
700000
Subsampled data
Function fitted to data

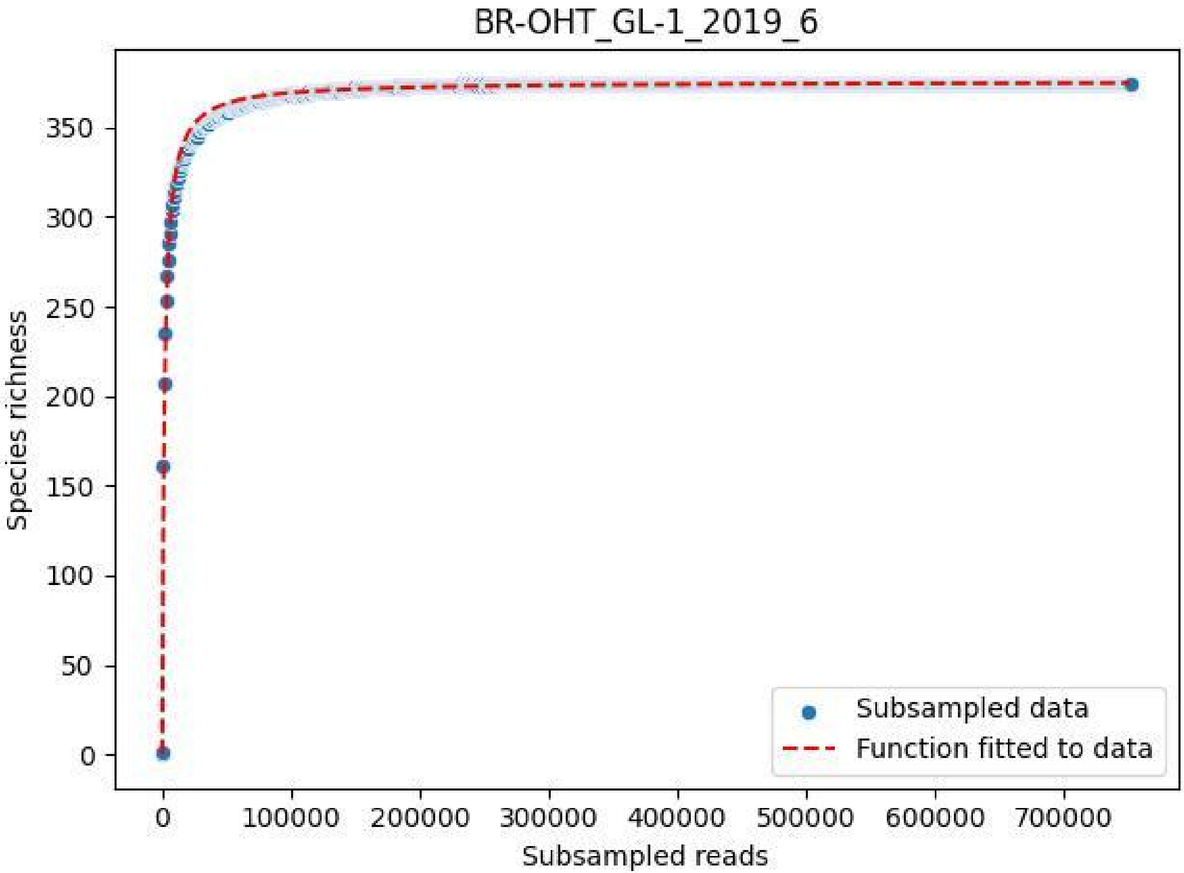
BR-OHT_GL-1_2019_6
Species richness
Subsampled reads
0
50
100
150
200
250
300
350
0
100000
200000
300000
400000
500000
600000
700000
Subsampled data
Function fitted to data

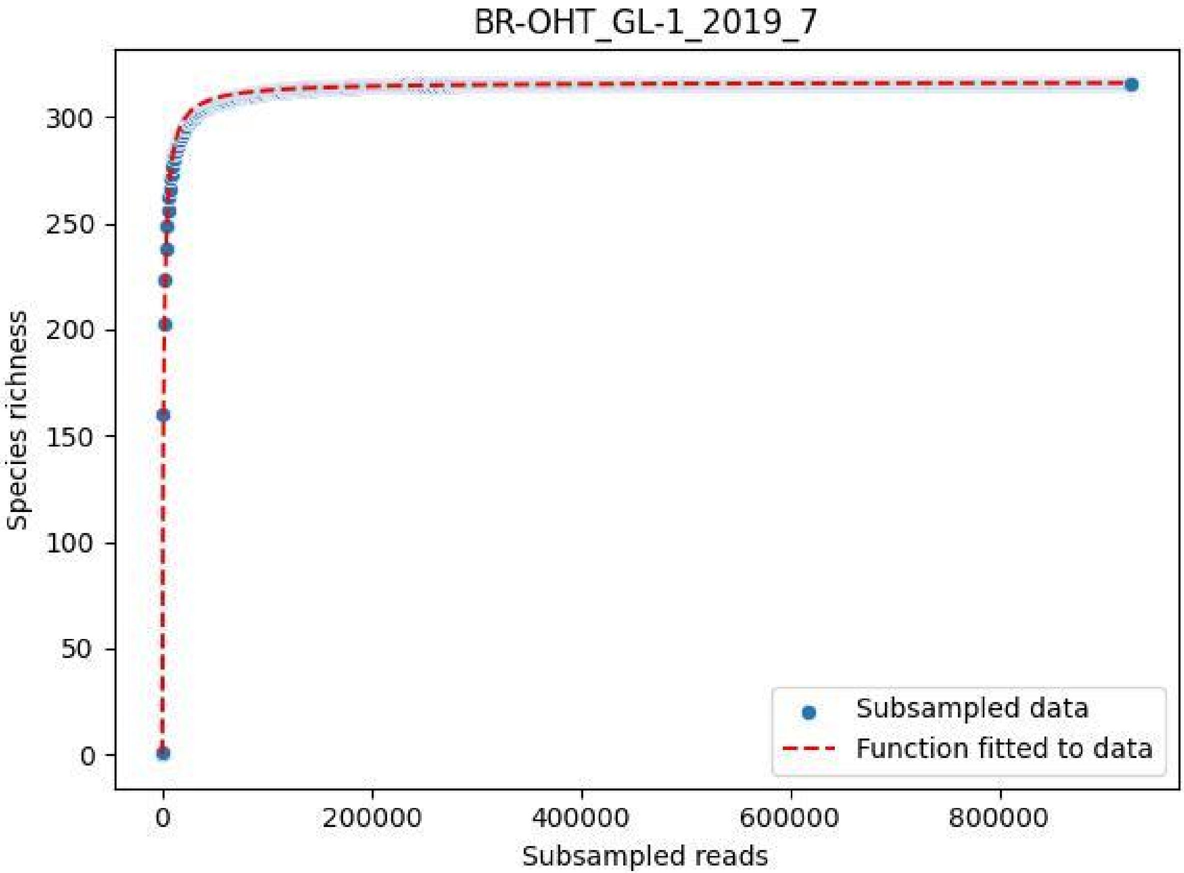

BR-OHT_GL-1_2019_7
Species richness
Subsampled reads
0
50
100
150
200
250
300
0
200000
400000
600000
800000
Subsampled data
Function fitted to data

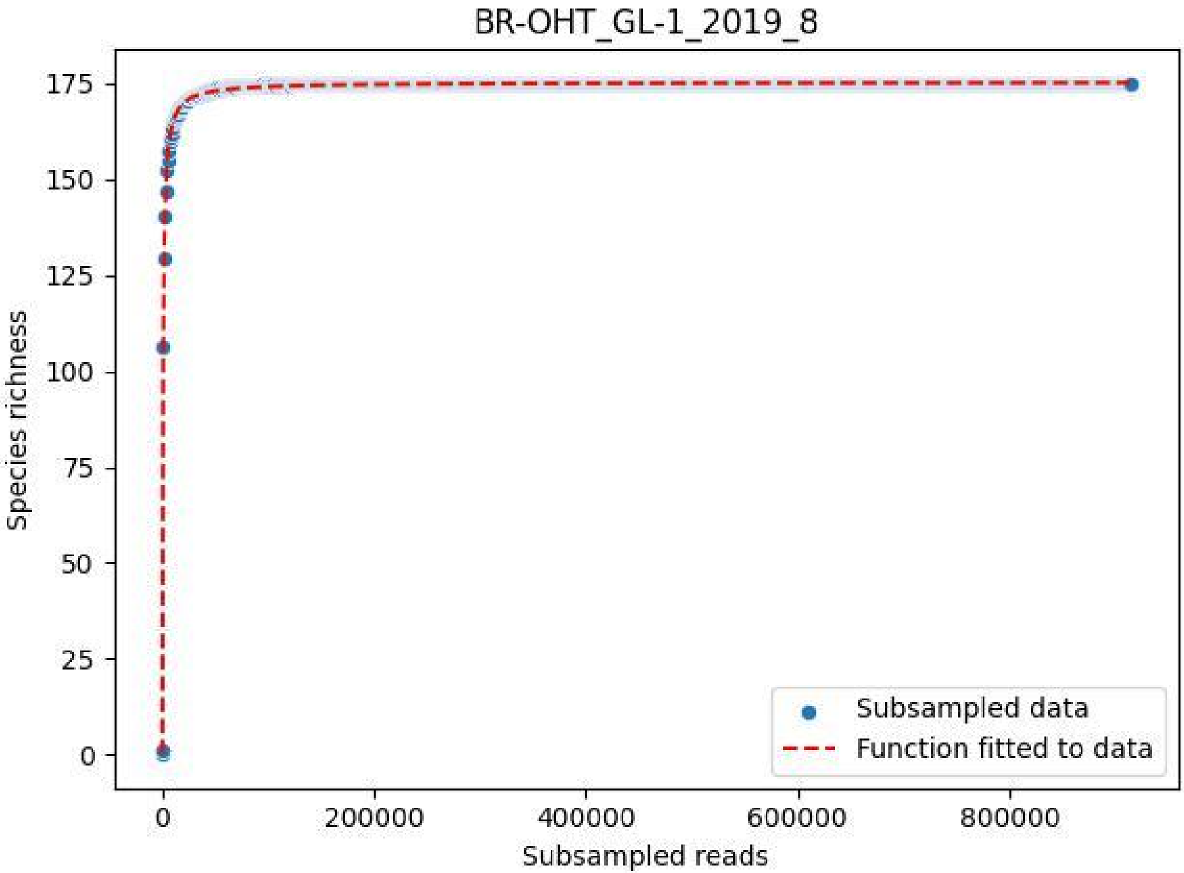
BR-OHT_GL-1_2019_8
Species richness
Subsampled reads
175
150
125
100
75
50
25
0
0
200000
400000
600000
800000
Subsampled data
Function fitted to data

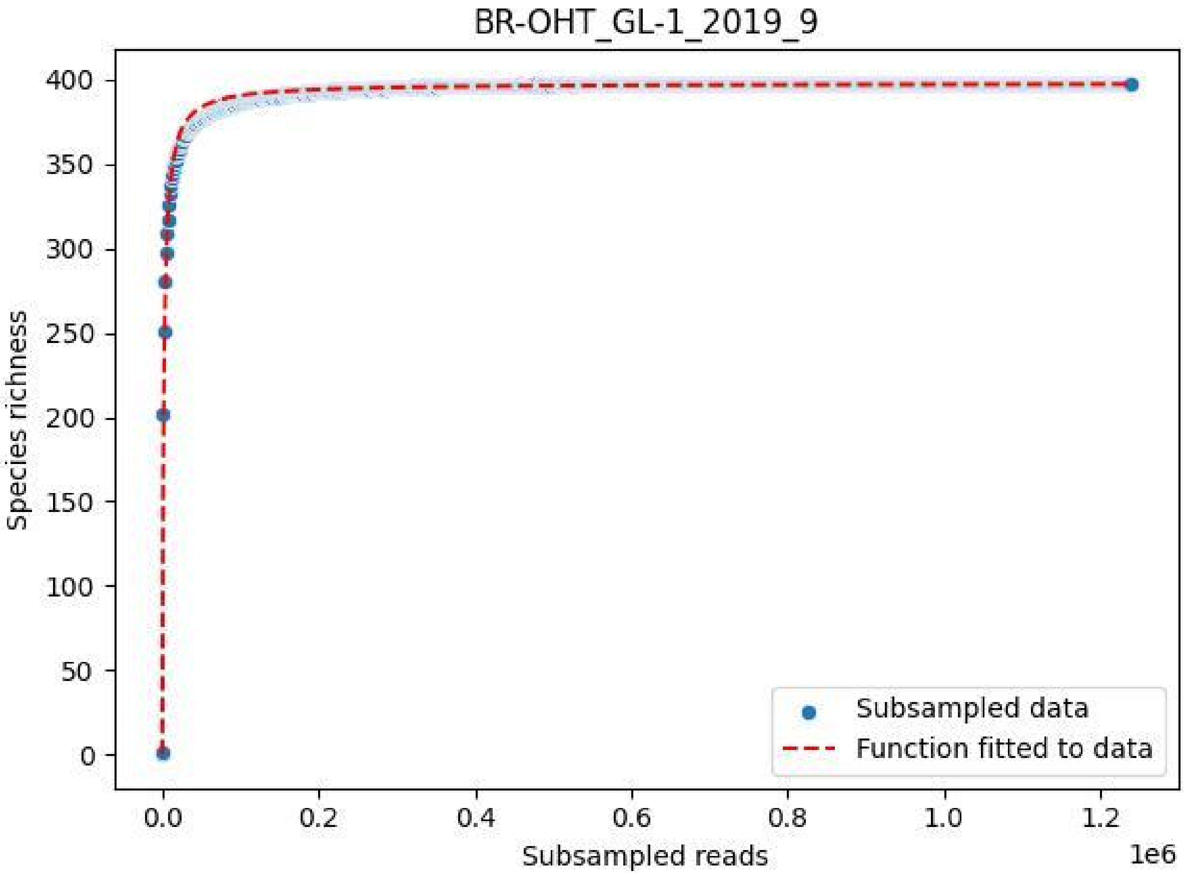
BR-OHT_GL-1_2019_9
Species richness
Subsampled reads
1e6
0
50
100
150
200
250
300
350
400
0.0
0.2
0.4
0.6
0.8
1.0
1.2
Subsampled data
Function fitted to data

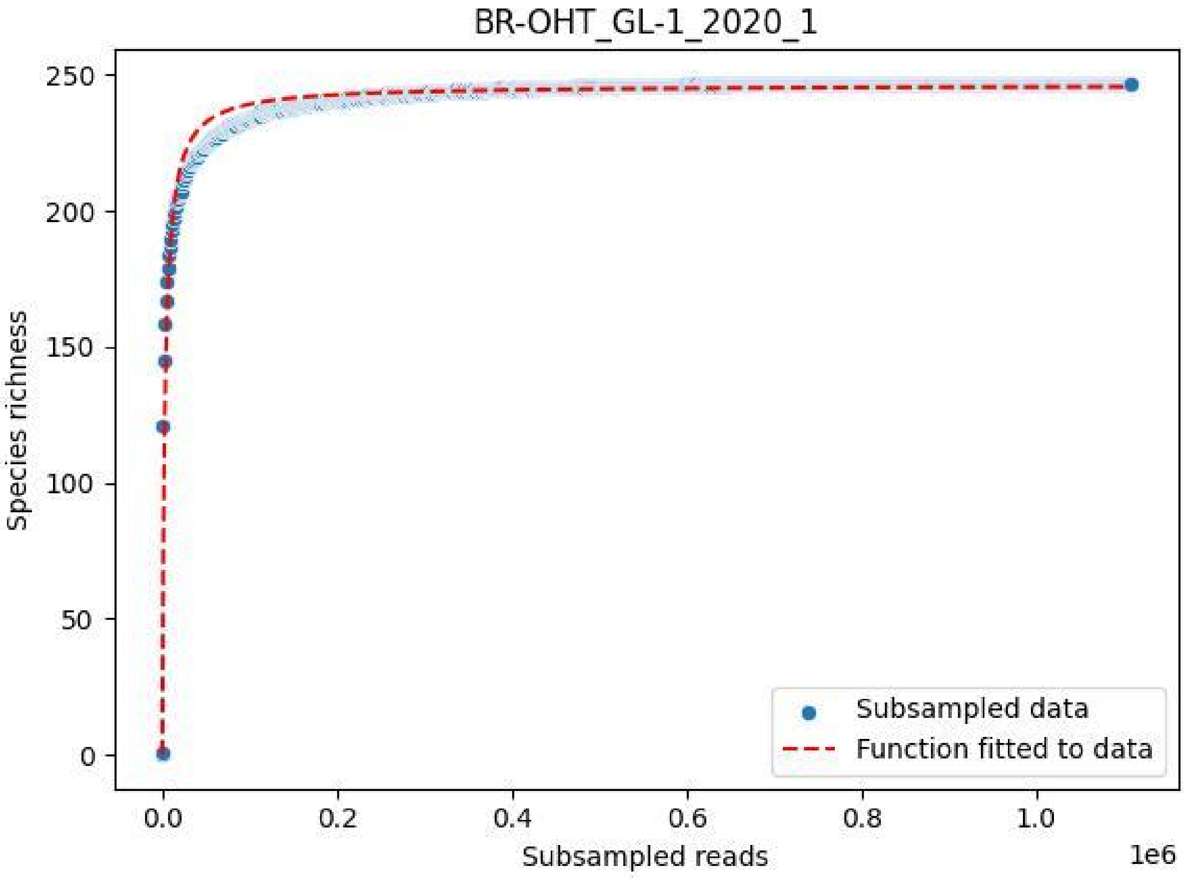
BR-OHT_GL-1_2020_1
250
200
150
100
50
0
Species richness
0.0
0.2
0.4
0.6
0.8
1.0
1e6
Subsampled reads
Subsampled data
Function fitted to data

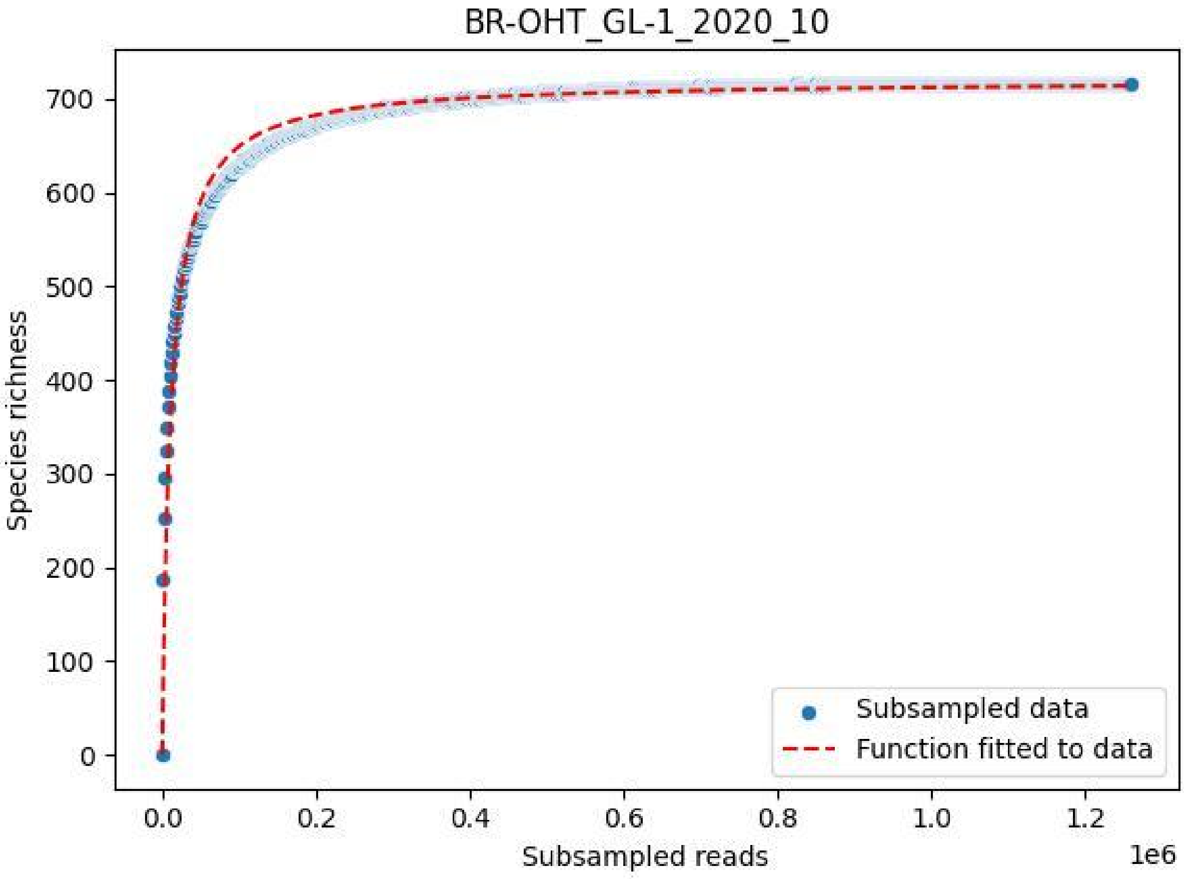
BR-OHT_GL-1_2020_10
Species richness
Subsampled reads
1e6
0
100
200
300
400
500
600
700
0.0
0.2
0.4
0.6
0.8
1.0
1.2
Subsampled data
Function fitted to data

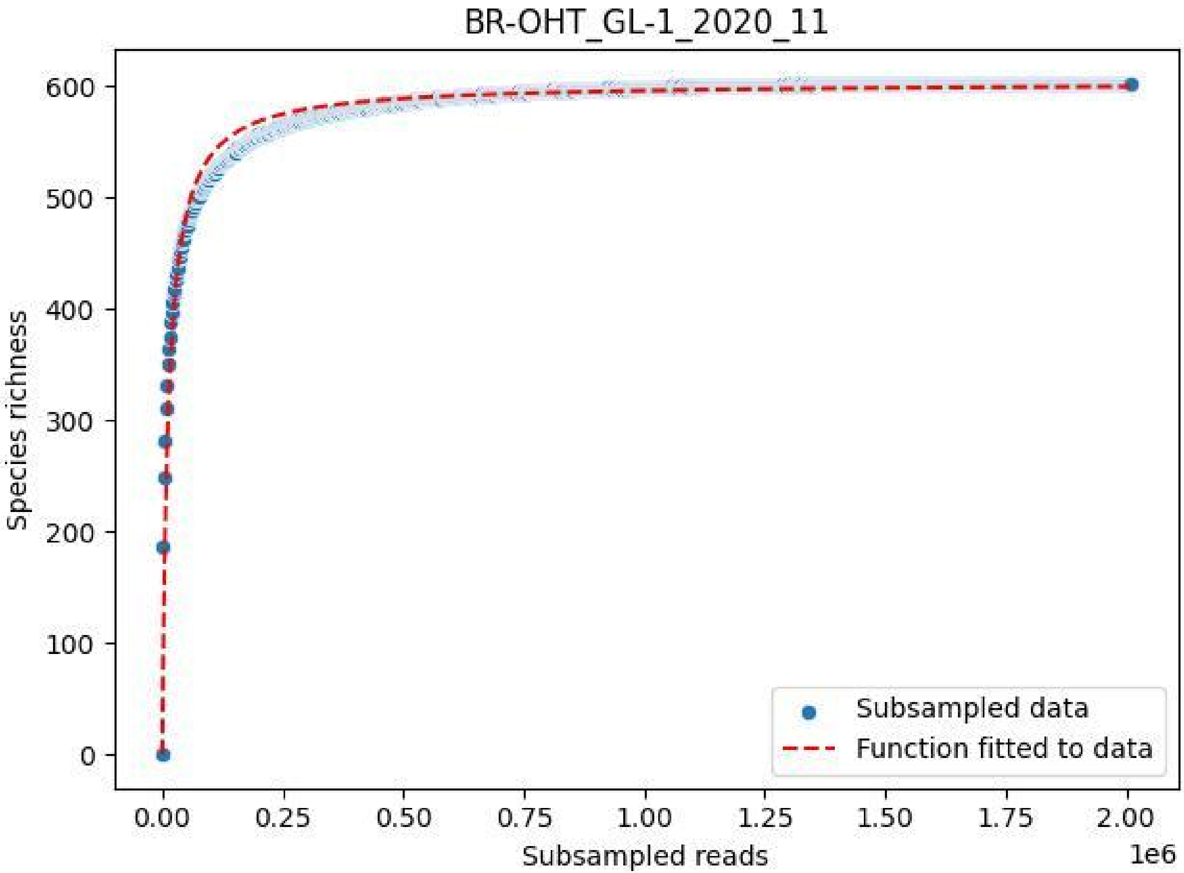

BR-OHT_GL-1_2020_11
Species richness
Subsampled reads
1e6
0
100
200
300
400
500
600
0.00
0.25
0.50
0.75
1.00
1.25
1.50
1.75
2.00
Subsampled data
Function fitted to data

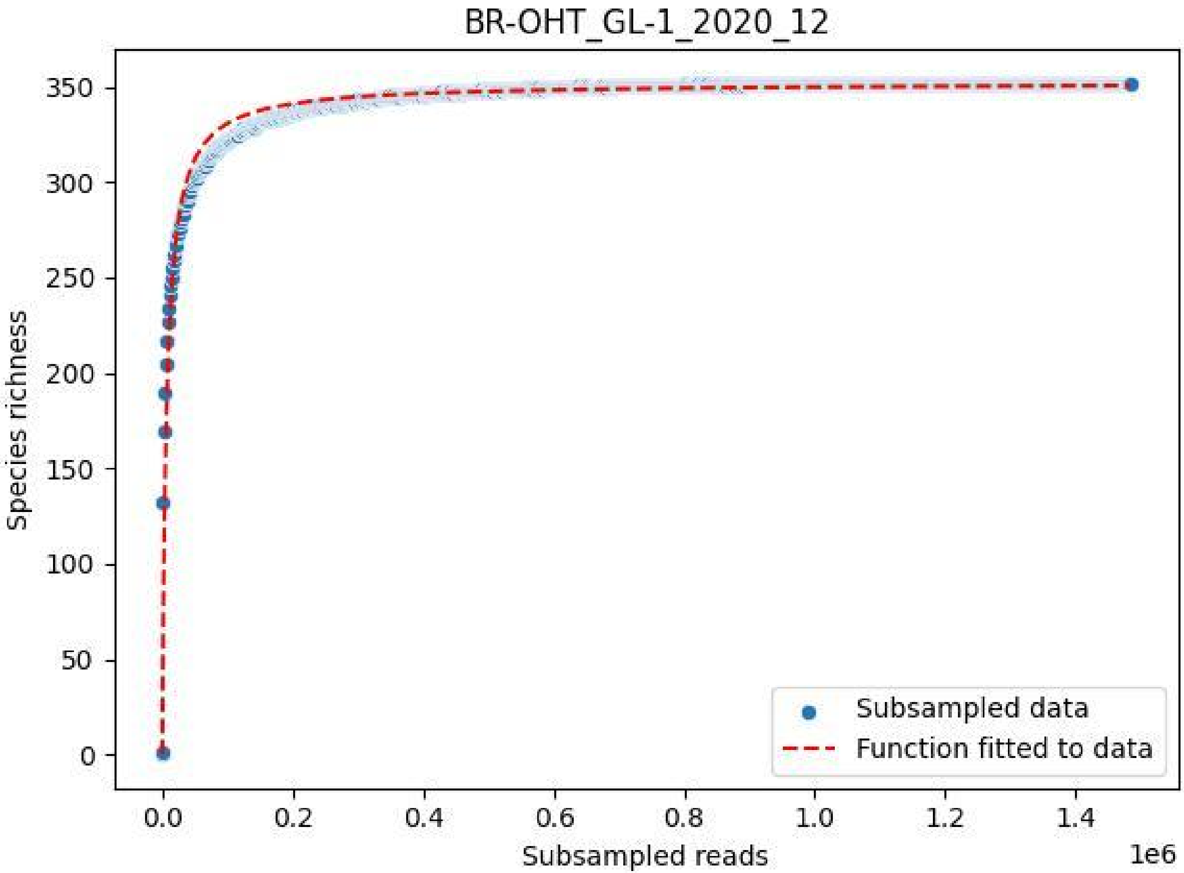
BR-OHT_GL-1_2020_12
Species richness
Subsampled reads
1e6
Subsampled data
Function fitted to data
0
50
100
150
200
250
300
350
0.0
0.2
0.4
0.6
0.8
1.0
1.2
1.4

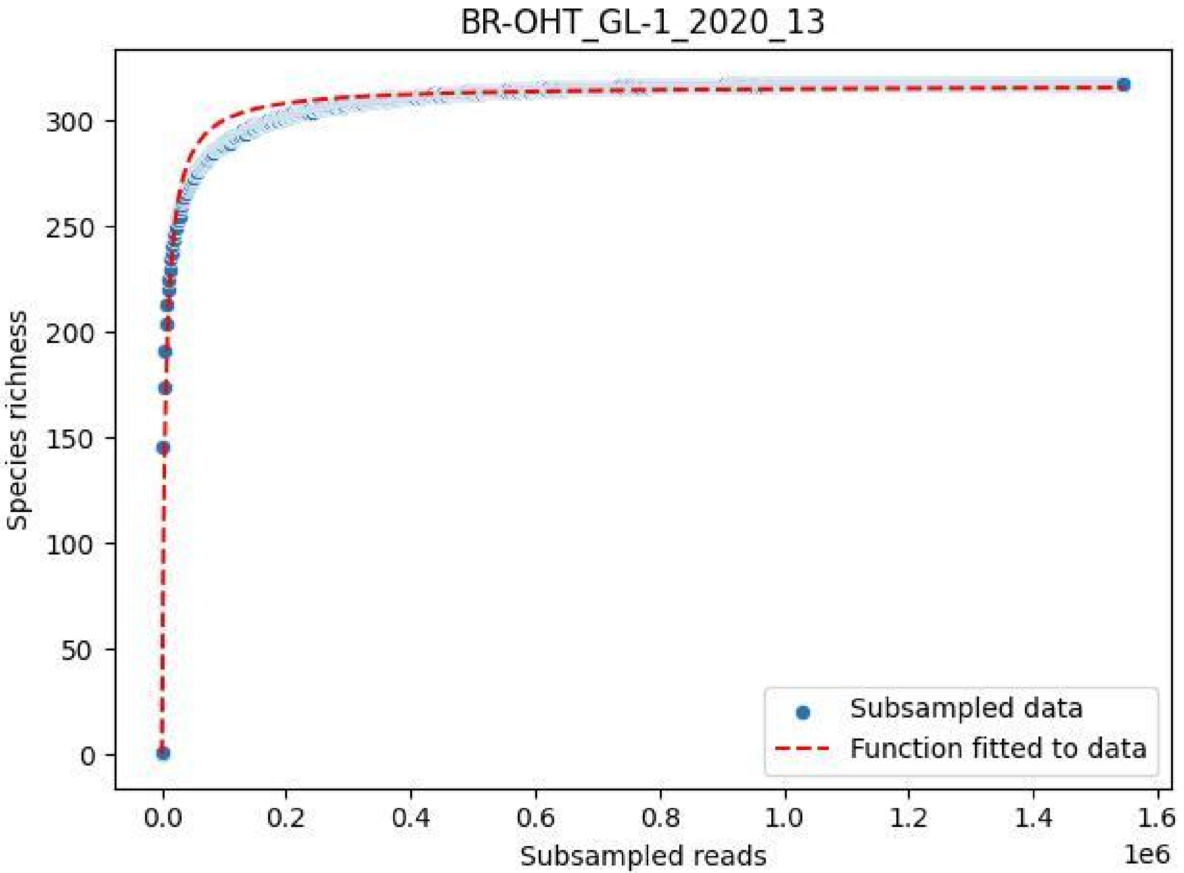

BR-OHT_GL-1_2020_13
Species richness
Subsampled reads
1e6
0
50
100
150
200
250
300
0.0
0.2
0.4
0.6
0.8
1.0
1.2
1.4
1.6
Subsampled data
Function fitted to data

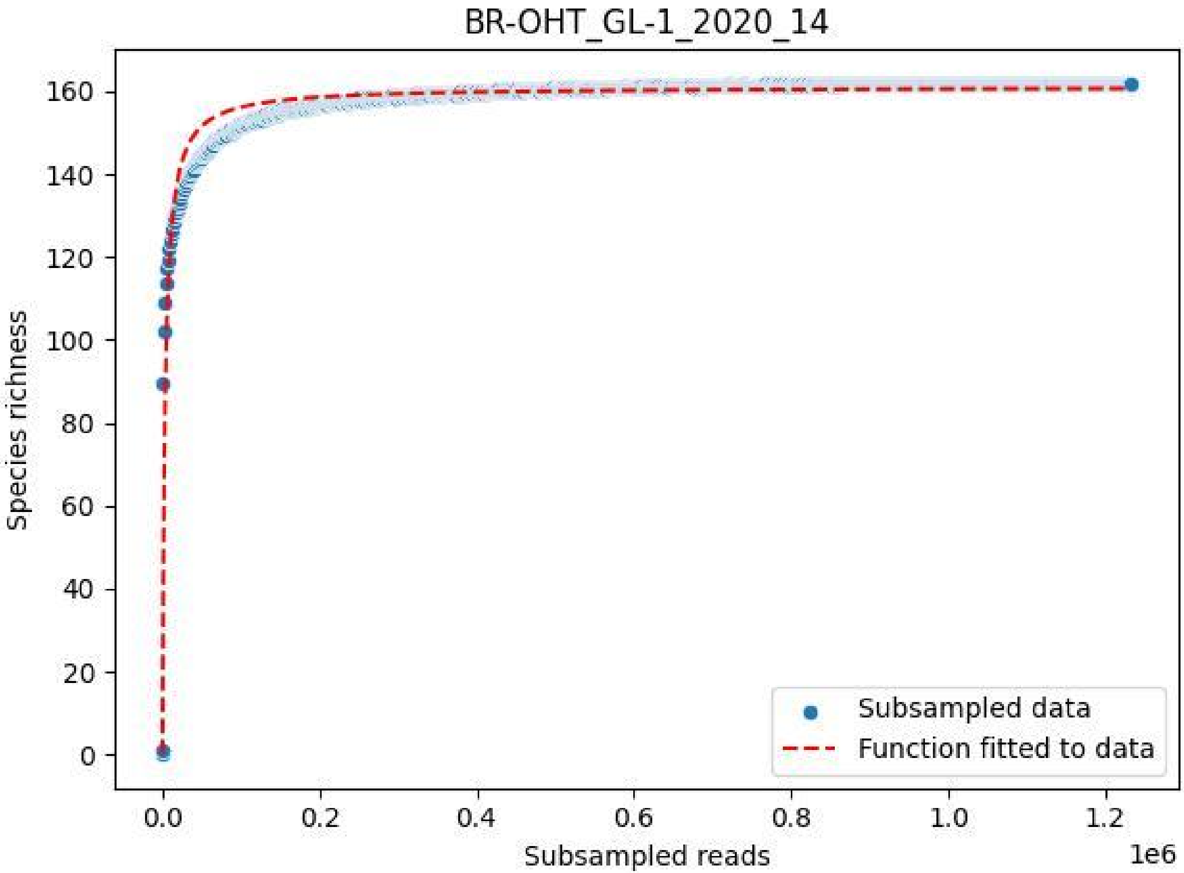
BR-OHT_GL-1_2020_14
Species richness
Subsampled reads
1e6
Subsampled data
Function fitted to data
0
20
40
60
80
100
120
140
160
0.0
0.2
0.4
0.6
0.8
1.0
1.2

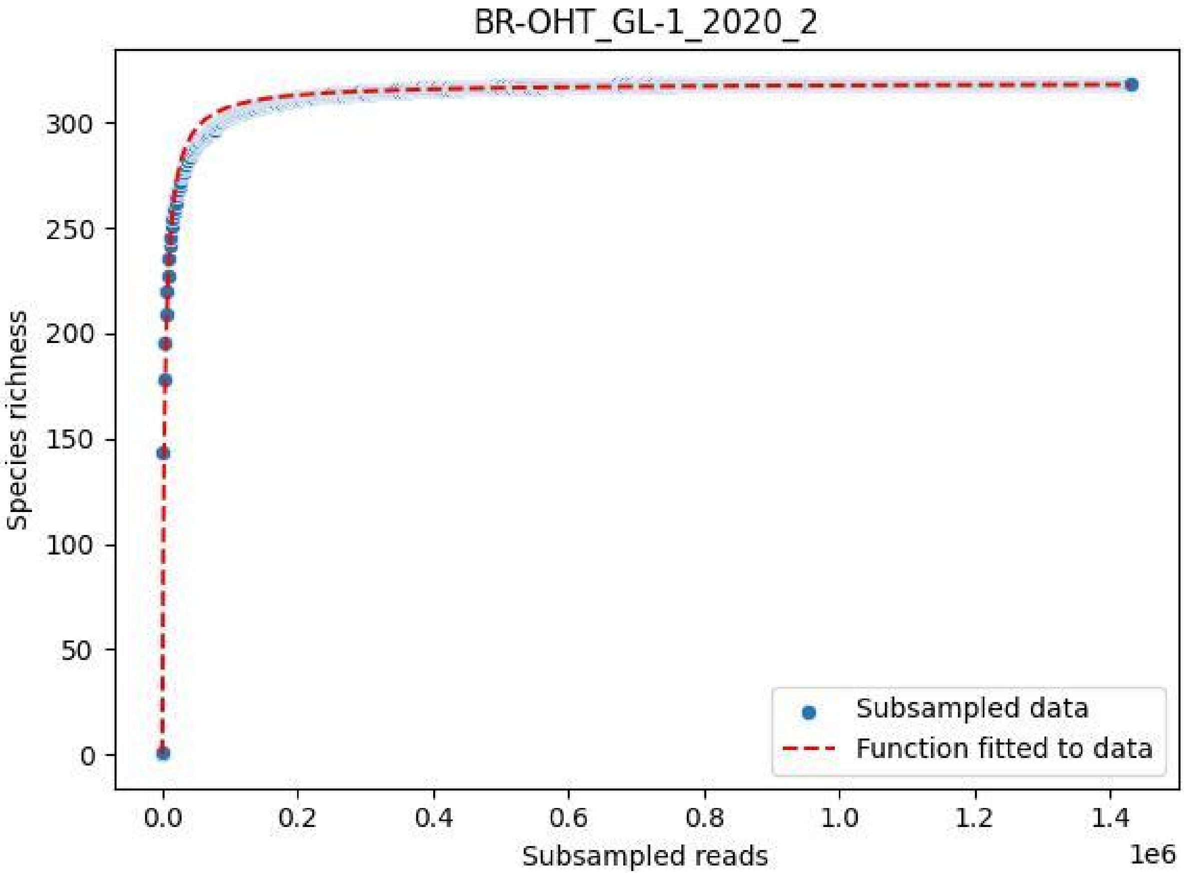
BR-OHT_GL-1_2020_2
Species richness
0
50
100
150
200
250
300
Subsampled reads
0.0
0.2
0.4
0.6
0.8
1.0
1.2
1.4
1e6
Subsampled data
Function fitted to data

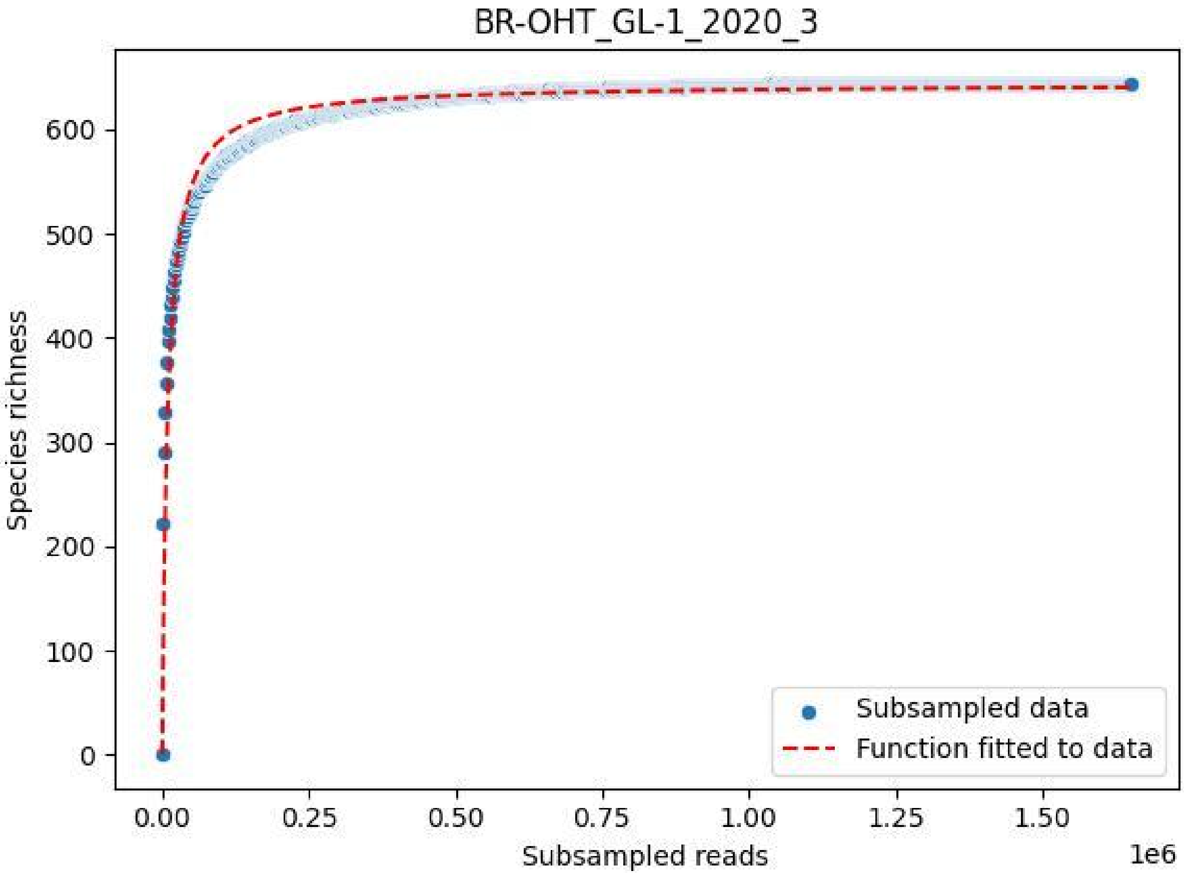
BR-OHT_GL-1_2020_3
Species richness
Subsampled reads
1e6
Subsampled data
Function fitted to data
0
100
200
300
400
500
600
0.00
0.25
0.50
0.75
1.00
1.25
1.50

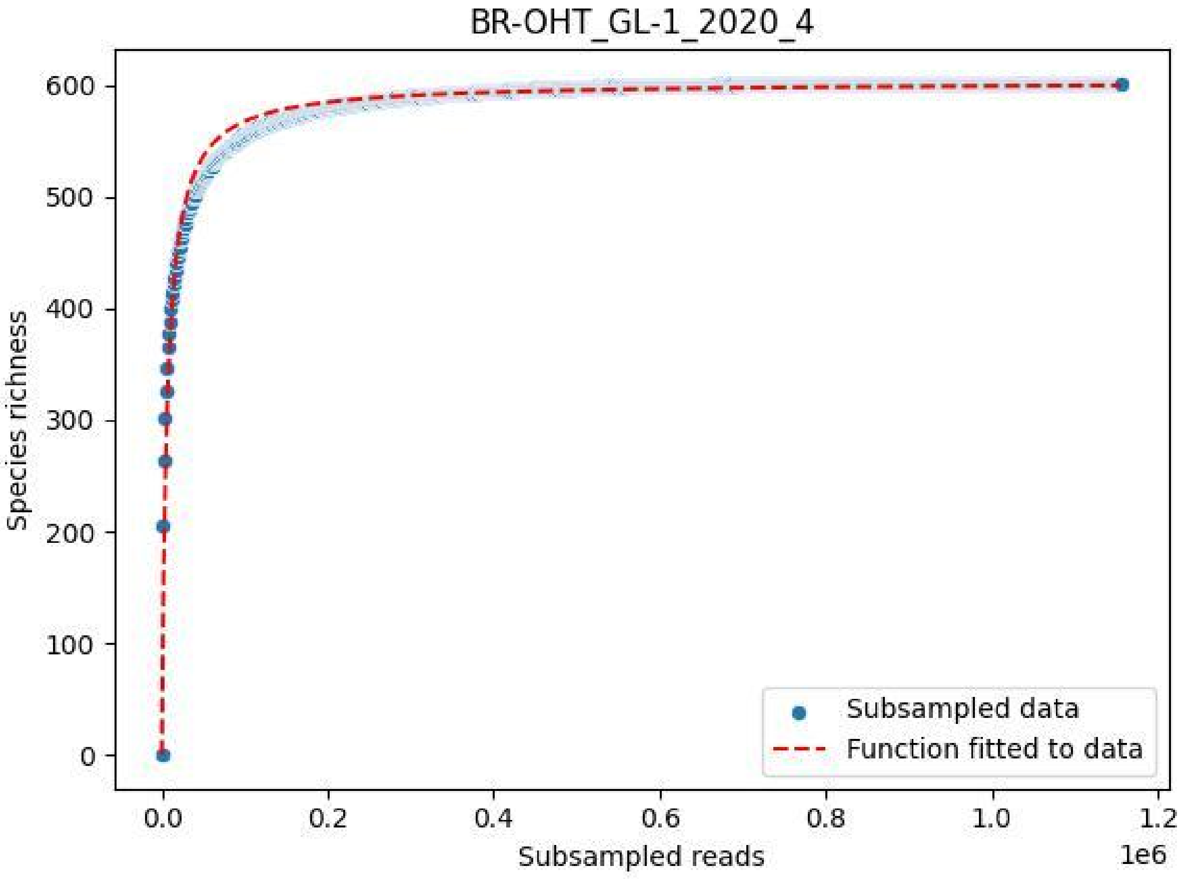
BR-OHT_GL-1_2020_4
Species richness
Subsampled reads
1e6
0
100
200
300
400
500
600
0.0
0.2
0.4
0.6
0.8
1.0
1.2
Subsampled data
Function fitted to data

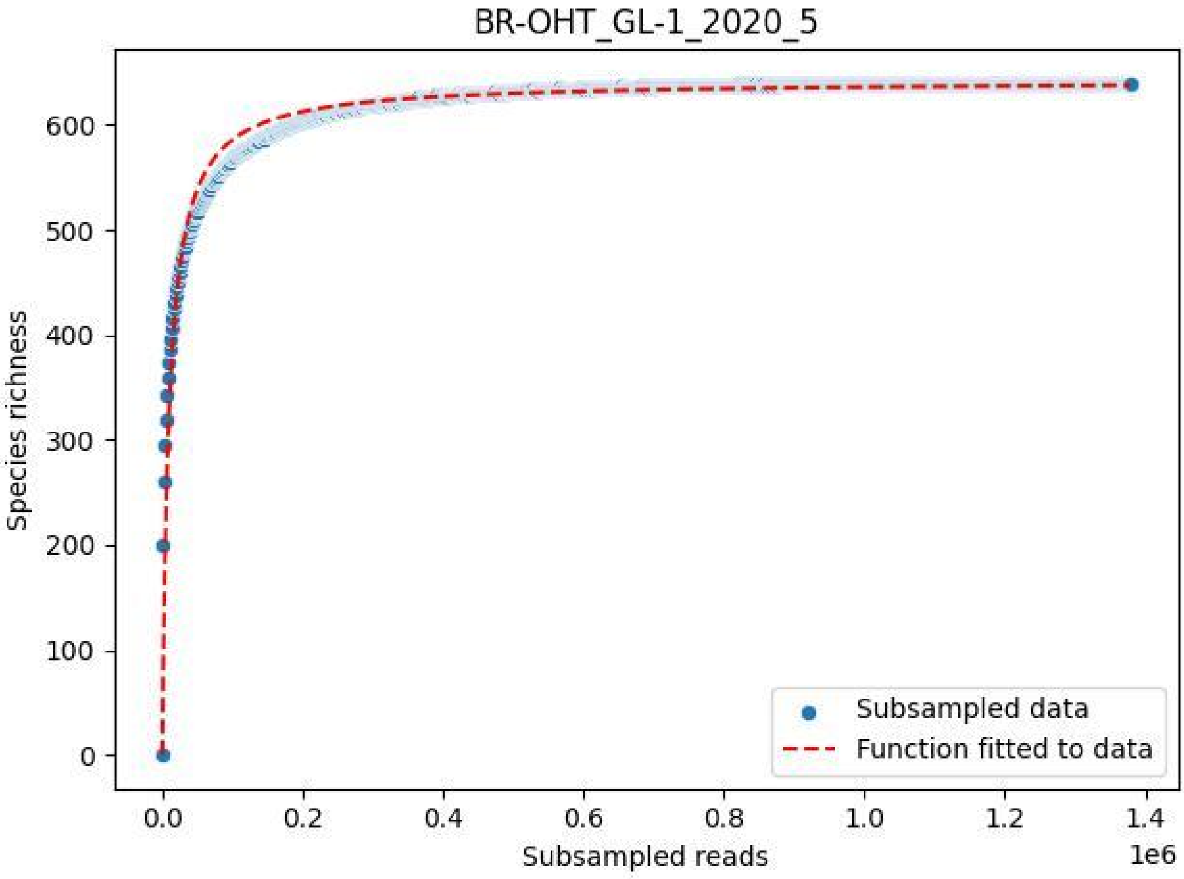
BR-OHT_GL-1_2020_5
Species richness
Subsampled reads
1e6
0
100
200
300
400
500
600
0.0
0.2
0.4
0.6
0.8
1.0
1.2
1.4
Subsampled data
Function fitted to data

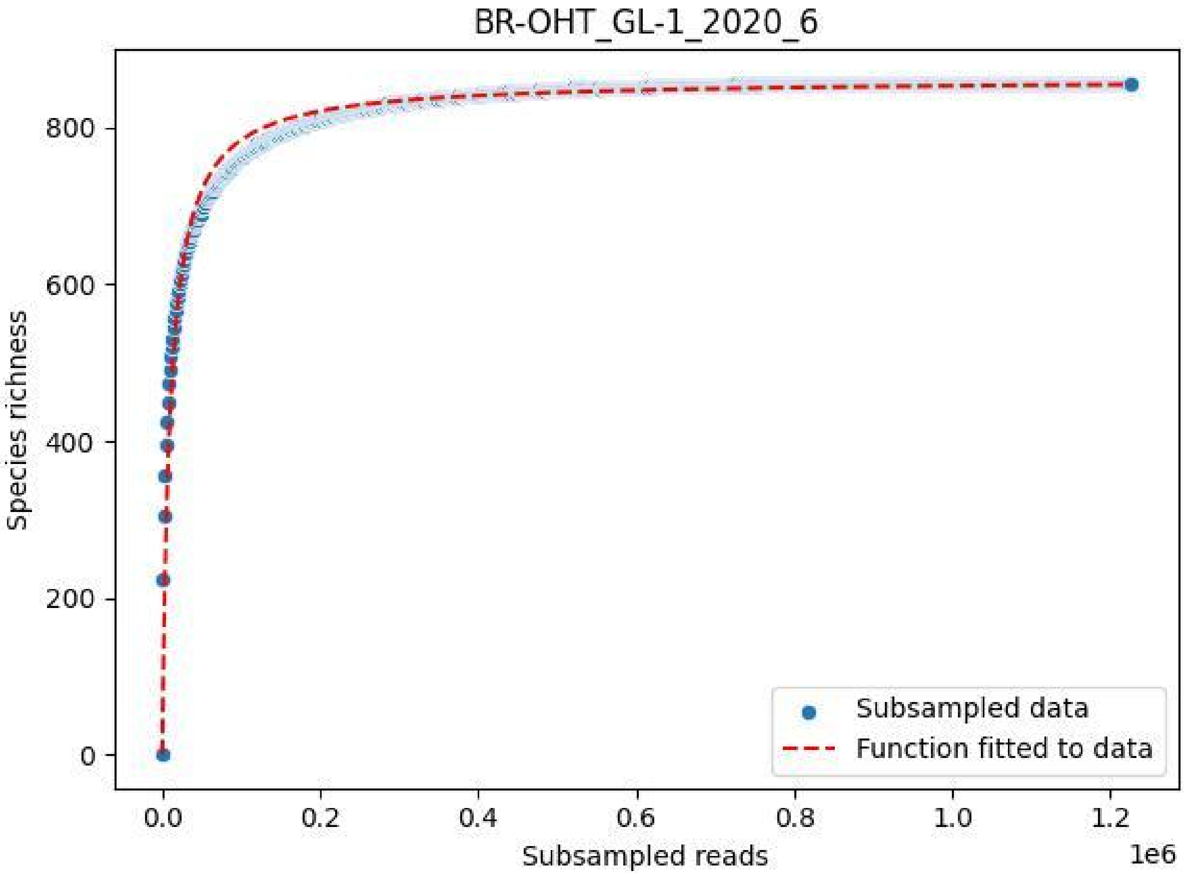
BR-OHT_GL-1_2020_6
Species richness
Subsampled reads
1e6
0
200
400
600
800
0.0
0.2
0.4
0.6
0.8
1.0
1.2
Subsampled data
Function fitted to data

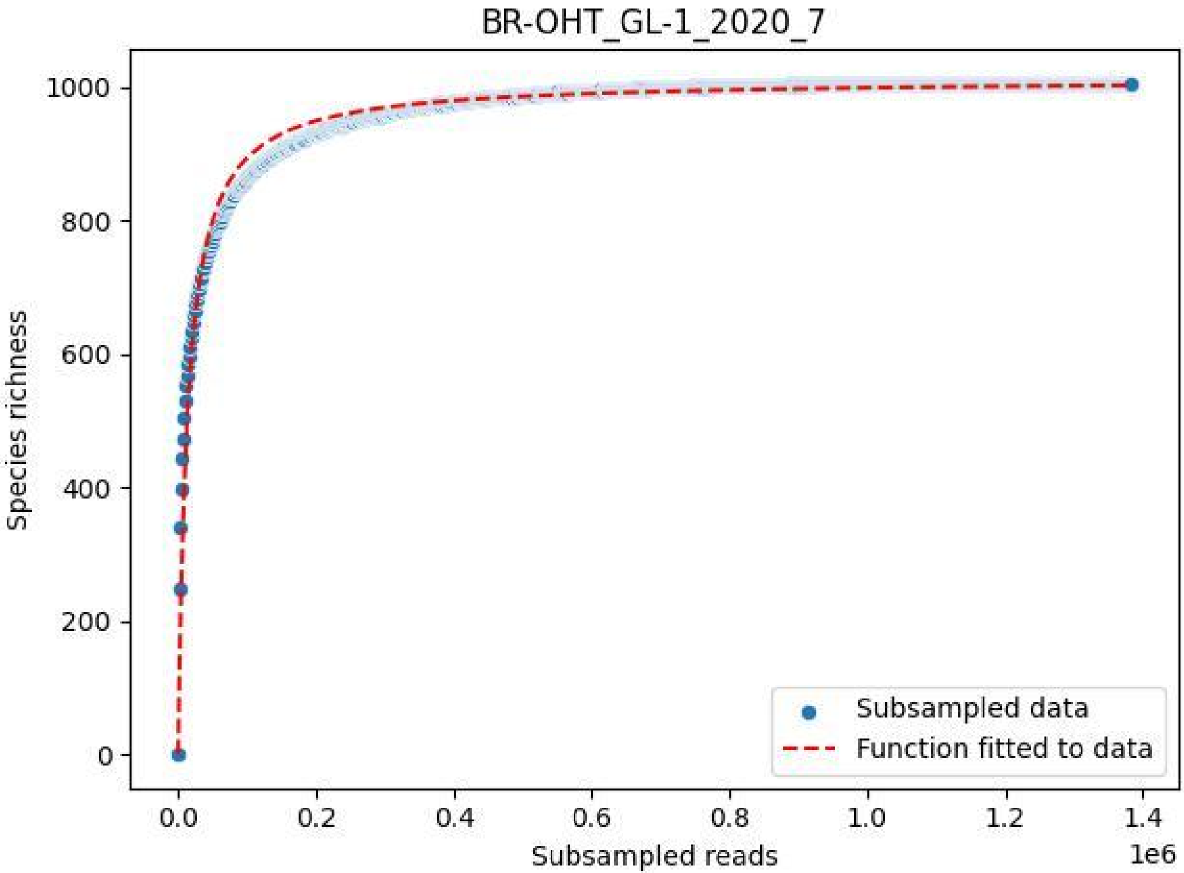

BR-OHT_GL-1_2020_7
Species richness
Subsampled reads
1e6
0
200
400
600
800
1000
0.0
0.2
0.4
0.6
0.8
1.0
1.2
1.4
Subsampled data
Function fitted to data

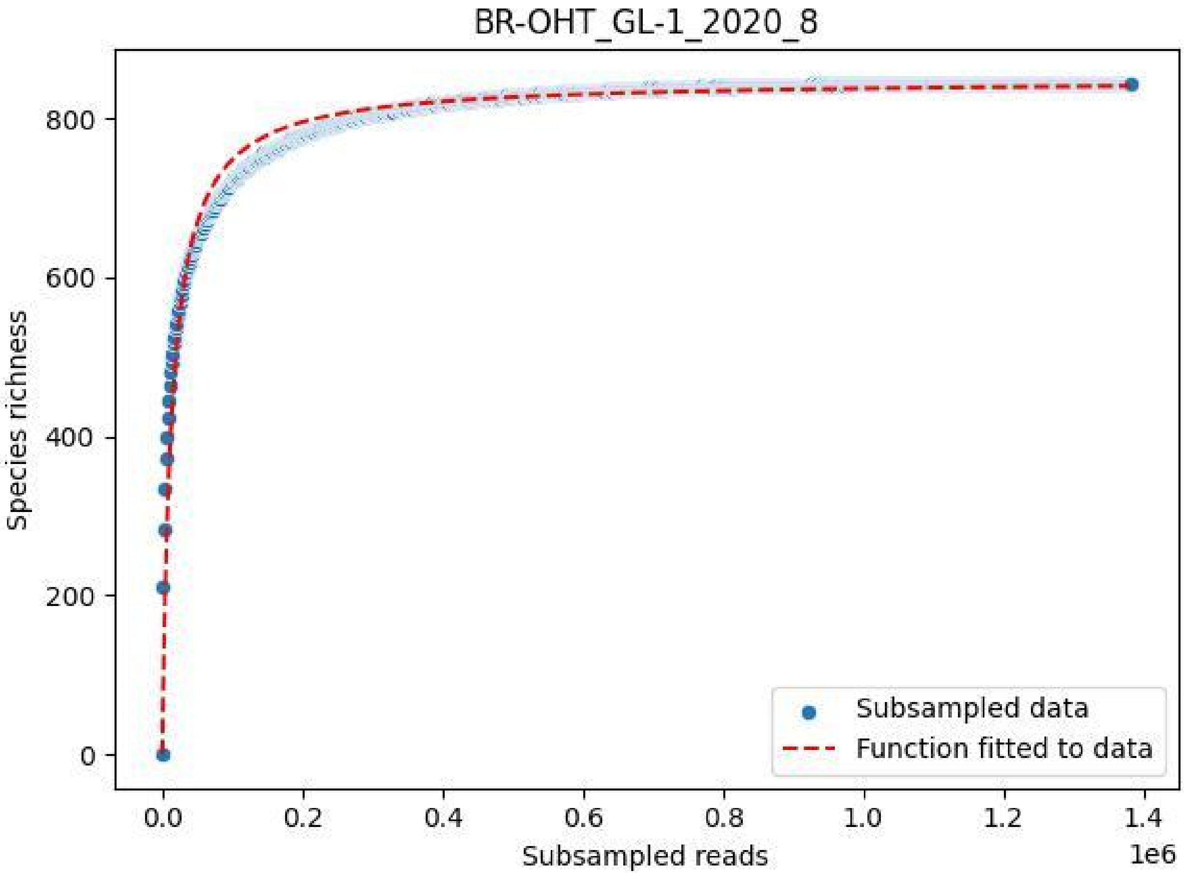
BR-OHT_GL-1_2020_8
Species richness
Subsampled reads
1e6
0
200
400
600
800
0.0
0.2
0.4
0.6
0.8
1.0
1.2
1.4
Subsampled data
Function fitted to data

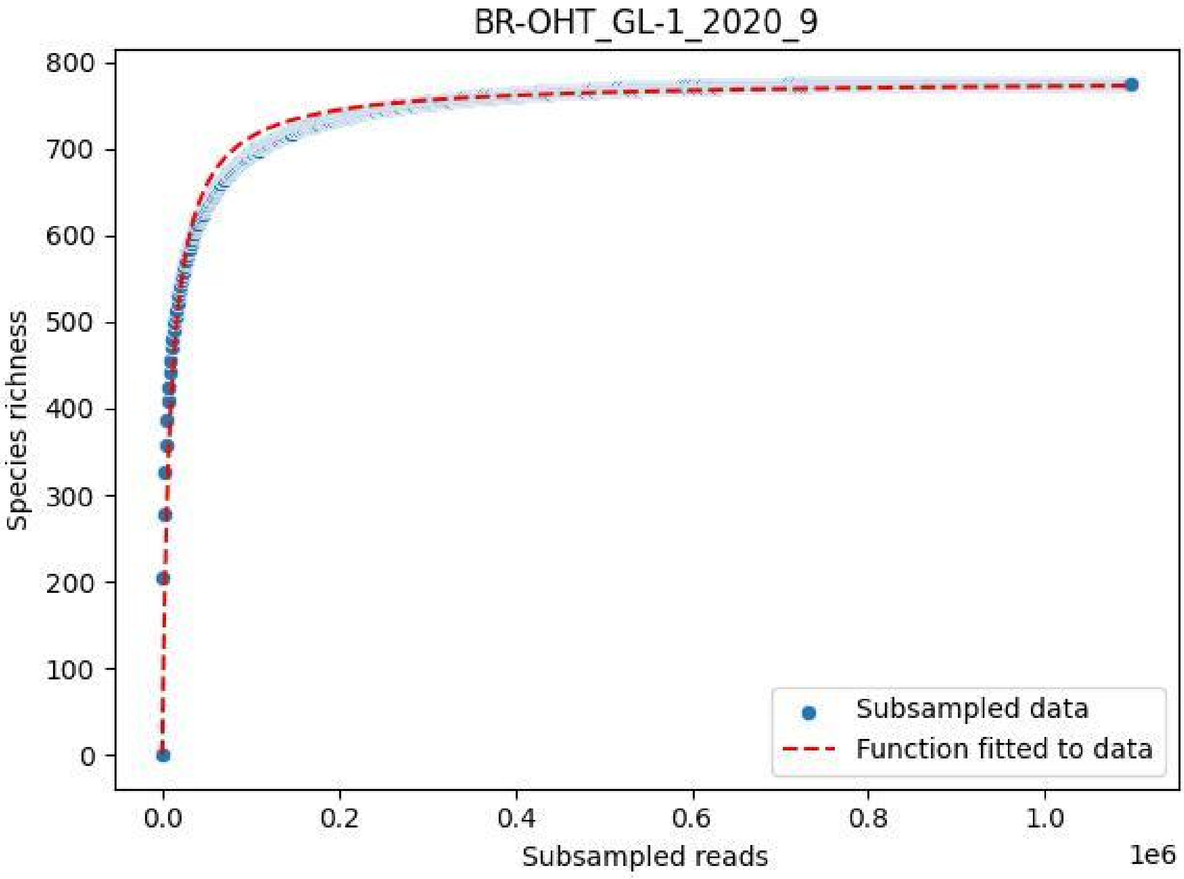
BR-OHT_GL-1_2020_9
Species richness
Subsampled reads
1e6
0
100
200
300
400
500
600
700
800
0.0
0.2
0.4
0.6
0.8
1.0
Subsampled data
Function fitted to data

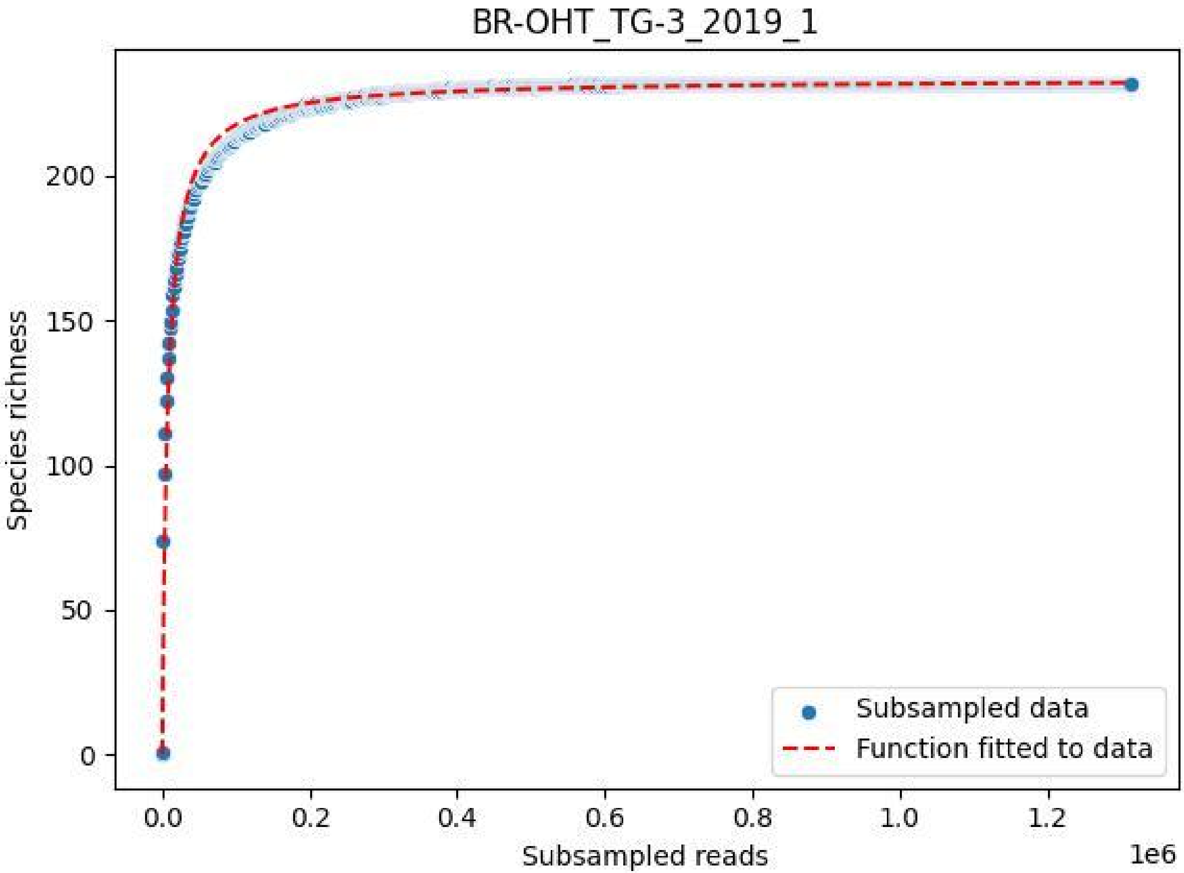
BR-OHT_TG-3_2019_1
Species richness
Subsampled reads
1e6
Subsampled data
Function fitted to data
0
50
100
150
200
0.0
0.2
0.4
0.6
0.8
1.0
1.2

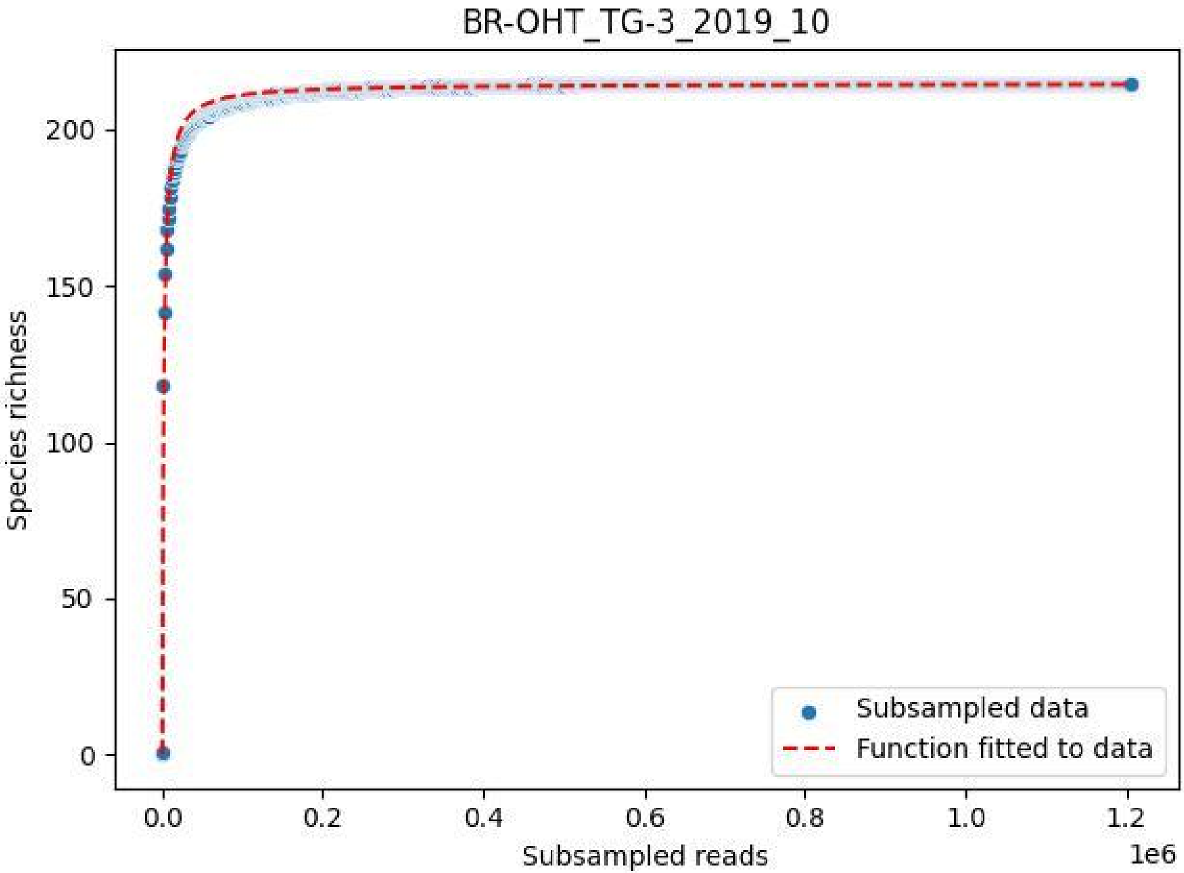
BR-OHT_TG-3_2019_10
Species richness
Subsampled reads
1e6
Subsampled data
Function fitted to data
0
50
100
150
200
0.0
0.2
0.4
0.6
0.8
1.0
1.2

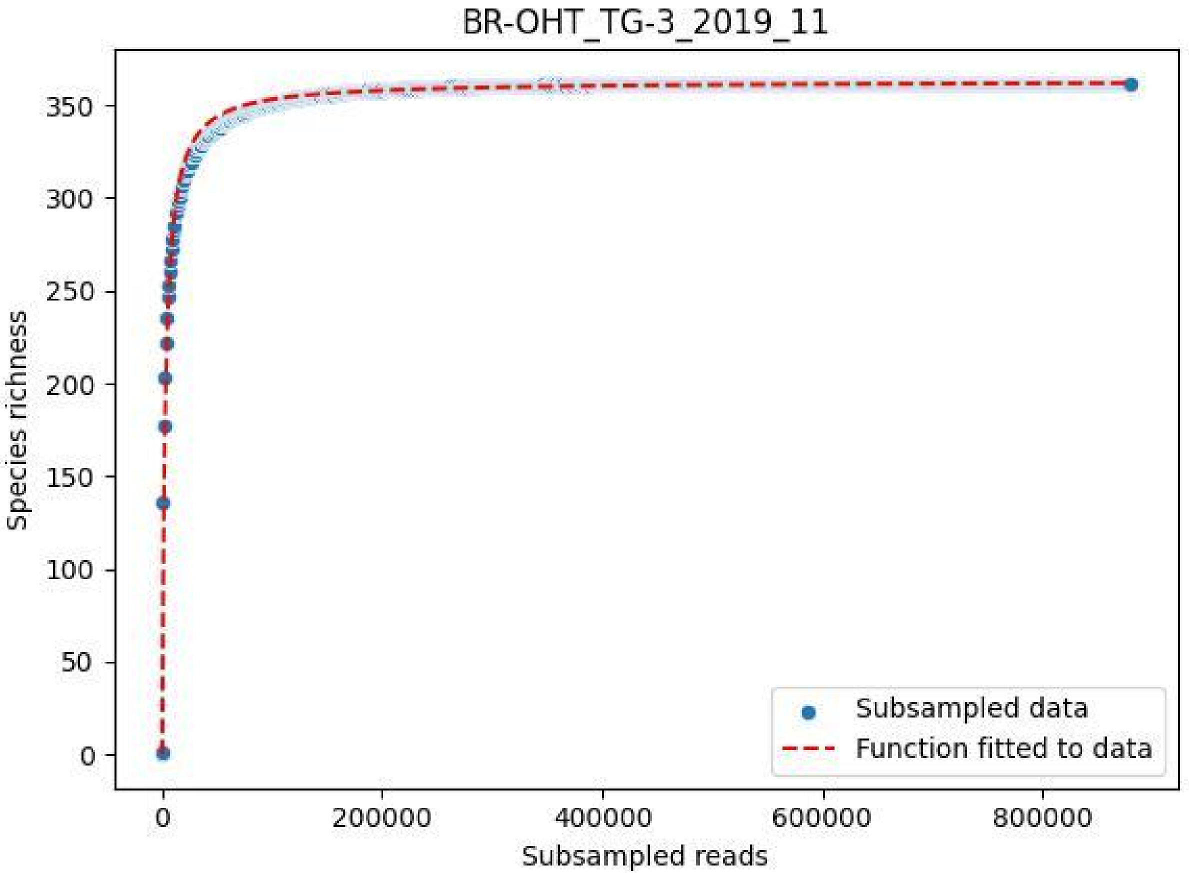
BR-OHT_TG-3_2019_11
Species richness
0
50
100
150
200
250
300
350
Subsampled reads
0
200000
400000
600000
800000
Subsampled data
Function fitted to data

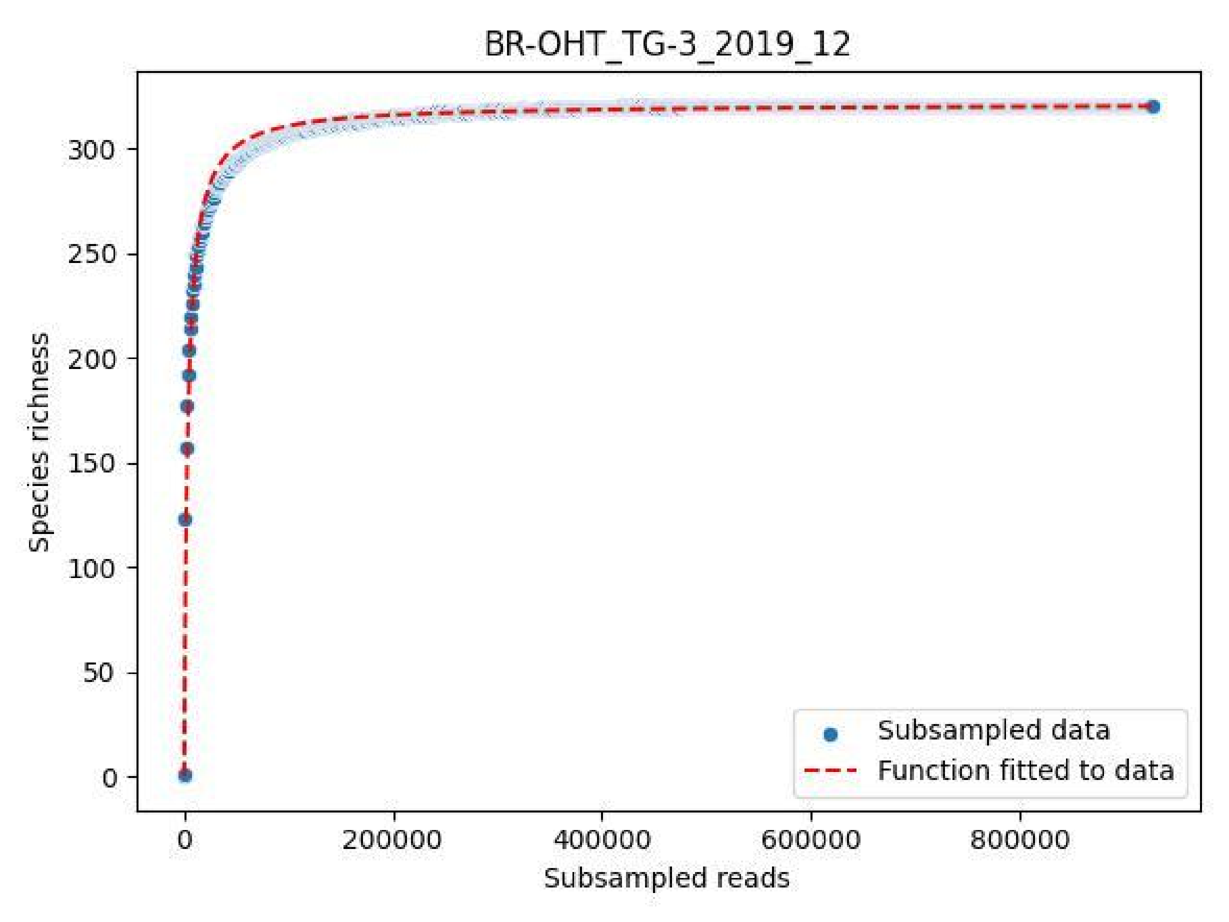

BR-OHT_TG-3_2019_12
Species richness
Subsampled reads
0
50
100
150
200
250
300
0
200000
400000
600000
800000
Subsampled data
Function fitted to data

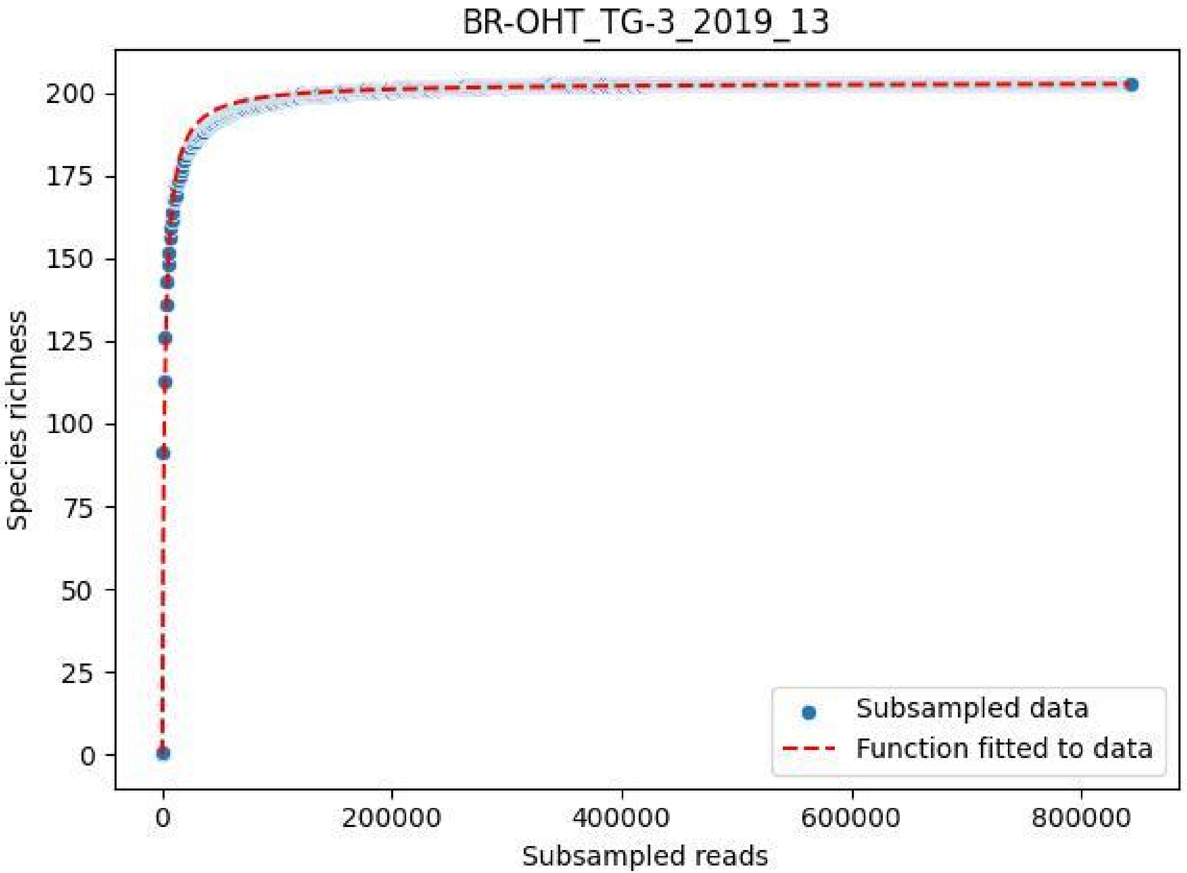
BR-OHT_TG-3_2019_13
Species richness
Subsampled reads
0
25
50
75
100
125
150
175
200
0
200000
400000
600000
800000
Subsampled data
Function fitted to data

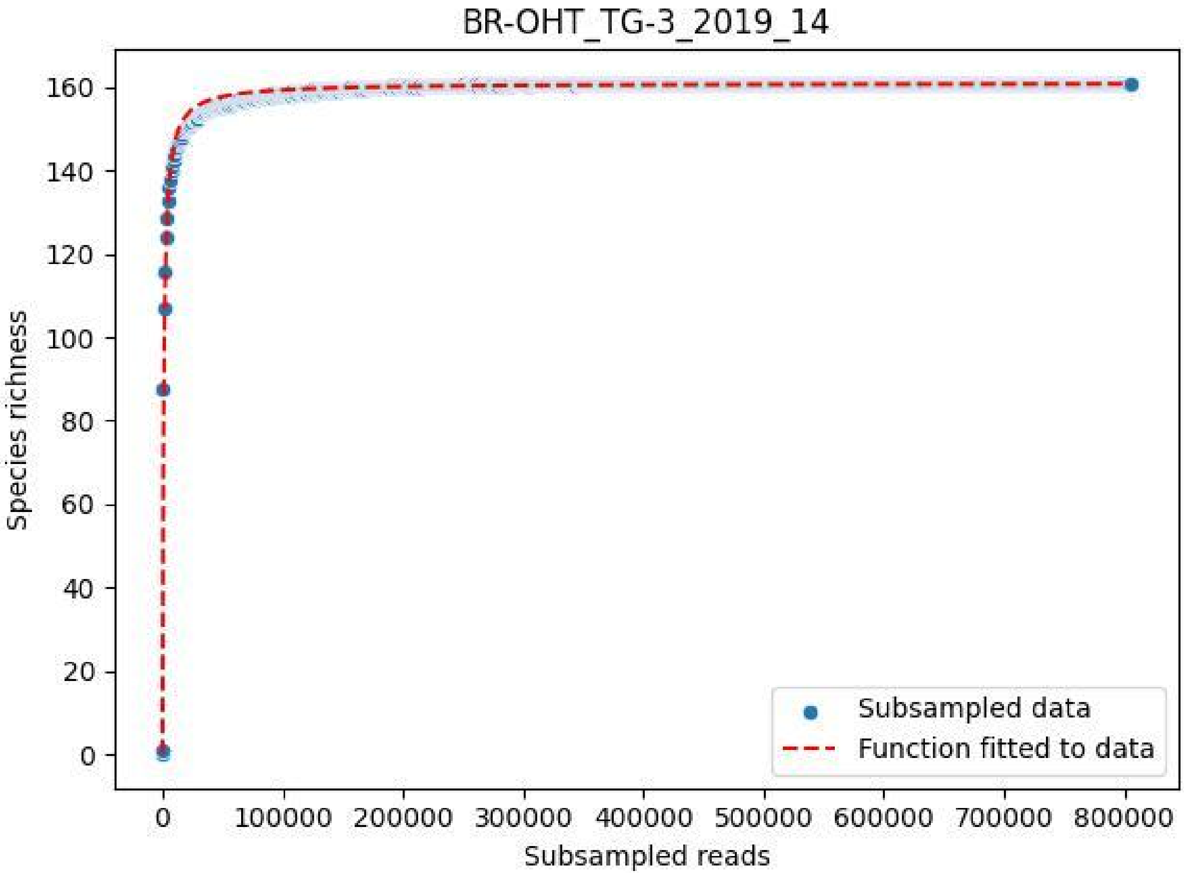

BR-OHT_TG-3_2019_14
Species richness
Subsampled reads
0
20
40
60
80
100
120
140
160
0
100000
200000
300000
400000
500000
600000
700000
800000
Subsampled data
Function fitted to data

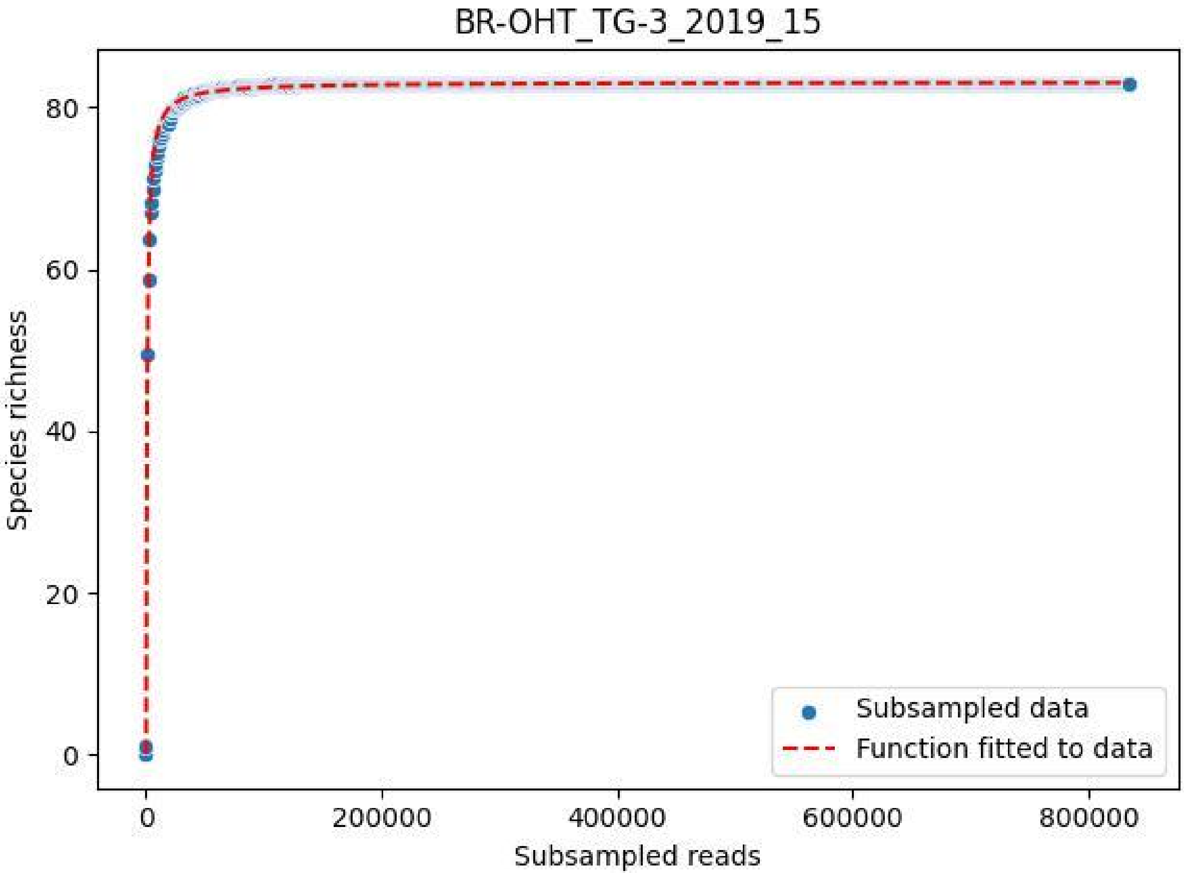
BR-OHT_TG-3_2019_15
Species richness
Subsampled reads
0
20
40
60
80
0
200000
400000
600000
800000
Subsampled data
Function fitted to data

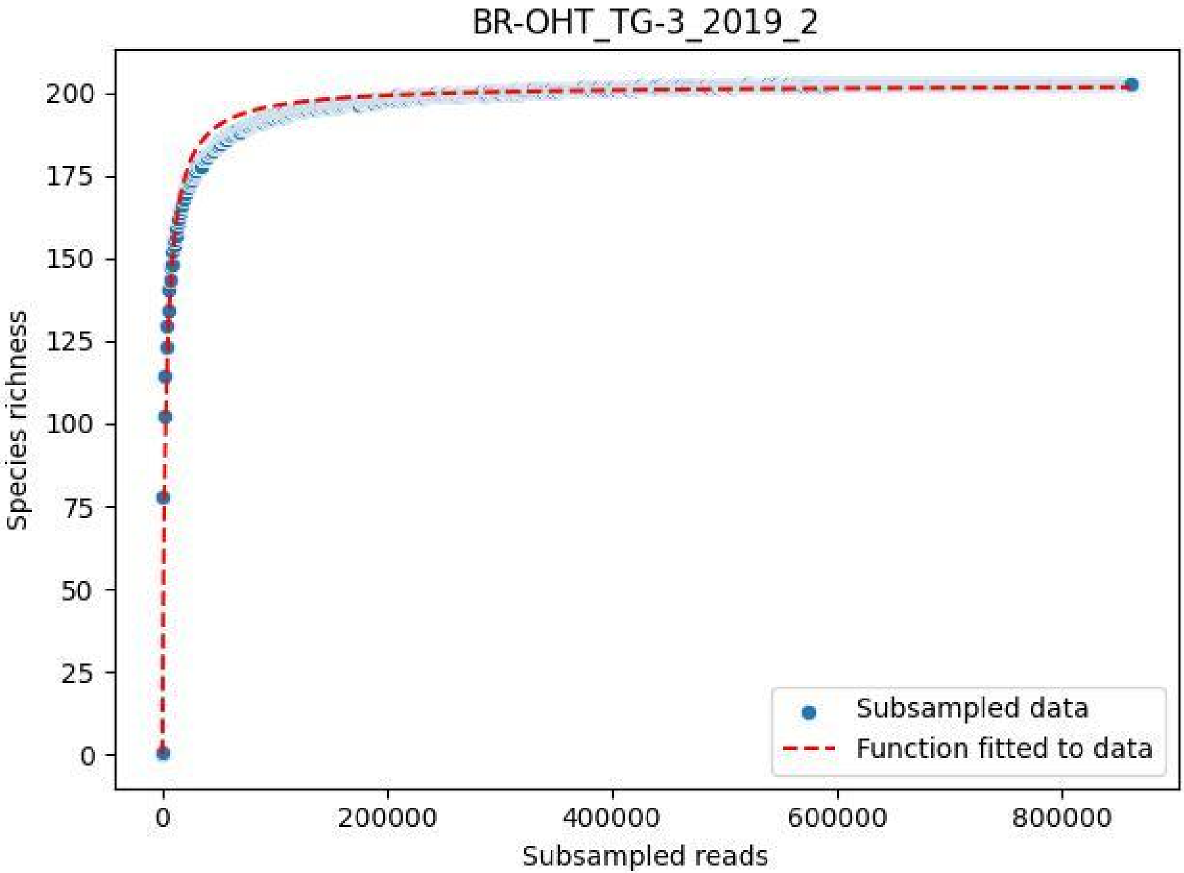
BR-OHT_TG-3_2019_2
Species richness
Subsampled reads
0
25
50
75
100
125
150
175
200
0
200000
400000
600000
800000
Subsampled data
Function fitted to data

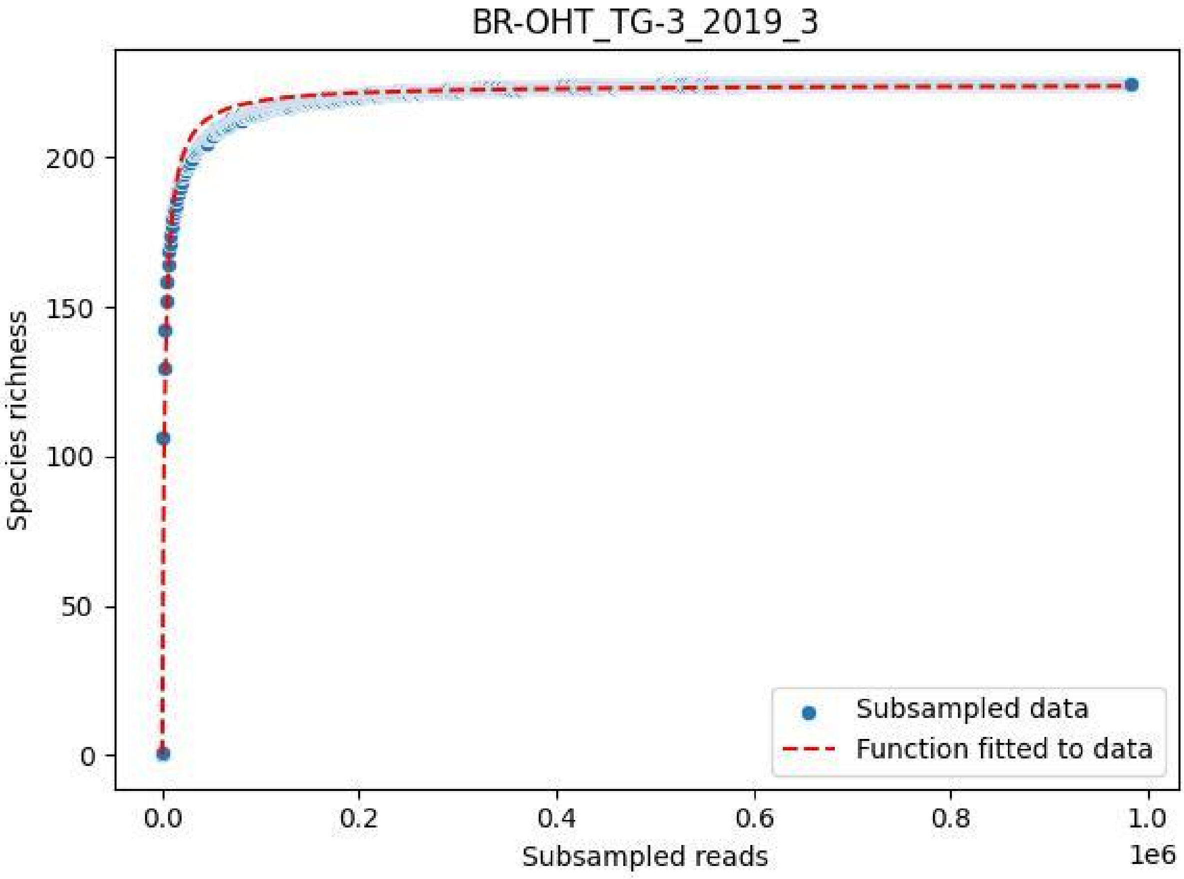
BR-OHT_TG-3_2019_3
Species richness
Subsampled reads
1e6
0
50
100
150
200
0.0
0.2
0.4
0.6
0.8
1.0
Subsampled data
Function fitted to data

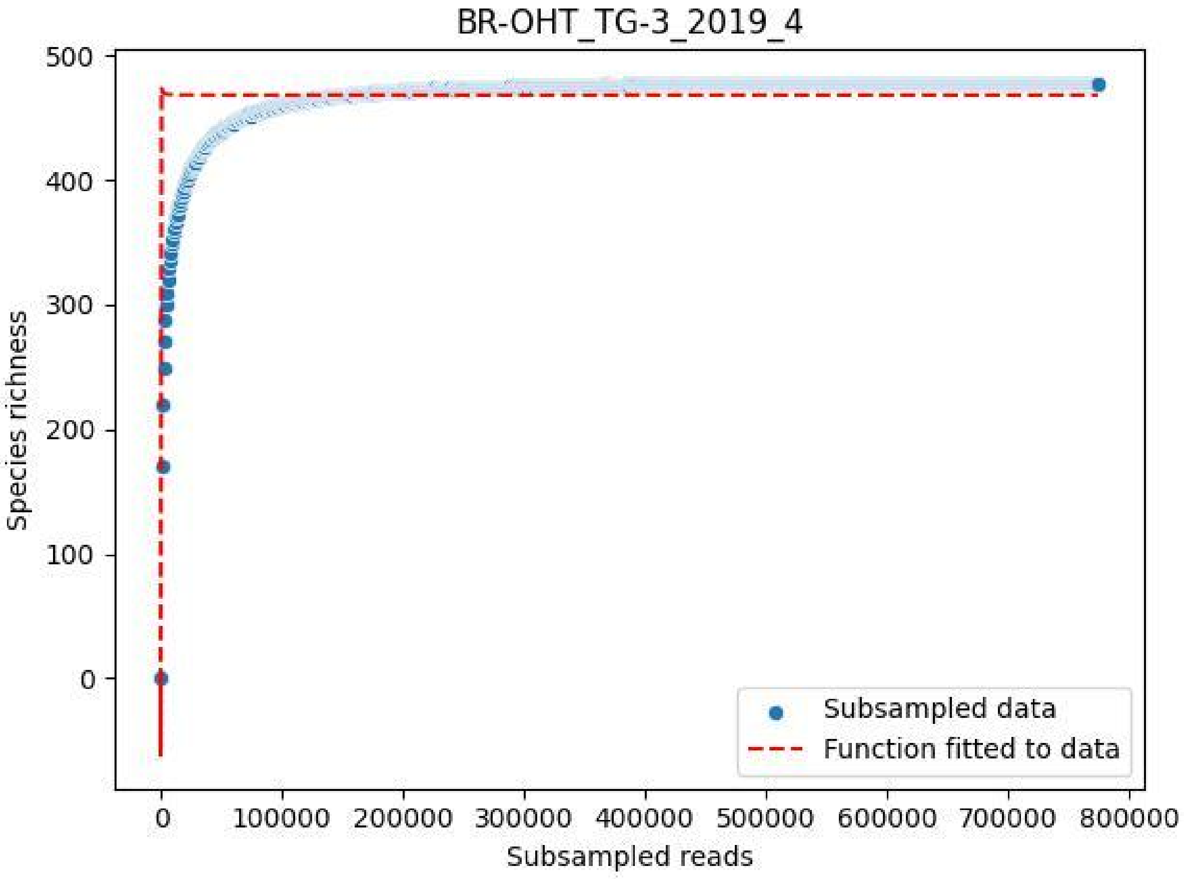
BR-OHT_TG-3_2019_4
500
400
300
200
100
0
Species richness
0
100000
200000
300000
400000
500000
600000
700000
800000
Subsampled reads
Subsampled data
Function fitted to data

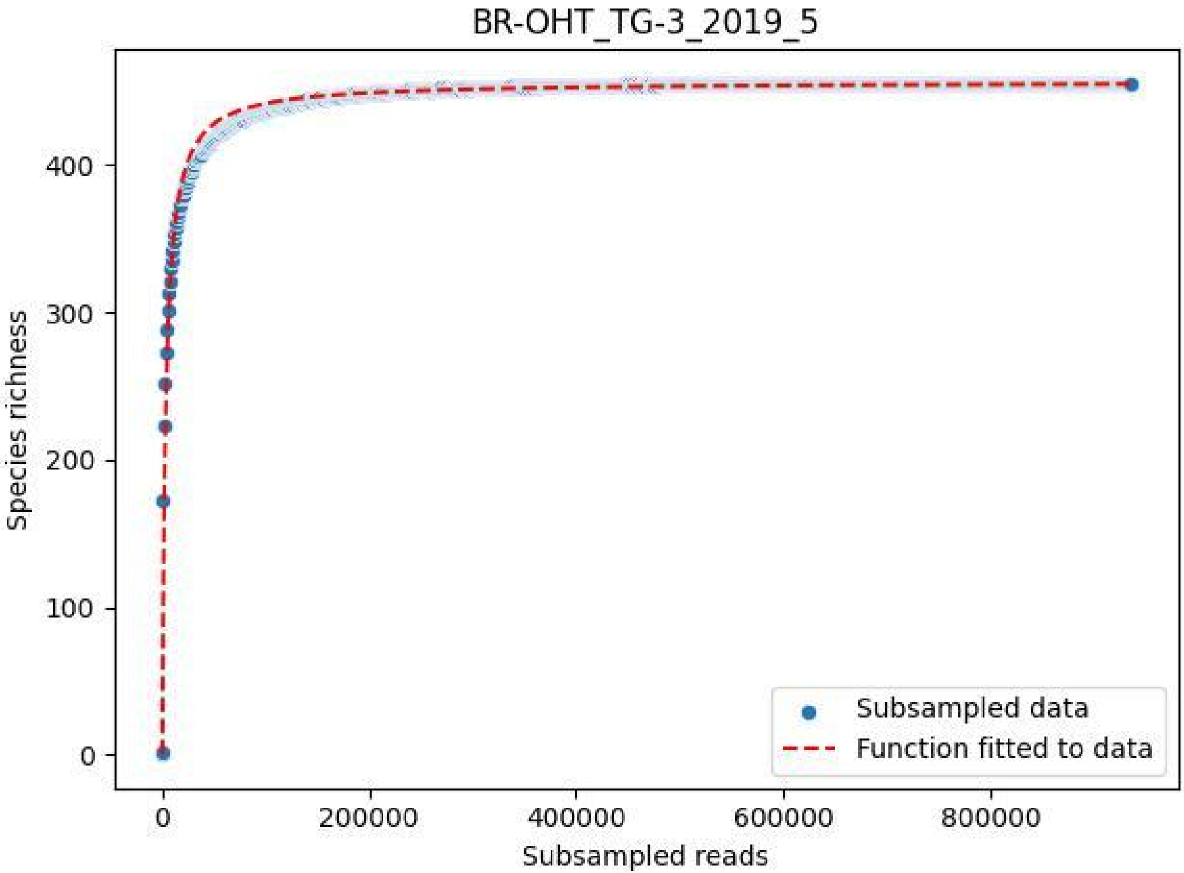

BR-OHT_TG-3_2019_5
Species richness
Subsampled reads
0
100
200
300
400
0
200000
400000
600000
800000
Subsampled data
Function fitted to data

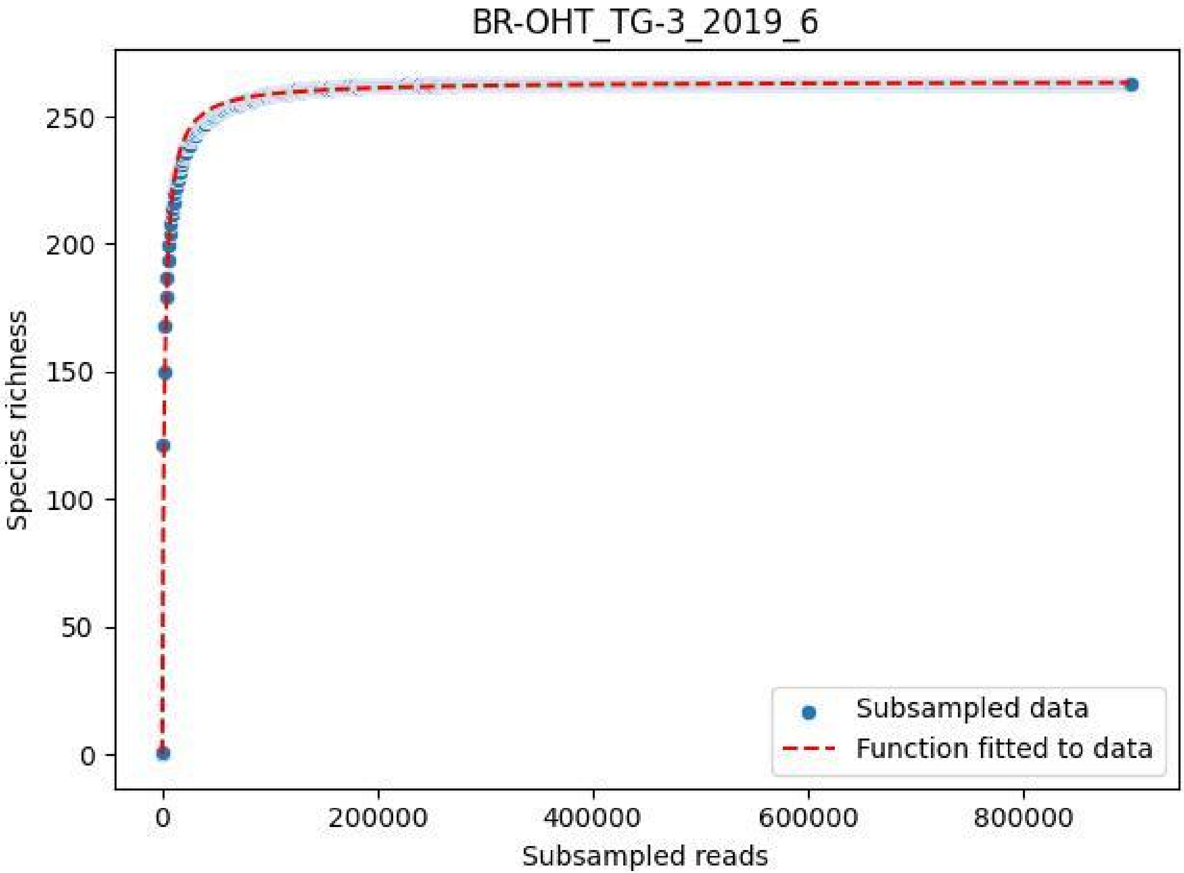

BR-OHT_TG-3_2019_6
Species richness
Subsampled reads
0
50
100
150
200
250
0
200000
400000
600000
800000
Subsampled data
Function fitted to data

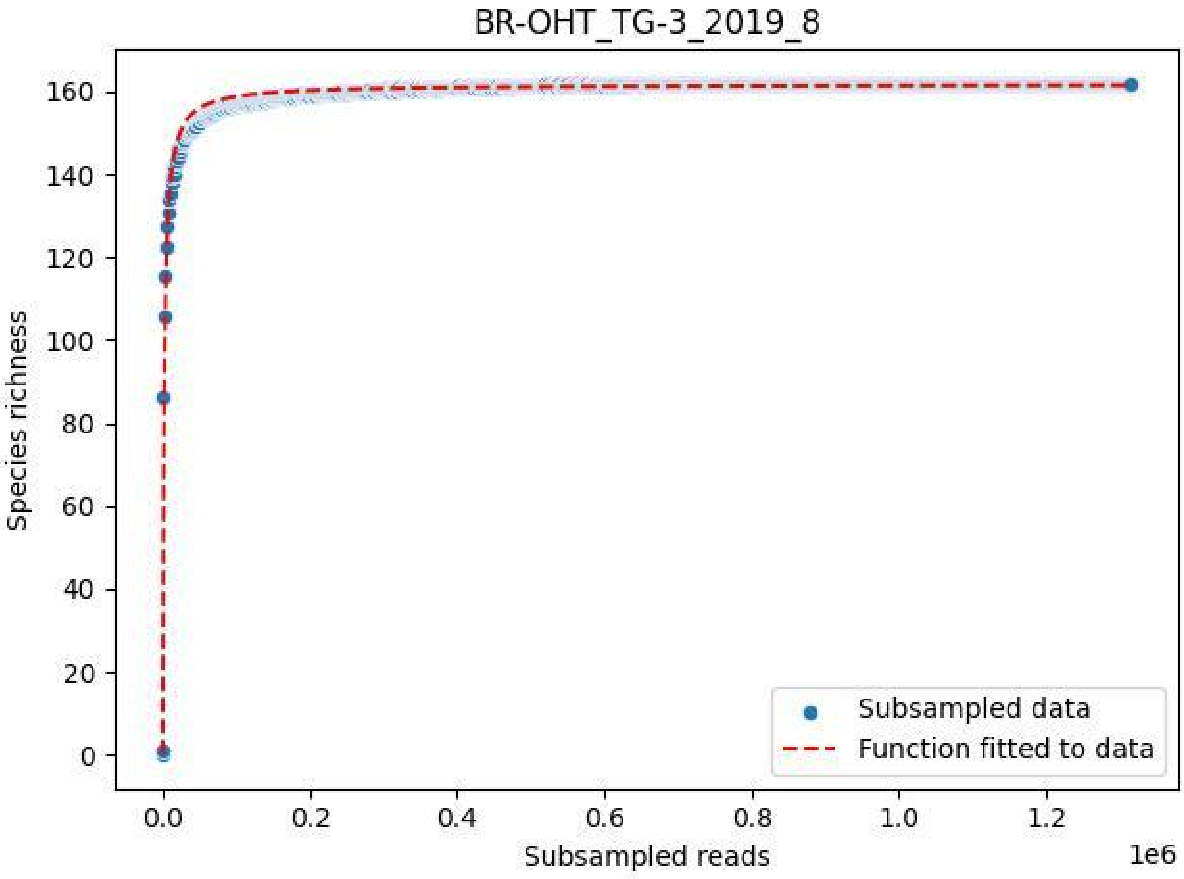
BR-OHT_TG-3_2019_8
Species richness
Subsampled reads
1e6
0
20
40
60
80
100
120
140
160
0.0
0.2
0.4
0.6
0.8
1.0
1.2
Subsampled data
Function fitted to data

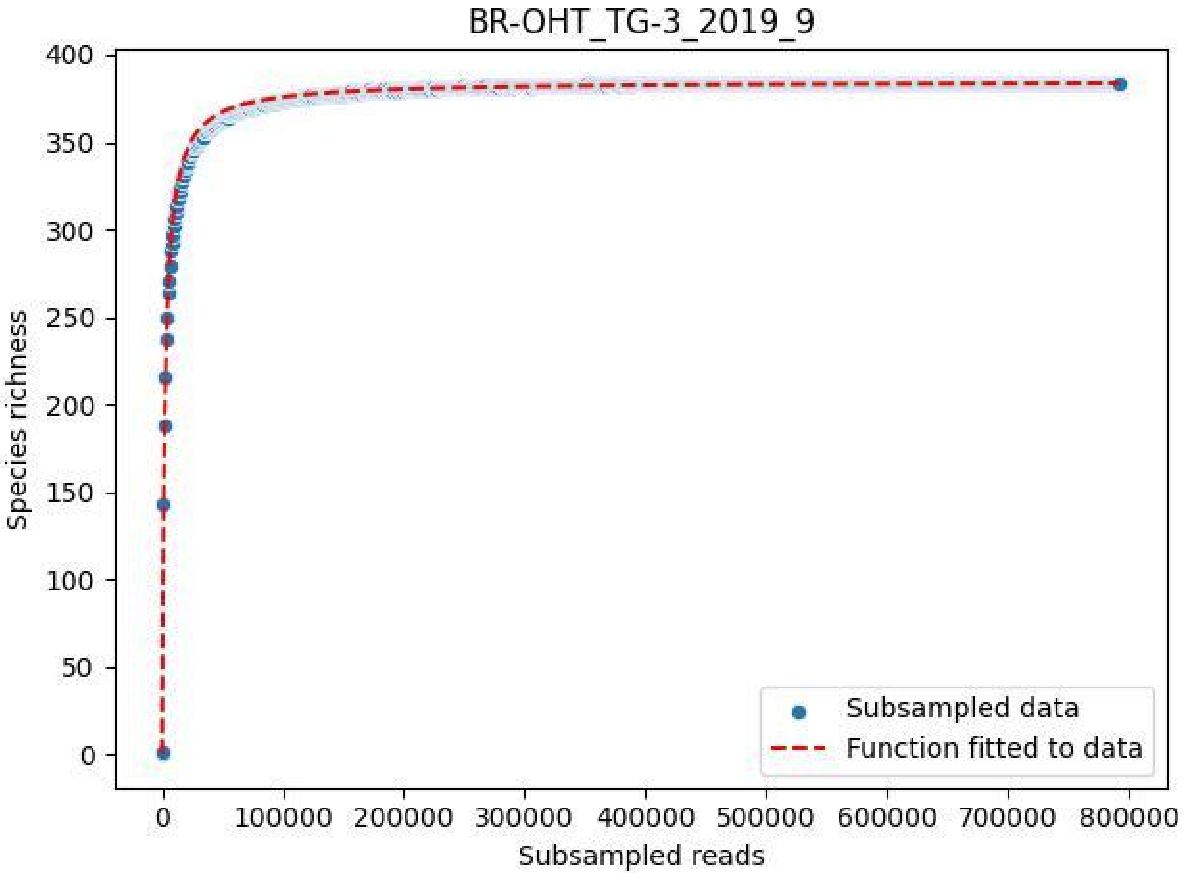
BR-OHT_TG-3_2019_9
Species richness
Subsampled reads
0
50
100
150
200
250
300
350
400
0
100000
200000
300000
400000
500000
600000
700000
800000
Subsampled data
Function fitted to data

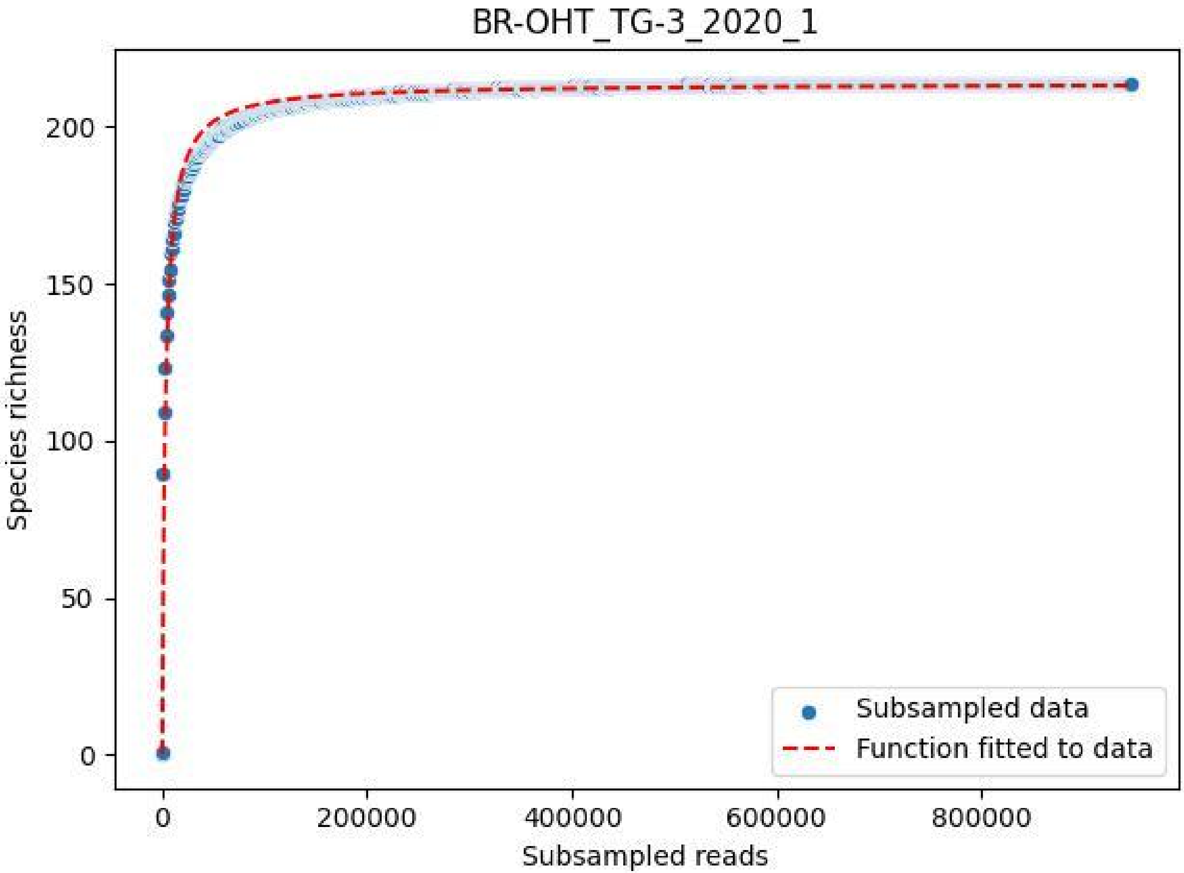
BR-OHT_TG-3_2020_1
Species richness
Subsampled reads
0
50
100
150
200
0
200000
400000
600000
800000
Subsampled data
Function fitted to data

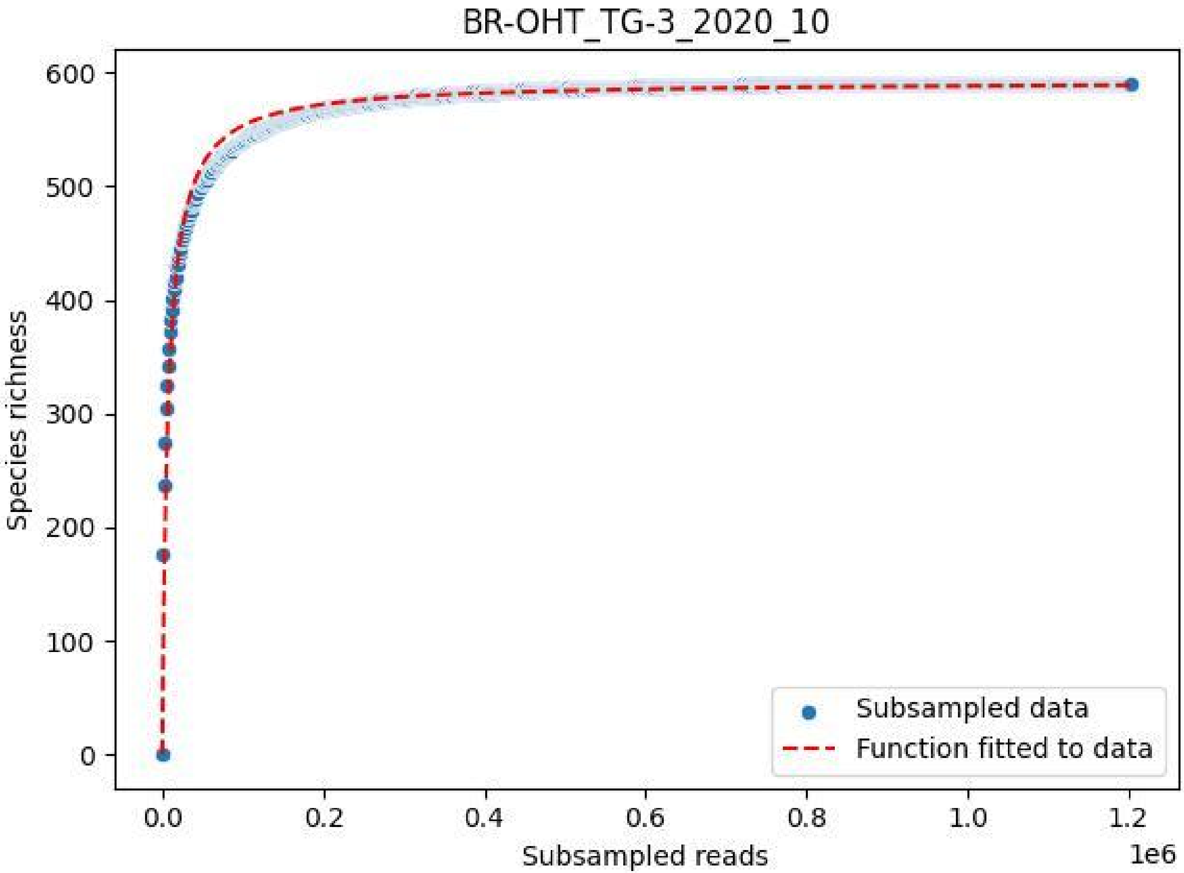

BR-OHT_TG-3_2020_10
600
500
400
300
200
100
0
Species richness
0.0
0.2
0.4
0.6
0.8
1.0
1.2
1e6
Subsampled reads
Subsampled data
Function fitted to data

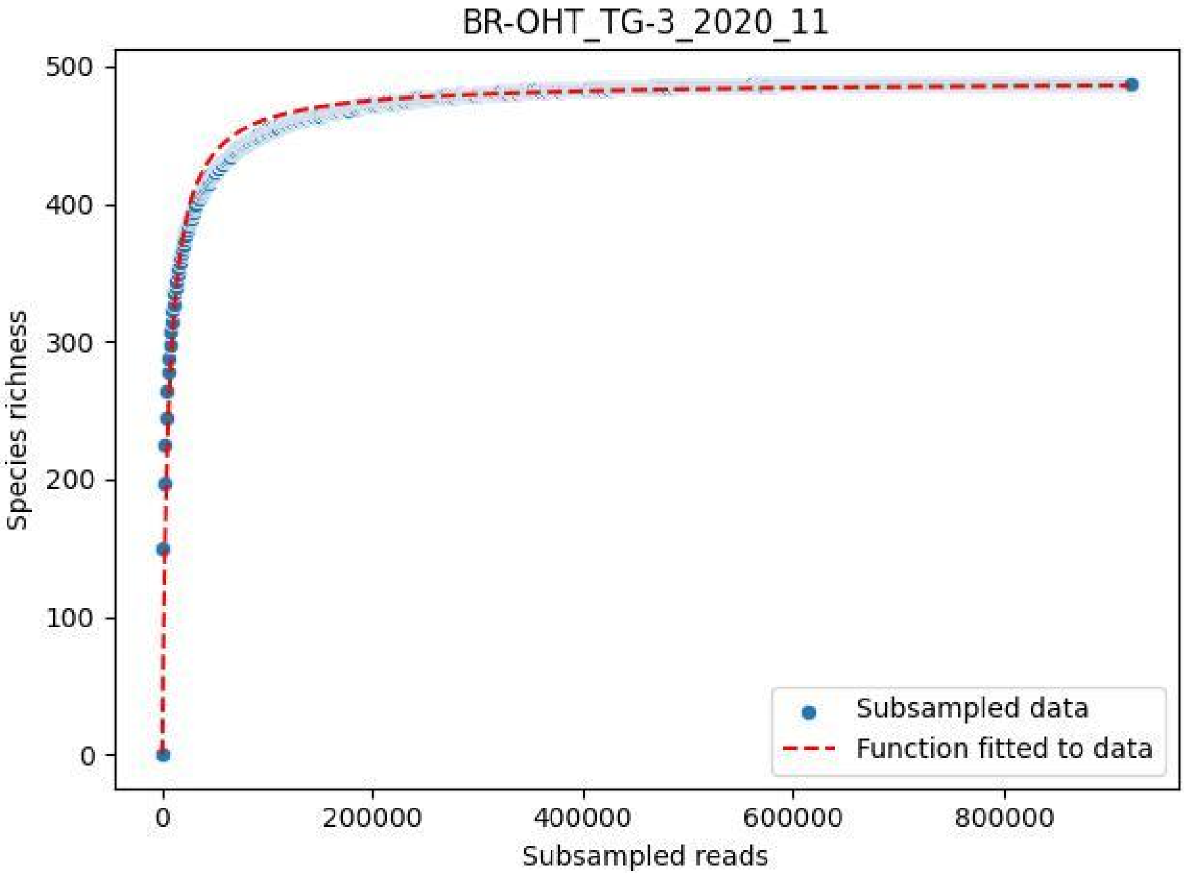

BR-OHT_TG-3_2020_11
Species richness
Subsampled reads
0
100
200
300
400
500
0
200000
400000
600000
800000
Subsampled data
Function fitted to data

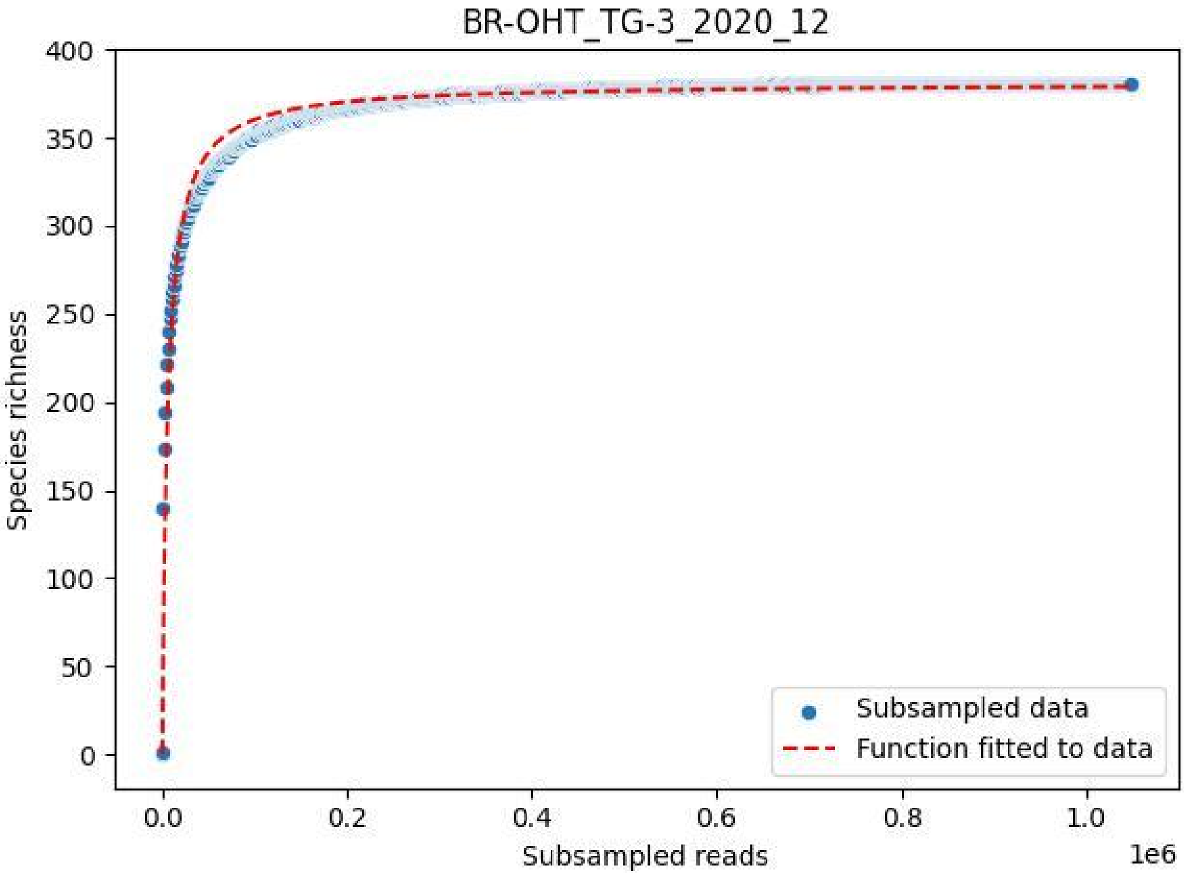
BR-OHT_TG-3_2020_12
Species richness
Subsampled reads
1e6
Subsampled data
Function fitted to data
0
50
100
150
200
250
300
350
400
0.0
0.2
0.4
0.6
0.8
1.0

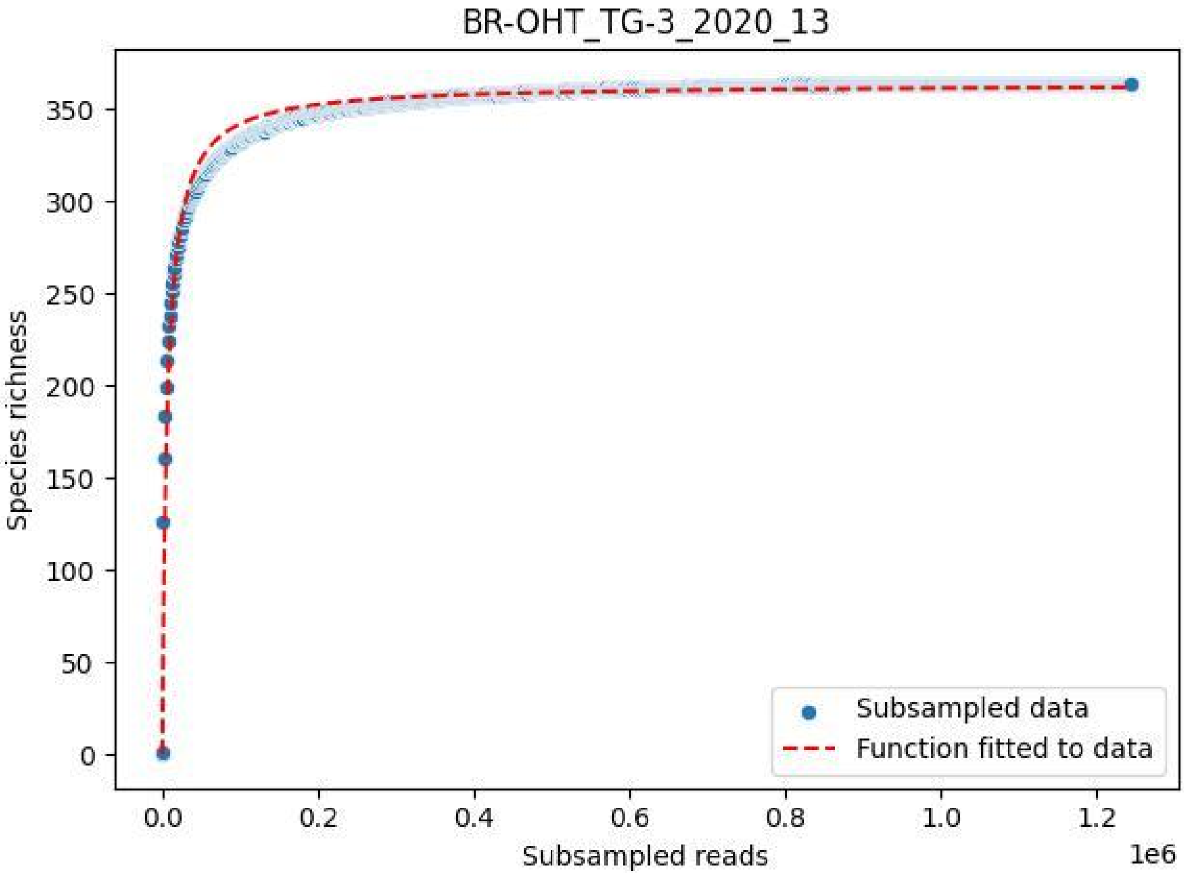
BR-OHT_TG-3_2020_13
Species richness
Subsampled reads
1e6
0
50
100
150
200
250
300
350
0.0
0.2
0.4
0.6
0.8
1.0
1.2
Subsampled data
Function fitted to data

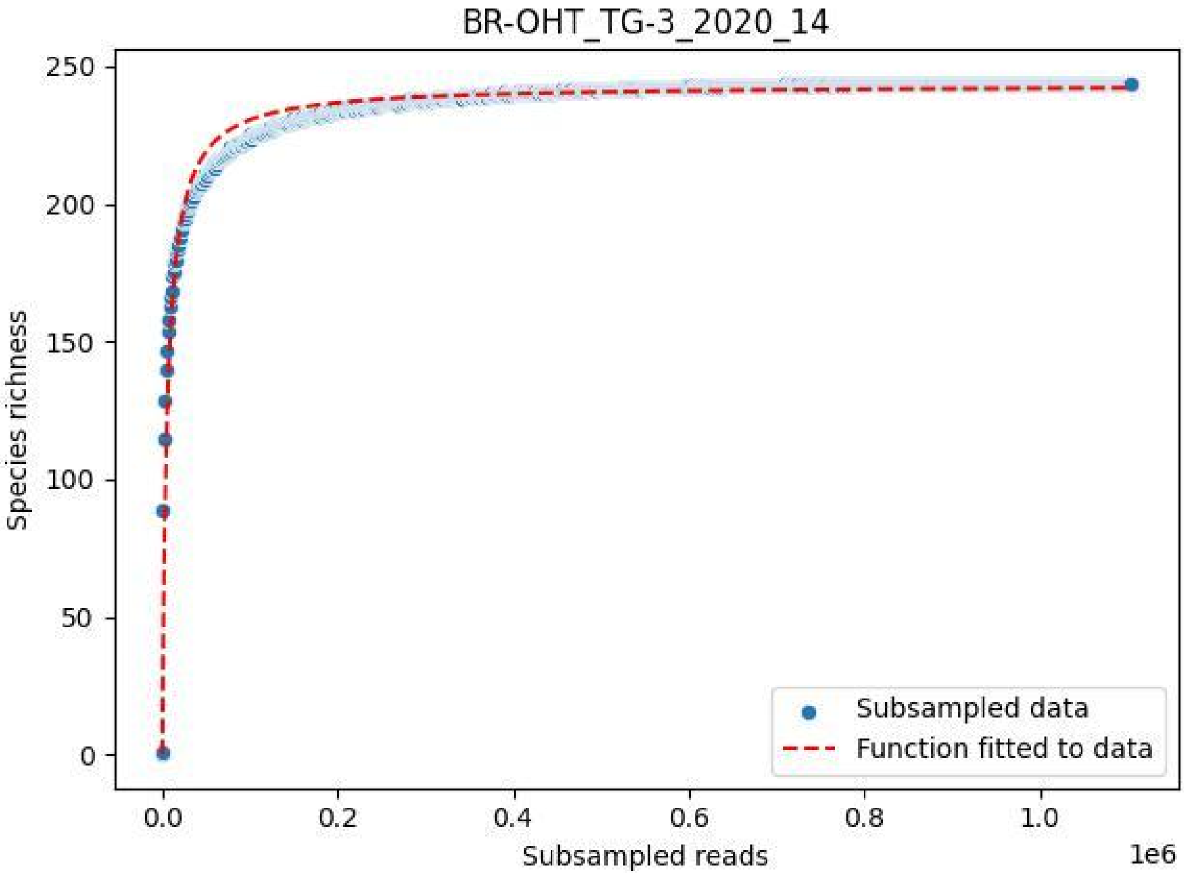
BR-OHT_TG-3_2020_14
Species richness
Subsampled reads
1e6
0
50
100
150
200
250
0.0
0.2
0.4
0.6
0.8
1.0
Subsampled data
Function fitted to data

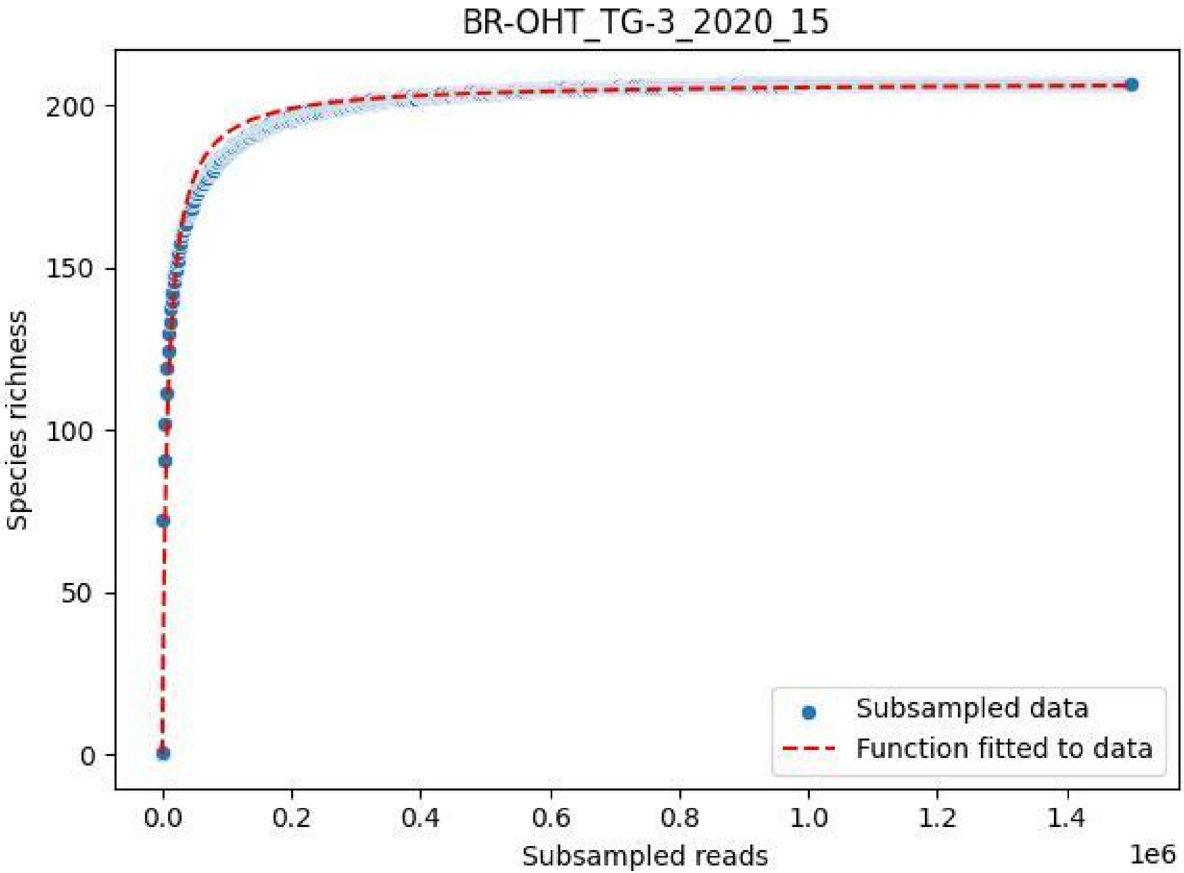

BR-OHT_TG-3_2020_15
Species richness
Subsampled reads
1e6
0
50
100
150
200
0.0
0.2
0.4
0.6
0.8
1.0
1.2
1.4
Subsampled data
Function fitted to data

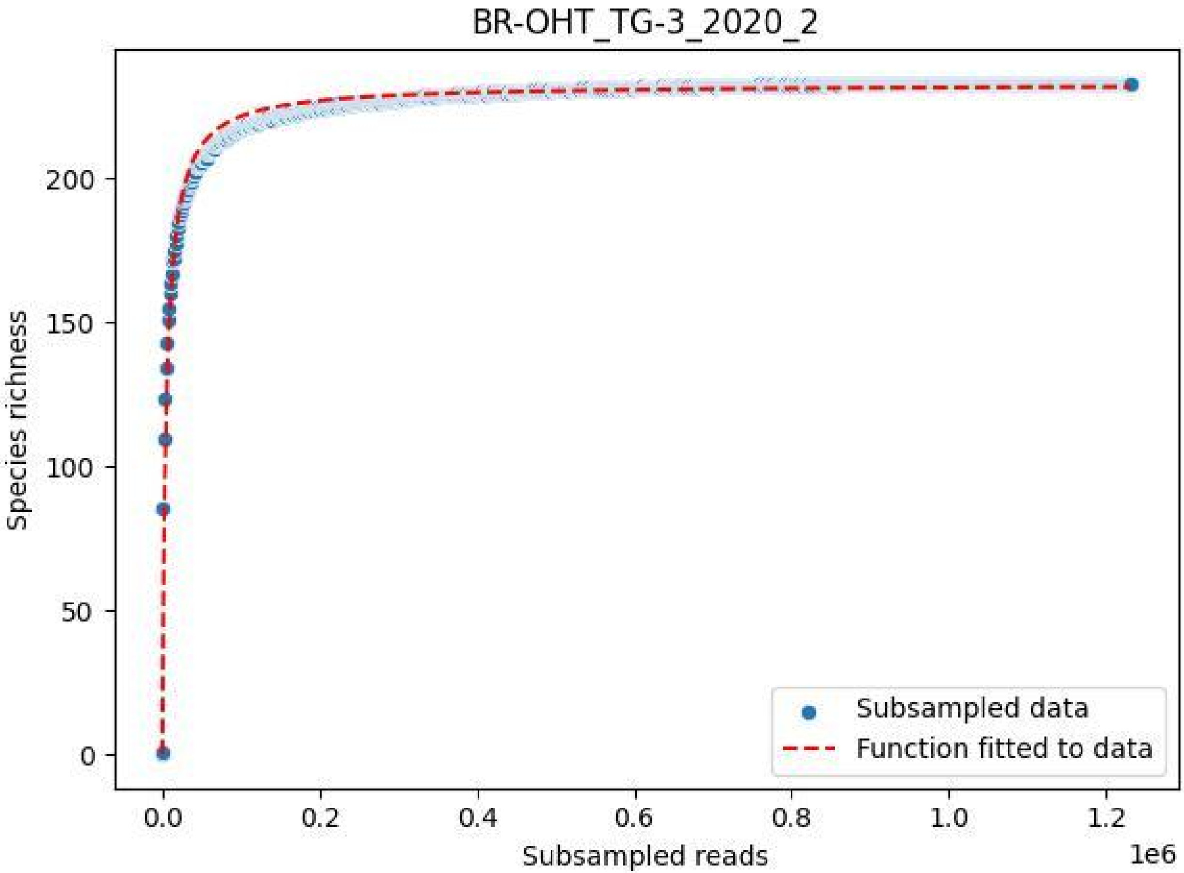
BR-OHT_TG-3_2020_2
Species richness
0
50
100
150
200
0.0
0.2
0.4
0.6
0.8
1.0
1.2
1e6
Subsampled reads
Subsampled data
Function fitted to data

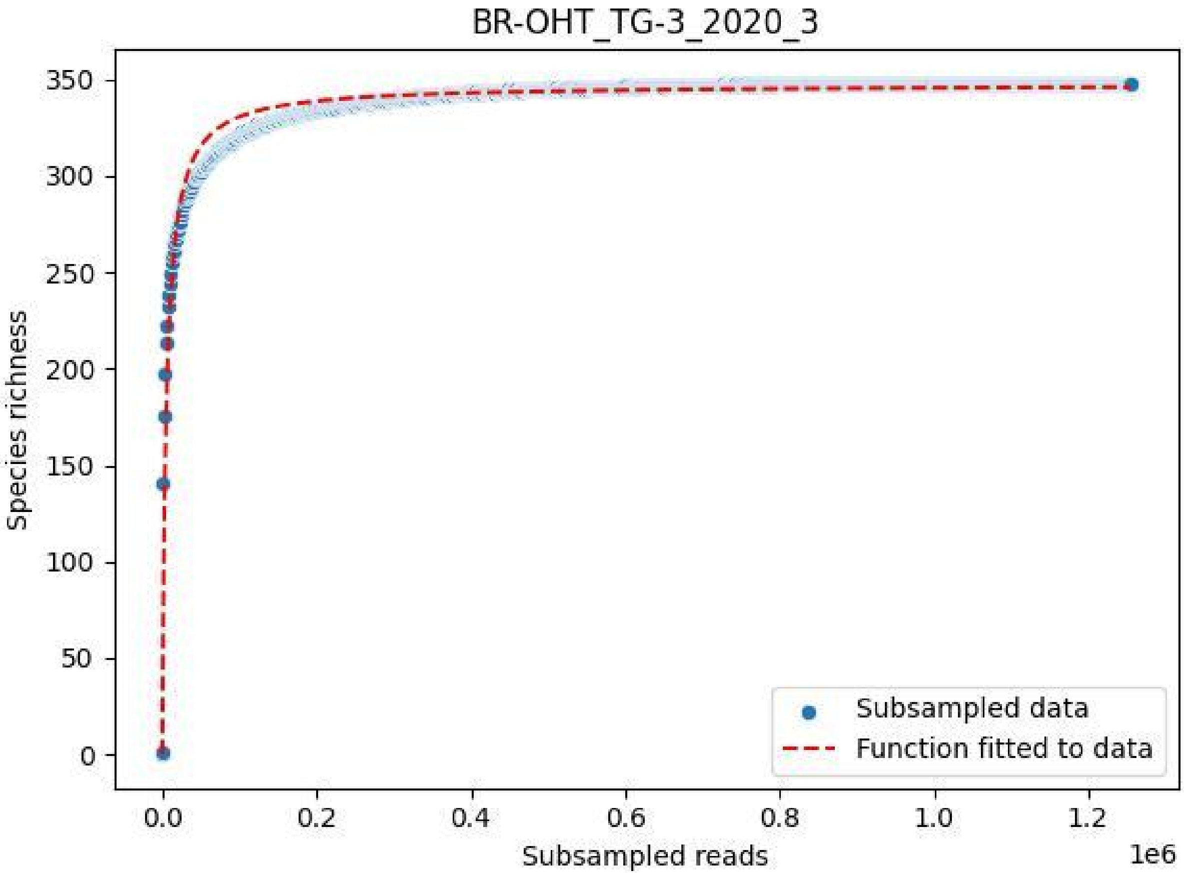

BR-OHT_TG-3_2020_3
Species richness
Subsampled reads
1e6
0
50
100
150
200
250
300
350
0.0
0.2
0.4
0.6
0.8
1.0
1.2
Subsampled data
Function fitted to data

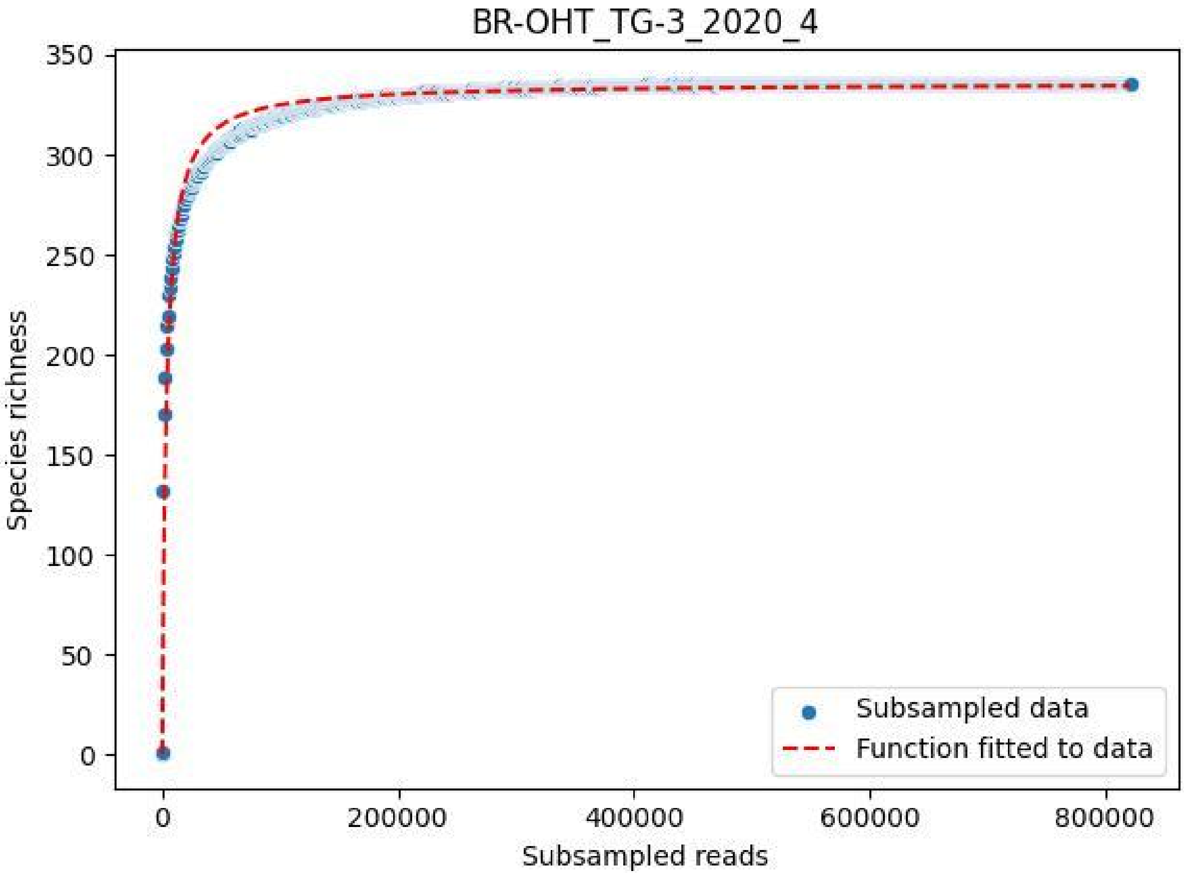

BR-OHT_TG-3_2020_4
Species richness
Subsampled reads
0
50
100
150
200
250
300
350
0
200000
400000
600000
800000
Subsampled data
Function fitted to data

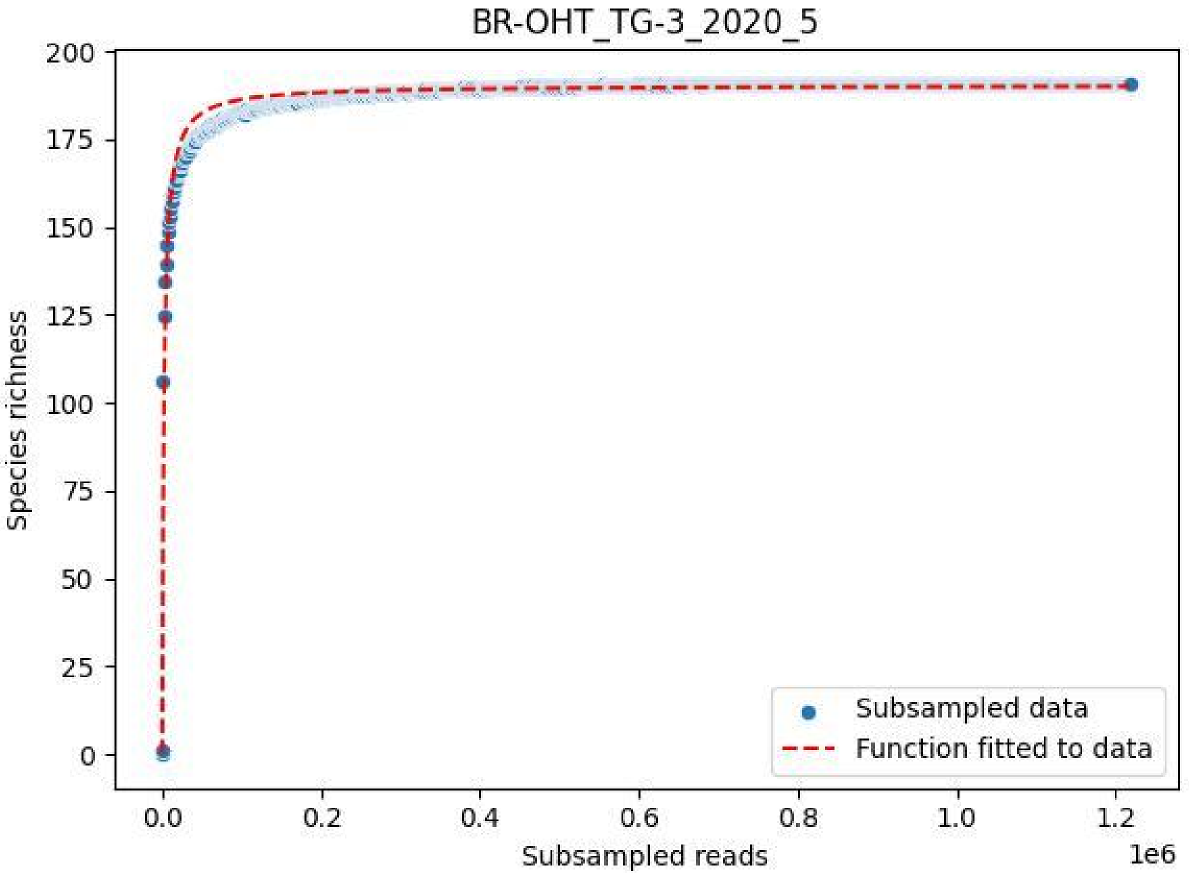
BR-OHT_TG-3_2020_5
200
175
150
125
100
75
50
25
0
Species richness
0.0
0.2
0.4
0.6
0.8
1.0
1.2
1e6
Subsampled reads
Subsampled data
Function fitted to data

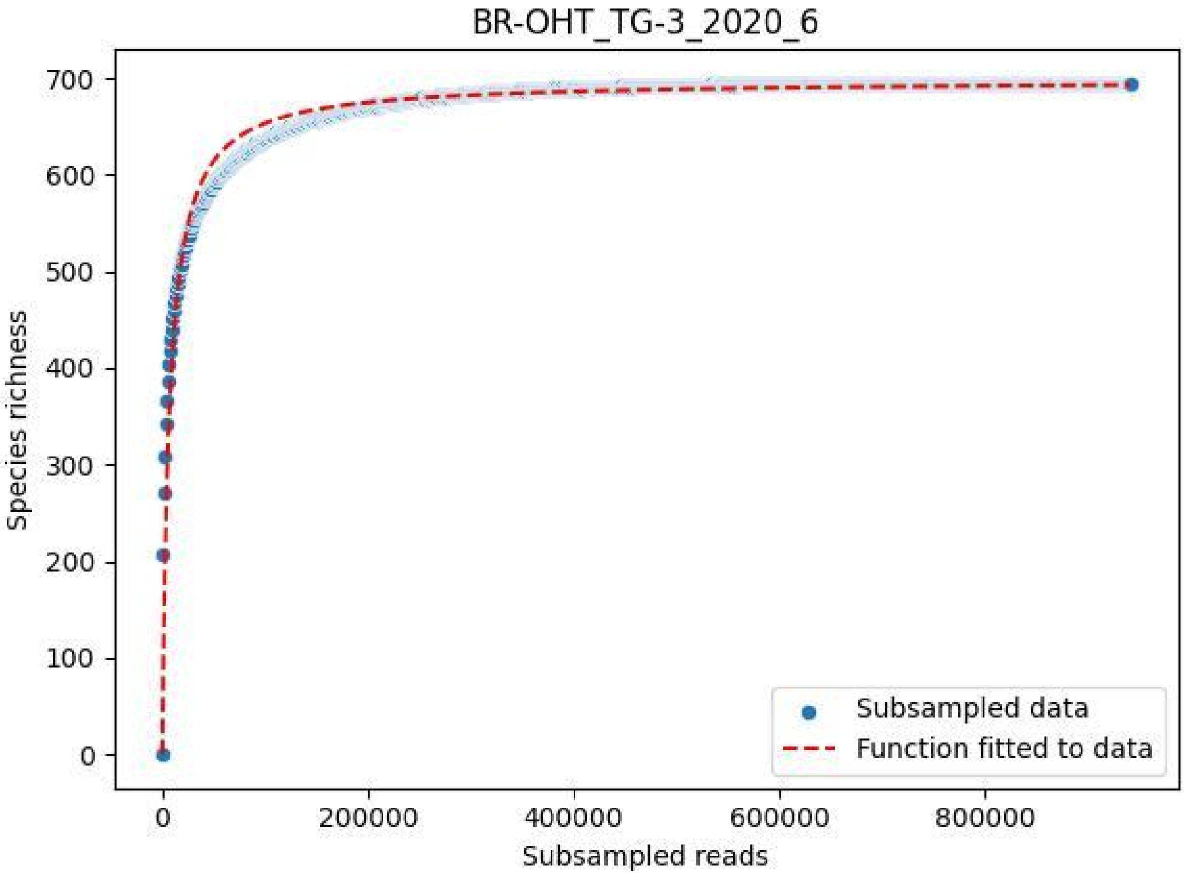

BR-OHT_TG-3_2020_6
Species richness
Subsampled reads
0
100
200
300
400
500
600
700
0
200000
400000
600000
800000
Subsampled data
Function fitted to data

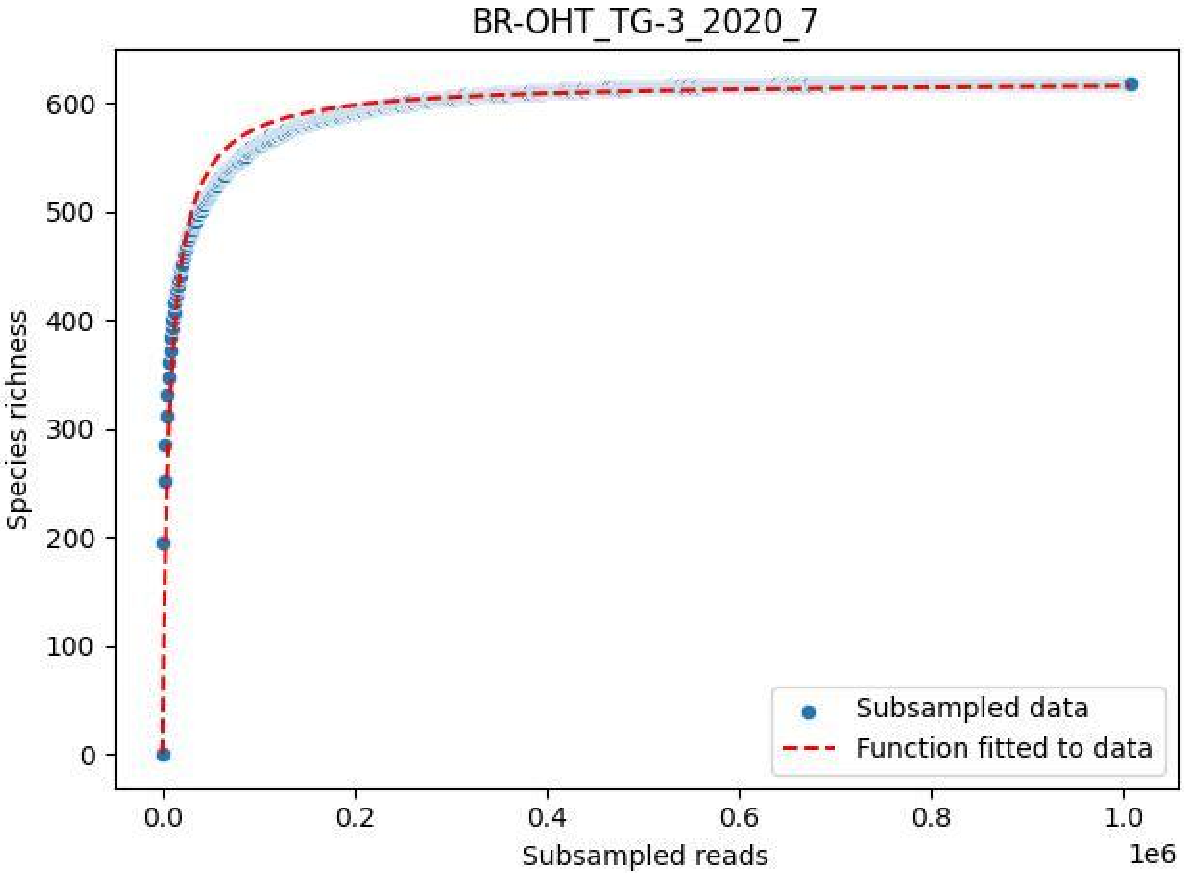
BR-OHT_TG-3_2020_7
Species richness
600
500
400
300
200
100
0
0.0
0.2
0.4
0.6
0.8
1.0
1e6
Subsampled reads
Subsampled data
Function fitted to data

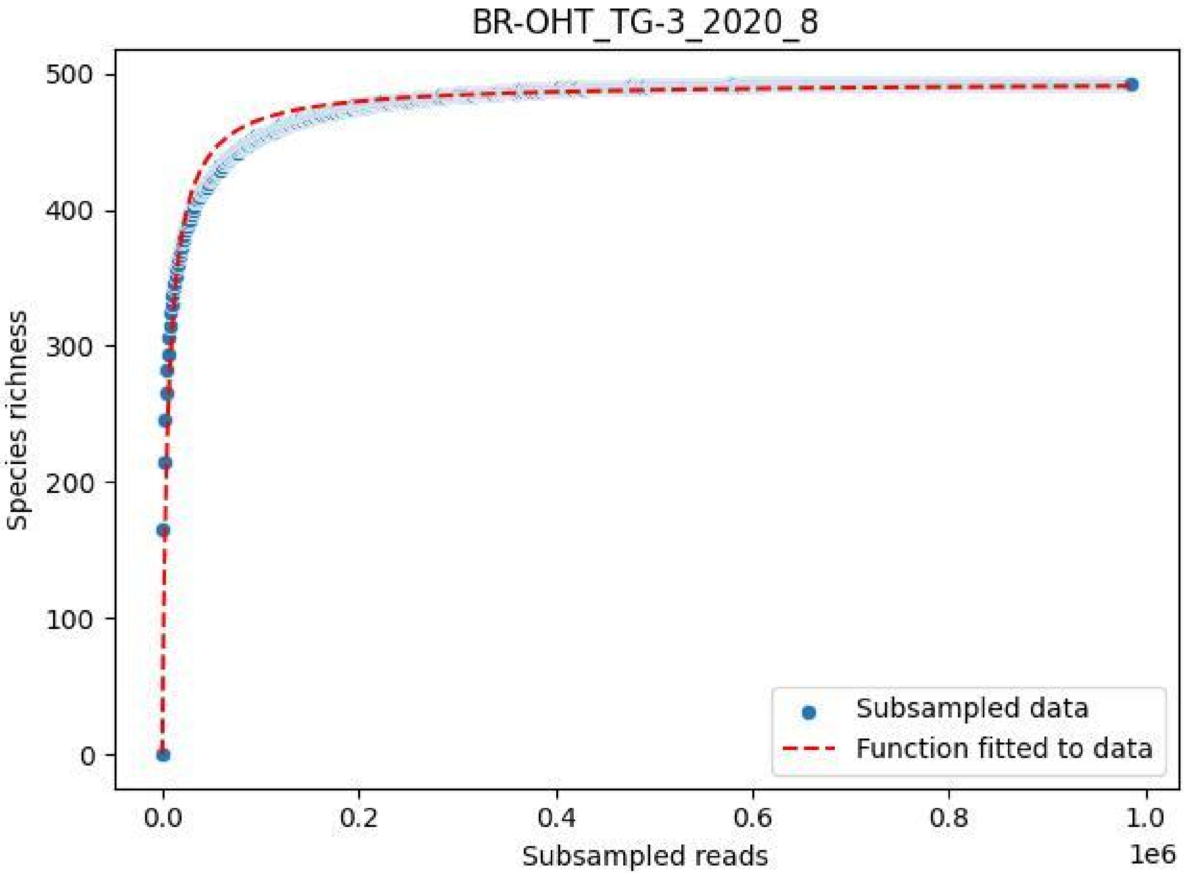
BR-OHT_TG-3_2020_8
Species richness
Subsampled reads
1e6
0
100
200
300
400
500
0.0
0.2
0.4
0.6
0.8
1.0
Subsampled data
Function fitted to data

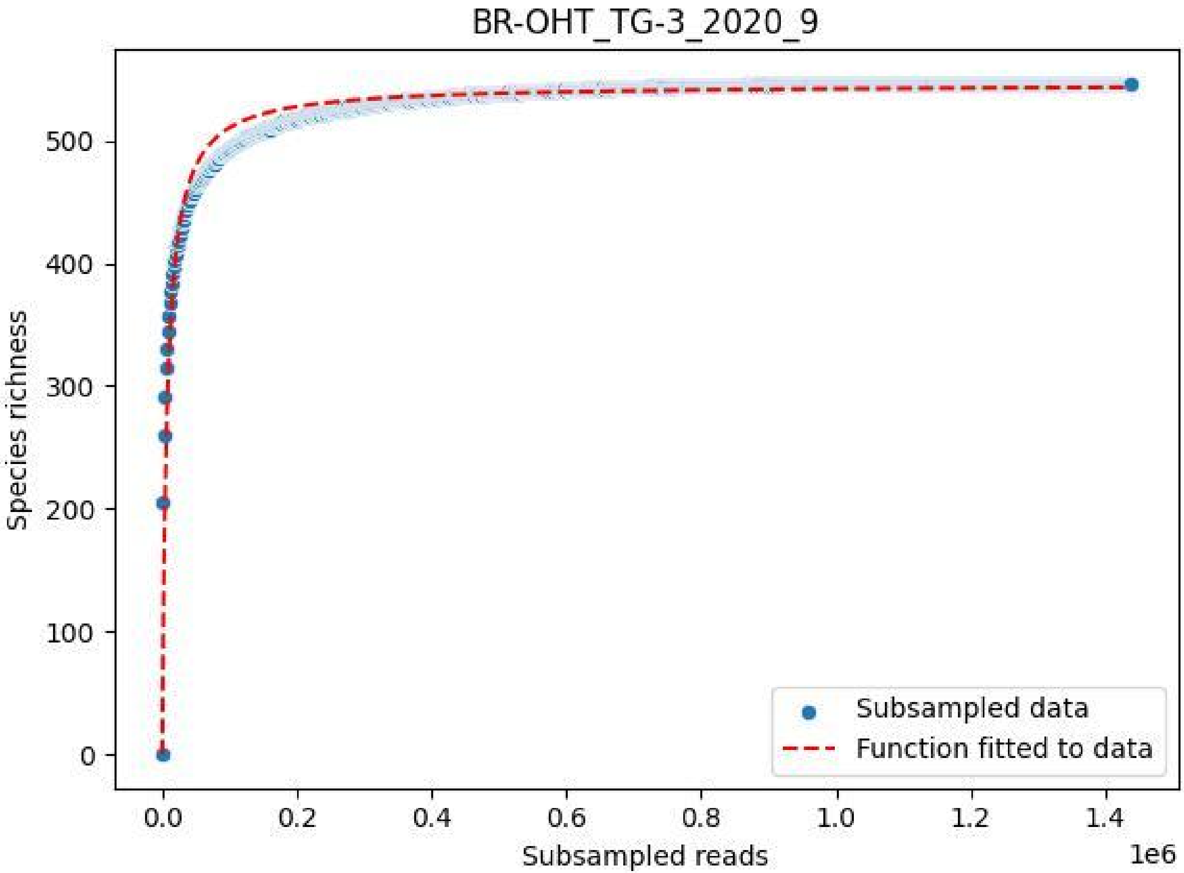
BR-OHT_TG-3_2020_9
Species richness
Subsampled reads
1e6
0
100
200
300
400
500
0.0
0.2
0.4
0.6
0.8
1.0
1.2
1.4
Subsampled data
Function fitted to data

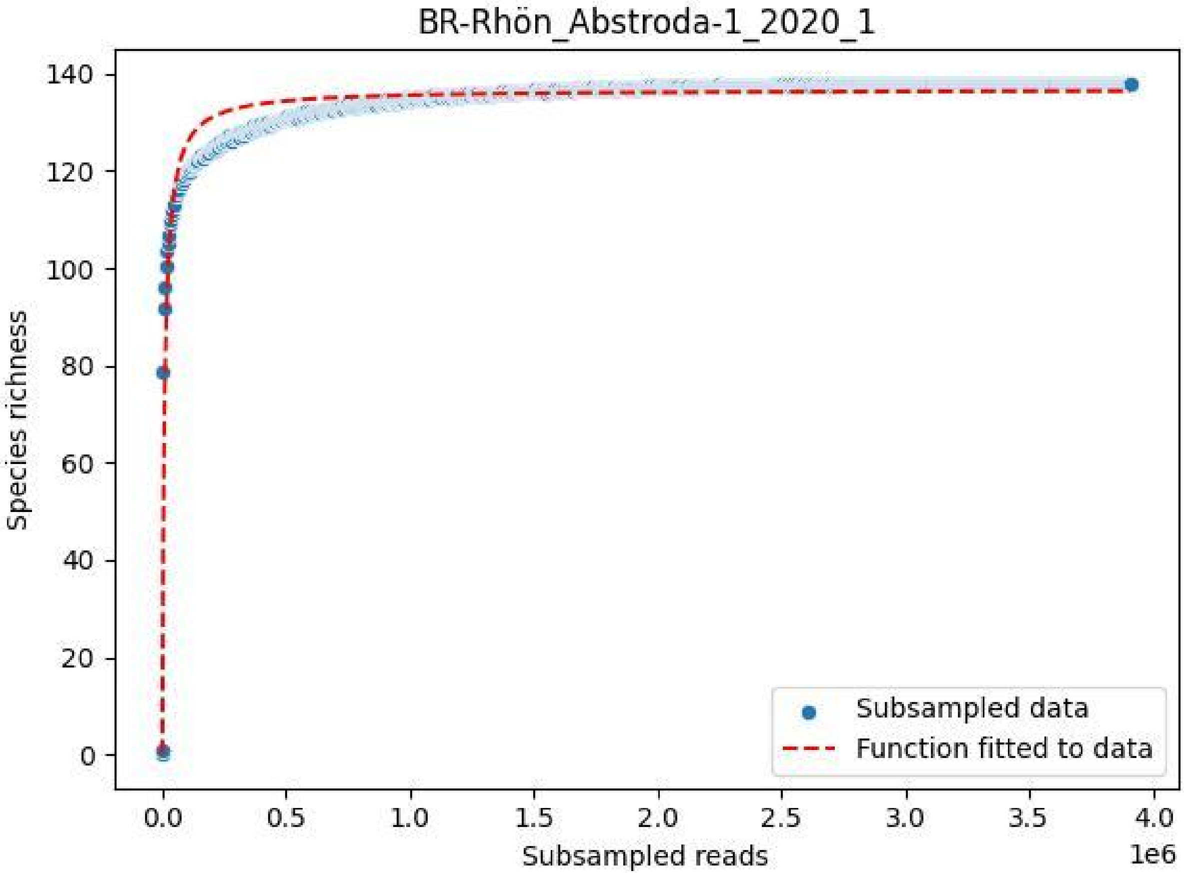

BR-Rhön_Abstroda-1_2020_1
Species richness
Subsampled reads
1e6
0
20
40
60
80
100
120
140
0.0
0.5
1.0
1.5
2.0
2.5
3.0
3.5
4.0
Subsampled data
Function fitted to data

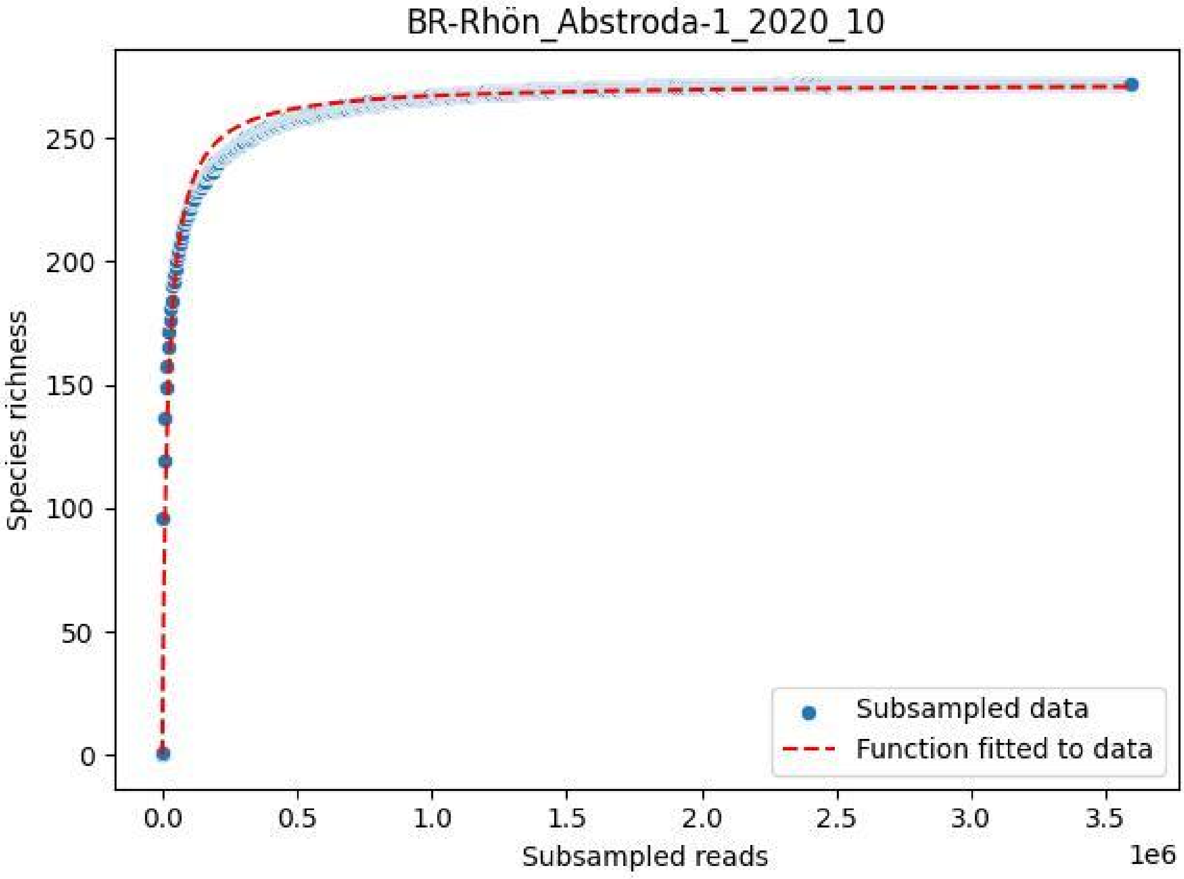
BR-Rhön_Abstroda-1_2020_10
Species richness
0
50
100
150
200
250
0.0
0.5
1.0
1.5
2.0
2.5
3.0
3.5
1e6
Subsampled reads
Subsampled data
Function fitted to data

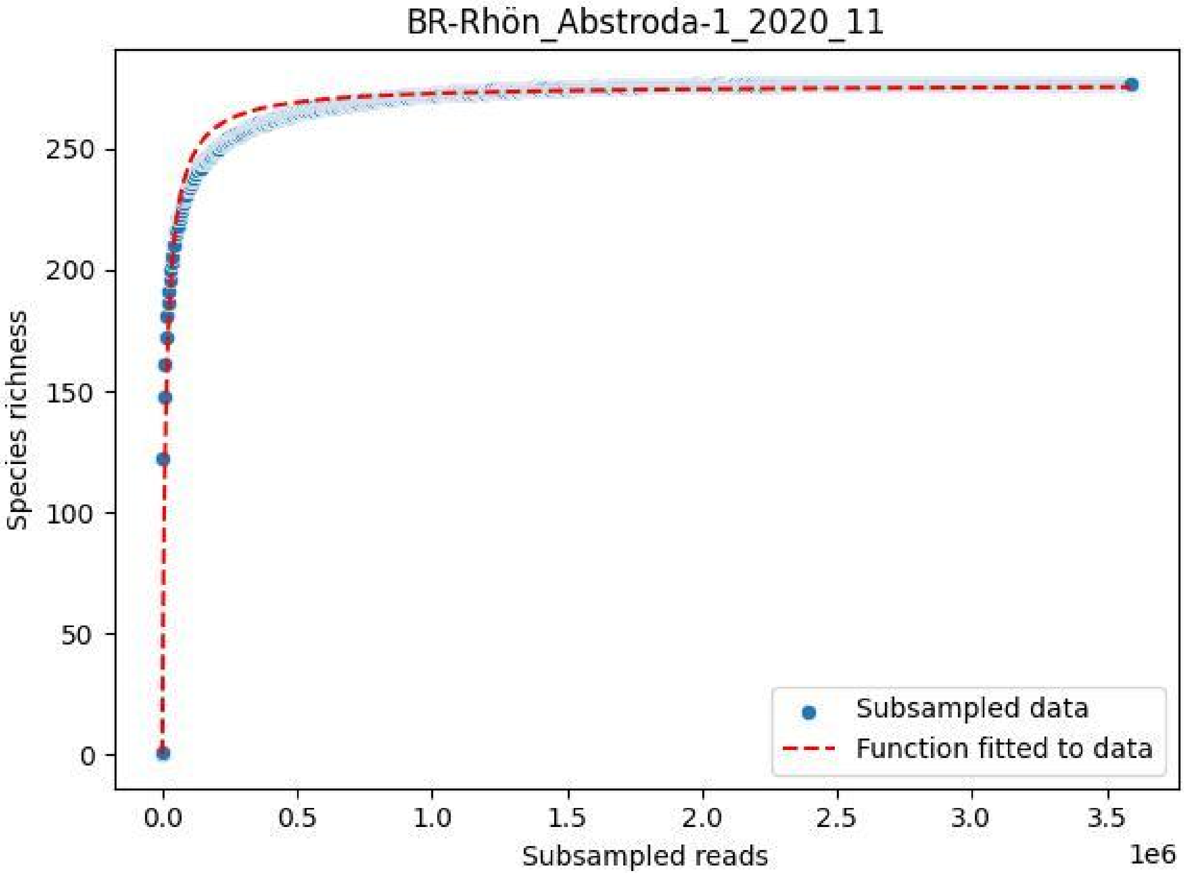
BR-Rhön_Abstroda-1_2020_11
Species richness
Subsampled reads
1e6
0
50
100
150
200
250
0.0
0.5
1.0
1.5
2.0
2.5
3.0
3.5
Subsampled data
Function fitted to data

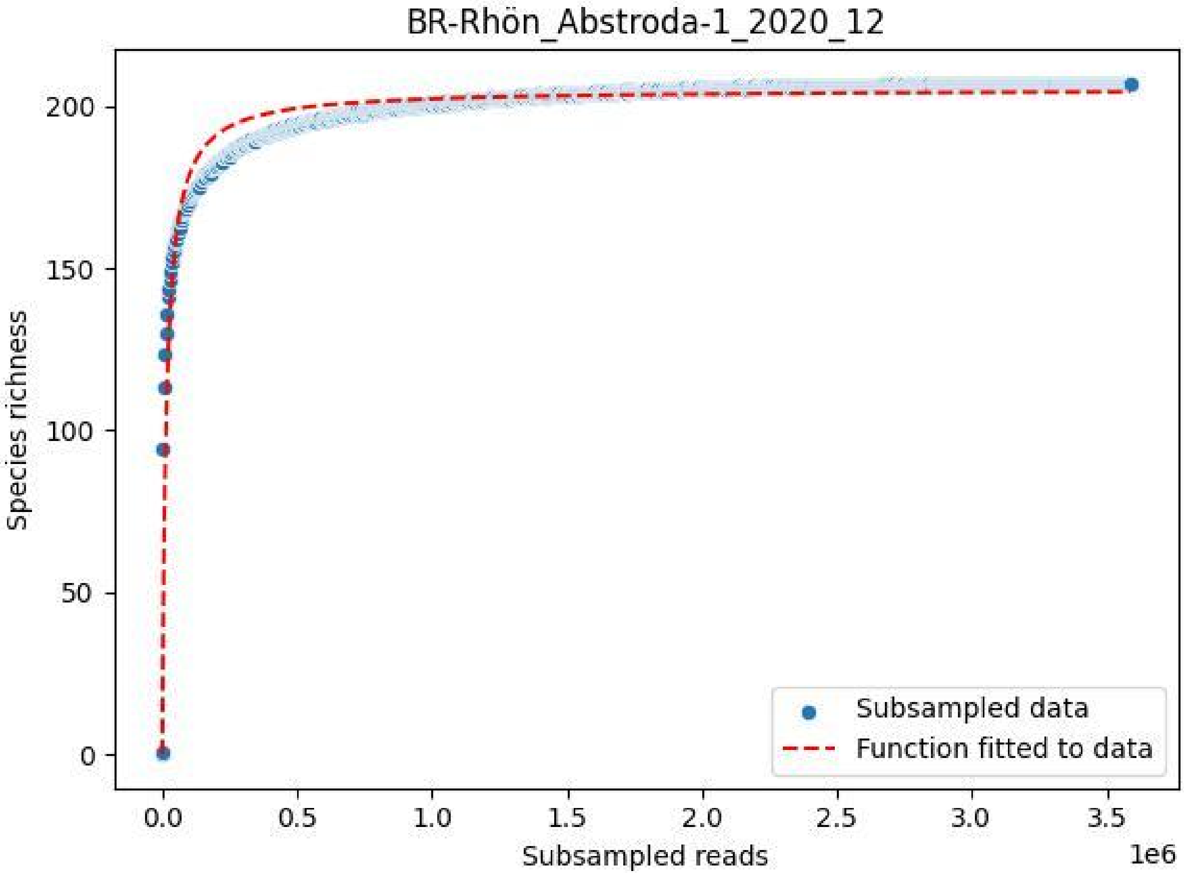

BR-Rhön_Abstroda-1_2020_12
Species richness
Subsampled reads
1e6
0
50
100
150
200
0.0
0.5
1.0
1.5
2.0
2.5
3.0
3.5
Subsampled data
Function fitted to data

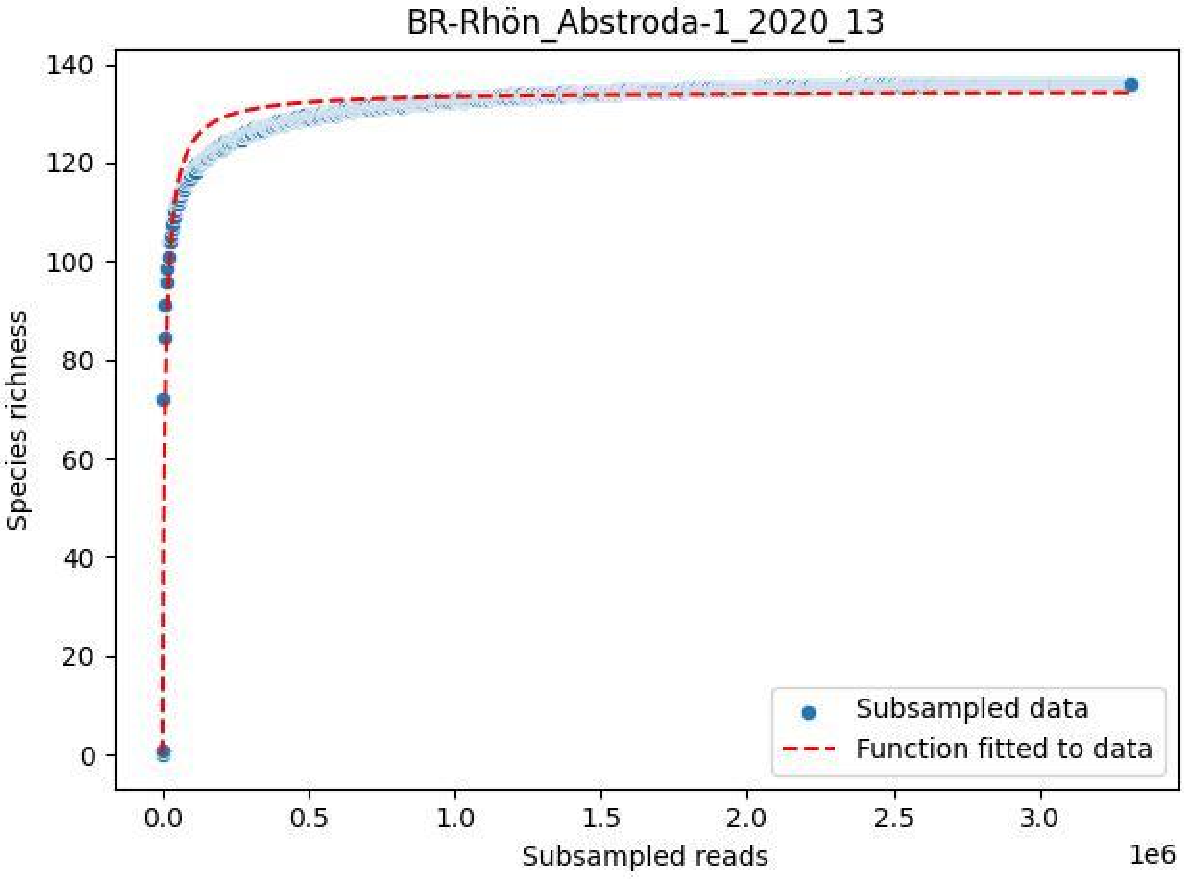

BR-Rhön_Abstroda-1_2020_13
Species richness
Subsampled reads
1e6
0
20
40
60
80
100
120
140
0.0
0.5
1.0
1.5
2.0
2.5
3.0
Subsampled data
Function fitted to data

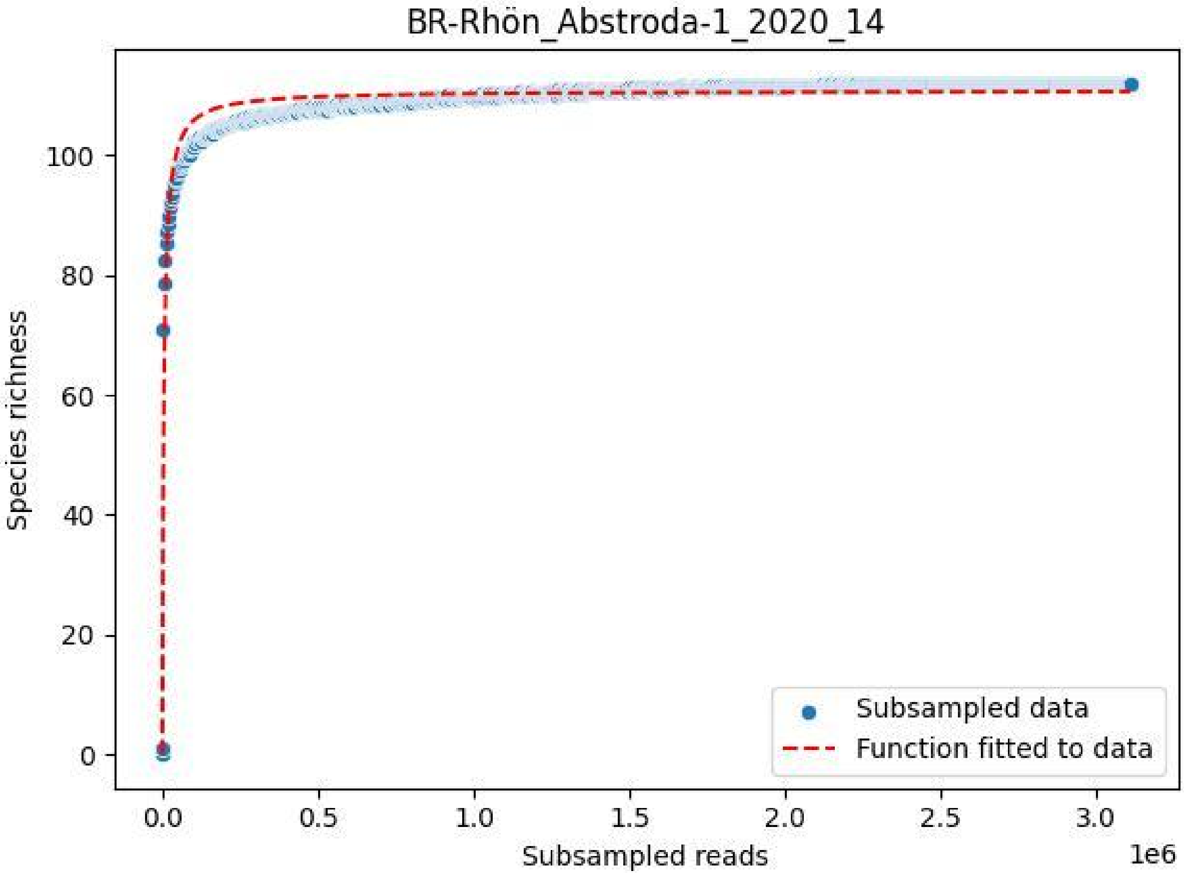
BR-Rhön_Abstroda-1_2020_14
Species richness
Subsampled reads
1e6
0
20
40
60
80
100
0.0
0.5
1.0
1.5
2.0
2.5
3.0
Subsampled data
Function fitted to data

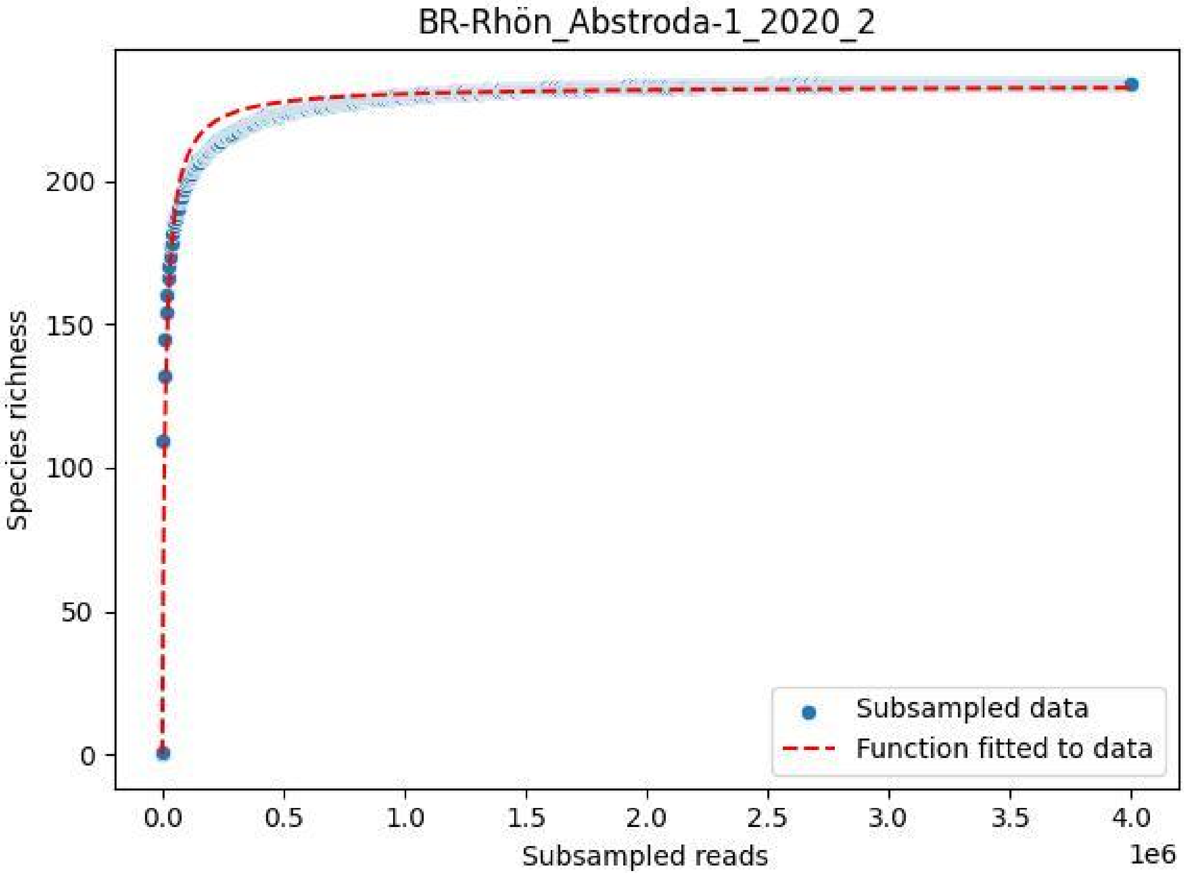
BR-Rhön_Abstroda-1_2020_2
Species richness
Subsampled reads
1e6
0
50
100
150
200
0.0
0.5
1.0
1.5
2.0
2.5
3.0
3.5
4.0
Subsampled data
Function fitted to data

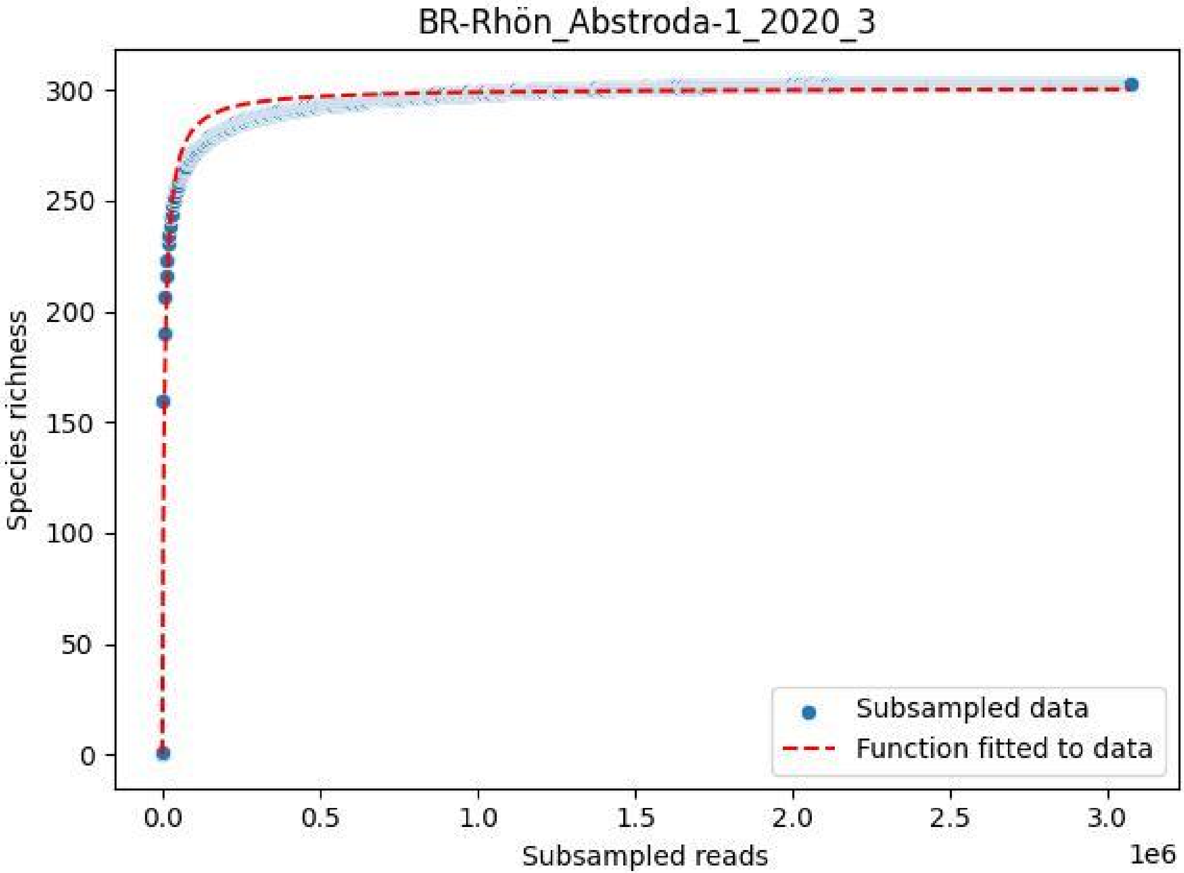
BR-Rhön_Abstroda-1_2020_3
Species richness
300
250
200
150
100
50
0
0.0
0.5
1.0
1.5
2.0
2.5
3.0
1e6
Subsampled reads
Subsampled data
Function fitted to data

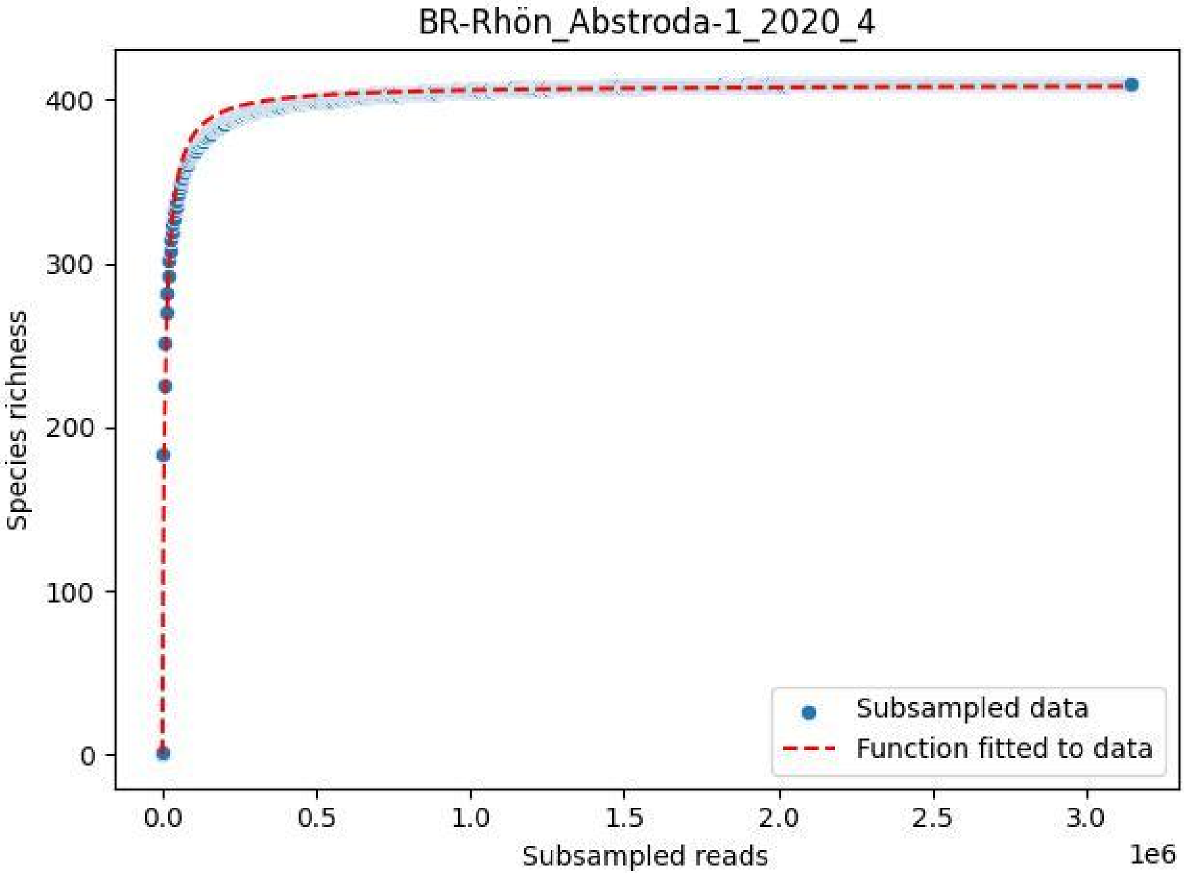
BR-Rhön_Abstroda-1_2020_4
Species richness
Subsampled reads
1e6
0
100
200
300
400
0.0
0.5
1.0
1.5
2.0
2.5
3.0
Subsampled data
Function fitted to data

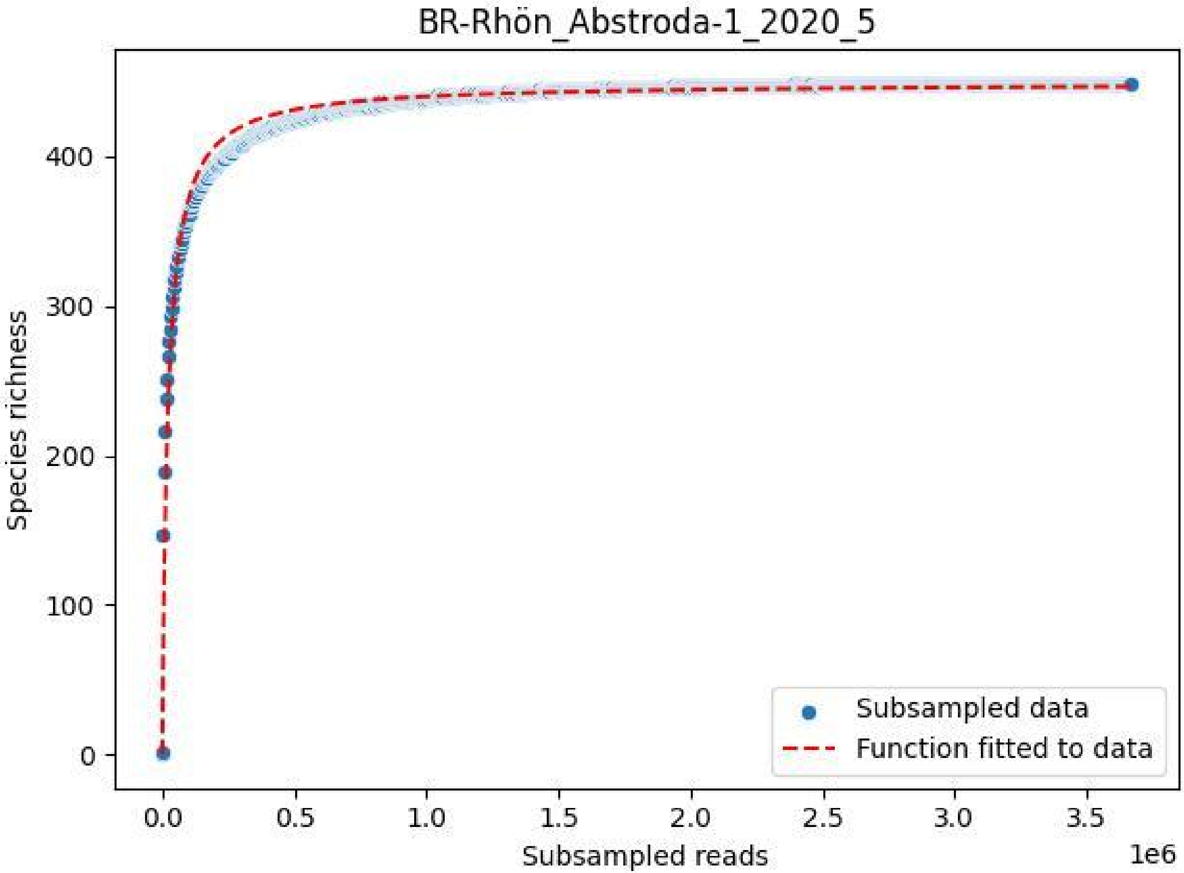

BR-Rhön_Abstroda-1_2020_5
Species richness
0
100
200
300
400
0.0
0.5
1.0
1.5
2.0
2.5
3.0
3.5
1e6
Subsampled reads
Subsampled data
Function fitted to data

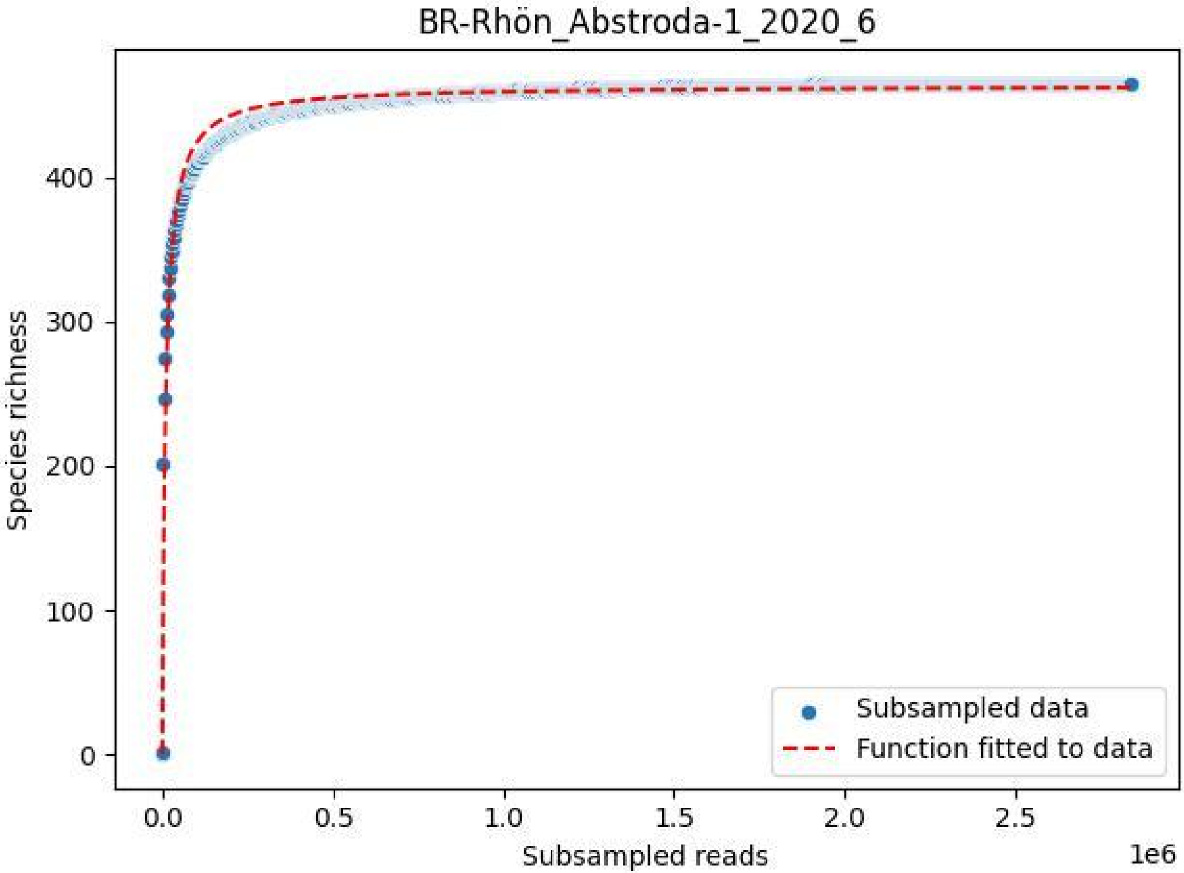
BR-Rhön_Abstroda-1_2020_6
Species richness
400
300
200
100
0
0.0
0.5
1.0
1.5
2.0
2.5
Subsampled reads
1e6
Subsampled data
Function fitted to data

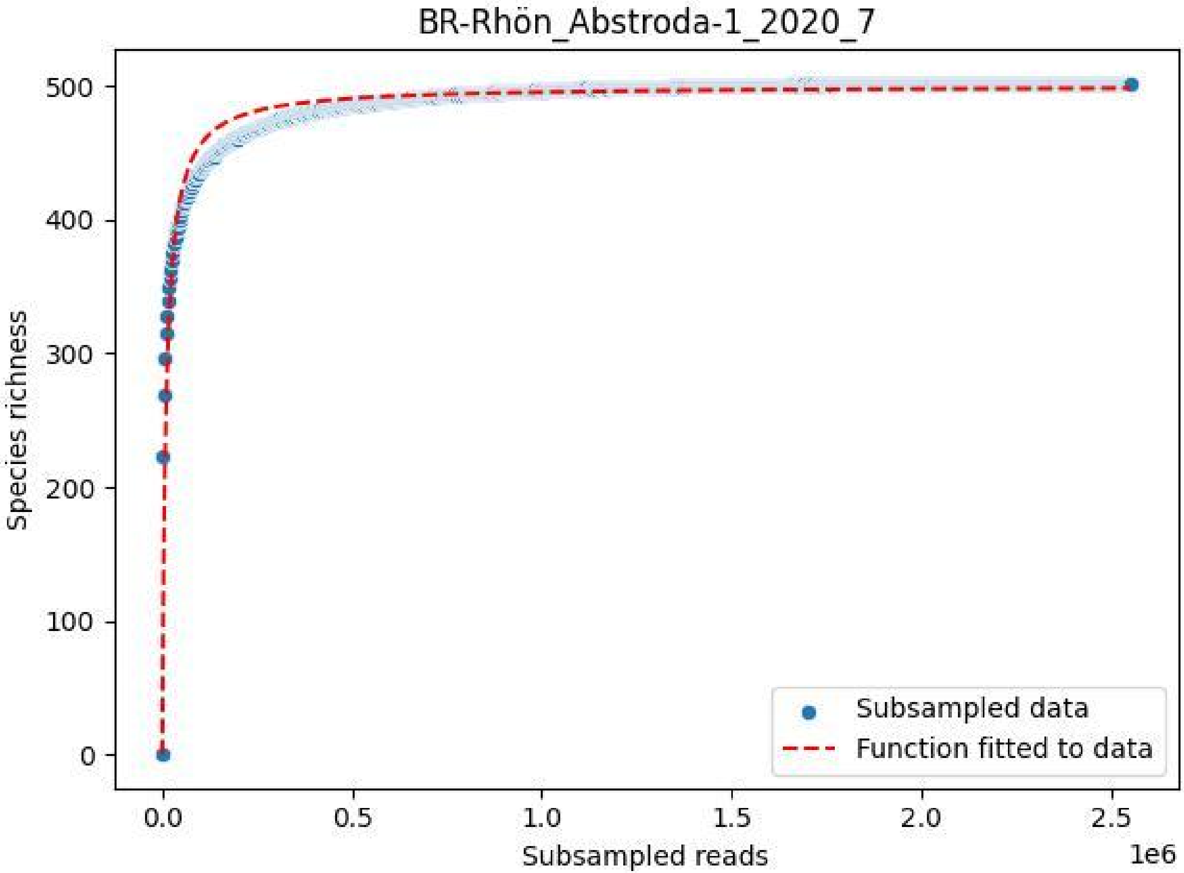
BR-Rhön_Abstroda-1_2020_7
Species richness
Subsampled reads
1e6
0
100
200
300
400
500
0.0
0.5
1.0
1.5
2.0
2.5
Subsampled data
Function fitted to data

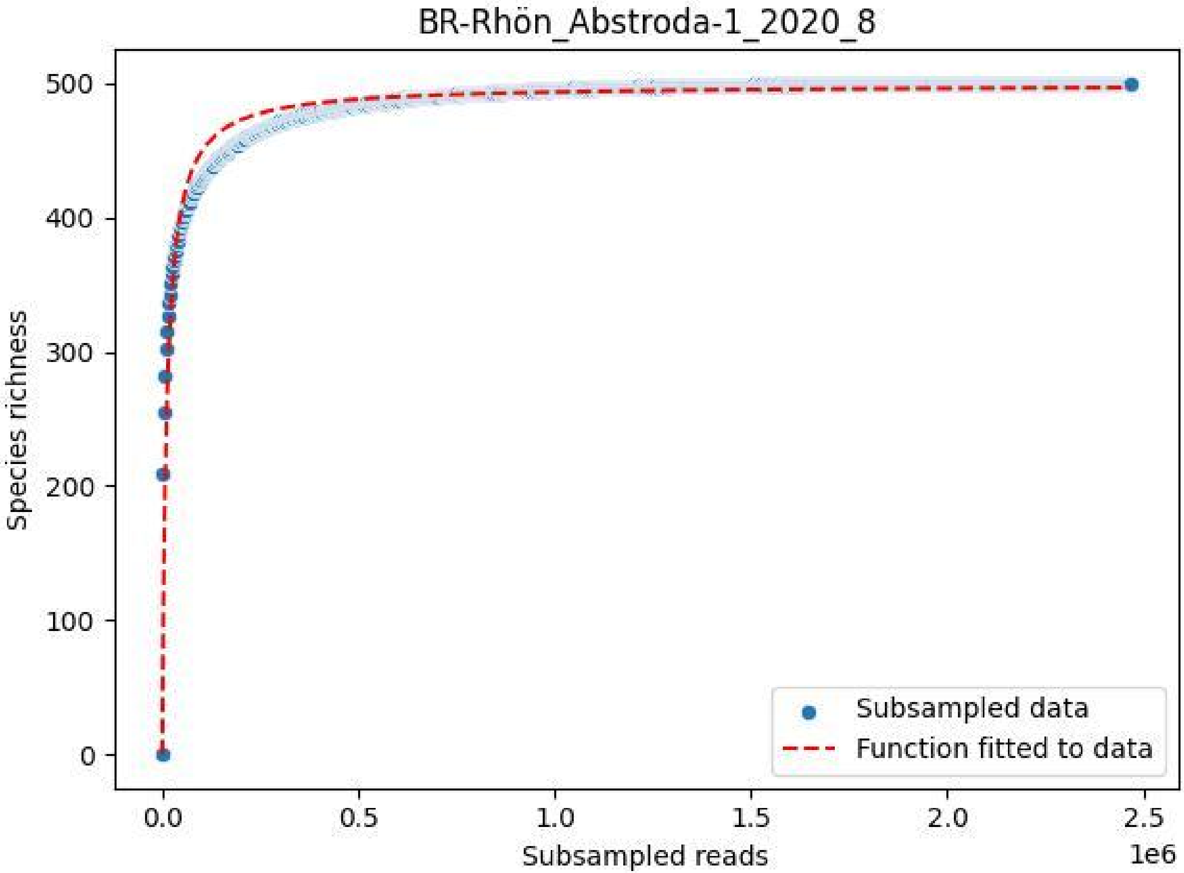
BR-Rhön_Abstroda-1_2020_8
Species richness
Subsampled reads
1e6
Subsampled data
Function fitted to data
0
100
200
300
400
500
0.0
0.5
1.0
1.5
2.0
2.5

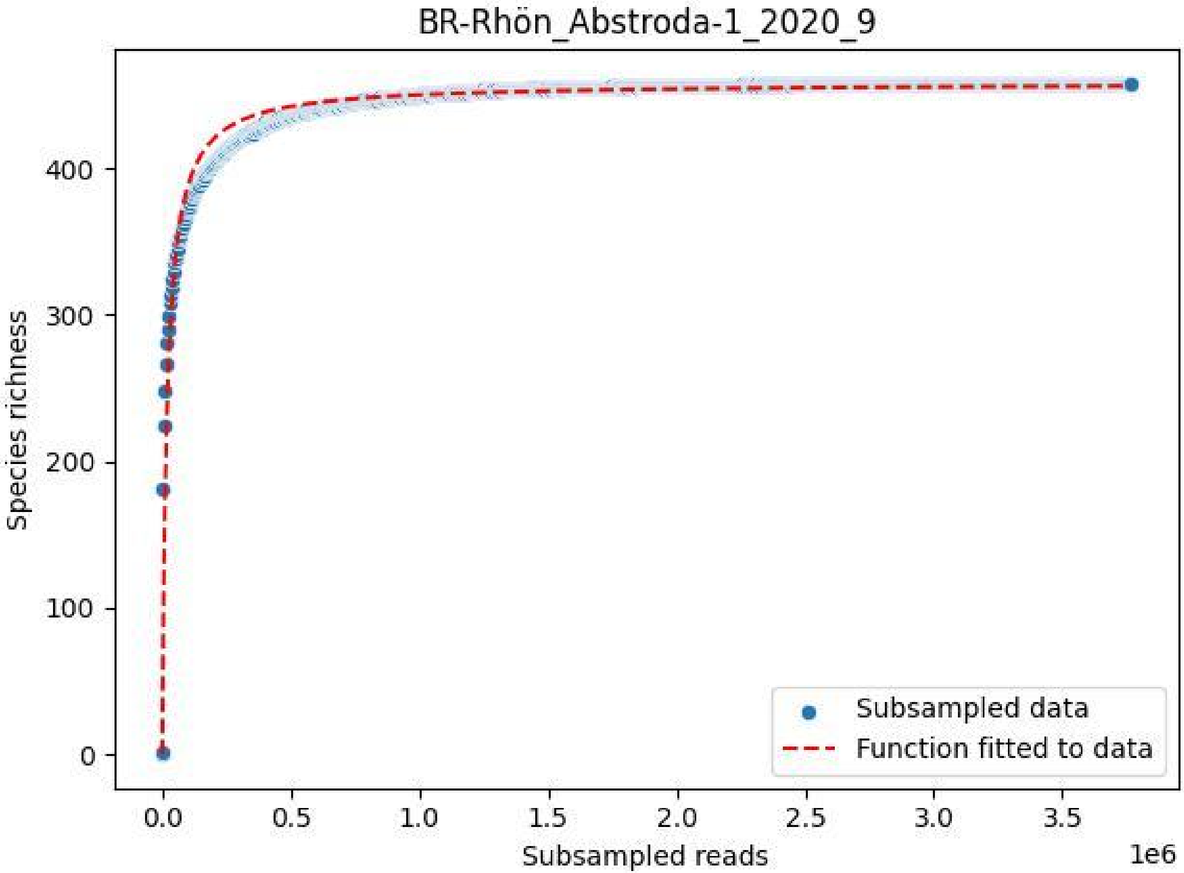
BR-Rhön_Abstroda-1_2020_9
Species richness
0
100
200
300
400
Subsampled reads
0.0
0.5
1.0
1.5
2.0
2.5
3.0
3.5
1e6
Subsampled data
Function fitted to data

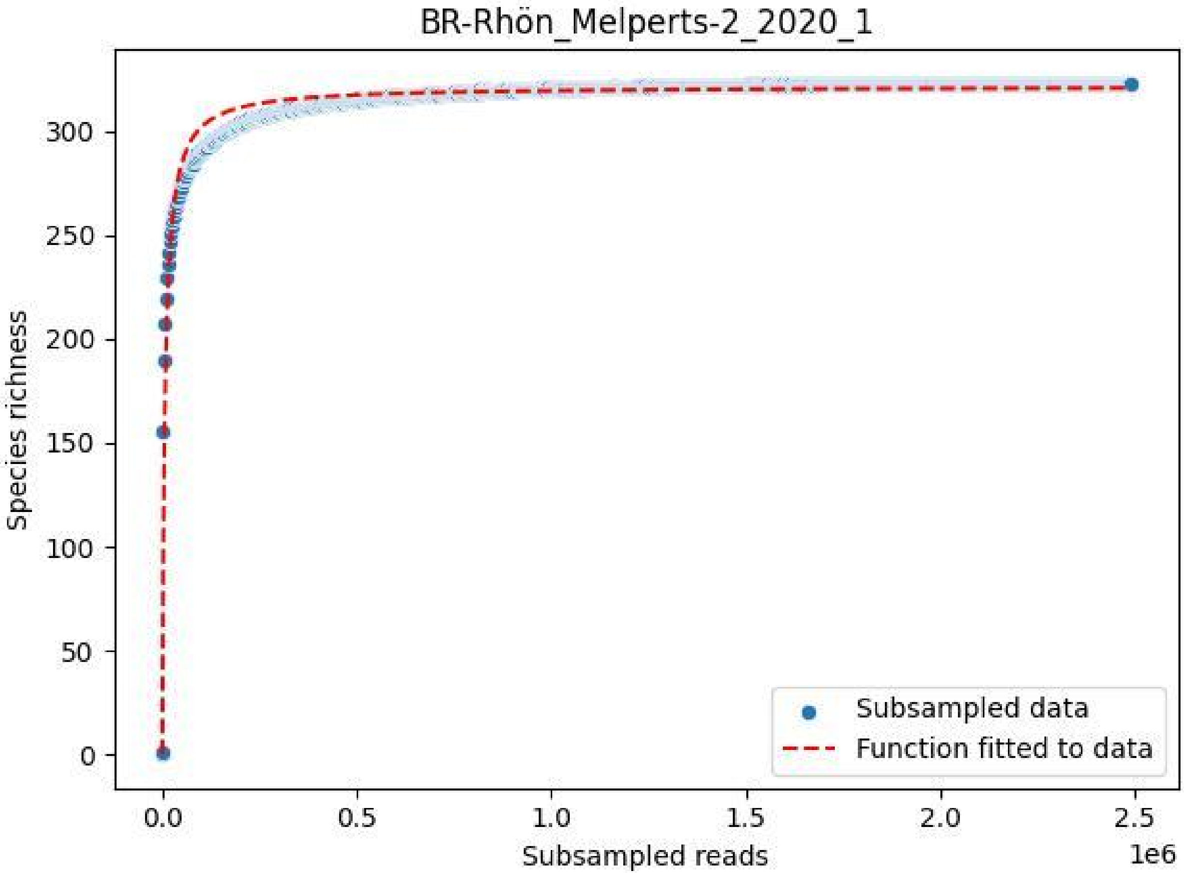
BR-Rhön_Melperts-2_2020_1
Species richness
0
50
100
150
200
250
300
0.0
0.5
1.0
1.5
2.0
2.5
1e6
Subsampled reads
Subsampled data
Function fitted to data

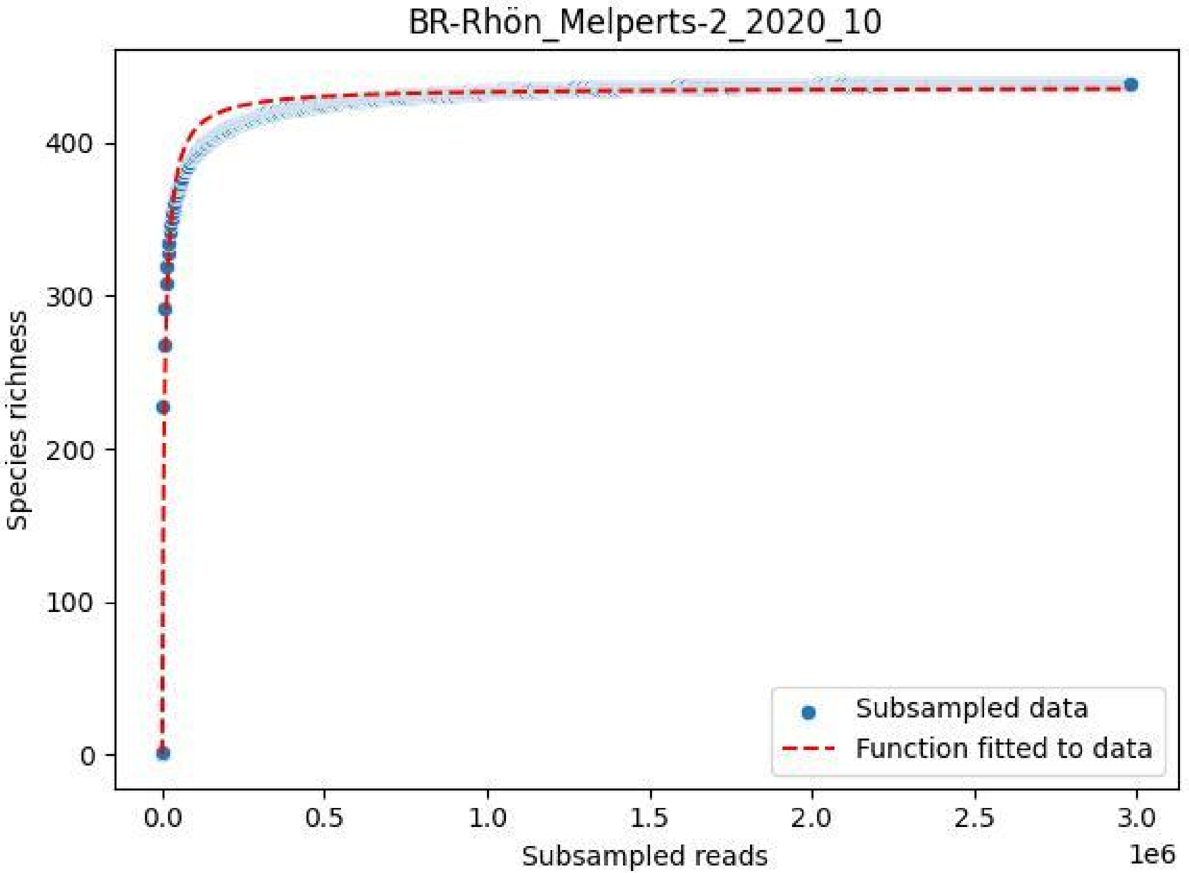

BR-Rhön_Melperts-2_2020_10
Species richness
Subsampled reads
1e6
400
300
200
100
0
0.0
0.5
1.0
1.5
2.0
2.5
3.0
Subsampled data
Function fitted to data

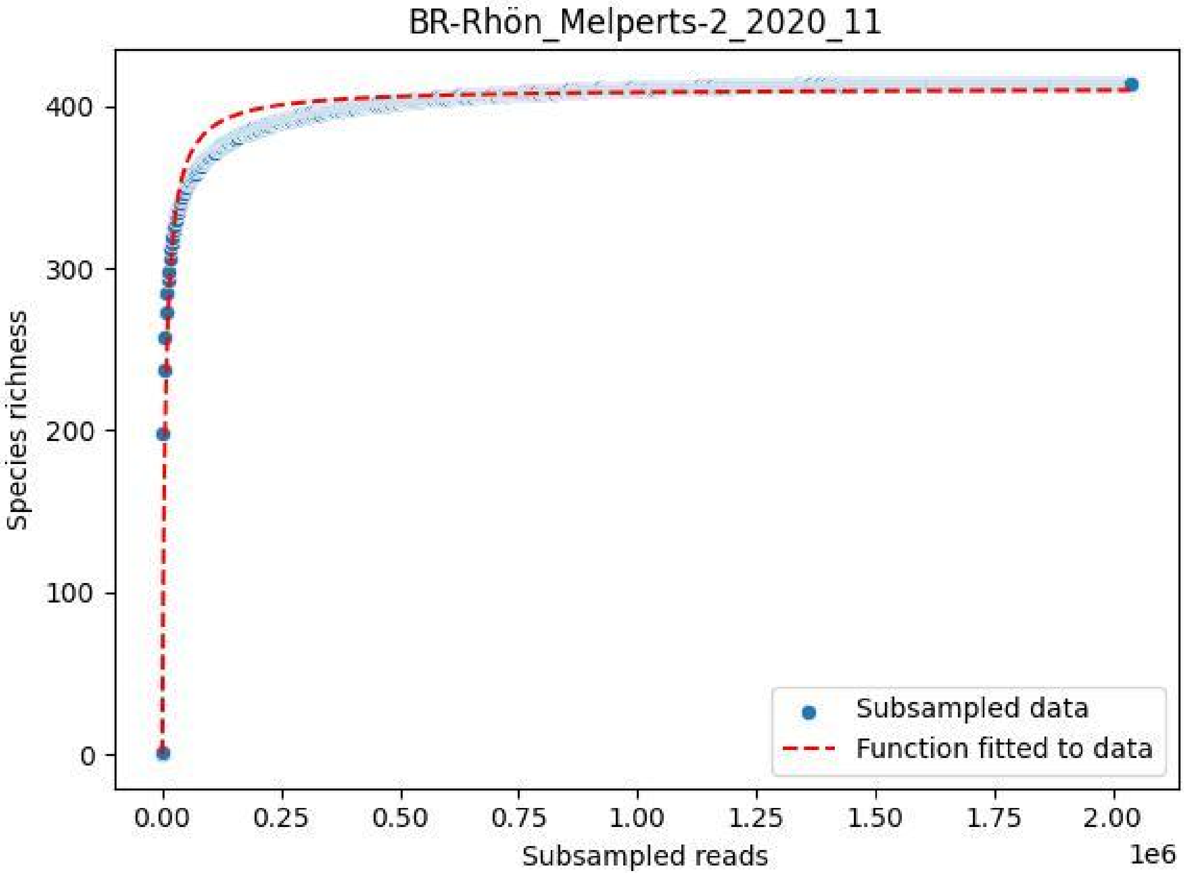
BR-Rhön_Melperts-2_2020_11
Species richness
Subsampled reads
1e6
0
100
200
300
400
0.00
0.25
0.50
0.75
1.00
1.25
1.50
1.75
2.00
Subsampled data
Function fitted to data

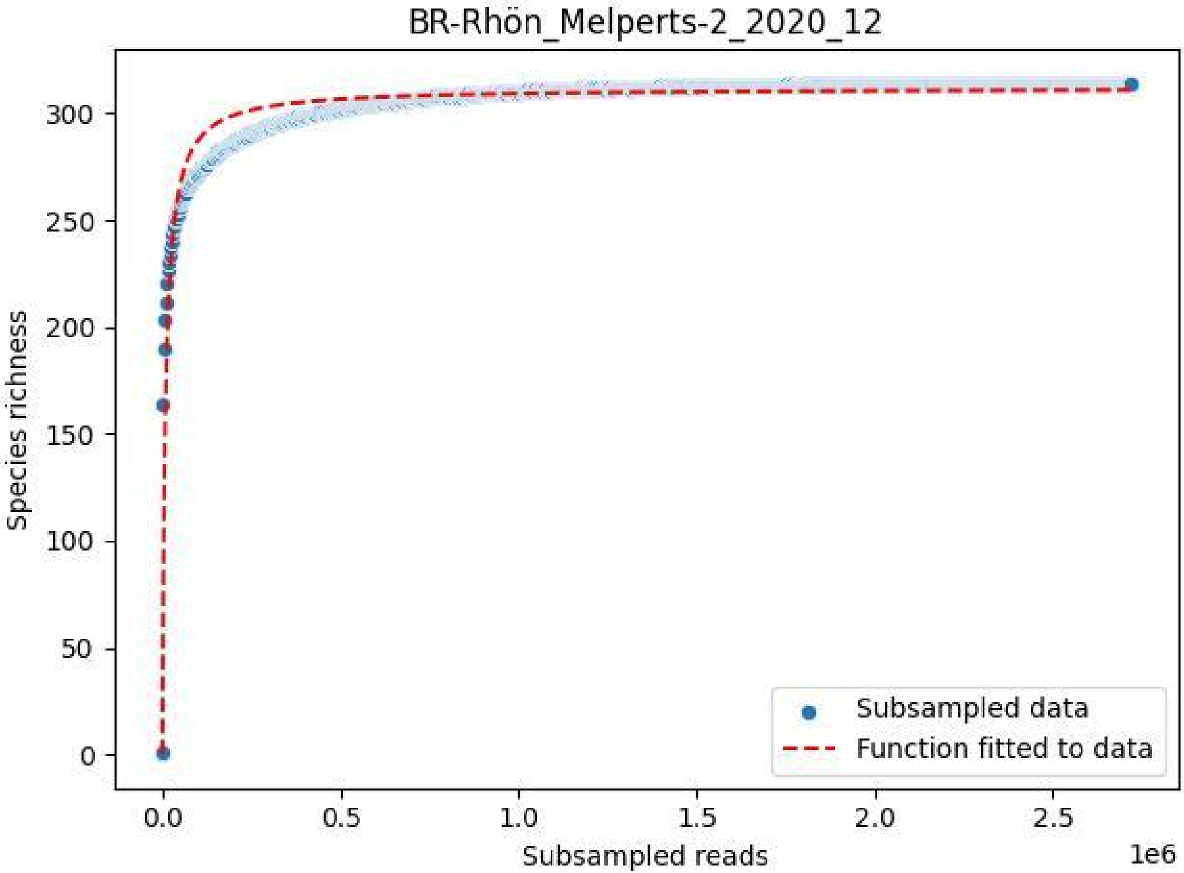
BR-Rhön_Melperts-2_2020_12
Species richness
300
250
200
150
100
50
0
0.0
0.5
1.0
1.5
2.0
2.5
1e6
Subsampled reads
Subsampled data
Function fitted to data

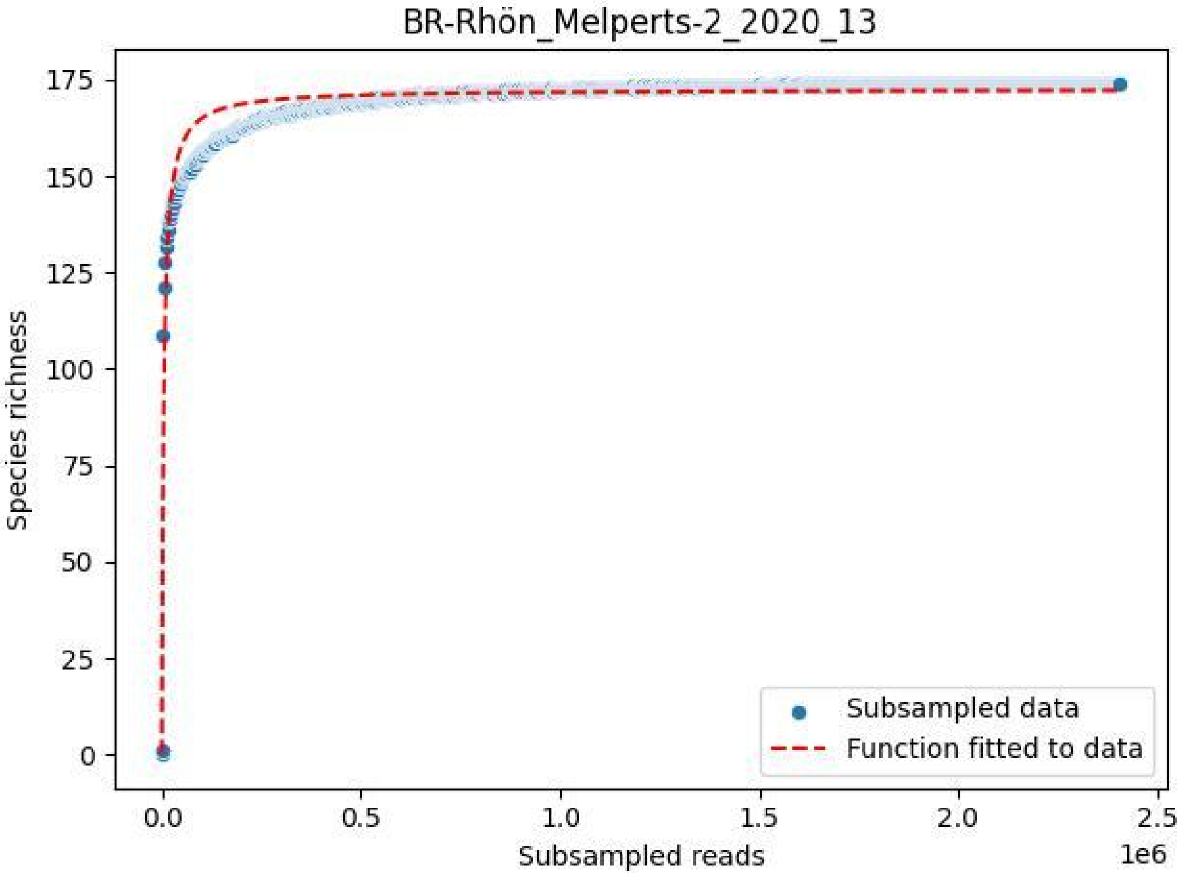
BR-Rhön_Melperts-2_2020_13
Species richness
Subsampled reads
1e6
175
150
125
100
75
50
25
0
0.0
0.5
1.0
1.5
2.0
2.5
Subsampled data
Function fitted to data

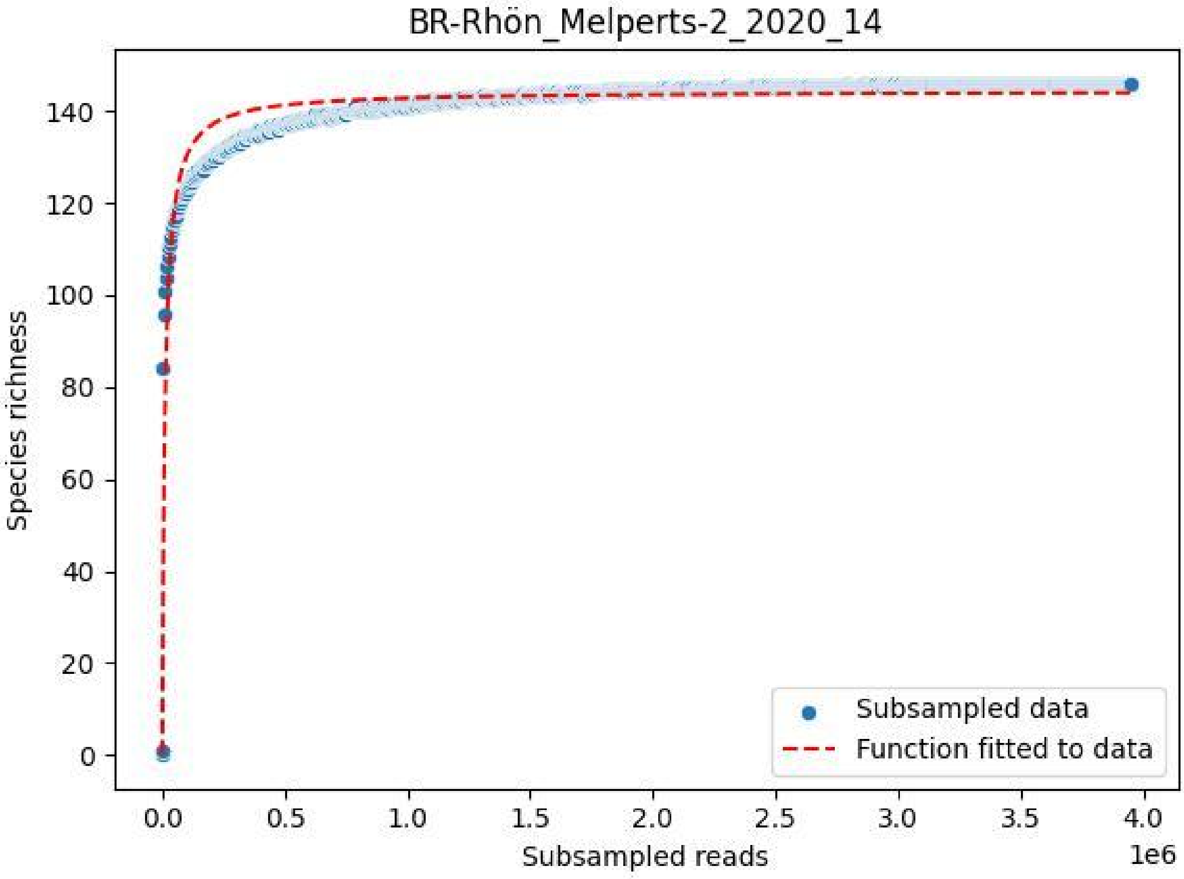
BR-Rhön_Melperts-2_2020_14
Species richness
Subsampled reads
1e6
0
20
40
60
80
100
120
140
0.0
0.5
1.0
1.5
2.0
2.5
3.0
3.5
4.0
Subsampled data
Function fitted to data

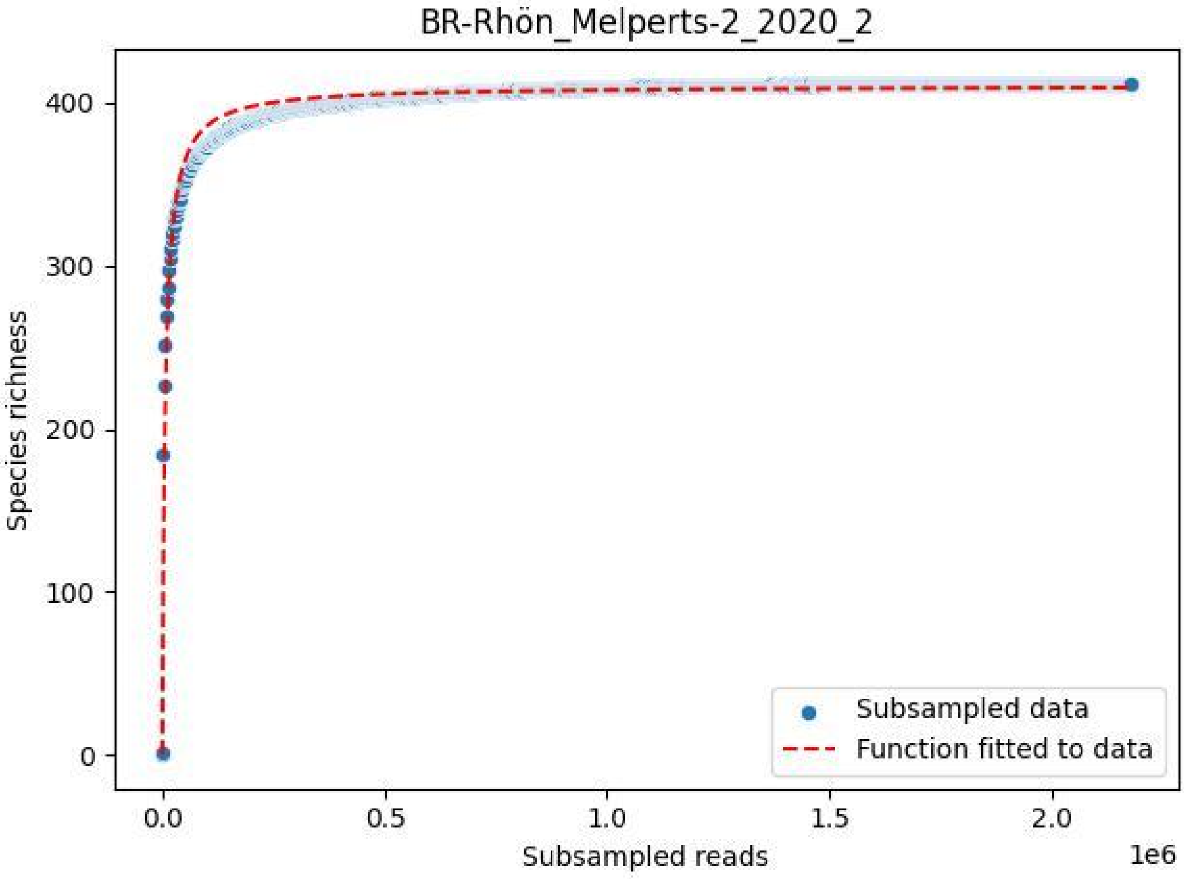

BR-Rhön_Melperts-2_2020_2
Species richness
400
300
200
100
0
0.0
0.5
1.0
1.5
2.0
1e6
Subsampled reads
Subsampled data
Function fitted to data

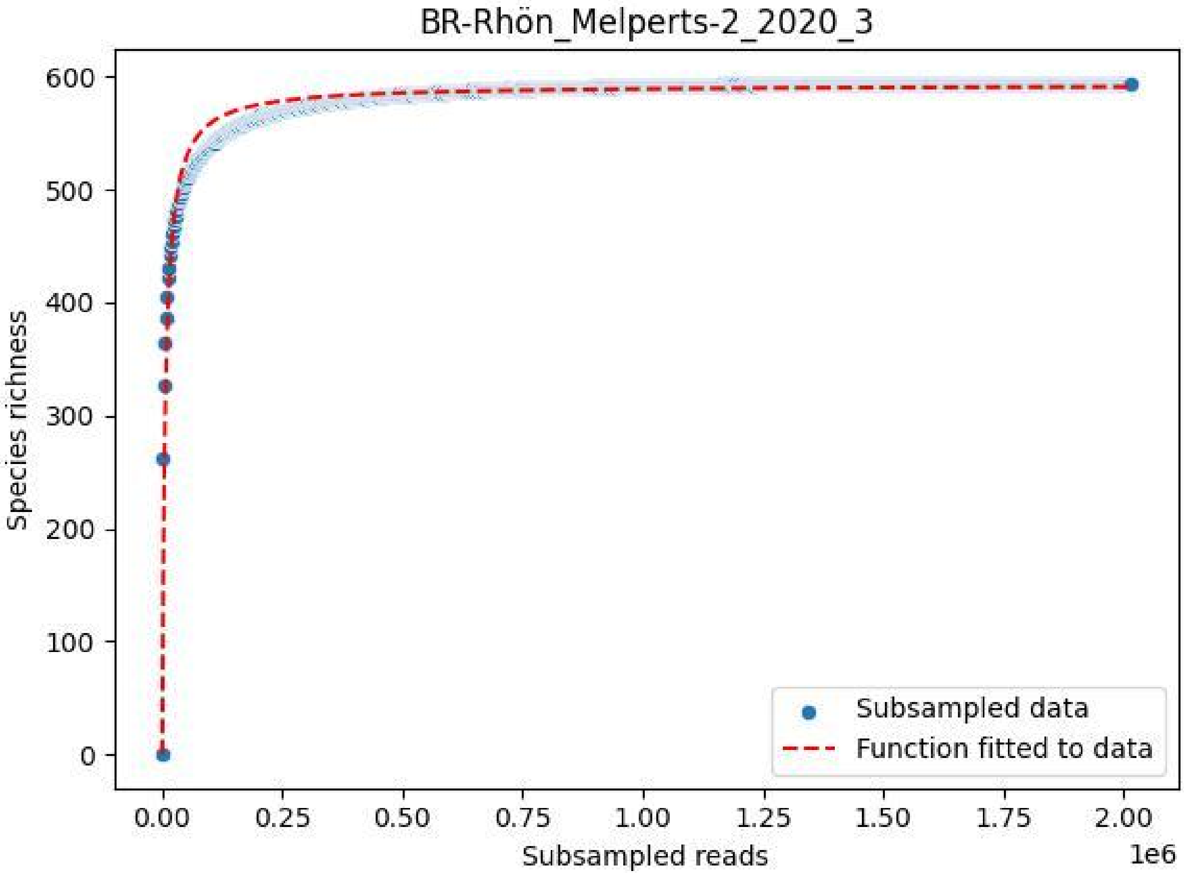
BR-Rhön_Melperts-2_2020_3
Species richness
600
500
400
300
200
100
0
0.00
0.25
0.50
0.75
1.00
1.25
1.50
1.75
2.00
1e6
Subsampled reads
Subsampled data
Function fitted to data

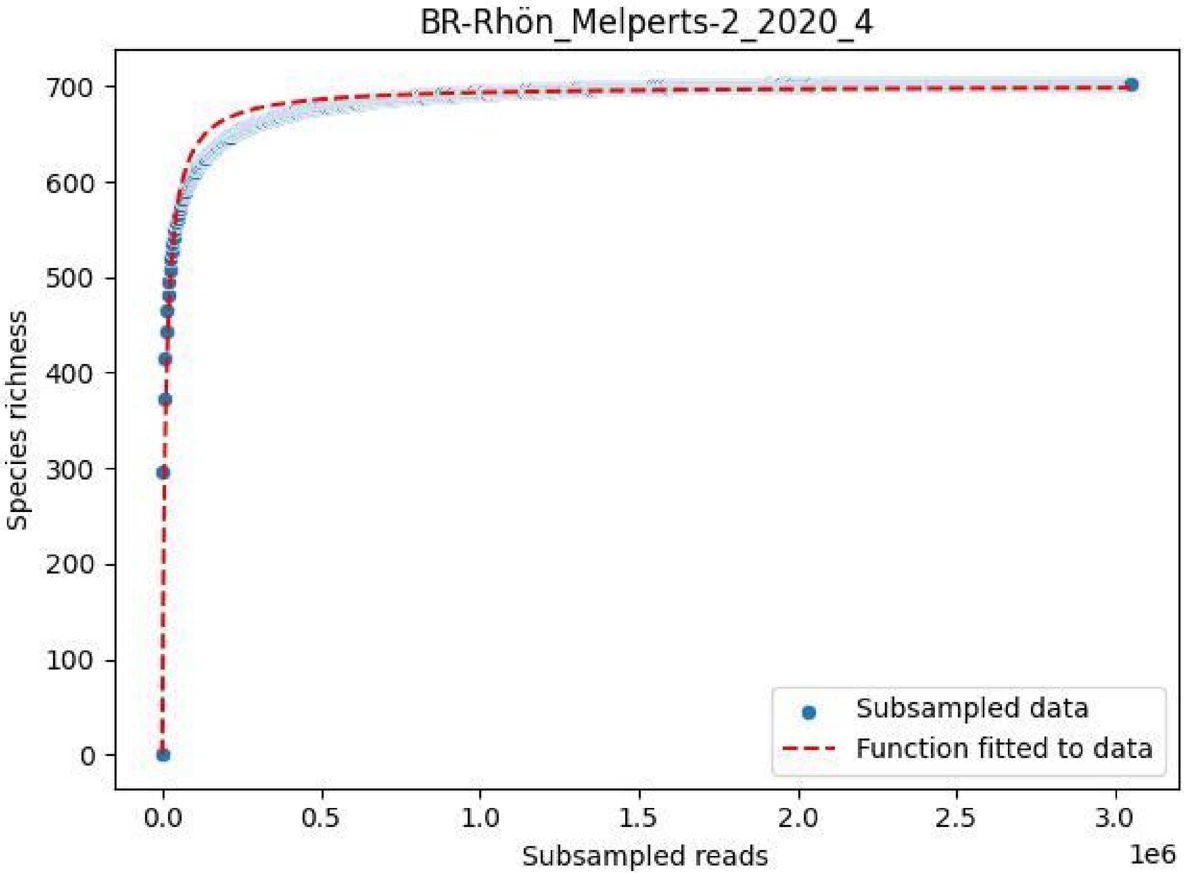

BR-Rhön_Melperts-2_2020_4
Species richness
Subsampled reads
1e6
0
100
200
300
400
500
600
700
0.0
0.5
1.0
1.5
2.0
2.5
3.0
Subsampled data
Function fitted to data

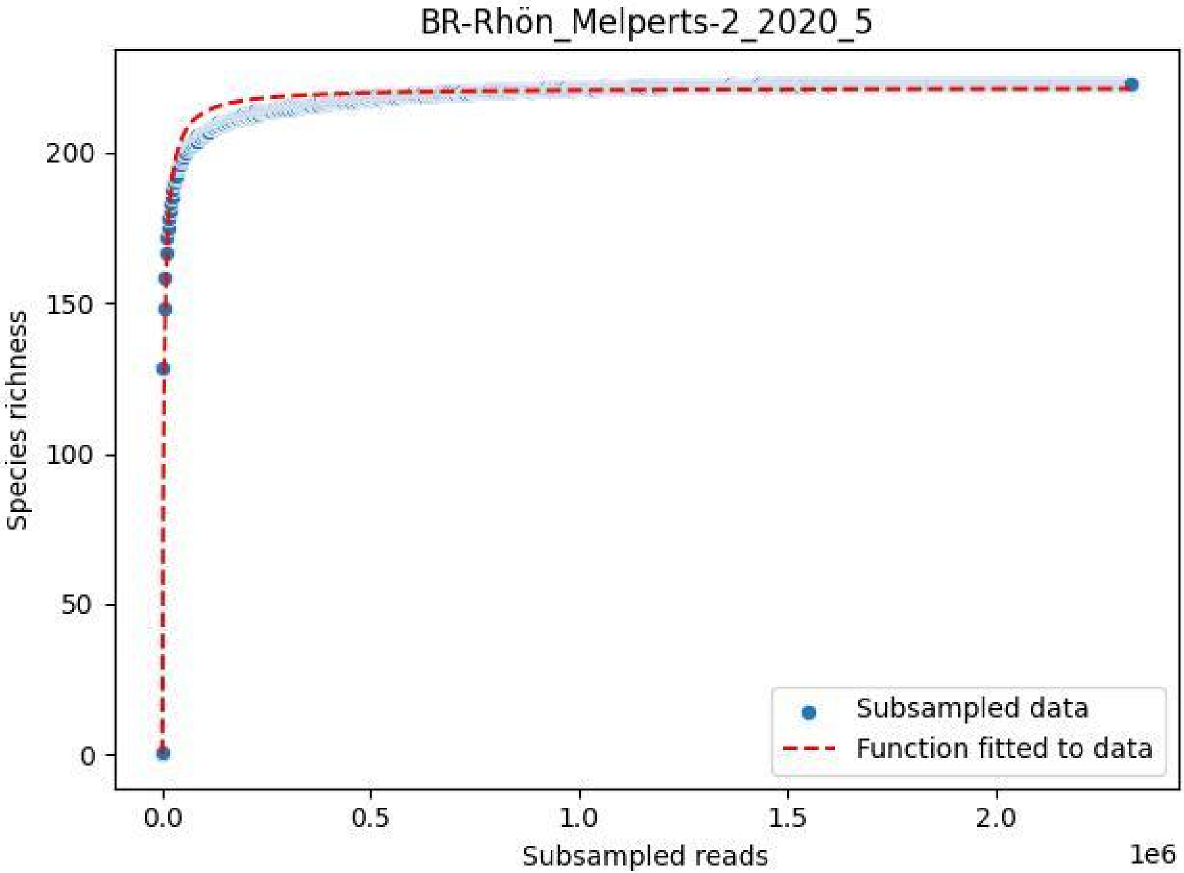

BR-Rhön_Melperts-2_2020_5
Species richness
Subsampled reads
1e6
0
50
100
150
200
0.0
0.5
1.0
1.5
2.0
Subsampled data
Function fitted to data

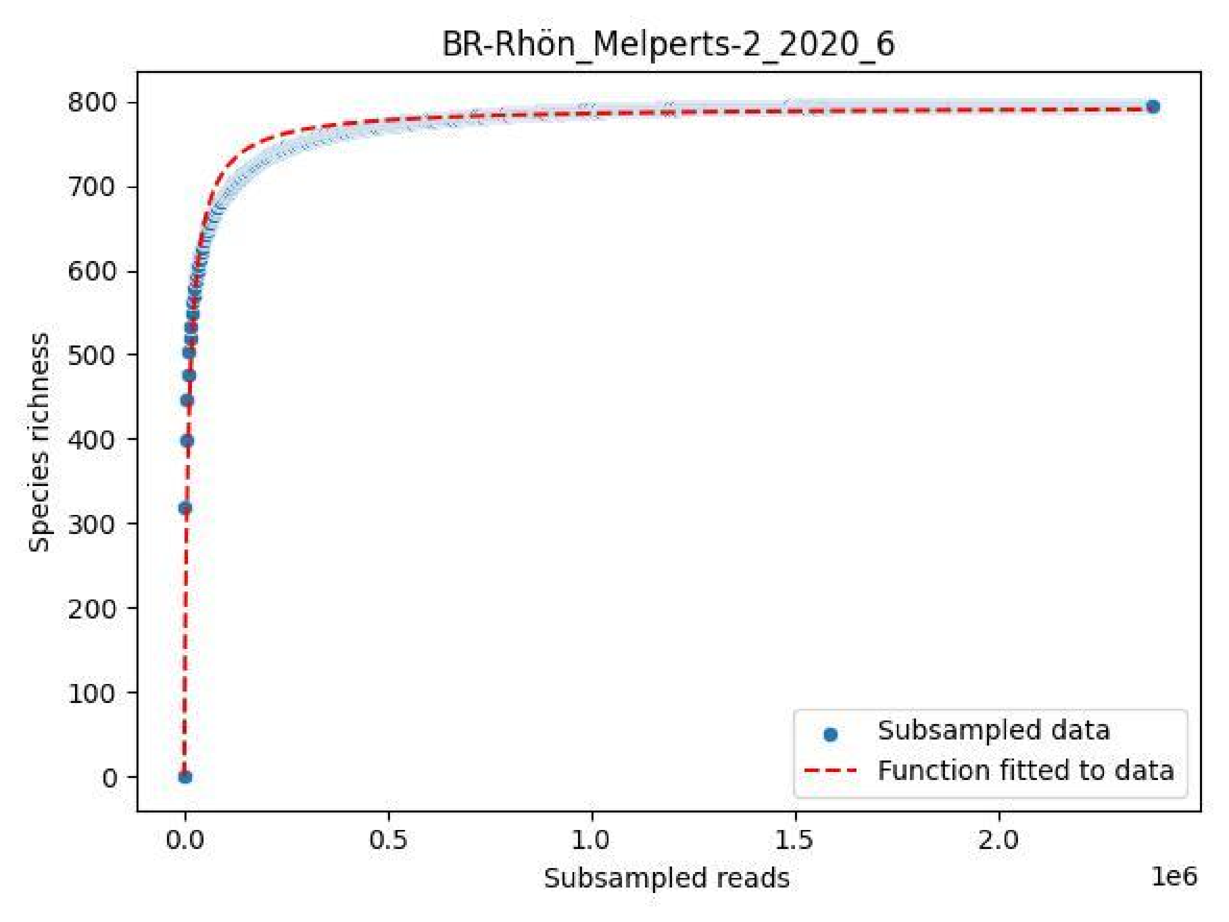
BR-Rhön_Melperts-2_2020_6
Species richness
800
700
600
500
400
300
200
100
0
0.0
0.5
1.0
1.5
2.0
1e6
Subsampled reads
Subsampled data
Function fitted to data

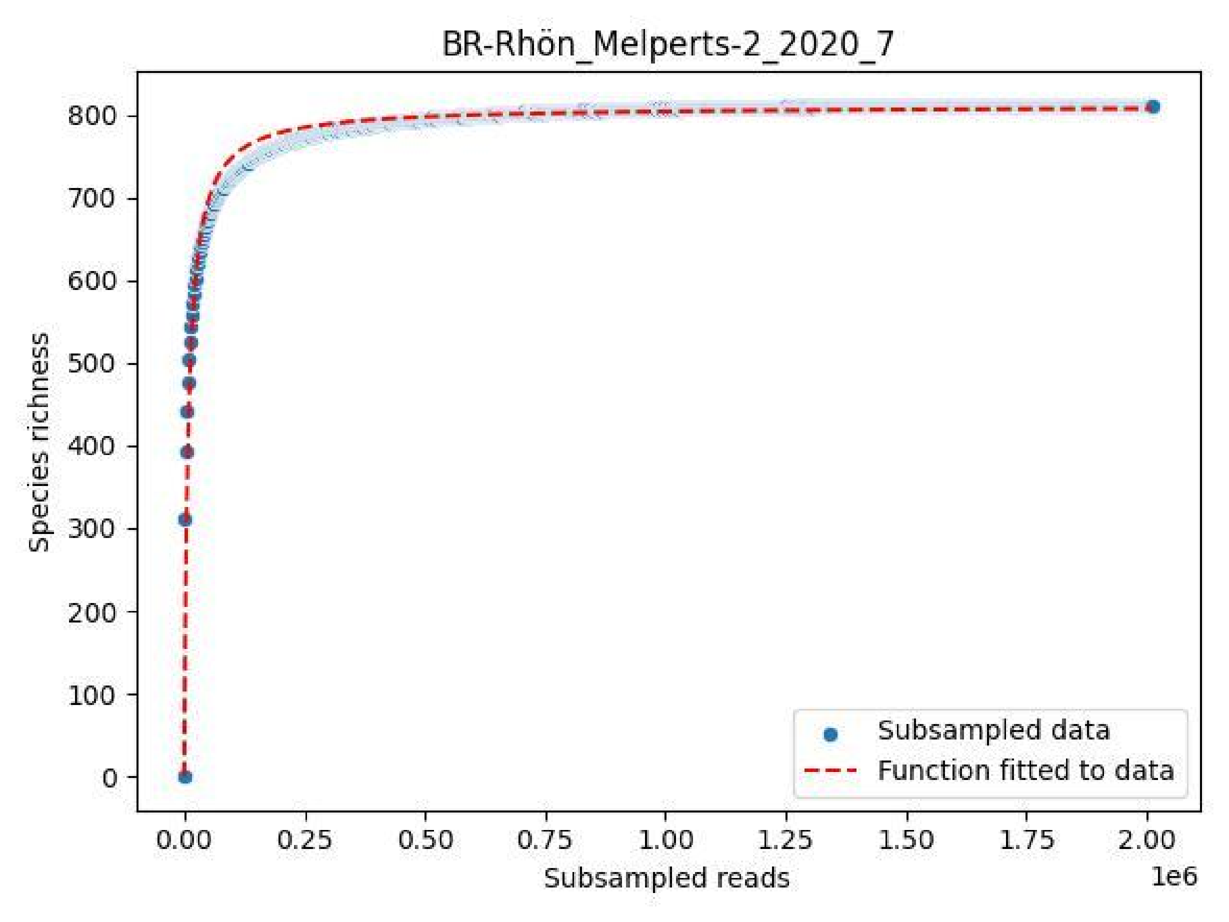
BR-Rhön_Melperts-2_2020_7
Species richness
0
100
200
300
400
500
600
700
800
0.00
0.25
0.50
0.75
1.00
1.25
1.50
1.75
2.00
1e6
Subsampled reads
Subsampled data
Function fitted to data

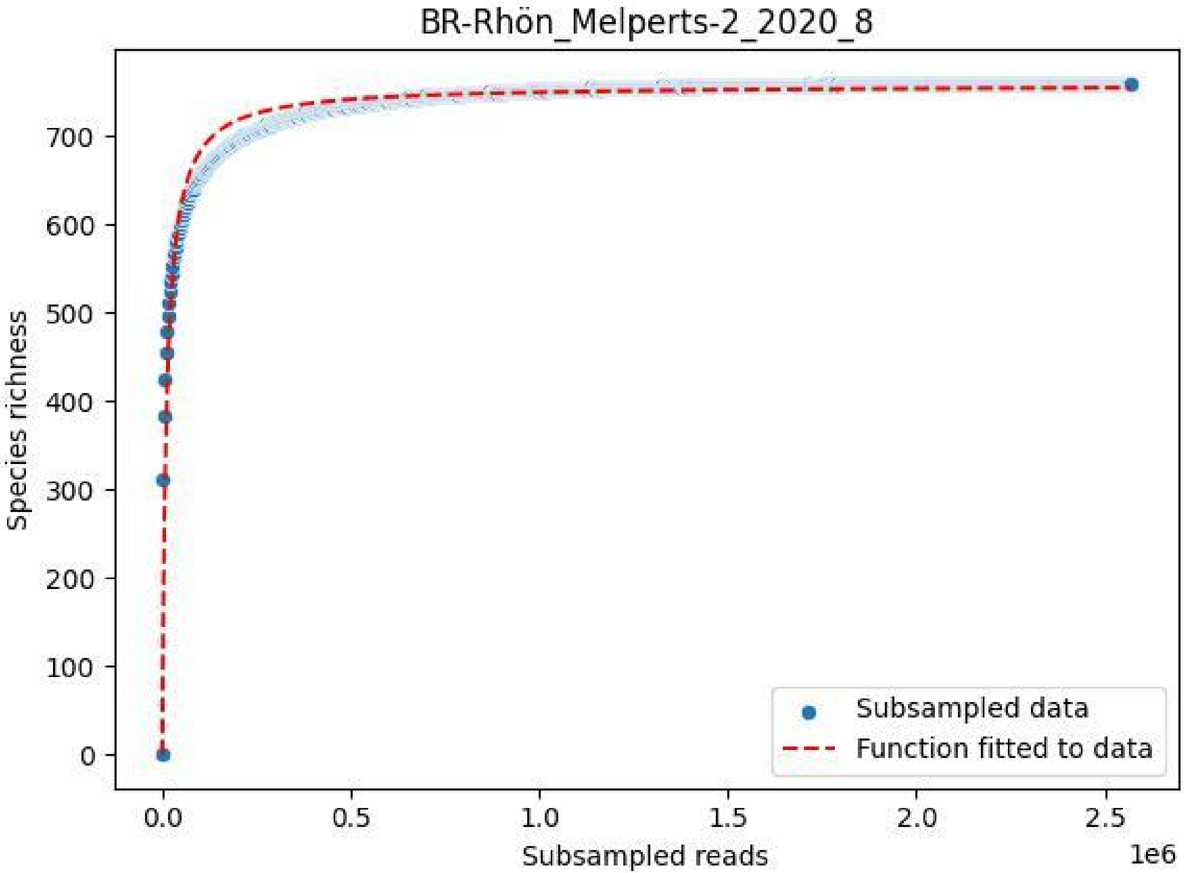
BR-Rhön_Melperts-2_2020_8
Species richness
Subsampled reads
1e6
0
100
200
300
400
500
600
700
0.0
0.5
1.0
1.5
2.0
2.5
Subsampled data
Function fitted to data

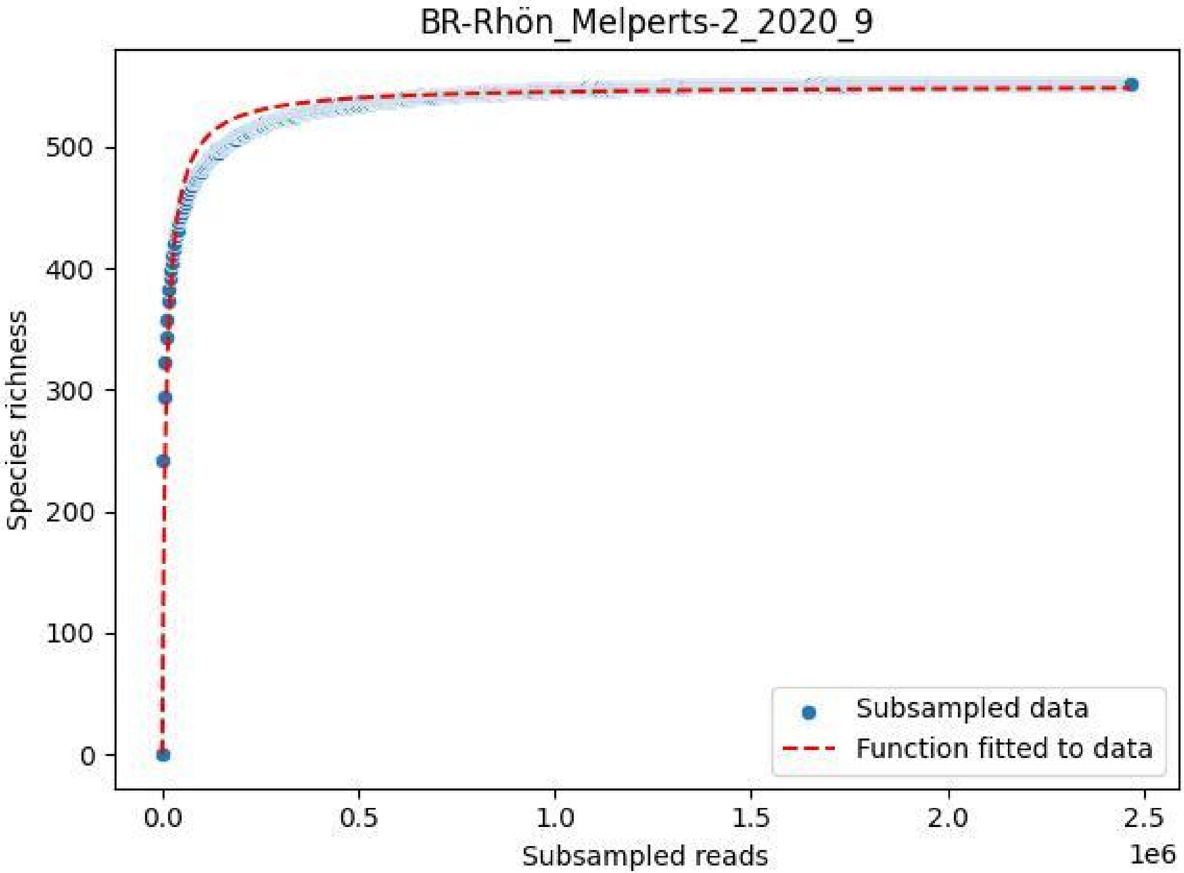
BR-Rhön_Melperts-2_2020_9
Species richness
Subsampled reads
1e6
0
100
200
300
400
500
0.0
0.5
1.0
1.5
2.0
2.5
Subsampled data
Function fitted to data

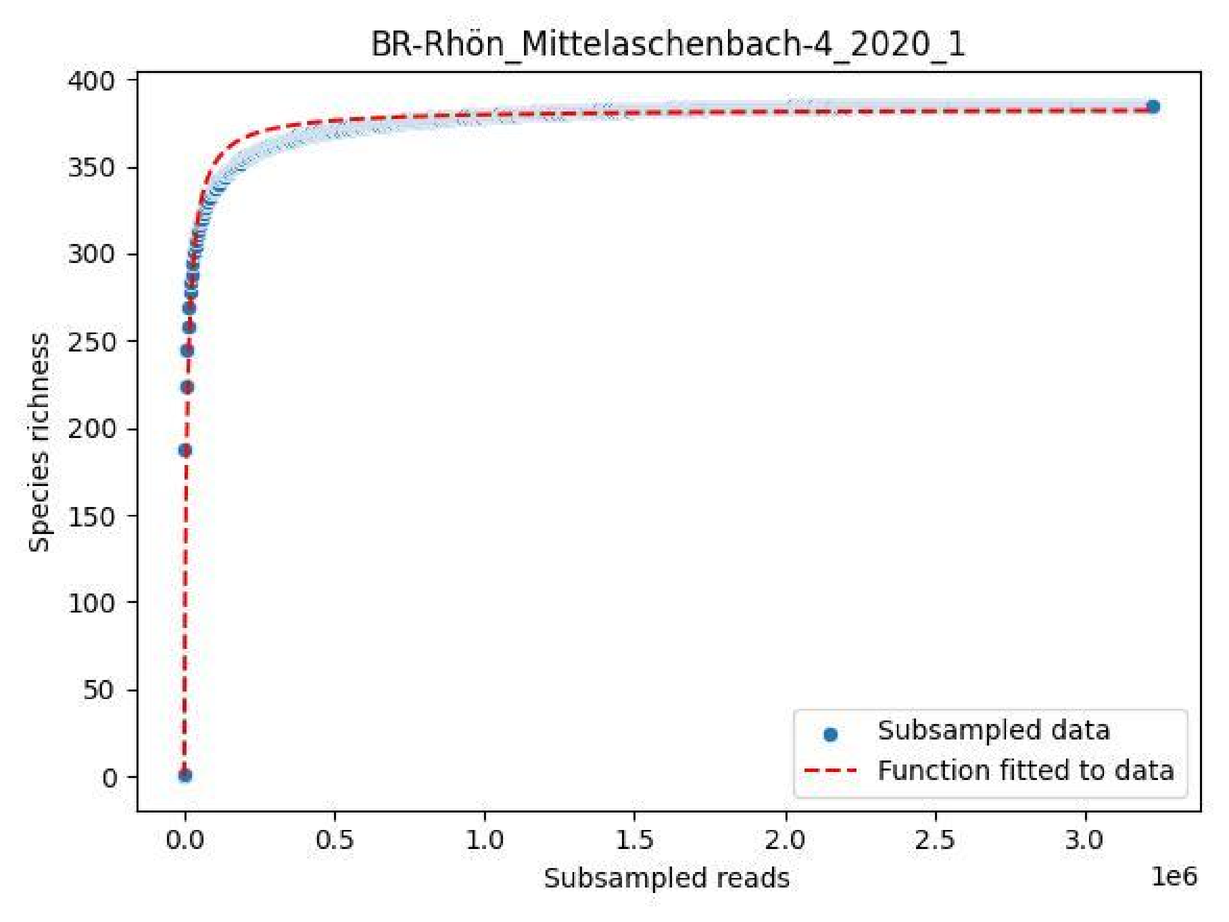

BR-Rhön_Mittelaschenbach-4_2020_1
Species richness
400
350
300
250
200
150
100
50
0
0.0
0.5
1.0
1.5
2.0
2.5
3.0
Subsampled reads
1e6
Subsampled data
Function fitted to data

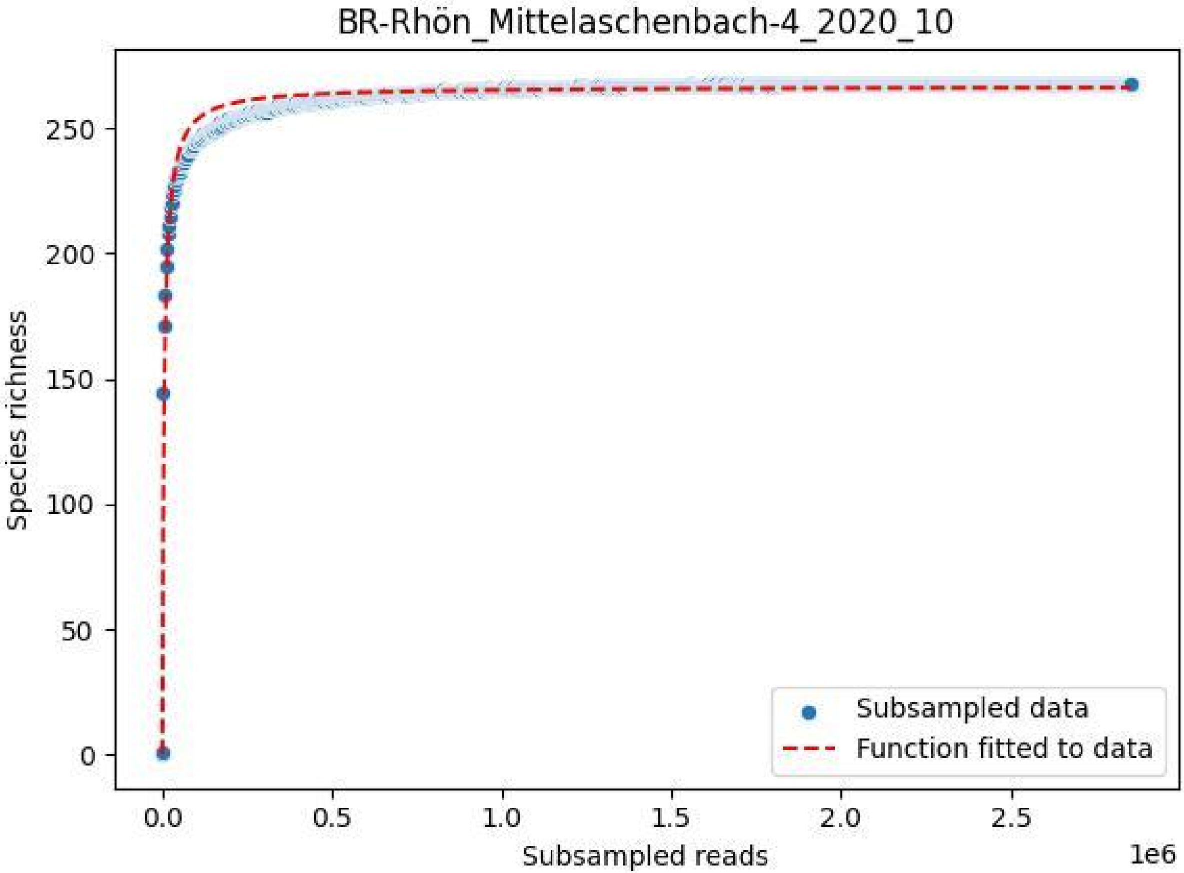

BR-Rhön_Mittelaschenbach-4_2020_10
Species richness
Subsampled reads
1e6
0
50
100
150
200
250
0.0
0.5
1.0
1.5
2.0
2.5
Subsampled data
Function fitted to data

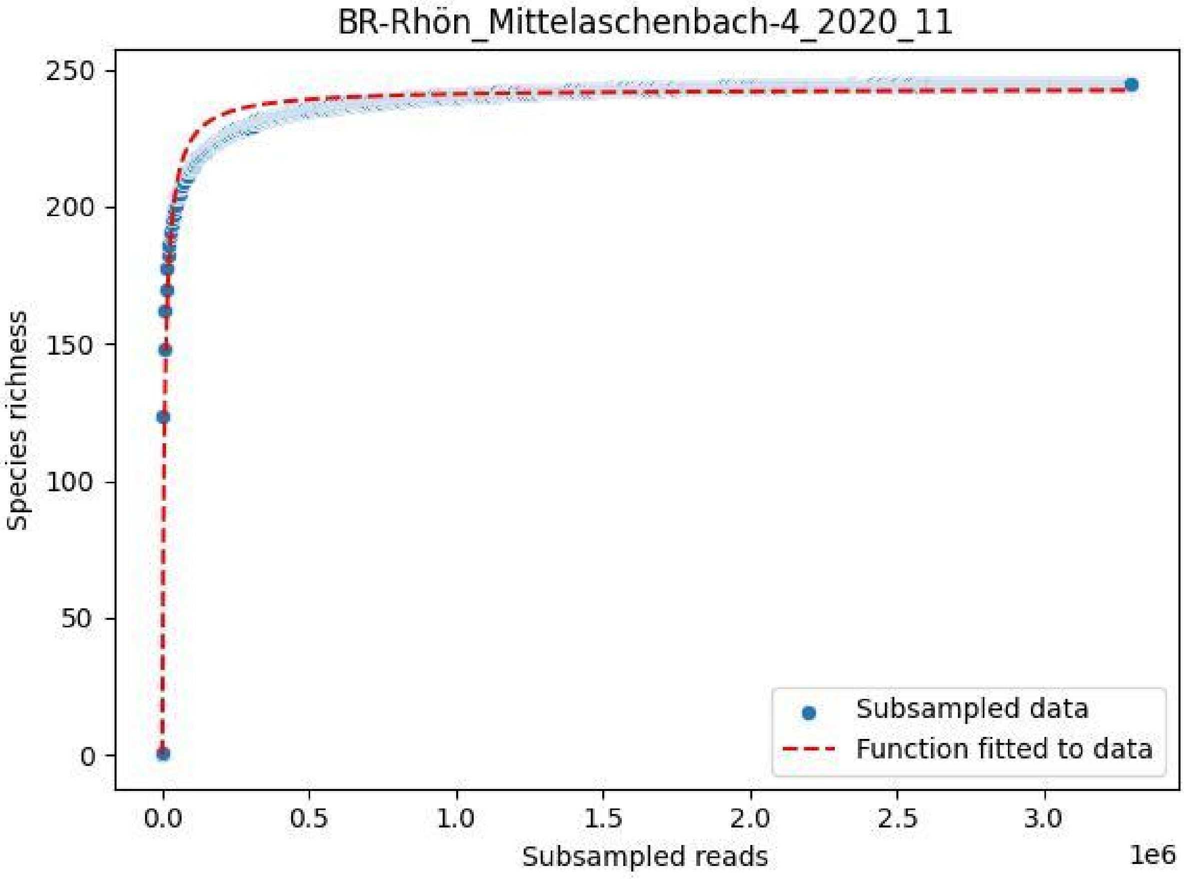
BR-Rhön_Mittelaschenbach-4_2020_11
Species richness
0
50
100
150
200
250
0.0
0.5
1.0
1.5
2.0
2.5
3.0
1e6
Subsampled reads
Subsampled data
Function fitted to data

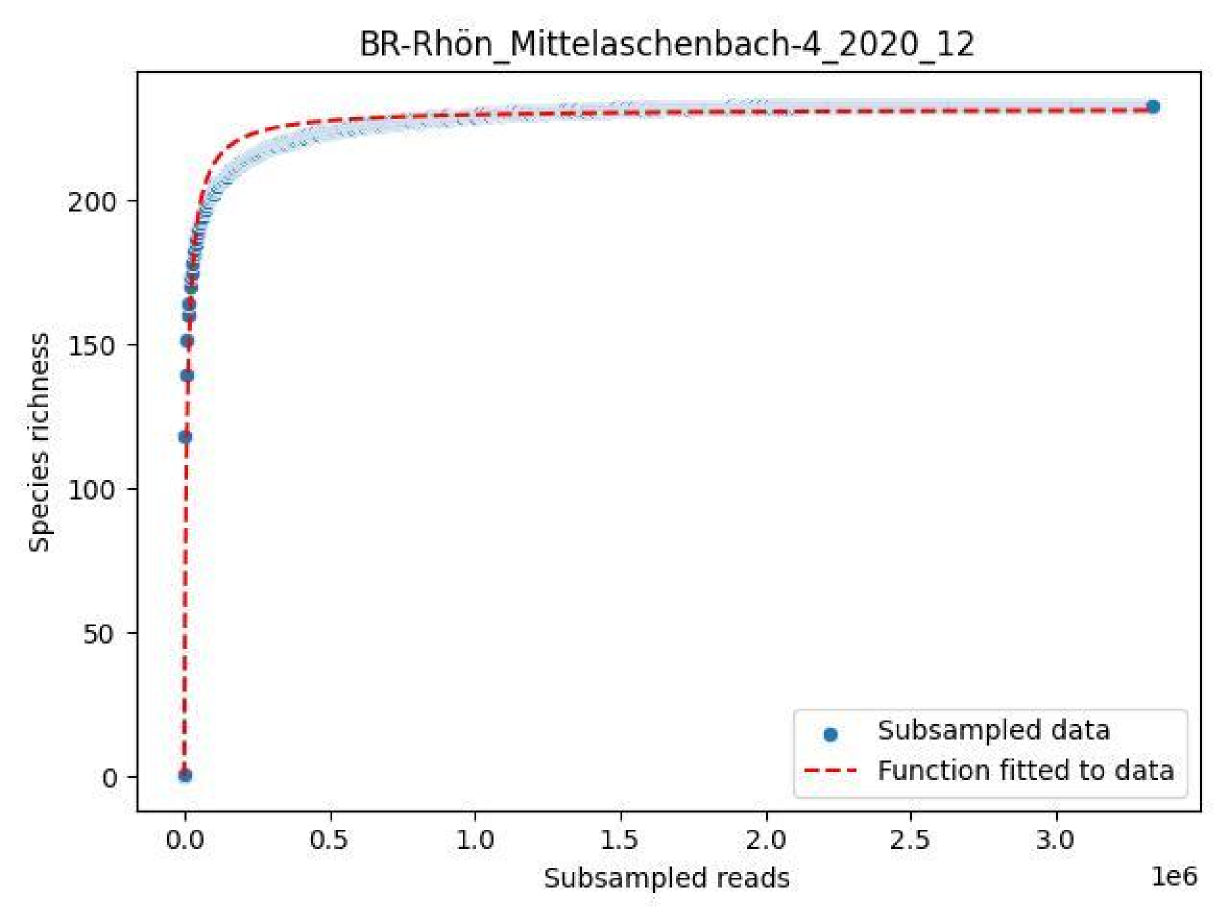
BR-Rhön_Mittelaschenbach-4_2020_12
Species richness
0
50
100
150
200
0.0
0.5
1.0
1.5
2.0
2.5
3.0
1e6
Subsampled reads
Subsampled data
Function fitted to data

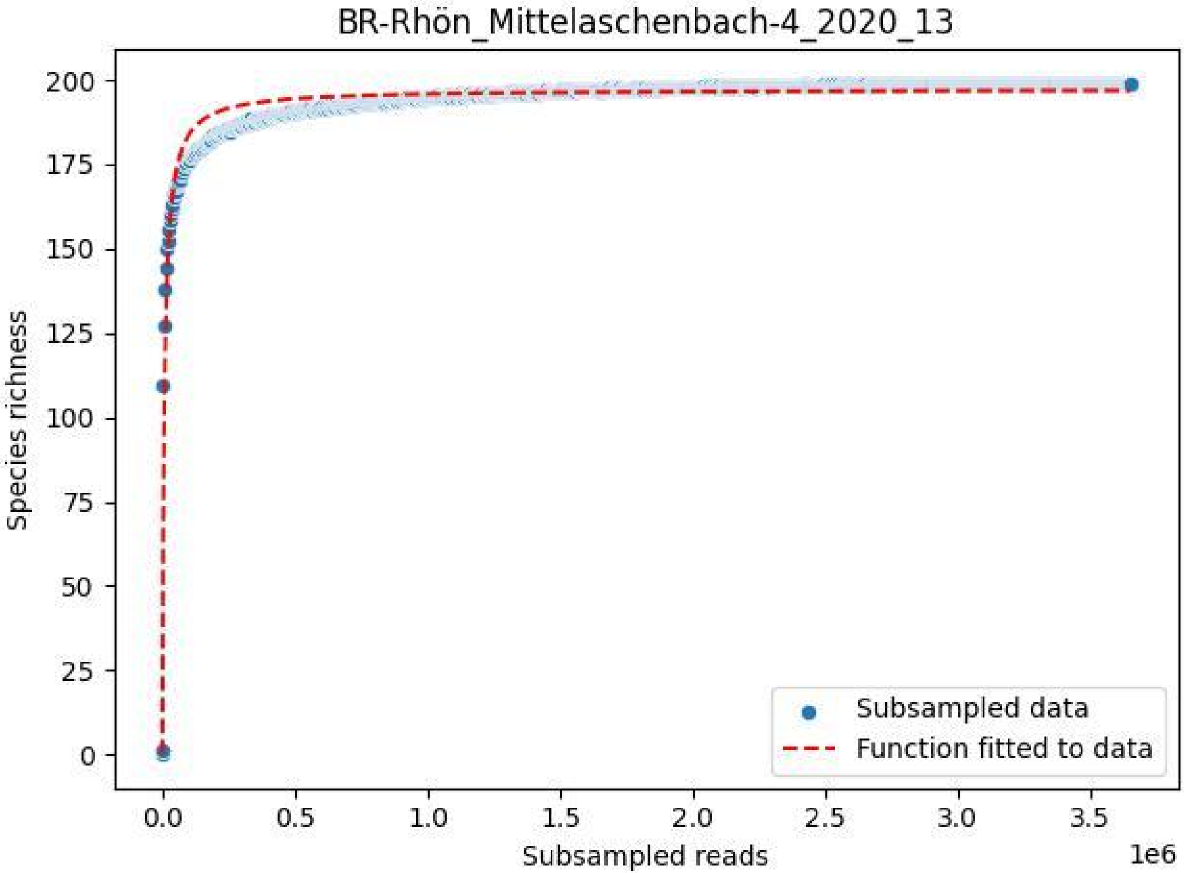

BR-Rhön_Mittelaschenbach-4_2020_13
Species richness
200
175
150
125
100
75
50
25
0
0.0
0.5
1.0
1.5
2.0
2.5
3.0
3.5
1e6
Subsampled reads
Subsampled data
Function fitted to data

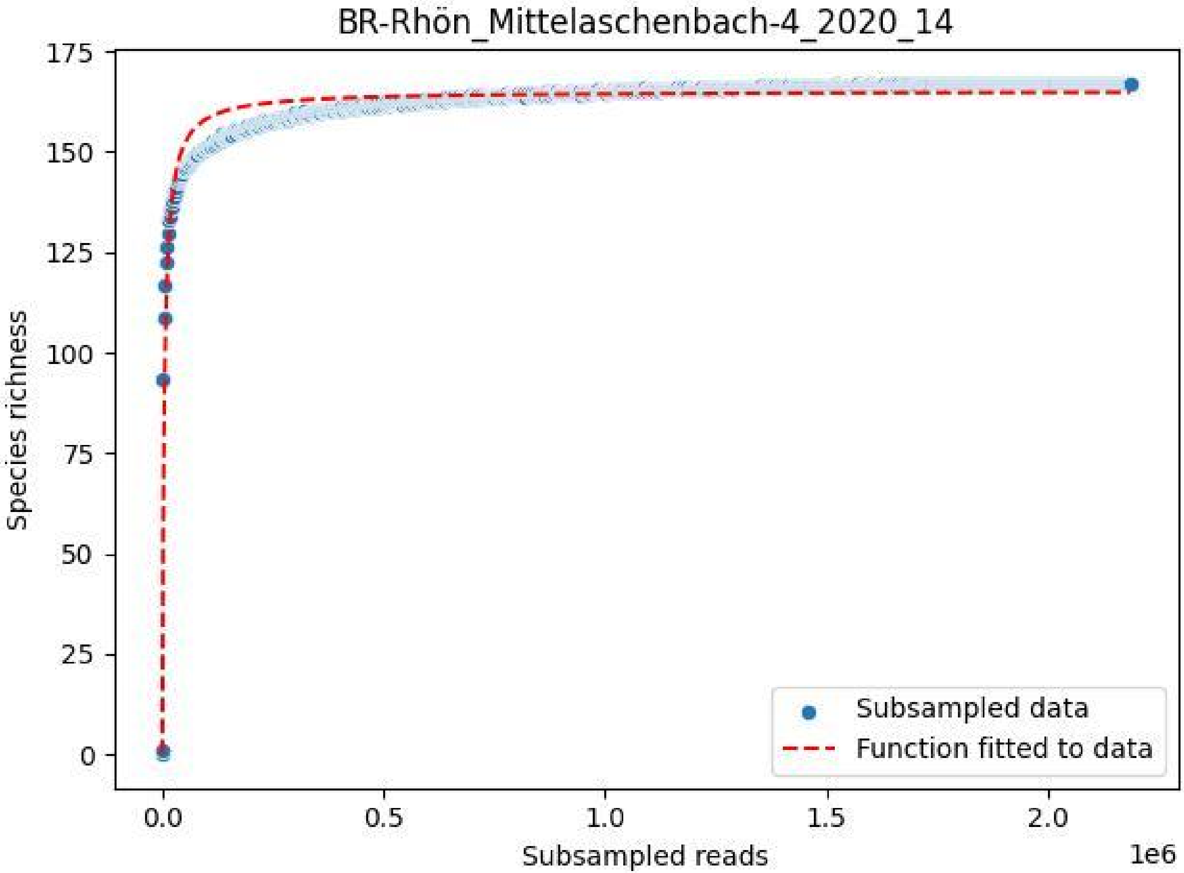
BR-Rhön_Mittelaschenbach-4_2020_14
Species richness
Subsampled reads
1e6
0
25
50
75
100
125
150
175
0.0
0.5
1.0
1.5
2.0
Subsampled data
Function fitted to data

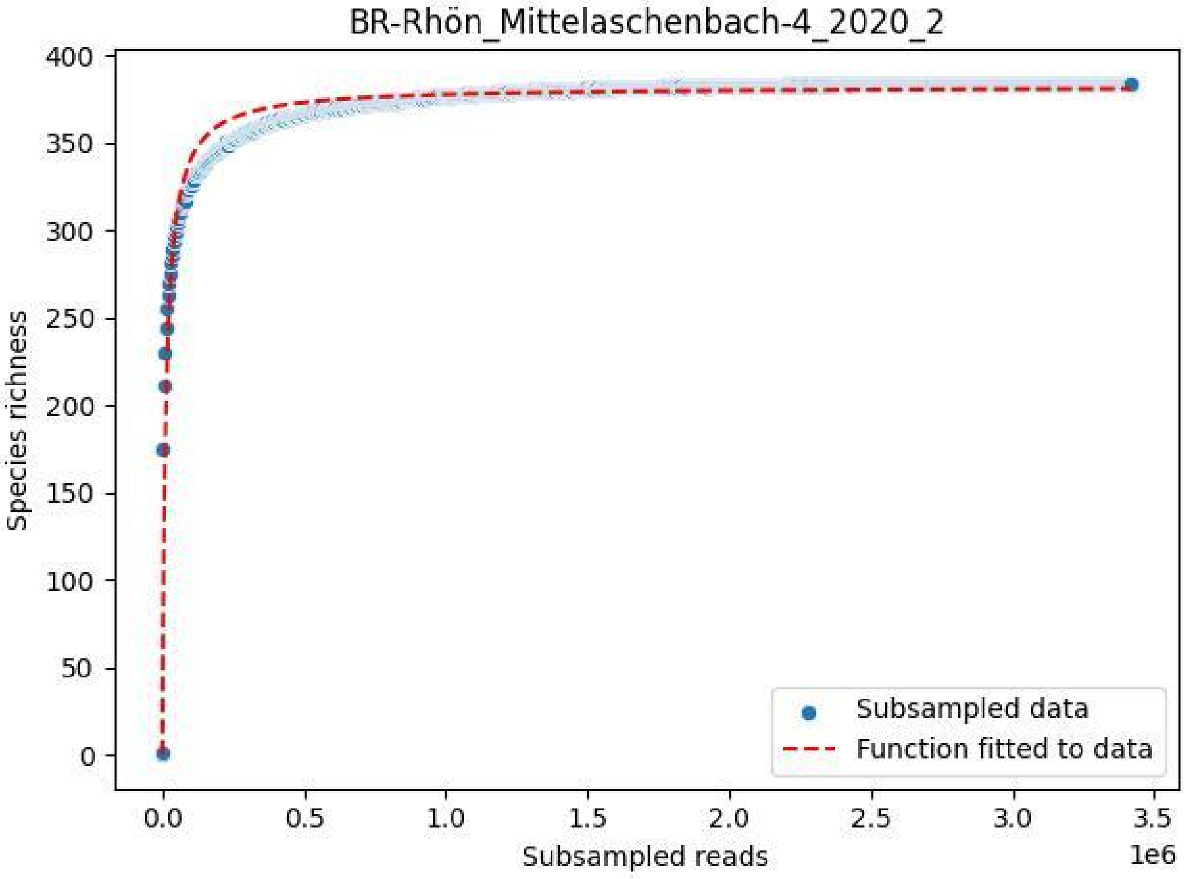
BR-Rhön_Mittelaschenbach-4_2020_2
Species richness
Subsampled reads
1e6
0
50
100
150
200
250
300
350
400
0.0
0.5
1.0
1.5
2.0
2.5
3.0
3.5
Subsampled data
Function fitted to data

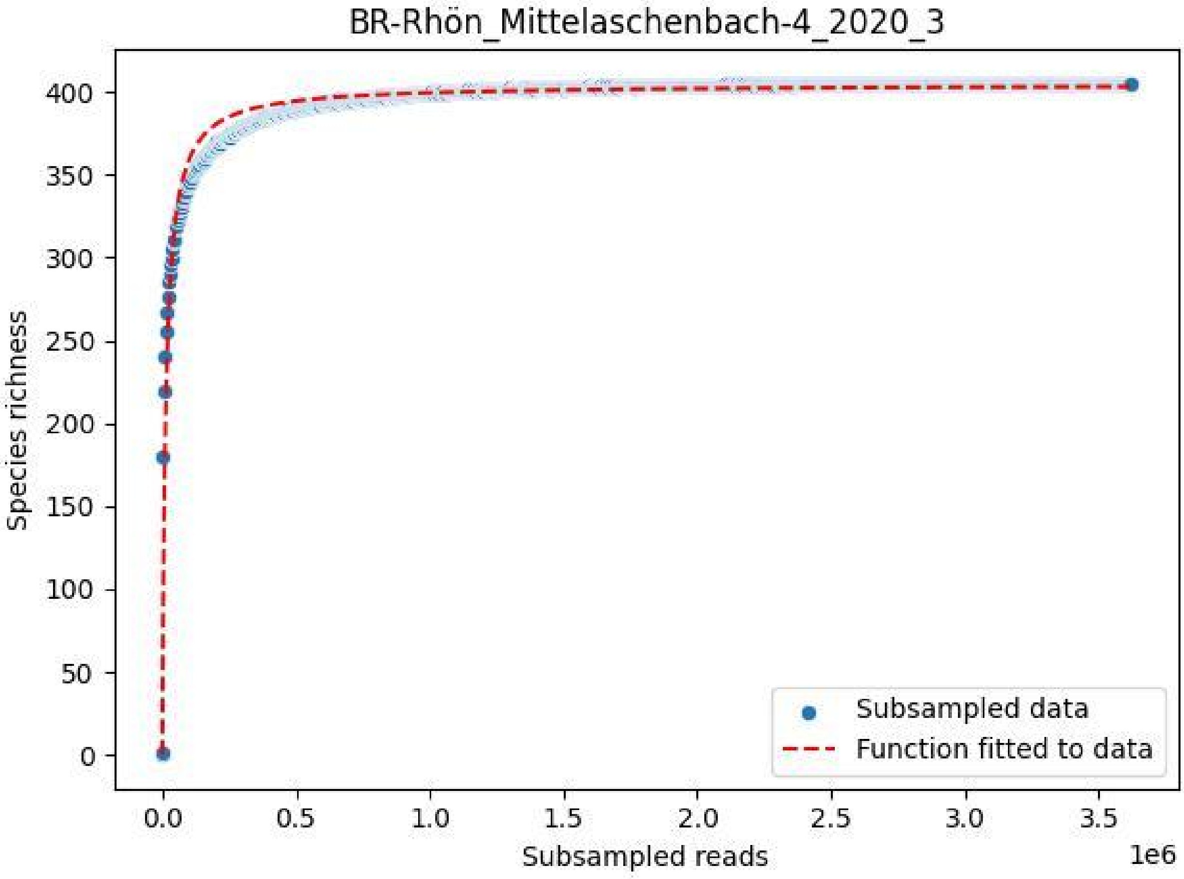

BR-Rhön_Mittelaschenbach-4_2020_3
Species richness
400
350
300
250
200
150
100
50
0
0.0
0.5
1.0
1.5
2.0
2.5
3.0
3.5
1e6
Subsampled reads
Subsampled data
Function fitted to data

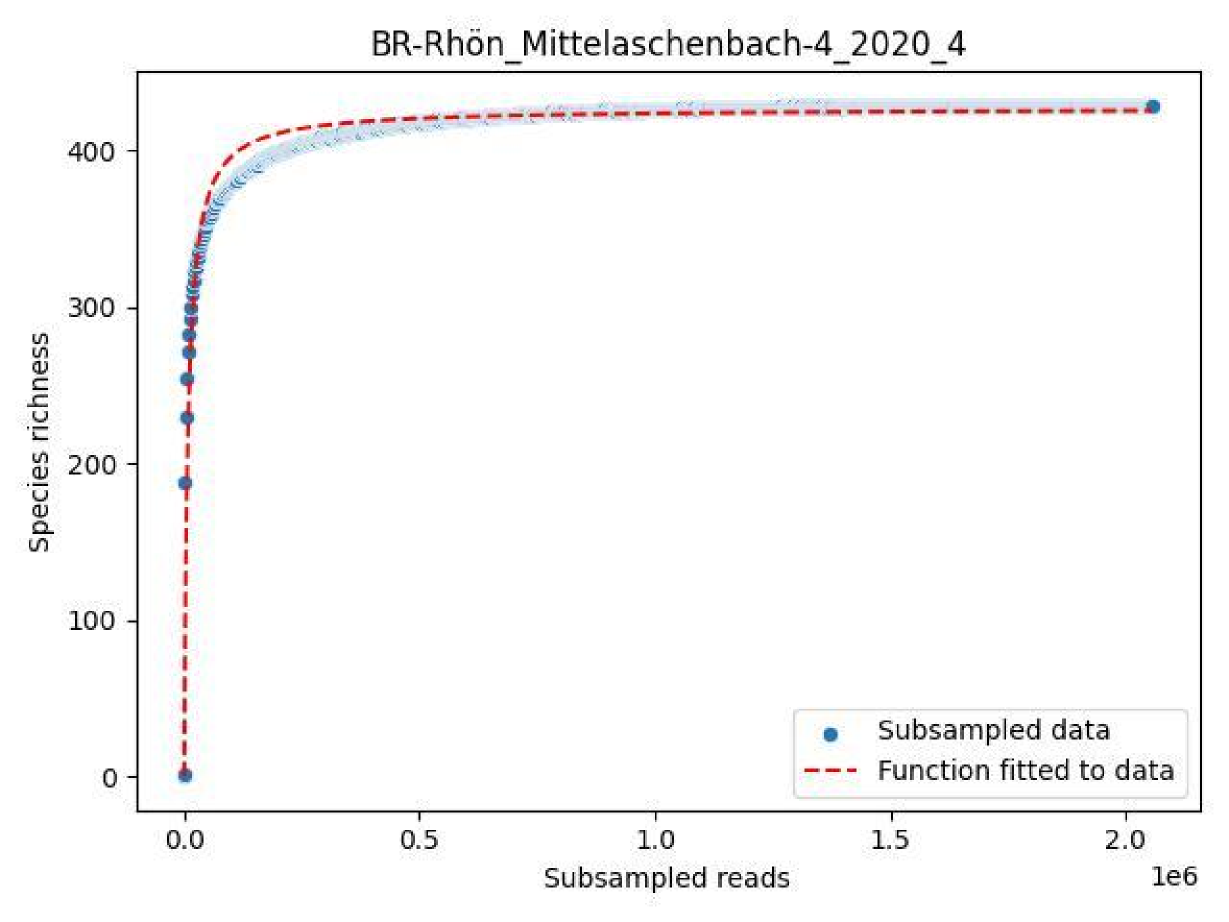
BR-Rhön_Mittelaschenbach-4_2020_4
Species richness
0
100
200
300
400
Subsampled reads
0.0
0.5
1.0
1.5
2.0
1e6
Subsampled data
Function fitted to data

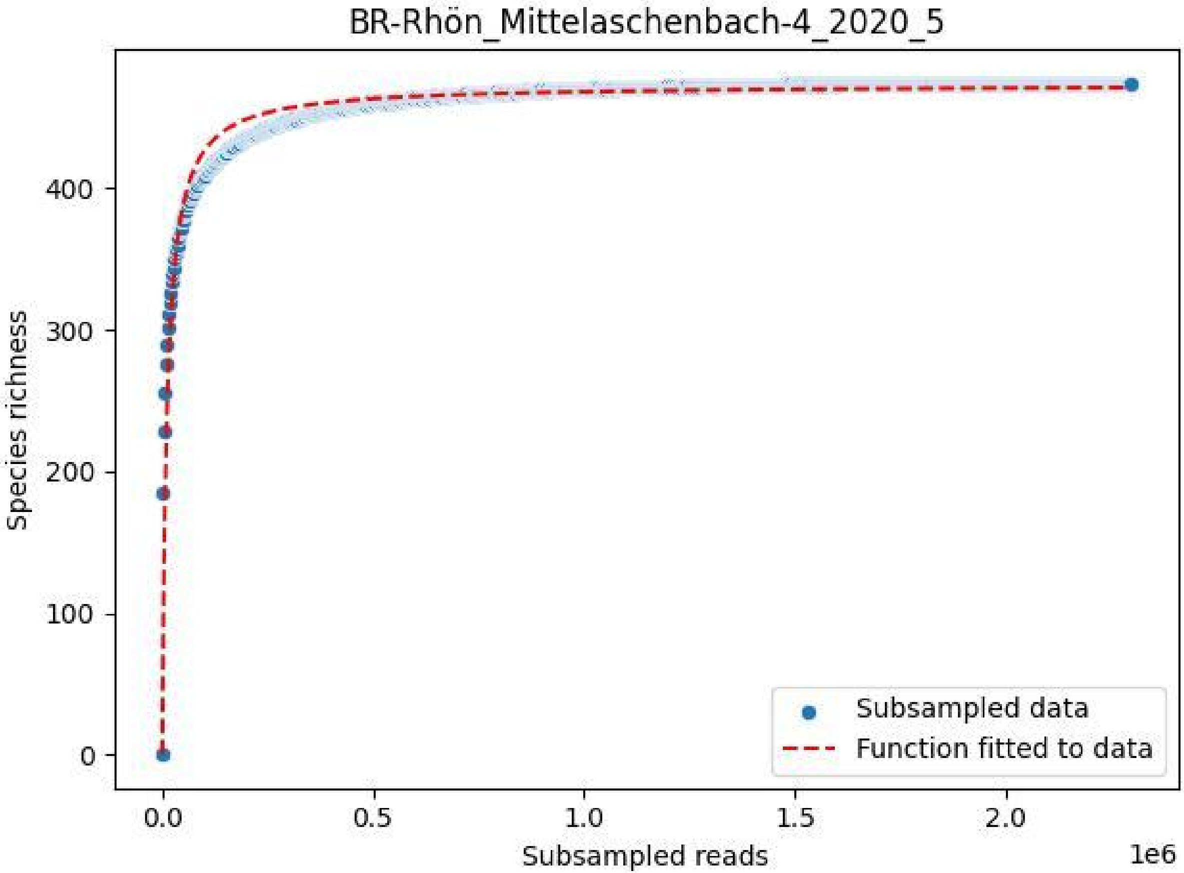

BR-Rhön_Mittelaschenbach-4_2020_5
Species richness
Subsampled reads
1e6
0
100
200
300
400
0.0
0.5
1.0
1.5
2.0
Subsampled data
Function fitted to data

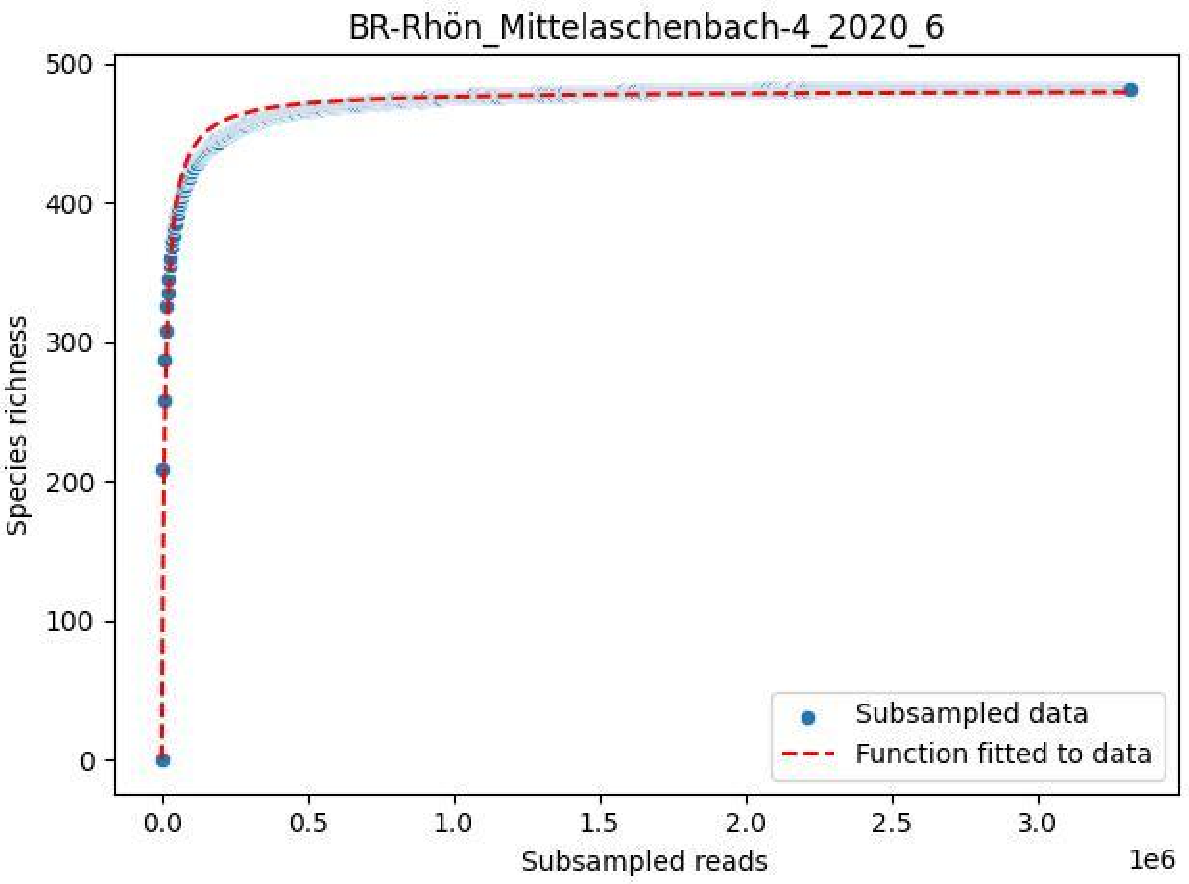

BR-Rhön_Mittelaschenbach-4_2020_6
Species richness
500
400
300
200
100
0
0.0
0.5
1.0
1.5
2.0
2.5
3.0
1e6
Subsampled reads
Subsampled data
Function fitted to data

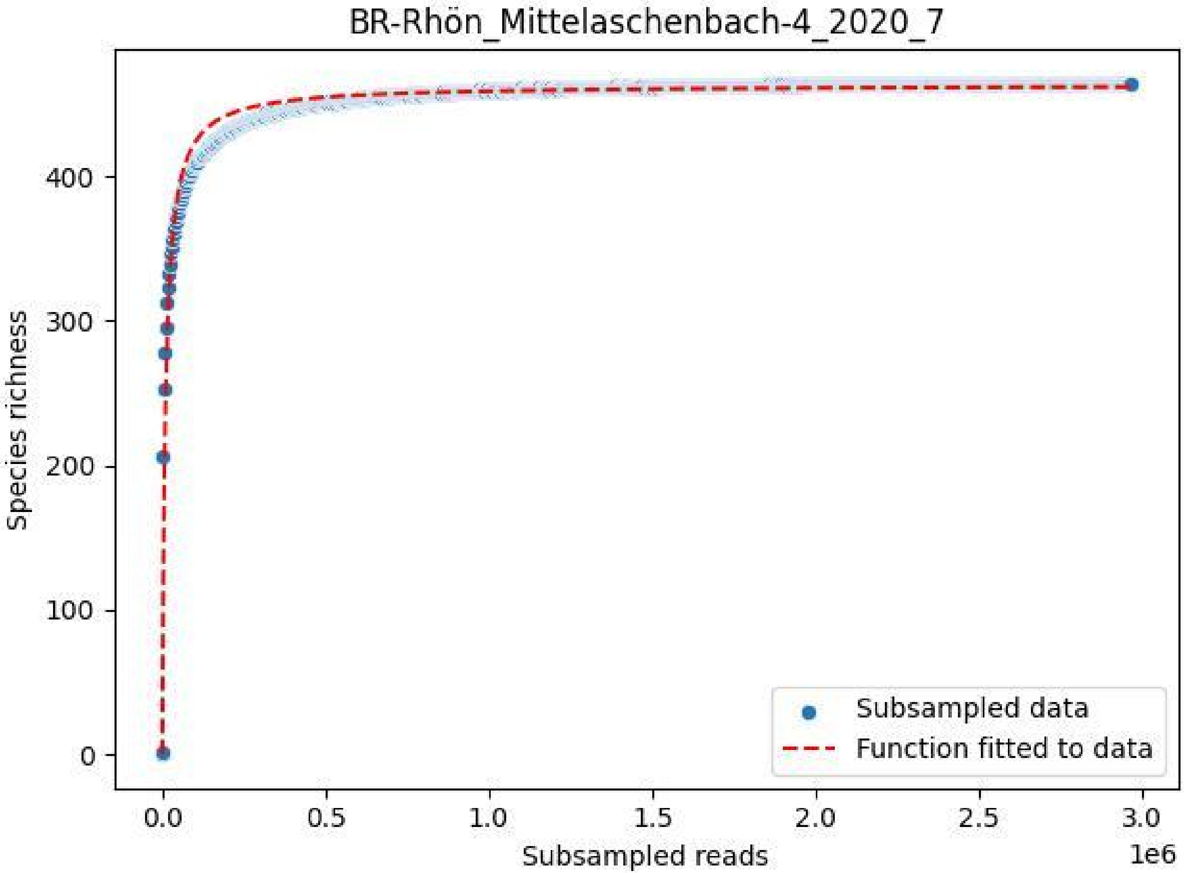
BR-Rhön_Mittelaschenbach-4_2020_7
Species richness
Subsampled reads
1e6
0
100
200
300
400
0.0
0.5
1.0
1.5
2.0
2.5
3.0
Subsampled data
Function fitted to data

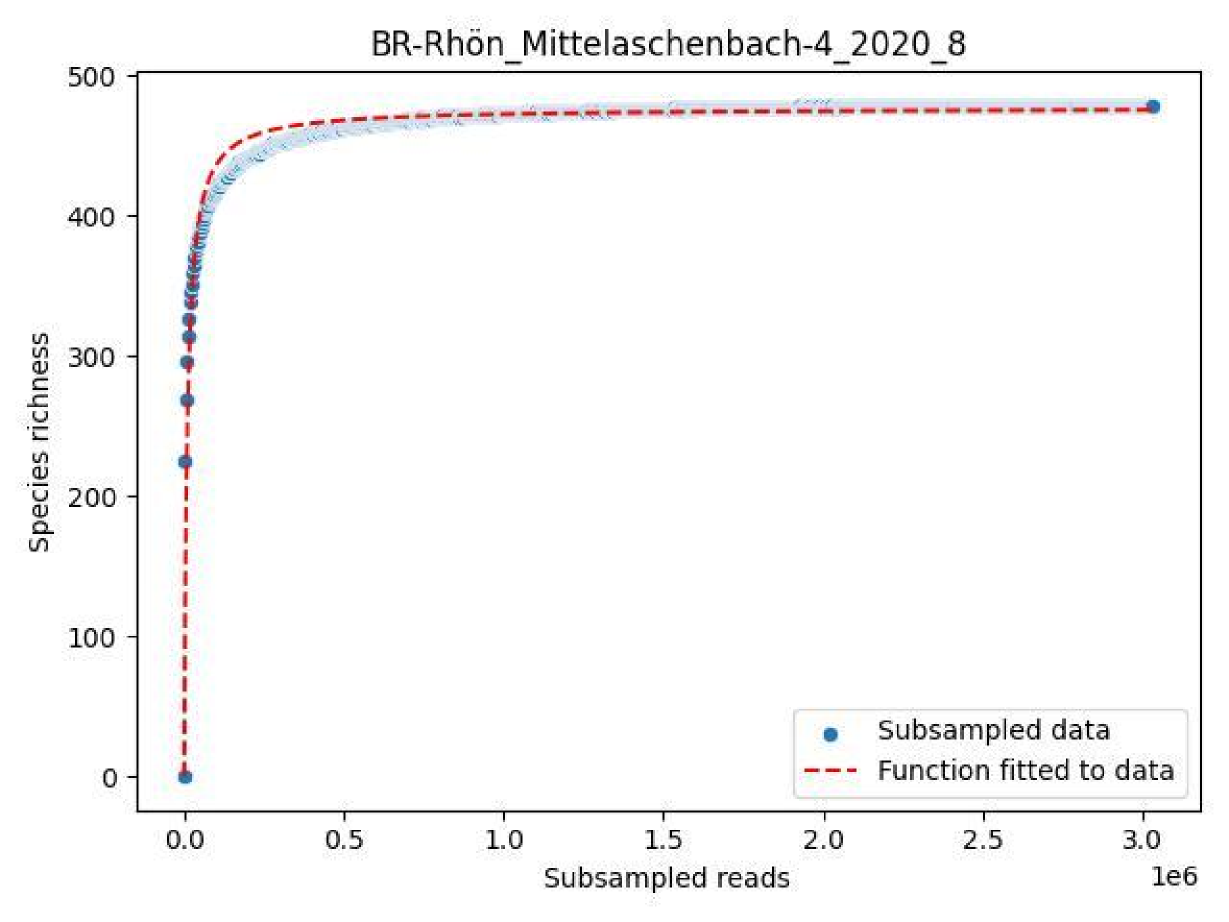

BR-Rhön_Mittelaschenbach-4_2020_8
Species richness
Subsampled reads
1e6
0
100
200
300
400
500
0.0
0.5
1.0
1.5
2.0
2.5
3.0
Subsampled data
Function fitted to data

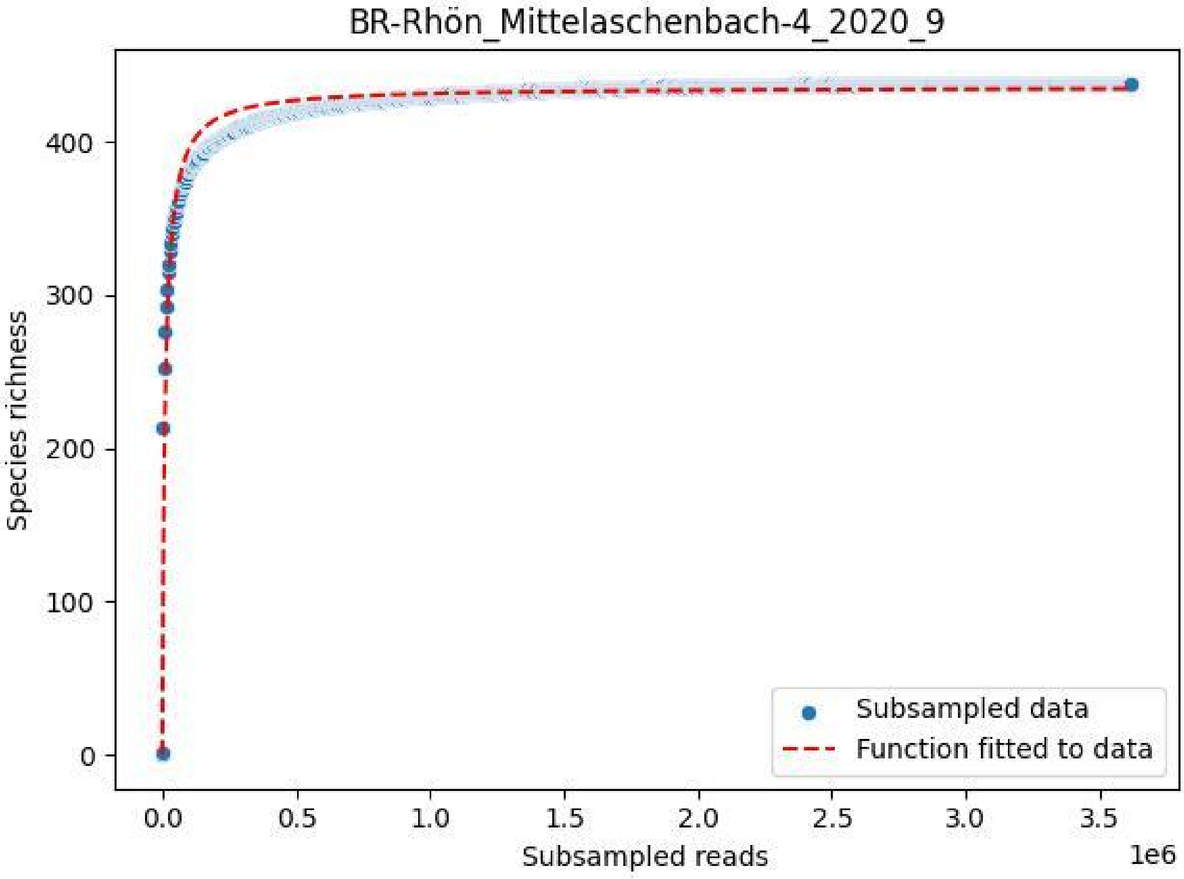
BR-Rhön_Mittelaschenbach-4_2020_9
Species richness
Subsampled reads
1e6
0
100
200
300
400
0.0
0.5
1.0
1.5
2.0
2.5
3.0
3.5
Subsampled data
Function fitted to data

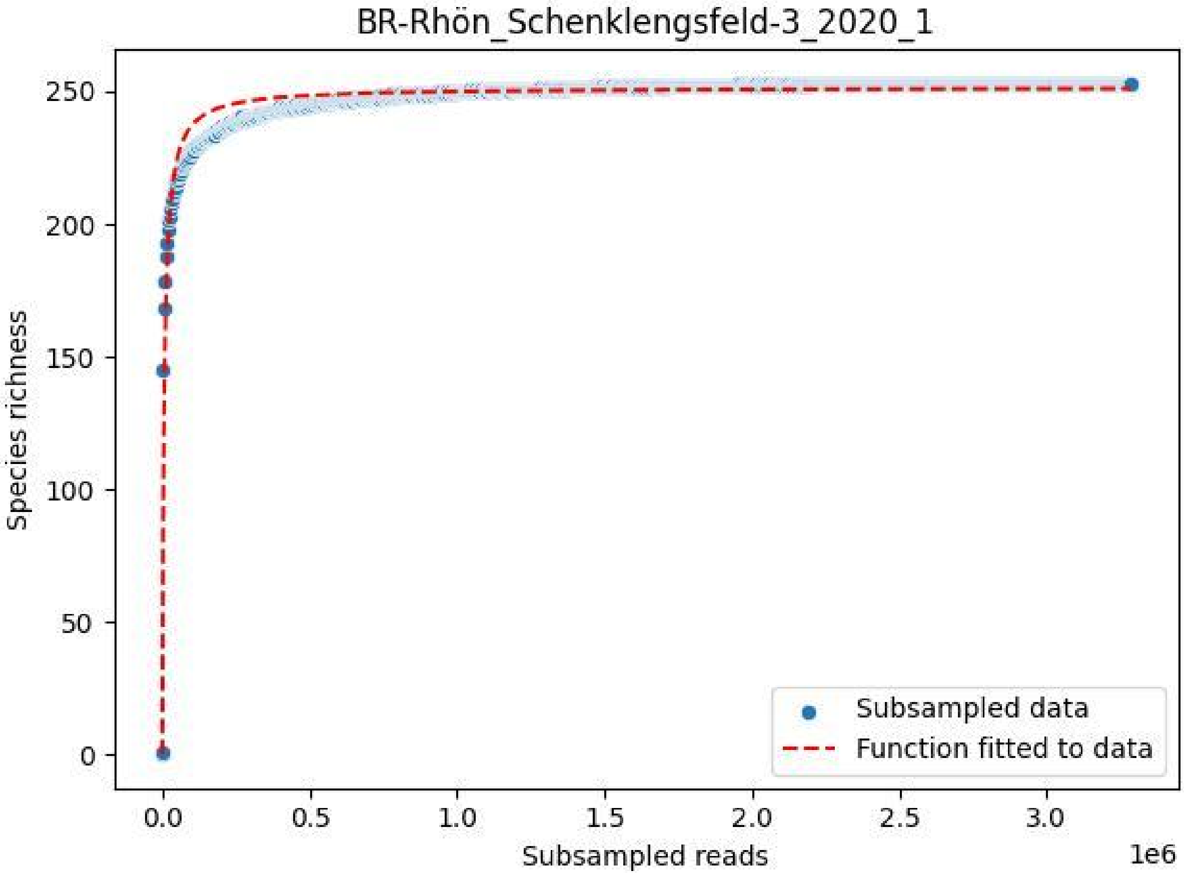
BR-Rhön_Schenklengsfeld-3_2020_1
Species richness
Subsampled reads
1e6
Subsampled data
Function fitted to data
0
50
100
150
200
250
0.0
0.5
1.0
1.5
2.0
2.5
3.0

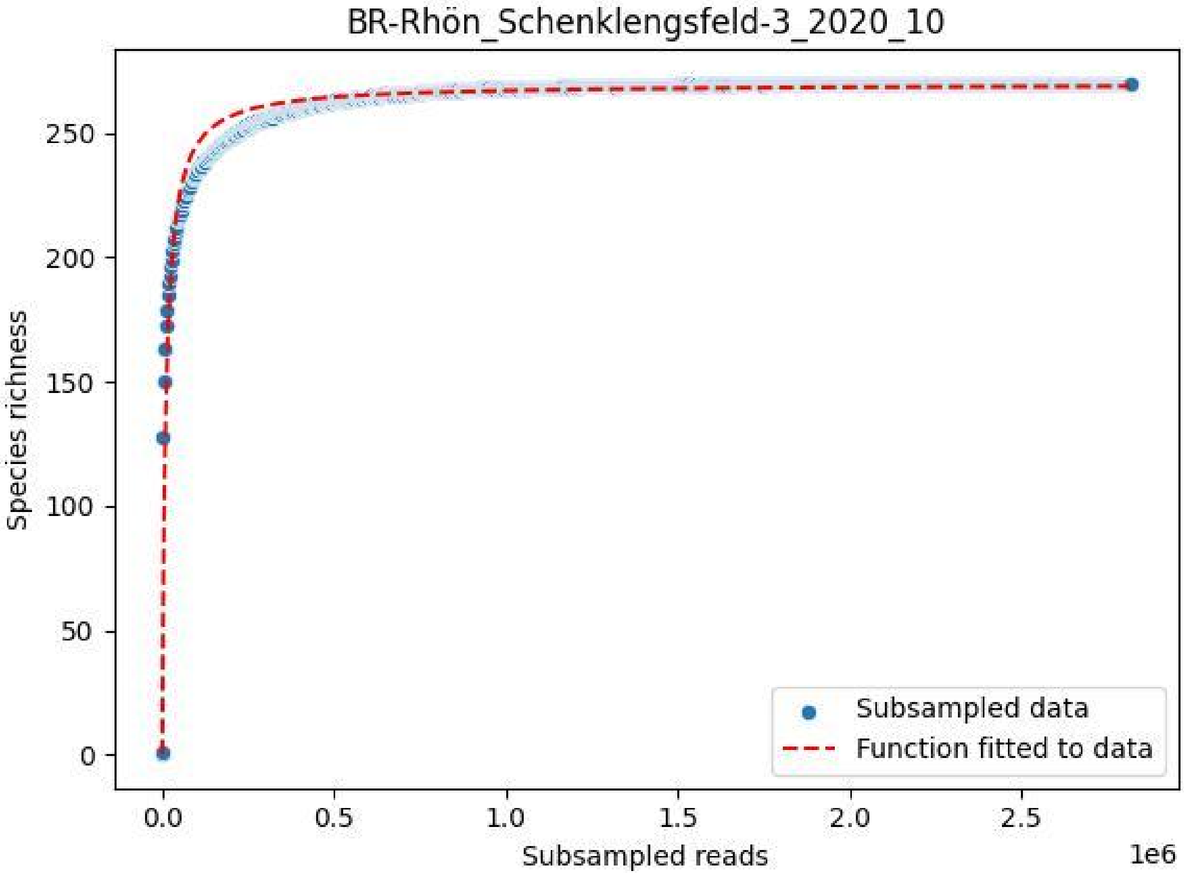

BR-Rhön_Schenklengsfeld-3_2020_10
Species richness
Subsampled reads
1e6
0
50
100
150
200
250
0.0
0.5
1.0
1.5
2.0
2.5
Subsampled data
Function fitted to data

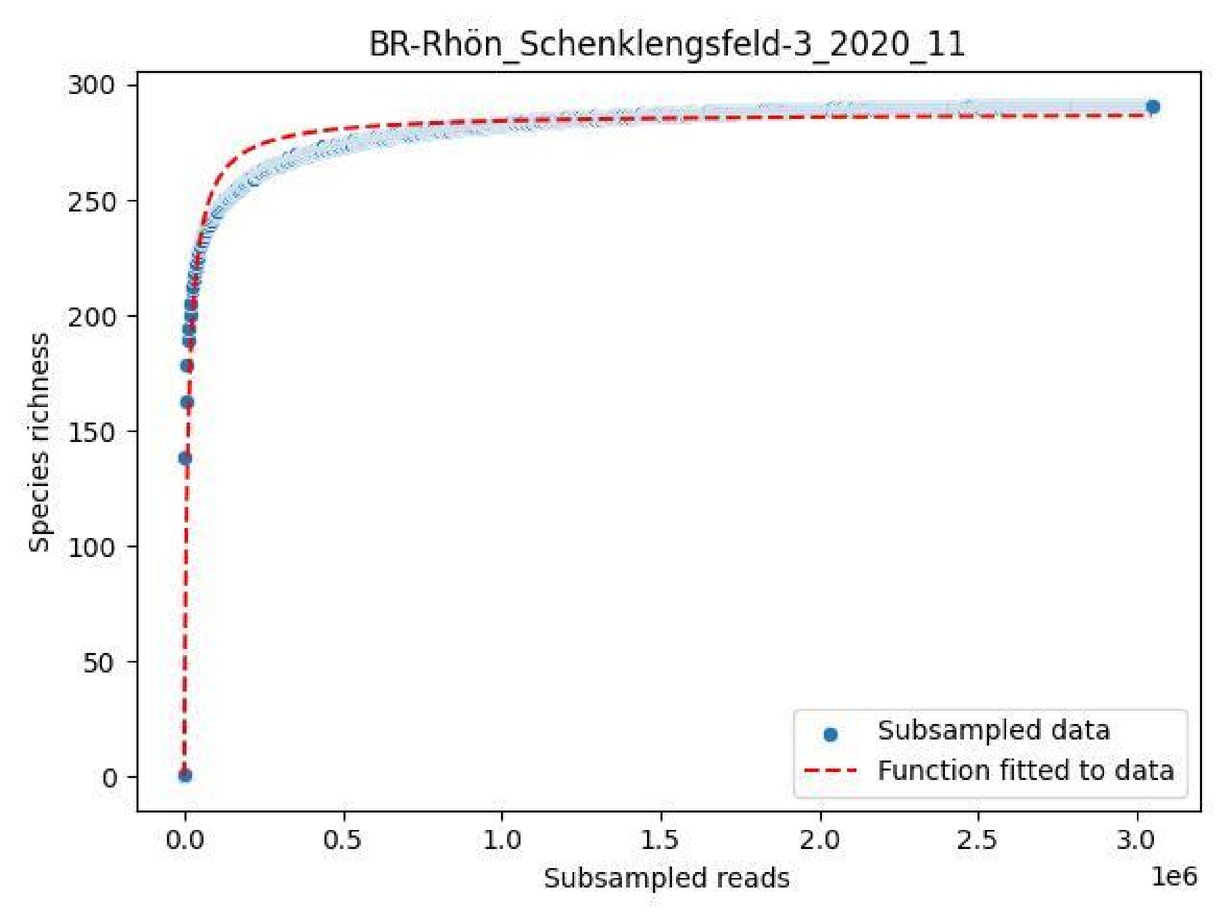

BR-Rhön_Schenklengsfeld-3_2020_11
Species richness
300
250
200
150
100
50
0
Subsampled reads
0.0
0.5
1.0
1.5
2.0
2.5
3.0
1e6
Subsampled data
Function fitted to data

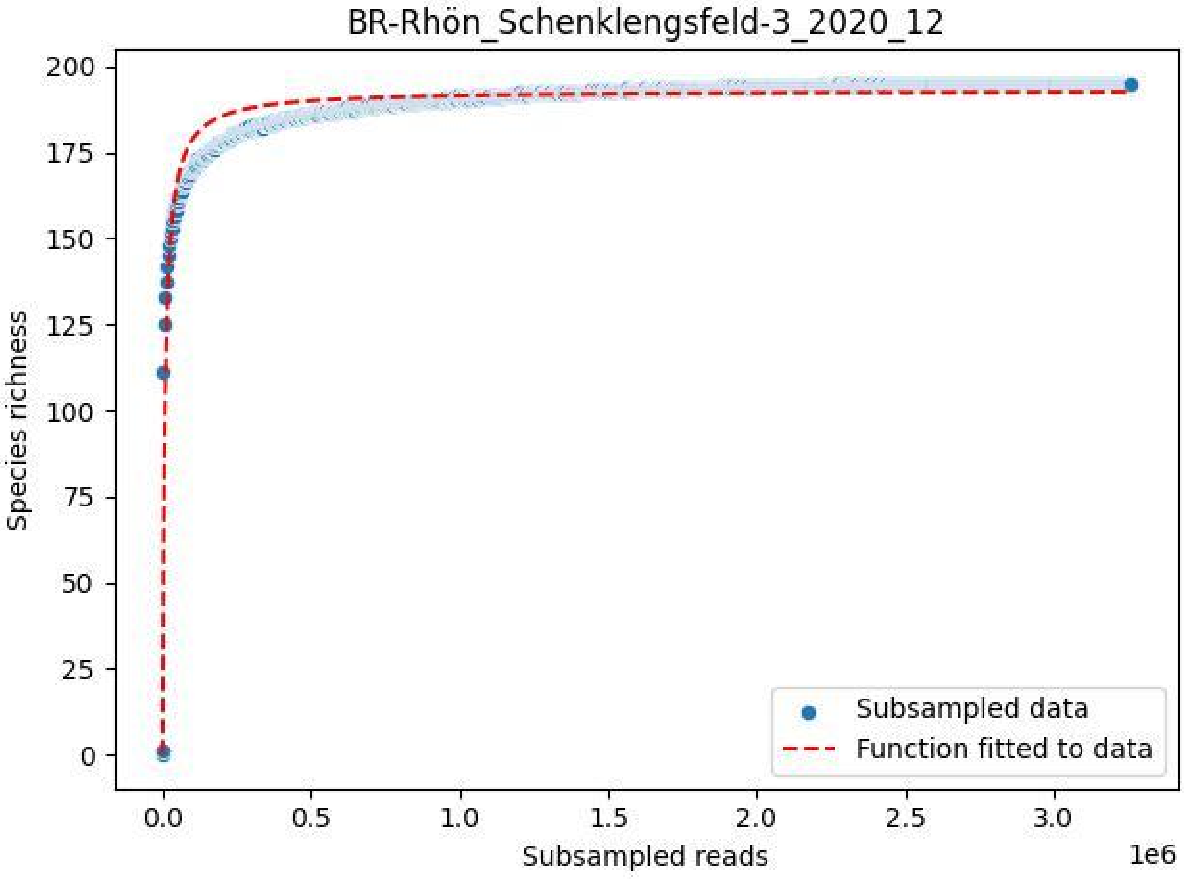

BR-Rhön_Schenklengsfeld-3_2020_12
Species richness
Subsampled reads
1e6
0
25
50
75
100
125
150
175
200
0.0
0.5
1.0
1.5
2.0
2.5
3.0
Subsampled data
Function fitted to data

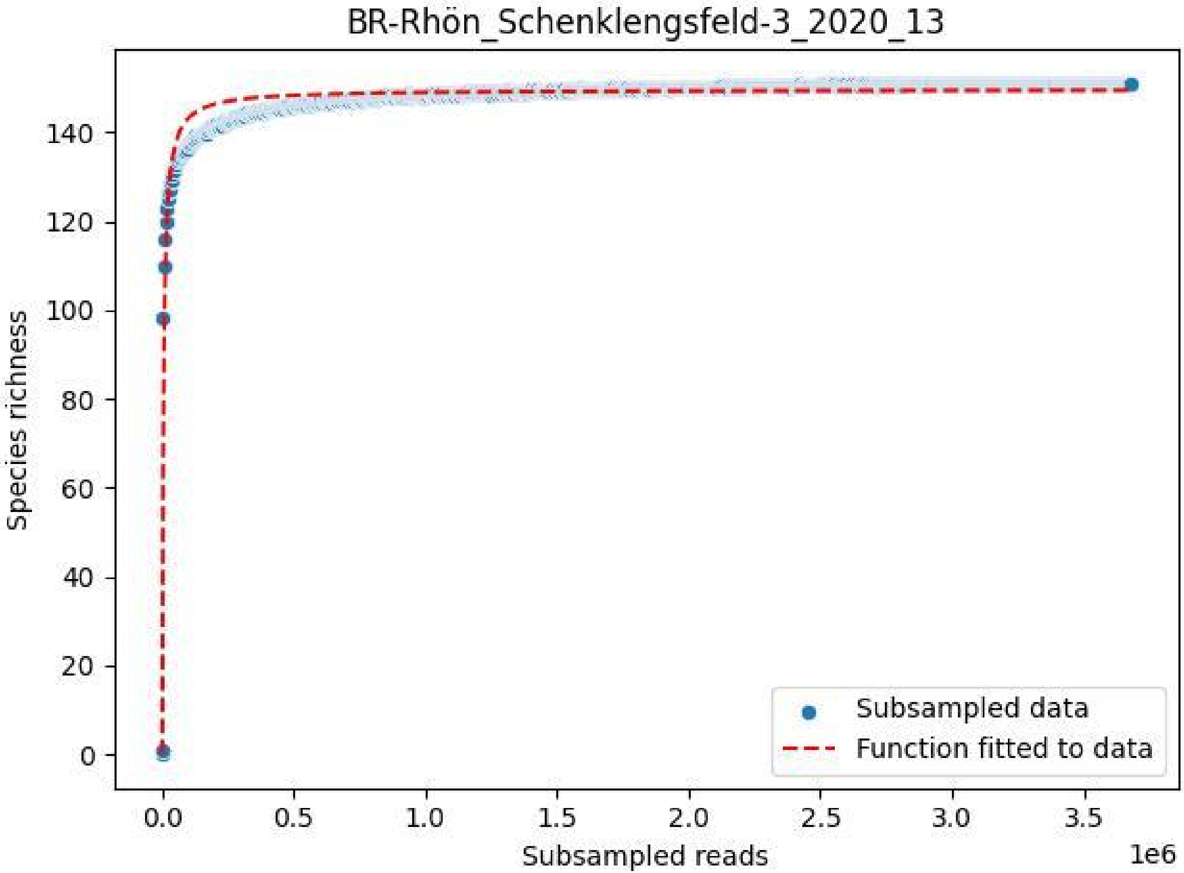
BR-Rhön_Schenklengsfeld-3_2020_13
Species richness
Subsampled reads
1e6
0
20
40
60
80
100
120
140
0.0
0.5
1.0
1.5
2.0
2.5
3.0
3.5
Subsampled data
Function fitted to data

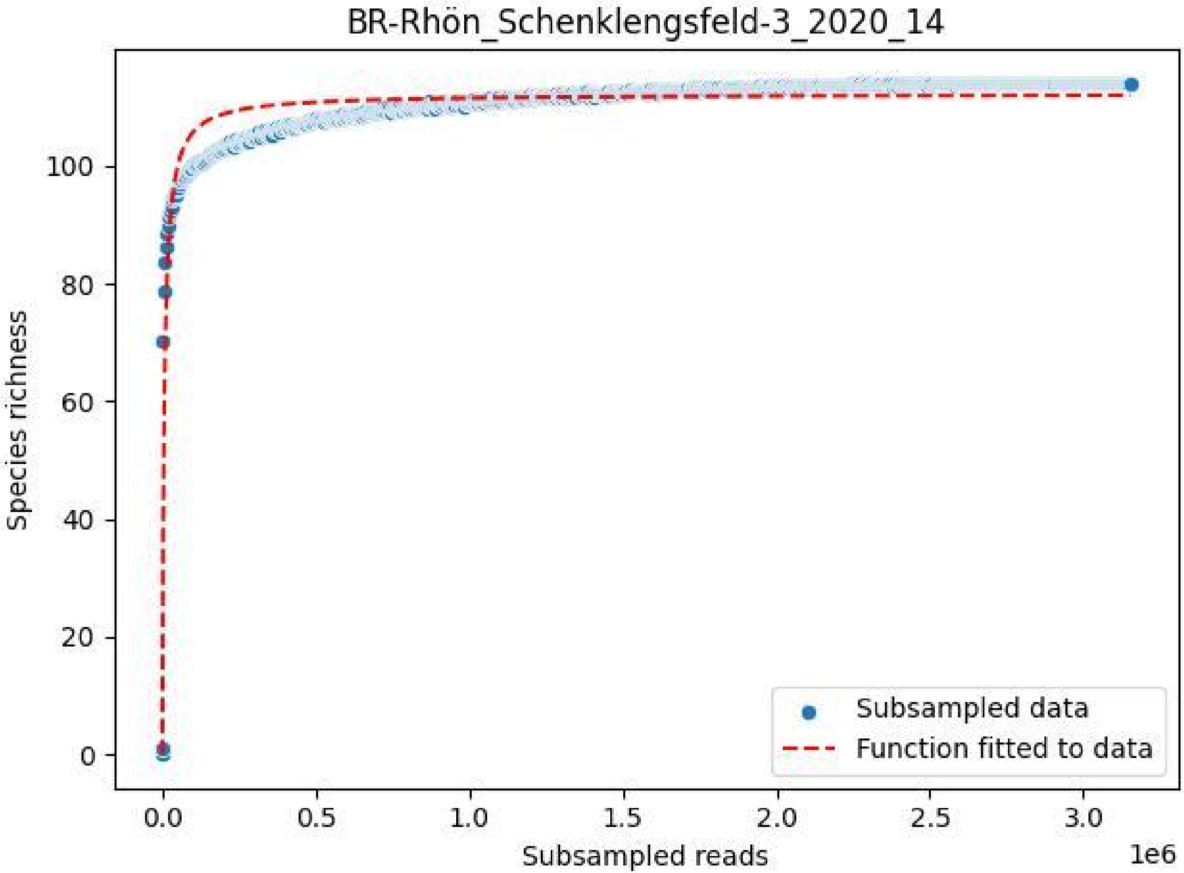
BR-Rhön_Schenklengsfeld-3_2020_14
Species richness
0
20
40
60
80
100
0.0
0.5
1.0
1.5
2.0
2.5
3.0
1e6
Subsampled reads
Subsampled data
Function fitted to data

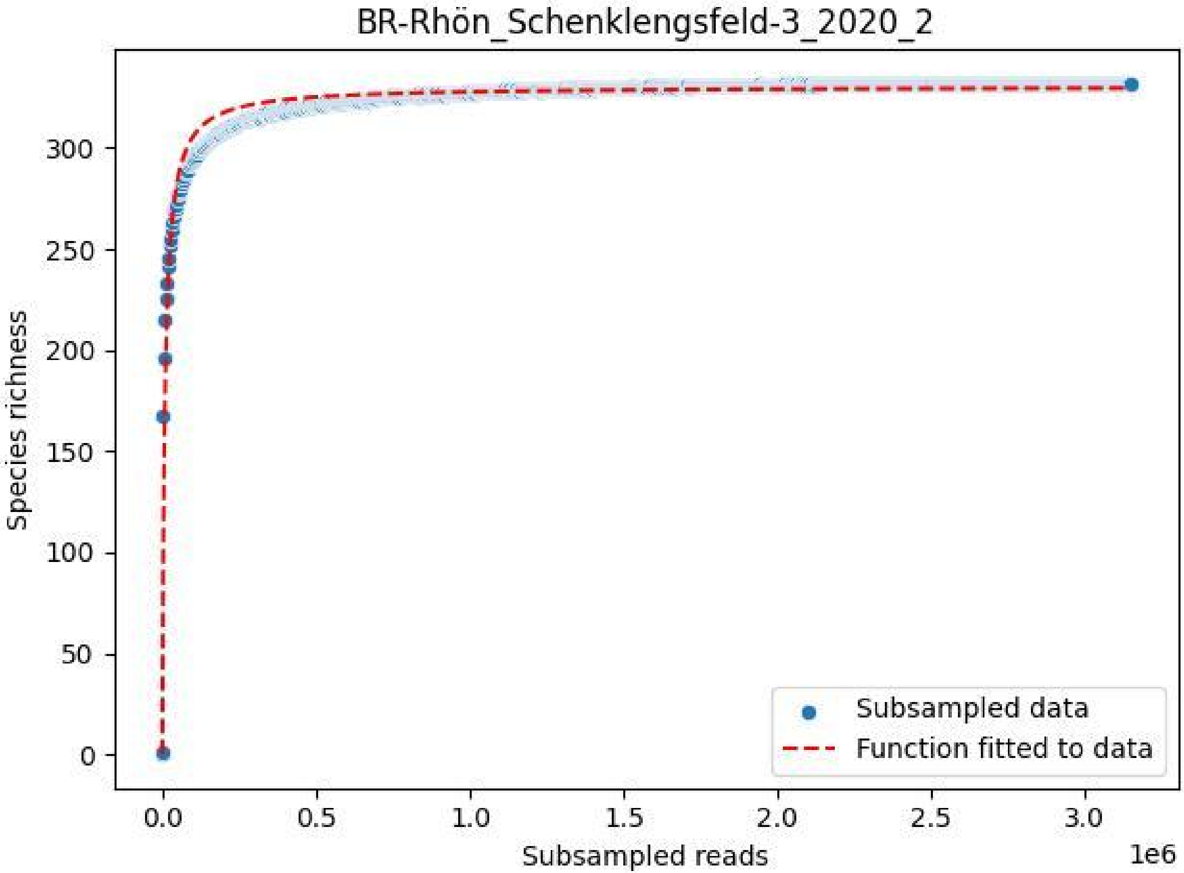
BR-Rhön_Schenklengsfeld-3_2020_2
Species richness
Subsampled reads
1e6
0
50
100
150
200
250
300
0.0
0.5
1.0
1.5
2.0
2.5
3.0
Subsampled data
Function fitted to data

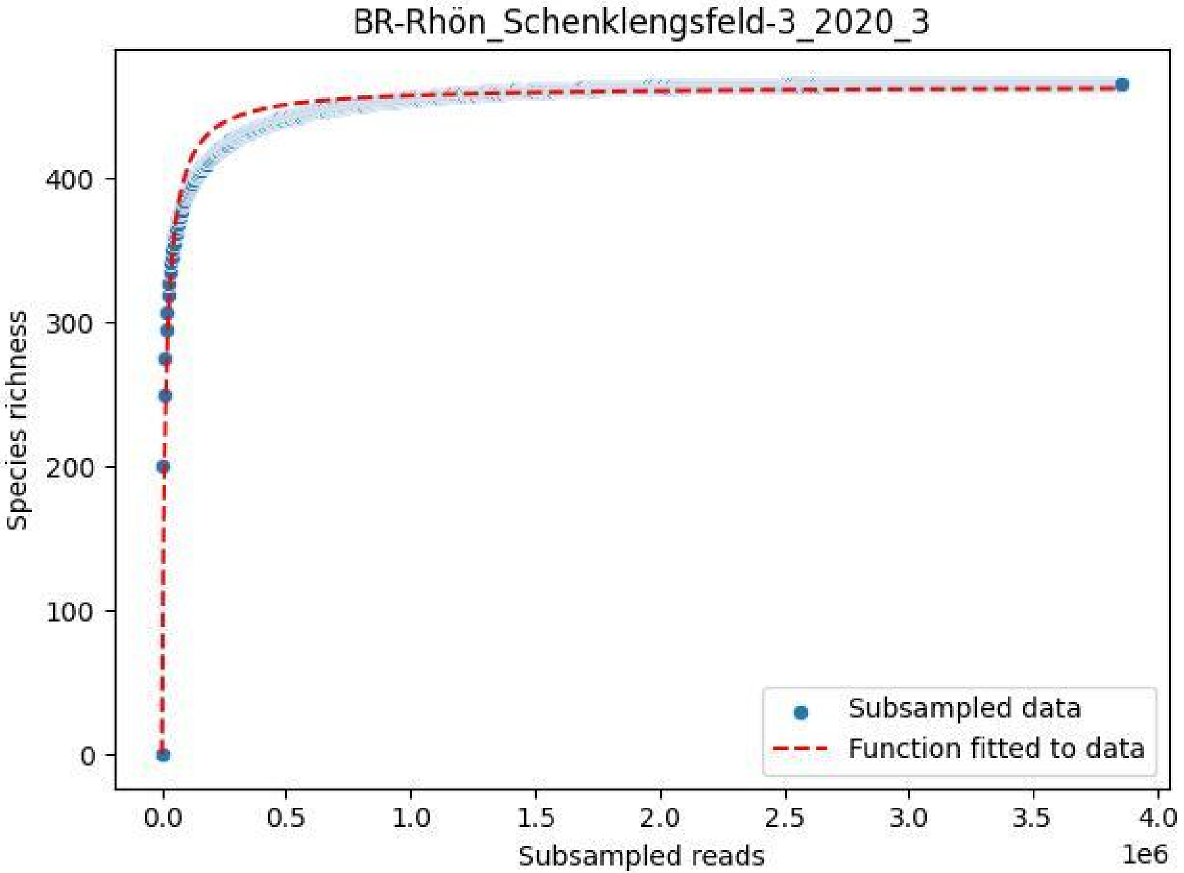
BR-Rhön_Schenklengsfeld-3_2020_3
Species richness
Subsampled reads
1e6
0
100
200
300
400
0.0
0.5
1.0
1.5
2.0
2.5
3.0
3.5
4.0
Subsampled data
Function fitted to data

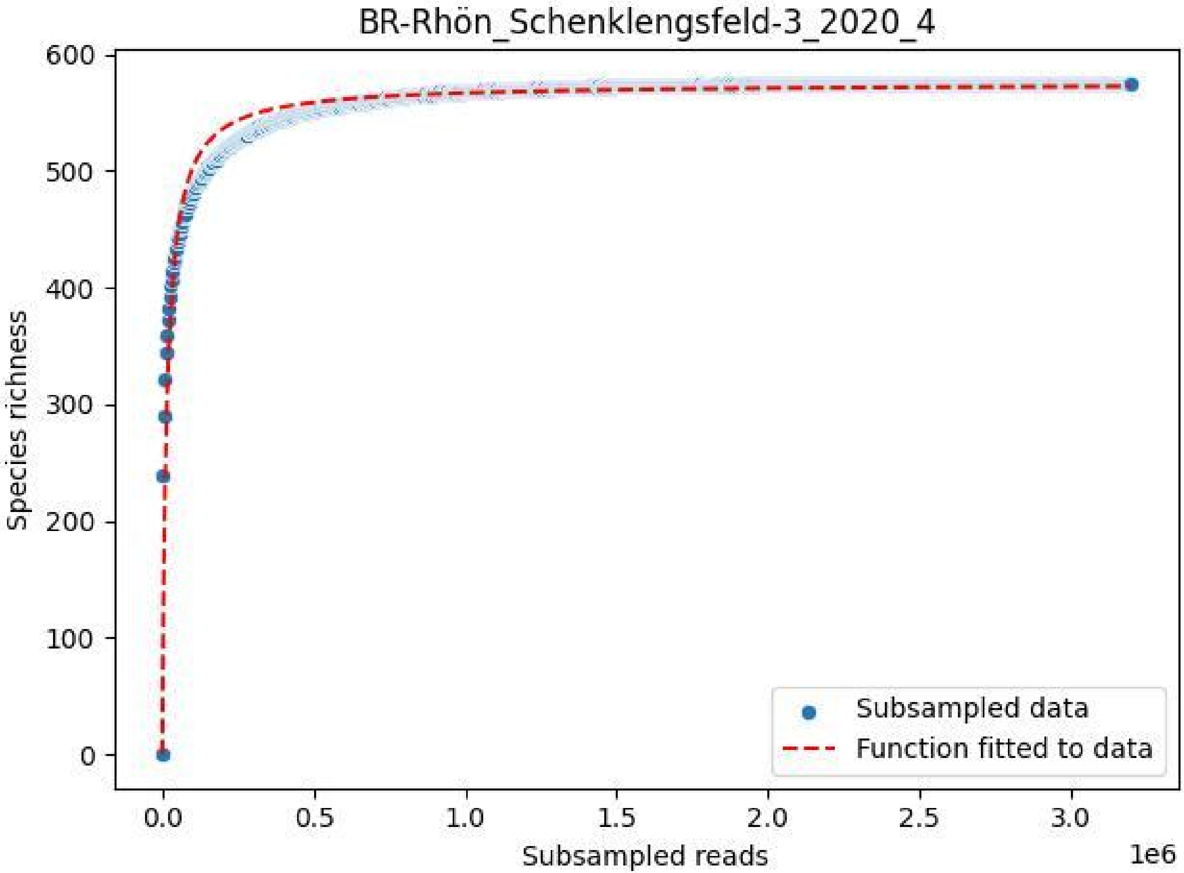
BR-Rhön_Schenklengsfeld-3_2020_4
Species richness
Subsampled reads
1e6
0
100
200
300
400
500
600
0.0
0.5
1.0
1.5
2.0
2.5
3.0
Subsampled data
Function fitted to data

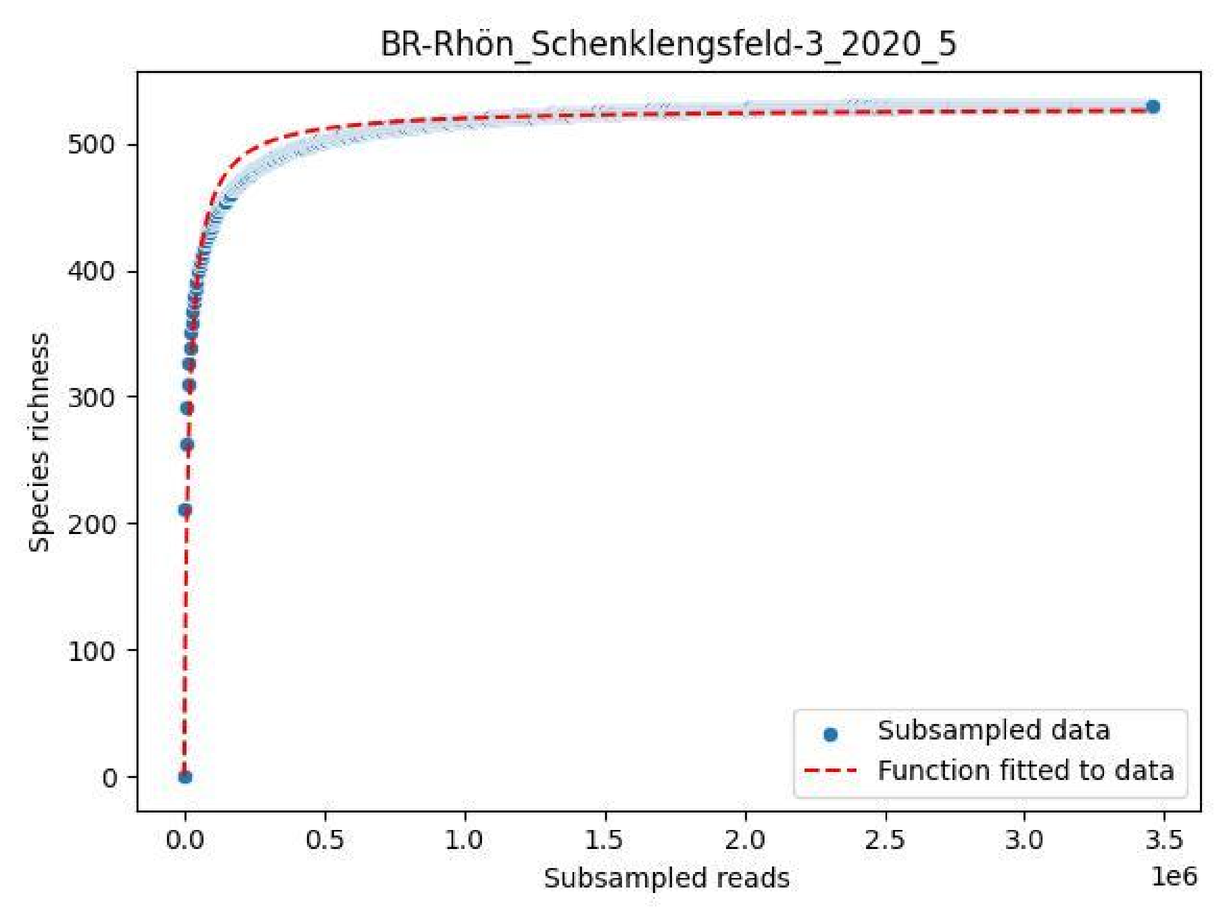
BR-Rhön_Schenklengsfeld-3_2020_5
Species richness
0
100
200
300
400
500
0.0
0.5
1.0
1.5
2.0
2.5
3.0
3.5
1e6
Subsampled reads
Subsampled data
Function fitted to data

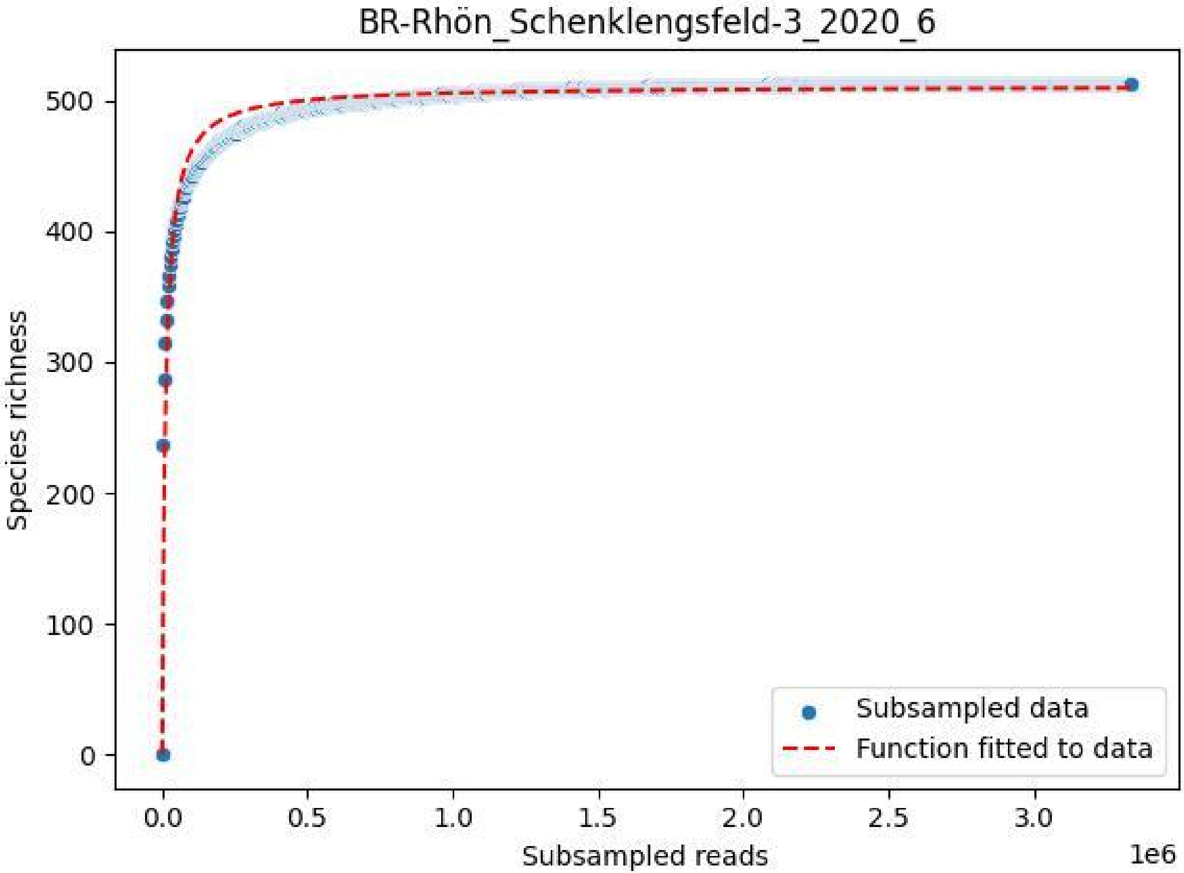
BR-Rhön_Schenklengsfeld-3_2020_6
Species richness
Subsampled reads
1e6
0
100
200
300
400
500
0.0
0.5
1.0
1.5
2.0
2.5
3.0
Subsampled data
Function fitted to data

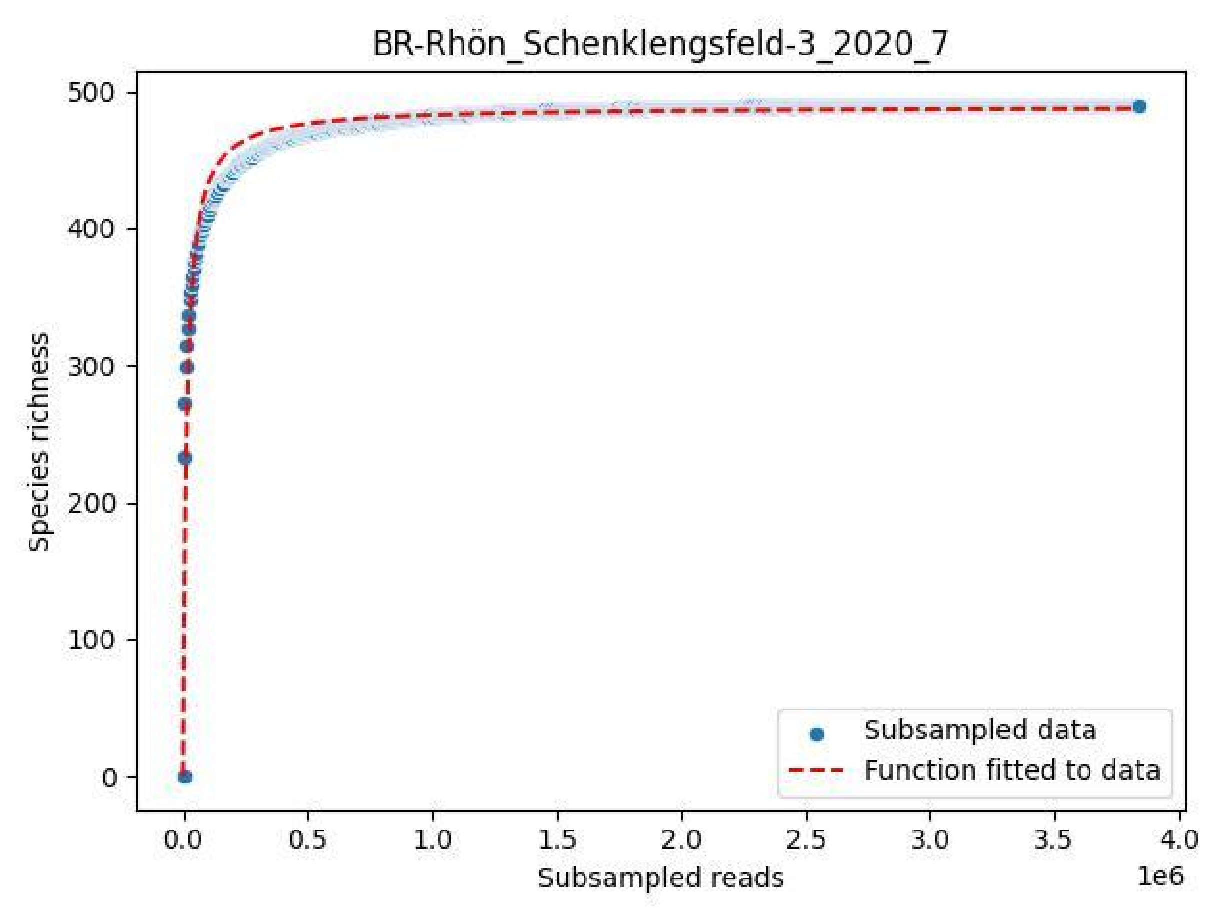
BR-Rhön_Schenklengsfeld-3_2020_7
Species richness
Subsampled reads
1e6
0
100
200
300
400
500
0.0
0.5
1.0
1.5
2.0
2.5
3.0
3.5
4.0
Subsampled data
Function fitted to data

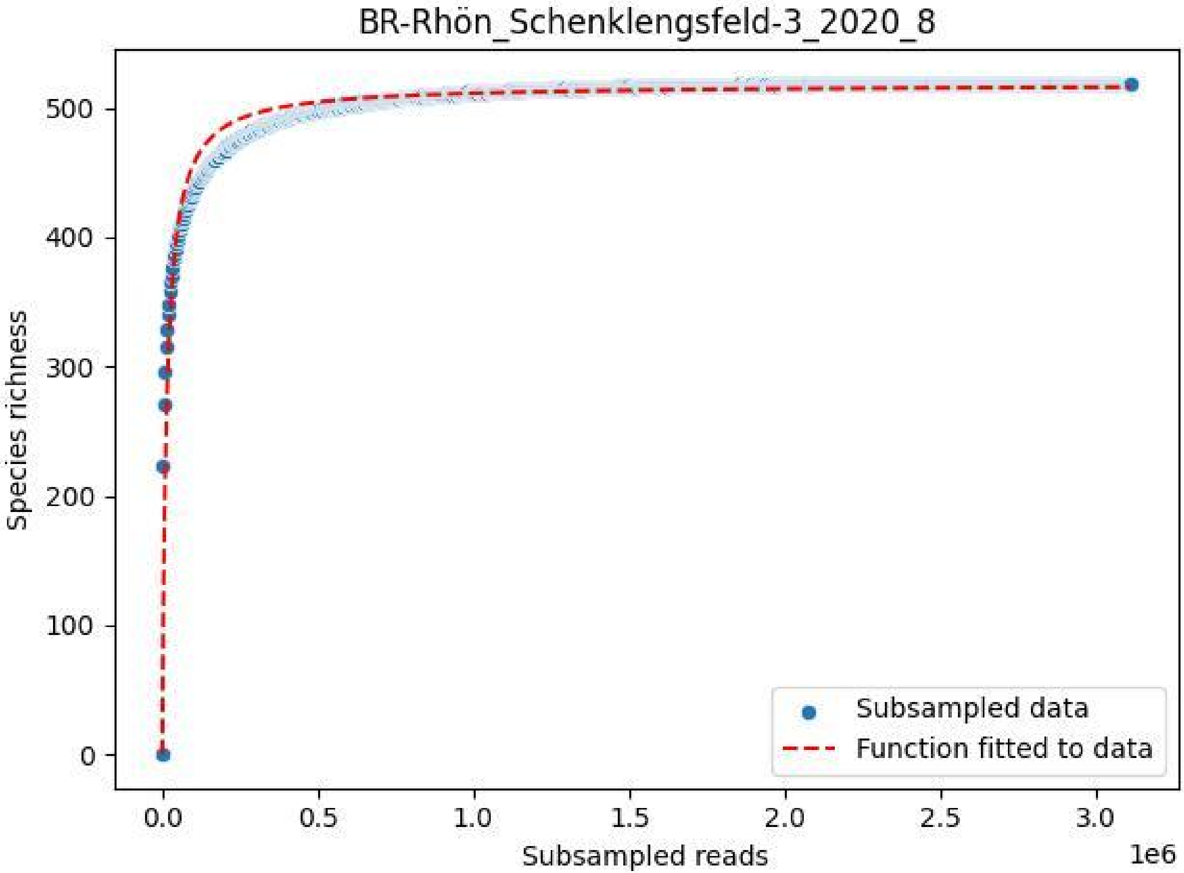
BR-Rhön_Schenklengsfeld-3_2020_8
Species richness
Subsampled reads
1e6
0
100
200
300
400
500
0.0
0.5
1.0
1.5
2.0
2.5
3.0
Subsampled data
Function fitted to data

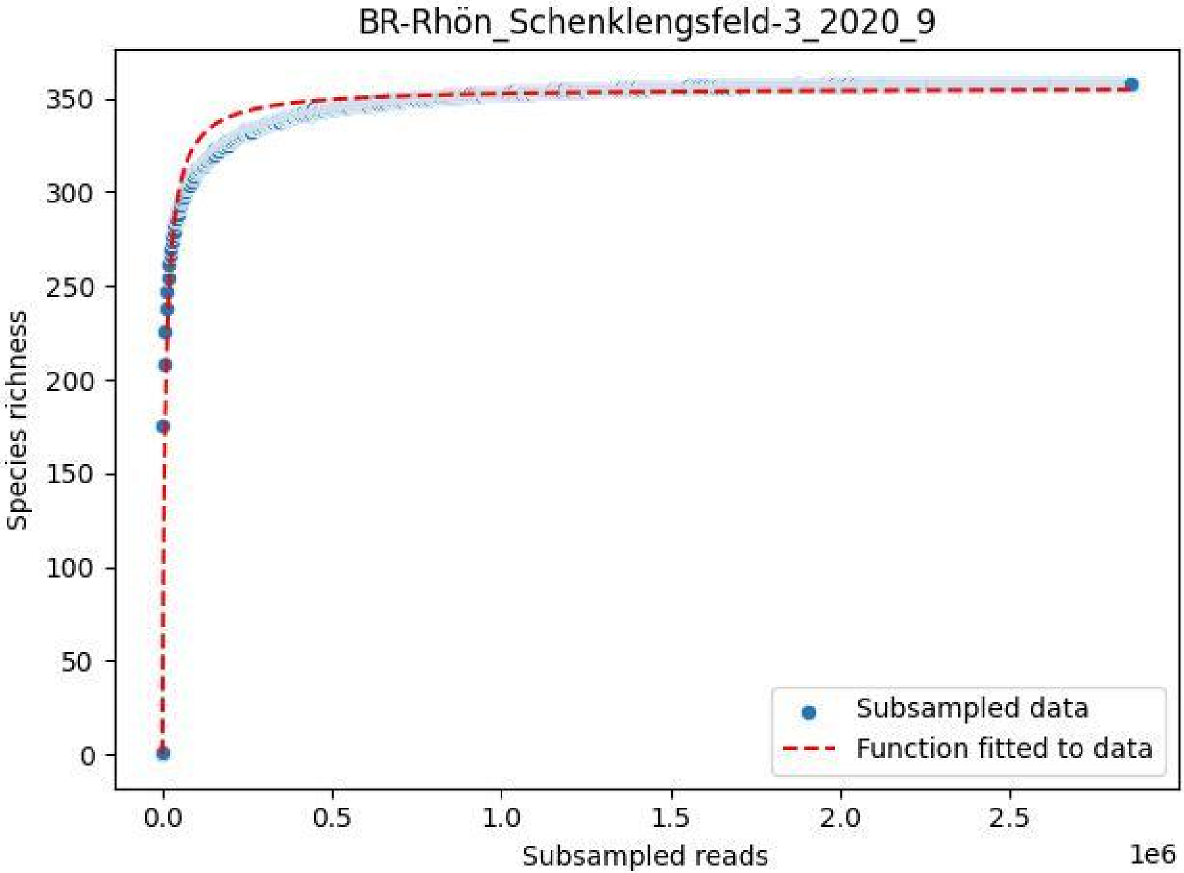
BR-Rhön_Schenklengsfeld-3_2020_9
Species richness
0
50
100
150
200
250
300
350
0.0
0.5
1.0
1.5
2.0
2.5
1e6
Subsampled reads
Subsampled data
Function fitted to data

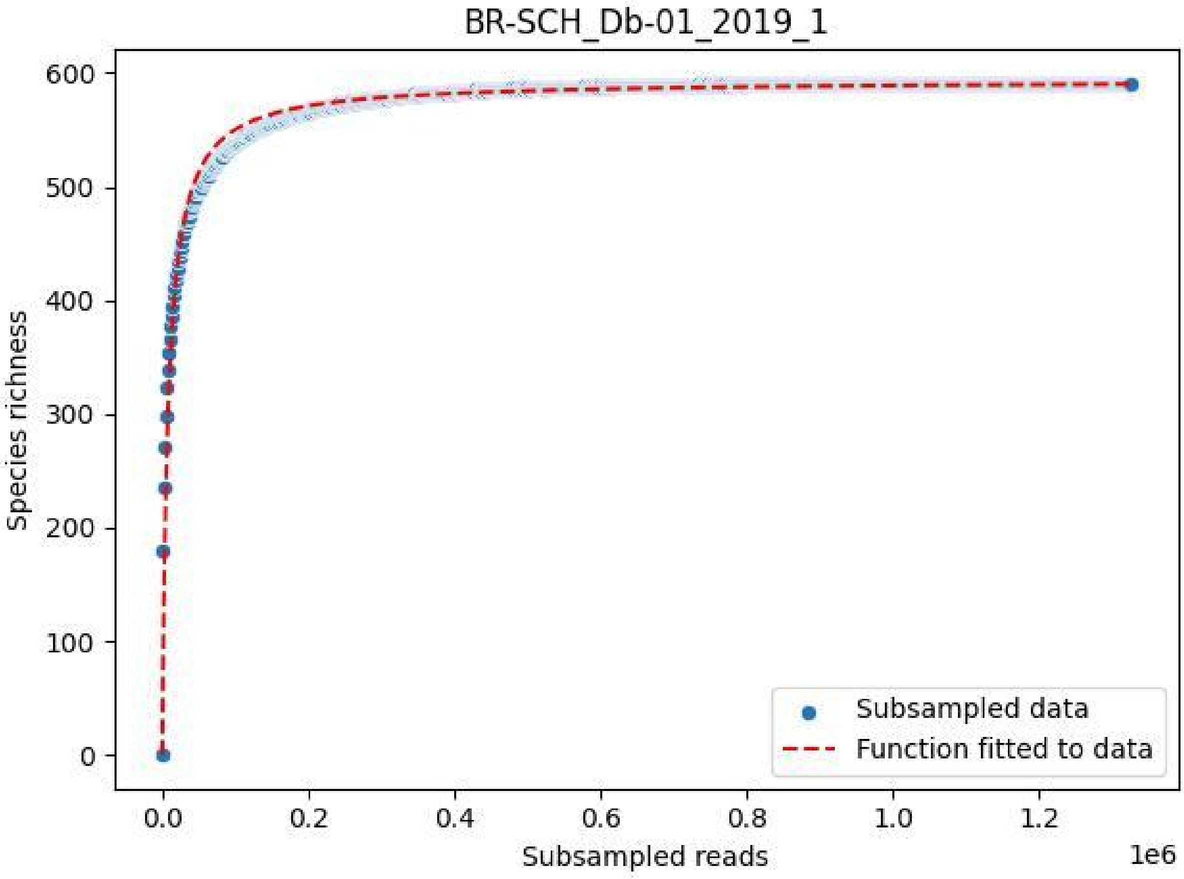

BR-SCH_Db-01_2019_1
Species richness
Subsampled reads
1e6
0
100
200
300
400
500
600
0.0
0.2
0.4
0.6
0.8
1.0
1.2
Subsampled data
Function fitted to data

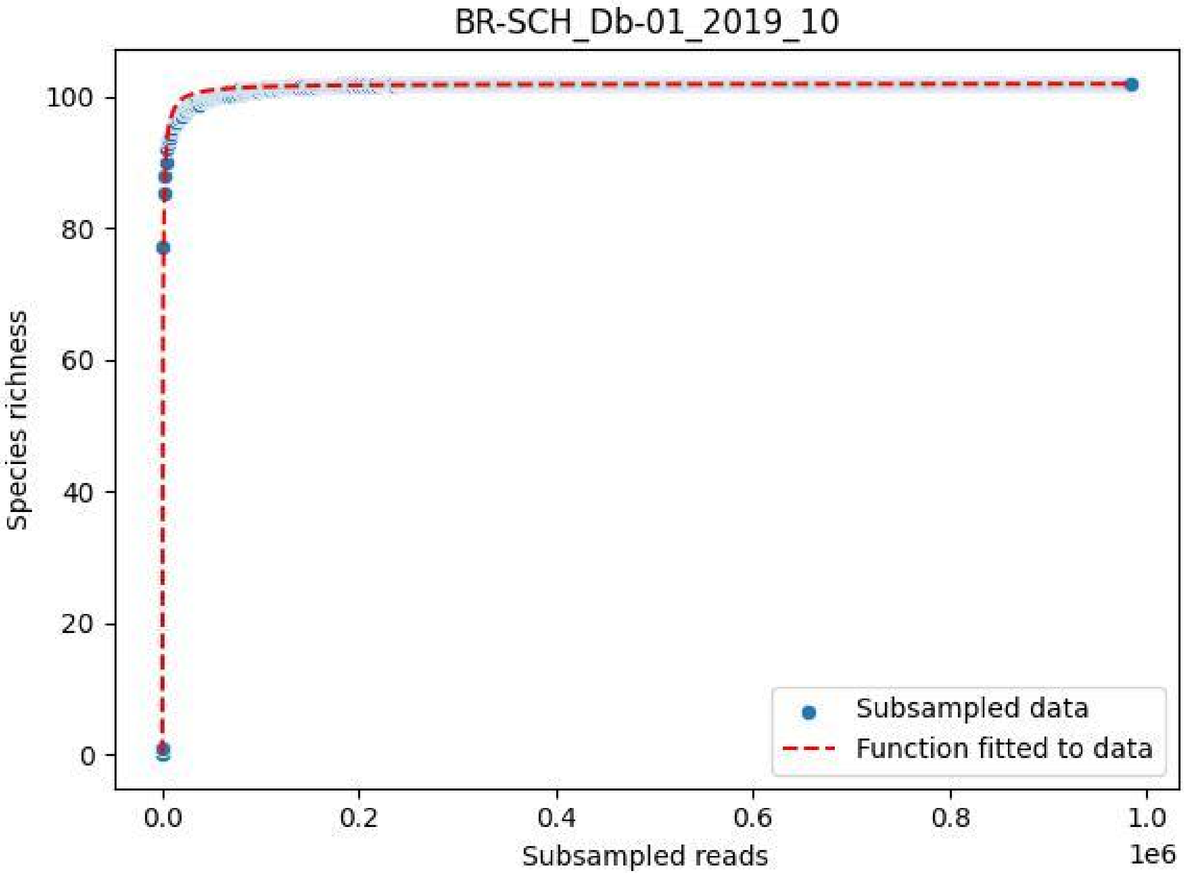
BR-SCH_Db-01_2019_10
Species richness
Subsampled reads
1e6
Subsampled data
Function fitted to data
0
20
40
60
80
100
0.0
0.2
0.4
0.6
0.8
1.0

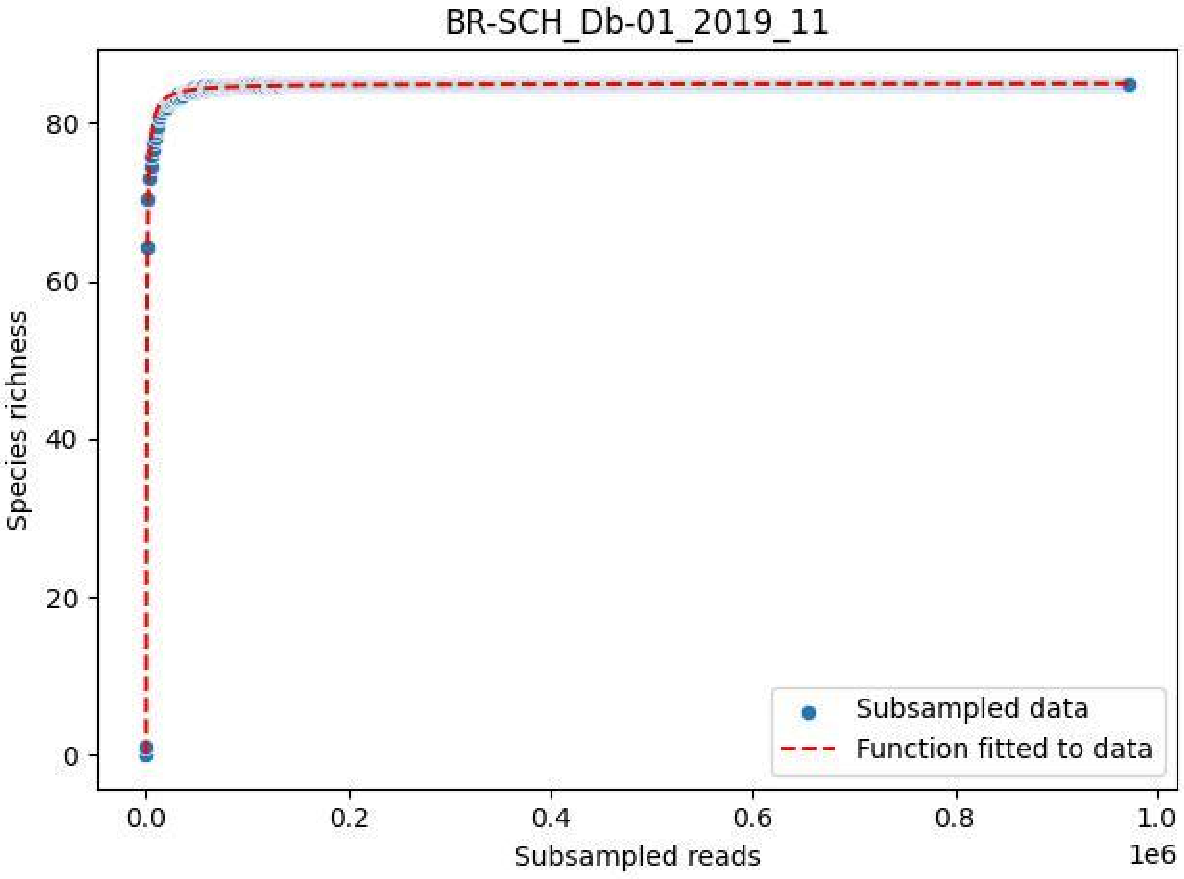
BR-SCH_Db-01_2019_11
Species richness
80
60
40
20
0
0.0
0.2
0.4
0.6
0.8
1.0
1e6
Subsampled reads
Subsampled data
Function fitted to data

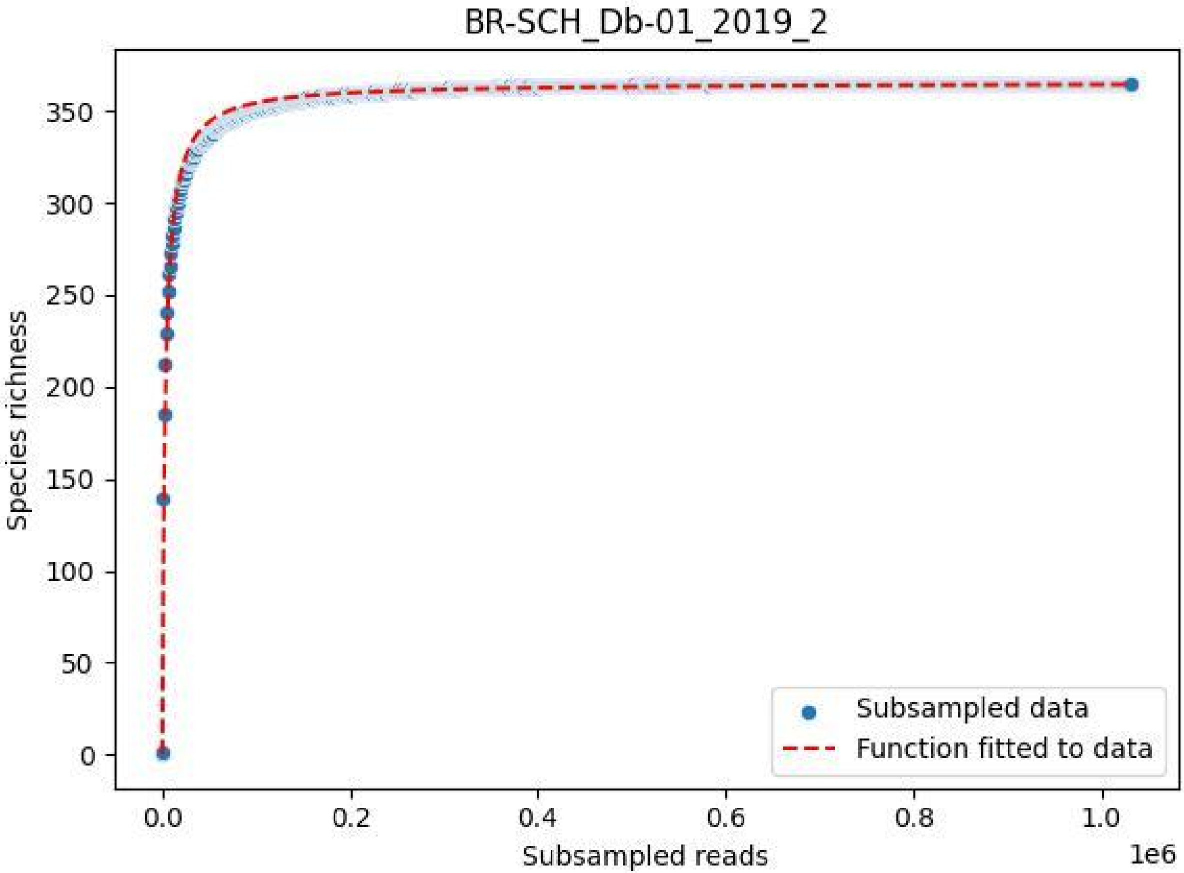

BR-SCH_Db-01_2019_2
Species richness
Subsampled reads
1e6
0
50
100
150
200
250
300
350
0.0
0.2
0.4
0.6
0.8
1.0
Subsampled data
Function fitted to data

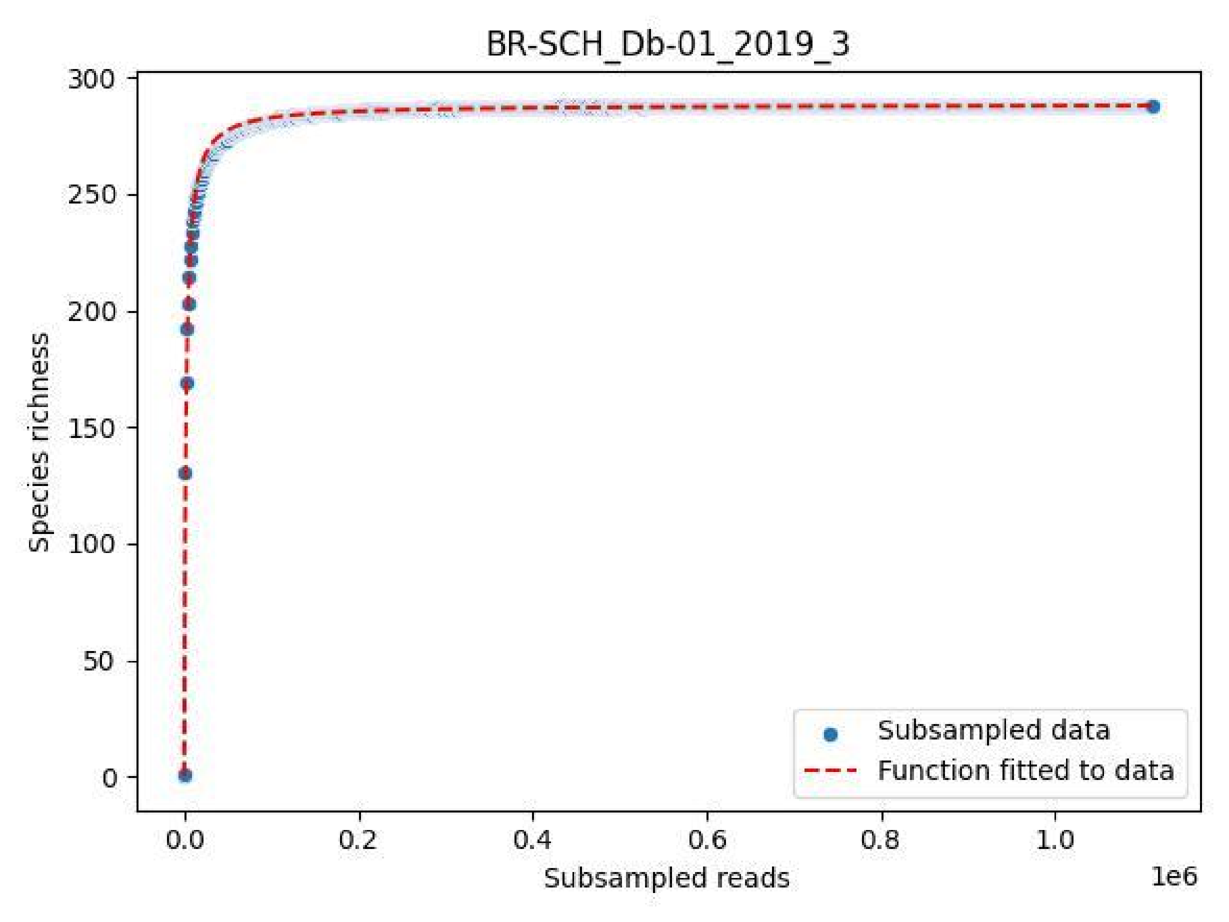
BR-SCH_Db-01_2019_3
Species richness
Subsampled reads
1e6
300
250
200
150
100
50
0
0.0
0.2
0.4
0.6
0.8
1.0
Subsampled data
Function fitted to data

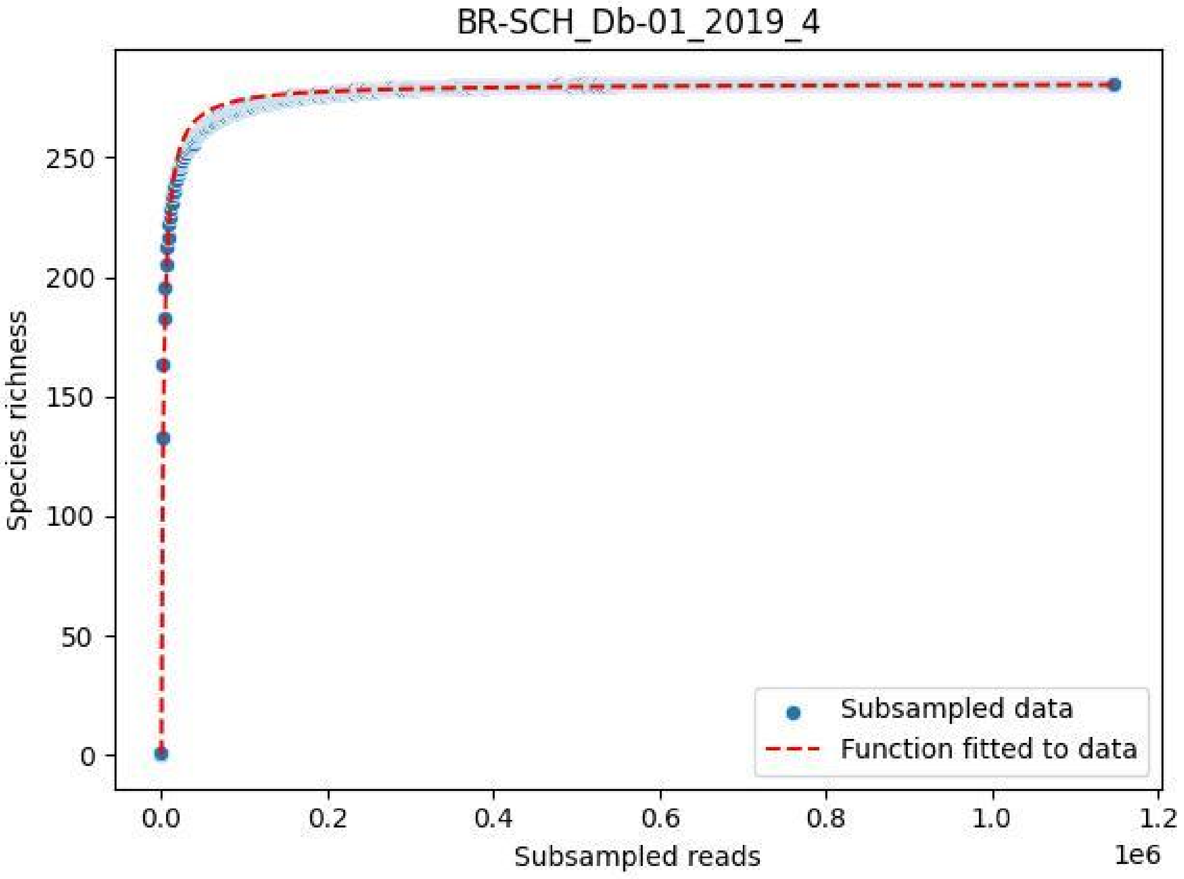

BR-SCH_Db-01_2019_4
Species richness
Subsampled reads
1e6
0
50
100
150
200
250
0.0
0.2
0.4
0.6
0.8
1.0
1.2
Subsampled data
Function fitted to data

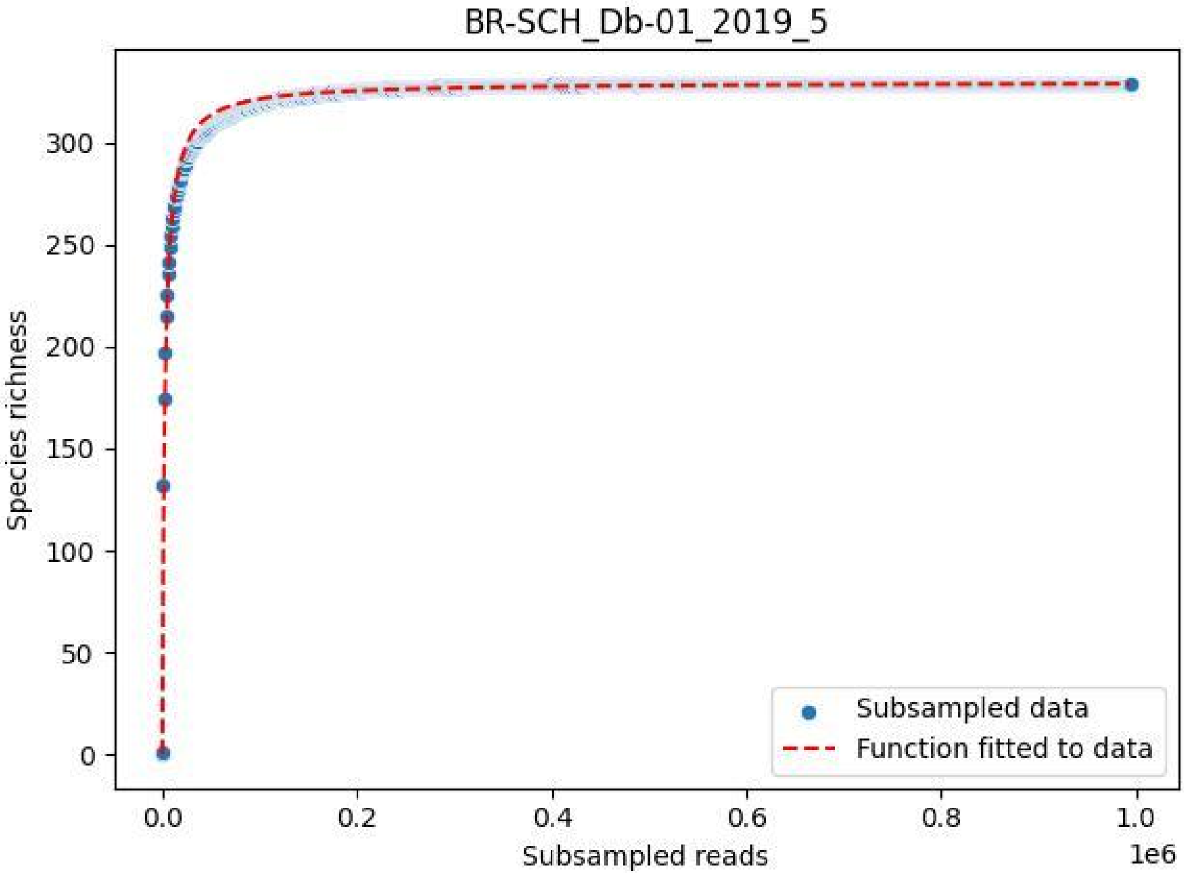

BR-SCH_Db-01_2019_5
Species richness
0
50
100
150
200
250
300
Subsampled reads
0.0
0.2
0.4
0.6
0.8
1.0
1e6
Subsampled data
Function fitted to data

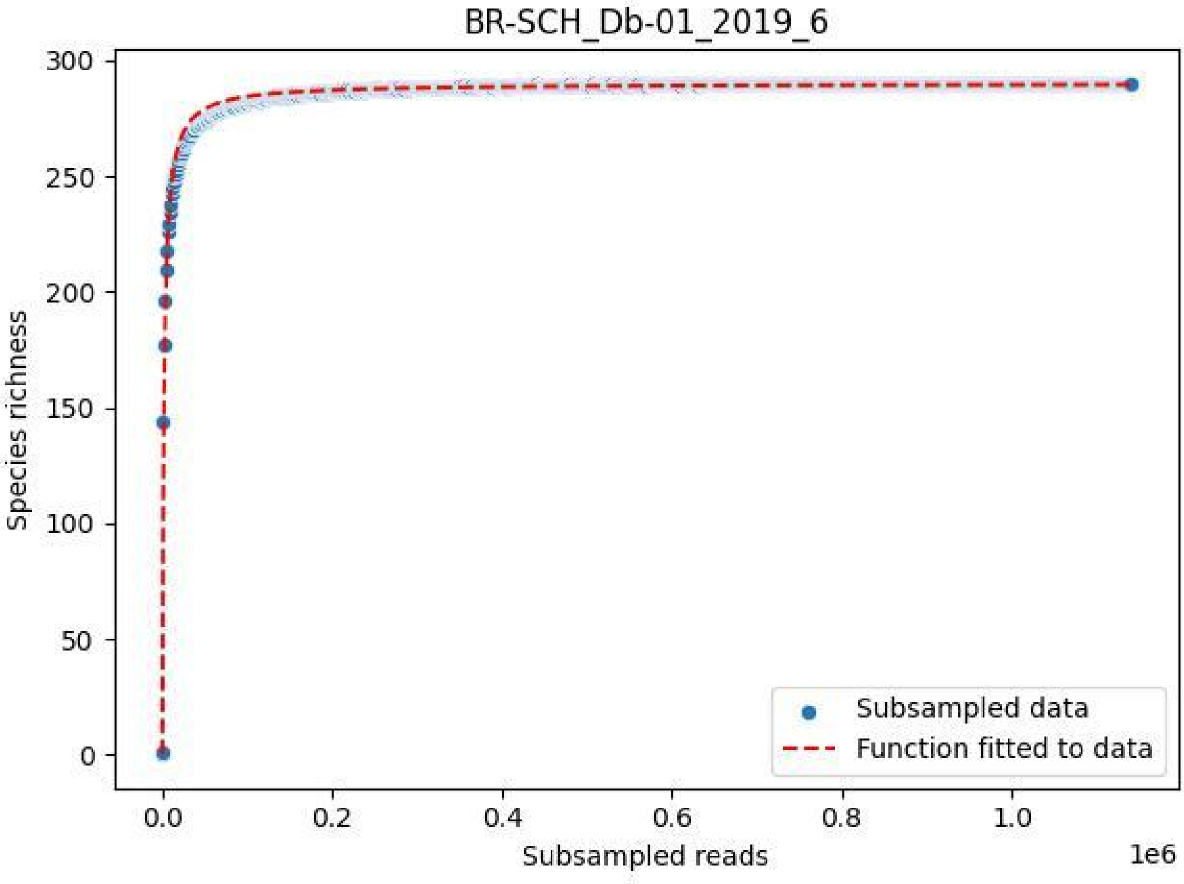
BR-SCH_Db-01_2019_6
Species richness
Subsampled reads
1e6
0
50
100
150
200
250
300
0.0
0.2
0.4
0.6
0.8
1.0
Subsampled data
Function fitted to data

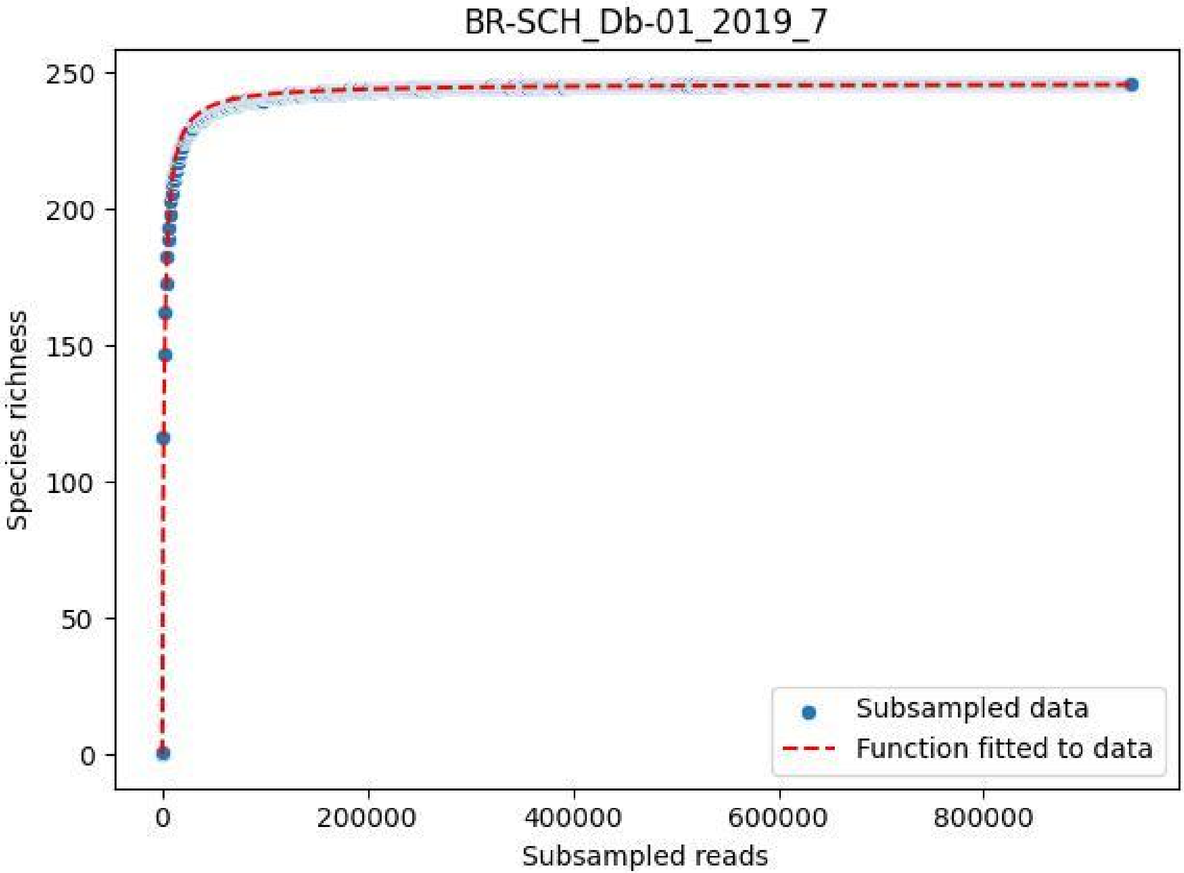
BR-SCH_Db-01_2019_7
Species richness
Subsampled reads
0
50
100
150
200
250
0
200000
400000
600000
800000
Subsampled data
Function fitted to data

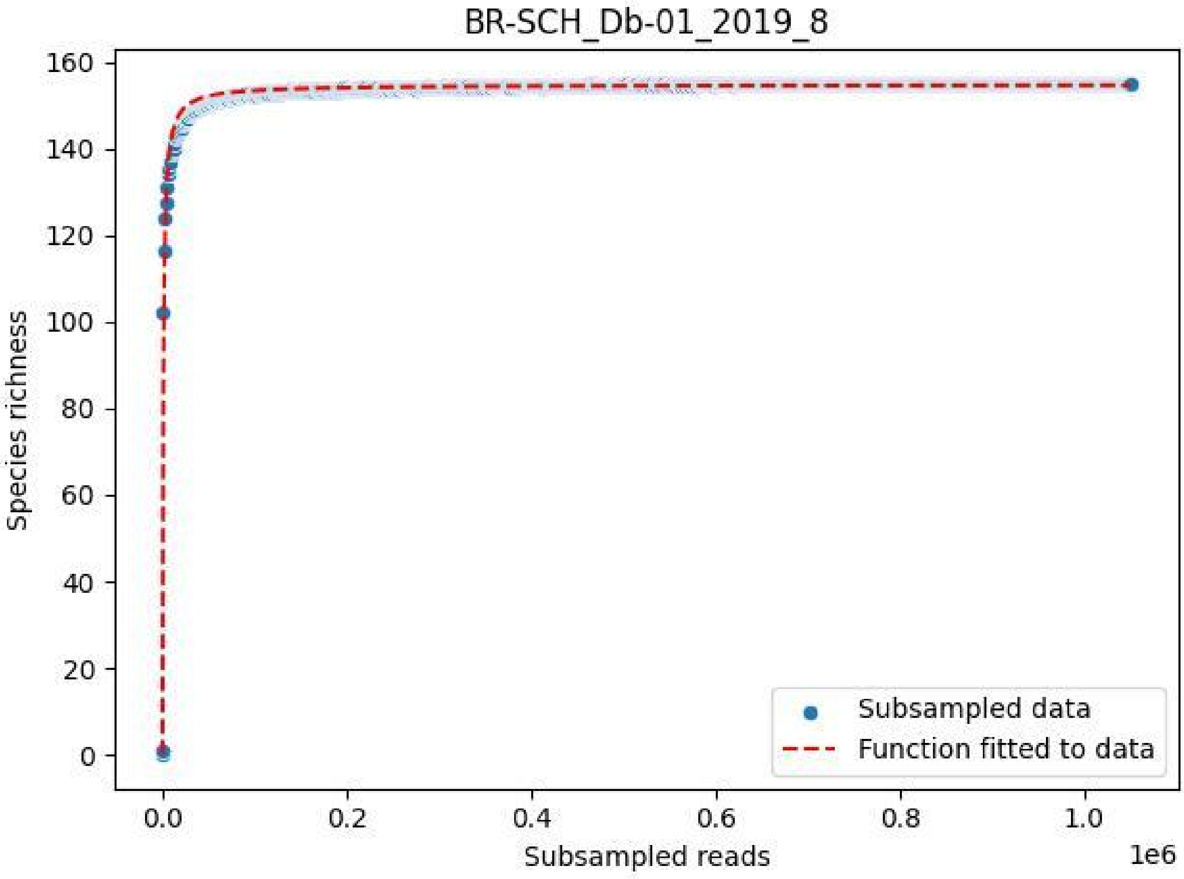

BR-SCH_Db-01_2019_8
Species richness
Subsampled reads
1e6
Subsampled data
Function fitted to data
0
20
40
60
80
100
120
140
160
0.0
0.2
0.4
0.6
0.8
1.0

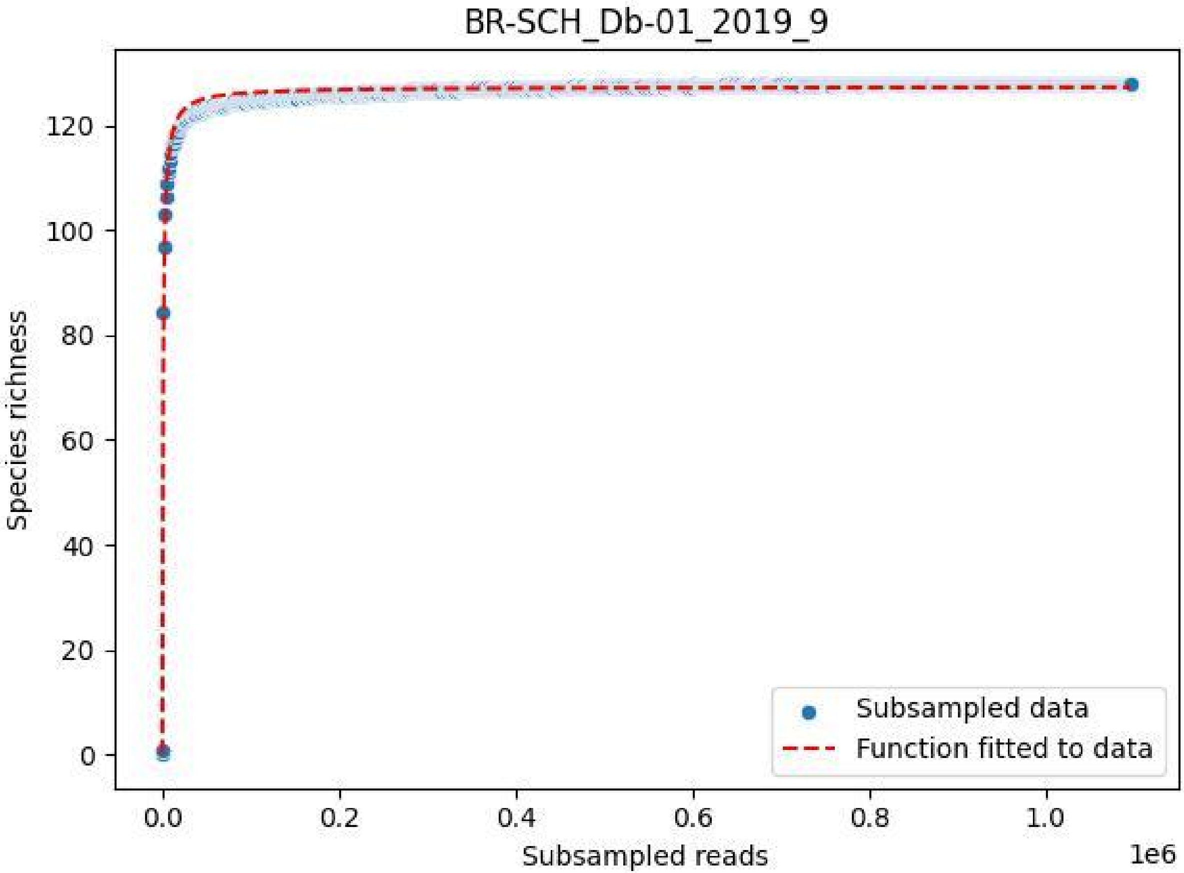

BR-SCH_Db-01_2019_9
Species richness
Subsampled reads
1e6
0.0
0.2
0.4
0.6
0.8
1.0
0
20
40
60
80
100
120
Subsampled data
Function fitted to data

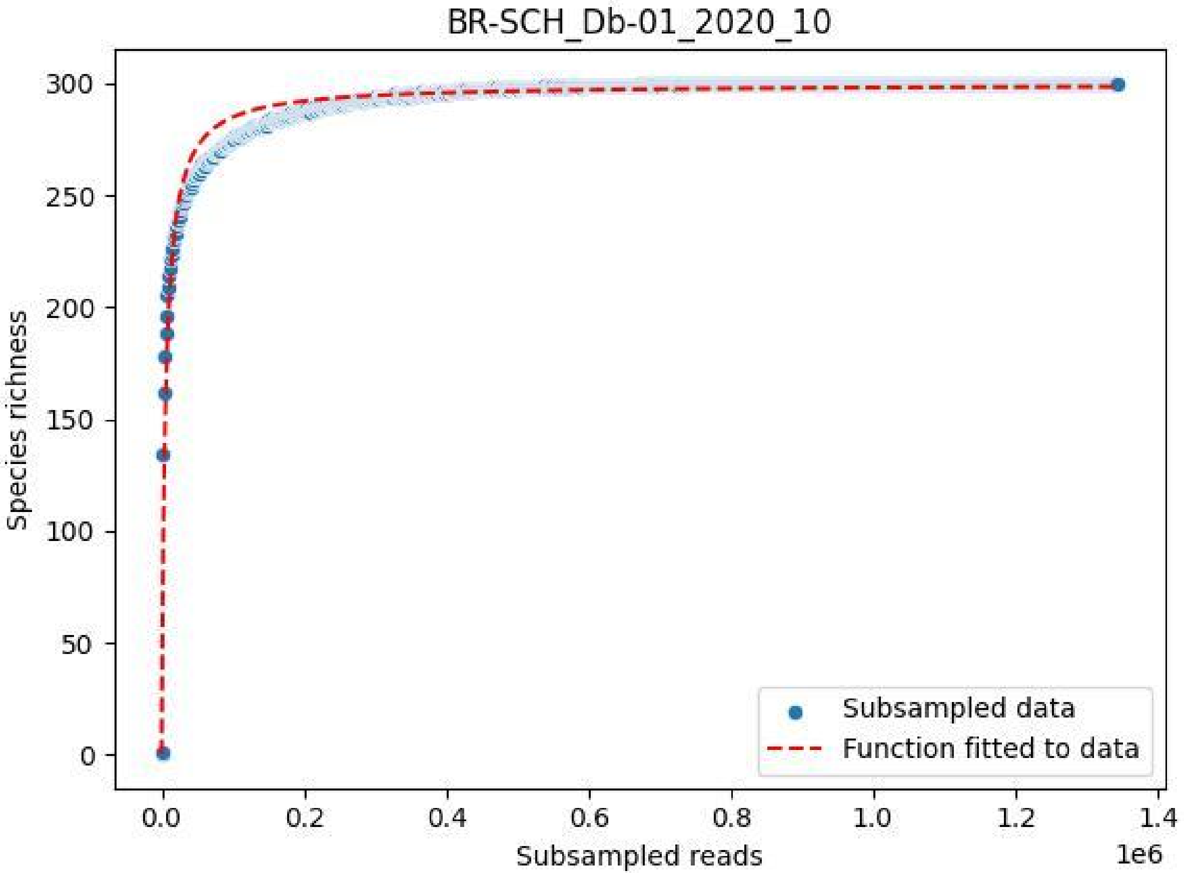
BR-SCH_Db-01_2020_10
Species richness
Subsampled reads
1e6
0
50
100
150
200
250
300
0.0
0.2
0.4
0.6
0.8
1.0
1.2
1.4
Subsampled data
Function fitted to data

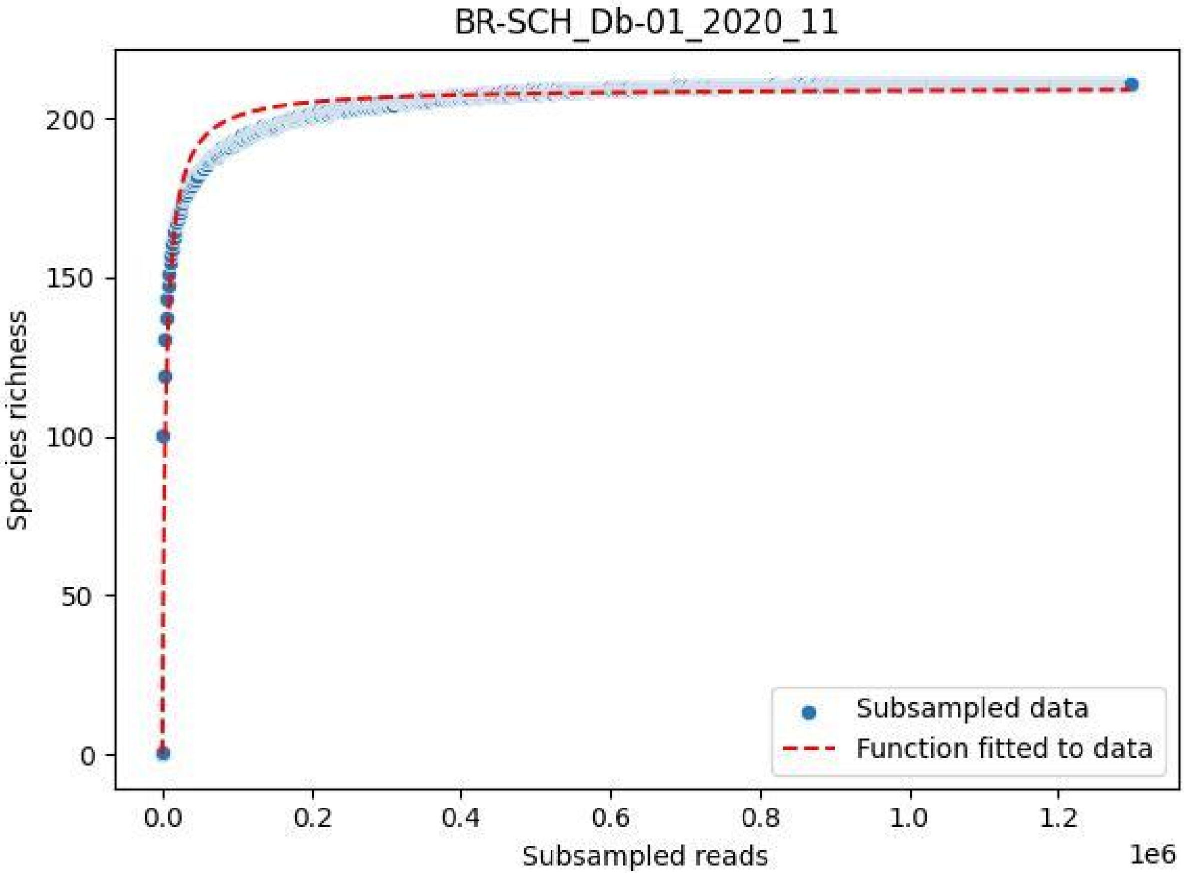

BR-SCH_Db-01_2020_11
Species richness
0
50
100
150
200
Subsampled reads
0.0
0.2
0.4
0.6
0.8
1.0
1.2
1e6
Subsampled data
Function fitted to data

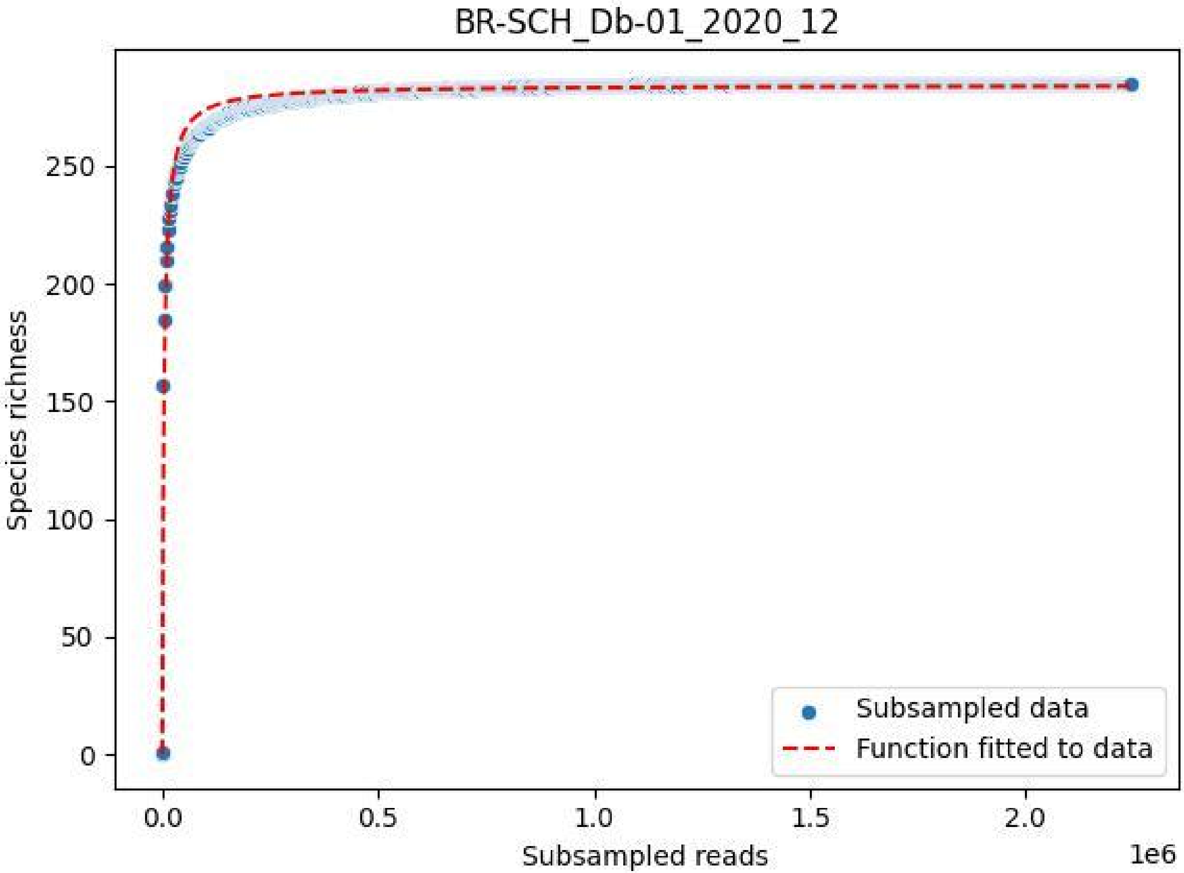
BR-SCH_Db-01_2020_12
Species richness
Subsampled reads
1e6
0
50
100
150
200
250
0.0
0.5
1.0
1.5
2.0
Subsampled data
Function fitted to data

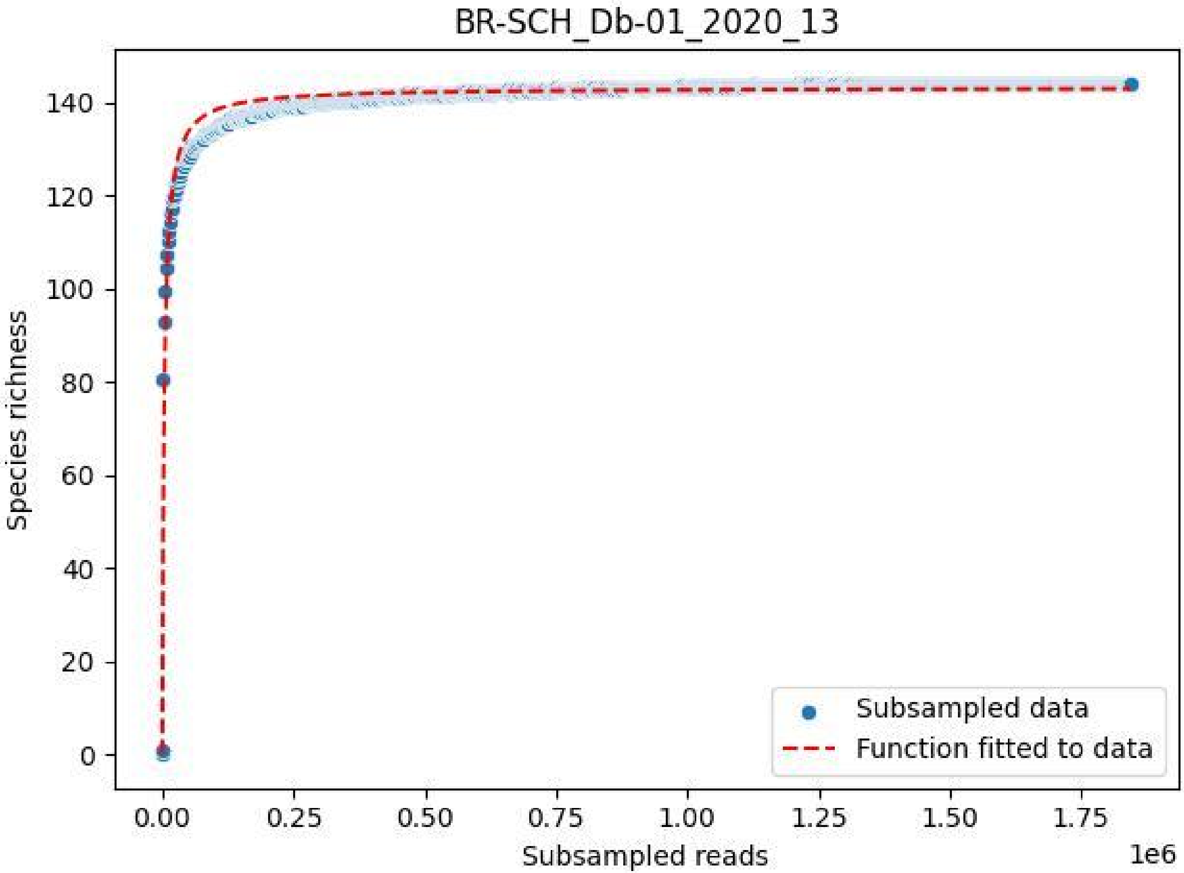

BR-SCH_Db-01_2020_13
Species richness
0
20
40
60
80
100
120
140
0.00
0.25
0.50
0.75
1.00
1.25
1.50
1.75
1e6
Subsampled reads
Subsampled data
Function fitted to data

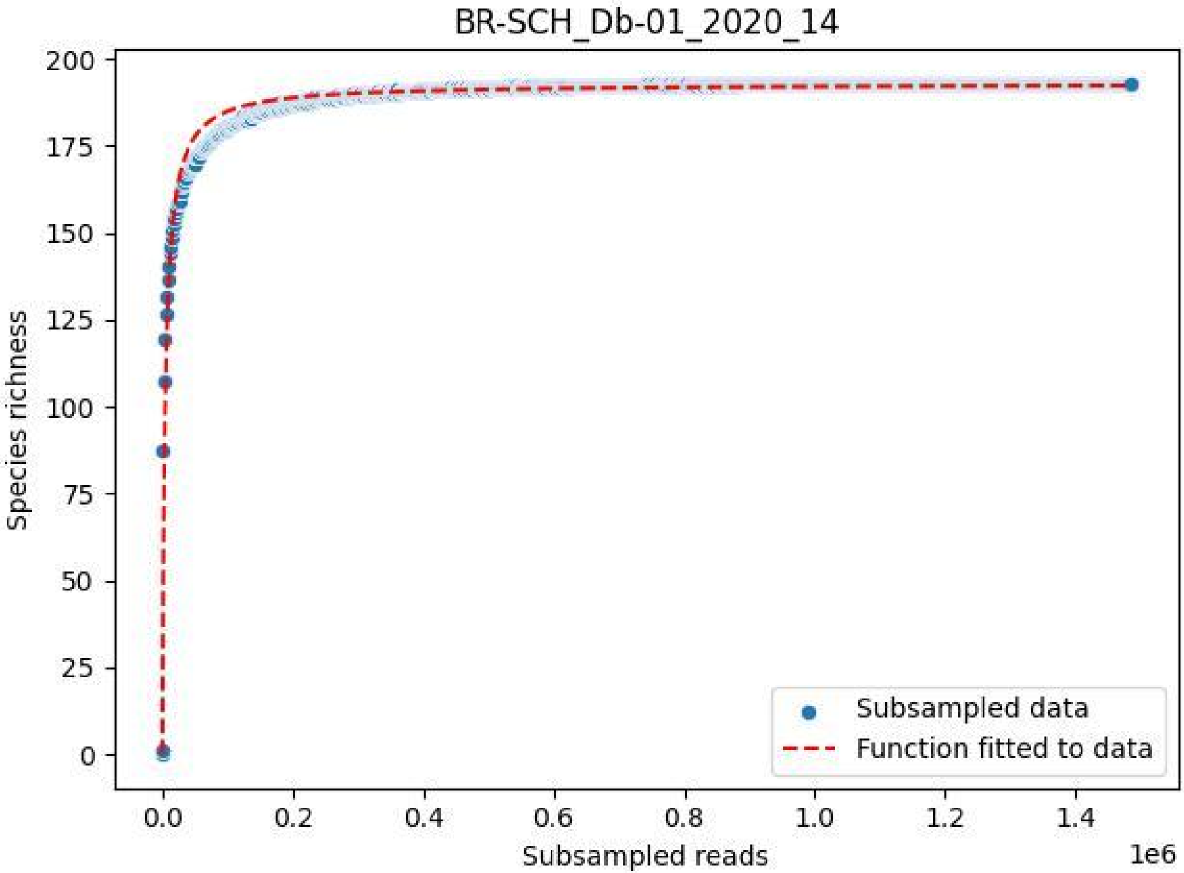
BR-SCH_Db-01_2020_14
Species richness
Subsampled reads
1e6
Subsampled data
Function fitted to data
200
175
150
125
100
75
50
25
0
0.0
0.2
0.4
0.6
0.8
1.0
1.2
1.4

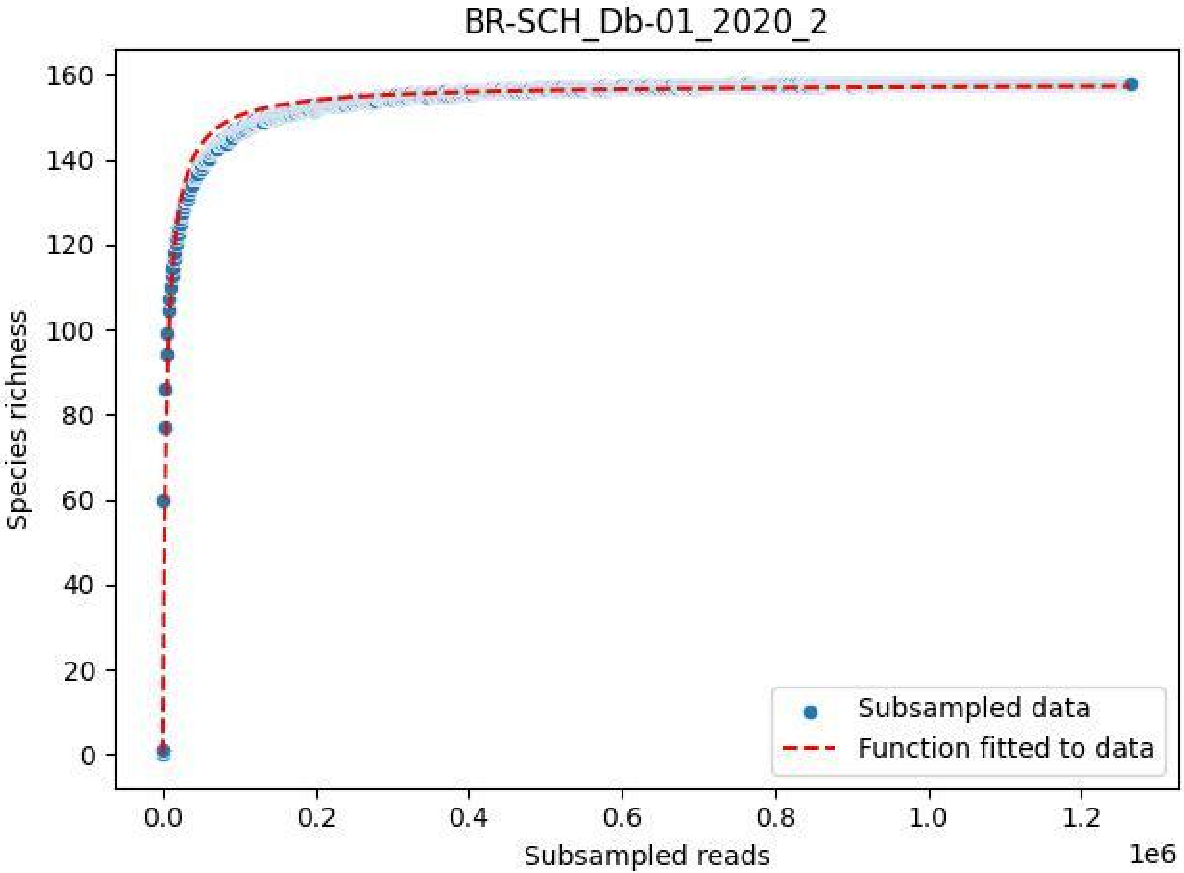
BR-SCH_Db-01_2020_2
Species richness
Subsampled reads
1e6
0
20
40
60
80
100
120
140
160
0.0
0.2
0.4
0.6
0.8
1.0
1.2
Subsampled data
Function fitted to data

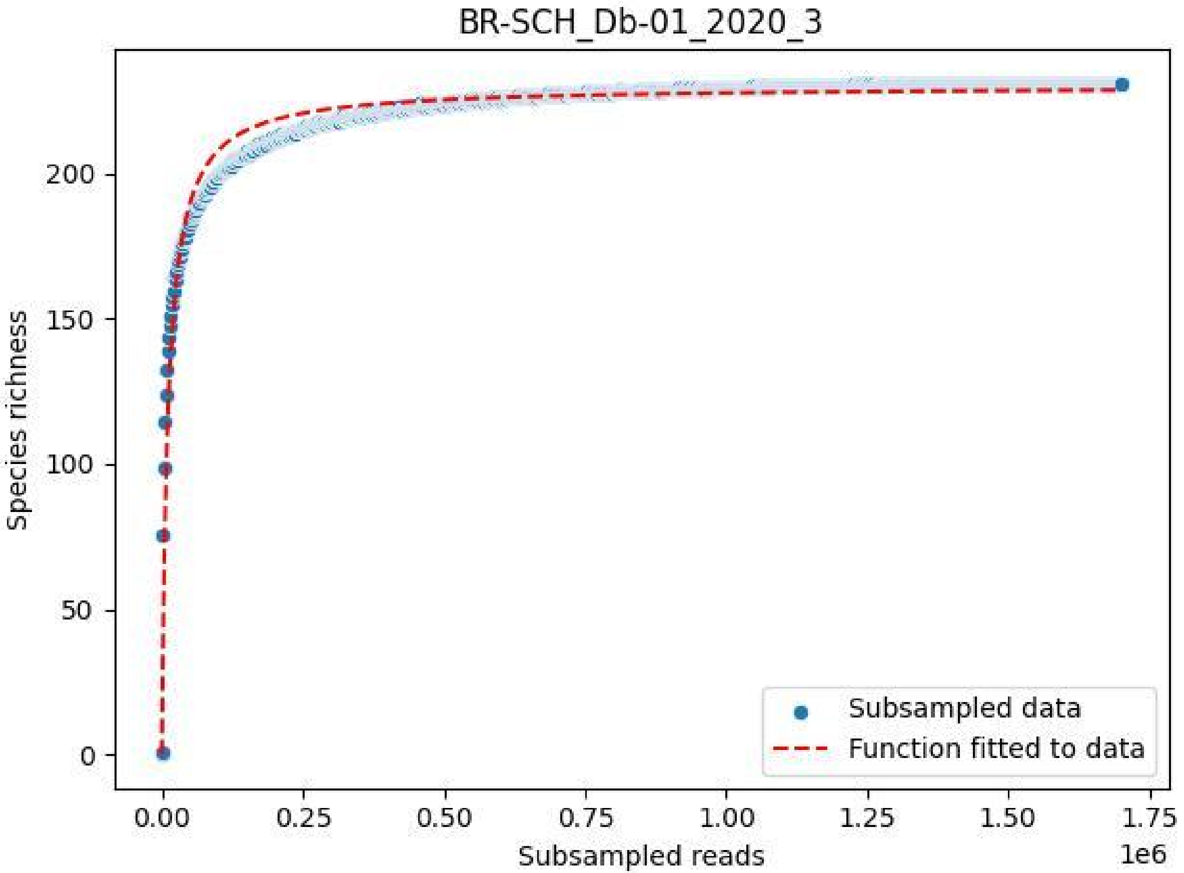
BR-SCH_Db-01_2020_3
Species richness
0
50
100
150
200
0.00
0.25
0.50
0.75
1.00
1.25
1.50
1.75
1e6
Subsampled reads
Subsampled data
Function fitted to data

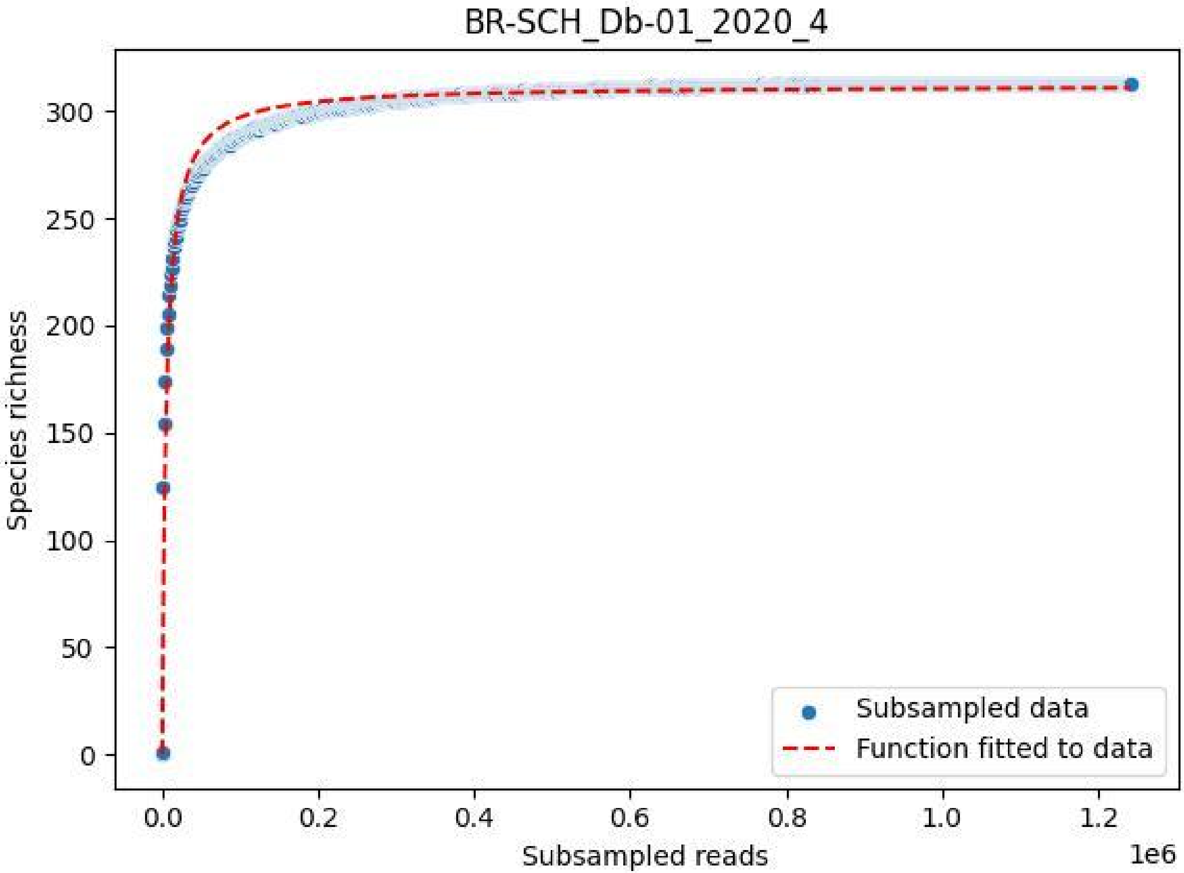
BR-SCH_Db-01_2020_4
Species richness
Subsampled reads
1e6
Subsampled data
Function fitted to data
0
50
100
150
200
250
300
0.0
0.2
0.4
0.6
0.8
1.0
1.2

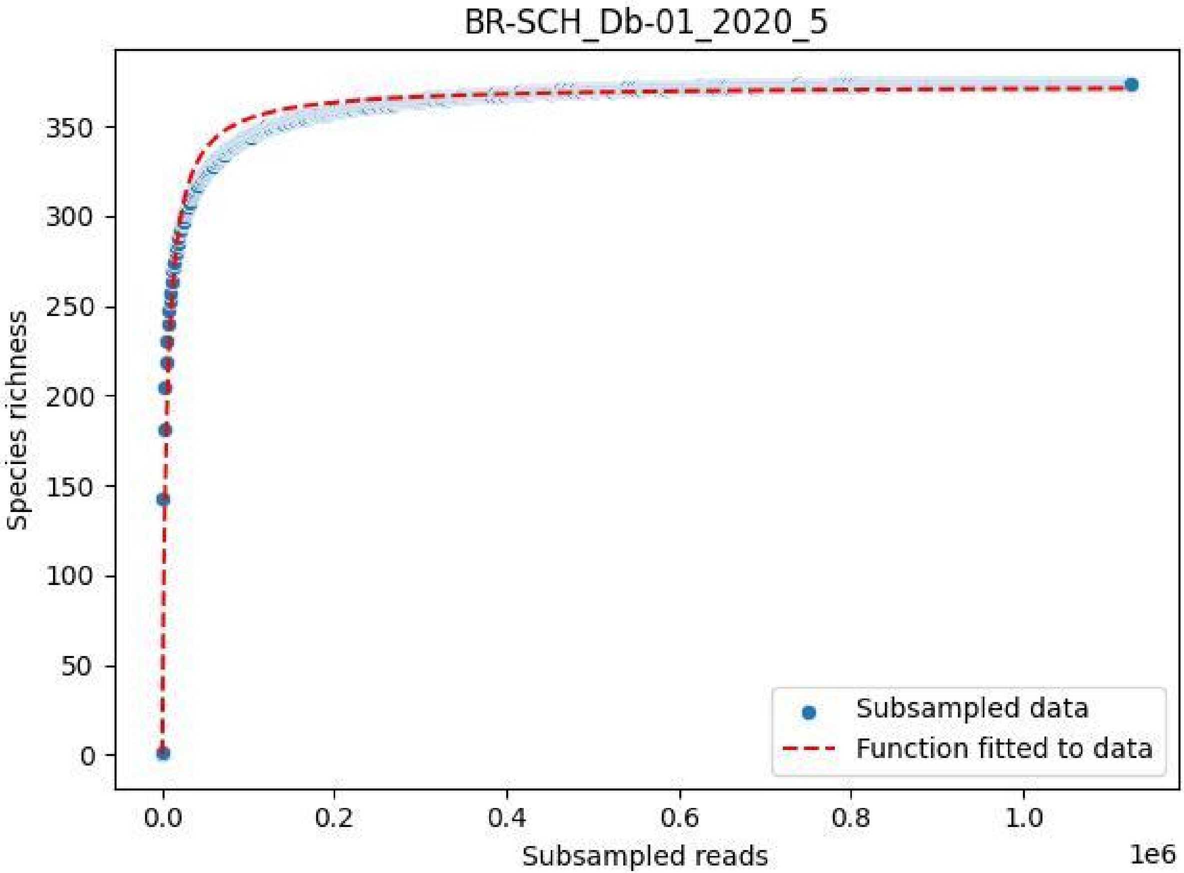

BR-SCH_Db-01_2020_5
Species richness
Subsampled reads
1e6
0
50
100
150
200
250
300
350
0.0
0.2
0.4
0.6
0.8
1.0
Subsampled data
Function fitted to data

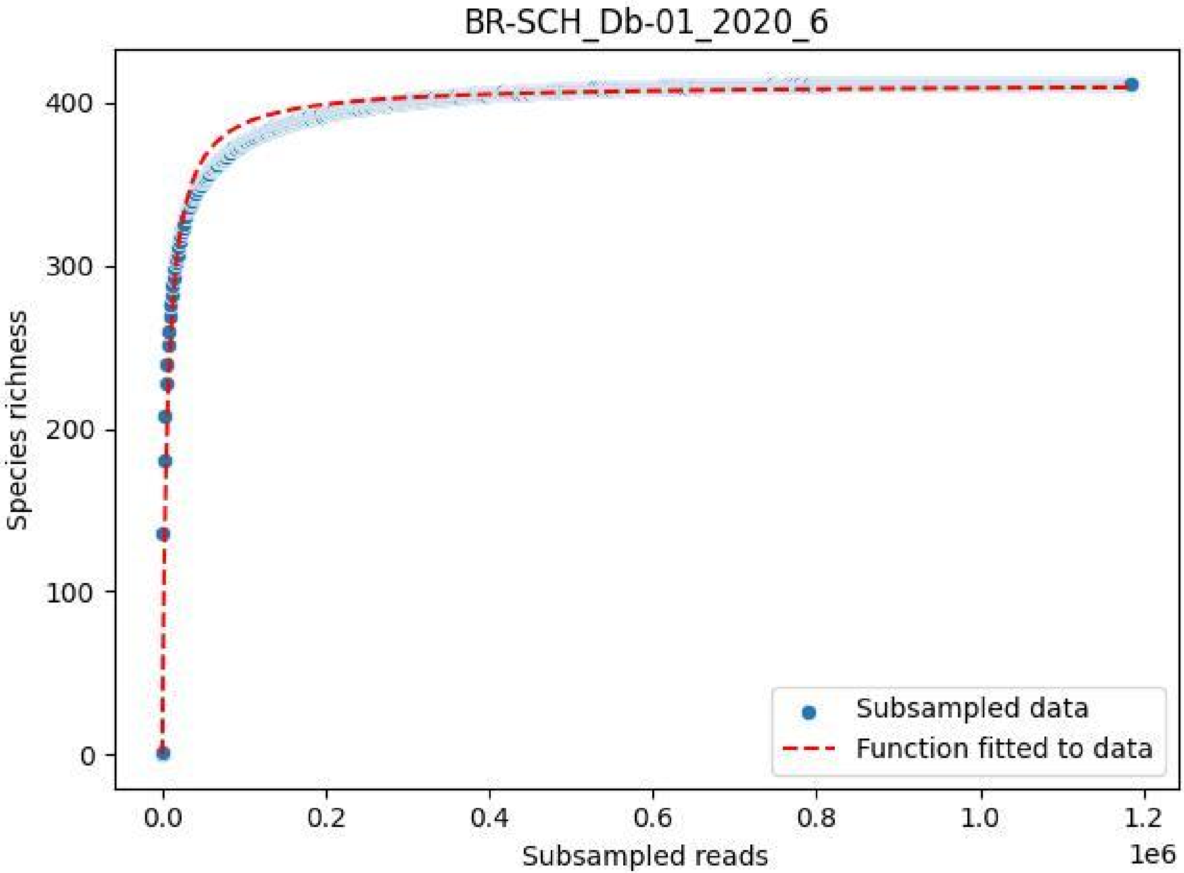
BR-SCH_Db-01_2020_6
Species richness
0
100
200
300
400
0.0
0.2
0.4
0.6
0.8
1.0
1.2
1e6
Subsampled reads
Subsampled data
Function fitted to data

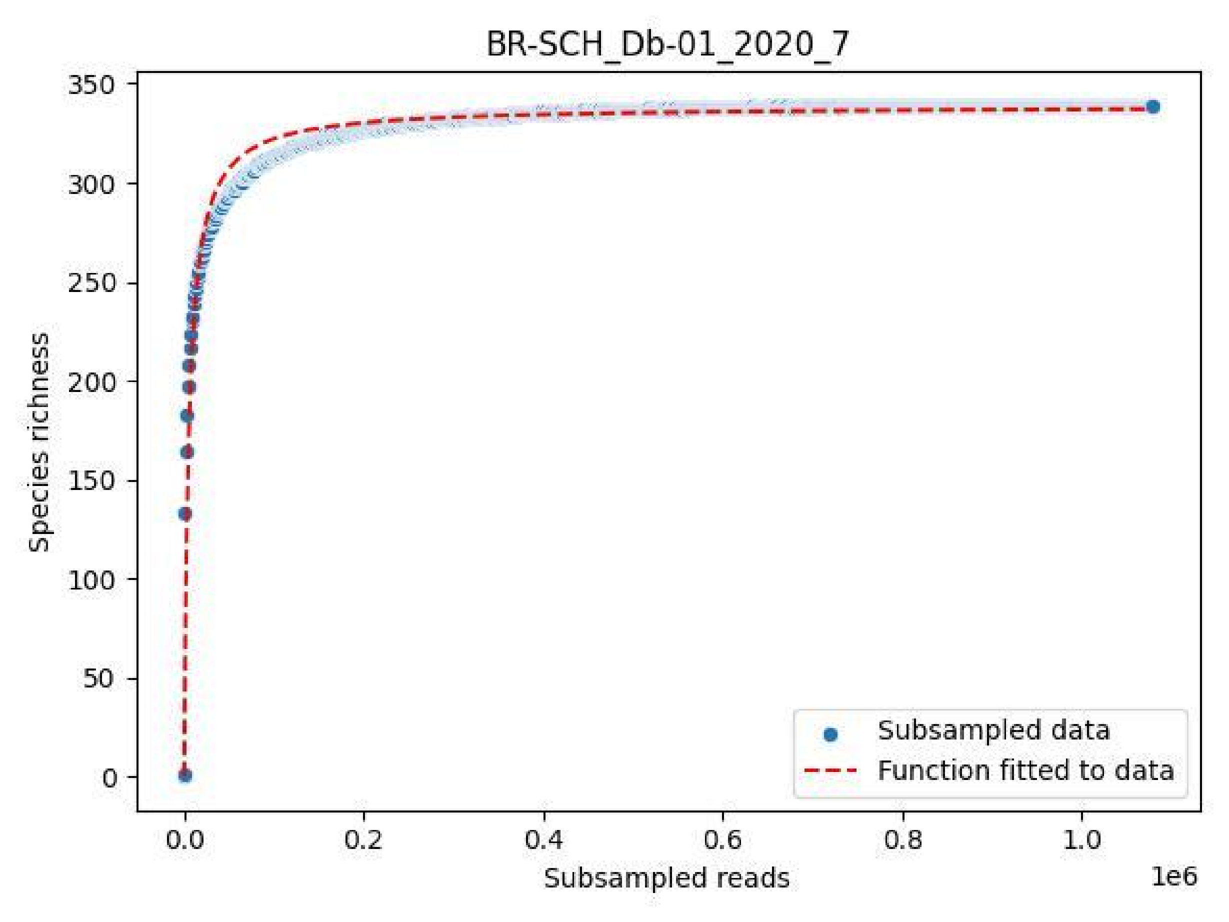
BR-SCH_Db-01_2020_7
Species richness
Subsampled reads
1e6
0
50
100
150
200
250
300
350
0.0
0.2
0.4
0.6
0.8
1.0
Subsampled data
Function fitted to data

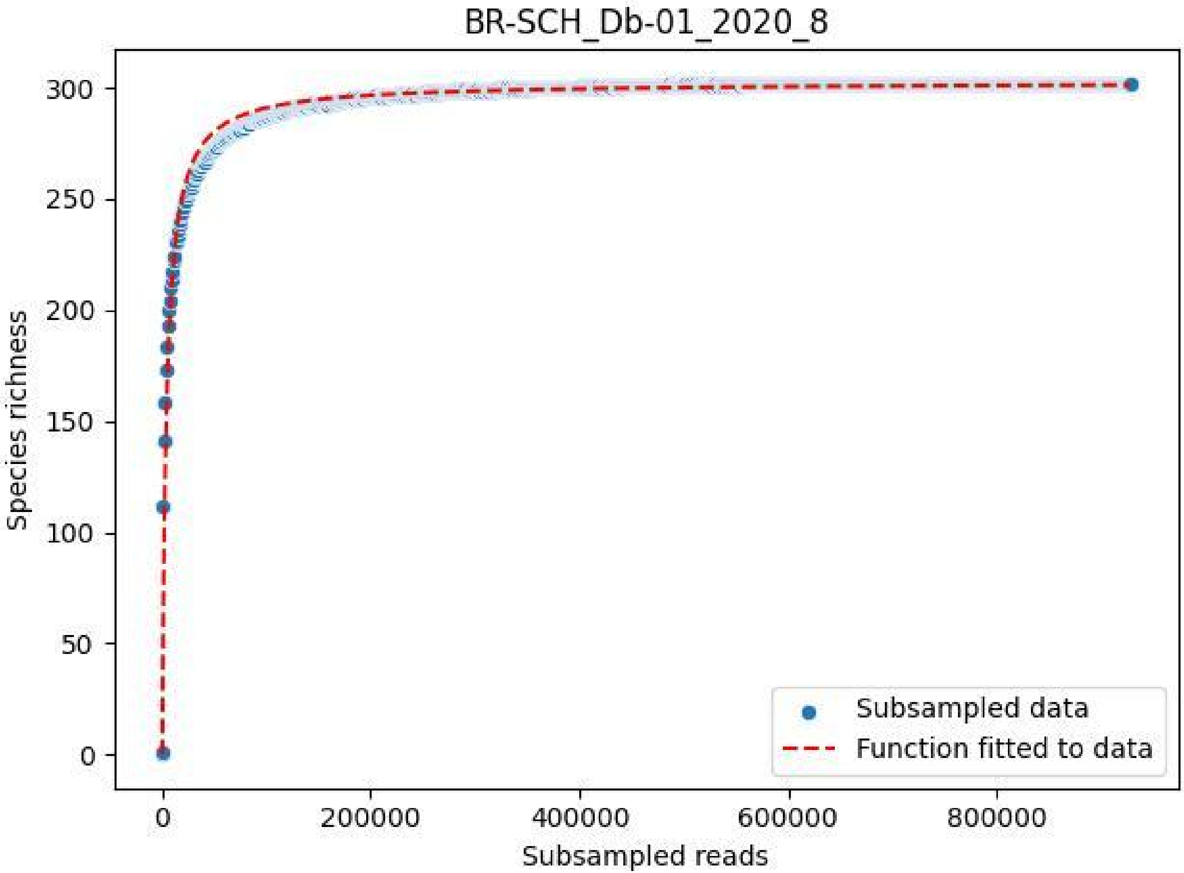
BR-SCH_Db-01_2020_8
Species richness
Subsampled reads
300
250
200
150
100
50
0
0
200000
400000
600000
800000
Subsampled data
Function fitted to data

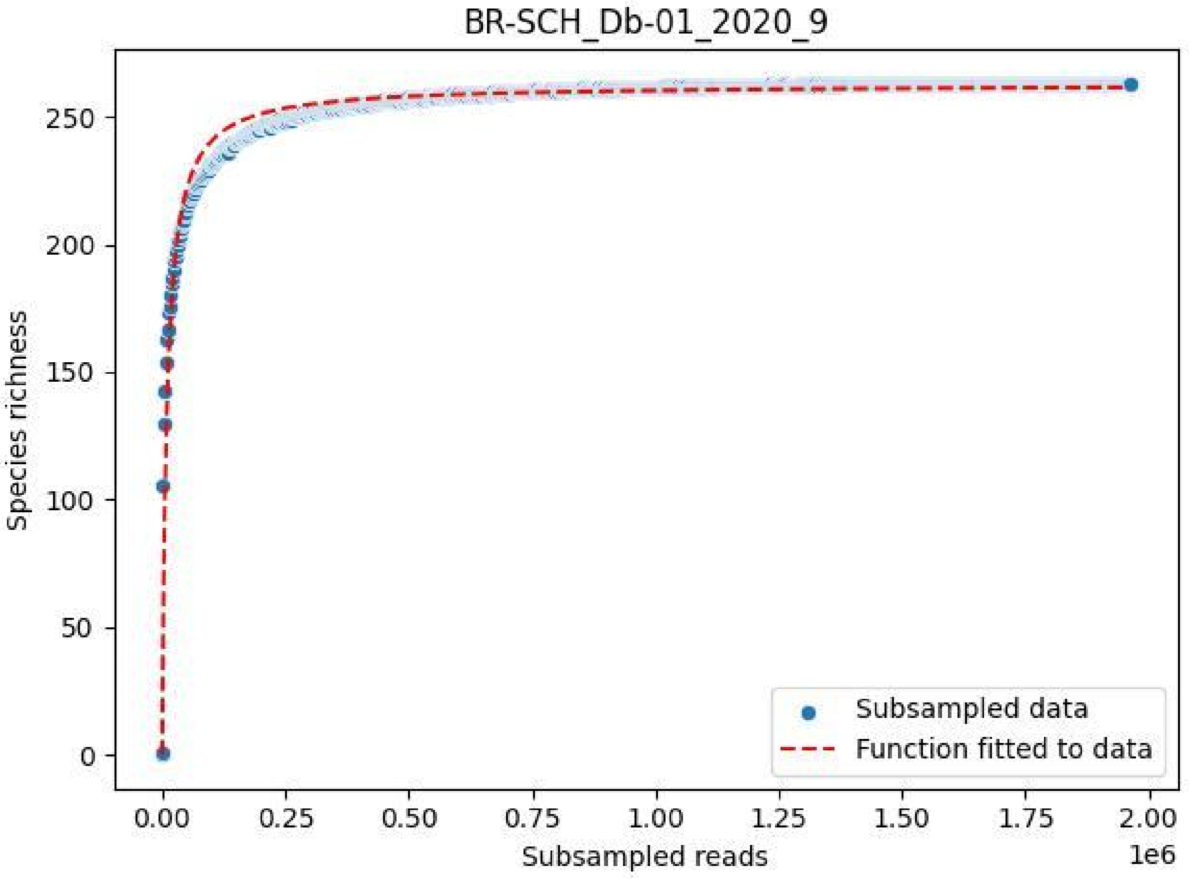
BR-SCH_Db-01_2020_9
Species richness
Subsampled reads
1e6
0
50
100
150
200
250
0.00
0.25
0.50
0.75
1.00
1.25
1.50
1.75
2.00
Subsampled data
Function fitted to data

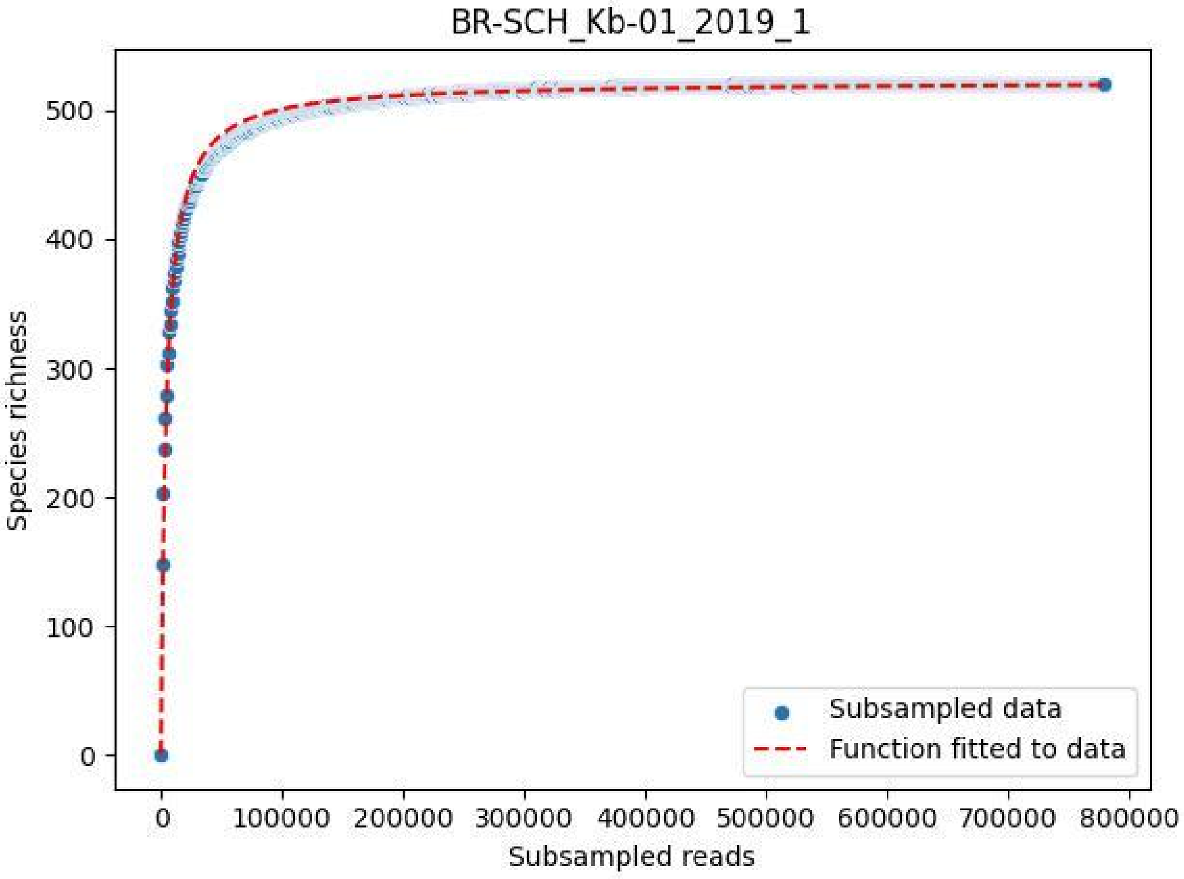
BR-SCH_Kb-01_2019_1
Species richness
Subsampled reads
0
100
200
300
400
500
0
100000
200000
300000
400000
500000
600000
700000
800000
Subsampled data
Function fitted to data

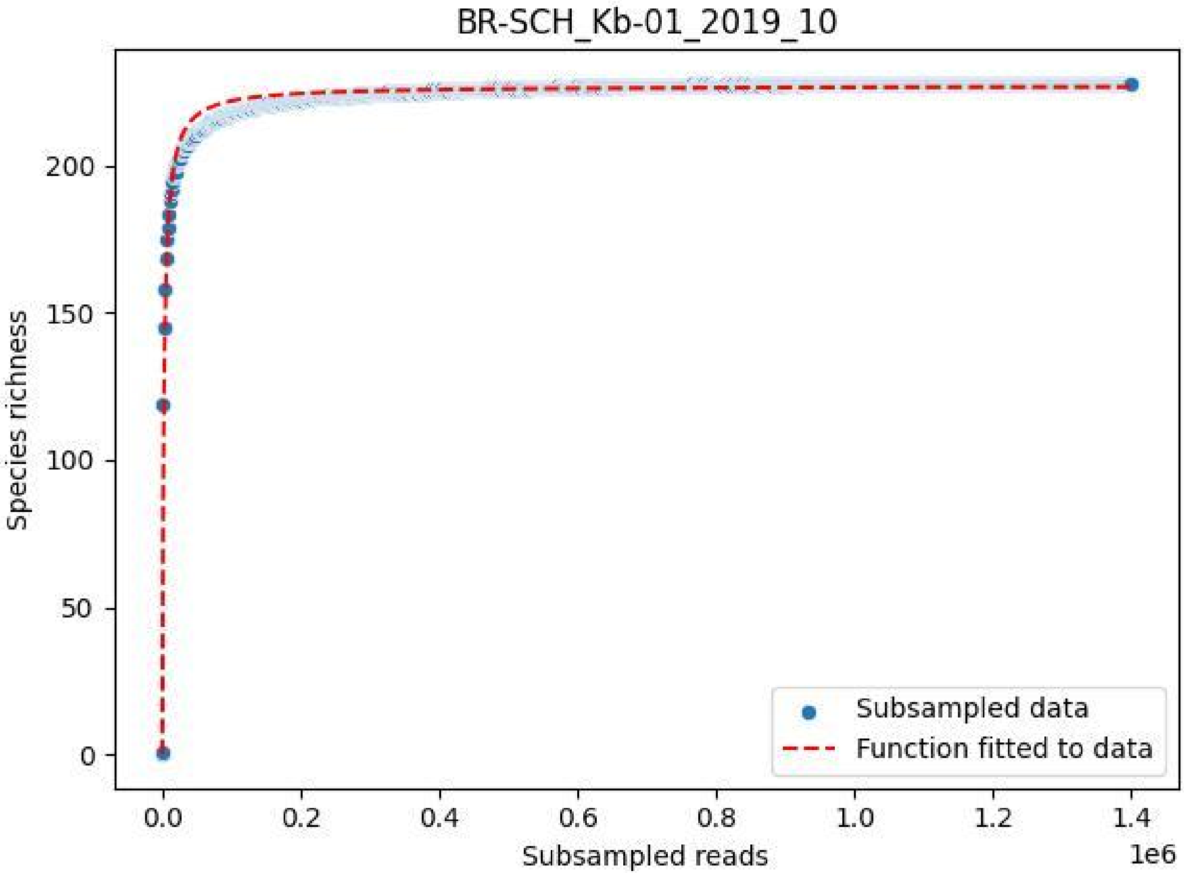
BR-SCH_Kb-01_2019_10
Species richness
200
150
100
50
0
0.0
0.2
0.4
0.6
0.8
1.0
1.2
1.4
Subsampled reads
1e6
Subsampled data
Function fitted to data

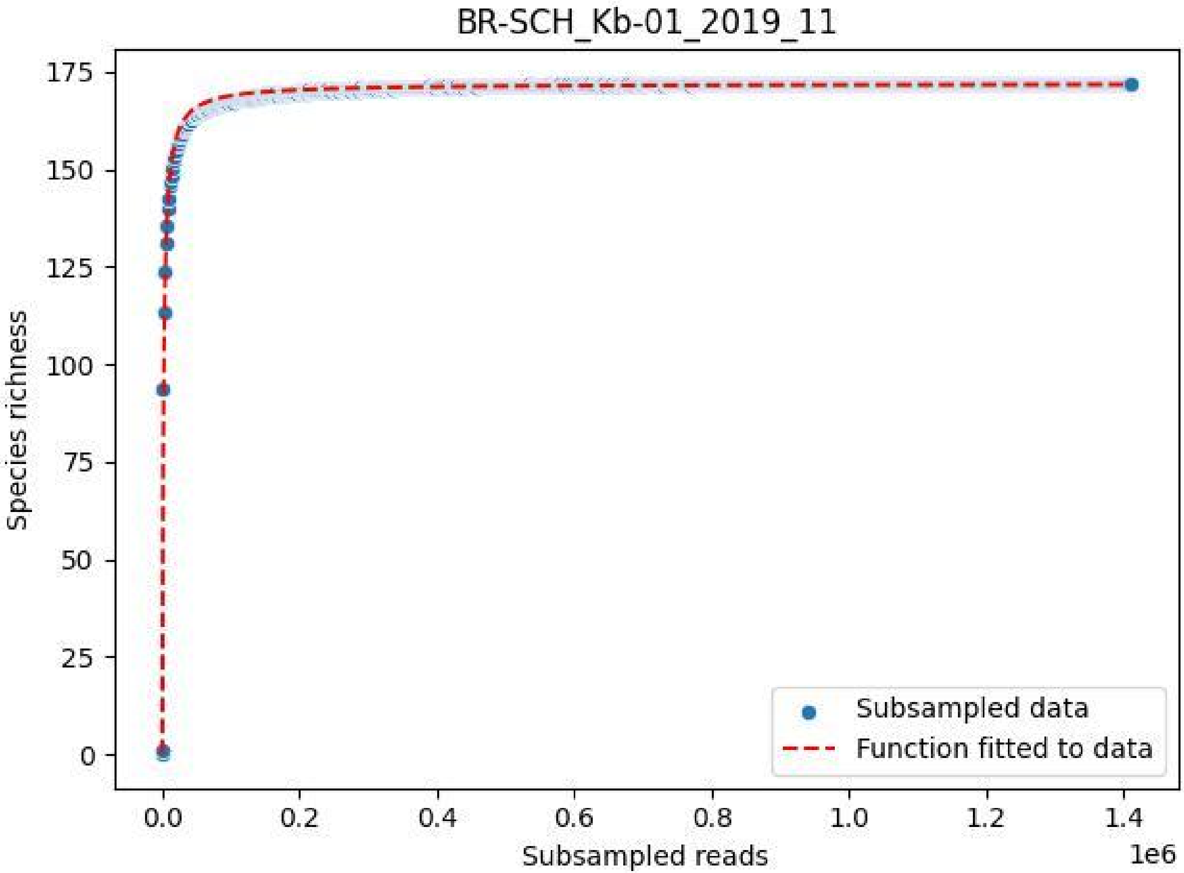
BR-SCH_Kb-01_2019_11
Species richness
Subsampled reads
1e6
0
25
50
75
100
125
150
175
0.0
0.2
0.4
0.6
0.8
1.0
1.2
1.4
Subsampled data
Function fitted to data

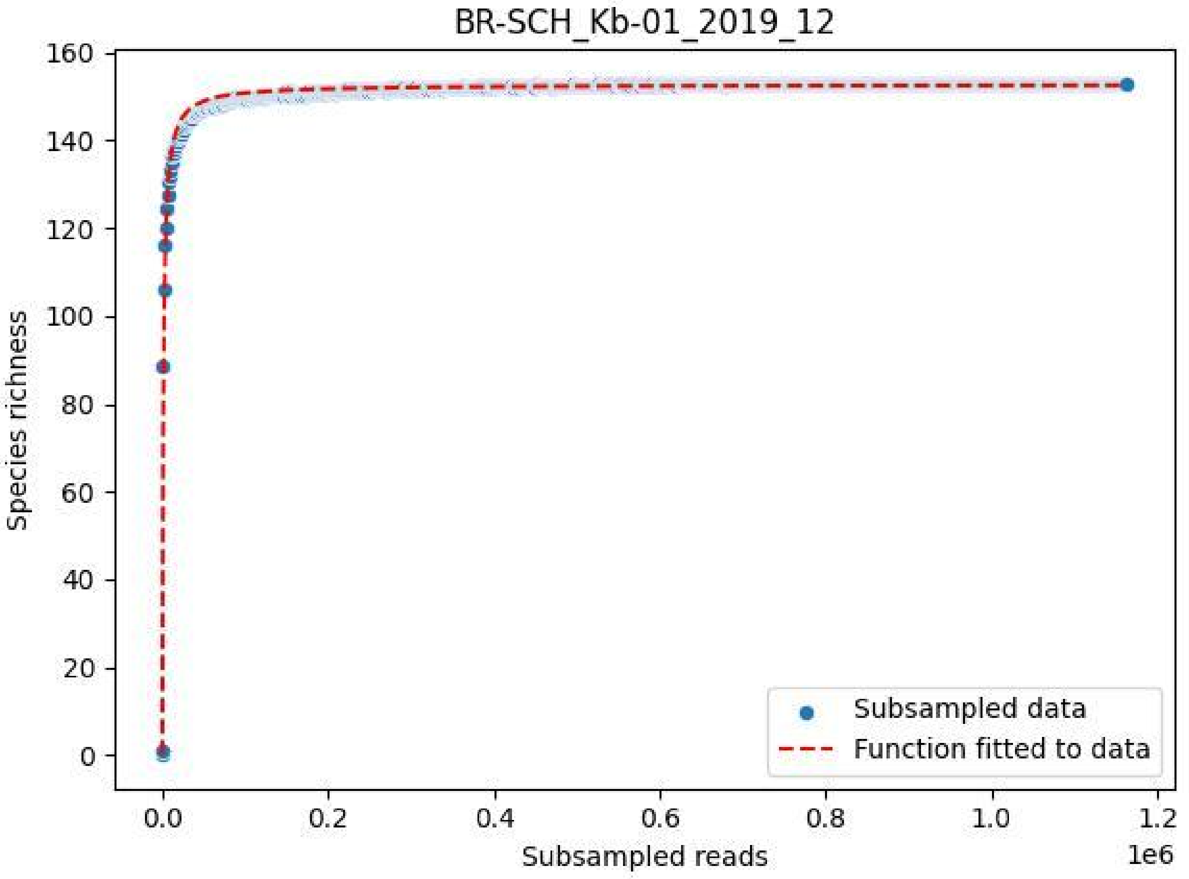

BR-SCH_Kb-01_2019_12
Species richness
Subsampled reads
1e6
0
20
40
60
80
100
120
140
160
0.0
0.2
0.4
0.6
0.8
1.0
1.2
Subsampled data
Function fitted to data

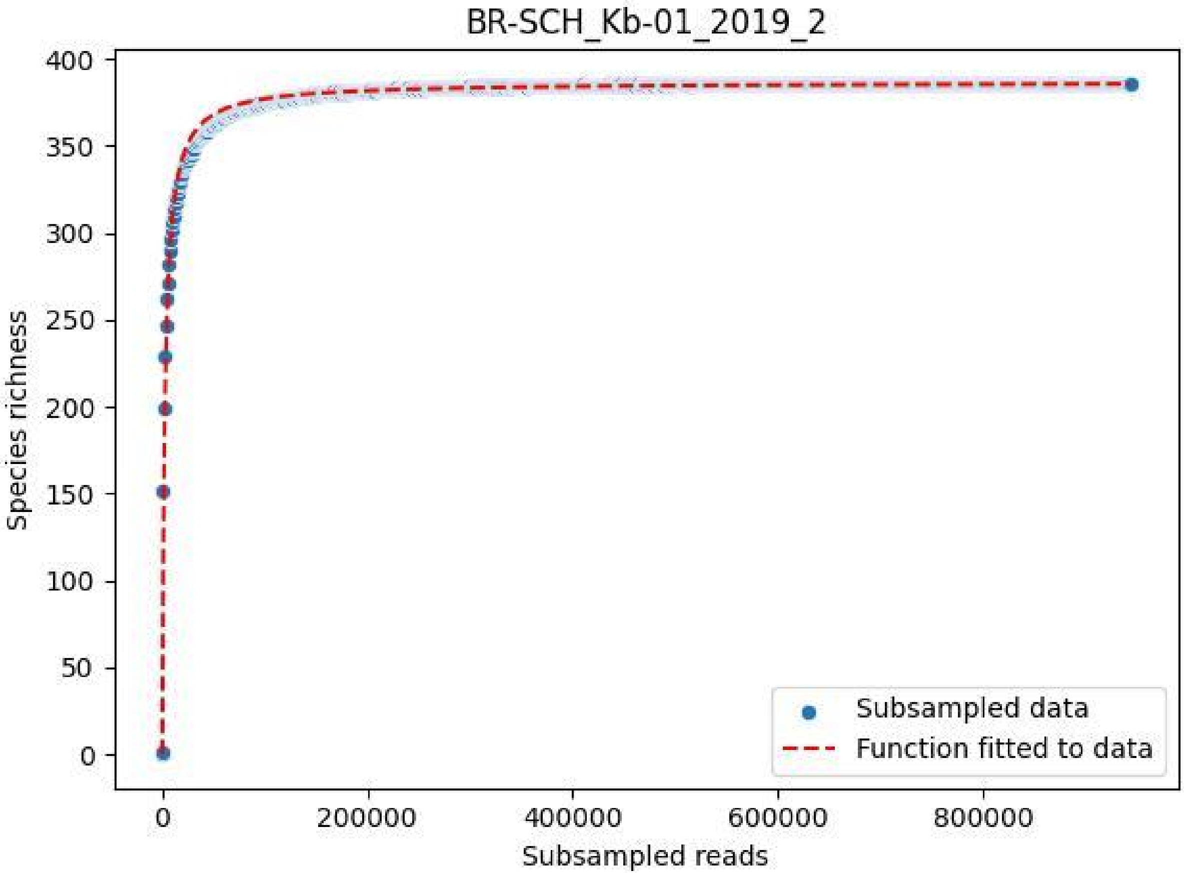
BR-SCH_Kb-01_2019_2
Species richness
Subsampled reads
400
350
300
250
200
150
100
50
0
0
200000
400000
600000
800000
Subsampled data
Function fitted to data

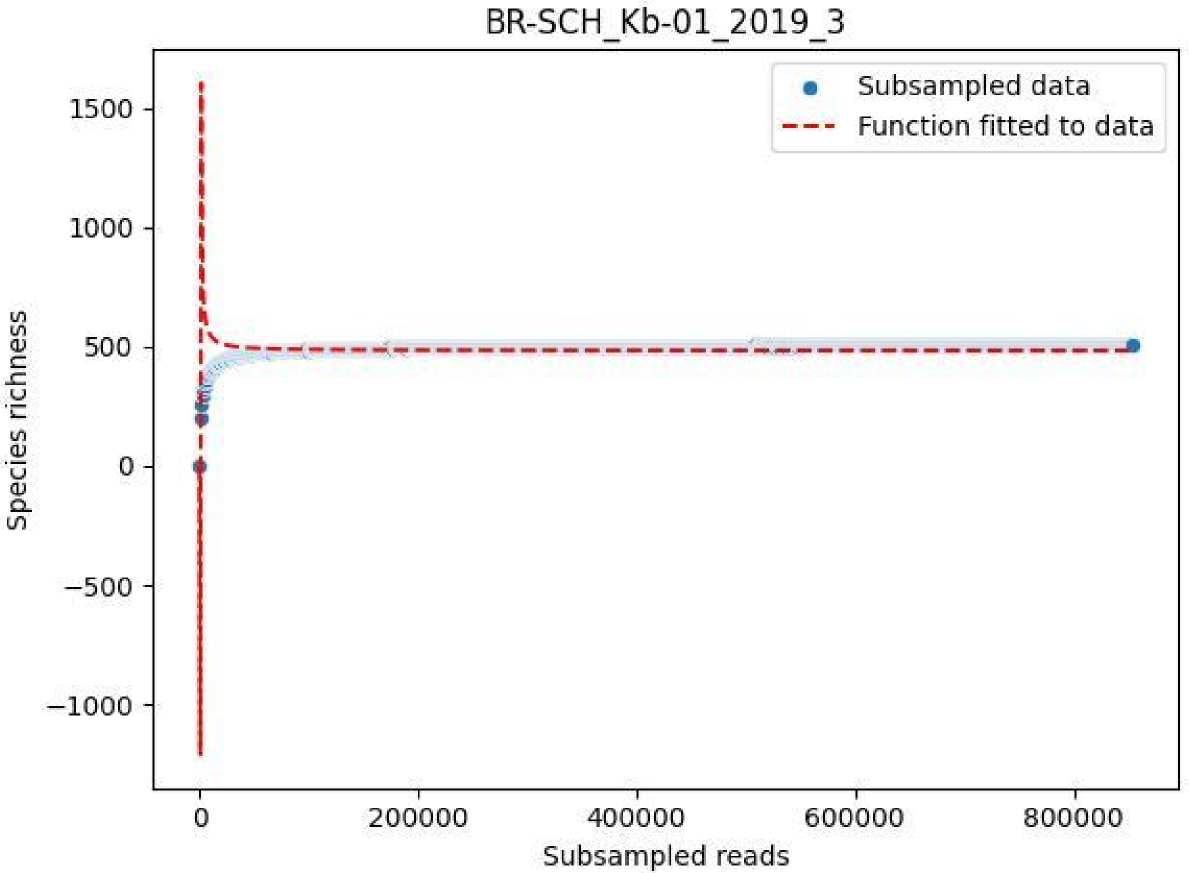

BR-SCH_Kb-01_2019_3
Subsampled data
Function fitted to data
1500
1000
500
0
−500
−1000
Species richness
0
200000
400000
600000
800000
Subsampled reads

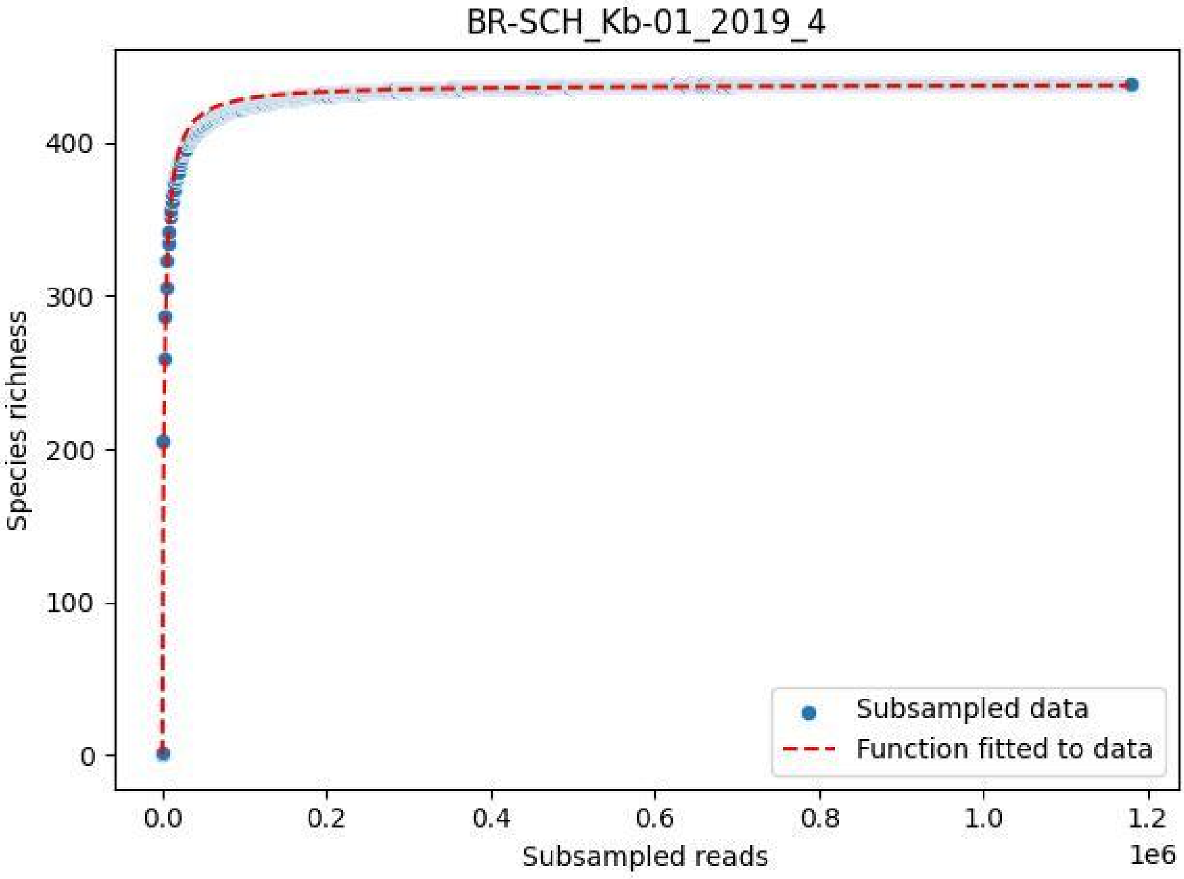
BR-SCH_Kb-01_2019_4
Species richness
Subsampled reads
1e6
0
100
200
300
400
0.0
0.2
0.4
0.6
0.8
1.0
1.2
Subsampled data
Function fitted to data

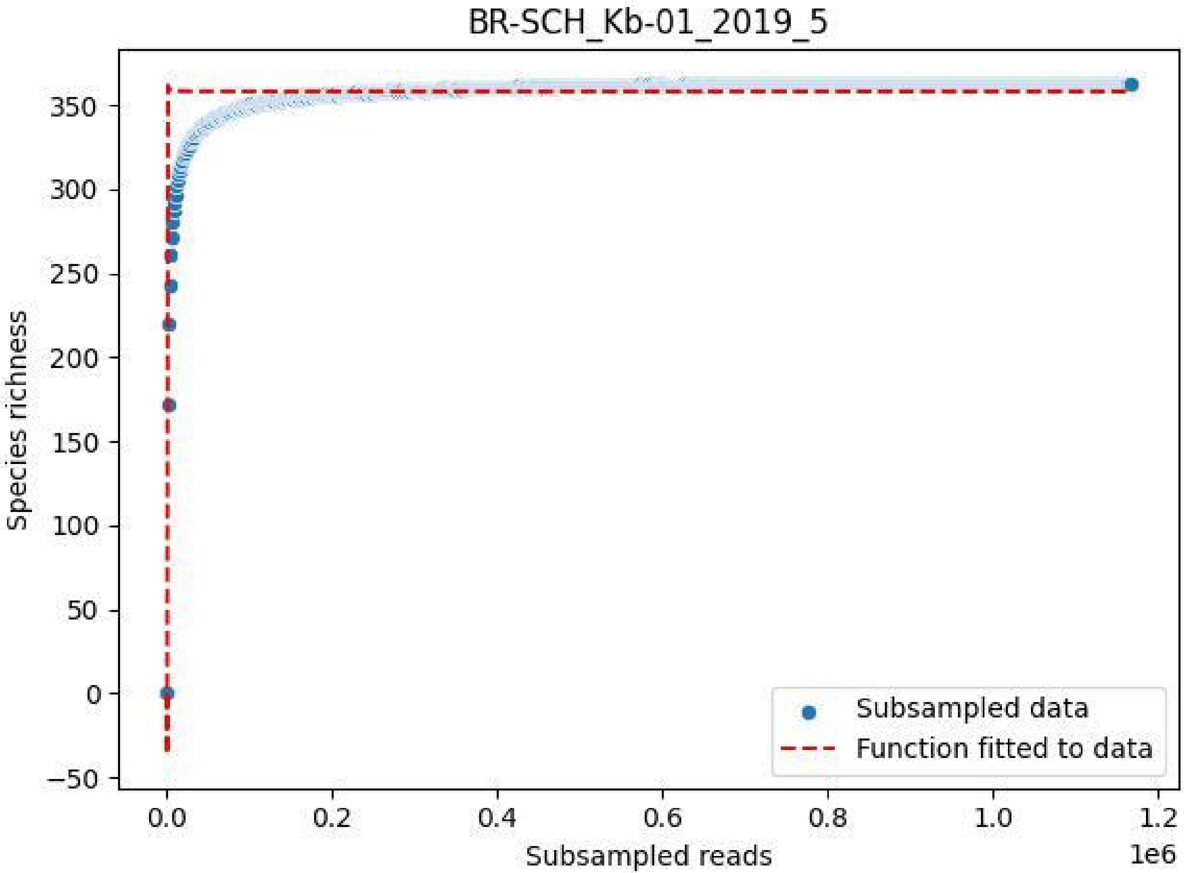

BR-SCH_Kb-01_2019_5
Species richness
Subsampled reads
1e6
350
300
250
200
150
100
50
0
−50
0.0
0.2
0.4
0.6
0.8
1.0
1.2
Subsampled data
Function fitted to data

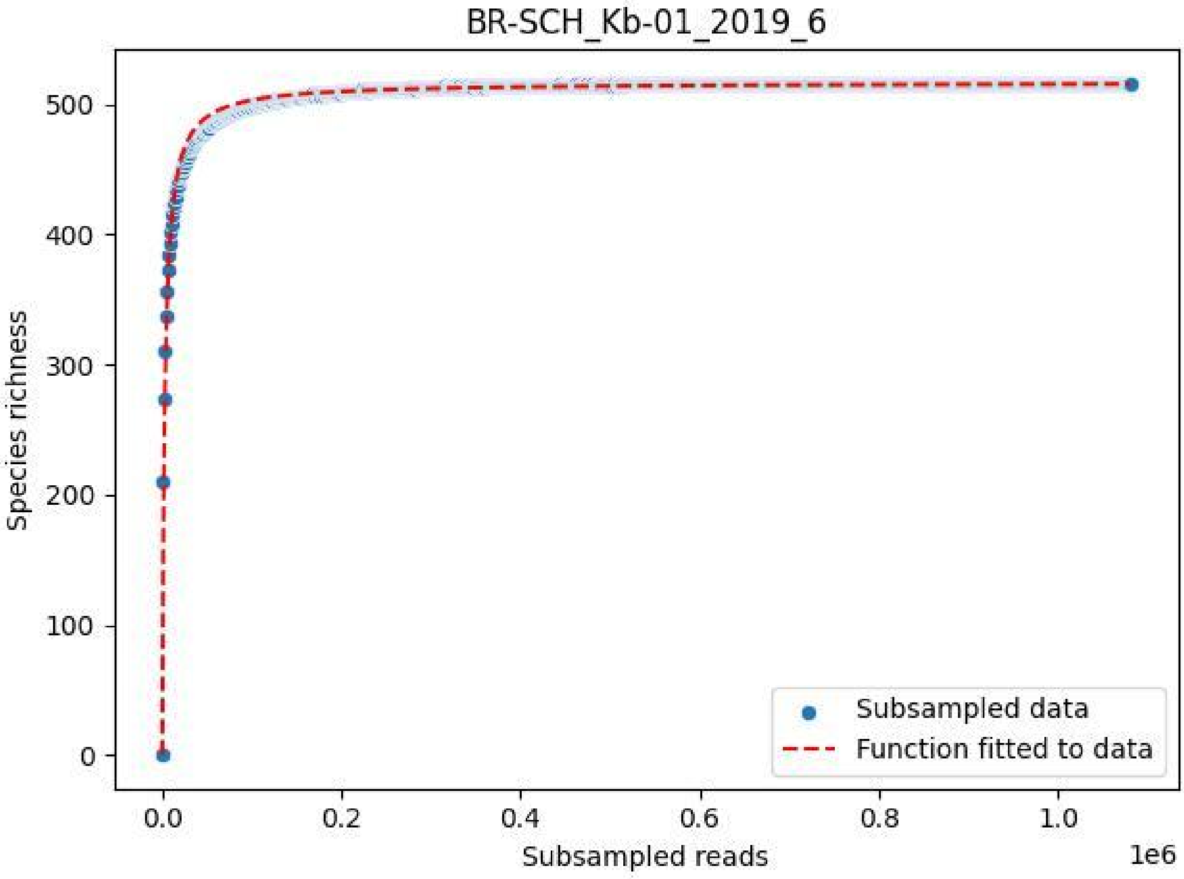
BR-SCH_Kb-01_2019_6
Species richness
Subsampled reads
1e6
Subsampled data
Function fitted to data
0
100
200
300
400
500
0.0
0.2
0.4
0.6
0.8
1.0

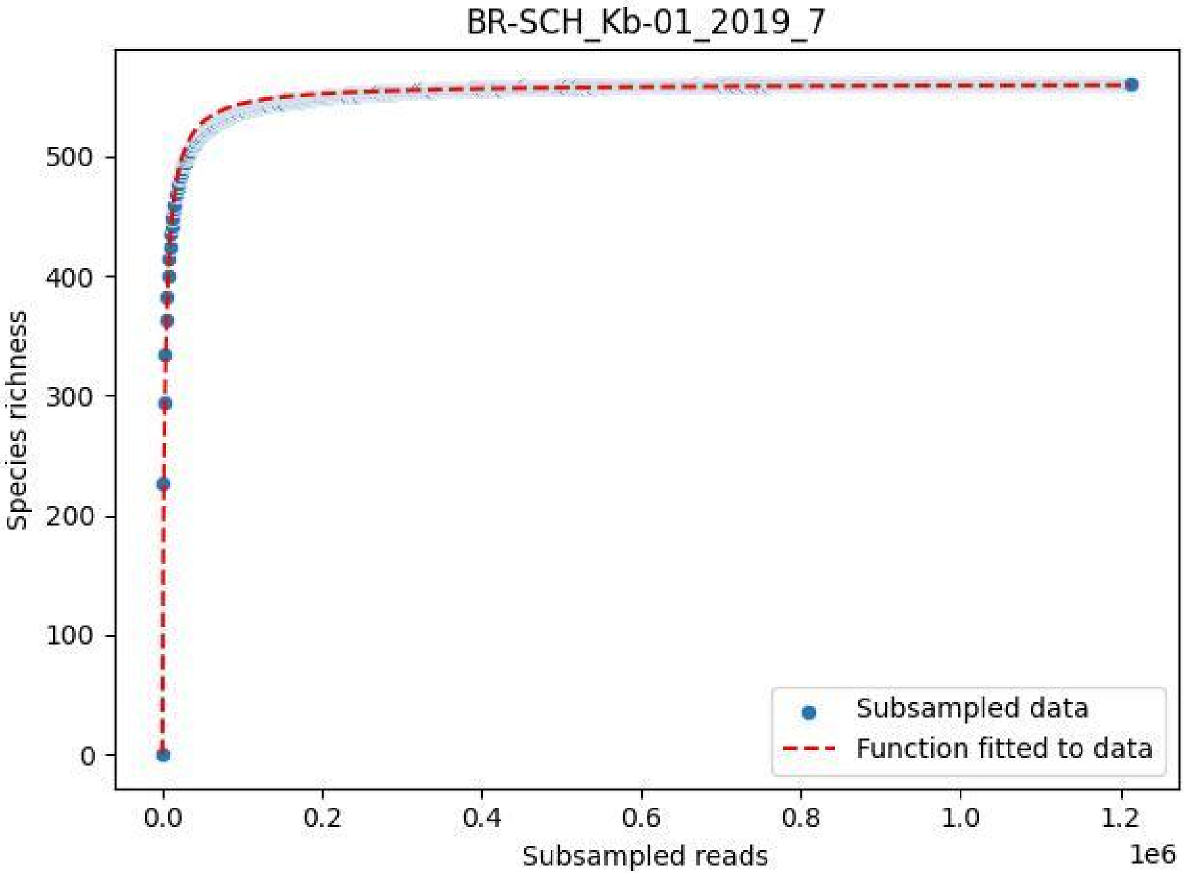
BR-SCH_Kb-01_2019_7
Species richness
500
400
300
200
100
0
0.0
0.2
0.4
0.6
0.8
1.0
1.2
1e6
Subsampled reads
Subsampled data
Function fitted to data

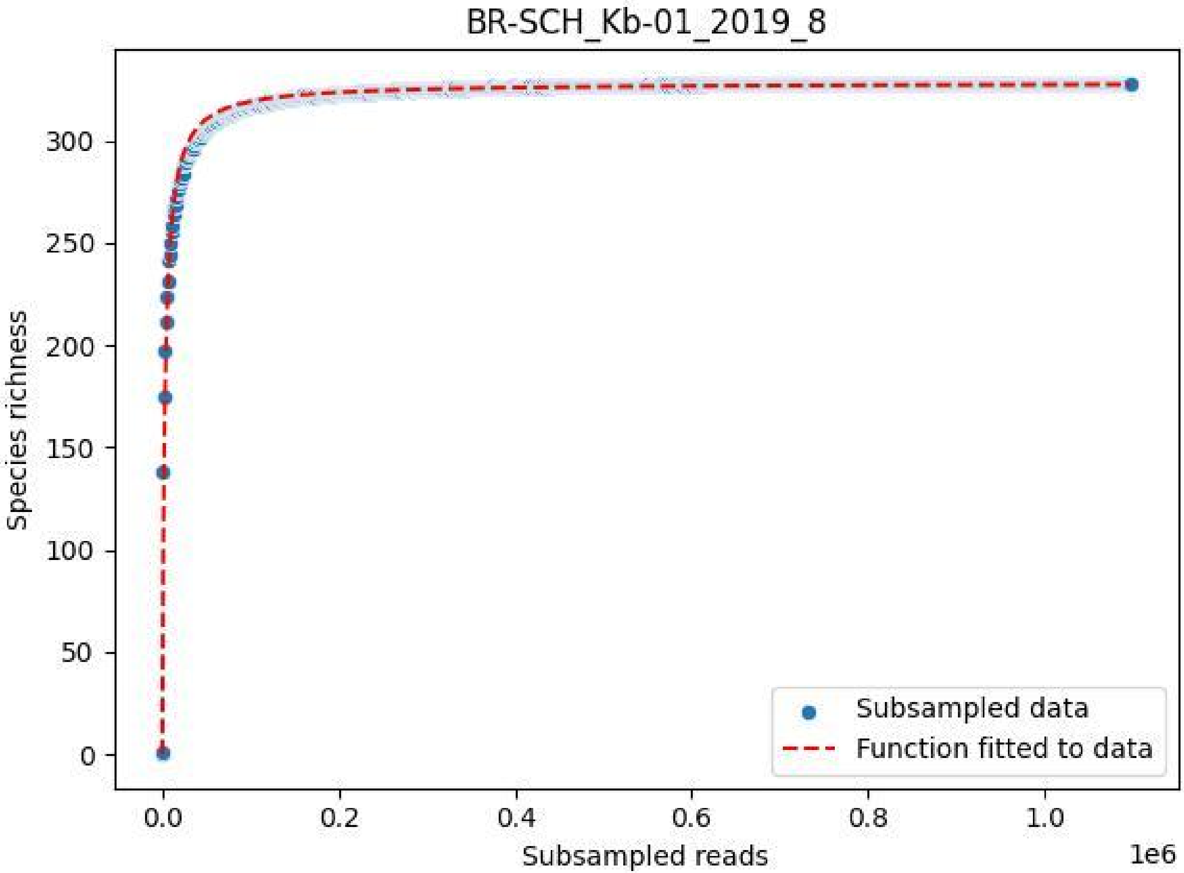
BR-SCH_Kb-01_2019_8
Species richness
Subsampled reads
1e6
0
50
100
150
200
250
300
0.0
0.2
0.4
0.6
0.8
1.0
Subsampled data
Function fitted to data

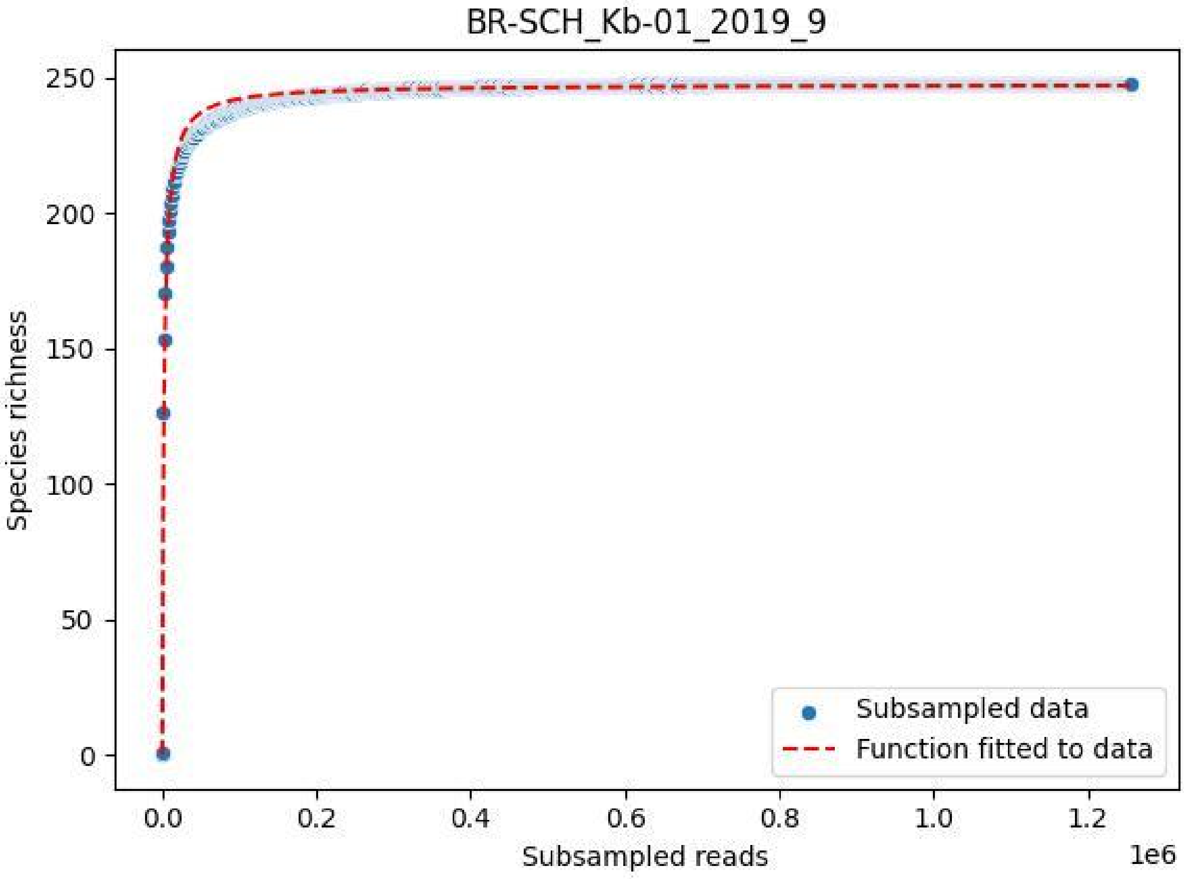
BR-SCH_Kb-01_2019_9
Species richness
Subsampled reads
1e6
Subsampled data
Function fitted to data
250
200
150
100
50
0
0.0
0.2
0.4
0.6
0.8
1.0
1.2

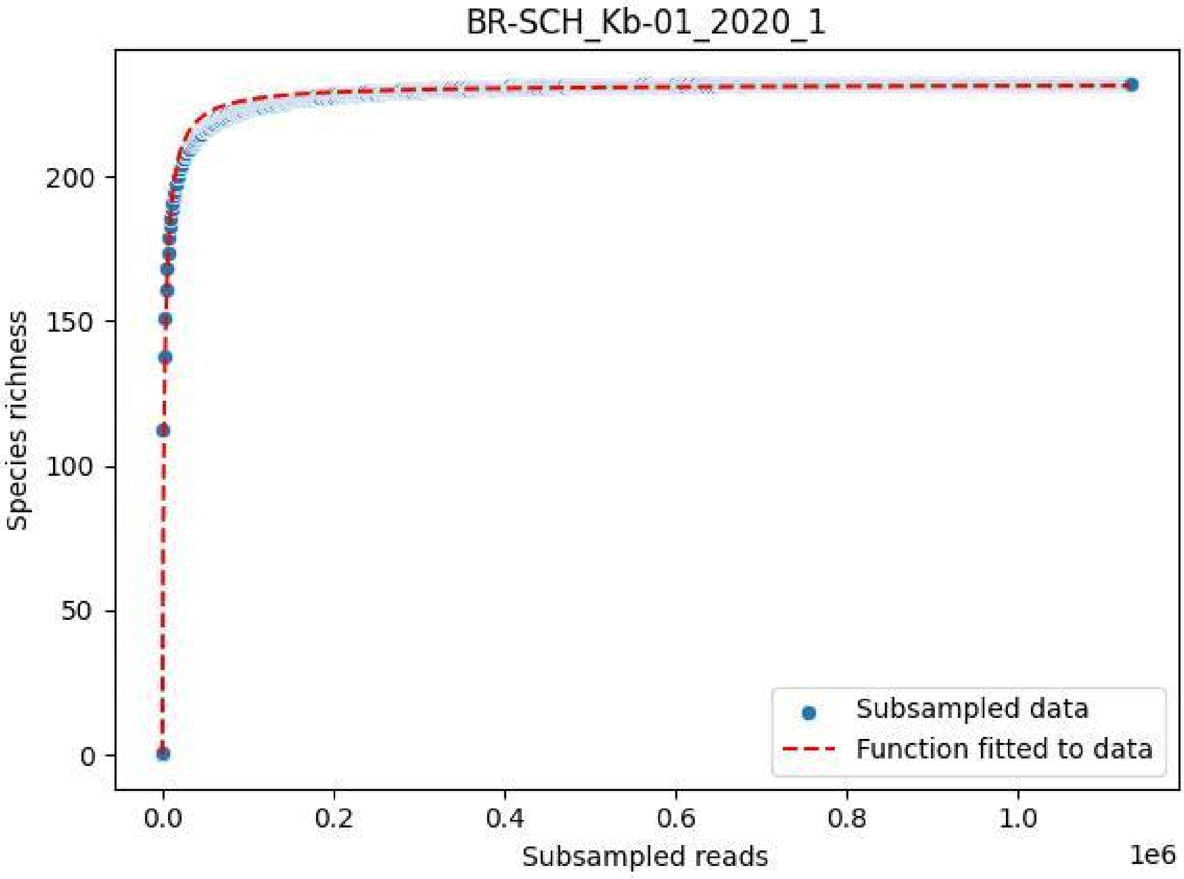

BR-SCH_Kb-01_2020_1
Species richness
Subsampled reads
1e6
0
50
100
150
200
0.0
0.2
0.4
0.6
0.8
1.0
Subsampled data
Function fitted to data

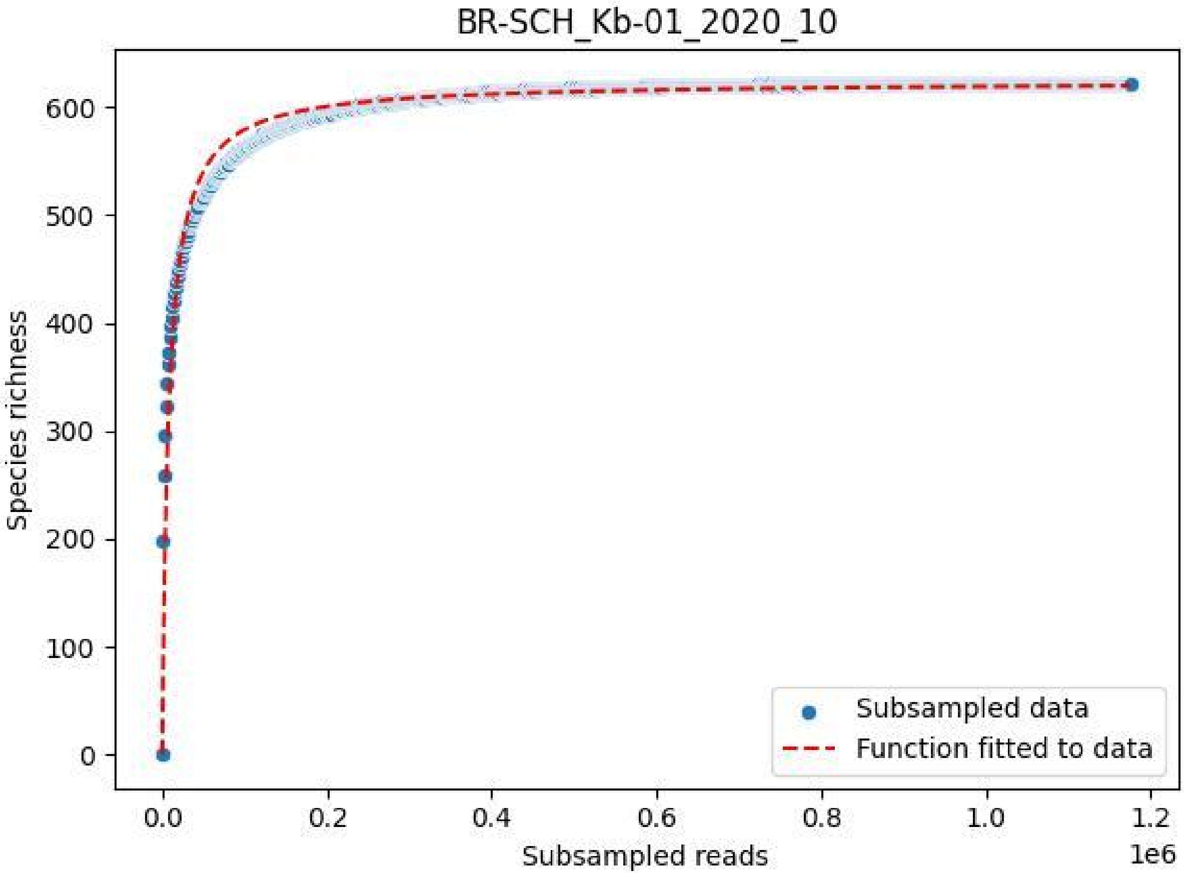
BR-SCH_Kb-01_2020_10
Species richness
600
500
400
300
200
100
0
0.0
0.2
0.4
0.6
0.8
1.0
1.2
1e6
Subsampled reads
Subsampled data
Function fitted to data

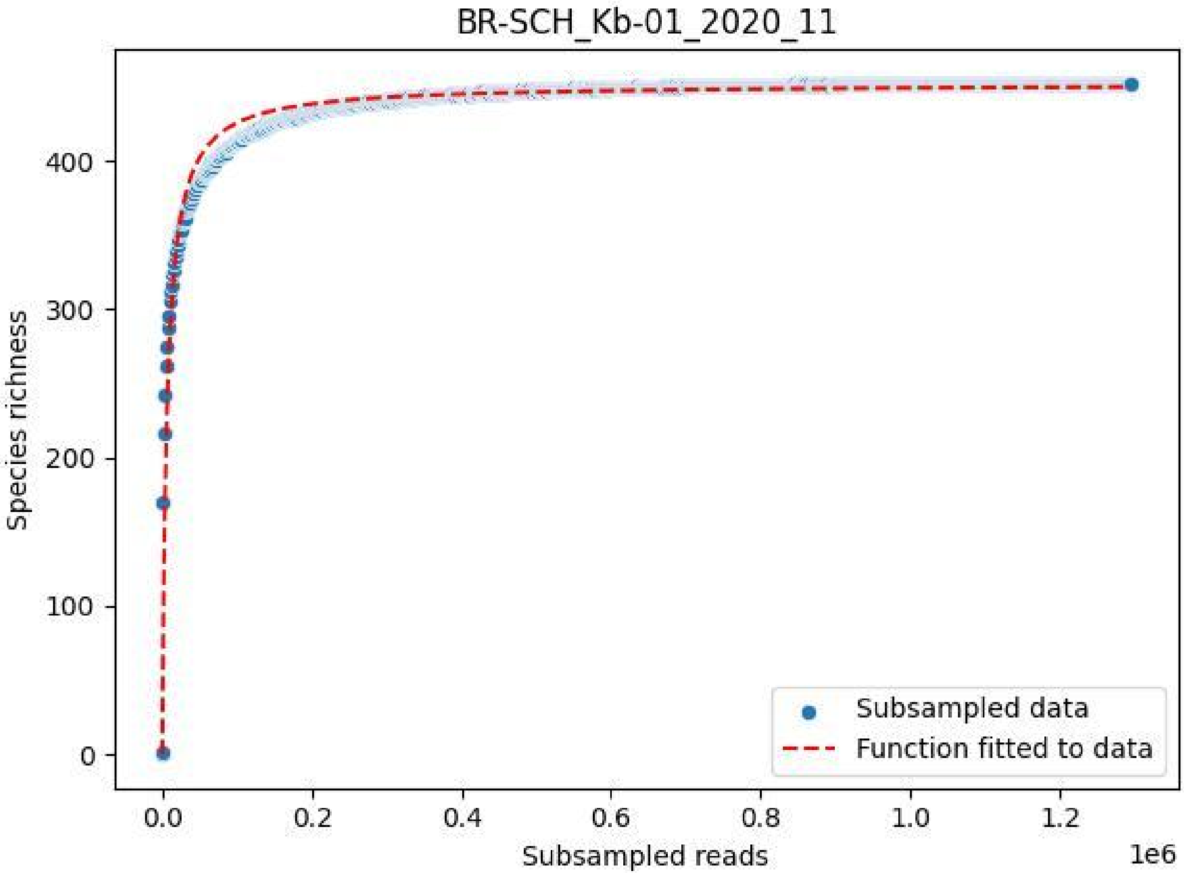
BR-SCH_Kb-01_2020_11
Species richness
0
100
200
300
400
Subsampled reads
0.0
0.2
0.4
0.6
0.8
1.0
1.2
1e6
Subsampled data
Function fitted to data

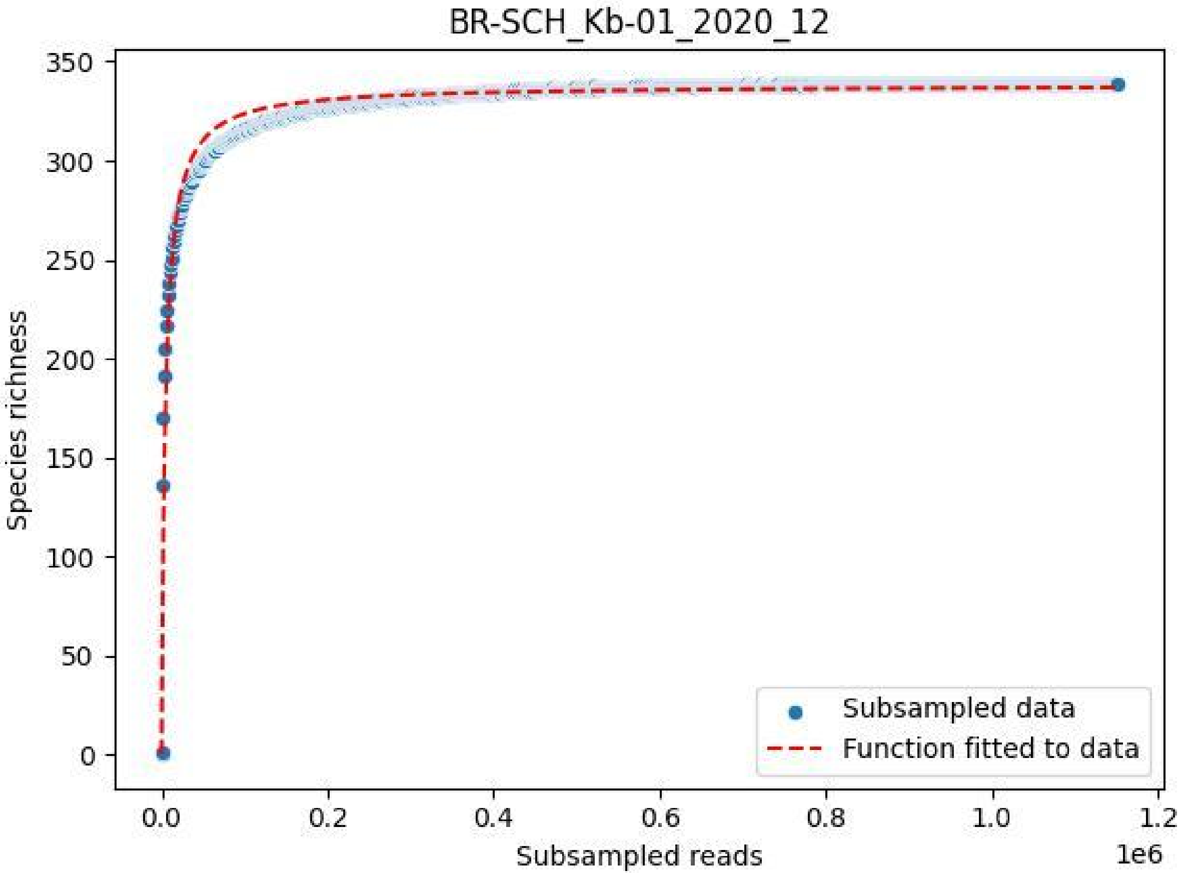
BR-SCH_Kb-01_2020_12
Species richness
Subsampled reads
1e6
0
50
100
150
200
250
300
350
0.0
0.2
0.4
0.6
0.8
1.0
1.2
Subsampled data
Function fitted to data

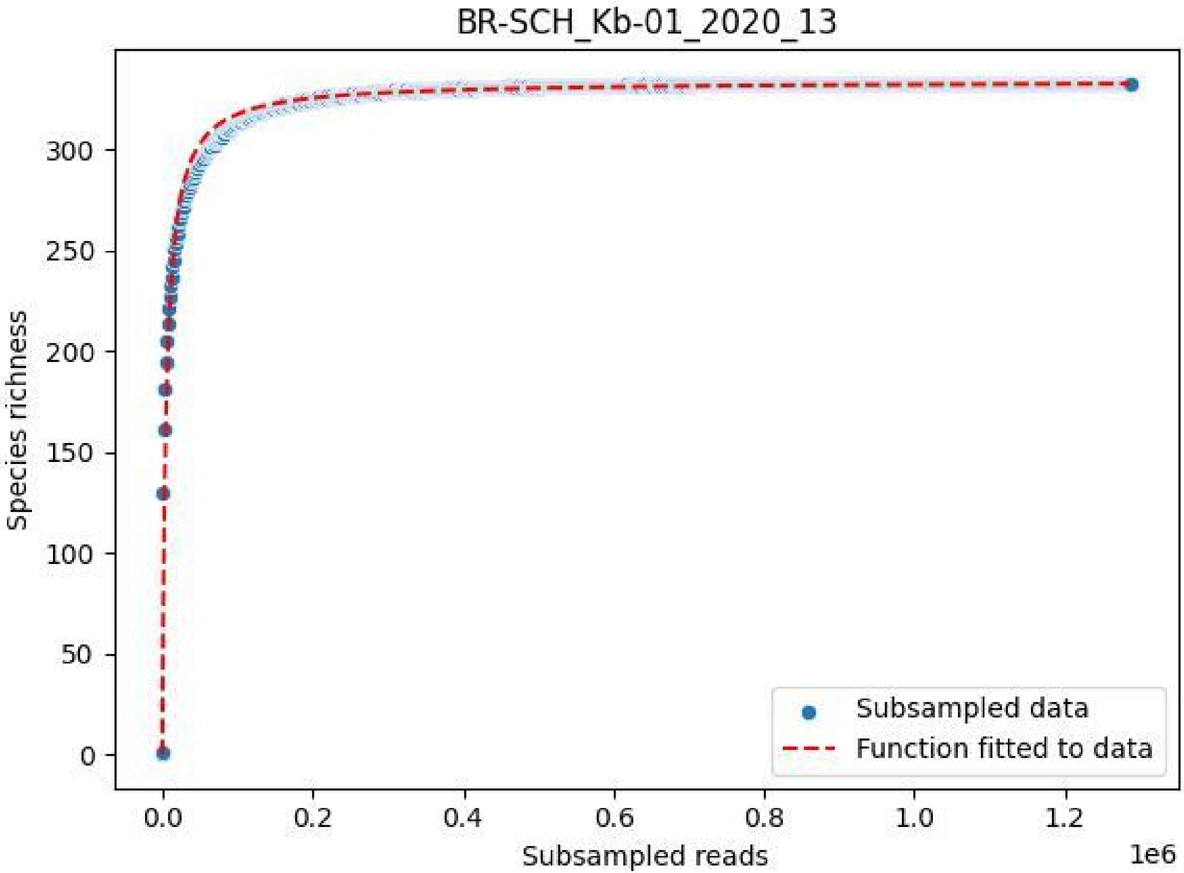
BR-SCH_Kb-01_2020_13
Species richness
300
250
200
150
100
50
0
0.0
0.2
0.4
0.6
0.8
1.0
1.2
1e6
Subsampled reads
Subsampled data
Function fitted to data

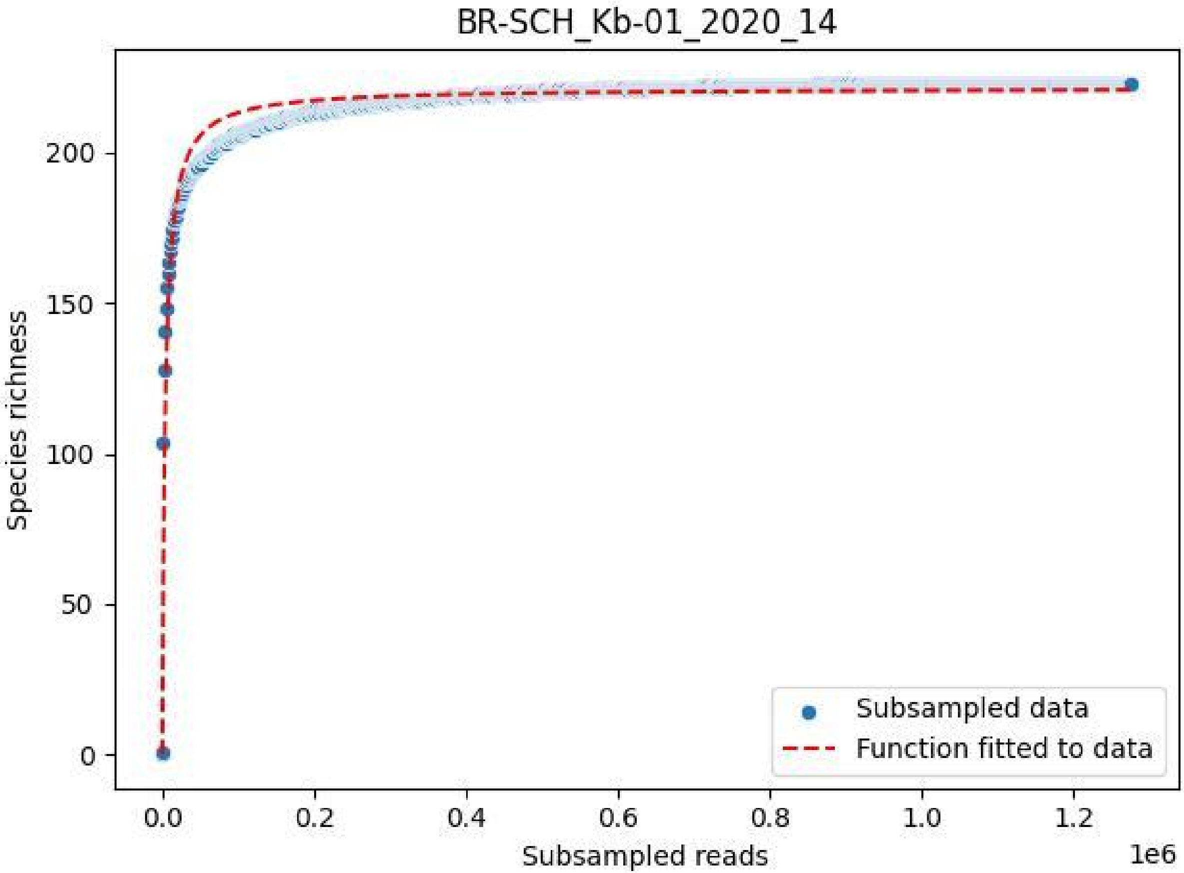
BR-SCH_Kb-01_2020_14
Species richness
0
50
100
150
200
0.0
0.2
0.4
0.6
0.8
1.0
1.2
1e6
Subsampled reads
Subsampled data
Function fitted to data

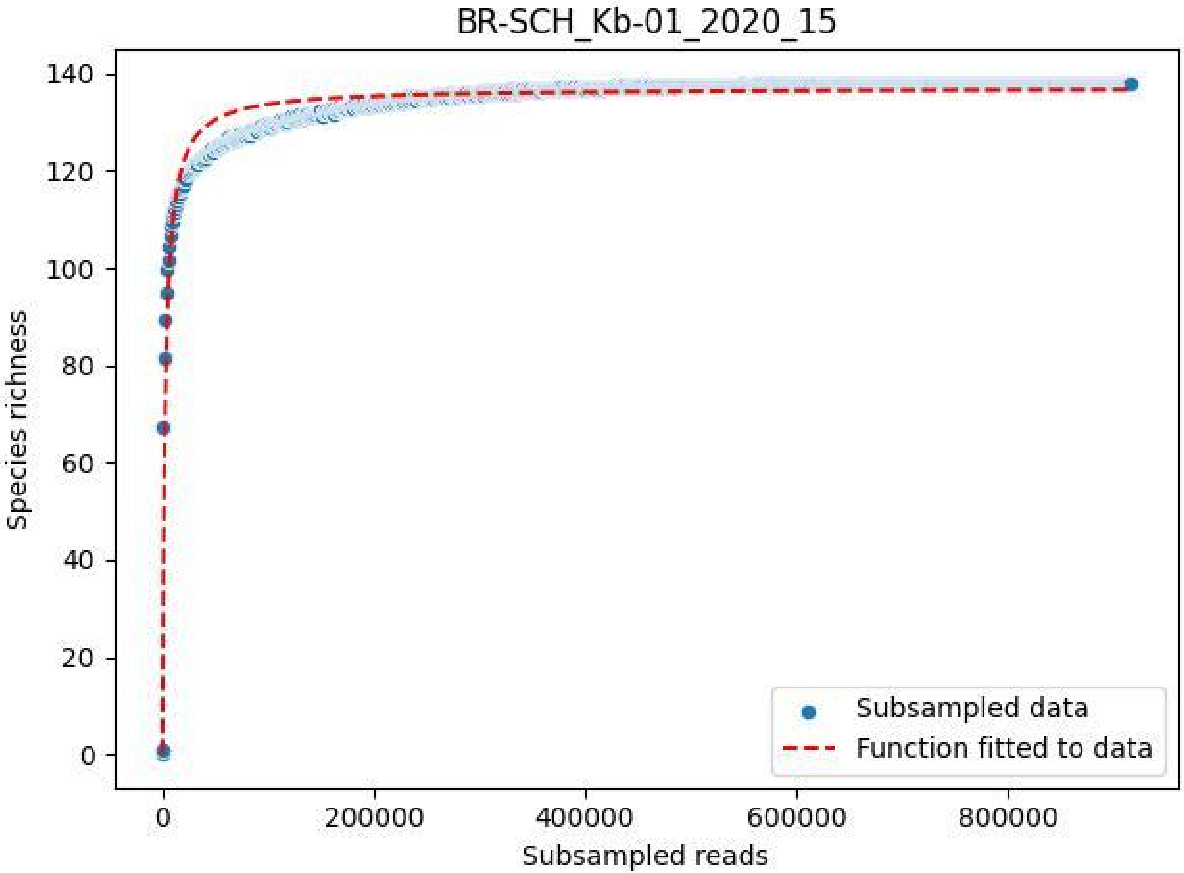
BR-SCH_Kb-01_2020_15
Species richness
Subsampled reads
Subsampled data
Function fitted to data
0
20
40
60
80
100
120
140
0
200000
400000
600000
800000

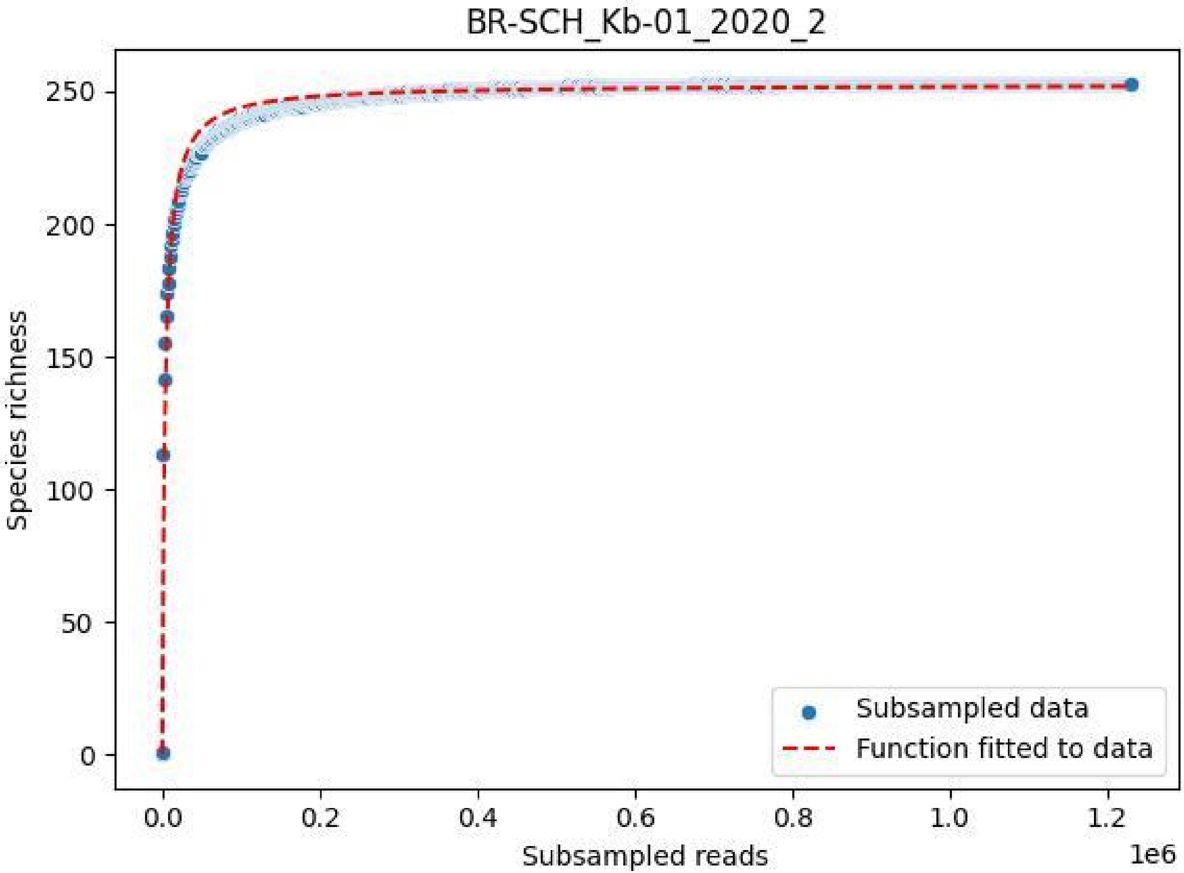
BR-SCH_Kb-01_2020_2
Species richness
Subsampled reads
1e6
250
200
150
100
50
0
0.0
0.2
0.4
0.6
0.8
1.0
1.2
Subsampled data
Function fitted to data

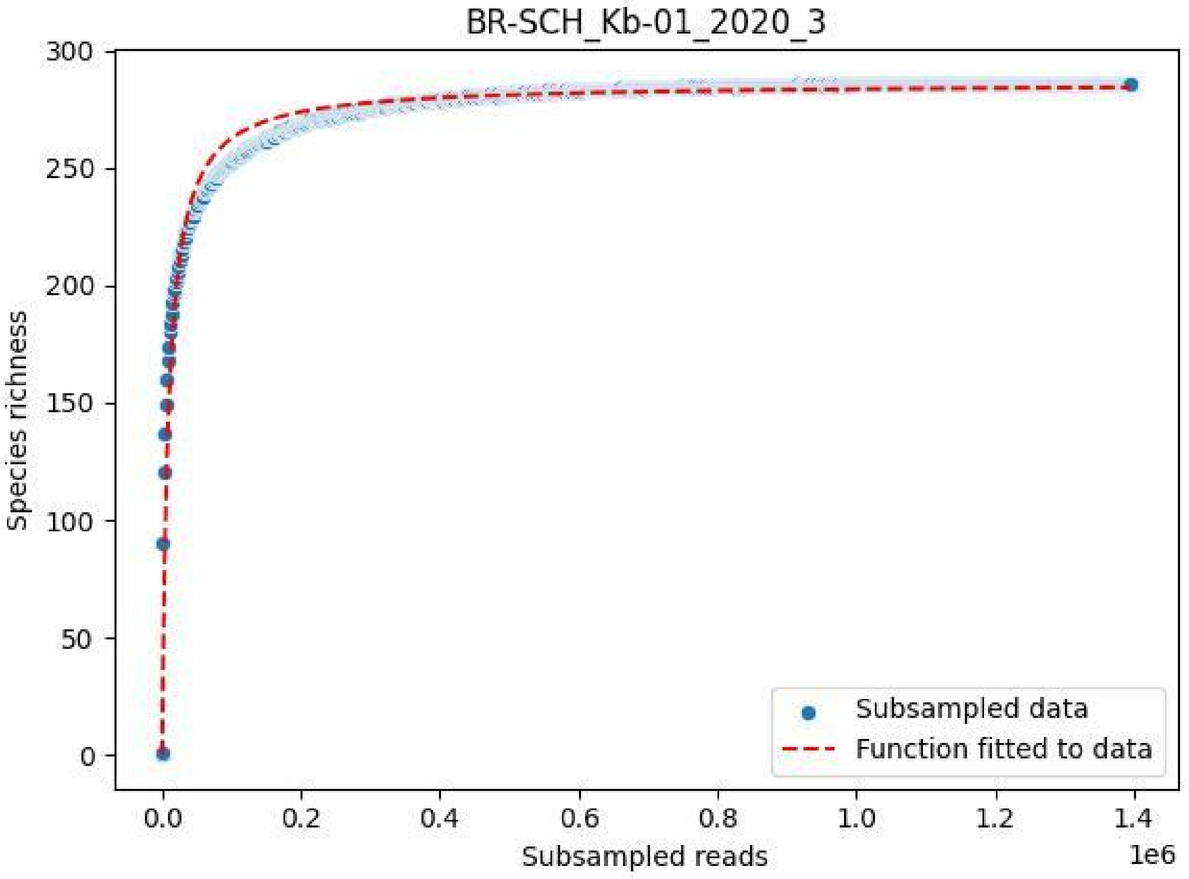
BR-SCH_Kb-01_2020_3
300
250
200
150
100
50
0
Species richness
0.0
0.2
0.4
0.6
0.8
1.0
1.2
1.4
1e6
Subsampled reads
Subsampled data
Function fitted to data

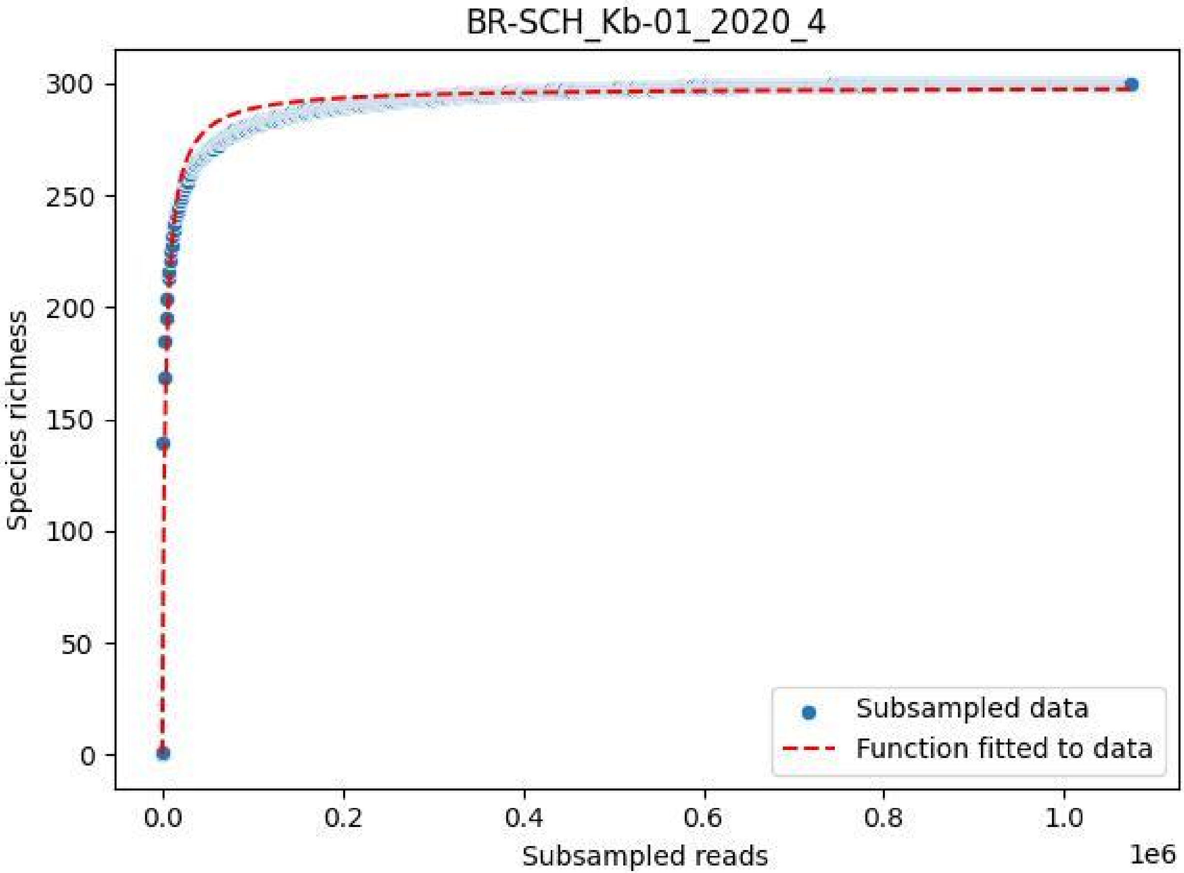
BR-SCH_Kb-01_2020_4
Species richness
300
250
200
150
100
50
0
0.0
0.2
0.4
0.6
0.8
1.0
1e6
Subsampled reads
Subsampled data
Function fitted to data

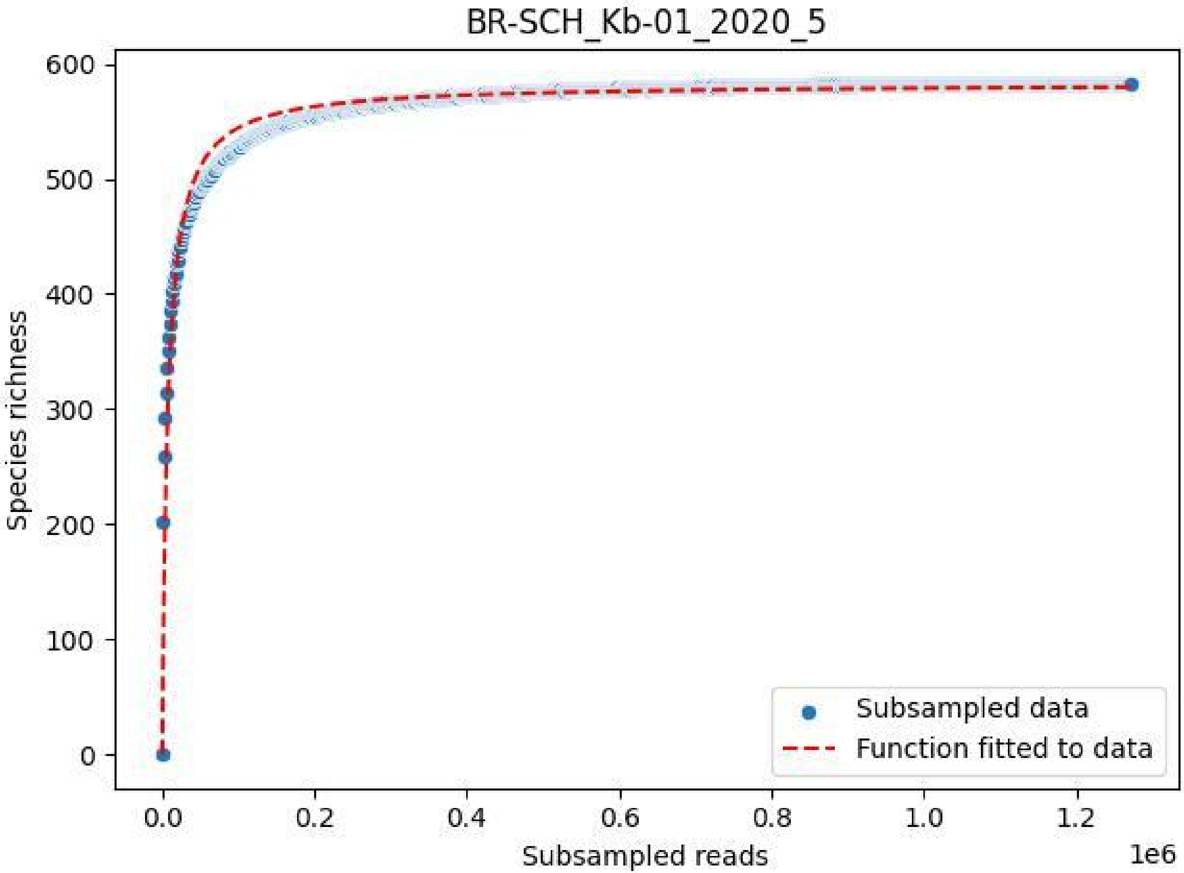
BR-SCH_Kb-01_2020_5
Species richness
Subsampled reads
1e6
Subsampled data
Function fitted to data
600
500
400
300
200
100
0
0.0
0.2
0.4
0.6
0.8
1.0
1.2

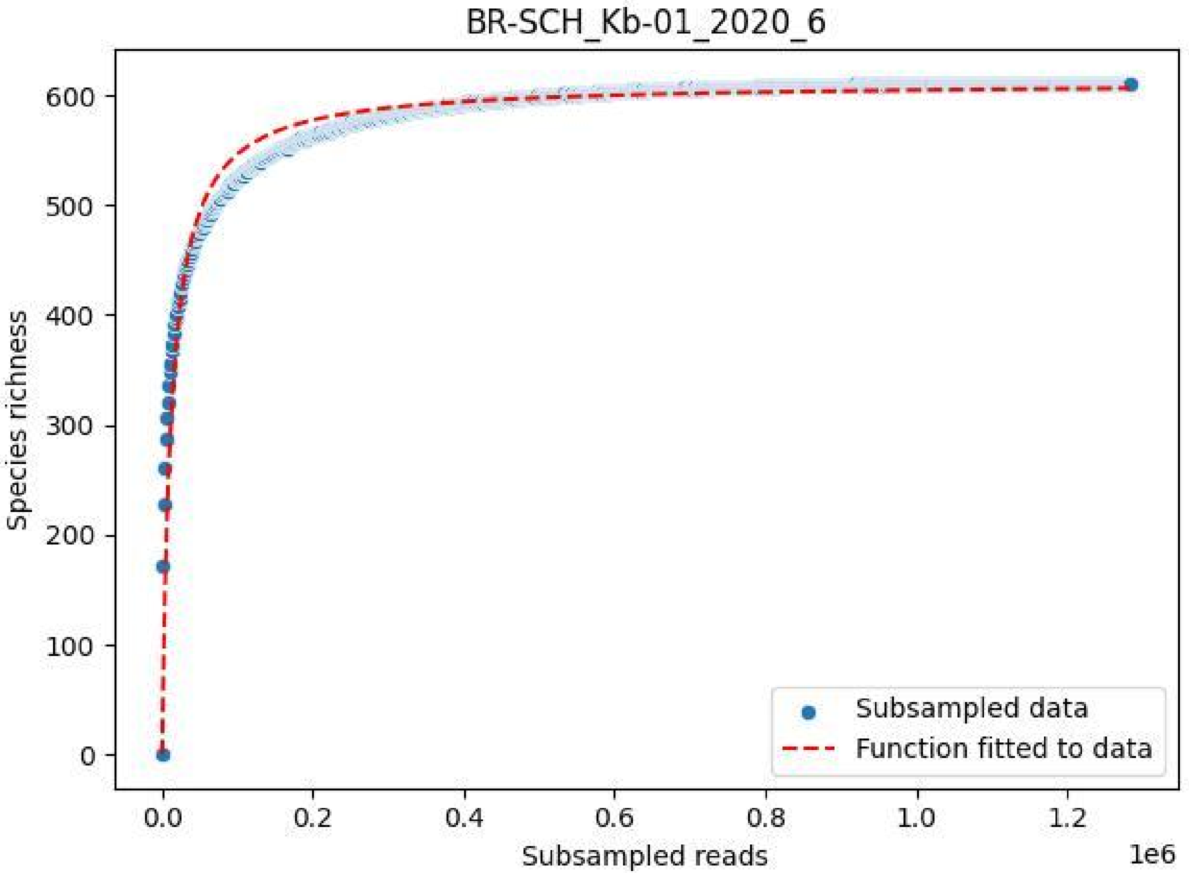
BR-SCH_Kb-01_2020_6
Species richness
Subsampled reads
1e6
0
100
200
300
400
500
600
0.0
0.2
0.4
0.6
0.8
1.0
1.2
Subsampled data
Function fitted to data

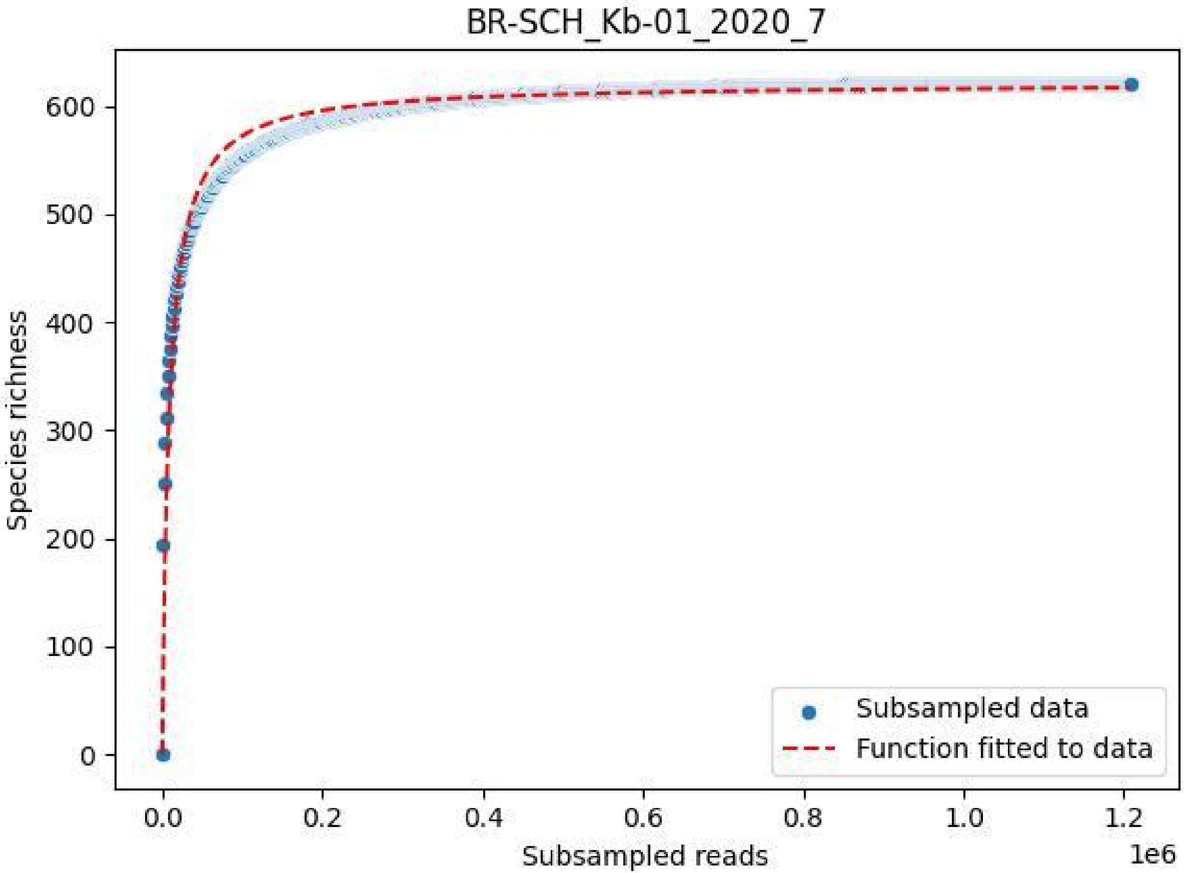
BR-SCH_Kb-01_2020_7
Species richness
Subsampled reads
1e6
0
100
200
300
400
500
600
0.0
0.2
0.4
0.6
0.8
1.0
1.2
Subsampled data
Function fitted to data

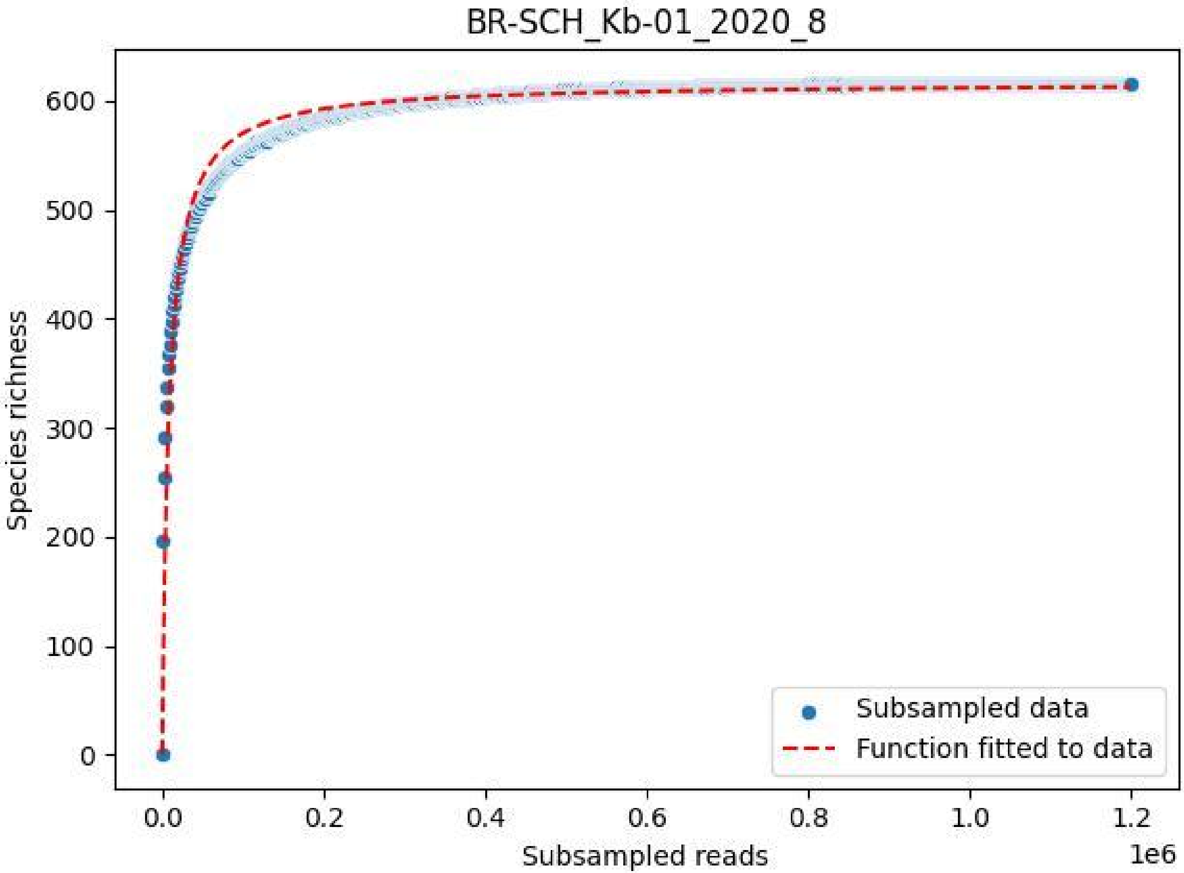

BR-SCH_Kb-01_2020_8
Species richness
Subsampled reads
1e6
0
100
200
300
400
500
600
0.0
0.2
0.4
0.6
0.8
1.0
1.2
Subsampled data
Function fitted to data

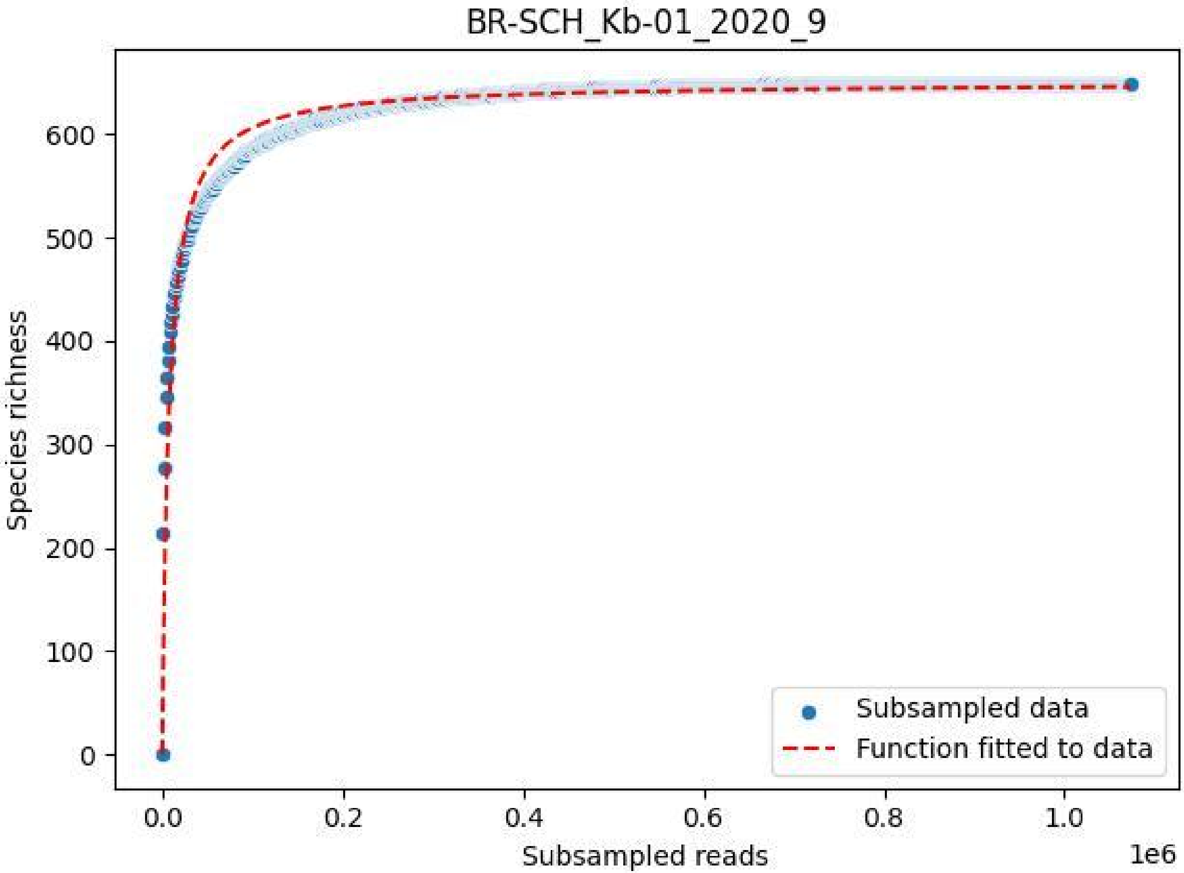
BR-SCH_Kb-01_2020_9
Species richness
600
500
400
300
200
100
0
0.0
0.2
0.4
0.6
0.8
1.0
1e6
Subsampled reads
Subsampled data
Function fitted to data

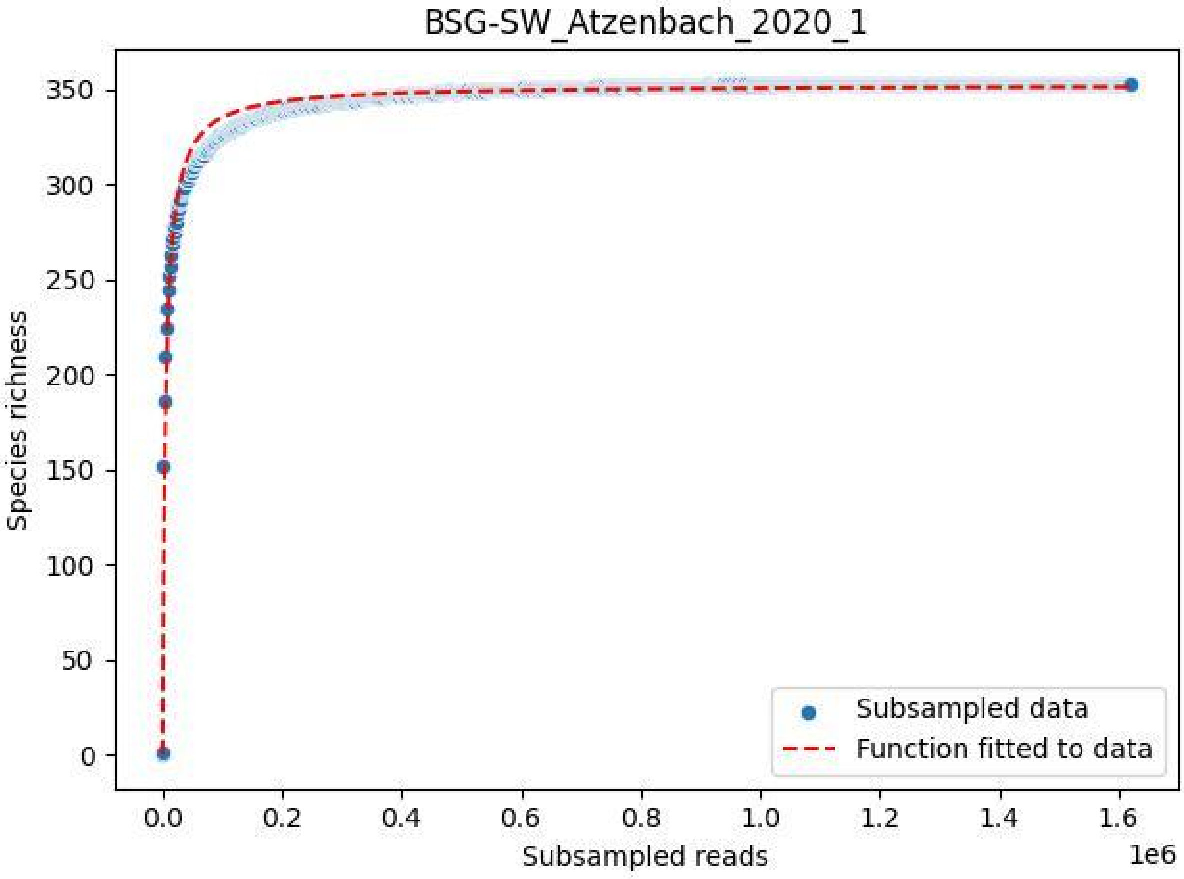
BSG-SW_Atzenbach_2020_1
Species richness
Subsampled reads
1e6
0
50
100
150
200
250
300
350
0.0
0.2
0.4
0.6
0.8
1.0
1.2
1.4
1.6
Subsampled data
Function fitted to data

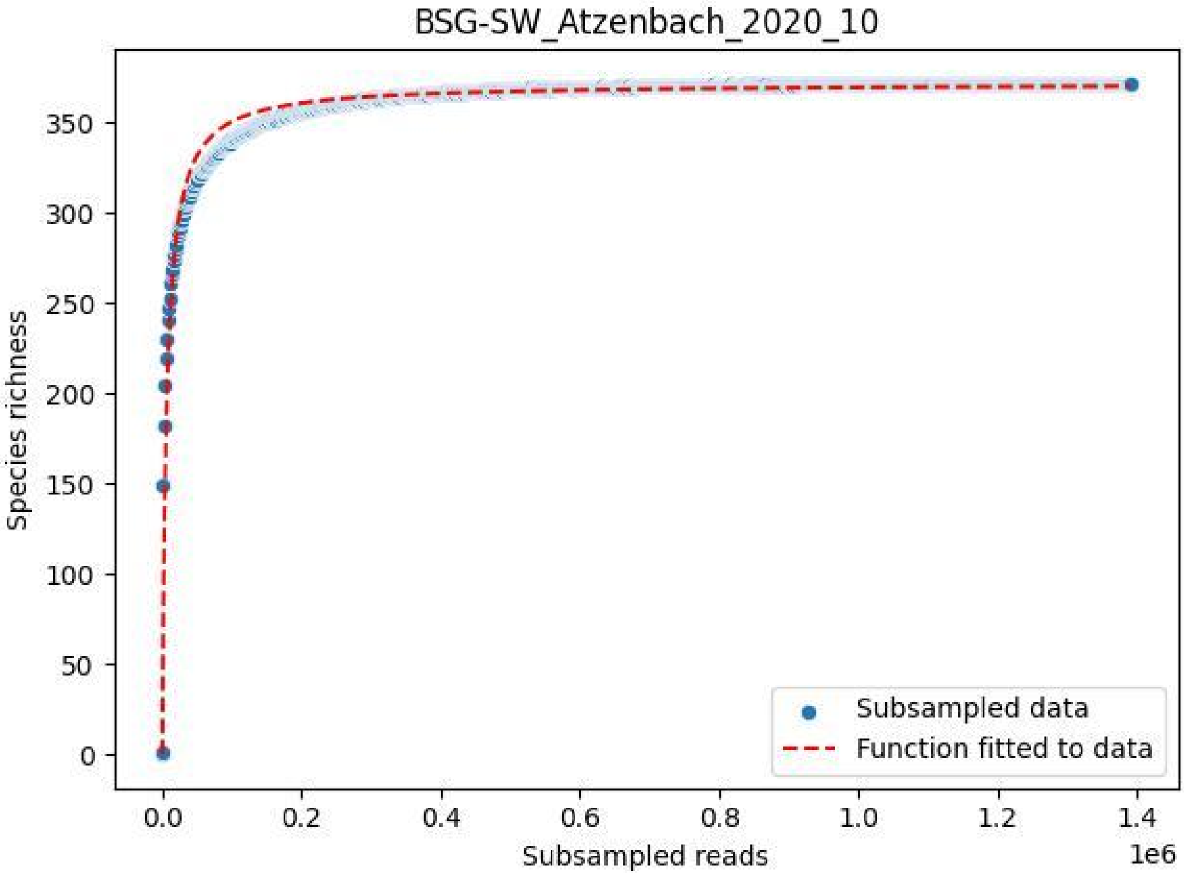
BSG-SW_Atzenbach_2020_10
Species richness
Subsampled reads
1e6
350
300
250
200
150
100
50
0
0.0
0.2
0.4
0.6
0.8
1.0
1.2
1.4
Subsampled data
Function fitted to data

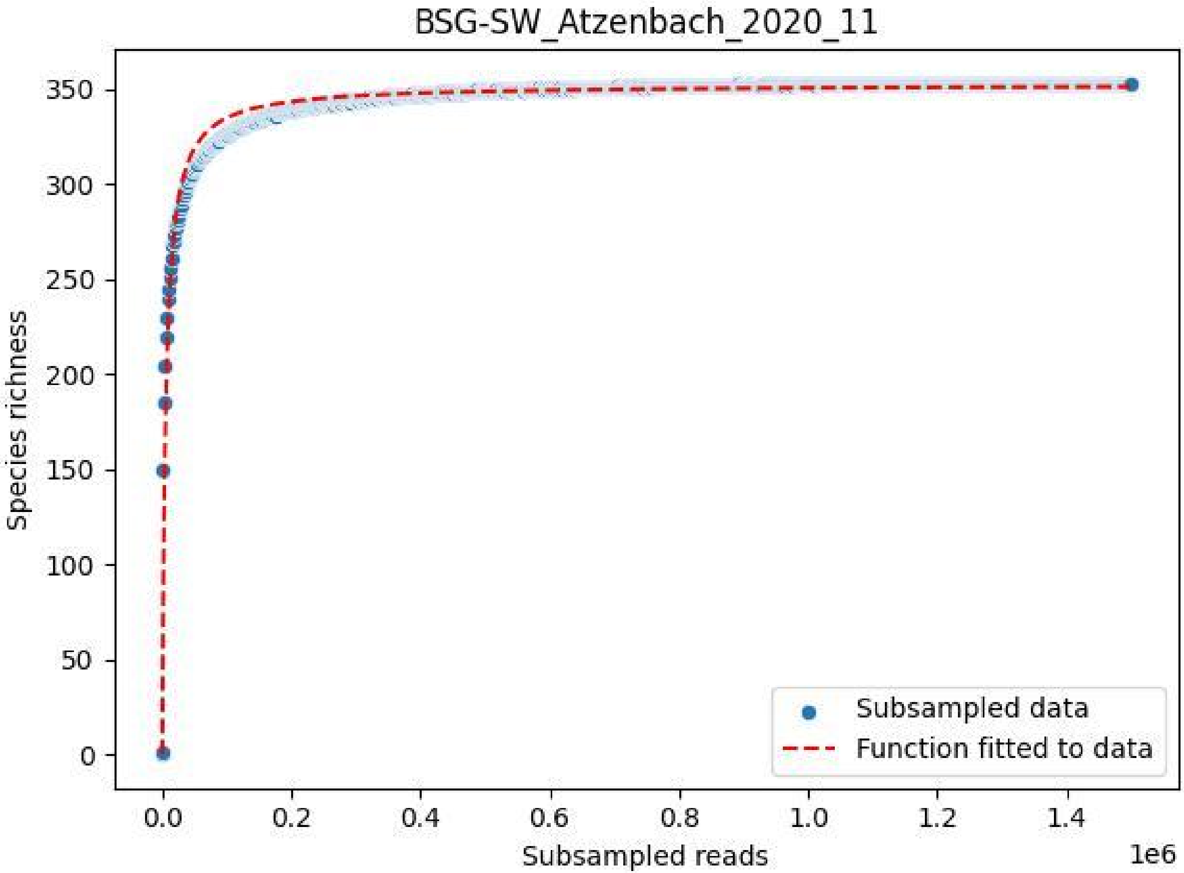

BSG-SW_Atzenbach_2020_11
Species richness
0
50
100
150
200
250
300
350
0.0
0.2
0.4
0.6
0.8
1.0
1.2
1.4
1e6
Subsampled reads
Subsampled data
Function fitted to data

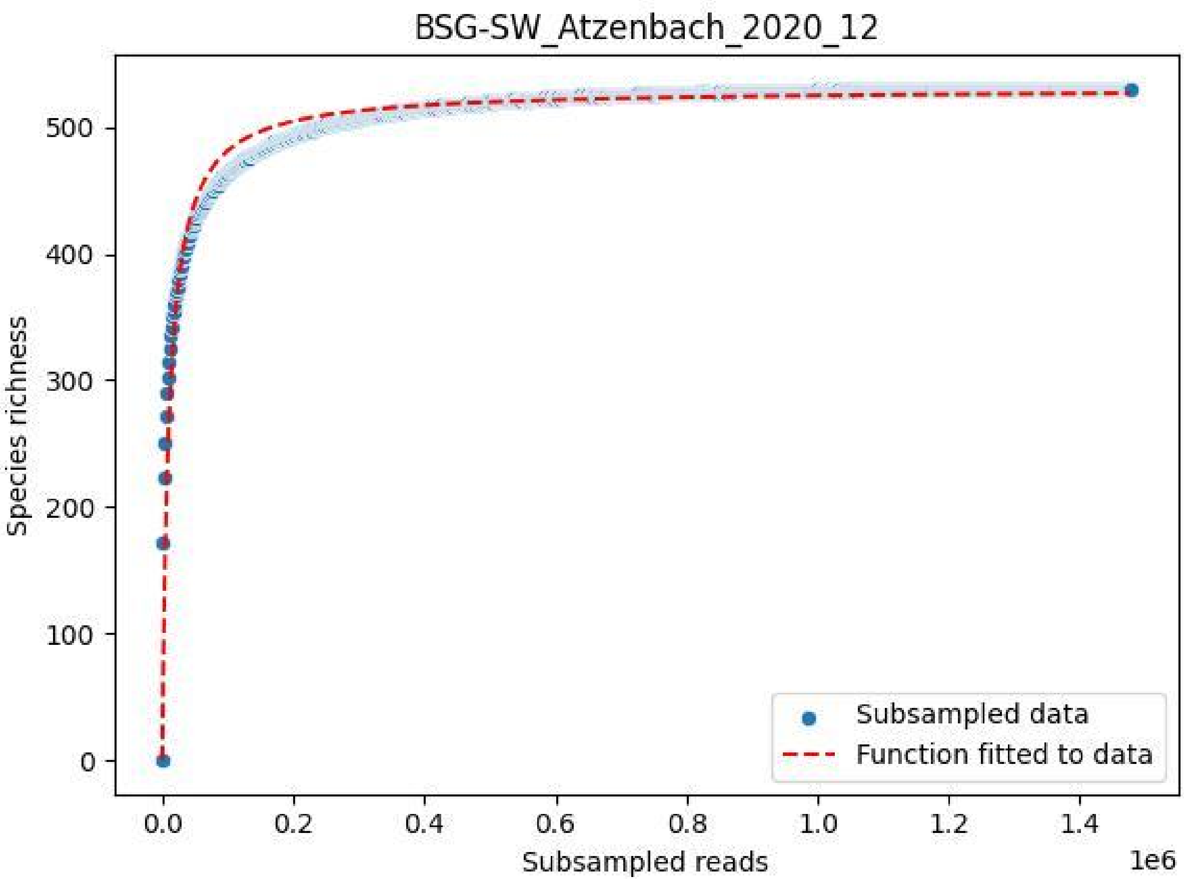
BSG-SW_Atzenbach_2020_12
Species richness
Subsampled reads
1e6
0
100
200
300
400
500
0.0
0.2
0.4
0.6
0.8
1.0
1.2
1.4
Subsampled data
Function fitted to data

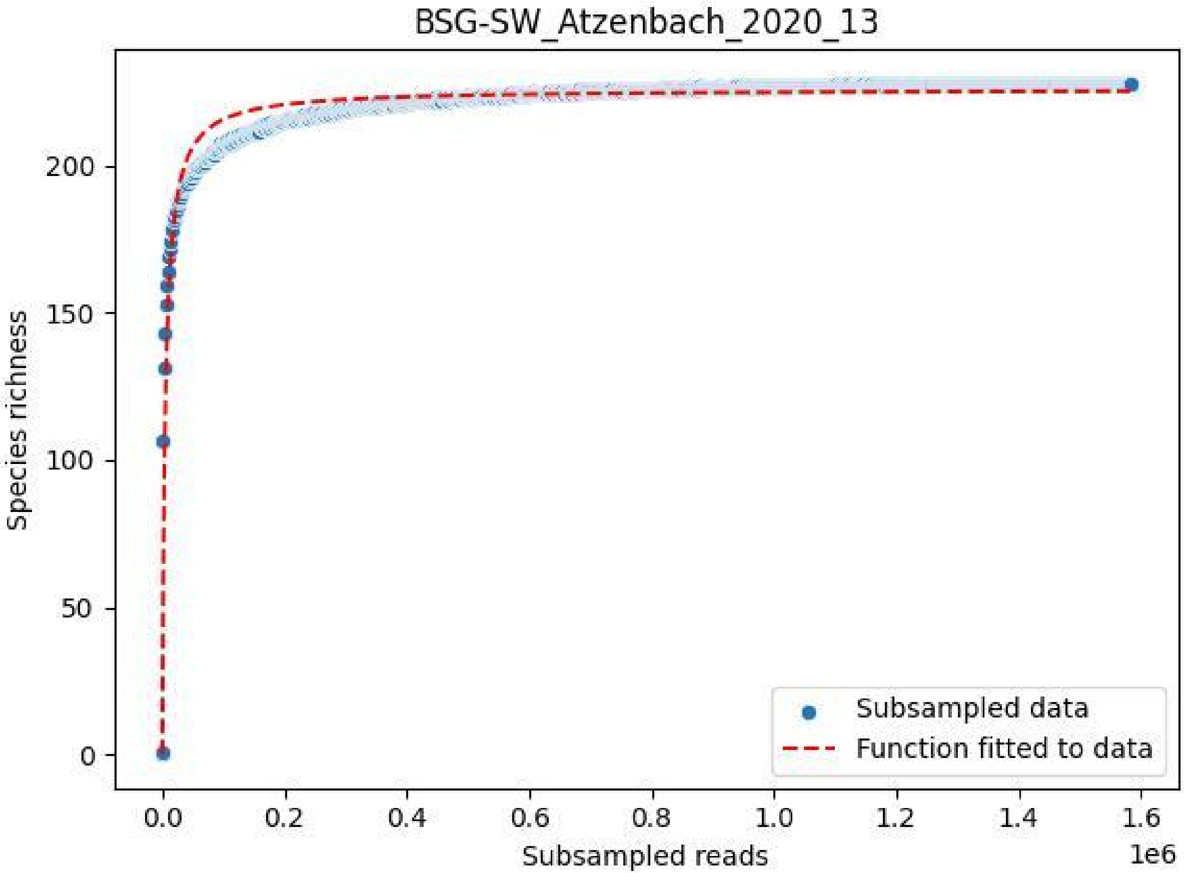
BSG-SW_Atzenbach_2020_13
Species richness
0
50
100
150
200
0.0
0.2
0.4
0.6
0.8
1.0
1.2
1.4
1.6
1e6
Subsampled reads
Subsampled data
Function fitted to data

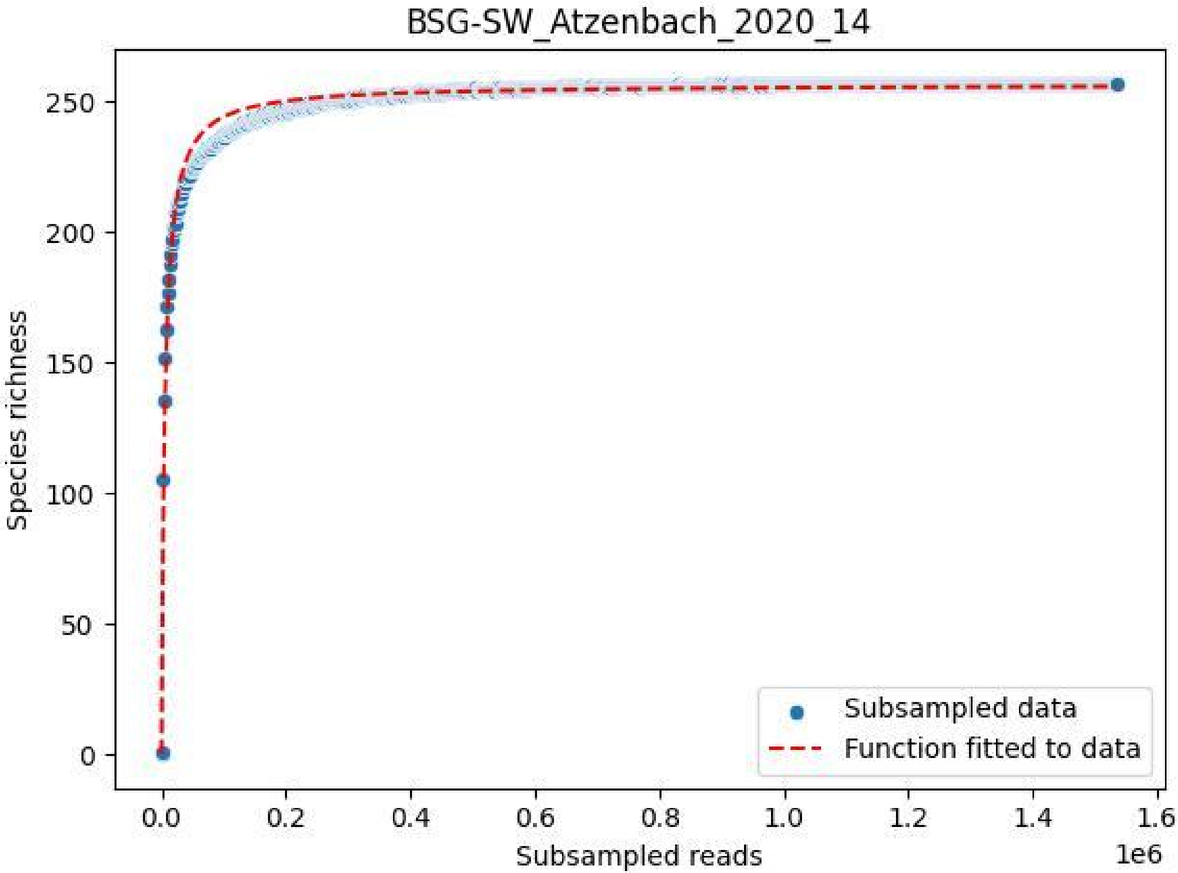
BSG-SW_Atzenbach_2020_14
Species richness
Subsampled reads
1e6
0
50
100
150
200
250
0.0
0.2
0.4
0.6
0.8
1.0
1.2
1.4
1.6
Subsampled data
Function fitted to data

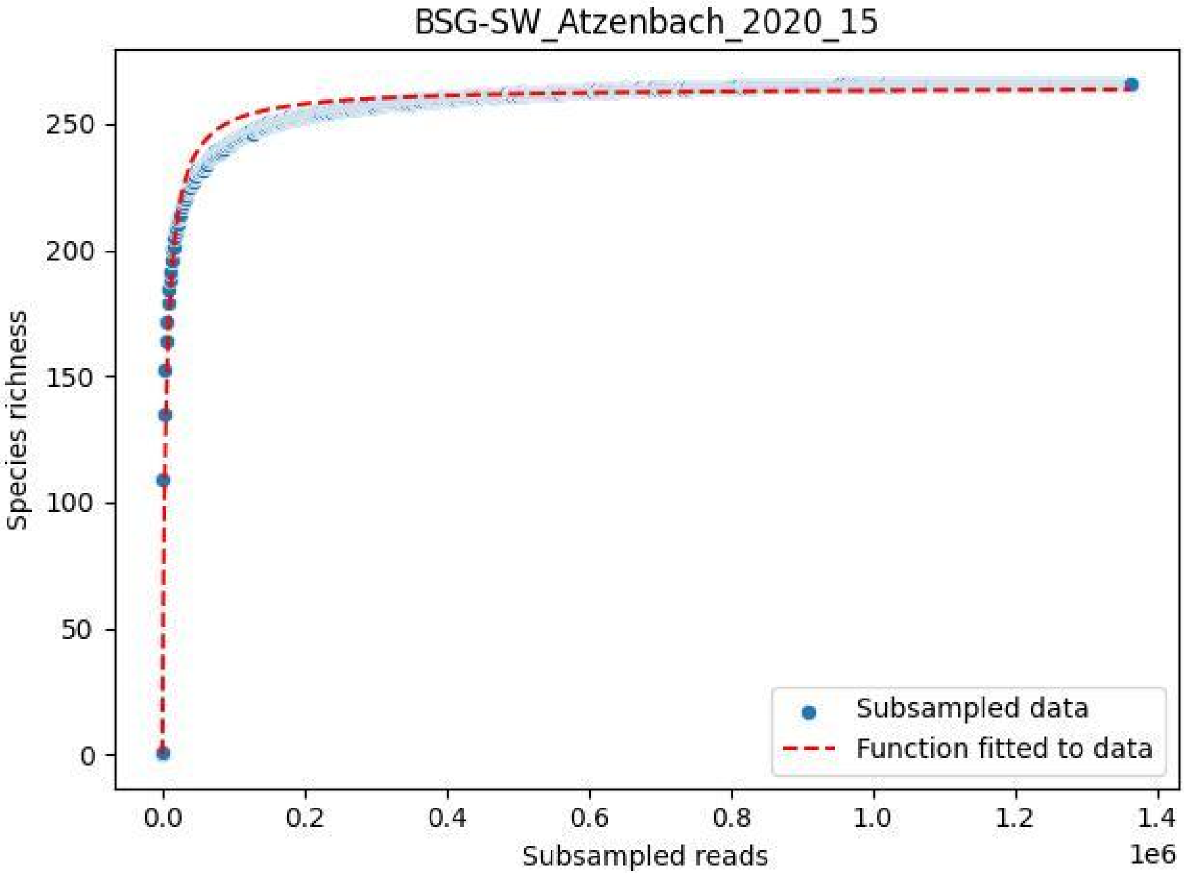
BSG-SW_Atzenbach_2020_15
Species richness
0
50
100
150
200
250
0.0
0.2
0.4
0.6
0.8
1.0
1.2
1.4
1e6
Subsampled reads
Subsampled data
Function fitted to data

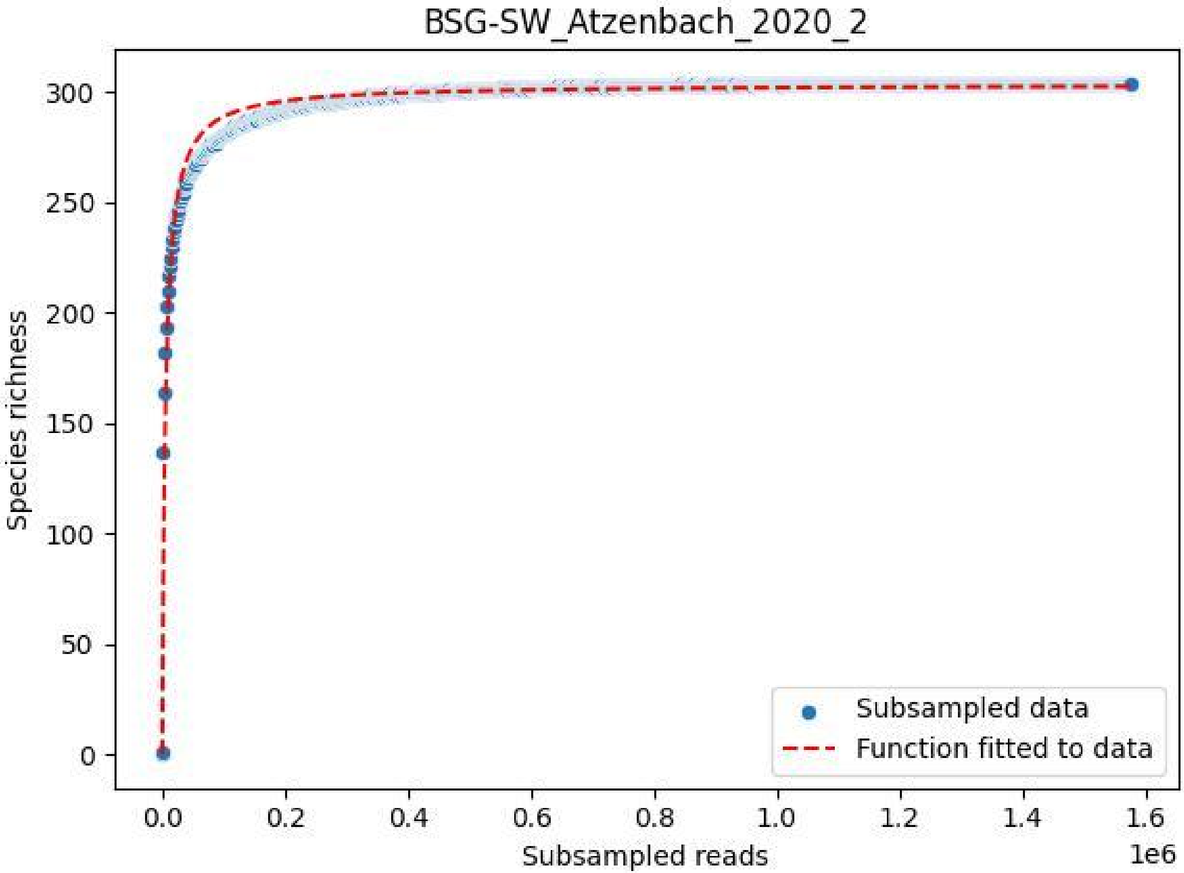
BSG-SW_Atzenbach_2020_2
Species richness
Subsampled reads
1e6
Subsampled data
Function fitted to data
0
50
100
150
200
250
300
0.0
0.2
0.4
0.6
0.8
1.0
1.2
1.4
1.6

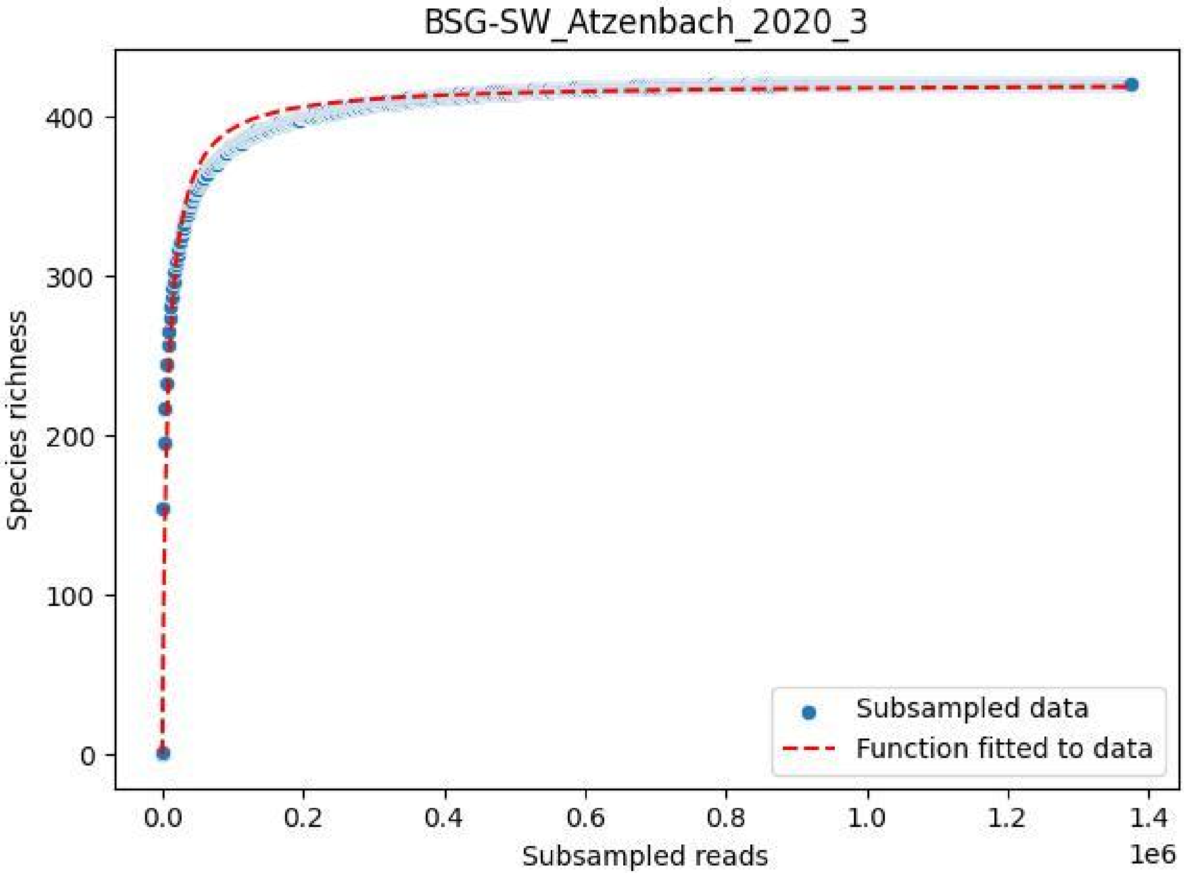

BSG-SW_Atzenbach_2020_3
Species richness
Subsampled reads
1e6
0
100
200
300
400
0.0
0.2
0.4
0.6
0.8
1.0
1.2
1.4
Subsampled data
Function fitted to data

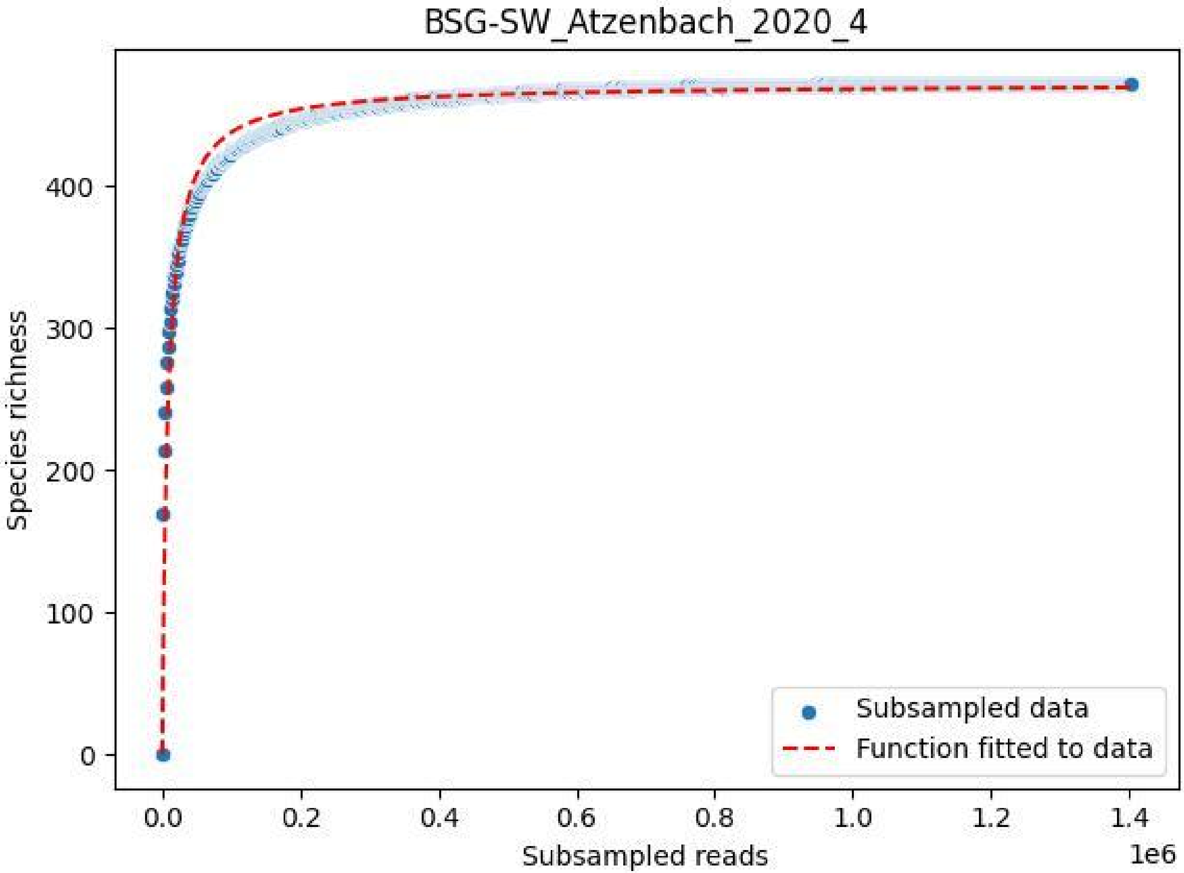
BSG-SW_Atzenbach_2020_4
Species richness
Subsampled reads
1e6
0
100
200
300
400
0.0
0.2
0.4
0.6
0.8
1.0
1.2
1.4
Subsampled data
Function fitted to data

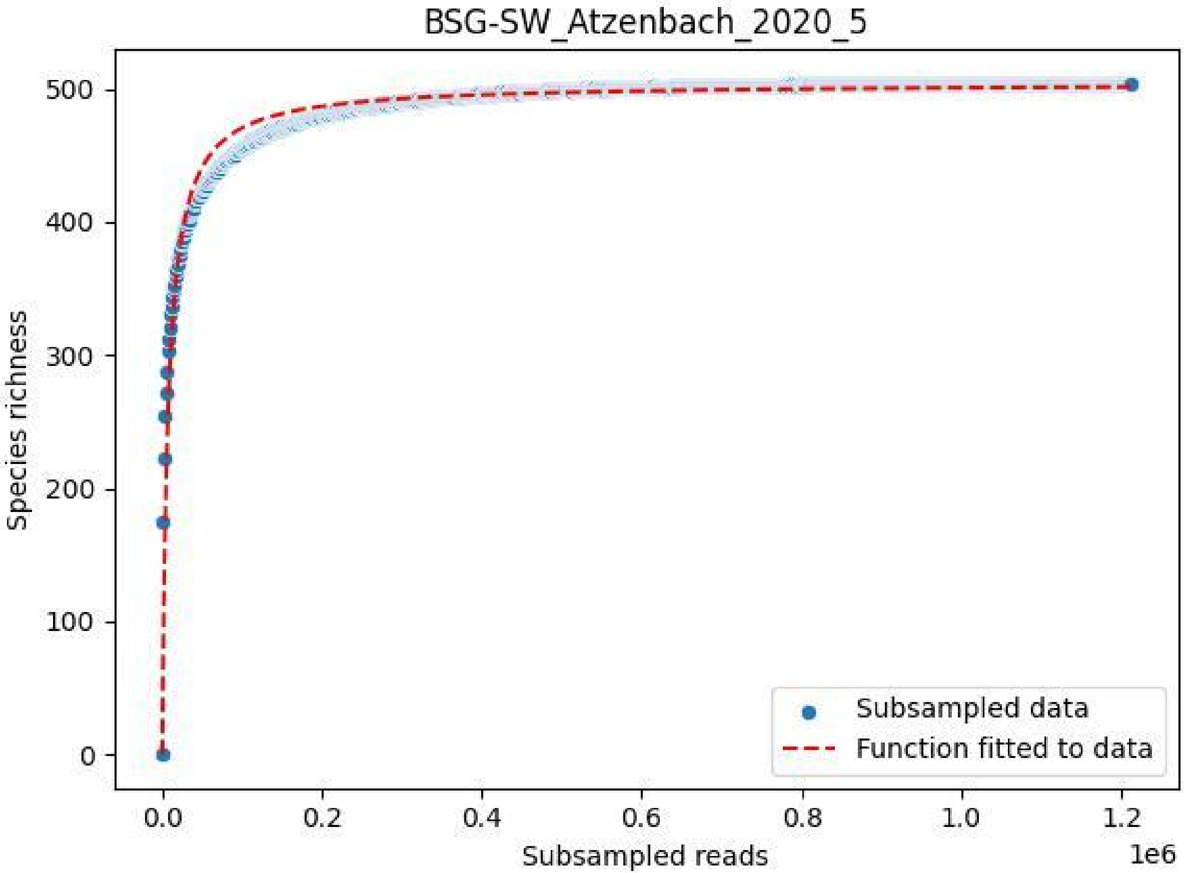
BSG-SW_Atzenbach_2020_5
Species richness
Subsampled reads
1e6
0
100
200
300
400
500
0.0
0.2
0.4
0.6
0.8
1.0
1.2
Subsampled data
Function fitted to data

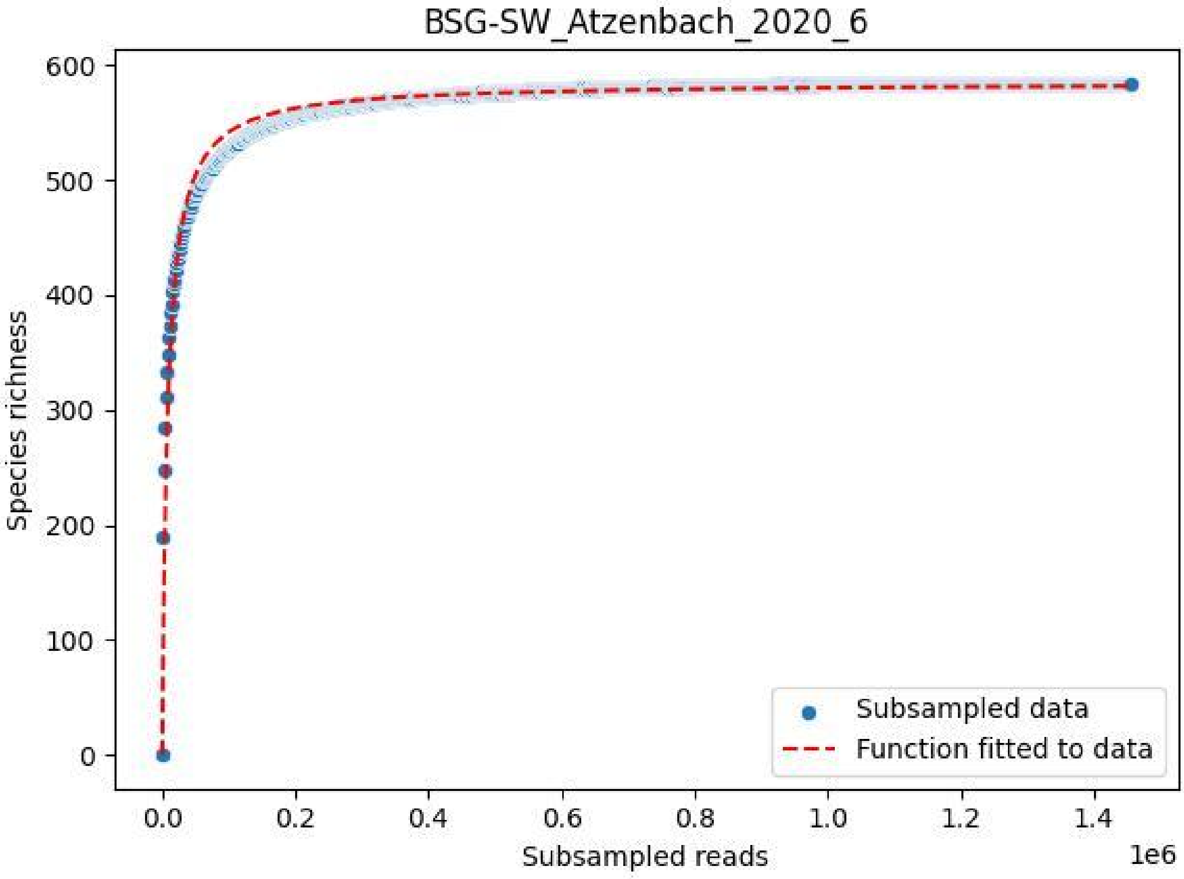
BSG-SW_Atzenbach_2020_6
Species richness
Subsampled reads
1e6
0
100
200
300
400
500
600
0.0
0.2
0.4
0.6
0.8
1.0
1.2
1.4
Subsampled data
Function fitted to data

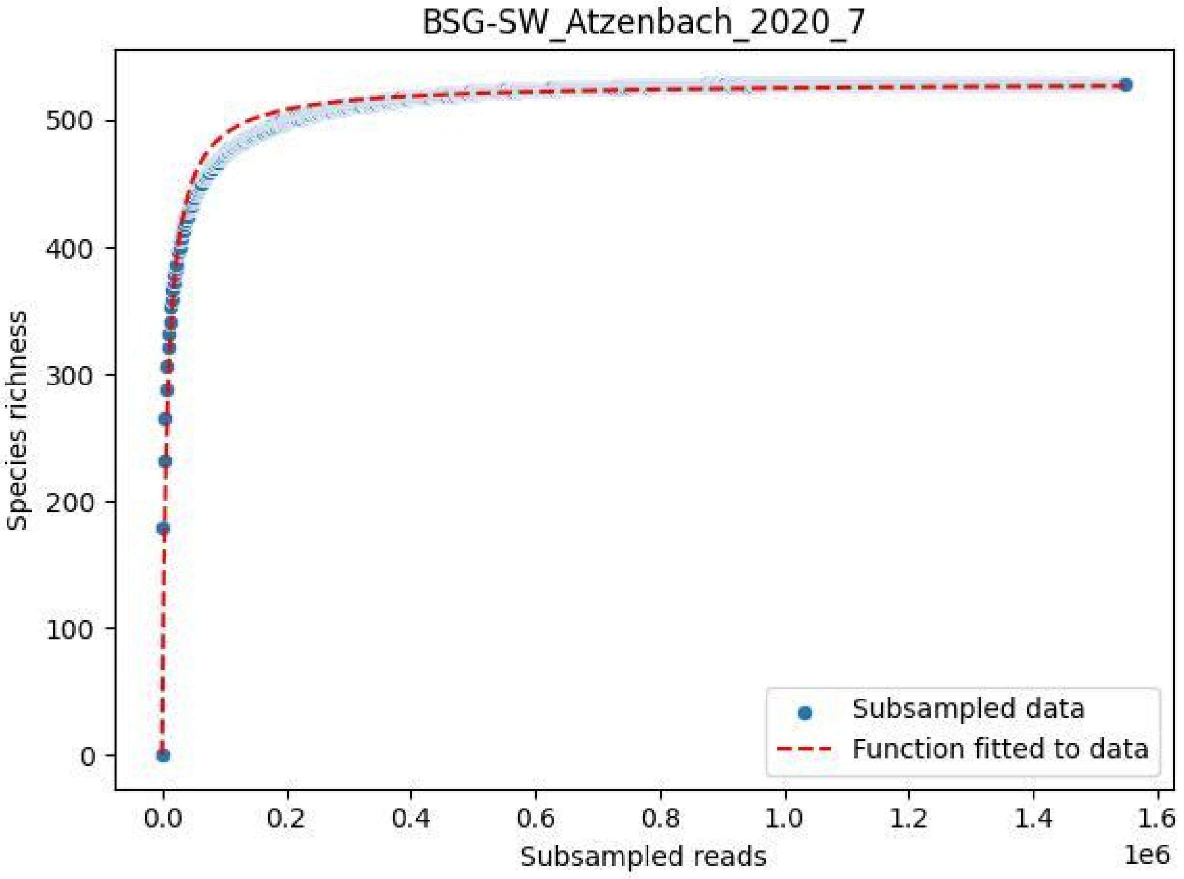
BSG-SW_Atzenbach_2020_7
Species richness
Subsampled reads
1e6
0
100
200
300
400
500
0.0
0.2
0.4
0.6
0.8
1.0
1.2
1.4
1.6
Subsampled data
Function fitted to data

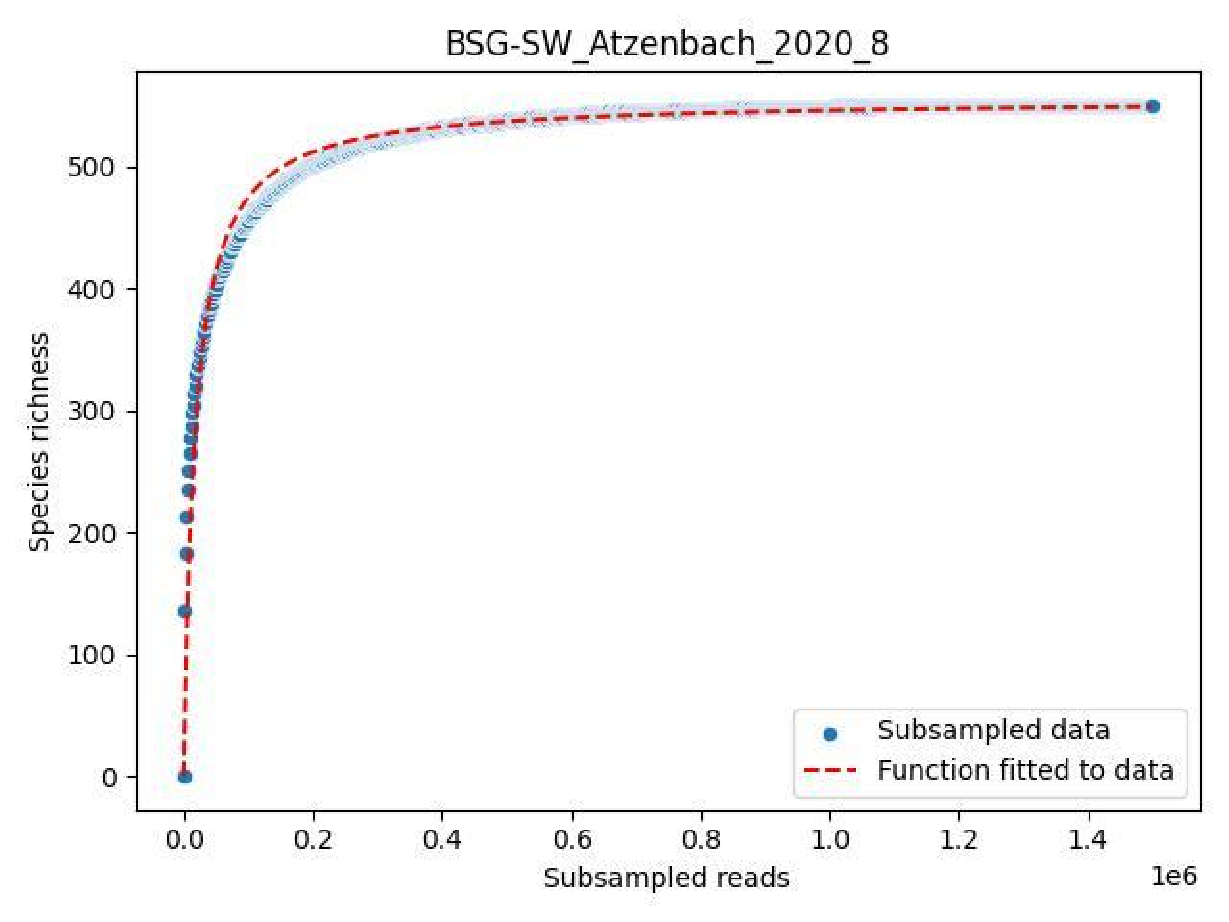

BSG-SW_Atzenbach_2020_8
Species richness
Subsampled reads
1e6
0
100
200
300
400
500
0.0
0.2
0.4
0.6
0.8
1.0
1.2
1.4
Subsampled data
Function fitted to data

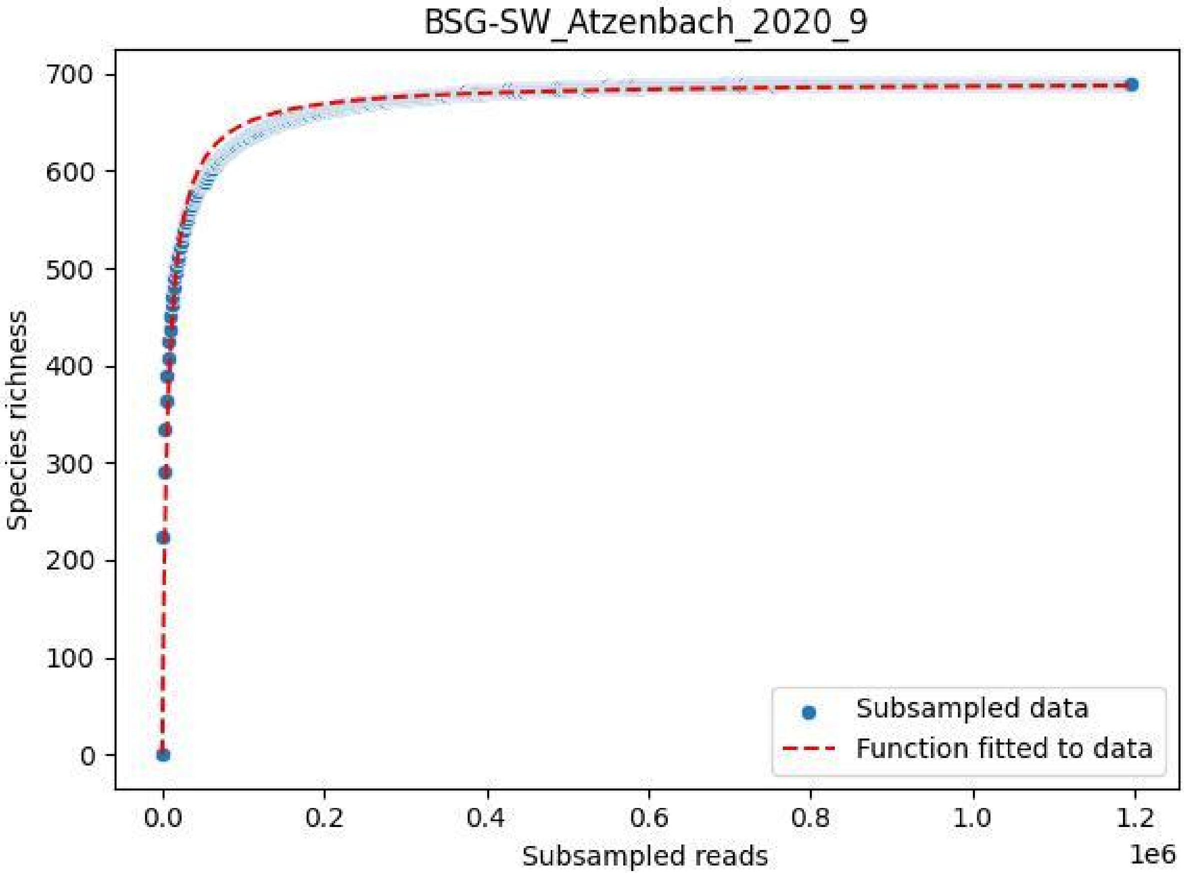

BSG-SW_Atzenbach_2020_9
Species richness
700
600
500
400
300
200
100
0
0.0
0.2
0.4
0.6
0.8
1.0
1.2
1e6
Subsampled reads
Subsampled data
Function fitted to data

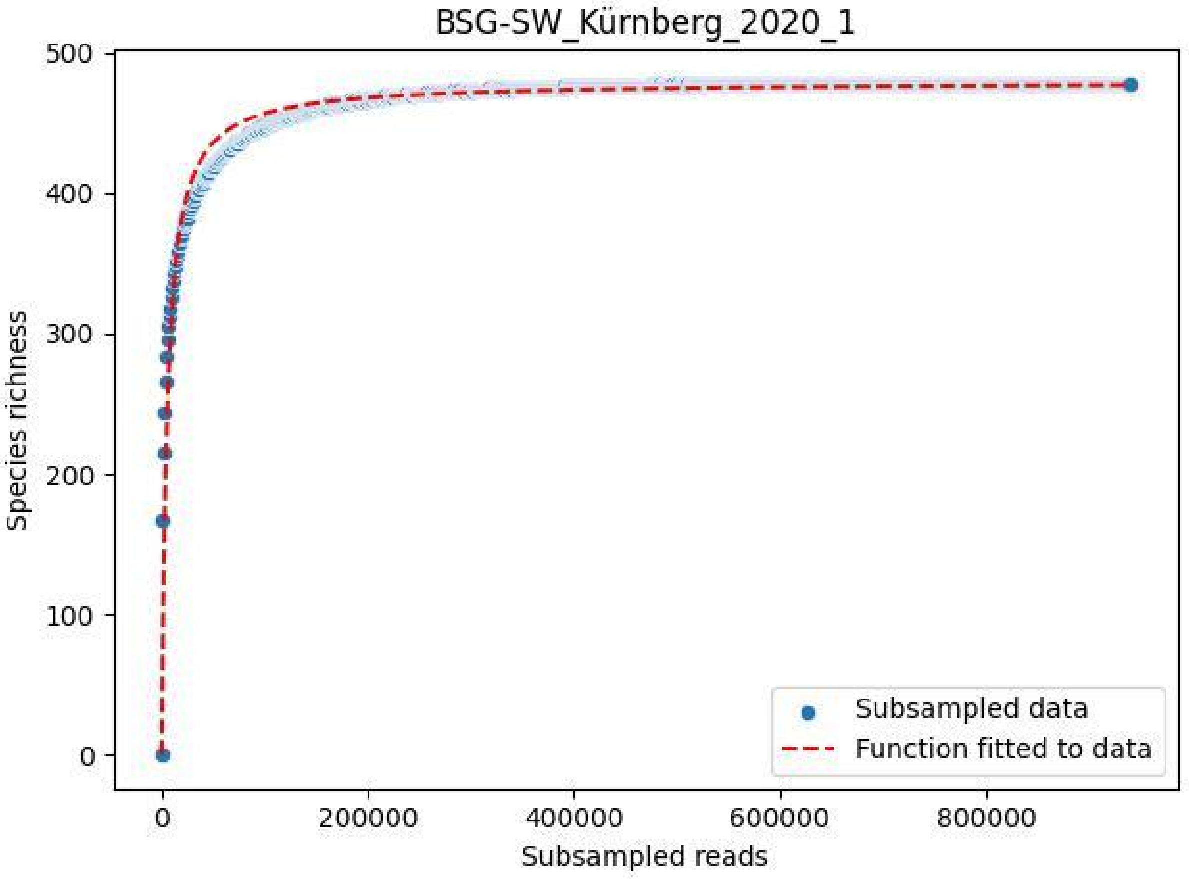
BSG-SW_Kürnberg_2020_1
Species richness
Subsampled reads
0
100
200
300
400
500
0
200000
400000
600000
800000
Subsampled data
Function fitted to data

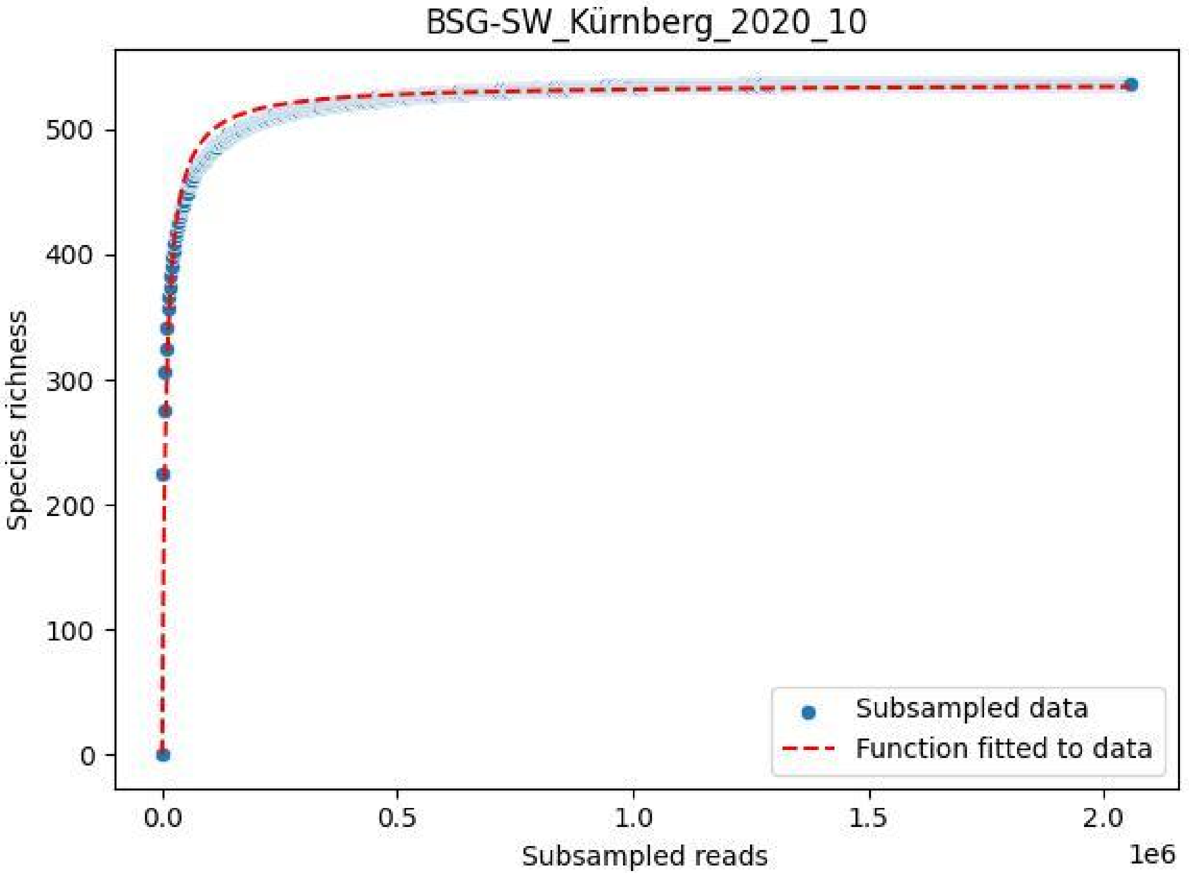
BSG-SW_Kürnberg_2020_10
Species richness
Subsampled reads
1e6
0
100
200
300
400
500
0.0
0.5
1.0
1.5
2.0
Subsampled data
Function fitted to data

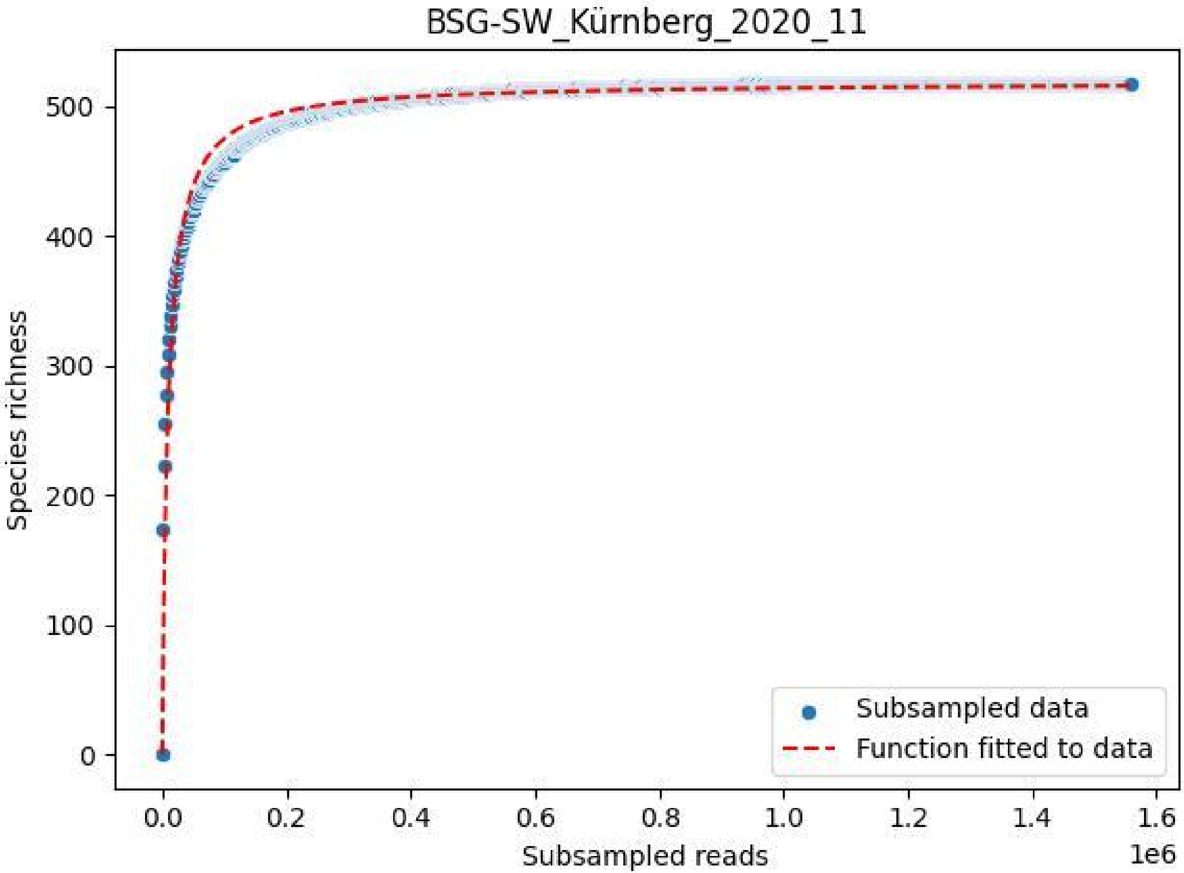
BSG-SW_Kürnberg_2020_11
Species richness
Subsampled reads
1e6
0
100
200
300
400
500
0.0
0.2
0.4
0.6
0.8
1.0
1.2
1.4
1.6
Subsampled data
Function fitted to data

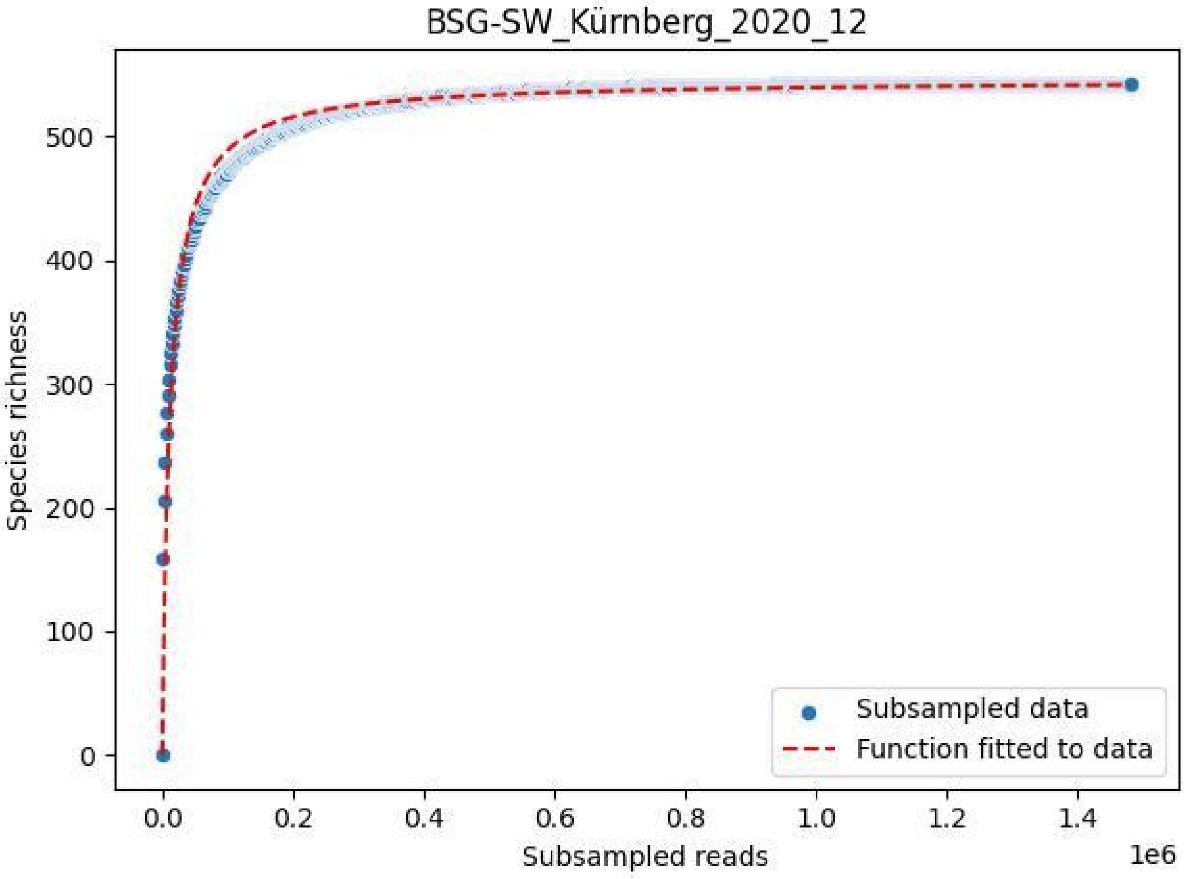
BSG-SW_Kürnberg_2020_12
Species richness
Subsampled reads
1e6
0
100
200
300
400
500
0.0
0.2
0.4
0.6
0.8
1.0
1.2
1.4
Subsampled data
Function fitted to data

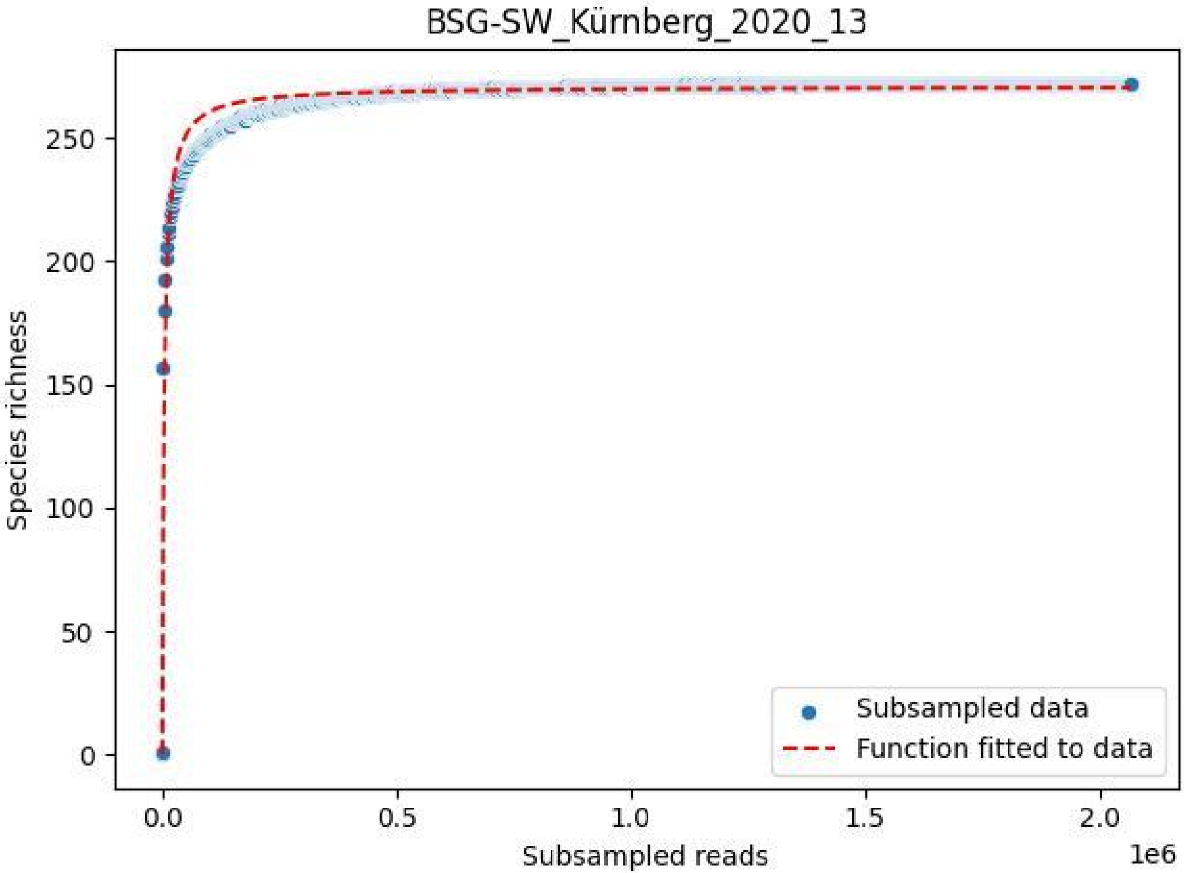
BSG-SW_Kürnberg_2020_13
Species richness
Subsampled reads
1e6
0
50
100
150
200
250
0.0
0.5
1.0
1.5
2.0
Subsampled data
Function fitted to data

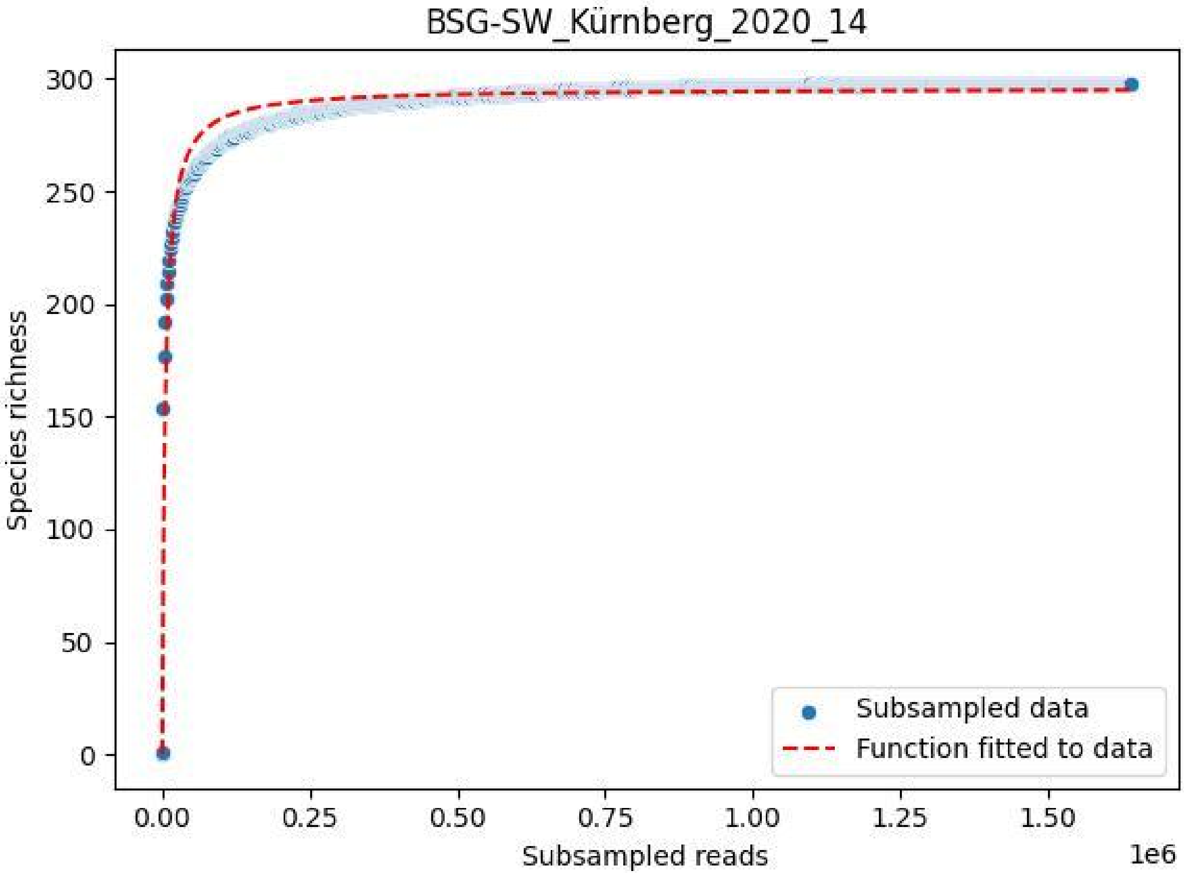
BSG-SW_Kürnberg_2020_14
Species richness
300
250
200
150
100
50
0
0.00
0.25
0.50
0.75
1.00
1.25
1.50
1e6
Subsampled reads
Subsampled data
Function fitted to data

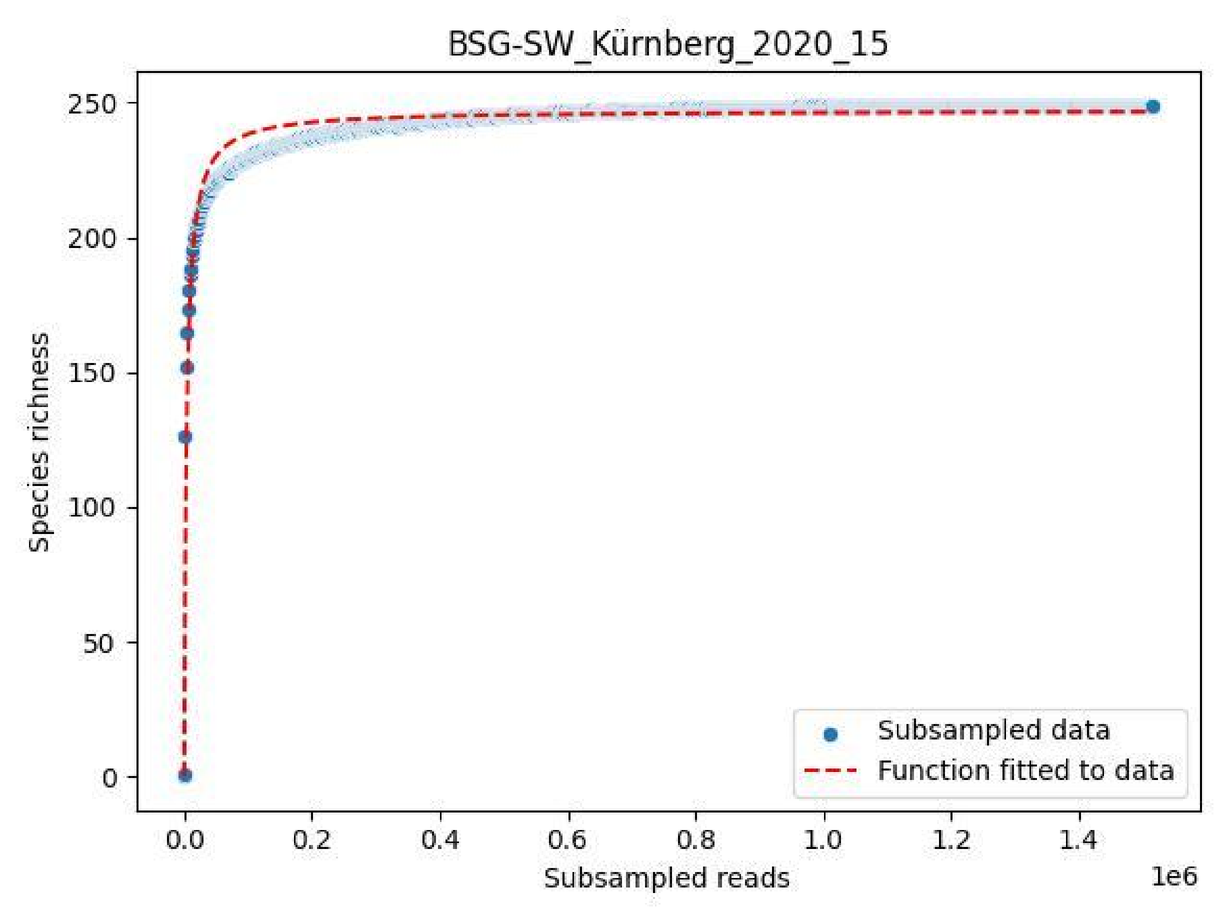
BSG-SW_Kürnberg_2020_15
Species richness
250
200
150
100
50
0
0.0
0.2
0.4
0.6
0.8
1.0
1.2
1.4
1e6
Subsampled reads
Subsampled data
Function fitted to data

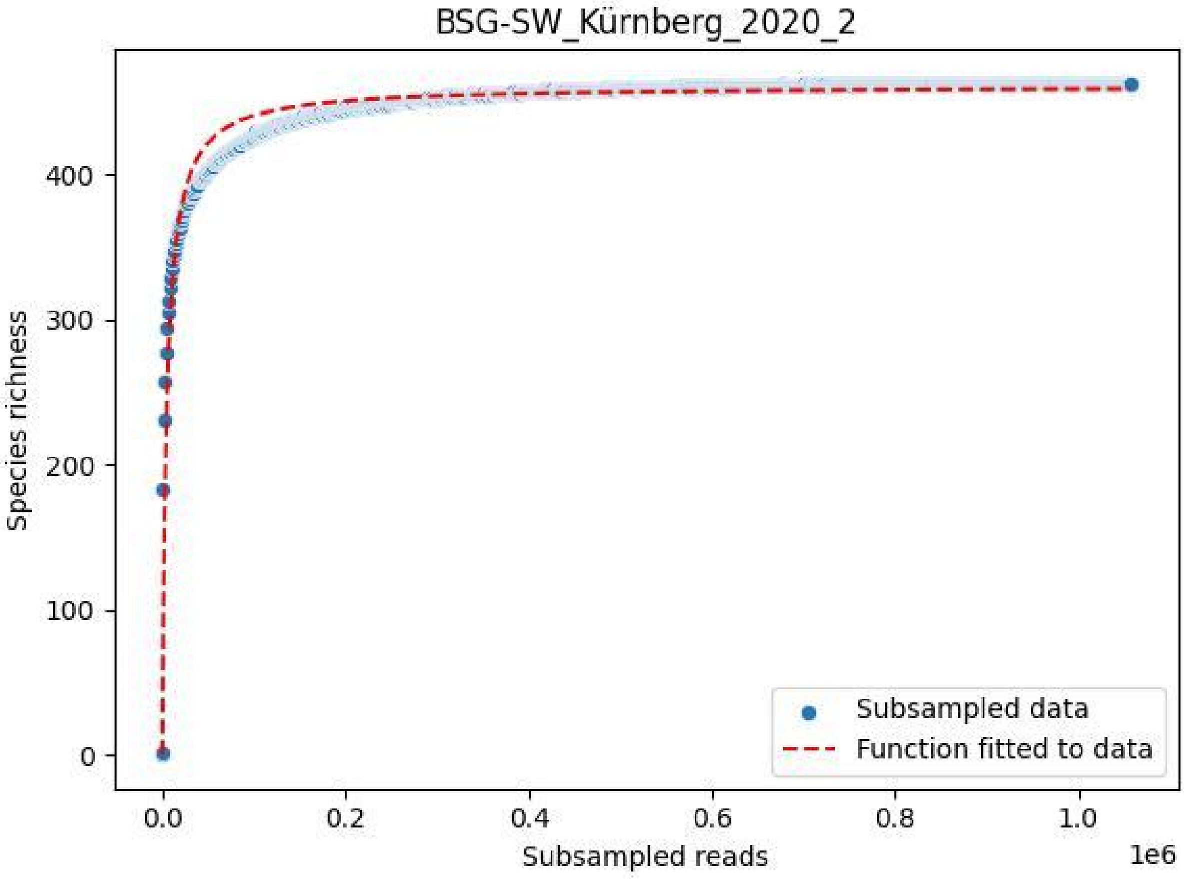
BSG-SW_Kürnberg_2020_2
Species richness
400
300
200
100
0
0.0
0.2
0.4
0.6
0.8
1.0
1e6
Subsampled reads
Subsampled data
Function fitted to data

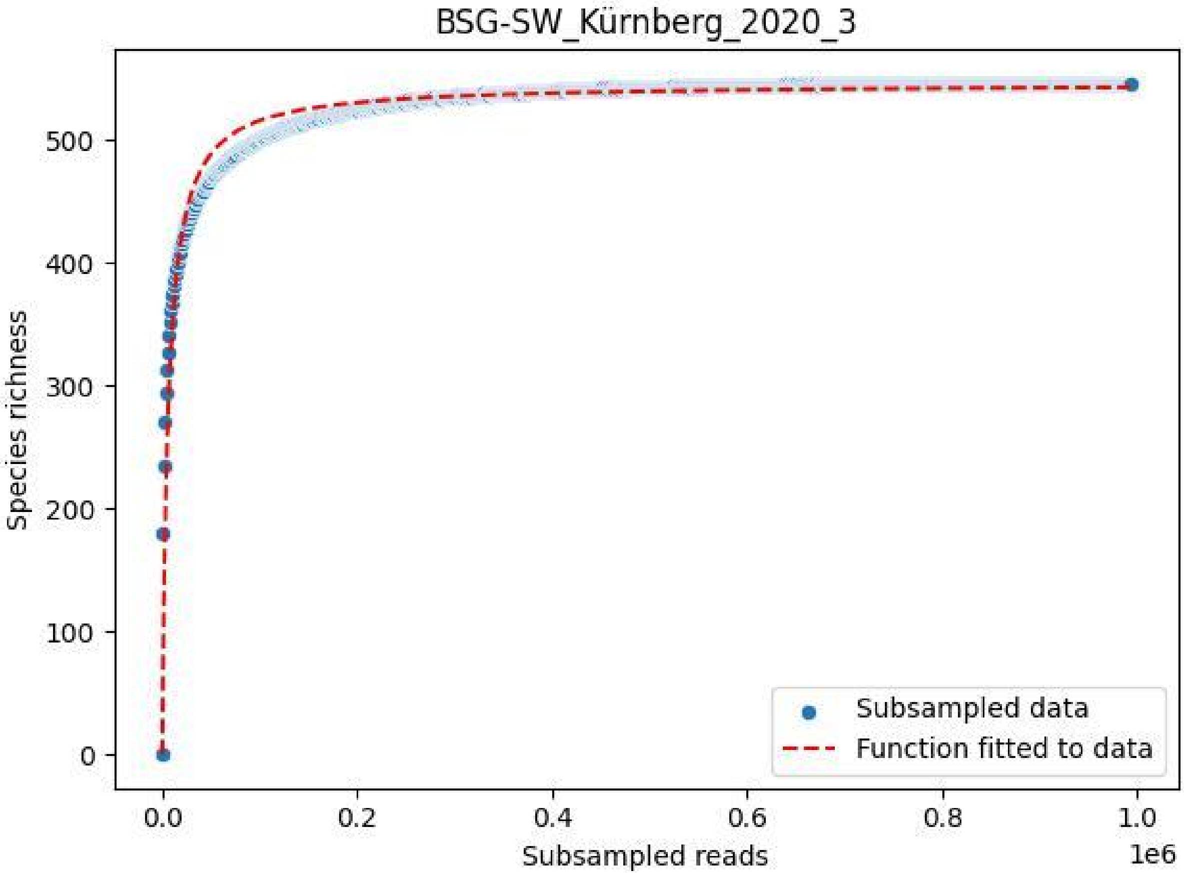

BSG-SW_Kürnberg_2020_3
Species richness
Subsampled reads
1e6
0
100
200
300
400
500
0.0
0.2
0.4
0.6
0.8
1.0
Subsampled data
Function fitted to data

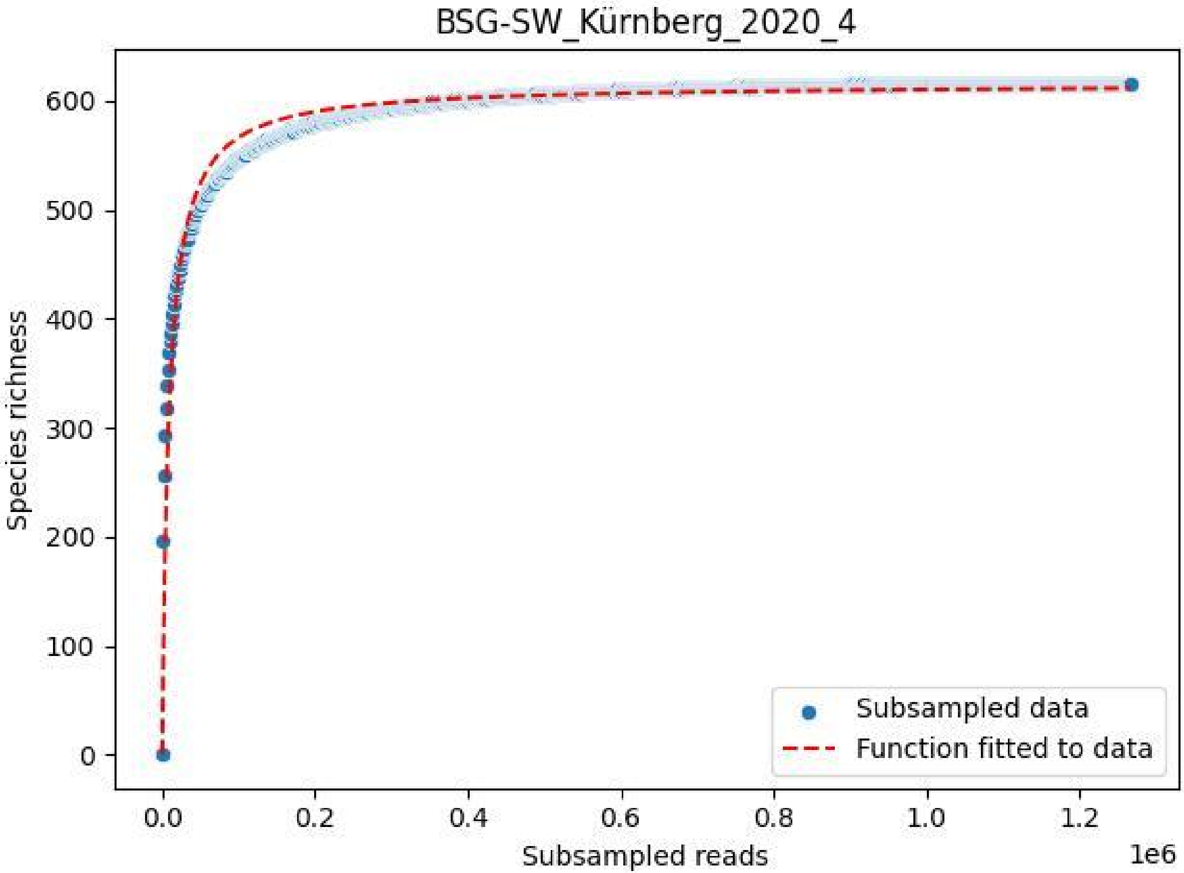

BSG-SW_Kürnberg_2020_4
Species richness
Subsampled reads
1e6
0
100
200
300
400
500
600
0.0
0.2
0.4
0.6
0.8
1.0
1.2
Subsampled data
Function fitted to data

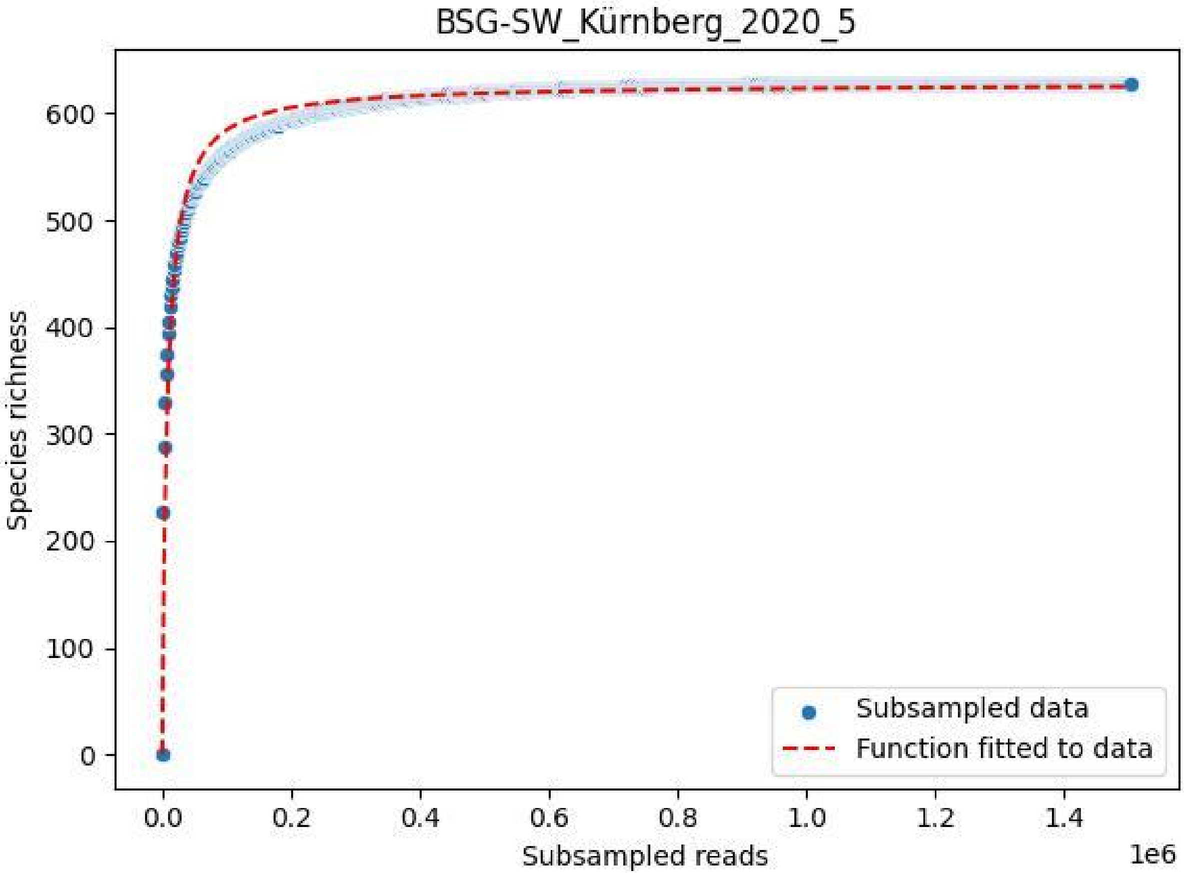

BSG-SW_Kürnberg_2020_5
Species richness
0
100
200
300
400
500
600
0.0
0.2
0.4
0.6
0.8
1.0
1.2
1.4
1e6
Subsampled reads
Subsampled data
Function fitted to data

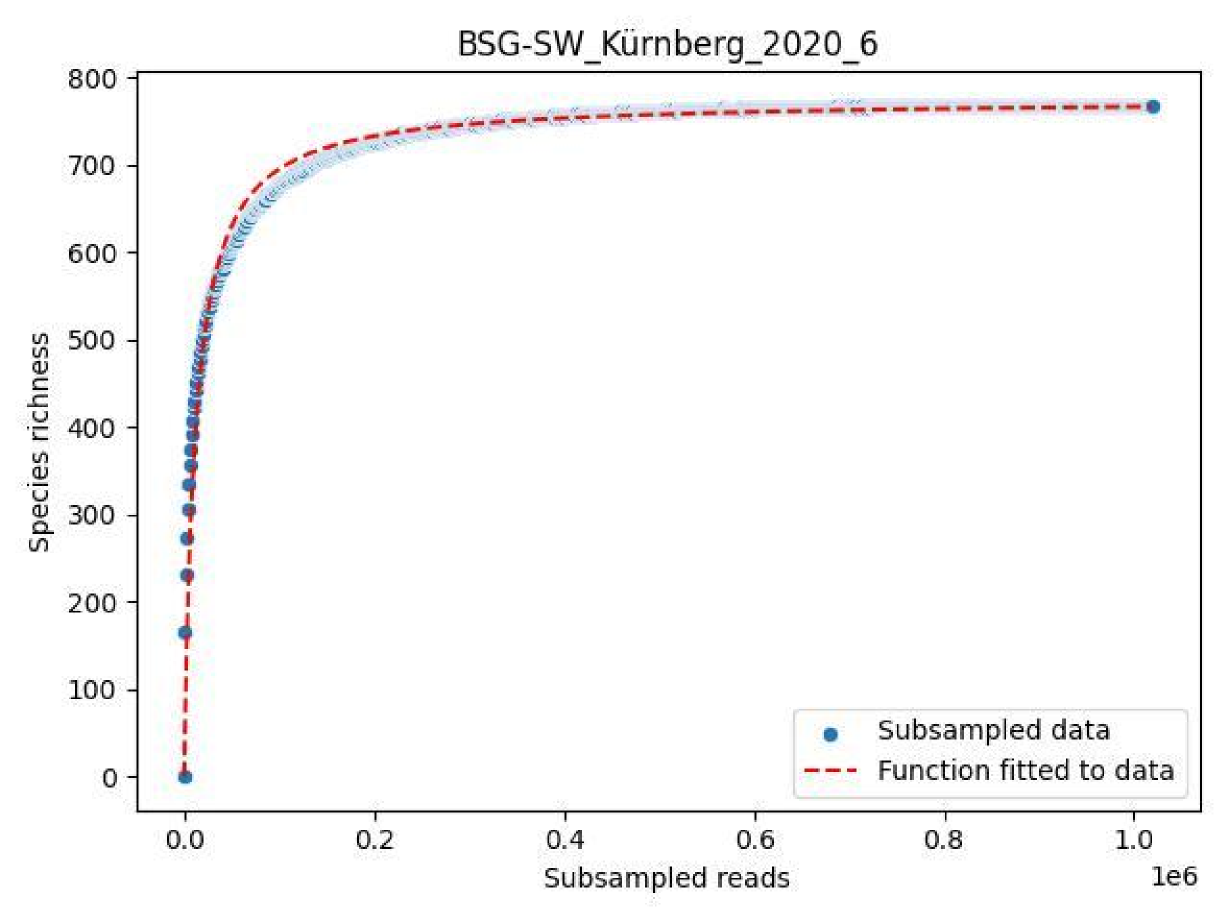
BSG-SW_Kürnberg_2020_6
Species richness
Subsampled reads
1e6
0
100
200
300
400
500
600
700
800
0.0
0.2
0.4
0.6
0.8
1.0
Subsampled data
Function fitted to data

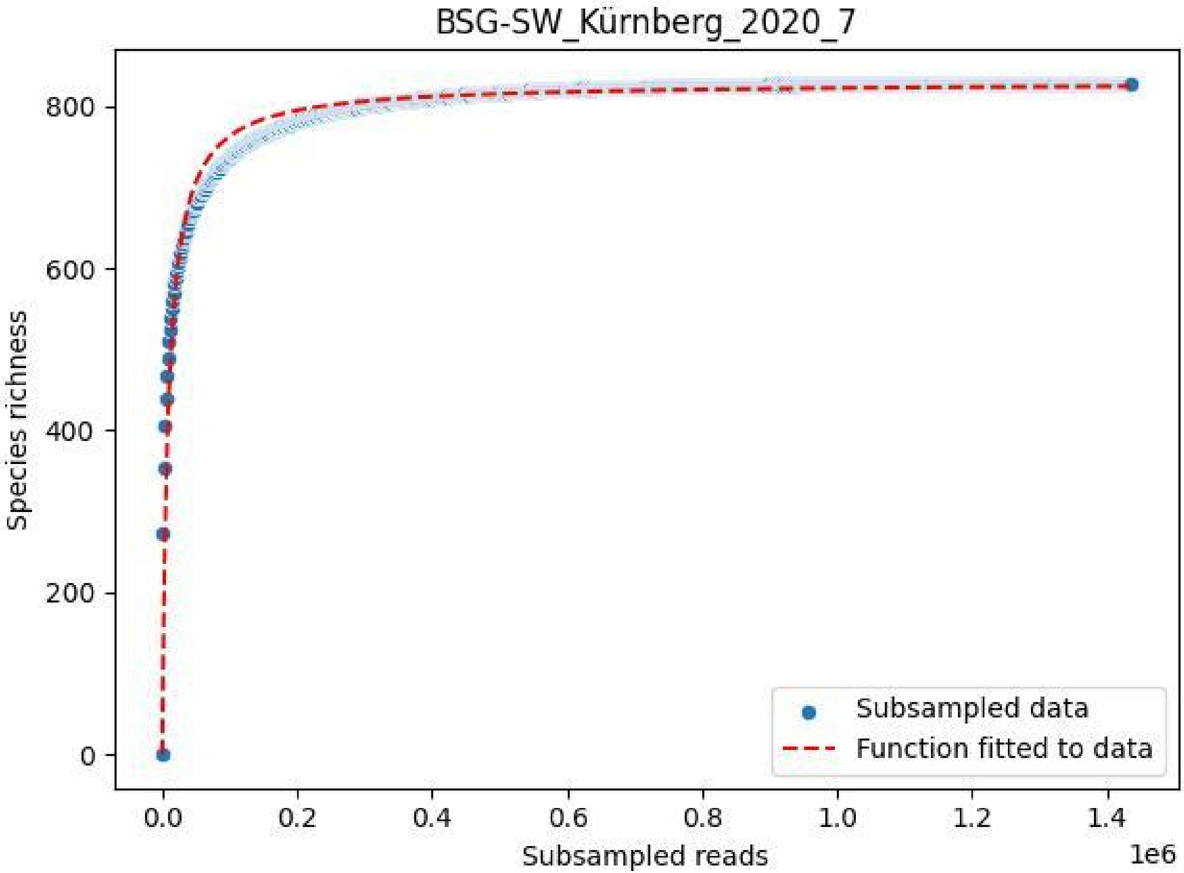
BSG-SW_Kürnberg_2020_7
Species richness
Subsampled reads
1e6
0
200
400
600
800
0.0
0.2
0.4
0.6
0.8
1.0
1.2
1.4
Subsampled data
Function fitted to data

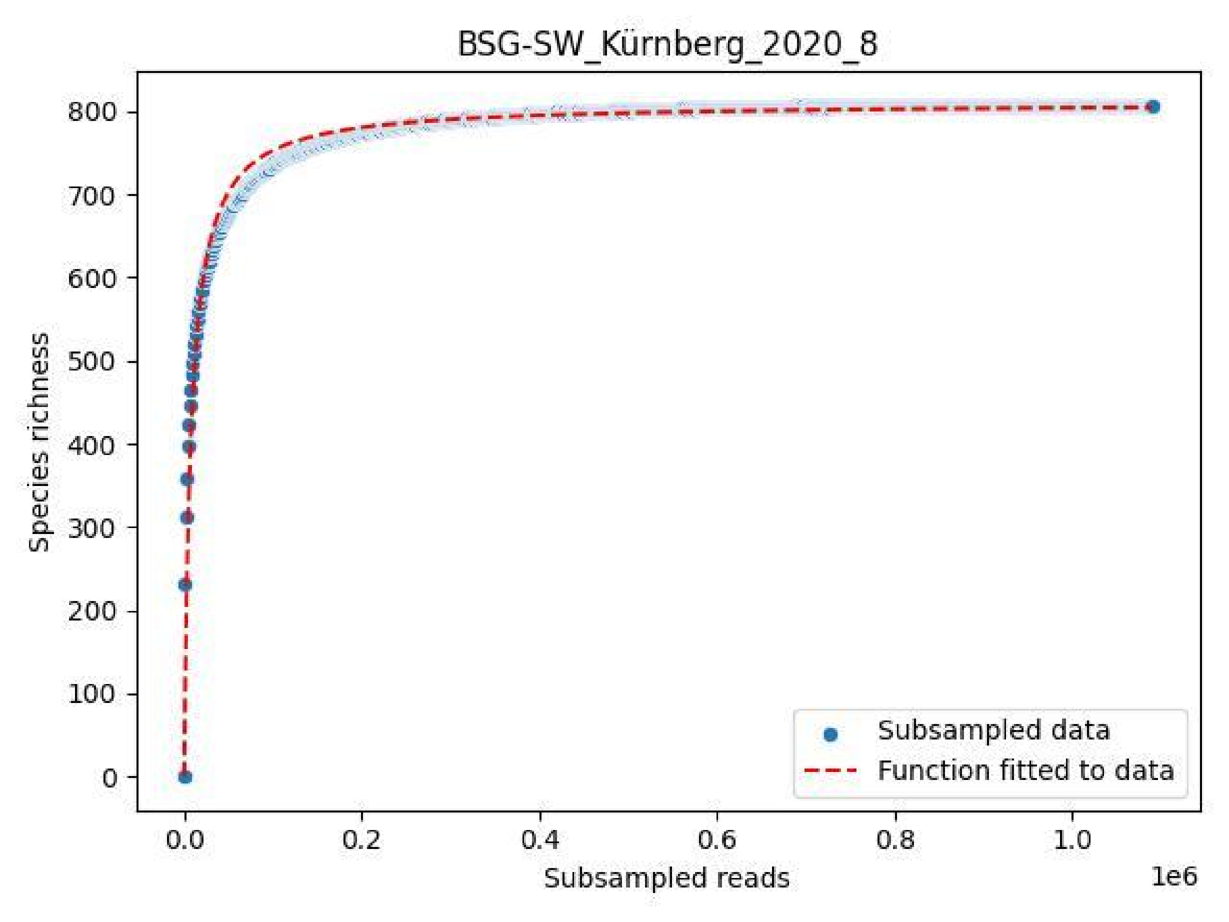

BSG-SW_Kürnberg_2020_8
Species richness
Subsampled reads
1e6
0
100
200
300
400
500
600
700
800
0.0
0.2
0.4
0.6
0.8
1.0
Subsampled data
Function fitted to data

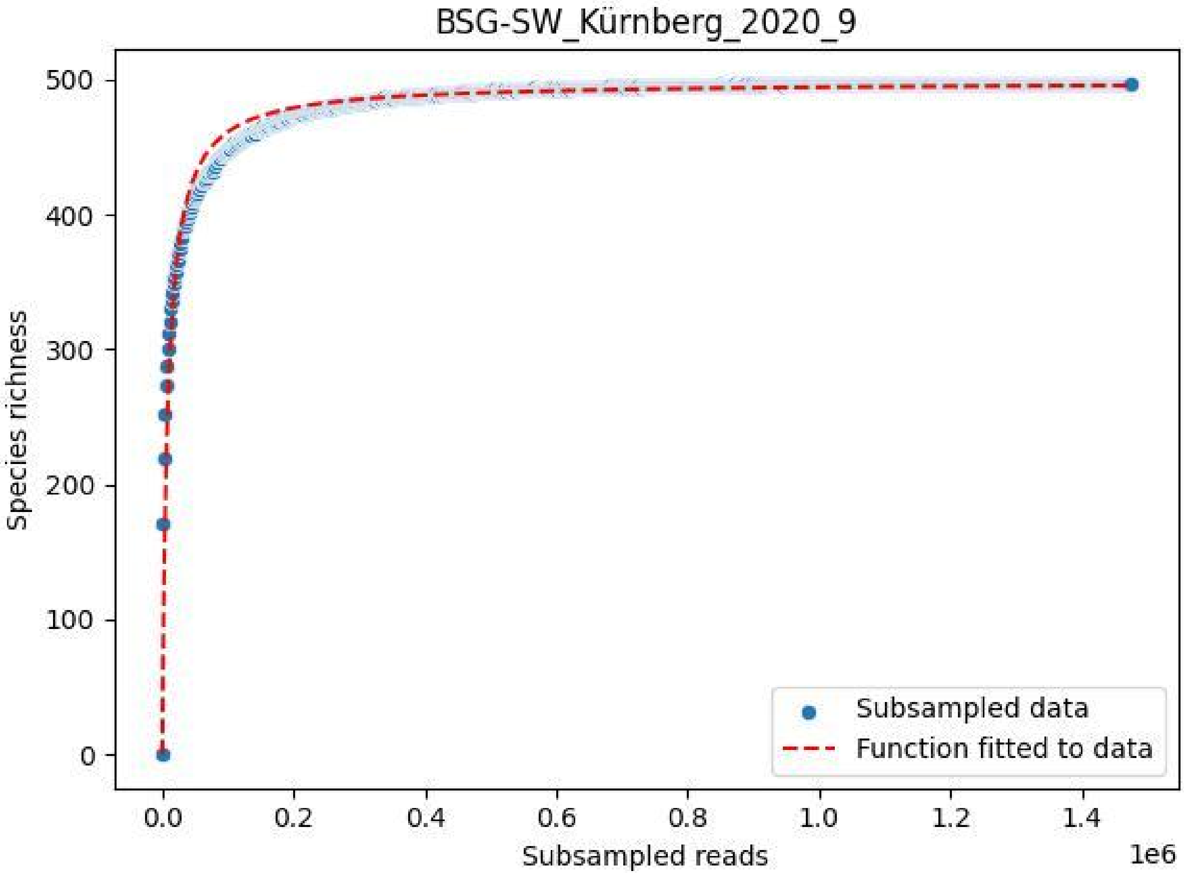
BSG-SW_Kürnberg_2020_9
Species richness
Subsampled reads
1e6
Subsampled data
Function fitted to data
0
100
200
300
400
500
0.0
0.2
0.4
0.6
0.8
1.0
1.2
1.4

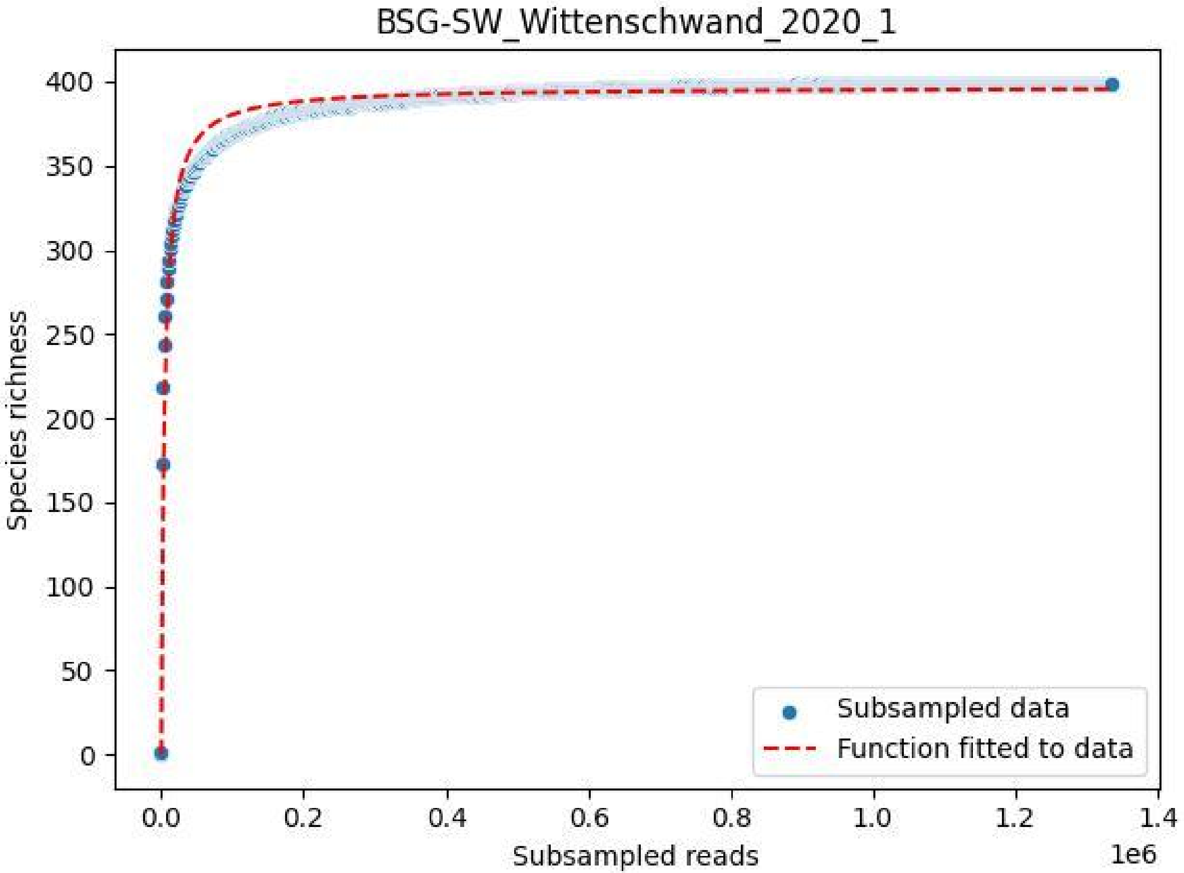
BSG-SW_Wittenschwand_2020_1
Species richness
Subsampled reads
1e6
0
50
100
150
200
250
300
350
400
0.0
0.2
0.4
0.6
0.8
1.0
1.2
1.4
Subsampled data
Function fitted to data

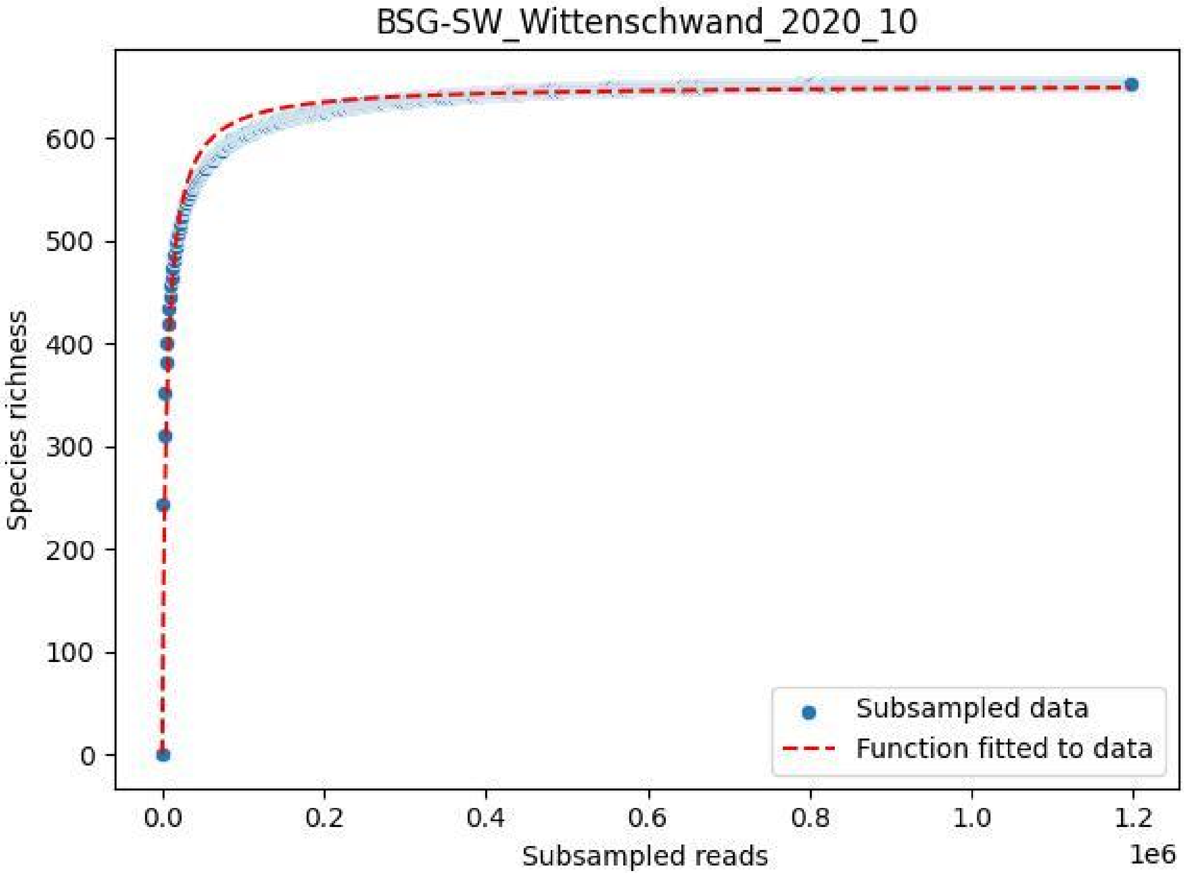

BSG-SW_Wittenschwand_2020_10
Species richness
Subsampled reads
1e6
0
100
200
300
400
500
600
0.0
0.2
0.4
0.6
0.8
1.0
1.2
Subsampled data
Function fitted to data

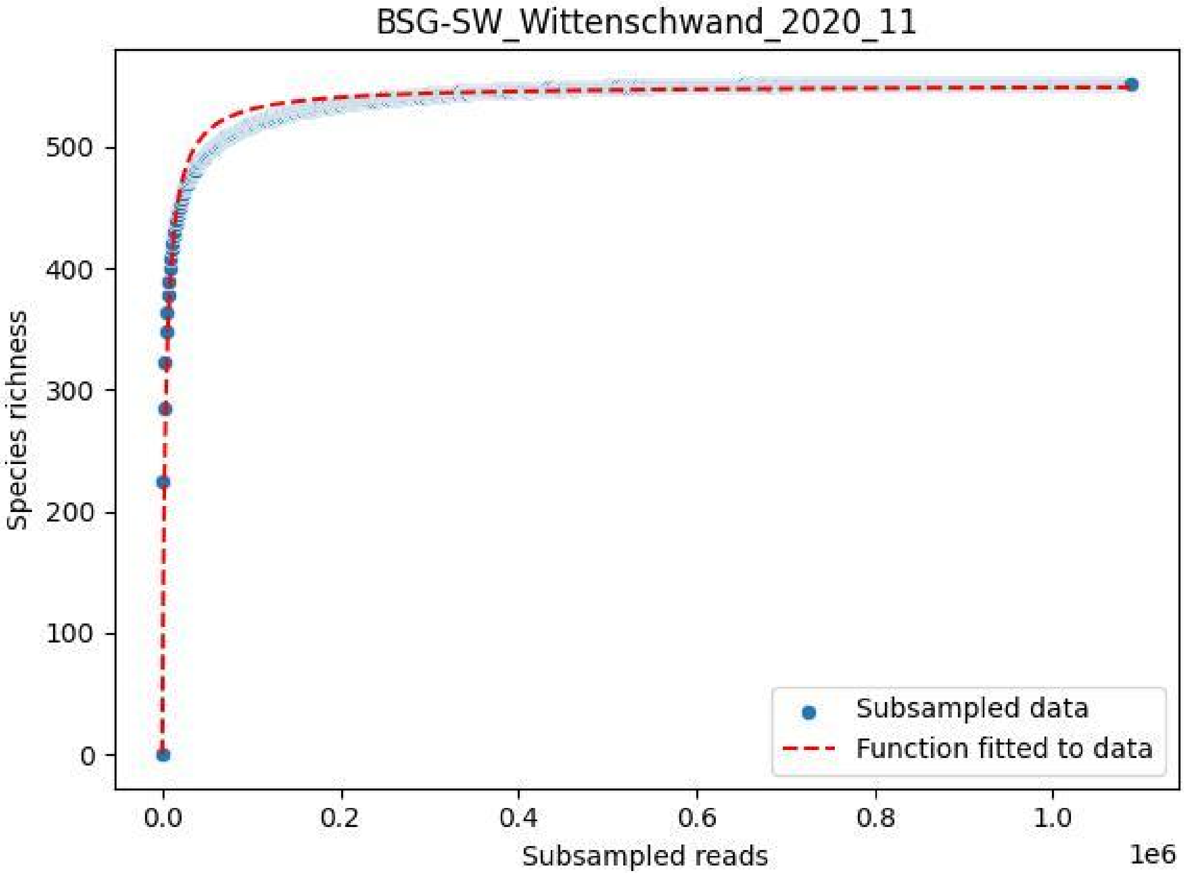
BSG-SW_Wittenschwand_2020_11
Species richness
Subsampled reads
1e6
0
100
200
300
400
500
0.0
0.2
0.4
0.6
0.8
1.0
Subsampled data
Function fitted to data

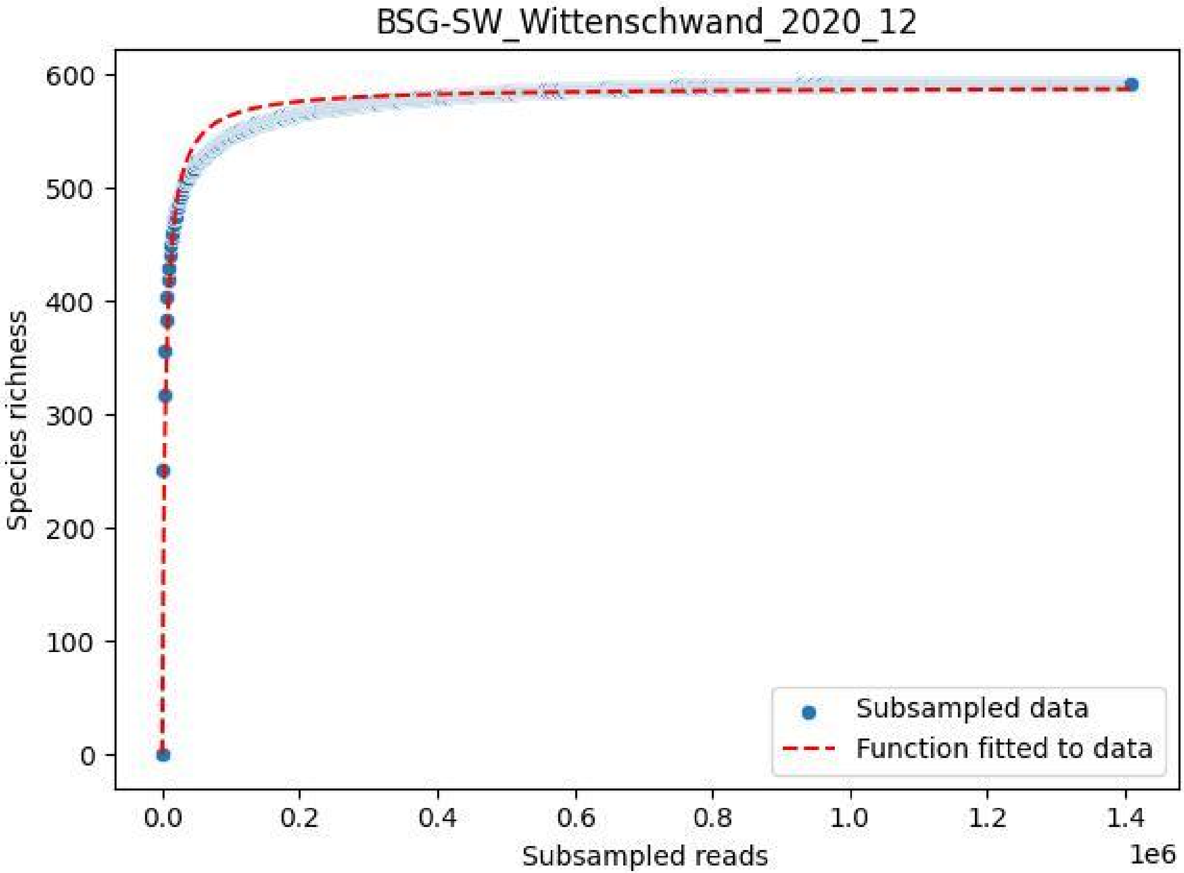
BSG-SW_Wittenschwand_2020_12
Species richness
0
100
200
300
400
500
600
0.0
0.2
0.4
0.6
0.8
1.0
1.2
1.4
Subsampled reads
1e6
Subsampled data
Function fitted to data

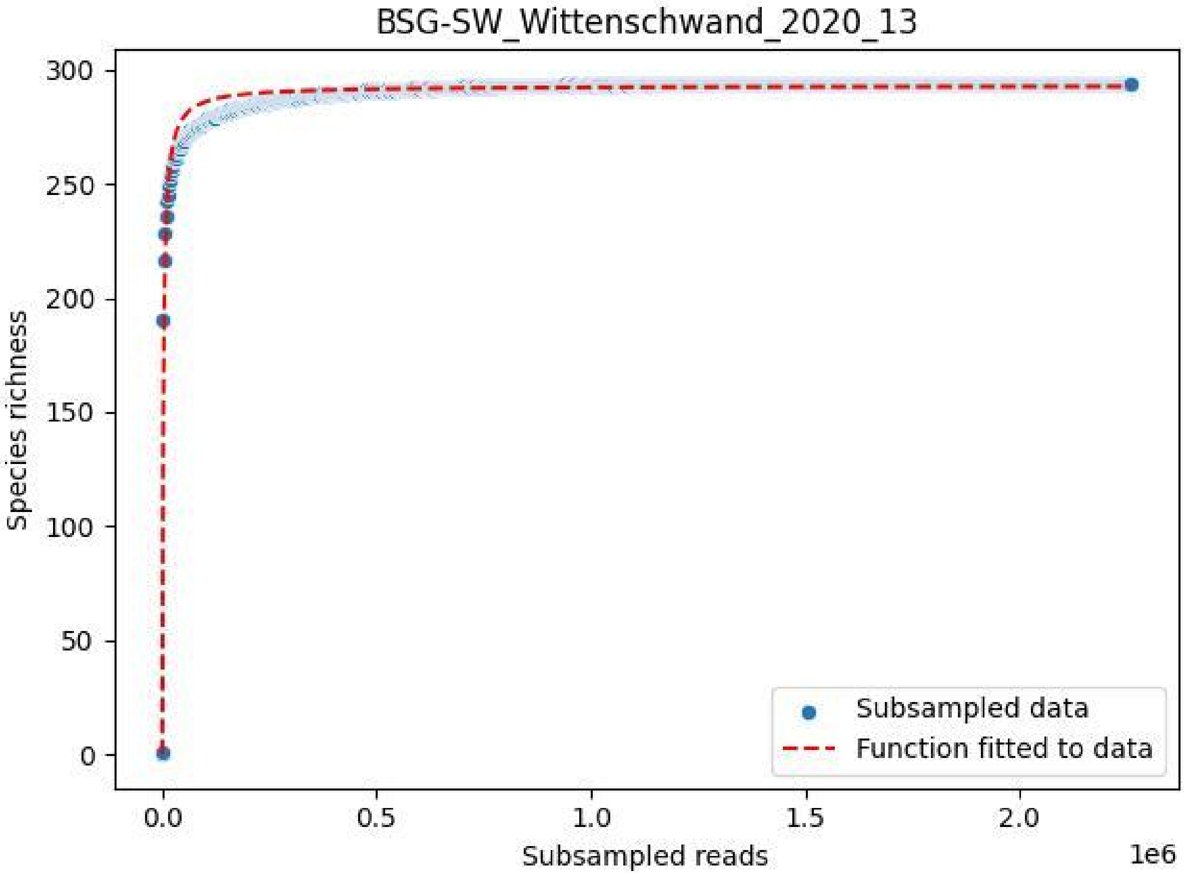
BSG-SW_Wittenschwand_2020_13
Species richness
Subsampled reads
1e6
0
50
100
150
200
250
300
0.0
0.5
1.0
1.5
2.0
Subsampled data
Function fitted to data

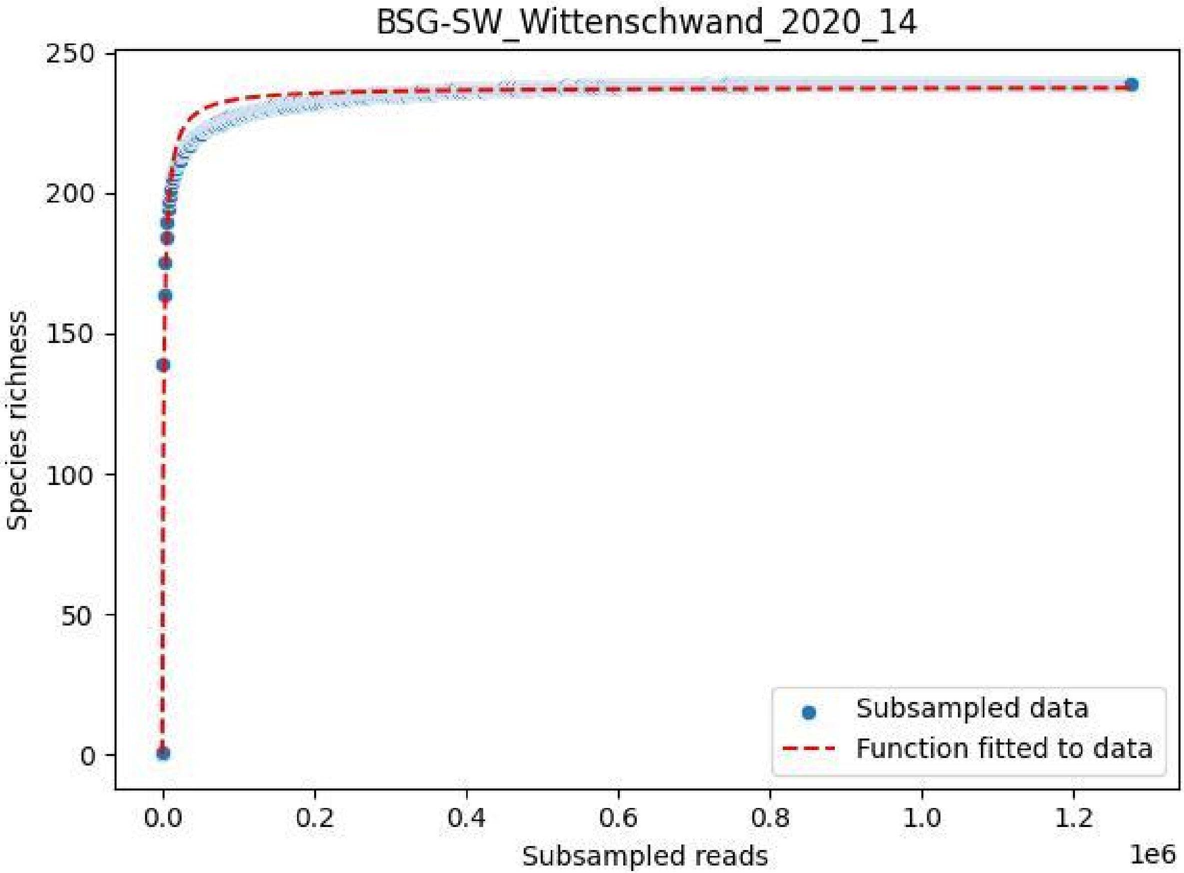

BSG-SW_Wittenschwand_2020_14
Species richness
Subsampled reads
1e6
0
50
100
150
200
250
0.0
0.2
0.4
0.6
0.8
1.0
1.2
Subsampled data
Function fitted to data

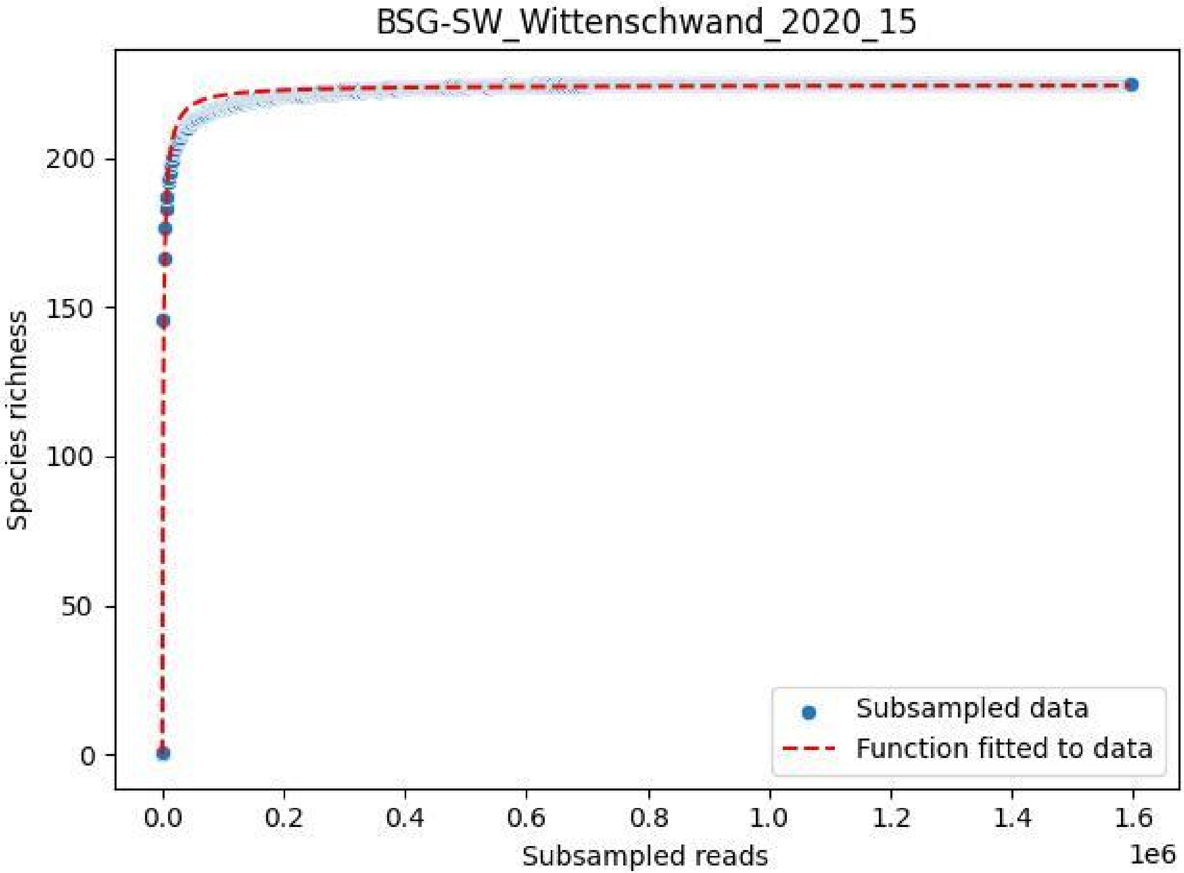
BSG-SW_Wittenschwand_2020_15
Species richness
Subsampled reads
1e6
0
50
100
150
200
0.0
0.2
0.4
0.6
0.8
1.0
1.2
1.4
1.6
Subsampled data
Function fitted to data

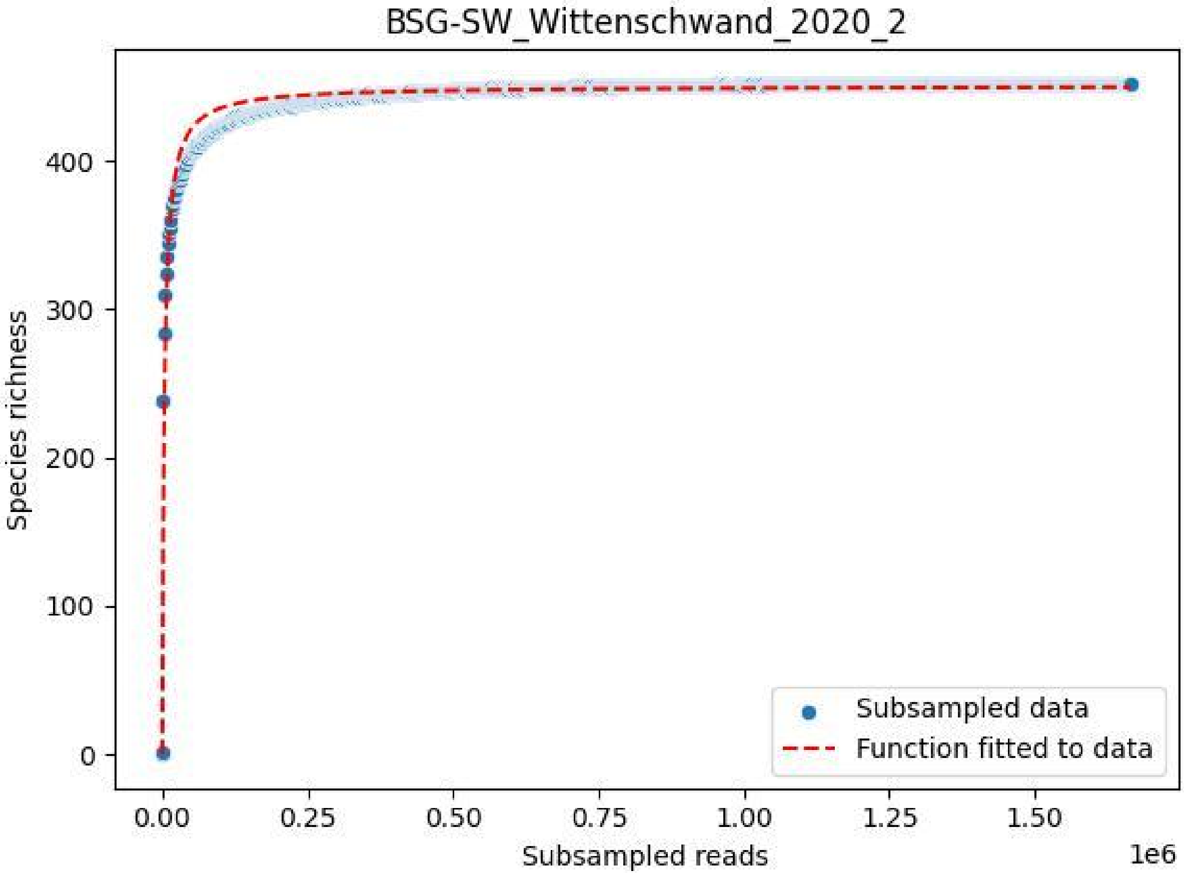
BSG-SW_Wittenschwand_2020_2
Species richness
400
300
200
100
0
0.00
0.25
0.50
0.75
1.00
1.25
1.50
Subsampled reads
1e6
Subsampled data
Function fitted to data

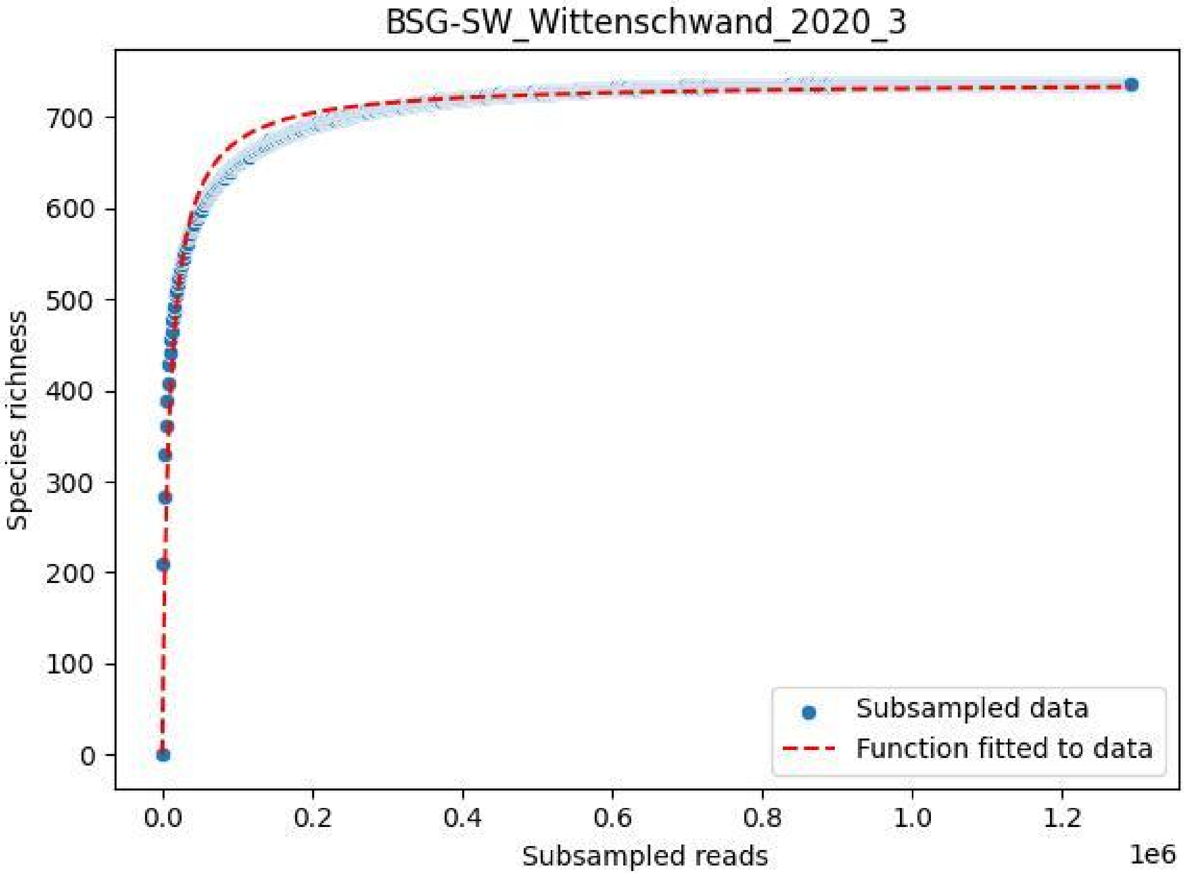

BSG-SW_Wittenschwand_2020_3
Species richness
Subsampled reads
1e6
0
100
200
300
400
500
600
700
0.0
0.2
0.4
0.6
0.8
1.0
1.2
Subsampled data
Function fitted to data

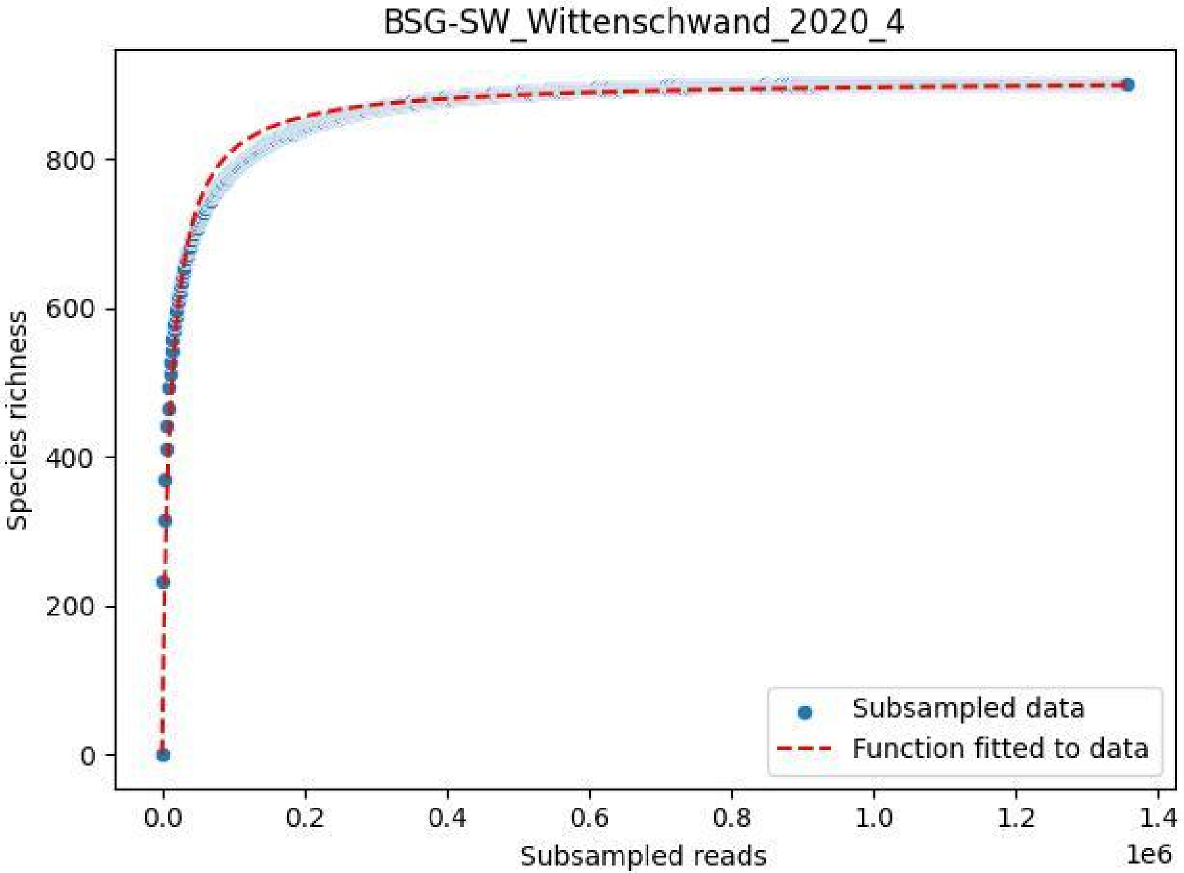

BSG-SW_Wittenschwand_2020_4
Species richness
Subsampled reads
1e6
0
200
400
600
800
0.0
0.2
0.4
0.6
0.8
1.0
1.2
1.4
Subsampled data
Function fitted to data

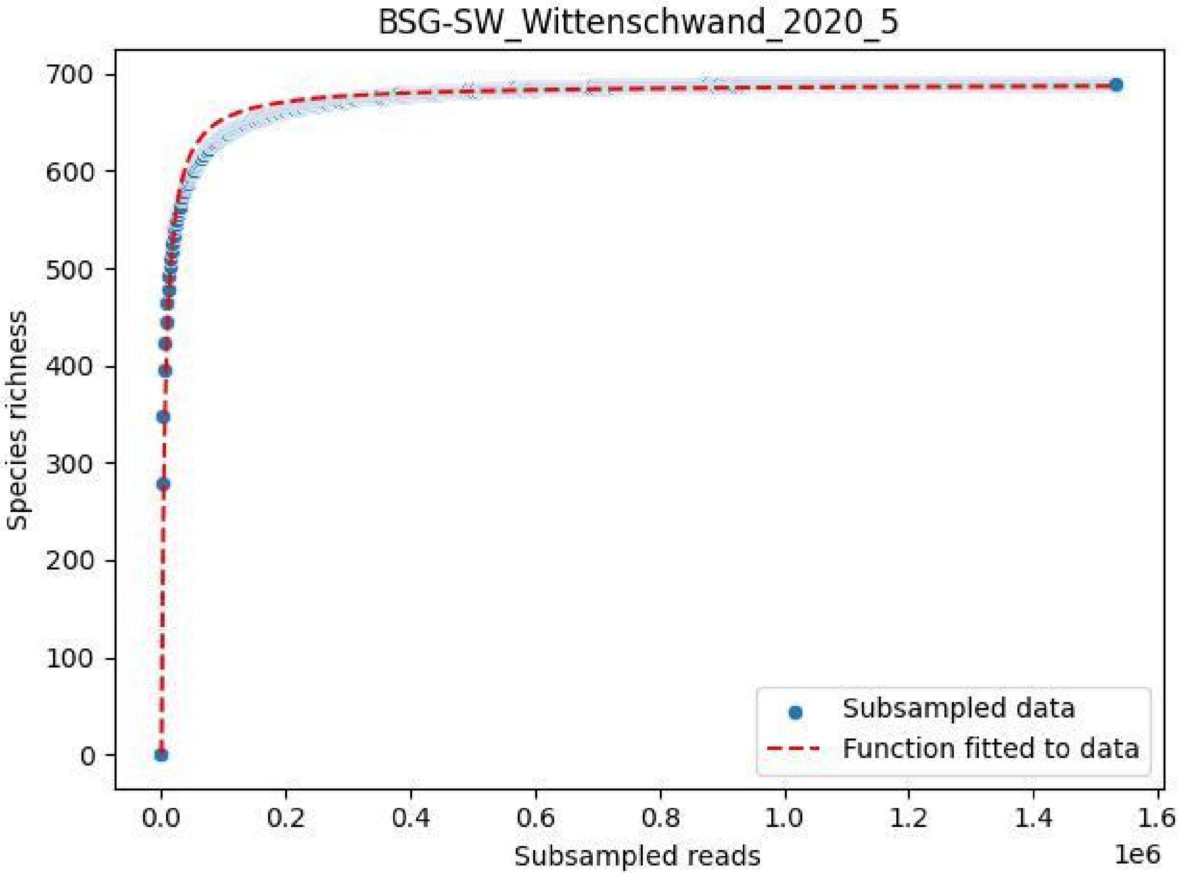
BSG-SW_Wittenschwand_2020_5
Species richness
Subsampled reads
1e6
0
100
200
300
400
500
600
700
0.0
0.2
0.4
0.6
0.8
1.0
1.2
1.4
1.6
Subsampled data
Function fitted to data

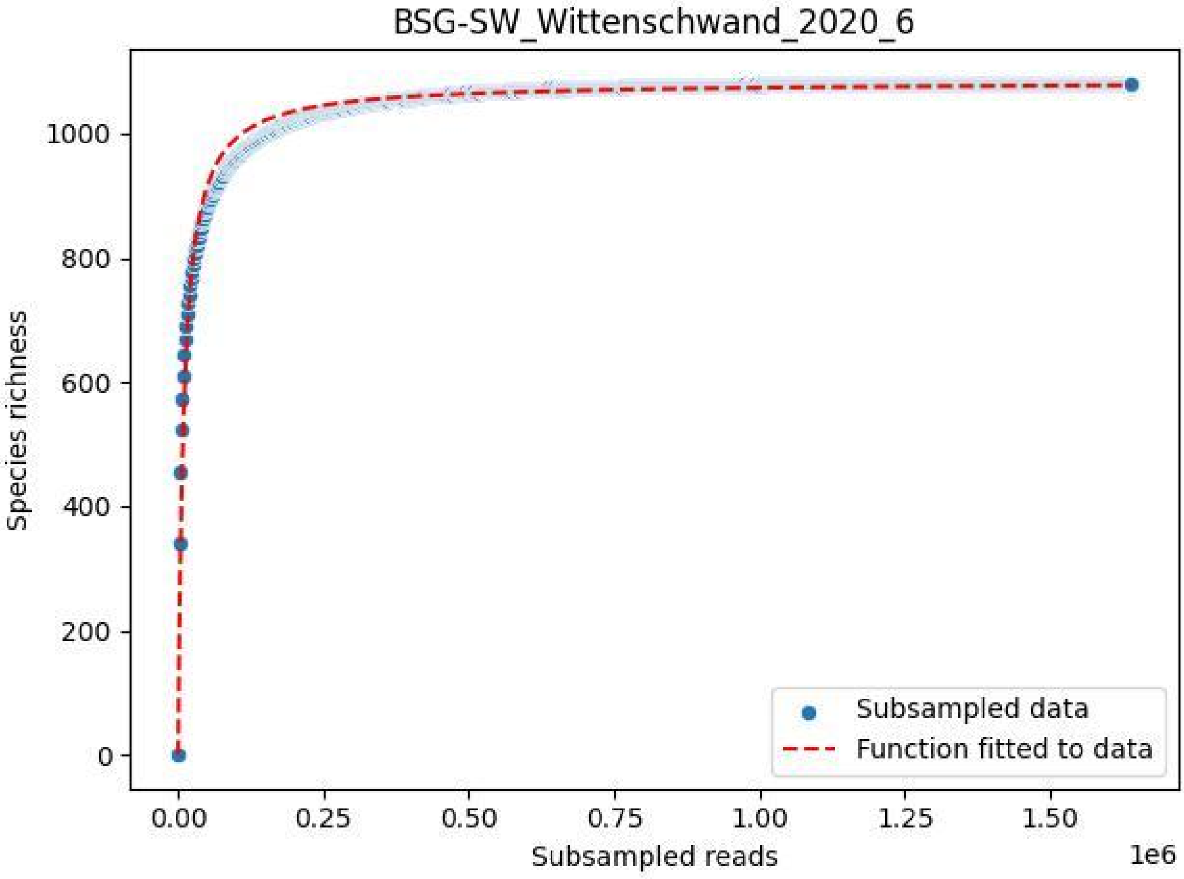
BSG-SW_Wittenschwand_2020_6
Species richness
Subsampled reads
1e6
0
200
400
600
800
1000
0.00
0.25
0.50
0.75
1.00
1.25
1.50
Subsampled data
Function fitted to data

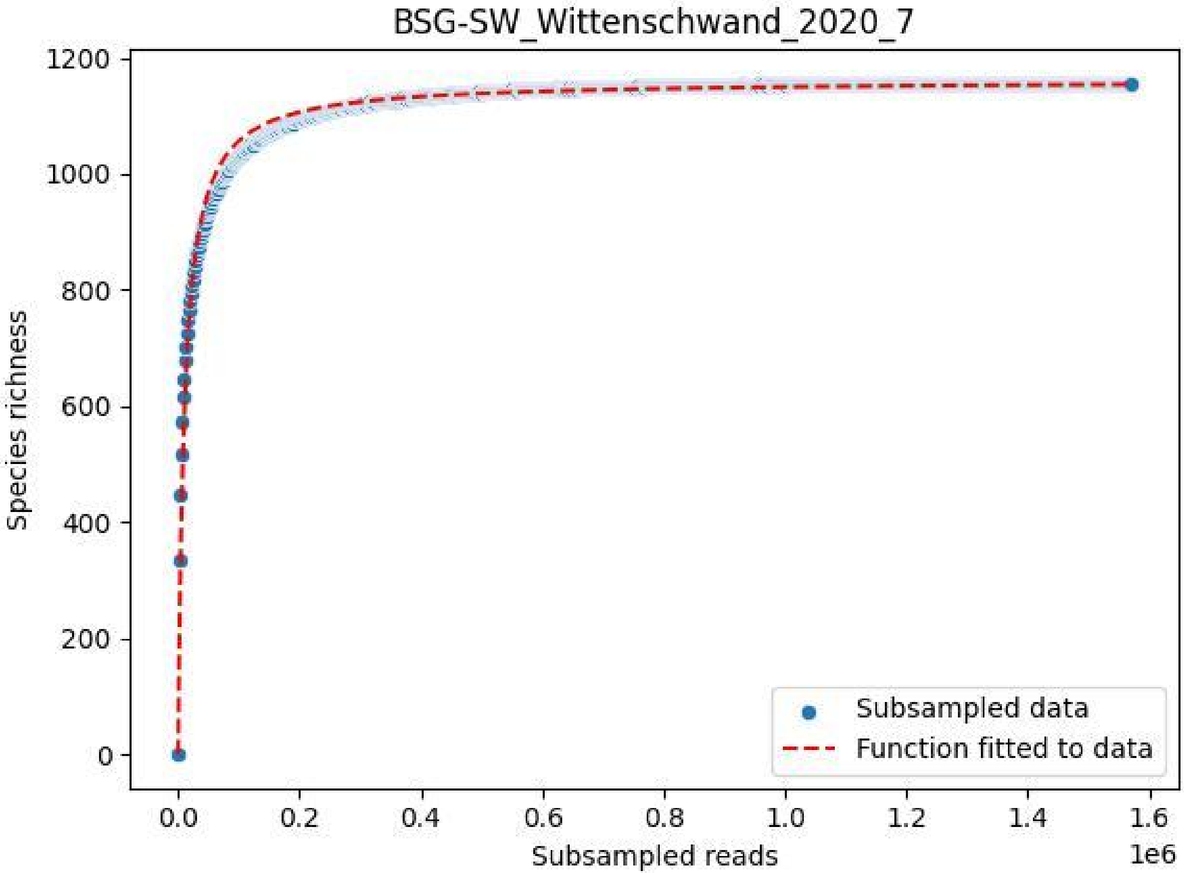
BSG-SW_Wittenschwand_2020_7
Species richness
Subsampled reads
1e6
0
200
400
600
800
1000
1200
0.0
0.2
0.4
0.6
0.8
1.0
1.2
1.4
1.6
Subsampled data
Function fitted to data

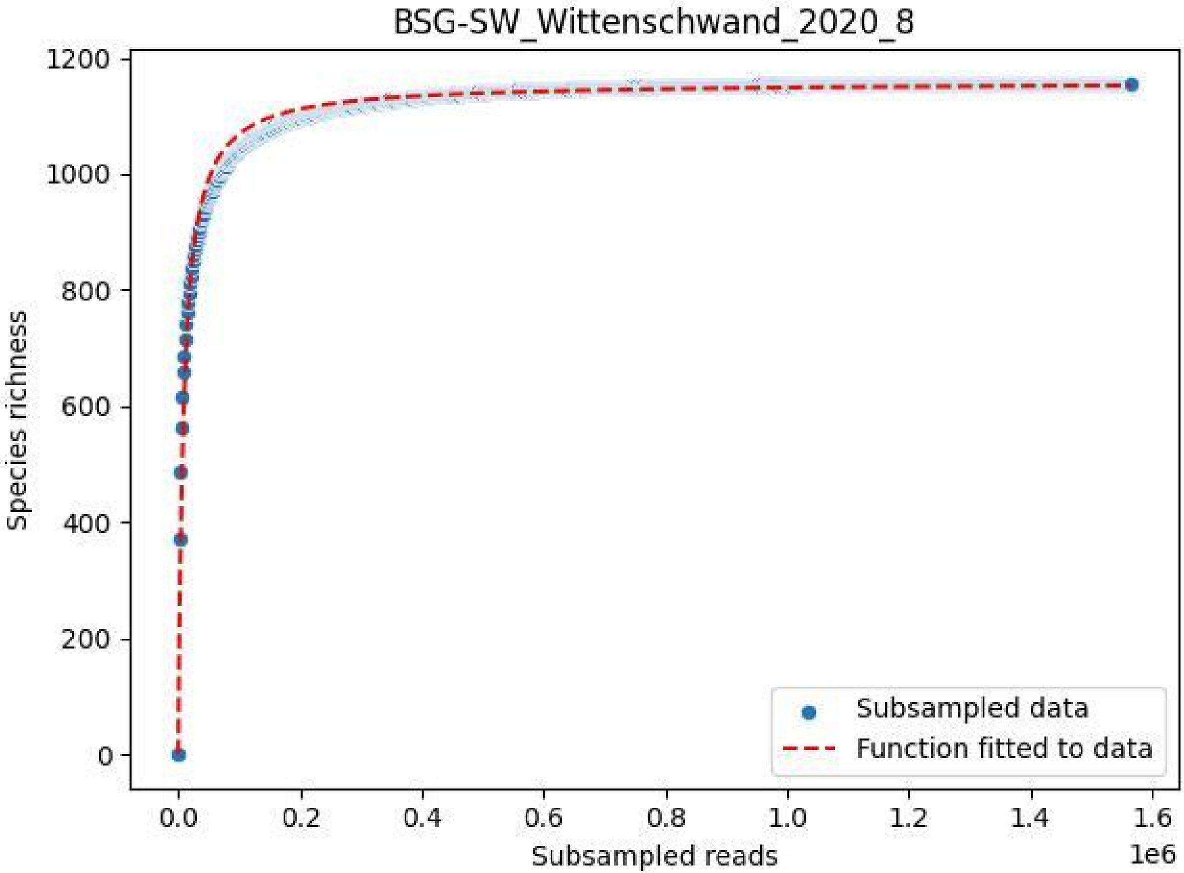
BSG-SW_Wittenschwand_2020_8
Species richness
Subsampled reads
1e6
0
200
400
600
800
1000
1200
0.0
0.2
0.4
0.6
0.8
1.0
1.2
1.4
1.6
Subsampled data
Function fitted to data

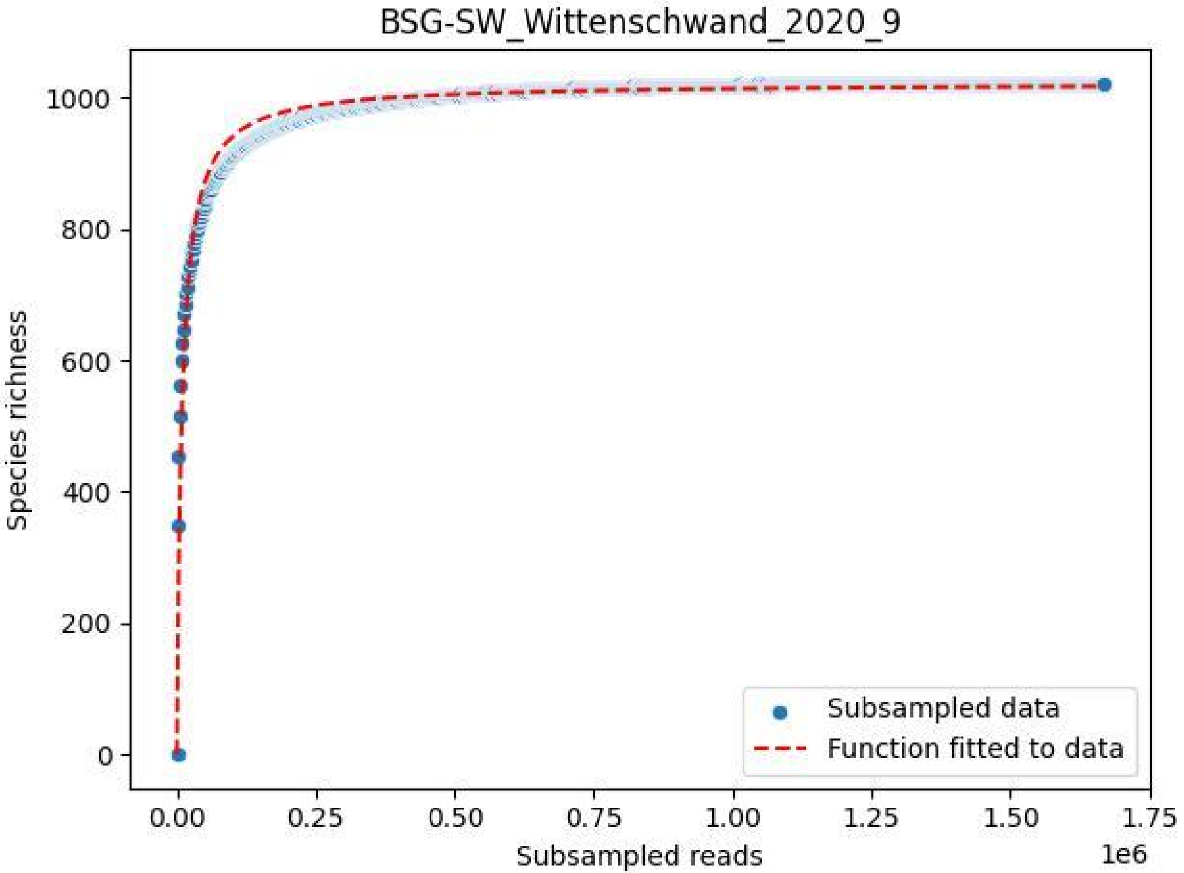
BSG-SW_Wittenschwand_2020_9
Species richness
Subsampled reads
1e6
0
200
400
600
800
1000
0.00
0.25
0.50
0.75
1.00
1.25
1.50
1.75
Subsampled data
Function fitted to data

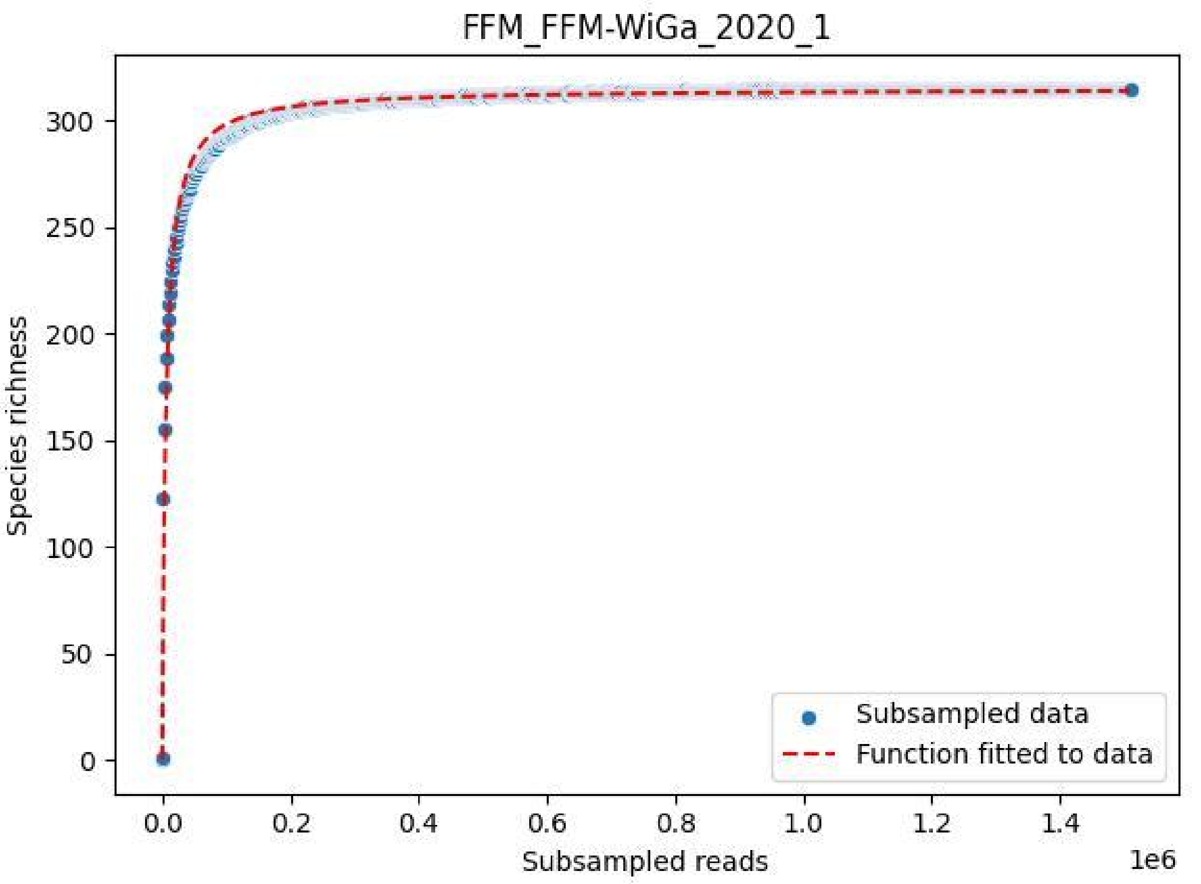

FFM_FFM-WiGa_2020_1
Species richness
Subsampled reads
1e6
0
50
100
150
200
250
300
0.0
0.2
0.4
0.6
0.8
1.0
1.2
1.4
Subsampled data
Function fitted to data

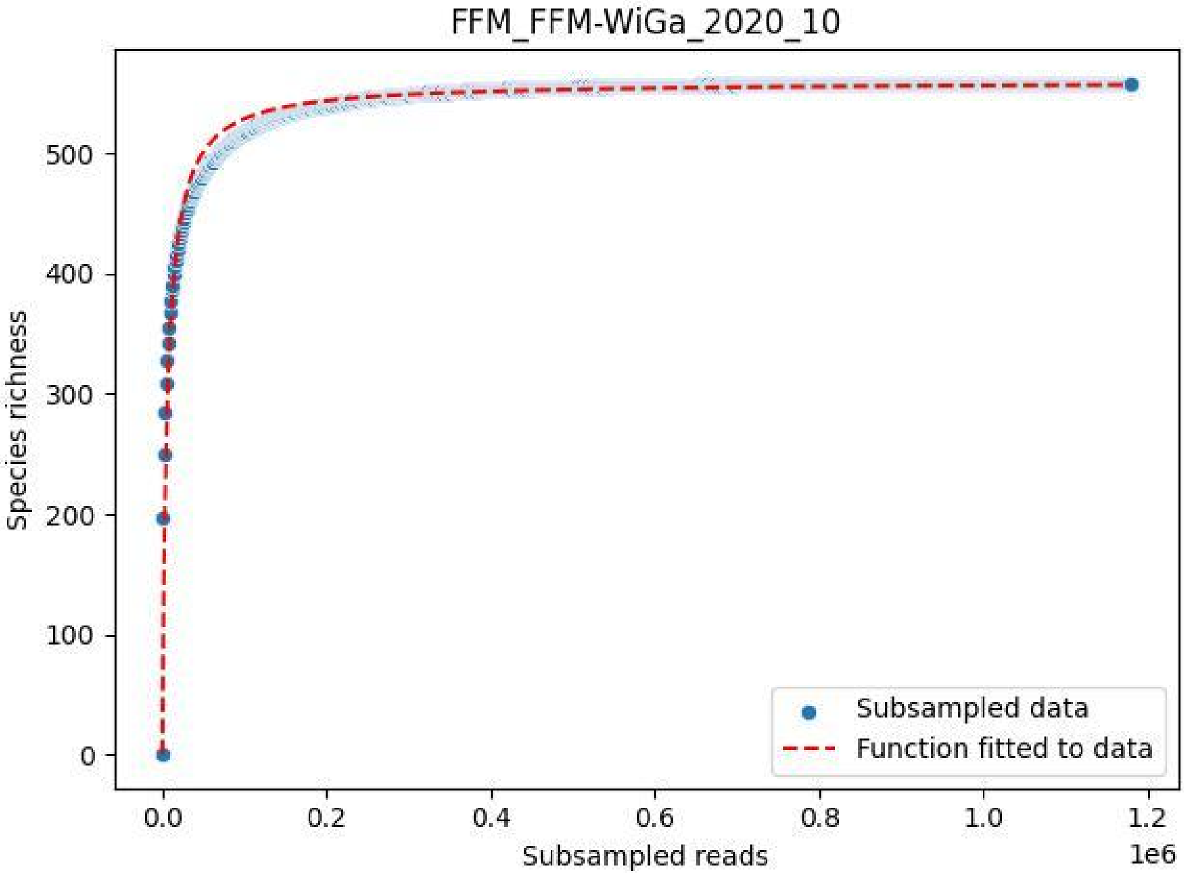

FFM_FFM-WiGa_2020_10
Species richness
Subsampled reads
1e6
0
100
200
300
400
500
0.0
0.2
0.4
0.6
0.8
1.0
1.2
Subsampled data
Function fitted to data

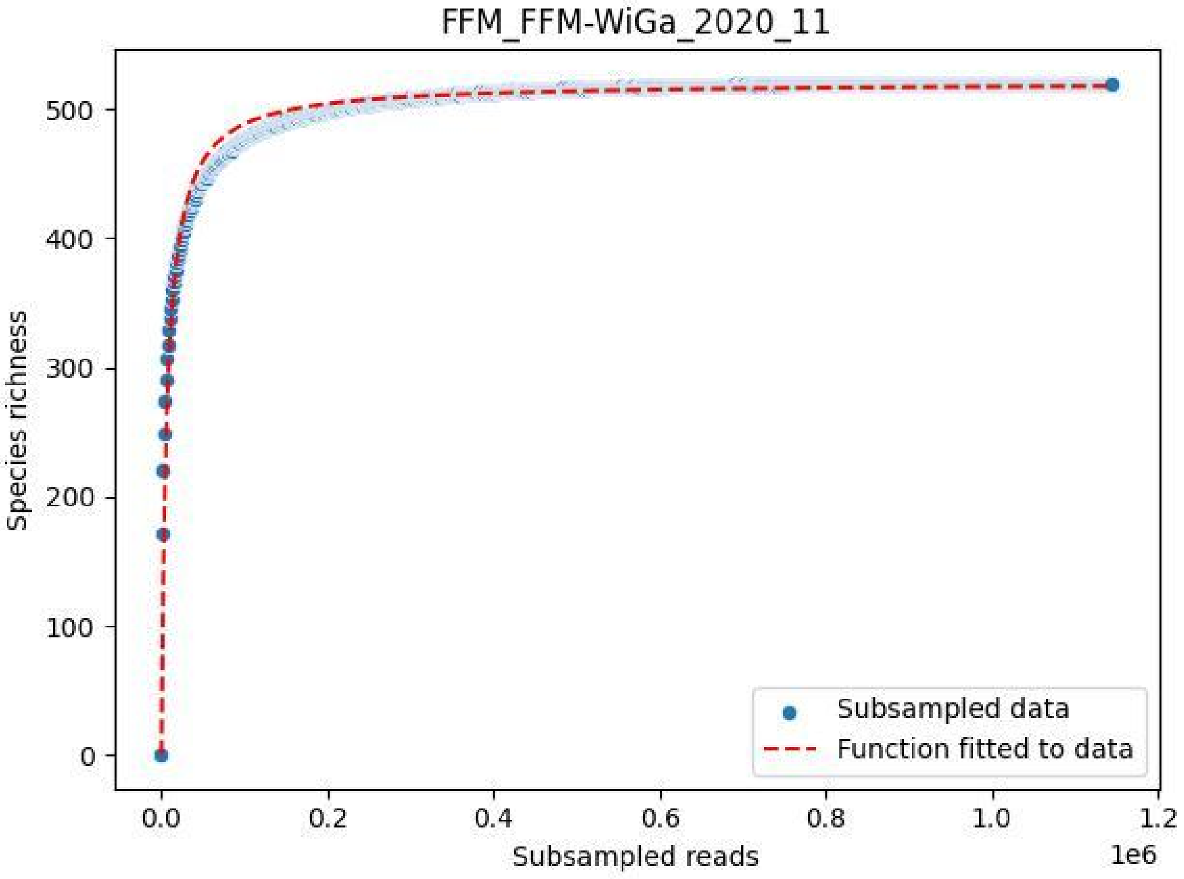
FFM_FFM-WiGa_2020_11
Species richness
Subsampled reads
1e6
0
100
200
300
400
500
0.0
0.2
0.4
0.6
0.8
1.0
1.2
Subsampled data
Function fitted to data

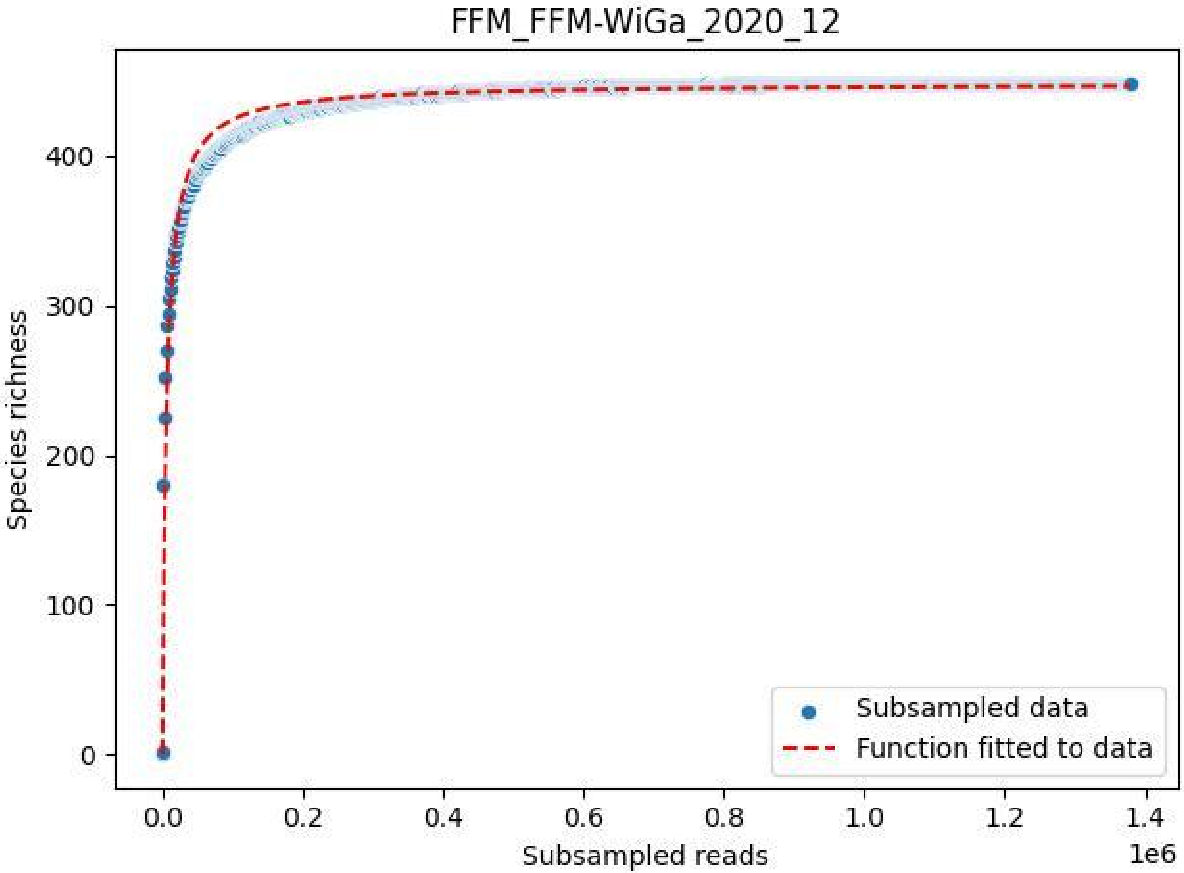
FFM_FFM-WiGa_2020_12
Species richness
Subsampled reads
1e6
0
100
200
300
400
0.0
0.2
0.4
0.6
0.8
1.0
1.2
1.4
Subsampled data
Function fitted to data

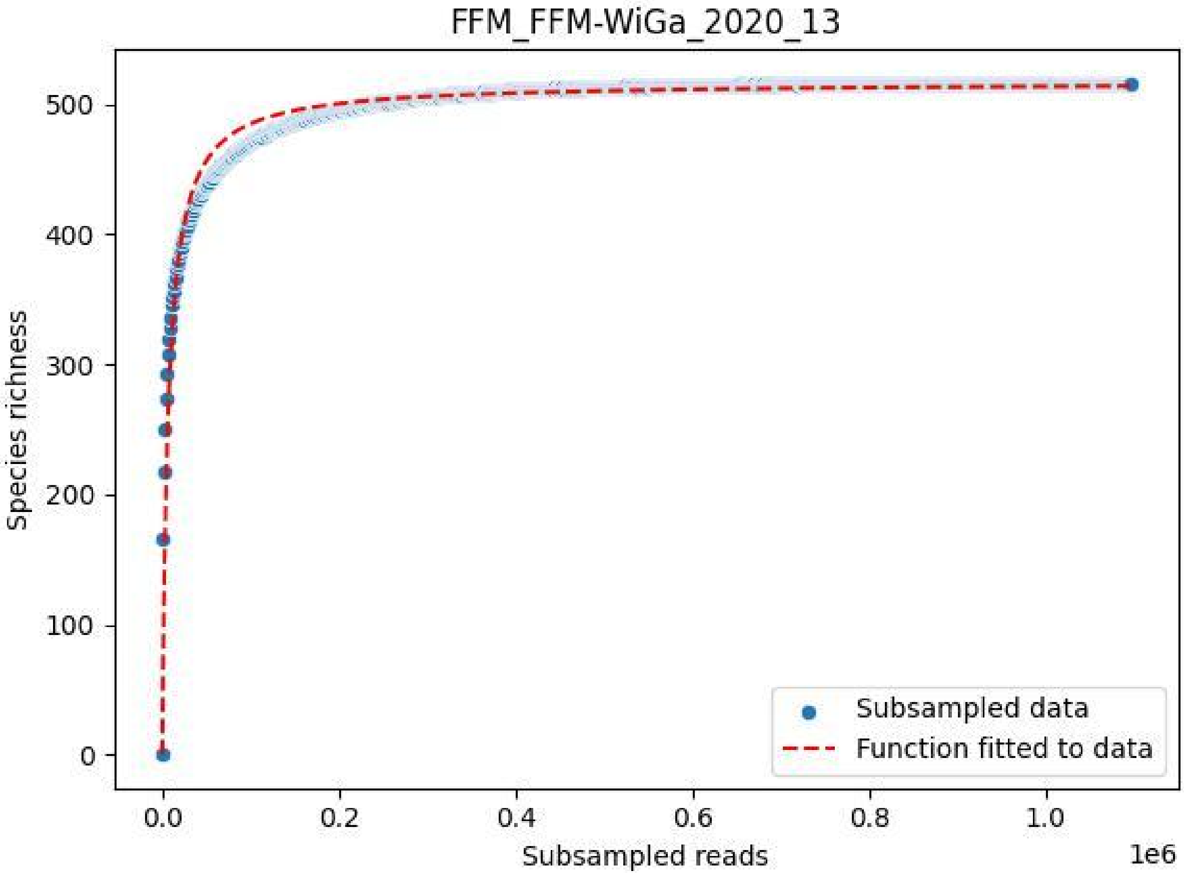
FFM_FFM-WiGa_2020_13
Species richness
Subsampled reads
1e6
0
100
200
300
400
500
0.0
0.2
0.4
0.6
0.8
1.0
Subsampled data
Function fitted to data

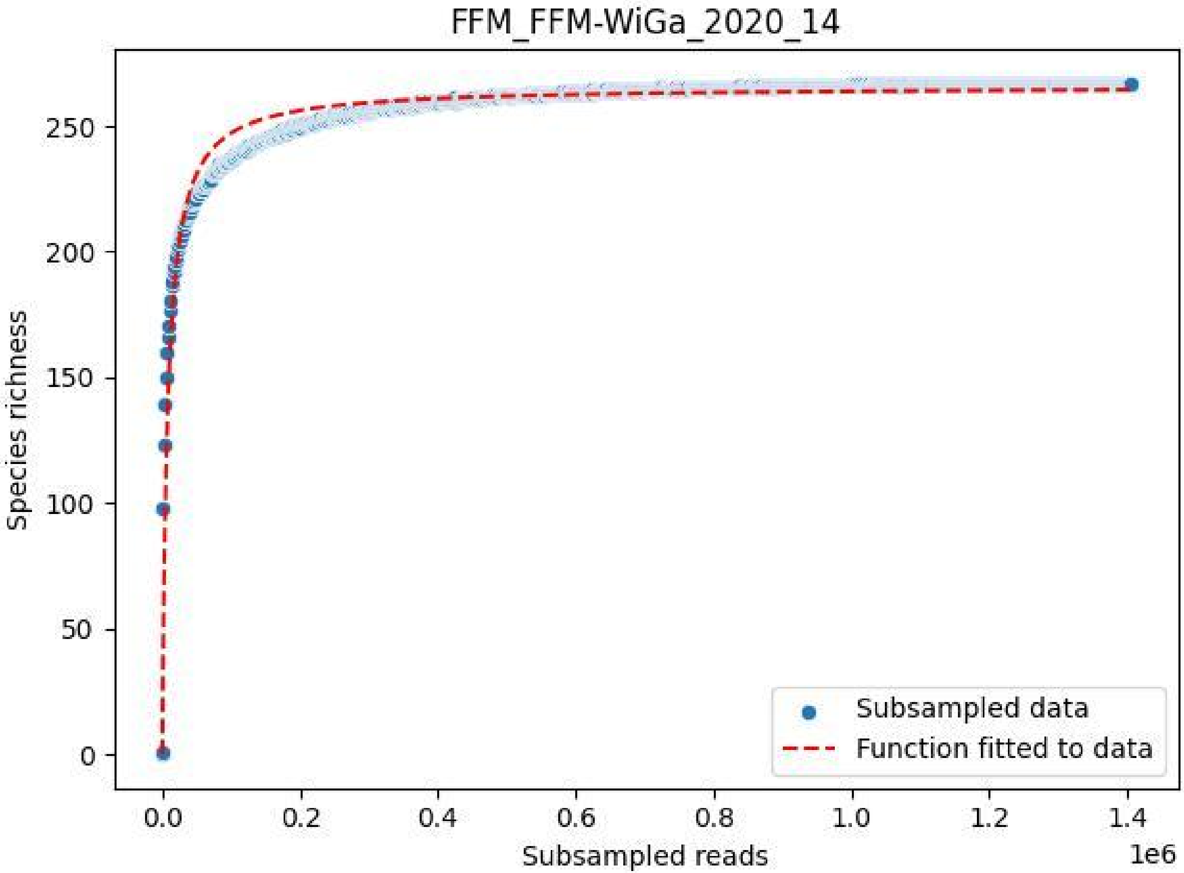
FFM_FFM-WiGa_2020_14
Species richness
0
50
100
150
200
250
0.0
0.2
0.4
0.6
0.8
1.0
1.2
1.4
1e6
Subsampled reads
Subsampled data
Function fitted to data

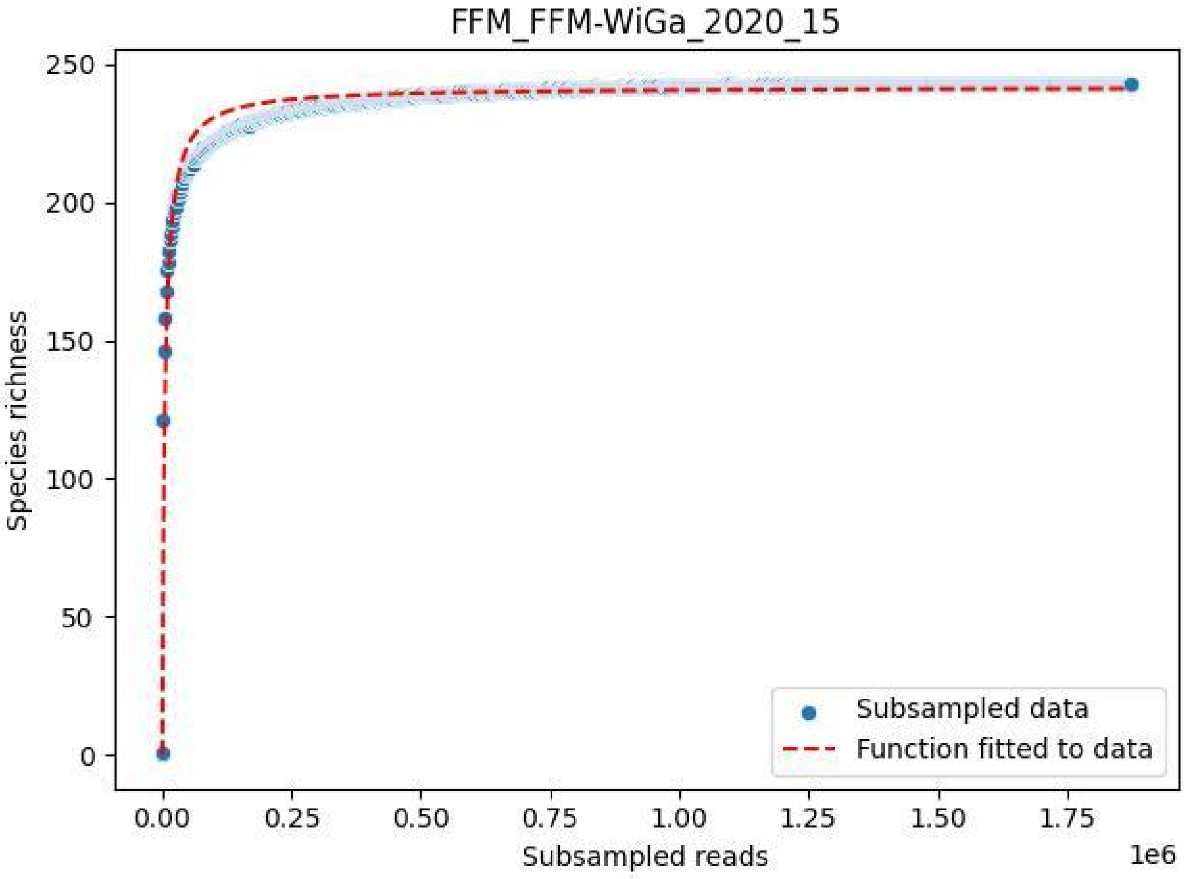

FFM_FFM-WiGa_2020_15
Species richness
Subsampled reads
1e6
0
50
100
150
200
250
0.00
0.25
0.50
0.75
1.00
1.25
1.50
1.75
Subsampled data
Function fitted to data

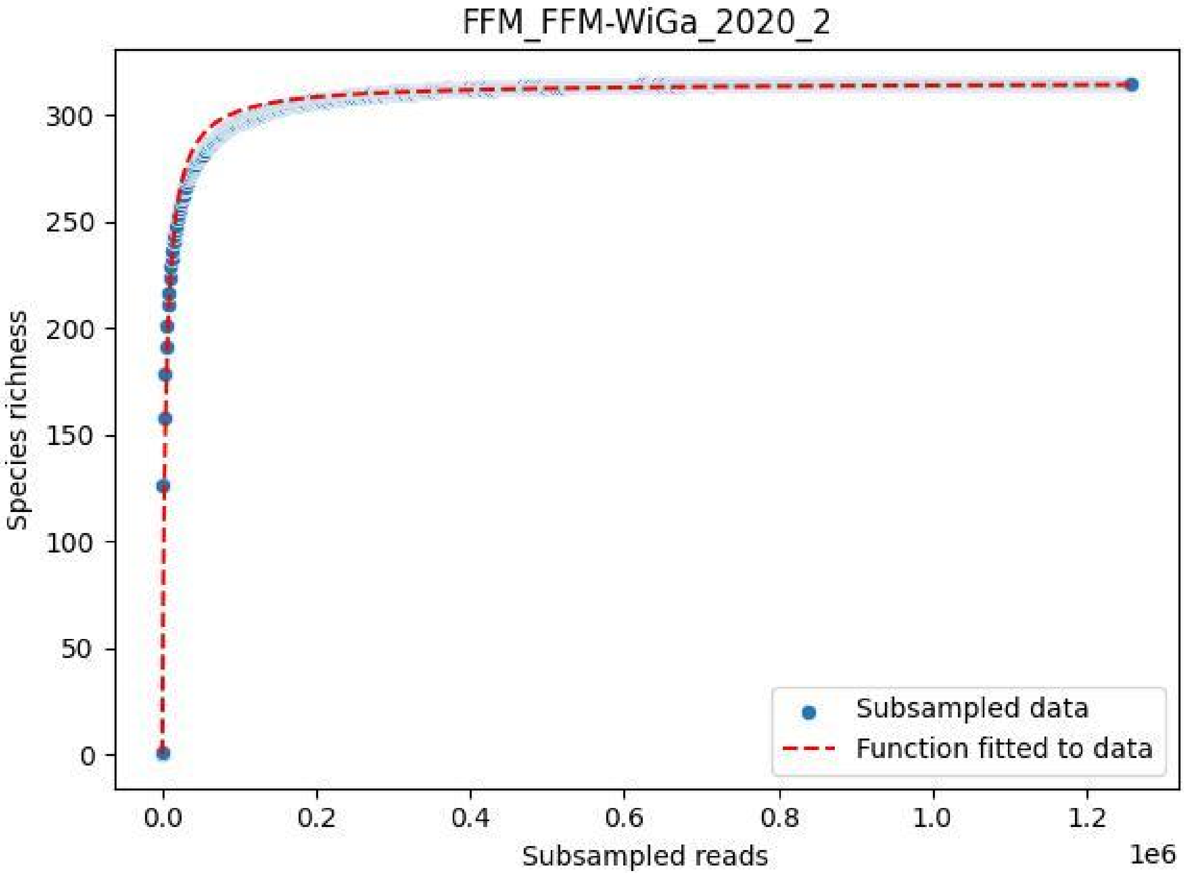
FFM_FFM-WiGa_2020_2
Species richness
Subsampled reads
1e6
0
50
100
150
200
250
300
0.0
0.2
0.4
0.6
0.8
1.0
1.2
Subsampled data
Function fitted to data

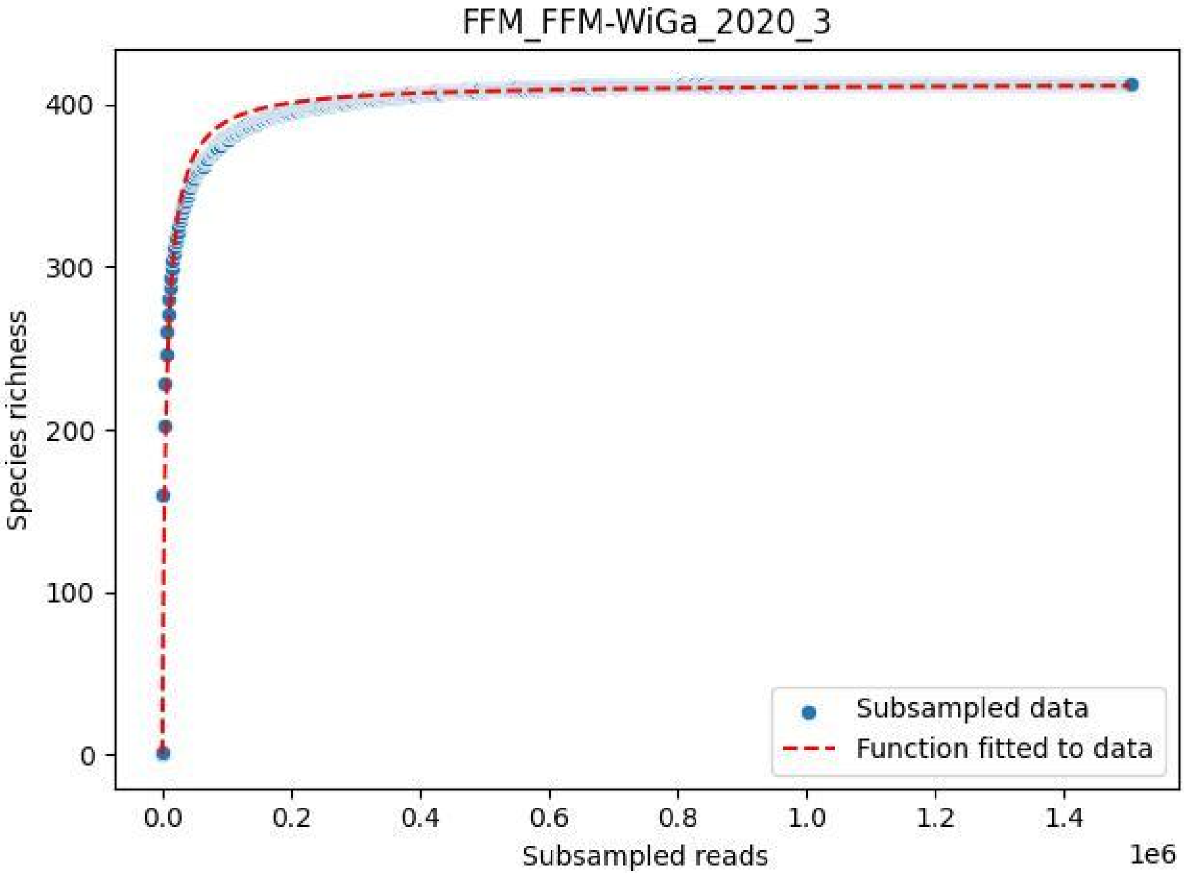
FFM_FFM-WiGa_2020_3
Species richness
0
100
200
300
400
0.0
0.2
0.4
0.6
0.8
1.0
1.2
1.4
1e6
Subsampled reads
Subsampled data
Function fitted to data

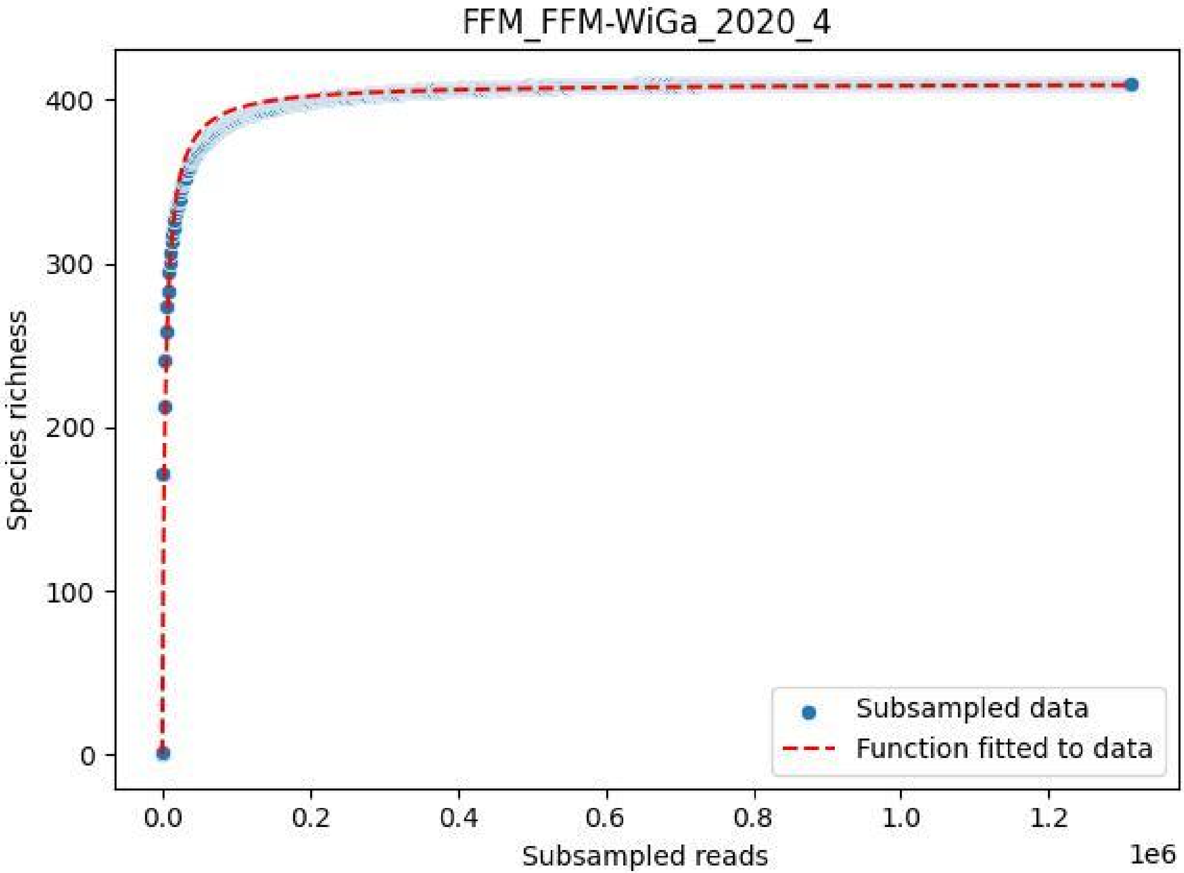

FFM_FFM-WiGa_2020_4
Species richness
Subsampled reads
1e6
0
100
200
300
400
0.0
0.2
0.4
0.6
0.8
1.0
1.2
Subsampled data
Function fitted to data

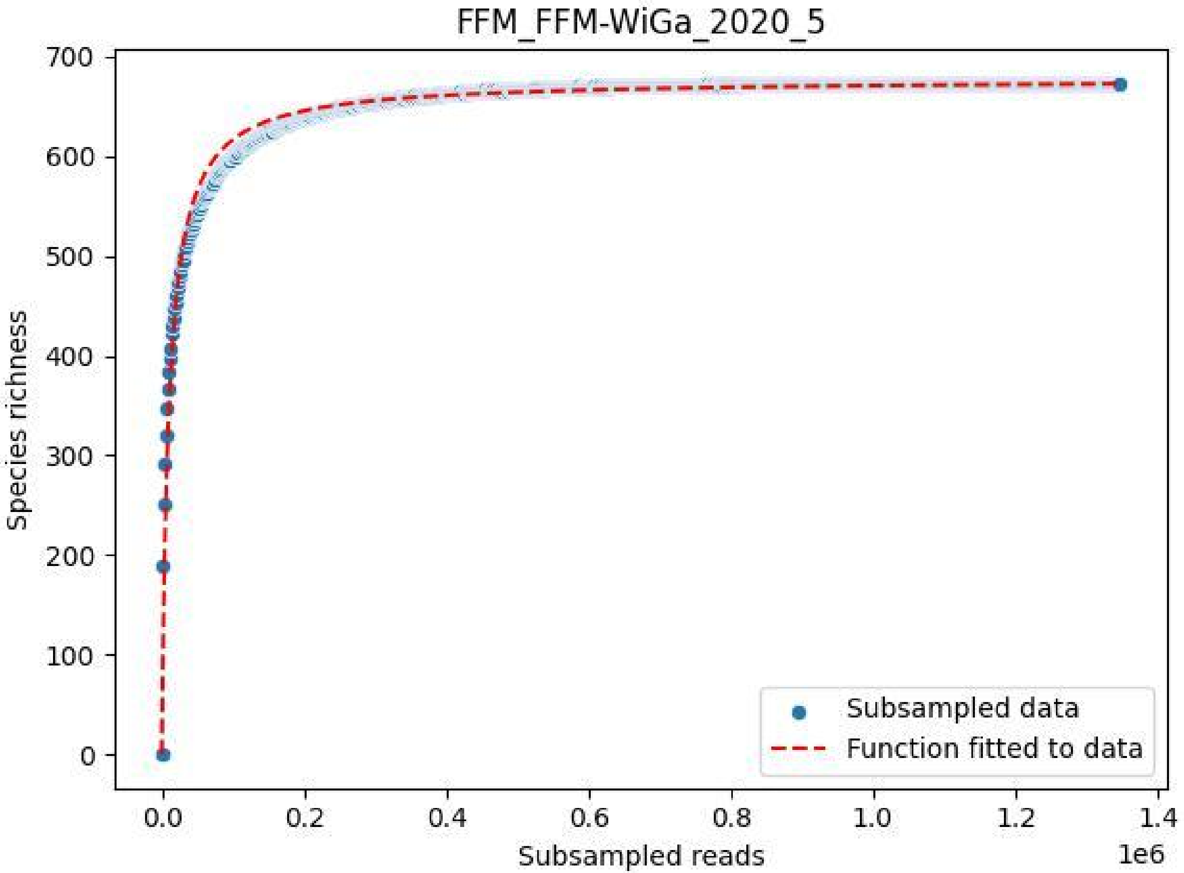

FFM_FFM-WiGa_2020_5
Species richness
Subsampled reads
1e6
Subsampled data
Function fitted to data
0
100
200
300
400
500
600
700
0.0
0.2
0.4
0.6
0.8
1.0
1.2
1.4

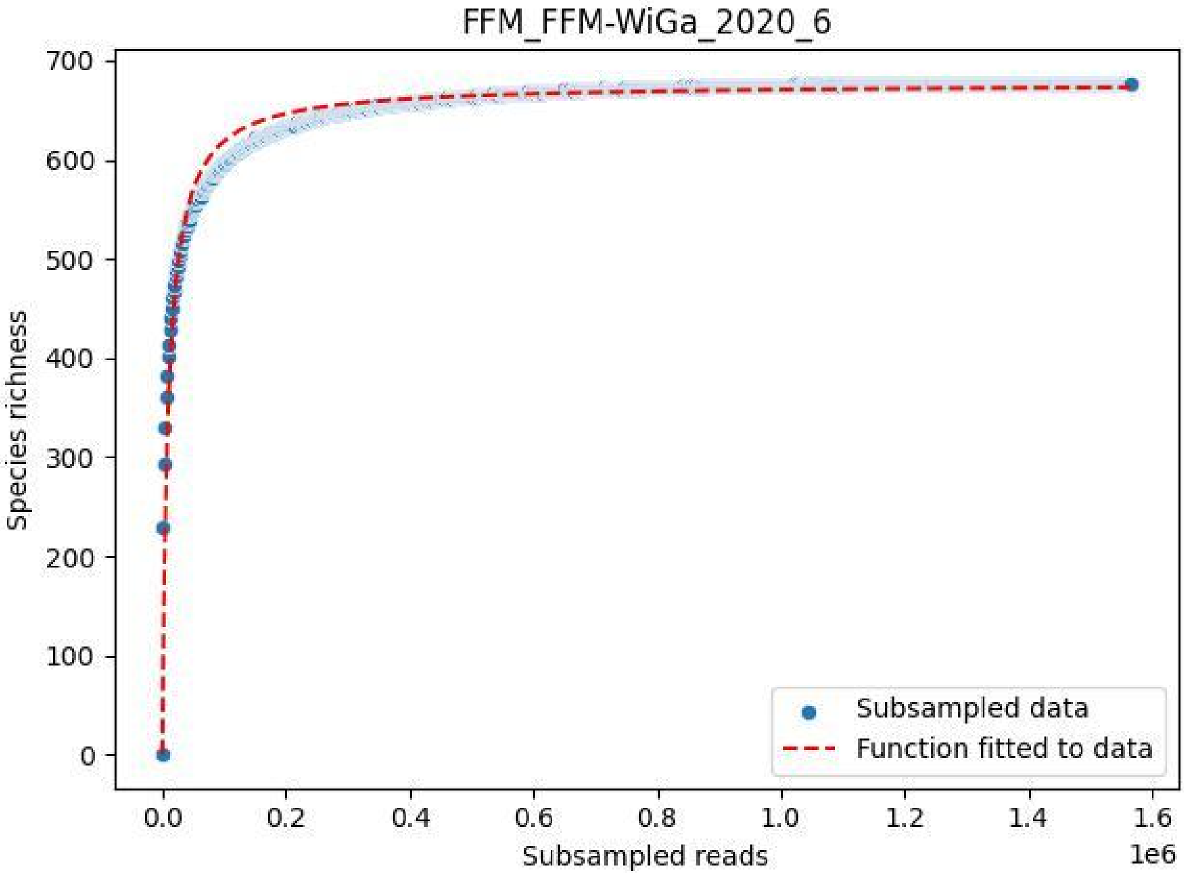
FFM_FFM-WiGa_2020_6
Species richness
Subsampled reads
1e6
0
100
200
300
400
500
600
700
0.0
0.2
0.4
0.6
0.8
1.0
1.2
1.4
1.6
Subsampled data
Function fitted to data

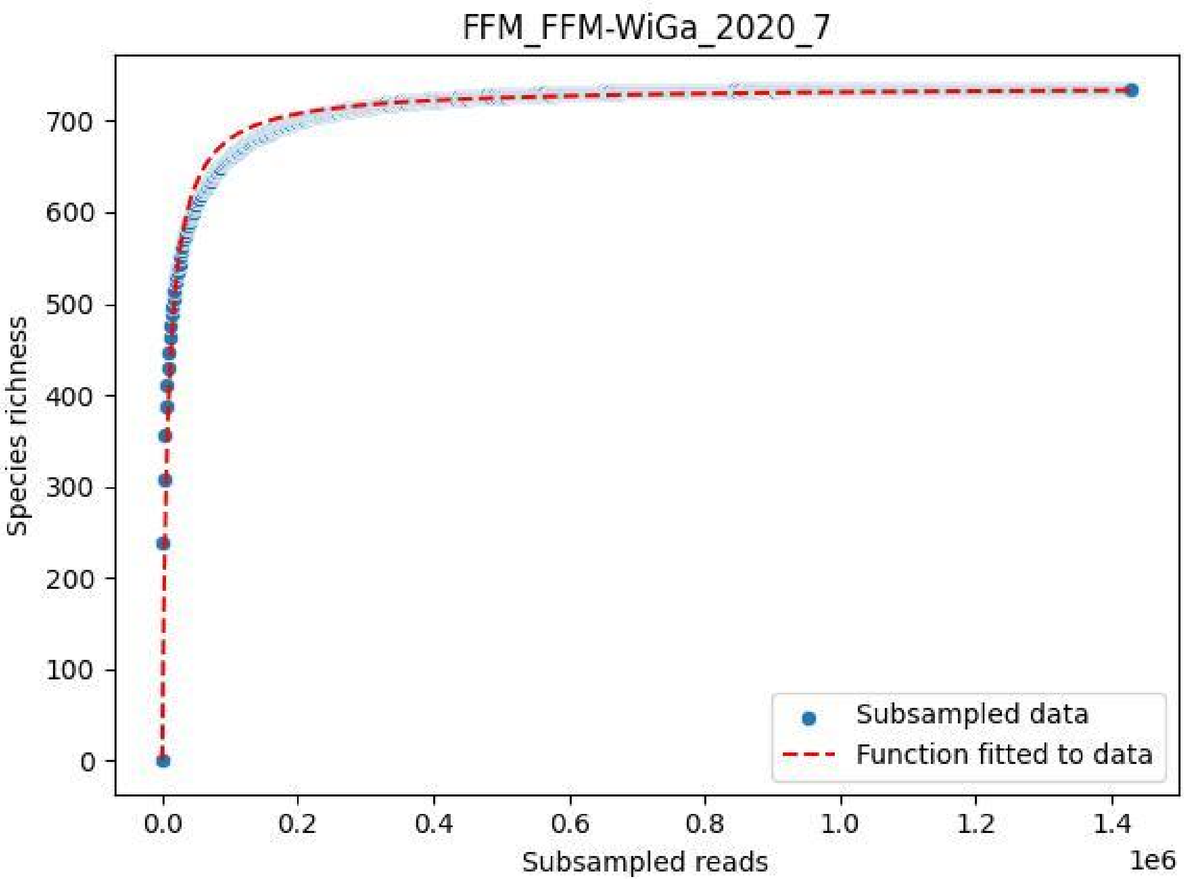
FFM_FFM-WiGa_2020_7
Species richness
Subsampled reads
1e6
0
100
200
300
400
500
600
700
0.0
0.2
0.4
0.6
0.8
1.0
1.2
1.4
Subsampled data
Function fitted to data

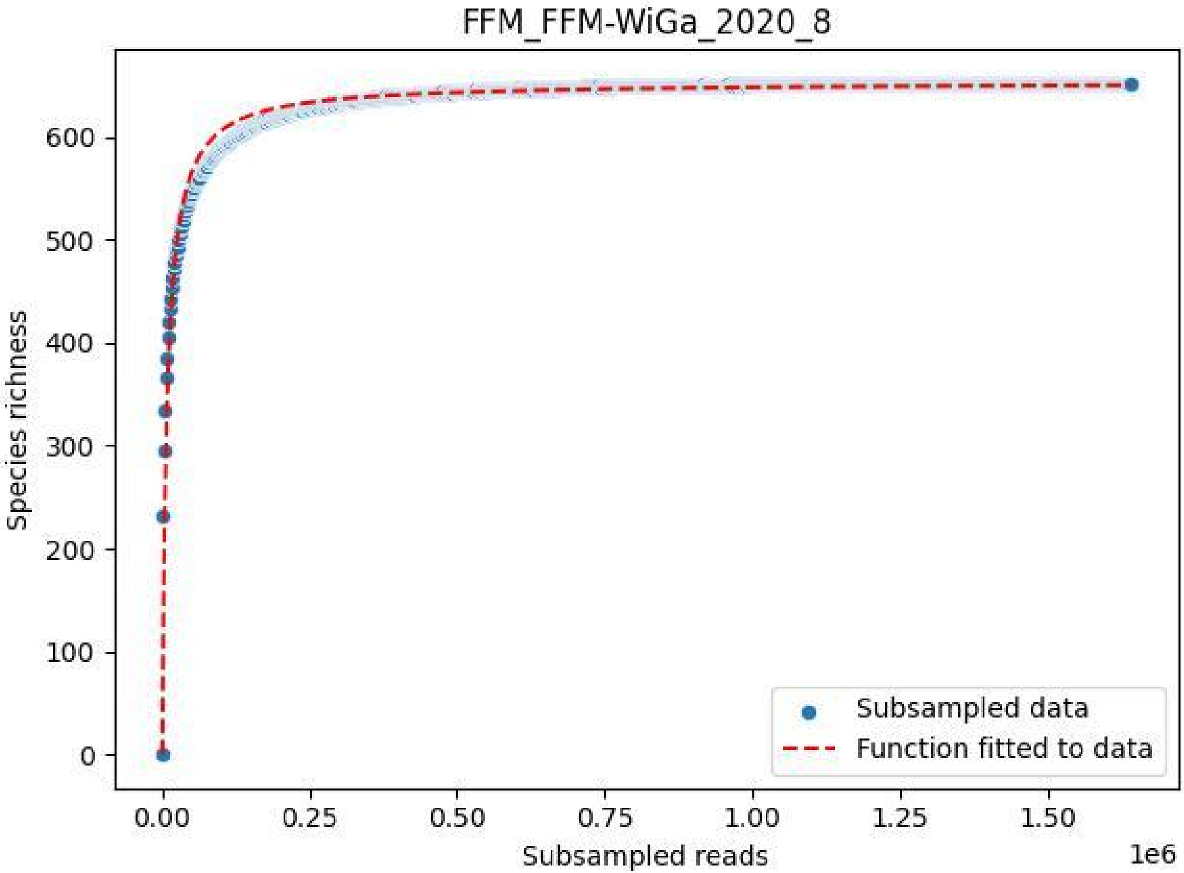

FFM_FFM-WiGa_2020_8
Species richness
Subsampled reads
1e6
0
100
200
300
400
500
600
0.00
0.25
0.50
0.75
1.00
1.25
1.50
Subsampled data
Function fitted to data

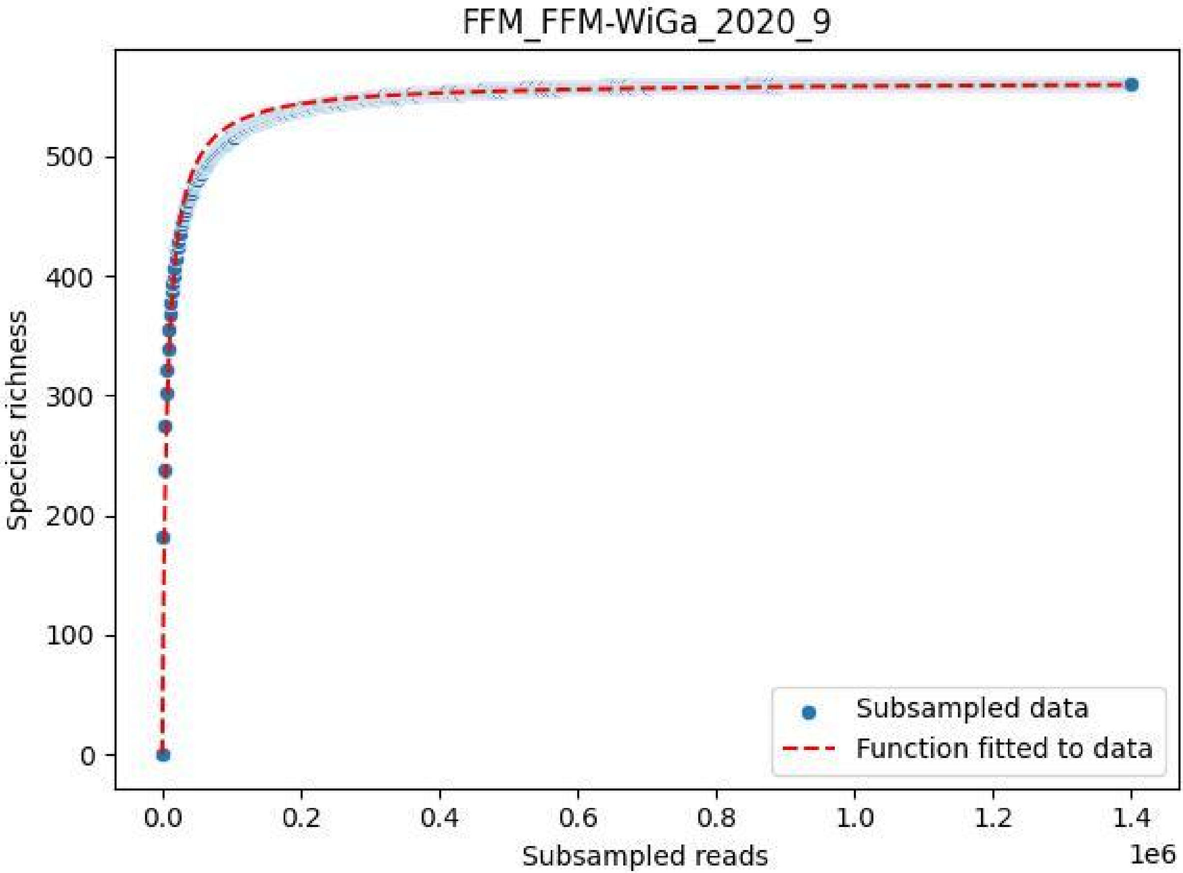
FFM_FFM-WiGa_2020_9
Species richness
Subsampled reads
1e6
0
100
200
300
400
500
0.0
0.2
0.4
0.6
0.8
1.0
1.2
1.4
Subsampled data
Function fitted to data

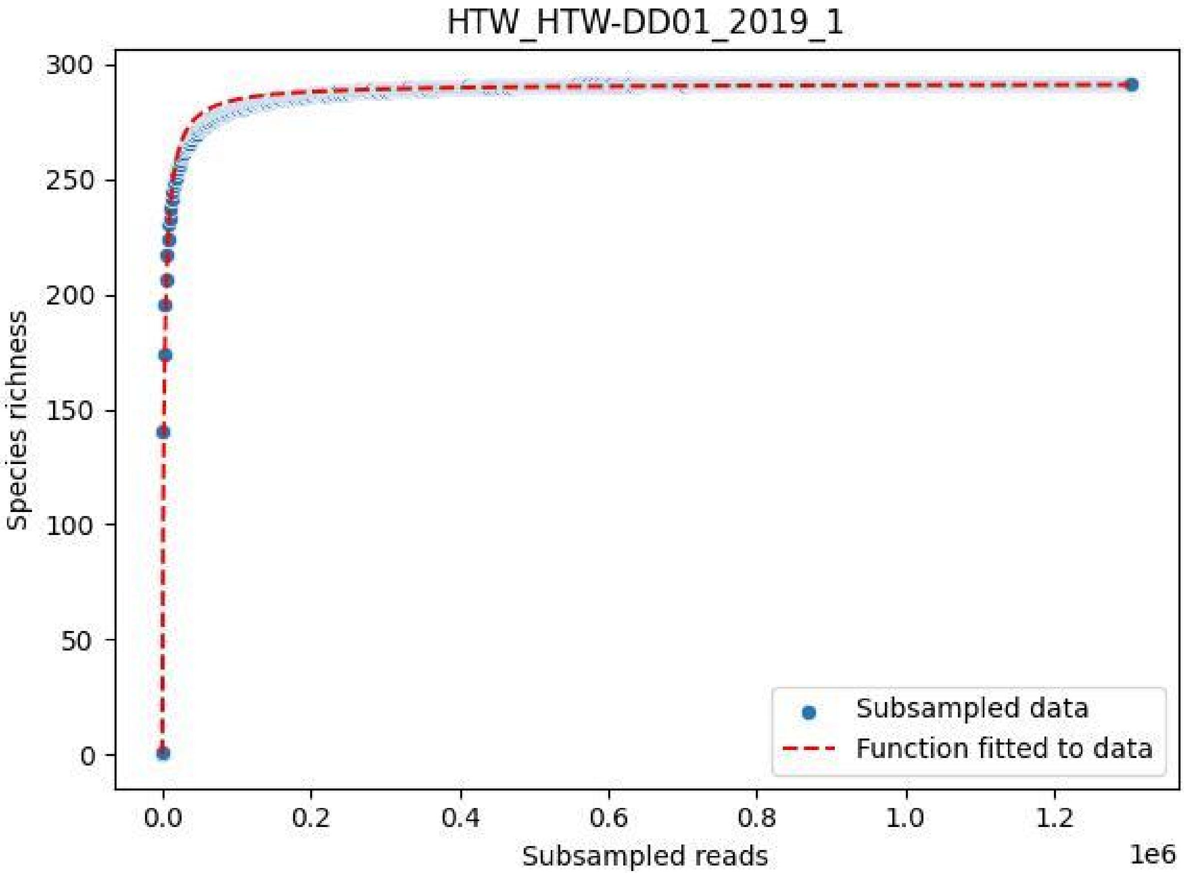
HTW_HTW-DD01_2019_1
Species richness
300
250
200
150
100
50
0
0.0
0.2
0.4
0.6
0.8
1.0
1.2
Subsampled reads
1e6
Subsampled data
Function fitted to data

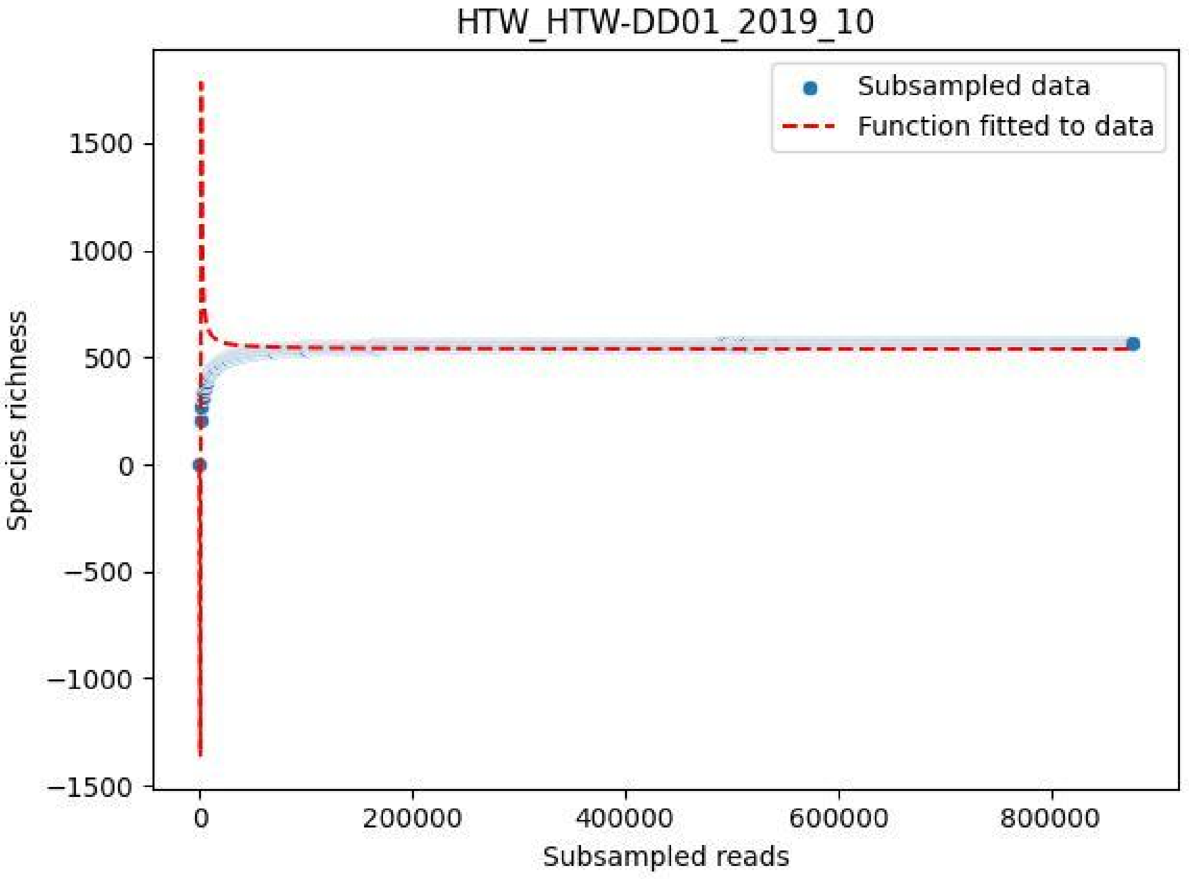
HTW_HTW-DD01_2019_10
Subsampled data
Function fitted to data
1500
1000
500
0
−500
−1000
−1500
Species richness
0
200000
400000
600000
800000
Subsampled reads

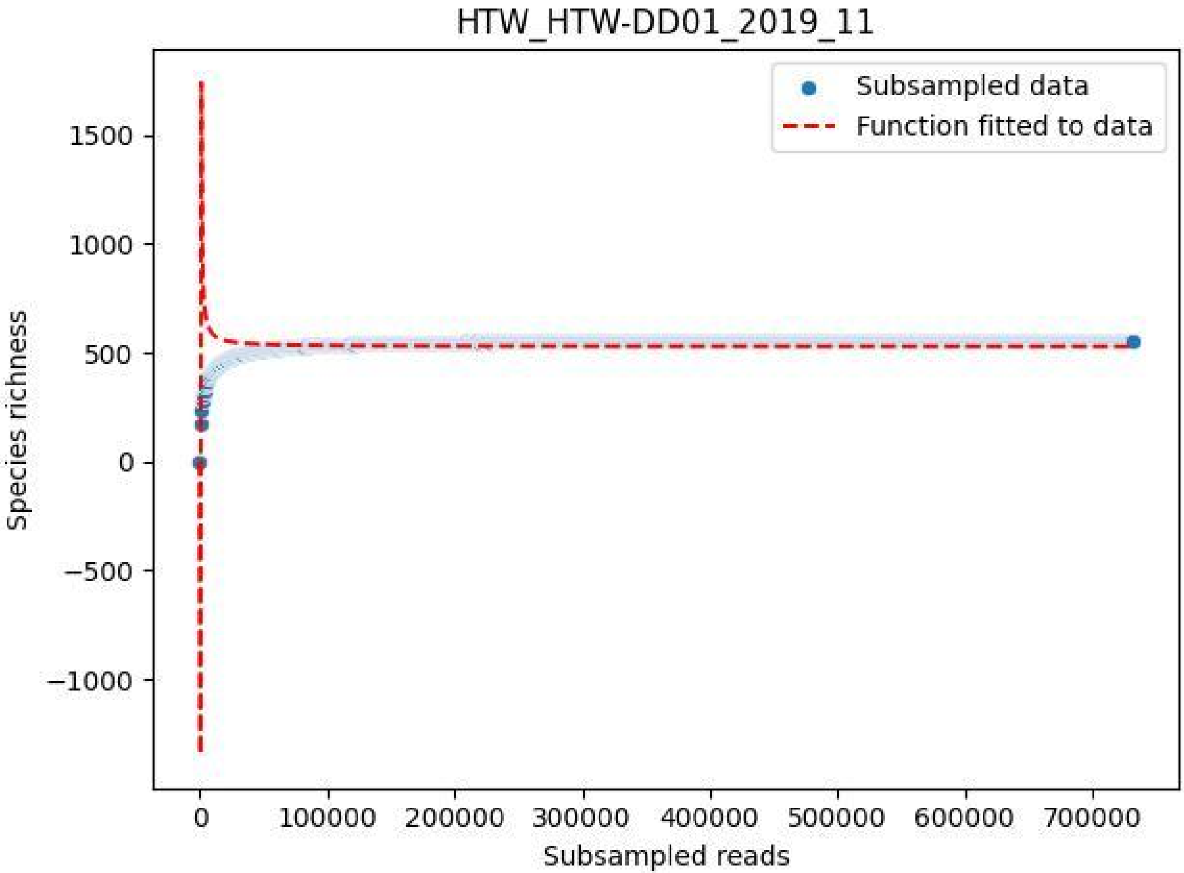
HTW_HTW-DD01_2019_11
Subsampled data
Function fitted to data
Species richness
Subsampled reads
1500
1000
500
0
−500
−1000
0
100000
200000
300000
400000
500000
600000
700000

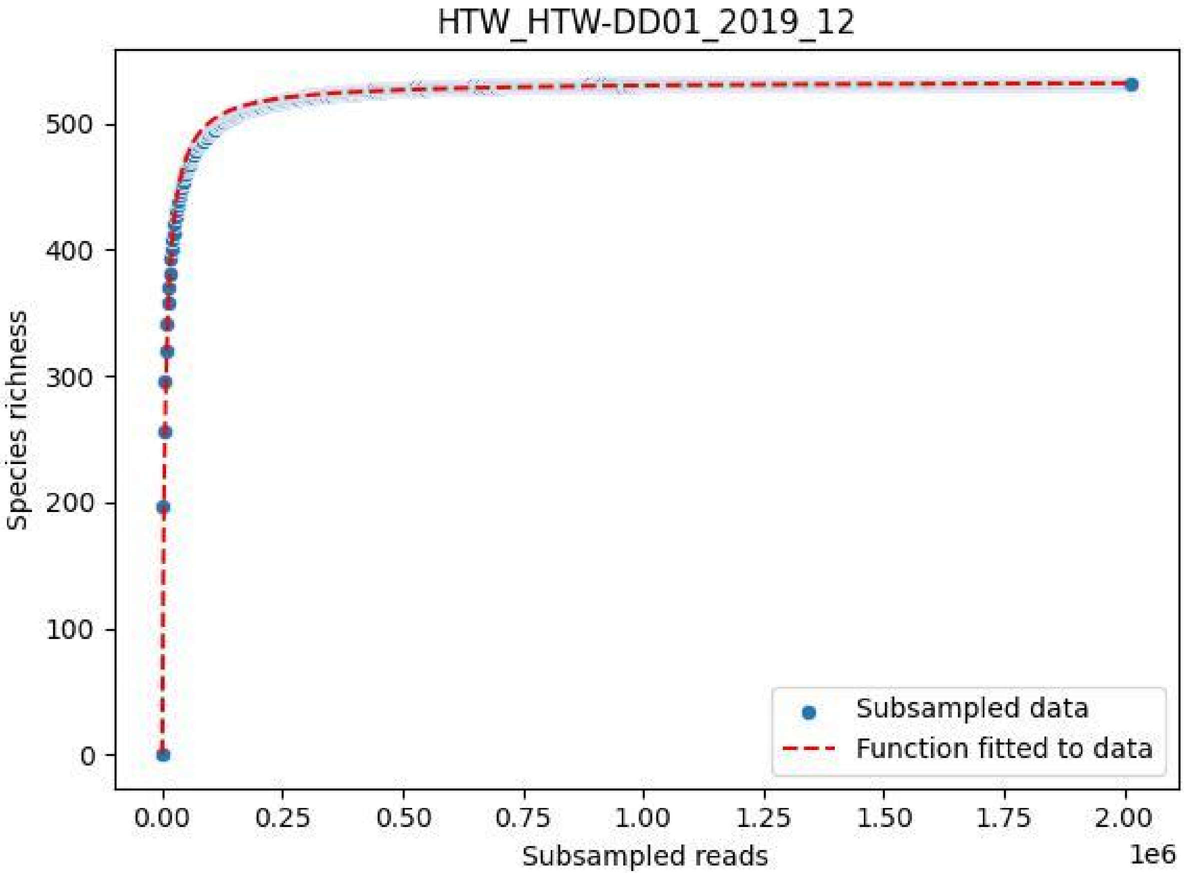
HTW_HTW-DD01_2019_12
Species richness
Subsampled reads
1e6
0
100
200
300
400
500
0.00
0.25
0.50
0.75
1.00
1.25
1.50
1.75
2.00
Subsampled data
Function fitted to data

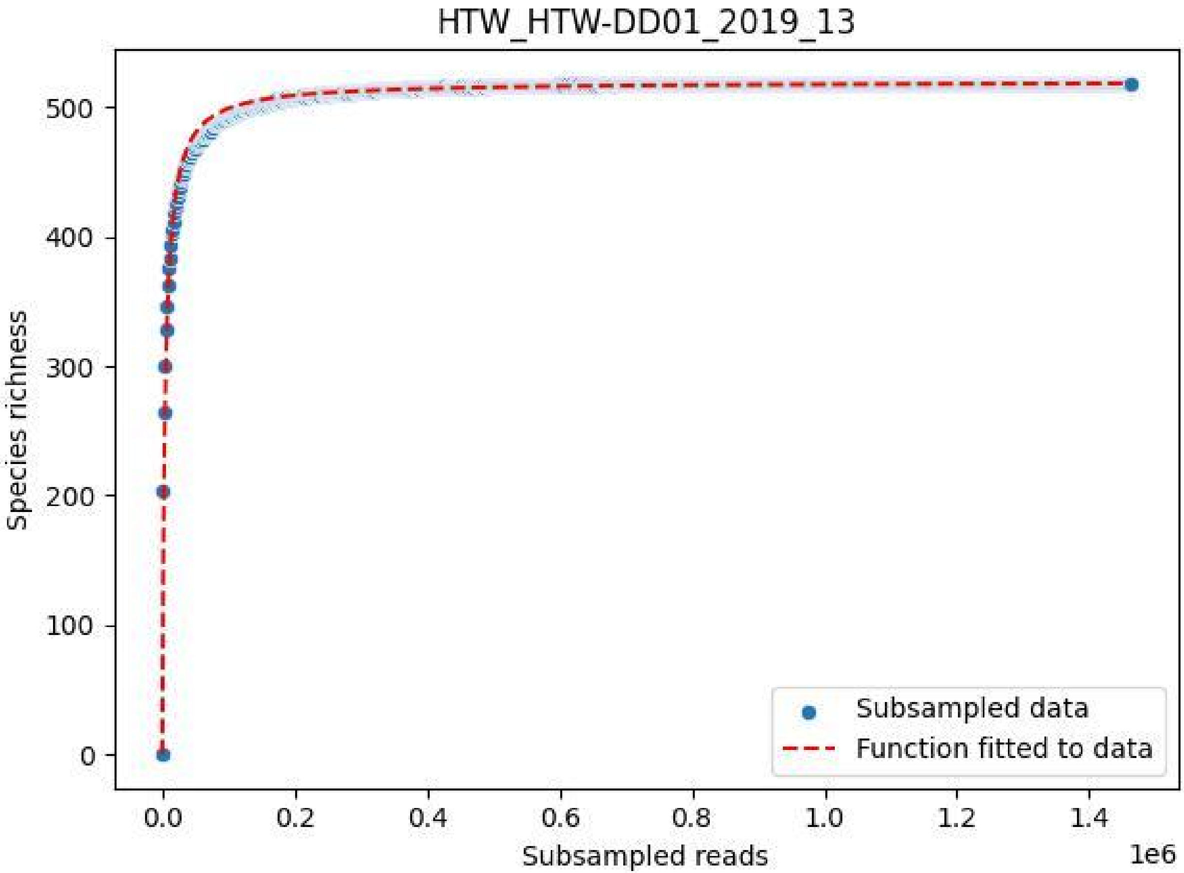

HTW_HTW-DD01_2019_13
Species richness
500
400
300
200
100
0
0.0
0.2
0.4
0.6
0.8
1.0
1.2
1.4
Subsampled reads
1e6
Subsampled data
Function fitted to data

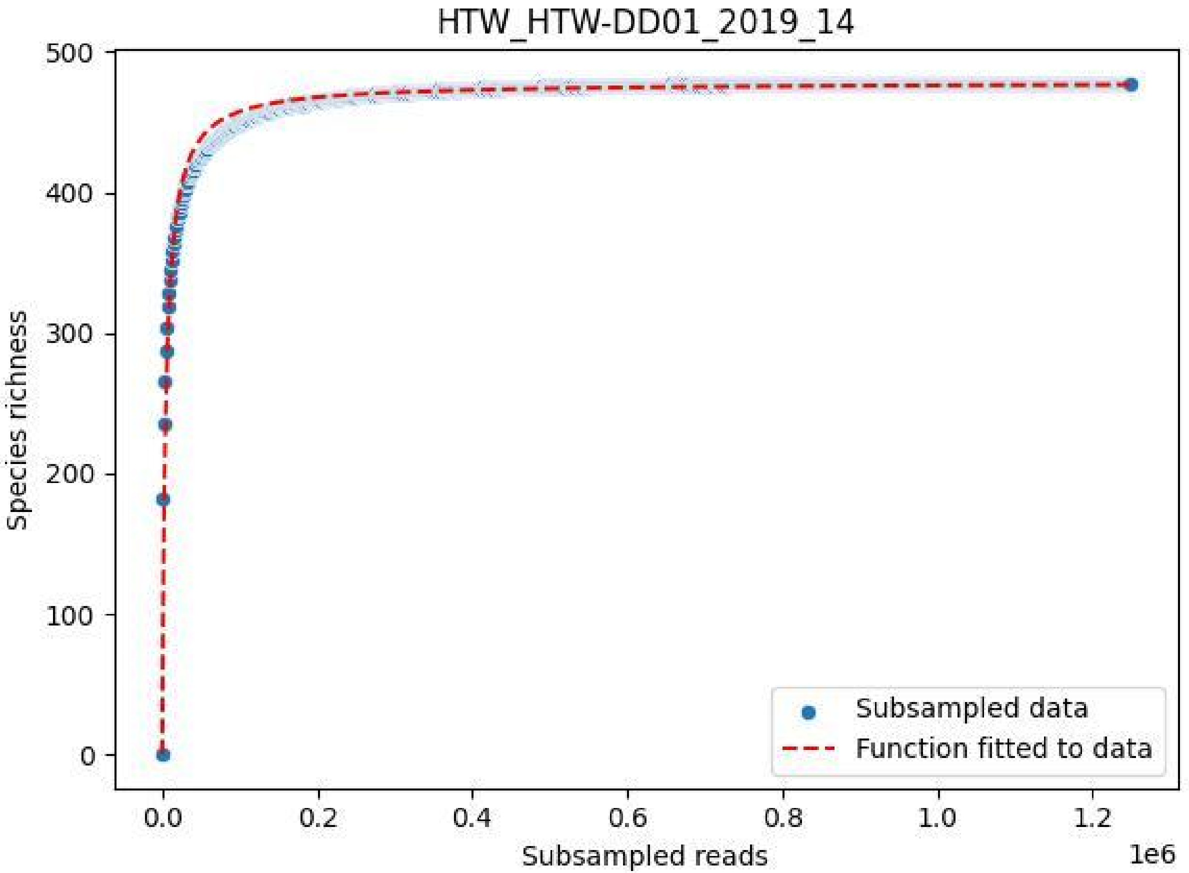

HTW_HTW-DD01_2019_14
Species richness
Subsampled reads
1e6
500
400
300
200
100
0
0.0
0.2
0.4
0.6
0.8
1.0
1.2
Subsampled data
Function fitted to data

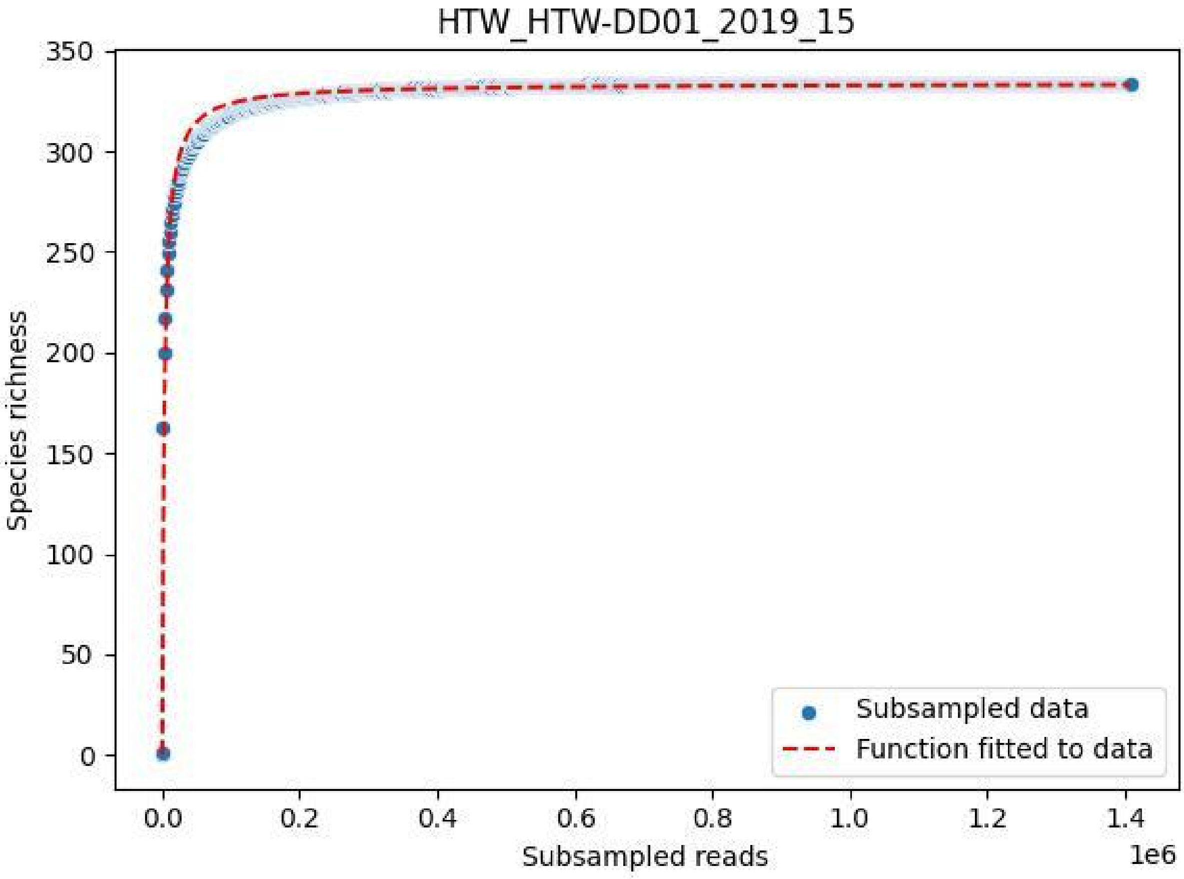
HTW_HTW-DD01_2019_15
Species richness
Subsampled reads
1e6
0.0
0.2
0.4
0.6
0.8
1.0
1.2
1.4
0
50
100
150
200
250
300
350
Subsampled data
Function fitted to data

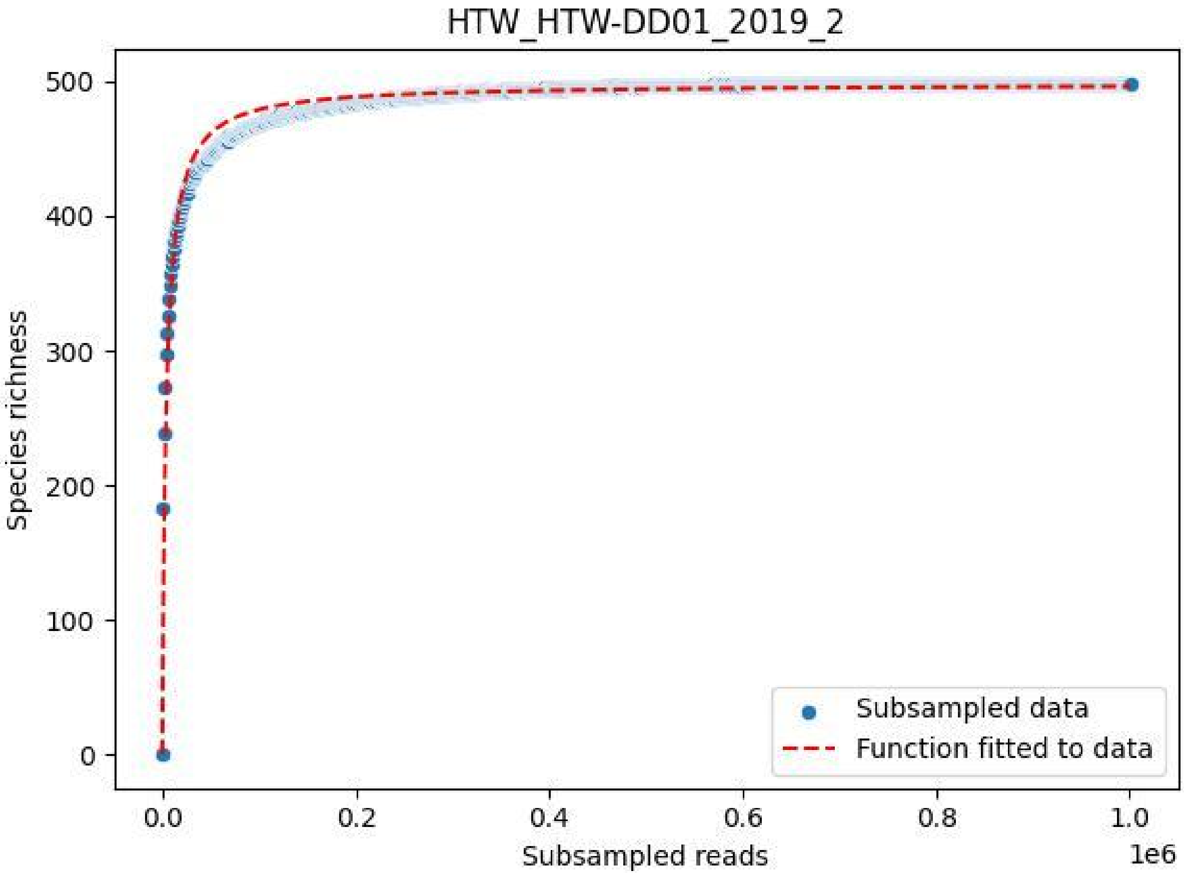
HTW_HTW-DD01_2019_2
Species richness
Subsampled reads
1e6
500
400
300
200
100
0
0.0
0.2
0.4
0.6
0.8
1.0
Subsampled data
Function fitted to data

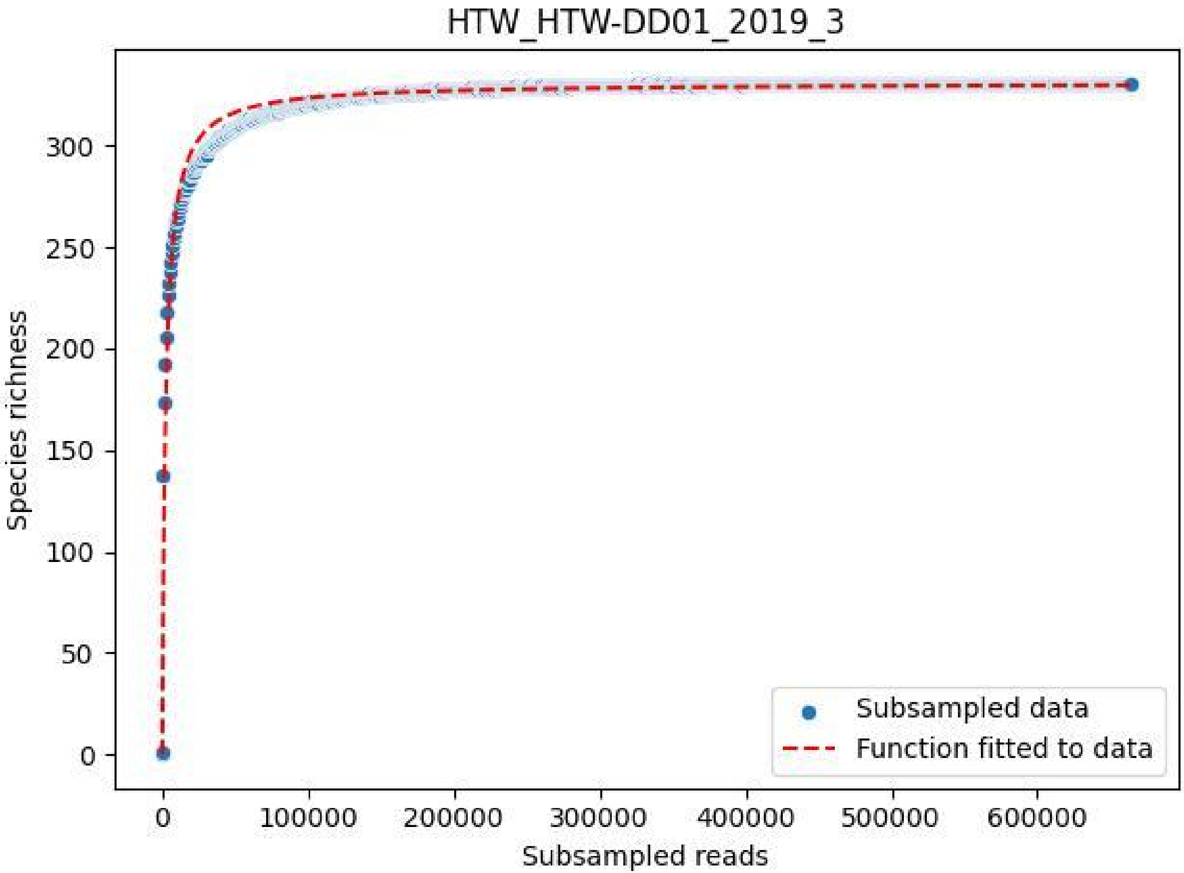
HTW_HTW-DD01_2019_3
Species richness
Subsampled reads
0
50
100
150
200
250
300
0
100000
200000
300000
400000
500000
600000
Subsampled data
Function fitted to data

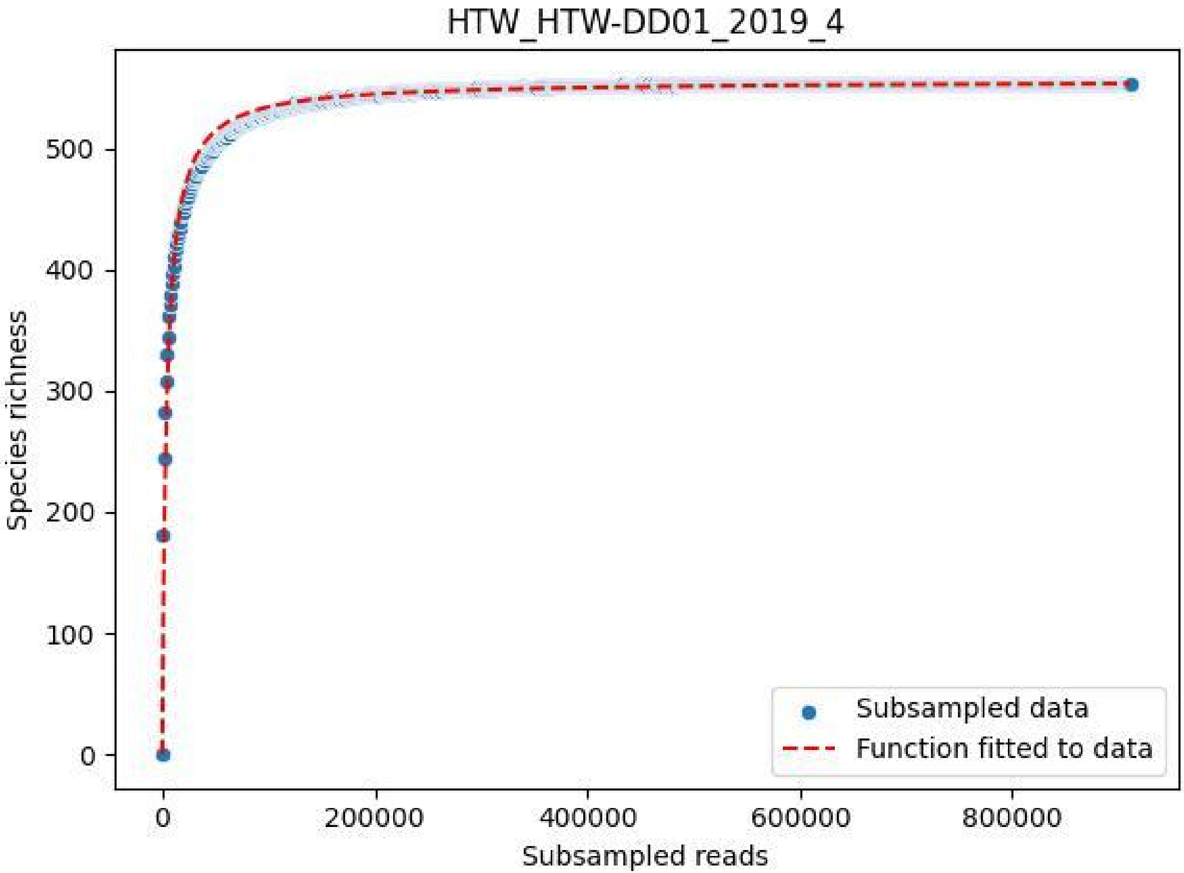
HTW_HTW-DD01_2019_4
Species richness
Subsampled reads
0
100
200
300
400
500
0
200000
400000
600000
800000
Subsampled data
Function fitted to data

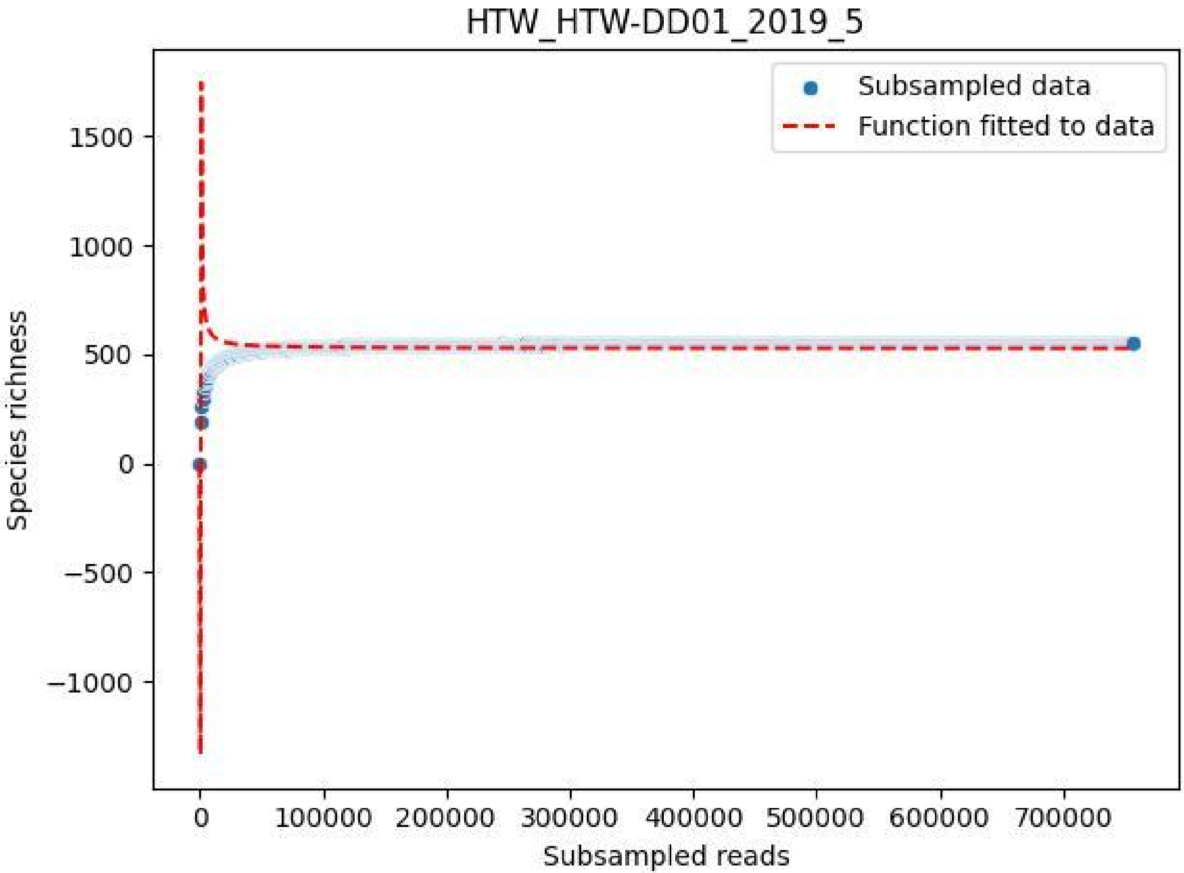
HTW_HTW-DD01_2019_5
Subsampled data
Function fitted to data
1500
1000
500
0
−500
−1000
Species richness
0
100000
200000
300000
400000
500000
600000
700000
Subsampled reads

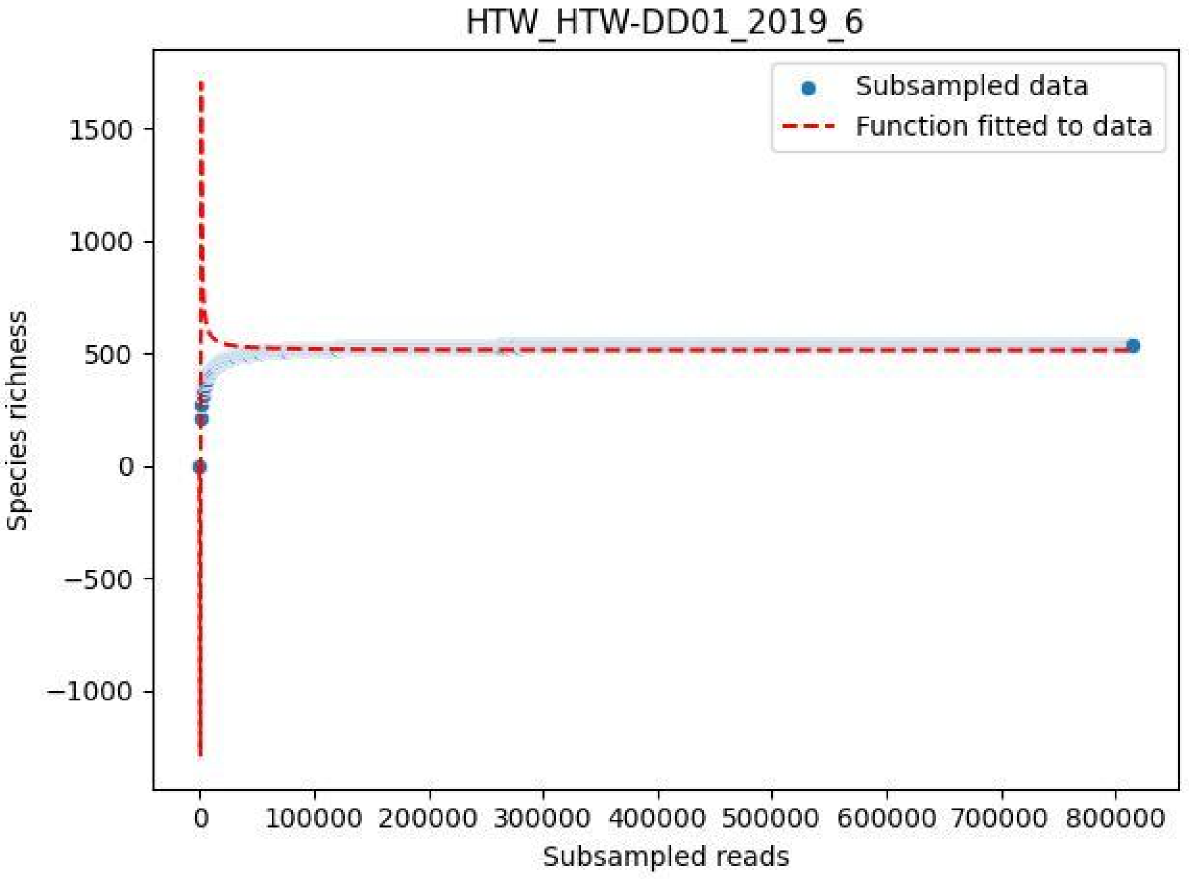
HTW_HTW-DD01_2019_6
Subsampled data
Function fitted to data
Species richness
1500
1000
500
0
−500
−1000
0
100000
200000
300000
400000
500000
600000
700000
800000
Subsampled reads

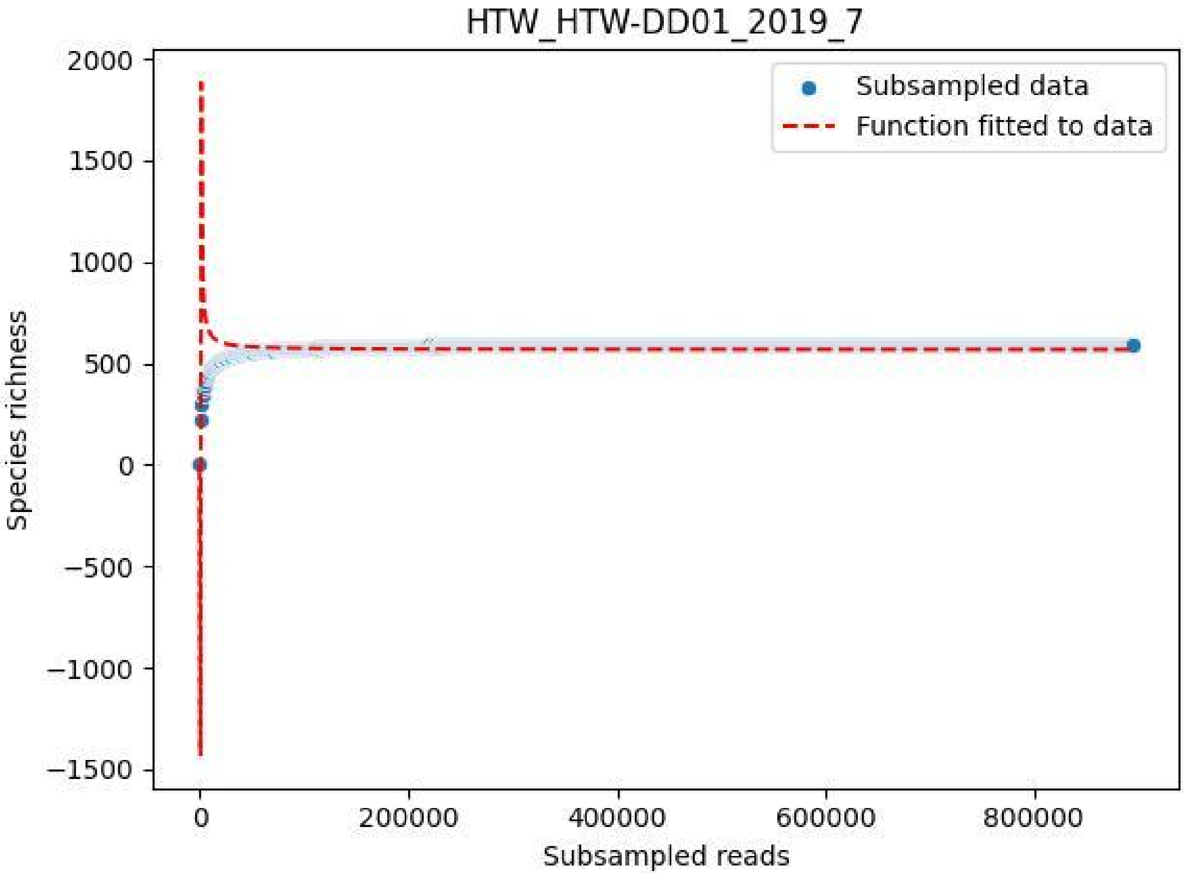
HTW_HTW-DD01_2019_7
Subsampled data
Function fitted to data
Species richness
Subsampled reads
2000
1500
1000
500
0
−500
−1000
−1500
0
200000
400000
600000
800000

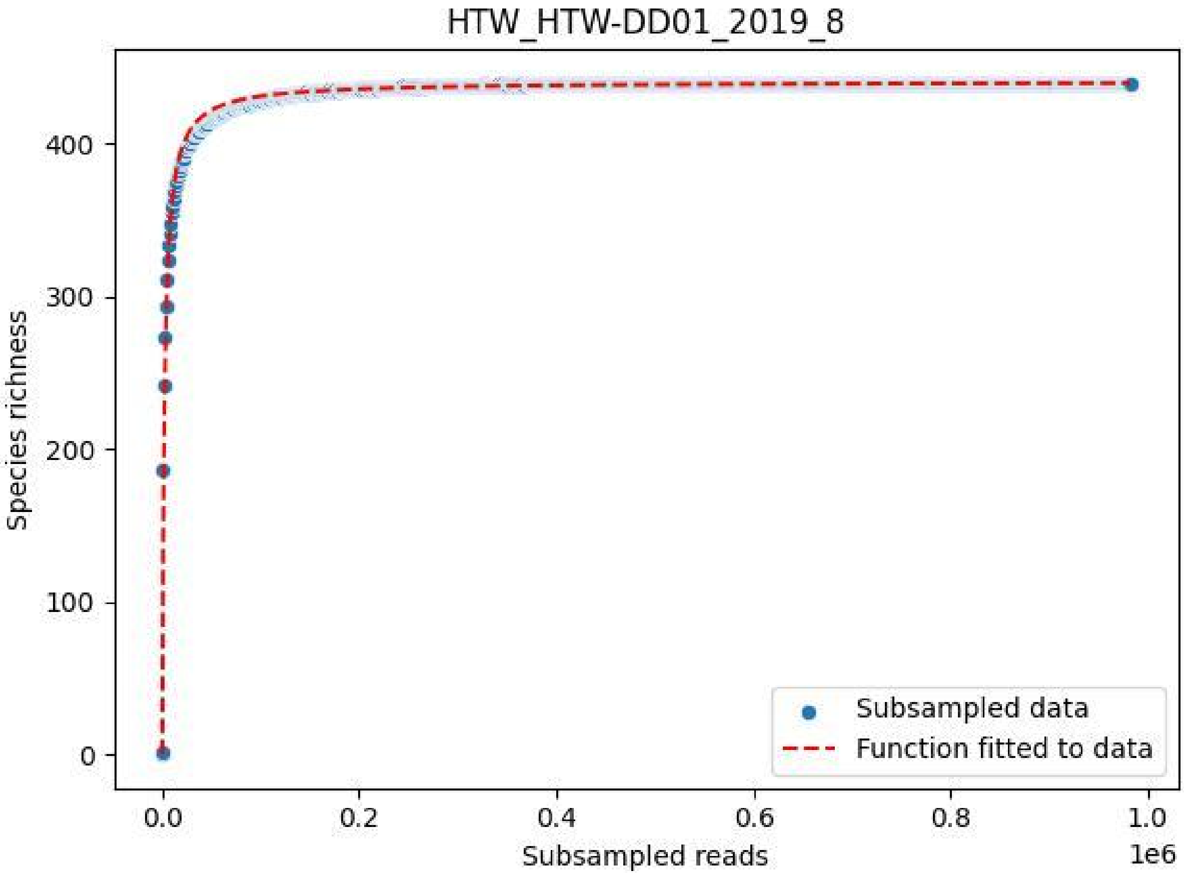
HTW_HTW-DD01_2019_8
Species richness
400
300
200
100
0
0.0
0.2
0.4
0.6
0.8
1.0
1e6
Subsampled reads
Subsampled data
Function fitted to data

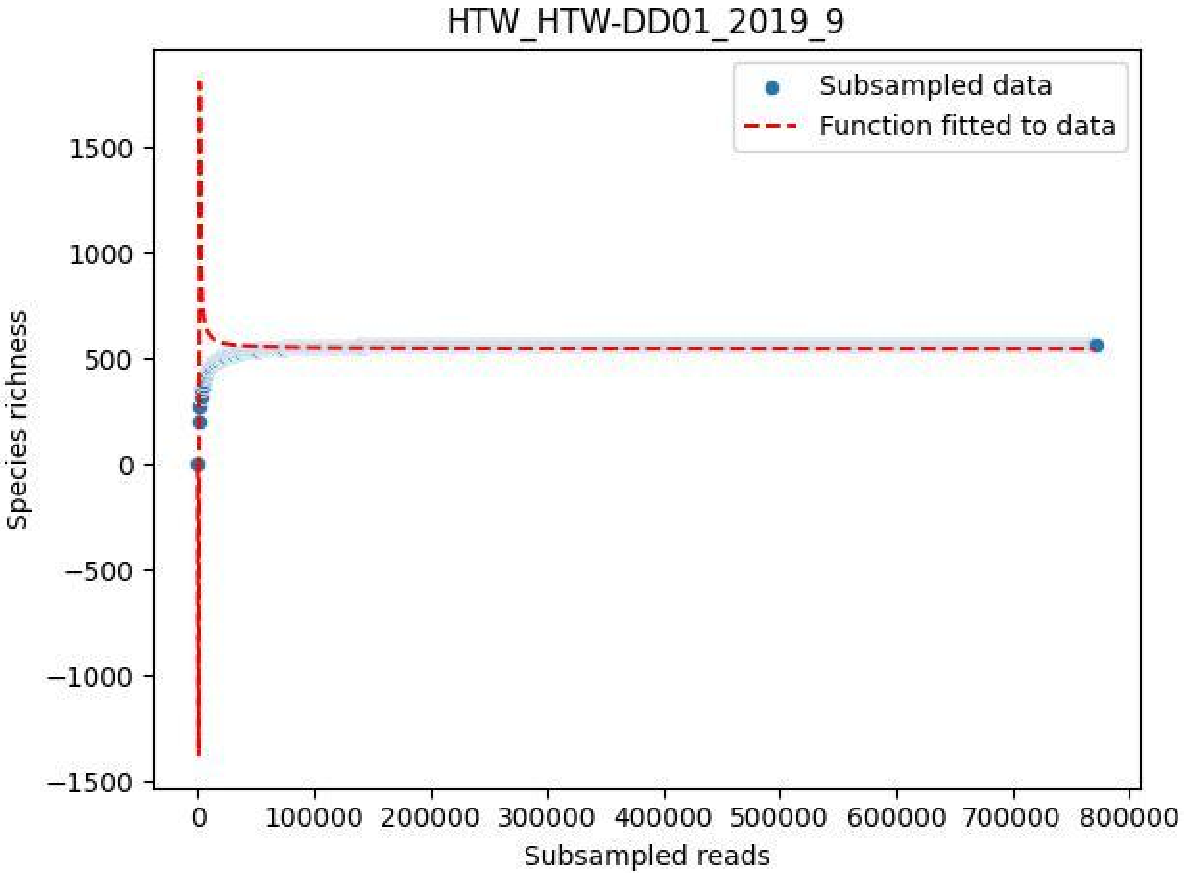
HTW_HTW-DD01_2019_9
Subsampled data
Function fitted to data
Species richness
Subsampled reads
1500
1000
500
0
−500
−1000
−1500
0
100000
200000
300000
400000
500000
600000
700000
800000

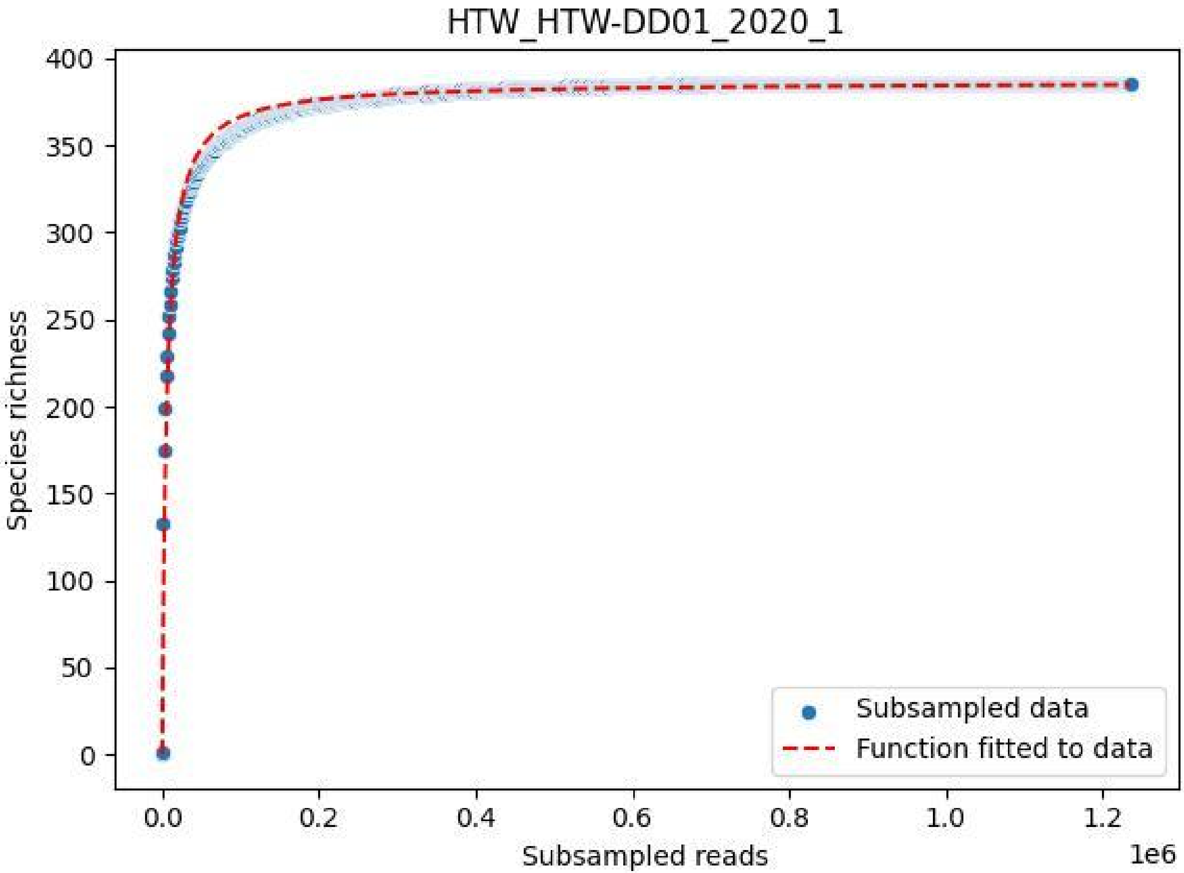
HTW_HTW-DD01_2020_1
Species richness
Subsampled reads
1e6
0
50
100
150
200
250
300
350
400
0.0
0.2
0.4
0.6
0.8
1.0
1.2
Subsampled data
Function fitted to data

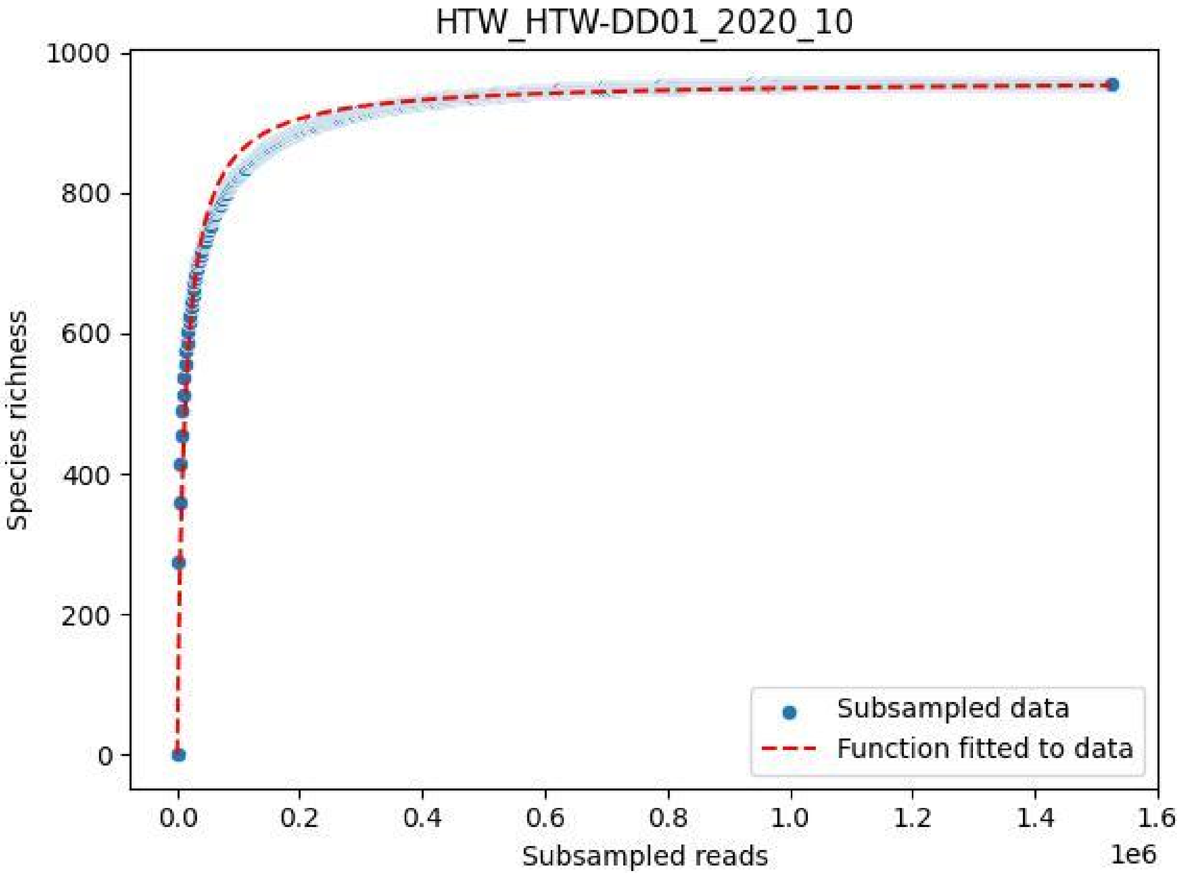
HTW_HTW-DD01_2020_10
Species richness
Subsampled reads
1e6
0
200
400
600
800
1000
0.0
0.2
0.4
0.6
0.8
1.0
1.2
1.4
1.6
Subsampled data
Function fitted to data

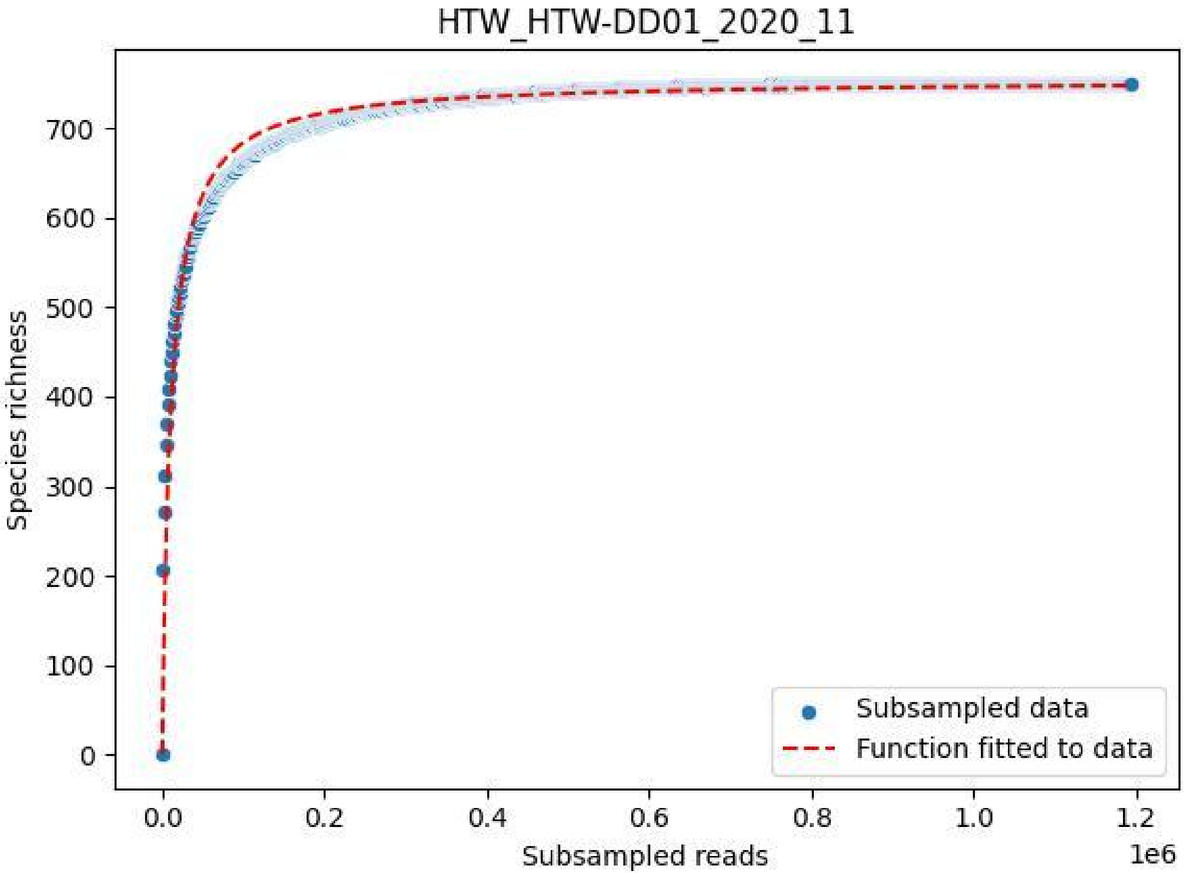
HTW_HTW-DD01_2020_11
Species richness
Subsampled reads
1e6
0
100
200
300
400
500
600
700
0.0
0.2
0.4
0.6
0.8
1.0
1.2
Subsampled data
Function fitted to data

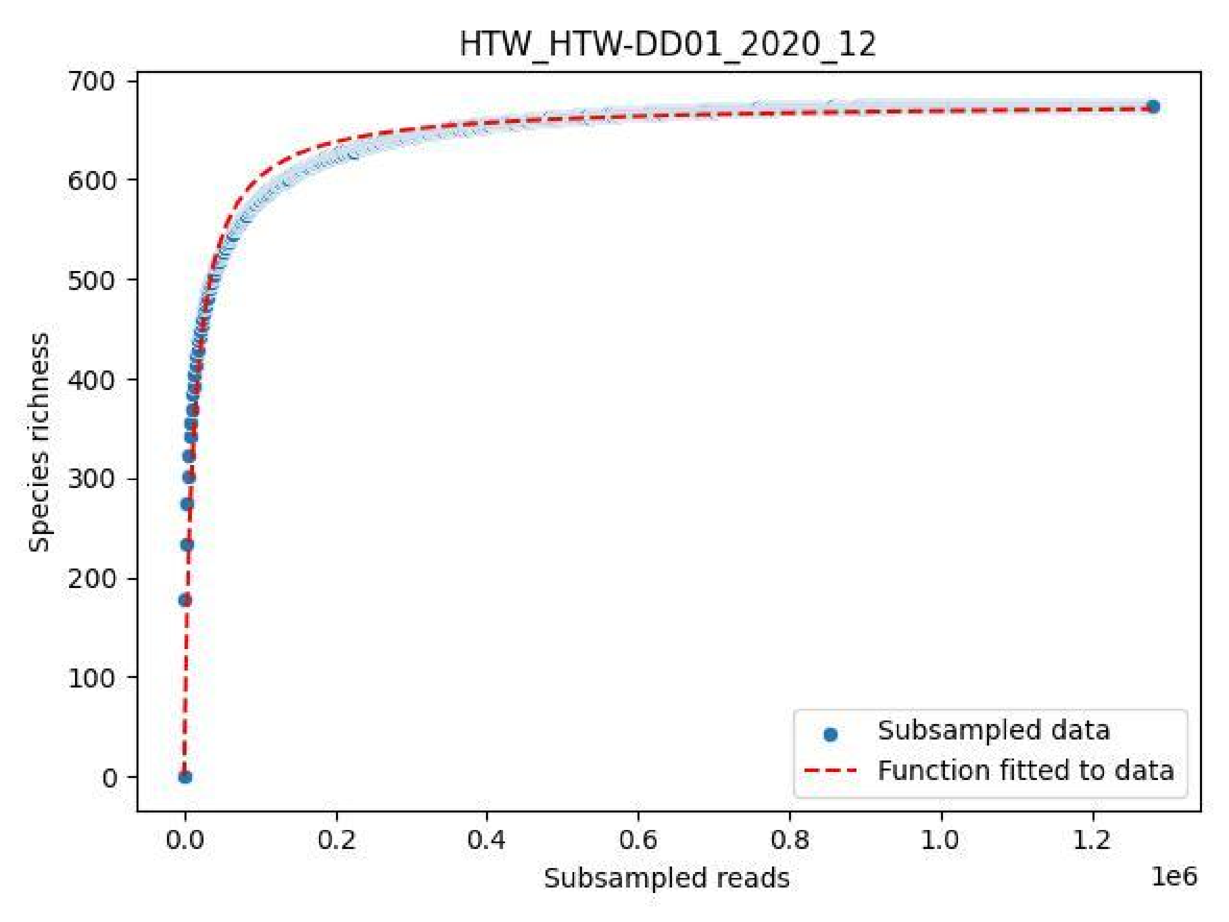
HTW_HTW-DD01_2020_12
Species richness
Subsampled reads
1e6
Subsampled data
Function fitted to data
0
100
200
300
400
500
600
700
0.0
0.2
0.4
0.6
0.8
1.0
1.2

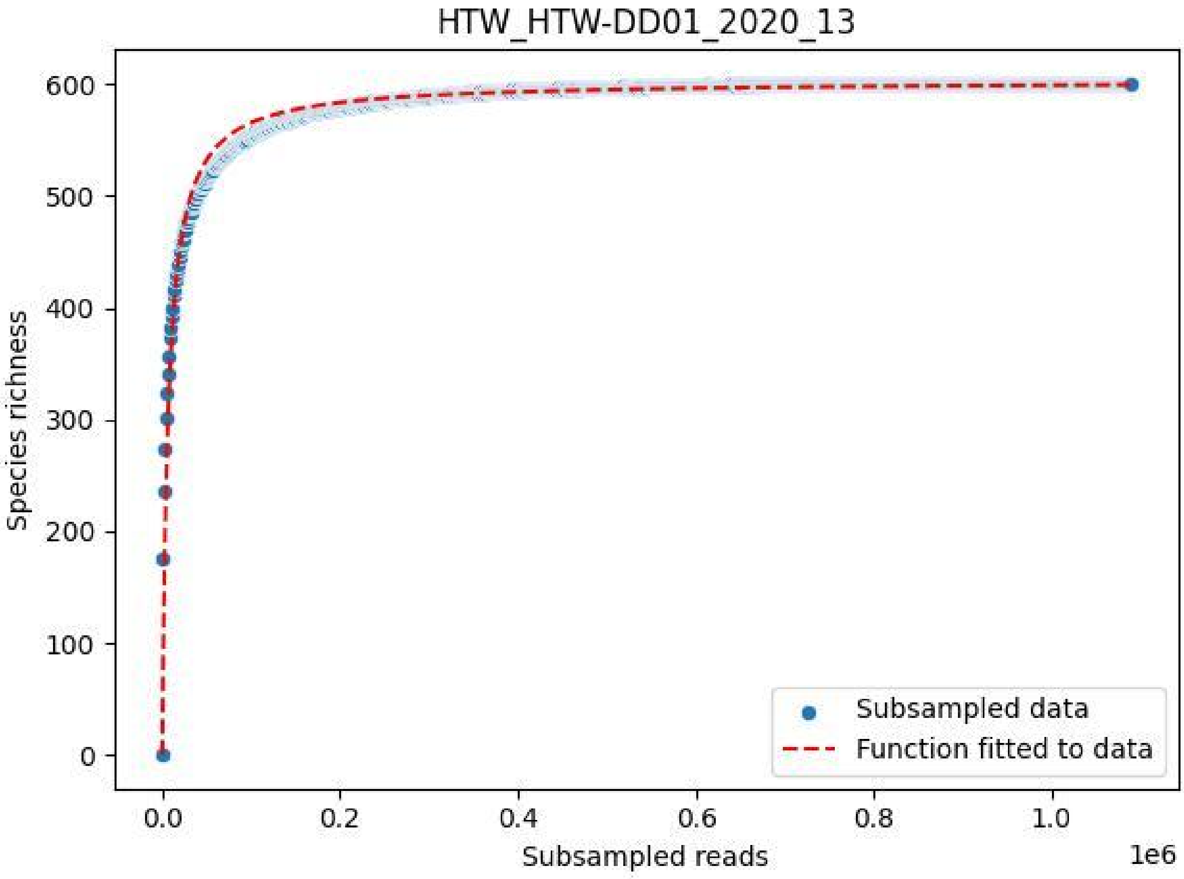
HTW_HTW-DD01_2020_13
Species richness
Subsampled reads
1e6
0
100
200
300
400
500
600
0.0
0.2
0.4
0.6
0.8
1.0
Subsampled data
Function fitted to data

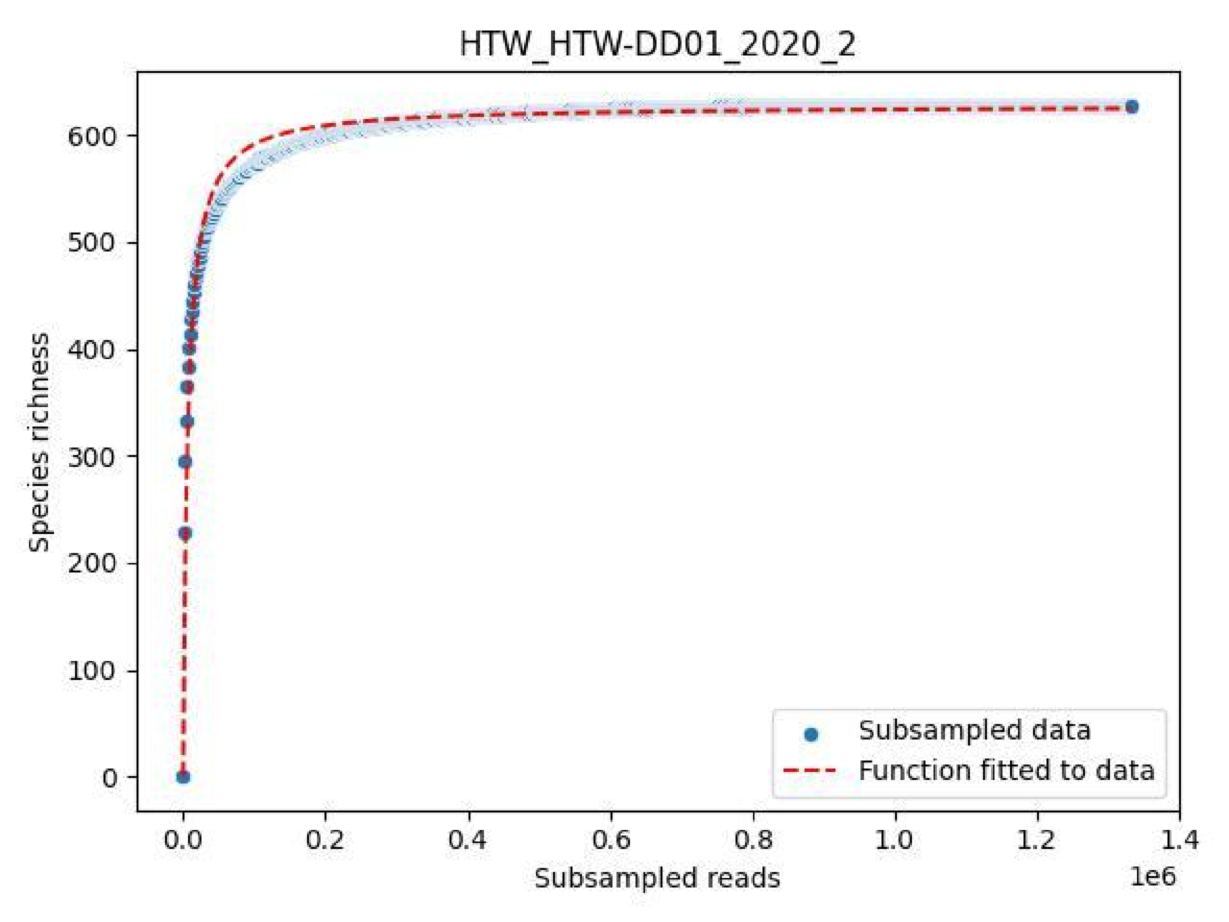
HTW_HTW-DD01_2020_2
Species richness
Subsampled reads
1e6
0
100
200
300
400
500
600
0.0
0.2
0.4
0.6
0.8
1.0
1.2
1.4
Subsampled data
Function fitted to data

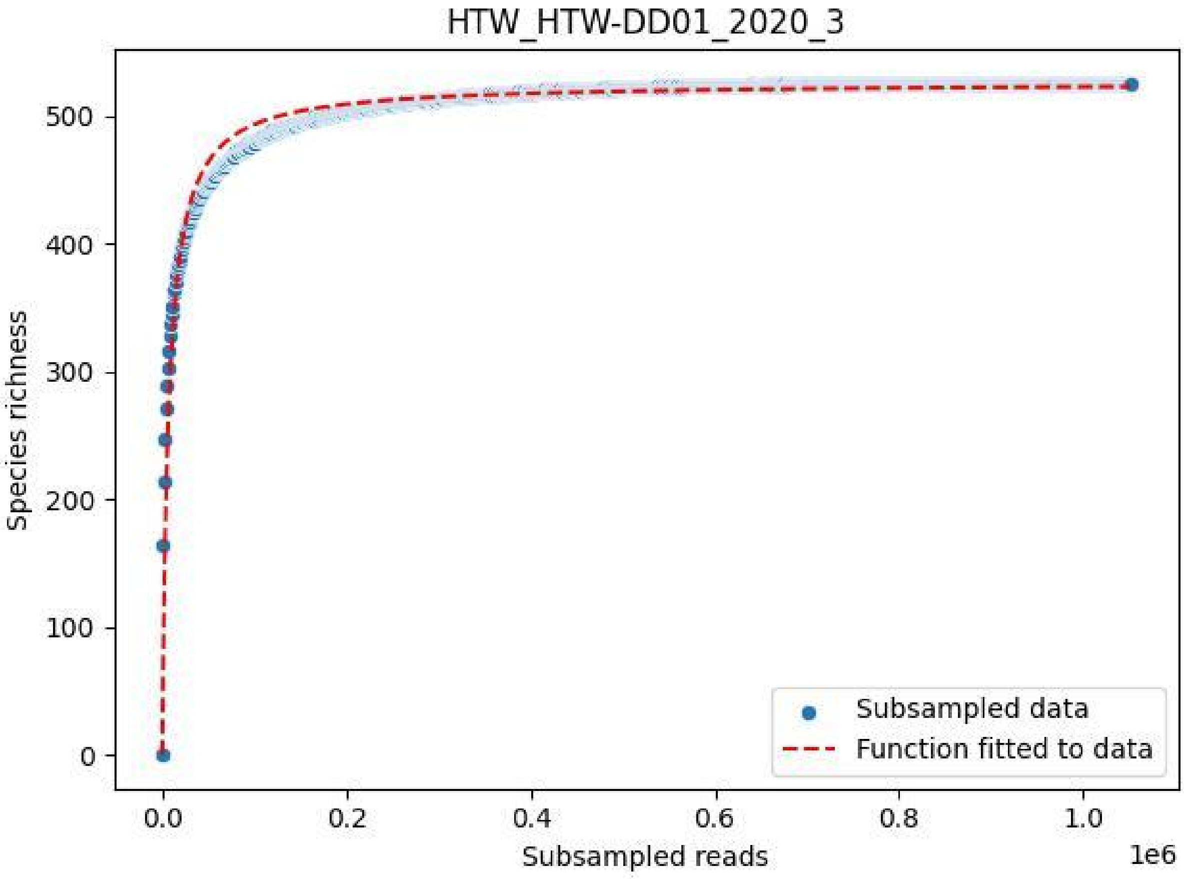

HTW_HTW-DD01_2020_3
Species richness
Subsampled reads
1e6
Subsampled data
Function fitted to data
0
100
200
300
400
500
0.0
0.2
0.4
0.6
0.8
1.0

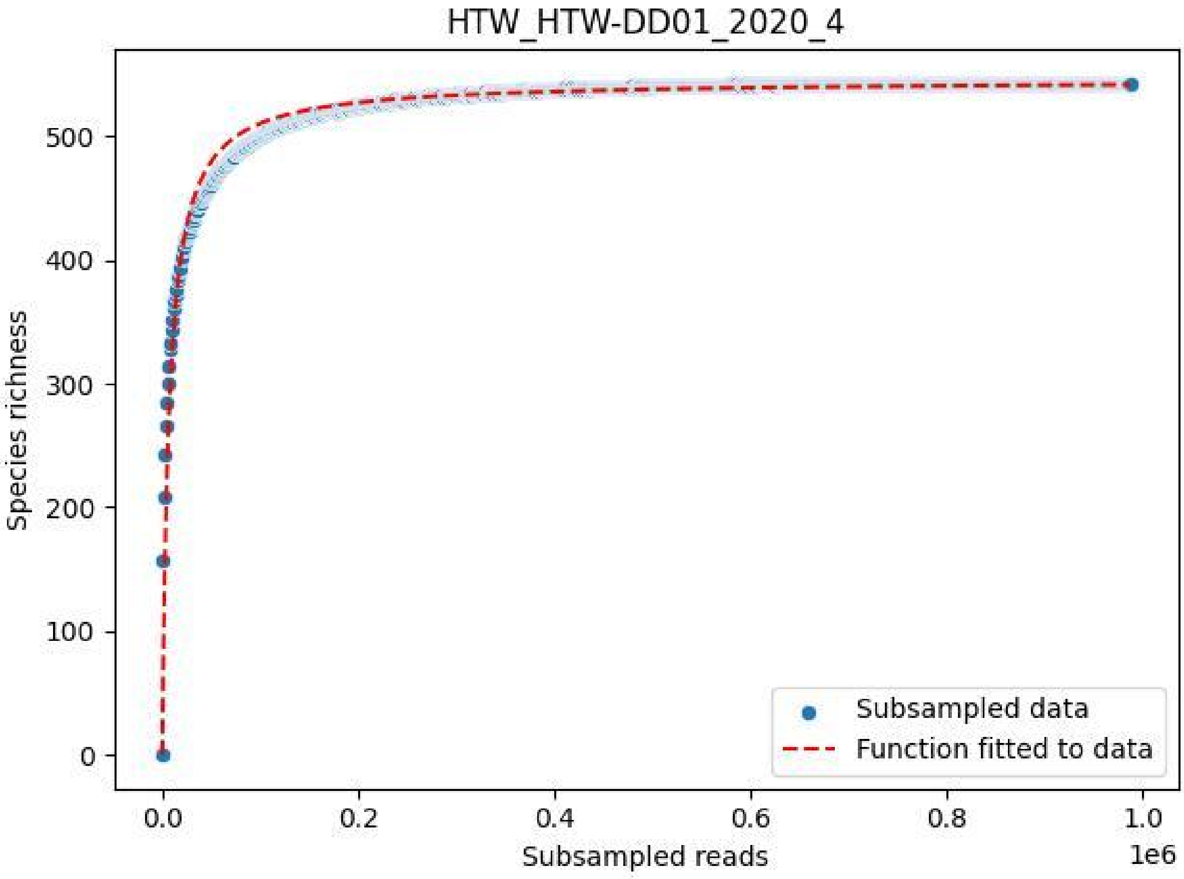
HTW_HTW-DD01_2020_4
Species richness
Subsampled reads
1e6
0
100
200
300
400
500
0.0
0.2
0.4
0.6
0.8
1.0
Subsampled data
Function fitted to data

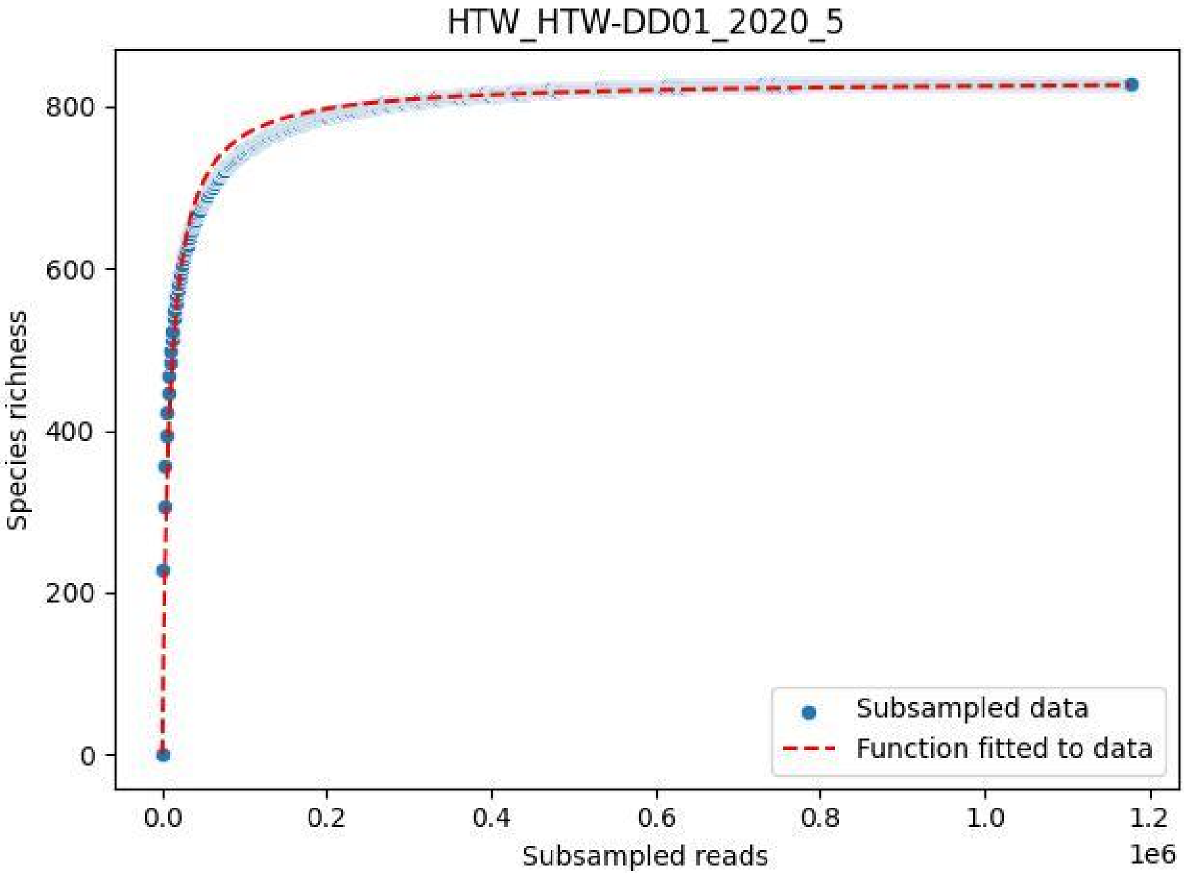
HTW_HTW-DD01_2020_5
Species richness
Subsampled reads
1e6
0
200
400
600
800
0.0
0.2
0.4
0.6
0.8
1.0
1.2
Subsampled data
Function fitted to data

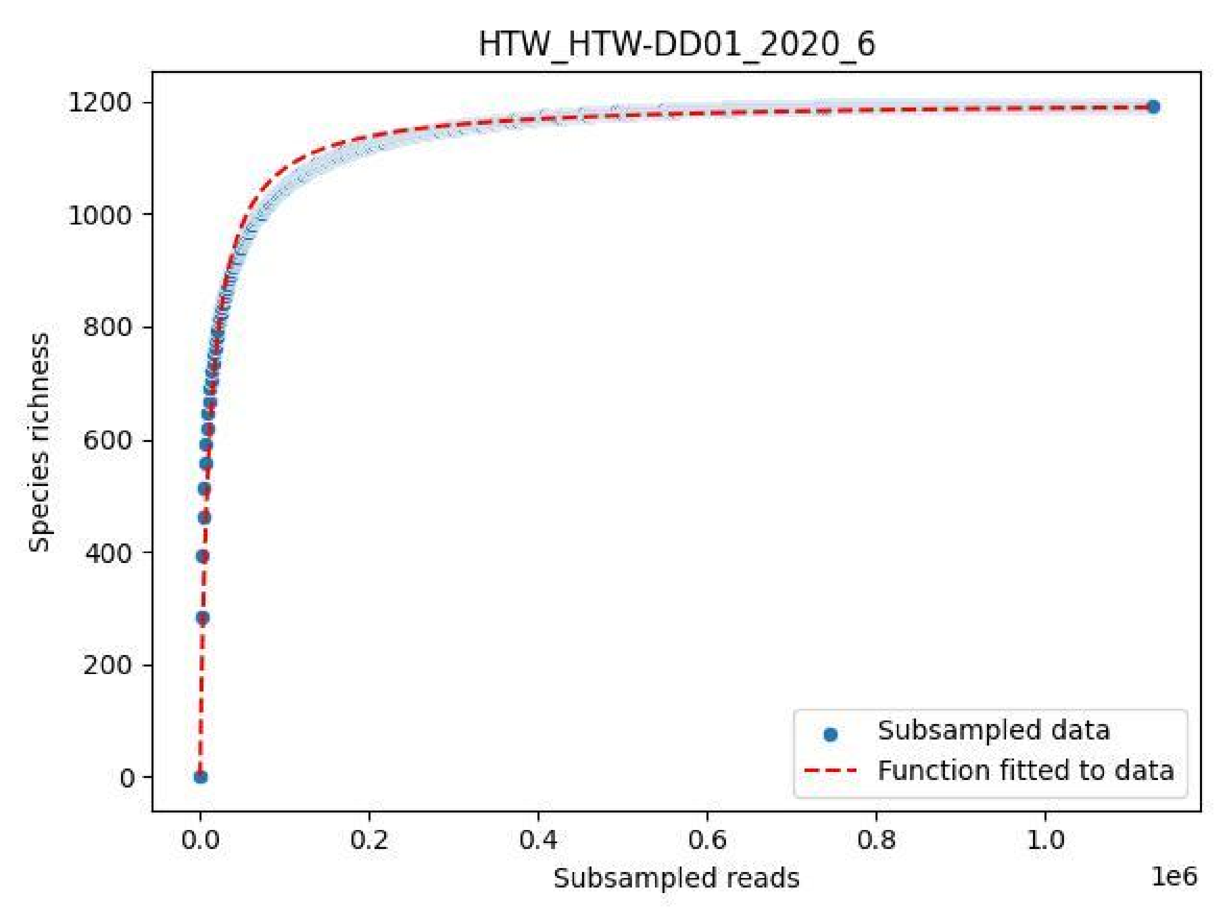

HTW_HTW-DD01_2020_6
Species richness
Subsampled reads
1e6
0
200
400
600
800
1000
1200
0.0
0.2
0.4
0.6
0.8
1.0
Subsampled data
Function fitted to data

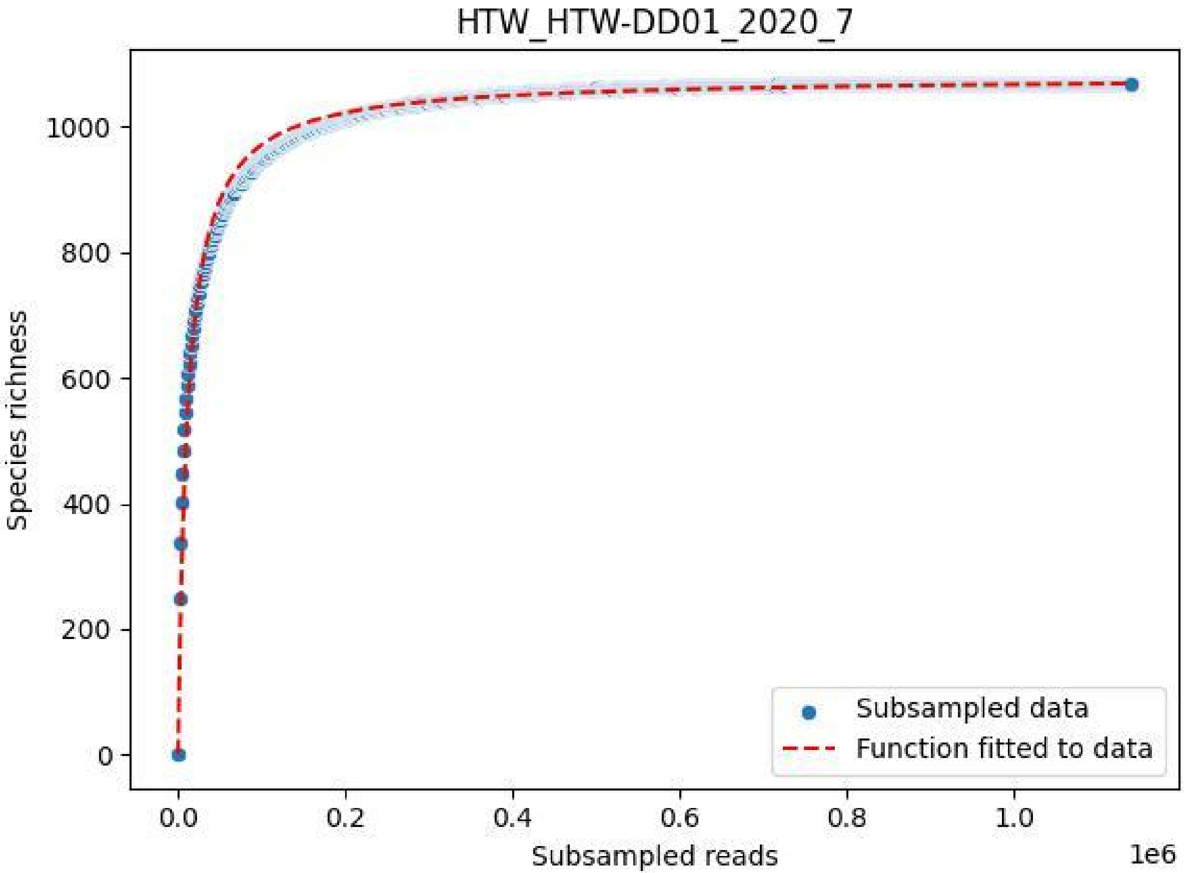
HTW_HTW-DD01_2020_7
Species richness
1000
800
600
400
200
0
0.0
0.2
0.4
0.6
0.8
1.0
1e6
Subsampled reads
Subsampled data
Function fitted to data

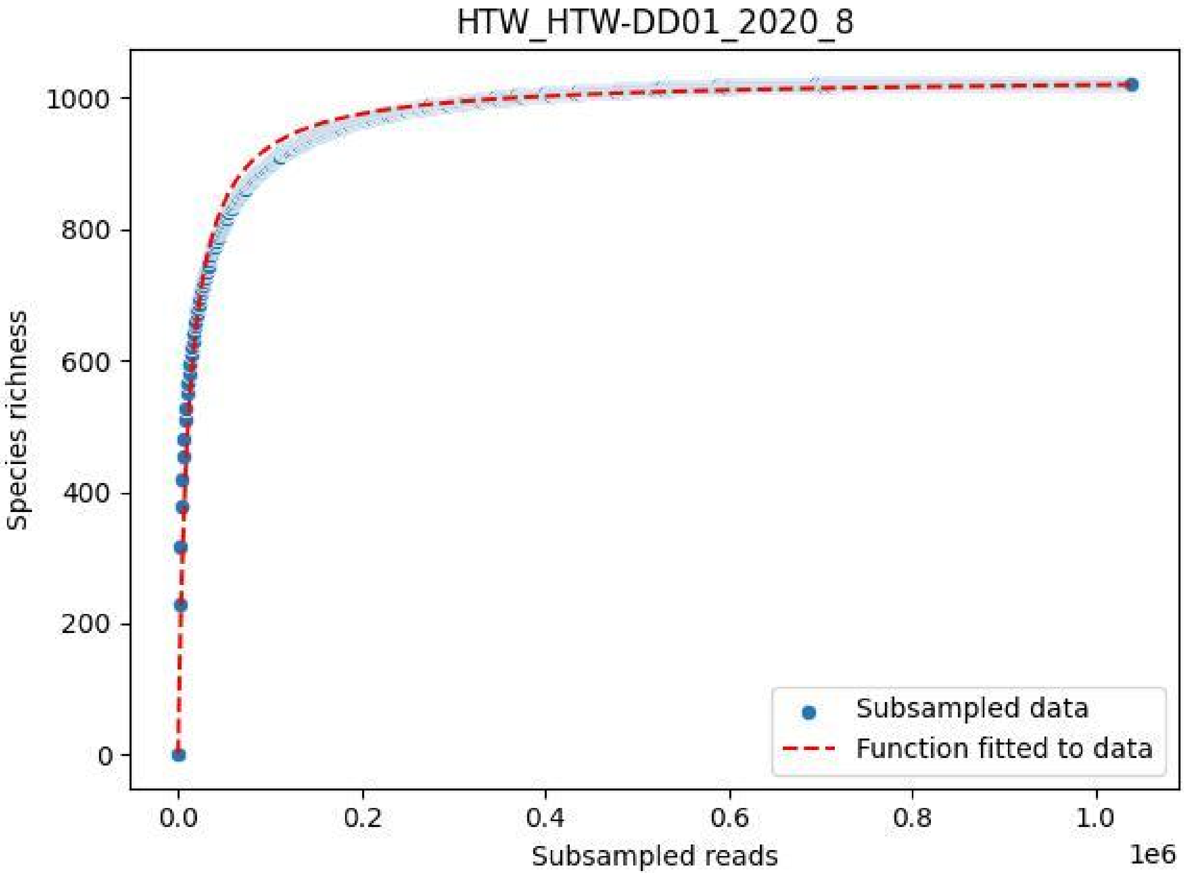
HTW_HTW-DD01_2020_8
Species richness
Subsampled reads
1e6
0
200
400
600
800
1000
0.0
0.2
0.4
0.6
0.8
1.0
Subsampled data
Function fitted to data

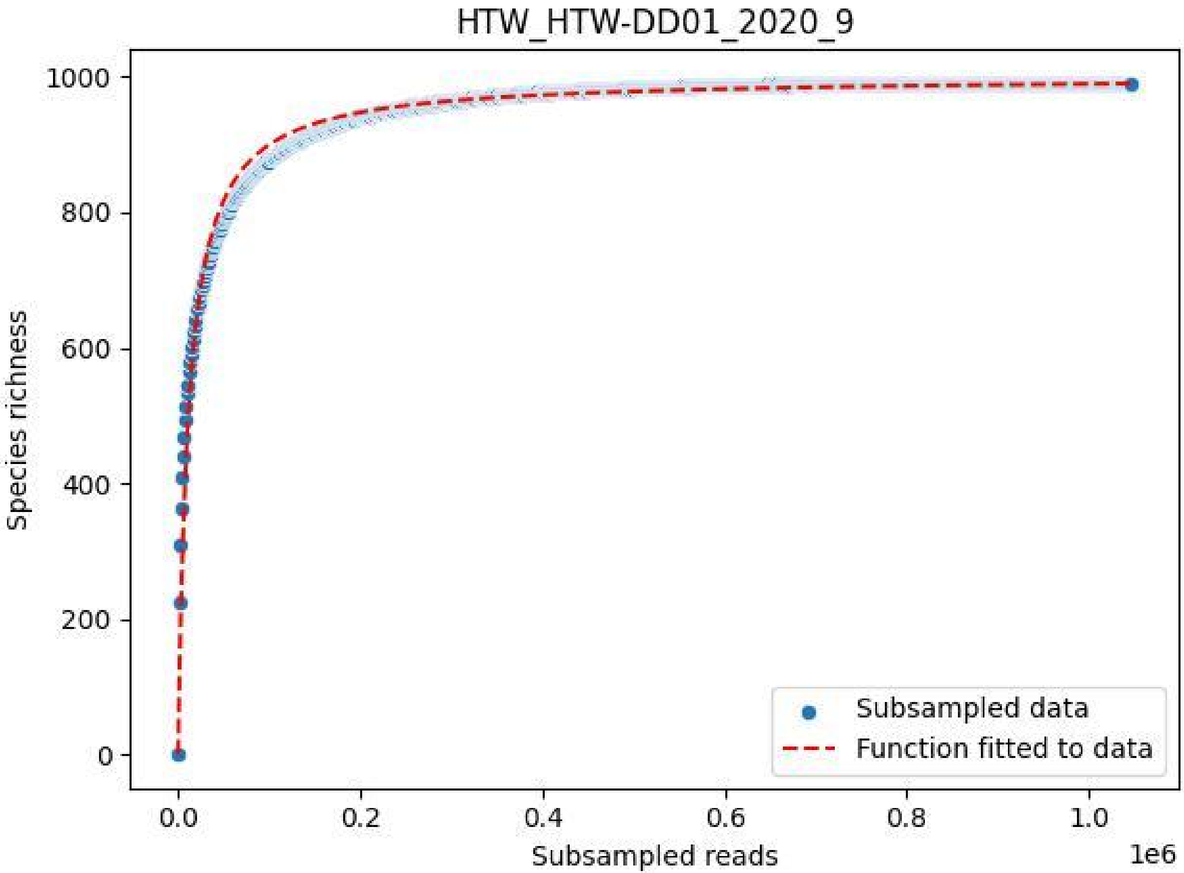
HTW_HTW-DD01_2020_9
Species richness
Subsampled reads
1e6
0
200
400
600
800
1000
0.0
0.2
0.4
0.6
0.8
1.0
Subsampled data
Function fitted to data

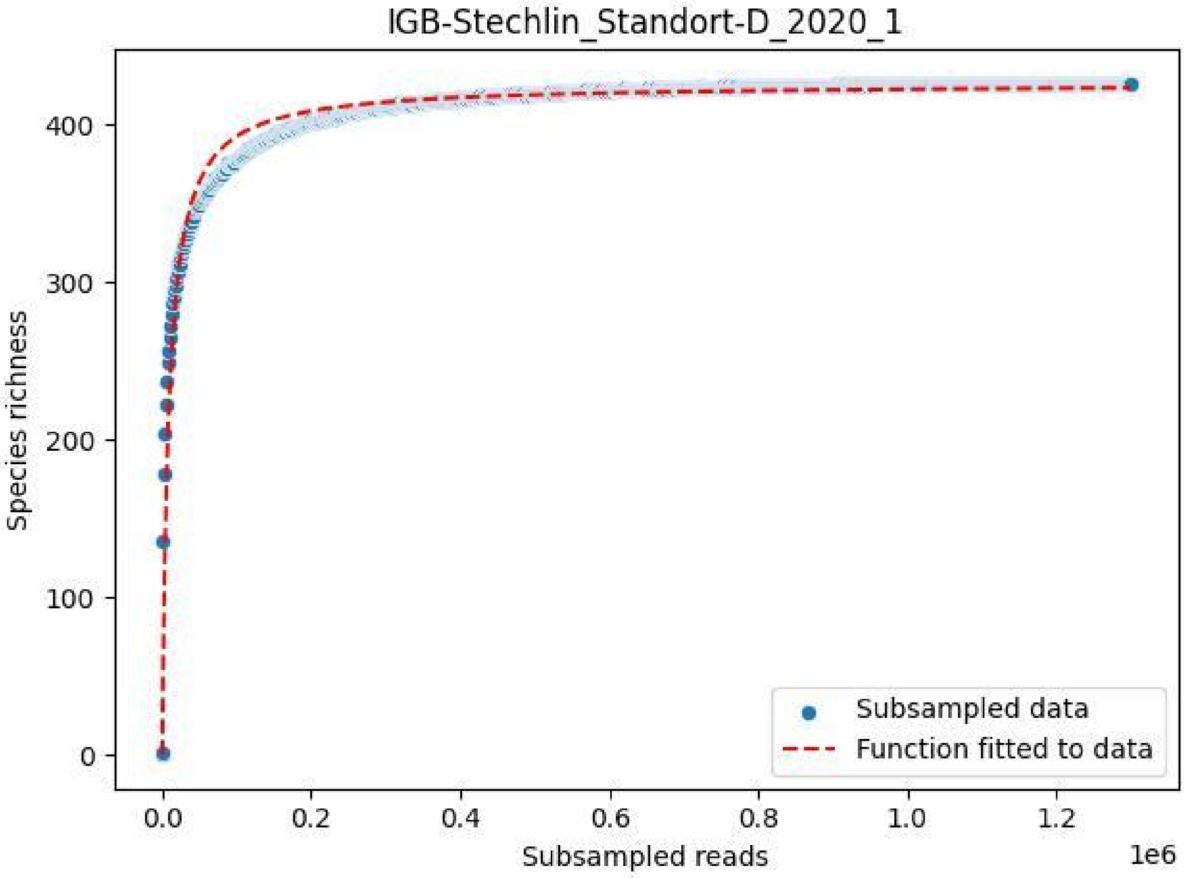
IGB-Stechlin_Standort-D_2020_1
Species richness
0
100
200
300
400
0.0
0.2
0.4
0.6
0.8
1.0
1.2
1e6
Subsampled reads
Subsampled data
Function fitted to data

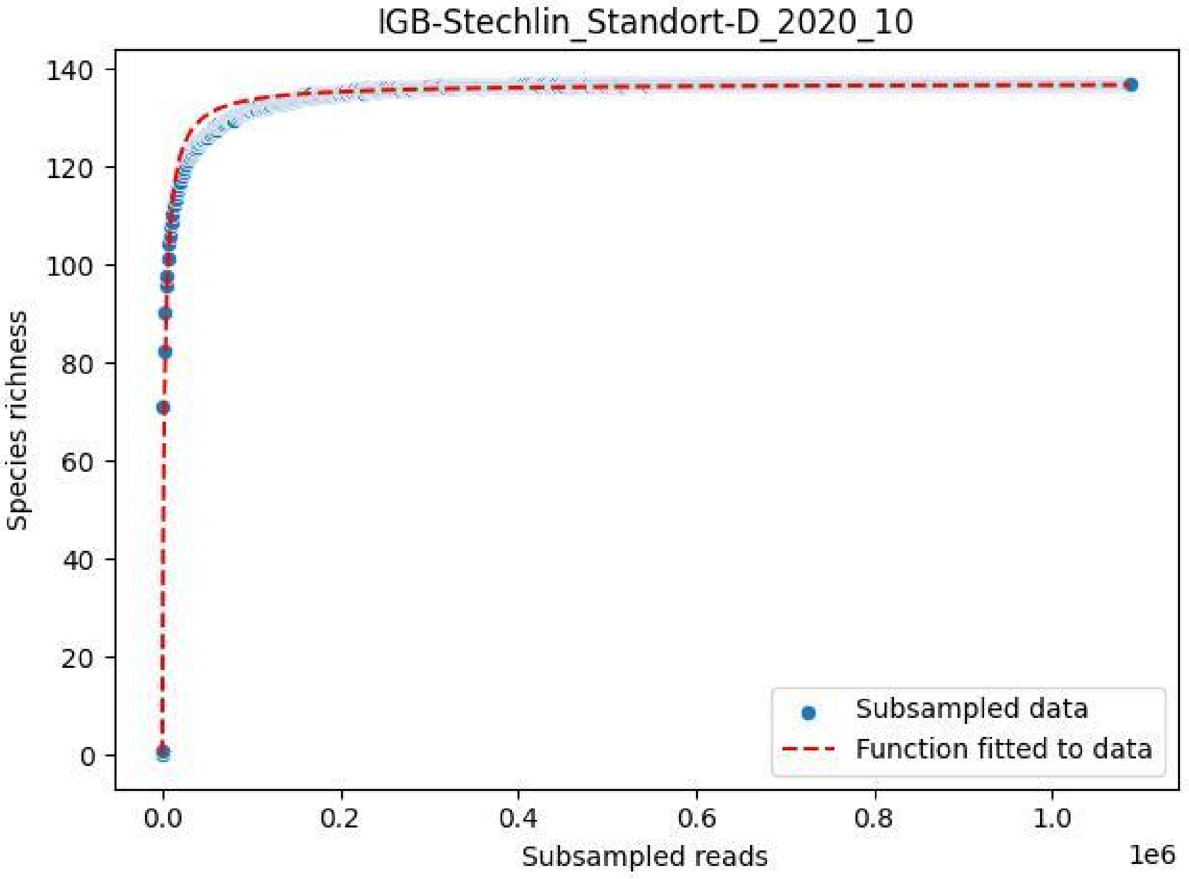

IGB-Stechlin_Standort-D_2020_10
140
120
100
80
60
40
20
0
Species richness
0.0
0.2
0.4
0.6
0.8
1.0
1e6
Subsampled reads
Subsampled data
Function fitted to data

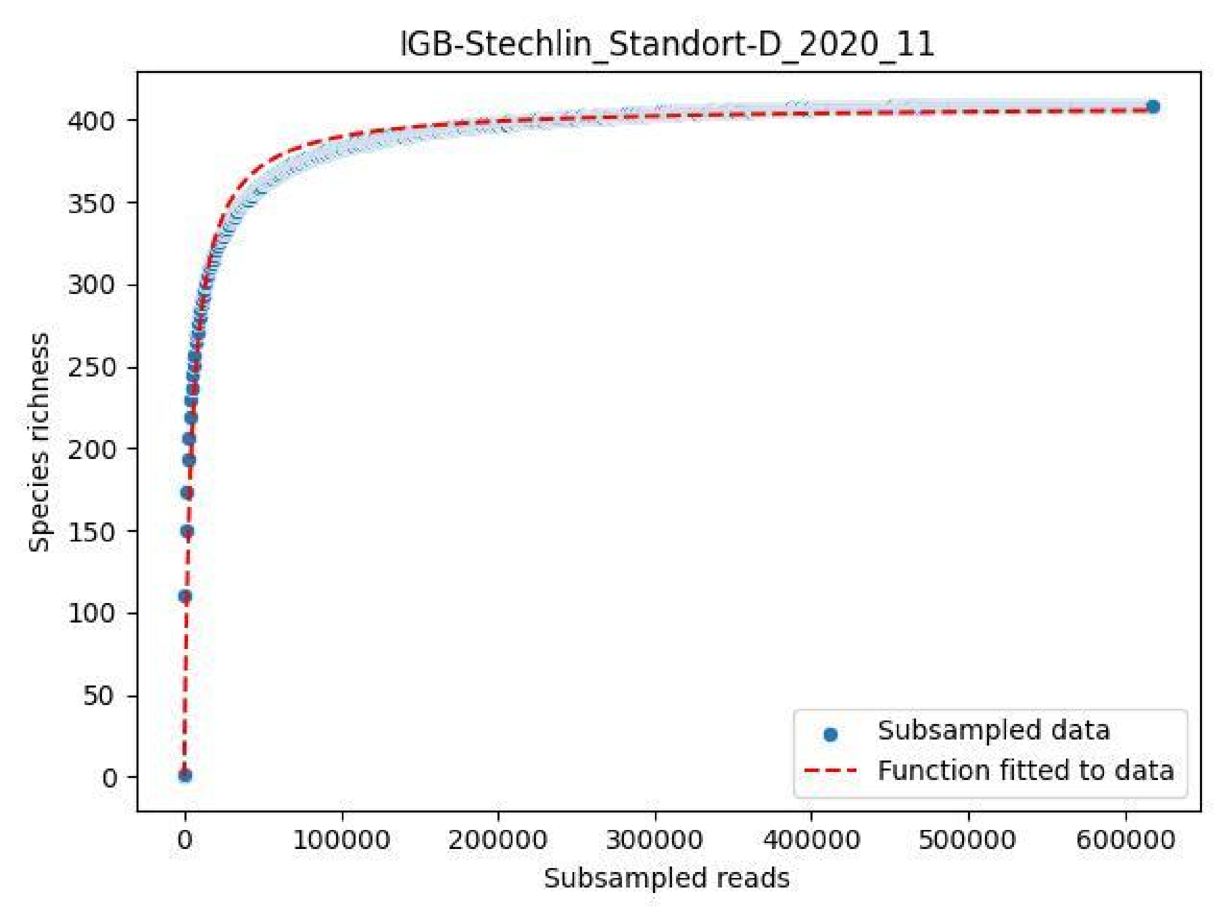
IGB-Stechlin_Standort-D_2020_11
Species richness
Subsampled reads
0
50
100
150
200
250
300
350
400
0
100000
200000
300000
400000
500000
600000
Subsampled data
Function fitted to data

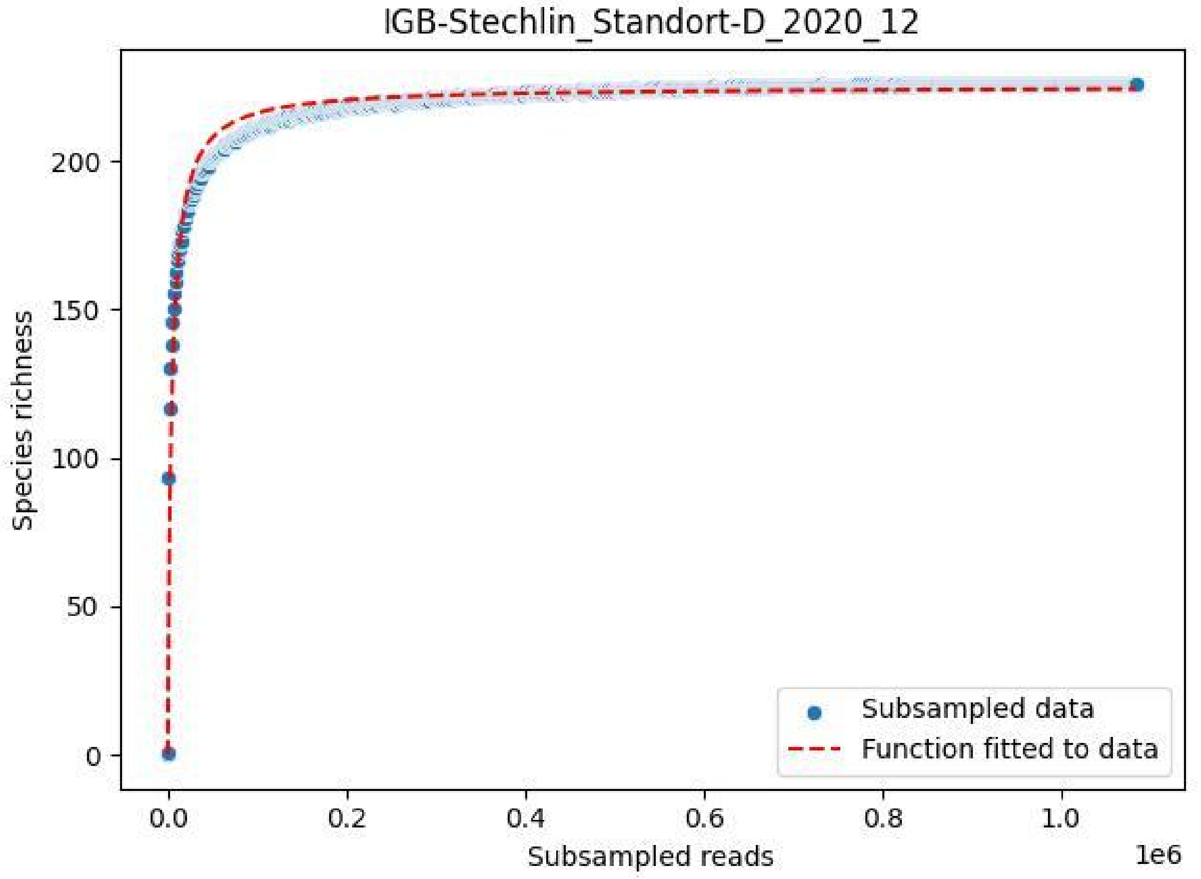
IGB-Stechlin_Standort-D_2020_12
Species richness
Subsampled reads
1e6
0
50
100
150
200
0.0
0.2
0.4
0.6
0.8
1.0
Subsampled data
Function fitted to data

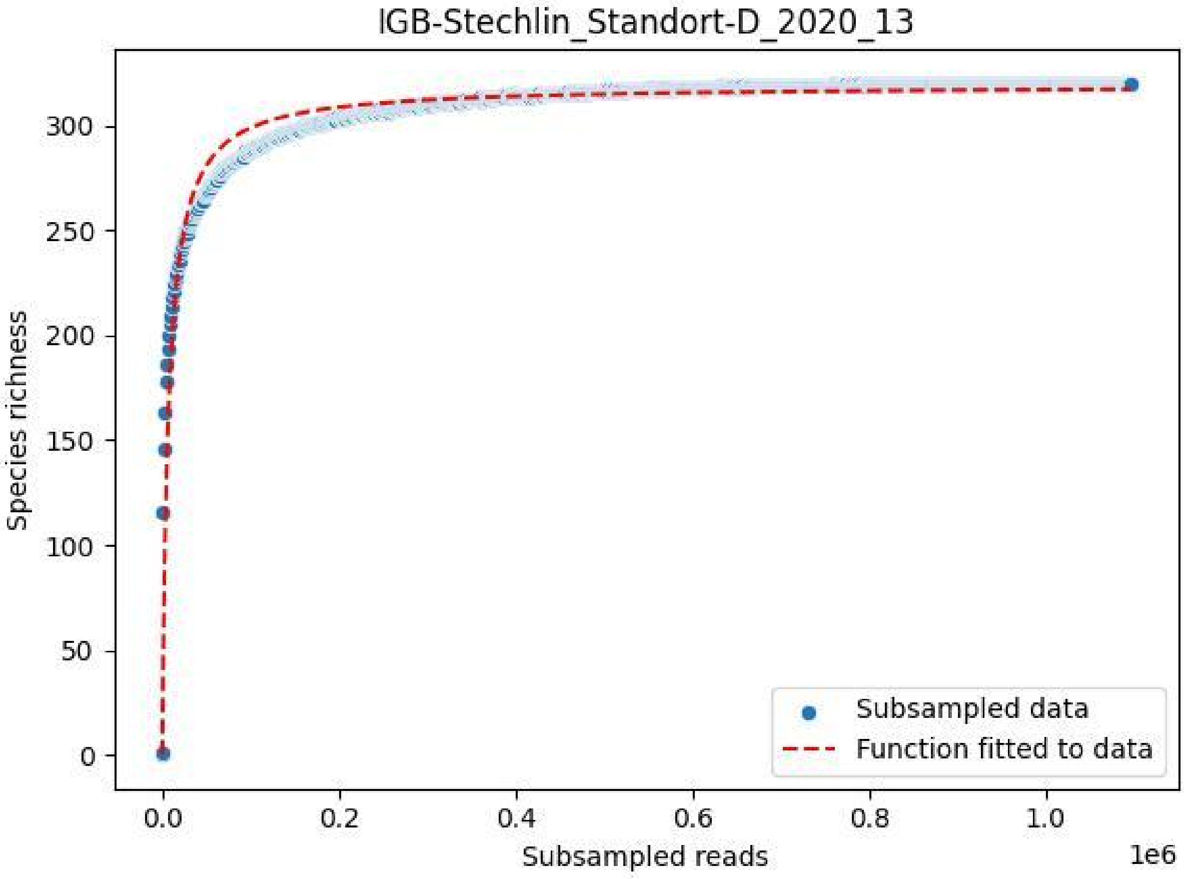

IGB-Stechlin_Standort-D_2020_13
Species richness
300
250
200
150
100
50
0
0.0
0.2
0.4
0.6
0.8
1.0
1e6
Subsampled reads
Subsampled data
Function fitted to data

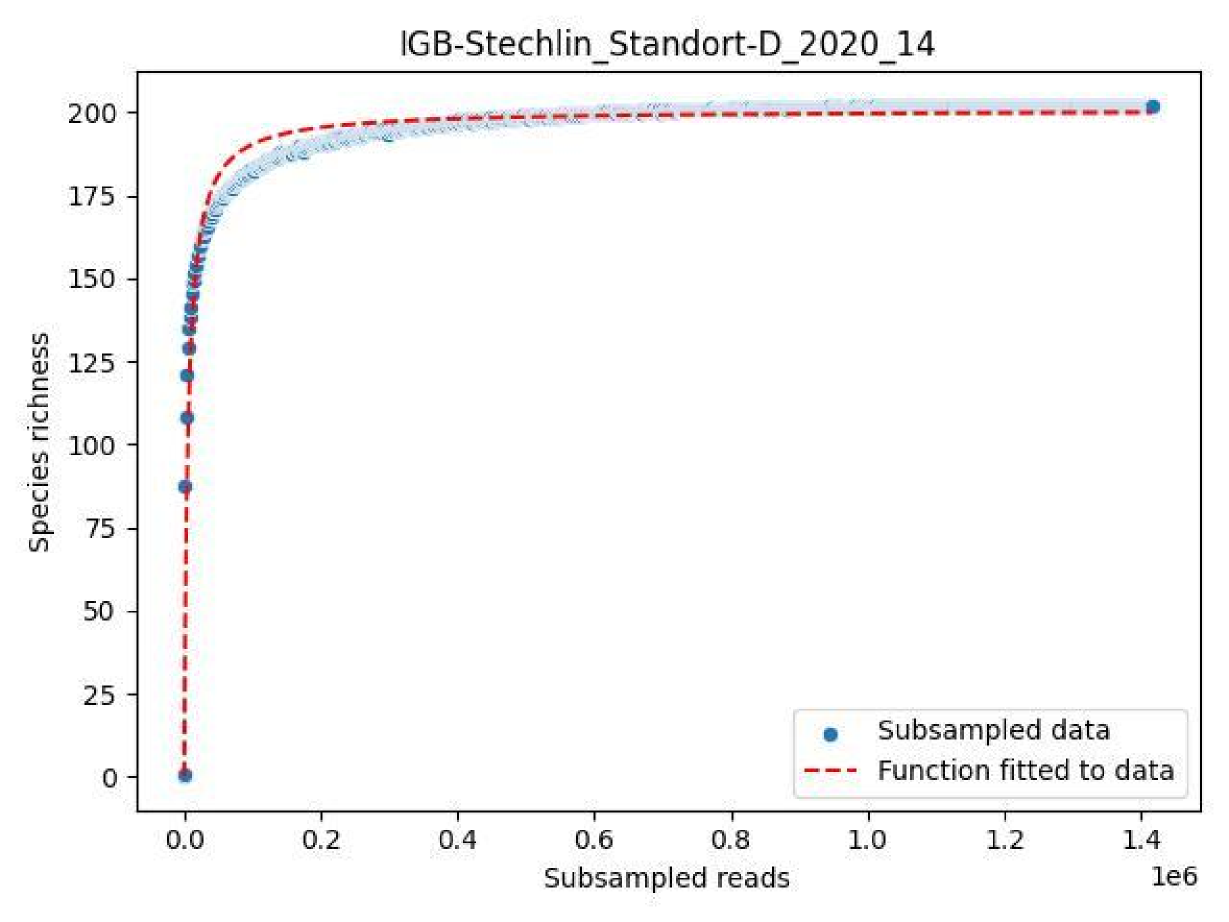
IGB-Stechlin_Standort-D_2020_14
Species richness
Subsampled reads
1e6
0
25
50
75
100
125
150
175
200
0.0
0.2
0.4
0.6
0.8
1.0
1.2
1.4
Subsampled data
Function fitted to data

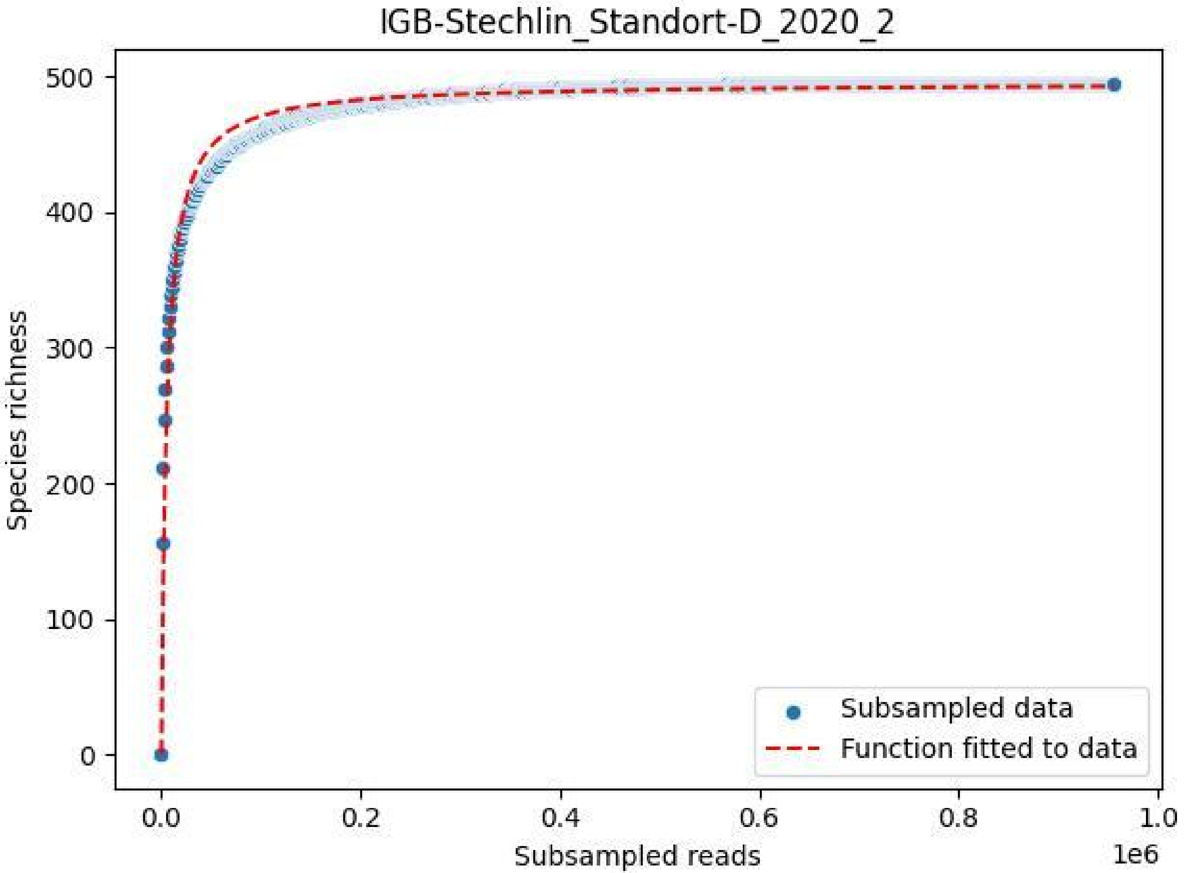
IGB-Stechlin_Standort-D_2020_2
500
400
300
200
100
0
Species richness
0.0
0.2
0.4
0.6
0.8
1.0
1e6
Subsampled reads
Subsampled data
Function fitted to data

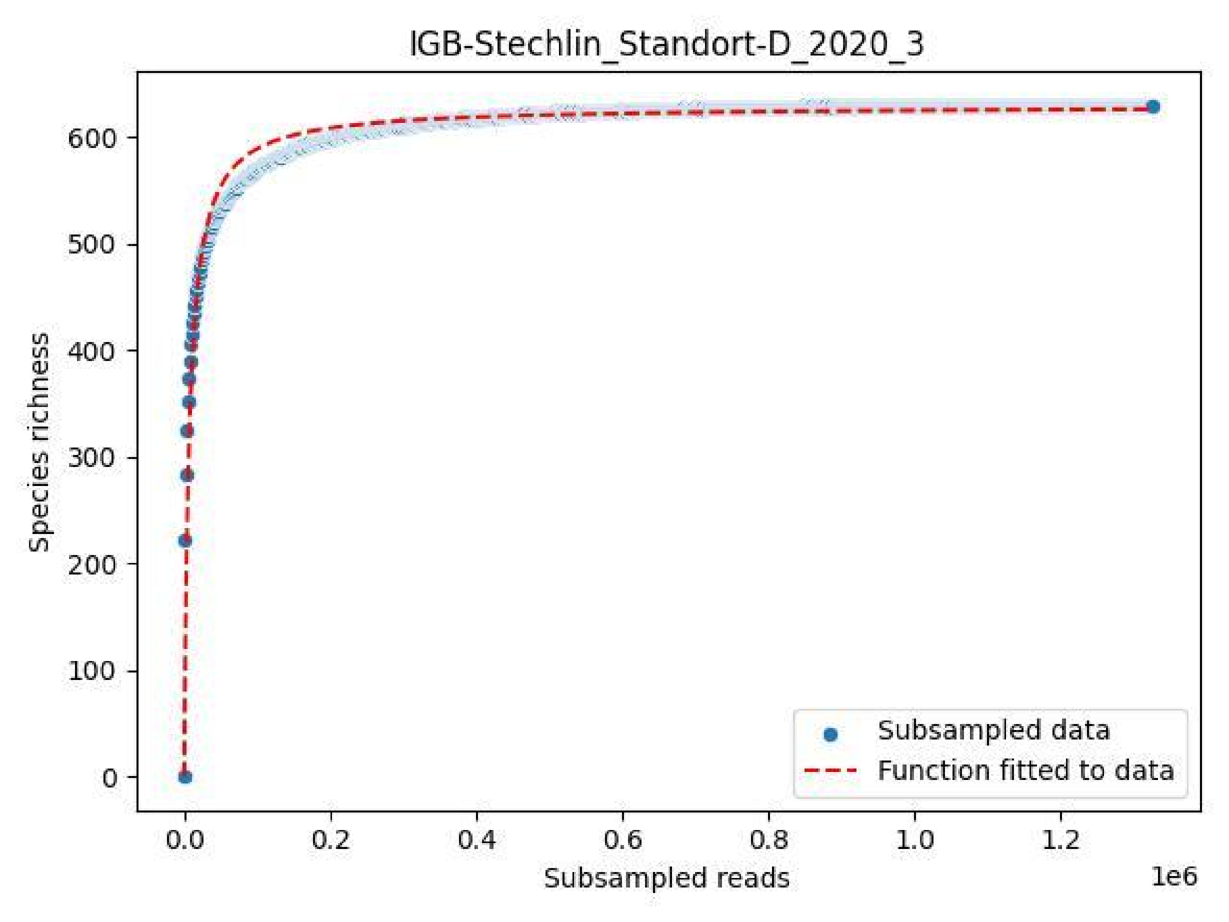
IGB-Stechlin_Standort-D_2020_3
Species richness
600
500
400
300
200
100
0
0.0
0.2
0.4
0.6
0.8
1.0
1.2
Subsampled reads
1e6
Subsampled data
Function fitted to data

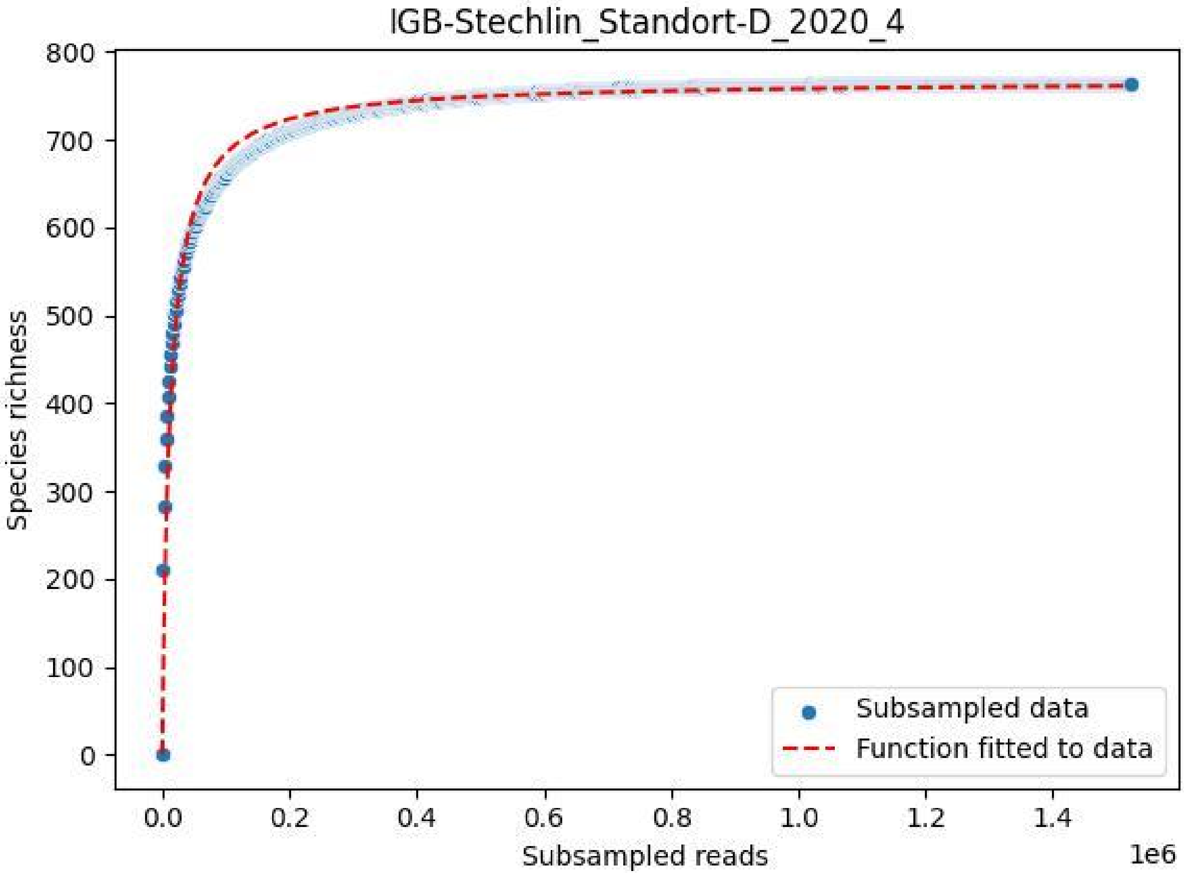
IGB-Stechlin_Standort-D_2020_4
Species richness
Subsampled reads
1e6
0
100
200
300
400
500
600
700
800
0.0
0.2
0.4
0.6
0.8
1.0
1.2
1.4
Subsampled data
Function fitted to data

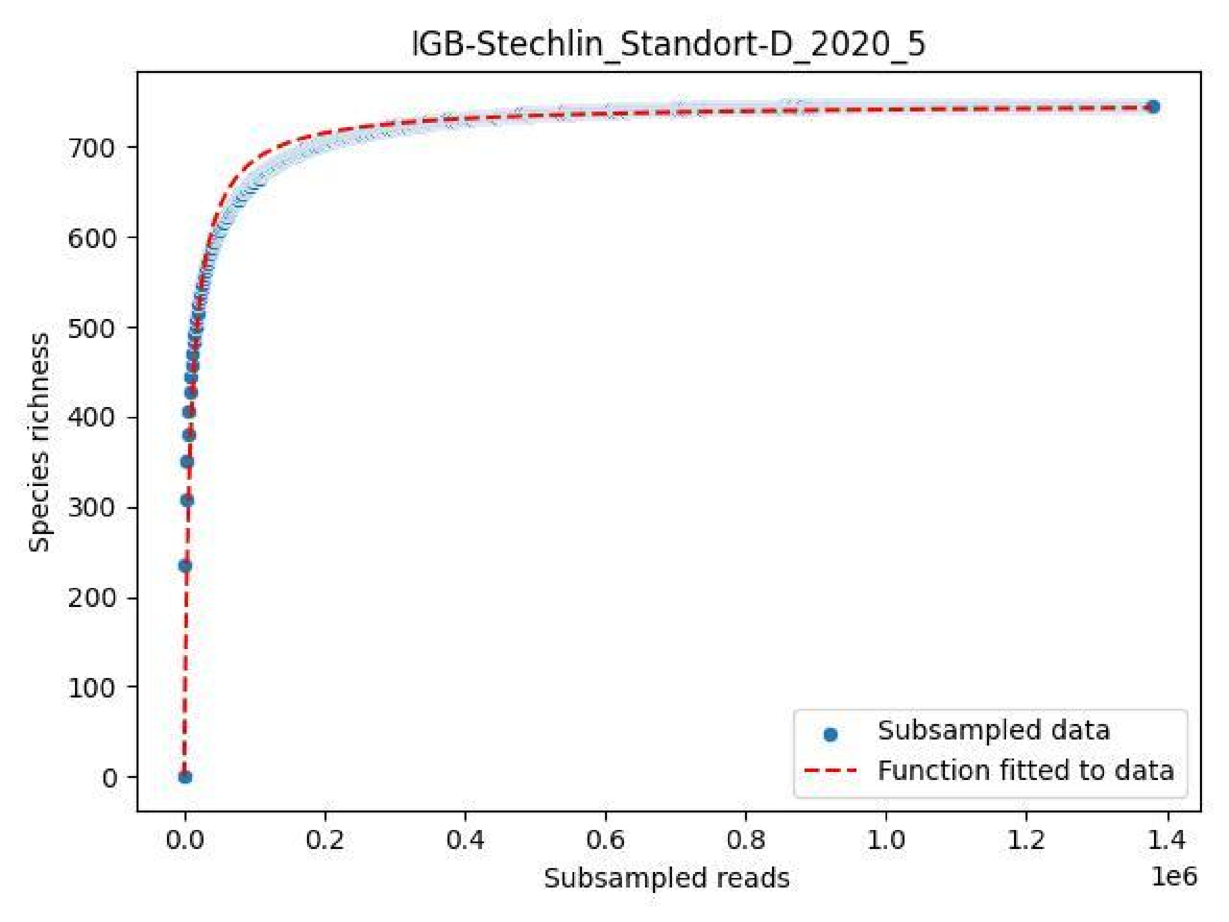
IGB-Stechlin_Standort-D_2020_5
Species richness
Subsampled reads
1e6
0
100
200
300
400
500
600
700
0.0
0.2
0.4
0.6
0.8
1.0
1.2
1.4
Subsampled data
Function fitted to data

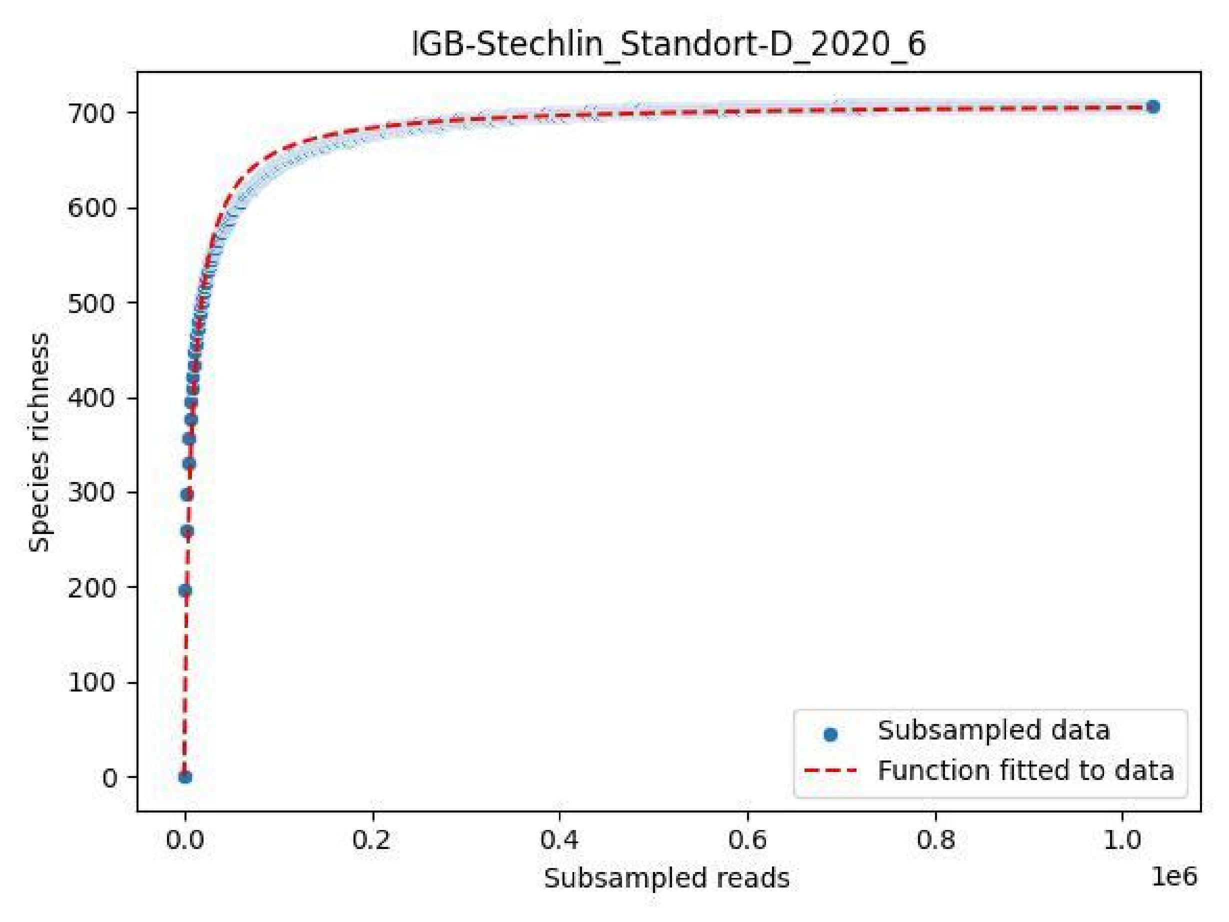
IGB-Stechlin_Standort-D_2020_6
Species richness
Subsampled reads
1e6
0.0
0.2
0.4
0.6
0.8
1.0
0
100
200
300
400
500
600
700
Subsampled data
Function fitted to data

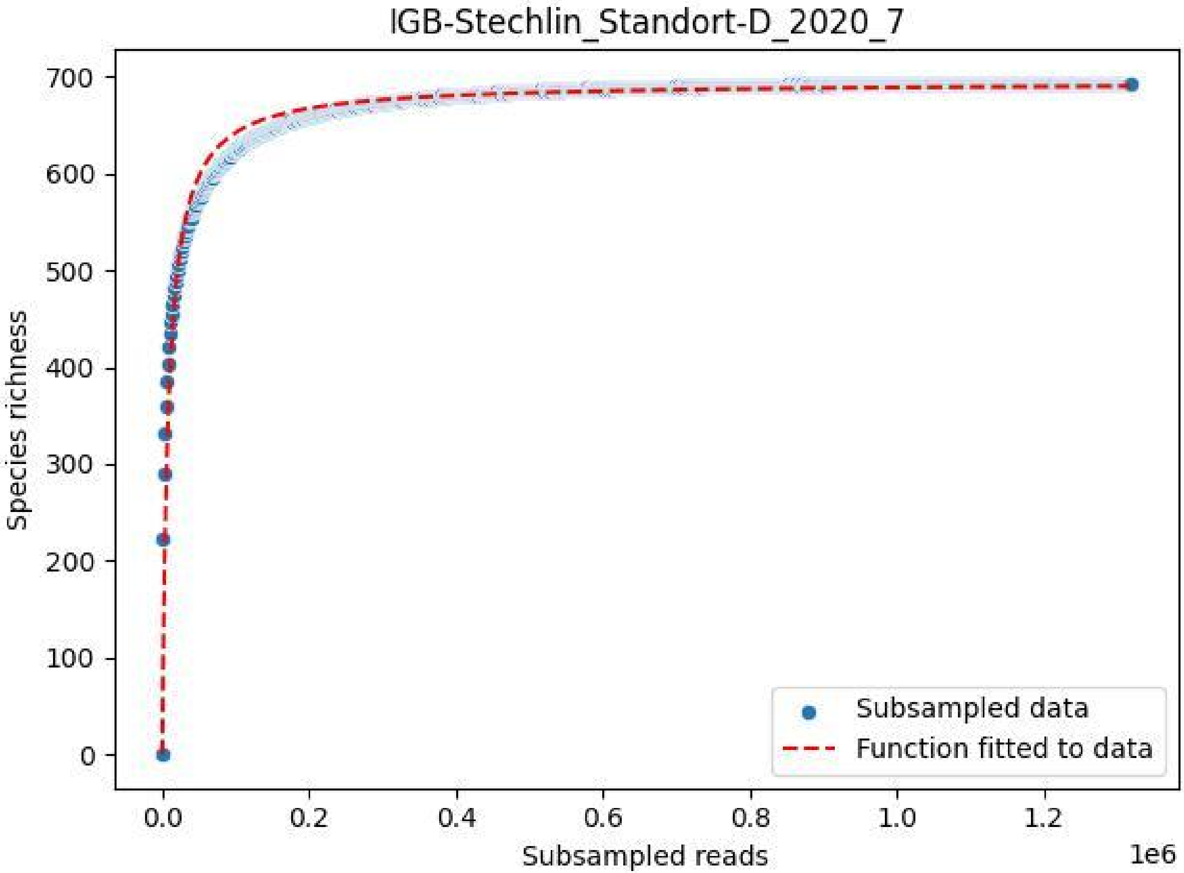

IGB-Stechlin_Standort-D_2020_7
Species richness
Subsampled reads
1e6
0
100
200
300
400
500
600
700
0.0
0.2
0.4
0.6
0.8
1.0
1.2
Subsampled data
Function fitted to data

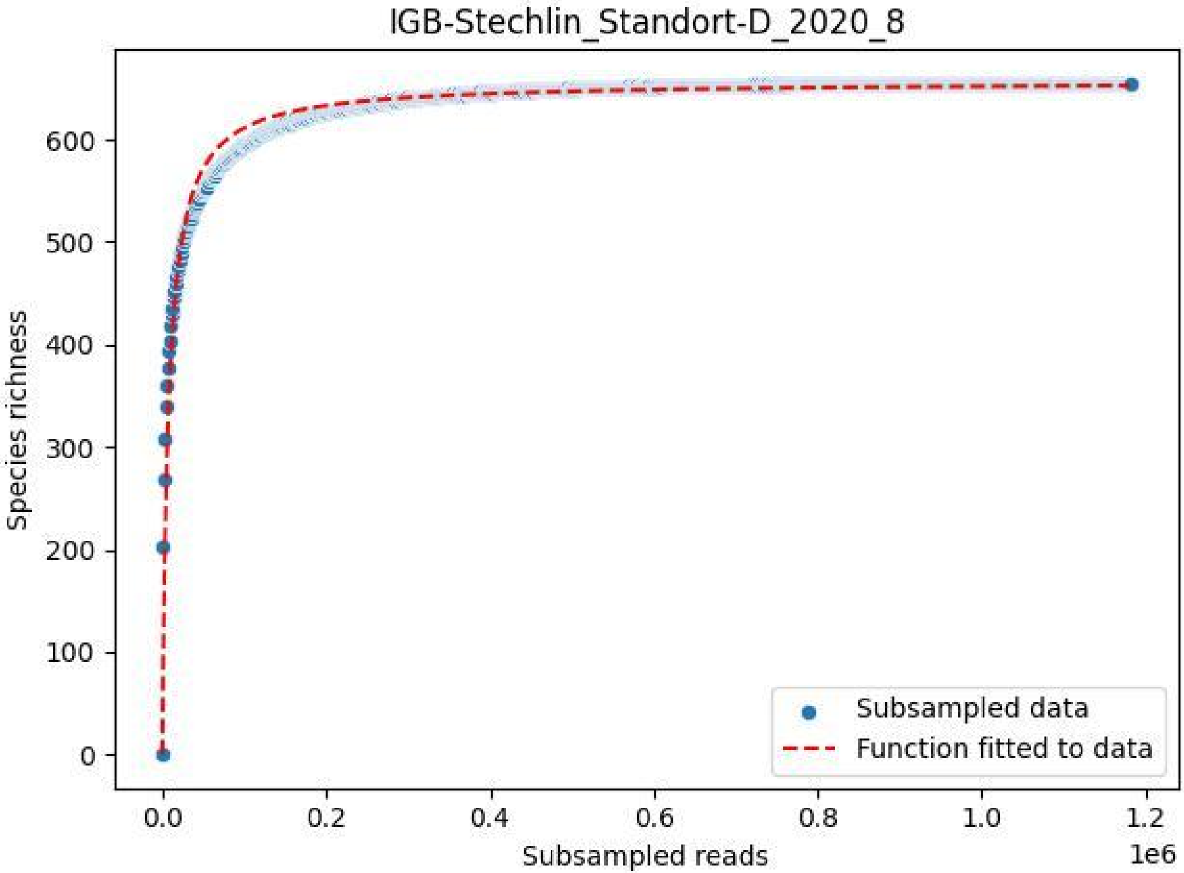
IGB-Stechlin_Standort-D_2020_8
Species richness
Subsampled reads
1e6
0
100
200
300
400
500
600
0.0
0.2
0.4
0.6
0.8
1.0
1.2
Subsampled data
Function fitted to data

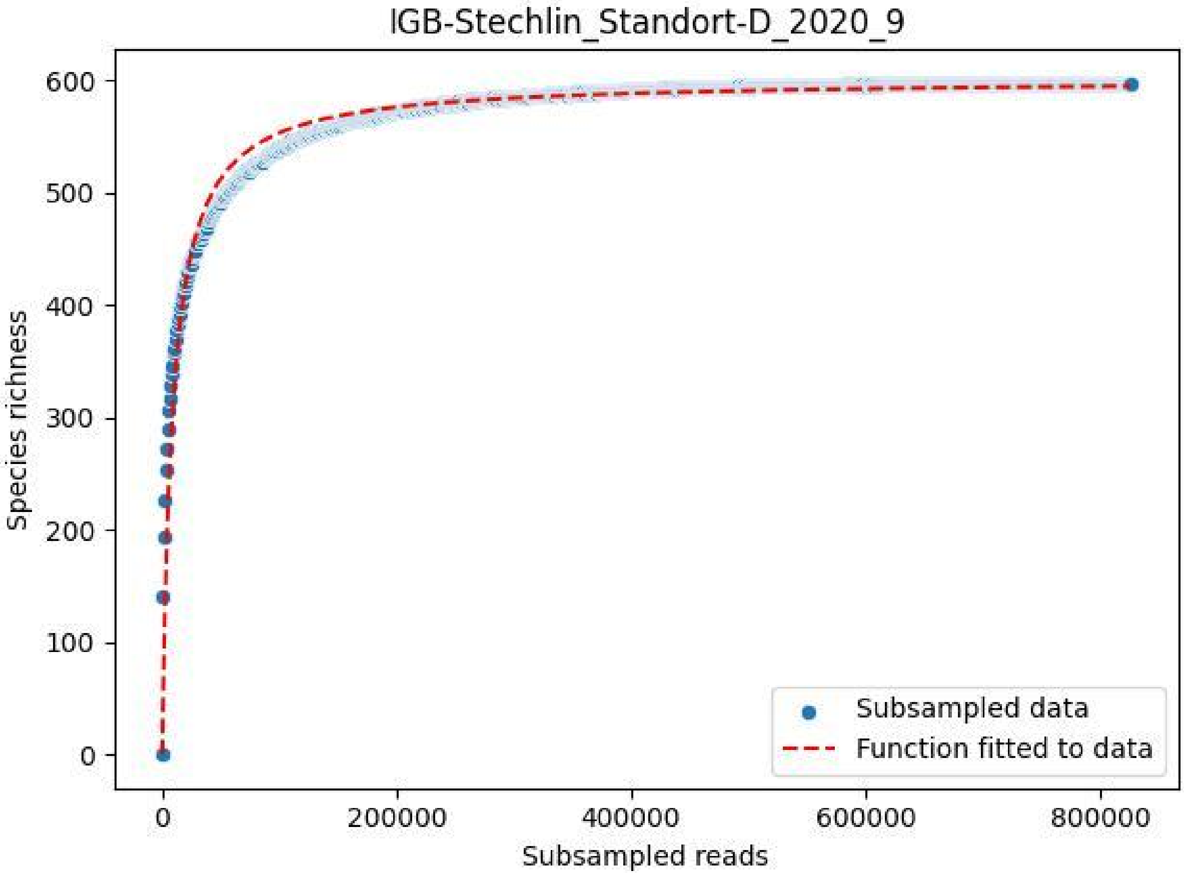

IGB-Stechlin_Standort-D_2020_9
Species richness
Subsampled reads
0
100
200
300
400
500
600
0
200000
400000
600000
800000
Subsampled data
Function fitted to data

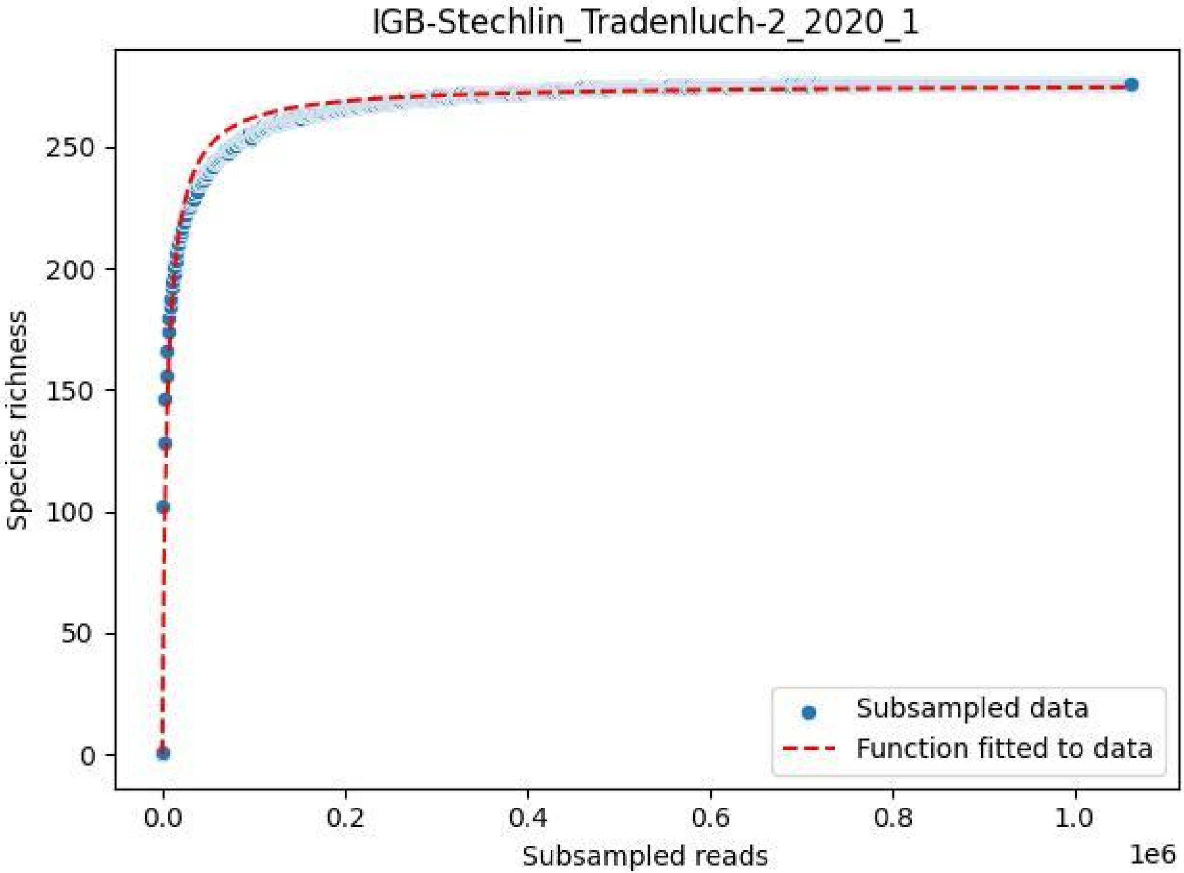
IGB-Stechlin_Tradenluch-2_2020_1
Species richness
0
50
100
150
200
250
0.0
0.2
0.4
0.6
0.8
1.0
1e6
Subsampled reads
Subsampled data
Function fitted to data

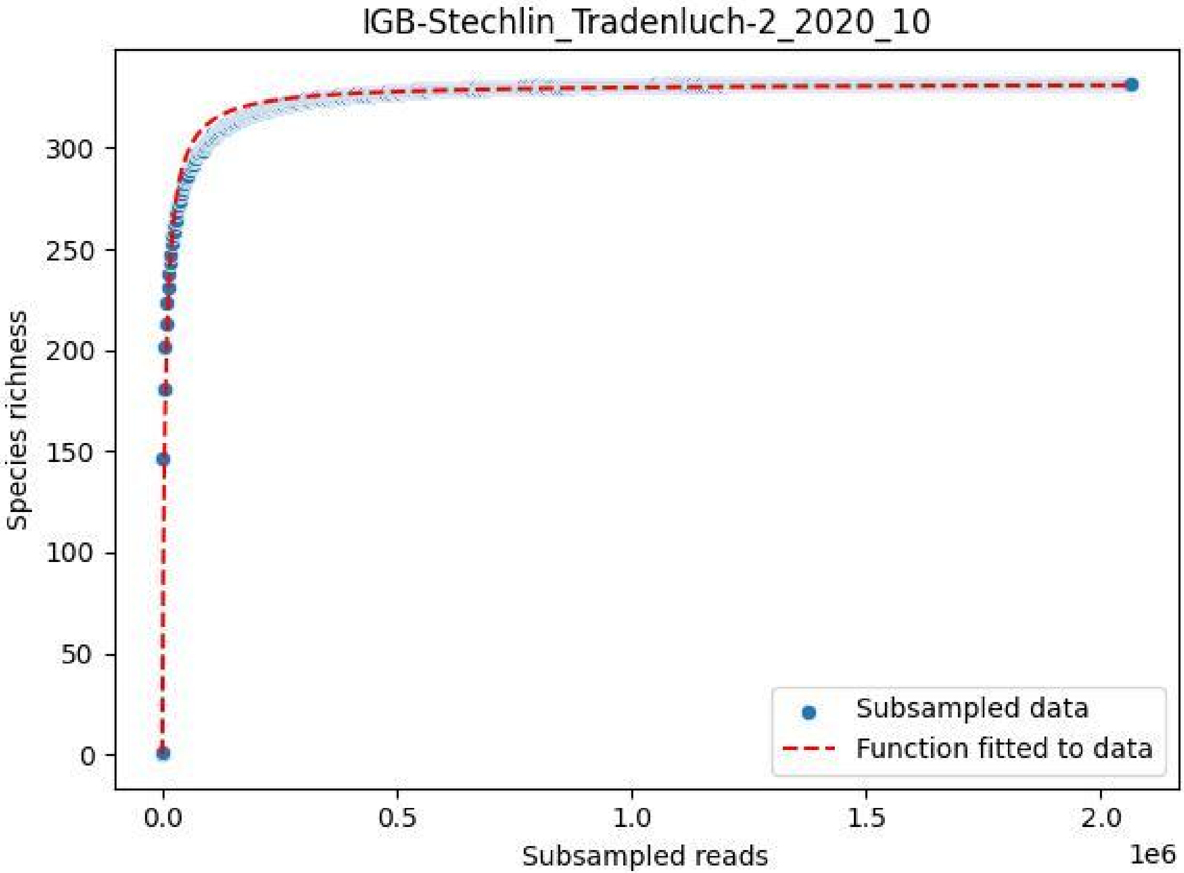

IGB-Stechlin_Tradenluch-2_2020_10
Species richness
300
250
200
150
100
50
0
0.0
0.5
1.0
1.5
2.0
Subsampled reads
1e6
Subsampled data
Function fitted to data

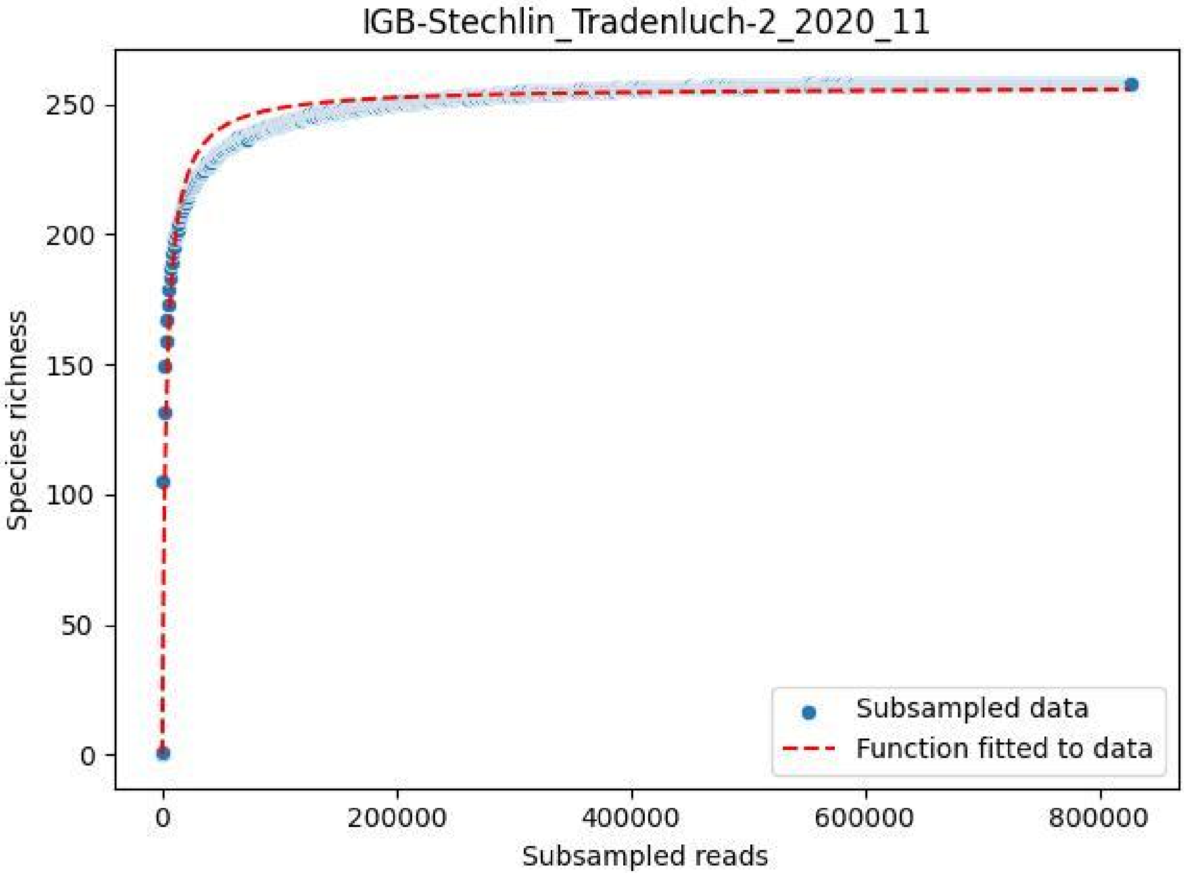
IGB-Stechlin_Tradenluch-2_2020_11
Species richness
Subsampled reads
0
50
100
150
200
250
0
200000
400000
600000
800000
Subsampled data
Function fitted to data

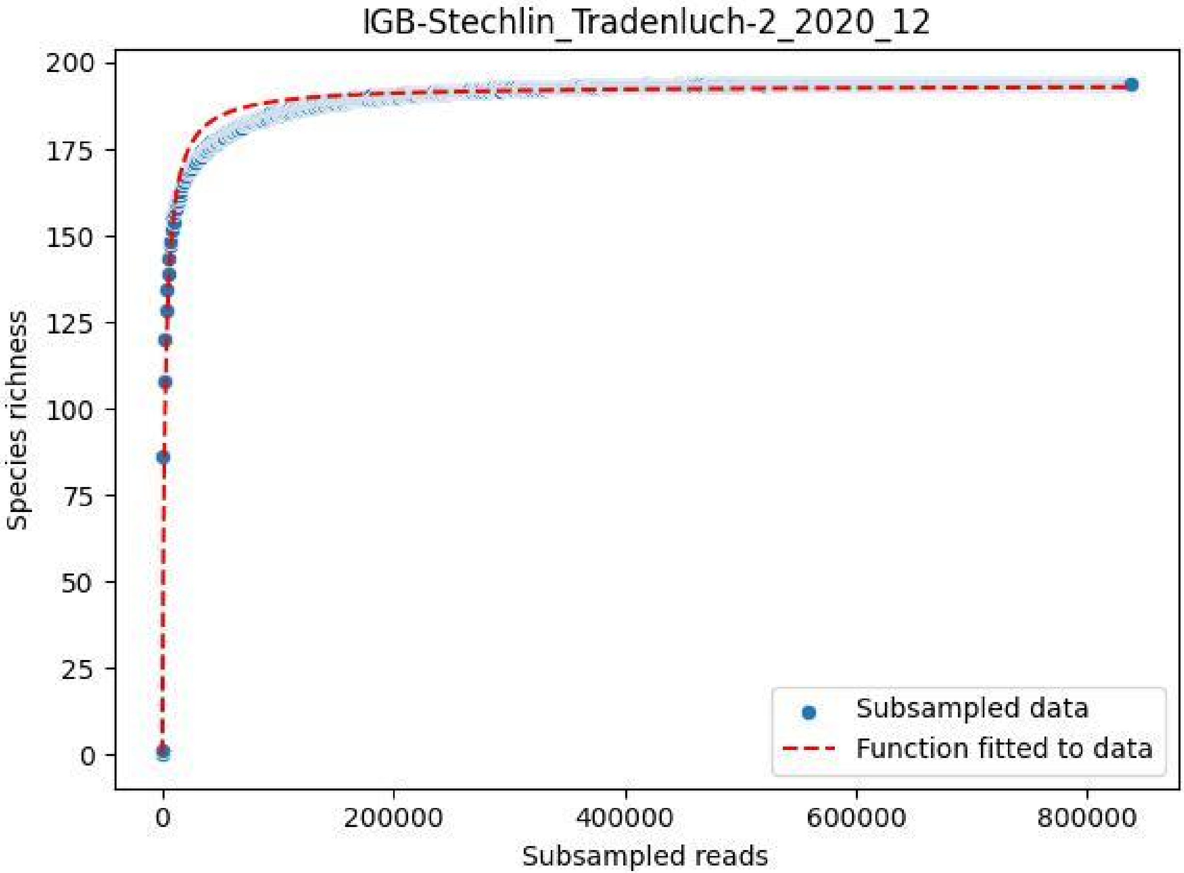
IGB-Stechlin_Tradenluch-2_2020_12
Species richness
Subsampled reads
0
25
50
75
100
125
150
175
200
0
200000
400000
600000
800000
Subsampled data
Function fitted to data

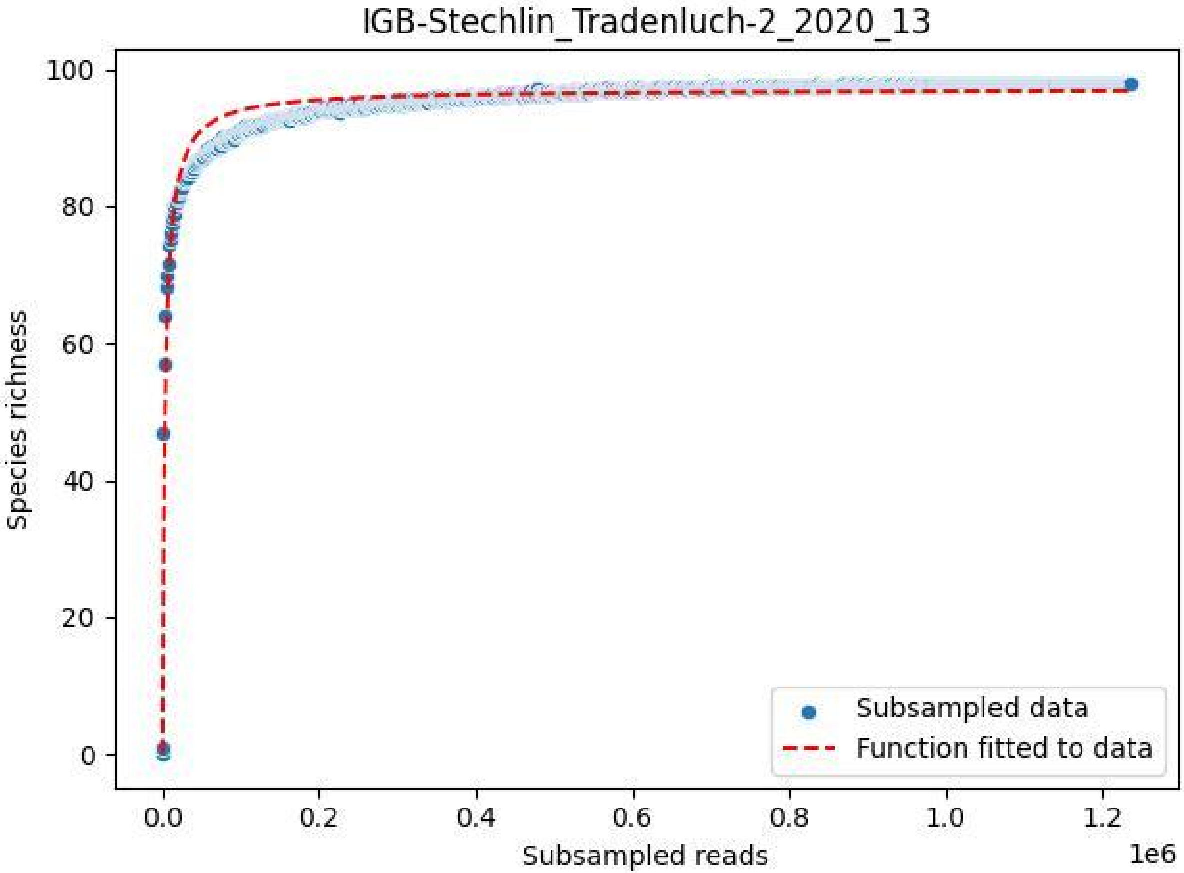
IGB-Stechlin_Tradenluch-2_2020_13
Species richness
Subsampled reads
1e6
0
20
40
60
80
100
0.0
0.2
0.4
0.6
0.8
1.0
1.2
Subsampled data
Function fitted to data

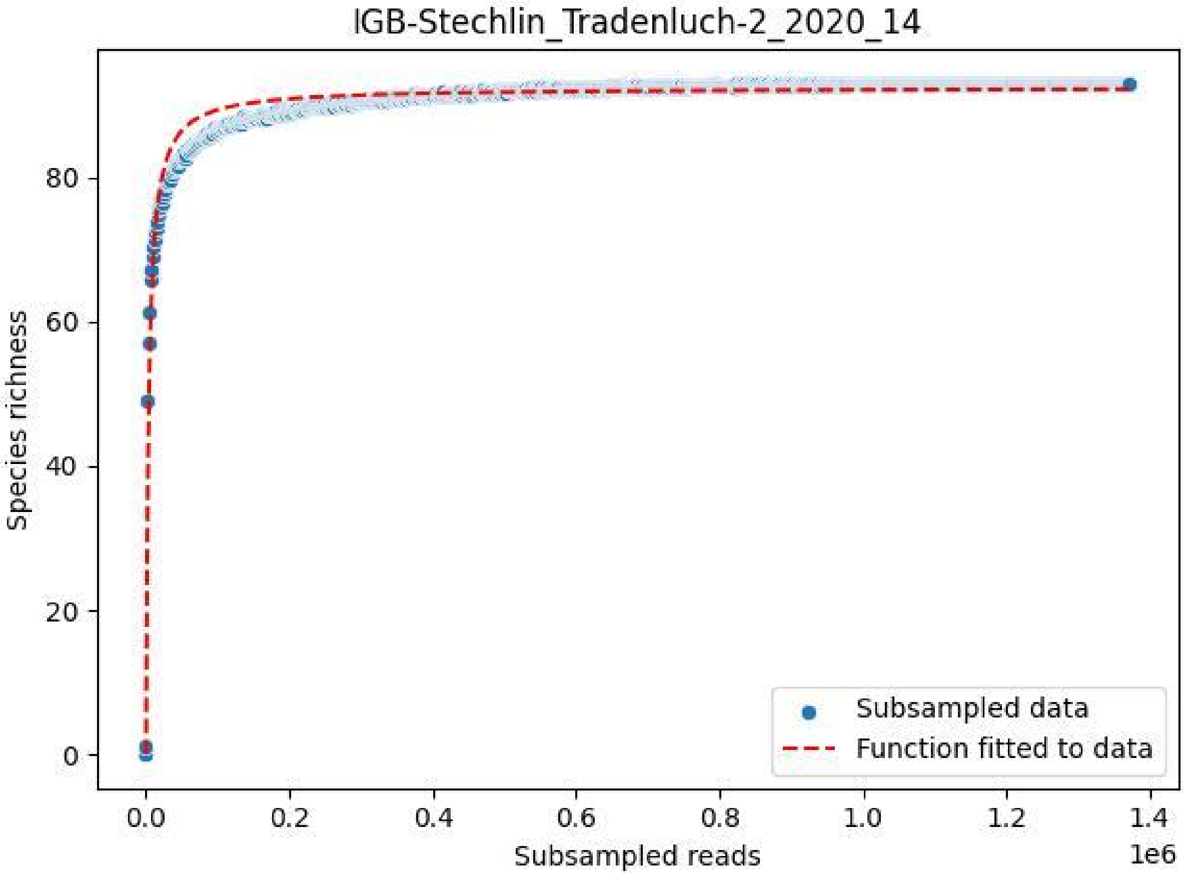

IGB-Stechlin_Tradenluch-2_2020_14
Species richness
0
20
40
60
80
Subsampled reads
0.0
0.2
0.4
0.6
0.8
1.0
1.2
1.4
1e6
Subsampled data
Function fitted to data

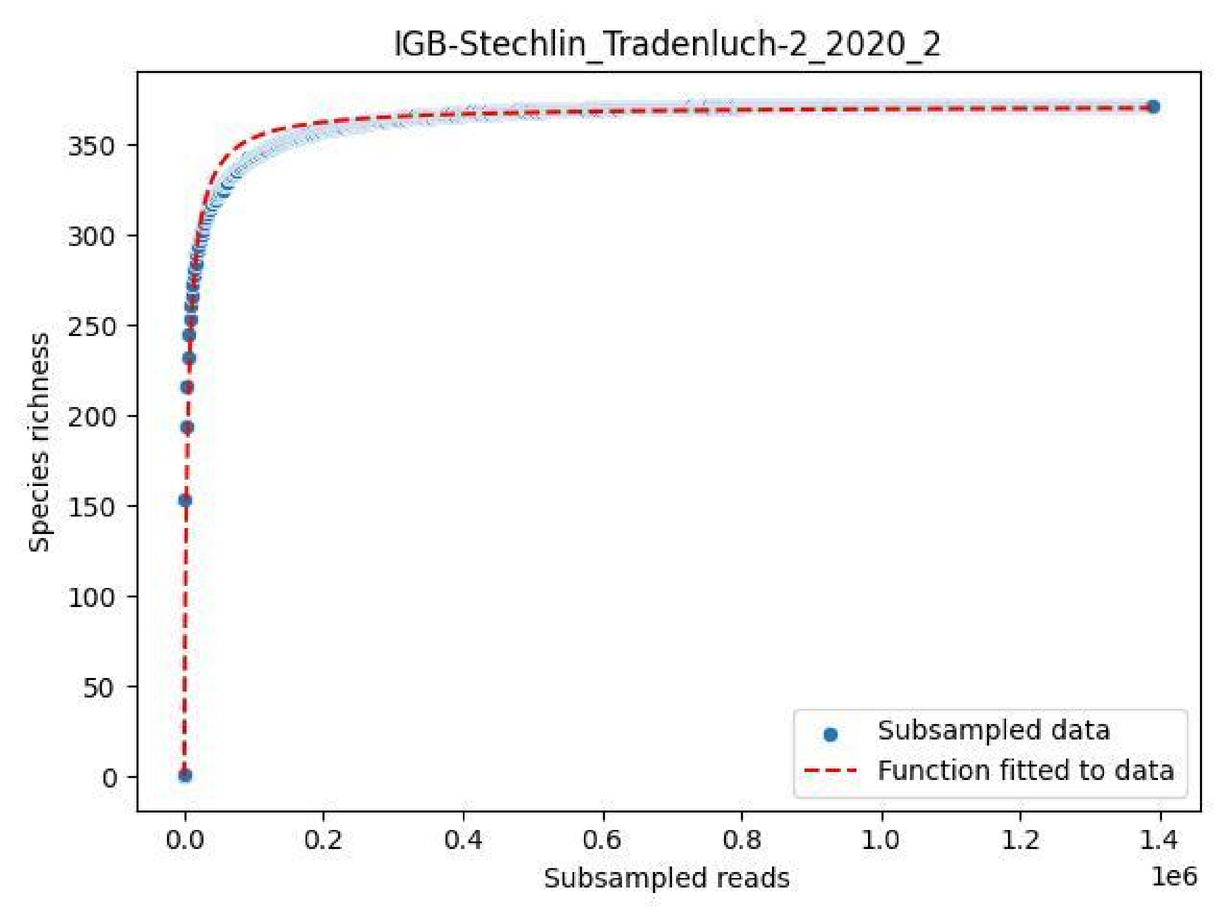
IGB-Stechlin_Tradenluch-2_2020_2
Species richness
Subsampled reads
1e6
0
50
100
150
200
250
300
350
0.0
0.2
0.4
0.6
0.8
1.0
1.2
1.4
Subsampled data
Function fitted to data

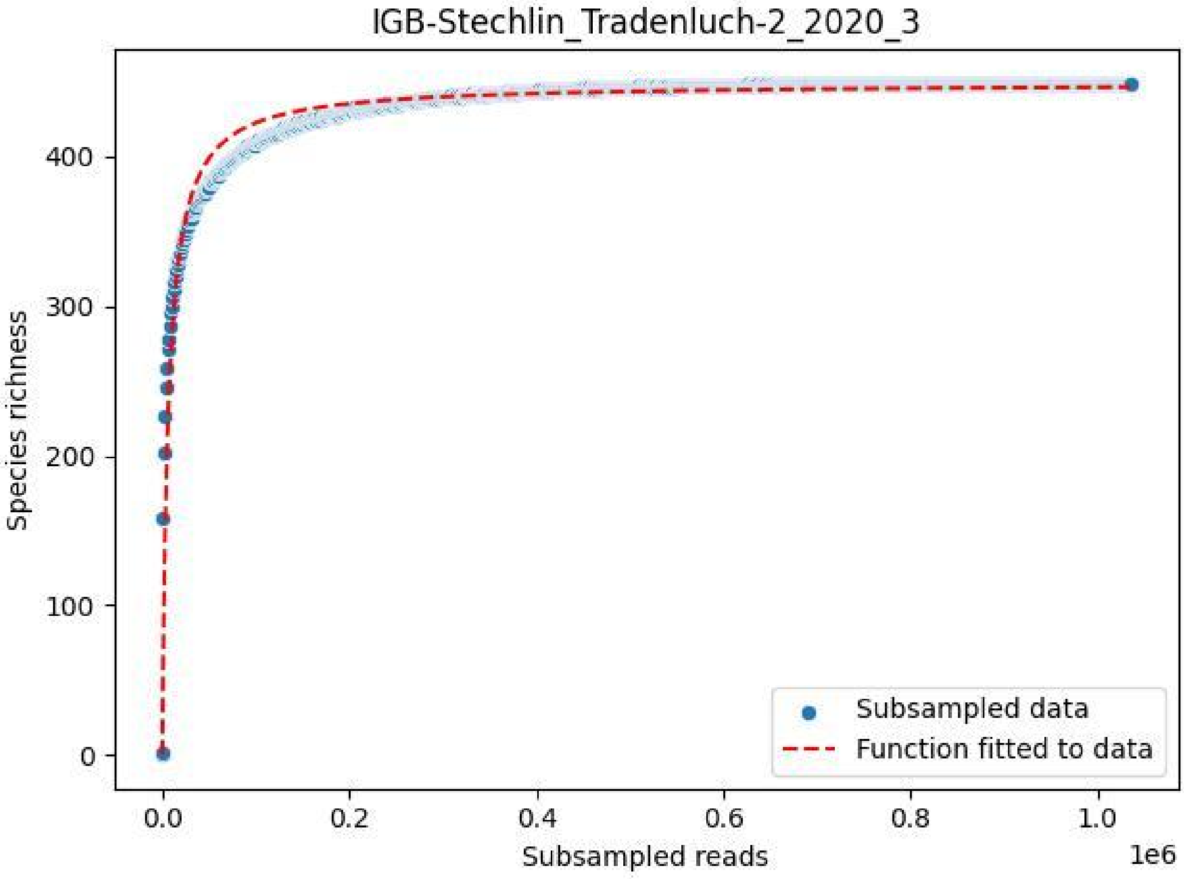
IGB-Stechlin_Tradenluch-2_2020_3
Species richness
Subsampled reads
0
100
200
300
400
0.0
0.2
0.4
0.6
0.8
1.0
1e6
Subsampled data
Function fitted to data

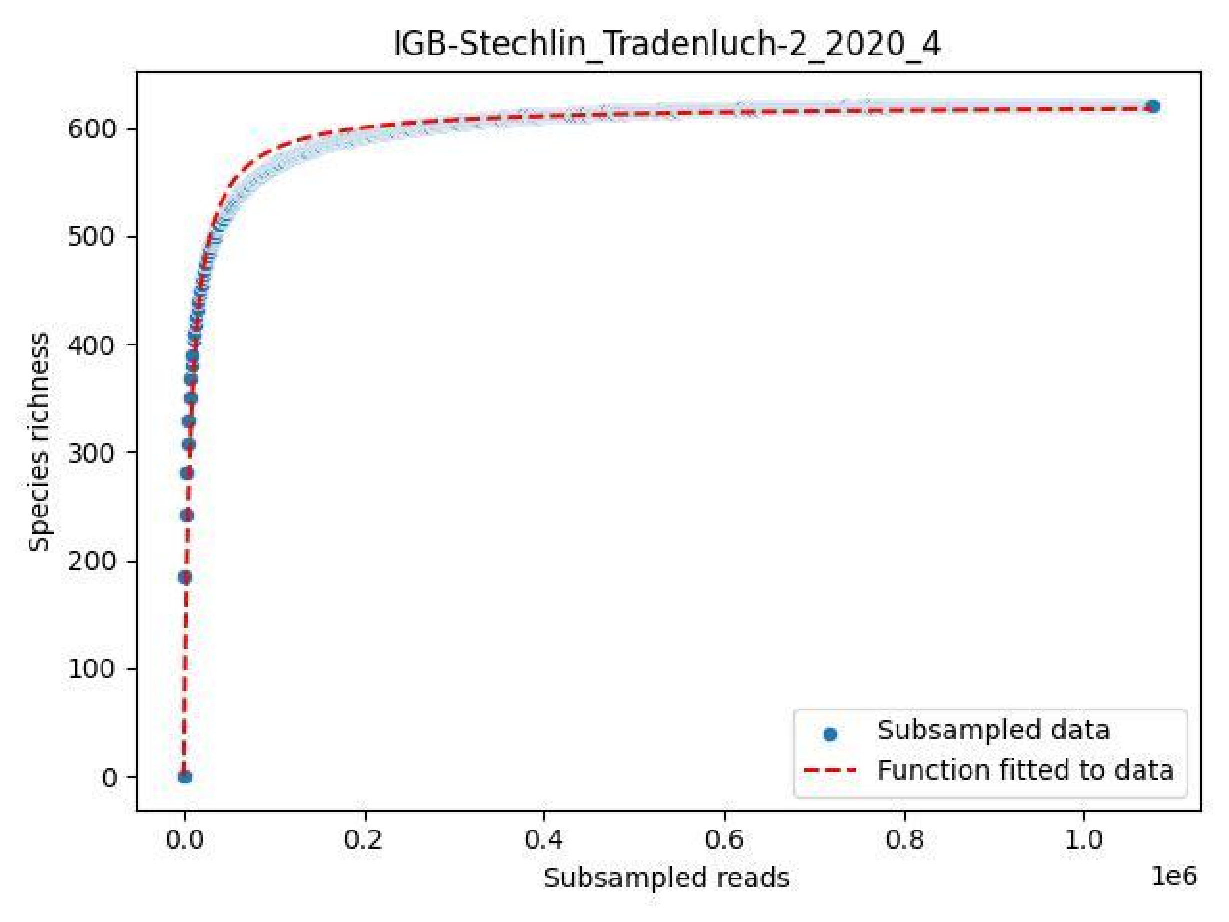

IGB-Stechlin_Tradenluch-2_2020_4
Species richness
Subsampled reads
1e6
0
100
200
300
400
500
600
0.0
0.2
0.4
0.6
0.8
1.0
Subsampled data
Function fitted to data

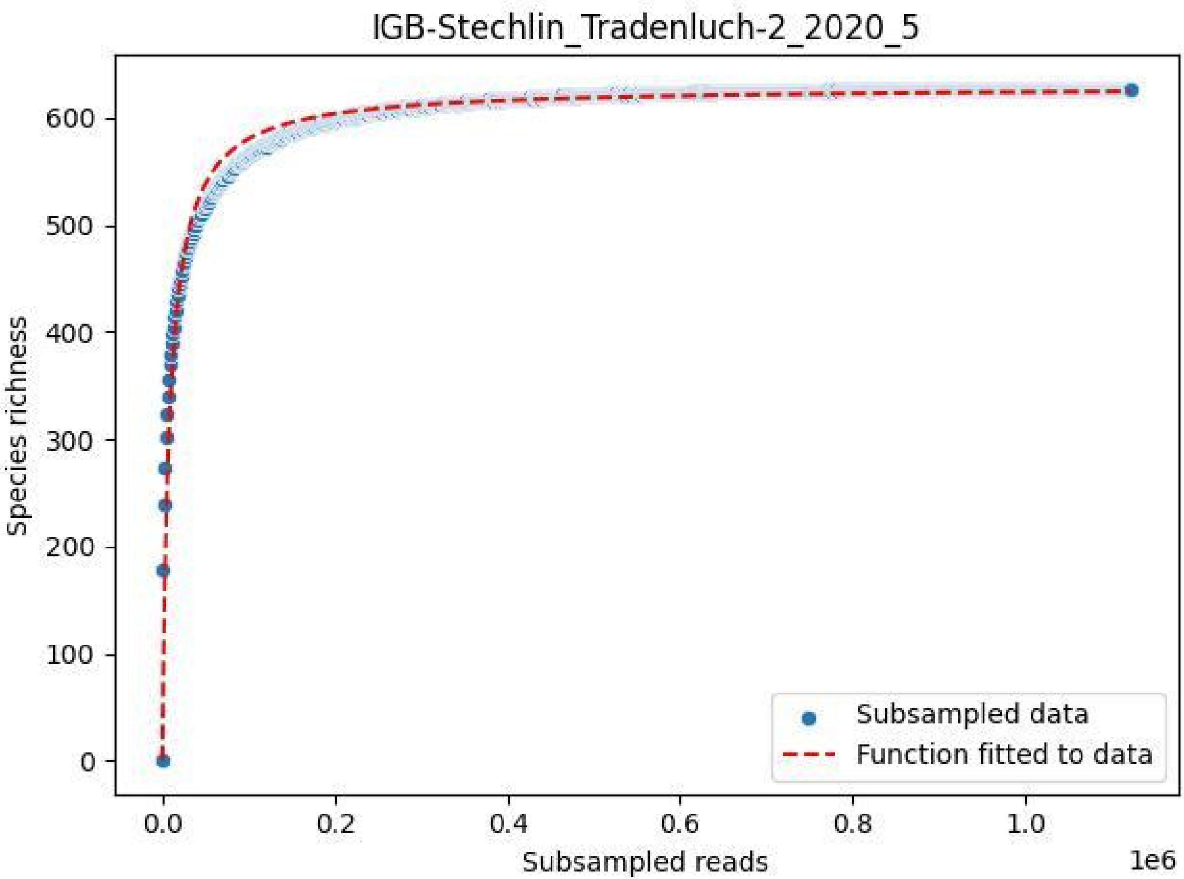
IGB-Stechlin_Tradenluch-2_2020_5
Species richness
Subsampled reads
1e6
0
100
200
300
400
500
600
0.0
0.2
0.4
0.6
0.8
1.0
Subsampled data
Function fitted to data

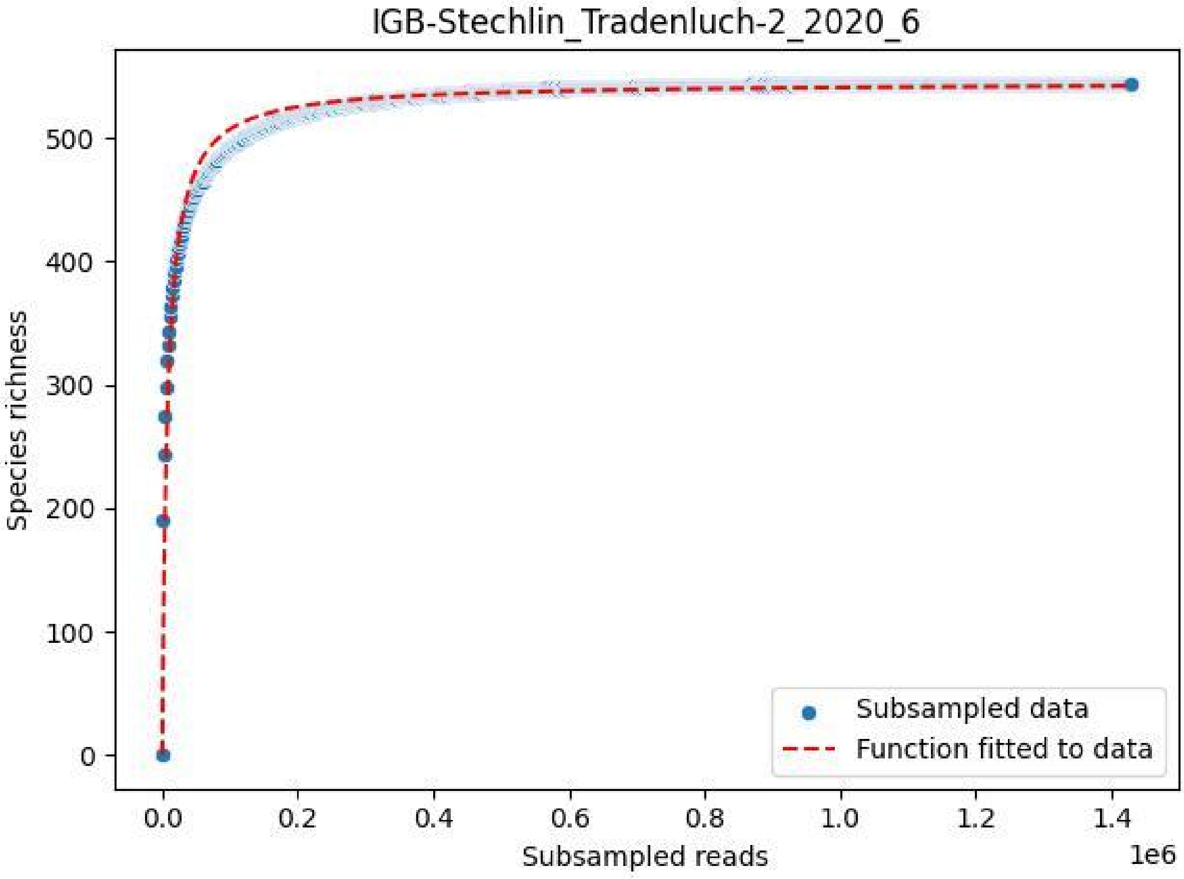
IGB-Stechlin_Tradenluch-2_2020_6
Species richness
Subsampled reads
1e6
0
100
200
300
400
500
0.0
0.2
0.4
0.6
0.8
1.0
1.2
1.4
Subsampled data
Function fitted to data

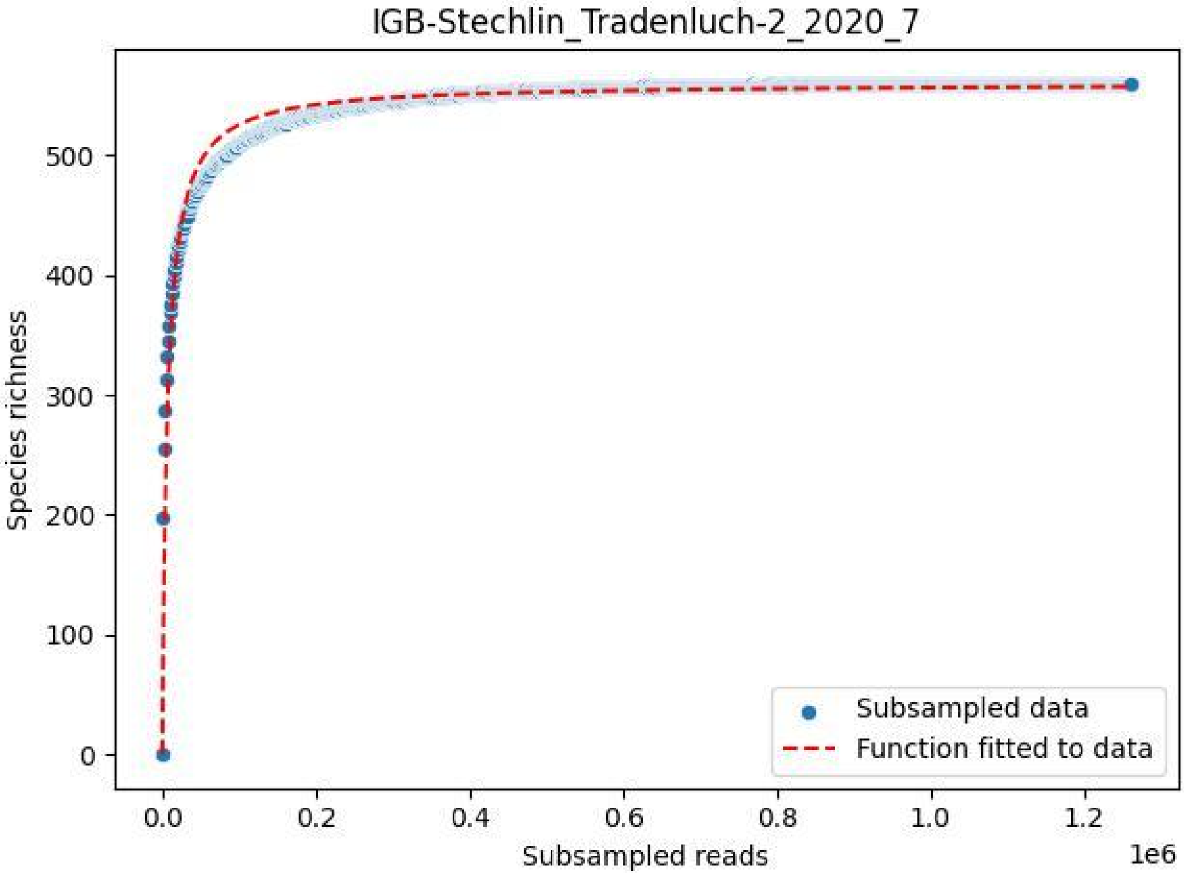
IGB-Stechlin_Tradenluch-2_2020_7
Species richness
Subsampled reads
1e6
0
100
200
300
400
500
0.0
0.2
0.4
0.6
0.8
1.0
1.2
Subsampled data
Function fitted to data

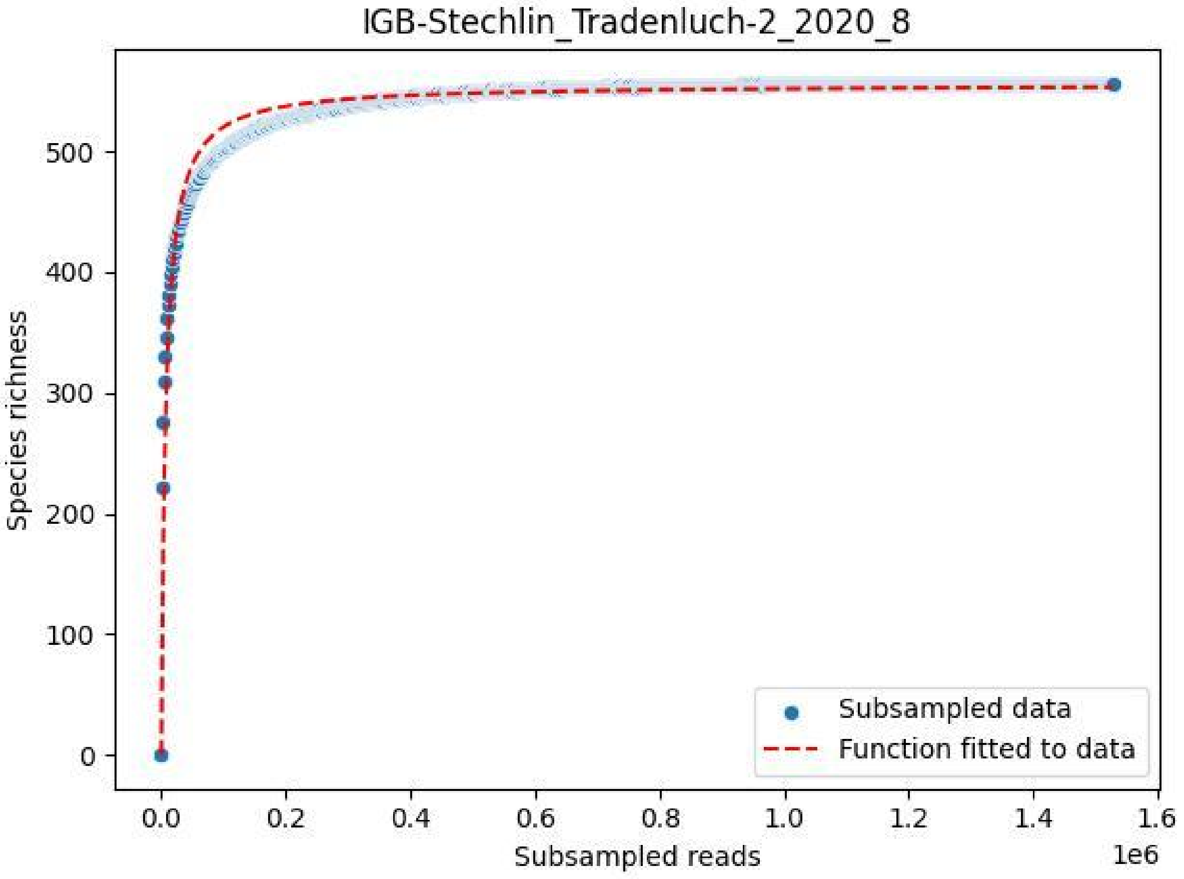
IGB-Stechlin_Tradenluch-2_2020_8
Species richness
Subsampled reads
1e6
Subsampled data
Function fitted to data
0
100
200
300
400
500
0.0
0.2
0.4
0.6
0.8
1.0
1.2
1.4
1.6

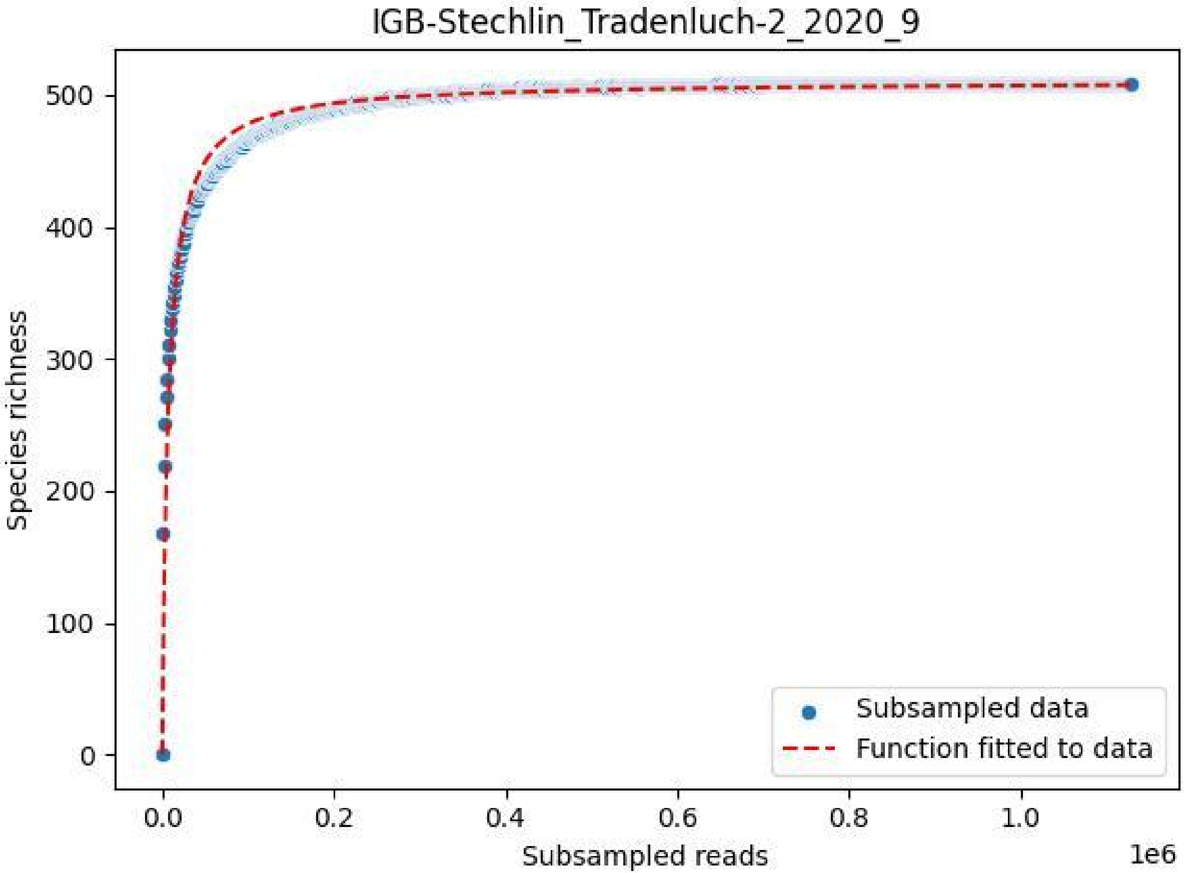

IGB-Stechlin_Tradenluch-2_2020_9
Species richness
500
400
300
200
100
0
0.0
0.2
0.4
0.6
0.8
1.0
1e6
Subsampled reads
Subsampled data
Function fitted to data

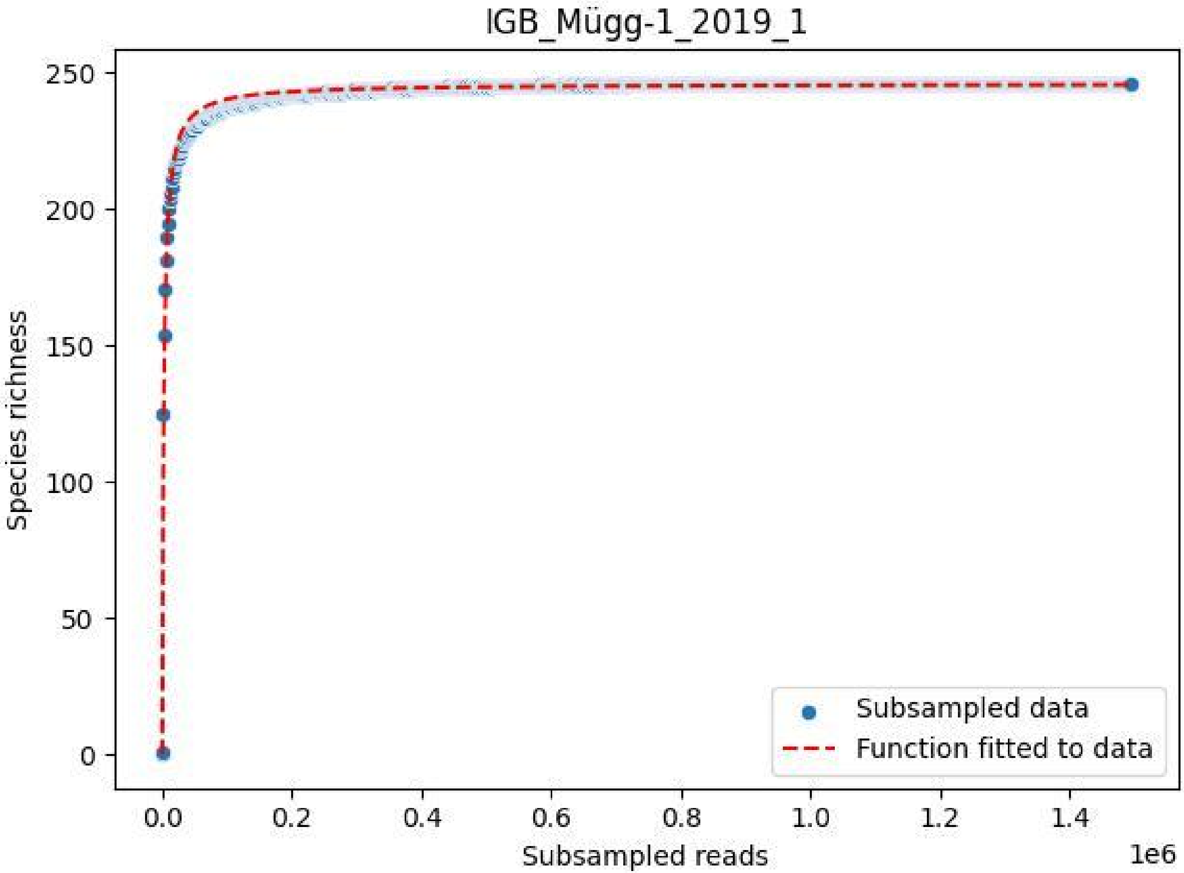
IGB_Mügg-1_2019_1
Species richness
Subsampled reads
1e6
0
50
100
150
200
250
0.0
0.2
0.4
0.6
0.8
1.0
1.2
1.4
Subsampled data
Function fitted to data

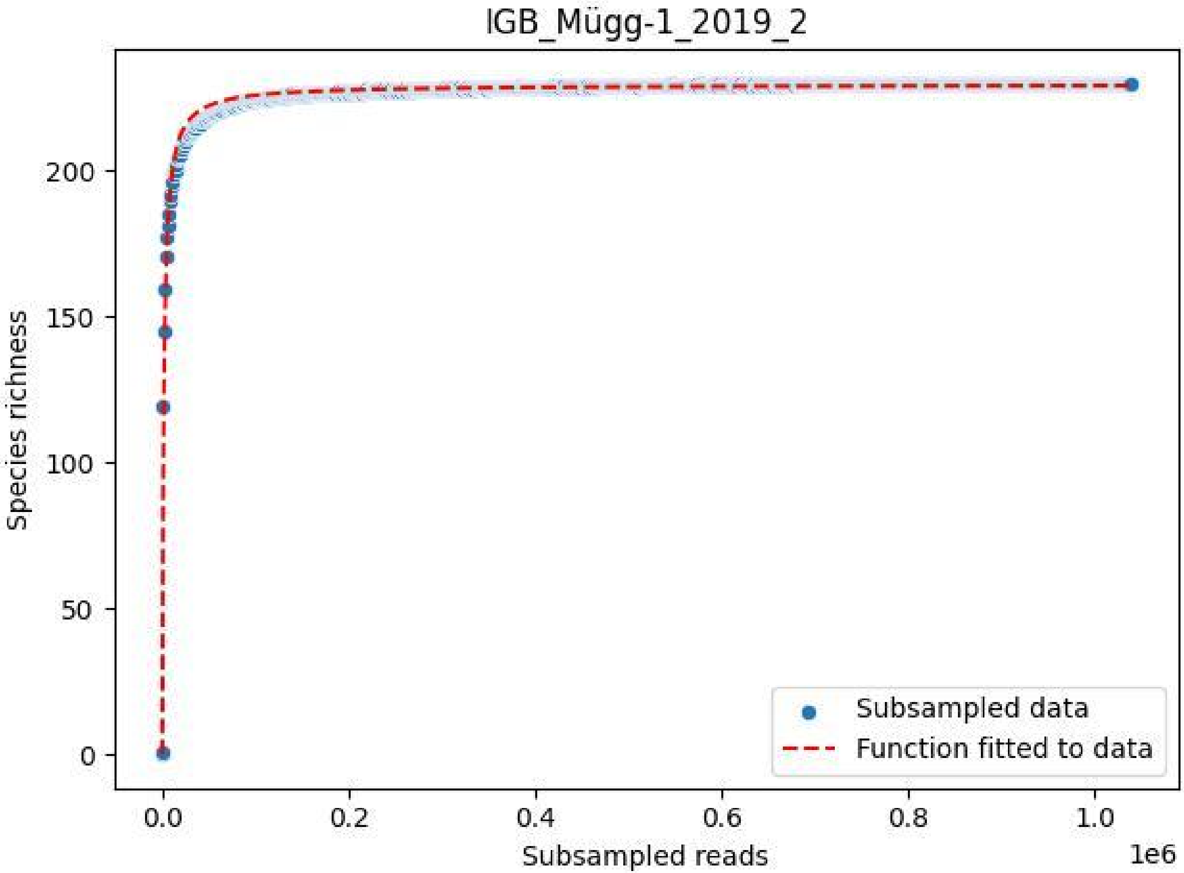
IGB_Mügg-1_2019_2
Species richness
0
50
100
150
200
0.0
0.2
0.4
0.6
0.8
1.0
1e6
Subsampled reads
Subsampled data
Function fitted to data

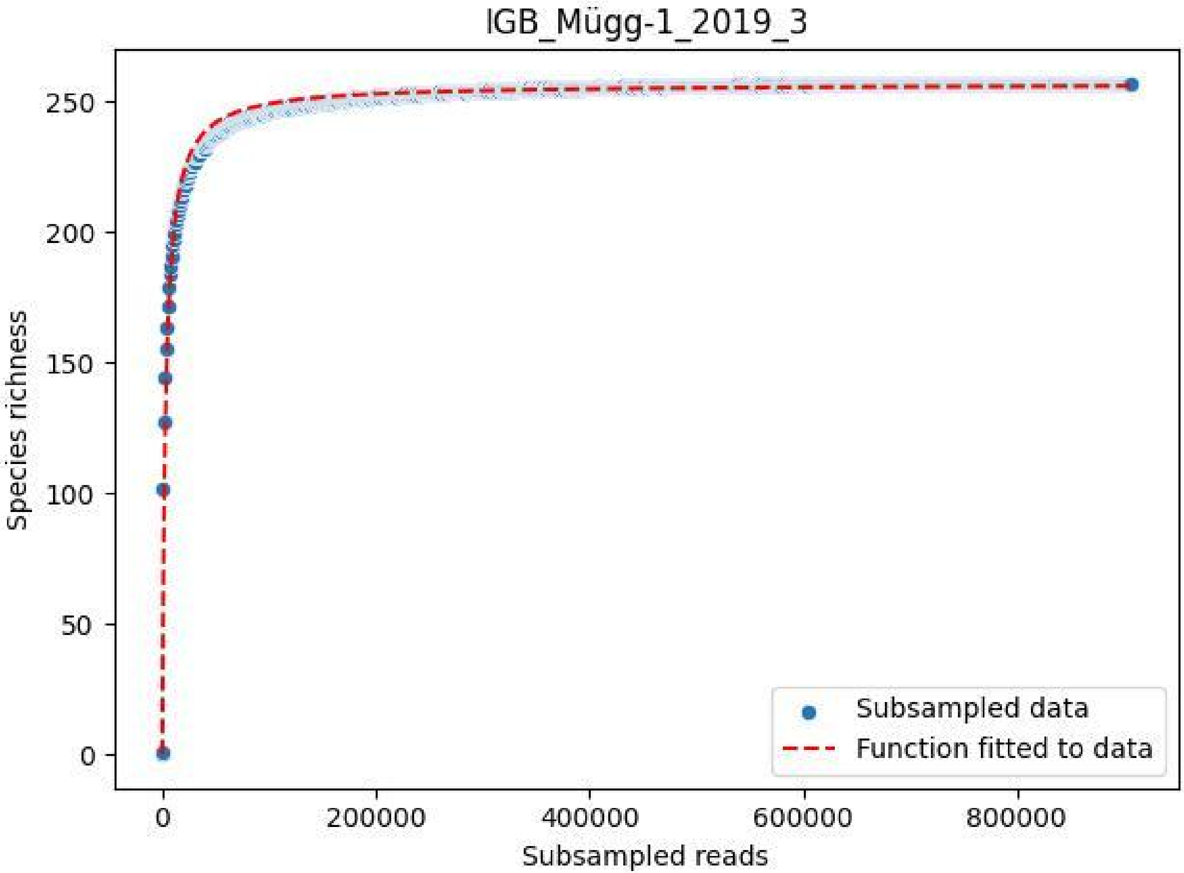

IGB_Mügg-1_2019_3
Species richness
0
50
100
150
200
250
Subsampled reads
0
200000
400000
600000
800000
Subsampled data
Function fitted to data

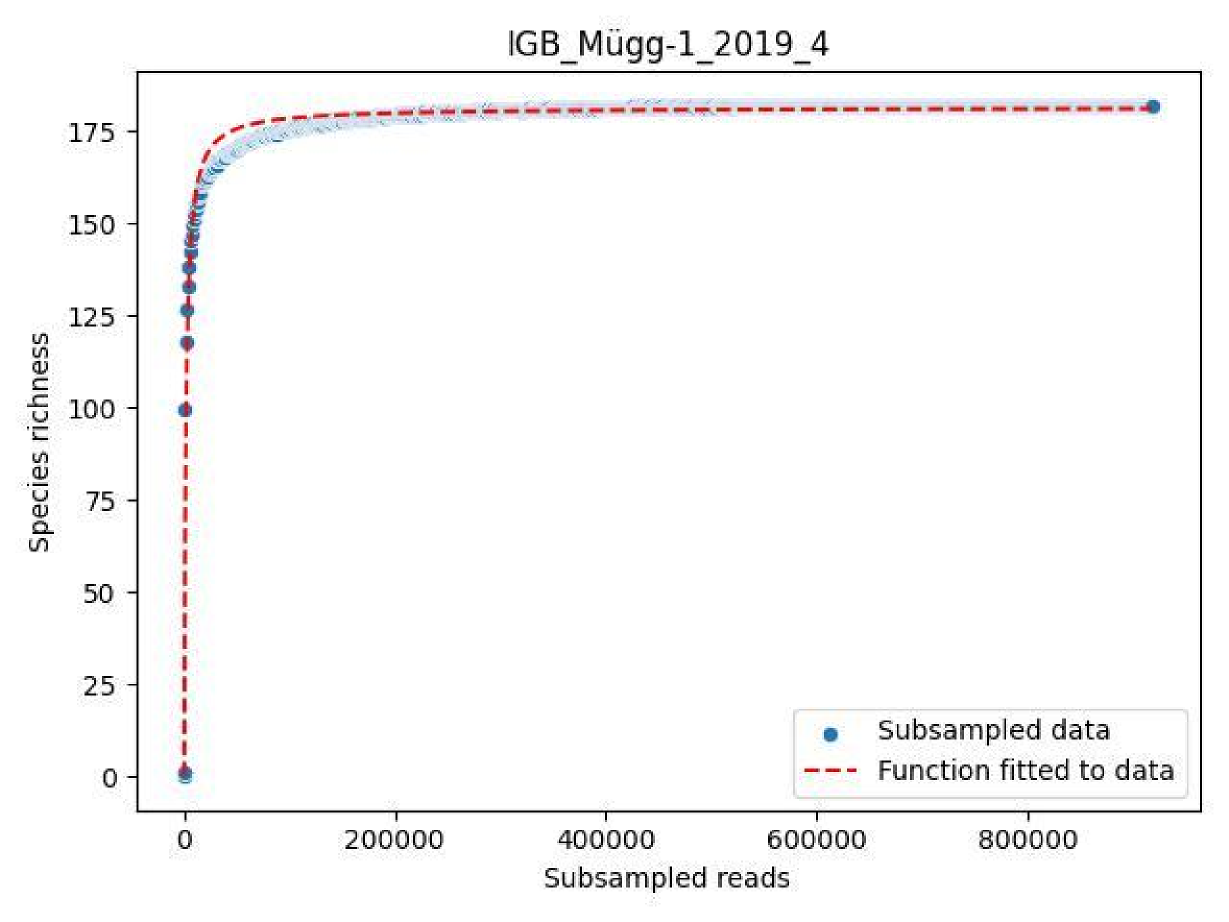
IGB_Mügg-1_2019_4
Species richness
0
25
50
75
100
125
150
175
0
200000
400000
600000
800000
Subsampled reads
Subsampled data
Function fitted to data

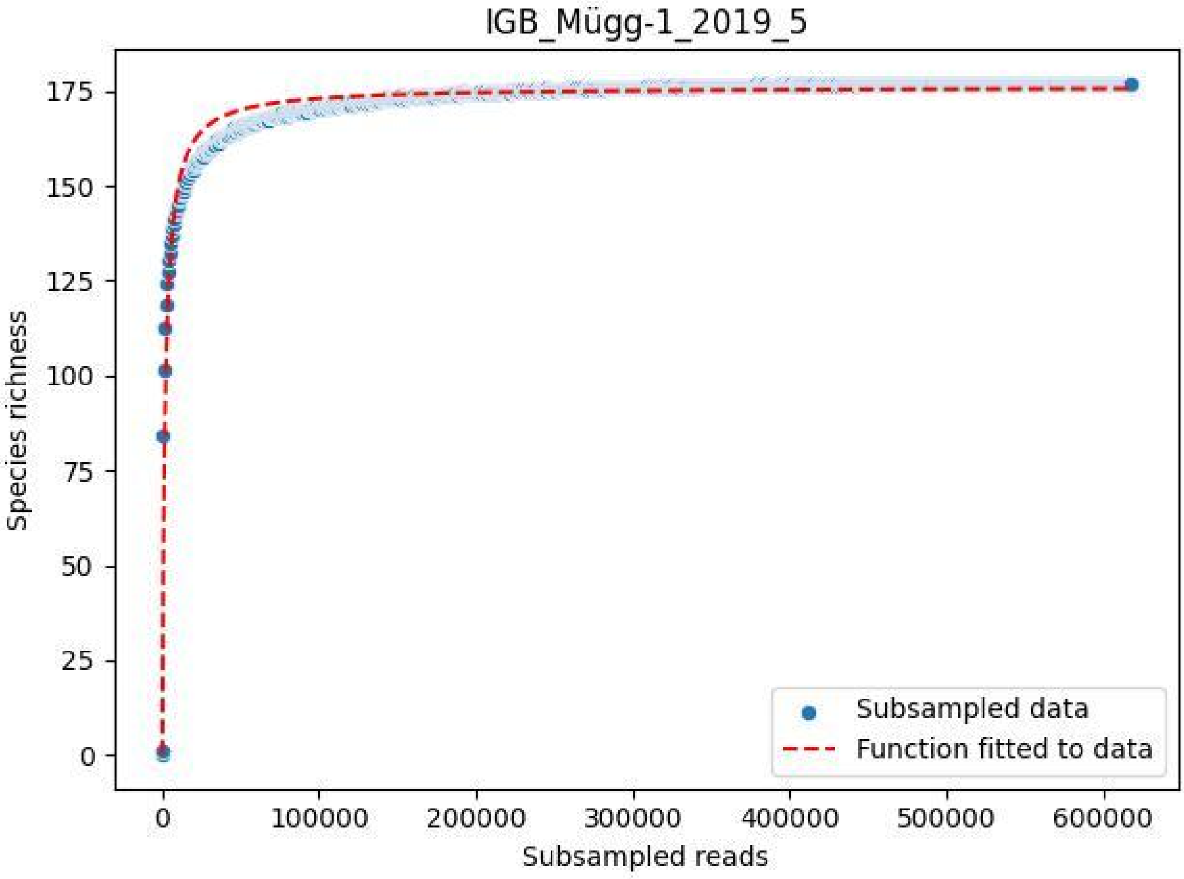

IGB_Mügg-1_2019_5
Species richness
0
25
50
75
100
125
150
175
Subsampled reads
0
100000
200000
300000
400000
500000
600000
Subsampled data
Function fitted to data

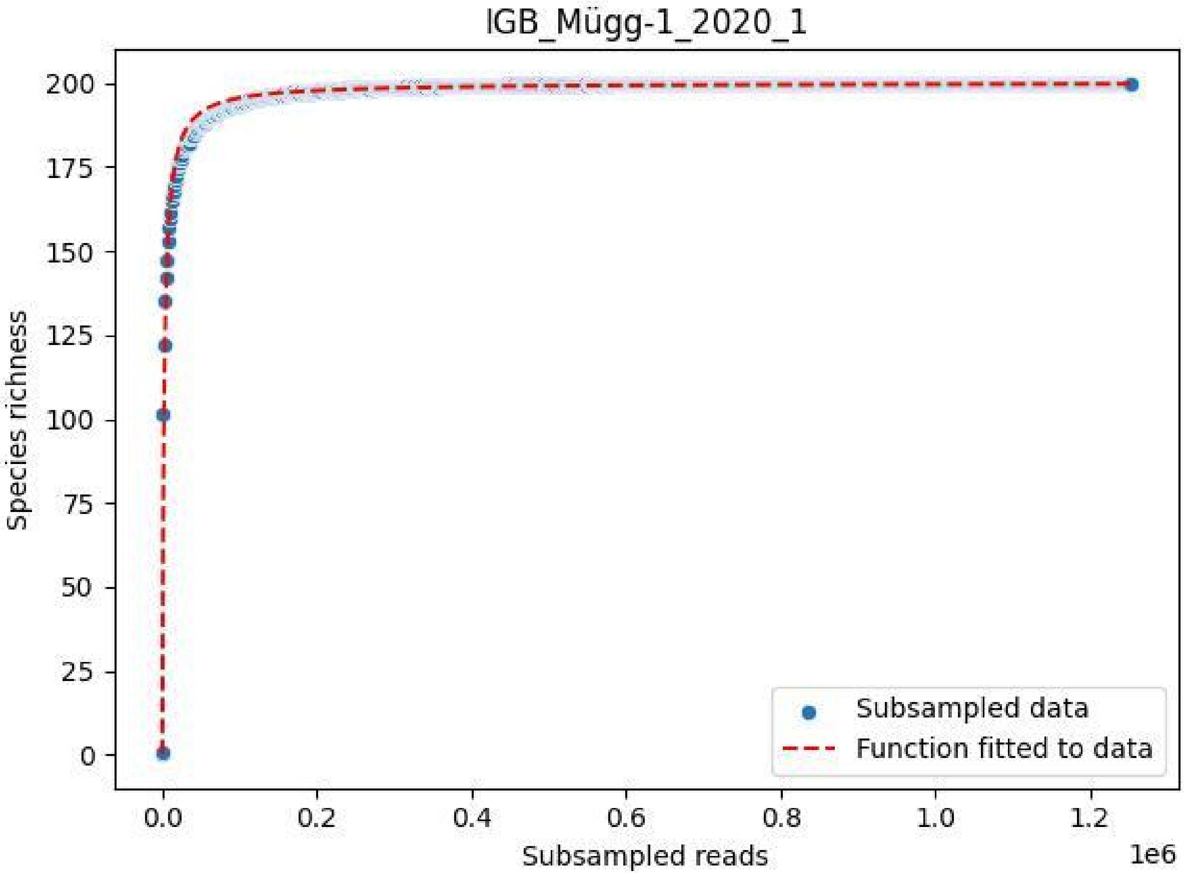
IGB_Mügg-1_2020_1
Species richness
Subsampled reads
1e6
Subsampled data
Function fitted to data
0
25
50
75
100
125
150
175
200
0.0
0.2
0.4
0.6
0.8
1.0
1.2

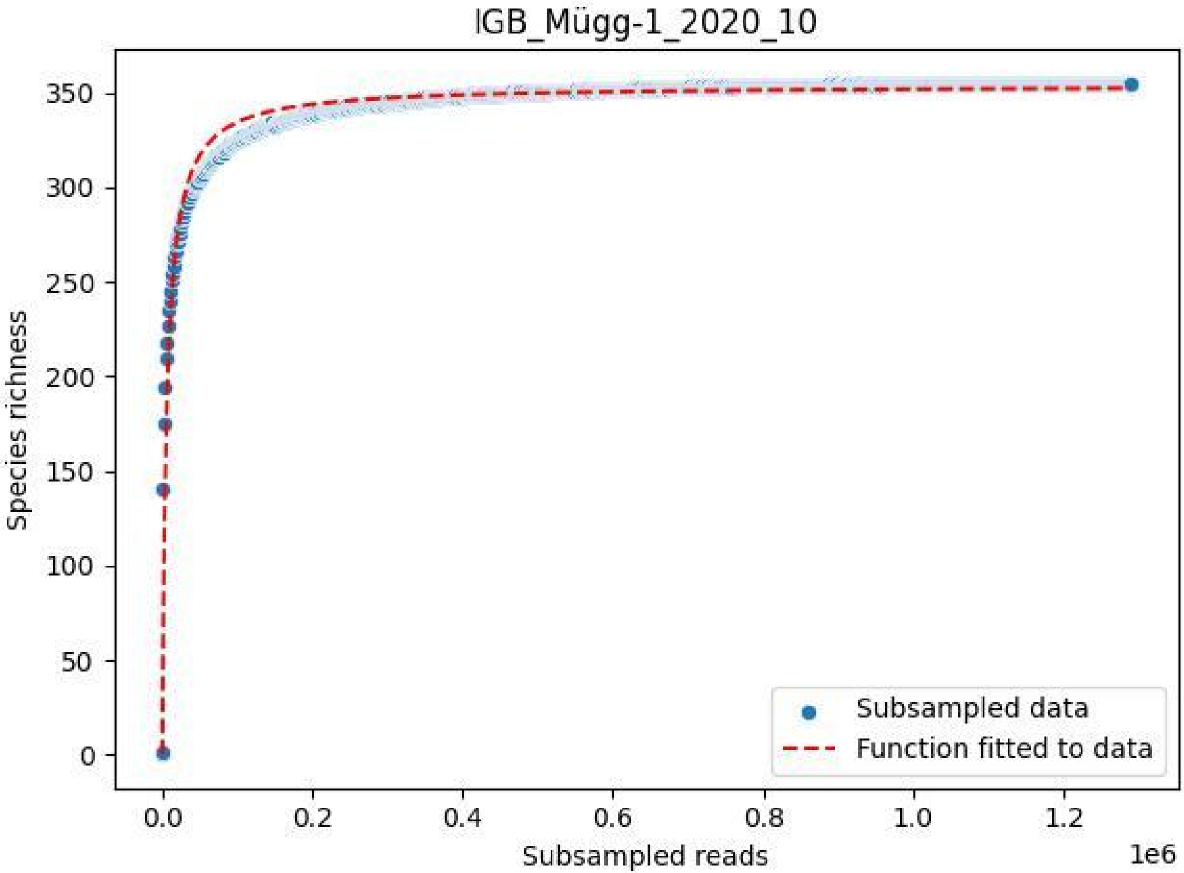
IGB_Mügg-1_2020_10
Species richness
Subsampled reads
1e6
0.0
0.2
0.4
0.6
0.8
1.0
1.2
0
50
100
150
200
250
300
350
Subsampled data
Function fitted to data

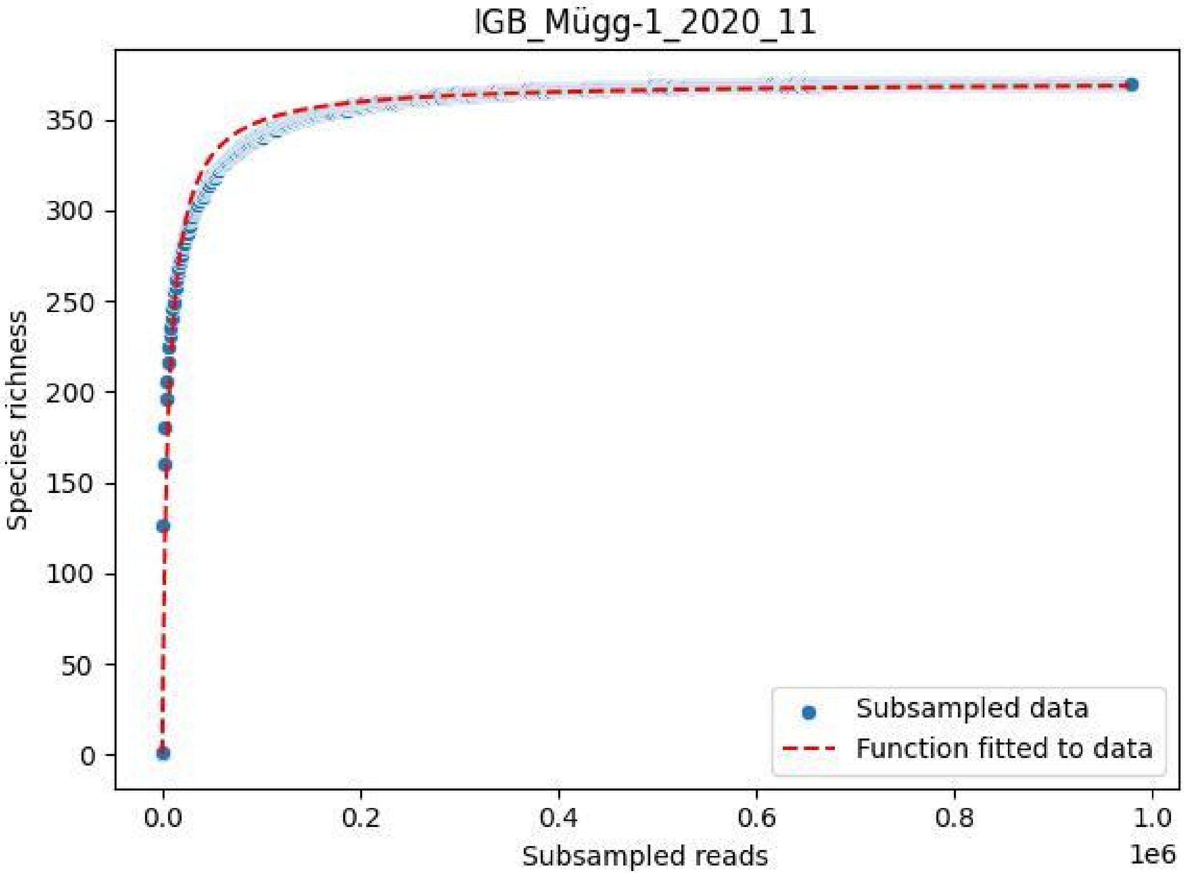

IGB_Mügg-1_2020_11
Species richness
0
50
100
150
200
250
300
350
Subsampled reads
0.0
0.2
0.4
0.6
0.8
1.0
1e6
Subsampled data
Function fitted to data

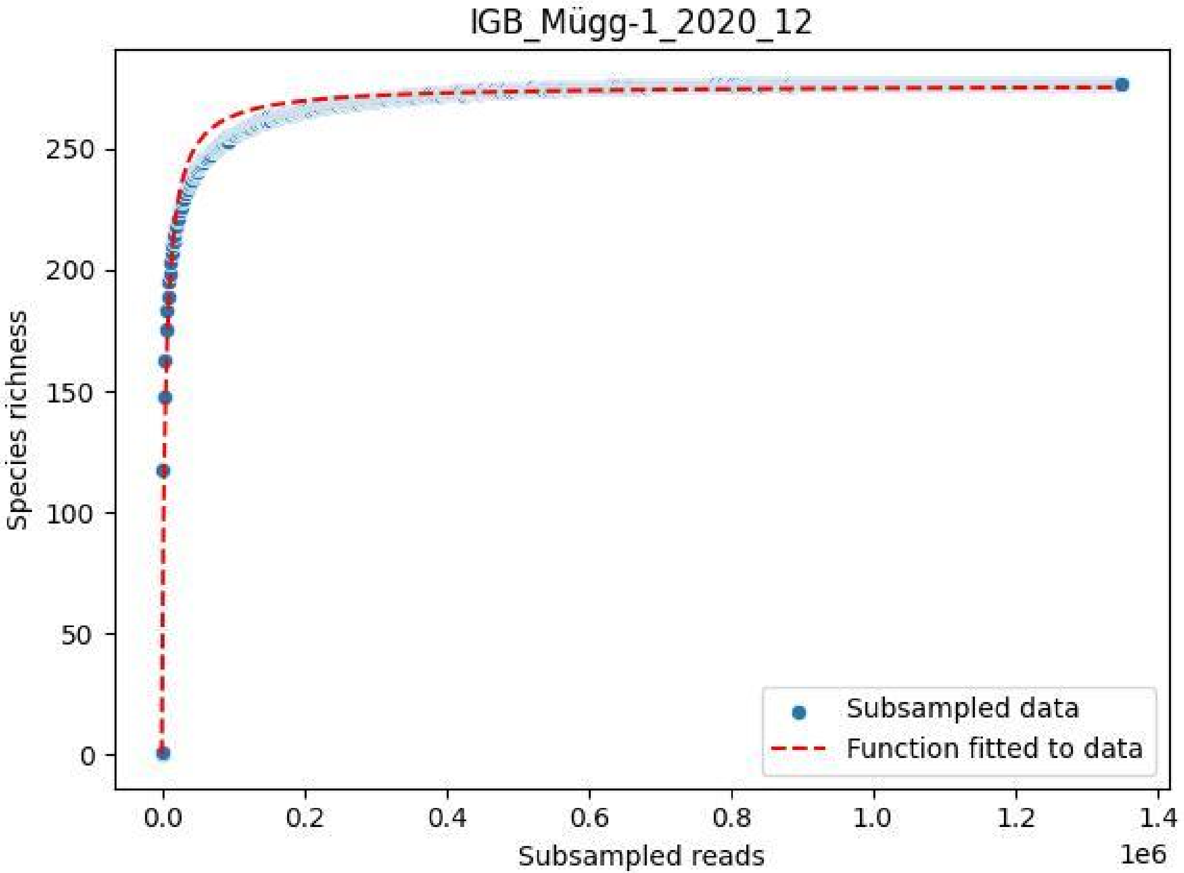
IGB_Mügg-1_2020_12
Species richness
Subsampled reads
1e6
Subsampled data
Function fitted to data
0
50
100
150
200
250
0.0
0.2
0.4
0.6
0.8
1.0
1.2
1.4

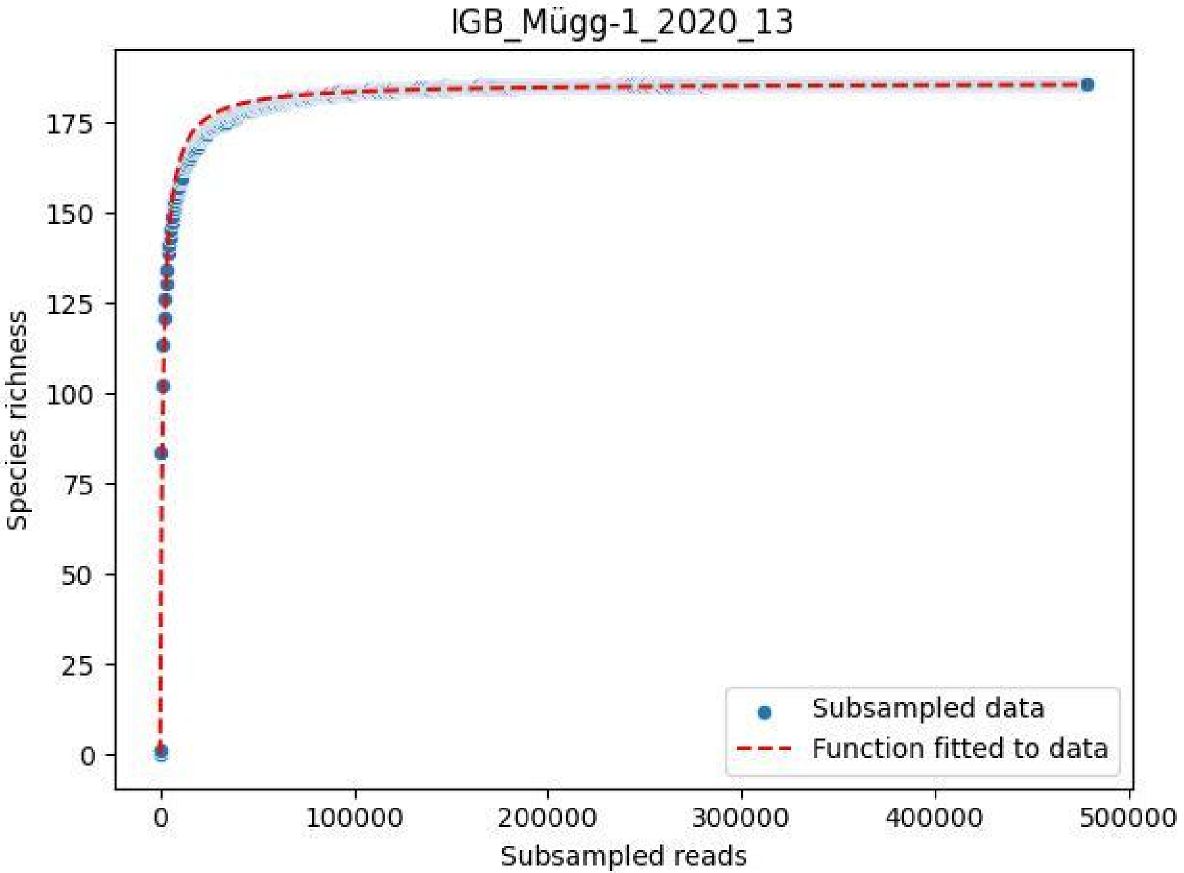

IGB_Mügg-1_2020_13
Species richness
175
150
125
100
75
50
25
0
Subsampled reads
0
100000
200000
300000
400000
500000
Subsampled data
Function fitted to data

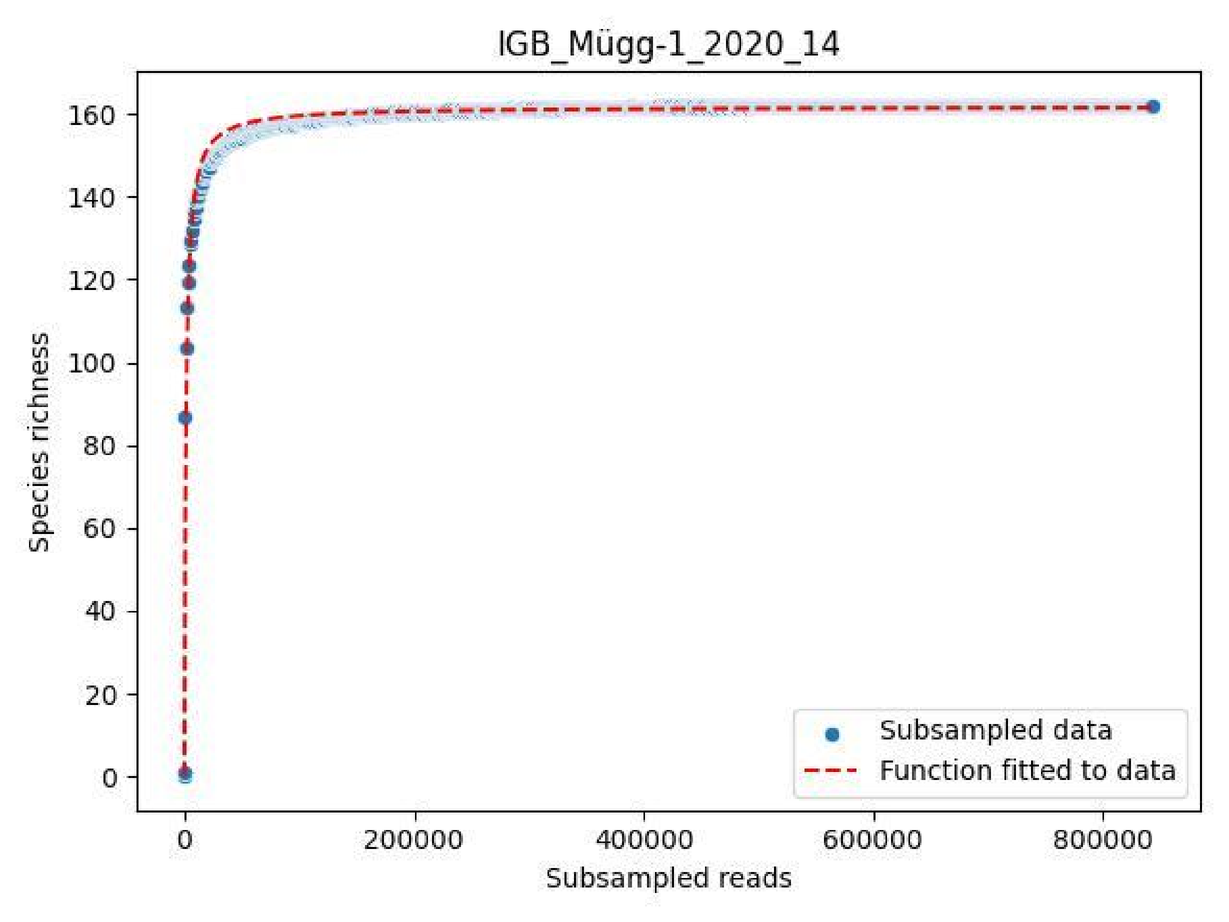
IGB_Mügg-1_2020_14
Species richness
Subsampled reads
0
20
40
60
80
100
120
140
160
0
200000
400000
600000
800000
Subsampled data
Function fitted to data

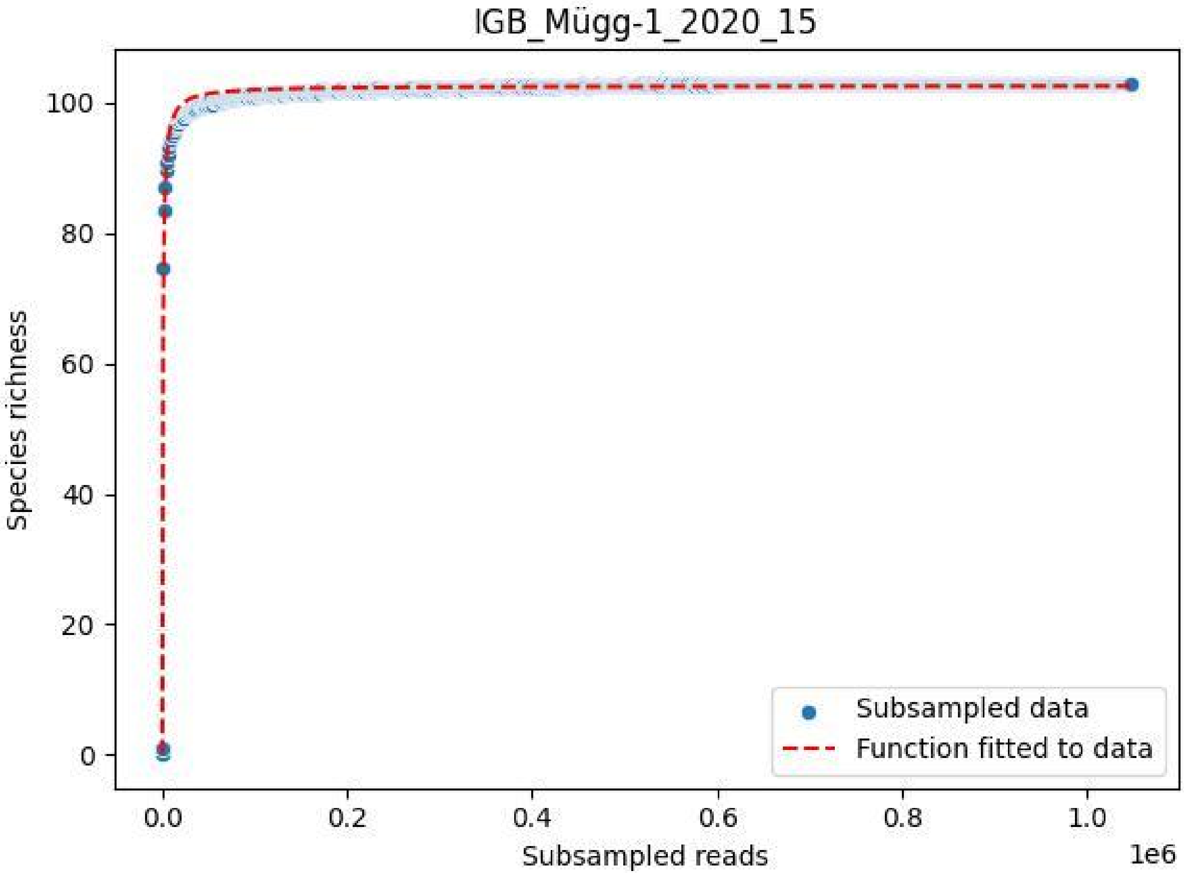
IGB_Mügg-1_2020_15
Species richness
100
80
60
40
20
0
0.0
0.2
0.4
0.6
0.8
1.0
1e6
Subsampled reads
Subsampled data
Function fitted to data

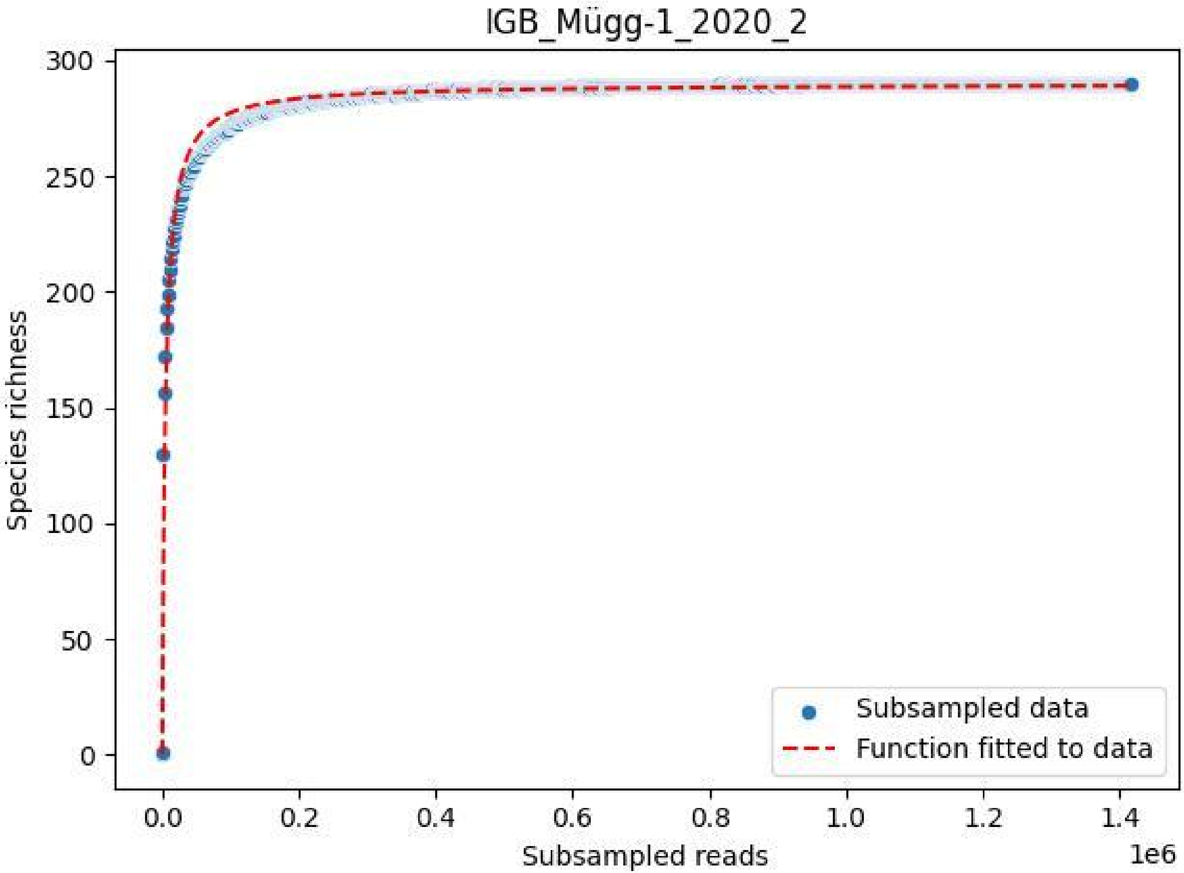
IGB_Mügg-1_2020_2
Species richness
Subsampled reads
1e6
Subsampled data
Function fitted to data
0
50
100
150
200
250
300
0.0
0.2
0.4
0.6
0.8
1.0
1.2
1.4

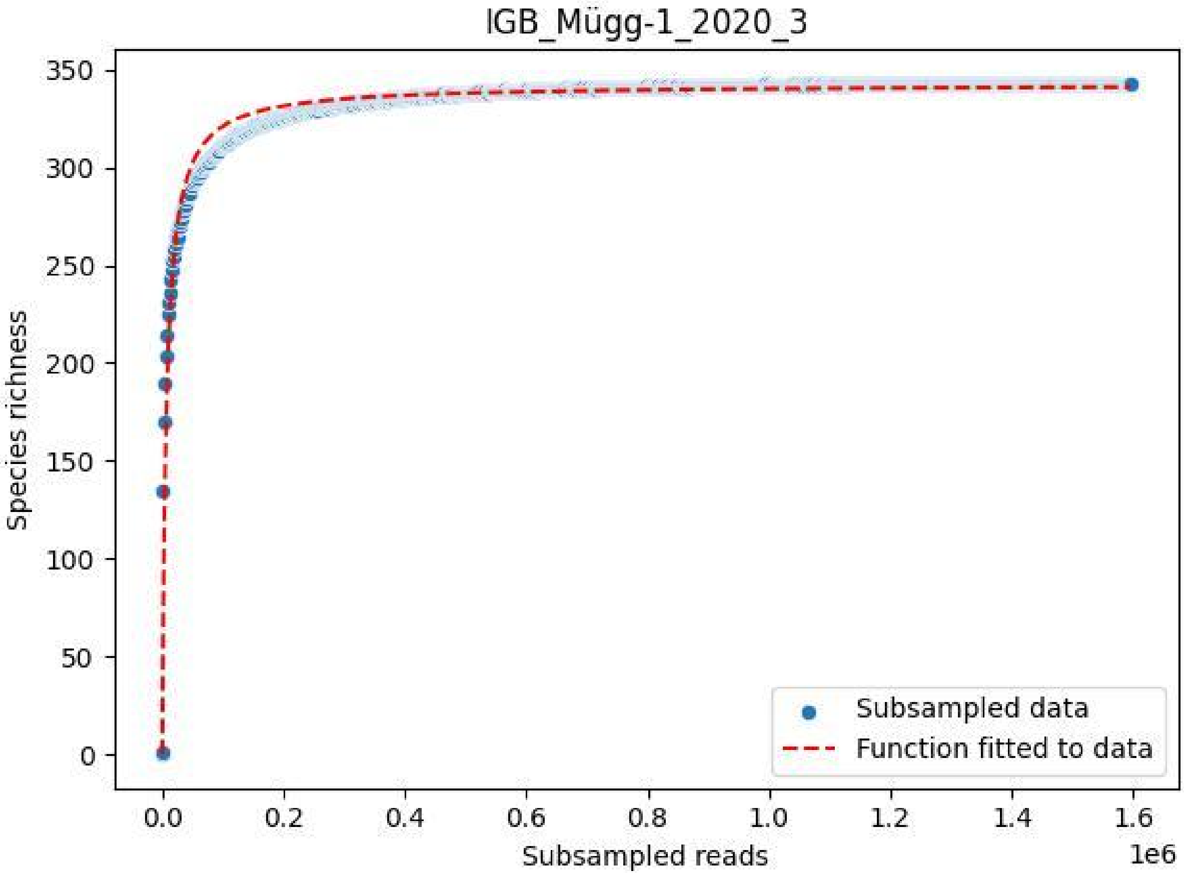
IGB_Mügg-1_2020_3
Species richness
Subsampled reads
1e6
0
50
100
150
200
250
300
350
0.0
0.2
0.4
0.6
0.8
1.0
1.2
1.4
1.6
Subsampled data
Function fitted to data

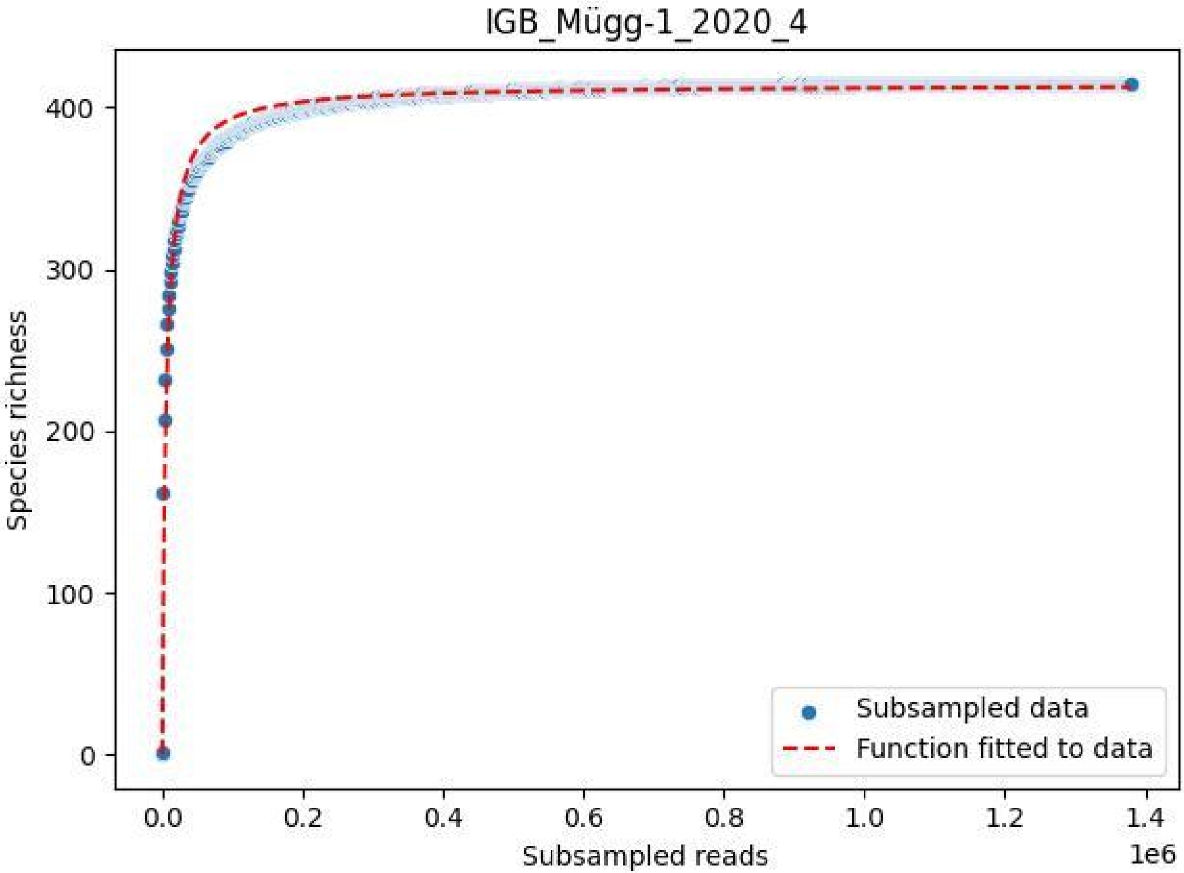

IGB_Mügg-1_2020_4
Species richness
400
300
200
100
0
0.0
0.2
0.4
0.6
0.8
1.0
1.2
1.4
1e6
Subsampled reads
Subsampled data
Function fitted to data

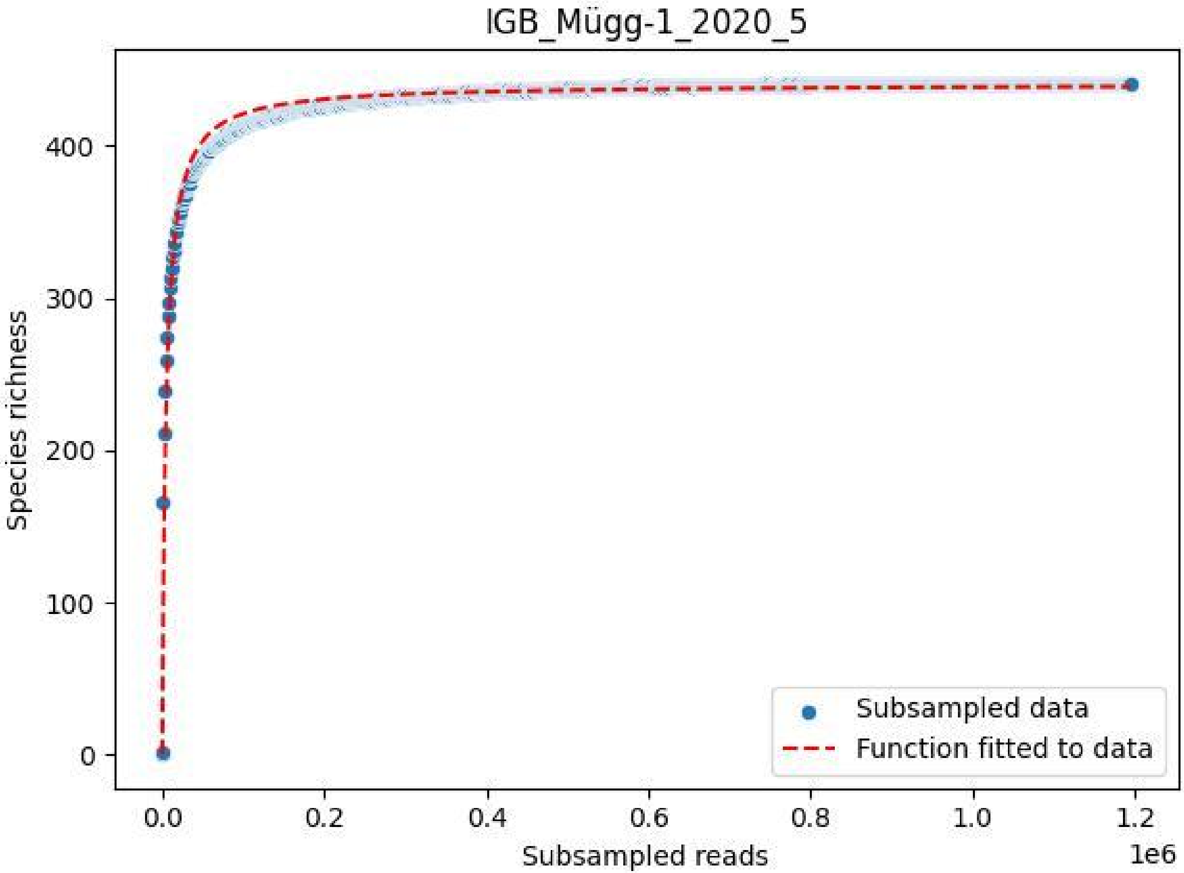
IGB_Mügg-1_2020_5
Species richness
400
300
200
100
0
0.0
0.2
0.4
0.6
0.8
1.0
1.2
Subsampled reads
1e6
Subsampled data
Function fitted to data

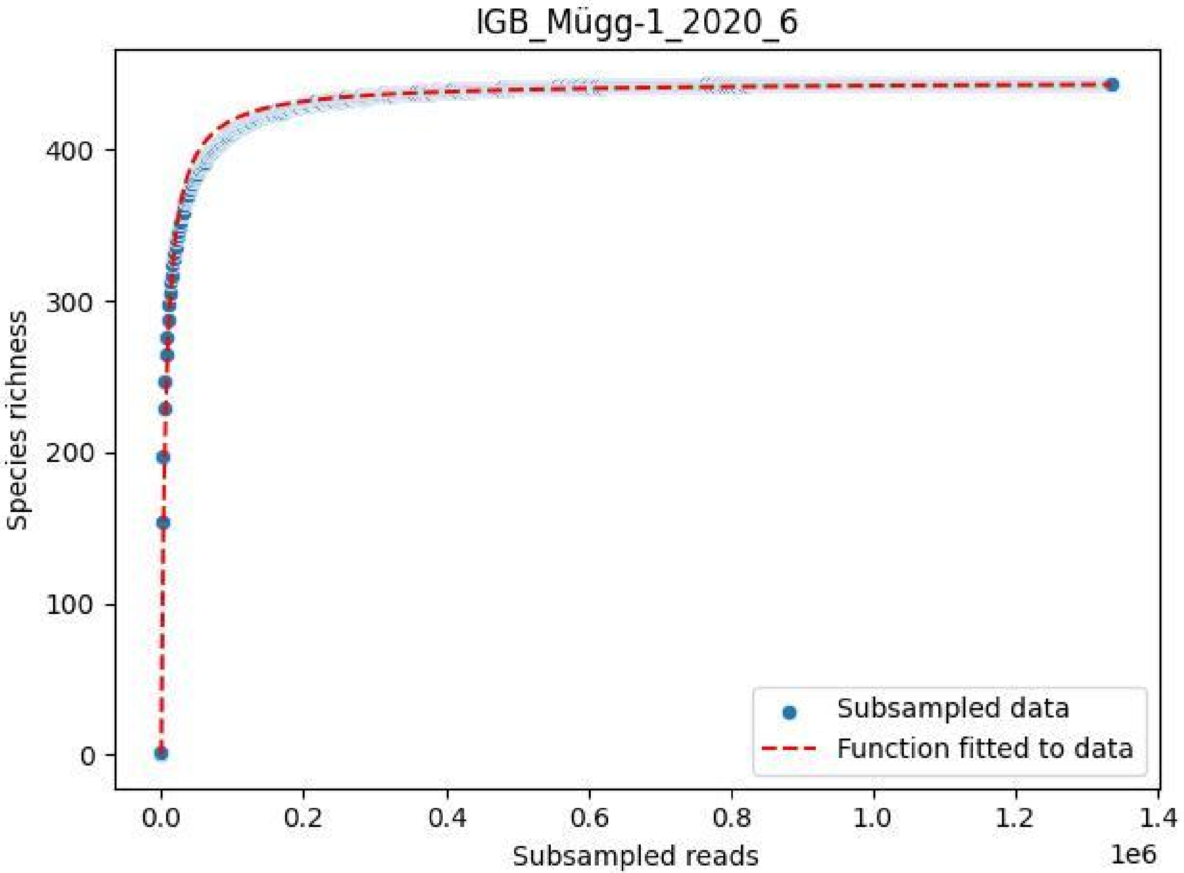

IGB_Mügg-1_2020_6
Species richness
Subsampled reads
1e6
0
100
200
300
400
0.0
0.2
0.4
0.6
0.8
1.0
1.2
1.4
Subsampled data
Function fitted to data

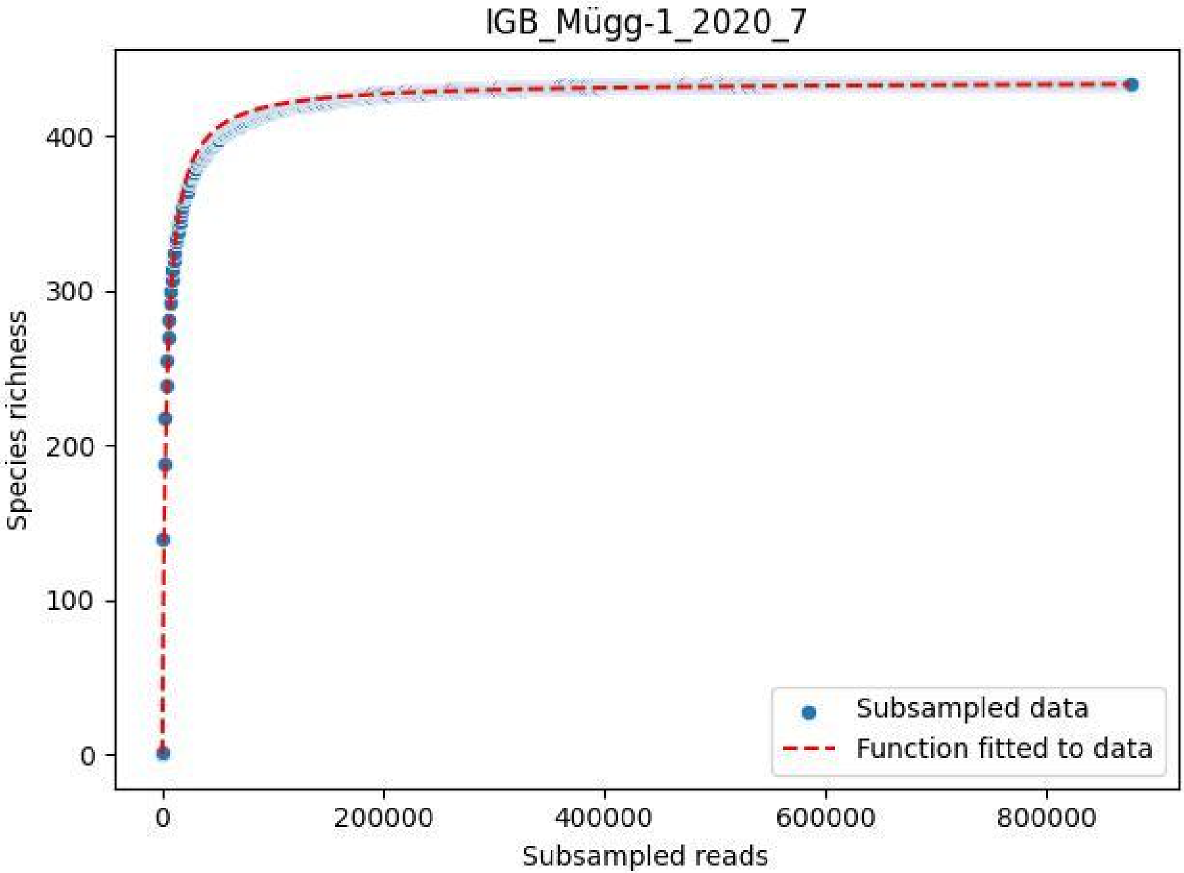
IGB_Mügg-1_2020_7
Species richness
Subsampled reads
0
100
200
300
400
0
200000
400000
600000
800000
Subsampled data
Function fitted to data

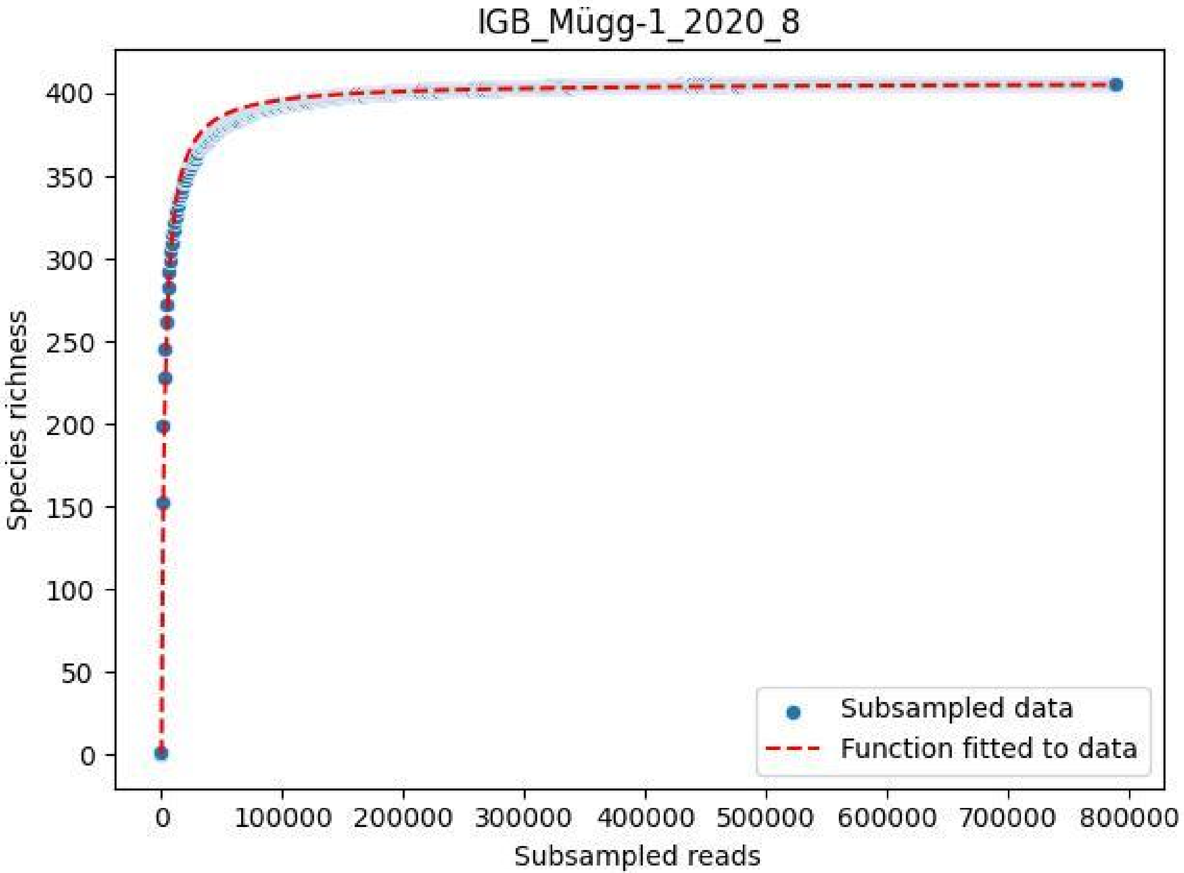
IGB_Mügg-1_2020_8
Species richness
Subsampled reads
Subsampled data
Function fitted to data
0
50
100
150
200
250
300
350
400
0
100000
200000
300000
400000
500000
600000
700000
800000

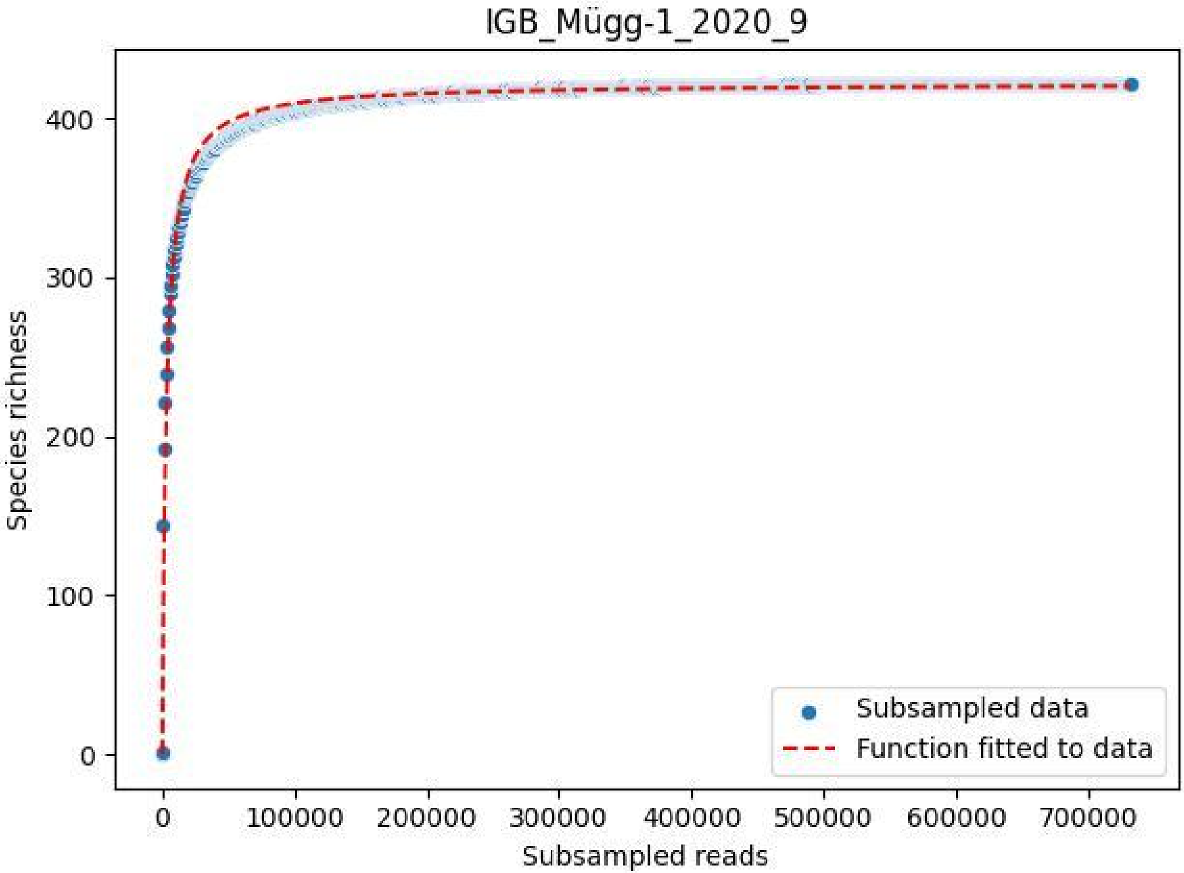
IGB_Mügg-1_2020_9
Species richness
Subsampled reads
0
100
200
300
400
0
100000
200000
300000
400000
500000
600000
700000
Subsampled data
Function fitted to data

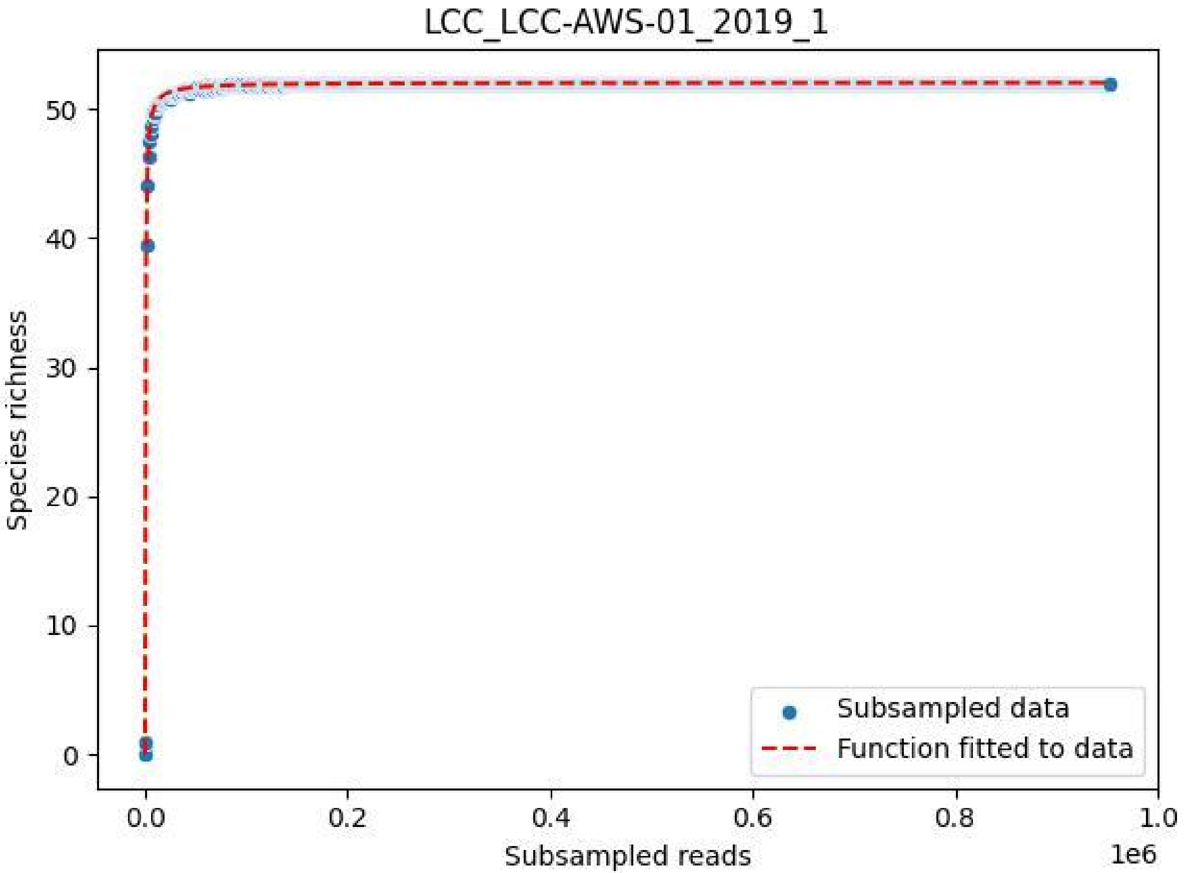

LCC_LCC-AWS-01_2019_1
Species richness
Subsampled reads
1e6
0
10
20
30
40
50
0.0
0.2
0.4
0.6
0.8
1.0
Subsampled data
Function fitted to data

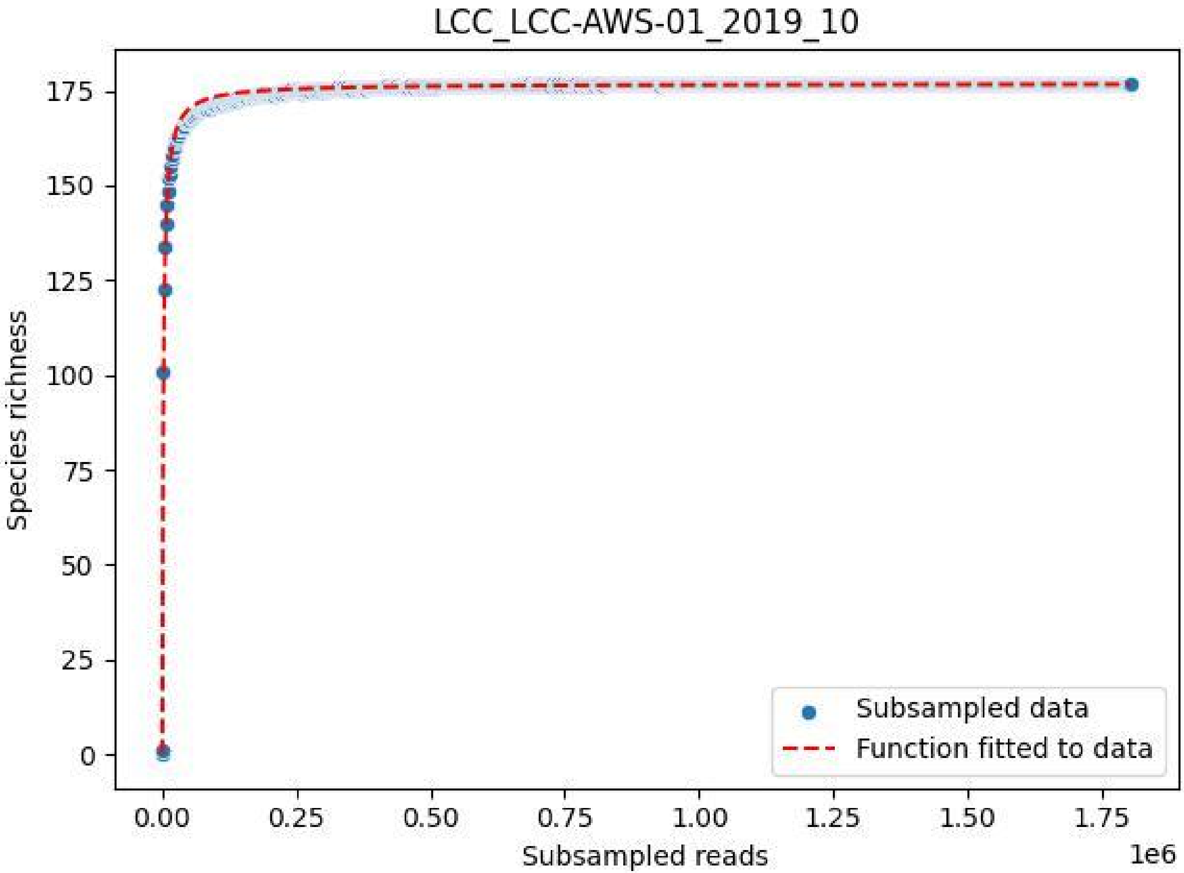
LCC_LCC-AWS-01_2019_10
Species richness
Subsampled reads
1e6
Subsampled data
Function fitted to data
0
25
50
75
100
125
150
175
0.00
0.25
0.50
0.75
1.00
1.25
1.50
1.75

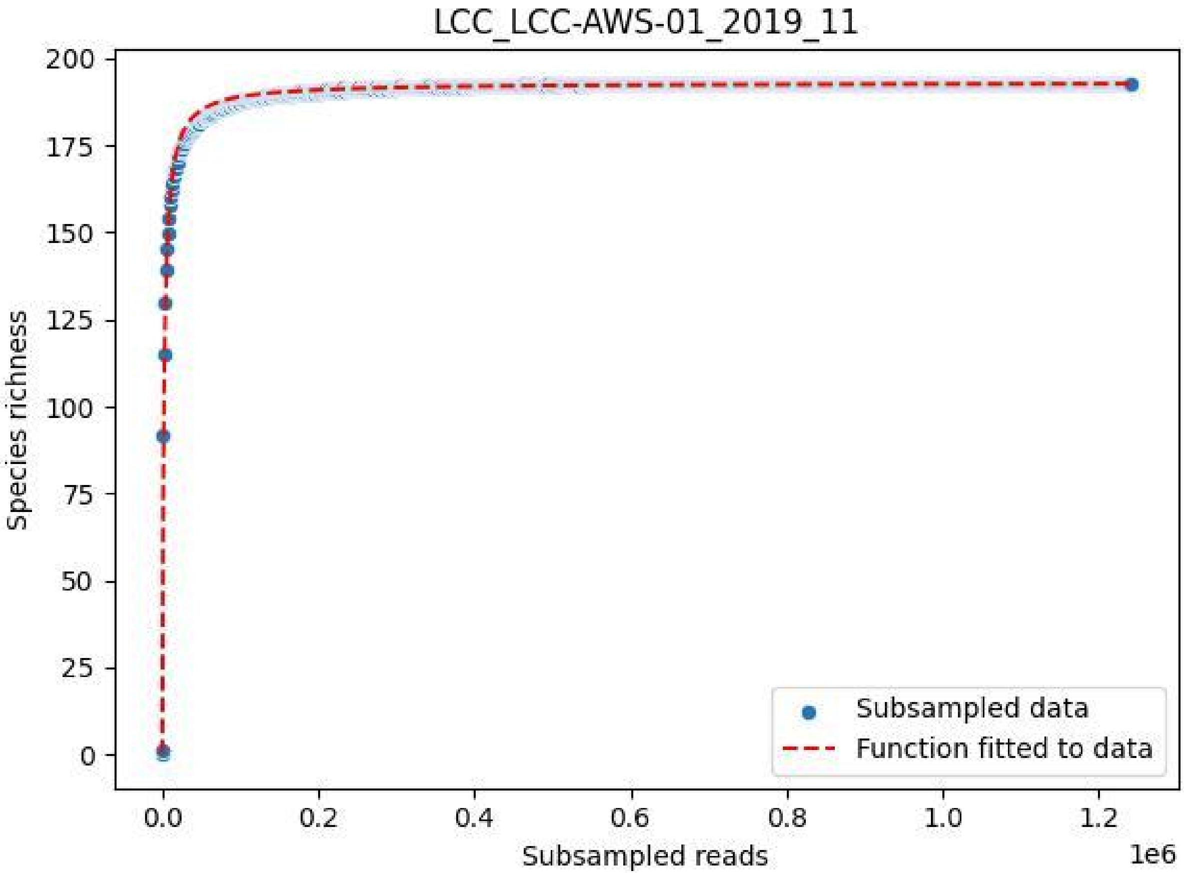
LCC_LCC-AWS-01_2019_11
Species richness
Subsampled reads
1e6
0
25
50
75
100
125
150
175
200
0.0
0.2
0.4
0.6
0.8
1.0
1.2
Subsampled data
Function fitted to data

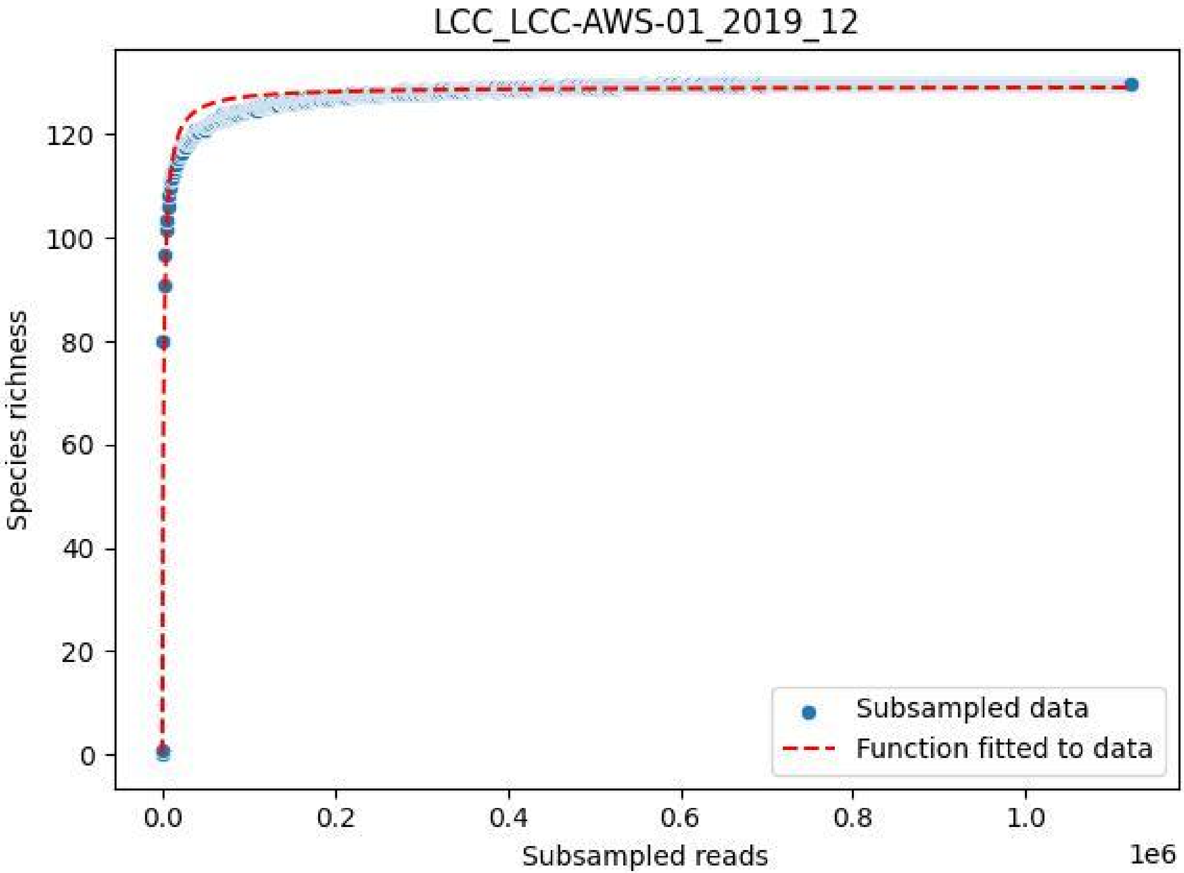

LCC_LCC-AWS-01_2019_12
Species richness
Subsampled reads
1e6
0.0
0.2
0.4
0.6
0.8
1.0
0
20
40
60
80
100
120
Subsampled data
Function fitted to data

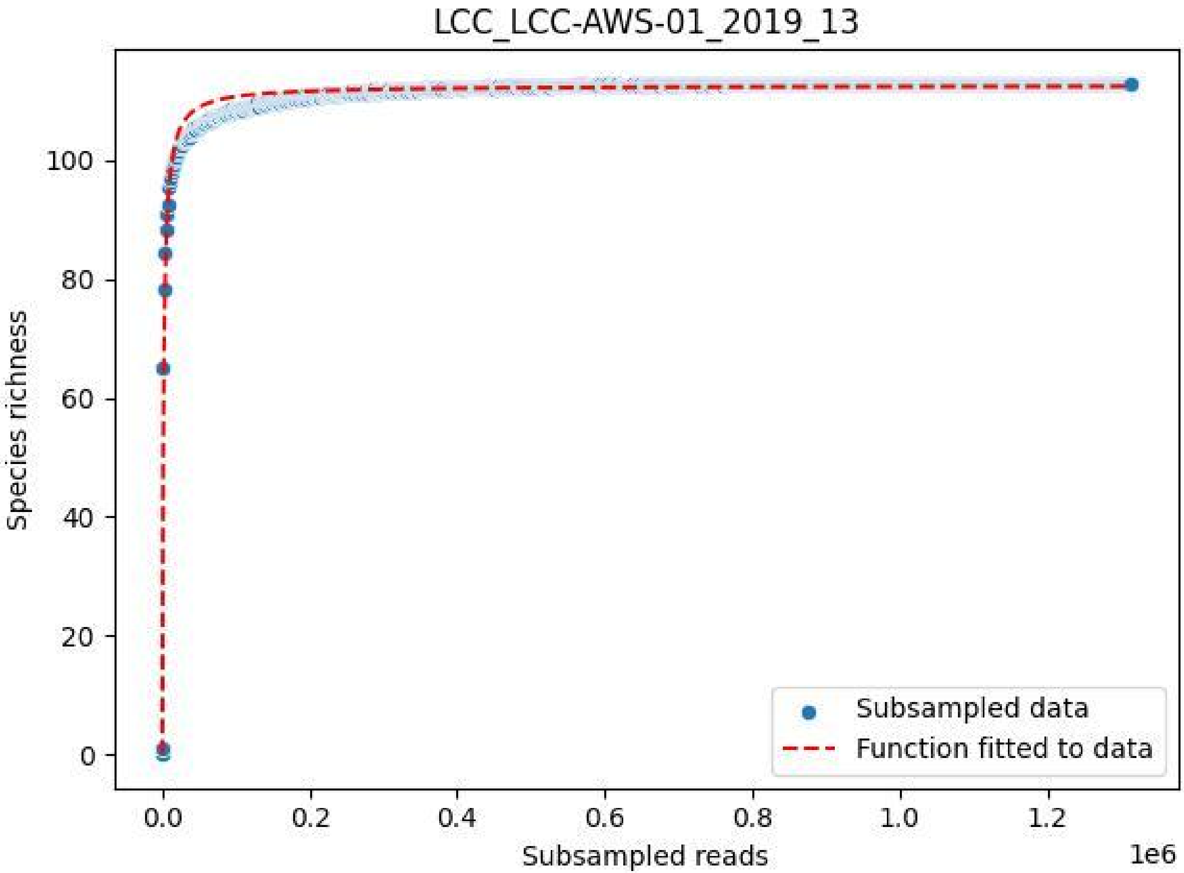
LCC_LCC-AWS-01_2019_13
Species richness
Subsampled reads
1e6
0
20
40
60
80
100
0.0
0.2
0.4
0.6
0.8
1.0
1.2
Subsampled data
Function fitted to data

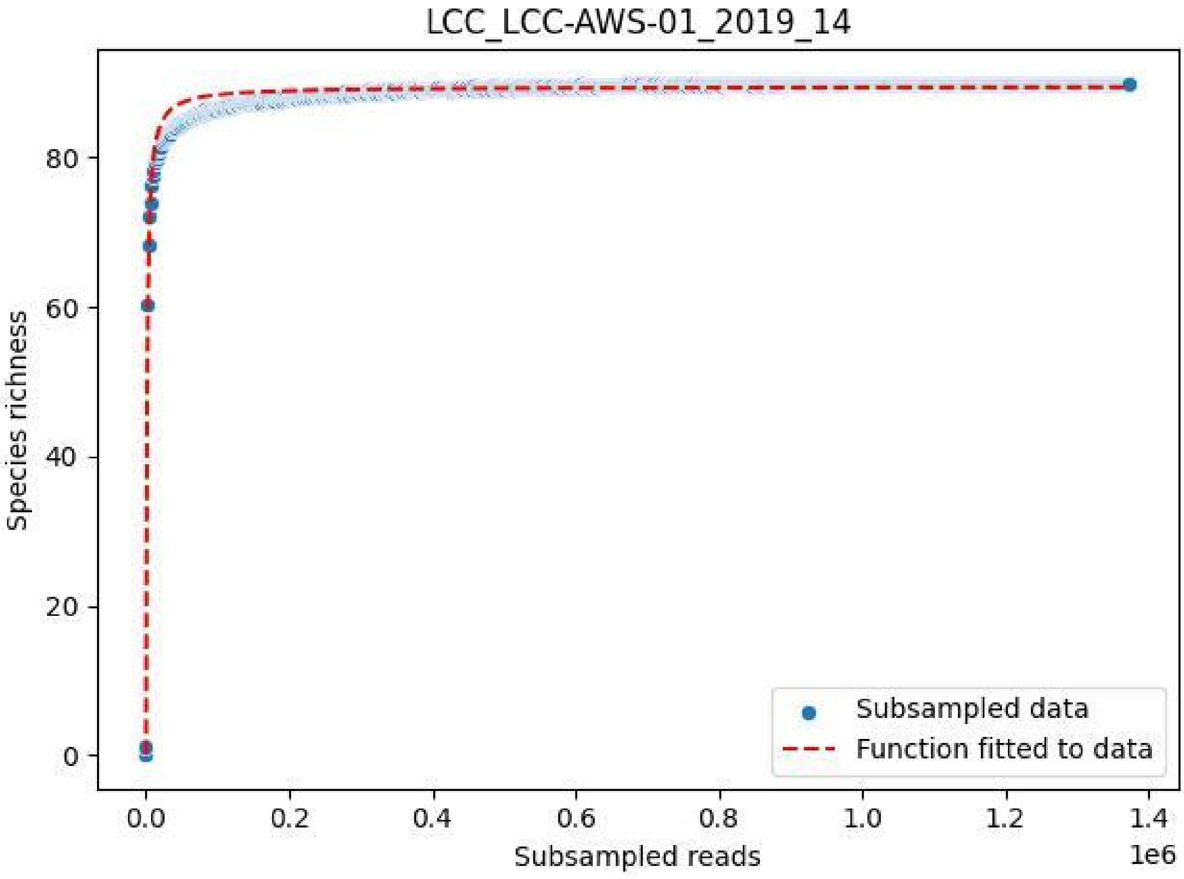
LCC_LCC-AWS-01_2019_14
Species richness
Subsampled reads
1e6
0
20
40
60
80
0.0
0.2
0.4
0.6
0.8
1.0
1.2
1.4
Subsampled data
Function fitted to data

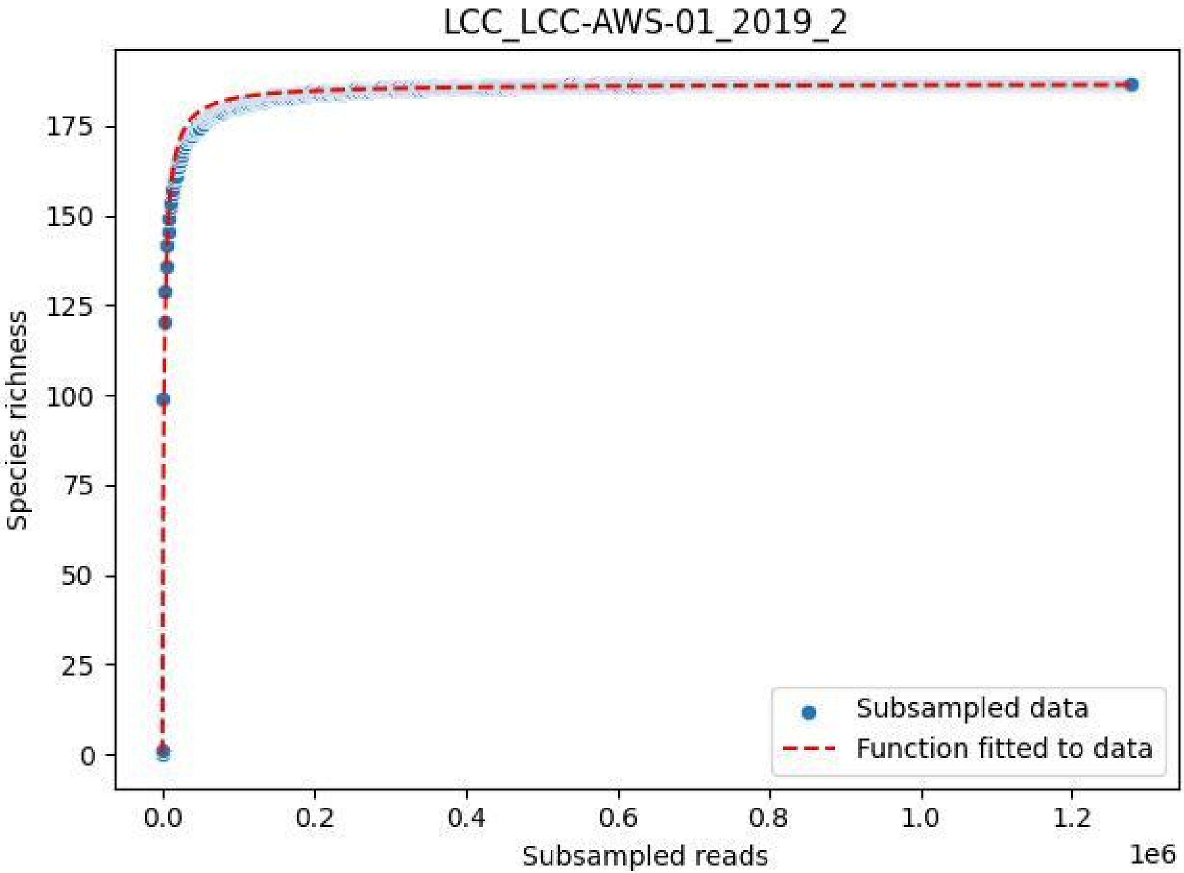
LCC_LCC-AWS-01_2019_2
Species richness
175
150
125
100
75
50
25
0
0.0
0.2
0.4
0.6
0.8
1.0
1.2
1e6
Subsampled reads
Subsampled data
Function fitted to data

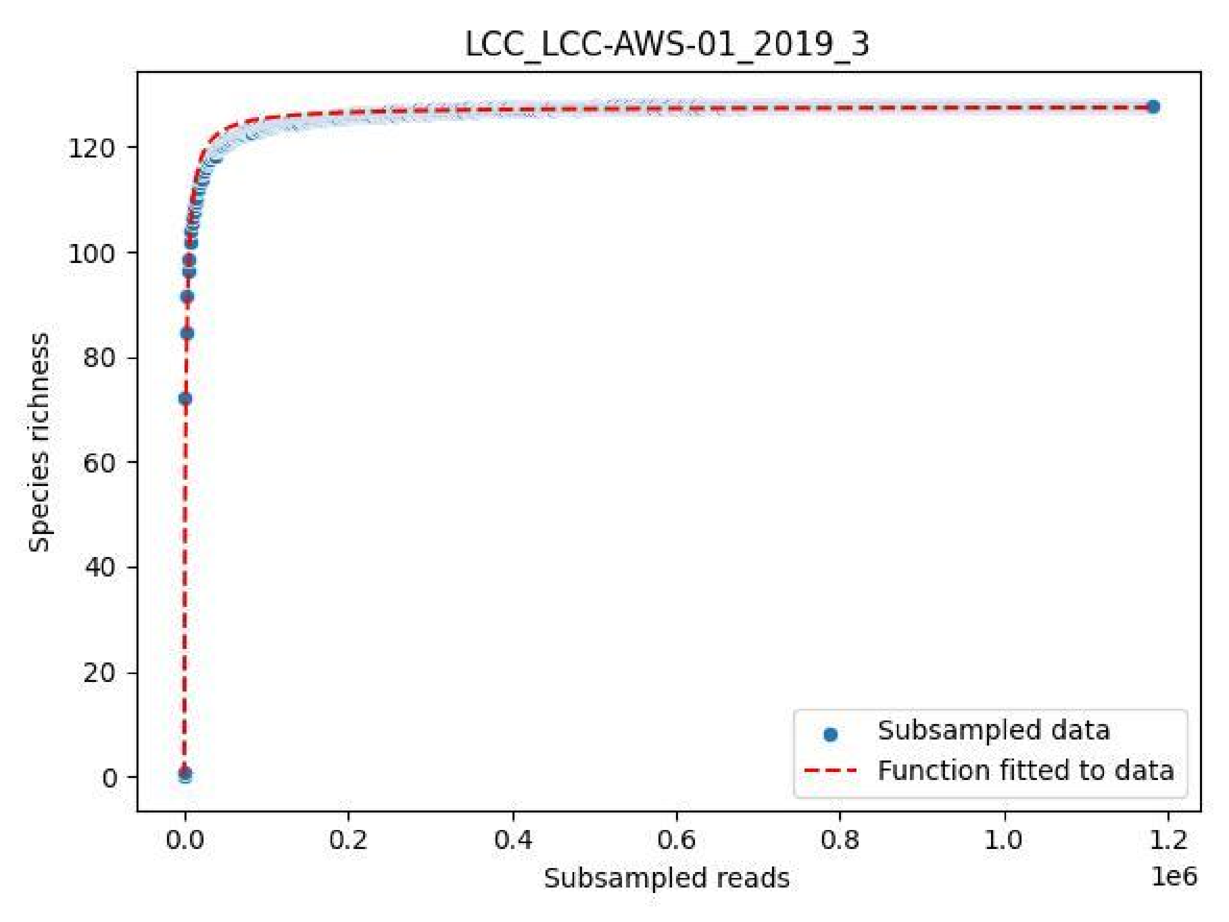

LCC_LCC-AWS-01_2019_3
Species richness
0
20
40
60
80
100
120
0.0
0.2
0.4
0.6
0.8
1.0
1.2
1e6
Subsampled reads
Subsampled data
Function fitted to data

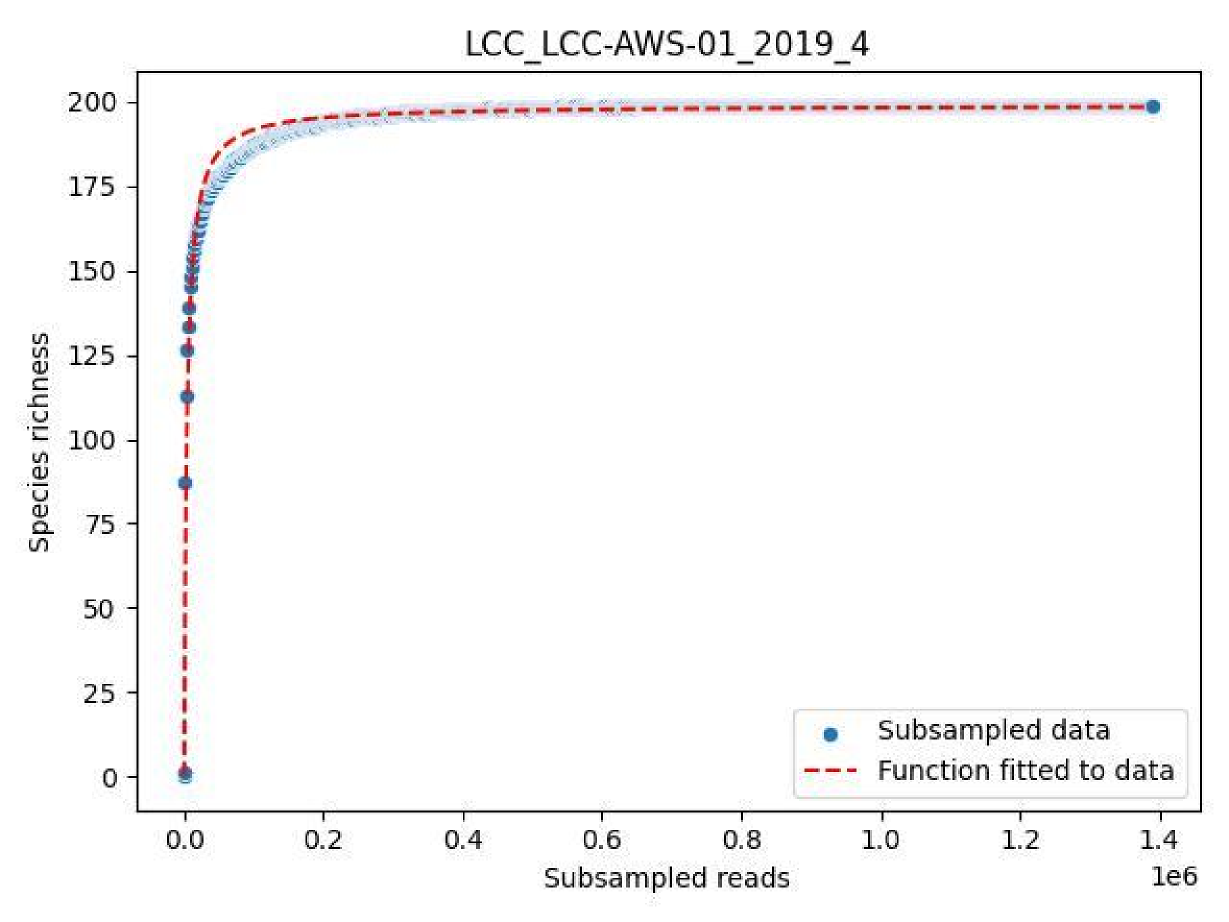

LCC_LCC-AWS-01_2019_4
Species richness
Subsampled reads
1e6
0
25
50
75
100
125
150
175
200
0.0
0.2
0.4
0.6
0.8
1.0
1.2
1.4
Subsampled data
Function fitted to data

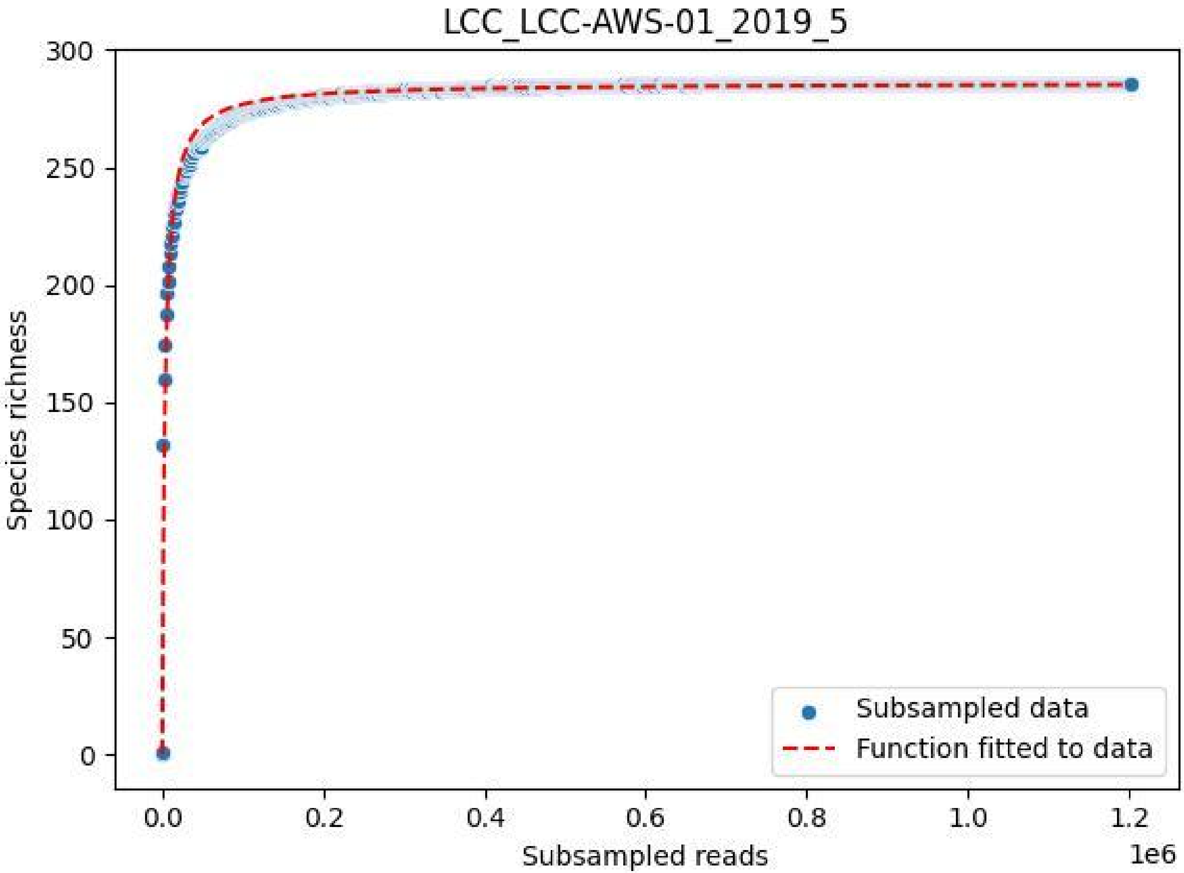

LCC_LCC-AWS-01_2019_5
Species richness
Subsampled reads
1e6
0
50
100
150
200
250
300
0.0
0.2
0.4
0.6
0.8
1.0
1.2
Subsampled data
Function fitted to data

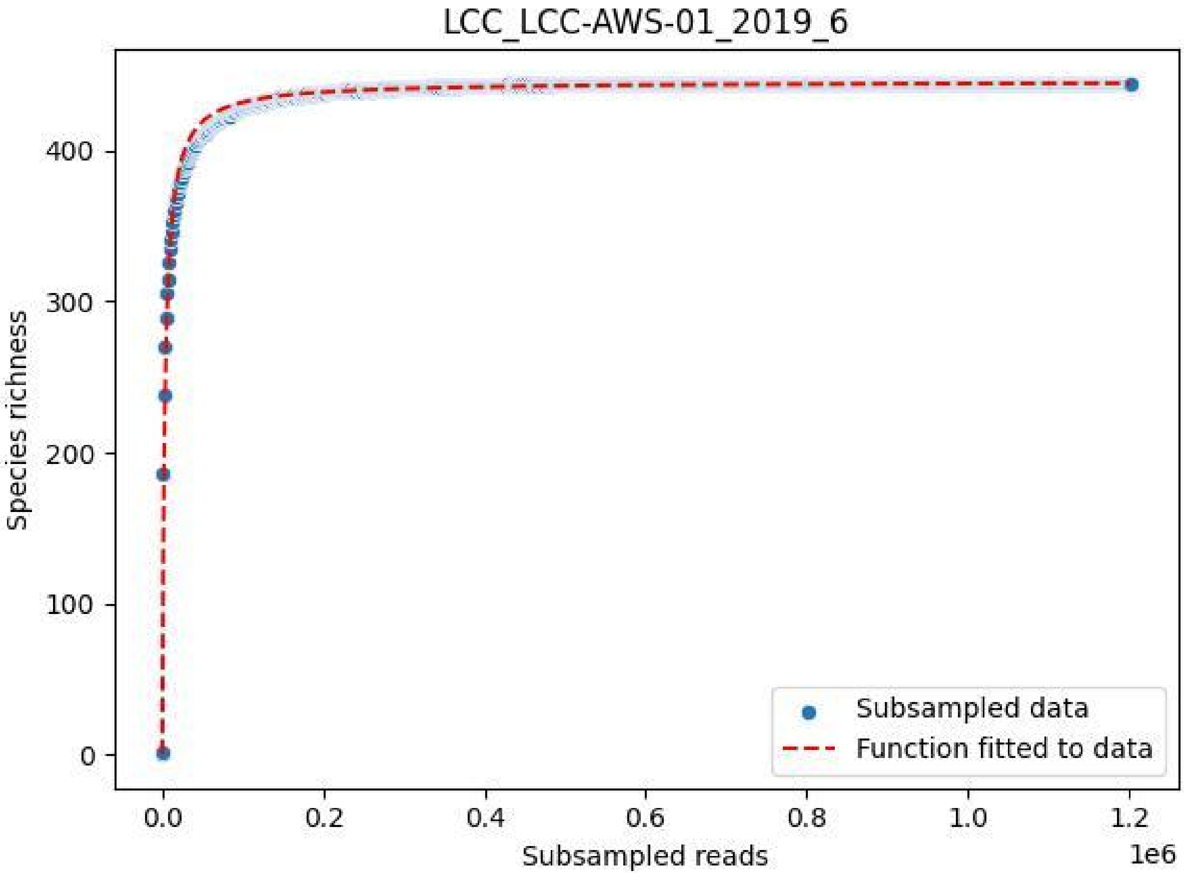

LCC_LCC-AWS-01_2019_6
Species richness
Subsampled reads
1e6
0
100
200
300
400
0.0
0.2
0.4
0.6
0.8
1.0
1.2
Subsampled data
Function fitted to data

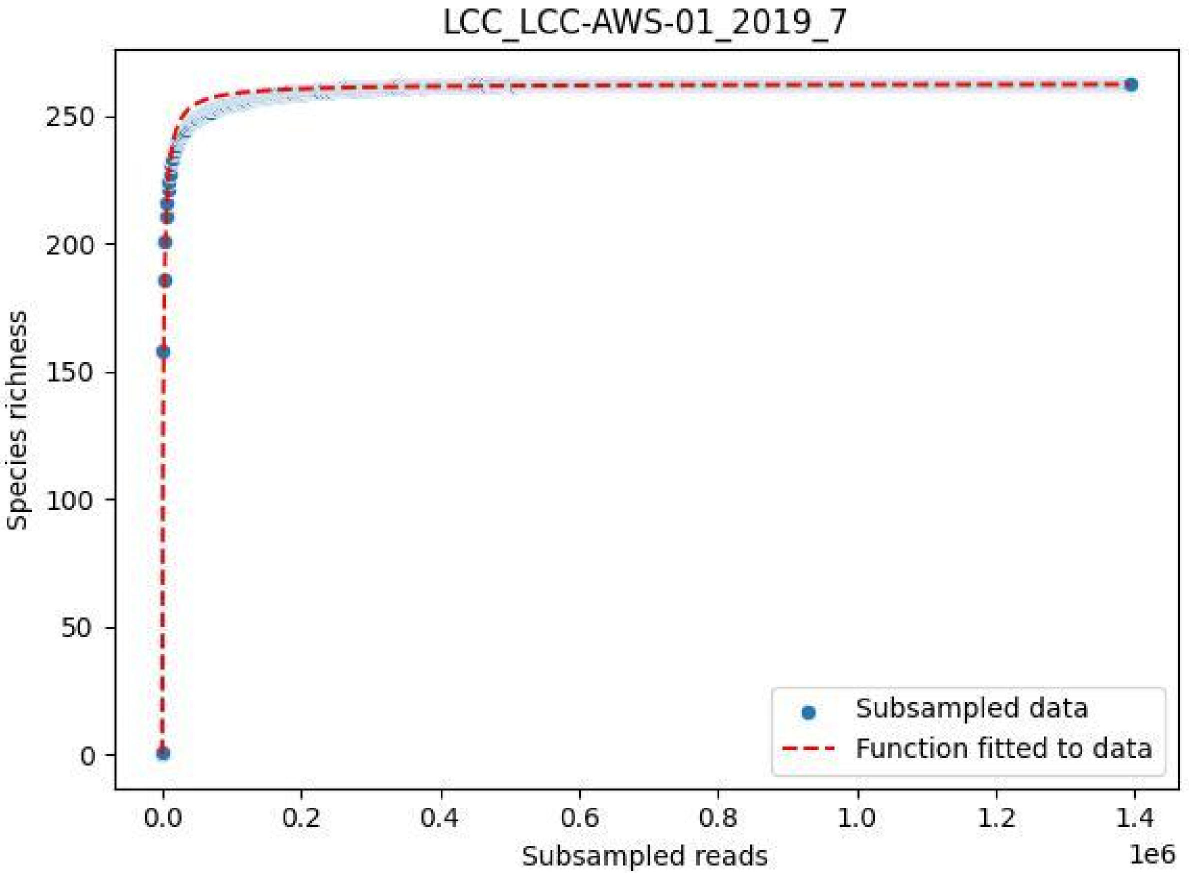

LCC_LCC-AWS-01_2019_7
Species richness
Subsampled reads
1e6
0
50
100
150
200
250
0.0
0.2
0.4
0.6
0.8
1.0
1.2
1.4
Subsampled data
Function fitted to data

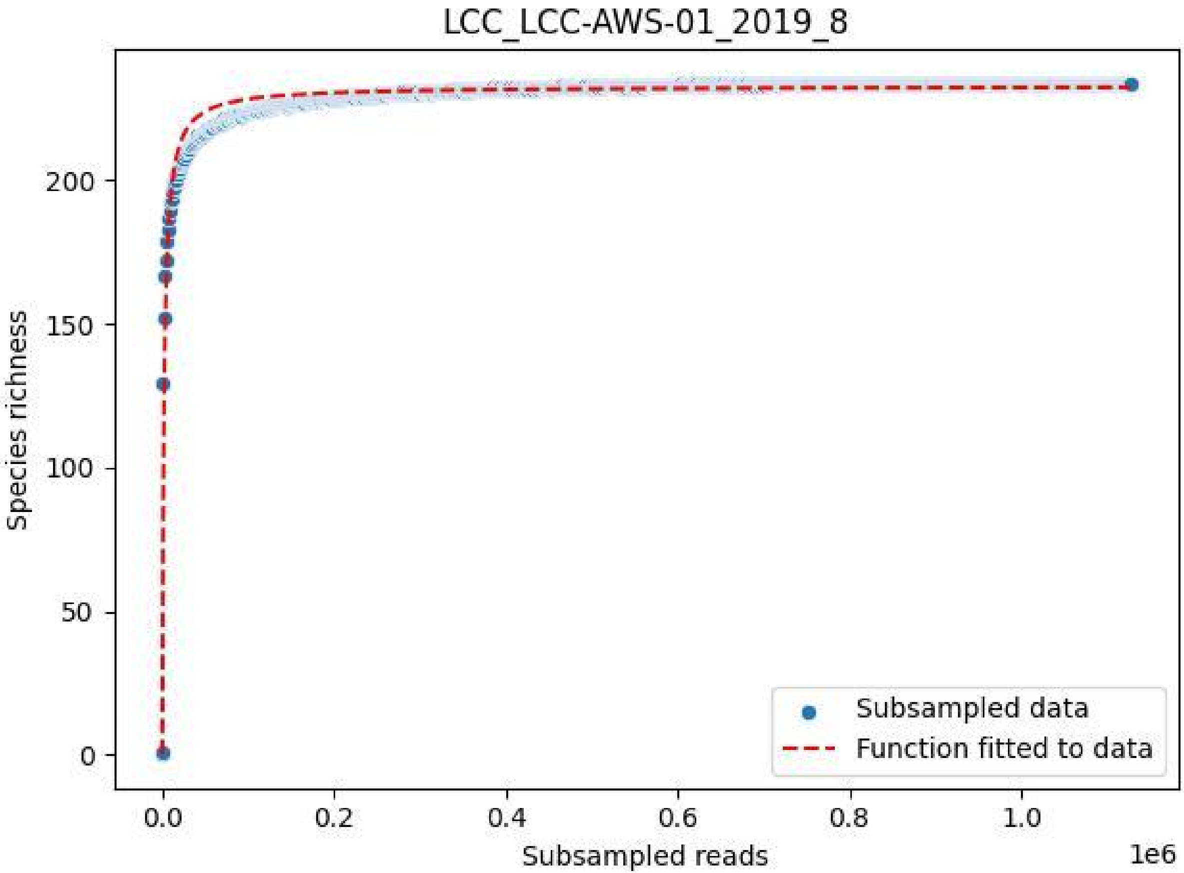

LCC_LCC-AWS-01_2019_8
Species richness
Subsampled reads
1e6
0
50
100
150
200
0.0
0.2
0.4
0.6
0.8
1.0
Subsampled data
Function fitted to data

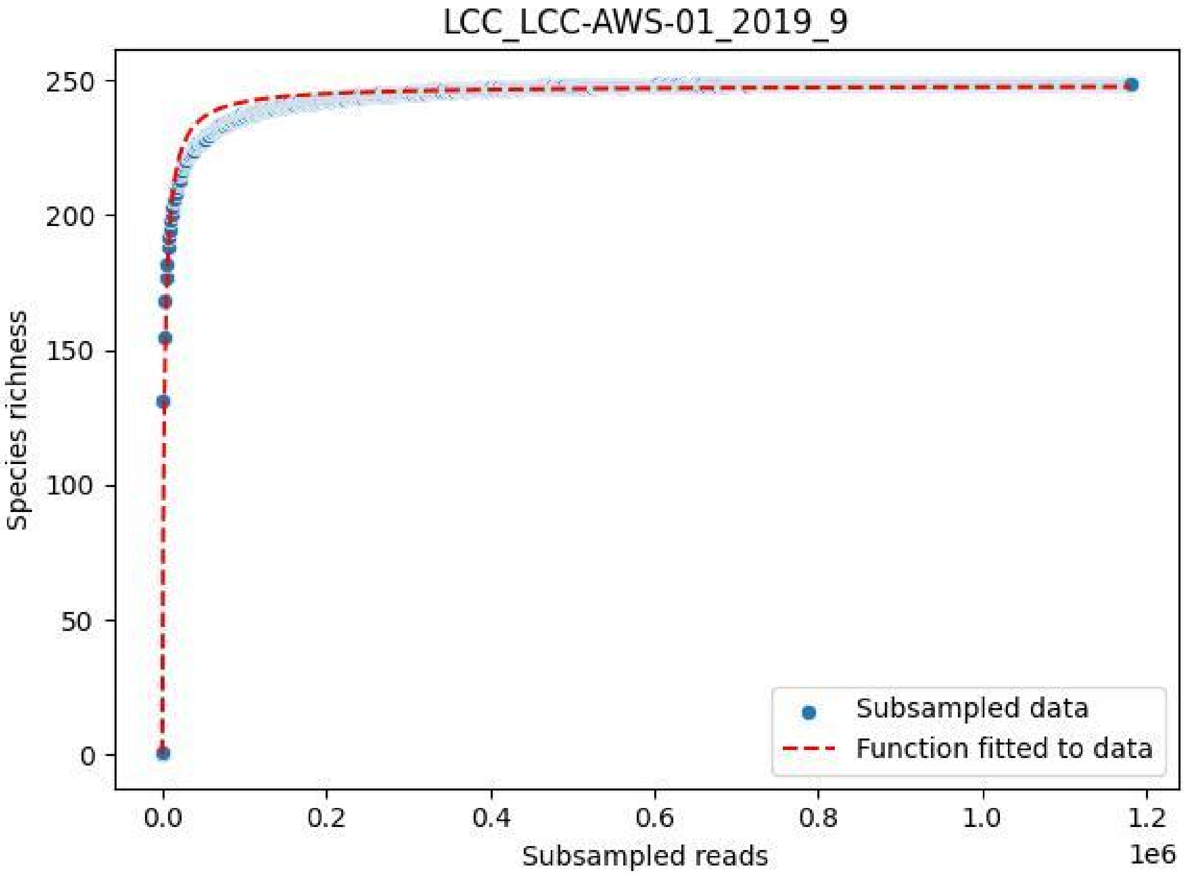
LCC_LCC-AWS-01_2019_9
Species richness
Subsampled reads
1e6
Subsampled data
Function fitted to data
0
50
100
150
200
250
0.0
0.2
0.4
0.6
0.8
1.0
1.2

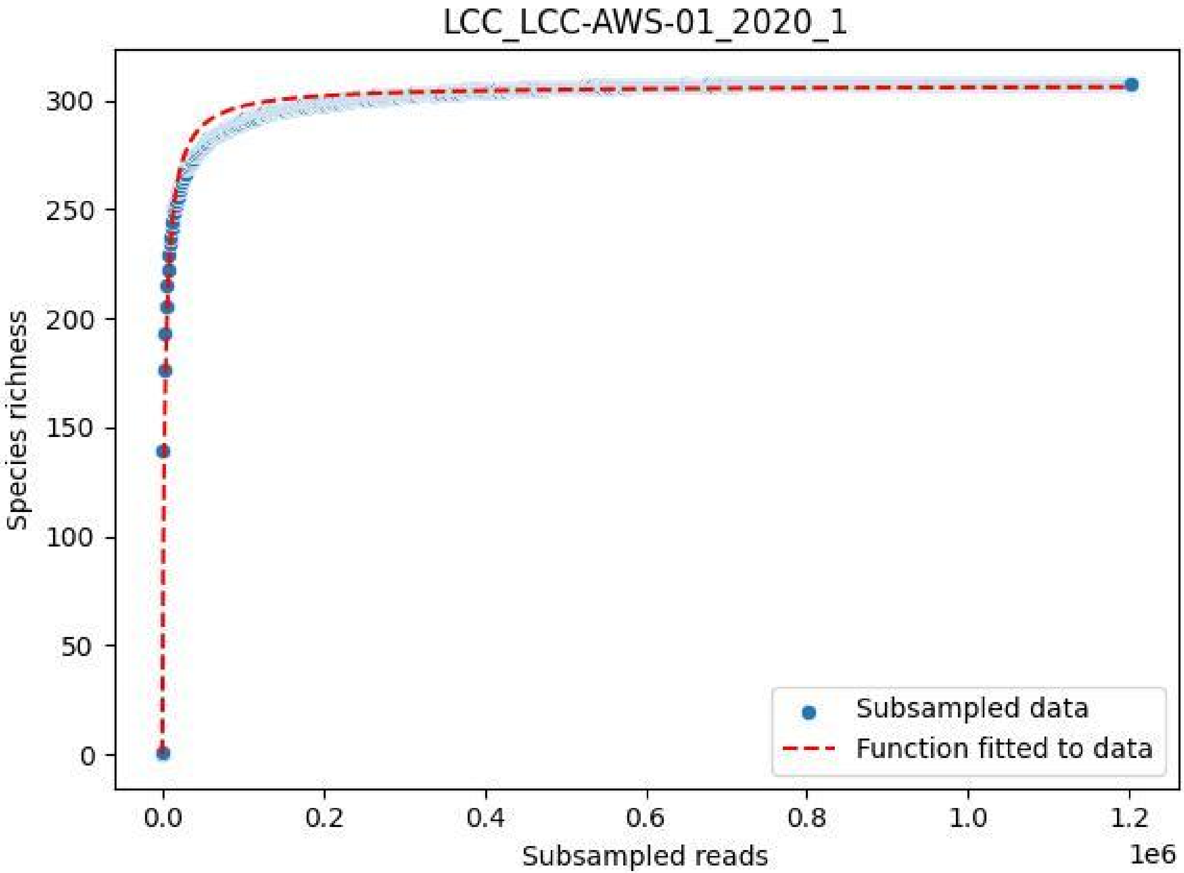
LCC_LCC-AWS-01_2020_1
Species richness
Subsampled reads
1e6
0
50
100
150
200
250
300
0.0
0.2
0.4
0.6
0.8
1.0
1.2
Subsampled data
Function fitted to data

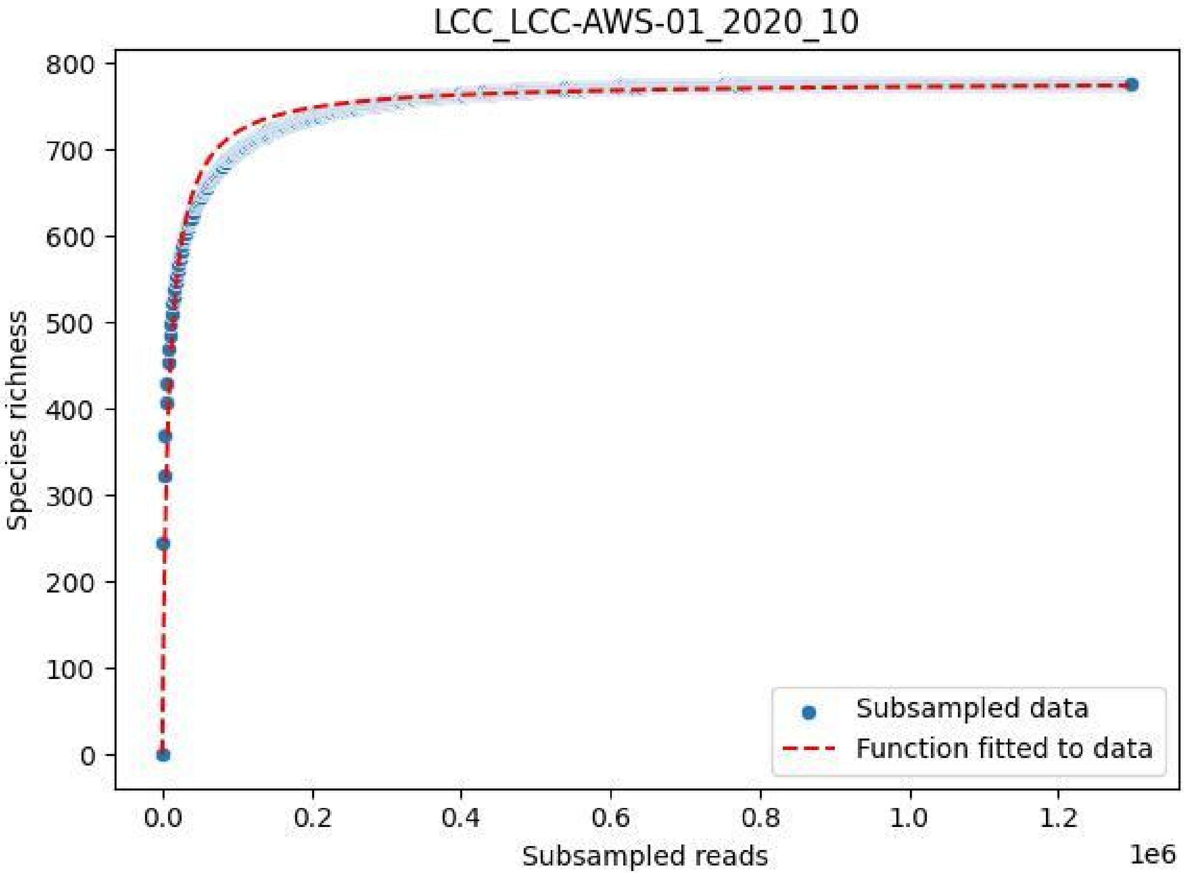
LCC_LCC-AWS-01_2020_10
Species richness
800
700
600
500
400
300
200
100
0
0.0
0.2
0.4
0.6
0.8
1.0
1.2
1e6
Subsampled reads
Subsampled data
Function fitted to data

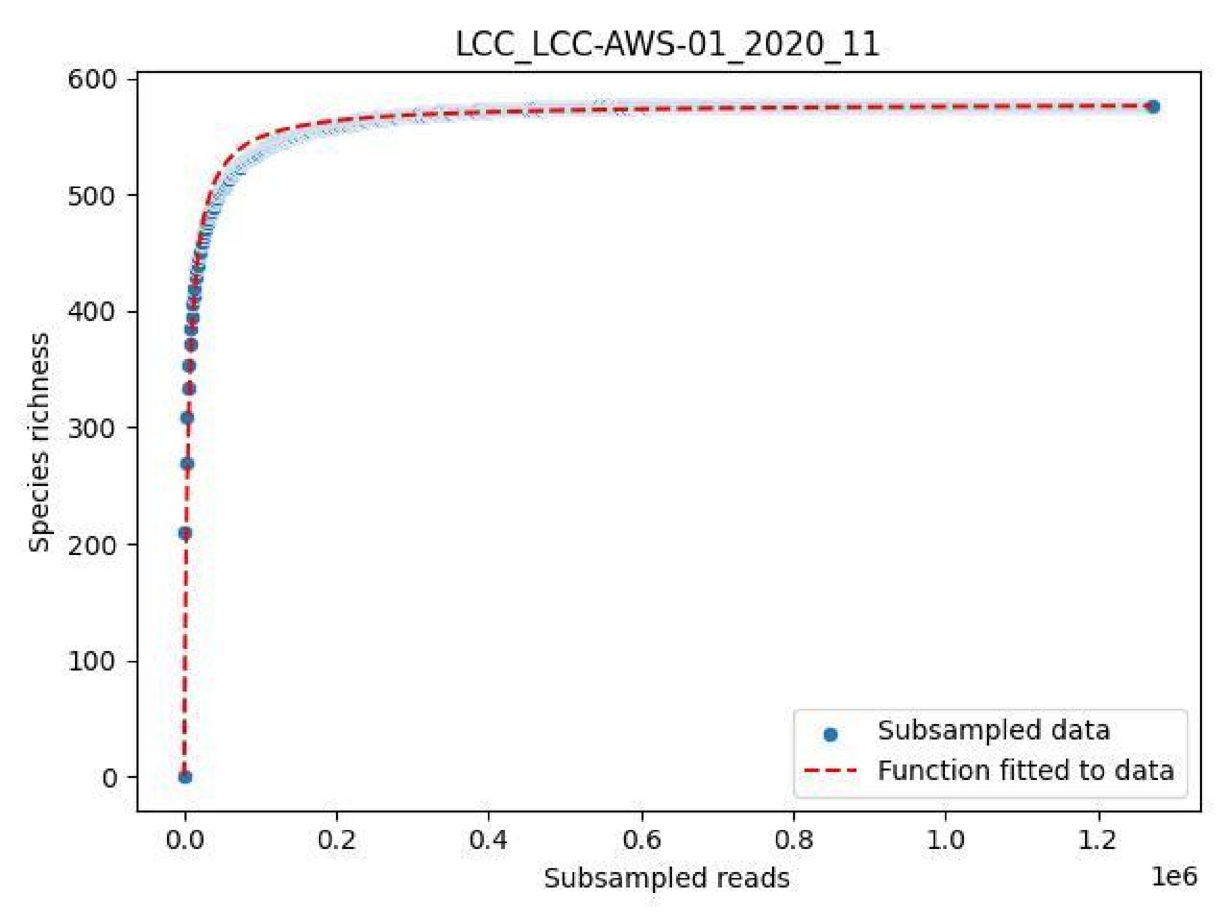
LCC_LCC-AWS-01_2020_11
Species richness
0
100
200
300
400
500
600
Subsampled reads
0.0
0.2
0.4
0.6
0.8
1.0
1.2
1e6
Subsampled data
Function fitted to data

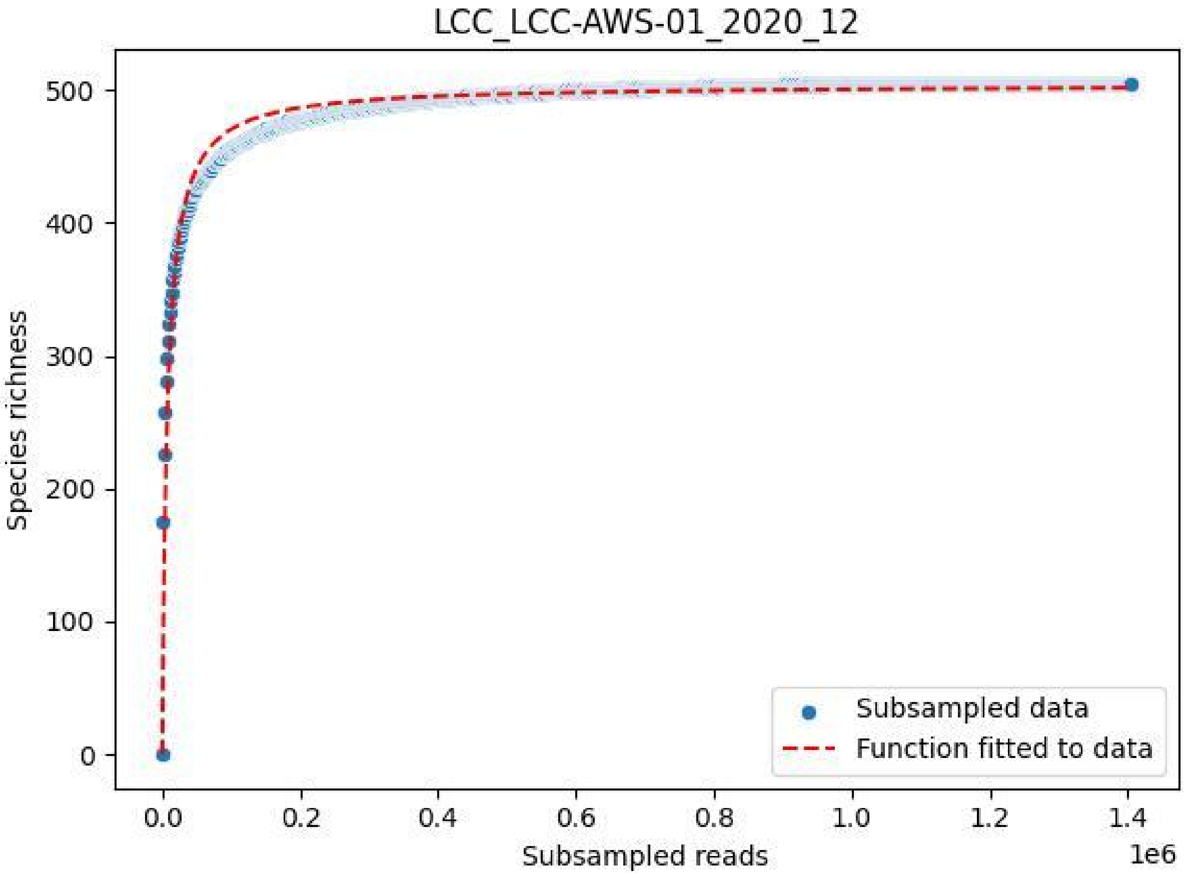
LCC_LCC-AWS-01_2020_12
Species richness
Subsampled reads
1e6
0
100
200
300
400
500
0.0
0.2
0.4
0.6
0.8
1.0
1.2
1.4
Subsampled data
Function fitted to data

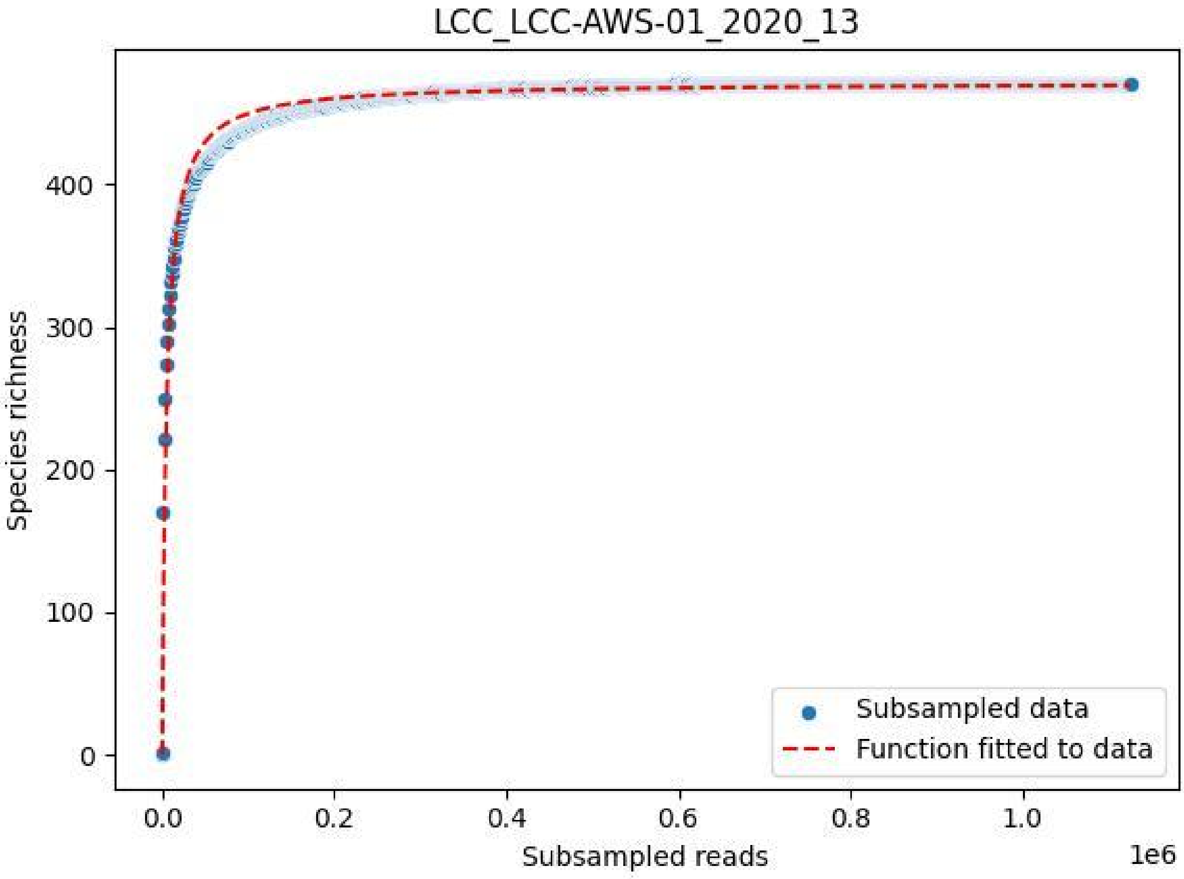

LCC_LCC-AWS-01_2020_13
Species richness
Subsampled reads
1e6
0
100
200
300
400
0.0
0.2
0.4
0.6
0.8
1.0
Subsampled data
Function fitted to data

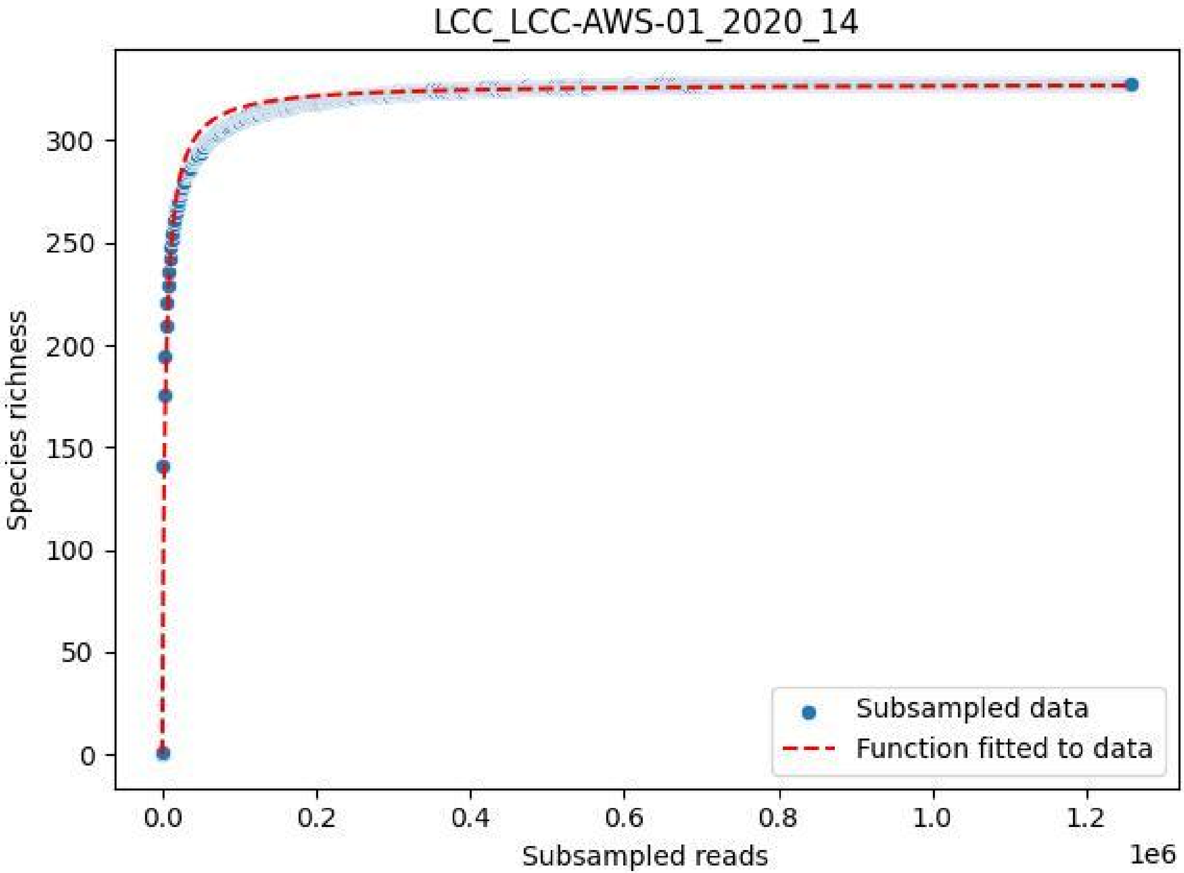
LCC_LCC-AWS-01_2020_14
Species richness
300
250
200
150
100
50
0
Subsampled reads
0.0
0.2
0.4
0.6
0.8
1.0
1.2
1e6
Subsampled data
Function fitted to data

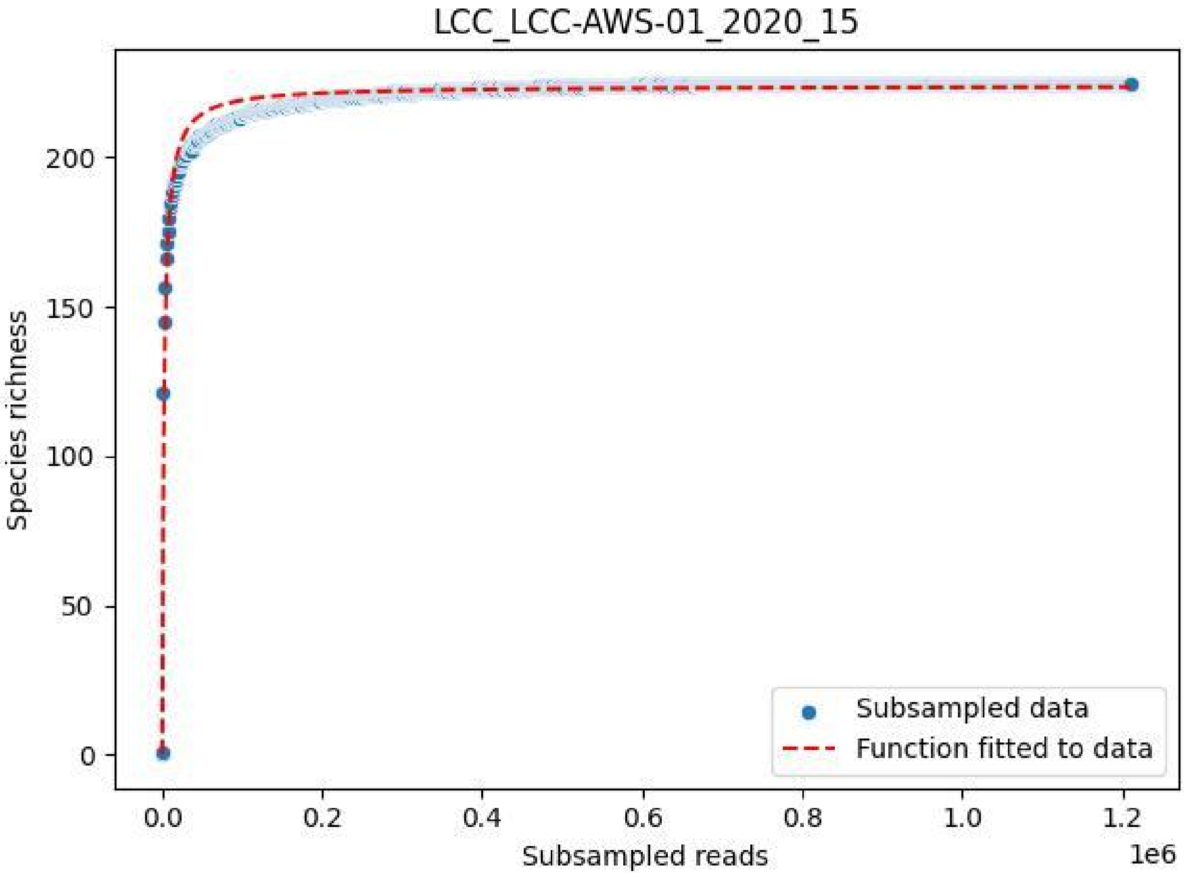
LCC_LCC-AWS-01_2020_15
Species richness
Subsampled reads
1e6
0
50
100
150
200
0.0
0.2
0.4
0.6
0.8
1.0
1.2
Subsampled data
Function fitted to data

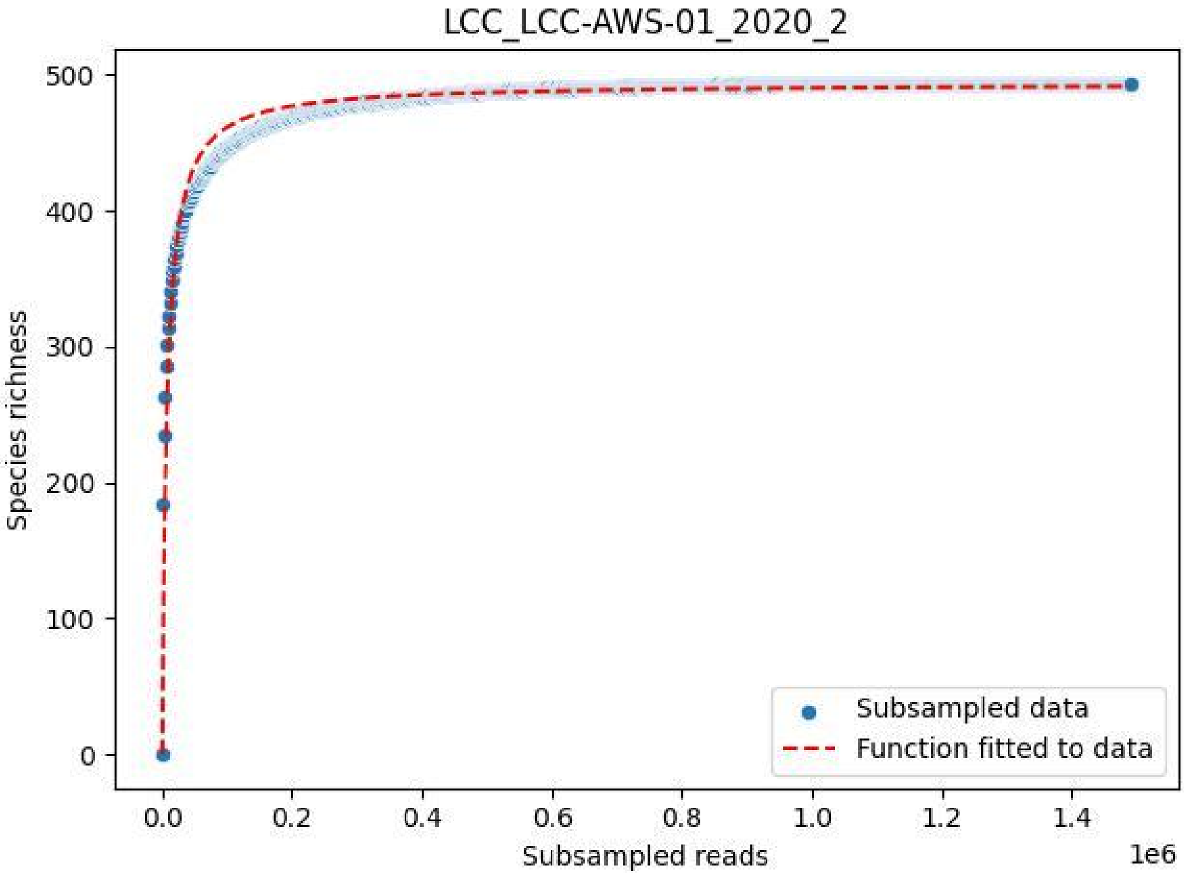

LCC_LCC-AWS-01_2020_2
Species richness
Subsampled reads
1e6
0
100
200
300
400
500
0.0
0.2
0.4
0.6
0.8
1.0
1.2
1.4
Subsampled data
Function fitted to data

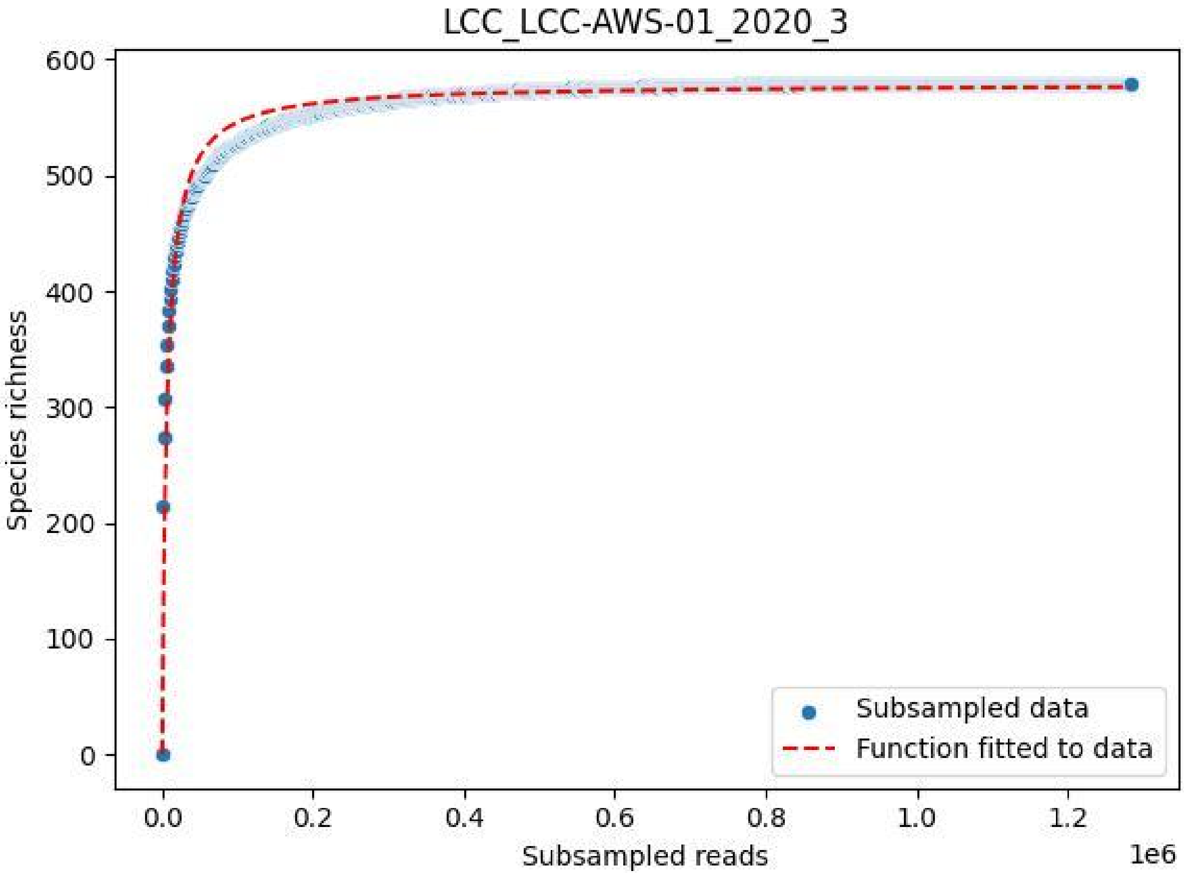
LCC_LCC-AWS-01_2020_3
Species richness
Subsampled reads
1e6
600
500
400
300
200
100
0
0.0
0.2
0.4
0.6
0.8
1.0
1.2
Subsampled data
Function fitted to data

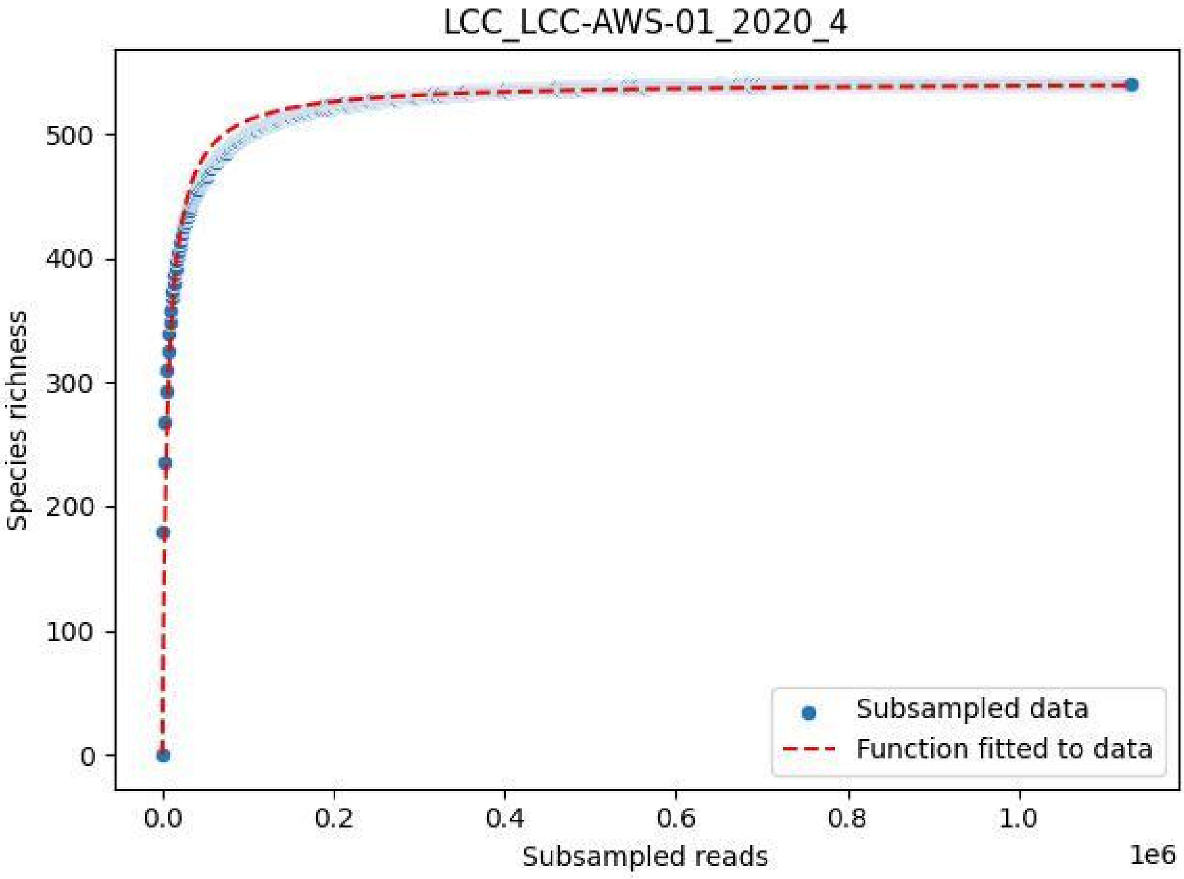
LCC_LCC-AWS-01_2020_4
Species richness
Subsampled reads
1e6
Subsampled data
Function fitted to data
0
100
200
300
400
500
0.0
0.2
0.4
0.6
0.8
1.0

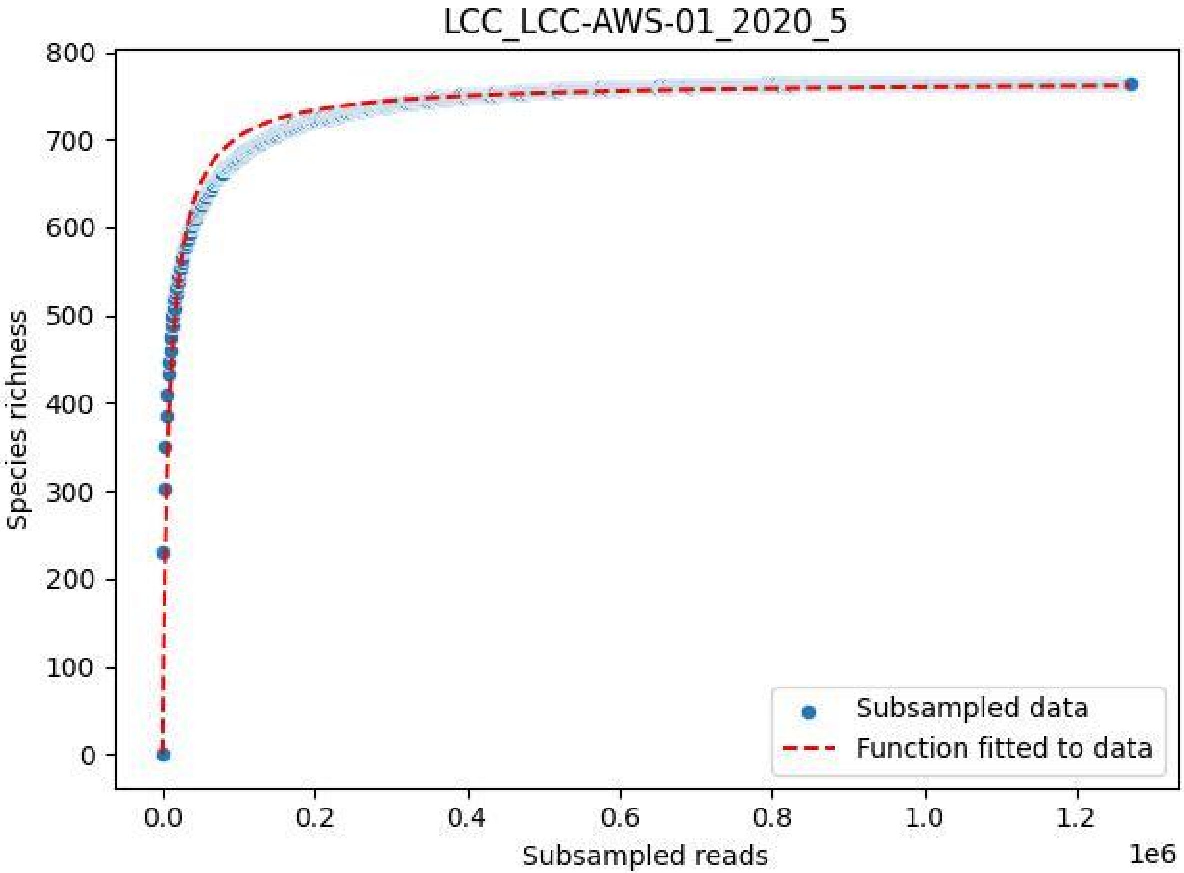
LCC_LCC-AWS-01_2020_5
Species richness
Subsampled reads
1e6
Subsampled data
Function fitted to data
0
100
200
300
400
500
600
700
800
0.0
0.2
0.4
0.6
0.8
1.0
1.2

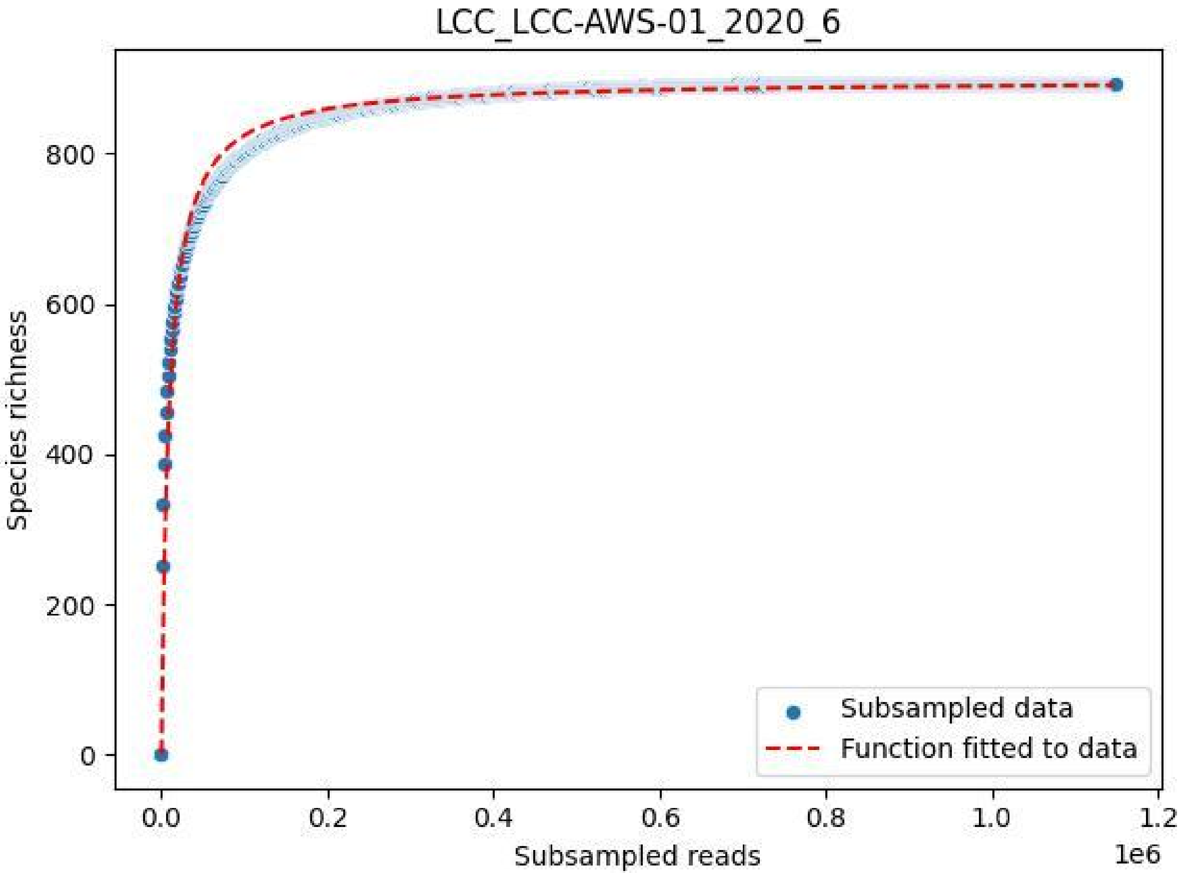
LCC_LCC-AWS-01_2020_6
Species richness
800
600
400
200
0
0.0
0.2
0.4
0.6
0.8
1.0
1.2
1e6
Subsampled reads
Subsampled data
Function fitted to data

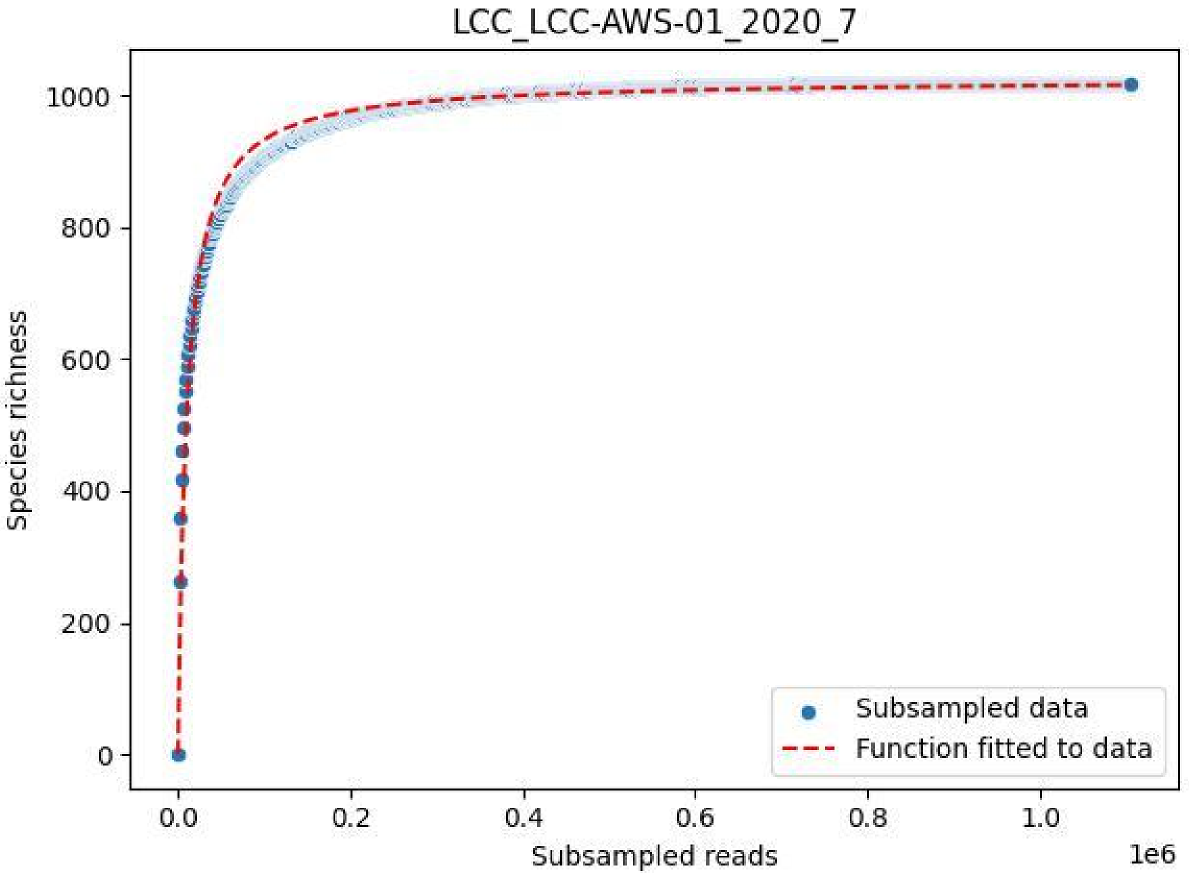

LCC_LCC-AWS-01_2020_7
Species richness
Subsampled reads
1e6
0
200
400
600
800
1000
0.0
0.2
0.4
0.6
0.8
1.0
Subsampled data
Function fitted to data

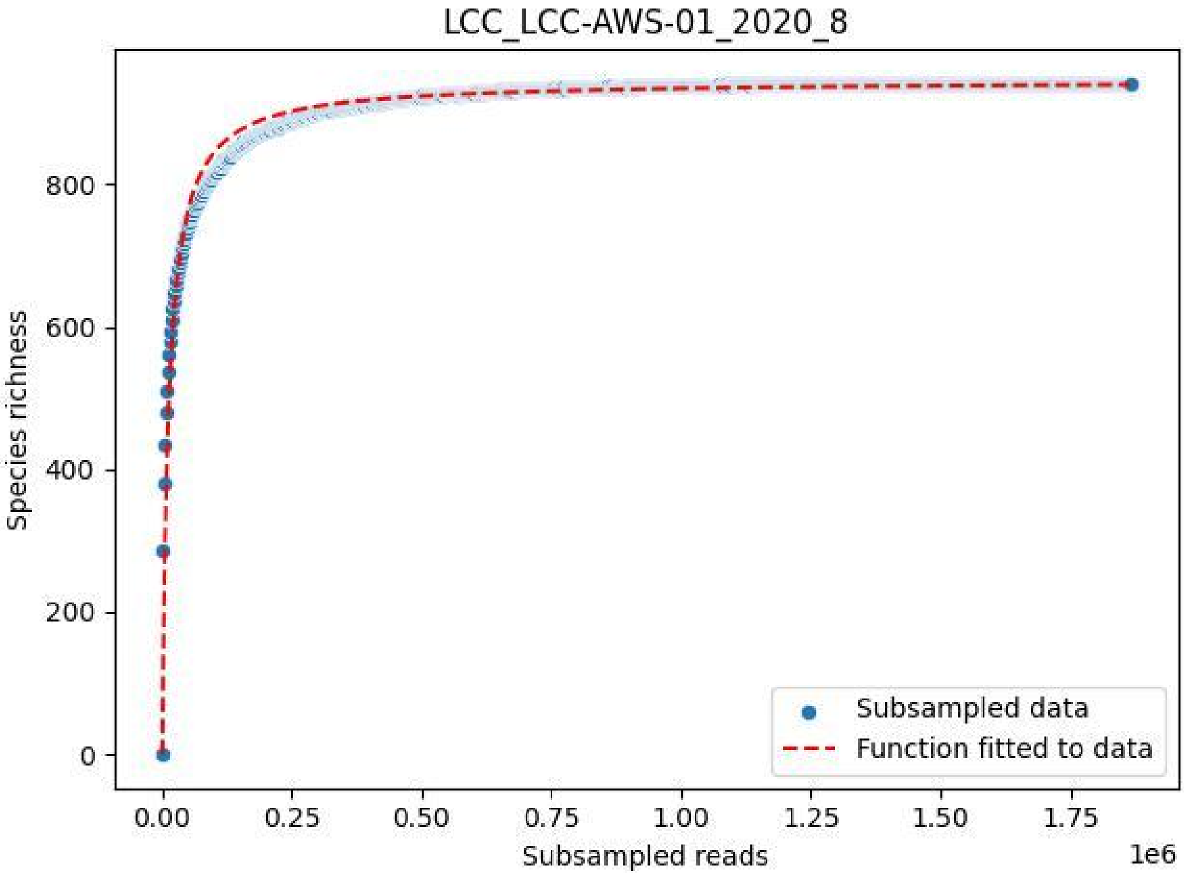

LCC_LCC-AWS-01_2020_8
Species richness
Subsampled reads
1e6
0
200
400
600
800
0.00
0.25
0.50
0.75
1.00
1.25
1.50
1.75
Subsampled data
Function fitted to data

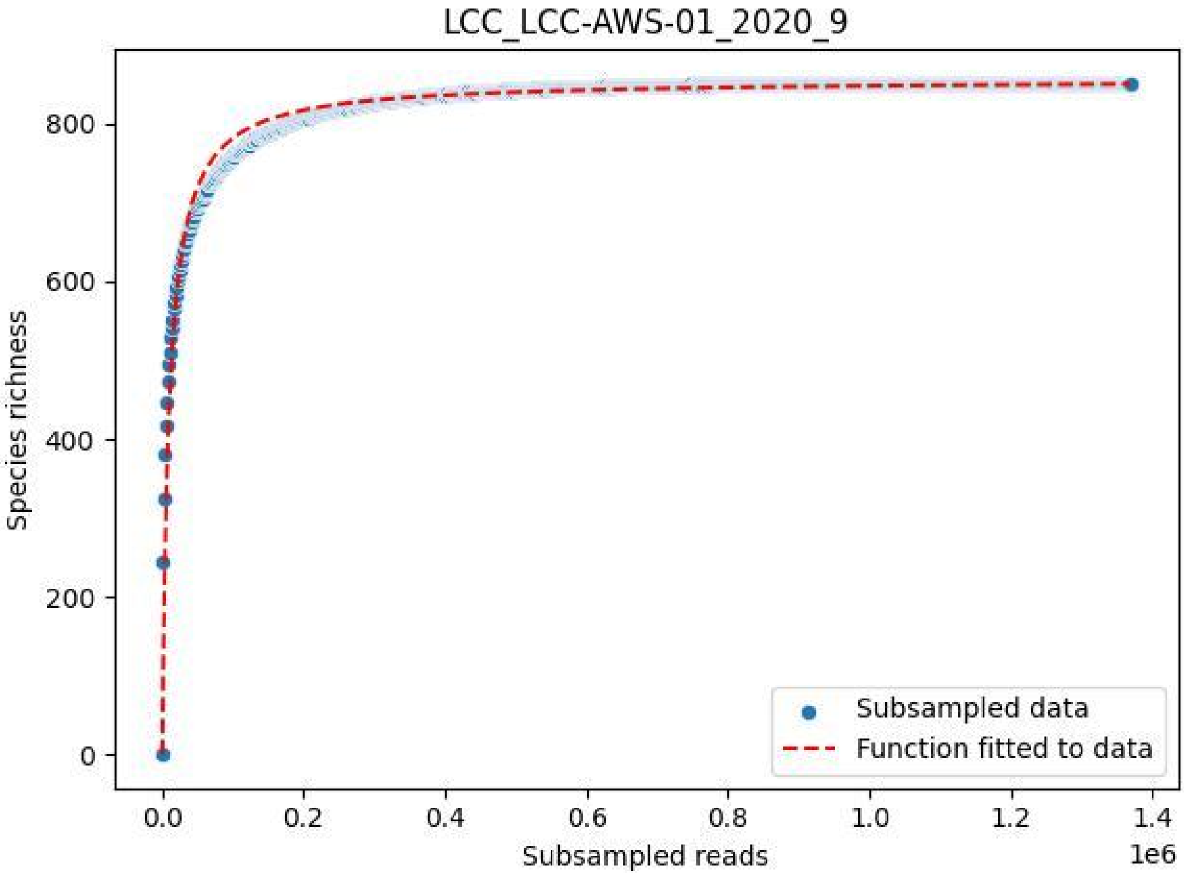
LCC_LCC-AWS-01_2020_9
Species richness
0
200
400
600
800
0.0
0.2
0.4
0.6
0.8
1.0
1.2
1.4
1e6
Subsampled reads
Subsampled data
Function fitted to data

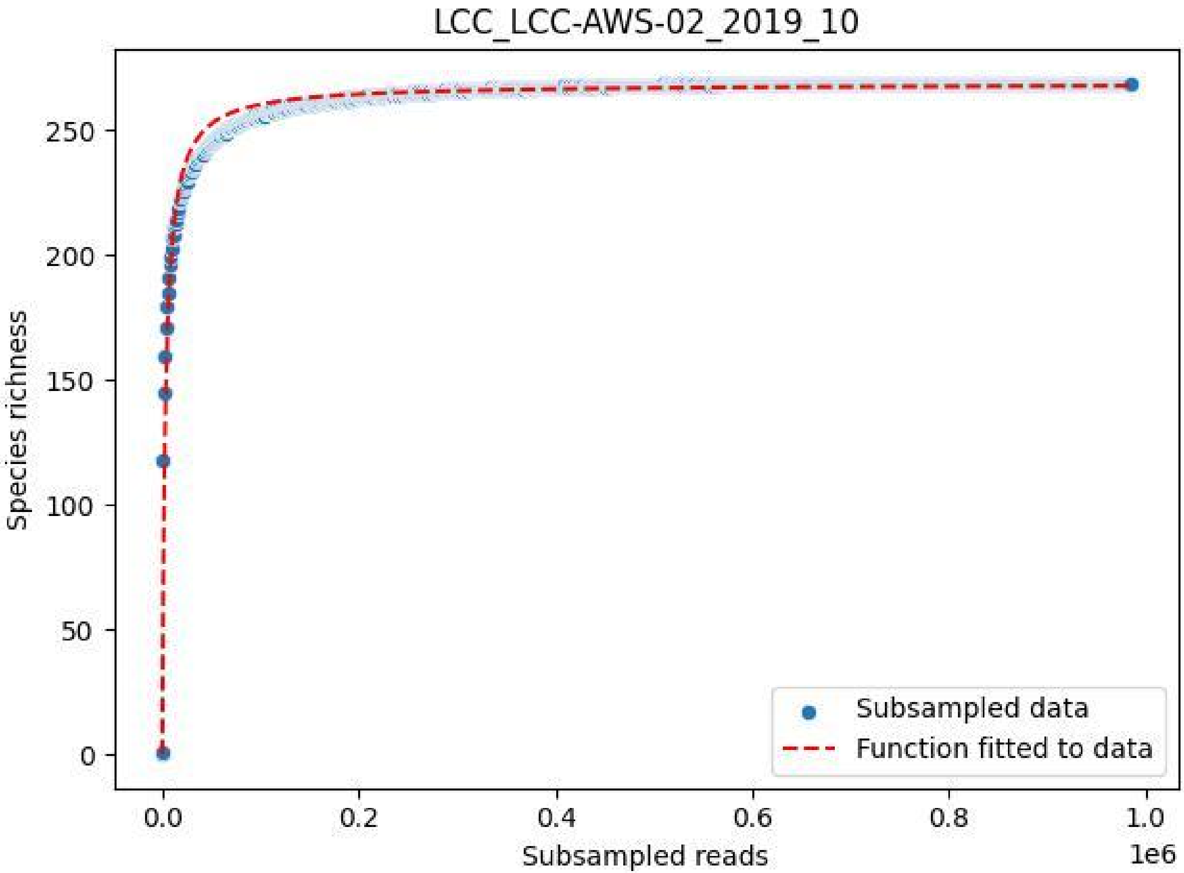
LCC_LCC-AWS-02_2019_10
Species richness
Subsampled reads
0
50
100
150
200
250
0.0
0.2
0.4
0.6
0.8
1.0
1e6
Subsampled data
Function fitted to data

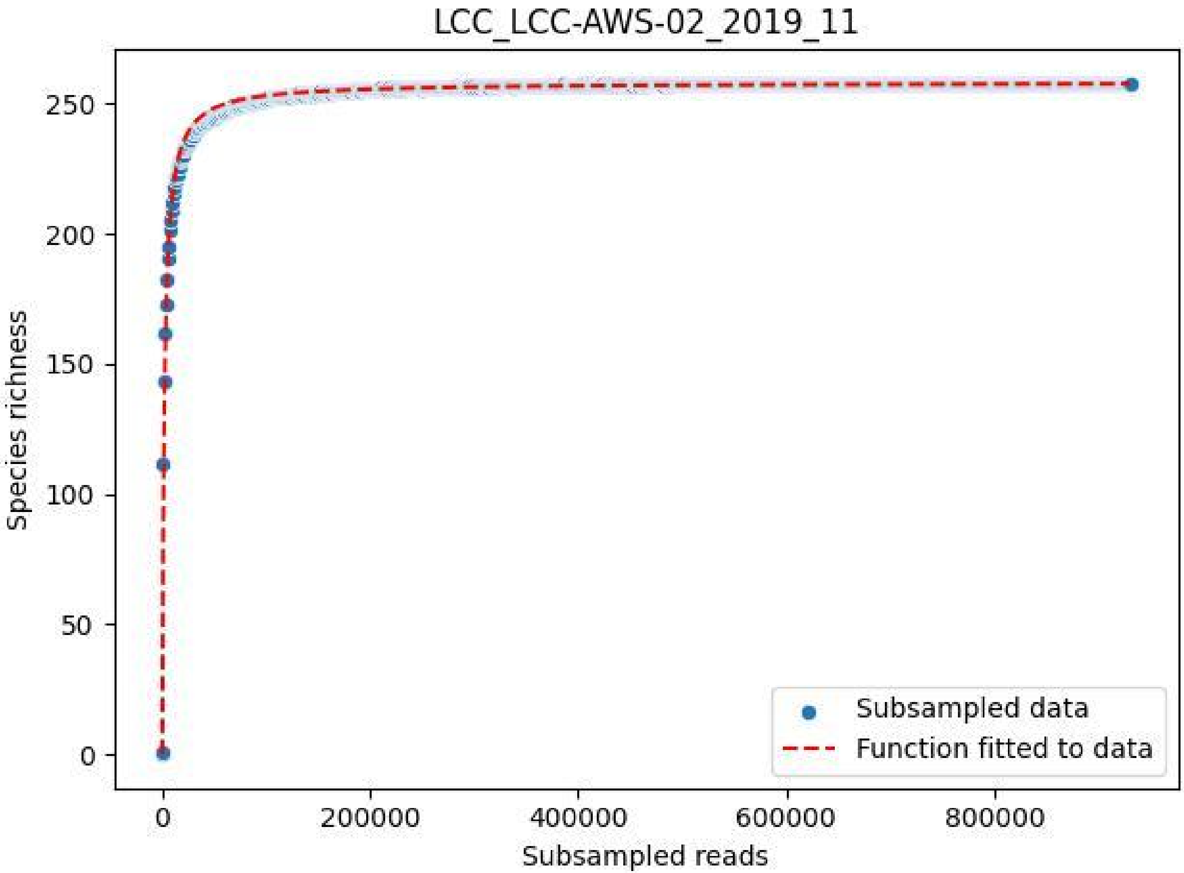

LCC_LCC-AWS-02_2019_11
Species richness
Subsampled reads
0
50
100
150
200
250
0
200000
400000
600000
800000
Subsampled data
Function fitted to data

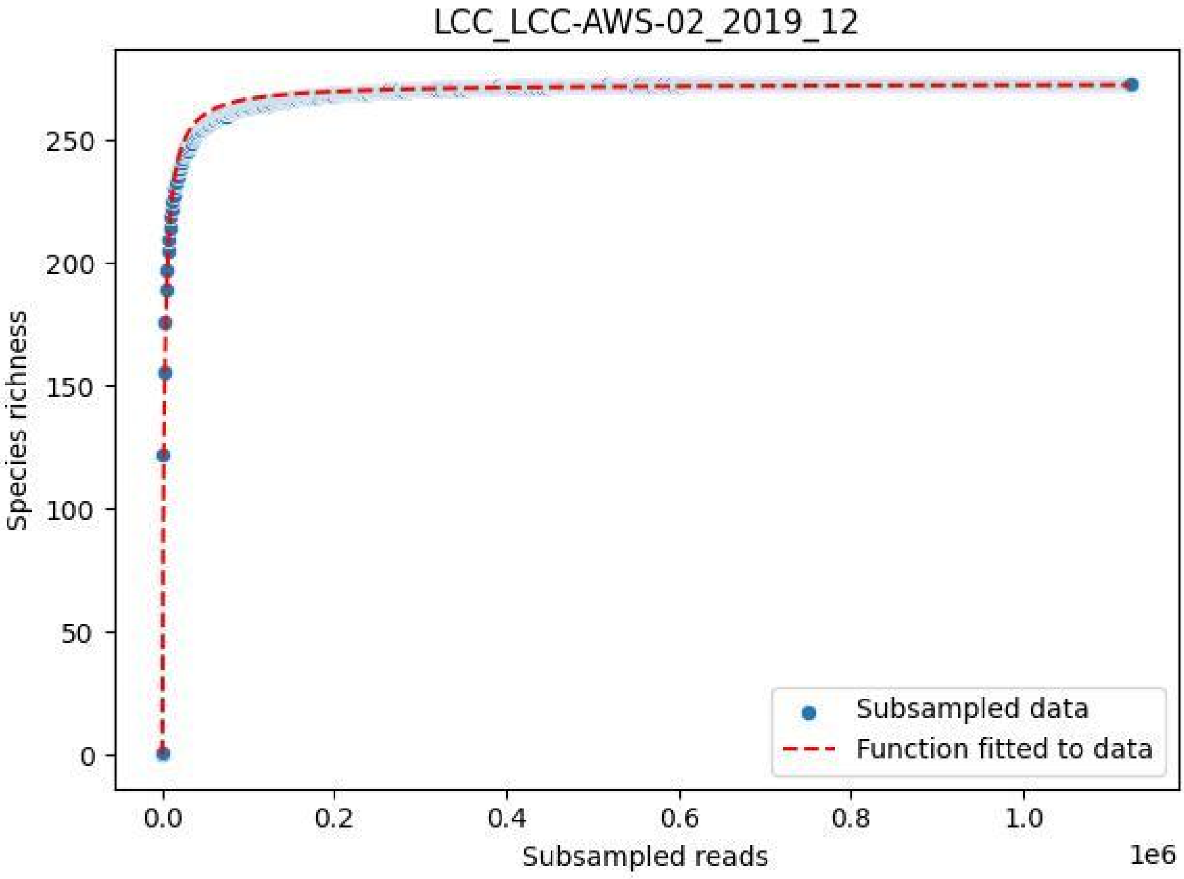

LCC_LCC-AWS-02_2019_12
Species richness
250
200
150
100
50
0
0.0
0.2
0.4
0.6
0.8
1.0
1e6
Subsampled reads
Subsampled data
Function fitted to data

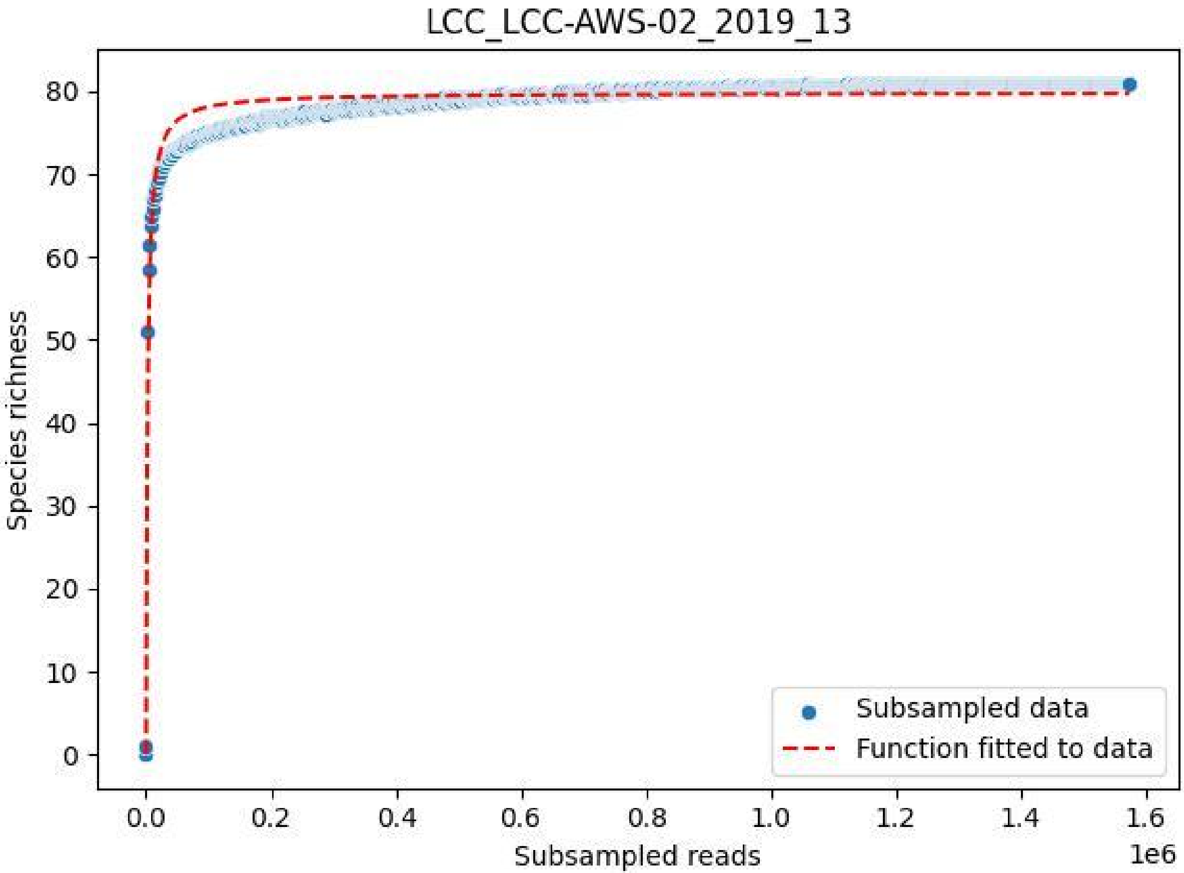

LCC_LCC-AWS-02_2019_13
Species richness
Subsampled reads
1e6
0
10
20
30
40
50
60
70
80
0.0
0.2
0.4
0.6
0.8
1.0
1.2
1.4
1.6
Subsampled data
Function fitted to data

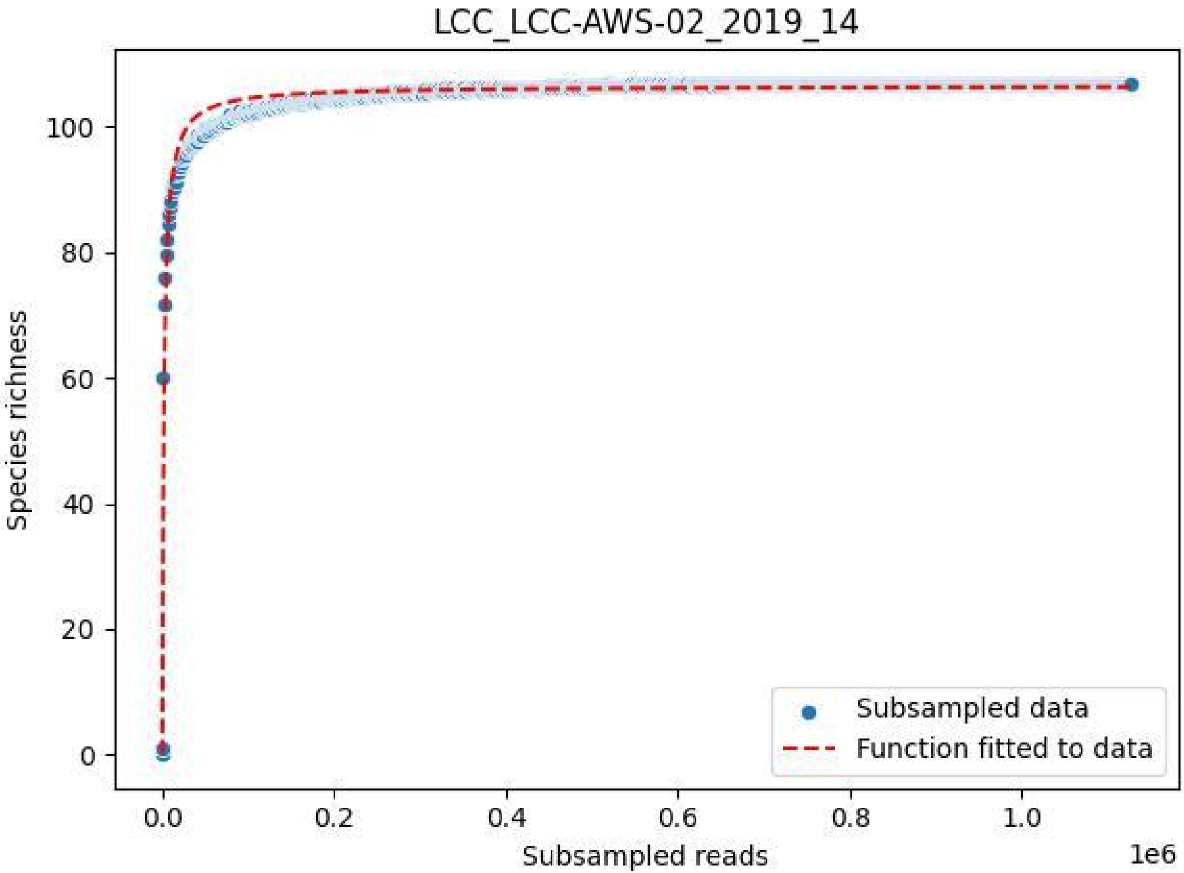
LCC_LCC-AWS-02_2019_14
Species richness
Subsampled reads
1e6
0
20
40
60
80
100
0.0
0.2
0.4
0.6
0.8
1.0
Subsampled data
Function fitted to data

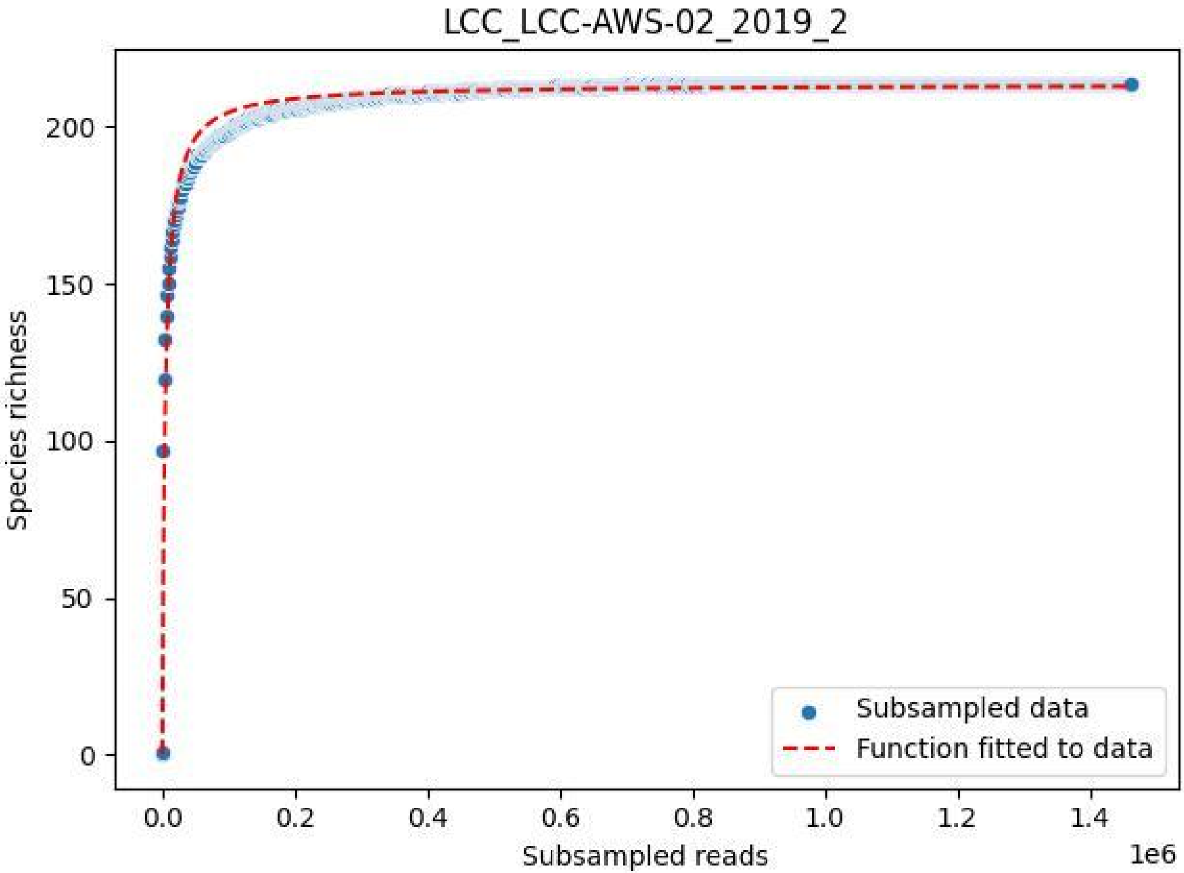
LCC_LCC-AWS-02_2019_2
Species richness
0
50
100
150
200
0.0
0.2
0.4
0.6
0.8
1.0
1.2
1.4
1e6
Subsampled reads
Subsampled data
Function fitted to data

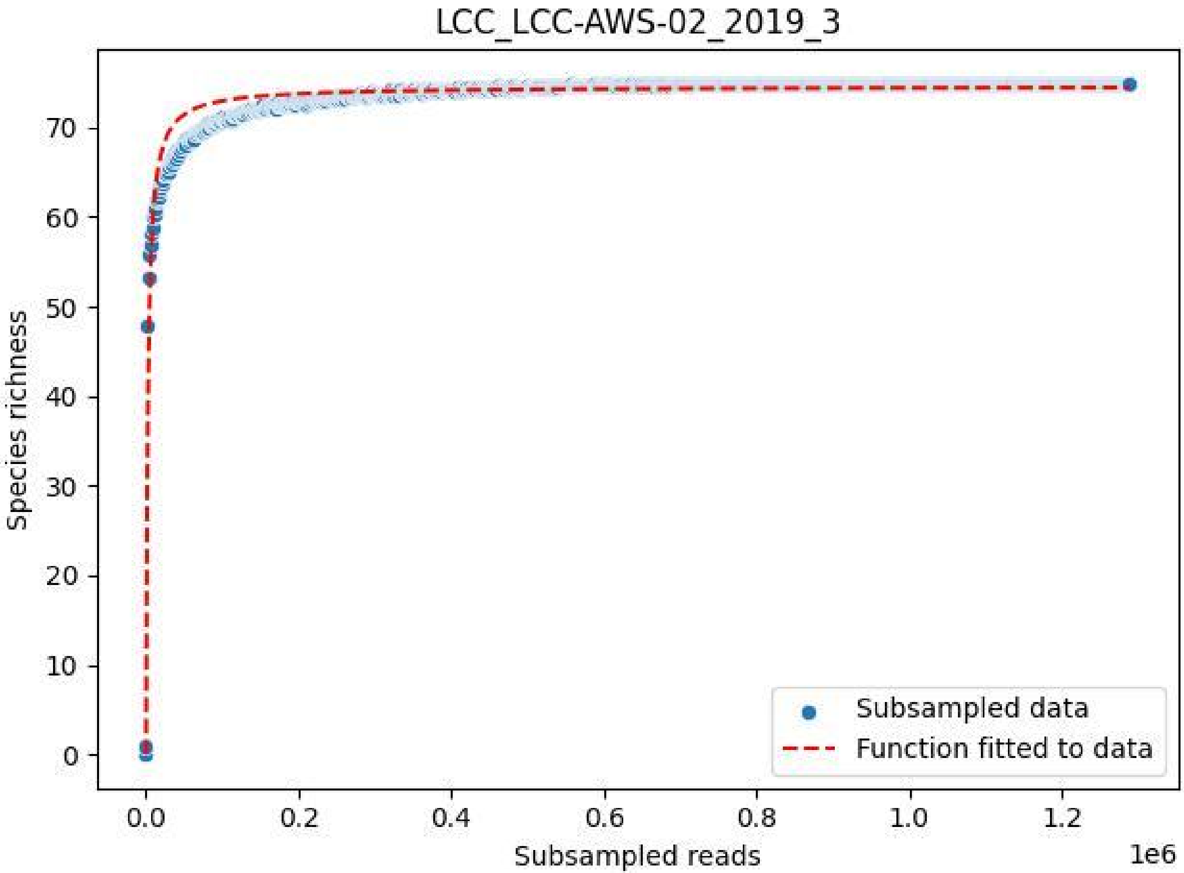
LCC_LCC-AWS-02_2019_3
Species richness
Subsampled reads
1e6
0
10
20
30
40
50
60
70
0.0
0.2
0.4
0.6
0.8
1.0
1.2
Subsampled data
Function fitted to data

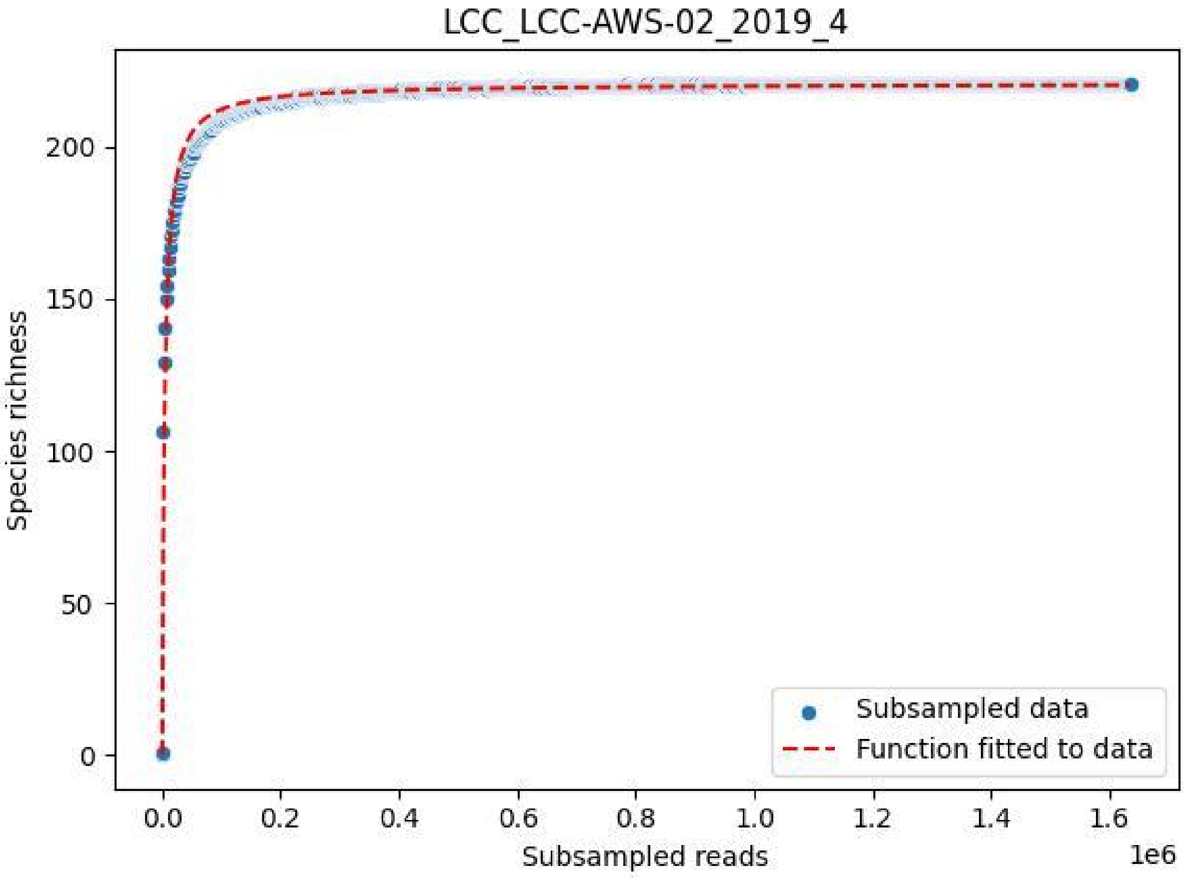
LCC_LCC-AWS-02_2019_4
Species richness
Subsampled reads
1e6
0
50
100
150
200
0.0
0.2
0.4
0.6
0.8
1.0
1.2
1.4
1.6
Subsampled data
Function fitted to data

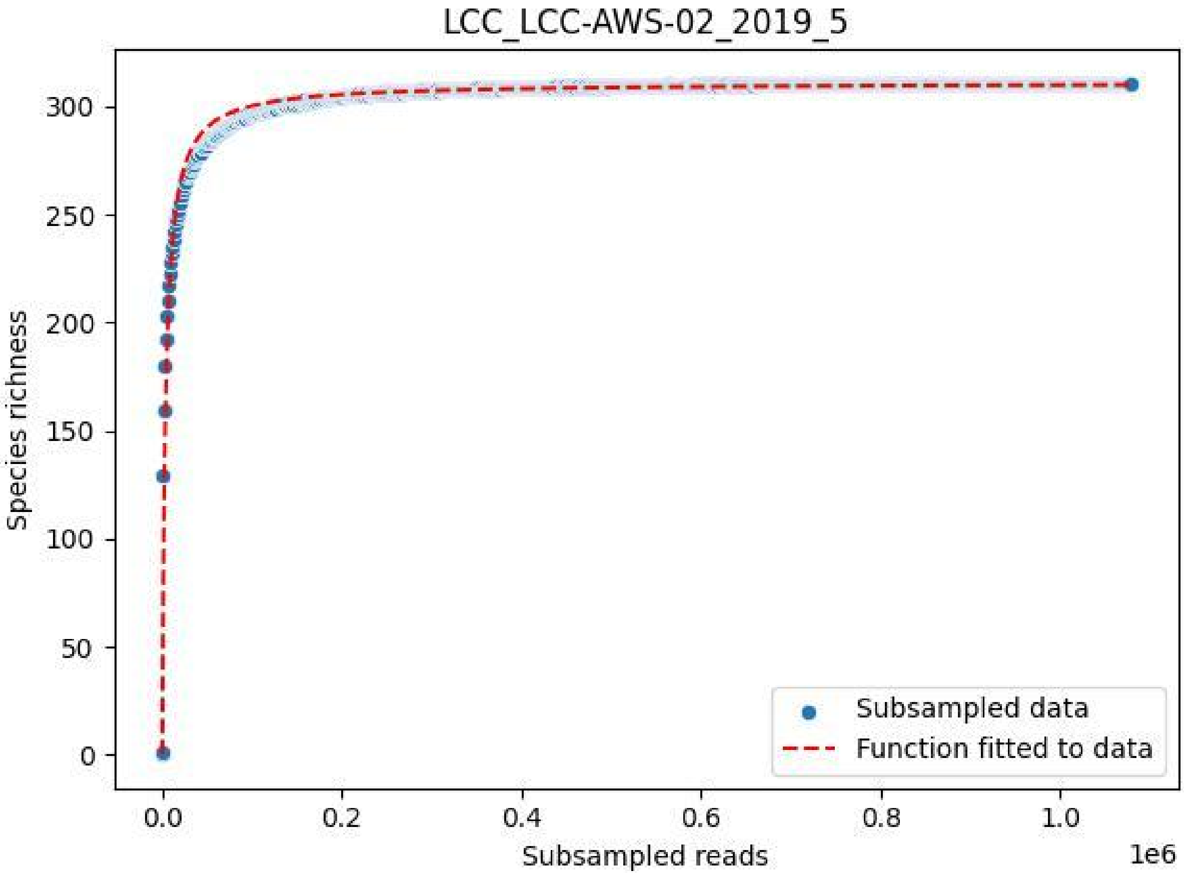

LCC_LCC-AWS-02_2019_5
Species richness
0
50
100
150
200
250
300
Subsampled reads
0.0
0.2
0.4
0.6
0.8
1.0
1e6
Subsampled data
Function fitted to data

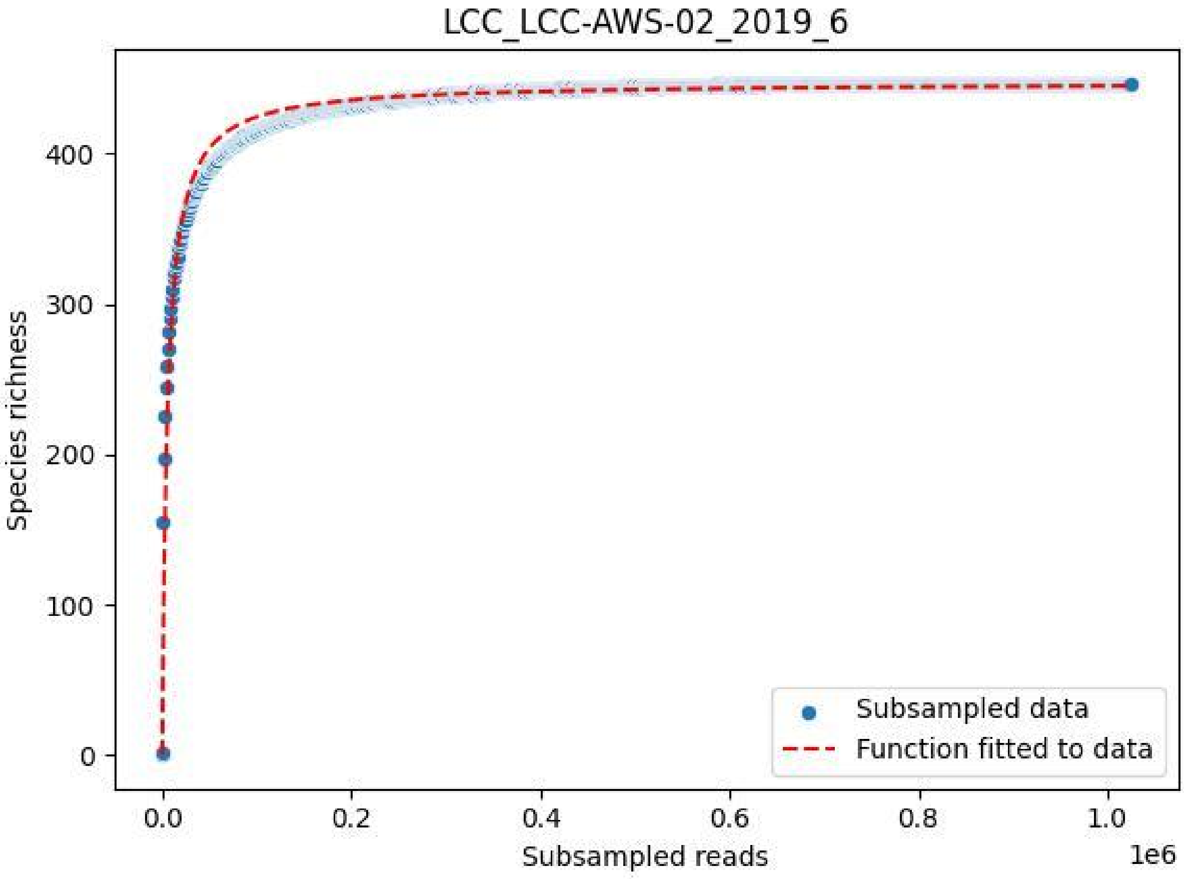
LCC_LCC-AWS-02_2019_6
Species richness
Subsampled reads
1e6
0
100
200
300
400
0.0
0.2
0.4
0.6
0.8
1.0
Subsampled data
Function fitted to data

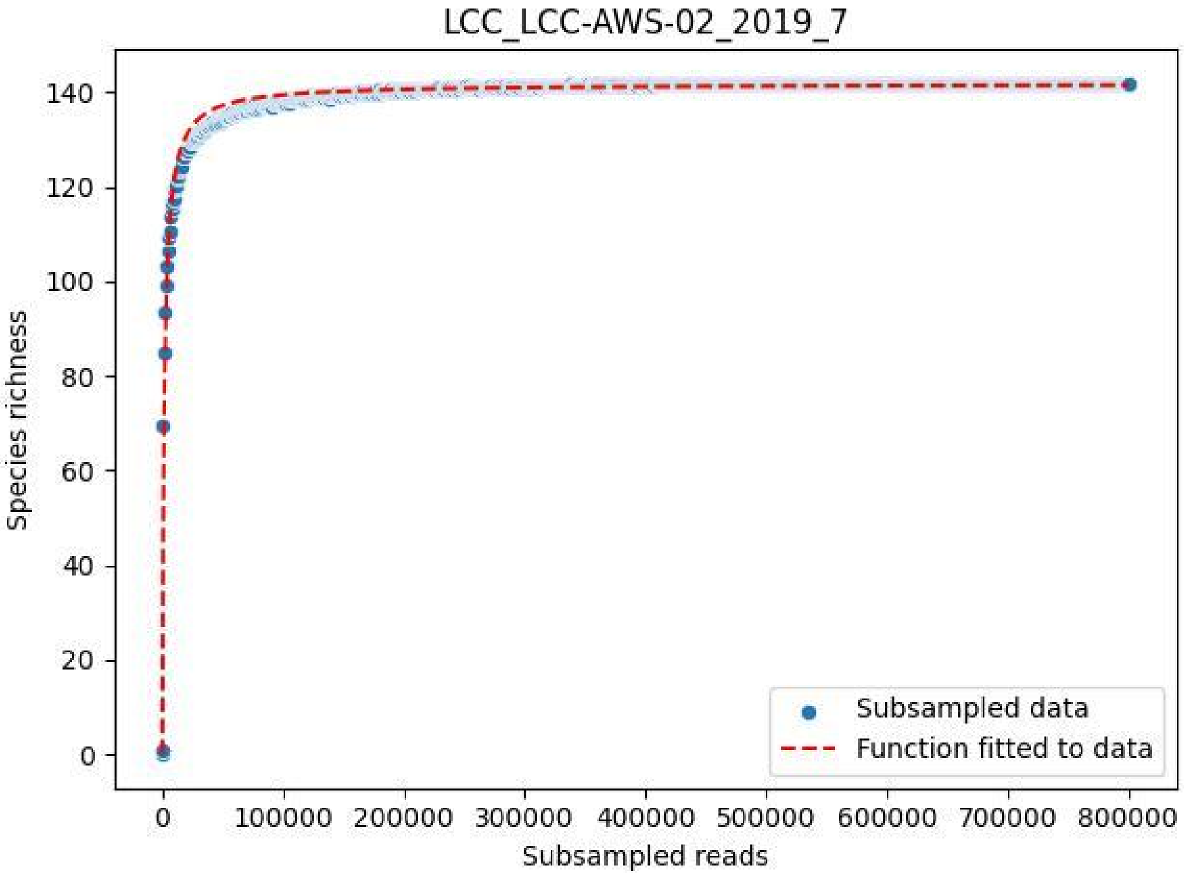

LCC_LCC-AWS-02_2019_7
Species richness
Subsampled reads
0
20
40
60
80
100
120
140
0
100000
200000
300000
400000
500000
600000
700000
800000
Subsampled data
Function fitted to data

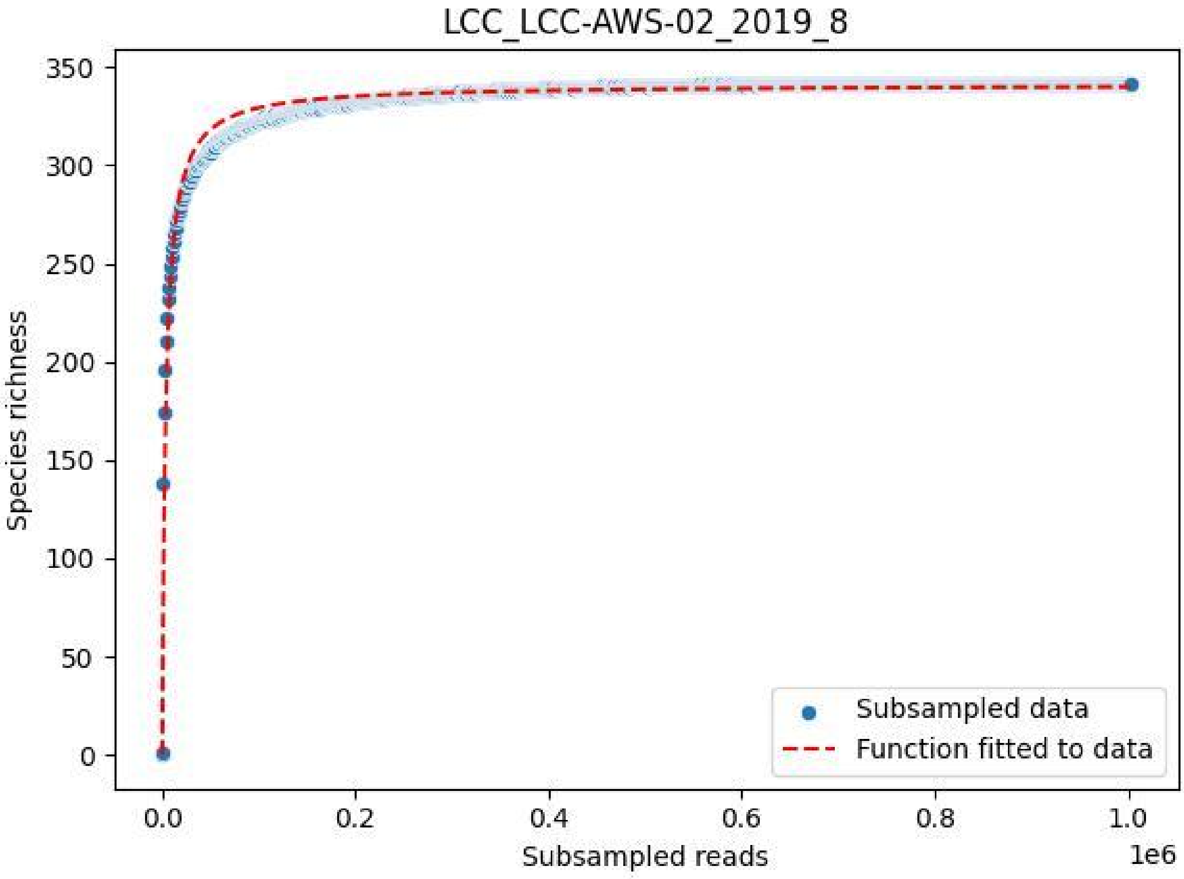
LCC_LCC-AWS-02_2019_8
Species richness
Subsampled reads
1e6
Subsampled data
Function fitted to data
0
50
100
150
200
250
300
350
0.0
0.2
0.4
0.6
0.8
1.0

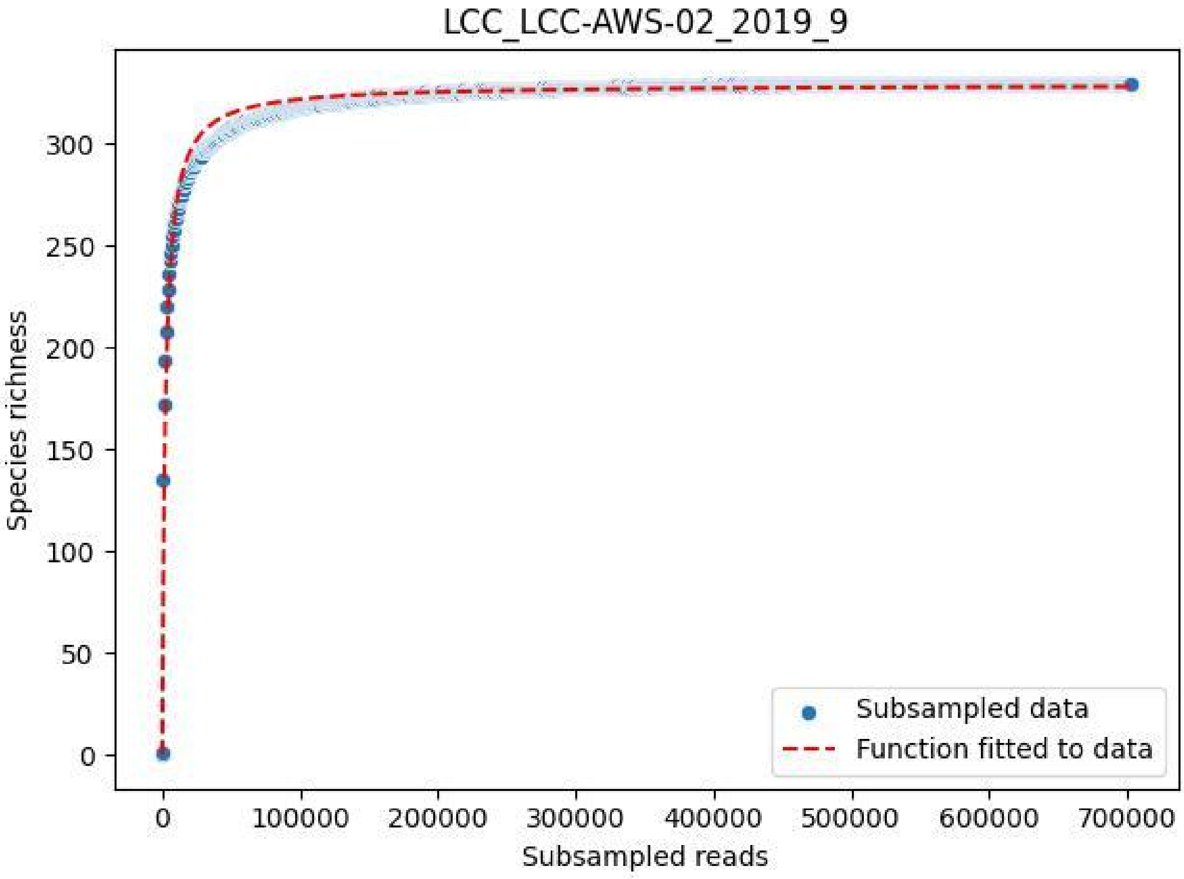

LCC_LCC-AWS-02_2019_9
Species richness
Subsampled reads
0
50
100
150
200
250
300
0
100000
200000
300000
400000
500000
600000
700000
Subsampled data
Function fitted to data

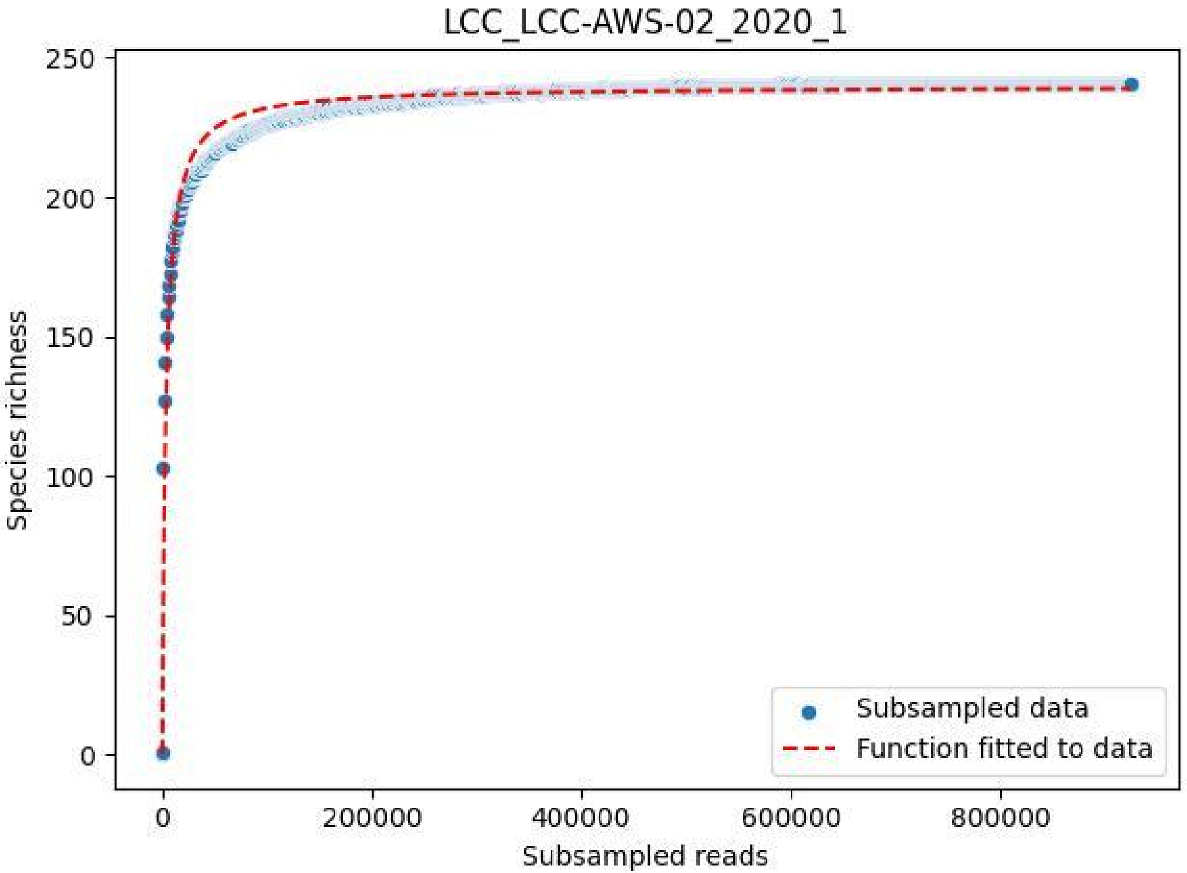
LCC_LCC-AWS-02_2020_1
Species richness
Subsampled reads
0
50
100
150
200
250
0
200000
400000
600000
800000
Subsampled data
Function fitted to data

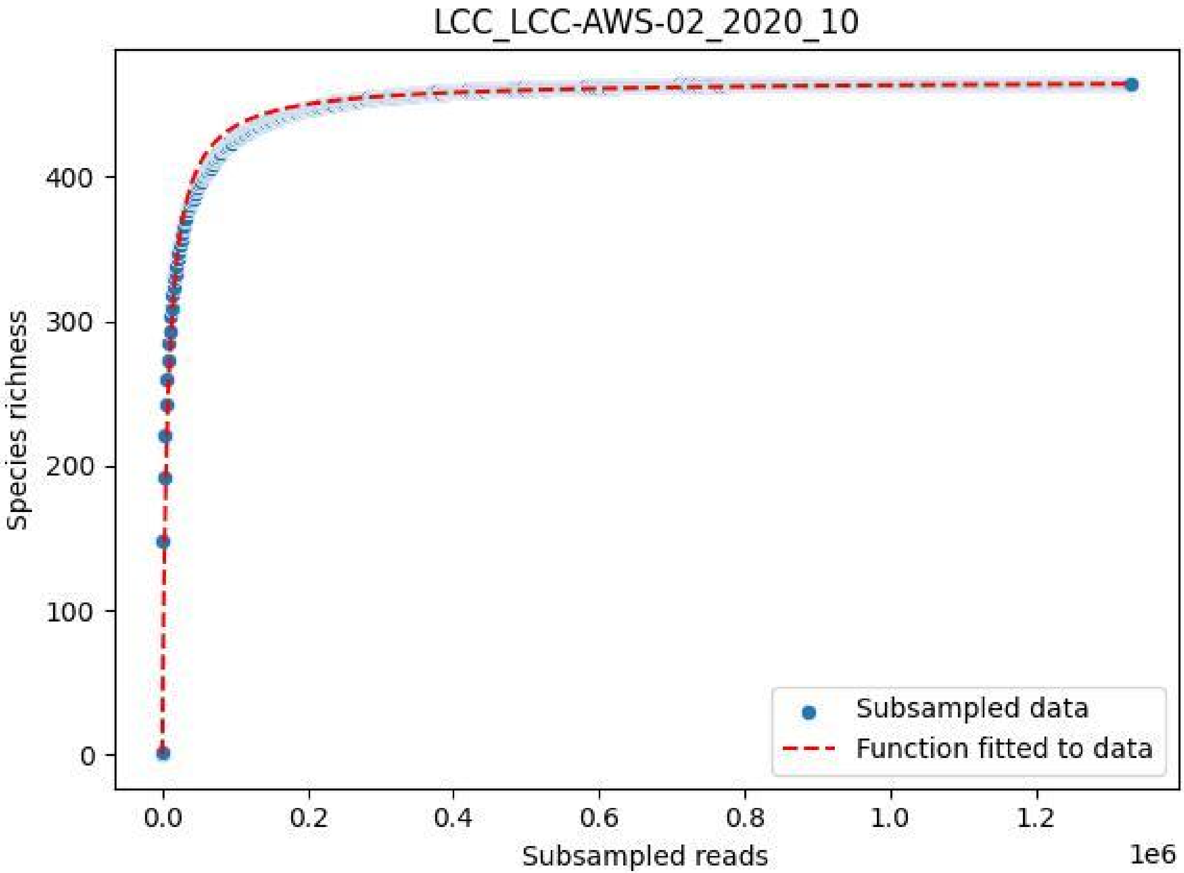
LCC_LCC-AWS-02_2020_10
Species richness
Subsampled reads
1e6
0
100
200
300
400
0.0
0.2
0.4
0.6
0.8
1.0
1.2
Subsampled data
Function fitted to data

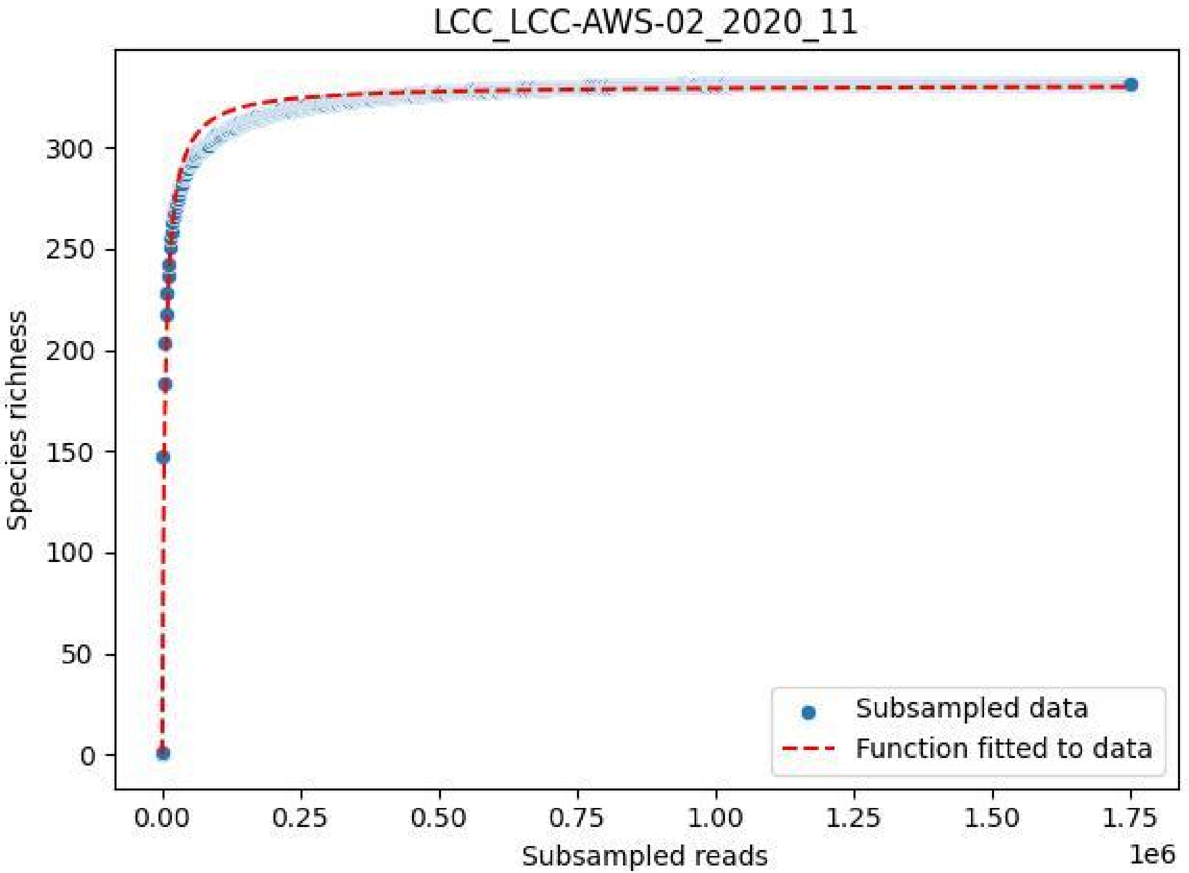
LCC_LCC-AWS-02_2020_11
Species richness
Subsampled reads
1e6
0
50
100
150
200
250
300
0.00
0.25
0.50
0.75
1.00
1.25
1.50
1.75
Subsampled data
Function fitted to data

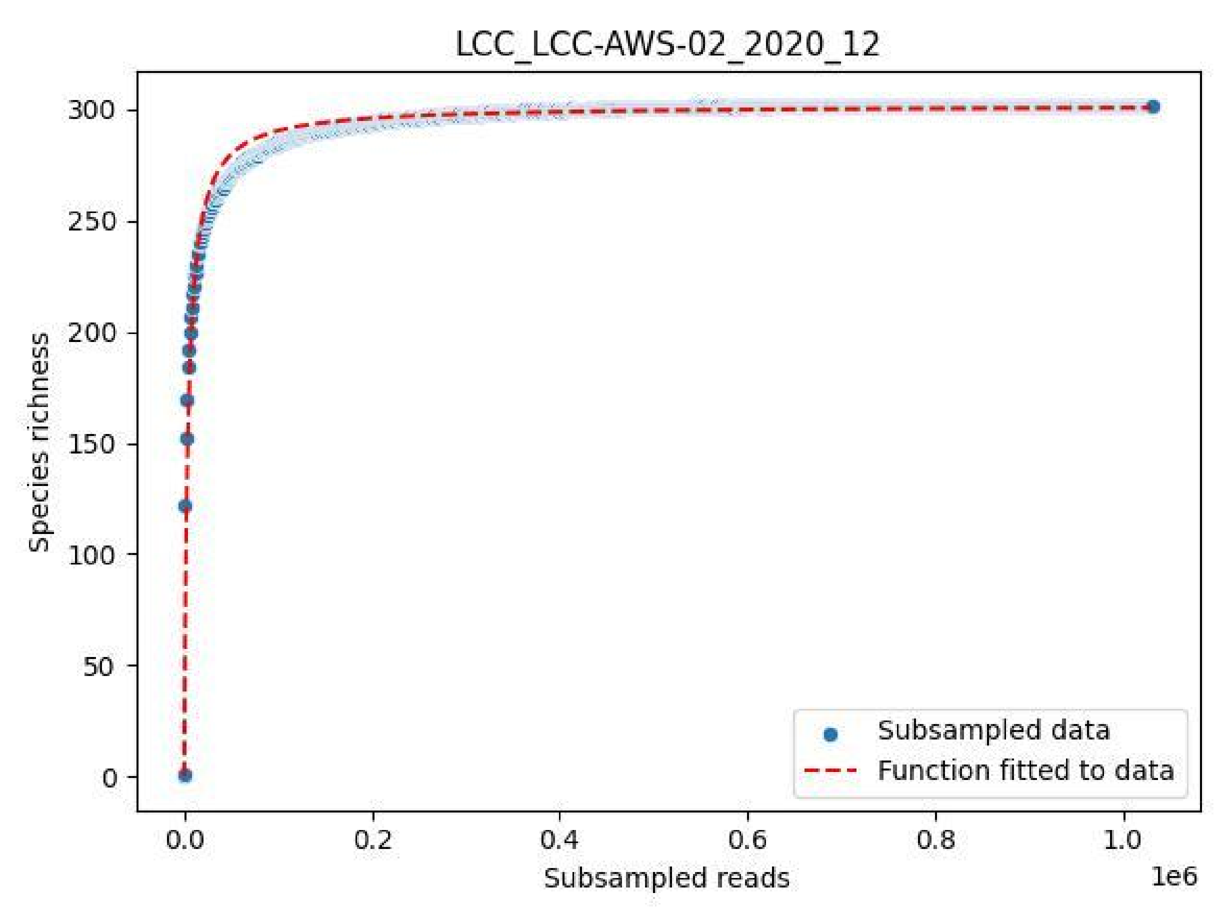

LCC_LCC-AWS-02_2020_12
Species richness
Subsampled reads
1e6
0
50
100
150
200
250
300
0.0
0.2
0.4
0.6
0.8
1.0
Subsampled data
Function fitted to data

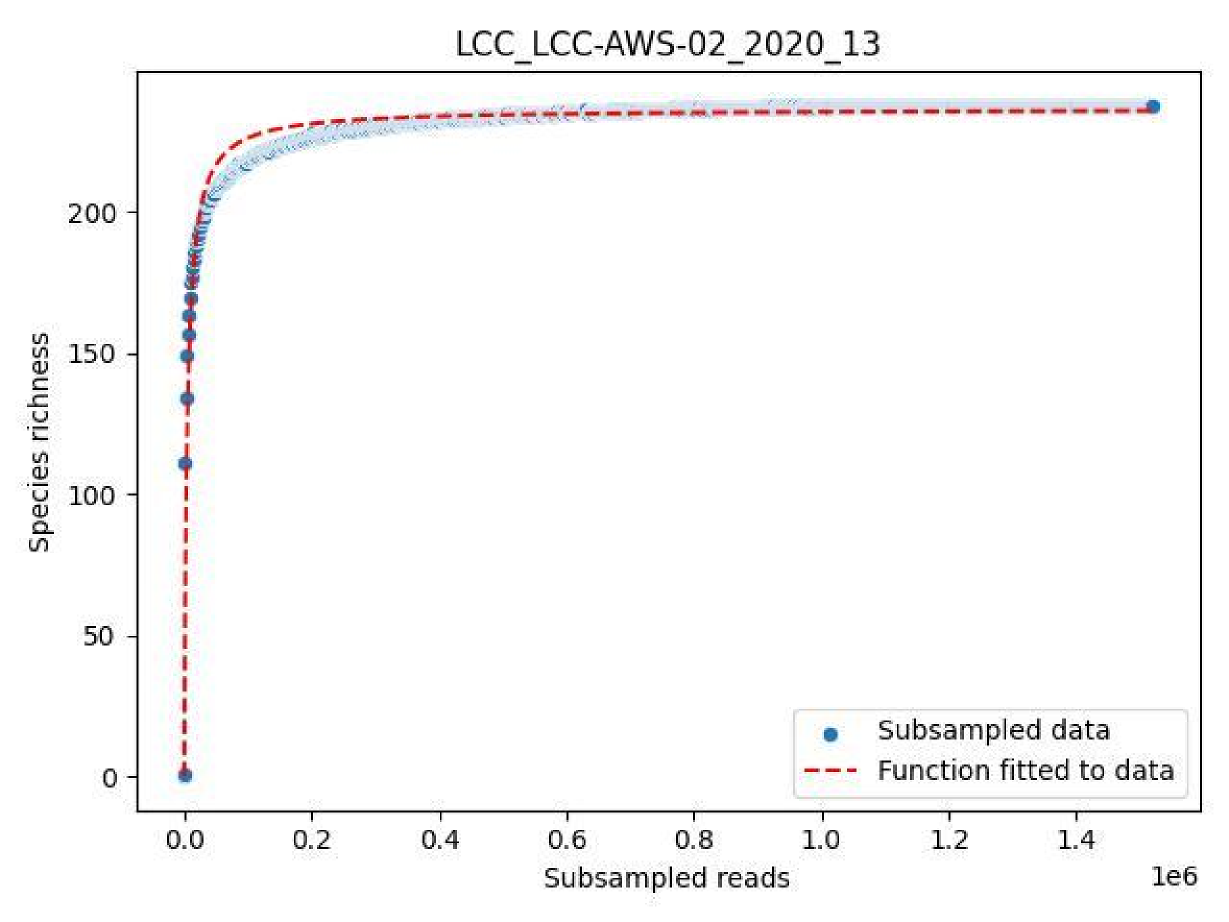

LCC_LCC-AWS-02_2020_13
Species richness
Subsampled reads
1e6
Subsampled data
Function fitted to data
0
50
100
150
200
0.0
0.2
0.4
0.6
0.8
1.0
1.2
1.4

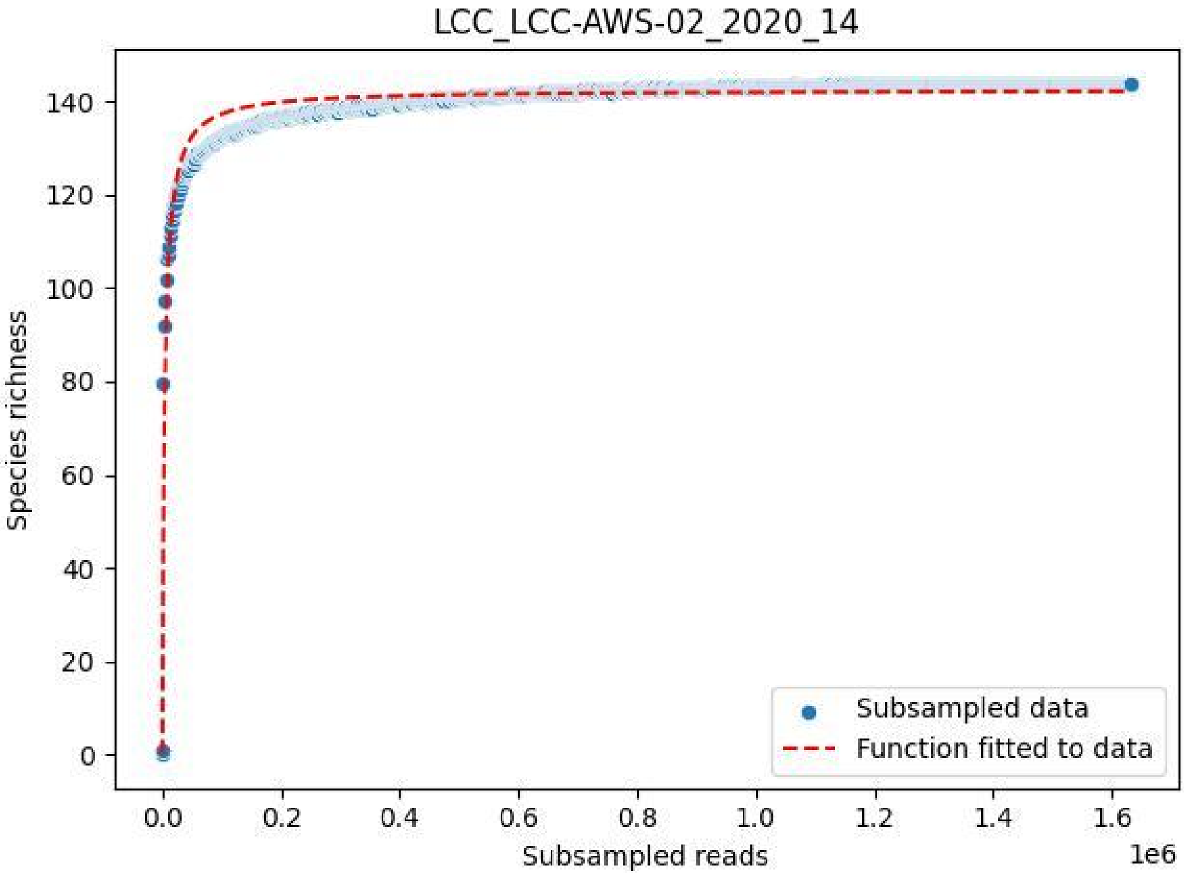
LCC_LCC-AWS-02_2020_14
Species richness
0
20
40
60
80
100
120
140
0.0
0.2
0.4
0.6
0.8
1.0
1.2
1.4
1.6
1e6
Subsampled reads
Subsampled data
Function fitted to data

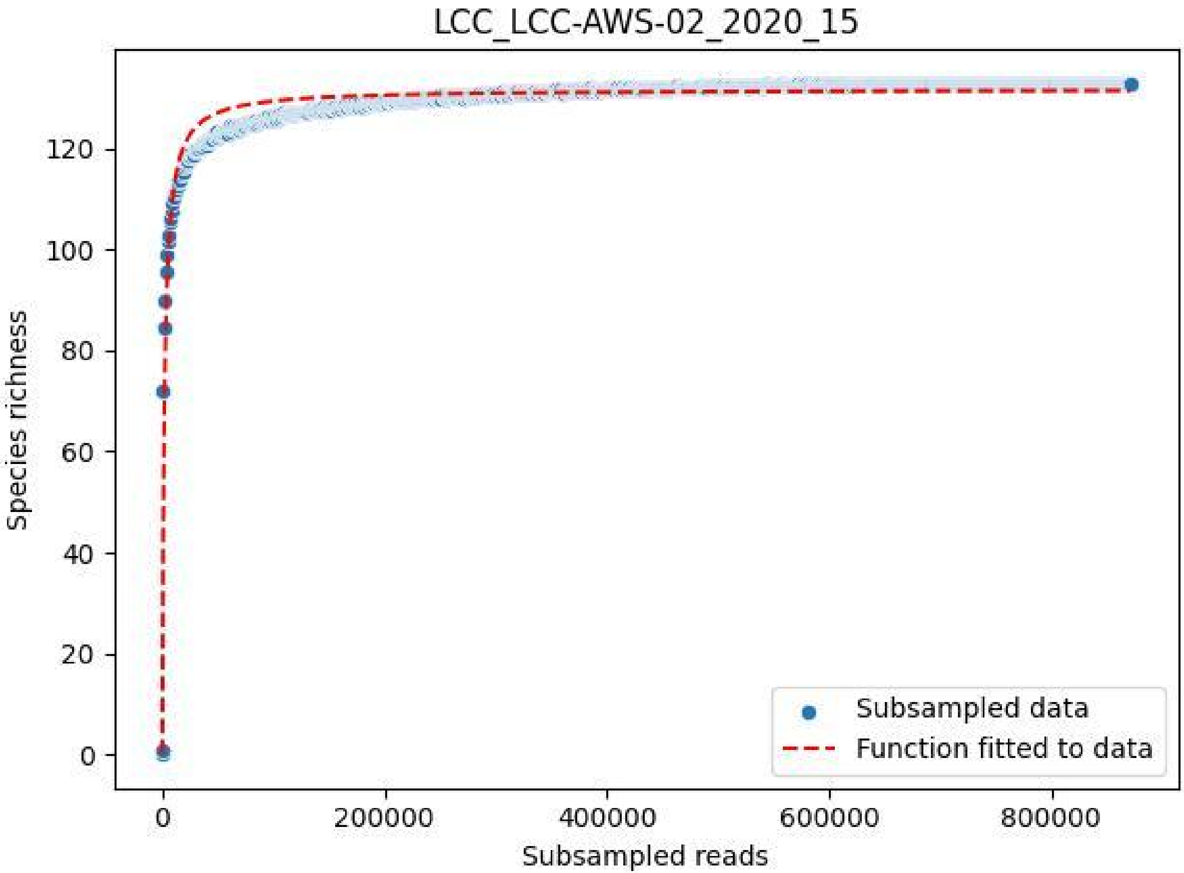
LCC_LCC-AWS-02_2020_15
Species richness
Subsampled reads
0
20
40
60
80
100
120
0
200000
400000
600000
800000
Subsampled data
Function fitted to data

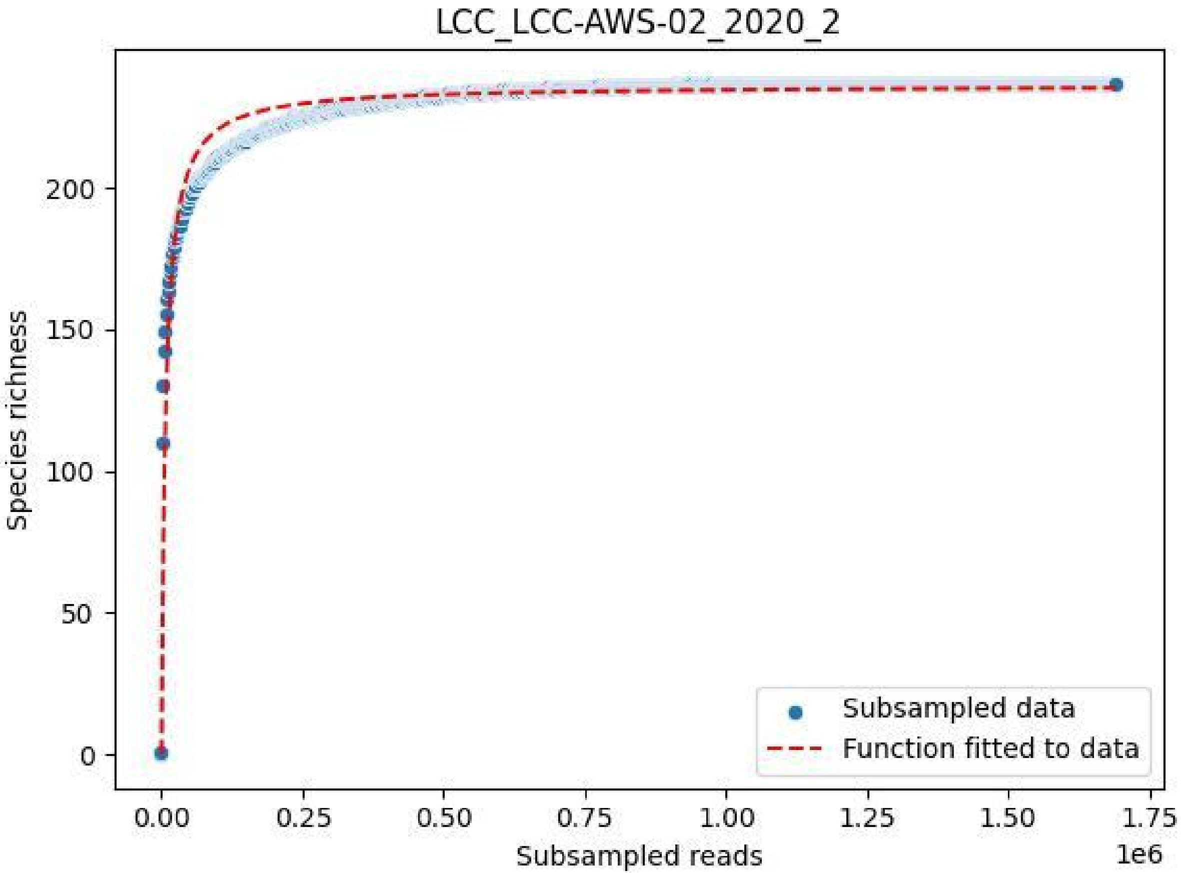
LCC_LCC-AWS-02_2020_2
Species richness
Subsampled reads
1e6
0
50
100
150
200
0.00
0.25
0.50
0.75
1.00
1.25
1.50
1.75
Subsampled data
Function fitted to data

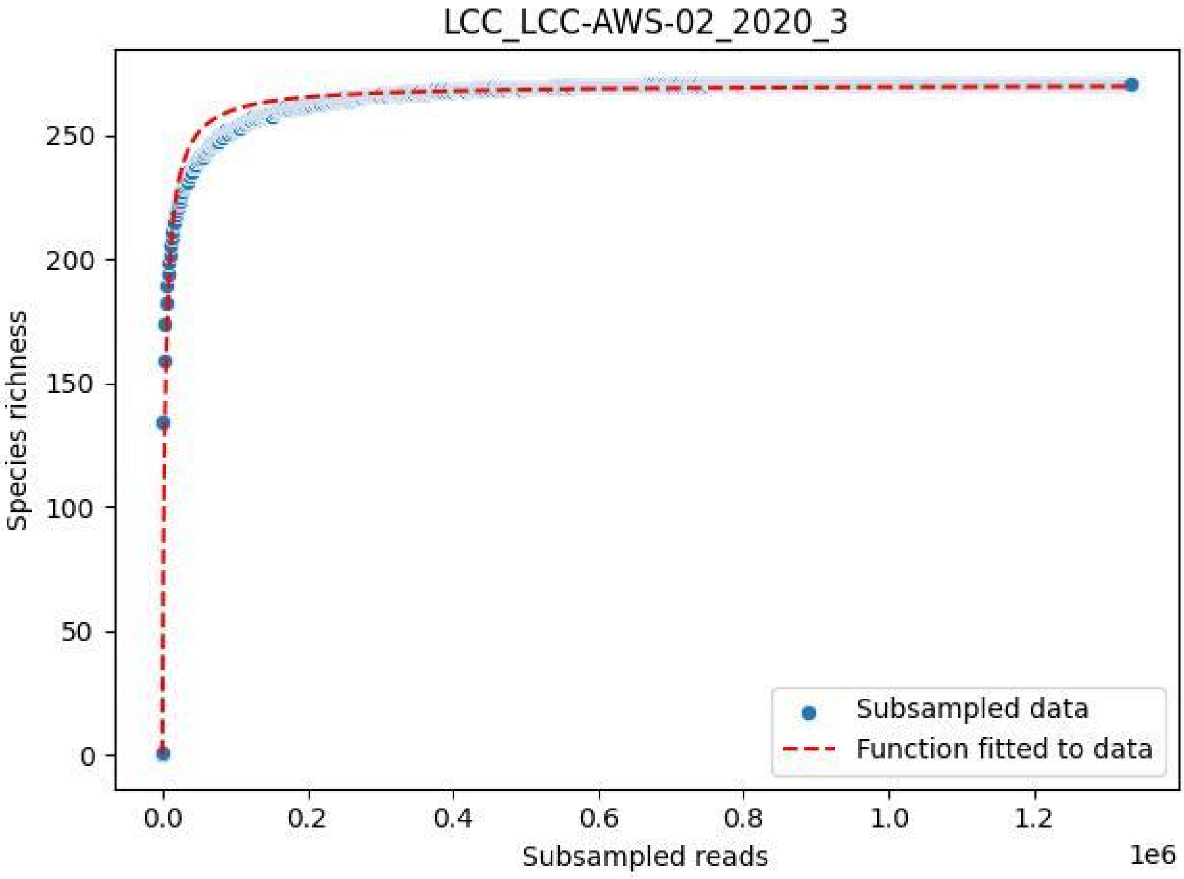
LCC_LCC-AWS-02_2020_3
Species richness
0
50
100
150
200
250
0.0
0.2
0.4
0.6
0.8
1.0
1.2
1e6
Subsampled reads
Subsampled data
Function fitted to data

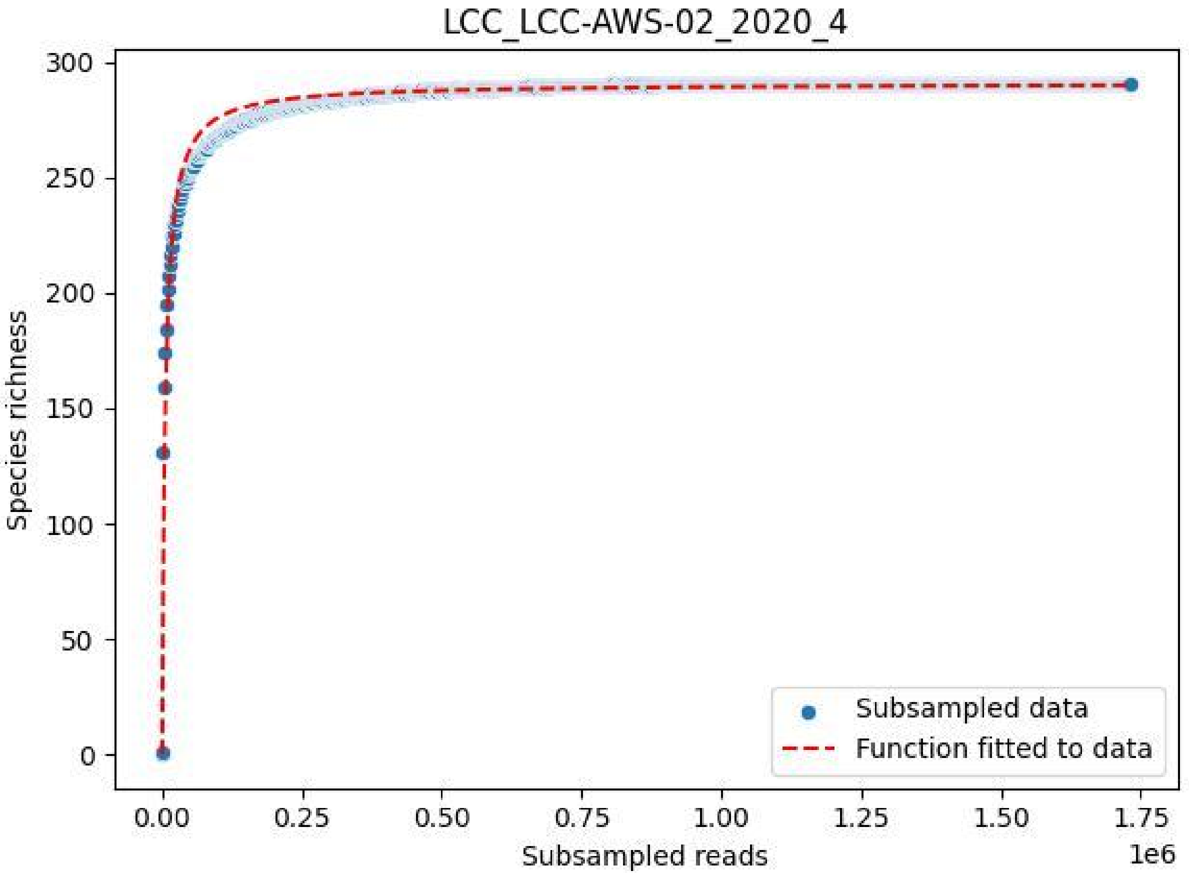
LCC_LCC-AWS-02_2020_4
Species richness
Subsampled reads
1e6
0
50
100
150
200
250
300
0.00
0.25
0.50
0.75
1.00
1.25
1.50
1.75
Subsampled data
Function fitted to data

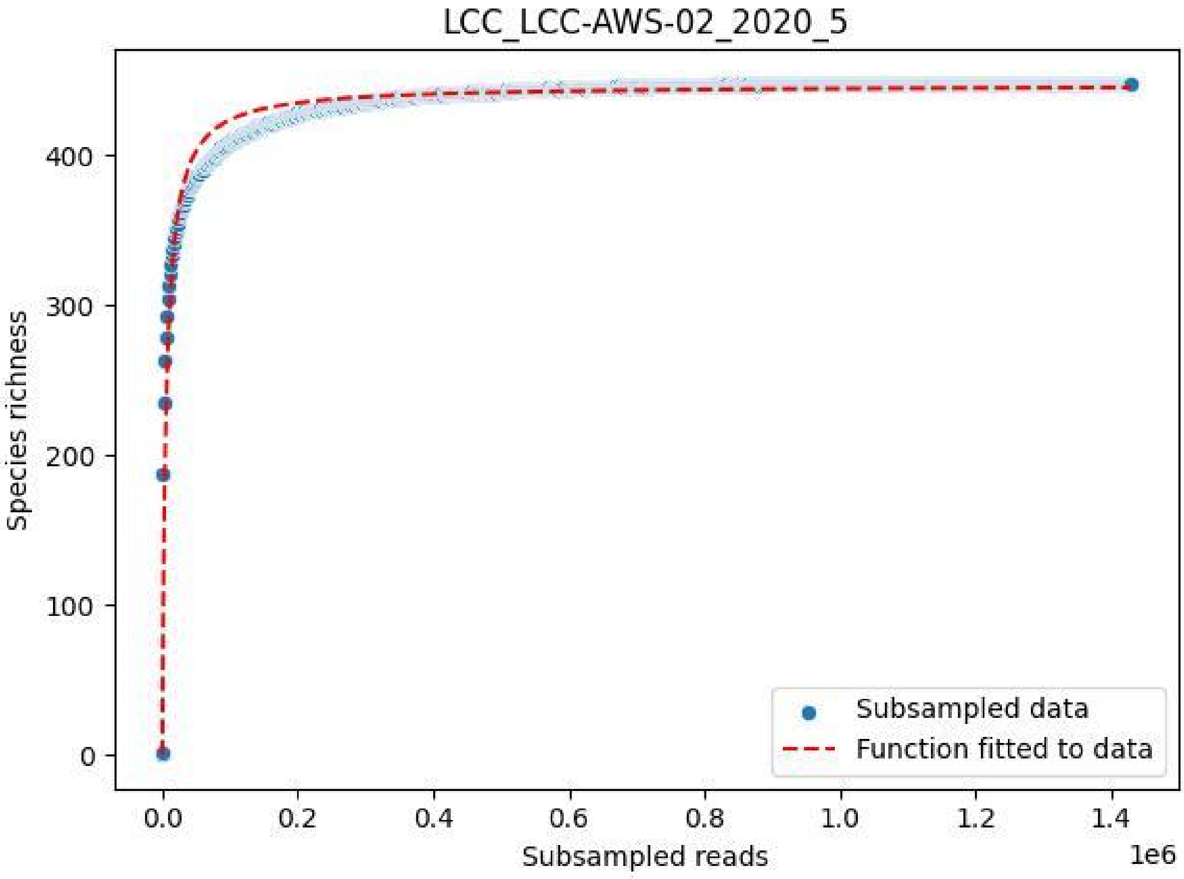
LCC_LCC-AWS-02_2020_5
Species richness
Subsampled reads
1e6
Subsampled data
Function fitted to data
0
100
200
300
400
0.0
0.2
0.4
0.6
0.8
1.0
1.2
1.4

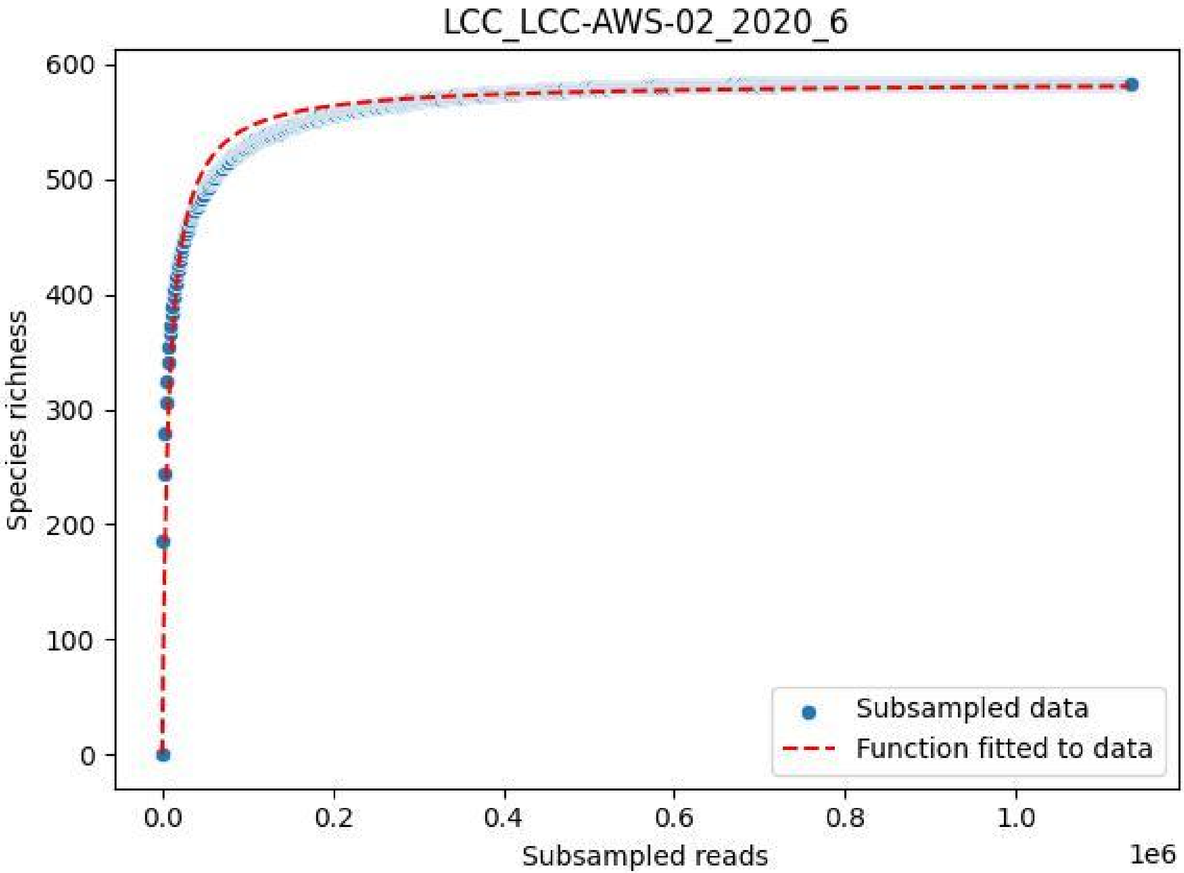
LCC_LCC-AWS-02_2020_6
Species richness
Subsampled reads
1e6
0
100
200
300
400
500
600
0.0
0.2
0.4
0.6
0.8
1.0
Subsampled data
Function fitted to data

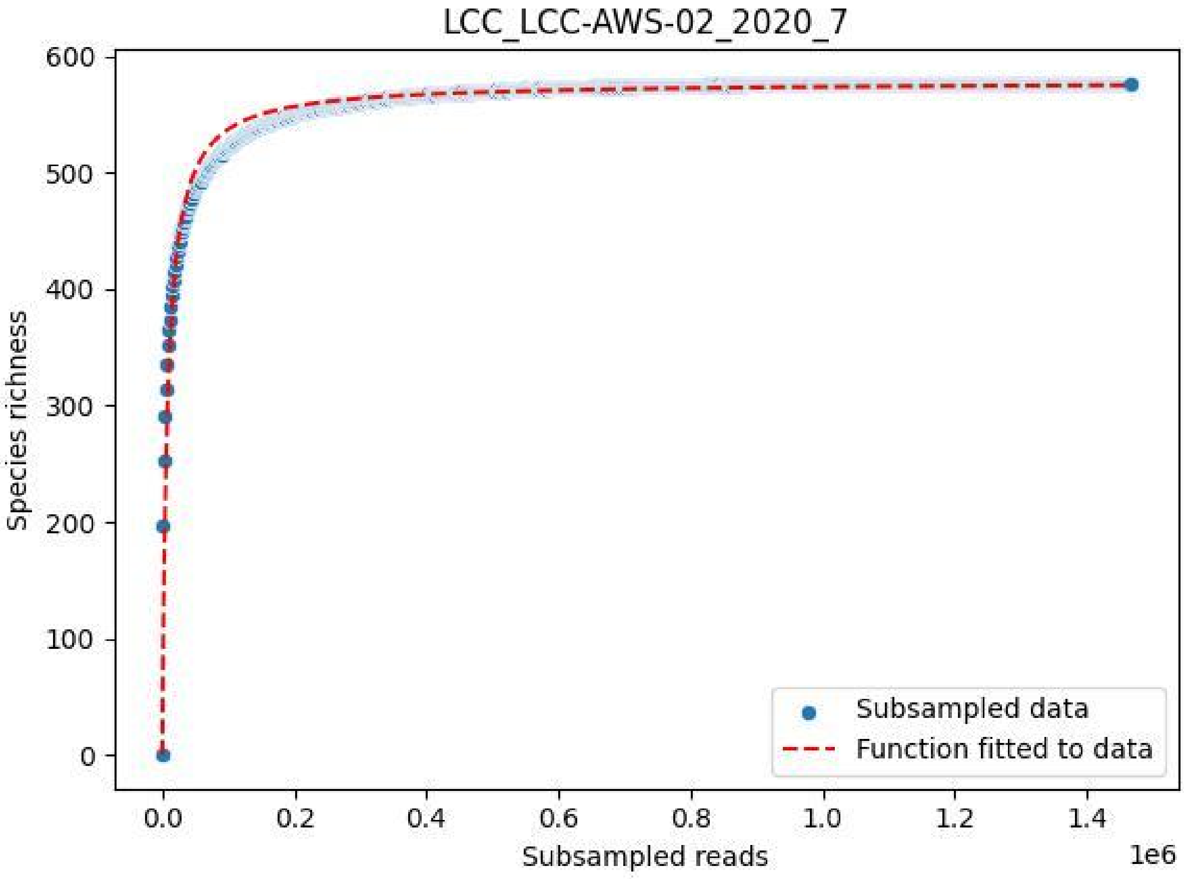
LCC_LCC-AWS-02_2020_7
Species richness
Subsampled reads
1e6
0
100
200
300
400
500
600
0.0
0.2
0.4
0.6
0.8
1.0
1.2
1.4
Subsampled data
Function fitted to data

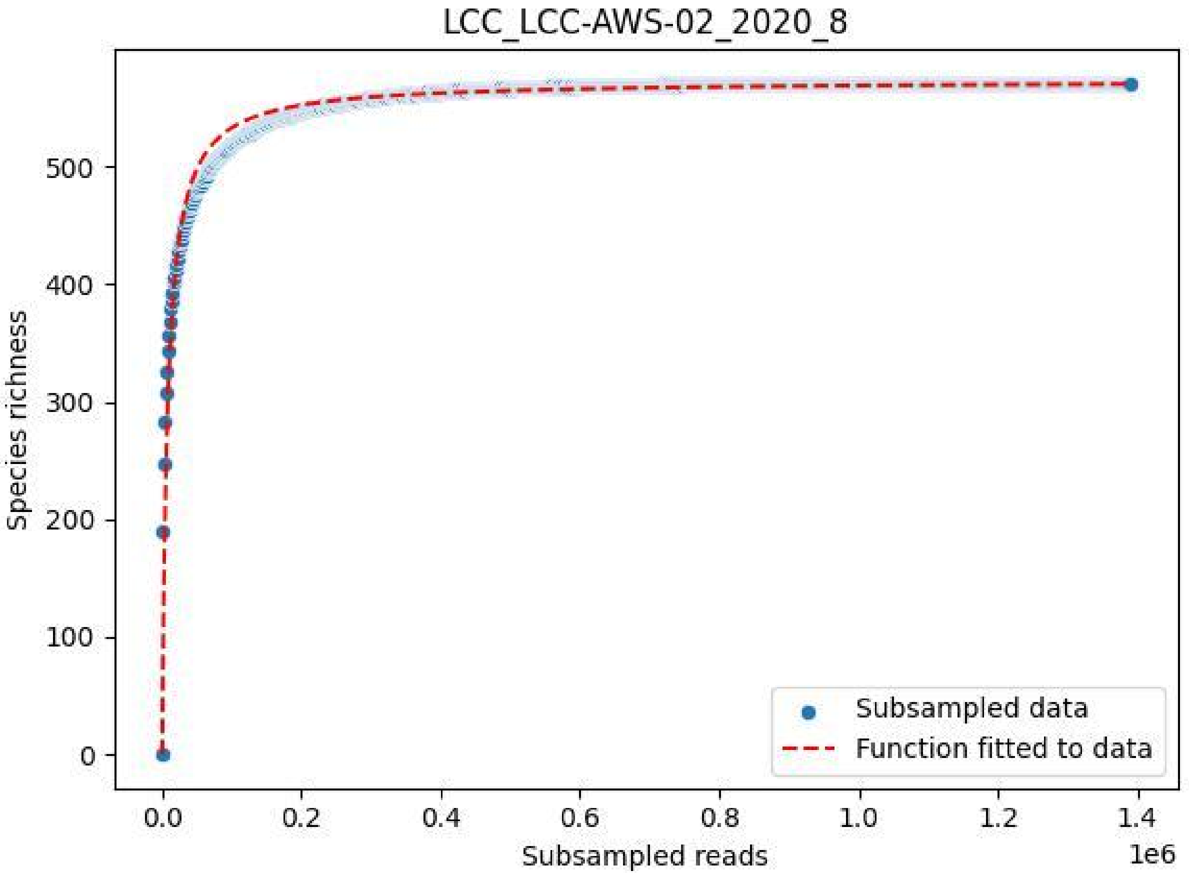
LCC_LCC-AWS-02_2020_8
Species richness
Subsampled reads
1e6
0
100
200
300
400
500
0.0
0.2
0.4
0.6
0.8
1.0
1.2
1.4
Subsampled data
Function fitted to data

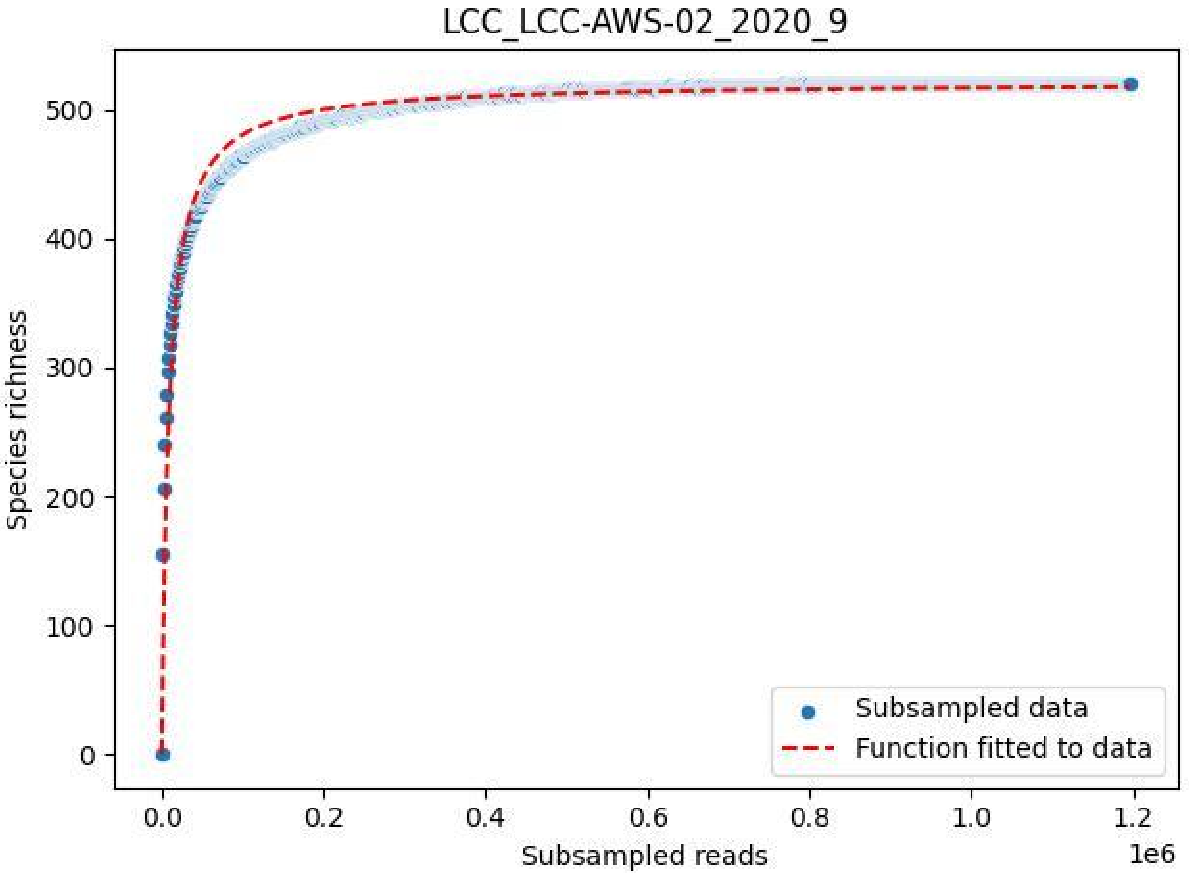

LCC_LCC-AWS-02_2020_9
Species richness
500
400
300
200
100
0
0.0
0.2
0.4
0.6
0.8
1.0
1.2
1e6
Subsampled reads
Subsampled data
Function fitted to data

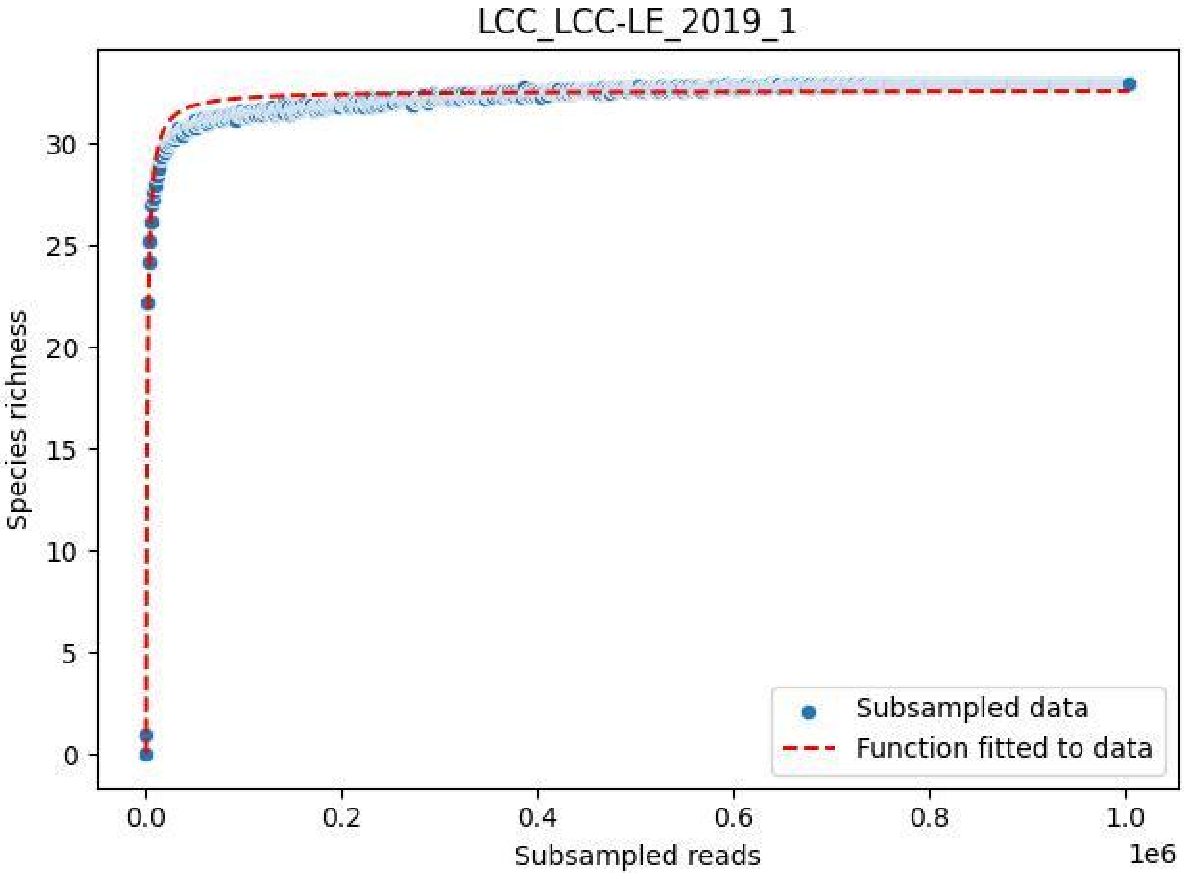
LCC_LCC-LE_2019_1
Species richness
Subsampled reads
1e6
0
5
10
15
20
25
30
0.0
0.2
0.4
0.6
0.8
1.0
Subsampled data
Function fitted to data

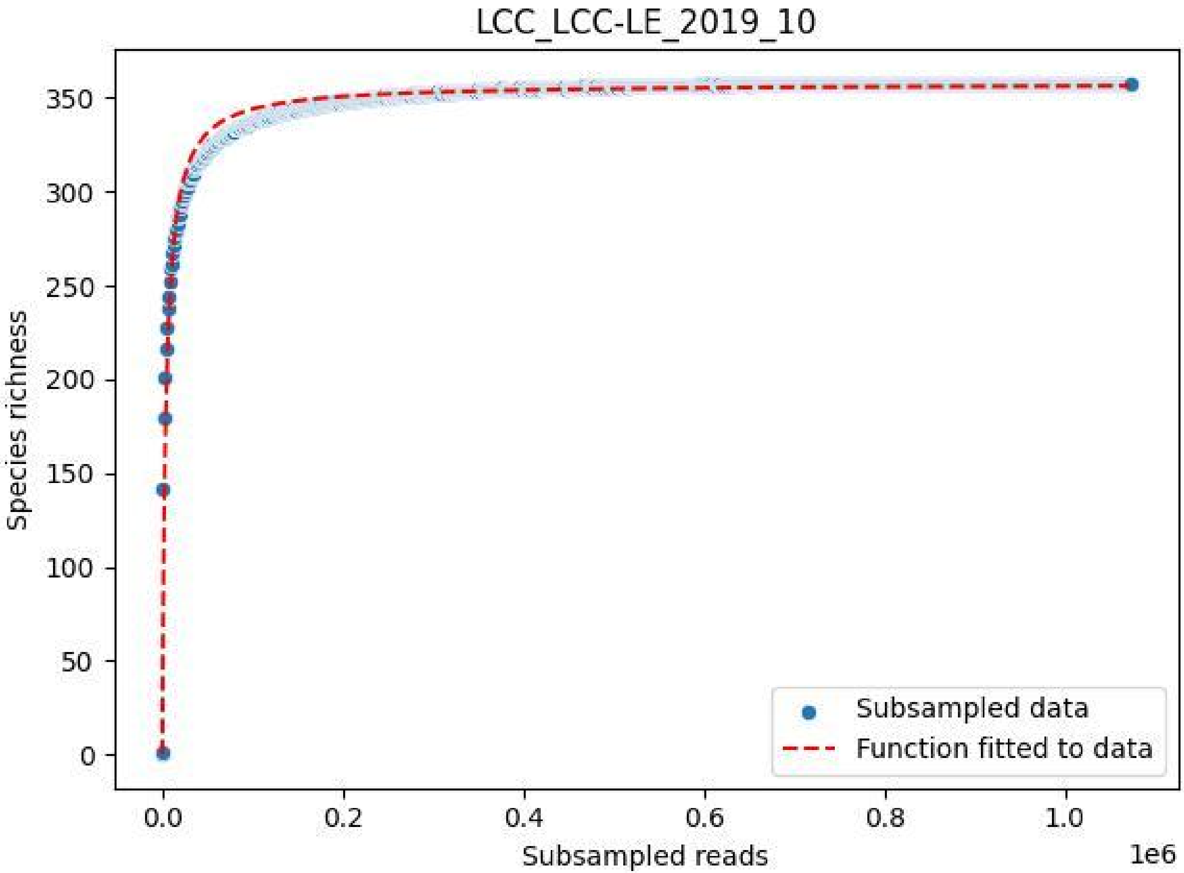

LCC_LCC-LE_2019_10
Species richness
Subsampled reads
1e6
0
50
100
150
200
250
300
350
0.0
0.2
0.4
0.6
0.8
1.0
Subsampled data
Function fitted to data

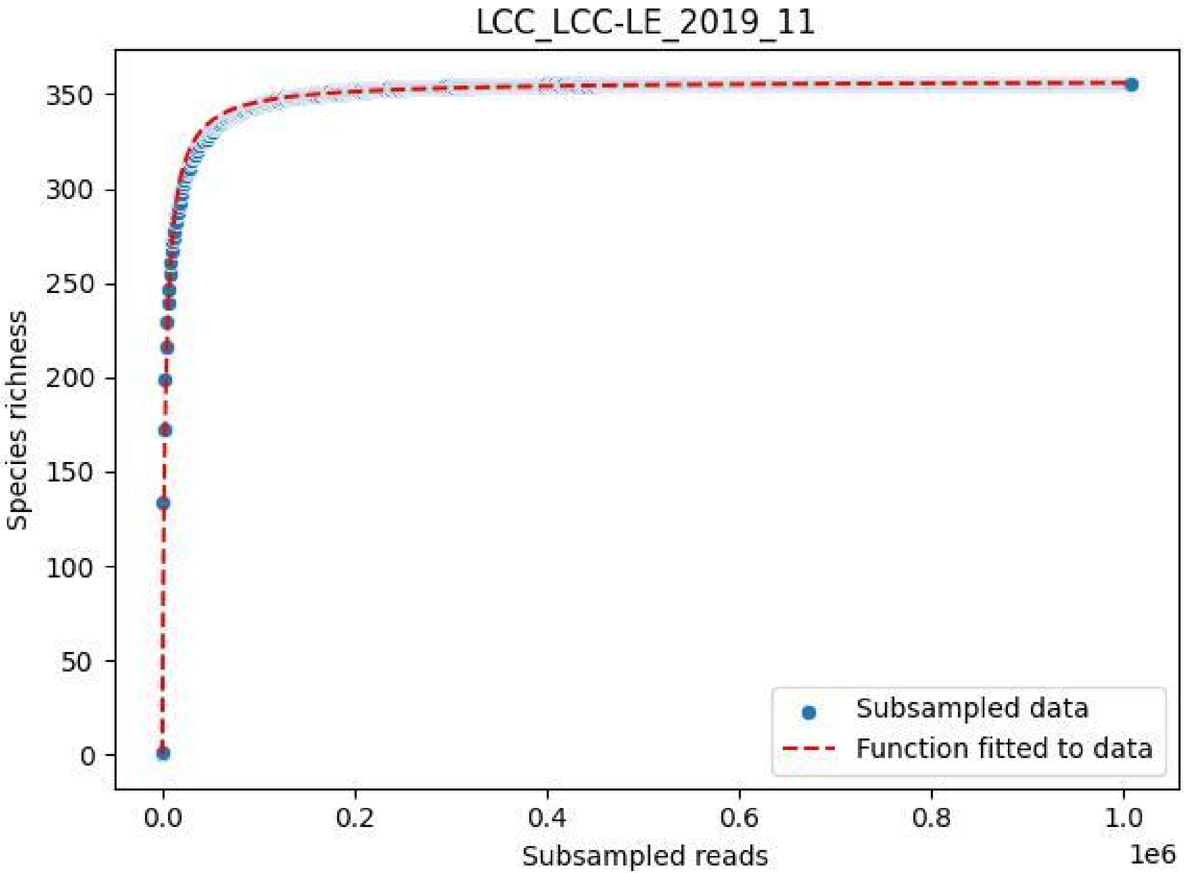
LCC_LCC-LE_2019_11
Species richness
Subsampled reads
1e6
0
50
100
150
200
250
300
350
0.0
0.2
0.4
0.6
0.8
1.0
Subsampled data
Function fitted to data

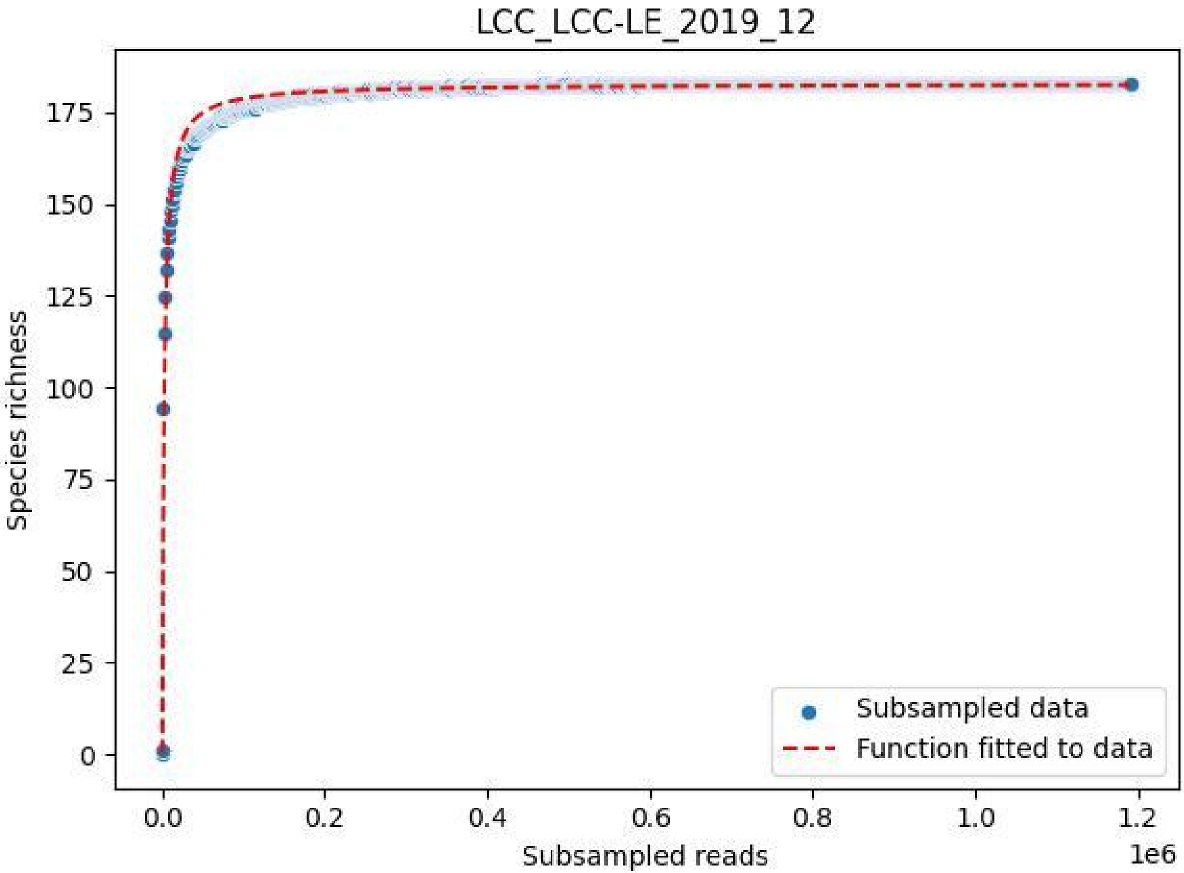
LCC_LCC-LE_2019_12
Species richness
Subsampled reads
1e6
0
25
50
75
100
125
150
175
0.0
0.2
0.4
0.6
0.8
1.0
1.2
Subsampled data
Function fitted to data

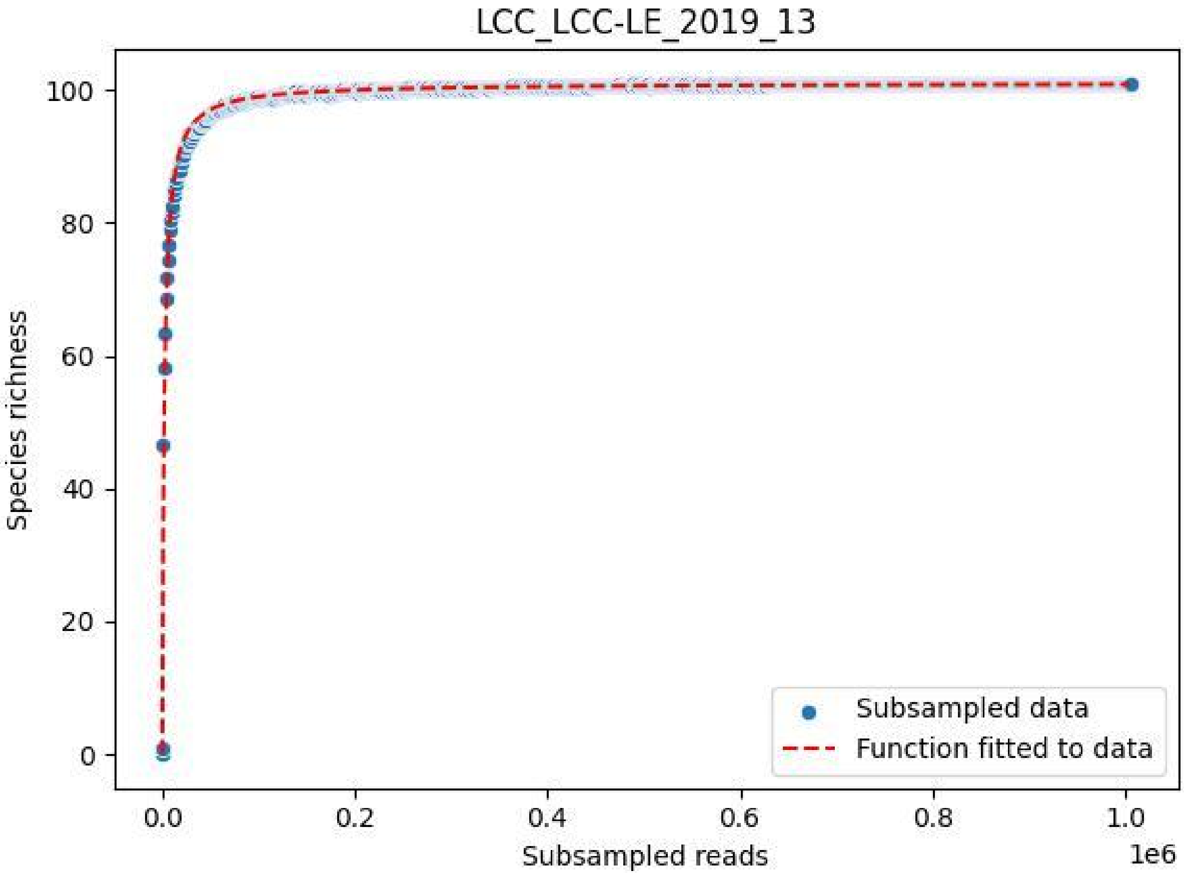
LCC_LCC-LE_2019_13
Species richness
Subsampled reads
1e6
0
20
40
60
80
100
0.0
0.2
0.4
0.6
0.8
1.0
Subsampled data
Function fitted to data

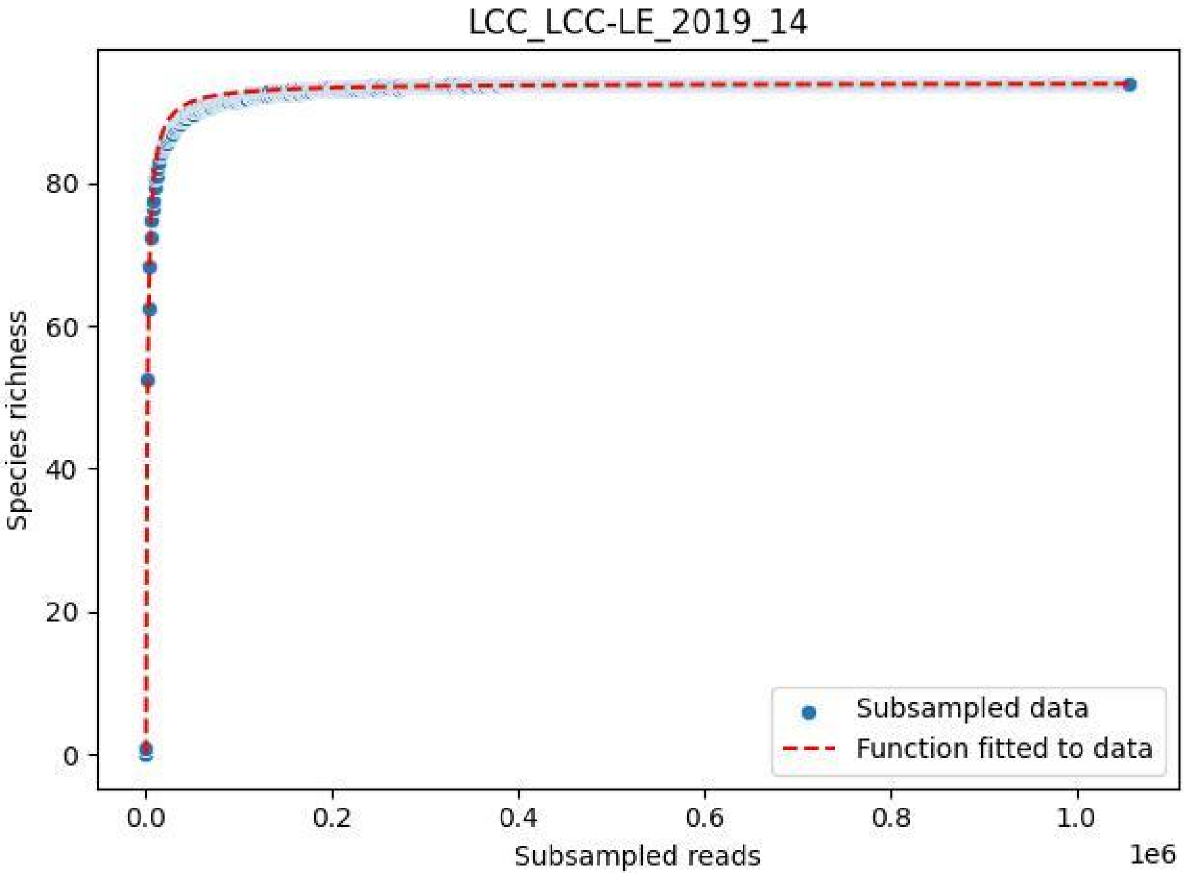

LCC_LCC-LE_2019_14
Species richness
0
20
40
60
80
0.0
0.2
0.4
0.6
0.8
1.0
1e6
Subsampled reads
Subsampled data
Function fitted to data

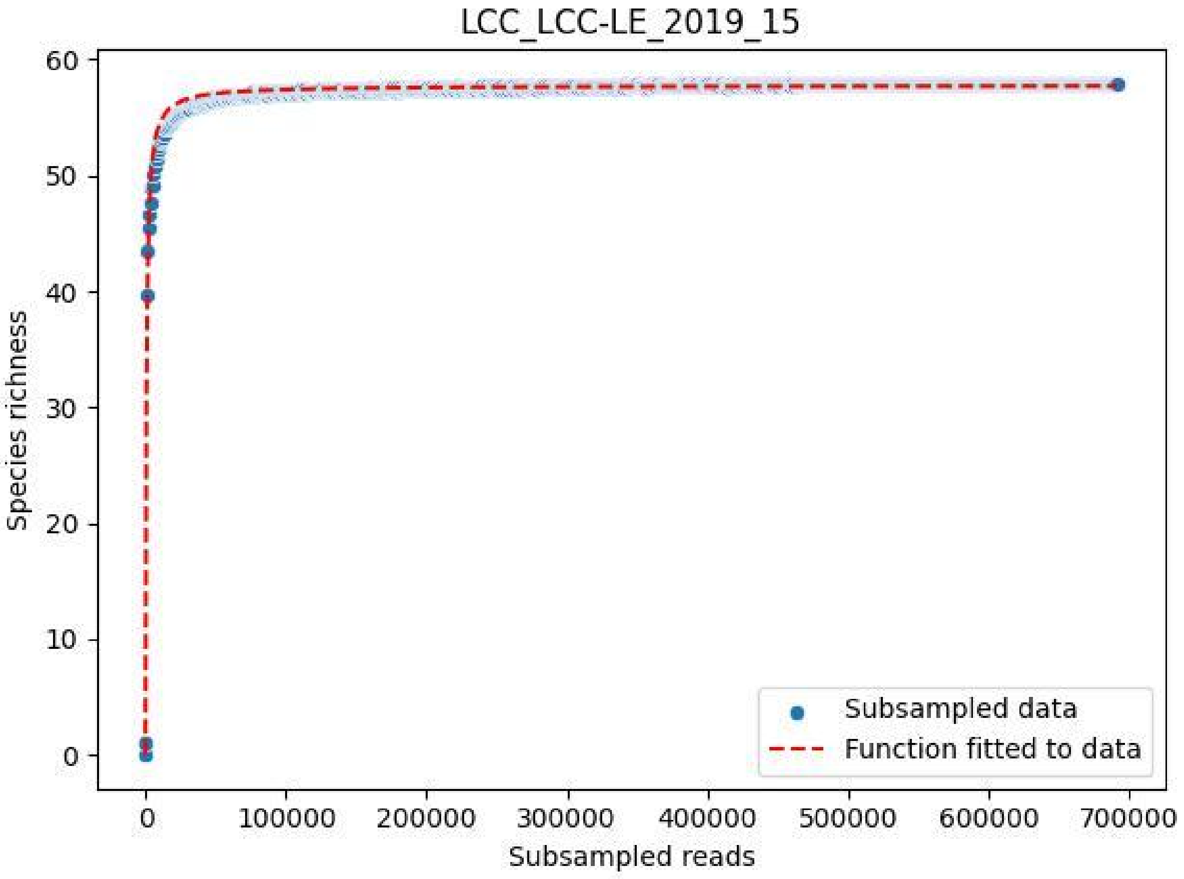

LCC_LCC-LE_2019_15
Species richness
Subsampled reads
0
10
20
30
40
50
60
0
100000
200000
300000
400000
500000
600000
700000
Subsampled data
Function fitted to data

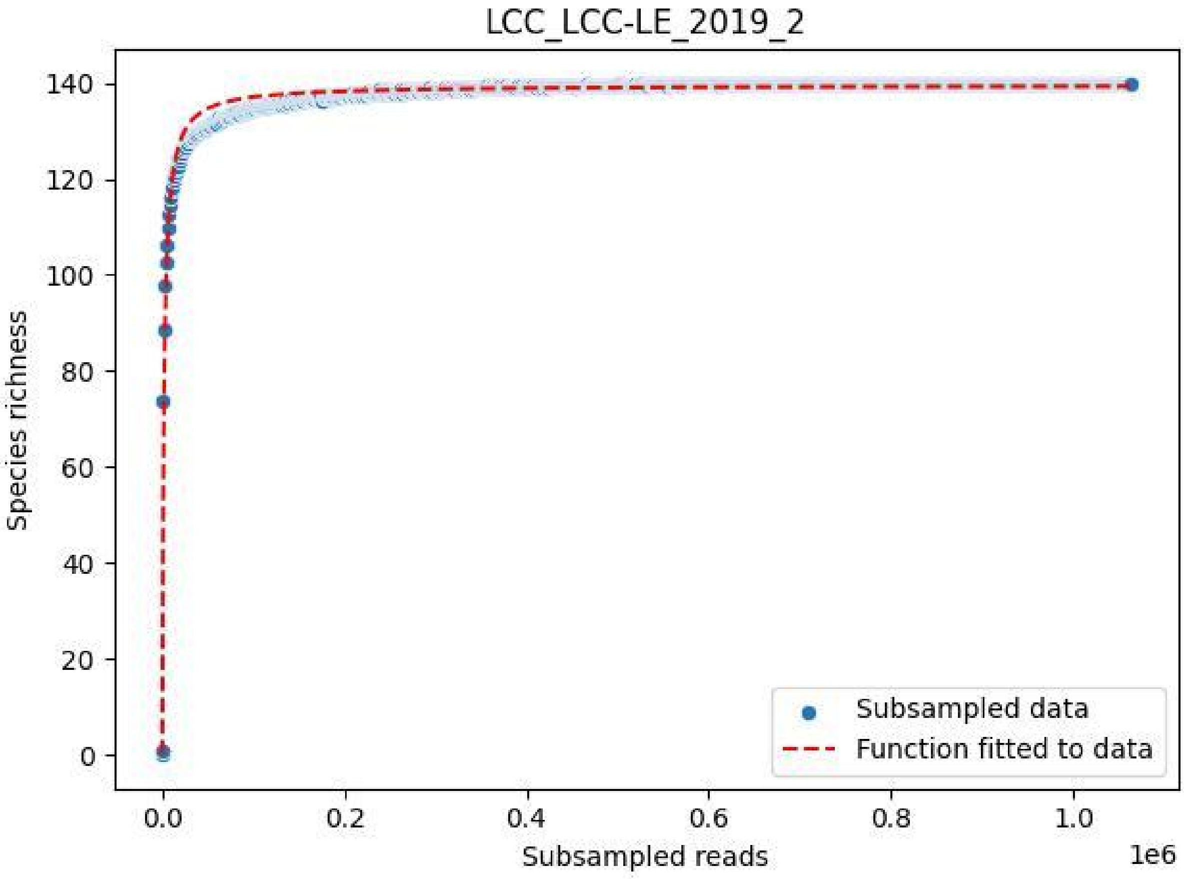
LCC_LCC-LE_2019_2
Species richness
Subsampled reads
1e6
0
20
40
60
80
100
120
140
0.0
0.2
0.4
0.6
0.8
1.0
Subsampled data
Function fitted to data

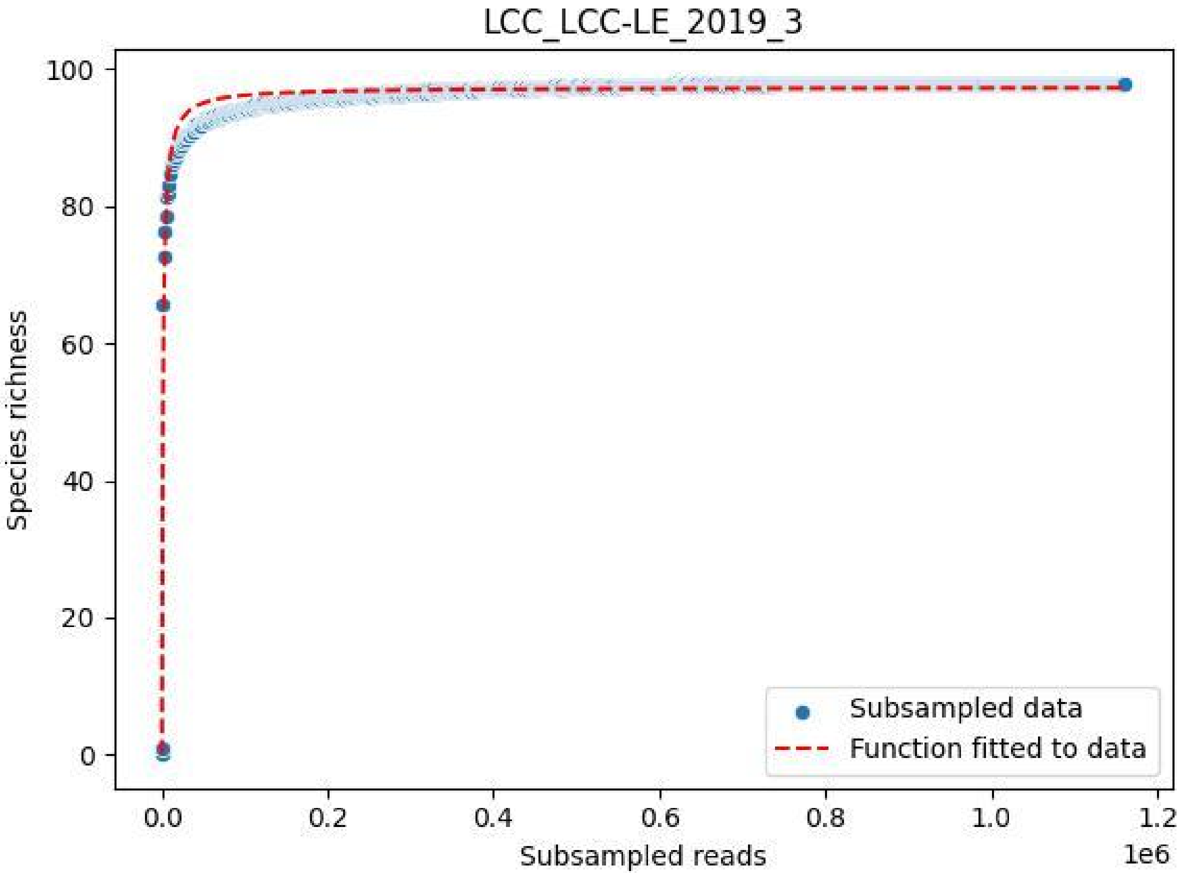

LCC_LCC-LE_2019_3
Species richness
Subsampled reads
1e6
0
20
40
60
80
100
0.0
0.2
0.4
0.6
0.8
1.0
1.2
Subsampled data
Function fitted to data

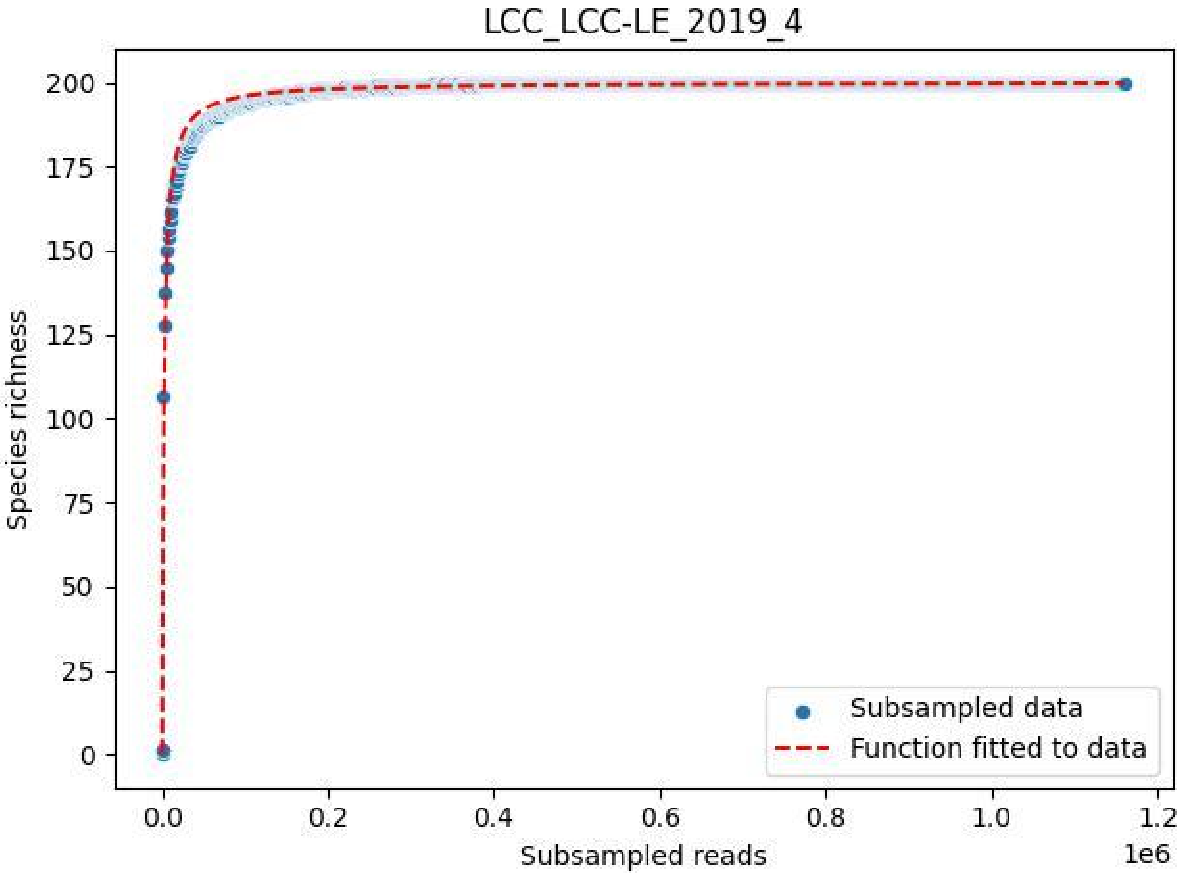
LCC_LCC-LE_2019_4
Species richness
Subsampled reads
1e6
0
25
50
75
100
125
150
175
200
0.0
0.2
0.4
0.6
0.8
1.0
1.2
Subsampled data
Function fitted to data

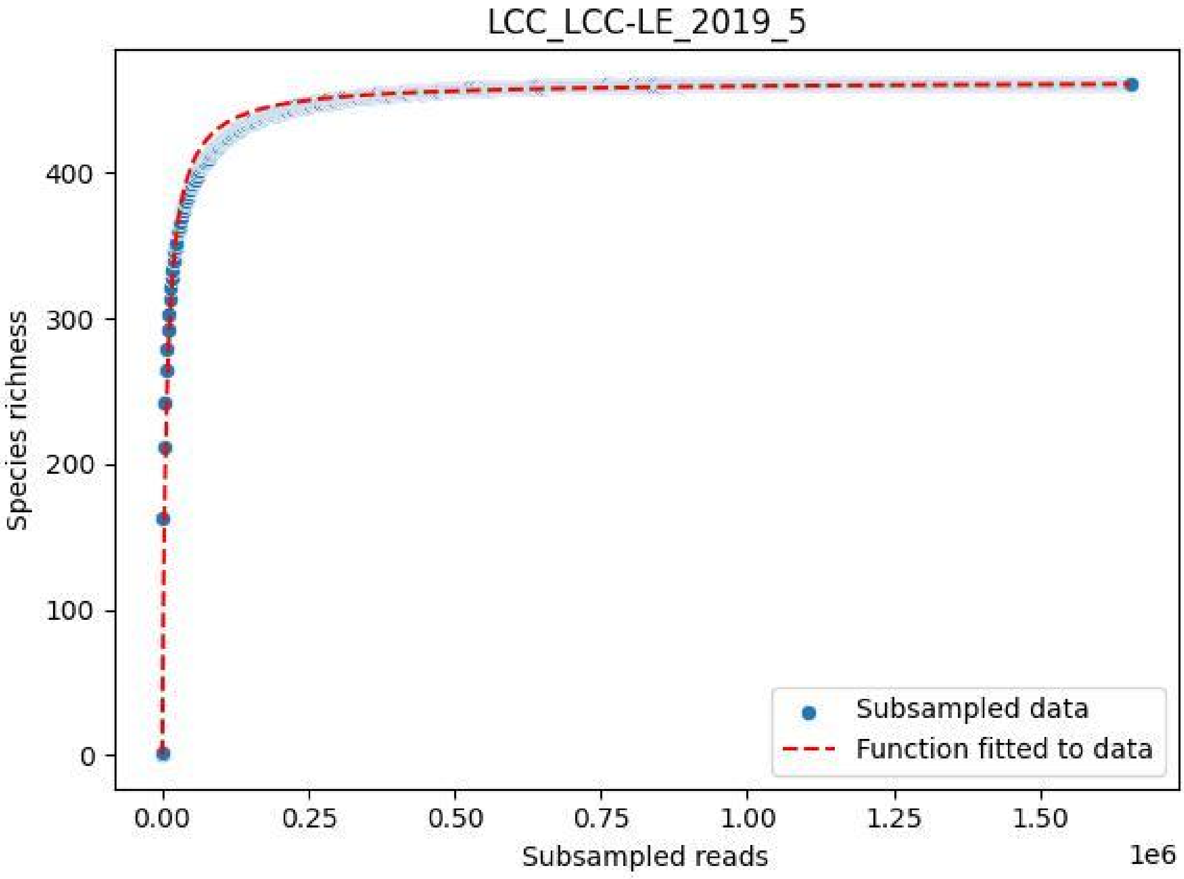

LCC_LCC-LE_2019_5
Species richness
0
100
200
300
400
Subsampled reads
0.00
0.25
0.50
0.75
1.00
1.25
1.50
1e6
Subsampled data
Function fitted to data

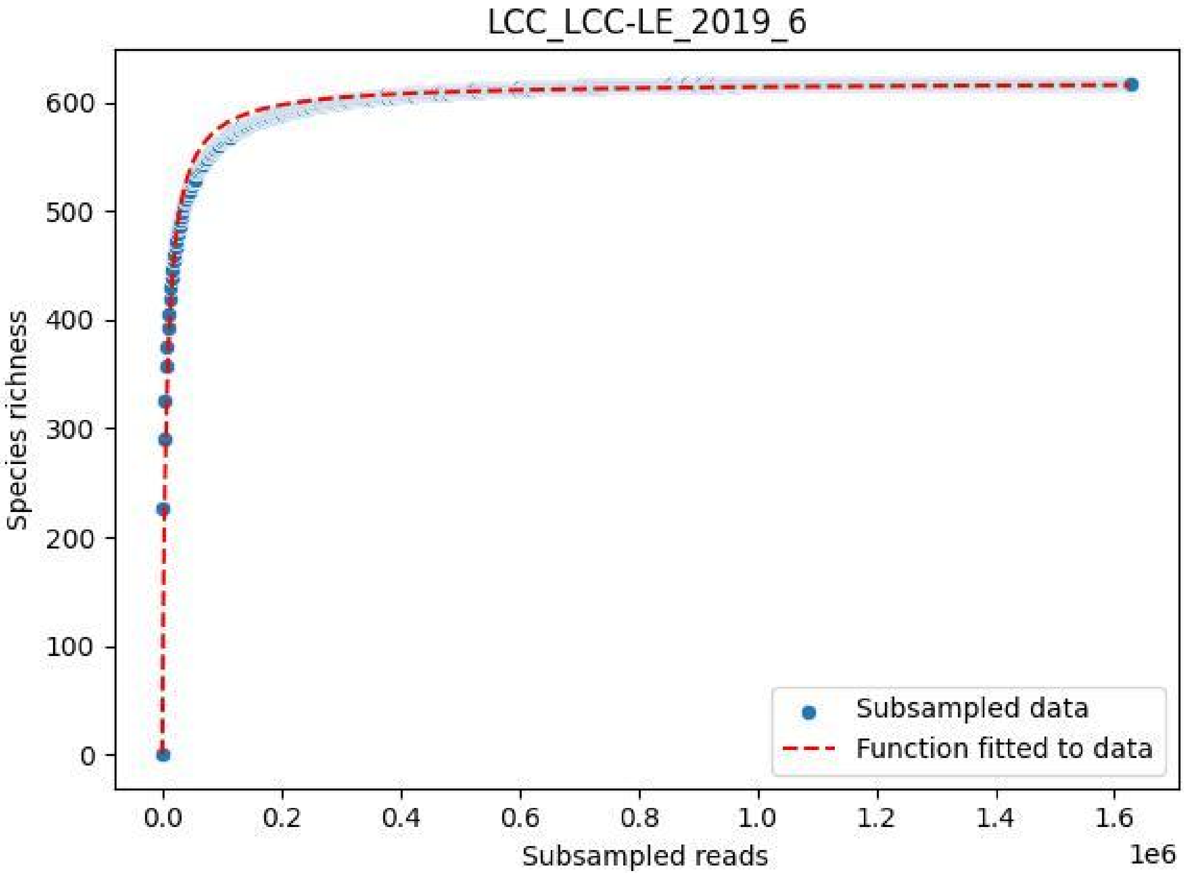

LCC_LCC-LE_2019_6
Species richness
0
100
200
300
400
500
600
0.0
0.2
0.4
0.6
0.8
1.0
1.2
1.4
1.6
1e6
Subsampled reads
Subsampled data
Function fitted to data

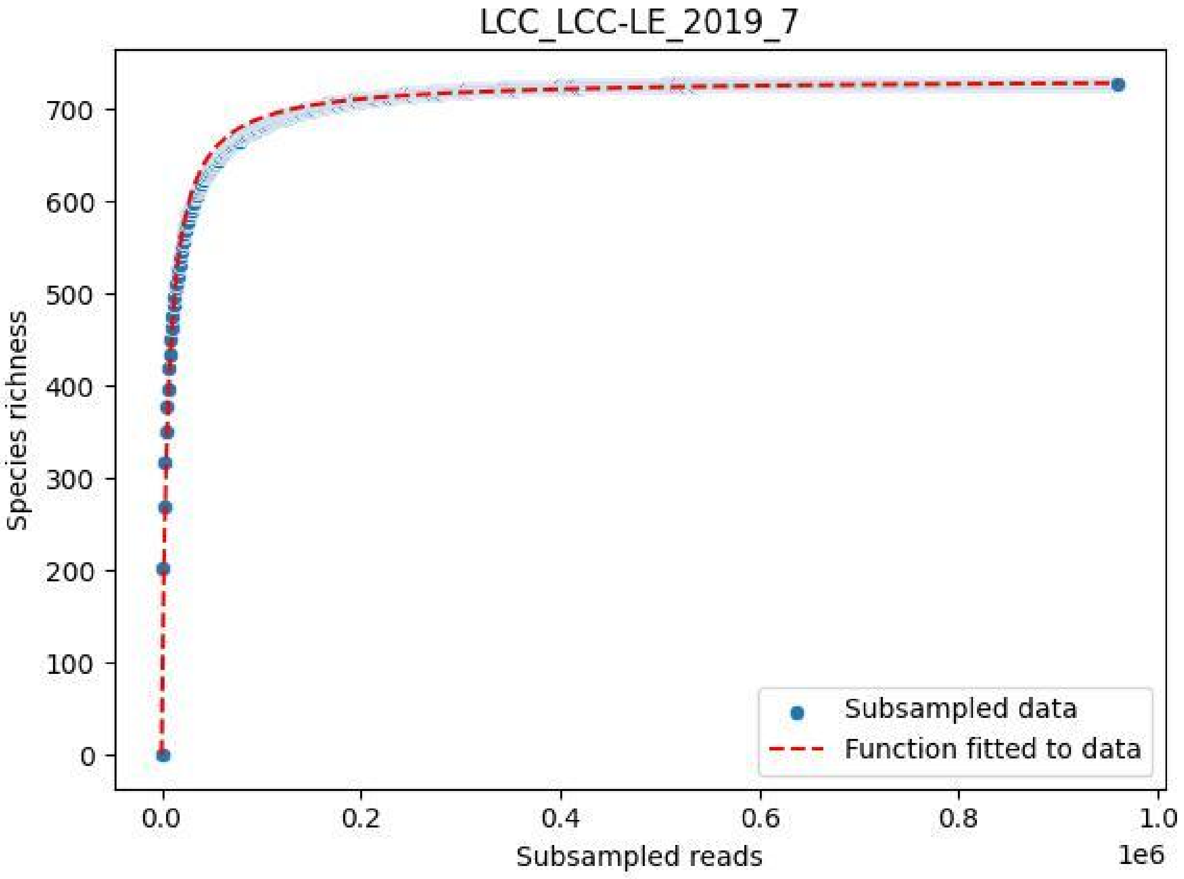

LCC_LCC-LE_2019_7
Species richness
700
600
500
400
300
200
100
0
0.0
0.2
0.4
0.6
0.8
1.0
1e6
Subsampled reads
Subsampled data
Function fitted to data

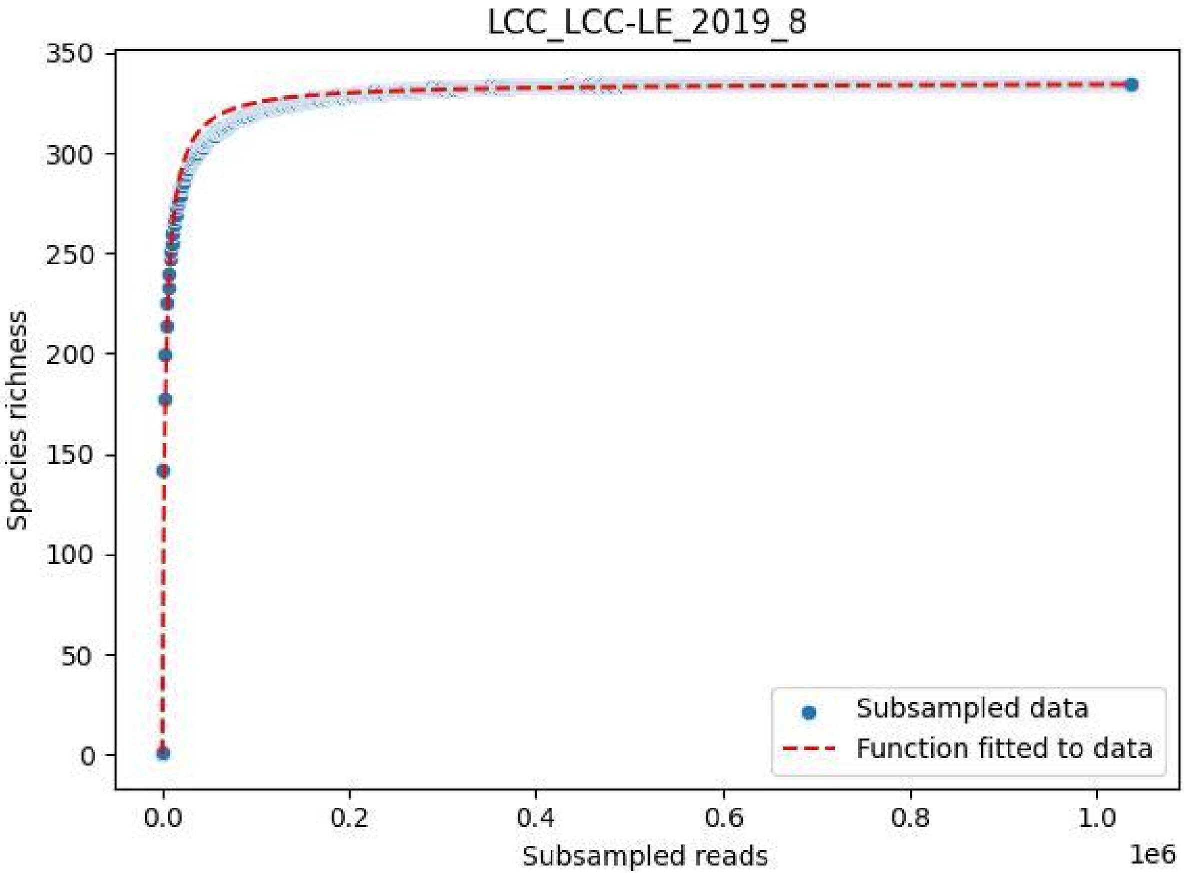
LCC_LCC-LE_2019_8
Species richness
Subsampled reads
1e6
0
50
100
150
200
250
300
350
0.0
0.2
0.4
0.6
0.8
1.0
Subsampled data
Function fitted to data

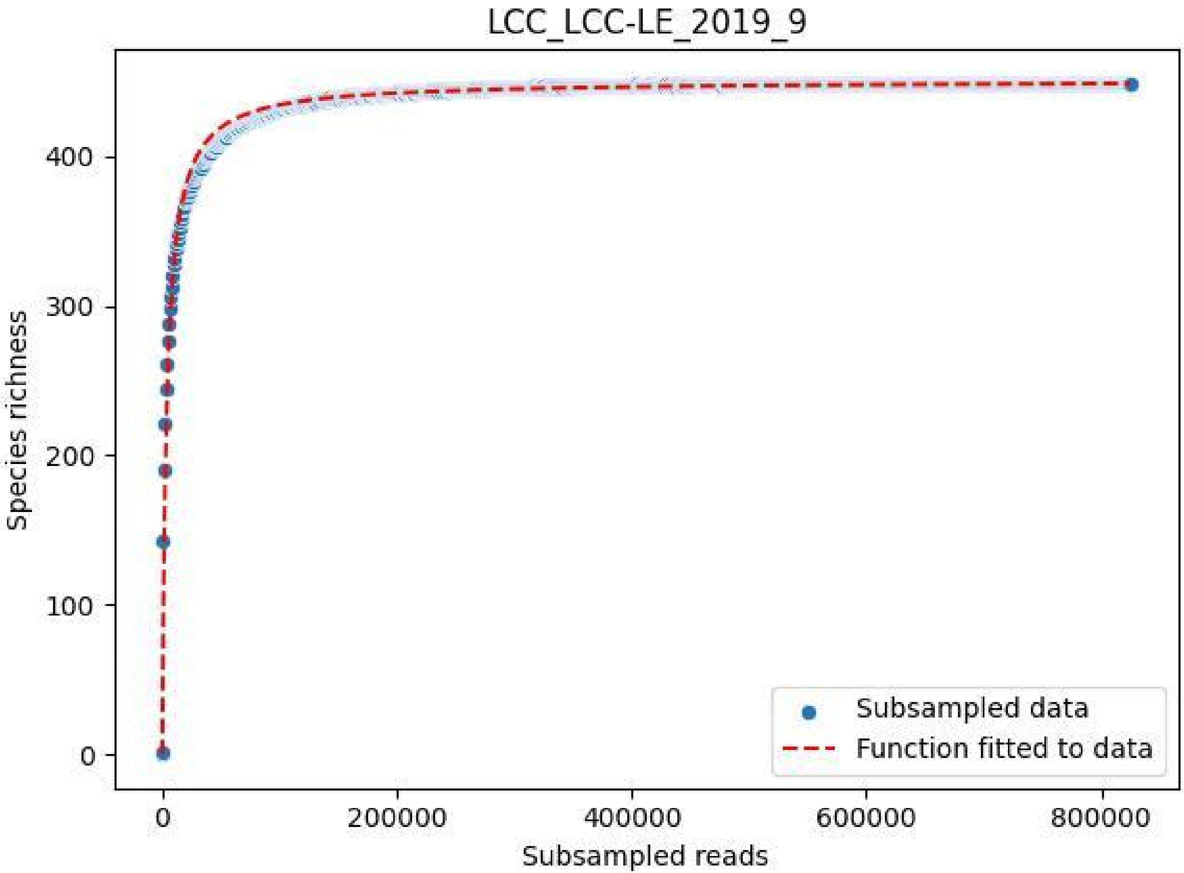
LCC_LCC-LE_2019_9
Species richness
Subsampled reads
0
100
200
300
400
0
200000
400000
600000
800000
Subsampled data
Function fitted to data

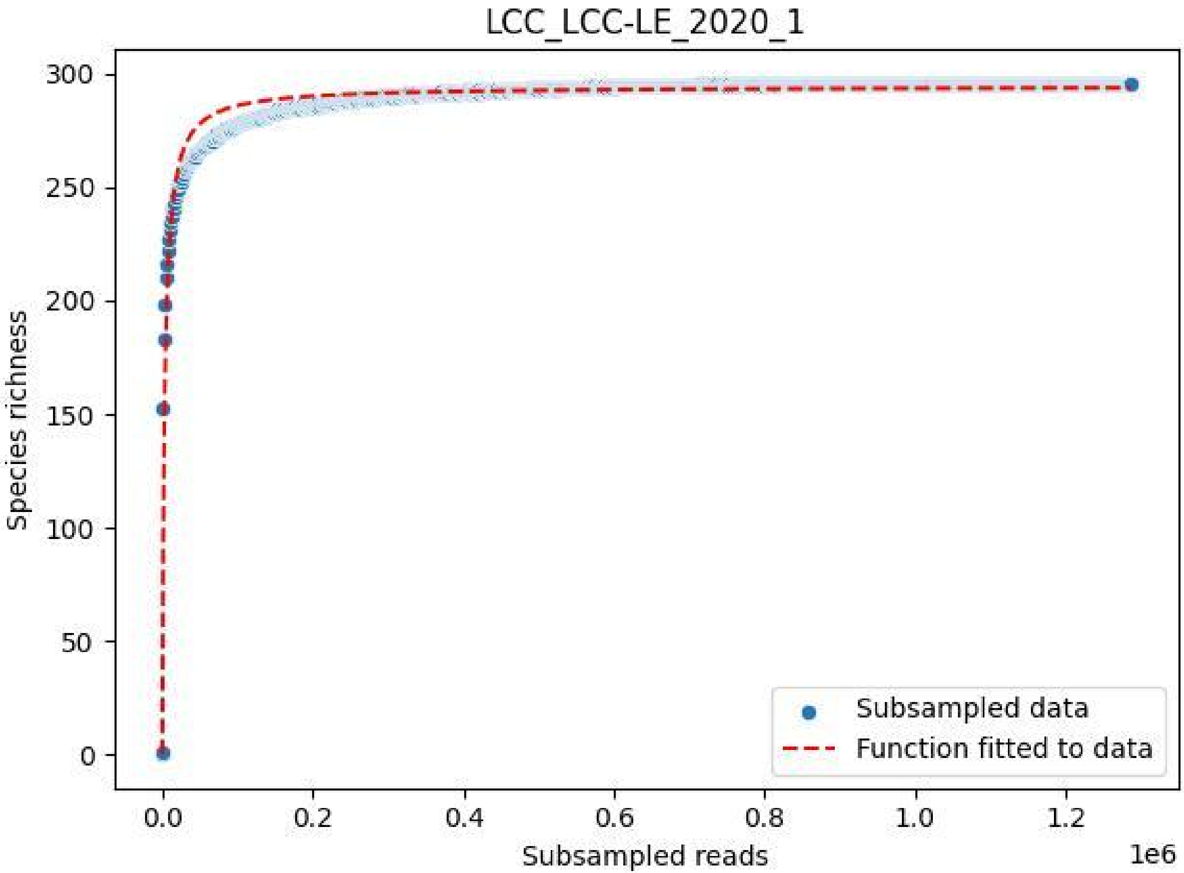
LCC_LCC-LE_2020_1
Species richness
300
250
200
150
100
50
0
0.0
0.2
0.4
0.6
0.8
1.0
1.2
1e6
Subsampled reads
Subsampled data
Function fitted to data

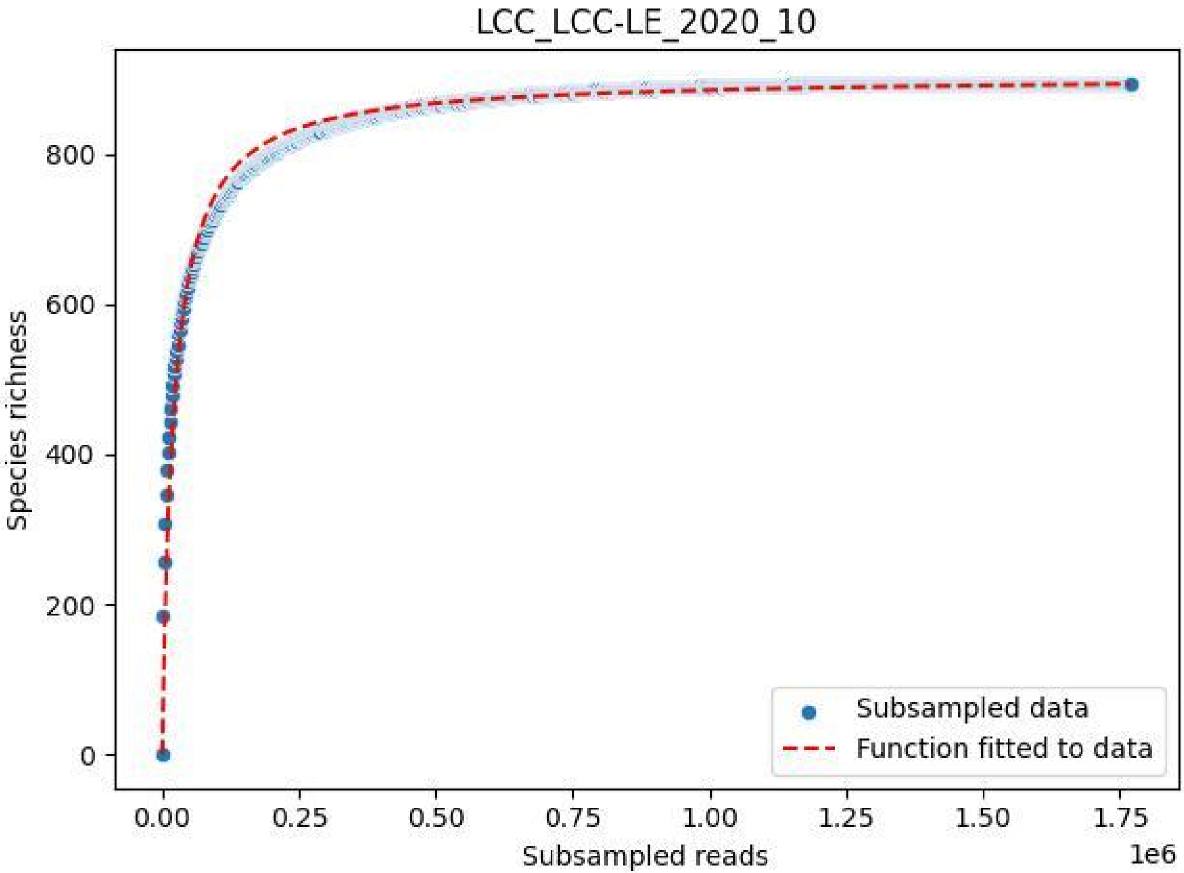
LCC_LCC-LE_2020_10
Species richness
Subsampled reads
1e6
0
200
400
600
800
0.00
0.25
0.50
0.75
1.00
1.25
1.50
1.75
Subsampled data
Function fitted to data

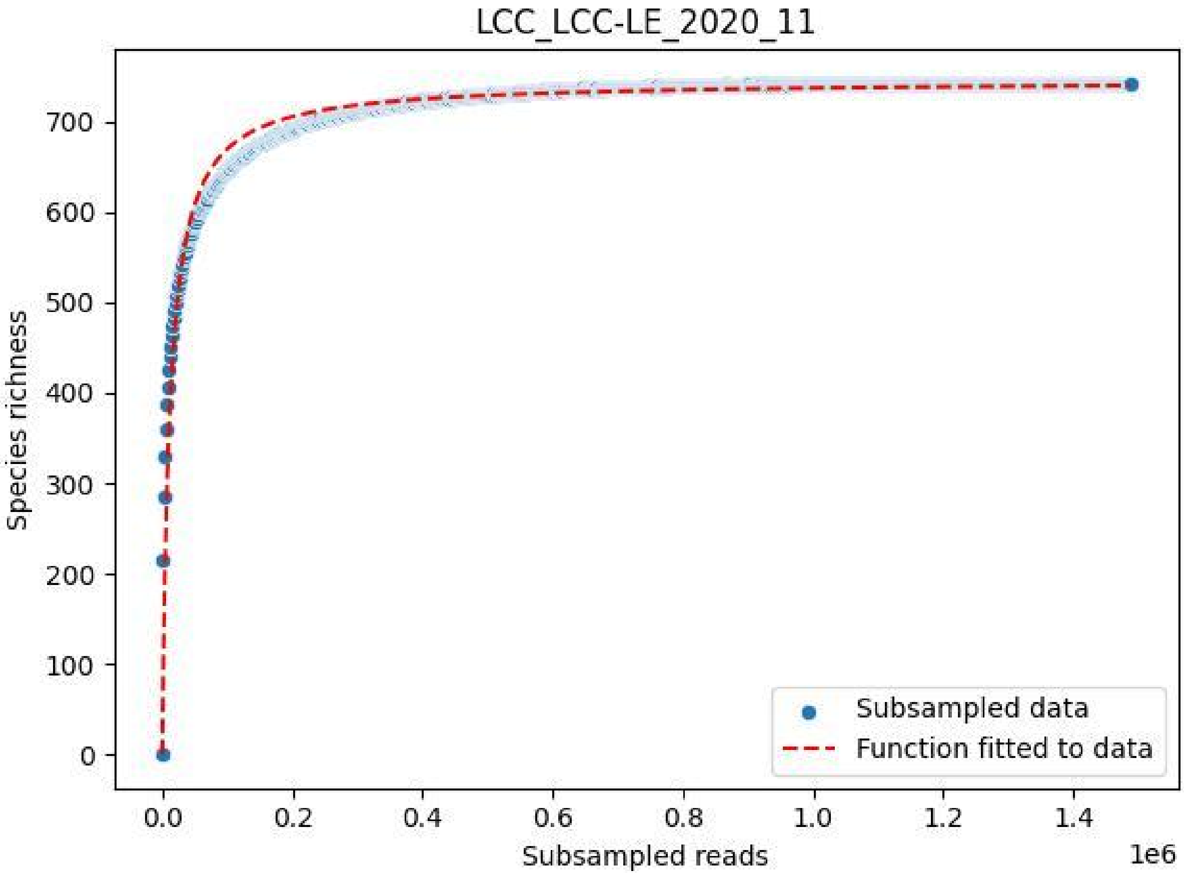

LCC_LCC-LE_2020_11
Species richness
0
100
200
300
400
500
600
700
Subsampled reads
0.0
0.2
0.4
0.6
0.8
1.0
1.2
1.4
1e6
Subsampled data
Function fitted to data

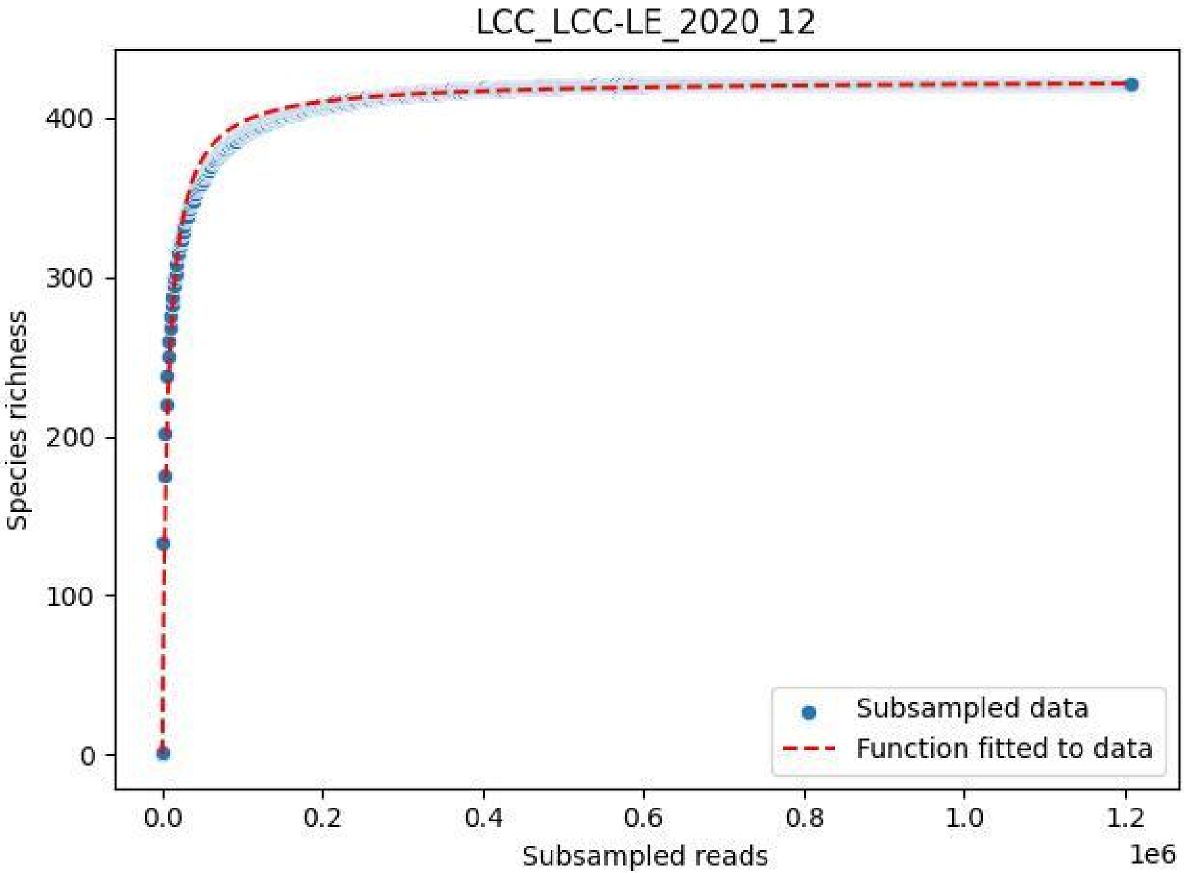
LCC_LCC-LE_2020_12
Species richness
0
100
200
300
400
Subsampled reads
0.0
0.2
0.4
0.6
0.8
1.0
1.2
1e6
Subsampled data
Function fitted to data

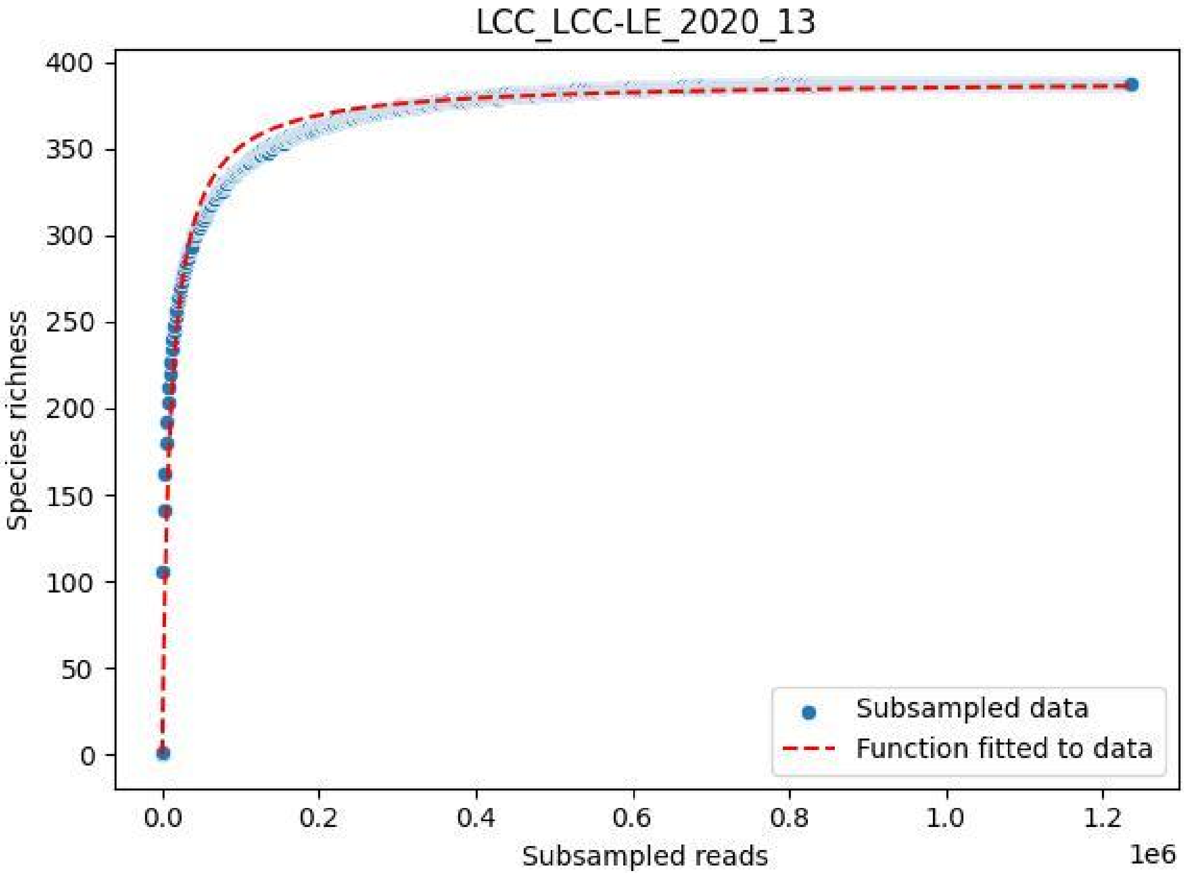

LCC_LCC-LE_2020_13
400
350
300
250
200
150
100
50
0
Species richness
0.0
0.2
0.4
0.6
0.8
1.0
1.2
Subsampled reads
1e6
Subsampled data
Function fitted to data

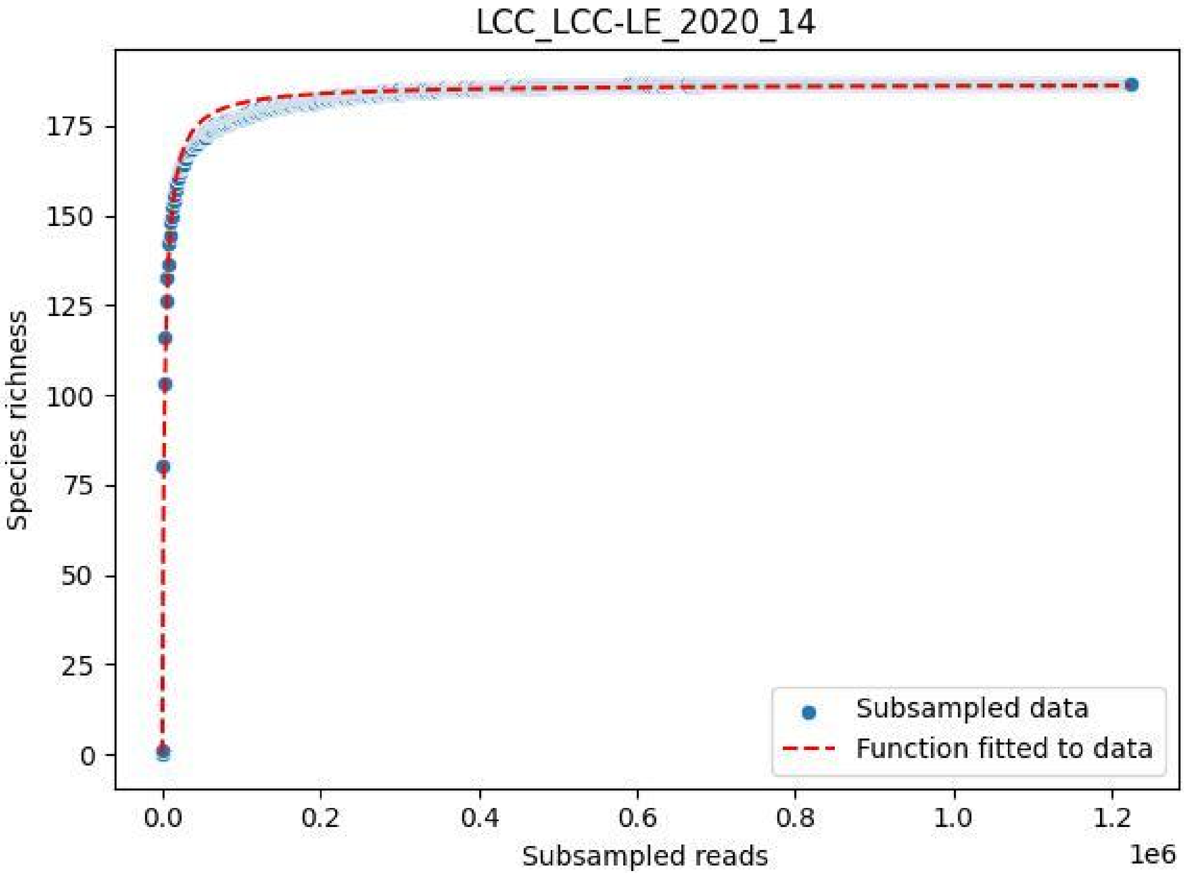

LCC_LCC-LE_2020_14
Species richness
Subsampled reads
1e6
0
25
50
75
100
125
150
175
0.0
0.2
0.4
0.6
0.8
1.0
1.2
Subsampled data
Function fitted to data

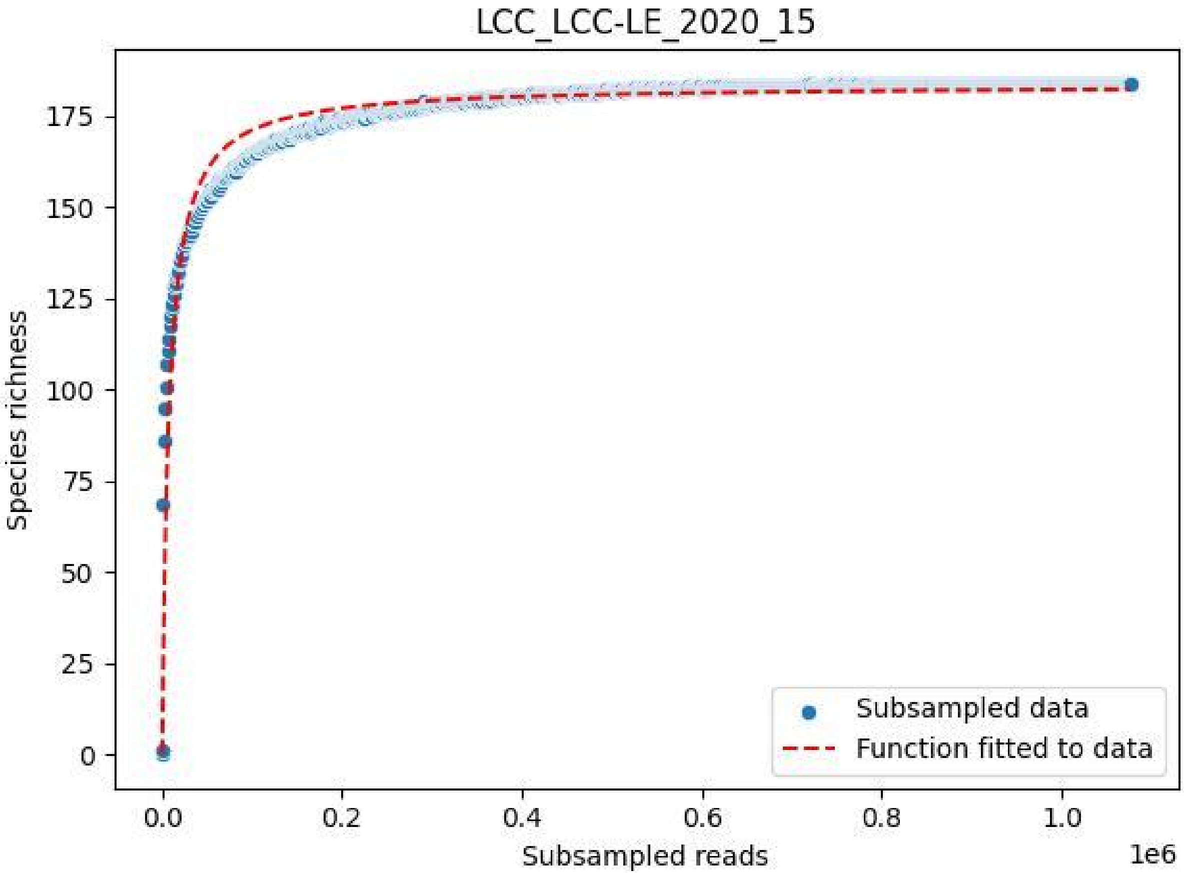

LCC_LCC-LE_2020_15
Species richness
Subsampled reads
1e6
0
25
50
75
100
125
150
175
0.0
0.2
0.4
0.6
0.8
1.0
Subsampled data
Function fitted to data

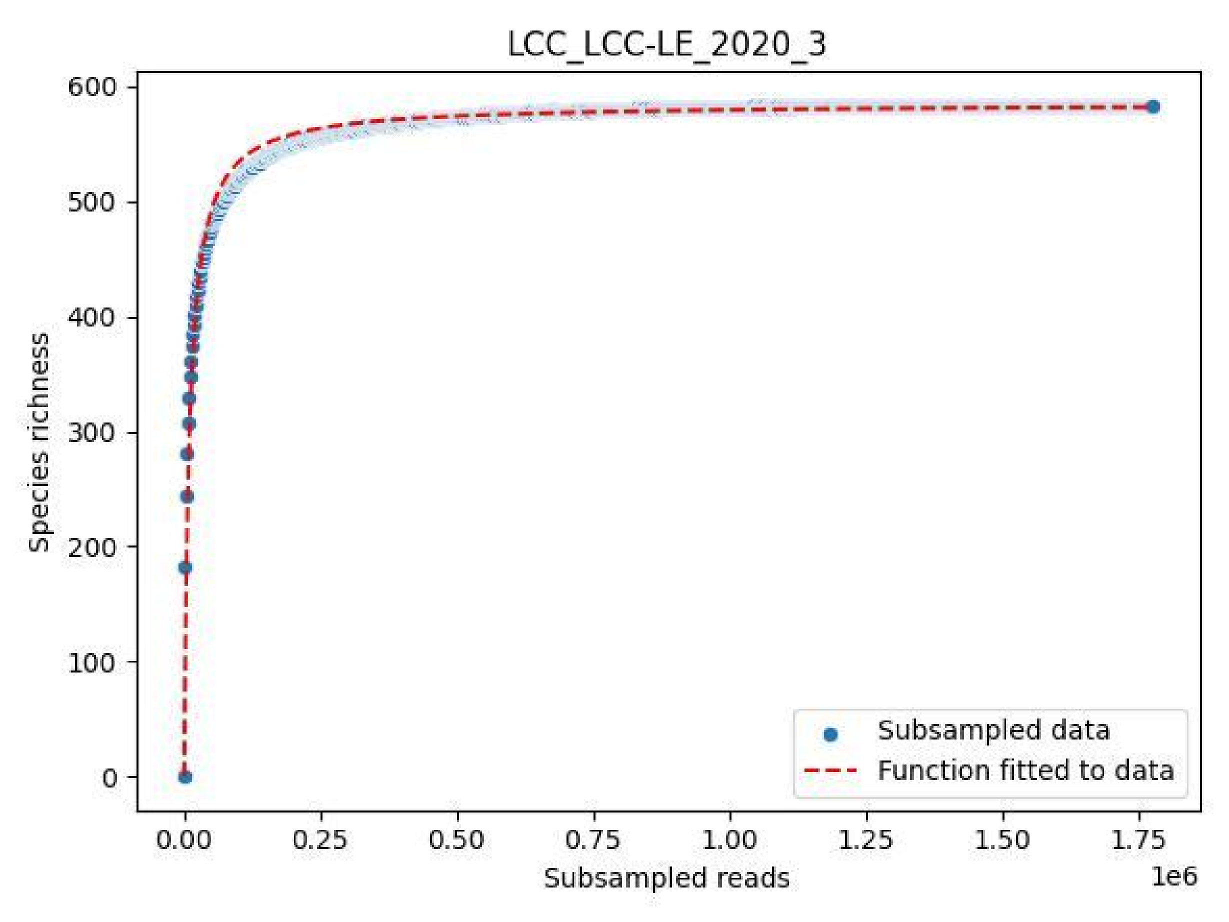

LCC_LCC-LE_2020_3
600
500
400
300
200
100
0
Species richness
0.00
0.25
0.50
0.75
1.00
1.25
1.50
1.75
1e6
Subsampled reads
Subsampled data
Function fitted to data

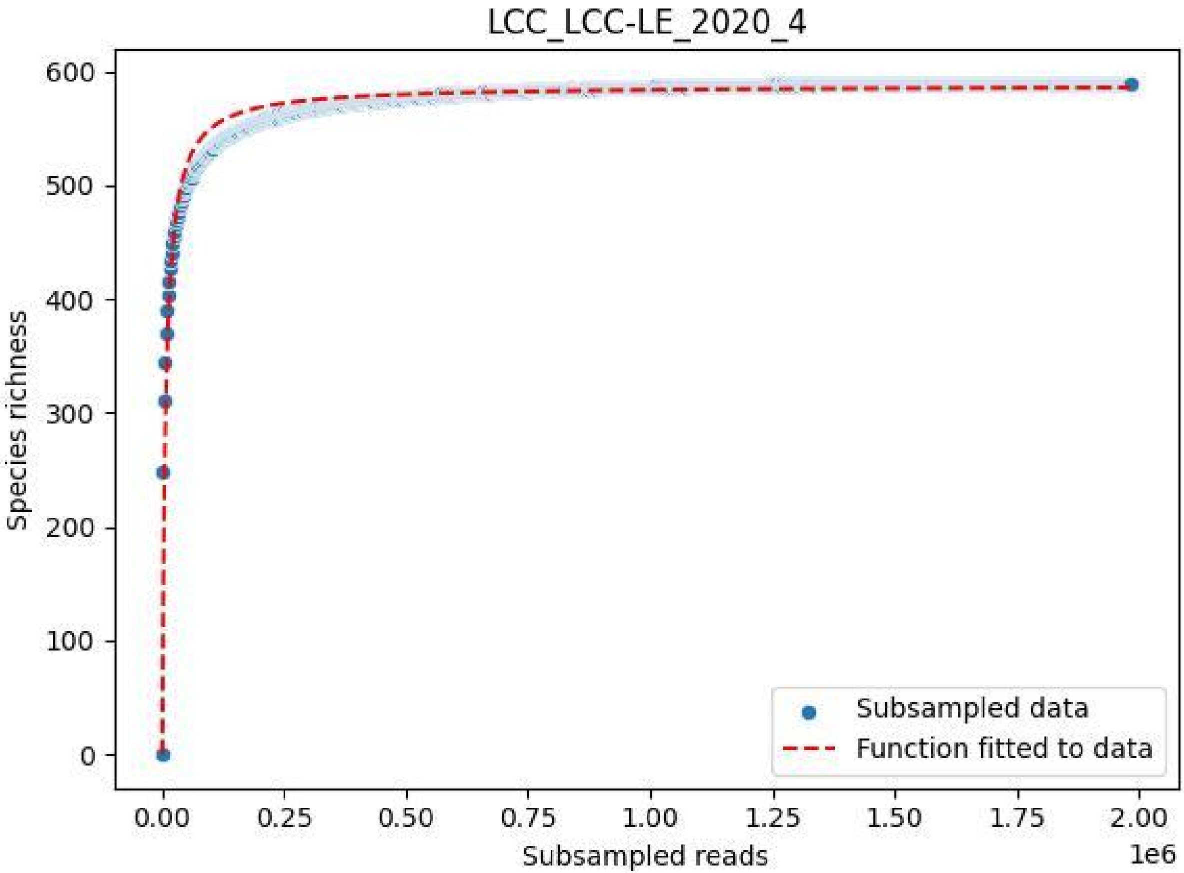

LCC_LCC-LE_2020_4
Species richness
Subsampled reads
1e6
0
100
200
300
400
500
600
0.00
0.25
0.50
0.75
1.00
1.25
1.50
1.75
2.00
Subsampled data
Function fitted to data

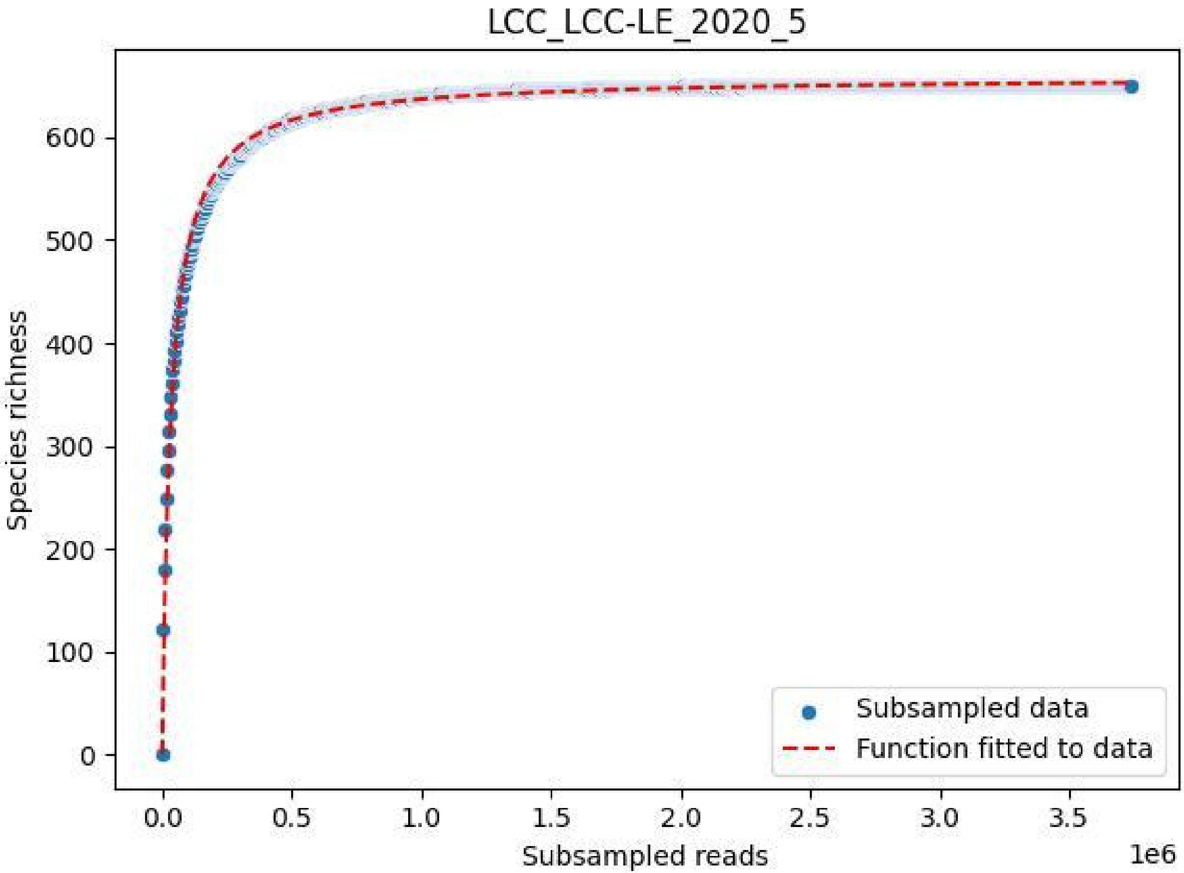
LCC_LCC-LE_2020_5
Species richness
0
100
200
300
400
500
600
0.0
0.5
1.0
1.5
2.0
2.5
3.0
3.5
1e6
Subsampled reads
Subsampled data
Function fitted to data

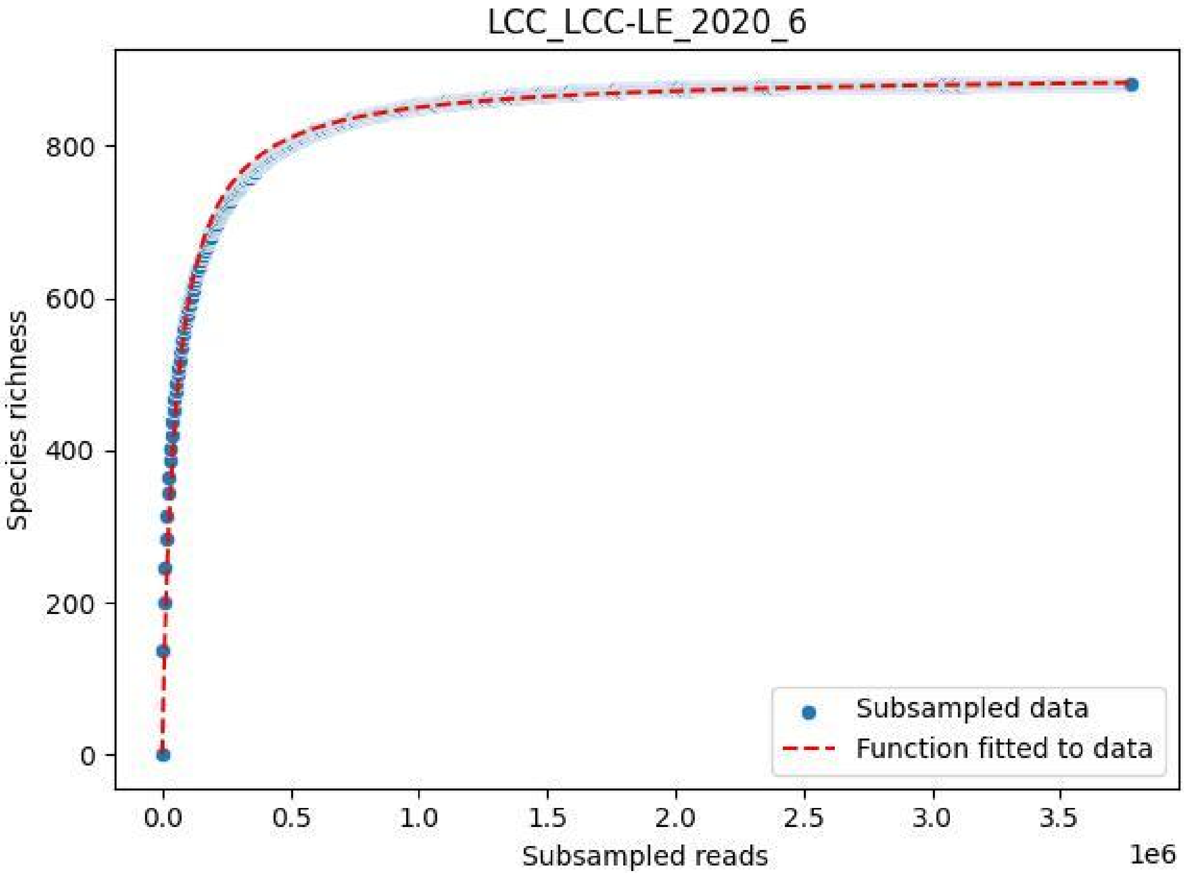
LCC_LCC-LE_2020_6
Species richness
Subsampled reads
1e6
Subsampled data
Function fitted to data
0
200
400
600
800
0.0
0.5
1.0
1.5
2.0
2.5
3.0
3.5

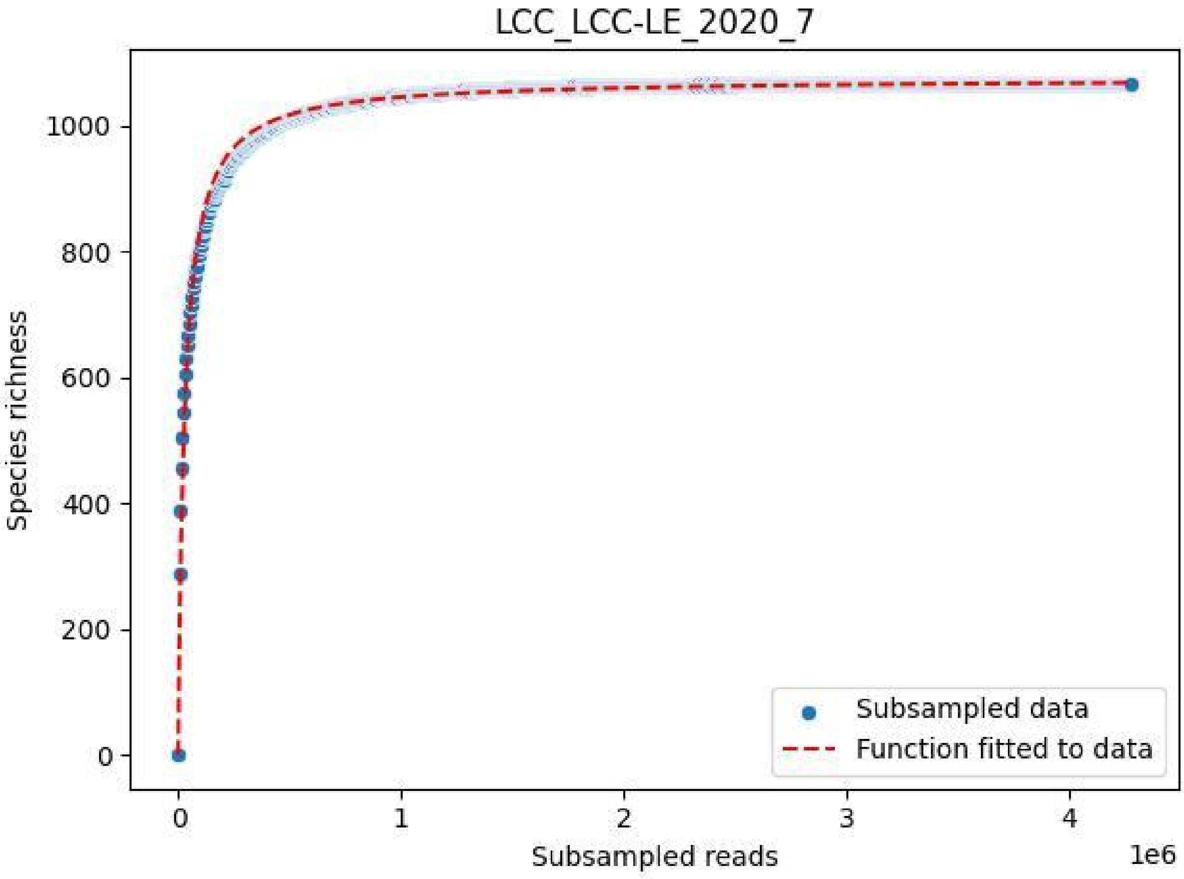
LCC_LCC-LE_2020_7
Species richness
0
200
400
600
800
1000
Subsampled reads
0
1
2
3
4
1e6
Subsampled data
Function fitted to data

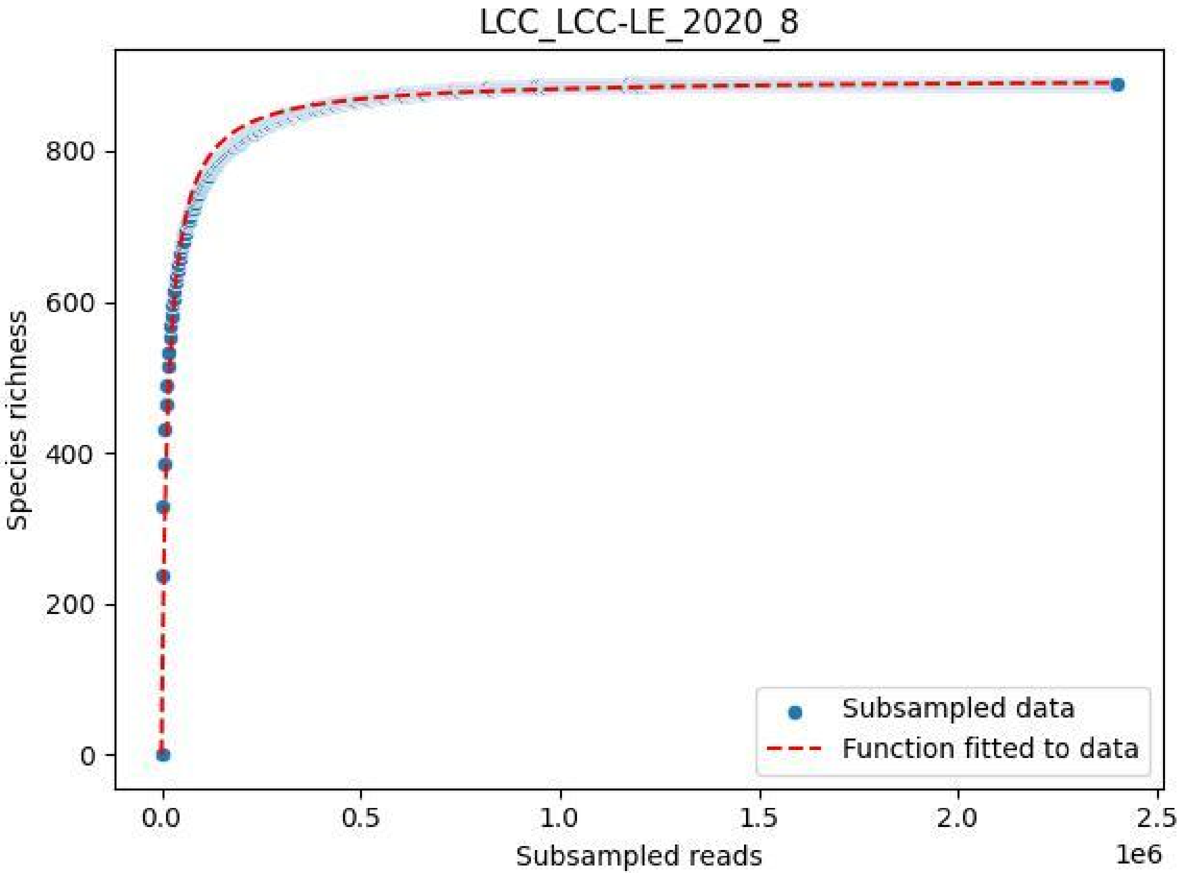

LCC_LCC-LE_2020_8
Species richness
Subsampled reads
1e6
0
200
400
600
800
0.0
0.5
1.0
1.5
2.0
2.5
Subsampled data
Function fitted to data

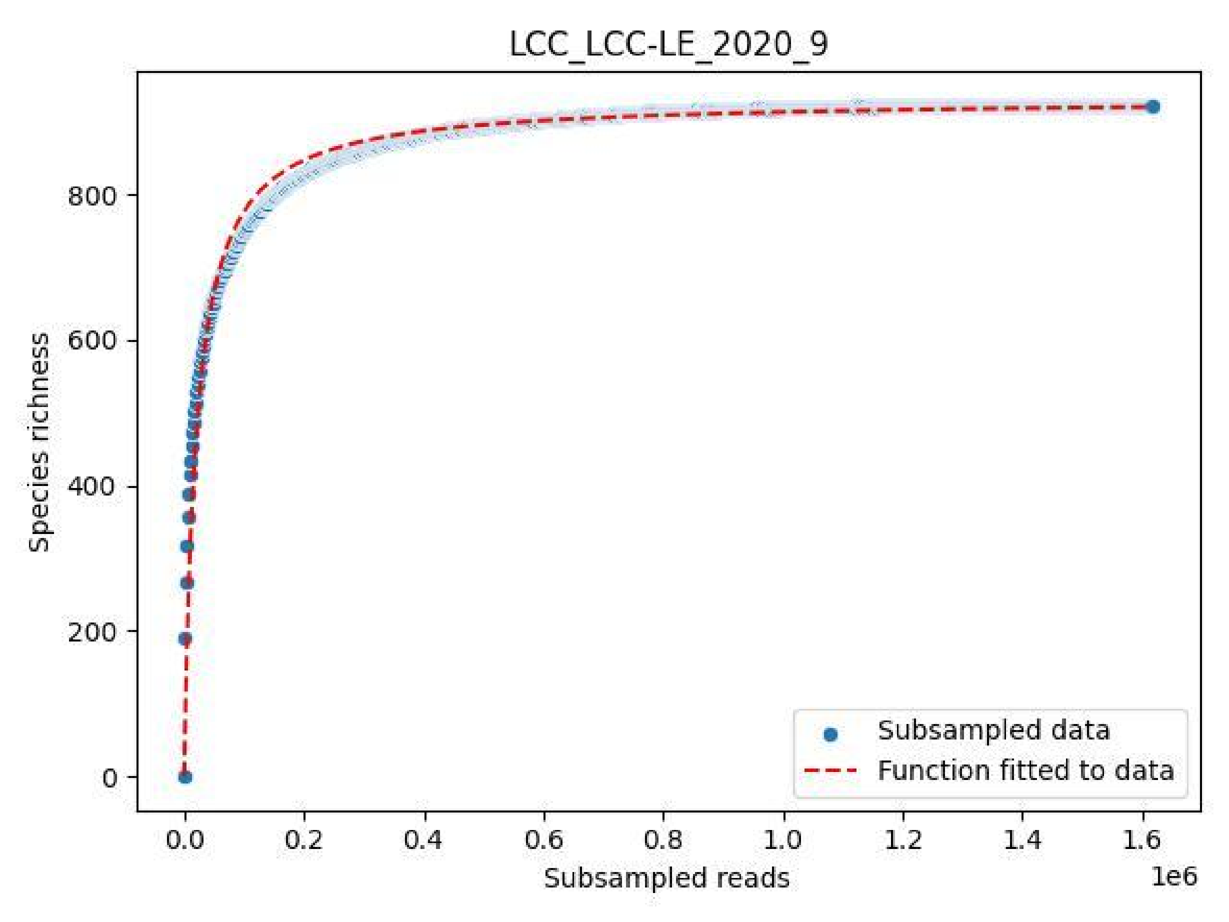

LCC_LCC-LE_2020_9
Species richness
Subsampled reads
1e6
0
200
400
600
800
0.0
0.2
0.4
0.6
0.8
1.0
1.2
1.4
1.6
Subsampled data
Function fitted to data

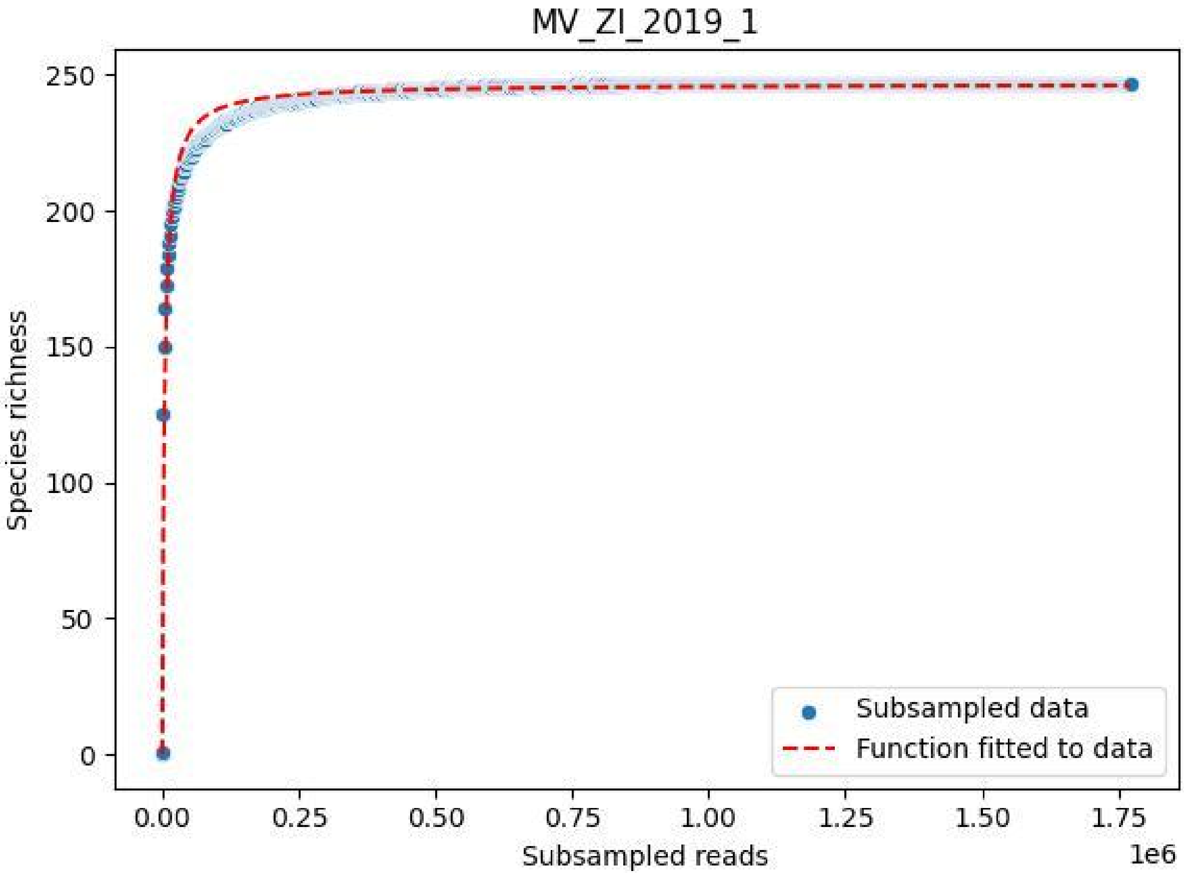
MV_ZI_2019_1
Species richness
Subsampled reads
1e6
0
50
100
150
200
250
0.00
0.25
0.50
0.75
1.00
1.25
1.50
1.75
Subsampled data
Function fitted to data

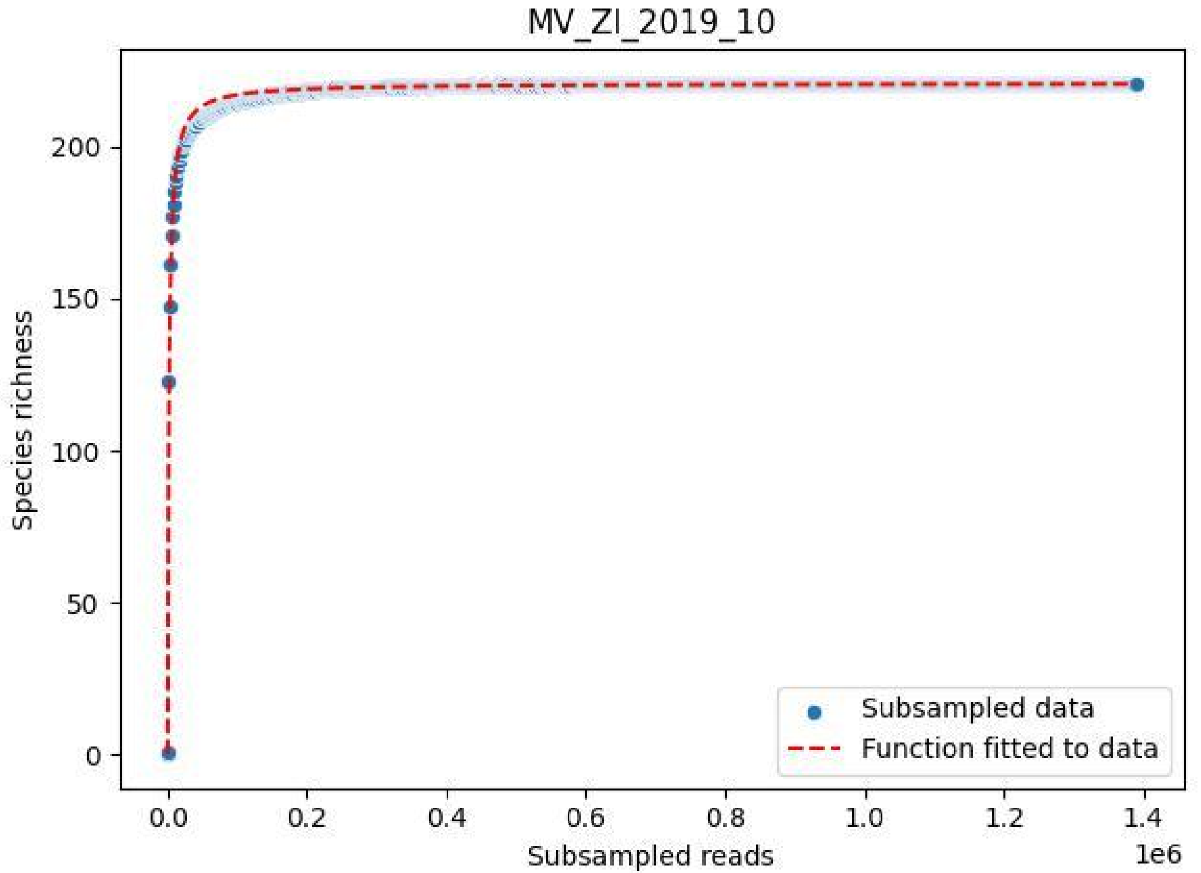

MV_ZI_2019_10
Species richness
Subsampled reads
1e6
0
50
100
150
200
0.0
0.2
0.4
0.6
0.8
1.0
1.2
1.4
Subsampled data
Function fitted to data

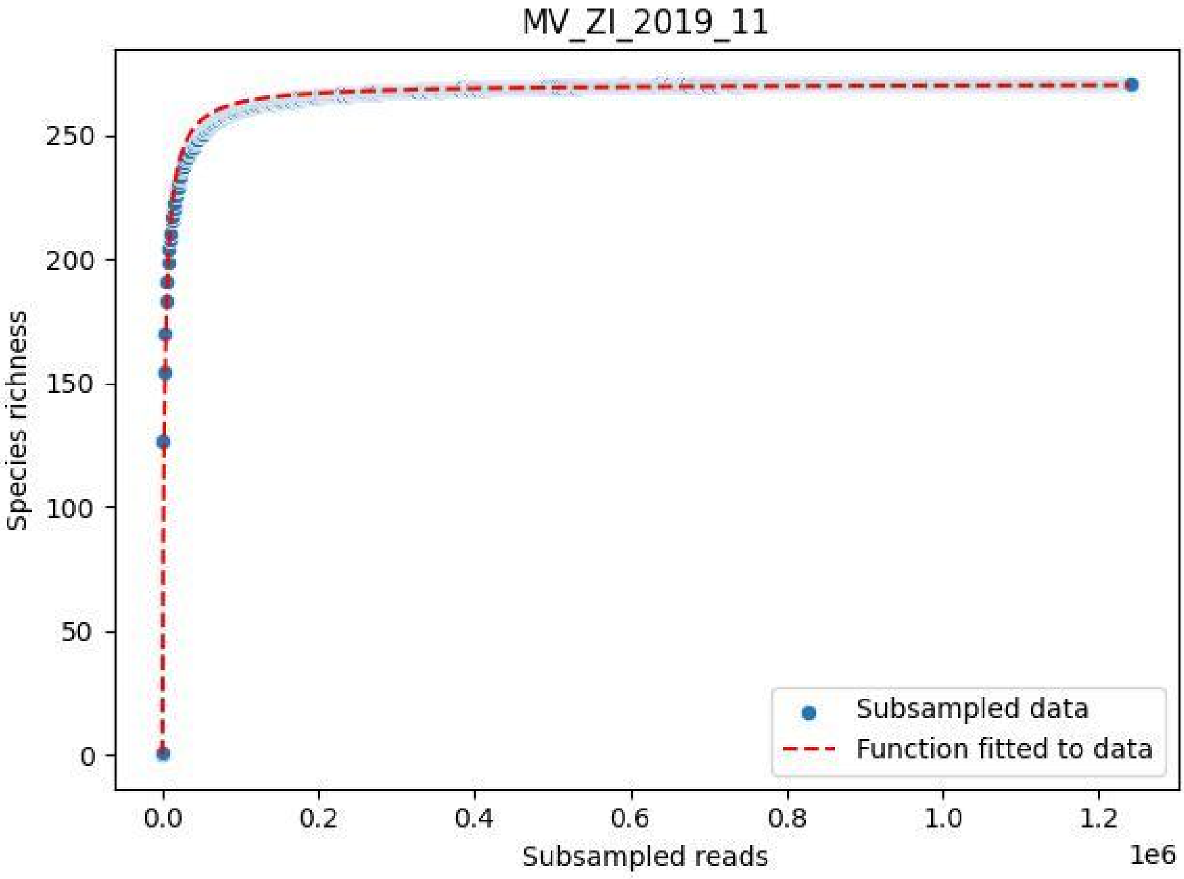
MV_ZI_2019_11
Species richness
Subsampled reads
1e6
Subsampled data
Function fitted to data
0
50
100
150
200
250
0.0
0.2
0.4
0.6
0.8
1.0
1.2

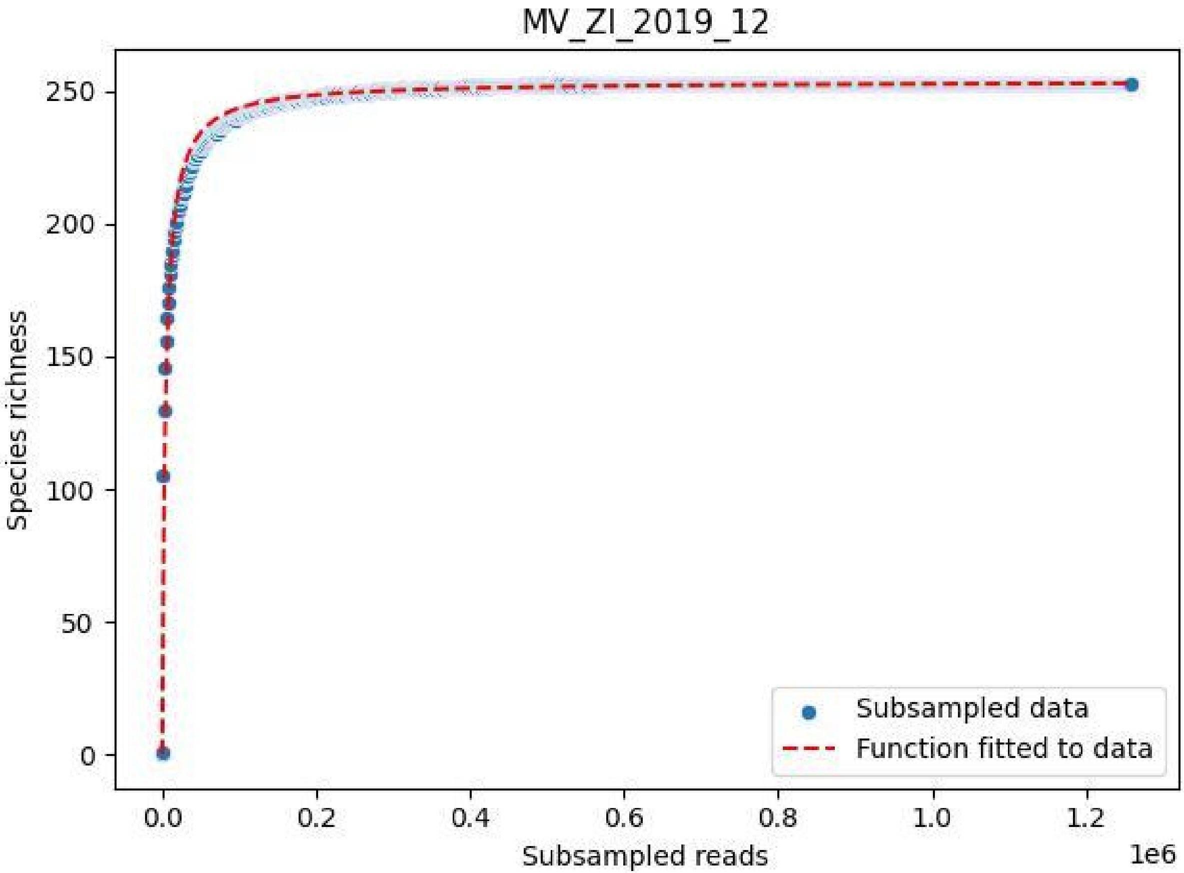
MV_ZI_2019_12
Species richness
Subsampled reads
1e6
0
50
100
150
200
250
0.0
0.2
0.4
0.6
0.8
1.0
1.2
Subsampled data
Function fitted to data

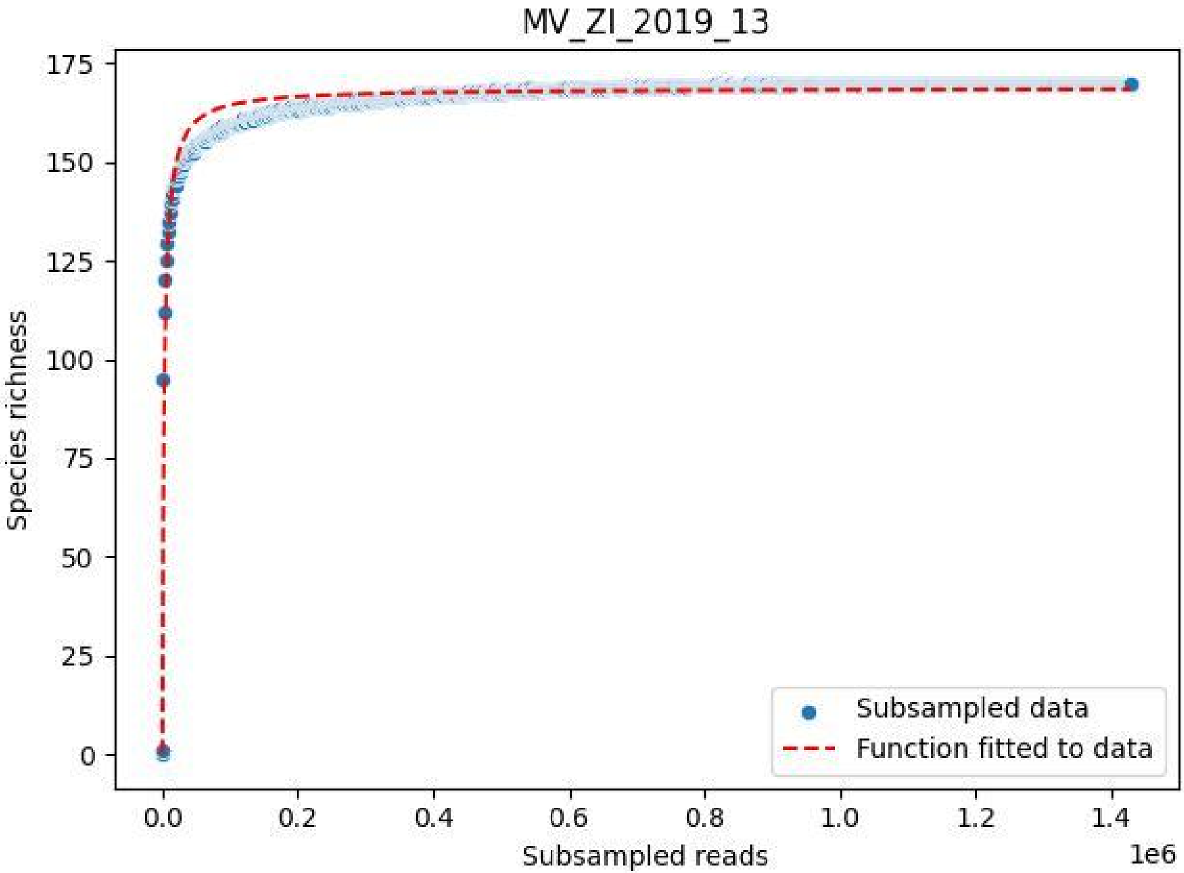
MV_ZI_2019_13
Species richness
175
150
125
100
75
50
25
0
0.0
0.2
0.4
0.6
0.8
1.0
1.2
1.4
1e6
Subsampled reads
Subsampled data
Function fitted to data

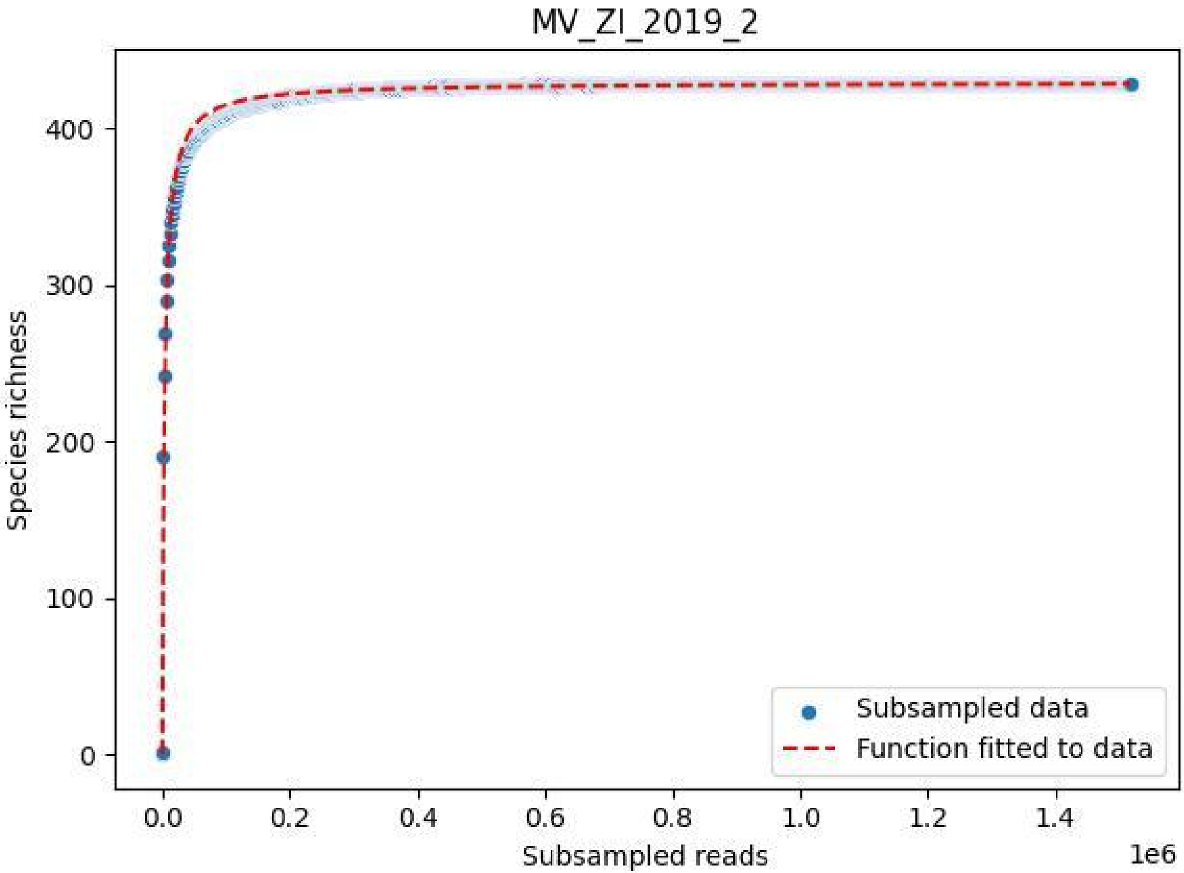

MV_ZI_2019_2
Species richness
400
300
200
100
0
0.0
0.2
0.4
0.6
0.8
1.0
1.2
1.4
Subsampled reads
1e6
Subsampled data
Function fitted to data

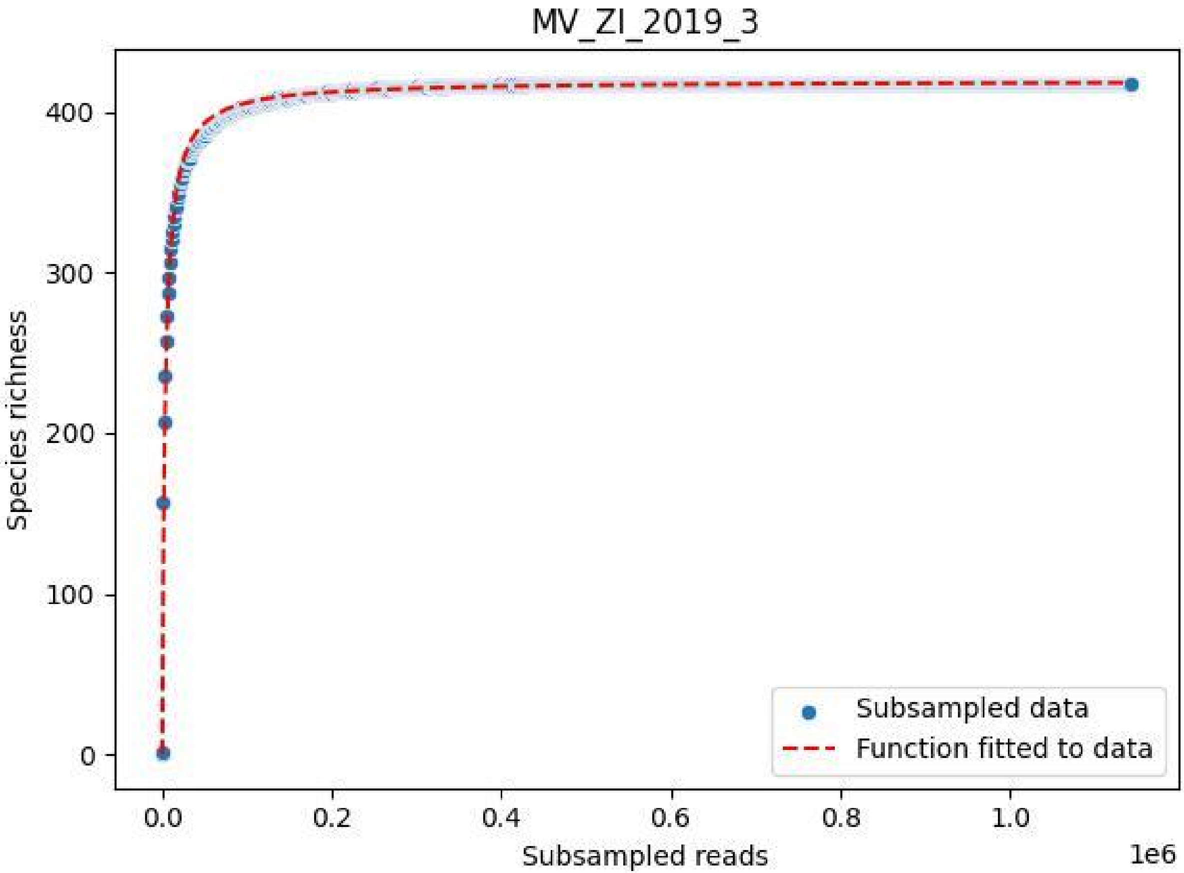
MV_ZI_2019_3
Species richness
Subsampled reads
1e6
0
100
200
300
400
0.0
0.2
0.4
0.6
0.8
1.0
Subsampled data
Function fitted to data

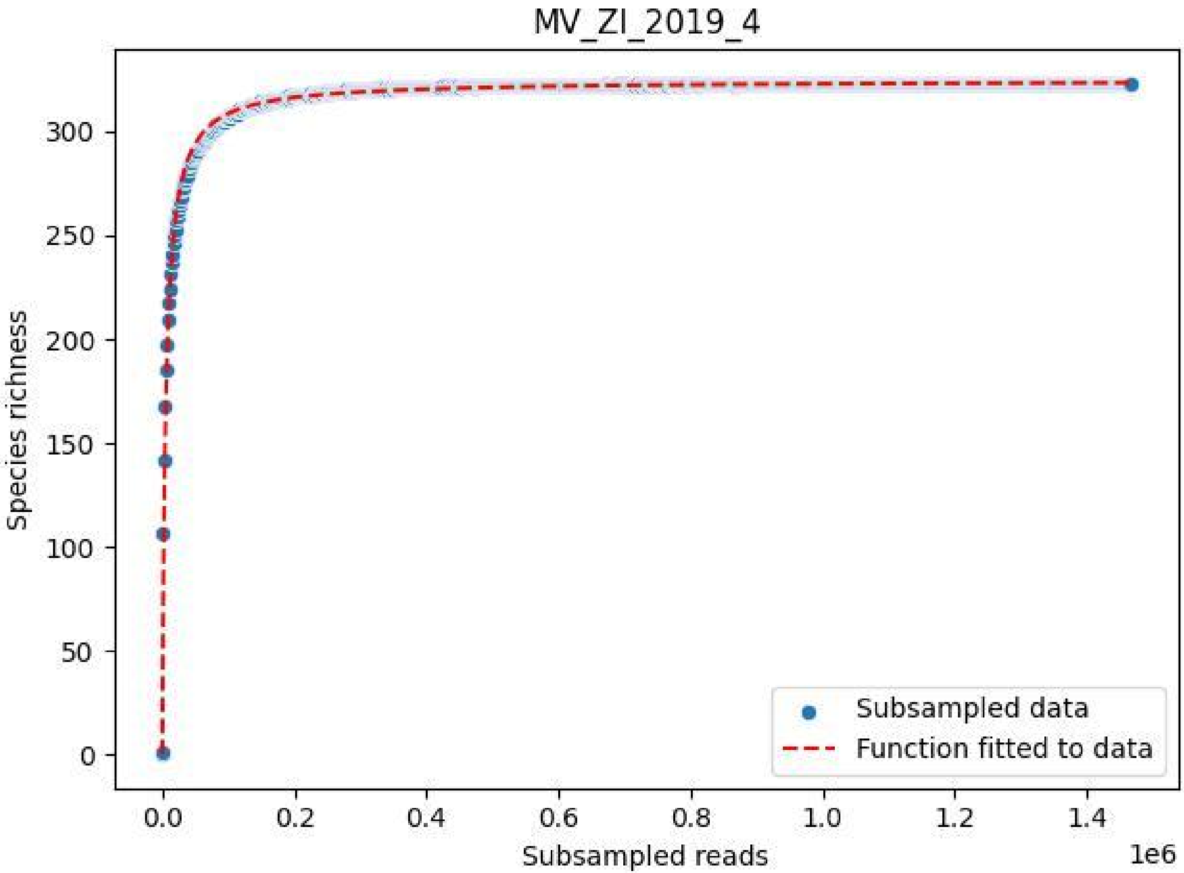
MV_ZI_2019_4
Species richness
300
250
200
150
100
50
0
0.0
0.2
0.4
0.6
0.8
1.0
1.2
1.4
Subsampled reads
1e6
Subsampled data
Function fitted to data

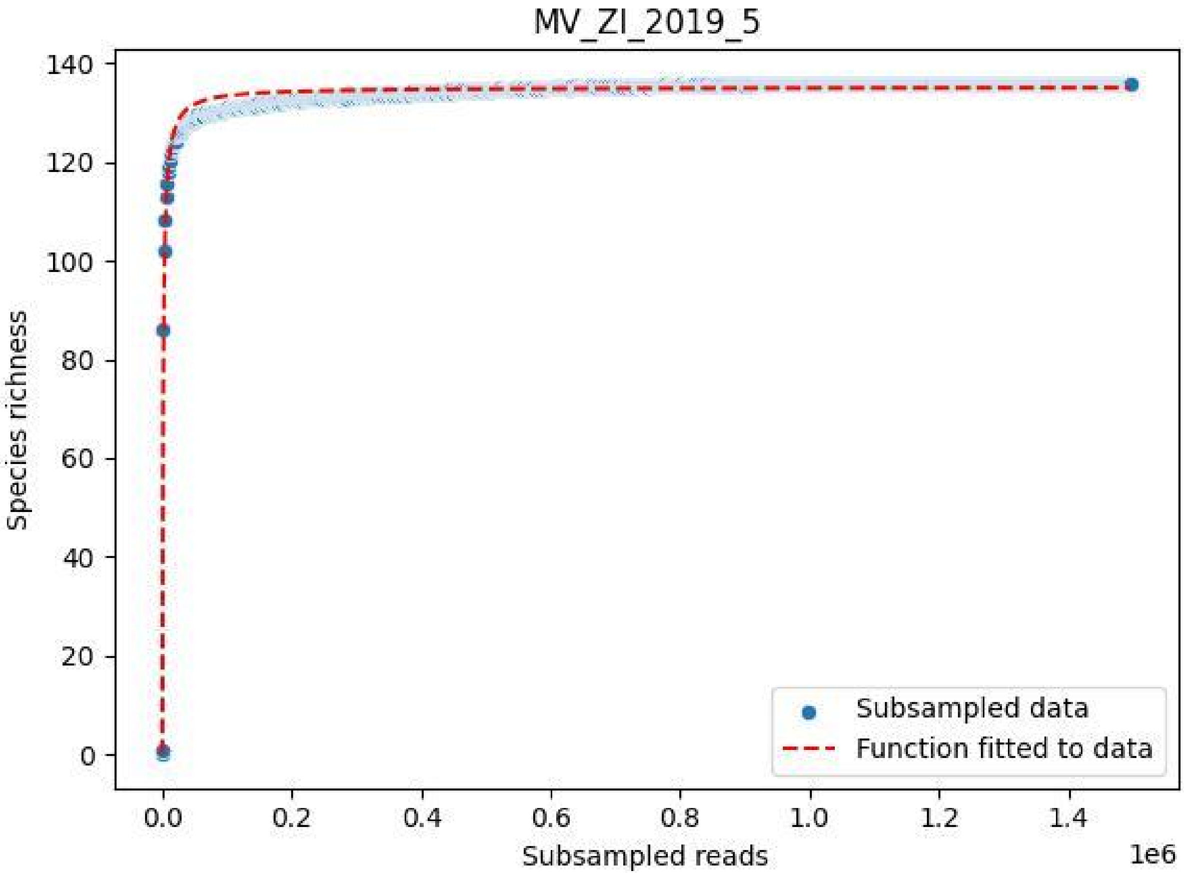
MV_ZI_2019_5
Species richness
Subsampled reads
1e6
0
20
40
60
80
100
120
140
0.0
0.2
0.4
0.6
0.8
1.0
1.2
1.4
Subsampled data
Function fitted to data

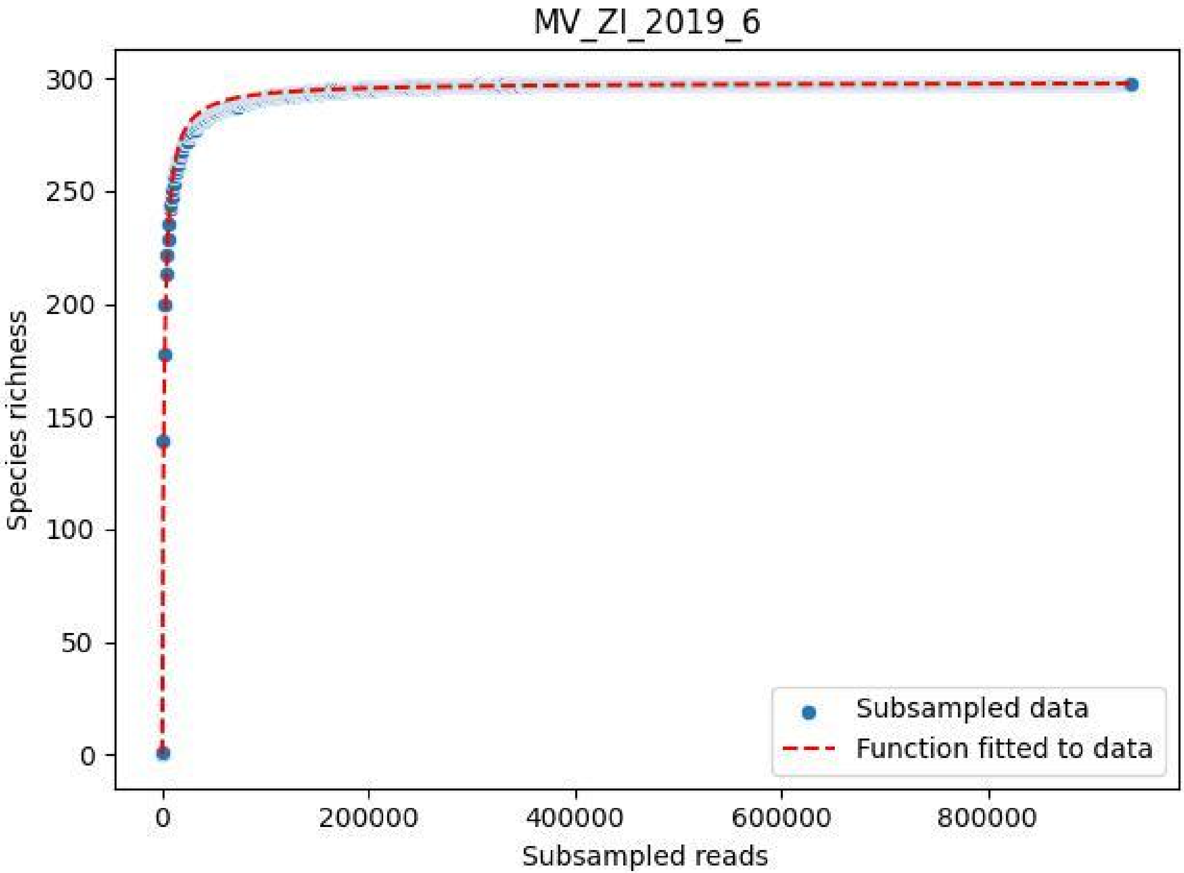
MV_ZI_2019_6
300
250
200
150
100
50
0
Species richness
0
200000
400000
600000
800000
Subsampled reads
Subsampled data
Function fitted to data

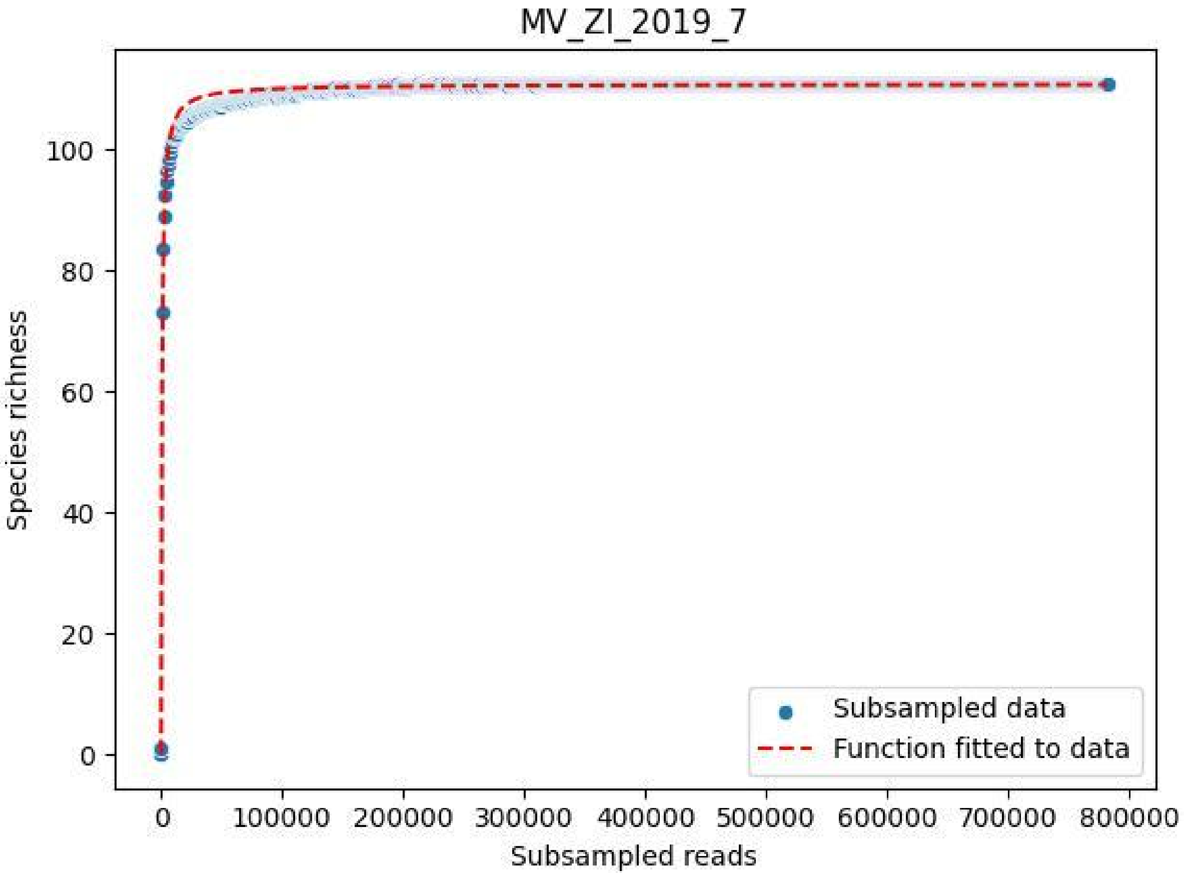

MV_ZI_2019_7
Species richness
Subsampled reads
0
20
40
60
80
100
0
100000
200000
300000
400000
500000
600000
700000
800000
Subsampled data
Function fitted to data

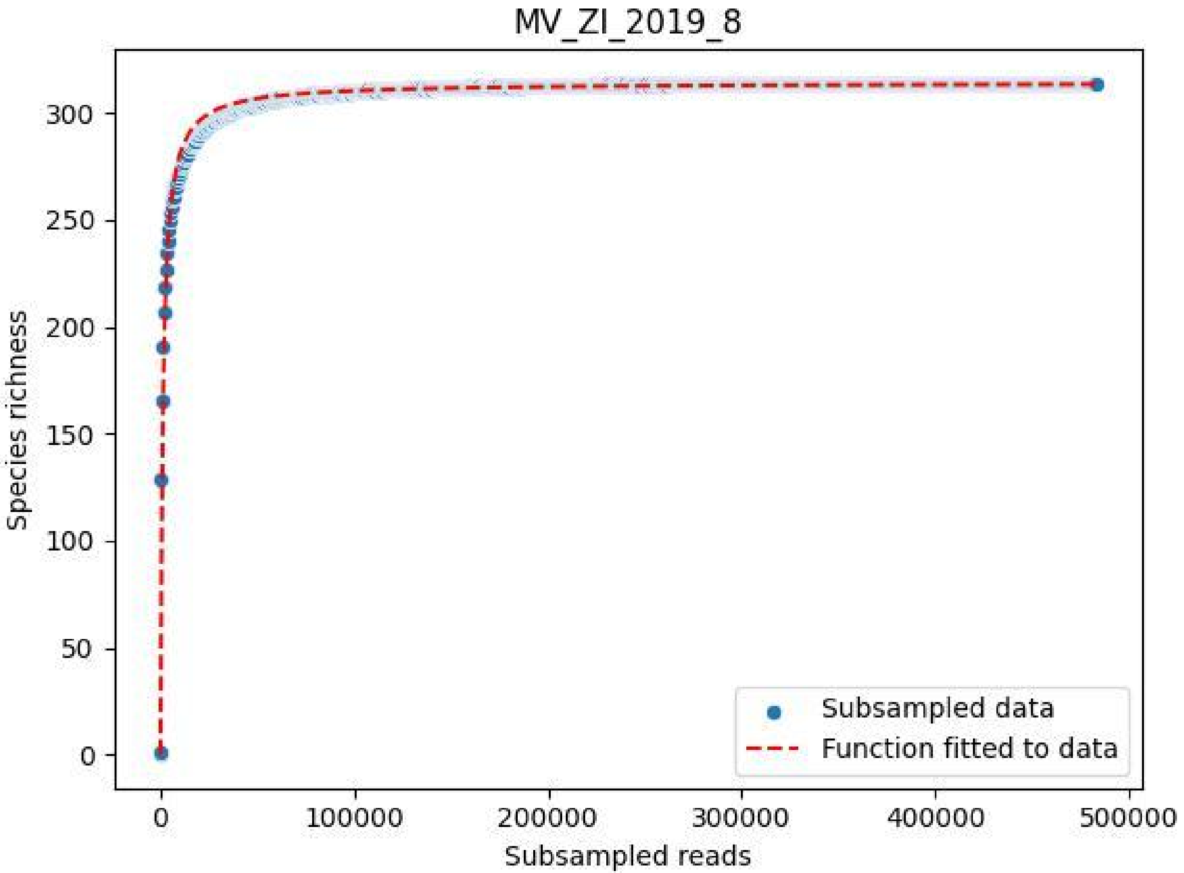

MV_ZI_2019_8
Species richness
Subsampled reads
0
50
100
150
200
250
300
0
100000
200000
300000
400000
500000
Subsampled data
Function fitted to data

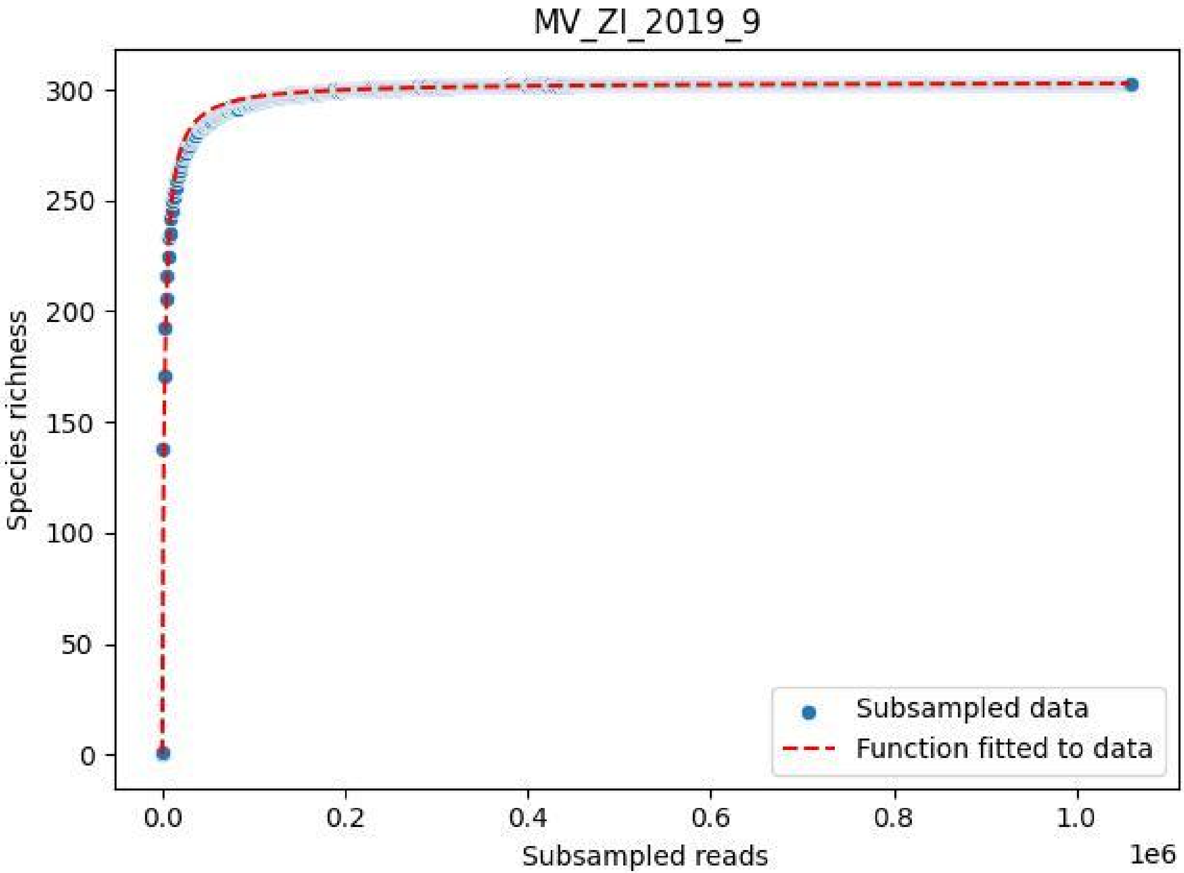

MV_ZI_2019_9
Species richness
Subsampled reads
1e6
0
50
100
150
200
250
300
0.0
0.2
0.4
0.6
0.8
1.0
Subsampled data
Function fitted to data

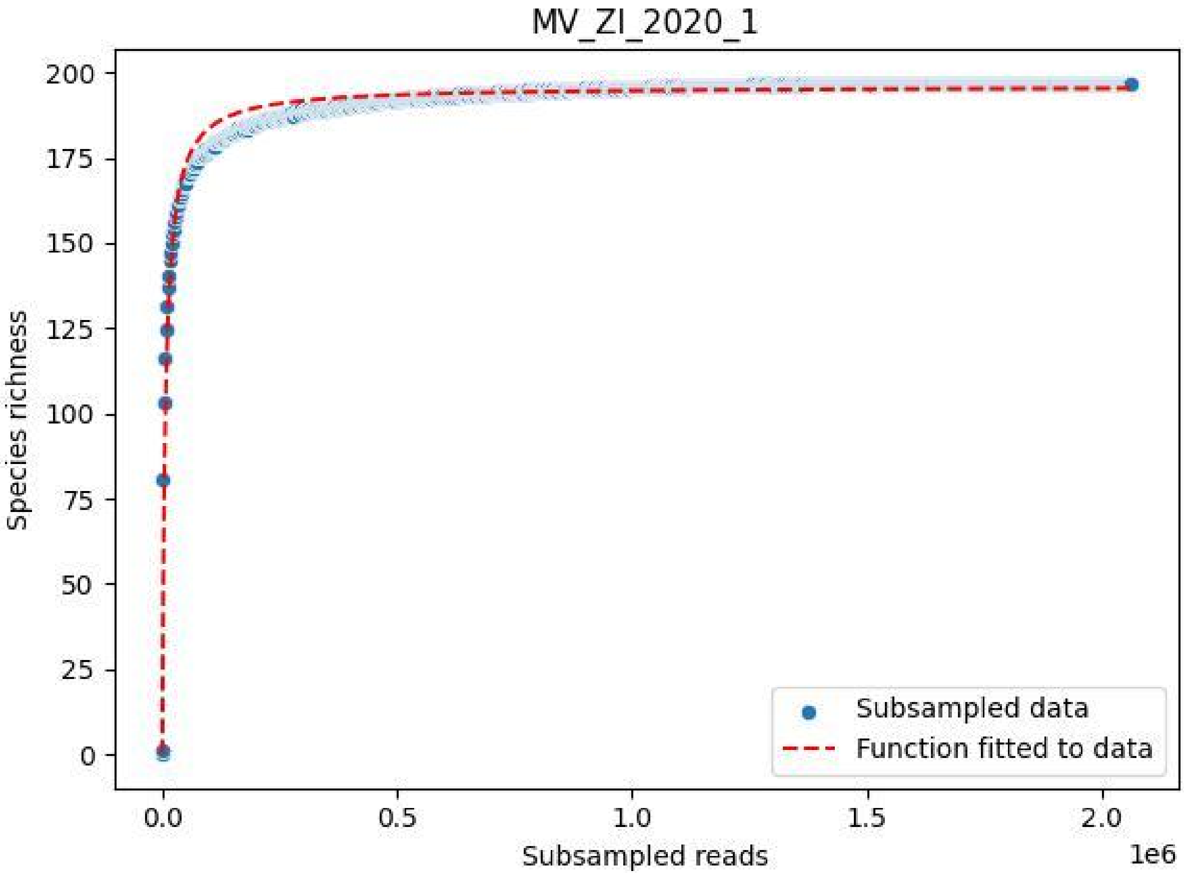
MV_ZI_2020_1
Species richness
Subsampled reads
1e6
0
25
50
75
100
125
150
175
200
0.0
0.5
1.0
1.5
2.0
Subsampled data
Function fitted to data

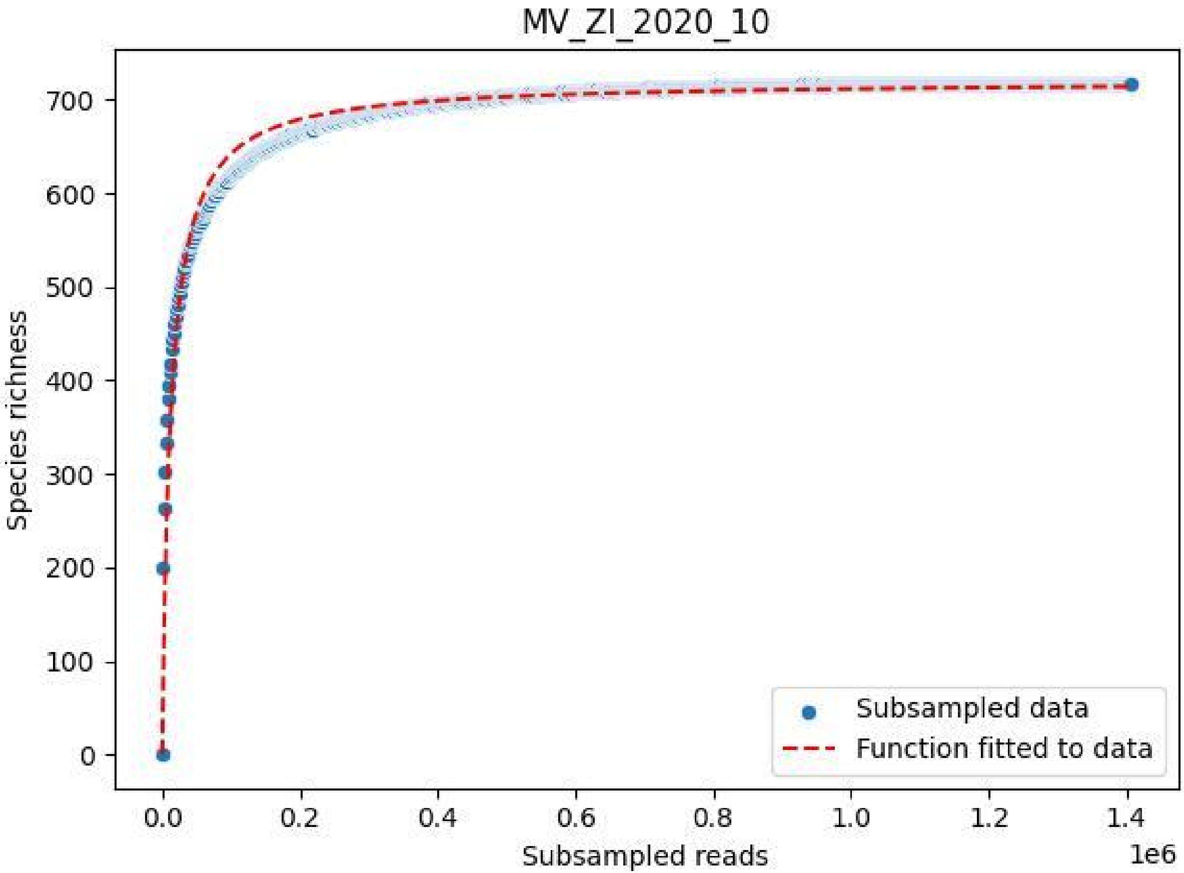

MV_ZI_2020_10
Species richness
Subsampled reads
1e6
0
100
200
300
400
500
600
700
0.0
0.2
0.4
0.6
0.8
1.0
1.2
1.4
Subsampled data
Function fitted to data

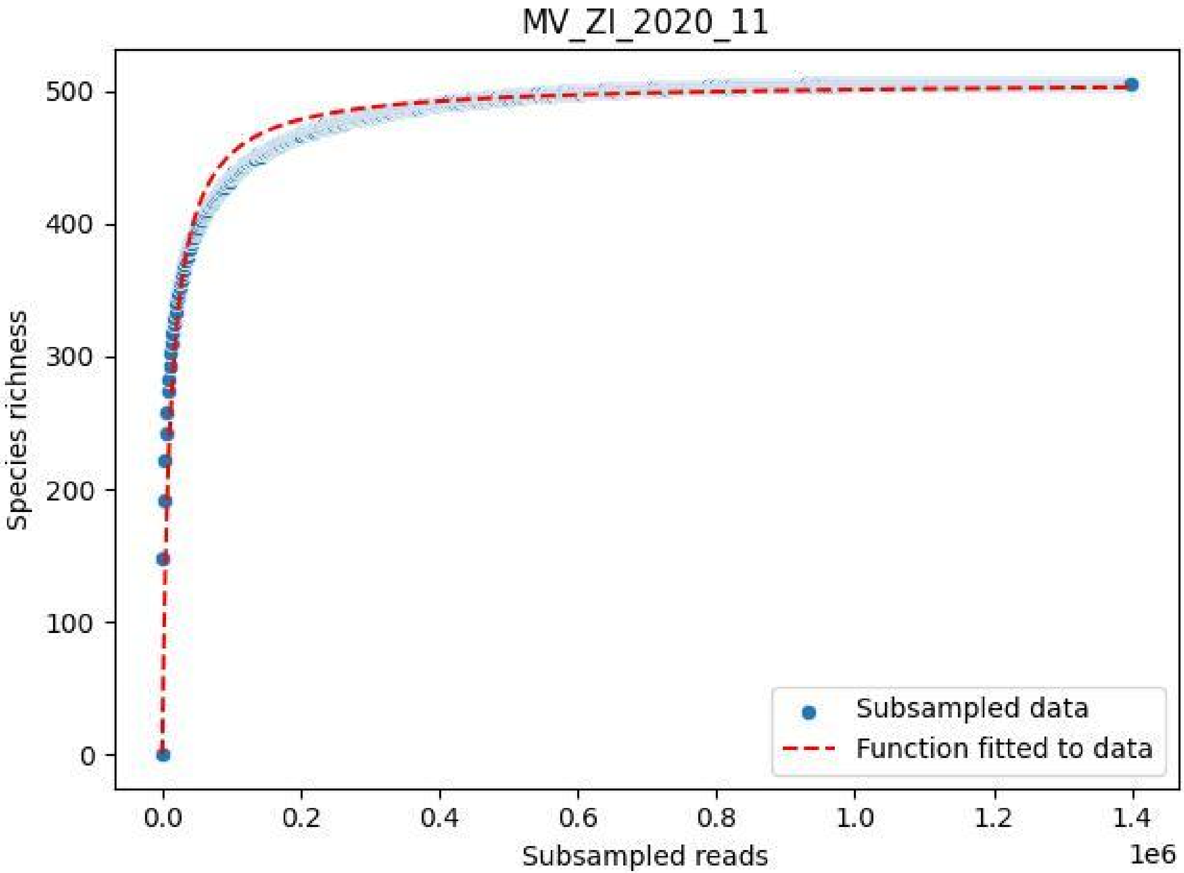

MV_ZI_2020_11
Species richness
Subsampled reads
1e6
0
100
200
300
400
500
0.0
0.2
0.4
0.6
0.8
1.0
1.2
1.4
Subsampled data
Function fitted to data

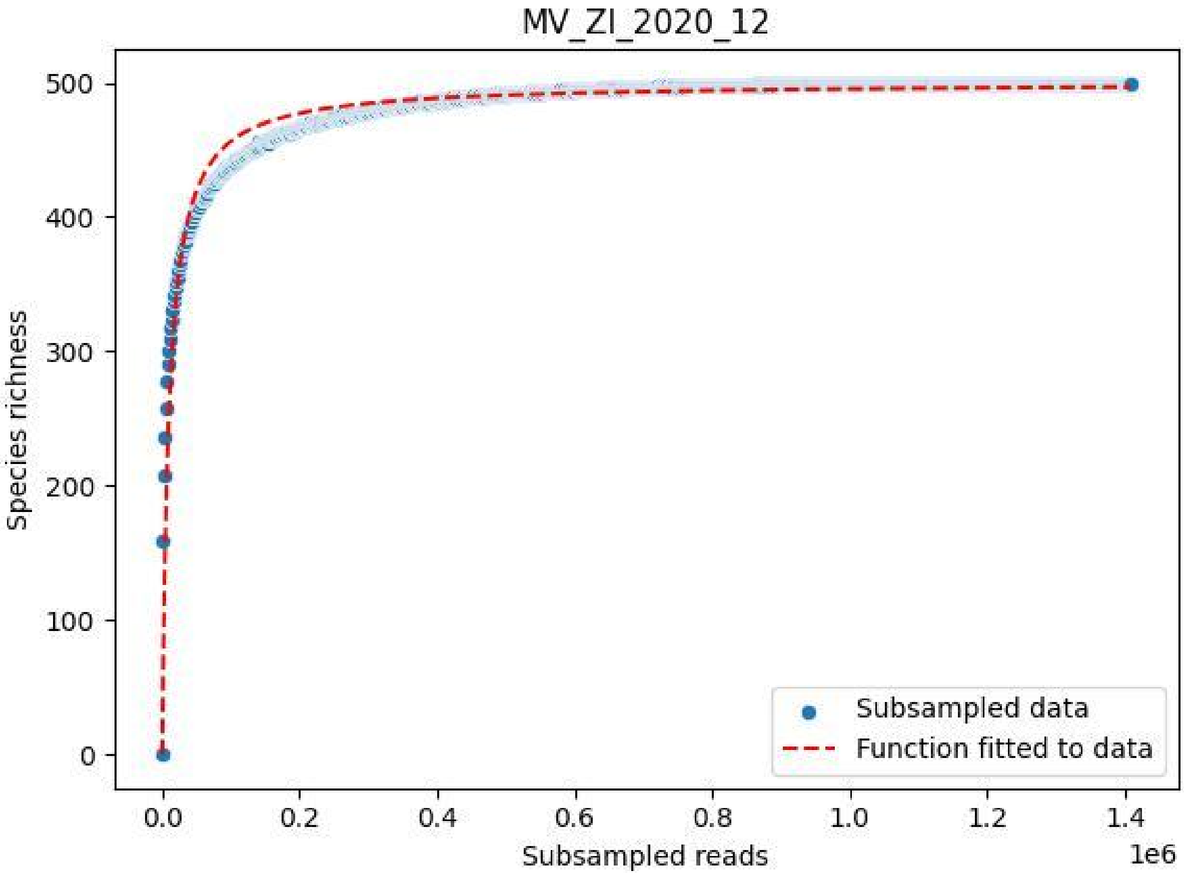
MV_ZI_2020_12
Species richness
Subsampled reads
1e6
Subsampled data
Function fitted to data
0
100
200
300
400
500
0.0
0.2
0.4
0.6
0.8
1.0
1.2
1.4

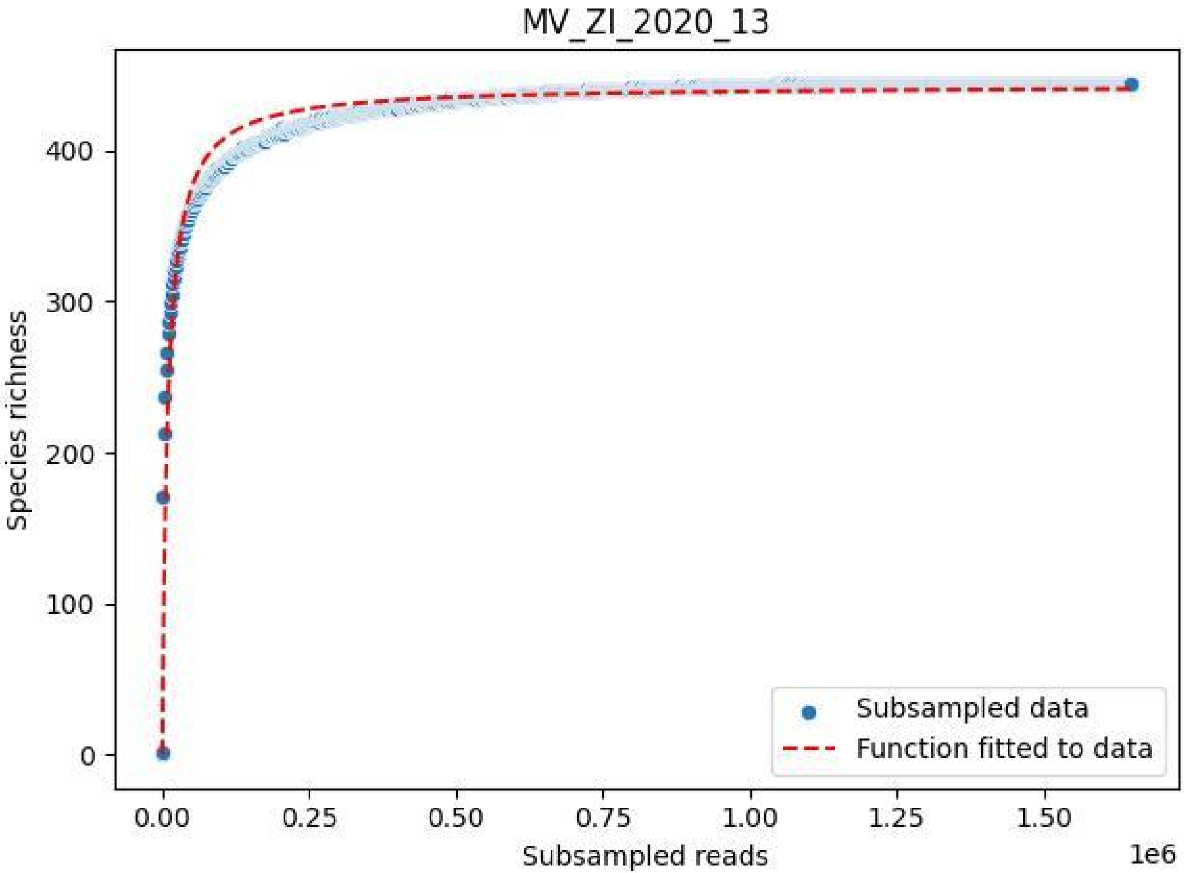
MV_ZI_2020_13
Species richness
400
300
200
100
0
0.00
0.25
0.50
0.75
1.00
1.25
1.50
Subsampled reads
1e6
Subsampled data
Function fitted to data

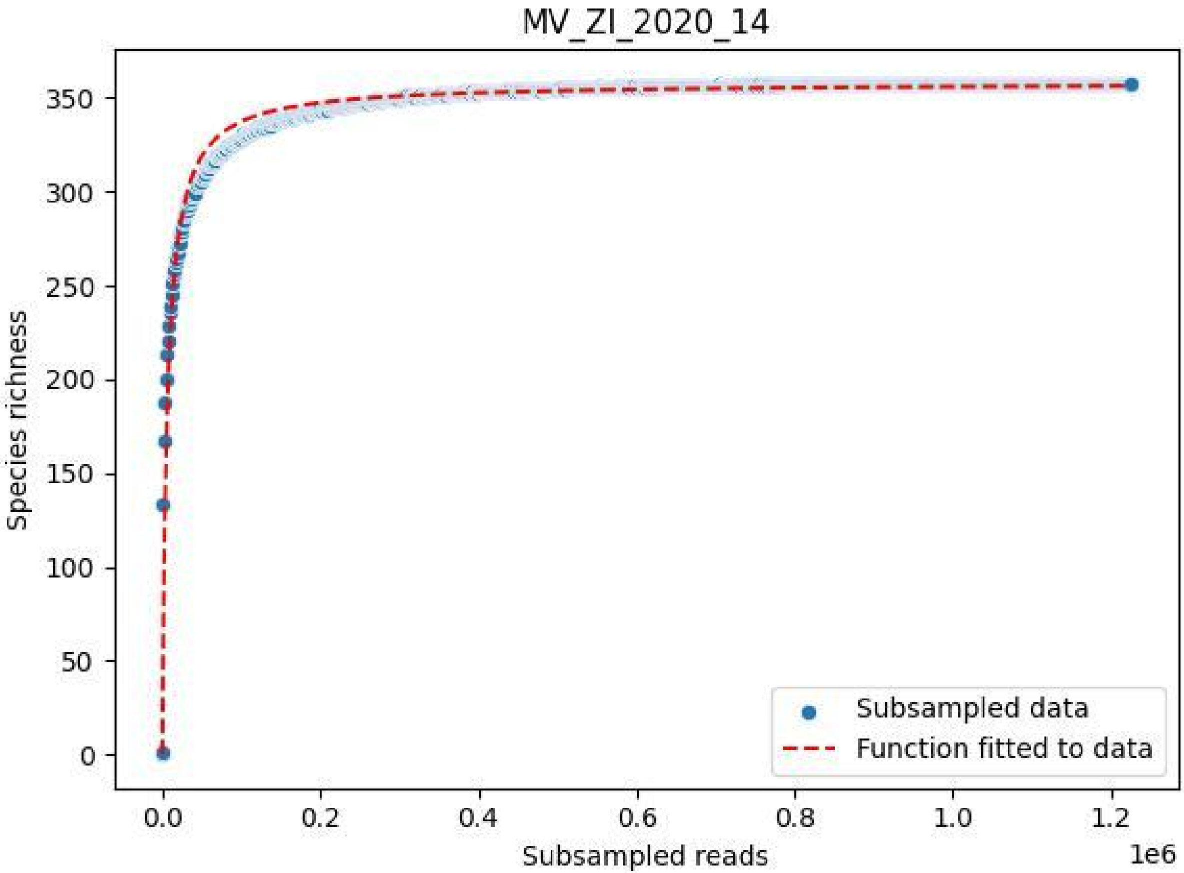

MV_ZI_2020_14
Species richness
350
300
250
200
150
100
50
0
0.0
0.2
0.4
0.6
0.8
1.0
1.2
1e6
Subsampled reads
Subsampled data
Function fitted to data

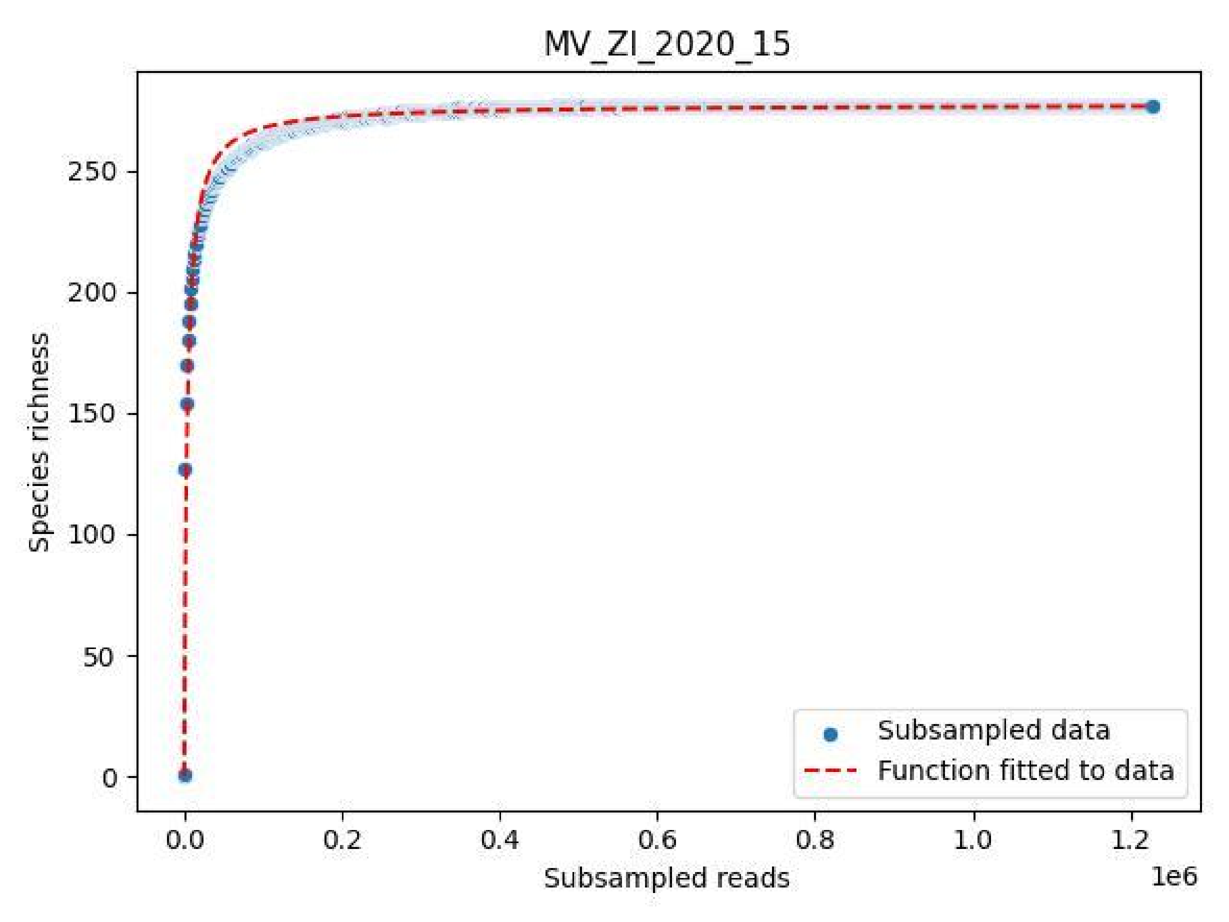

MV_ZI_2020_15
Species richness
0
50
100
150
200
250
0.0
0.2
0.4
0.6
0.8
1.0
1.2
1e6
Subsampled reads
Subsampled data
Function fitted to data

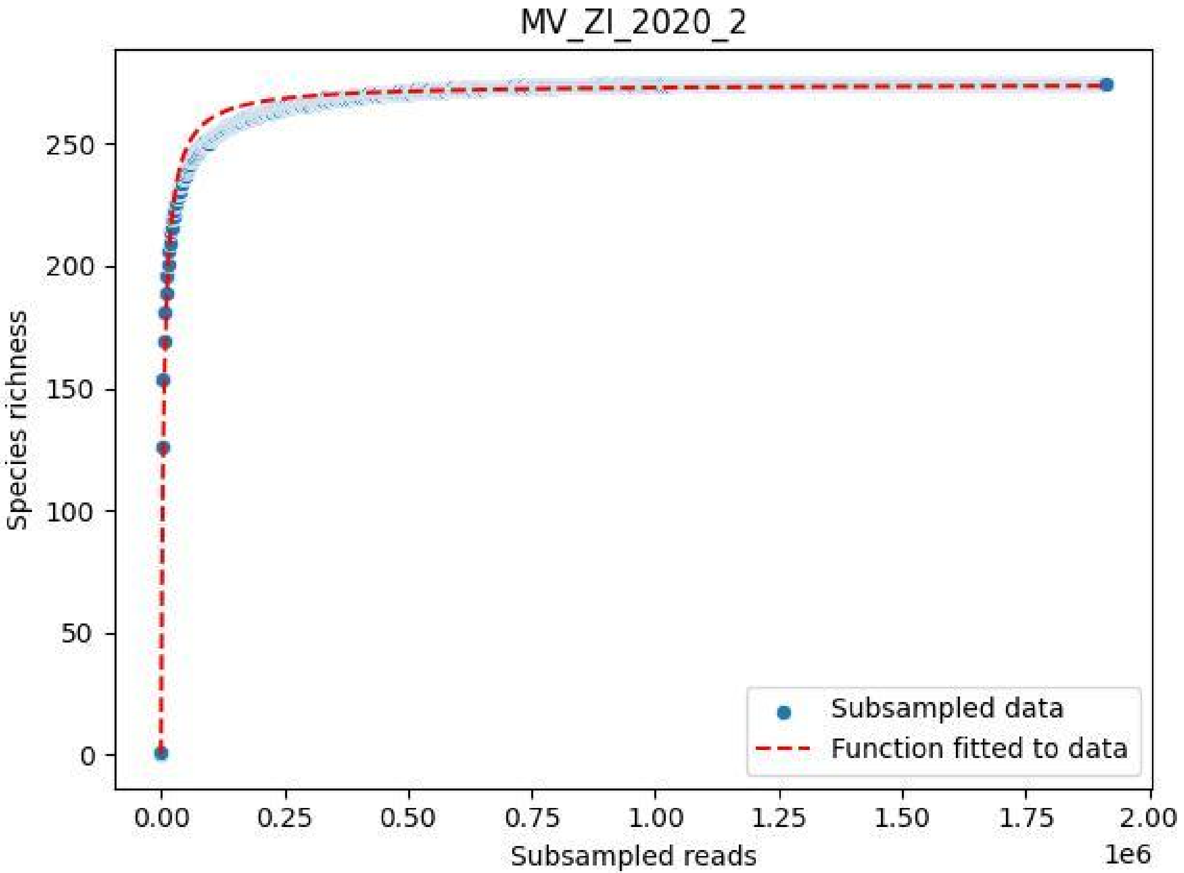
MV_ZI_2020_2
Species richness
Subsampled reads
1e6
0
50
100
150
200
250
0.00
0.25
0.50
0.75
1.00
1.25
1.50
1.75
2.00
Subsampled data
Function fitted to data

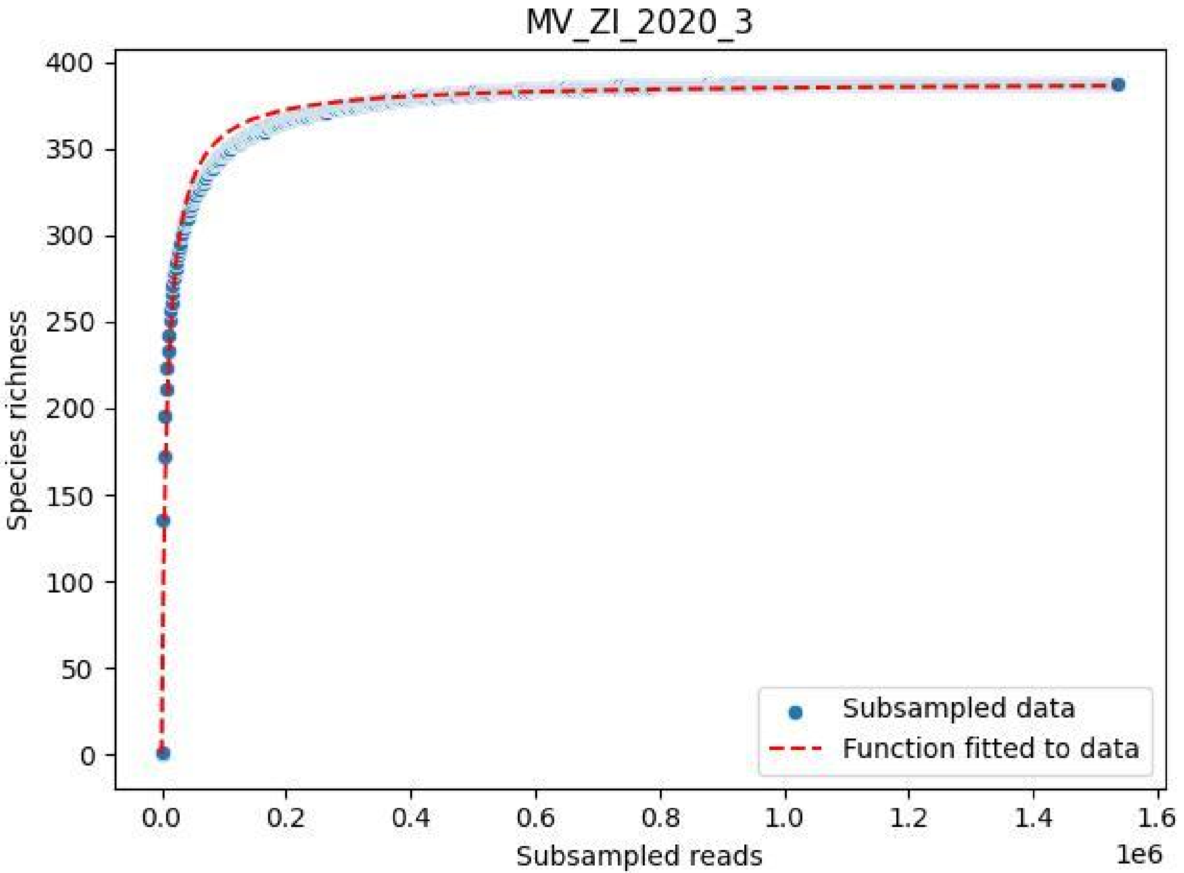
MV_ZI_2020_3
Species richness
Subsampled reads
1e6
Subsampled data
Function fitted to data
0
50
100
150
200
250
300
350
400
0.0
0.2
0.4
0.6
0.8
1.0
1.2
1.4
1.6

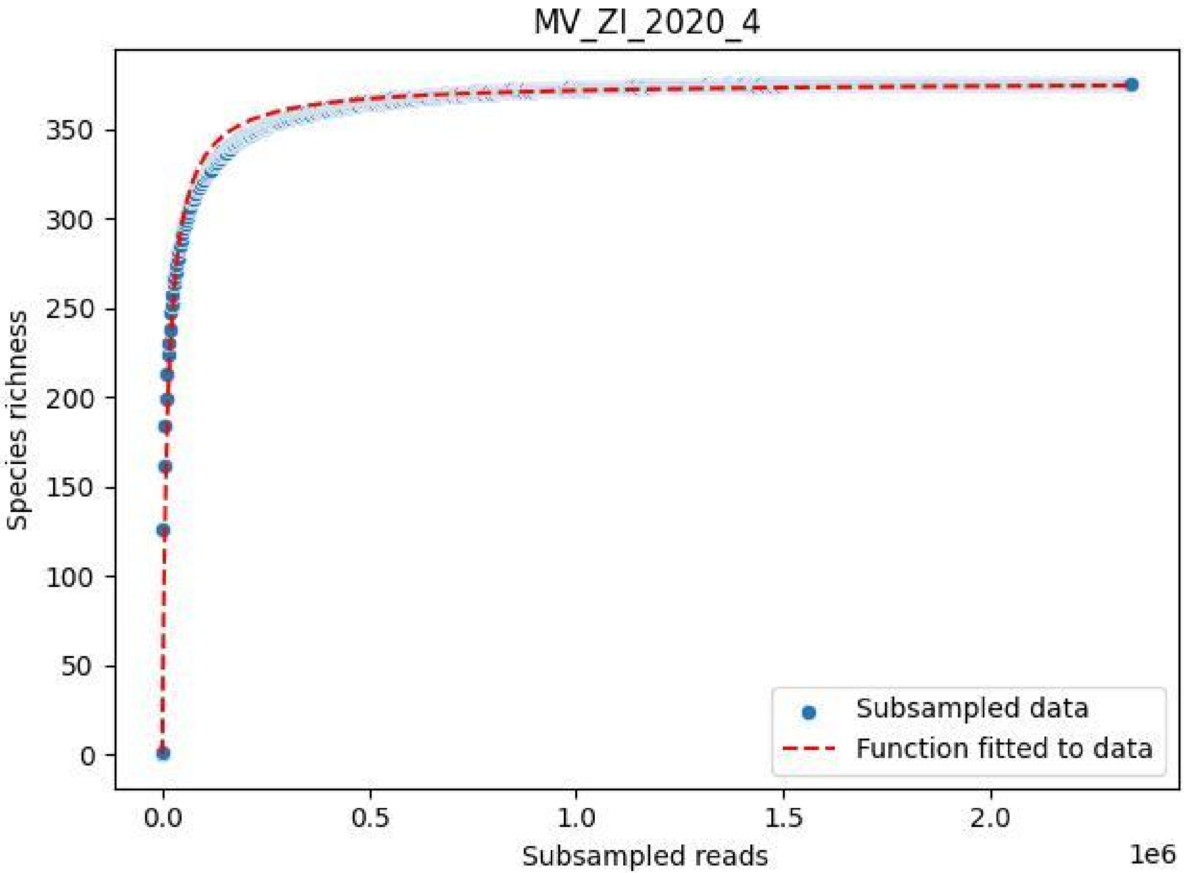

MV_ZI_2020_4
Species richness
Subsampled reads
1e6
0
50
100
150
200
250
300
350
0.0
0.5
1.0
1.5
2.0
Subsampled data
Function fitted to data

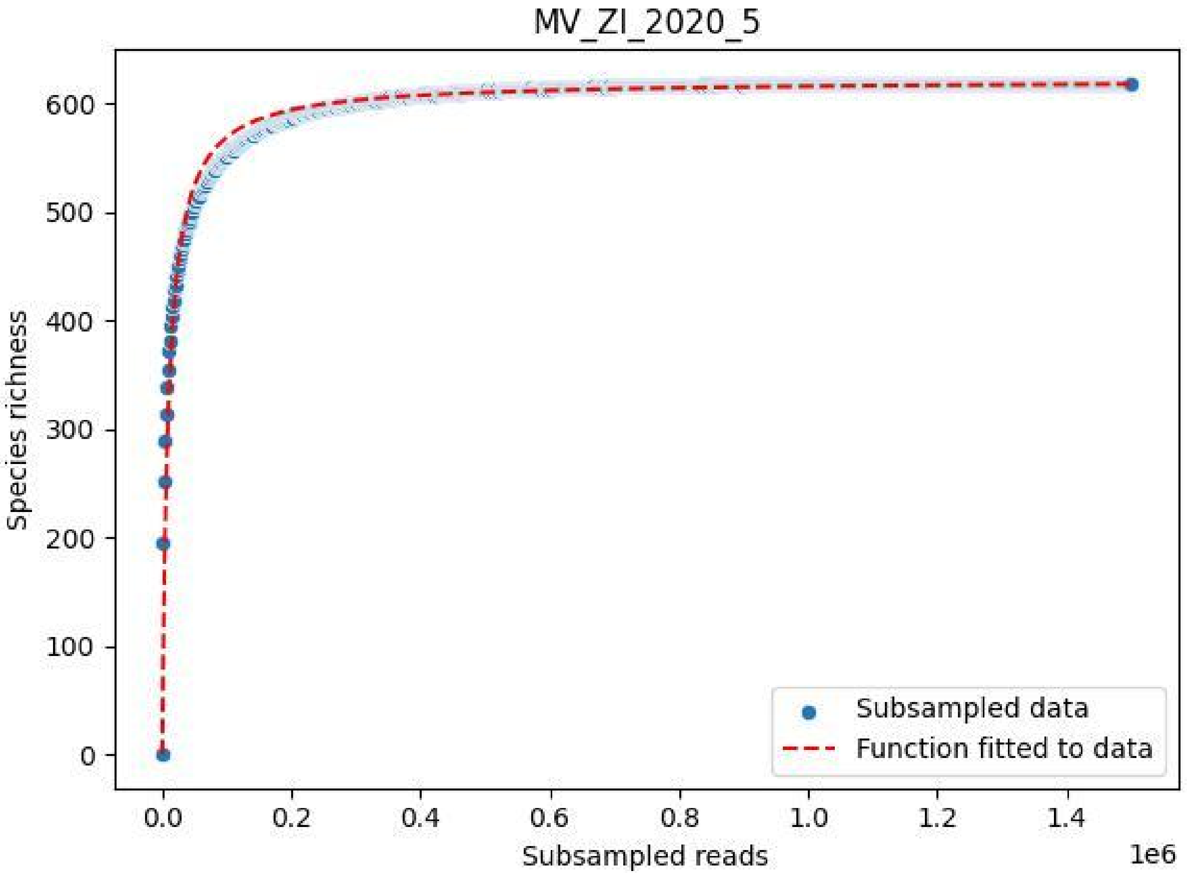

MV_ZI_2020_5
Species richness
Subsampled reads
1e6
0
100
200
300
400
500
600
0.0
0.2
0.4
0.6
0.8
1.0
1.2
1.4
Subsampled data
Function fitted to data

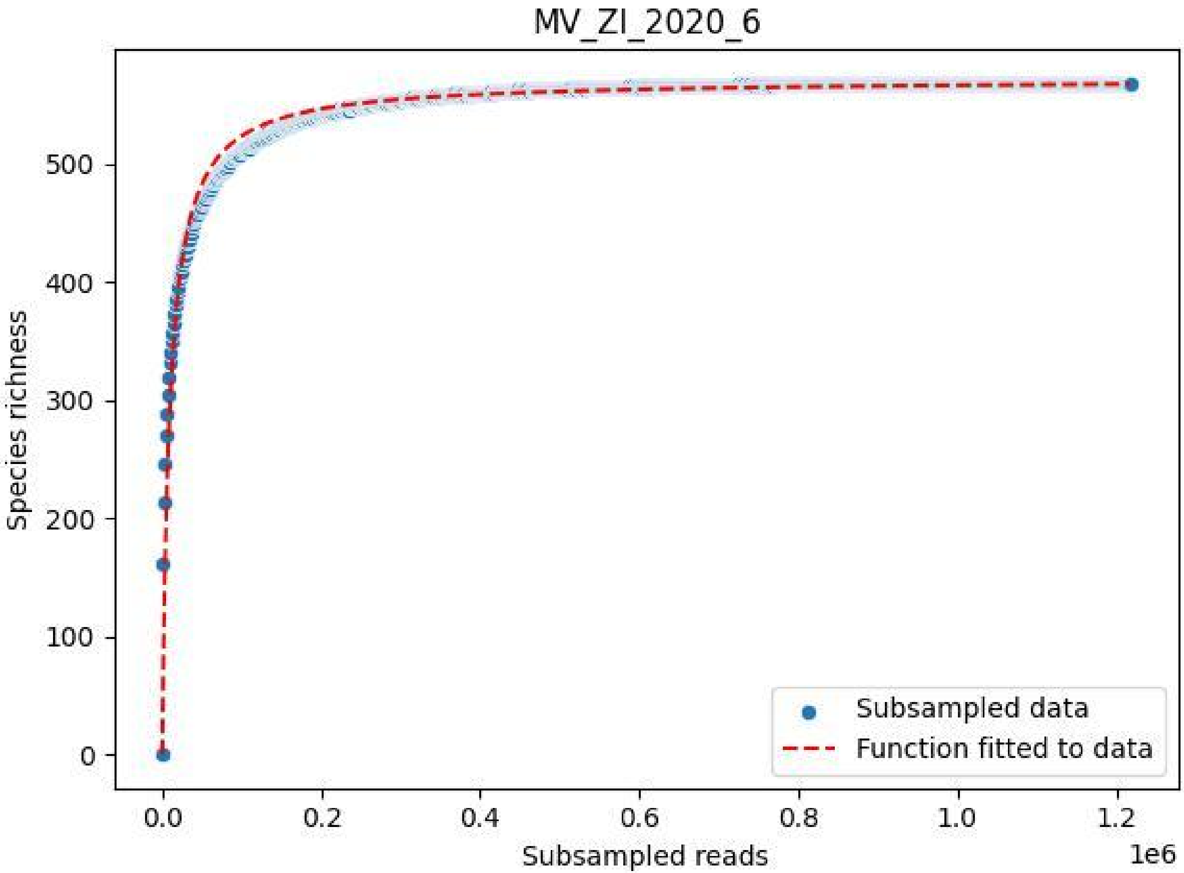
MV_ZI_2020_6
Species richness
Subsampled reads
1e6
0
100
200
300
400
500
0.0
0.2
0.4
0.6
0.8
1.0
1.2
Subsampled data
Function fitted to data

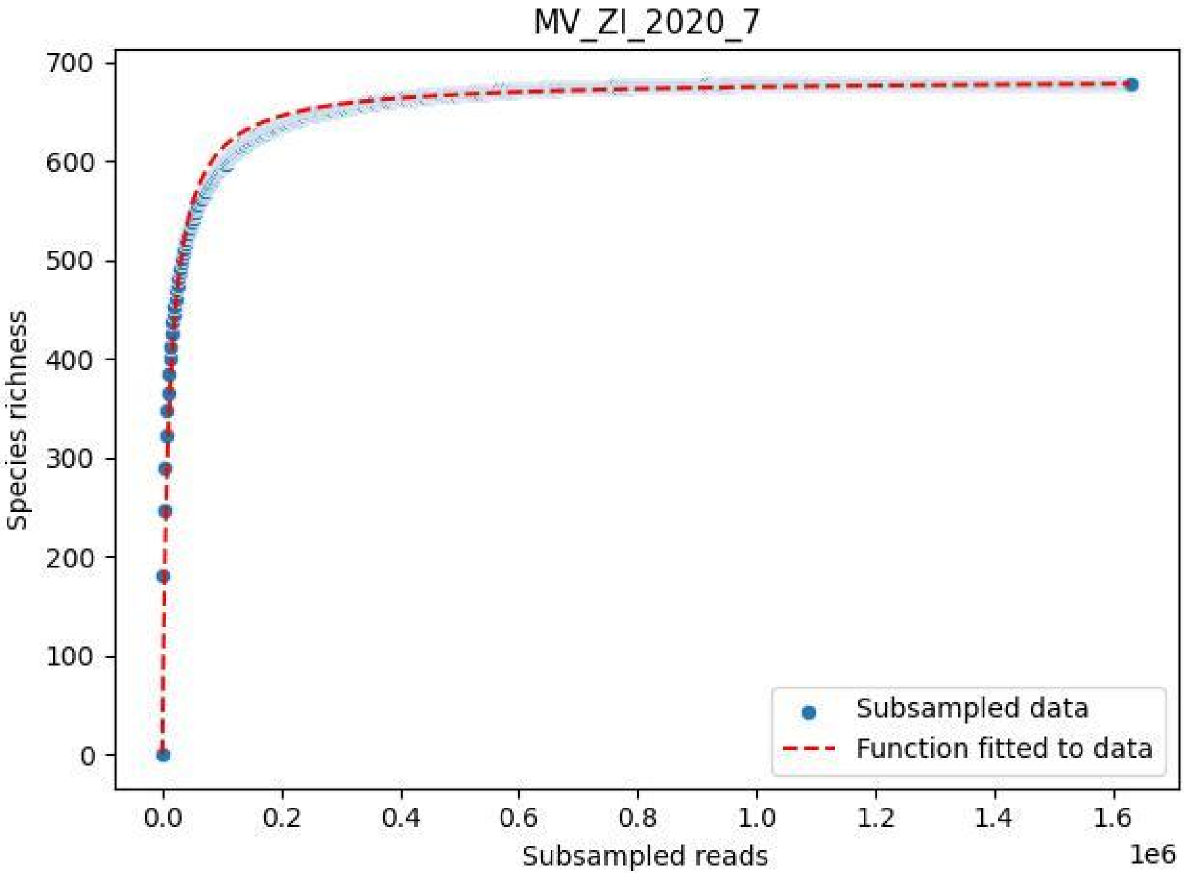

MV_ZI_2020_7
Species richness
Subsampled reads
1e6
Subsampled data
Function fitted to data

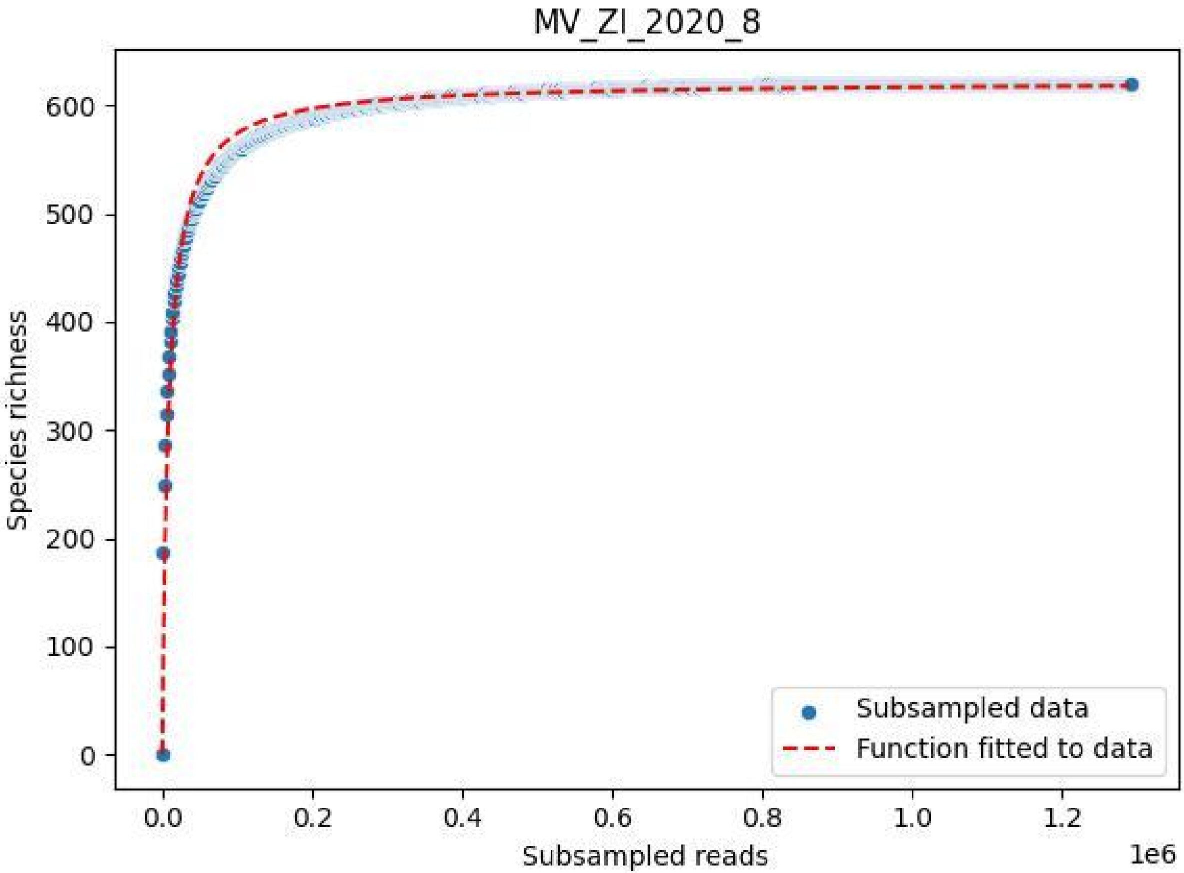

MV_ZI_2020_8
Species richness
Subsampled reads
1e6
0
100
200
300
400
500
600
0.0
0.2
0.4
0.6
0.8
1.0
1.2
Subsampled data
Function fitted to data

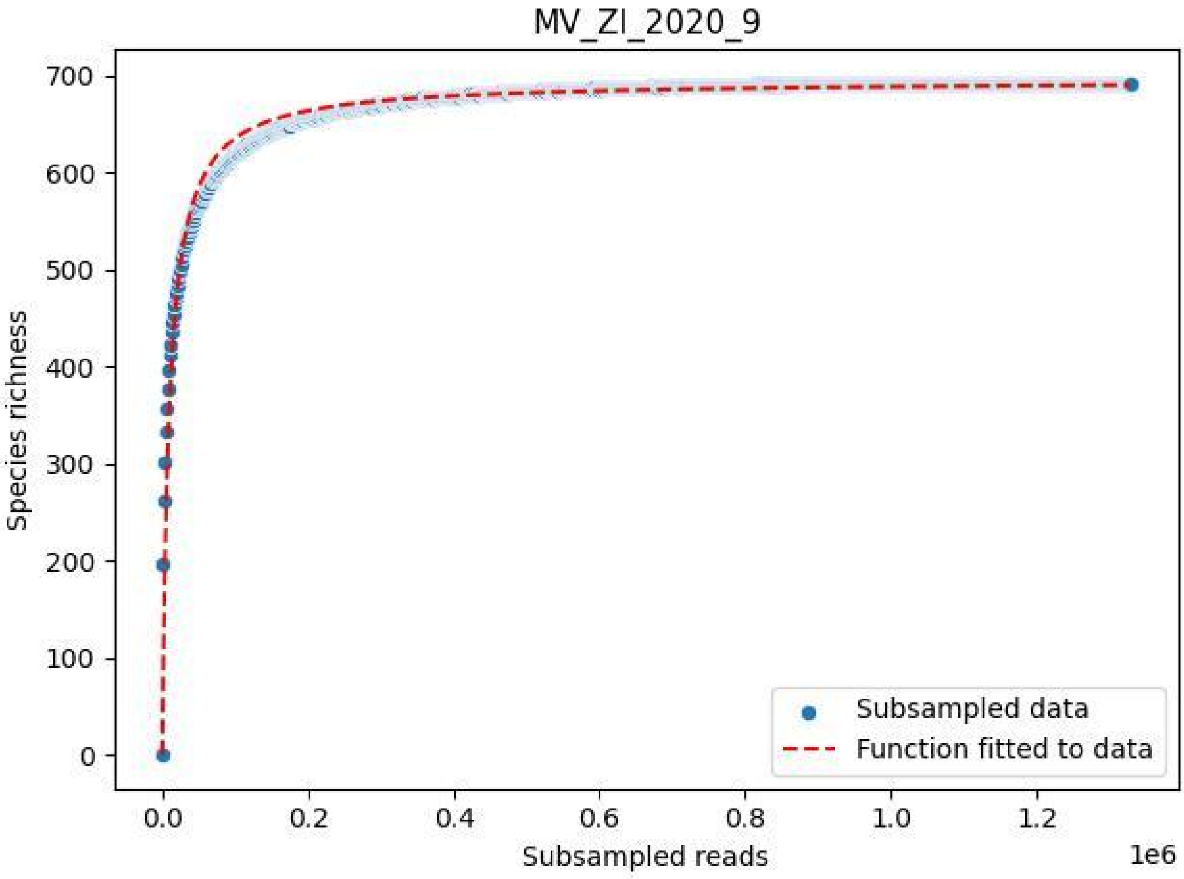
MV_ZI_2020_9
700
600
500
400
300
200
100
0
Species richness
Subsampled data
Function fitted to data
0.0
0.2
0.4
0.6
0.8
1.0
1.2
Subsampled reads
1e6

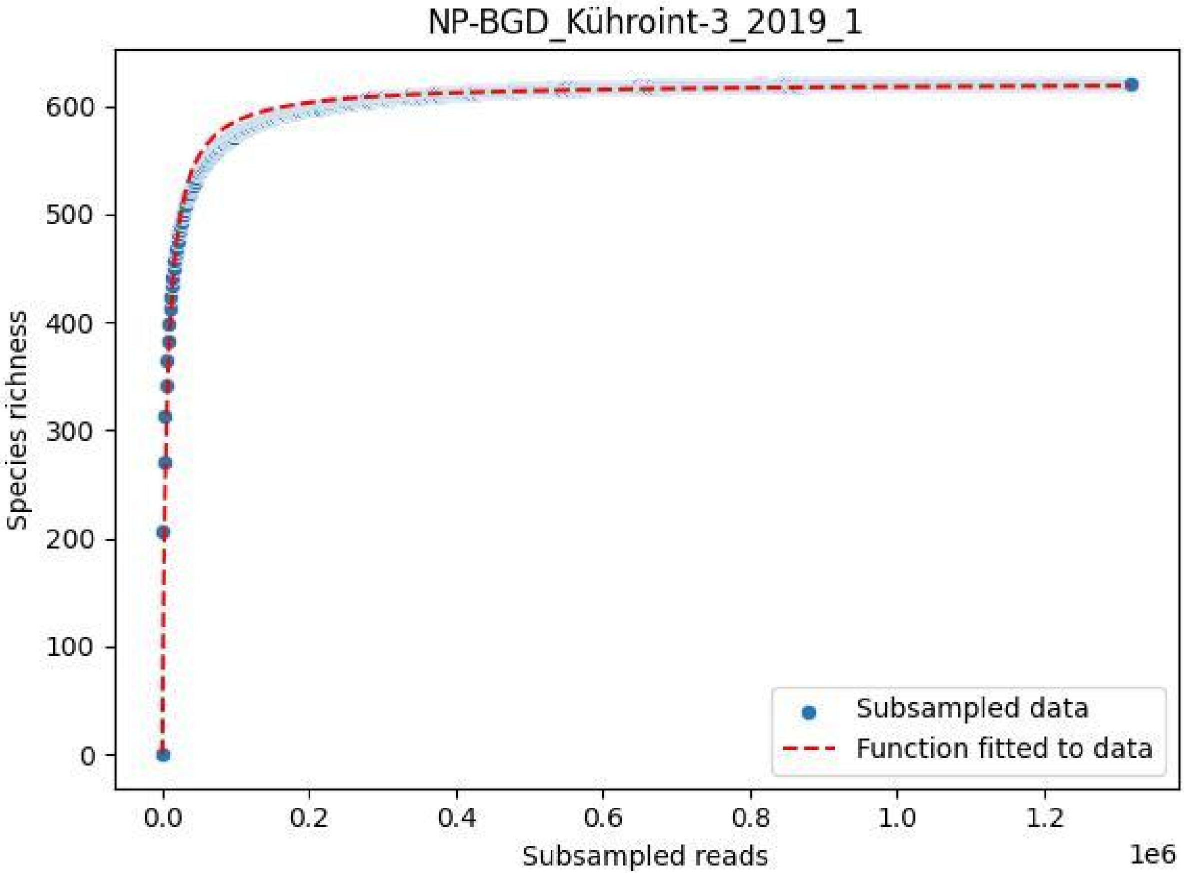
NP-BGD_Kühroint-3_2019_1
Species richness
0
100
200
300
400
500
600
0.0
0.2
0.4
0.6
0.8
1.0
1.2
1e6
Subsampled reads
Subsampled data
Function fitted to data

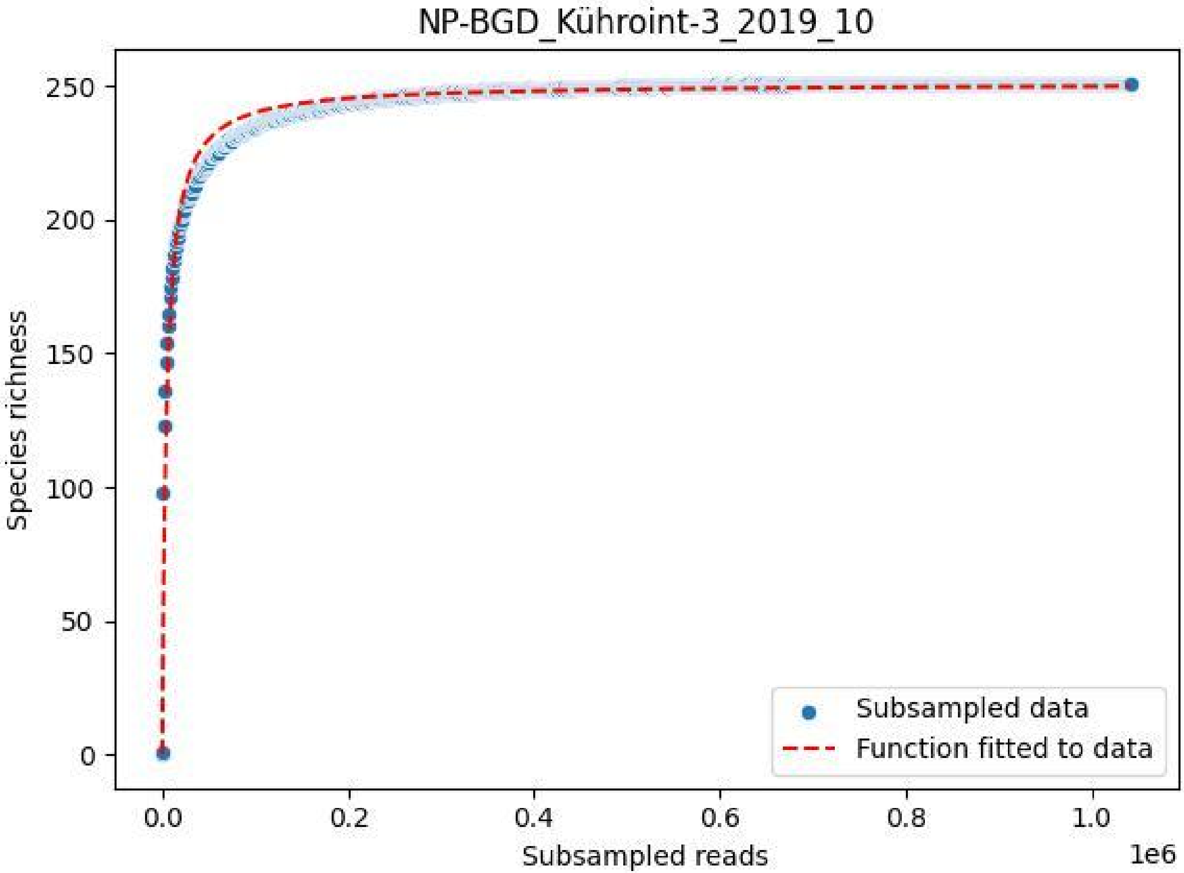

NP-BGD_Kühroint-3_2019_10
Species richness
Subsampled reads
1e6
0
50
100
150
200
250
0.0
0.2
0.4
0.6
0.8
1.0
Subsampled data
Function fitted to data

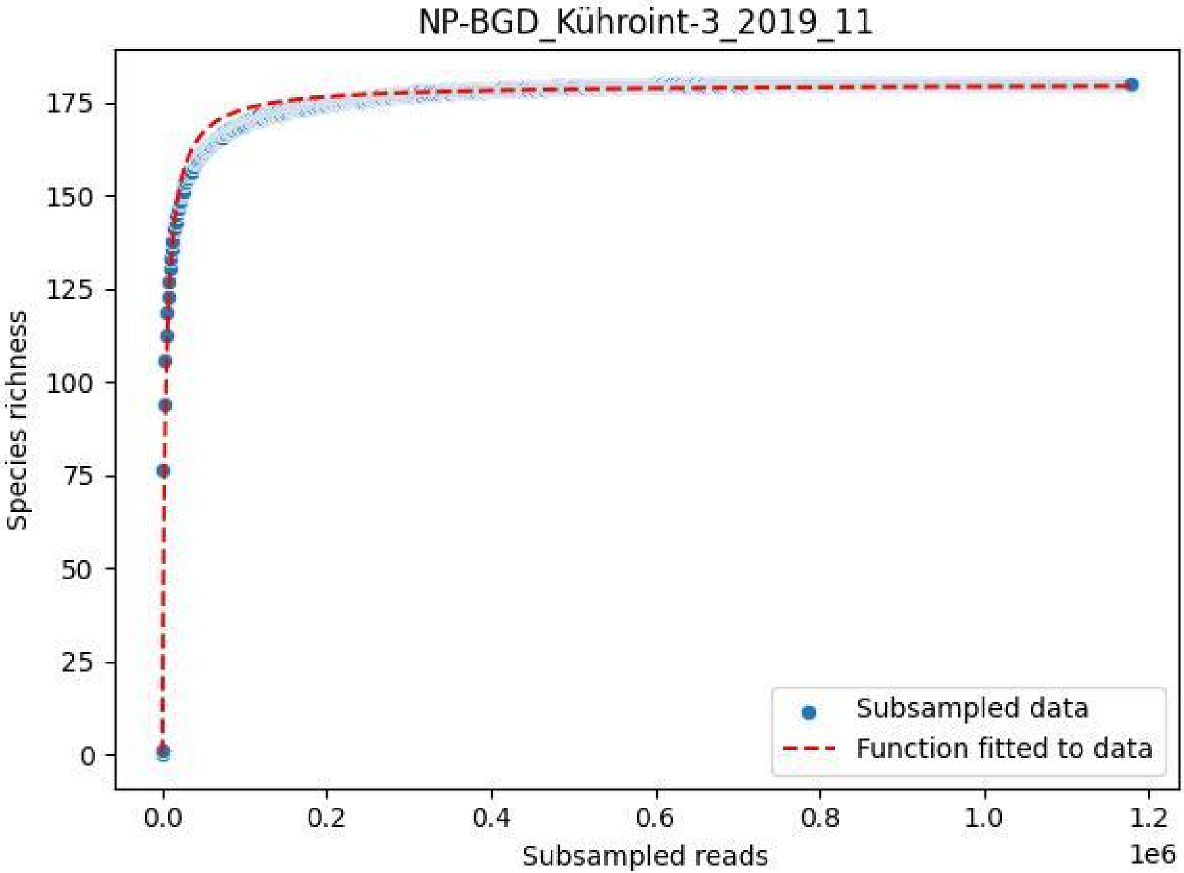
NP-BGD_Kühroint-3_2019_11
Species richness
175
150
125
100
75
50
25
0
0.0
0.2
0.4
0.6
0.8
1.0
1.2
1e6
Subsampled reads
Subsampled data
Function fitted to data

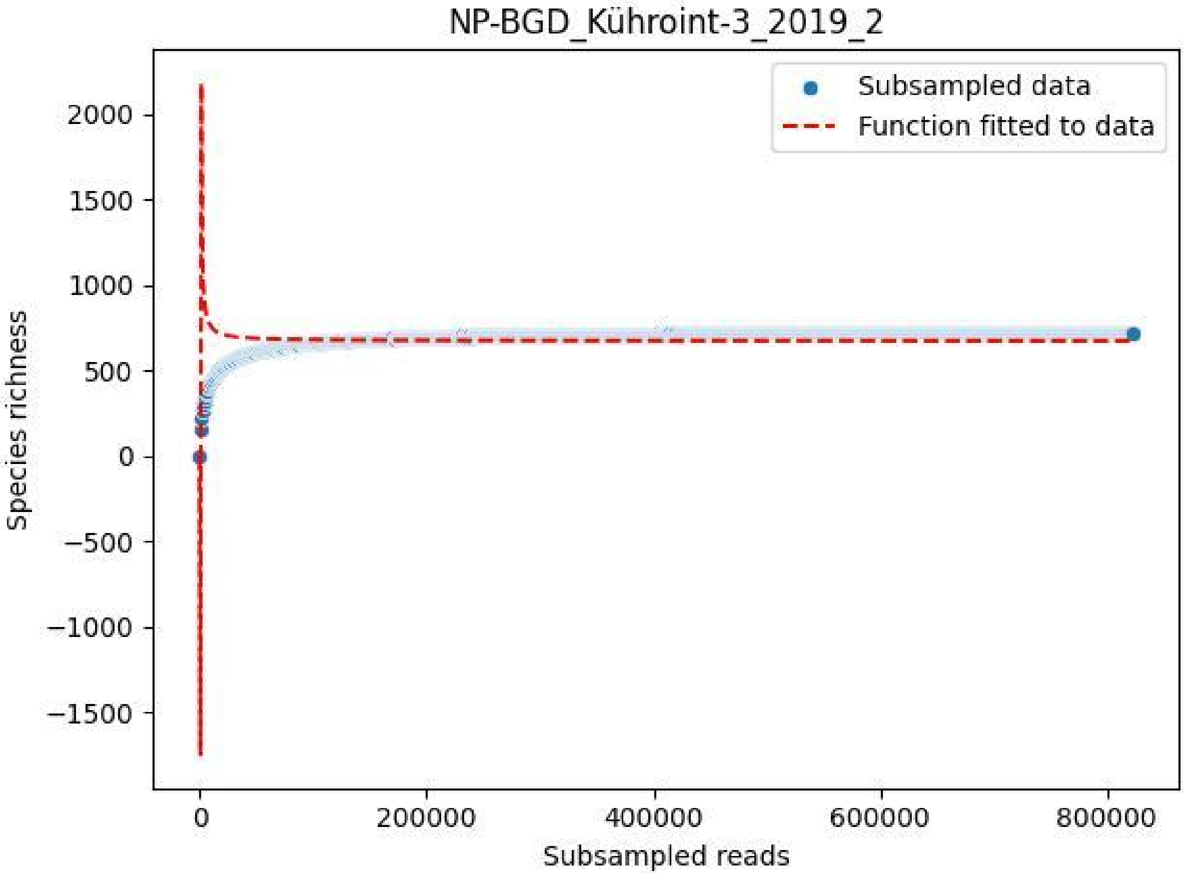
NP-BGD_Kühroint-3_2019_2
Subsampled data
Function fitted to data
Species richness
Subsampled reads
2000
1500
1000
500
0
−500
−1000
−1500
0
200000
400000
600000
800000

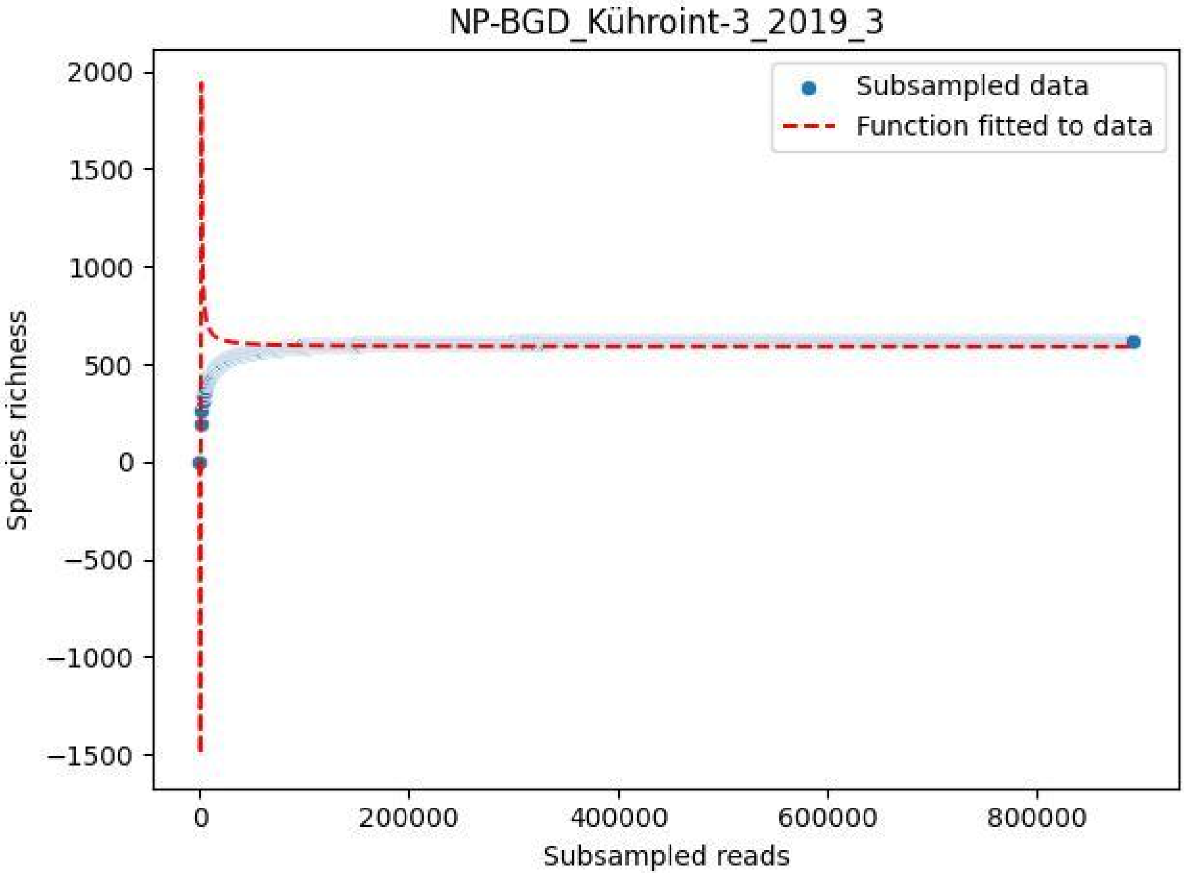
NP-BGD_Kühroint-3_2019_3
Subsampled data
Function fitted to data
Species richness
Subsampled reads
2000
1500
1000
500
0
−500
−1000
−1500
0
200000
400000
600000
800000

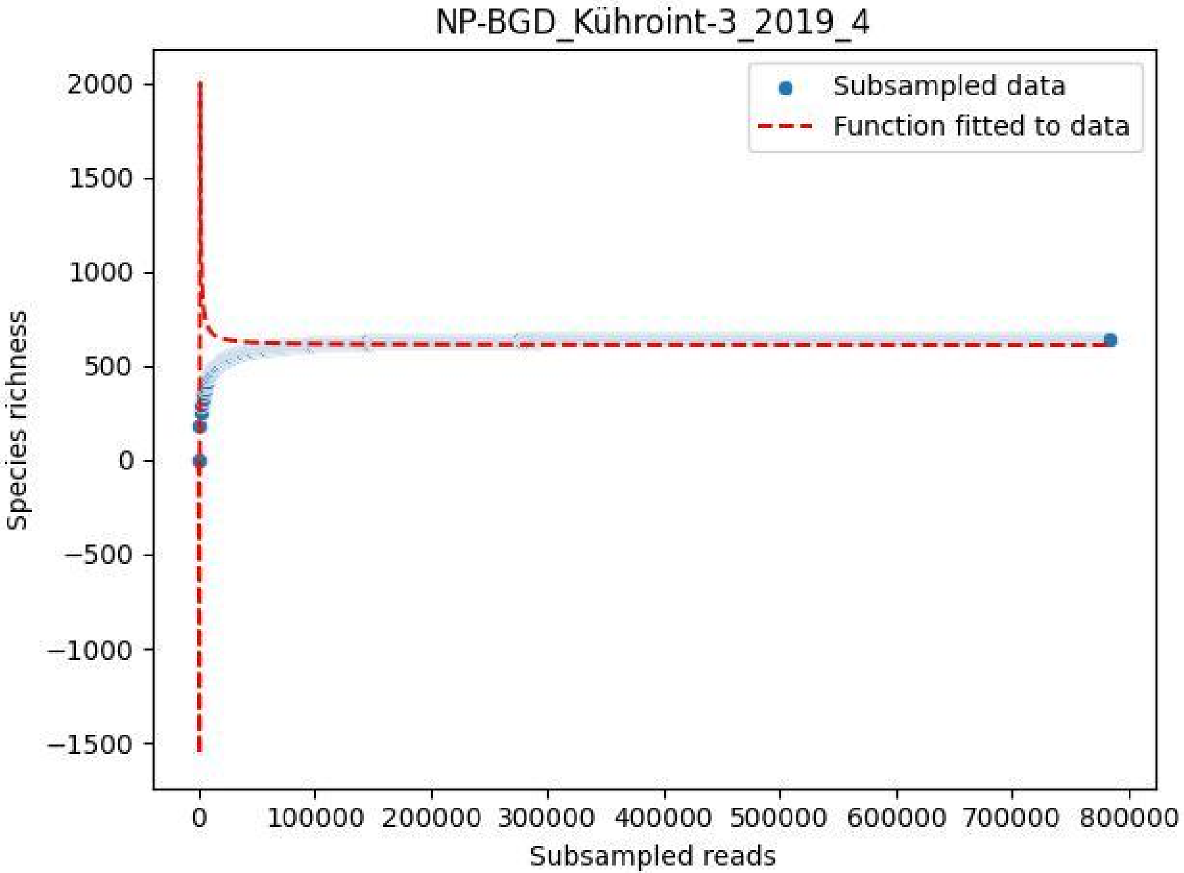

NP-BGD_Kühroint-3_2019_4
Subsampled data
Function fitted to data
Species richness
Subsampled reads
2000
1500
1000
500
0
−500
−1000
−1500
0
100000
200000
300000
400000
500000
600000
700000
800000

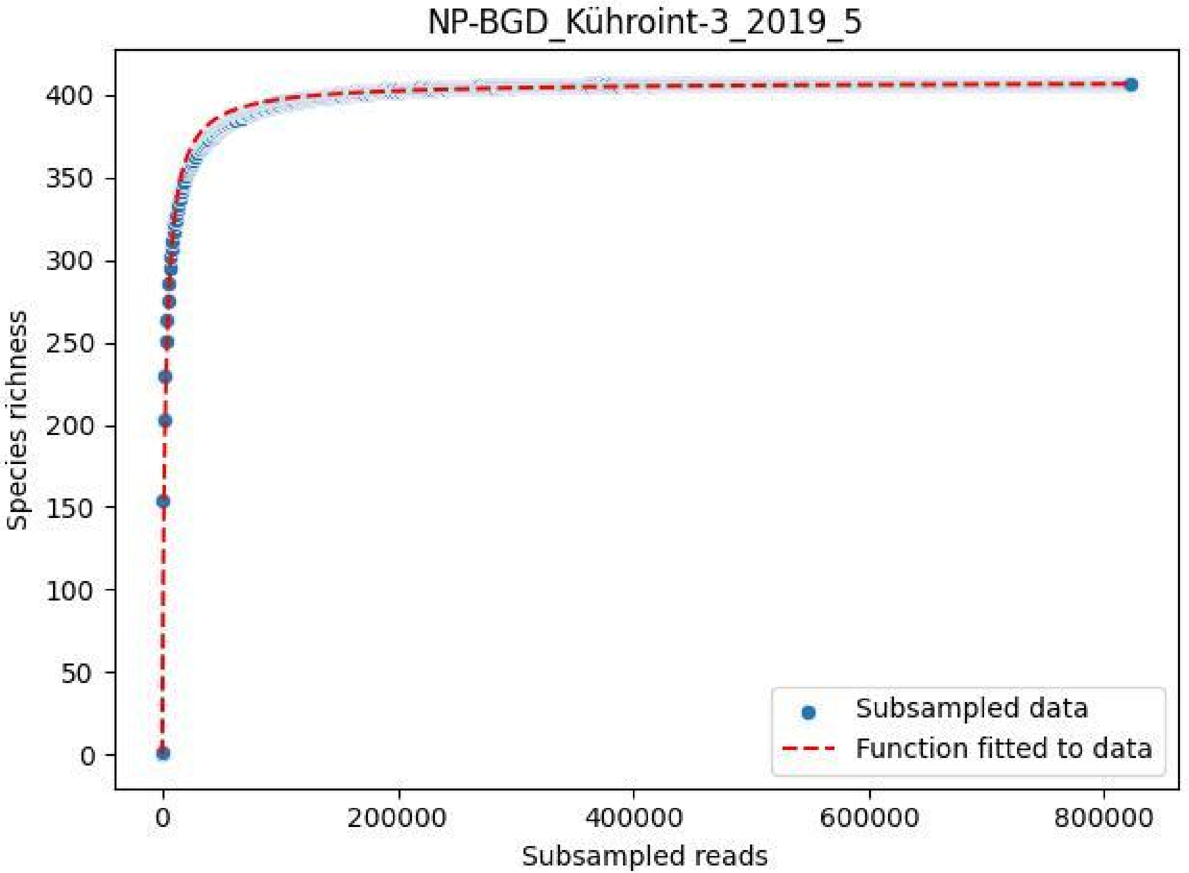
NP-BGD_Kühroint-3_2019_5
Species richness
Subsampled reads
0
50
100
150
200
250
300
350
400
0
200000
400000
600000
800000
Subsampled data
Function fitted to data

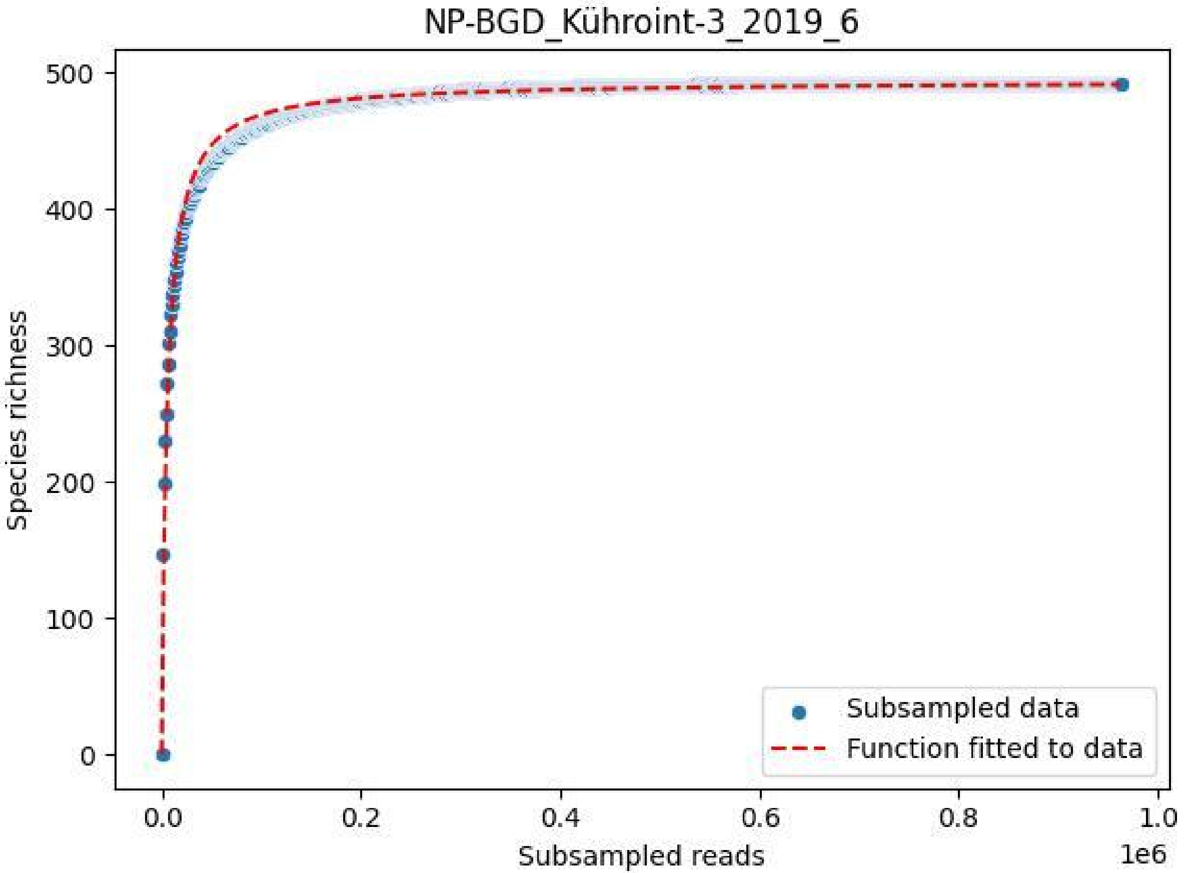
NP-BGD_Kühroint-3_2019_6
Species richness
Subsampled reads
1e6
500
400
300
200
100
0
0.0
0.2
0.4
0.6
0.8
1.0
Subsampled data
Function fitted to data

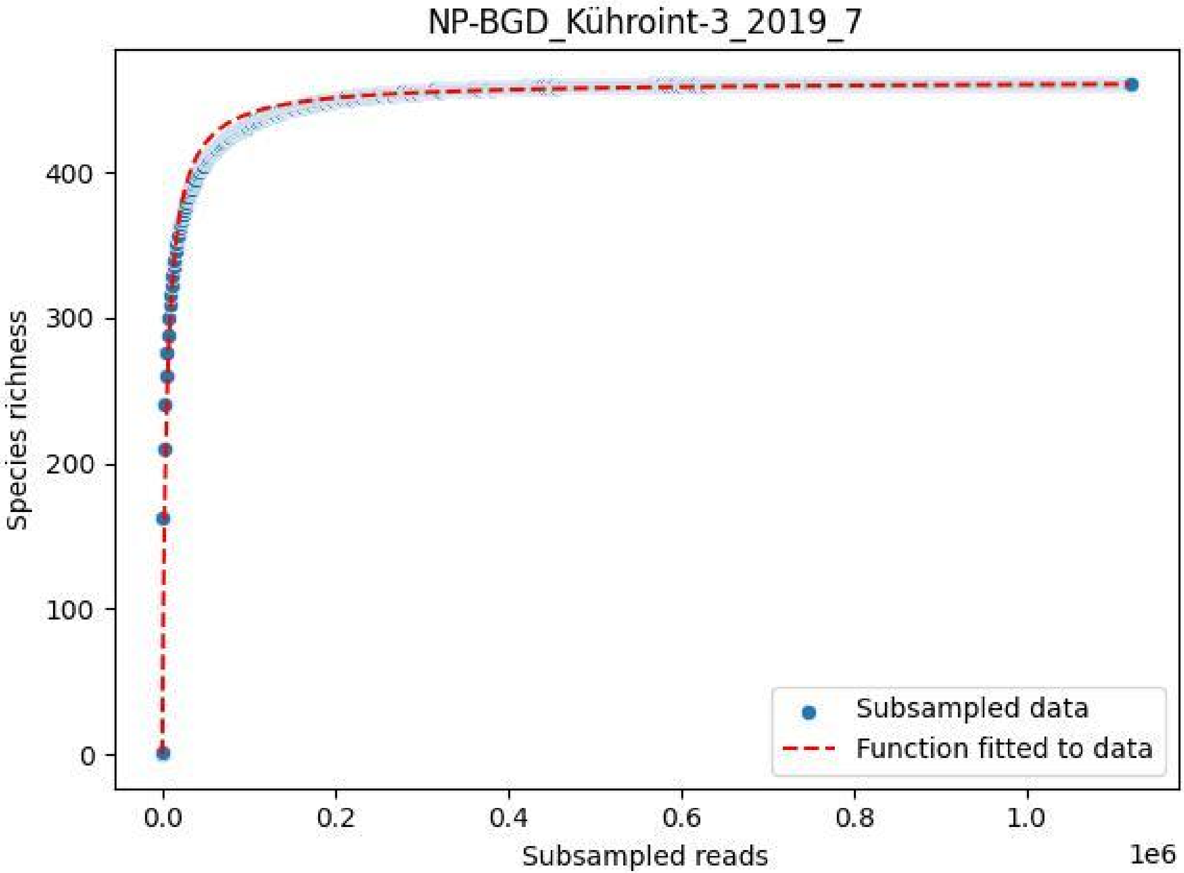
NP-BGD_Kühroint-3_2019_7
Species richness
0
100
200
300
400
Subsampled reads
0.0
0.2
0.4
0.6
0.8
1.0
1e6
Subsampled data
Function fitted to data

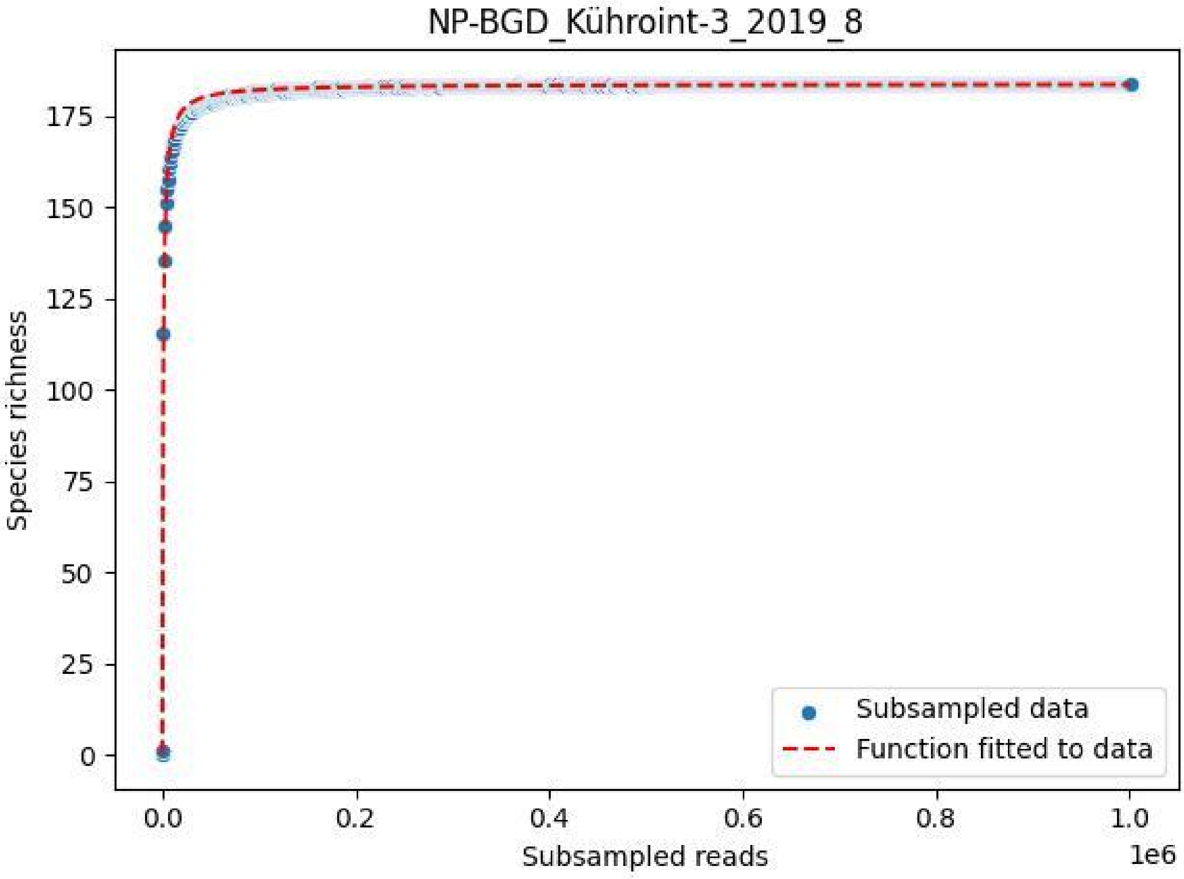
NP-BGD_Kühroint-3_2019_8
Species richness
Subsampled reads
1e6
Subsampled data
Function fitted to data
0
25
50
75
100
125
150
175
0.0
0.2
0.4
0.6
0.8
1.0

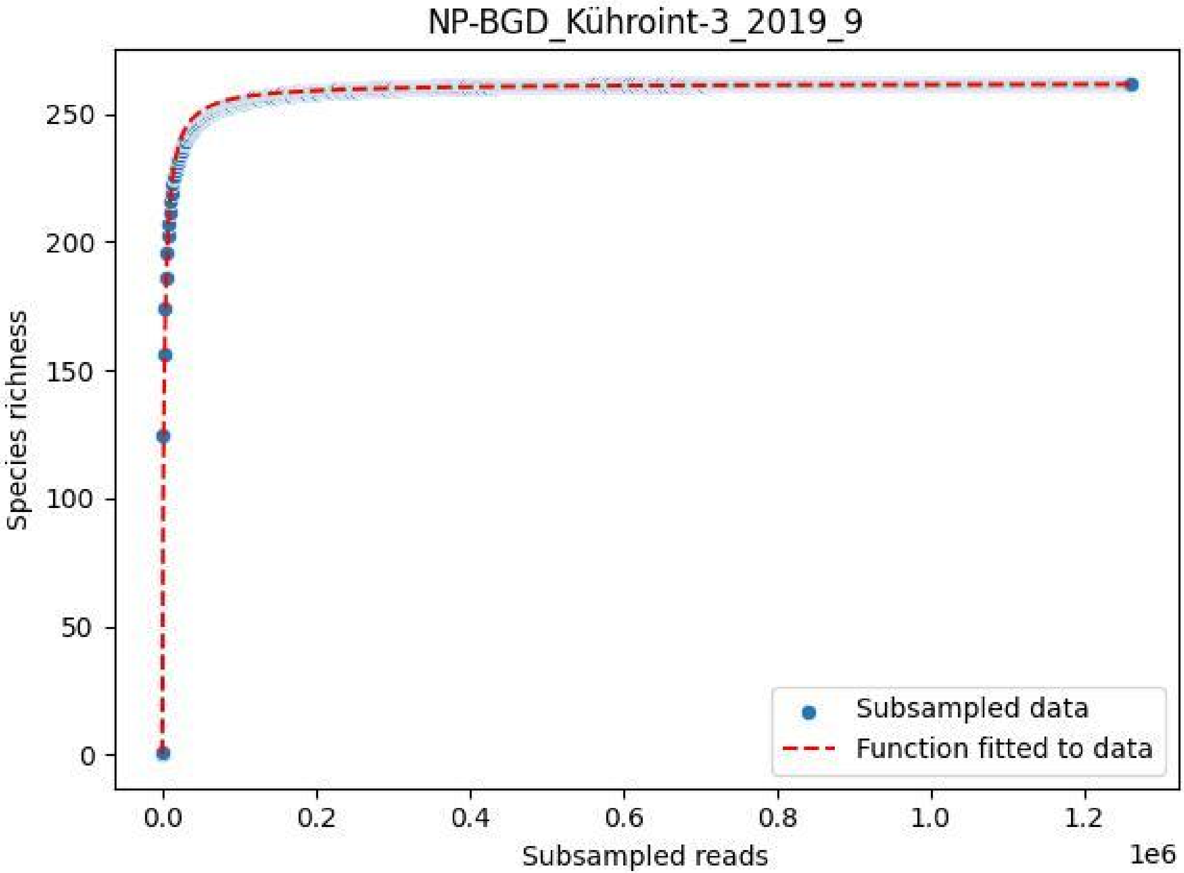
NP-BGD_Kühroint-3_2019_9
Species richness
0
50
100
150
200
250
0.0
0.2
0.4
0.6
0.8
1.0
1.2
1e6
Subsampled reads
Subsampled data
Function fitted to data

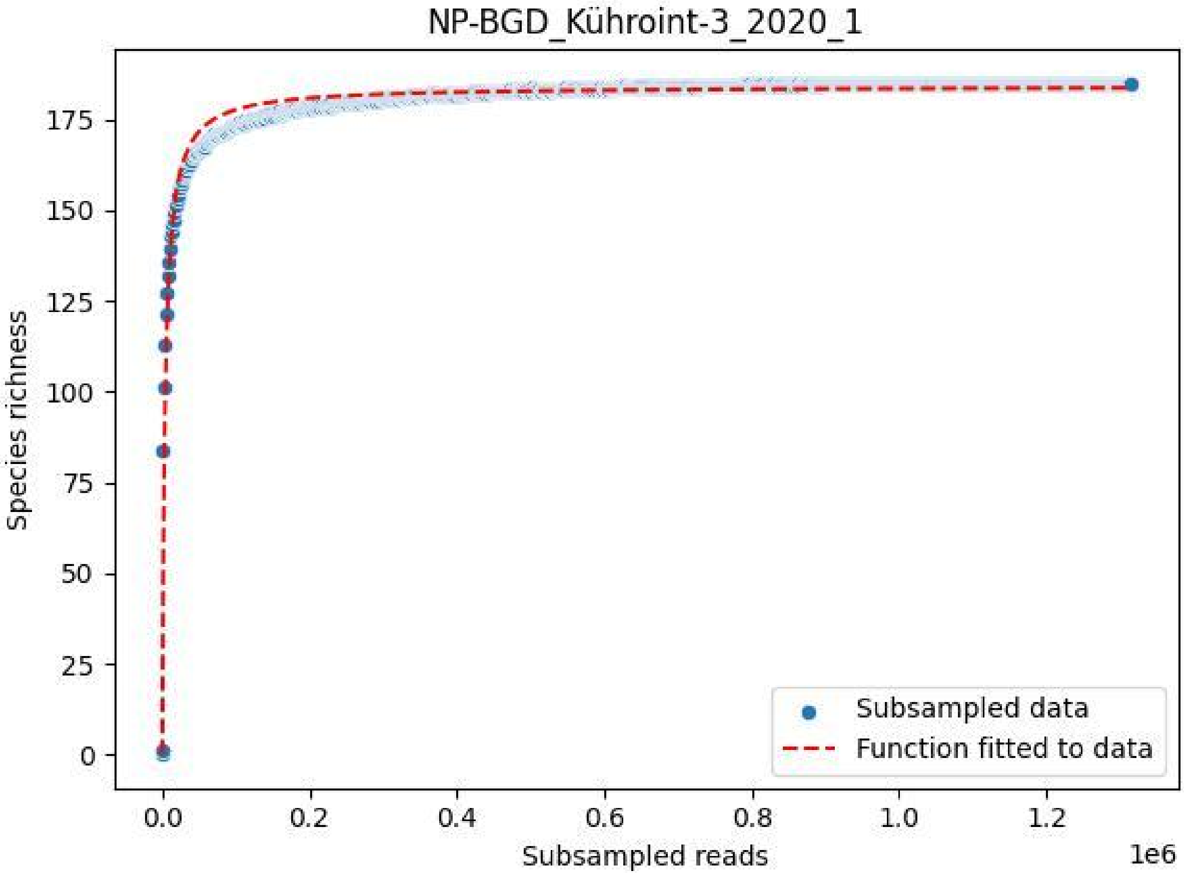
NP-BGD_Kühroint-3_2020_1
Species richness
Subsampled reads
1e6
0
25
50
75
100
125
150
175
0.0
0.2
0.4
0.6
0.8
1.0
1.2
Subsampled data
Function fitted to data

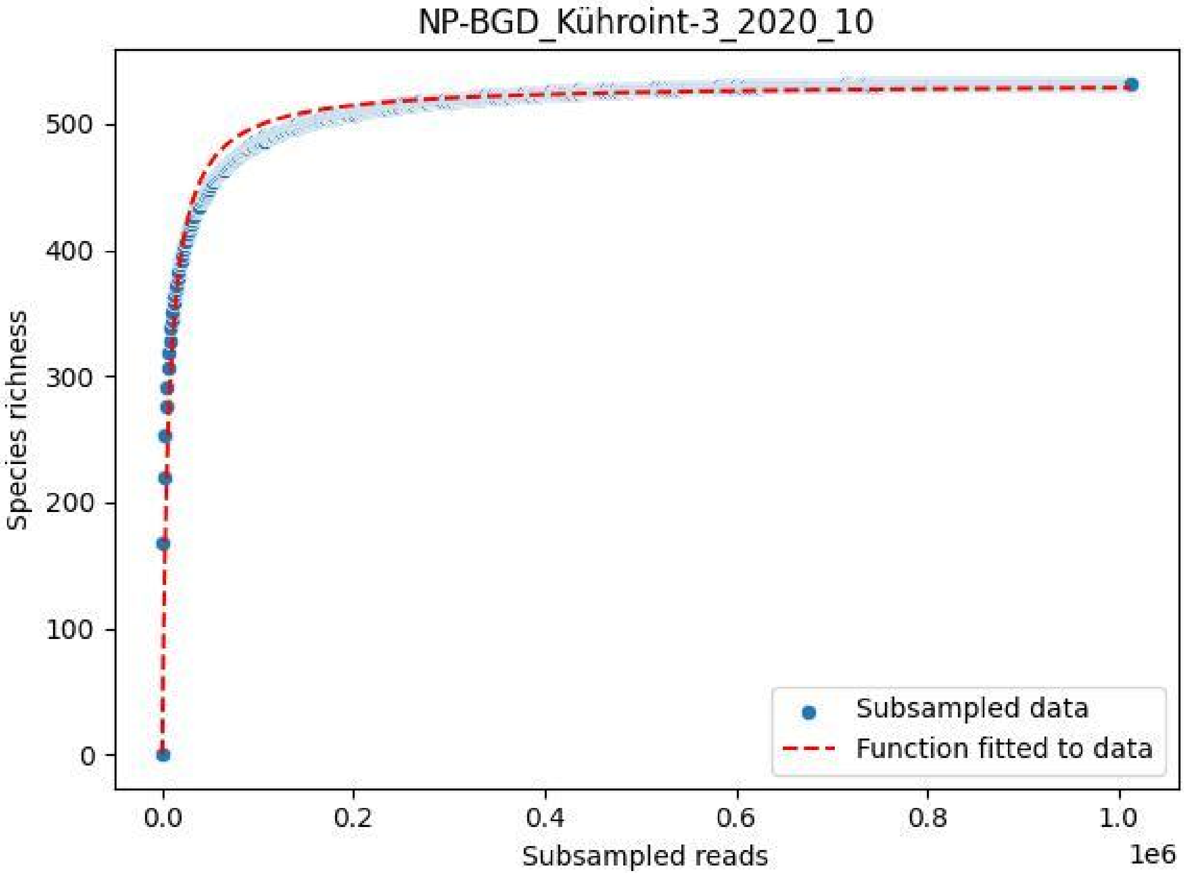
NP-BGD_Kühroint-3_2020_10
Species richness
Subsampled reads
1e6
0
100
200
300
400
500
0.0
0.2
0.4
0.6
0.8
1.0
Subsampled data
Function fitted to data

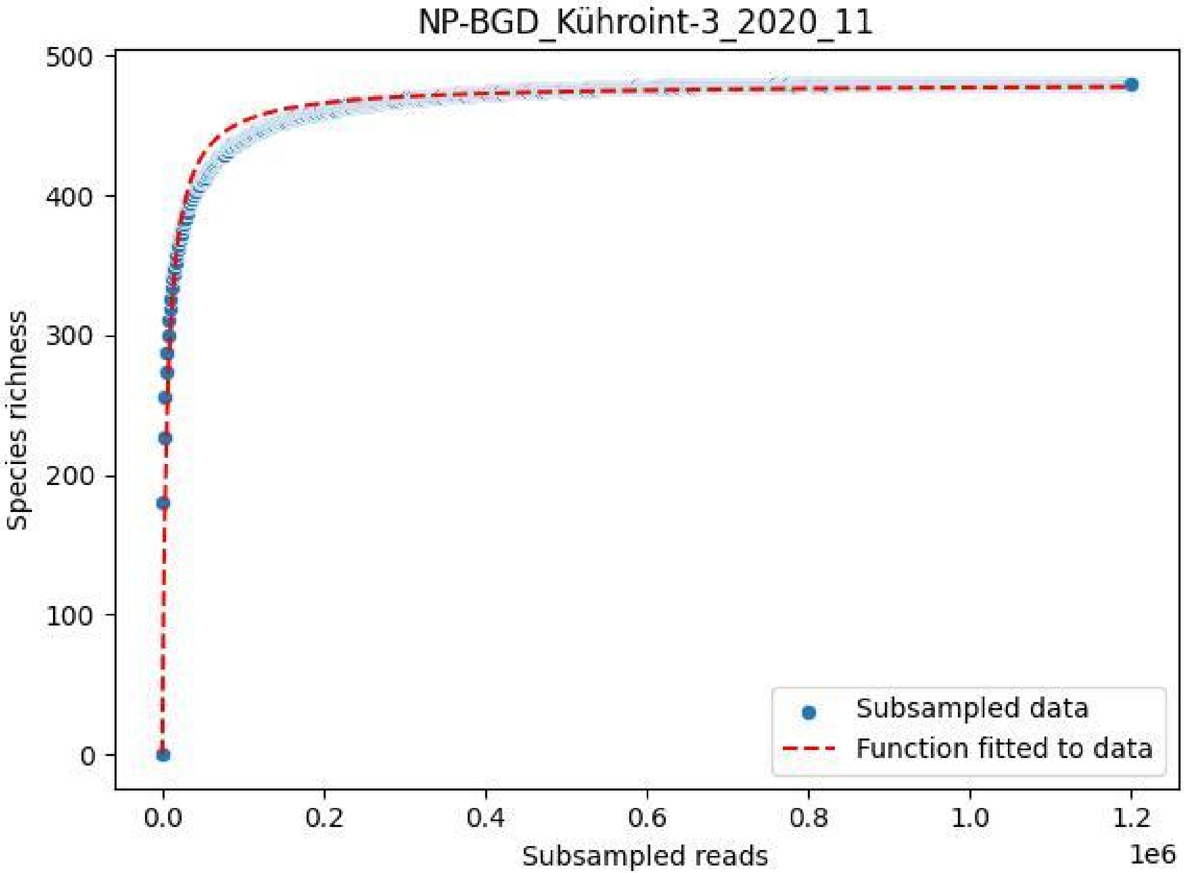

NP-BGD_Kühroint-3_2020_11
Species richness
Subsampled reads
1e6
0
100
200
300
400
500
0.0
0.2
0.4
0.6
0.8
1.0
1.2
Subsampled data
Function fitted to data

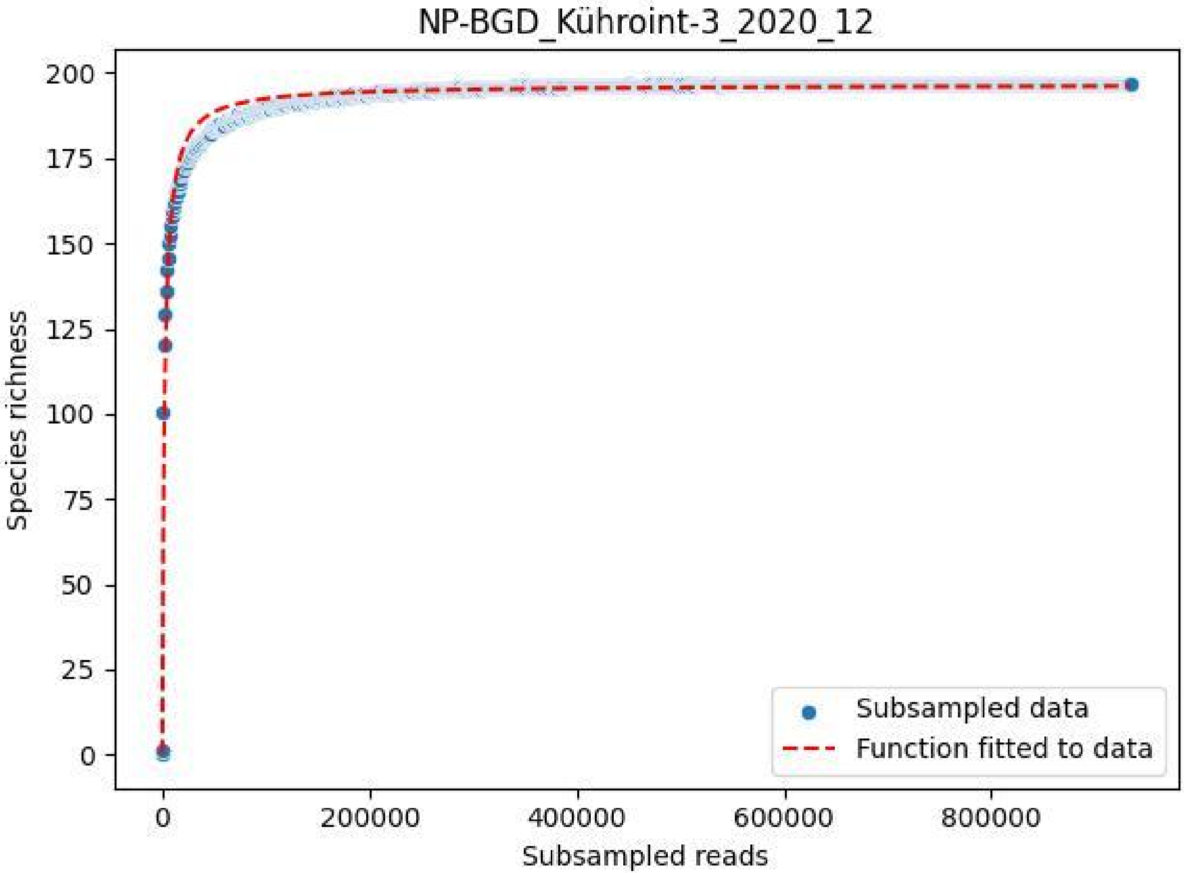

NP-BGD_Kühroint-3_2020_12
Species richness
200
175
150
125
100
75
50
25
0
0
200000
400000
600000
800000
Subsampled reads
Subsampled data
Function fitted to data

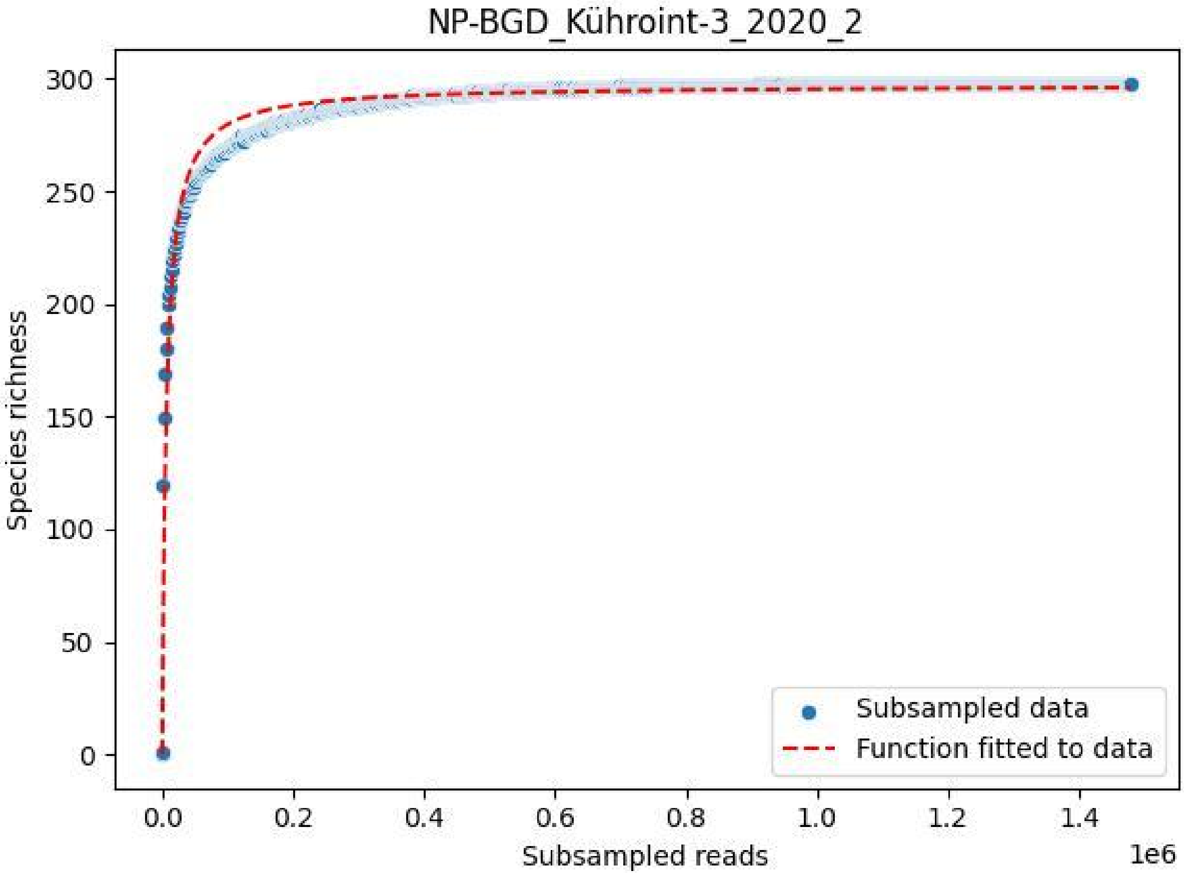
NP-BGD_Kühroint-3_2020_2
Species richness
Subsampled reads
1e6
0
50
100
150
200
250
300
0.0
0.2
0.4
0.6
0.8
1.0
1.2
1.4
Subsampled data
Function fitted to data

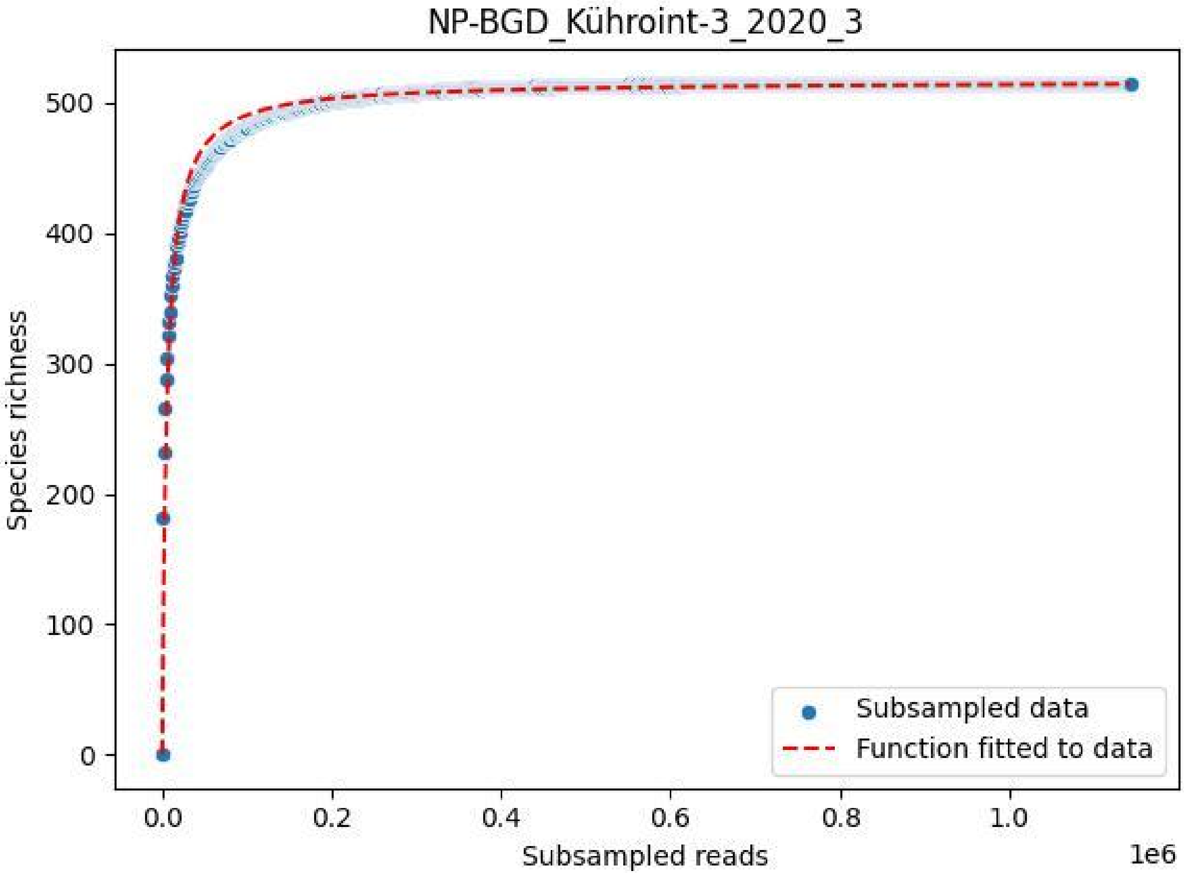
NP-BGD_Kühroint-3_2020_3
Species richness
Subsampled reads
1e6
0
100
200
300
400
500
0.0
0.2
0.4
0.6
0.8
1.0
Subsampled data
Function fitted to data

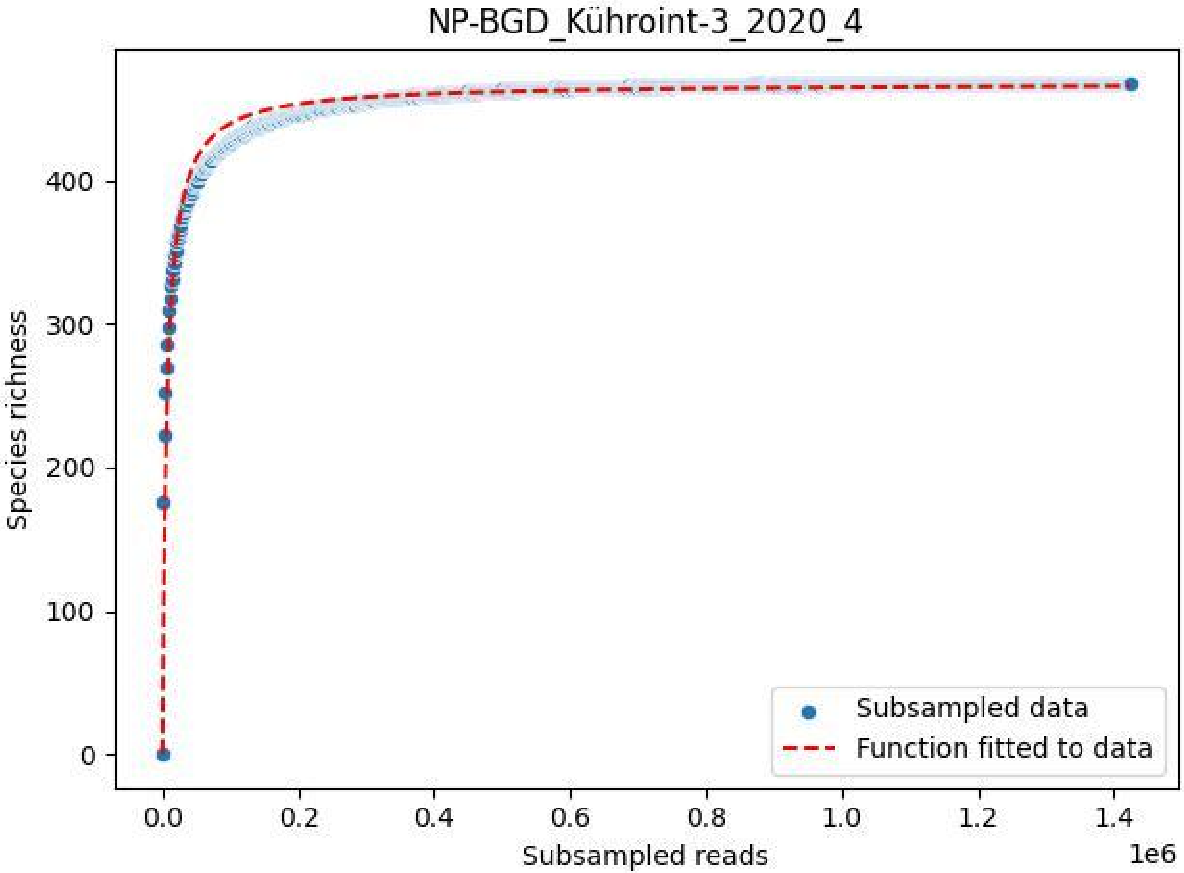
NP-BGD_Kühroint-3_2020_4
Species richness
Subsampled reads
1e6
0
100
200
300
400
0.0
0.2
0.4
0.6
0.8
1.0
1.2
1.4
Subsampled data
Function fitted to data

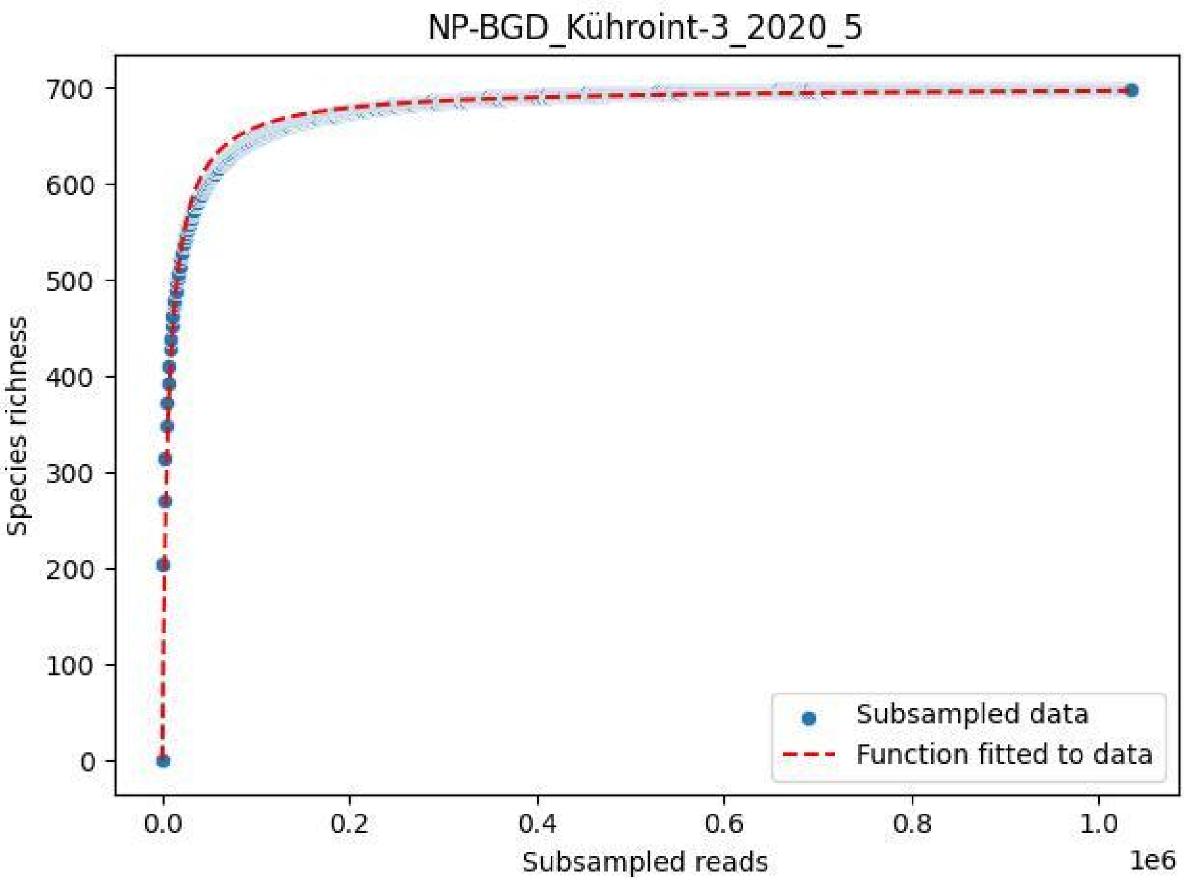
NP-BGD_Kühroint-3_2020_5
Species richness
Subsampled reads
1e6
0
100
200
300
400
500
600
700
0.0
0.2
0.4
0.6
0.8
1.0
Subsampled data
Function fitted to data

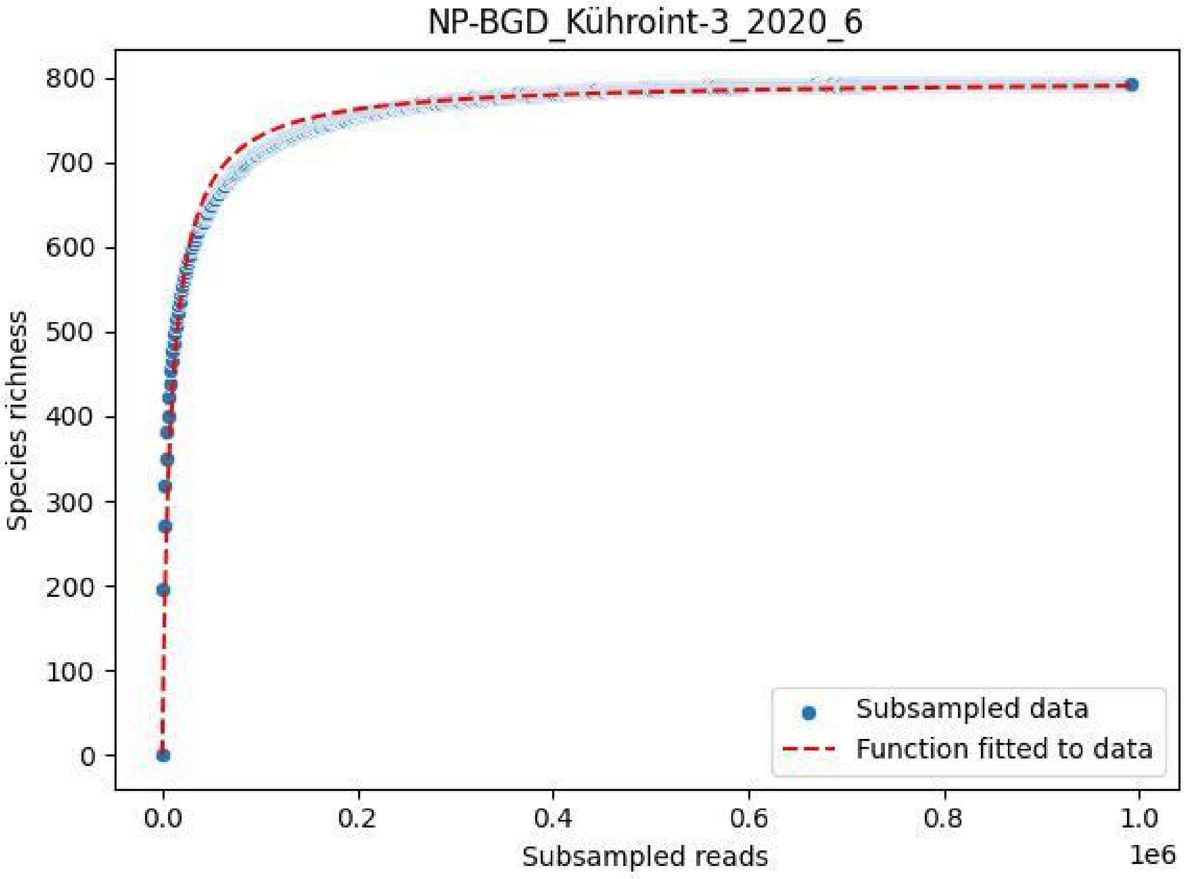
NP-BGD_Kühroint-3_2020_6
Species richness
0
100
200
300
400
500
600
700
800
0.0
0.2
0.4
0.6
0.8
1.0
1e6
Subsampled reads
Subsampled data
Function fitted to data

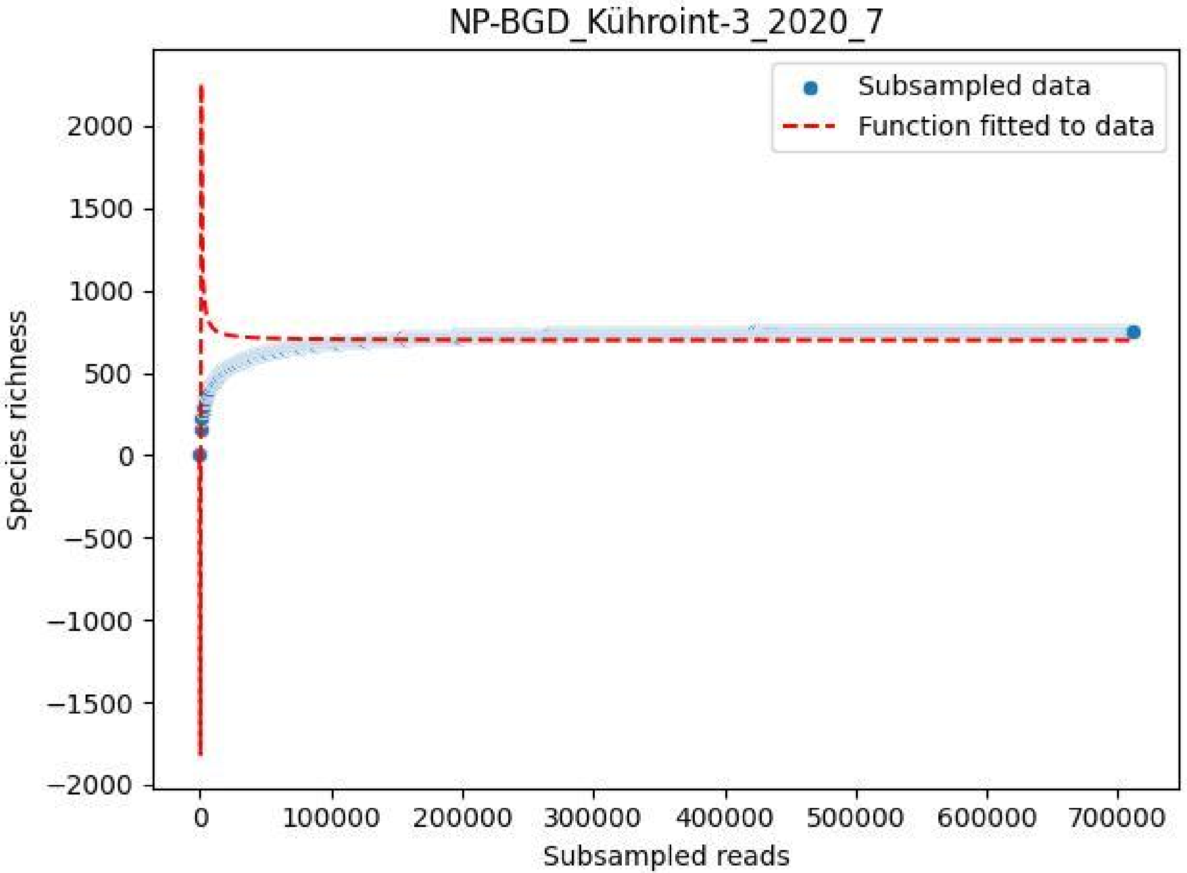
NP-BGD_Kühroint-3_2020_7
Subsampled data
Function fitted to data
Species richness
Subsampled reads
2000
1500
1000
500
0
−500
−1000
−1500
−2000
0
100000
200000
300000
400000
500000
600000
700000

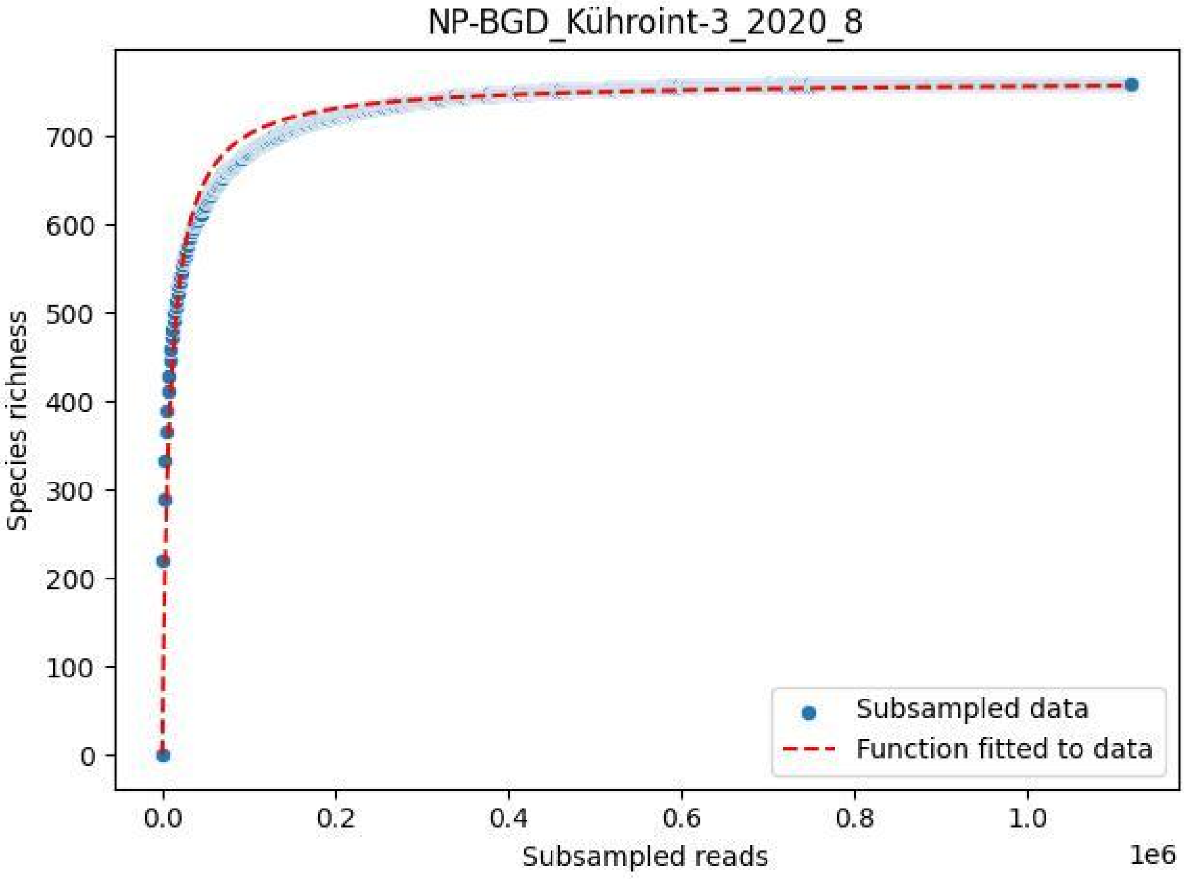
NP-BGD_Kühroint-3_2020_8
Species richness
0
100
200
300
400
500
600
700
0.0
0.2
0.4
0.6
0.8
1.0
1e6
Subsampled reads
Subsampled data
Function fitted to data

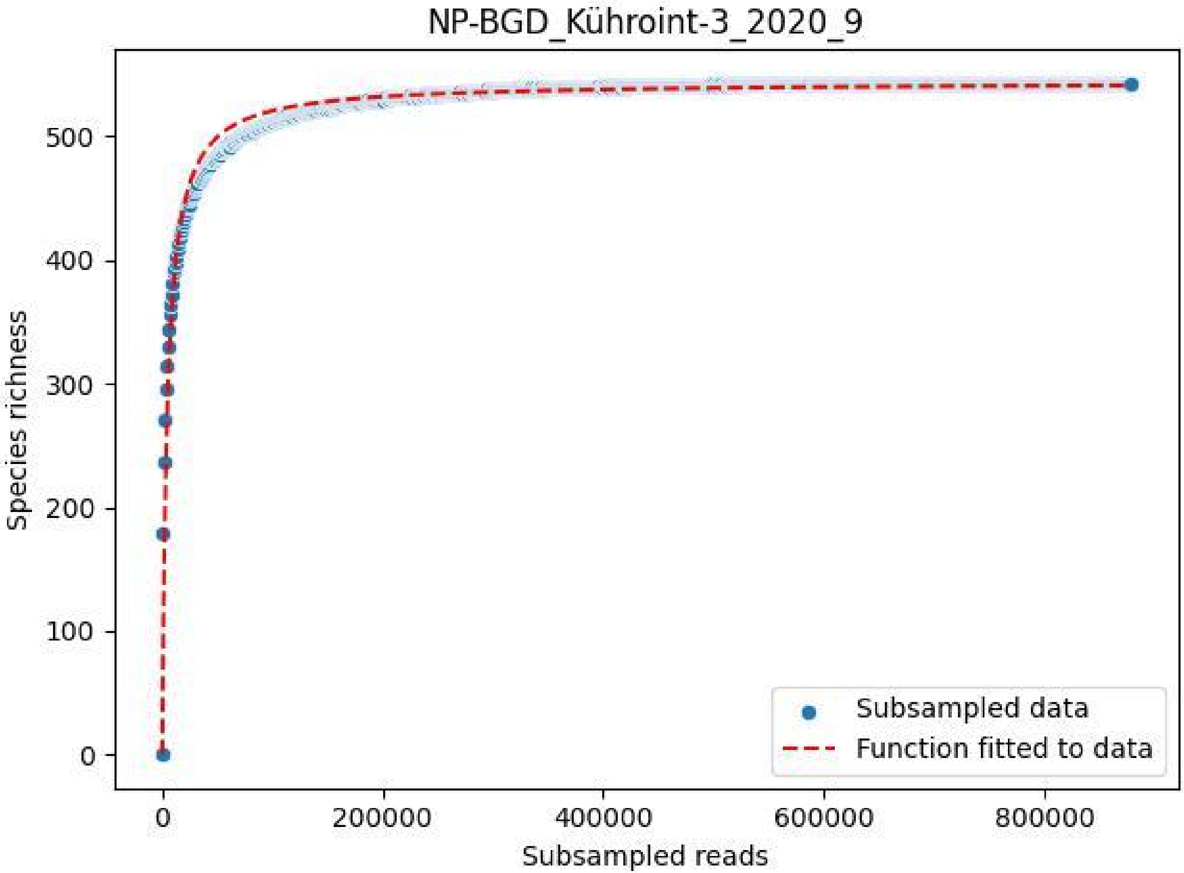
NP-BGD_Kühroint-3_2020_9
Species richness
0
100
200
300
400
500
0
200000
400000
600000
800000
Subsampled reads
Subsampled data
Function fitted to data

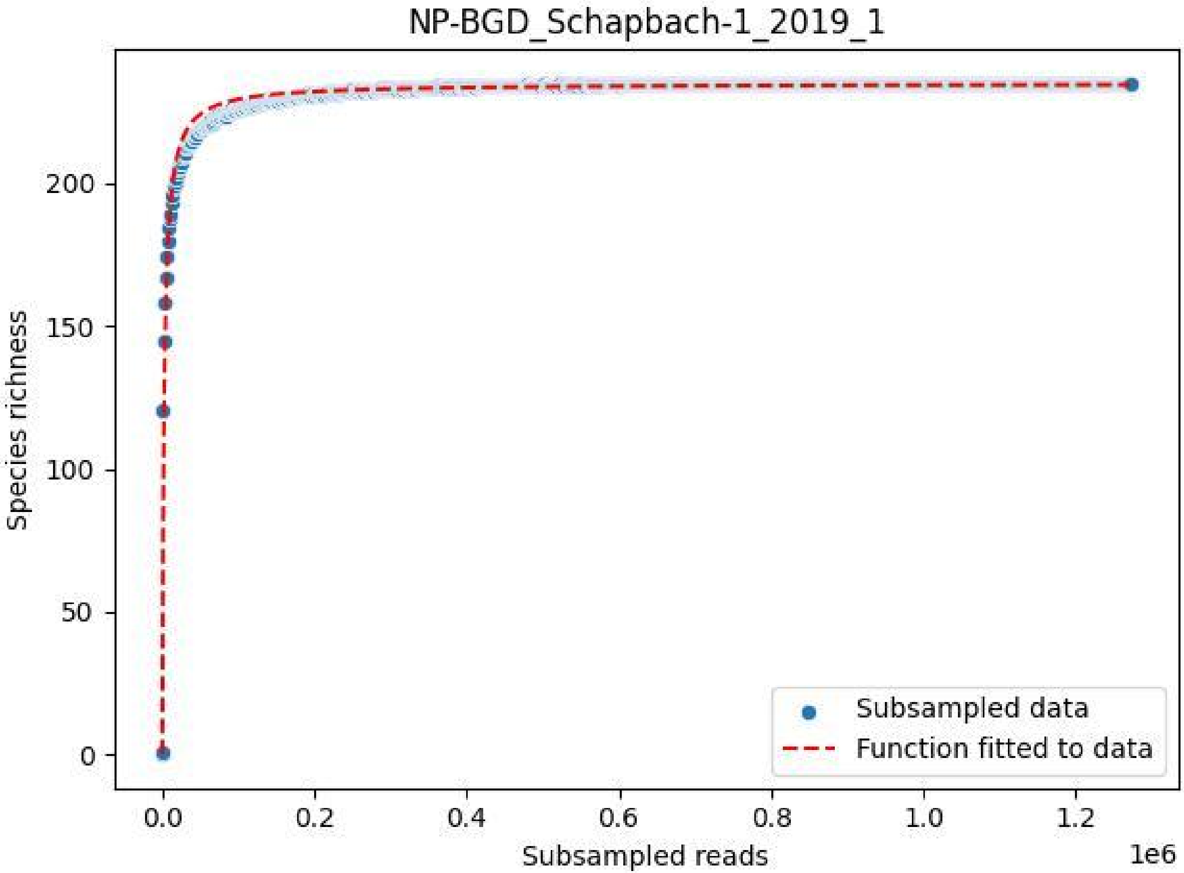
NP-BGD_Schapbach-1_2019_1
Species richness
Subsampled reads
1e6
0
50
100
150
200
0.0
0.2
0.4
0.6
0.8
1.0
1.2
Subsampled data
Function fitted to data

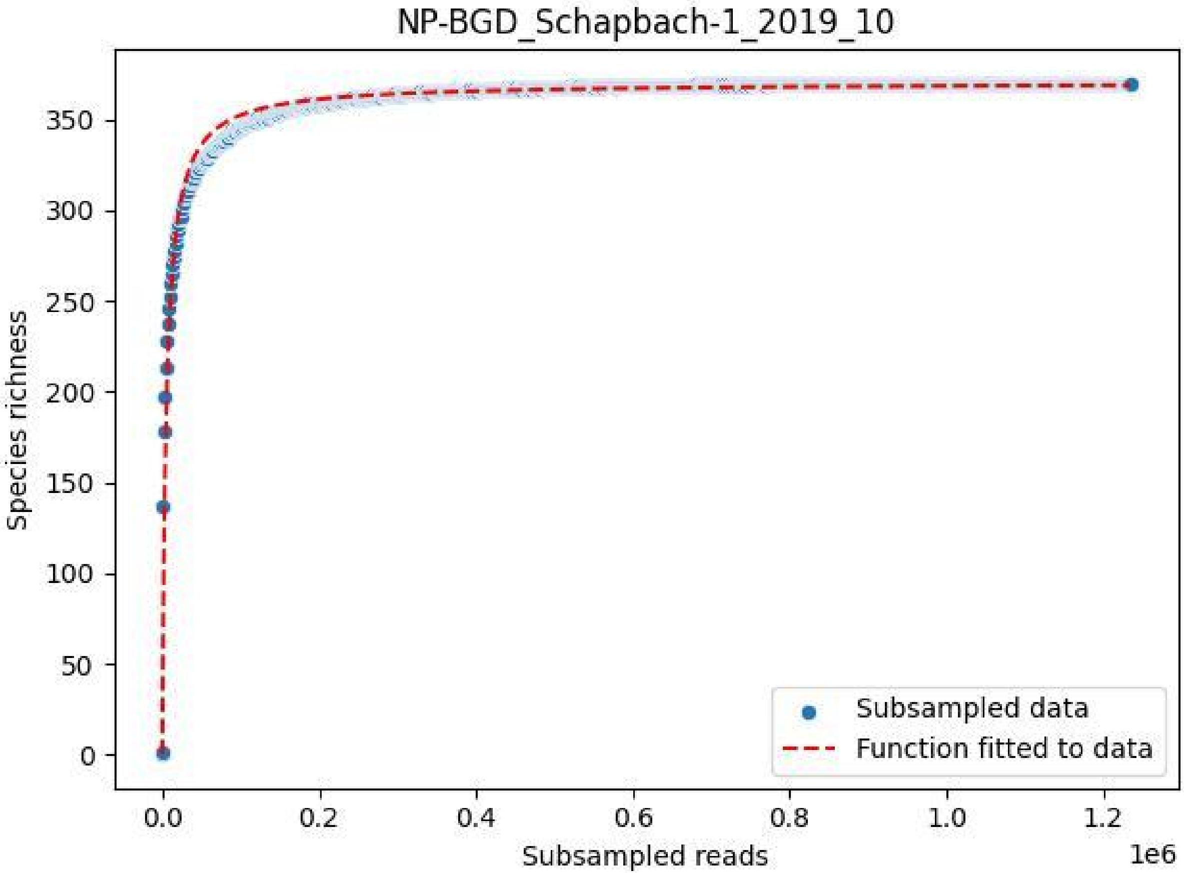

NP-BGD_Schapbach-1_2019_10
Species richness
Subsampled reads
1e6
0
50
100
150
200
250
300
350
0.0
0.2
0.4
0.6
0.8
1.0
1.2
Subsampled data
Function fitted to data

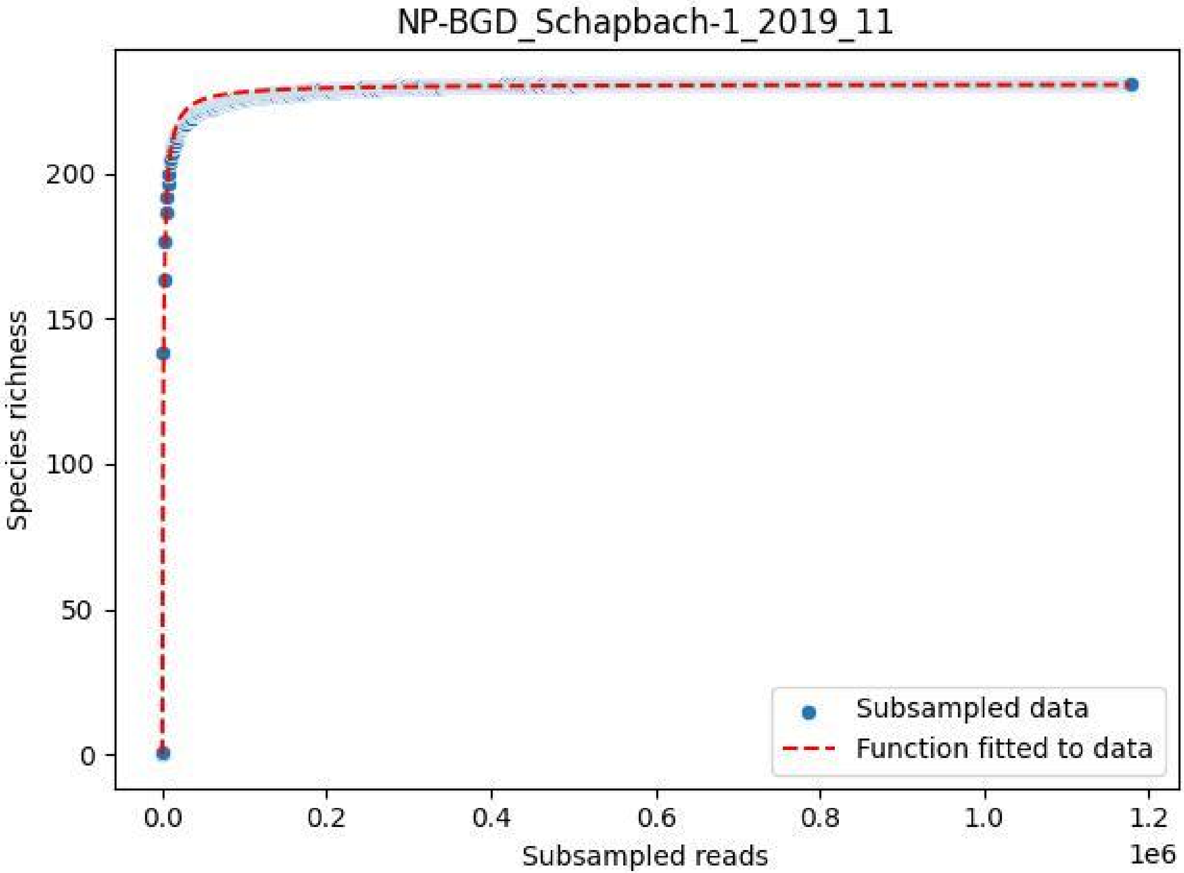
NP-BGD_Schapbach-1_2019_11
Species richness
Subsampled reads
1e6
0
50
100
150
200
0.0
0.2
0.4
0.6
0.8
1.0
1.2
Subsampled data
Function fitted to data

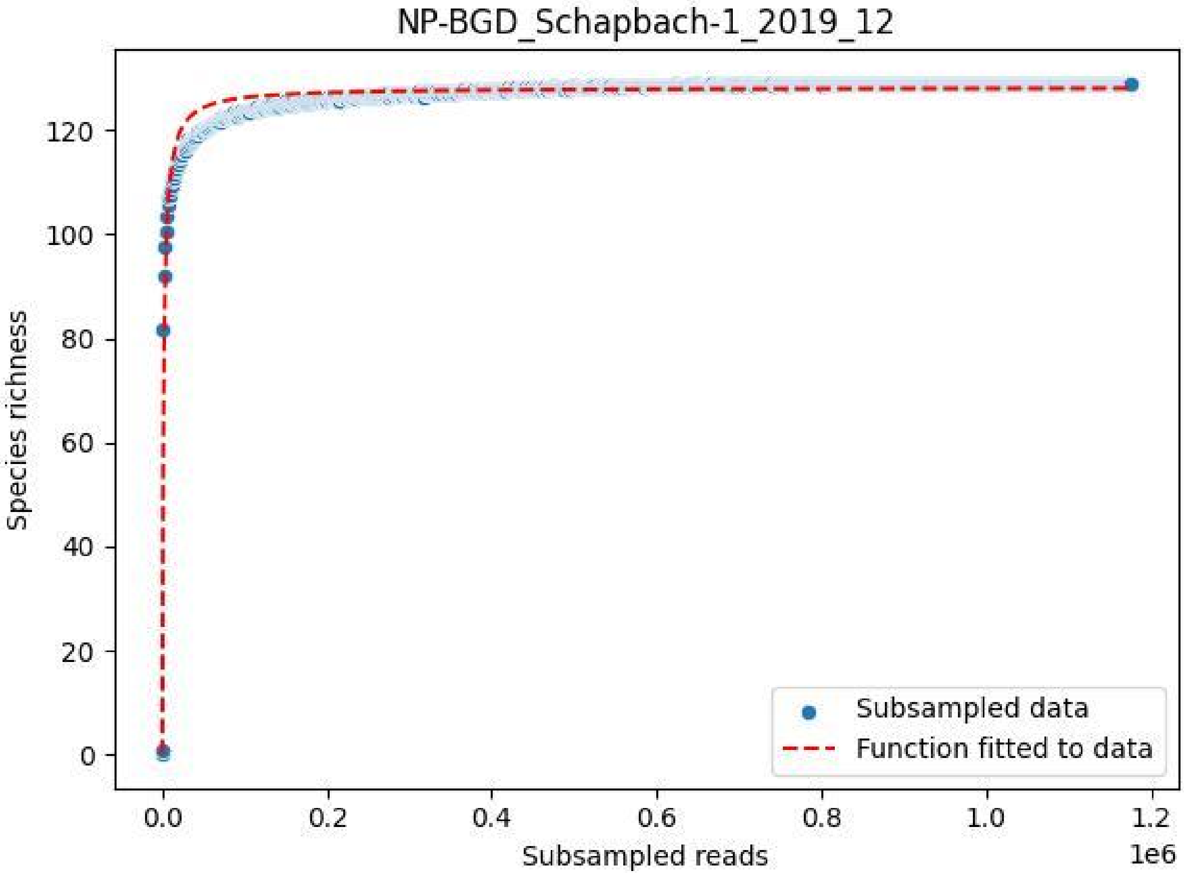
NP-BGD_Schapbach-1_2019_12
Species richness
Subsampled reads
1e6
0
20
40
60
80
100
120
0.0
0.2
0.4
0.6
0.8
1.0
1.2
Subsampled data
Function fitted to data

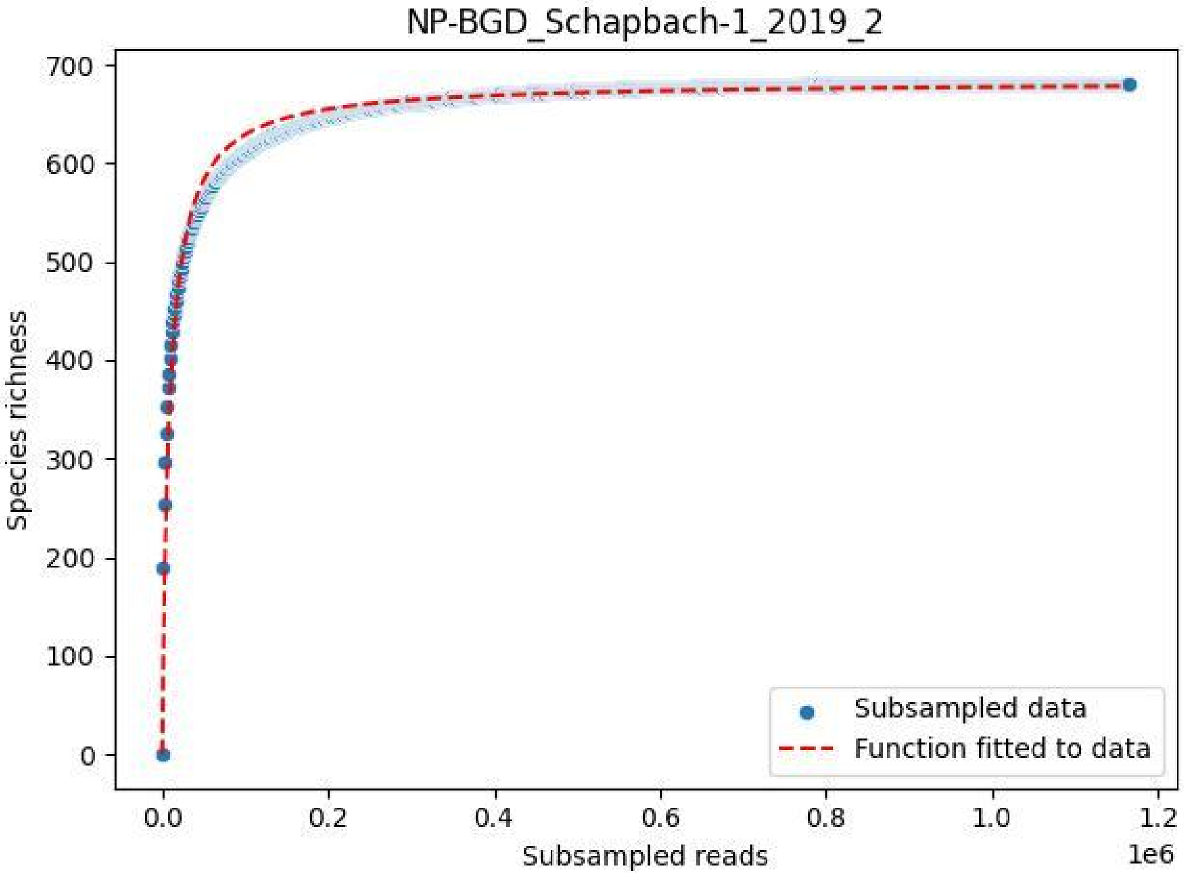
NP-BGD_Schapbach-1_2019_2
Species richness
Subsampled reads
1e6
0
100
200
300
400
500
600
700
0.0
0.2
0.4
0.6
0.8
1.0
1.2
Subsampled data
Function fitted to data

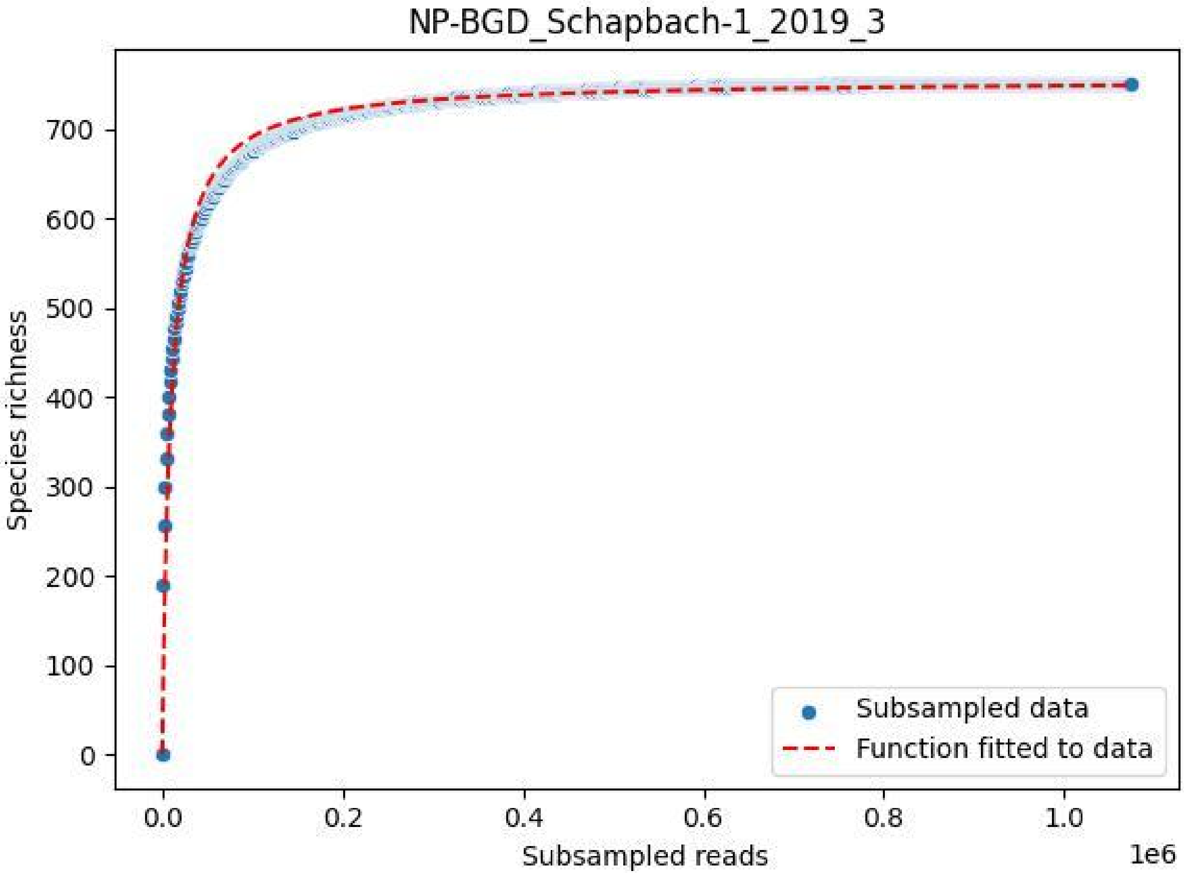
NP-BGD_Schapbach-1_2019_3
Species richness
Subsampled reads
1e6
Subsampled data
Function fitted to data
0
100
200
300
400
500
600
700
0.0
0.2
0.4
0.6
0.8
1.0

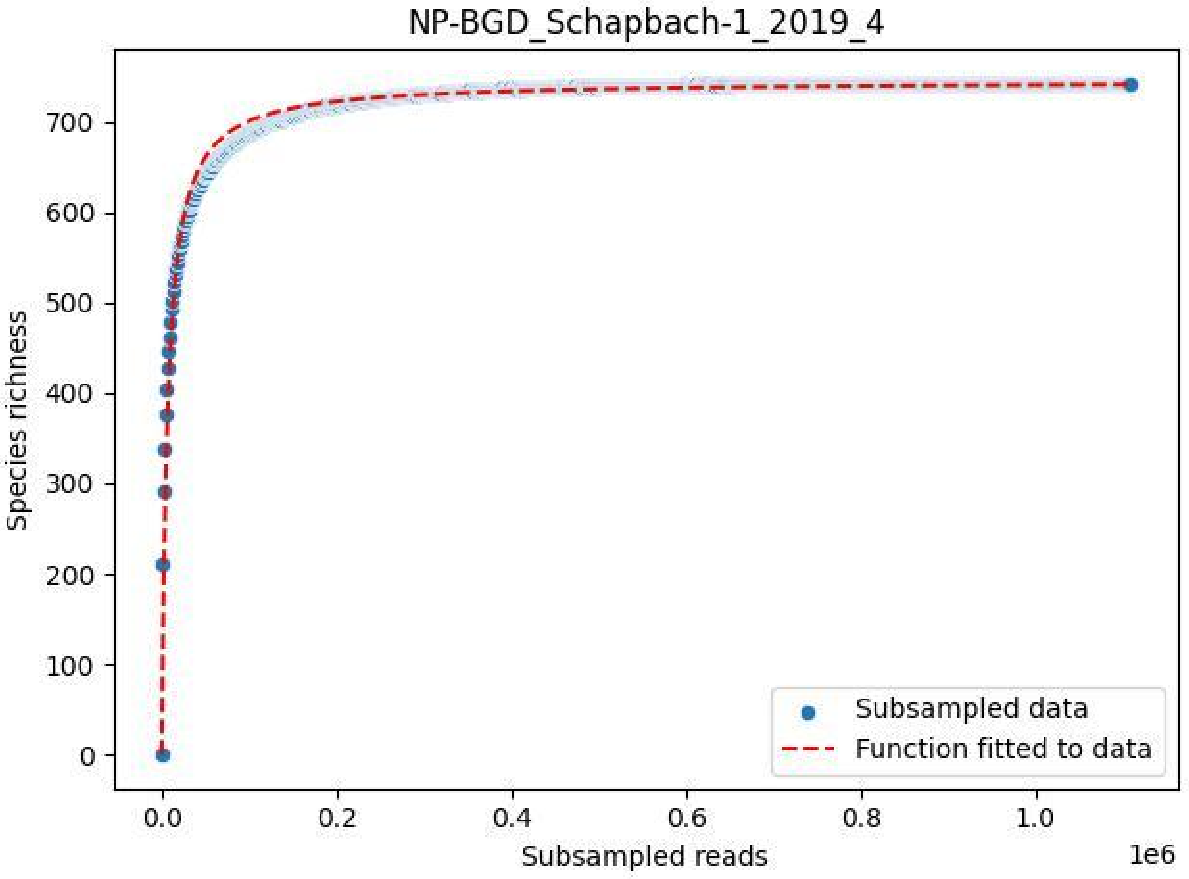
NP-BGD_Schapbach-1_2019_4
Species richness
Subsampled reads
1e6
0
100
200
300
400
500
600
700
0.0
0.2
0.4
0.6
0.8
1.0
Subsampled data
Function fitted to data

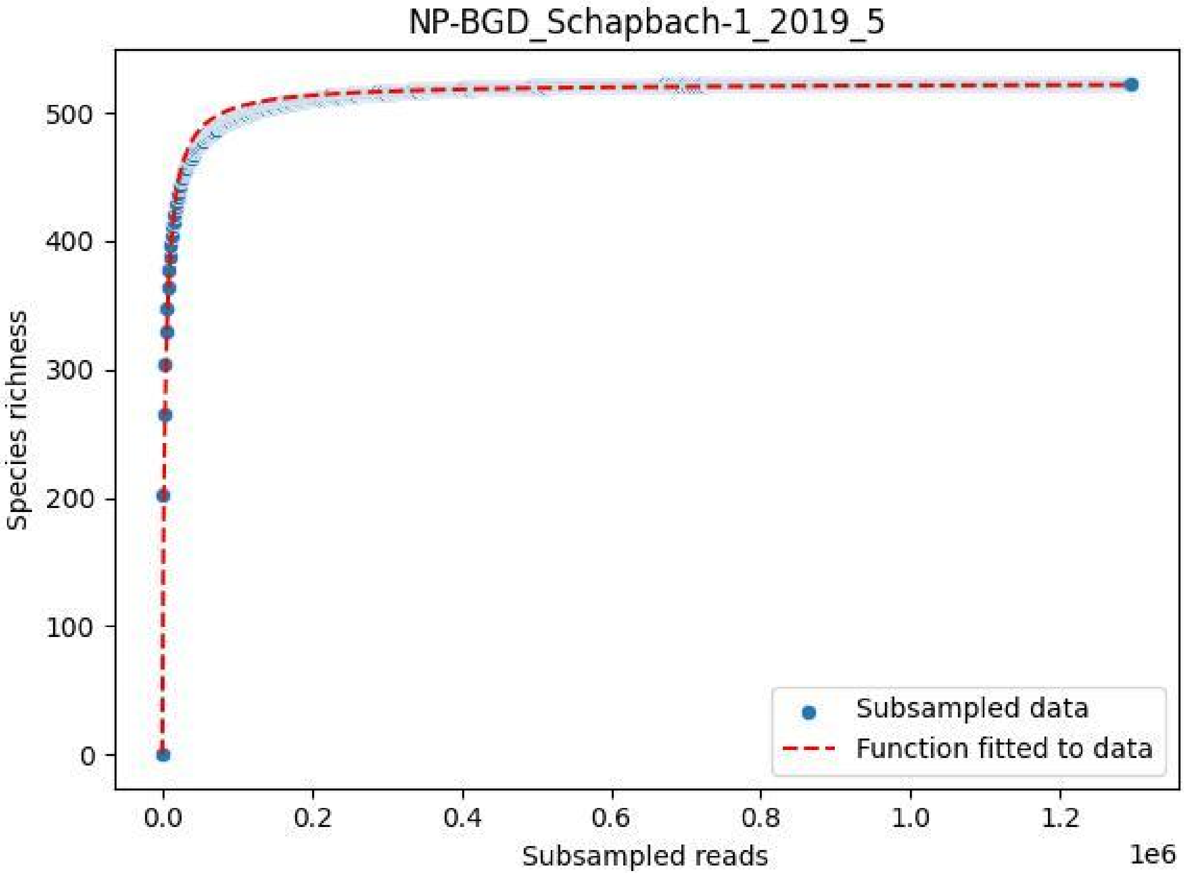

NP-BGD_Schapbach-1_2019_5
Species richness
Subsampled reads
1e6
0
100
200
300
400
500
0.0
0.2
0.4
0.6
0.8
1.0
1.2
Subsampled data
Function fitted to data

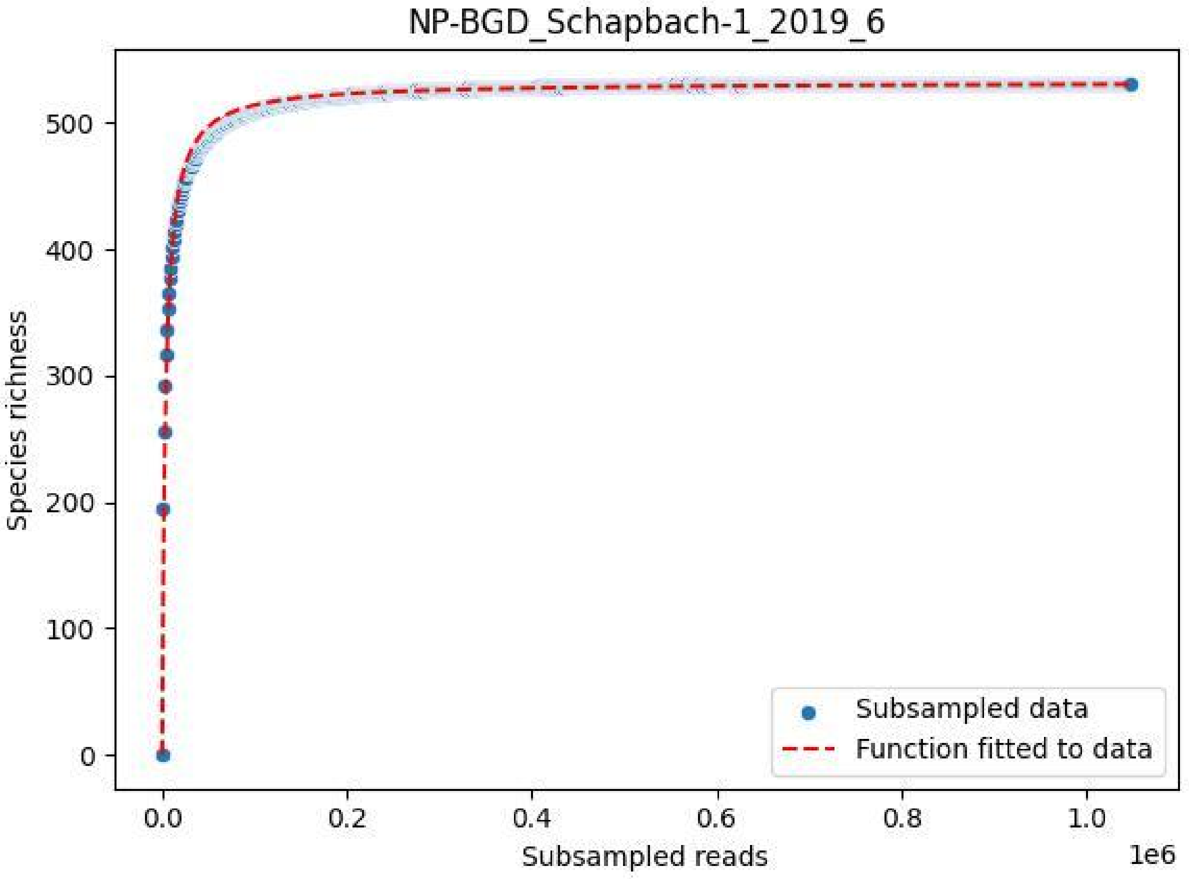
NP-BGD_Schapbach-1_2019_6
Species richness
Subsampled reads
1e6
0
100
200
300
400
500
0.0
0.2
0.4
0.6
0.8
1.0
Subsampled data
Function fitted to data

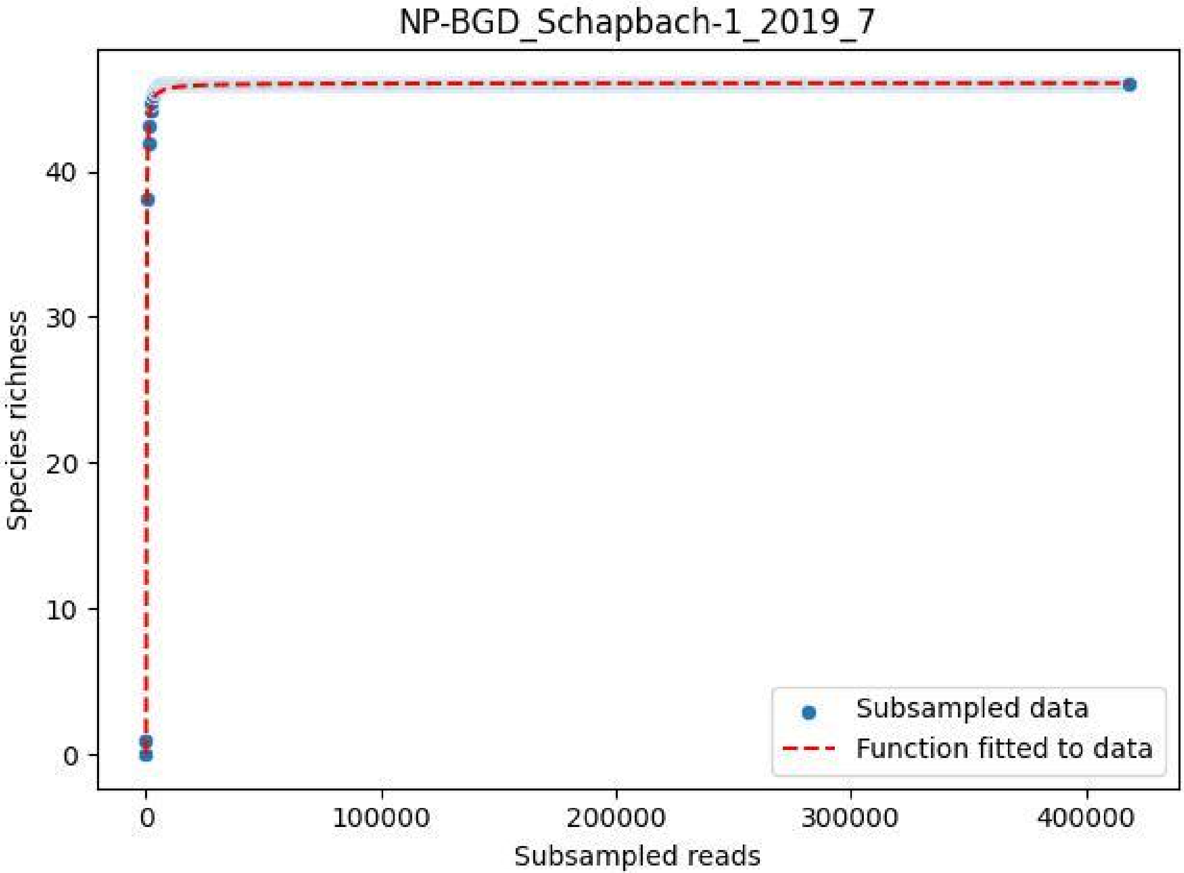
NP-BGD_Schapbach-1_2019_7
Species richness
Subsampled reads
0
10
20
30
40
0
100000
200000
300000
400000
Subsampled data
Function fitted to data

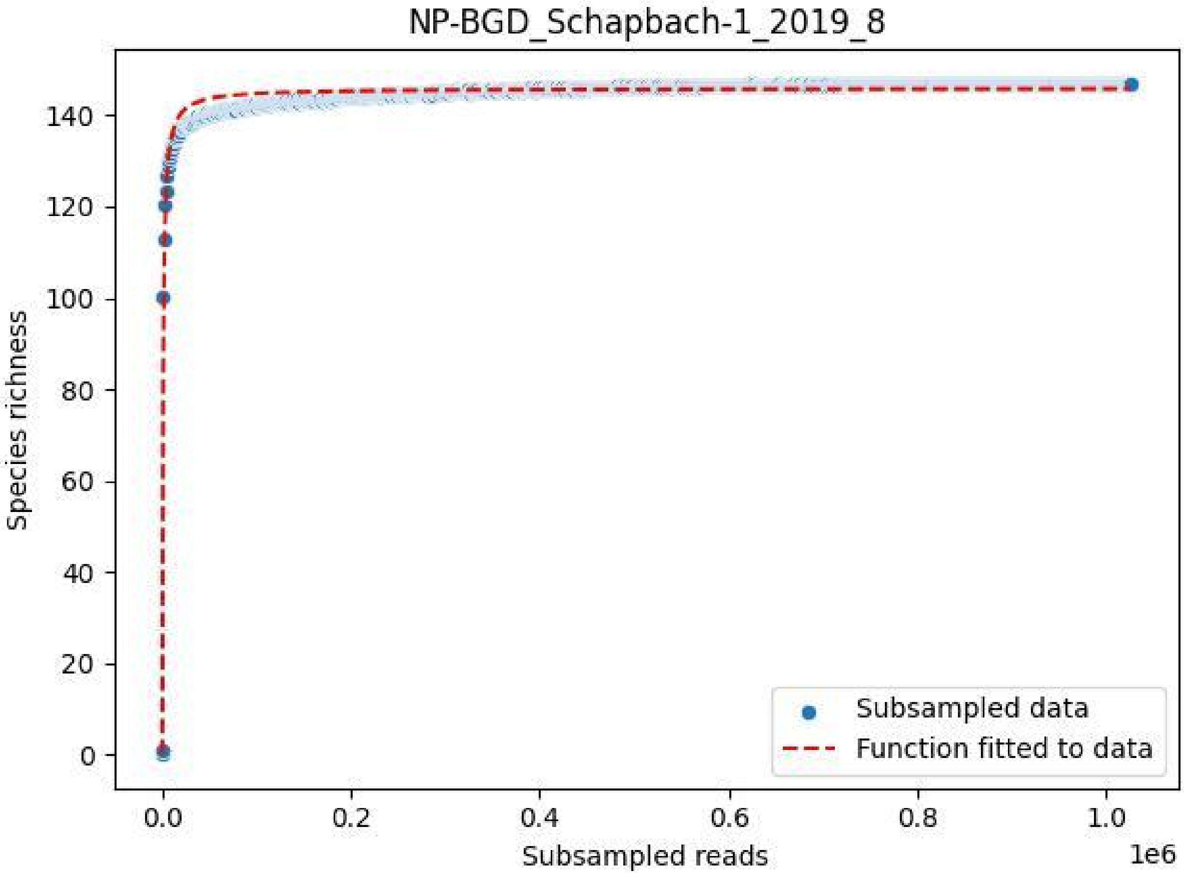
NP-BGD_Schapbach-1_2019_8
Species richness
Subsampled reads
1e6
0
20
40
60
80
100
120
140
0.0
0.2
0.4
0.6
0.8
1.0
Subsampled data
Function fitted to data

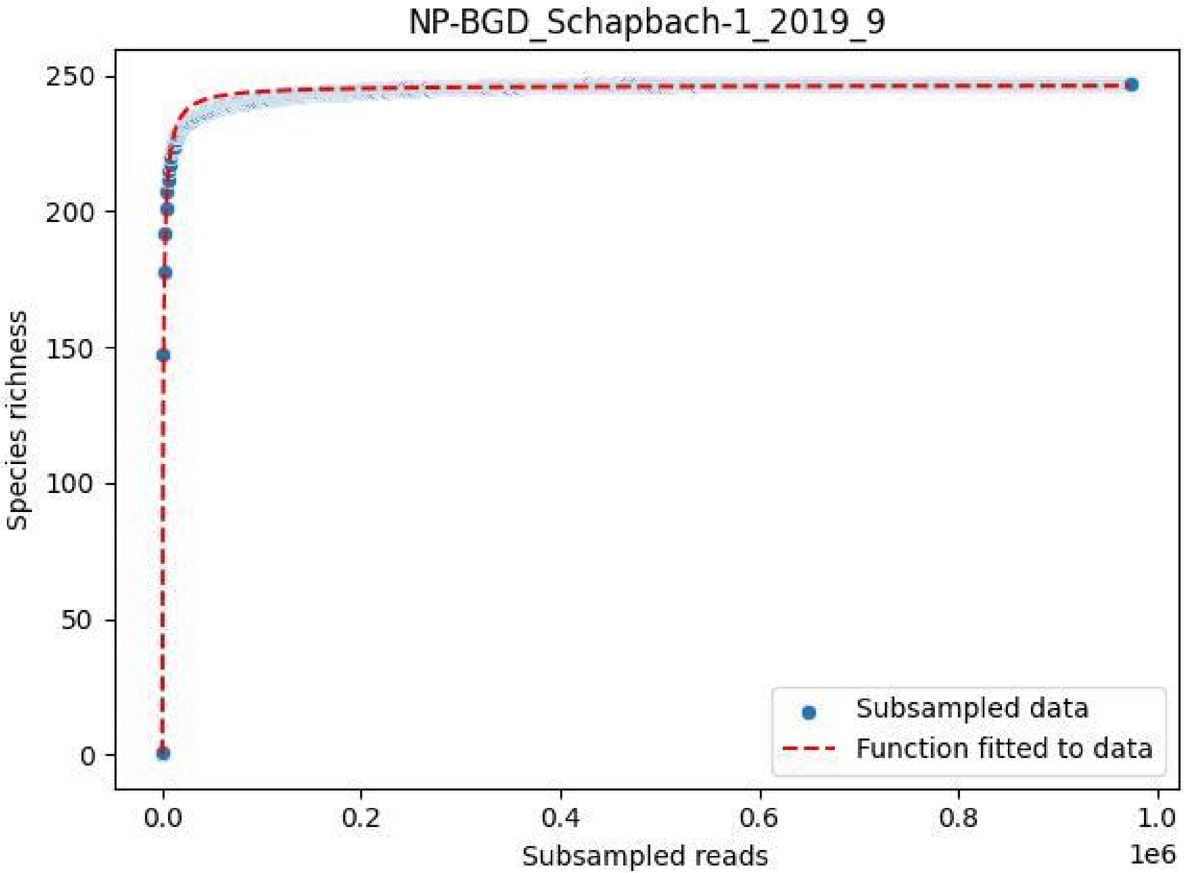
NP-BGD_Schapbach-1_2019_9
Species richness
250
200
150
100
50
0
0.0
0.2
0.4
0.6
0.8
1.0
1e6
Subsampled reads
Subsampled data
Function fitted to data

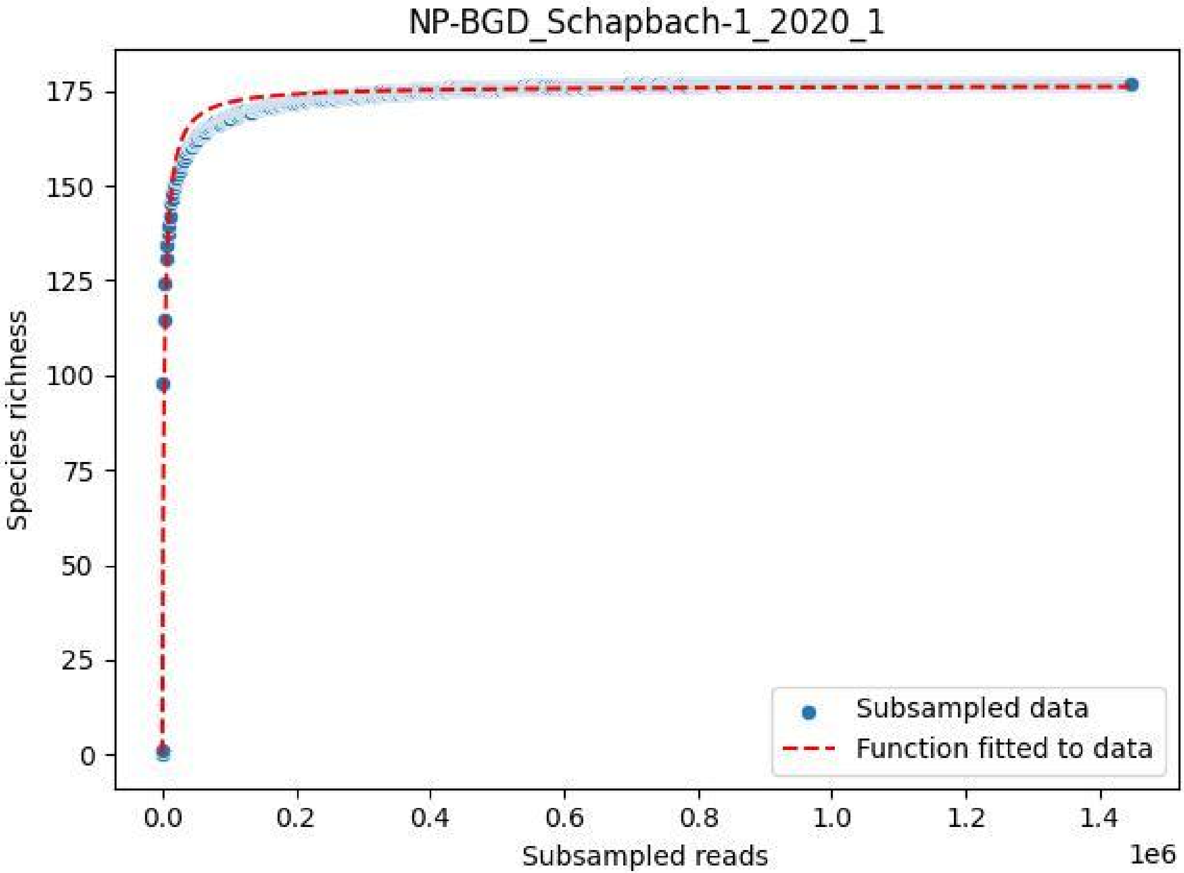

NP-BGD_Schapbach-1_2020_1
Species richness
Subsampled reads
1e6
0
25
50
75
100
125
150
175
0.0
0.2
0.4
0.6
0.8
1.0
1.2
1.4
Subsampled data
Function fitted to data

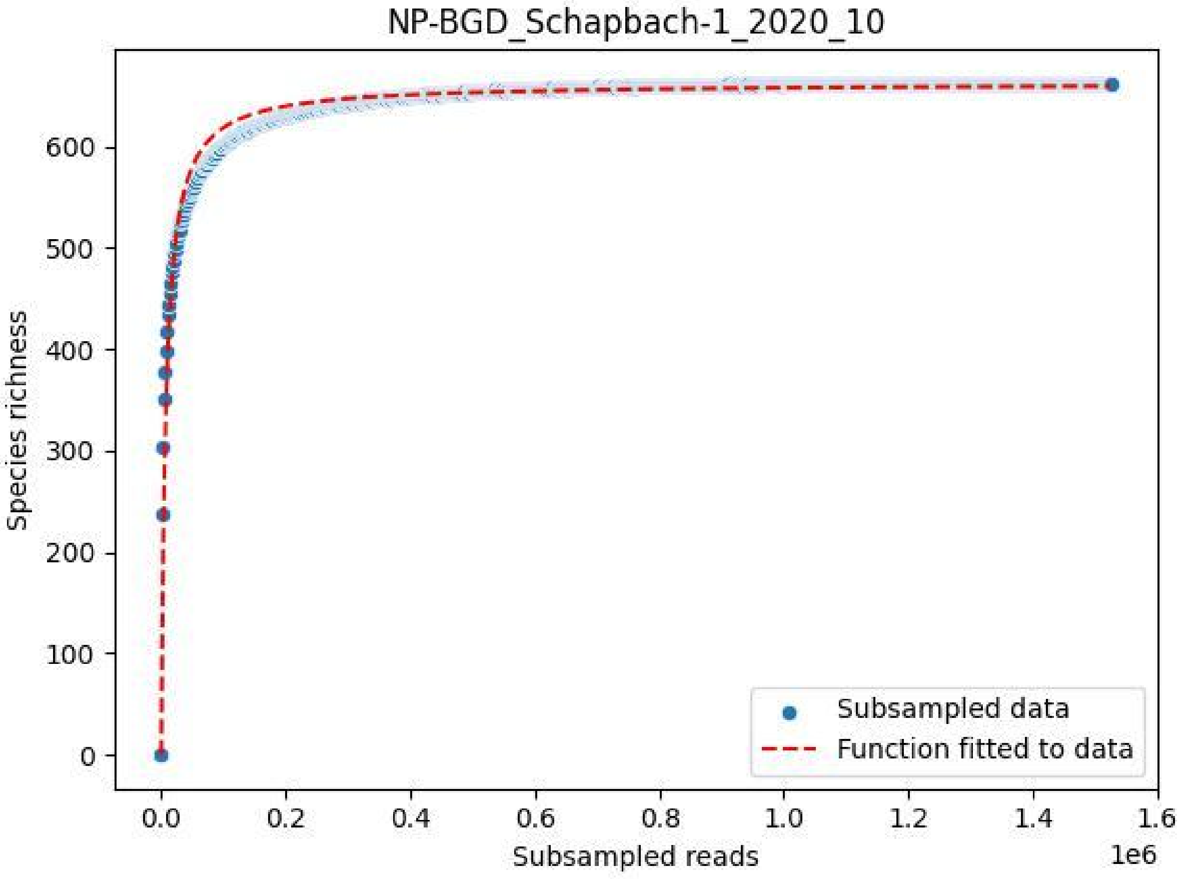
NP-BGD_Schapbach-1_2020_10
Species richness
Subsampled reads
1e6
0
100
200
300
400
500
600
0.0
0.2
0.4
0.6
0.8
1.0
1.2
1.4
1.6
Subsampled data
Function fitted to data

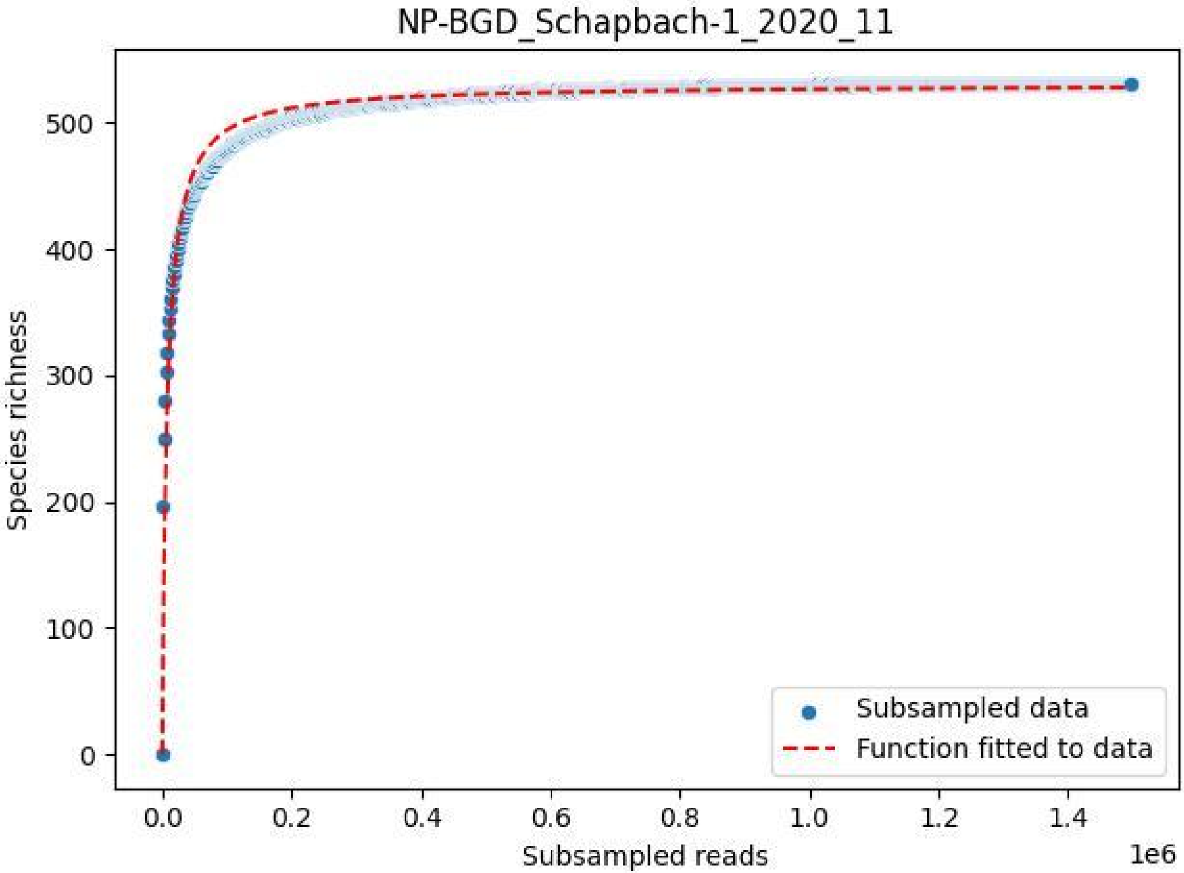
NP-BGD_Schapbach-1_2020_11
Species richness
Subsampled reads
1e6
Subsampled data
Function fitted to data
0
100
200
300
400
500
0.0
0.2
0.4
0.6
0.8
1.0
1.2
1.4

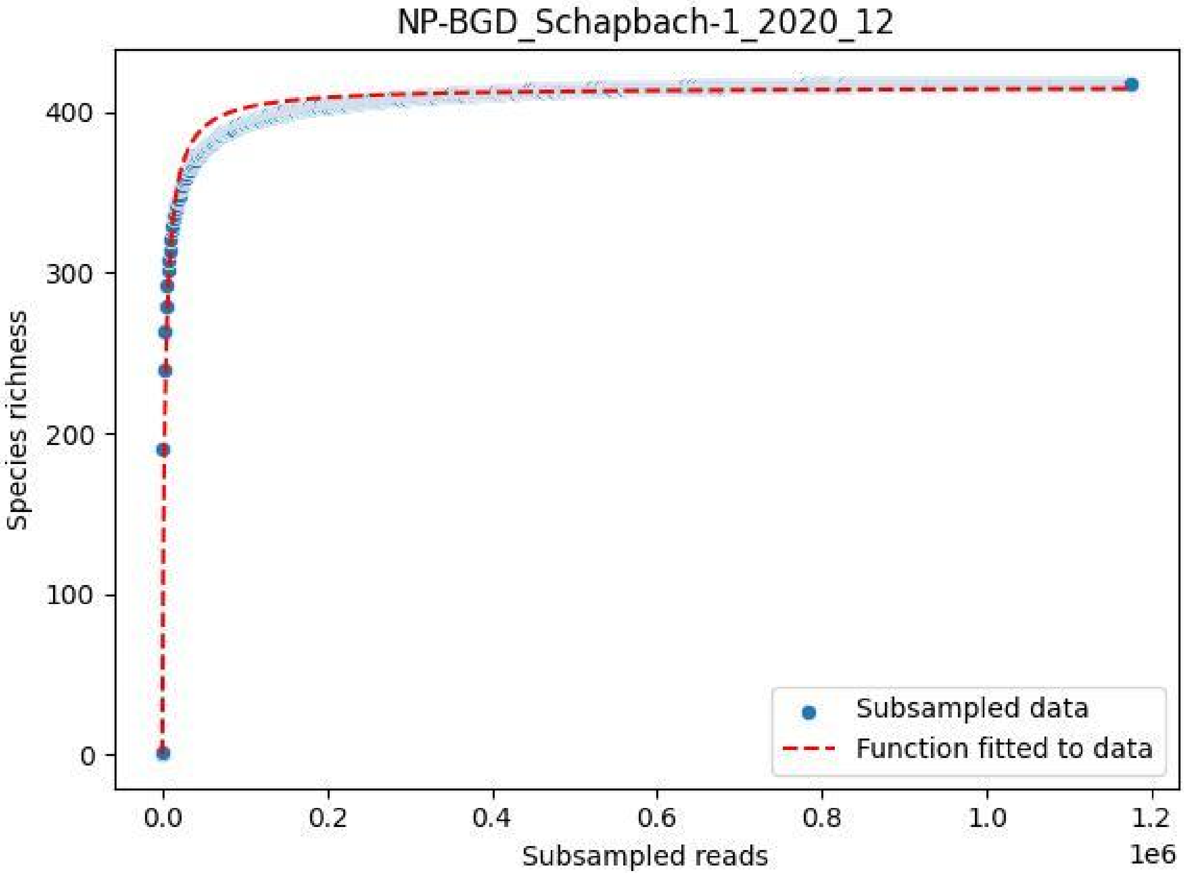
NP-BGD_Schapbach-1_2020_12
Species richness
Subsampled reads
1e6
0
100
200
300
400
0.0
0.2
0.4
0.6
0.8
1.0
1.2
Subsampled data
Function fitted to data

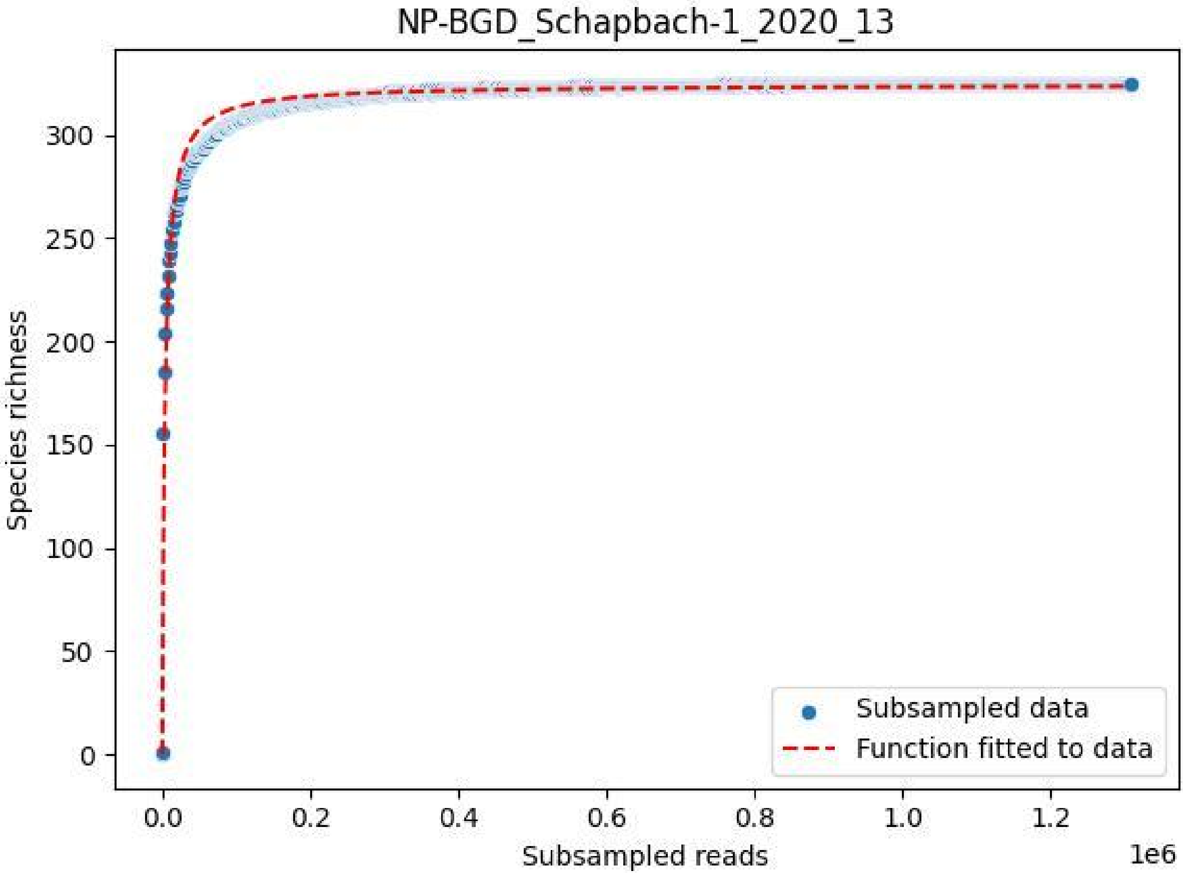
NP-BGD_Schapbach-1_2020_13
Species richness
Subsampled reads
1e6
0.0
0.2
0.4
0.6
0.8
1.0
1.2
0
50
100
150
200
250
300
Subsampled data
Function fitted to data

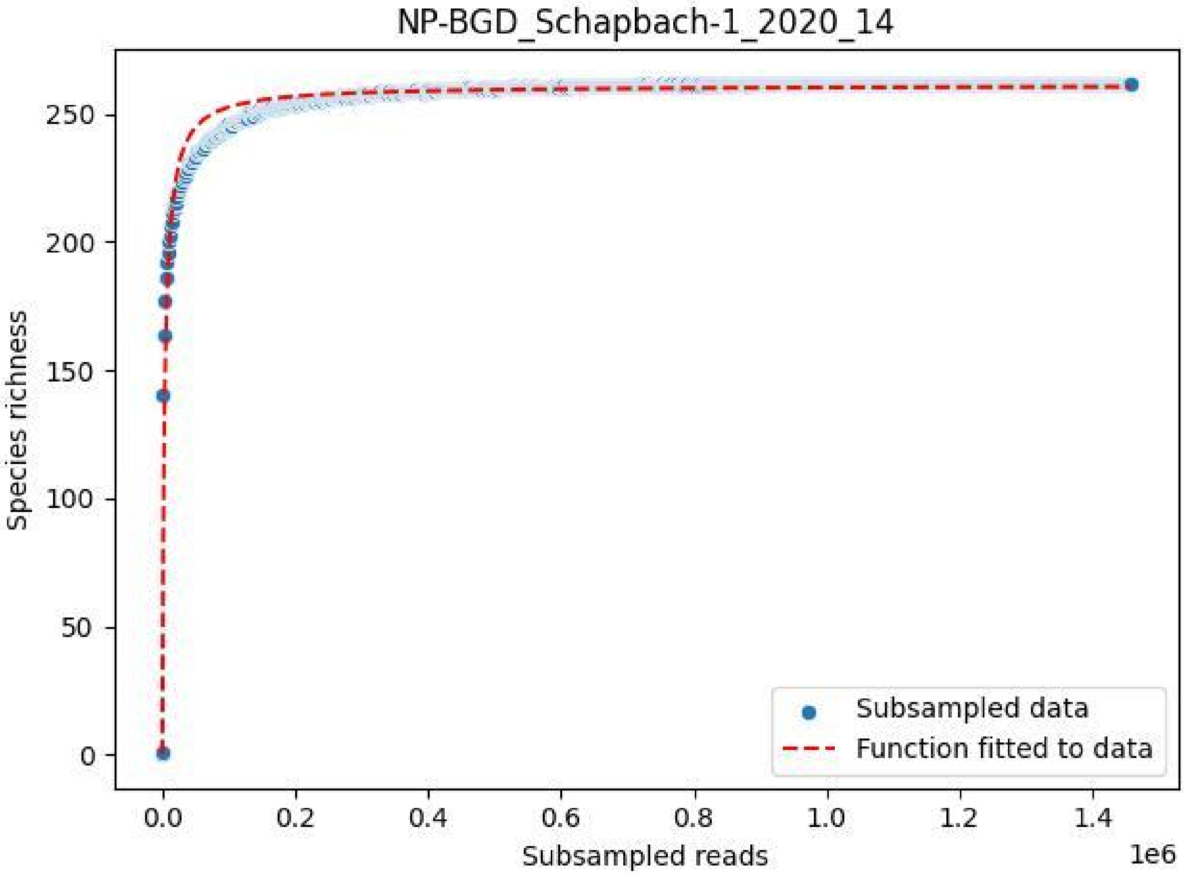
NP-BGD_Schapbach-1_2020_14
Species richness
Subsampled reads
1e6
0
50
100
150
200
250
0.0
0.2
0.4
0.6
0.8
1.0
1.2
1.4
Subsampled data
Function fitted to data

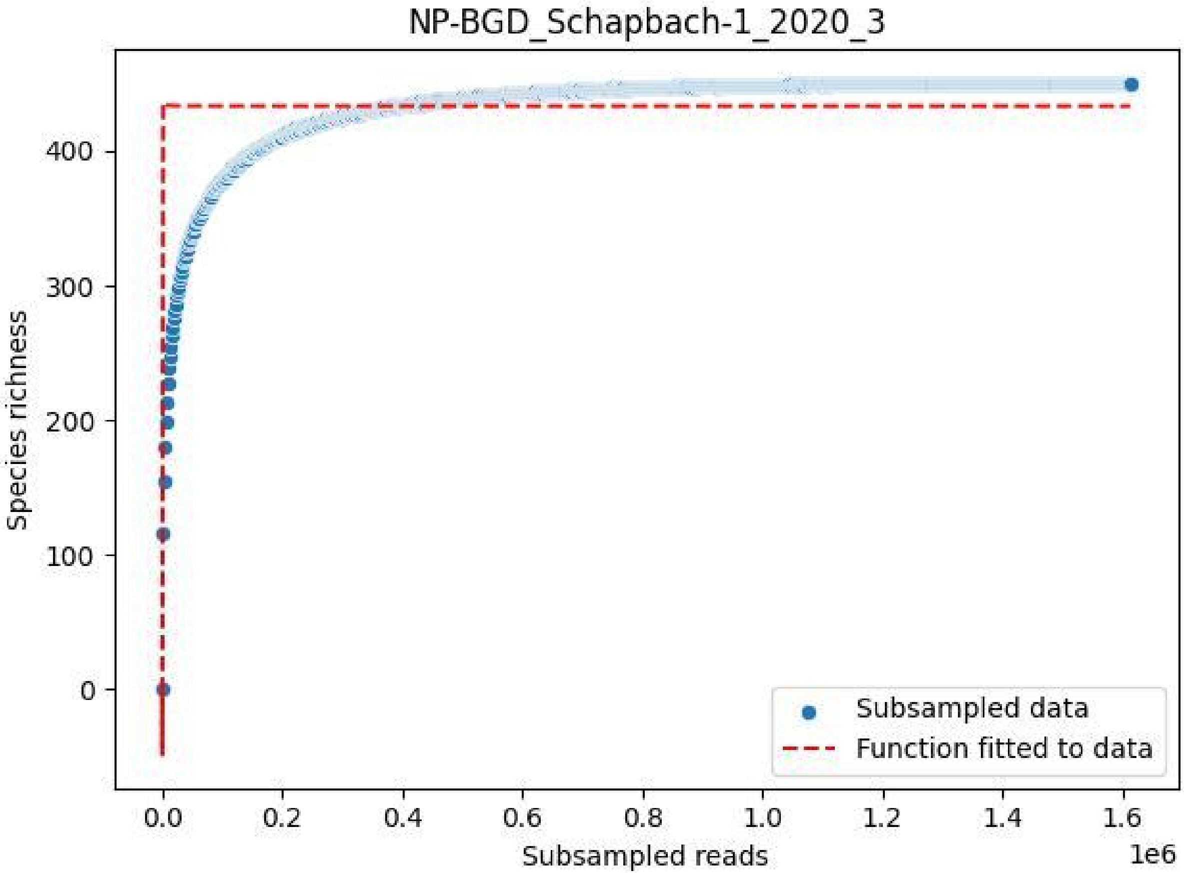

NP-BGD_Schapbach-1_2020_3
Species richness
Subsampled reads
1e6
0
100
200
300
400
0.0
0.2
0.4
0.6
0.8
1.0
1.2
1.4
1.6
Subsampled data
Function fitted to data

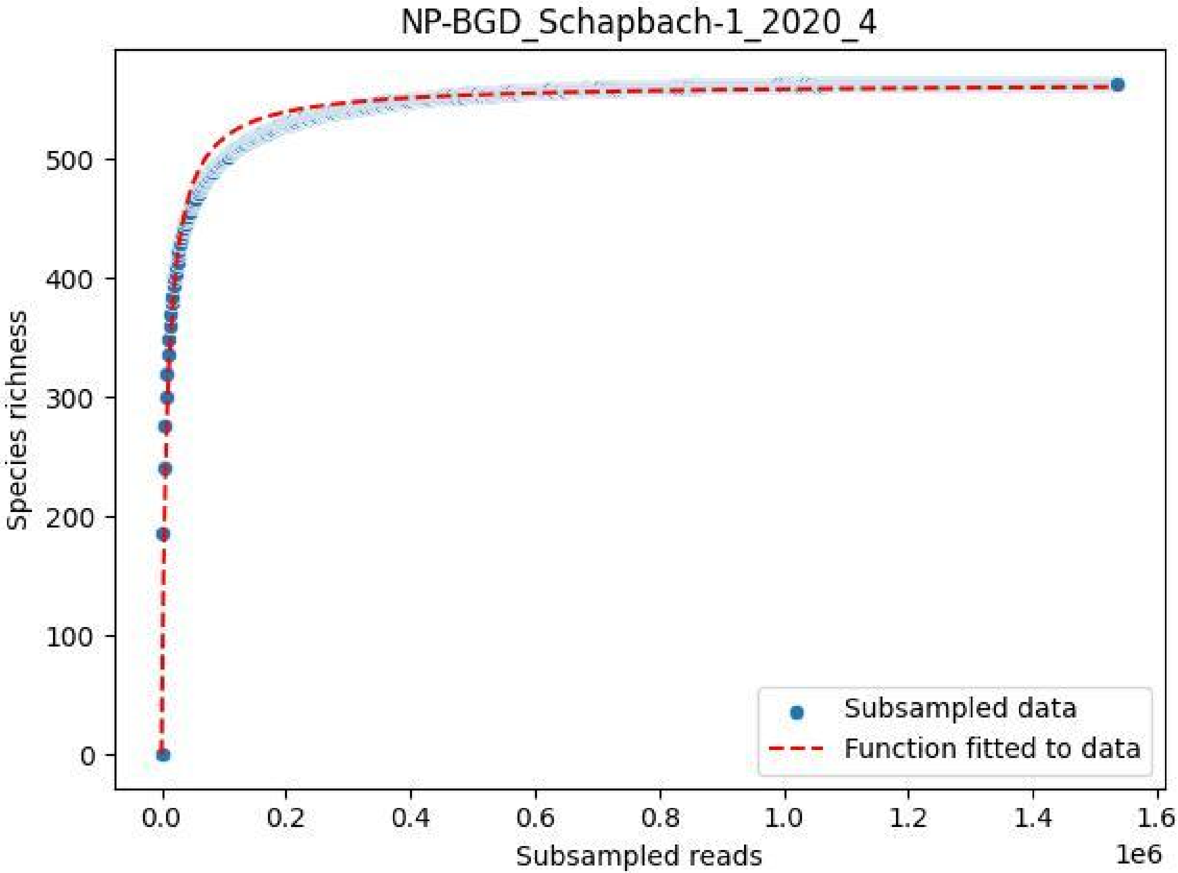
NP-BGD_Schapbach-1_2020_4
Species richness
Subsampled reads
1e6
0
100
200
300
400
500
0.0
0.2
0.4
0.6
0.8
1.0
1.2
1.4
1.6
Subsampled data
Function fitted to data

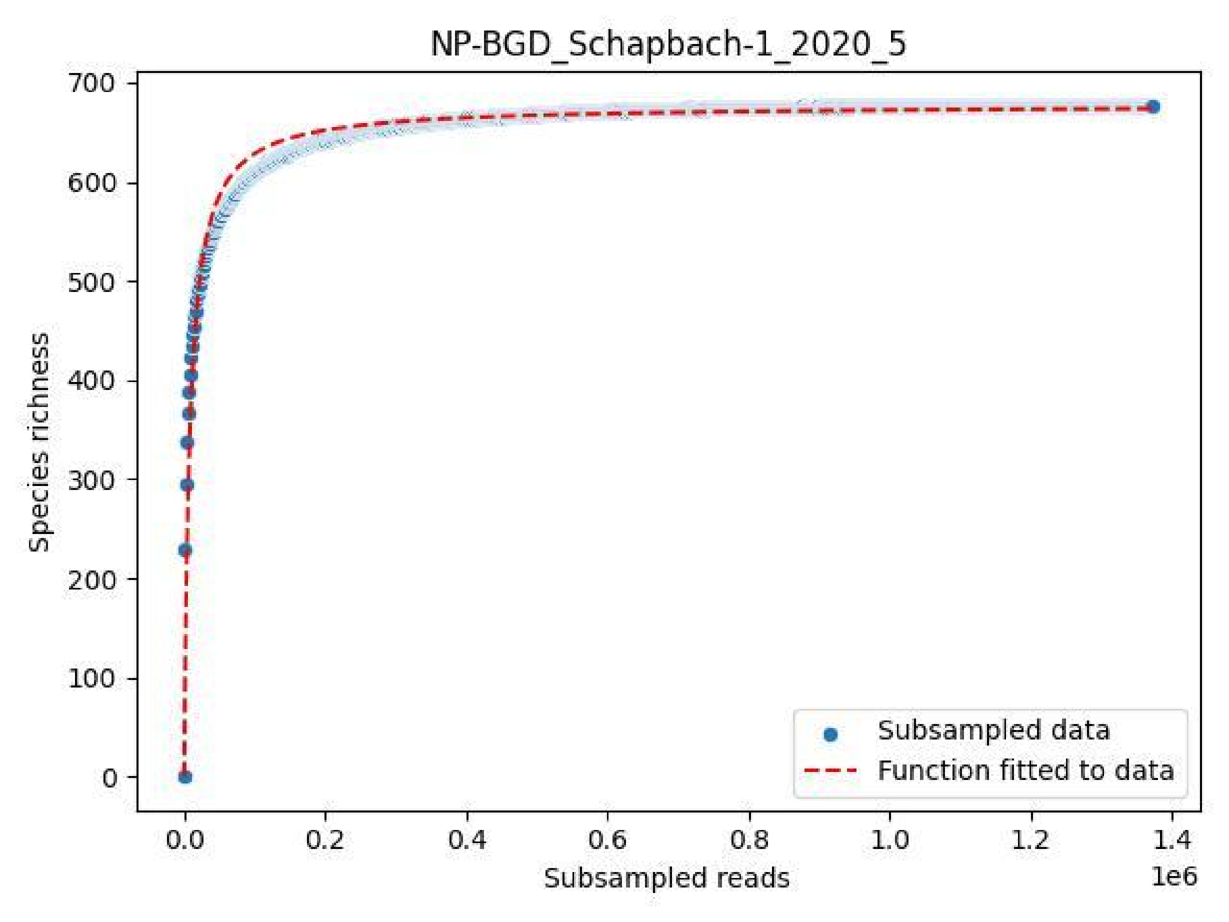
NP-BGD_Schapbach-1_2020_5
Species richness
700
600
500
400
300
200
100
0
0.0
0.2
0.4
0.6
0.8
1.0
1.2
1.4
1e6
Subsampled reads
Subsampled data
Function fitted to data

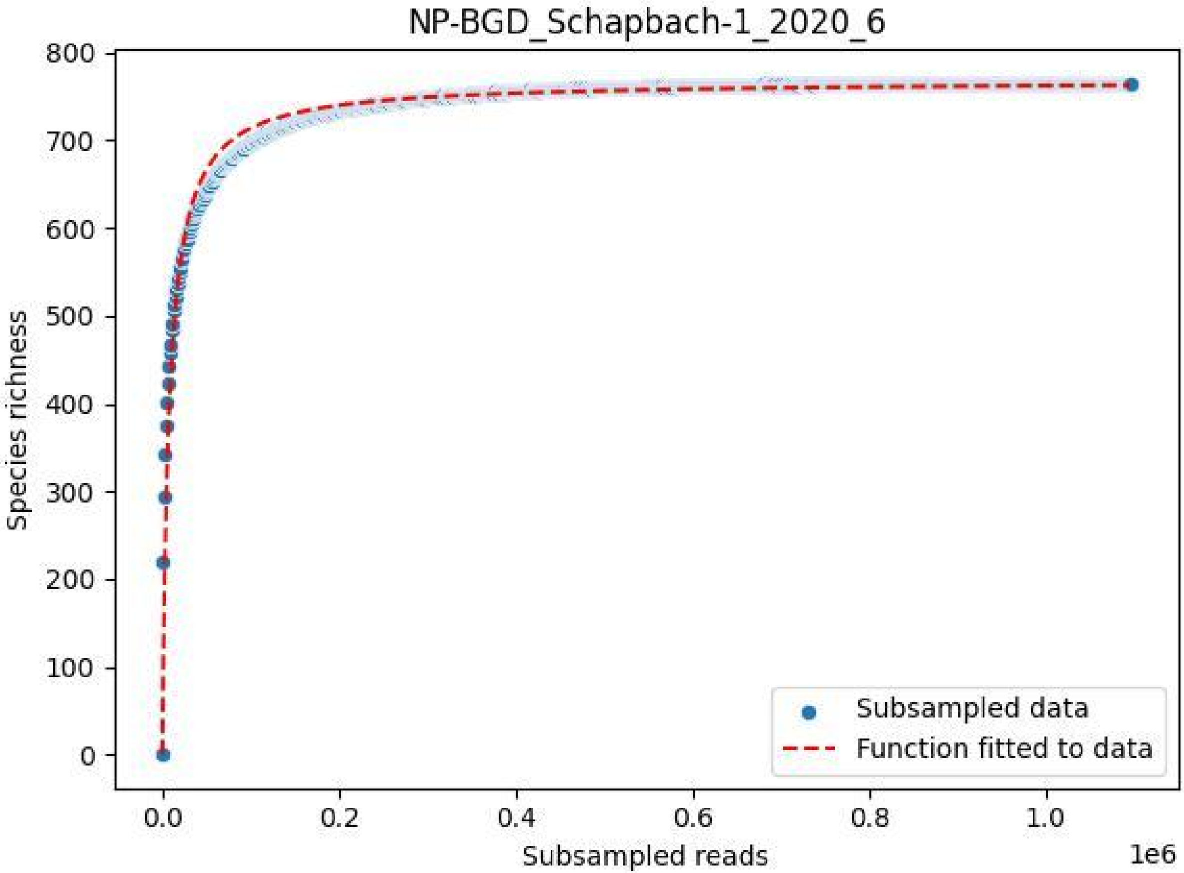
NP-BGD_Schapbach-1_2020_6
Species richness
Subsampled reads
1e6
0
100
200
300
400
500
600
700
800
0.0
0.2
0.4
0.6
0.8
1.0
Subsampled data
Function fitted to data

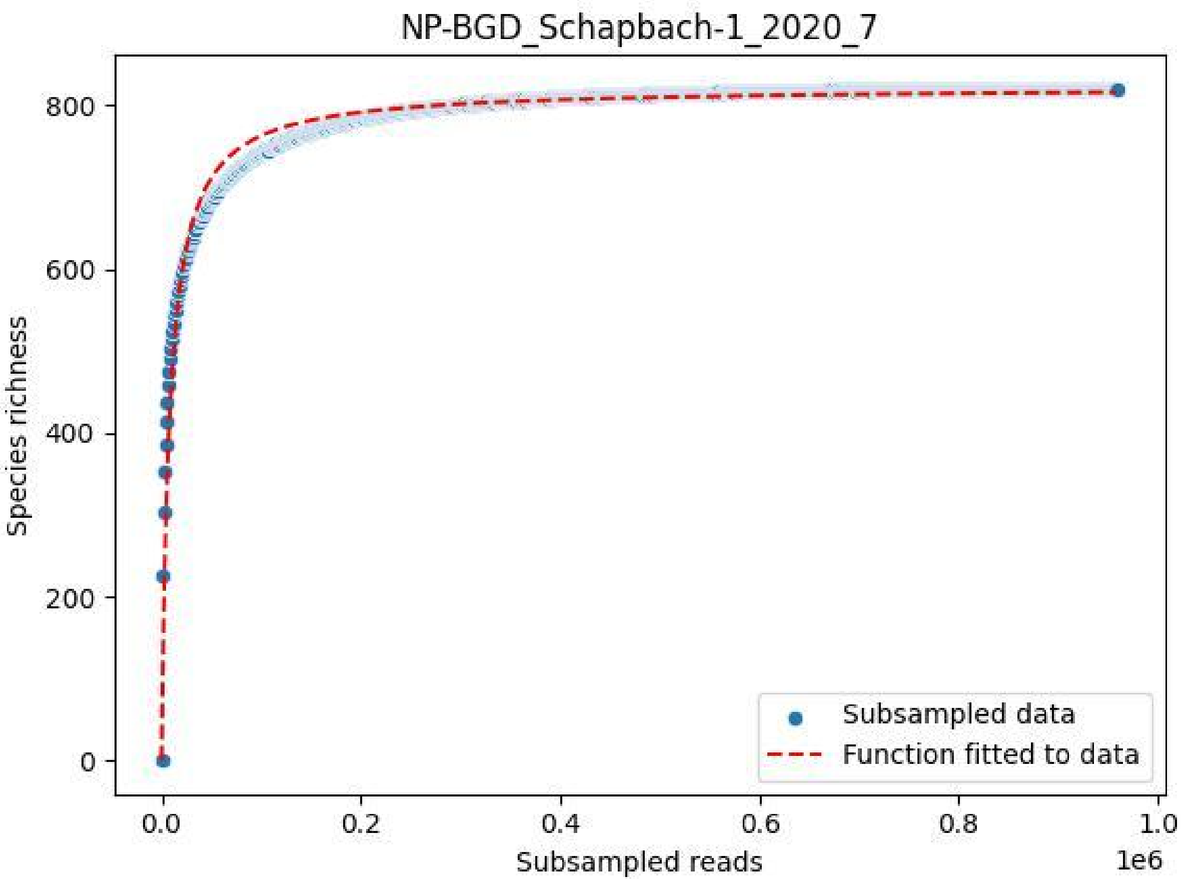

NP-BGD_Schapbach-1_2020_7
Species richness
0
200
400
600
800
Subsampled reads
0.0
0.2
0.4
0.6
0.8
1.0
1e6
Subsampled data
Function fitted to data

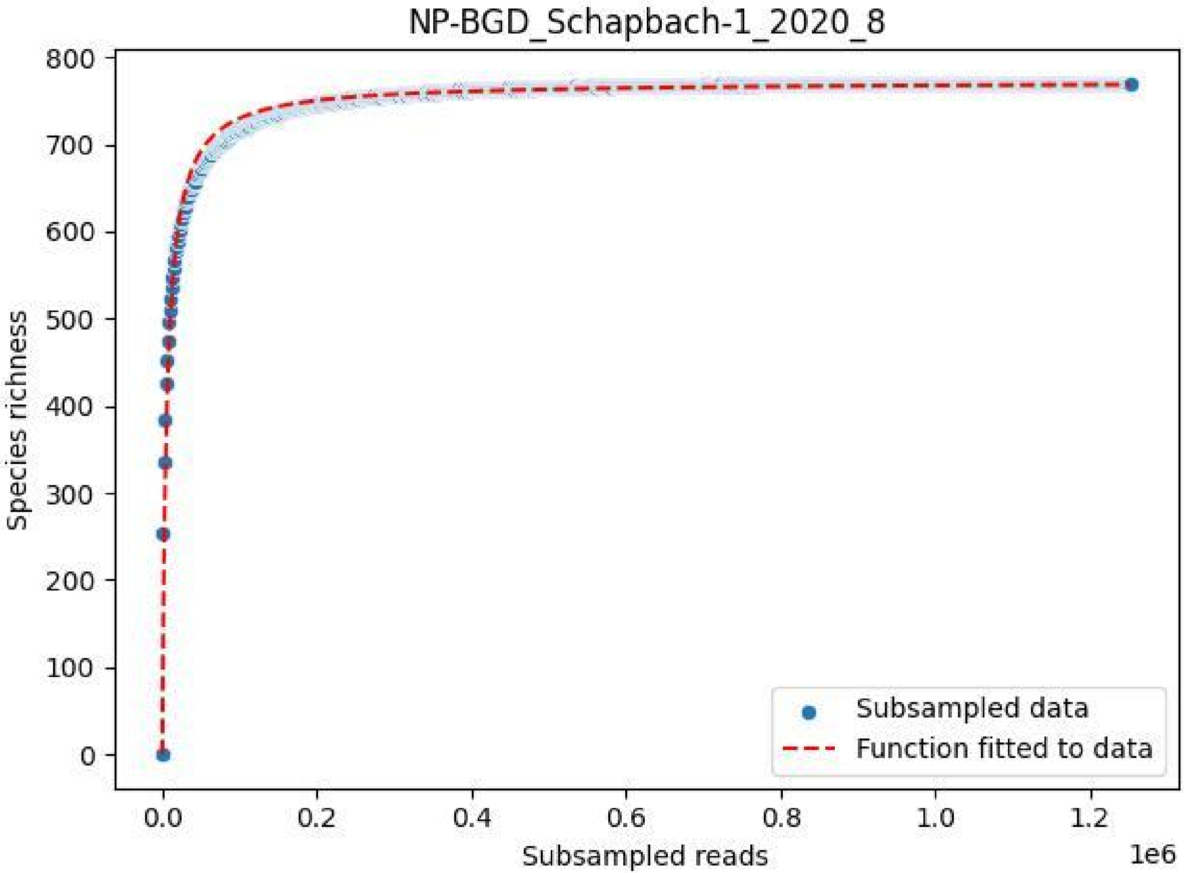
NP-BGD_Schapbach-1_2020_8
Species richness
Subsampled reads
1e6
0
100
200
300
400
500
600
700
800
0.0
0.2
0.4
0.6
0.8
1.0
1.2
Subsampled data
Function fitted to data

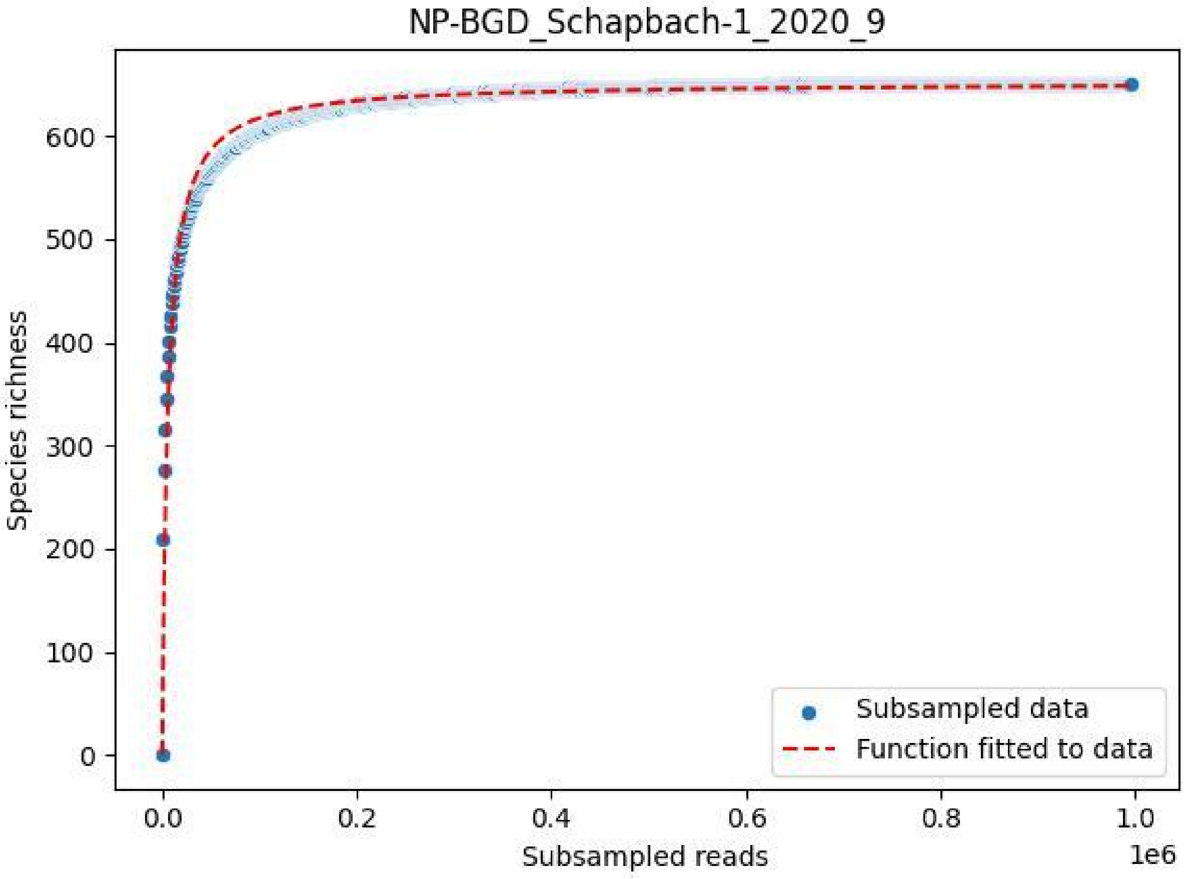

NP-BGD_Schapbach-1_2020_9
Species richness
Subsampled reads
1e6
0
100
200
300
400
500
600
0.0
0.2
0.4
0.6
0.8
1.0
Subsampled data
Function fitted to data

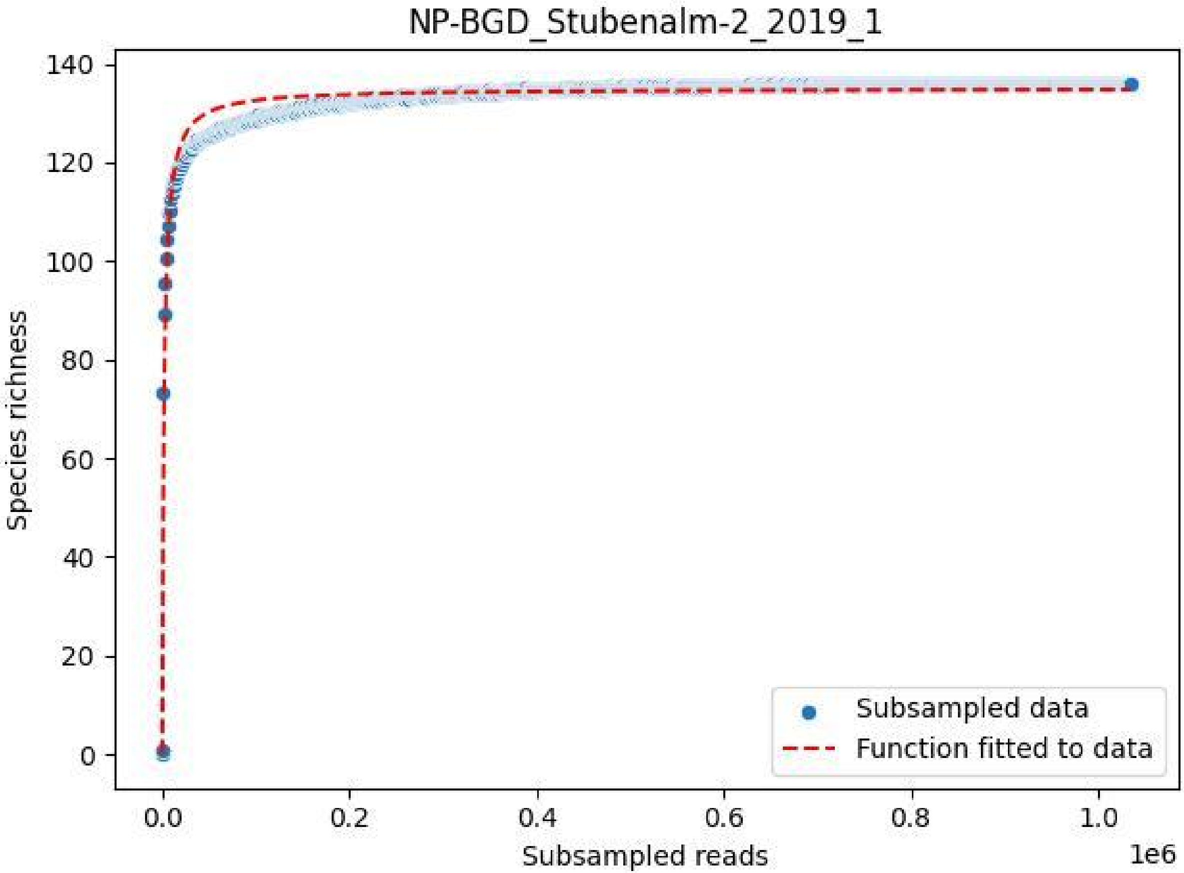
NP-BGD_Stubenalm-2_2019_1
Species richness
Subsampled reads
1e6
0
20
40
60
80
100
120
140
0.0
0.2
0.4
0.6
0.8
1.0
Subsampled data
Function fitted to data

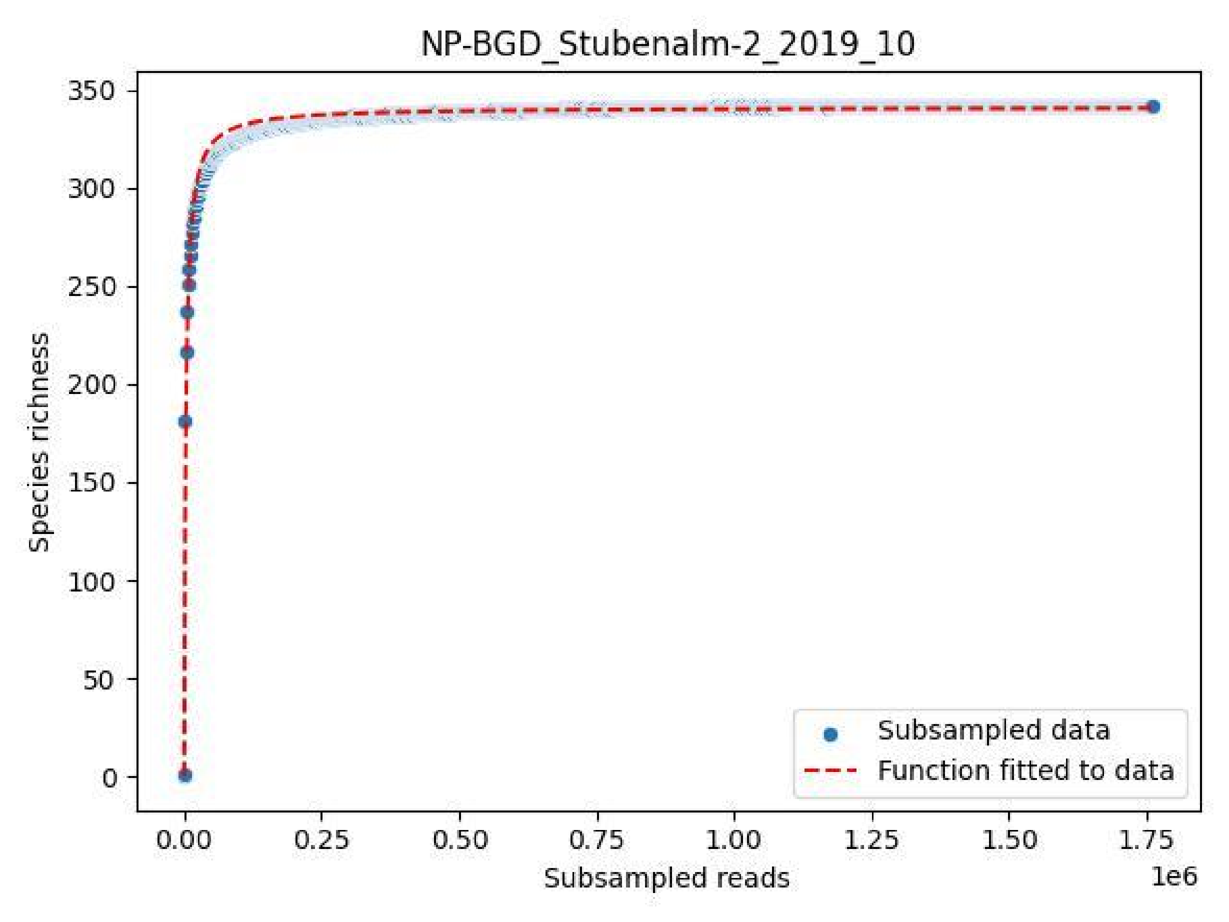
NP-BGD_Stubenalm-2_2019_10
Species richness
Subsampled reads
1e6
0
50
100
150
200
250
300
350
0.00
0.25
0.50
0.75
1.00
1.25
1.50
1.75
Subsampled data
Function fitted to data

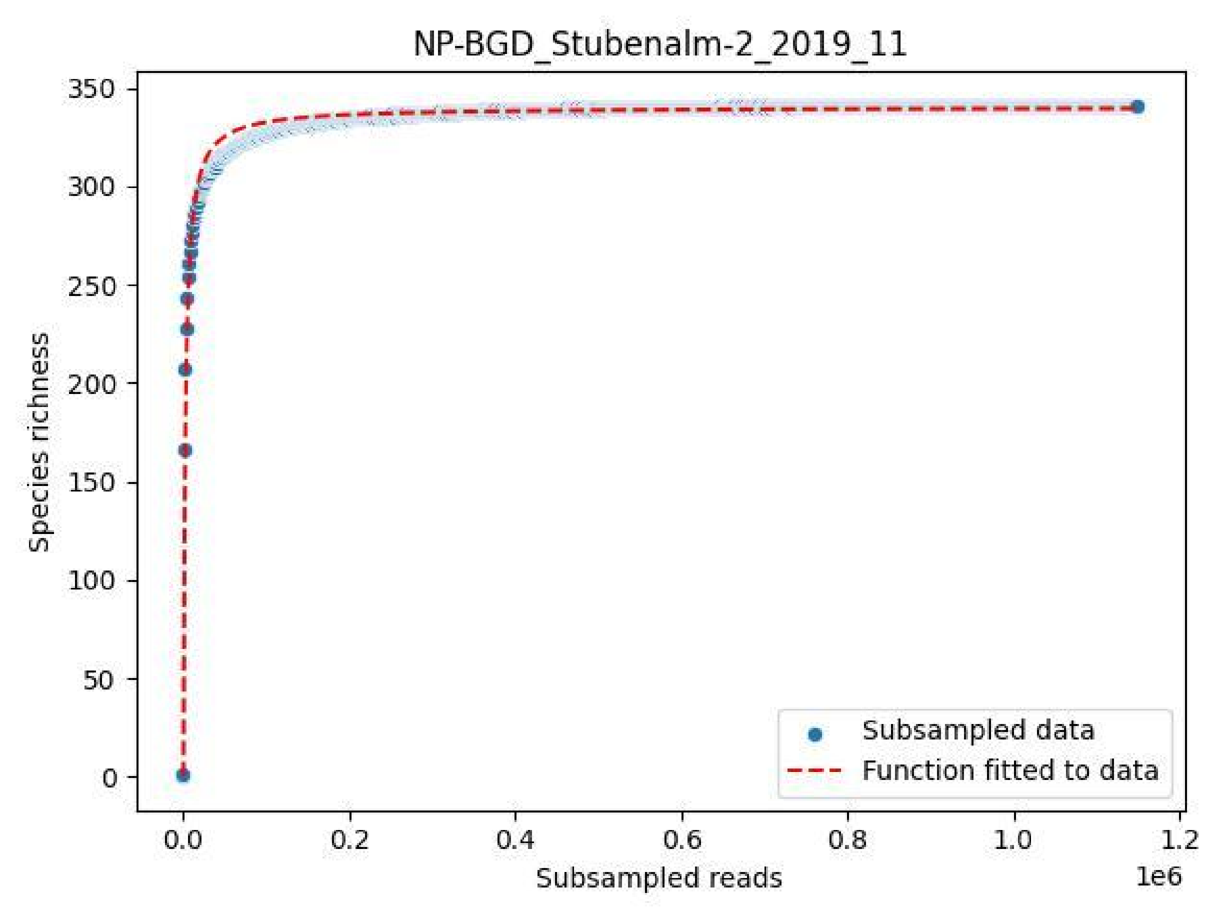
NP-BGD_Stubenalm-2_2019_11
Species richness
Subsampled reads
1e6
Subsampled data
Function fitted to data
0
50
100
150
200
250
300
350
0.0
0.2
0.4
0.6
0.8
1.0
1.2

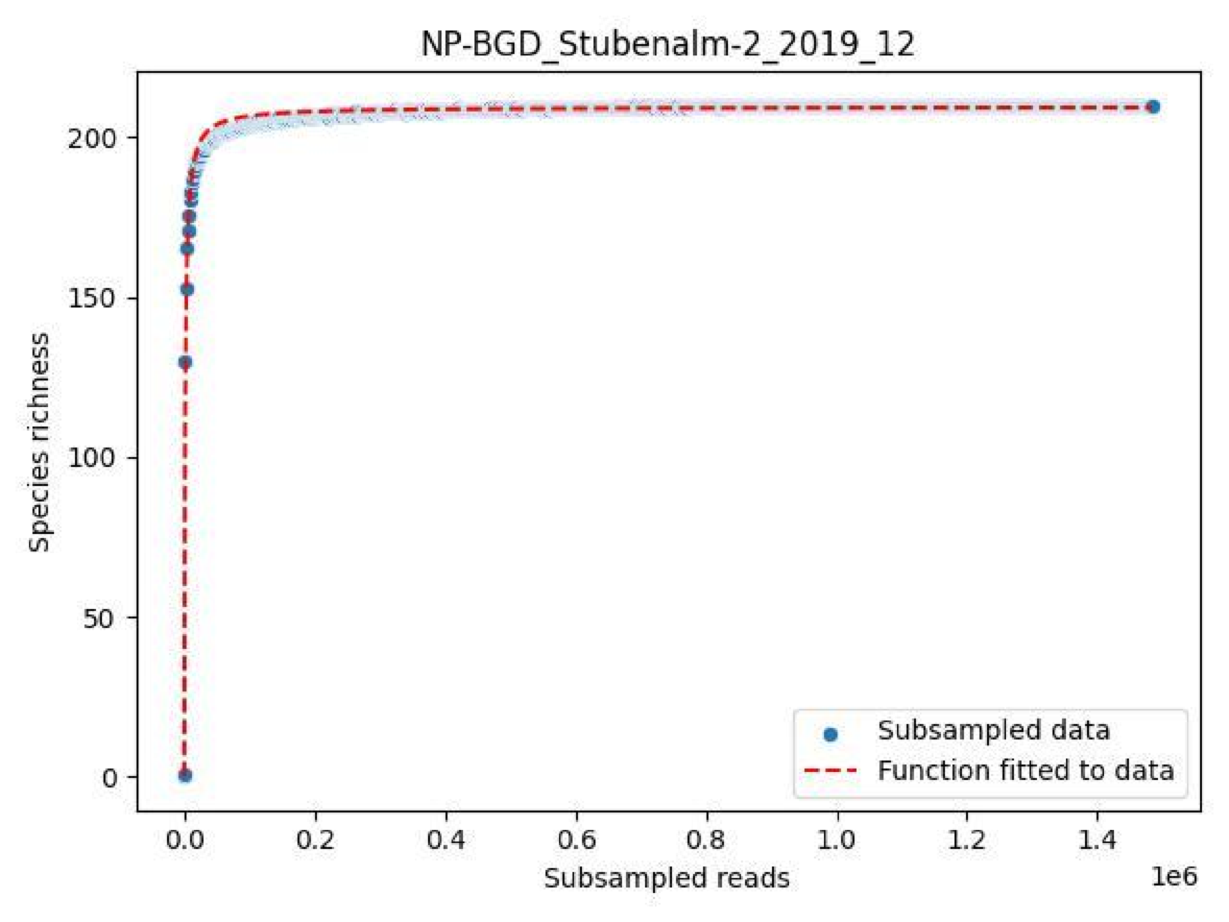

NP-BGD_Stubenalm-2_2019_12
Species richness
Subsampled reads
1e6
0
50
100
150
200
0.0
0.2
0.4
0.6
0.8
1.0
1.2
1.4
Subsampled data
Function fitted to data

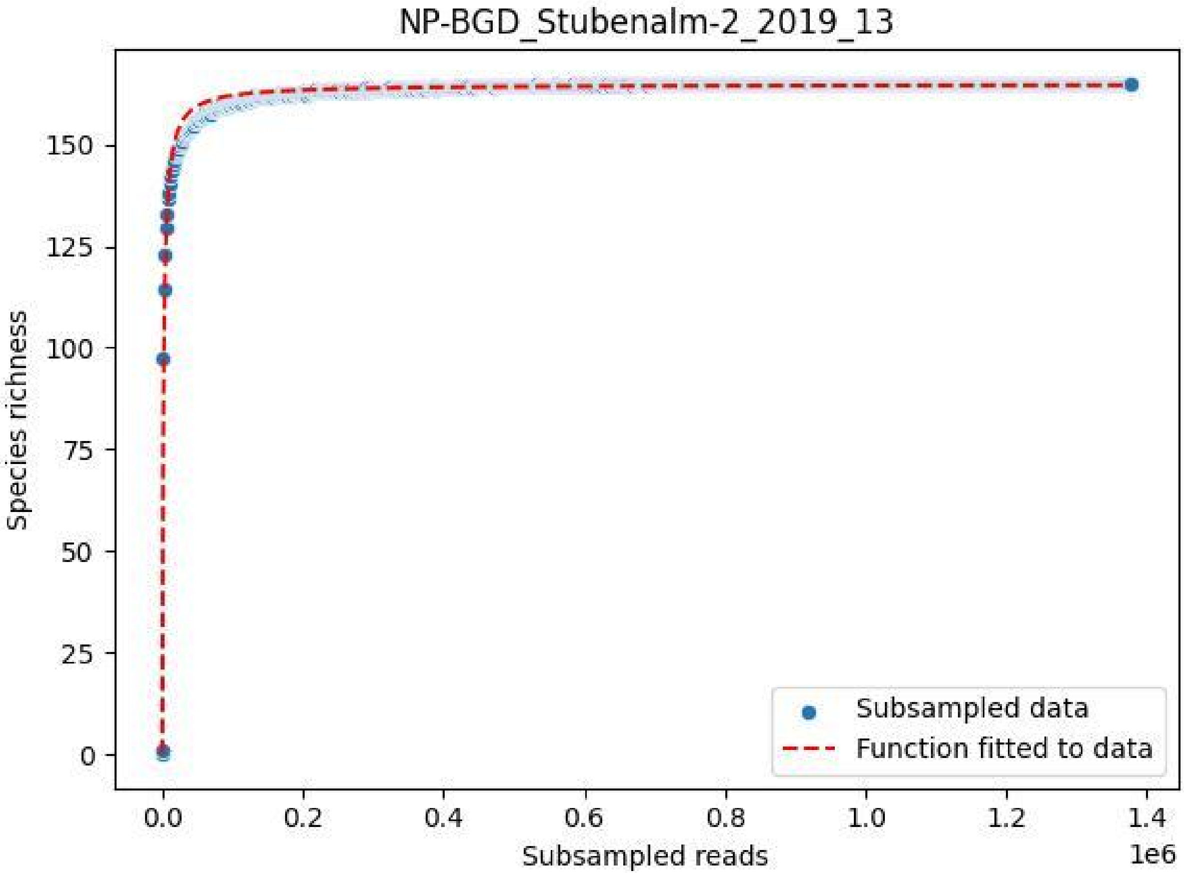
NP-BGD_Stubenalm-2_2019_13
Species richness
Subsampled reads
1e6
0
25
50
75
100
125
150
0.0
0.2
0.4
0.6
0.8
1.0
1.2
1.4
Subsampled data
Function fitted to data

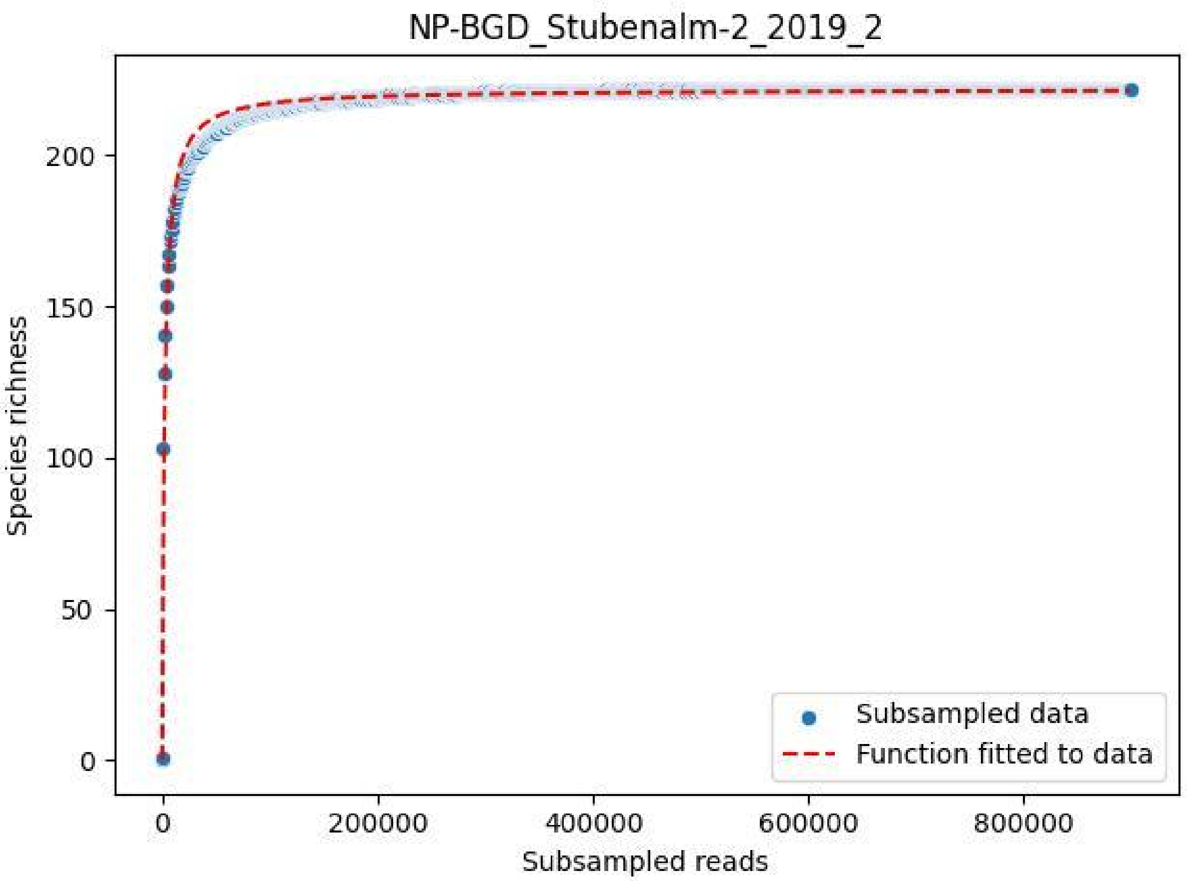

NP-BGD_Stubenalm-2_2019_2
Species richness
Subsampled reads
0
50
100
150
200
200000
400000
600000
800000
Subsampled data
Function fitted to data

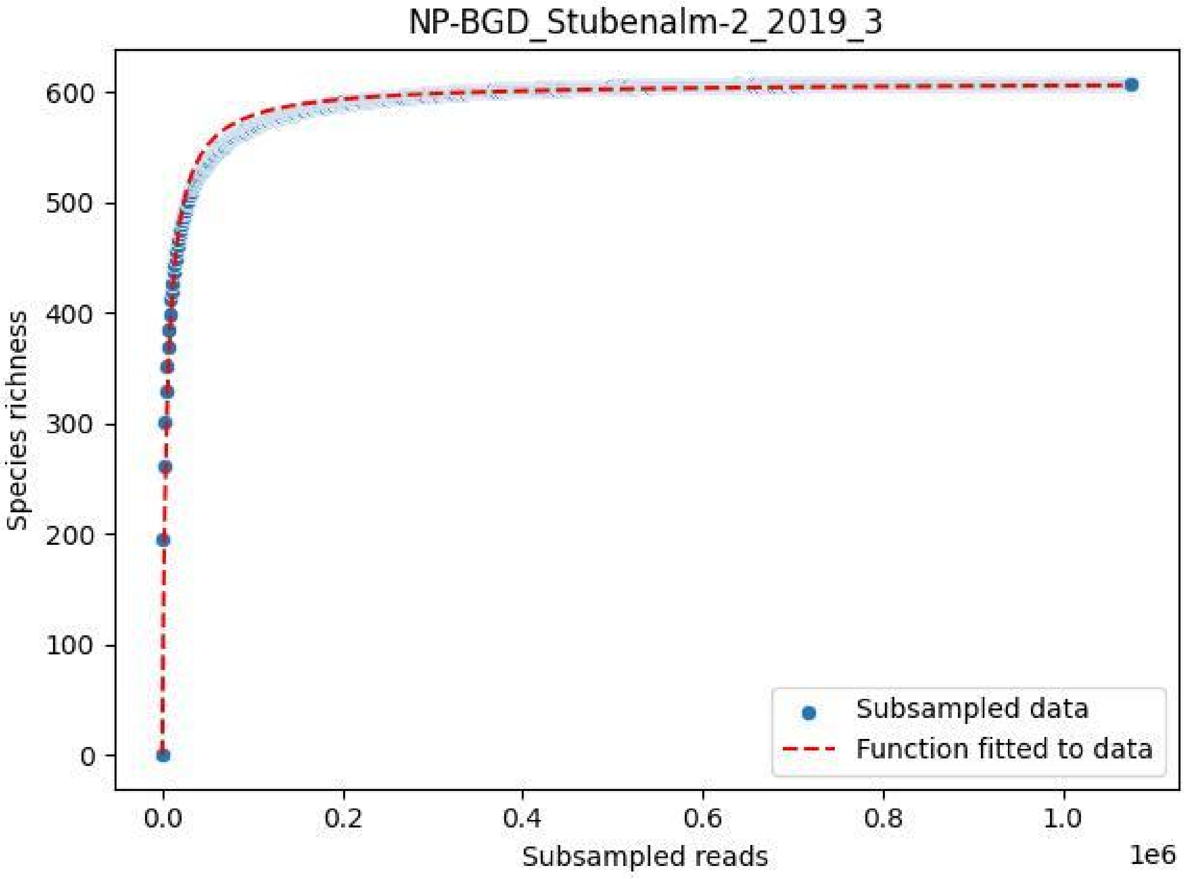
NP-BGD_Stubenalm-2_2019_3
Species richness
Subsampled reads
1e6
Subsampled data
Function fitted to data
0
100
200
300
400
500
600
0.0
0.2
0.4
0.6
0.8
1.0

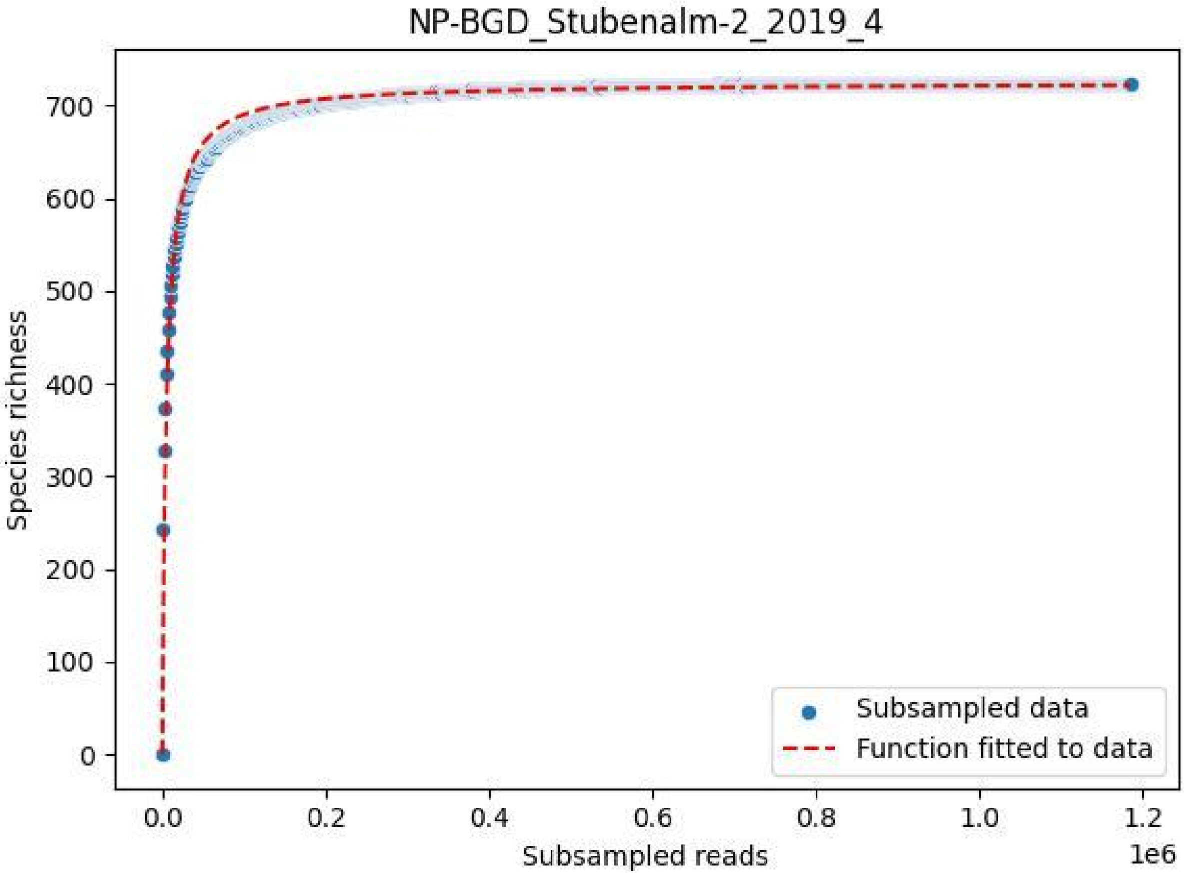
NP-BGD_Stubenalm-2_2019_4
Species richness
Subsampled reads
1e6
0
100
200
300
400
500
600
700
0.0
0.2
0.4
0.6
0.8
1.0
1.2
Subsampled data
Function fitted to data

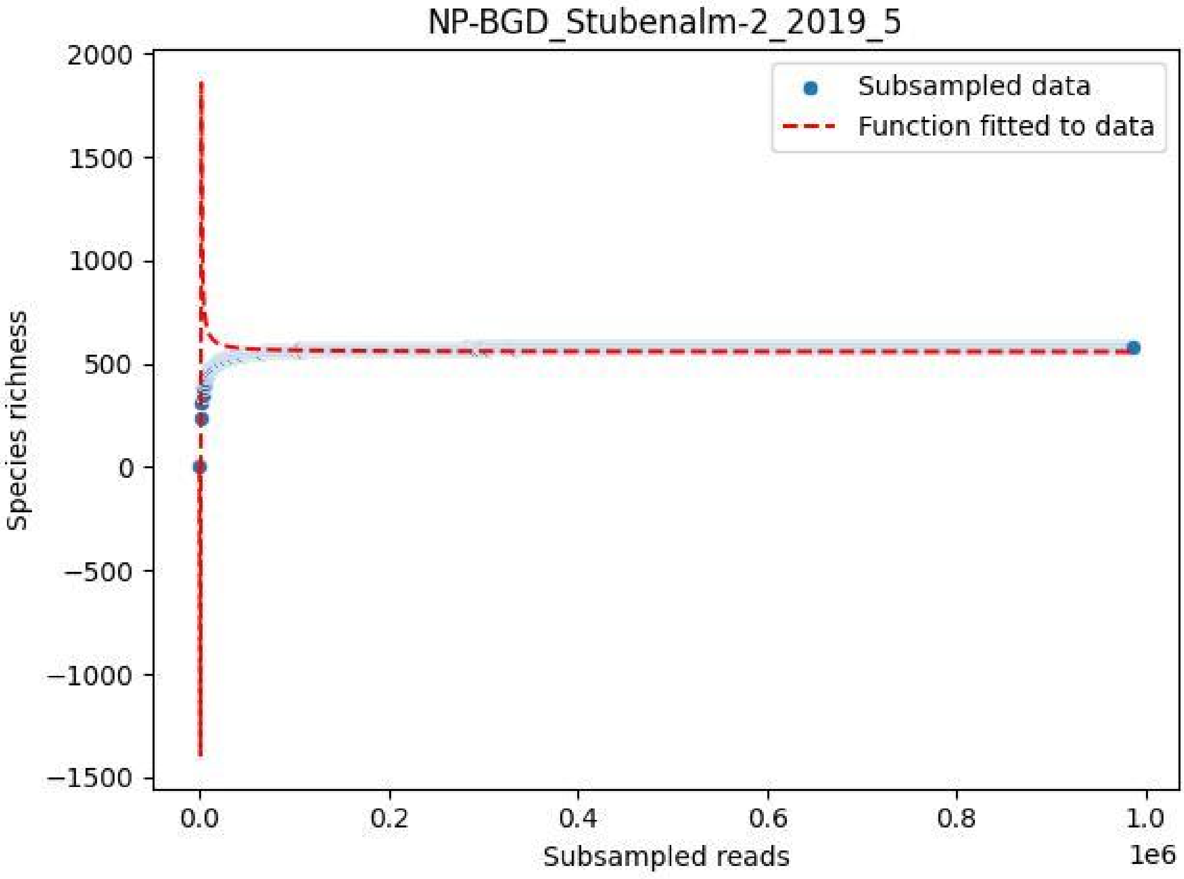
NP-BGD_Stubenalm-2_2019_5
Subsampled data
Function fitted to data
Species richness
Subsampled reads
1e6
2000
1500
1000
500
0
−500
−1000
−1500
0.0
0.2
0.4
0.6
0.8
1.0

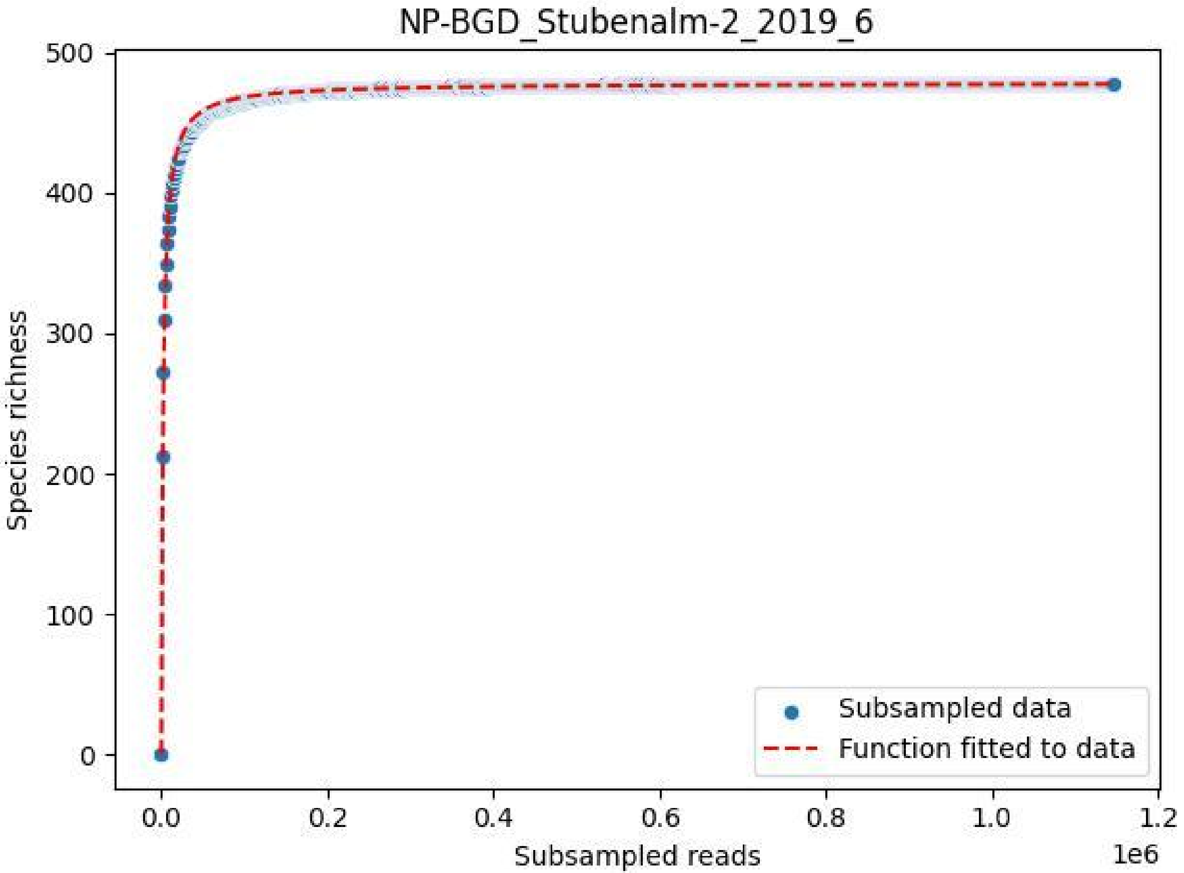
NP-BGD_Stubenalm-2_2019_6
Species richness
Subsampled reads
1e6
Subsampled data
Function fitted to data
0
100
200
300
400
500
0.0
0.2
0.4
0.6
0.8
1.0
1.2

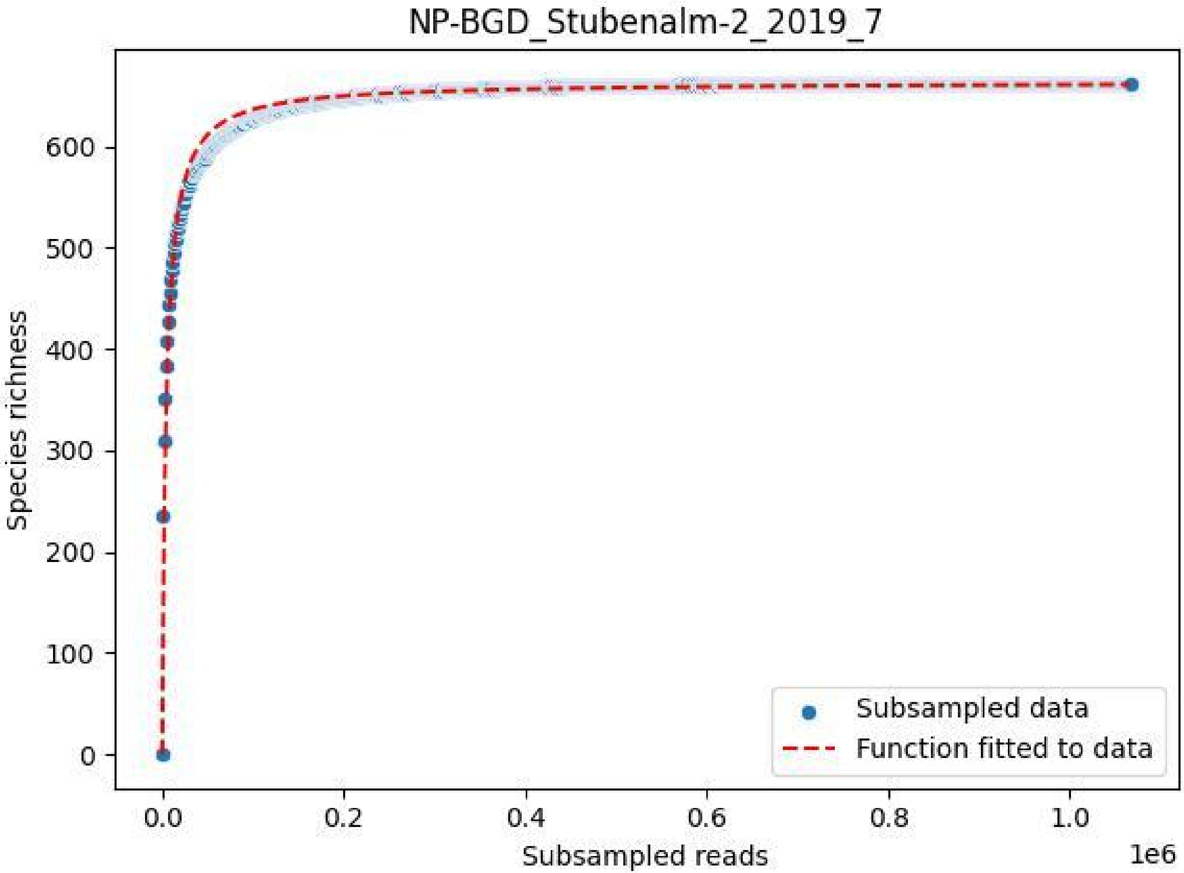
NP-BGD_Stubenalm-2_2019_7
Species richness
0
100
200
300
400
500
600
0.0
0.2
0.4
0.6
0.8
1.0
1e6
Subsampled reads
Subsampled data
Function fitted to data

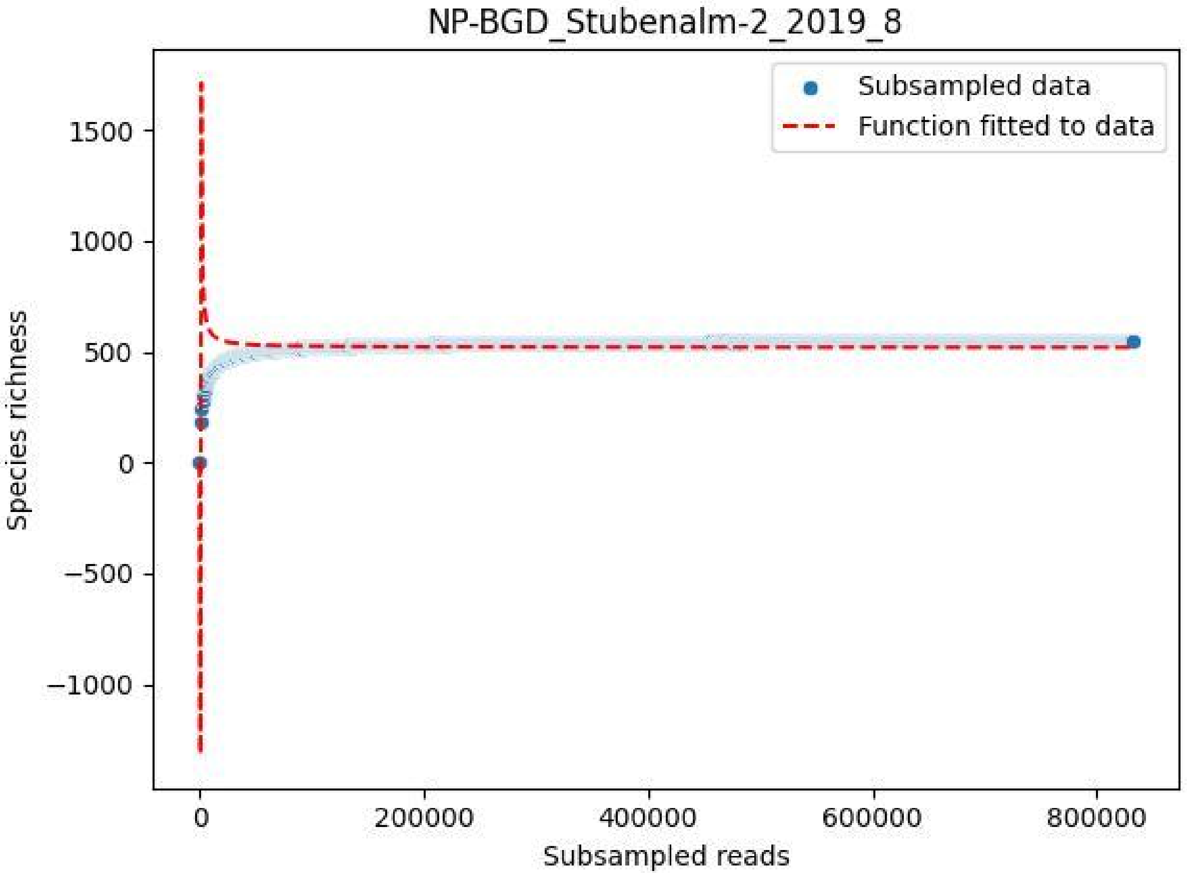
NP-BGD_Stubenalm-2_2019_8
Subsampled data
Function fitted to data
1500
1000
500
0
−500
−1000
Species richness
0
200000
400000
600000
800000
Subsampled reads

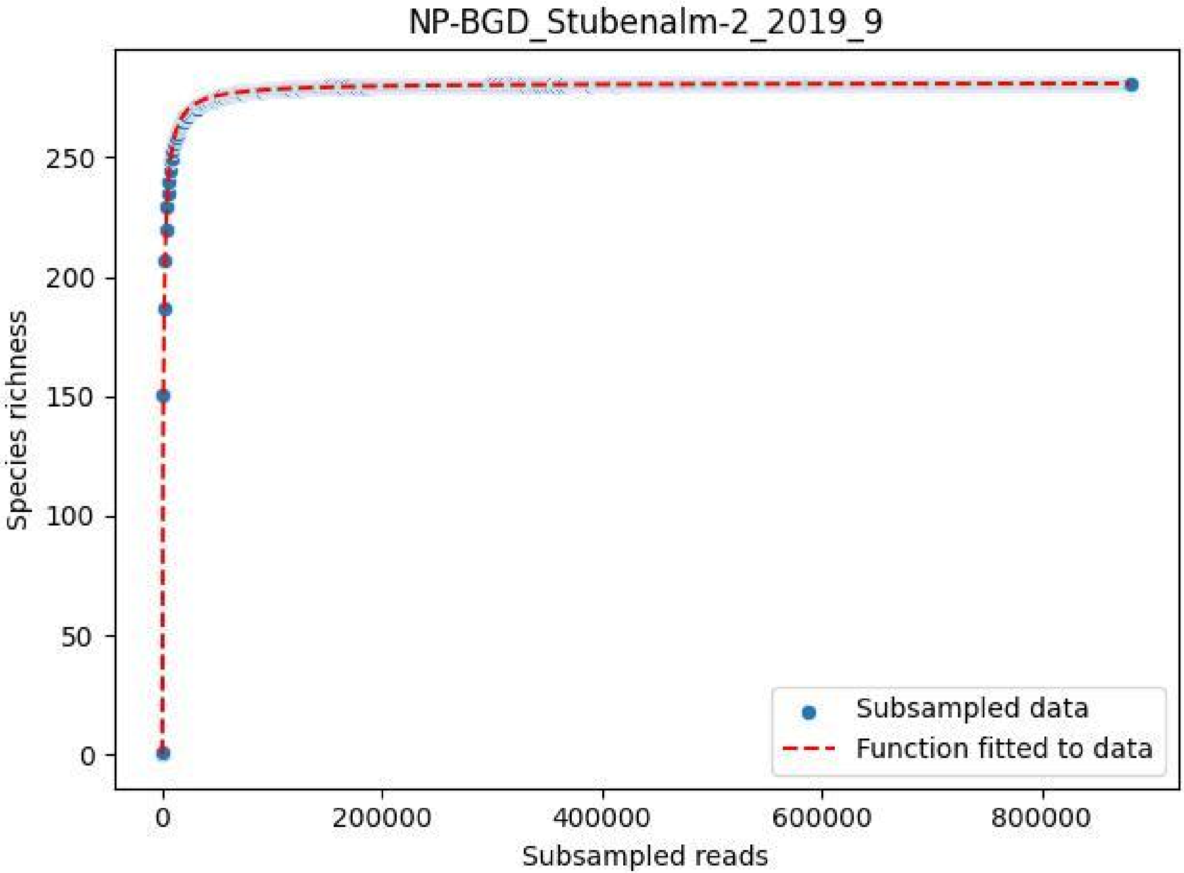
NP-BGD_Stubenalm-2_2019_9
Species richness
Subsampled reads
0
50
100
150
200
250
0
200000
400000
600000
800000
Subsampled data
Function fitted to data

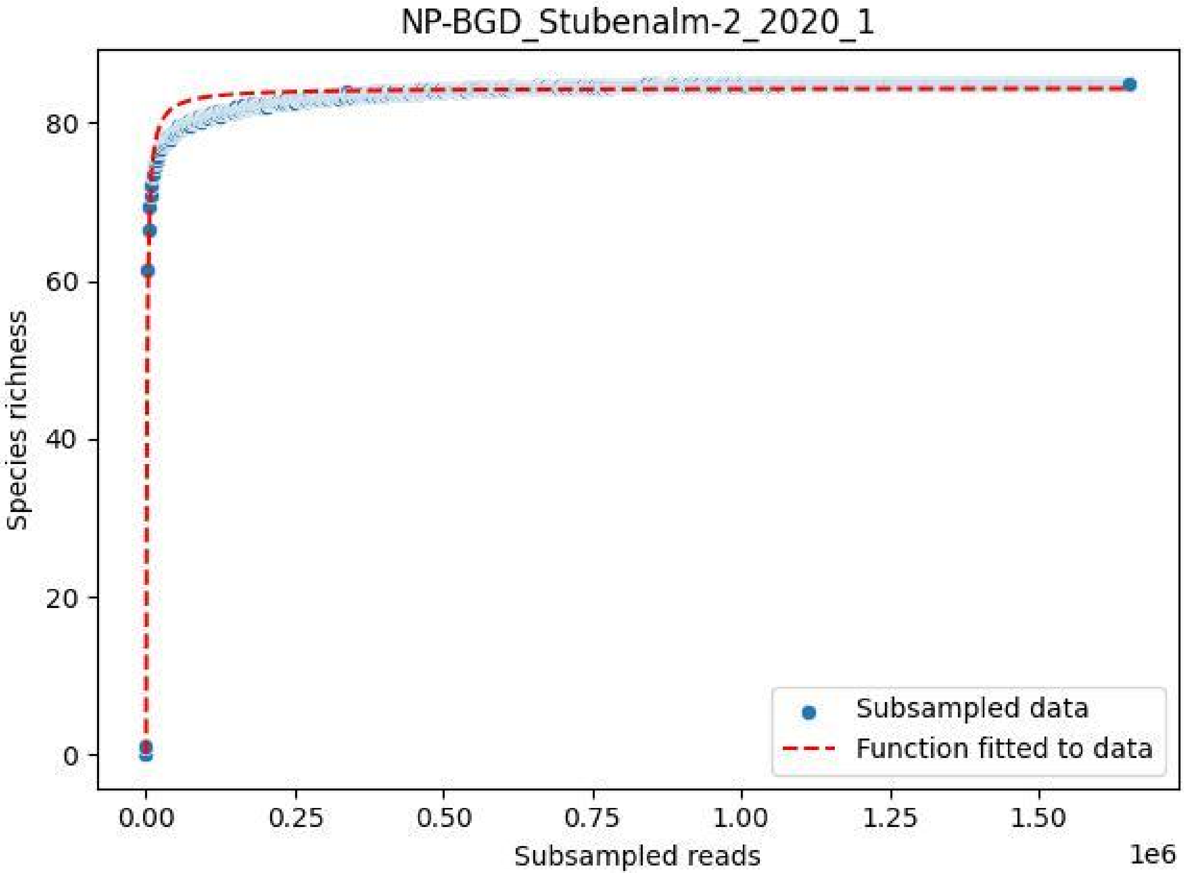

NP-BGD_Stubenalm-2_2020_1
Species richness
Subsampled reads
1e6
0
20
40
60
80
0.00
0.25
0.50
0.75
1.00
1.25
1.50
Subsampled data
Function fitted to data

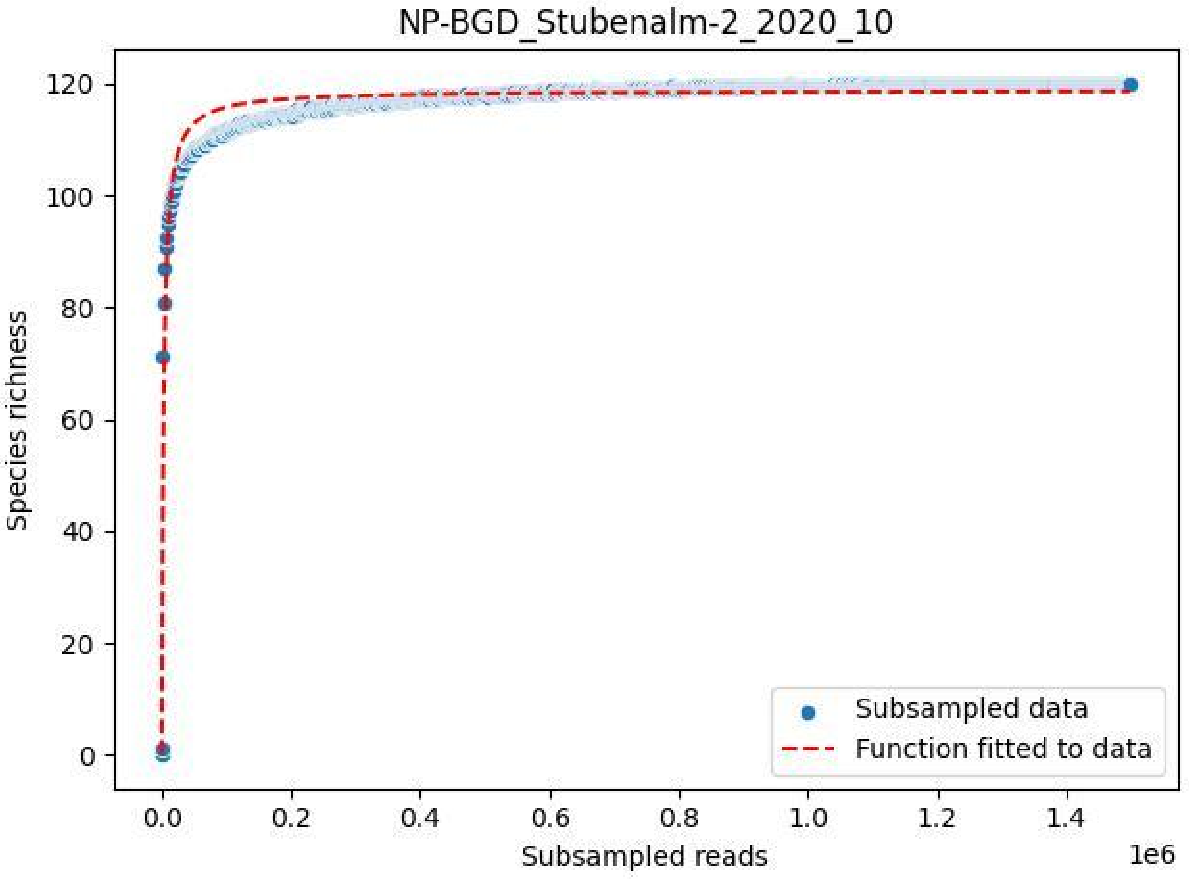
NP-BGD_Stubenalm-2_2020_10
Species richness
Subsampled reads
1e6
0
20
40
60
80
100
120
0.0
0.2
0.4
0.6
0.8
1.0
1.2
1.4
Subsampled data
Function fitted to data

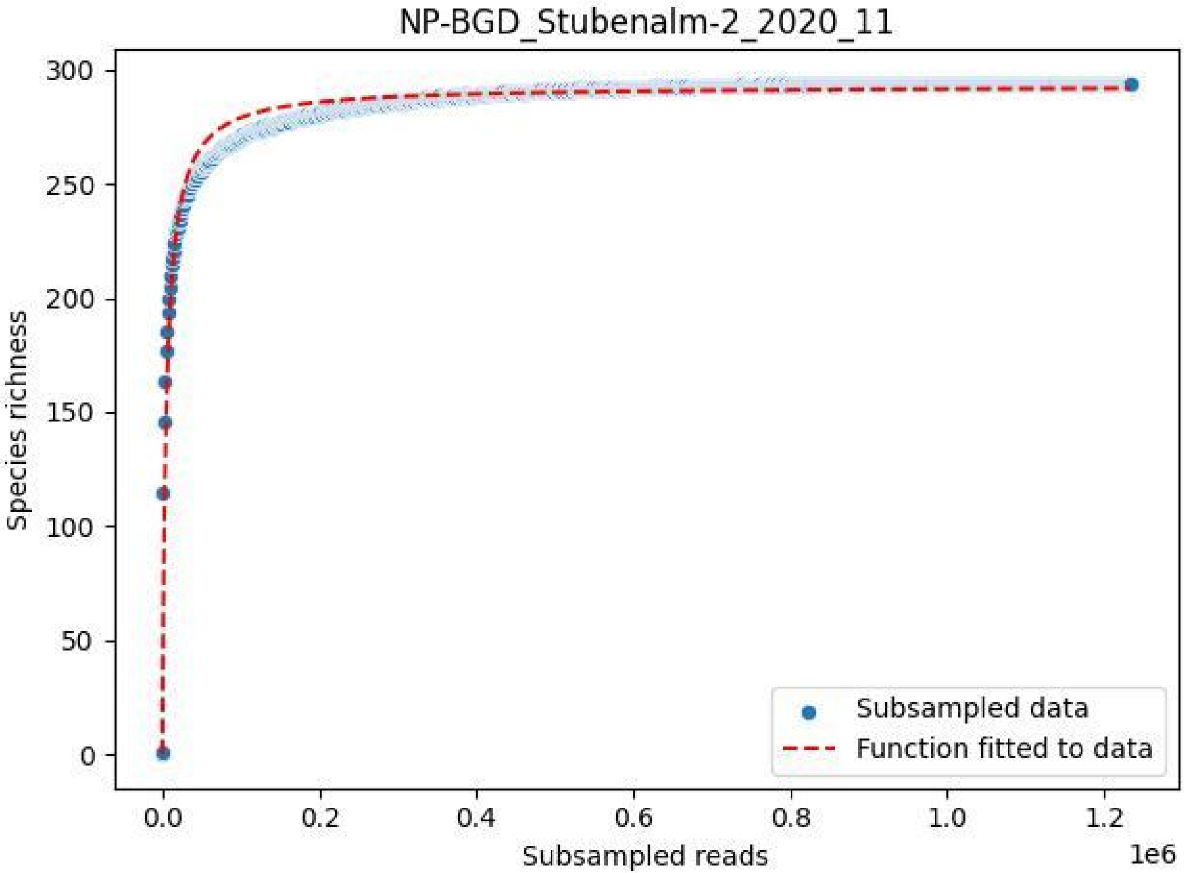
NP-BGD_Stubenalm-2_2020_11
Species richness
Subsampled reads
1e6
0.0
0.2
0.4
0.6
0.8
1.0
1.2
0
50
100
150
200
250
300
Subsampled data
Function fitted to data

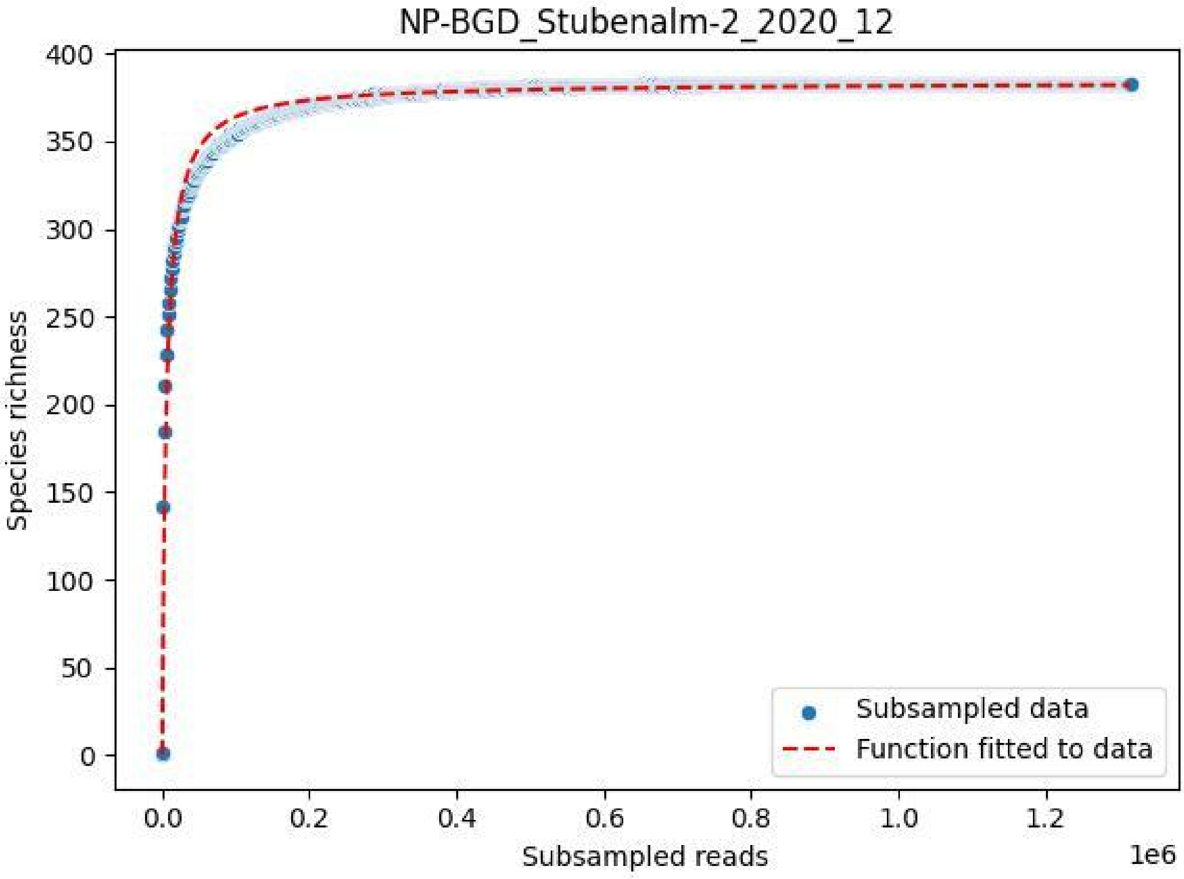
NP-BGD_Stubenalm-2_2020_12
Species richness
Subsampled reads
1e6
0
50
100
150
200
250
300
350
400
0.0
0.2
0.4
0.6
0.8
1.0
1.2
Subsampled data
Function fitted to data

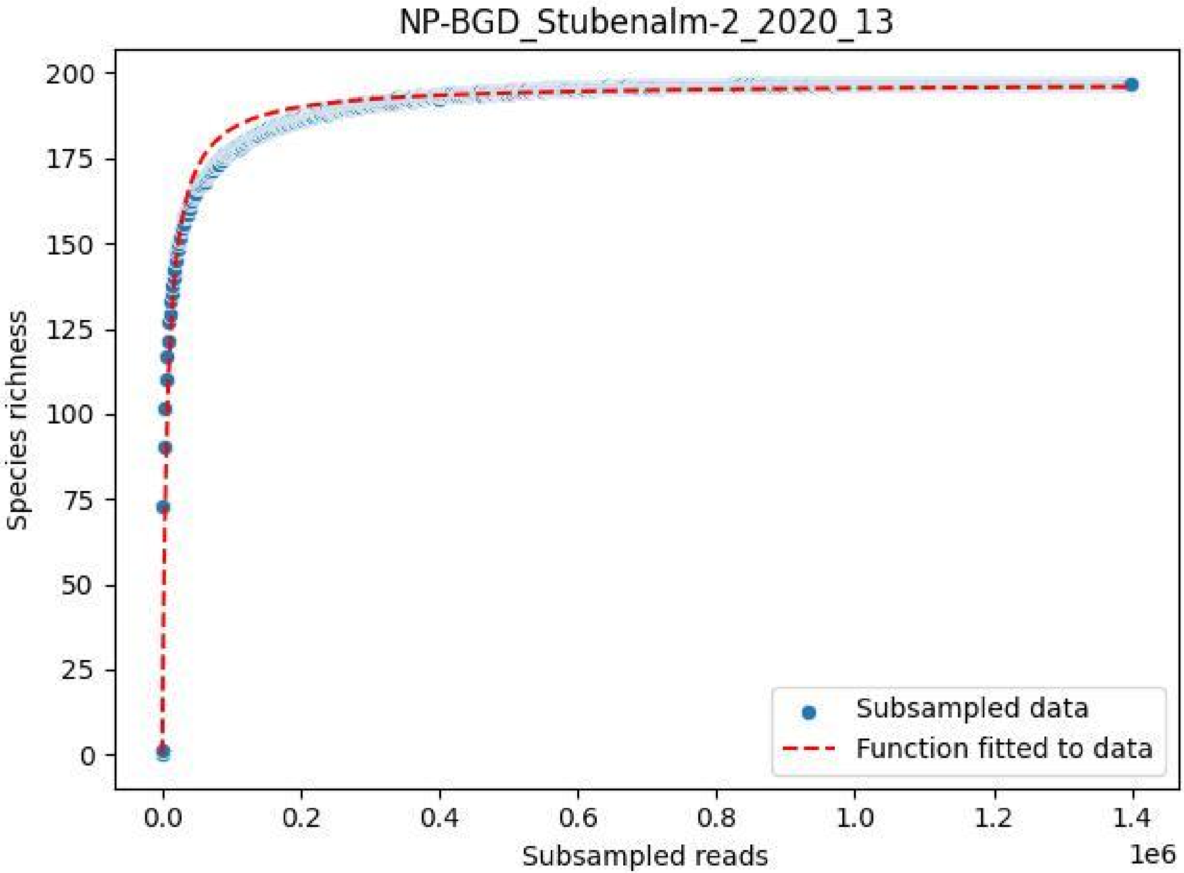
NP-BGD_Stubenalm-2_2020_13
Species richness
200
175
150
125
100
75
50
25
0
0.0
0.2
0.4
0.6
0.8
1.0
1.2
1.4
1e6
Subsampled reads
Subsampled data
Function fitted to data

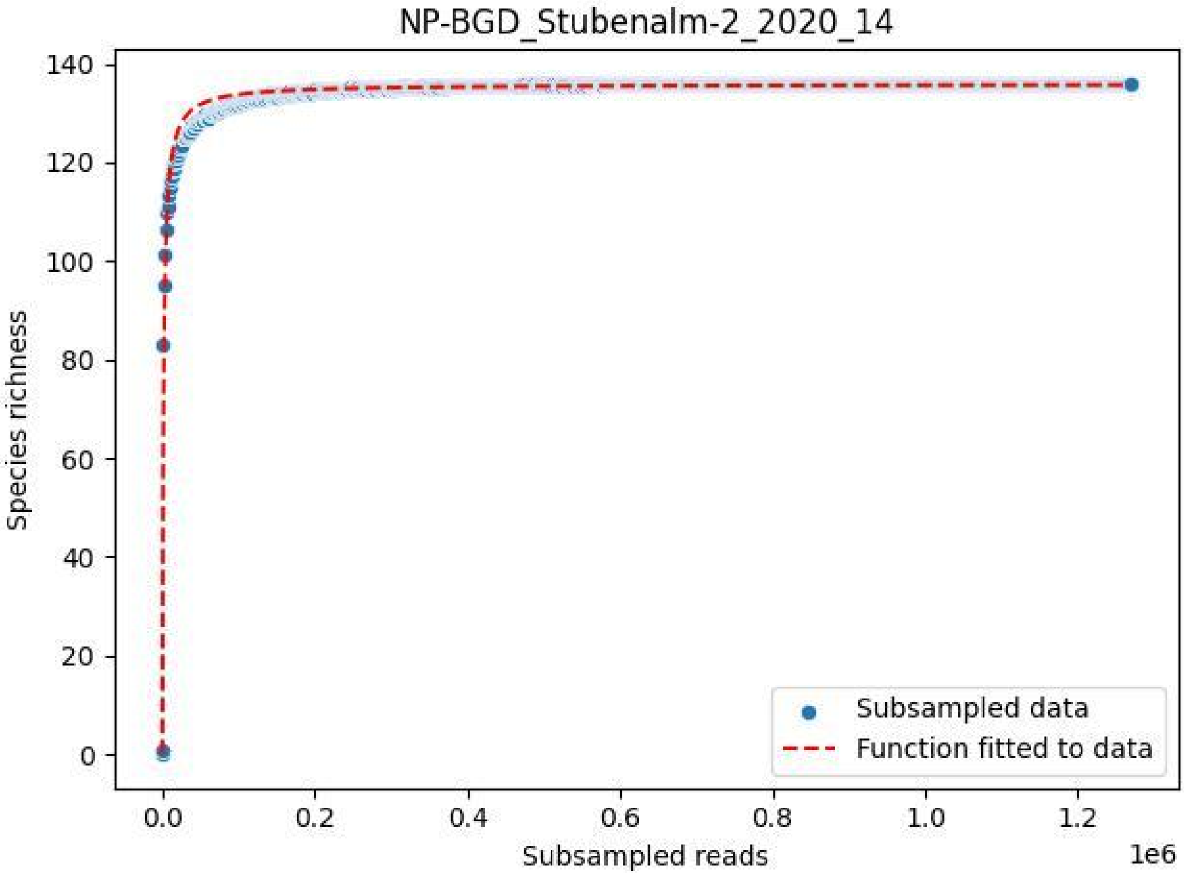
NP-BGD_Stubenalm-2_2020_14
Species richness
0
20
40
60
80
100
120
140
0.0
0.2
0.4
0.6
0.8
1.0
1.2
1e6
Subsampled reads
Subsampled data
Function fitted to data

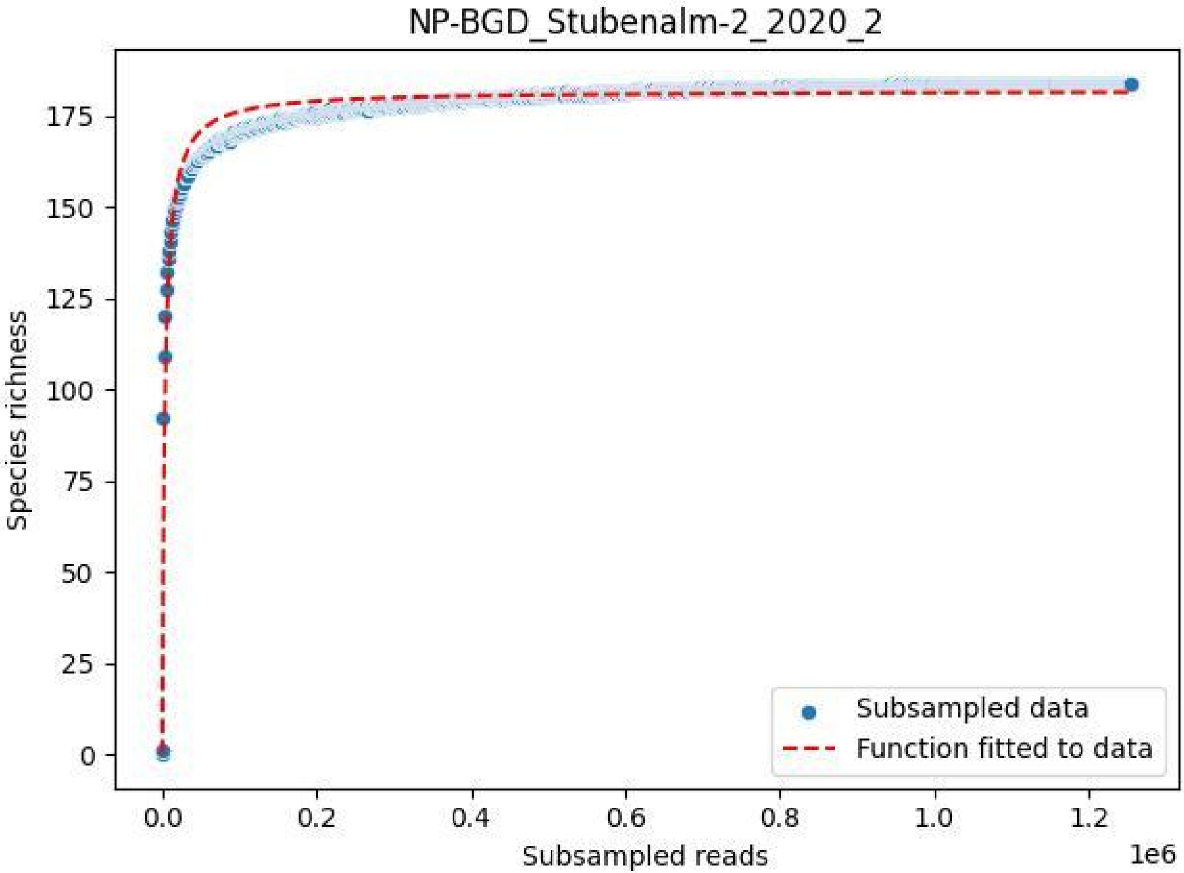

NP-BGD_Stubenalm-2_2020_2
Species richness
Subsampled reads
1e6
0.0
0.2
0.4
0.6
0.8
1.0
1.2
0
25
50
75
100
125
150
175
Subsampled data
Function fitted to data

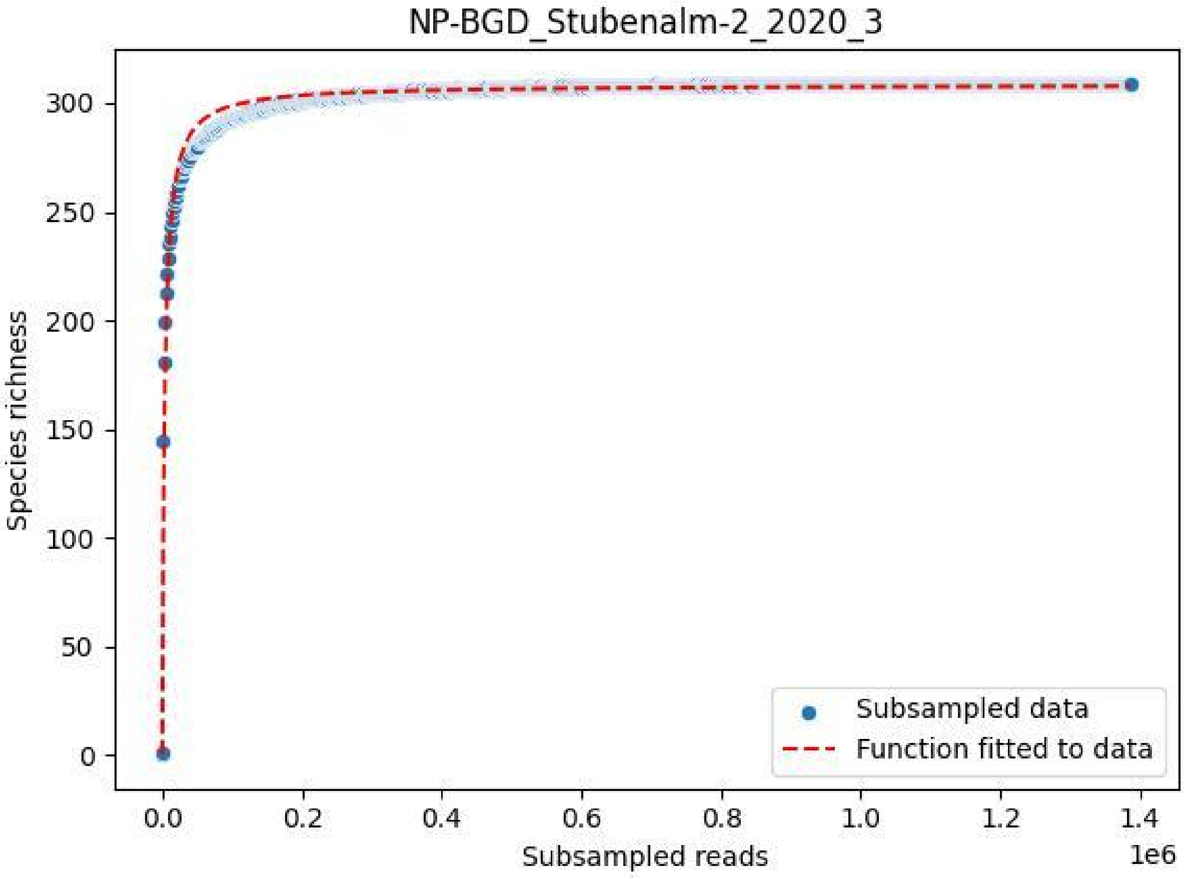
NP-BGD_Stubenalm-2_2020_3
Species richness
0
50
100
150
200
250
300
0.0
0.2
0.4
0.6
0.8
1.0
1.2
1.4
1e6
Subsampled reads
Subsampled data
Function fitted to data

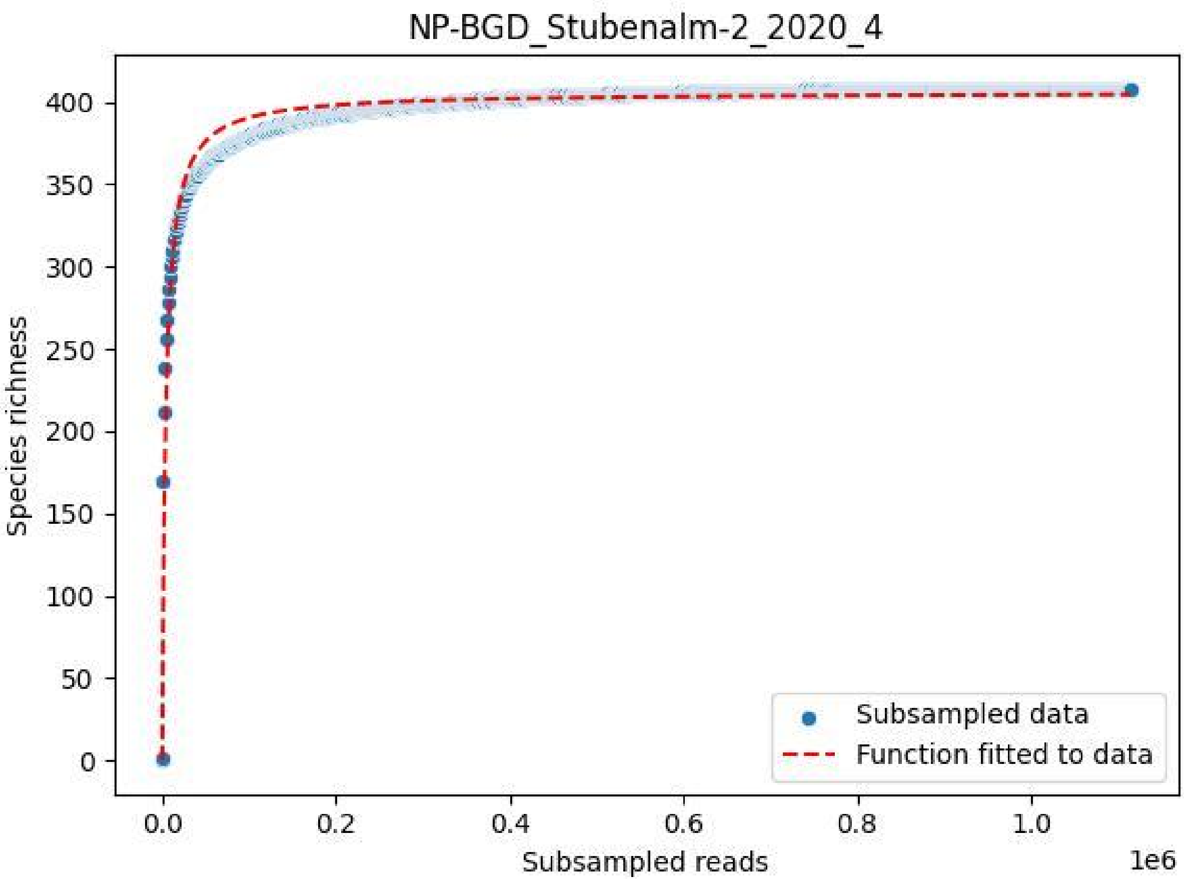

NP-BGD_Stubenalm-2_2020_4
Species richness
Subsampled reads
1e6
0
50
100
150
200
250
300
350
400
0.0
0.2
0.4
0.6
0.8
1.0
Subsampled data
Function fitted to data

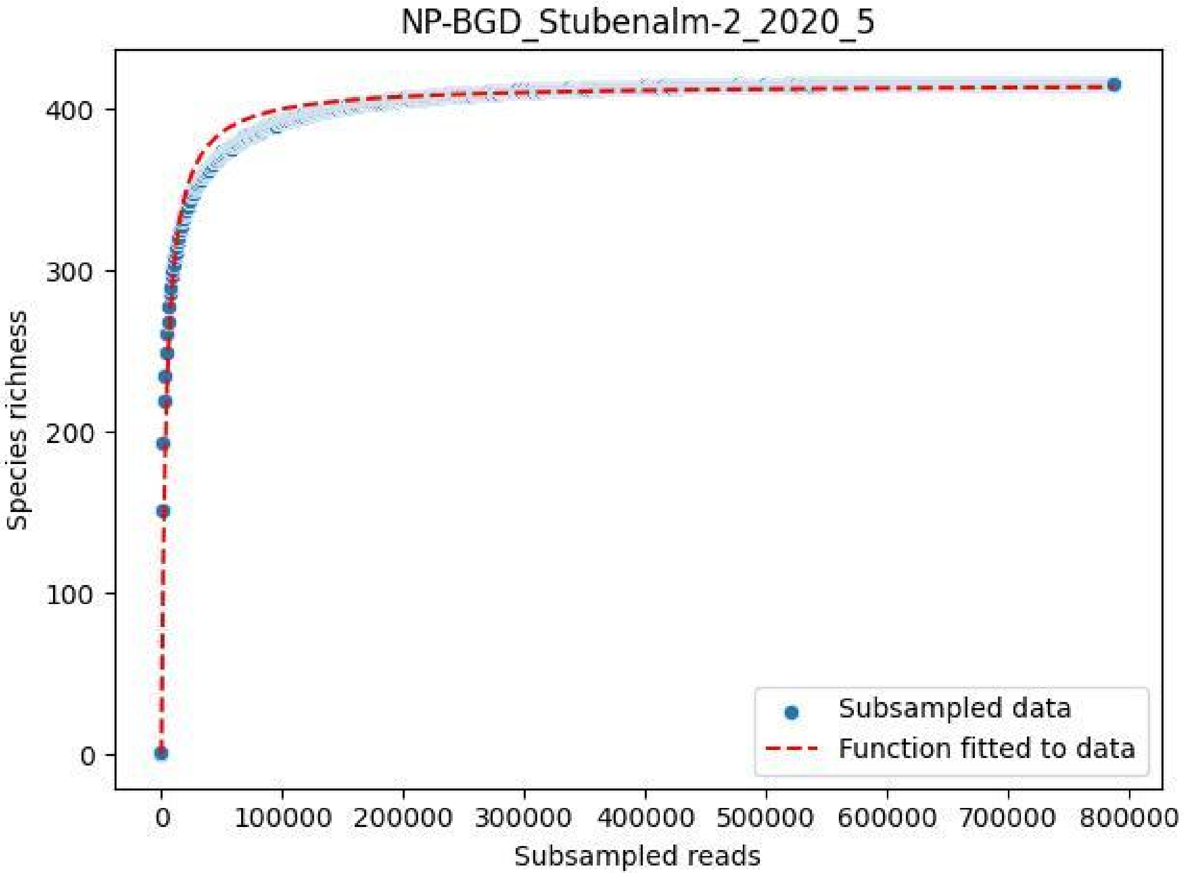

NP-BGD_Stubenalm-2_2020_5
Species richness
0
100
200
300
400
0
100000
200000
300000
400000
500000
600000
700000
800000
Subsampled reads
Subsampled data
Function fitted to data

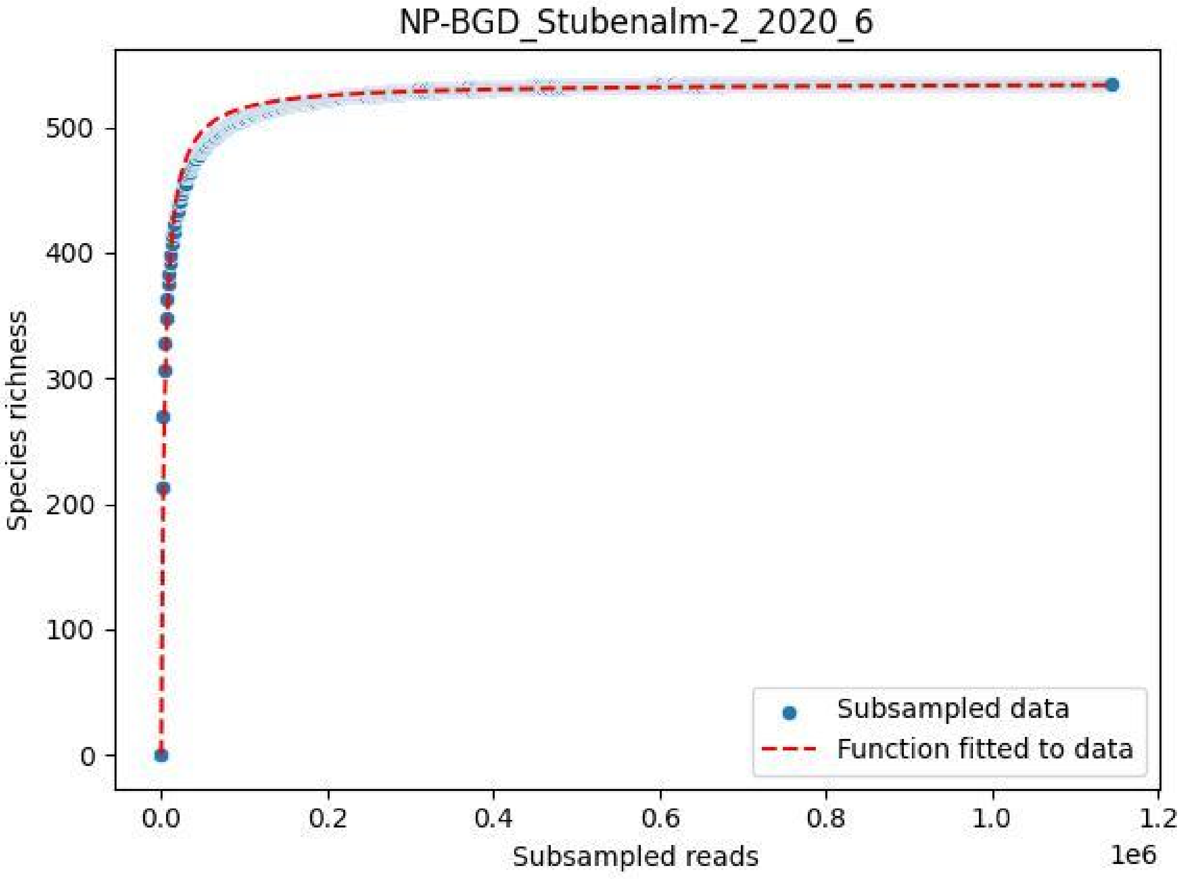
NP-BGD_Stubenalm-2_2020_6
Species richness
Subsampled reads
1e6
Subsampled data
Function fitted to data
0
100
200
300
400
500
0.0
0.2
0.4
0.6
0.8
1.0
1.2

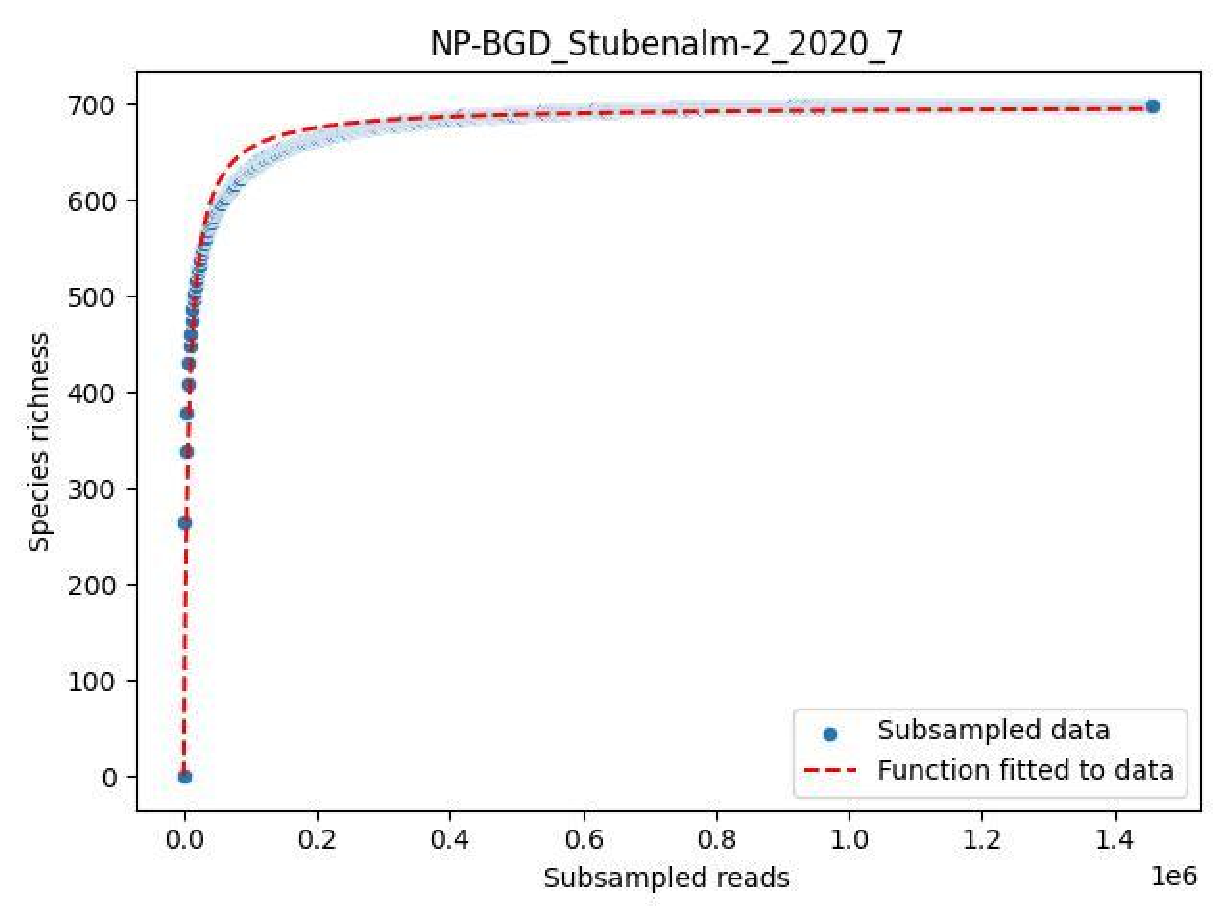
NP-BGD_Stubenalm-2_2020_7
Species richness
0
100
200
300
400
500
600
700
0.0
0.2
0.4
0.6
0.8
1.0
1.2
1.4
1e6
Subsampled reads
Subsampled data
Function fitted to data

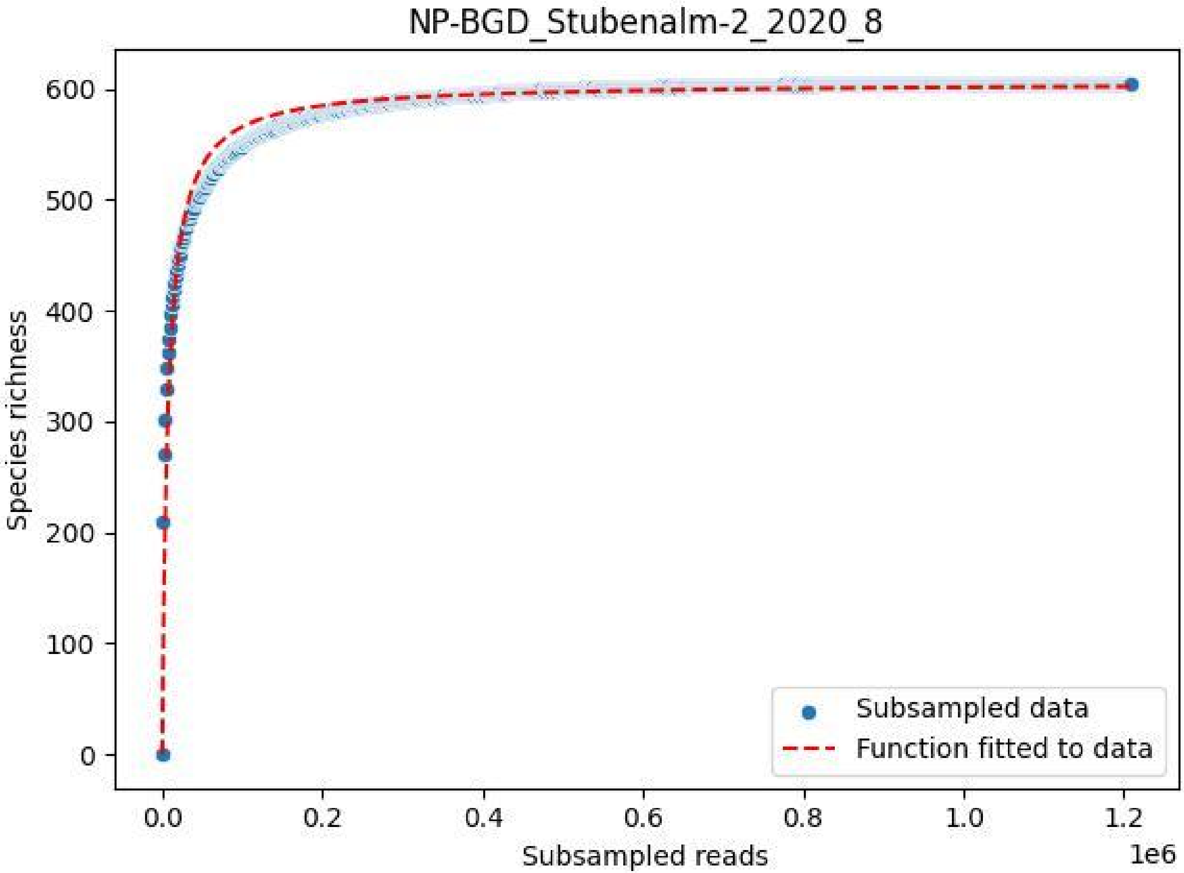
NP-BGD_Stubenalm-2_2020_8
Species richness
Subsampled reads
1e6
600
500
400
300
200
100
0
0.0
0.2
0.4
0.6
0.8
1.0
1.2
Subsampled data
Function fitted to data

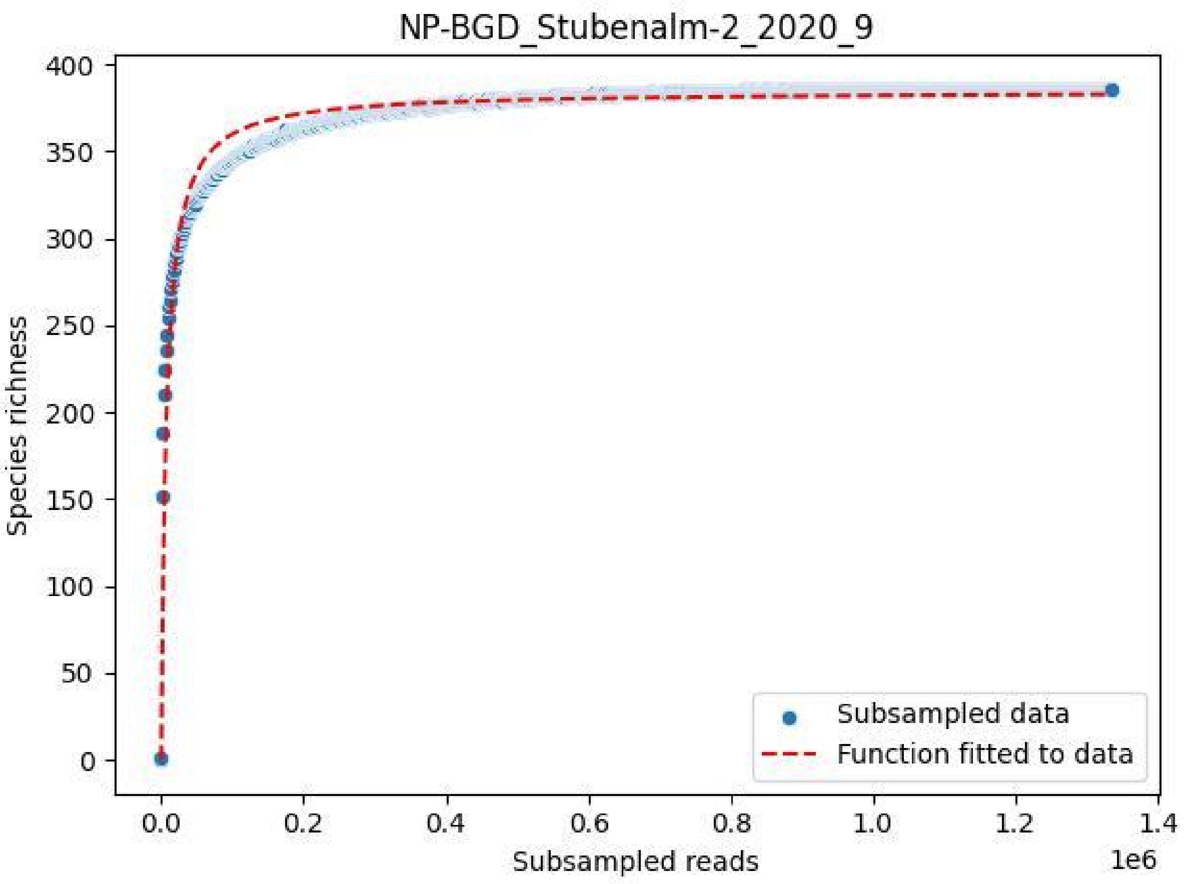

NP-BGD_Stubenalm-2_2020_9
Species richness
Subsampled reads
1e6
0
50
100
150
200
250
300
350
400
0.0
0.2
0.4
0.6
0.8
1.0
1.2
1.4
Subsampled data
Function fitted to data

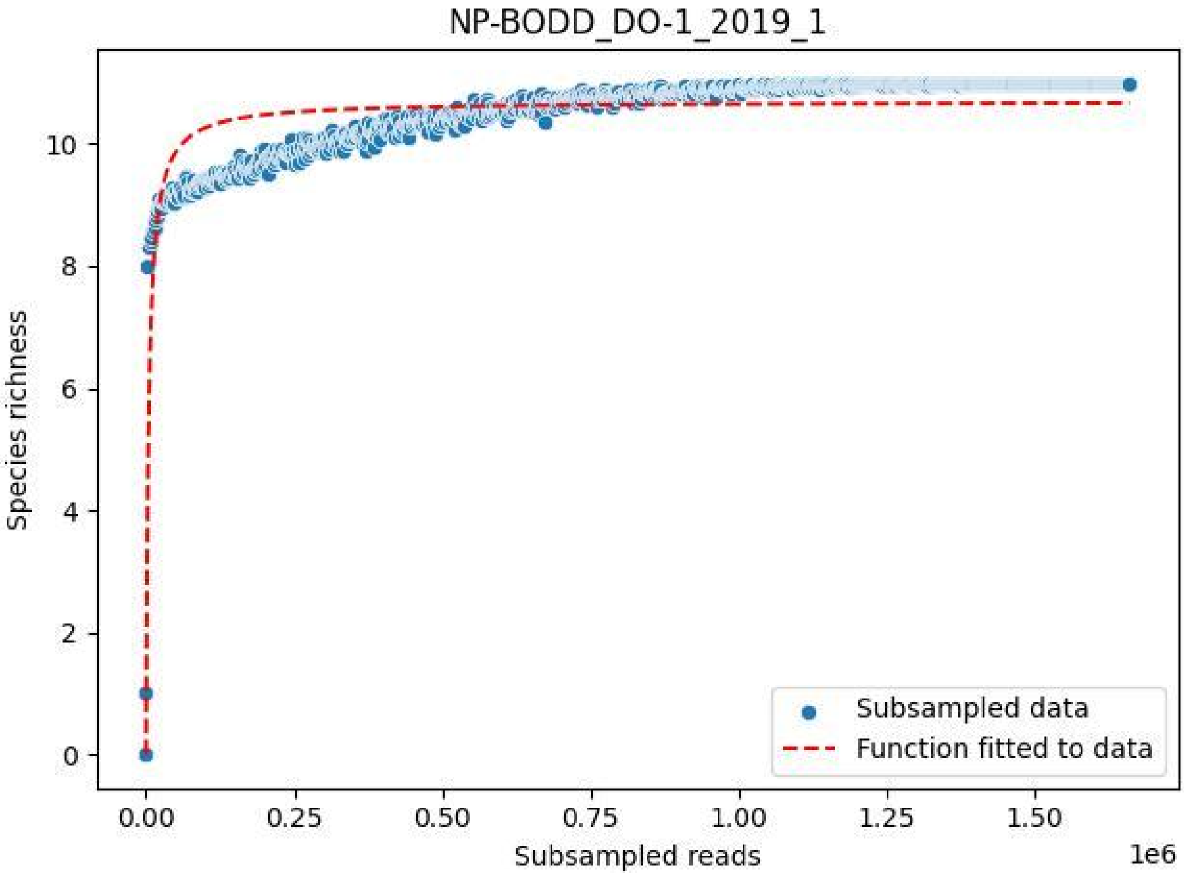
NP-BODD_DO-1_2019_1
Species richness
Subsampled reads
1e6
0
2
4
6
8
10
0.00
0.25
0.50
0.75
1.00
1.25
1.50
Subsampled data
Function fitted to data

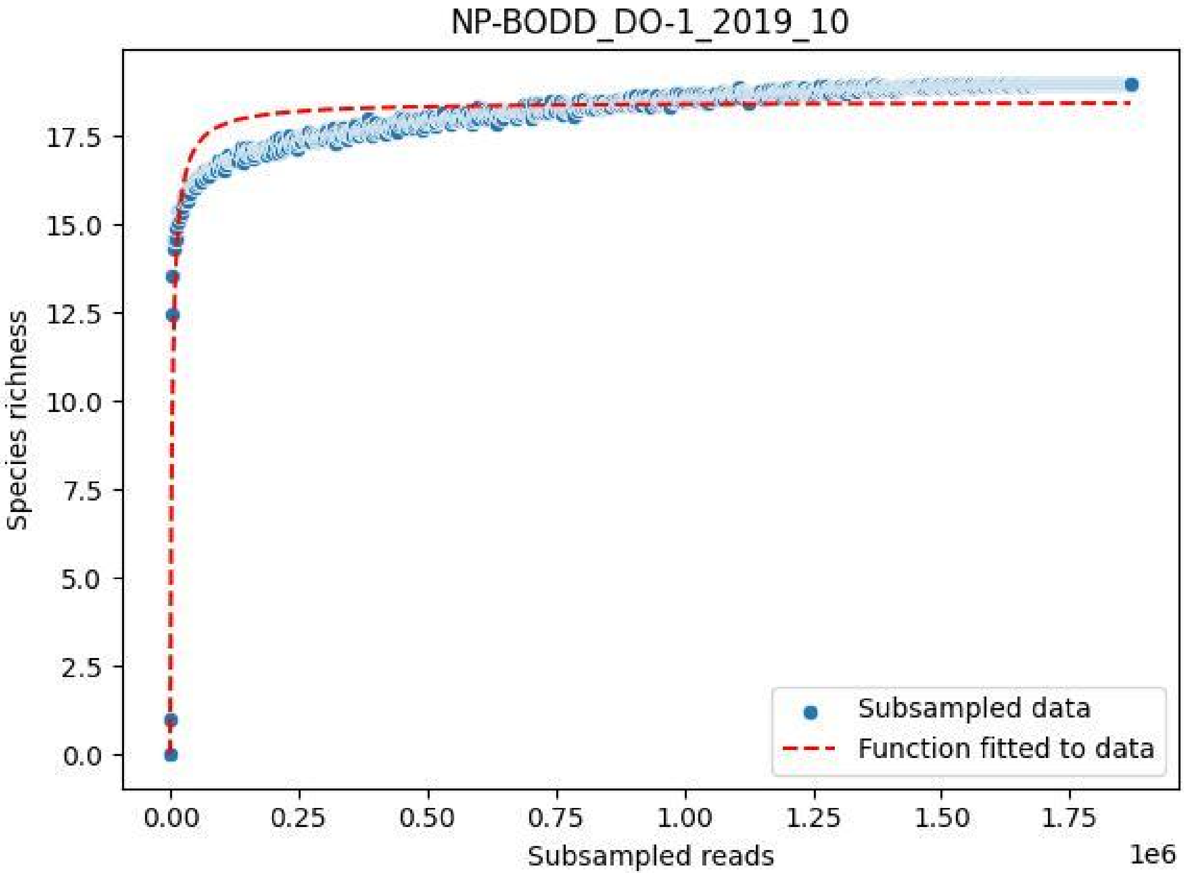
NP-BODD_DO-1_2019_10
Species richness
0.0
2.5
5.0
7.5
10.0
12.5
15.0
17.5
0.00
0.25
0.50
0.75
1.00
1.25
1.50
1.75
1e6
Subsampled reads
Subsampled data
Function fitted to data

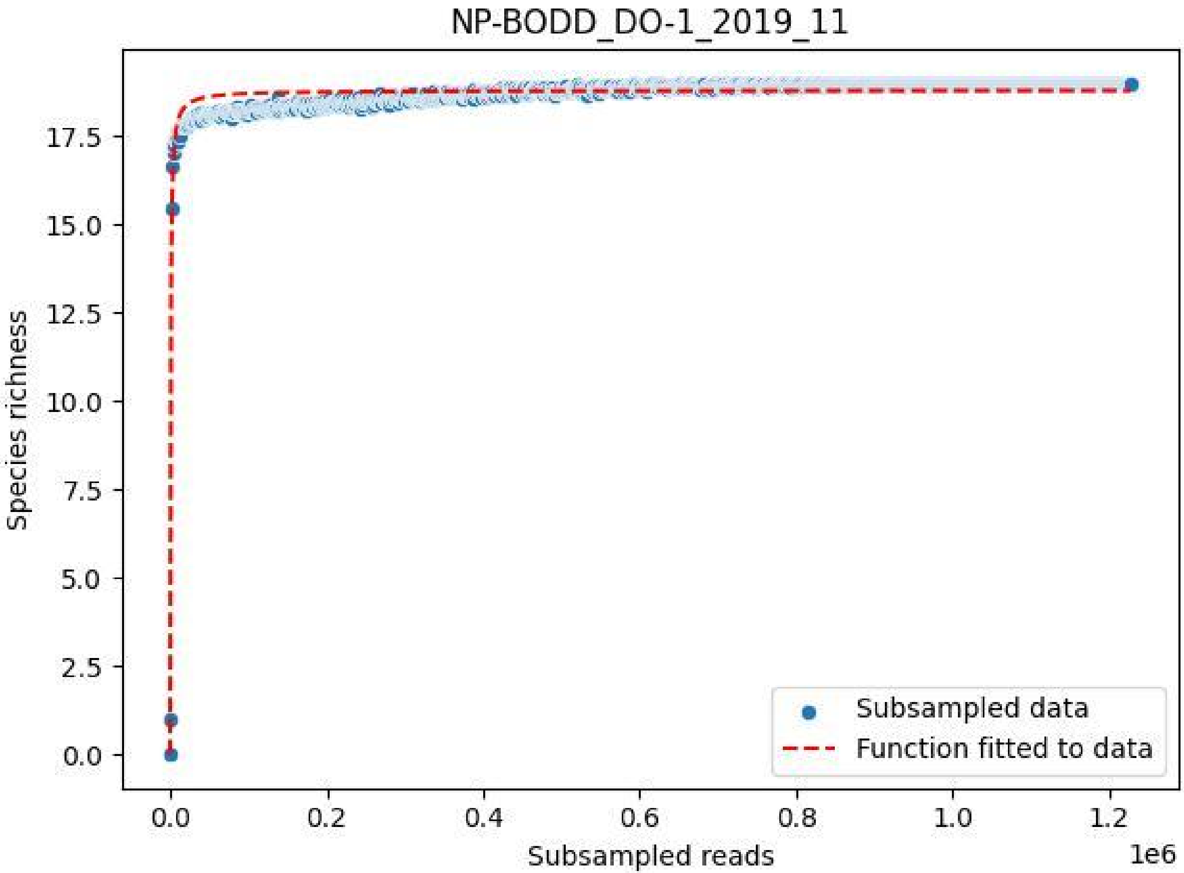

NP-BODD_DO-1_2019_11
Species richness
Subsampled reads
1e6
0.0
2.5
5.0
7.5
10.0
12.5
15.0
17.5
0.0
0.2
0.4
0.6
0.8
1.0
1.2
Subsampled data
Function fitted to data

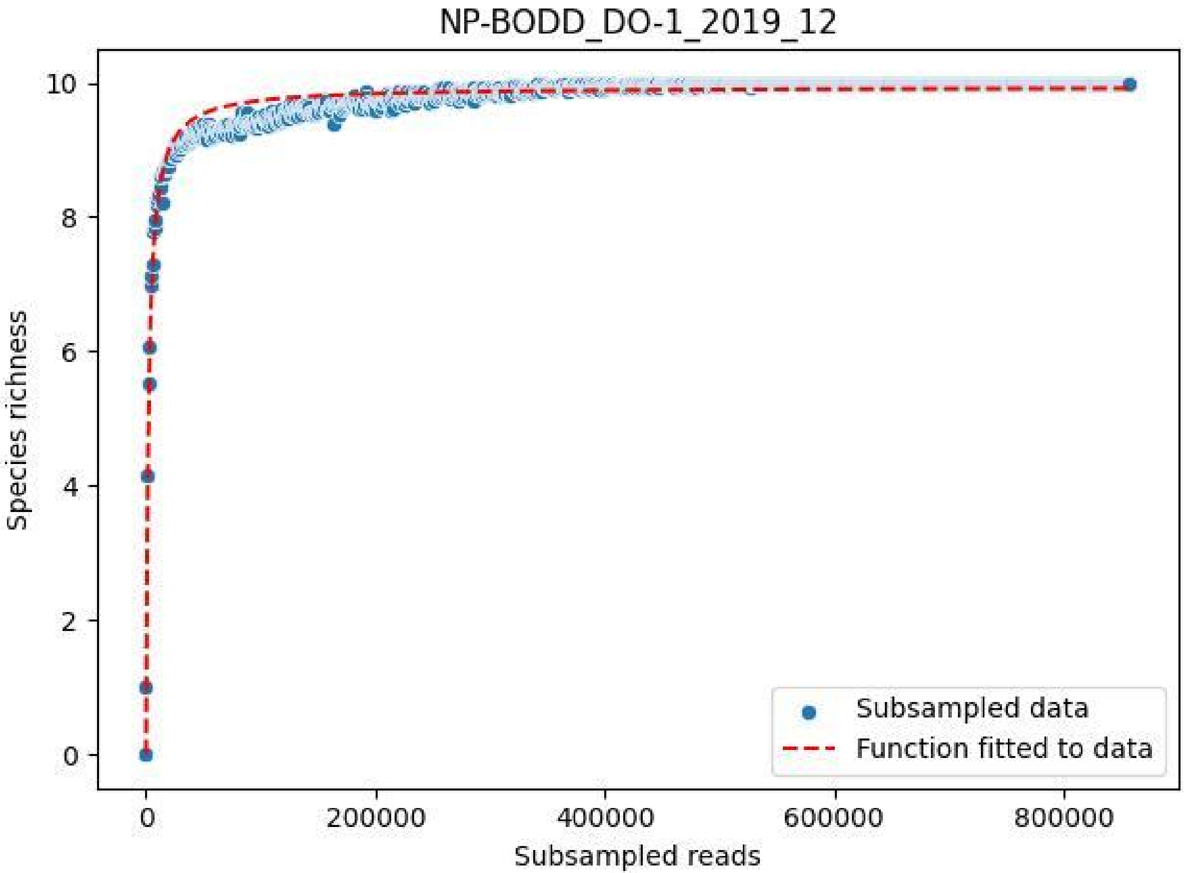
NP-BODD_DO-1_2019_12
Species richness
Subsampled reads
0
2
4
6
8
10
0
200000
400000
600000
800000
Subsampled data
Function fitted to data

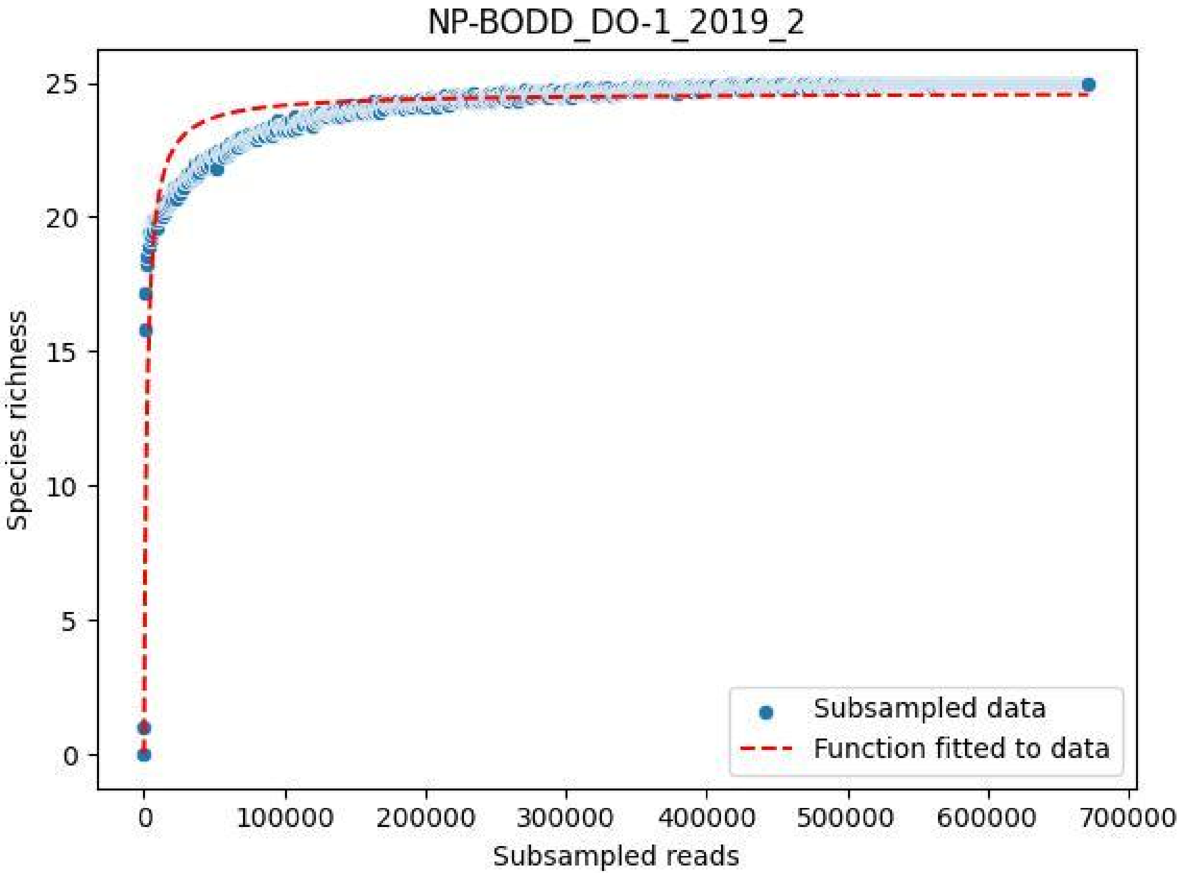
NP-BODD_DO-1_2019_2
Species richness
Subsampled reads
0
5
10
15
20
25
0
100000
200000
300000
400000
500000
600000
700000
Subsampled data
Function fitted to data

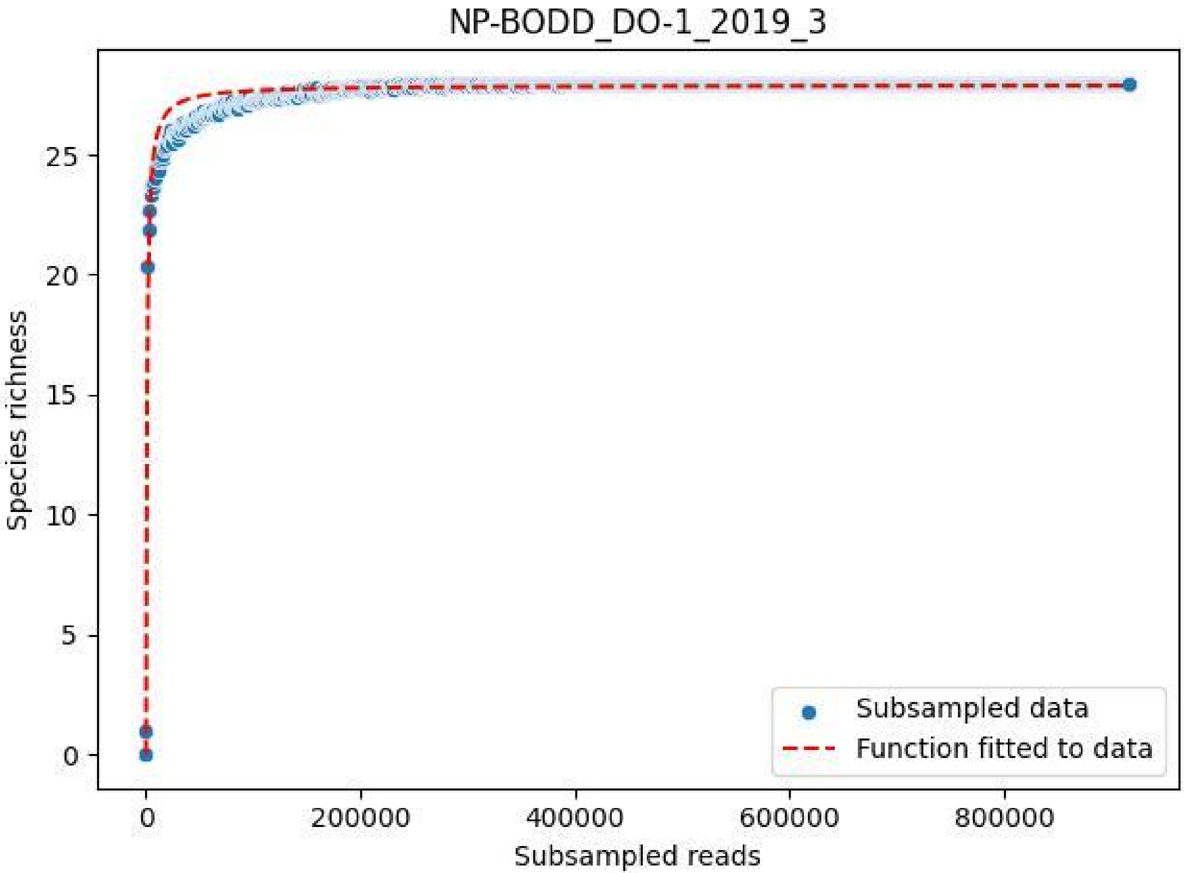
NP-BODD_DO-1_2019_3
Species richness
Subsampled reads
0
5
10
15
20
25
200000
400000
600000
800000
Subsampled data
Function fitted to data

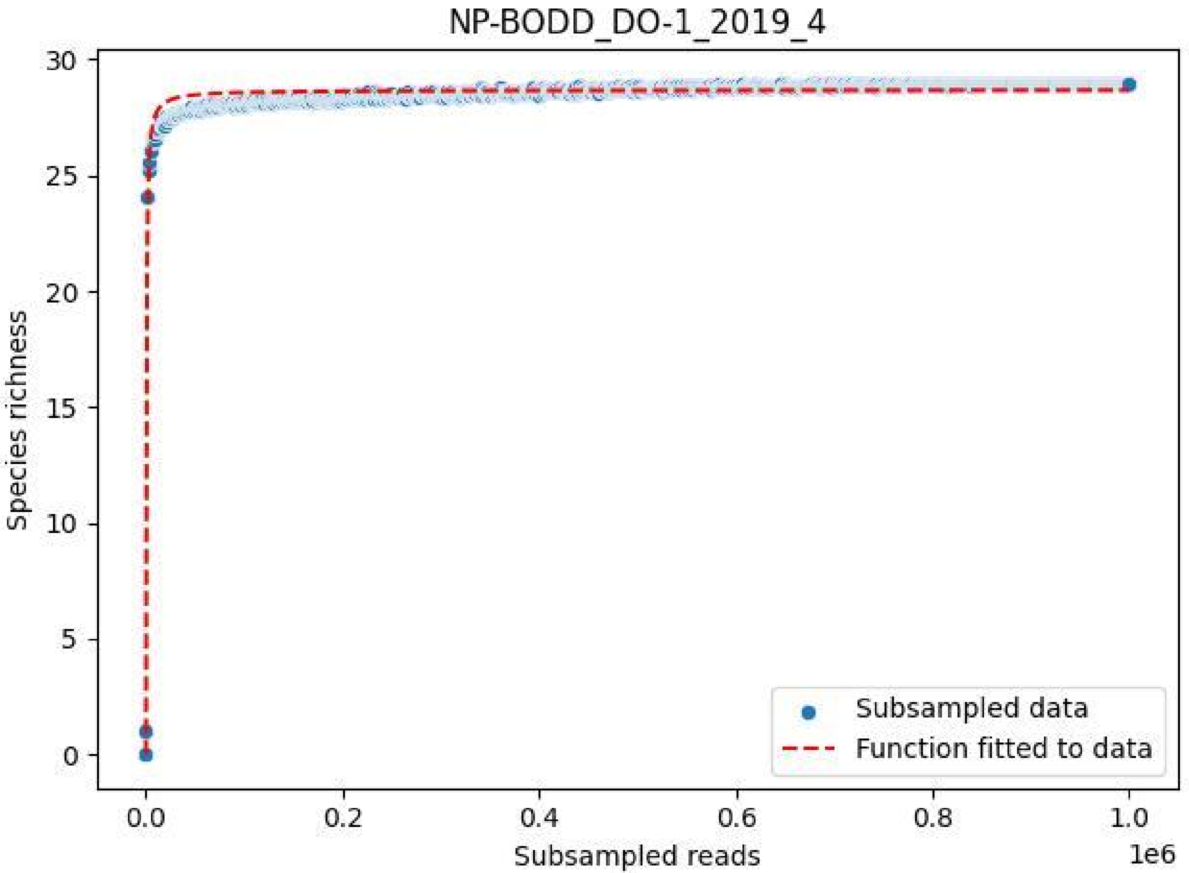
NP-BODD_DO-1_2019_4
Species richness
Subsampled reads
1e6
0
5
10
15
20
25
30
0.0
0.2
0.4
0.6
0.8
1.0
Subsampled data
Function fitted to data

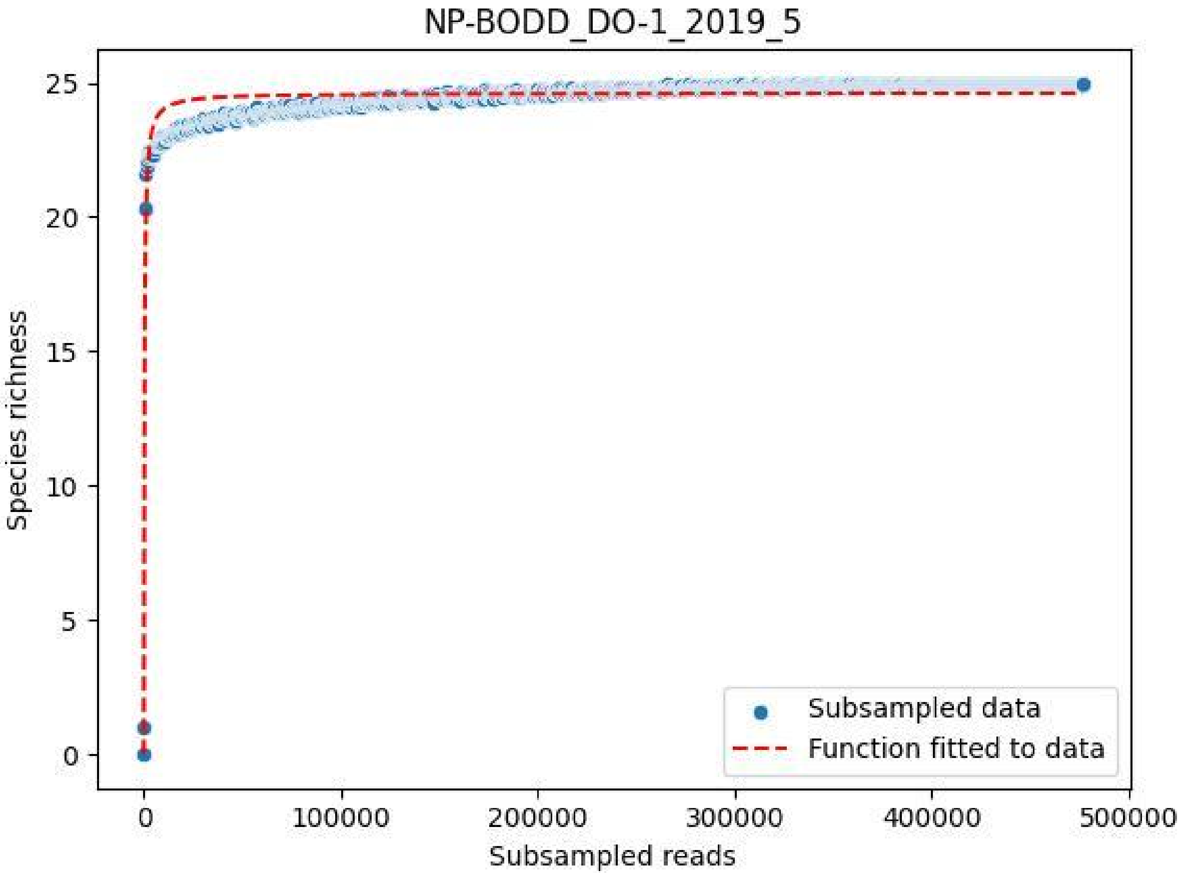
NP-BODD_DO-1_2019_5
Species richness
Subsampled reads
0
5
10
15
20
25
0
100000
200000
300000
400000
500000
Subsampled data
Function fitted to data

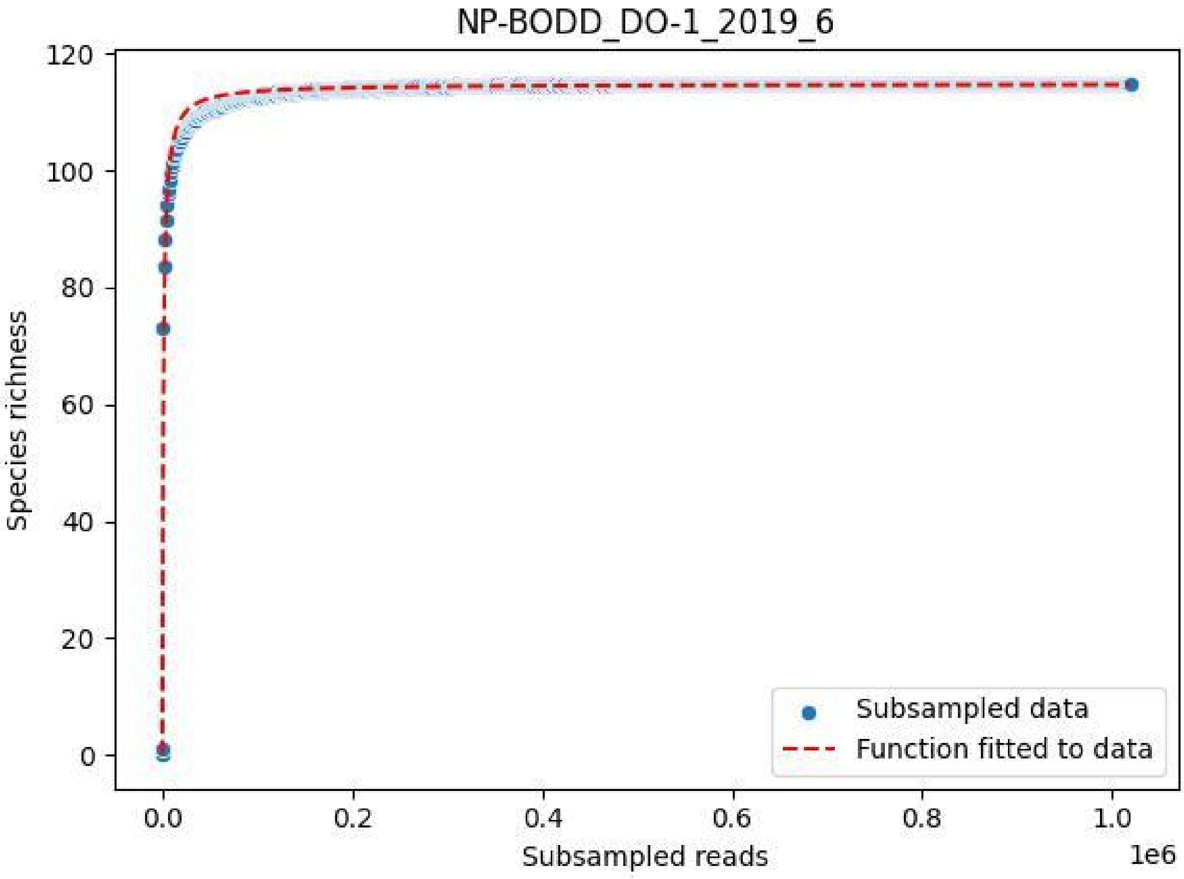
NP-BODD_DO-1_2019_6
Species richness
Subsampled reads
1e6
0
20
40
60
80
100
120
0.0
0.2
0.4
0.6
0.8
1.0
Subsampled data
Function fitted to data

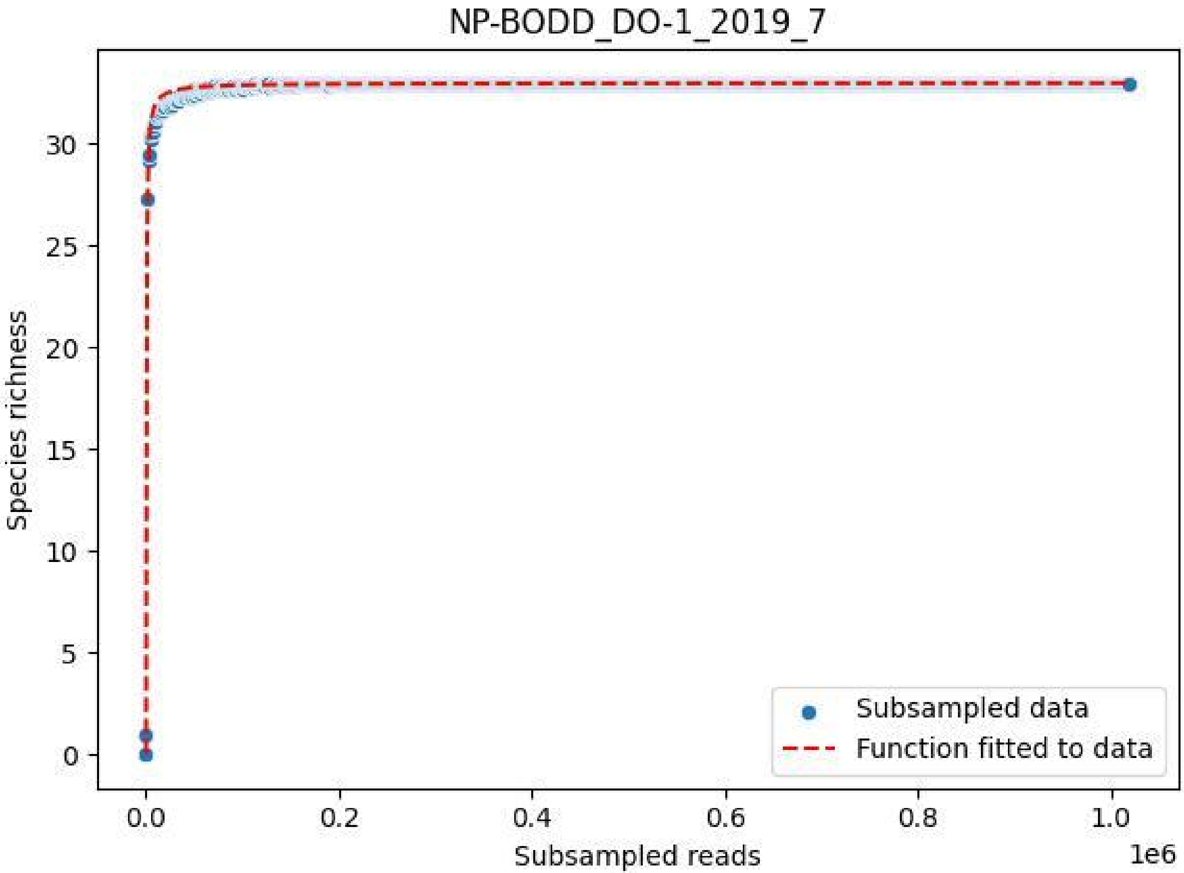
NP-BODD_DO-1_2019_7
Species richness
Subsampled reads
1e6
0
5
10
15
20
25
30
0.0
0.2
0.4
0.6
0.8
1.0
Subsampled data
Function fitted to data

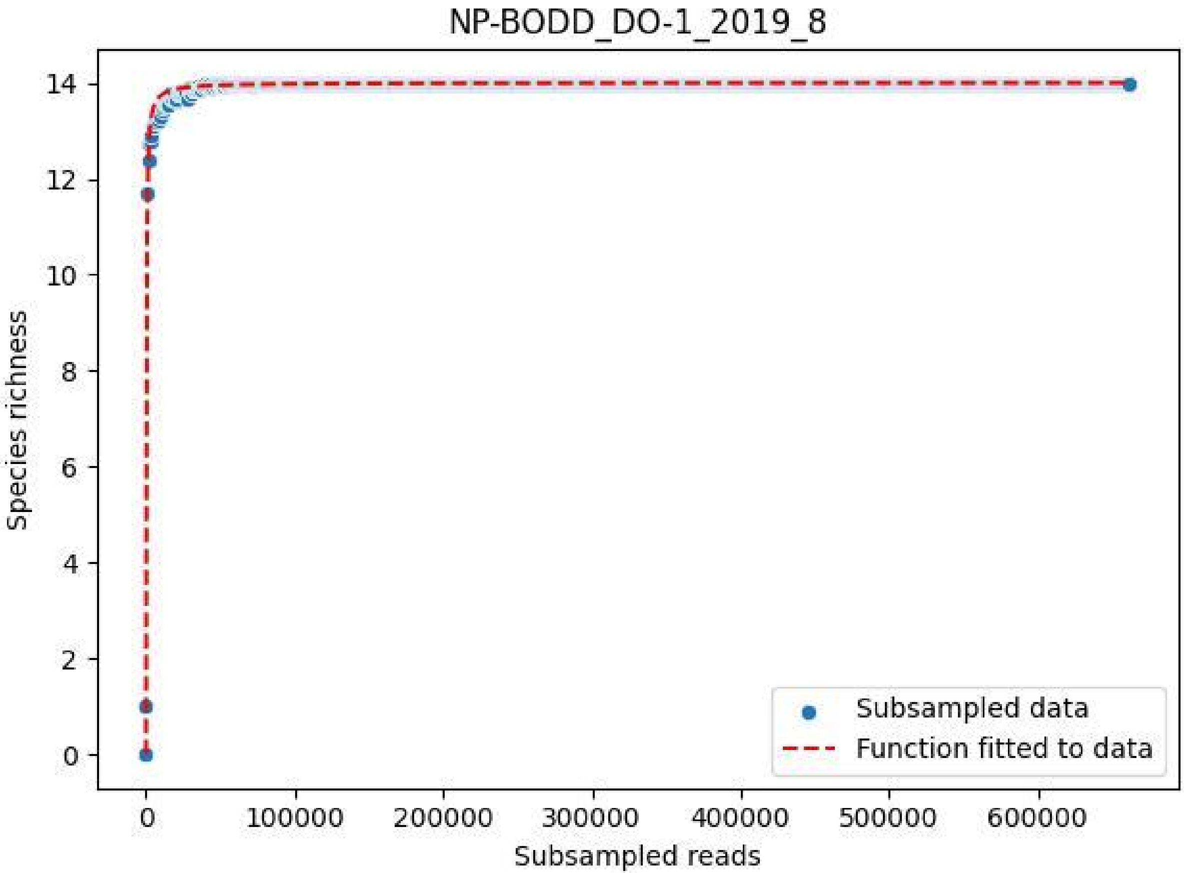
NP-BODD_DO-1_2019_8
Species richness
0
2
4
6
8
10
12
14
Subsampled reads
0
100000
200000
300000
400000
500000
600000
Subsampled data
Function fitted to data

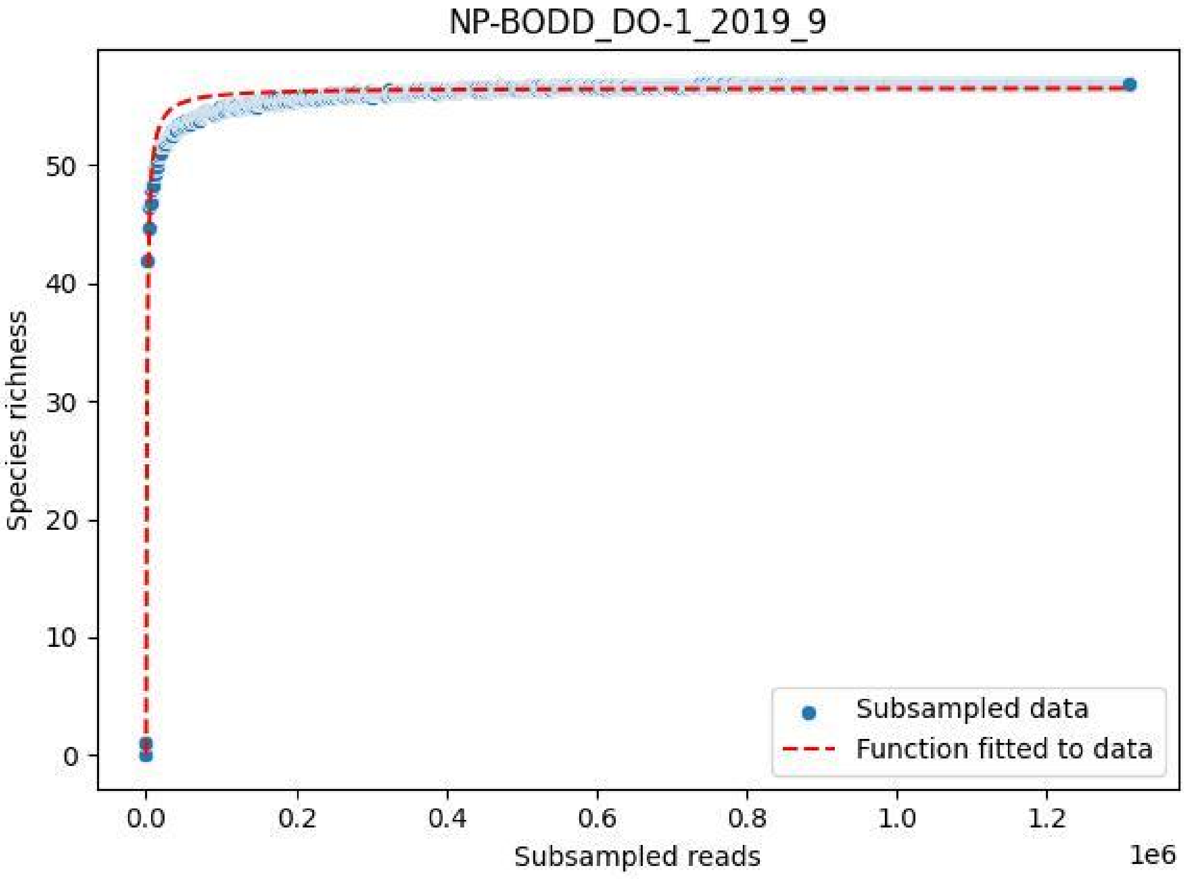

NP-BODD_DO-1_2019_9
Species richness
Subsampled reads
1e6
0
10
20
30
40
50
0.0
0.2
0.4
0.6
0.8
1.0
1.2
Subsampled data
Function fitted to data

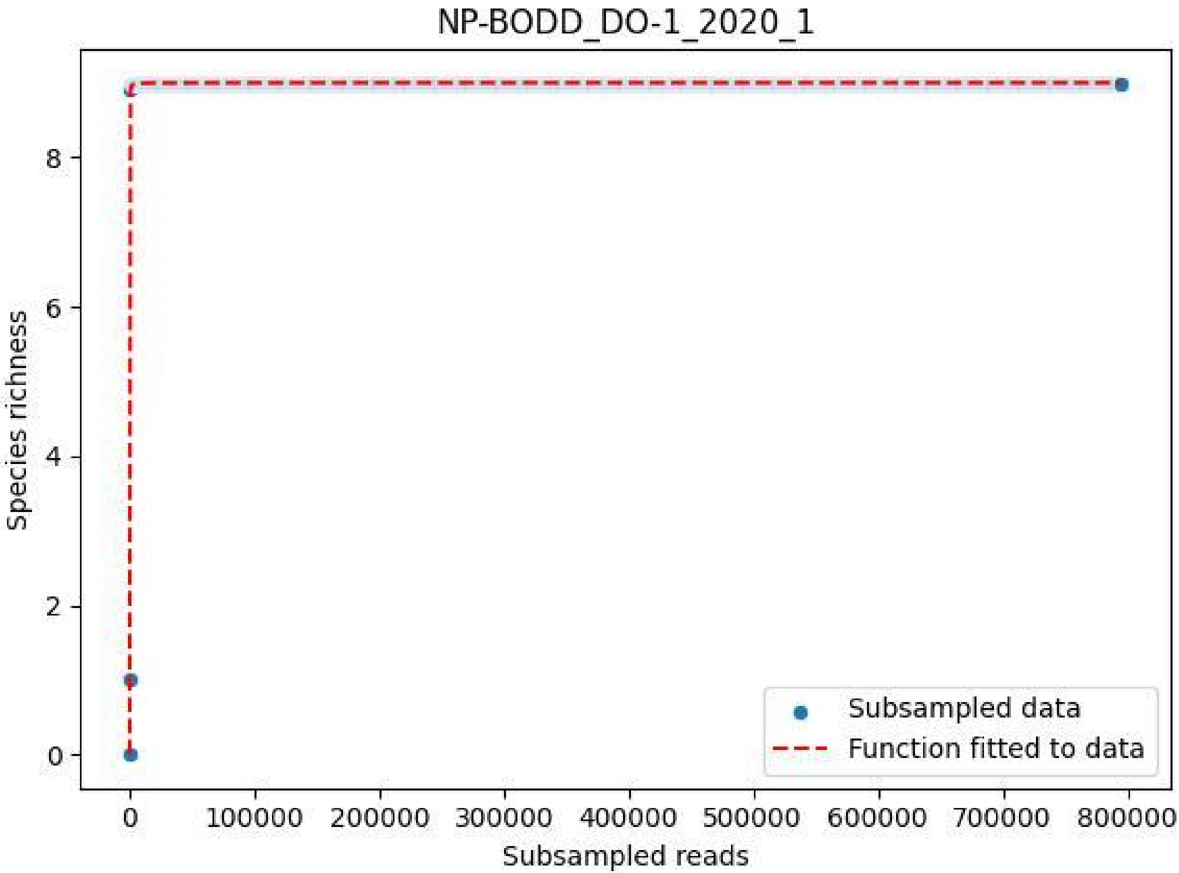
NP-BODD_DO-1_2020_1
Species richness
Subsampled reads
0
2
4
6
8
0
100000
200000
300000
400000
500000
600000
700000
800000
Subsampled data
Function fitted to data

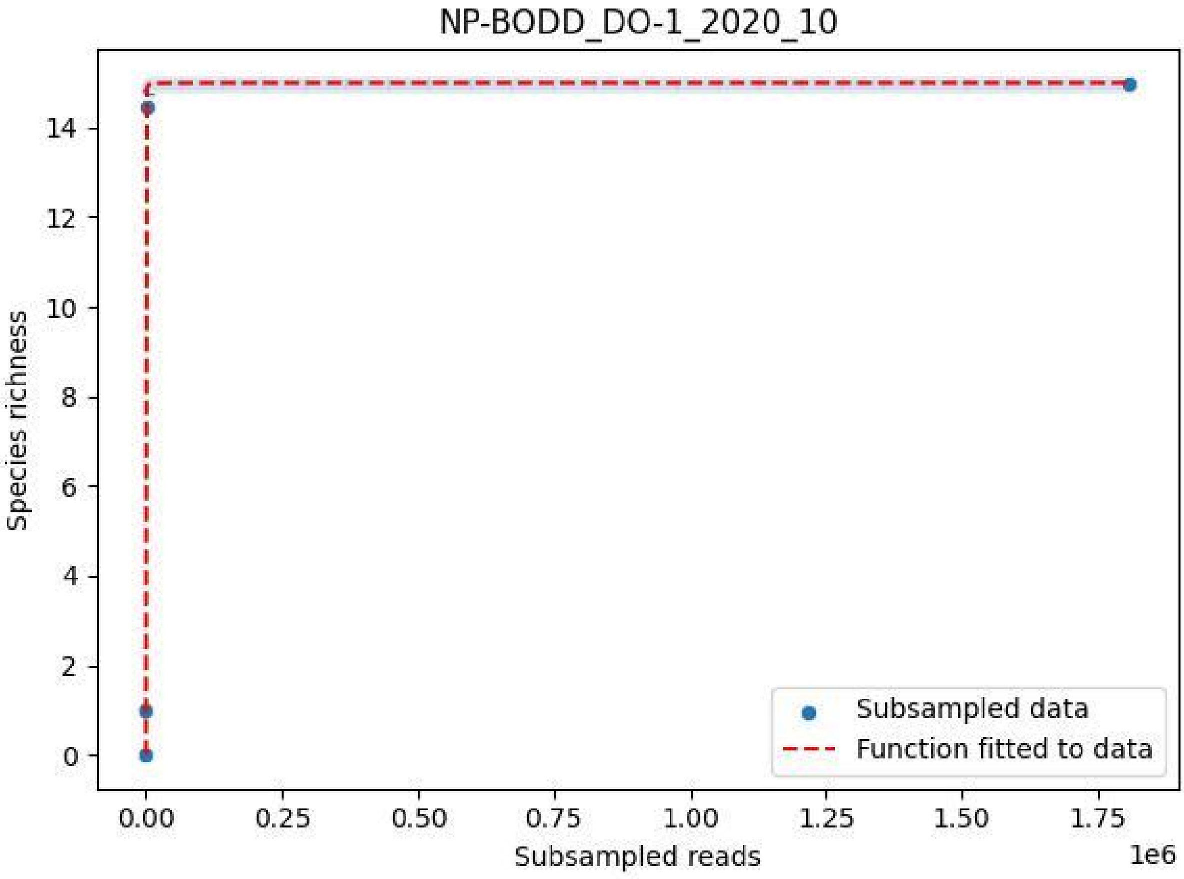
NP-BODD_DO-1_2020_10
Species richness
0
2
4
6
8
10
12
14
Subsampled data
Function fitted to data
0.00
0.25
0.50
0.75
1.00
1.25
1.50
1.75
1e6
Subsampled reads

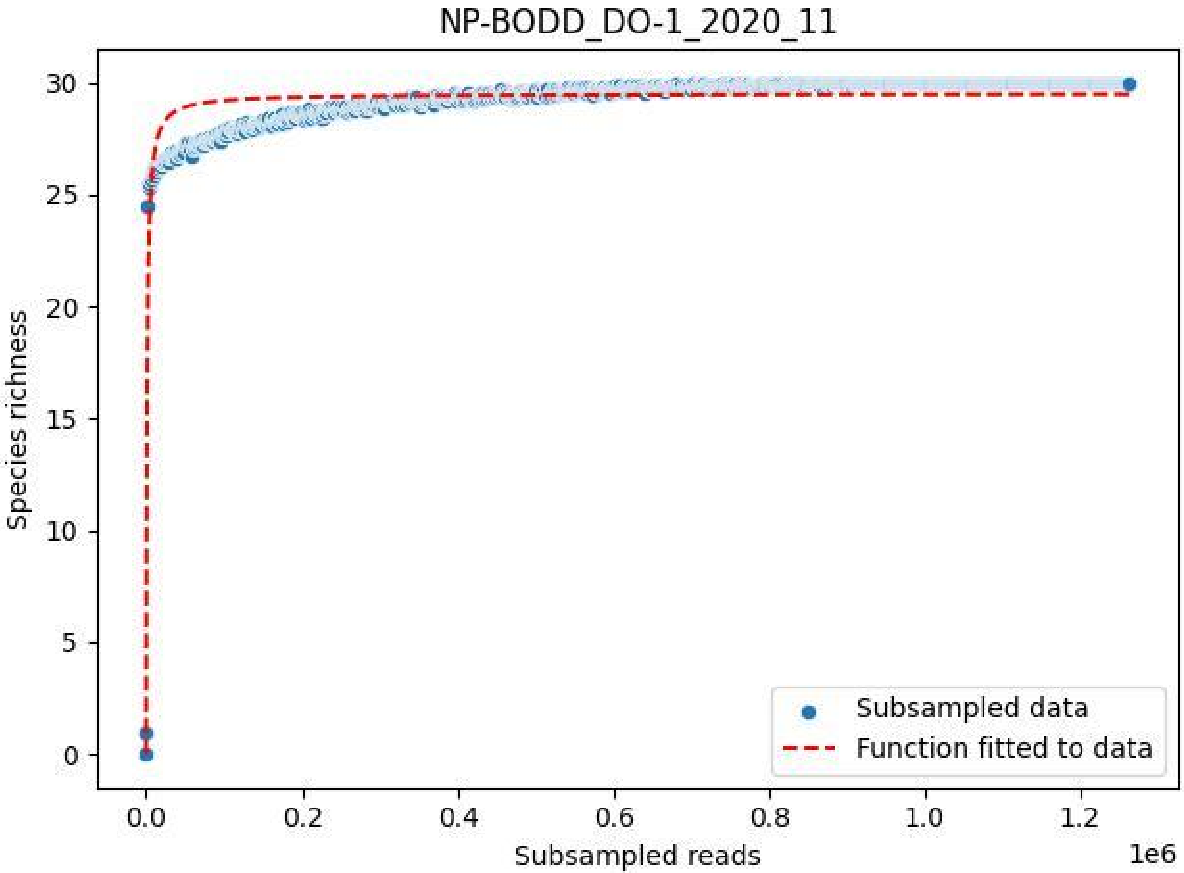
NP-BODD_DO-1_2020_11
Species richness
Subsampled reads
1e6
0.0
0.2
0.4
0.6
0.8
1.0
1.2
0
5
10
15
20
25
30
Subsampled data
Function fitted to data

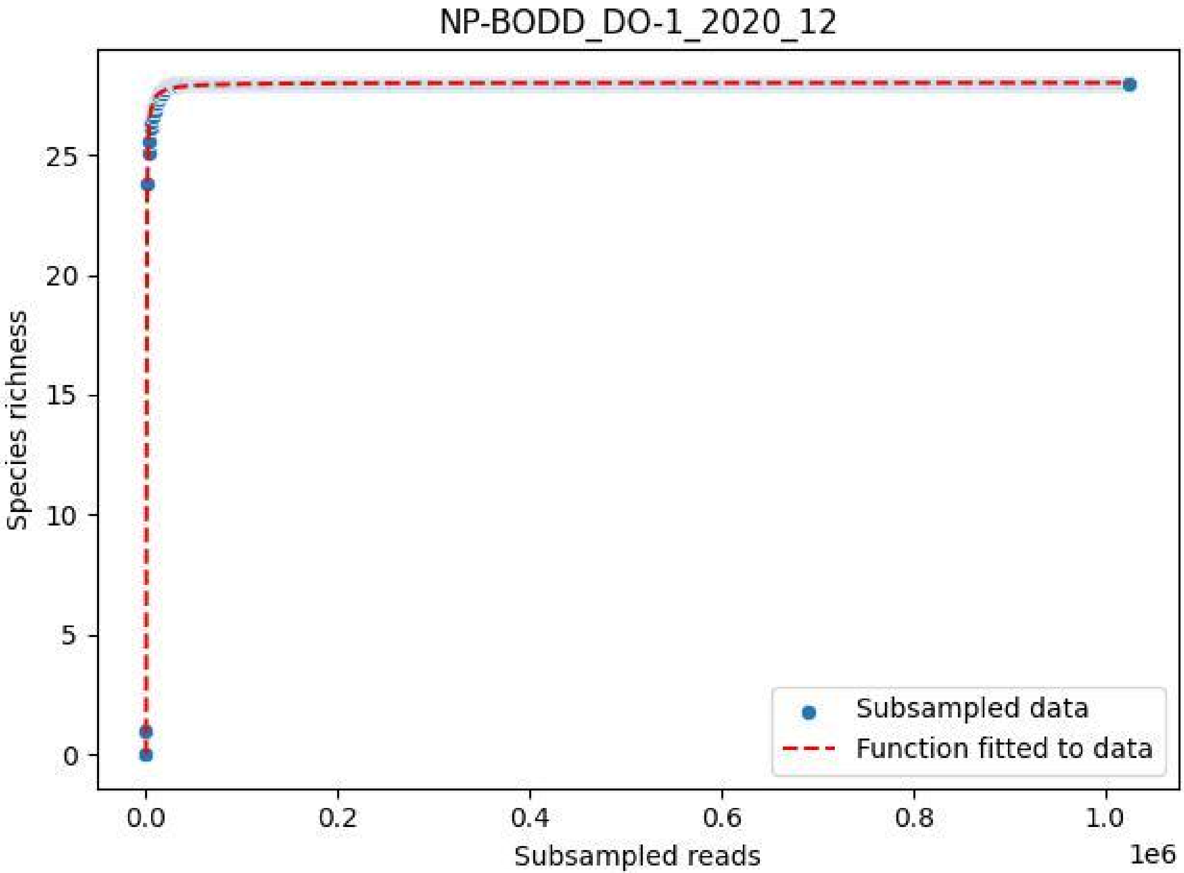
NP-BODD_DO-1_2020_12
Species richness
Subsampled reads
0
5
10
15
20
25
0.0
0.2
0.4
0.6
0.8
1.0
1e6
Subsampled data
Function fitted to data

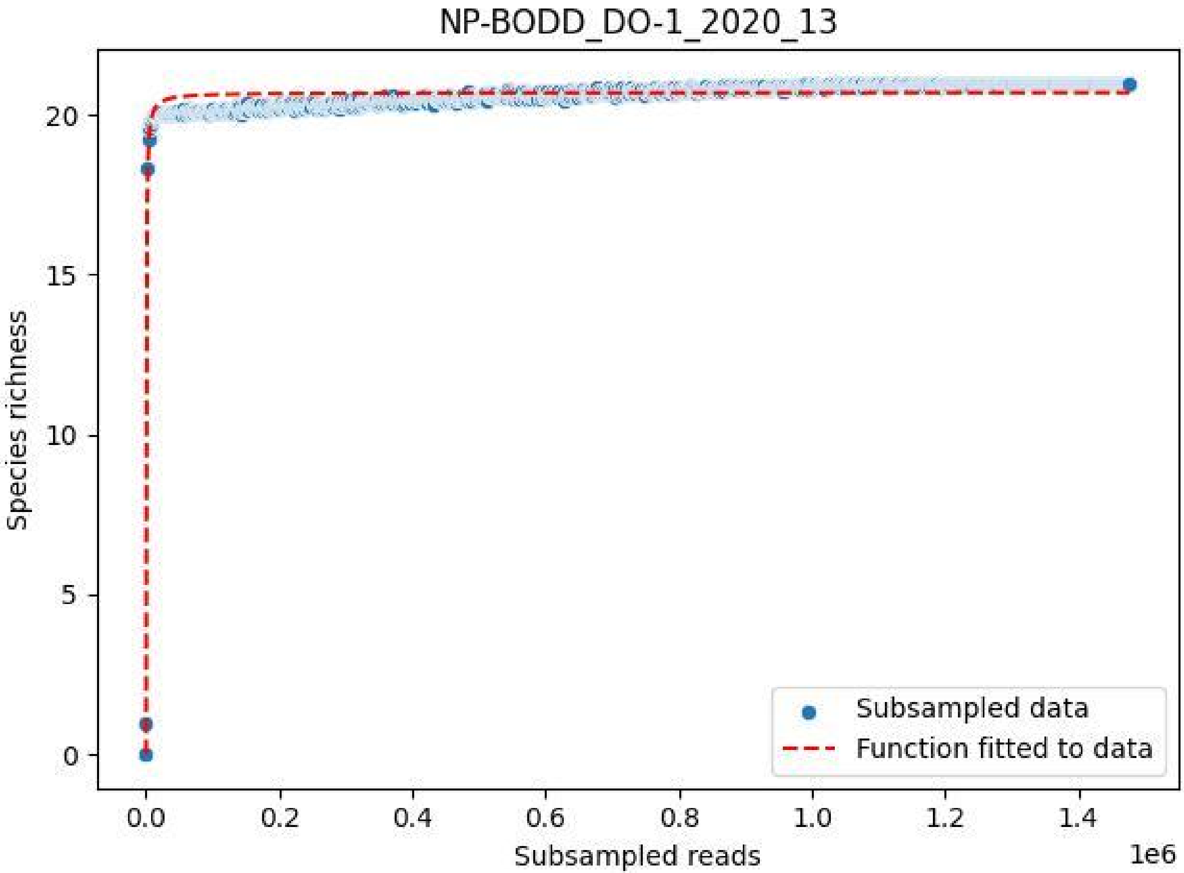
NP-BODD_DO-1_2020_13
Species richness
Subsampled reads
1e6
0.0
0.2
0.4
0.6
0.8
1.0
1.2
1.4
0
5
10
15
20
Subsampled data
Function fitted to data

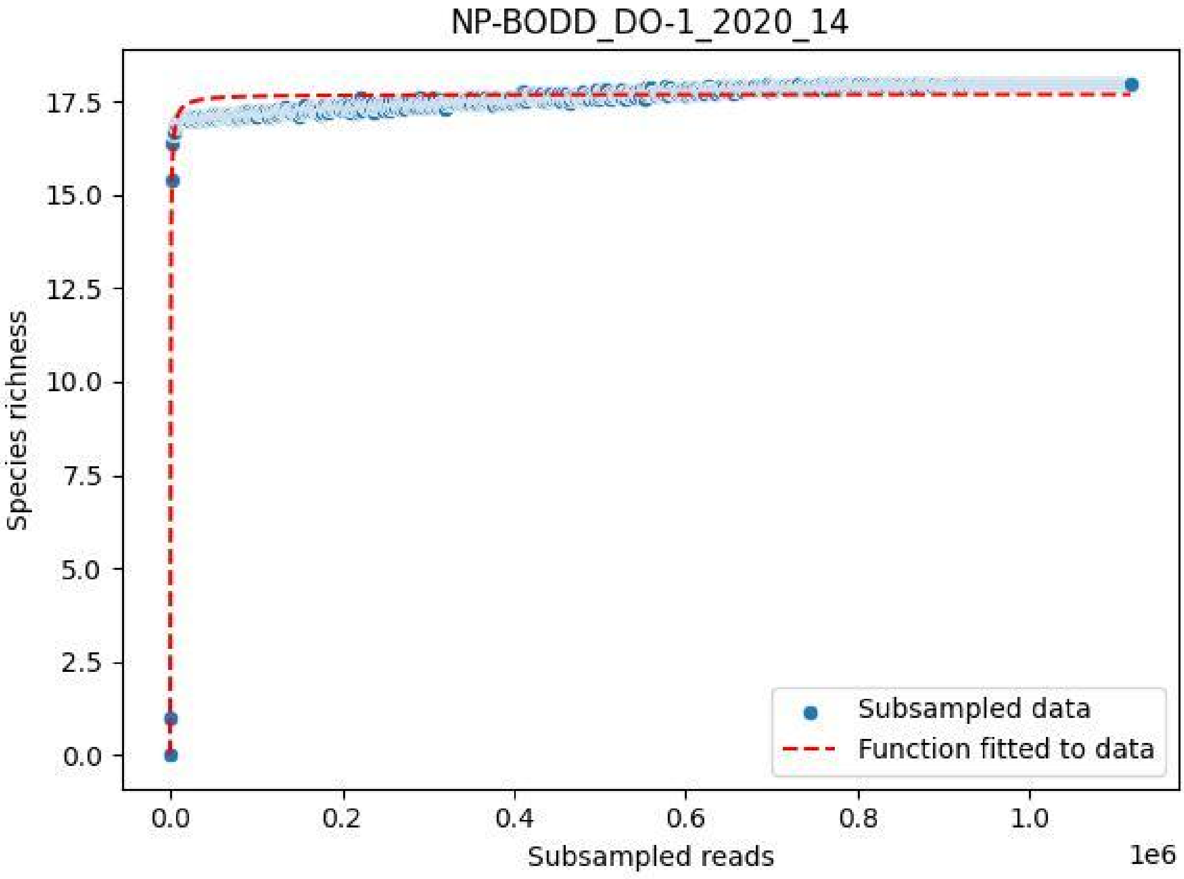
NP-BODD_DO-1_2020_14
Species richness
17.5
15.0
12.5
10.0
7.5
5.0
2.5
0.0
0.0
0.2
0.4
0.6
0.8
1.0
1e6
Subsampled reads
Subsampled data
Function fitted to data

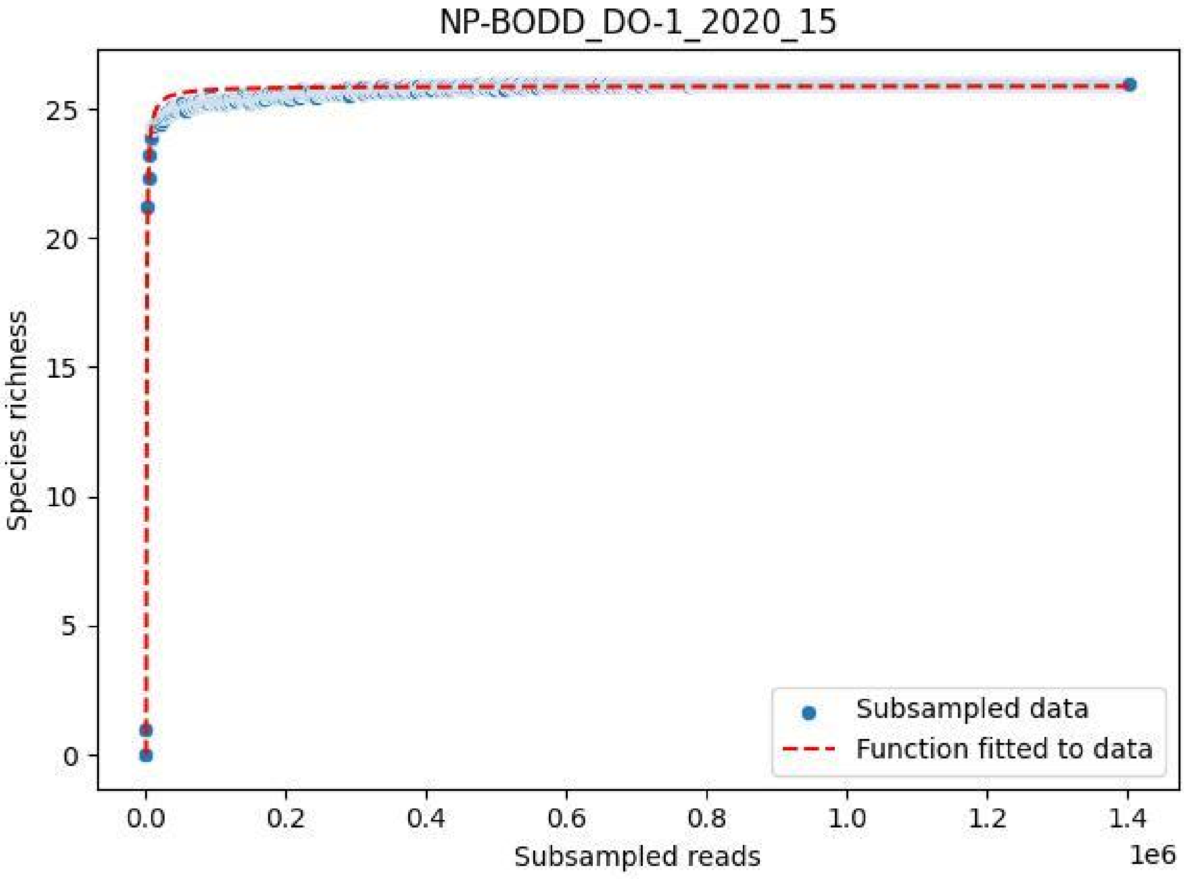
NP-BODD_DO-1_2020_15
Species richness
Subsampled reads
1e6
0
5
10
15
20
25
0.0
0.2
0.4
0.6
0.8
1.0
1.2
1.4
Subsampled data
Function fitted to data

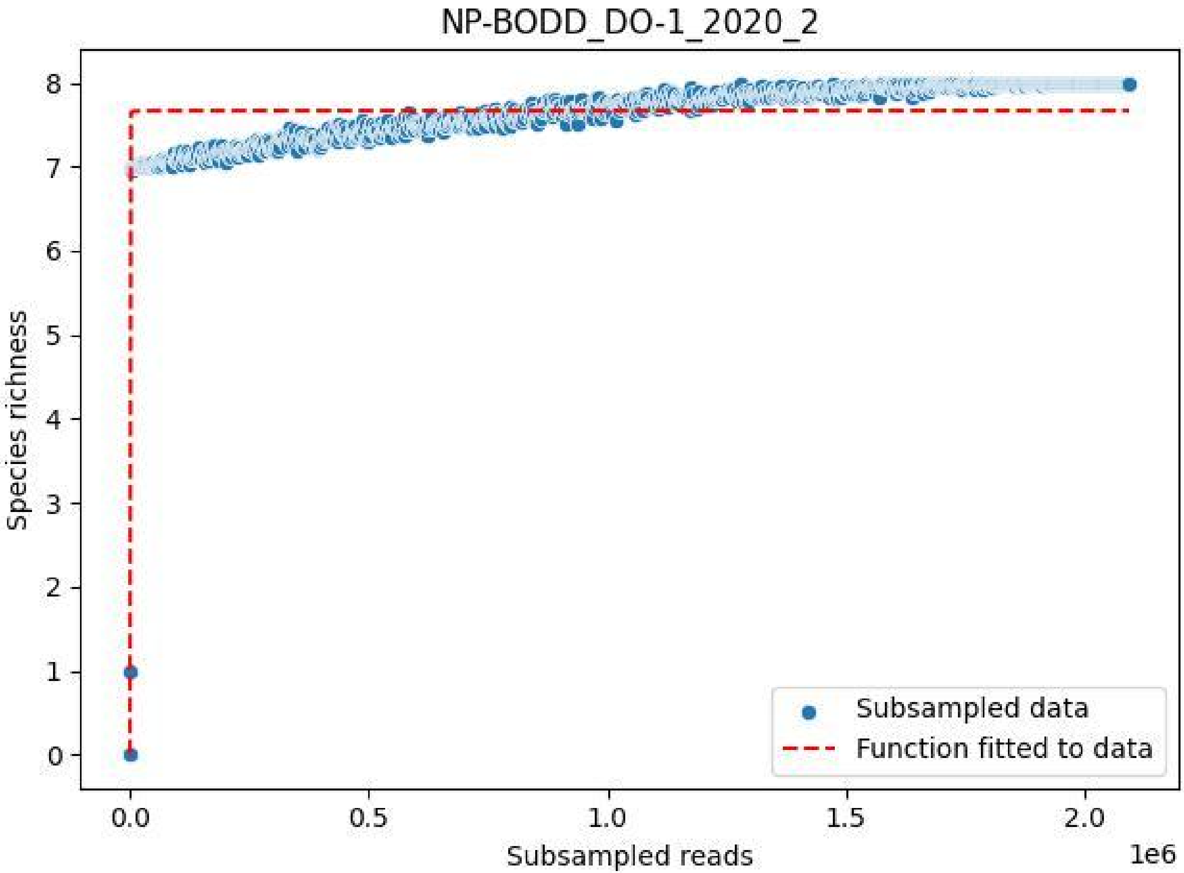
NP-BODD_DO-1_2020_2
Species richness
Subsampled reads
1e6
0.0
0.5
1.0
1.5
2.0
0
1
2
3
4
5
6
7
8
Subsampled data
Function fitted to data

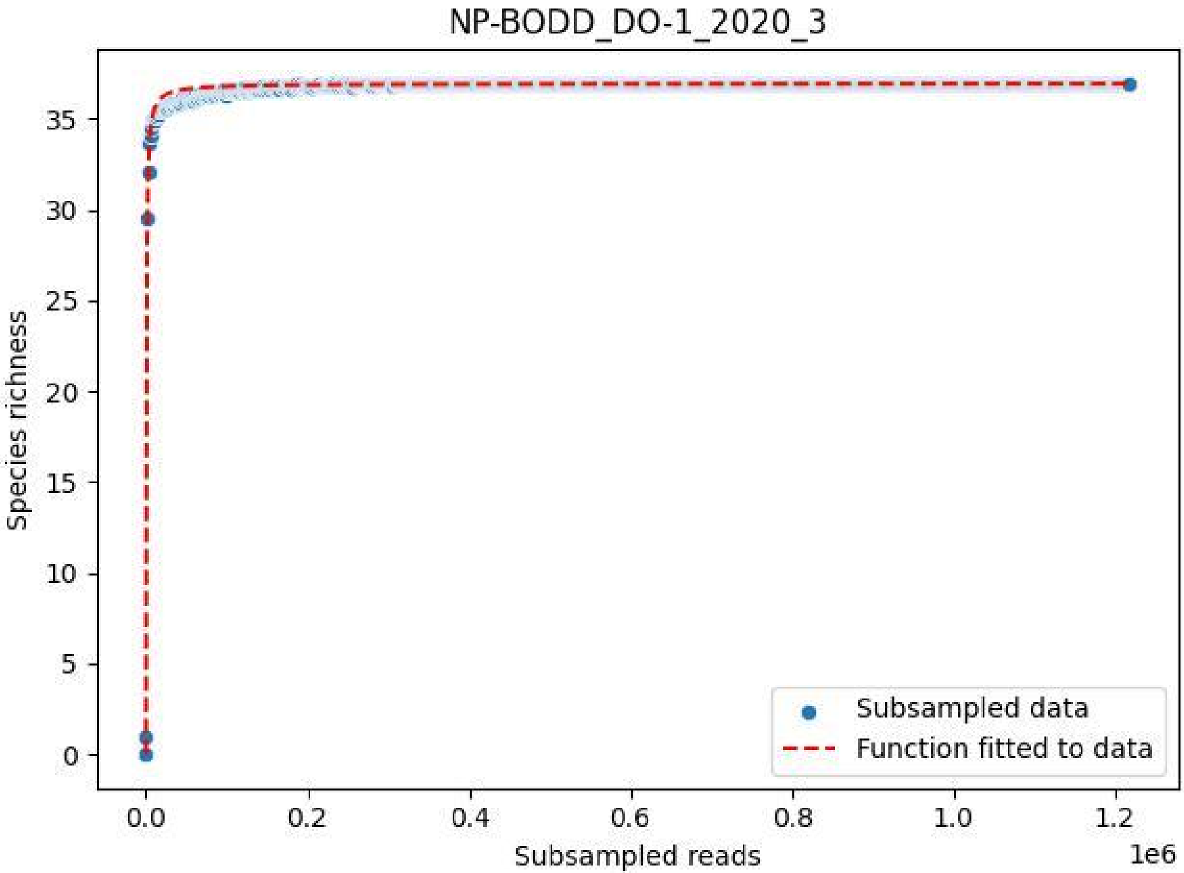
NP-BODD_DO-1_2020_3
Species richness
Subsampled reads
1e6
Subsampled data
Function fitted to data
0
5
10
15
20
25
30
35
0.0
0.2
0.4
0.6
0.8
1.0
1.2

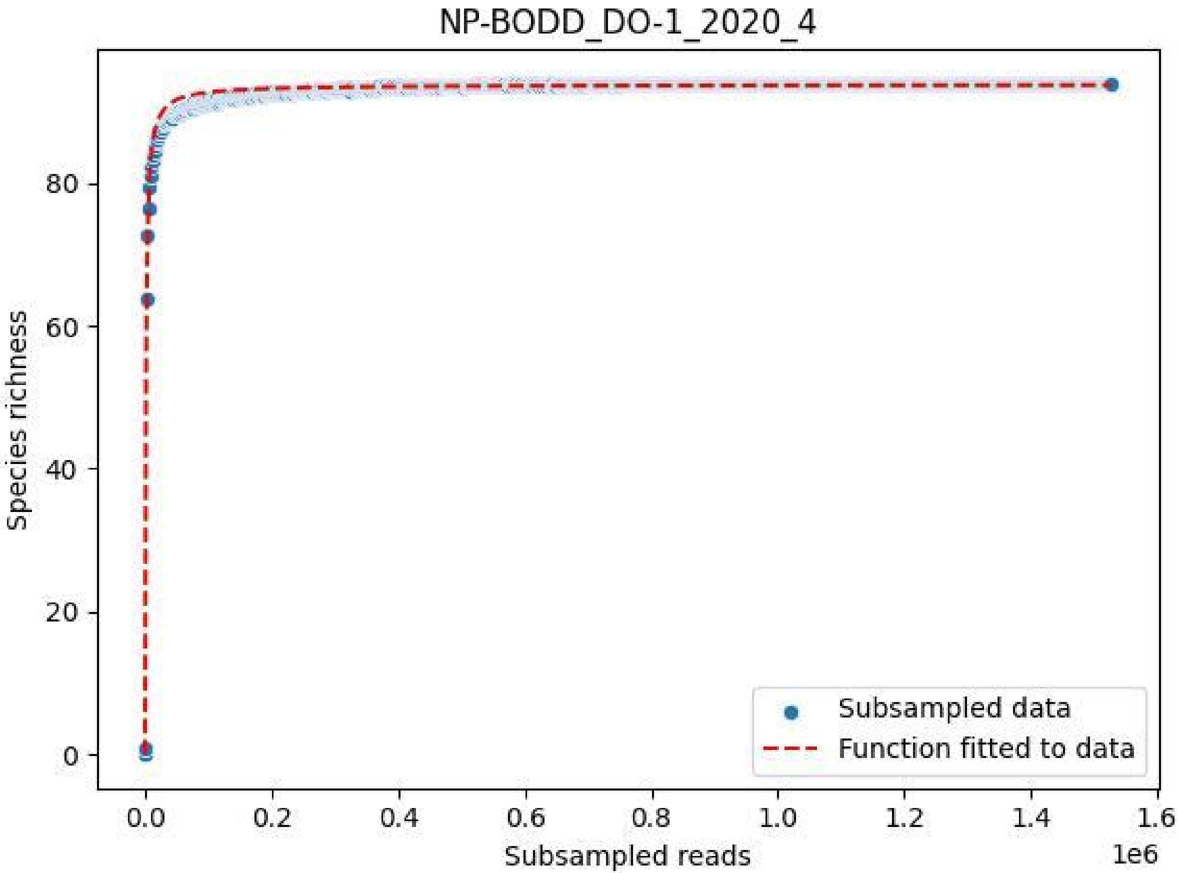
NP-BODD_DO-1_2020_4
Species richness
Subsampled reads
1e6
0.0
0.2
0.4
0.6
0.8
1.0
1.2
1.4
1.6
0
20
40
60
80
Subsampled data
Function fitted to data

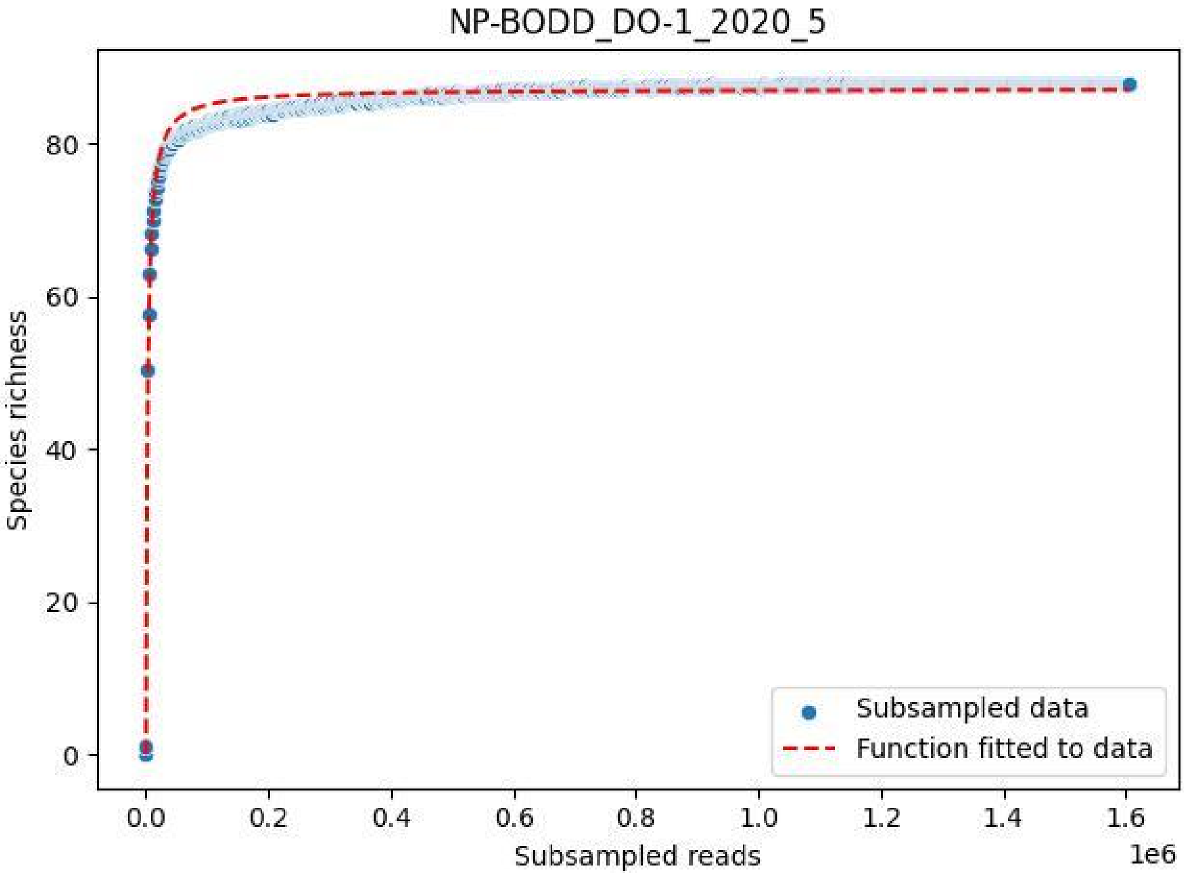

NP-BODD_DO-1_2020_5
Species richness
Subsampled reads
1e6
0
20
40
60
80
0.0
0.2
0.4
0.6
0.8
1.0
1.2
1.4
1.6
Subsampled data
Function fitted to data

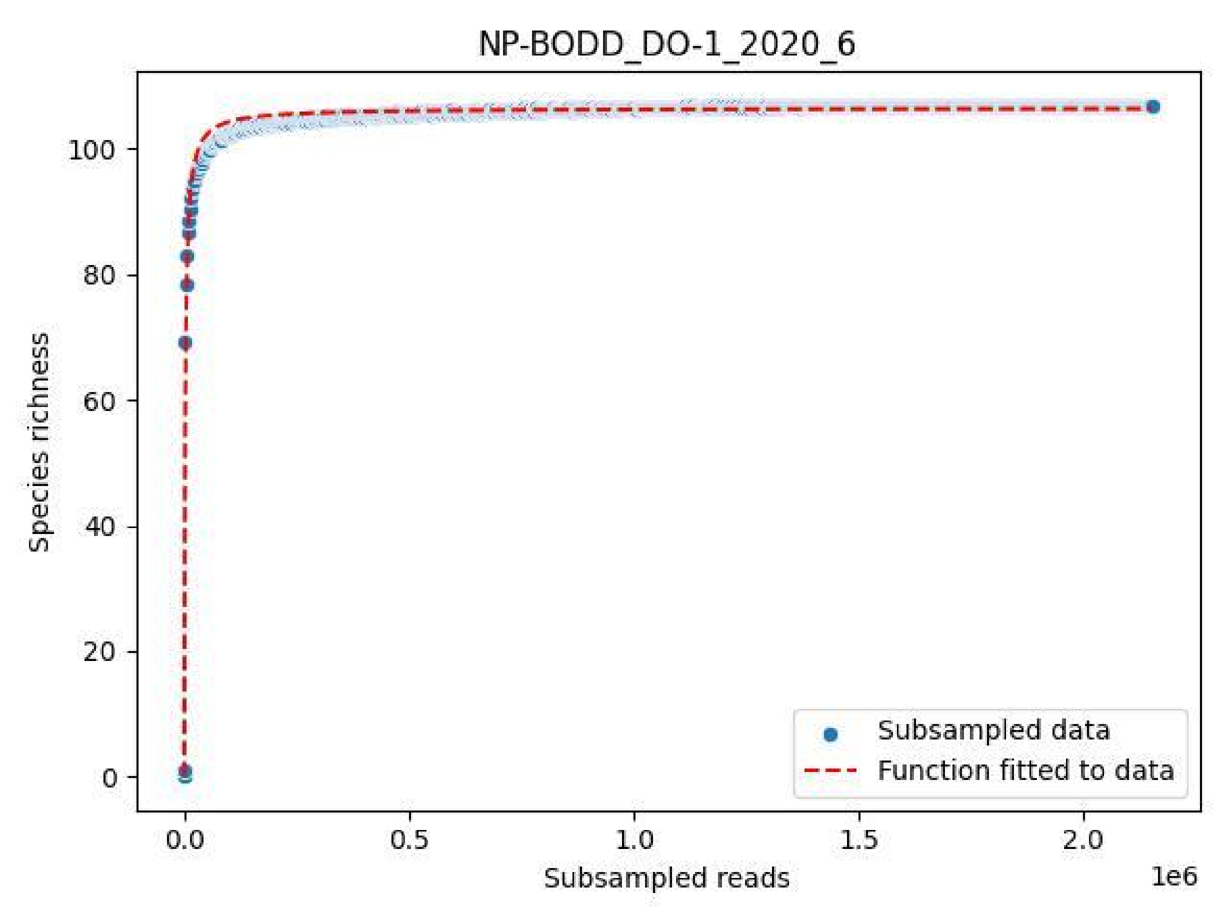

NP-BODD_DO-1_2020_6
Species richness
Subsampled reads
1e6
0
20
40
60
80
100
0.0
0.5
1.0
1.5
2.0
Subsampled data
Function fitted to data

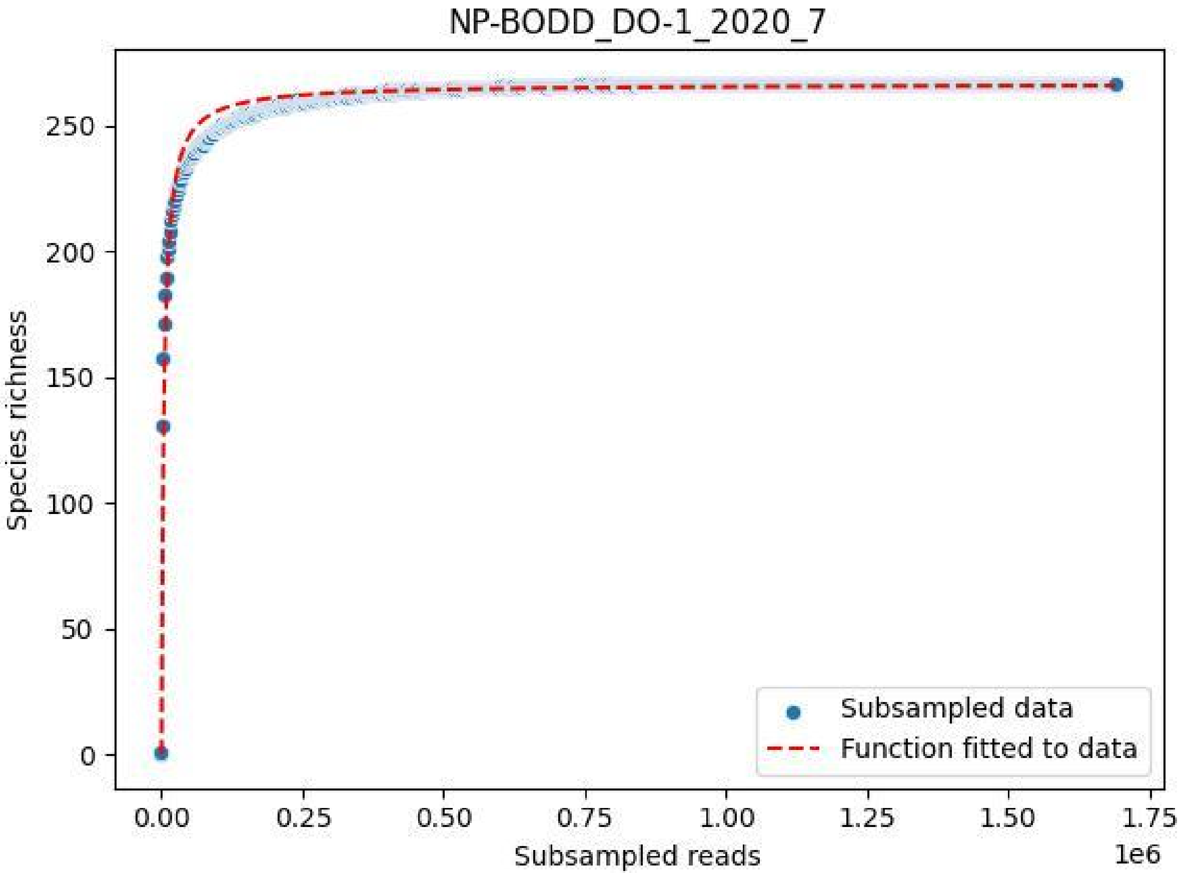
NP-BODD_DO-1_2020_7
Species richness
Subsampled reads
1e6
0
50
100
150
200
250
0.00
0.25
0.50
0.75
1.00
1.25
1.50
1.75
Subsampled data
Function fitted to data

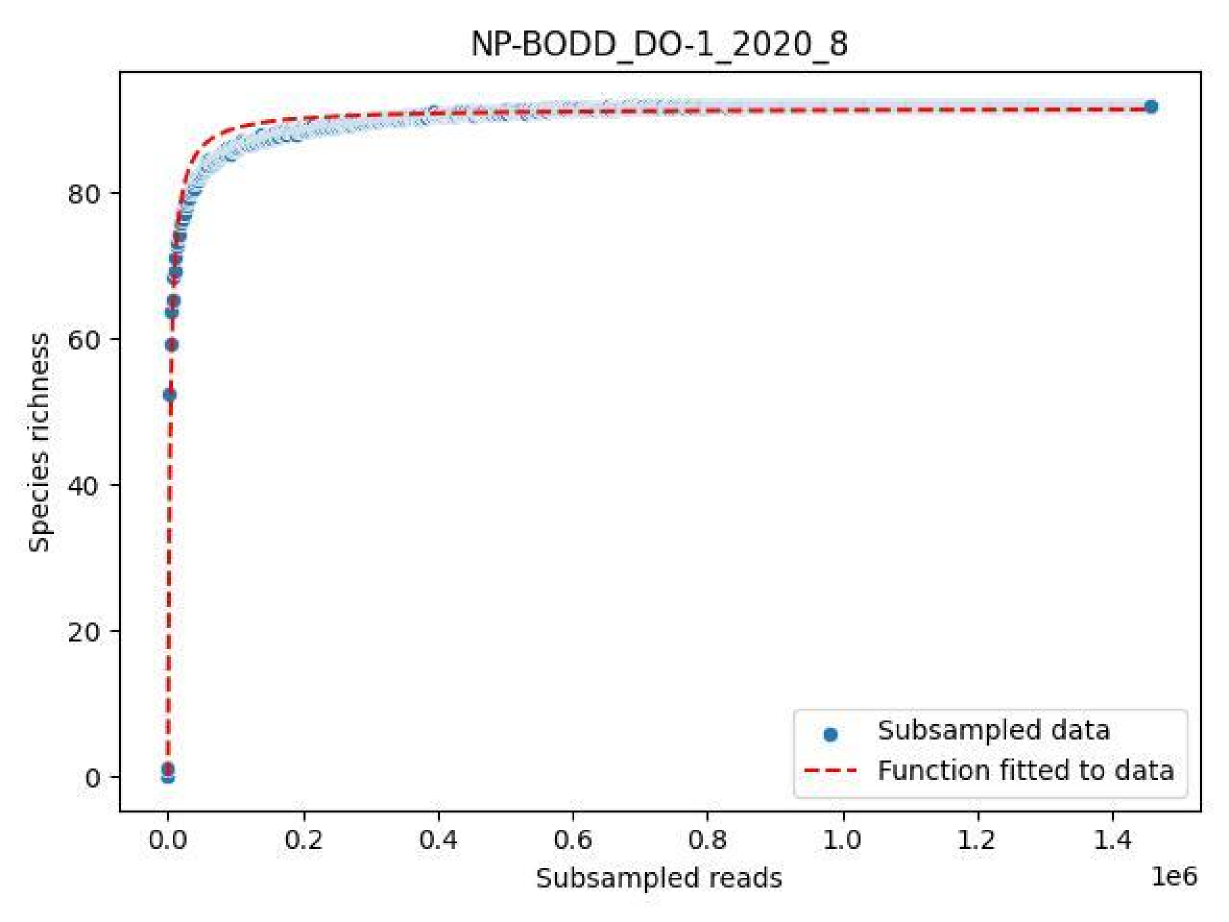
NP-BODD_DO-1_2020_8
Species richness
0
20
40
60
80
0.0
0.2
0.4
0.6
0.8
1.0
1.2
1.4
1e6
Subsampled reads
Subsampled data
Function fitted to data

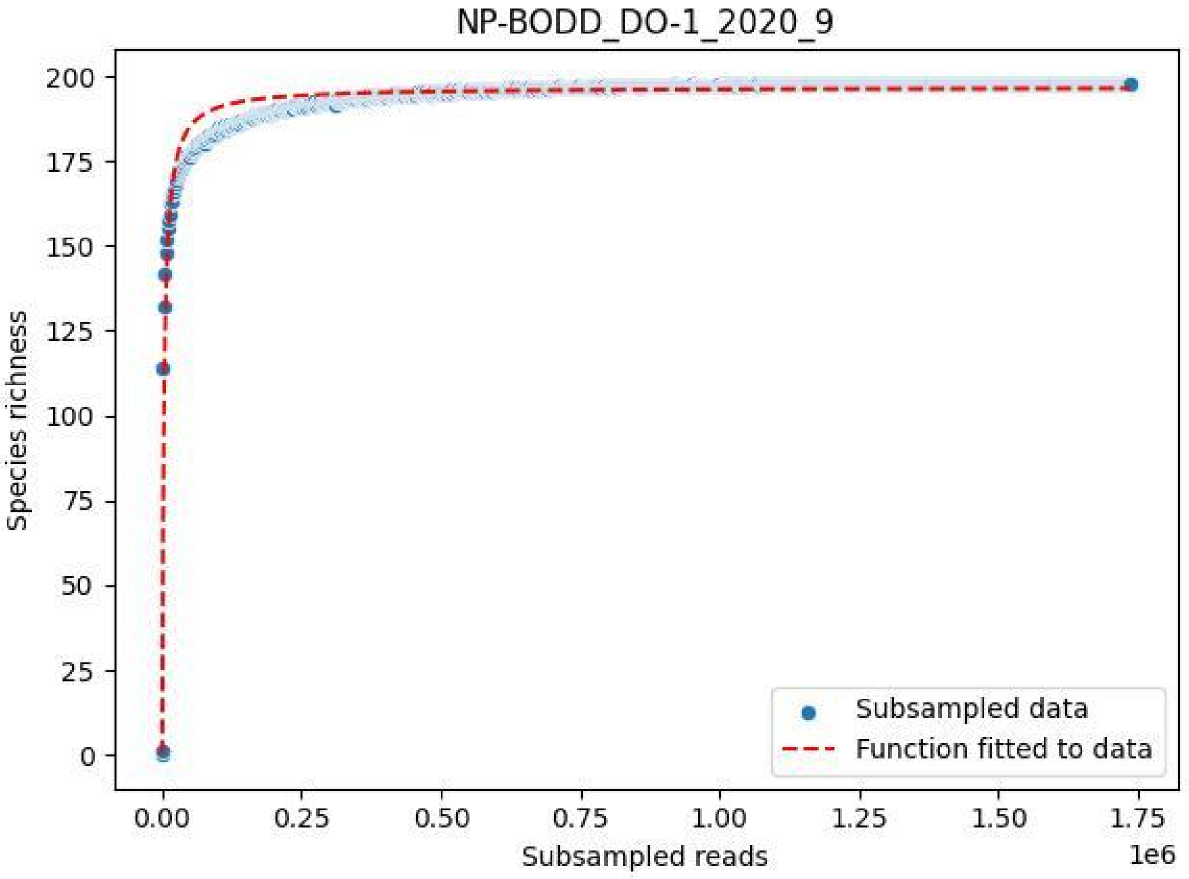
NP-BODD_DO-1_2020_9
Species richness
Subsampled reads
1e6
0
25
50
75
100
125
150
175
200
0.00
0.25
0.50
0.75
1.00
1.25
1.50
1.75
Subsampled data
Function fitted to data

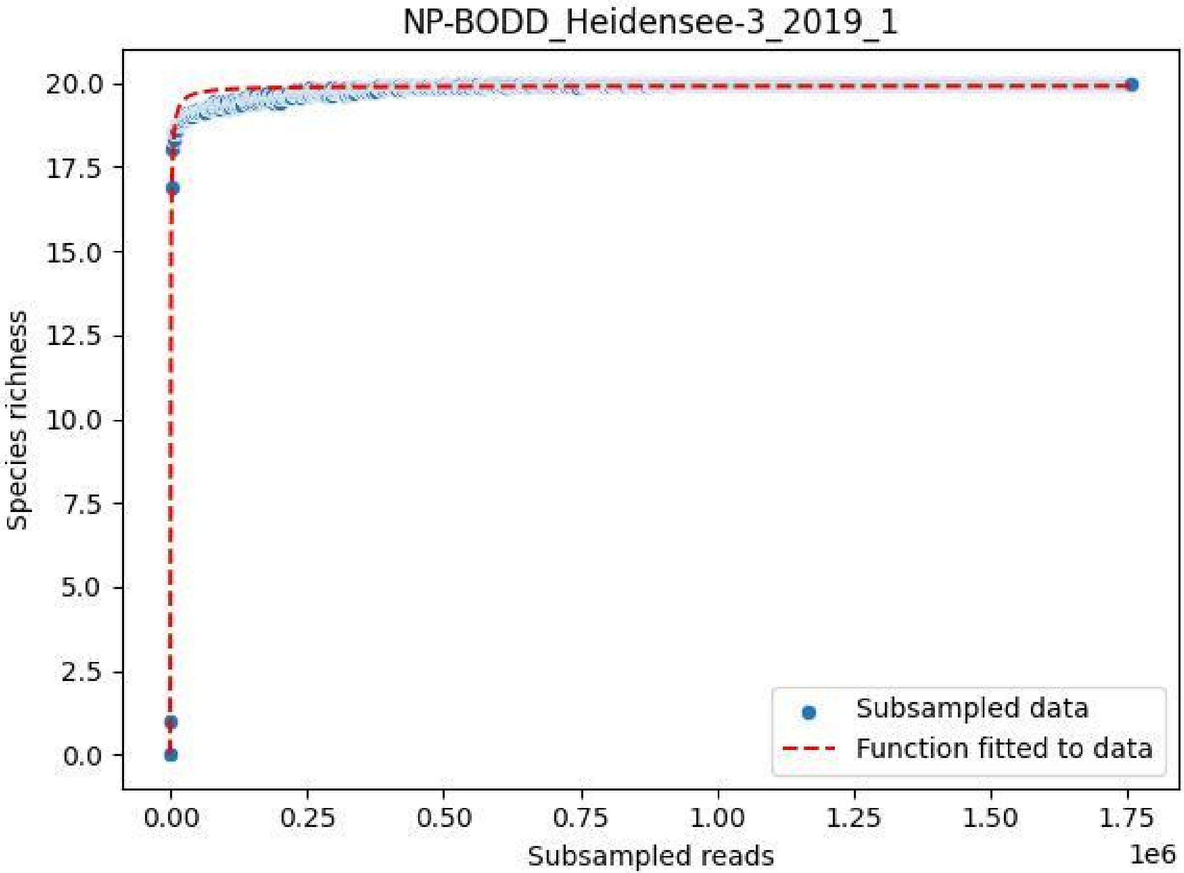

NP-BODD_Heidensee-3_2019_1
Species richness
Subsampled reads
1e6
0.0
2.5
5.0
7.5
10.0
12.5
15.0
17.5
20.0
0.00
0.25
0.50
0.75
1.00
1.25
1.50
1.75
Subsampled data
Function fitted to data

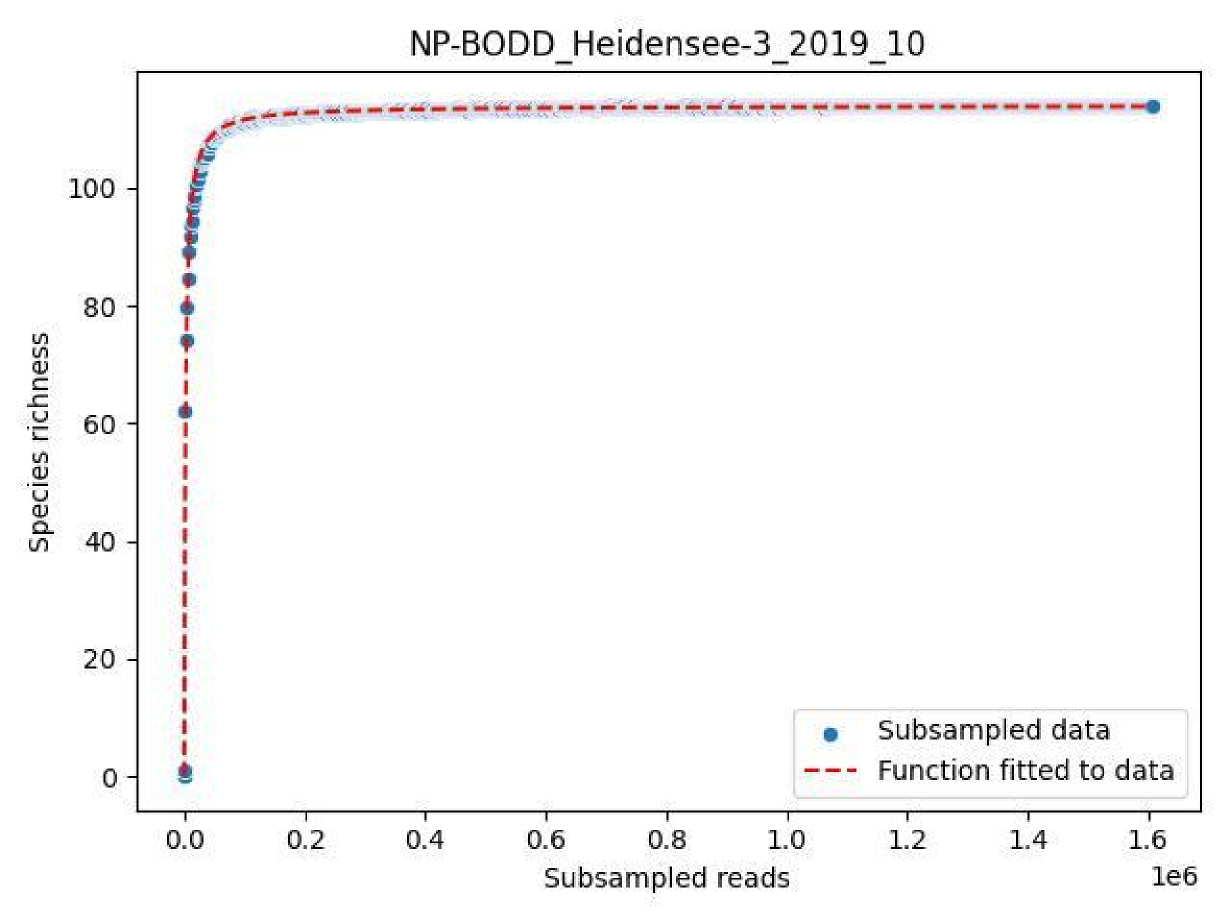

NP-BODD_Heidensee-3_2019_10
Species richness
Subsampled reads
1e6
0
20
40
60
80
100
0.0
0.2
0.4
0.6
0.8
1.0
1.2
1.4
1.6
Subsampled data
Function fitted to data

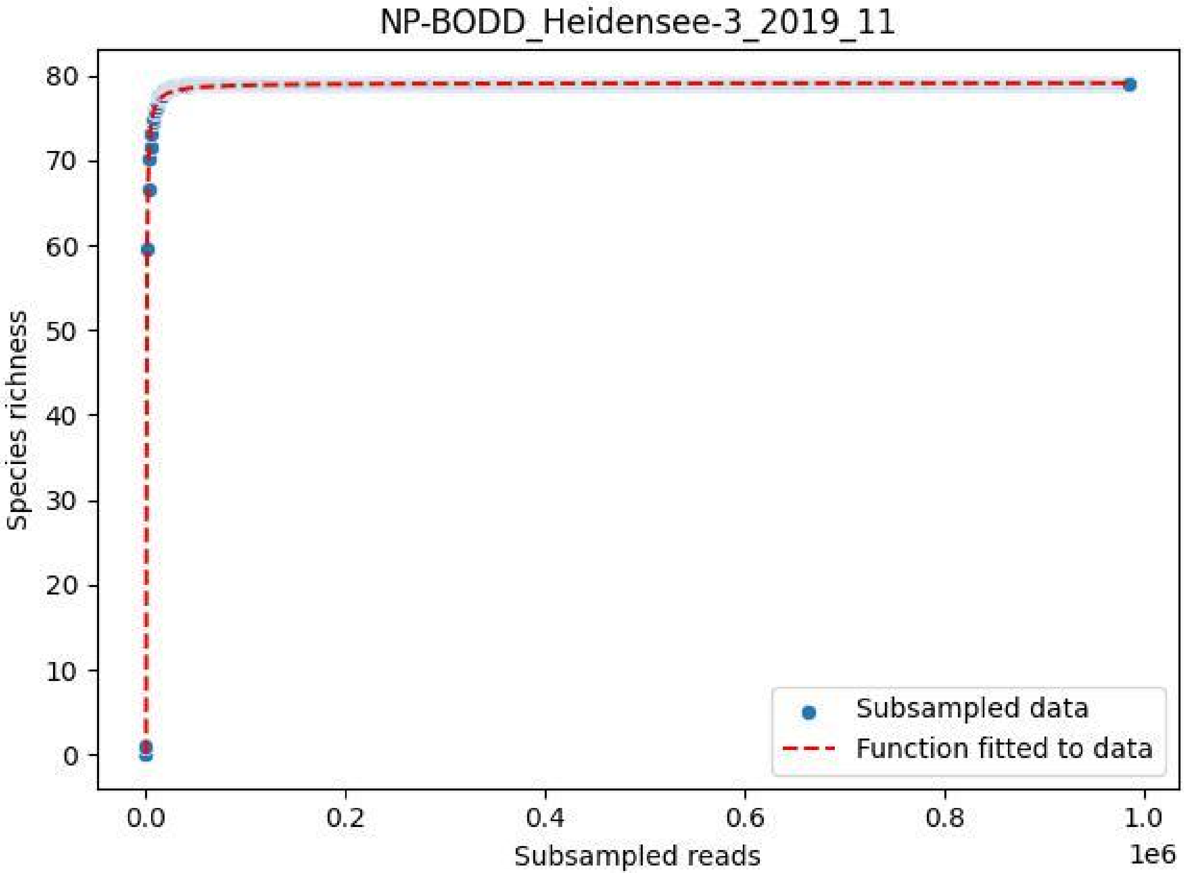
NP-BODD_Heidensee-3_2019_11
Species richness
Subsampled reads
1e6
Subsampled data
Function fitted to data
0
10
20
30
40
50
60
70
80
0.0
0.2
0.4
0.6
0.8
1.0

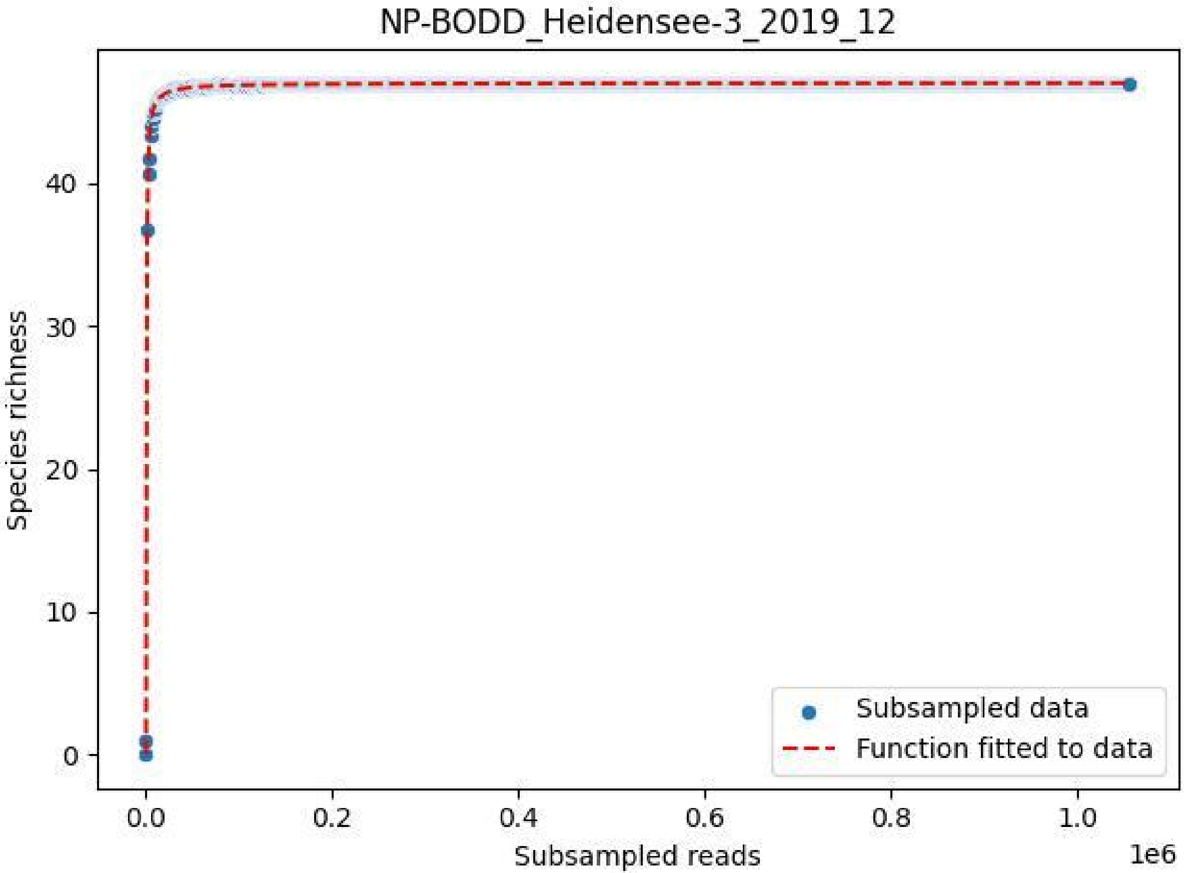

NP-BODD_Heidensee-3_2019_12
Species richness
Subsampled reads
1e6
0
10
20
30
40
0.0
0.2
0.4
0.6
0.8
1.0
Subsampled data
Function fitted to data

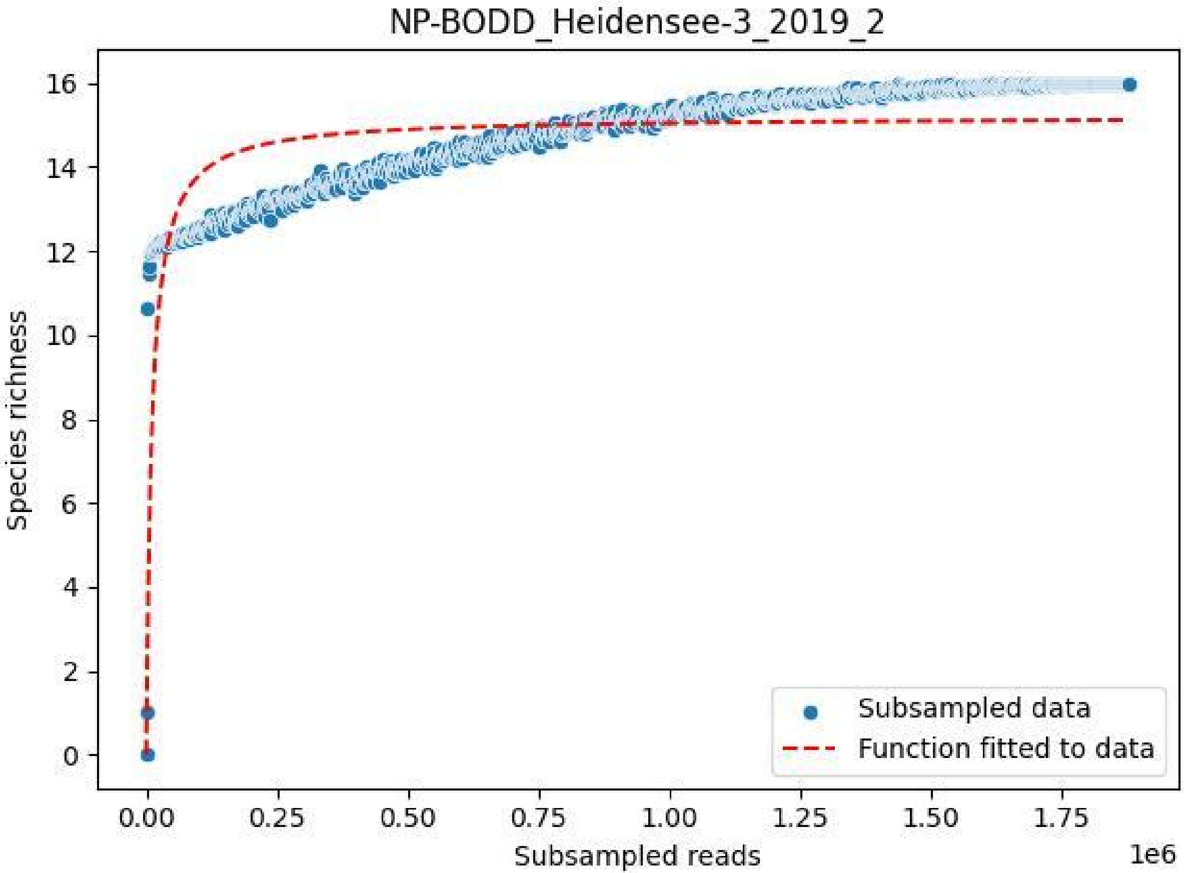
NP-BODD_Heidensee-3_2019_2
Species richness
Subsampled reads
1e6
0
2
4
6
8
10
12
14
16
0.00
0.25
0.50
0.75
1.00
1.25
1.50
1.75
Subsampled data
Function fitted to data

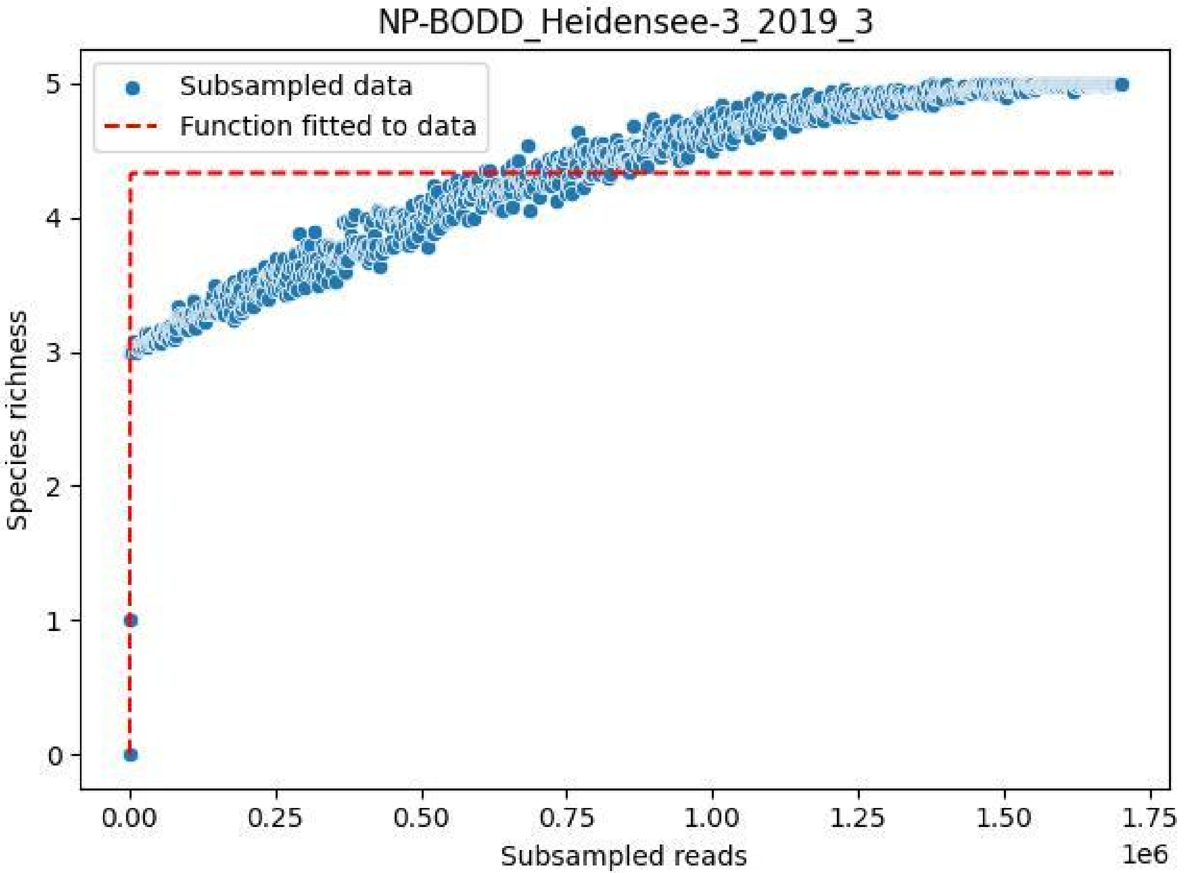
NP-BODD_Heidensee-3_2019_3
Subsampled data
Function fitted to data
Species richness
Subsampled reads
1e6
0
1
2
3
4
5
0.00
0.25
0.50
0.75
1.00
1.25
1.50
1.75

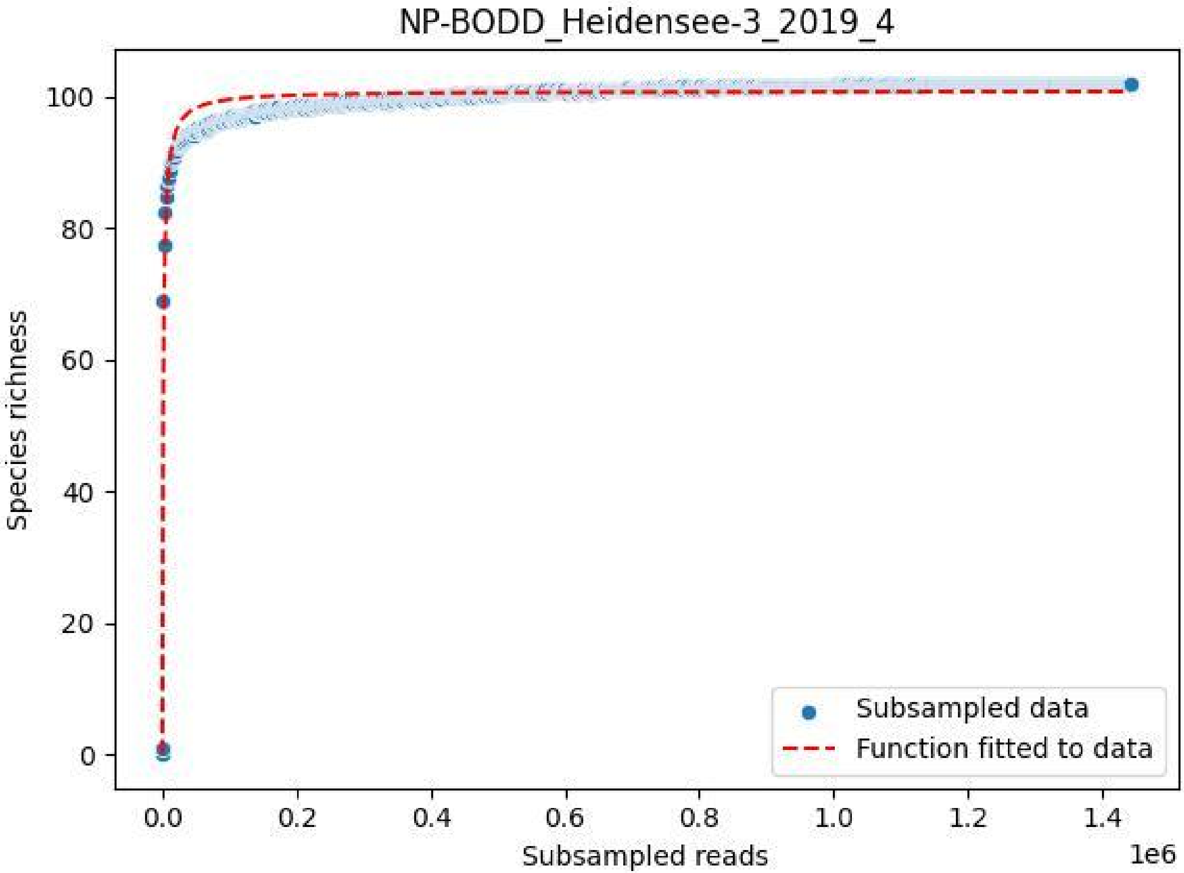
NP-BODD_Heidensee-3_2019_4
Species richness
Subsampled reads
1e6
0
20
40
60
80
100
0.0
0.2
0.4
0.6
0.8
1.0
1.2
1.4
Subsampled data
Function fitted to data

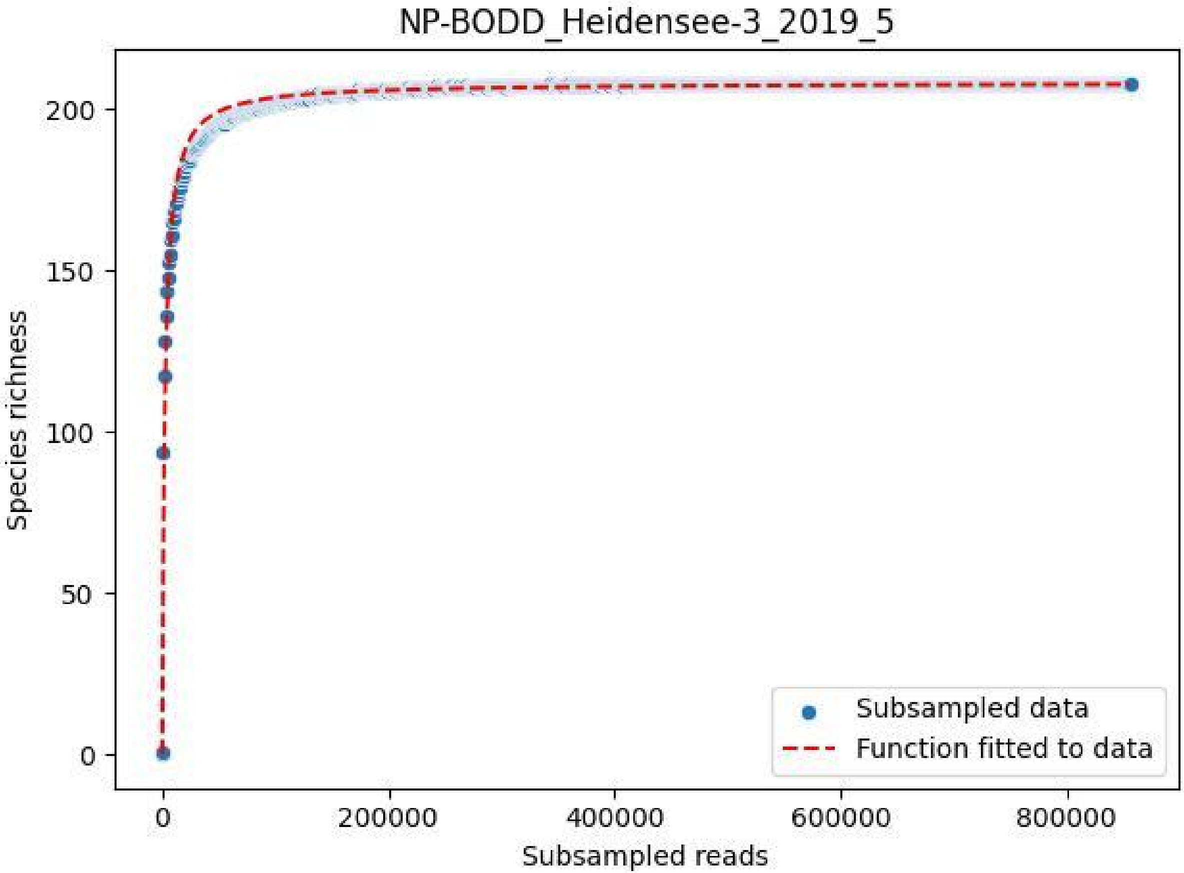

NP-BODD_Heidensee-3_2019_5
Species richness
Subsampled reads
0
50
100
150
200
0
200000
400000
600000
800000
Subsampled data
Function fitted to data

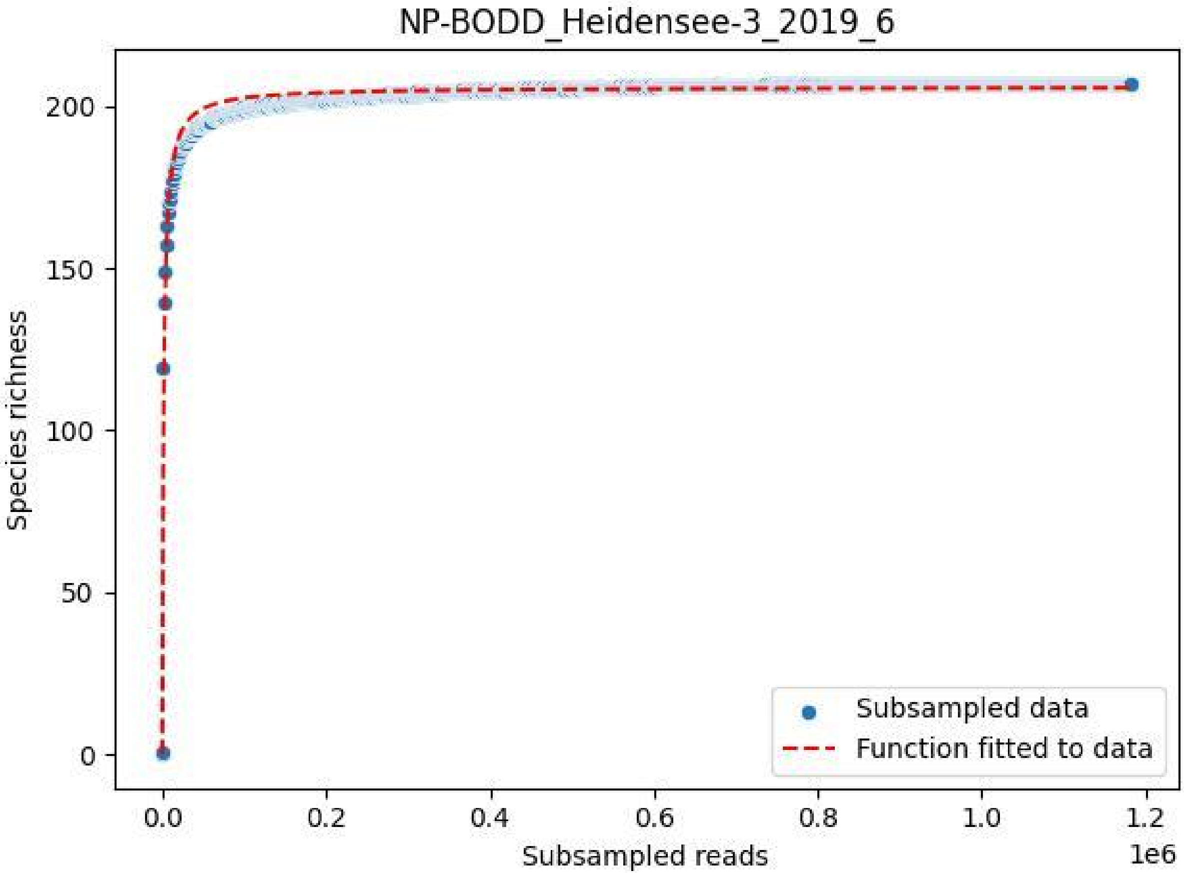

NP-BODD_Heidensee-3_2019_6
Species richness
Subsampled reads
1e6
0
50
100
150
200
0.0
0.2
0.4
0.6
0.8
1.0
1.2
Subsampled data
Function fitted to data

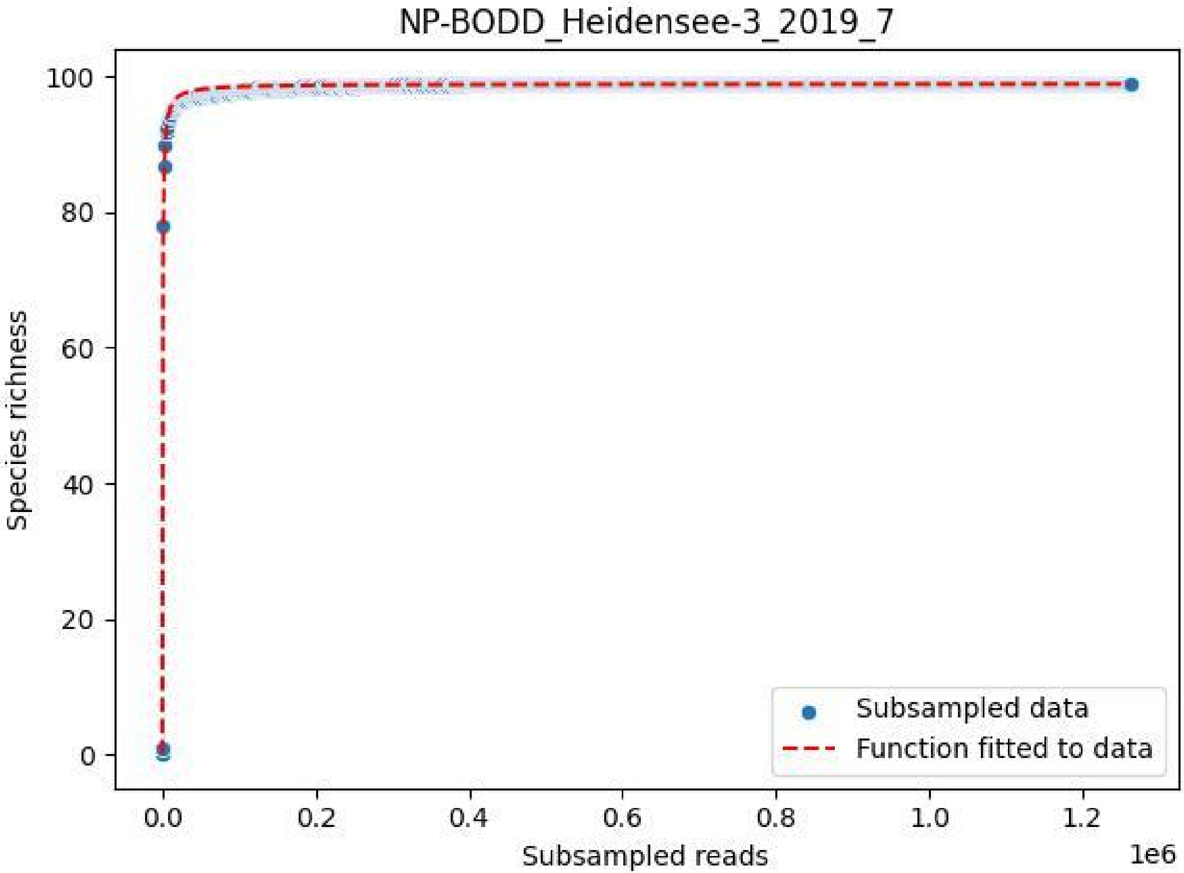

NP-BODD_Heidensee-3_2019_7
Species richness
Subsampled reads
1e6
0
20
40
60
80
100
0.0
0.2
0.4
0.6
0.8
1.0
1.2
Subsampled data
Function fitted to data

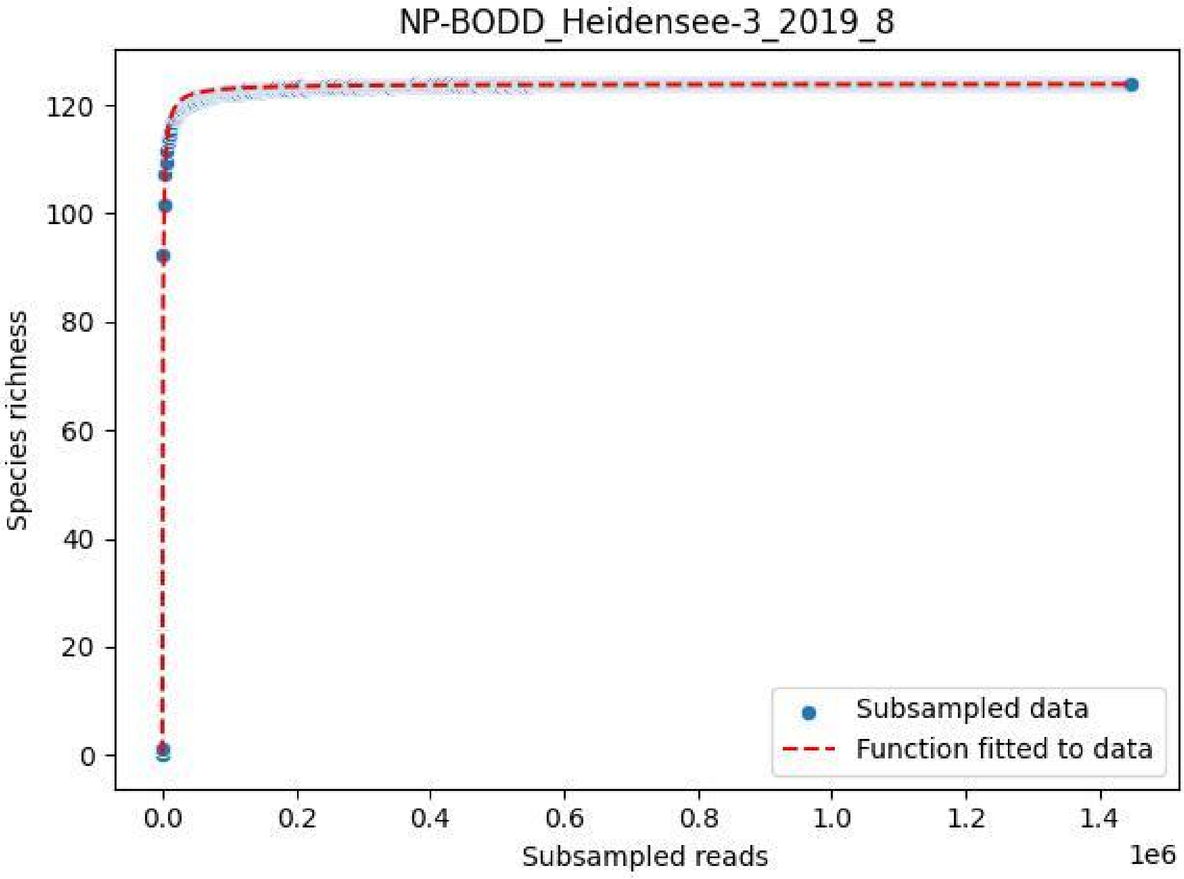
NP-BODD_Heidensee-3_2019_8
Species richness
Subsampled reads
1e6
0
20
40
60
80
100
120
0.0
0.2
0.4
0.6
0.8
1.0
1.2
1.4
Subsampled data
Function fitted to data

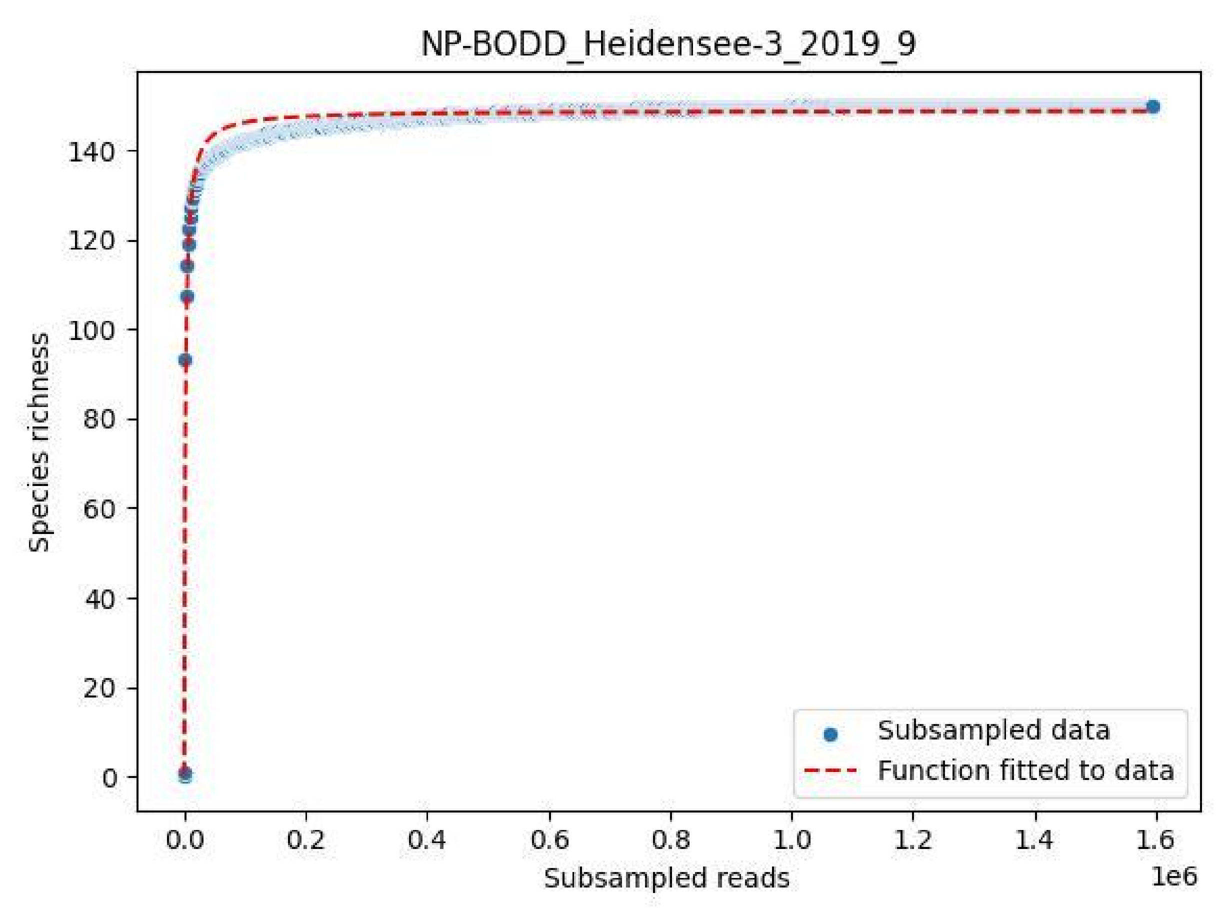
NP-BODD_Heidensee-3_2019_9
Species richness
0
20
40
60
80
100
120
140
0.0
0.2
0.4
0.6
0.8
1.0
1.2
1.4
1.6
1e6
Subsampled reads
Subsampled data
Function fitted to data

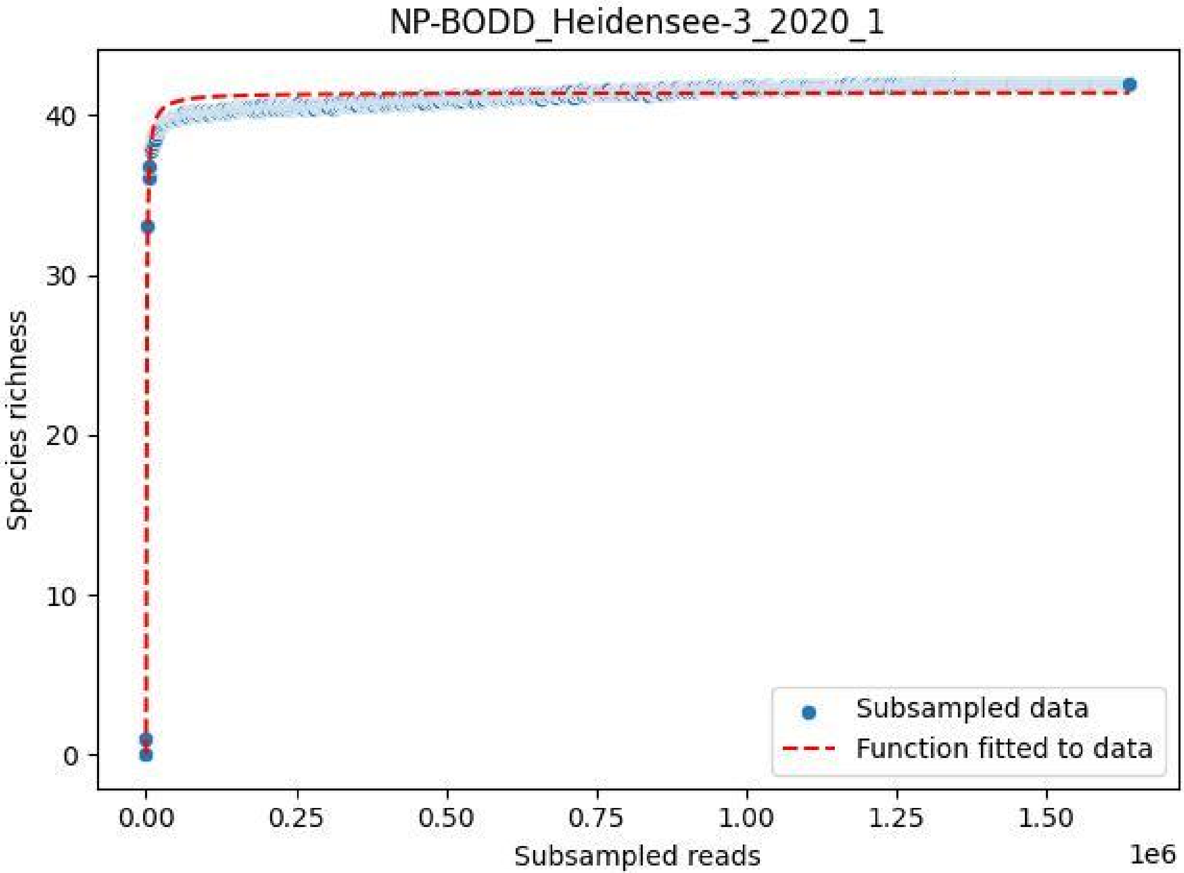

NP-BODD_Heidensee-3_2020_1
Species richness
Subsampled reads
1e6
0
10
20
30
40
0.00
0.25
0.50
0.75
1.00
1.25
1.50
Subsampled data
Function fitted to data

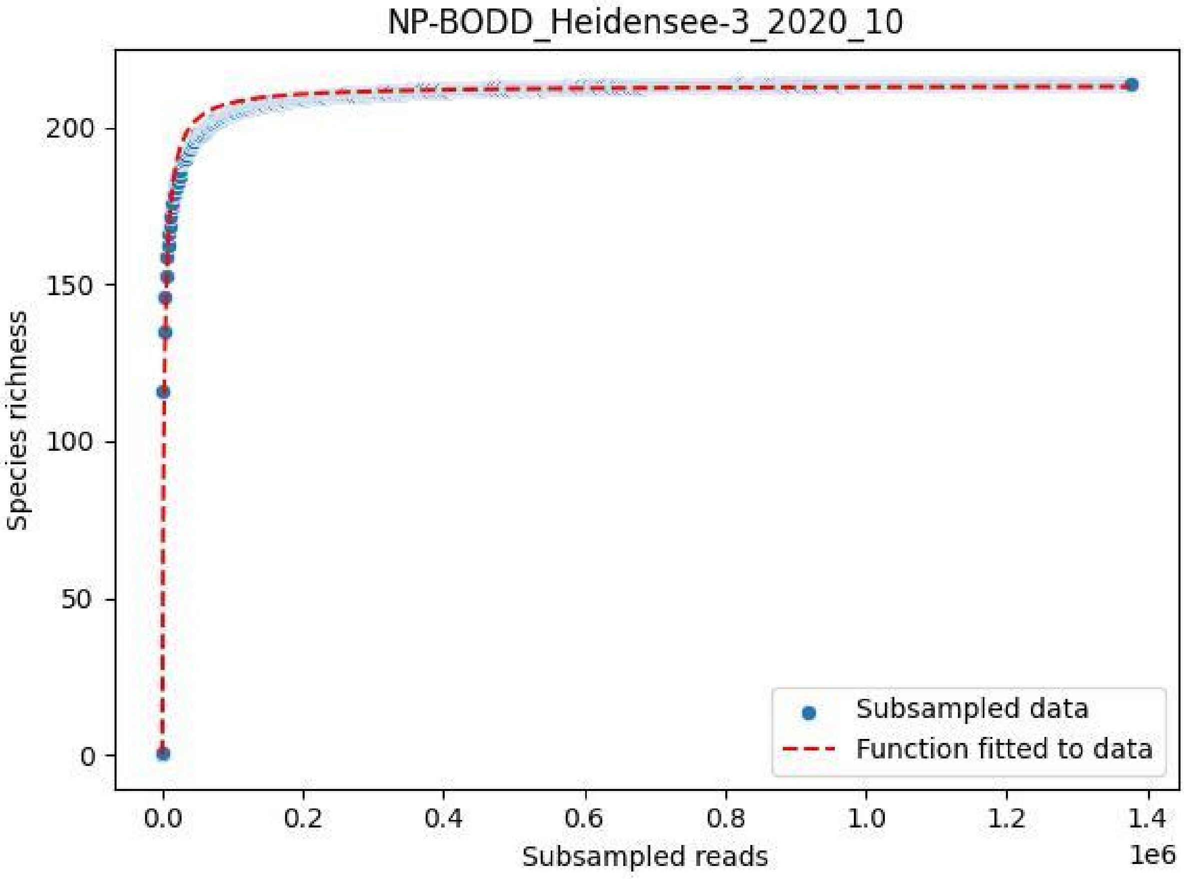
NP-BODD_Heidensee-3_2020_10
Species richness
Subsampled reads
1e6
0
50
100
150
200
0.0
0.2
0.4
0.6
0.8
1.0
1.2
1.4
Subsampled data
Function fitted to data

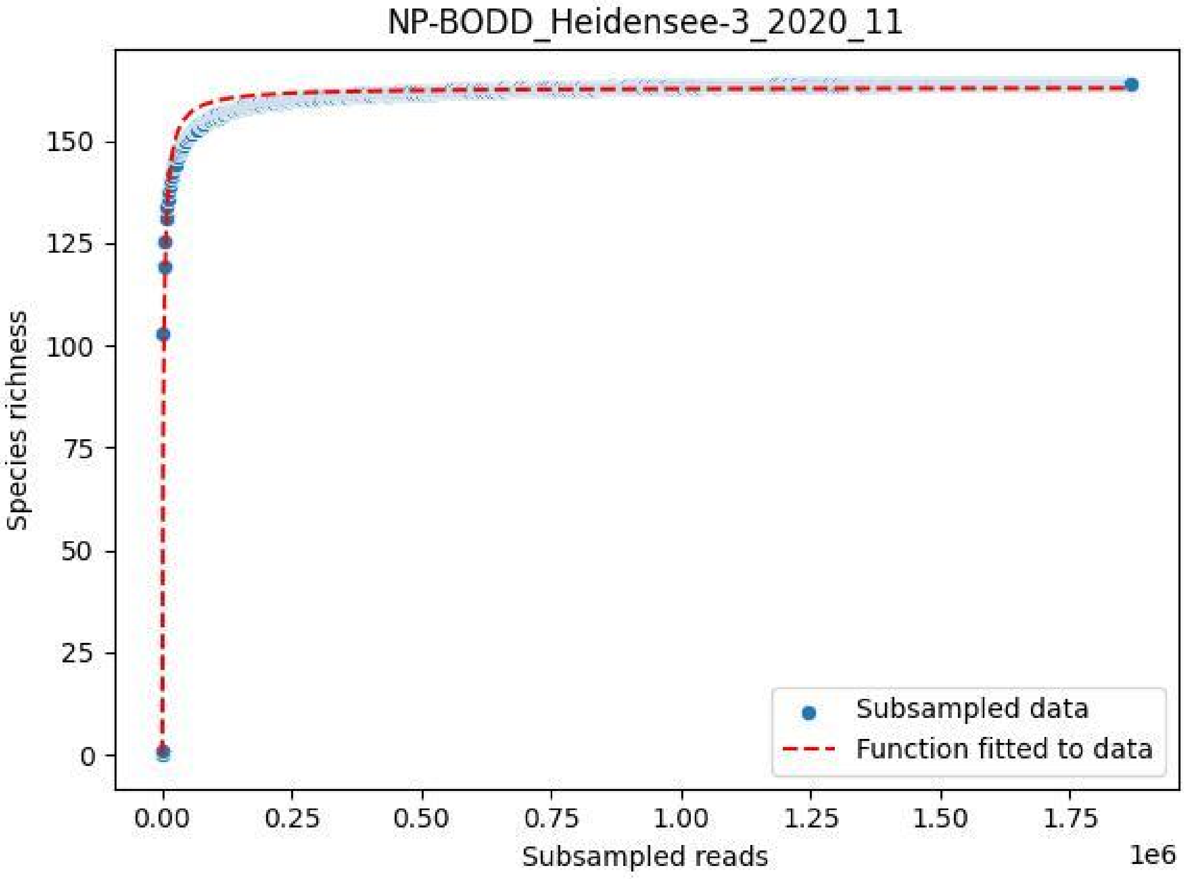
NP-BODD_Heidensee-3_2020_11
Species richness
150
125
100
75
50
25
0
0.00
0.25
0.50
0.75
1.00
1.25
1.50
1.75
Subsampled reads
1e6
Subsampled data
Function fitted to data

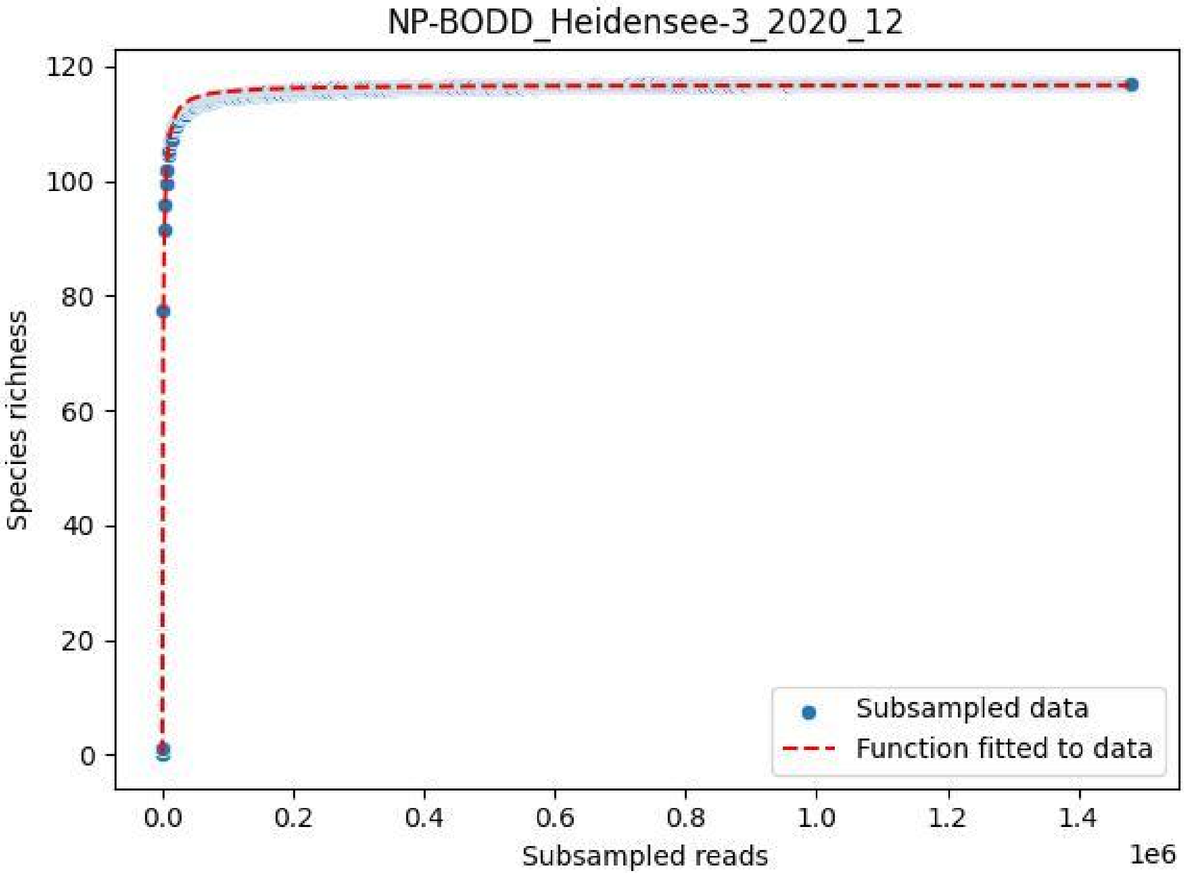

NP-BODD_Heidensee-3_2020_12
Species richness
Subsampled reads
1e6
0
20
40
60
80
100
120
0.0
0.2
0.4
0.6
0.8
1.0
1.2
1.4
Subsampled data
Function fitted to data

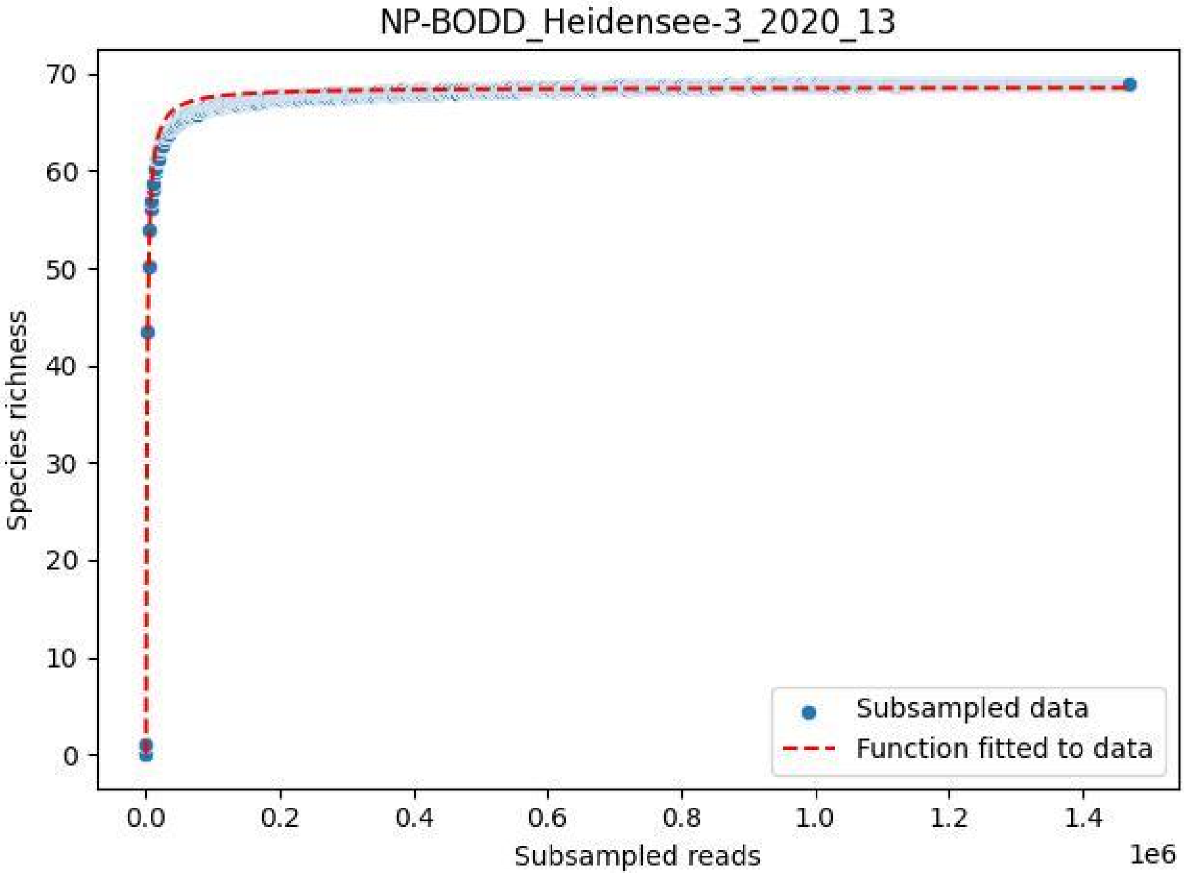
NP-BODD_Heidensee-3_2020_13
Species richness
Subsampled reads
1e6
0
10
20
30
40
50
60
70
0.0
0.2
0.4
0.6
0.8
1.0
1.2
1.4
Subsampled data
Function fitted to data

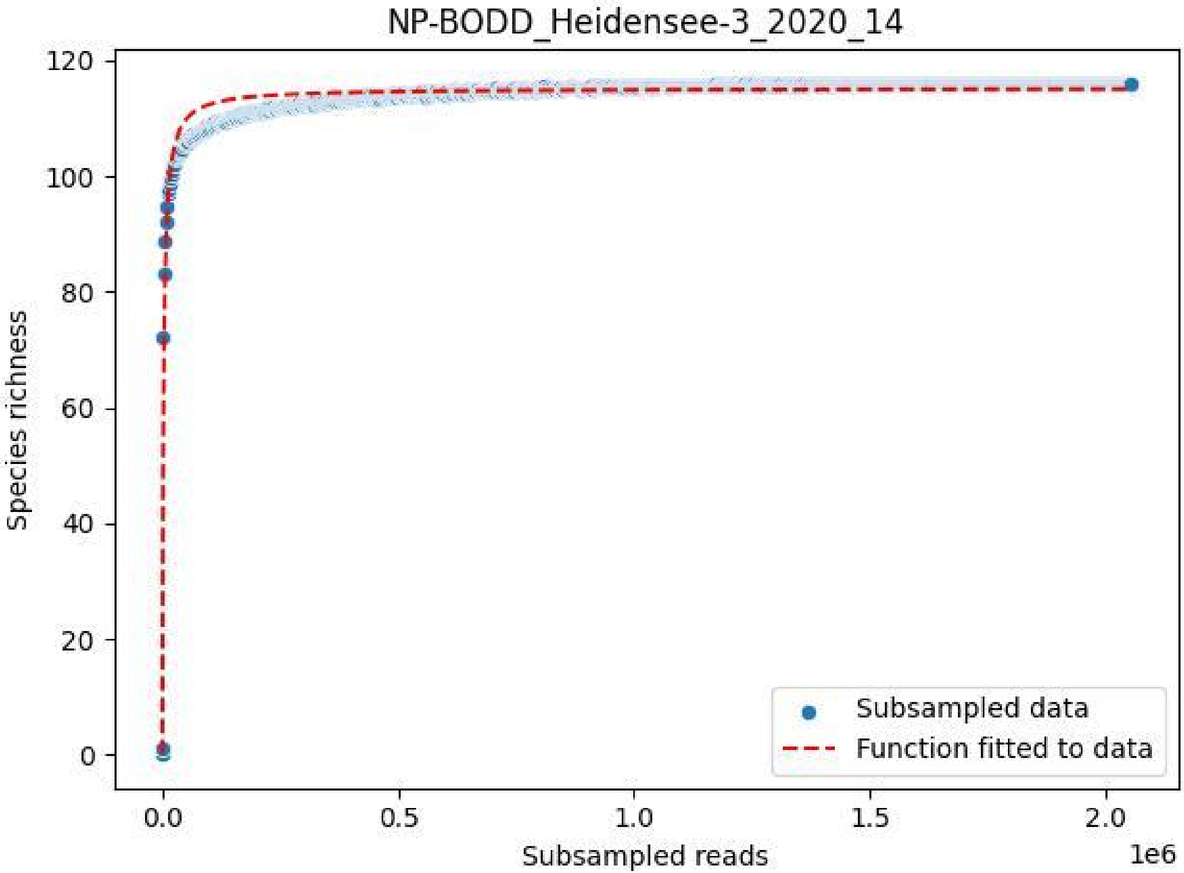
NP-BODD_Heidensee-3_2020_14
Species richness
Subsampled reads
1e6
0
20
40
60
80
100
120
0.0
0.5
1.0
1.5
2.0
Subsampled data
Function fitted to data

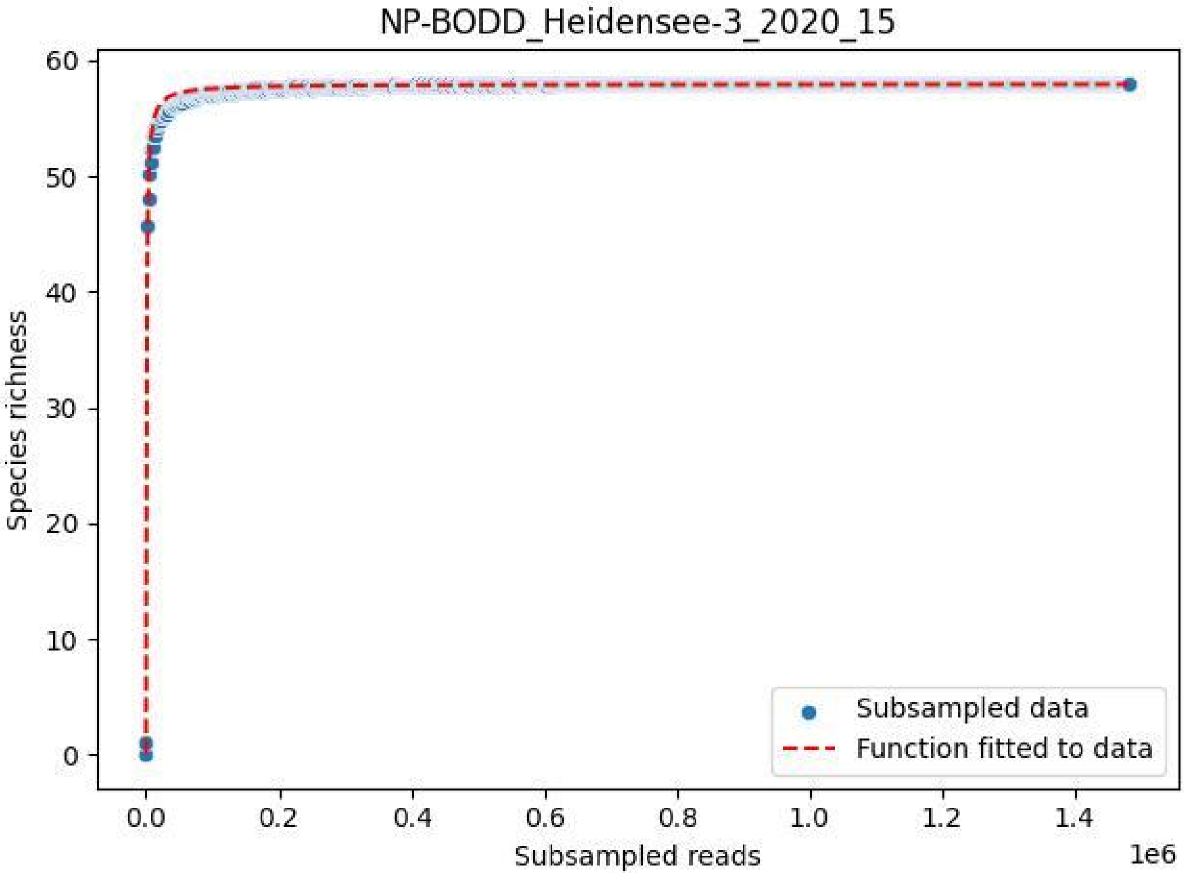

NP-BODD_Heidensee-3_2020_15
Species richness
Subsampled reads
1e6
Subsampled data
Function fitted to data
0
10
20
30
40
50
60
0.0
0.2
0.4
0.6
0.8
1.0
1.2
1.4

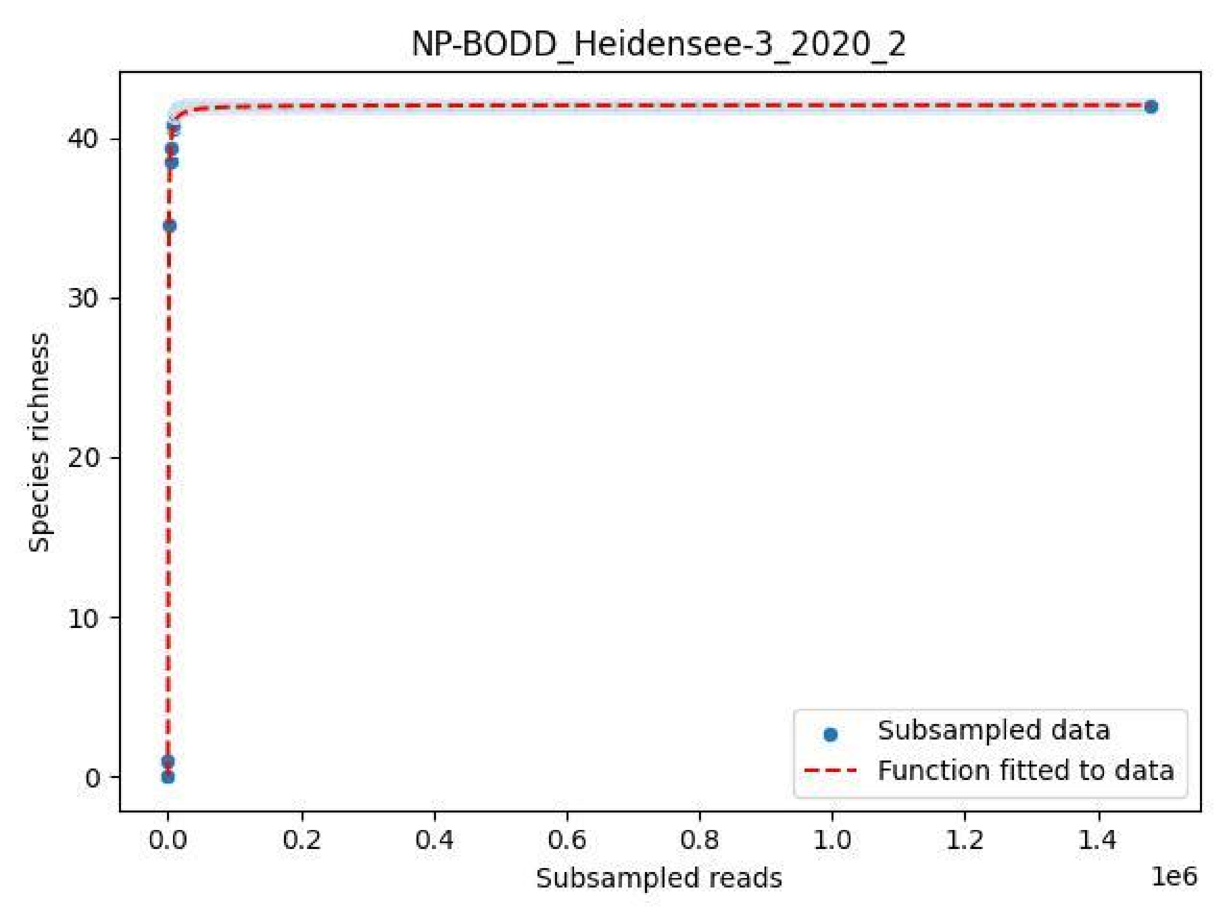
NP-BODD_Heidensee-3_2020_2
Species richness
Subsampled reads
1e6
Subsampled data
Function fitted to data
0
10
20
30
40
0.0
0.2
0.4
0.6
0.8
1.0
1.2
1.4

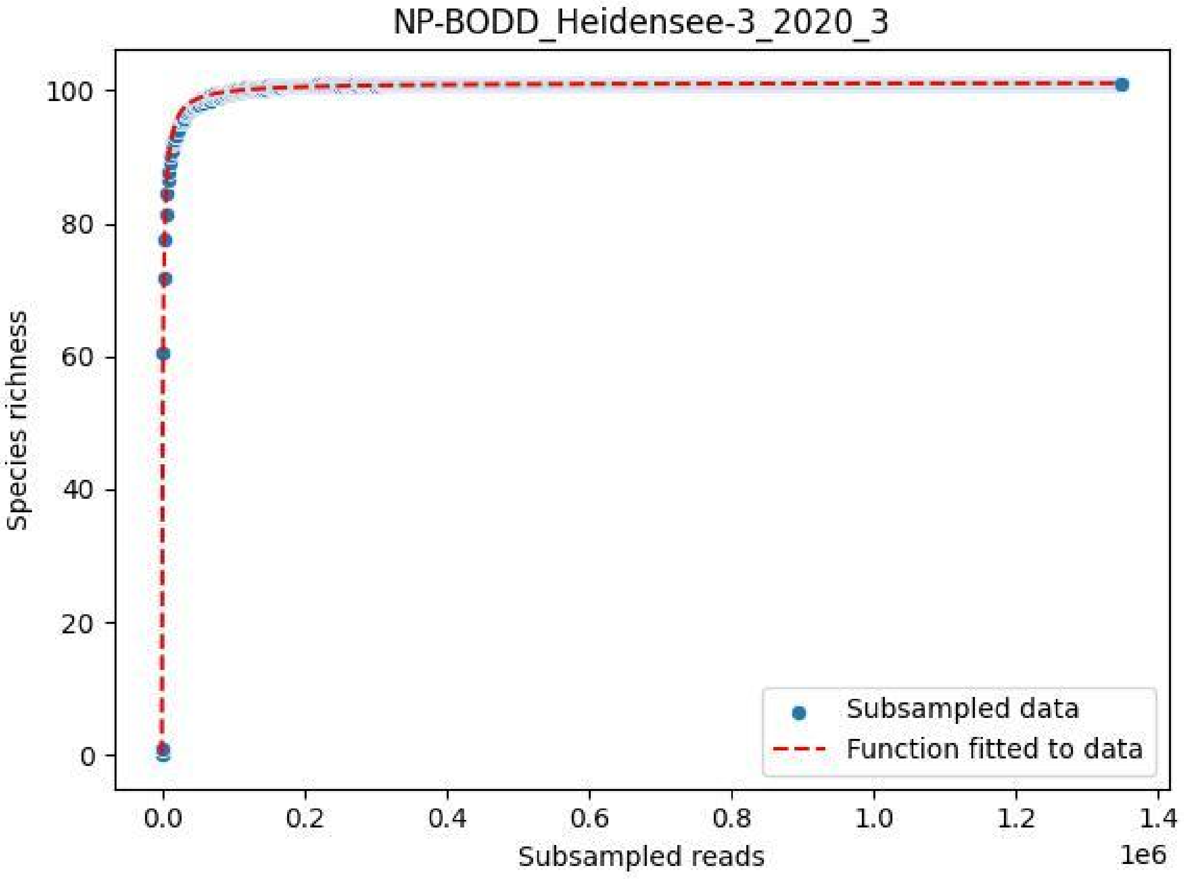
NP-BODD_Heidensee-3_2020_3
Species richness
Subsampled reads
1e6
0
20
40
60
80
100
0.0
0.2
0.4
0.6
0.8
1.0
1.2
1.4
Subsampled data
Function fitted to data

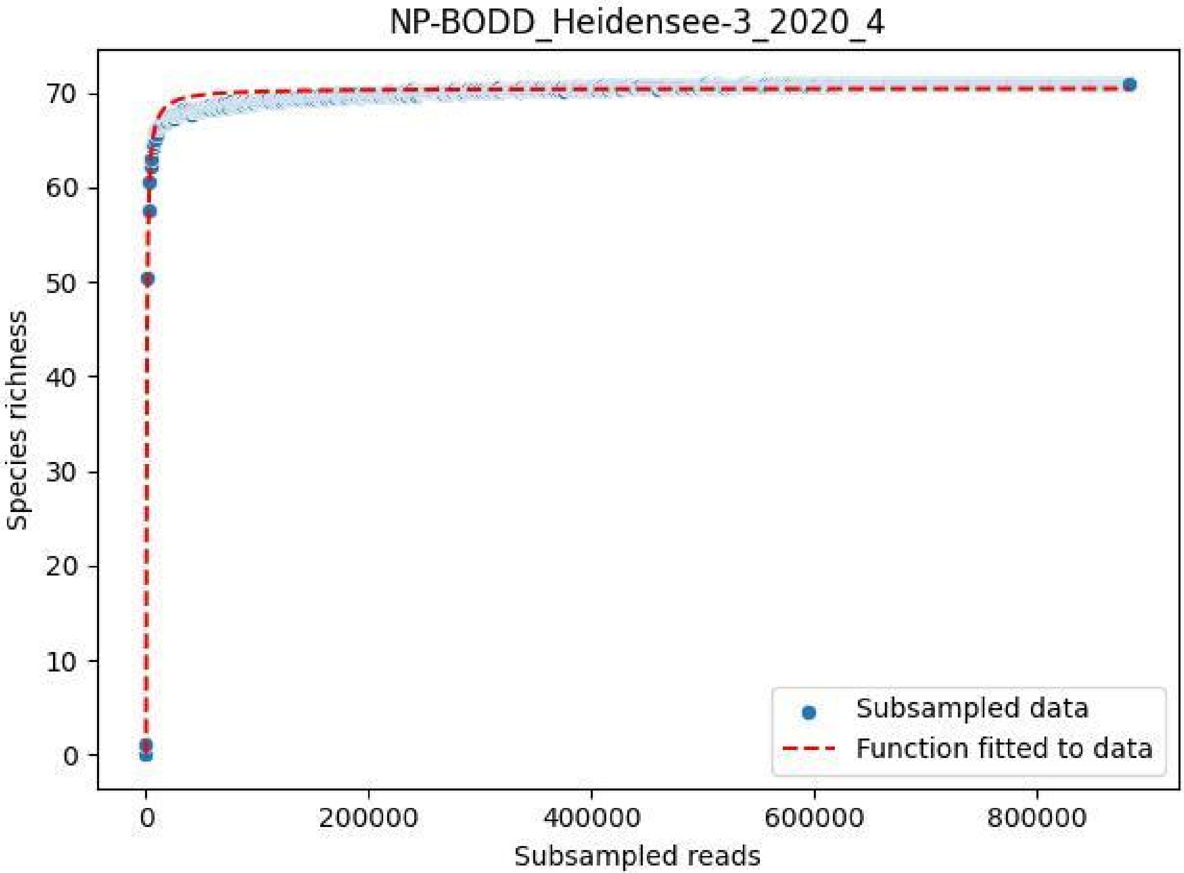

NP-BODD_Heidensee-3_2020_4
Species richness
Subsampled reads
0
10
20
30
40
50
60
70
0
200000
400000
600000
800000
Subsampled data
Function fitted to data

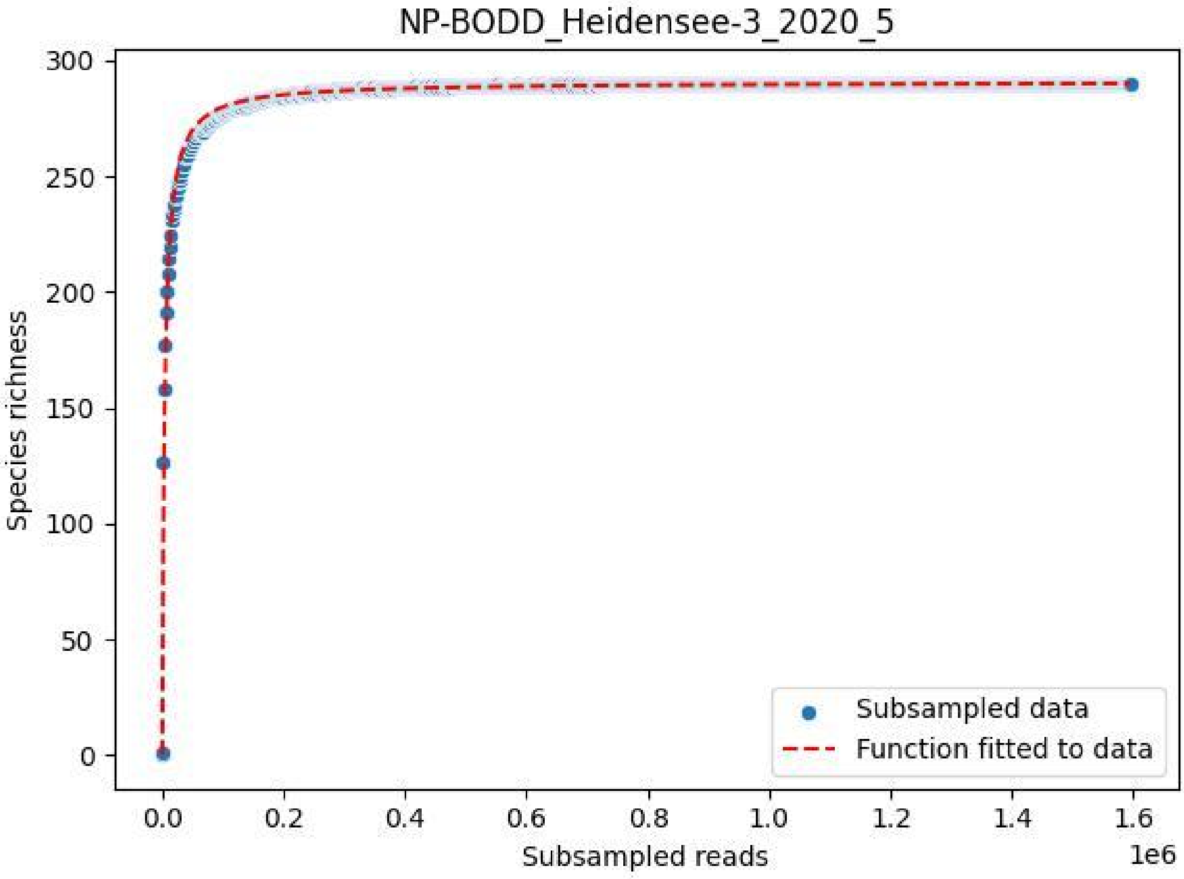

NP-BODD_Heidensee-3_2020_5
Species richness
Subsampled reads
1e6
300
250
200
150
100
50
0
0.0
0.2
0.4
0.6
0.8
1.0
1.2
1.4
1.6
Subsampled data
Function fitted to data

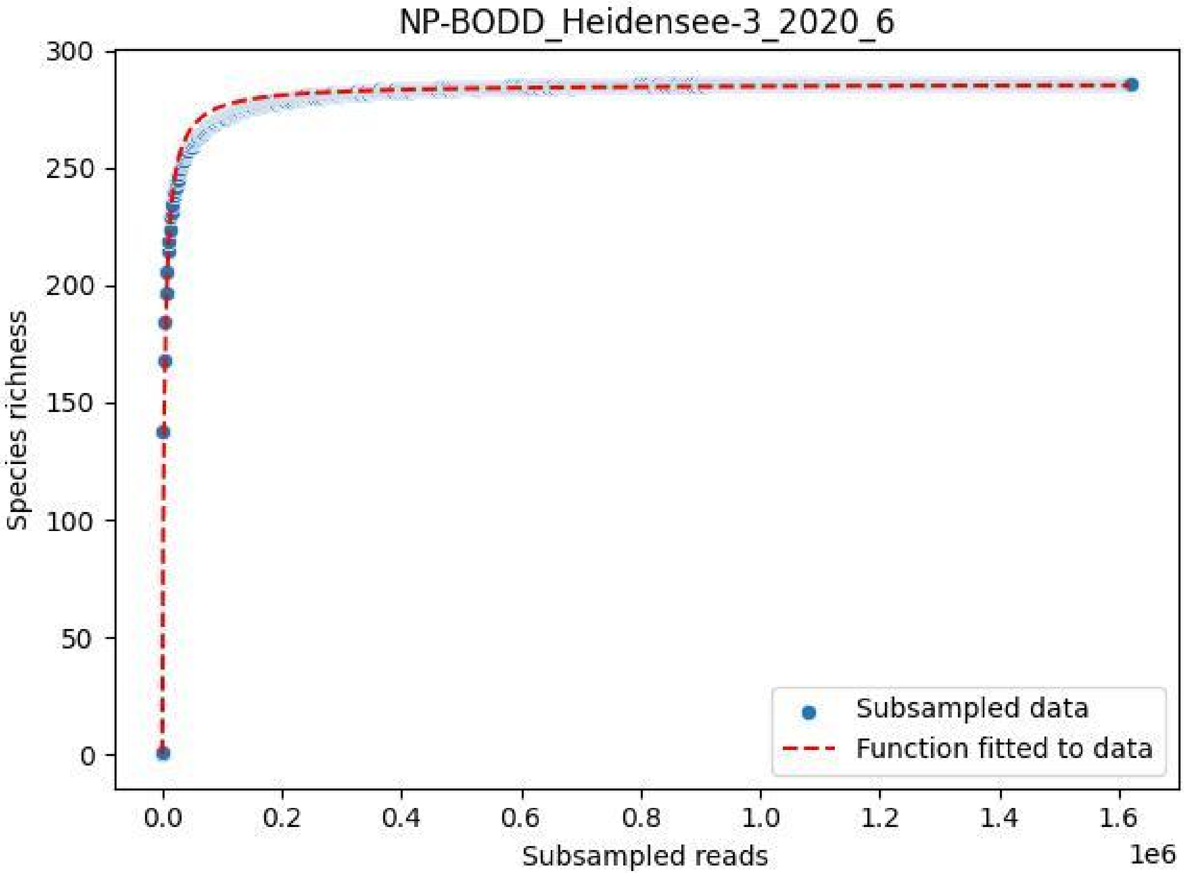
NP-BODD_Heidensee-3_2020_6
Species richness
Subsampled reads
1e6
Subsampled data
Function fitted to data
0
50
100
150
200
250
300
0.0
0.2
0.4
0.6
0.8
1.0
1.2
1.4
1.6

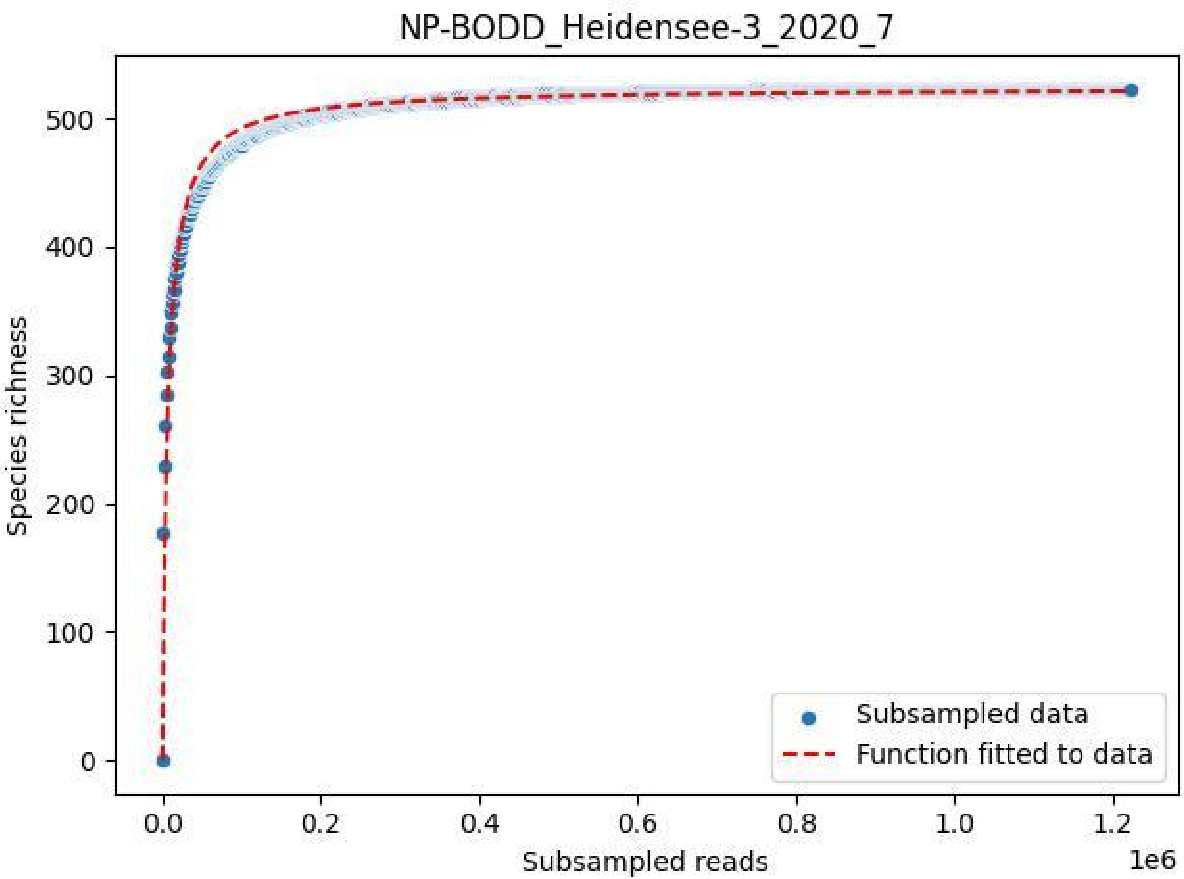
NP-BODD_Heidensee-3_2020_7
Species richness
Subsampled reads
1e6
0
100
200
300
400
500
0.0
0.2
0.4
0.6
0.8
1.0
1.2
Subsampled data
Function fitted to data

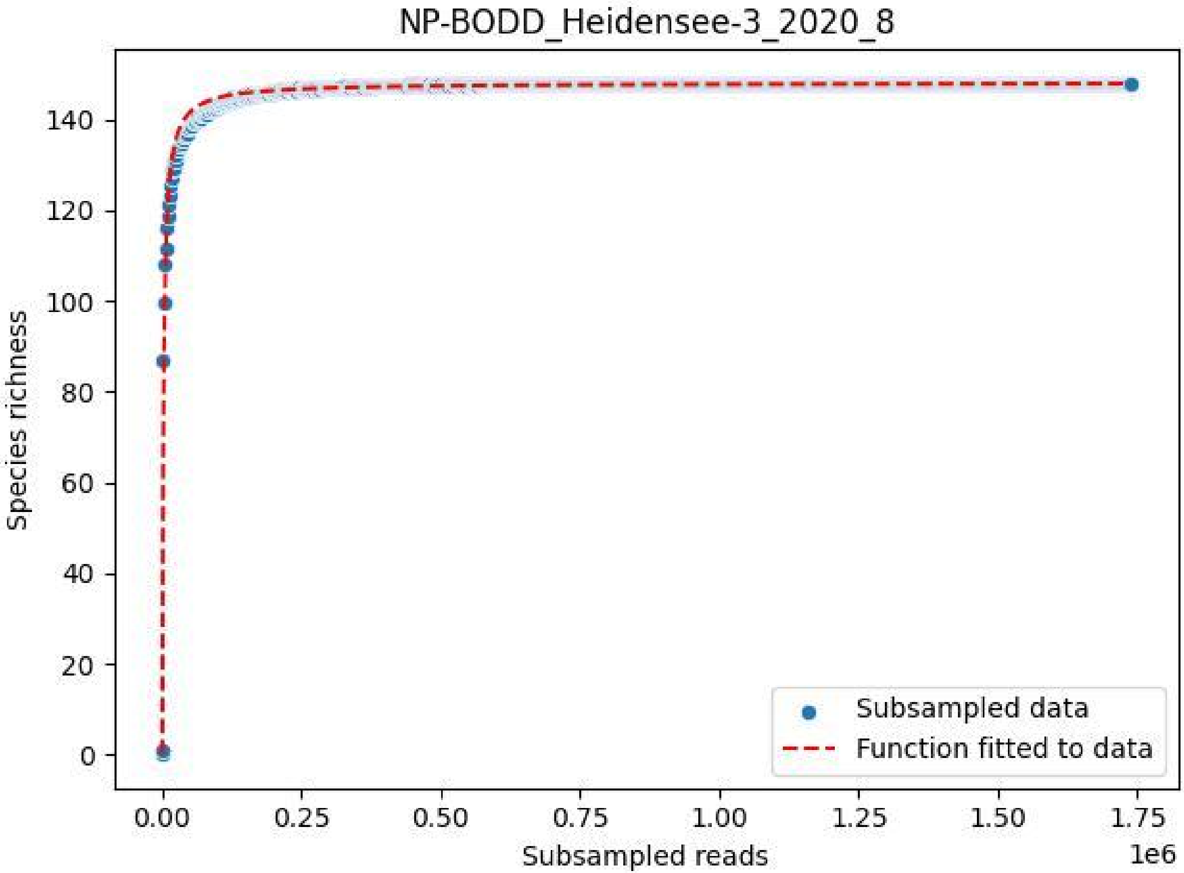
NP-BODD_Heidensee-3_2020_8
Species richness
0
20
40
60
80
100
120
140
Subsampled reads
0.00
0.25
0.50
0.75
1.00
1.25
1.50
1.75
1e6
Subsampled data
Function fitted to data

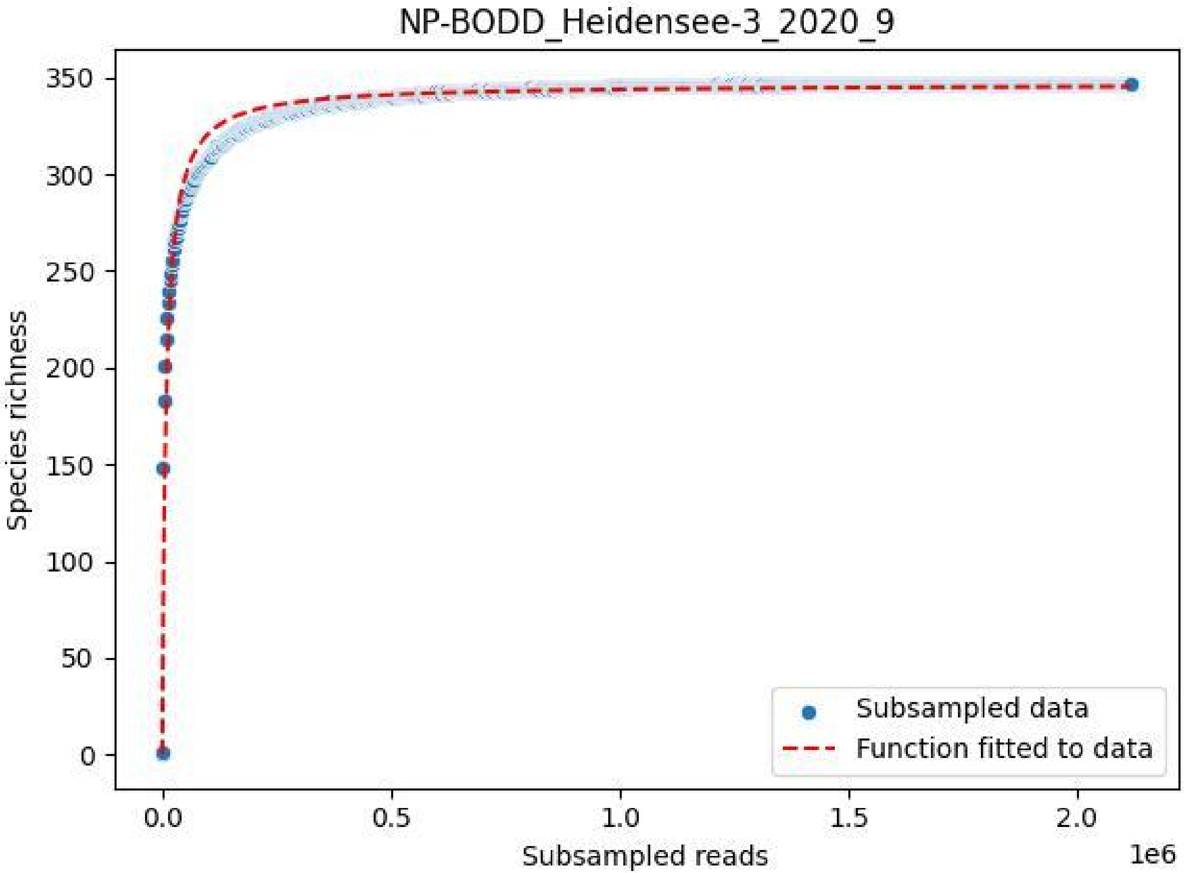
NP-BODD_Heidensee-3_2020_9
Species richness
Subsampled reads
1e6
0.0
0.5
1.0
1.5
2.0
0
50
100
150
200
250
300
350
Subsampled data
Function fitted to data

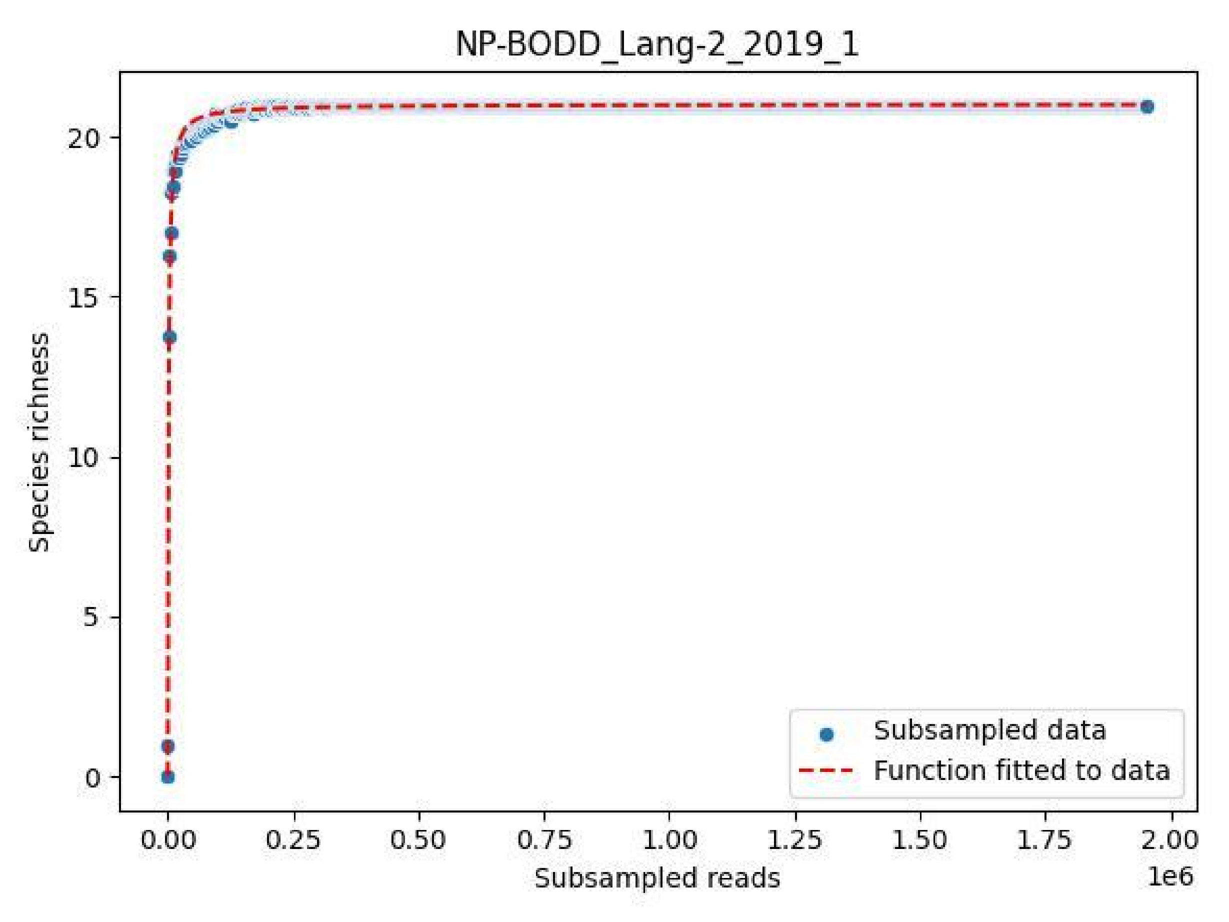
NP-BODD_Lang-2_2019_1
Species richness
20
15
10
5
0
0.00
0.25
0.50
0.75
1.00
1.25
1.50
1.75
2.00
1e6
Subsampled reads
Subsampled data
Function fitted to data

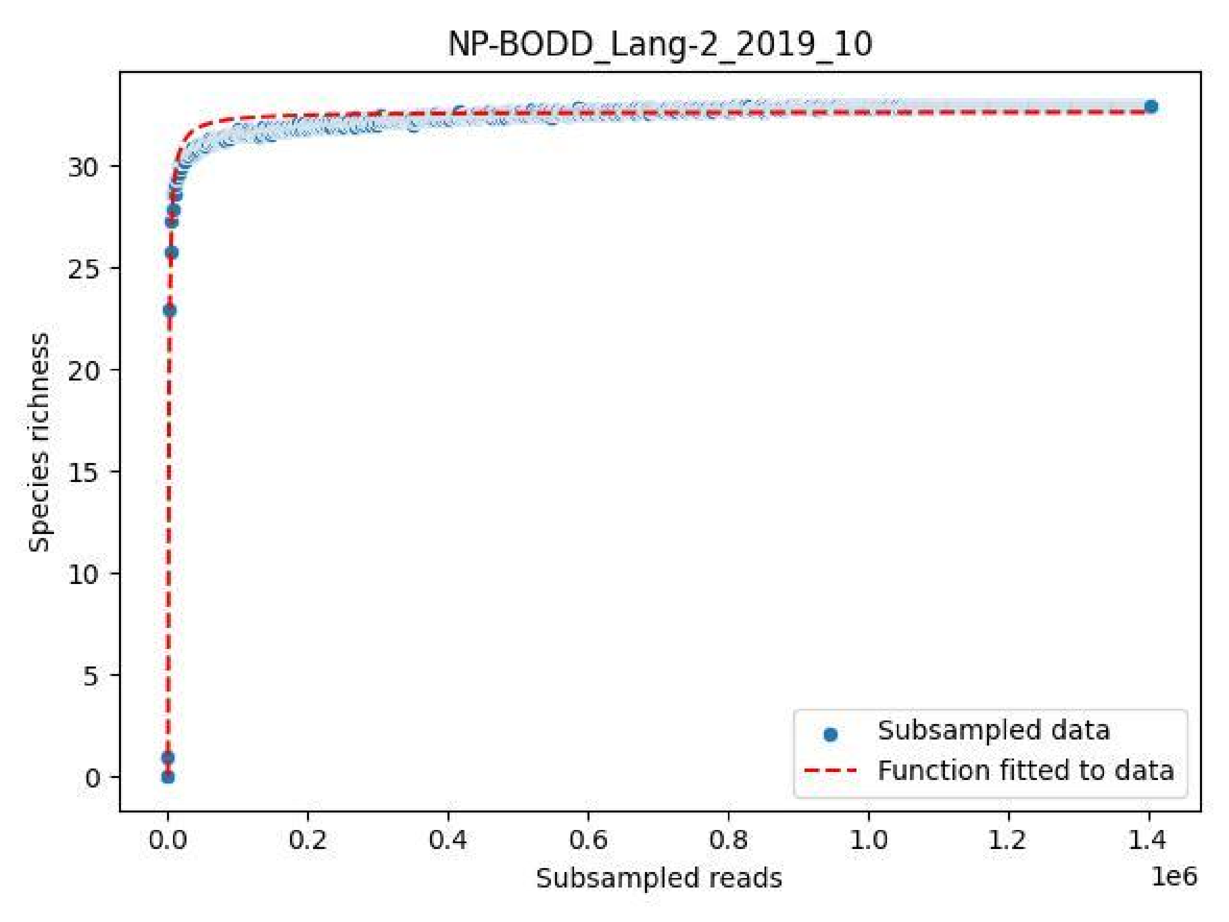
NP-BODD_Lang-2_2019_10
Species richness
Subsampled reads
1e6
0
5
10
15
20
25
30
0.0
0.2
0.4
0.6
0.8
1.0
1.2
1.4
Subsampled data
Function fitted to data

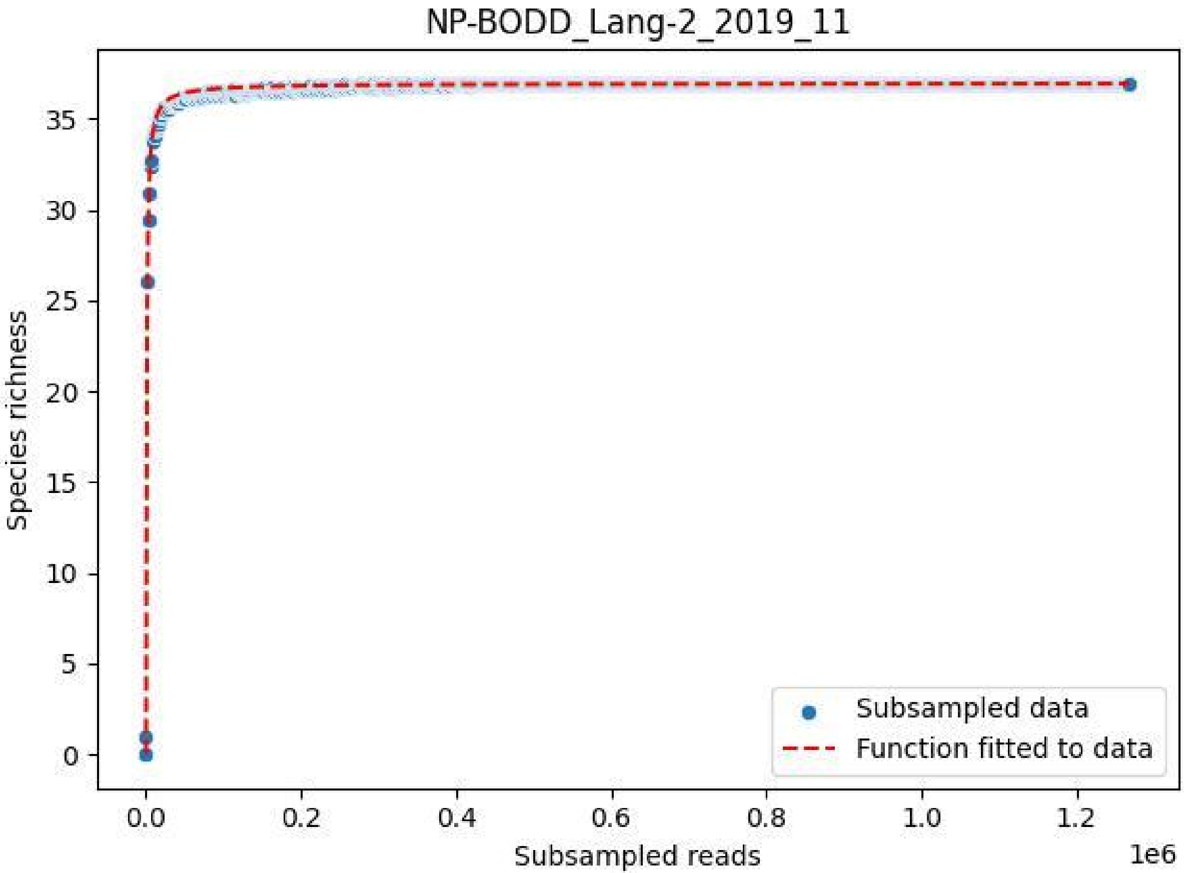
NP-BODD_Lang-2_2019_11
Species richness
Subsampled reads
1e6
0
5
10
15
20
25
30
35
0.0
0.2
0.4
0.6
0.8
1.0
1.2
Subsampled data
Function fitted to data

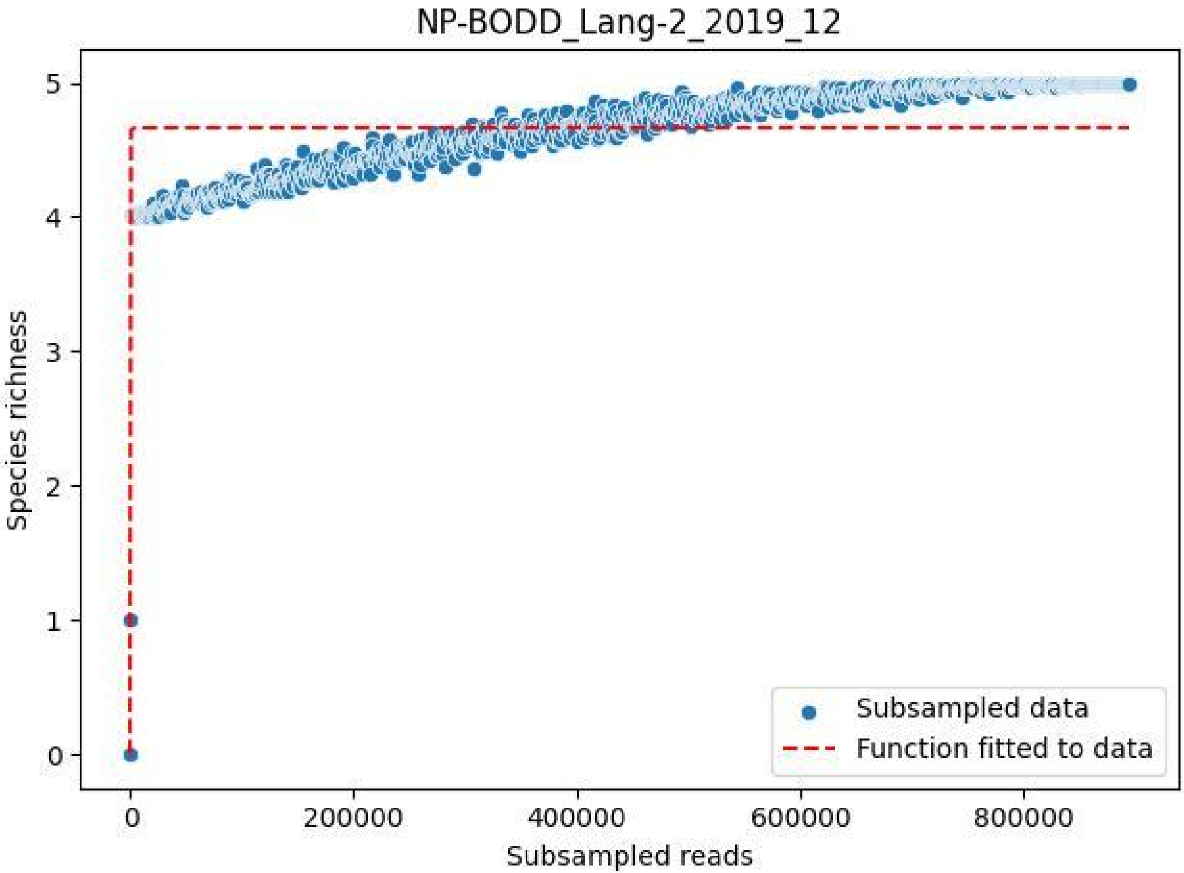

NP-BODD_Lang-2_2019_12
Species richness
Subsampled reads
0
1
2
3
4
5
0
200000
400000
600000
800000
Subsampled data
Function fitted to data

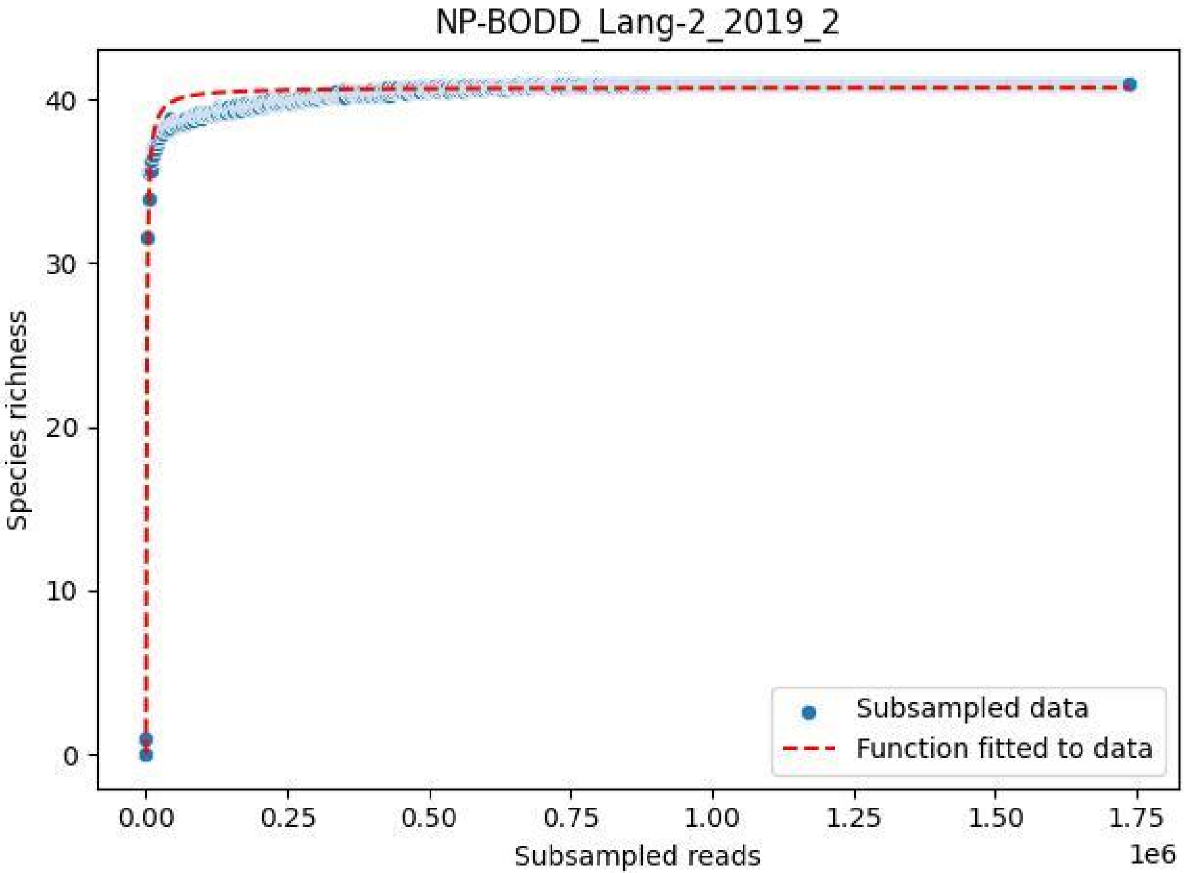
NP-BODD_Lang-2_2019_2
Species richness
Subsampled reads
1e6
0
10
20
30
40
0.00
0.25
0.50
0.75
1.00
1.25
1.50
1.75
Subsampled data
Function fitted to data

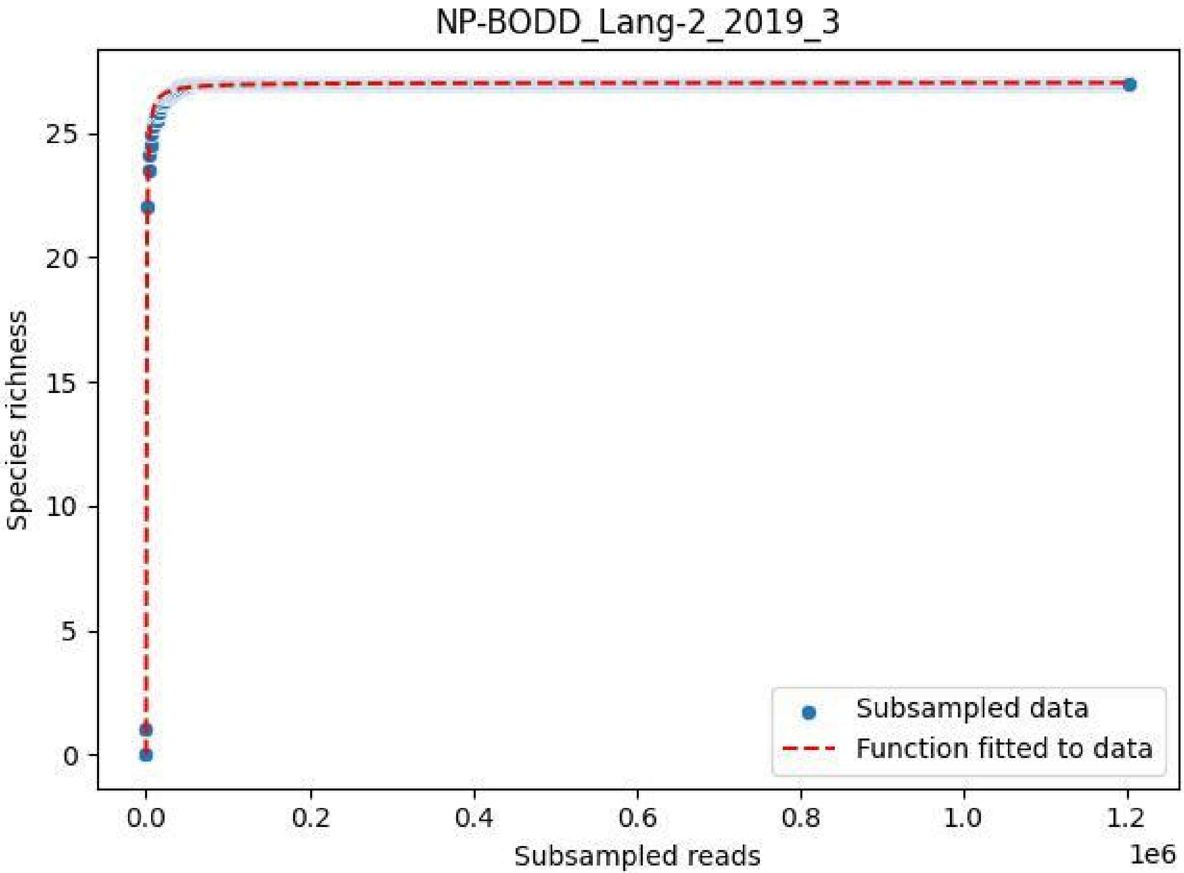
NP-BODD_Lang-2_2019_3
Species richness
Subsampled reads
1e6
0
5
10
15
20
25
0.0
0.2
0.4
0.6
0.8
1.0
1.2
Subsampled data
Function fitted to data

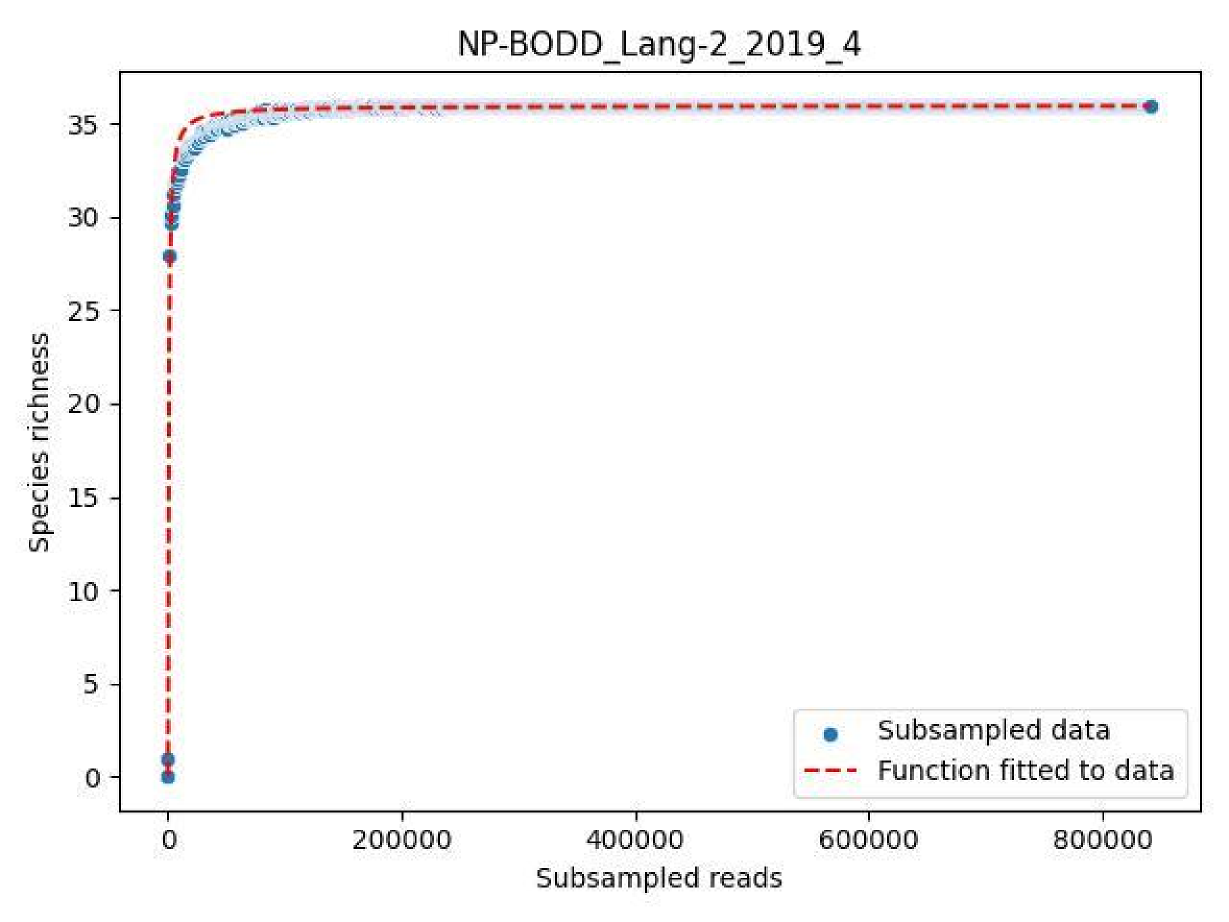
NP-BODD_Lang-2_2019_4
Species richness
0
5
10
15
20
25
30
35
Subsampled reads
0
200000
400000
600000
800000
Subsampled data
Function fitted to data

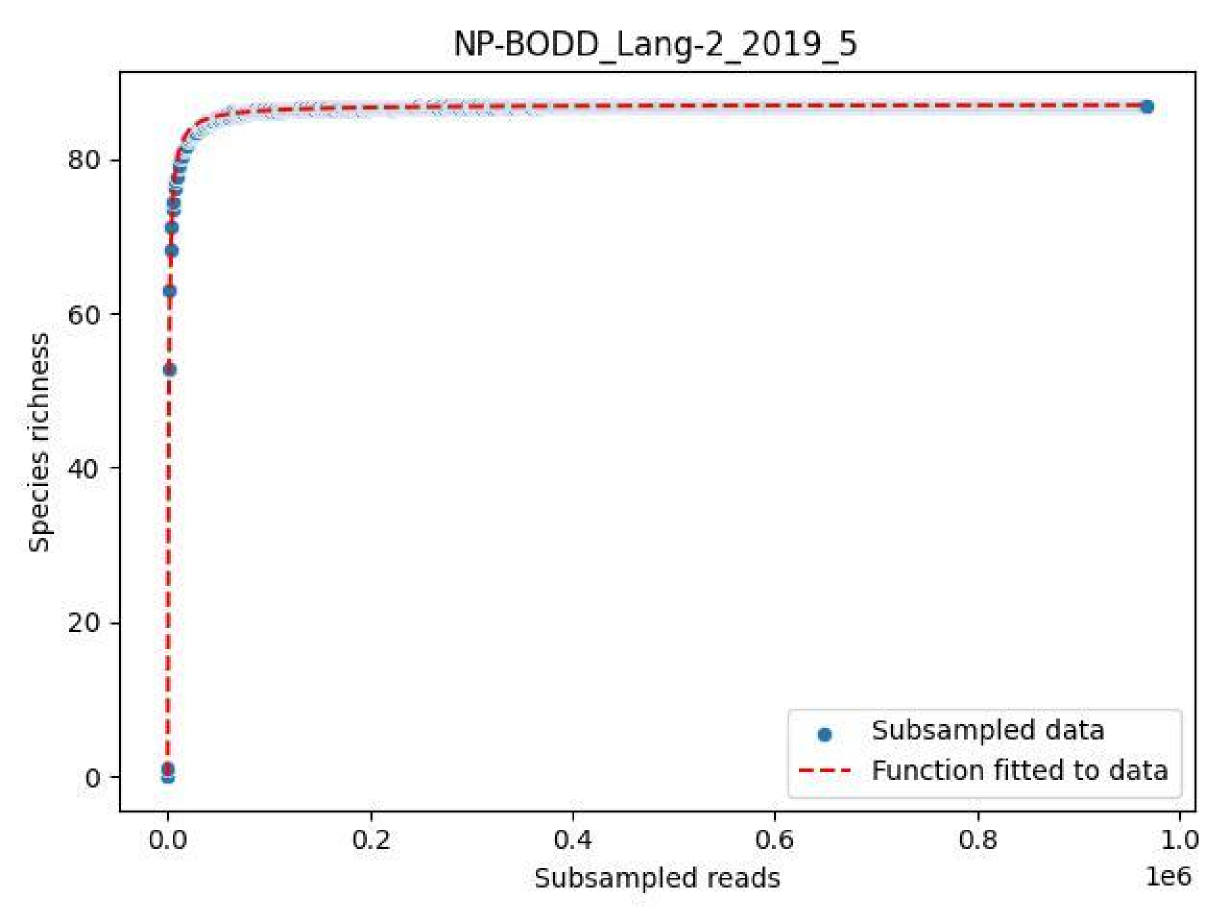
NP-BODD_Lang-2_2019_5
Species richness
0
20
40
60
80
0.0
0.2
0.4
0.6
0.8
1.0
1e6
Subsampled reads
Subsampled data
Function fitted to data

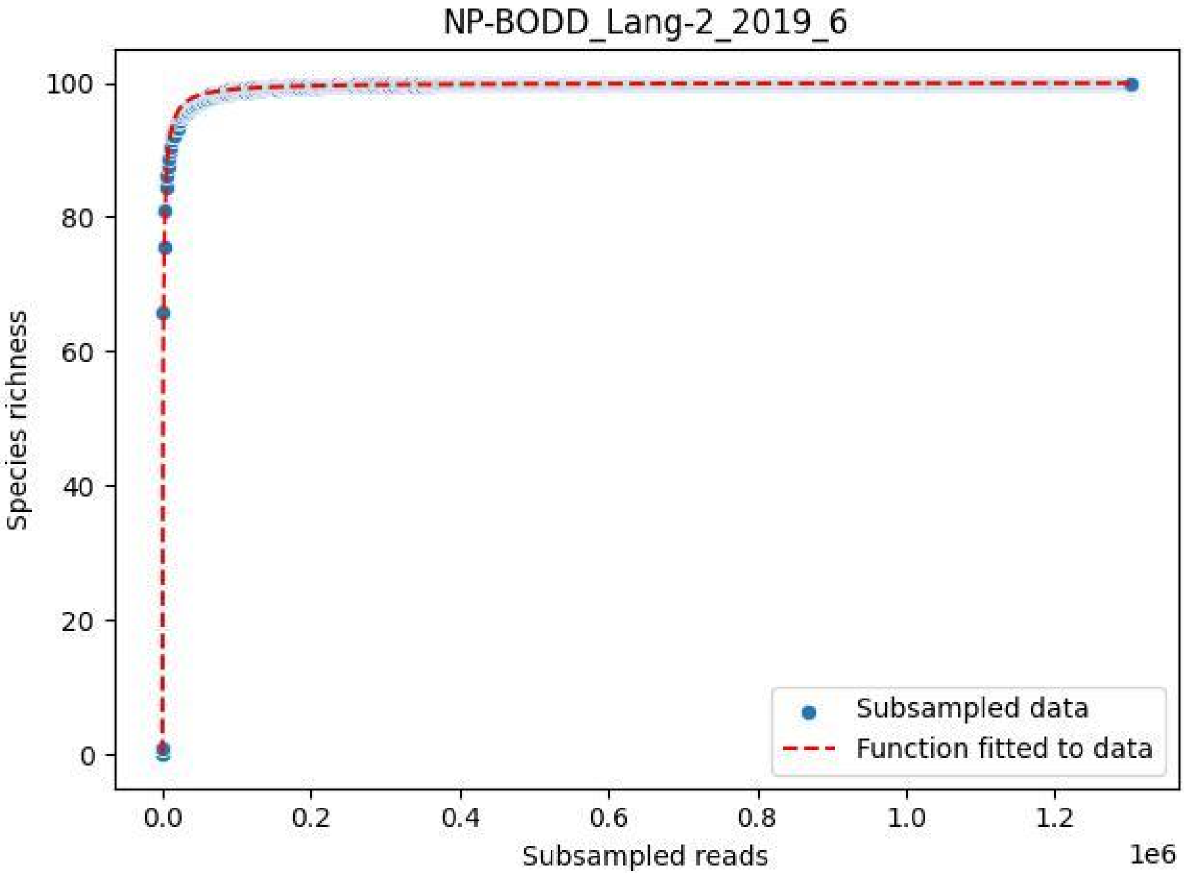
NP-BODD_Lang-2_2019_6
Species richness
100
80
60
40
20
0
0.0
0.2
0.4
0.6
0.8
1.0
1.2
Subsampled reads
1e6
Subsampled data
Function fitted to data

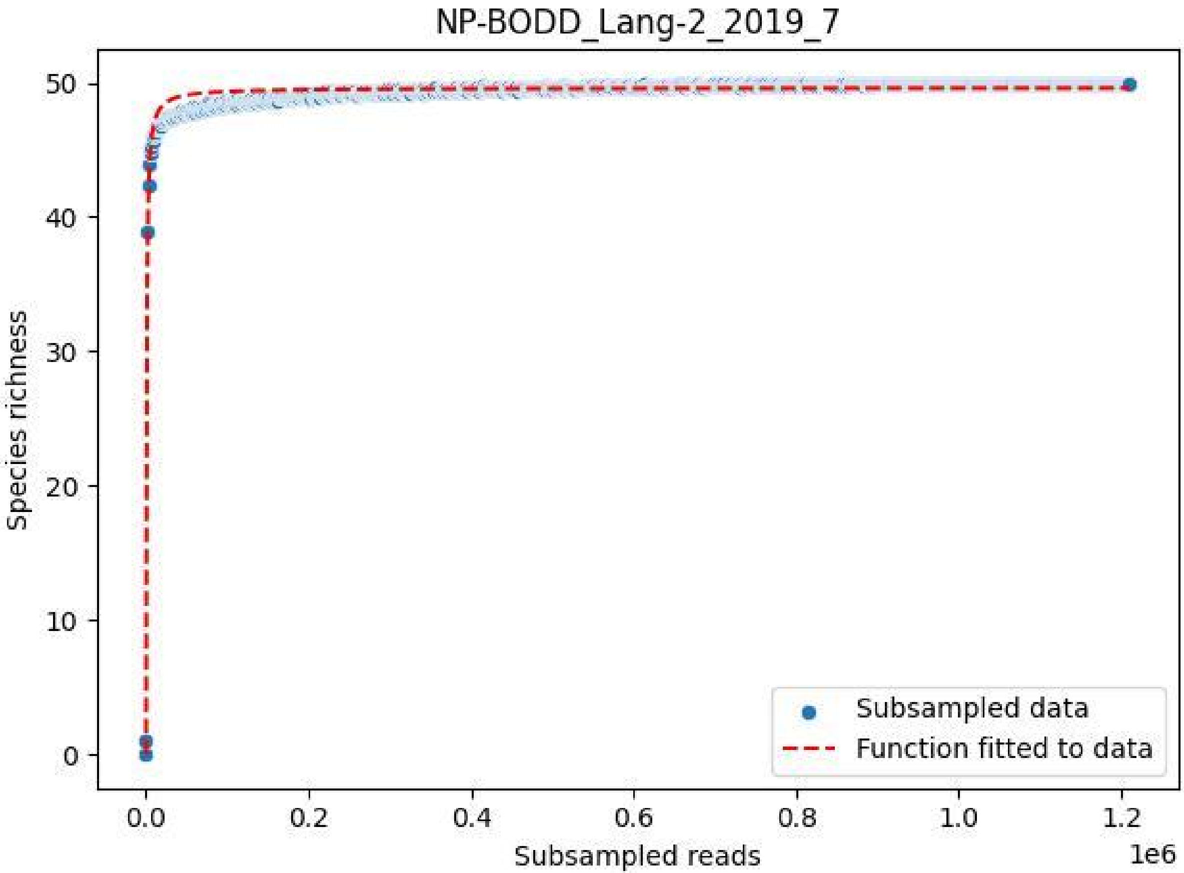
NP-BODD_Lang-2_2019_7
Species richness
Subsampled reads
1e6
0
10
20
30
40
50
0.0
0.2
0.4
0.6
0.8
1.0
1.2
Subsampled data
Function fitted to data

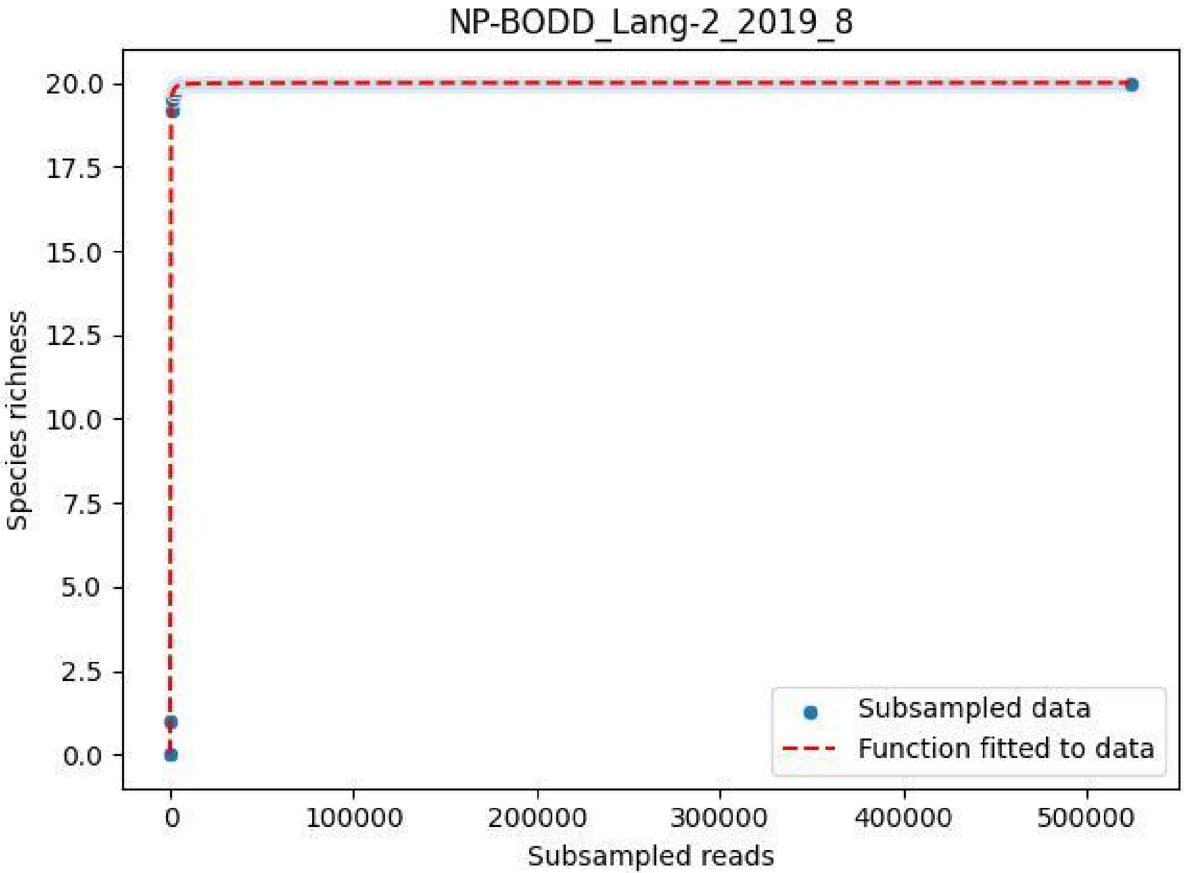

NP-BODD_Lang-2_2019_8
Species richness
Subsampled reads
20.0
17.5
15.0
12.5
10.0
7.5
5.0
2.5
0.0
0
100000
200000
300000
400000
500000
Subsampled data
Function fitted to data

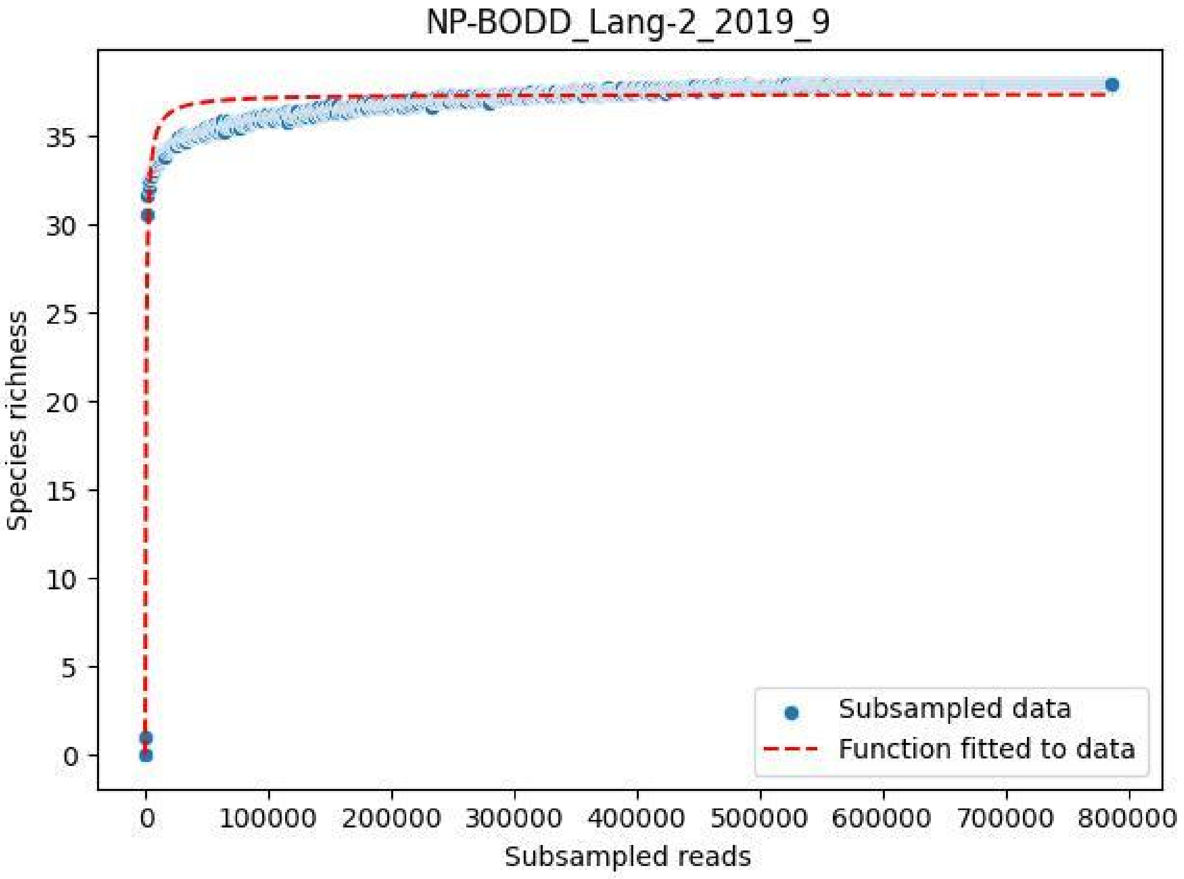

NP-BODD_Lang-2_2019_9
Species richness
0
5
10
15
20
25
30
35
0
100000
200000
300000
400000
500000
600000
700000
800000
Subsampled reads
Subsampled data
Function fitted to data

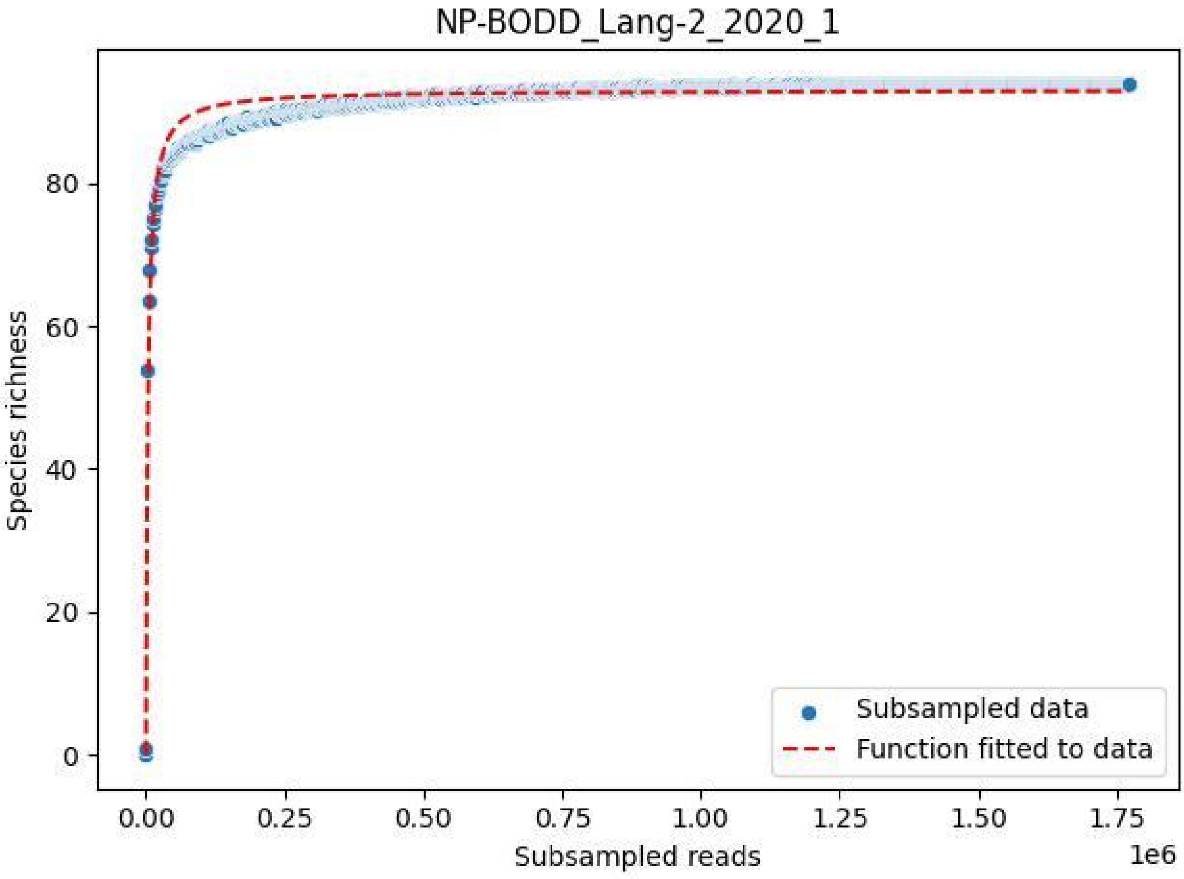
NP-BODD_Lang-2_2020_1
Species richness
0
20
40
60
80
0.00
0.25
0.50
0.75
1.00
1.25
1.50
1.75
1e6
Subsampled reads
Subsampled data
Function fitted to data

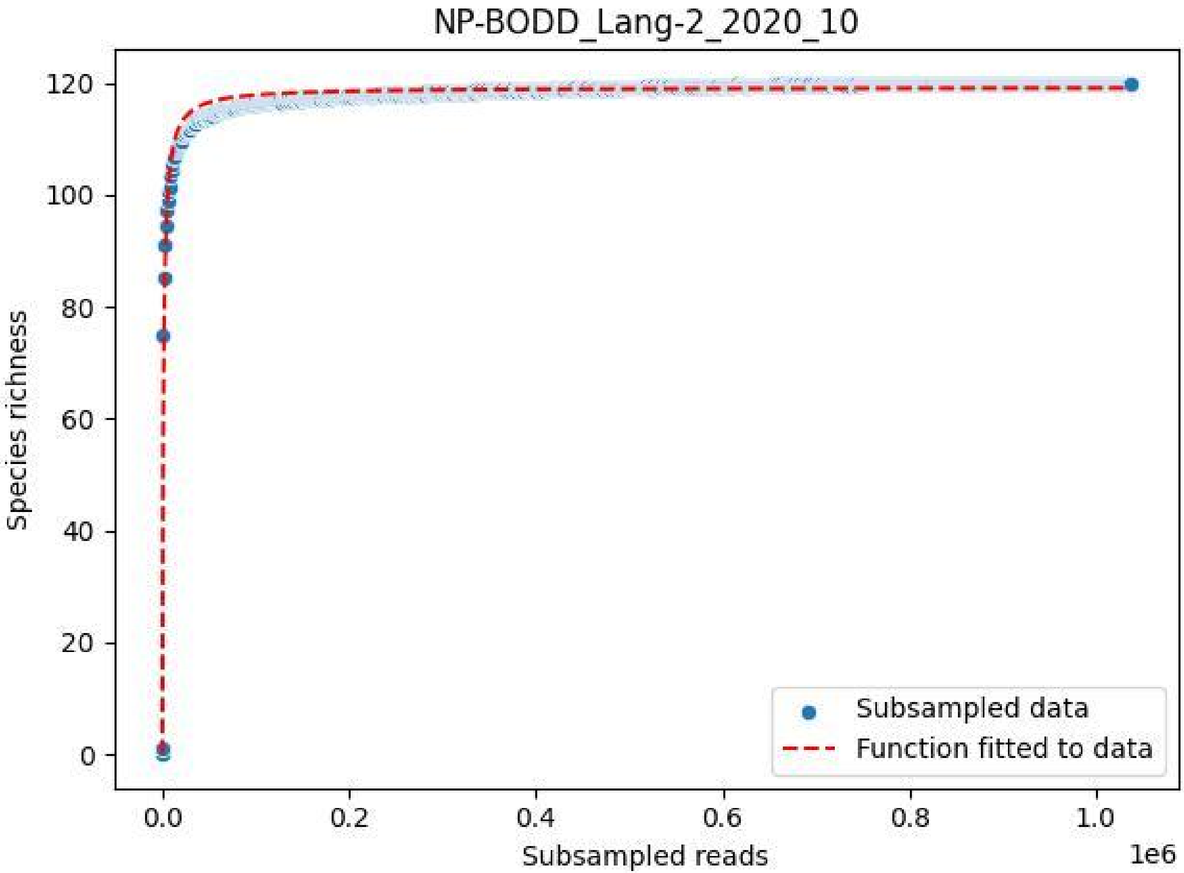
NP-BODD_Lang-2_2020_10
Species richness
Subsampled reads
1e6
0
20
40
60
80
100
120
0.0
0.2
0.4
0.6
0.8
1.0
Subsampled data
Function fitted to data

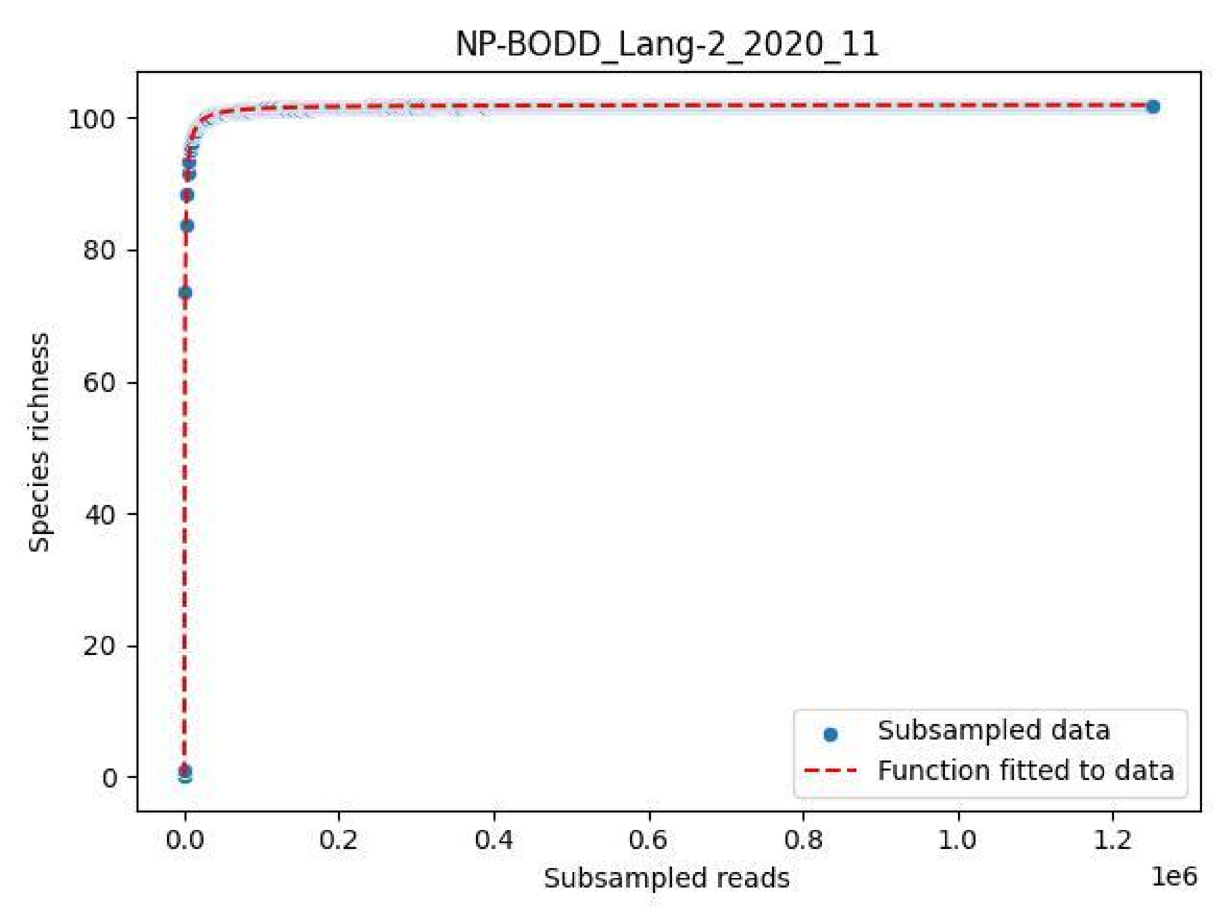
NP-BODD_Lang-2_2020_11
Species richness
Subsampled reads
1e6
0
20
40
60
80
100
0.0
0.2
0.4
0.6
0.8
1.0
1.2
Subsampled data
Function fitted to data

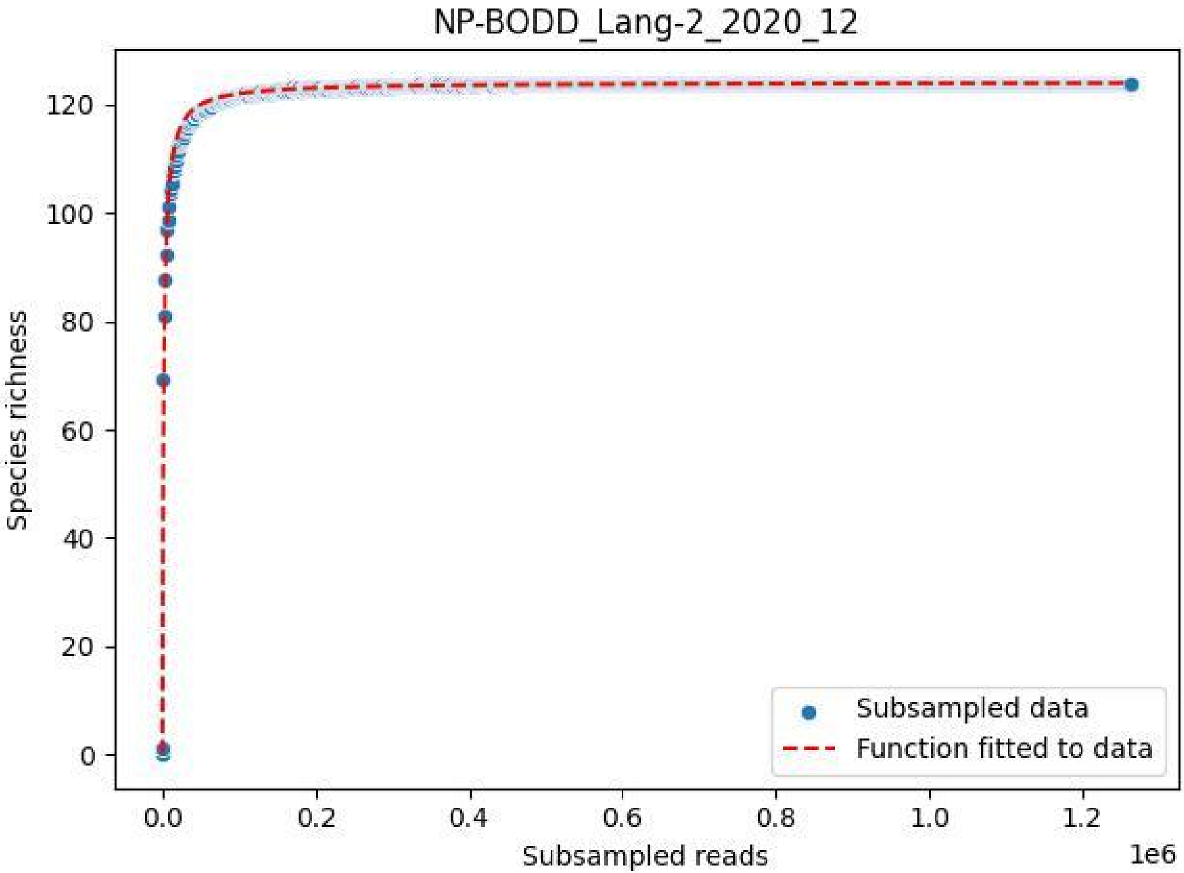

NP-BODD_Lang-2_2020_12
Species richness
Subsampled reads
1e6
0
20
40
60
80
100
120
0.0
0.2
0.4
0.6
0.8
1.0
1.2
Subsampled data
Function fitted to data

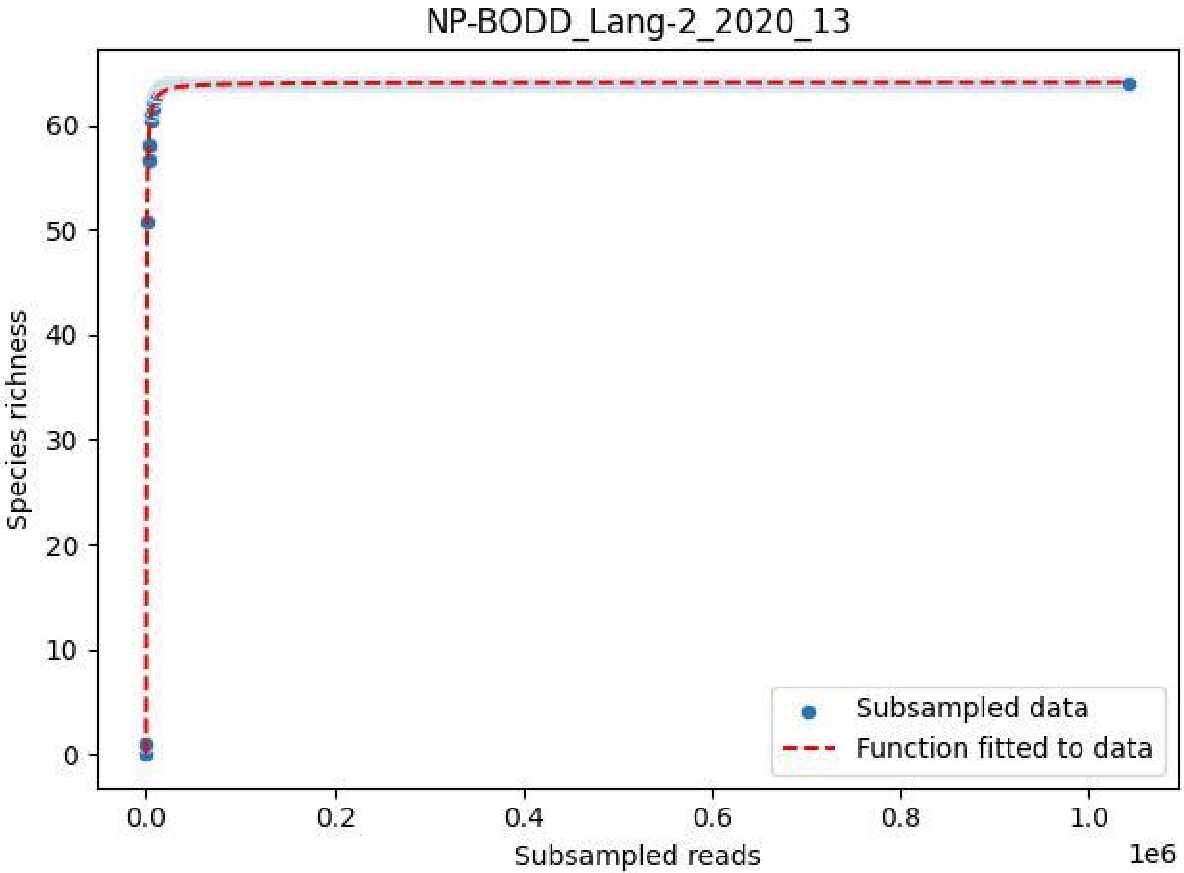
NP-BODD_Lang-2_2020_13
Species richness
Subsampled reads
1e6
0
10
20
30
40
50
60
0.0
0.2
0.4
0.6
0.8
1.0
Subsampled data
Function fitted to data

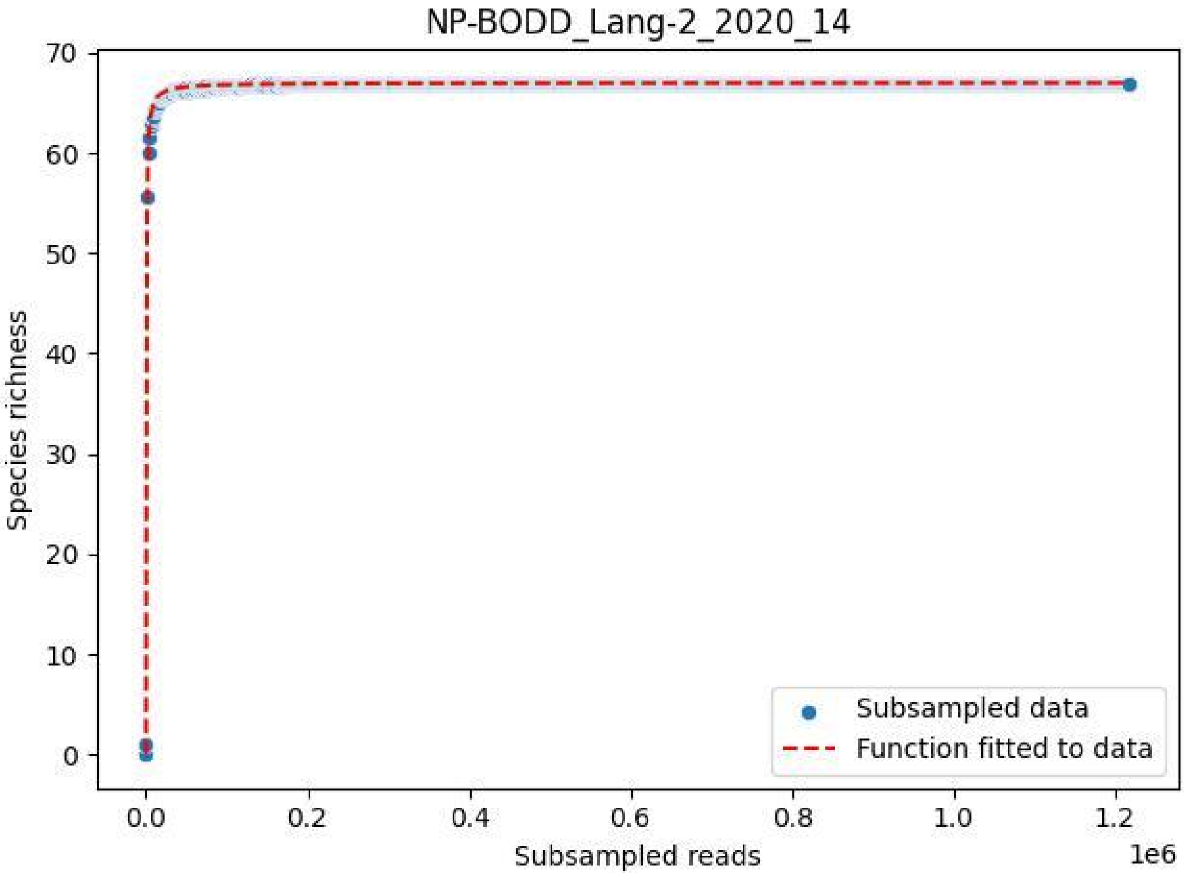
NP-BODD_Lang-2_2020_14
Species richness
Subsampled reads
1e6
0
10
20
30
40
50
60
70
0.0
0.2
0.4
0.6
0.8
1.0
1.2
Subsampled data
Function fitted to data

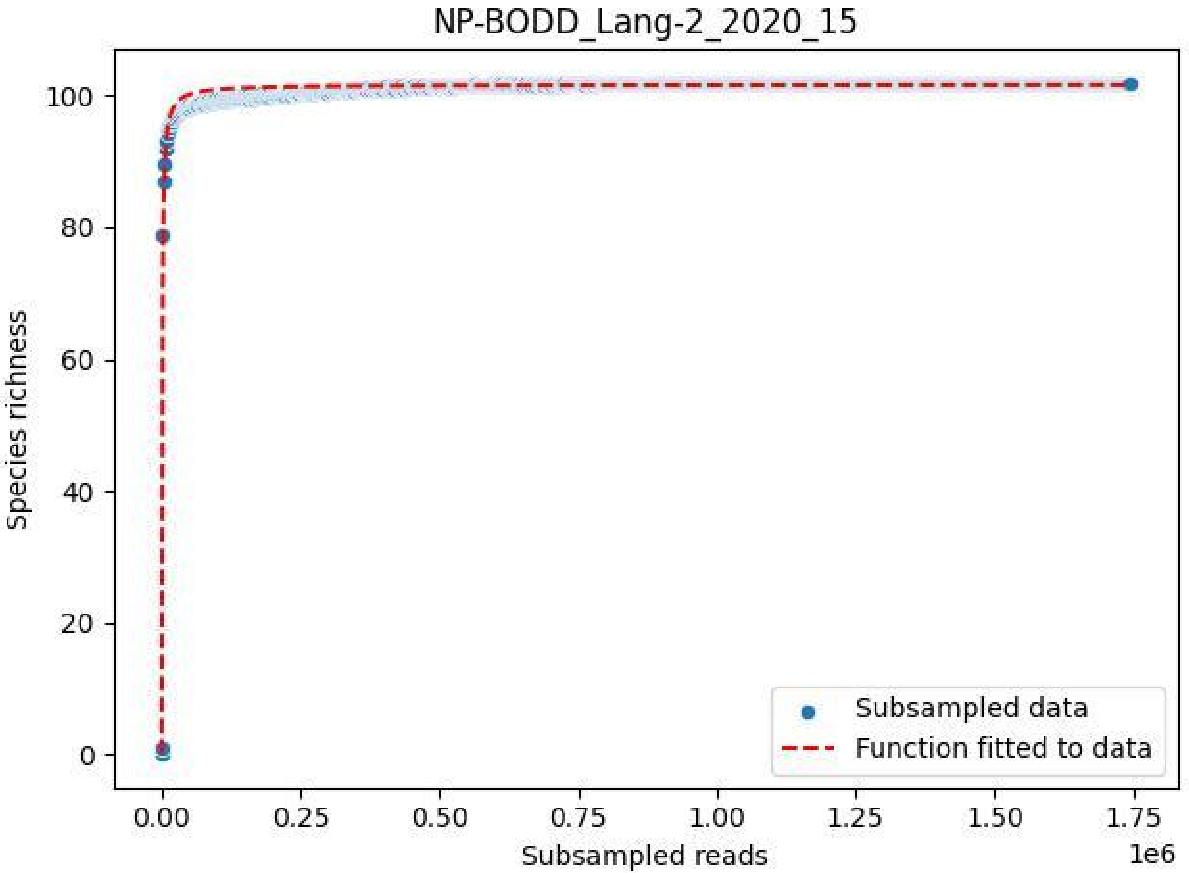

NP-BODD_Lang-2_2020_15
Species richness
Subsampled reads
1e6
0
20
40
60
80
100
0.00
0.25
0.50
0.75
1.00
1.25
1.50
1.75
Subsampled data
Function fitted to data

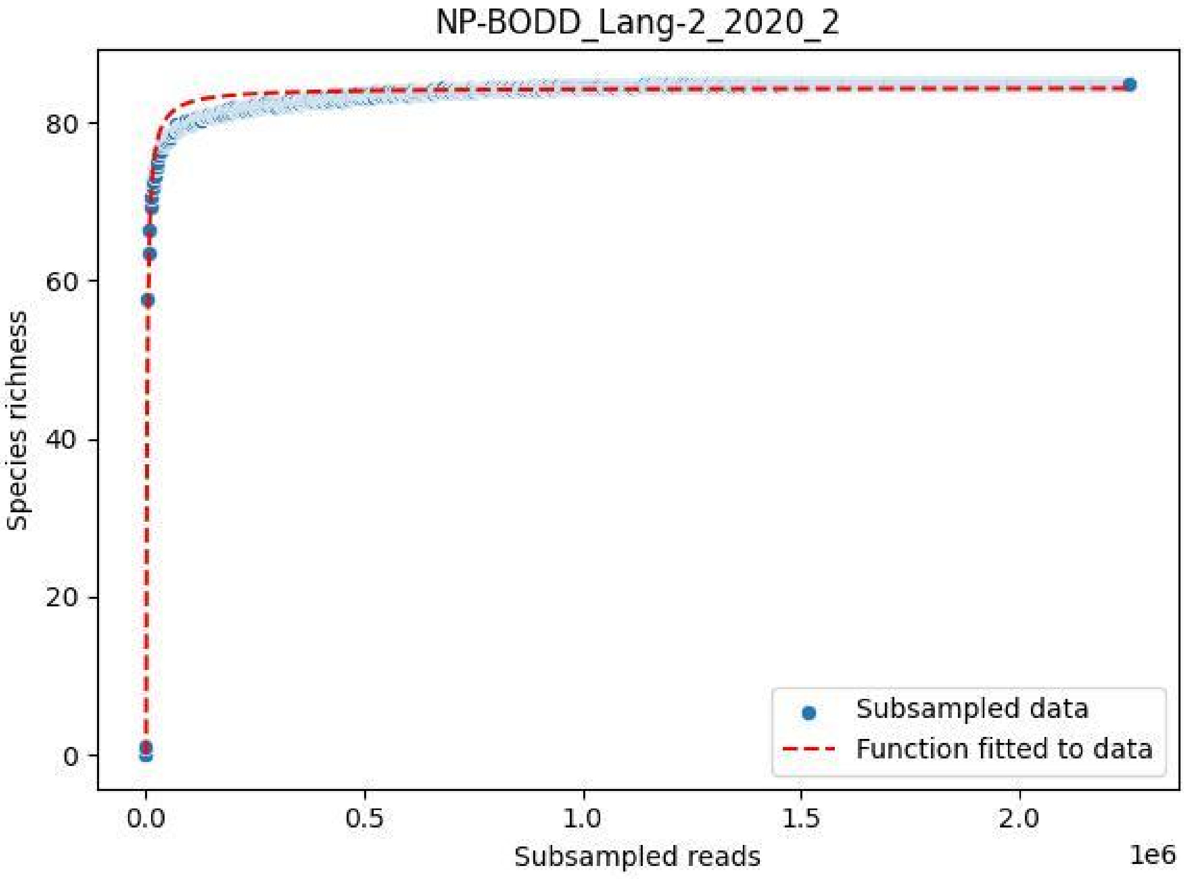
NP-BODD_Lang-2_2020_2
Species richness
Subsampled reads
1e6
0
20
40
60
80
0.0
0.5
1.0
1.5
2.0
Subsampled data
Function fitted to data

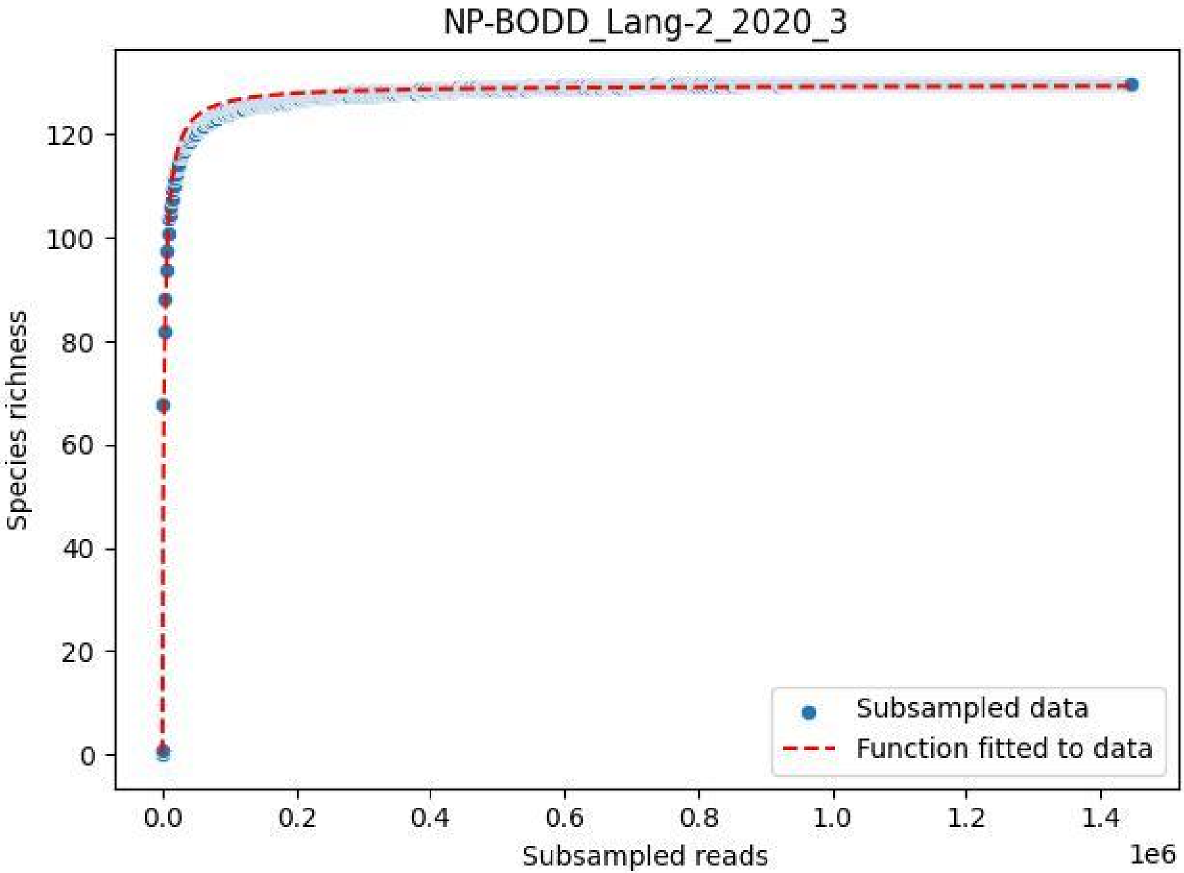
NP-BODD_Lang-2_2020_3
Species richness
Subsampled reads
1e6
0
20
40
60
80
100
120
0.0
0.2
0.4
0.6
0.8
1.0
1.2
1.4
Subsampled data
Function fitted to data

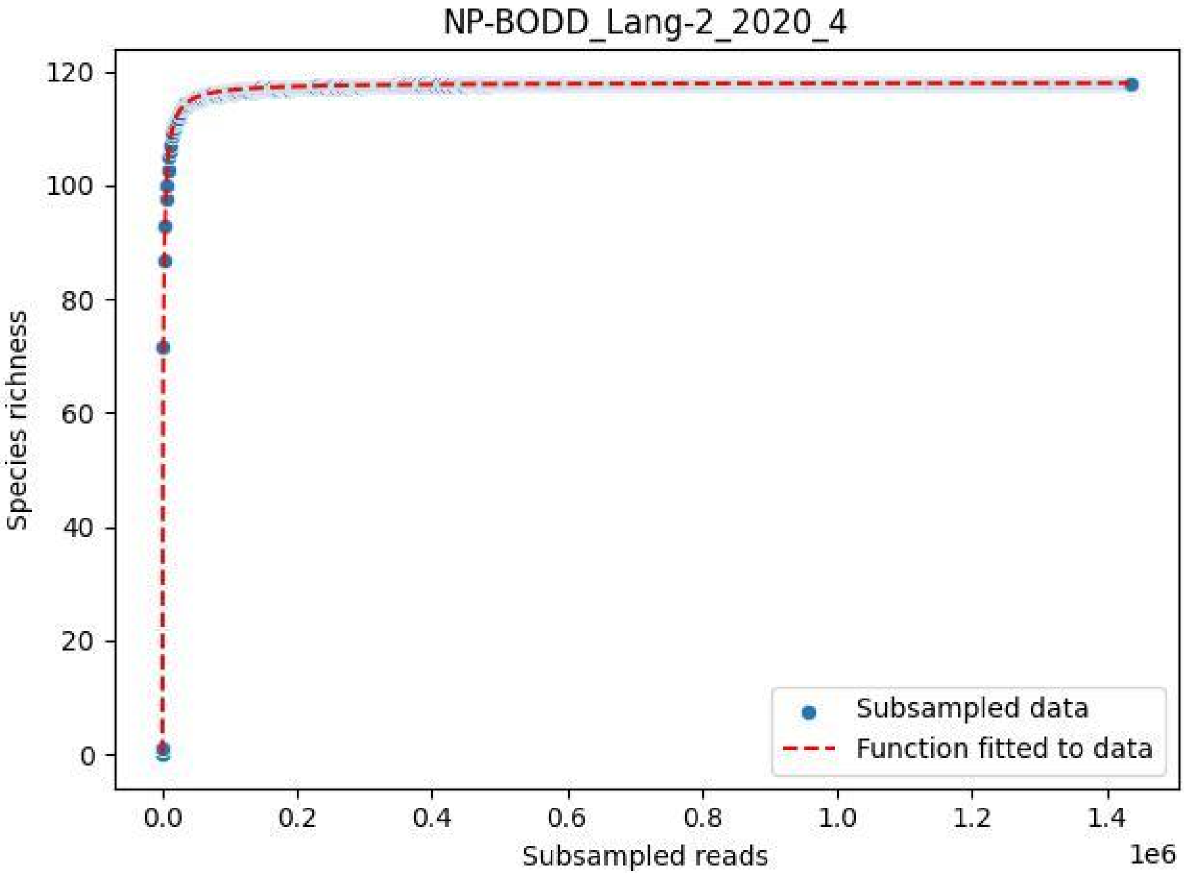
NP-BODD_Lang-2_2020_4
Species richness
Subsampled reads
1e6
0.0
0.2
0.4
0.6
0.8
1.0
1.2
1.4
0
20
40
60
80
100
120
Subsampled data
Function fitted to data

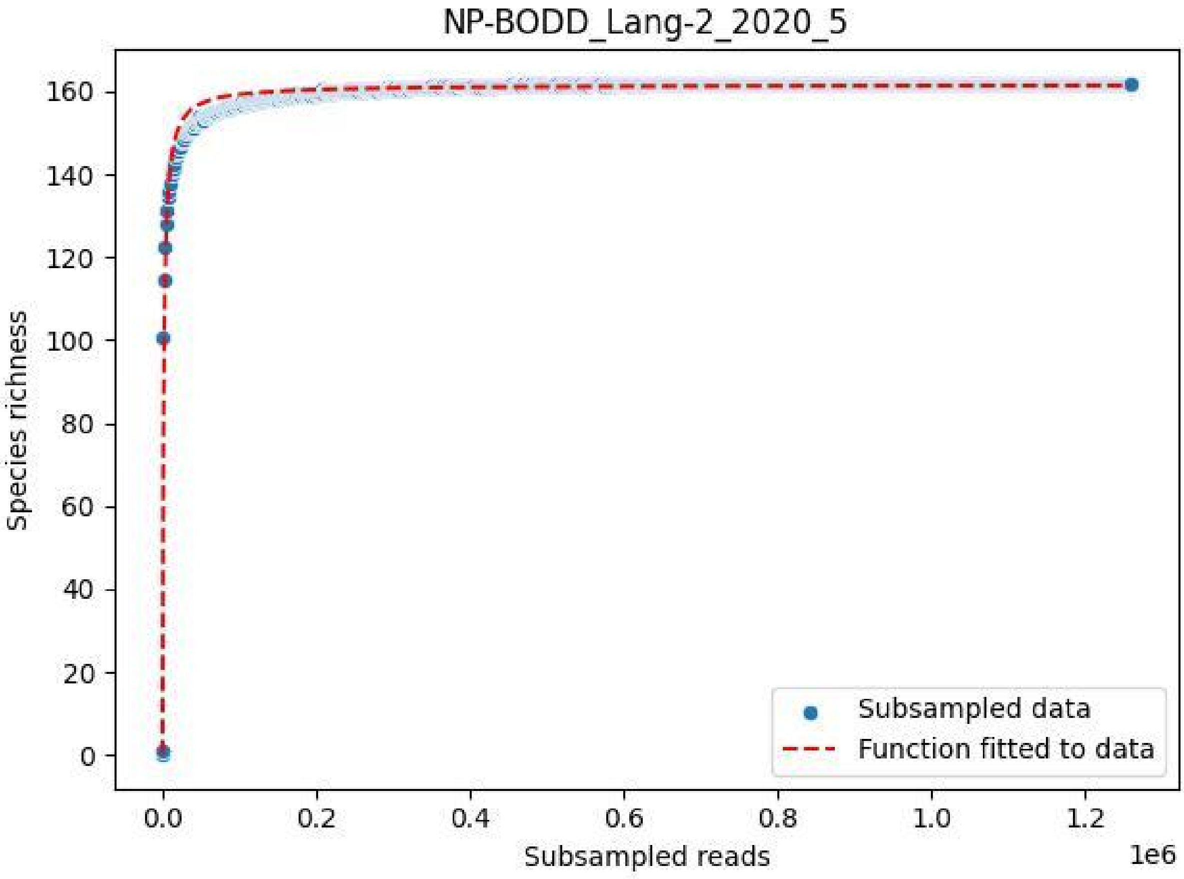

NP-BODD_Lang-2_2020_5
Species richness
Subsampled reads
1e6
Subsampled data
Function fitted to data
0
20
40
60
80
100
120
140
160
0.0
0.2
0.4
0.6
0.8
1.0
1.2

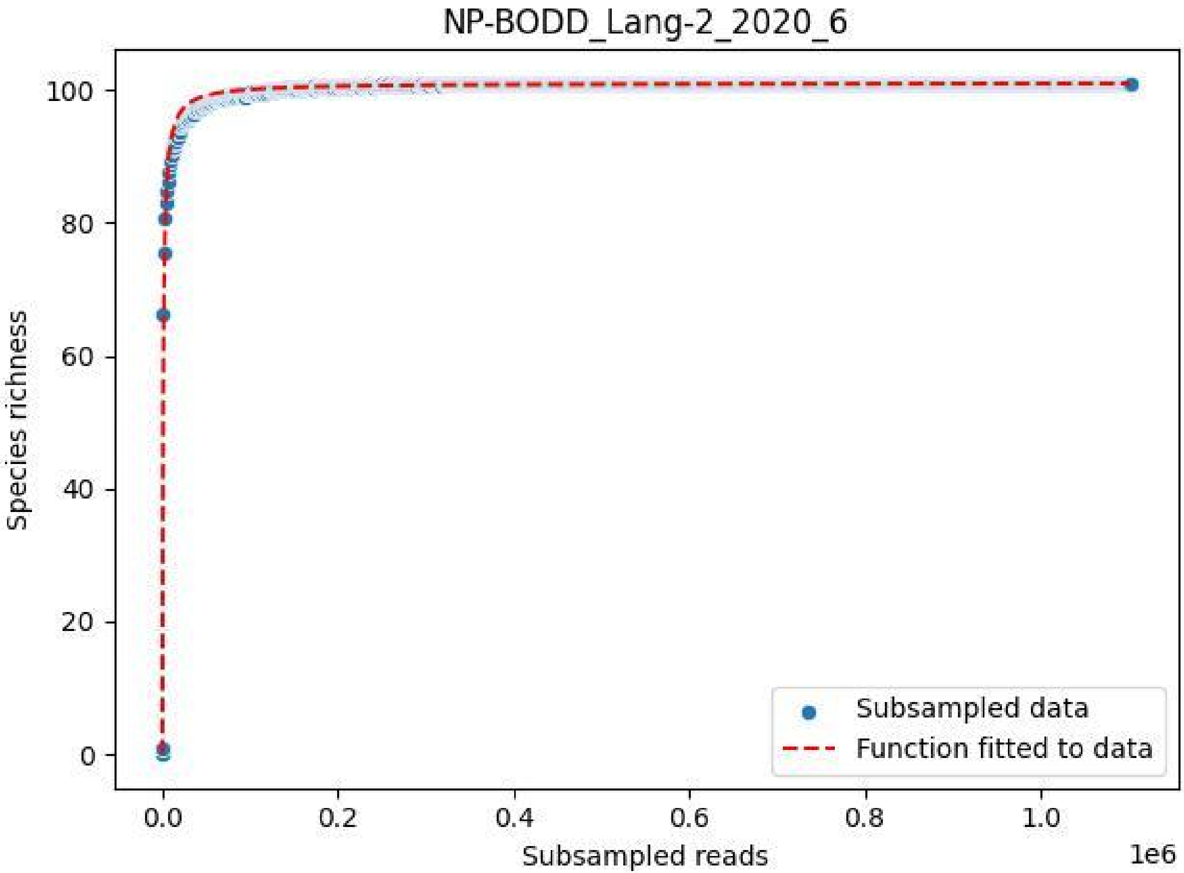

NP-BODD_Lang-2_2020_6
Species richness
Subsampled reads
1e6
0
20
40
60
80
100
0.0
0.2
0.4
0.6
0.8
1.0
Subsampled data
Function fitted to data

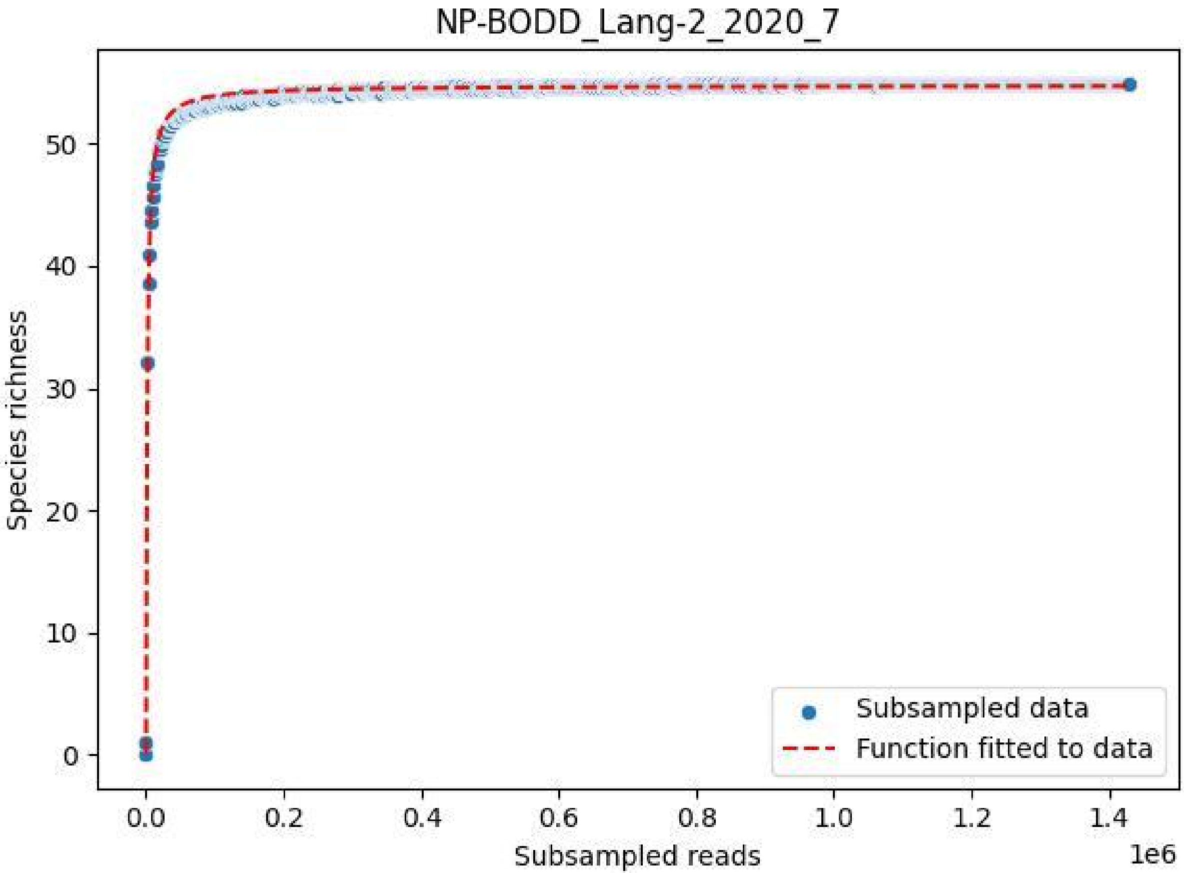

NP-BODD_Lang-2_2020_7
Species richness
Subsampled reads
1e6
0
10
20
30
40
50
0.0
0.2
0.4
0.6
0.8
1.0
1.2
1.4
Subsampled data
Function fitted to data

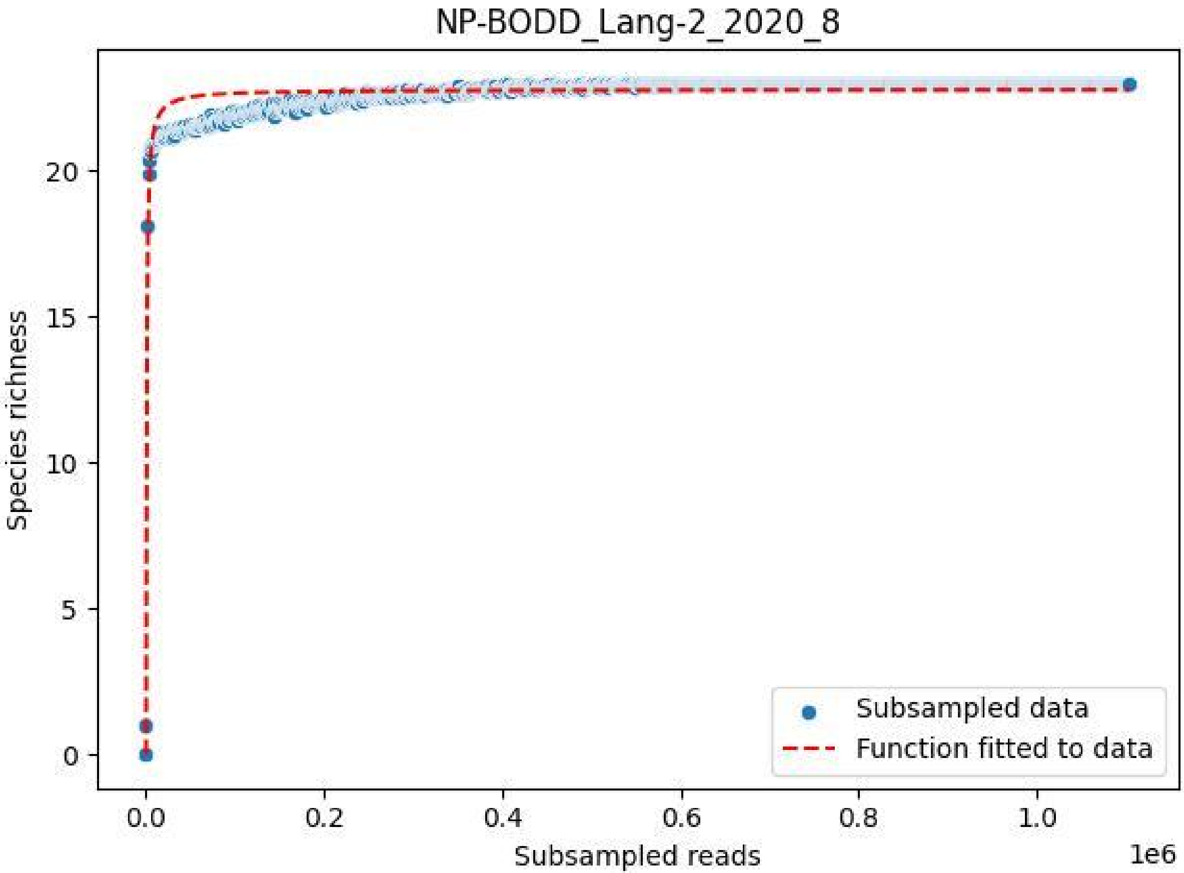

NP-BODD_Lang-2_2020_8
Species richness
0
5
10
15
20
0.0
0.2
0.4
0.6
0.8
1.0
1e6
Subsampled reads
Subsampled data
Function fitted to data

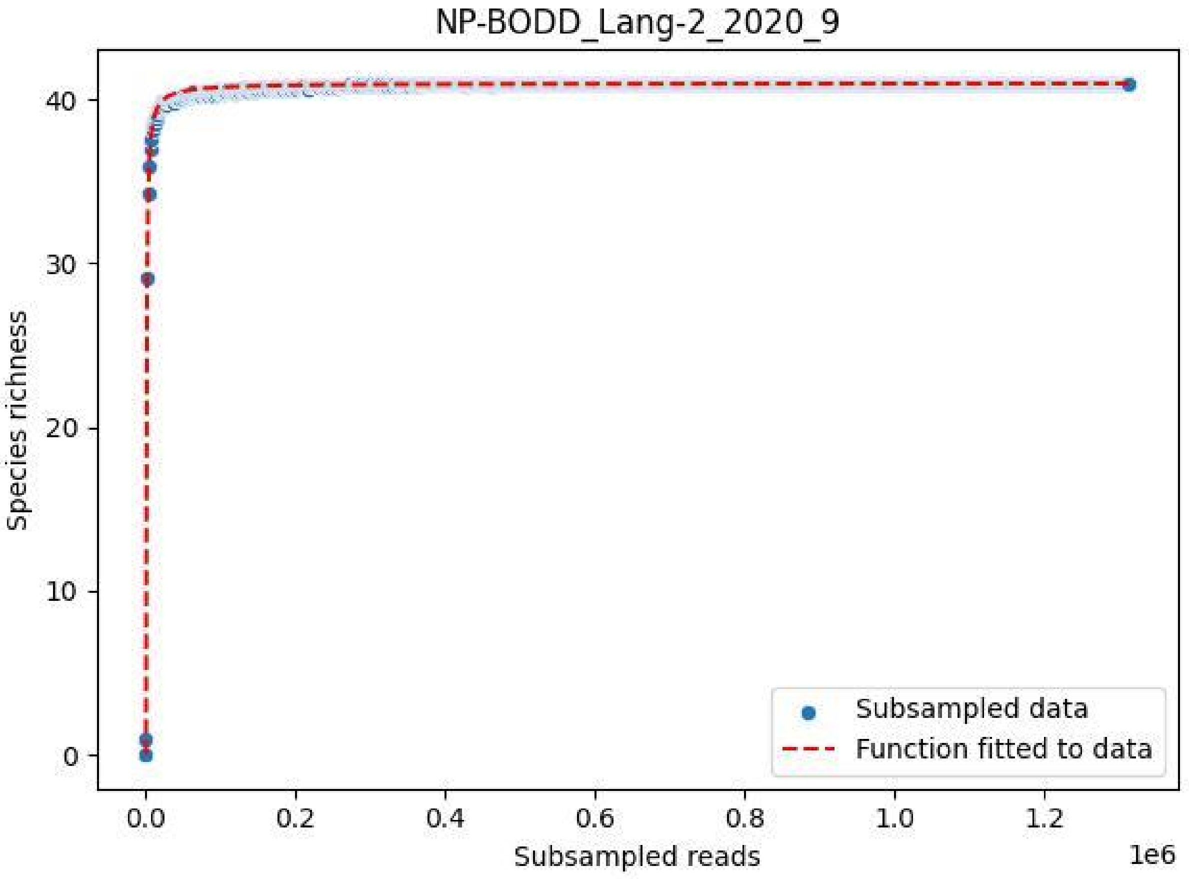
NP-BODD_Lang-2_2020_9
Species richness
Subsampled reads
1e6
0
10
20
30
40
0.0
0.2
0.4
0.6
0.8
1.0
1.2
Subsampled data
Function fitted to data

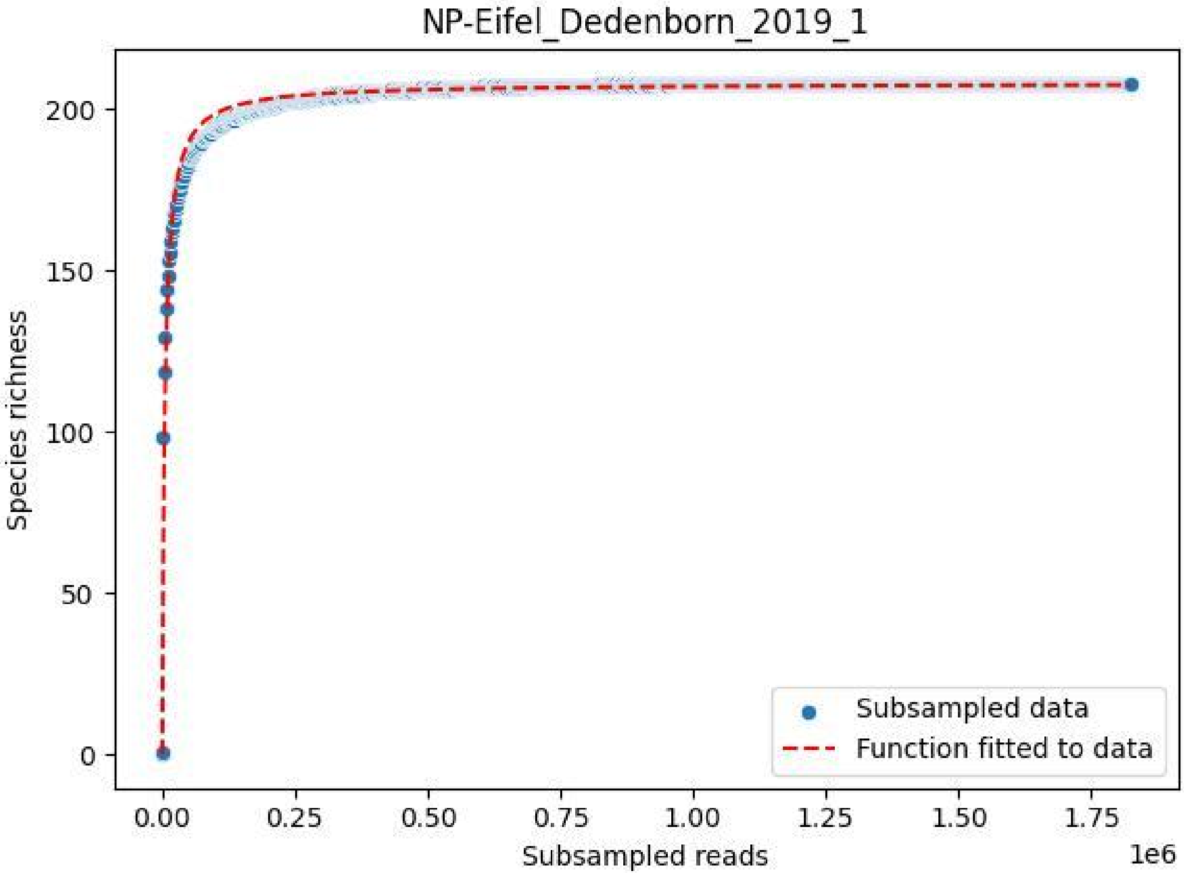
NP-Eifel_Dedenborn_2019_1
Species richness
Subsampled reads
1e6
0
50
100
150
200
0.00
0.25
0.50
0.75
1.00
1.25
1.50
1.75
Subsampled data
Function fitted to data

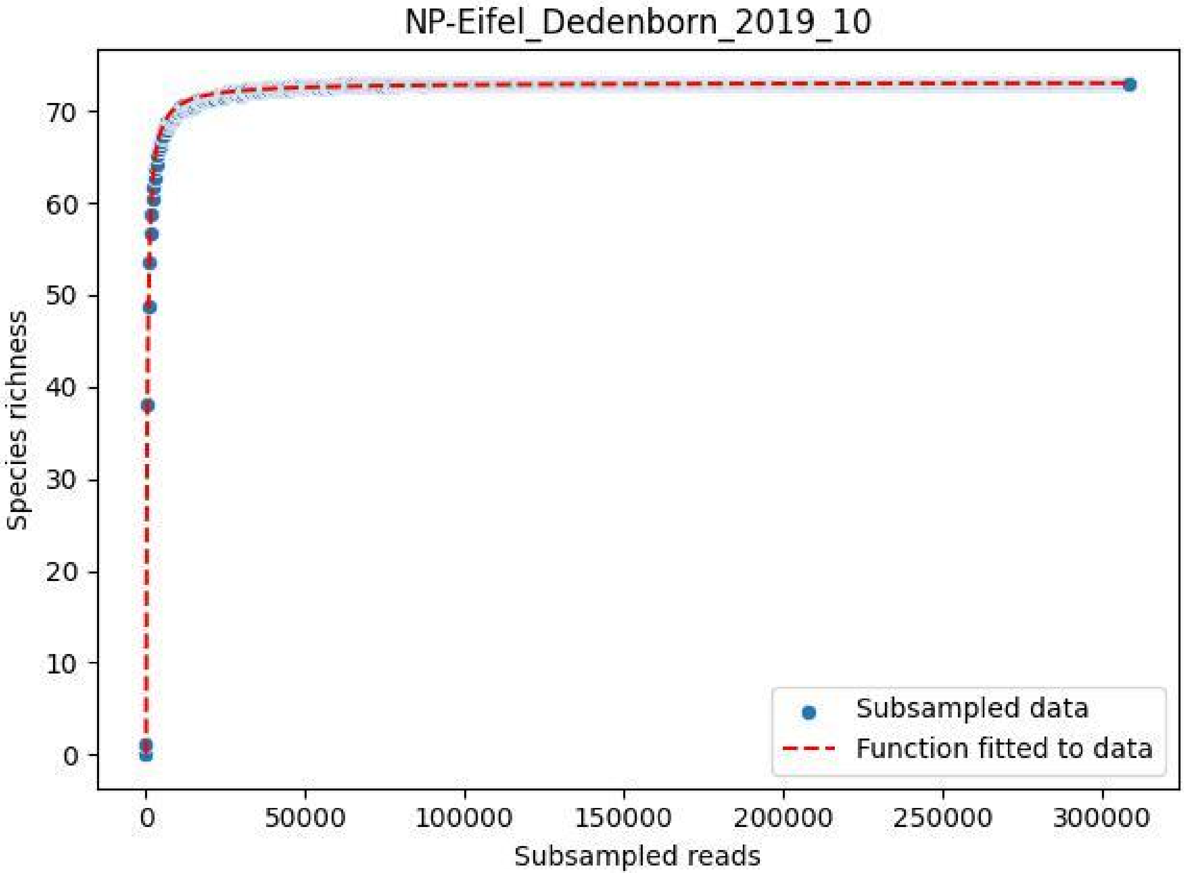
NP-Eifel_Dedenborn_2019_10
Species richness
0
10
20
30
40
50
60
70
0
50000
100000
150000
200000
250000
300000
Subsampled reads
Subsampled data
Function fitted to data

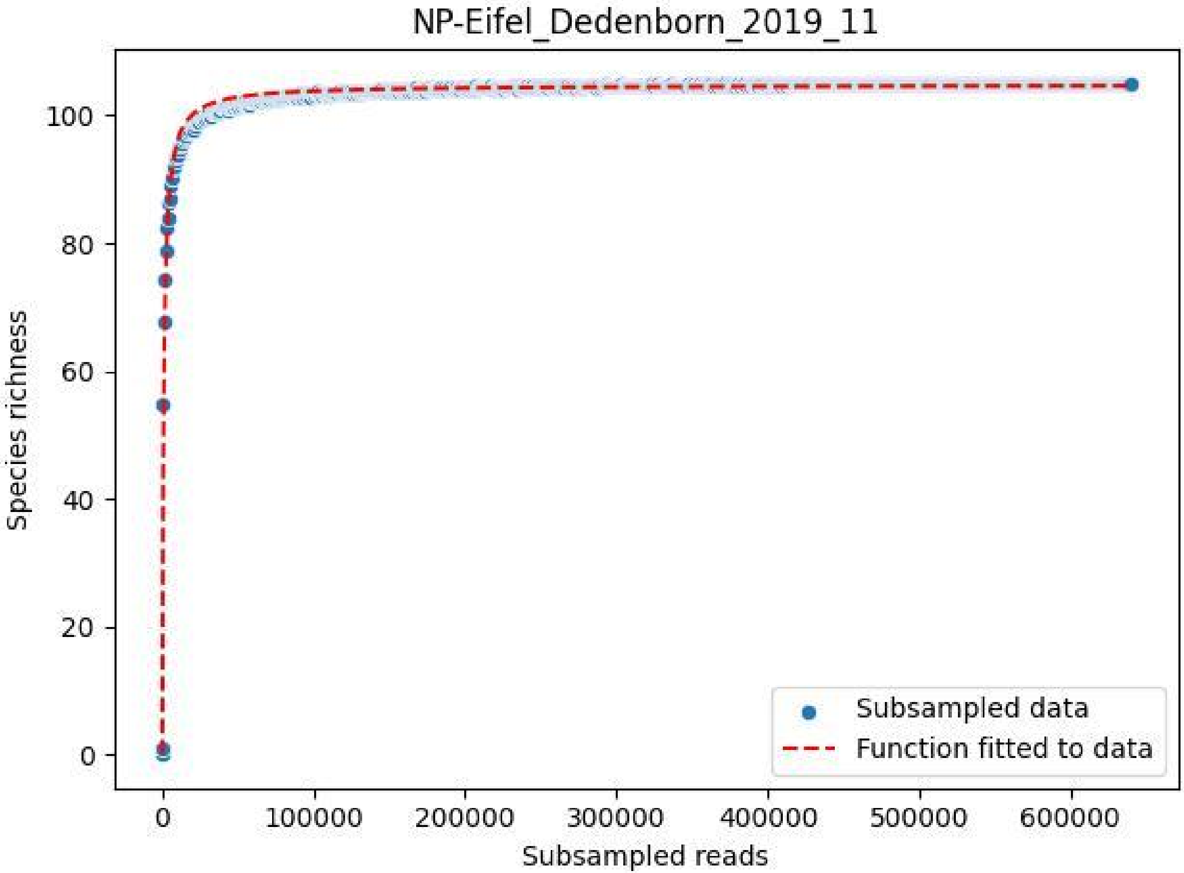
NP-Eifel_Dedenborn_2019_11
Species richness
0
20
40
60
80
100
0
100000
200000
300000
400000
500000
600000
Subsampled reads
Subsampled data
Function fitted to data

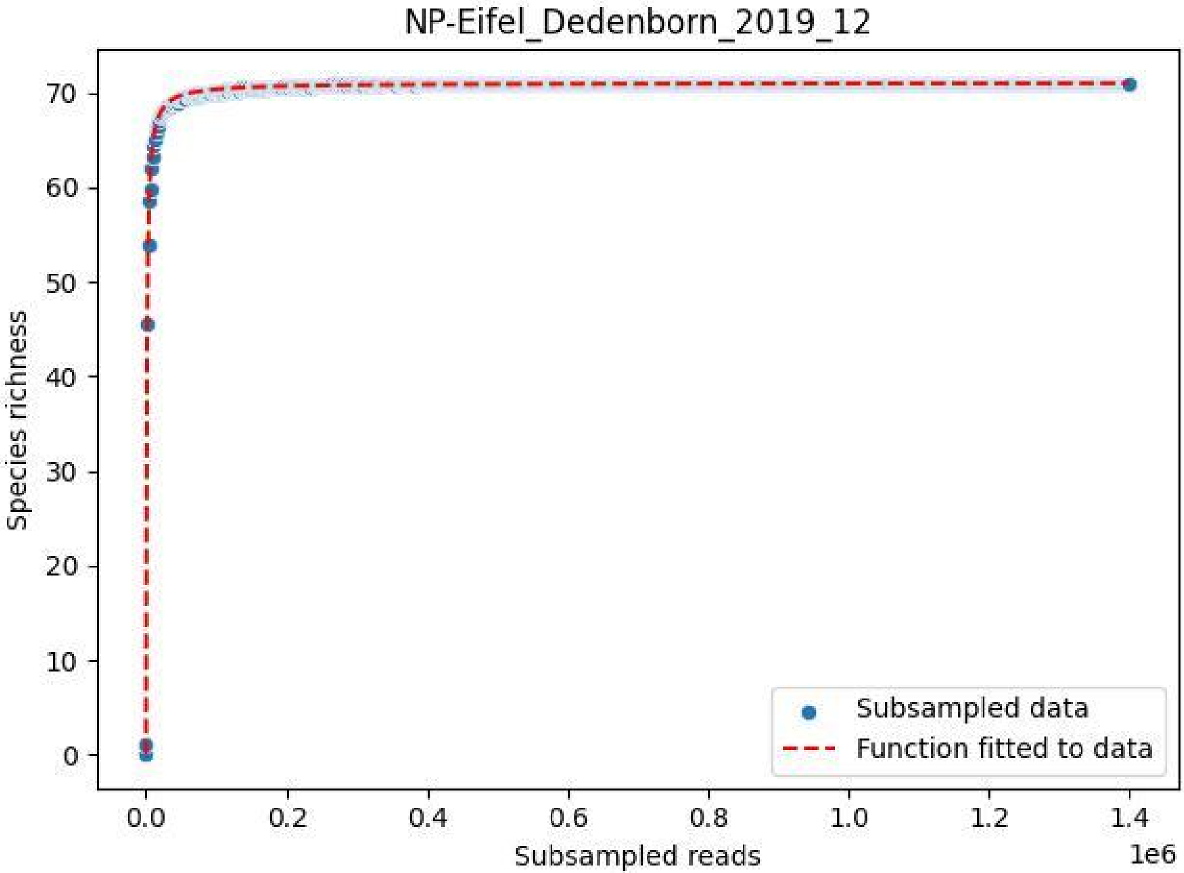
NP-Eifel_Dedenborn_2019_12
Species richness
Subsampled reads
1e6
0
10
20
30
40
50
60
70
0.0
0.2
0.4
0.6
0.8
1.0
1.2
1.4
Subsampled data
Function fitted to data

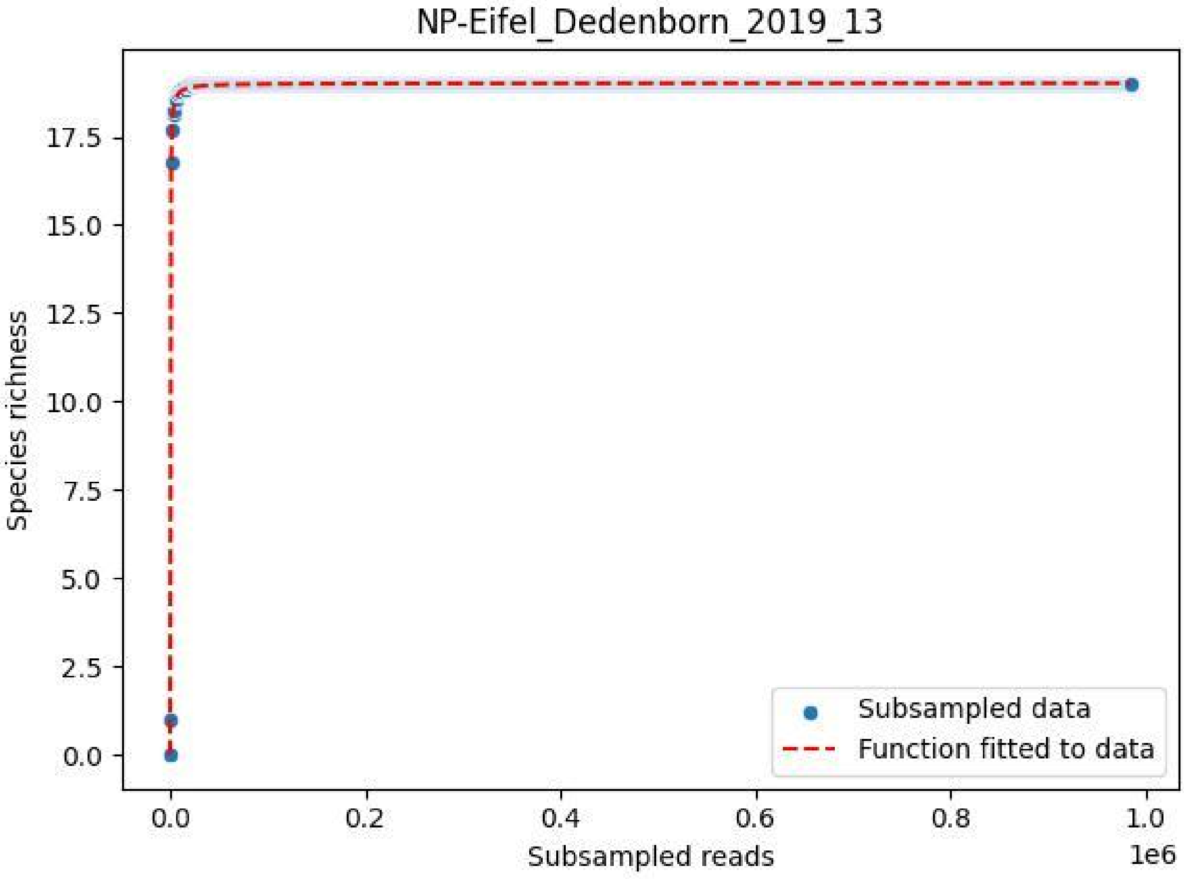
NP-Eifel_Dedenborn_2019_13
Species richness
17.5
15.0
12.5
10.0
7.5
5.0
2.5
0.0
0.0
0.2
0.4
0.6
0.8
1.0
1e6
Subsampled reads
Subsampled data
Function fitted to data

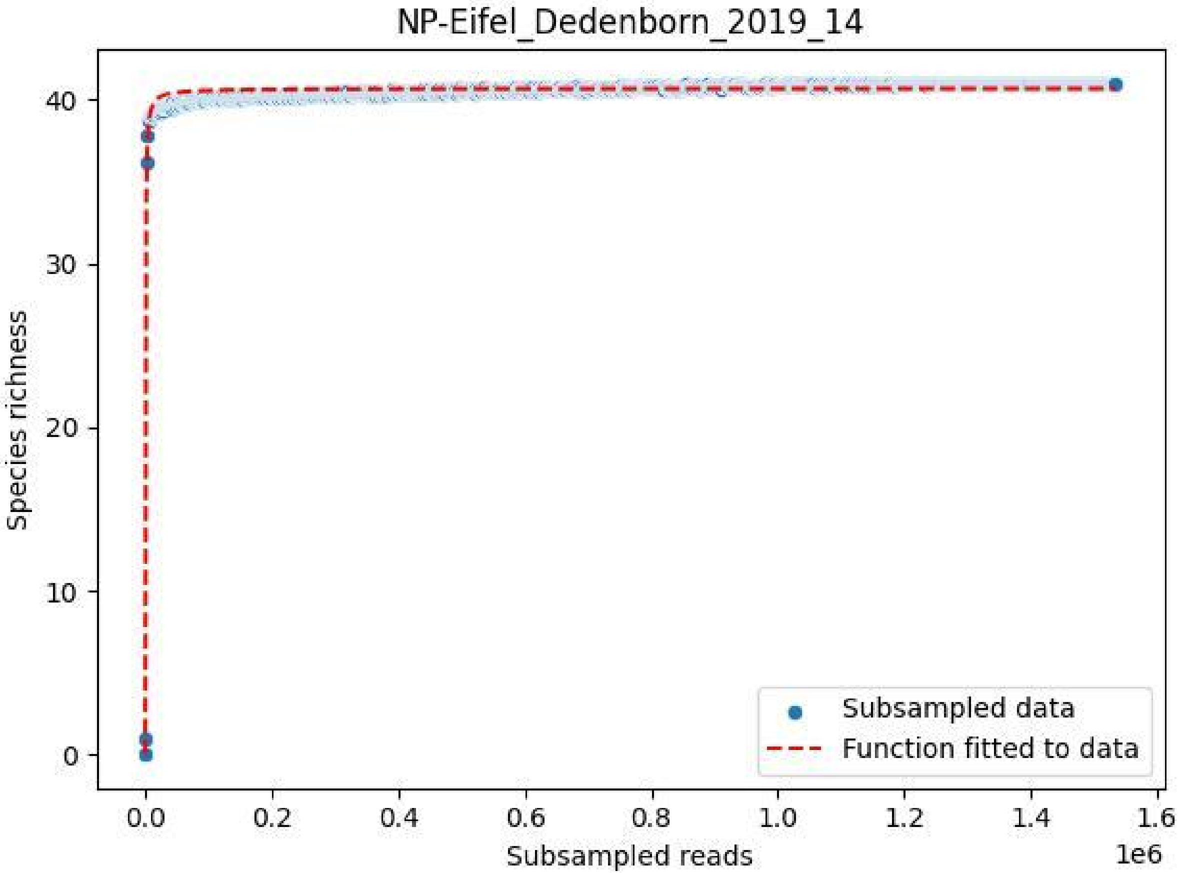
NP-Eifel_Dedenborn_2019_14
Species richness
Subsampled reads
1e6
0
10
20
30
40
0.0
0.2
0.4
0.6
0.8
1.0
1.2
1.4
1.6
Subsampled data
Function fitted to data

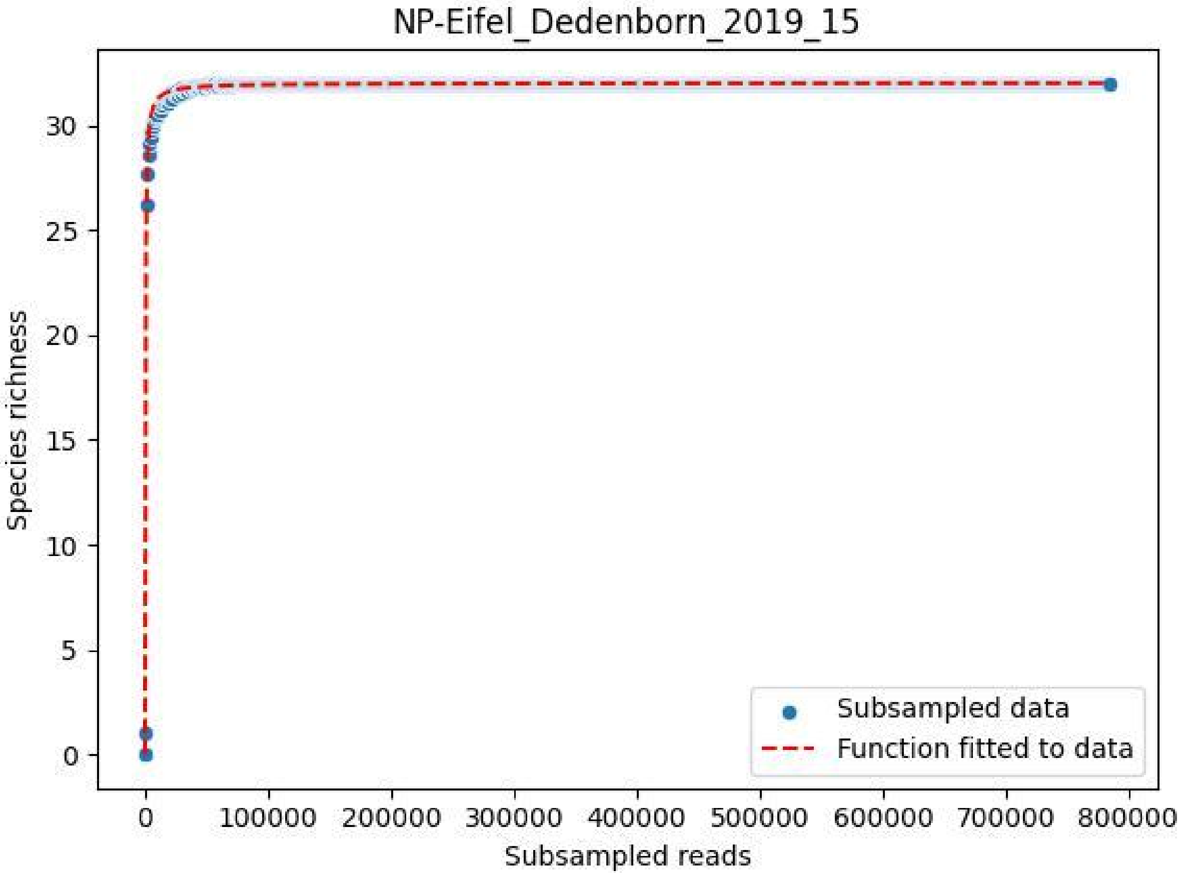
NP-Eifel_Dedenborn_2019_15
Species richness
0
5
10
15
20
25
30
0
100000
200000
300000
400000
500000
600000
700000
800000
Subsampled reads
Subsampled data
Function fitted to data

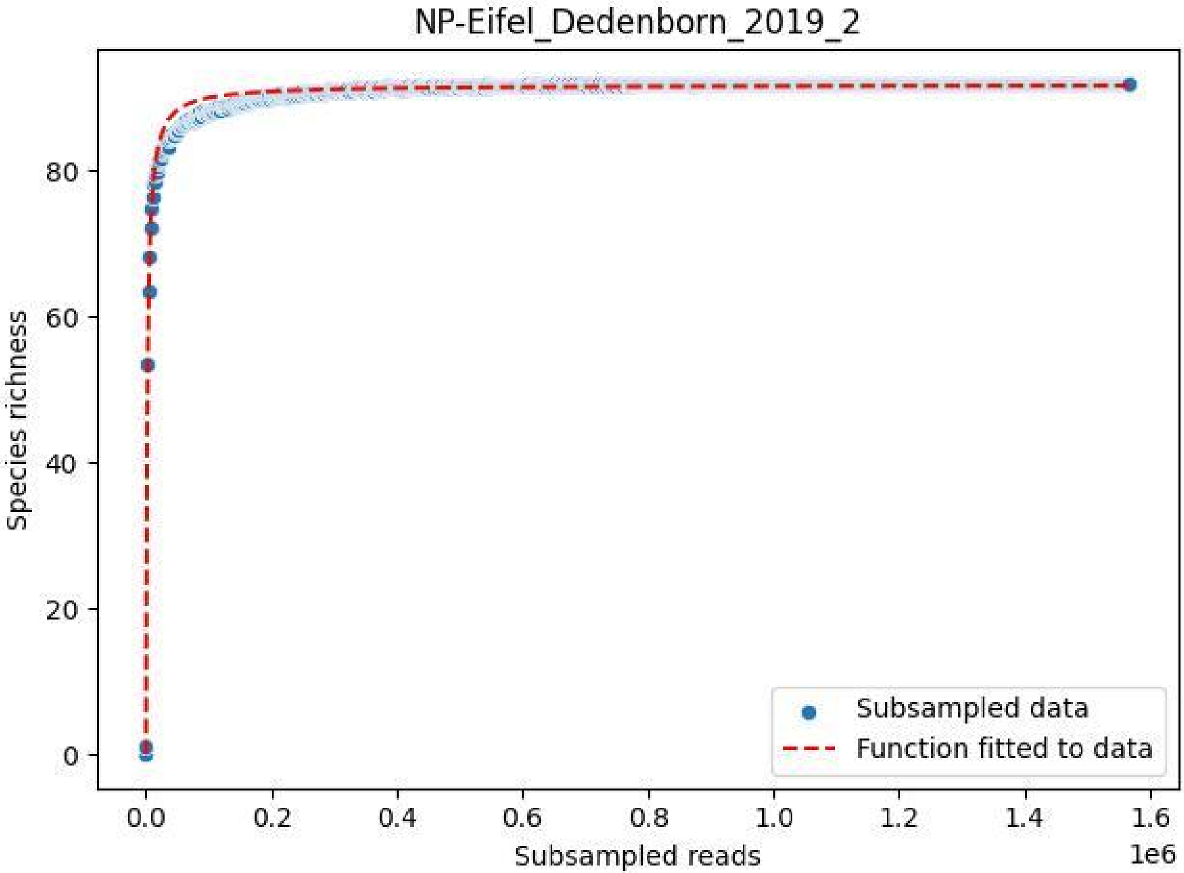

NP-Eifel_Dedenborn_2019_2
Species richness
Subsampled reads
1e6
0
20
40
60
80
0.0
0.2
0.4
0.6
0.8
1.0
1.2
1.4
1.6
Subsampled data
Function fitted to data

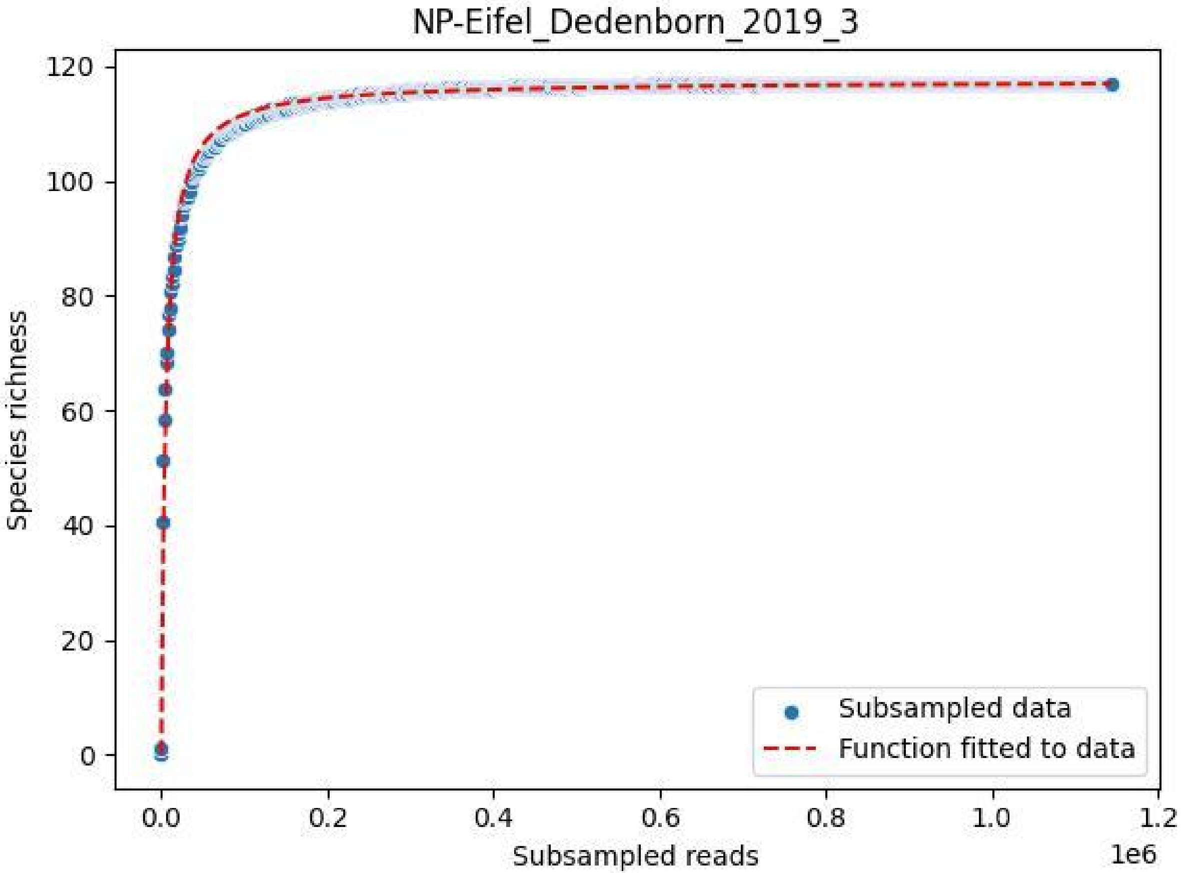

NP-Eifel_Dedenborn_2019_3
Species richness
Subsampled reads
1e6
0
20
40
60
80
100
120
0.0
0.2
0.4
0.6
0.8
1.0
1.2
Subsampled data
Function fitted to data

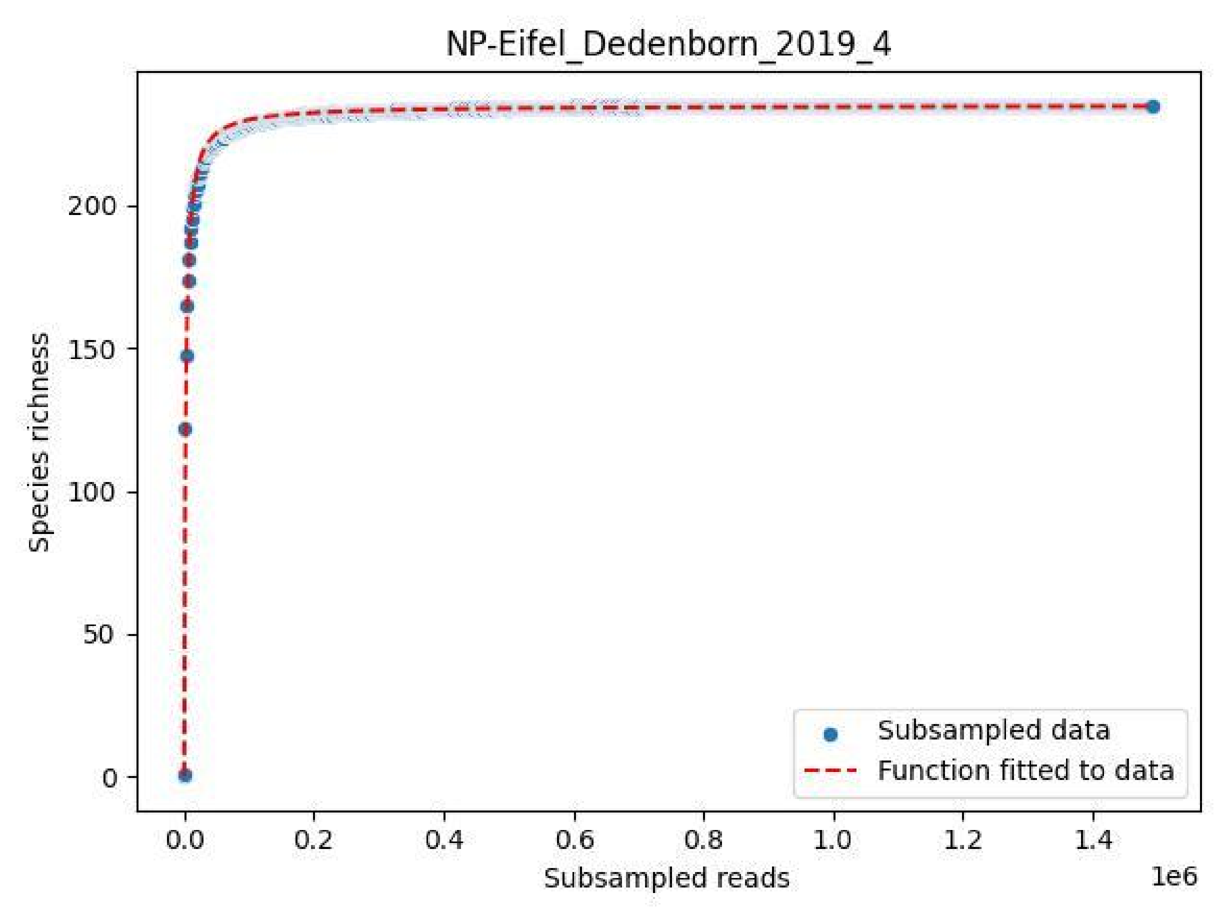

NP-Eifel_Dedenborn_2019_4
Species richness
Subsampled reads
1e6
0
50
100
150
200
0.0
0.2
0.4
0.6
0.8
1.0
1.2
1.4
Subsampled data
Function fitted to data

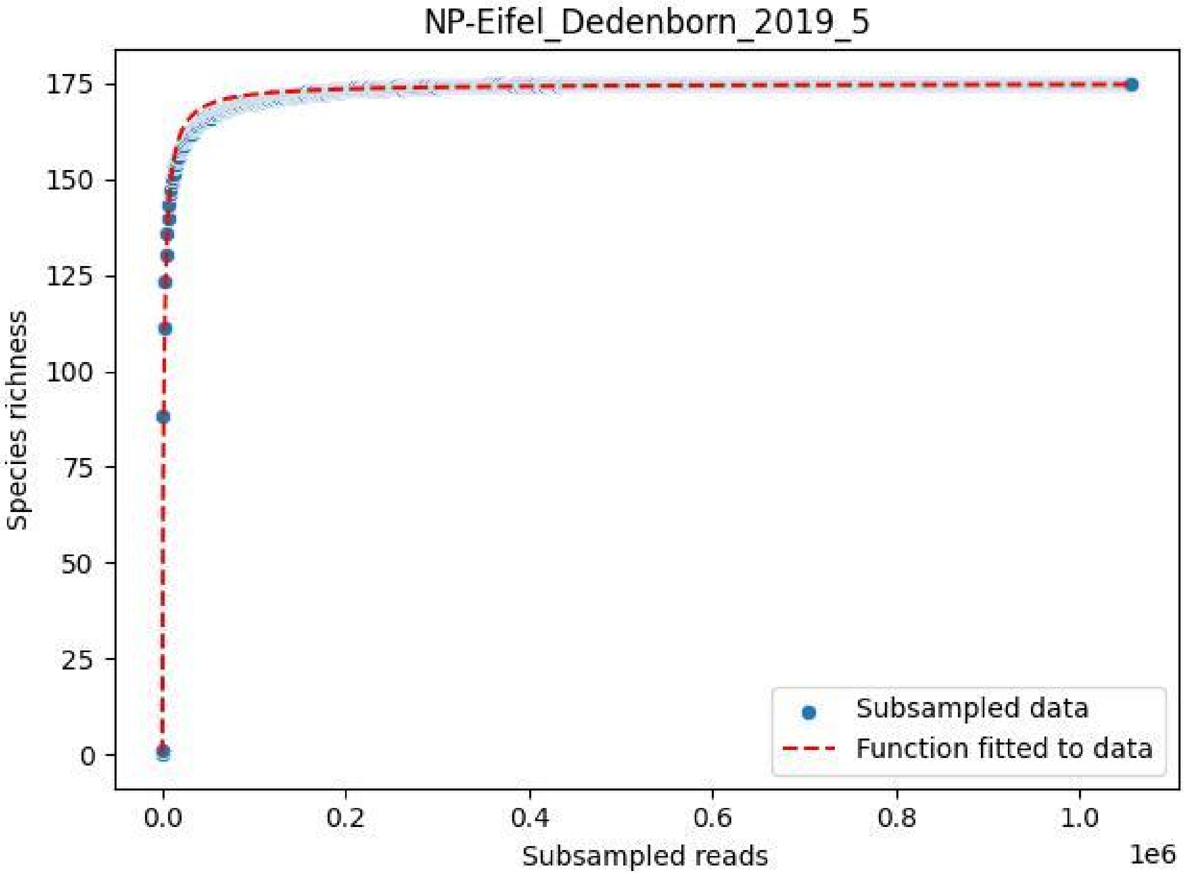
NP-Eifel_Dedenborn_2019_5
Species richness
Subsampled reads
1e6
0
25
50
75
100
125
150
175
0.0
0.2
0.4
0.6
0.8
1.0
Subsampled data
Function fitted to data

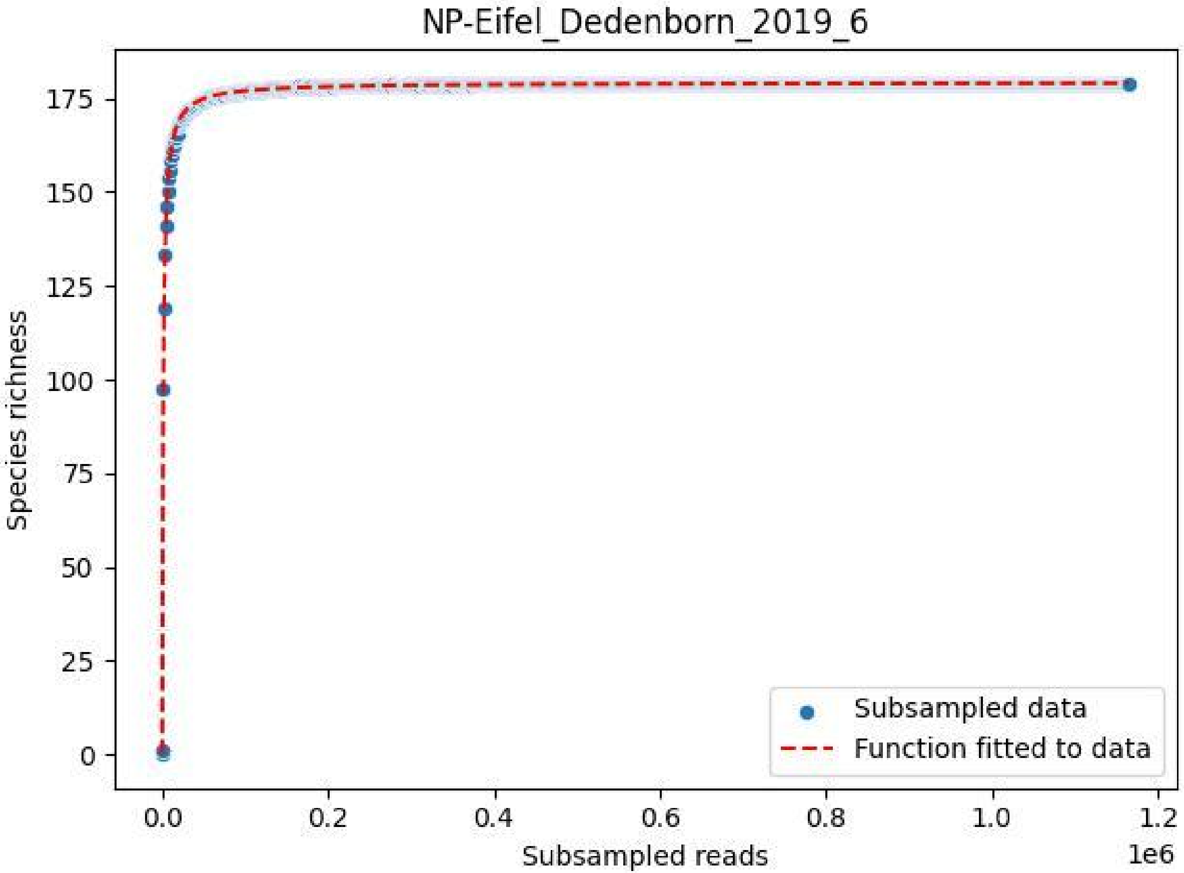

NP-Eifel_Dedenborn_2019_6
Species richness
Subsampled reads
1e6
Subsampled data
Function fitted to data
0
25
50
75
100
125
150
175
0.0
0.2
0.4
0.6
0.8
1.0
1.2

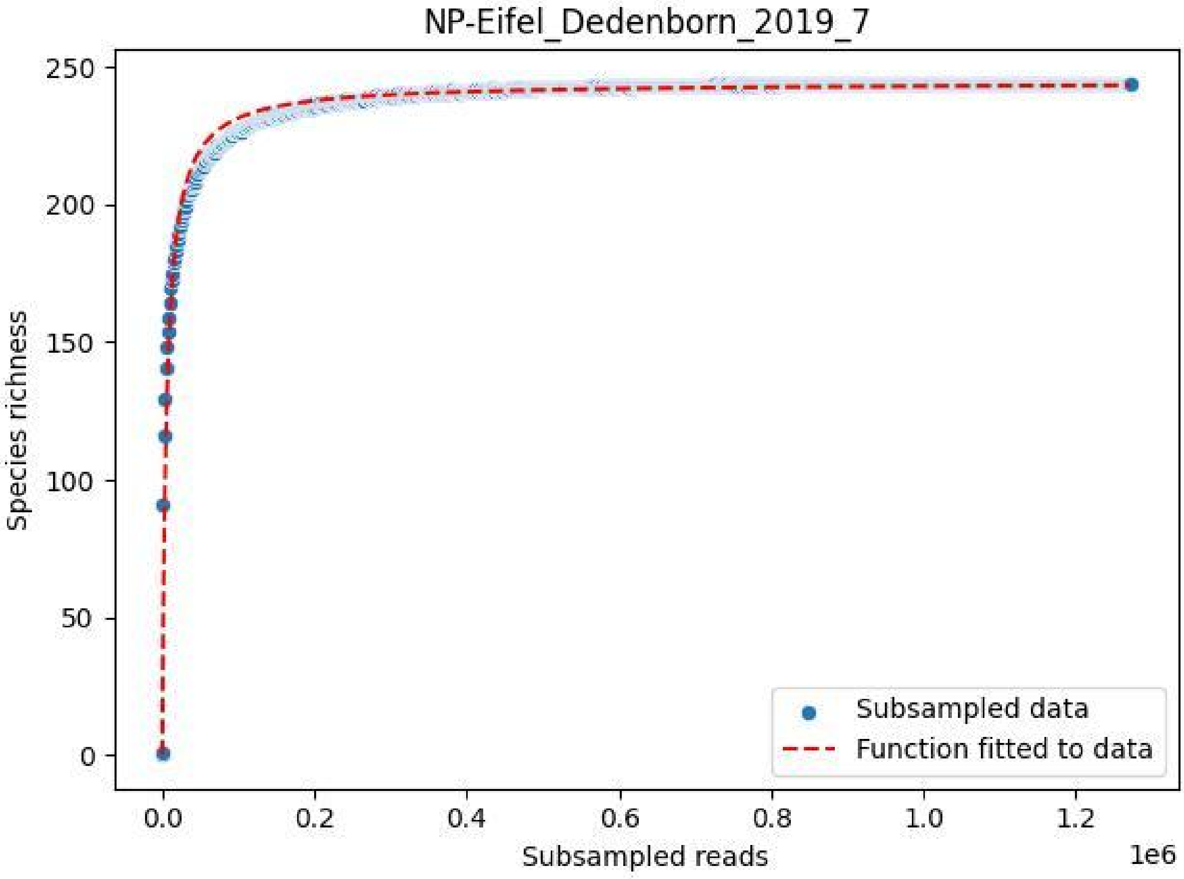

NP-Eifel_Dedenborn_2019_7
Species richness
Subsampled reads
1e6
0
50
100
150
200
250
0.0
0.2
0.4
0.6
0.8
1.0
1.2
Subsampled data
Function fitted to data

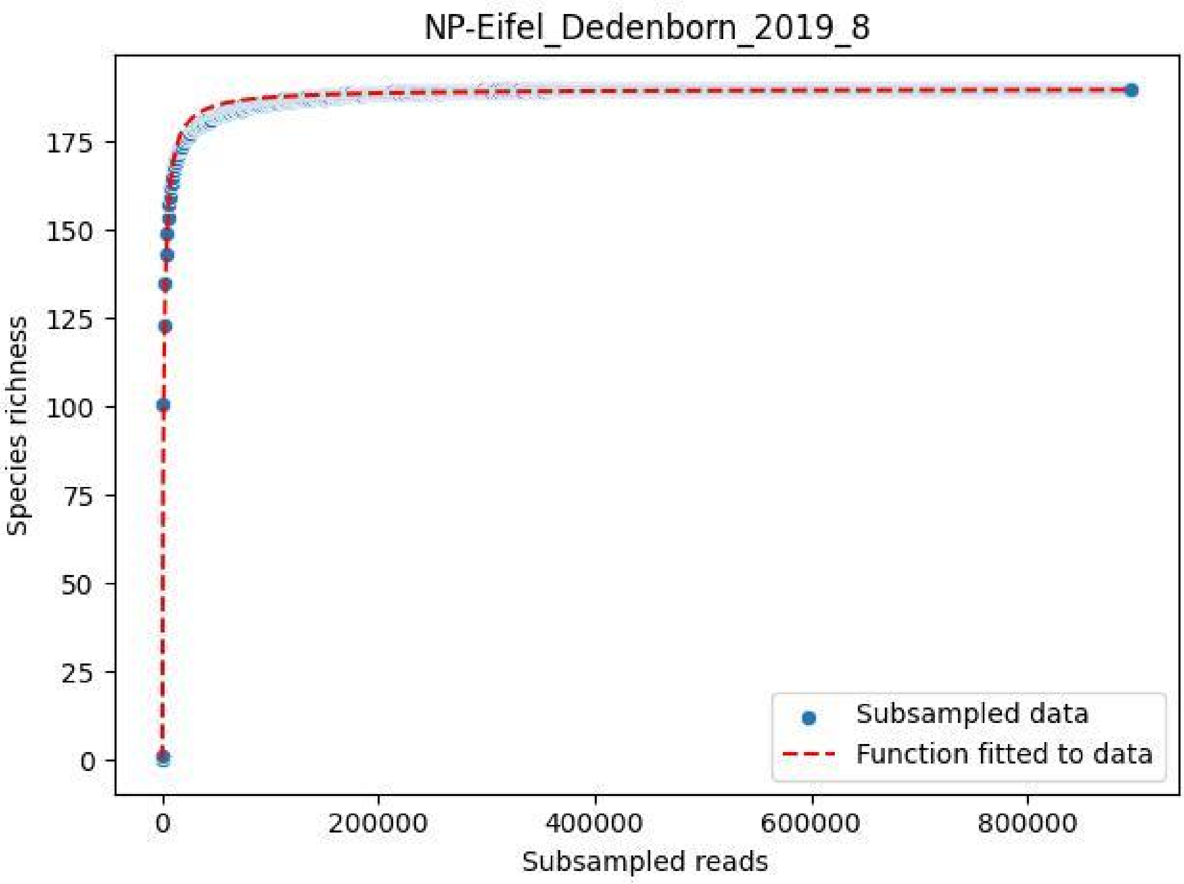
NP-Eifel_Dedenborn_2019_8
Species richness
Subsampled reads
0
25
50
75
100
125
150
175
0
200000
400000
600000
800000
Subsampled data
Function fitted to data

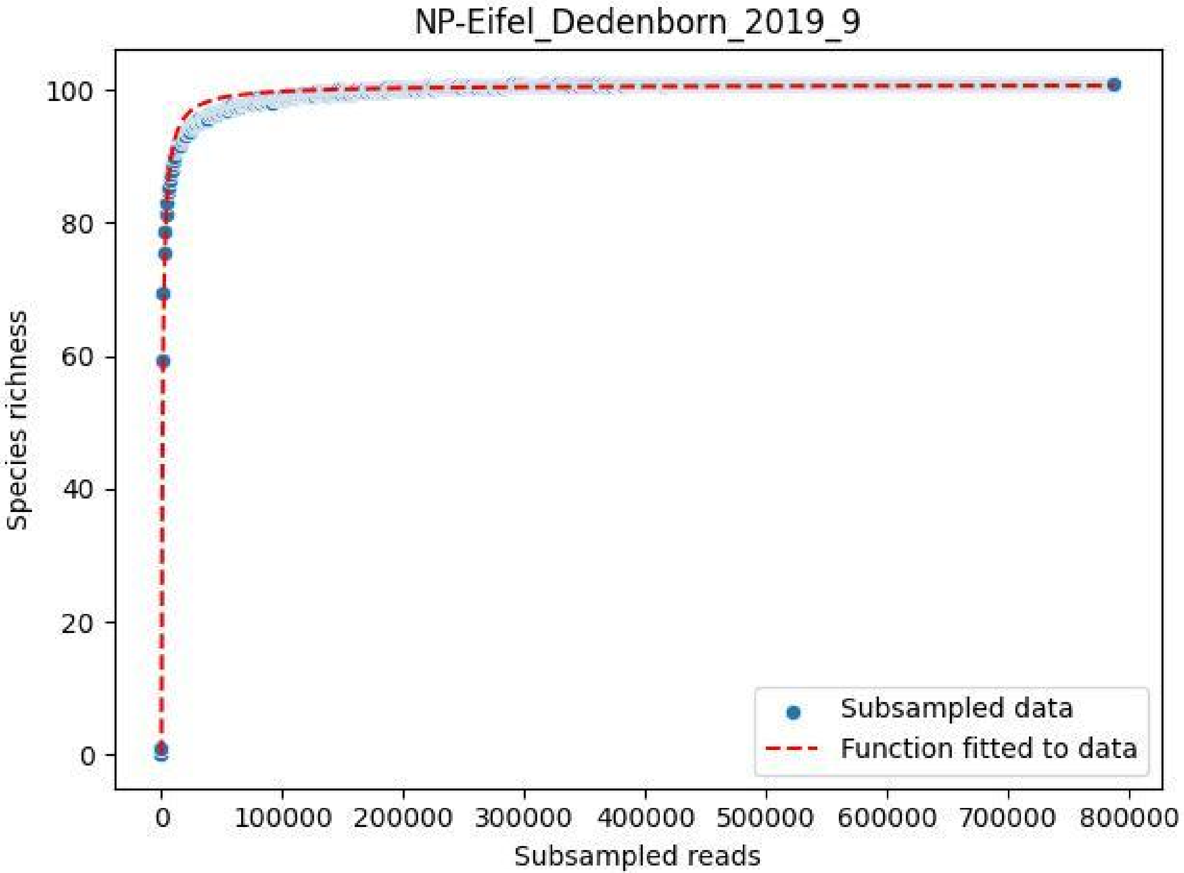
NP-Eifel_Dedenborn_2019_9
Species richness
Subsampled reads
0
20
40
60
80
100
0
100000
200000
300000
400000
500000
600000
700000
800000
Subsampled data
Function fitted to data

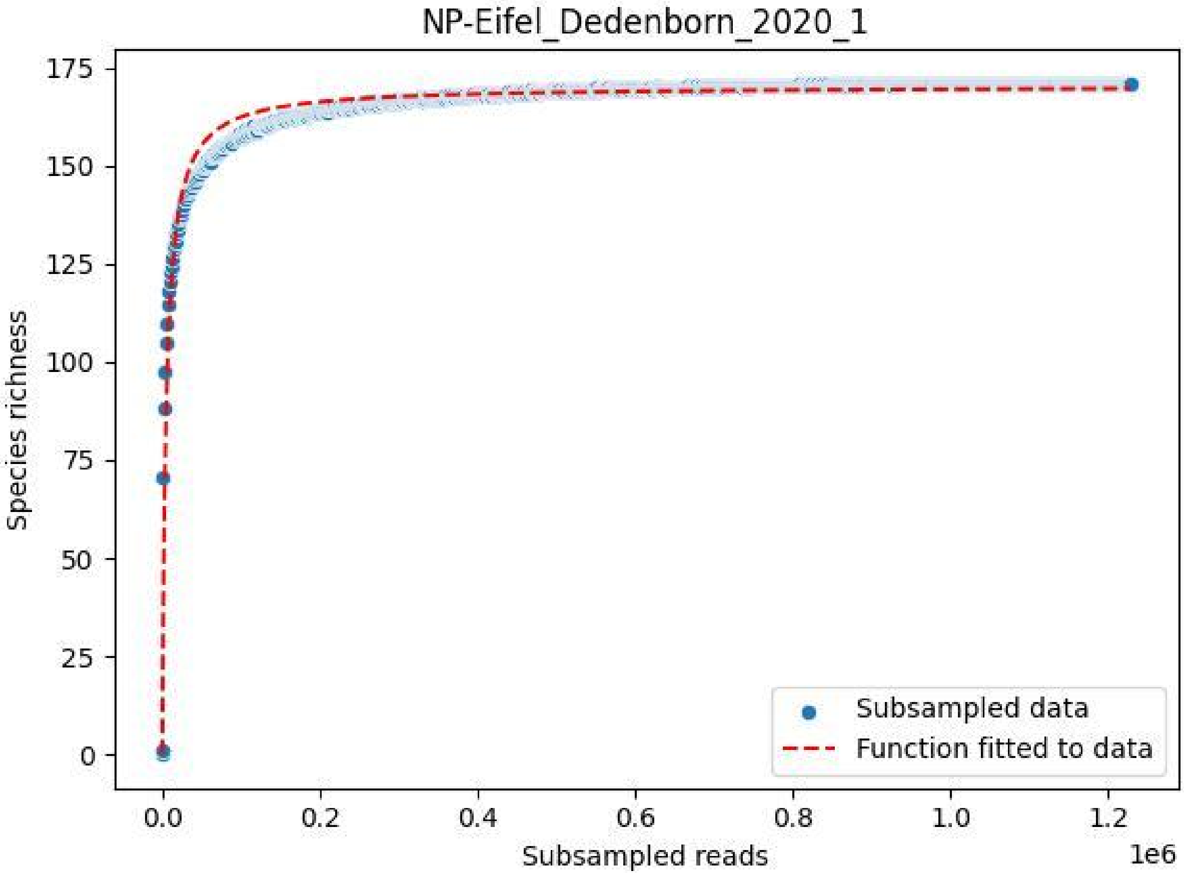
NP-Eifel_Dedenborn_2020_1
Species richness
Subsampled reads
1e6
0
25
50
75
100
125
150
175
0.0
0.2
0.4
0.6
0.8
1.0
1.2
Subsampled data
Function fitted to data

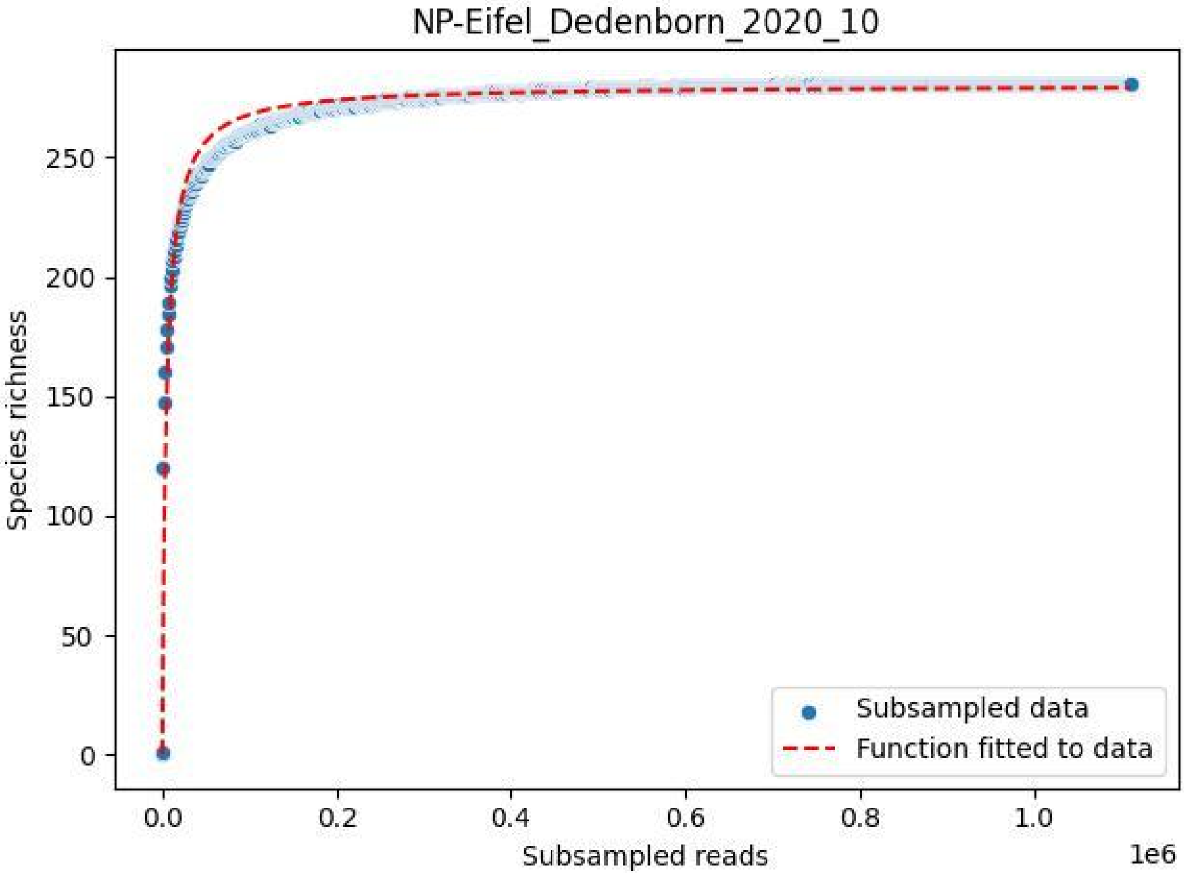

NP-Eifel_Dedenborn_2020_10
Species richness
0
50
100
150
200
250
0.0
0.2
0.4
0.6
0.8
1.0
1e6
Subsampled reads
Subsampled data
Function fitted to data

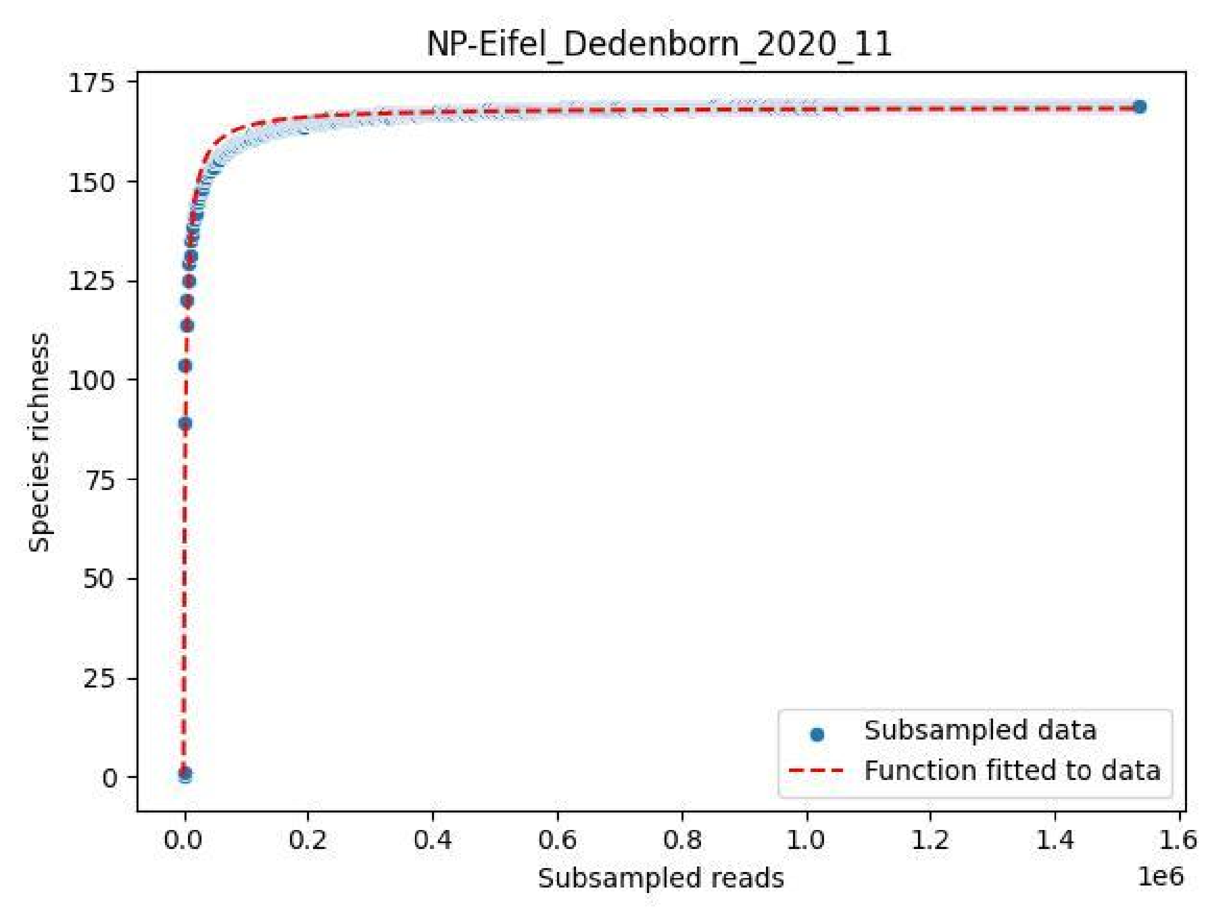

NP-Eifel_Dedenborn_2020_11
Species richness
Subsampled reads
1e6
Subsampled data
Function fitted to data
175
150
125
100
75
50
25
0
0.0
0.2
0.4
0.6
0.8
1.0
1.2
1.4
1.6

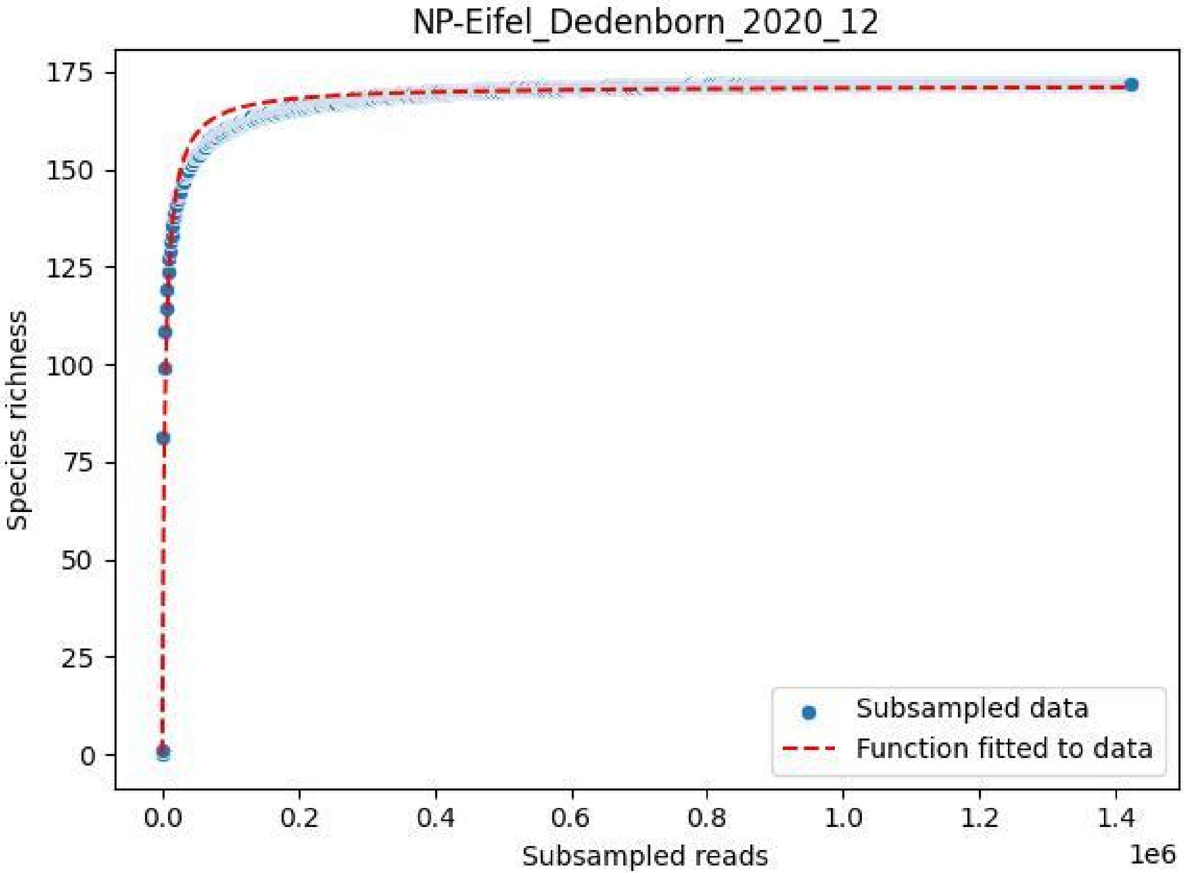
NP-Eifel_Dedenborn_2020_12
175
150
125
100
75
50
25
0
Species richness
0.0
0.2
0.4
0.6
0.8
1.0
1.2
1.4
1e6
Subsampled reads
Subsampled data
Function fitted to data

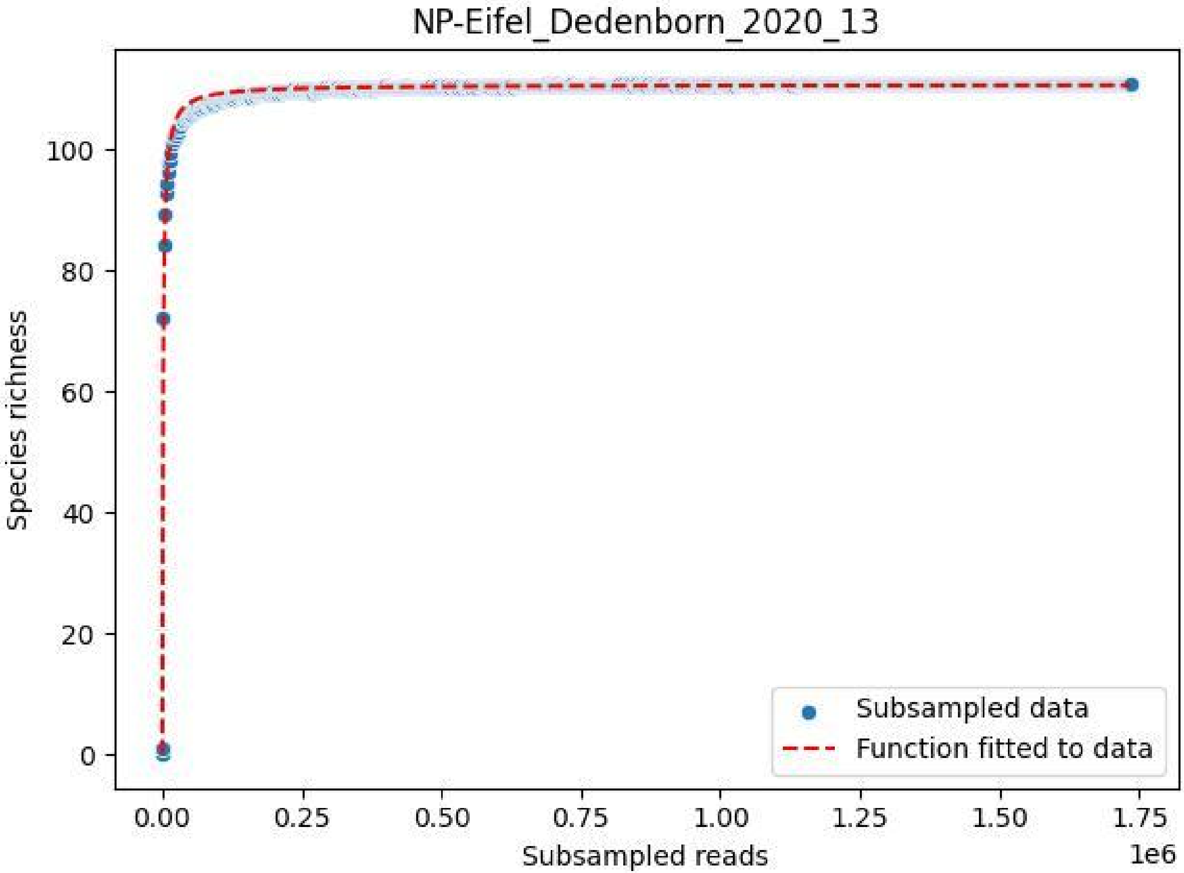
NP-Eifel_Dedenborn_2020_13
Species richness
Subsampled reads
1e6
0
20
40
60
80
100
0.00
0.25
0.50
0.75
1.00
1.25
1.50
1.75
Subsampled data
Function fitted to data

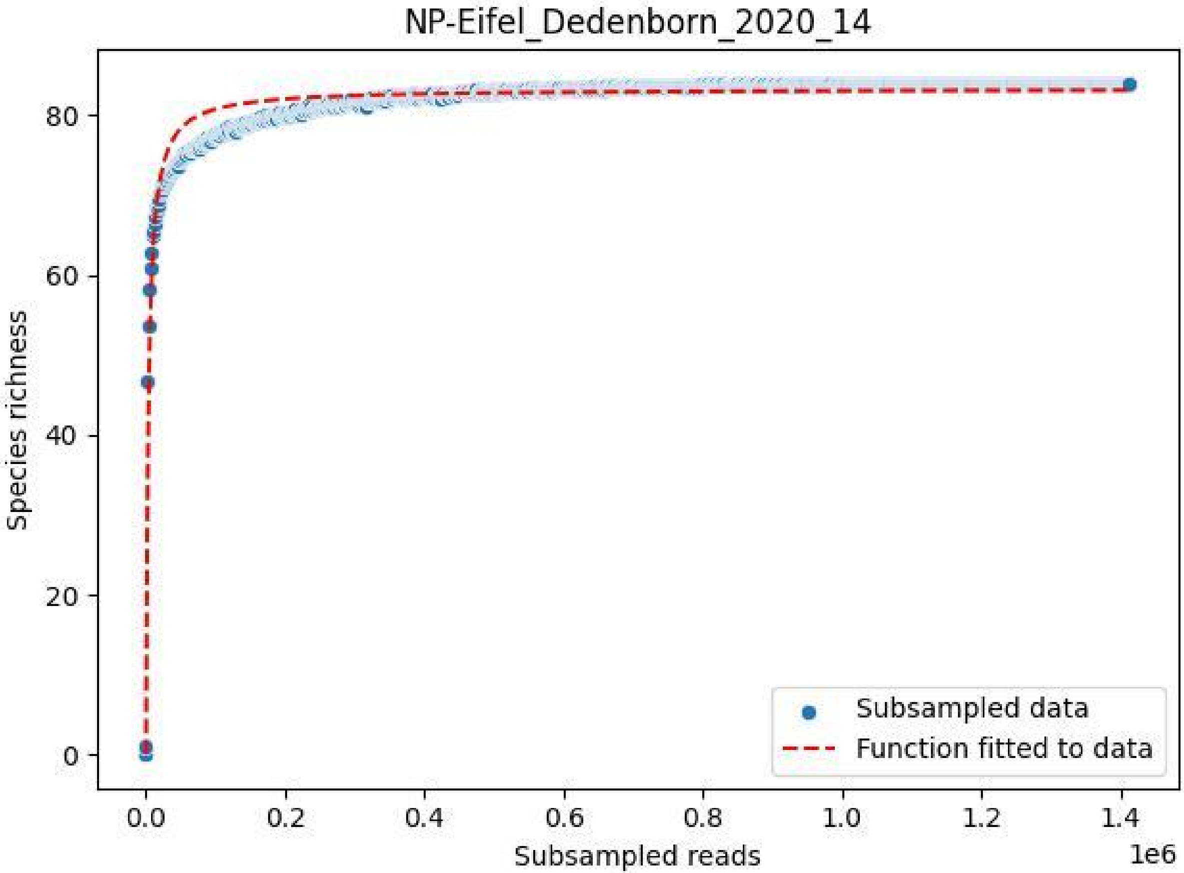
NP-Eifel_Dedenborn_2020_14
Species richness
Subsampled reads
1e6
0
20
40
60
80
0.0
0.2
0.4
0.6
0.8
1.0
1.2
1.4
Subsampled data
Function fitted to data

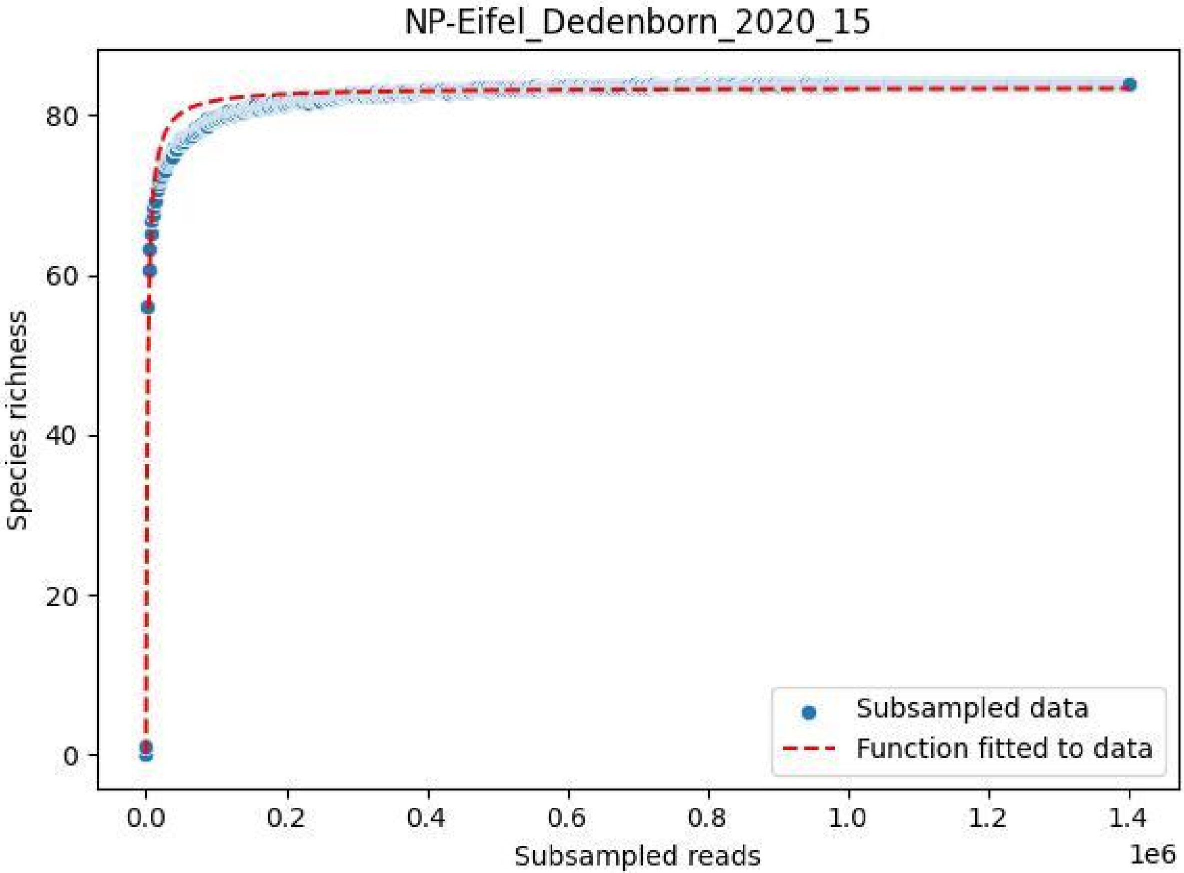
NP-Eifel_Dedenborn_2020_15
Species richness
Subsampled reads
1e6
0
20
40
60
80
0.0
0.2
0.4
0.6
0.8
1.0
1.2
1.4
Subsampled data
Function fitted to data

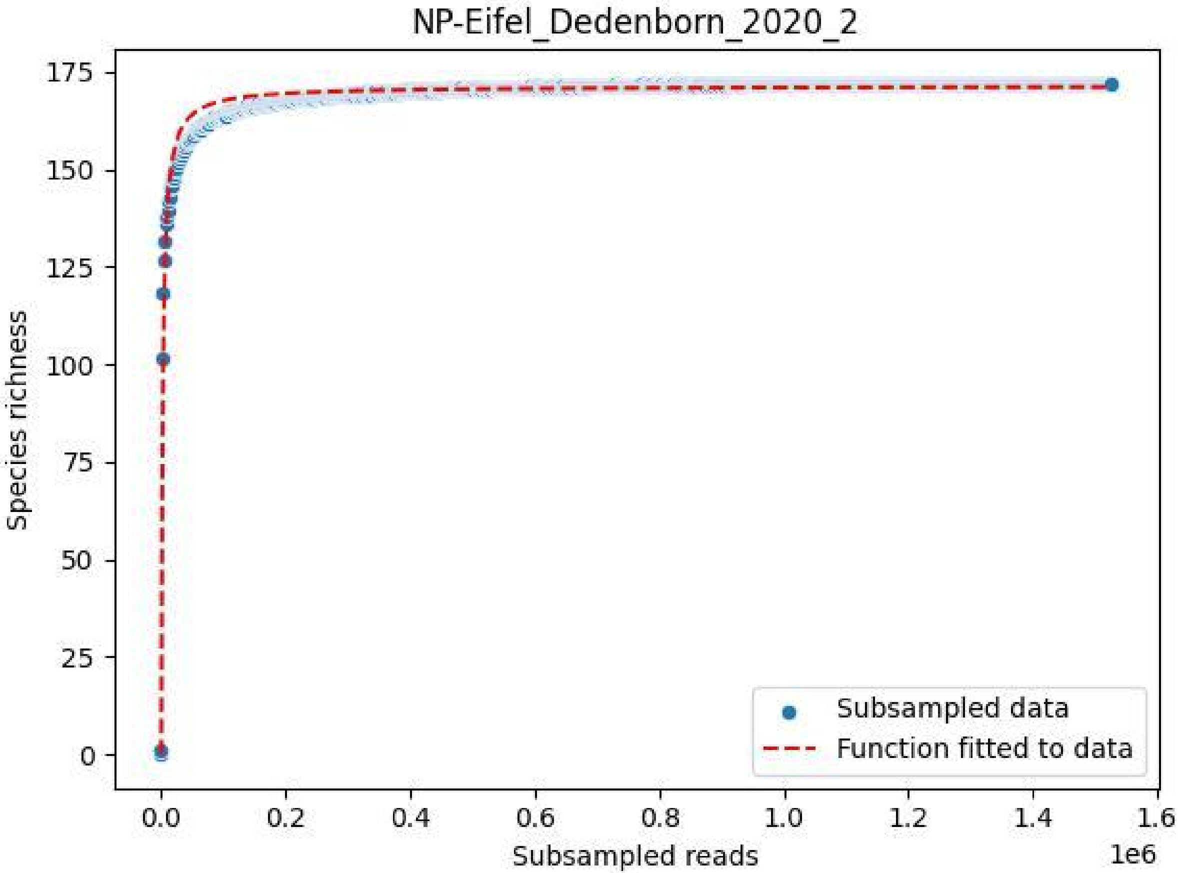

NP-Eifel_Dedenborn_2020_2
Species richness
Subsampled reads
1e6
0.0
0.2
0.4
0.6
0.8
1.0
1.2
1.4
1.6
0
25
50
75
100
125
150
175
Subsampled data
Function fitted to data

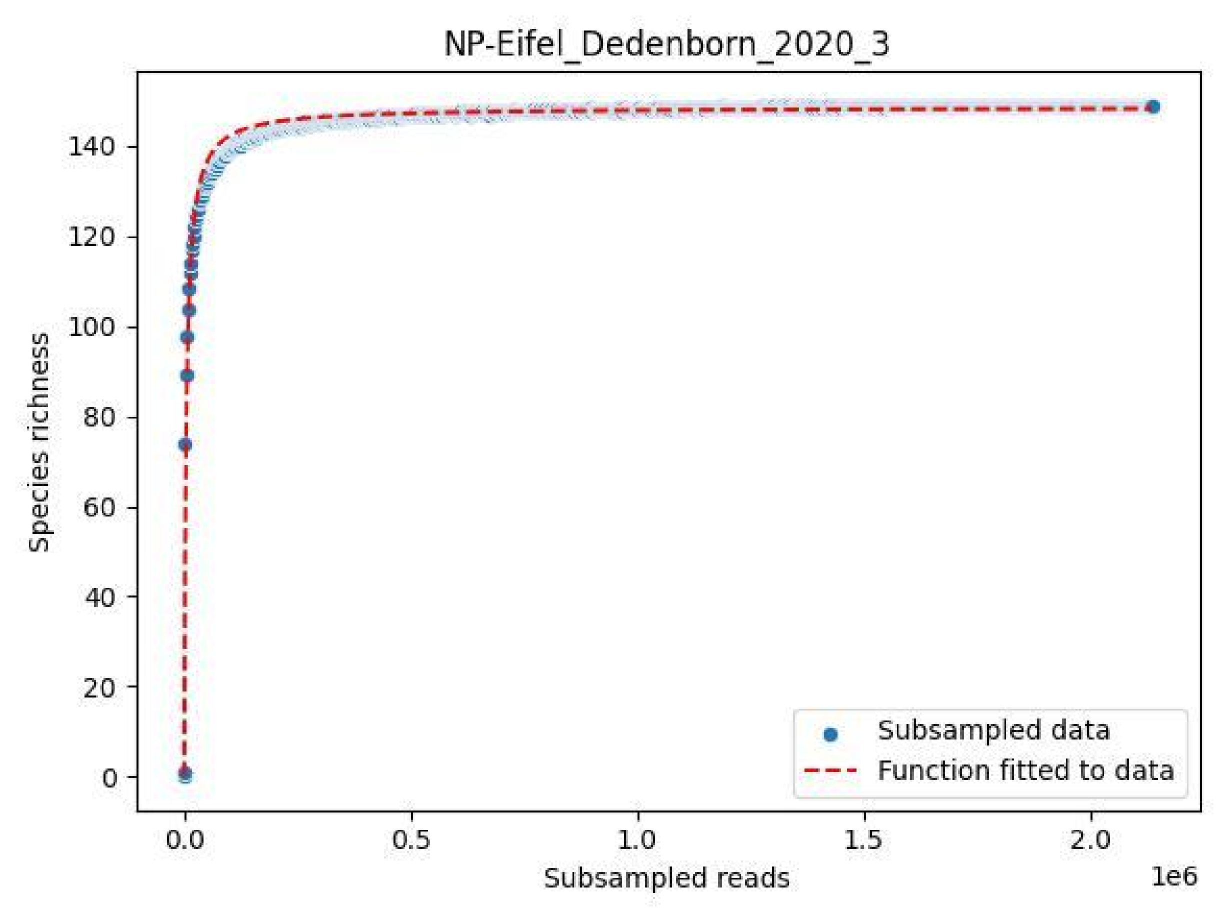

NP-Eifel_Dedenborn_2020_3
Species richness
Subsampled reads
1e6
0
20
40
60
80
100
120
140
0.0
0.5
1.0
1.5
2.0
Subsampled data
Function fitted to data

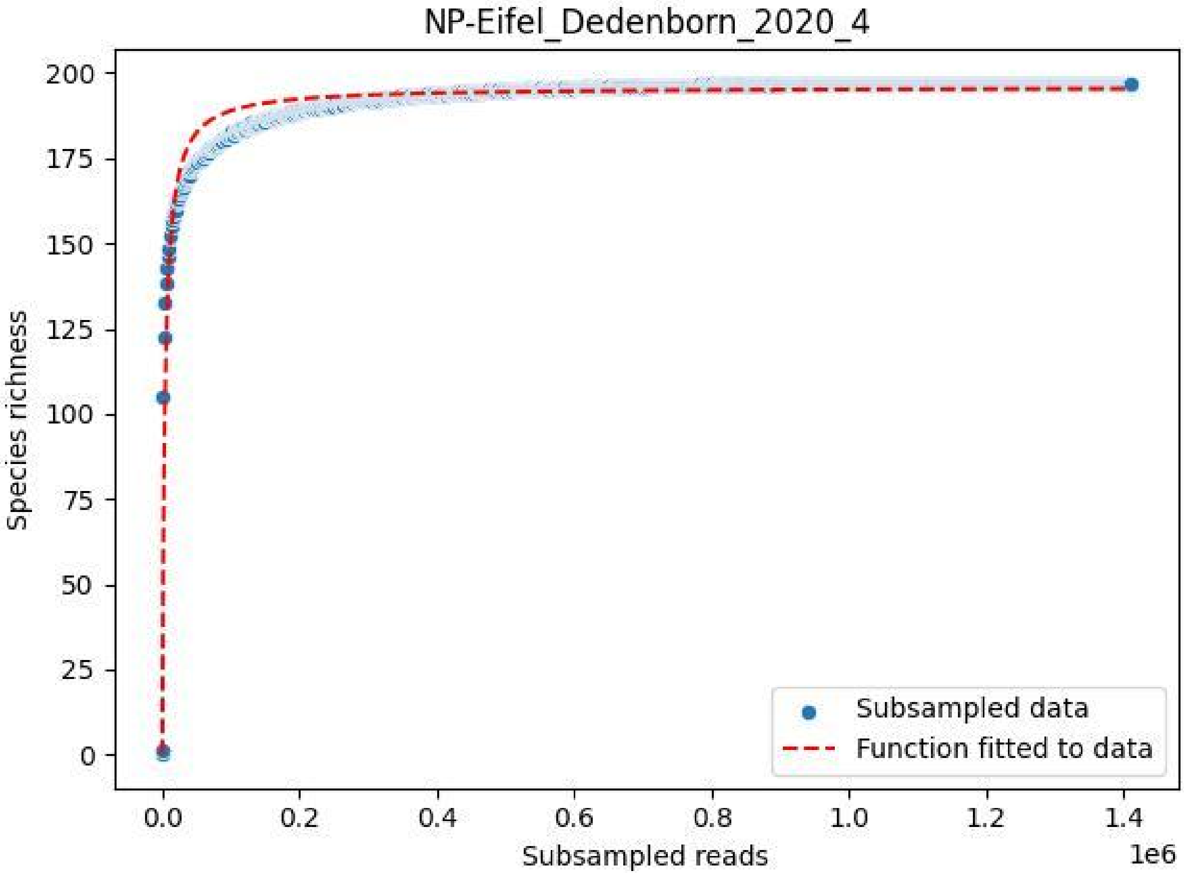

NP-Eifel_Dedenborn_2020_4
Species richness
Subsampled reads
1e6
0
25
50
75
100
125
150
175
200
0.0
0.2
0.4
0.6
0.8
1.0
1.2
1.4
Subsampled data
Function fitted to data

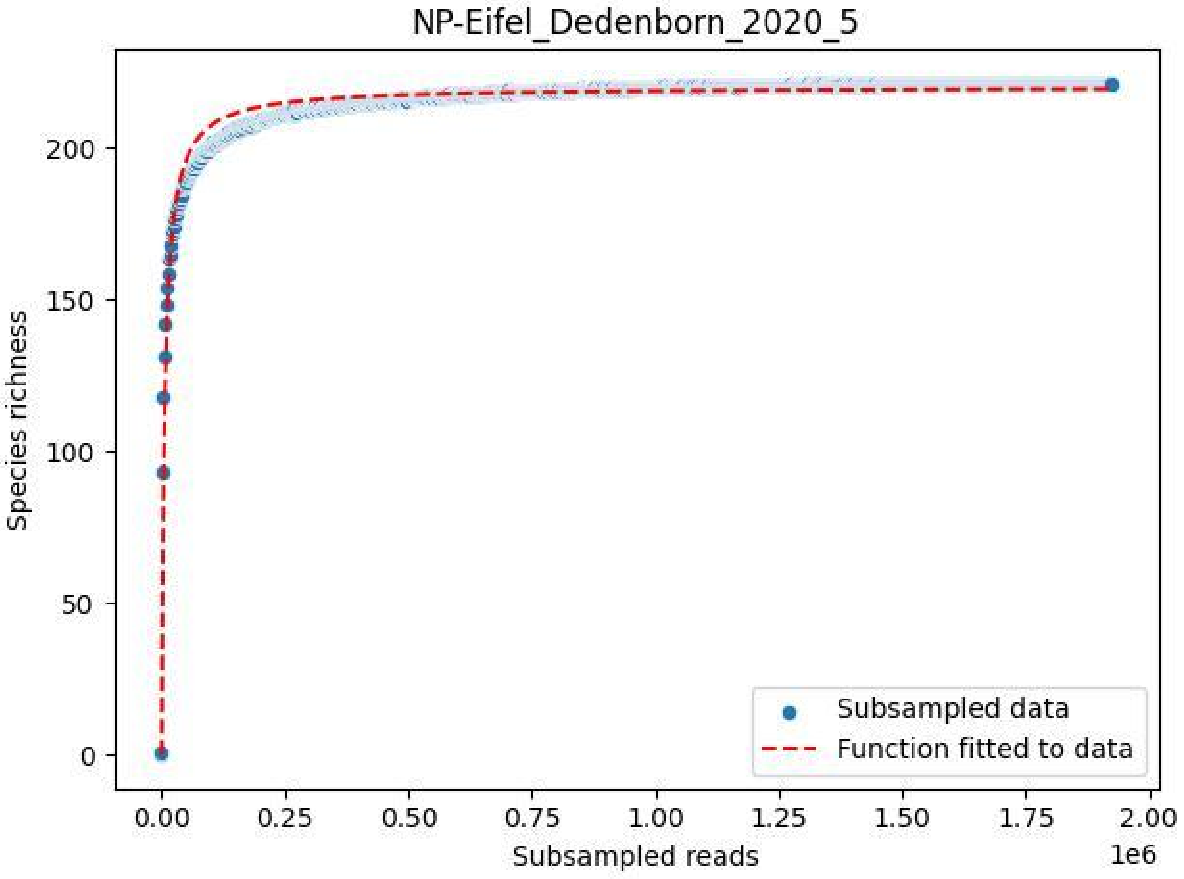
NP-Eifel_Dedenborn_2020_5
Species richness
0
50
100
150
200
0.00
0.25
0.50
0.75
1.00
1.25
1.50
1.75
2.00
1e6
Subsampled reads
Subsampled data
Function fitted to data

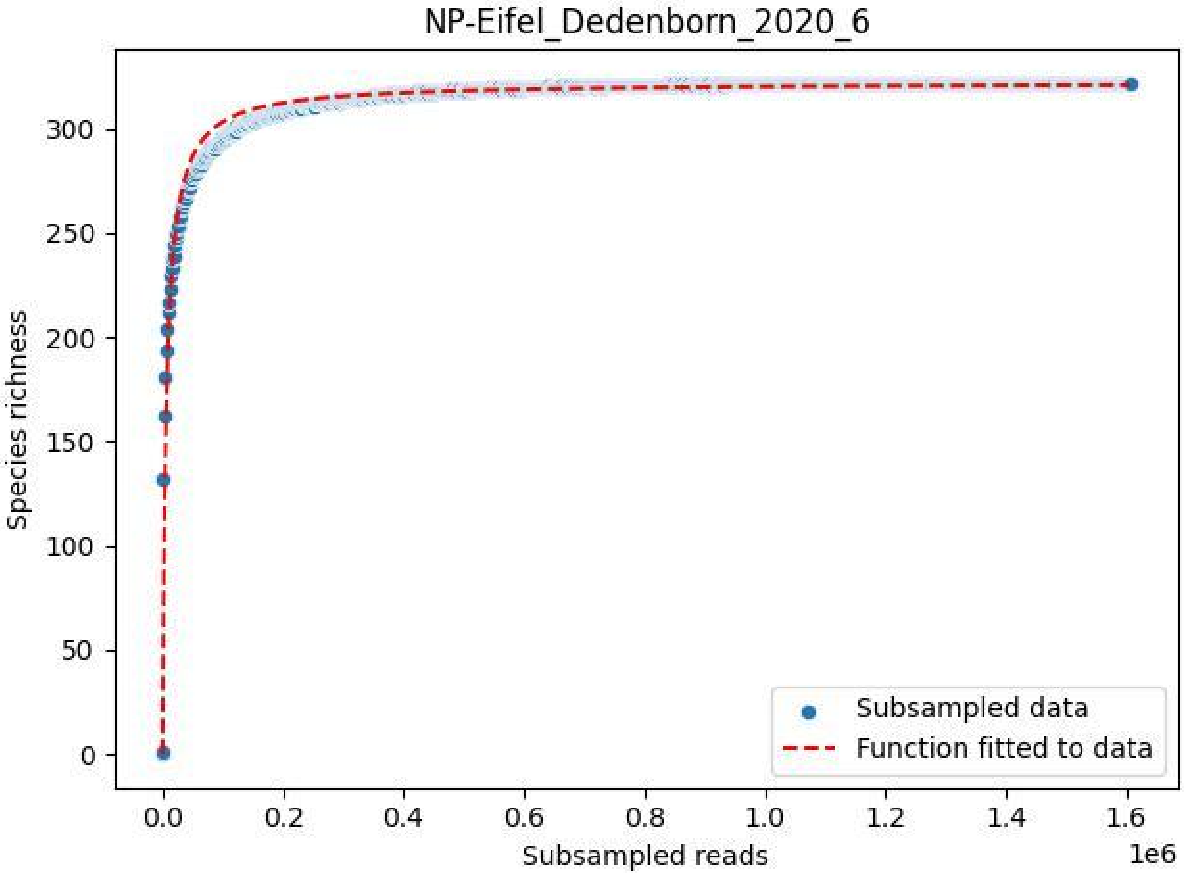
NP-Eifel_Dedenborn_2020_6
Species richness
Subsampled reads
1e6
0
50
100
150
200
250
300
0.0
0.2
0.4
0.6
0.8
1.0
1.2
1.4
1.6
Subsampled data
Function fitted to data

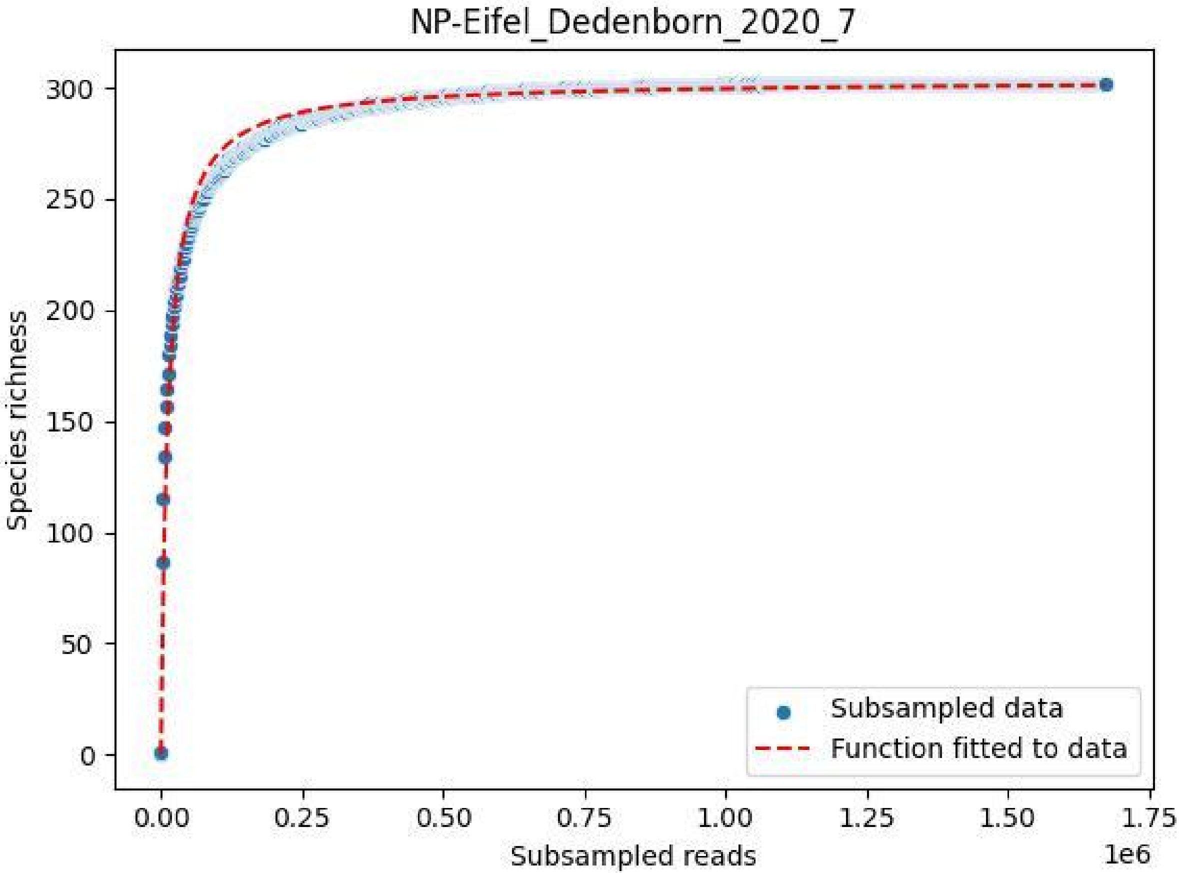
NP-Eifel_Dedenborn_2020_7
Species richness
0
50
100
150
200
250
300
0.00
0.25
0.50
0.75
1.00
1.25
1.50
1.75
1e6
Subsampled reads
Subsampled data
Function fitted to data

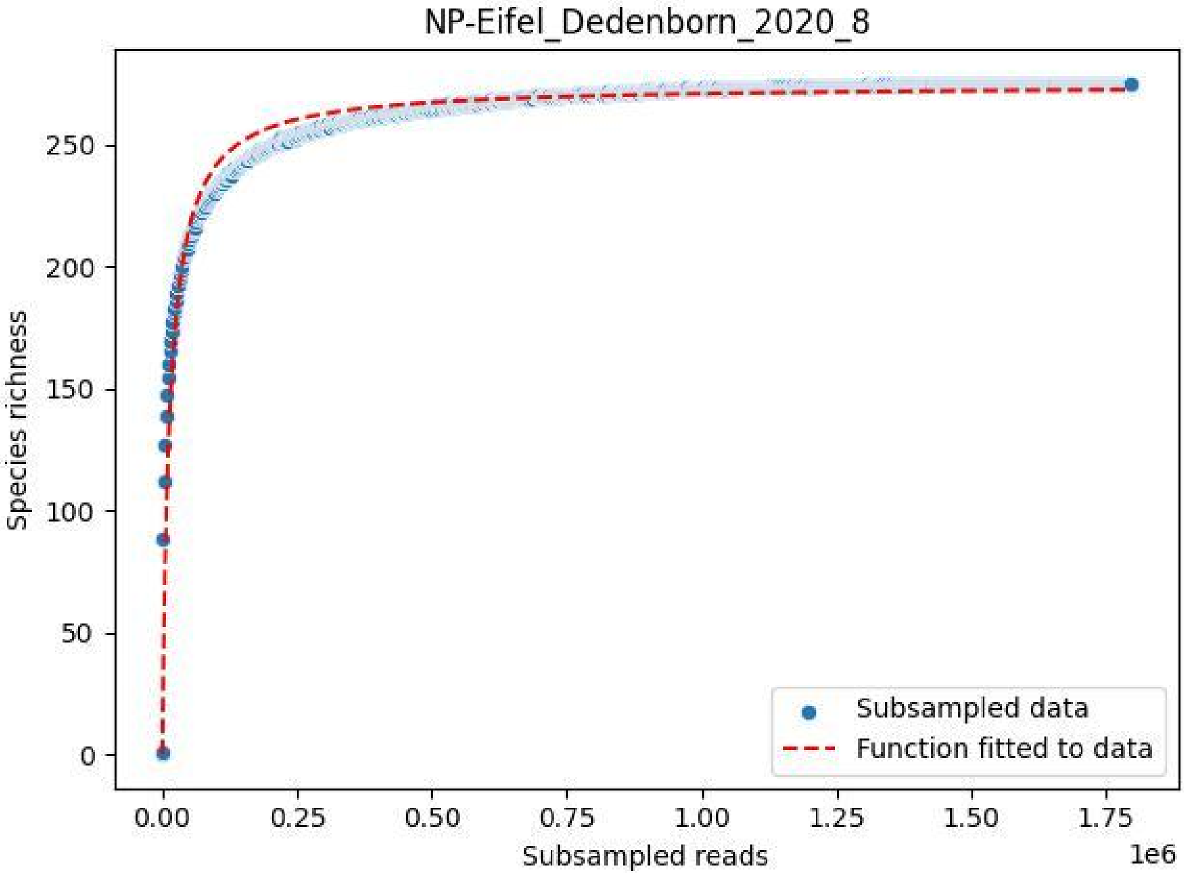
NP-Eifel_Dedenborn_2020_8
Species richness
Subsampled reads
1e6
Subsampled data
Function fitted to data
0
50
100
150
200
250
0.00
0.25
0.50
0.75
1.00
1.25
1.50
1.75

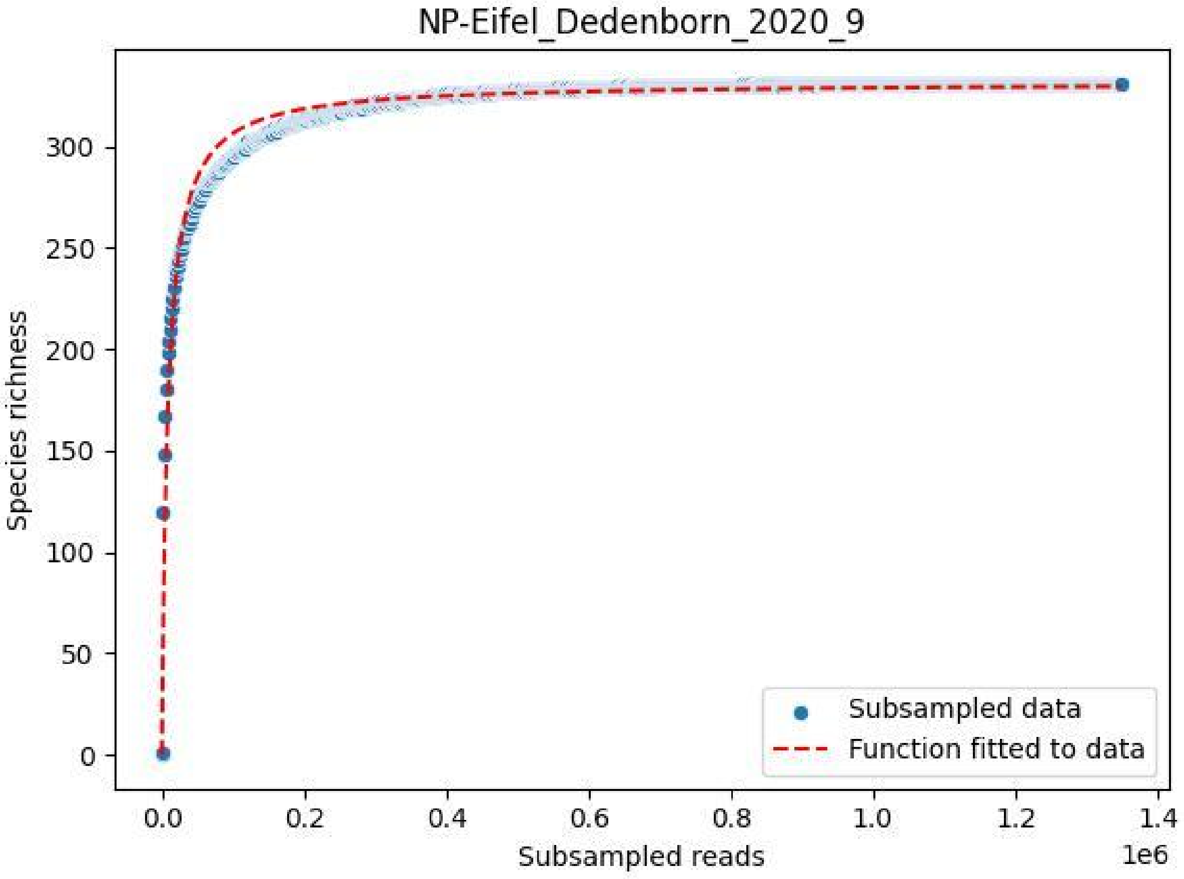
NP-Eifel_Dedenborn_2020_9
Species richness
0
50
100
150
200
250
300
0.0
0.2
0.4
0.6
0.8
1.0
1.2
1.4
1e6
Subsampled reads
Subsampled data
Function fitted to data

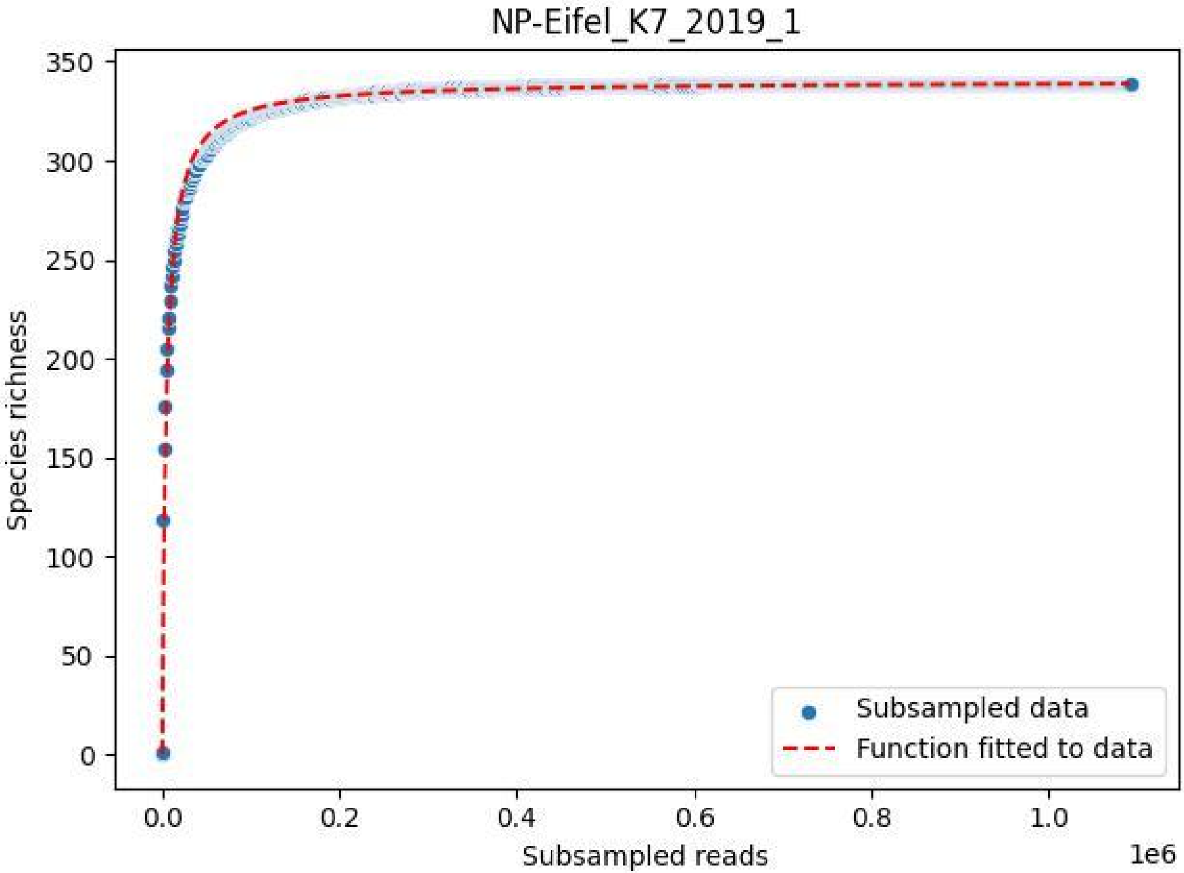

NP-Eifel_K7_2019_1
Species richness
Subsampled reads
1e6
0
50
100
150
200
250
300
350
0.0
0.2
0.4
0.6
0.8
1.0
Subsampled data
Function fitted to data

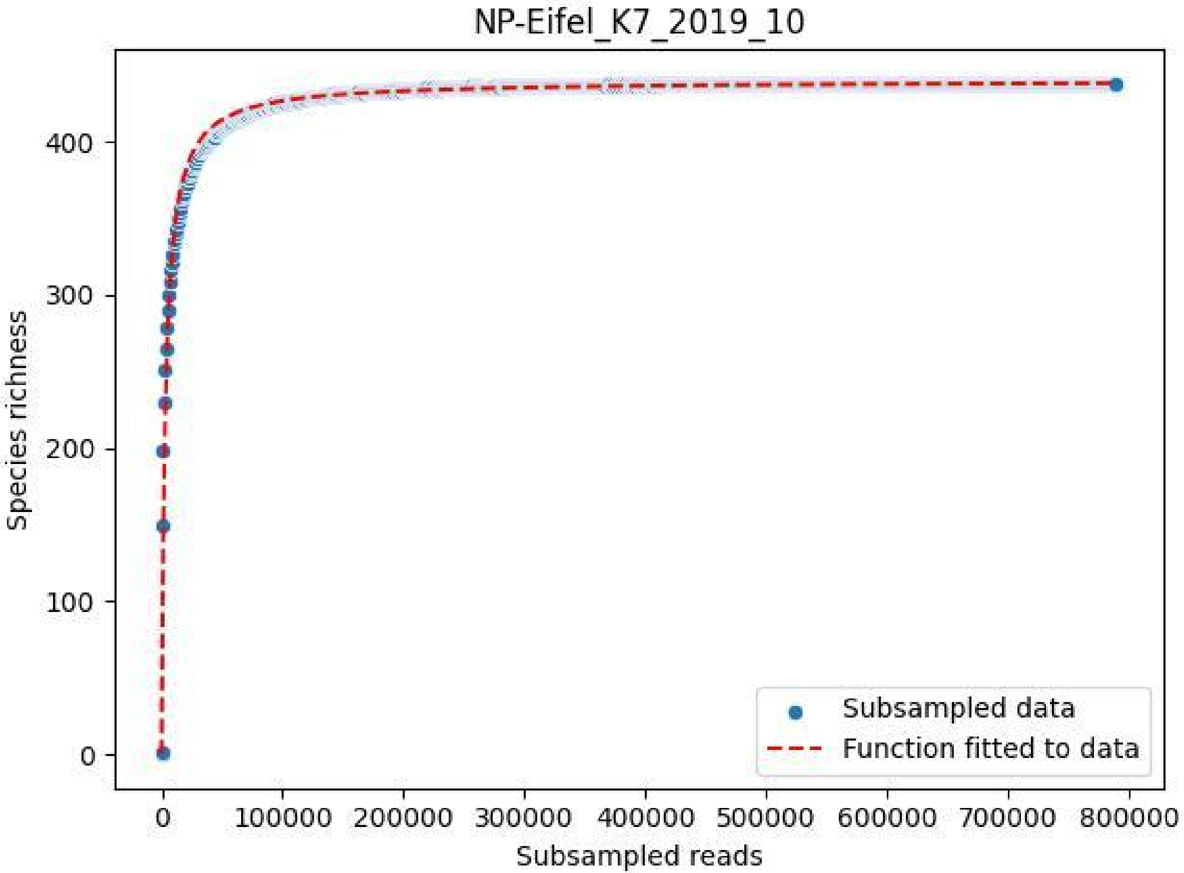
NP-Eifel_K7_2019_10
Species richness
0
100
200
300
400
Subsampled reads
0
100000
200000
300000
400000
500000
600000
700000
800000
Subsampled data
Function fitted to data

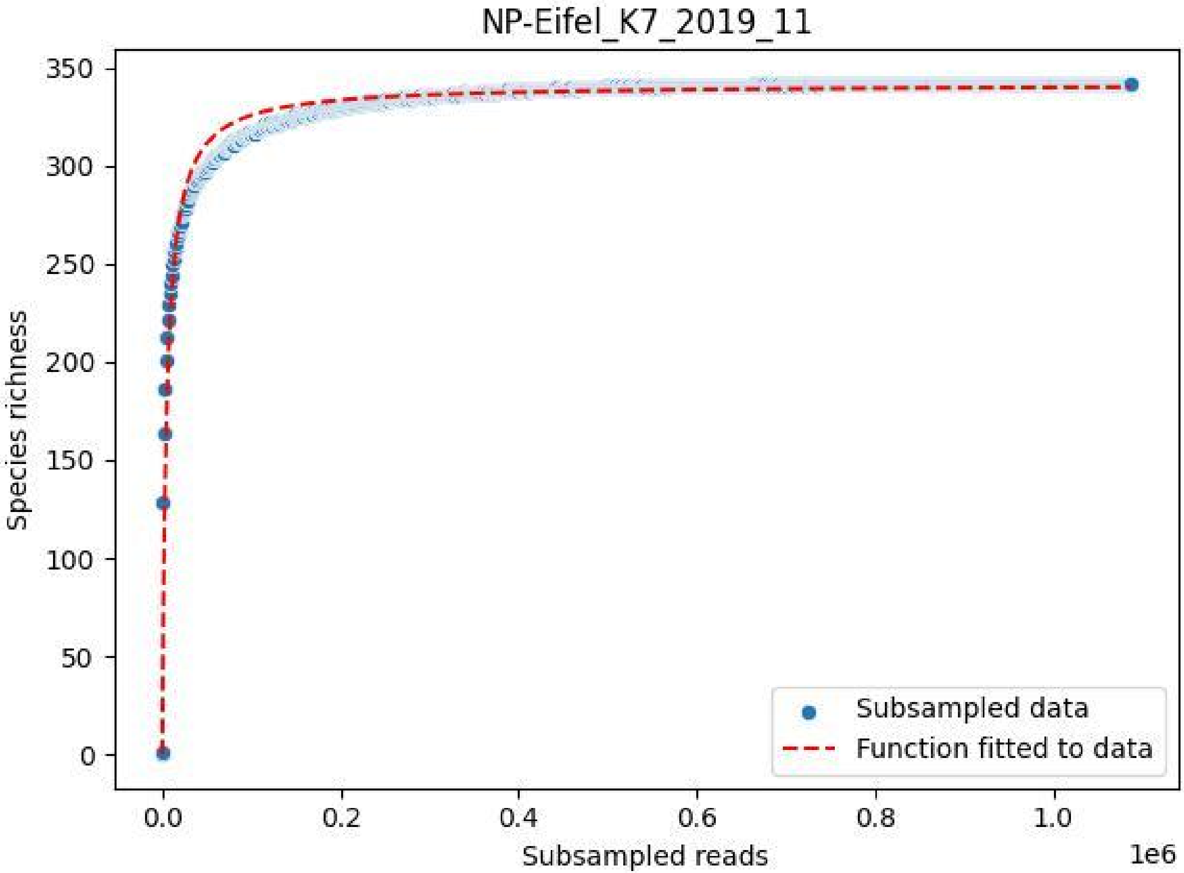

NP-Eifel_K7_2019_11
Species richness
Subsampled reads
1e6
0
50
100
150
200
250
300
350
0.0
0.2
0.4
0.6
0.8
1.0
Subsampled data
Function fitted to data

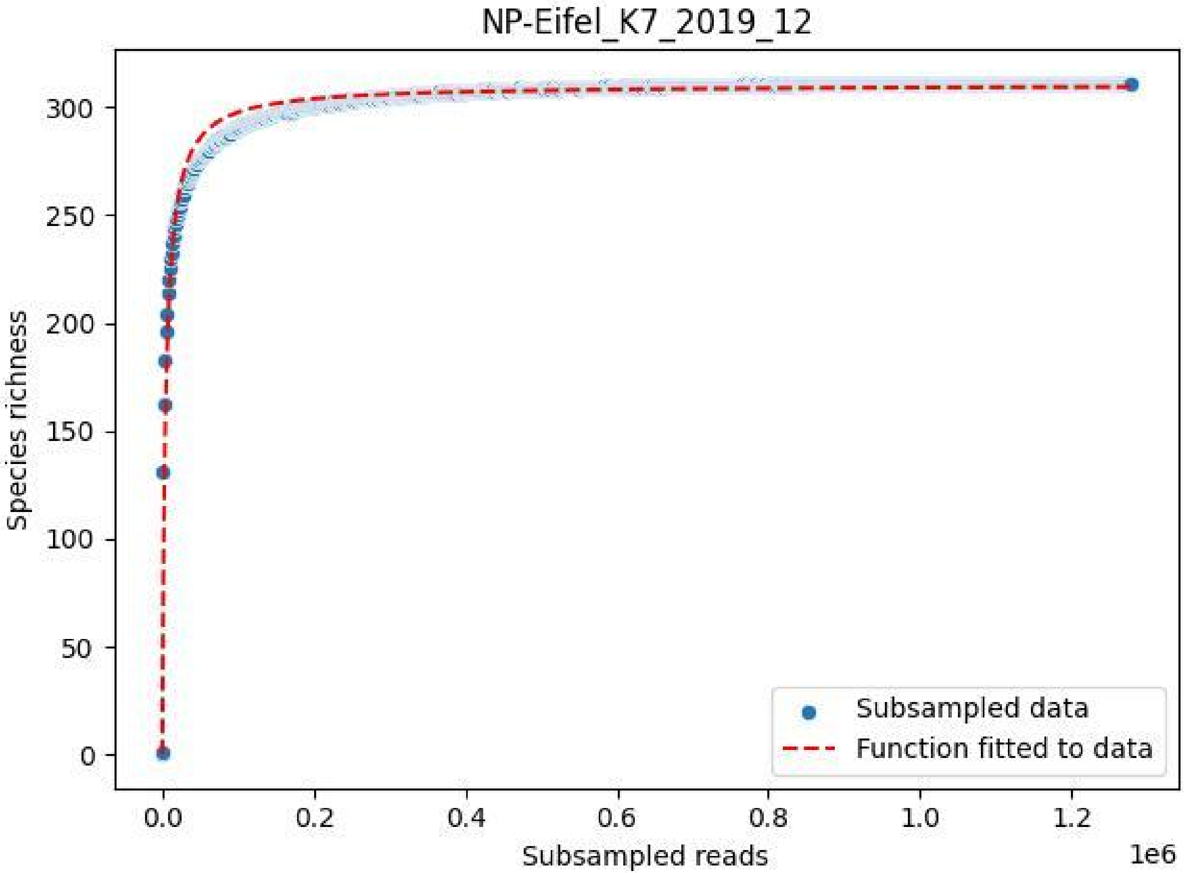
NP-Eifel_K7_2019_12
Species richness
Subsampled reads
1e6
300
250
200
150
100
50
0
0.0
0.2
0.4
0.6
0.8
1.0
1.2
Subsampled data
Function fitted to data

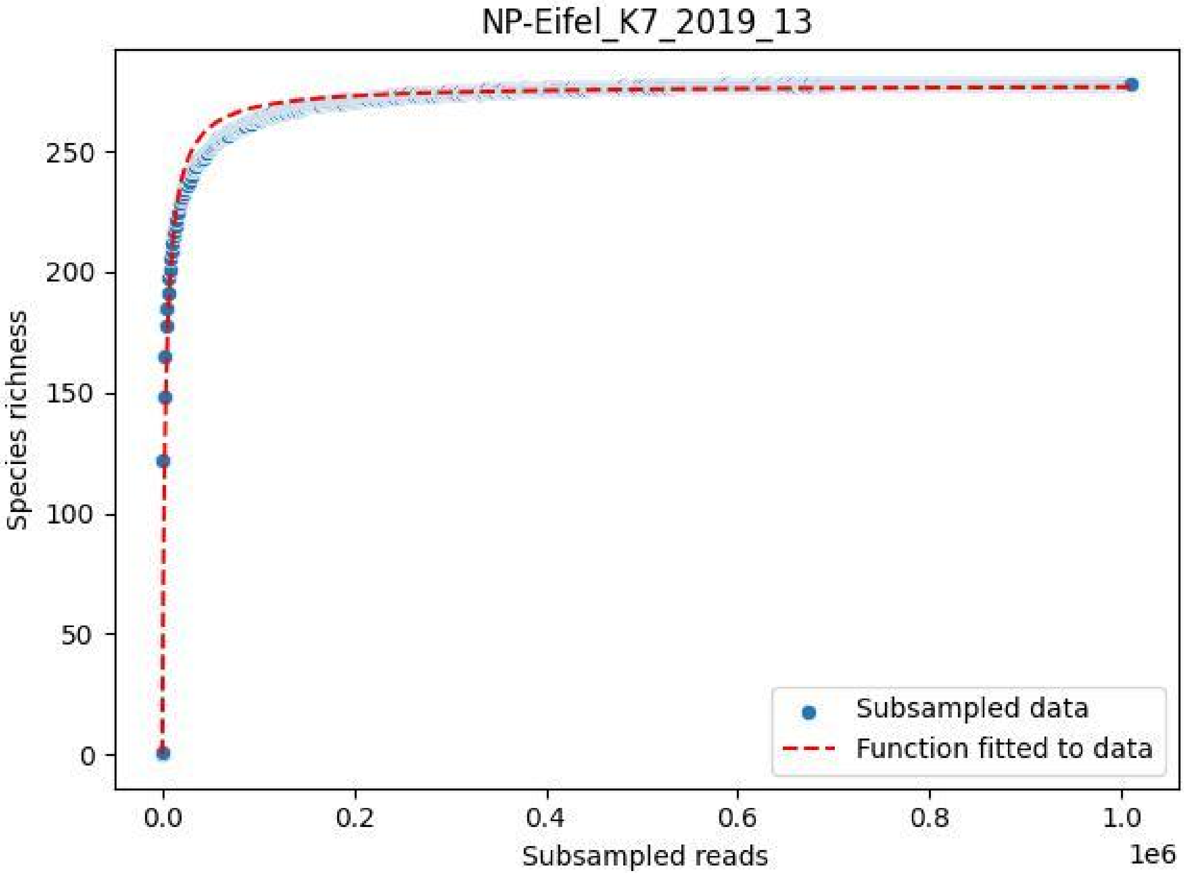
NP-Eifel_K7_2019_13
Species richness
Subsampled reads
1e6
0
50
100
150
200
250
0.0
0.2
0.4
0.6
0.8
1.0
Subsampled data
Function fitted to data

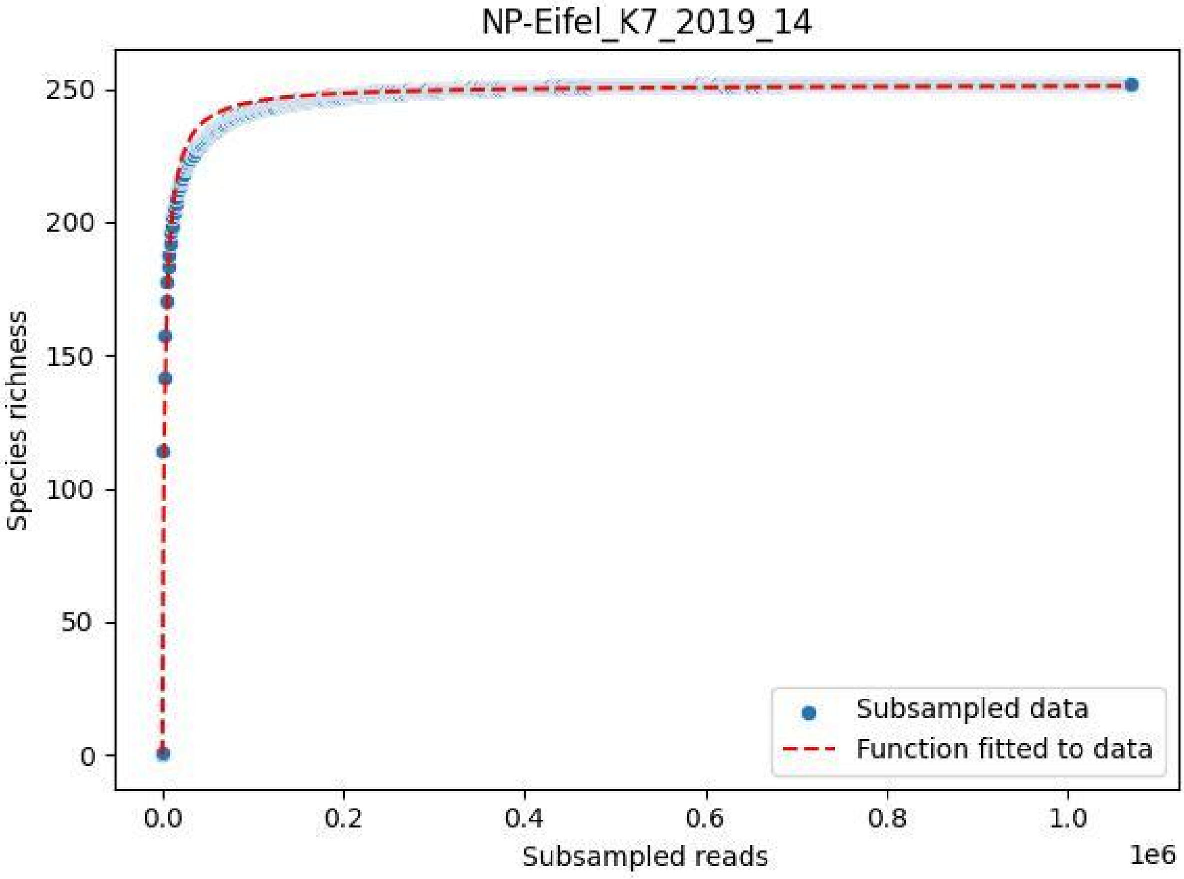
NP-Eifel_K7_2019_14
Species richness
Subsampled reads
1e6
0
50
100
150
200
250
0.0
0.2
0.4
0.6
0.8
1.0
Subsampled data
Function fitted to data

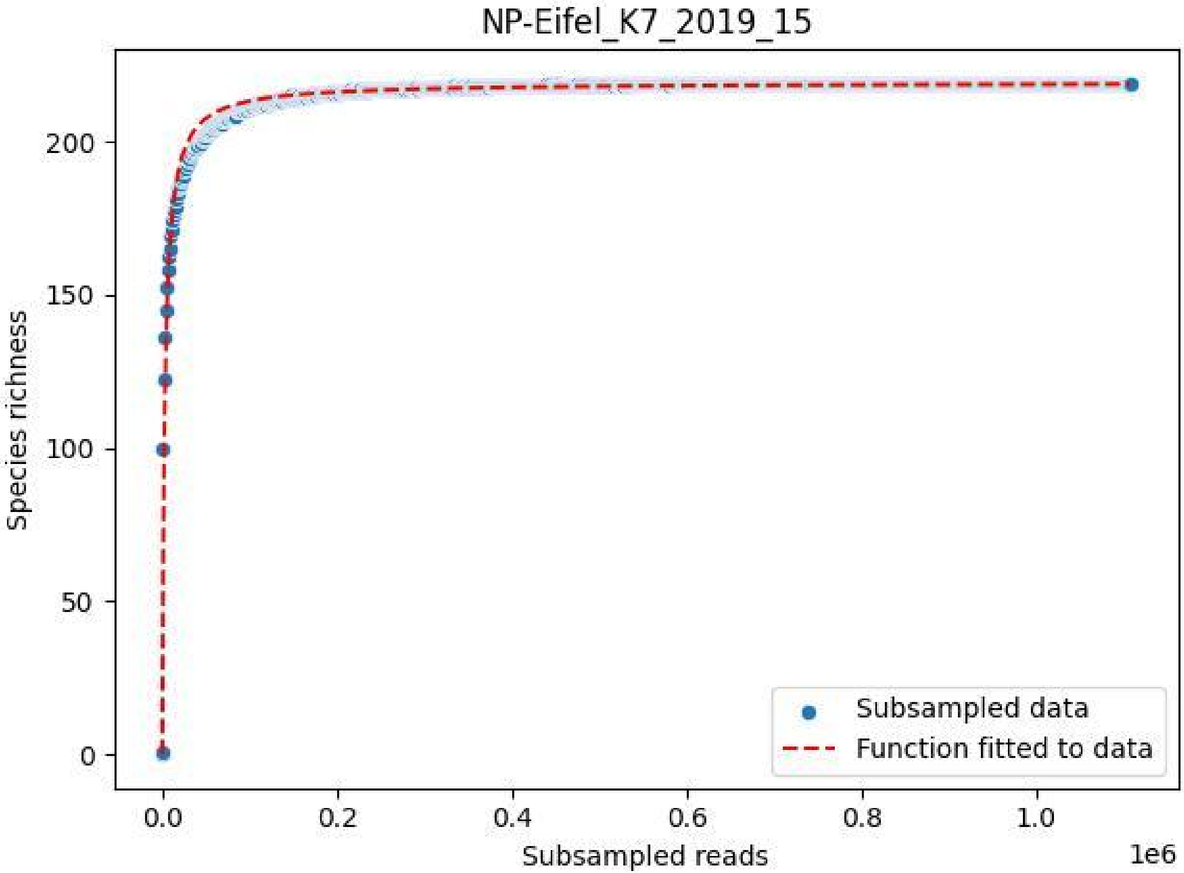
NP-Eifel_K7_2019_15
Species richness
Subsampled reads
1e6
0
50
100
150
200
0.0
0.2
0.4
0.6
0.8
1.0
Subsampled data
Function fitted to data

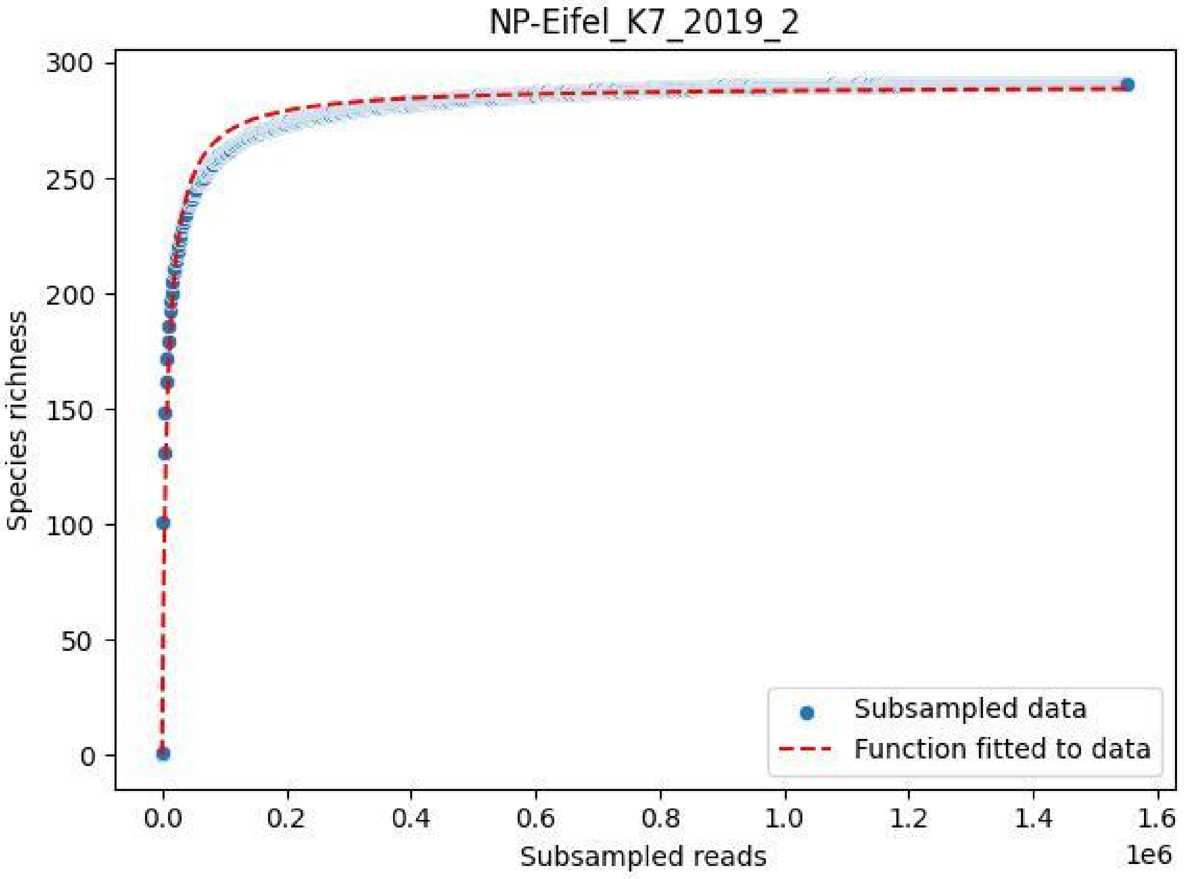

NP-Eifel_K7_2019_2
Species richness
Subsampled reads
1e6
300
250
200
150
100
50
0
0.0
0.2
0.4
0.6
0.8
1.0
1.2
1.4
1.6
Subsampled data
Function fitted to data

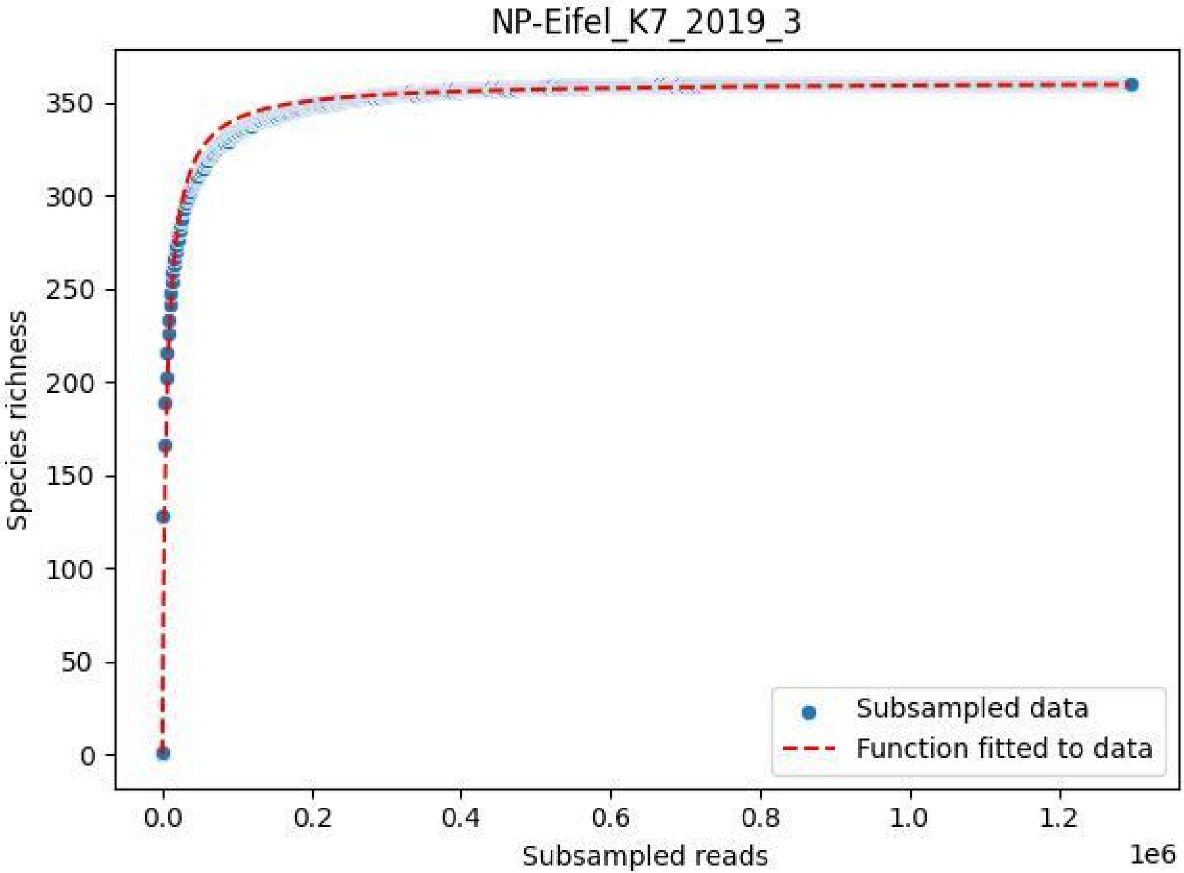

NP-Eifel_K7_2019_3
Species richness
350
300
250
200
150
100
50
0
0.0
0.2
0.4
0.6
0.8
1.0
1.2
1e6
Subsampled reads
Subsampled data
Function fitted to data

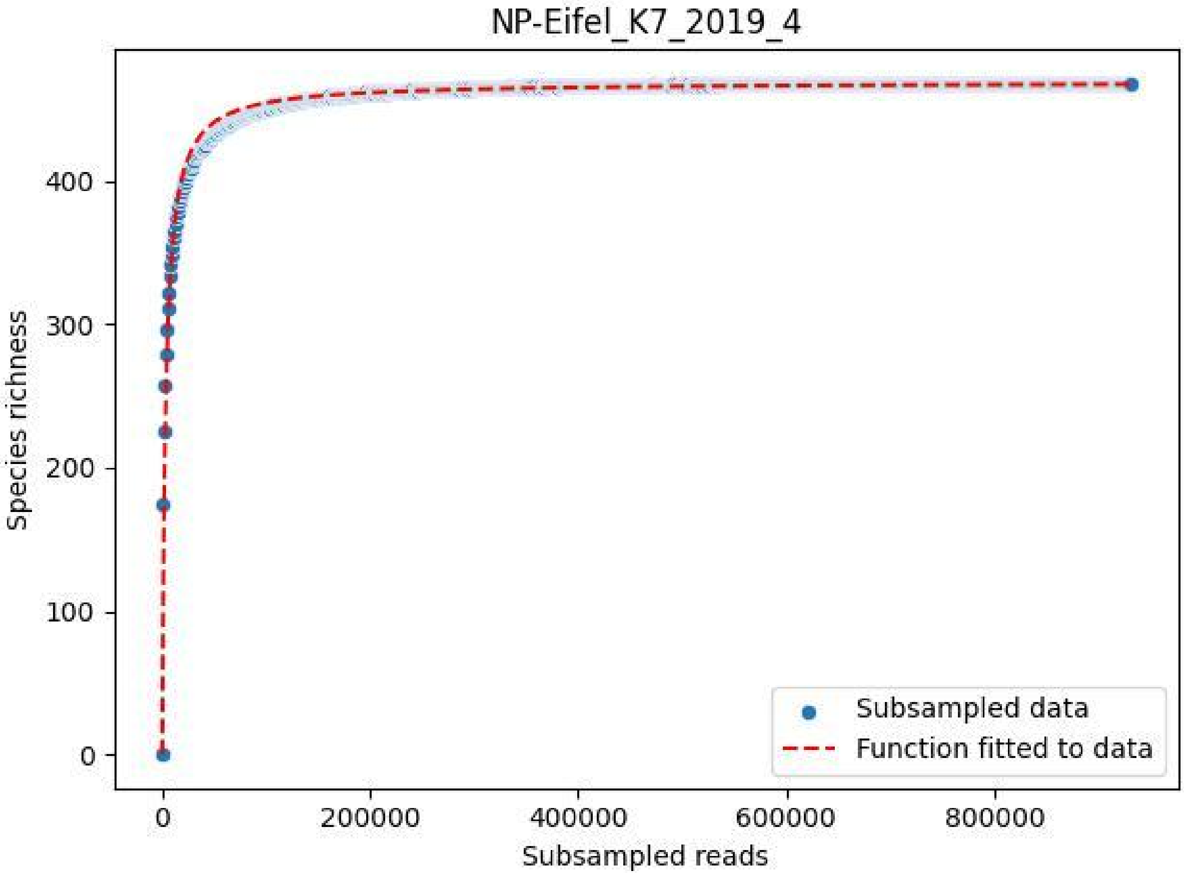
NP-Eifel_K7_2019_4
Species richness
Subsampled reads
0
100
200
300
400
0
200000
400000
600000
800000
Subsampled data
Function fitted to data

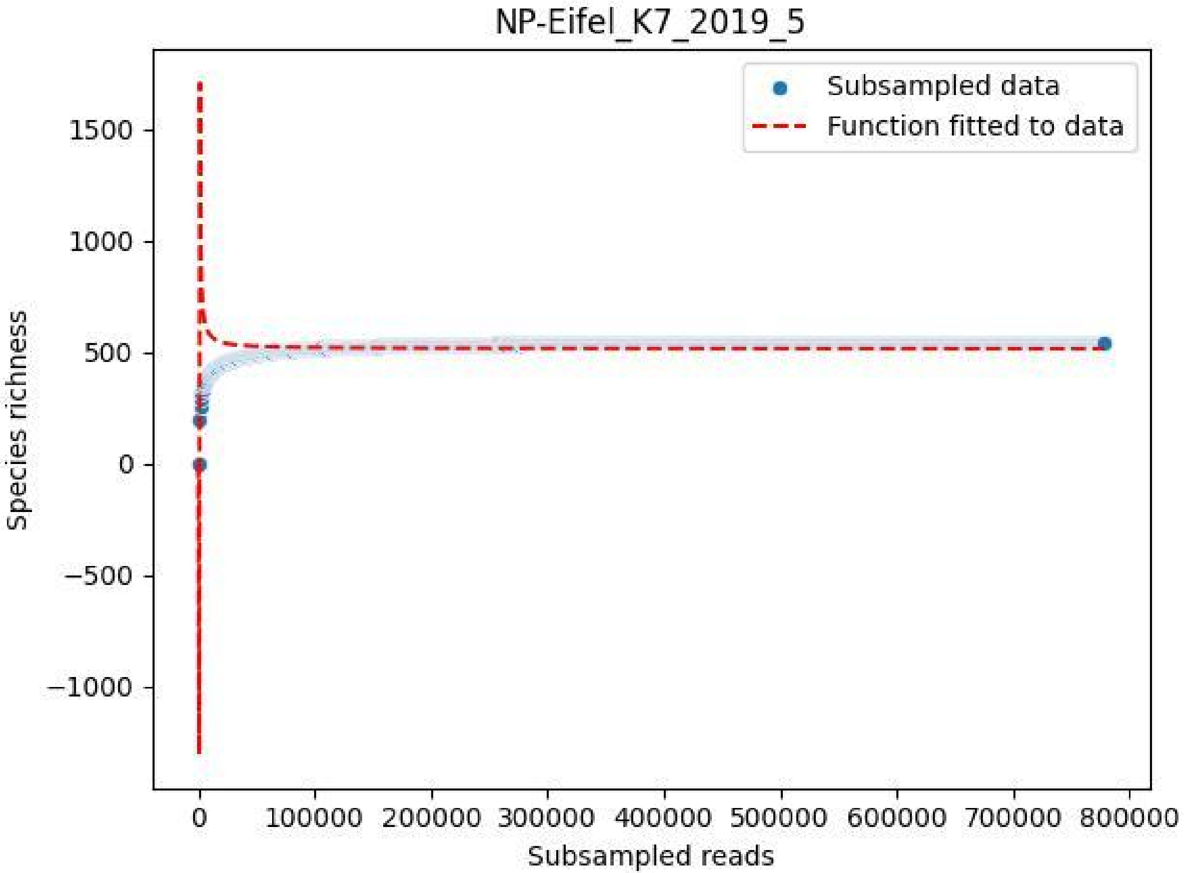
NP-Eifel_K7_2019_5
Subsampled data
Function fitted to data
Species richness
1500
1000
500
0
−500
−1000
0
100000
200000
300000
400000
500000
600000
700000
800000
Subsampled reads

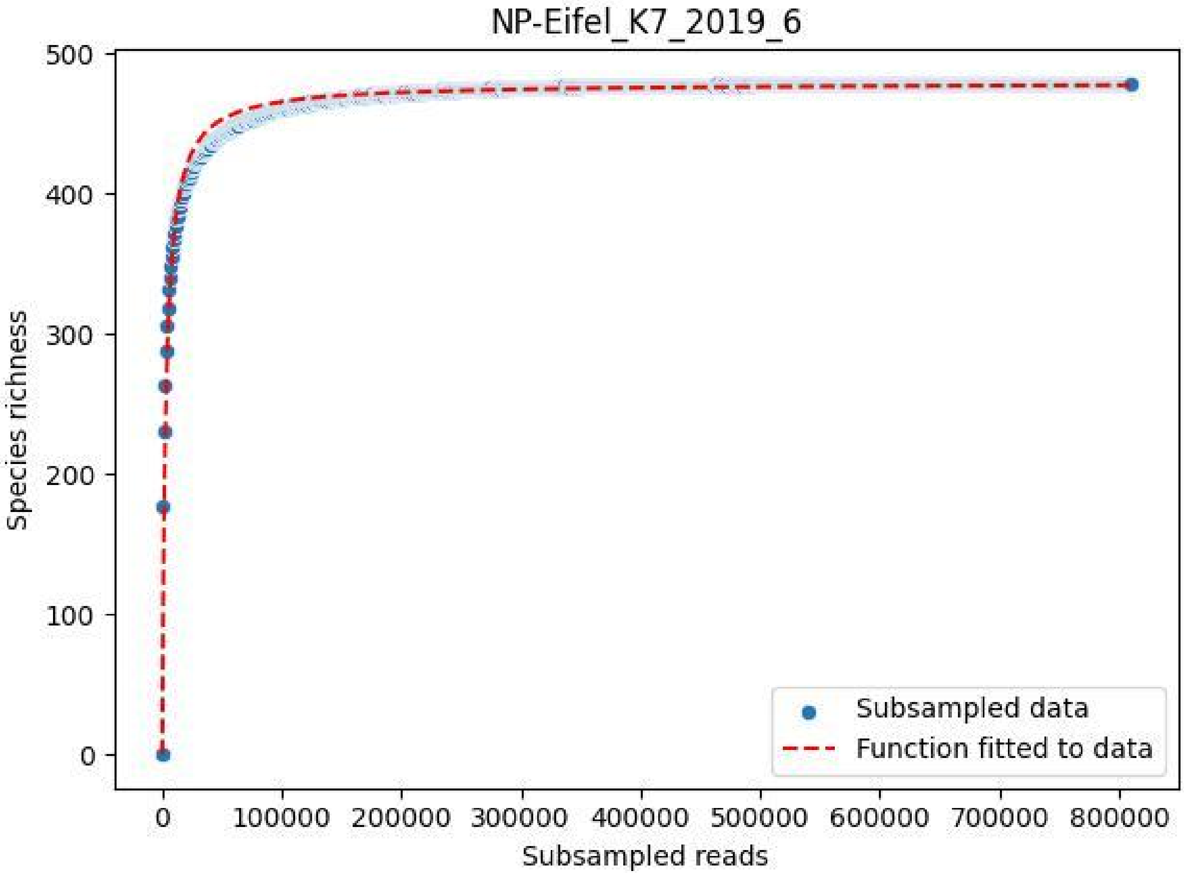
NP-Eifel_K7_2019_6
Species richness
Subsampled reads
0
100
200
300
400
500
0
100000
200000
300000
400000
500000
600000
700000
800000
Subsampled data
Function fitted to data

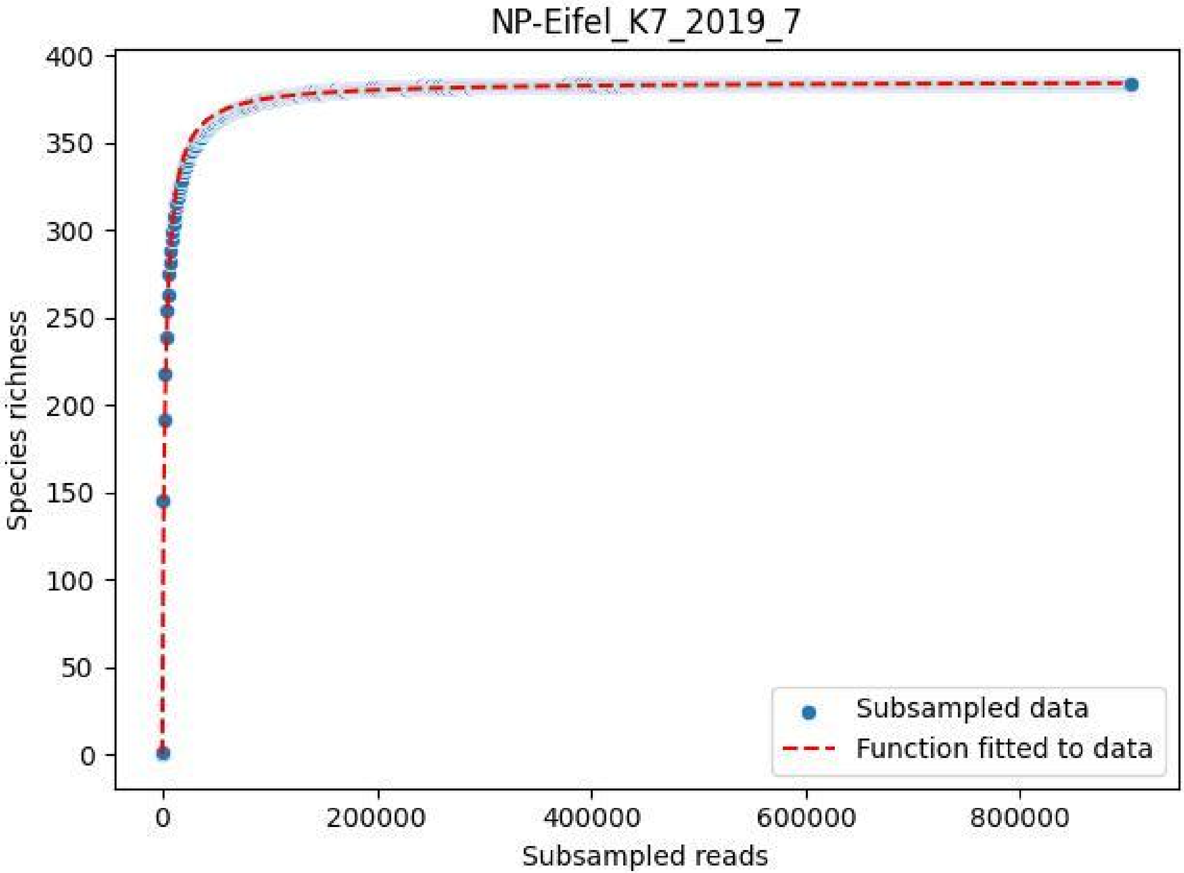

NP-Eifel_K7_2019_7
Species richness
Subsampled reads
0
50
100
150
200
250
300
350
400
0
200000
400000
600000
800000
Subsampled data
Function fitted to data

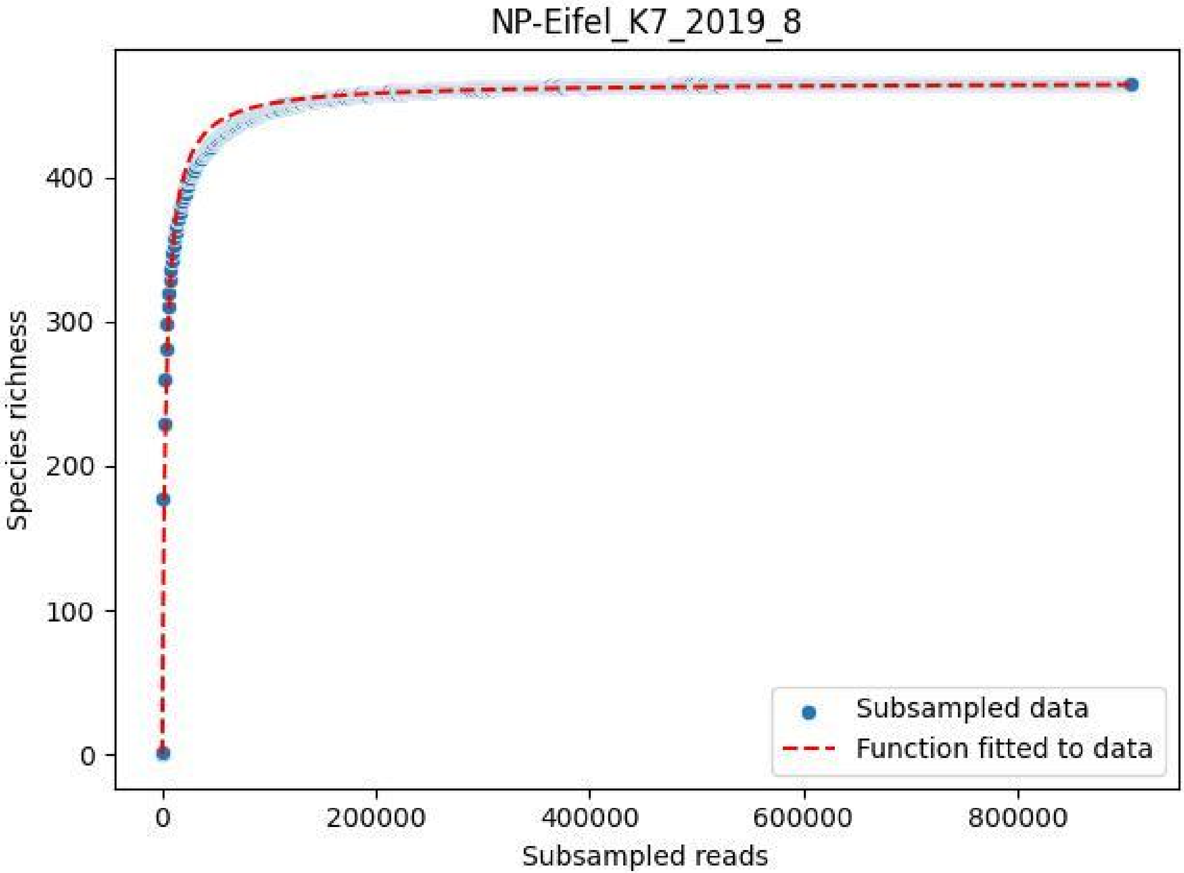
NP-Eifel_K7_2019_8
Species richness
Subsampled reads
0
100
200
300
400
0
200000
400000
600000
800000
Subsampled data
Function fitted to data

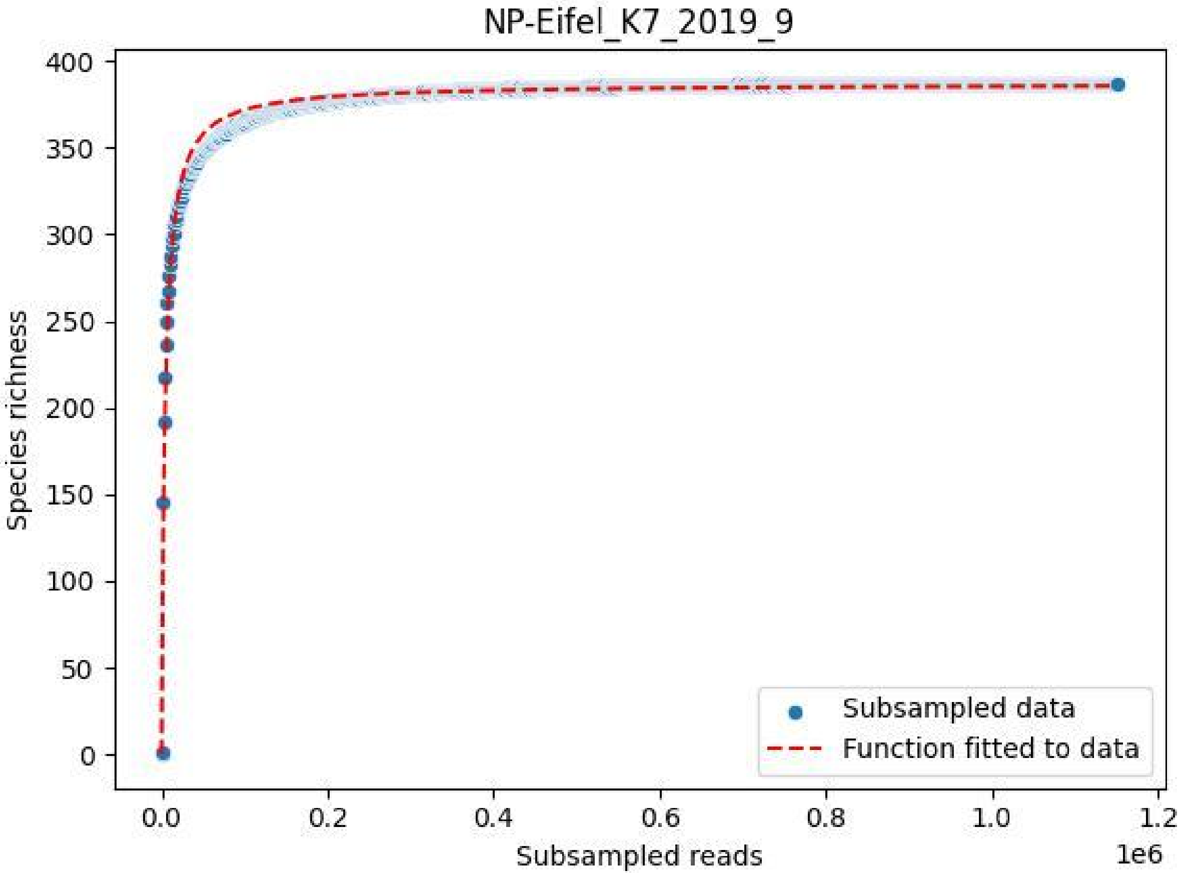
NP-Eifel_K7_2019_9
Species richness
400
350
300
250
200
150
100
50
0
0.0
0.2
0.4
0.6
0.8
1.0
1.2
Subsampled reads
1e6
Subsampled data
Function fitted to data

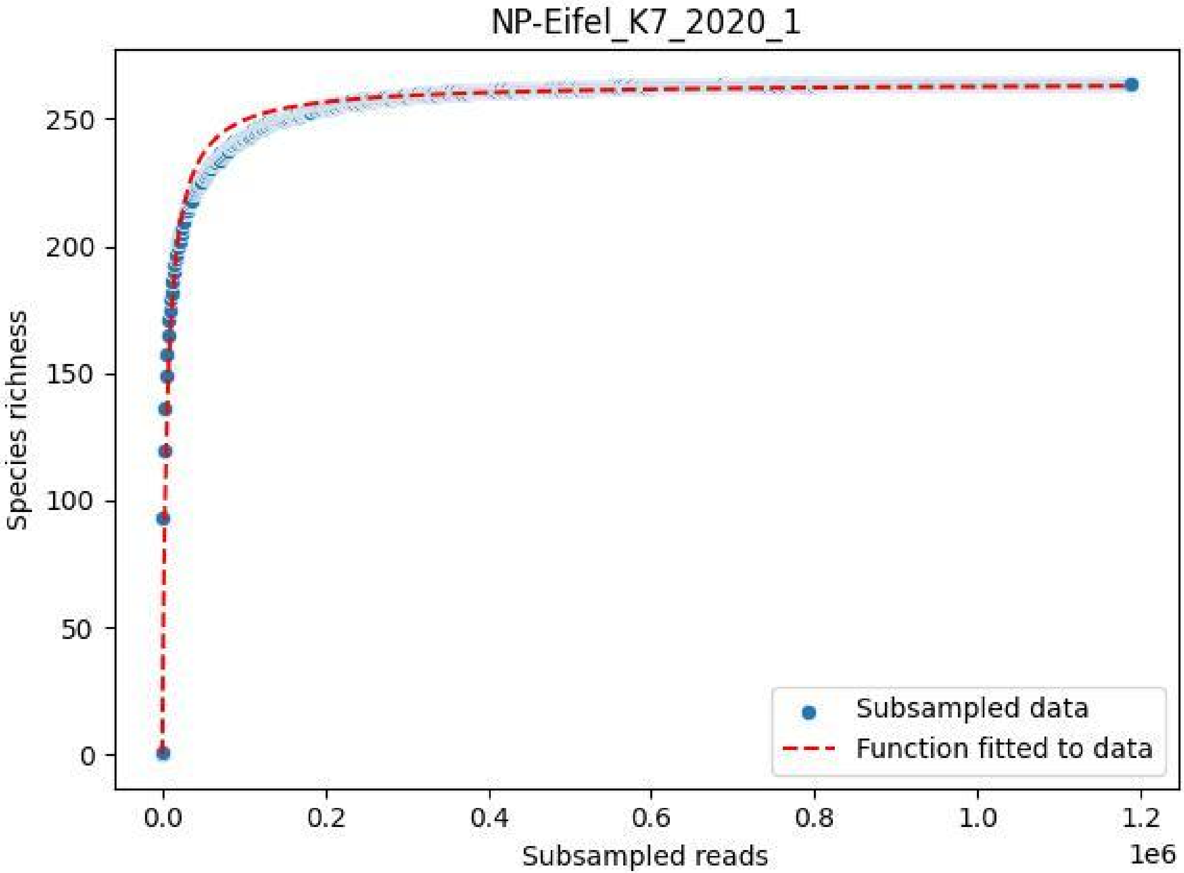
NP-Eifel_K7_2020_1
Species richness
Subsampled reads
1e6
0
50
100
150
200
250
0.0
0.2
0.4
0.6
0.8
1.0
1.2
Subsampled data
Function fitted to data

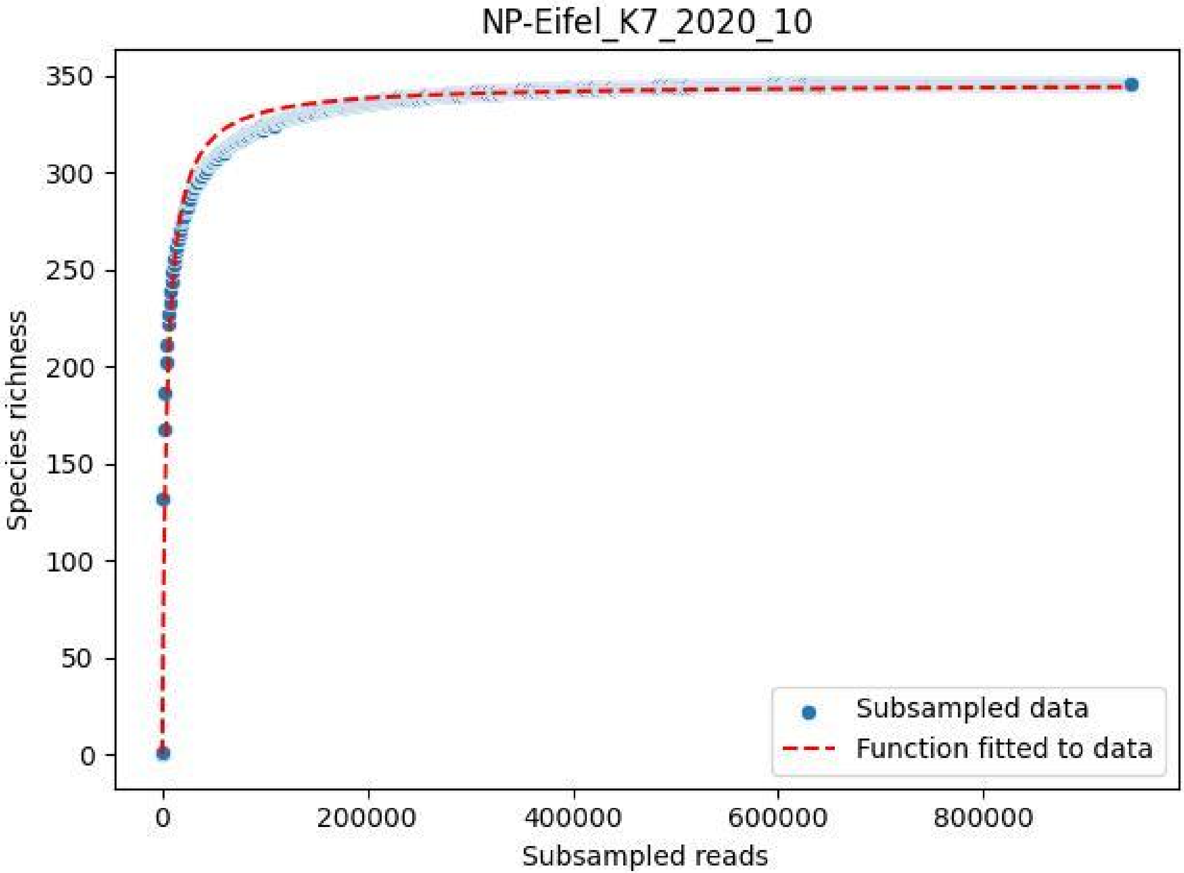

NP-Eifel_K7_2020_10
Species richness
Subsampled reads
0
50
100
150
200
250
300
350
0
200000
400000
600000
800000
Subsampled data
Function fitted to data

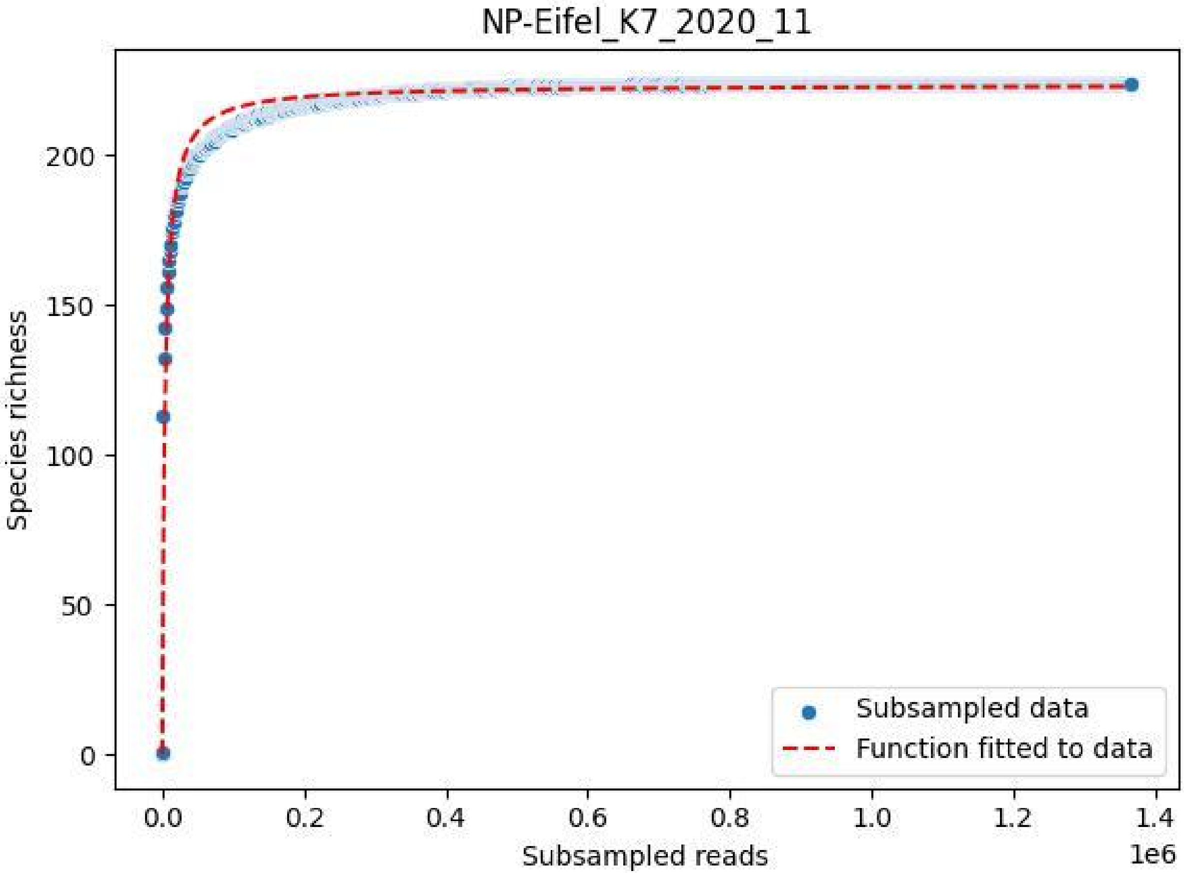
NP-Eifel_K7_2020_11
Species richness
Subsampled reads
1e6
0
50
100
150
200
0.0
0.2
0.4
0.6
0.8
1.0
1.2
1.4
Subsampled data
Function fitted to data

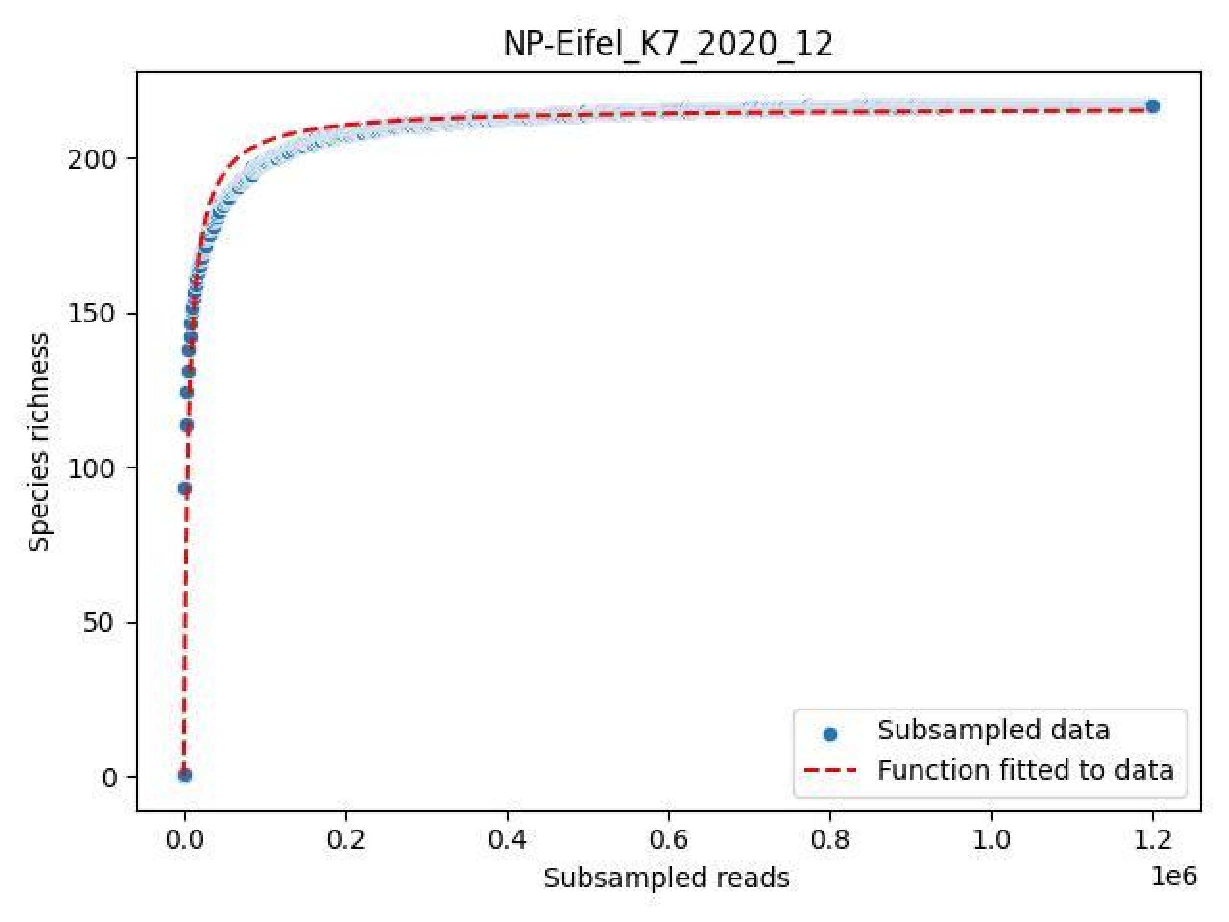
NP-Eifel_K7_2020_12
Species richness
Subsampled reads
1e6
0
50
100
150
200
0.0
0.2
0.4
0.6
0.8
1.0
1.2
Subsampled data
Function fitted to data

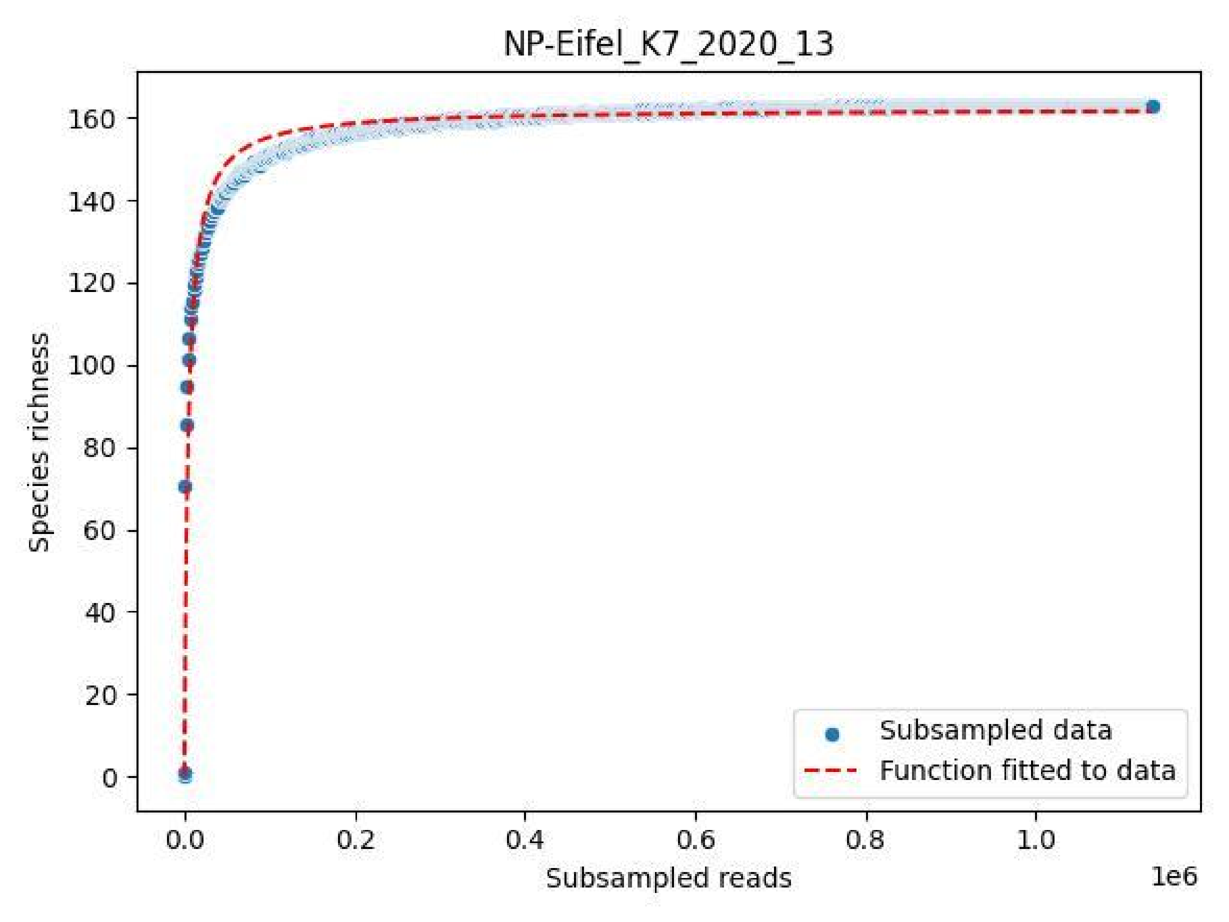
NP-Eifel_K7_2020_13
Species richness
0
20
40
60
80
100
120
140
160
0.0
0.2
0.4
0.6
0.8
1.0
1e6
Subsampled reads
Subsampled data
Function fitted to data

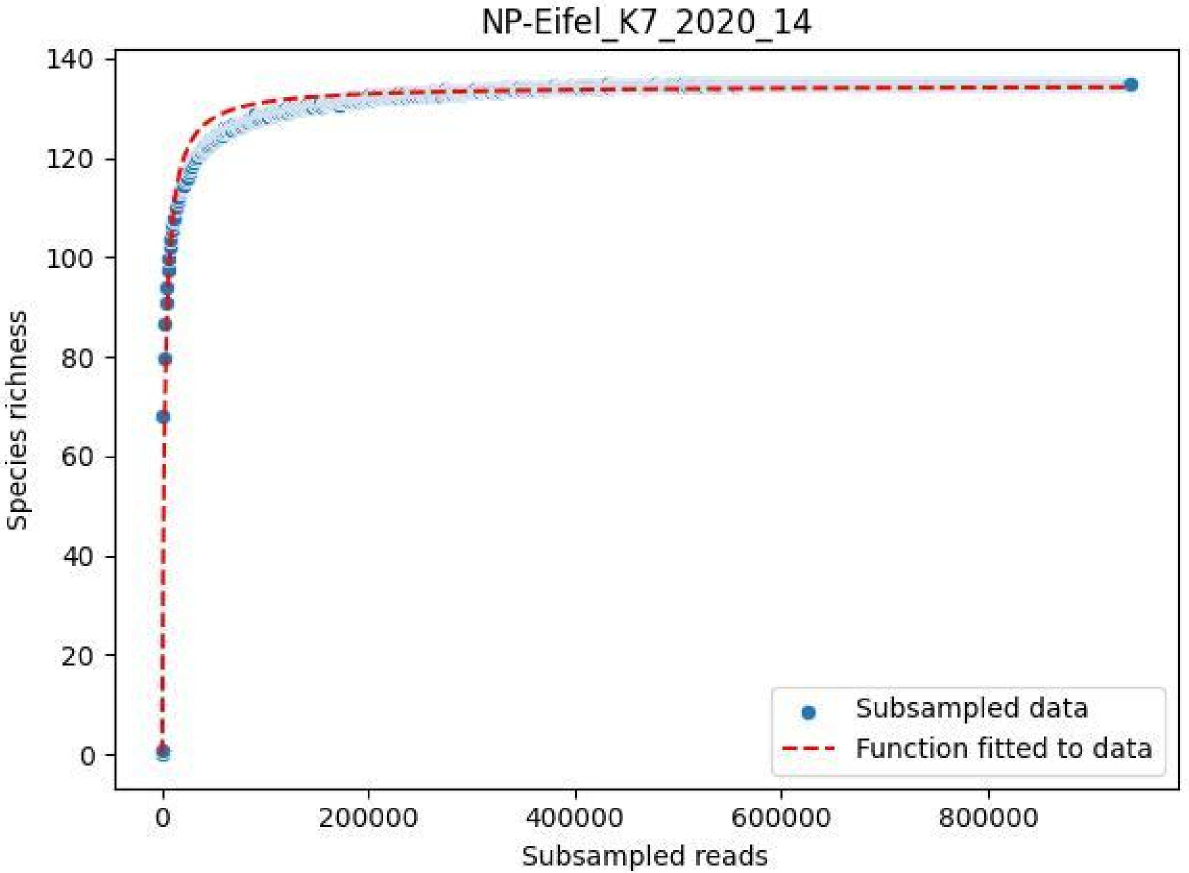
NP-Eifel_K7_2020_14
Species richness
Subsampled reads
0
20
40
60
80
100
120
140
0
200000
400000
600000
800000
Subsampled data
Function fitted to data

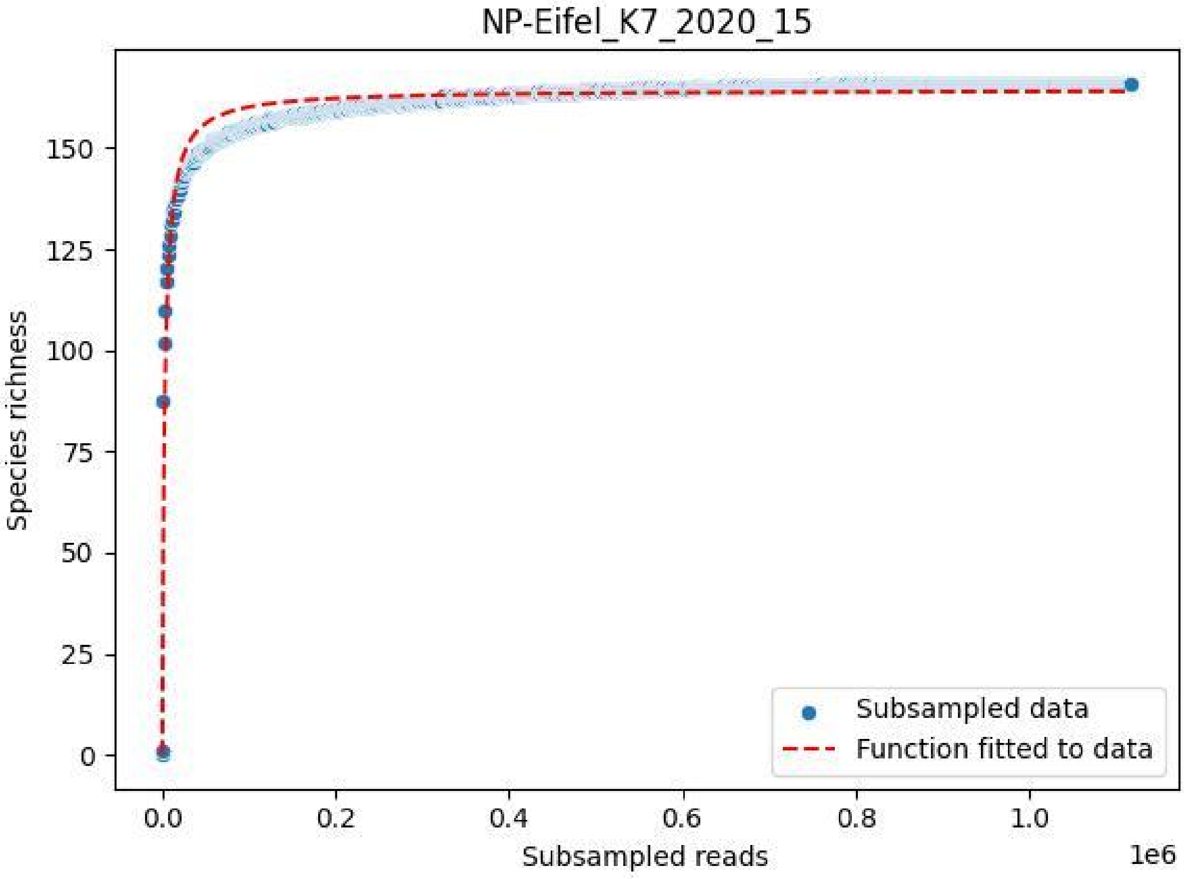
NP-Eifel_K7_2020_15
Species richness
0
25
50
75
100
125
150
0.0
0.2
0.4
0.6
0.8
1.0
1e6
Subsampled reads
Subsampled data
Function fitted to data

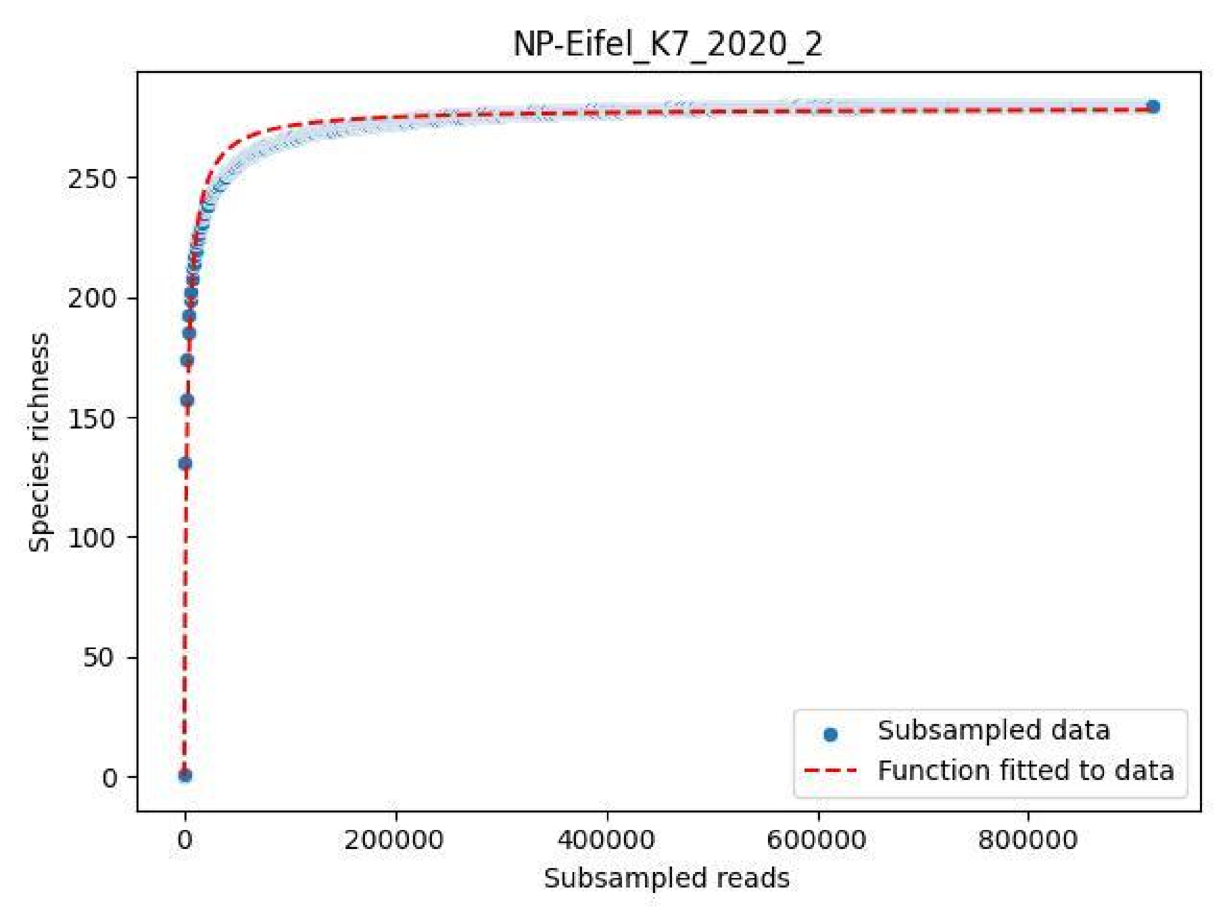
NP-Eifel_K7_2020_2
Species richness
Subsampled reads
0
50
100
150
200
250
0
200000
400000
600000
800000
Subsampled data
Function fitted to data

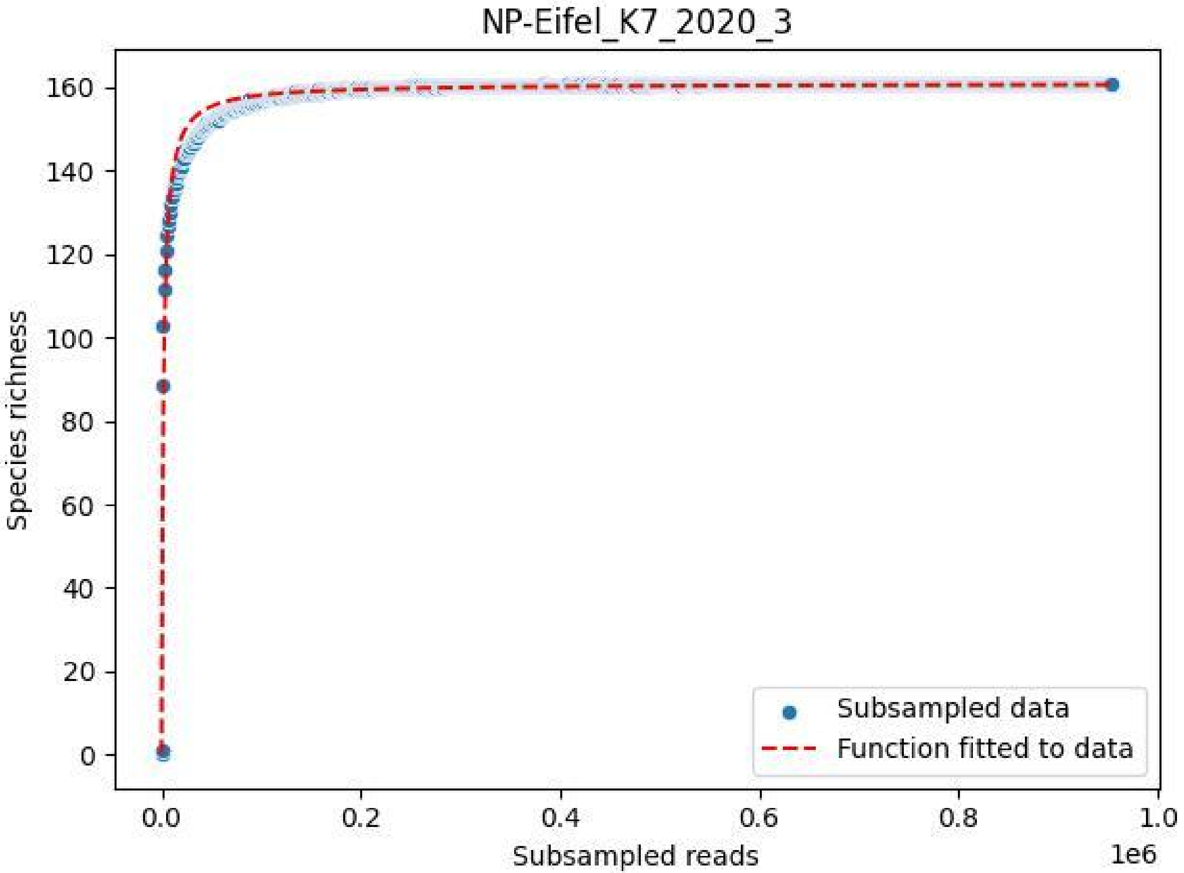
NP-Eifel_K7_2020_3
Species richness
Subsampled reads
1e6
Subsampled data
Function fitted to data
0
20
40
60
80
100
120
140
160
0.0
0.2
0.4
0.6
0.8
1.0

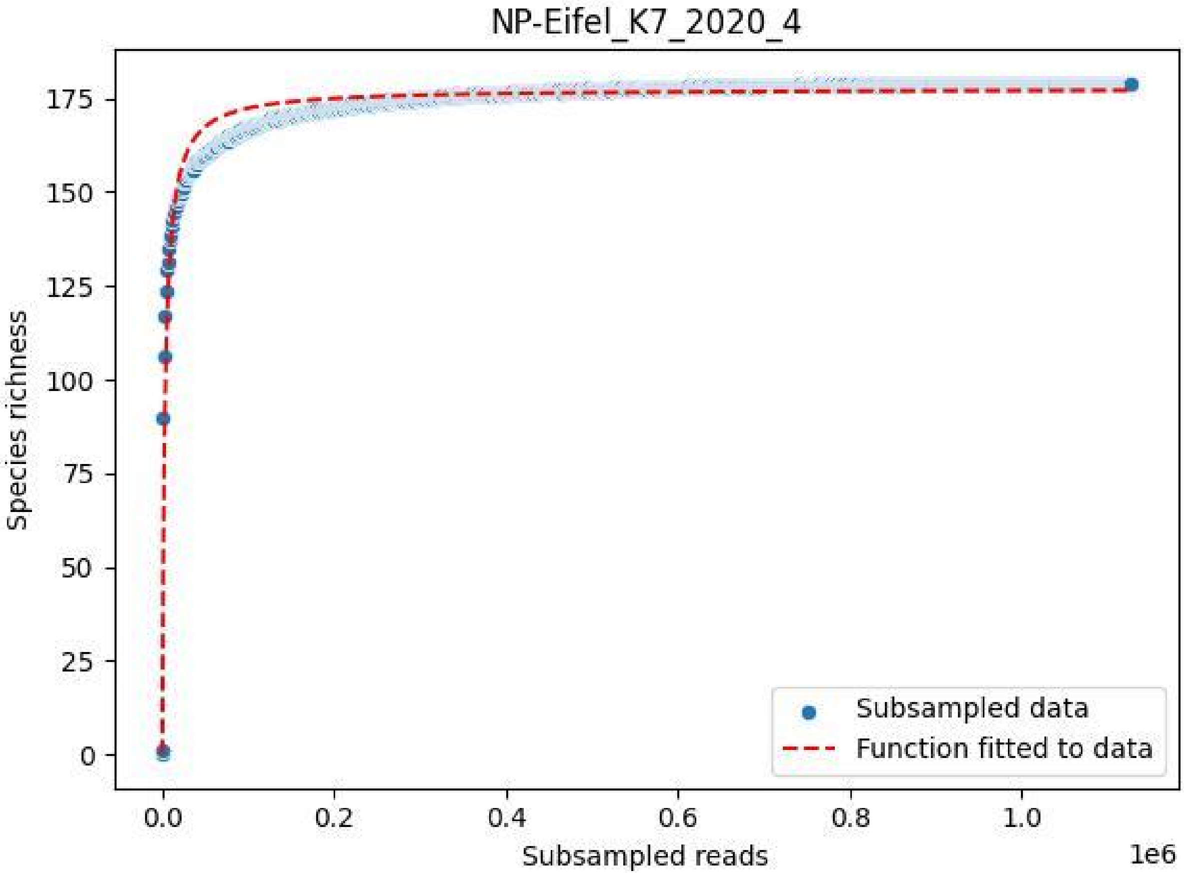

NP-Eifel_K7_2020_4
Species richness
Subsampled reads
1e6
0
25
50
75
100
125
150
175
0.0
0.2
0.4
0.6
0.8
1.0
Subsampled data
Function fitted to data

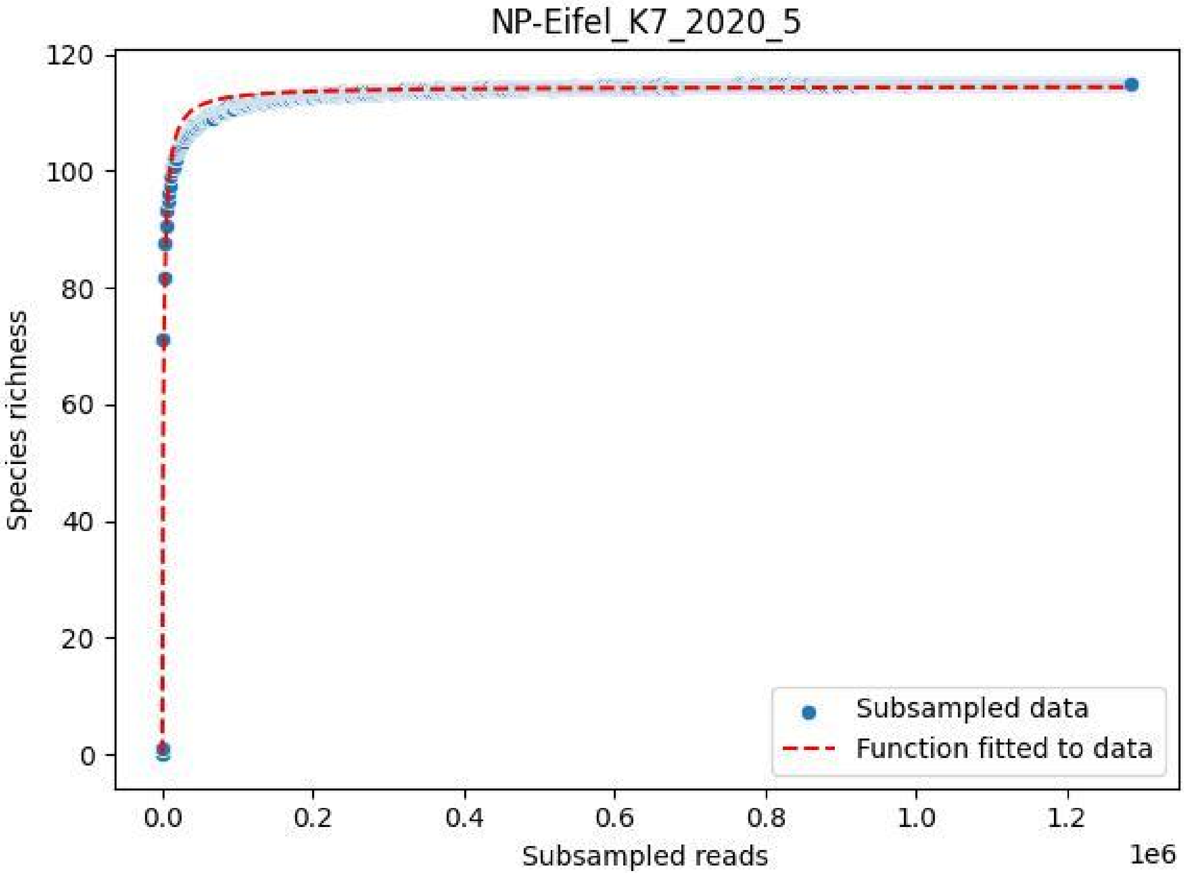
NP-Eifel_K7_2020_5
Species richness
Subsampled reads
1e6
0
20
40
60
80
100
120
0.0
0.2
0.4
0.6
0.8
1.0
1.2
Subsampled data
Function fitted to data

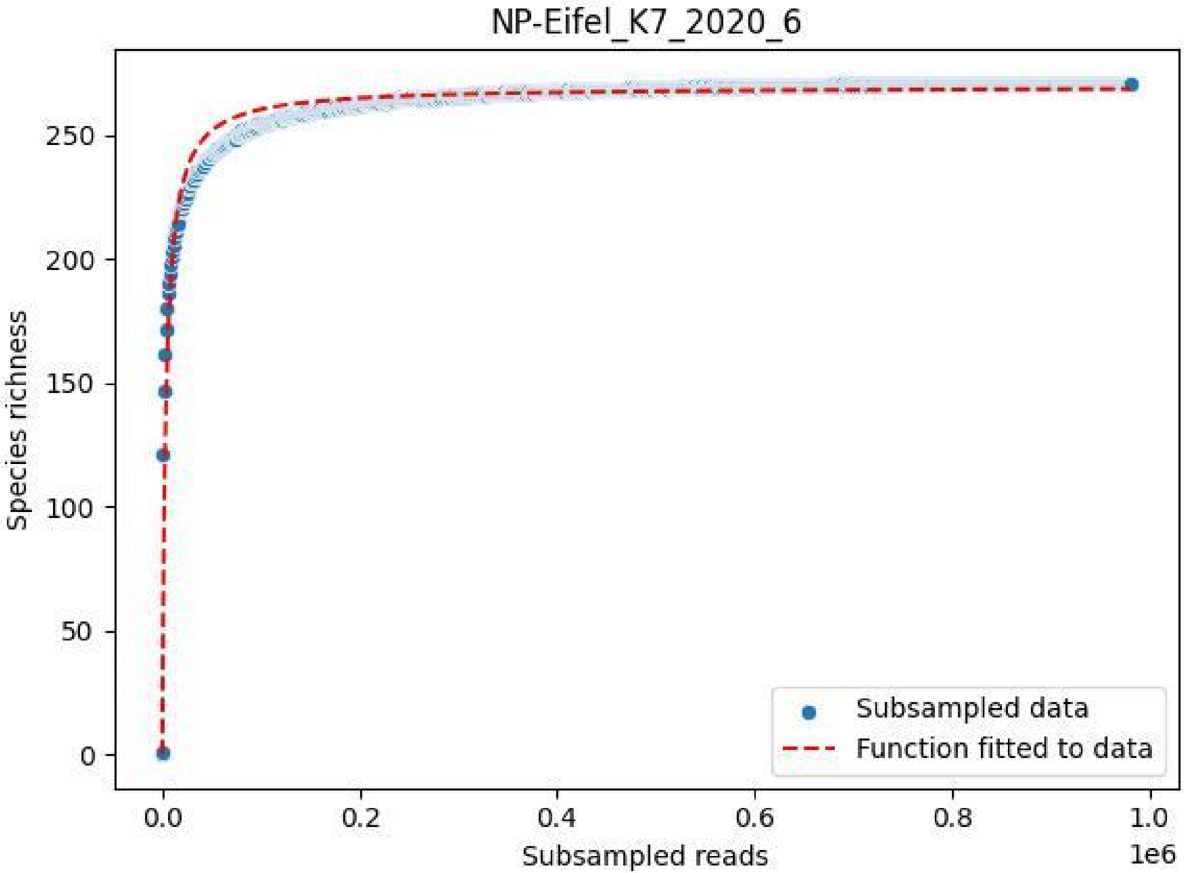
NP-Eifel_K7_2020_6
Species richness
Subsampled reads
1e6
0
50
100
150
200
250
0.0
0.2
0.4
0.6
0.8
1.0
Subsampled data
Function fitted to data

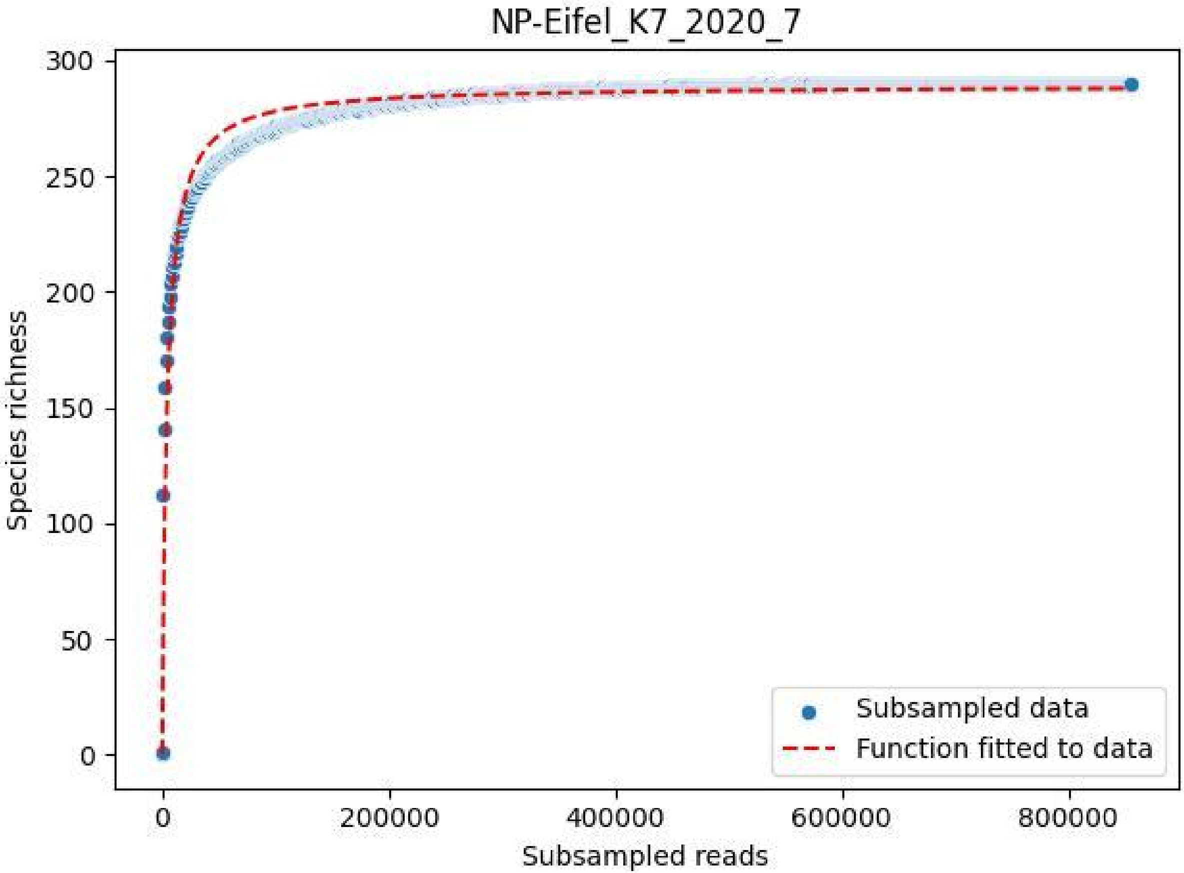
NP-Eifel_K7_2020_7
Species richness
Subsampled reads
0
50
100
150
200
250
300
0
200000
400000
600000
800000
Subsampled data
Function fitted to data

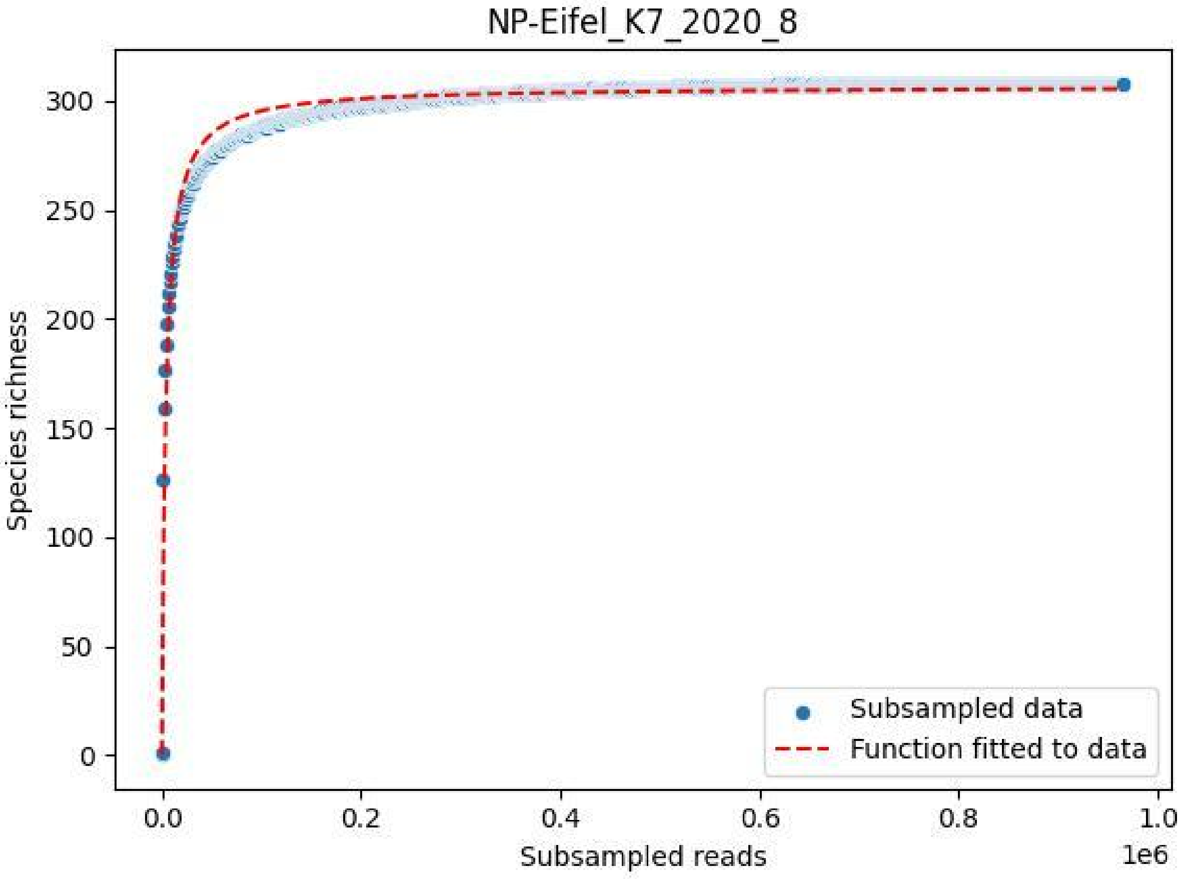

NP-Eifel_K7_2020_8
Species richness
300
250
200
150
100
50
0
0.0
0.2
0.4
0.6
0.8
1.0
1e6
Subsampled reads
Subsampled data
Function fitted to data

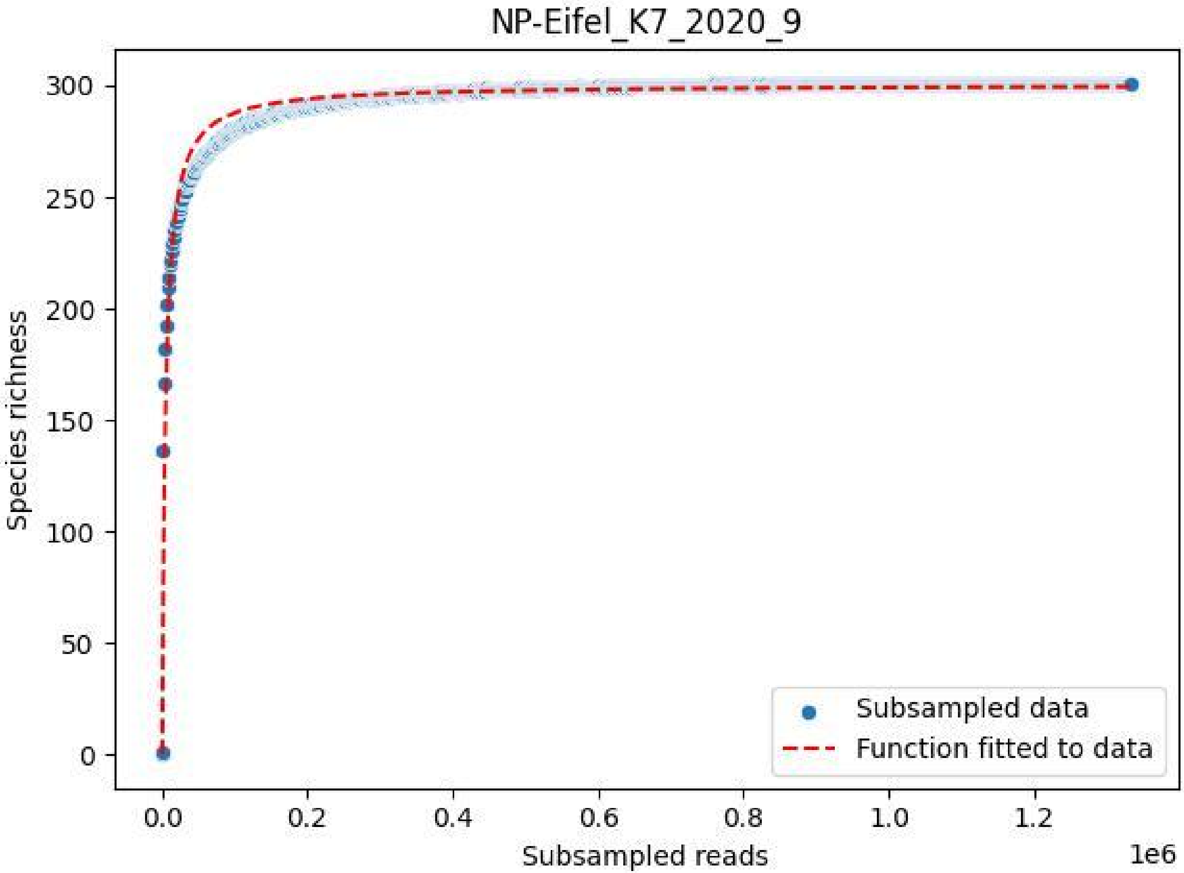
NP-Eifel_K7_2020_9
Species richness
Subsampled reads
1e6
300
250
200
150
100
50
0
0.0
0.2
0.4
0.6
0.8
1.0
1.2
Subsampled data
Function fitted to data

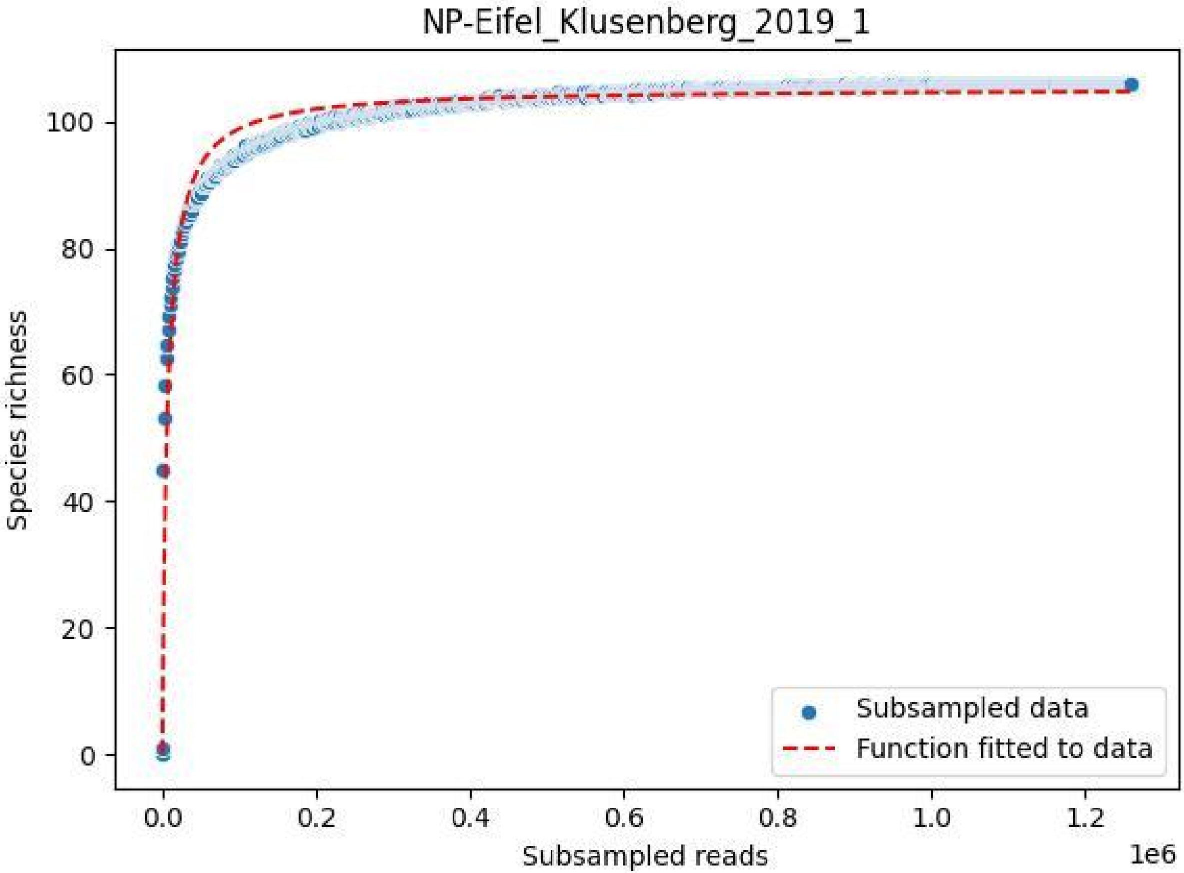
NP-Eifel_Klusenberg_2019_1
Species richness
0
20
40
60
80
100
0.0
0.2
0.4
0.6
0.8
1.0
1.2
1e6
Subsampled reads
Subsampled data
Function fitted to data

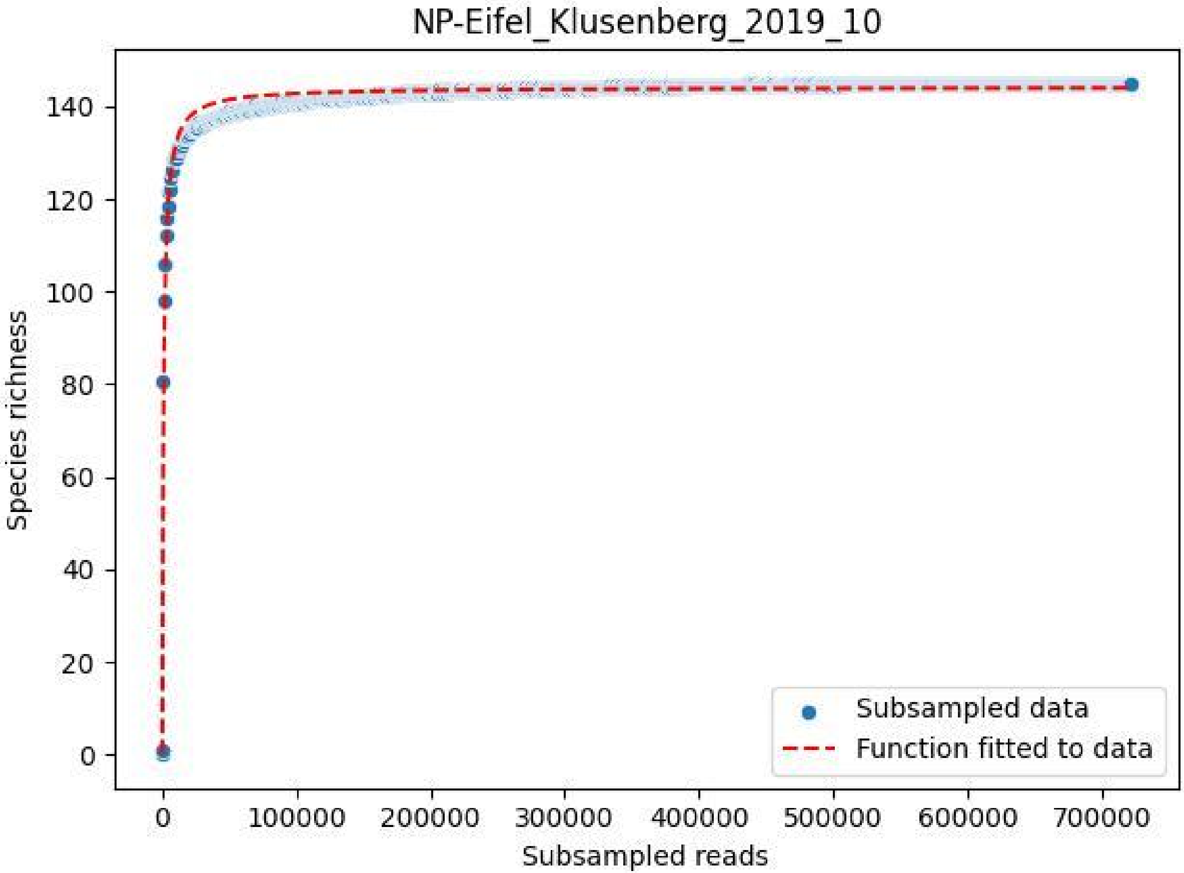
NP-Eifel_Klusenberg_2019_10
Species richness
Subsampled reads
0
20
40
60
80
100
120
140
0
100000
200000
300000
400000
500000
600000
700000
Subsampled data
Function fitted to data

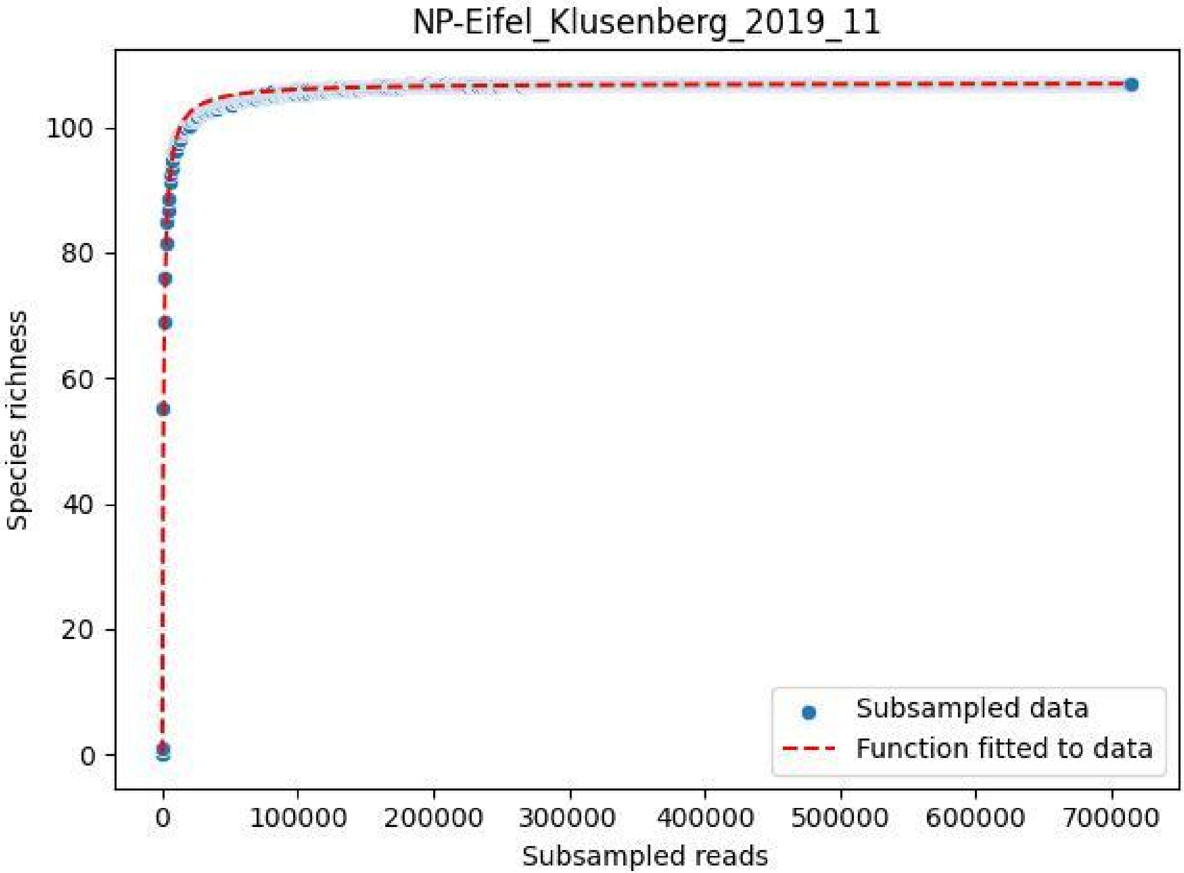
NP-Eifel_Klusenberg_2019_11
Species richness
Subsampled reads
0
20
40
60
80
100
0
100000
200000
300000
400000
500000
600000
700000
Subsampled data
Function fitted to data

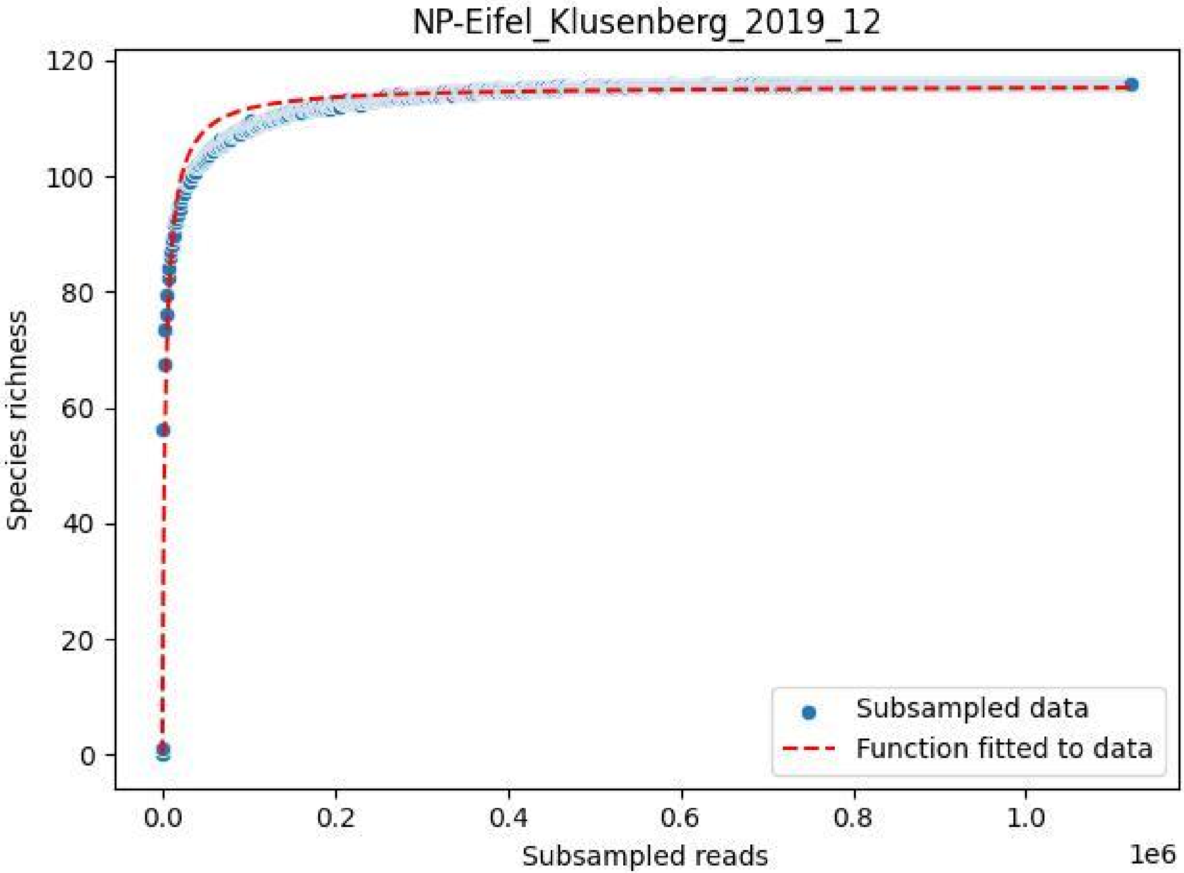
NP-Eifel_Klusenberg_2019_12
Species richness
Subsampled reads
1e6
0
20
40
60
80
100
120
0.0
0.2
0.4
0.6
0.8
1.0
Subsampled data
Function fitted to data

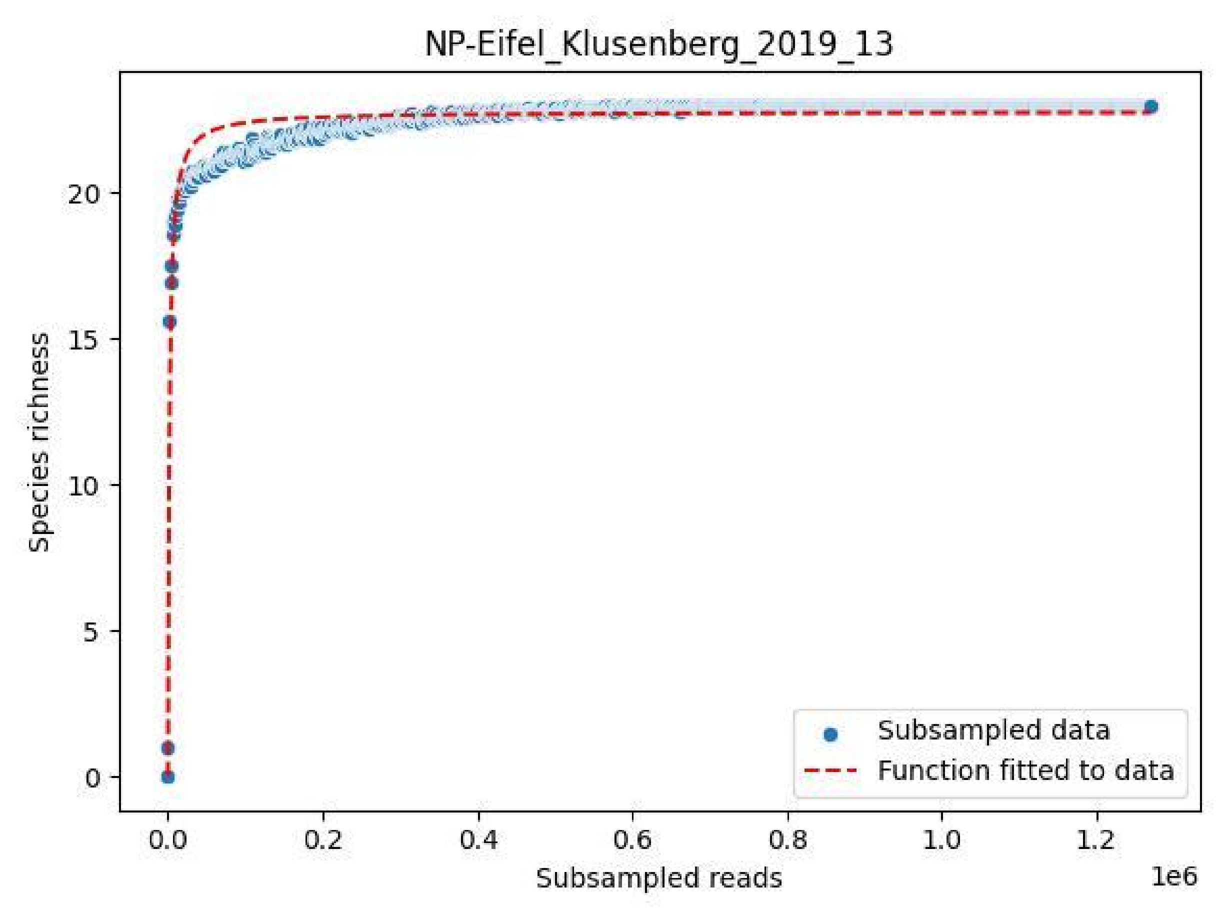

NP-Eifel_Klusenberg_2019_13
Species richness
Subsampled reads
1e6
0
5
10
15
20
0.0
0.2
0.4
0.6
0.8
1.0
1.2
Subsampled data
Function fitted to data

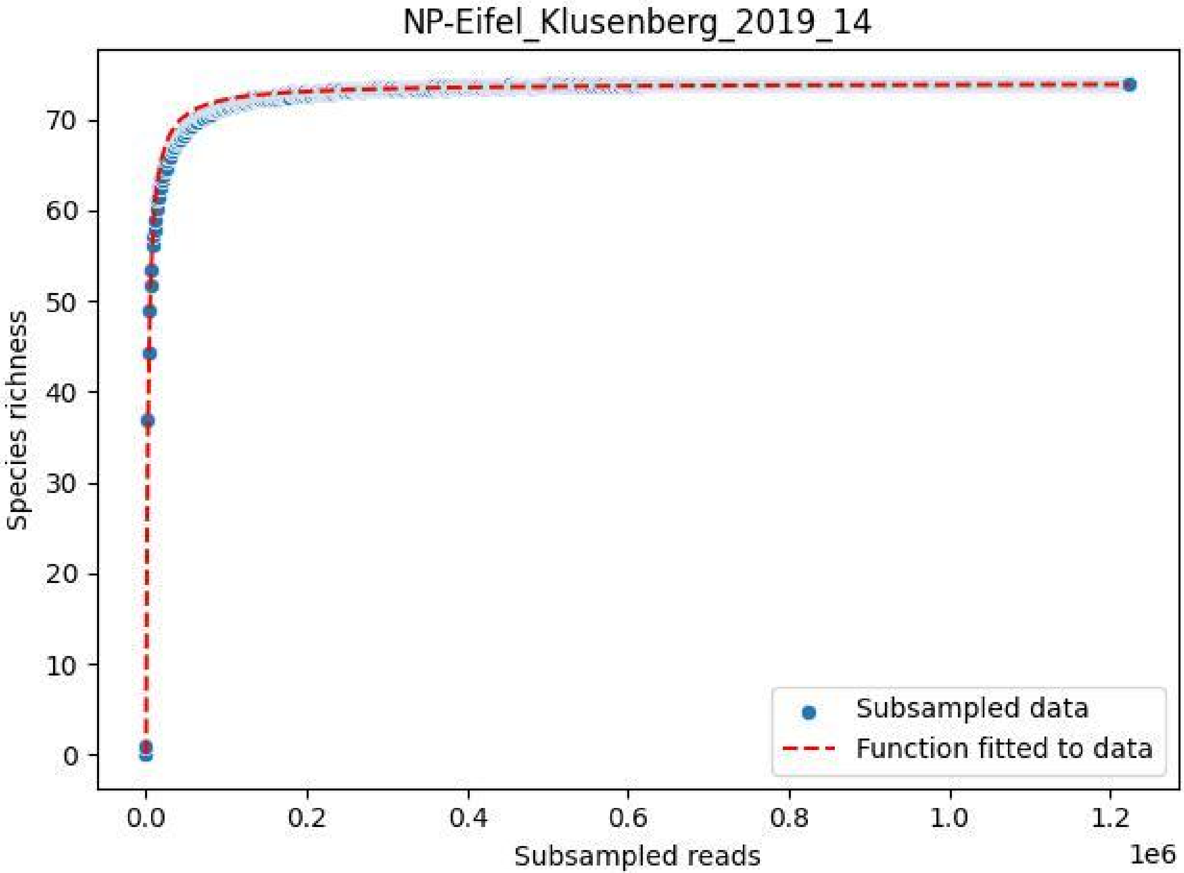
NP-Eifel_Klusenberg_2019_14
Species richness
Subsampled reads
1e6
Subsampled data
Function fitted to data
0
10
20
30
40
50
60
70
0.0
0.2
0.4
0.6
0.8
1.0
1.2

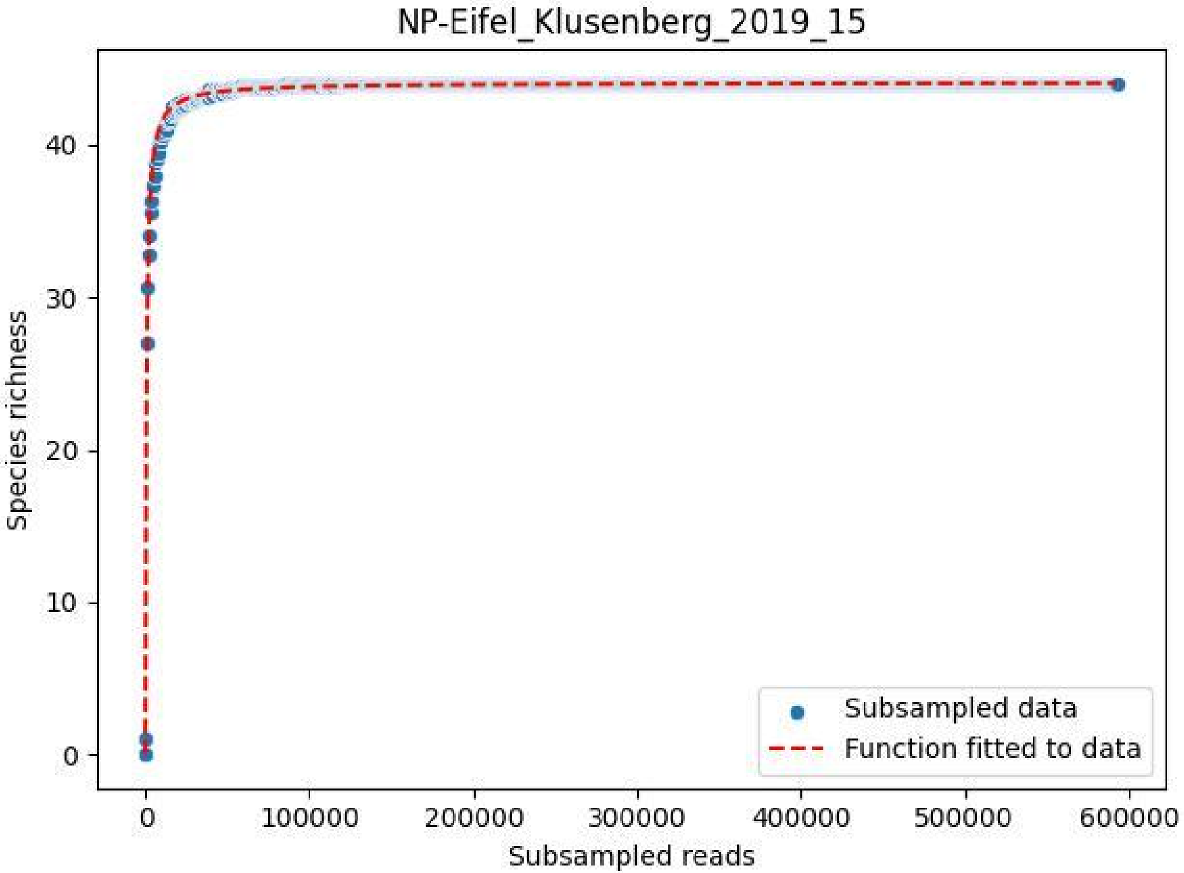
NP-Eifel_Klusenberg_2019_15
Species richness
Subsampled reads
0
10
20
30
40
0
100000
200000
300000
400000
500000
600000
Subsampled data
Function fitted to data

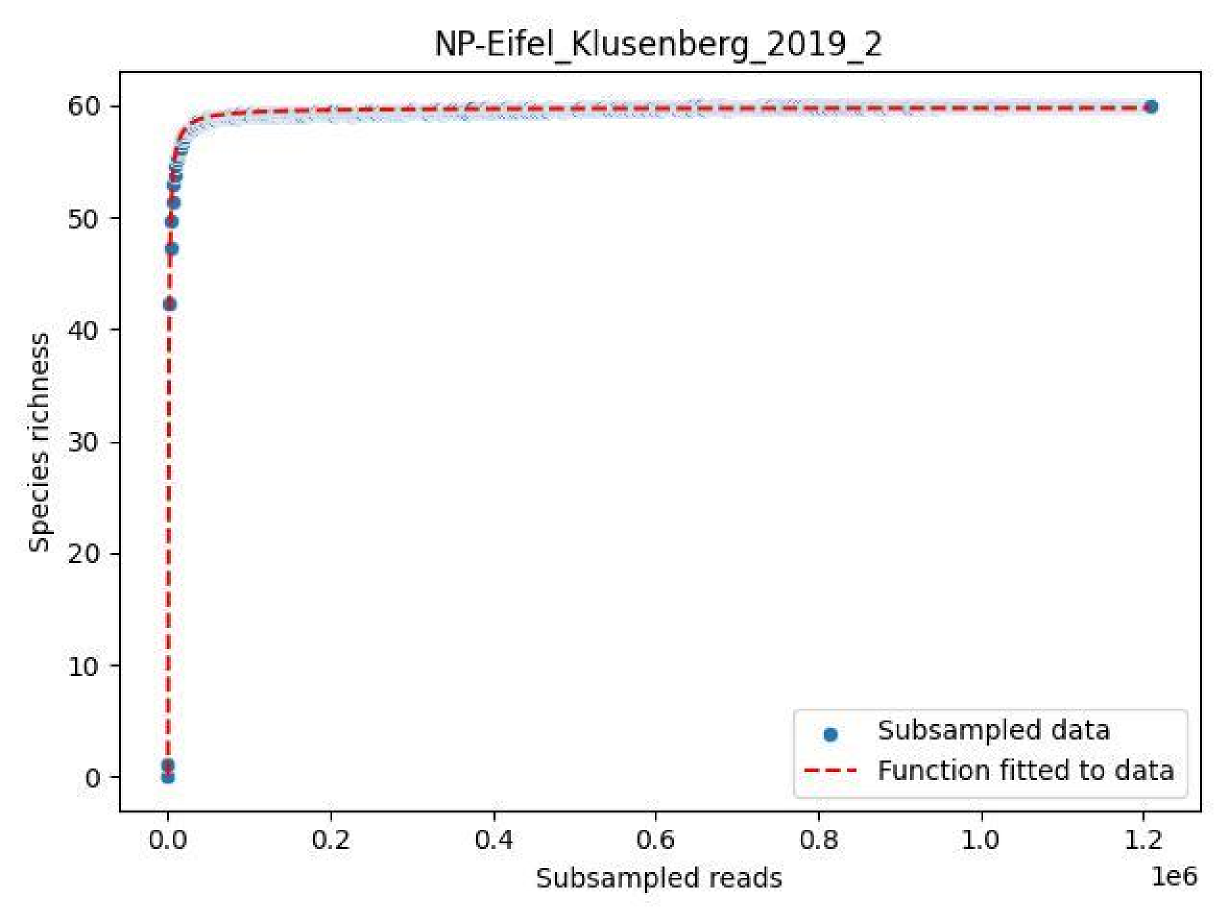
NP-Eifel_Klusenberg_2019_2
Species richness
Subsampled reads
1e6
Subsampled data
Function fitted to data
0
10
20
30
40
50
60
0.0
0.2
0.4
0.6
0.8
1.0
1.2

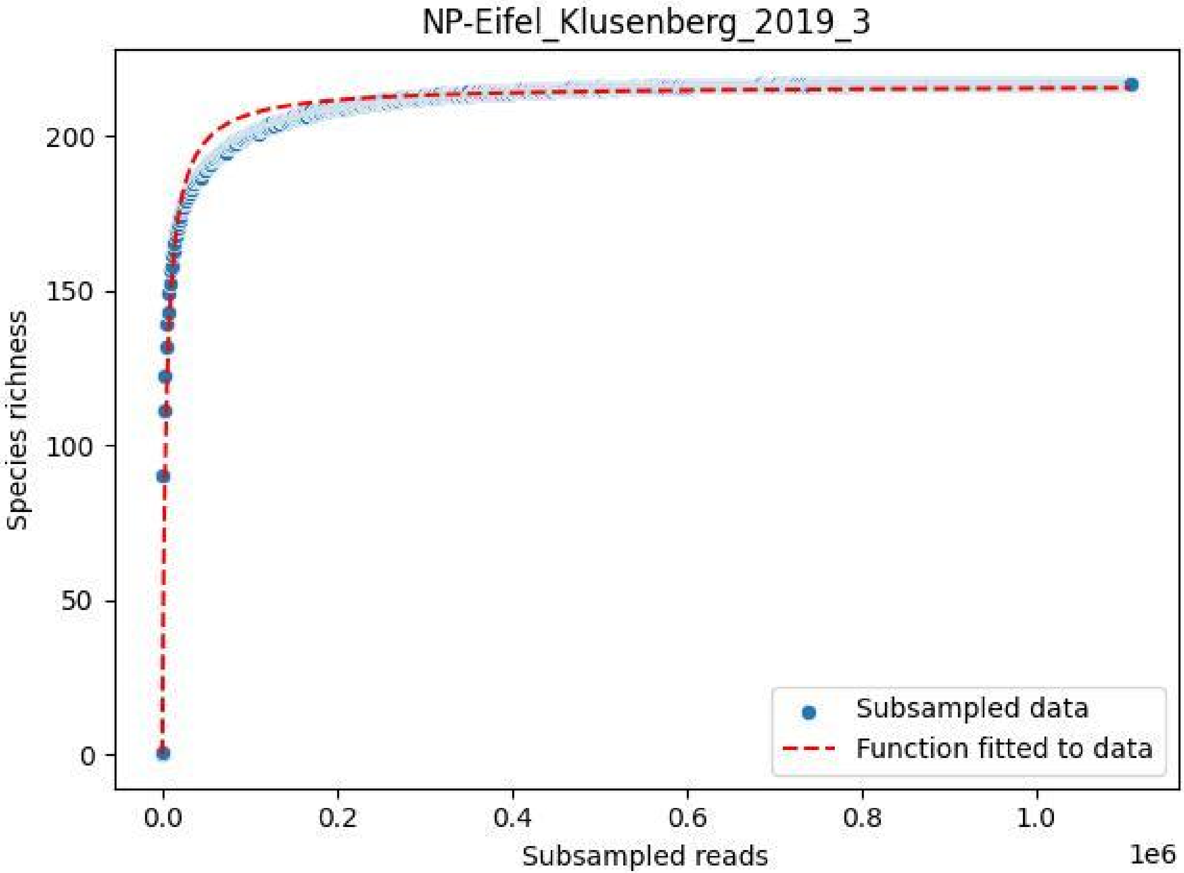
NP-Eifel_Klusenberg_2019_3
Species richness
Subsampled reads
1e6
Subsampled data
Function fitted to data
0
50
100
150
200
0.0
0.2
0.4
0.6
0.8
1.0

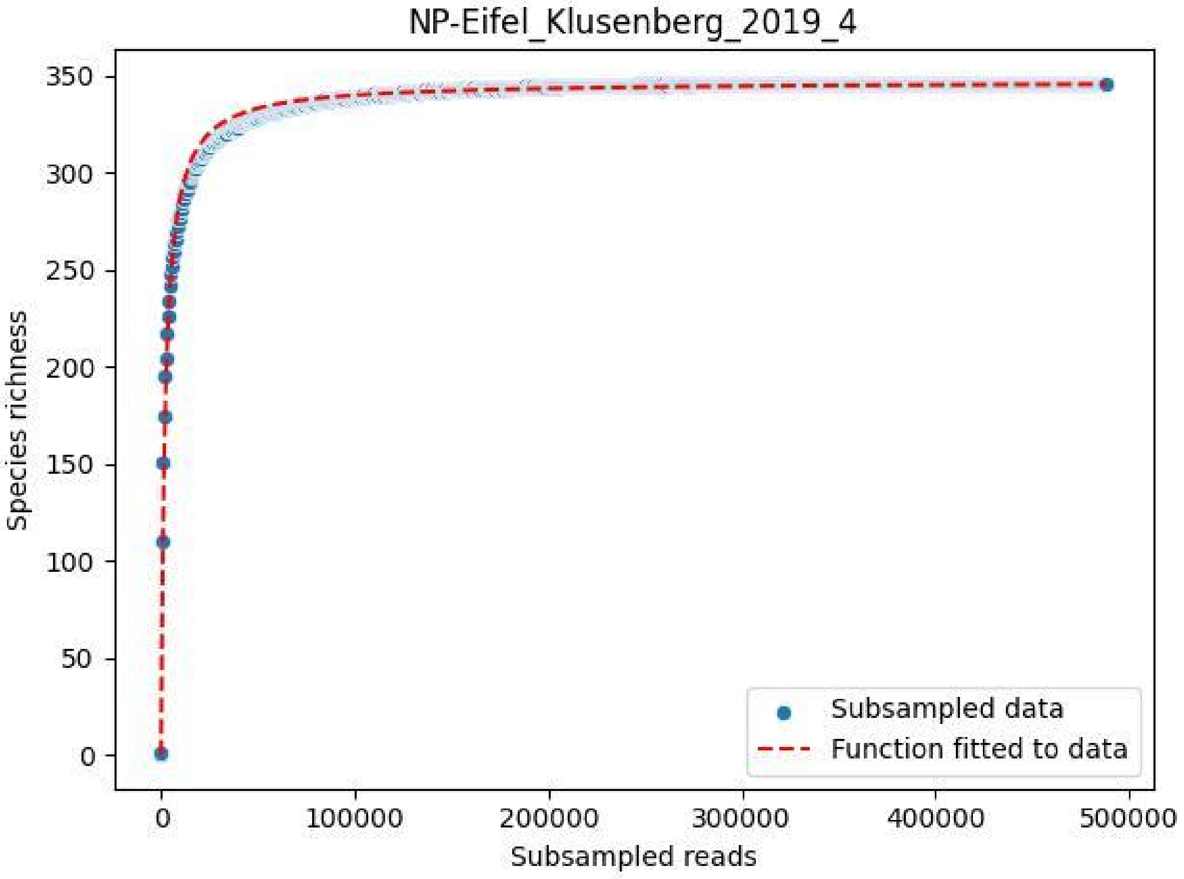
NP-Eifel_Klusenberg_2019_4
Species richness
Subsampled reads
0
50
100
150
200
250
300
350
0
100000
200000
300000
400000
500000
Subsampled data
Function fitted to data

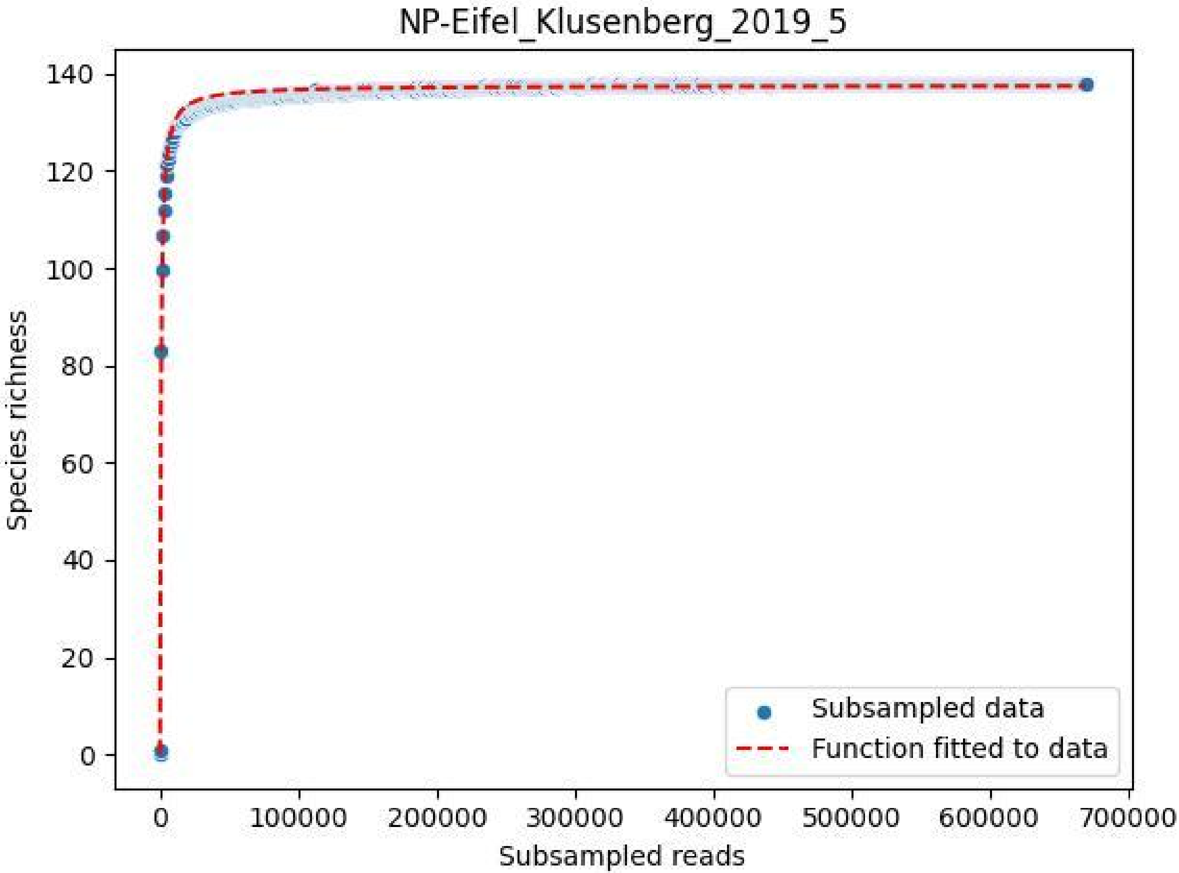
NP-Eifel_Klusenberg_2019_5
Species richness
Subsampled reads
0
20
40
60
80
100
120
140
0
100000
200000
300000
400000
500000
600000
700000
Subsampled data
Function fitted to data

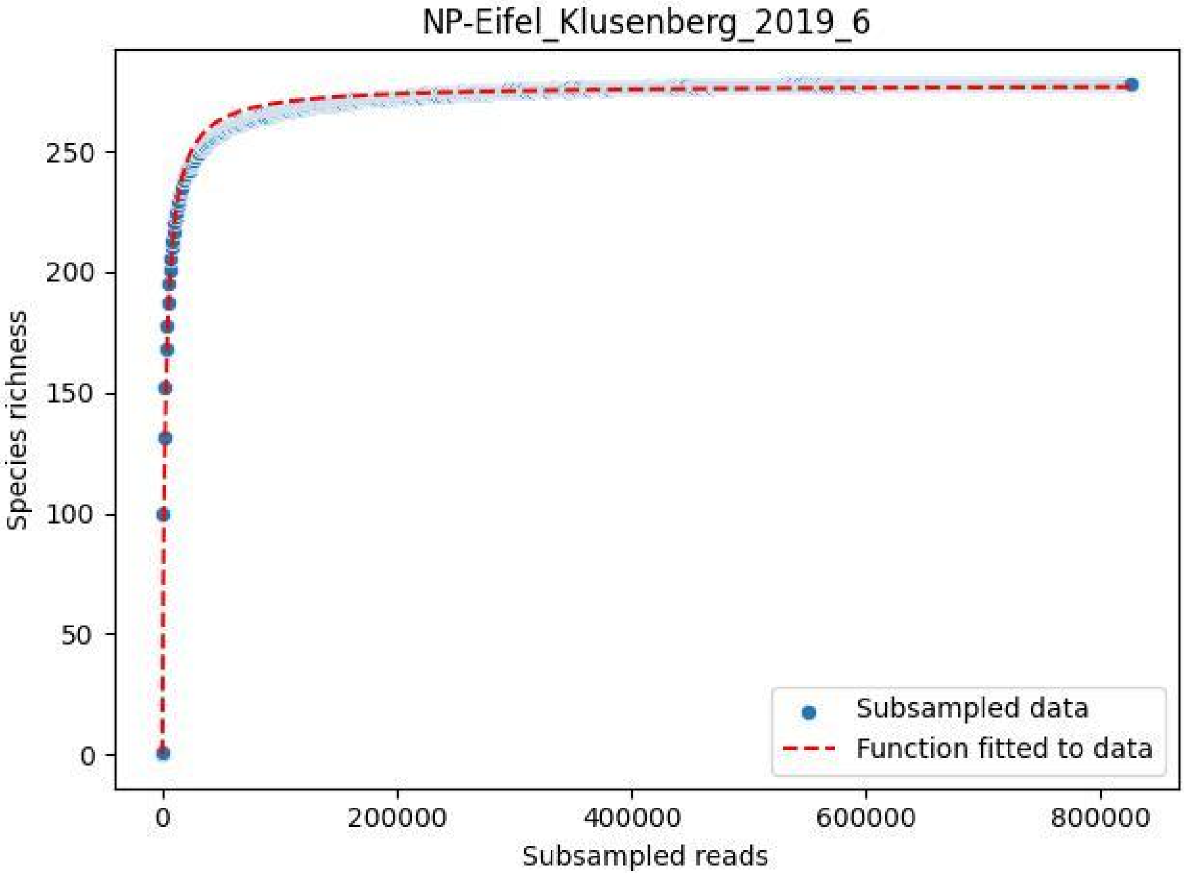
NP-Eifel_Klusenberg_2019_6
Species richness
Subsampled reads
0
50
100
150
200
250
0
200000
400000
600000
800000
Subsampled data
Function fitted to data

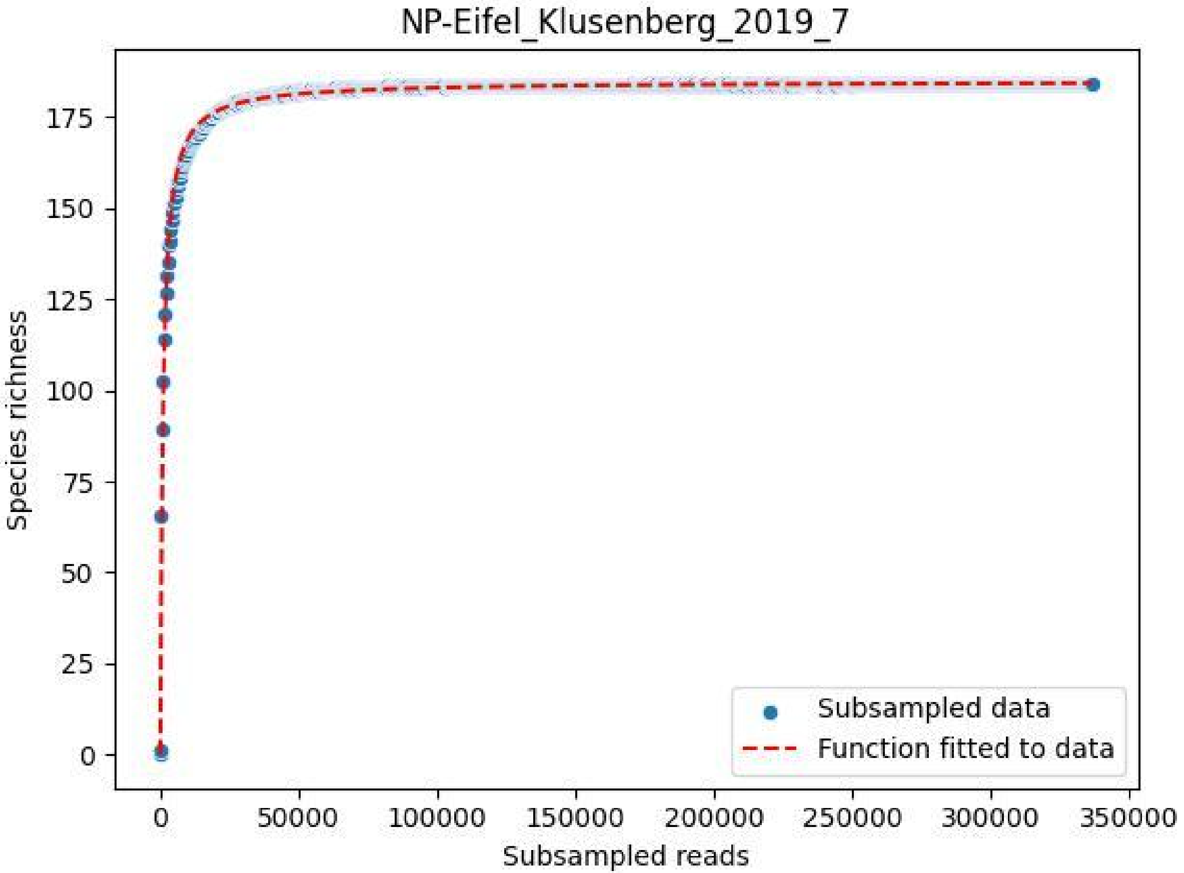

NP-Eifel_Klusenberg_2019_7
Species richness
0
25
50
75
100
125
150
175
Subsampled reads
0
50000
100000
150000
200000
250000
300000
350000
Subsampled data
Function fitted to data

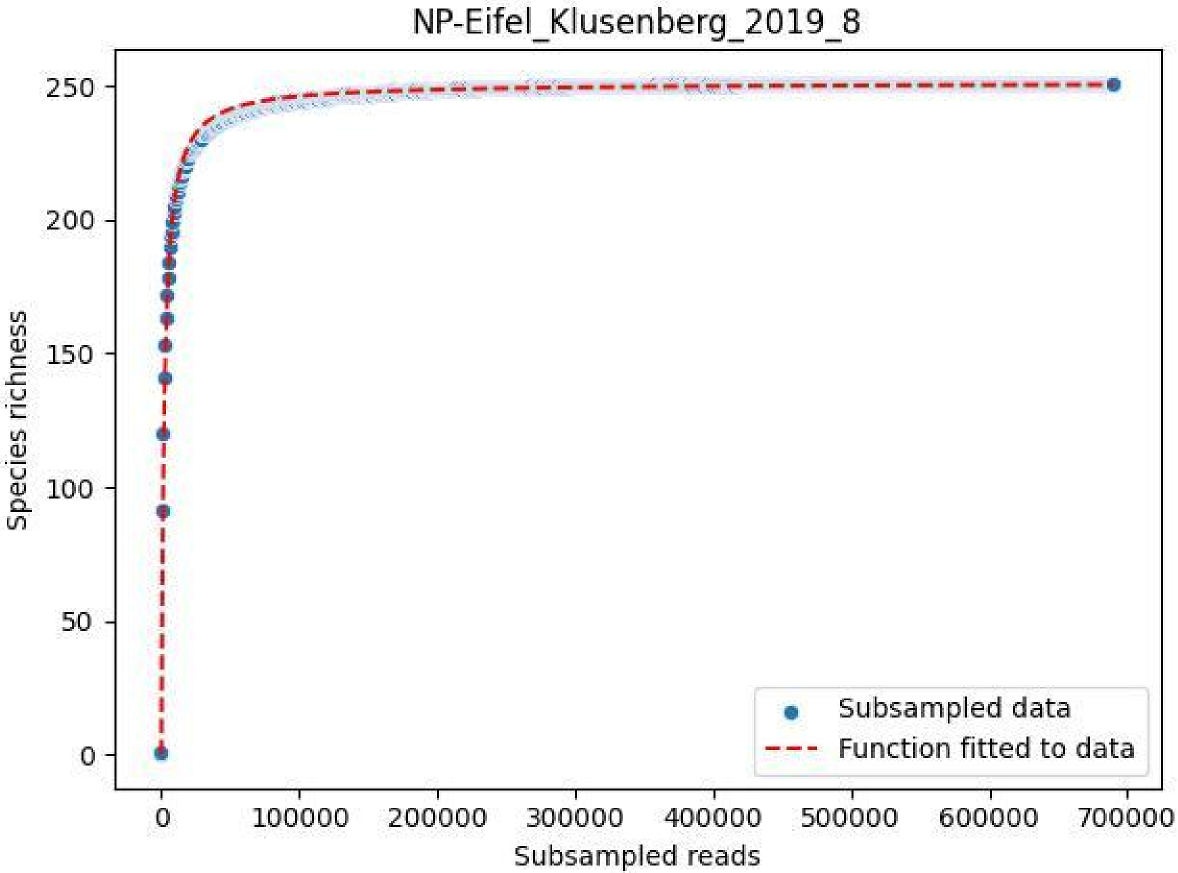

NP-Eifel_Klusenberg_2019_8
Species richness
Subsampled reads
0
50
100
150
200
250
0
100000
200000
300000
400000
500000
600000
700000
Subsampled data
Function fitted to data

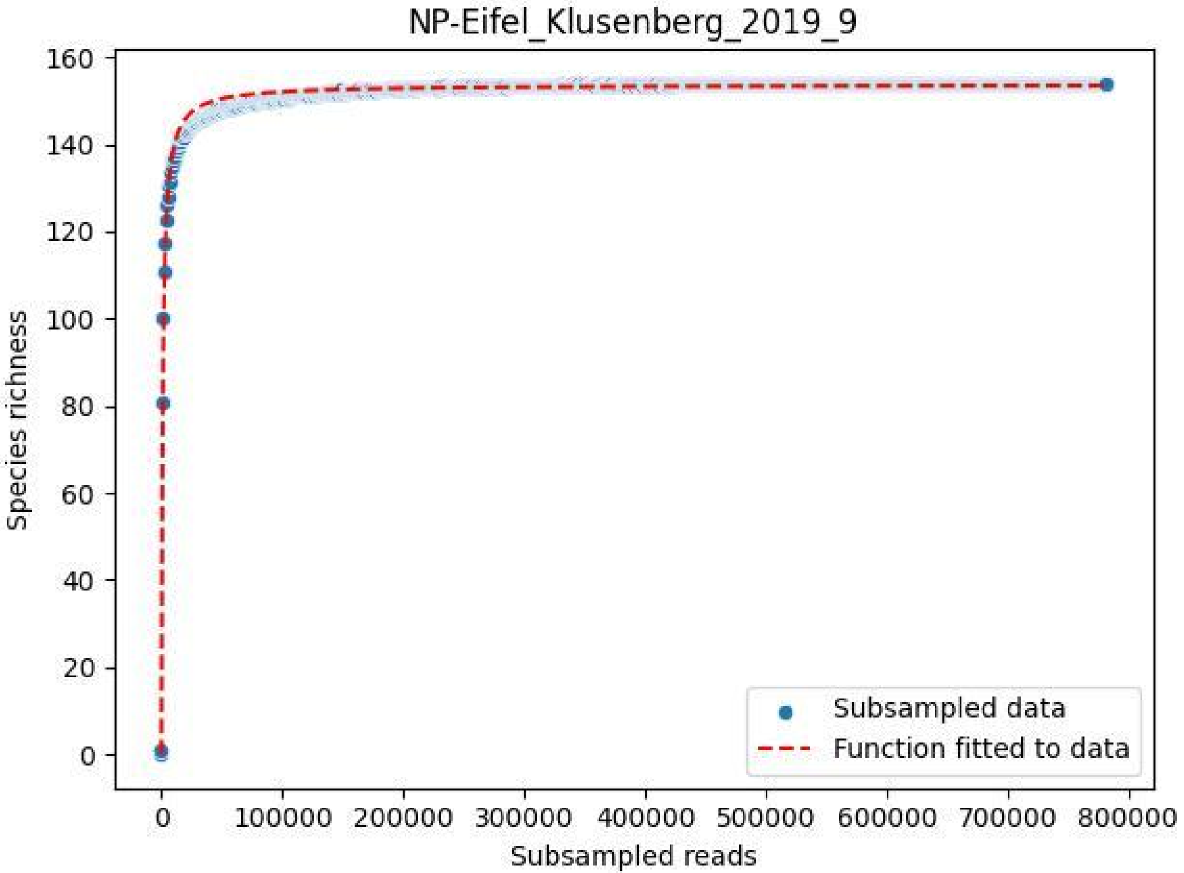
NP-Eifel_Klusenberg_2019_9
Species richness
Subsampled reads
0
20
40
60
80
100
120
140
160
0
100000
200000
300000
400000
500000
600000
700000
800000
Subsampled data
Function fitted to data

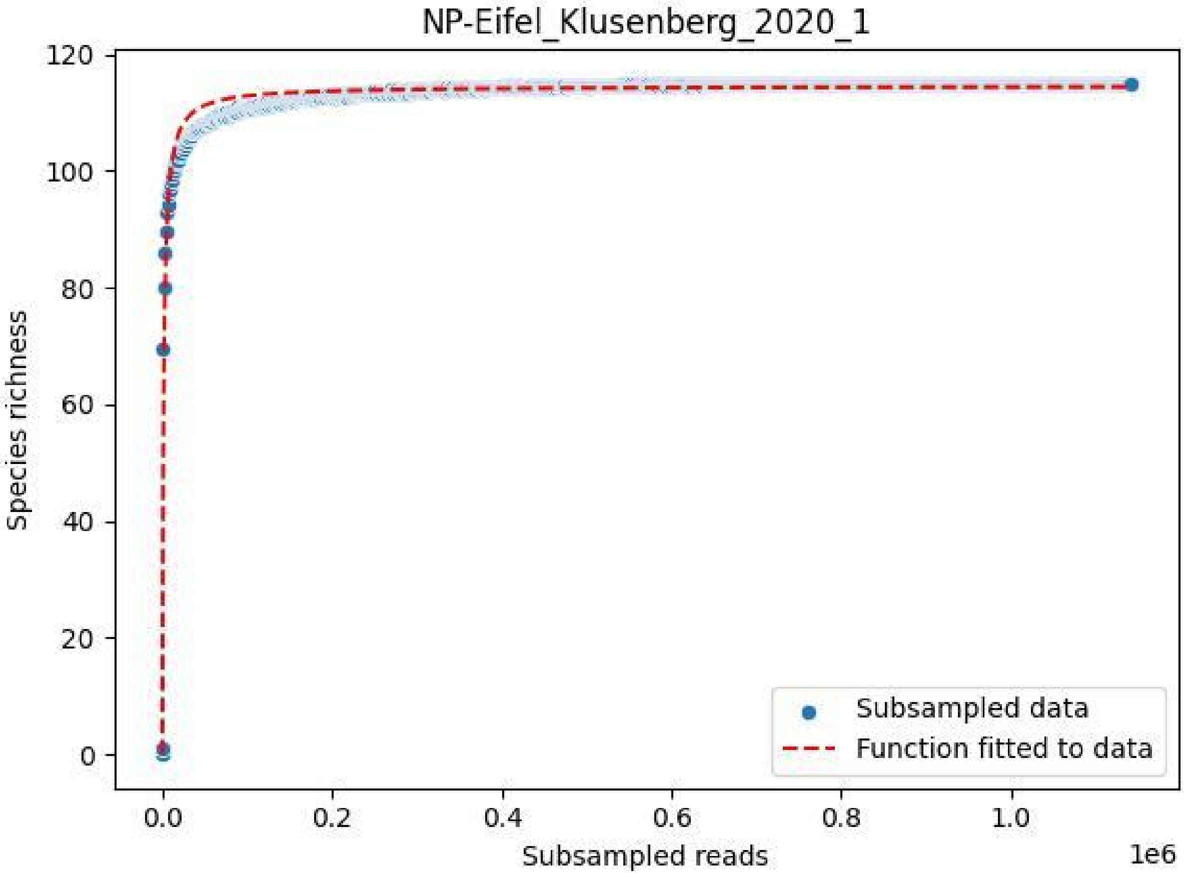
NP-Eifel_Klusenberg_2020_1
Species richness
Subsampled reads
1e6
0
20
40
60
80
100
120
0.0
0.2
0.4
0.6
0.8
1.0
Subsampled data
Function fitted to data

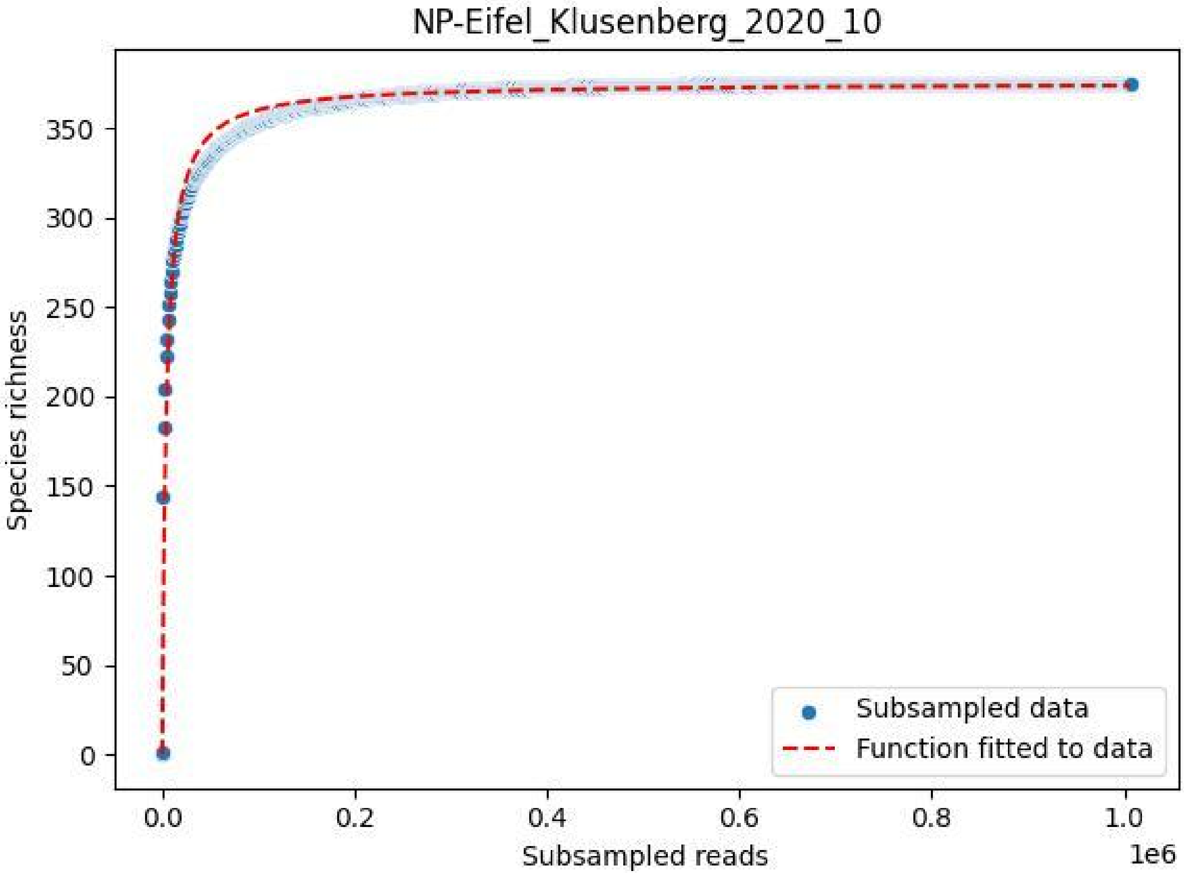
NP-Eifel_Klusenberg_2020_10
Species richness
350
300
250
200
150
100
50
0
0.0
0.2
0.4
0.6
0.8
1.0
1e6
Subsampled reads
Subsampled data
Function fitted to data

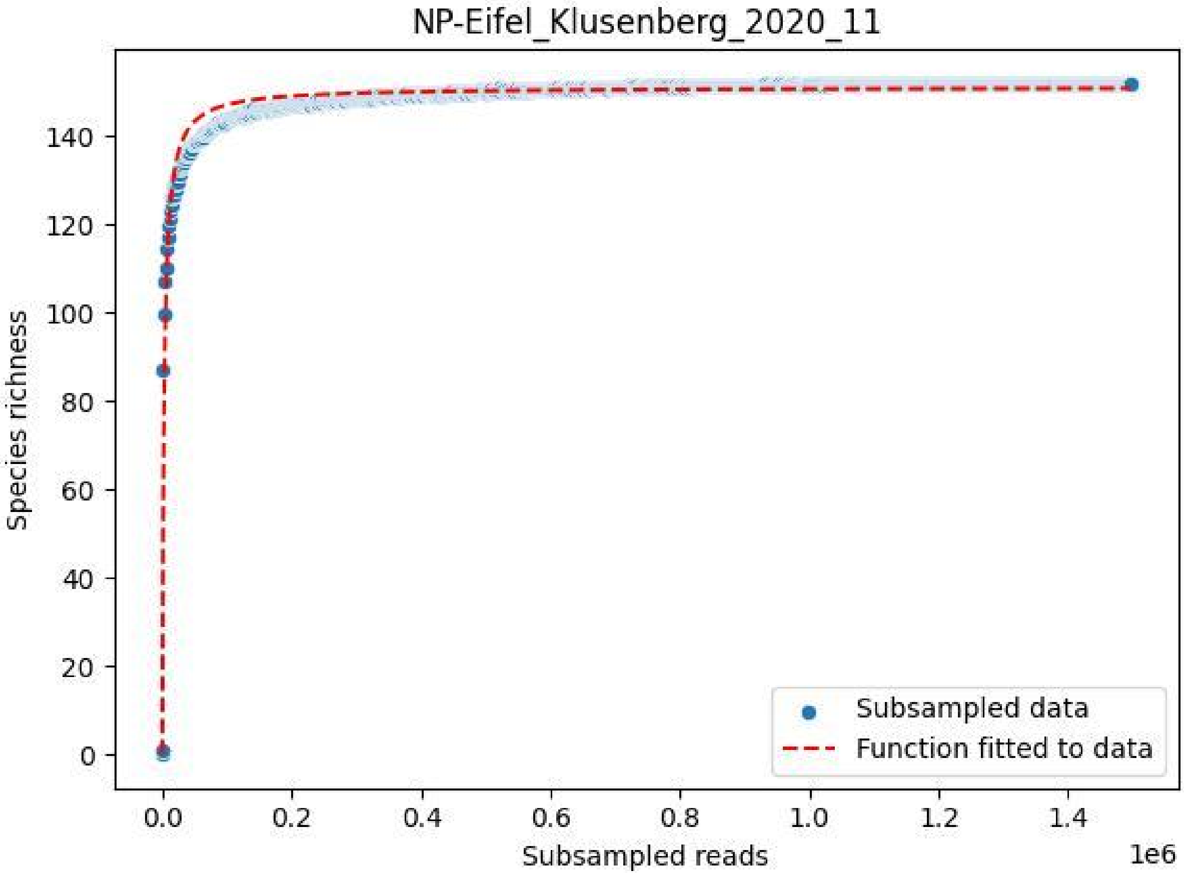
NP-Eifel_Klusenberg_2020_11
Species richness
0
20
40
60
80
100
120
140
0.0
0.2
0.4
0.6
0.8
1.0
1.2
1.4
1e6
Subsampled reads
Subsampled data
Function fitted to data

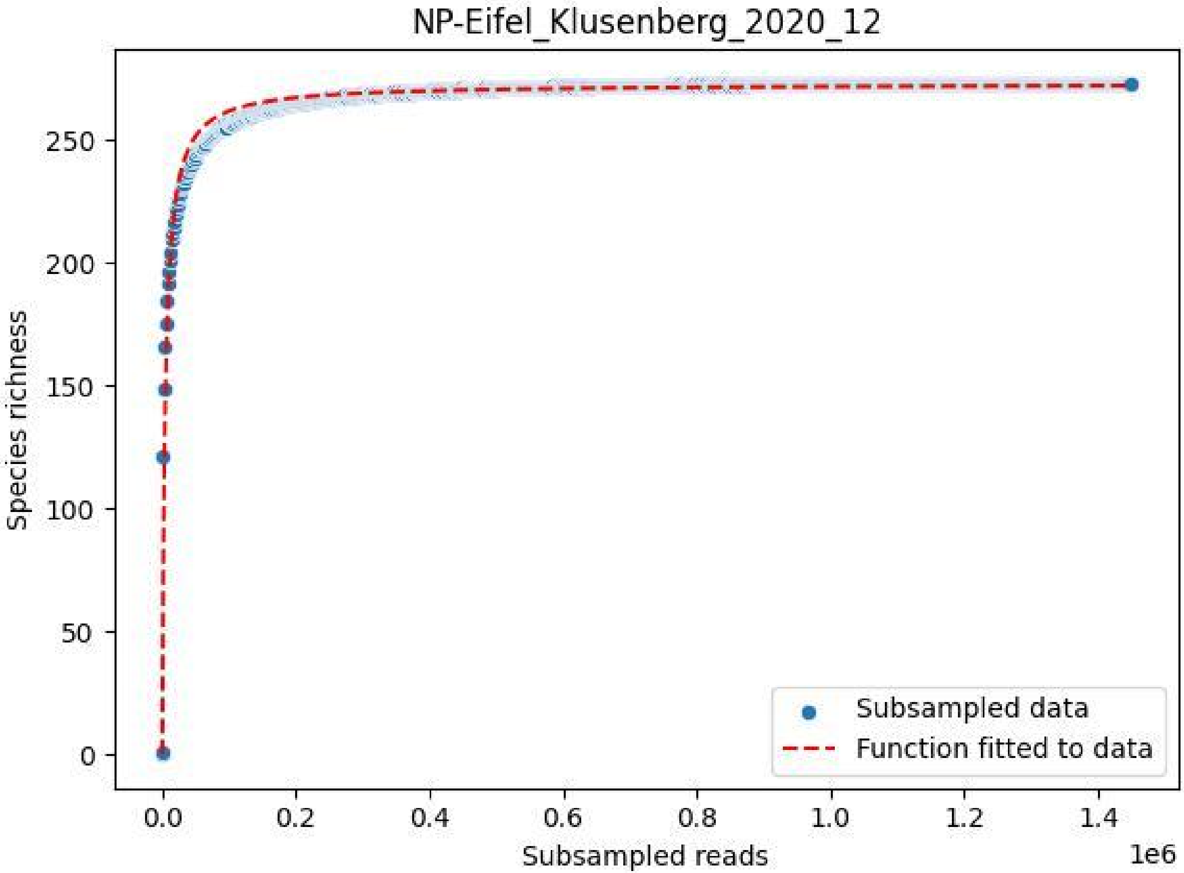
NP-Eifel_Klusenberg_2020_12
Species richness
Subsampled reads
1e6
0
50
100
150
200
250
0.0
0.2
0.4
0.6
0.8
1.0
1.2
1.4
Subsampled data
Function fitted to data

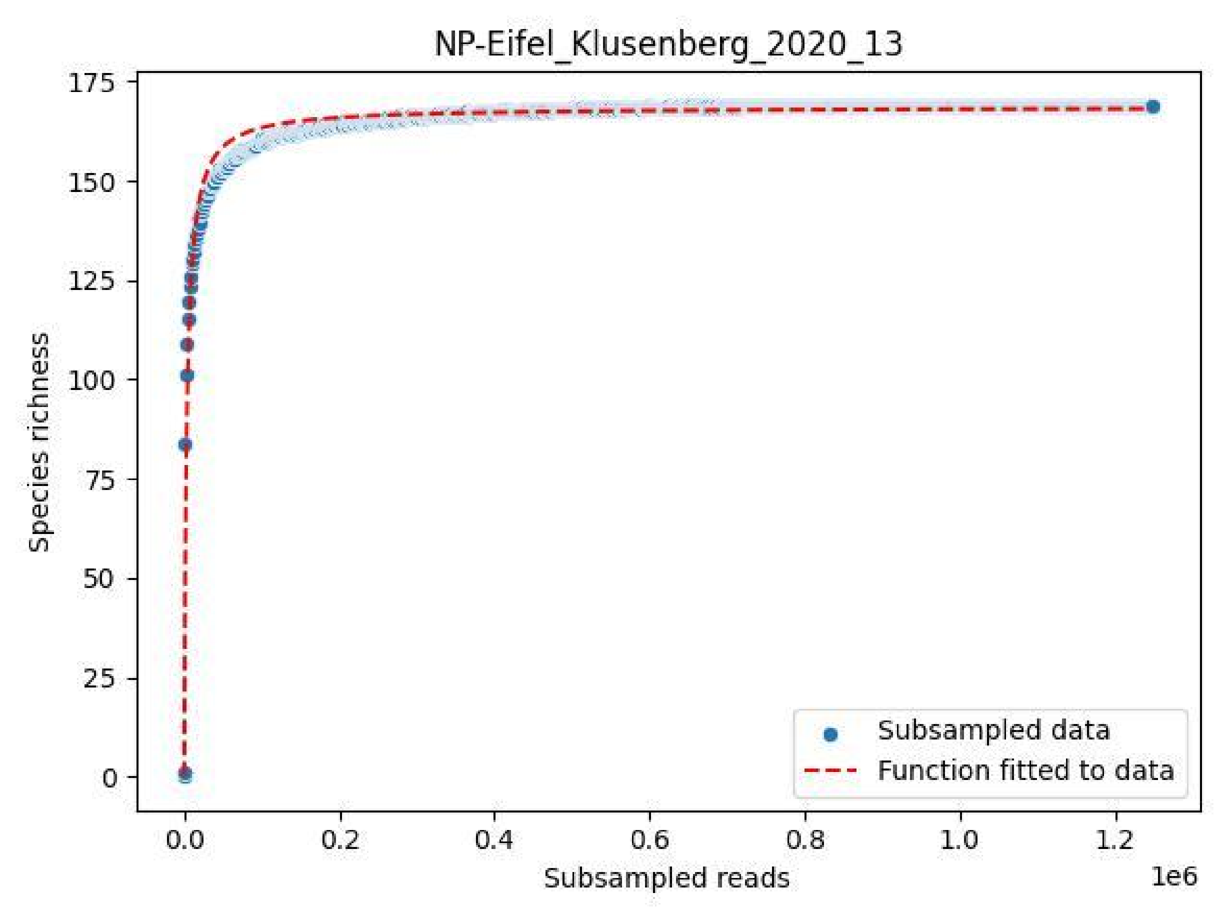

NP-Eifel_Klusenberg_2020_13
Species richness
Subsampled reads
1e6
0
25
50
75
100
125
150
175
0.0
0.2
0.4
0.6
0.8
1.0
1.2
Subsampled data
Function fitted to data

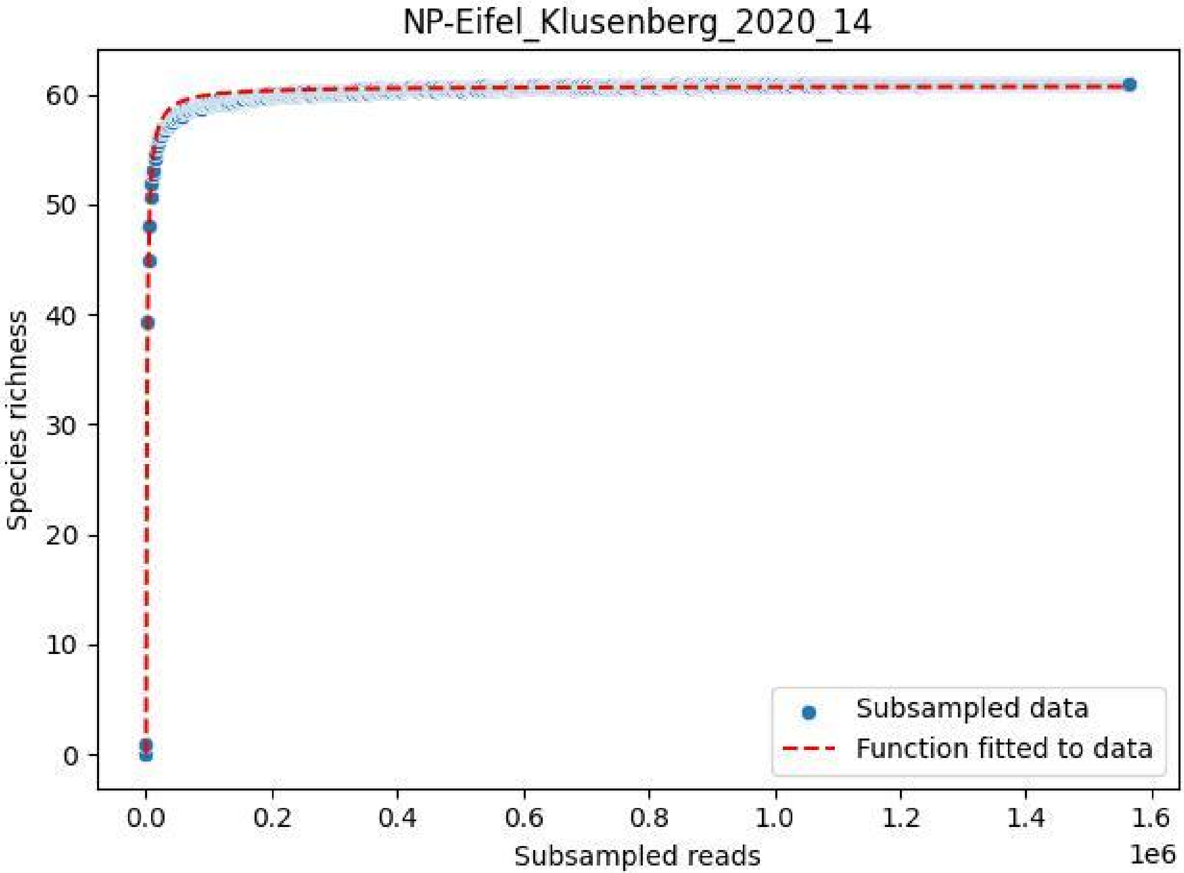

NP-Eifel_Klusenberg_2020_14
Species richness
Subsampled reads
1e6
0
10
20
30
40
50
60
0.0
0.2
0.4
0.6
0.8
1.0
1.2
1.4
1.6
Subsampled data
Function fitted to data

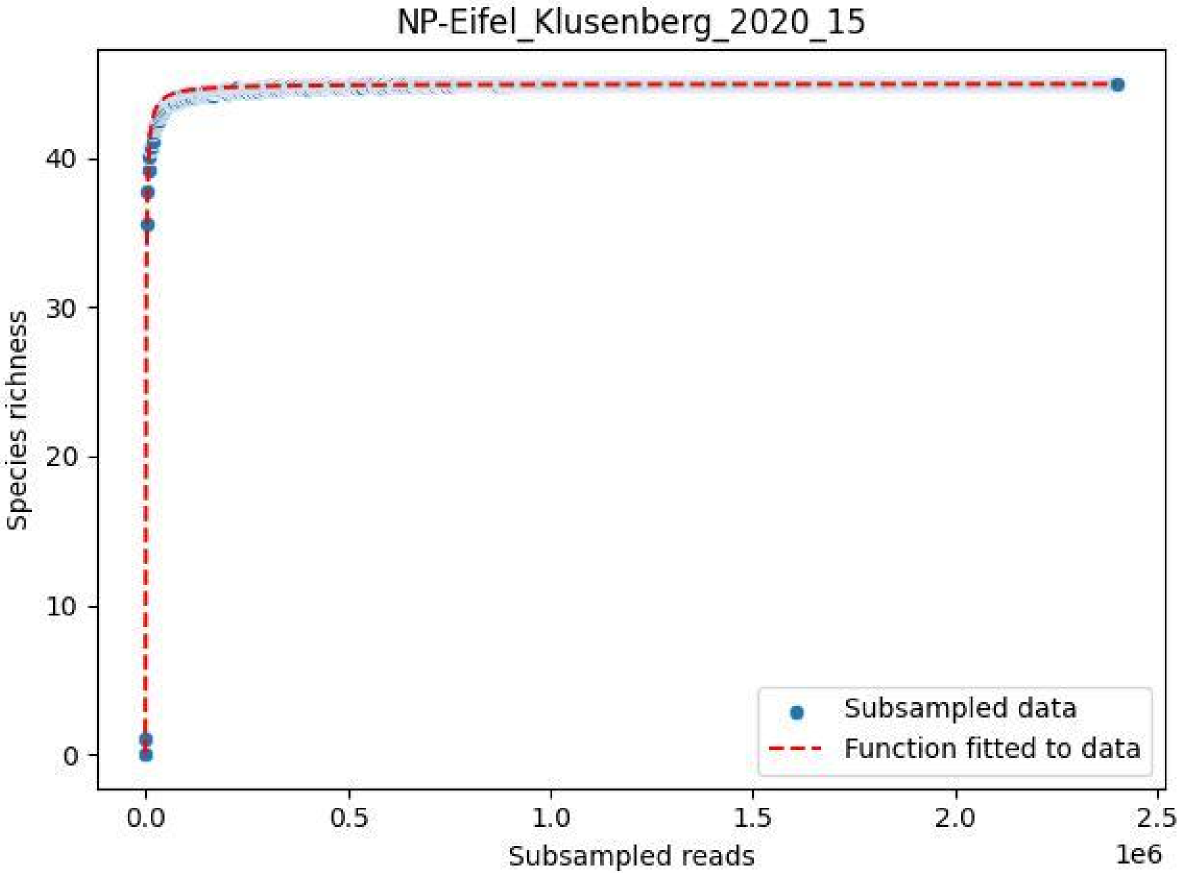

NP-Eifel_Klusenberg_2020_15
Species richness
Subsampled reads
1e6
0
10
20
30
40
0.0
0.5
1.0
1.5
2.0
2.5
Subsampled data
Function fitted to data

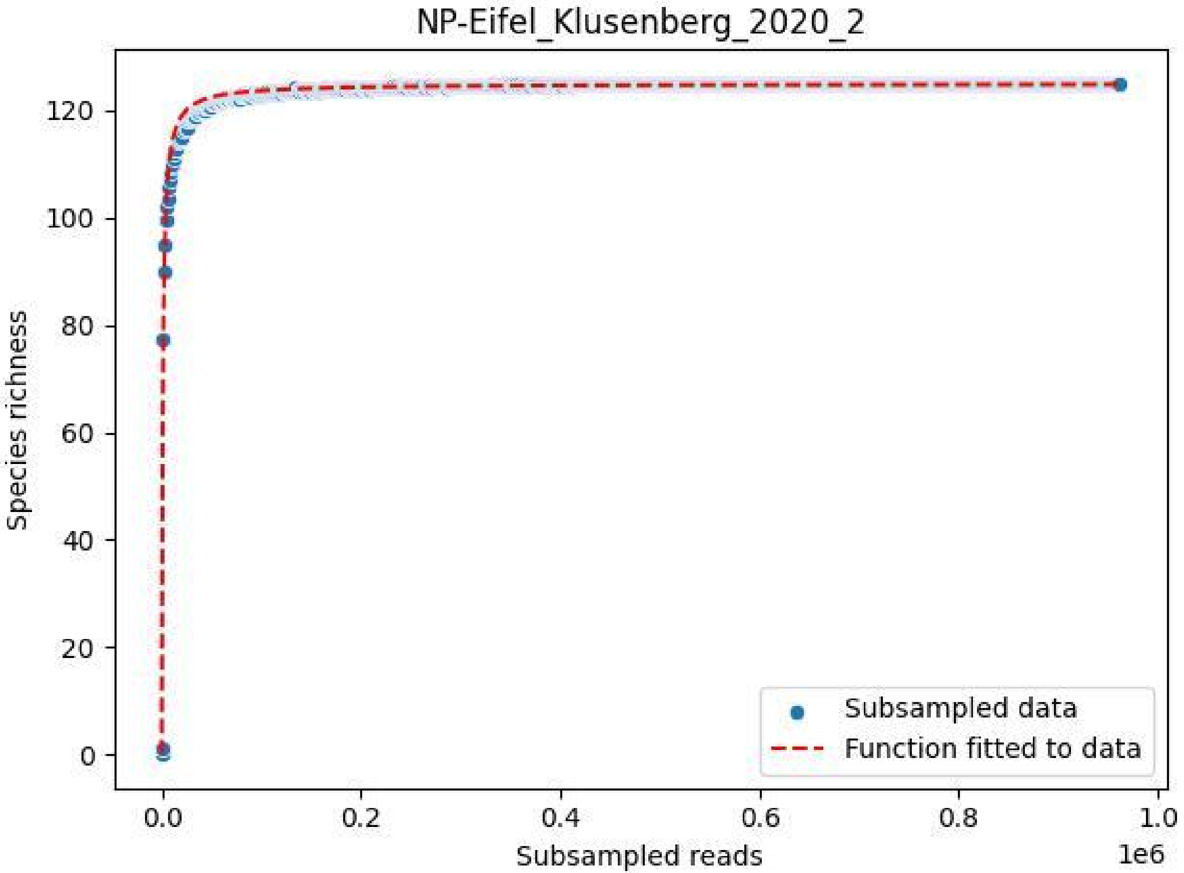
NP-Eifel_Klusenberg_2020_2
Species richness
Subsampled reads
120
100
80
60
40
20
0
0.0
0.2
0.4
0.6
0.8
1.0
1e6
Subsampled data
Function fitted to data

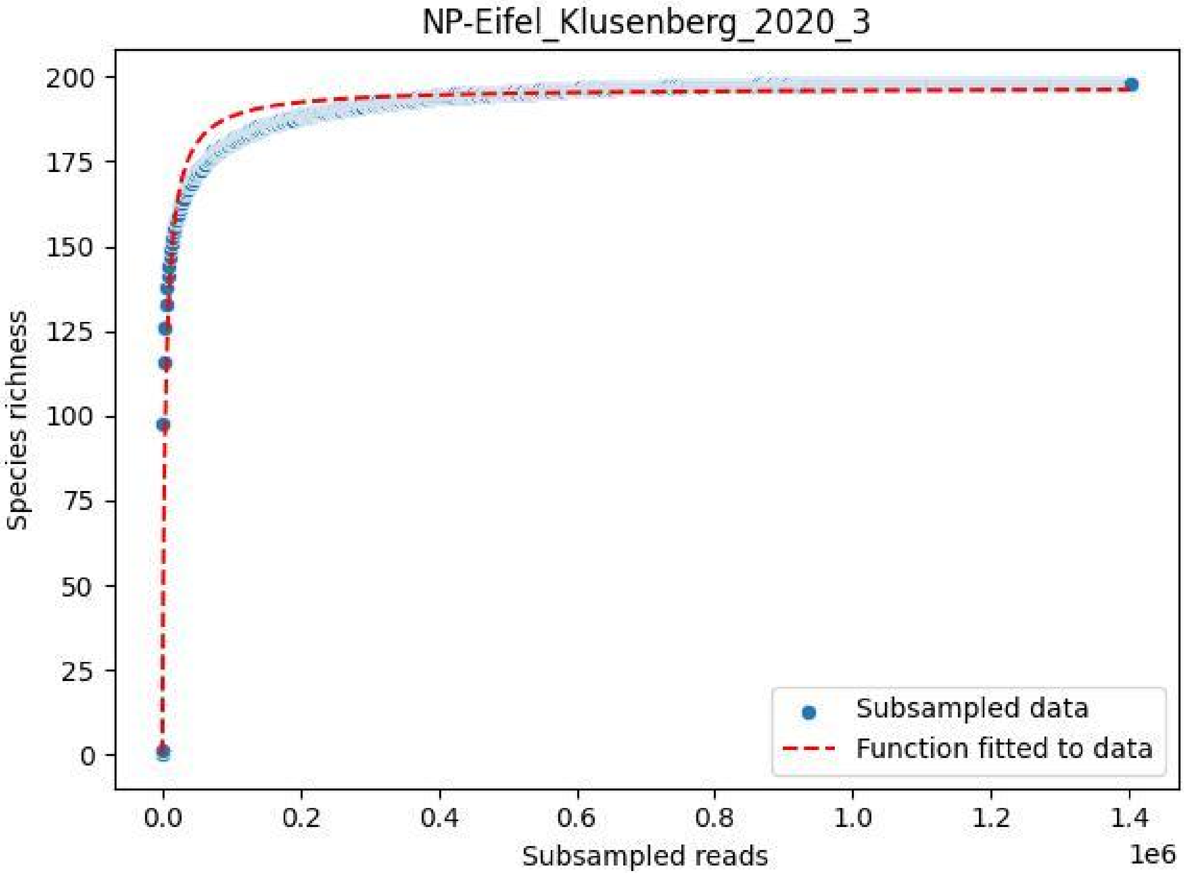
NP-Eifel_Klusenberg_2020_3
200
175
150
125
100
75
50
25
0
Species richness
0.0
0.2
0.4
0.6
0.8
1.0
1.2
1.4
1e6
Subsampled reads
Subsampled data
Function fitted to data

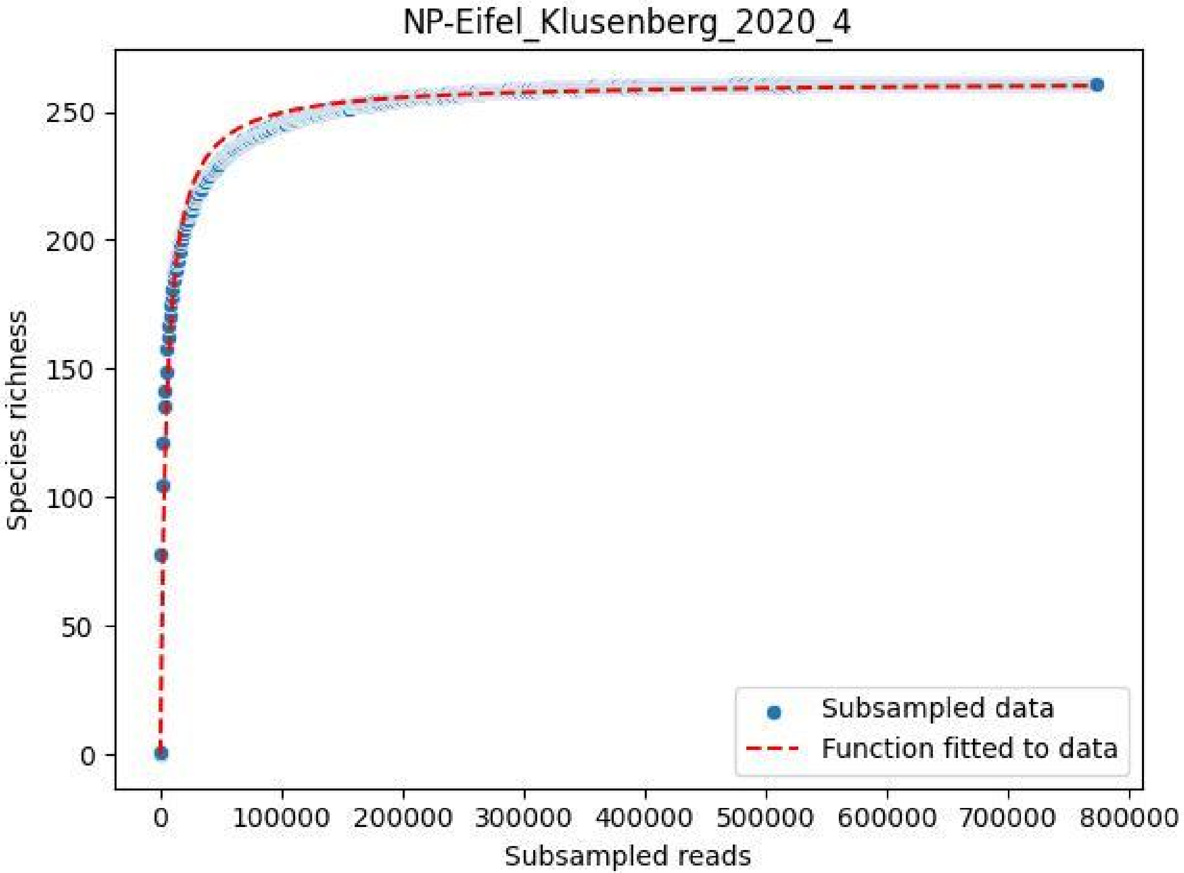
NP-Eifel_Klusenberg_2020_4
Species richness
0
50
100
150
200
250
Subsampled reads
0
100000
200000
300000
400000
500000
600000
700000
800000
Subsampled data
Function fitted to data

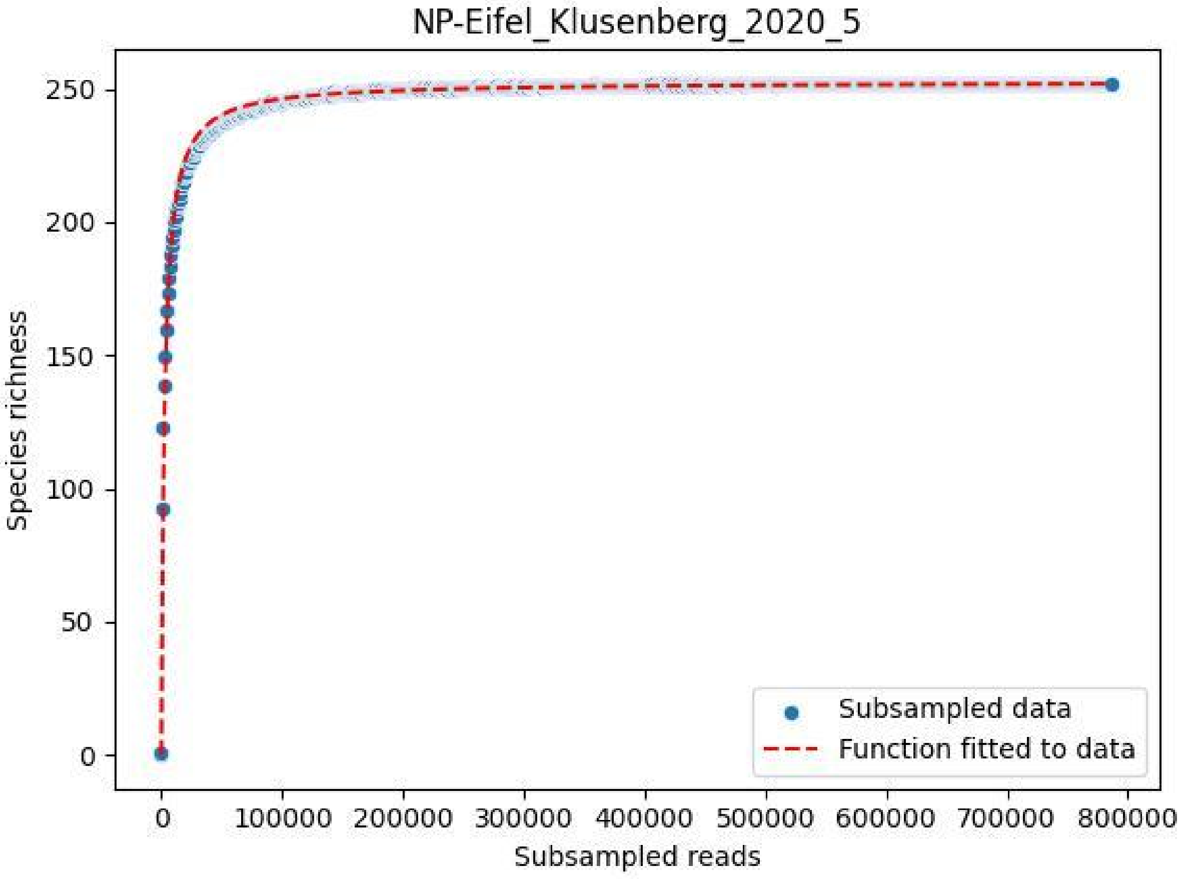
NP-Eifel_Klusenberg_2020_5
Species richness
Subsampled reads
0
50
100
150
200
250
0
100000
200000
300000
400000
500000
600000
700000
800000
Subsampled data
Function fitted to data

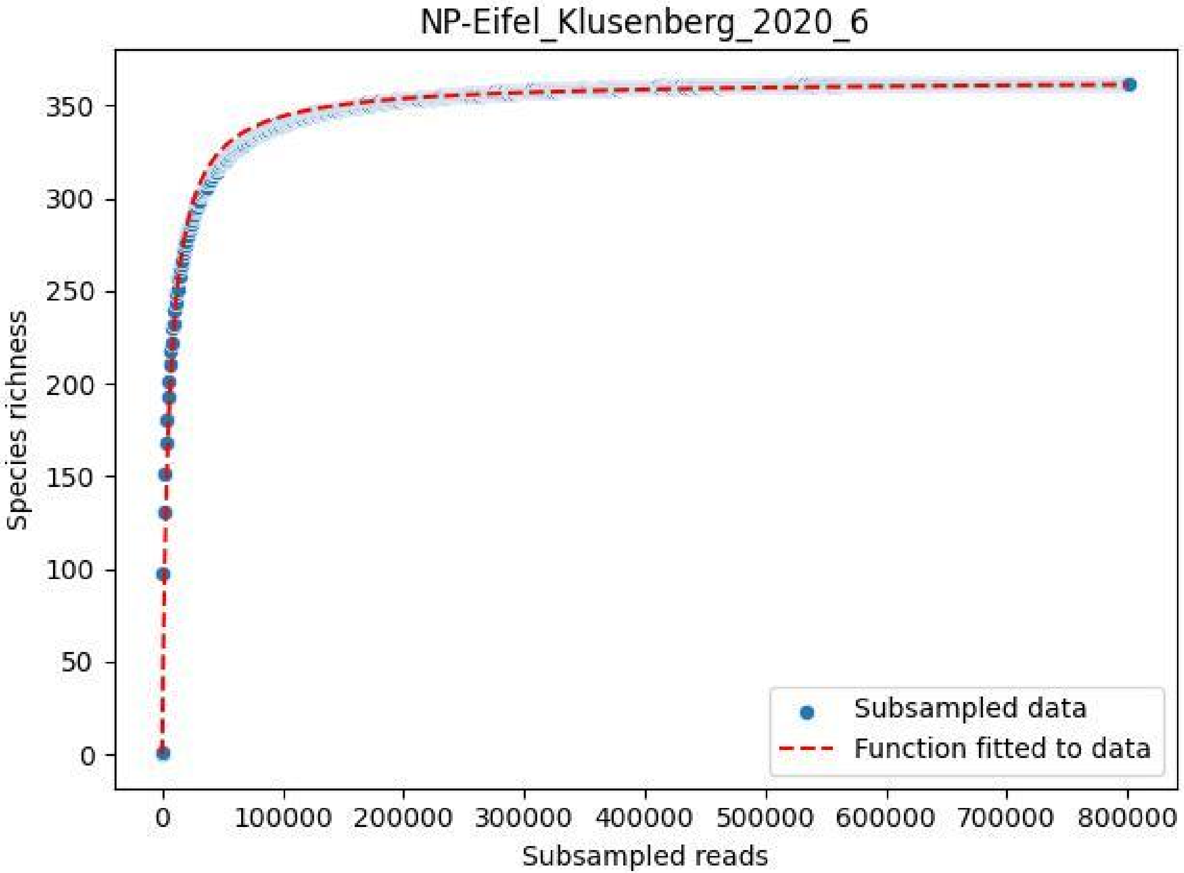
NP-Eifel_Klusenberg_2020_6
Species richness
Subsampled reads
0
50
100
150
200
250
300
350
0
100000
200000
300000
400000
500000
600000
700000
800000
Subsampled data
Function fitted to data

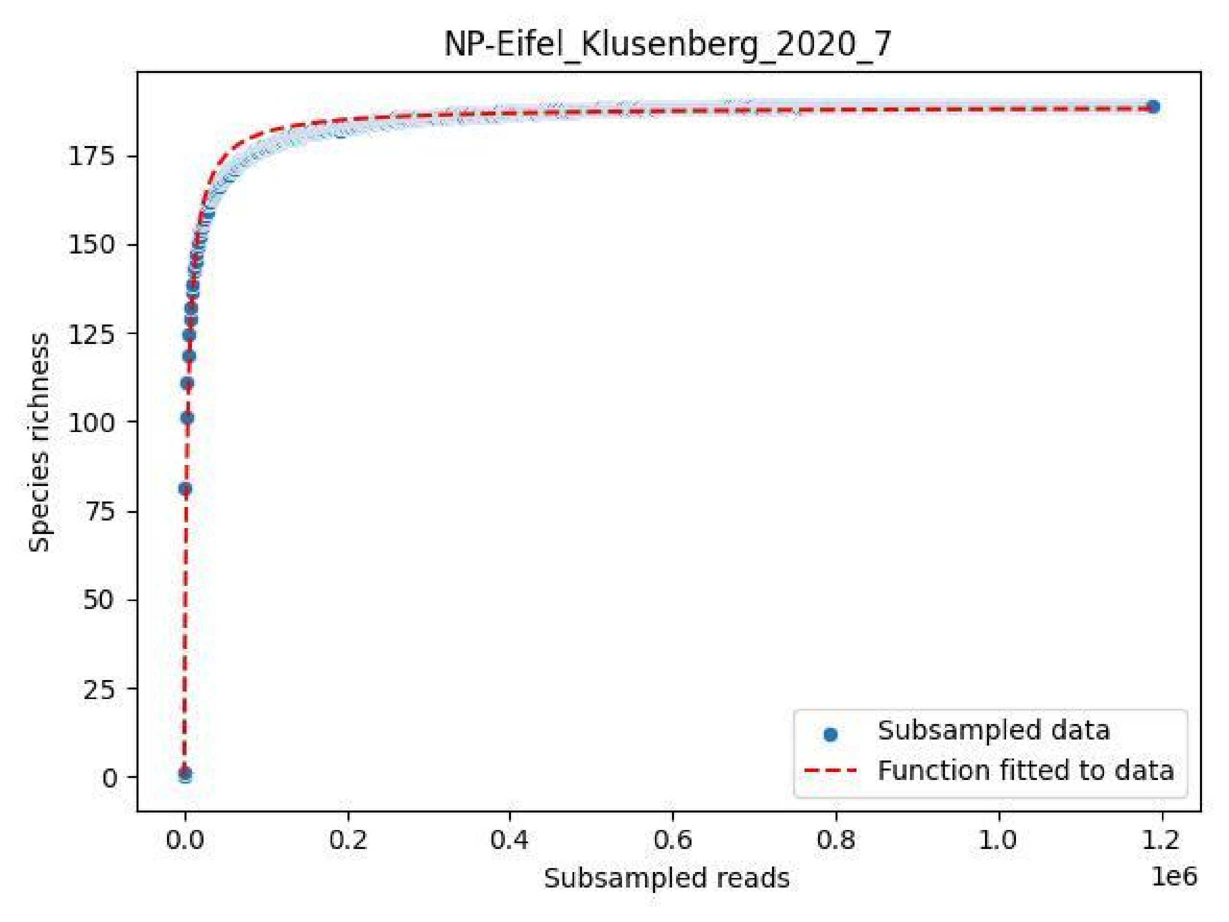
NP-Eifel_Klusenberg_2020_7
Species richness
Subsampled reads
1e6
0
25
50
75
100
125
150
175
0.0
0.2
0.4
0.6
0.8
1.0
1.2
Subsampled data
Function fitted to data

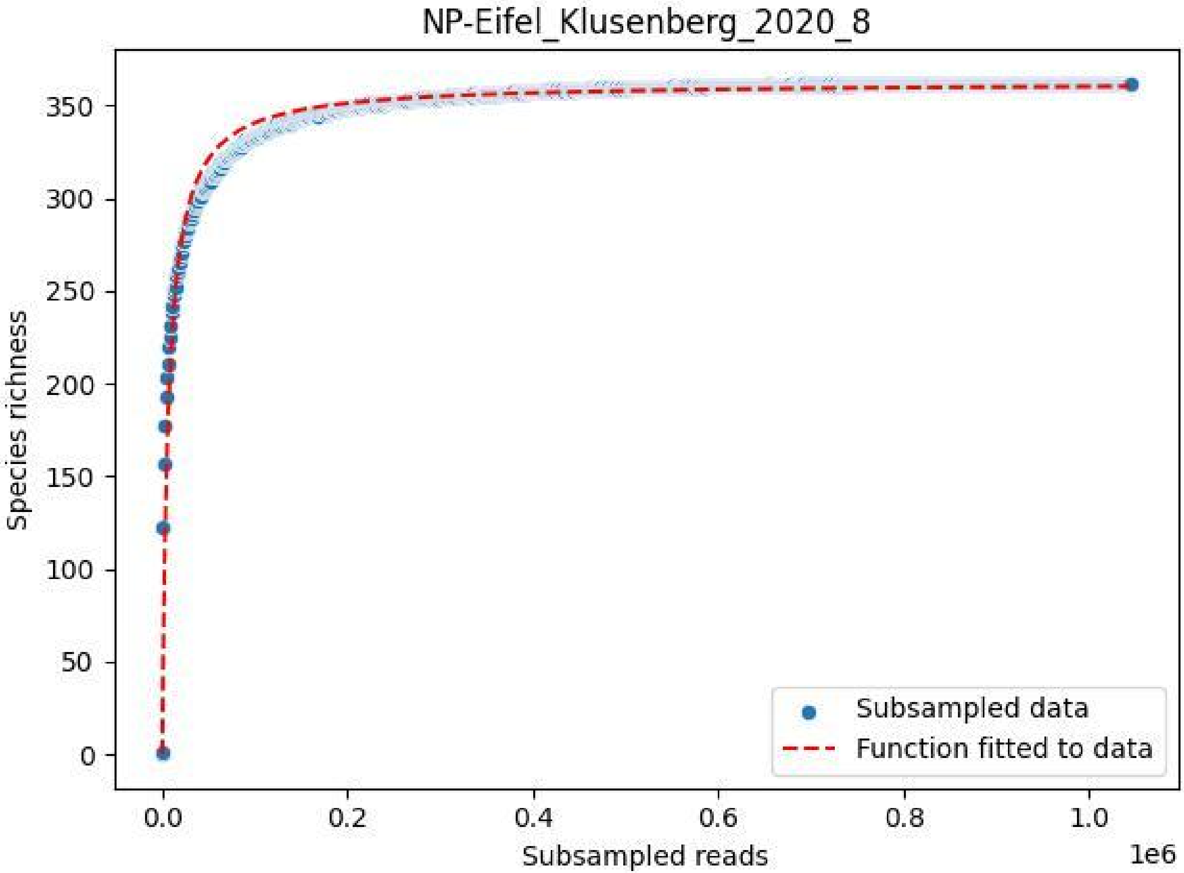
NP-Eifel_Klusenberg_2020_8
Species richness
Subsampled reads
1e6
0
50
100
150
200
250
300
350
0.0
0.2
0.4
0.6
0.8
1.0
Subsampled data
Function fitted to data

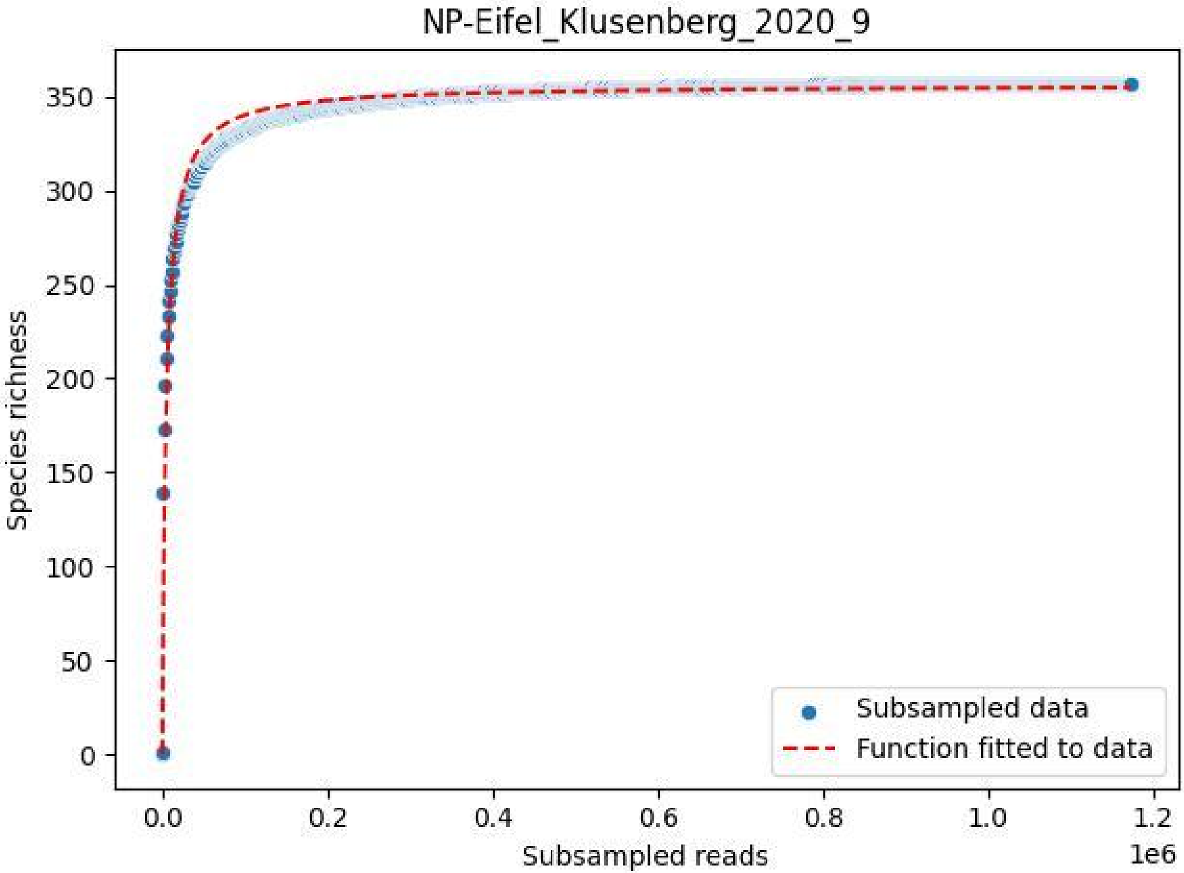

NP-Eifel_Klusenberg_2020_9
Species richness
Subsampled reads
1e6
0
50
100
150
200
250
300
350
0.0
0.2
0.4
0.6
0.8
1.0
1.2
Subsampled data
Function fitted to data

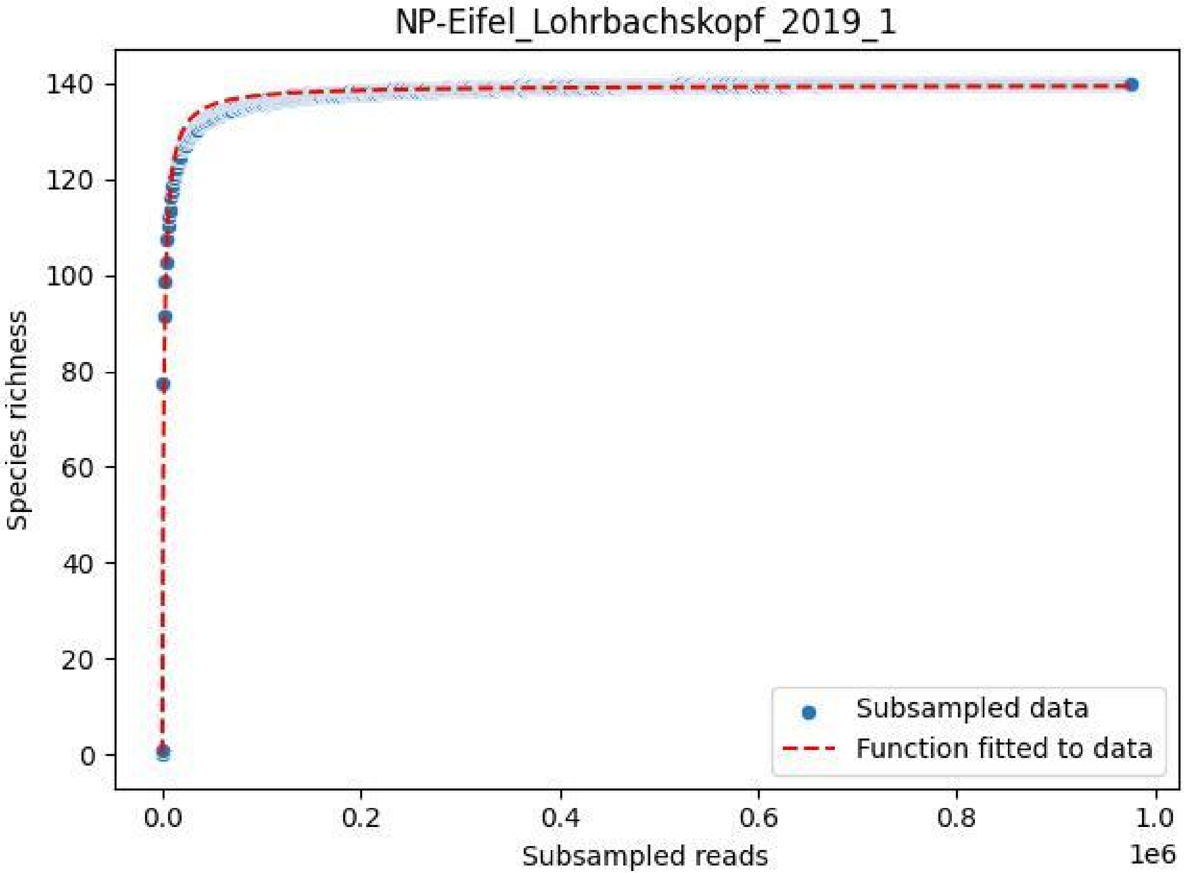
NP-Eifel_Lohrbachskopf_2019_1
Species richness
140
120
100
80
60
40
20
0
0.0
0.2
0.4
0.6
0.8
1.0
1e6
Subsampled reads
Subsampled data
Function fitted to data

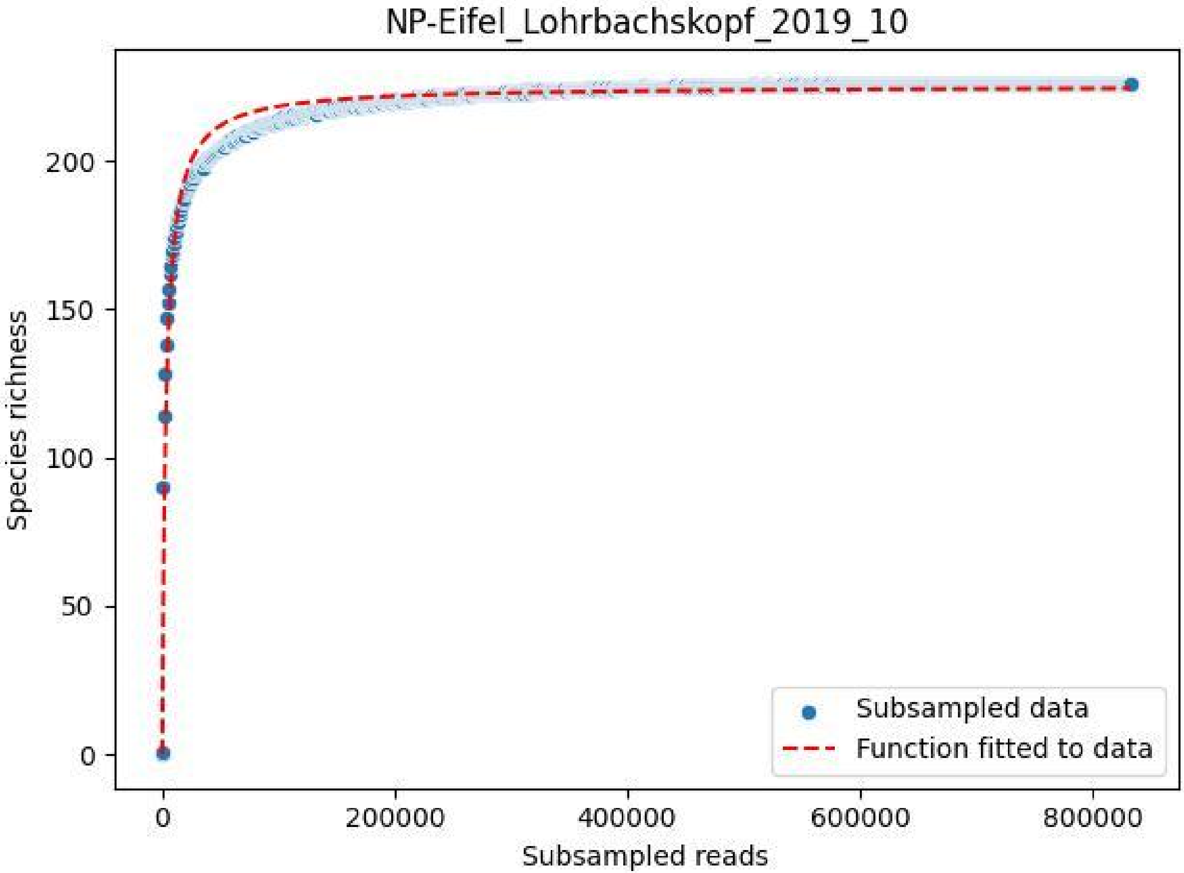
NP-Eifel_Lohrbachskopf_2019_10
Species richness
Subsampled reads
0
50
100
150
200
0
200000
400000
600000
800000
Subsampled data
Function fitted to data

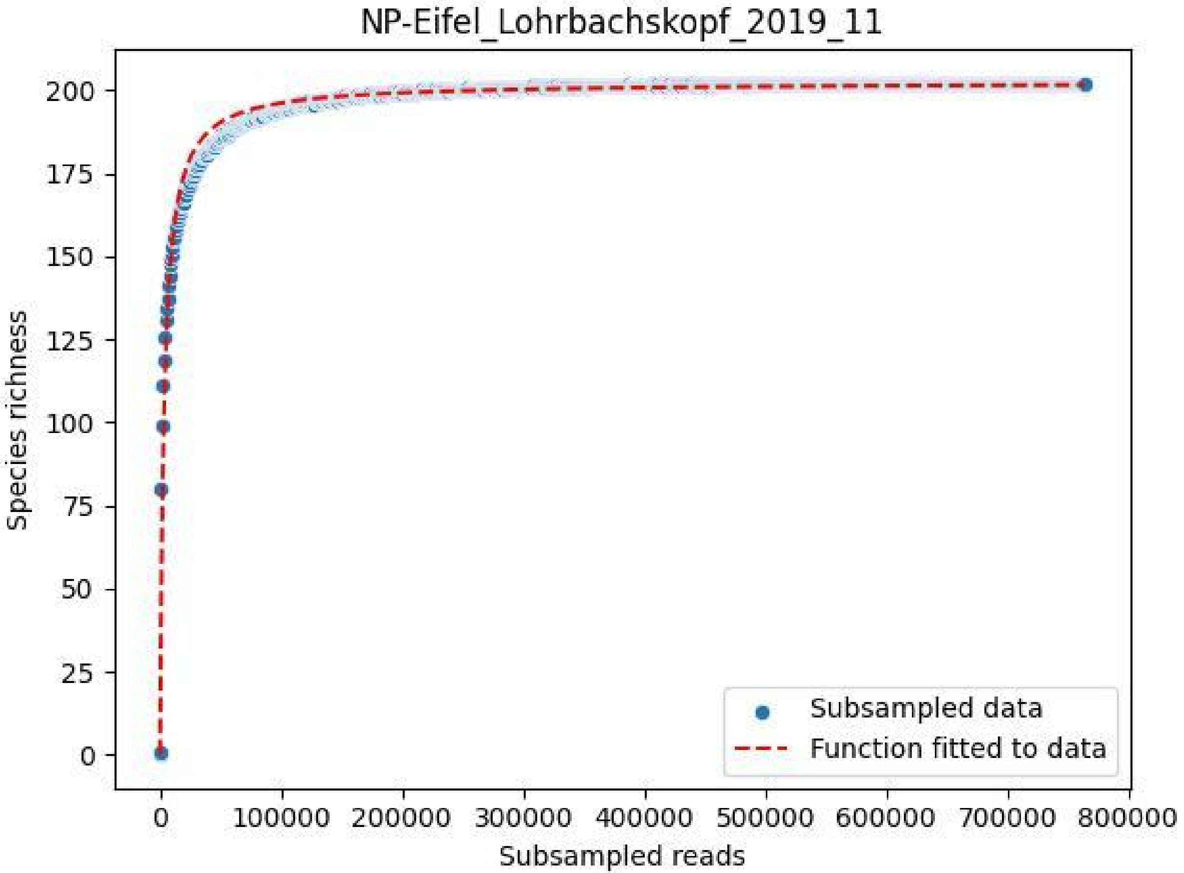

NP-Eifel_Lohrbachskopf_2019_11
Species richness
0
25
50
75
100
125
150
175
200
Subsampled reads
0
100000
200000
300000
400000
500000
600000
700000
800000
Subsampled data
Function fitted to data

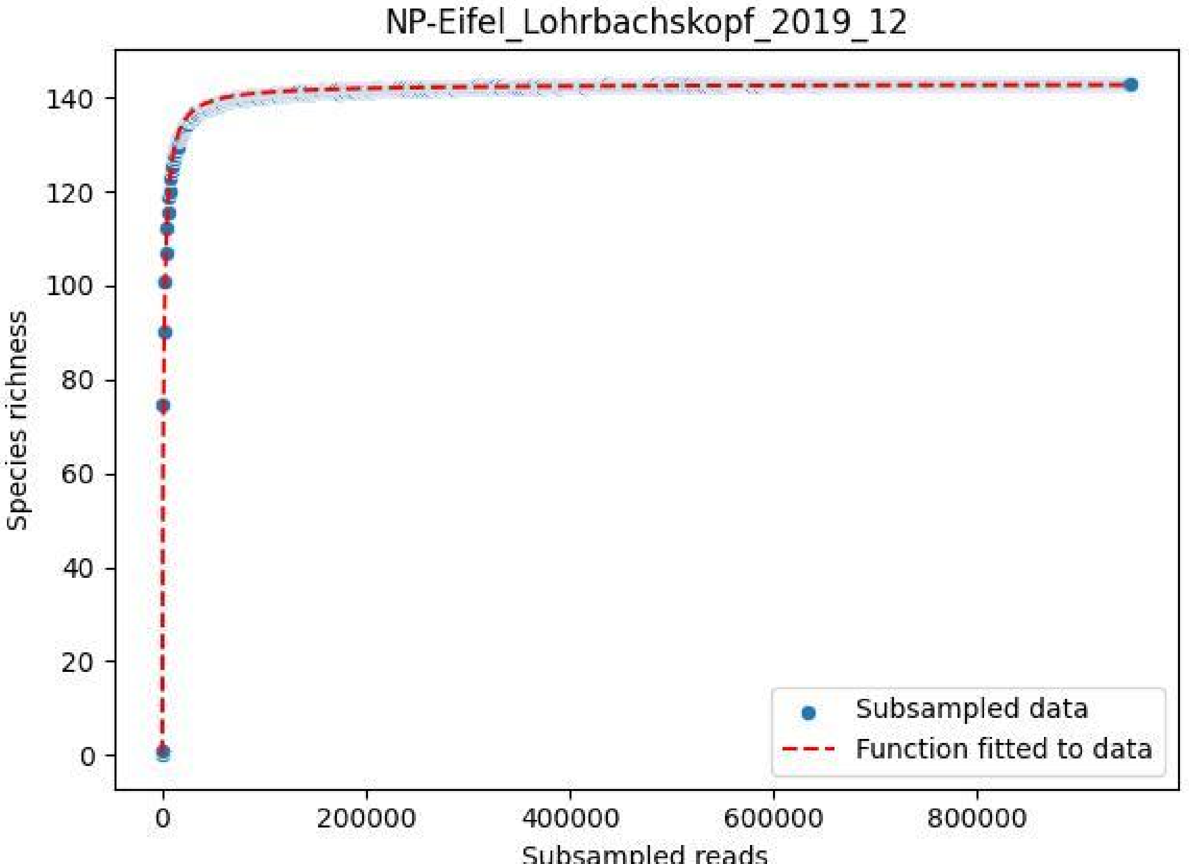
NP-Eifel_Lohrbachskopf_2019_12
Species richness
Subsampled reads
0
20
40
60
80
100
120
140
0
200000
400000
600000
800000
Subsampled data
Function fitted to data

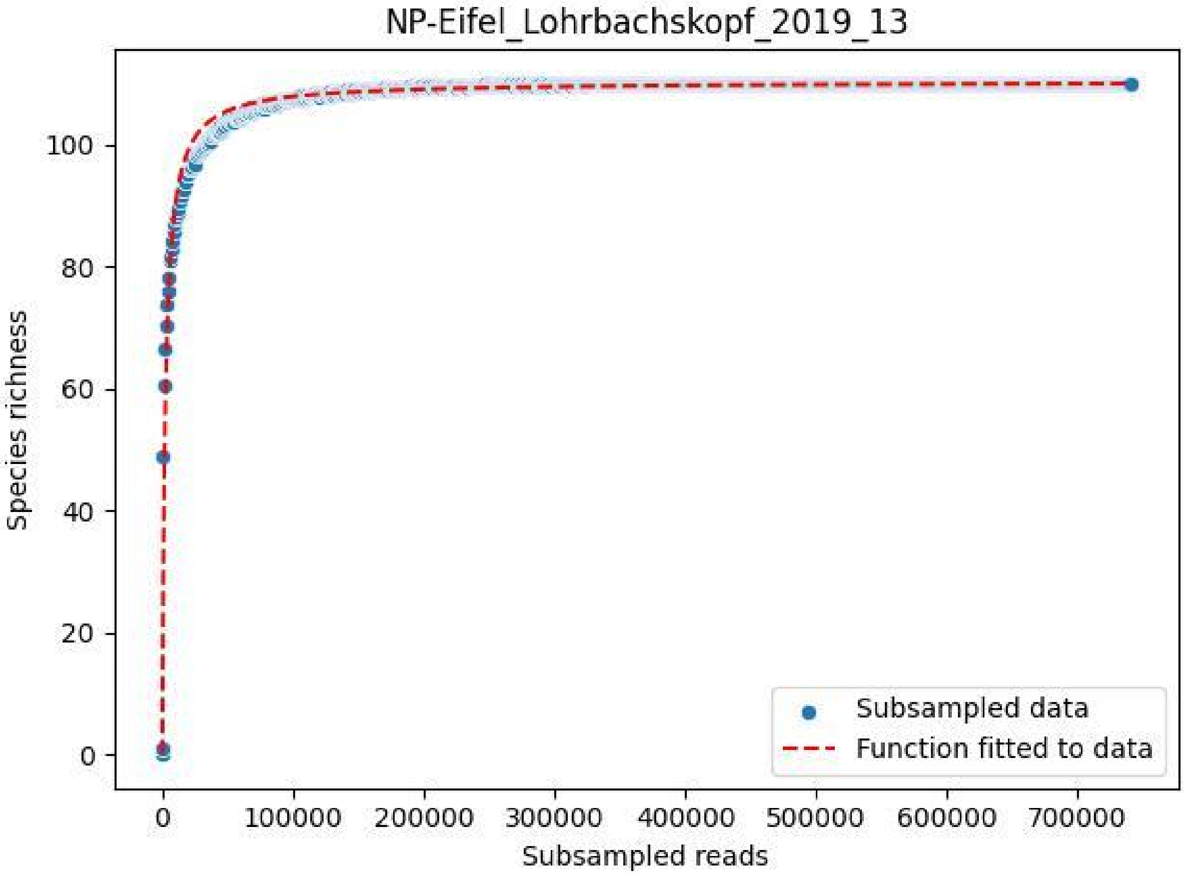
NP-Eifel_Lohrbachskopf_2019_13
Species richness
Subsampled reads
0
20
40
60
80
100
0
100000
200000
300000
400000
500000
600000
700000
Subsampled data
Function fitted to data

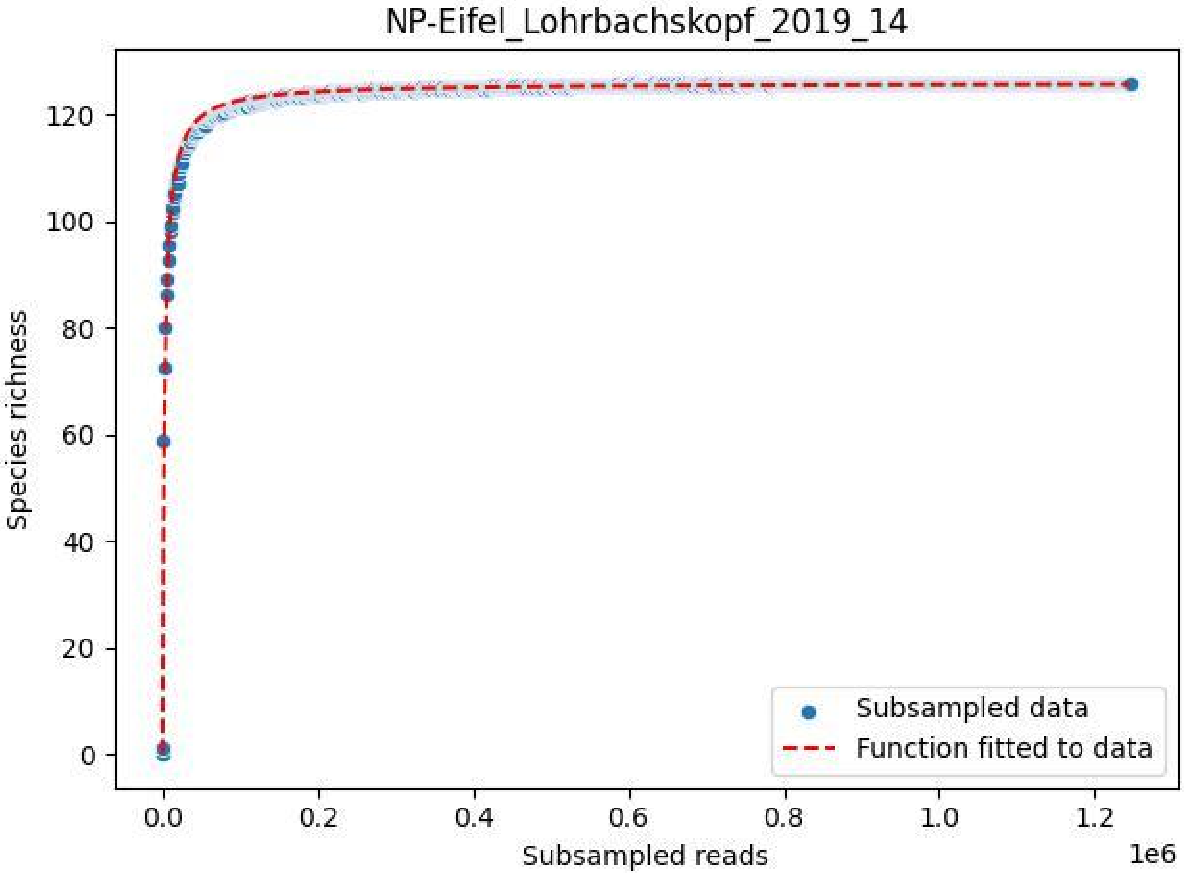

NP-Eifel_Lohrbachskopf_2019_14
Species richness
Subsampled reads
1e6
0
20
40
60
80
100
120
0.0
0.2
0.4
0.6
0.8
1.0
1.2
Subsampled data
Function fitted to data

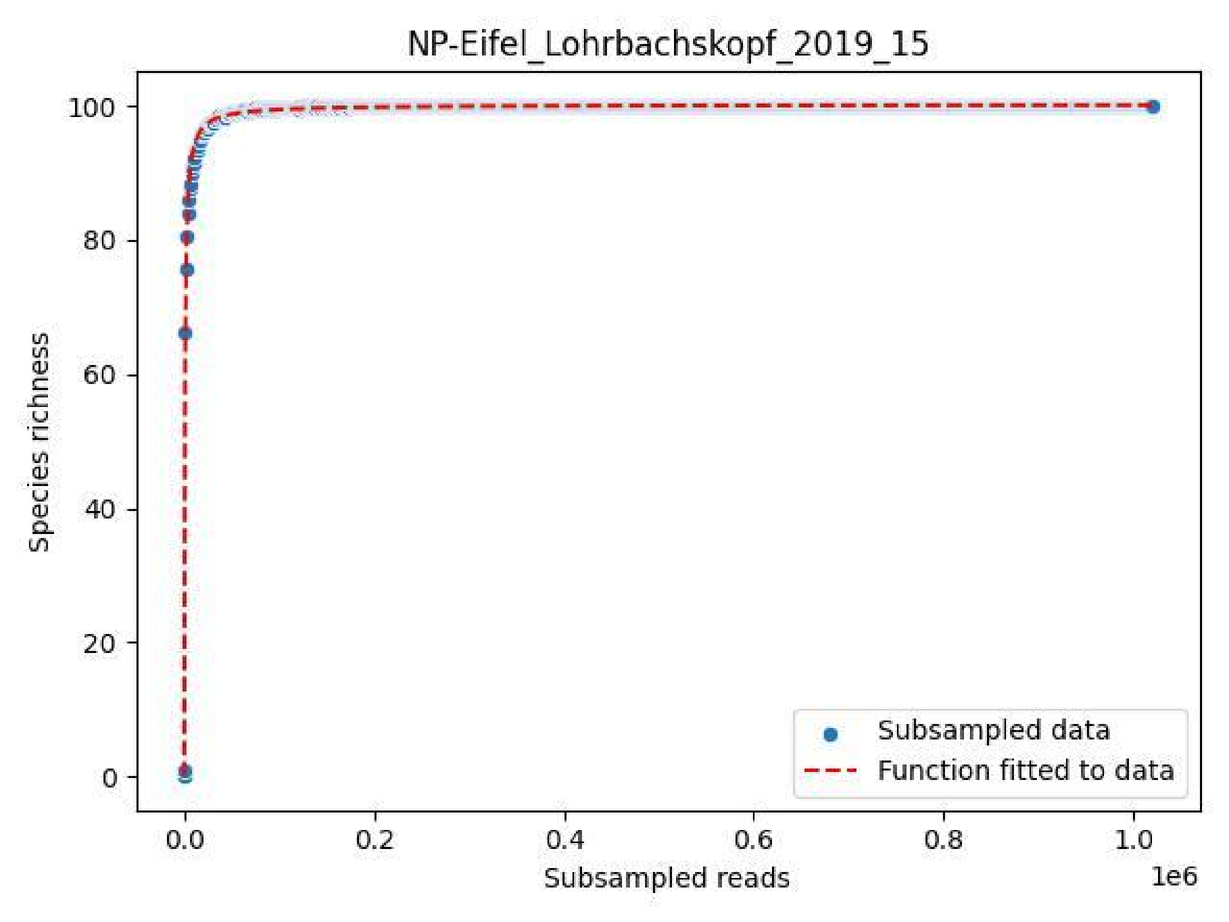
NP-Eifel_Lohrbachskopf_2019_15
Species richness
Subsampled reads
1e6
Subsampled data
Function fitted to data
0
20
40
60
80
100
0.0
0.2
0.4
0.6
0.8
1.0

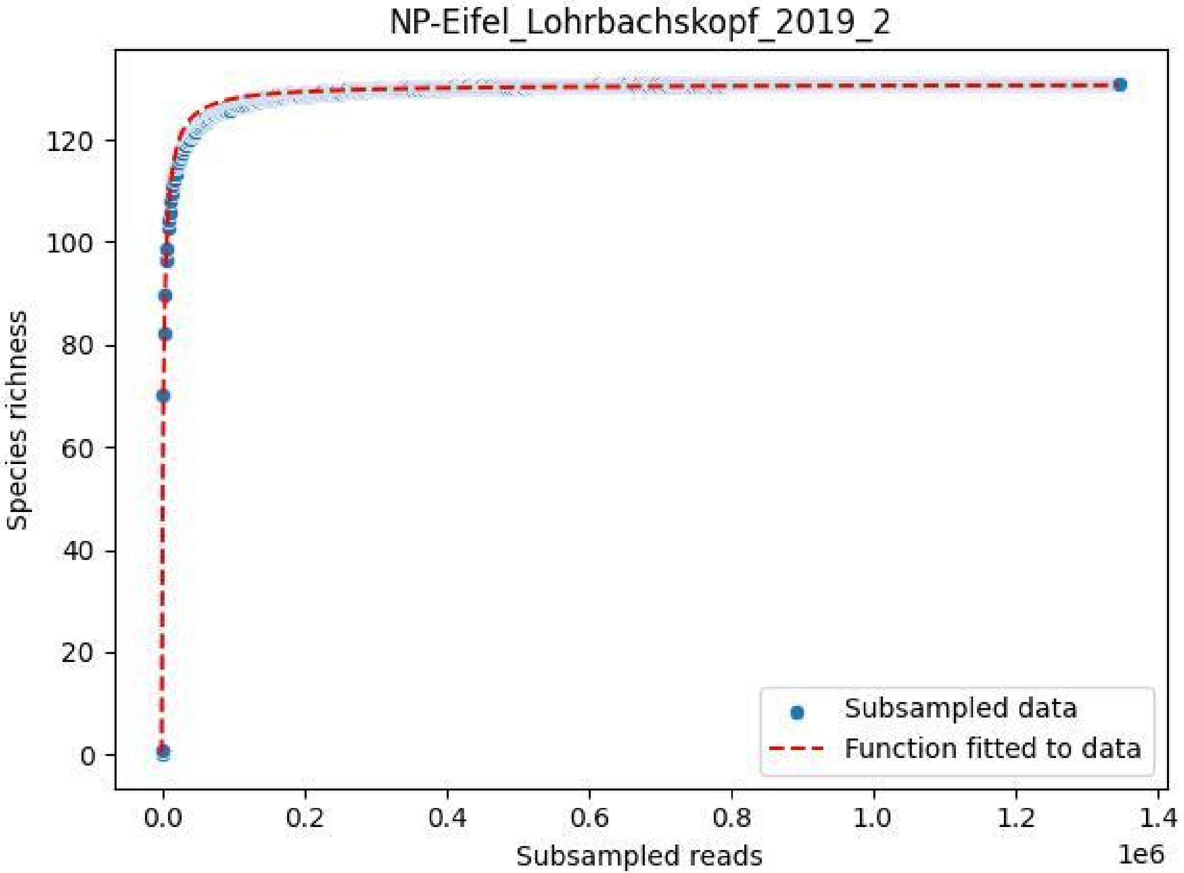
NP-Eifel_Lohrbachskopf_2019_2
Species richness
Subsampled reads
1e6
0
20
40
60
80
100
120
0.0
0.2
0.4
0.6
0.8
1.0
1.2
1.4
Subsampled data
Function fitted to data

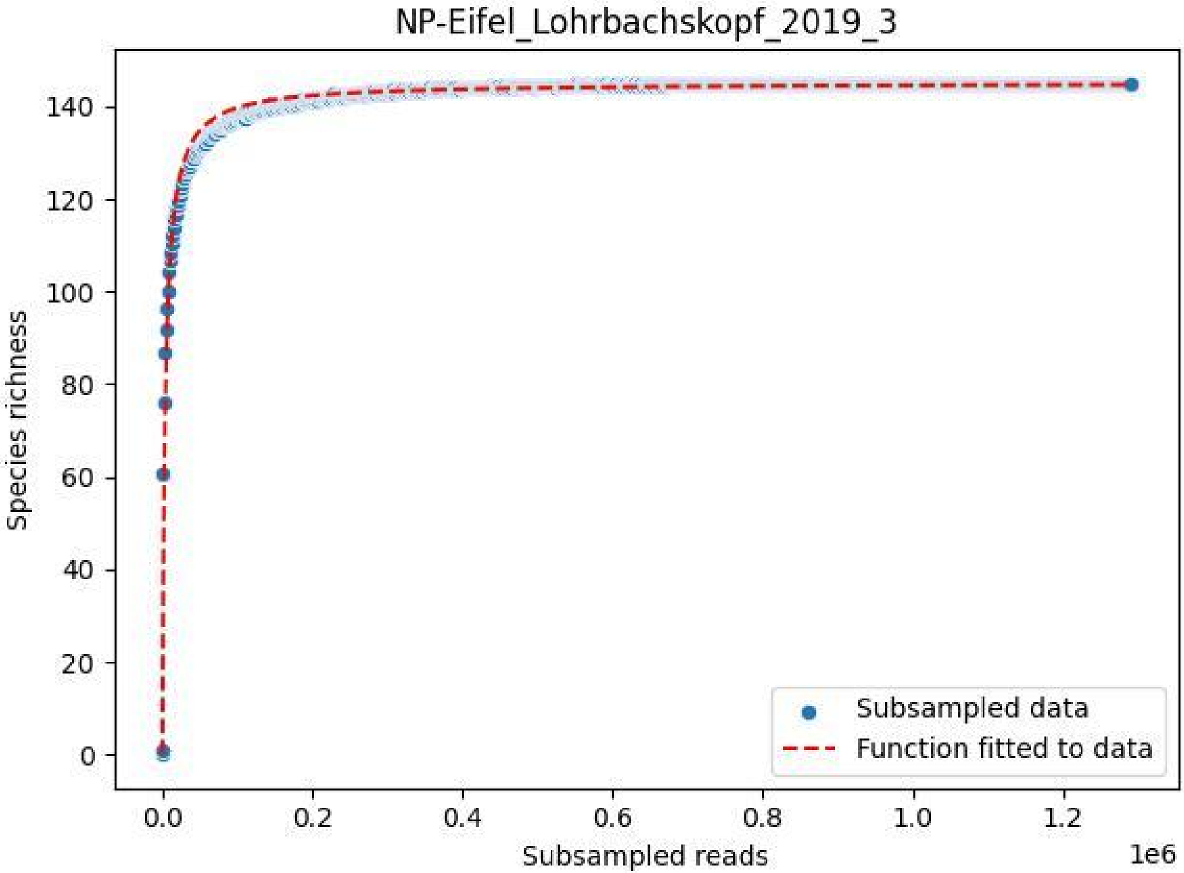
NP-Eifel_Lohrbachskopf_2019_3
Species richness
Subsampled reads
1e6
0
20
40
60
80
100
120
140
0.0
0.2
0.4
0.6
0.8
1.0
1.2
Subsampled data
Function fitted to data

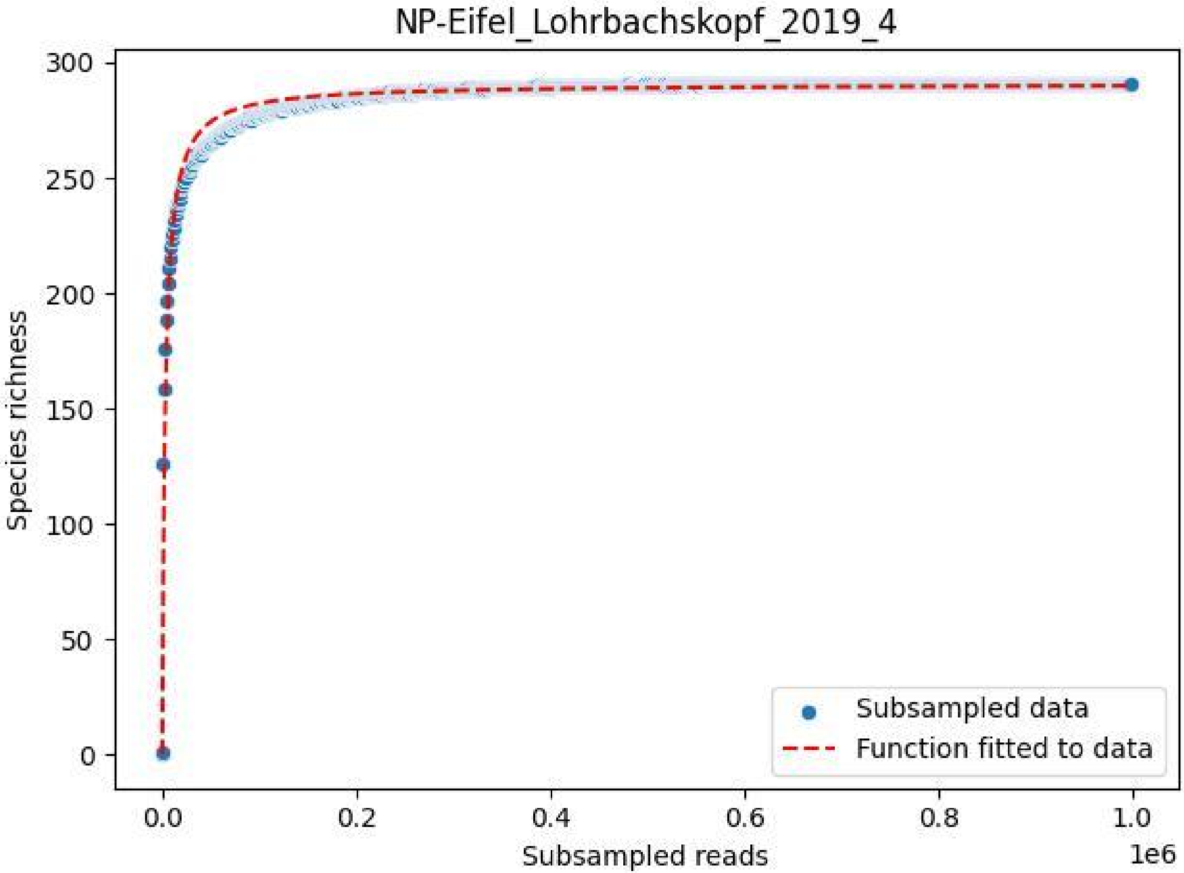
NP-Eifel_Lohrbachskopf_2019_4
Species richness
Subsampled reads
1e6
Subsampled data
Function fitted to data
0
50
100
150
200
250
300
0.0
0.2
0.4
0.6
0.8
1.0

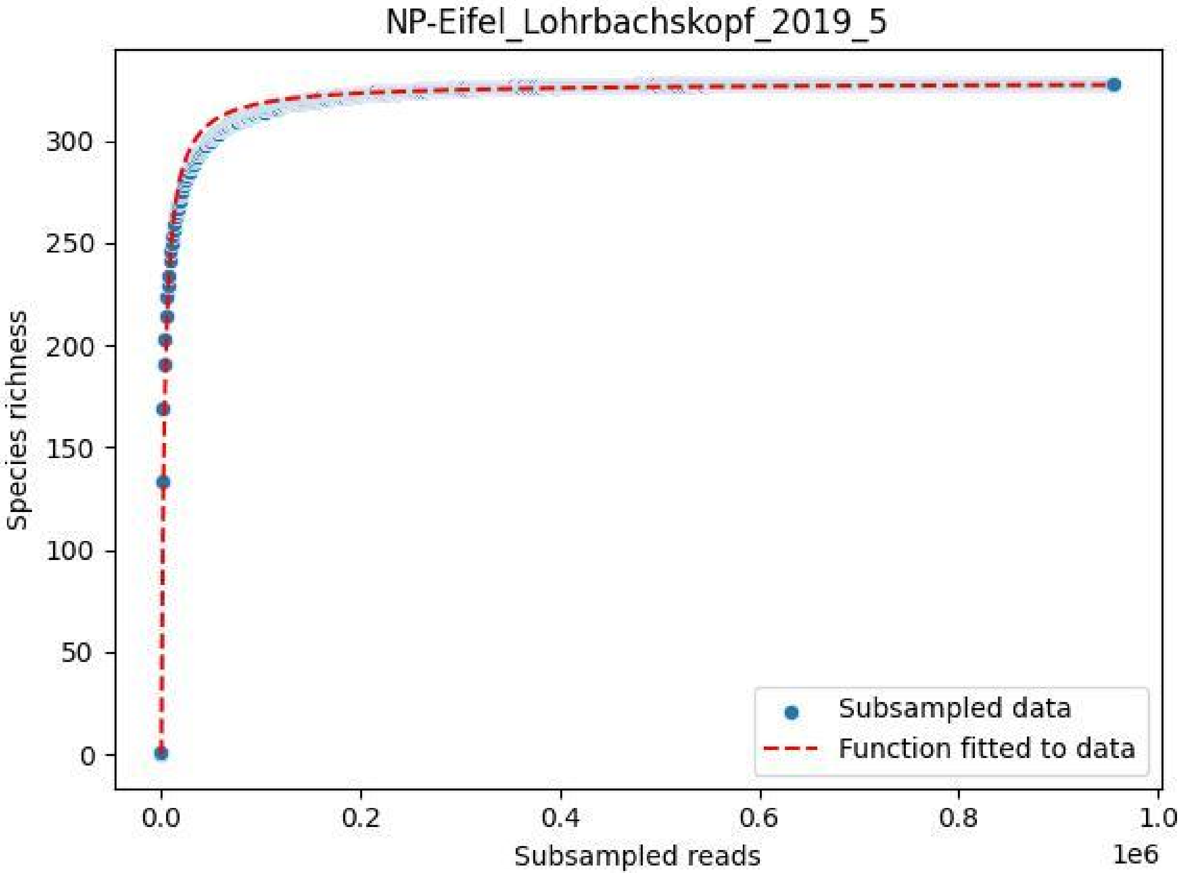
NP-Eifel_Lohrbachskopf_2019_5
Species richness
Subsampled reads
1e6
0
50
100
150
200
250
300
0.0
0.2
0.4
0.6
0.8
1.0
Subsampled data
Function fitted to data

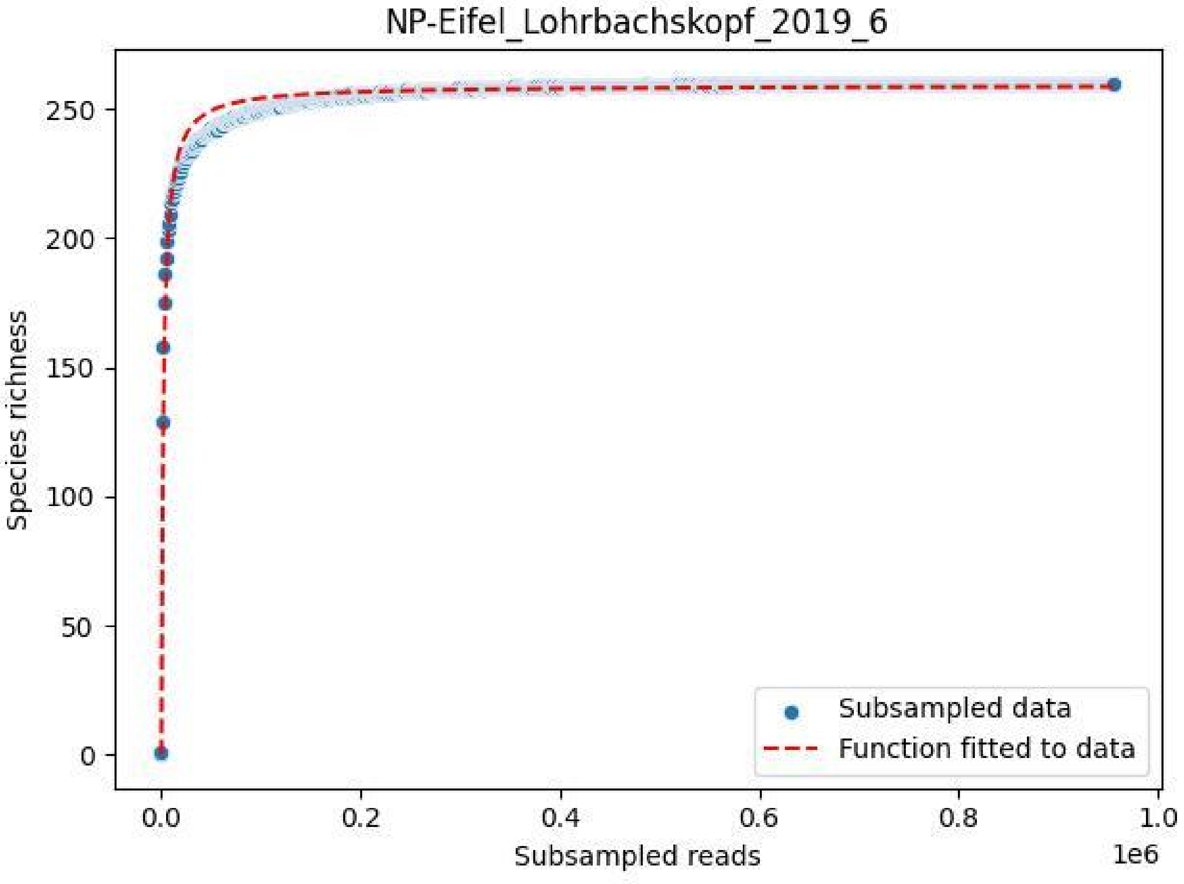
NP-Eifel_Lohrbachskopf_2019_6
Species richness
250
200
150
100
50
0
0.0
0.2
0.4
0.6
0.8
1.0
1e6
Subsampled reads
Subsampled data
Function fitted to data

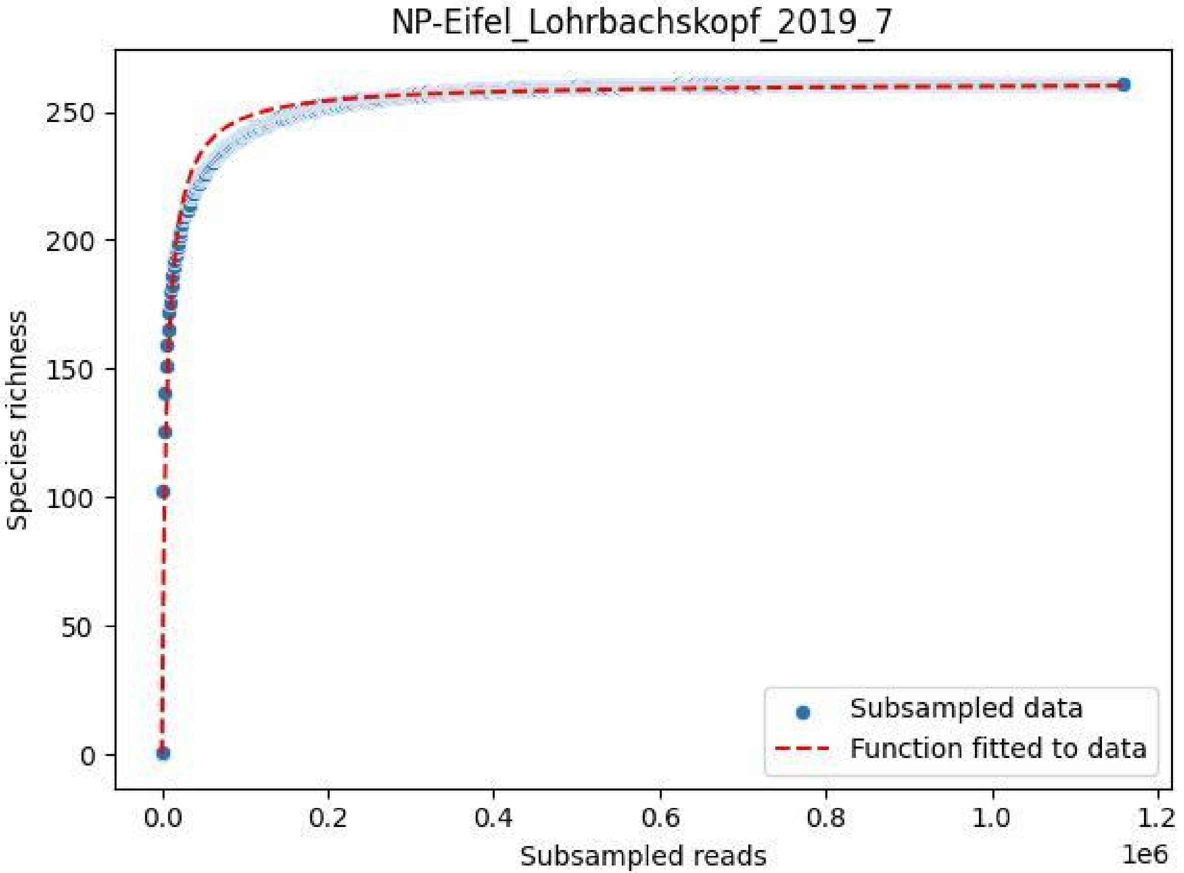
NP-Eifel_Lohrbachskopf_2019_7
Species richness
Subsampled reads
1e6
0
50
100
150
200
250
0.0
0.2
0.4
0.6
0.8
1.0
1.2
Subsampled data
Function fitted to data

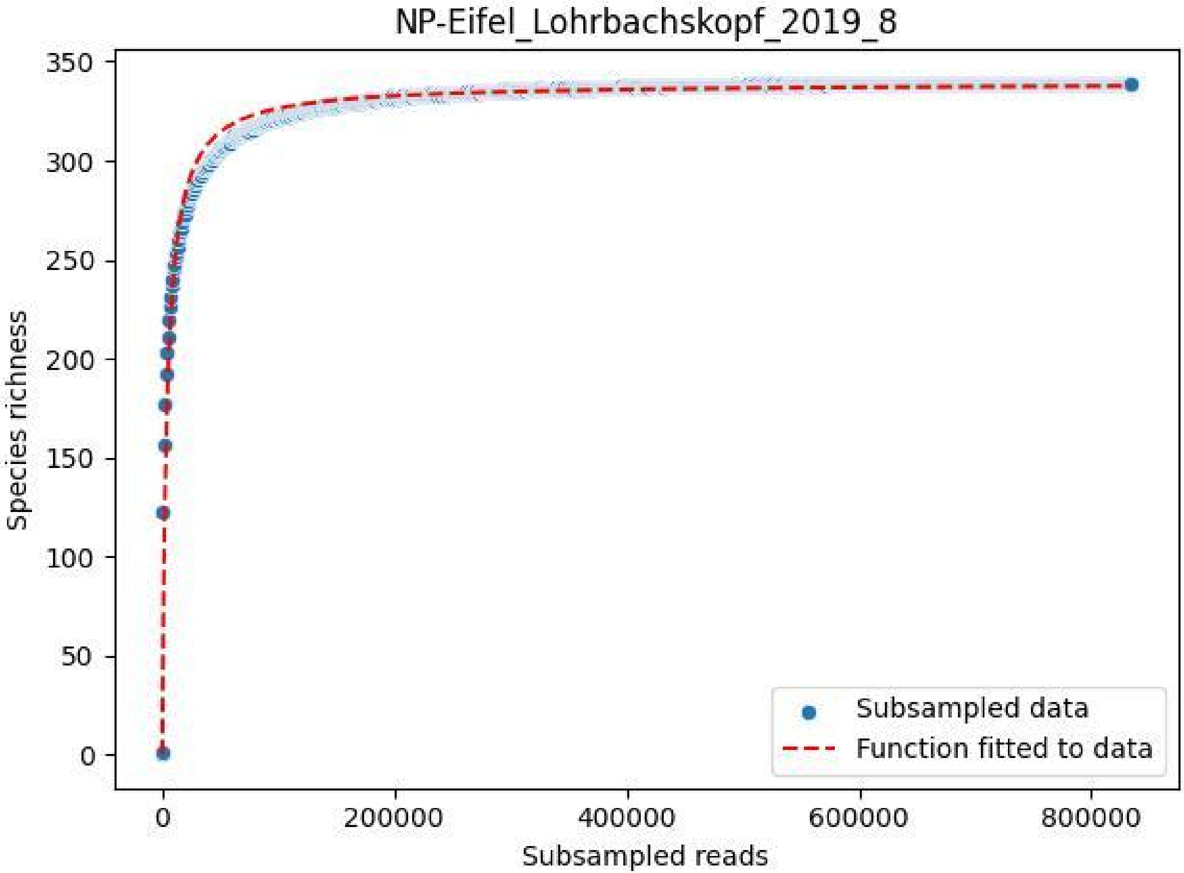
NP-Eifel_Lohrbachskopf_2019_8
Species richness
Subsampled reads
0
50
100
150
200
250
300
350
0
200000
400000
600000
800000
Subsampled data
Function fitted to data

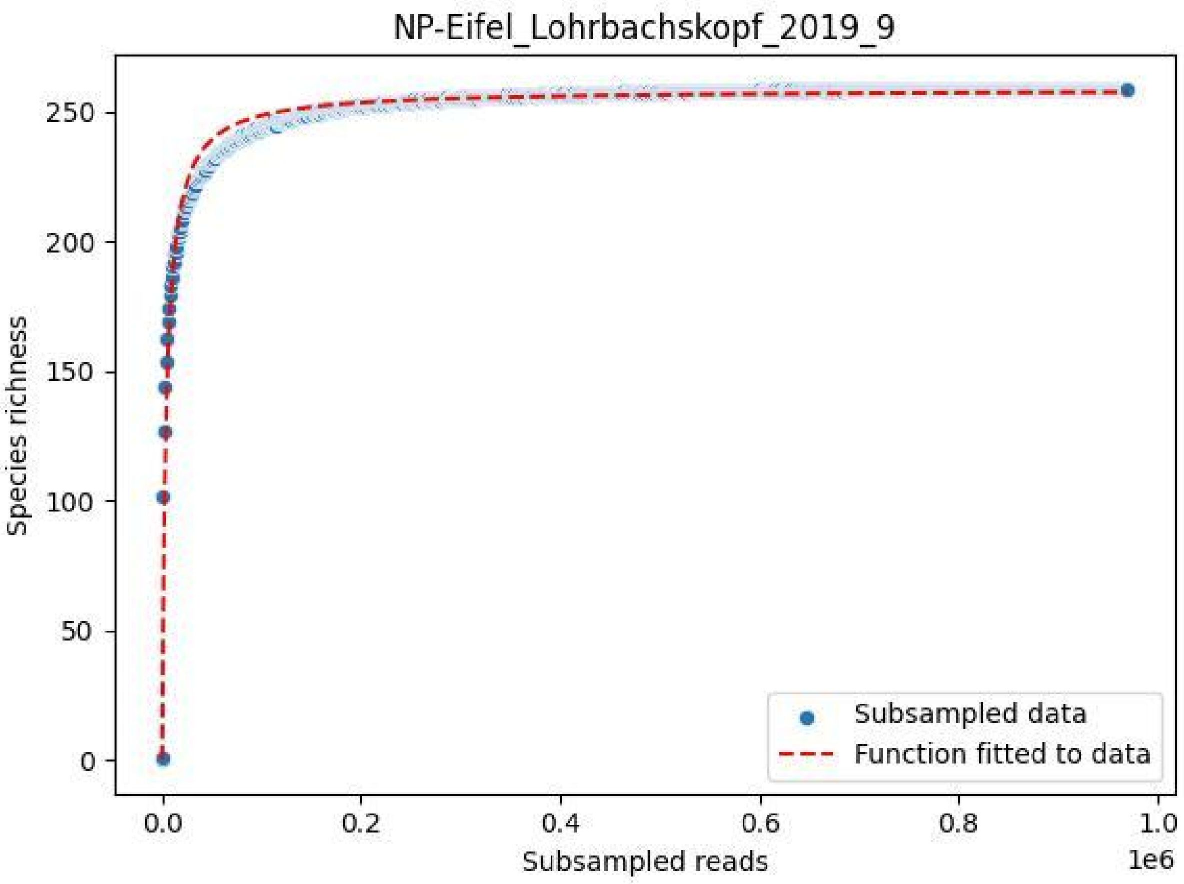
NP-Eifel_Lohrbachskopf_2019_9
Species richness
0
50
100
150
200
250
0.0
0.2
0.4
0.6
0.8
1.0
1e6
Subsampled reads
Subsampled data
Function fitted to data

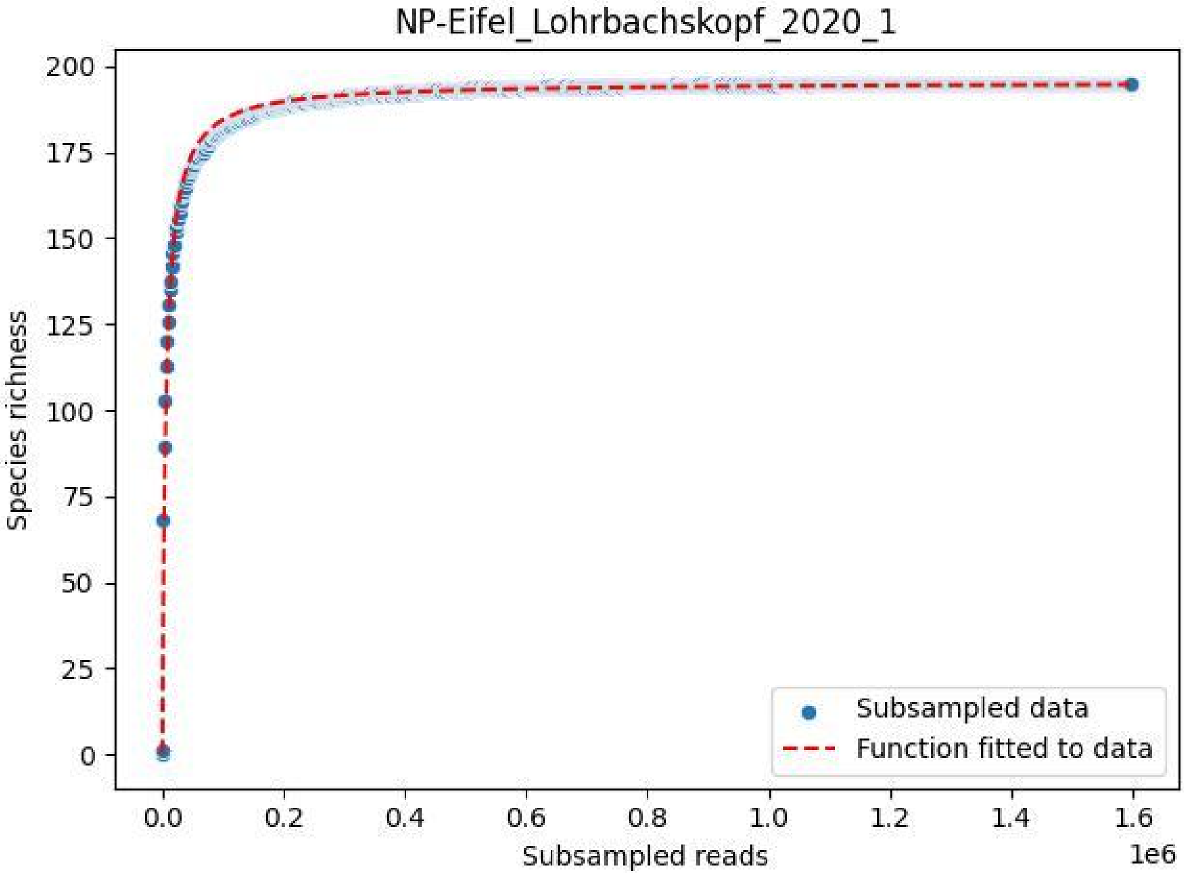
NP-Eifel_Lohrbachskopf_2020_1
Species richness
Subsampled reads
1e6
0
25
50
75
100
125
150
175
200
0.0
0.2
0.4
0.6
0.8
1.0
1.2
1.4
1.6
Subsampled data
Function fitted to data

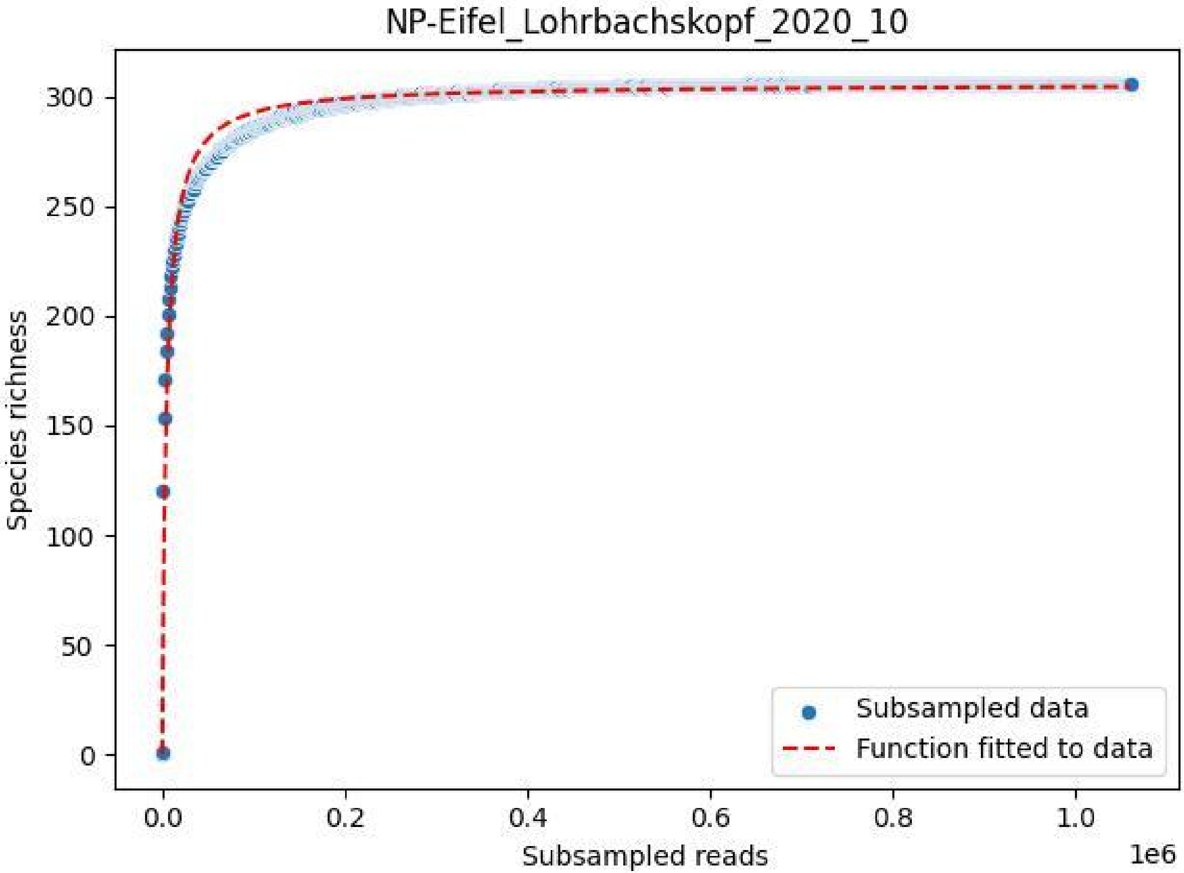
NP-Eifel_Lohrbachskopf_2020_10
Species richness
Subsampled reads
1e6
0
50
100
150
200
250
300
0.0
0.2
0.4
0.6
0.8
1.0
Subsampled data
Function fitted to data

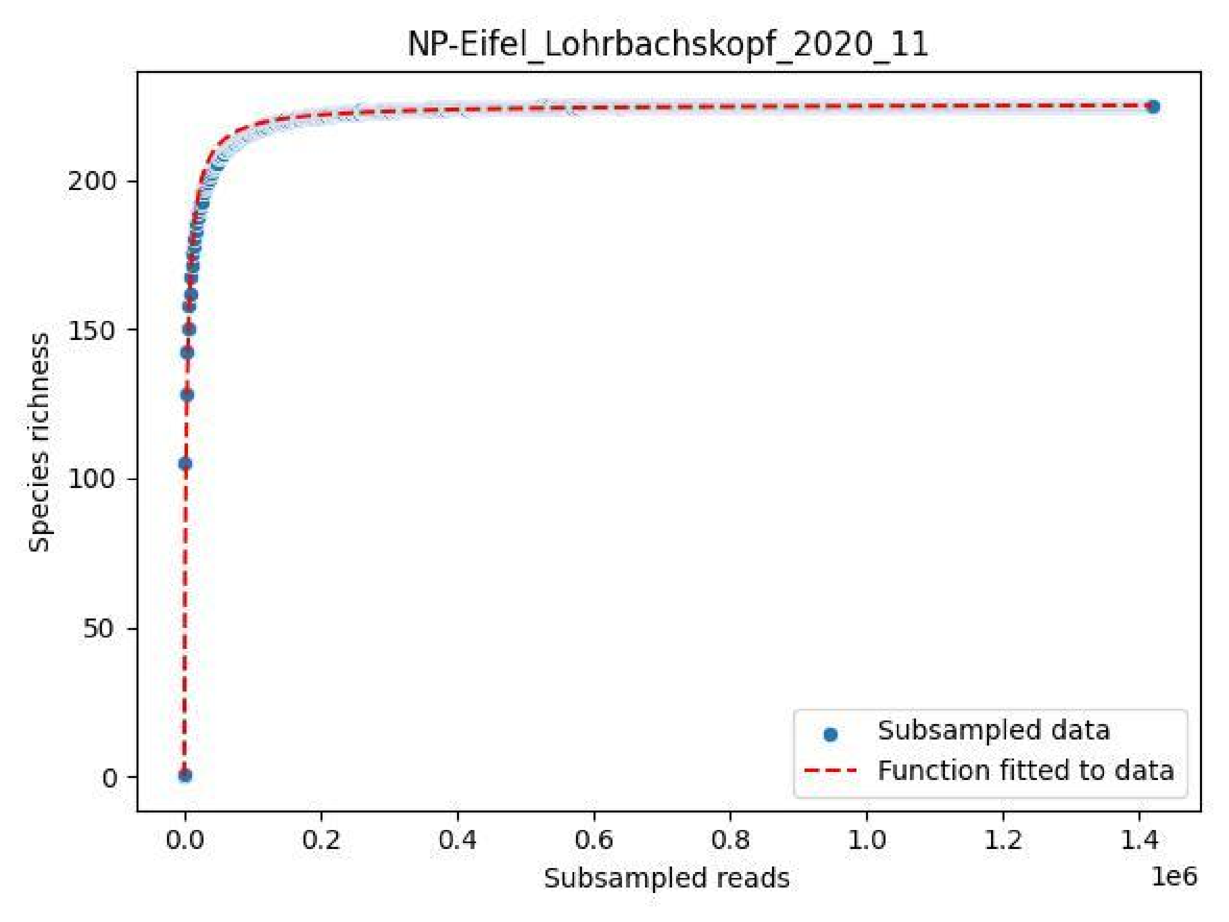

NP-Eifel_Lohrbachskopf_2020_11
Species richness
Subsampled reads
1e6
0
50
100
150
200
0.0
0.2
0.4
0.6
0.8
1.0
1.2
1.4
Subsampled data
Function fitted to data

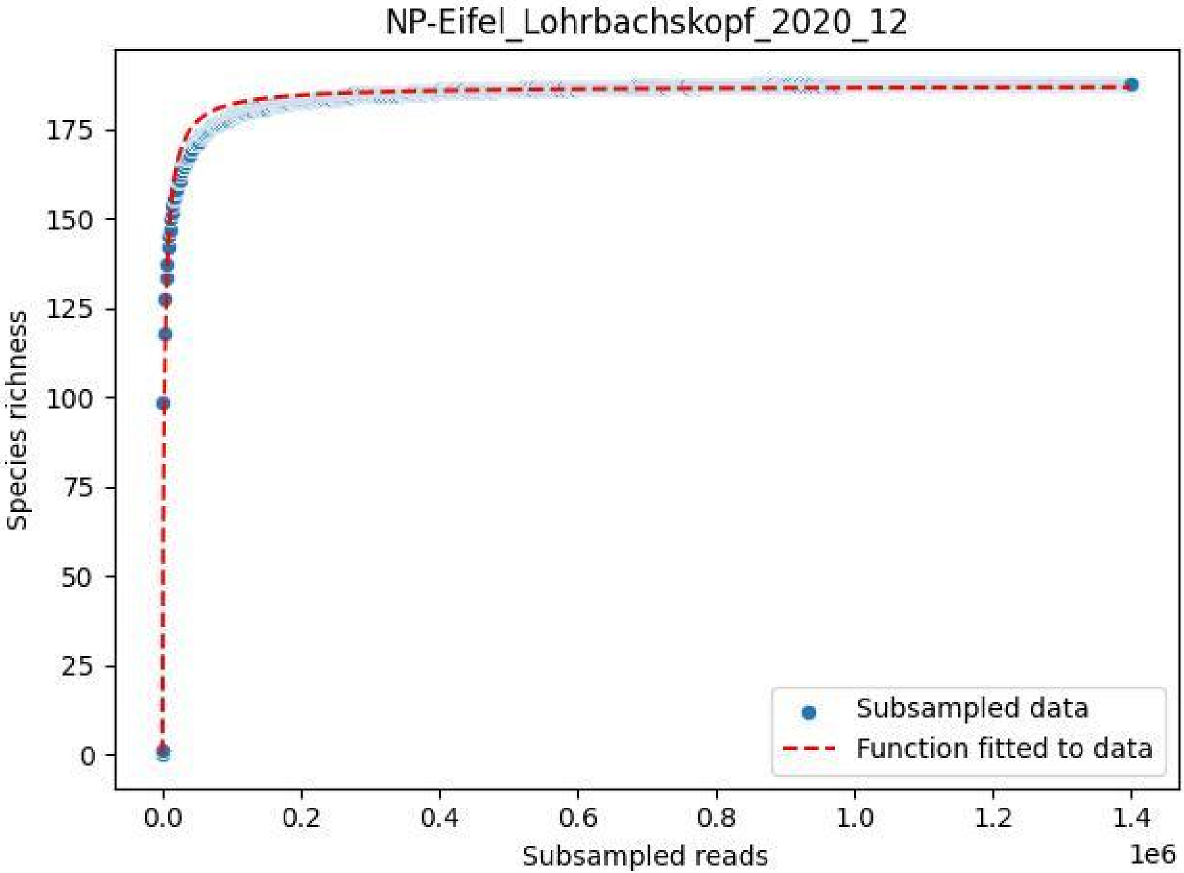
NP-Eifel_Lohrbachskopf_2020_12
Species richness
Subsampled reads
1e6
0
25
50
75
100
125
150
175
0.0
0.2
0.4
0.6
0.8
1.0
1.2
1.4
Subsampled data
Function fitted to data

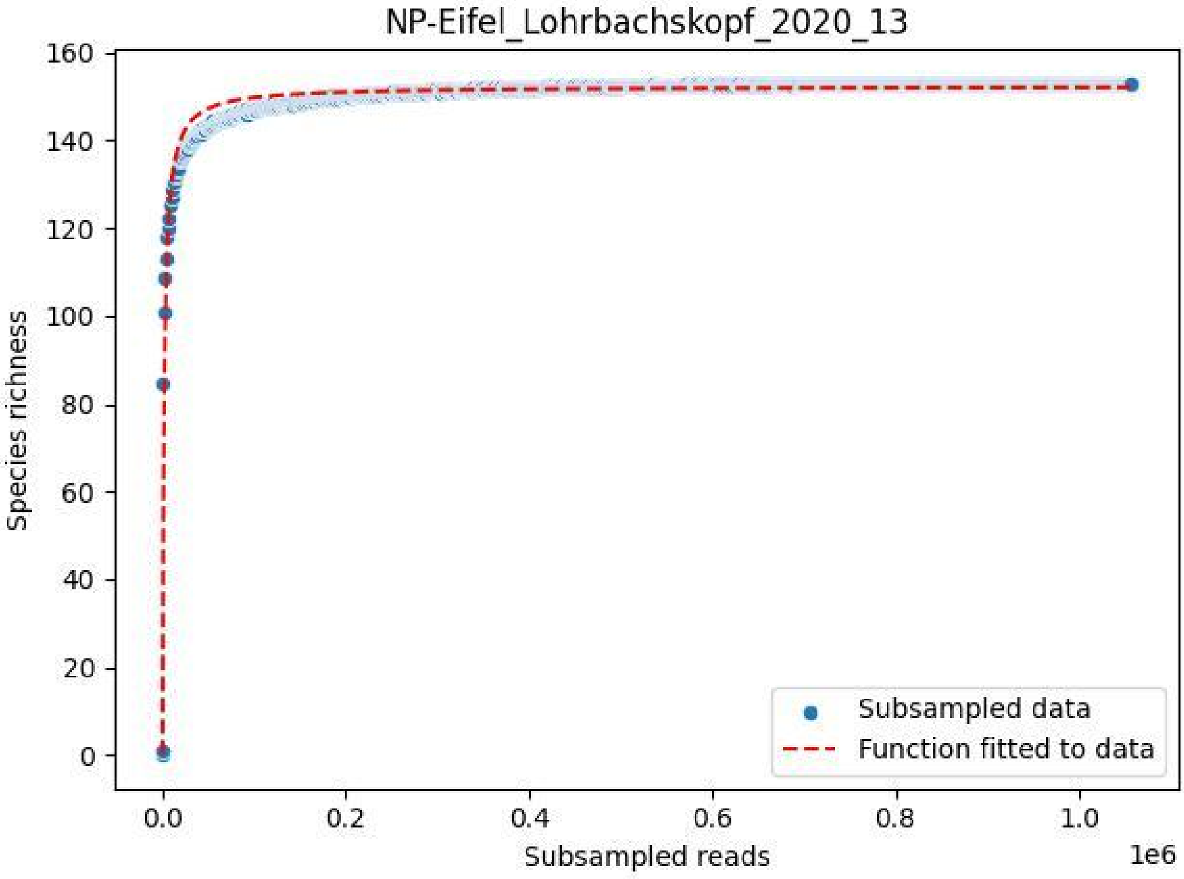

NP-Eifel_Lohrbachskopf_2020_13
160
140
120
100
80
60
40
20
0
Species richness
0.0
0.2
0.4
0.6
0.8
1.0
1e6
Subsampled reads
Subsampled data
Function fitted to data

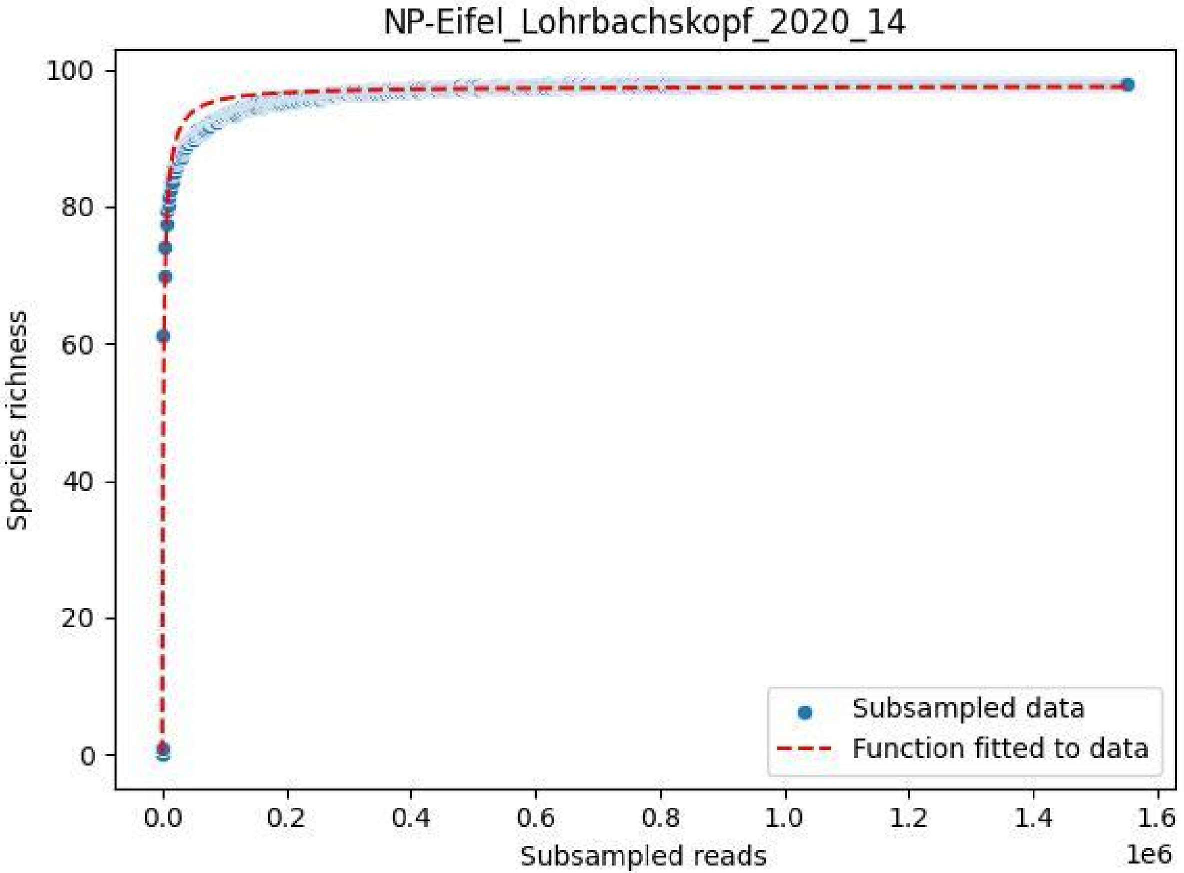

NP-Eifel_Lohrbachskopf_2020_14
Species richness
Subsampled reads
1e6
0
20
40
60
80
100
0.0
0.2
0.4
0.6
0.8
1.0
1.2
1.4
1.6
Subsampled data
Function fitted to data

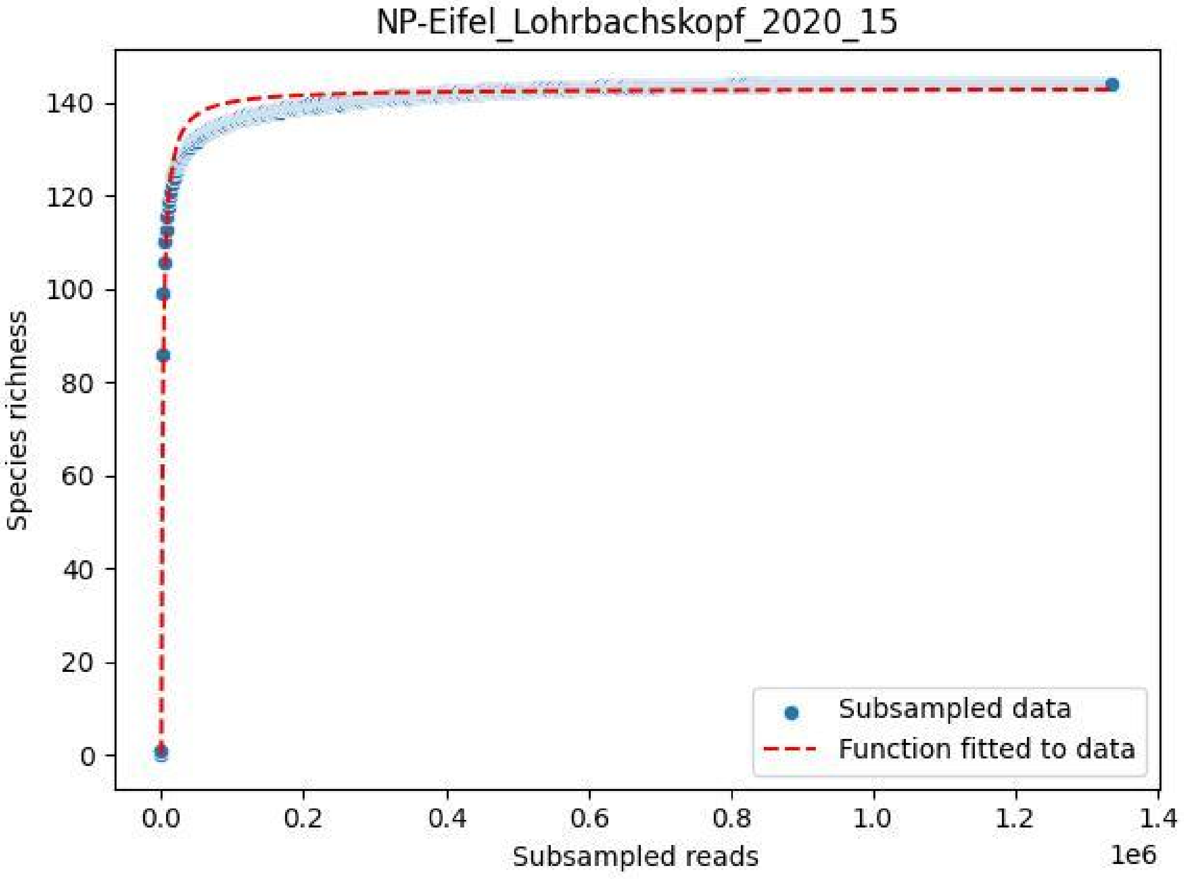
NP-Eifel_Lohrbachskopf_2020_15
Species richness
Subsampled reads
1e6
Subsampled data
Function fitted to data
0
20
40
60
80
100
120
140
0.0
0.2
0.4
0.6
0.8
1.0
1.2
1.4

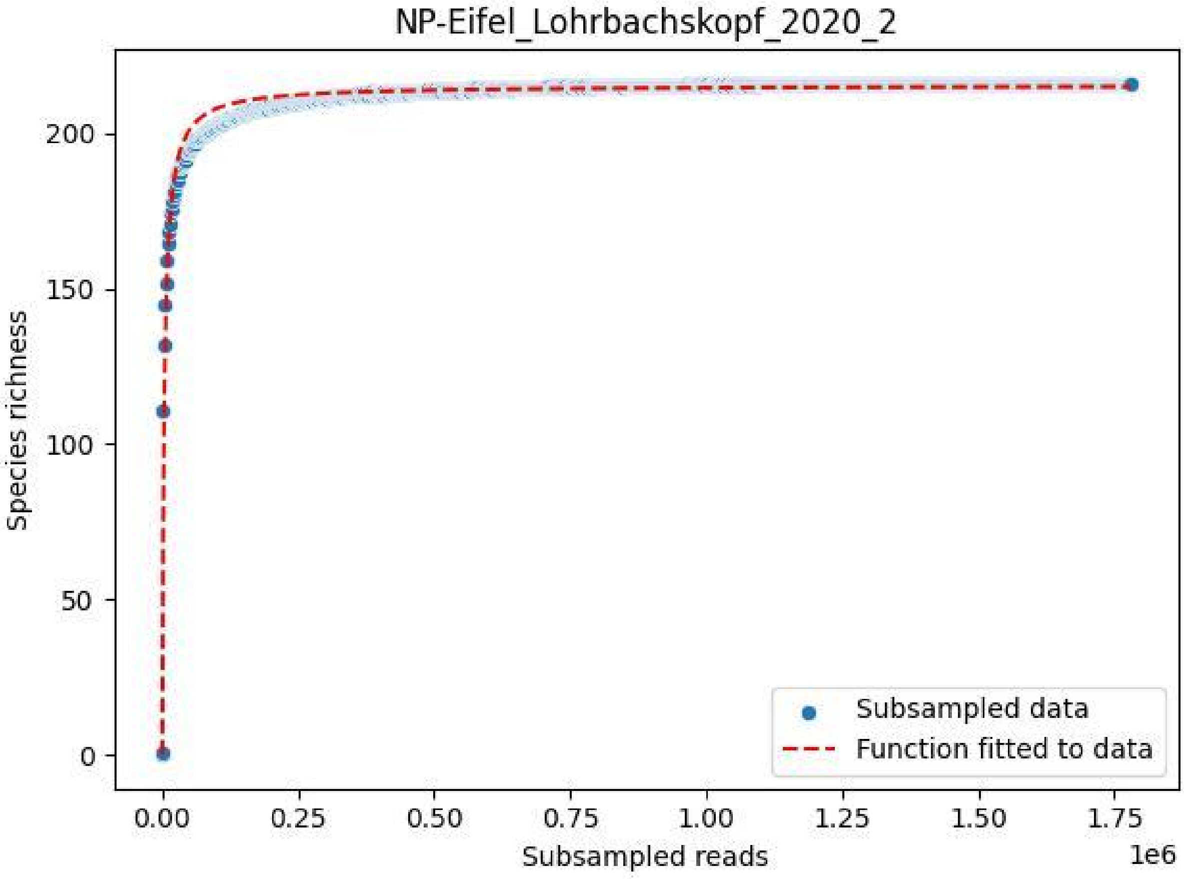
NP-Eifel_Lohrbachskopf_2020_2
Species richness
Subsampled reads
1e6
0
50
100
150
200
0.00
0.25
0.50
0.75
1.00
1.25
1.50
1.75
Subsampled data
Function fitted to data

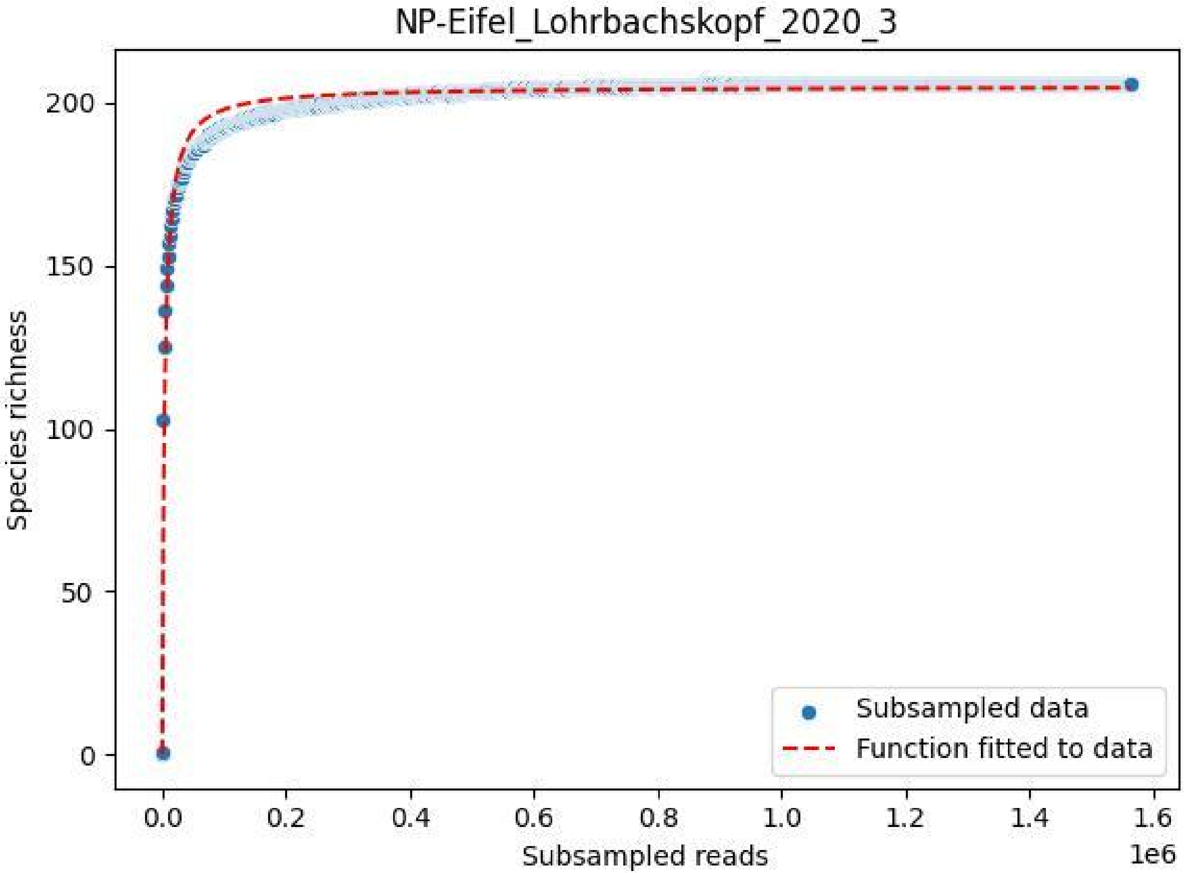
NP-Eifel_Lohrbachskopf_2020_3
Species richness
Subsampled reads
1e6
0
50
100
150
200
0.0
0.2
0.4
0.6
0.8
1.0
1.2
1.4
1.6
Subsampled data
Function fitted to data

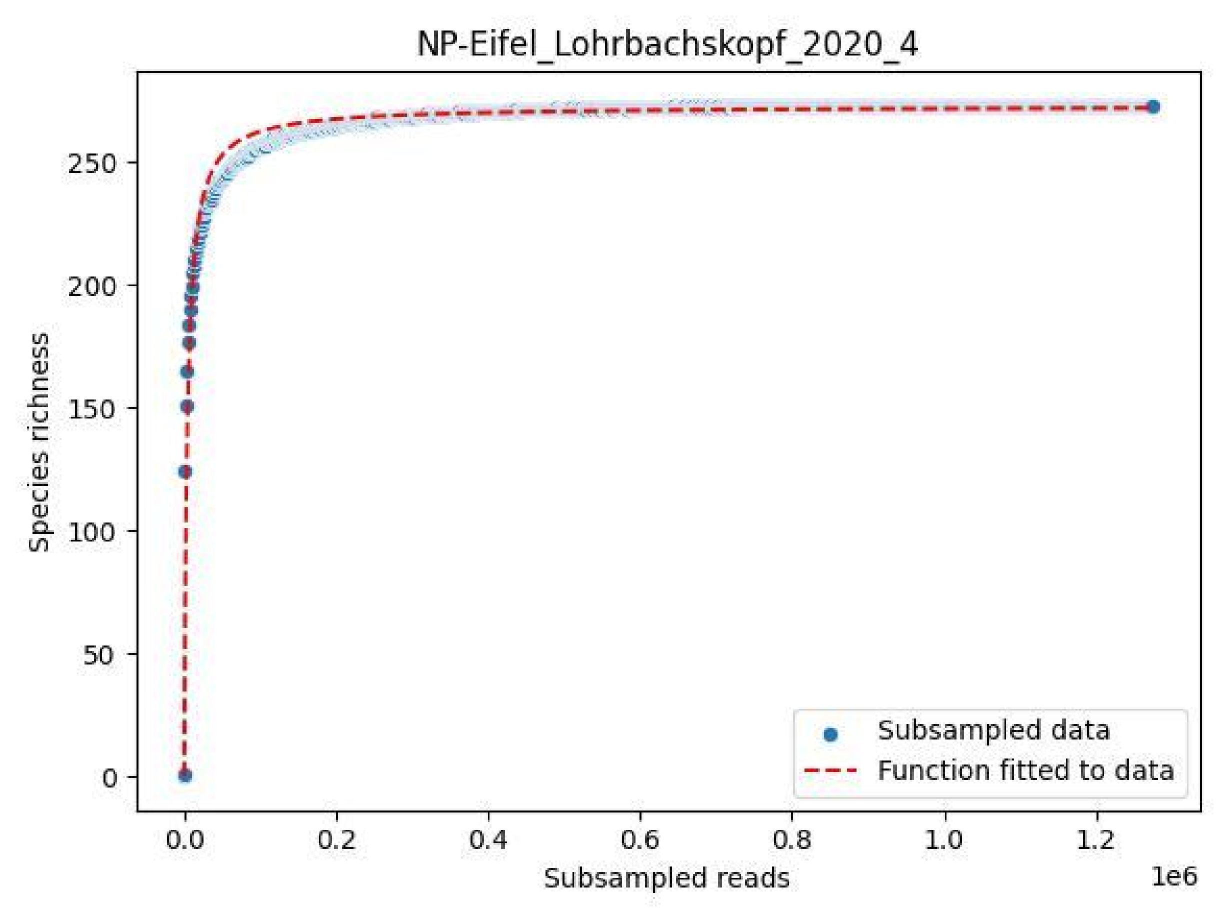
NP-Eifel_Lohrbachskopf_2020_4
Species richness
Subsampled reads
1e6
0
50
100
150
200
250
0.0
0.2
0.4
0.6
0.8
1.0
1.2
Subsampled data
Function fitted to data

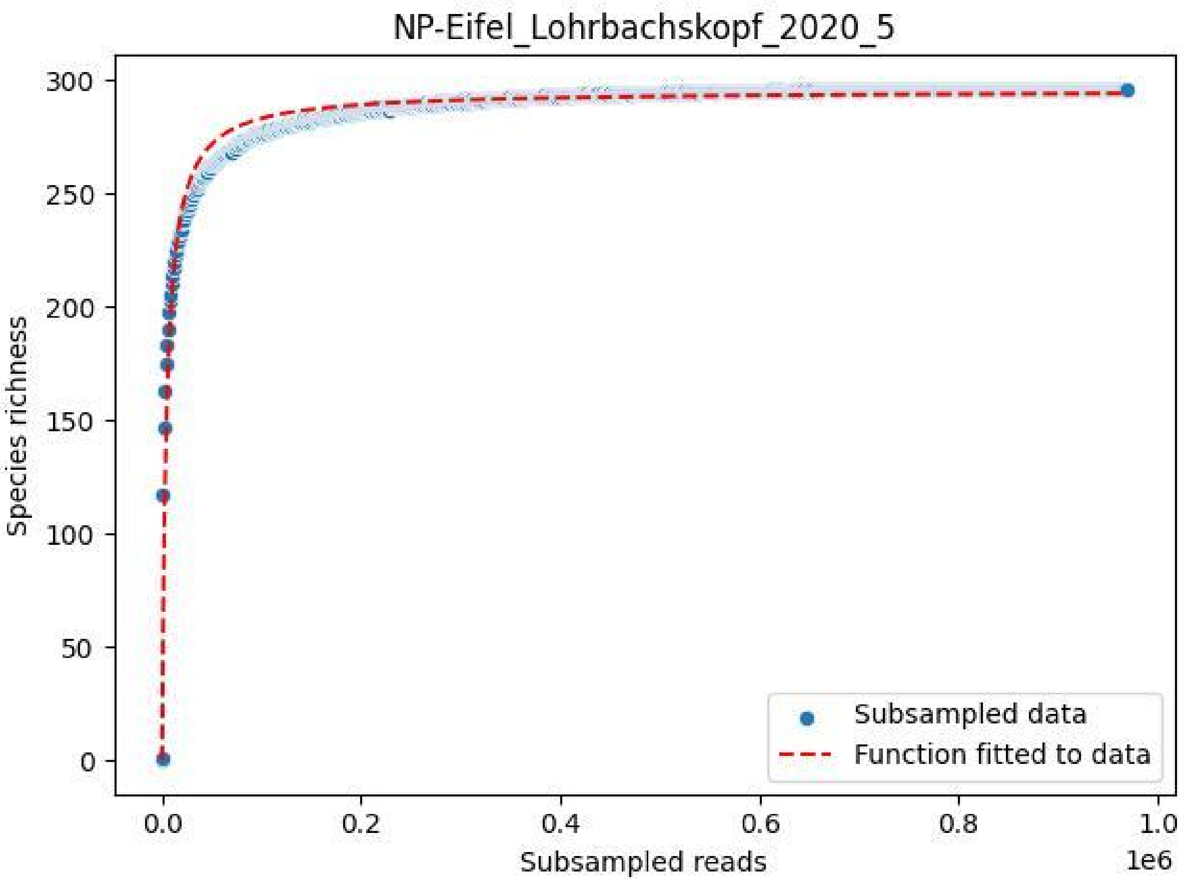
NP-Eifel_Lohrbachskopf_2020_5
Species richness
Subsampled reads
1e6
Subsampled data
Function fitted to data
0
50
100
150
200
250
300
0.0
0.2
0.4
0.6
0.8
1.0

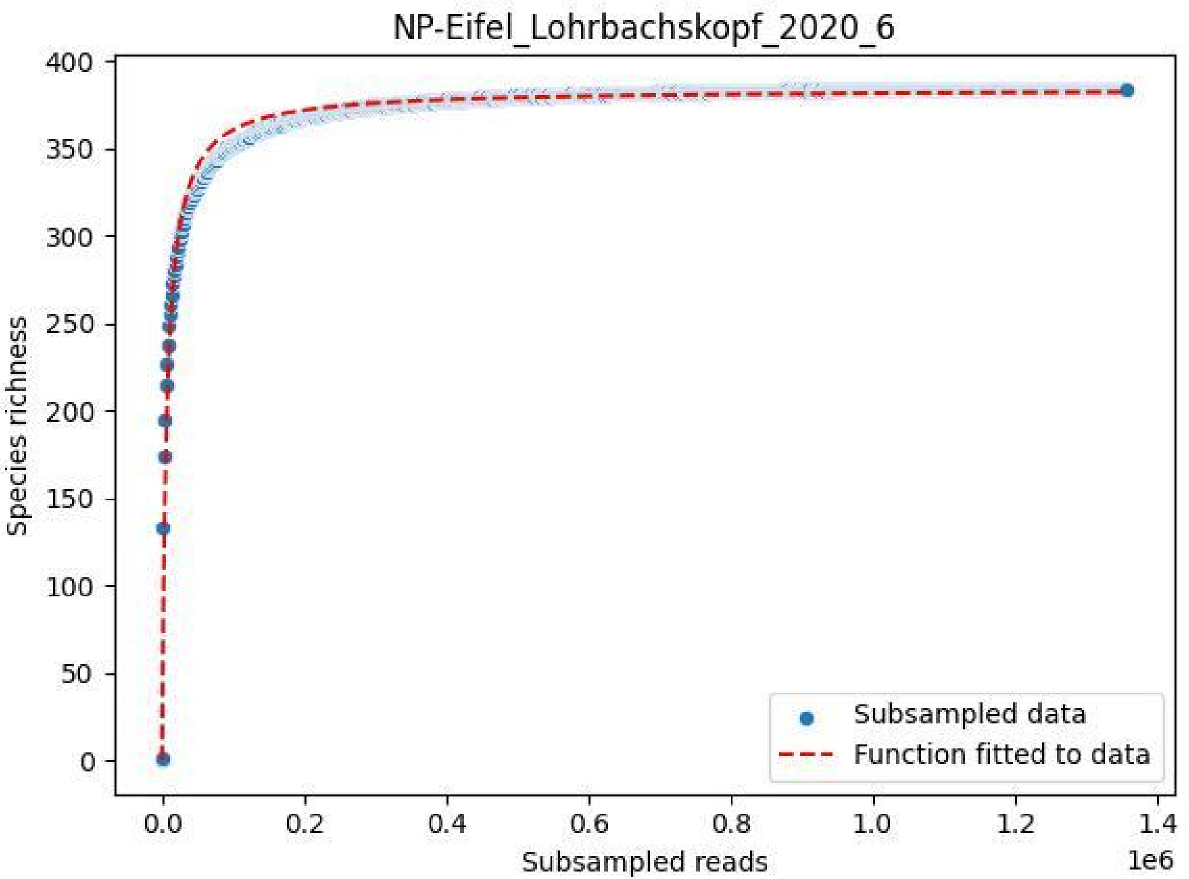
NP-Eifel_Lohrbachskopf_2020_6
Species richness
Subsampled reads
1e6
0.0
0.2
0.4
0.6
0.8
1.0
1.2
1.4
0
50
100
150
200
250
300
350
400
Subsampled data
Function fitted to data

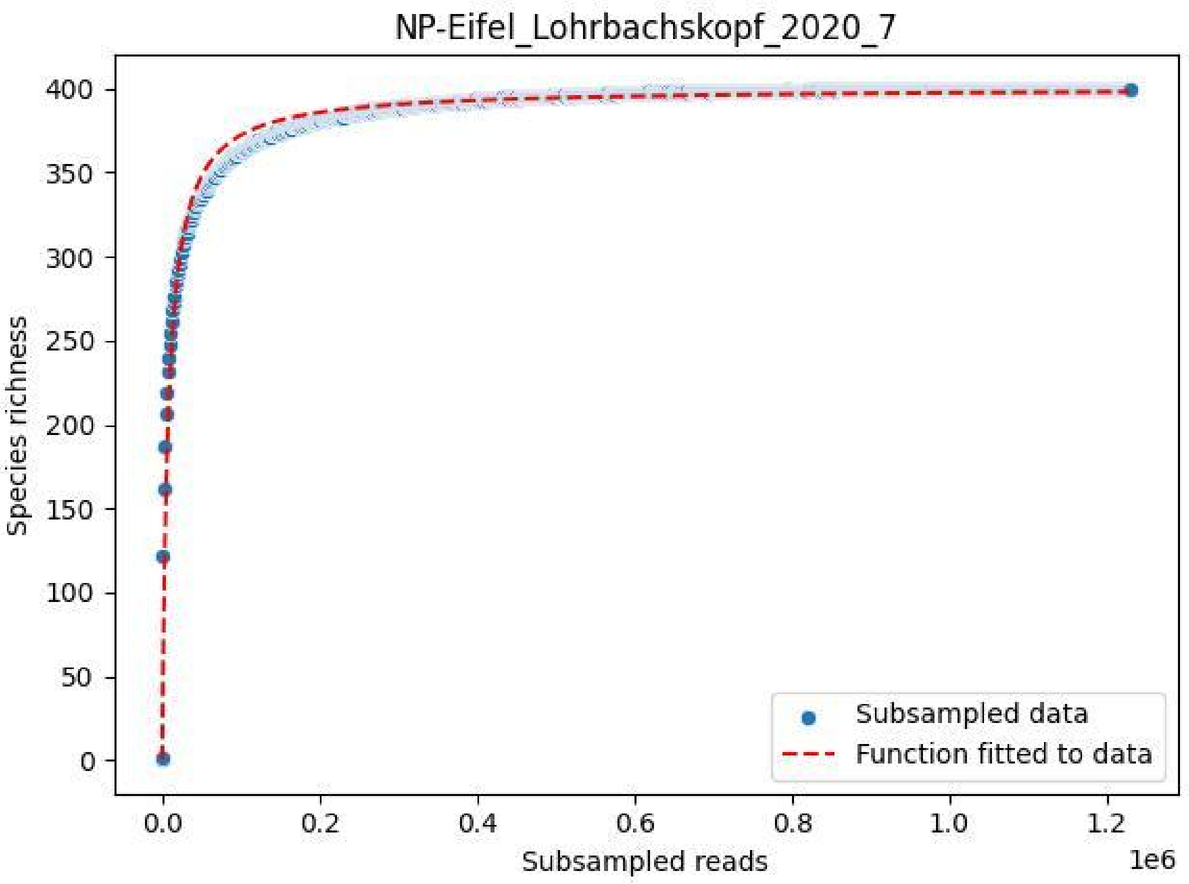
NP-Eifel_Lohrbachskopf_2020_7
Species richness
Subsampled reads
1e6
0
50
100
150
200
250
300
350
400
0.0
0.2
0.4
0.6
0.8
1.0
1.2
Subsampled data
Function fitted to data

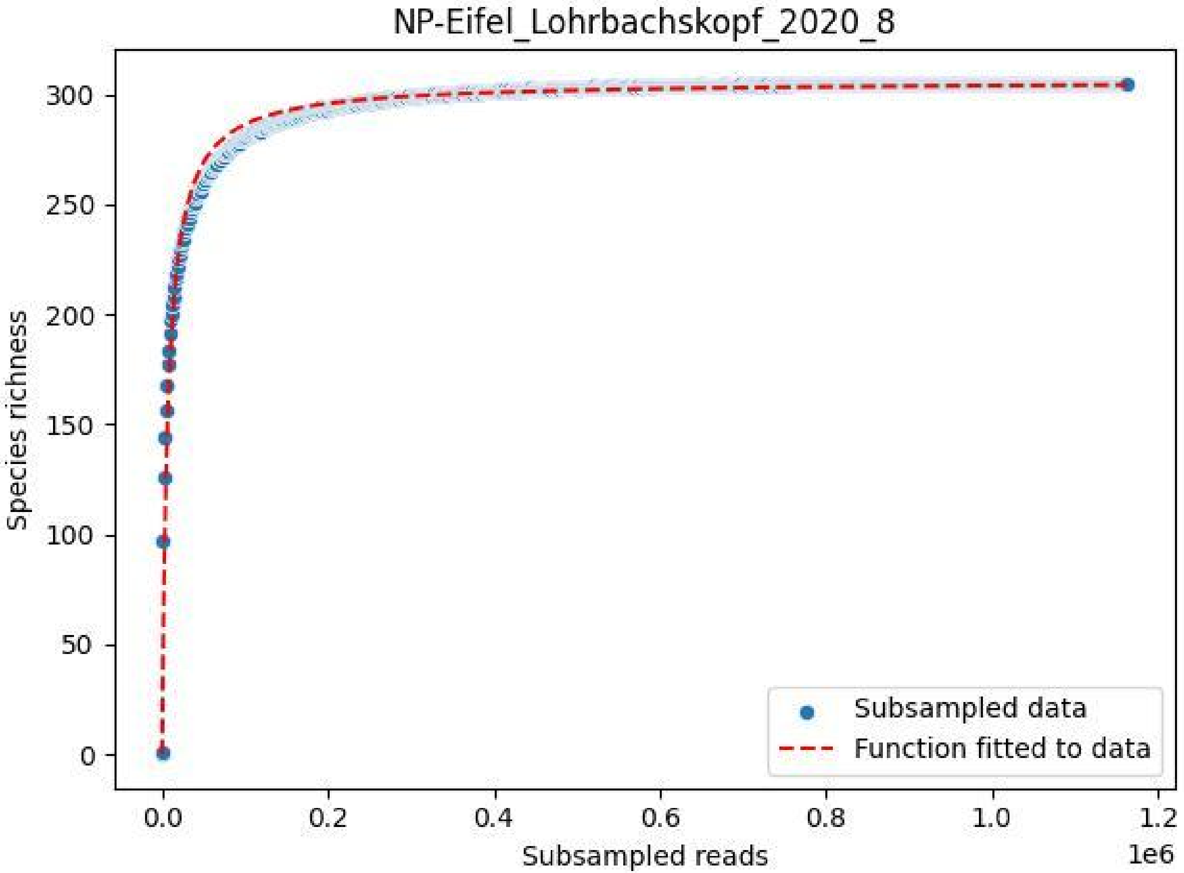

NP-Eifel_Lohrbachskopf_2020_8
Species richness
Subsampled reads
1e6
0
50
100
150
200
250
300
0.0
0.2
0.4
0.6
0.8
1.0
1.2
Subsampled data
Function fitted to data

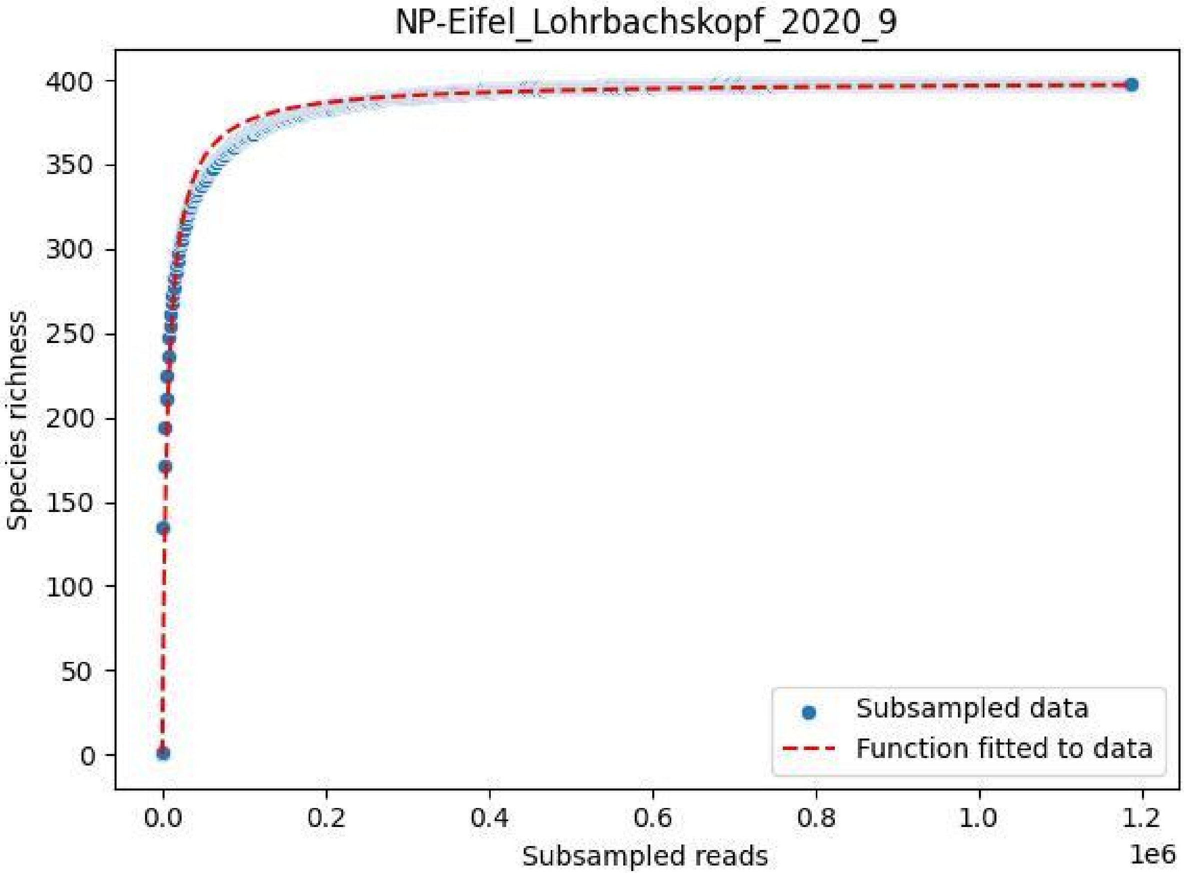
NP-Eifel_Lohrbachskopf_2020_9
Species richness
Subsampled reads
1e6
0
50
100
150
200
250
300
350
400
0.0
0.2
0.4
0.6
0.8
1.0
1.2
Subsampled data
Function fitted to data

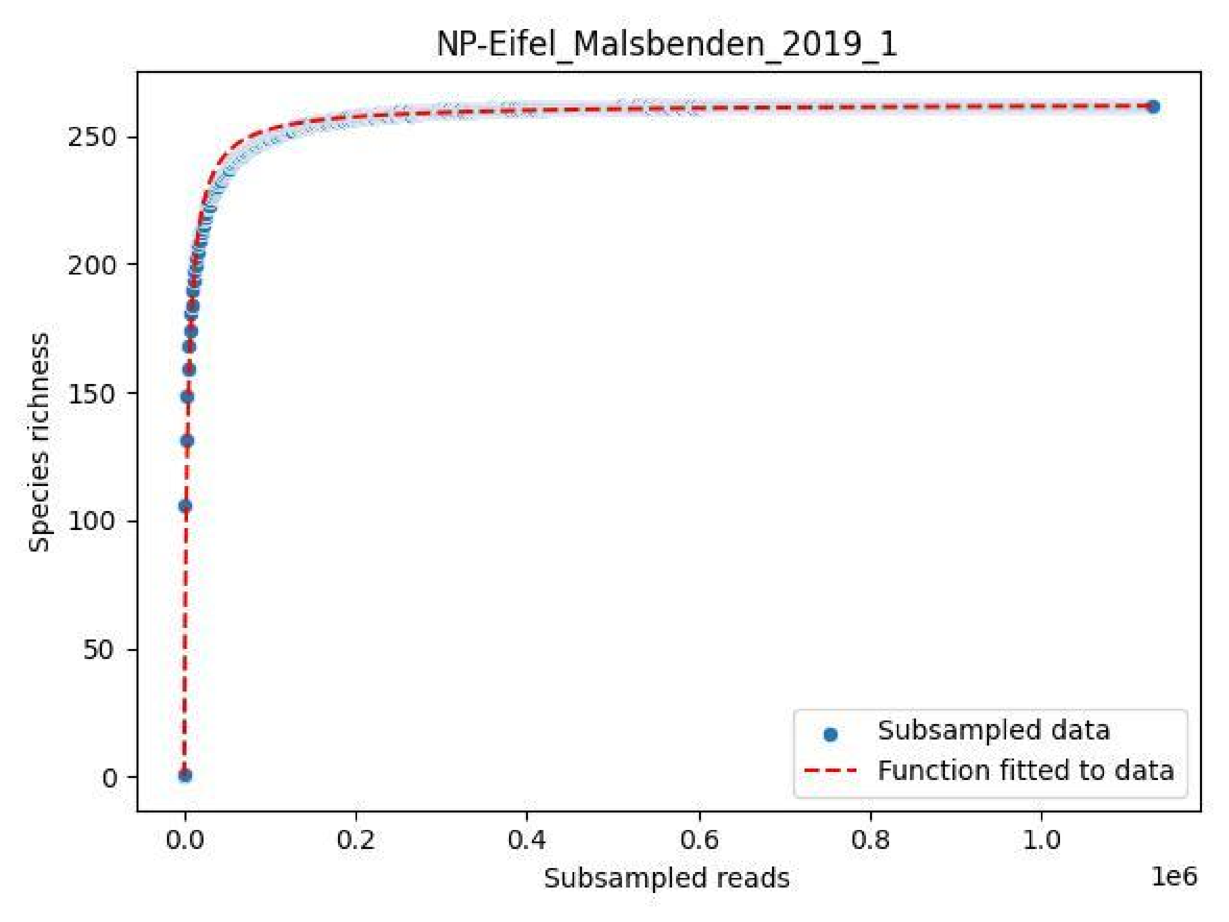
NP-Eifel_Malsbenden_2019_1
Species richness
250
200
150
100
50
0
0.0
0.2
0.4
0.6
0.8
1.0
1e6
Subsampled reads
Subsampled data
Function fitted to data

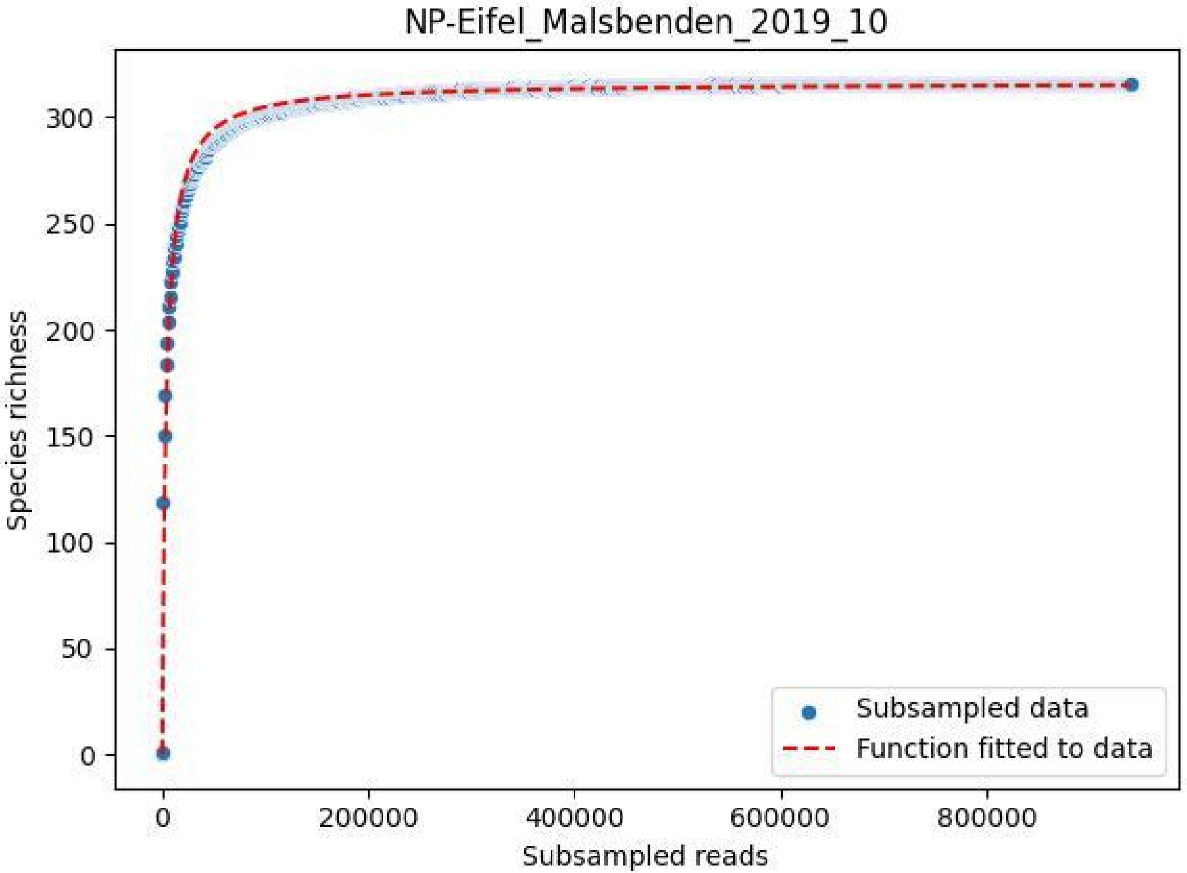
NP-Eifel_Malsbenden_2019_10
Species richness
Subsampled reads
0
50
100
150
200
250
300
0
200000
400000
600000
800000
Subsampled data
Function fitted to data

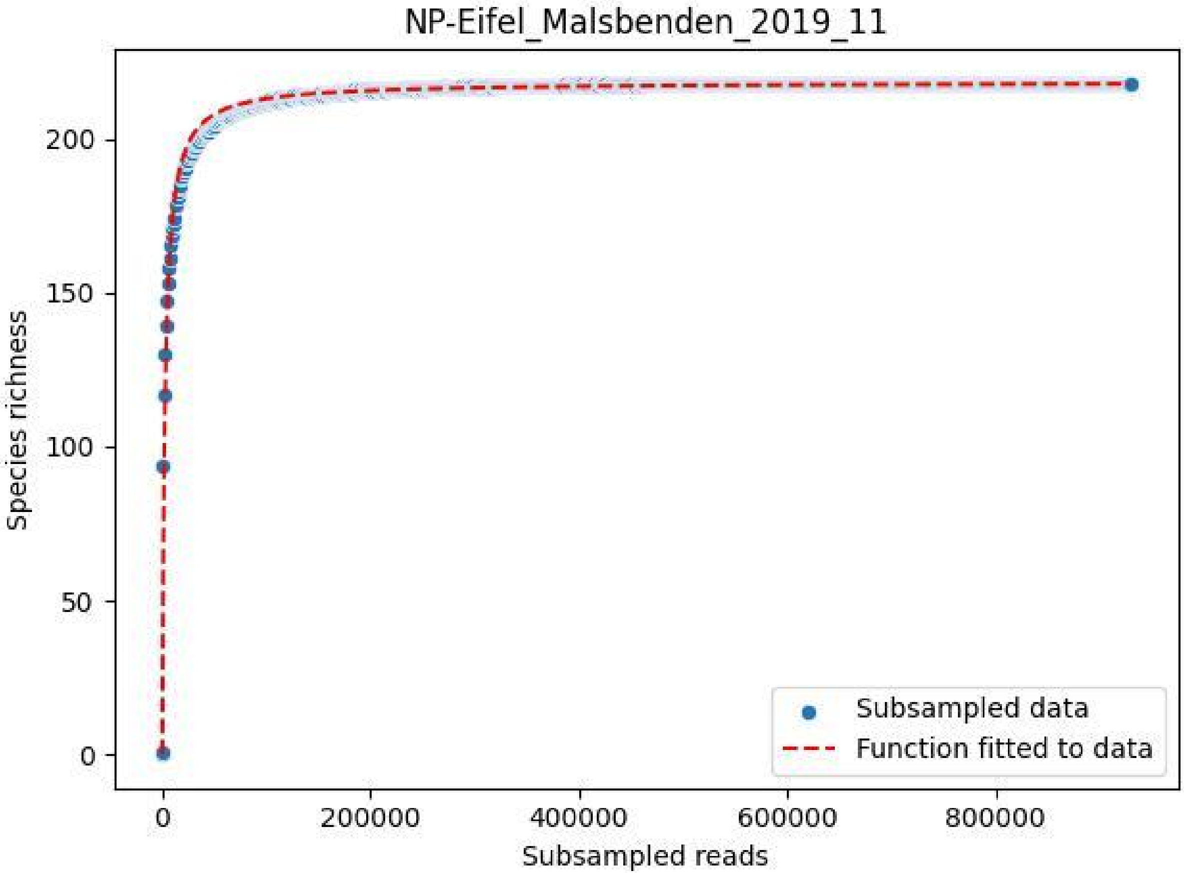

NP-Eifel_Malsbenden_2019_11
Species richness
Subsampled reads
0
50
100
150
200
0
200000
400000
600000
800000
Subsampled data
Function fitted to data

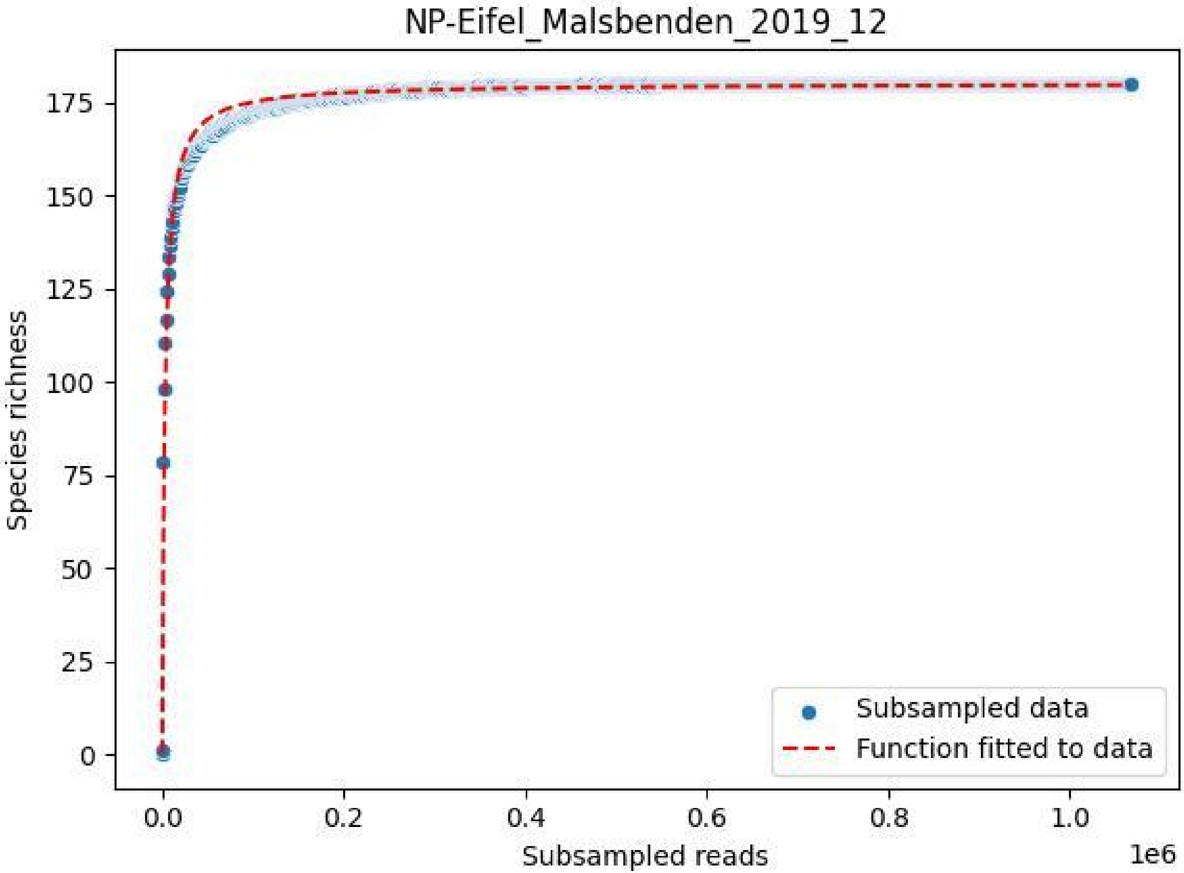

NP-Eifel_Malsbenden_2019_12
Species richness
Subsampled reads
1e6
0
25
50
75
100
125
150
175
0.0
0.2
0.4
0.6
0.8
1.0
Subsampled data
Function fitted to data

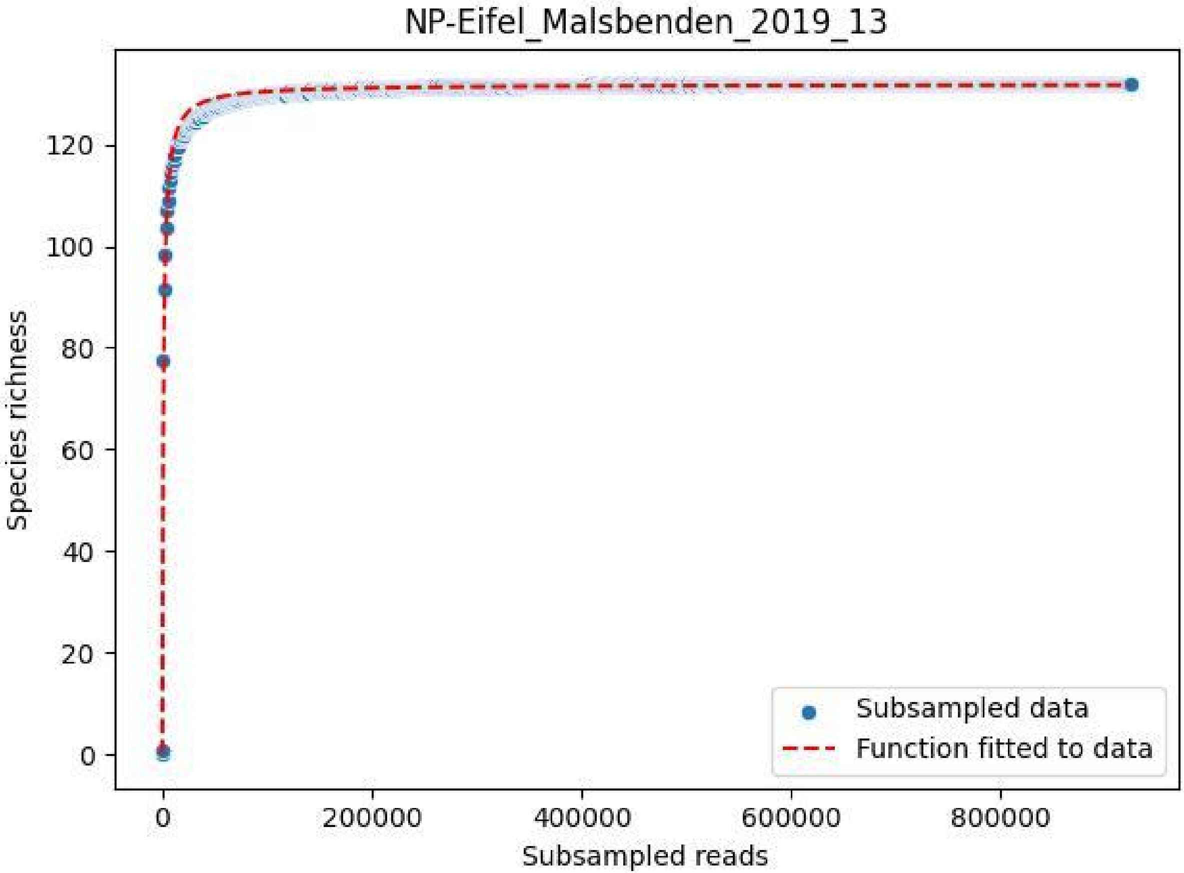
NP-Eifel_Malsbenden_2019_13
Species richness
Subsampled reads
0
20
40
60
80
100
120
0
200000
400000
600000
800000
Subsampled data
Function fitted to data

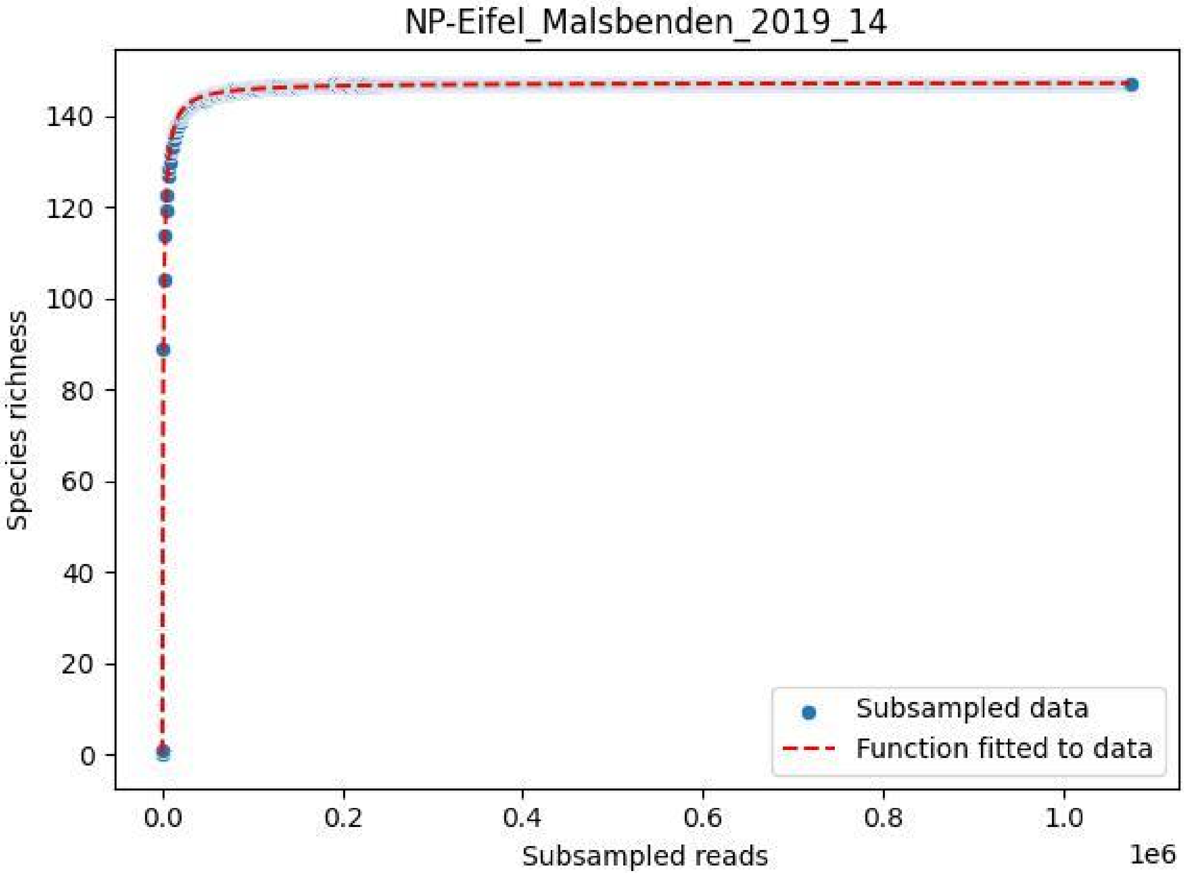
NP-Eifel_Malsbenden_2019_14
Species richness
Subsampled reads
1e6
Subsampled data
Function fitted to data
0
20
40
60
80
100
120
140
0.0
0.2
0.4
0.6
0.8
1.0

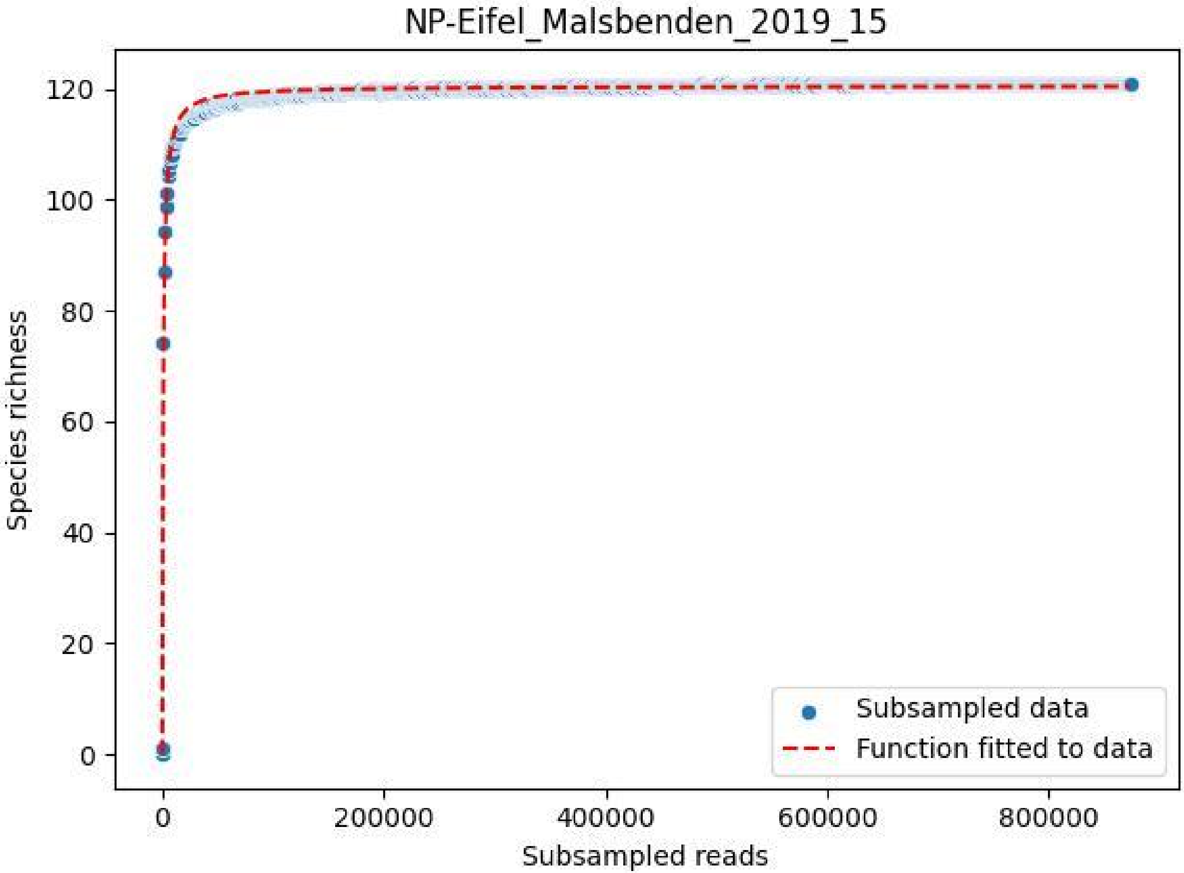
NP-Eifel_Malsbenden_2019_15
Species richness
Subsampled reads
0
20
40
60
80
100
120
0
200000
400000
600000
800000
Subsampled data
Function fitted to data

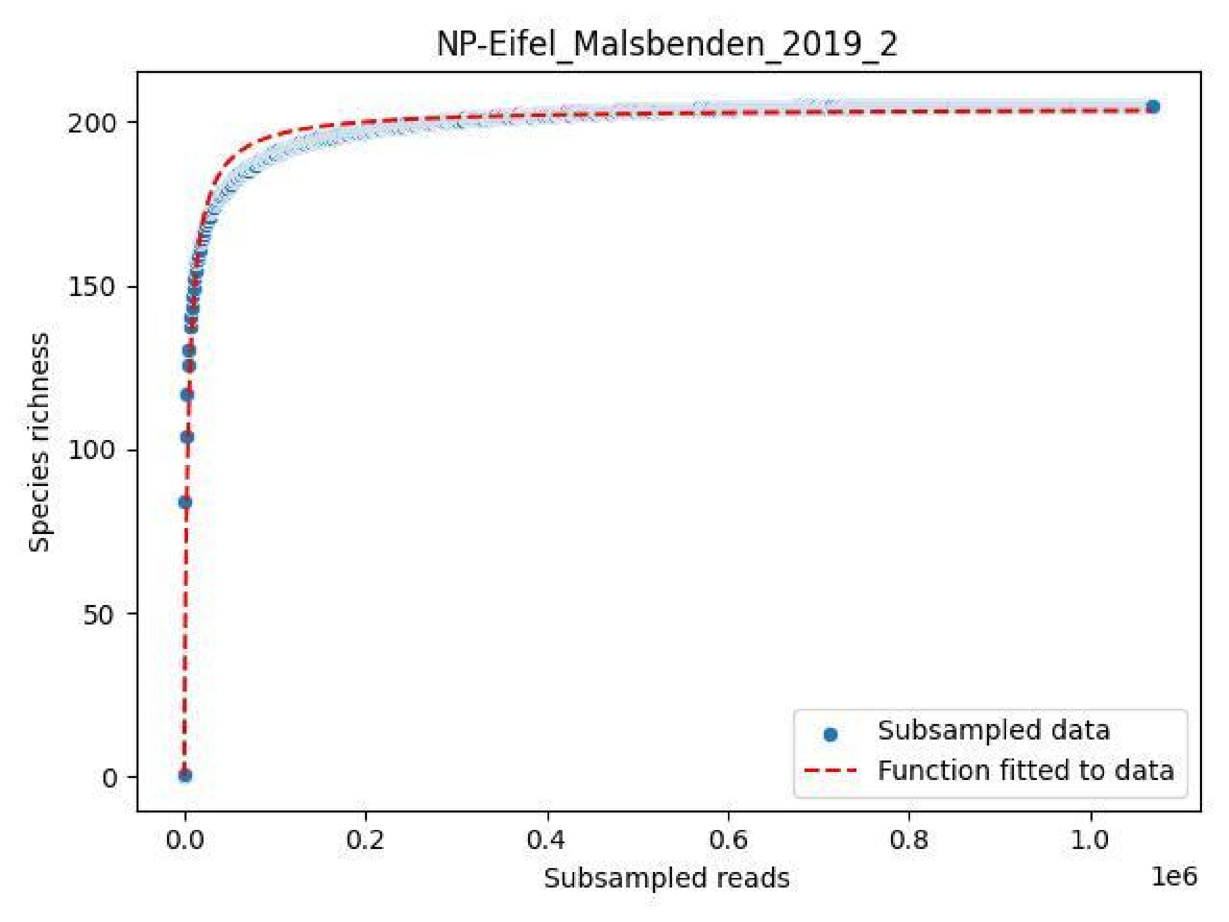
NP-Eifel_Malsbenden_2019_2
Species richness
0
50
100
150
200
0.0
0.2
0.4
0.6
0.8
1.0
1e6
Subsampled reads
Subsampled data
Function fitted to data

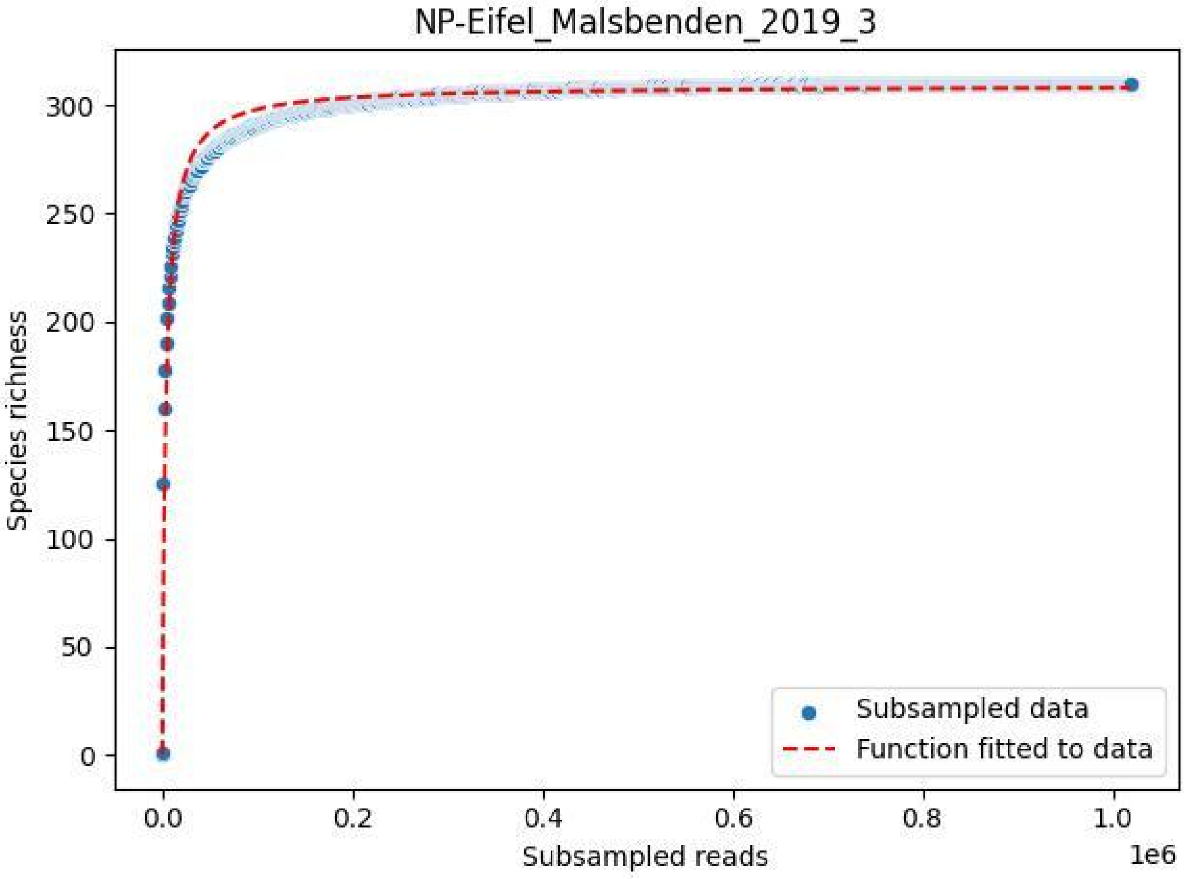
NP-Eifel_Malsbenden_2019_3
Species richness
300
250
200
150
100
50
0
0.0
0.2
0.4
0.6
0.8
1.0
1e6
Subsampled reads
Subsampled data
Function fitted to data

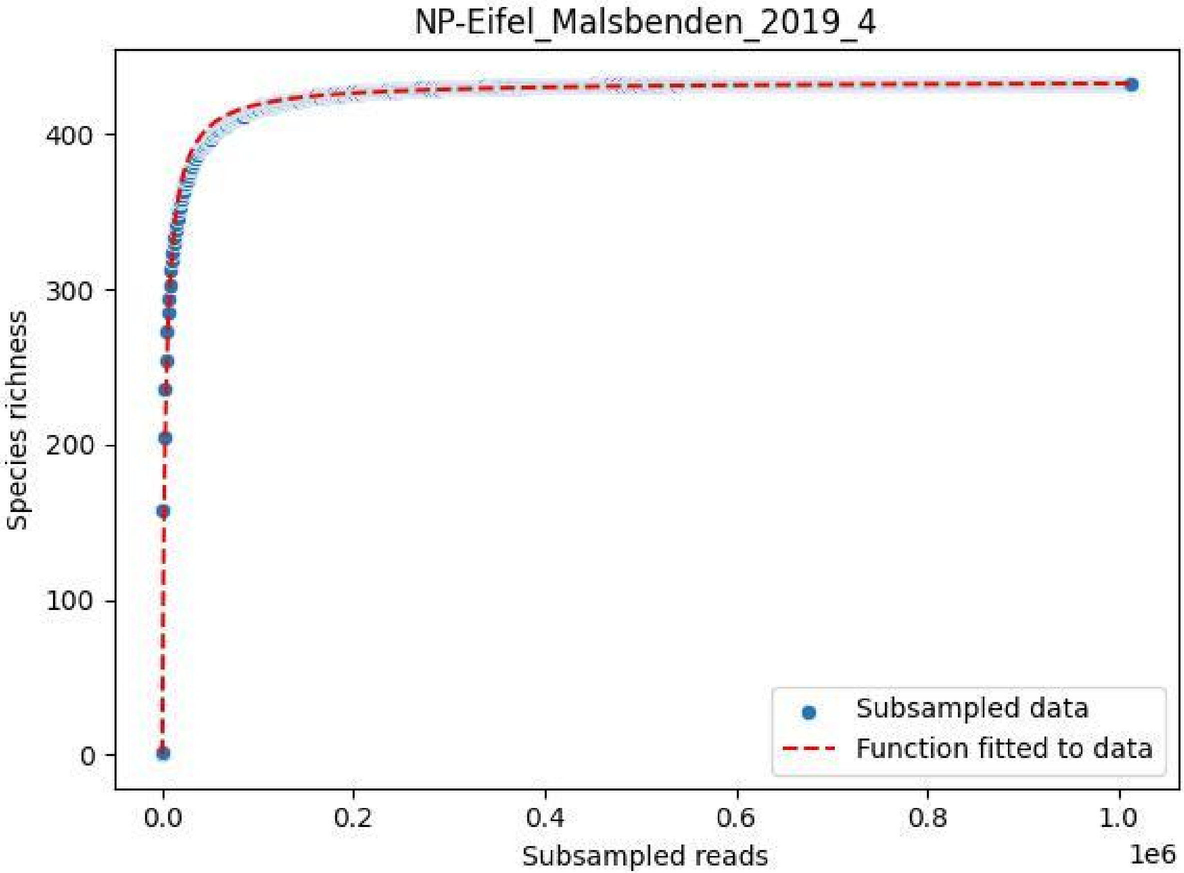

NP-Eifel_Malsbenden_2019_4
Species richness
Subsampled reads
1e6
0
100
200
300
400
0.0
0.2
0.4
0.6
0.8
1.0
Subsampled data
Function fitted to data

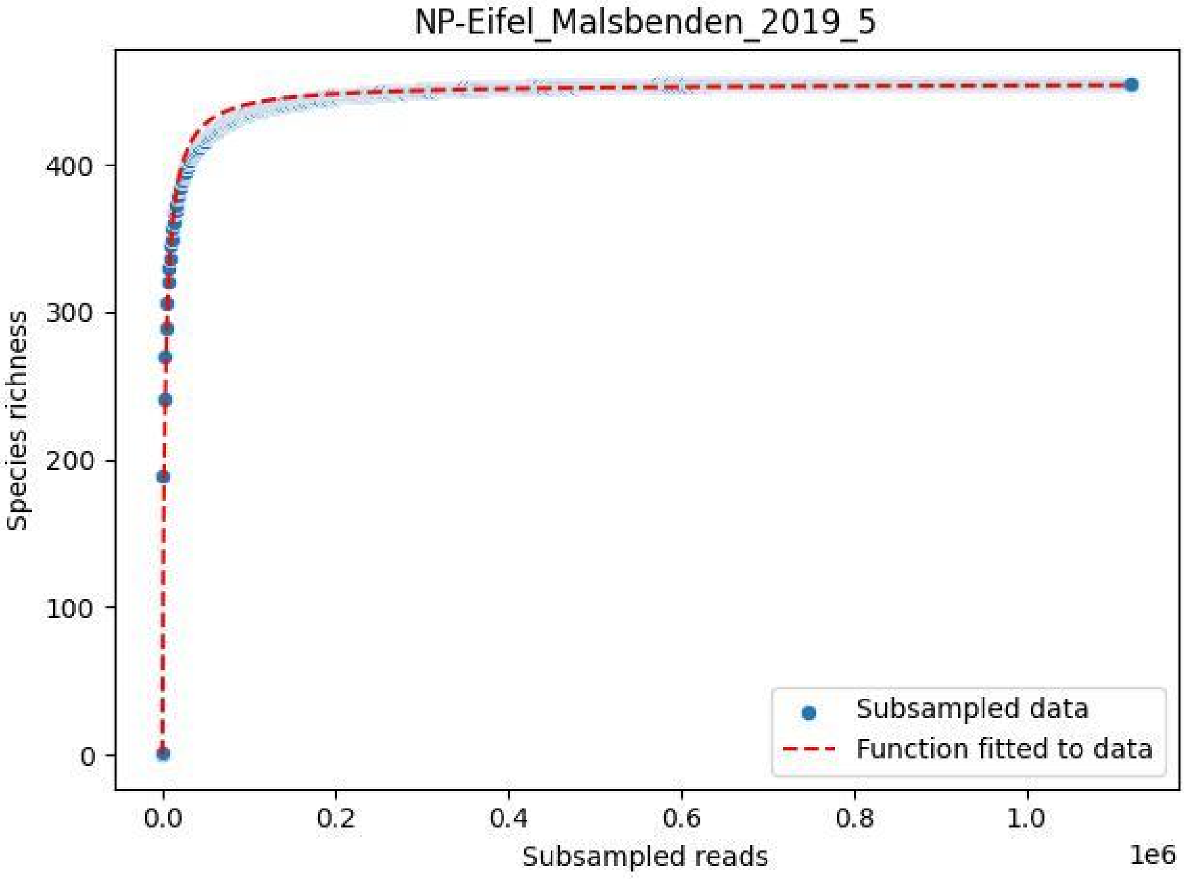
NP-Eifel_Malsbenden_2019_5
Species richness
400
300
200
100
0
0.0
0.2
0.4
0.6
0.8
1.0
1e6
Subsampled reads
Subsampled data
Function fitted to data

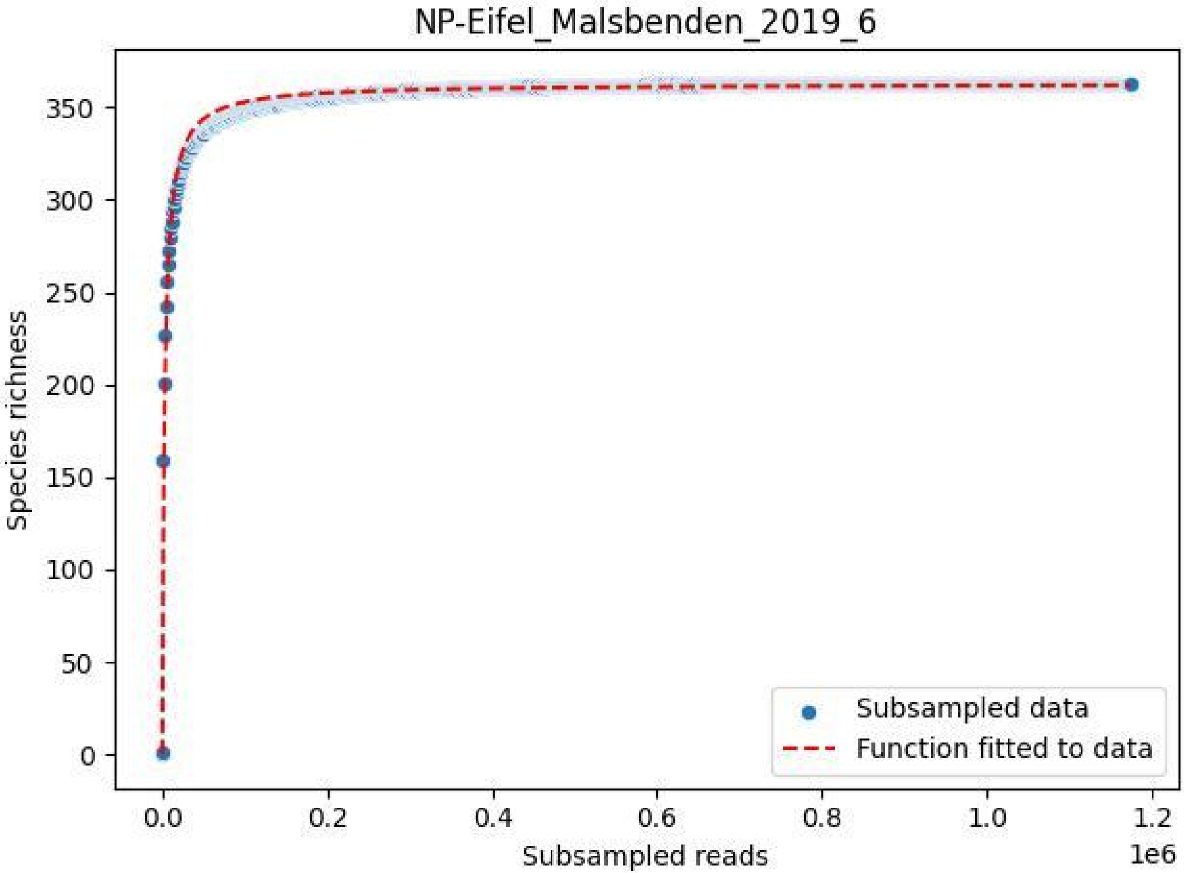
NP-Eifel_Malsbenden_2019_6
Species richness
Subsampled reads
1e6
0
50
100
150
200
250
300
350
0.0
0.2
0.4
0.6
0.8
1.0
1.2
Subsampled data
Function fitted to data

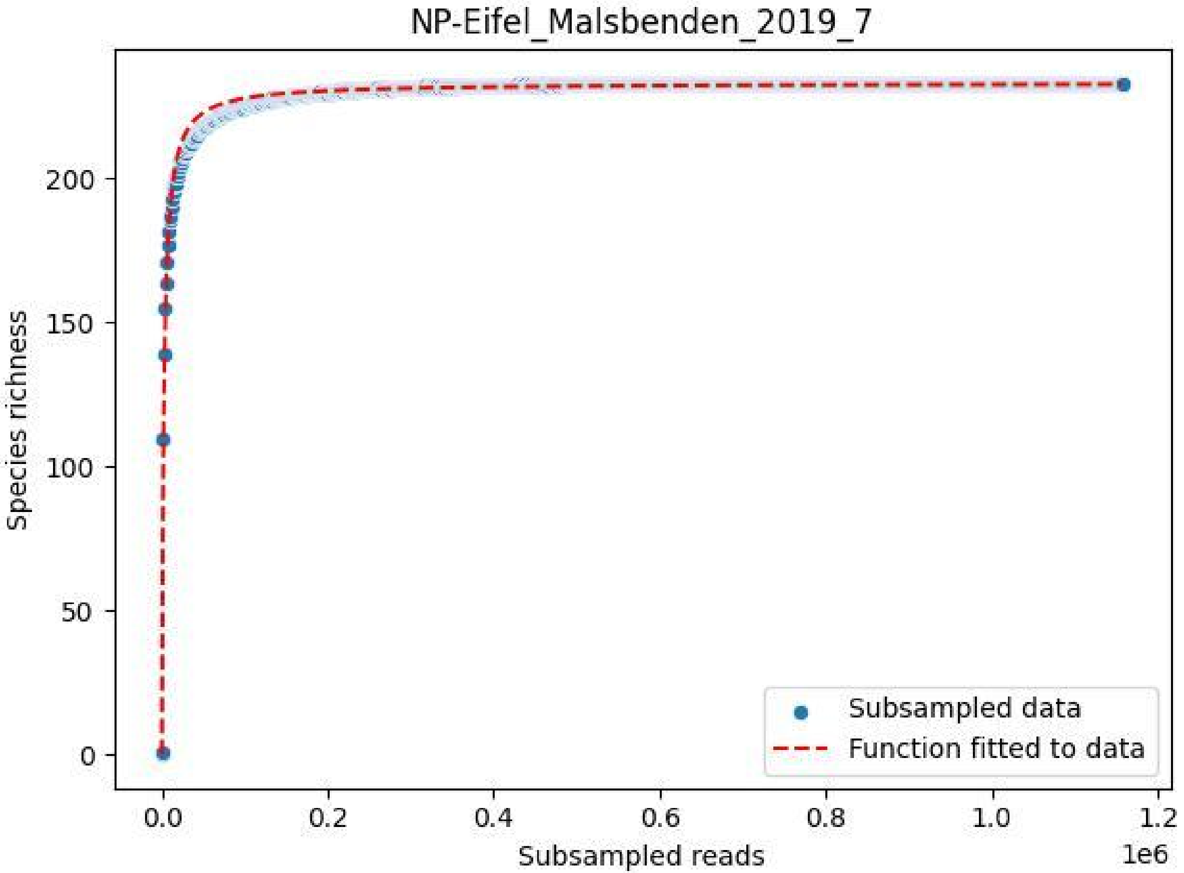
NP-Eifel_Malsbenden_2019_7
Species richness
Subsampled reads
1e6
Subsampled data
Function fitted to data
0
50
100
150
200
0.0
0.2
0.4
0.6
0.8
1.0
1.2

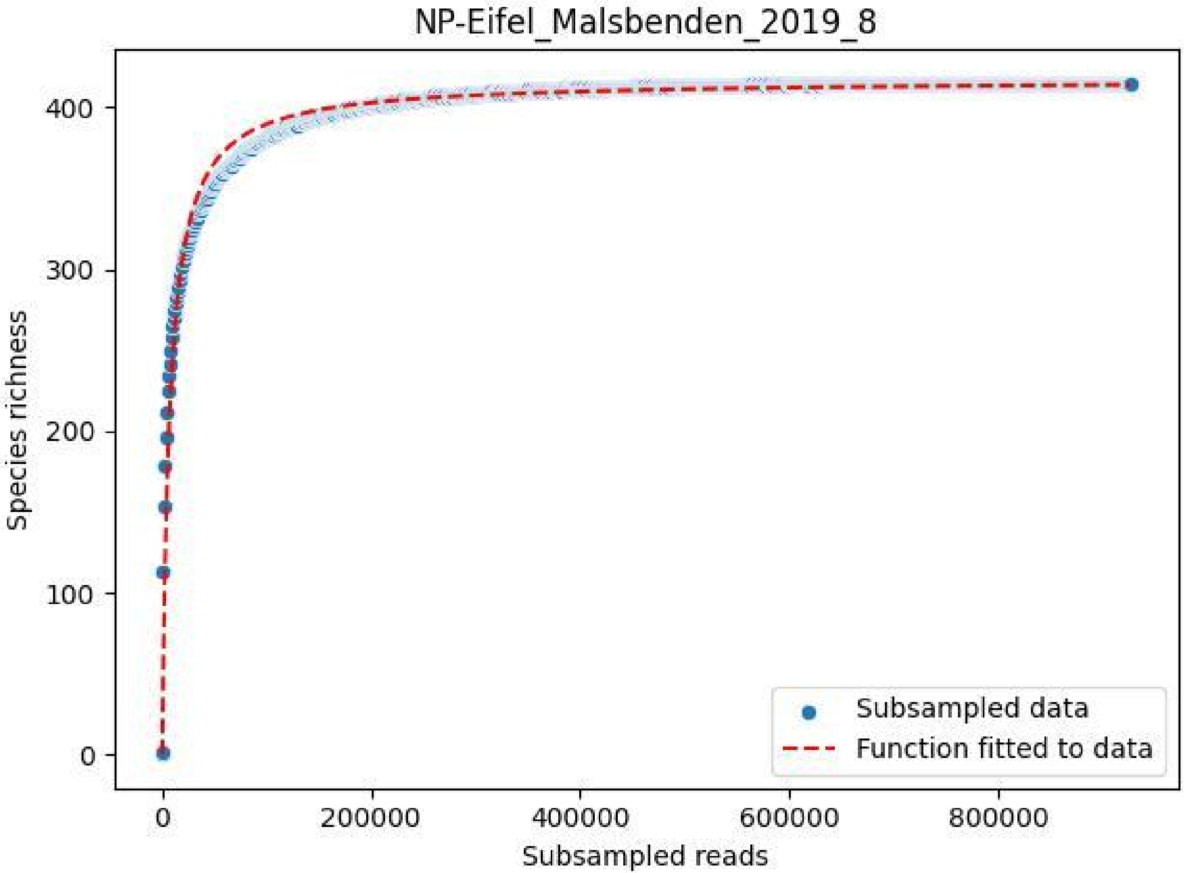
NP-Eifel_Malsbenden_2019_8
Species richness
Subsampled reads
0
100
200
300
400
0
200000
400000
600000
800000
Subsampled data
Function fitted to data

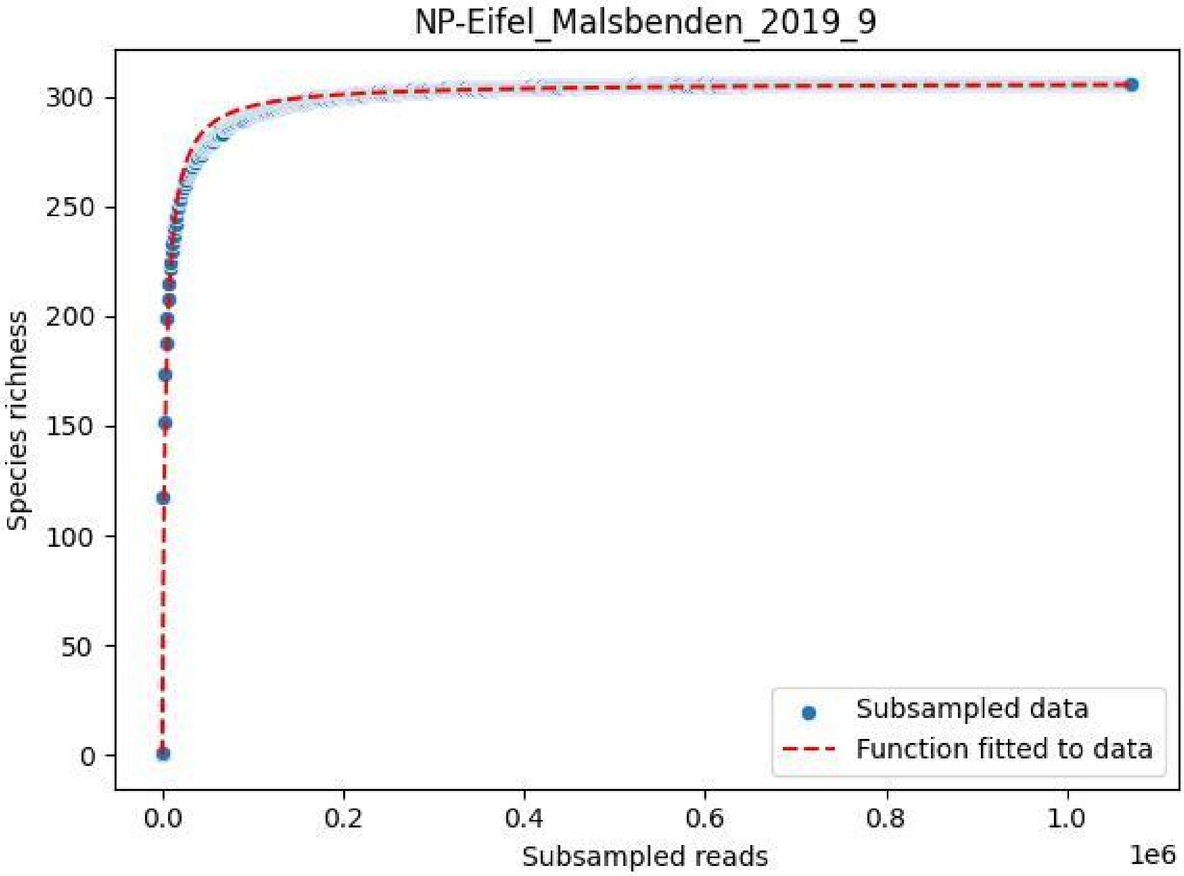
NP-Eifel_Malsbenden_2019_9
Species richness
Subsampled reads
1e6
Subsampled data
Function fitted to data
0
50
100
150
200
250
300
0.0
0.2
0.4
0.6
0.8
1.0

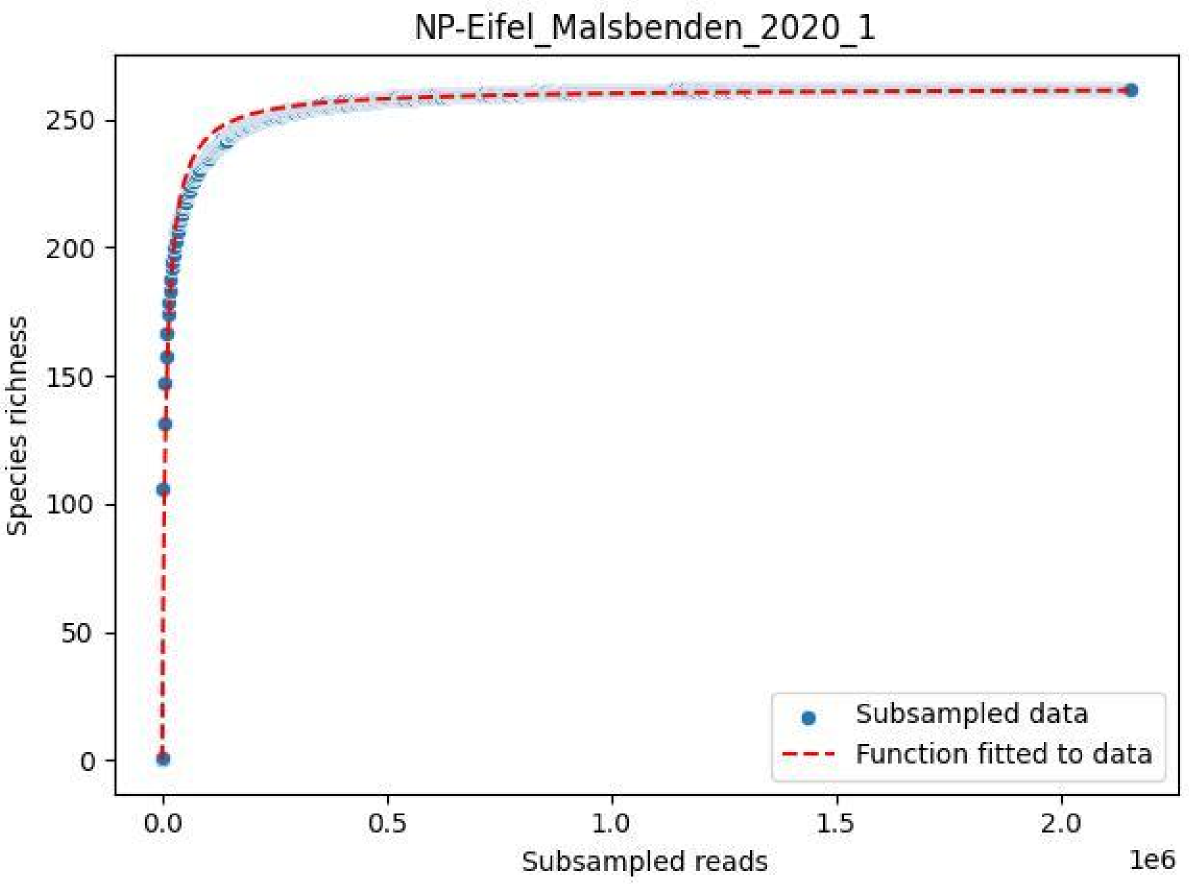
NP-Eifel_Malsbenden_2020_1
Species richness
Subsampled reads
1e6
Subsampled data
Function fitted to data
0
50
100
150
200
250
0.0
0.5
1.0
1.5
2.0

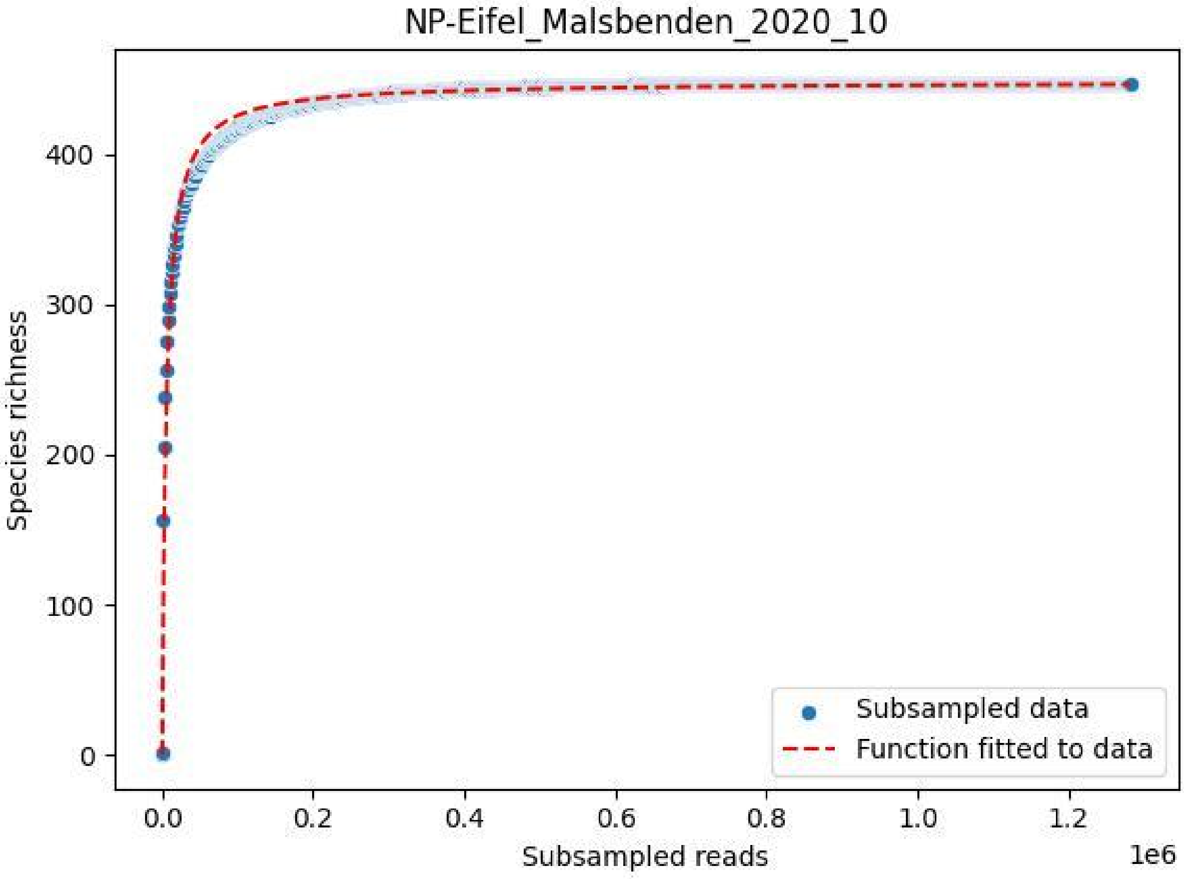
NP-Eifel_Malsbenden_2020_10
Species richness
400
300
200
100
0
0.0
0.2
0.4
0.6
0.8
1.0
1.2
1e6
Subsampled reads
Subsampled data
Function fitted to data

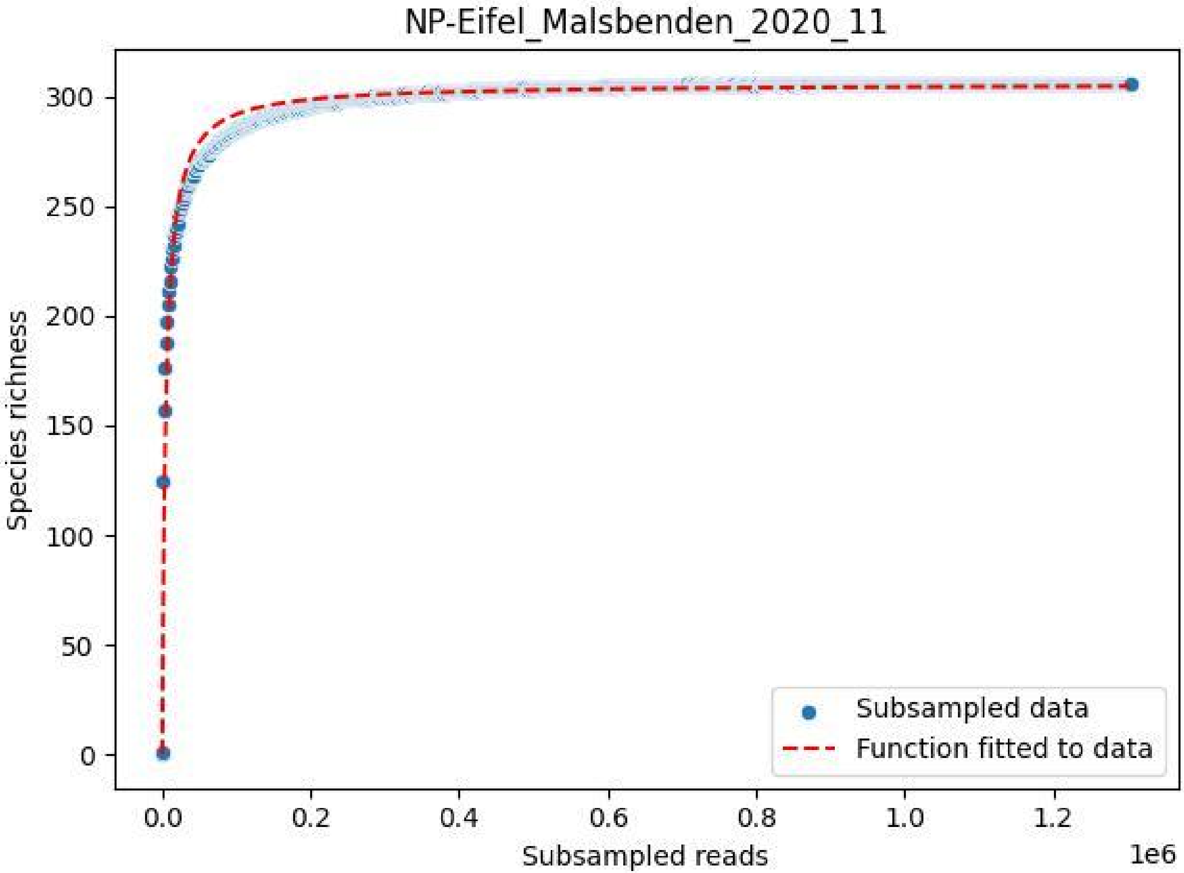
NP-Eifel_Malsbenden_2020_11
Species richness
Subsampled reads
1e6
Subsampled data
Function fitted to data
300
250
200
150
100
50
0
0.0
0.2
0.4
0.6
0.8
1.0
1.2

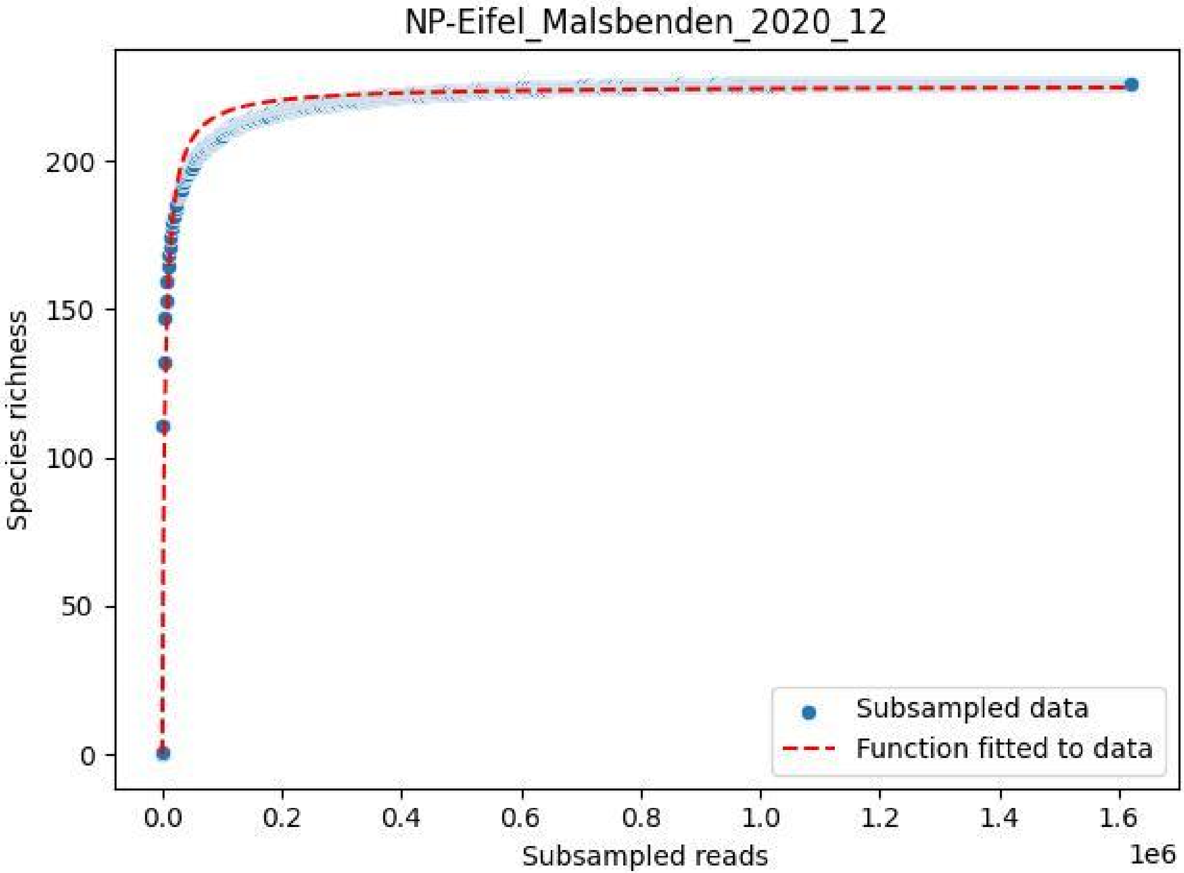
NP-Eifel_Malsbenden_2020_12
Species richness
Subsampled reads
1e6
Subsampled data
Function fitted to data
0
50
100
150
200
0.0
0.2
0.4
0.6
0.8
1.0
1.2
1.4
1.6

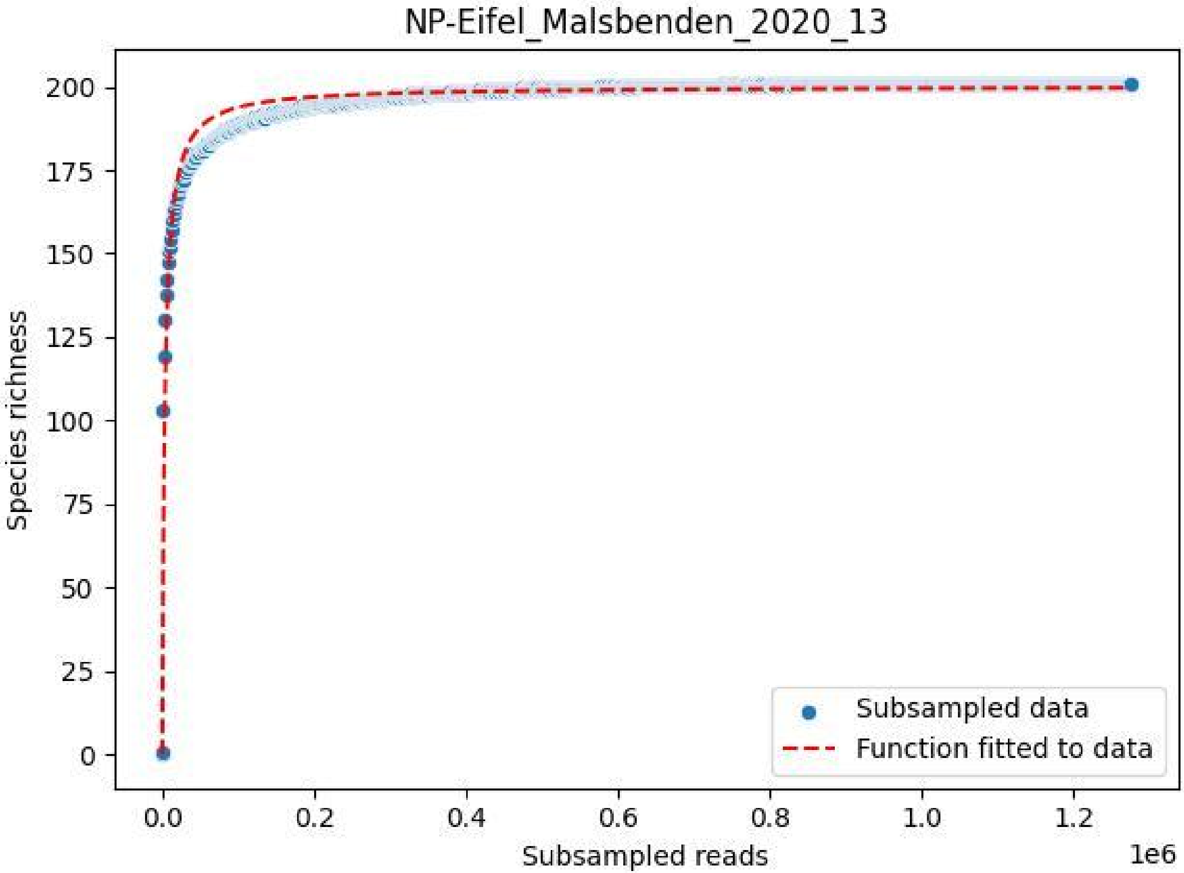

NP-Eifel_Malsbenden_2020_13
Species richness
Subsampled reads
1e6
0
25
50
75
100
125
150
175
200
0.0
0.2
0.4
0.6
0.8
1.0
1.2
Subsampled data
Function fitted to data

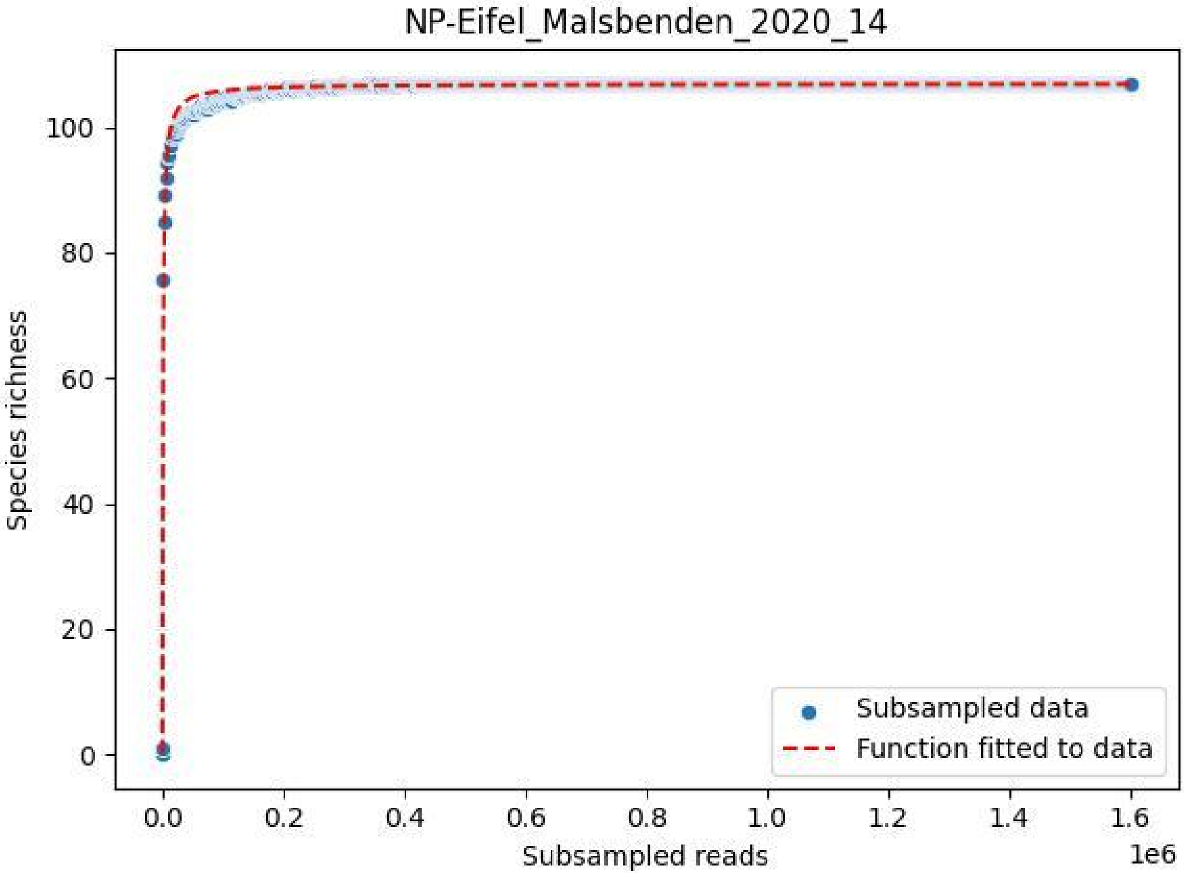
NP-Eifel_Malsbenden_2020_14
Species richness
0
20
40
60
80
100
0.0
0.2
0.4
0.6
0.8
1.0
1.2
1.4
1.6
1e6
Subsampled reads
Subsampled data
Function fitted to data

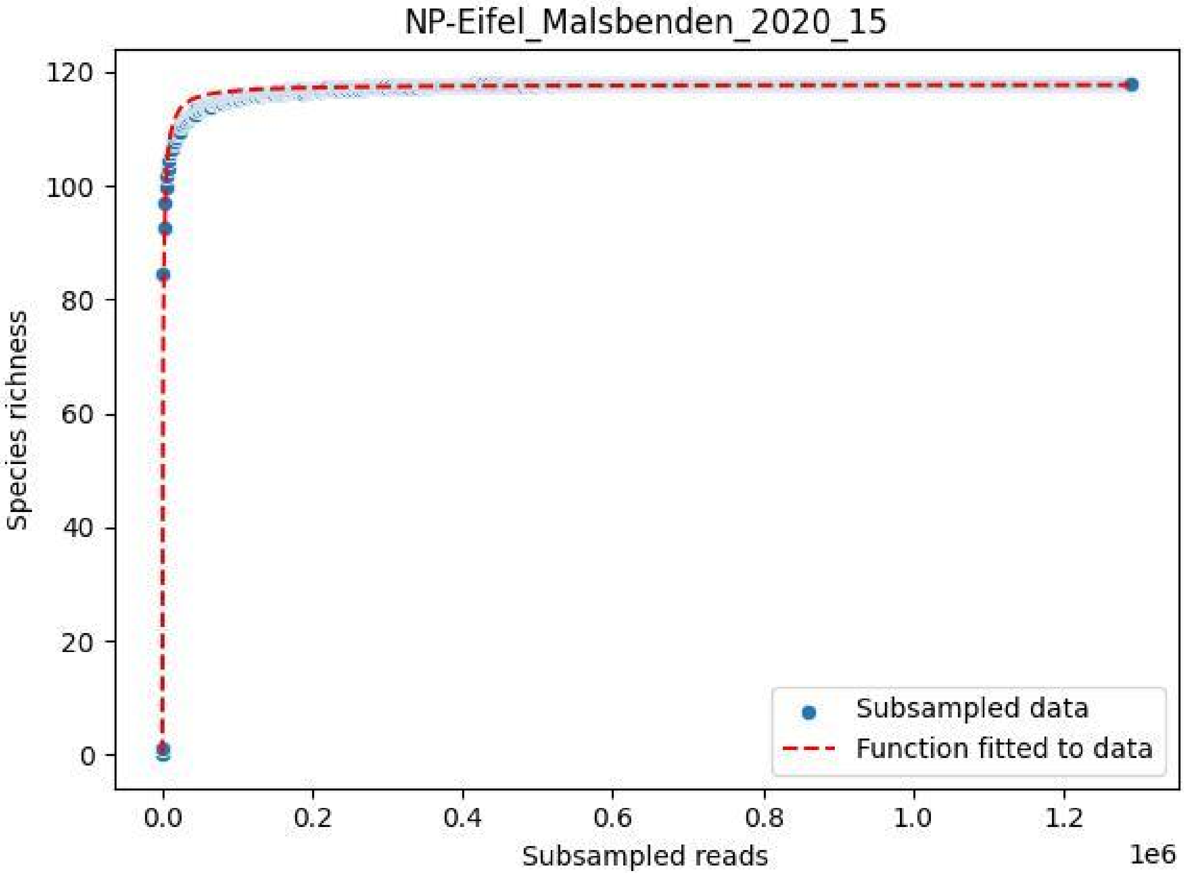
NP-Eifel_Malsbenden_2020_15
Species richness
Subsampled reads
1e6
0
20
40
60
80
100
120
0.0
0.2
0.4
0.6
0.8
1.0
1.2
Subsampled data
Function fitted to data

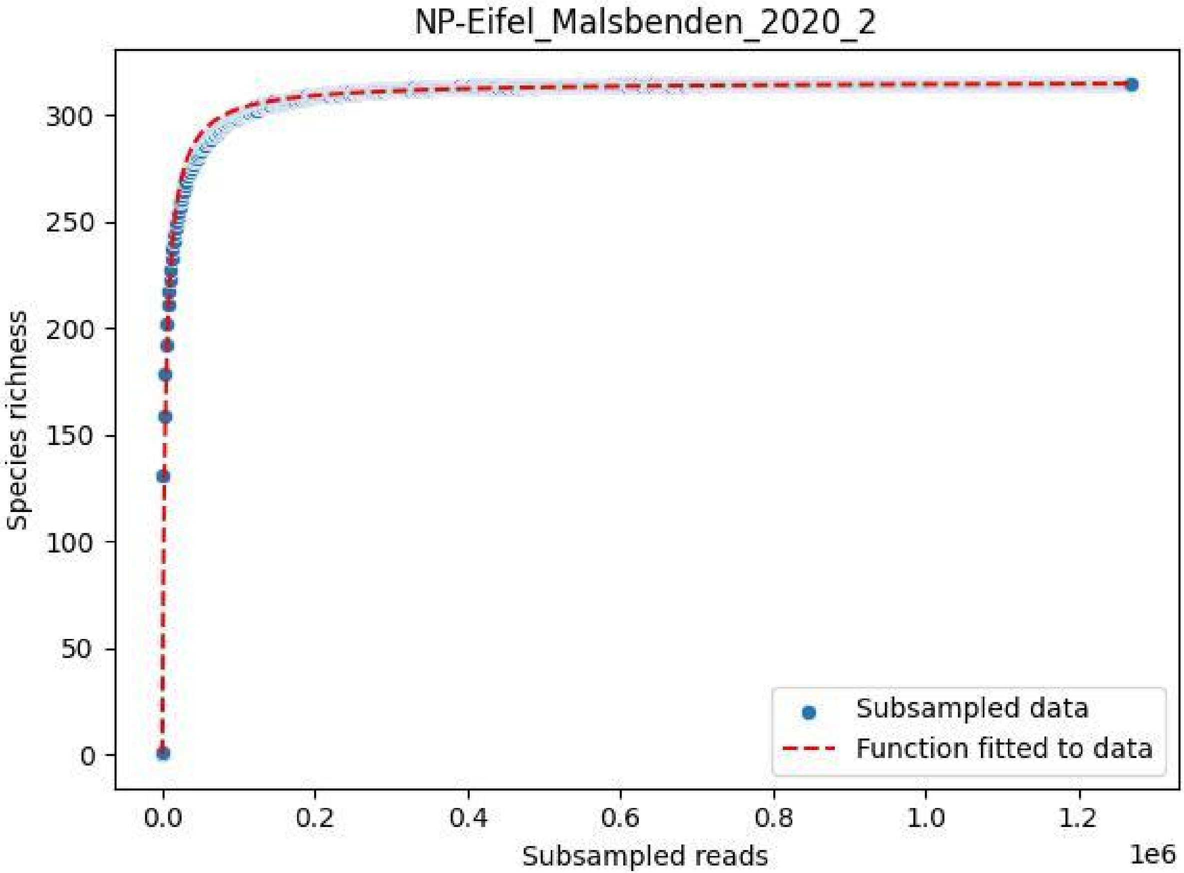

NP-Eifel_Malsbenden_2020_2
Species richness
Subsampled reads
1e6
0
50
100
150
200
250
300
0.0
0.2
0.4
0.6
0.8
1.0
1.2
Subsampled data
Function fitted to data

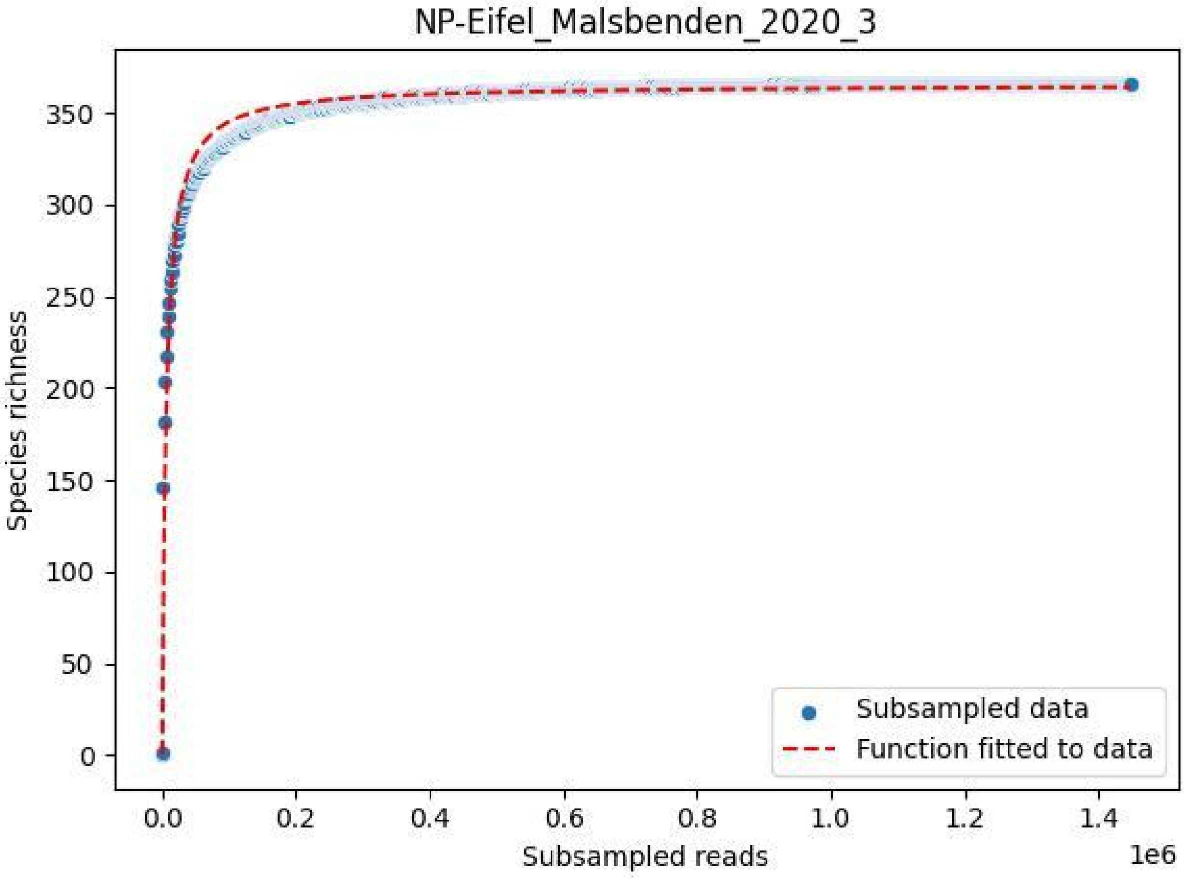
NP-Eifel_Malsbenden_2020_3
Species richness
Subsampled reads
1e6
Subsampled data
Function fitted to data
0
50
100
150
200
250
300
350
0.0
0.2
0.4
0.6
0.8
1.0
1.2
1.4

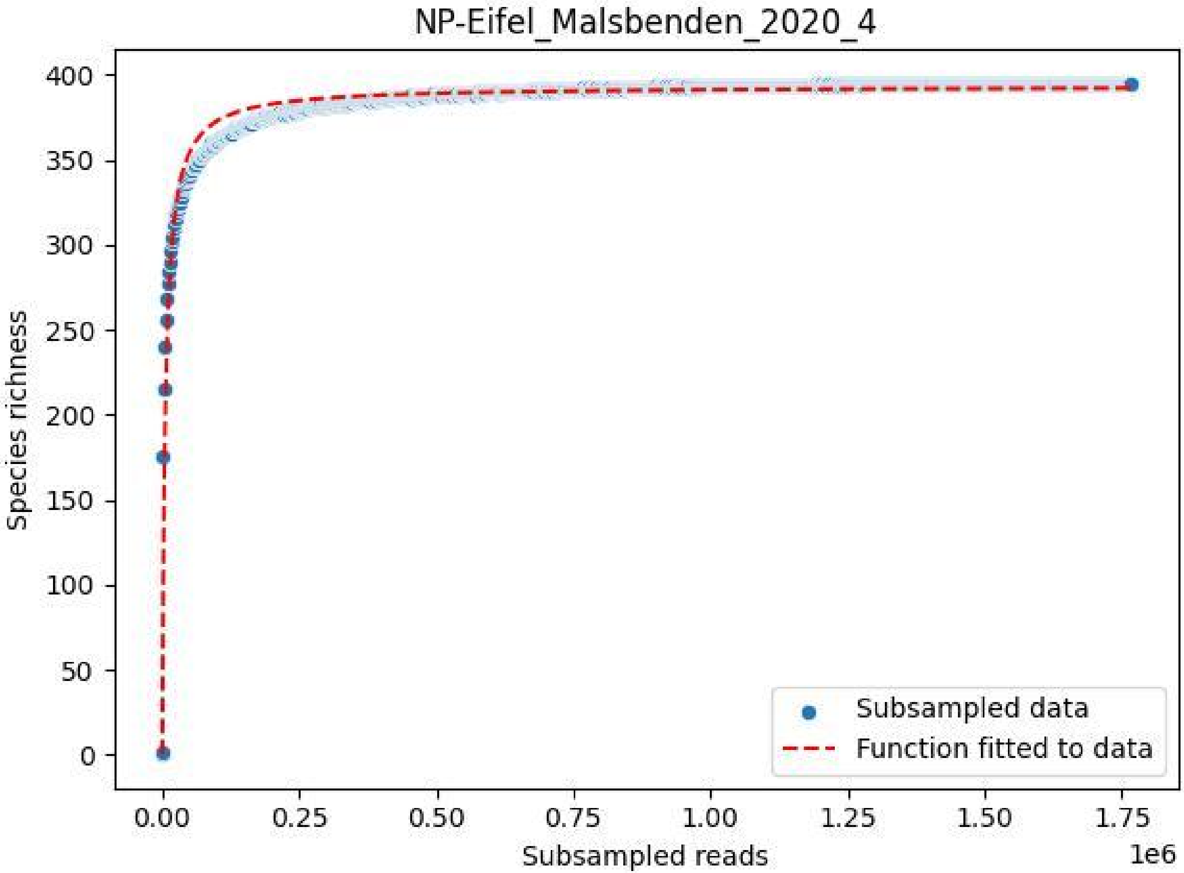
NP-Eifel_Malsbenden_2020_4
Species richness
Subsampled reads
1e6
0
50
100
150
200
250
300
350
400
0.00
0.25
0.50
0.75
1.00
1.25
1.50
1.75
Subsampled data
Function fitted to data

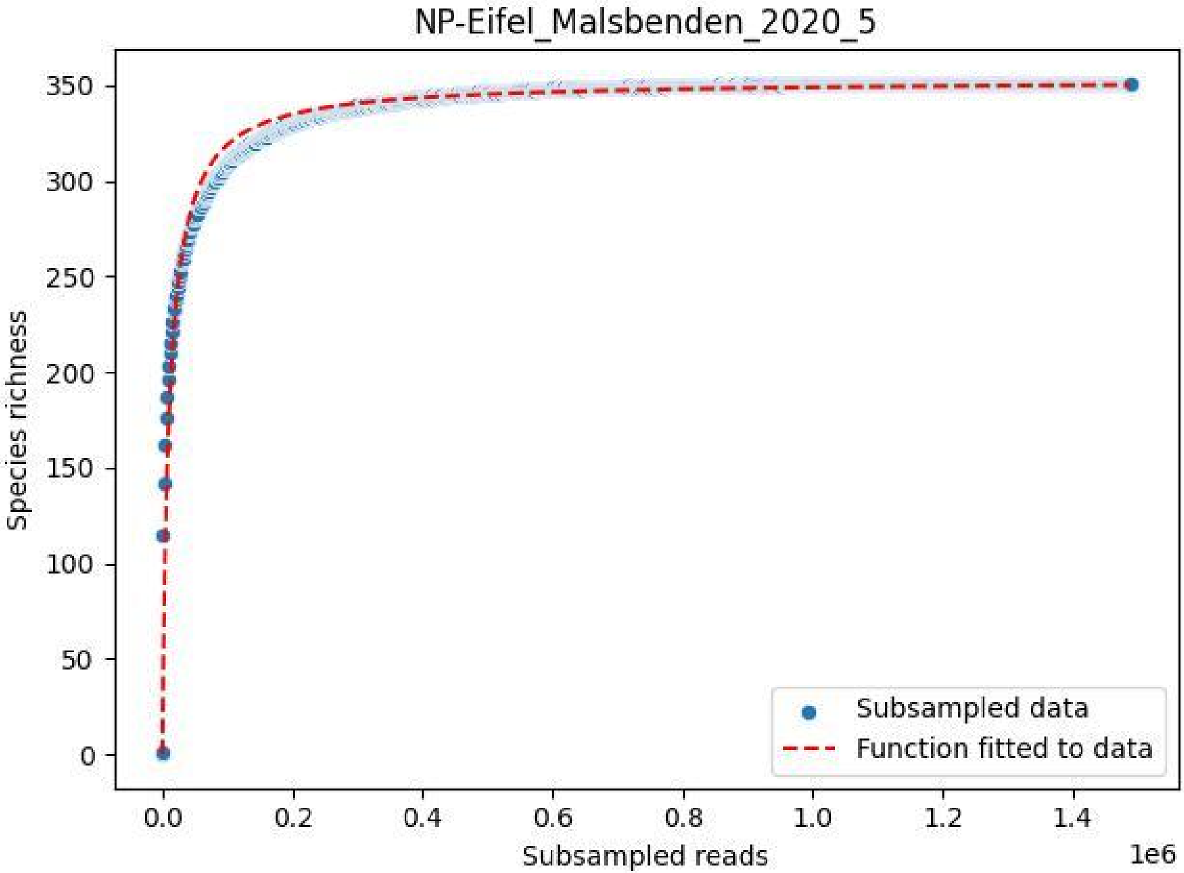

NP-Eifel_Malsbenden_2020_5
Species richness
Subsampled reads
1e6
0
50
100
150
200
250
300
350
0.0
0.2
0.4
0.6
0.8
1.0
1.2
1.4
Subsampled data
Function fitted to data

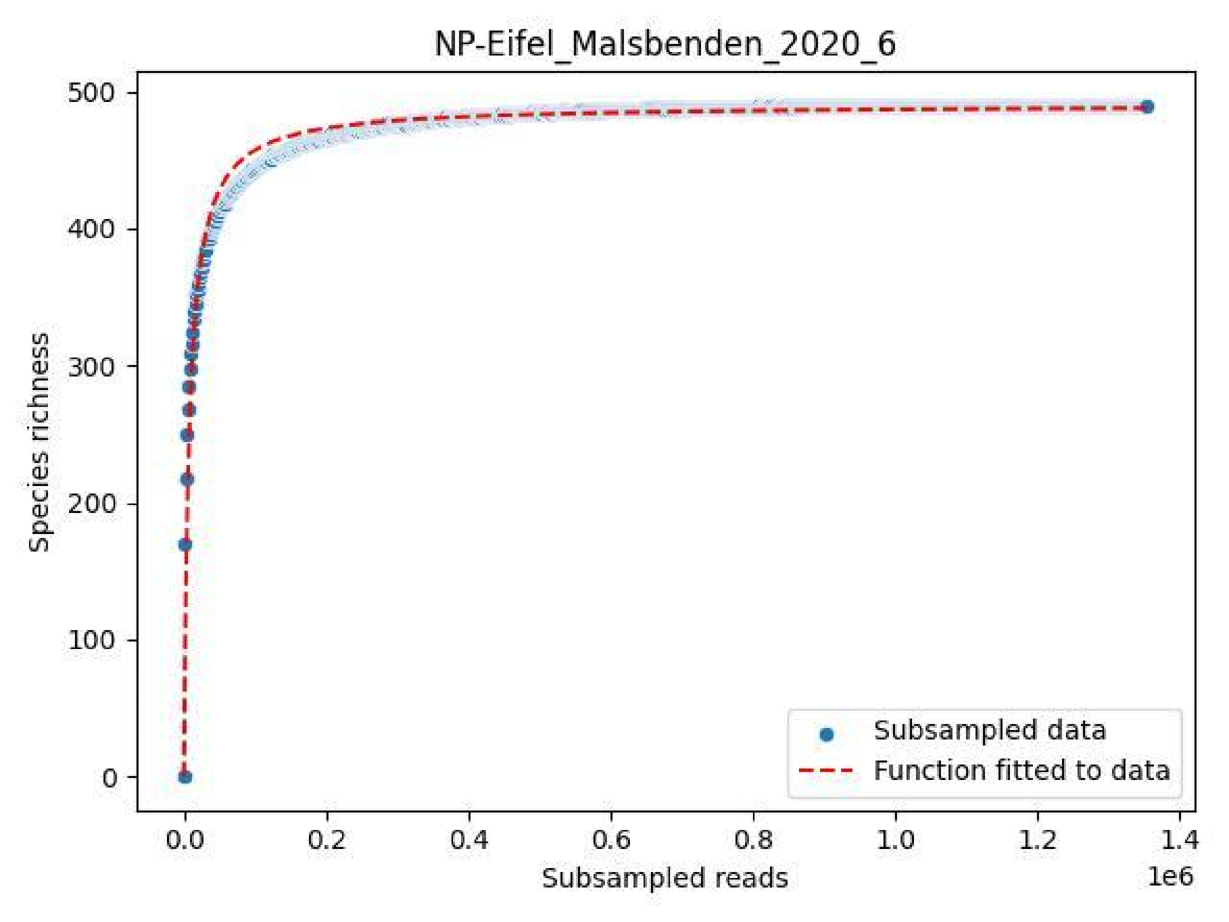

NP-Eifel_Malsbenden_2020_6
Species richness
500
400
300
200
100
0
0.0
0.2
0.4
0.6
0.8
1.0
1.2
1.4
1e6
Subsampled reads
Subsampled data
Function fitted to data

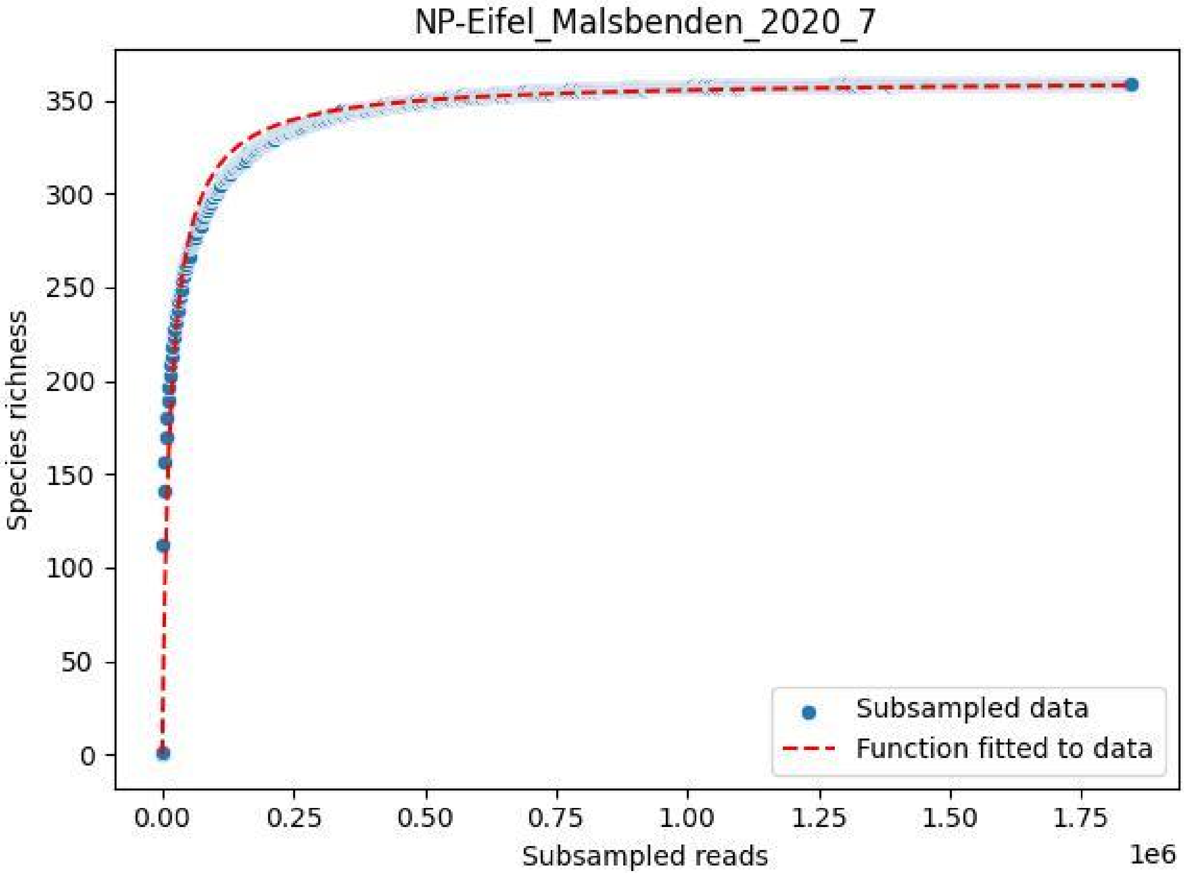
NP-Eifel_Malsbenden_2020_7
Species richness
Subsampled reads
1e6
0
50
100
150
200
250
300
350
0.00
0.25
0.50
0.75
1.00
1.25
1.50
1.75
Subsampled data
Function fitted to data

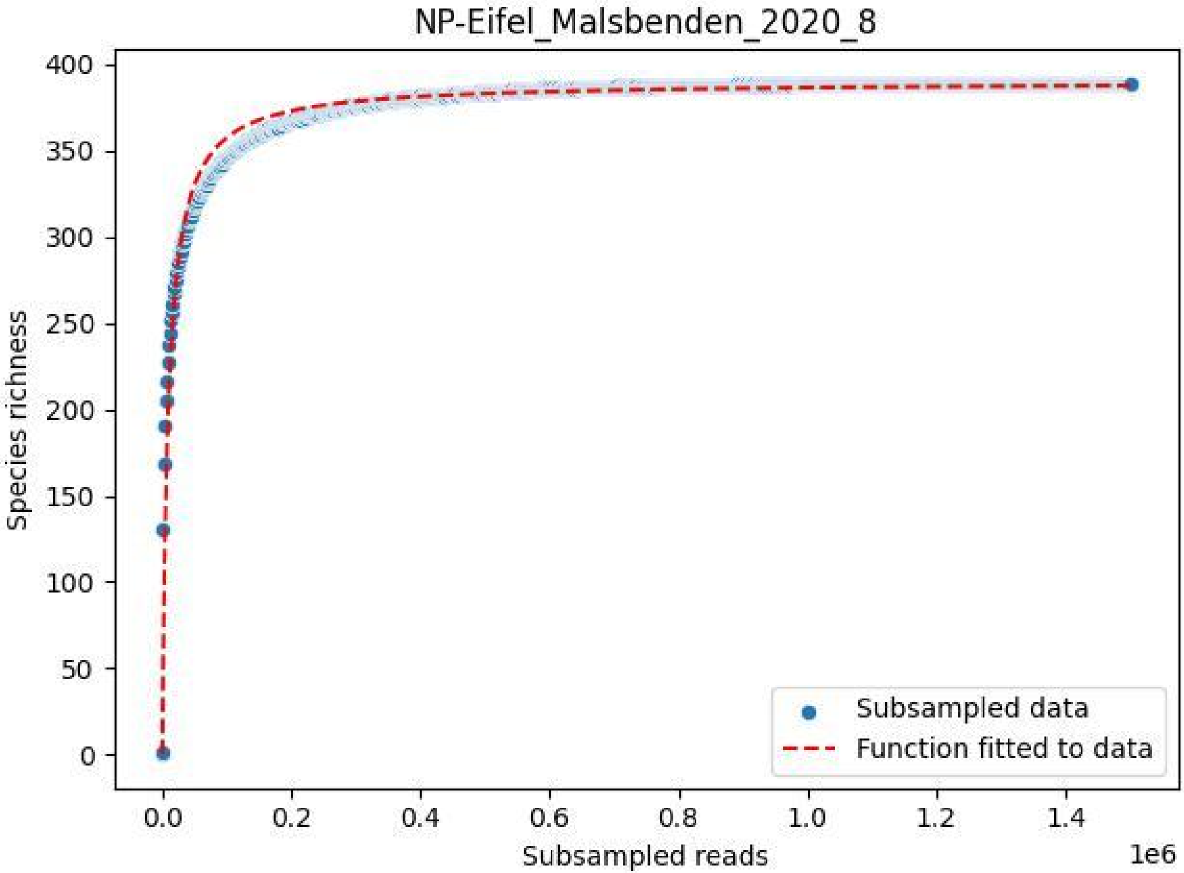

NP-Eifel_Malsbenden_2020_8
Species richness
Subsampled reads
1e6
Subsampled data
Function fitted to data
0
50
100
150
200
250
300
350
400
0.0
0.2
0.4
0.6
0.8
1.0
1.2
1.4

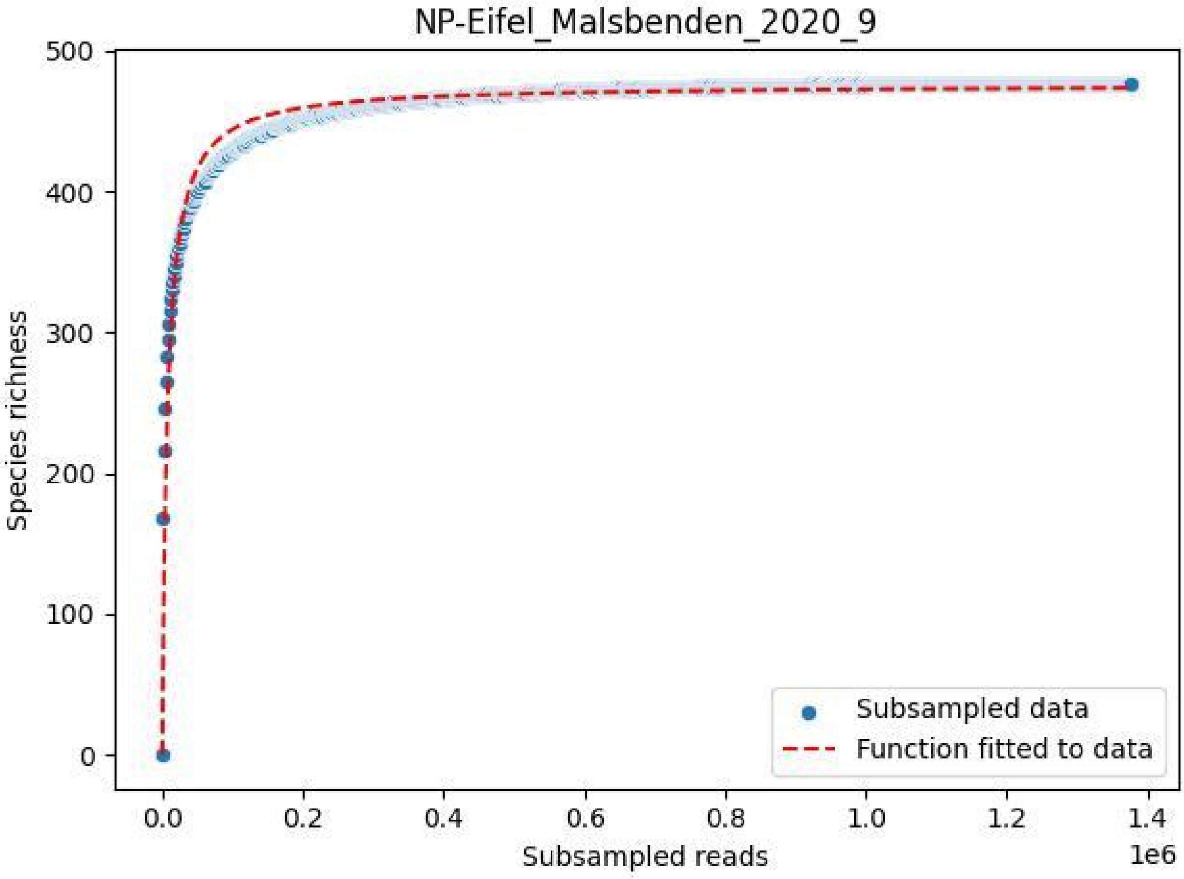
NP-Eifel_Malsbenden_2020_9
Species richness
Subsampled reads
1e6
0
100
200
300
400
500
0.0
0.2
0.4
0.6
0.8
1.0
1.2
1.4
Subsampled data
Function fitted to data

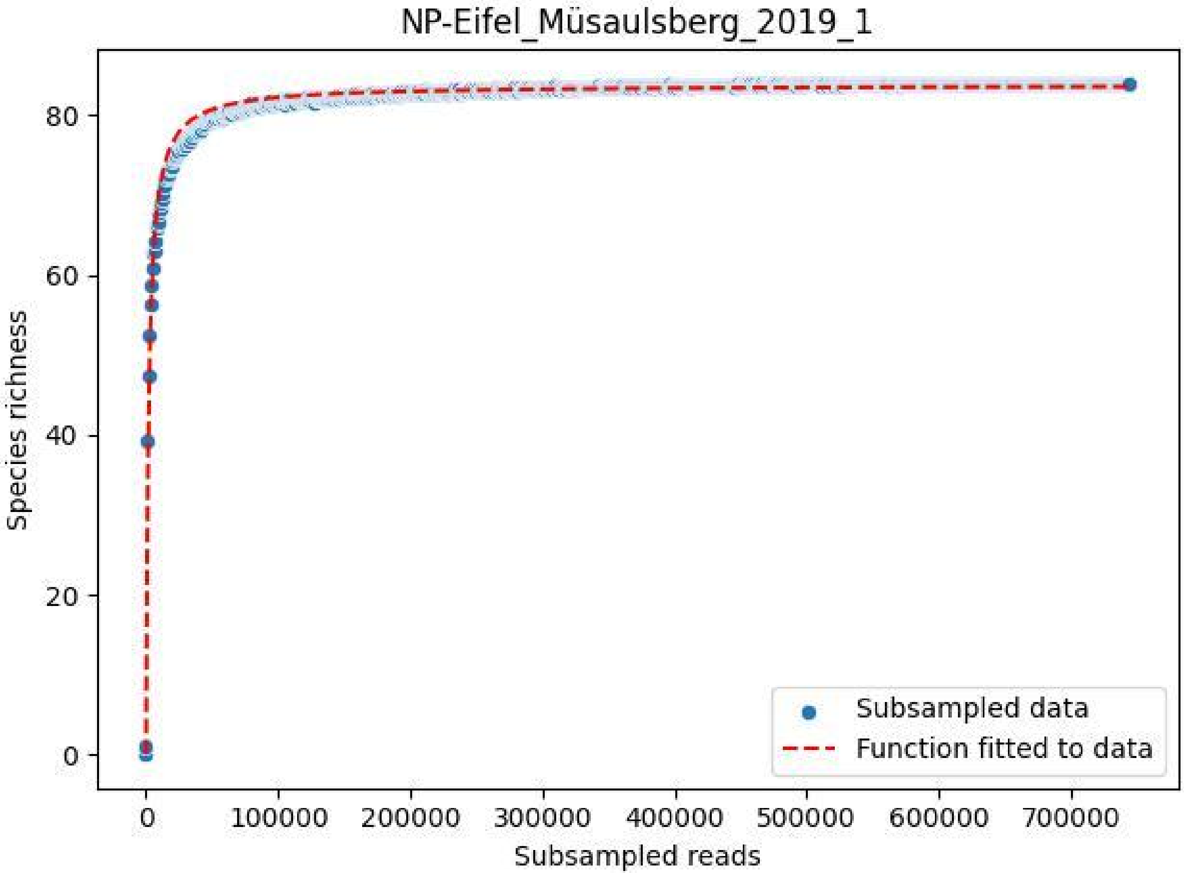

NP-Eifel_Müsaulsberg_2019_1
Species richness
Subsampled reads
0
20
40
60
80
0
100000
200000
300000
400000
500000
600000
700000
Subsampled data
Function fitted to data

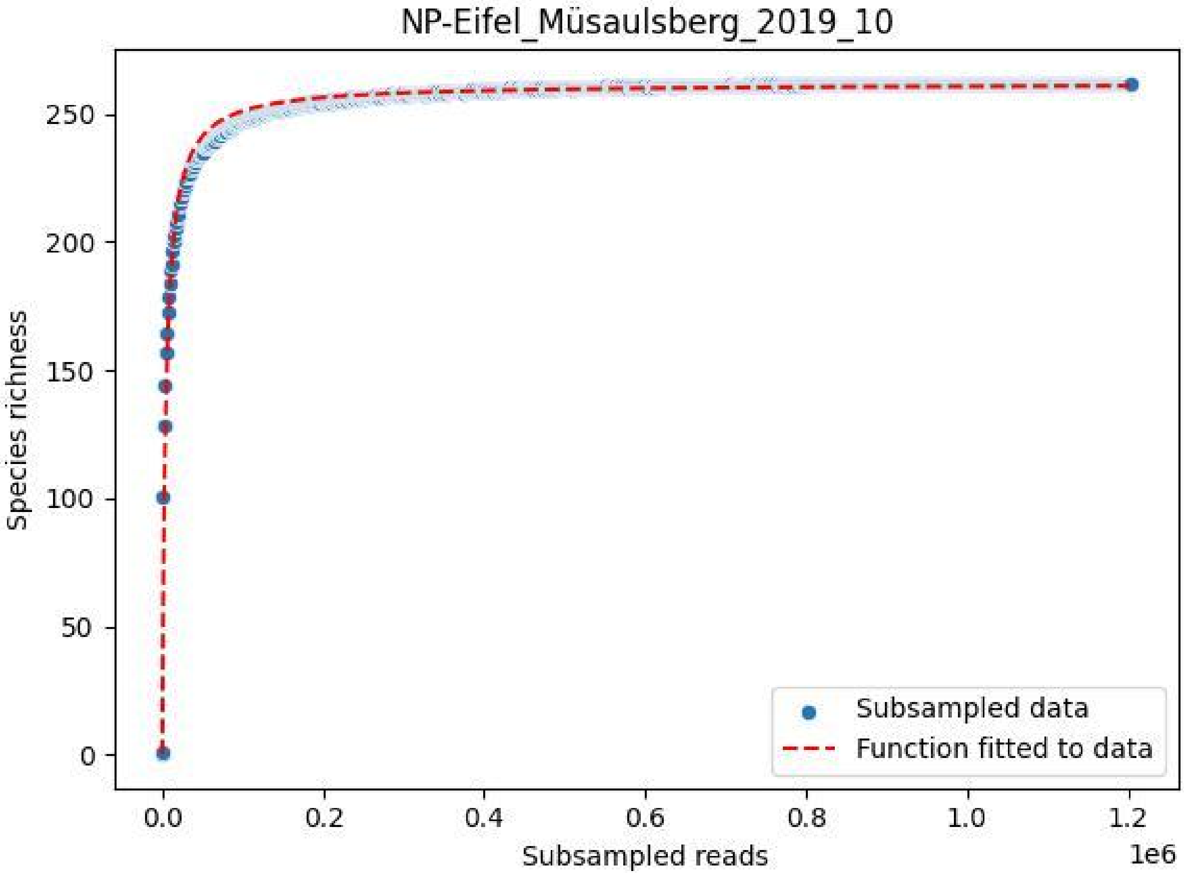
NP-Eifel_Müsaulsberg_2019_10
Species richness
Subsampled reads
1e6
0
50
100
150
200
250
0.0
0.2
0.4
0.6
0.8
1.0
1.2
Subsampled data
Function fitted to data

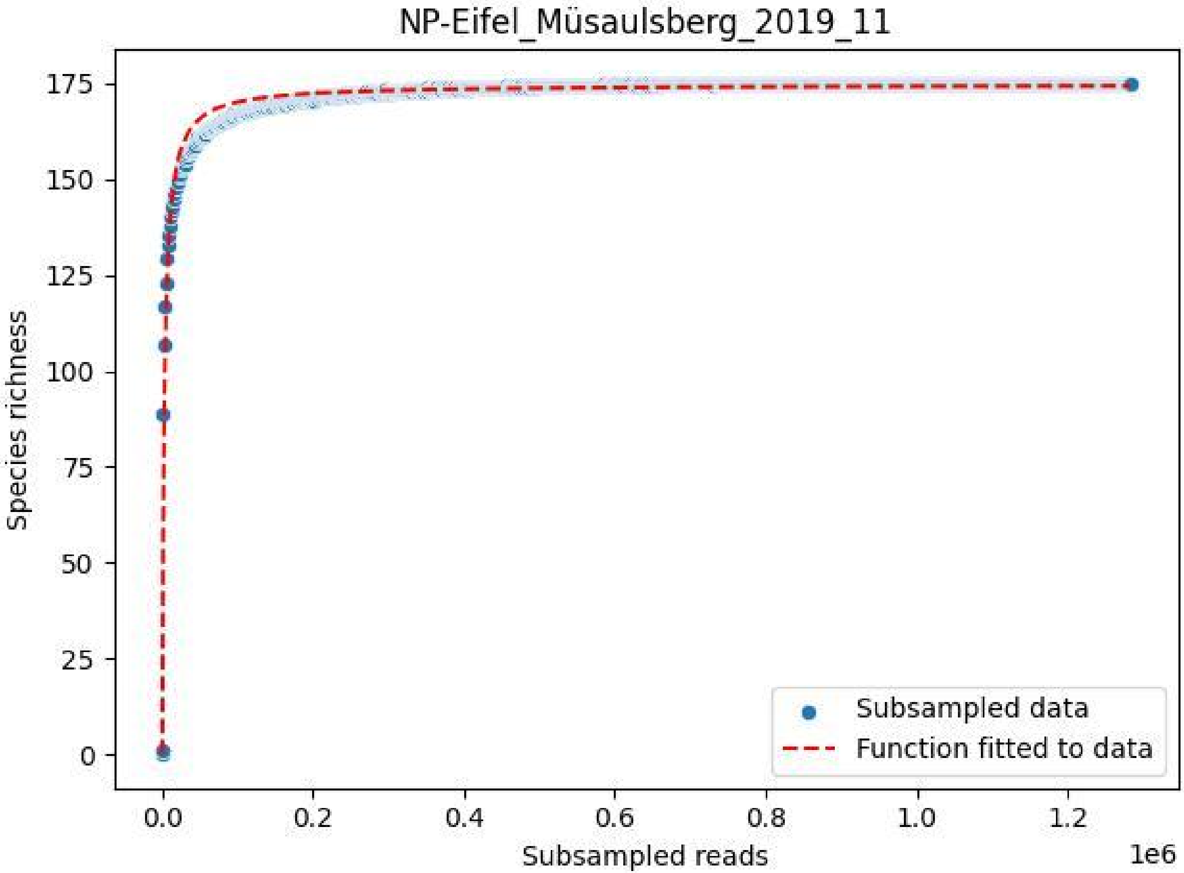

NP-Eifel_Müsaulsberg_2019_11
Species richness
175
150
125
100
75
50
25
0
0.0
0.2
0.4
0.6
0.8
1.0
1.2
1e6
Subsampled reads
Subsampled data
Function fitted to data

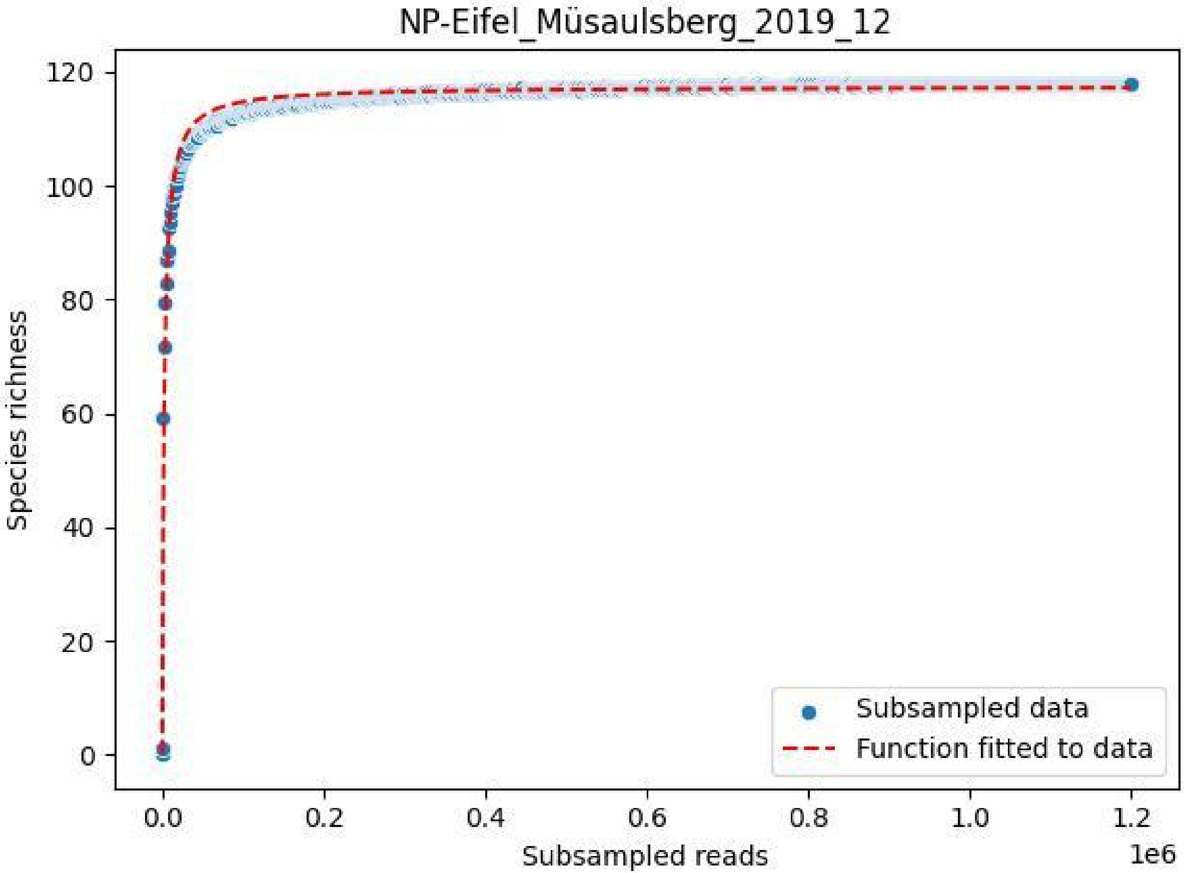
NP-Eifel_Müsaulsberg_2019_12
Species richness
Subsampled reads
1e6
0
20
40
60
80
100
120
0.0
0.2
0.4
0.6
0.8
1.0
1.2
Subsampled data
Function fitted to data

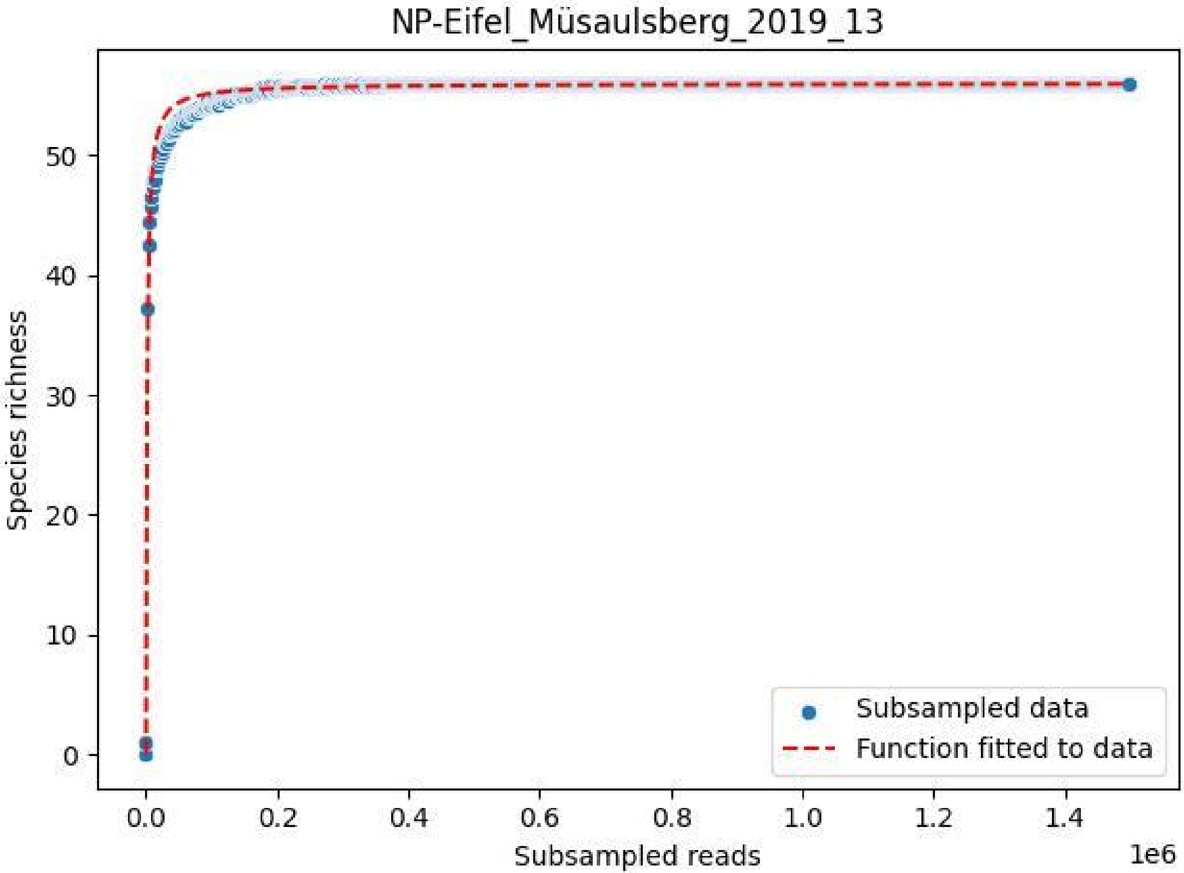
NP-Eifel_Müsaulsberg_2019_13
Species richness
Subsampled reads
1e6
0
10
20
30
40
50
0.0
0.2
0.4
0.6
0.8
1.0
1.2
1.4
Subsampled data
Function fitted to data

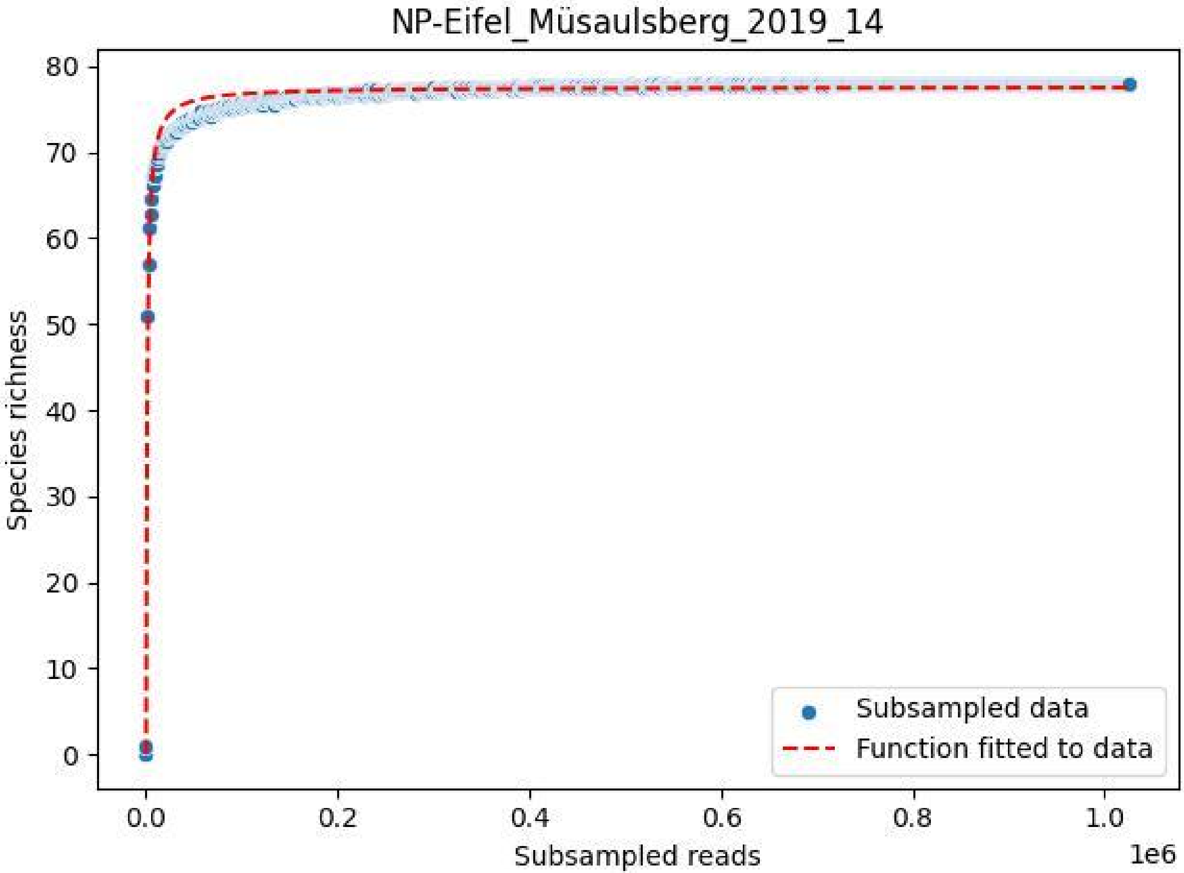
NP-Eifel_Müsaulsberg_2019_14
Species richness
Subsampled reads
1e6
Subsampled data
Function fitted to data
0
10
20
30
40
50
60
70
80
0.0
0.2
0.4
0.6
0.8
1.0

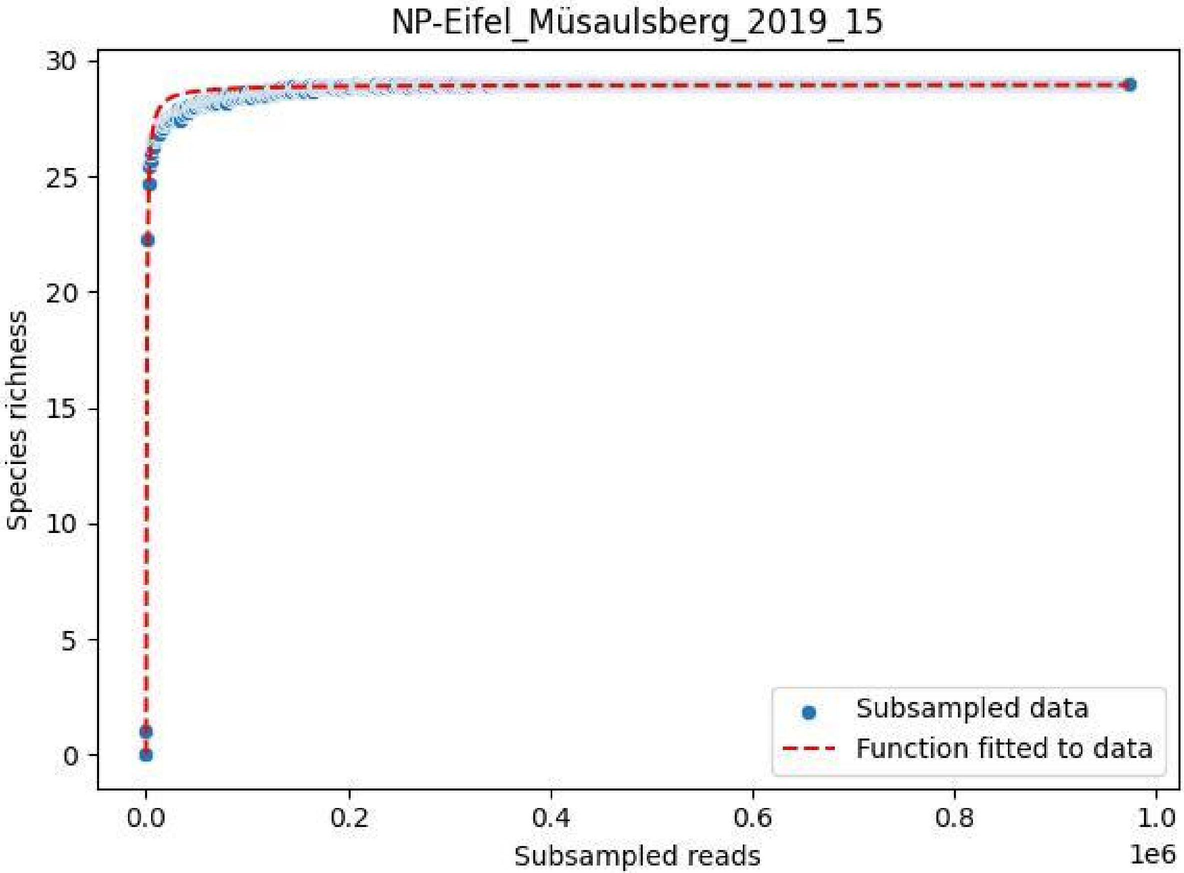
NP-Eifel_Müsaulsberg_2019_15
Species richness
Subsampled reads
1e6
0
5
10
15
20
25
30
0.0
0.2
0.4
0.6
0.8
1.0
Subsampled data
Function fitted to data

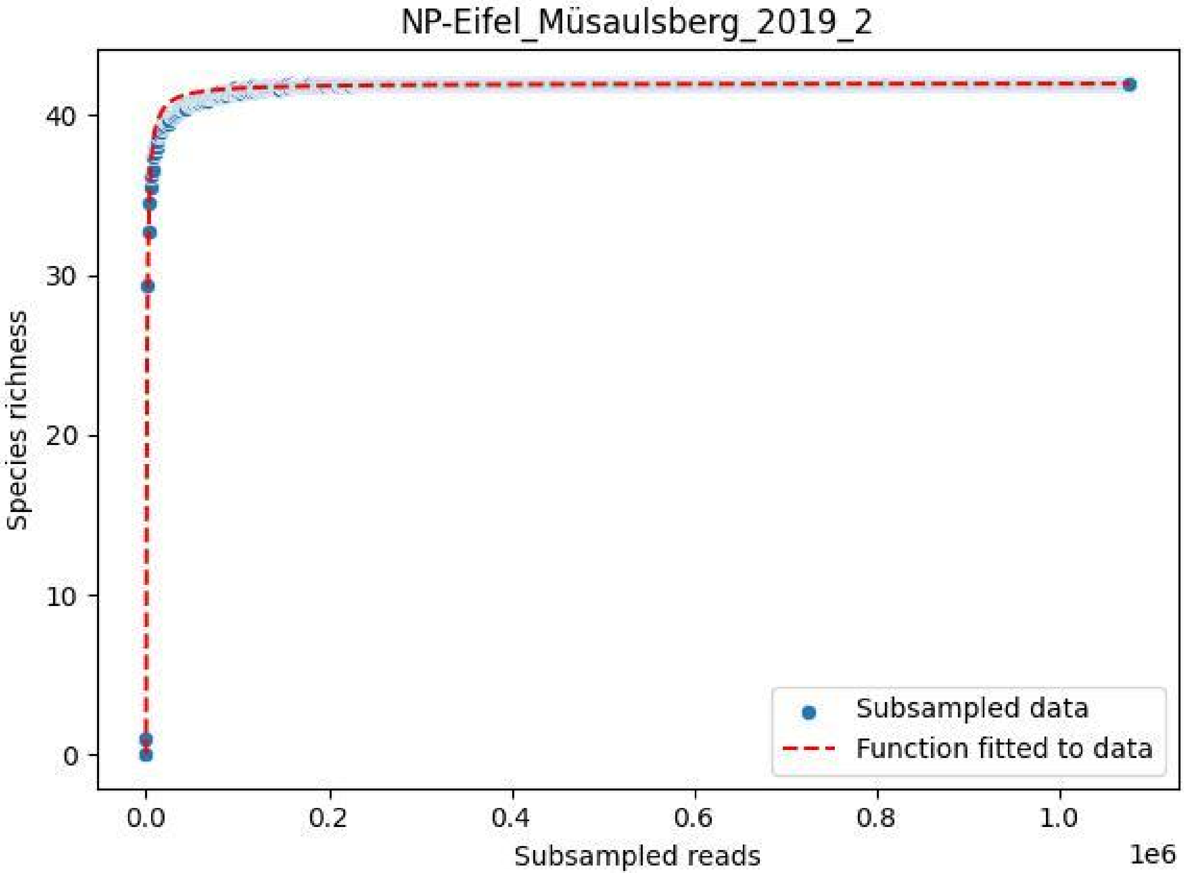

NP-Eifel_Müsaulsberg_2019_2
Species richness
Subsampled reads
1e6
0
10
20
30
40
0.0
0.2
0.4
0.6
0.8
1.0
Subsampled data
Function fitted to data

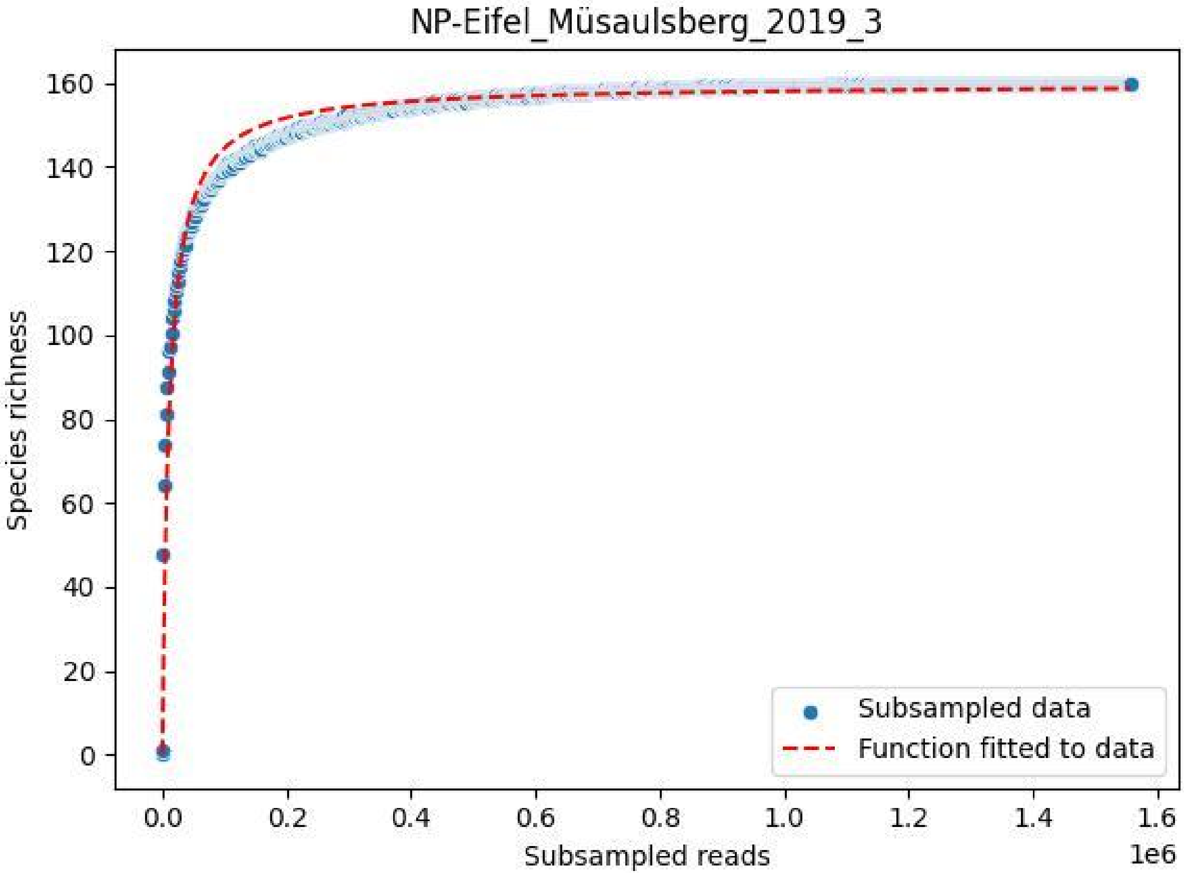
NP-Eifel_Müsaulsberg_2019_3
Species richness
Subsampled reads
1e6
0
20
40
60
80
100
120
140
160
0.0
0.2
0.4
0.6
0.8
1.0
1.2
1.4
1.6
Subsampled data
Function fitted to data

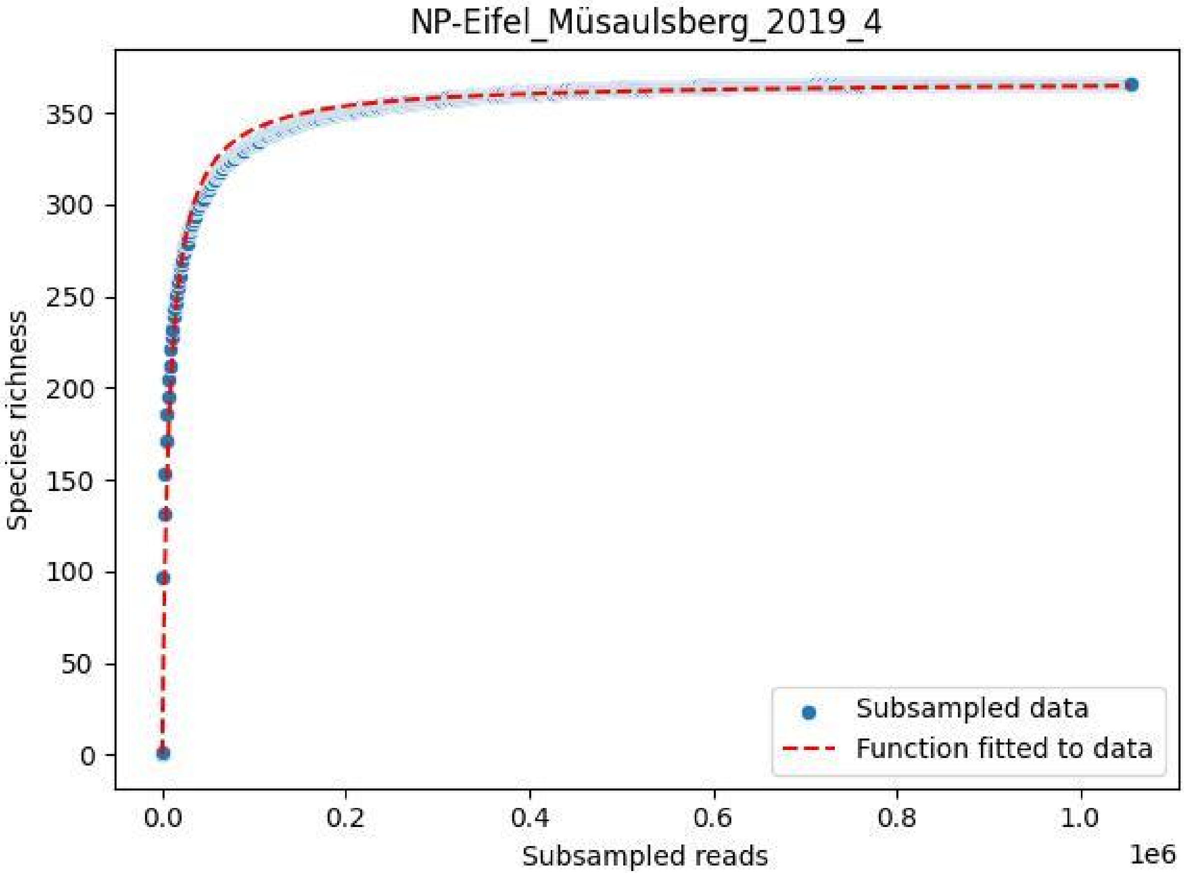

NP-Eifel_Müsaulsberg_2019_4
Species richness
Subsampled reads
1e6
Subsampled data
Function fitted to data
0
50
100
150
200
250
300
350
0.0
0.2
0.4
0.6
0.8
1.0

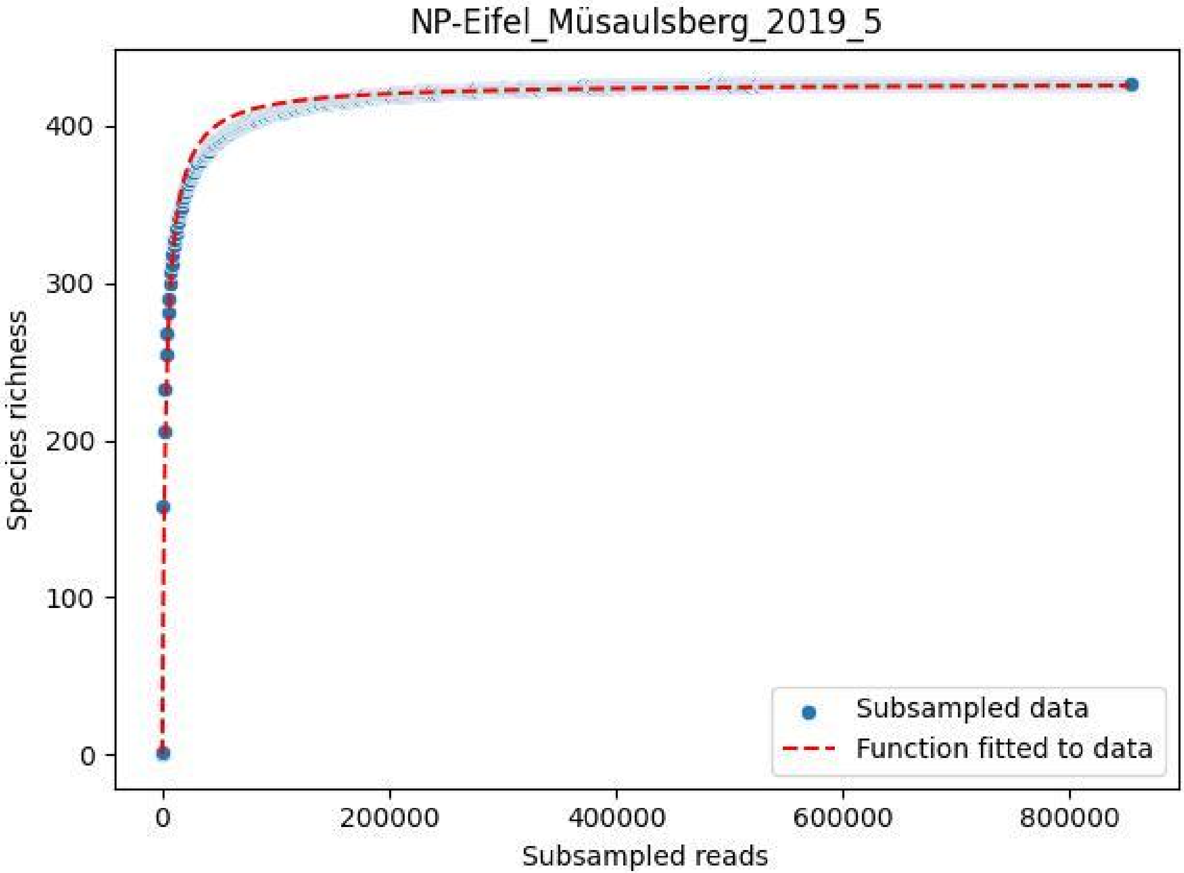
NP-Eifel_Müsaulsberg_2019_5
Species richness
0
100
200
300
400
Subsampled reads
0
200000
400000
600000
800000
Subsampled data
Function fitted to data

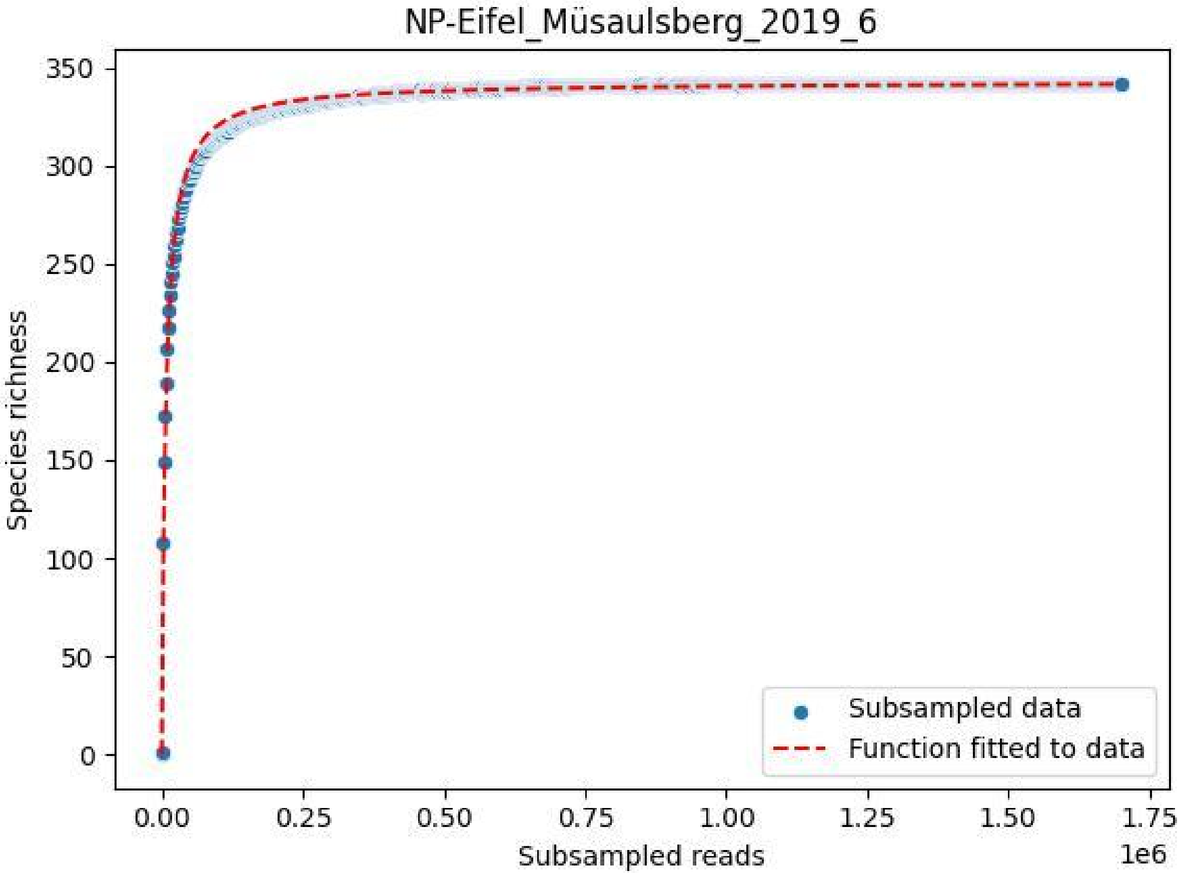

NP-Eifel_Müsaulsberg_2019_6
Species richness
Subsampled reads
1e6
0
50
100
150
200
250
300
350
0.00
0.25
0.50
0.75
1.00
1.25
1.50
1.75
Subsampled data
Function fitted to data

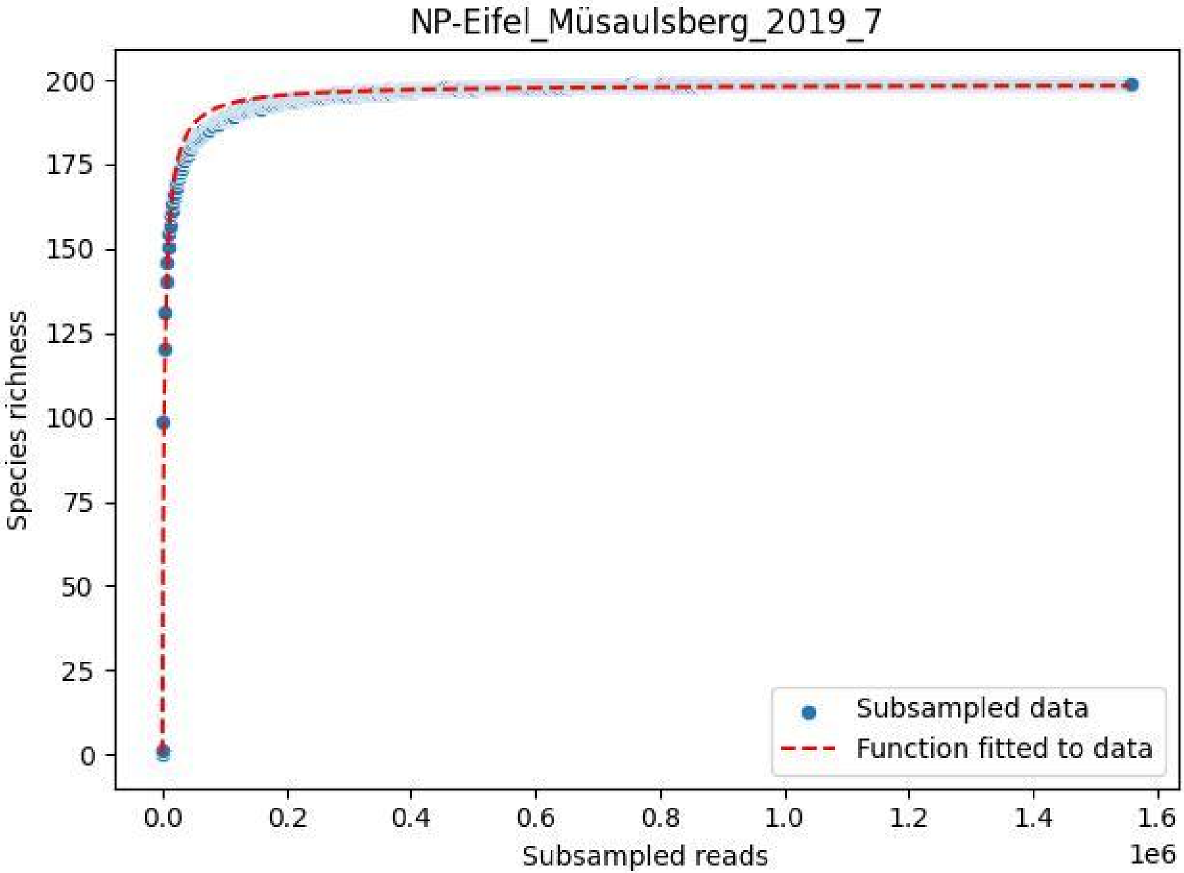
NP-Eifel_Müsaulsberg_2019_7
Species richness
Subsampled reads
1e6
0
25
50
75
100
125
150
175
200
0.0
0.2
0.4
0.6
0.8
1.0
1.2
1.4
1.6
Subsampled data
Function fitted to data

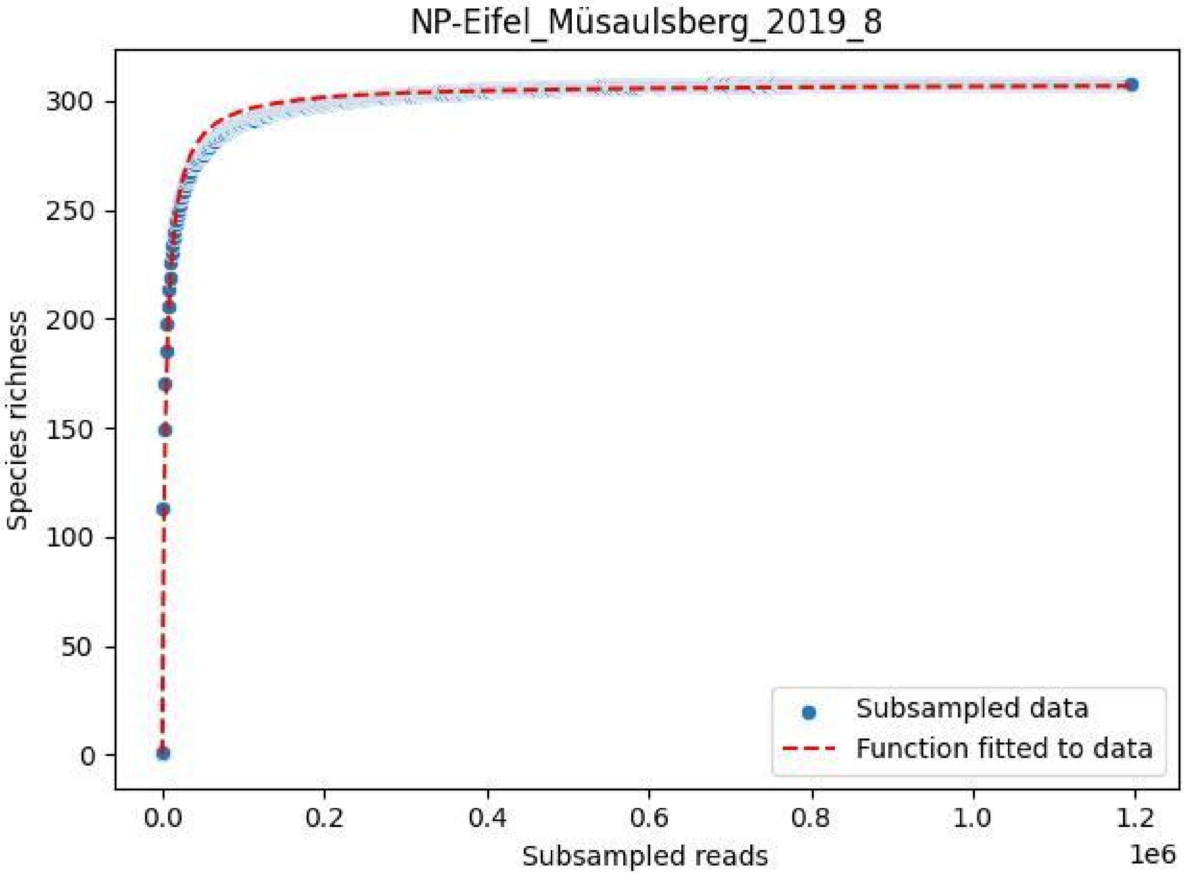
NP-Eifel_Müsaulsberg_2019_8
Species richness
Subsampled reads
1e6
0.0
0.2
0.4
0.6
0.8
1.0
1.2
0
50
100
150
200
250
300
Subsampled data
Function fitted to data

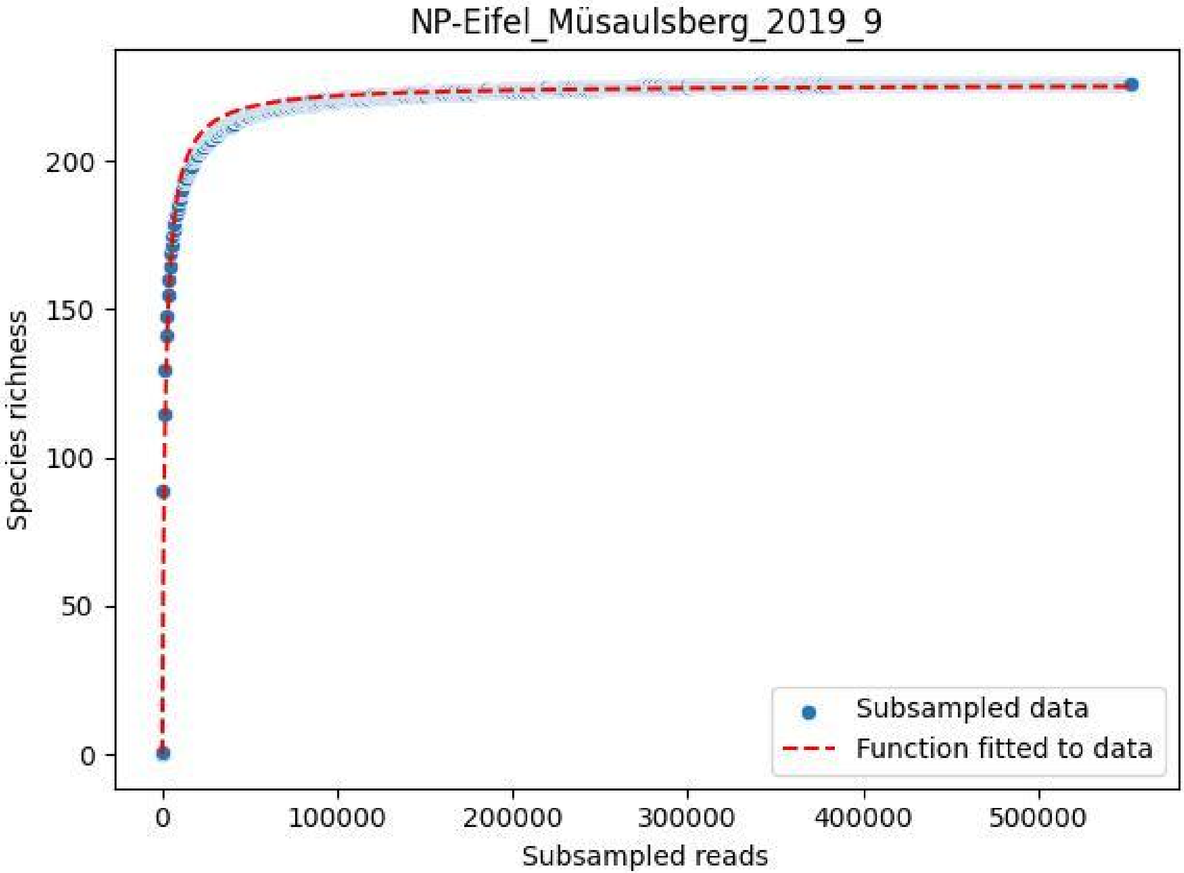
NP-Eifel_Müsaulsberg_2019_9
Species richness
Subsampled reads
0
50
100
150
200
0
100000
200000
300000
400000
500000
Subsampled data
Function fitted to data

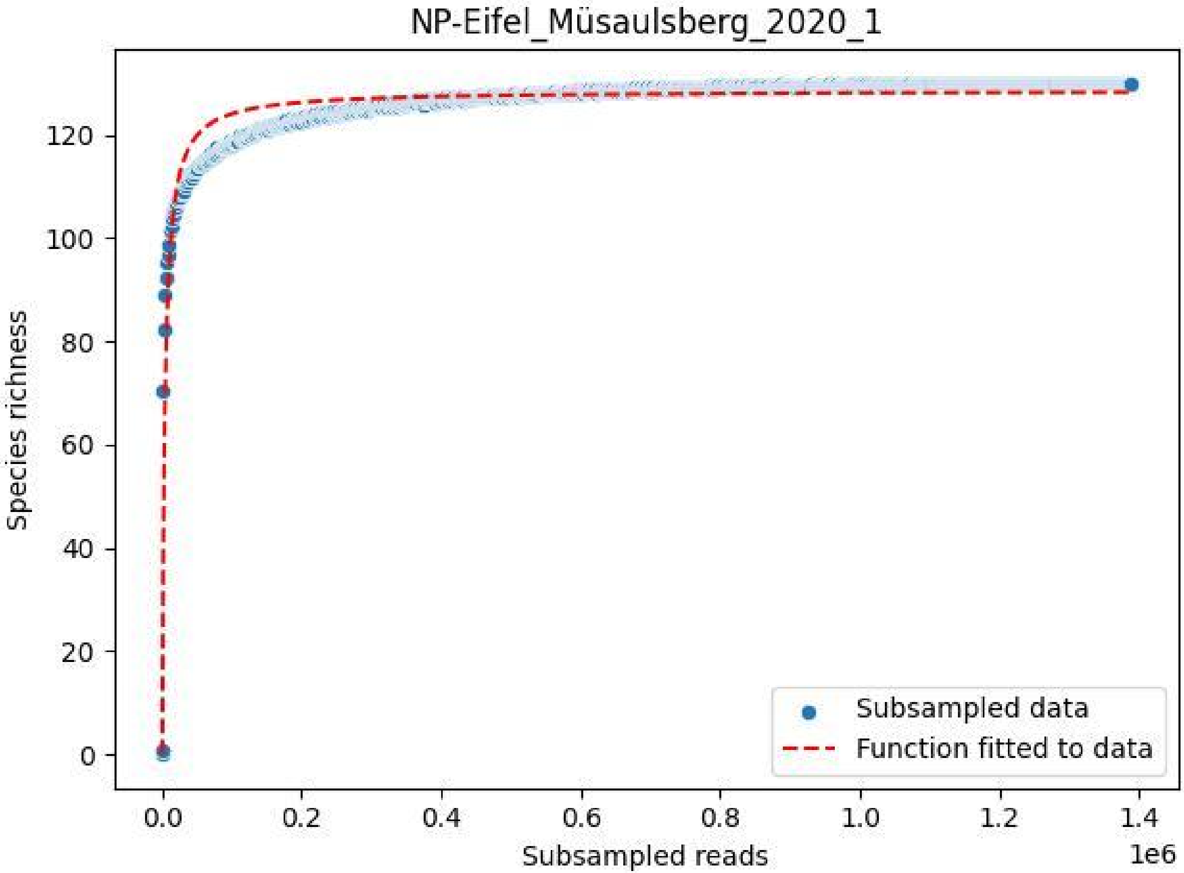

NP-Eifel_Müsaulsberg_2020_1
Species richness
Subsampled reads
1e6
Subsampled data
Function fitted to data
0
20
40
60
80
100
120
0.0
0.2
0.4
0.6
0.8
1.0
1.2
1.4

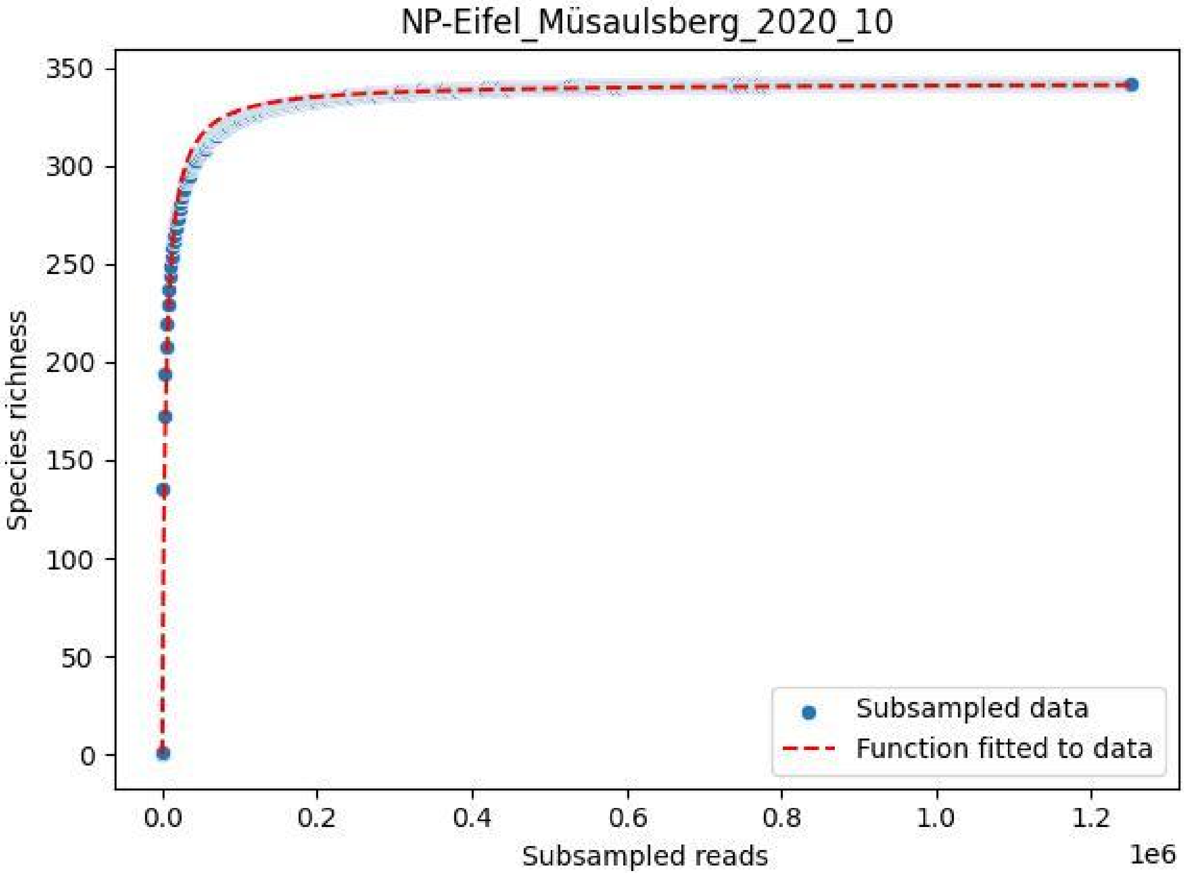

NP-Eifel_Müsaulsberg_2020_10
Species richness
Subsampled reads
1e6
Subsampled data
Function fitted to data
0
50
100
150
200
250
300
350
0.0
0.2
0.4
0.6
0.8
1.0
1.2

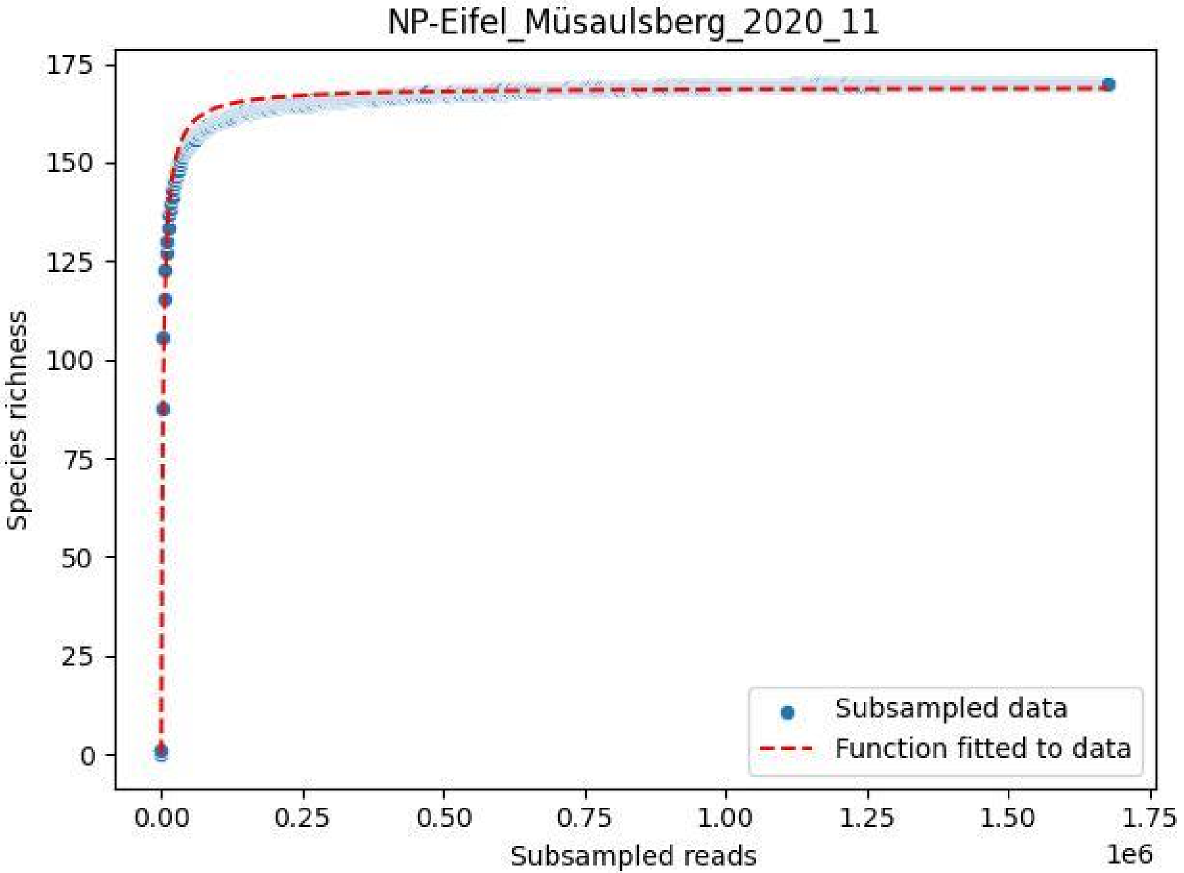

NP-Eifel_Müsaulsberg_2020_11
Species richness
Subsampled reads
1e6
175
150
125
100
75
50
25
0
0.00
0.25
0.50
0.75
1.00
1.25
1.50
1.75
Subsampled data
Function fitted to data

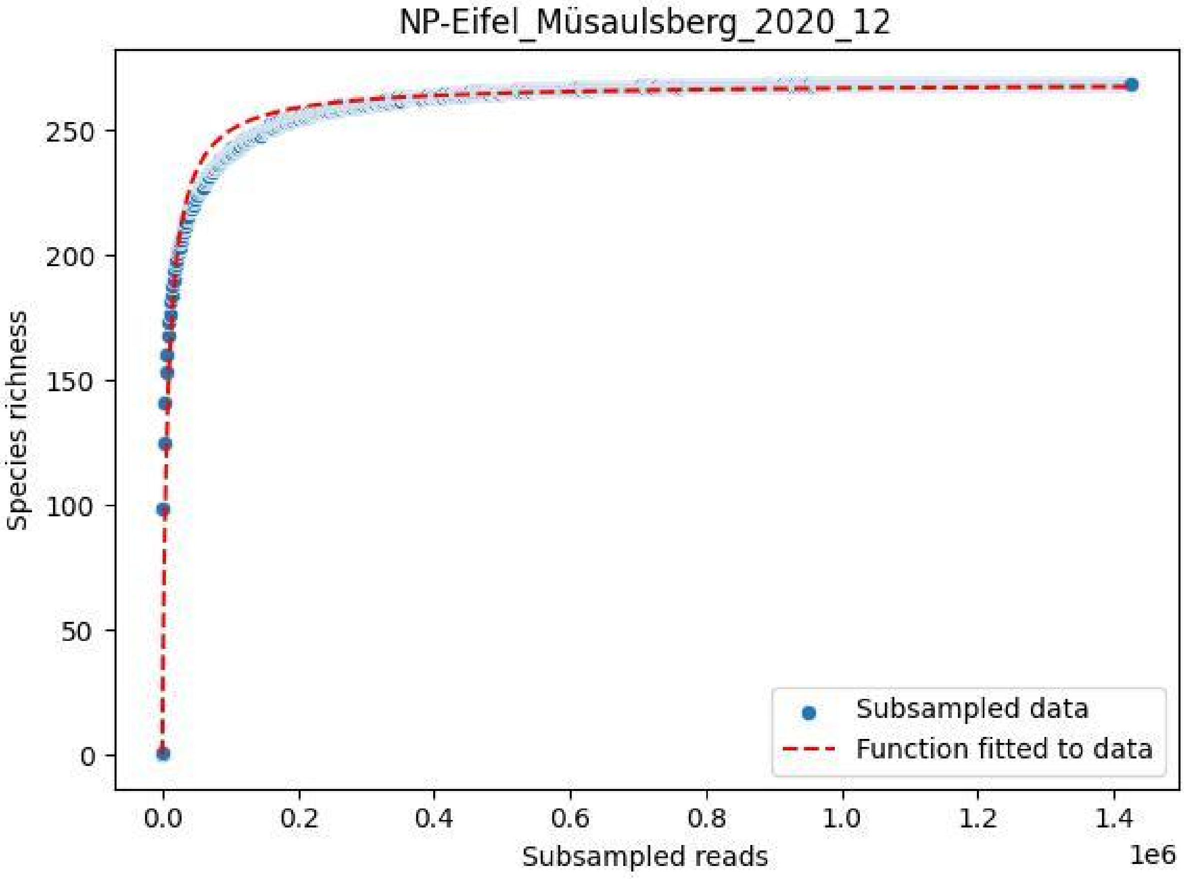
NP-Eifel_Müsaulsberg_2020_12
Species richness
Subsampled reads
1e6
0
50
100
150
200
250
0.0
0.2
0.4
0.6
0.8
1.0
1.2
1.4
Subsampled data
Function fitted to data

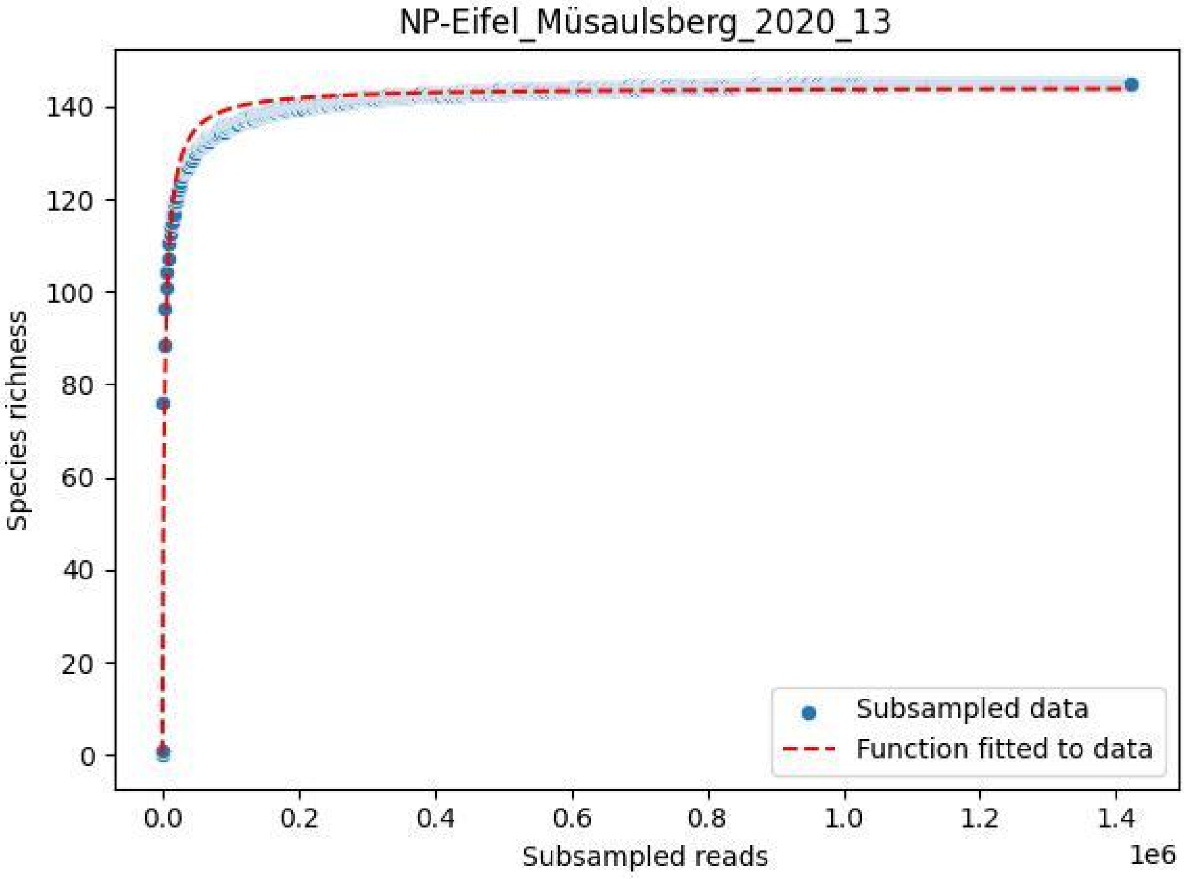
NP-Eifel_Müsaulsberg_2020_13
Species richness
Subsampled reads
1e6
Subsampled data
Function fitted to data
0
20
40
60
80
100
120
140
0.0
0.2
0.4
0.6
0.8
1.0
1.2
1.4

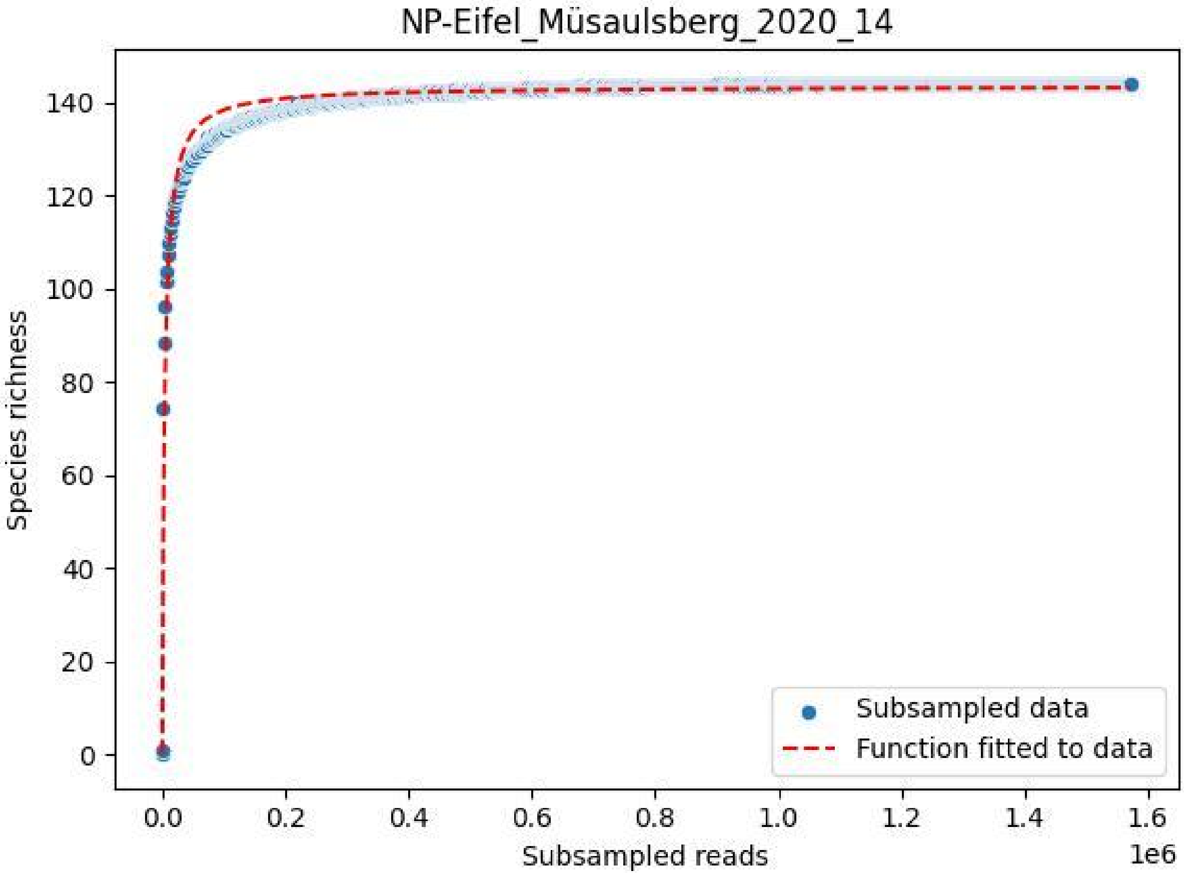
NP-Eifel_Müsaulsberg_2020_14
Species richness
Subsampled reads
1e6
0
20
40
60
80
100
120
140
0.0
0.2
0.4
0.6
0.8
1.0
1.2
1.4
1.6
Subsampled data
Function fitted to data

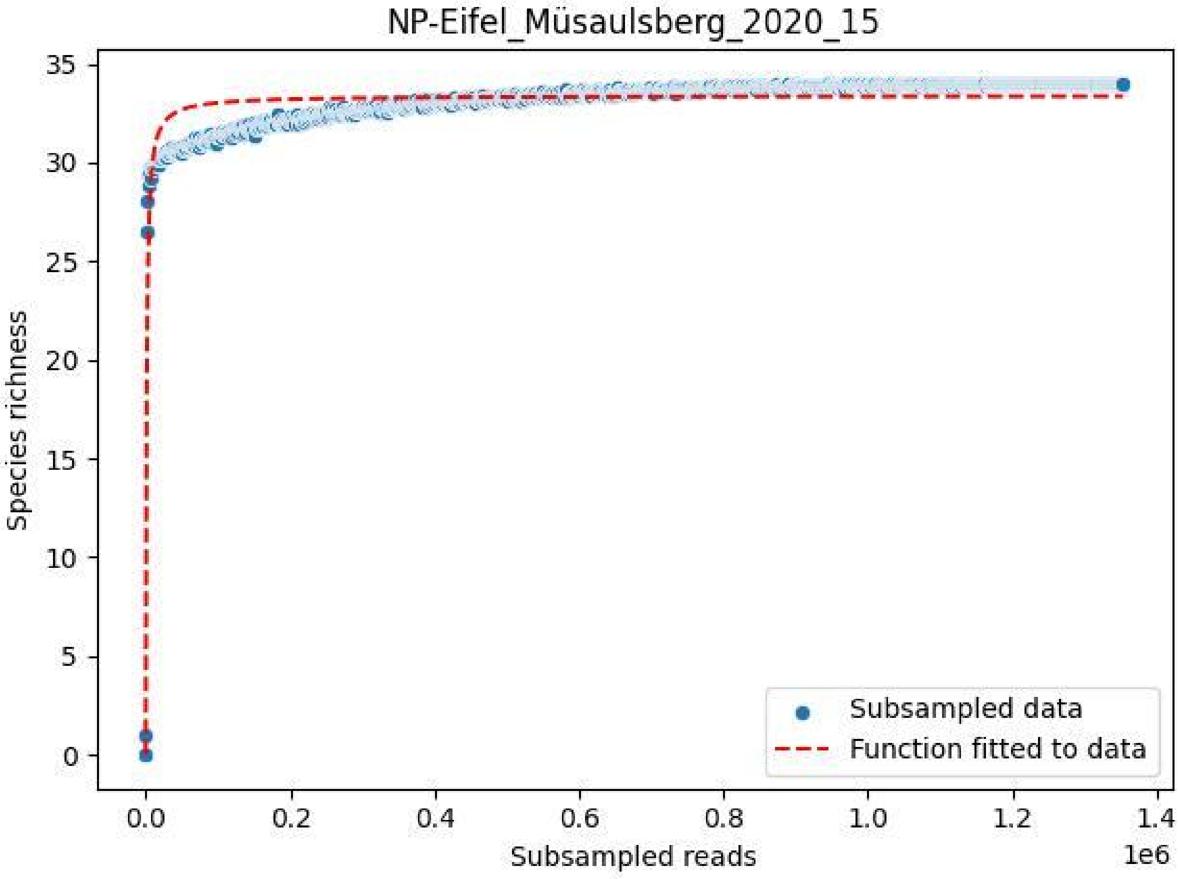

NP-Eifel_Müsaulsberg_2020_15
Species richness
Subsampled reads
1e6
Subsampled data
Function fitted to data
0
5
10
15
20
25
30
35
0.0
0.2
0.4
0.6
0.8
1.0
1.2
1.4

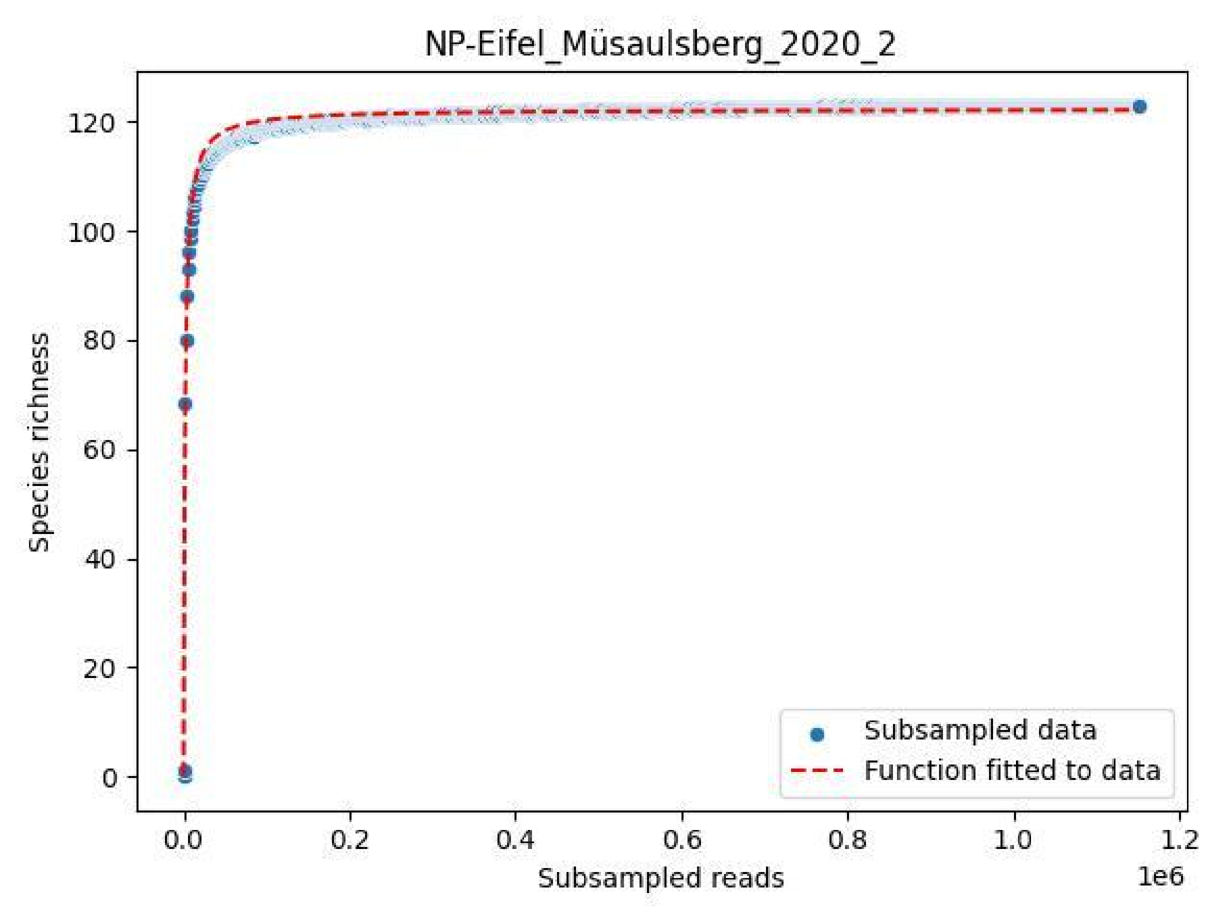
NP-Eifel_Müsaulsberg_2020_2
Species richness
0
20
40
60
80
100
120
Subsampled reads
0.0
0.2
0.4
0.6
0.8
1.0
1.2
1e6
Subsampled data
Function fitted to data

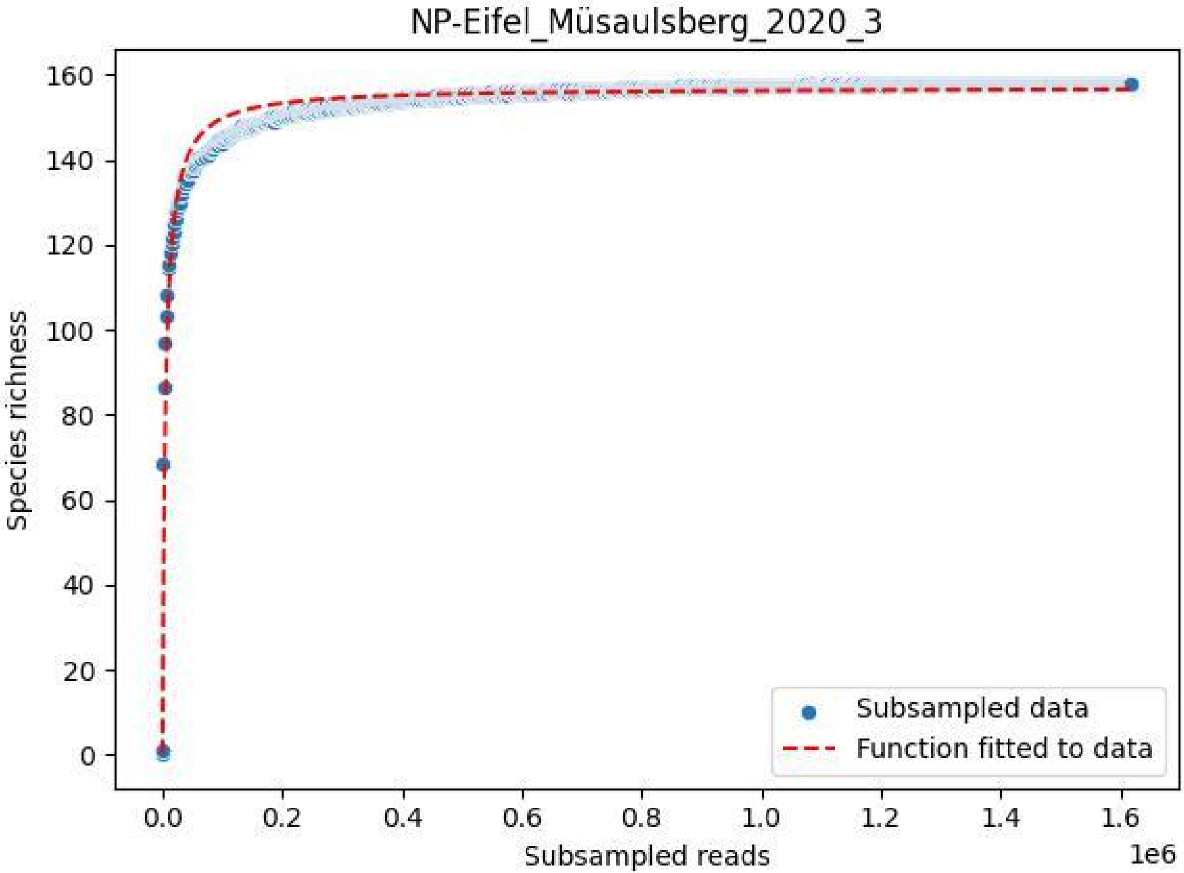

NP-Eifel_Müsaulsberg_2020_3
Species richness
Subsampled reads
1e6
Subsampled data
Function fitted to data
0
20
40
60
80
100
120
140
160
0.0
0.2
0.4
0.6
0.8
1.0
1.2
1.4
1.6

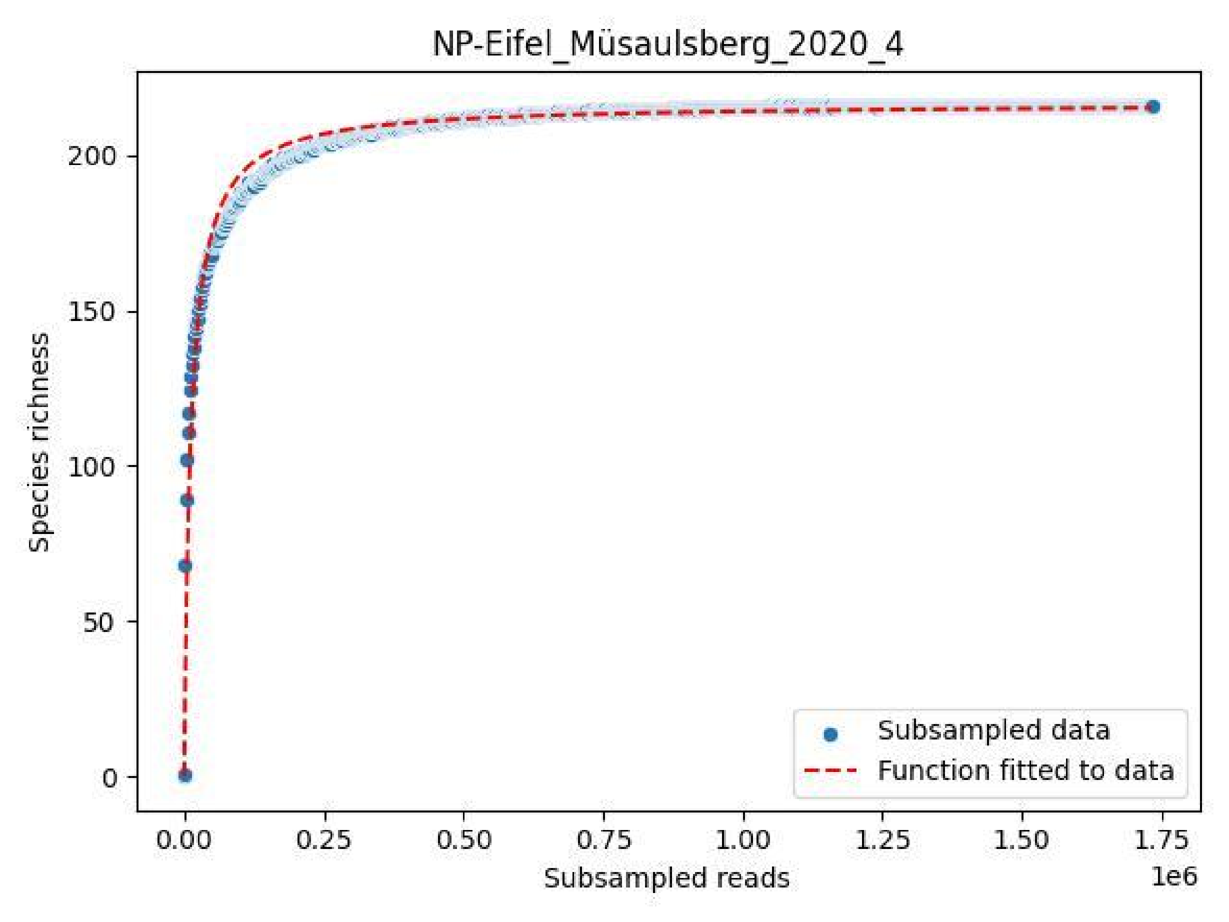
NP-Eifel_Müsaulsberg_2020_4
Species richness
Subsampled reads
1e6
0
50
100
150
200
0.00
0.25
0.50
0.75
1.00
1.25
1.50
1.75
Subsampled data
Function fitted to data

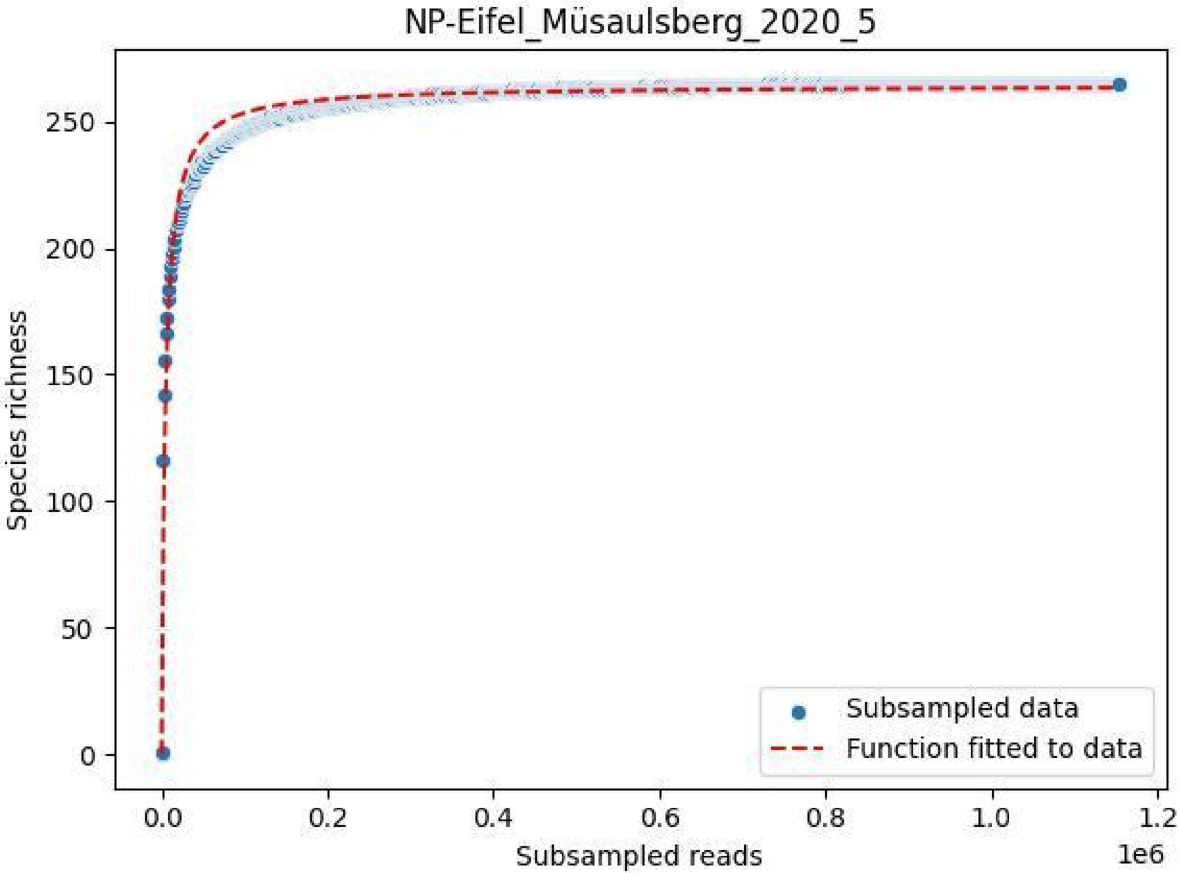

NP-Eifel_Müsaulsberg_2020_5
Species richness
Subsampled reads
1e6
0
50
100
150
200
250
0.0
0.2
0.4
0.6
0.8
1.0
1.2
Subsampled data
Function fitted to data

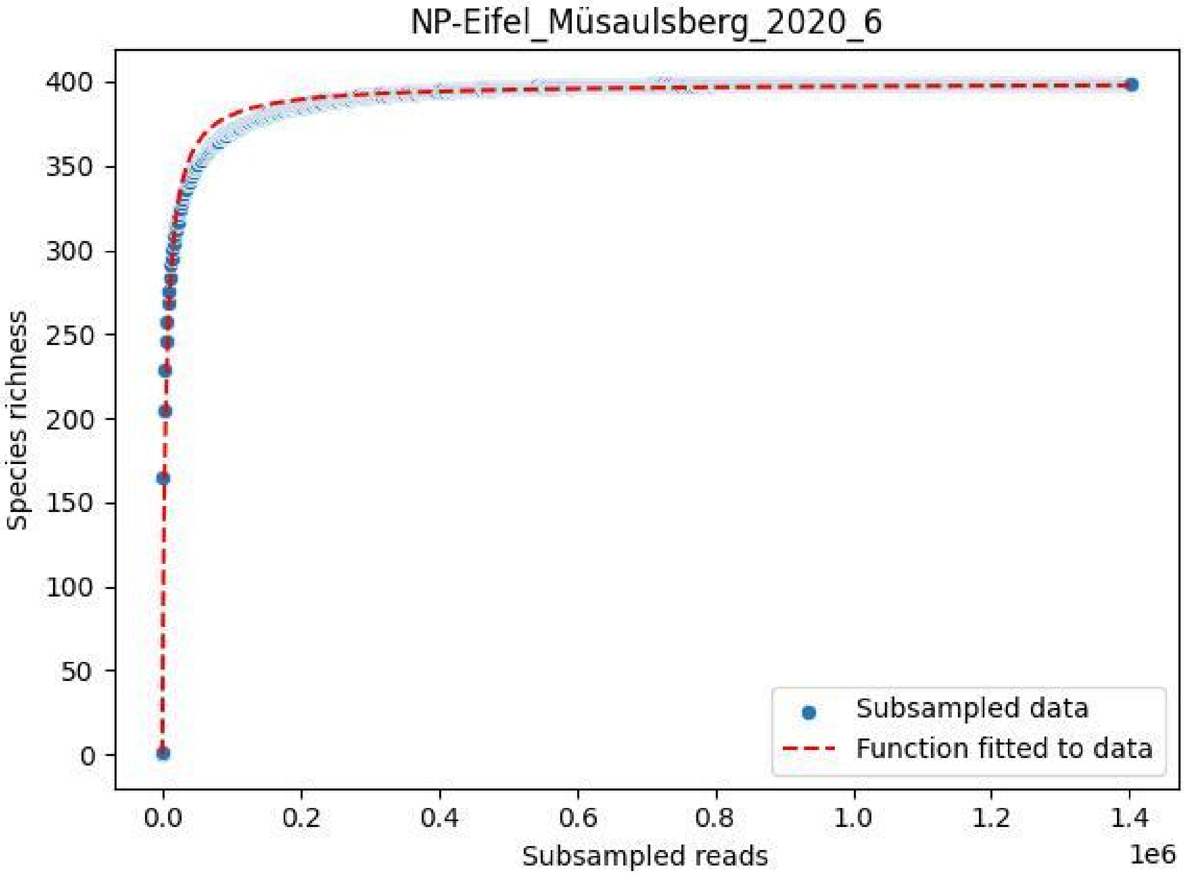

NP-Eifel_Müsaulsberg_2020_6
Species richness
Subsampled reads
1e6
0
50
100
150
200
250
300
350
400
0.0
0.2
0.4
0.6
0.8
1.0
1.2
1.4
Subsampled data
Function fitted to data

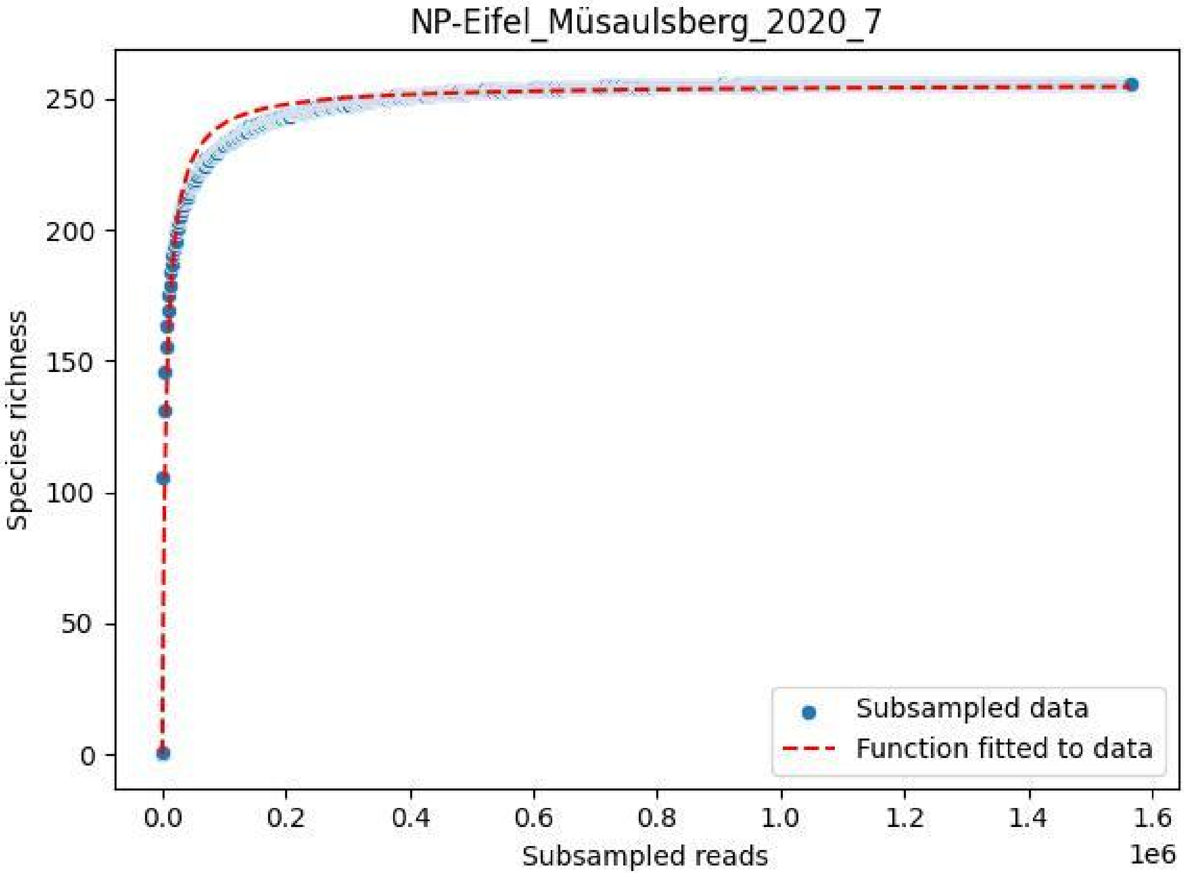
NP-Eifel_Müsaulsberg_2020_7
Species richness
0
50
100
150
200
250
Subsampled reads
0.0
0.2
0.4
0.6
0.8
1.0
1.2
1.4
1.6
1e6
Subsampled data
Function fitted to data

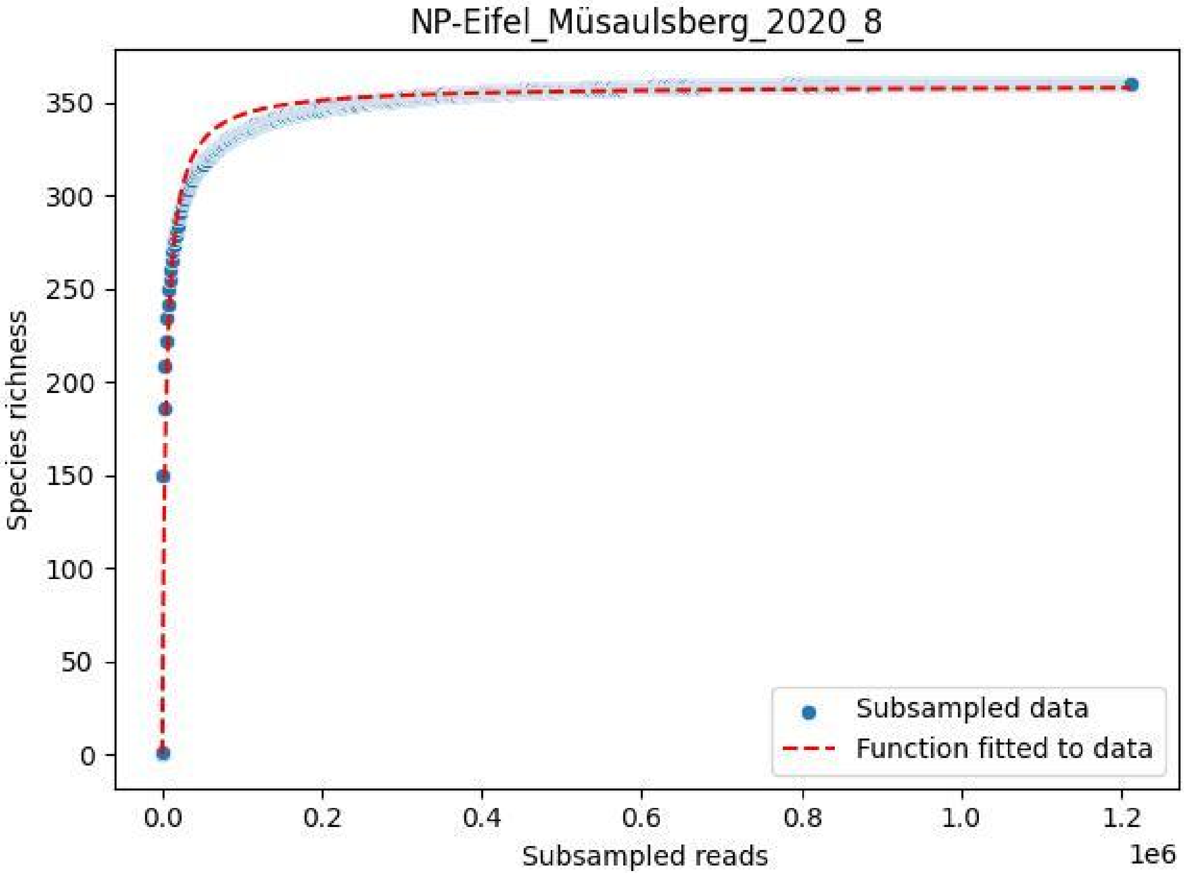
NP-Eifel_Müsaulsberg_2020_8
Species richness
0
50
100
150
200
250
300
350
0.0
0.2
0.4
0.6
0.8
1.0
1.2
1e6
Subsampled reads
Subsampled data
Function fitted to data

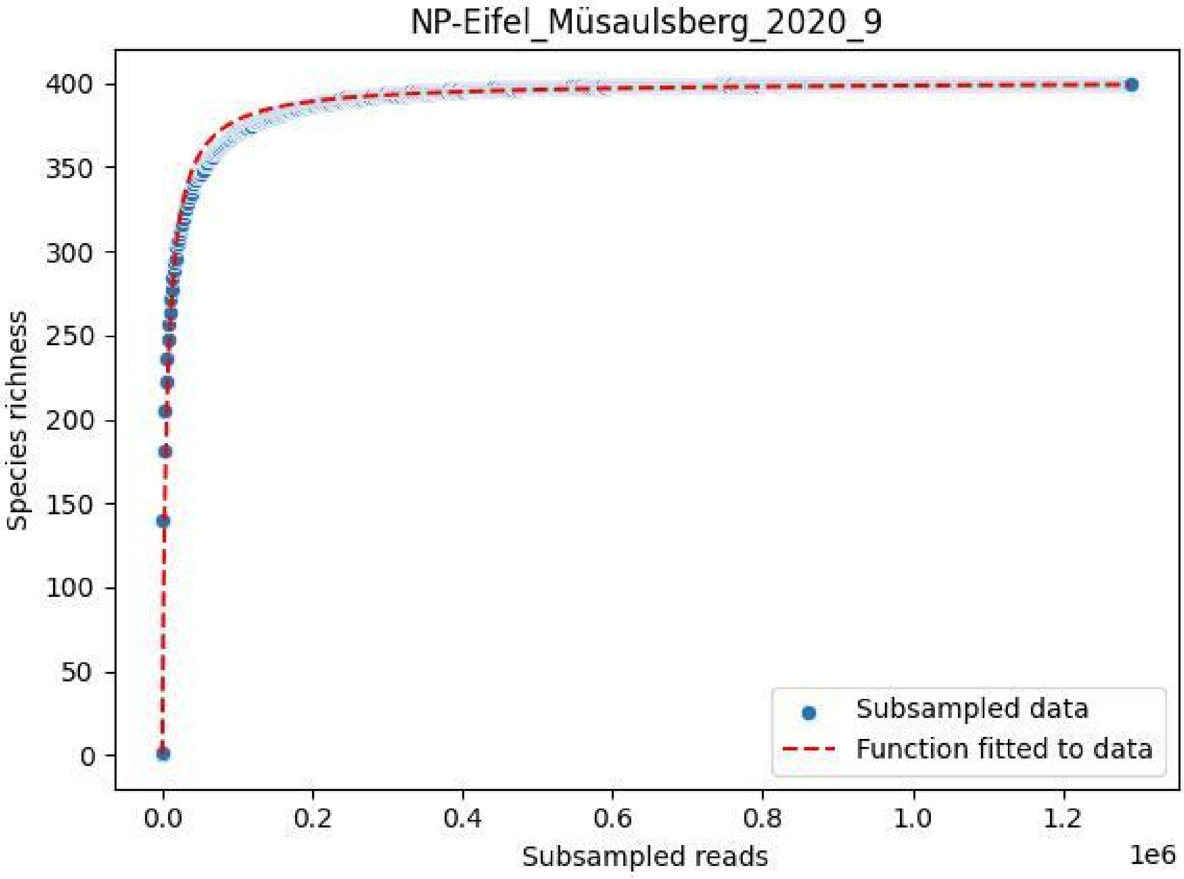
NP-Eifel_Müsaulsberg_2020_9
Species richness
Subsampled reads
1e6
Subsampled data
Function fitted to data
0
50
100
150
200
250
300
350
400
0.0
0.2
0.4
0.6
0.8
1.0
1.2

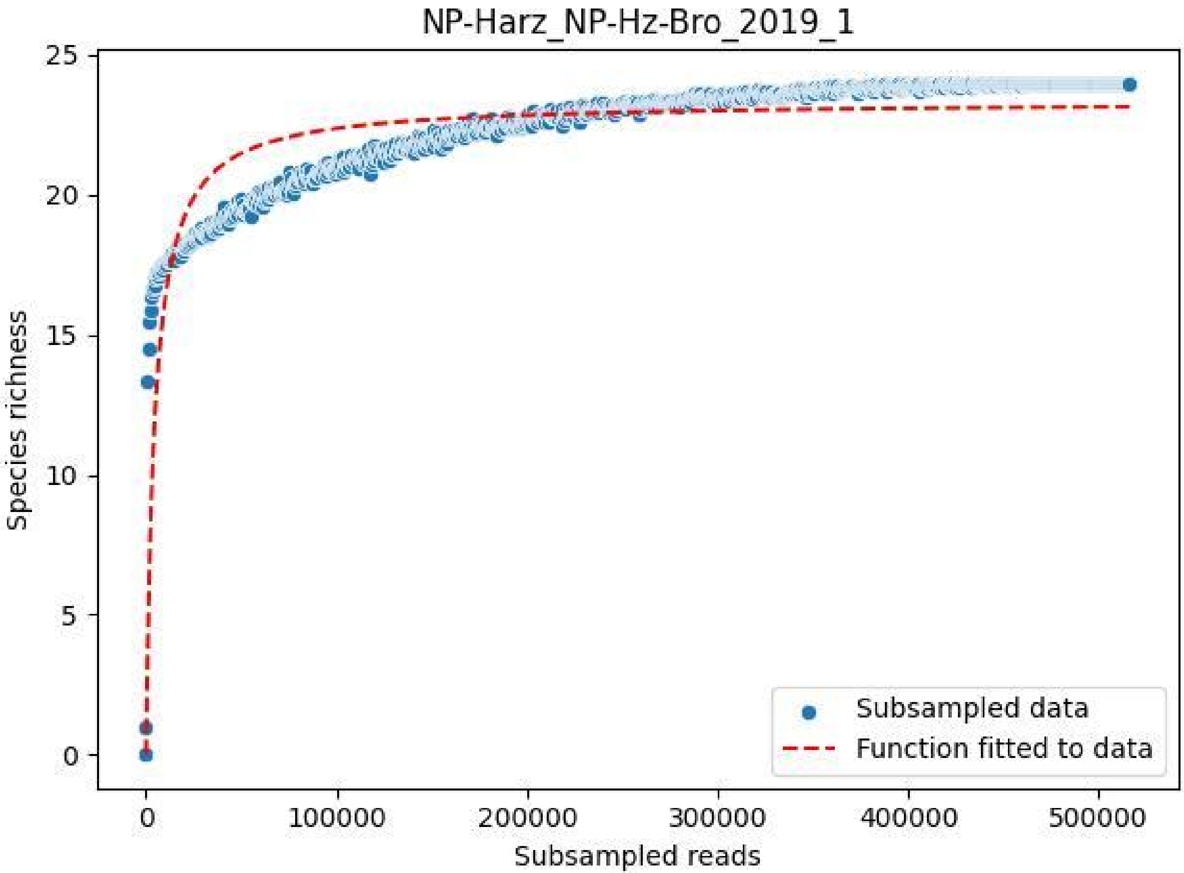
NP-Harz_NP-Hz-Bro_2019_1
Species richness
Subsampled reads
0
5
10
15
20
25
0
100000
200000
300000
400000
500000
Subsampled data
Function fitted to data

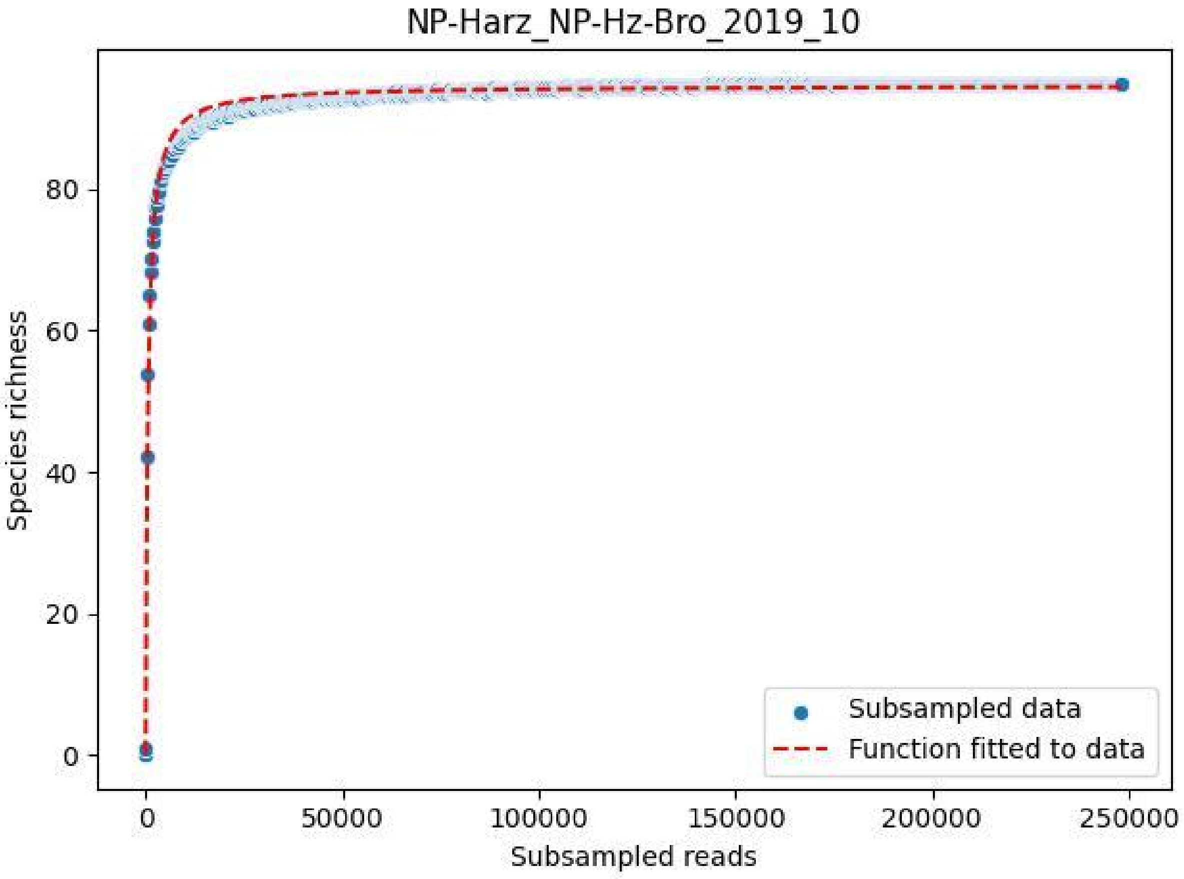

NP-Harz_NP-Hz-Bro_2019_10
Species richness
Subsampled reads
0
20
40
60
80
0
50000
100000
150000
200000
250000
Subsampled data
Function fitted to data

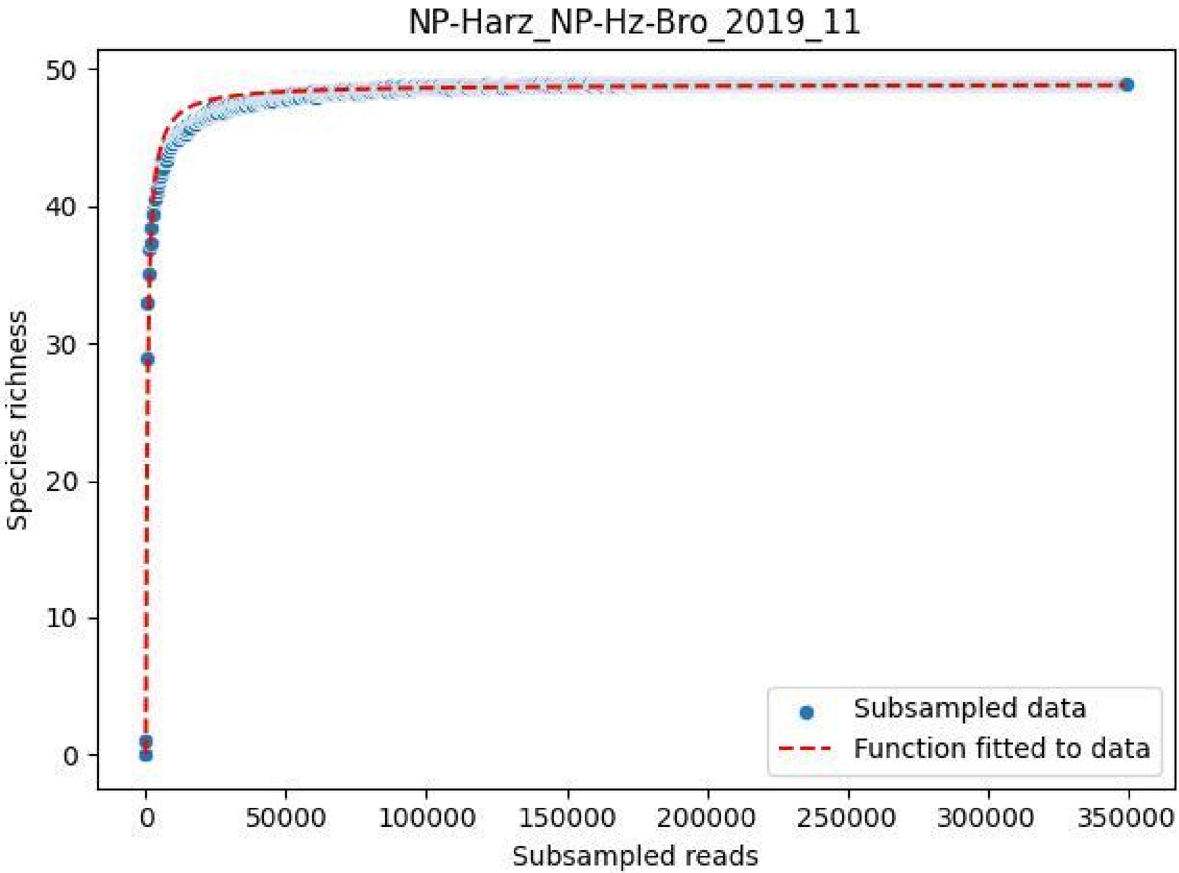
NP-Harz_NP-Hz-Bro_2019_11
Species richness
Subsampled reads
0
10
20
30
40
50
0
50000
100000
150000
200000
250000
300000
350000
Subsampled data
Function fitted to data

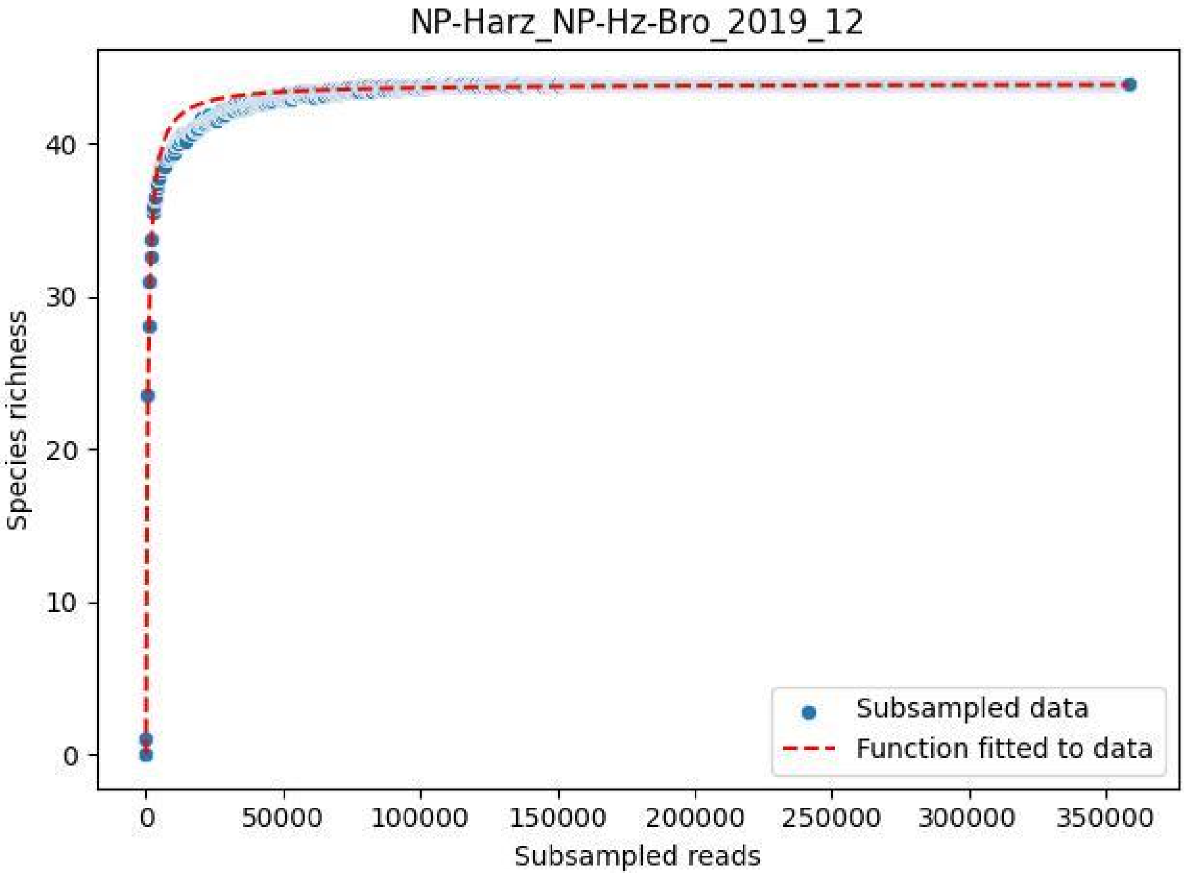
NP-Harz_NP-Hz-Bro_2019_12
Species richness
Subsampled reads
Subsampled data
Function fitted to data
0
10
20
30
40
0
50000
100000
150000
200000
250000
300000
350000

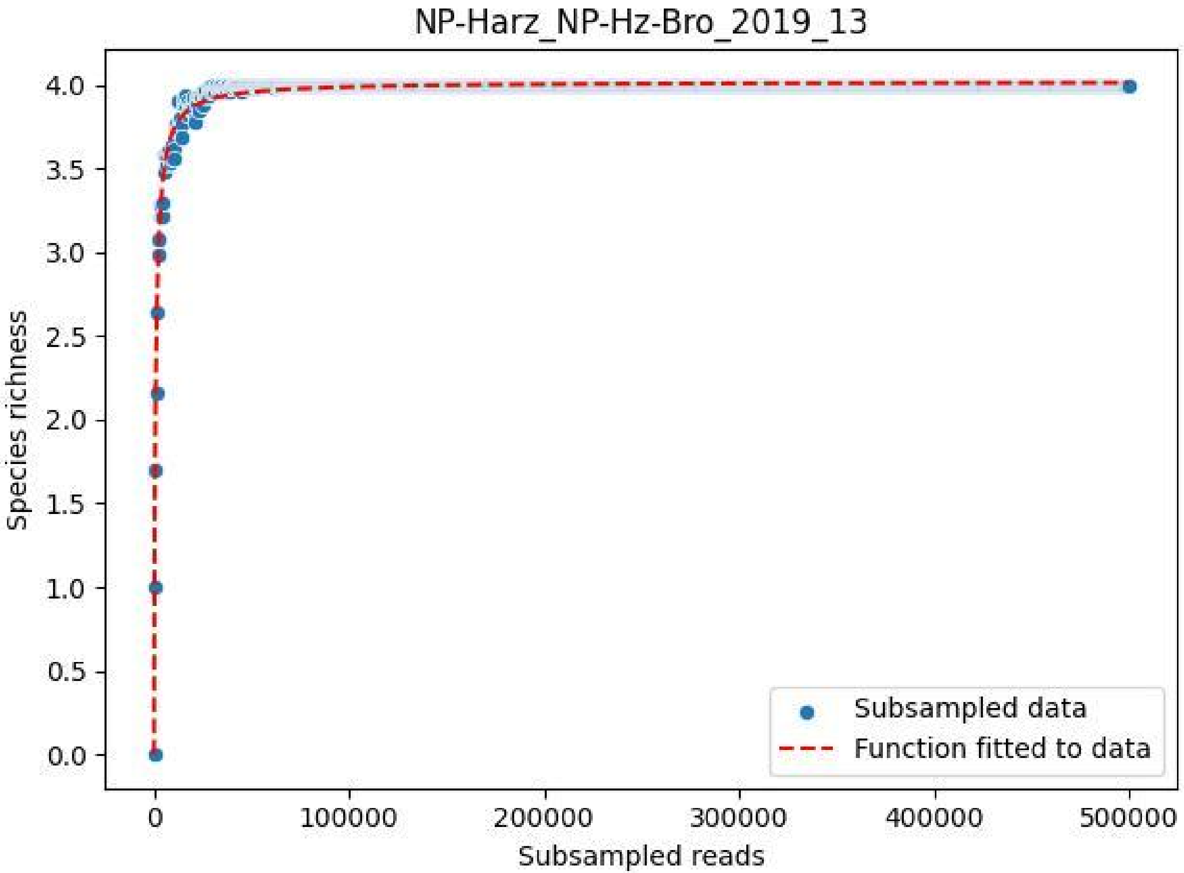
NP-Harz_NP-Hz-Bro_2019_13
Species richness
Subsampled reads
0.0
0.5
1.0
1.5
2.0
2.5
3.0
3.5
4.0
0
100000
200000
300000
400000
500000
Subsampled data
Function fitted to data

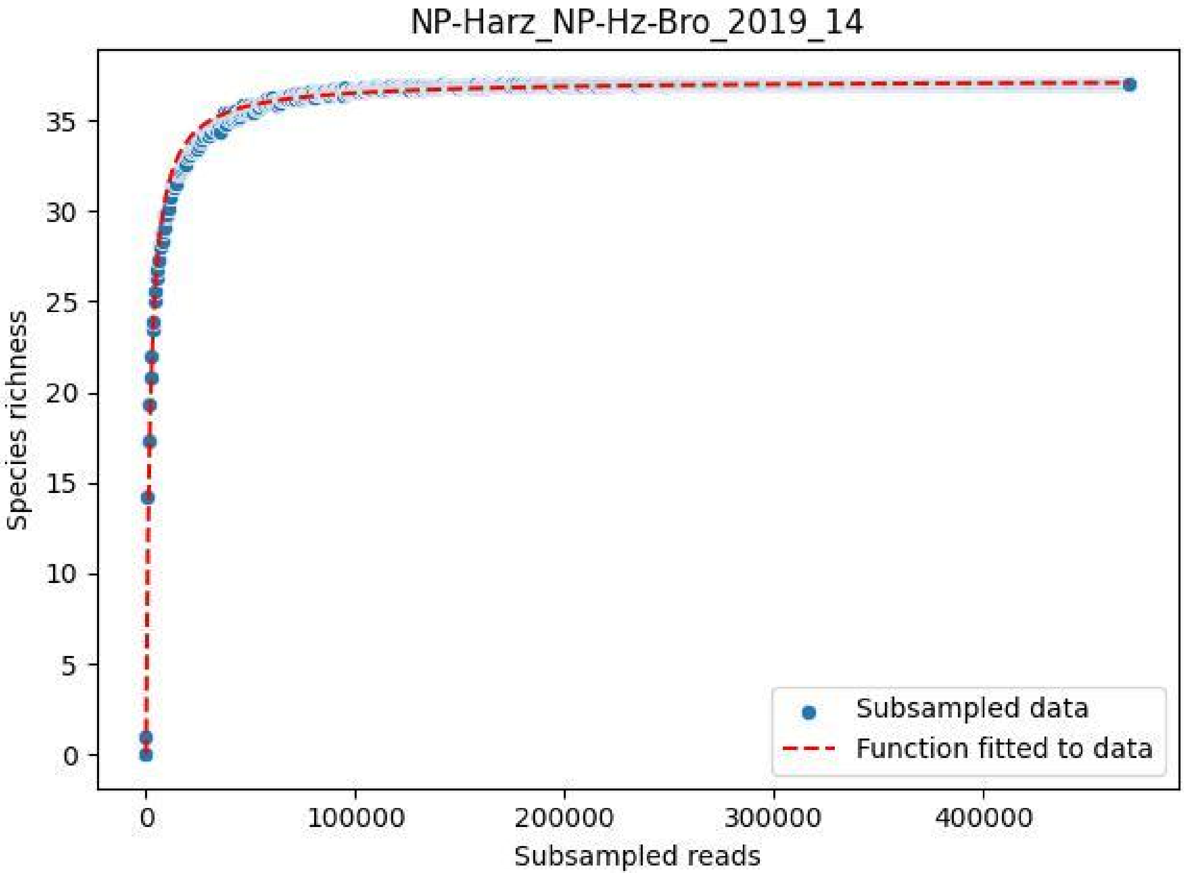

NP-Harz_NP-Hz-Bro_2019_14
Species richness
Subsampled reads
0
5
10
15
20
25
30
35
0
100000
200000
300000
400000
Subsampled data
Function fitted to data

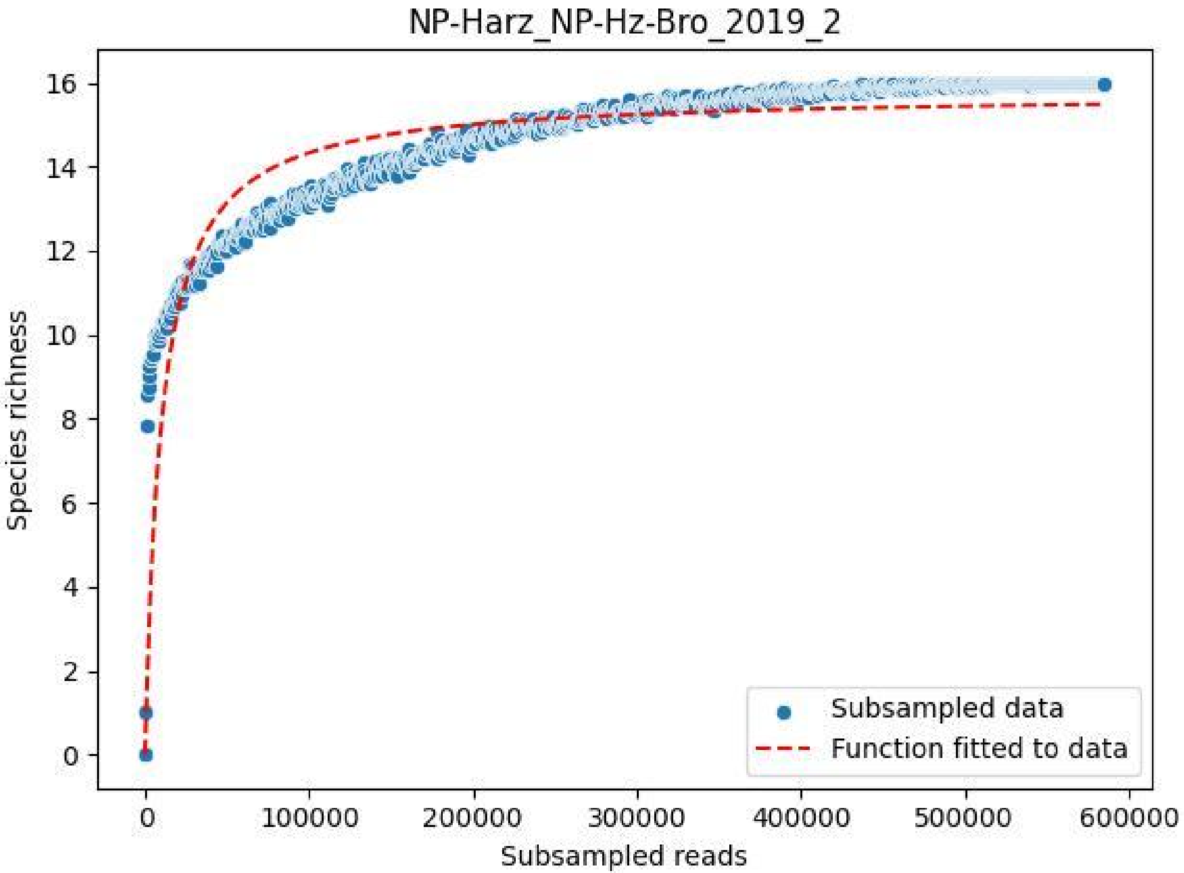
NP-Harz_NP-Hz-Bro_2019_2
Species richness
Subsampled reads
0
2
4
6
8
10
12
14
16
0
100000
200000
300000
400000
500000
600000
Subsampled data
Function fitted to data

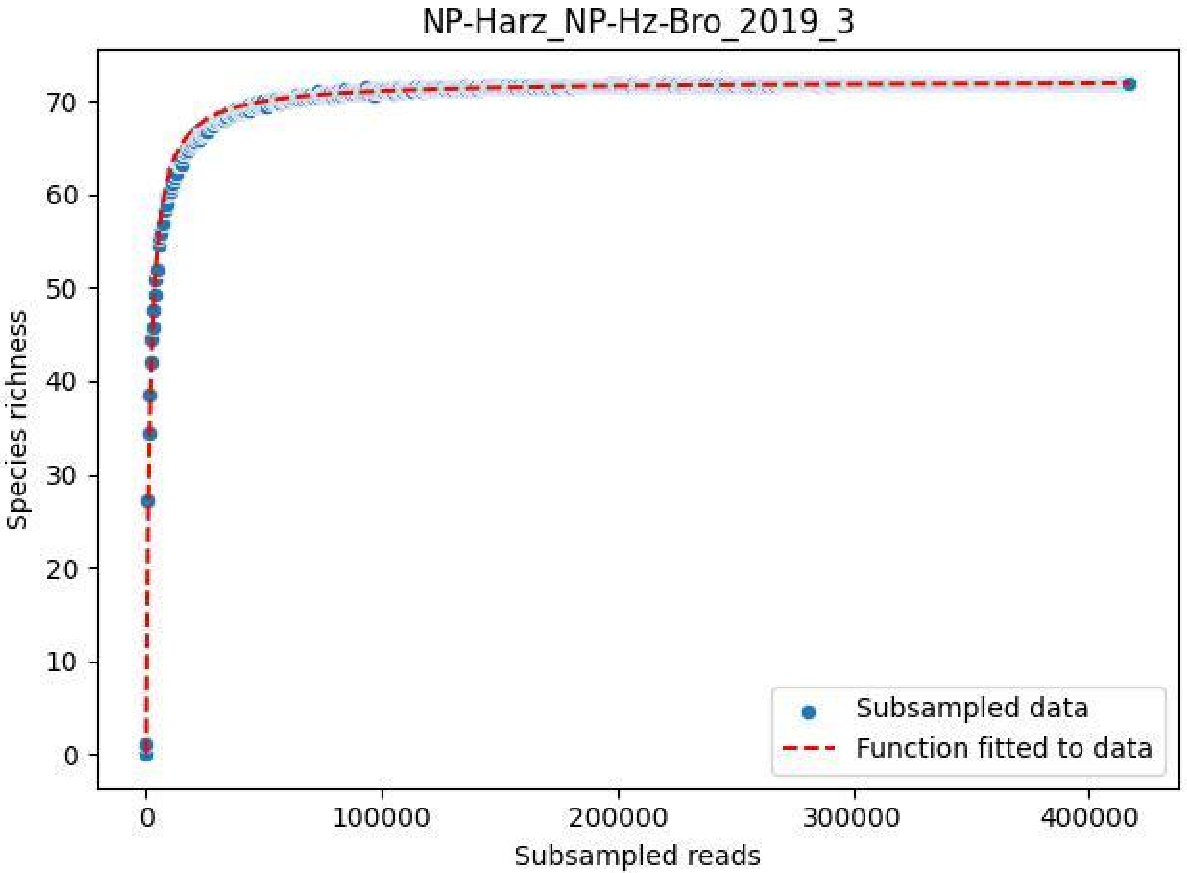
NP-Harz_NP-Hz-Bro_2019_3
Species richness
Subsampled reads
0
10
20
30
40
50
60
70
0
100000
200000
300000
400000
Subsampled data
Function fitted to data

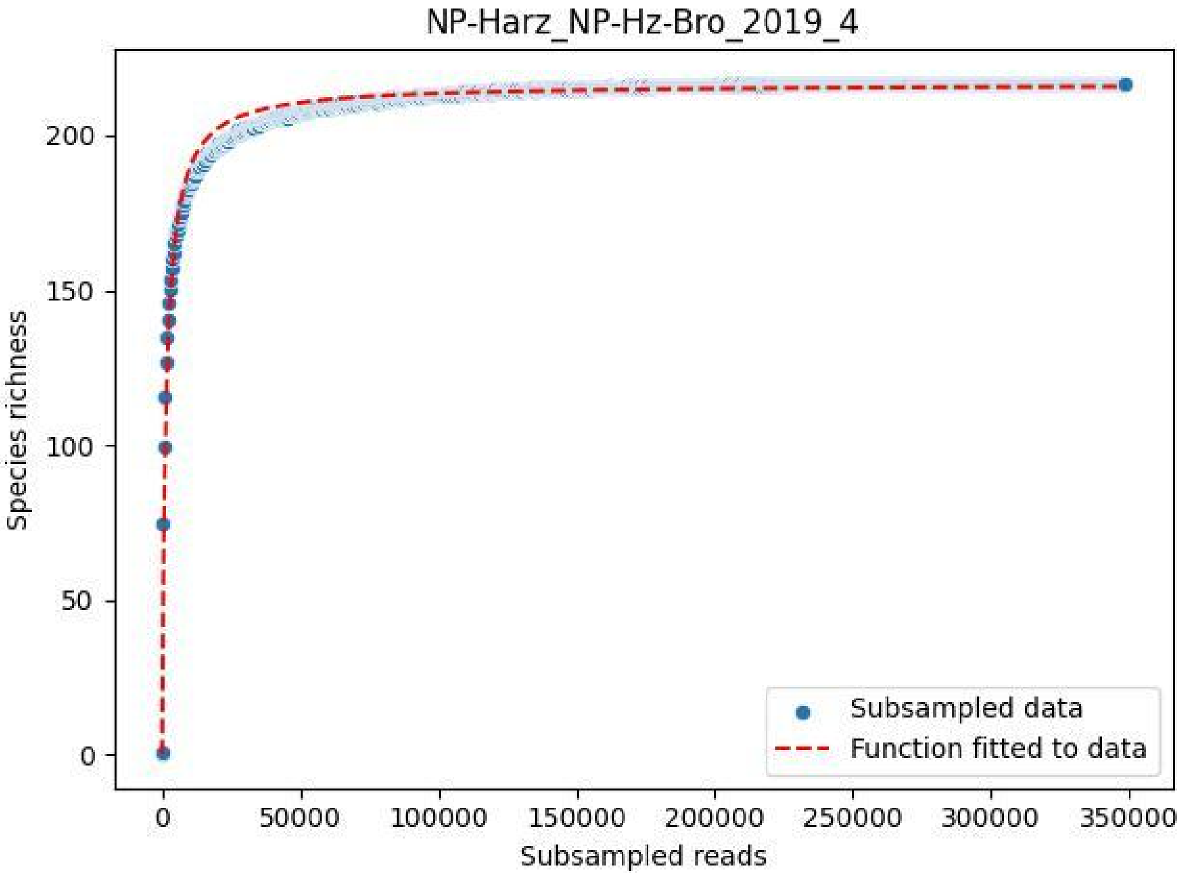
NP-Harz_NP-Hz-Bro_2019_4
Species richness
Subsampled reads
0
50
100
150
200
0
50000
100000
150000
200000
250000
300000
350000
Subsampled data
Function fitted to data

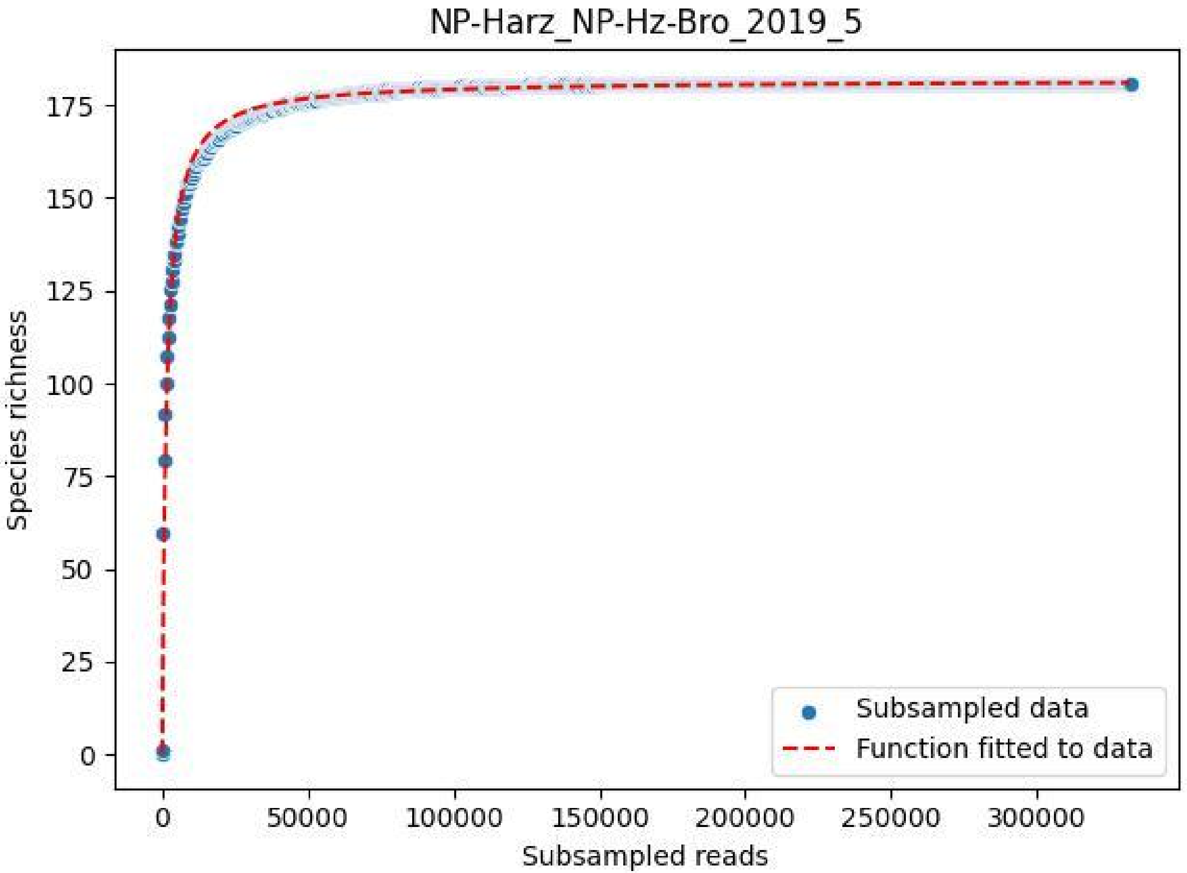
NP-Harz_NP-Hz-Bro_2019_5
Species richness
Subsampled reads
0
25
50
75
100
125
150
175
0
50000
100000
150000
200000
250000
300000
Subsampled data
Function fitted to data

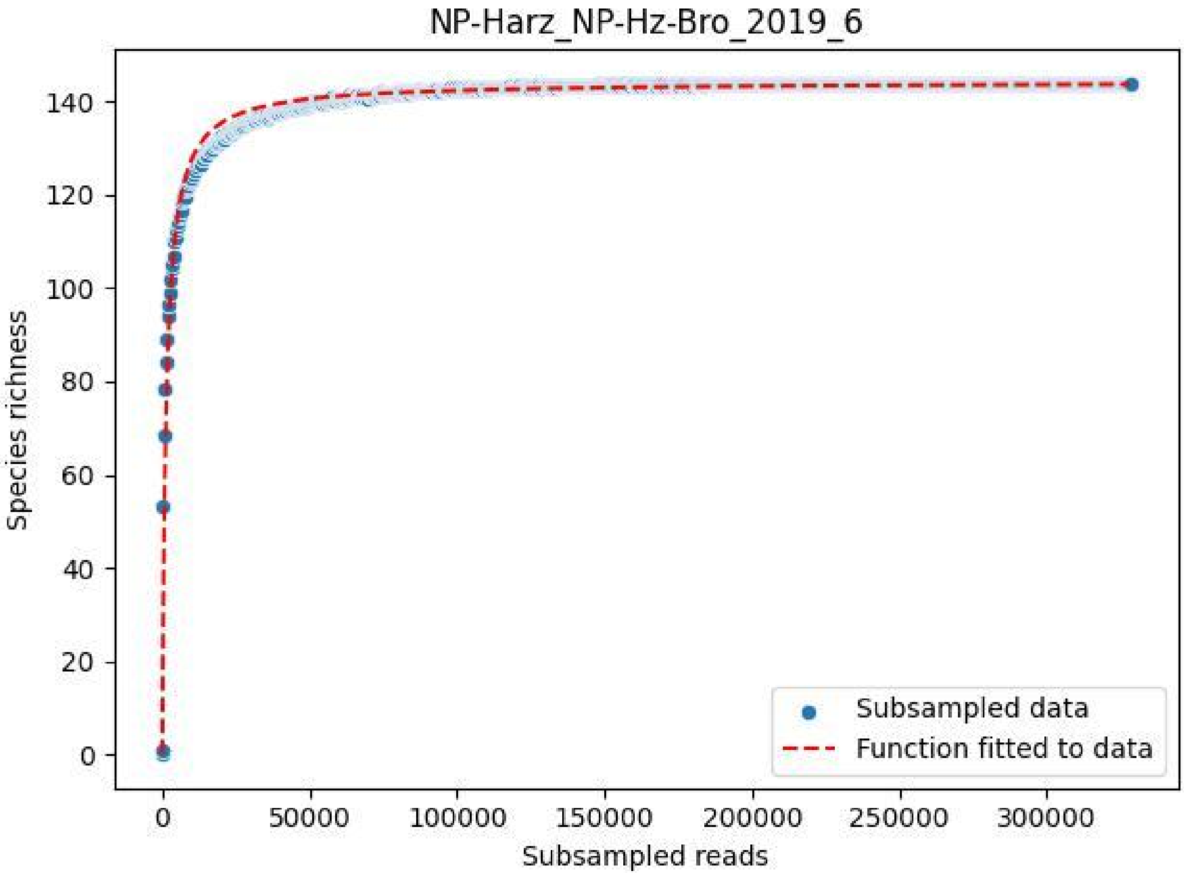
NP-Harz_NP-Hz-Bro_2019_6
Species richness
Subsampled reads
0
20
40
60
80
100
120
140
0
50000
100000
150000
200000
250000
300000
Subsampled data
Function fitted to data

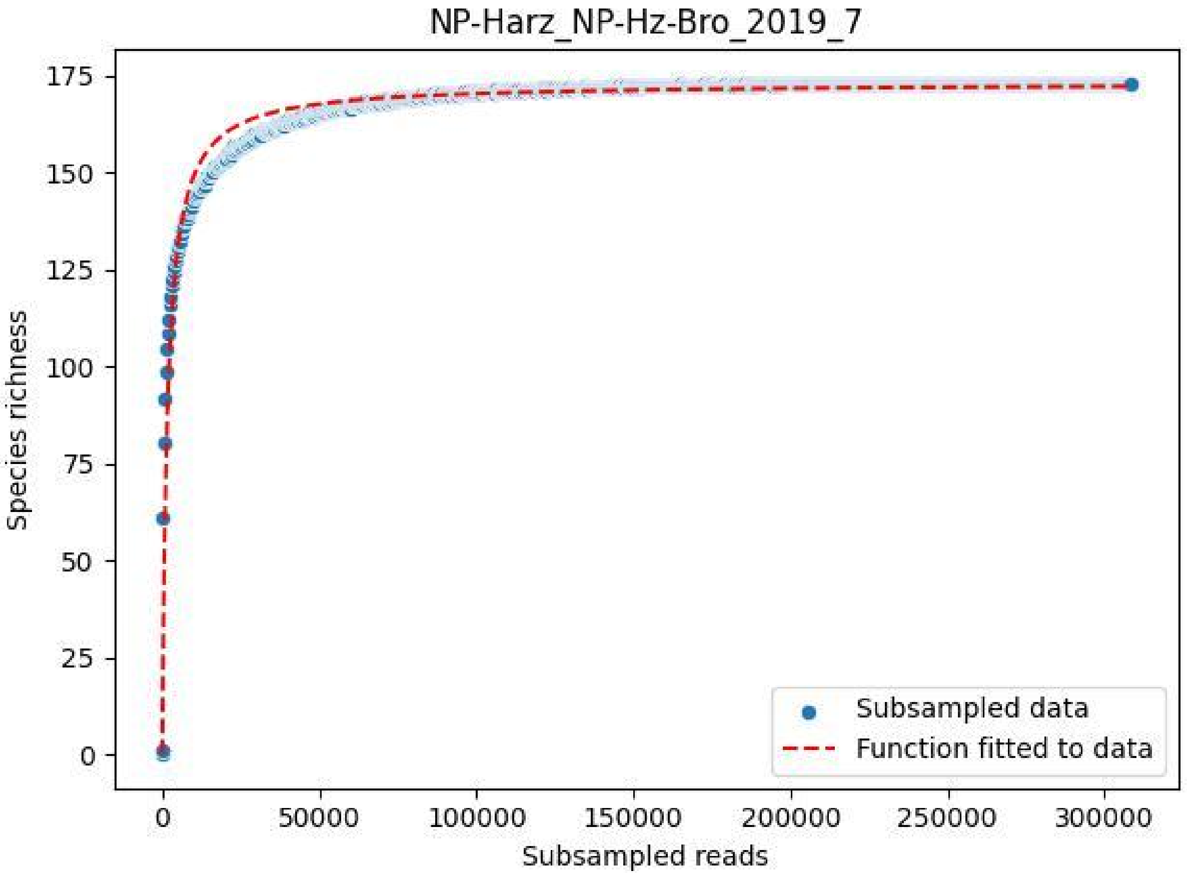
NP-Harz_NP-Hz-Bro_2019_7
Species richness
Subsampled reads
0
25
50
75
100
125
150
175
0
50000
100000
150000
200000
250000
300000
Subsampled data
Function fitted to data

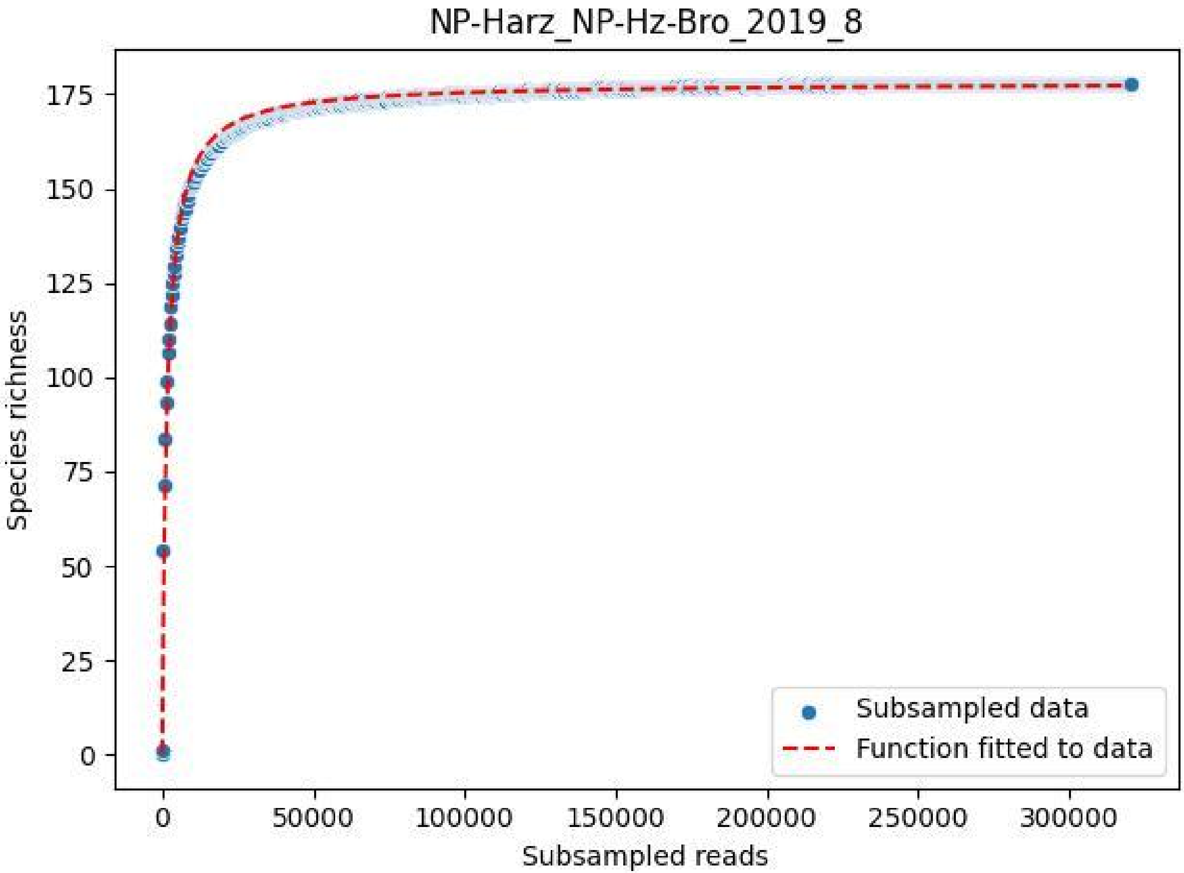
NP-Harz_NP-Hz-Bro_2019_8
Species richness
Subsampled reads
0
25
50
75
100
125
150
175
0
50000
100000
150000
200000
250000
300000
Subsampled data
Function fitted to data

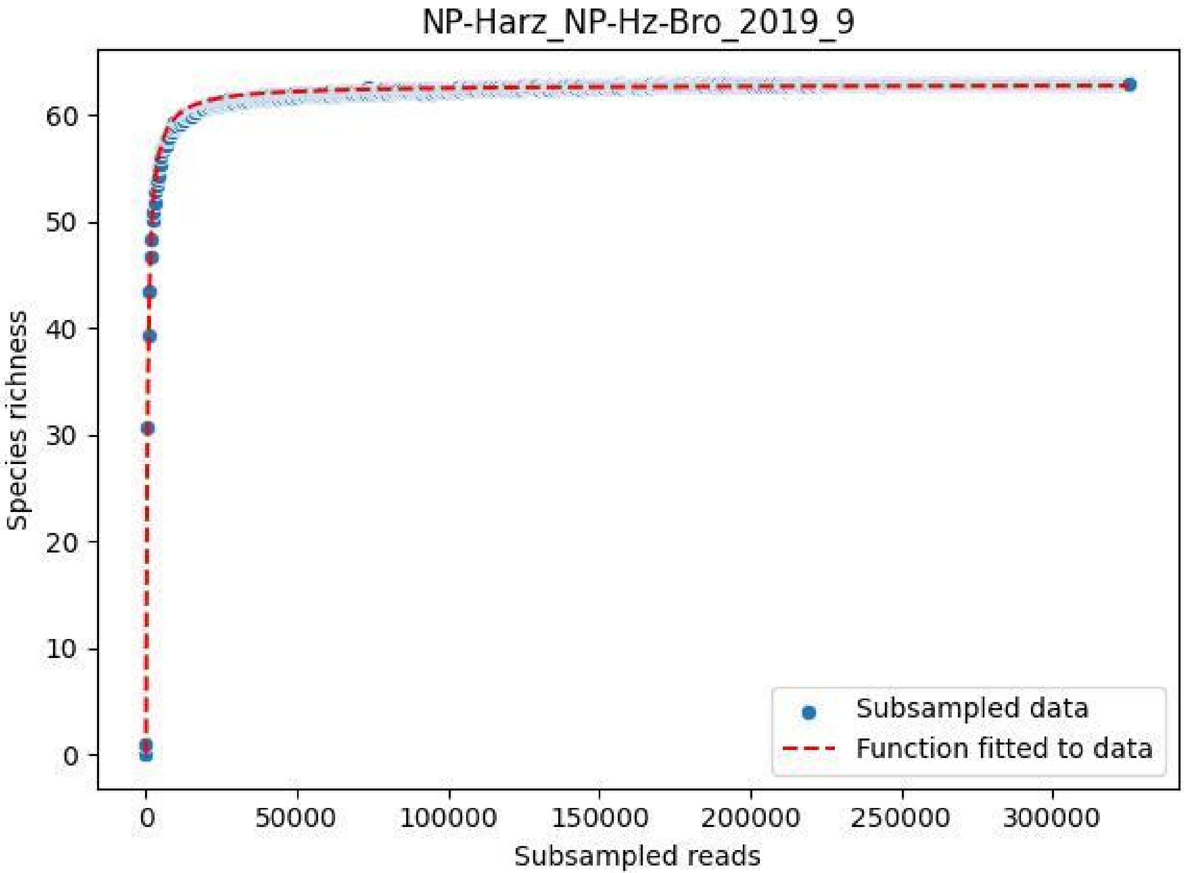
NP-Harz_NP-Hz-Bro_2019_9
Species richness
Subsampled reads
Subsampled data
Function fitted to data
0
10
20
30
40
50
60
0
50000
100000
150000
200000
250000
300000

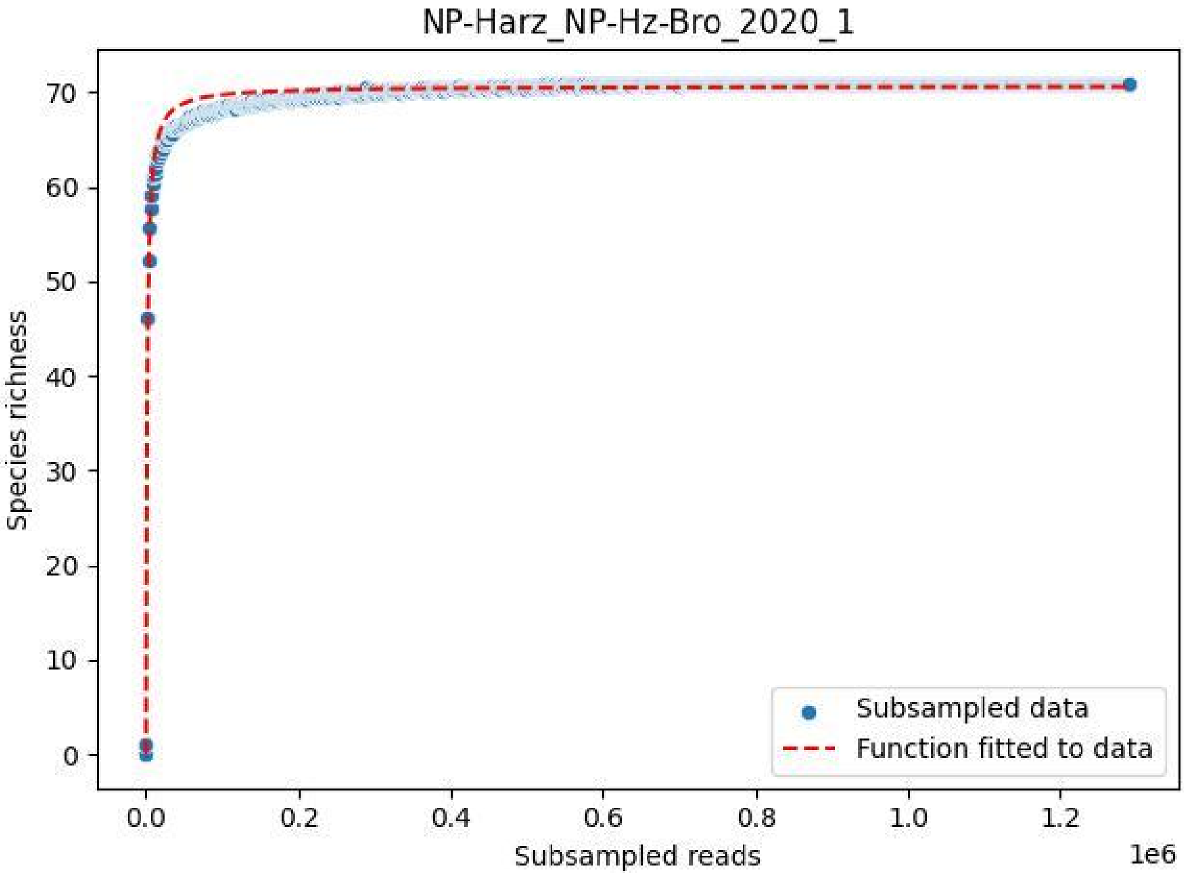
NP-Harz_NP-Hz-Bro_2020_1
Species richness
Subsampled reads
1e6
0
10
20
30
40
50
60
70
0.0
0.2
0.4
0.6
0.8
1.0
1.2
Subsampled data
Function fitted to data

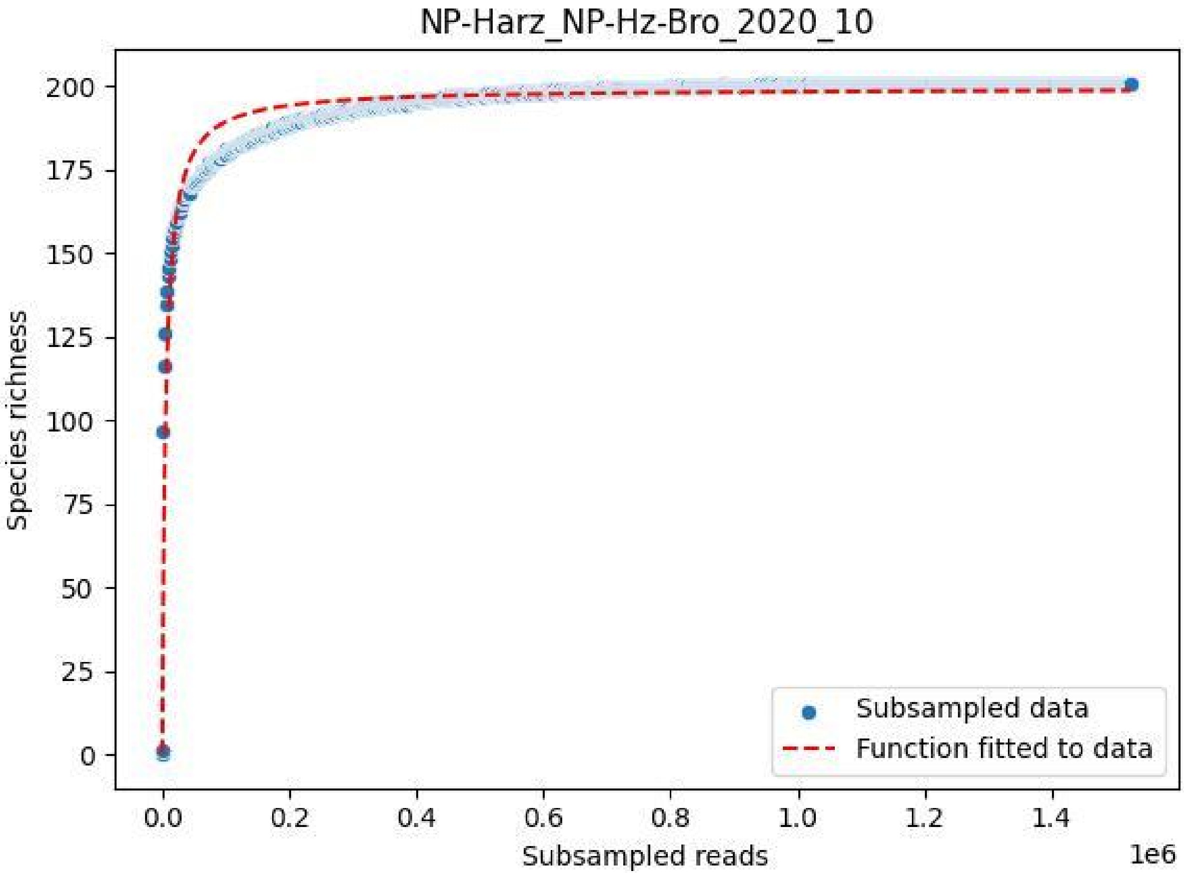

NP-Harz_NP-Hz-Bro_2020_10
Species richness
Subsampled reads
1e6
200
175
150
125
100
75
50
25
0
0.0
0.2
0.4
0.6
0.8
1.0
1.2
1.4
Subsampled data
Function fitted to data

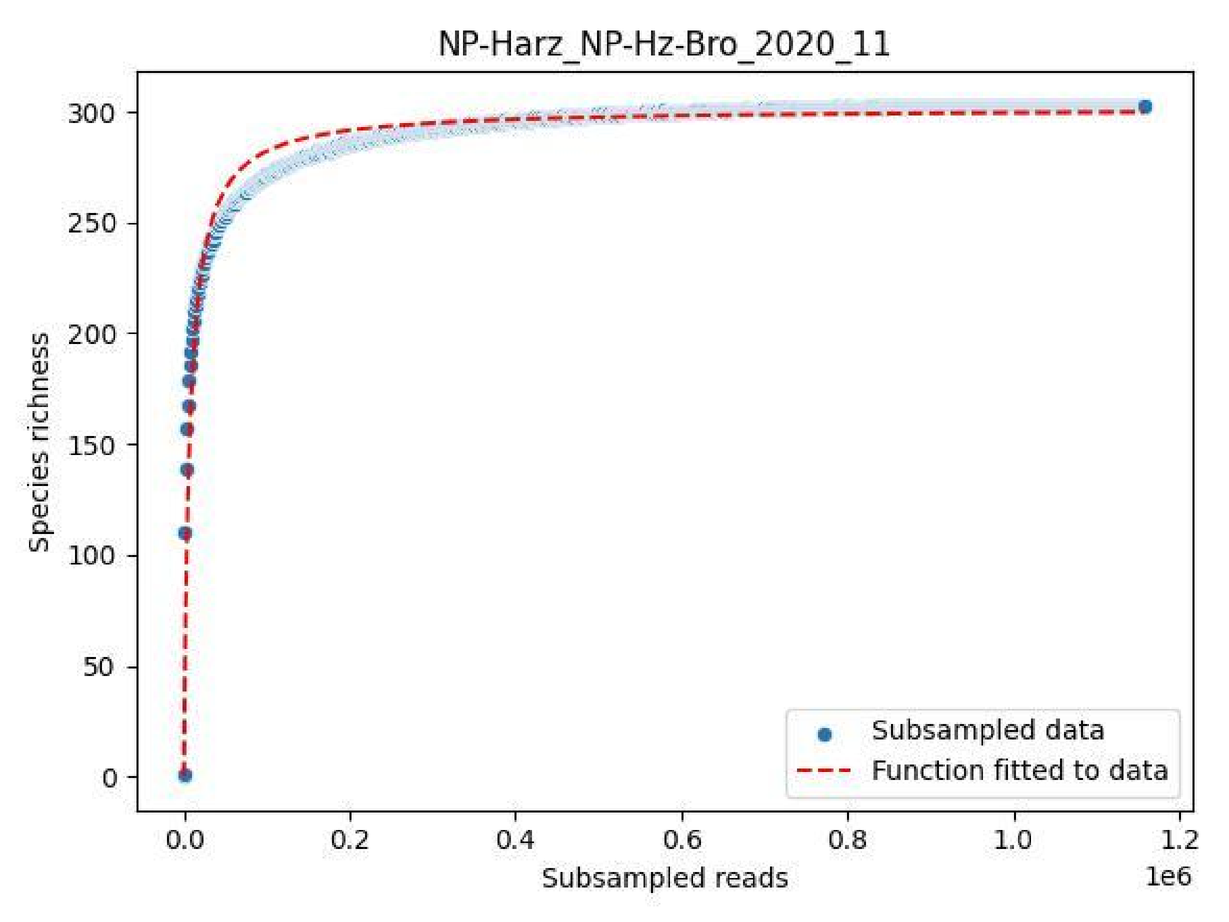

NP-Harz_NP-Hz-Bro_2020_11
Species richness
Subsampled reads
1e6
0
50
100
150
200
250
300
0.0
0.2
0.4
0.6
0.8
1.0
1.2
Subsampled data
Function fitted to data

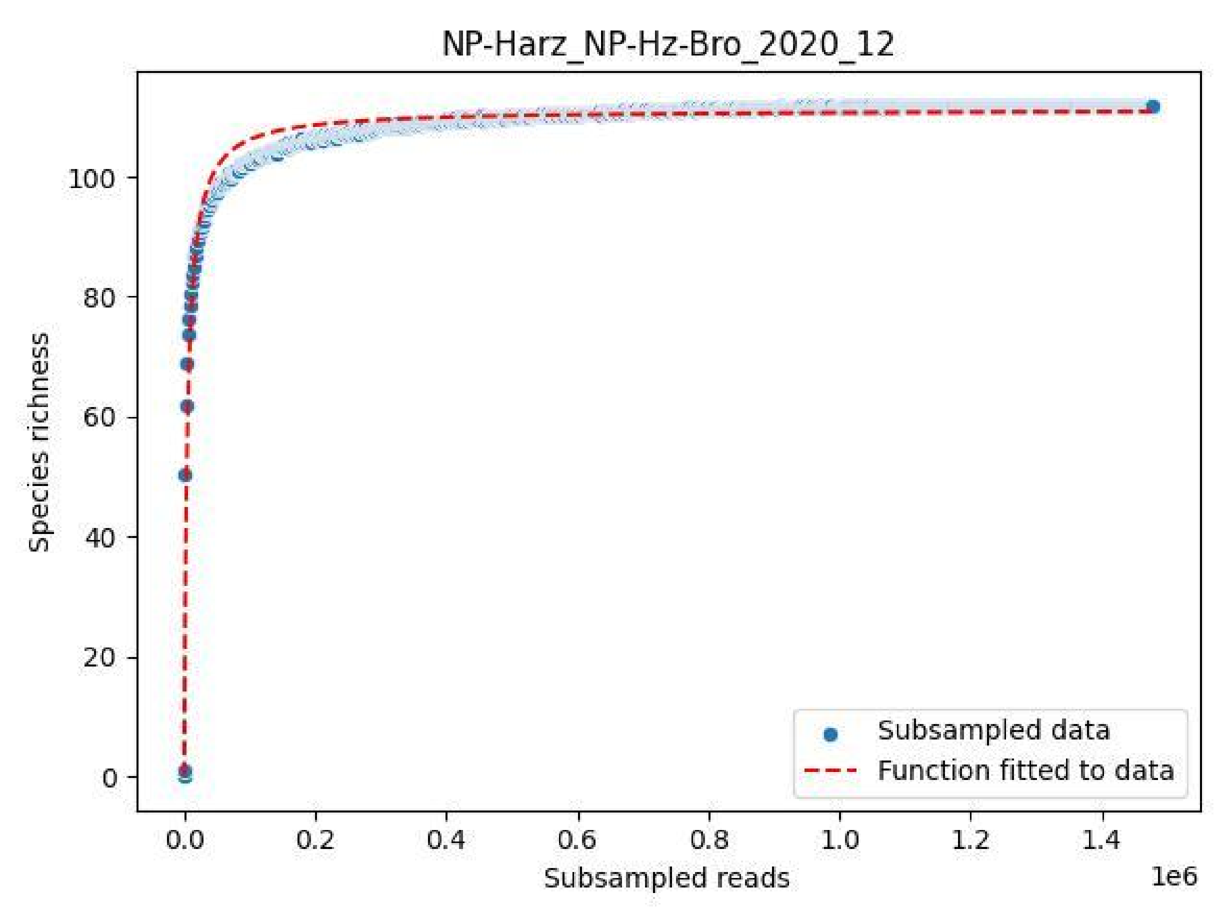
NP-Harz_NP-Hz-Bro_2020_12
Species richness
Subsampled reads
1e6
Subsampled data
Function fitted to data
0
20
40
60
80
100
0.0
0.2
0.4
0.6
0.8
1.0
1.2
1.4

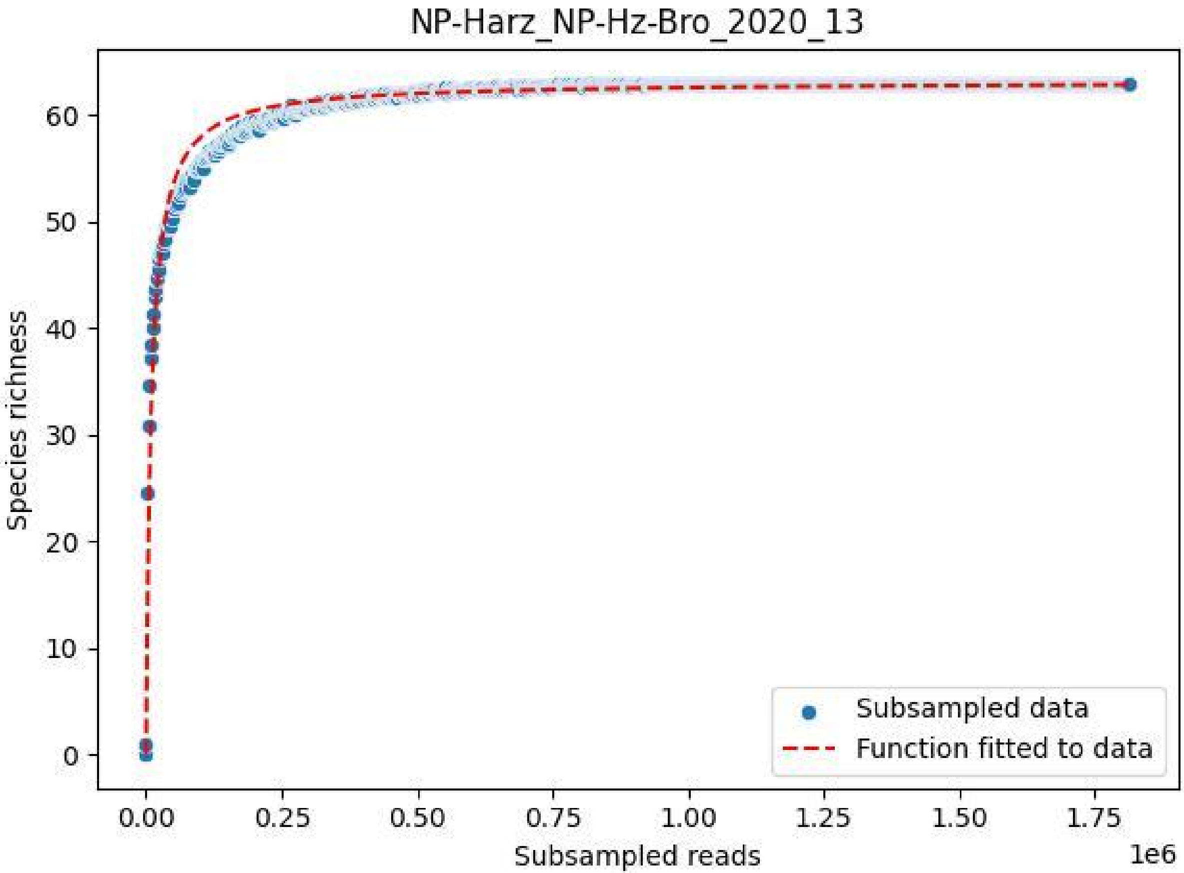
NP-Harz_NP-Hz-Bro_2020_13
Species richness
Subsampled reads
1e6
0
10
20
30
40
50
60
0.00
0.25
0.50
0.75
1.00
1.25
1.50
1.75
Subsampled data
Function fitted to data

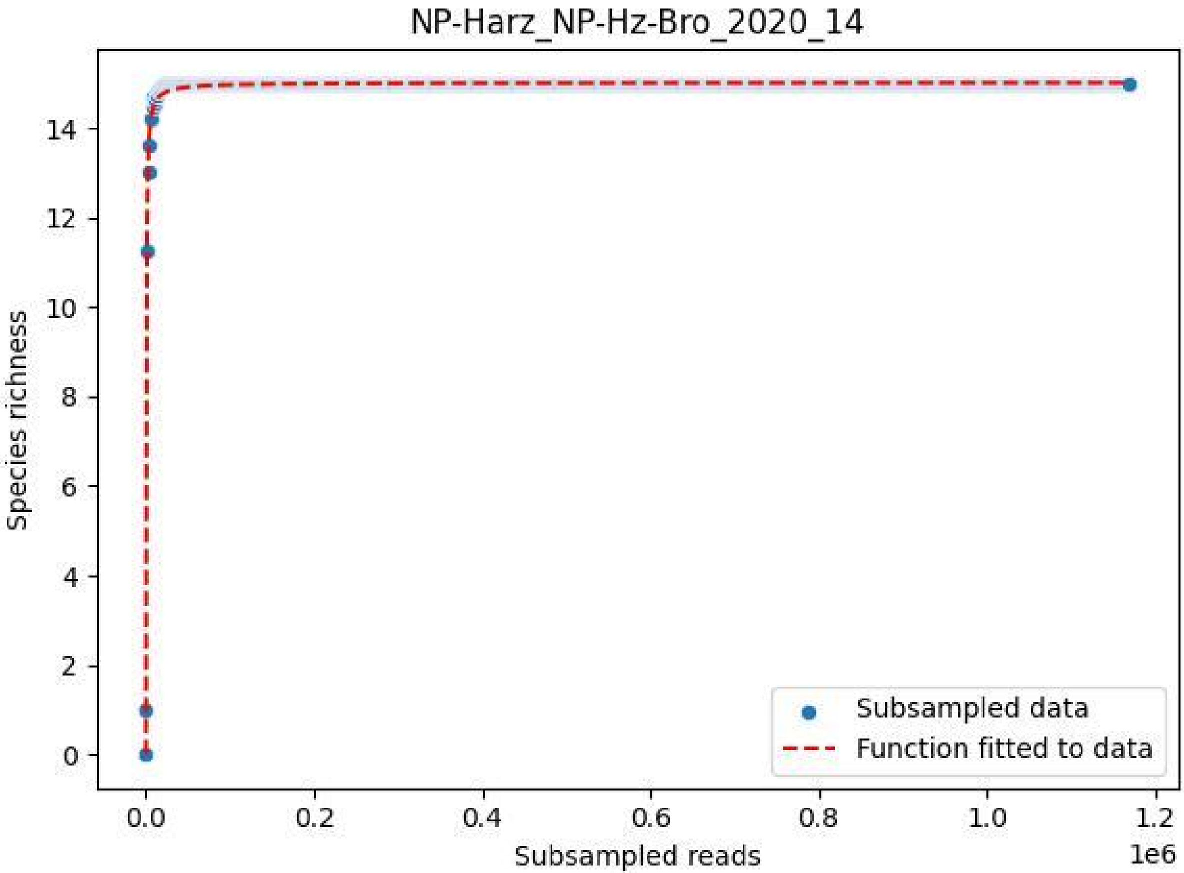
NP-Harz_NP-Hz-Bro_2020_14
Species richness
Subsampled reads
1e6
0.0
0.2
0.4
0.6
0.8
1.0
1.2
0
2
4
6
8
10
12
14
Subsampled data
Function fitted to data

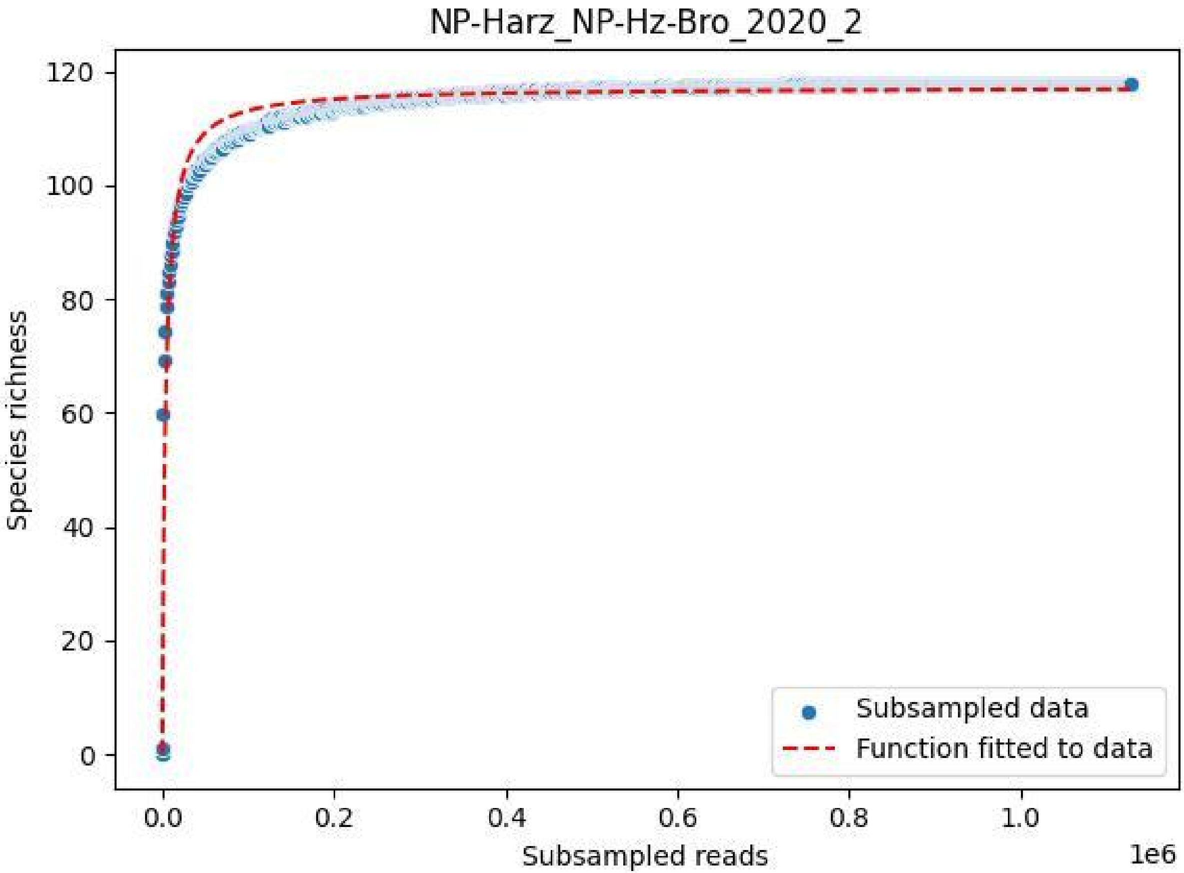
NP-Harz_NP-Hz-Bro_2020_2
Species richness
Subsampled reads
1e6
0.0
0.2
0.4
0.6
0.8
1.0
0
20
40
60
80
100
120
Subsampled data
Function fitted to data

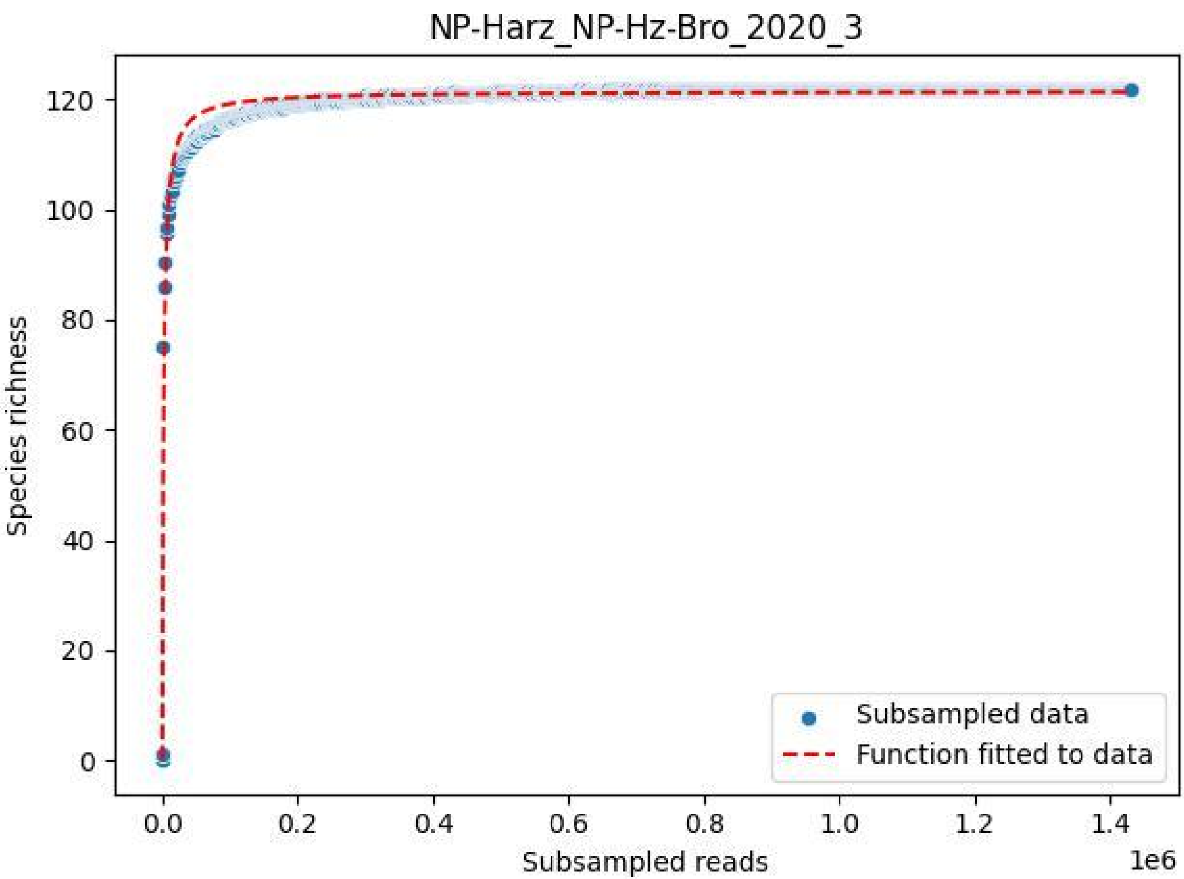
NP-Harz_NP-Hz-Bro_2020_3
Species richness
0
20
40
60
80
100
120
0.0
0.2
0.4
0.6
0.8
1.0
1.2
1.4
1e6
Subsampled reads
Subsampled data
Function fitted to data

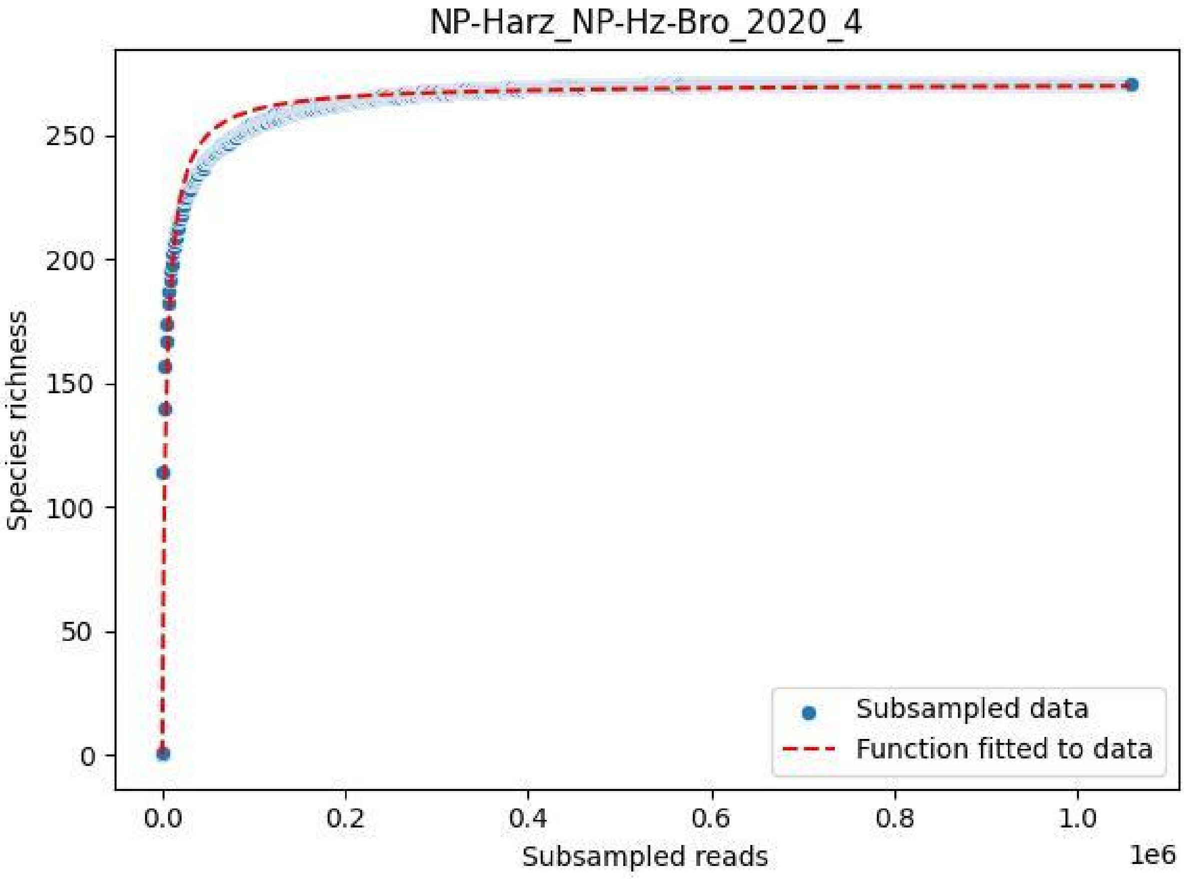
NP-Harz_NP-Hz-Bro_2020_4
Species richness
Subsampled reads
1e6
0
50
100
150
200
250
0.0
0.2
0.4
0.6
0.8
1.0
Subsampled data
Function fitted to data

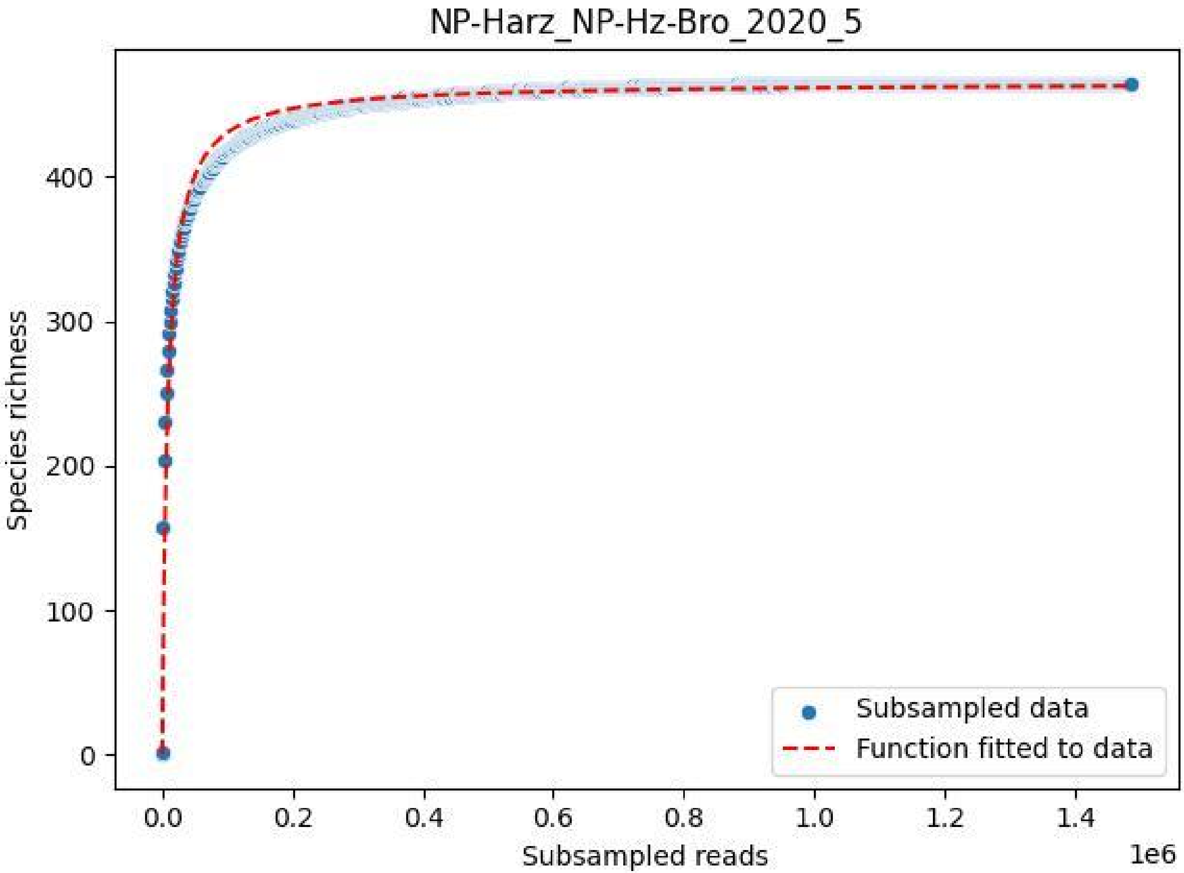

NP-Harz_NP-Hz-Bro_2020_5
Species richness
Subsampled reads
1e6
0
100
200
300
400
0.0
0.2
0.4
0.6
0.8
1.0
1.2
1.4
Subsampled data
Function fitted to data

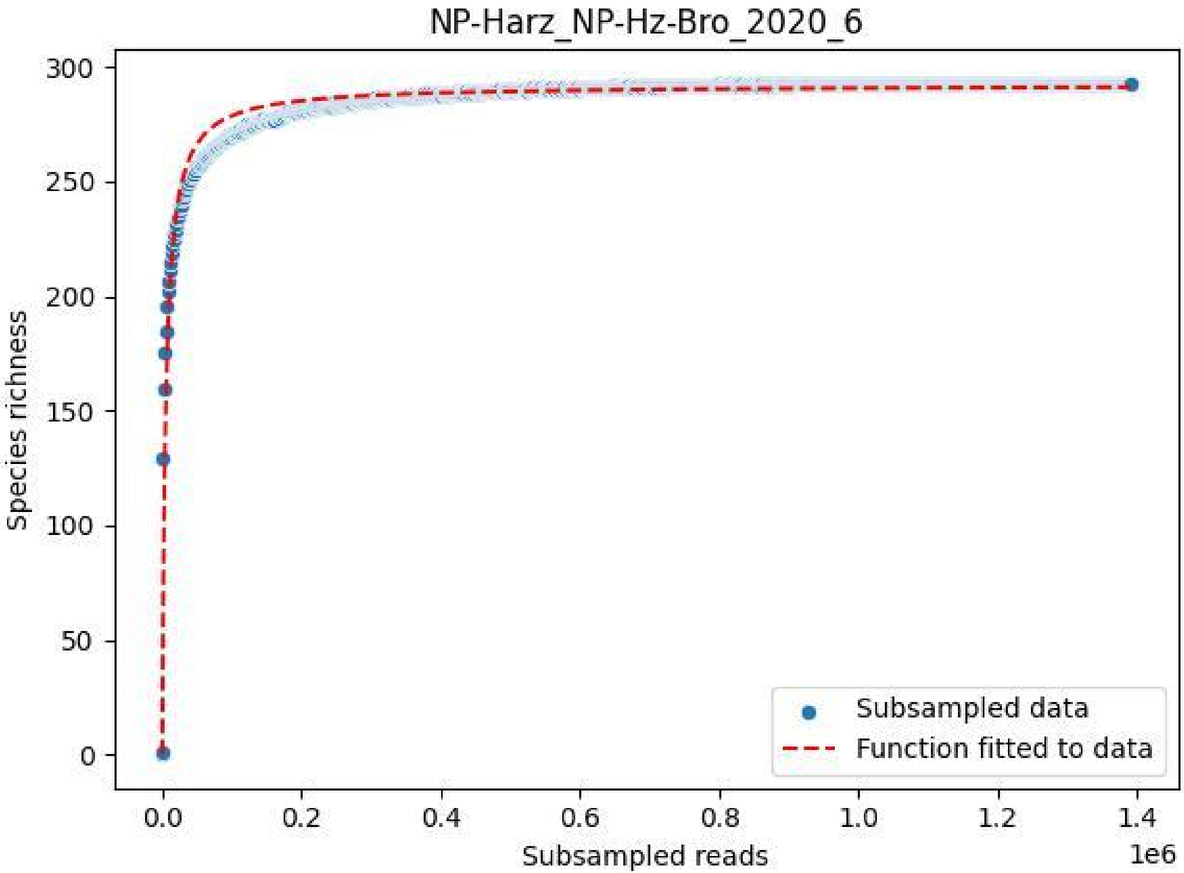
NP-Harz_NP-Hz-Bro_2020_6
Species richness
Subsampled reads
1e6
300
250
200
150
100
50
0
0.0
0.2
0.4
0.6
0.8
1.0
1.2
1.4
Subsampled data
Function fitted to data

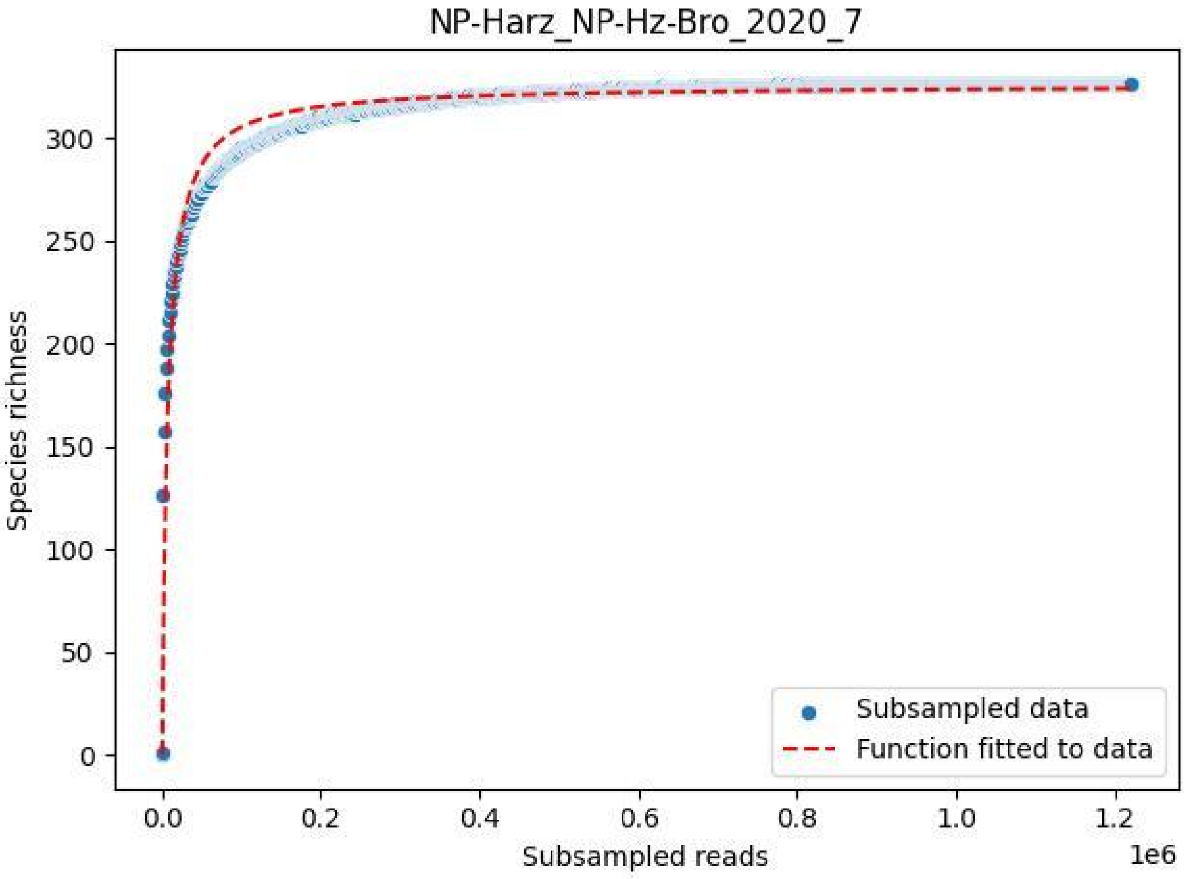

NP-Harz_NP-Hz-Bro_2020_7
Species richness
Subsampled reads
1e6
0
50
100
150
200
250
300
0.0
0.2
0.4
0.6
0.8
1.0
1.2
Subsampled data
Function fitted to data

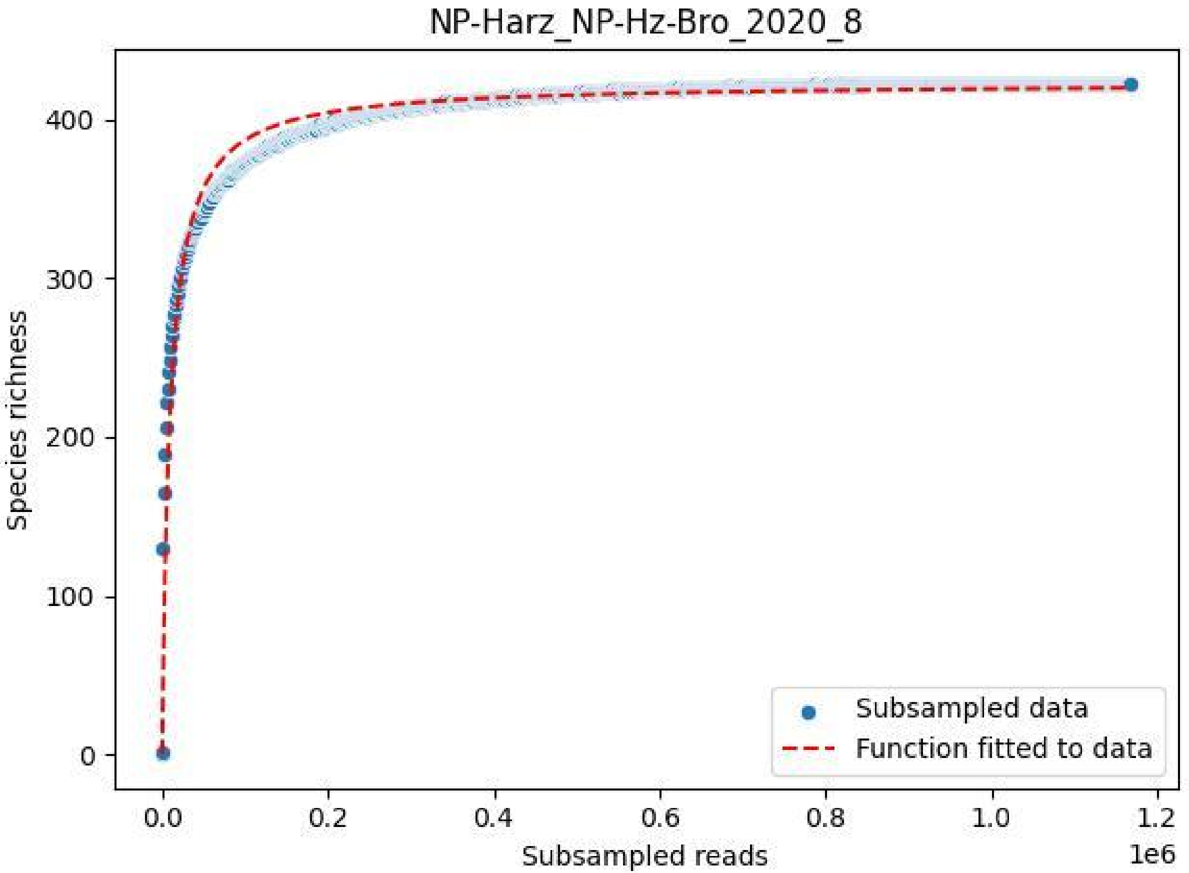

NP-Harz_NP-Hz-Bro_2020_8
Species richness
Subsampled reads
1e6
0
100
200
300
400
0.0
0.2
0.4
0.6
0.8
1.0
1.2
Subsampled data
Function fitted to data

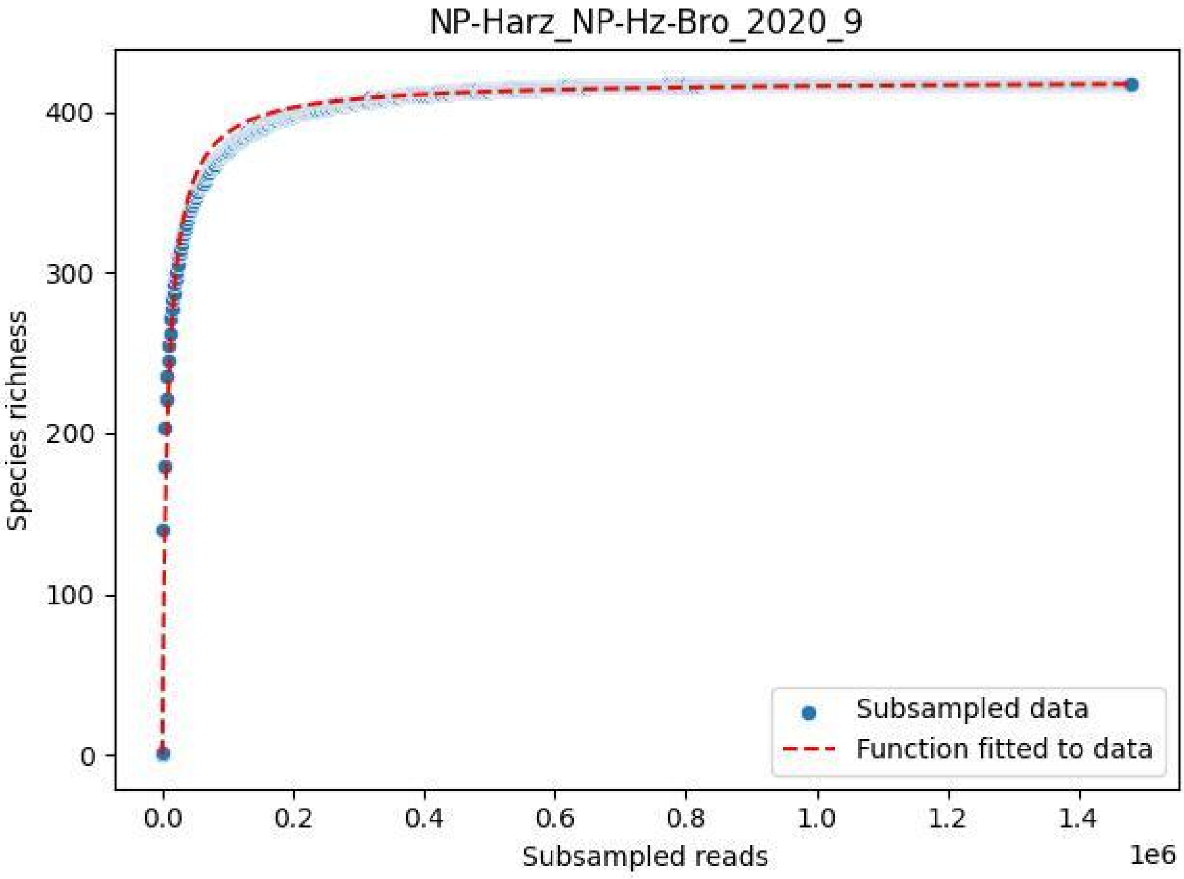
NP-Harz_NP-Hz-Bro_2020_9
Species richness
Subsampled reads
1e6
0
100
200
300
400
0.0
0.2
0.4
0.6
0.8
1.0
1.2
1.4
Subsampled data
Function fitted to data

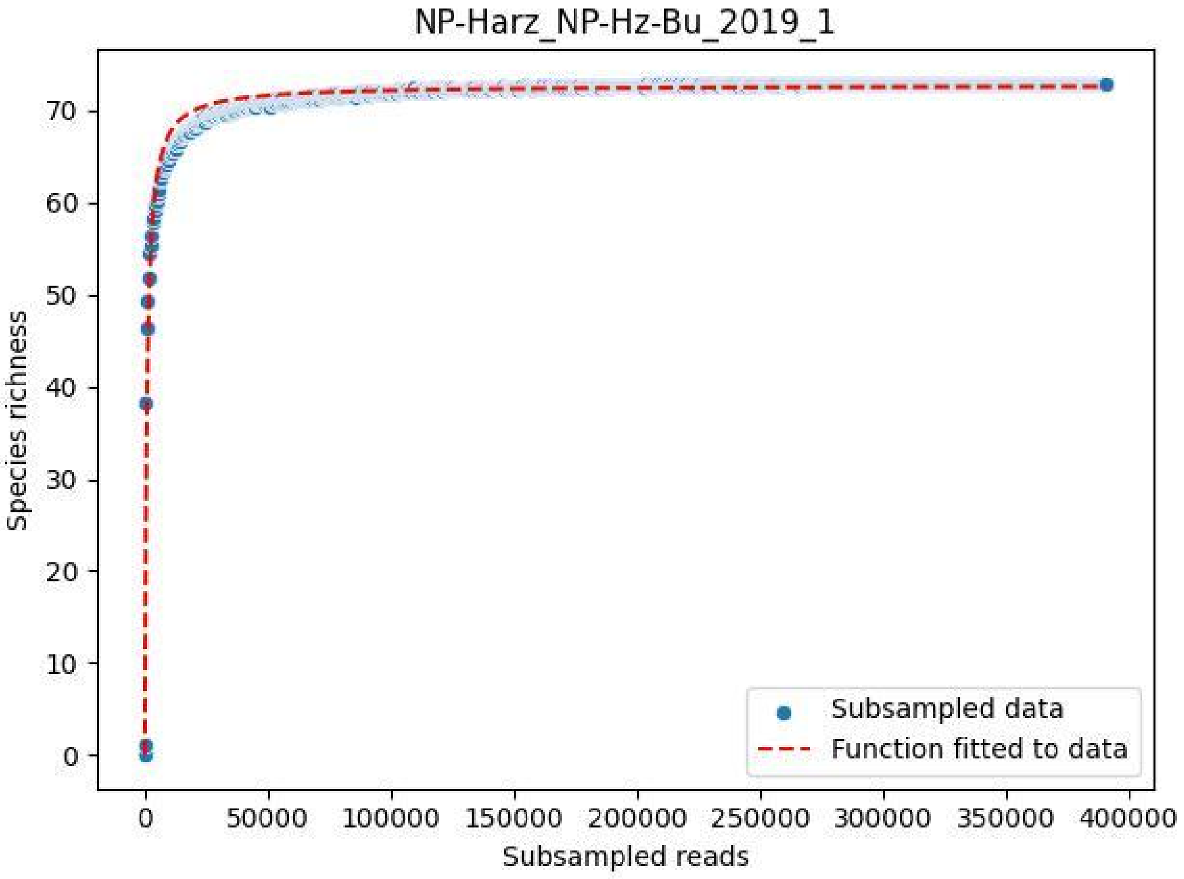
NP-Harz_NP-Hz-Bu_2019_1
Species richness
Subsampled reads
0
10
20
30
40
50
60
70
0
50000
100000
150000
200000
250000
300000
350000
400000
Subsampled data
Function fitted to data

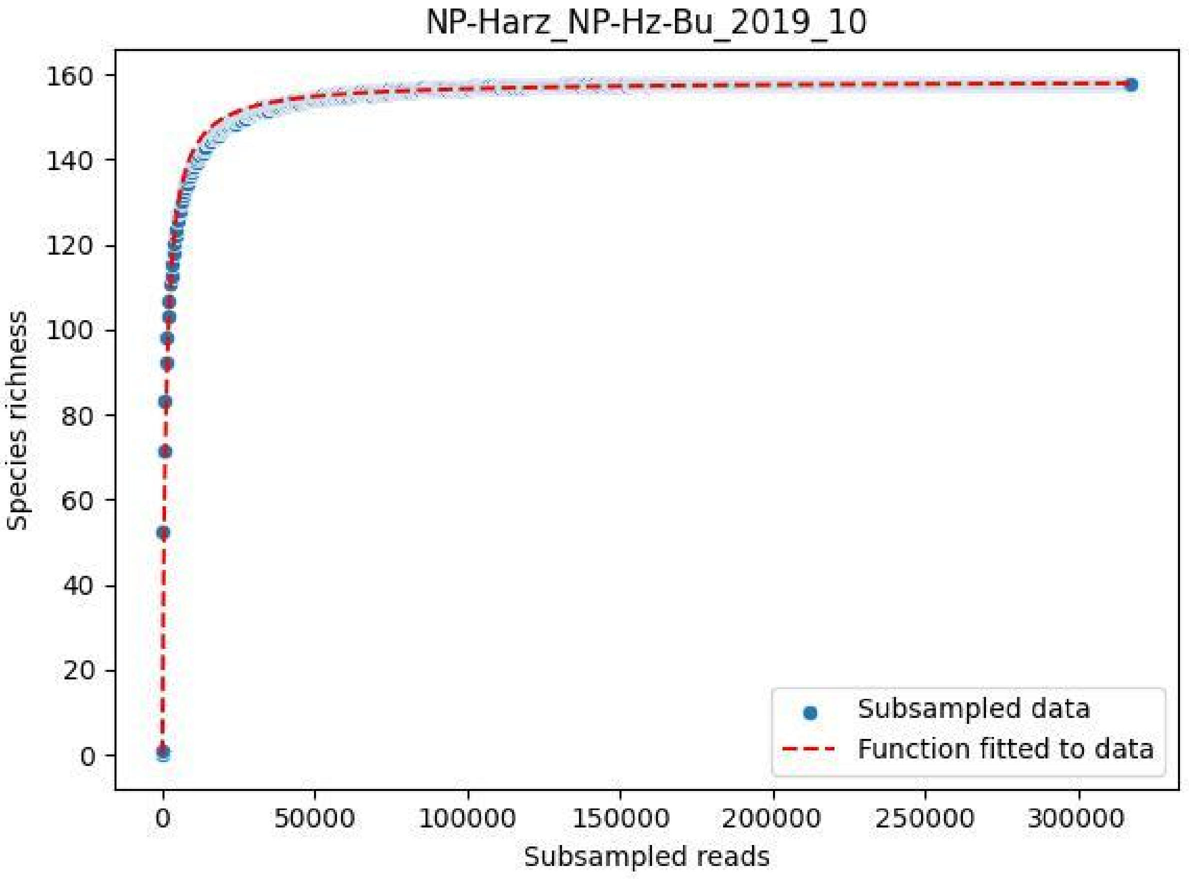

NP-Harz_NP-Hz-Bu_2019_10
Species richness
Subsampled reads
0
20
40
60
80
100
120
140
160
0
50000
100000
150000
200000
250000
300000
Subsampled data
Function fitted to data

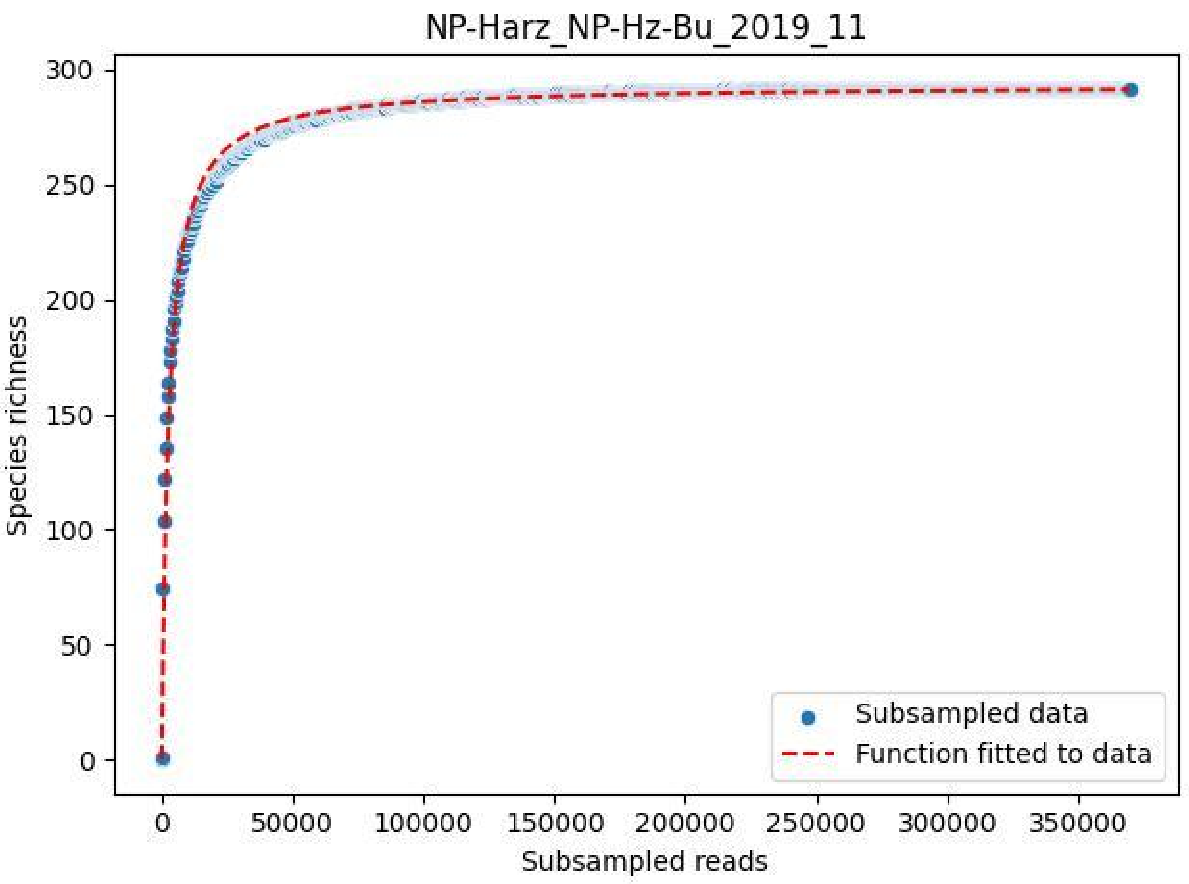

NP-Harz_NP-Hz-Bu_2019_11
Species richness
Subsampled reads
0
50
100
150
200
250
300
0
50000
100000
150000
200000
250000
300000
350000
Subsampled data
Function fitted to data

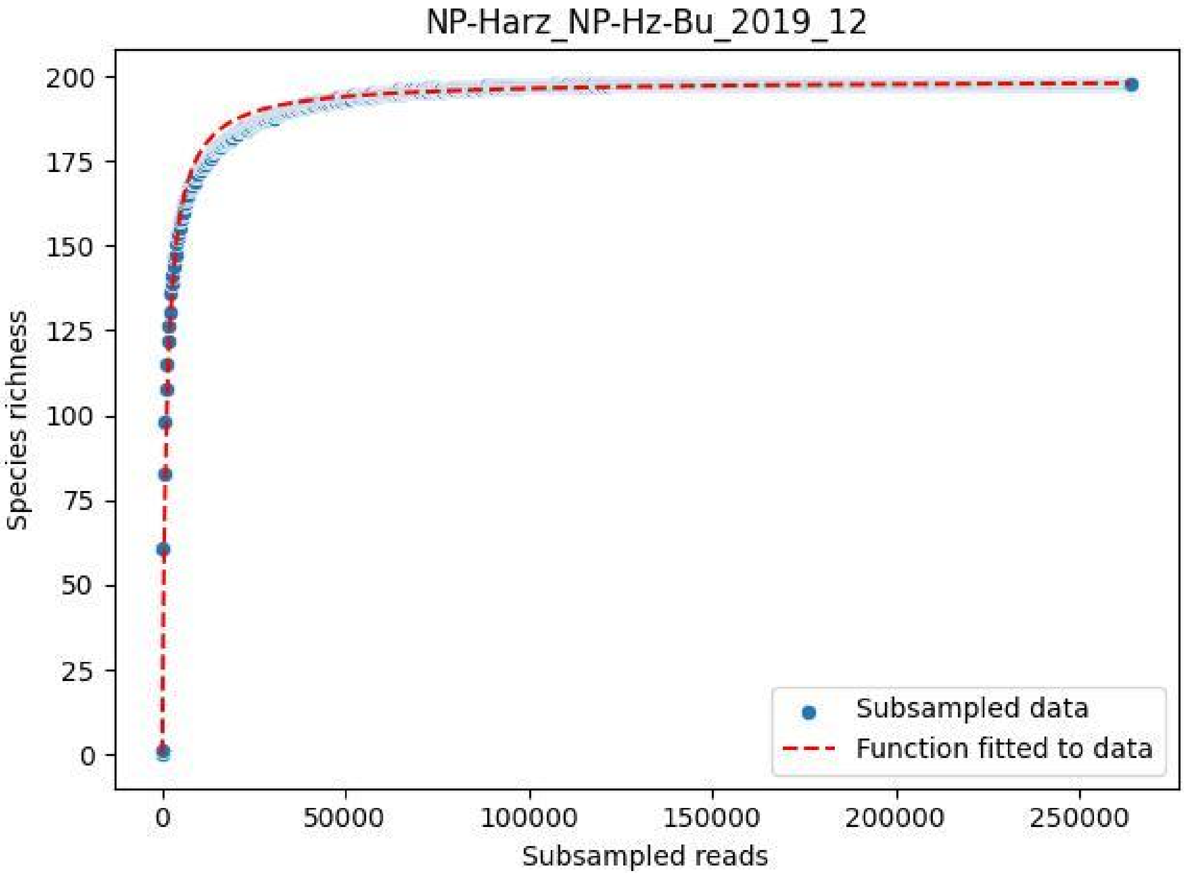
NP-Harz_NP-Hz-Bu_2019_12
Species richness
Subsampled reads
0
25
50
75
100
125
150
175
200
0
50000
100000
150000
200000
250000
Subsampled data
Function fitted to data

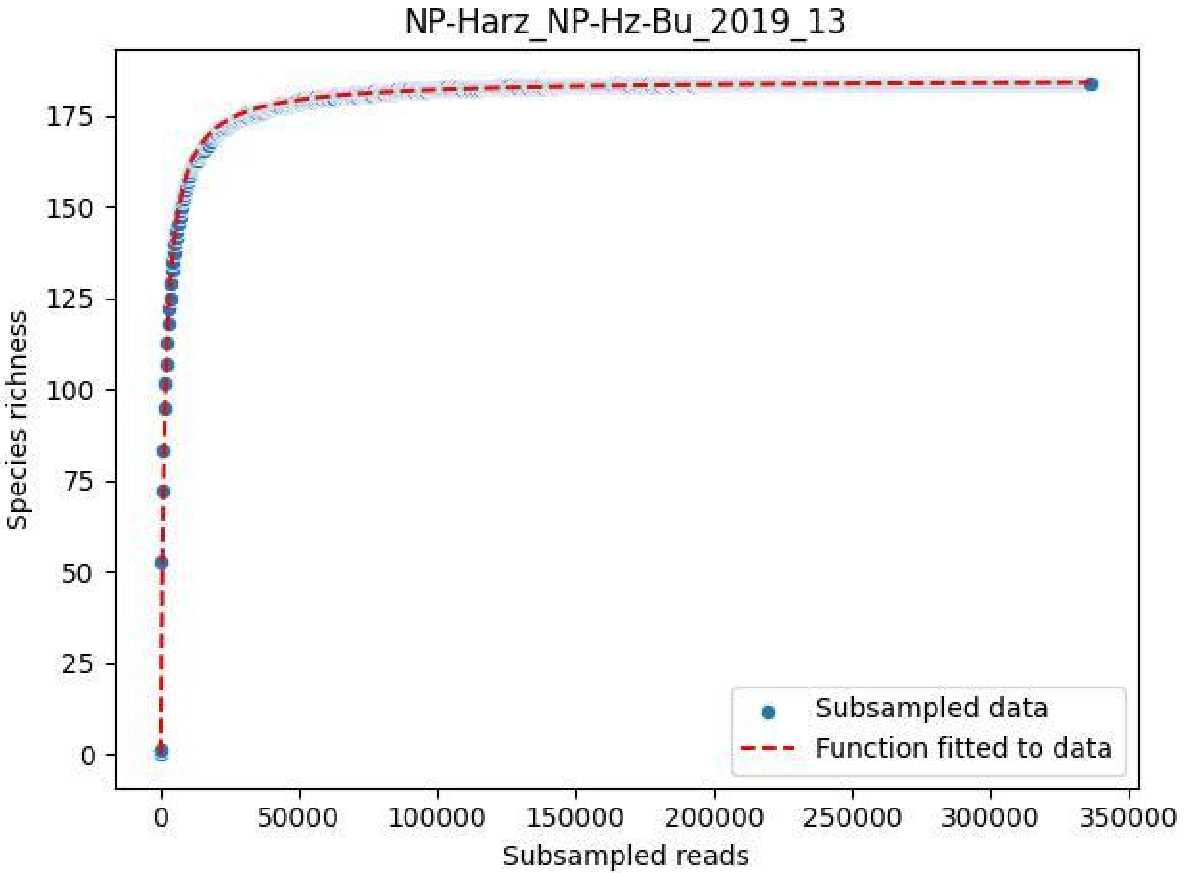

NP-Harz_NP-Hz-Bu_2019_13
Species richness
Subsampled reads
0
25
50
75
100
125
150
175
0
50000
100000
150000
200000
250000
300000
350000
Subsampled data
Function fitted to data

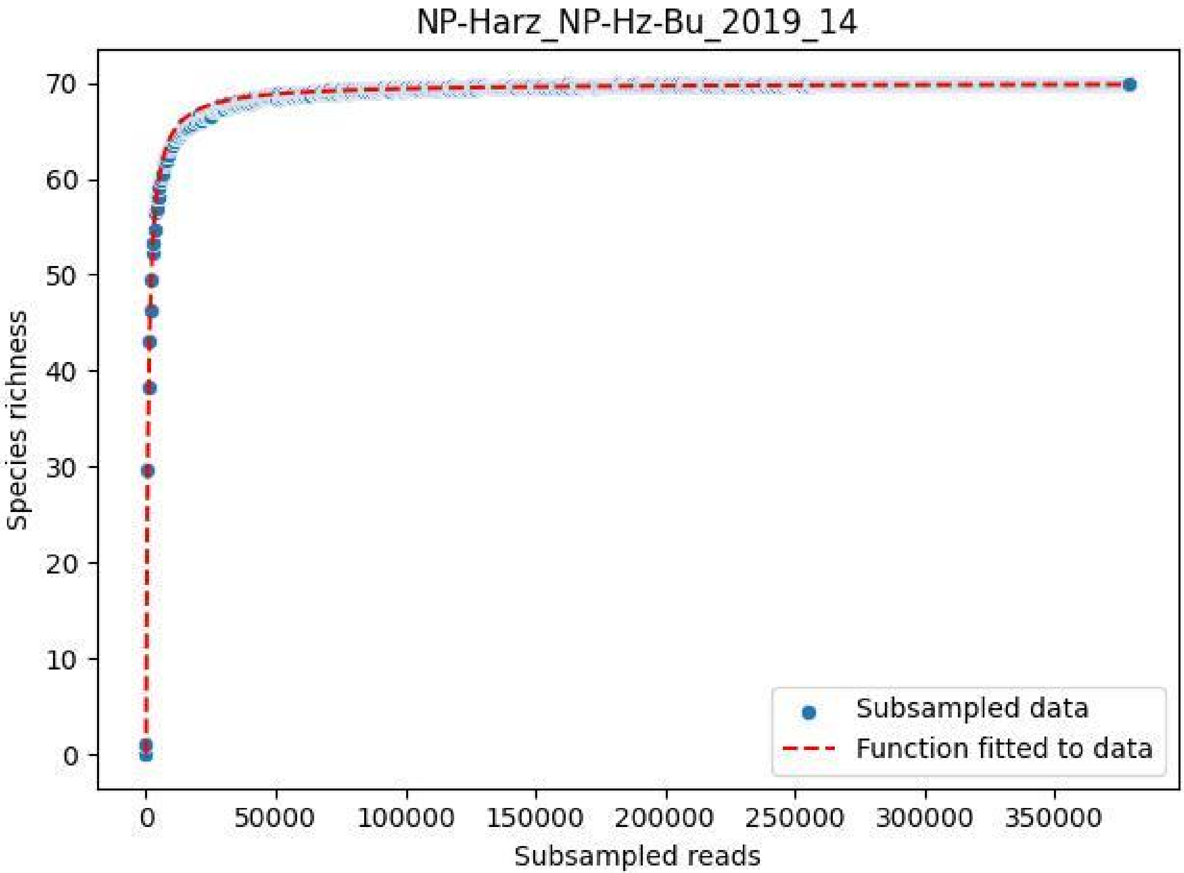

NP-Harz_NP-Hz-Bu_2019_14
Species richness
Subsampled reads
0
10
20
30
40
50
60
70
0
50000
100000
150000
200000
250000
300000
350000
Subsampled data
Function fitted to data

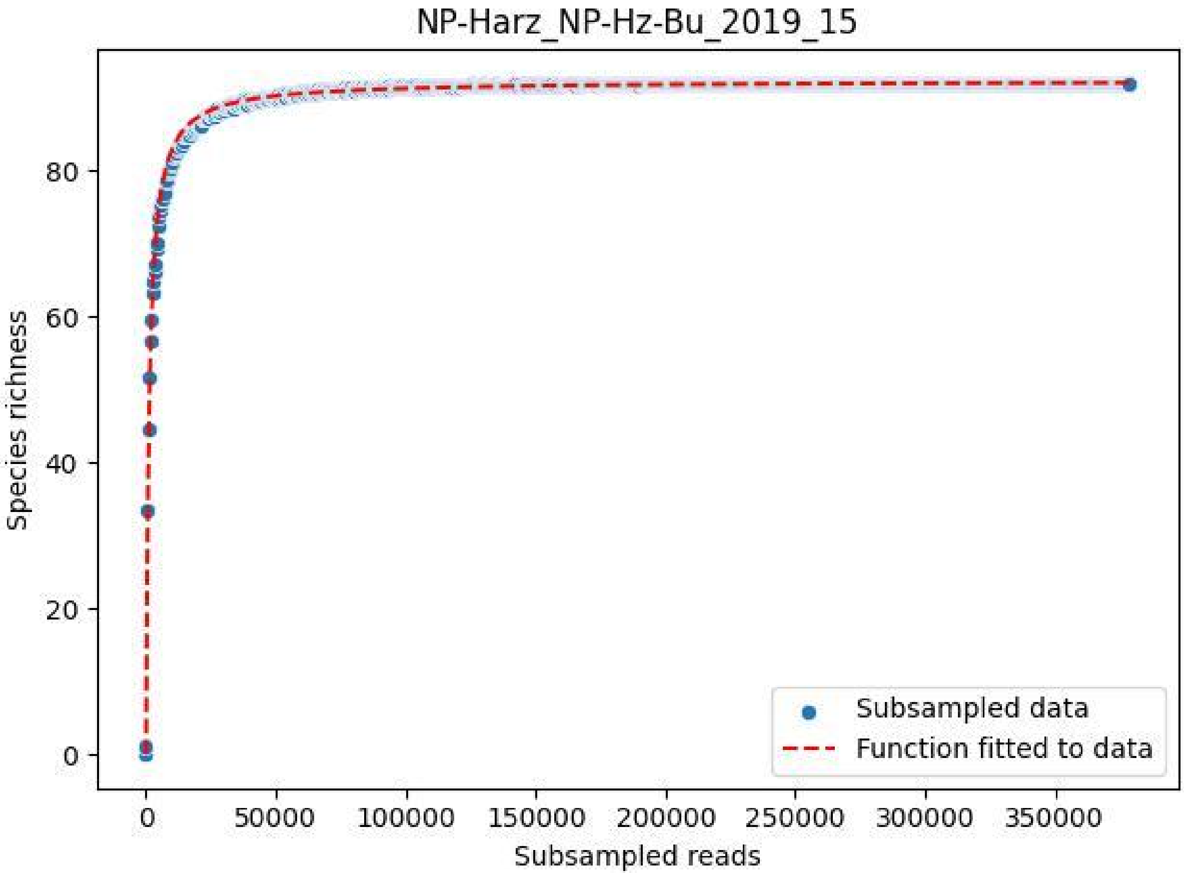
NP-Harz_NP-Hz-Bu_2019_15
Species richness
0
20
40
60
80
Subsampled reads
0
50000
100000
150000
200000
250000
300000
350000
Subsampled data
Function fitted to data

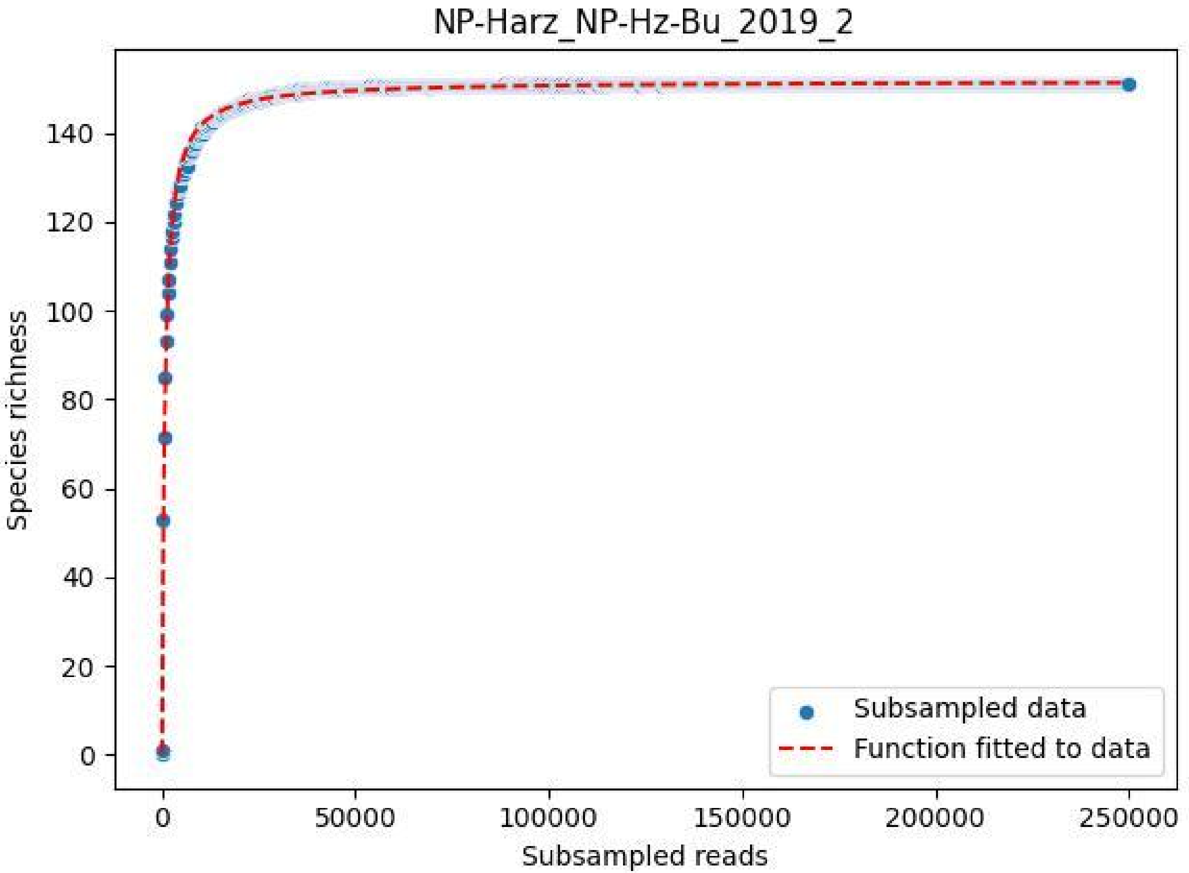

NP-Harz_NP-Hz-Bu_2019_2
Species richness
Subsampled reads
0
20
40
60
80
100
120
140
0
50000
100000
150000
200000
250000
Subsampled data
Function fitted to data

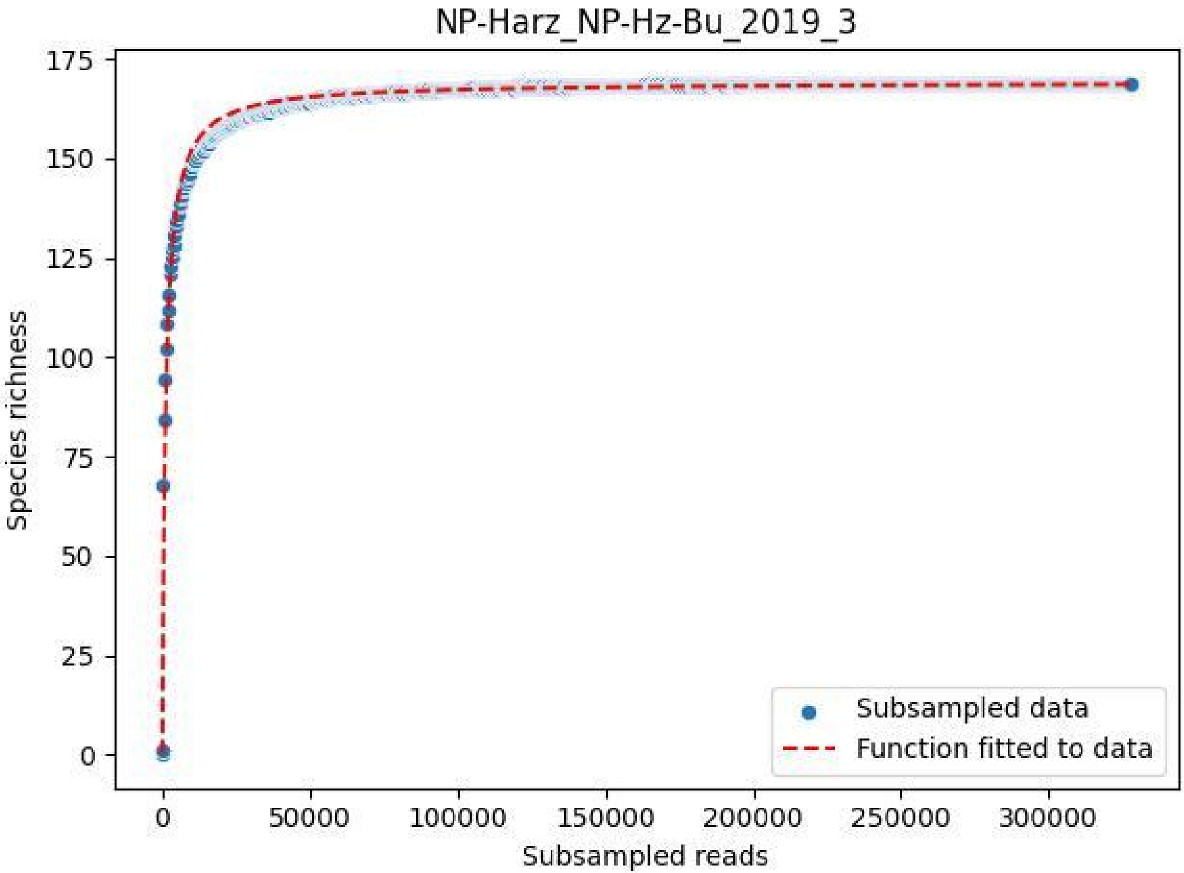

NP-Harz_NP-Hz-Bu_2019_3
Species richness
Subsampled reads
0
25
50
75
100
125
150
175
0
50000
100000
150000
200000
250000
300000
Subsampled data
Function fitted to data

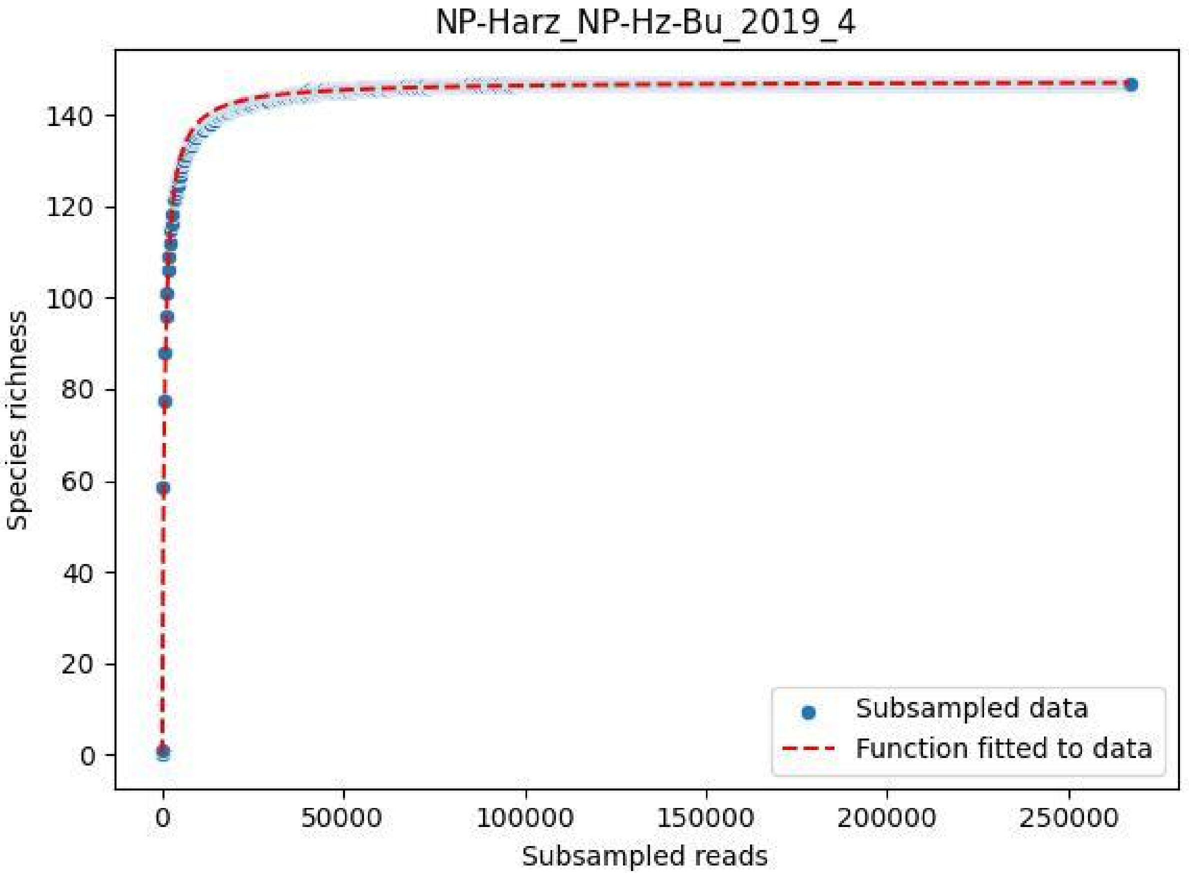
NP-Harz_NP-Hz-Bu_2019_4
Species richness
Subsampled reads
0
20
40
60
80
100
120
140
0
50000
100000
150000
200000
250000
Subsampled data
Function fitted to data

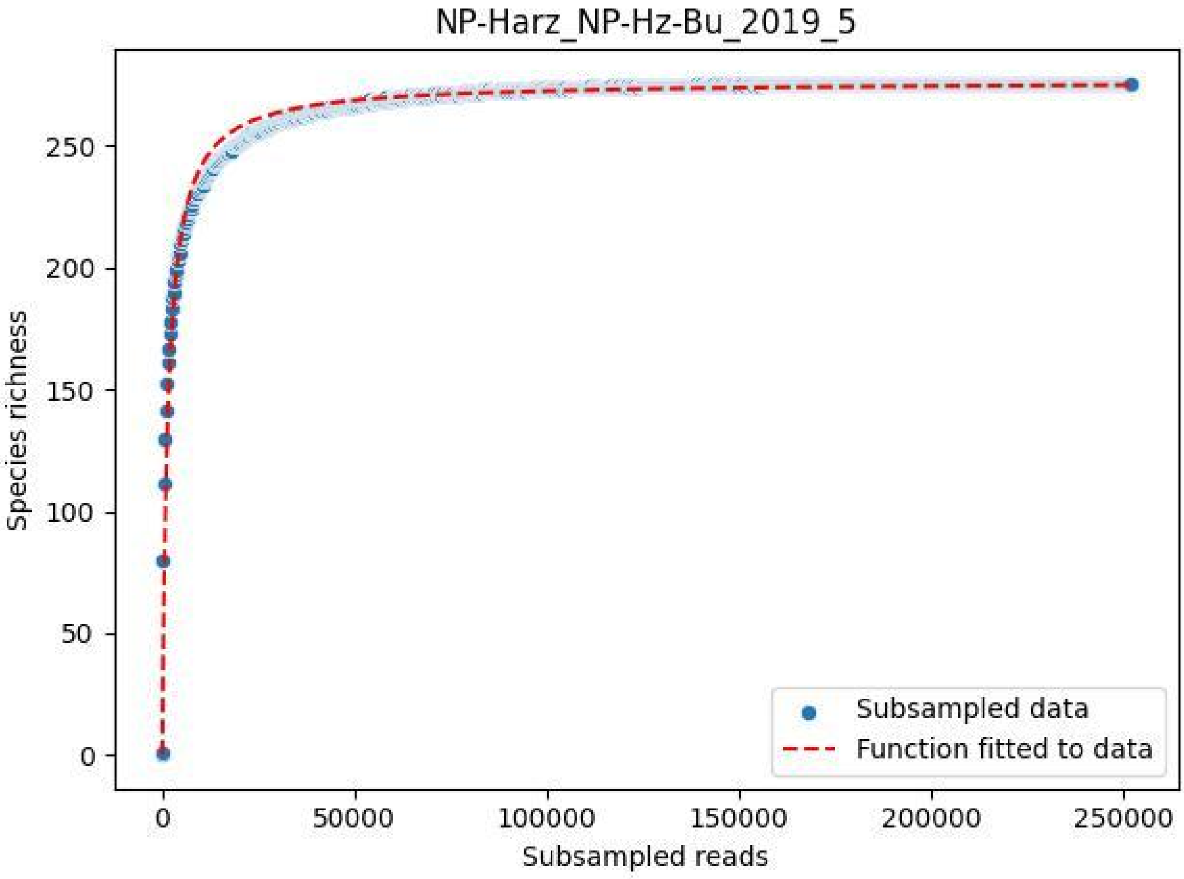
NP-Harz_NP-Hz-Bu_2019_5
Species richness
Subsampled reads
0
50
100
150
200
250
0
50000
100000
150000
200000
250000
Subsampled data
Function fitted to data

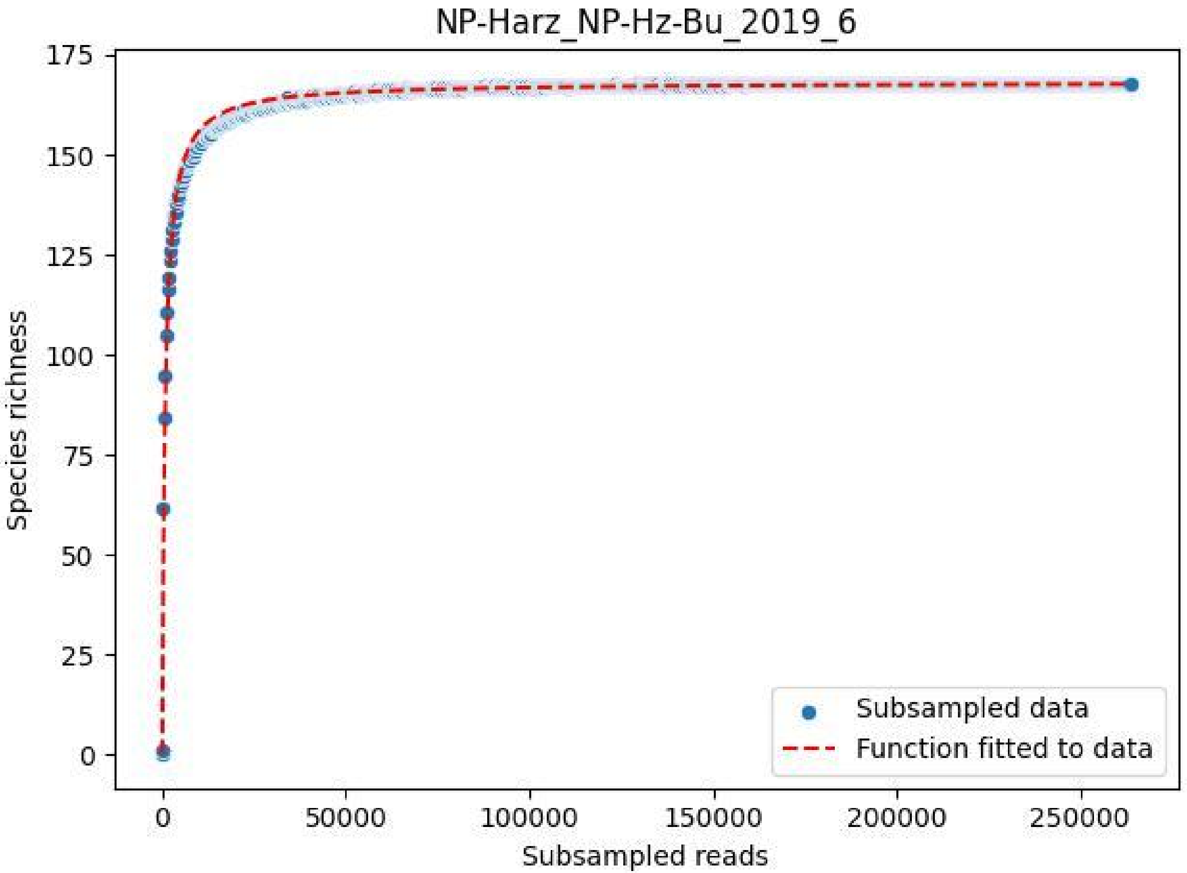
NP-Harz_NP-Hz-Bu_2019_6
Species richness
Subsampled reads
175
150
125
100
75
50
25
0
0
50000
100000
150000
200000
250000
Subsampled data
Function fitted to data

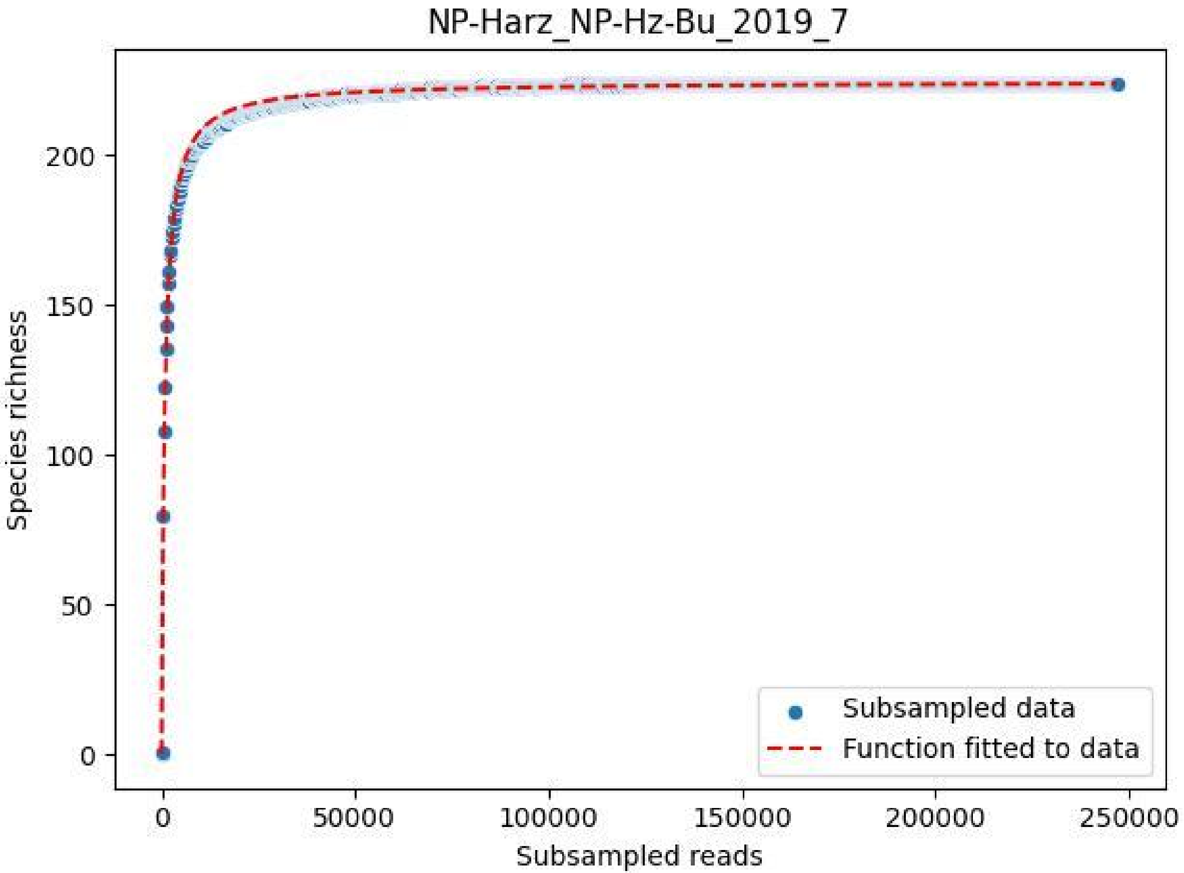
NP-Harz_NP-Hz-Bu_2019_7
Species richness
0
50
100
150
200
0
50000
100000
150000
200000
250000
Subsampled reads
Subsampled data
Function fitted to data

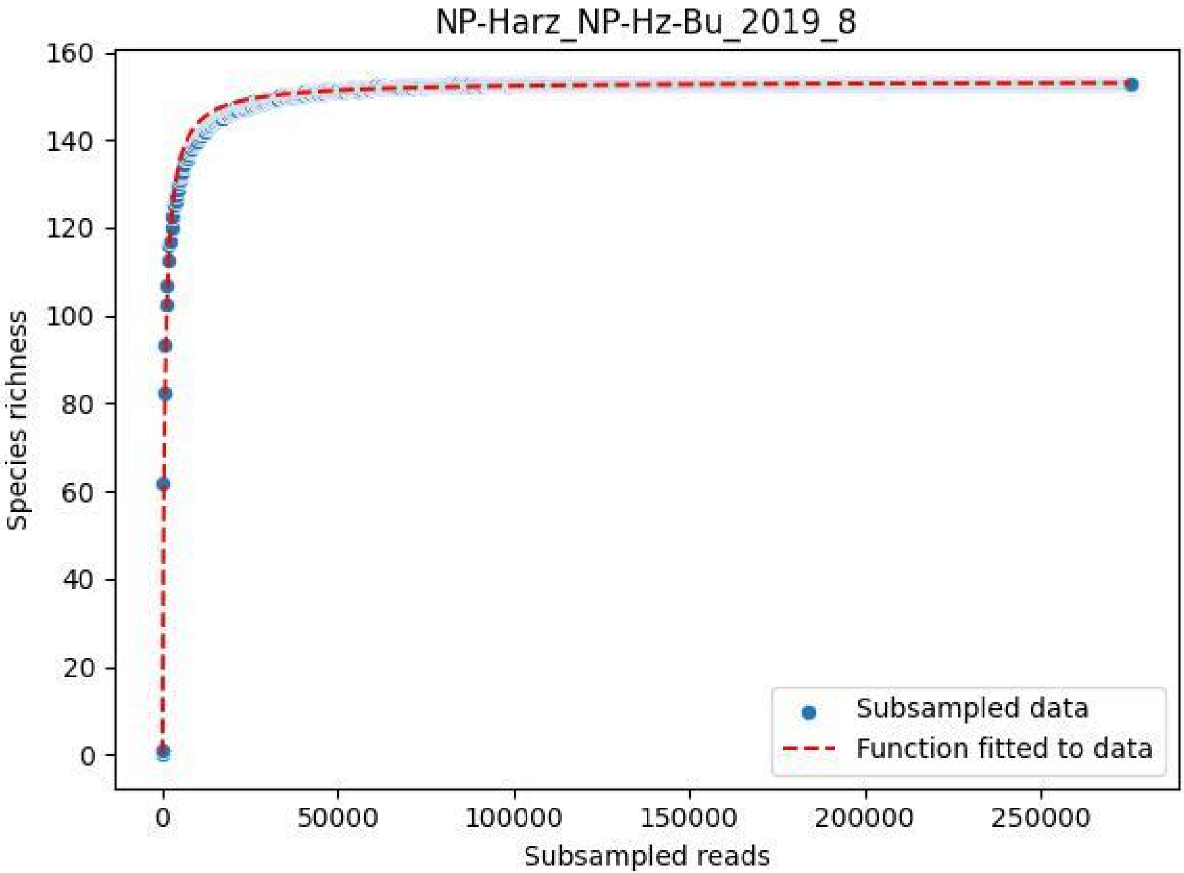

NP-Harz_NP-Hz-Bu_2019_8
Species richness
Subsampled reads
0
20
40
60
80
100
120
140
160
0
50000
100000
150000
200000
250000
Subsampled data
Function fitted to data

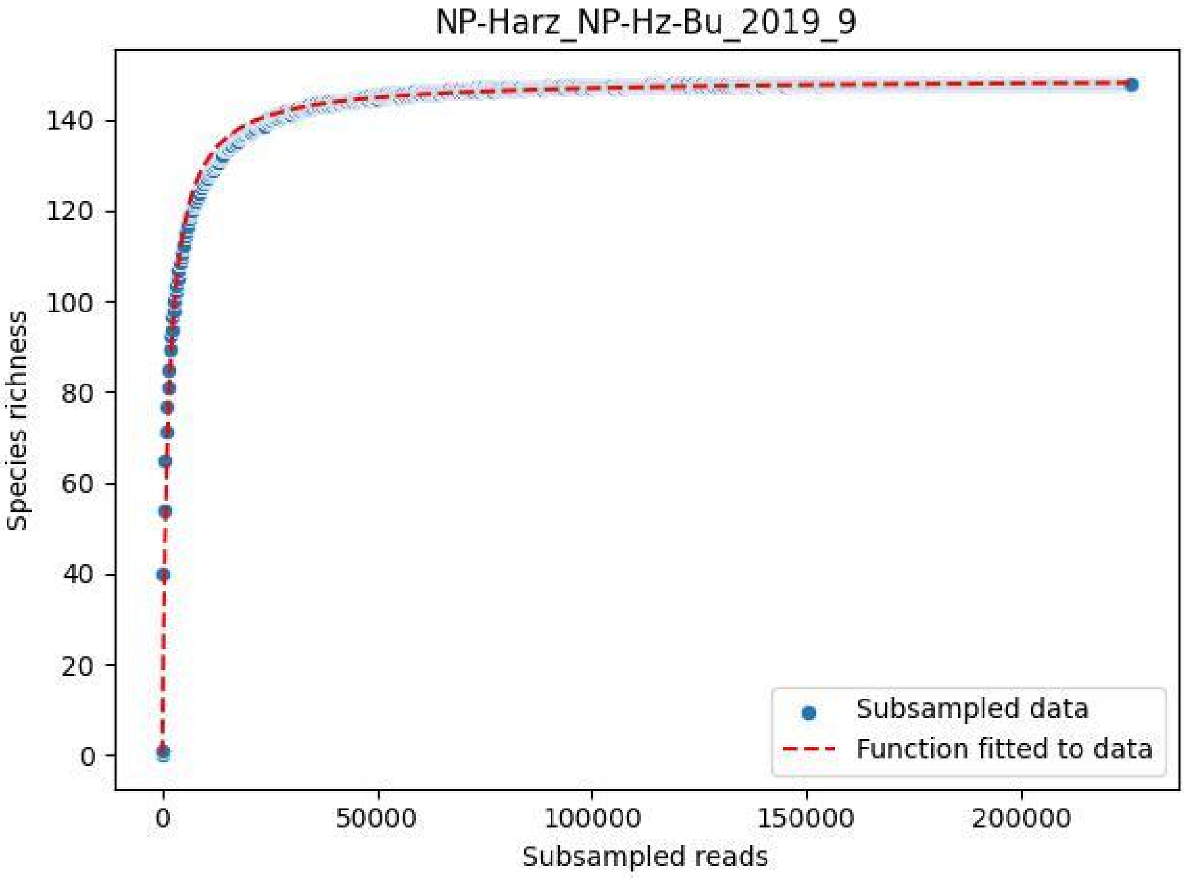
NP-Harz_NP-Hz-Bu_2019_9
Species richness
Subsampled reads
0
20
40
60
80
100
120
140
0
50000
100000
150000
200000
Subsampled data
Function fitted to data

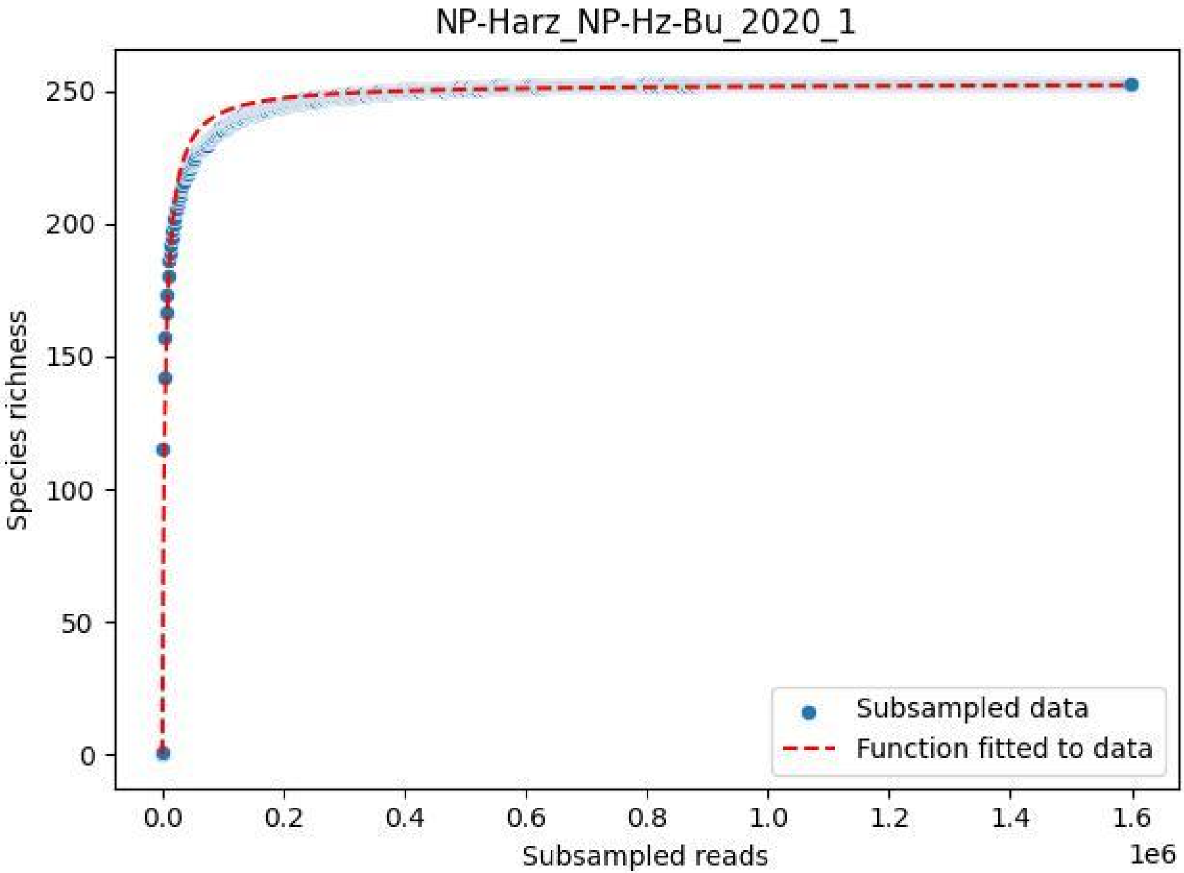
NP-Harz_NP-Hz-Bu_2020_1
Species richness
Subsampled reads
1e6
0.0
0.2
0.4
0.6
0.8
1.0
1.2
1.4
1.6
0
50
100
150
200
250
Subsampled data
Function fitted to data

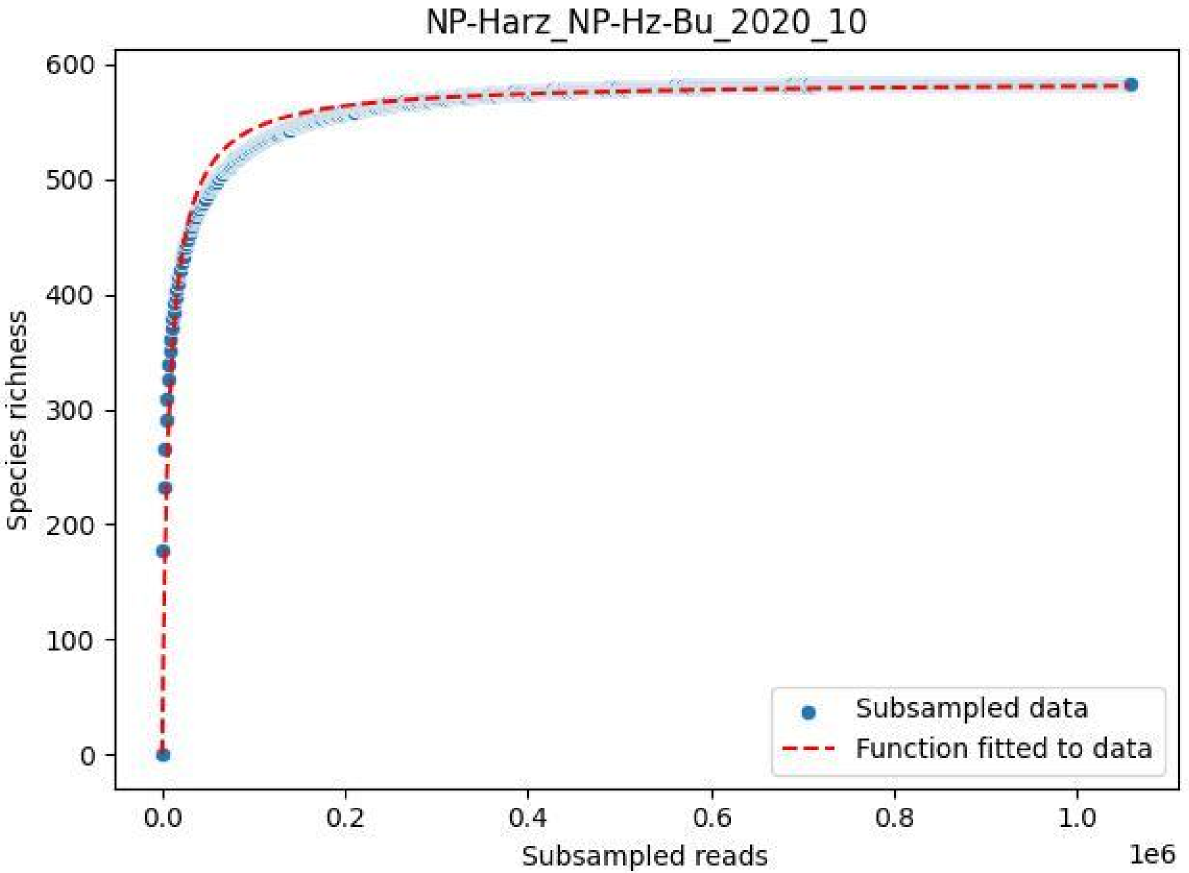

NP-Harz_NP-Hz-Bu_2020_10
Species richness
0
100
200
300
400
500
600
Subsampled reads
0.0
0.2
0.4
0.6
0.8
1.0
1e6
Subsampled data
Function fitted to data

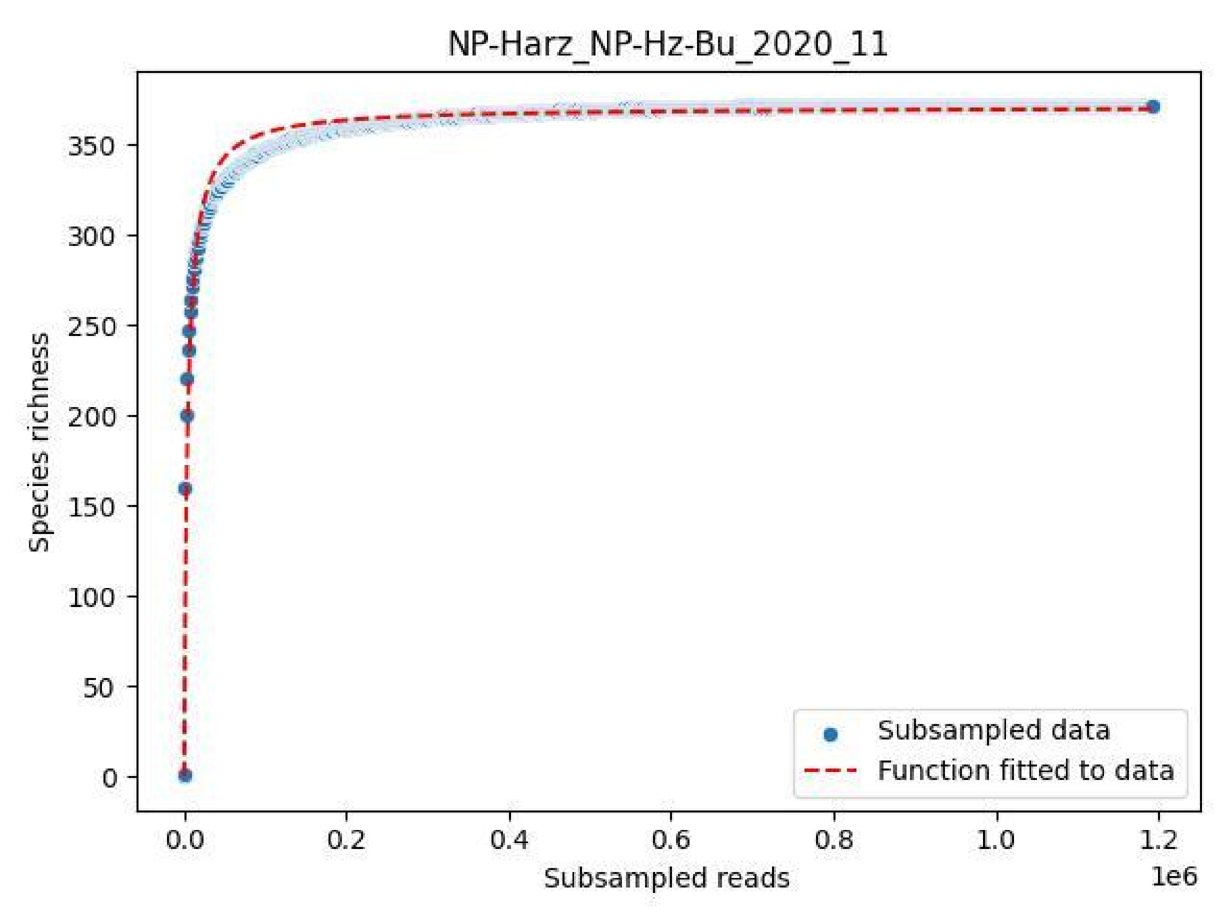

NP-Harz_NP-Hz-Bu_2020_11
Species richness
Subsampled reads
1e6
0
50
100
150
200
250
300
350
0.0
0.2
0.4
0.6
0.8
1.0
1.2
Subsampled data
Function fitted to data

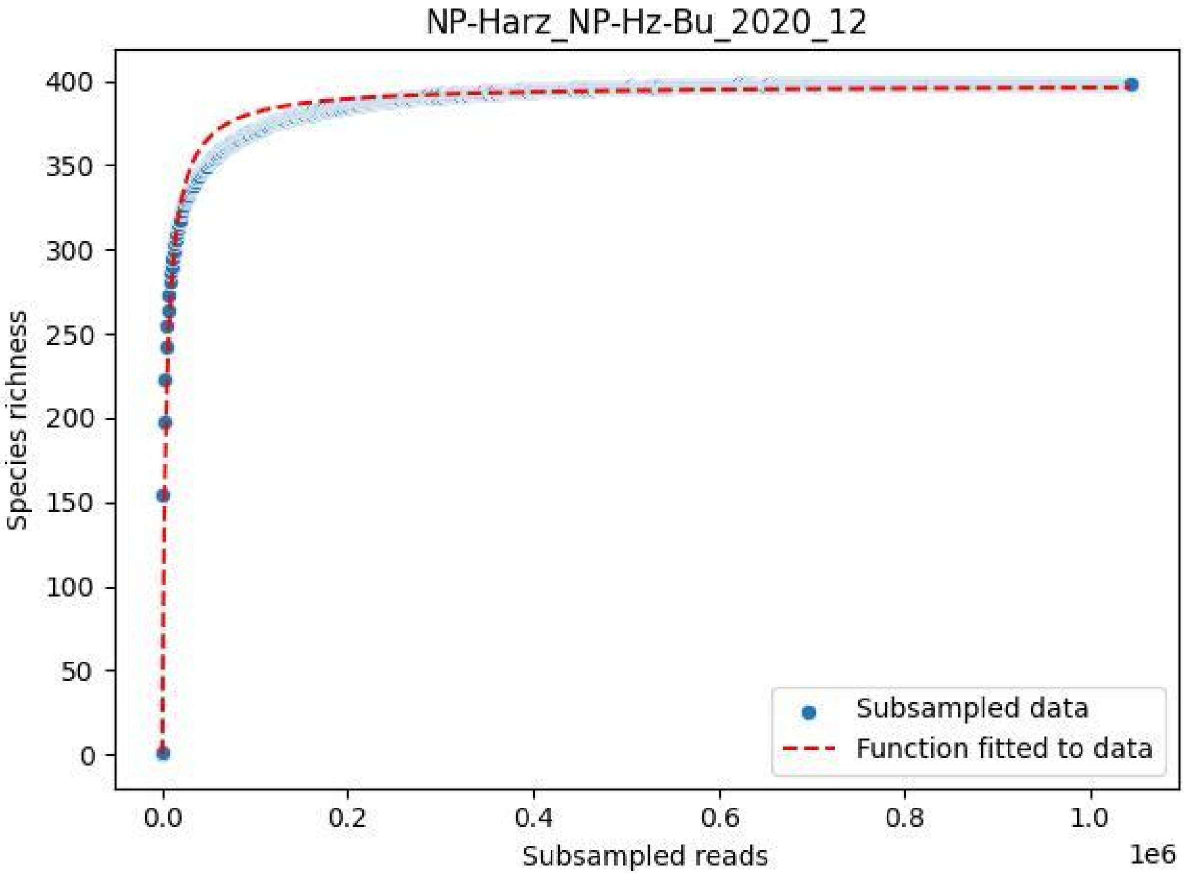
NP-Harz_NP-Hz-Bu_2020_12
Species richness
Subsampled reads
1e6
0
50
100
150
200
250
300
350
400
0.0
0.2
0.4
0.6
0.8
1.0
Subsampled data
Function fitted to data

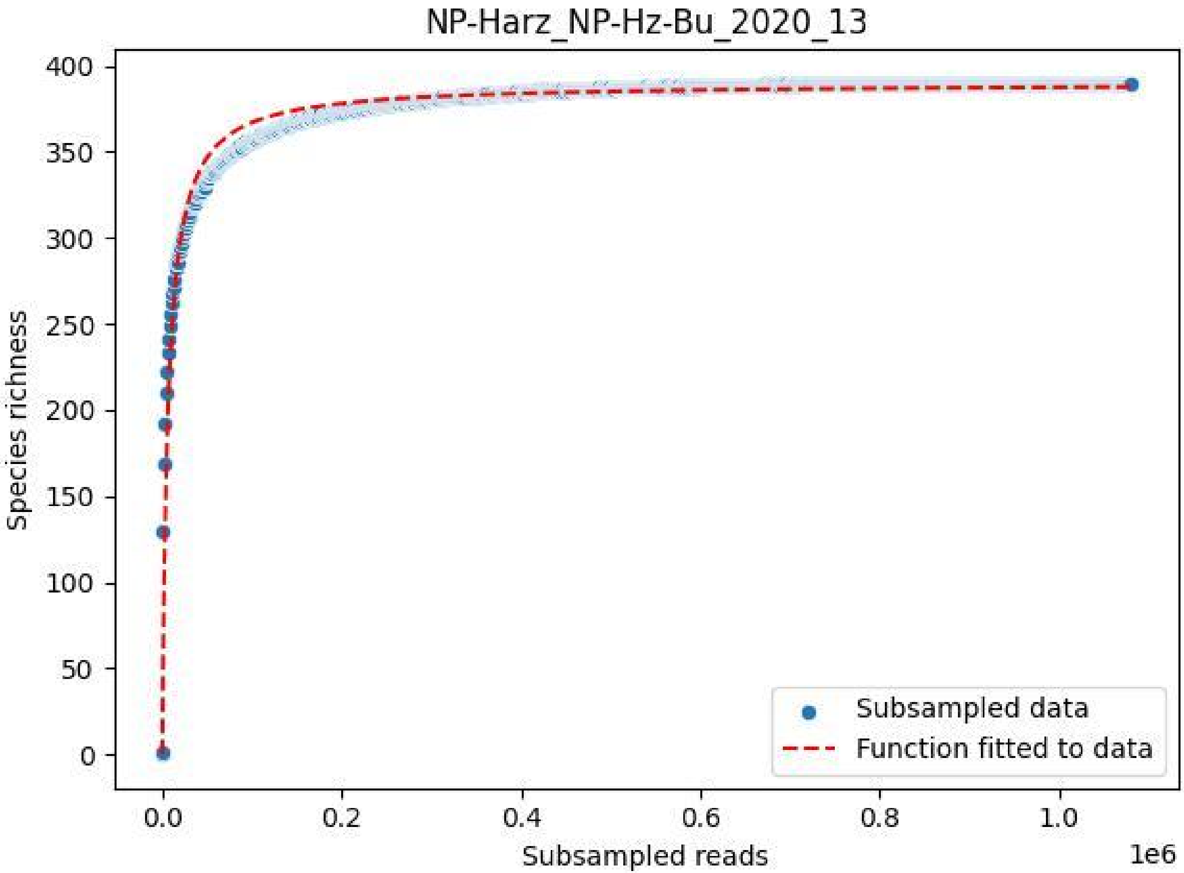
NP-Harz_NP-Hz-Bu_2020_13
Species richness
400
350
300
250
200
150
100
50
0
0.0
0.2
0.4
0.6
0.8
1.0
1e6
Subsampled reads
Subsampled data
Function fitted to data

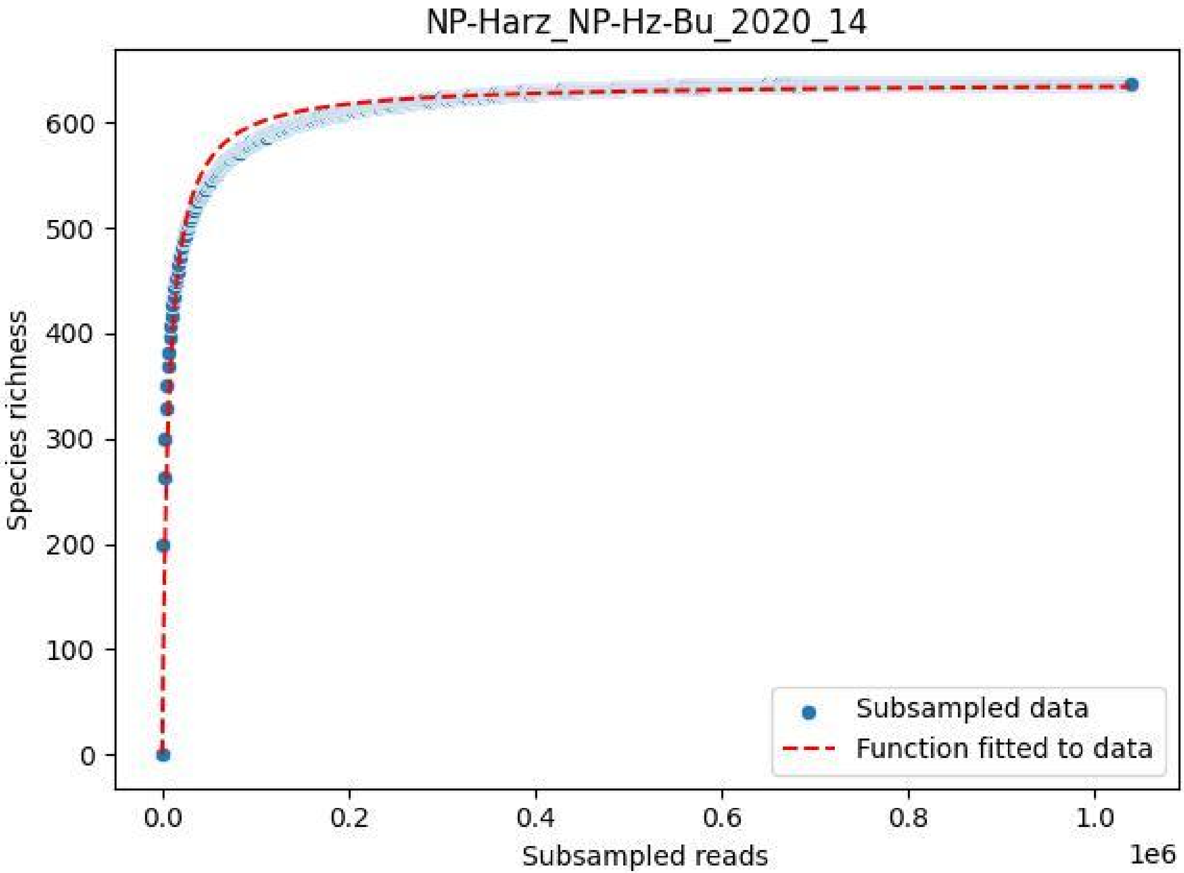
NP-Harz_NP-Hz-Bu_2020_14
Species richness
Subsampled reads
1e6
Subsampled data
Function fitted to data
0
100
200
300
400
500
600
0.0
0.2
0.4
0.6
0.8
1.0

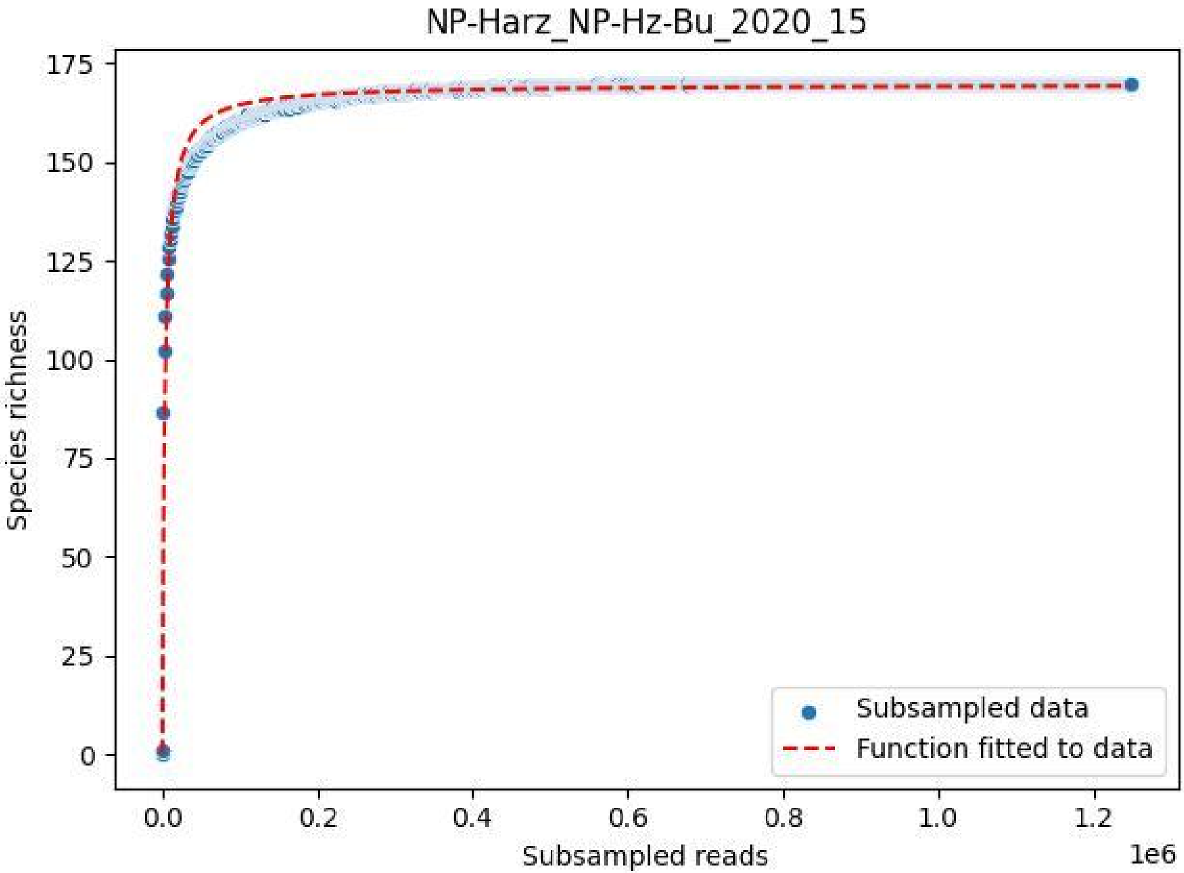
NP-Harz_NP-Hz-Bu_2020_15
Species richness
Subsampled reads
1e6
0.0
0.2
0.4
0.6
0.8
1.0
1.2
0
25
50
75
100
125
150
175
Subsampled data
Function fitted to data

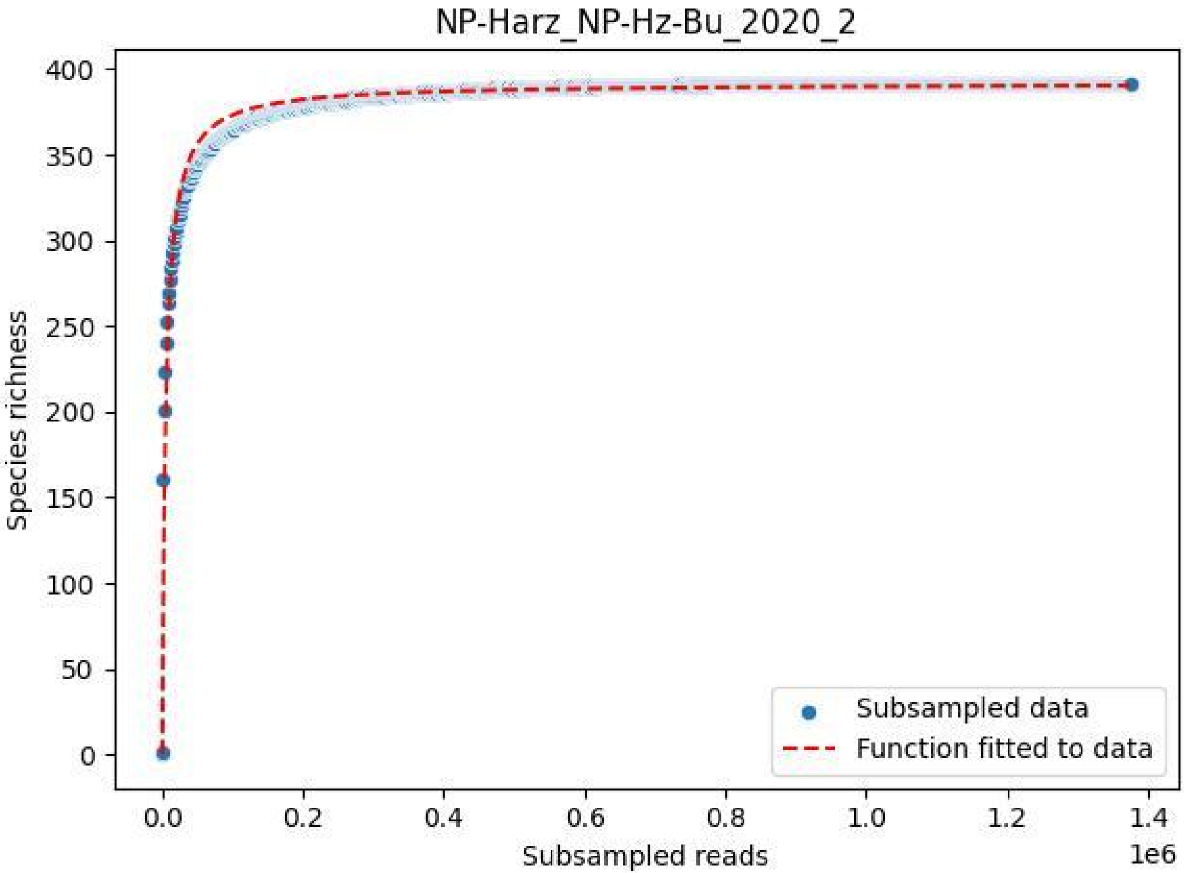

NP-Harz_NP-Hz-Bu_2020_2
Species richness
Subsampled reads
1e6
0
50
100
150
200
250
300
350
400
0.0
0.2
0.4
0.6
0.8
1.0
1.2
1.4
Subsampled data
Function fitted to data

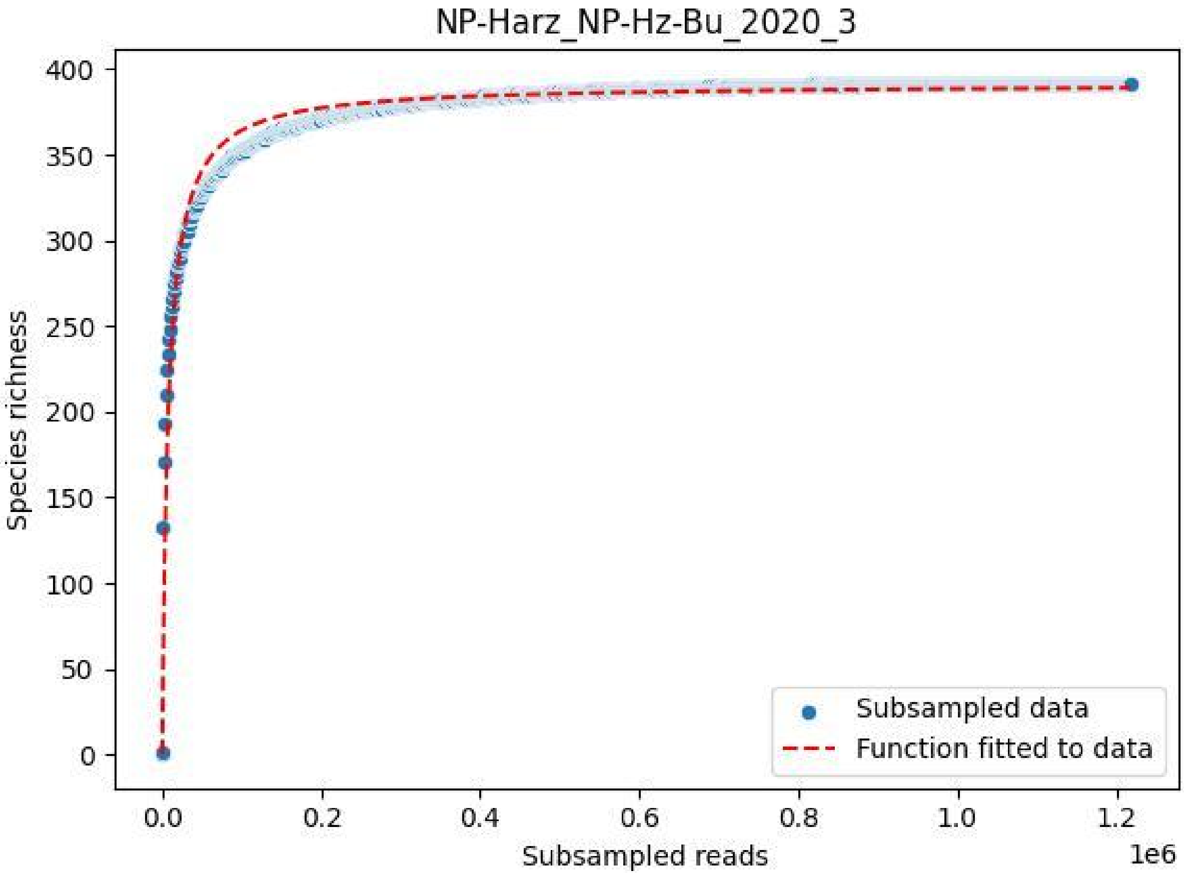
NP-Harz_NP-Hz-Bu_2020_3
Species richness
Subsampled reads
1e6
0
50
100
150
200
250
300
350
400
0.0
0.2
0.4
0.6
0.8
1.0
1.2
Subsampled data
Function fitted to data

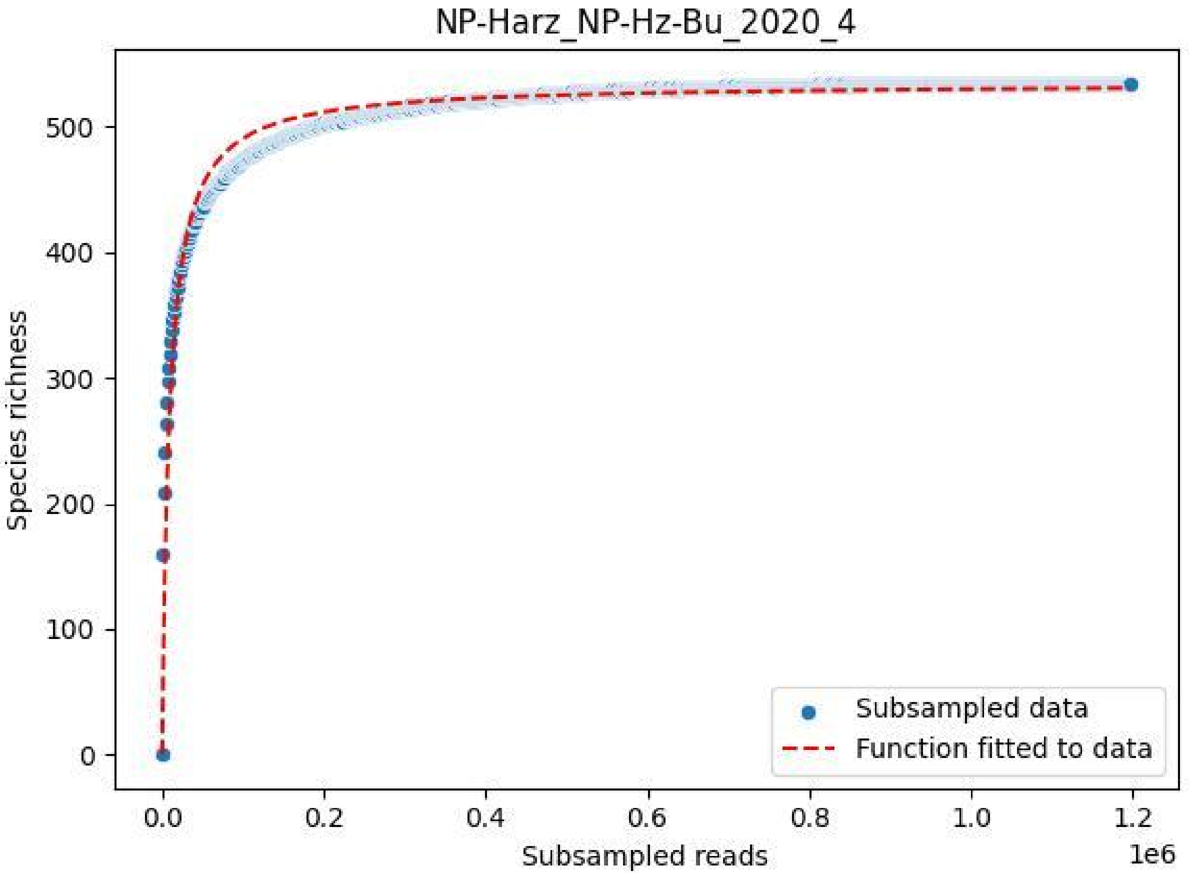
NP-Harz_NP-Hz-Bu_2020_4
Species richness
0
100
200
300
400
500
0.0
0.2
0.4
0.6
0.8
1.0
1.2
1e6
Subsampled reads
Subsampled data
Function fitted to data

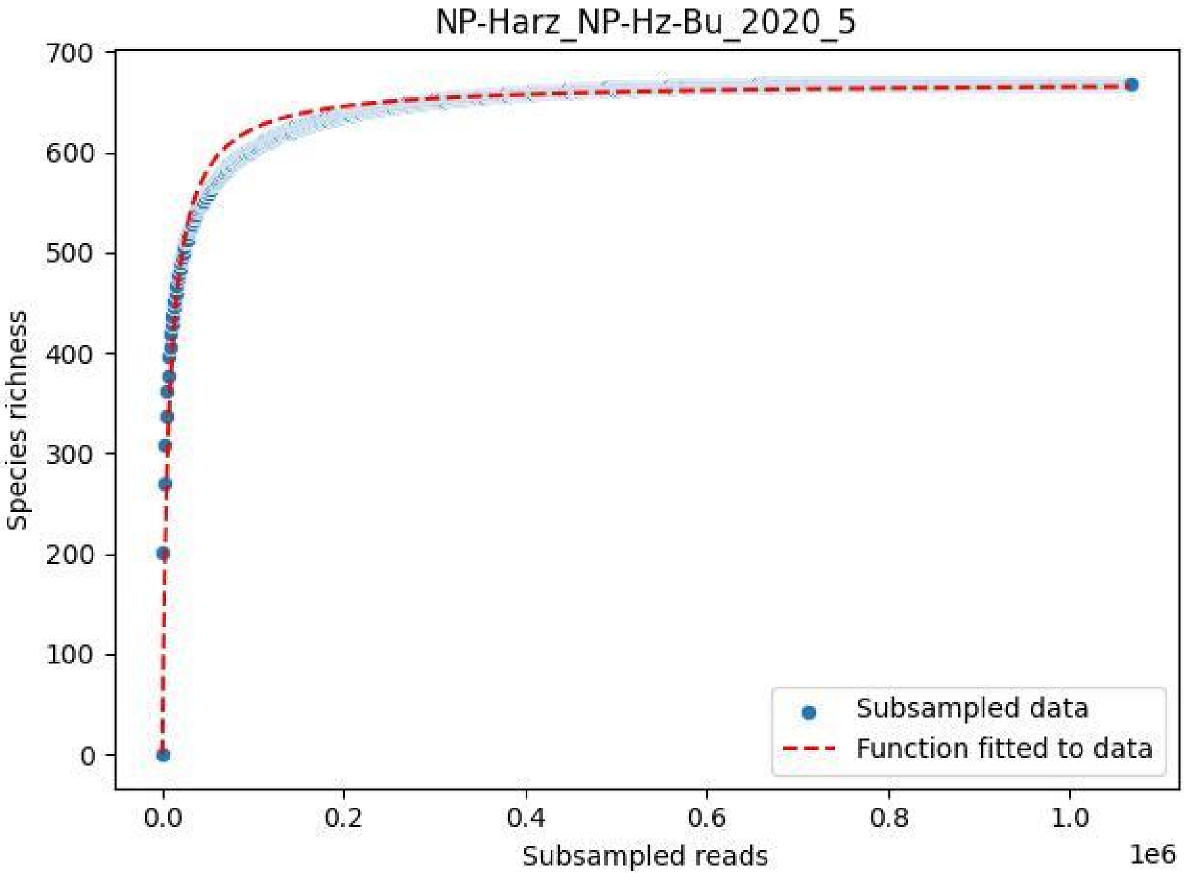
NP-Harz_NP-Hz-Bu_2020_5
Species richness
Subsampled reads
1e6
0
100
200
300
400
500
600
700
0.0
0.2
0.4
0.6
0.8
1.0
Subsampled data
Function fitted to data

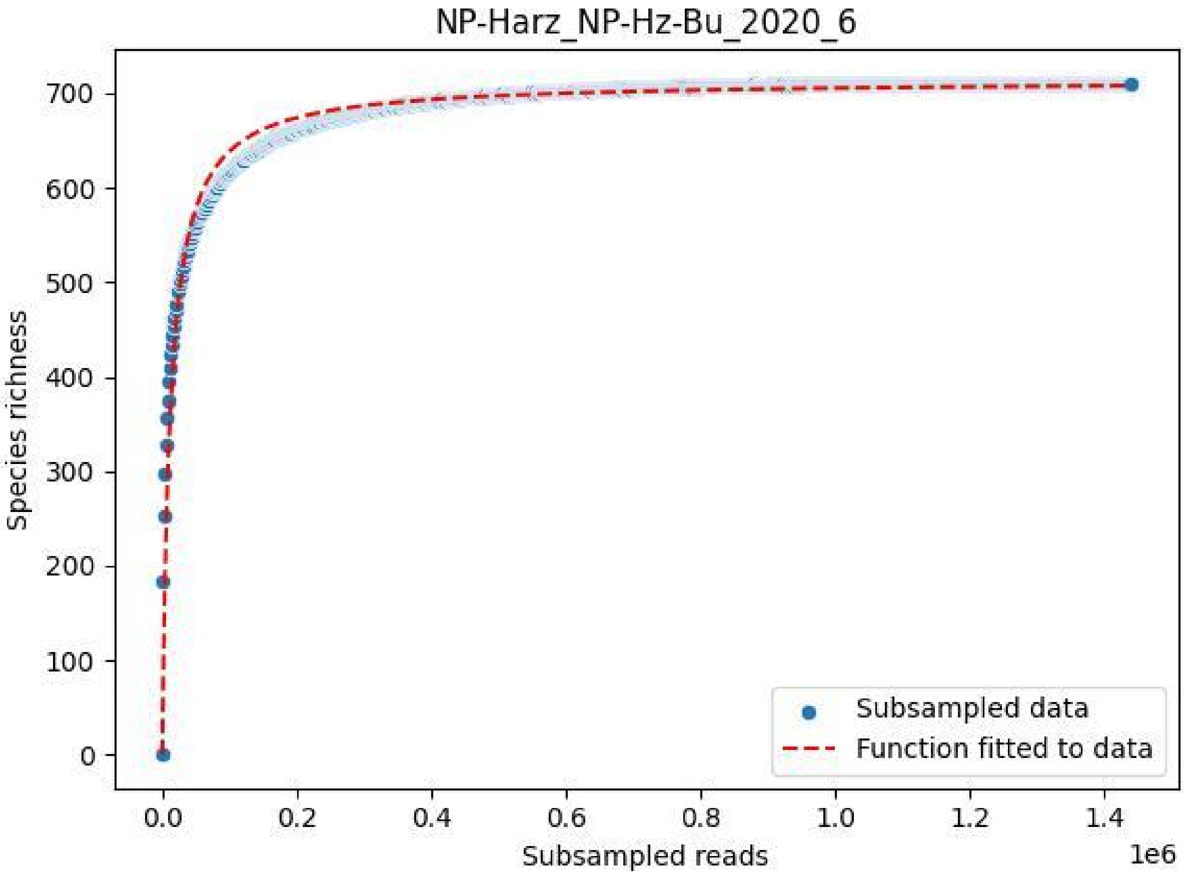
NP-Harz_NP-Hz-Bu_2020_6
Species richness
Subsampled reads
1e6
0
100
200
300
400
500
600
700
0.0
0.2
0.4
0.6
0.8
1.0
1.2
1.4
Subsampled data
Function fitted to data

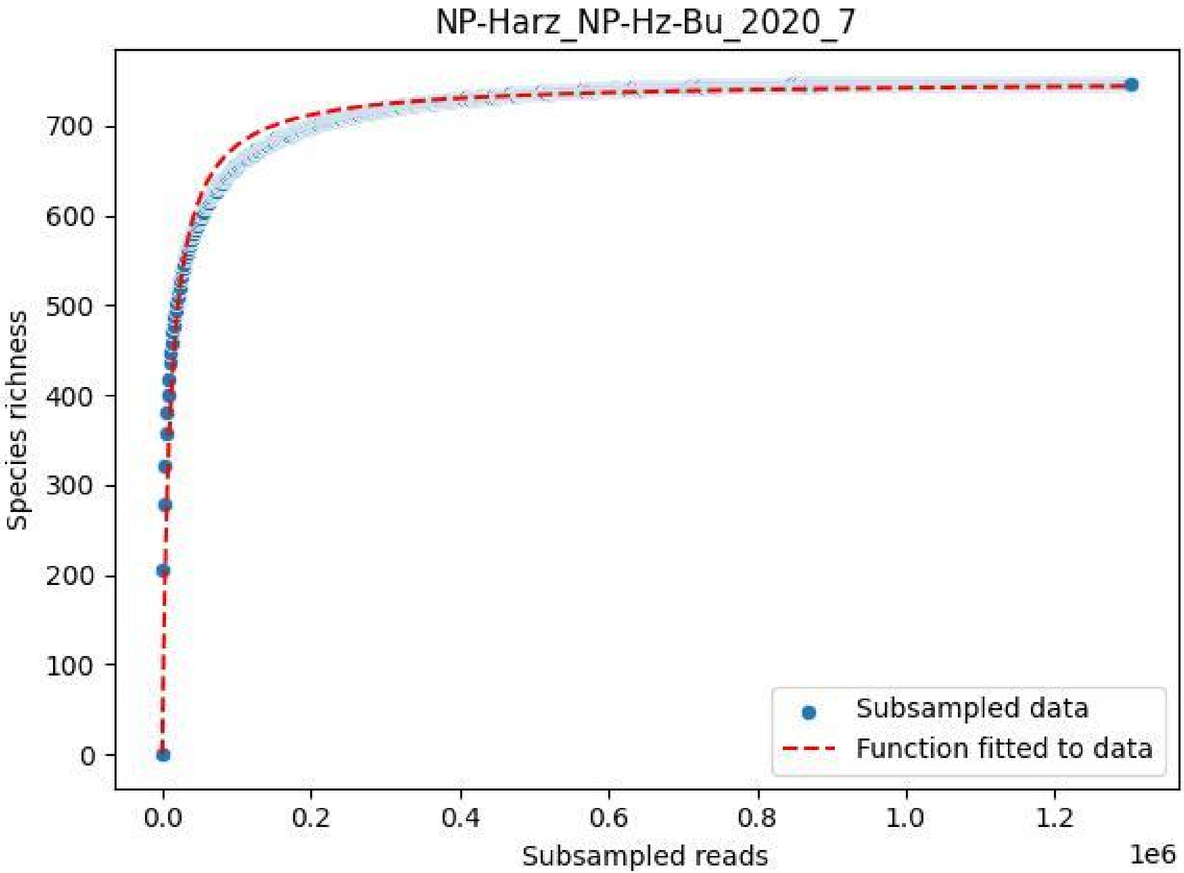
NP-Harz_NP-Hz-Bu_2020_7
Species richness
Subsampled reads
1e6
0
100
200
300
400
500
600
700
0.0
0.2
0.4
0.6
0.8
1.0
1.2
Subsampled data
Function fitted to data

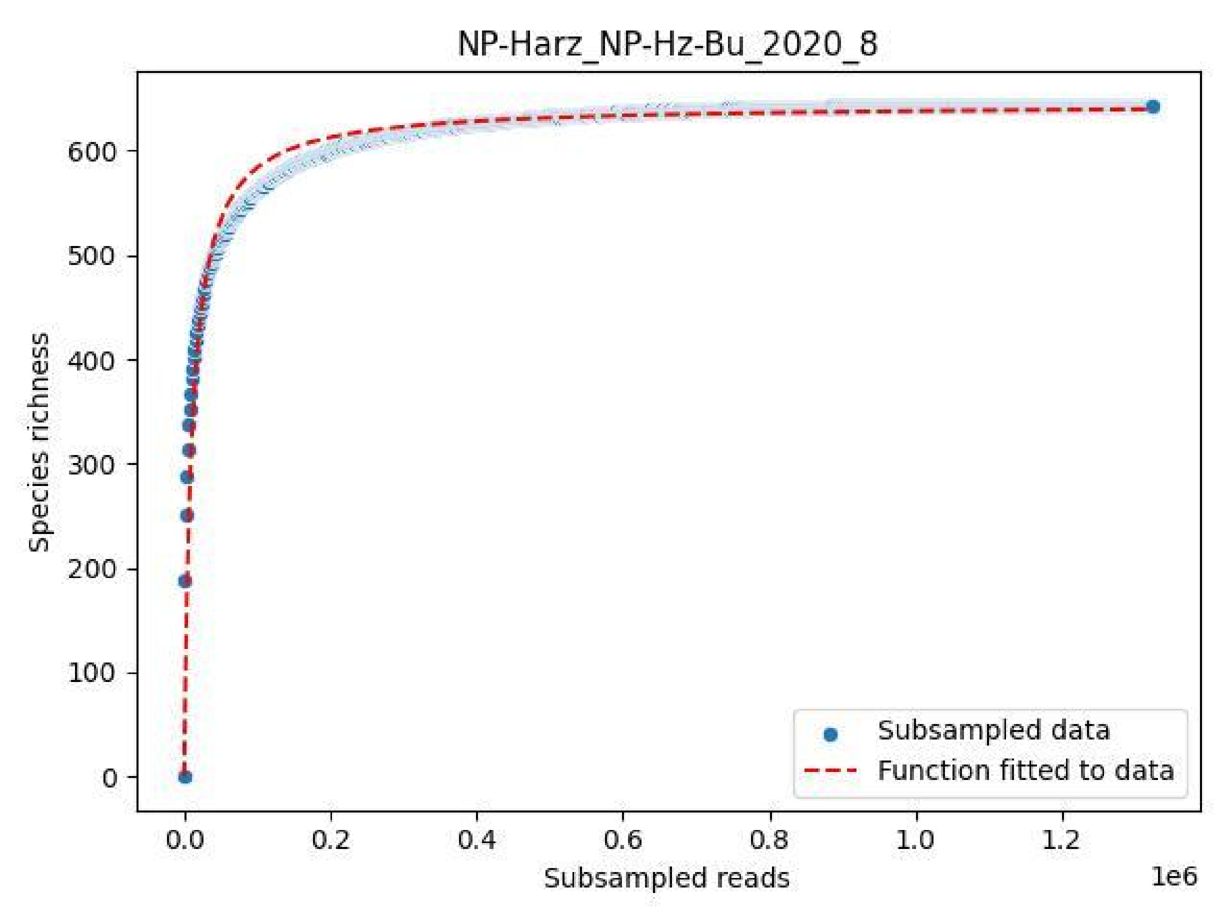
NP-Harz_NP-Hz-Bu_2020_8
Species richness
Subsampled reads
1e6
0
100
200
300
400
500
600
0.0
0.2
0.4
0.6
0.8
1.0
1.2
Subsampled data
Function fitted to data

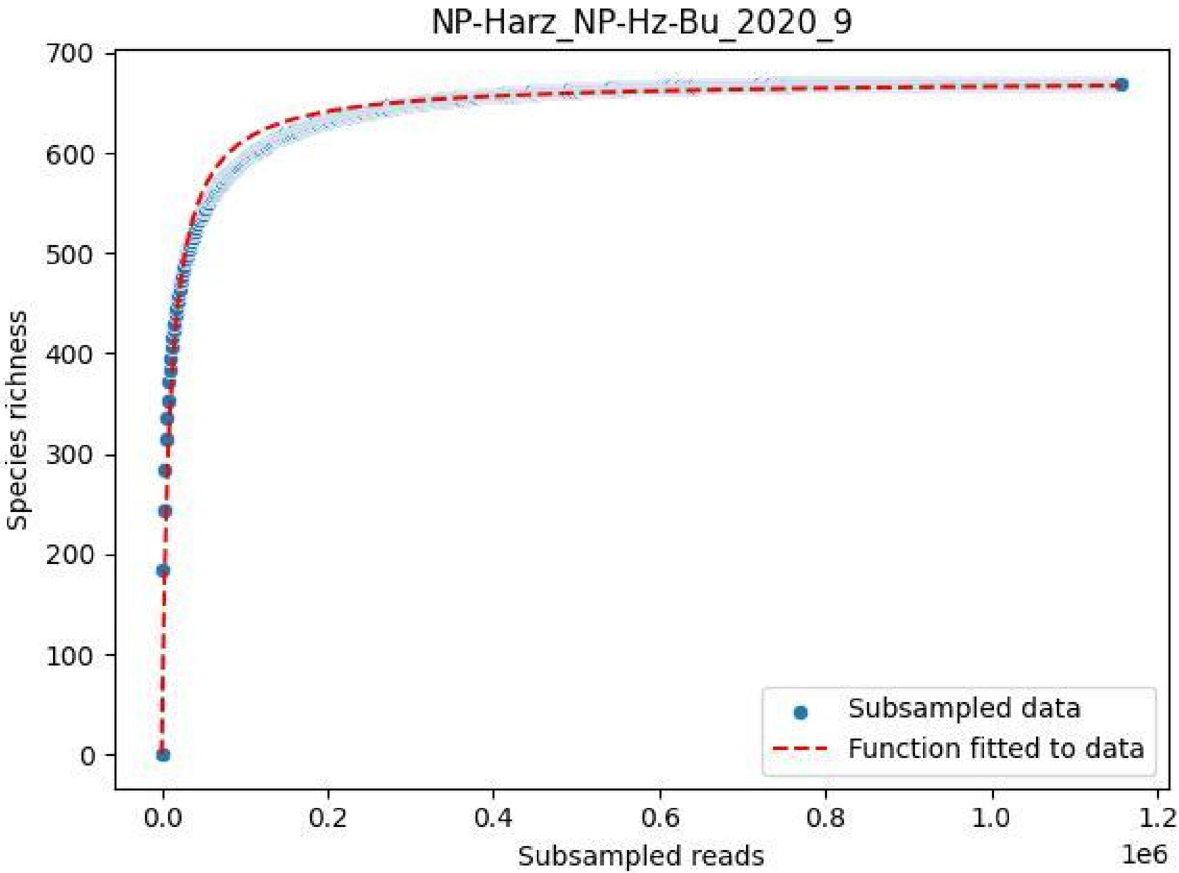
NP-Harz_NP-Hz-Bu_2020_9
Species richness
Subsampled reads
1e6
0
100
200
300
400
500
600
700
0.0
0.2
0.4
0.6
0.8
1.0
1.2
Subsampled data
Function fitted to data

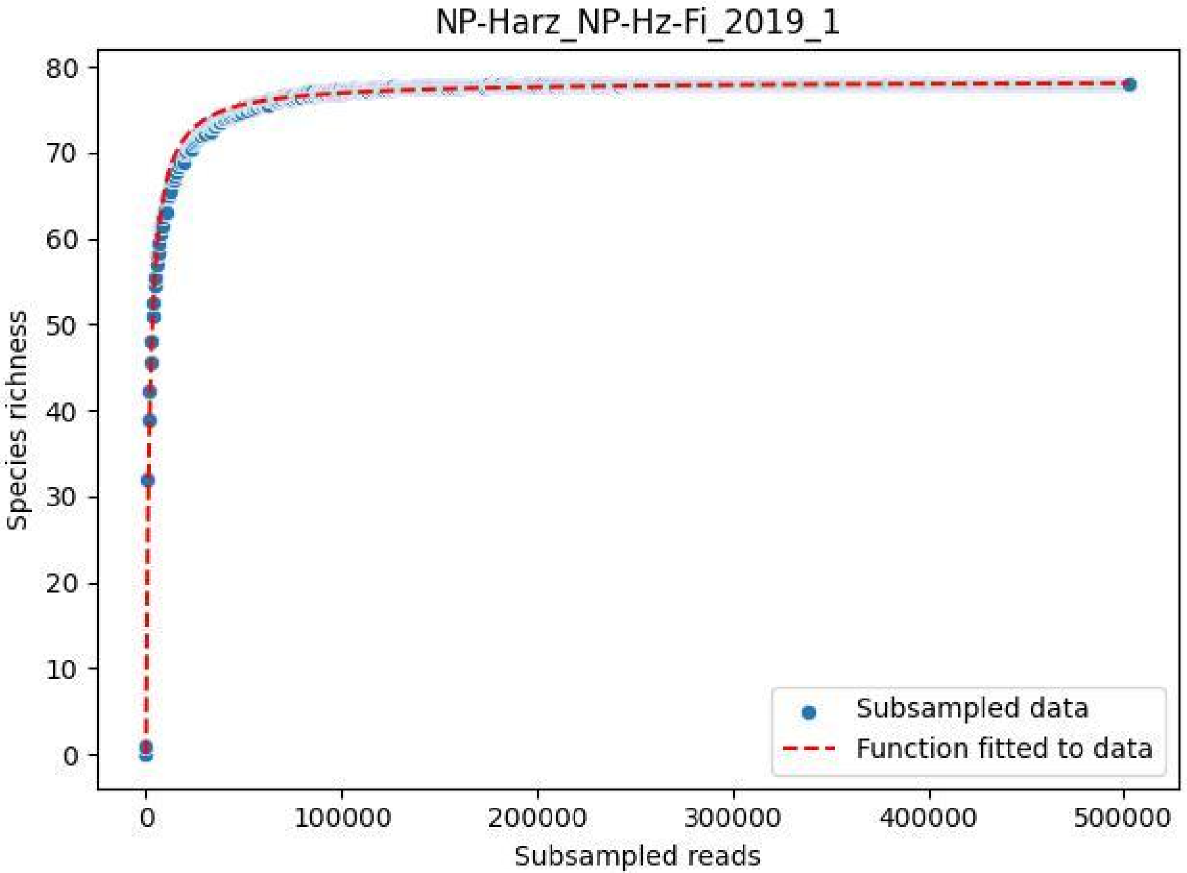
NP-Harz_NP-Hz-Fi_2019_1
Species richness
Subsampled reads
0
10
20
30
40
50
60
70
80
0
100000
200000
300000
400000
500000
Subsampled data
Function fitted to data

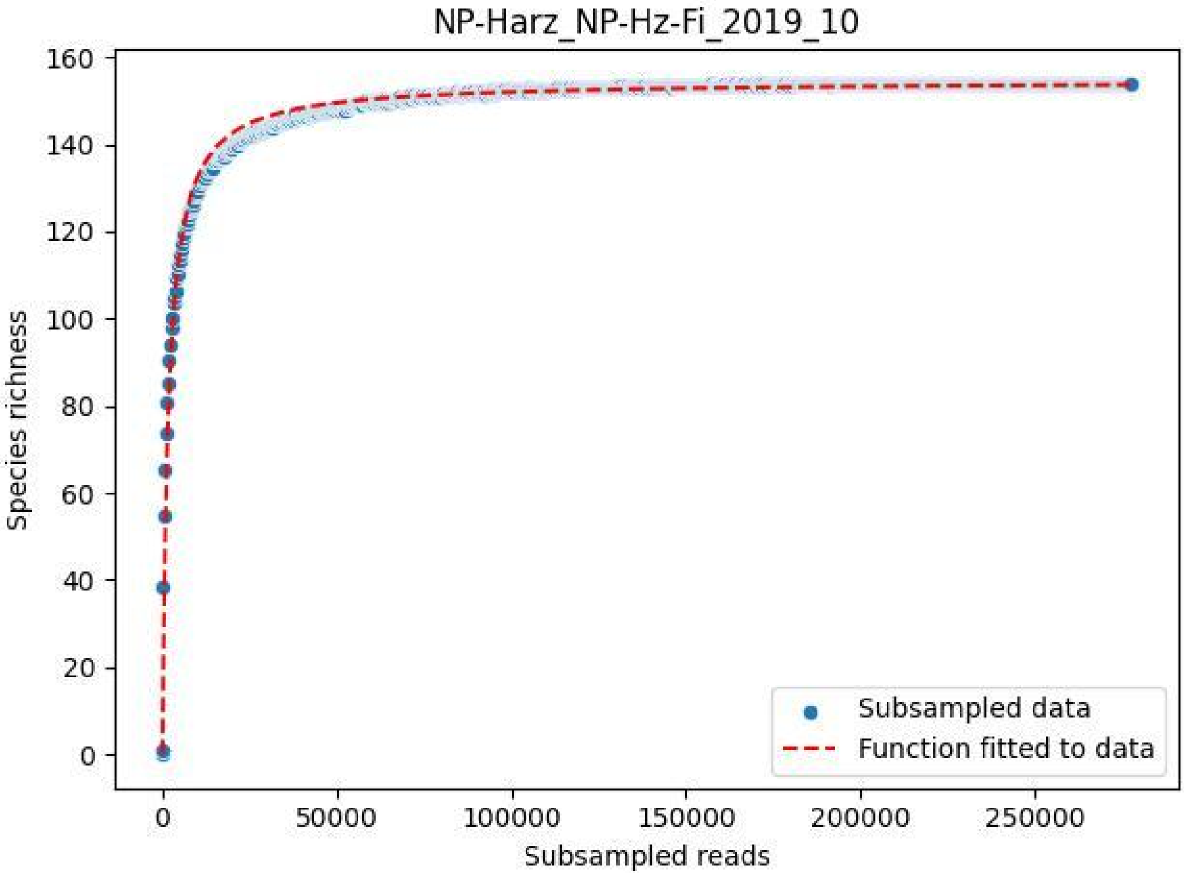
NP-Harz_NP-Hz-Fi_2019_10
Species richness
Subsampled reads
0
20
40
60
80
100
120
140
160
0
50000
100000
150000
200000
250000
Subsampled data
Function fitted to data

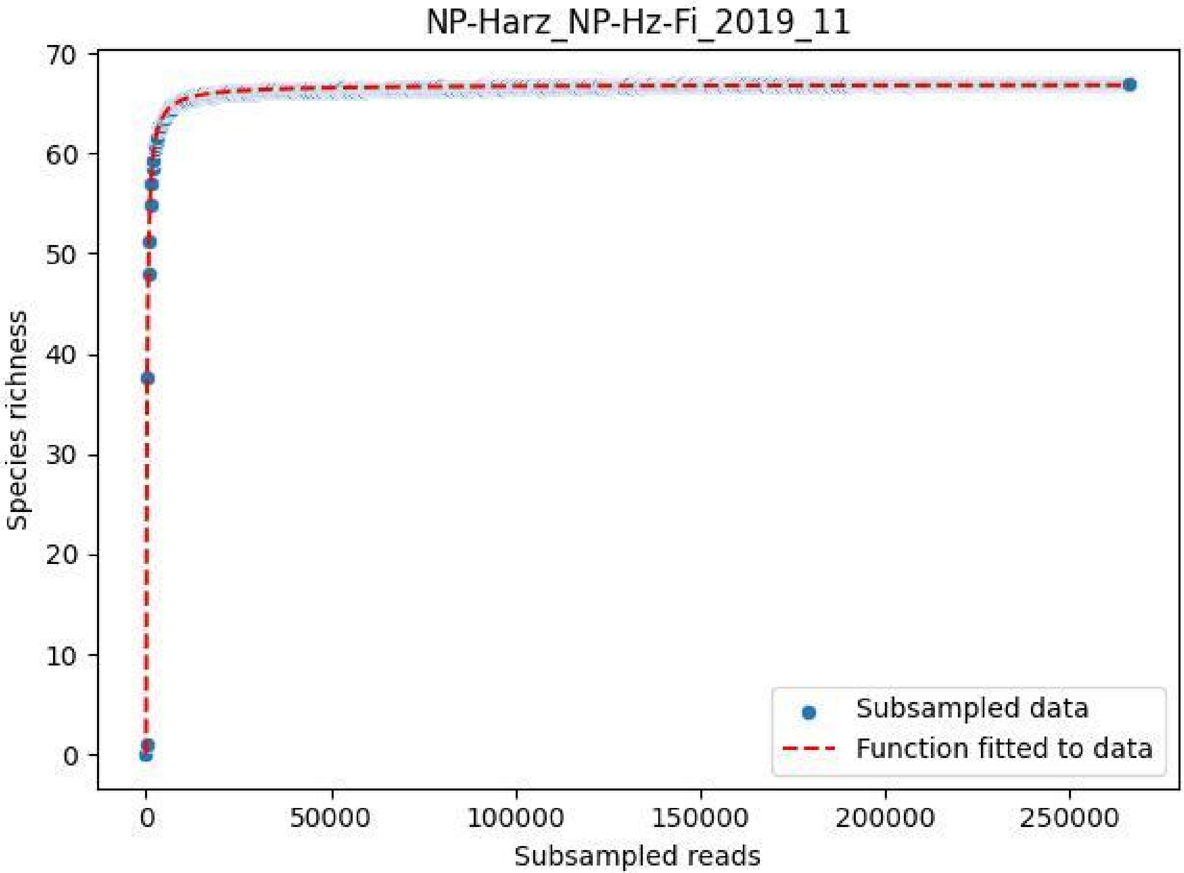

NP-Harz_NP-Hz-Fi_2019_11
Species richness
Subsampled reads
0
10
20
30
40
50
60
70
0
50000
100000
150000
200000
250000
Subsampled data
Function fitted to data

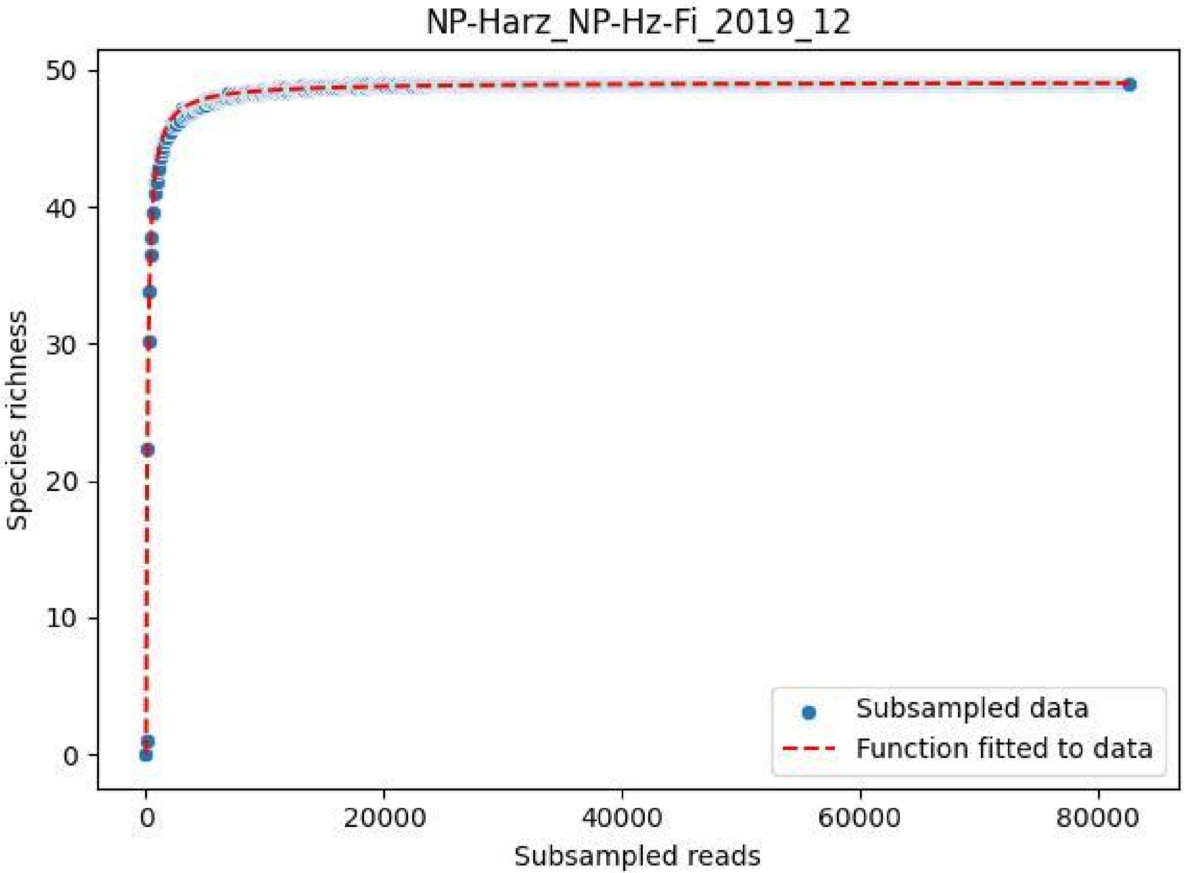
NP-Harz_NP-Hz-Fi_2019_12
Species richness
Subsampled reads
0
10
20
30
40
50
0
20000
40000
60000
80000
Subsampled data
Function fitted to data

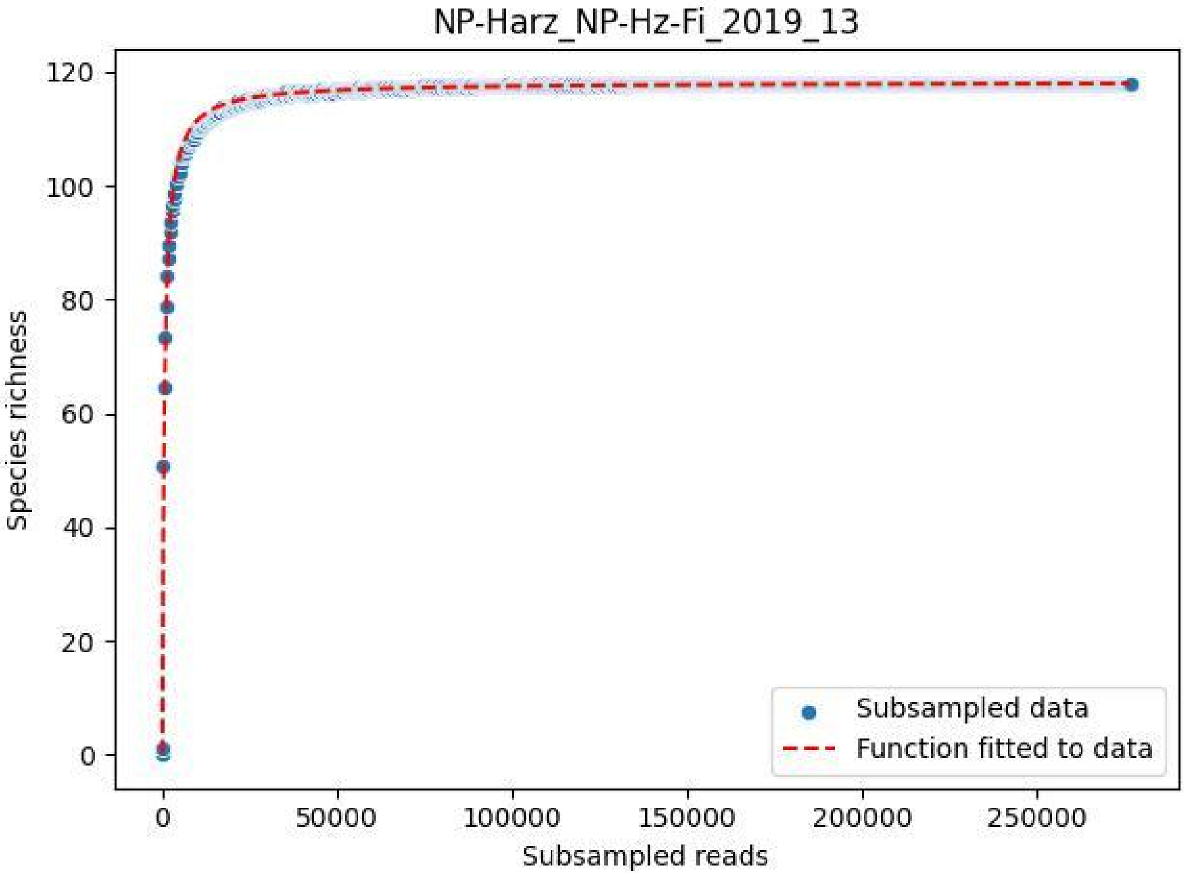
NP-Harz_NP-Hz-Fi_2019_13
Species richness
Subsampled reads
0
20
40
60
80
100
120
0
50000
100000
150000
200000
250000
Subsampled data
Function fitted to data

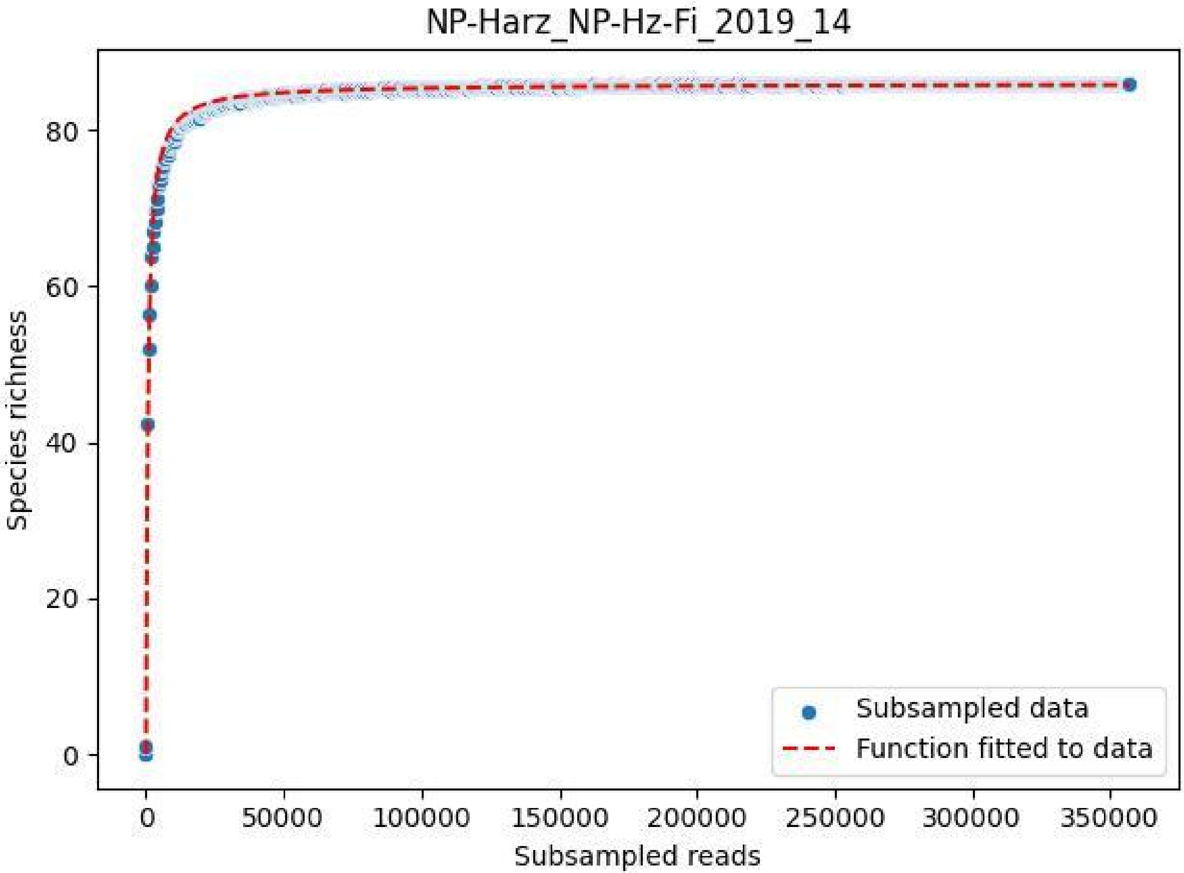
NP-Harz_NP-Hz-Fi_2019_14
Species richness
0
20
40
60
80
0
50000
100000
150000
200000
250000
300000
350000
Subsampled reads
Subsampled data
Function fitted to data

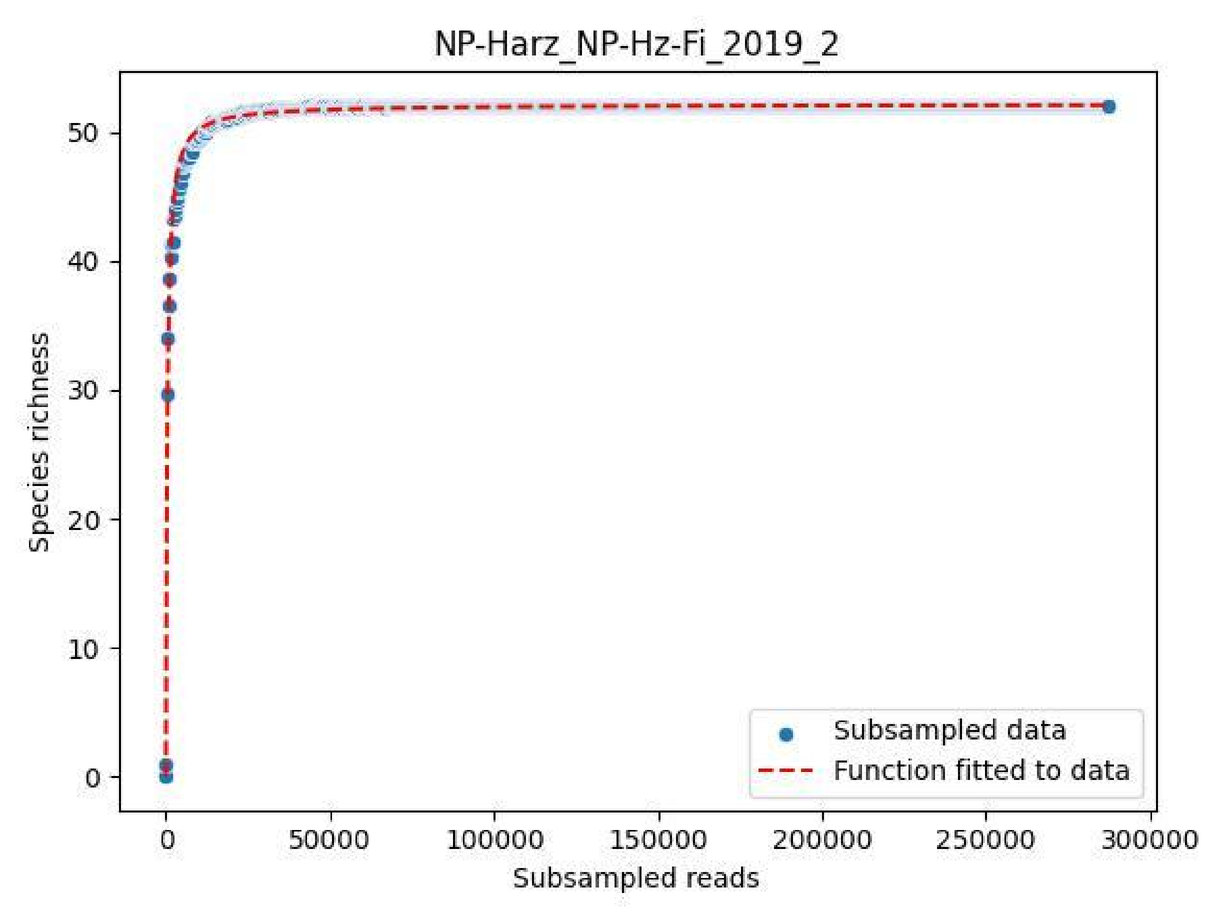
NP-Harz_NP-Hz-Fi_2019_2
Species richness
Subsampled reads
0
10
20
30
40
50
0
50000
100000
150000
200000
250000
300000
Subsampled data
Function fitted to data

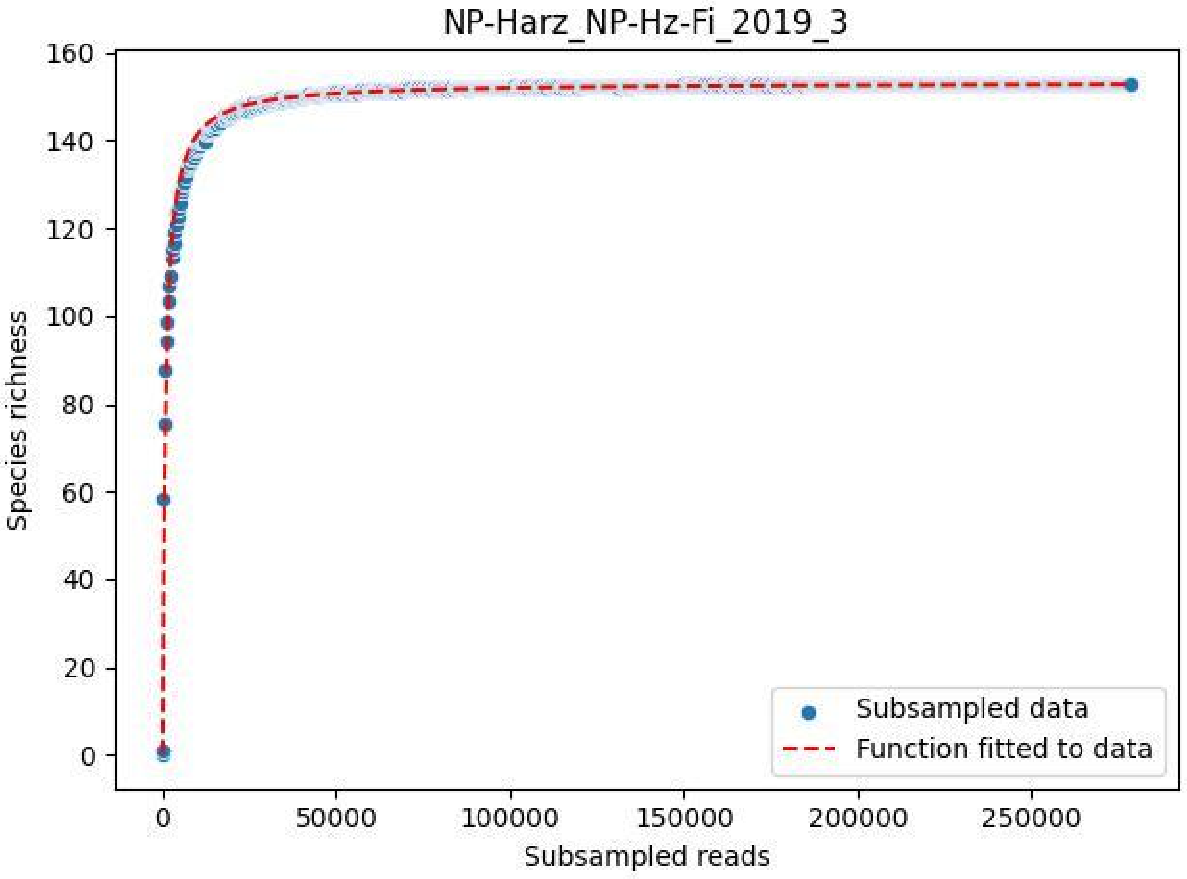

NP-Harz_NP-Hz-Fi_2019_3
Species richness
Subsampled reads
0
20
40
60
80
100
120
140
160
0
50000
100000
150000
200000
250000
Subsampled data
Function fitted to data

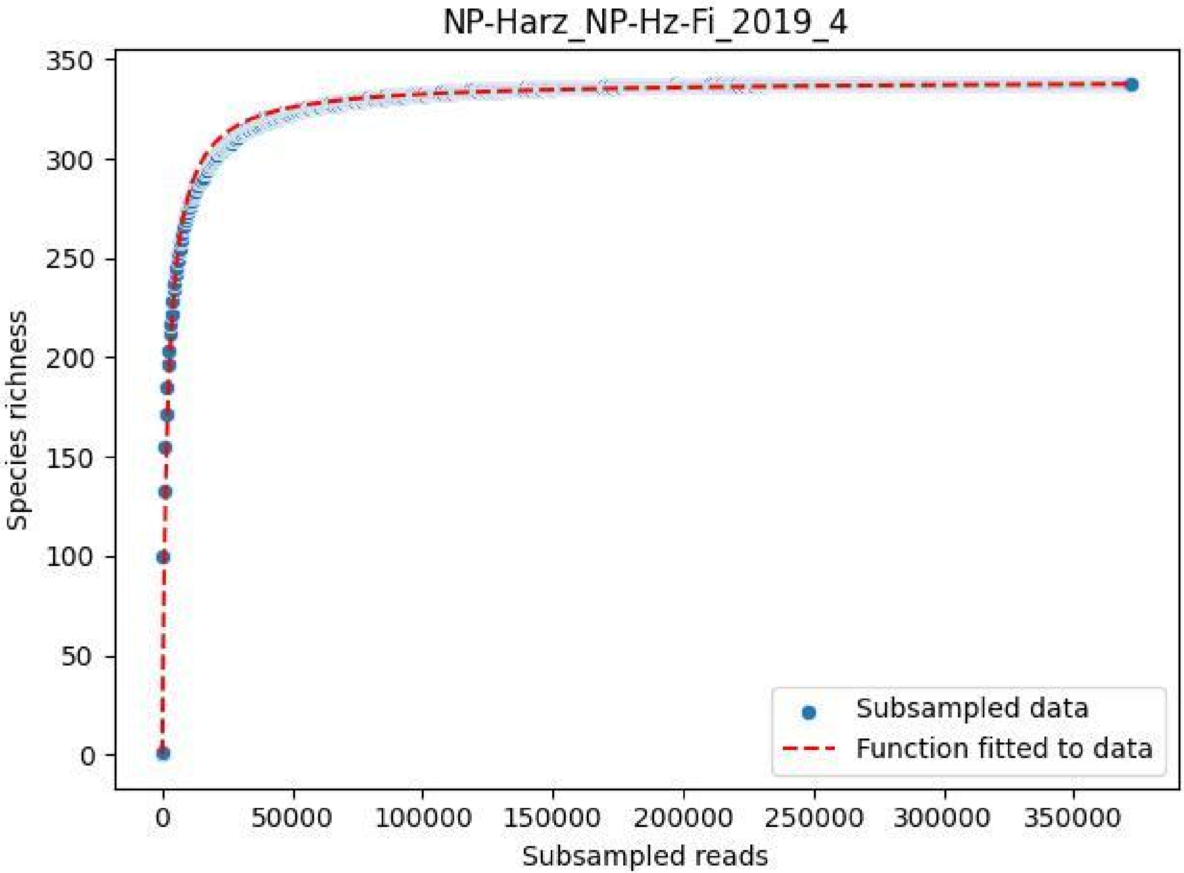

NP-Harz_NP-Hz-Fi_2019_4
Species richness
Subsampled reads
0
50
100
150
200
250
300
350
0
50000
100000
150000
200000
250000
300000
350000
Subsampled data
Function fitted to data

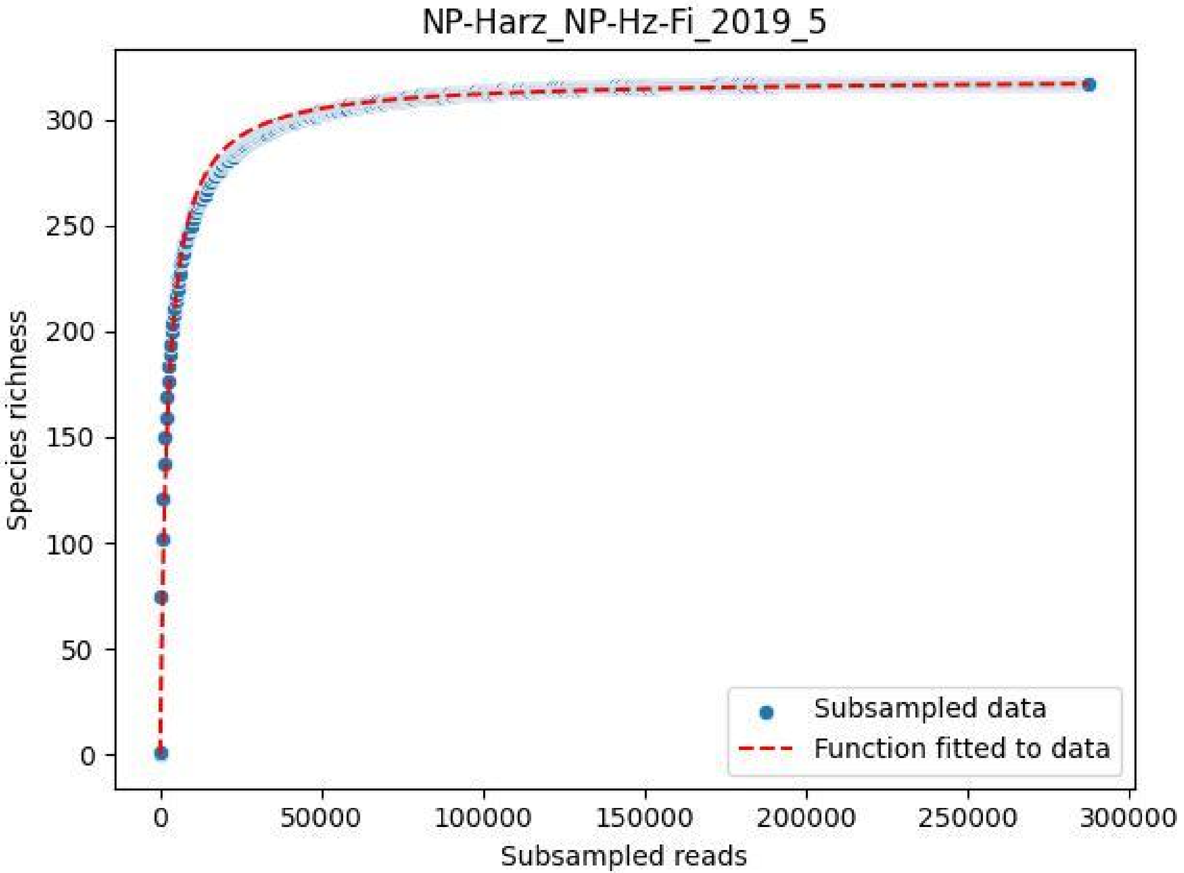
NP-Harz_NP-Hz-Fi_2019_5
Species richness
Subsampled reads
0
50
100
150
200
250
300
0
50000
100000
150000
200000
250000
300000
Subsampled data
Function fitted to data

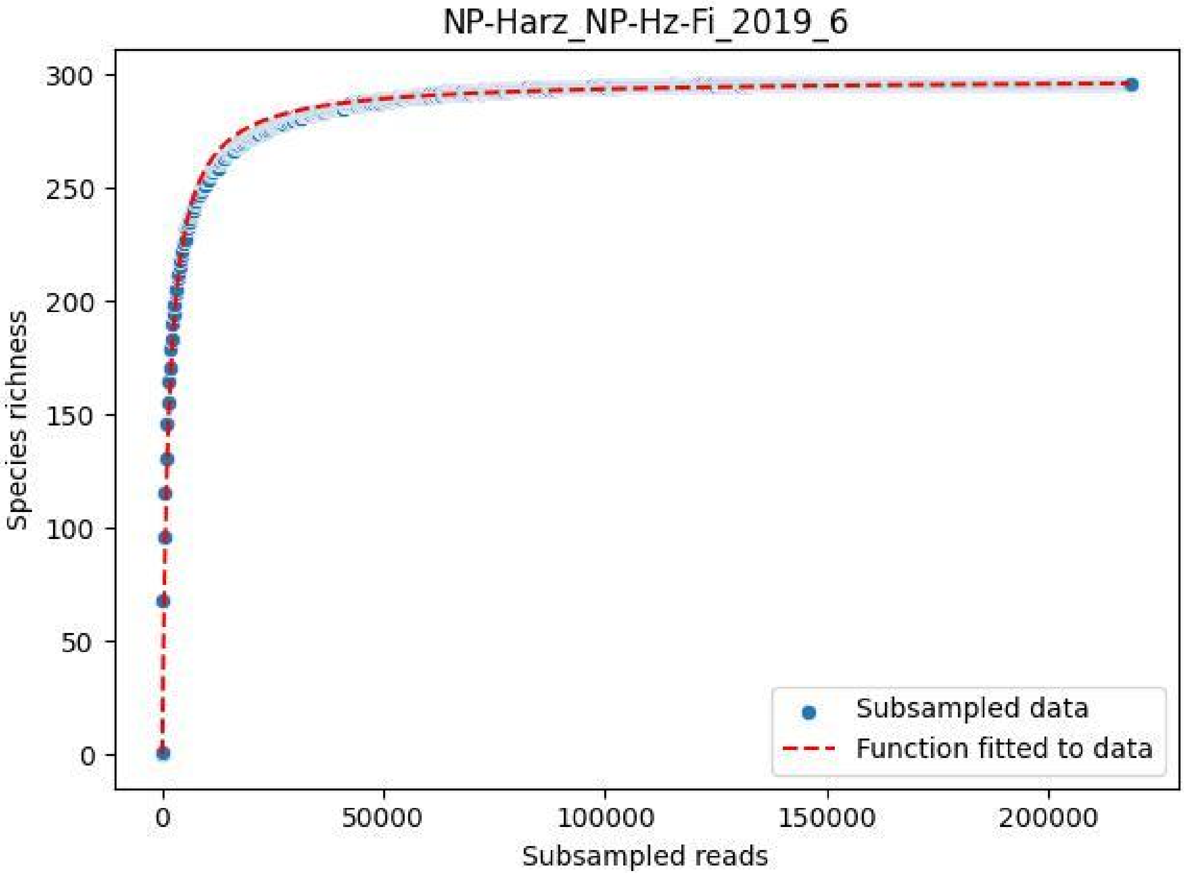
NP-Harz_NP-Hz-Fi_2019_6
Species richness
Subsampled reads
0
50
100
150
200
250
300
0
50000
100000
150000
200000
Subsampled data
Function fitted to data

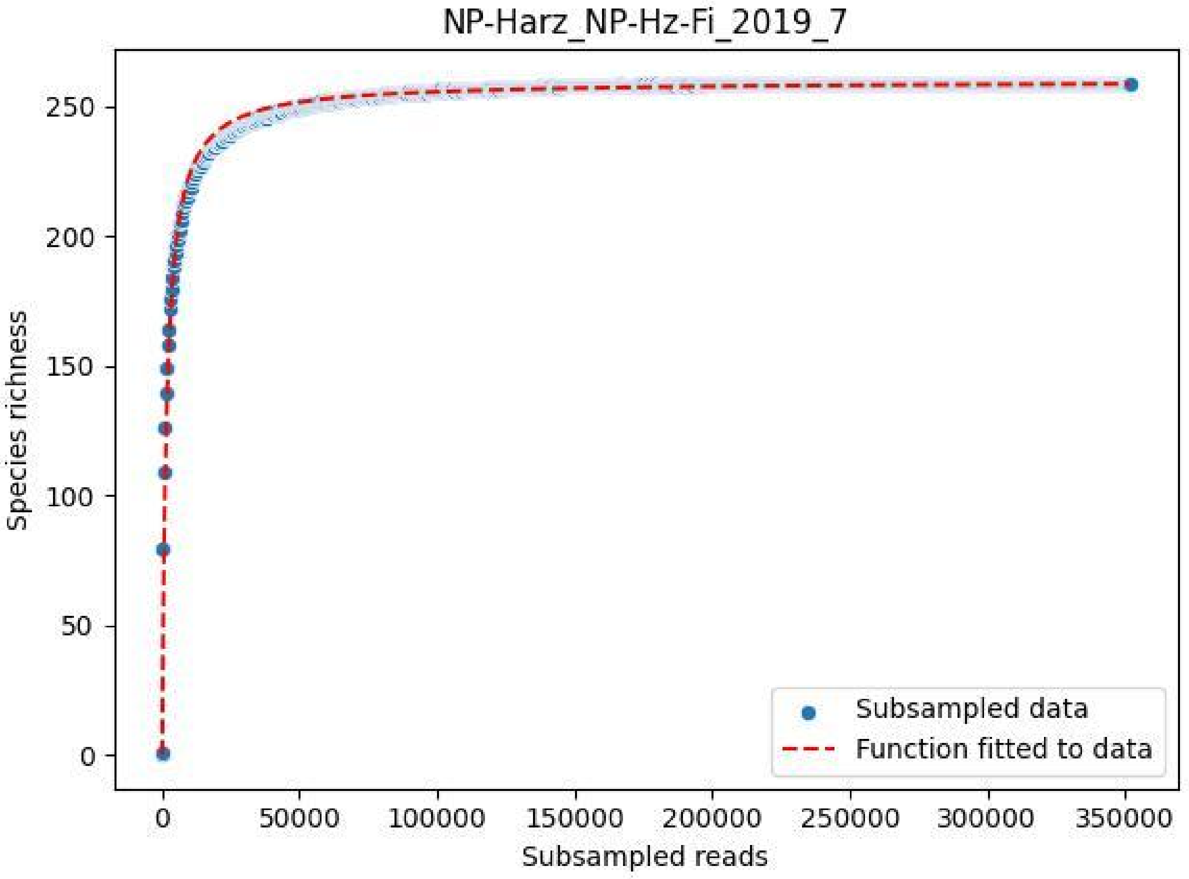

NP-Harz_NP-Hz-Fi_2019_7
Species richness
Subsampled reads
0
50
100
150
200
250
0
50000
100000
150000
200000
250000
300000
350000
Subsampled data
Function fitted to data

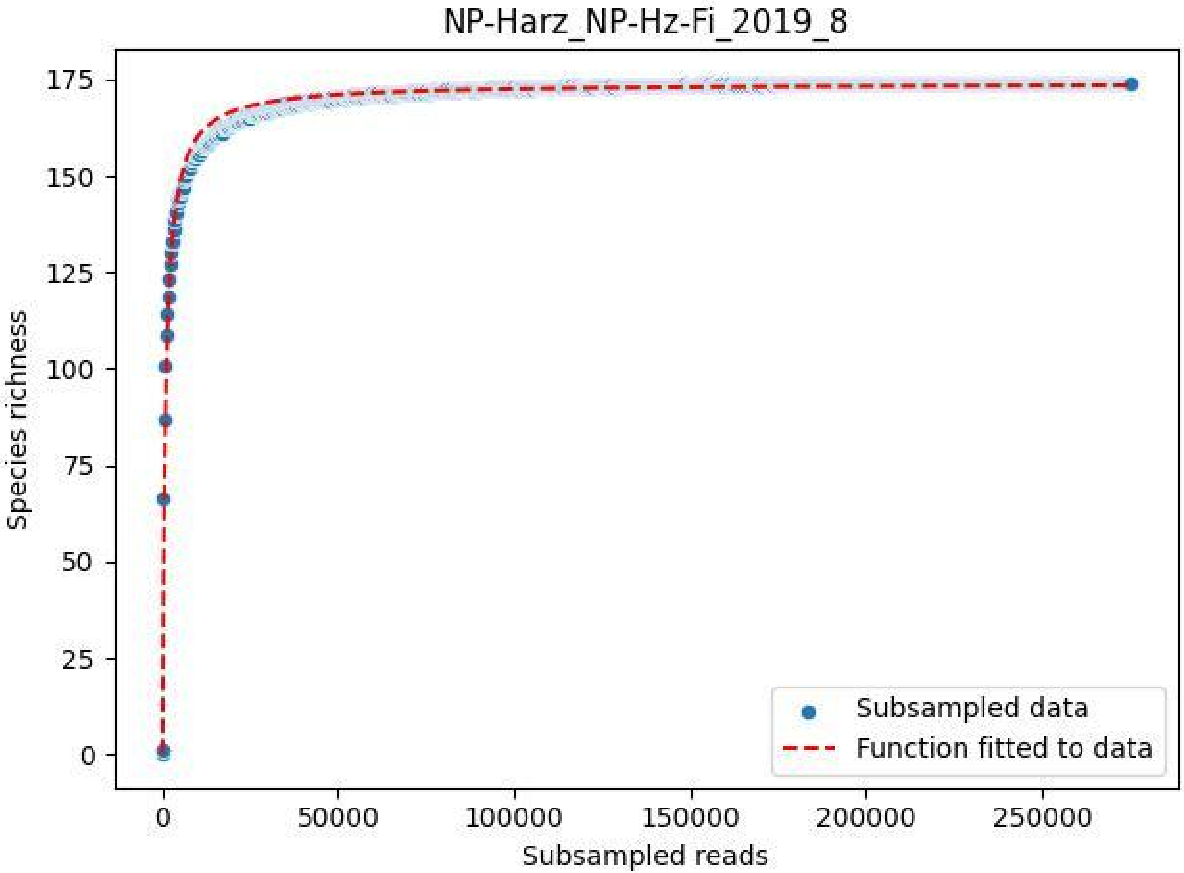
NP-Harz_NP-Hz-Fi_2019_8
Species richness
Subsampled reads
0
25
50
75
100
125
150
175
0
50000
100000
150000
200000
250000
Subsampled data
Function fitted to data

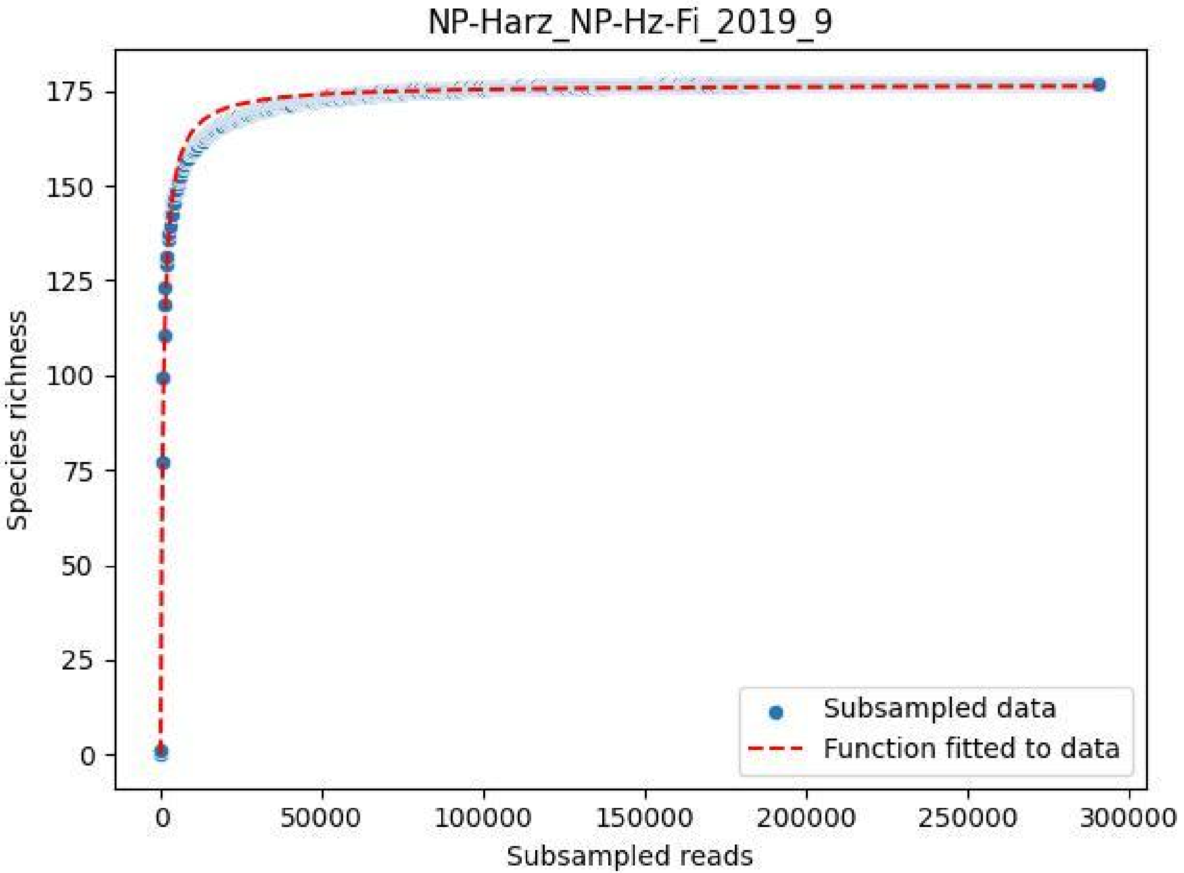

NP-Harz_NP-Hz-Fi_2019_9
Species richness
Subsampled reads
0
25
50
75
100
125
150
175
0
50000
100000
150000
200000
250000
300000
Subsampled data
Function fitted to data

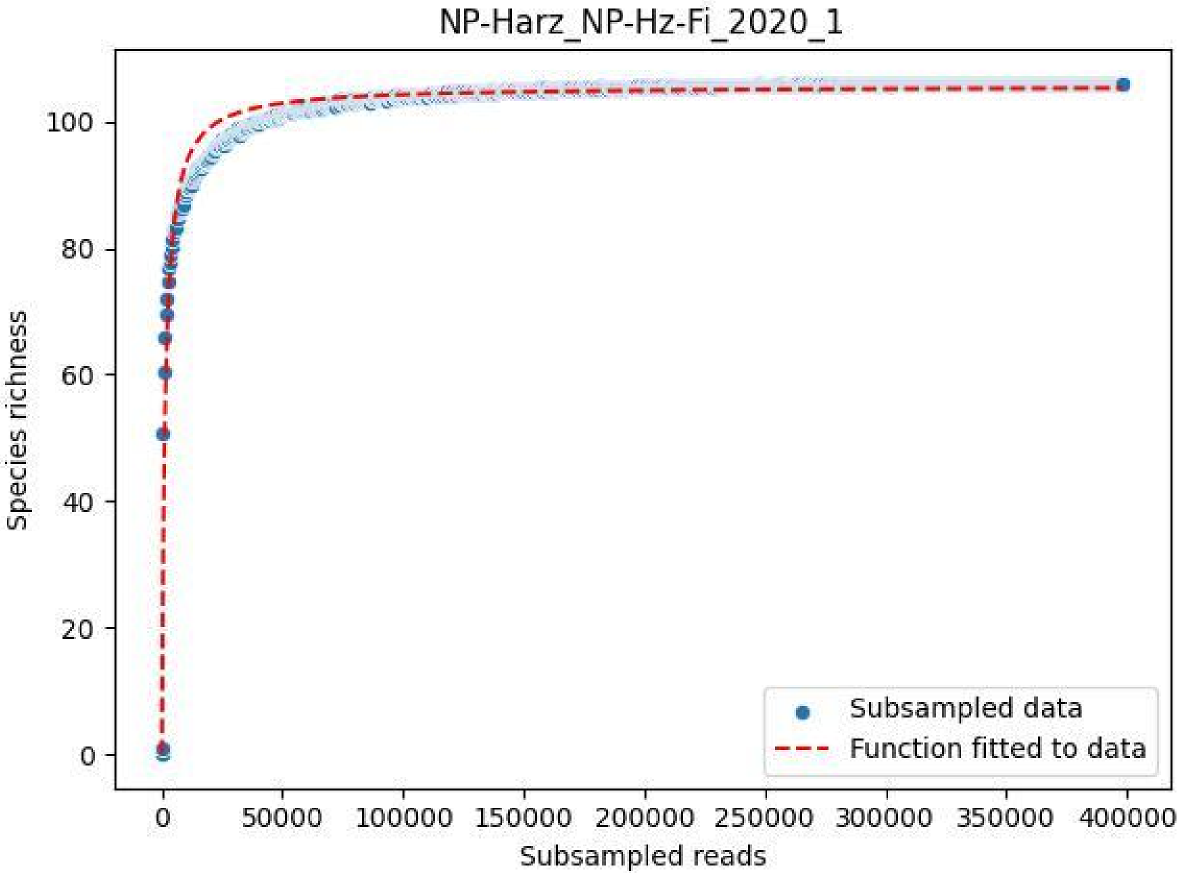
NP-Harz_NP-Hz-Fi_2020_1
Species richness
Subsampled reads
0
20
40
60
80
100
0
50000
100000
150000
200000
250000
300000
350000
400000
Subsampled data
Function fitted to data

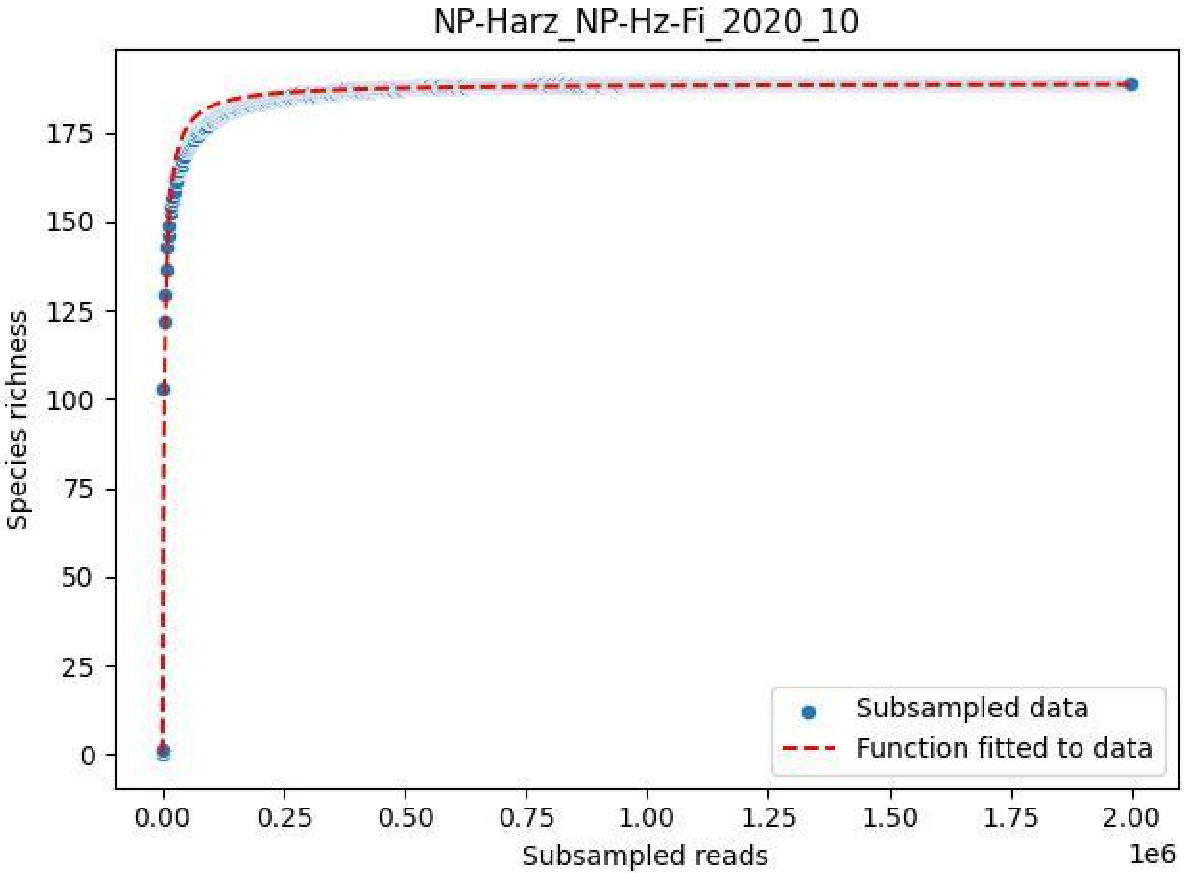

NP-Harz_NP-Hz-Fi_2020_10
Species richness
0
25
50
75
100
125
150
175
0.00
0.25
0.50
0.75
1.00
1.25
1.50
1.75
2.00
1e6
Subsampled reads
Subsampled data
Function fitted to data

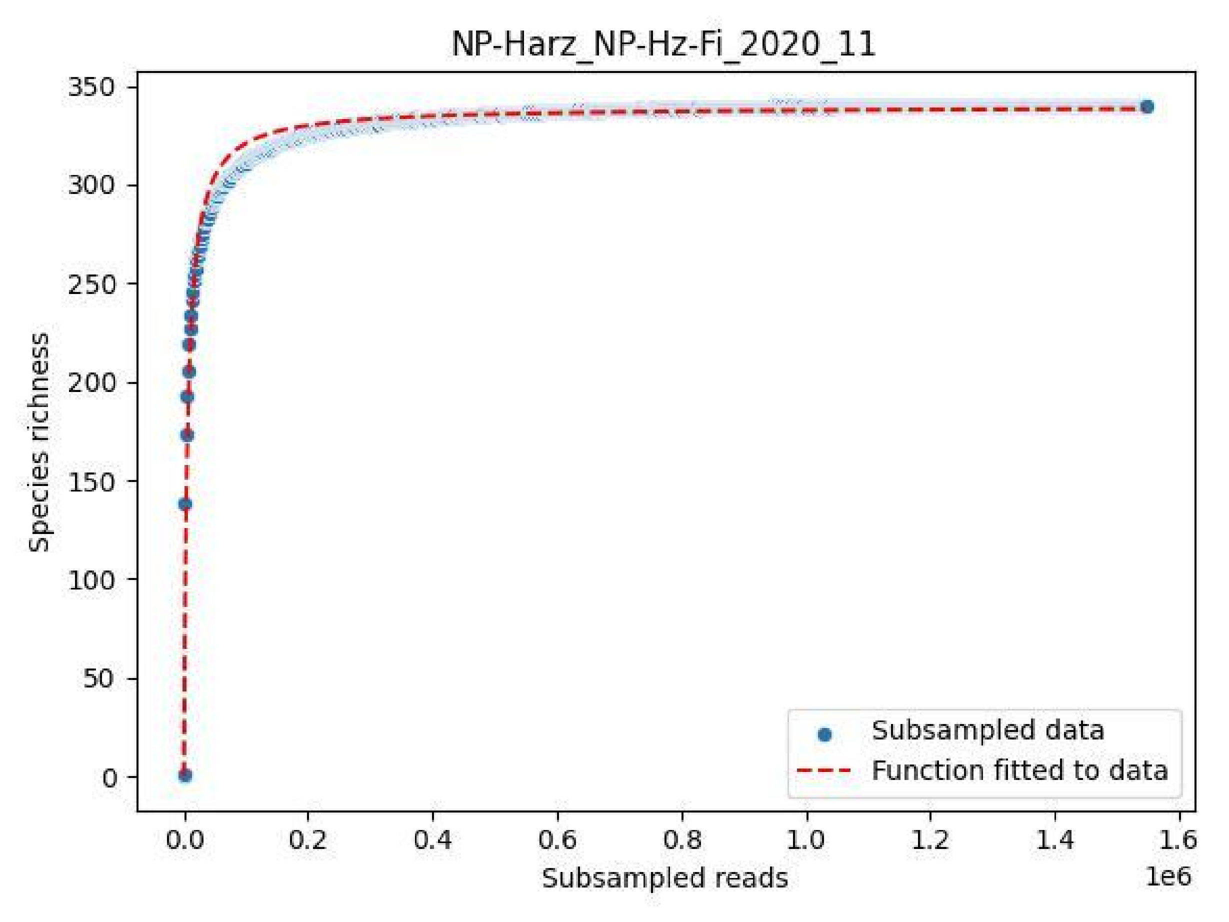

NP-Harz_NP-Hz-Fi_2020_11
Species richness
Subsampled reads
1e6
0
50
100
150
200
250
300
350
0.0
0.2
0.4
0.6
0.8
1.0
1.2
1.4
1.6
Subsampled data
Function fitted to data

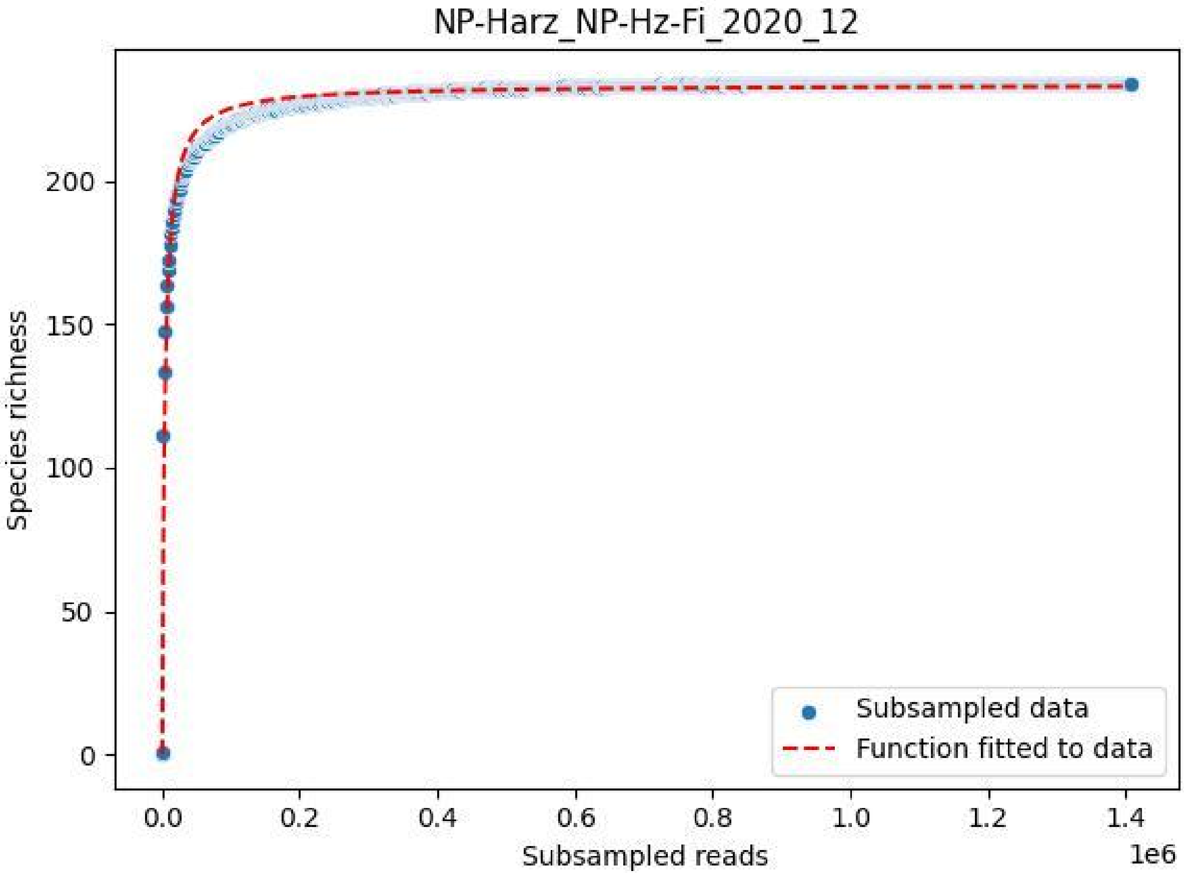
NP-Harz_NP-Hz-Fi_2020_12
Species richness
Subsampled reads
1e6
Subsampled data
Function fitted to data
0
50
100
150
200
0.0
0.2
0.4
0.6
0.8
1.0
1.2
1.4

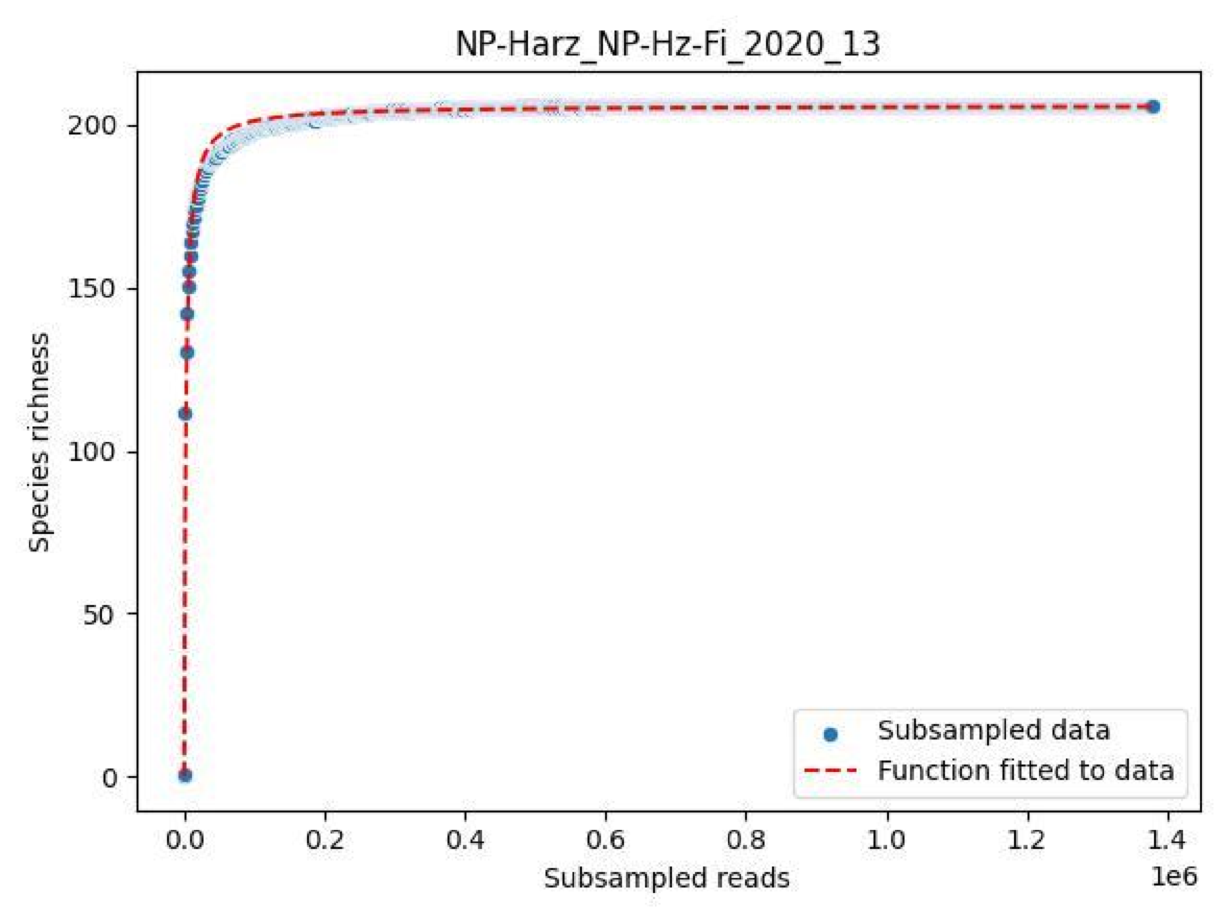

NP-Harz_NP-Hz-Fi_2020_13
Species richness
200
150
100
50
0
0.0
0.2
0.4
0.6
0.8
1.0
1.2
1.4
Subsampled reads
1e6
Subsampled data
Function fitted to data

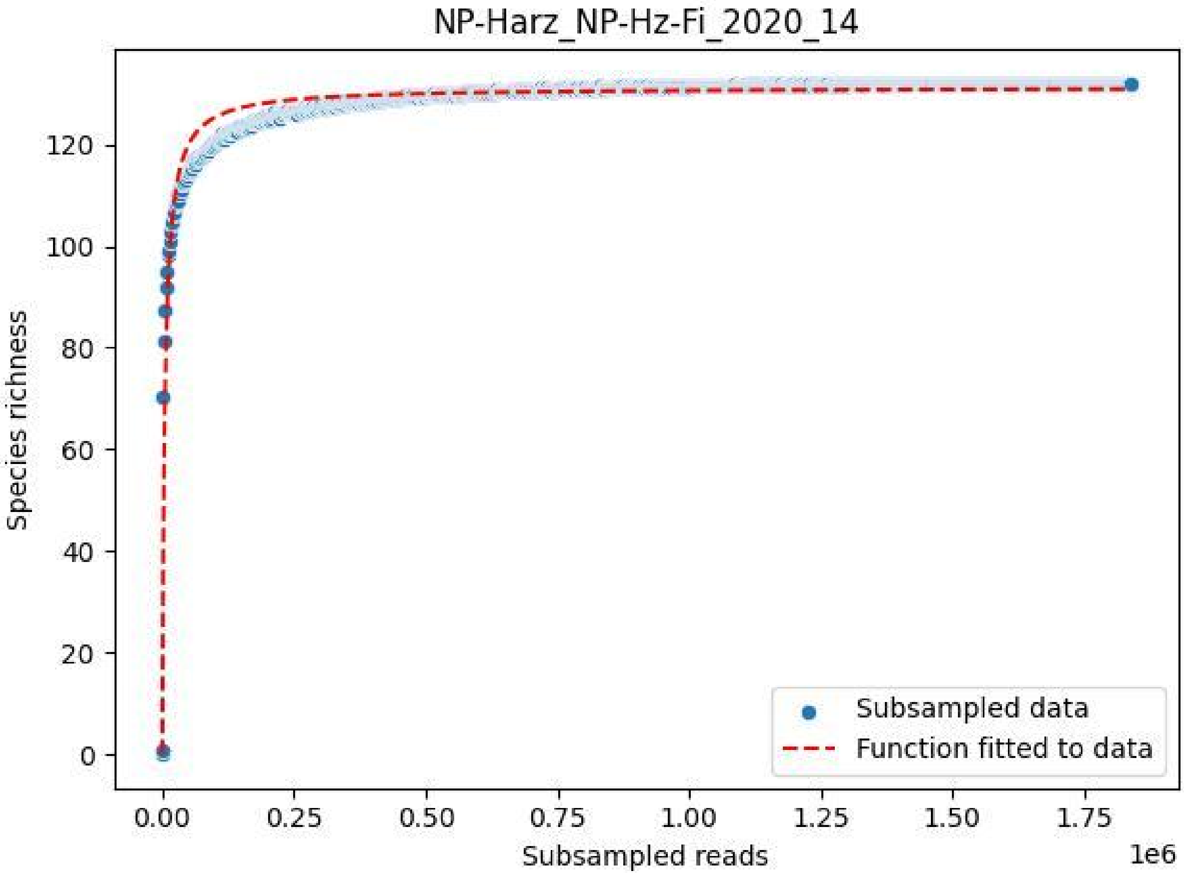
NP-Harz_NP-Hz-Fi_2020_14
Species richness
0
20
40
60
80
100
120
0.00
0.25
0.50
0.75
1.00
1.25
1.50
1.75
1e6
Subsampled reads
Subsampled data
Function fitted to data

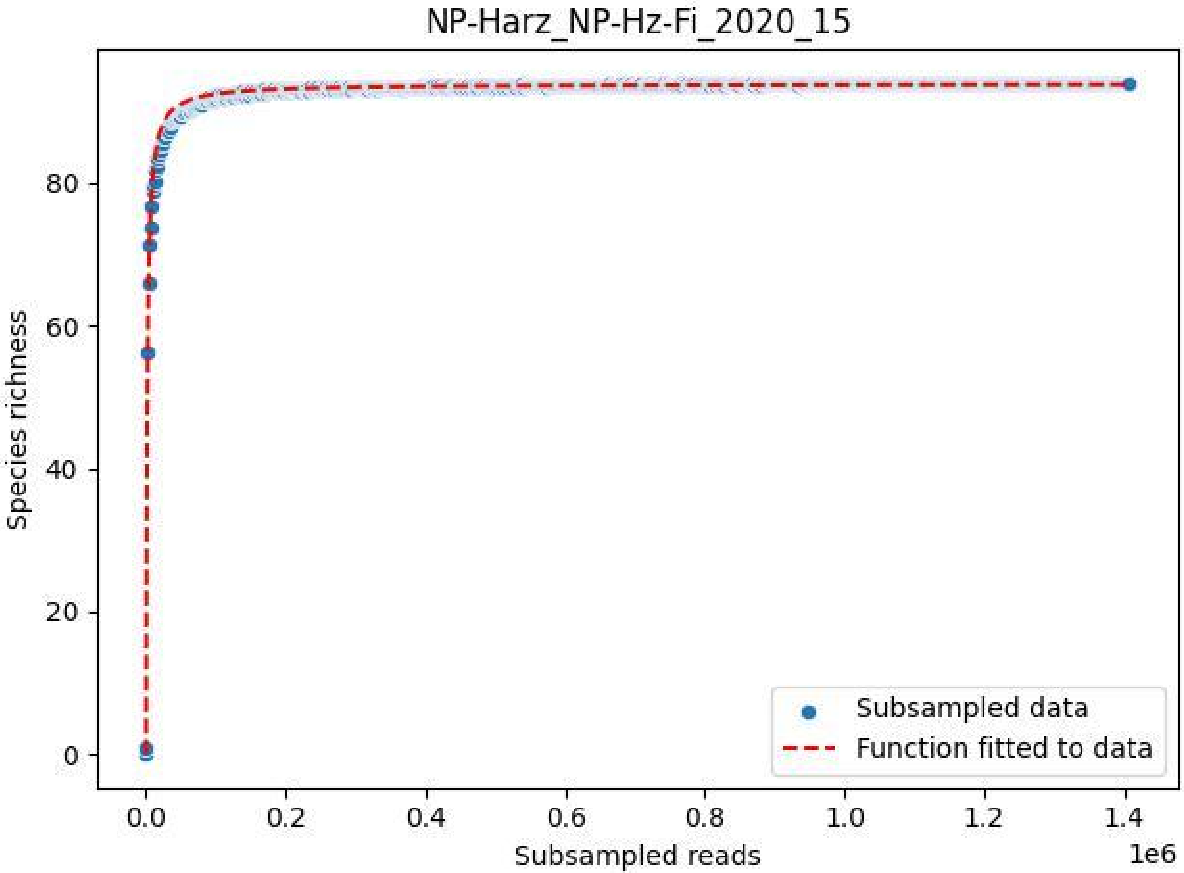
NP-Harz_NP-Hz-Fi_2020_15
Species richness
Subsampled reads
1e6
0
20
40
60
80
0.0
0.2
0.4
0.6
0.8
1.0
1.2
1.4
Subsampled data
Function fitted to data

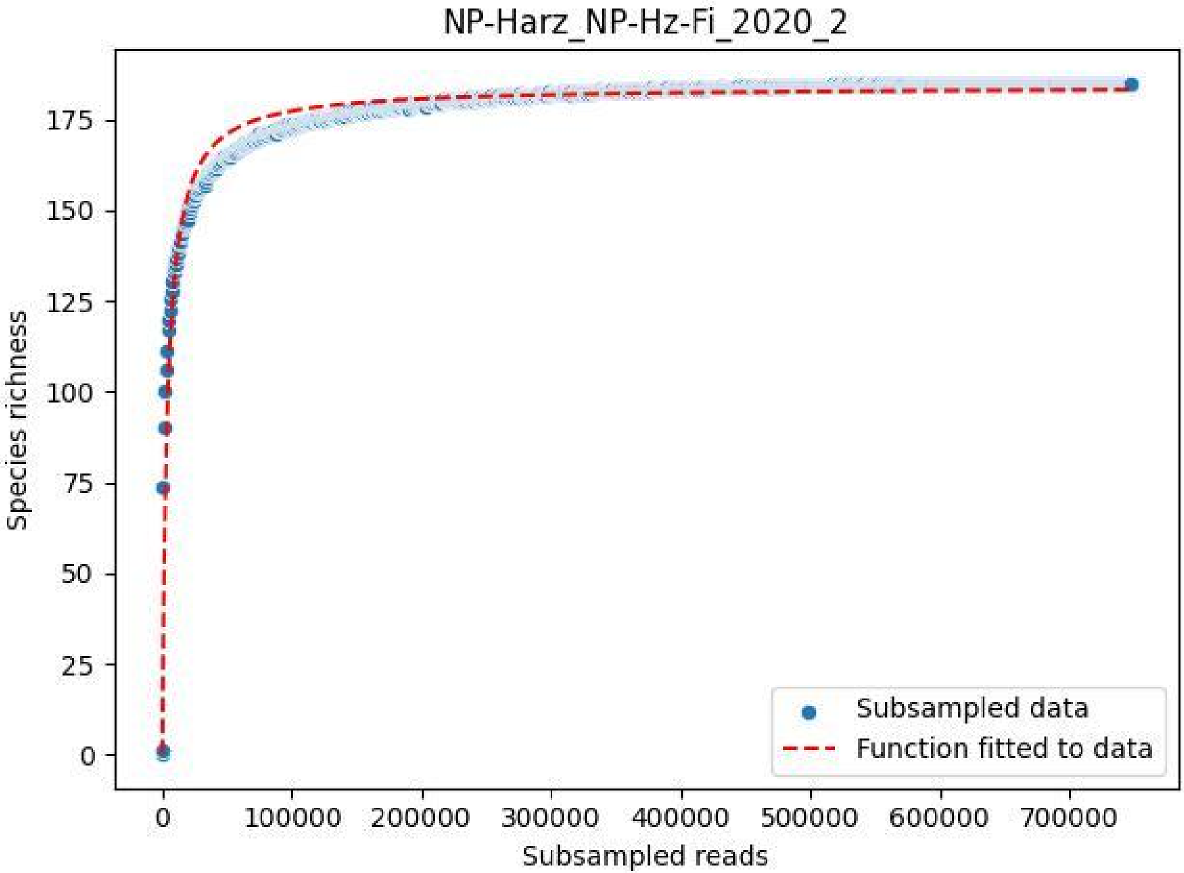
NP-Harz_NP-Hz-Fi_2020_2
Species richness
Subsampled reads
0
25
50
75
100
125
150
175
0
100000
200000
300000
400000
500000
600000
700000
Subsampled data
Function fitted to data

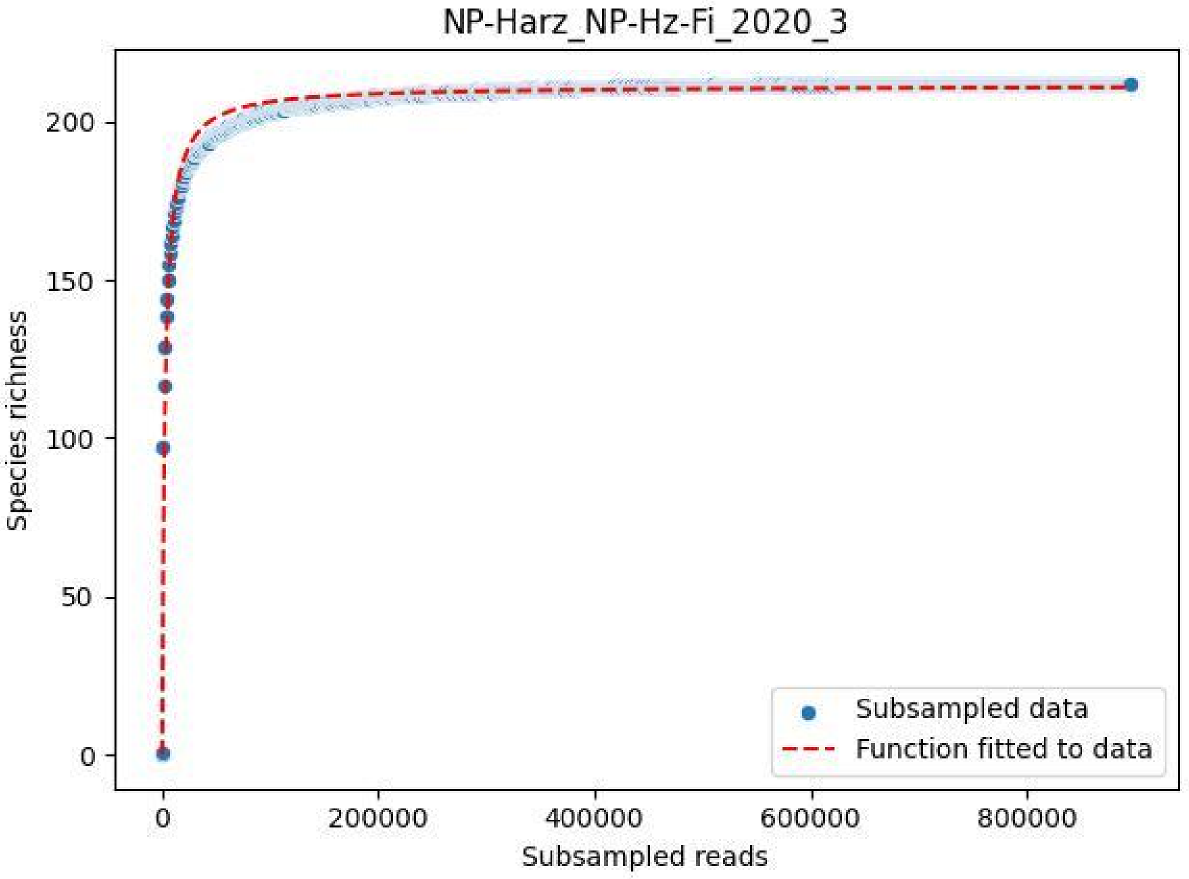
NP-Harz_NP-Hz-Fi_2020_3
Species richness
Subsampled reads
0
50
100
150
200
0
200000
400000
600000
800000
Subsampled data
Function fitted to data

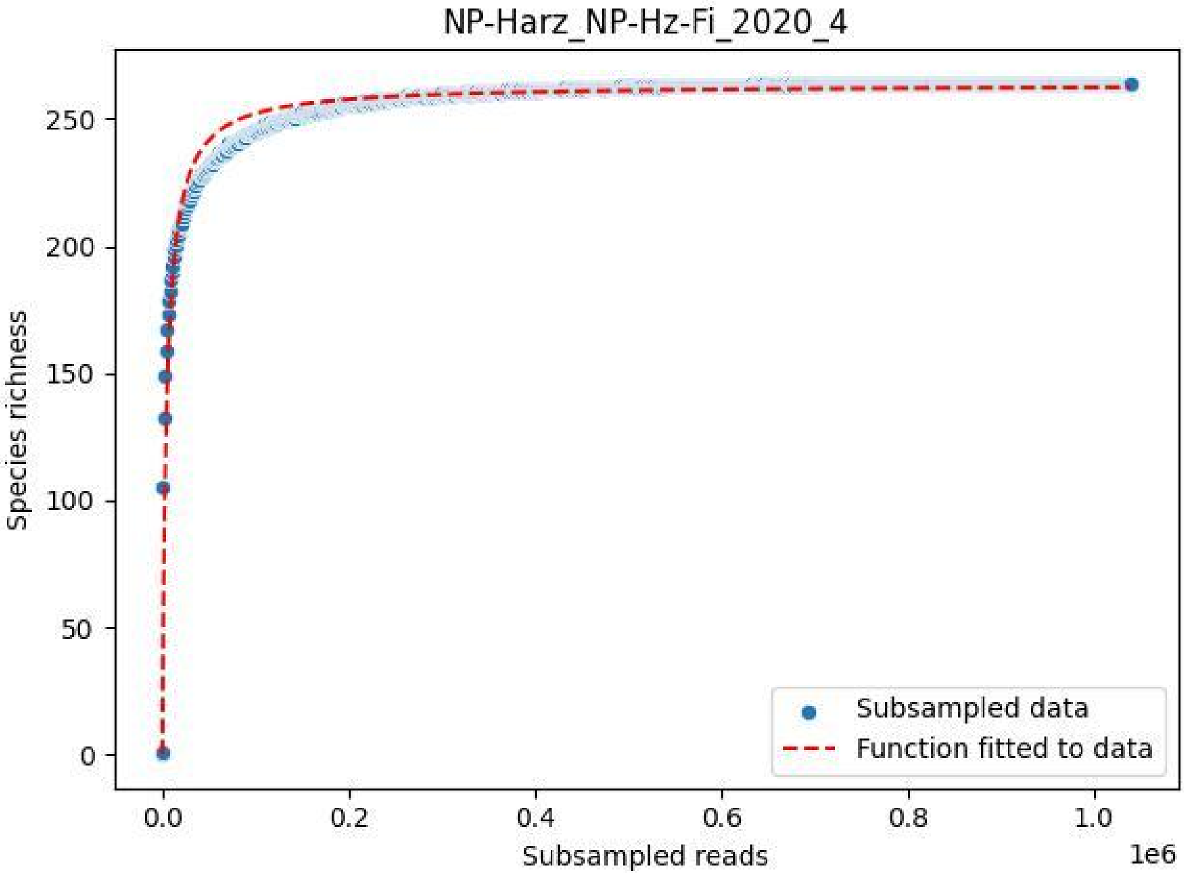
NP-Harz_NP-Hz-Fi_2020_4
Species richness
Subsampled reads
1e6
0
50
100
150
200
250
0.0
0.2
0.4
0.6
0.8
1.0
Subsampled data
Function fitted to data

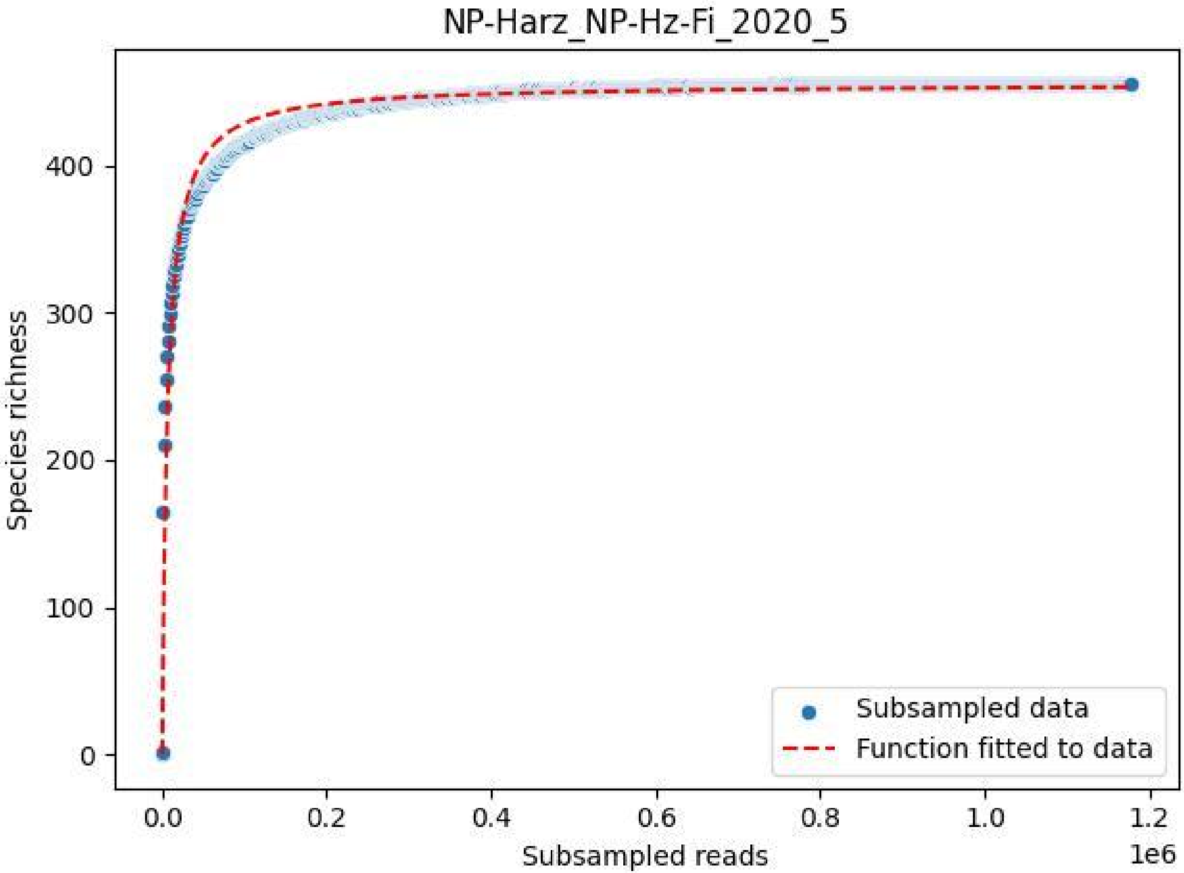
NP-Harz_NP-Hz-Fi_2020_5
Species richness
Subsampled reads
0
100
200
300
400
0.0
0.2
0.4
0.6
0.8
1.0
1.2
1e6
Subsampled data
Function fitted to data

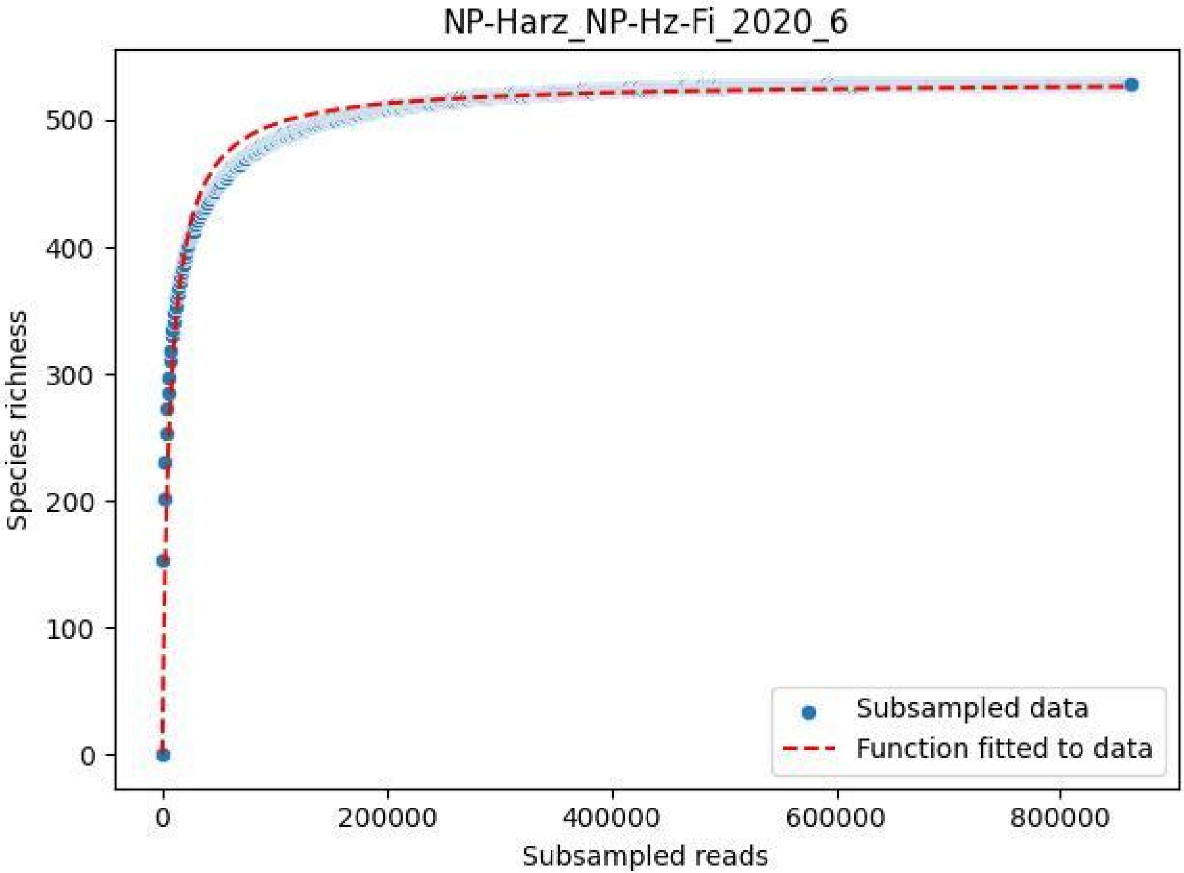
NP-Harz_NP-Hz-Fi_2020_6
Species richness
Subsampled reads
0
100
200
300
400
500
0
200000
400000
600000
800000
Subsampled data
Function fitted to data

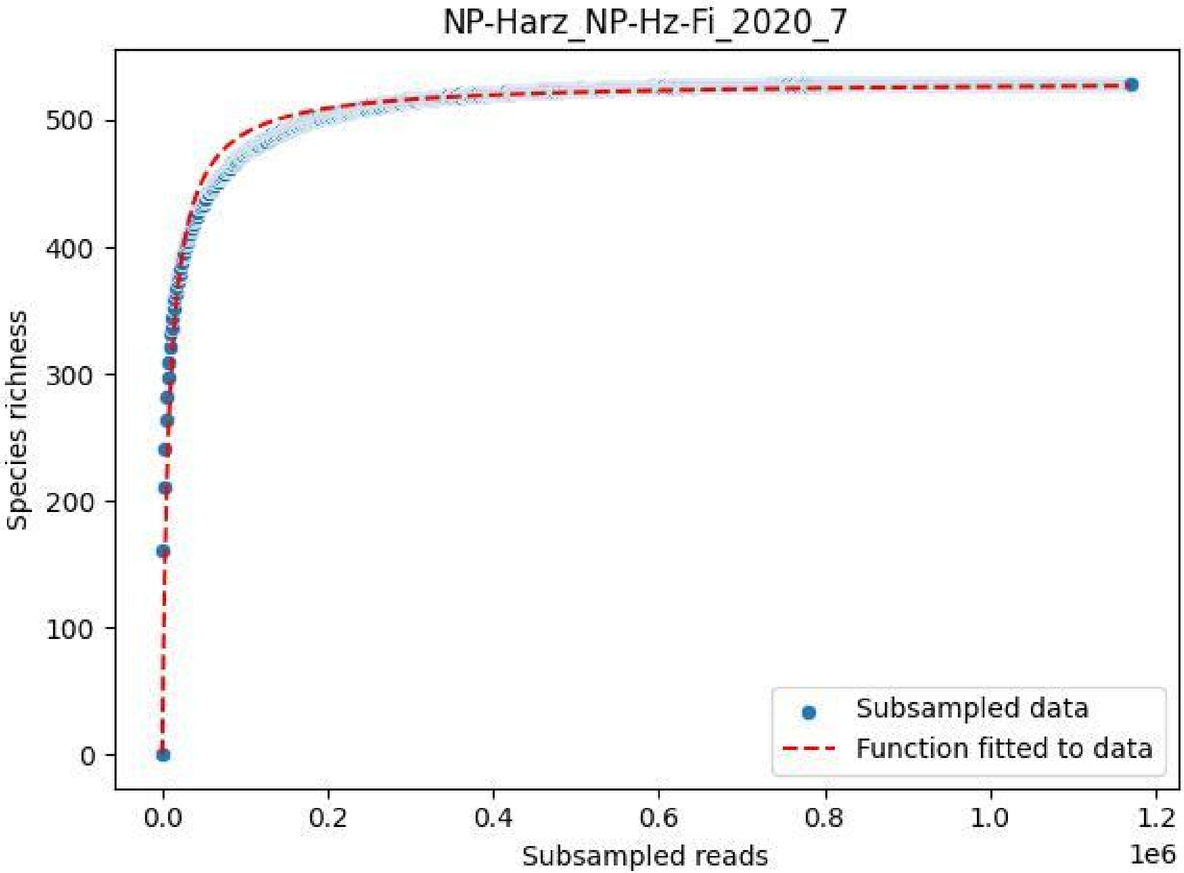
NP-Harz_NP-Hz-Fi_2020_7
Species richness
Subsampled reads
1e6
0
100
200
300
400
500
0.0
0.2
0.4
0.6
0.8
1.0
1.2
Subsampled data
Function fitted to data

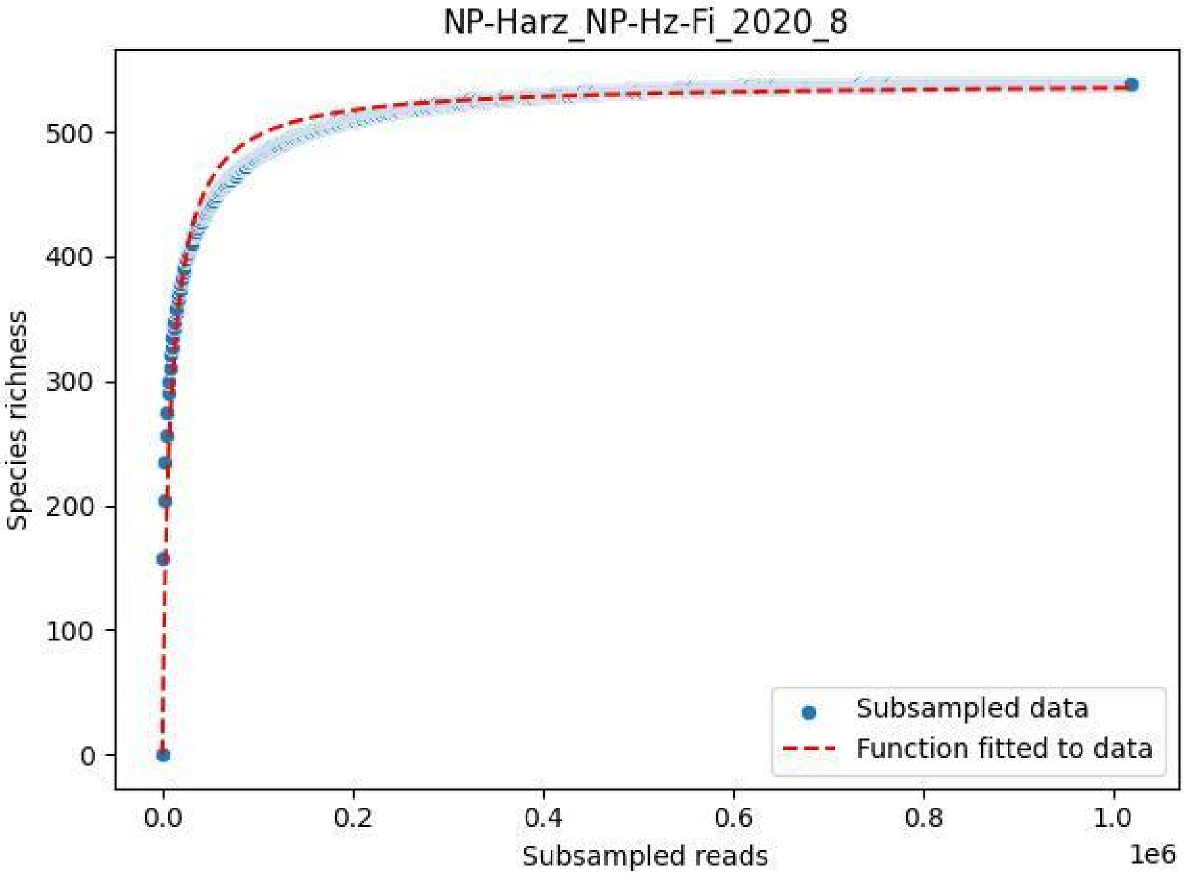
NP-Harz_NP-Hz-Fi_2020_8
Species richness
Subsampled reads
1e6
0
100
200
300
400
500
0.0
0.2
0.4
0.6
0.8
1.0
Subsampled data
Function fitted to data

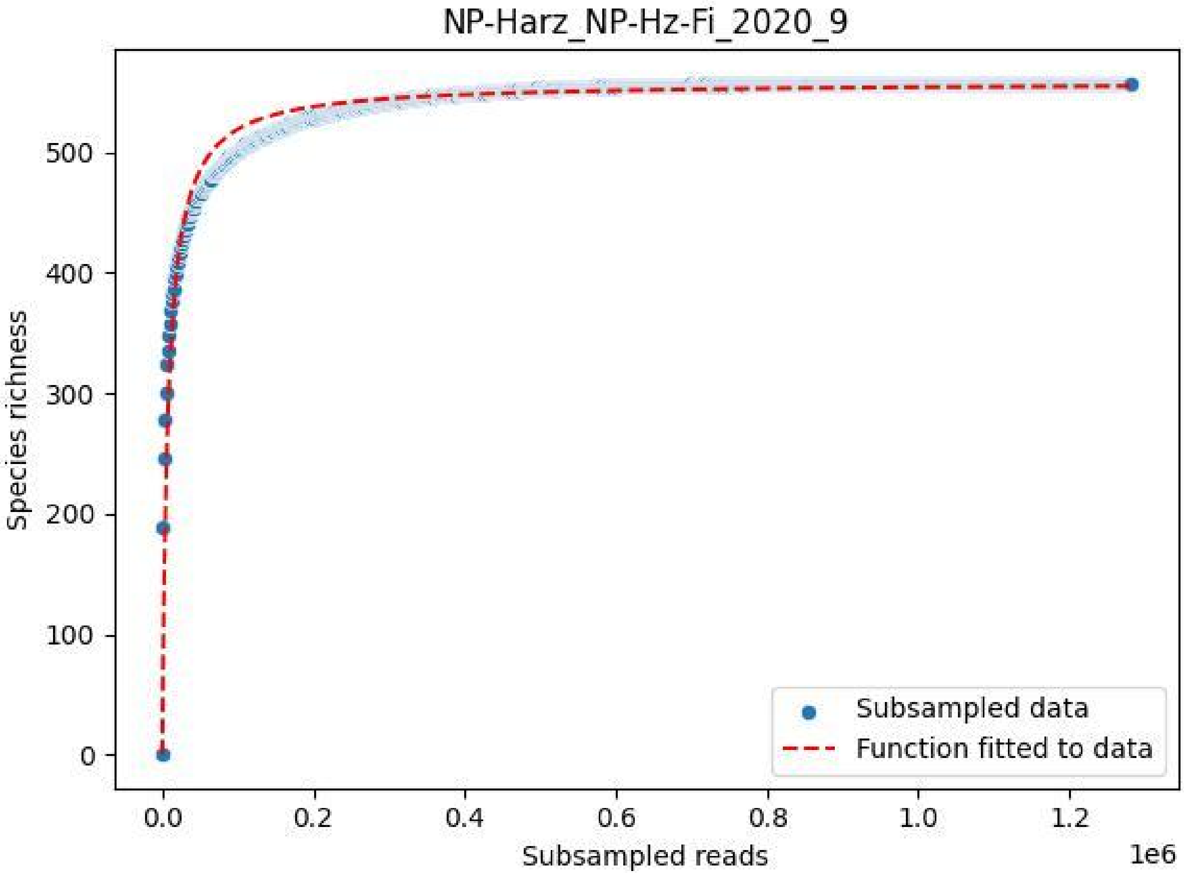
NP-Harz_NP-Hz-Fi_2020_9
Species richness
Subsampled reads
1e6
0
100
200
300
400
500
0.0
0.2
0.4
0.6
0.8
1.0
1.2
Subsampled data
Function fitted to data

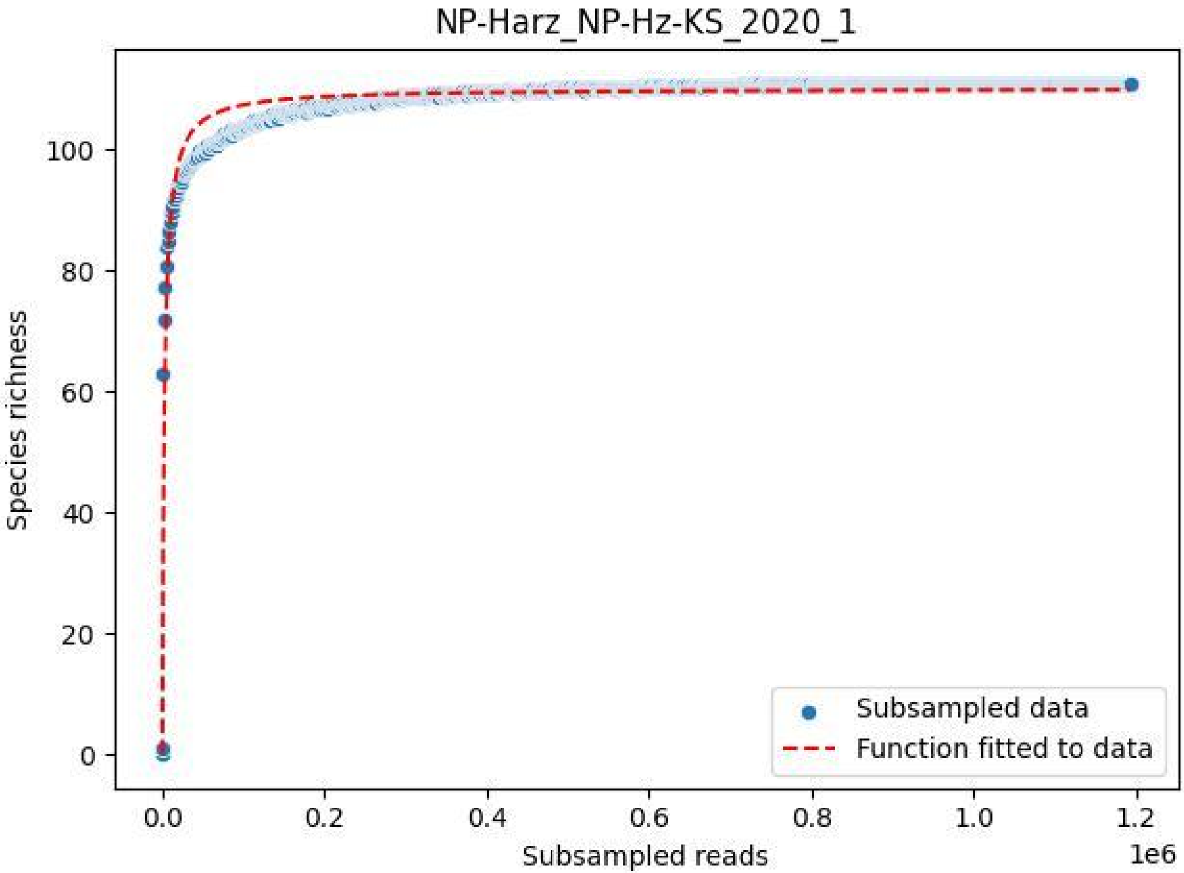
NP-Harz_NP-Hz-KS_2020_1
Species richness
Subsampled reads
1e6
0
20
40
60
80
100
0.0
0.2
0.4
0.6
0.8
1.0
1.2
Subsampled data
Function fitted to data

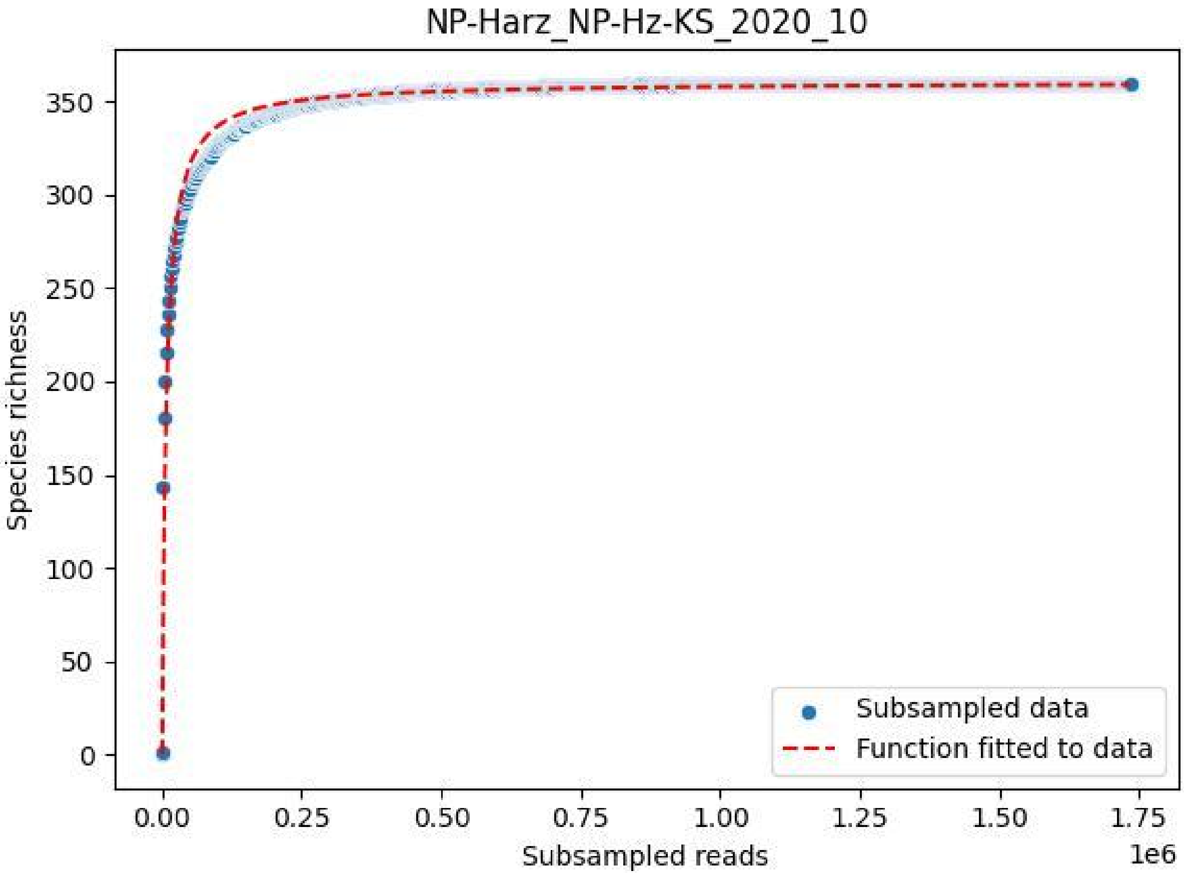

NP-Harz_NP-Hz-KS_2020_10
Species richness
Subsampled reads
1e6
0
50
100
150
200
250
300
350
0.00
0.25
0.50
0.75
1.00
1.25
1.50
1.75
Subsampled data
Function fitted to data

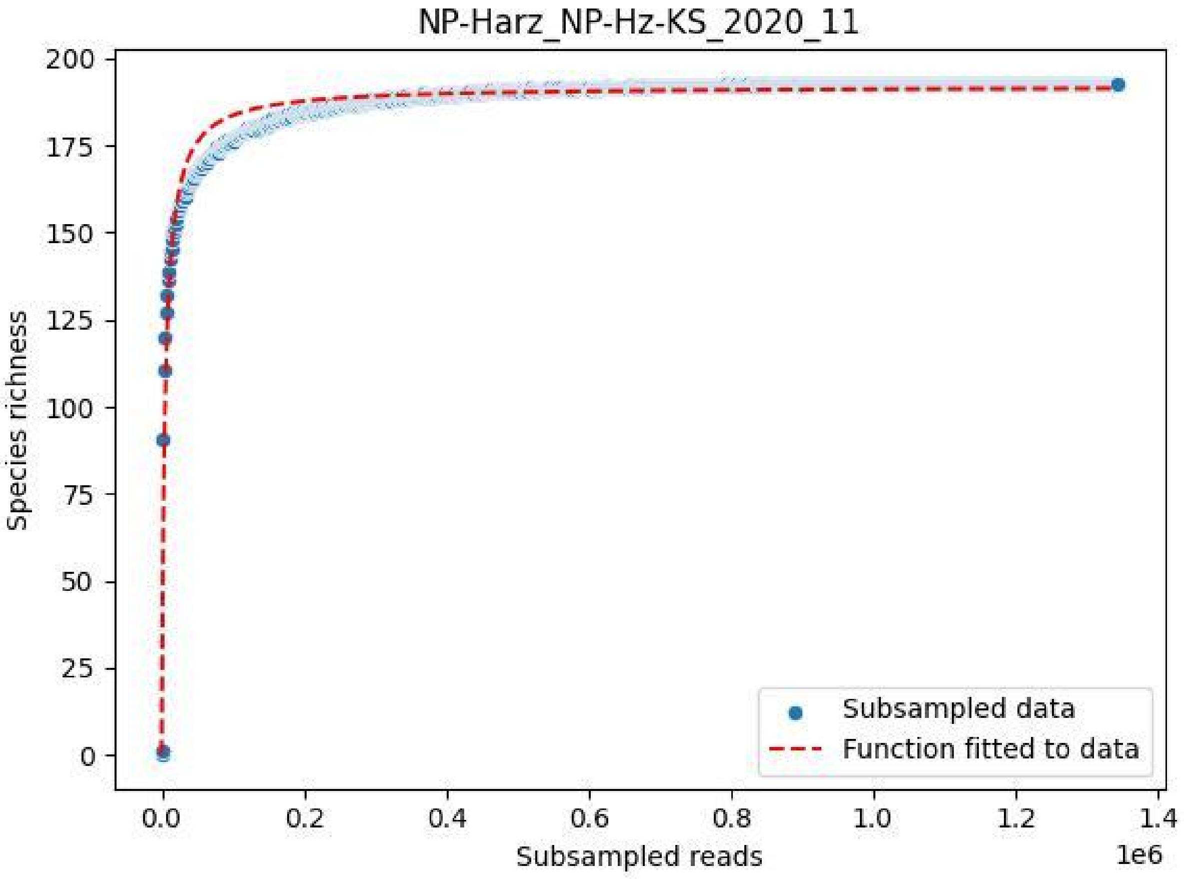
NP-Harz_NP-Hz-KS_2020_11
Species richness
Subsampled reads
1e6
0
25
50
75
100
125
150
175
200
0.0
0.2
0.4
0.6
0.8
1.0
1.2
1.4
Subsampled data
Function fitted to data

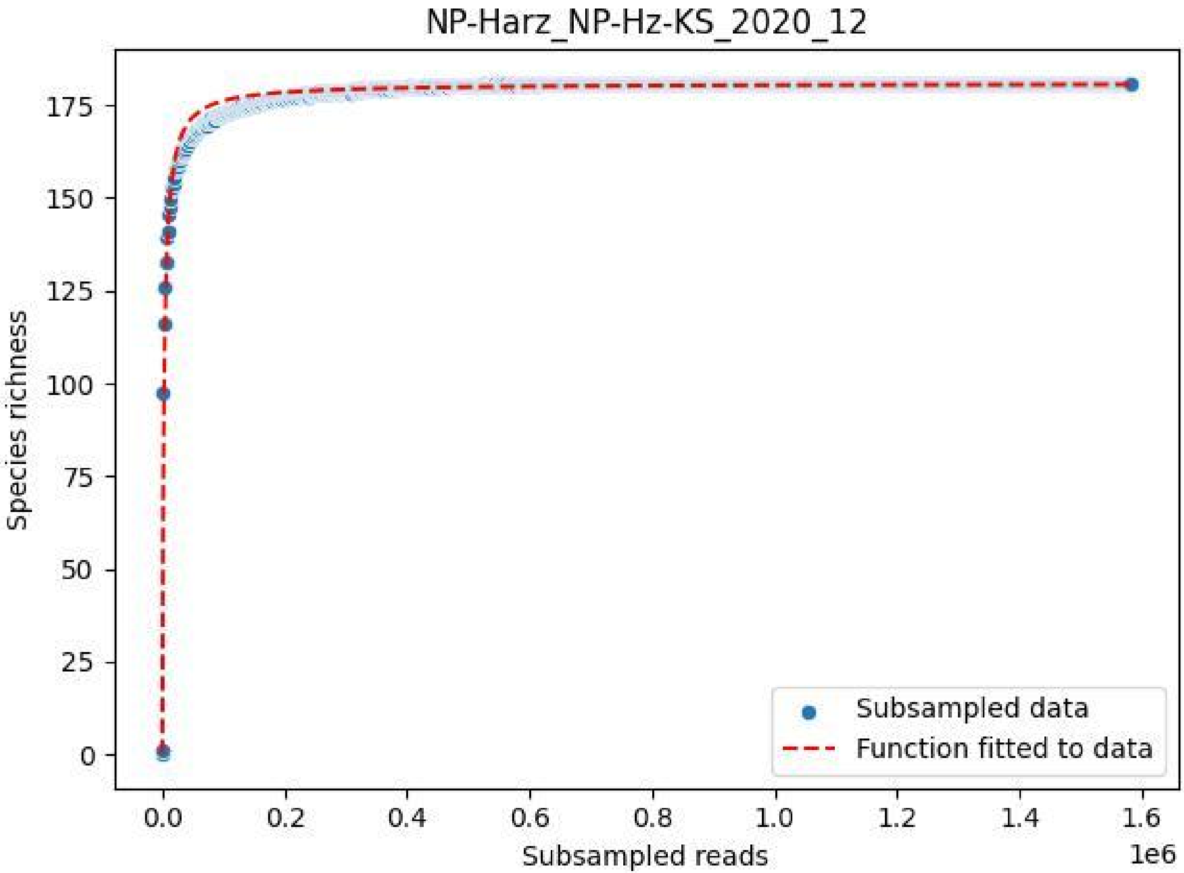

NP-Harz_NP-Hz-KS_2020_12
Species richness
Subsampled reads
1e6
0
25
50
75
100
125
150
175
0.0
0.2
0.4
0.6
0.8
1.0
1.2
1.4
1.6
Subsampled data
Function fitted to data

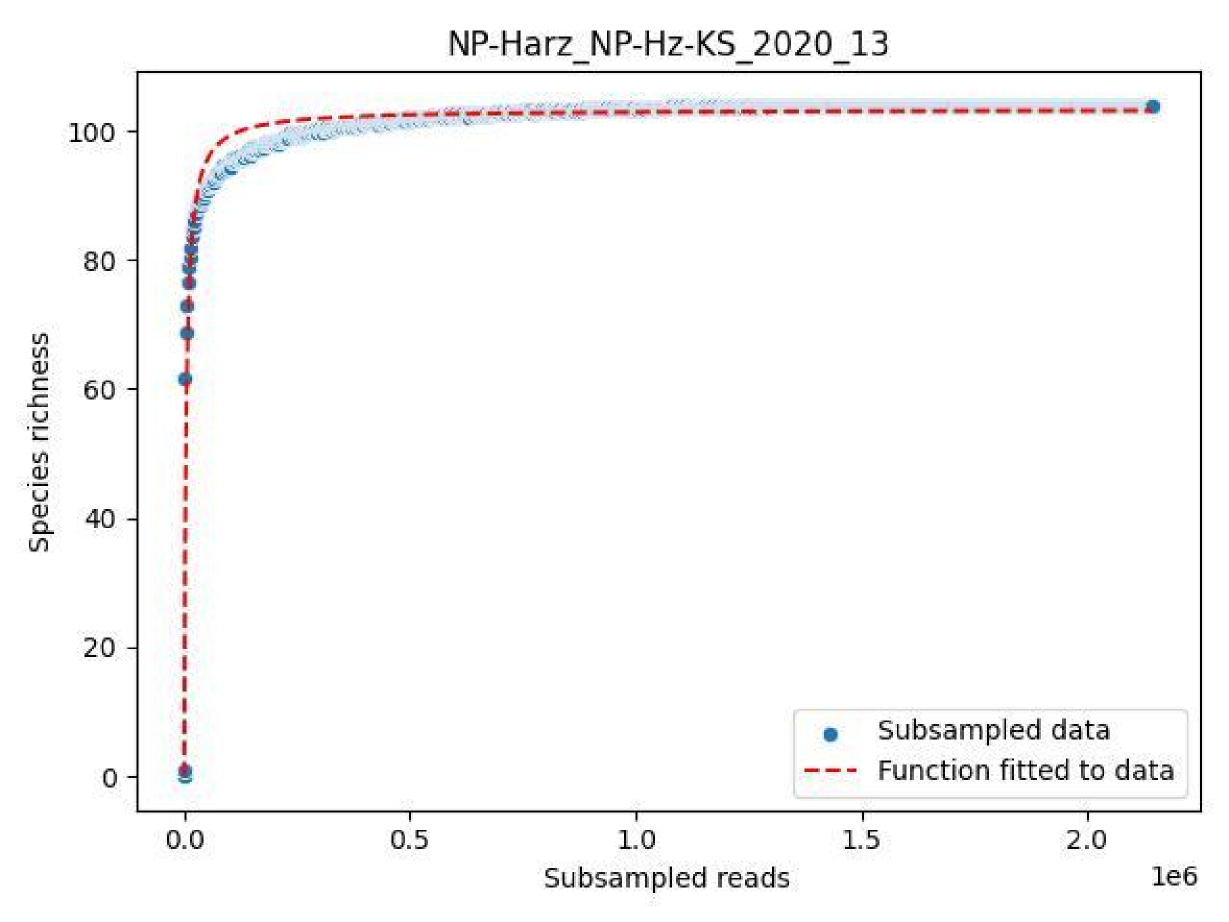
NP-Harz_NP-Hz-KS_2020_13
Species richness
Subsampled reads
1e6
0.0
0.5
1.0
1.5
2.0
0
20
40
60
80
100
Subsampled data
Function fitted to data

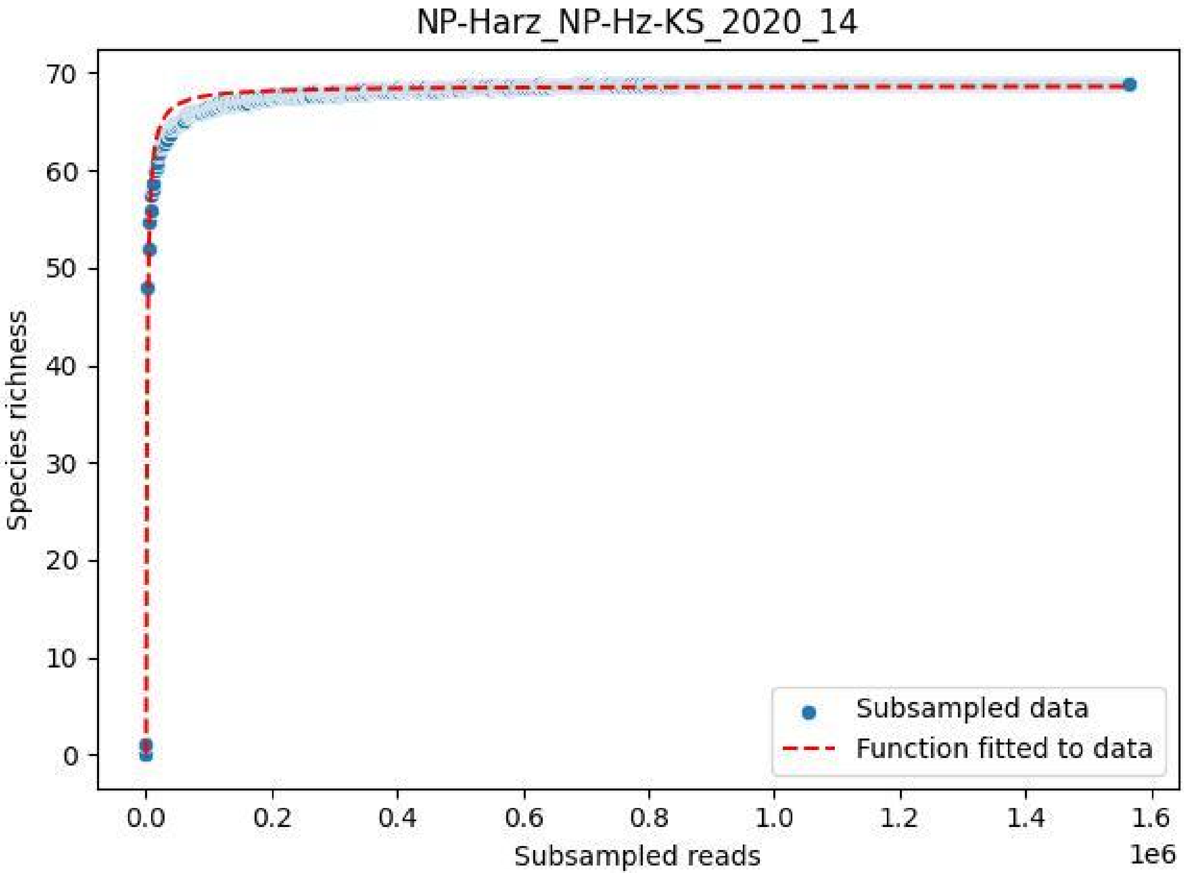
NP-Harz_NP-Hz-KS_2020_14
Species richness
Subsampled reads
1e6
0
10
20
30
40
50
60
70
0.0
0.2
0.4
0.6
0.8
1.0
1.2
1.4
1.6
Subsampled data
Function fitted to data

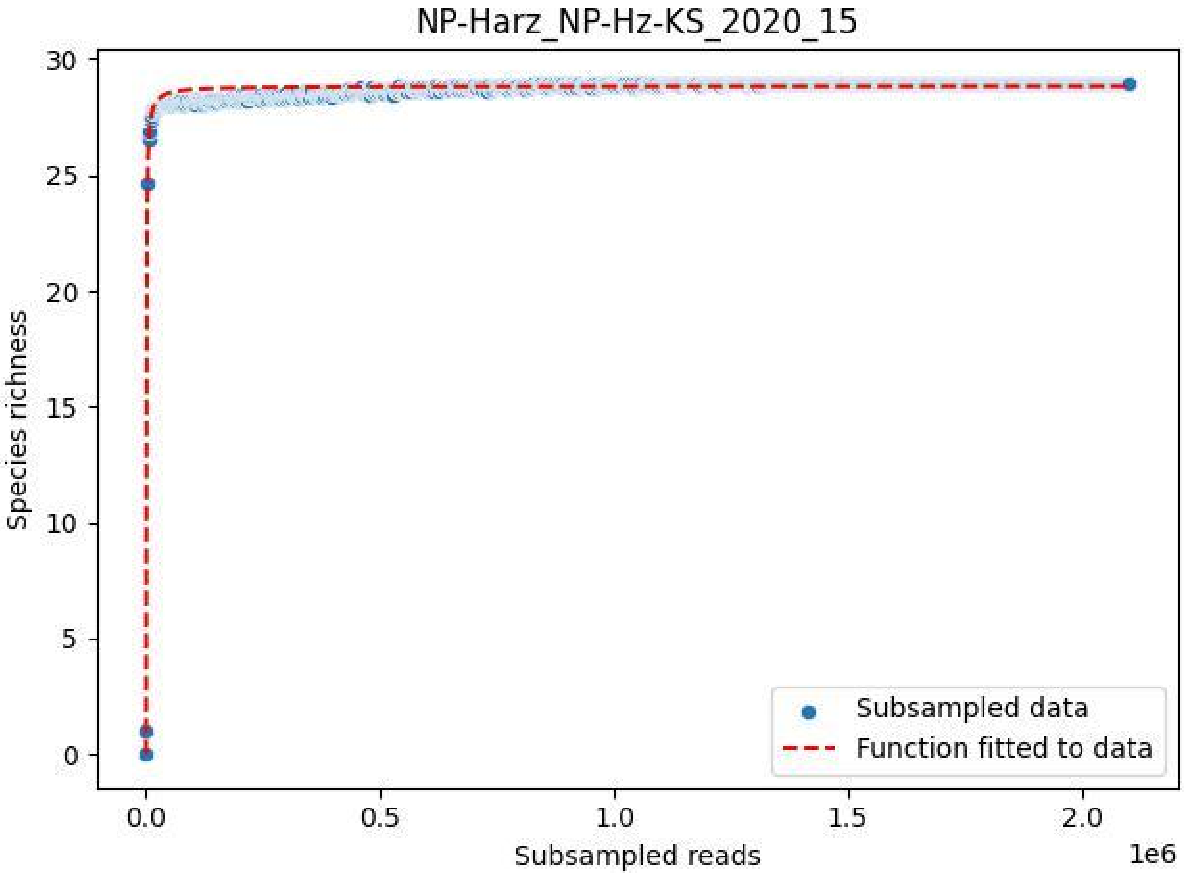
NP-Harz_NP-Hz-KS_2020_15
Species richness
Subsampled reads
1e6
Subsampled data
Function fitted to data
30
25
20
15
10
5
0
0.0
0.5
1.0
1.5
2.0

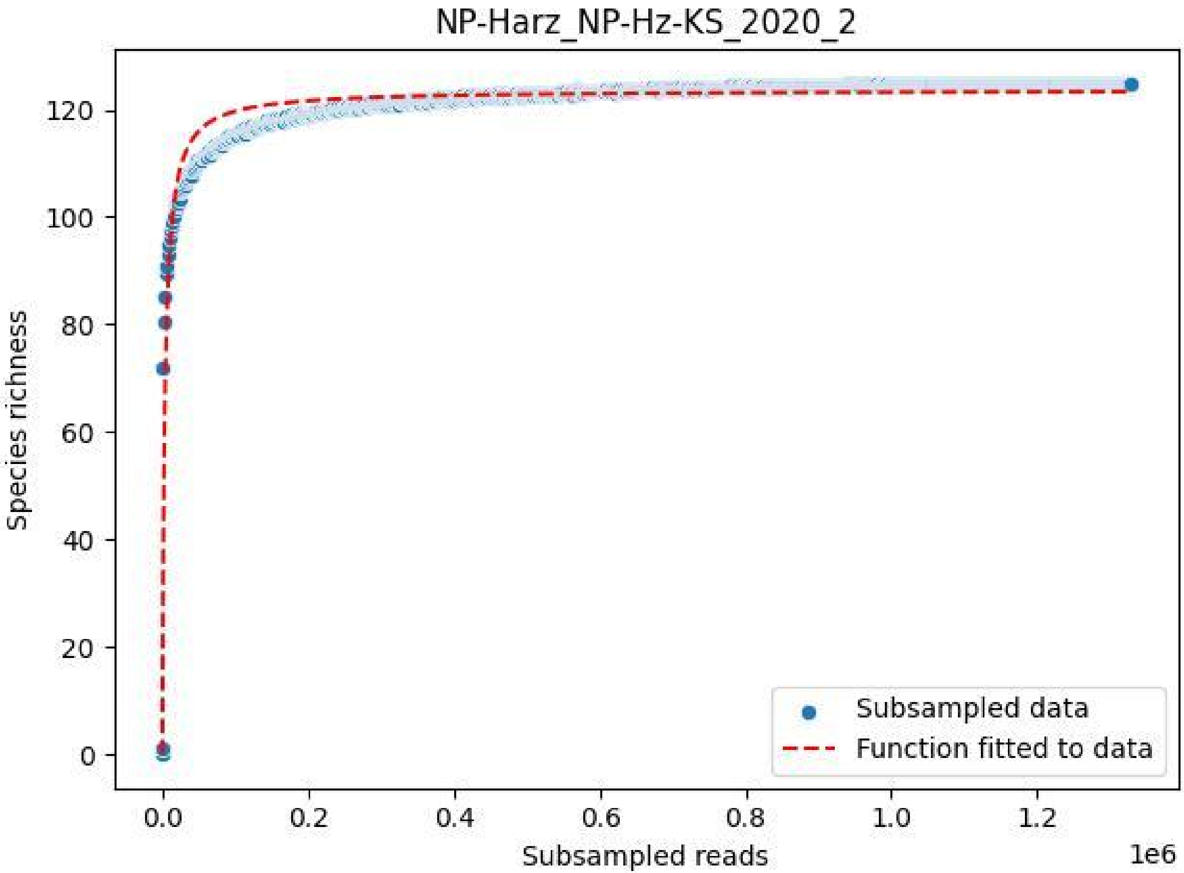

NP-Harz_NP-Hz-KS_2020_2
Species richness
0
20
40
60
80
100
120
0.0
0.2
0.4
0.6
0.8
1.0
1.2
1e6
Subsampled reads
Subsampled data
Function fitted to data

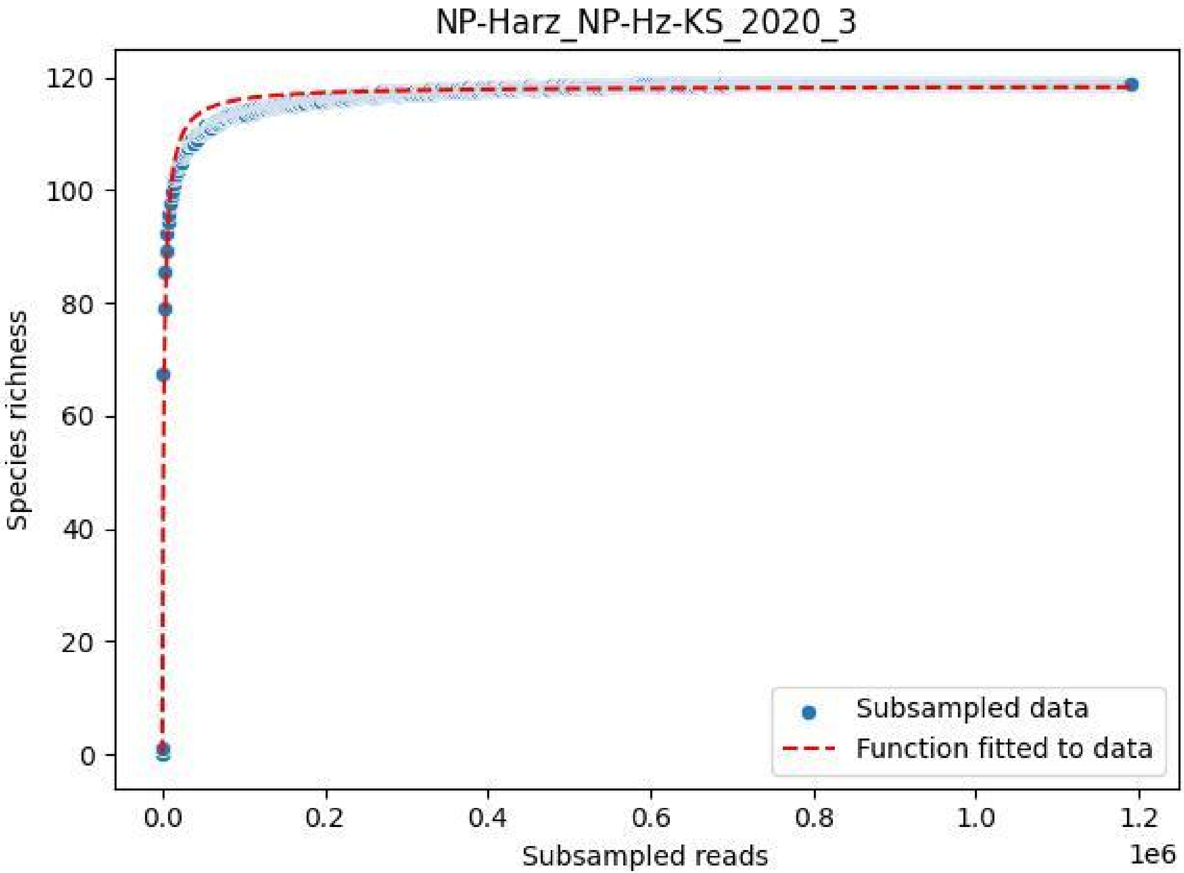
NP-Harz_NP-Hz-KS_2020_3
Species richness
Subsampled reads
1e6
0
20
40
60
80
100
120
0.0
0.2
0.4
0.6
0.8
1.0
1.2
Subsampled data
Function fitted to data

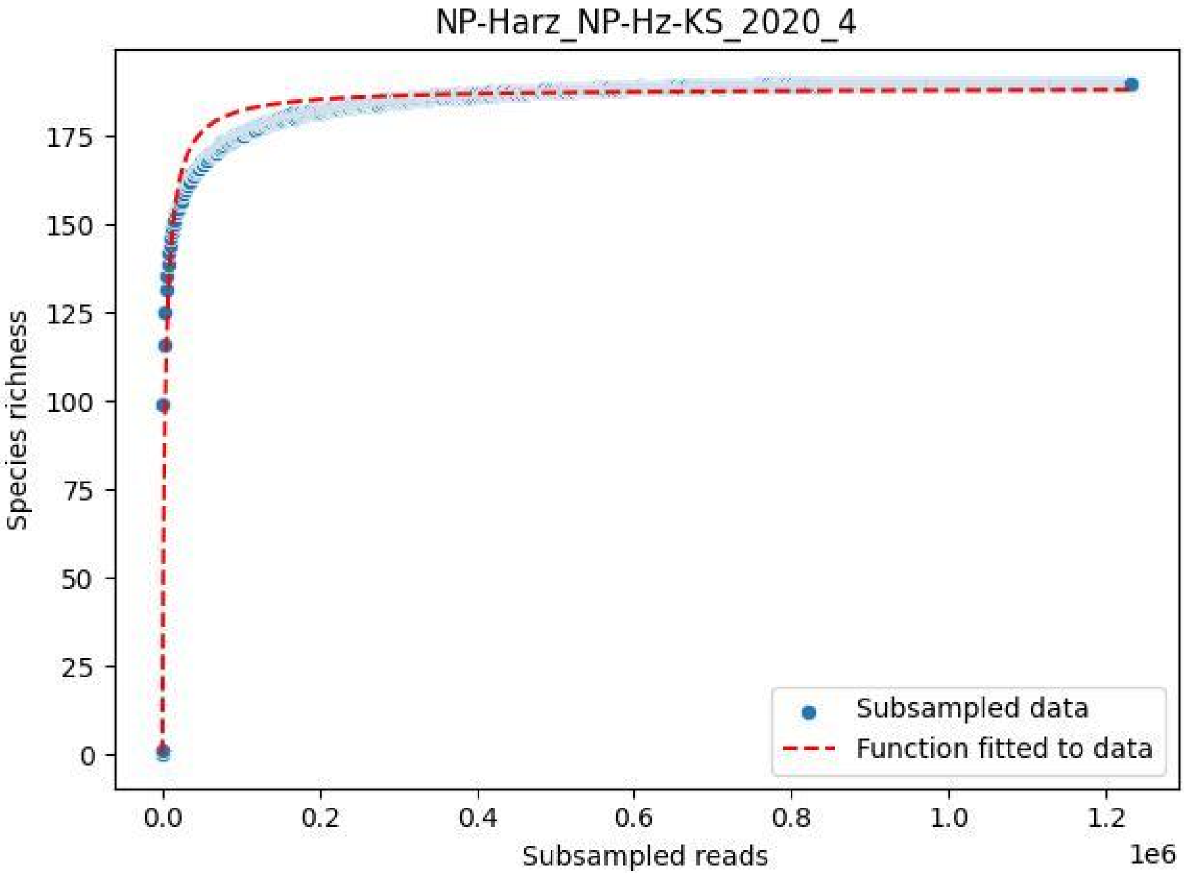
NP-Harz_NP-Hz-KS_2020_4
Species richness
0
25
50
75
100
125
150
175
0.0
0.2
0.4
0.6
0.8
1.0
1.2
1e6
Subsampled reads
Subsampled data
Function fitted to data

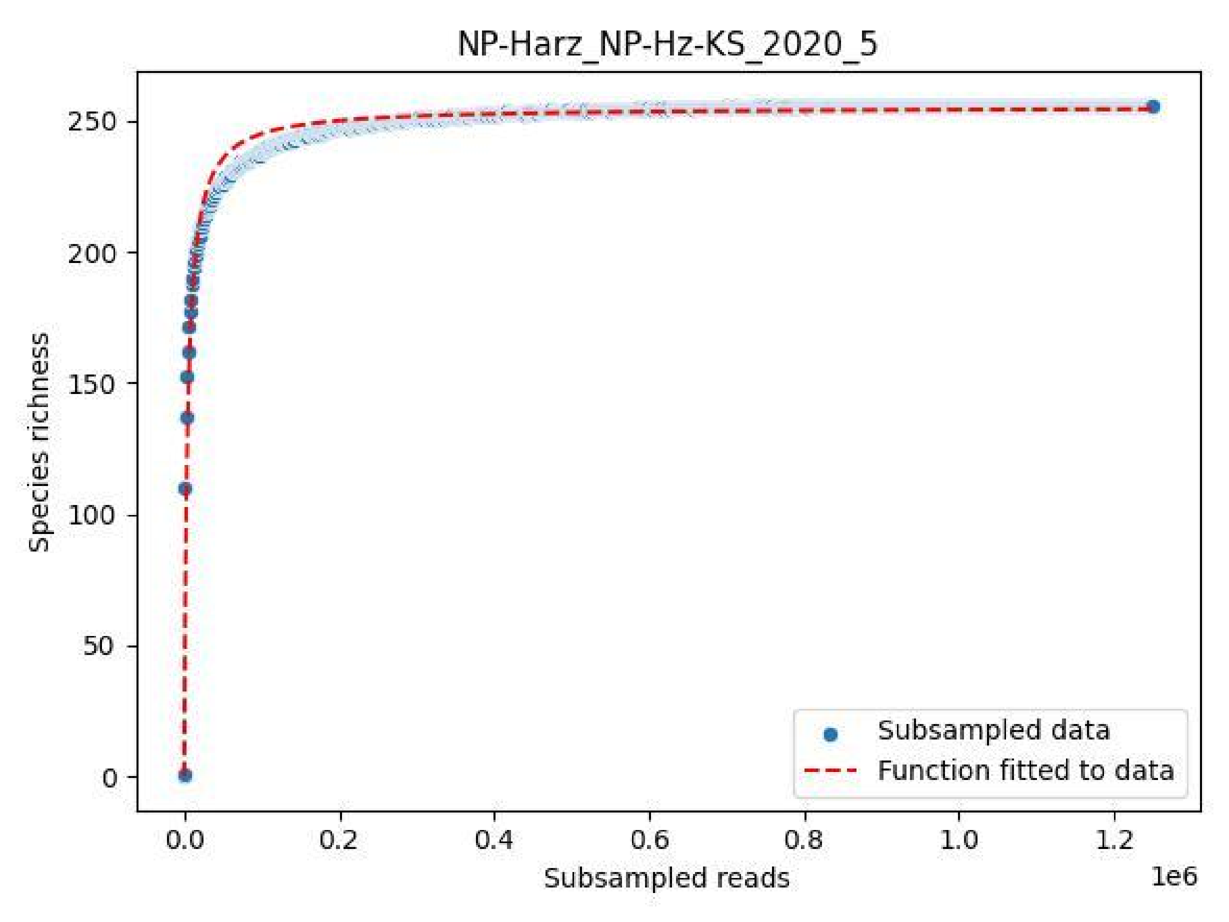
NP-Harz_NP-Hz-KS_2020_5
Species richness
Subsampled reads
1e6
0
50
100
150
200
250
0.0
0.2
0.4
0.6
0.8
1.0
1.2
Subsampled data
Function fitted to data

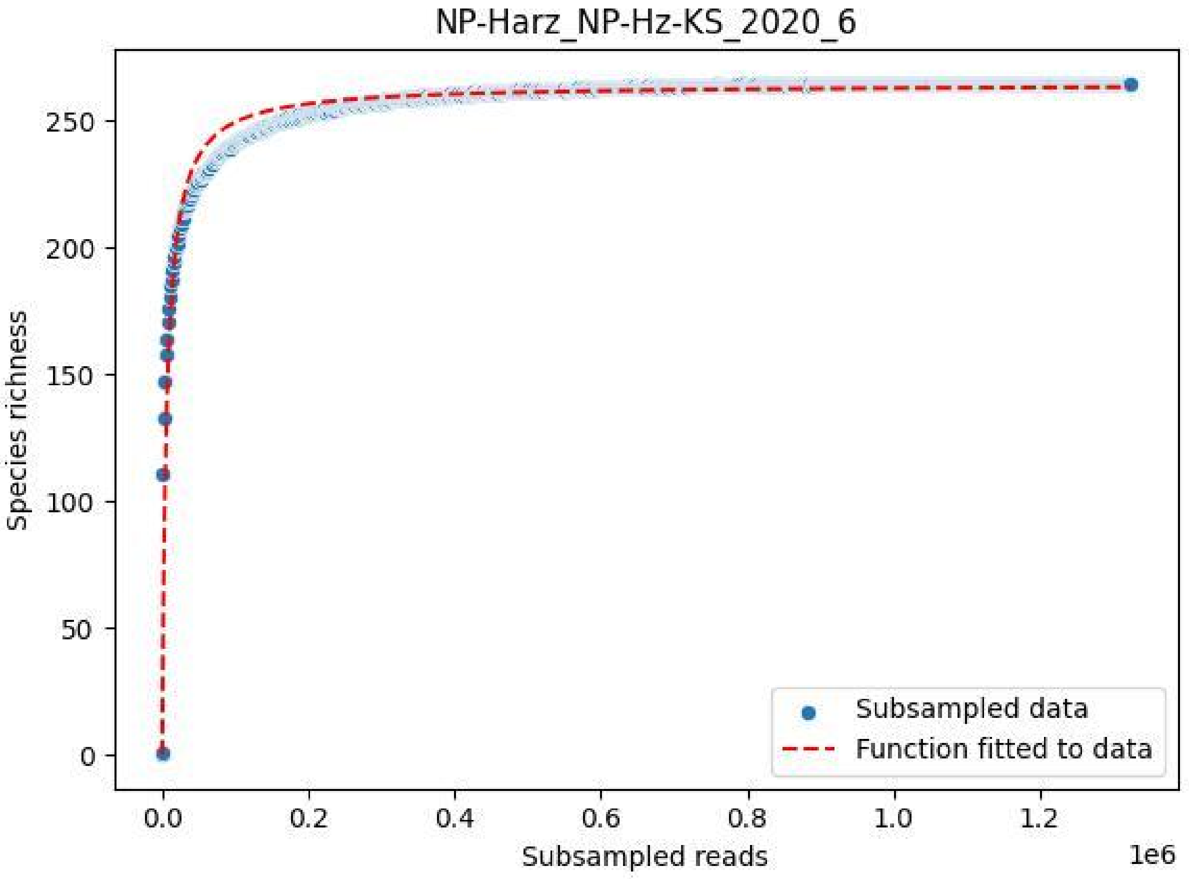
NP-Harz_NP-Hz-KS_2020_6
Species richness
Subsampled reads
0
50
100
150
200
250
0.0
0.2
0.4
0.6
0.8
1.0
1.2
1e6
Subsampled data
Function fitted to data

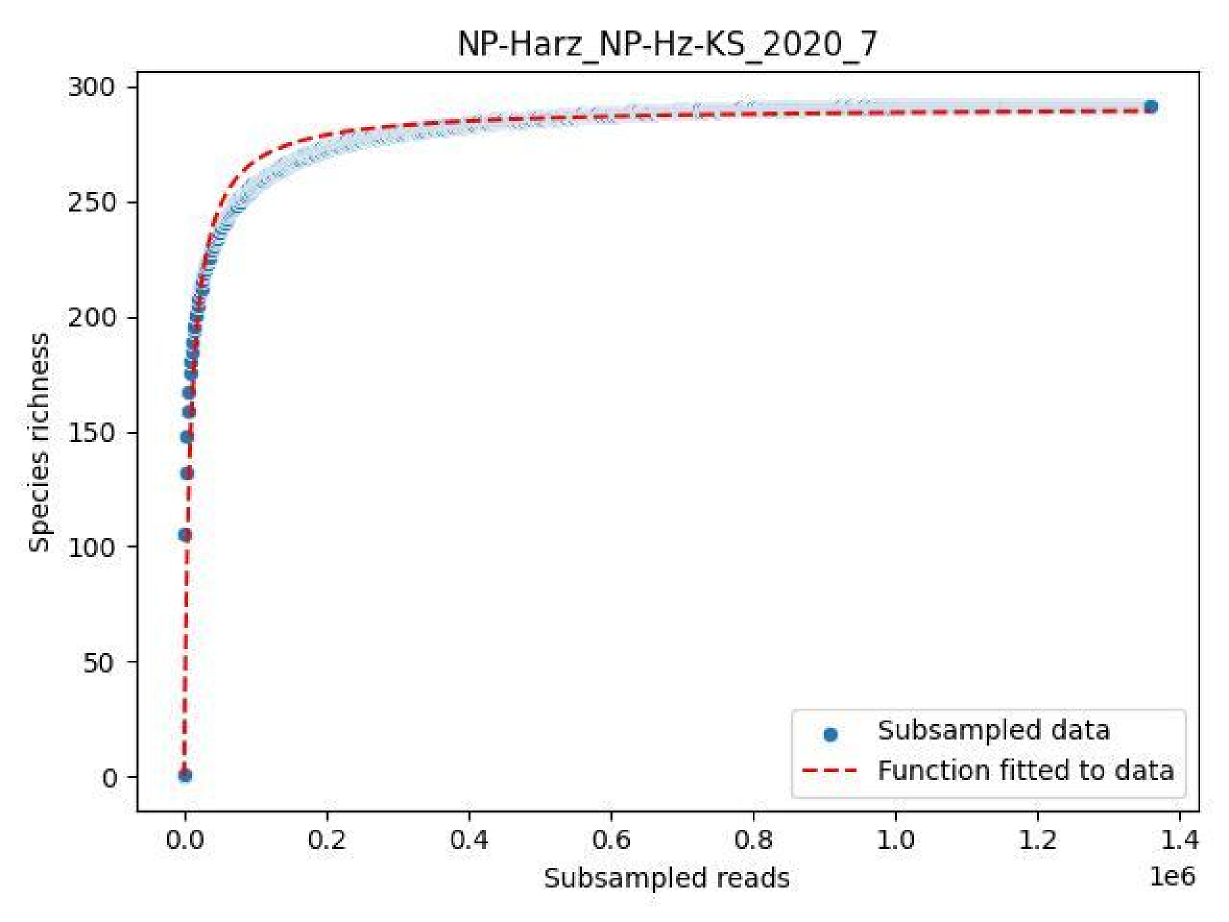

NP-Harz_NP-Hz-KS_2020_7
Species richness
Subsampled reads
1e6
0
50
100
150
200
250
300
0.0
0.2
0.4
0.6
0.8
1.0
1.2
1.4
Subsampled data
Function fitted to data

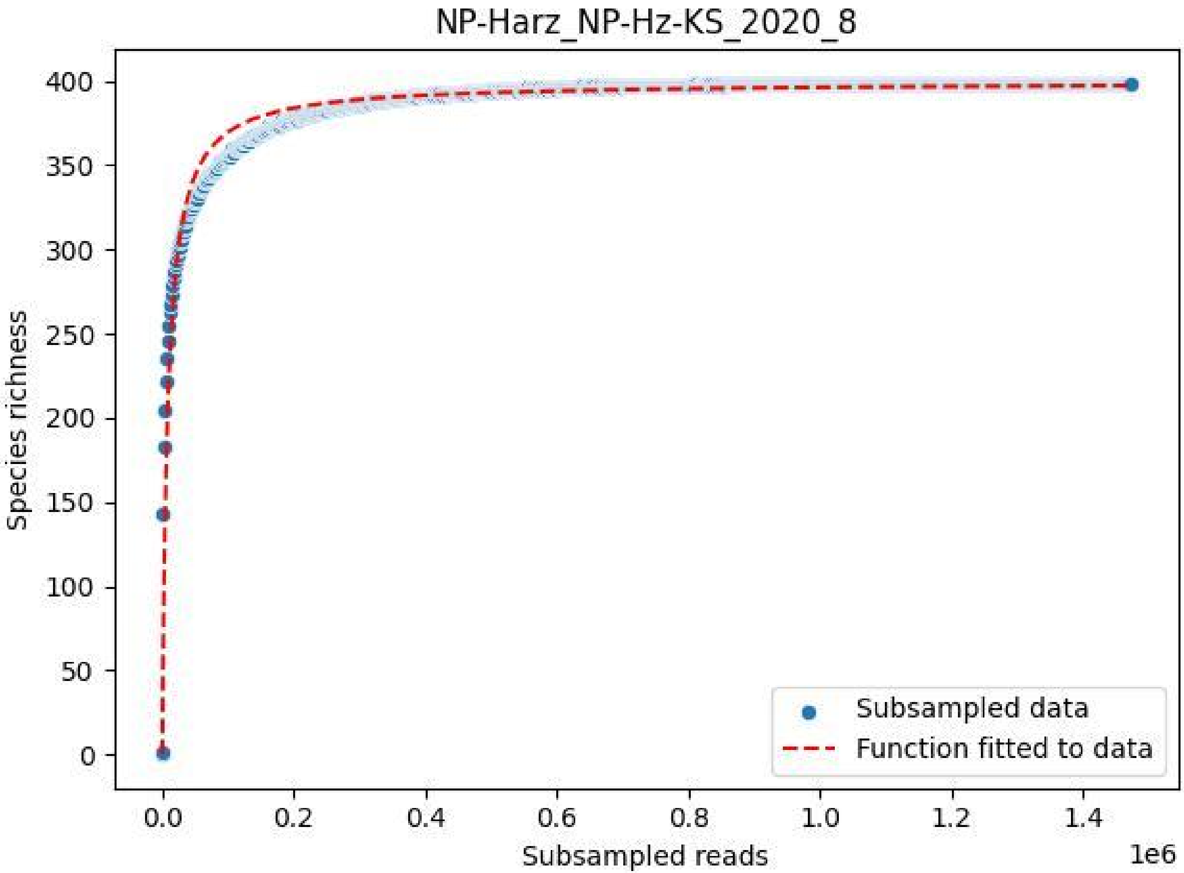
NP-Harz_NP-Hz-KS_2020_8
Species richness
0
50
100
150
200
250
300
350
400
Subsampled reads
0.0
0.2
0.4
0.6
0.8
1.0
1.2
1.4
1e6
Subsampled data
Function fitted to data

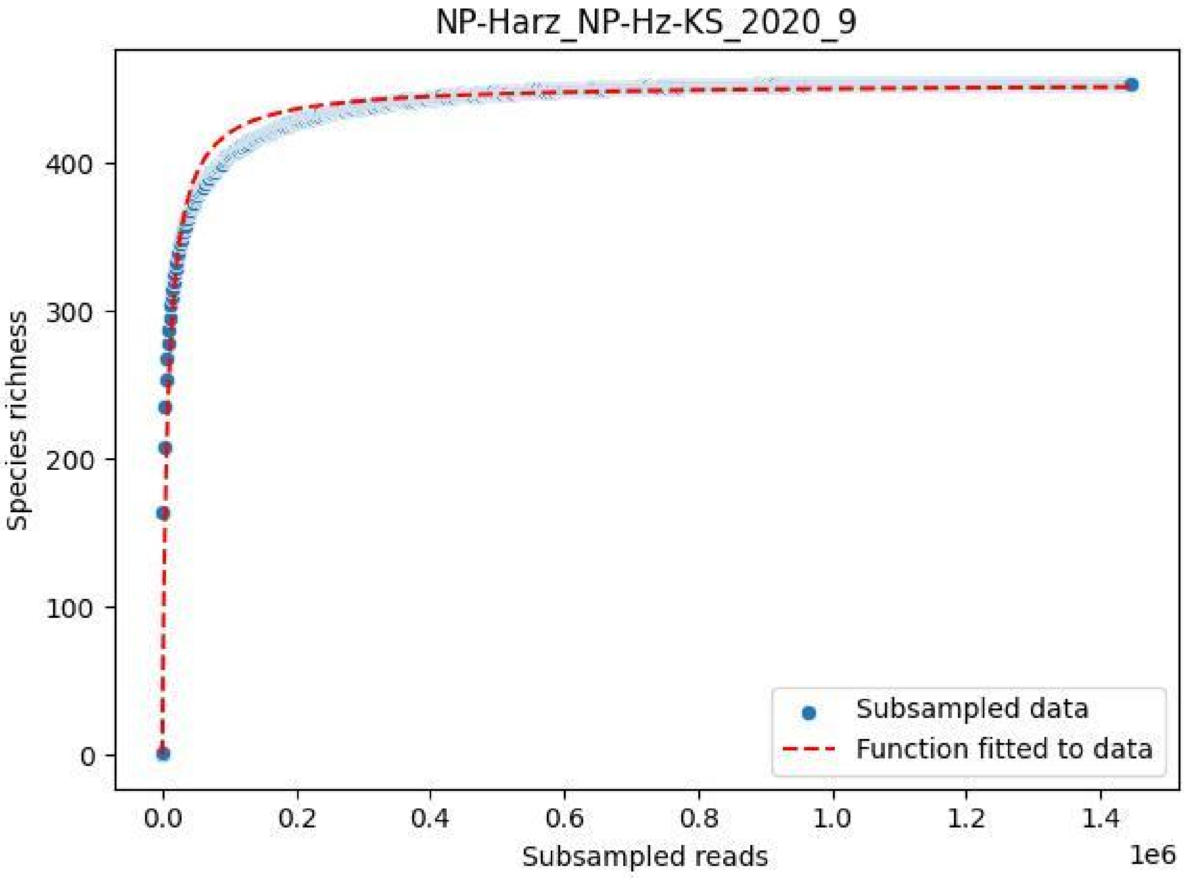

NP-Harz_NP-Hz-KS_2020_9
Species richness
Subsampled reads
1e6
0
100
200
300
400
0.0
0.2
0.4
0.6
0.8
1.0
1.2
1.4
Subsampled data
Function fitted to data

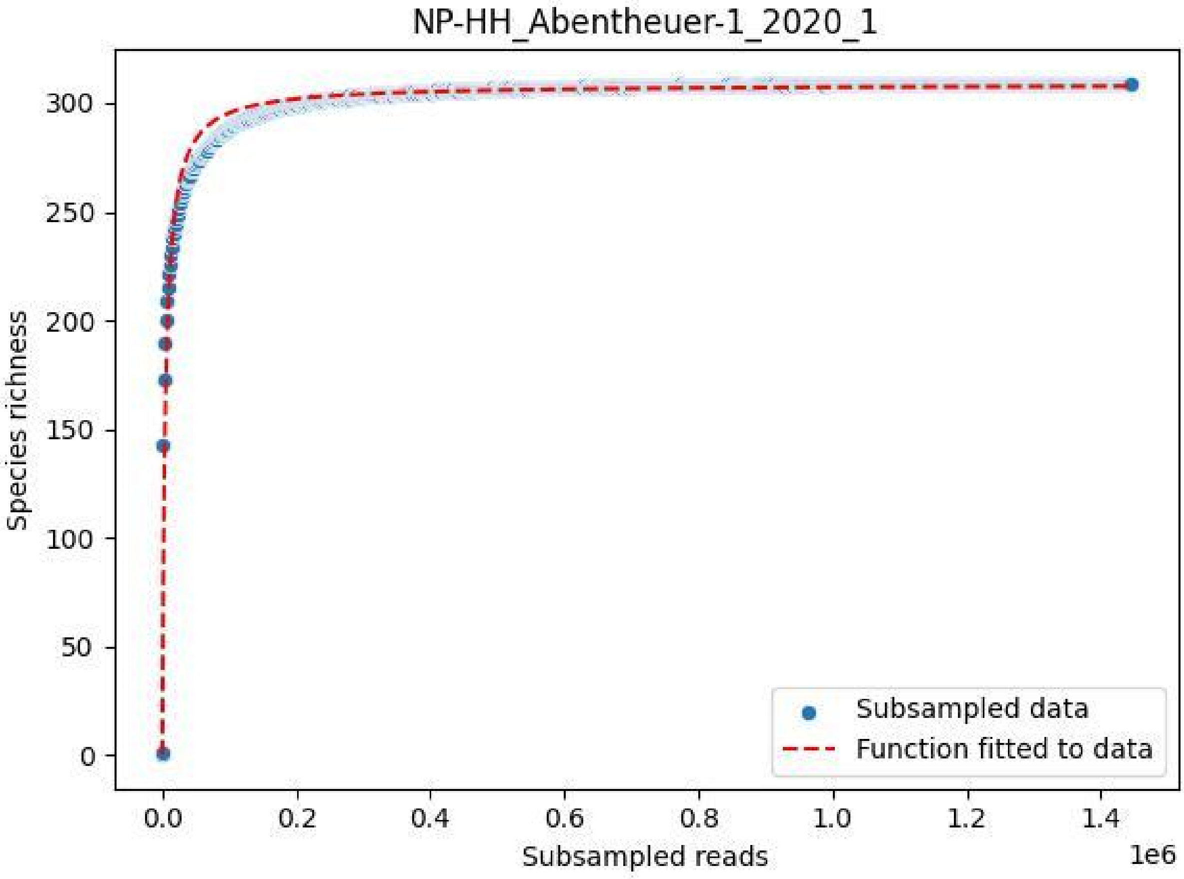
NP-HH_Abentheuer-1_2020_1
Species richness
0
50
100
150
200
250
300
0.0
0.2
0.4
0.6
0.8
1.0
1.2
1.4
1e6
Subsampled reads
Subsampled data
Function fitted to data

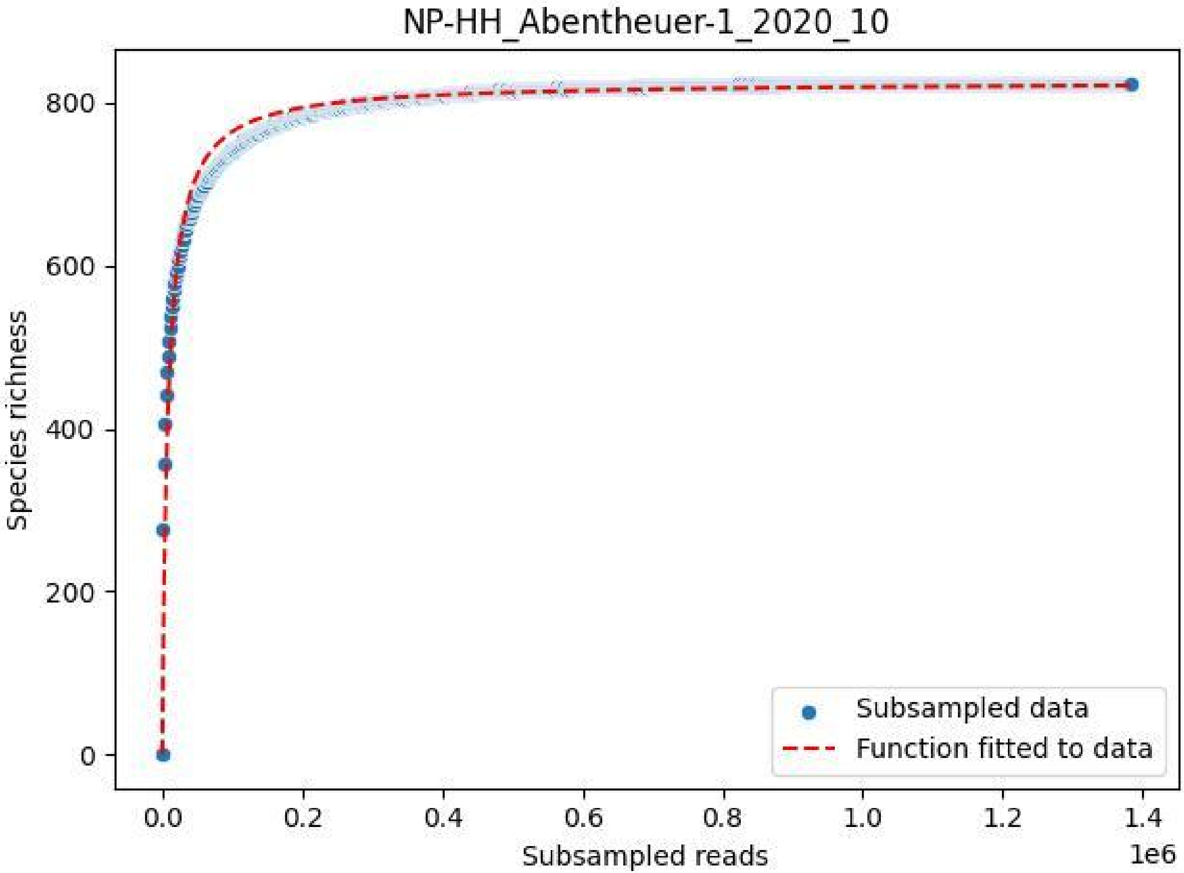

NP-HH_Abentheuer-1_2020_10
Species richness
Subsampled reads
1e6
0
200
400
600
800
0.0
0.2
0.4
0.6
0.8
1.0
1.2
1.4
Subsampled data
Function fitted to data

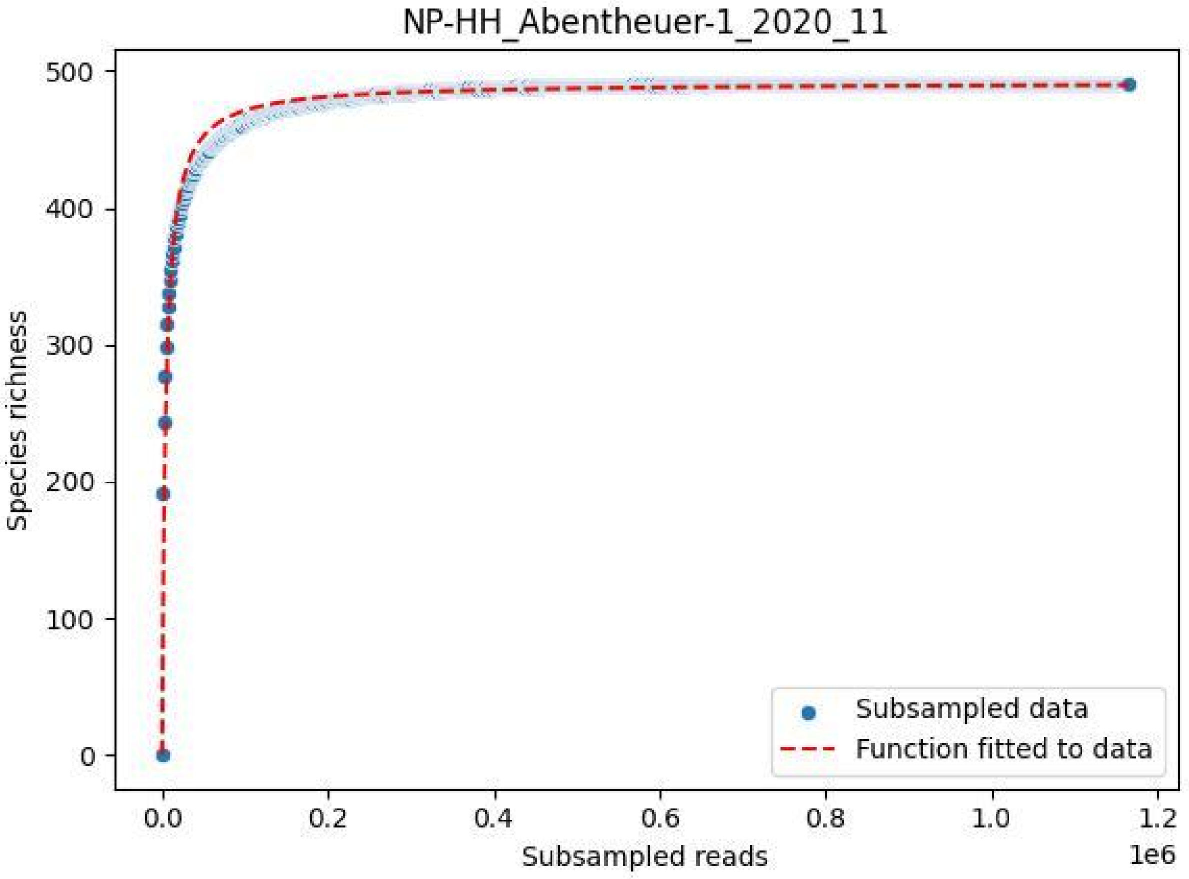

NP-HH_Abentheuer-1_2020_11
Species richness
Subsampled reads
1e6
0
100
200
300
400
500
0.0
0.2
0.4
0.6
0.8
1.0
1.2
Subsampled data
Function fitted to data

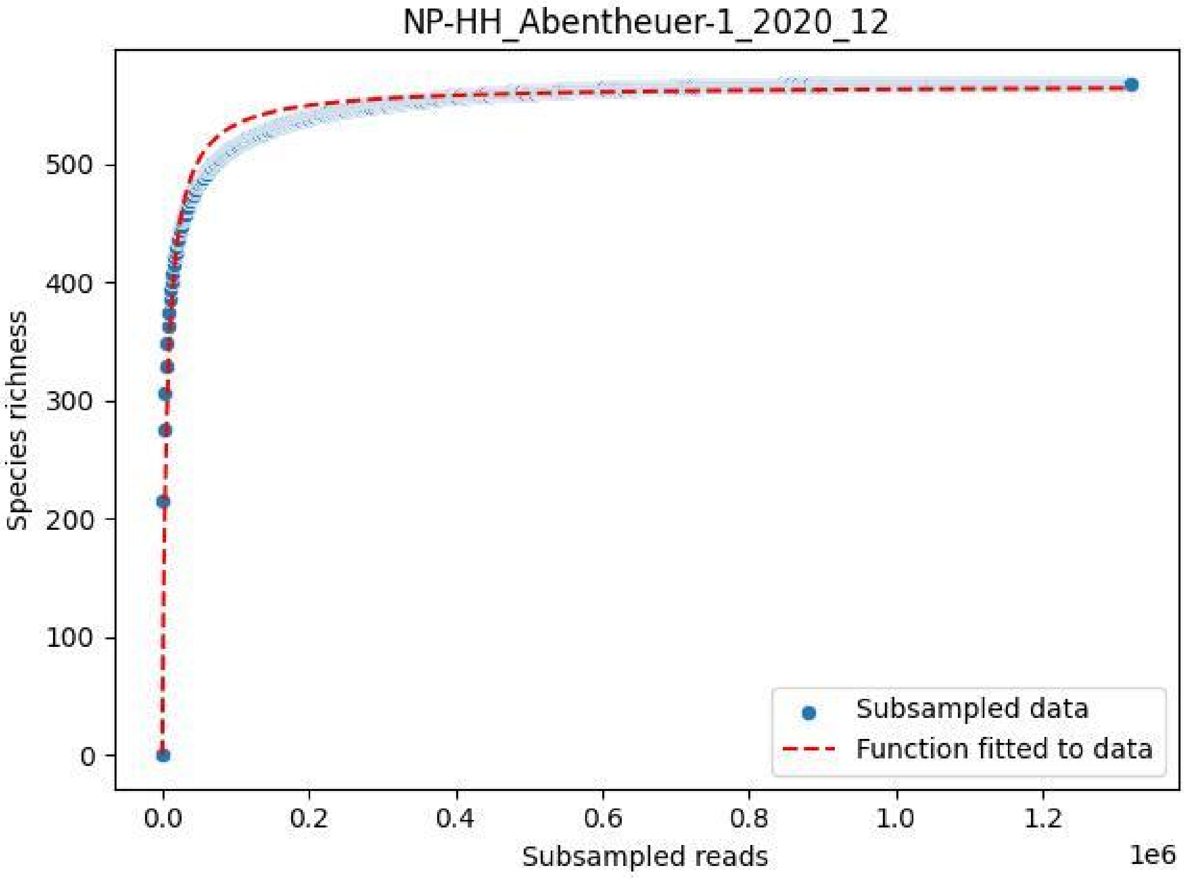
NP-HH_Abentheuer-1_2020_12
Species richness
0
100
200
300
400
500
0.0
0.2
0.4
0.6
0.8
1.0
1.2
1e6
Subsampled reads
Subsampled data
Function fitted to data

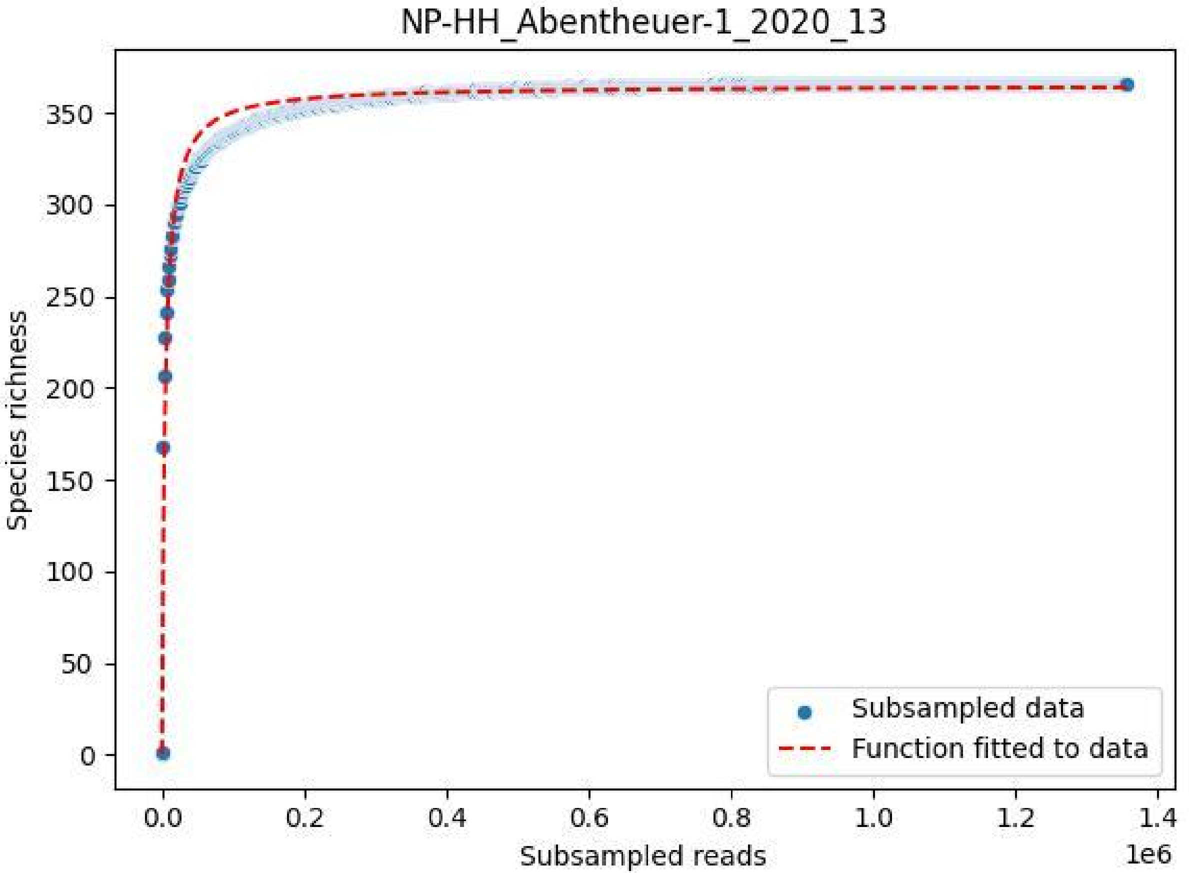

NP-HH_Abentheuer-1_2020_13
Species richness
Subsampled reads
1e6
0
50
100
150
200
250
300
350
0.0
0.2
0.4
0.6
0.8
1.0
1.2
1.4
Subsampled data
Function fitted to data

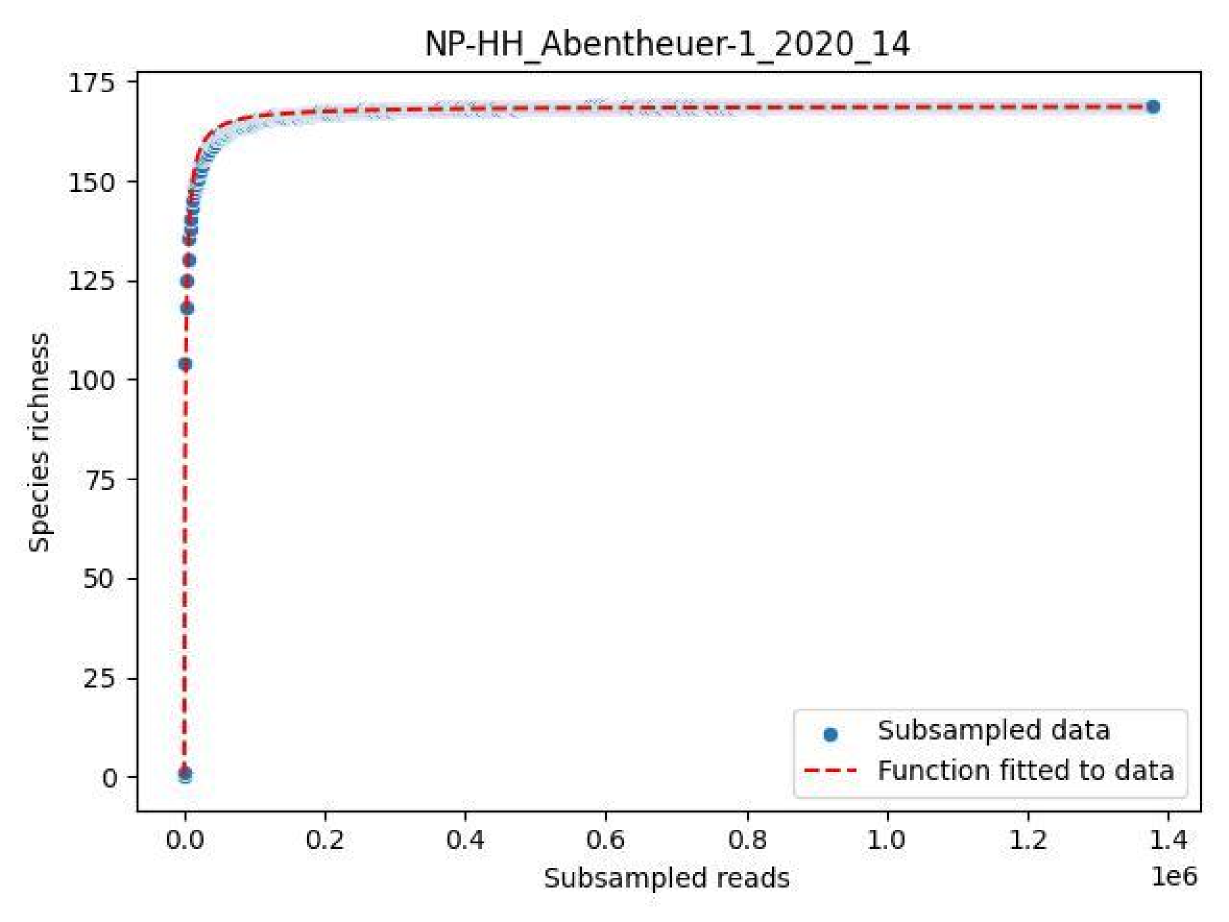

NP-HH_Abentheuer-1_2020_14
Species richness
Subsampled reads
1e6
0
25
50
75
100
125
150
175
0.0
0.2
0.4
0.6
0.8
1.0
1.2
1.4
Subsampled data
Function fitted to data

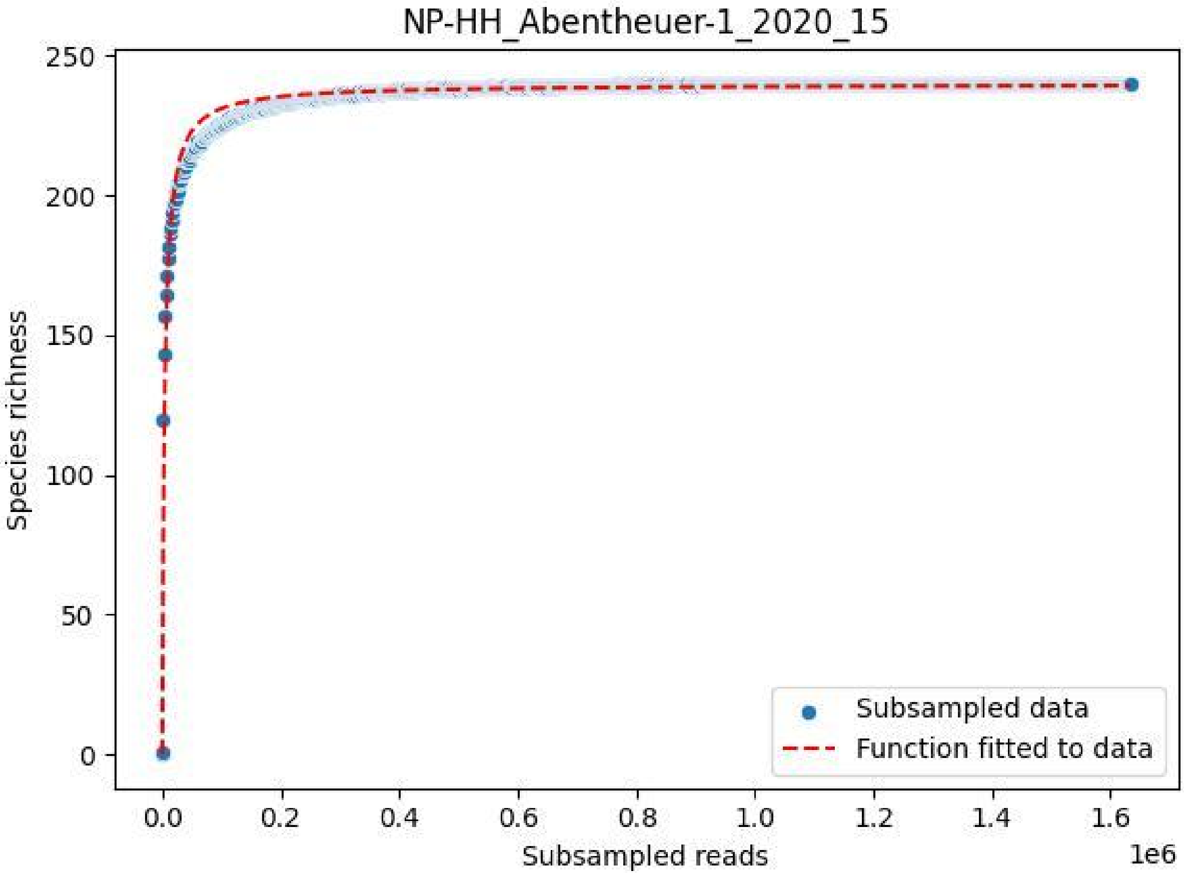

NP-HH_Abentheuer-1_2020_15
Species richness
Subsampled reads
1e6
0
50
100
150
200
250
0.0
0.2
0.4
0.6
0.8
1.0
1.2
1.4
1.6
Subsampled data
Function fitted to data

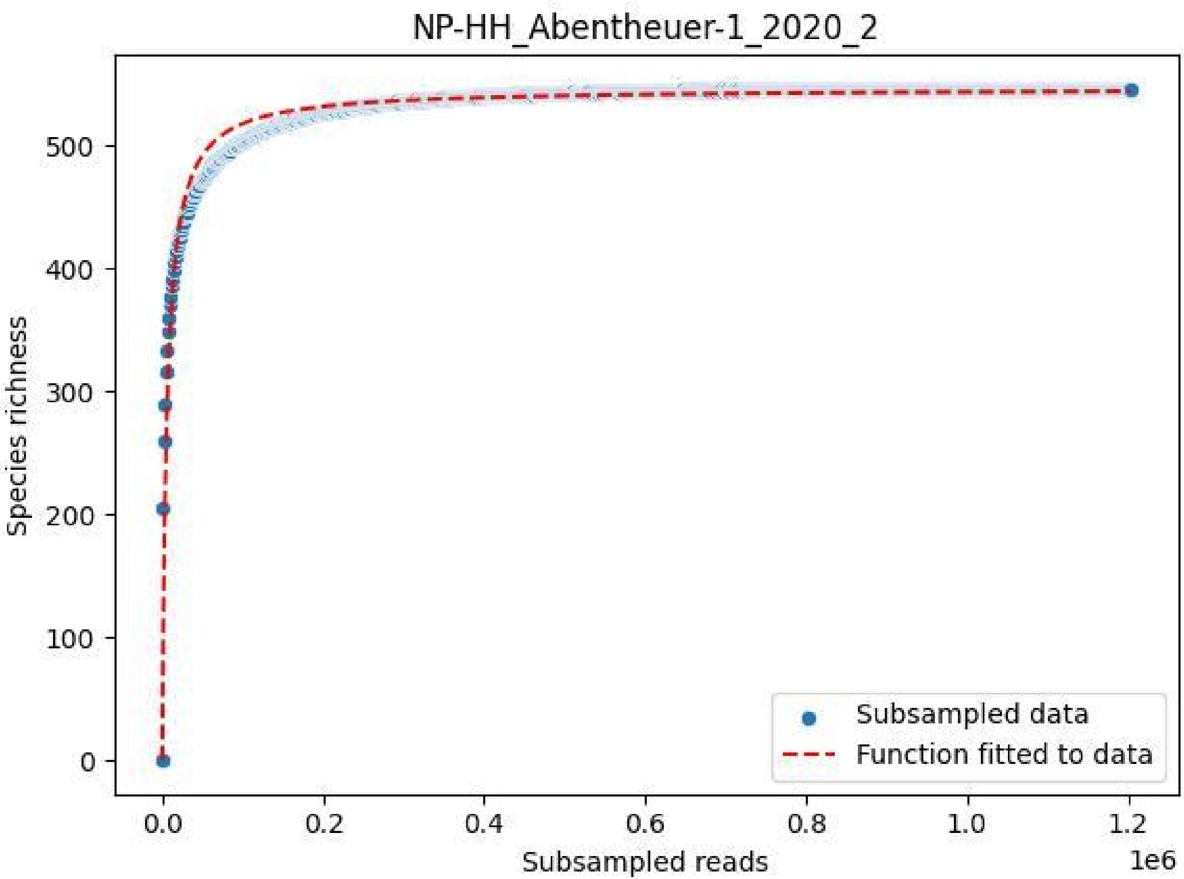
NP-HH_Abentheuer-1_2020_2
Species richness
Subsampled reads
1e6
0
100
200
300
400
500
0.0
0.2
0.4
0.6
0.8
1.0
1.2
Subsampled data
Function fitted to data

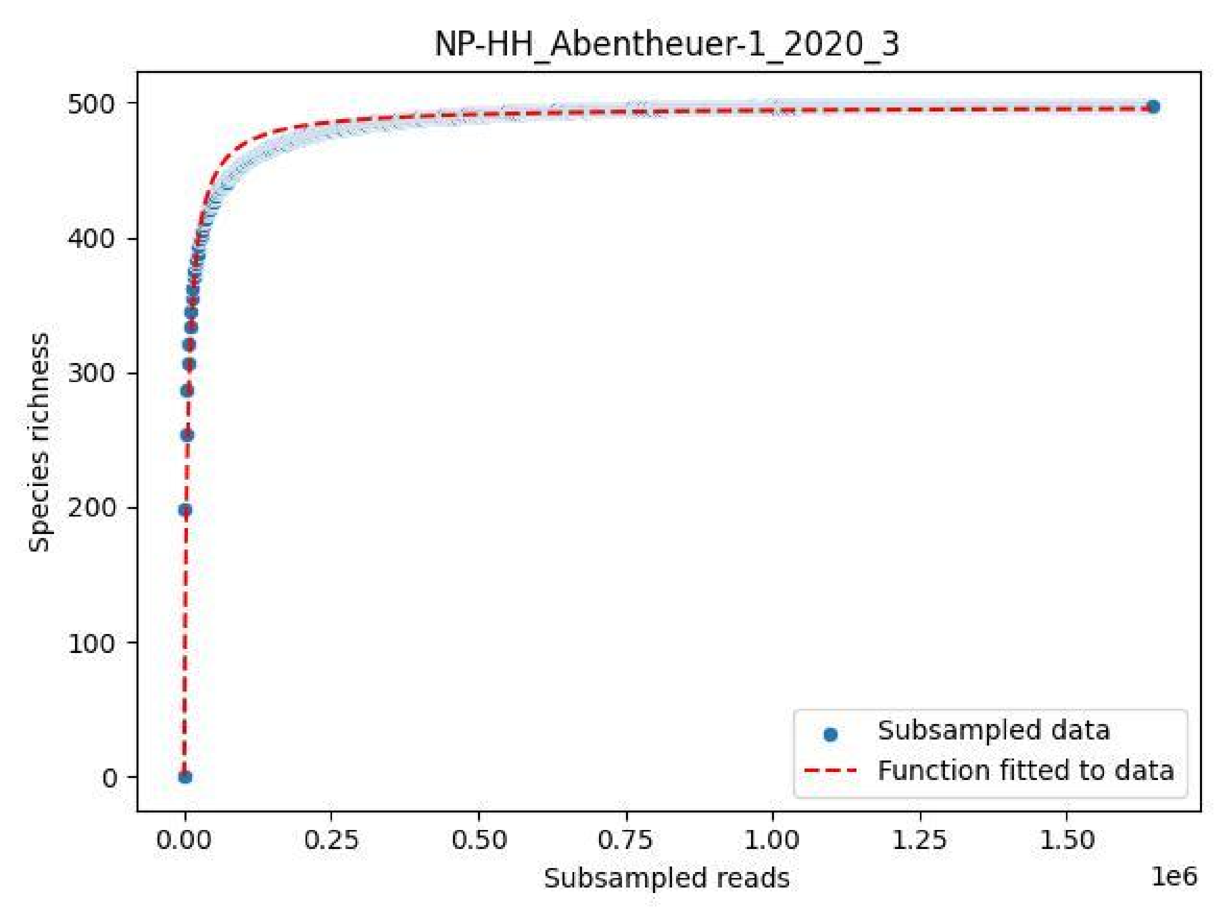
NP-HH_Abentheuer-1_2020_3
Species richness
Subsampled reads
1e6
0
100
200
300
400
500
0.00
0.25
0.50
0.75
1.00
1.25
1.50
Subsampled data
Function fitted to data

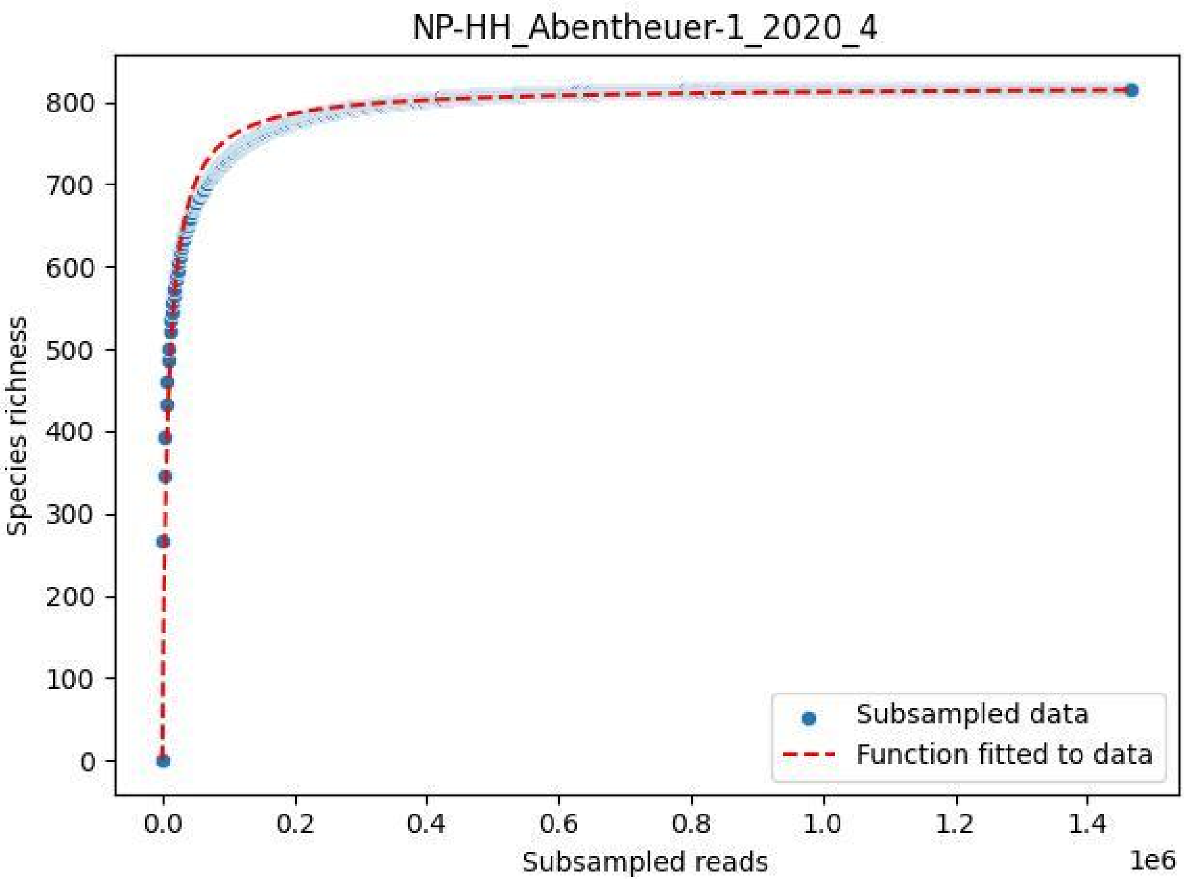
NP-HH_Abentheuer-1_2020_4
Species richness
0
100
200
300
400
500
600
700
800
Subsampled reads
0.0
0.2
0.4
0.6
0.8
1.0
1.2
1.4
1e6
Subsampled data
Function fitted to data

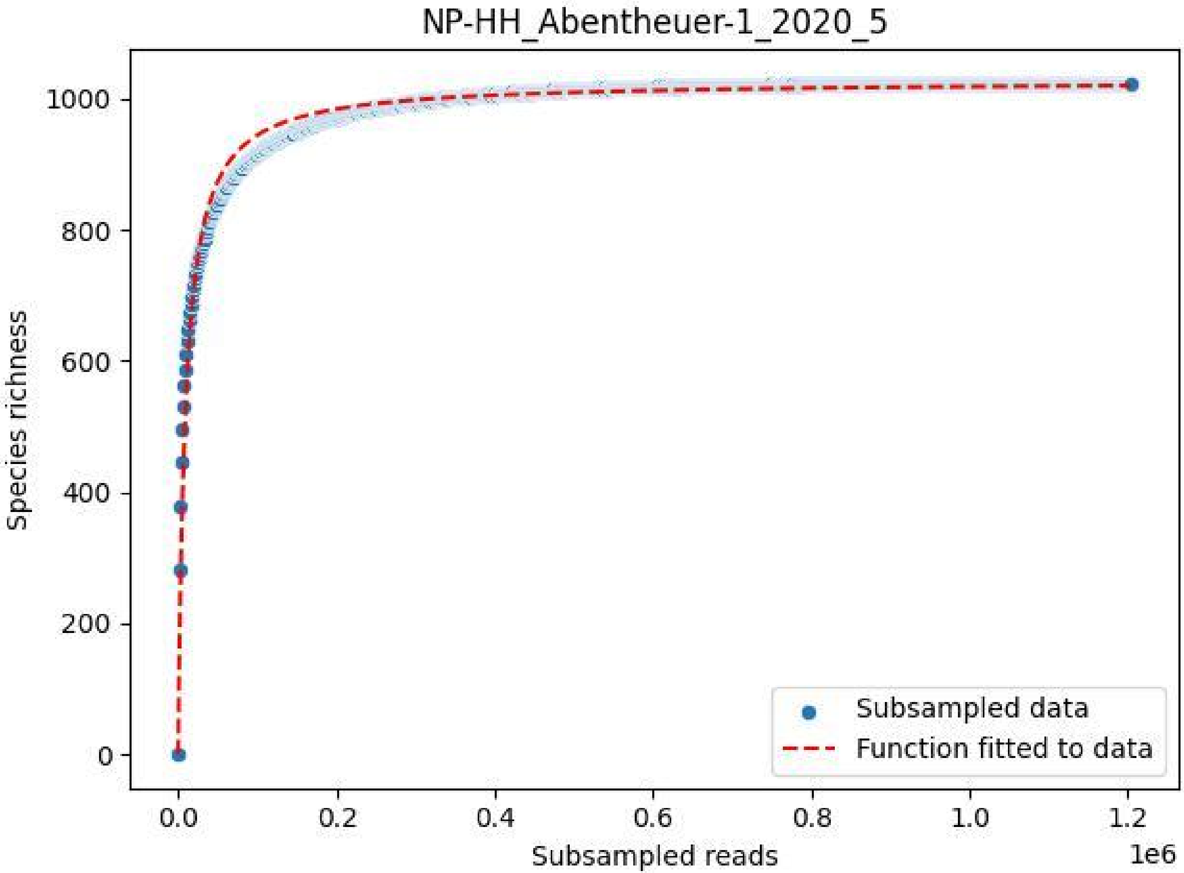
NP-HH_Abentheuer-1_2020_5
Species richness
1000
800
600
400
200
0
0.0
0.2
0.4
0.6
0.8
1.0
1.2
1e6
Subsampled reads
Subsampled data
Function fitted to data

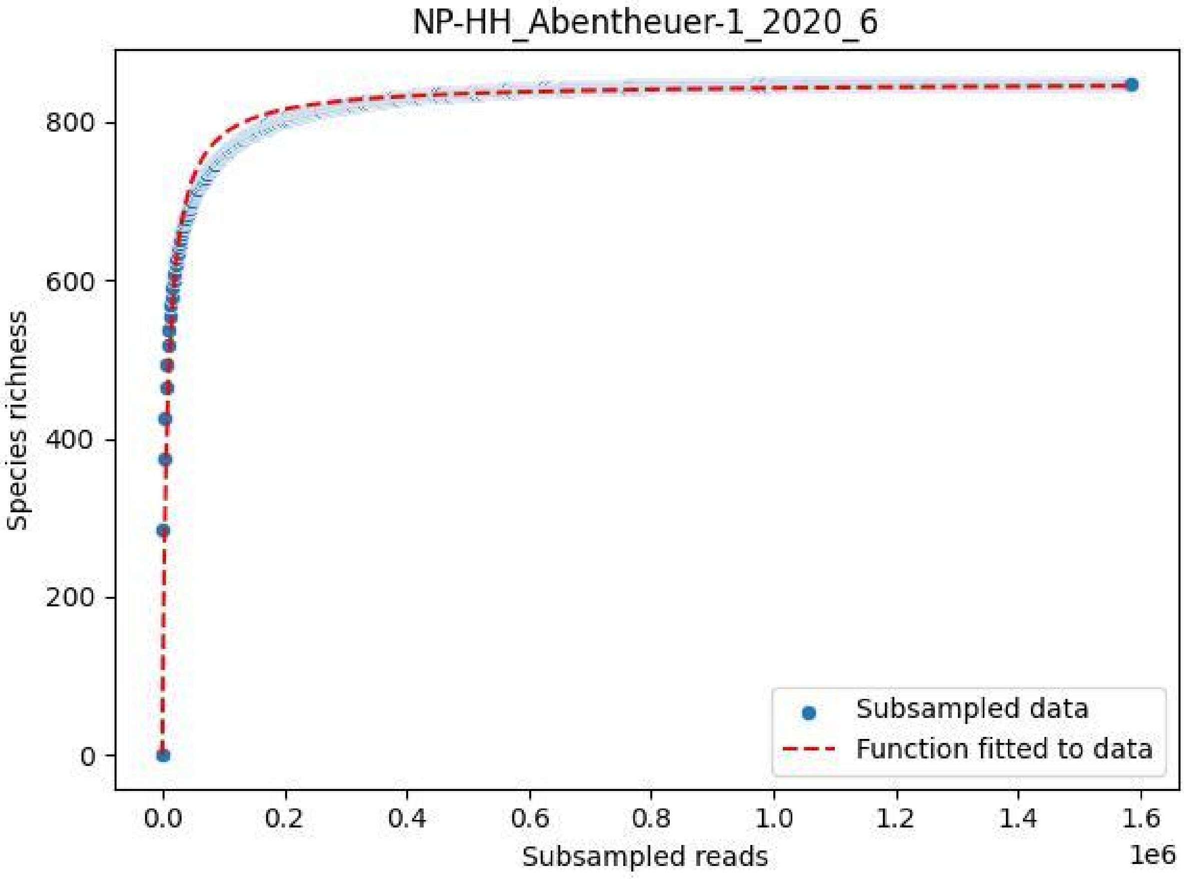
NP-HH_Abentheuer-1_2020_6
Species richness
Subsampled reads
1e6
Subsampled data
Function fitted to data
0
200
400
600
800
0.0
0.2
0.4
0.6
0.8
1.0
1.2
1.4
1.6

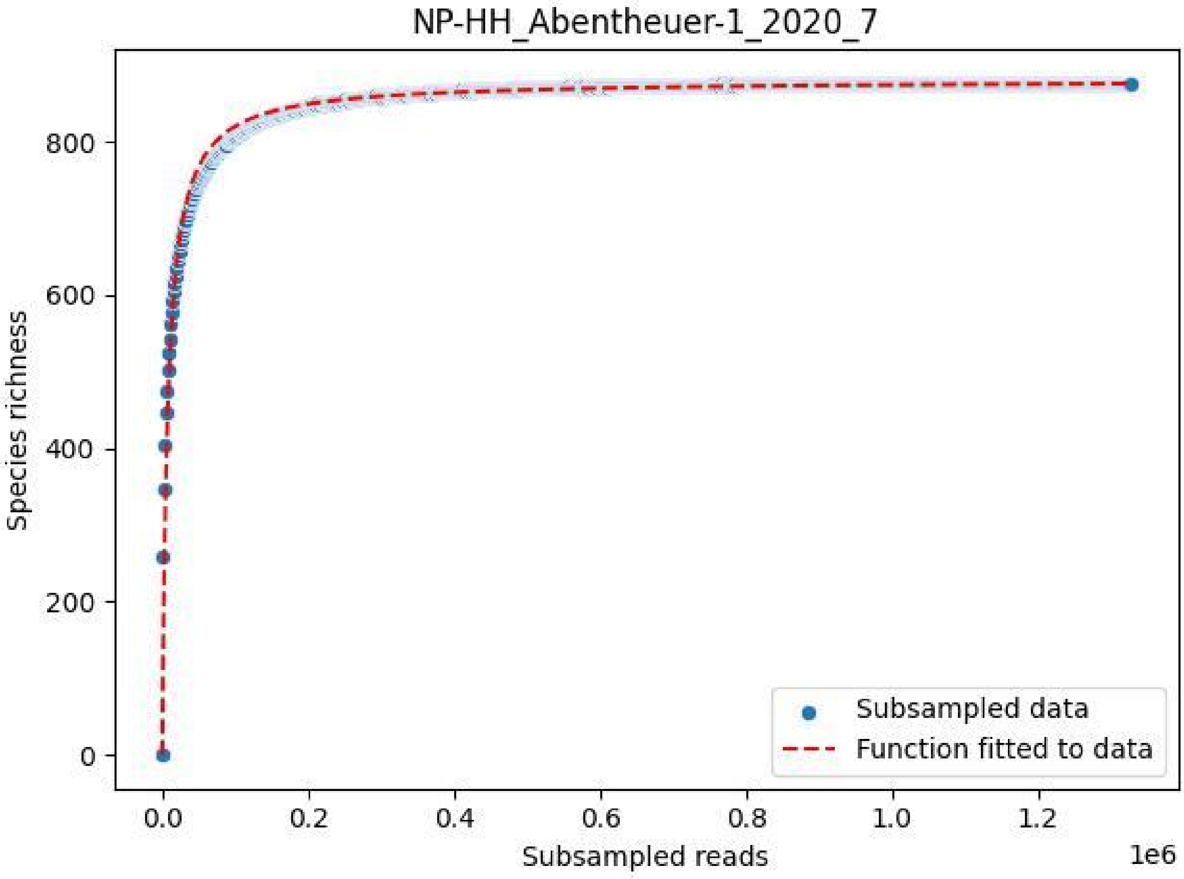

NP-HH_Abentheuer-1_2020_7
Species richness
Subsampled reads
1e6
0
200
400
600
800
0.0
0.2
0.4
0.6
0.8
1.0
1.2
Subsampled data
Function fitted to data

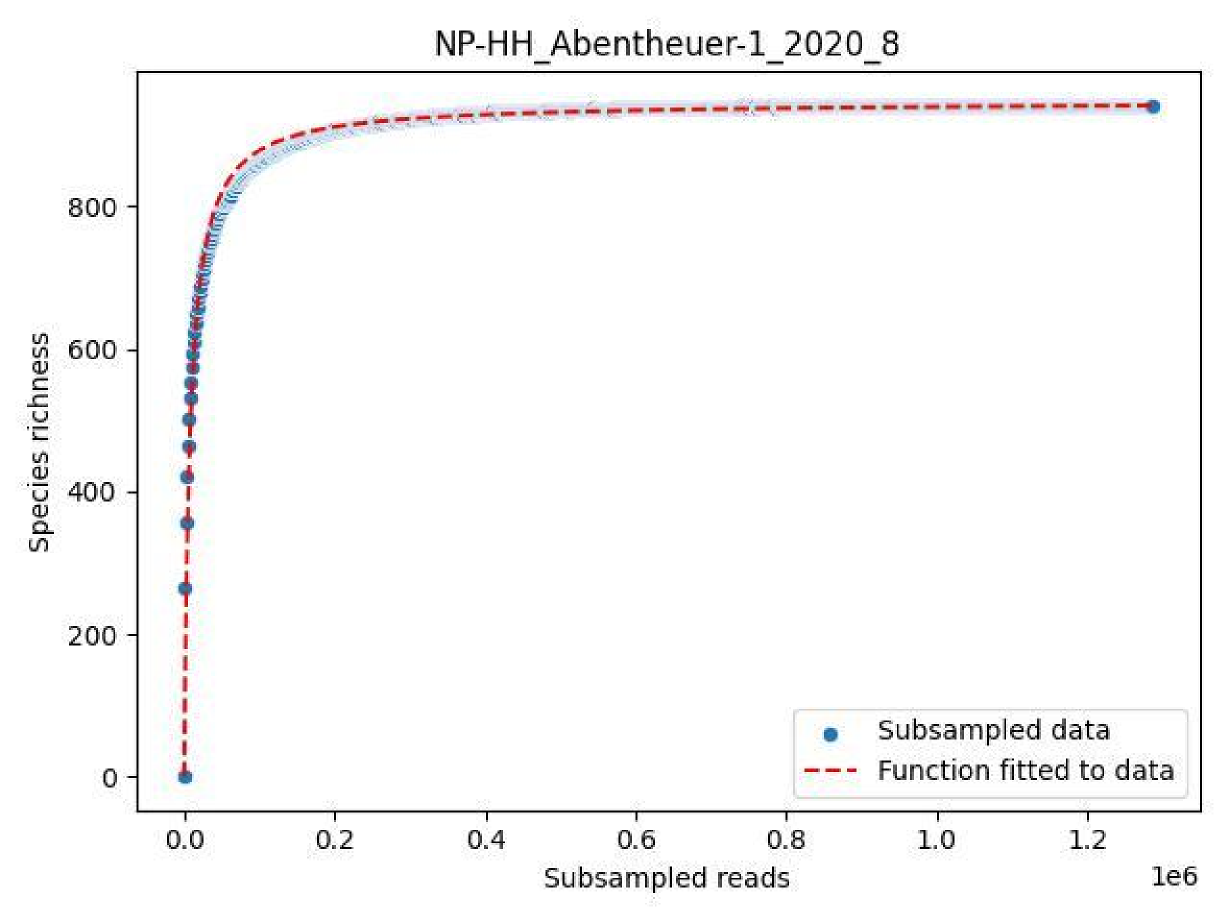

NP-HH_Abentheuer-1_2020_8
Species richness
0
200
400
600
800
0.0
0.2
0.4
0.6
0.8
1.0
1.2
1e6
Subsampled reads
Subsampled data
Function fitted to data

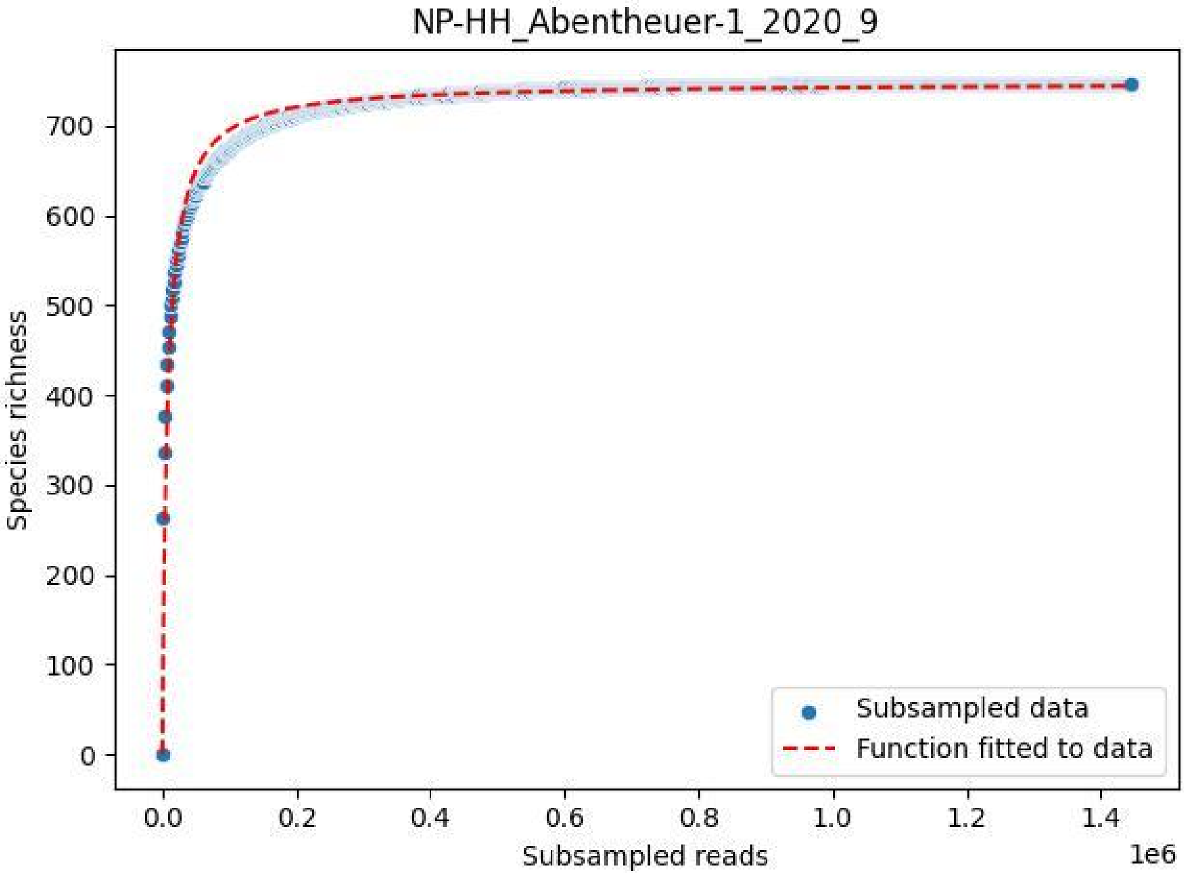
NP-HH_Abentheuer-1_2020_9
Species richness
Subsampled reads
1e6
0
100
200
300
400
500
600
700
0.0
0.2
0.4
0.6
0.8
1.0
1.2
1.4
Subsampled data
Function fitted to data

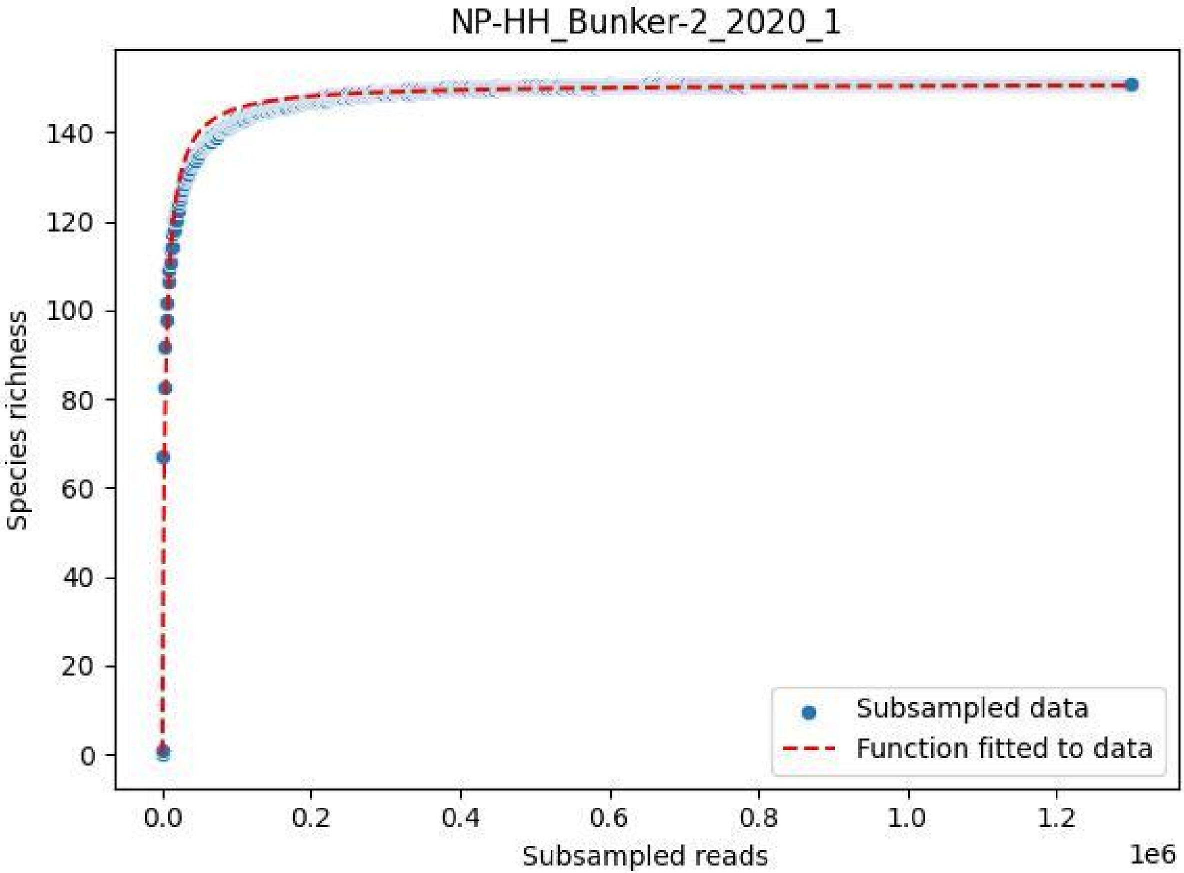
NP-HH_Bunker-2_2020_1
Species richness
0
20
40
60
80
100
120
140
0.0
0.2
0.4
0.6
0.8
1.0
1.2
1e6
Subsampled reads
Subsampled data
Function fitted to data

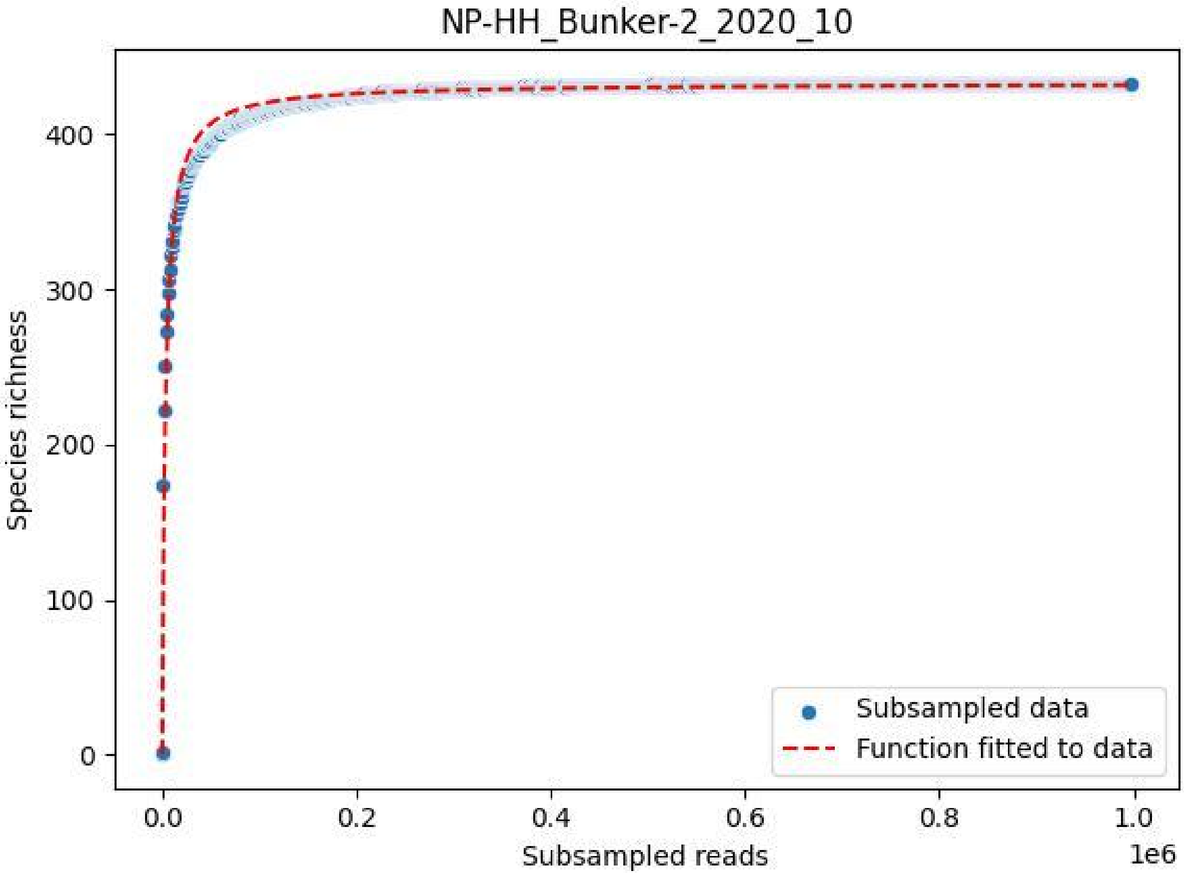
NP-HH_Bunker-2_2020_10
Species richness
Subsampled reads
1e6
0
100
200
300
400
0.0
0.2
0.4
0.6
0.8
1.0
Subsampled data
Function fitted to data

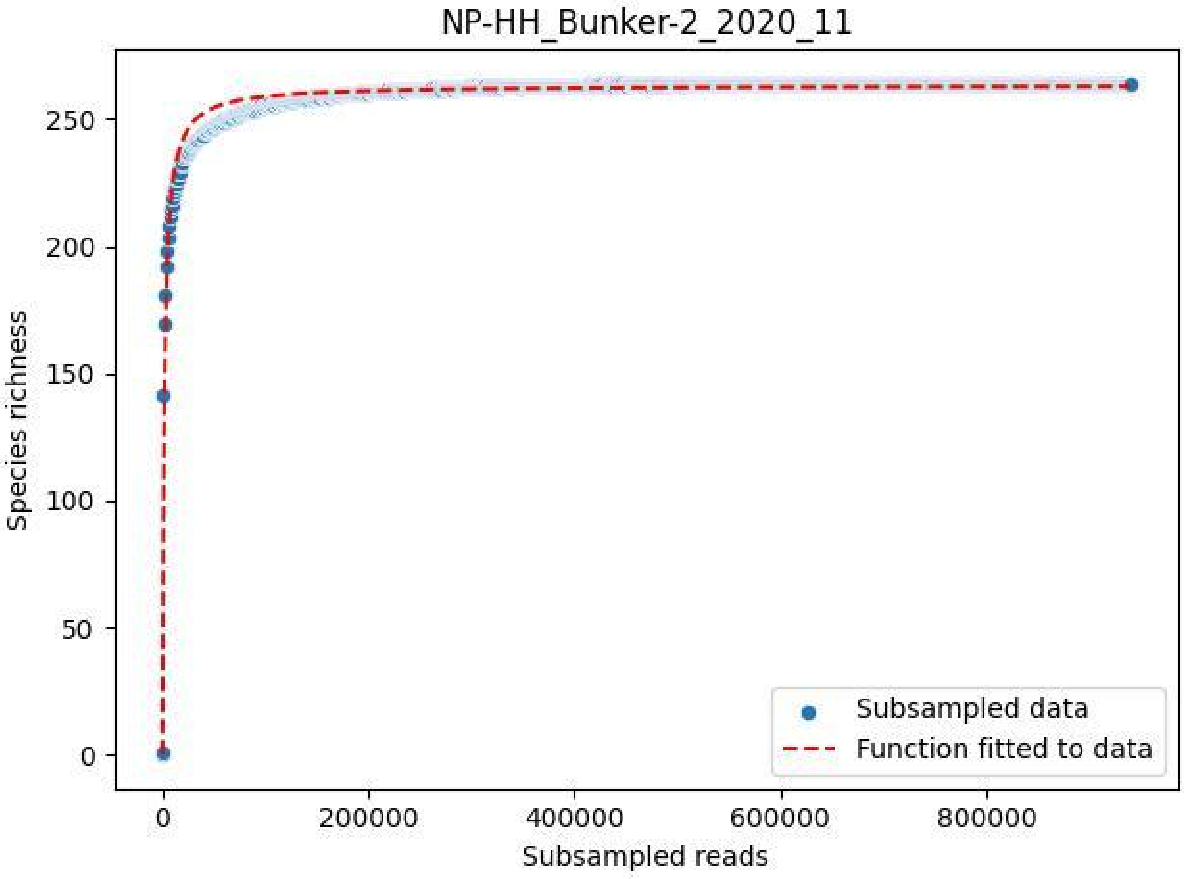

NP-HH_Bunker-2_2020_11
Species richness
0
50
100
150
200
250
Subsampled reads
0
200000
400000
600000
800000
Subsampled data
Function fitted to data

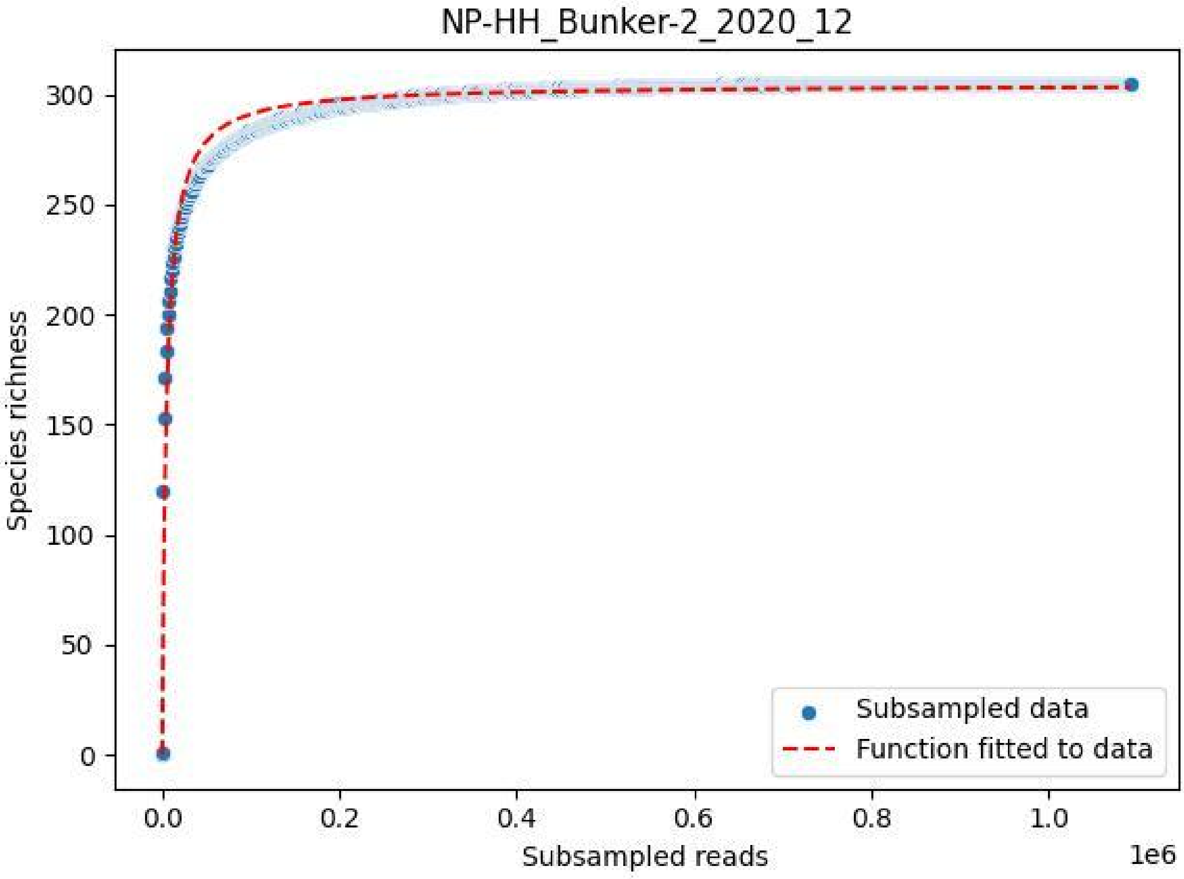
NP-HH_Bunker-2_2020_12
Species richness
300
250
200
150
100
50
0
0.0
0.2
0.4
0.6
0.8
1.0
1e6
Subsampled reads
Subsampled data
Function fitted to data

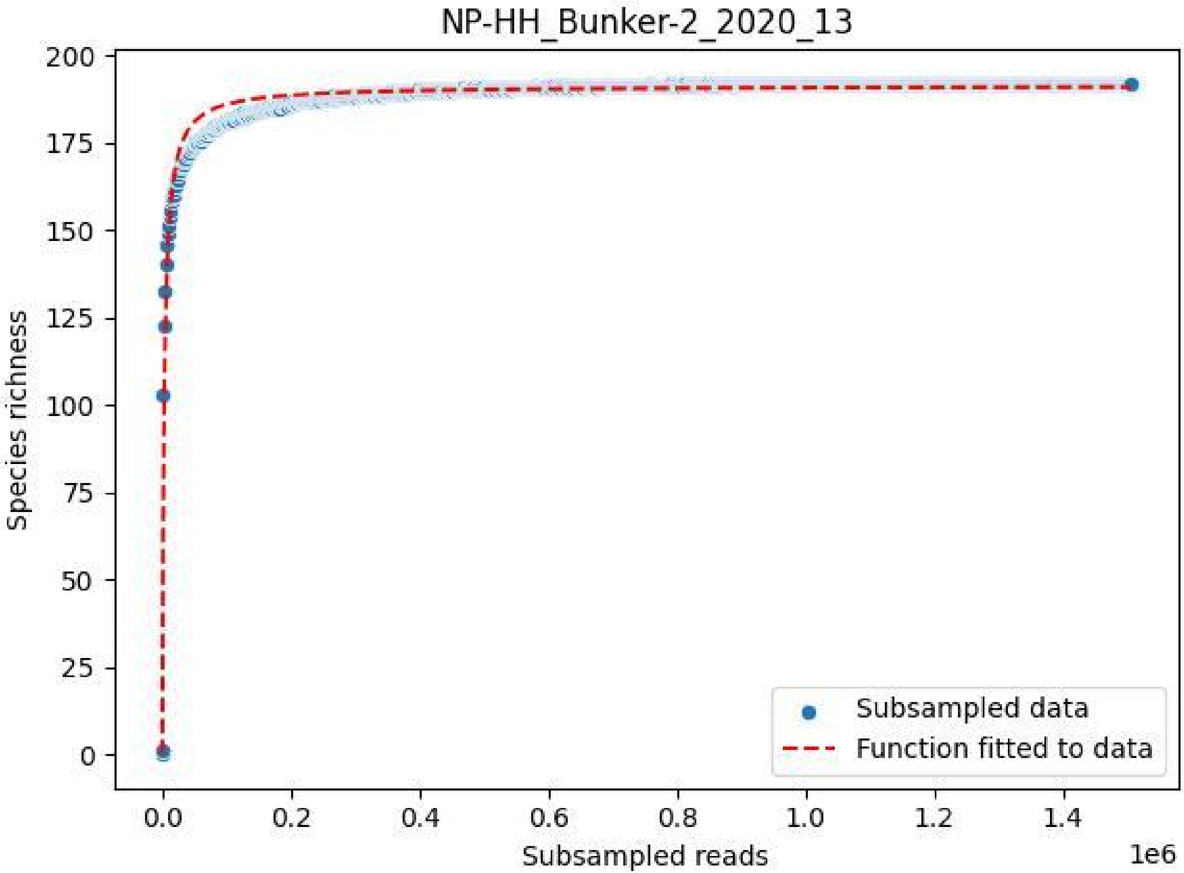

NP-HH_Bunker-2_2020_13
Species richness
200
175
150
125
100
75
50
25
0
0.0
0.2
0.4
0.6
0.8
1.0
1.2
1.4
1e6
Subsampled reads
Subsampled data
Function fitted to data

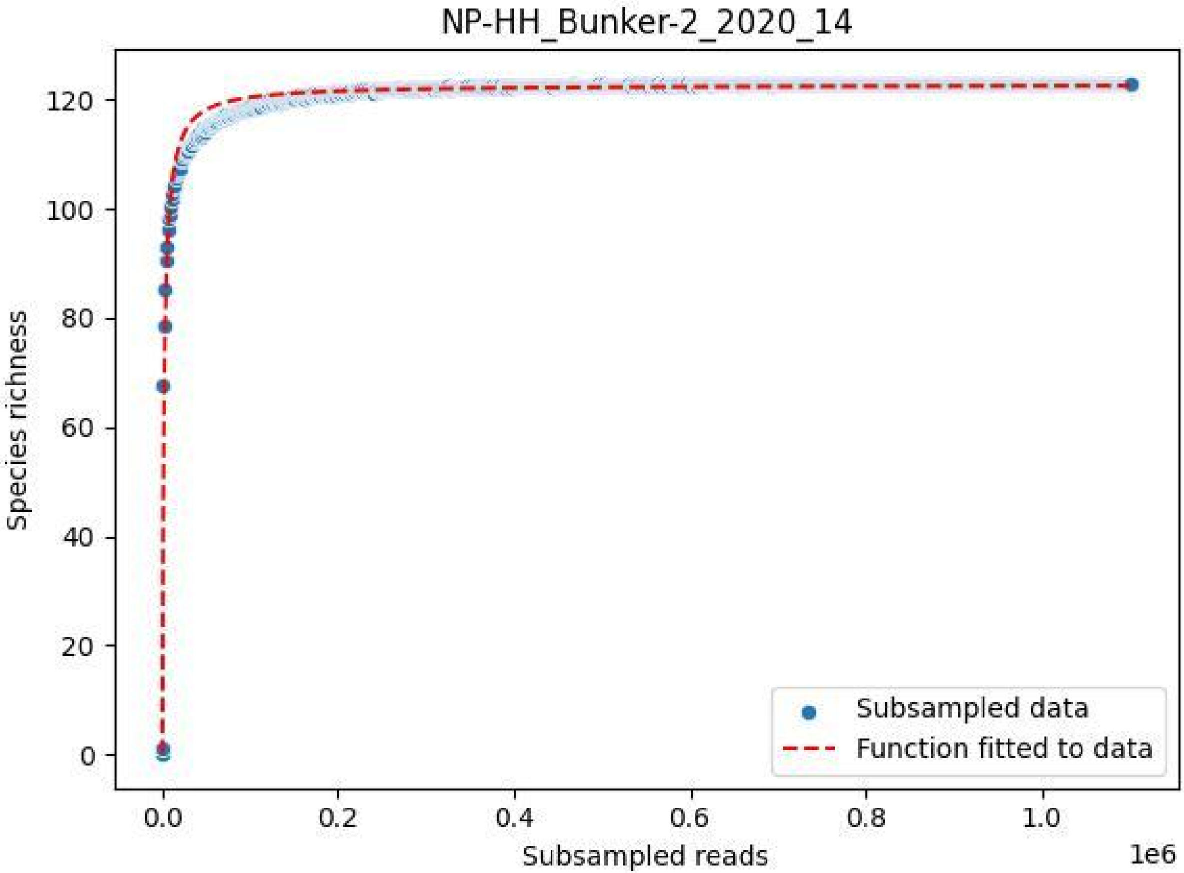
NP-HH_Bunker-2_2020_14
Species richness
120
100
80
60
40
20
0
0.0
0.2
0.4
0.6
0.8
1.0
1e6
Subsampled reads
Subsampled data
Function fitted to data

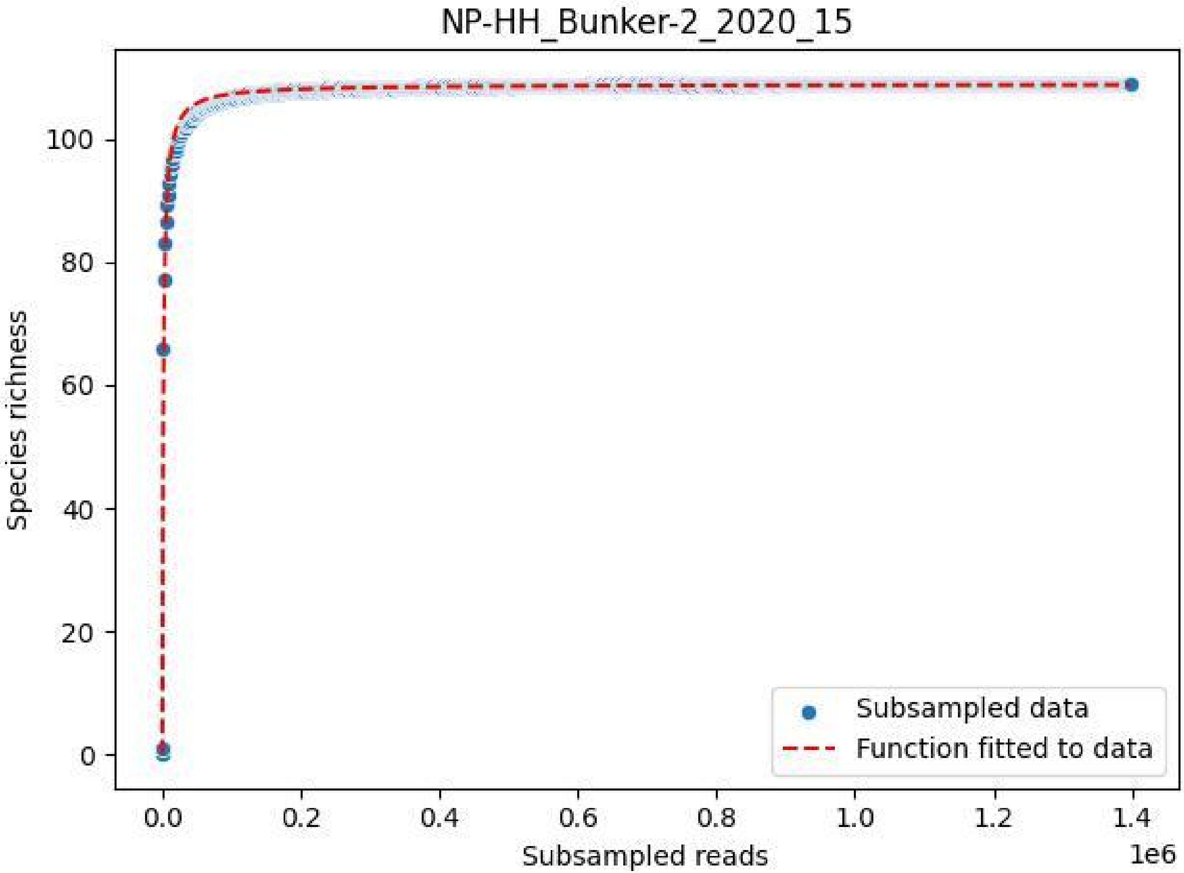

NP-HH_Bunker-2_2020_15
Species richness
Subsampled reads
1e6
0
20
40
60
80
100
0.0
0.2
0.4
0.6
0.8
1.0
1.2
1.4
Subsampled data
Function fitted to data

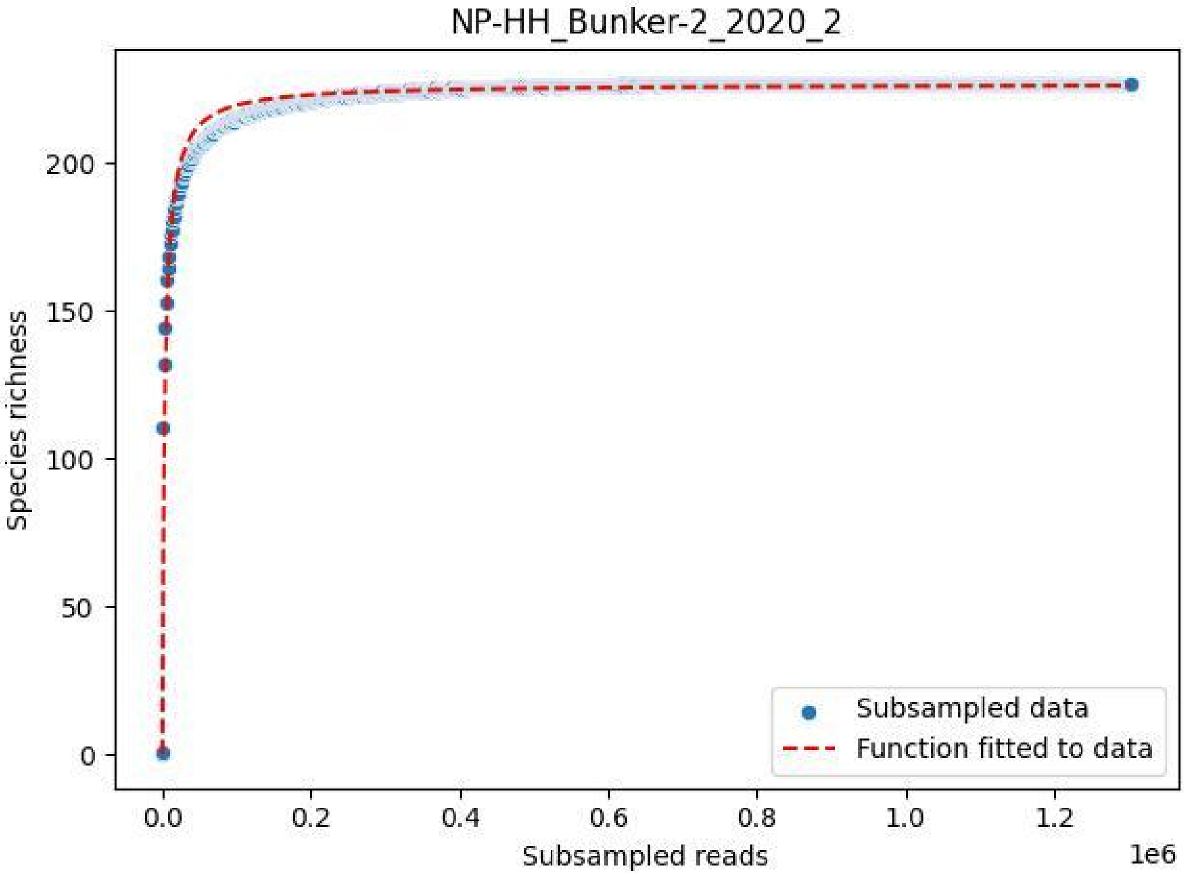

NP-HH_Bunker-2_2020_2
Species richness
Subsampled reads
1e6
0
50
100
150
200
0.0
0.2
0.4
0.6
0.8
1.0
1.2
Subsampled data
Function fitted to data

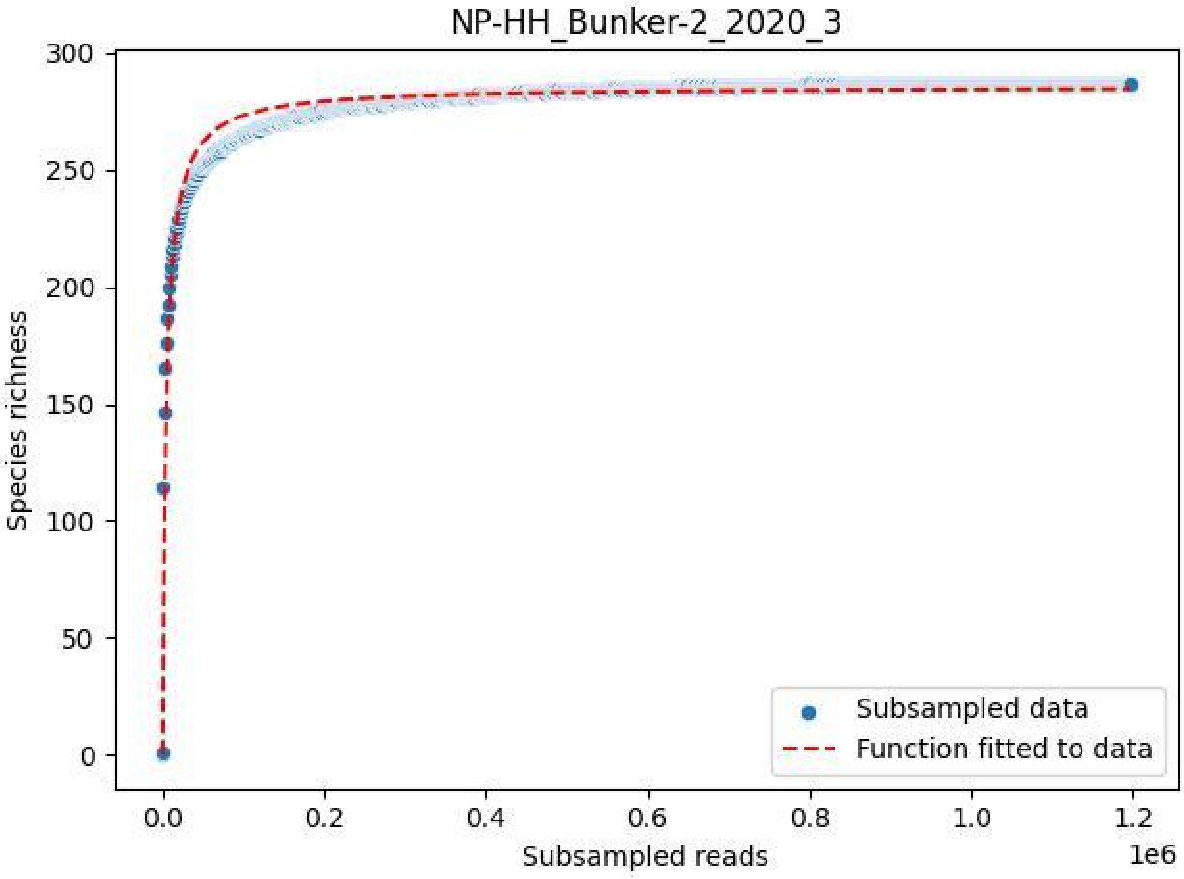
NP-HH_Bunker-2_2020_3
Species richness
Subsampled reads
1e6
Subsampled data
Function fitted to data
0
50
100
150
200
250
300
0.0
0.2
0.4
0.6
0.8
1.0
1.2

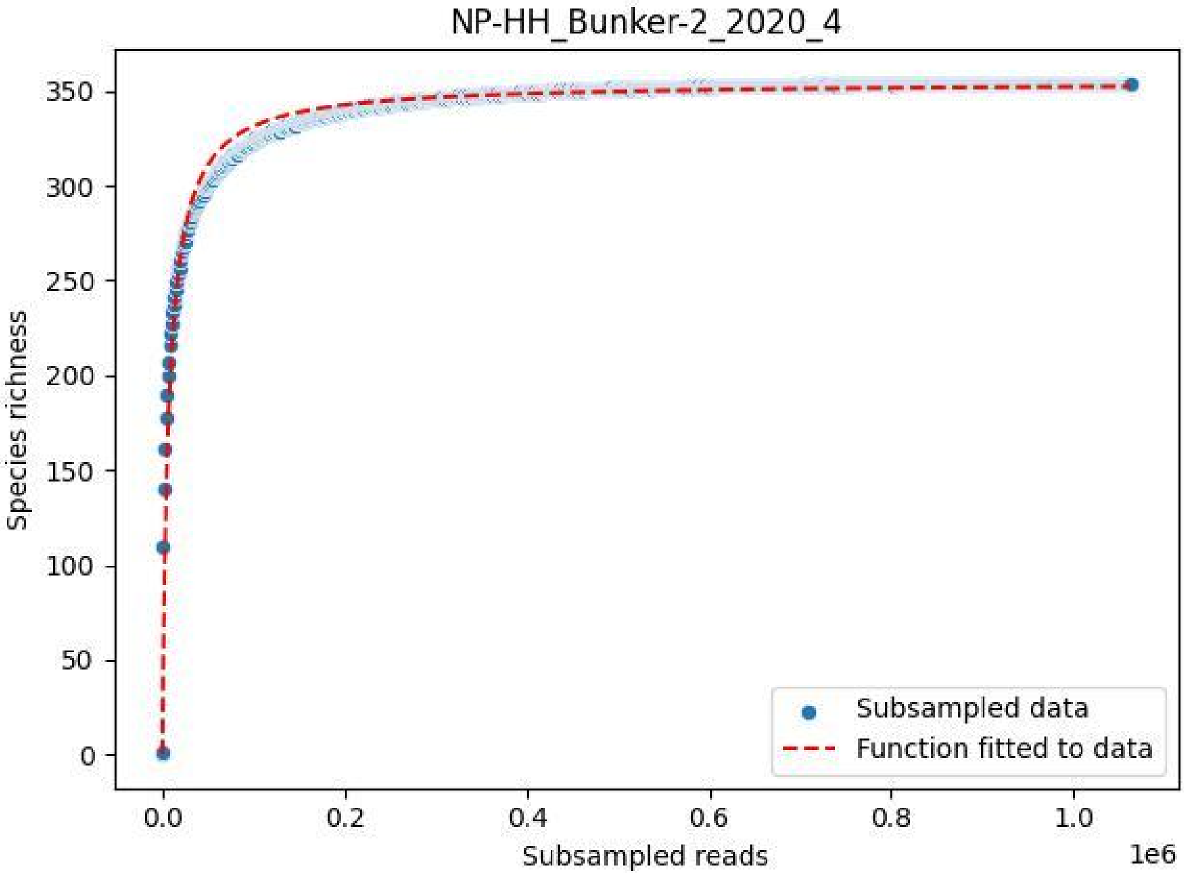
NP-HH_Bunker-2_2020_4
Species richness
Subsampled reads
1e6
0
50
100
150
200
250
300
350
0.0
0.2
0.4
0.6
0.8
1.0
Subsampled data
Function fitted to data

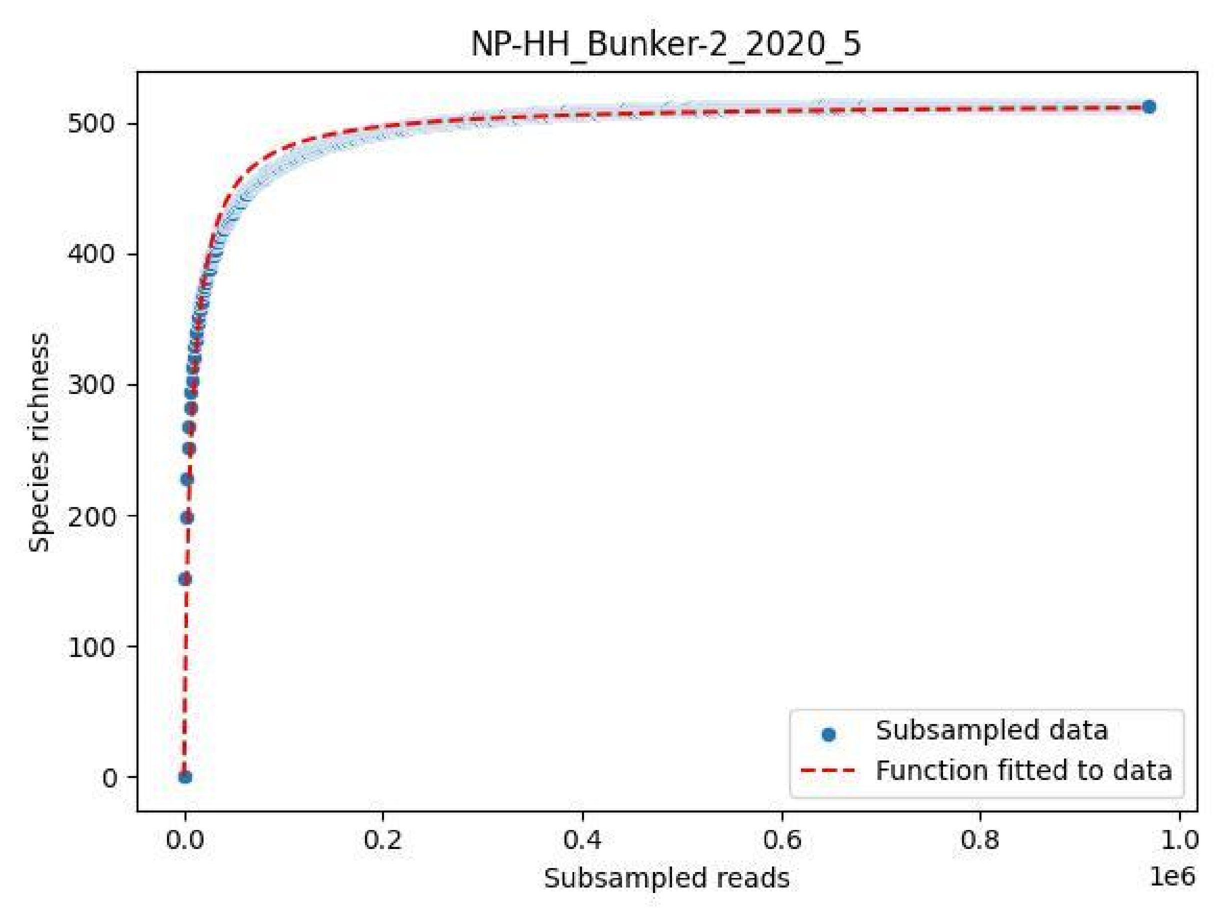
NP-HH_Bunker-2_2020_5
Species richness
Subsampled reads
1e6
0
100
200
300
400
500
0.0
0.2
0.4
0.6
0.8
1.0
Subsampled data
Function fitted to data

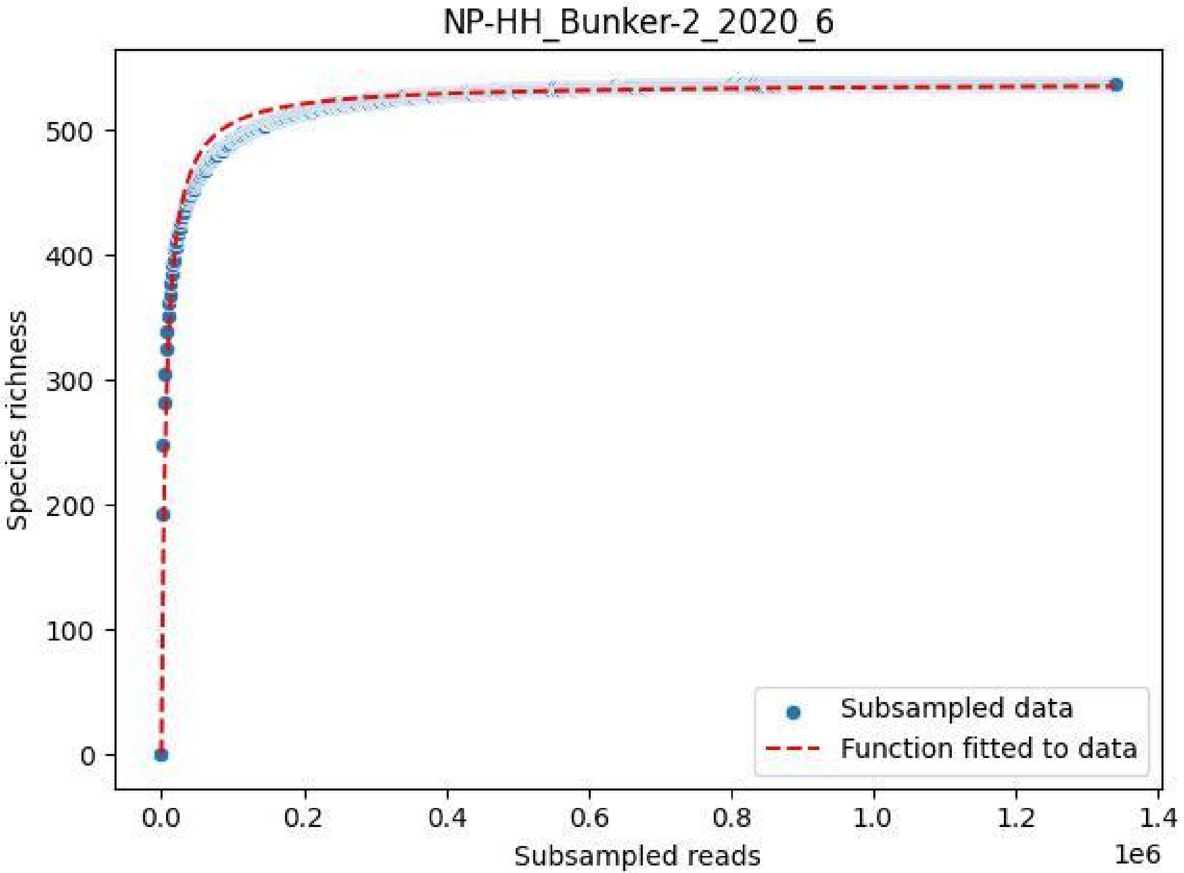

NP-HH_Bunker-2_2020_6
Species richness
0
100
200
300
400
500
0.0
0.2
0.4
0.6
0.8
1.0
1.2
1.4
1e6
Subsampled reads
Subsampled data
Function fitted to data

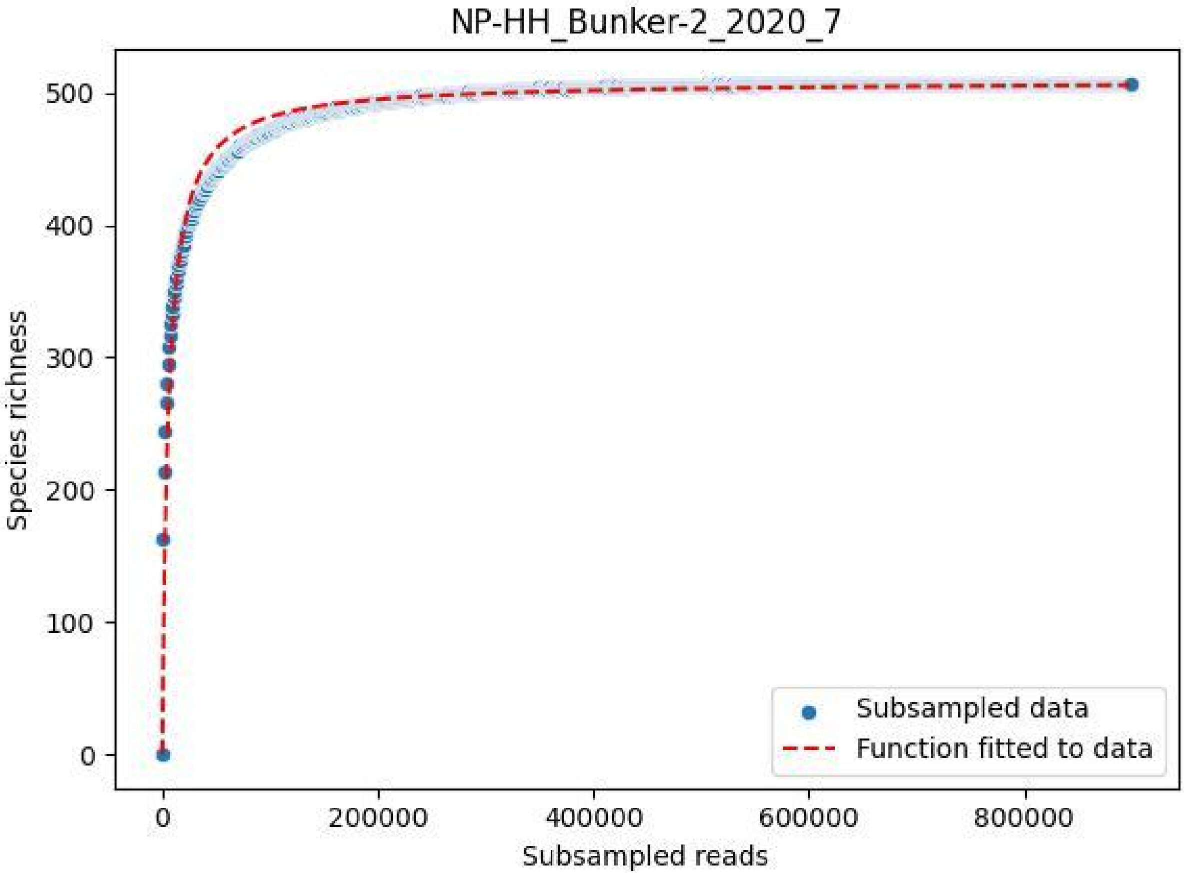

NP-HH_Bunker-2_2020_7
Species richness
Subsampled reads
0
100
200
300
400
500
0
200000
400000
600000
800000
Subsampled data
Function fitted to data

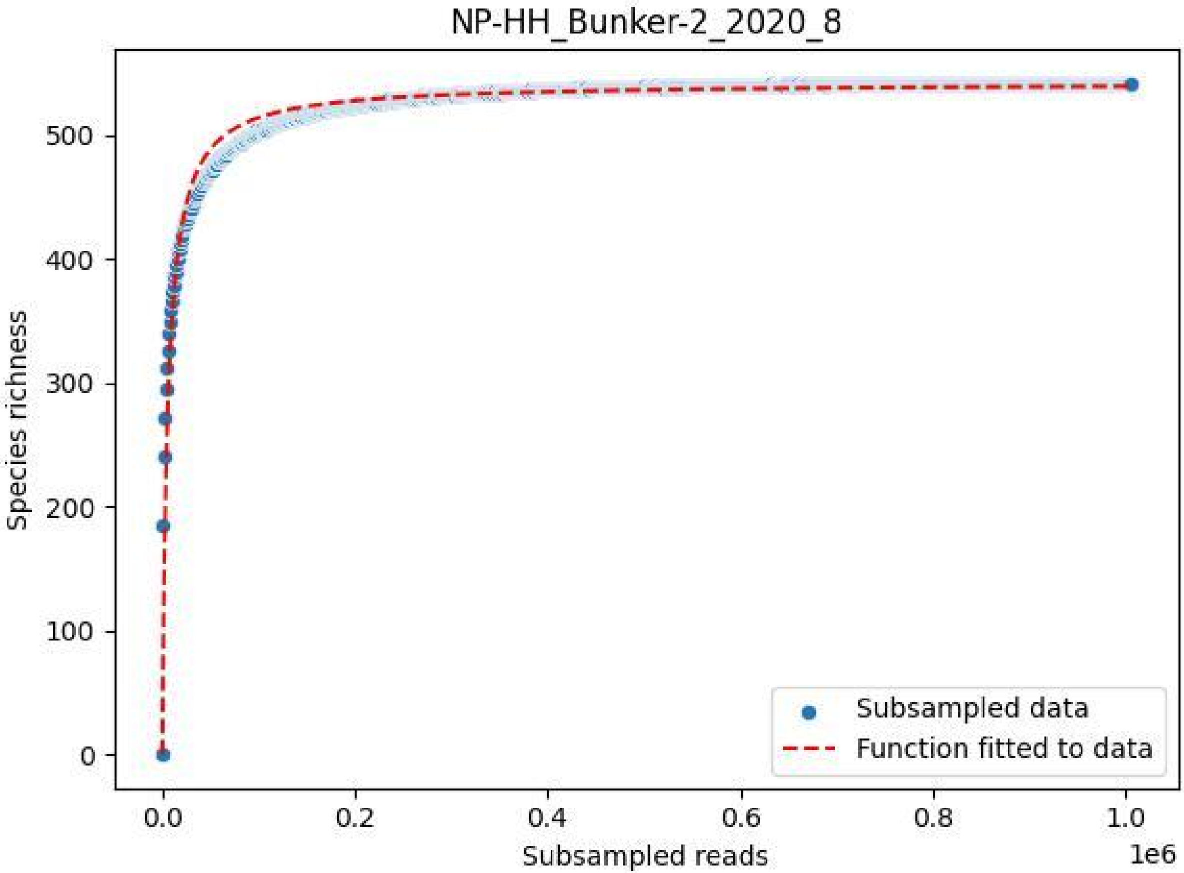
NP-HH_Bunker-2_2020_8
Species richness
0
100
200
300
400
500
0.0
0.2
0.4
0.6
0.8
1.0
1e6
Subsampled reads
Subsampled data
Function fitted to data

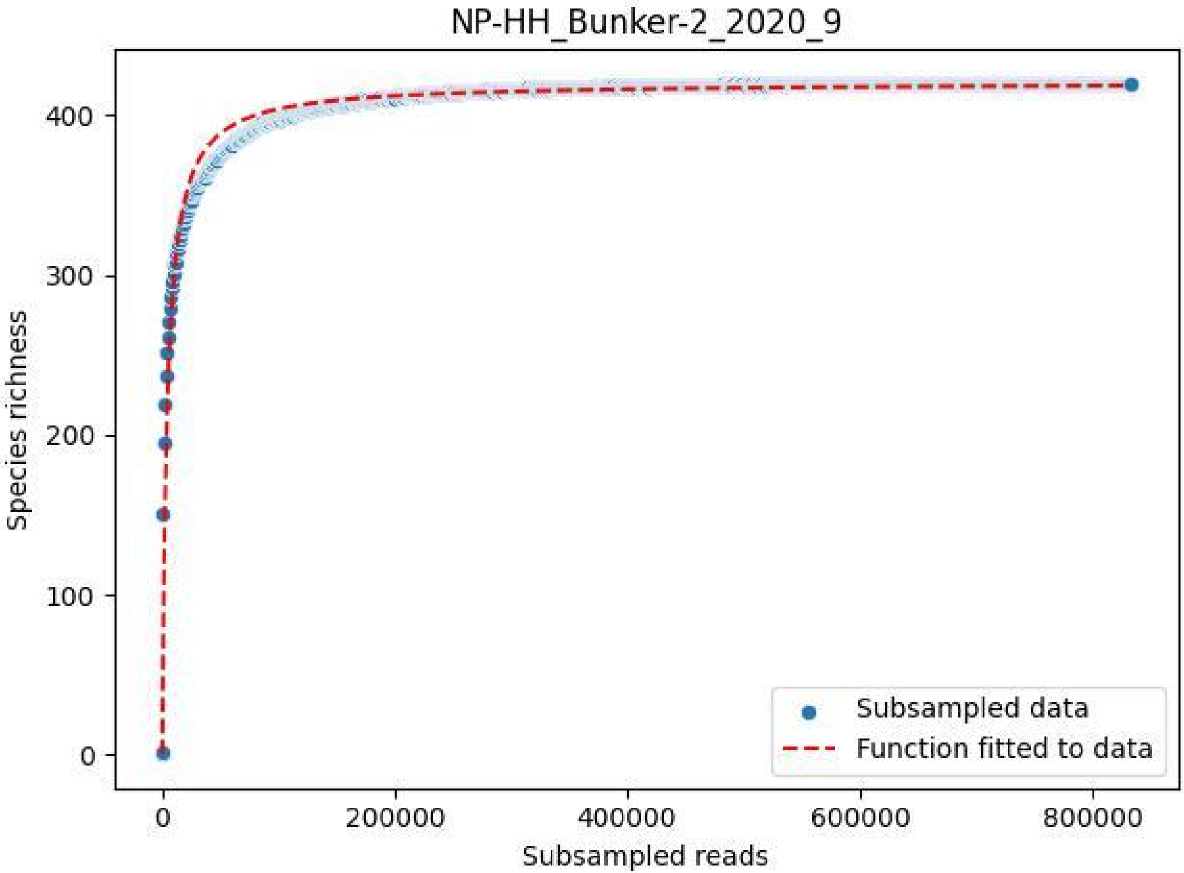

NP-HH_Bunker-2_2020_9
Species richness
Subsampled reads
0
100
200
300
400
0
200000
400000
600000
800000
Subsampled data
Function fitted to data

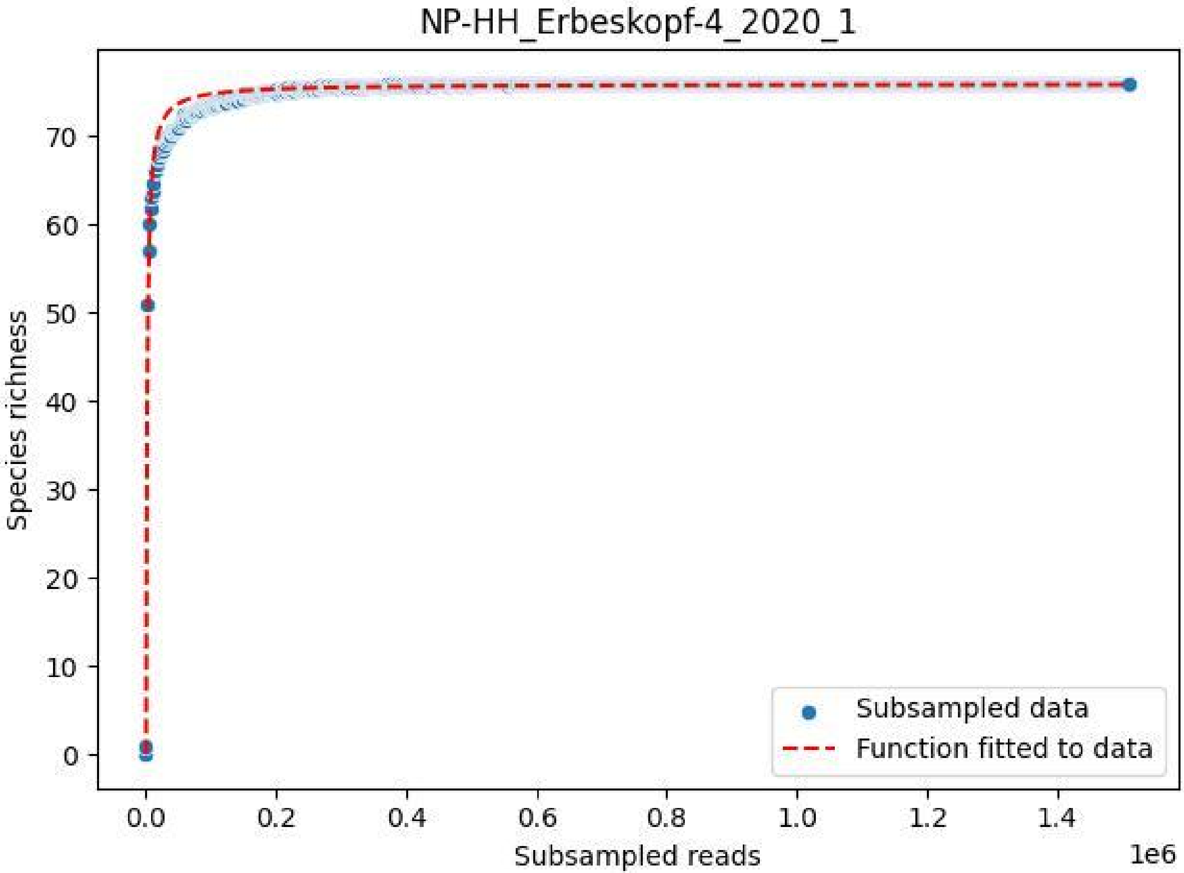

NP-HH_Erbeskopf-4_2020_1
Species richness
Subsampled reads
1e6
0
10
20
30
40
50
60
70
0.0
0.2
0.4
0.6
0.8
1.0
1.2
1.4
Subsampled data
Function fitted to data

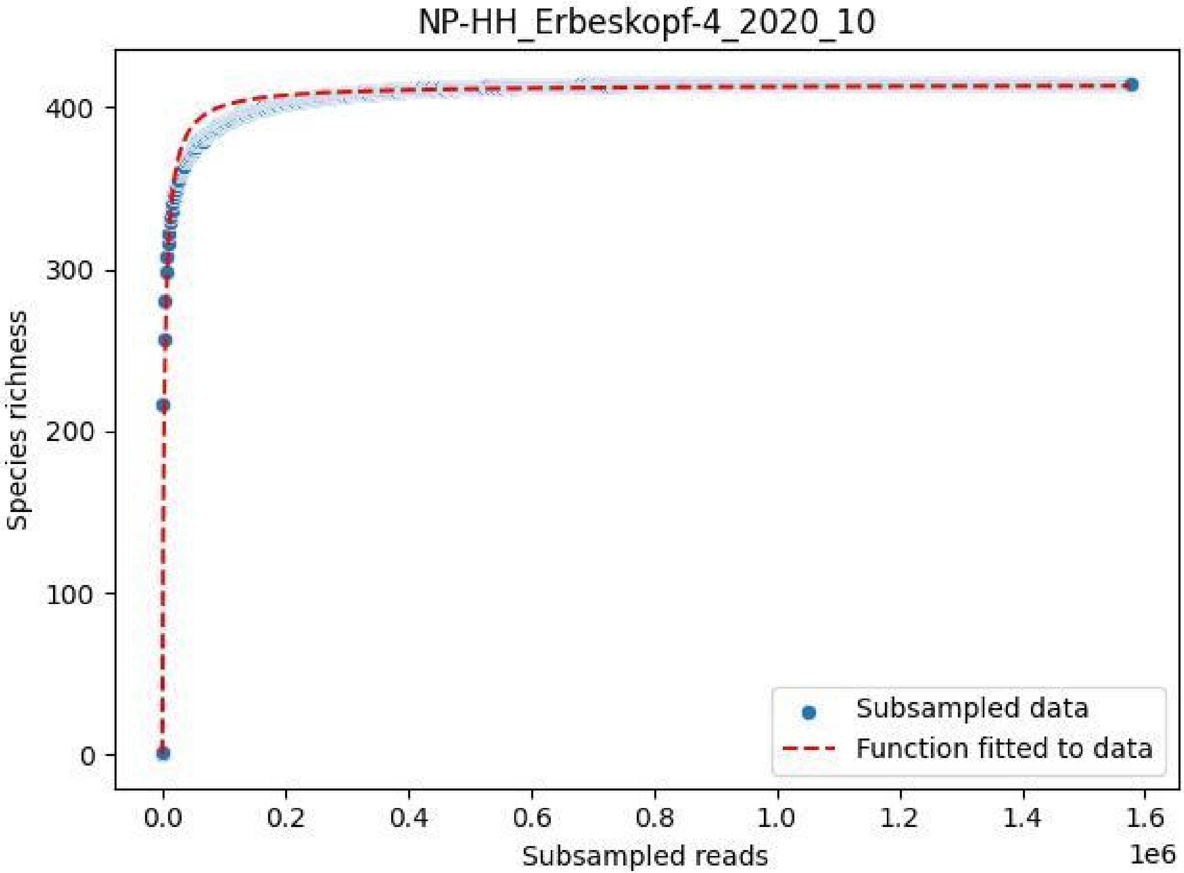

NP-HH_Erbeskopf-4_2020_10
Species richness
Subsampled reads
1e6
0
100
200
300
400
0.0
0.2
0.4
0.6
0.8
1.0
1.2
1.4
1.6
Subsampled data
Function fitted to data

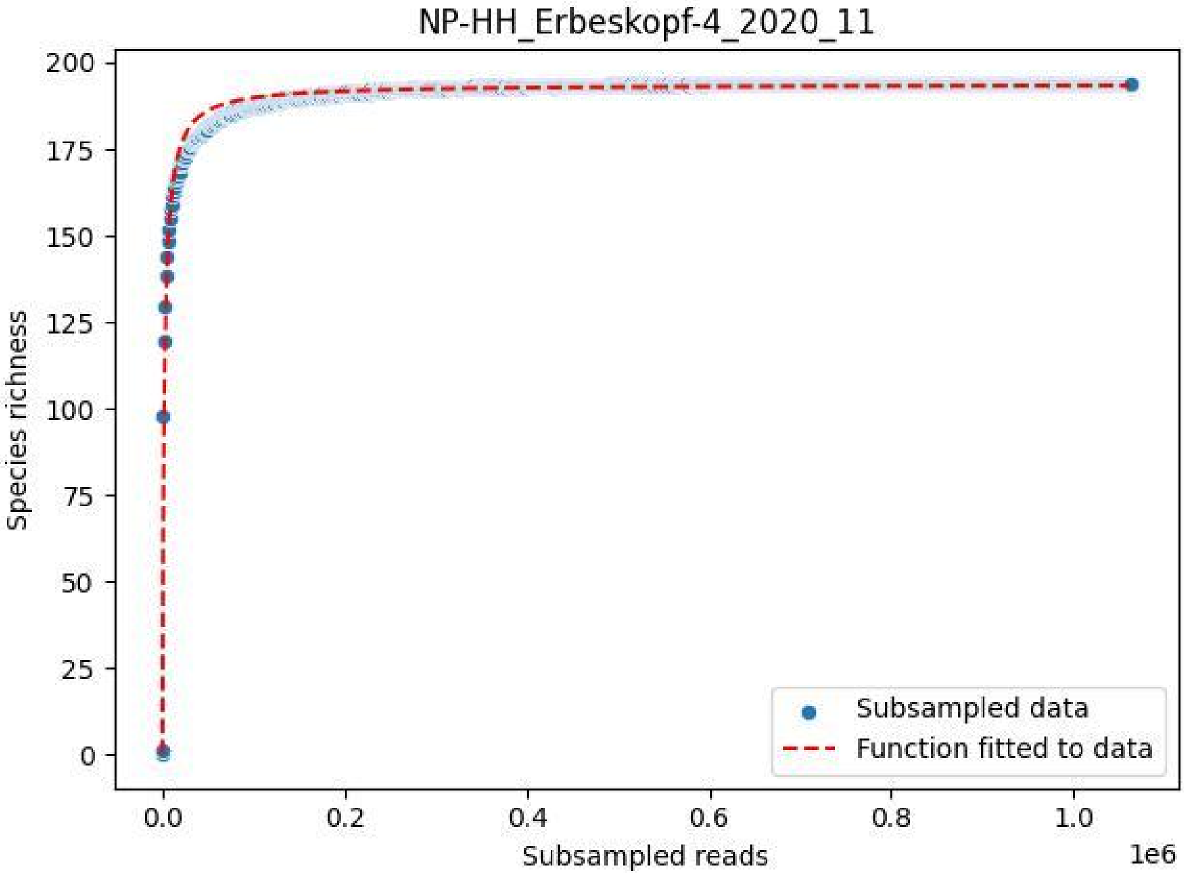

NP-HH_Erbeskopf-4_2020_11
Species richness
Subsampled reads
1e6
0.0
0.2
0.4
0.6
0.8
1.0
0
25
50
75
100
125
150
175
200
Subsampled data
Function fitted to data

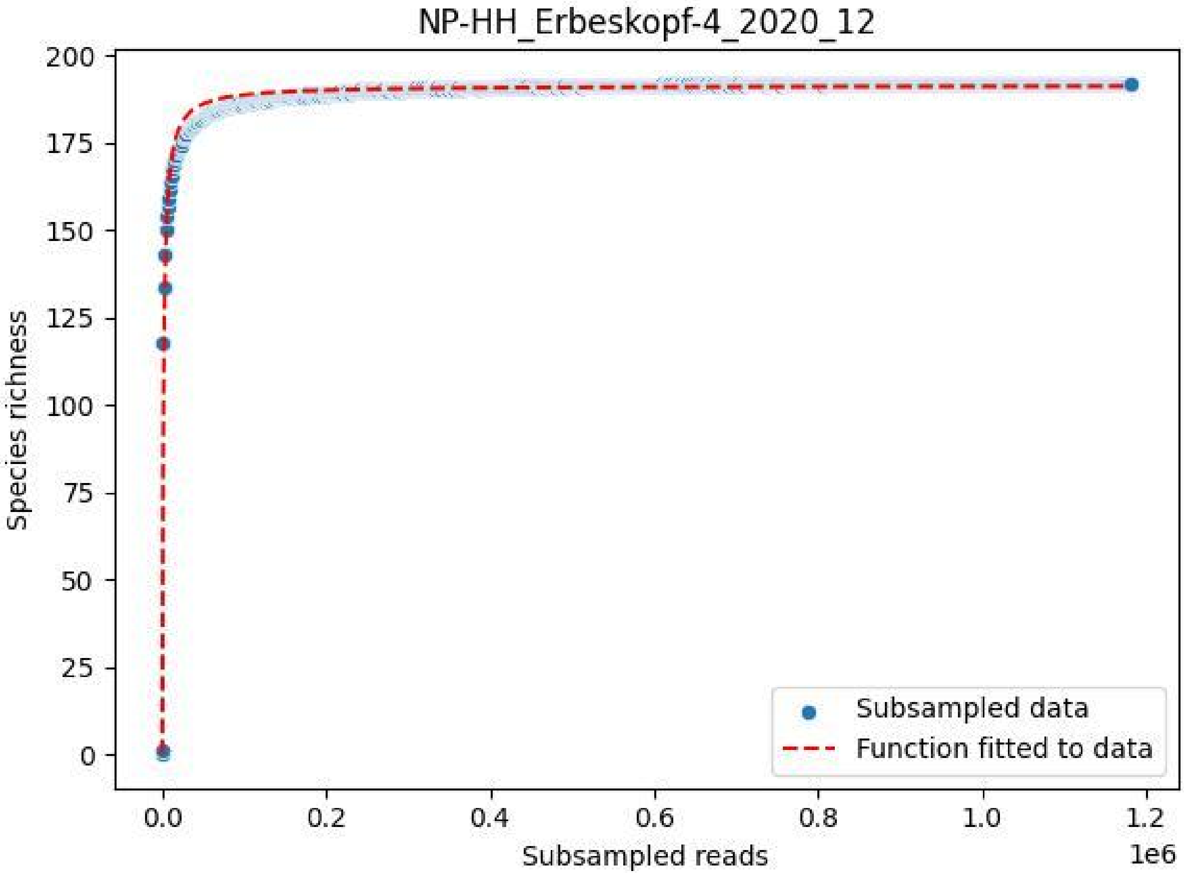
NP-HH_Erbeskopf-4_2020_12
200
175
150
125
100
75
50
25
0
Species richness
0.0
0.2
0.4
0.6
0.8
1.0
1.2
1e6
Subsampled reads
Subsampled data
Function fitted to data

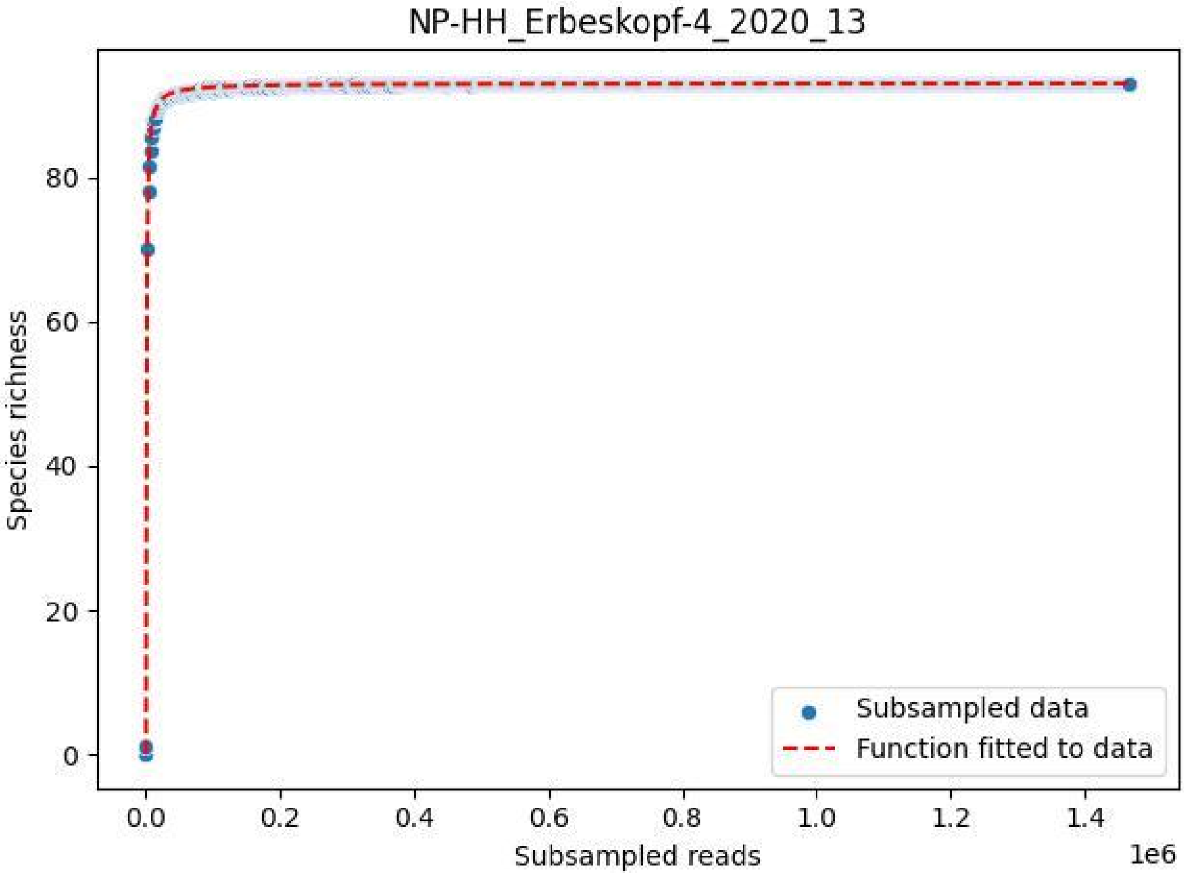
NP-HH_Erbeskopf-4_2020_13
Species richness
Subsampled reads
1e6
0
20
40
60
80
0.0
0.2
0.4
0.6
0.8
1.0
1.2
1.4
Subsampled data
Function fitted to data

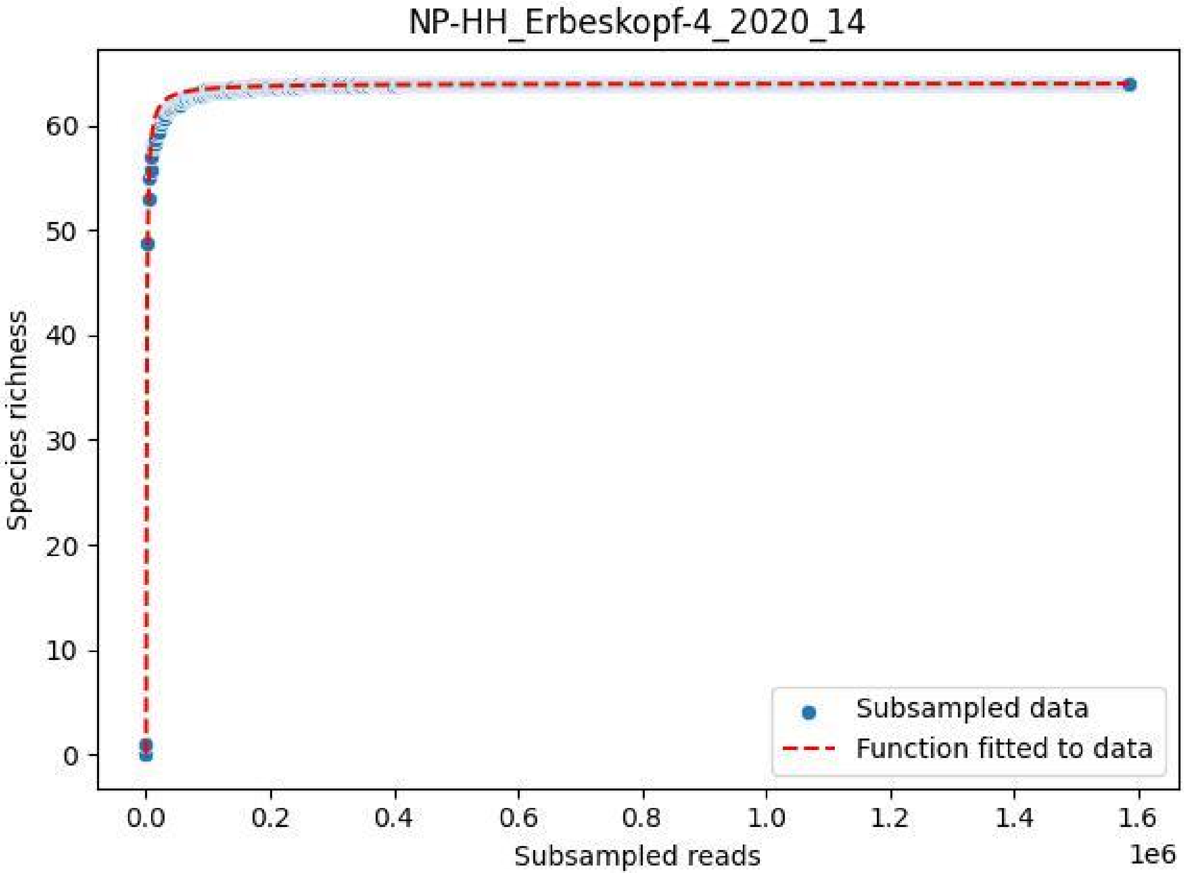

NP-HH_Erbeskopf-4_2020_14
Species richness
Subsampled reads
1e6
0
10
20
30
40
50
60
0.0
0.2
0.4
0.6
0.8
1.0
1.2
1.4
1.6
Subsampled data
Function fitted to data

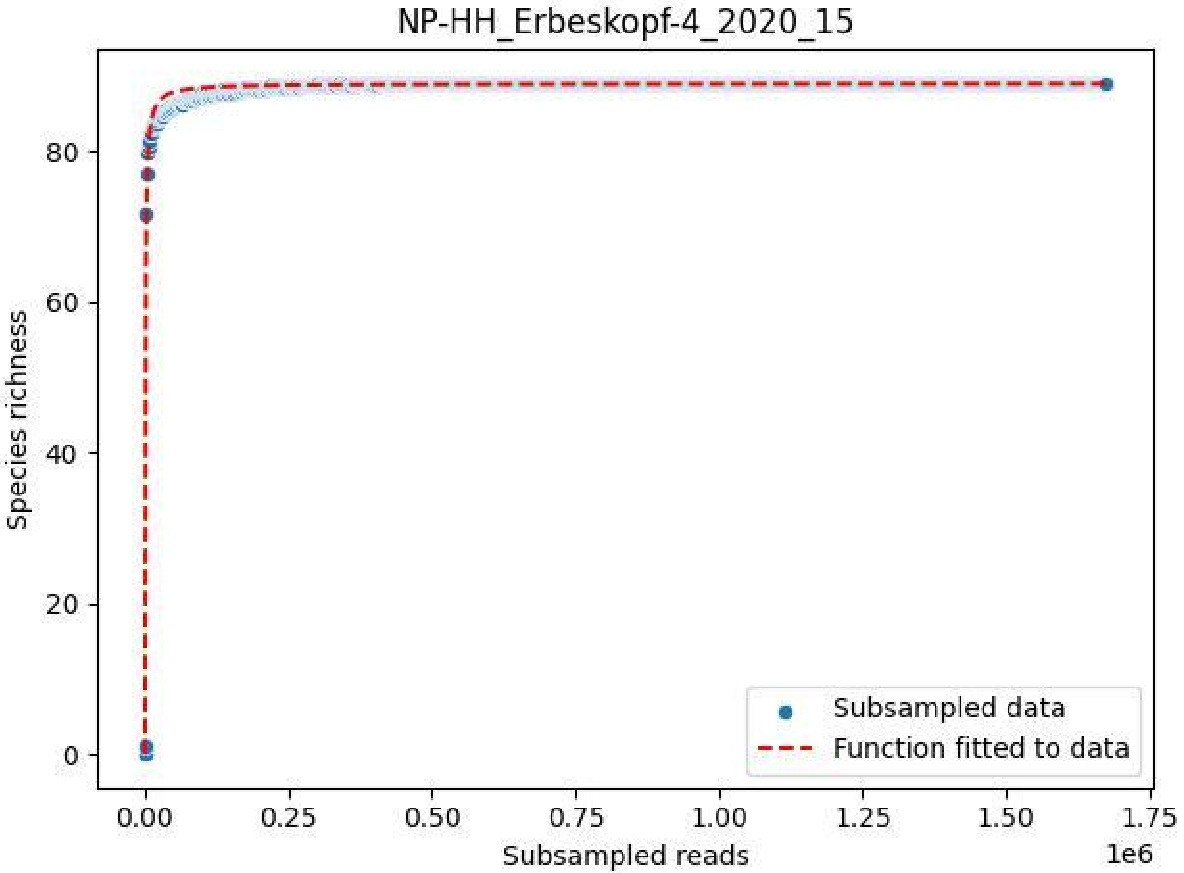

NP-HH_Erbeskopf-4_2020_15
Species richness
0
20
40
60
80
Subsampled reads
0.00
0.25
0.50
0.75
1.00
1.25
1.50
1.75
1e6
Subsampled data
Function fitted to data

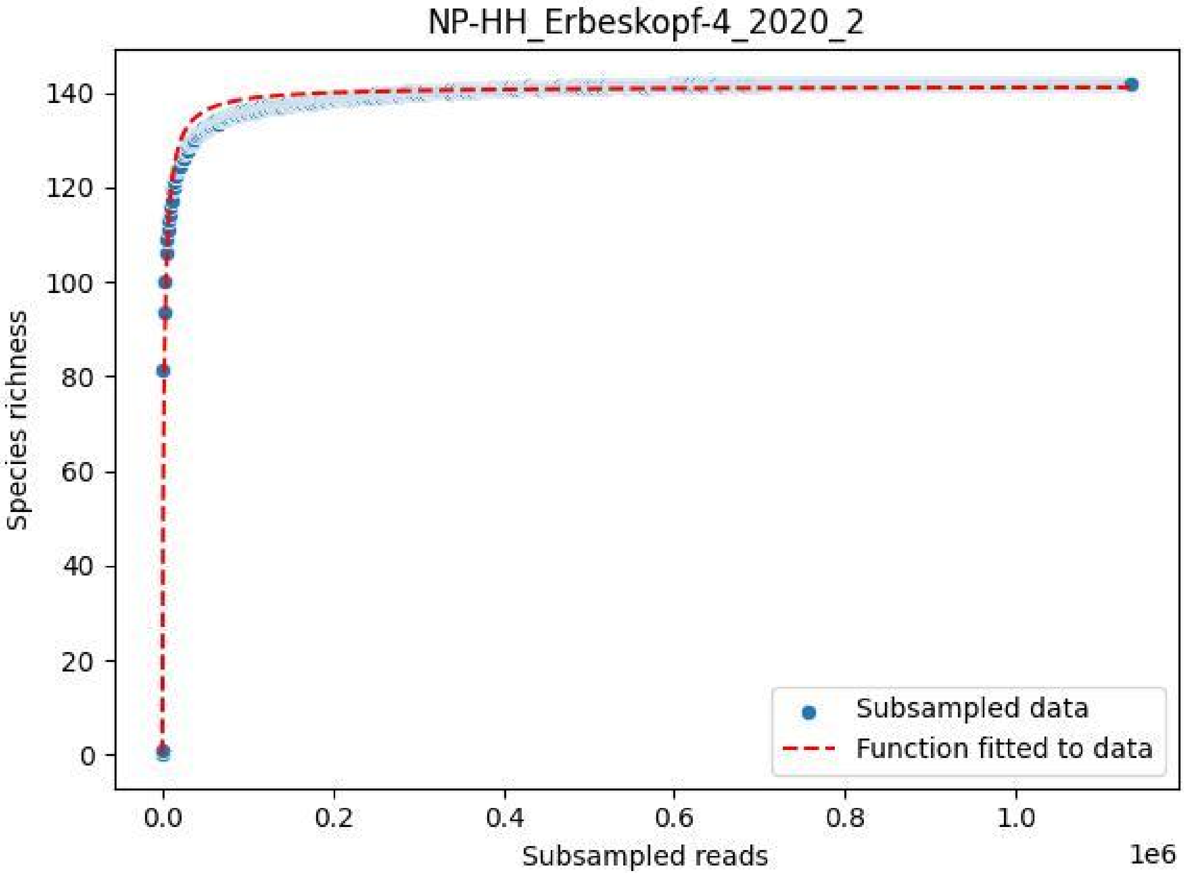
NP-HH_Erbeskopf-4_2020_2
Species richness
Subsampled reads
1e6
0
20
40
60
80
100
120
140
0.0
0.2
0.4
0.6
0.8
1.0
Subsampled data
Function fitted to data

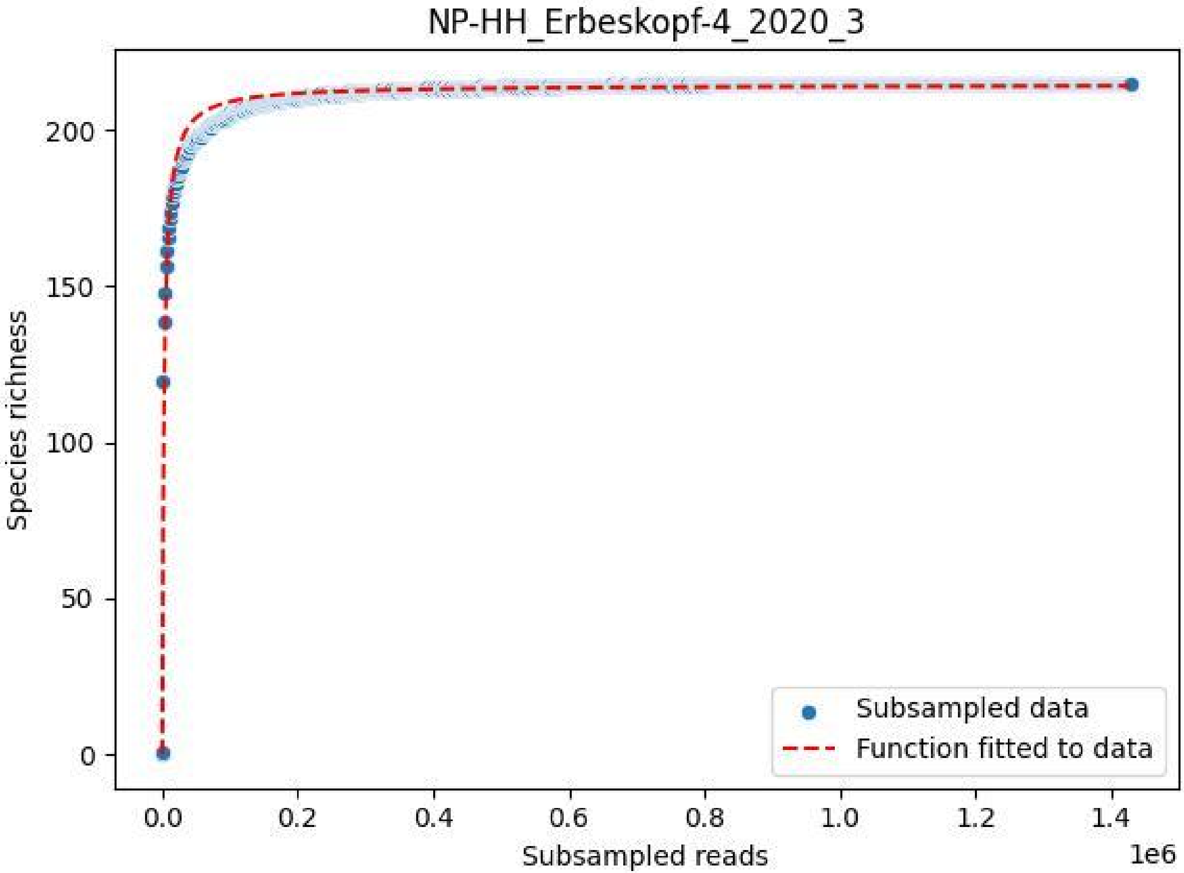

NP-HH_Erbeskopf-4_2020_3
Species richness
Subsampled reads
1e6
0
50
100
150
200
0.0
0.2
0.4
0.6
0.8
1.0
1.2
1.4
Subsampled data
Function fitted to data

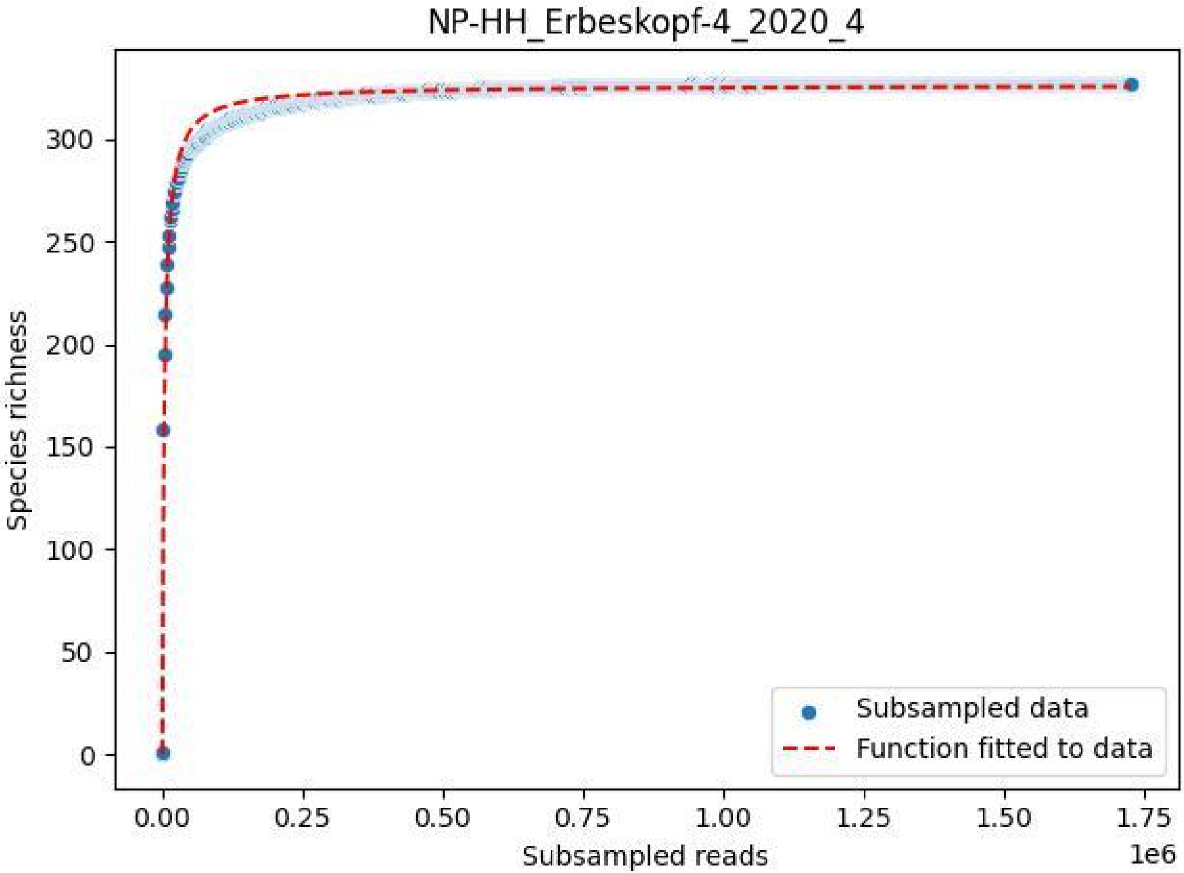

NP-HH_Erbeskopf-4_2020_4
Species richness
Subsampled reads
1e6
0
50
100
150
200
250
300
0.00
0.25
0.50
0.75
1.00
1.25
1.50
1.75
Subsampled data
Function fitted to data

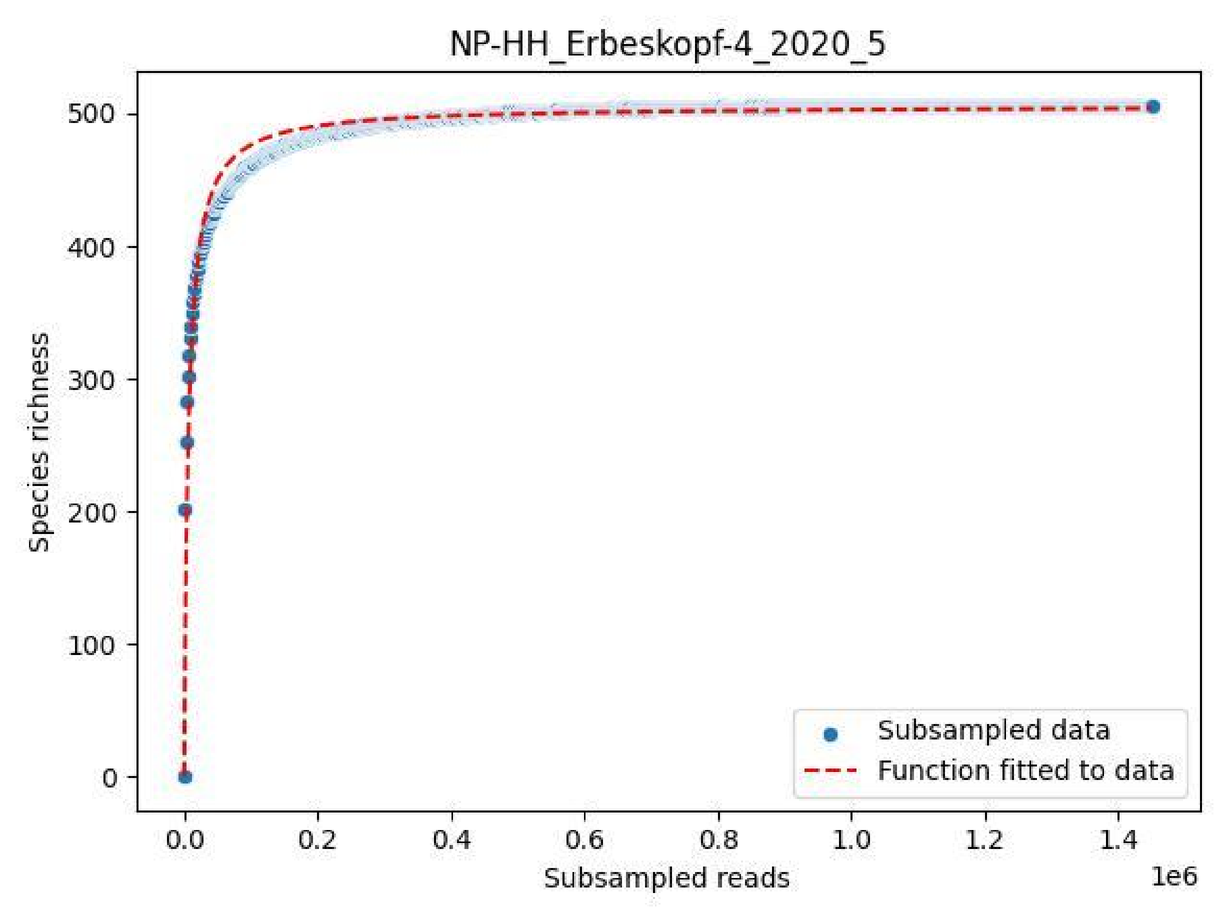
NP-HH_Erbeskopf-4_2020_5
Species richness
Subsampled reads
1e6
Subsampled data
Function fitted to data
0
100
200
300
400
500
0.0
0.2
0.4
0.6
0.8
1.0
1.2
1.4

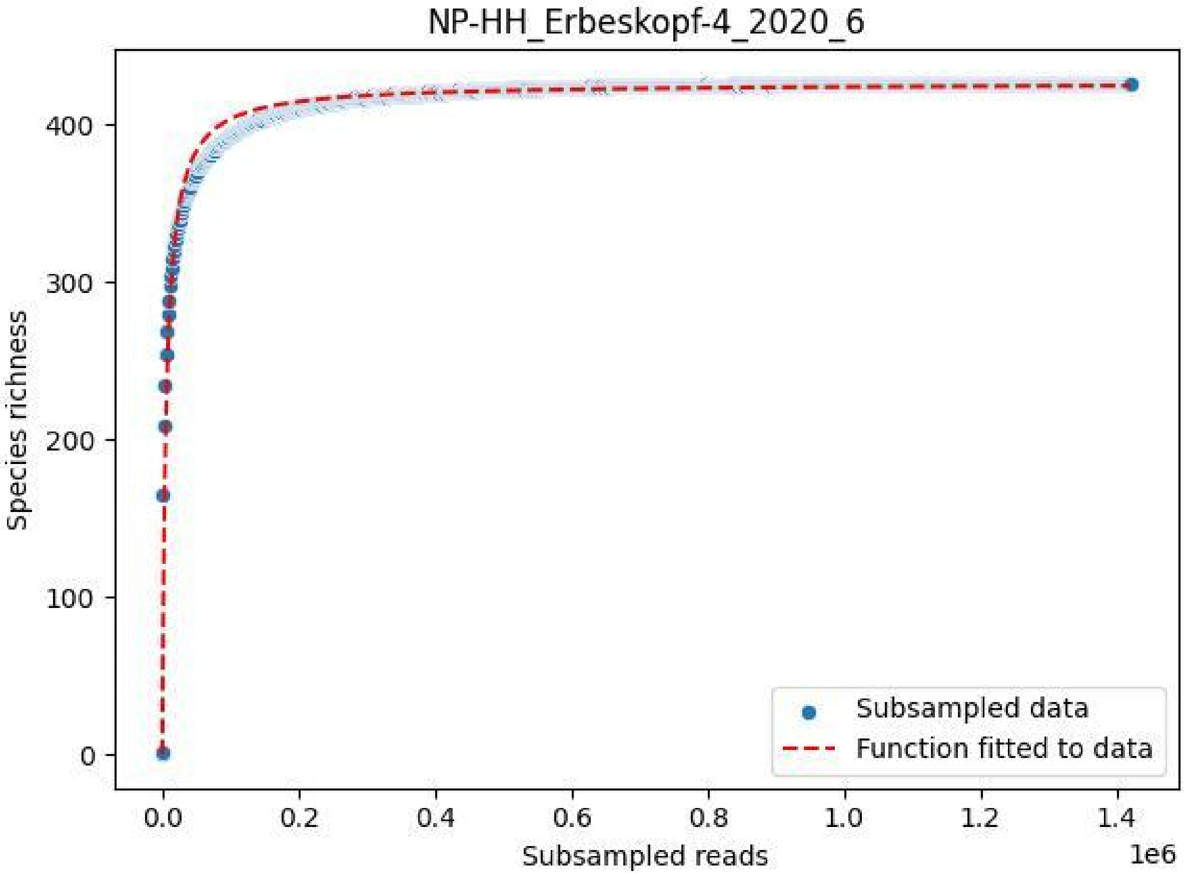
NP-HH_Erbeskopf-4_2020_6
Species richness
400
300
200
100
0
0.0
0.2
0.4
0.6
0.8
1.0
1.2
1.4
Subsampled reads
1e6
Subsampled data
Function fitted to data

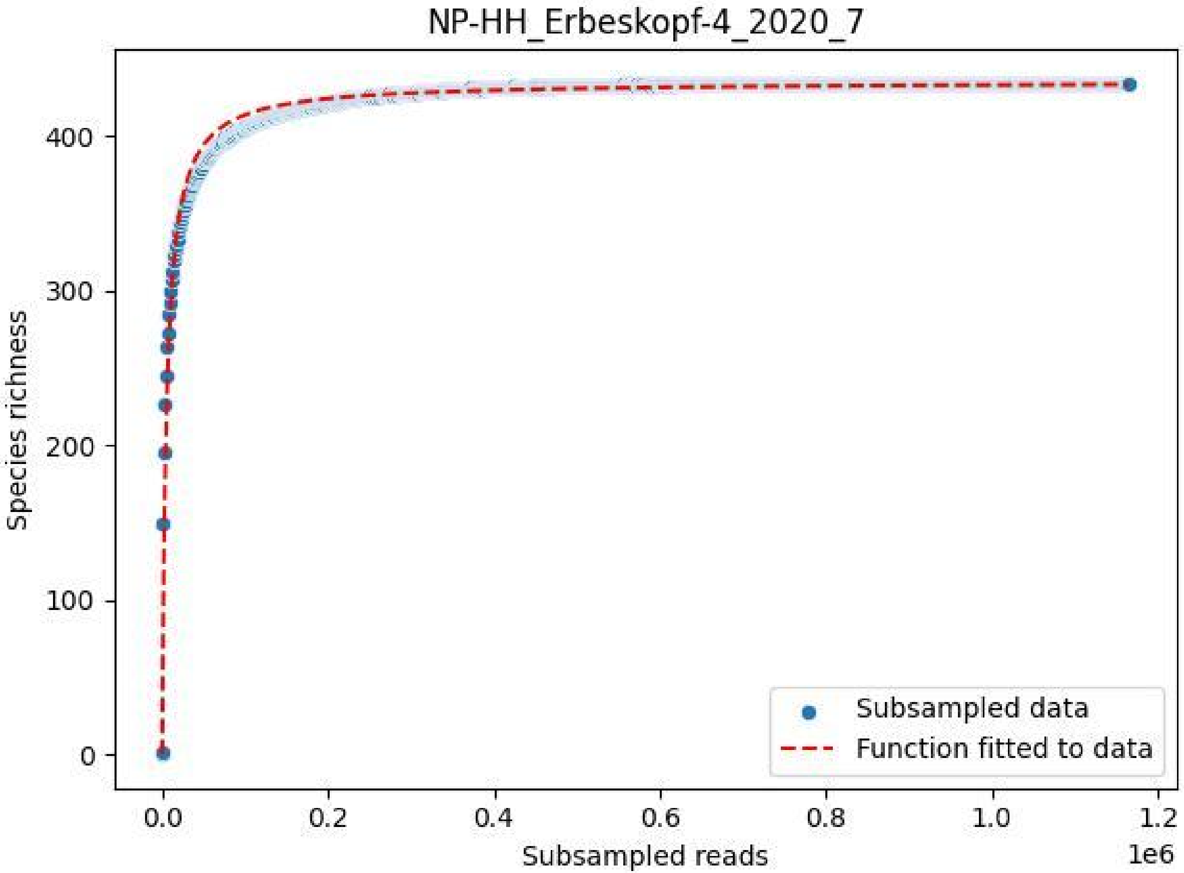

NP-HH_Erbeskopf-4_2020_7
Species richness
Subsampled reads
1e6
0
100
200
300
400
0.0
0.2
0.4
0.6
0.8
1.0
1.2
Subsampled data
Function fitted to data

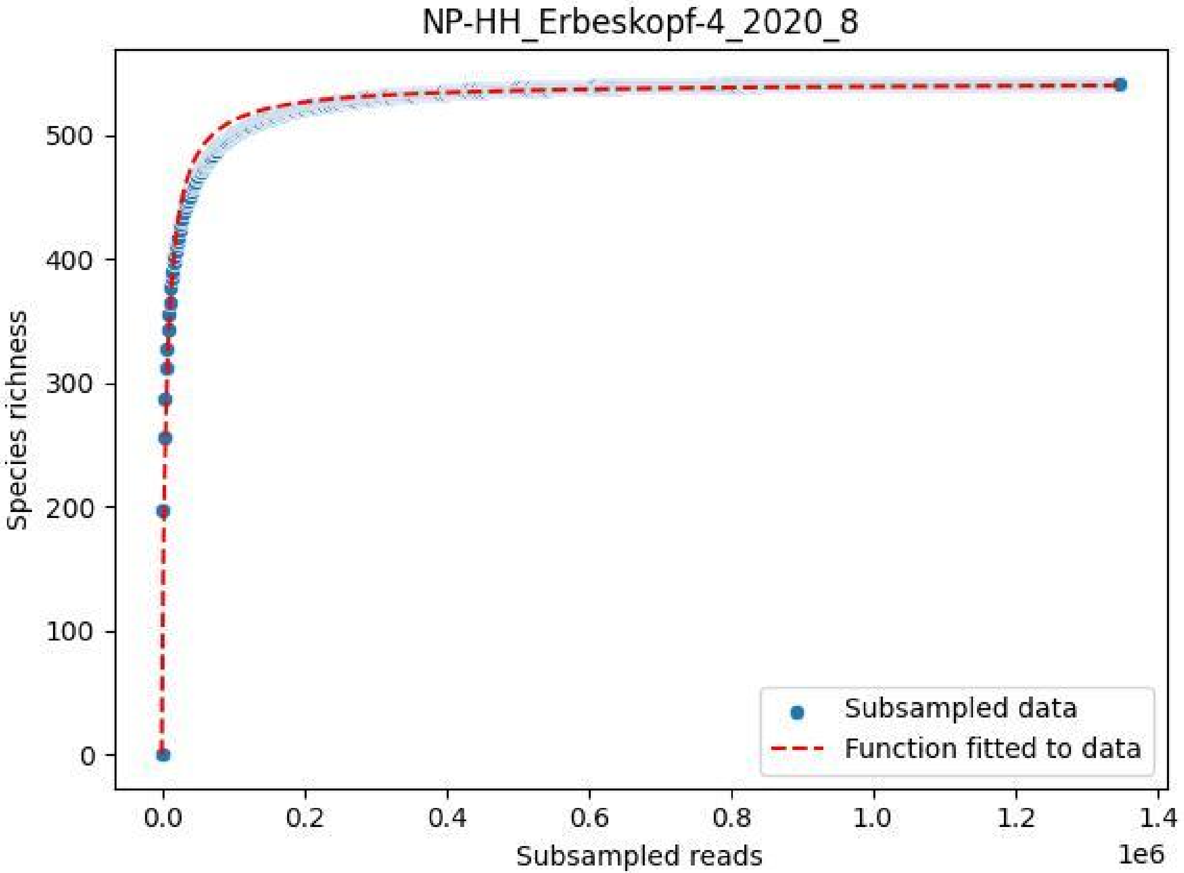
NP-HH_Erbeskopf-4_2020_8
Species richness
Subsampled reads
1e6
0
100
200
300
400
500
0.0
0.2
0.4
0.6
0.8
1.0
1.2
1.4
Subsampled data
Function fitted to data

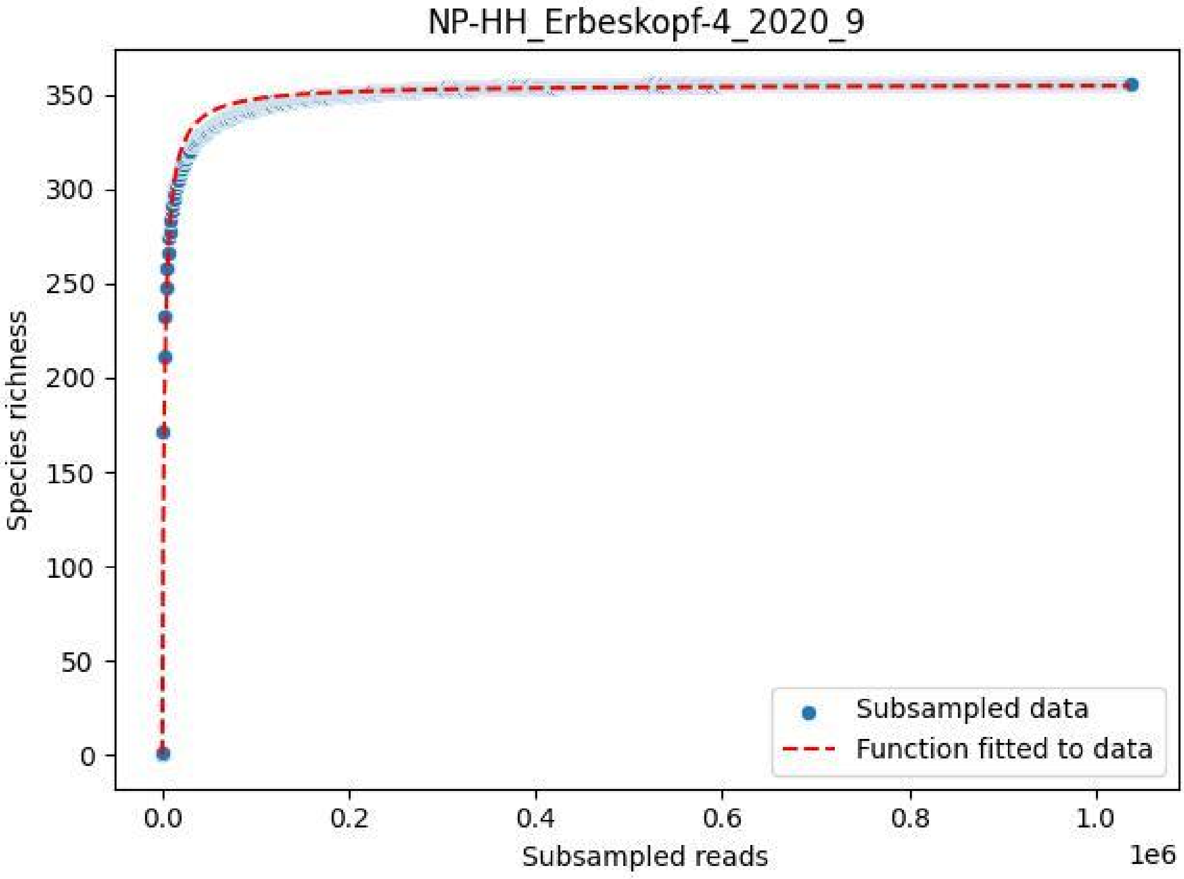
NP-HH_Erbeskopf-4_2020_9
Species richness
Subsampled reads
1e6
0
50
100
150
200
250
300
350
0.0
0.2
0.4
0.6
0.8
1.0
Subsampled data
Function fitted to data

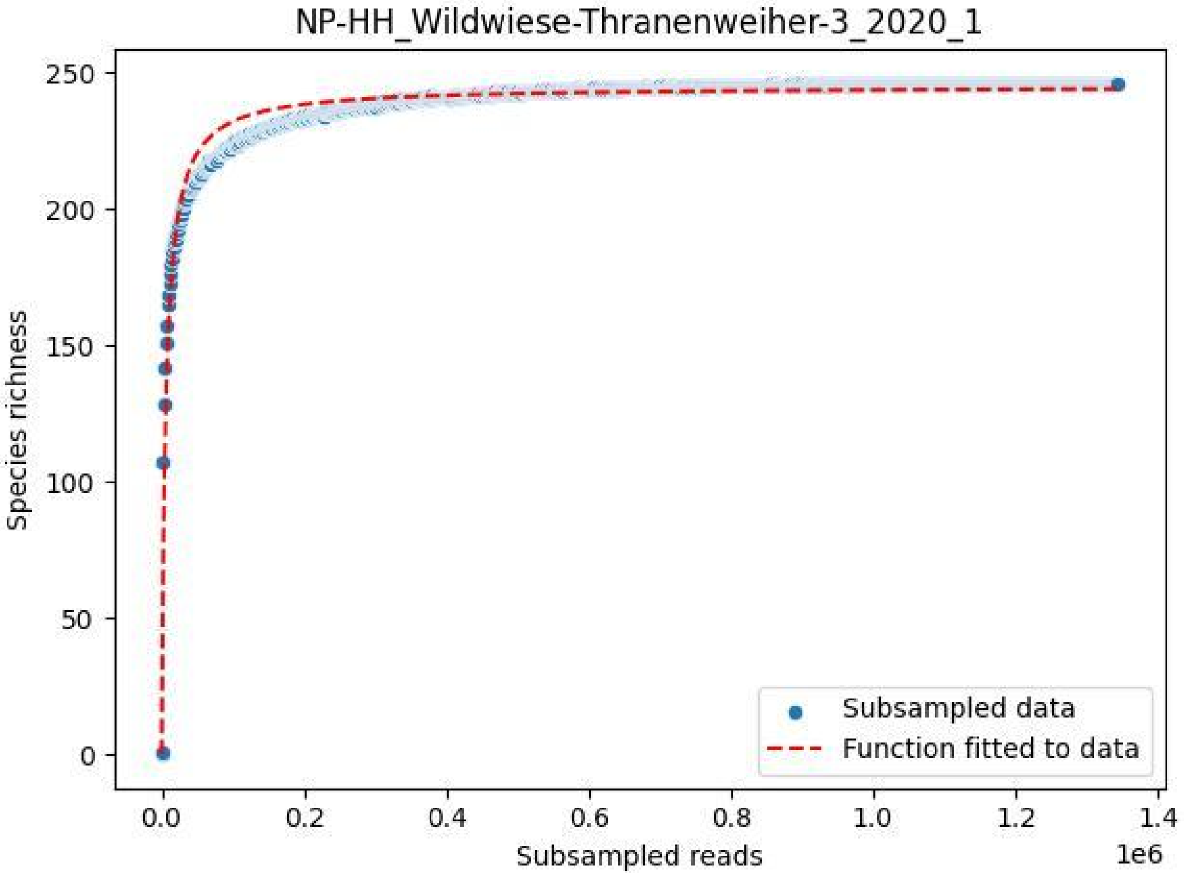

NP-HH_Wildwiese-Thranenweiher-3_2020_1
Species richness
Subsampled reads
1e6
0
50
100
150
200
250
0.0
0.2
0.4
0.6
0.8
1.0
1.2
1.4
Subsampled data
Function fitted to data

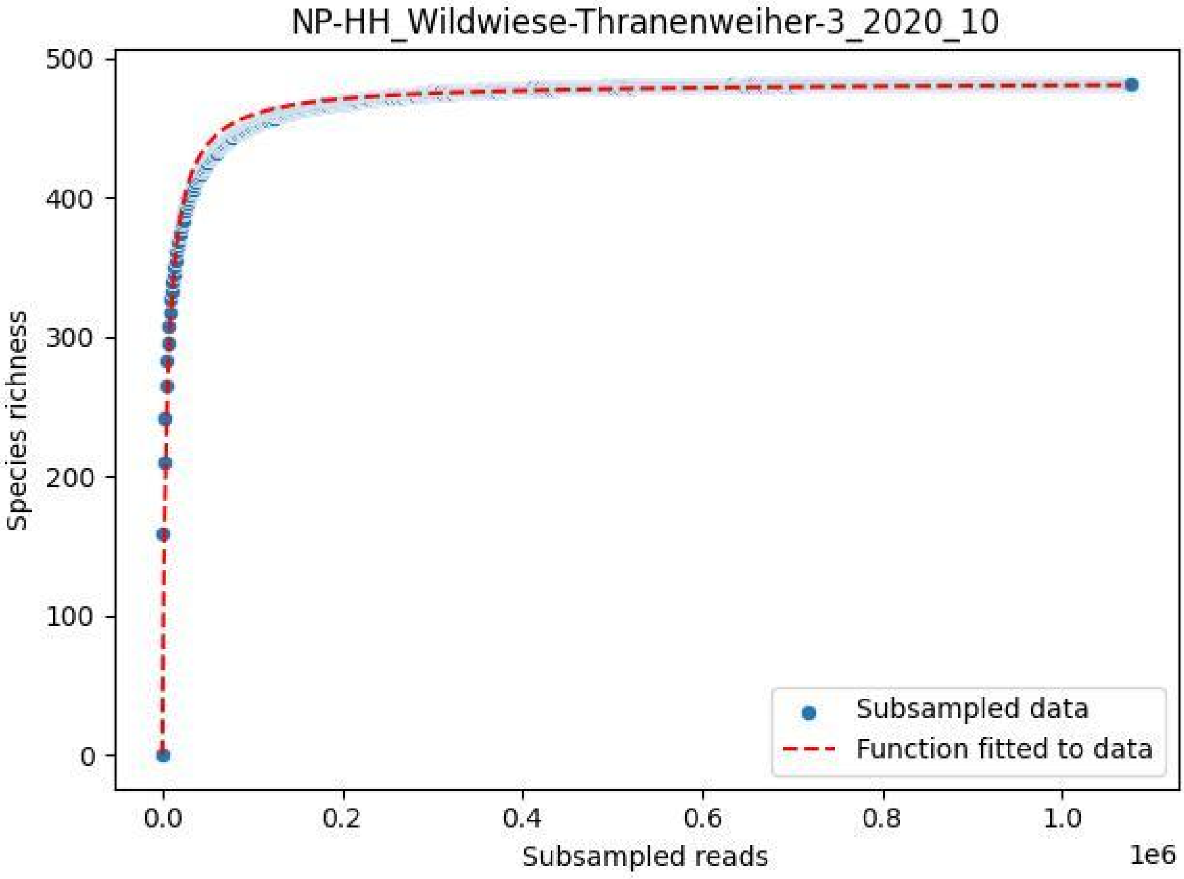
NP-HH_Wildwiese-Thranenweiher-3_2020_10
Species richness
Subsampled reads
1e6
Subsampled data
Function fitted to data
0
100
200
300
400
500
0.0
0.2
0.4
0.6
0.8
1.0

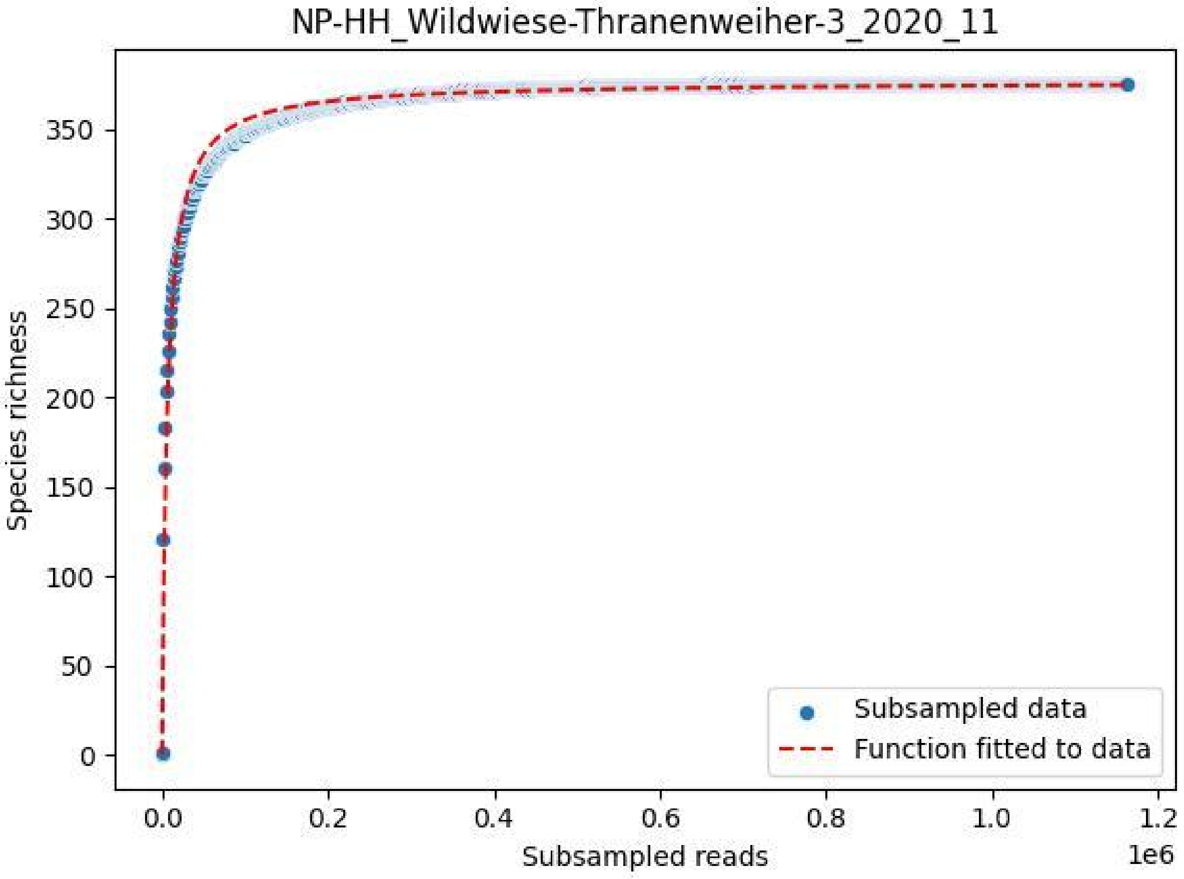
NP-HH_Wildwiese-Thranenweiher-3_2020_11
Species richness
Subsampled reads
1e6
0
50
100
150
200
250
300
350
0.0
0.2
0.4
0.6
0.8
1.0
1.2
Subsampled data
Function fitted to data

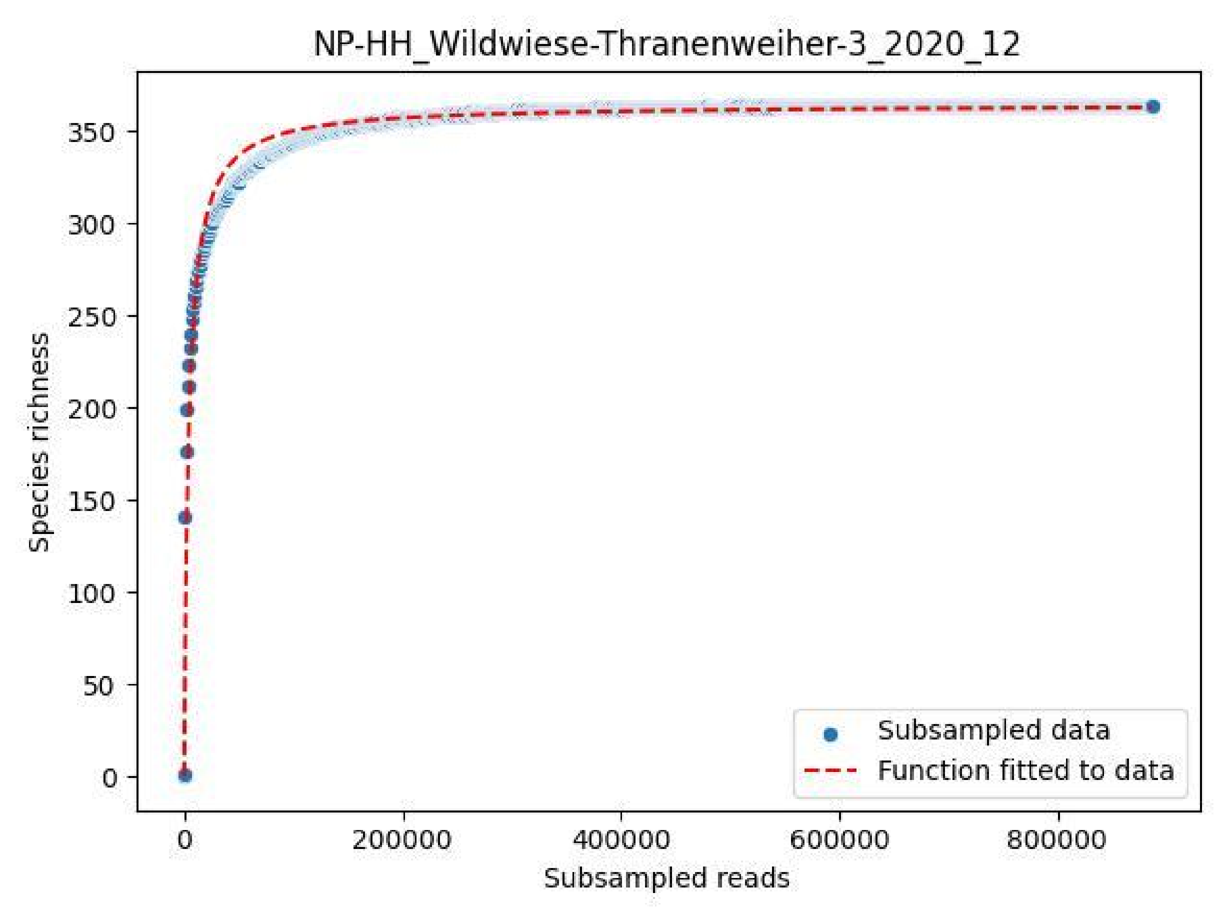

NP-HH_Wildwiese-Thranenweiher-3_2020_12
Species richness
Subsampled reads
0
50
100
150
200
250
300
350
0
200000
400000
600000
800000
Subsampled data
Function fitted to data

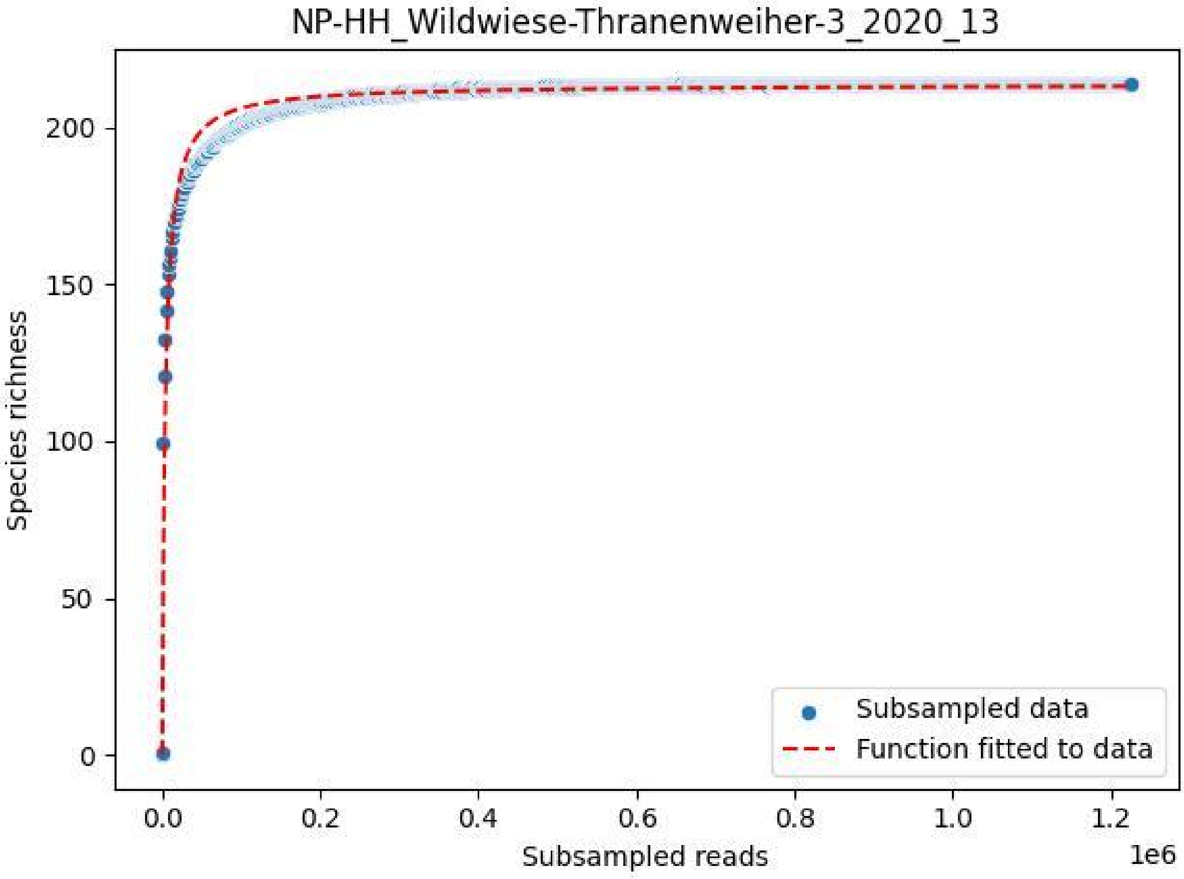

NP-HH_Wildwiese-Thranenweiher-3_2020_13
Species richness
Subsampled reads
1e6
0
50
100
150
200
0.0
0.2
0.4
0.6
0.8
1.0
1.2
Subsampled data
Function fitted to data

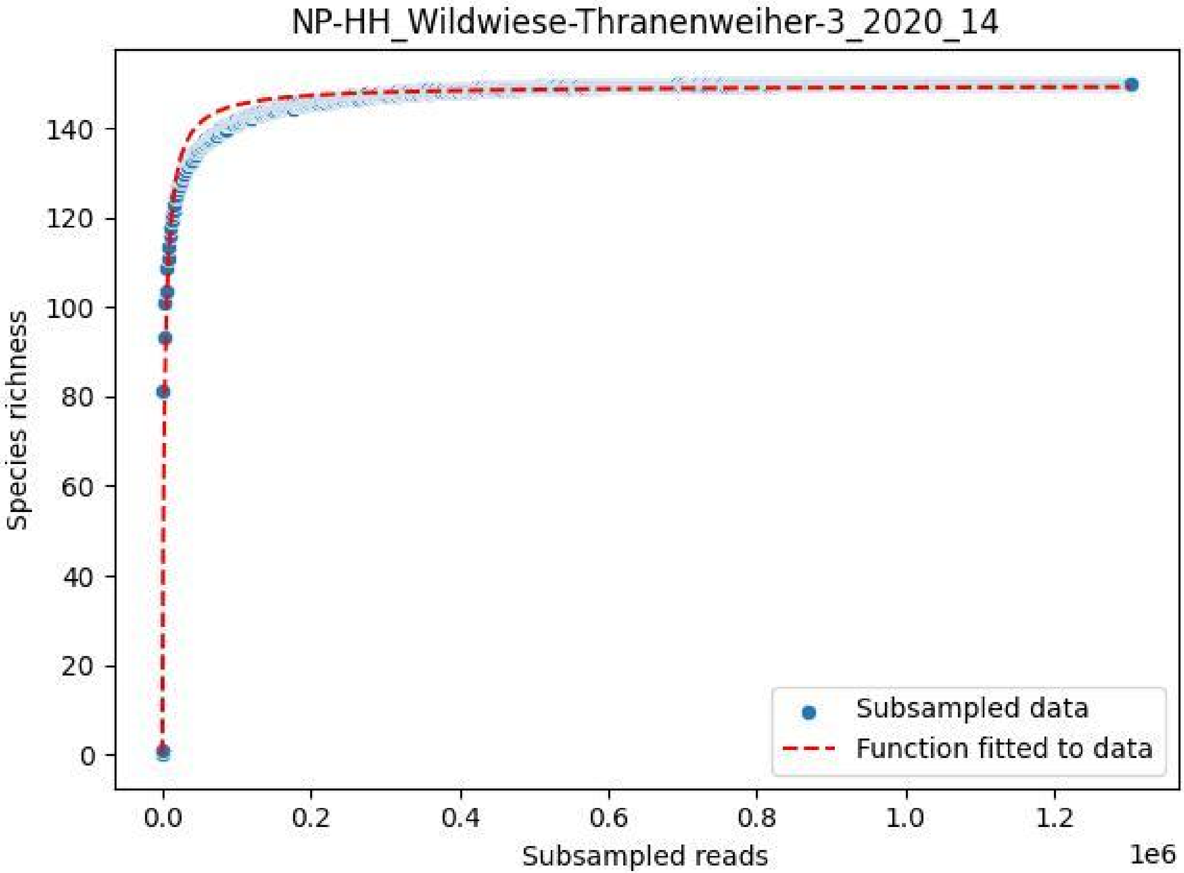

NP-HH_Wildwiese-Thranenweiher-3_2020_14
Species richness
Subsampled reads
1e6
0
20
40
60
80
100
120
140
0.0
0.2
0.4
0.6
0.8
1.0
1.2
Subsampled data
Function fitted to data

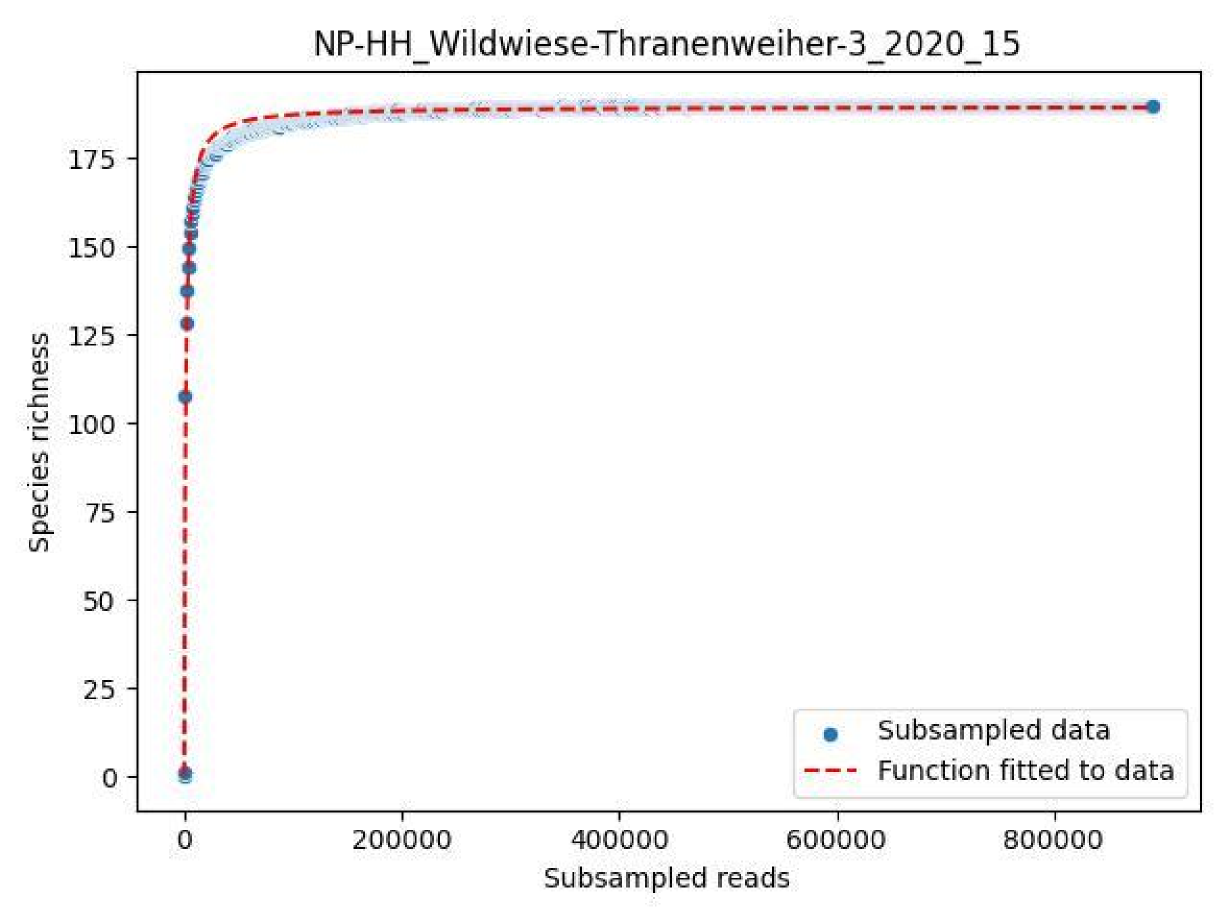

NP-HH_Wildwiese-Thranenweiher-3_2020_15
Species richness
Subsampled reads
0
25
50
75
100
125
150
175
0
200000
400000
600000
800000
Subsampled data
Function fitted to data

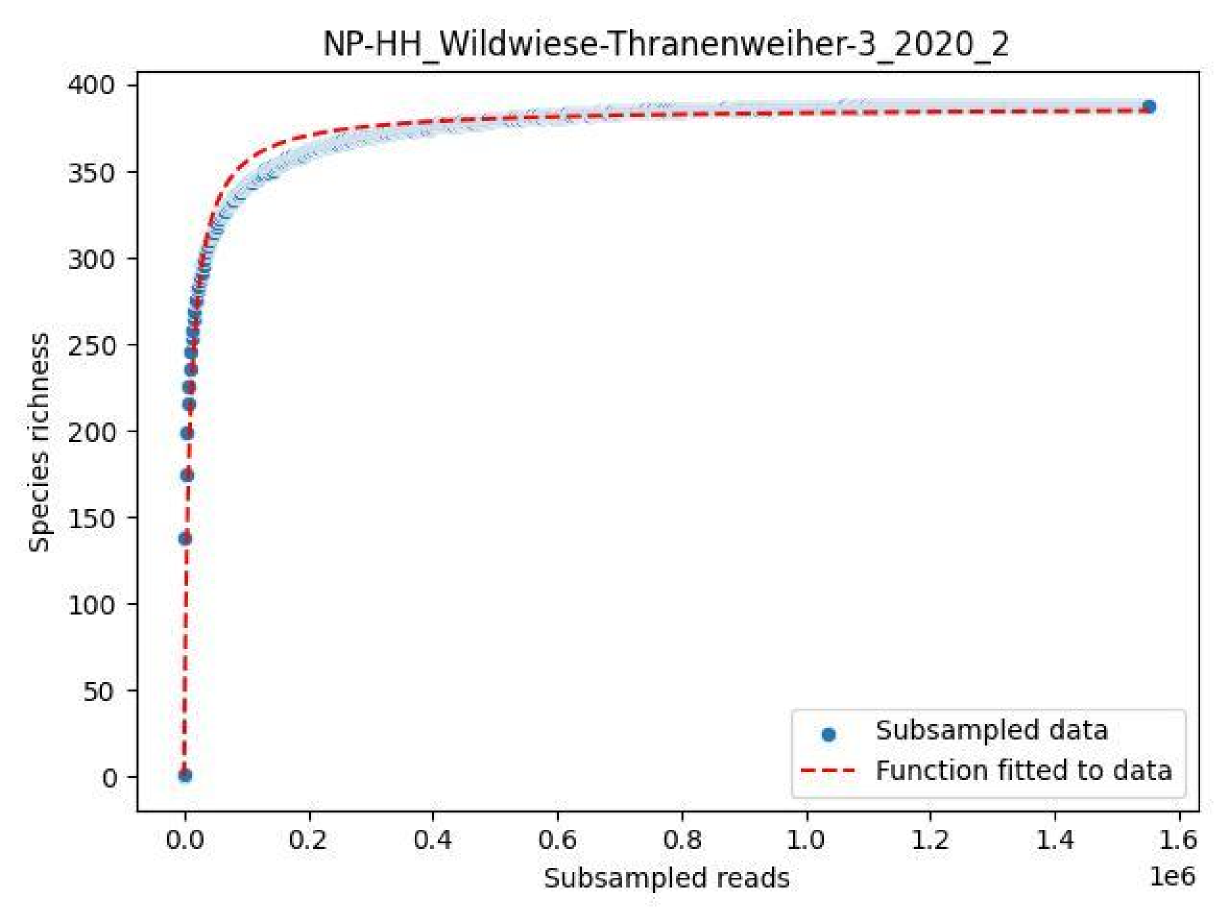
NP-HH_Wildwiese-Thranenweiher-3_2020_2
Species richness
400
350
300
250
200
150
100
50
0
0.0
0.2
0.4
0.6
0.8
1.0
1.2
1.4
1.6
1e6
Subsampled reads
Subsampled data
Function fitted to data

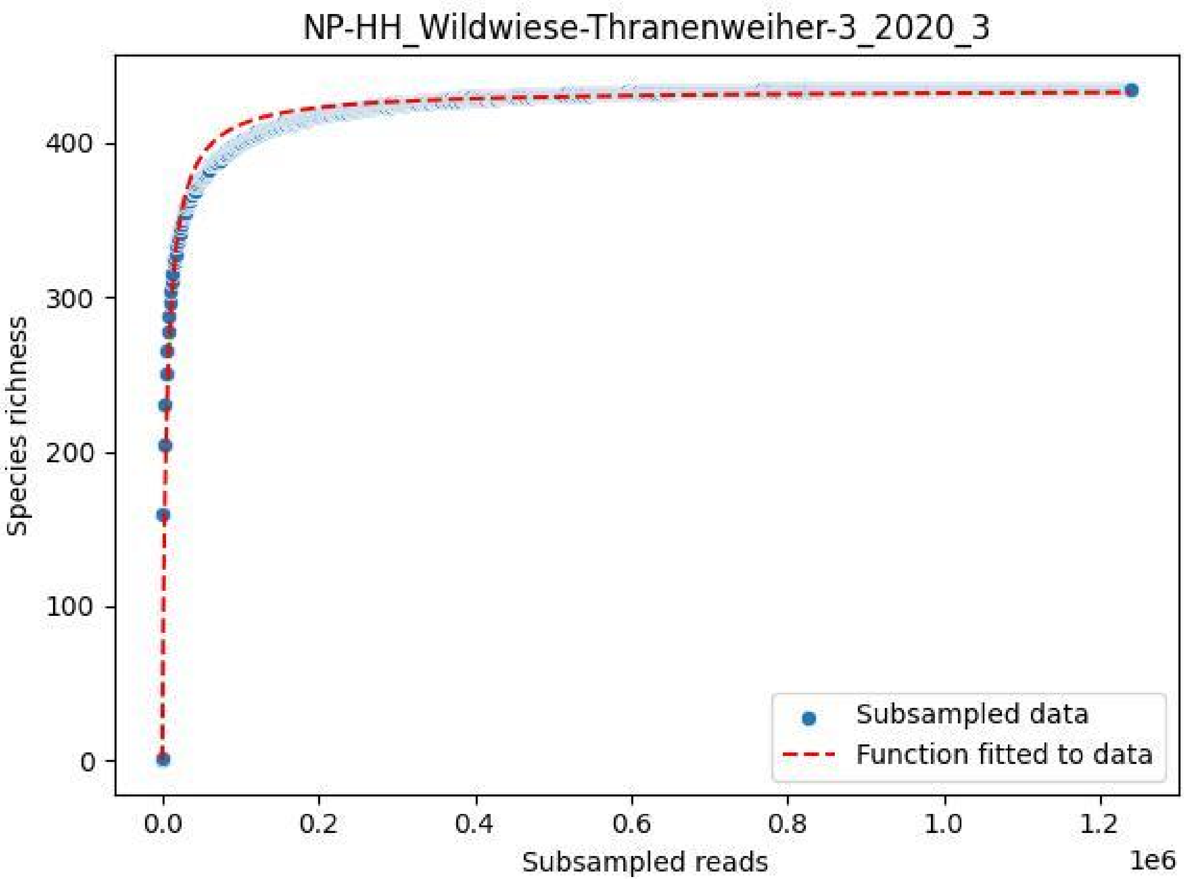
NP-HH_Wildwiese-Thranenweiher-3_2020_3
Species richness
Subsampled reads
1e6
0
100
200
300
400
0.0
0.2
0.4
0.6
0.8
1.0
1.2
Subsampled data
Function fitted to data

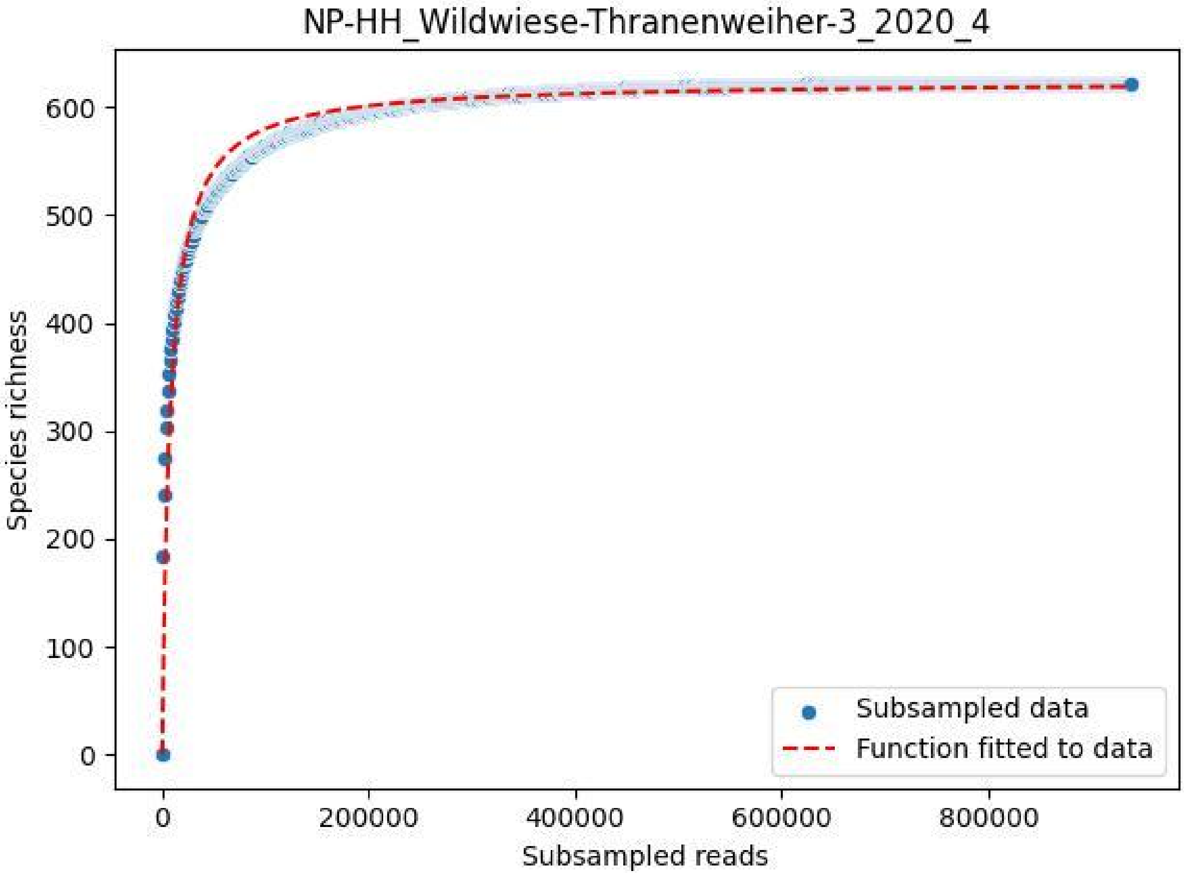
NP-HH_Wildwiese-Thranenweiher-3_2020_4
Species richness
Subsampled reads
0
100
200
300
400
500
600
0
200000
400000
600000
800000
Subsampled data
Function fitted to data

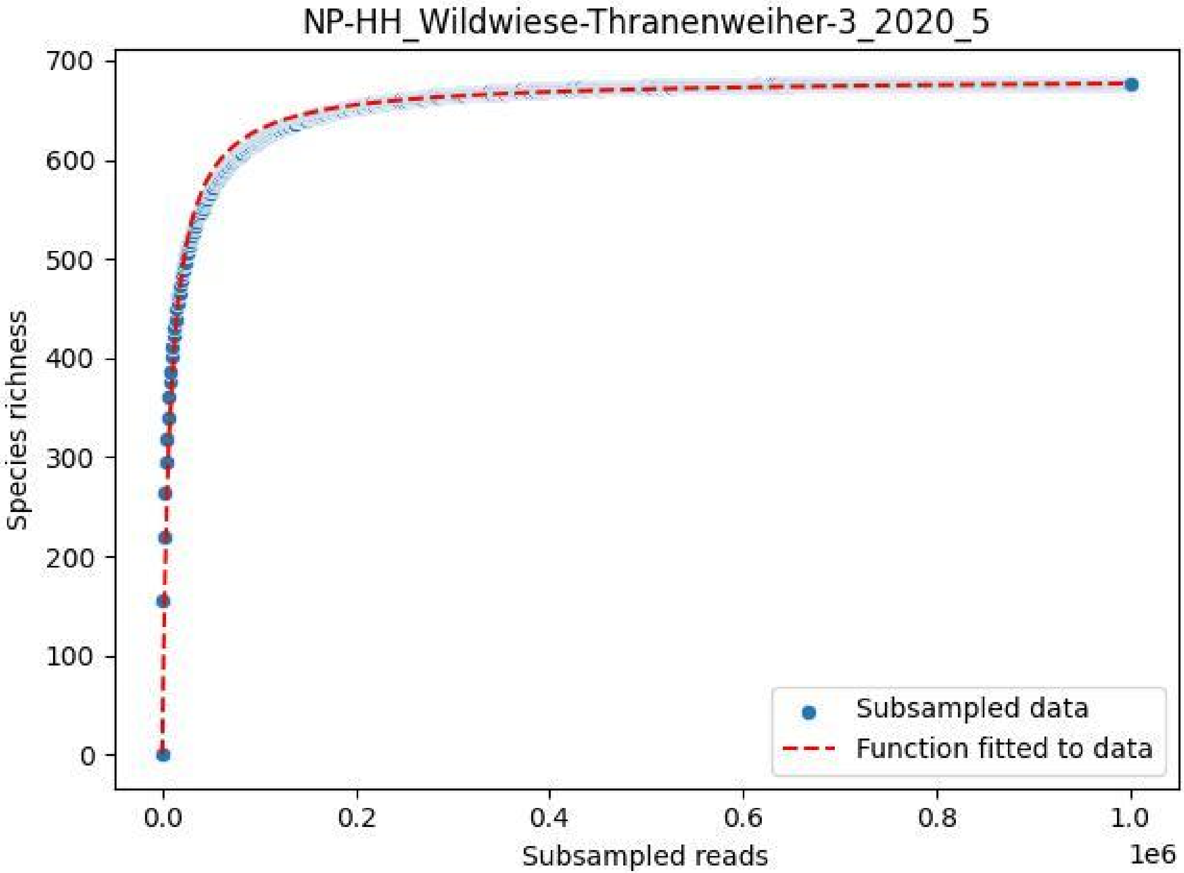
NP-HH_Wildwiese-Thranenweiher-3_2020_5
Species richness
Subsampled reads
1e6
Subsampled data
Function fitted to data
0
100
200
300
400
500
600
700
0.0
0.2
0.4
0.6
0.8
1.0

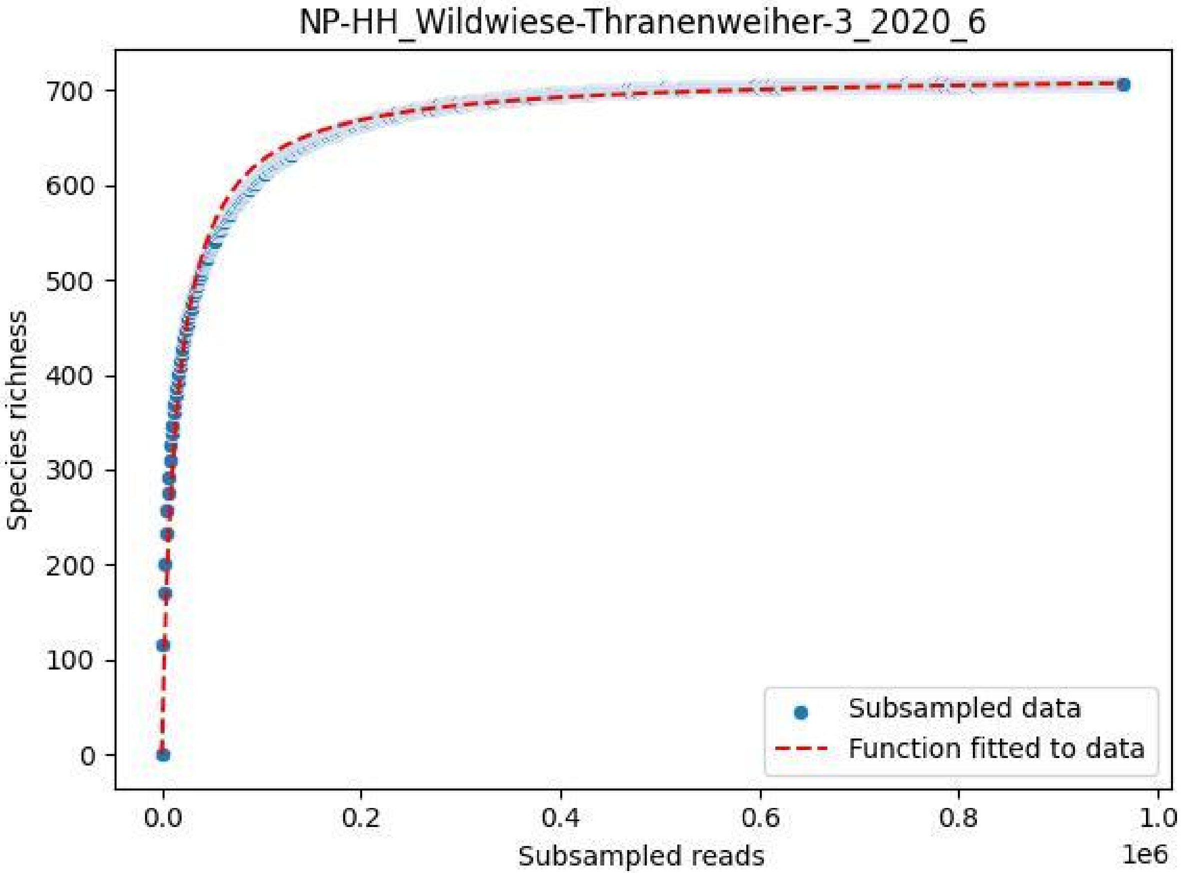

NP-HH_Wildwiese-Thranenweiher-3_2020_6
Species richness
Subsampled reads
1e6
0
100
200
300
400
500
600
700
0.0
0.2
0.4
0.6
0.8
1.0
Subsampled data
Function fitted to data

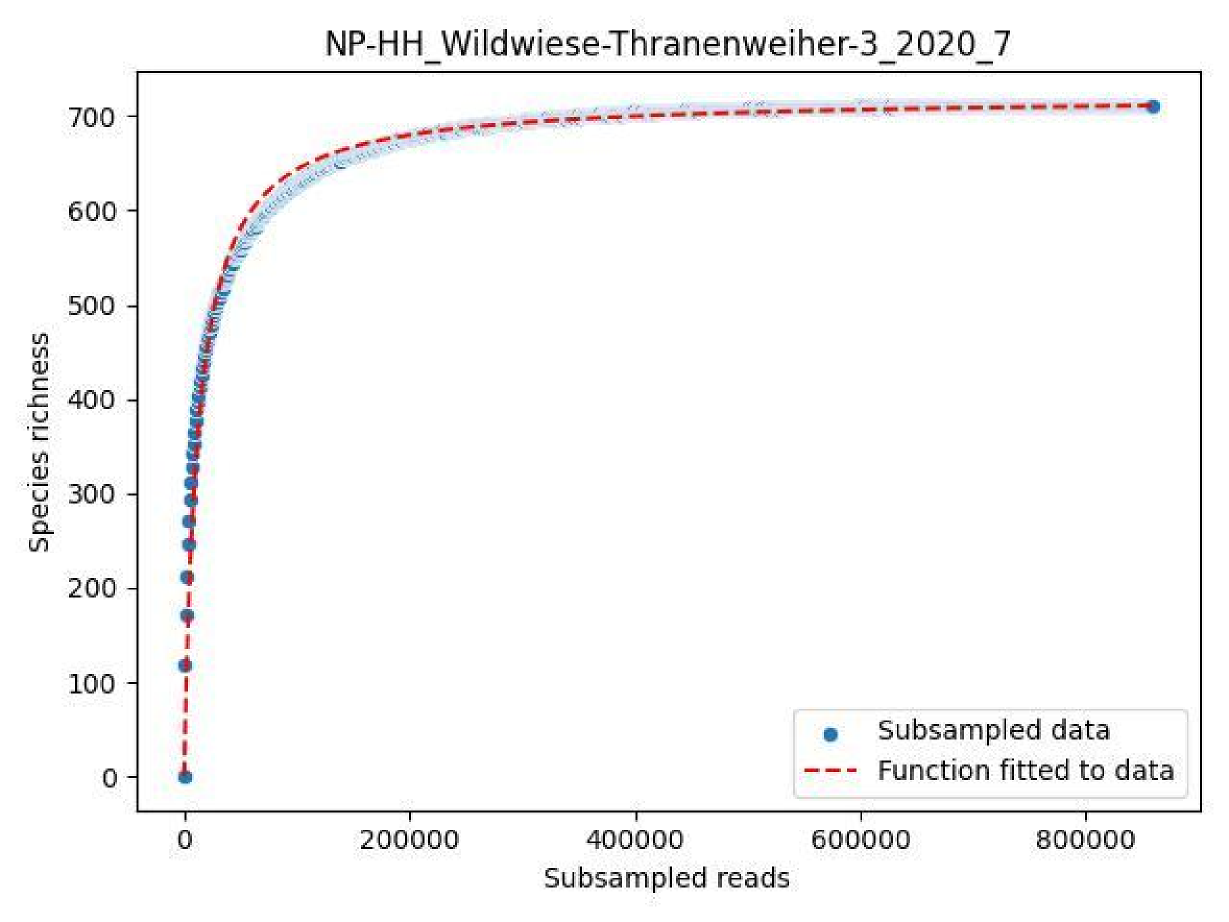

NP-HH_Wildwiese-Thranenweiher-3_2020_7
Species richness
Subsampled reads
0
100
200
300
400
500
600
700
0
200000
400000
600000
800000
Subsampled data
Function fitted to data

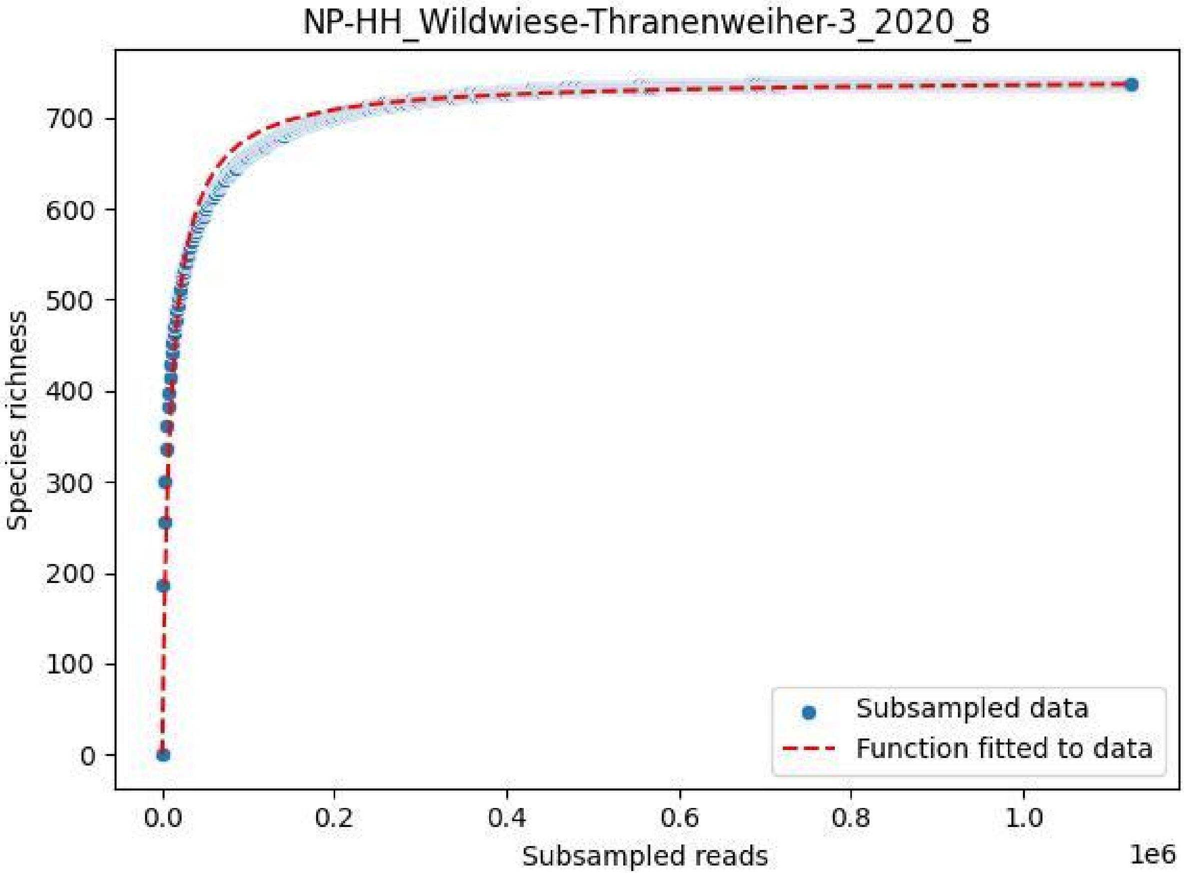

NP-HH_Wildwiese-Thranenweiher-3_2020_8
Species richness
0
100
200
300
400
500
600
700
0.0
0.2
0.4
0.6
0.8
1.0
1e6
Subsampled reads
Subsampled data
Function fitted to data

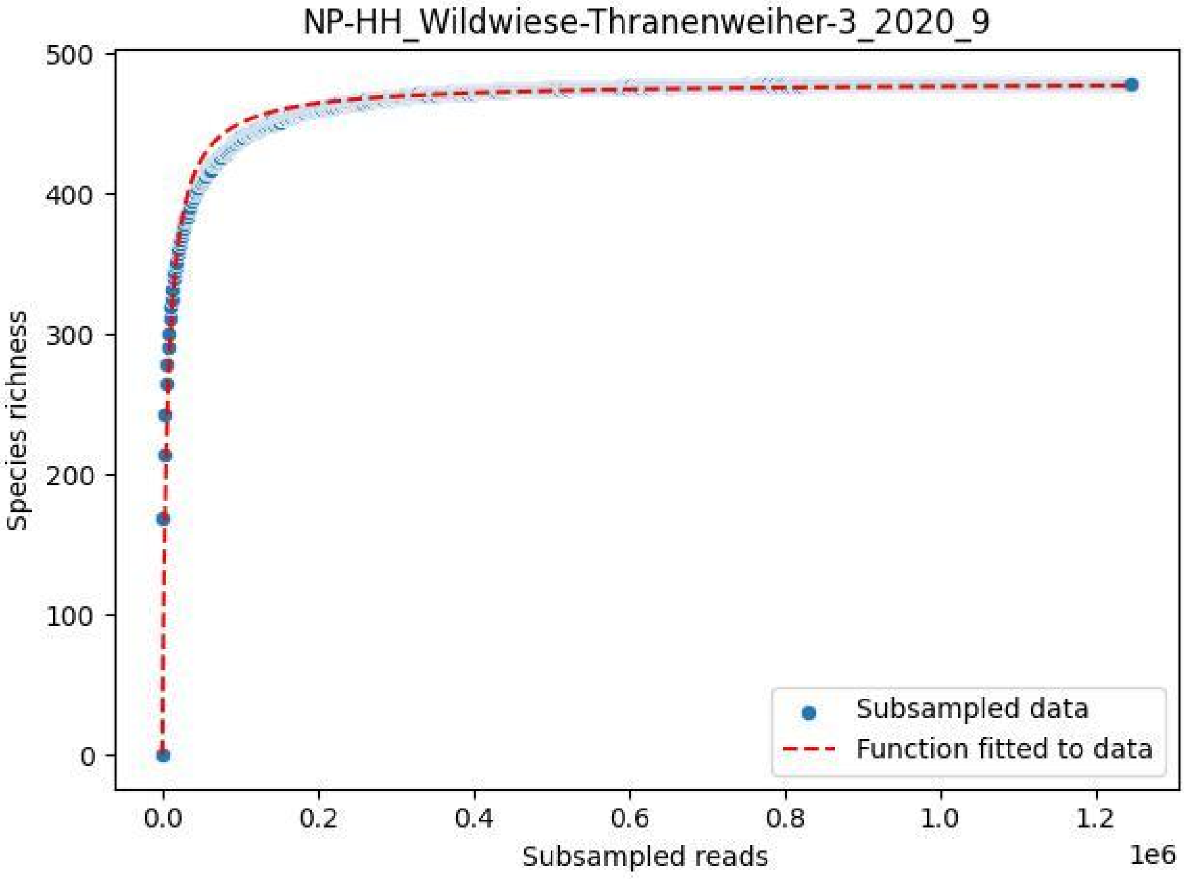

NP-HH_Wildwiese-Thranenweiher-3_2020_9
Species richness
Subsampled reads
1e6
0
100
200
300
400
500
0.0
0.2
0.4
0.6
0.8
1.0
1.2
Subsampled data
Function fitted to data

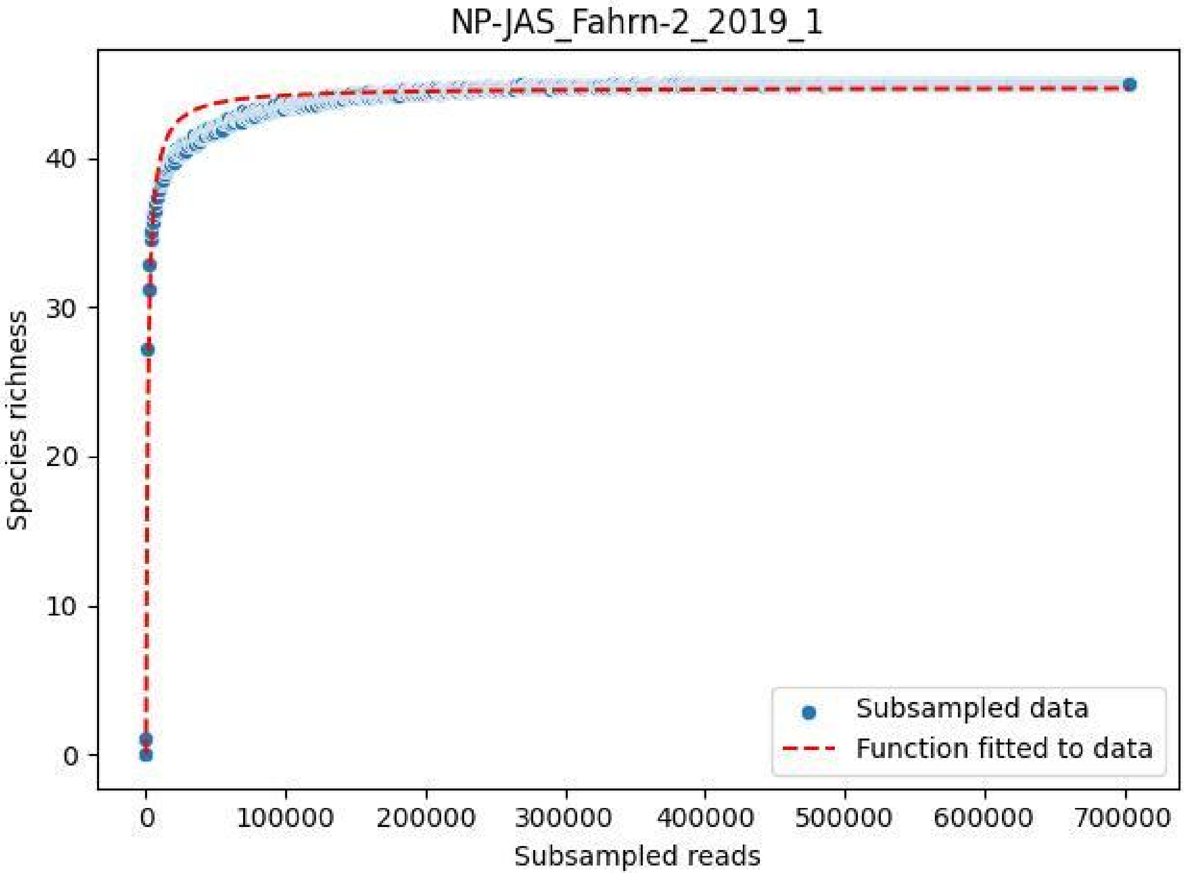

NP-JAS_Fahrn-2_2019_1
Species richness
Subsampled reads
0
10
20
30
40
0
100000
200000
300000
400000
500000
600000
700000
Subsampled data
Function fitted to data

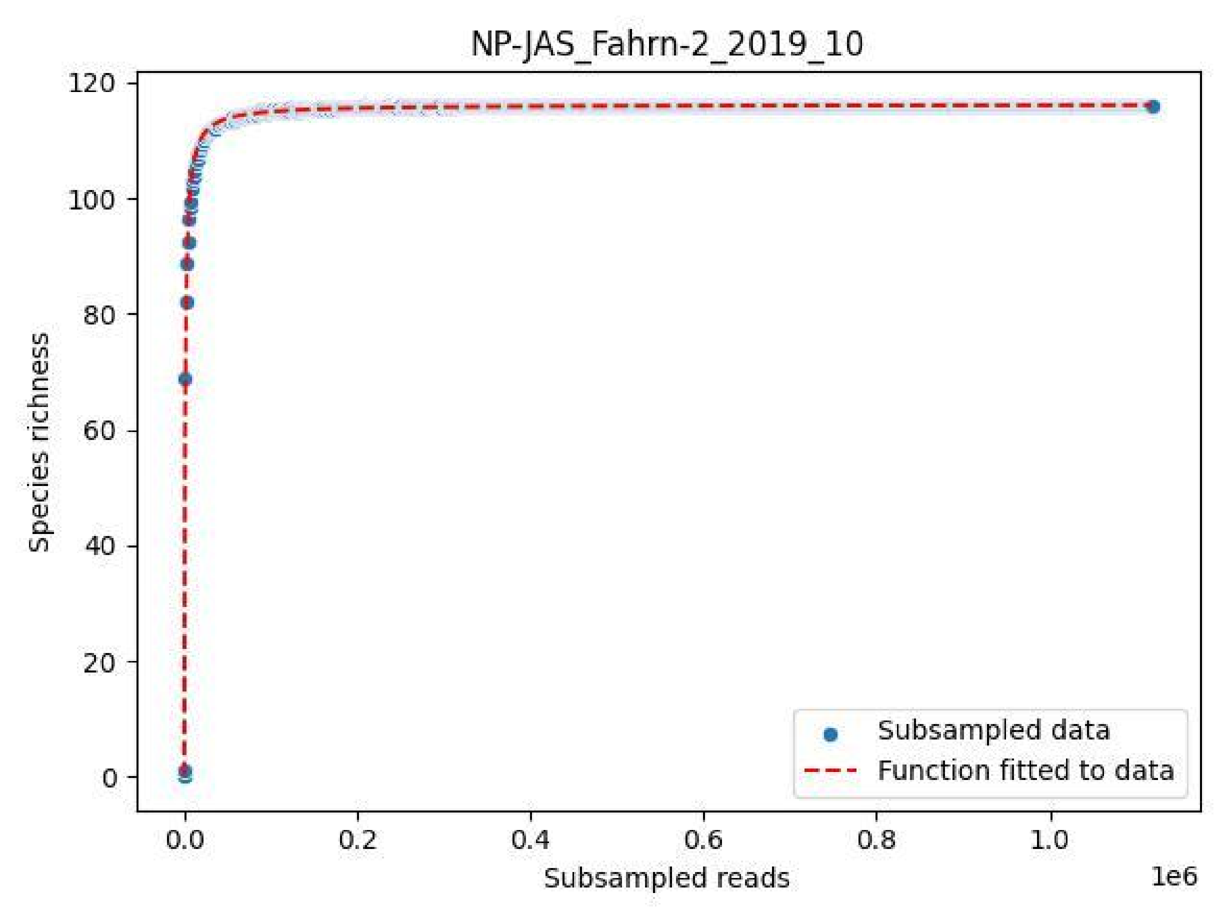
NP-JAS_Fahrn-2_2019_10
Species richness
Subsampled reads
1e6
0
20
40
60
80
100
120
0.0
0.2
0.4
0.6
0.8
1.0
Subsampled data
Function fitted to data

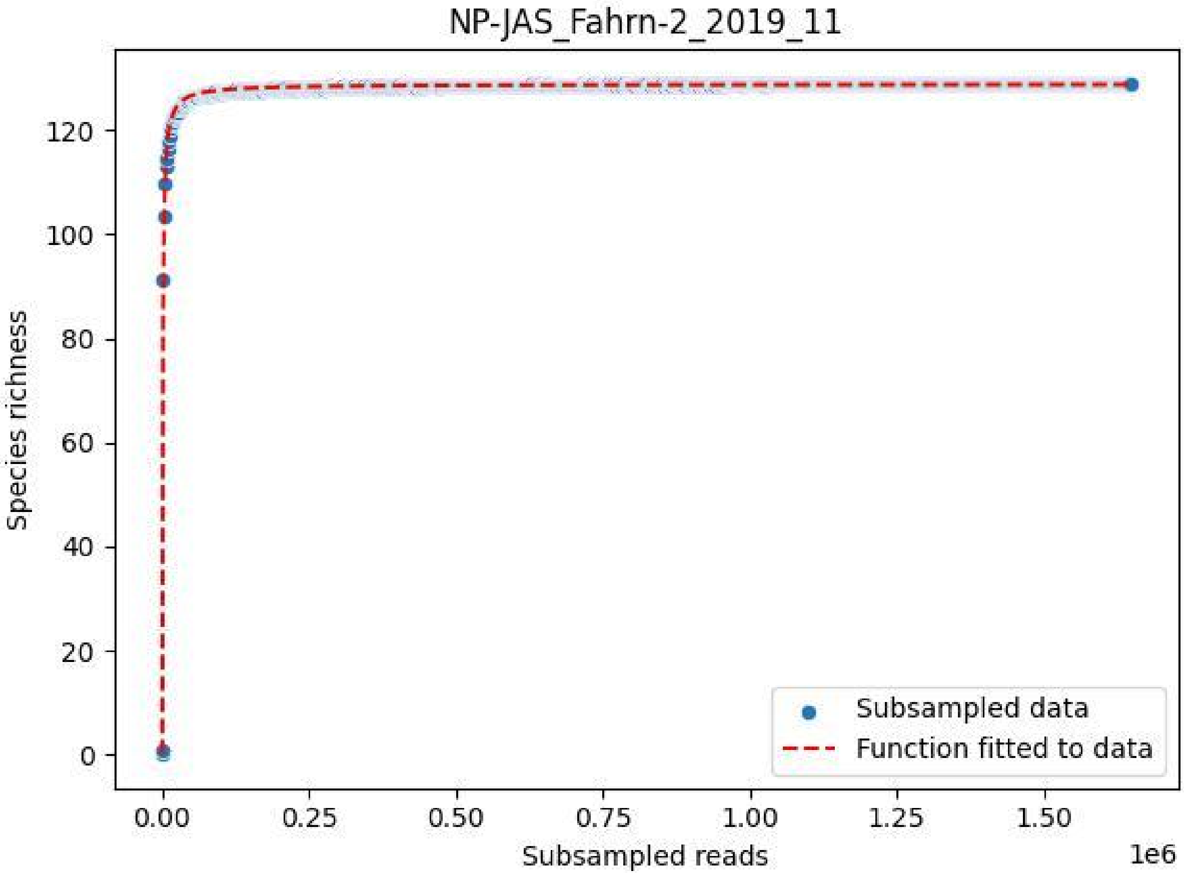

NP-JAS_Fahrn-2_2019_11
Species richness
Subsampled reads
1e6
0
20
40
60
80
100
120
0.00
0.25
0.50
0.75
1.00
1.25
1.50
Subsampled data
Function fitted to data

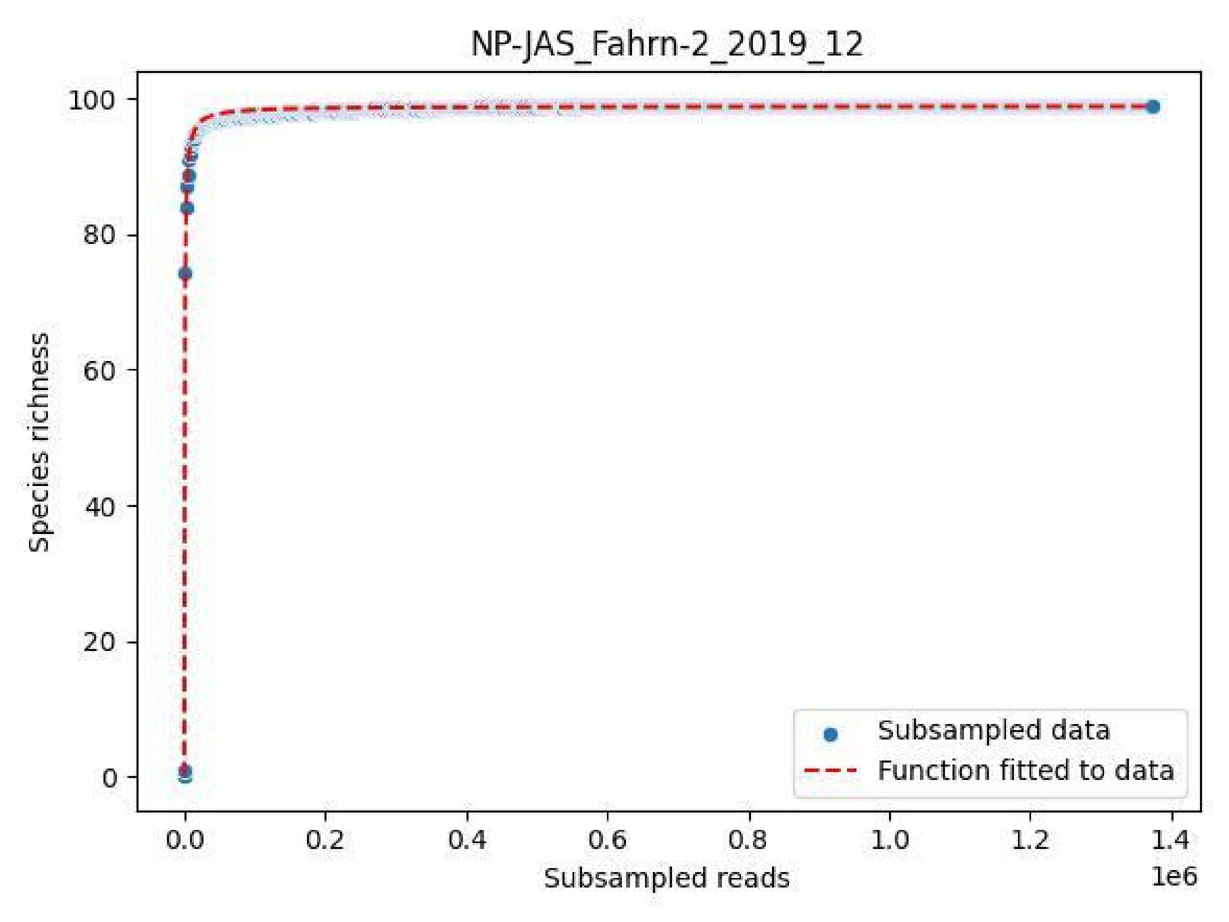

NP-JAS_Fahrn-2_2019_12
Species richness
Subsampled reads
1e6
0
20
40
60
80
100
0.0
0.2
0.4
0.6
0.8
1.0
1.2
1.4
Subsampled data
Function fitted to data

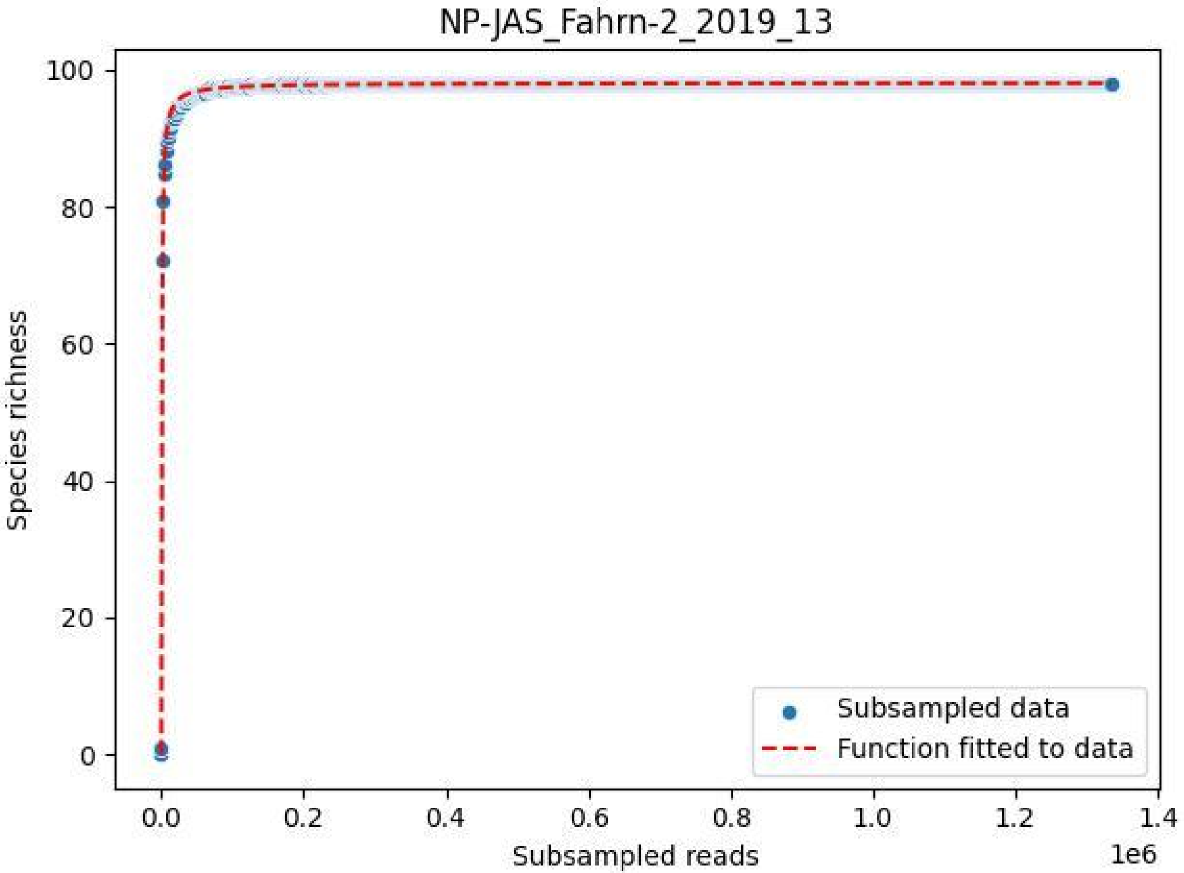

NP-JAS_Fahrn-2_2019_13
100
80
60
40
20
0
Species richness
0.0
0.2
0.4
0.6
0.8
1.0
1.2
1.4
1e6
Subsampled reads
Subsampled data
Function fitted to data

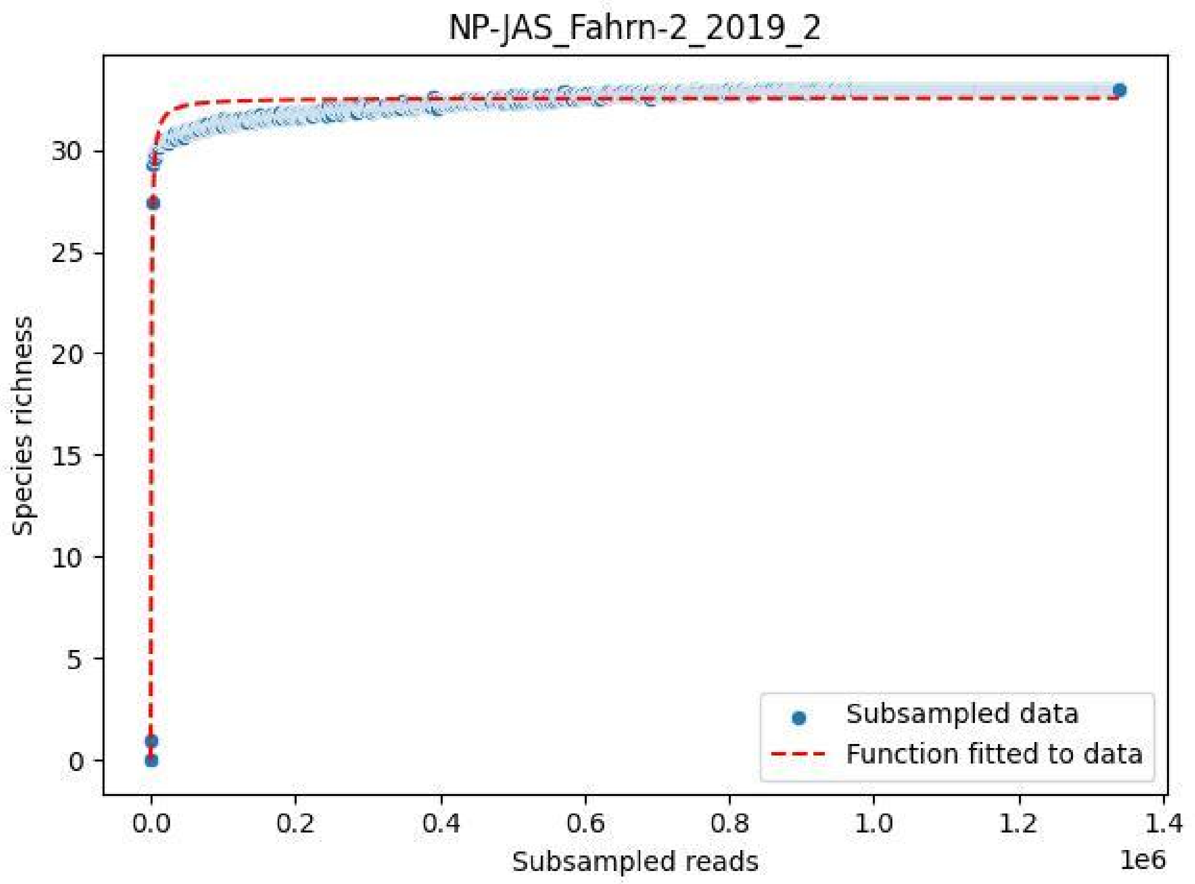
NP-JAS_Fahrn-2_2019_2
Species richness
Subsampled reads
1e6
0
5
10
15
20
25
30
0.0
0.2
0.4
0.6
0.8
1.0
1.2
1.4
Subsampled data
Function fitted to data

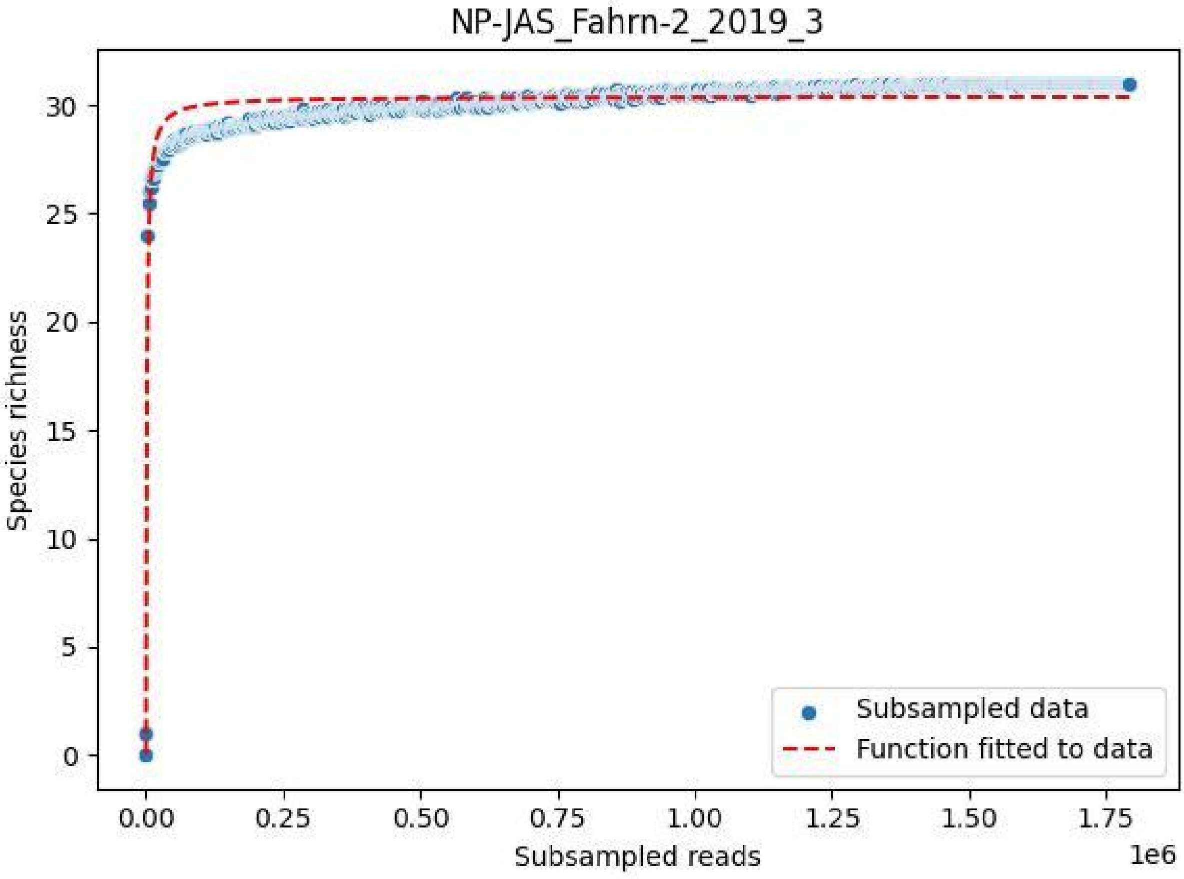

NP-JAS_Fahrn-2_2019_3
Species richness
0
5
10
15
20
25
30
0.00
0.25
0.50
0.75
1.00
1.25
1.50
1.75
1e6
Subsampled reads
Subsampled data
Function fitted to data

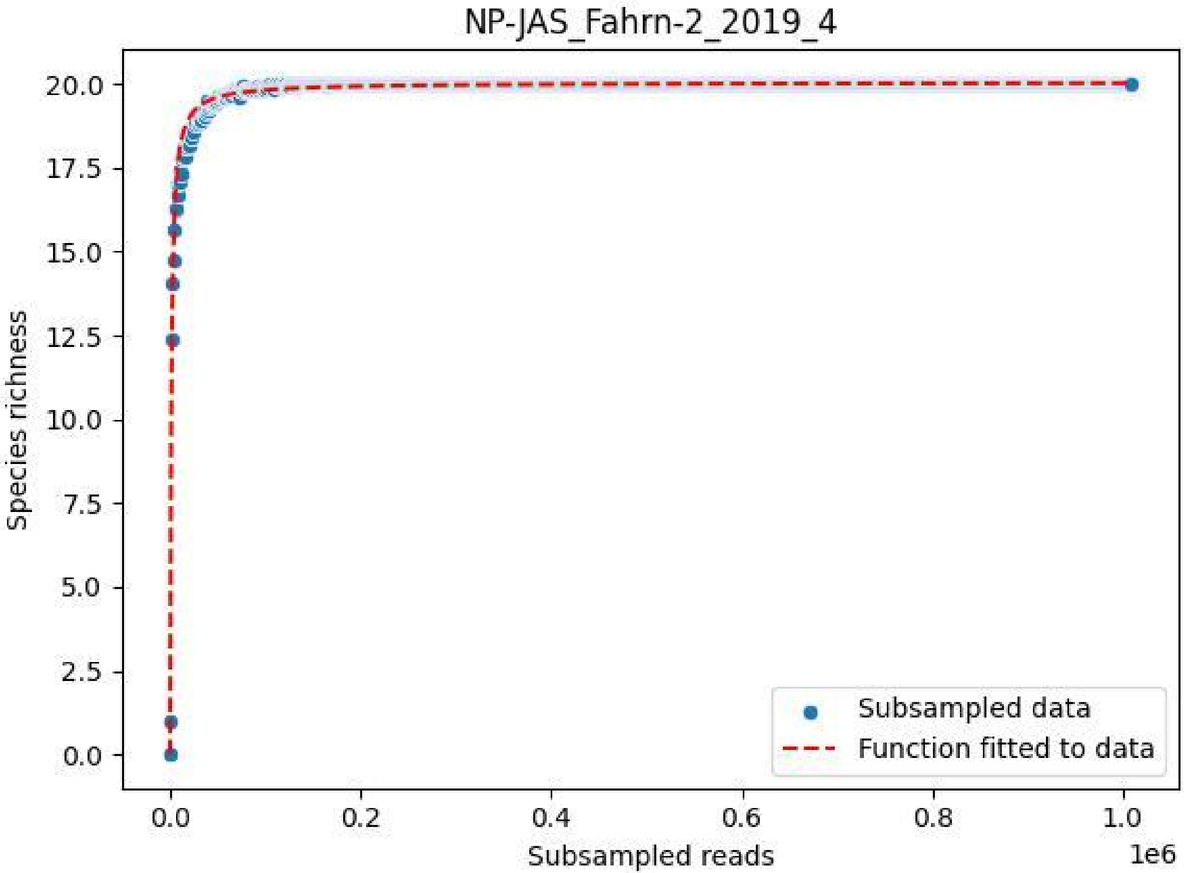

NP-JAS_Fahrn-2_2019_4
Species richness
Subsampled reads
1e6
Subsampled data
Function fitted to data
0.0
2.5
5.0
7.5
10.0
12.5
15.0
17.5
20.0
0.0
0.2
0.4
0.6
0.8
1.0

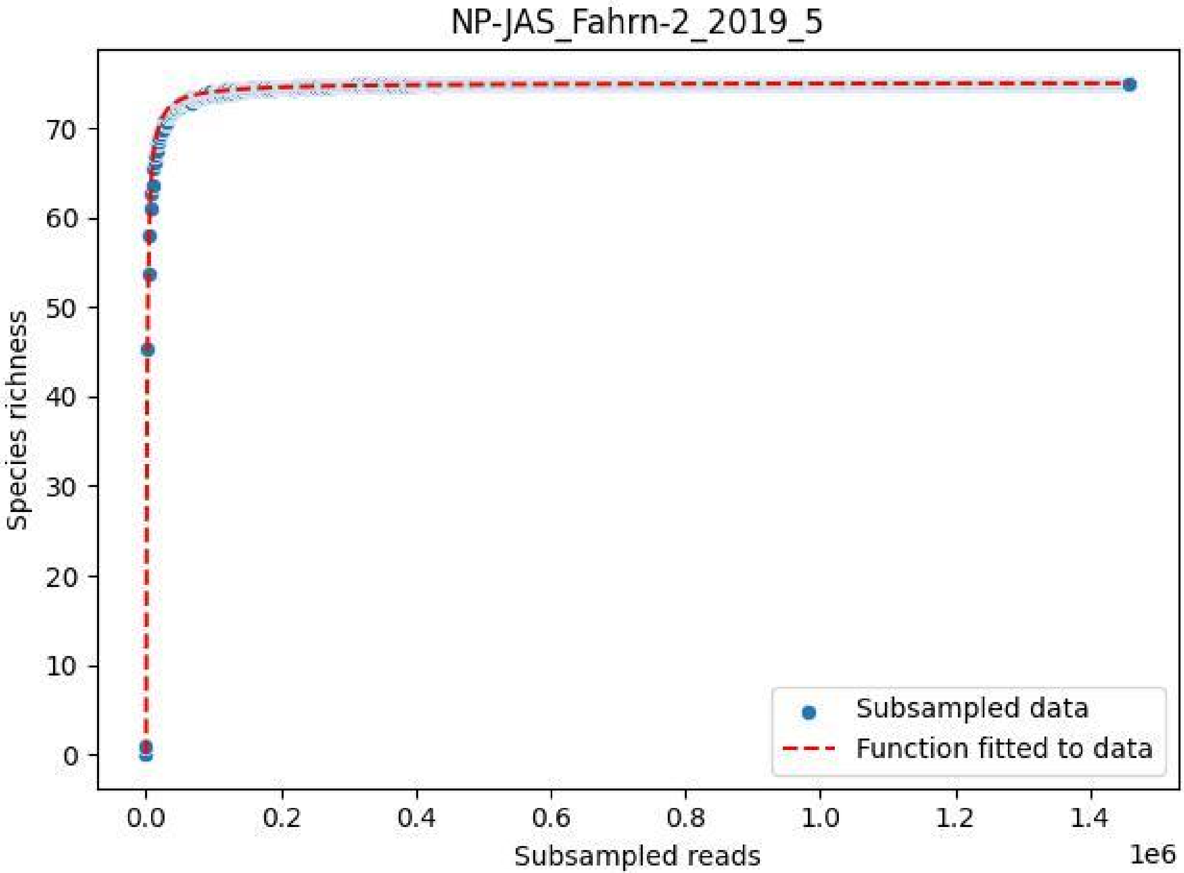
NP-JAS_Fahrn-2_2019_5
Species richness
Subsampled reads
1e6
0
10
20
30
40
50
60
70
0.0
0.2
0.4
0.6
0.8
1.0
1.2
1.4
Subsampled data
Function fitted to data

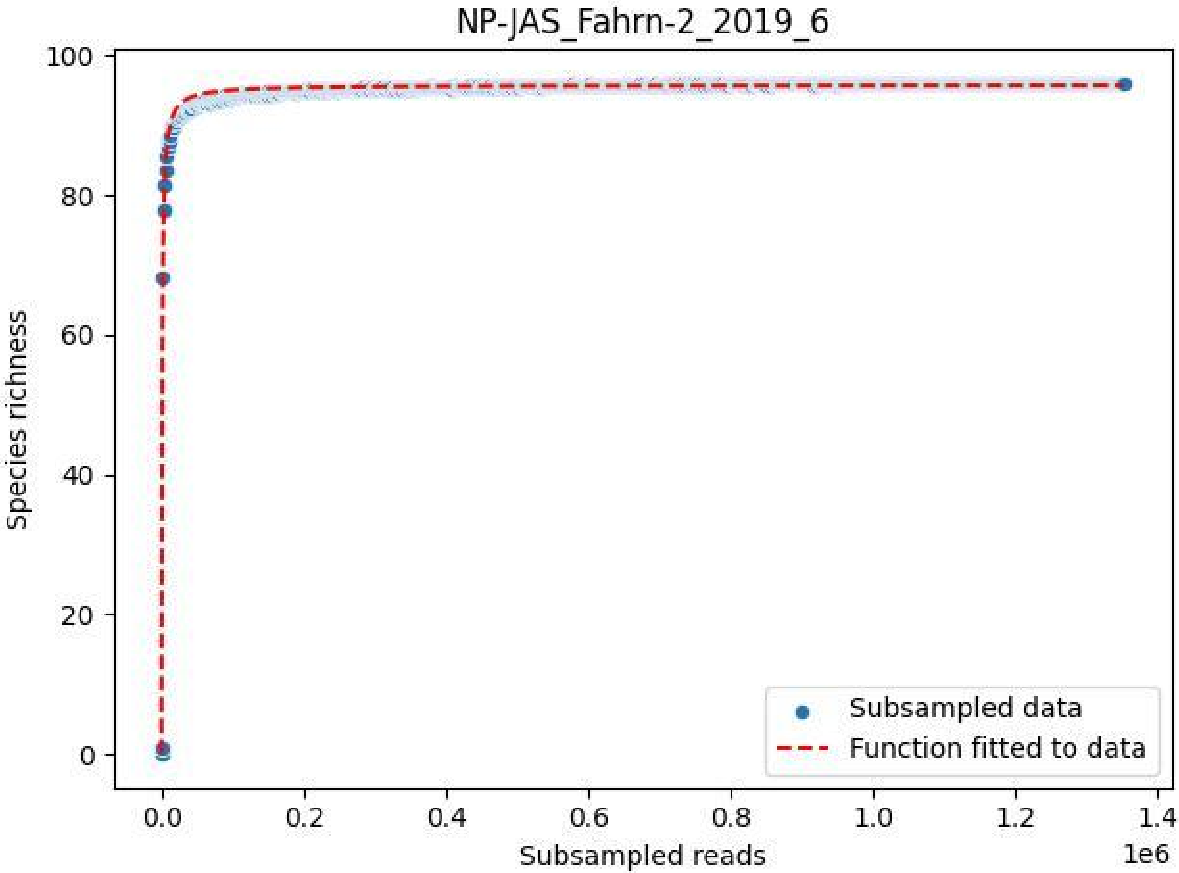

NP-JAS_Fahrn-2_2019_6
Species richness
Subsampled reads
1e6
0
20
40
60
80
100
0.0
0.2
0.4
0.6
0.8
1.0
1.2
1.4
Subsampled data
Function fitted to data

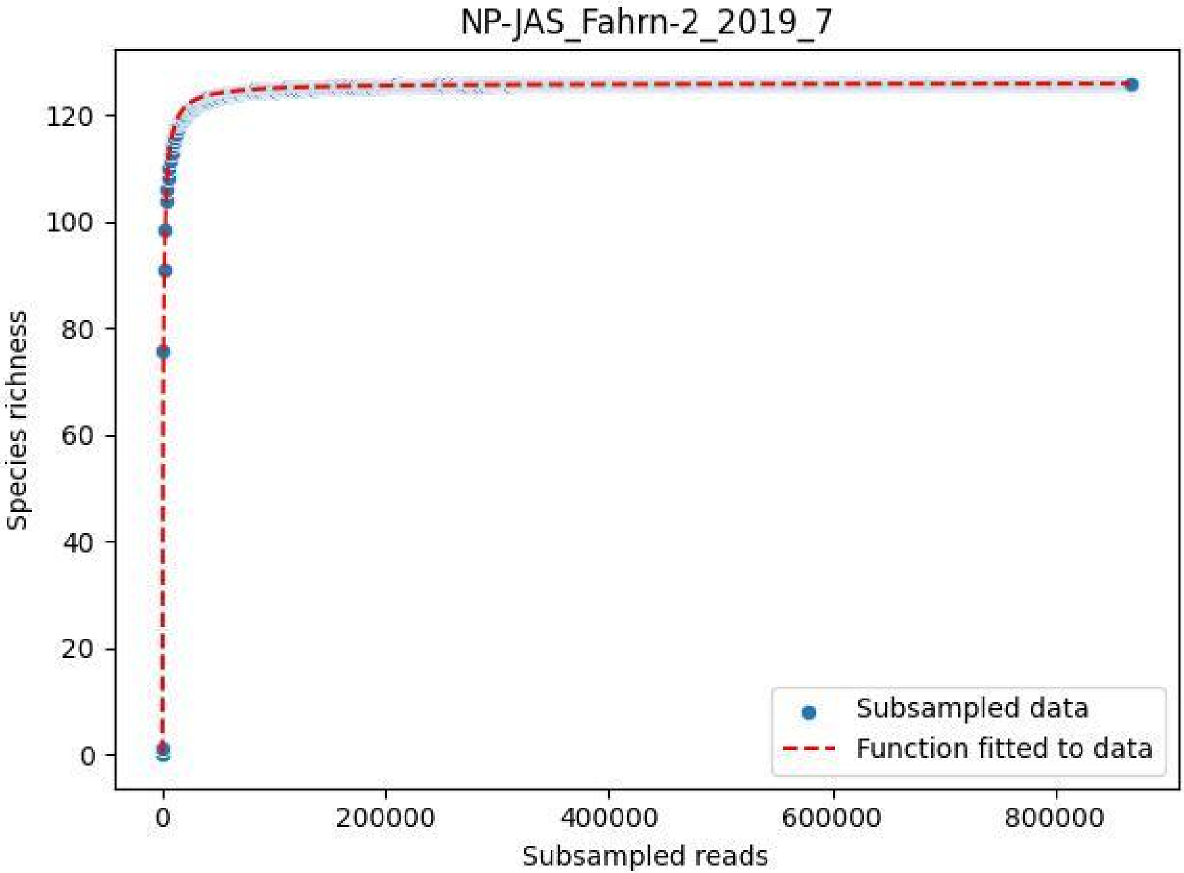

NP-JAS_Fahrn-2_2019_7
Species richness
Subsampled reads
0
20
40
60
80
100
120
0
200000
400000
600000
800000
Subsampled data
Function fitted to data

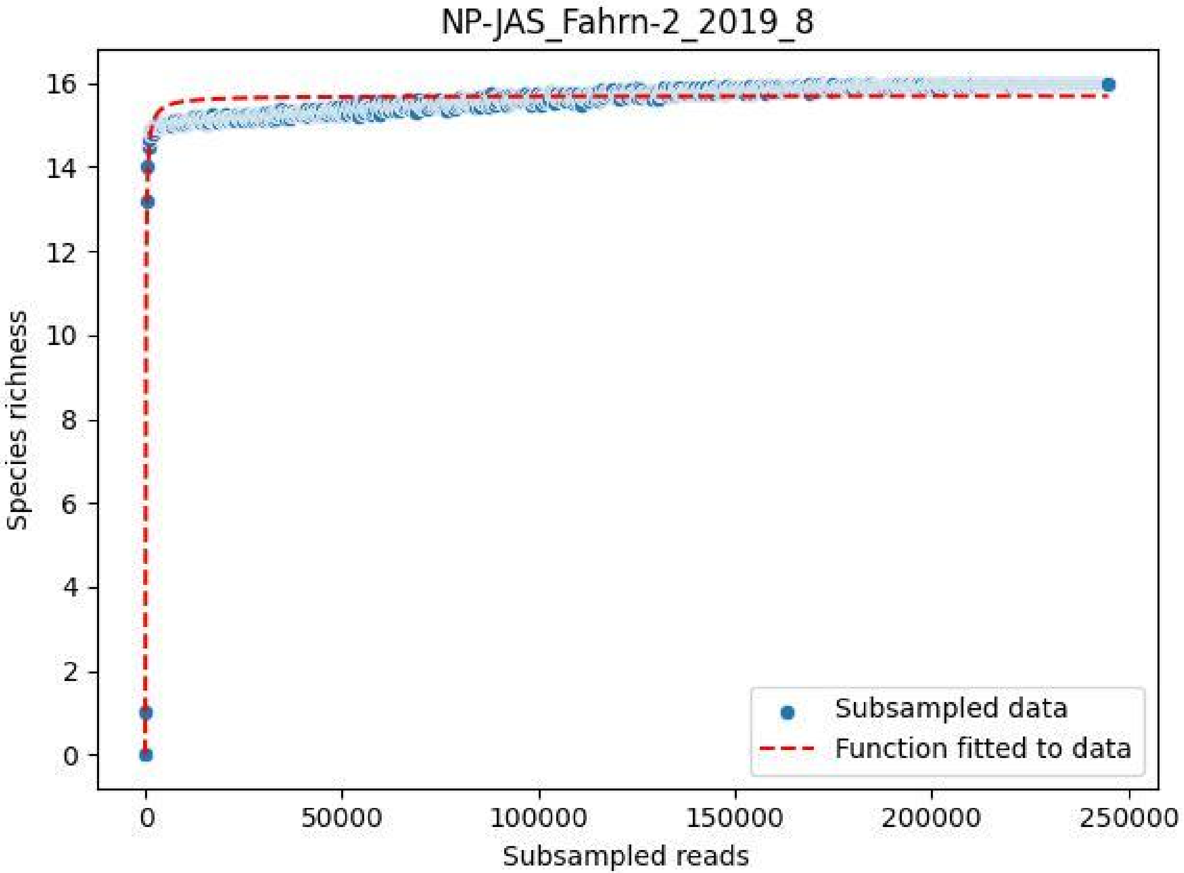
NP-JAS_Fahrn-2_2019_8
Species richness
Subsampled reads
0
2
4
6
8
10
12
14
16
0
50000
100000
150000
200000
250000
Subsampled data
Function fitted to data

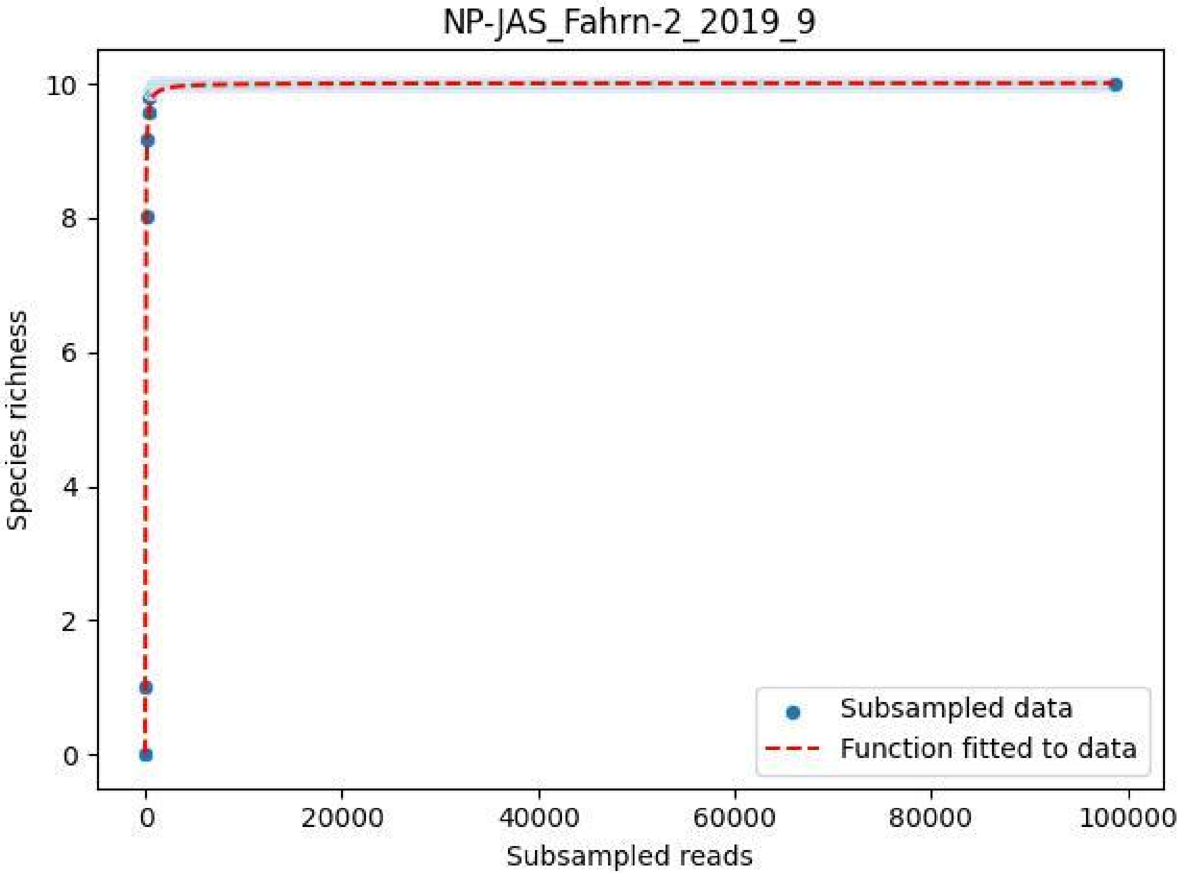

NP-JAS_Fahrn-2_2019_9
Species richness
Subsampled reads
0
2
4
6
8
10
0
20000
40000
60000
80000
100000
Subsampled data
Function fitted to data

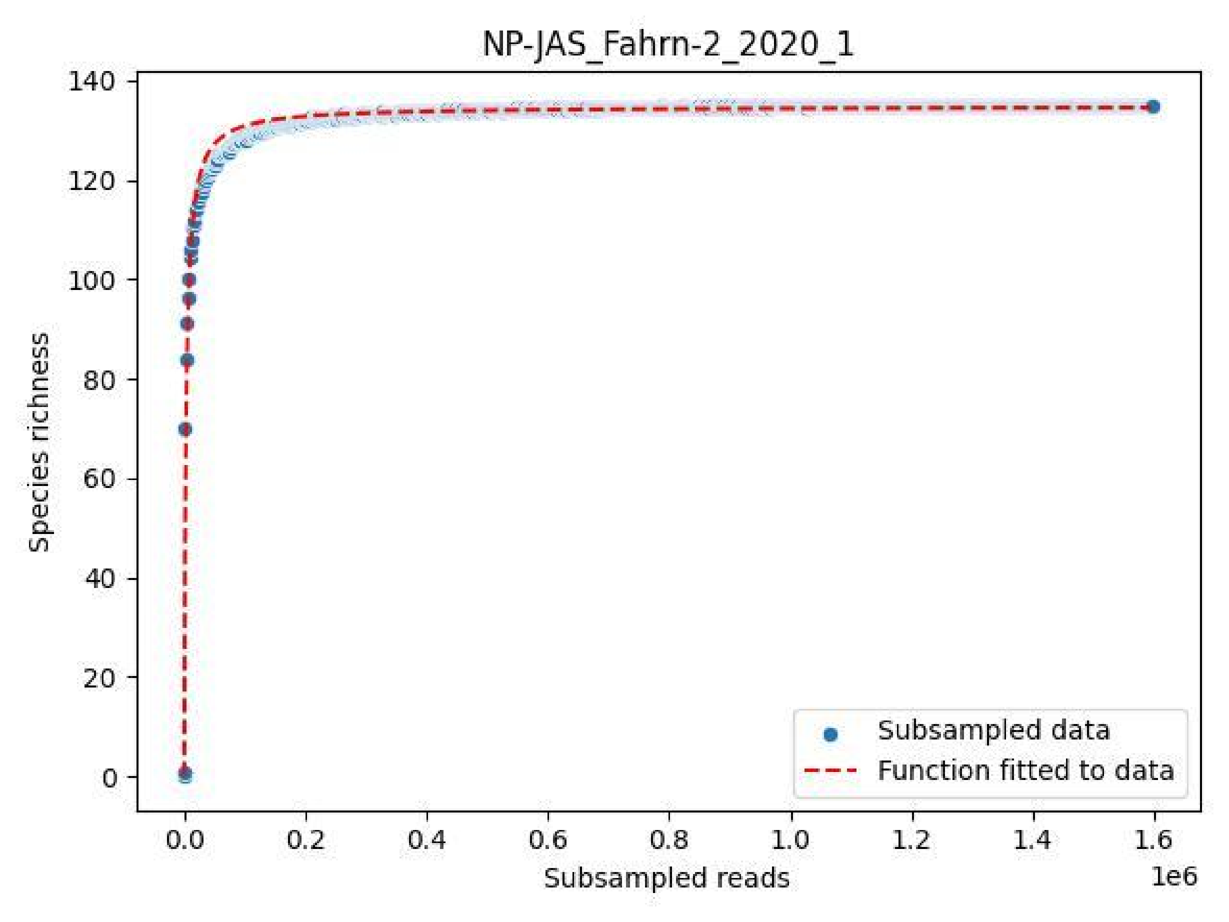
NP-JAS_Fahrn-2_2020_1
Species richness
Subsampled reads
1e6
0
20
40
60
80
100
120
140
0.0
0.2
0.4
0.6
0.8
1.0
1.2
1.4
1.6
Subsampled data
Function fitted to data

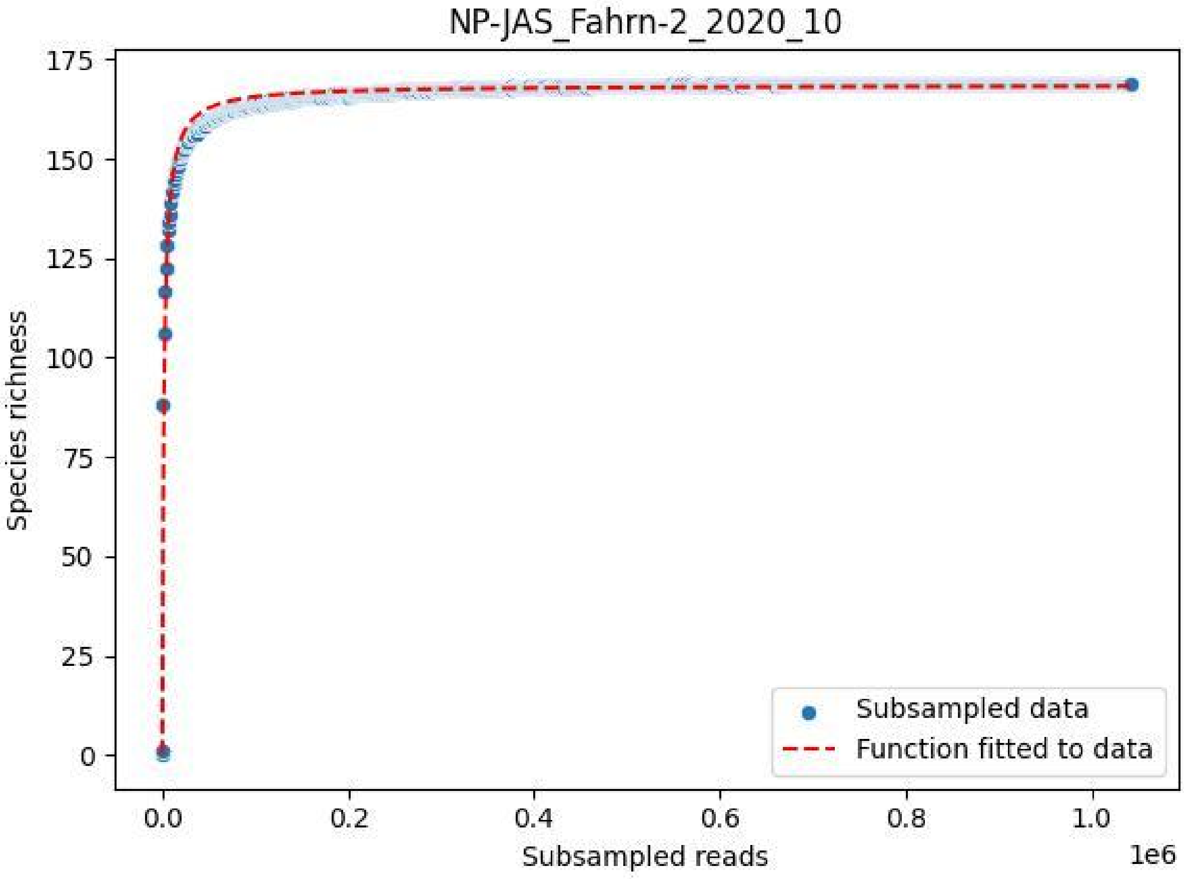

NP-JAS_Fahrn-2_2020_10
175
150
125
100
75
50
25
0
Species richness
0.0
0.2
0.4
0.6
0.8
1.0
1e6
Subsampled reads
Subsampled data
Function fitted to data

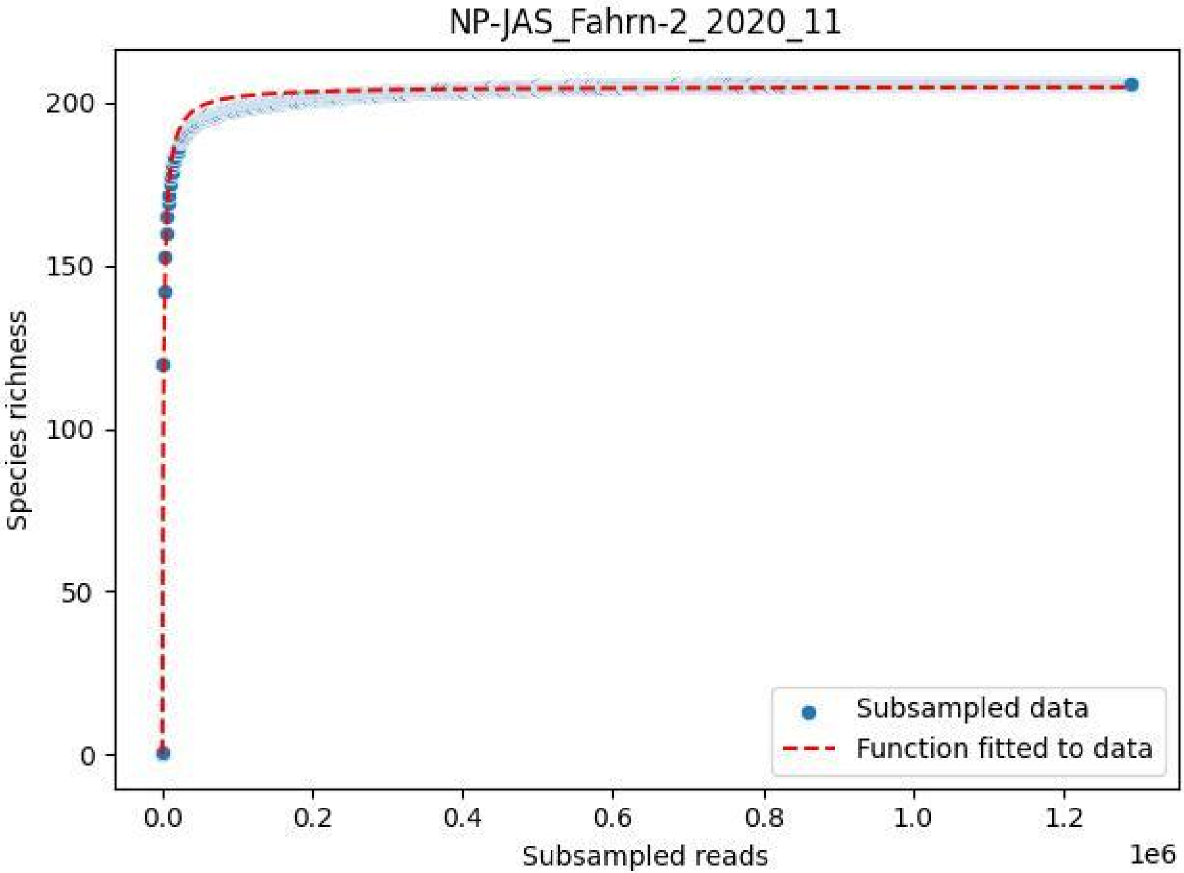

NP-JAS_Fahrn-2_2020_11
Species richness
0
50
100
150
200
0.0
0.2
0.4
0.6
0.8
1.0
1.2
1e6
Subsampled reads
Subsampled data
Function fitted to data

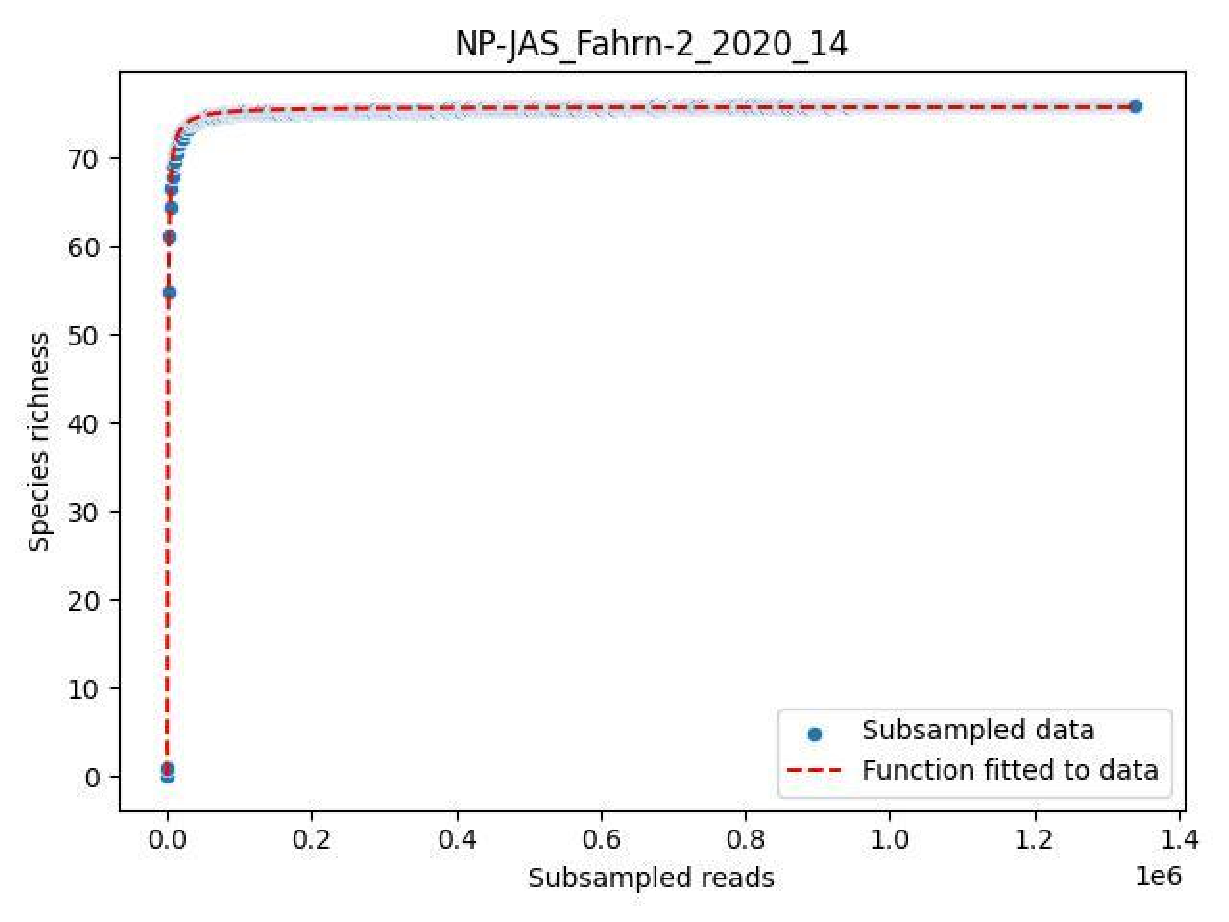

NP-JAS_Fahrn-2_2020_14
Species richness
Subsampled reads
1e6
0
10
20
30
40
50
60
70
0.0
0.2
0.4
0.6
0.8
1.0
1.2
1.4
Subsampled data
Function fitted to data

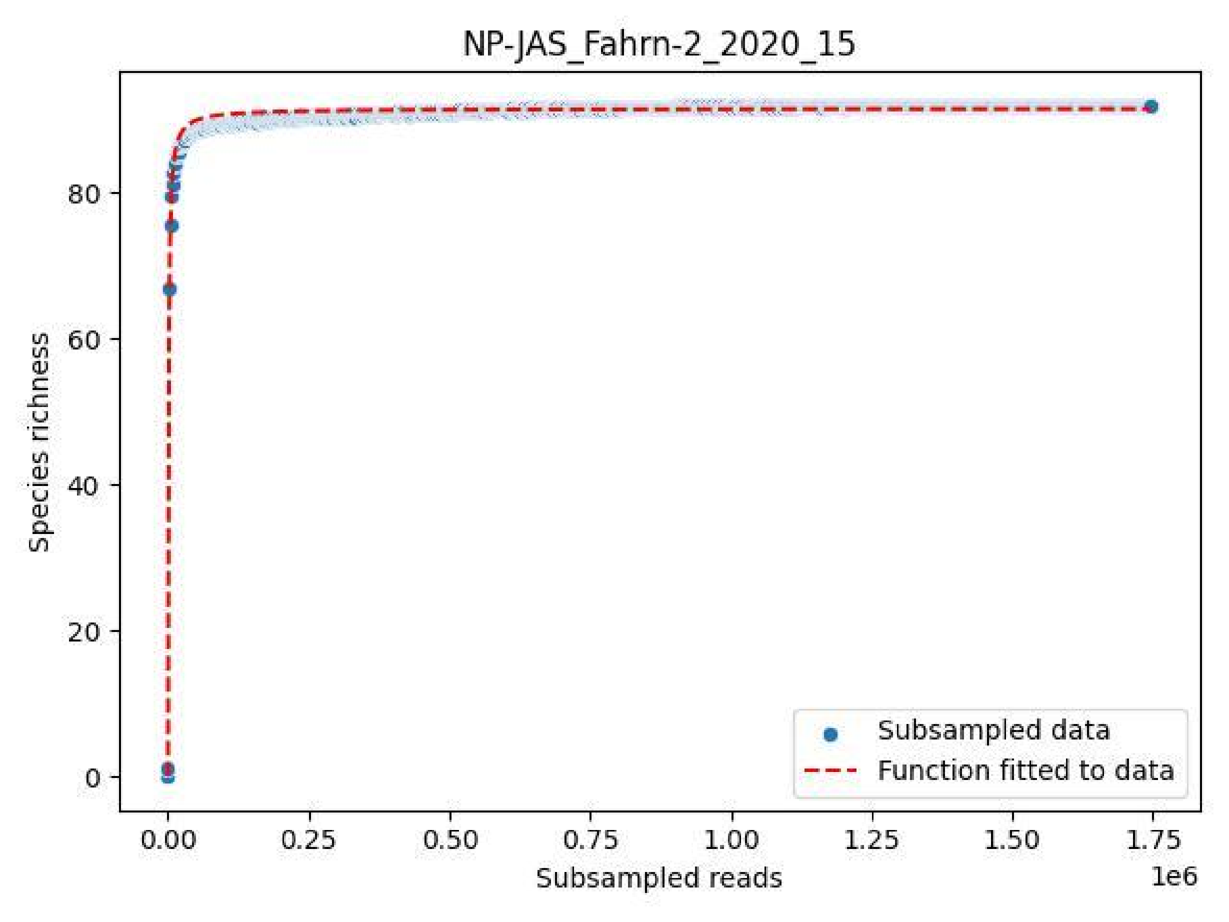

NP-JAS_Fahrn-2_2020_15
Species richness
0
20
40
60
80
0.00
0.25
0.50
0.75
1.00
1.25
1.50
1.75
1e6
Subsampled reads
Subsampled data
Function fitted to data

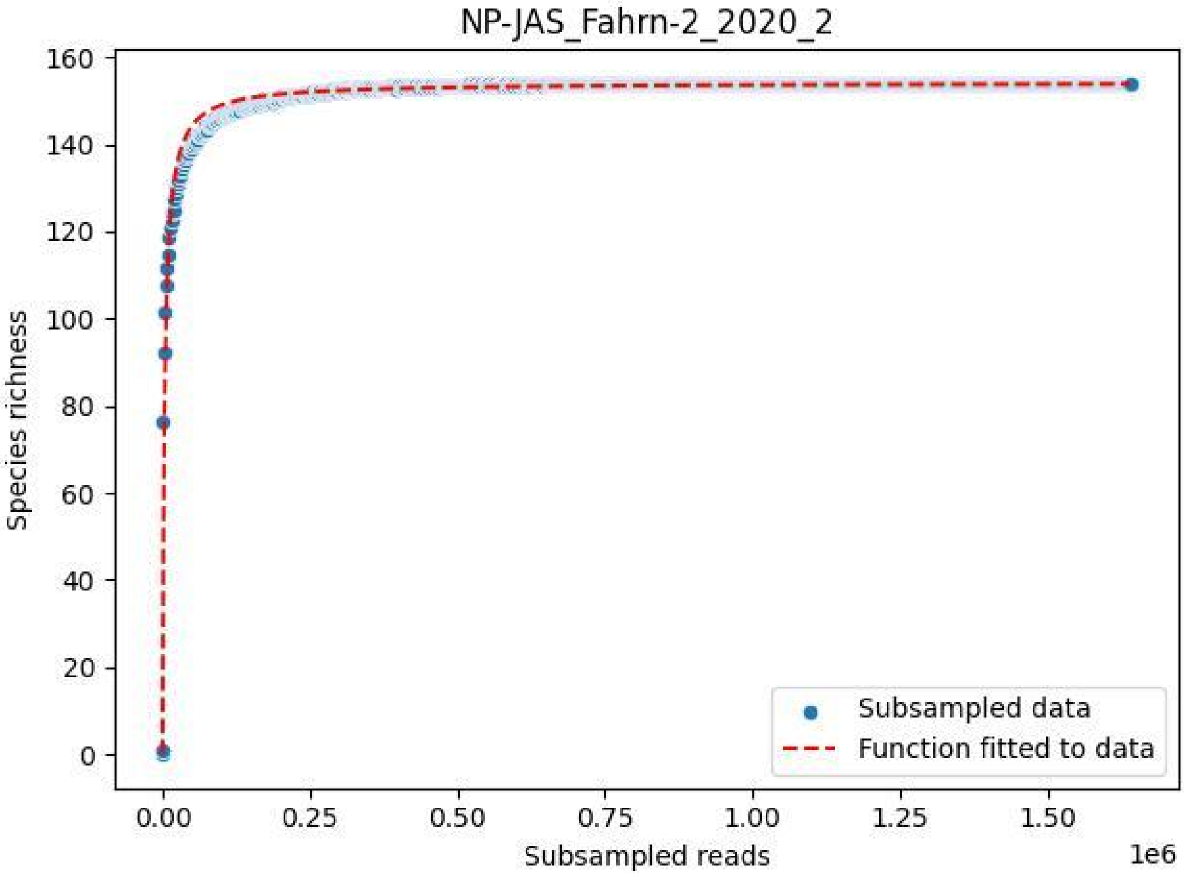

NP-JAS_Fahrn-2_2020_2
Species richness
0
20
40
60
80
100
120
140
160
Subsampled reads
0.00
0.25
0.50
0.75
1.00
1.25
1.50
1e6
Subsampled data
Function fitted to data

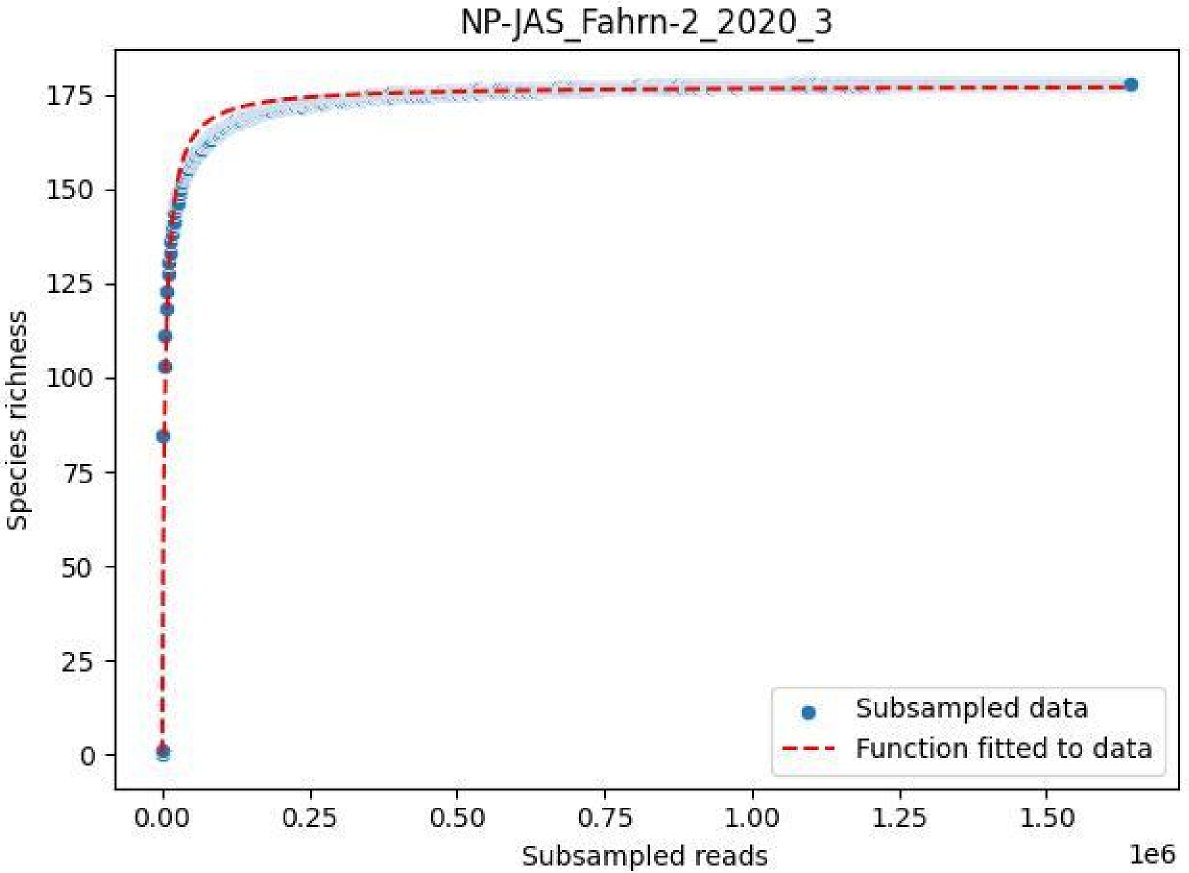
NP-JAS_Fahrn-2_2020_3
Species richness
Subsampled reads
1e6
0
25
50
75
100
125
150
175
0.00
0.25
0.50
0.75
1.00
1.25
1.50
Subsampled data
Function fitted to data

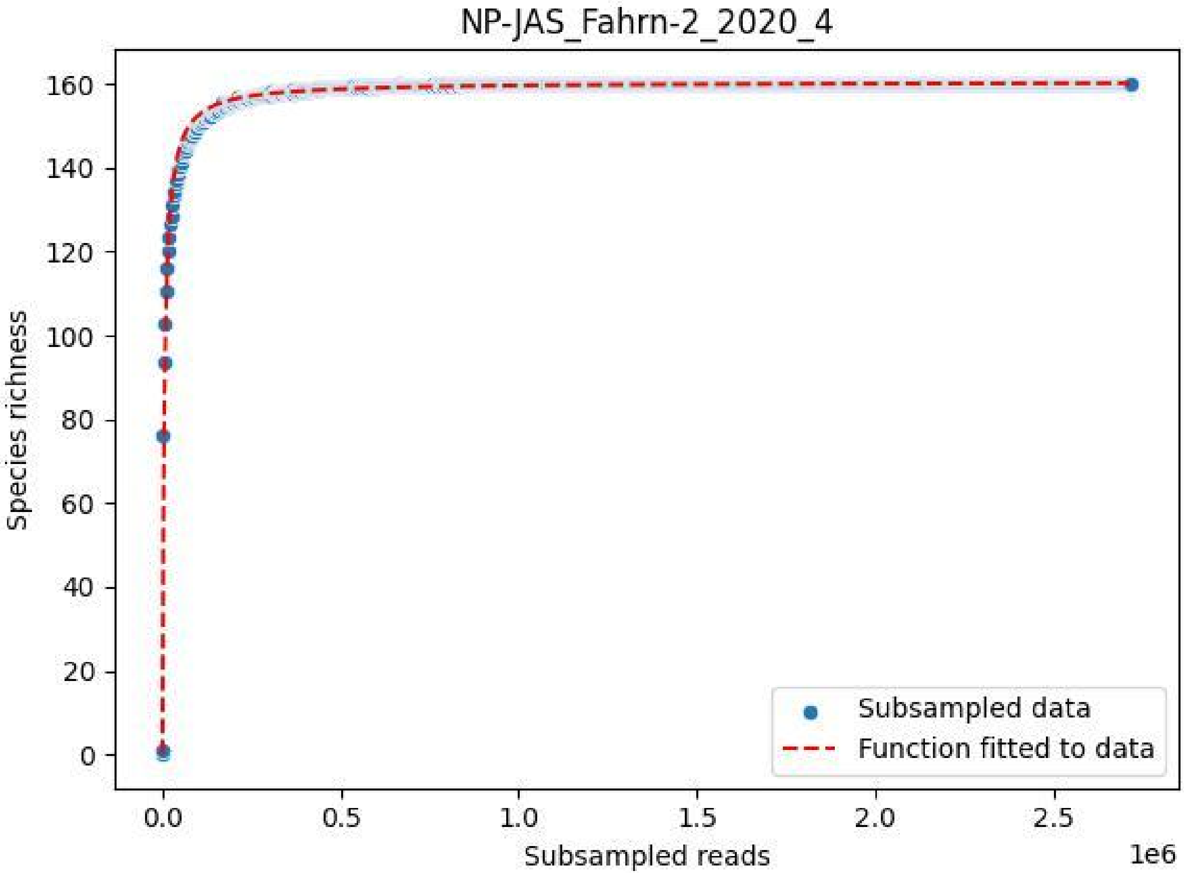
NP-JAS_Fahrn-2_2020_4
Species richness
Subsampled reads
1e6
0
20
40
60
80
100
120
140
160
0.0
0.5
1.0
1.5
2.0
2.5
Subsampled data
Function fitted to data

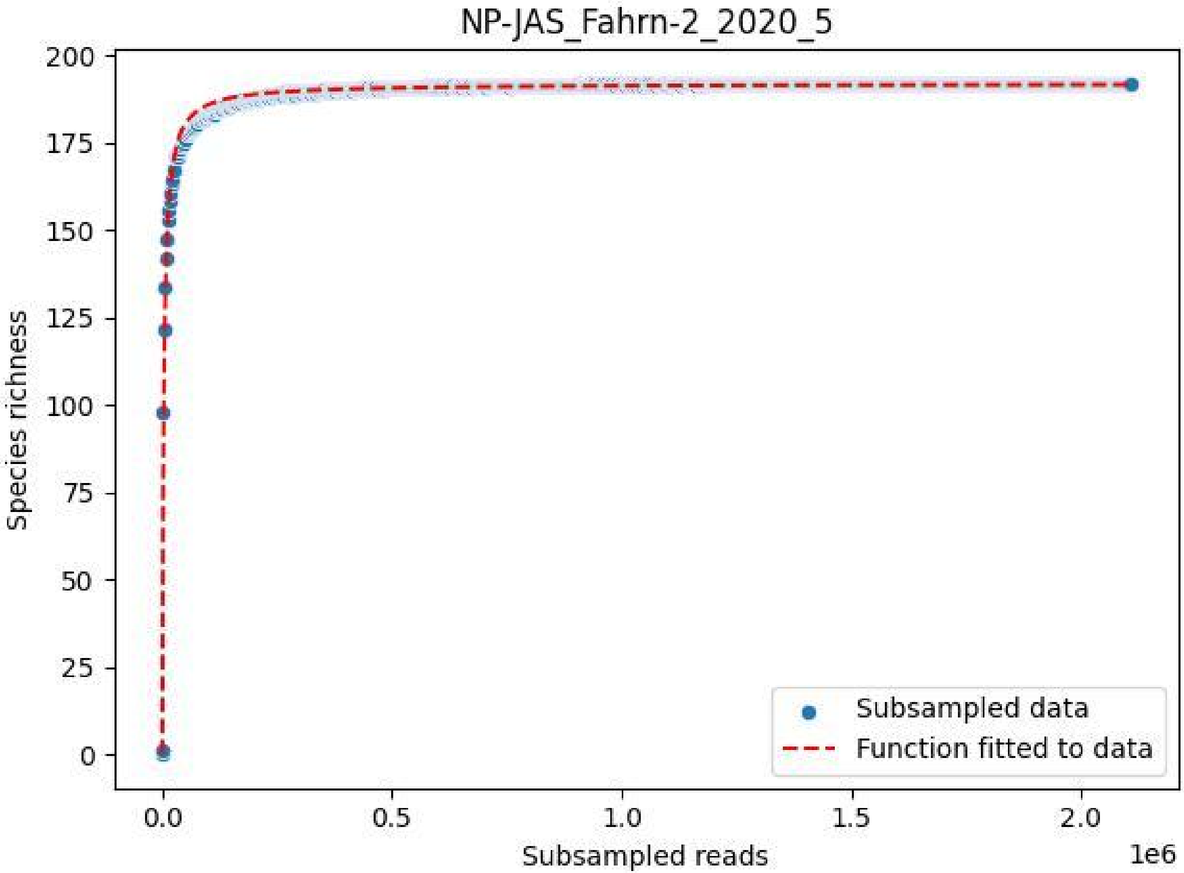

NP-JAS_Fahrn-2_2020_5
Species richness
Subsampled reads
1e6
0
25
50
75
100
125
150
175
200
0.0
0.5
1.0
1.5
2.0
Subsampled data
Function fitted to data

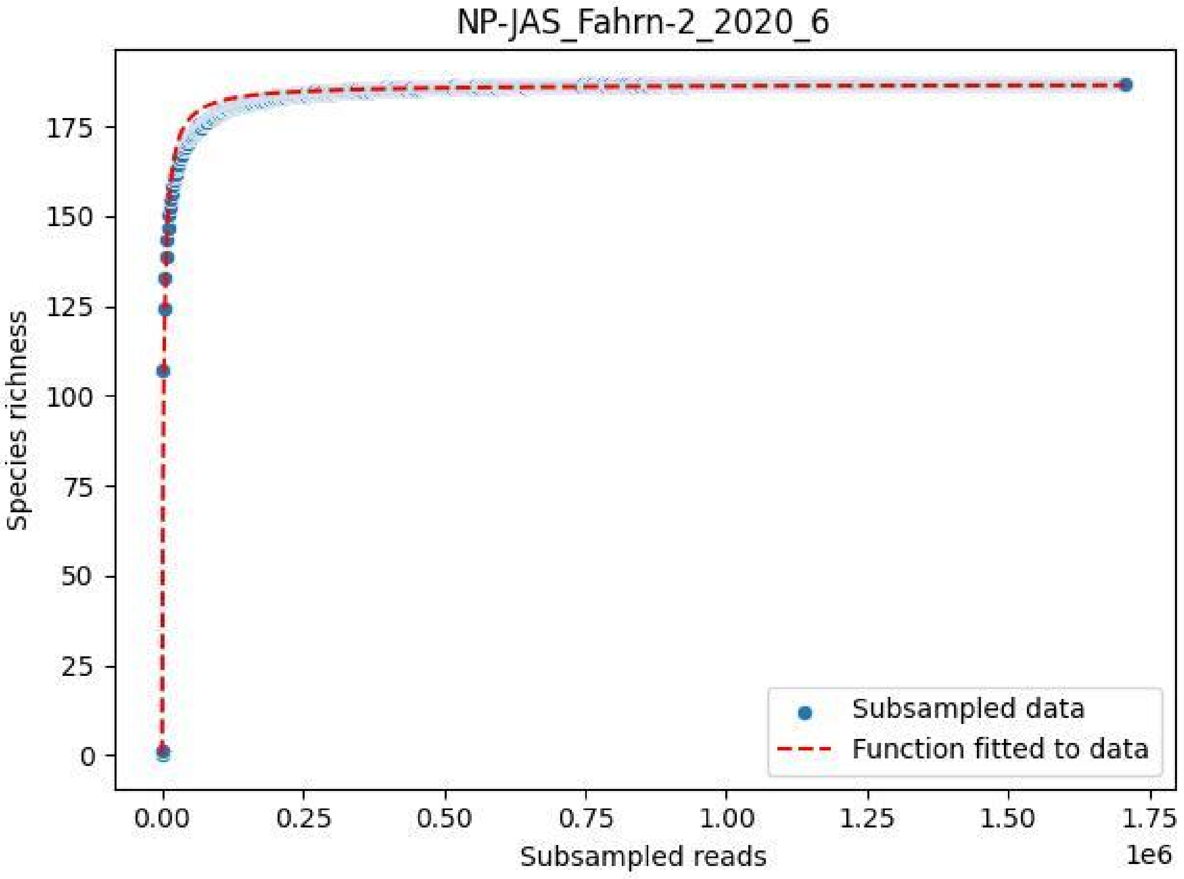

NP-JAS_Fahrn-2_2020_6
Species richness
Subsampled reads
1e6
0
25
50
75
100
125
150
175
0.00
0.25
0.50
0.75
1.00
1.25
1.50
1.75
Subsampled data
Function fitted to data

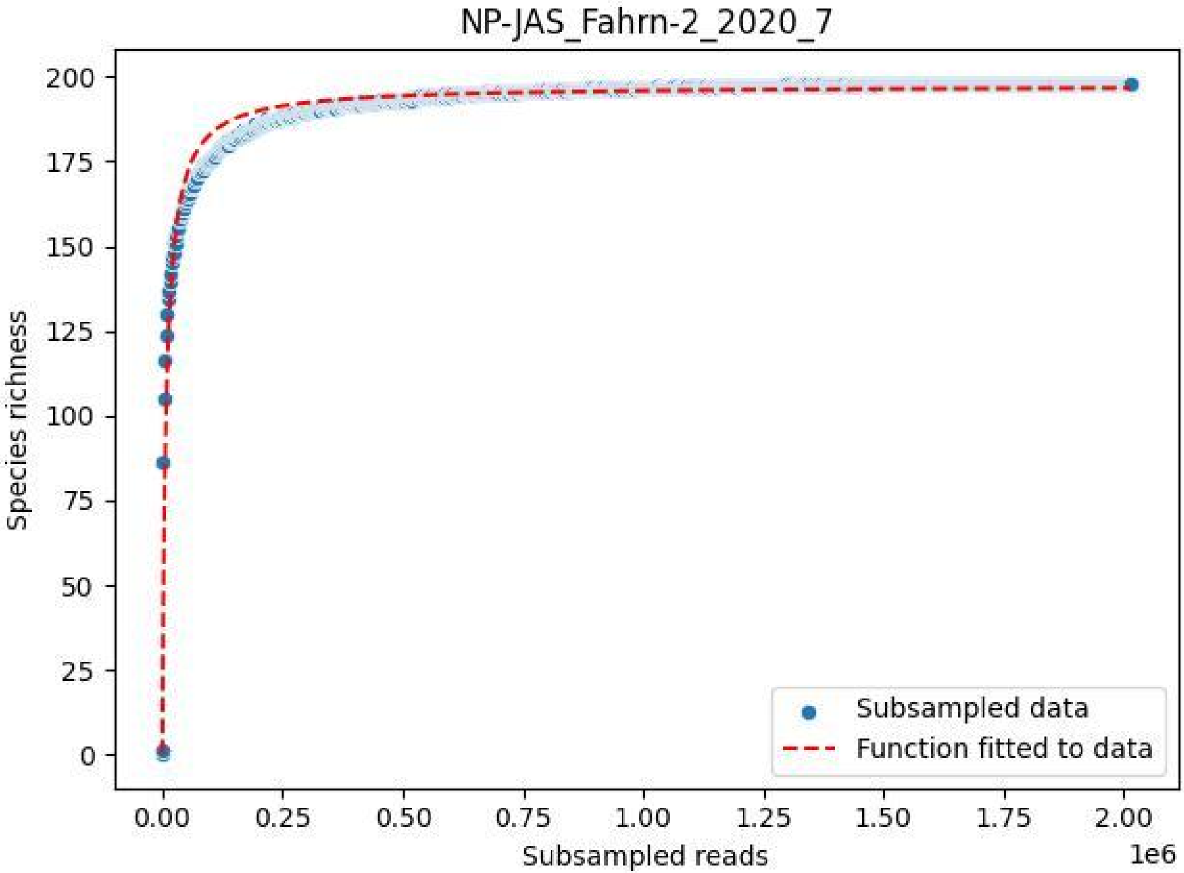
NP-JAS_Fahrn-2_2020_7
Species richness
200
175
150
125
100
75
50
25
0
0.00
0.25
0.50
0.75
1.00
1.25
1.50
1.75
2.00
1e6
Subsampled reads
Subsampled data
Function fitted to data

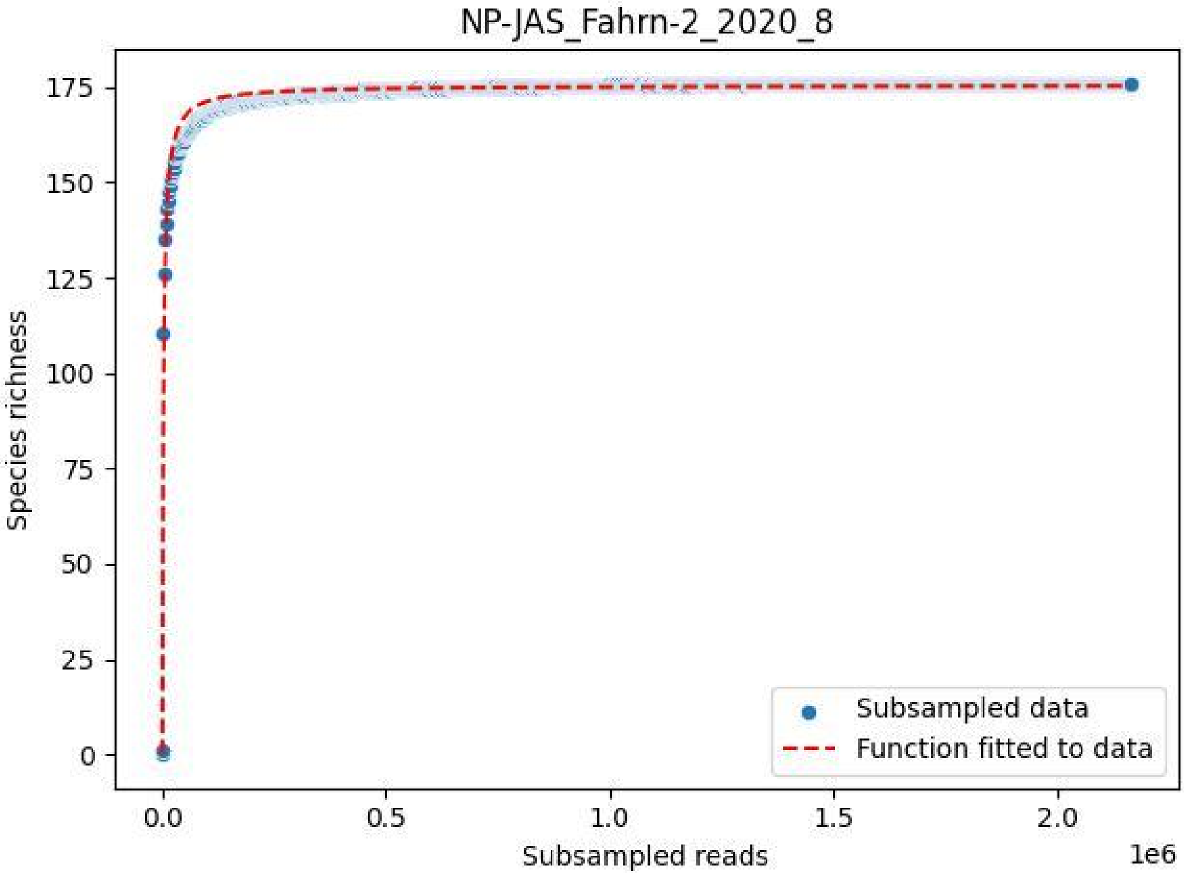
NP-JAS_Fahrn-2_2020_8
Species richness
Subsampled reads
1e6
0
25
50
75
100
125
150
175
0.0
0.5
1.0
1.5
2.0
Subsampled data
Function fitted to data

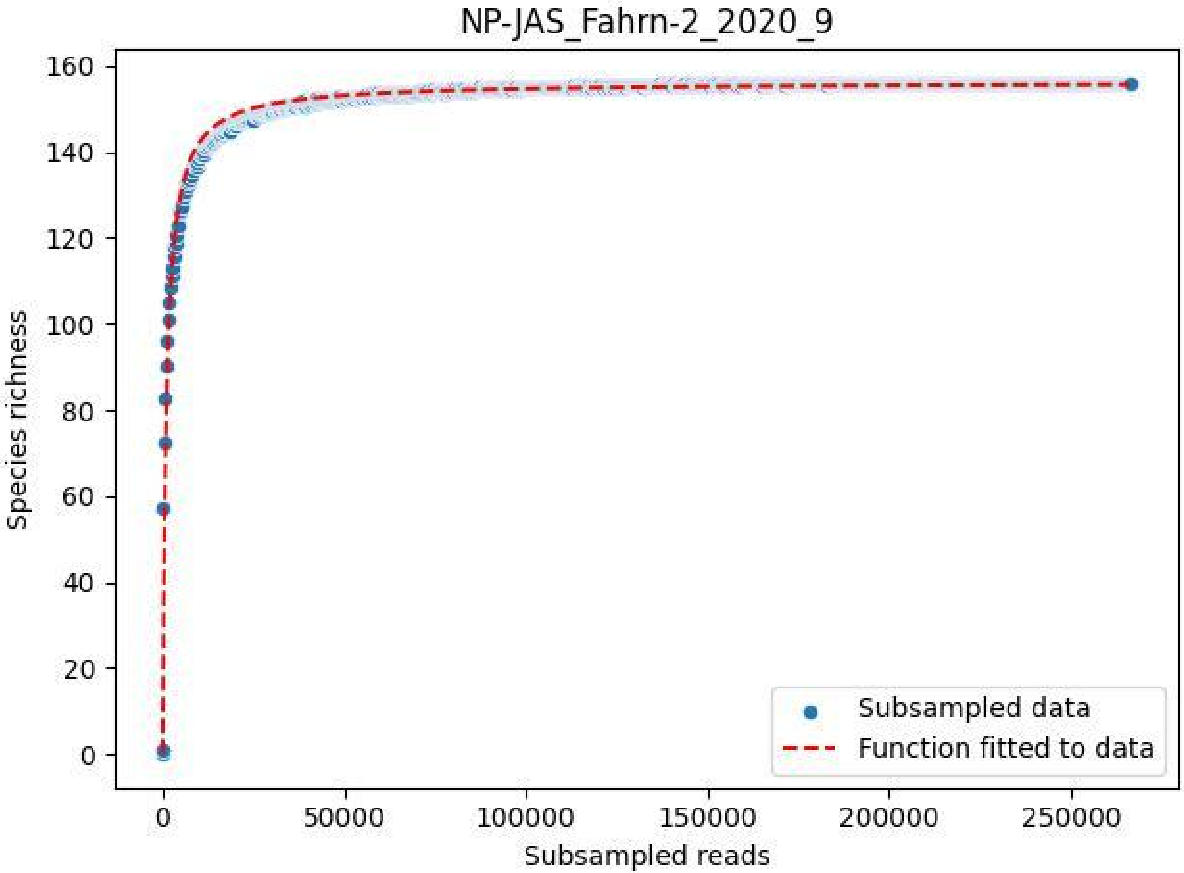

NP-JAS_Fahrn-2_2020_9
Species richness
Subsampled reads
0
20
40
60
80
100
120
140
160
0
50000
100000
150000
200000
250000
Subsampled data
Function fitted to data

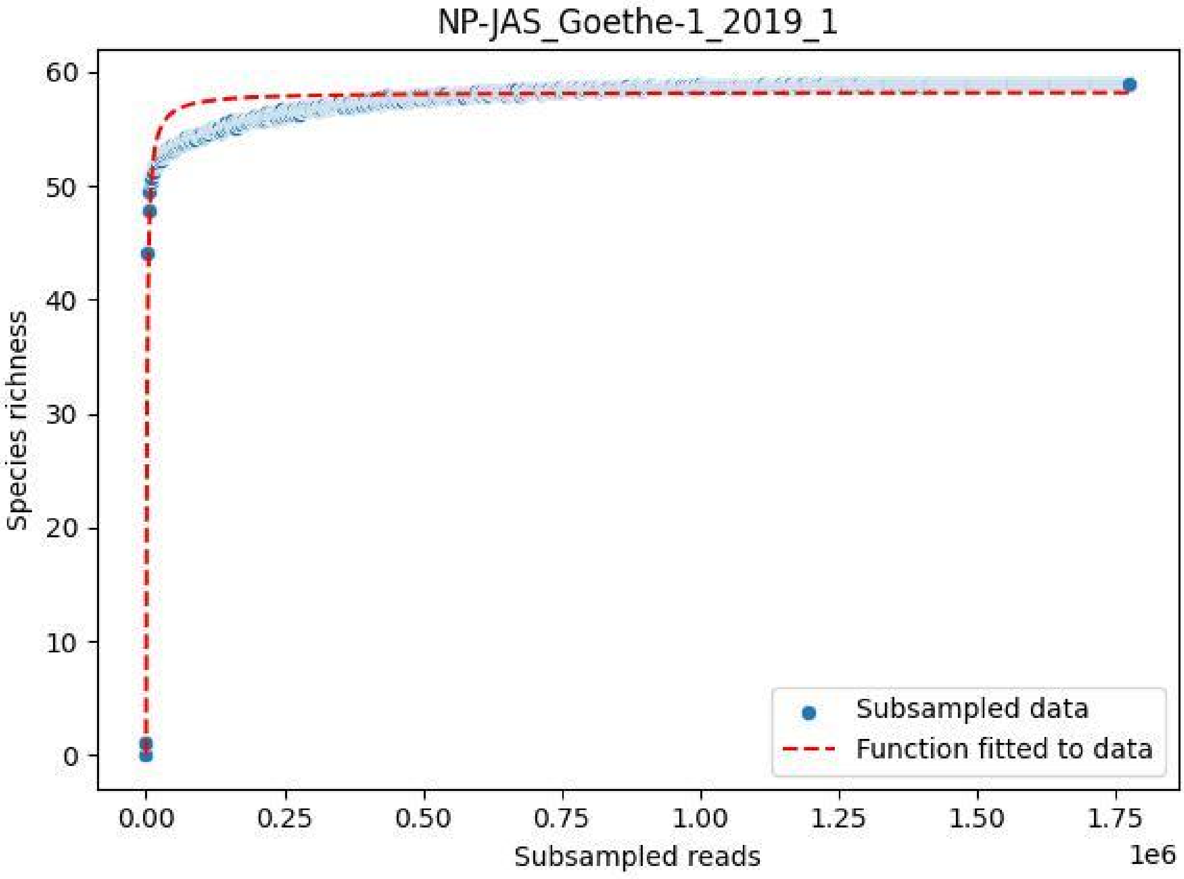
NP-JAS_Goethe-1_2019_1
Species richness
Subsampled reads
1e6
0
10
20
30
40
50
60
0.00
0.25
0.50
0.75
1.00
1.25
1.50
1.75
Subsampled data
Function fitted to data

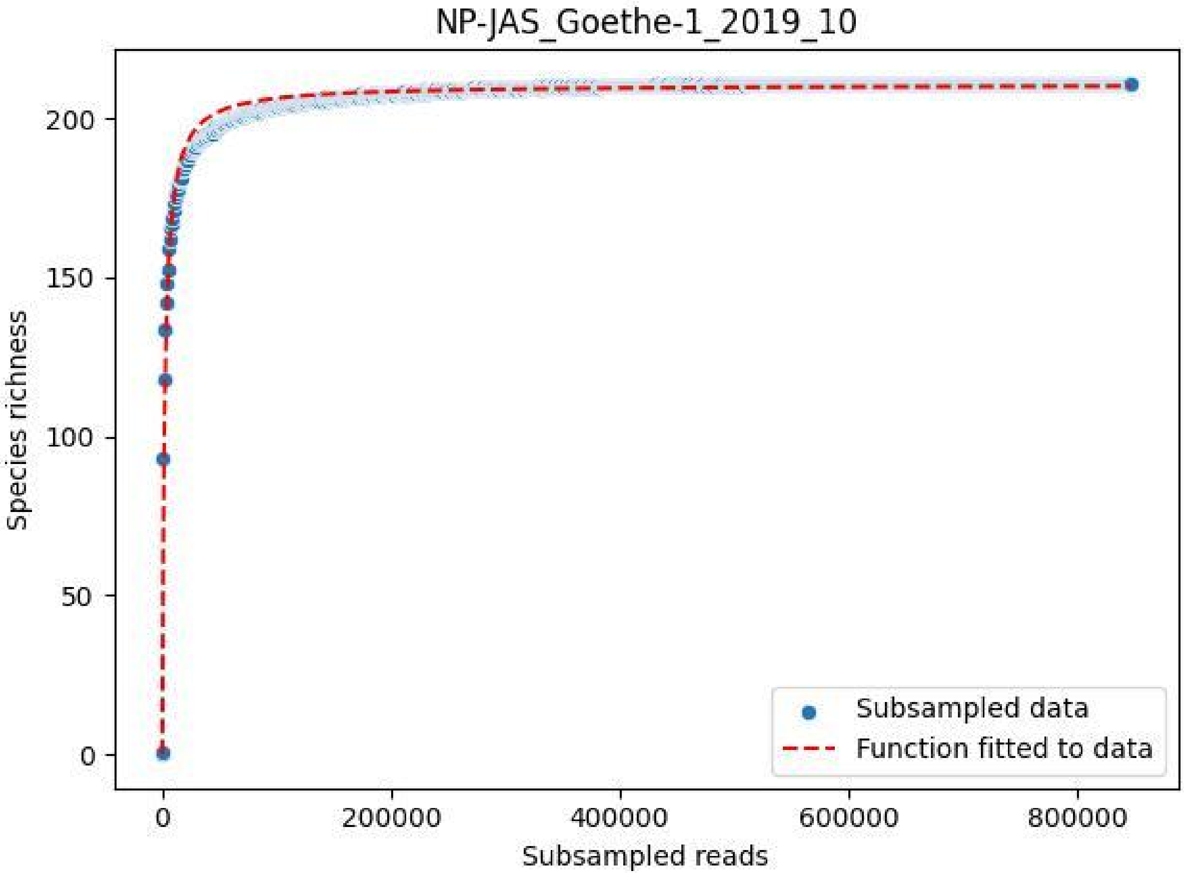
NP-JAS_Goethe-1_2019_10
Species richness
Subsampled reads
0
50
100
150
200
0
200000
400000
600000
800000
Subsampled data
Function fitted to data

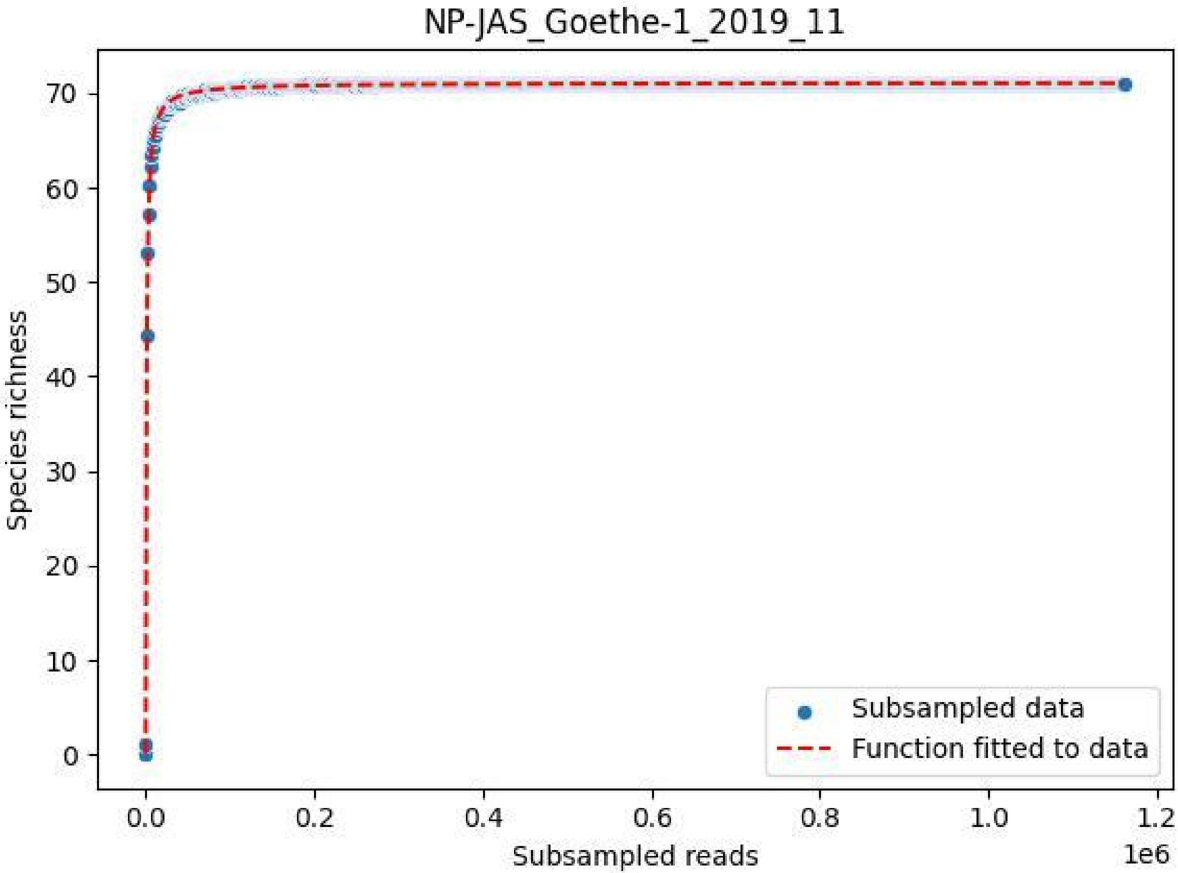

NP-JAS_Goethe-1_2019_11
Species richness
Subsampled reads
1e6
0
10
20
30
40
50
60
70
0.0
0.2
0.4
0.6
0.8
1.0
1.2
Subsampled data
Function fitted to data

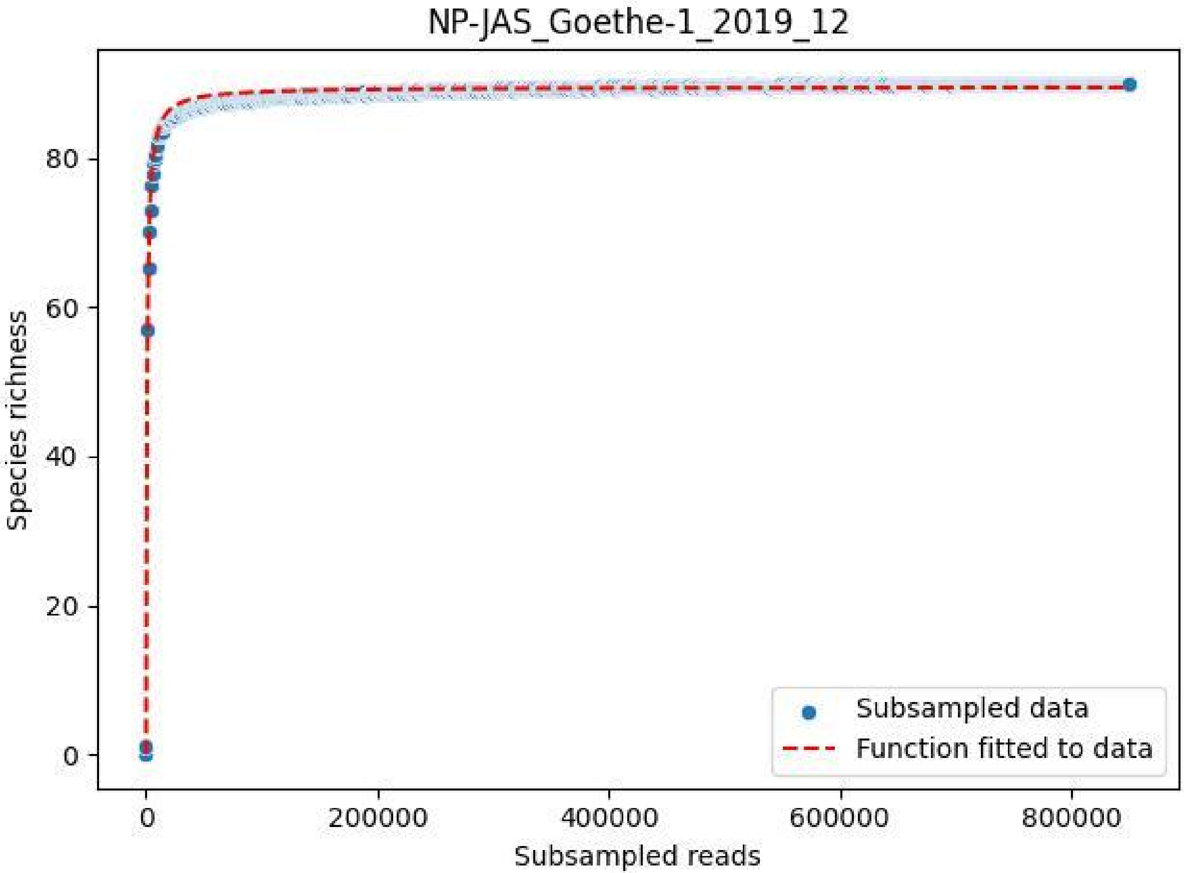

NP-JAS_Goethe-1_2019_12
Species richness
Subsampled reads
0
20
40
60
80
0
200000
400000
600000
800000
Subsampled data
Function fitted to data

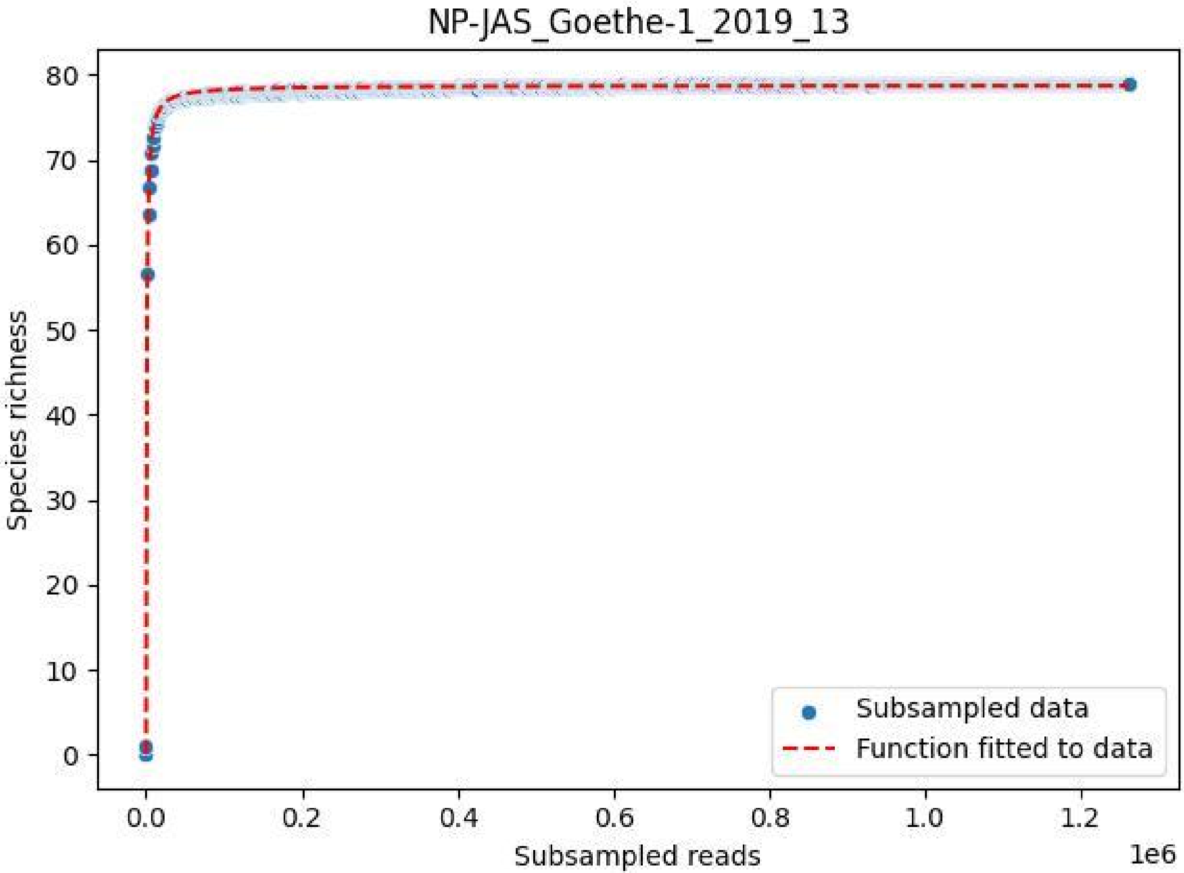

NP-JAS_Goethe-1_2019_13
Species richness
Subsampled reads
1e6
Subsampled data
Function fitted to data
0
10
20
30
40
50
60
70
80
0.0
0.2
0.4
0.6
0.8
1.0
1.2

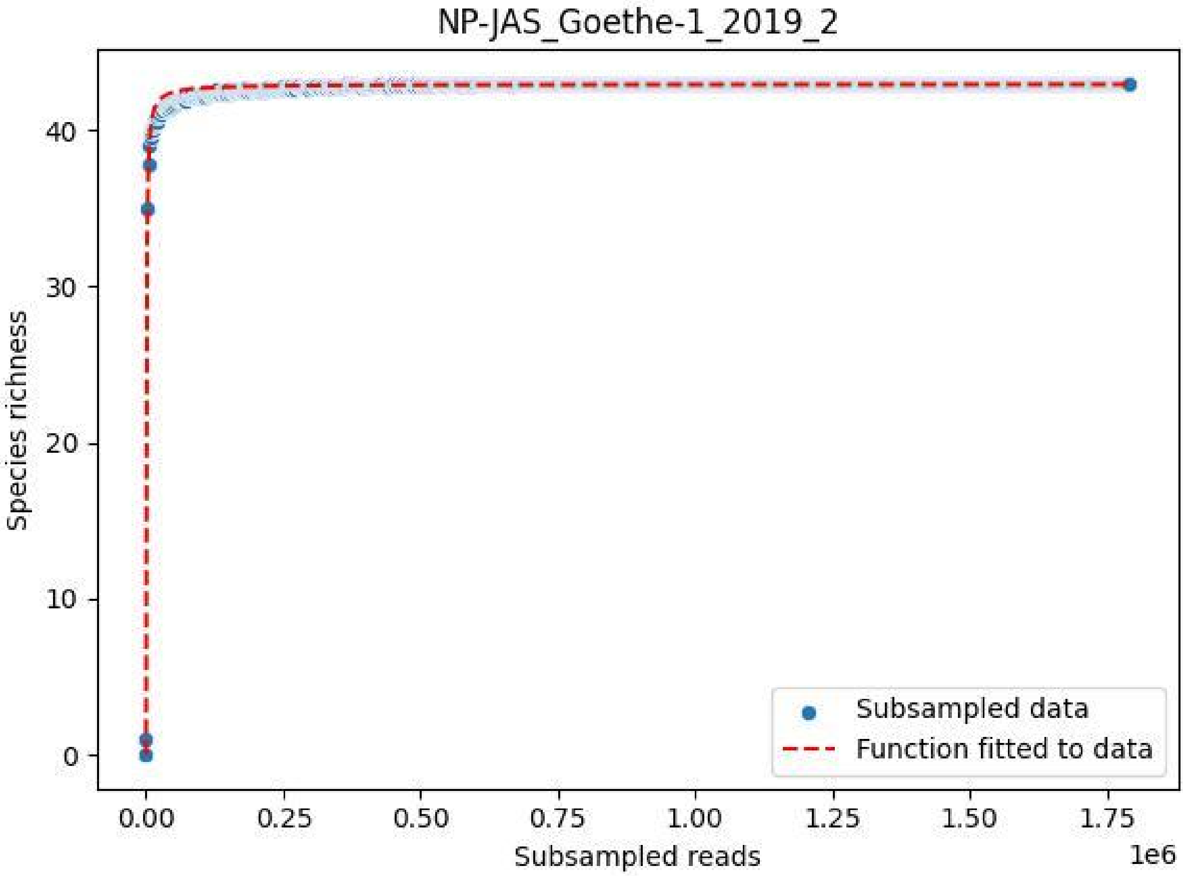
NP-JAS_Goethe-1_2019_2
Species richness
Subsampled reads
1e6
0
10
20
30
40
0.00
0.25
0.50
0.75
1.00
1.25
1.50
1.75
Subsampled data
Function fitted to data

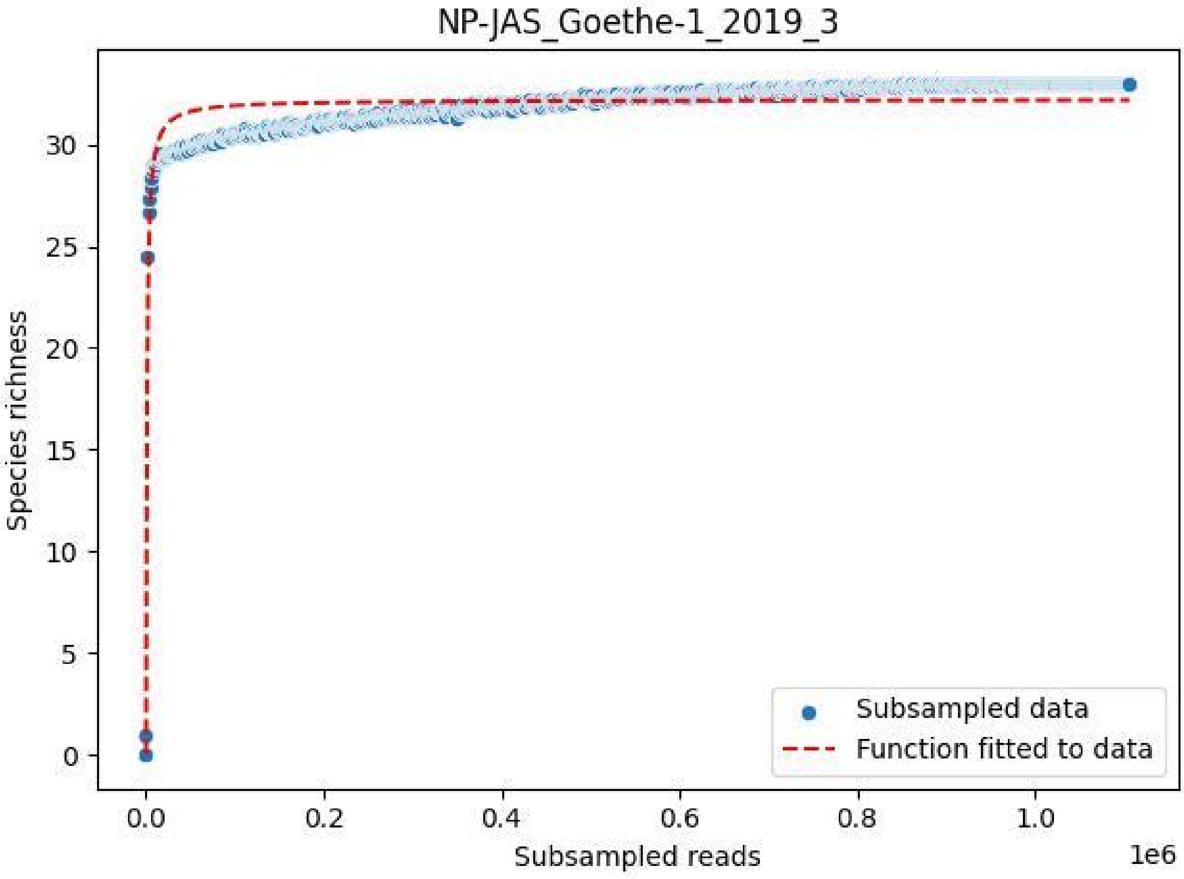
NP-JAS_Goethe-1_2019_3
Species richness
Subsampled reads
1e6
0
5
10
15
20
25
30
0.0
0.2
0.4
0.6
0.8
1.0
Subsampled data
Function fitted to data

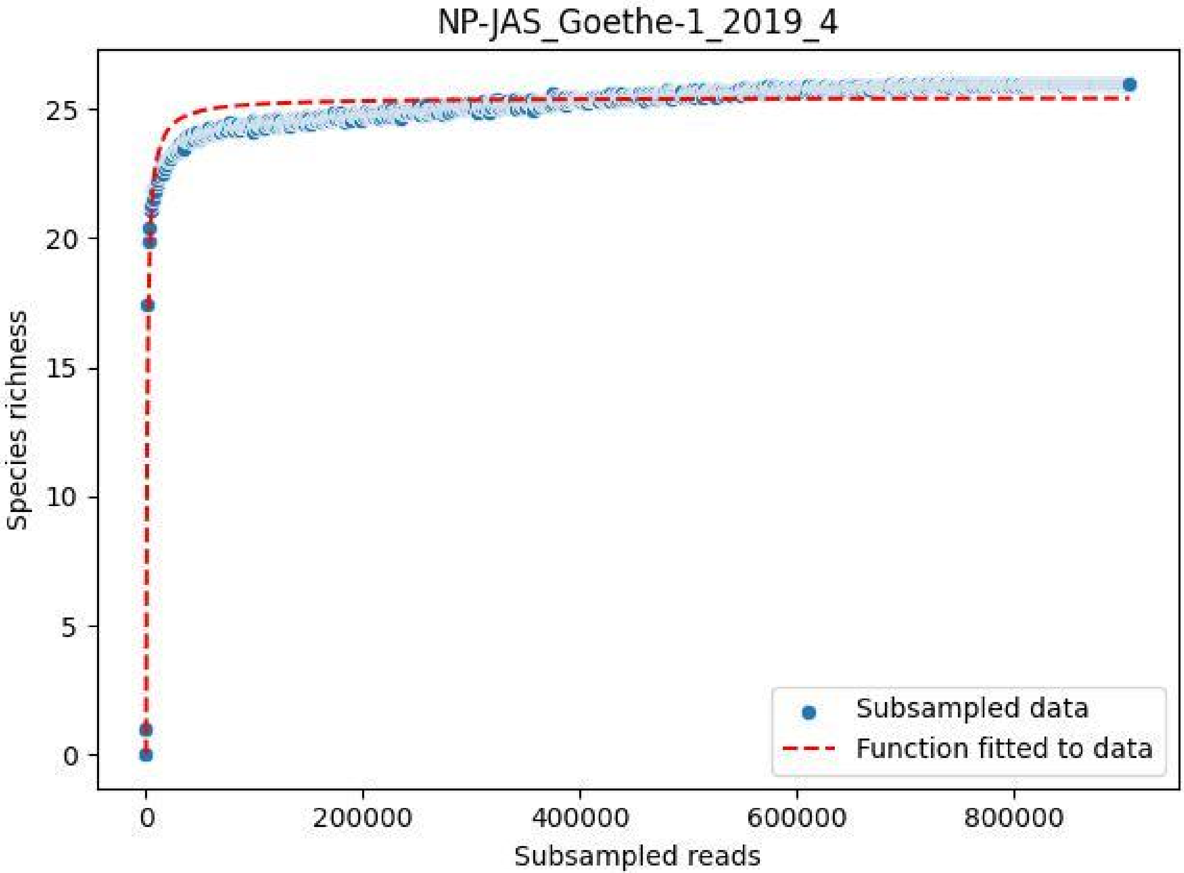

NP-JAS_Goethe-1_2019_4
Species richness
Subsampled reads
0
5
10
15
20
25
0
200000
400000
600000
800000
Subsampled data
Function fitted to data

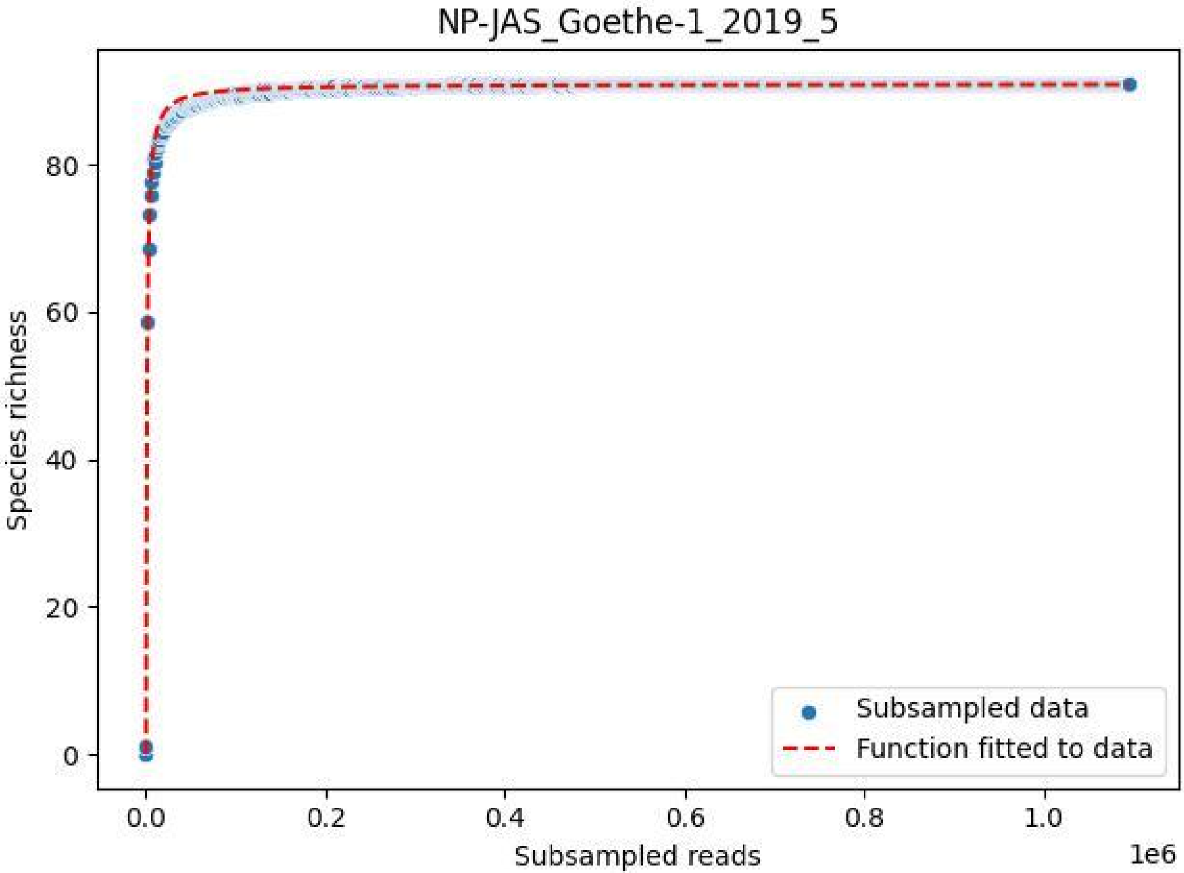

NP-JAS_Goethe-1_2019_5
Species richness
0
20
40
60
80
0.0
0.2
0.4
0.6
0.8
1.0
1e6
Subsampled reads
Subsampled data
Function fitted to data

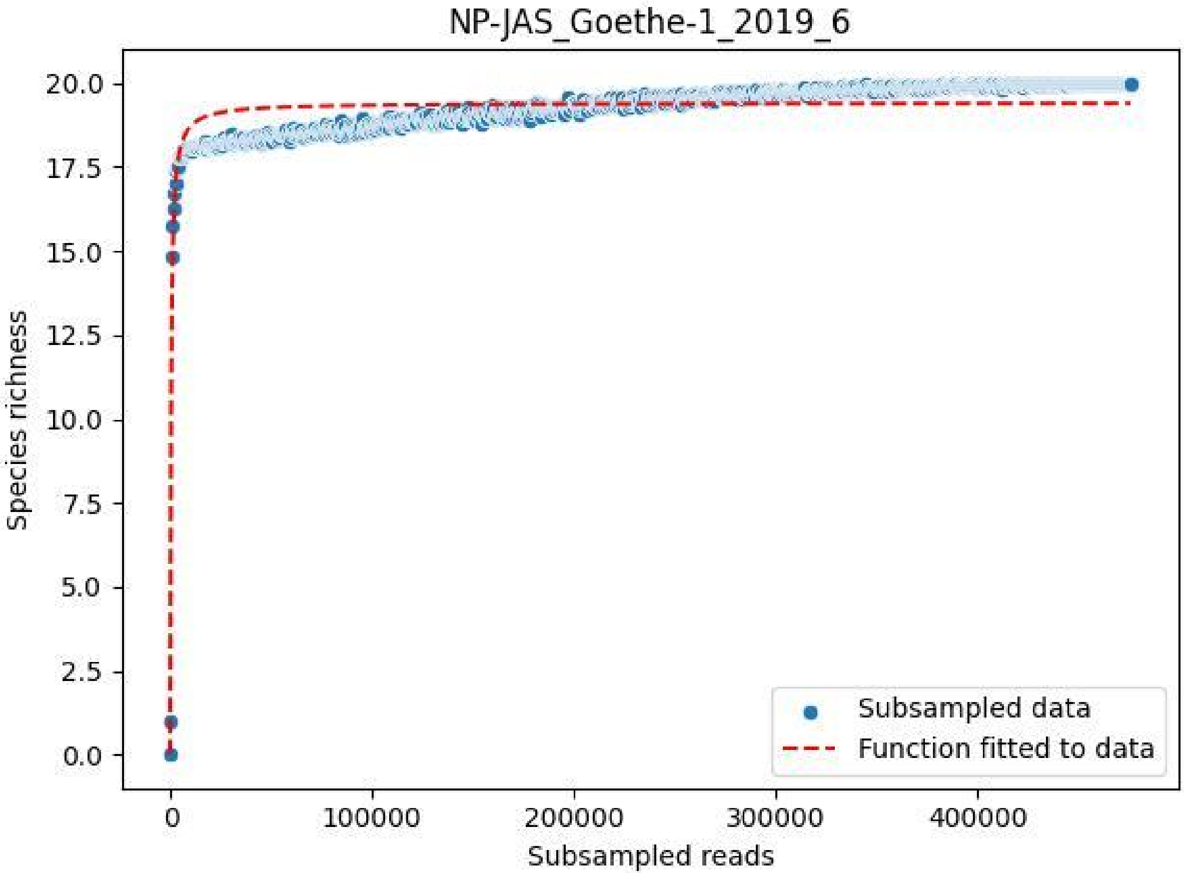

NP-JAS_Goethe-1_2019_6
Species richness
Subsampled reads
0.0
2.5
5.0
7.5
10.0
12.5
15.0
17.5
20.0
0
100000
200000
300000
400000
Subsampled data
Function fitted to data

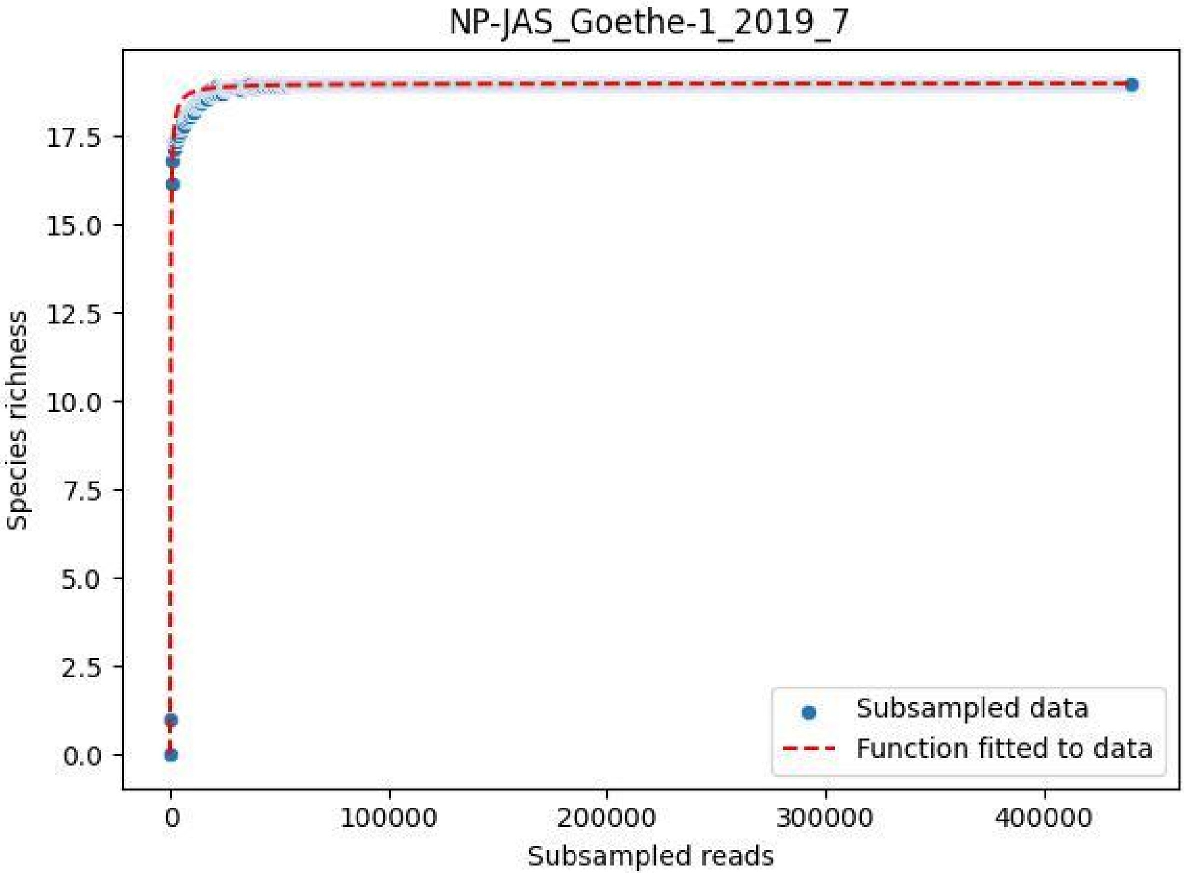

NP-JAS_Goethe-1_2019_7
Species richness
Subsampled reads
0.0
2.5
5.0
7.5
10.0
12.5
15.0
17.5
0
100000
200000
300000
400000
Subsampled data
Function fitted to data

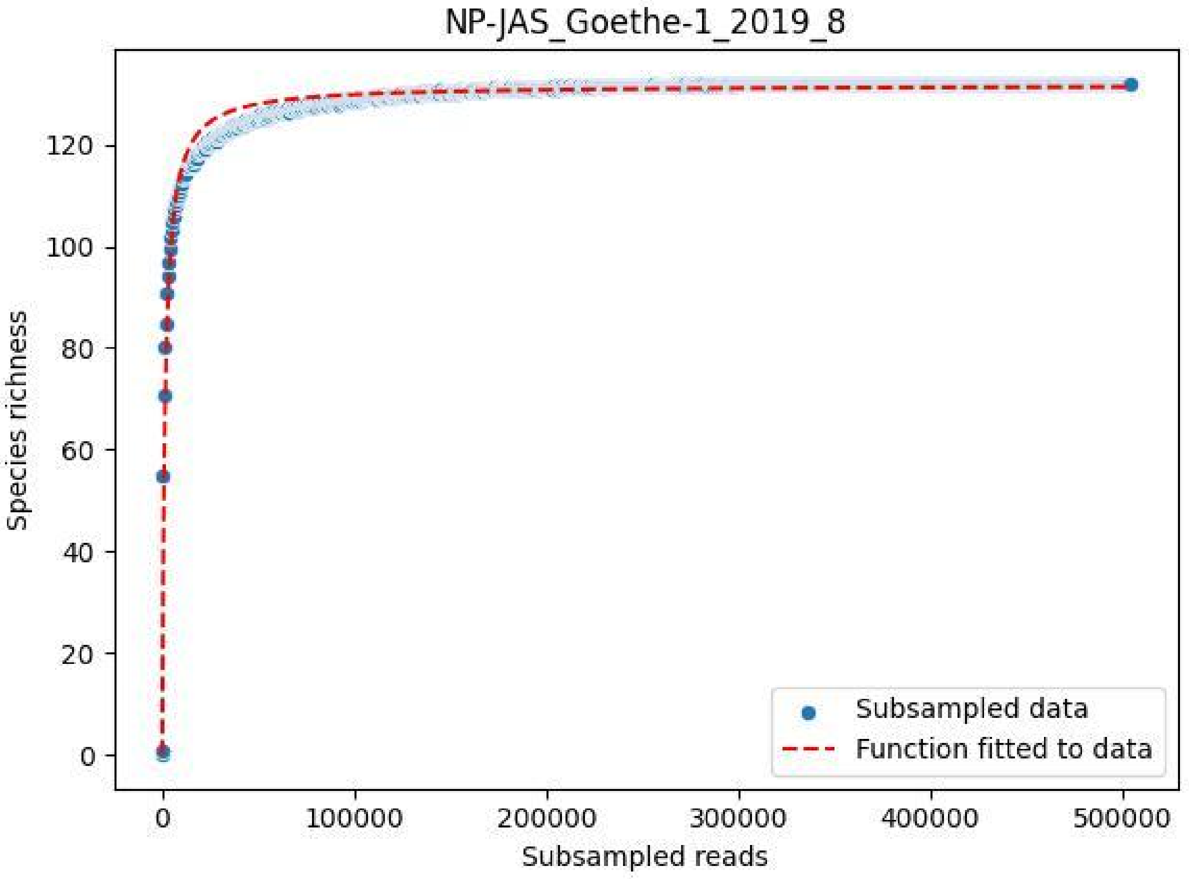
NP-JAS_Goethe-1_2019_8
Species richness
0
20
40
60
80
100
120
0
100000
200000
300000
400000
500000
Subsampled reads
Subsampled data
Function fitted to data

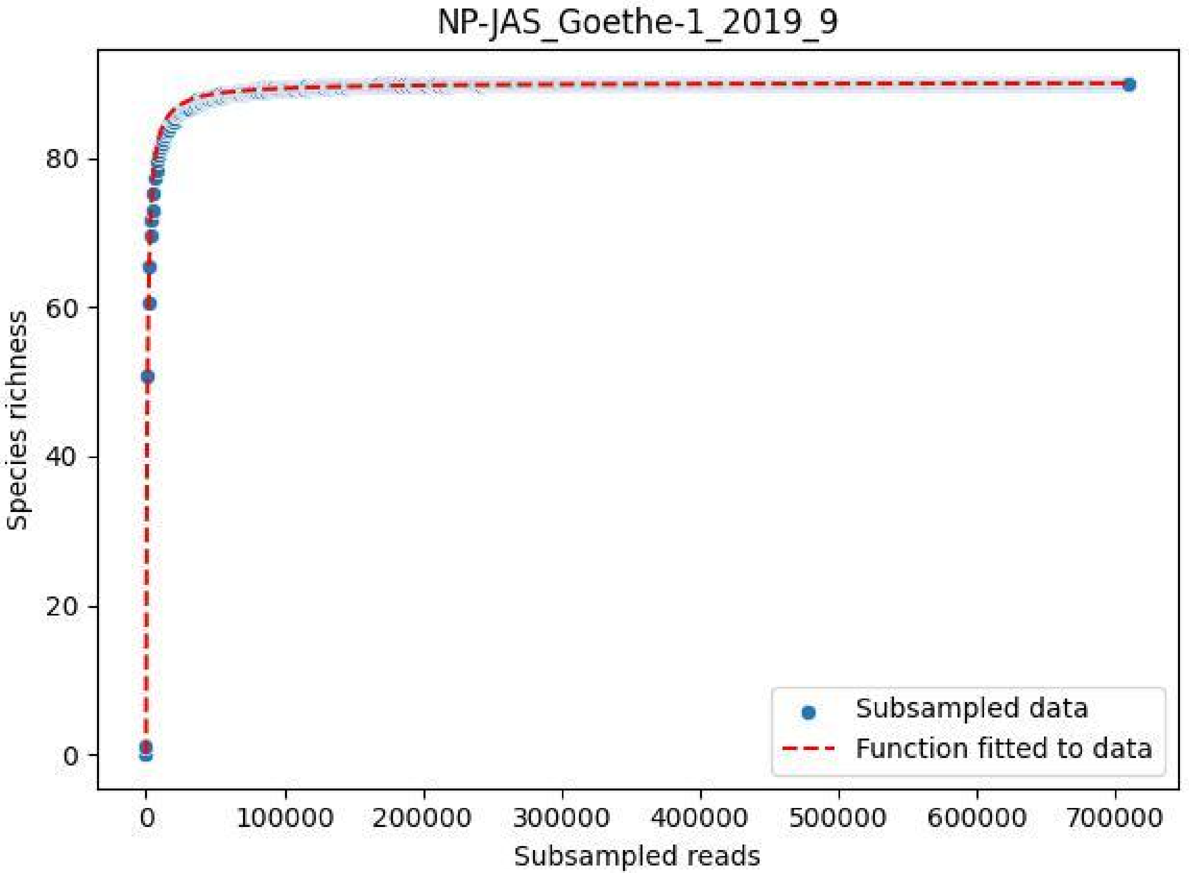

NP-JAS_Goethe-1_2019_9
Species richness
Subsampled reads
0
20
40
60
80
0
100000
200000
300000
400000
500000
600000
700000
Subsampled data
Function fitted to data

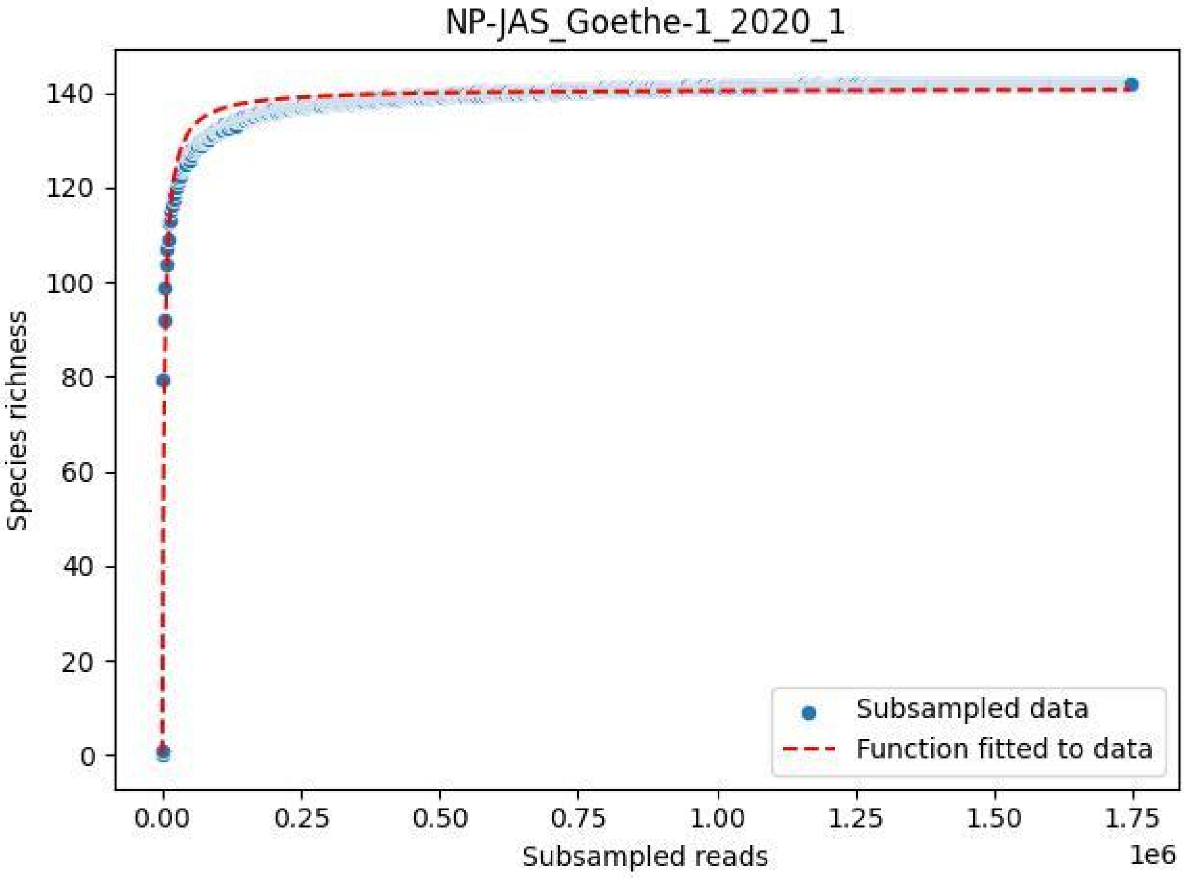
NP-JAS_Goethe-1_2020_1
Species richness
Subsampled reads
1e6
0
20
40
60
80
100
120
140
0.00
0.25
0.50
0.75
1.00
1.25
1.50
1.75
Subsampled data
Function fitted to data

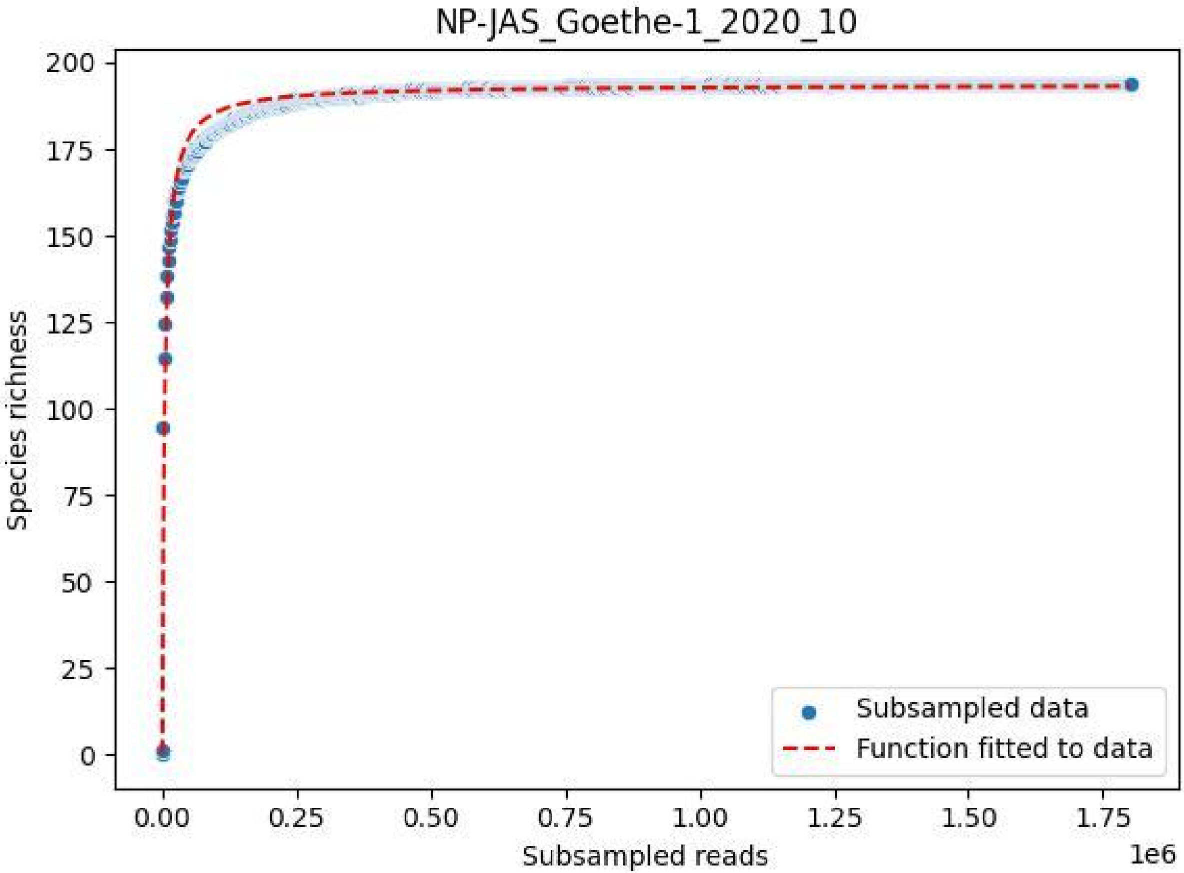

NP-JAS_Goethe-1_2020_10
Species richness
Subsampled reads
1e6
0
25
50
75
100
125
150
175
200
0.00
0.25
0.50
0.75
1.00
1.25
1.50
1.75
Subsampled data
Function fitted to data

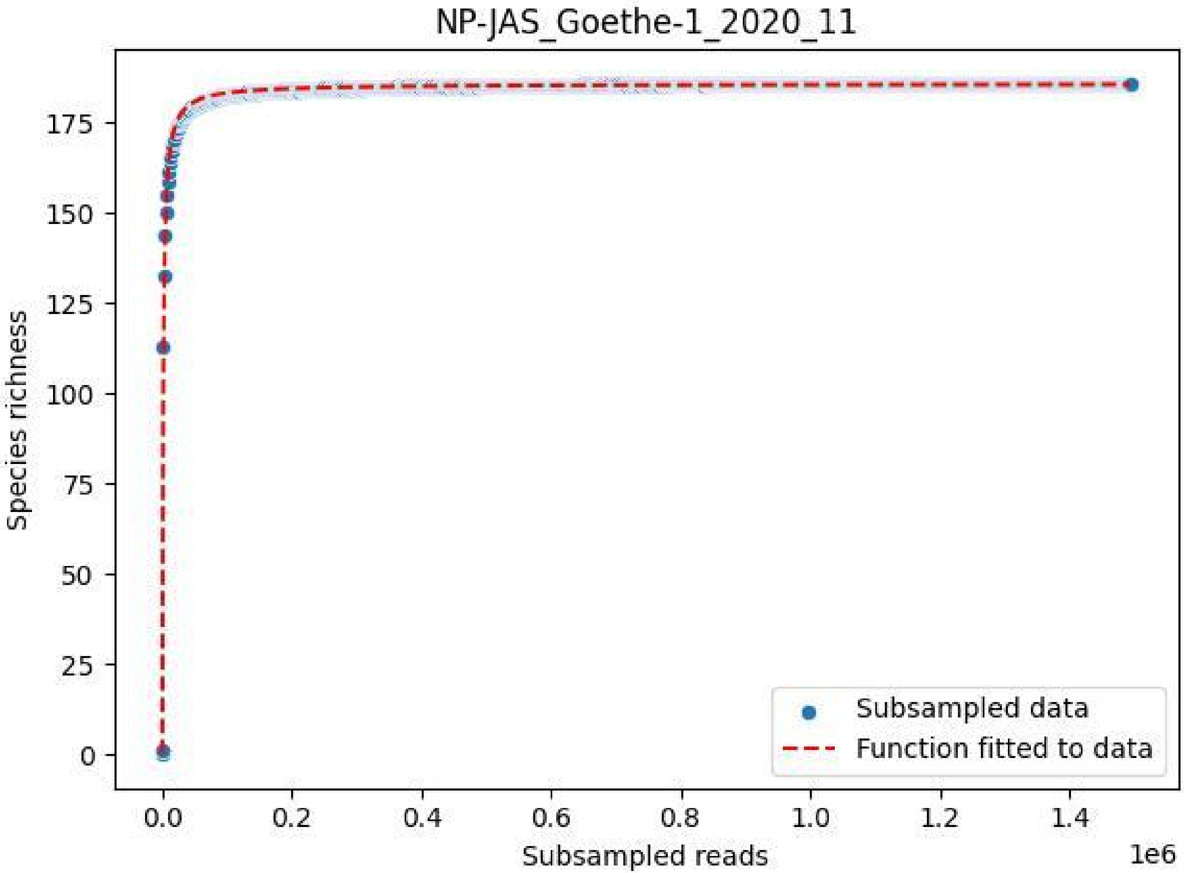

NP-JAS_Goethe-1_2020_11
Species richness
Subsampled reads
1e6
0
25
50
75
100
125
150
175
0.0
0.2
0.4
0.6
0.8
1.0
1.2
1.4
Subsampled data
Function fitted to data

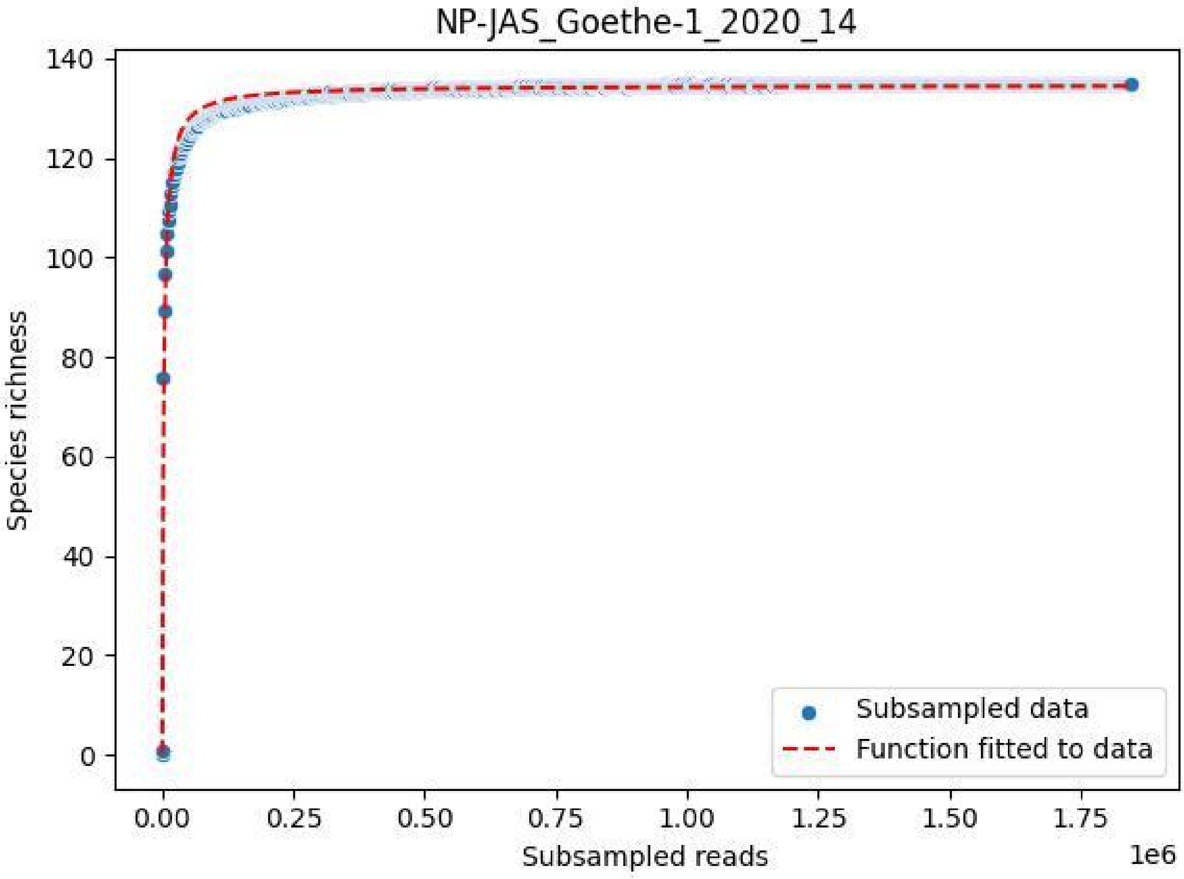

NP-JAS_Goethe-1_2020_14
Species richness
Subsampled reads
1e6
0
20
40
60
80
100
120
140
0.00
0.25
0.50
0.75
1.00
1.25
1.50
1.75
Subsampled data
Function fitted to data

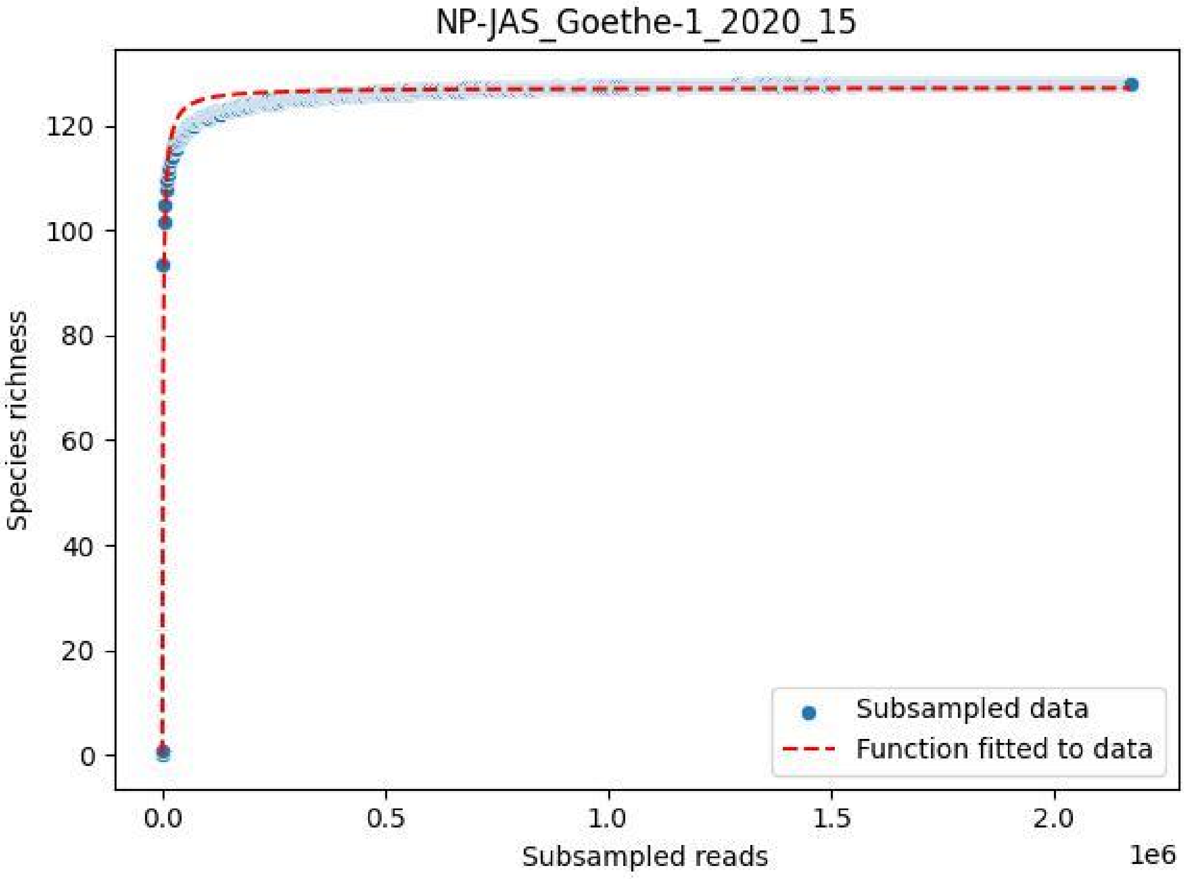
NP-JAS_Goethe-1_2020_15
Species richness
Subsampled reads
1e6
0
20
40
60
80
100
120
0.0
0.5
1.0
1.5
2.0
Subsampled data
Function fitted to data

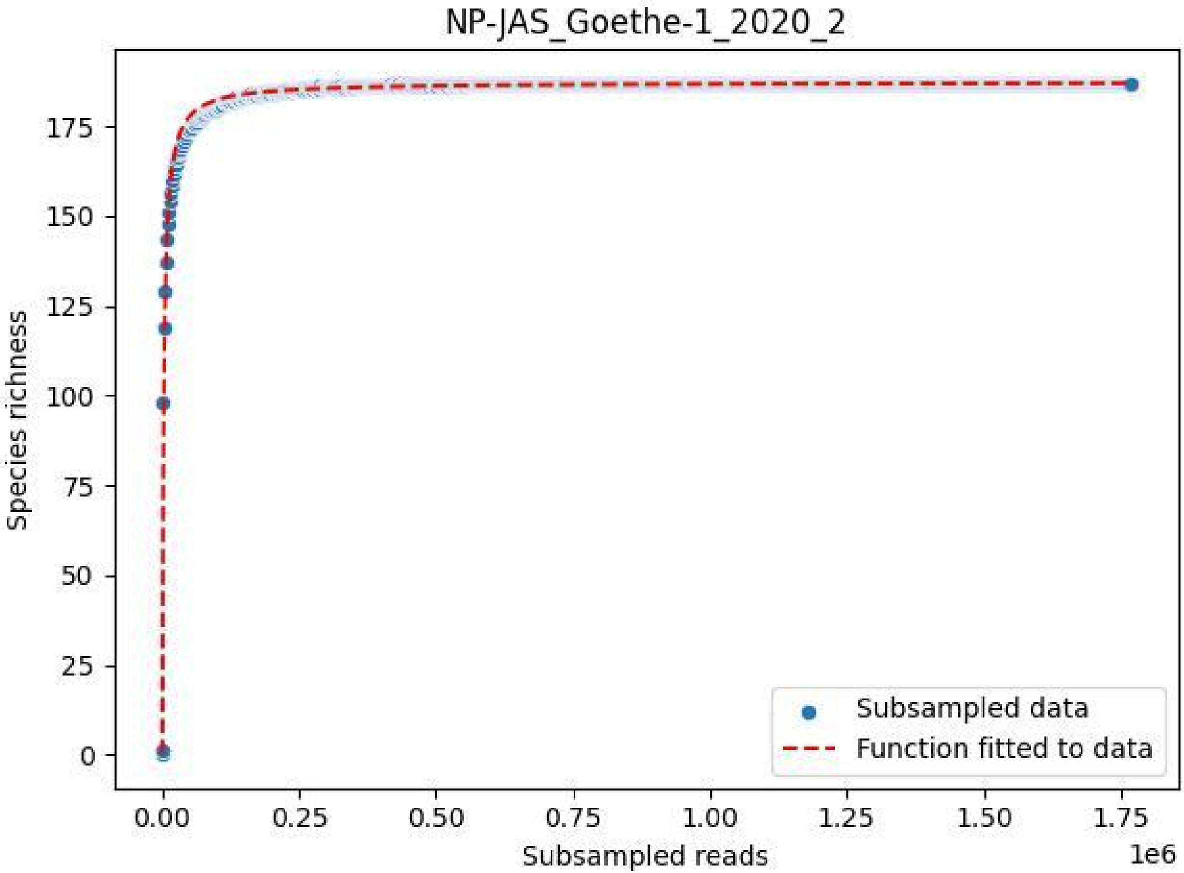
NP-JAS_Goethe-1_2020_2
Species richness
Subsampled reads
1e6
0
25
50
75
100
125
150
175
0.00
0.25
0.50
0.75
1.00
1.25
1.50
1.75
Subsampled data
Function fitted to data

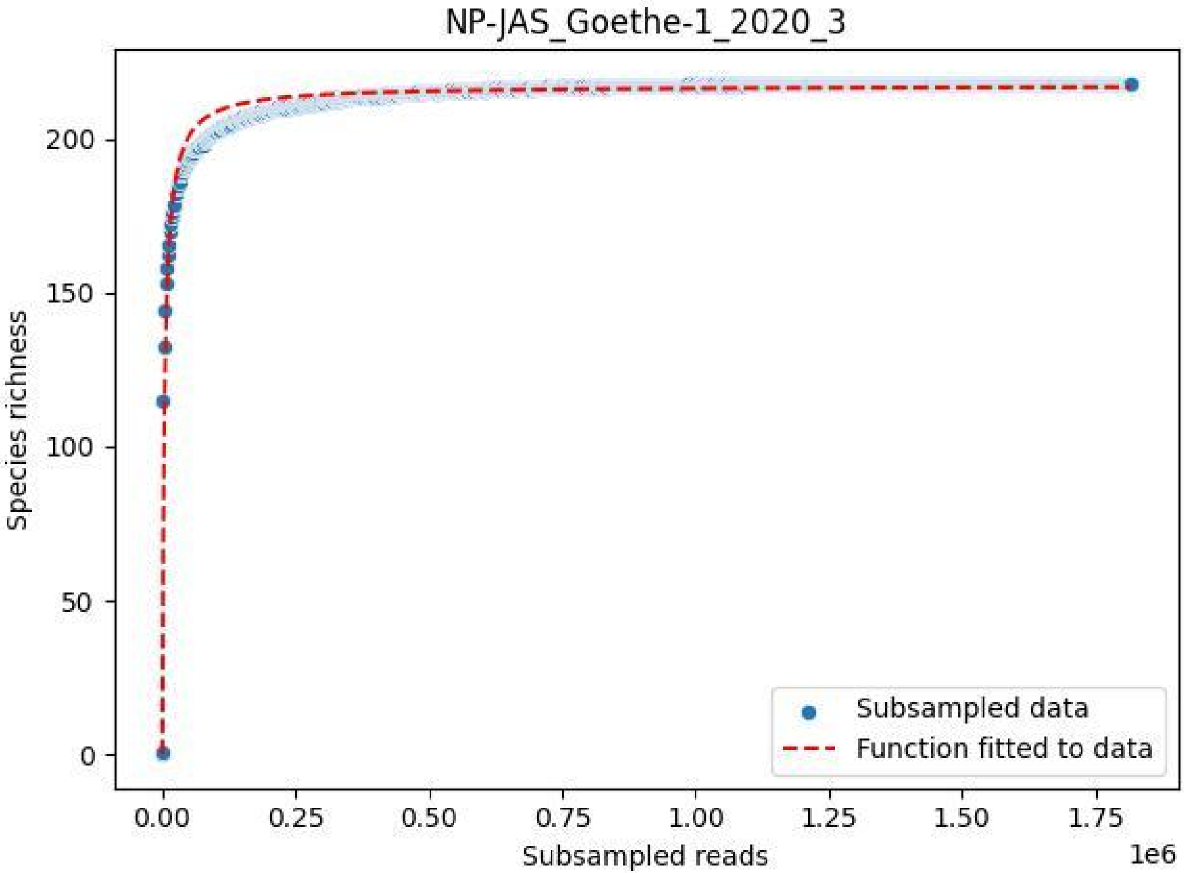

NP-JAS_Goethe-1_2020_3
Species richness
Subsampled reads
1e6
0
50
100
150
200
0.00
0.25
0.50
0.75
1.00
1.25
1.50
1.75
Subsampled data
Function fitted to data

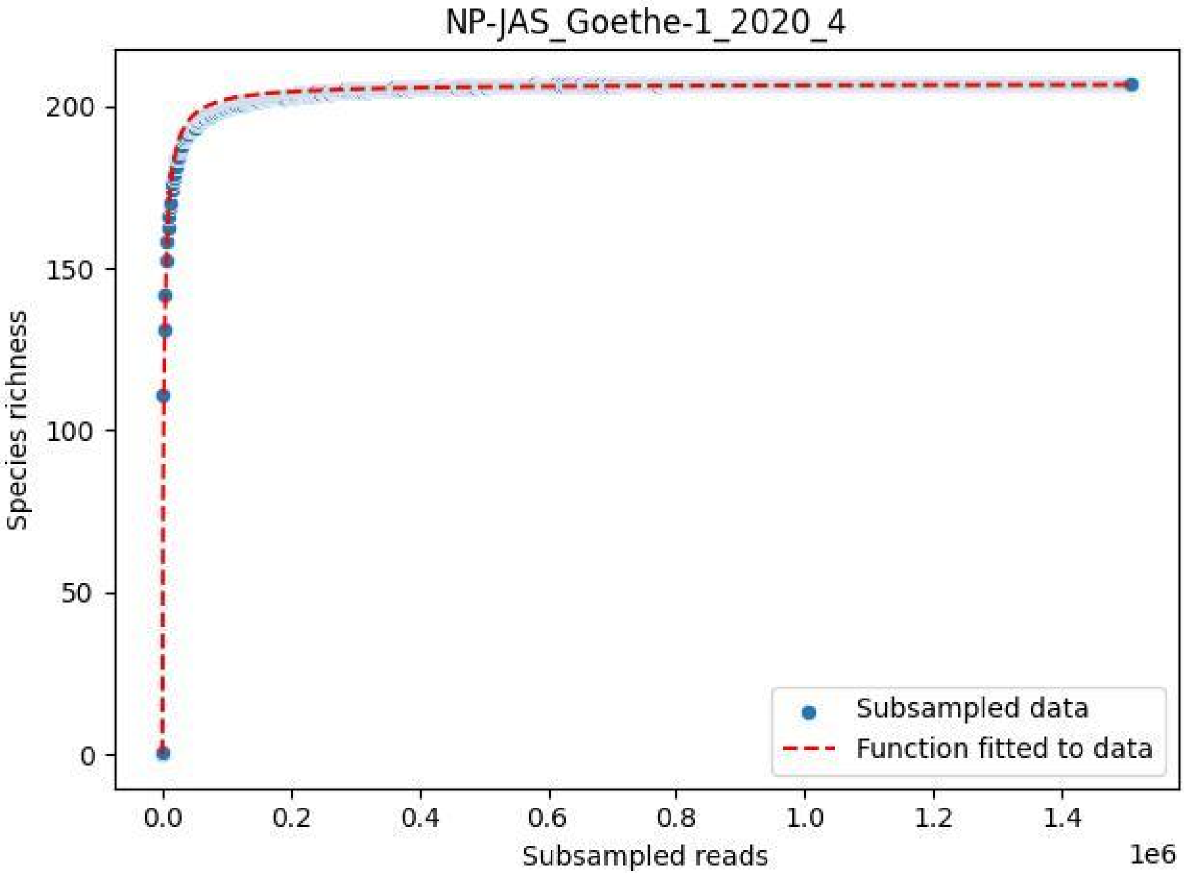

NP-JAS_Goethe-1_2020_4
Species richness
Subsampled reads
1e6
0
50
100
150
200
0.0
0.2
0.4
0.6
0.8
1.0
1.2
1.4
Subsampled data
Function fitted to data

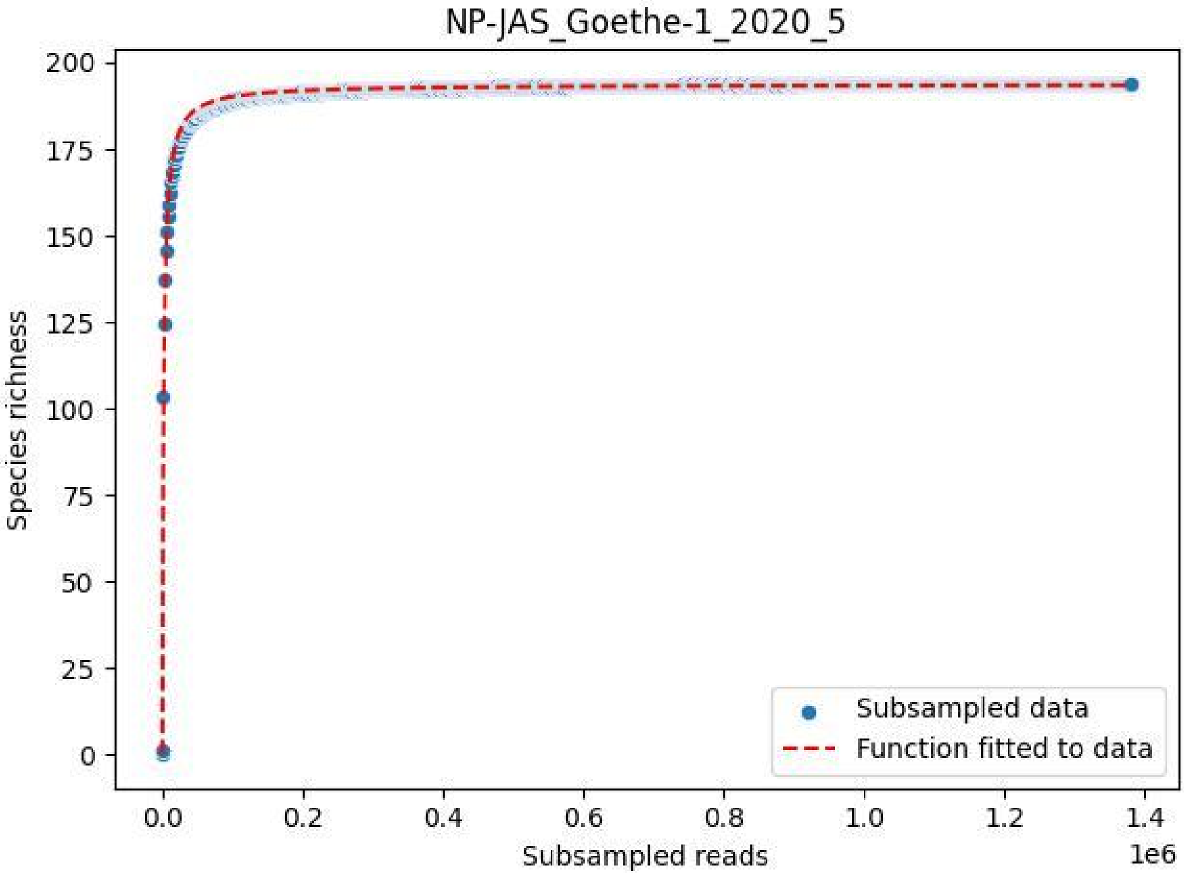
NP-JAS_Goethe-1_2020_5
Species richness
Subsampled reads
1e6
Subsampled data
Function fitted to data
0
25
50
75
100
125
150
175
200
0.0
0.2
0.4
0.6
0.8
1.0
1.2
1.4

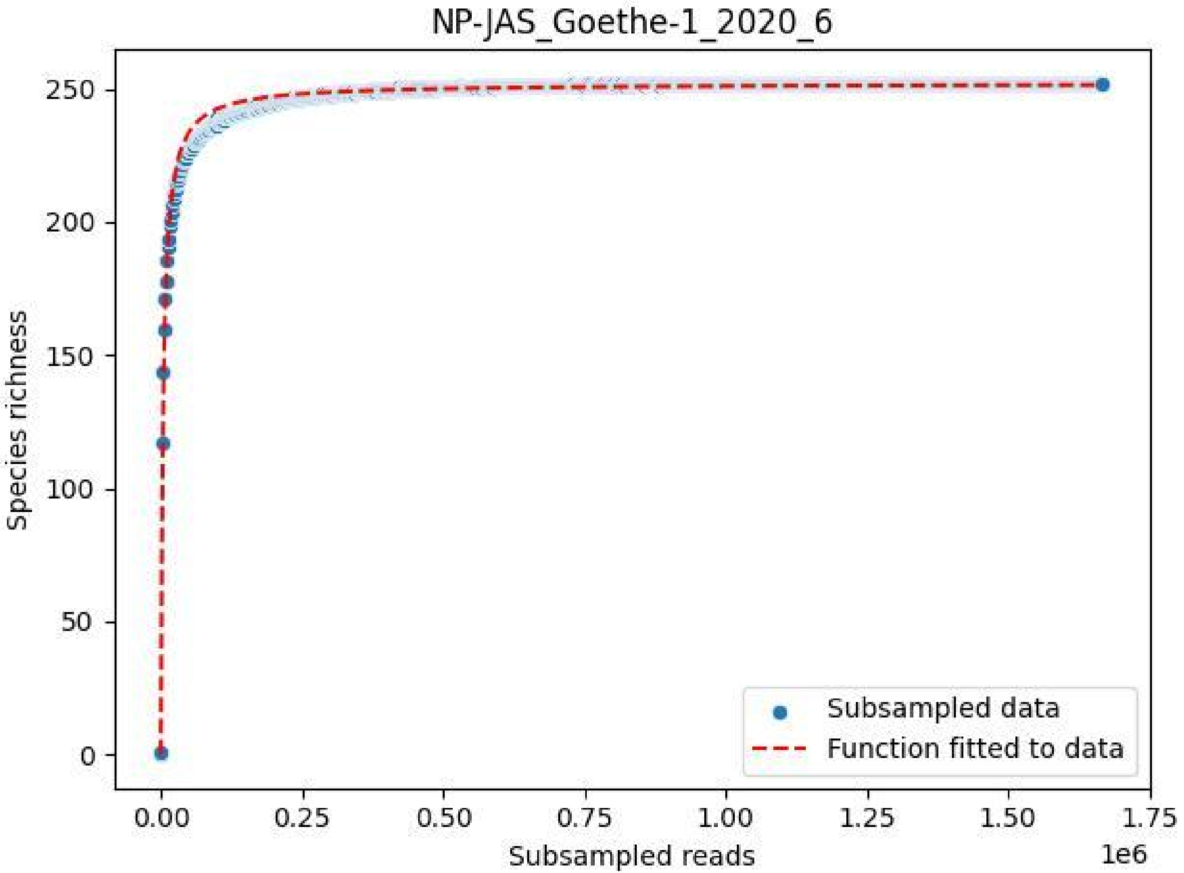
NP-JAS_Goethe-1_2020_6
Species richness
Subsampled reads
1e6
0
50
100
150
200
250
0.00
0.25
0.50
0.75
1.00
1.25
1.50
1.75
Subsampled data
Function fitted to data

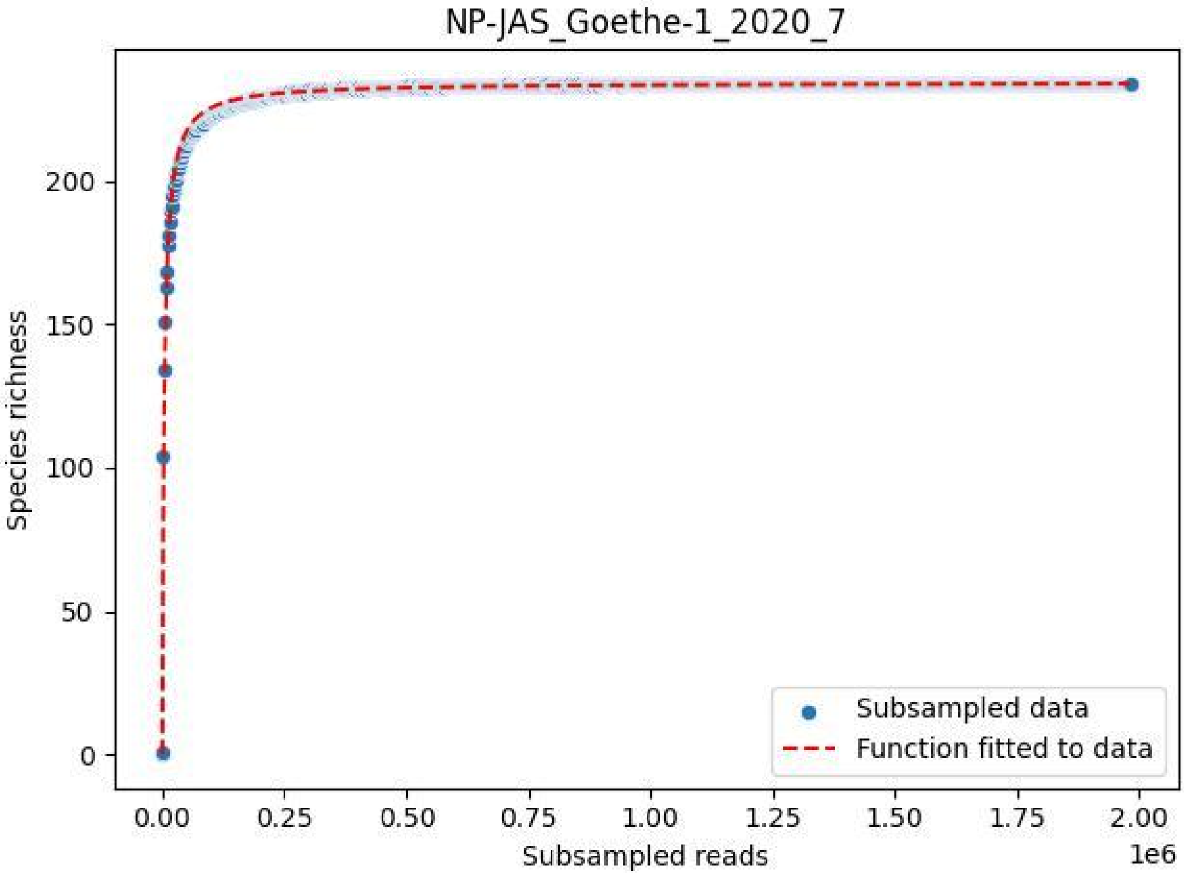
NP-JAS_Goethe-1_2020_7
Species richness
Subsampled reads
1e6
0
50
100
150
200
0.00
0.25
0.50
0.75
1.00
1.25
1.50
1.75
2.00
Subsampled data
Function fitted to data

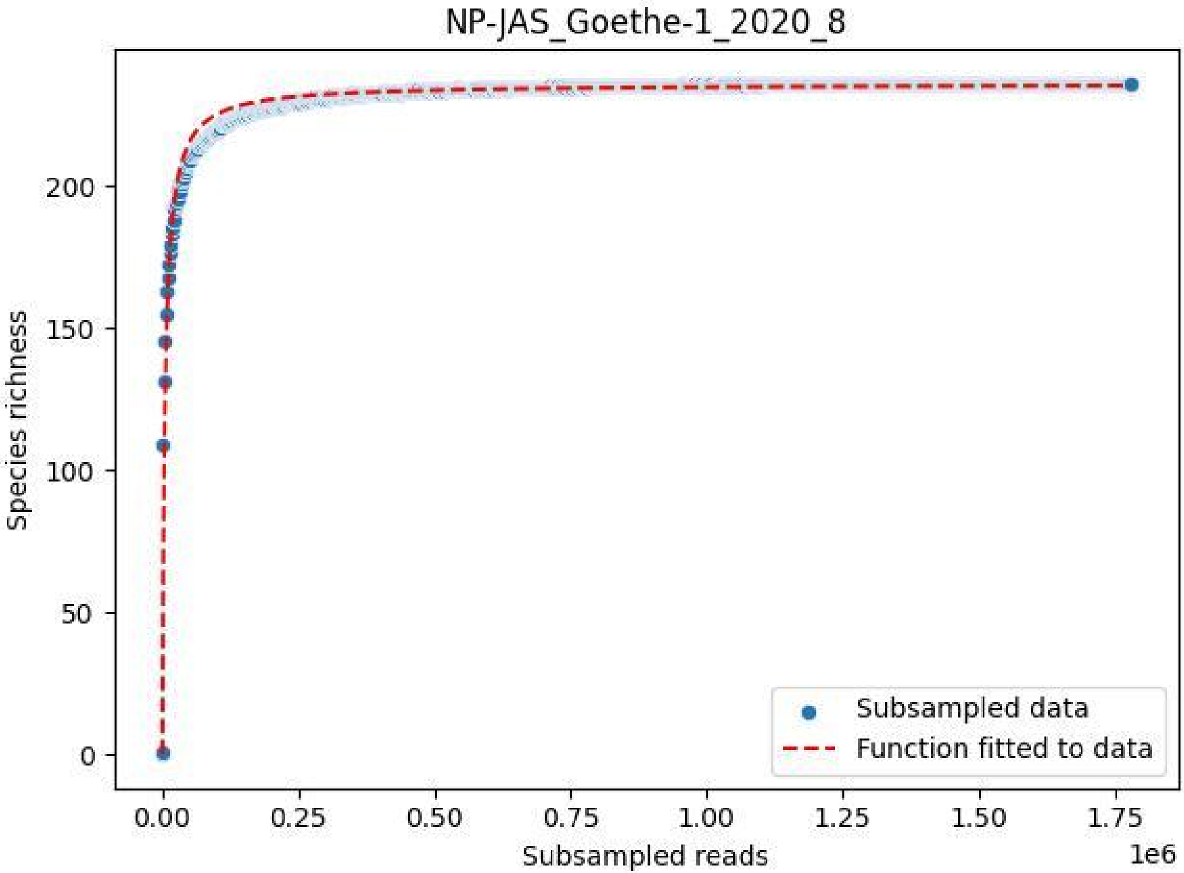

NP-JAS_Goethe-1_2020_8
Species richness
Subsampled reads
1e6
0
50
100
150
200
0.00
0.25
0.50
0.75
1.00
1.25
1.50
1.75
Subsampled data
Function fitted to data

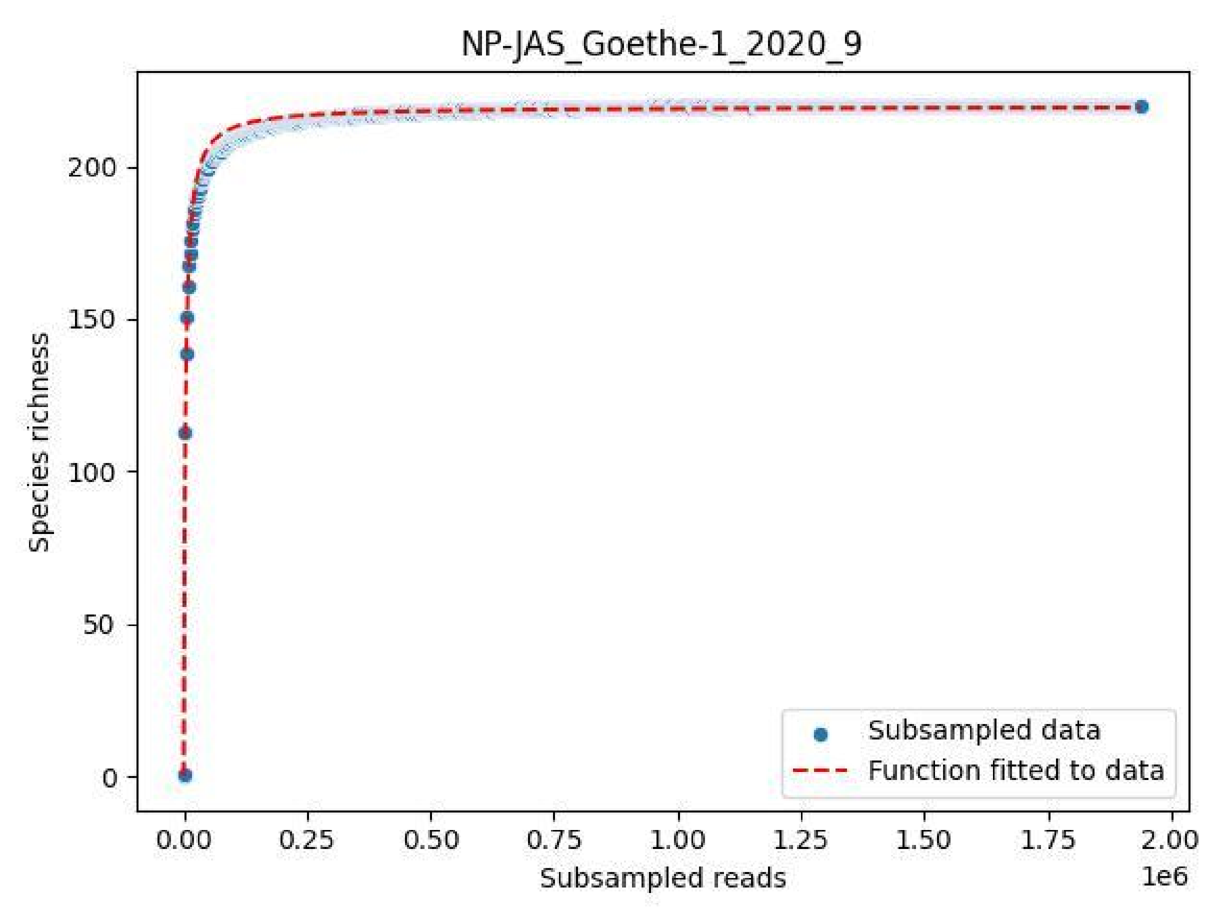
NP-JAS_Goethe-1_2020_9
Species richness
Subsampled reads
1e6
0
50
100
150
200
0.00
0.25
0.50
0.75
1.00
1.25
1.50
1.75
2.00
Subsampled data
Function fitted to data

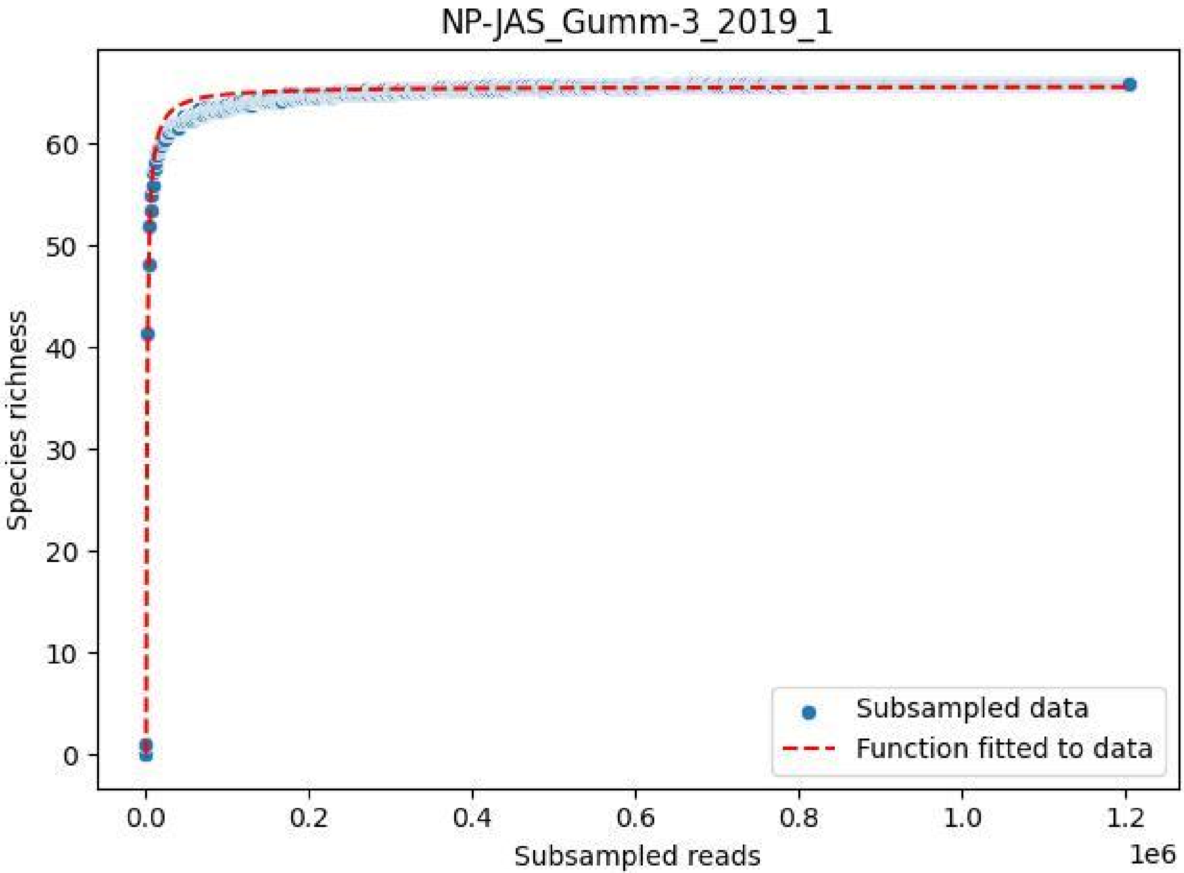

NP-JAS_Gumm-3_2019_1
Species richness
Subsampled reads
1e6
0
10
20
30
40
50
60
0.0
0.2
0.4
0.6
0.8
1.0
1.2
Subsampled data
Function fitted to data

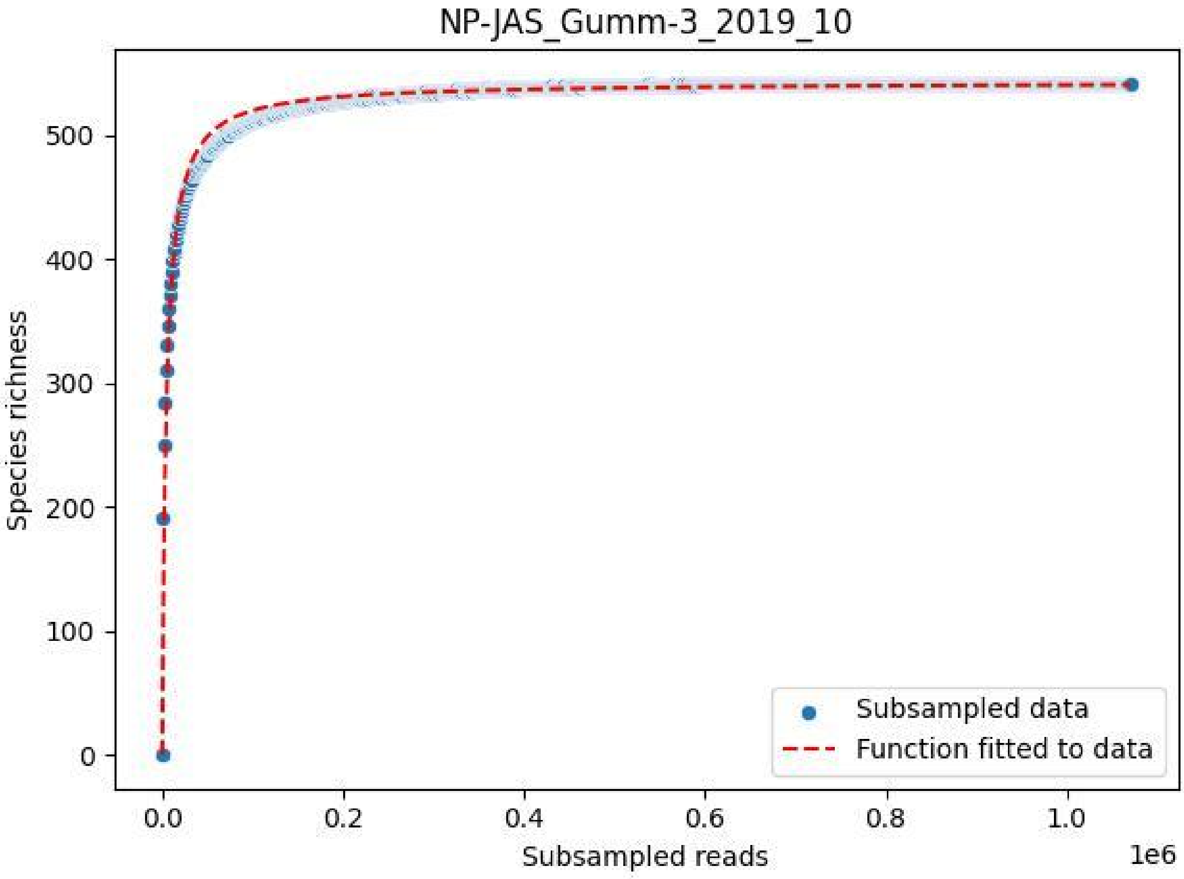

NP-JAS_Gumm-3_2019_10
Species richness
Subsampled reads
1e6
0
100
200
300
400
500
0.0
0.2
0.4
0.6
0.8
1.0
Subsampled data
Function fitted to data

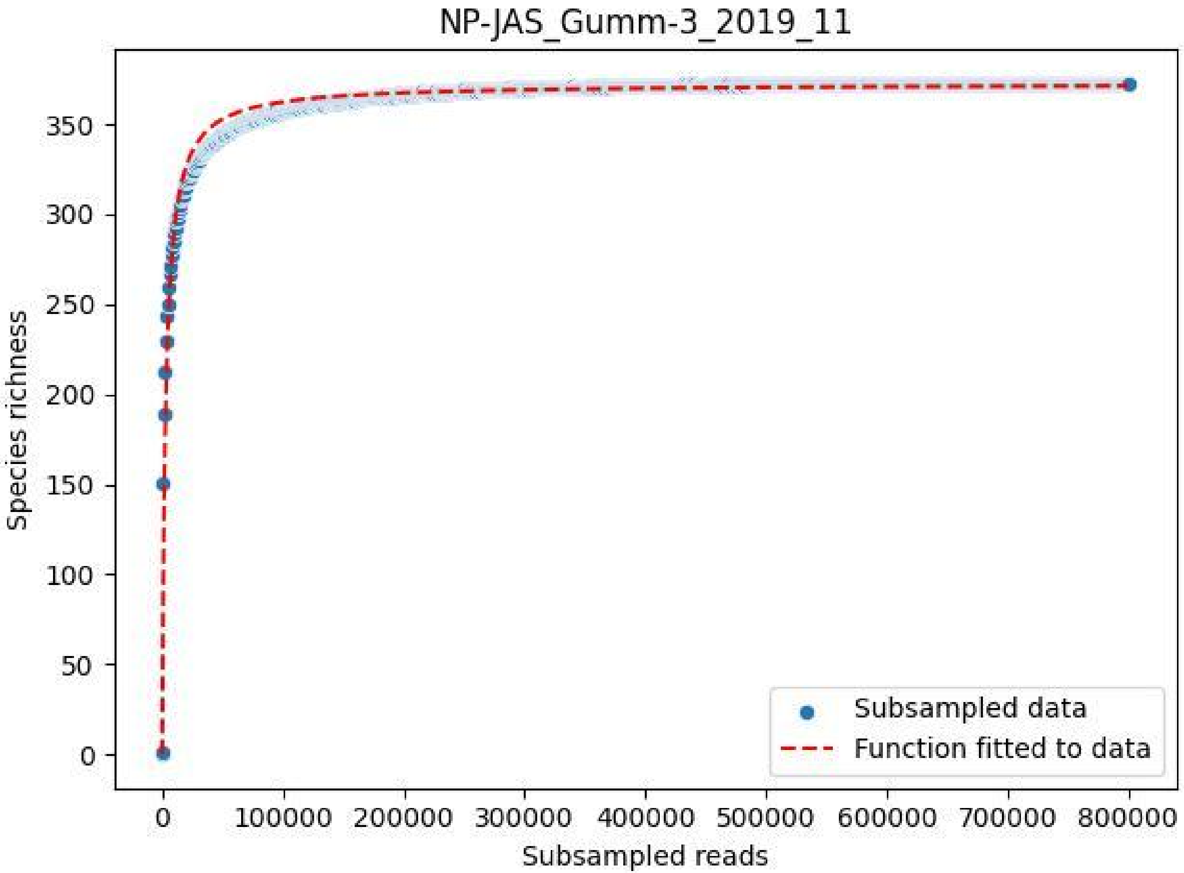

NP-JAS_Gumm-3_2019_11
Species richness
Subsampled reads
0
50
100
150
200
250
300
350
0
100000
200000
300000
400000
500000
600000
700000
800000
Subsampled data
Function fitted to data

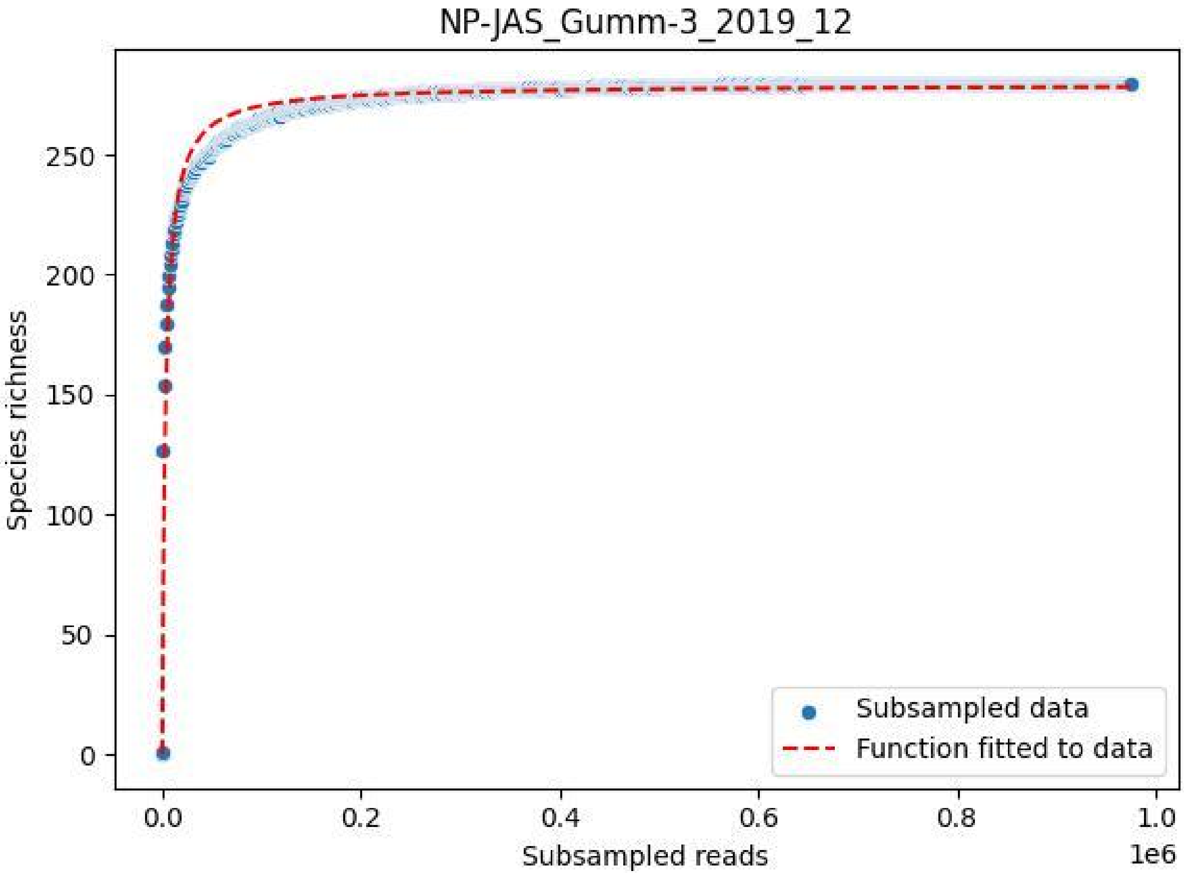
NP-JAS_Gumm-3_2019_12
Species richness
Subsampled reads
1e6
0
50
100
150
200
250
0.0
0.2
0.4
0.6
0.8
1.0
Subsampled data
Function fitted to data

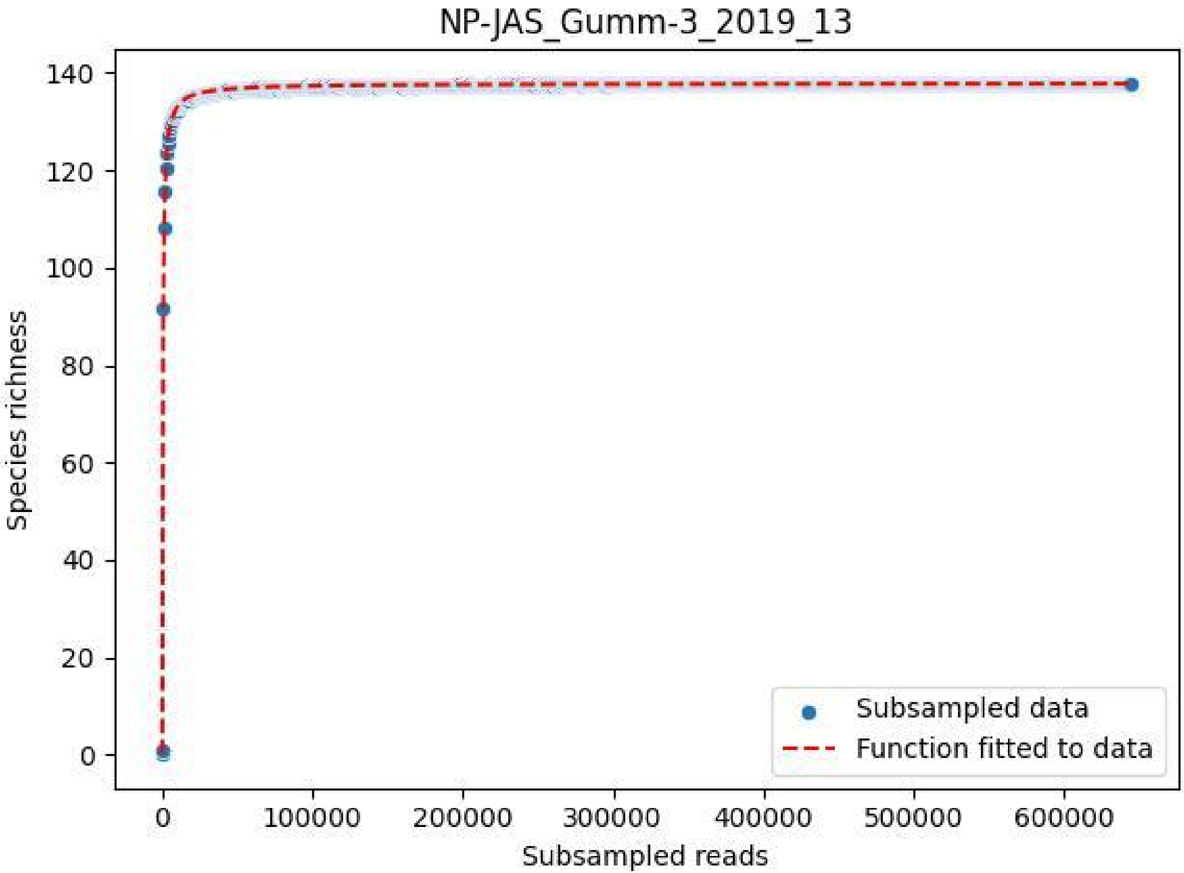
NP-JAS_Gumm-3_2019_13
Species richness
Subsampled reads
0
20
40
60
80
100
120
140
0
100000
200000
300000
400000
500000
600000
Subsampled data
Function fitted to data

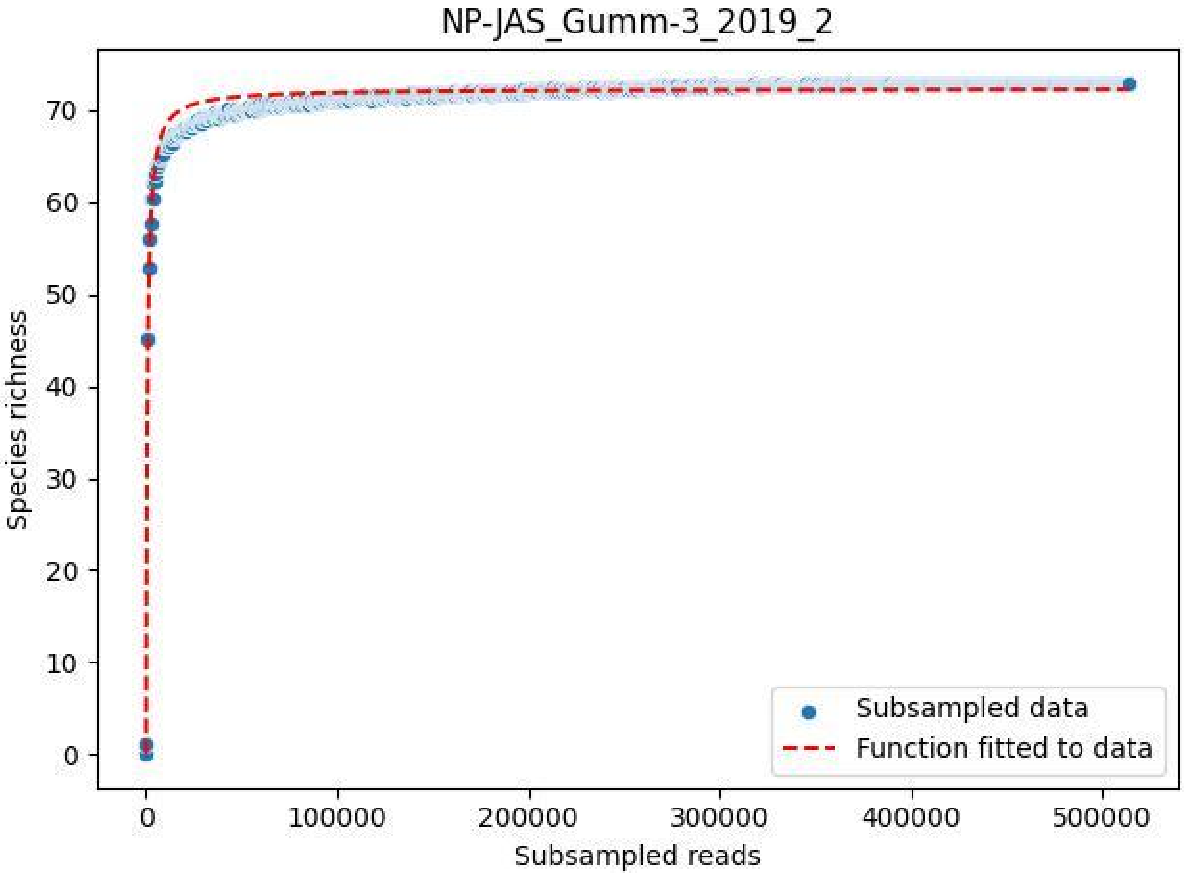

NP-JAS_Gumm-3_2019_2
Species richness
Subsampled reads
0
10
20
30
40
50
60
70
0
100000
200000
300000
400000
500000
Subsampled data
Function fitted to data

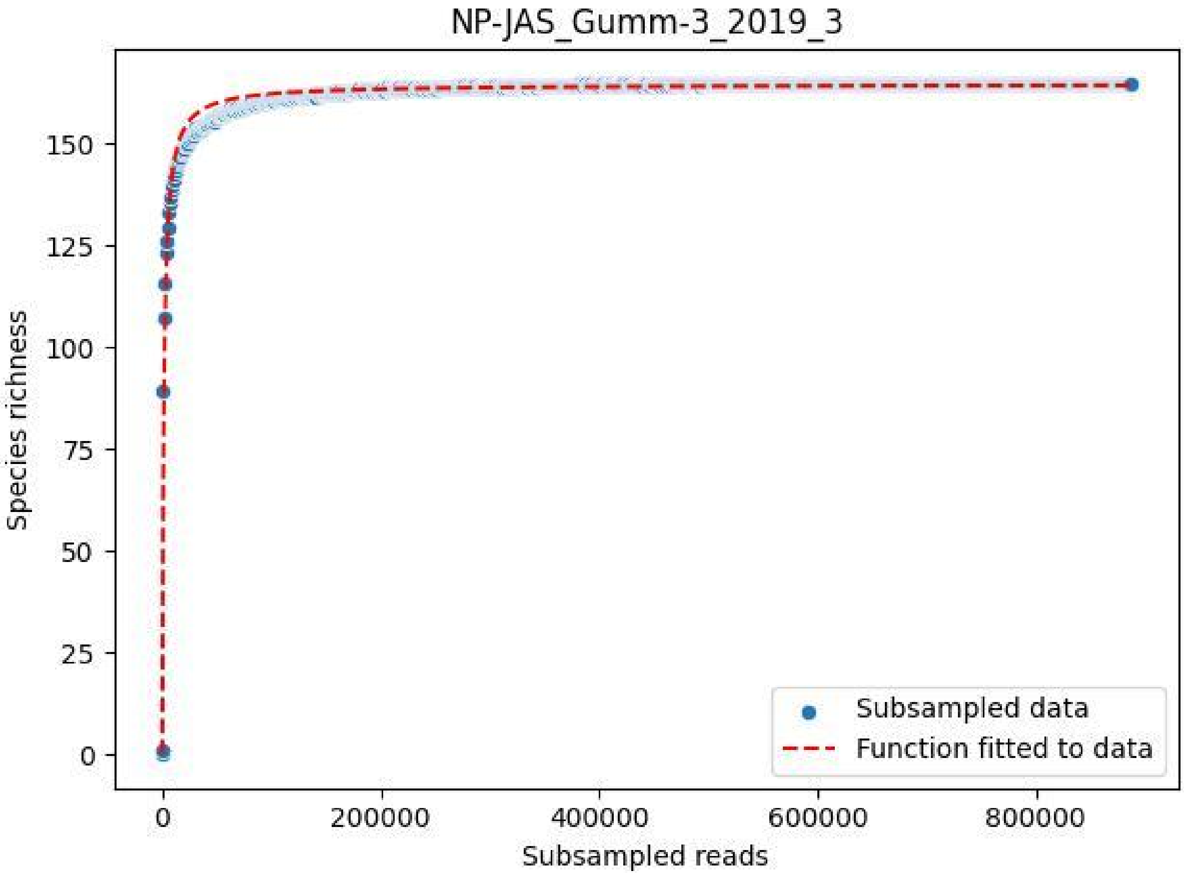

NP-JAS_Gumm-3_2019_3
Species richness
Subsampled reads
0
25
50
75
100
125
150
200000
400000
600000
800000
Subsampled data
Function fitted to data

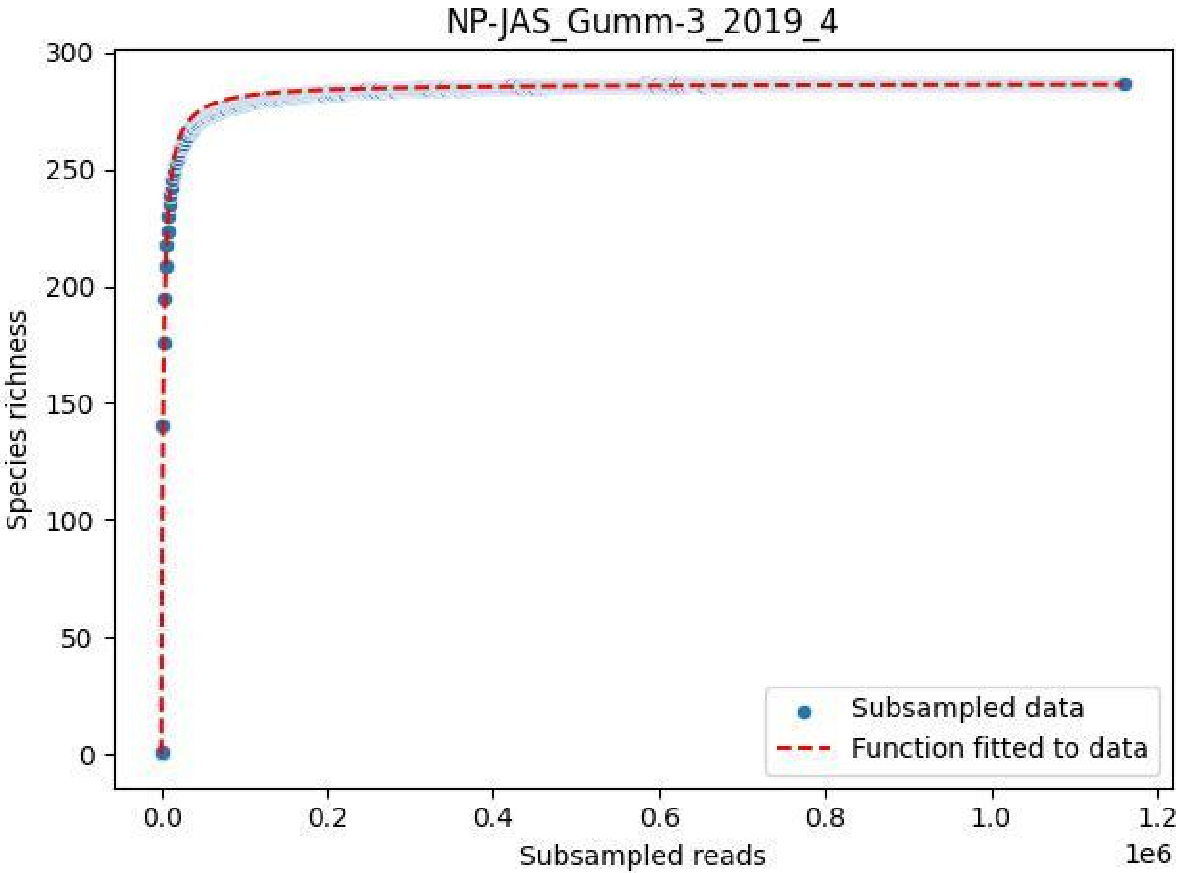

NP-JAS_Gumm-3_2019_4
Species richness
Subsampled reads
1e6
0
50
100
150
200
250
300
0.0
0.2
0.4
0.6
0.8
1.0
1.2
Subsampled data
Function fitted to data

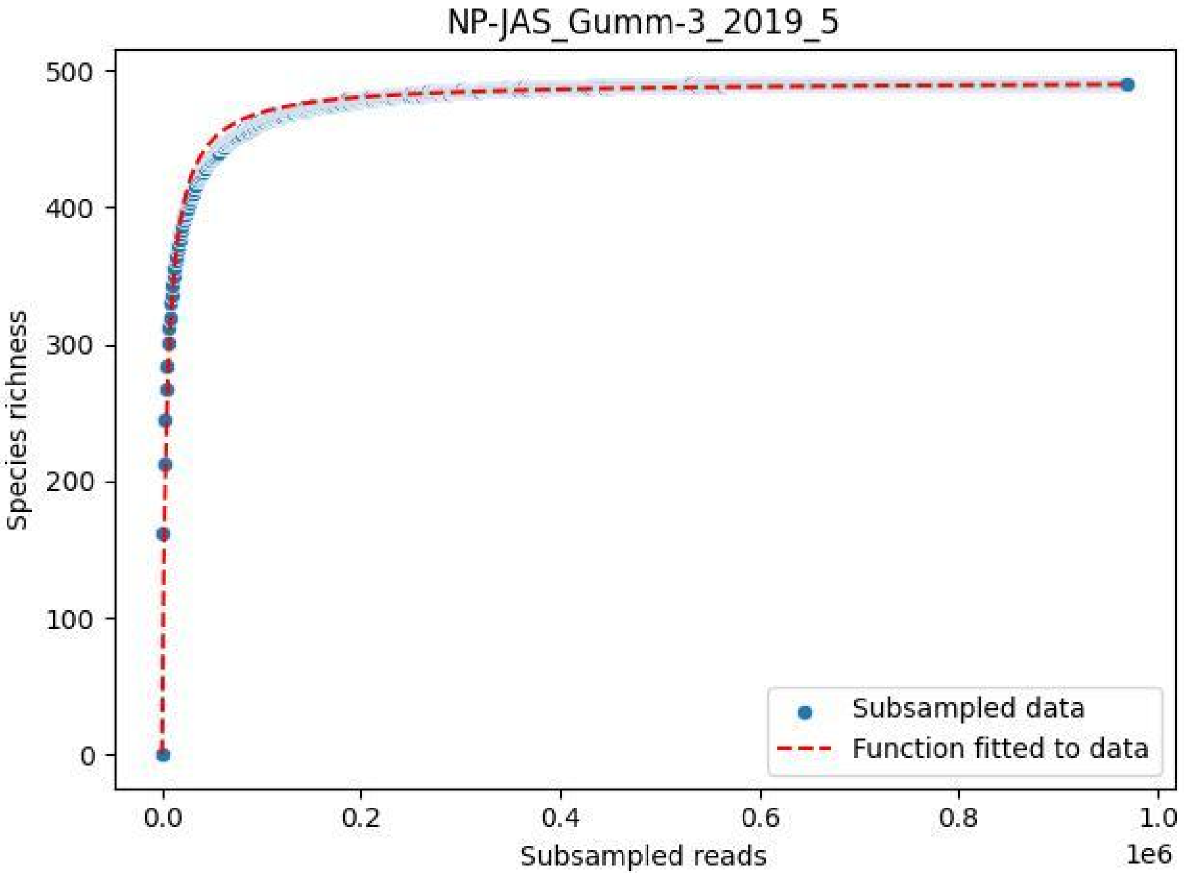
NP-JAS_Gumm-3_2019_5
Species richness
Subsampled reads
1e6
0
100
200
300
400
500
0.0
0.2
0.4
0.6
0.8
1.0
Subsampled data
Function fitted to data

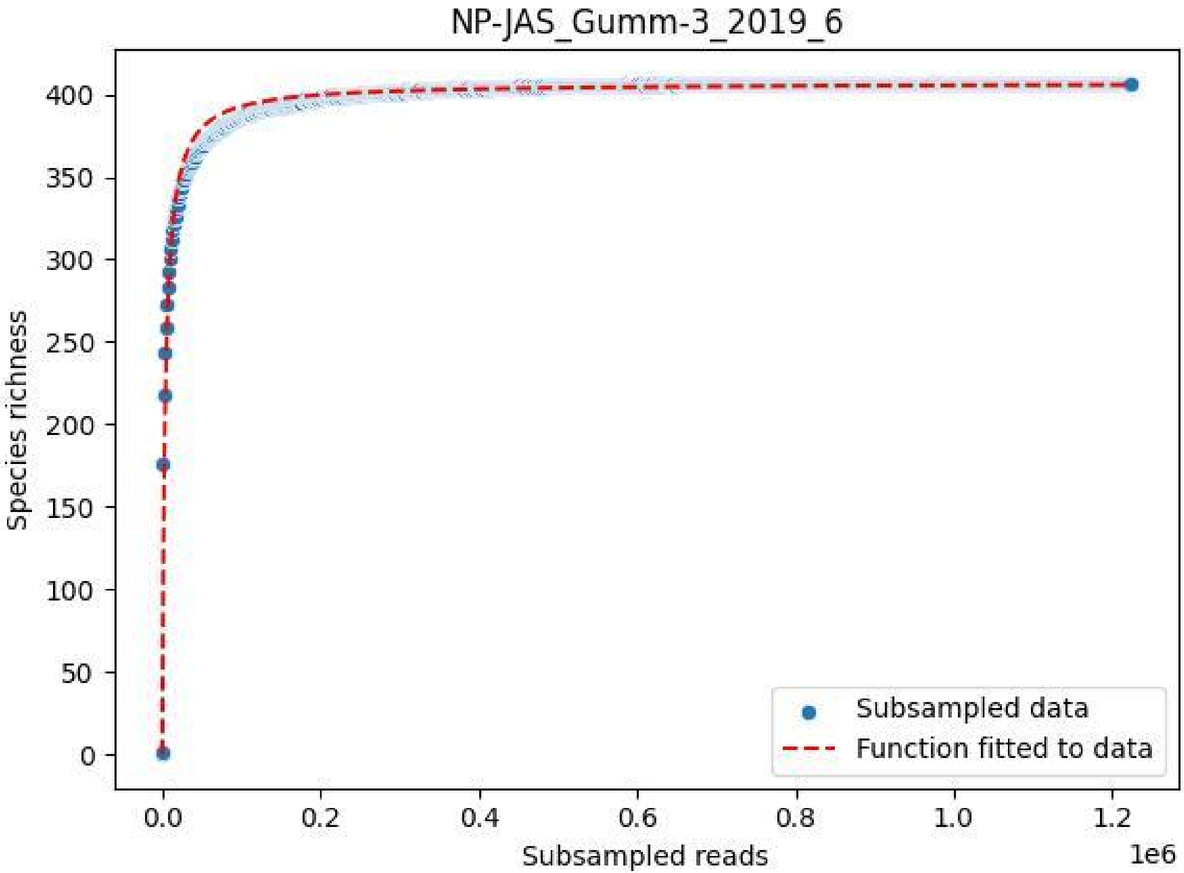

NP-JAS_Gumm-3_2019_6
Species richness
Subsampled reads
1e6
0
50
100
150
200
250
300
350
400
0.0
0.2
0.4
0.6
0.8
1.0
1.2
Subsampled data
Function fitted to data

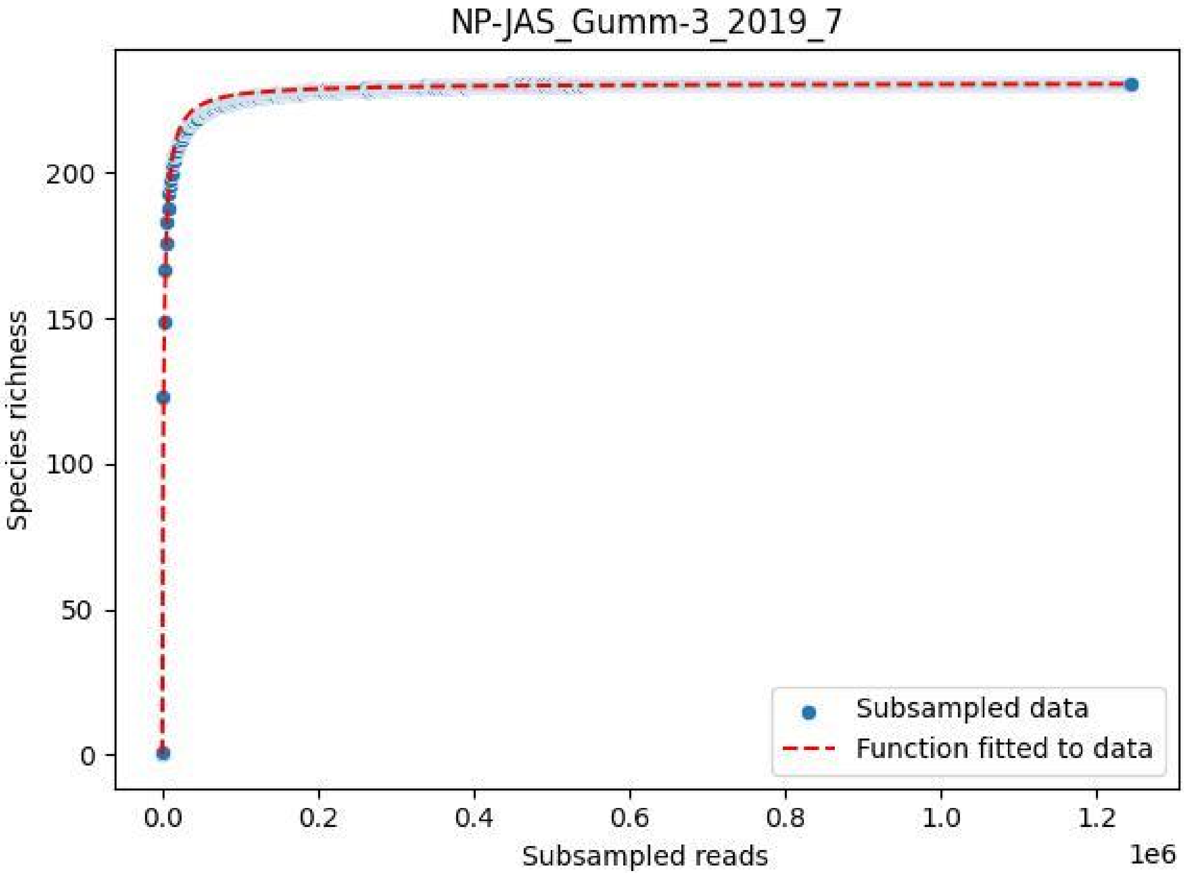
NP-JAS_Gumm-3_2019_7
Species richness
Subsampled reads
1e6
0
50
100
150
200
0.0
0.2
0.4
0.6
0.8
1.0
1.2
Subsampled data
Function fitted to data

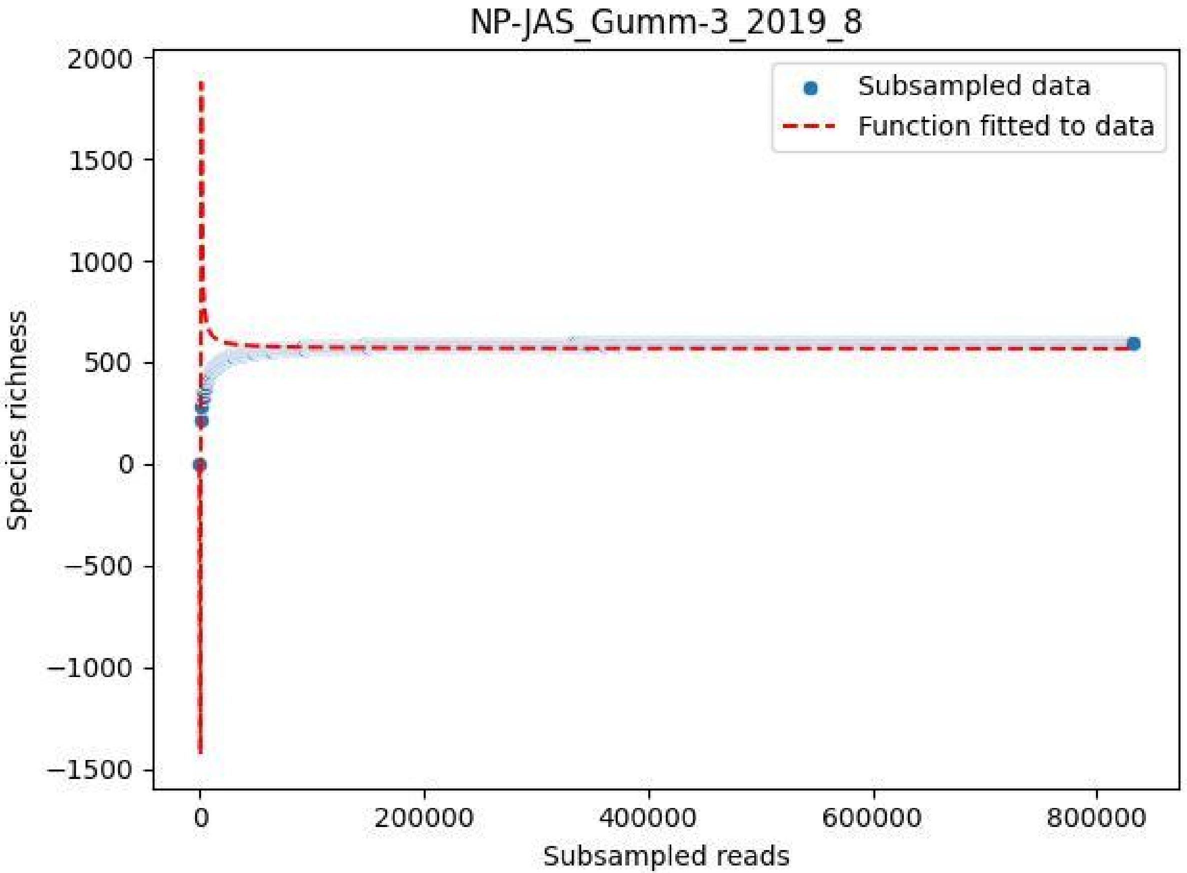

NP-JAS_Gumm-3_2019_8
Subsampled data
Function fitted to data
Species richness
Subsampled reads
2000
1500
1000
500
0
−500
−1000
−1500
0
200000
400000
600000
800000

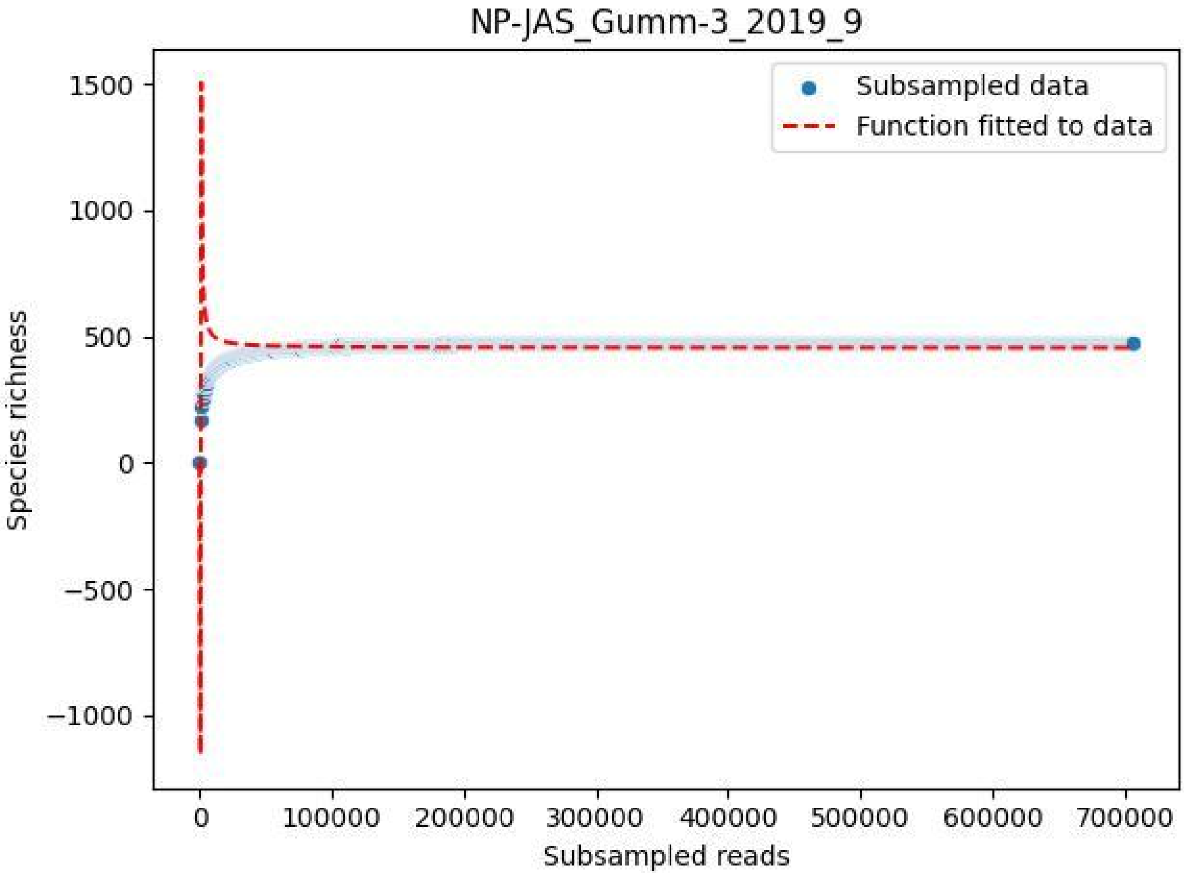

NP-JAS_Gumm-3_2019_9
Subsampled data
Function fitted to data
1500
1000
500
0
−500
−1000
Species richness
0
100000
200000
300000
400000
500000
600000
700000
Subsampled reads

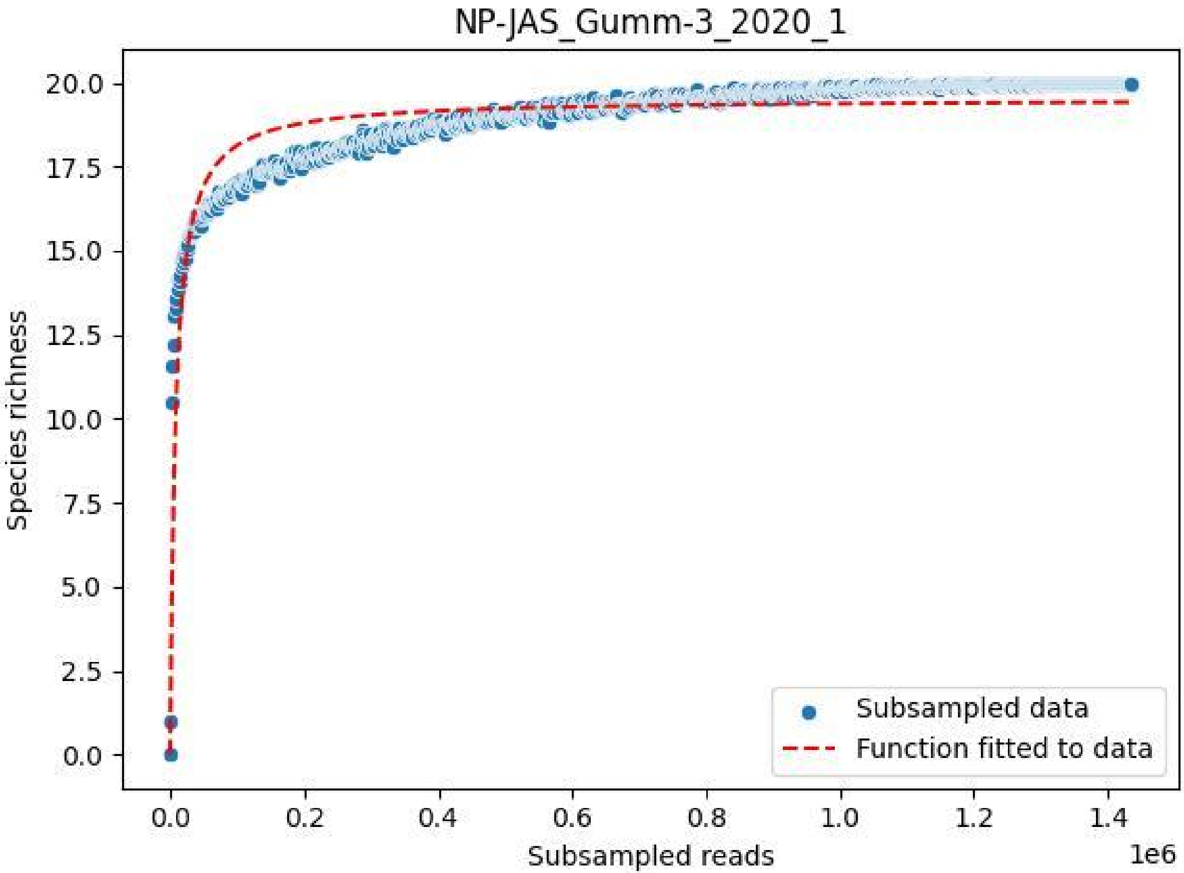
NP-JAS_Gumm-3_2020_1
Species richness
20.0
17.5
15.0
12.5
10.0
7.5
5.0
2.5
0.0
0.0
0.2
0.4
0.6
0.8
1.0
1.2
1.4
1e6
Subsampled reads
Subsampled data
Function fitted to data

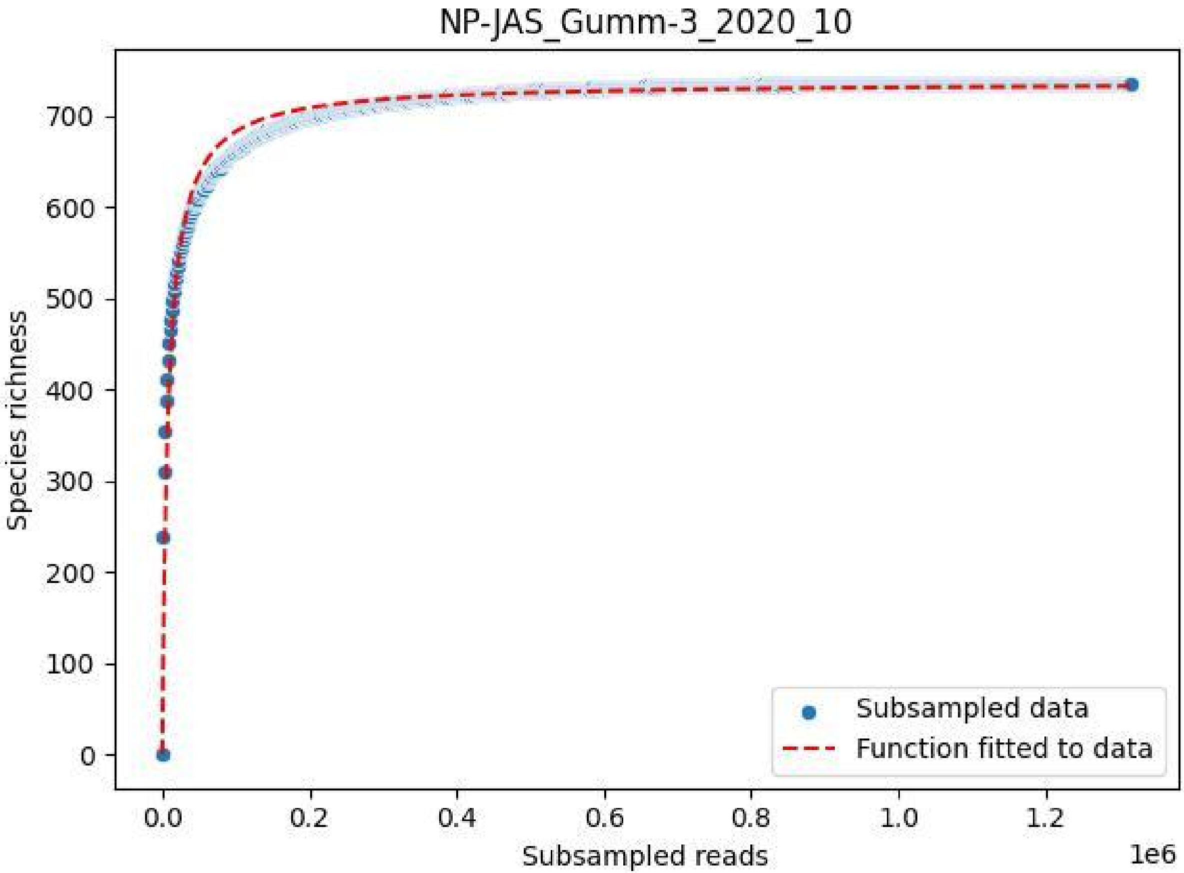
NP-JAS_Gumm-3_2020_10
Species richness
0
100
200
300
400
500
600
700
0.0
0.2
0.4
0.6
0.8
1.0
1.2
1e6
Subsampled reads
Subsampled data
Function fitted to data

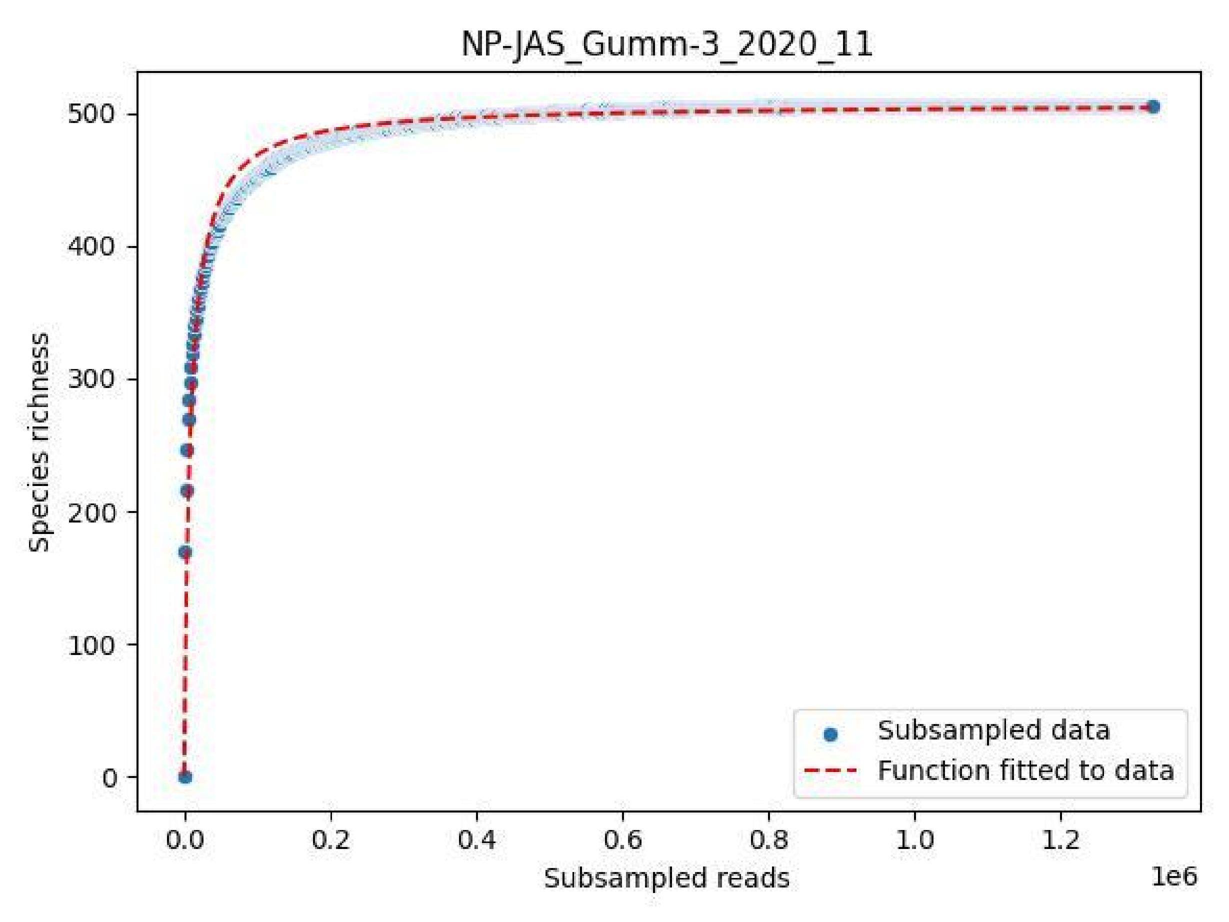

NP-JAS_Gumm-3_2020_11
Species richness
500
400
300
200
100
0
0.0
0.2
0.4
0.6
0.8
1.0
1.2
1e6
Subsampled reads
Subsampled data
Function fitted to data

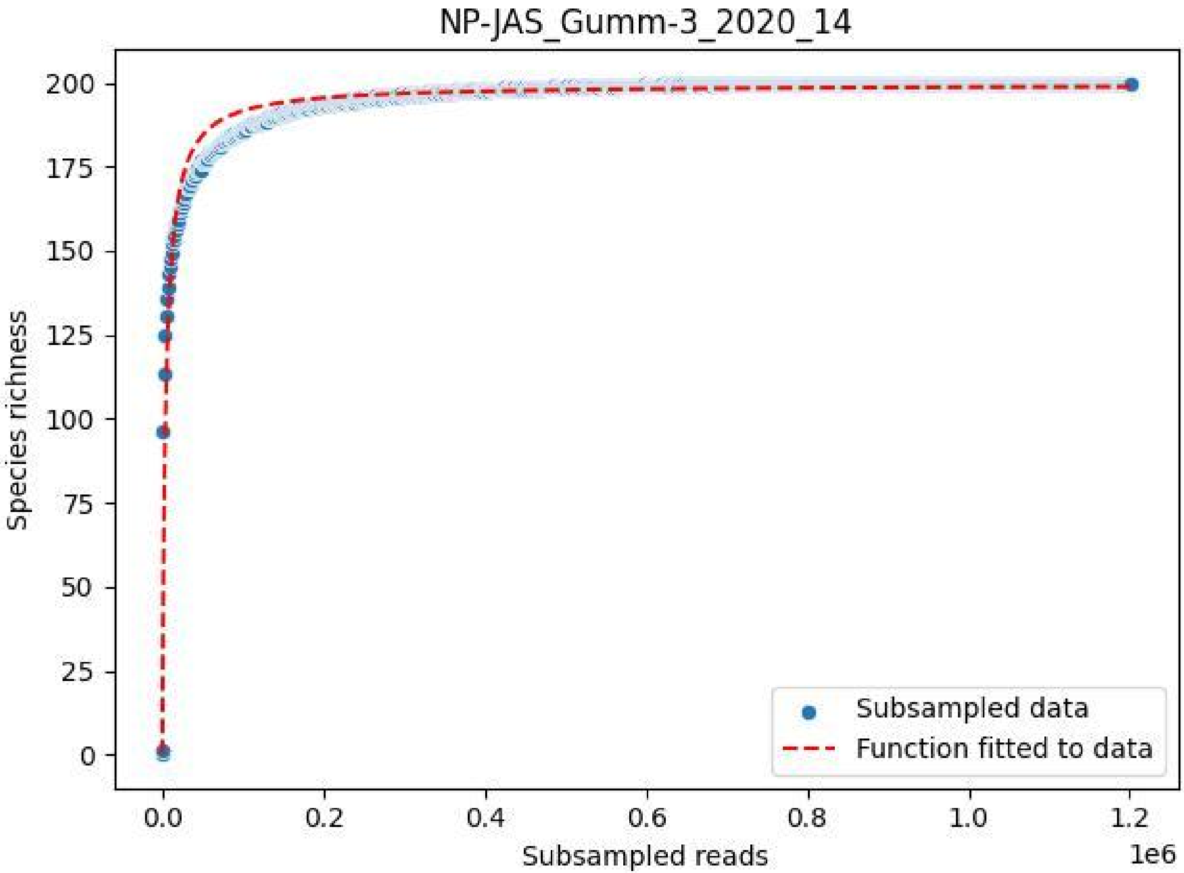

NP-JAS_Gumm-3_2020_14
Species richness
Subsampled reads
1e6
Subsampled data
Function fitted to data
0
25
50
75
100
125
150
175
200
0.0
0.2
0.4
0.6
0.8
1.0
1.2

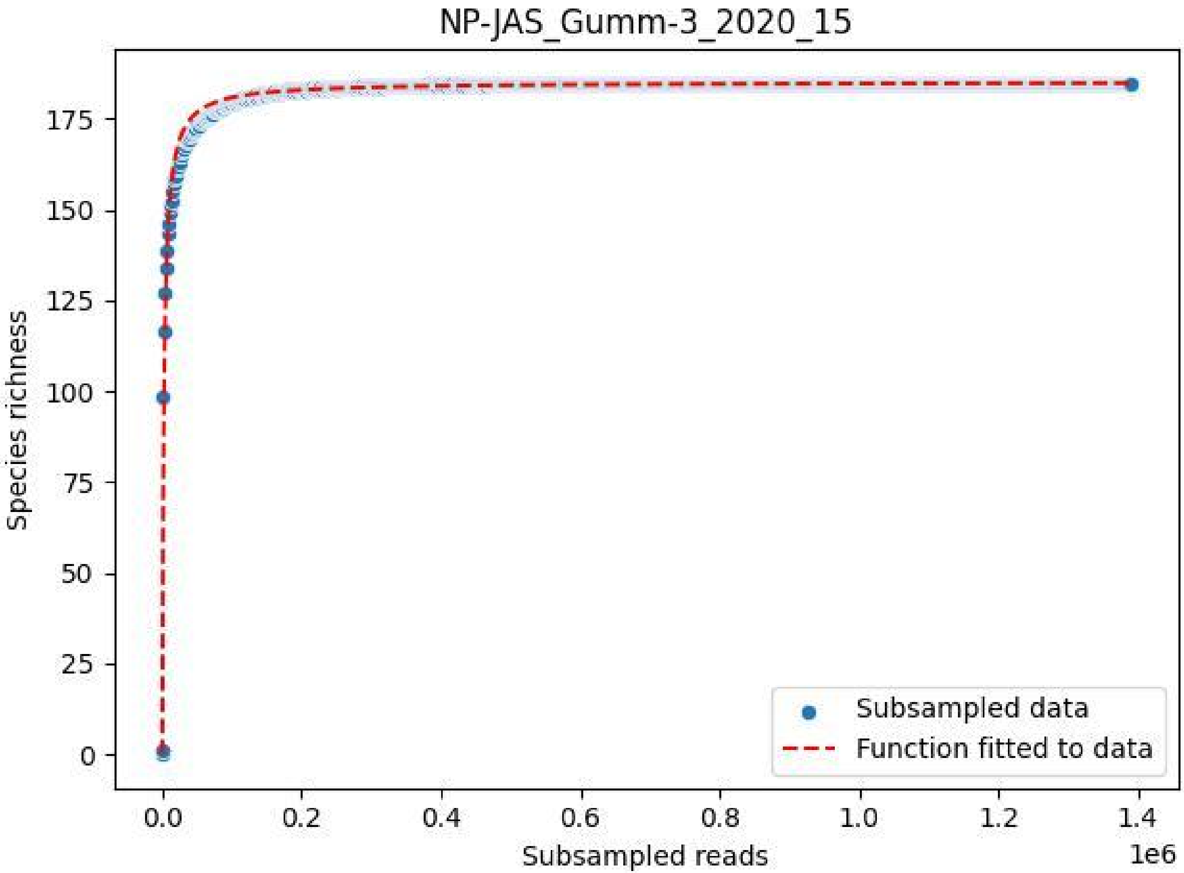
NP-JAS_Gumm-3_2020_15
Species richness
Subsampled reads
1e6
0
25
50
75
100
125
150
175
0.0
0.2
0.4
0.6
0.8
1.0
1.2
1.4
Subsampled data
Function fitted to data

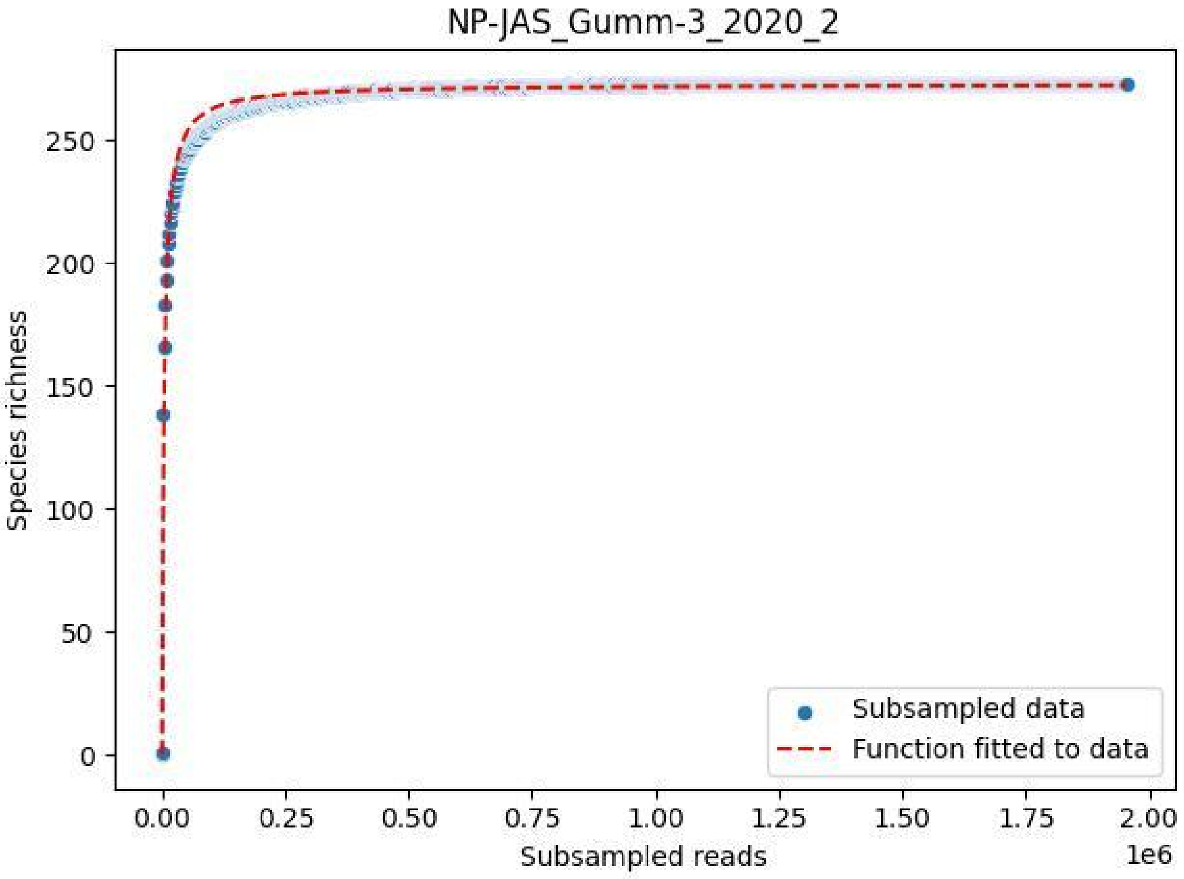

NP-JAS_Gumm-3_2020_2
Species richness
Subsampled reads
1e6
0
50
100
150
200
250
0.00
0.25
0.50
0.75
1.00
1.25
1.50
1.75
2.00
Subsampled data
Function fitted to data

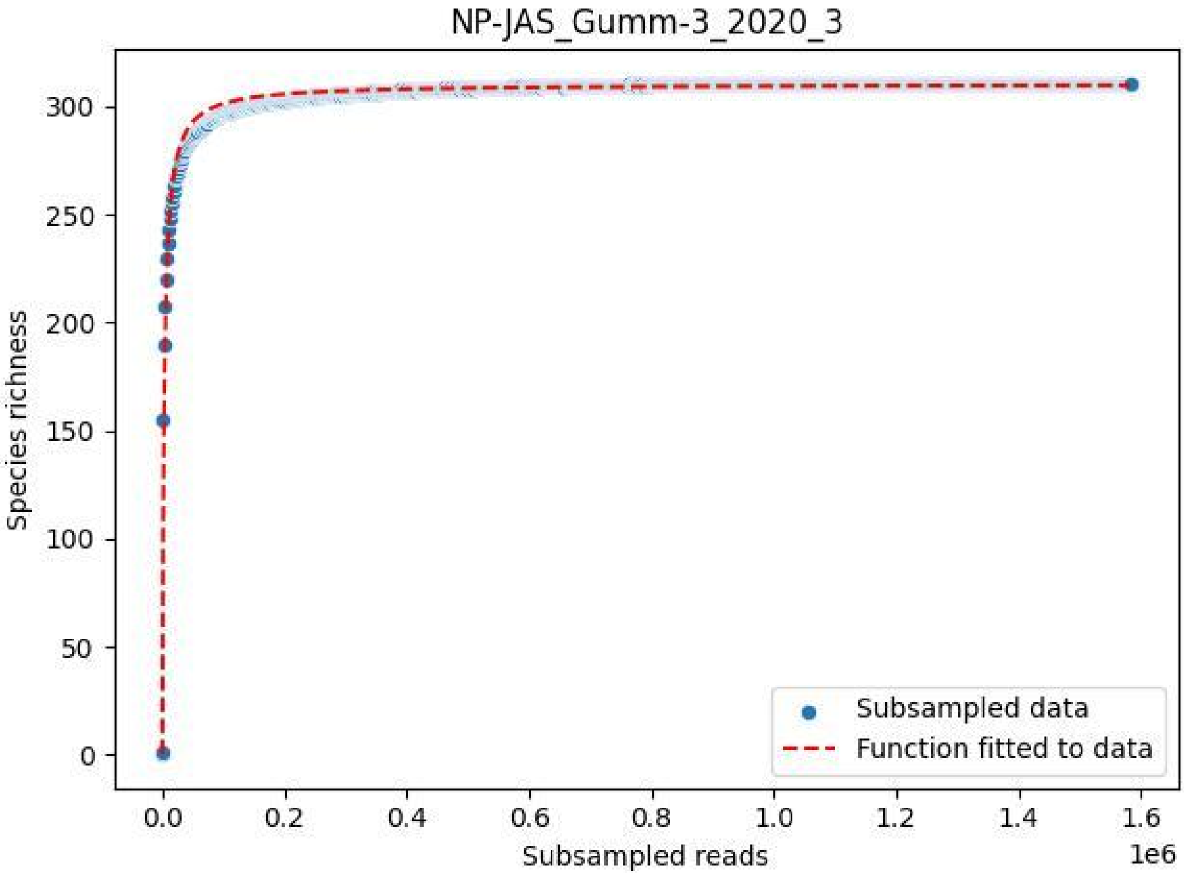
NP-JAS_Gumm-3_2020_3
Species richness
Subsampled reads
1e6
0
50
100
150
200
250
300
0.0
0.2
0.4
0.6
0.8
1.0
1.2
1.4
1.6
Subsampled data
Function fitted to data

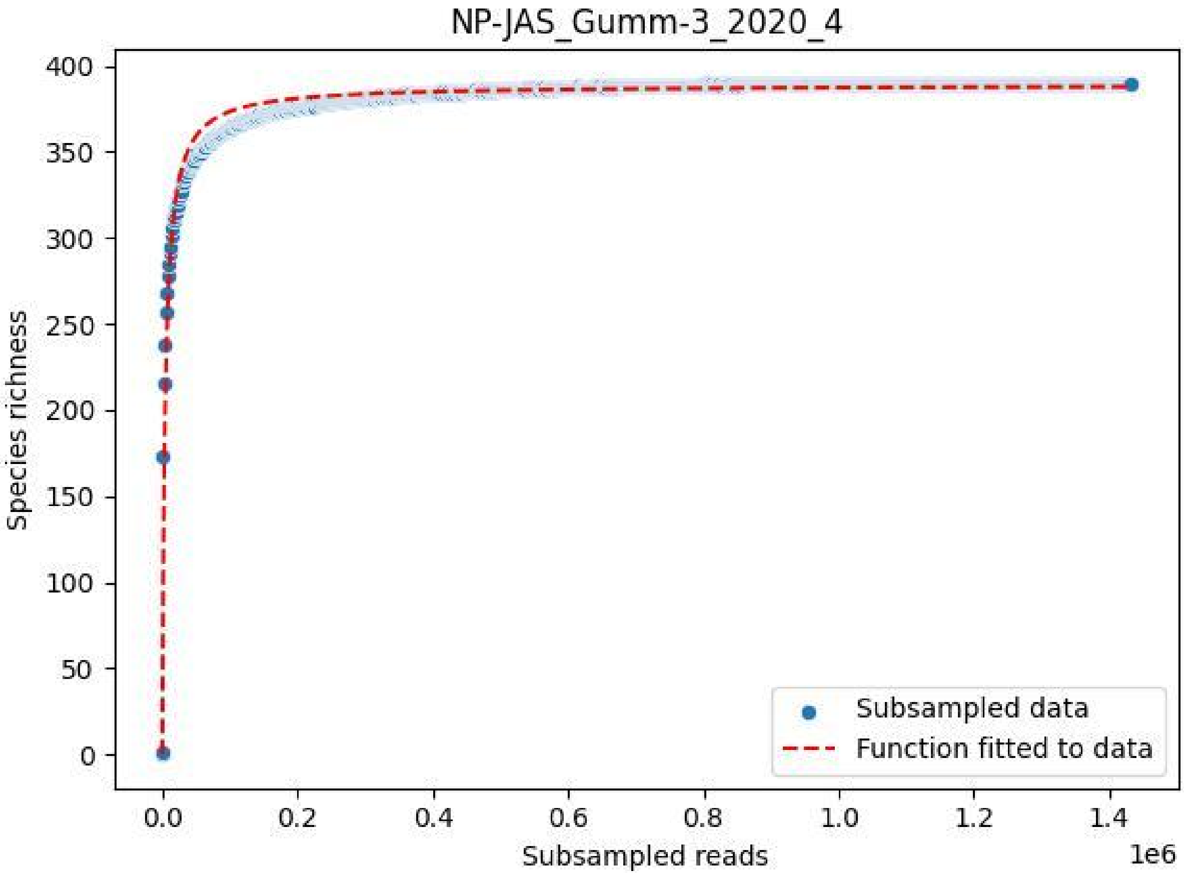

NP-JAS_Gumm-3_2020_4
Species richness
Subsampled reads
1e6
0
50
100
150
200
250
300
350
400
0.0
0.2
0.4
0.6
0.8
1.0
1.2
1.4
Subsampled data
Function fitted to data

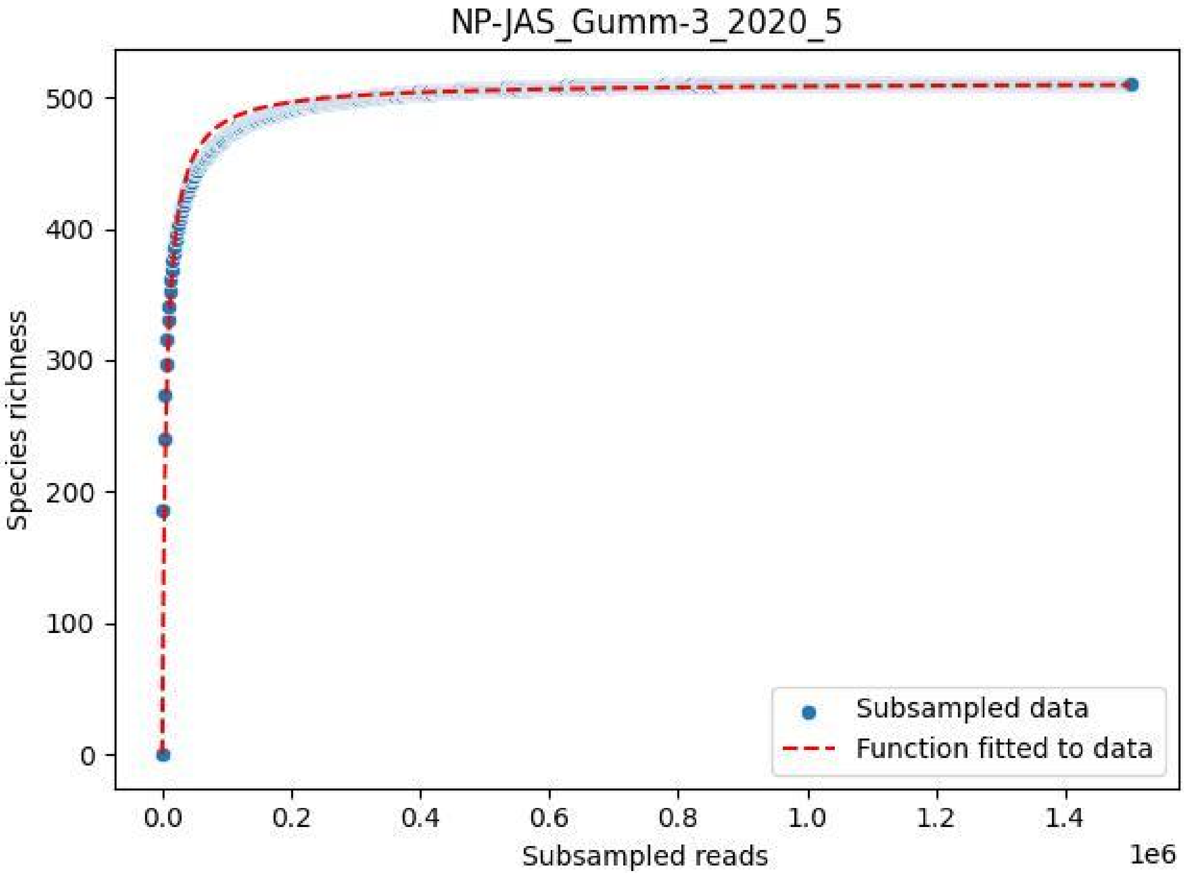

NP-JAS_Gumm-3_2020_5
Species richness
Subsampled reads
1e6
0
100
200
300
400
500
0.0
0.2
0.4
0.6
0.8
1.0
1.2
1.4
Subsampled data
Function fitted to data

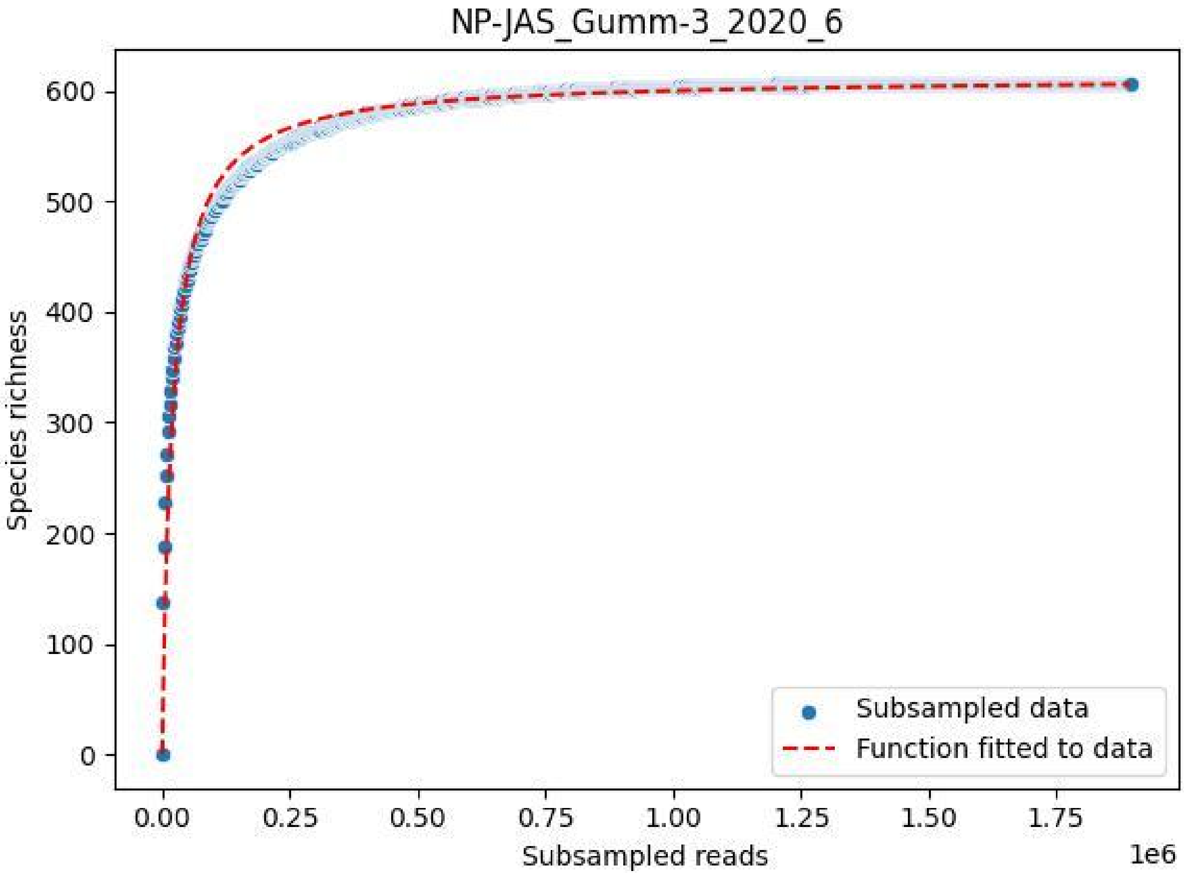

NP-JAS_Gumm-3_2020_6
Species richness
600
500
400
300
200
100
0
0.00
0.25
0.50
0.75
1.00
1.25
1.50
1.75
1e6
Subsampled reads
Subsampled data
Function fitted to data

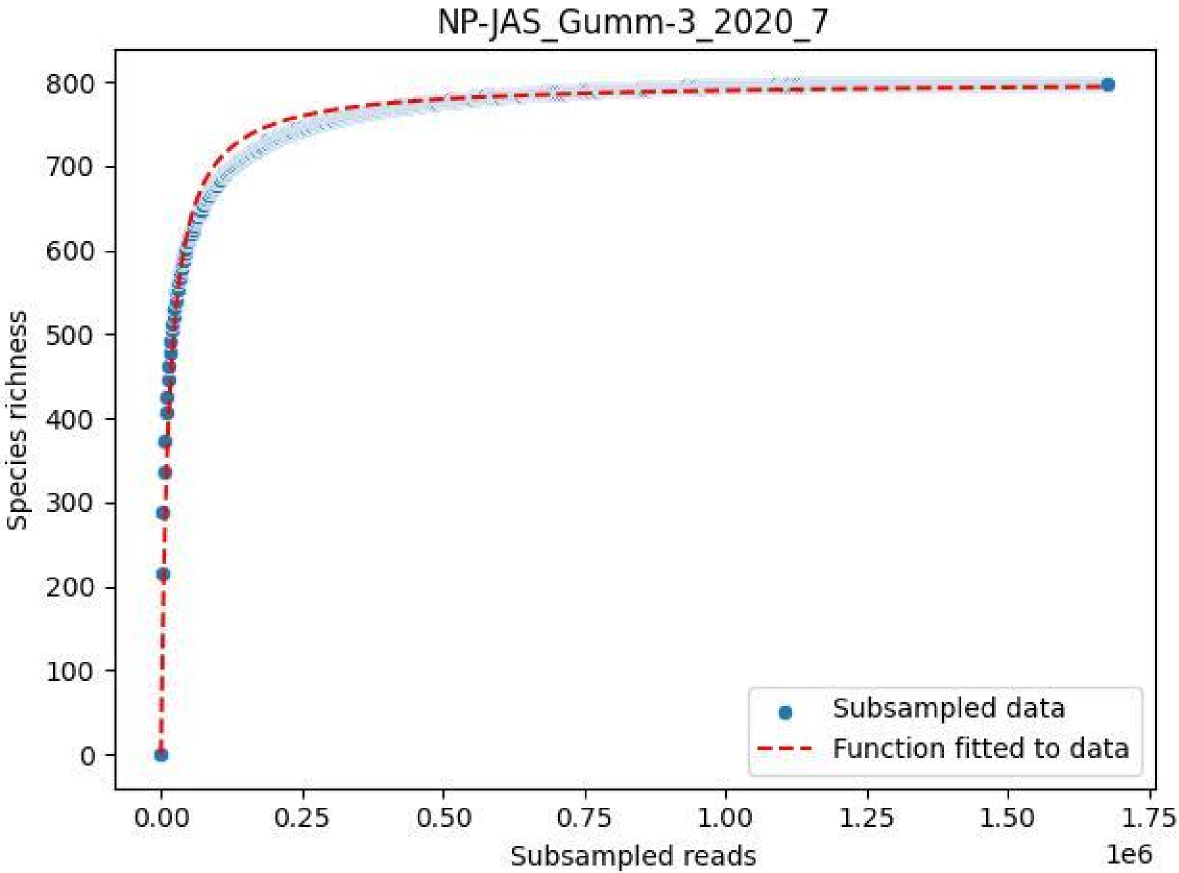

NP-JAS_Gumm-3_2020_7
800
700
600
500
400
300
200
100
0
Species richness
0.00
0.25
0.50
0.75
1.00
1.25
1.50
1.75
1e6
Subsampled reads
Subsampled data
Function fitted to data

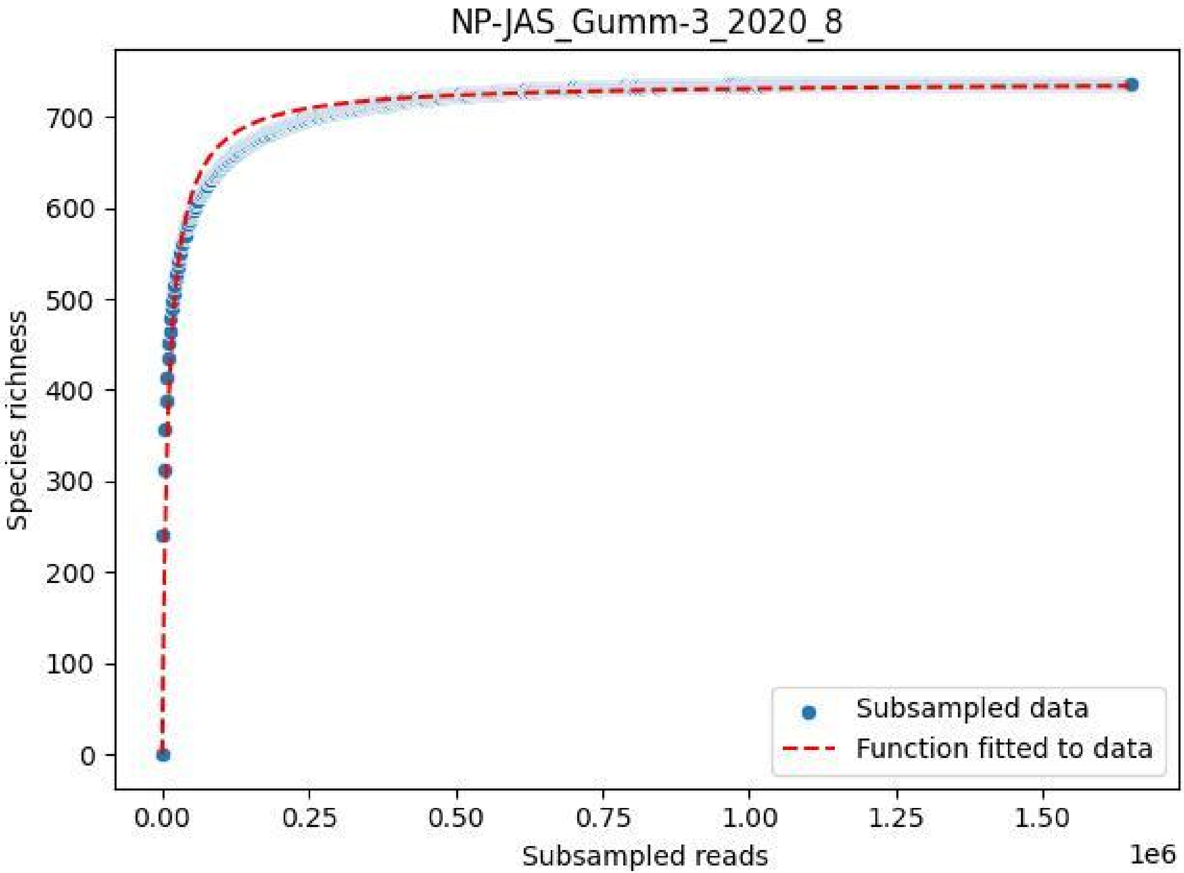
NP-JAS_Gumm-3_2020_8
Species richness
Subsampled reads
1e6
0.00
0.25
0.50
0.75
1.00
1.25
1.50
0
100
200
300
400
500
600
700
Subsampled data
Function fitted to data

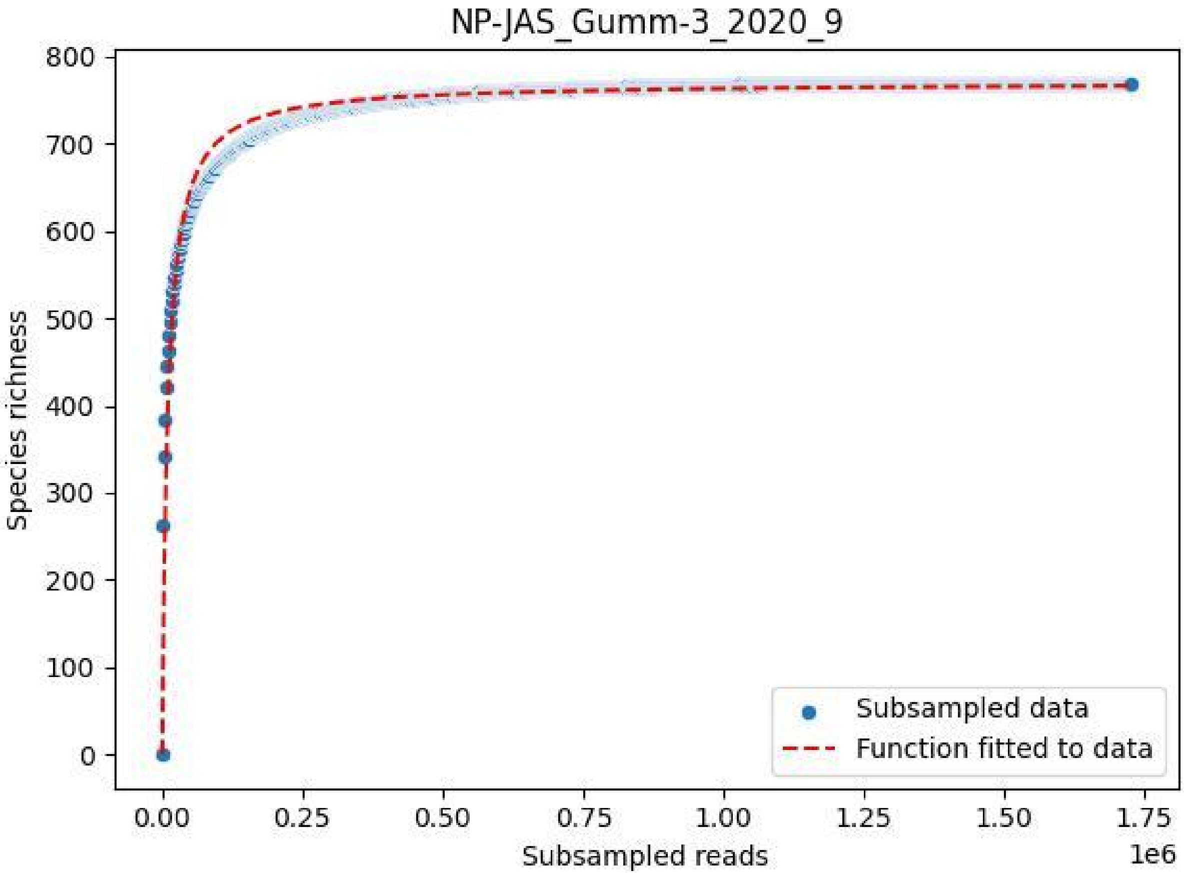
NP-JAS_Gumm-3_2020_9
Species richness
Subsampled reads
1e6
0
100
200
300
400
500
600
700
800
0.00
0.25
0.50
0.75
1.00
1.25
1.50
1.75
Subsampled data
Function fitted to data

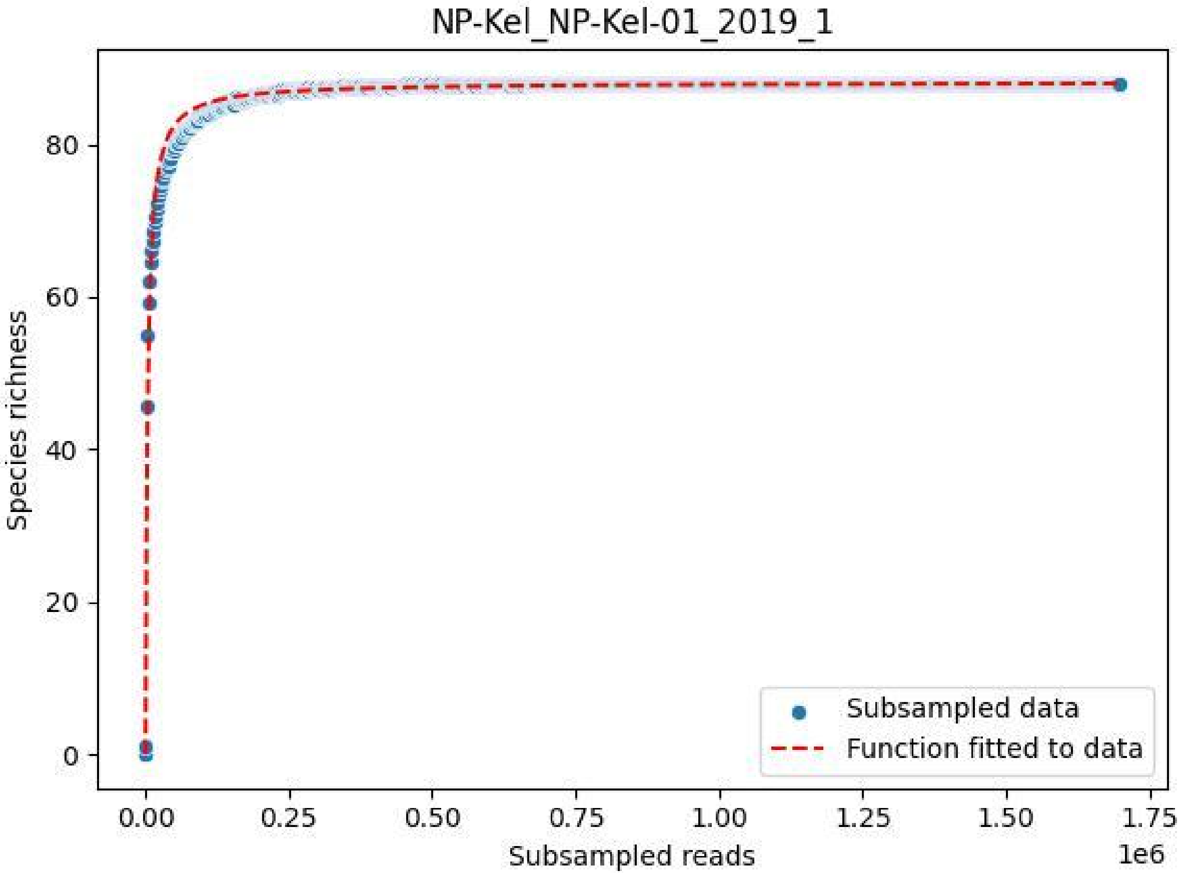
NP-Kel_NP-Kel-01_2019_1
Species richness
Subsampled reads
1e6
0
20
40
60
80
0.00
0.25
0.50
0.75
1.00
1.25
1.50
1.75
Subsampled data
Function fitted to data

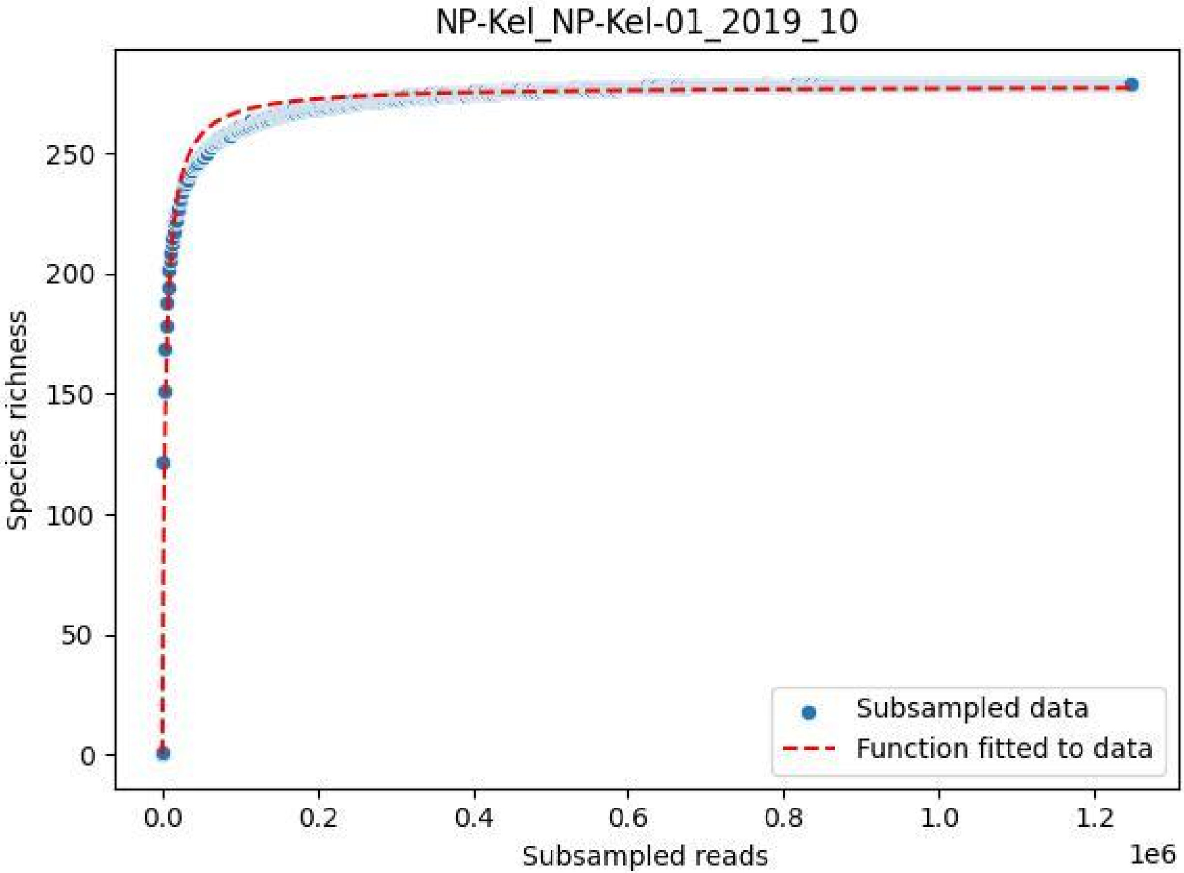
NP-Kel_NP-Kel-01_2019_10
Species richness
250
200
150
100
50
0
0.0
0.2
0.4
0.6
0.8
1.0
1.2
1e6
Subsampled reads
Subsampled data
Function fitted to data

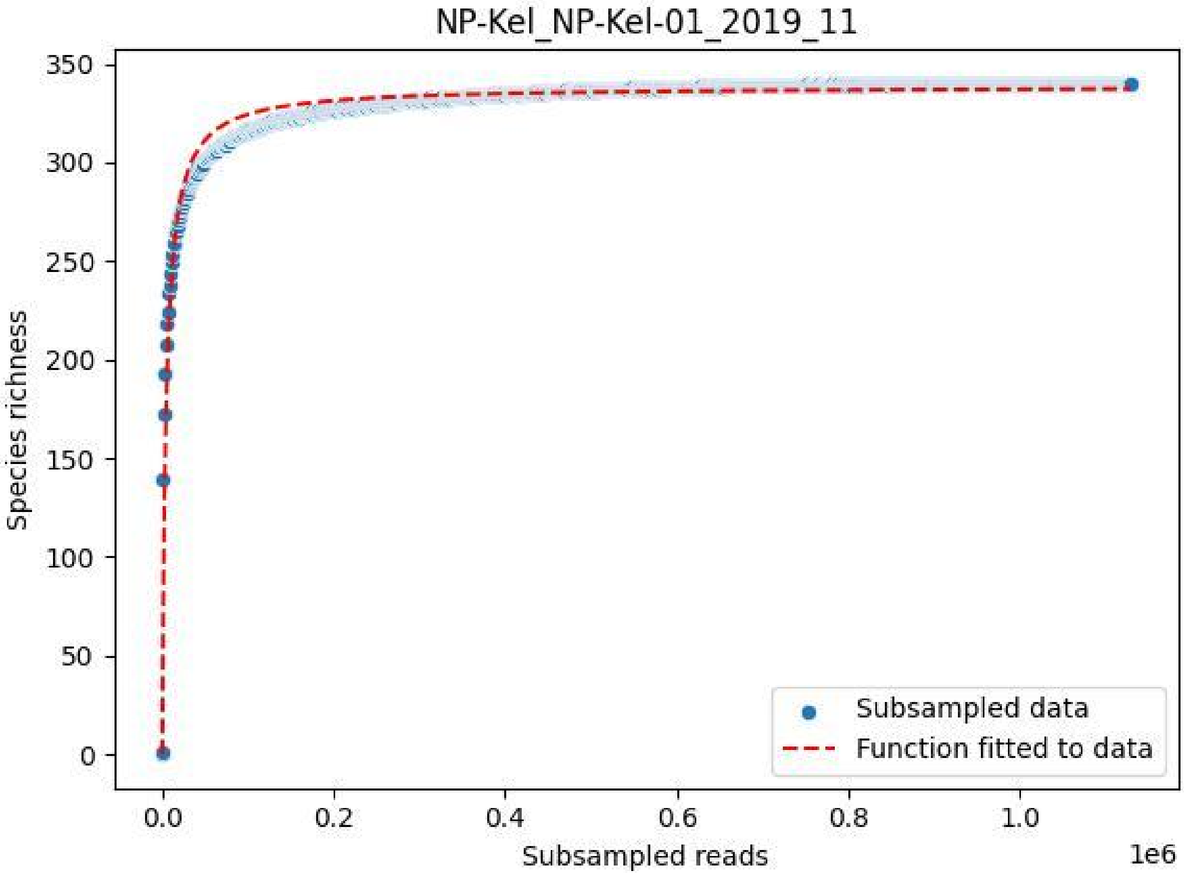
NP-Kel_NP-Kel-01_2019_11
Species richness
Subsampled reads
1e6
0
50
100
150
200
250
300
350
0.0
0.2
0.4
0.6
0.8
1.0
Subsampled data
Function fitted to data

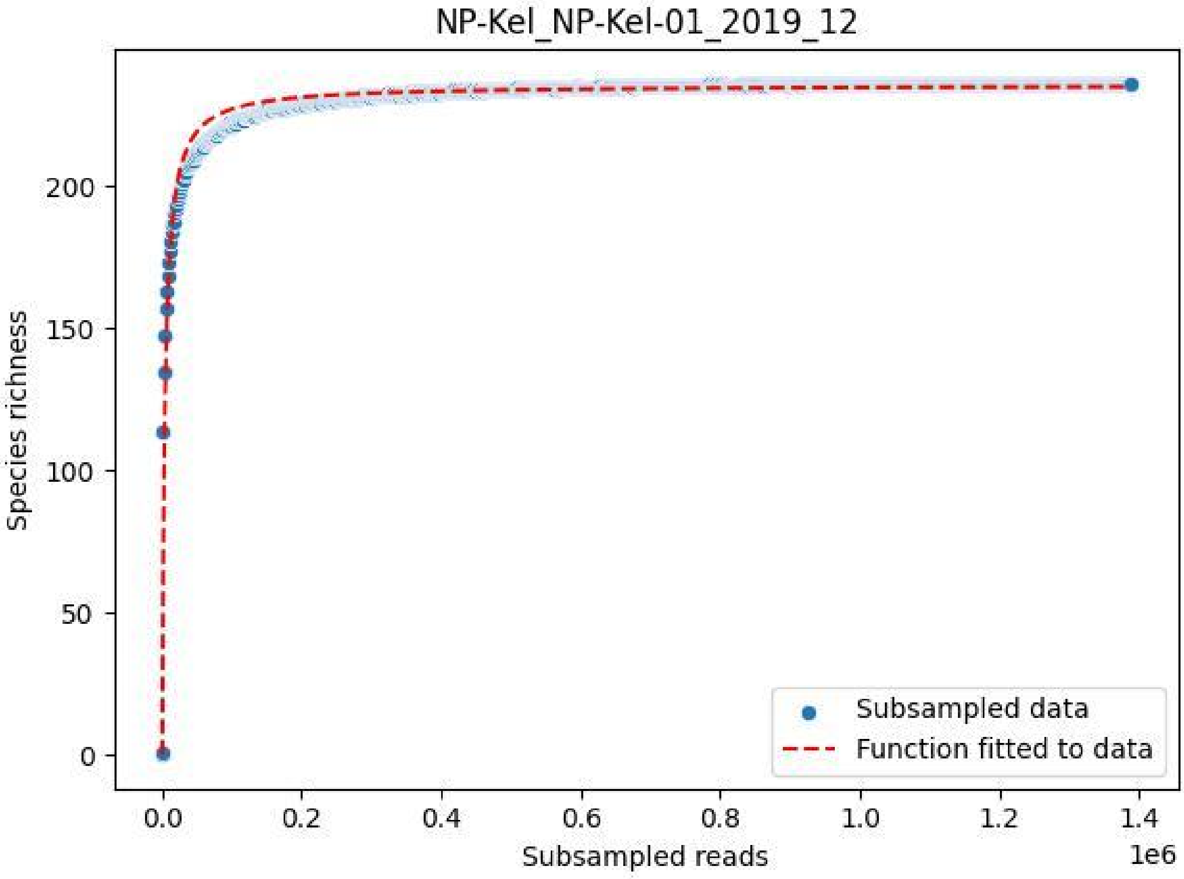

NP-Kel_NP-Kel-01_2019_12
Species richness
Subsampled reads
1e6
0
50
100
150
200
0.0
0.2
0.4
0.6
0.8
1.0
1.2
1.4
Subsampled data
Function fitted to data

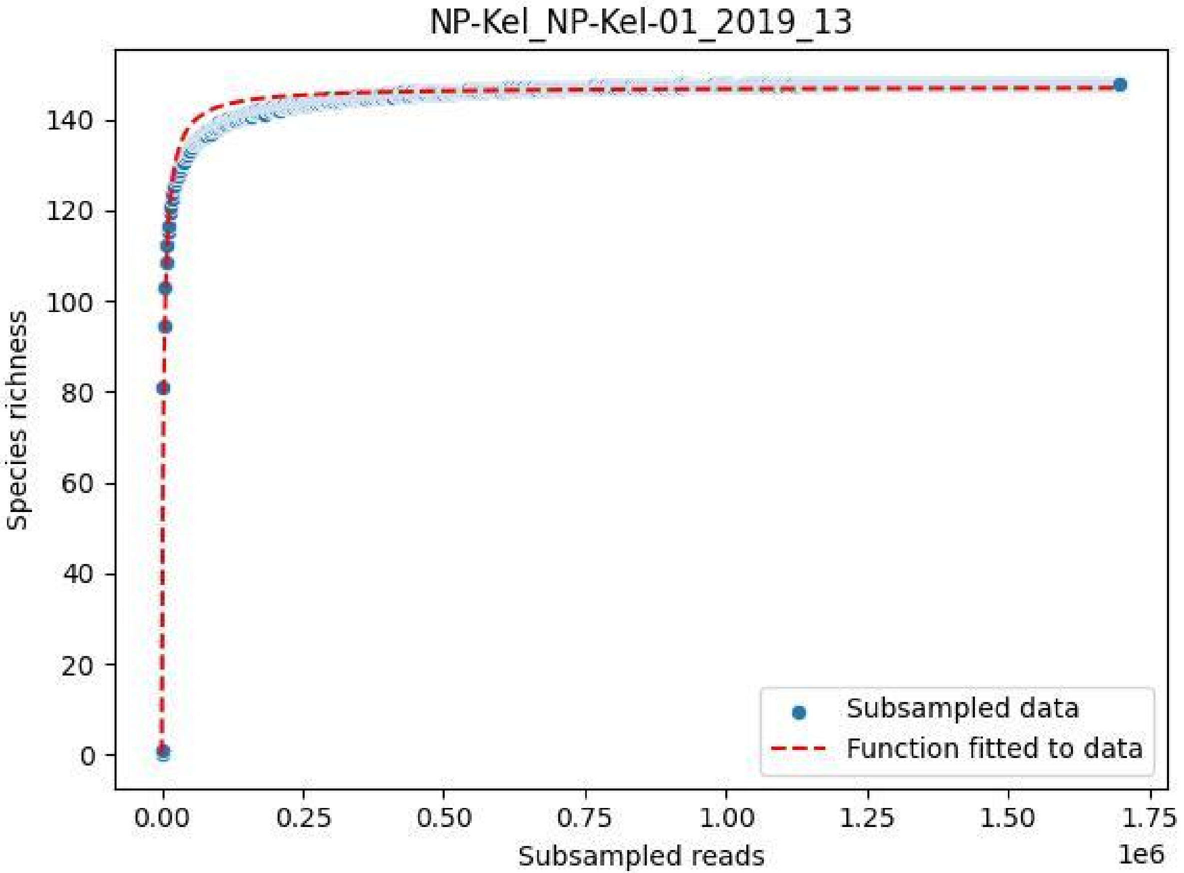
NP-Kel_NP-Kel-01_2019_13
Species richness
Subsampled reads
1e6
0
20
40
60
80
100
120
140
0.00
0.25
0.50
0.75
1.00
1.25
1.50
1.75
Subsampled data
Function fitted to data

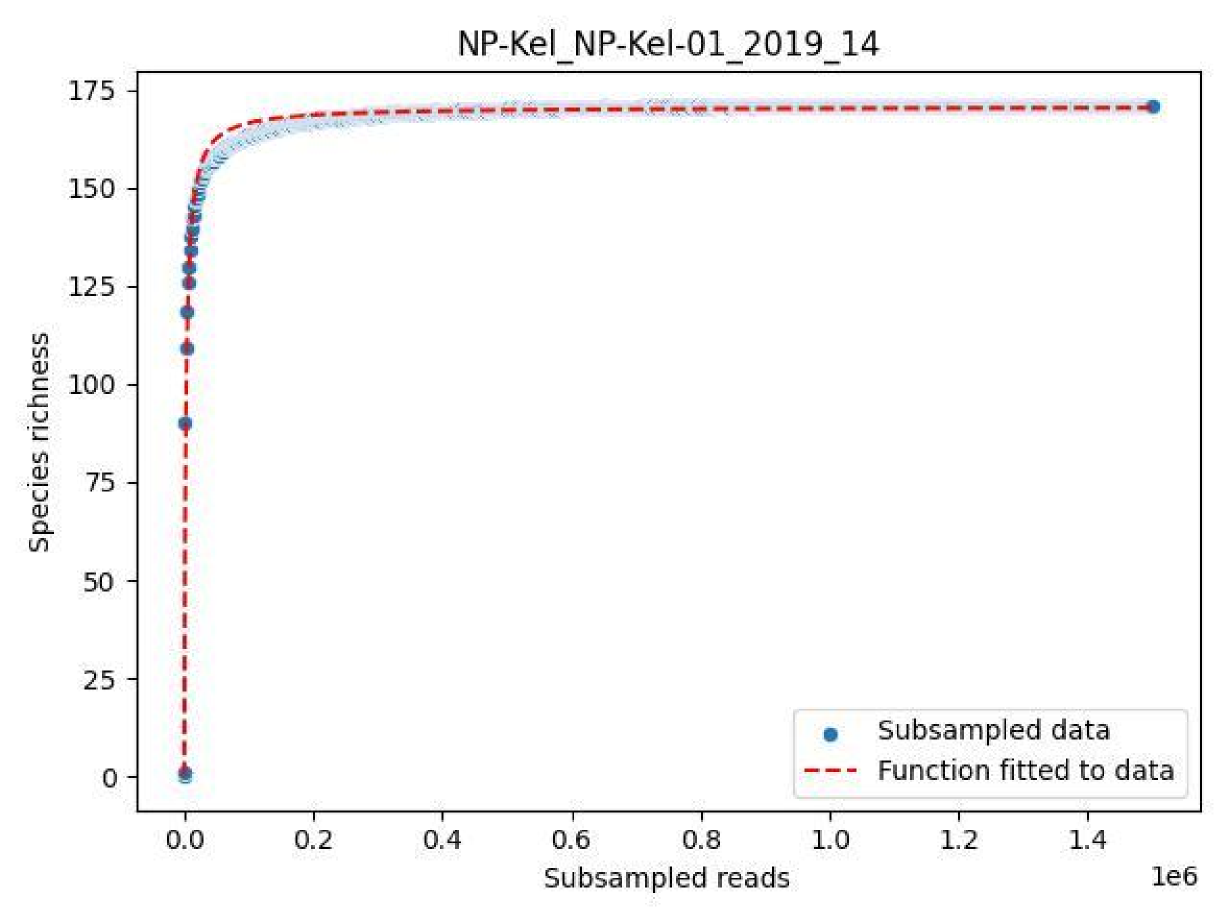

NP-Kel_NP-Kel-01_2019_14
Species richness
Subsampled reads
1e6
0
25
50
75
100
125
150
175
0.0
0.2
0.4
0.6
0.8
1.0
1.2
1.4
Subsampled data
Function fitted to data

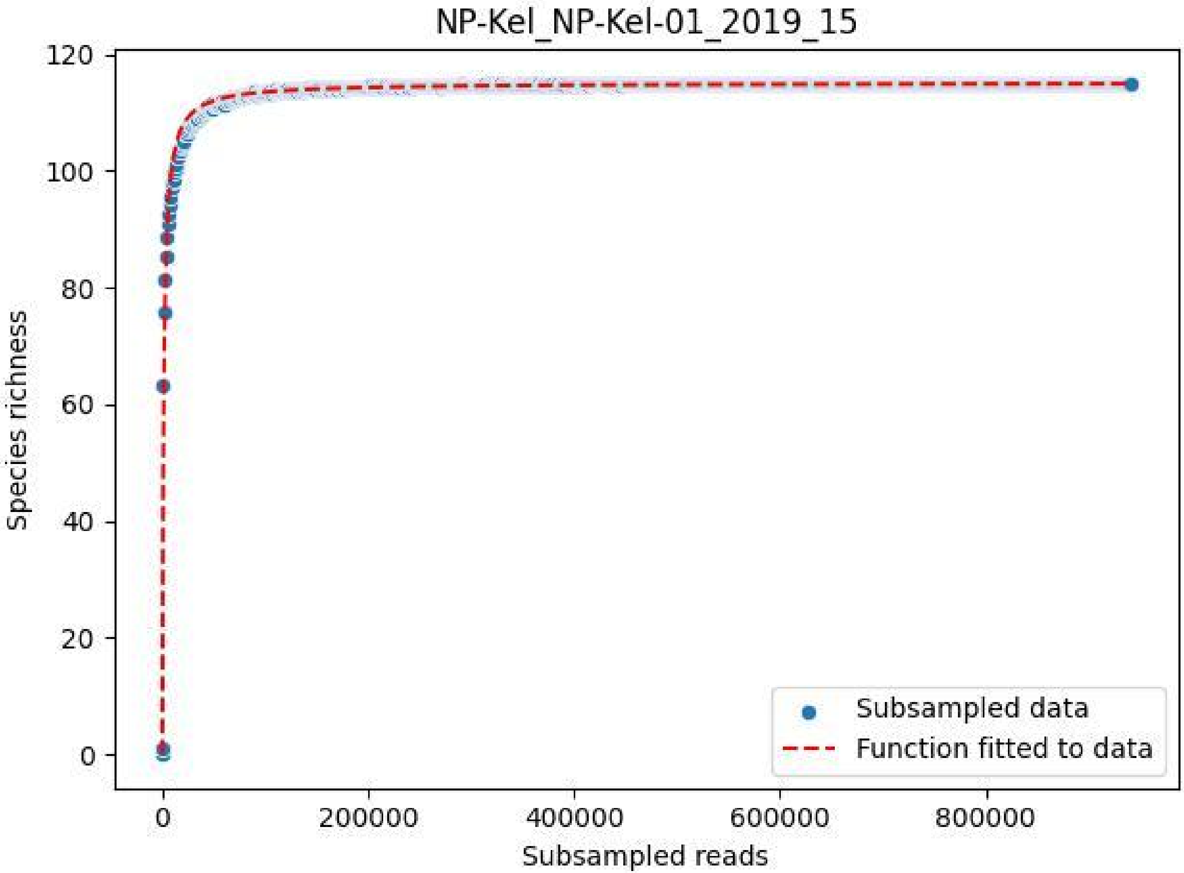
NP-Kel_NP-Kel-01_2019_15
Species richness
Subsampled reads
0
20
40
60
80
100
120
0
200000
400000
600000
800000
Subsampled data
Function fitted to data

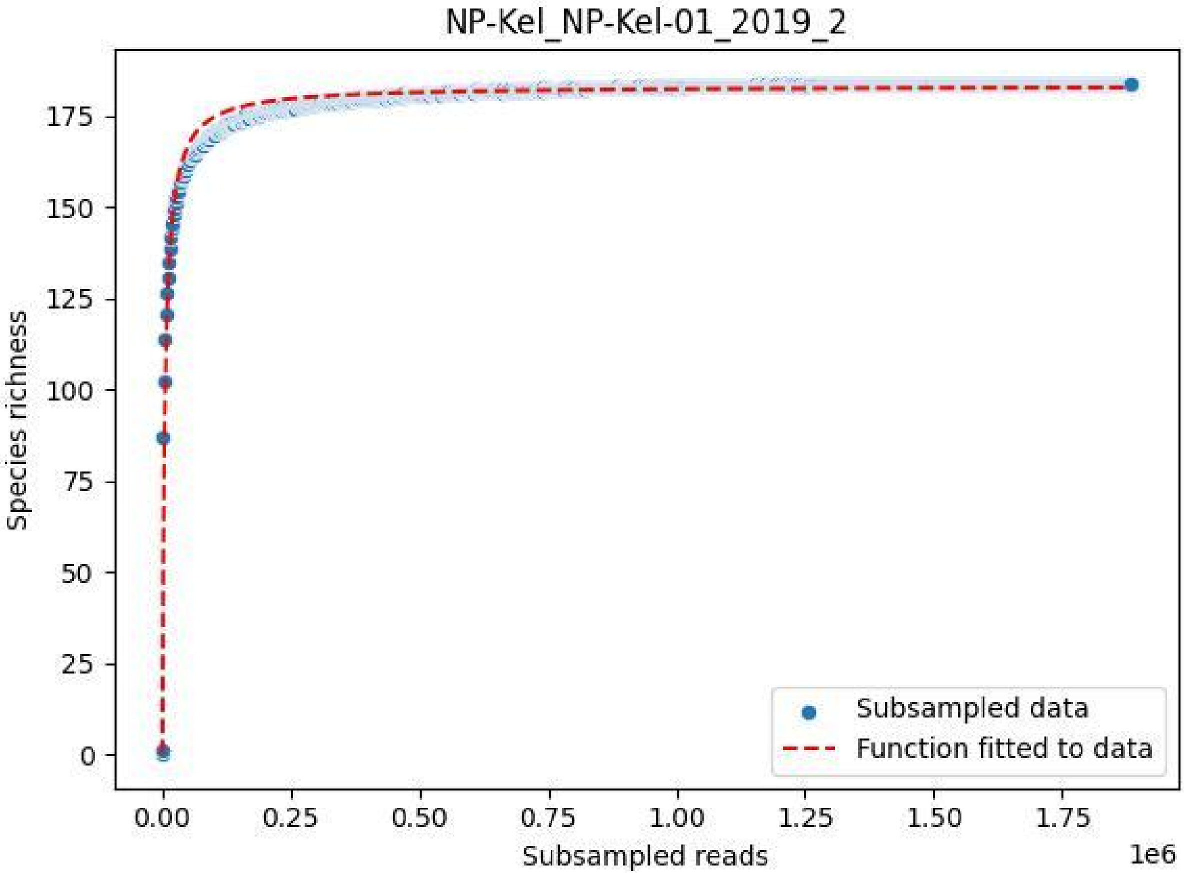

NP-Kel_NP-Kel-01_2019_2
Species richness
Subsampled reads
1e6
0
25
50
75
100
125
150
175
0.00
0.25
0.50
0.75
1.00
1.25
1.50
1.75
Subsampled data
Function fitted to data

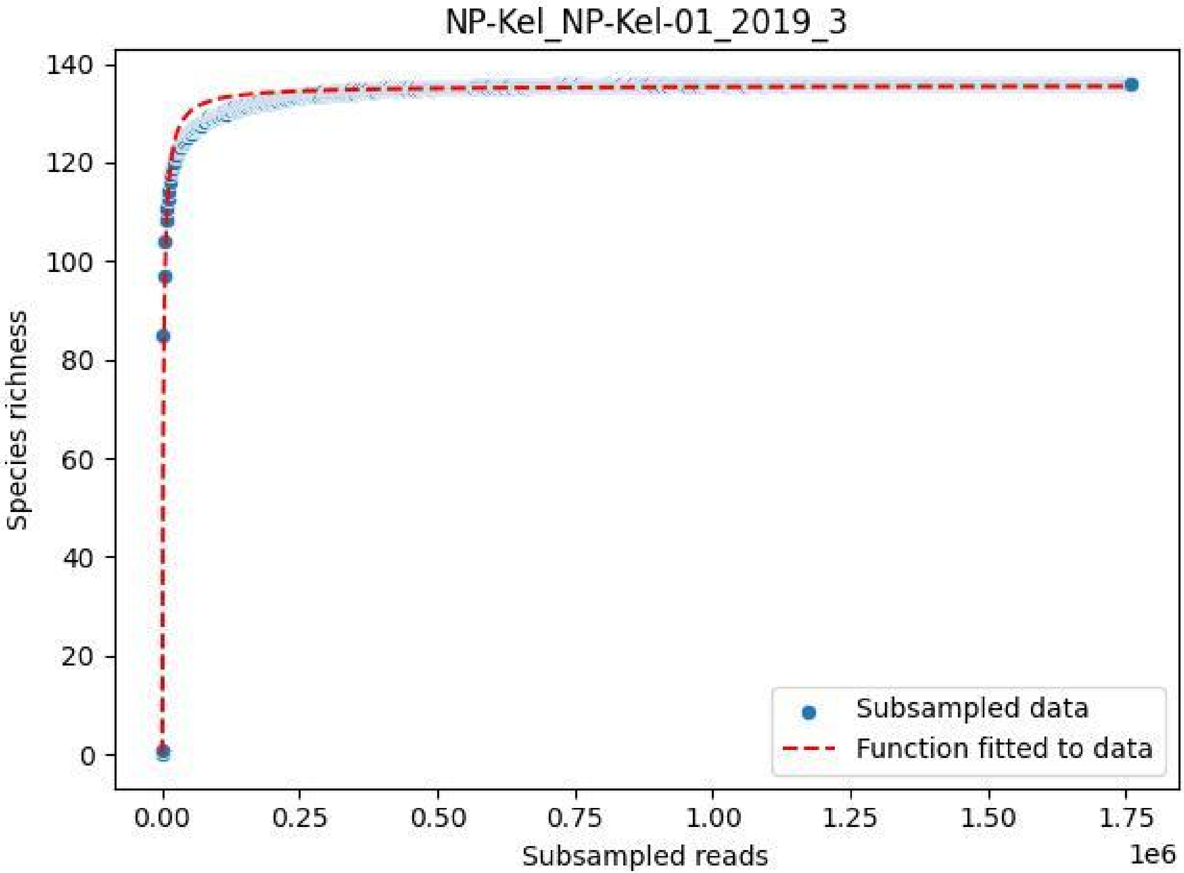

NP-Kel_NP-Kel-01_2019_3
Species richness
Subsampled reads
1e6
0.00
0.25
0.50
0.75
1.00
1.25
1.50
1.75
0
20
40
60
80
100
120
140
Subsampled data
Function fitted to data

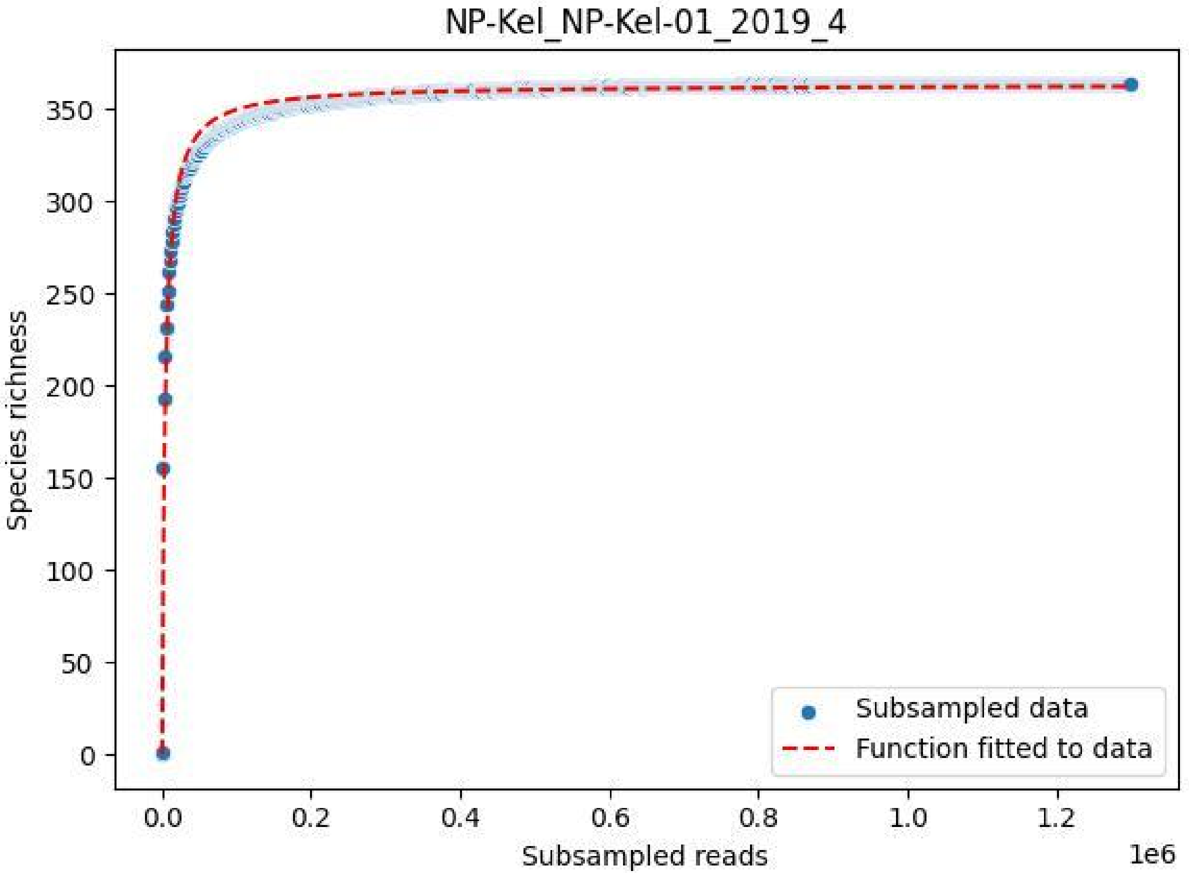
NP-Kel_NP-Kel-01_2019_4
Species richness
Subsampled reads
1e6
0
50
100
150
200
250
300
350
0.0
0.2
0.4
0.6
0.8
1.0
1.2
Subsampled data
Function fitted to data

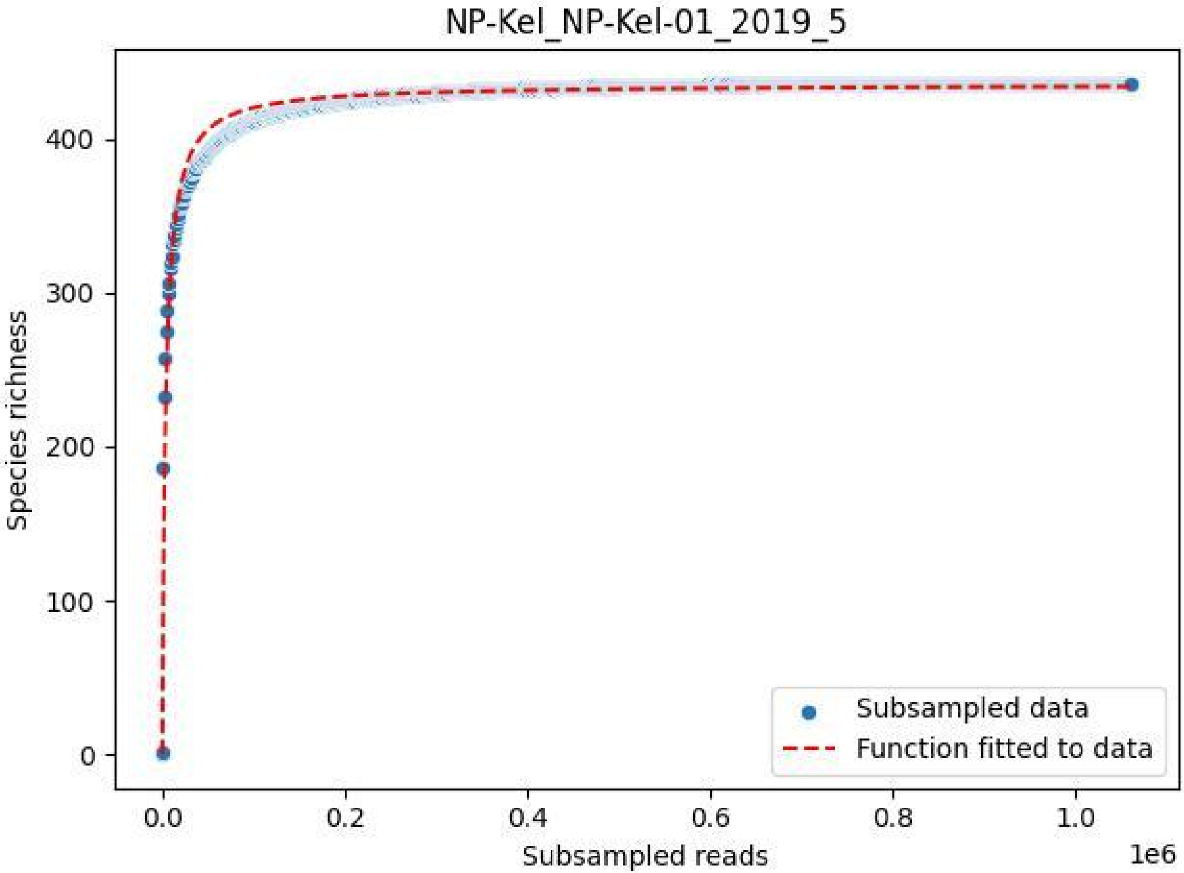

NP-Kel_NP-Kel-01_2019_5
Species richness
400
300
200
100
0
0.0
0.2
0.4
0.6
0.8
1.0
1e6
Subsampled reads
Subsampled data
Function fitted to data

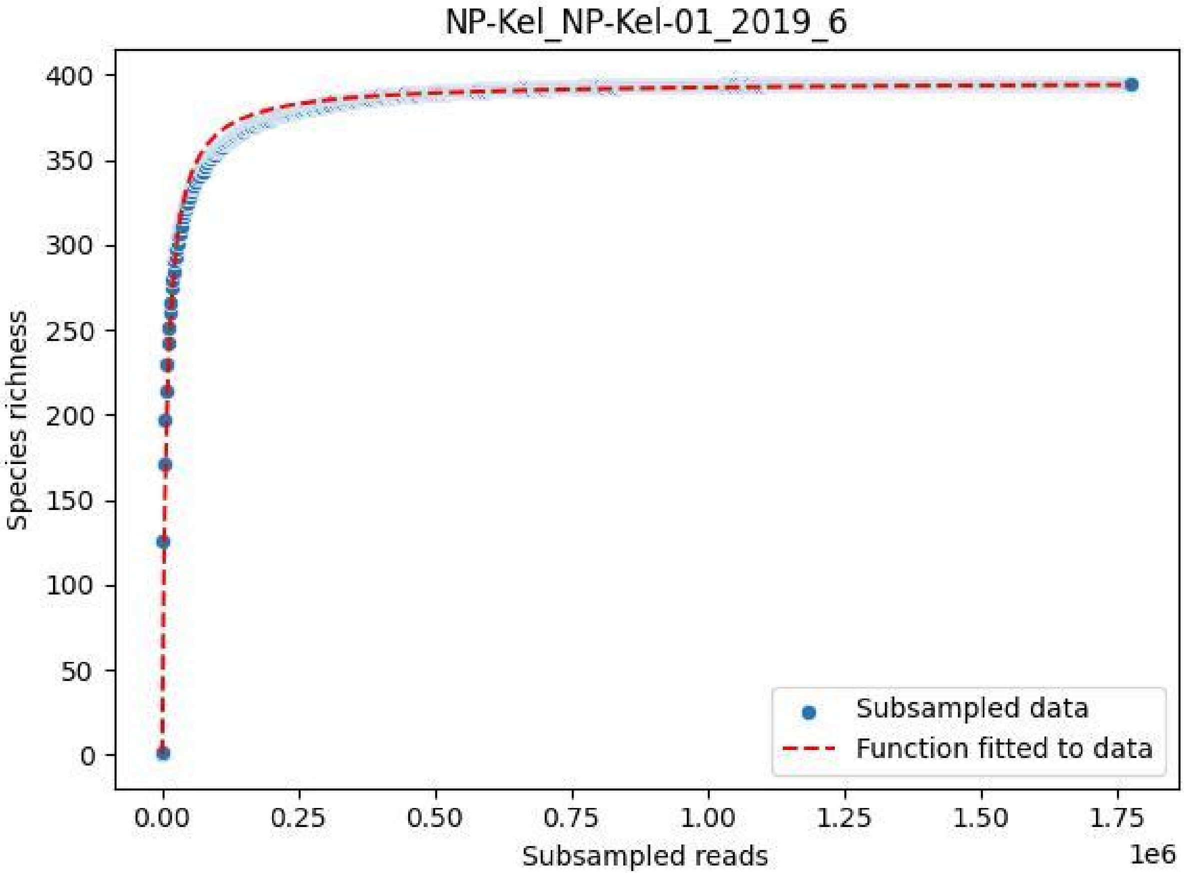

NP-Kel_NP-Kel-01_2019_6
Species richness
Subsampled reads
1e6
0
50
100
150
200
250
300
350
400
0.00
0.25
0.50
0.75
1.00
1.25
1.50
1.75
Subsampled data
Function fitted to data

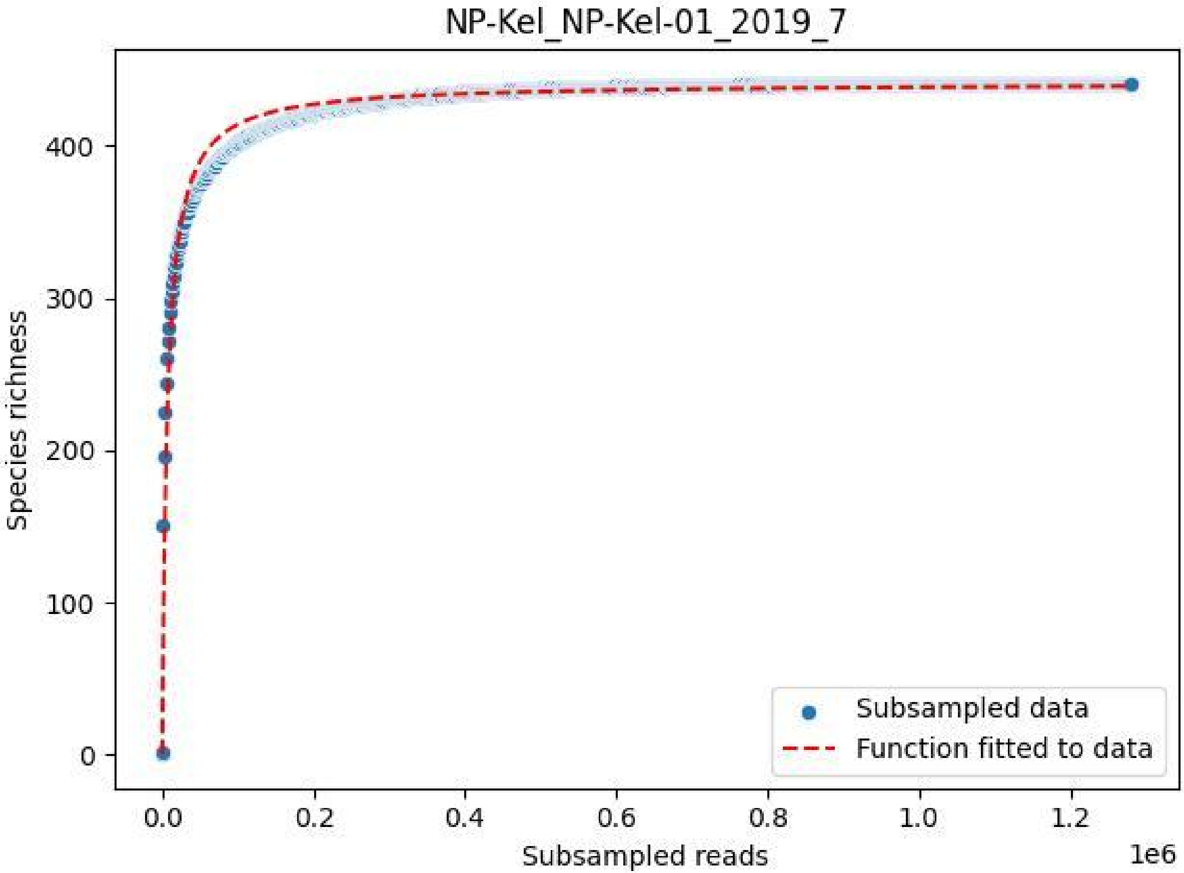

NP-Kel_NP-Kel-01_2019_7
Species richness
Subsampled reads
1e6
0
100
200
300
400
0.0
0.2
0.4
0.6
0.8
1.0
1.2
Subsampled data
Function fitted to data

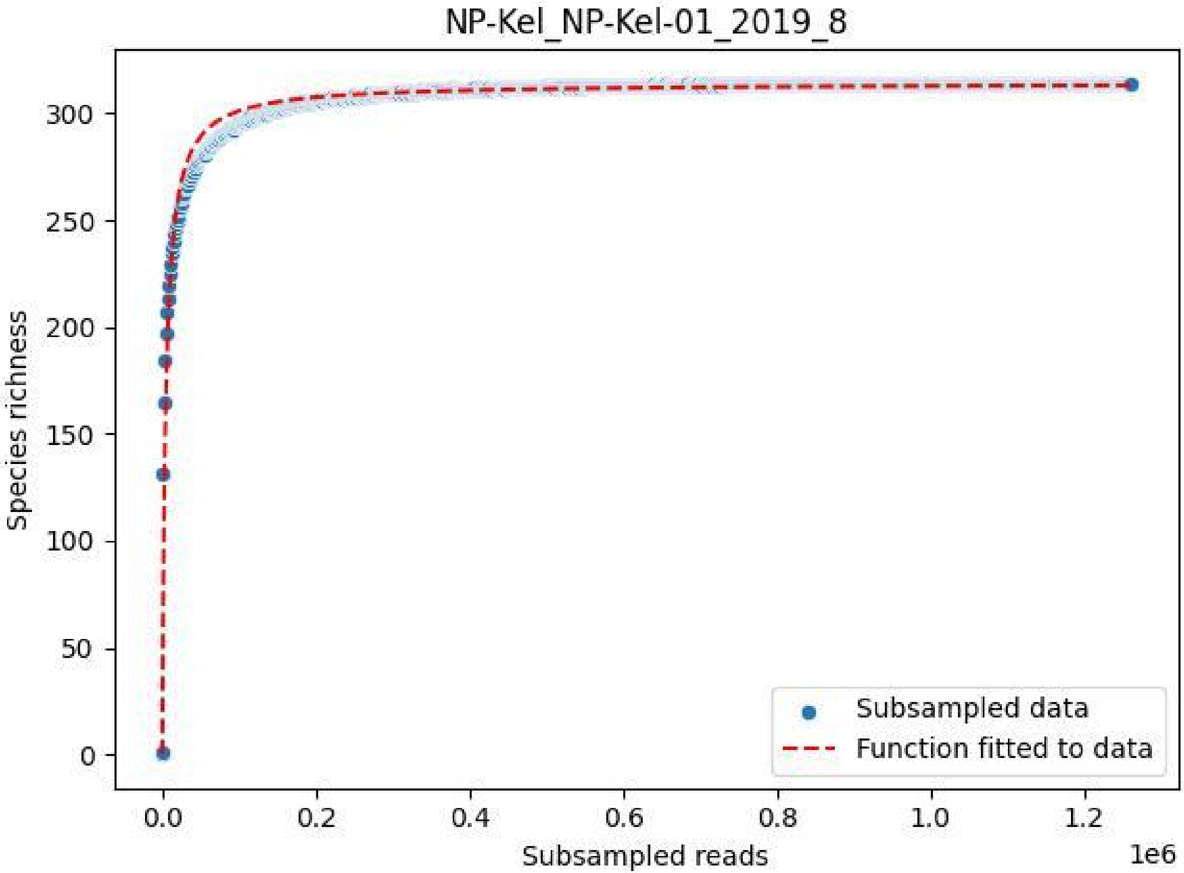

NP-Kel_NP-Kel-01_2019_8
Species richness
Subsampled reads
1e6
0
50
100
150
200
250
300
0.0
0.2
0.4
0.6
0.8
1.0
1.2
Subsampled data
Function fitted to data

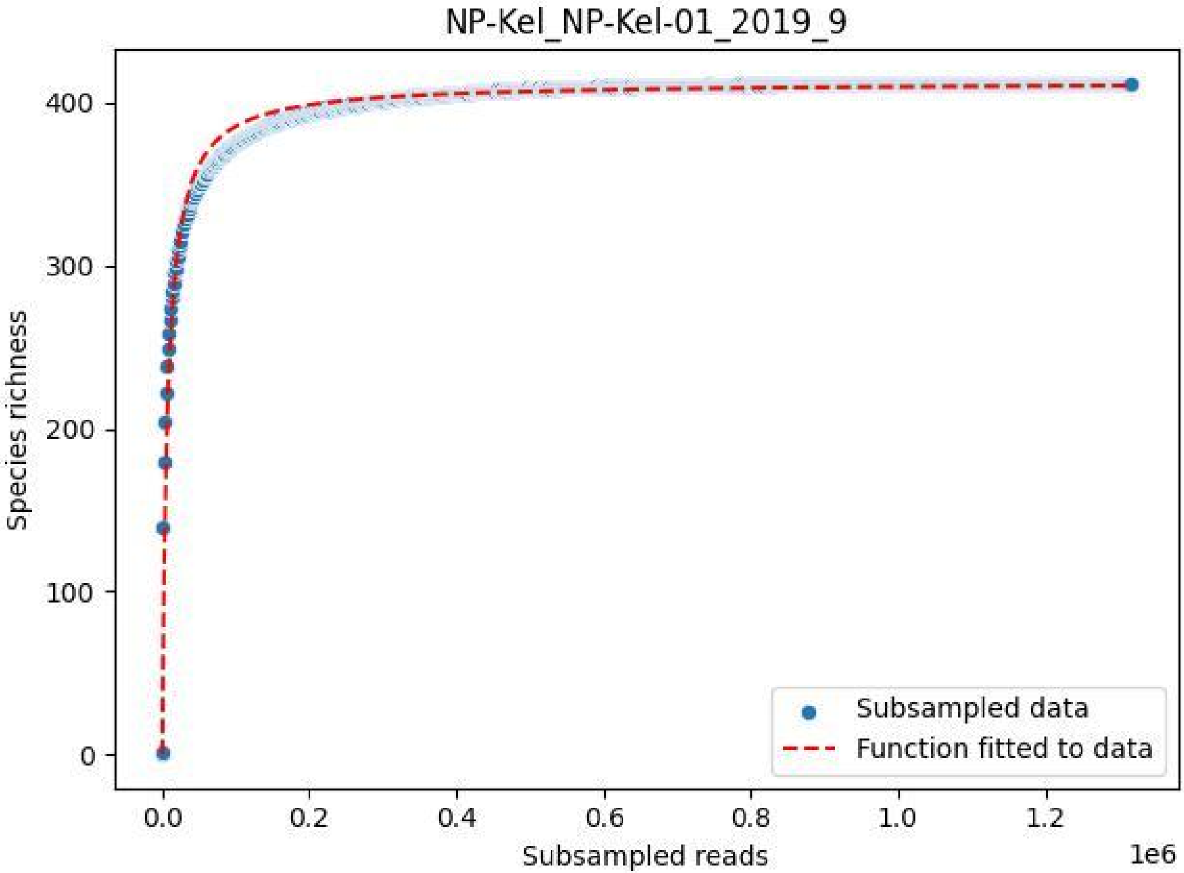
NP-Kel_NP-Kel-01_2019_9
Species richness
Subsampled reads
1e6
0
100
200
300
400
0.0
0.2
0.4
0.6
0.8
1.0
1.2
Subsampled data
Function fitted to data

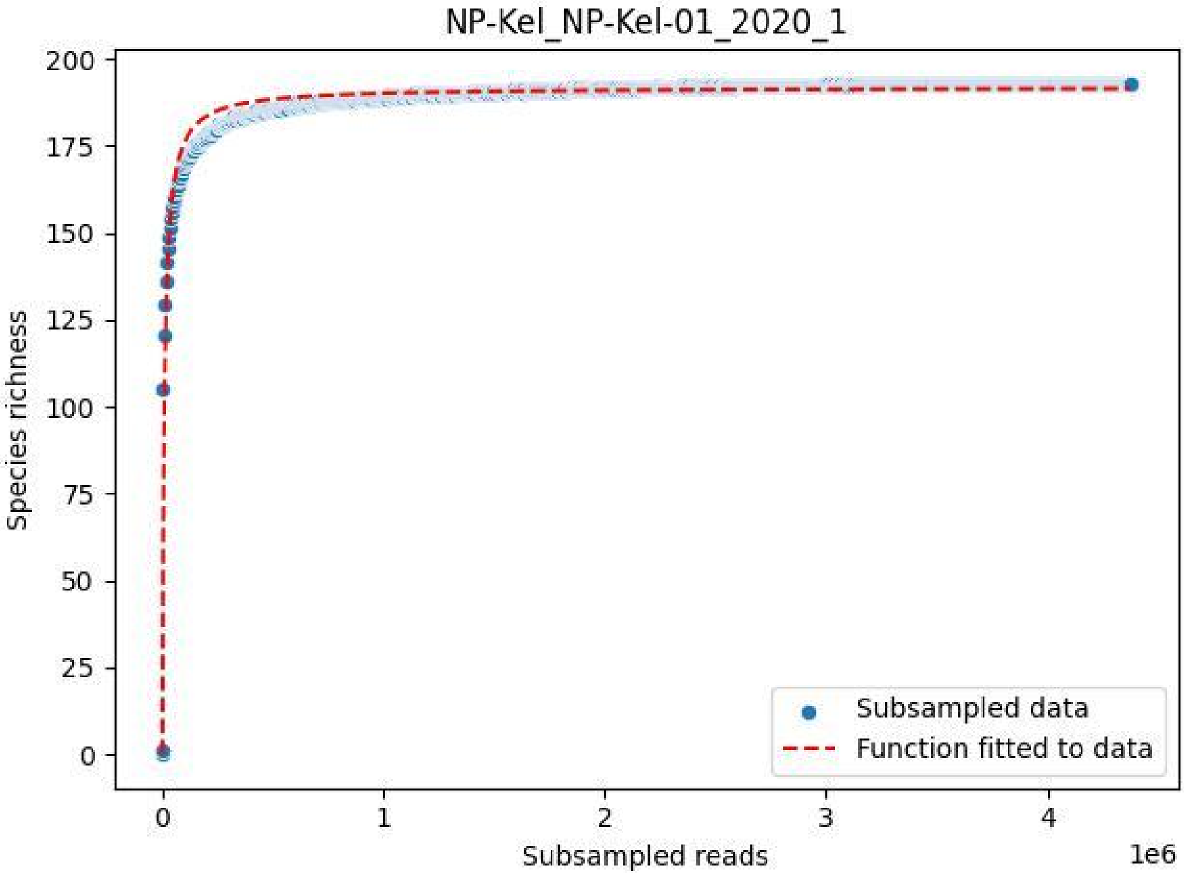
NP-Kel_NP-Kel-01_2020_1
Species richness
Subsampled reads
1e6
0
25
50
75
100
125
150
175
200
0
1
2
3
4
Subsampled data
Function fitted to data

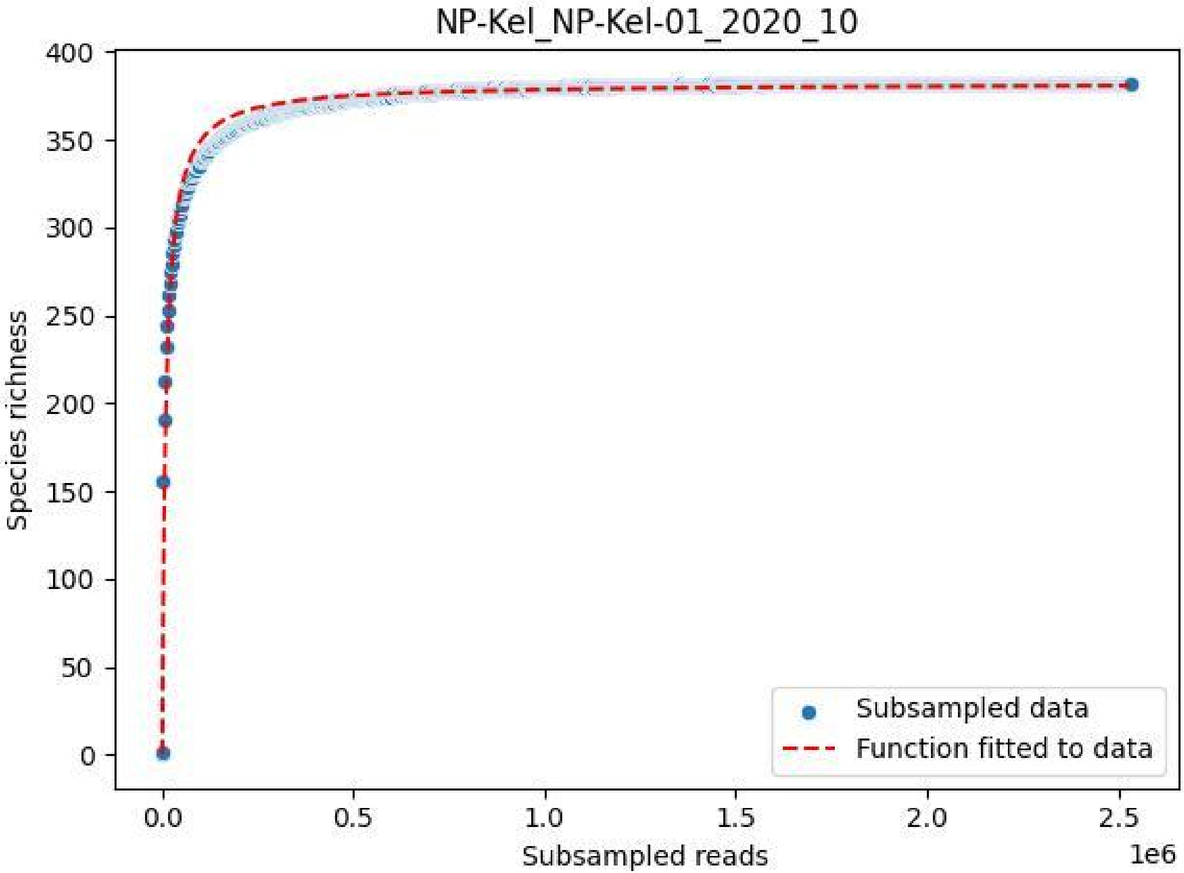

NP-Kel_NP-Kel-01_2020_10
Species richness
Subsampled reads
1e6
0
50
100
150
200
250
300
350
400
0.0
0.5
1.0
1.5
2.0
2.5
Subsampled data
Function fitted to data

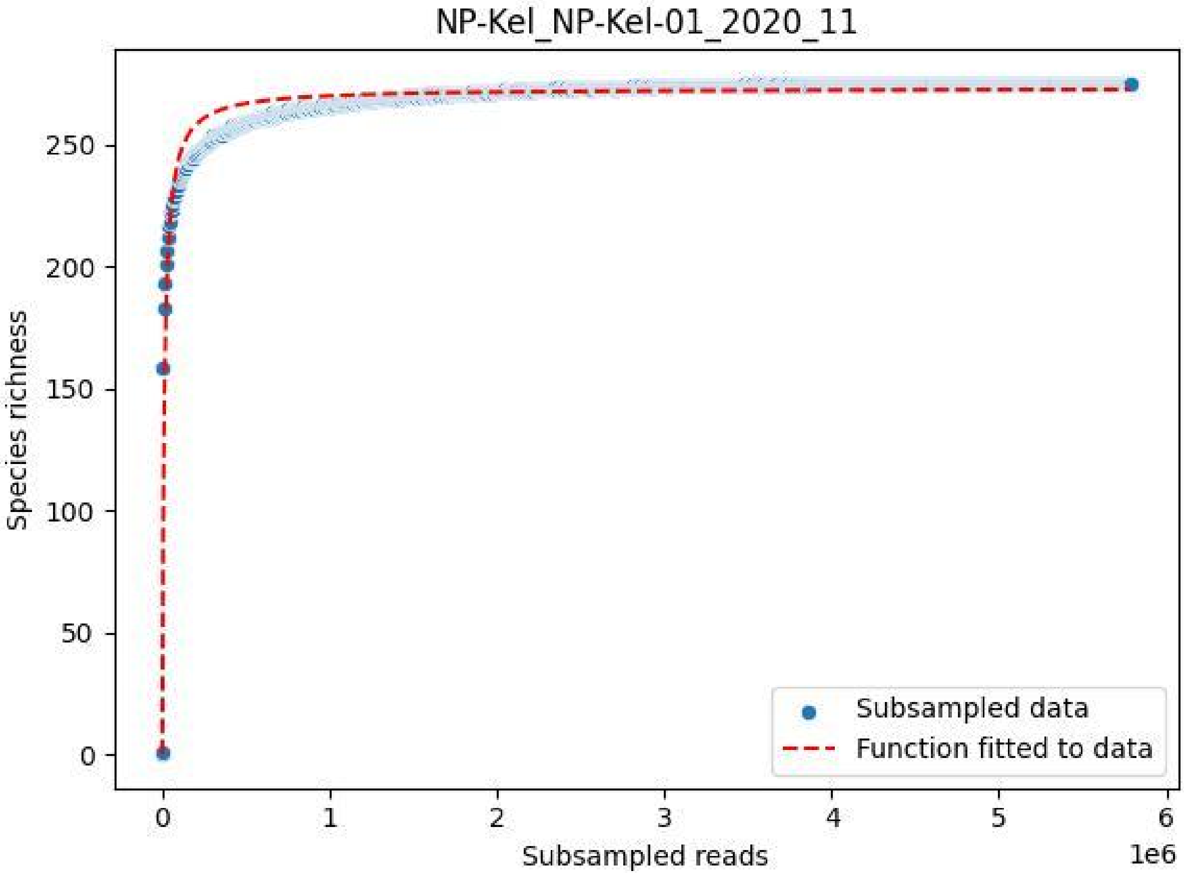
NP-Kel_NP-Kel-01_2020_11
Species richness
0
50
100
150
200
250
0
1
2
3
4
5
6
1e6
Subsampled reads
Subsampled data
Function fitted to data

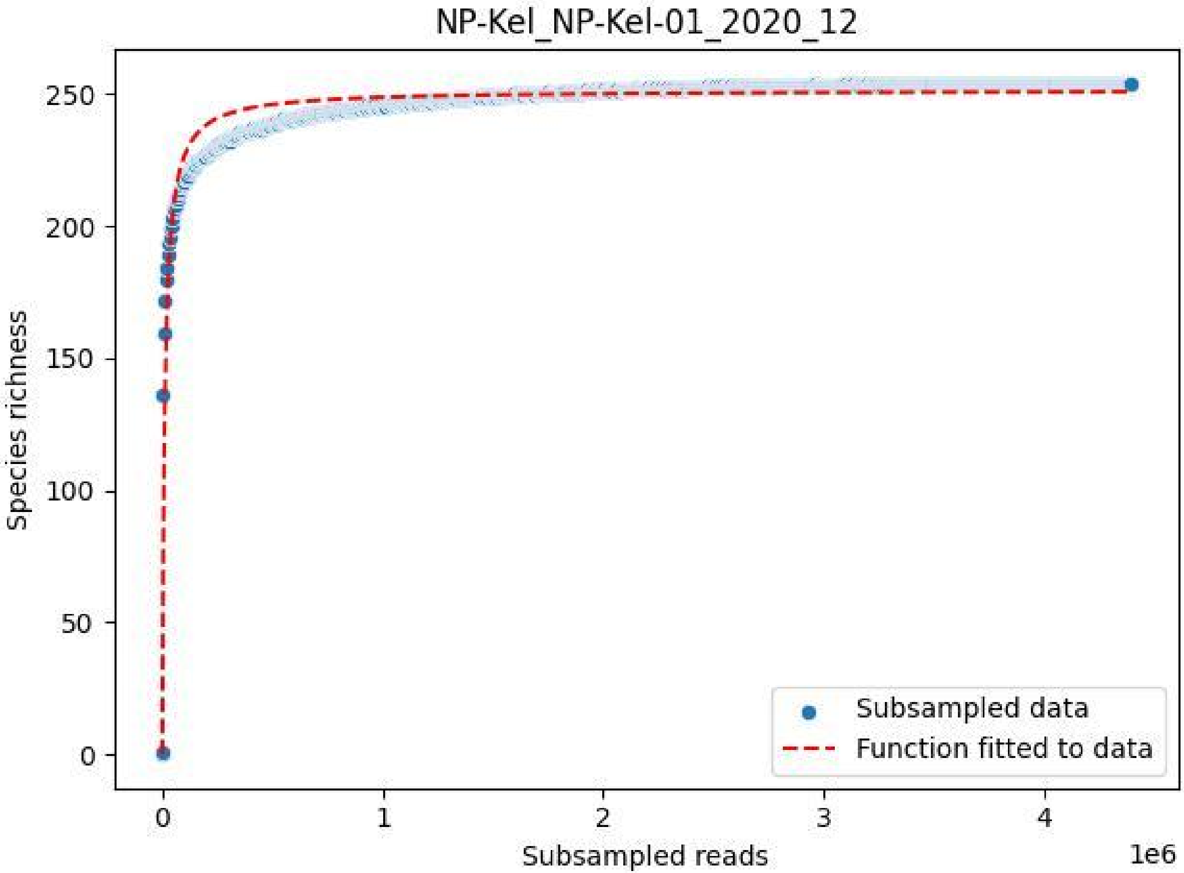

NP-Kel_NP-Kel-01_2020_12
Species richness
250
200
150
100
50
0
0
1
2
3
4
1e6
Subsampled reads
Subsampled data
Function fitted to data

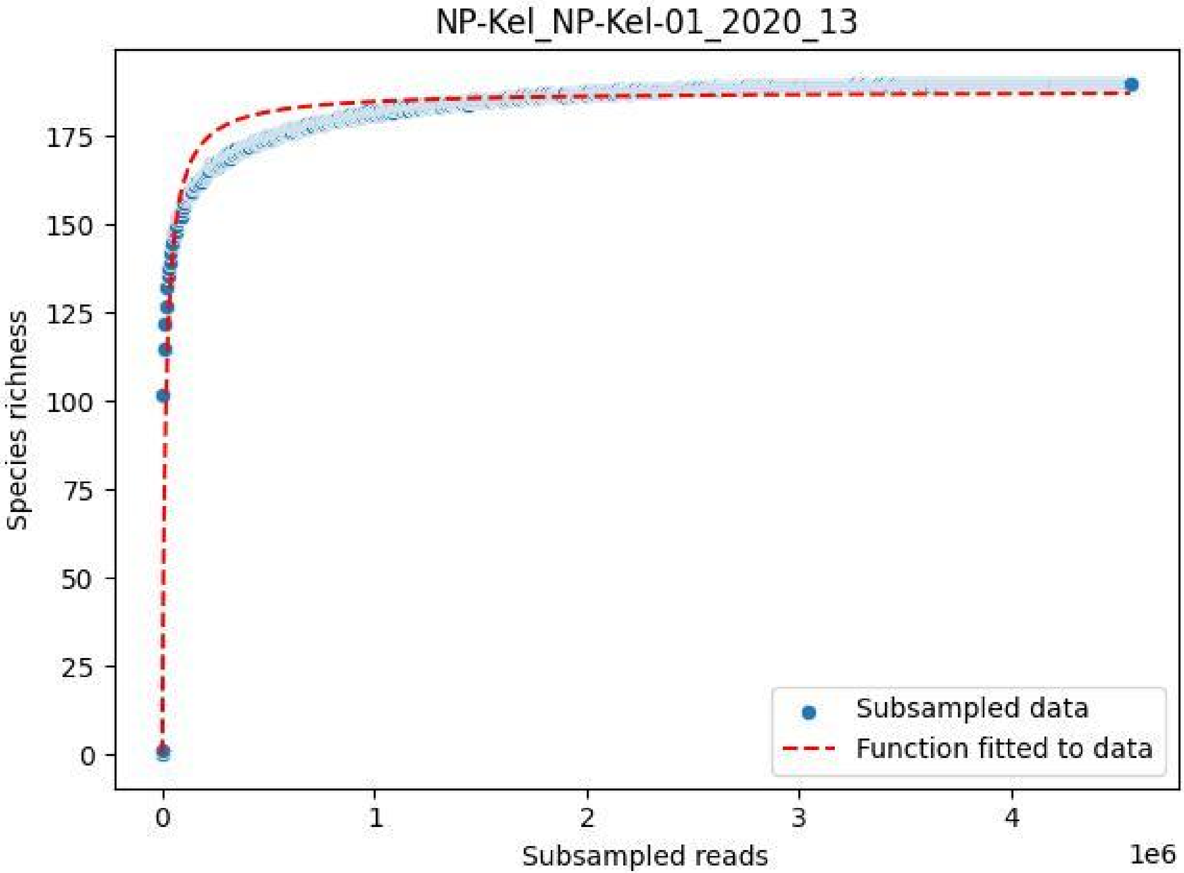
NP-Kel_NP-Kel-01_2020_13
Species richness
Subsampled reads
1e6
0
25
50
75
100
125
150
175
0
1
2
3
4
Subsampled data
Function fitted to data

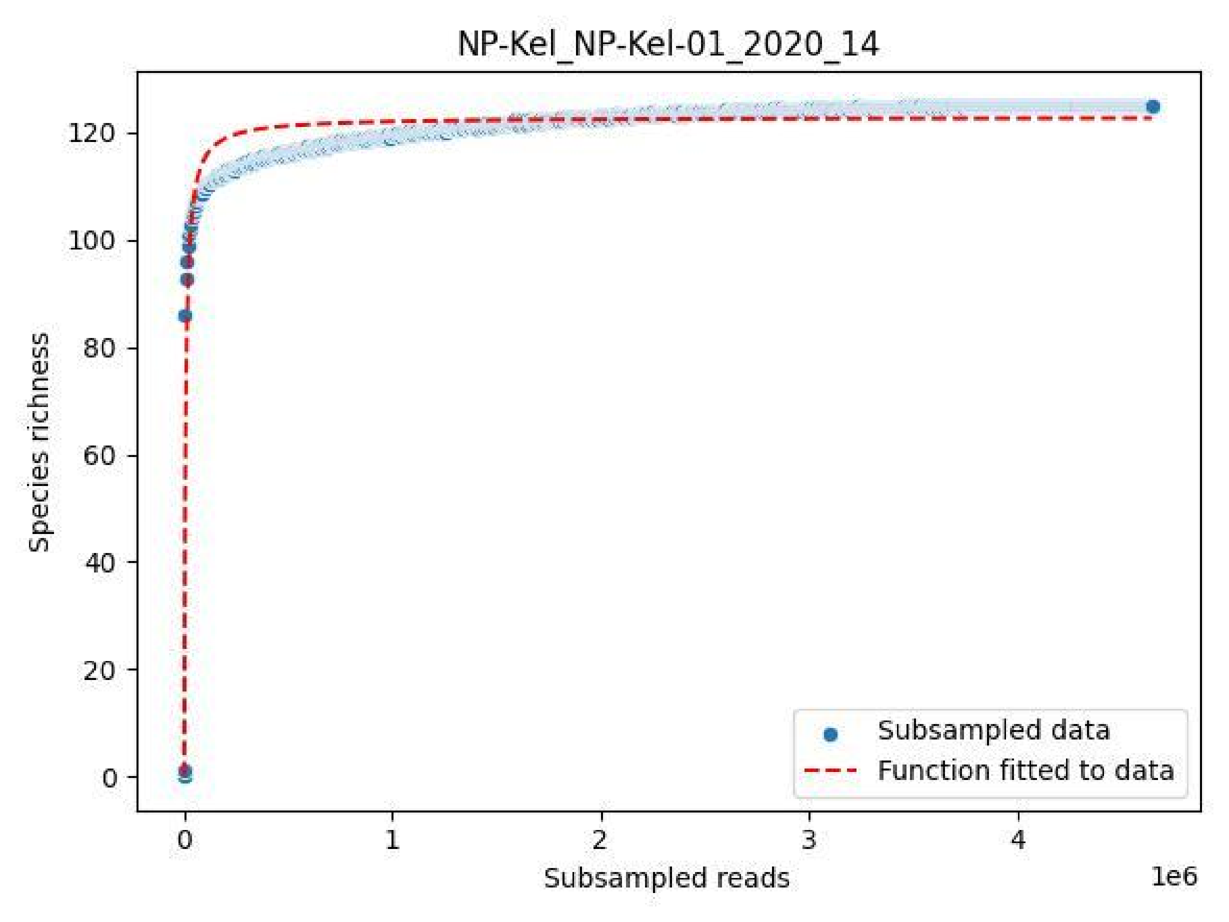
NP-Kel_NP-Kel-01_2020_14
Species richness
Subsampled reads
1e6
0
20
40
60
80
100
120
0
1
2
3
4
Subsampled data
Function fitted to data

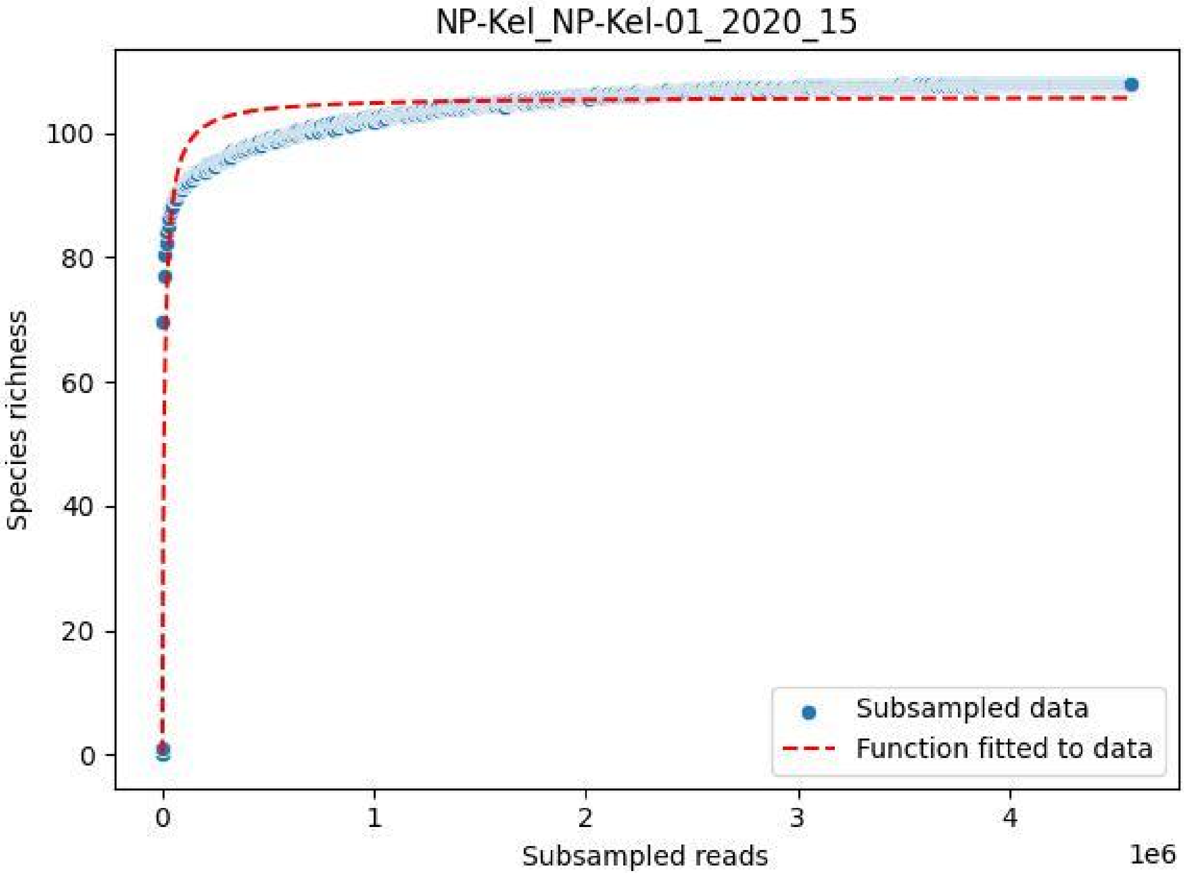
NP-Kel_NP-Kel-01_2020_15
Species richness
0
20
40
60
80
100
0
1
2
3
4
1e6
Subsampled reads
Subsampled data
Function fitted to data

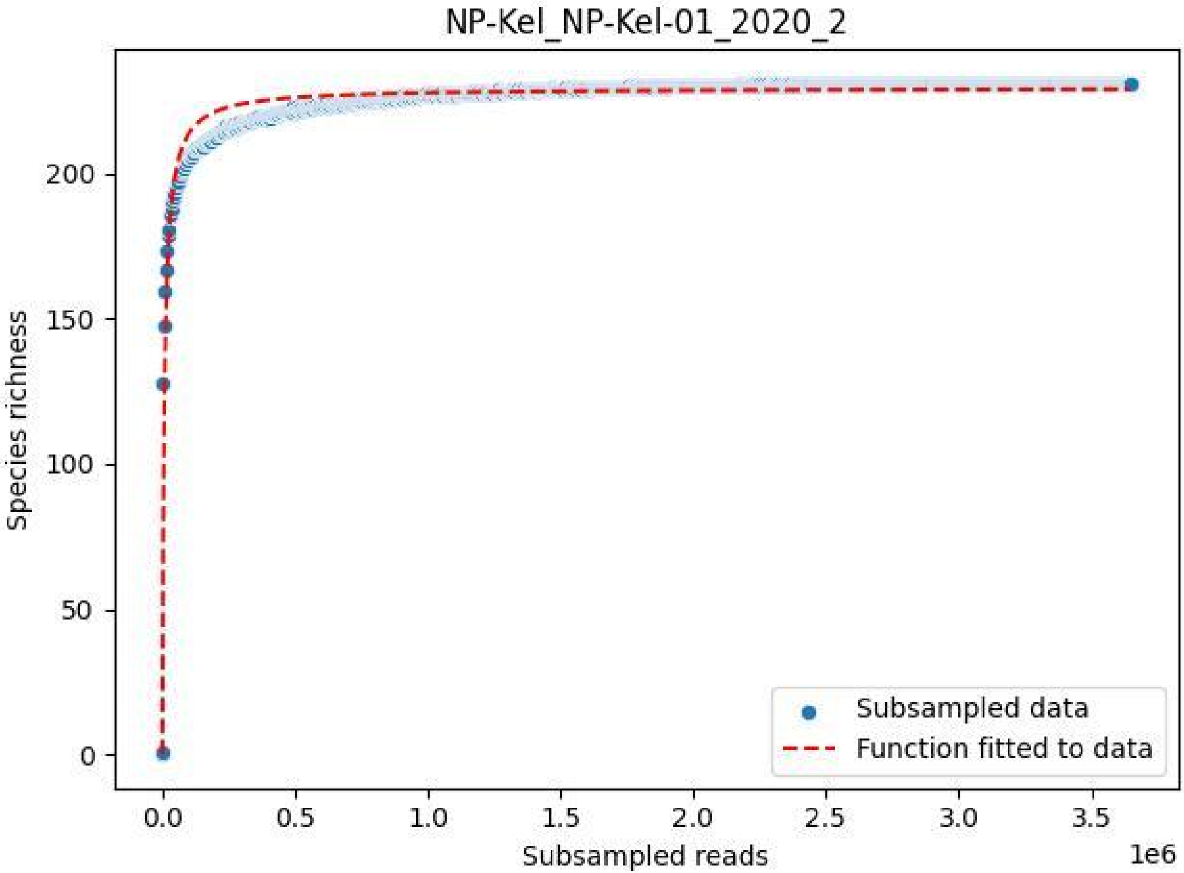

NP-Kel_NP-Kel-01_2020_2
Species richness
0
50
100
150
200
0.0
0.5
1.0
1.5
2.0
2.5
3.0
3.5
1e6
Subsampled reads
Subsampled data
Function fitted to data

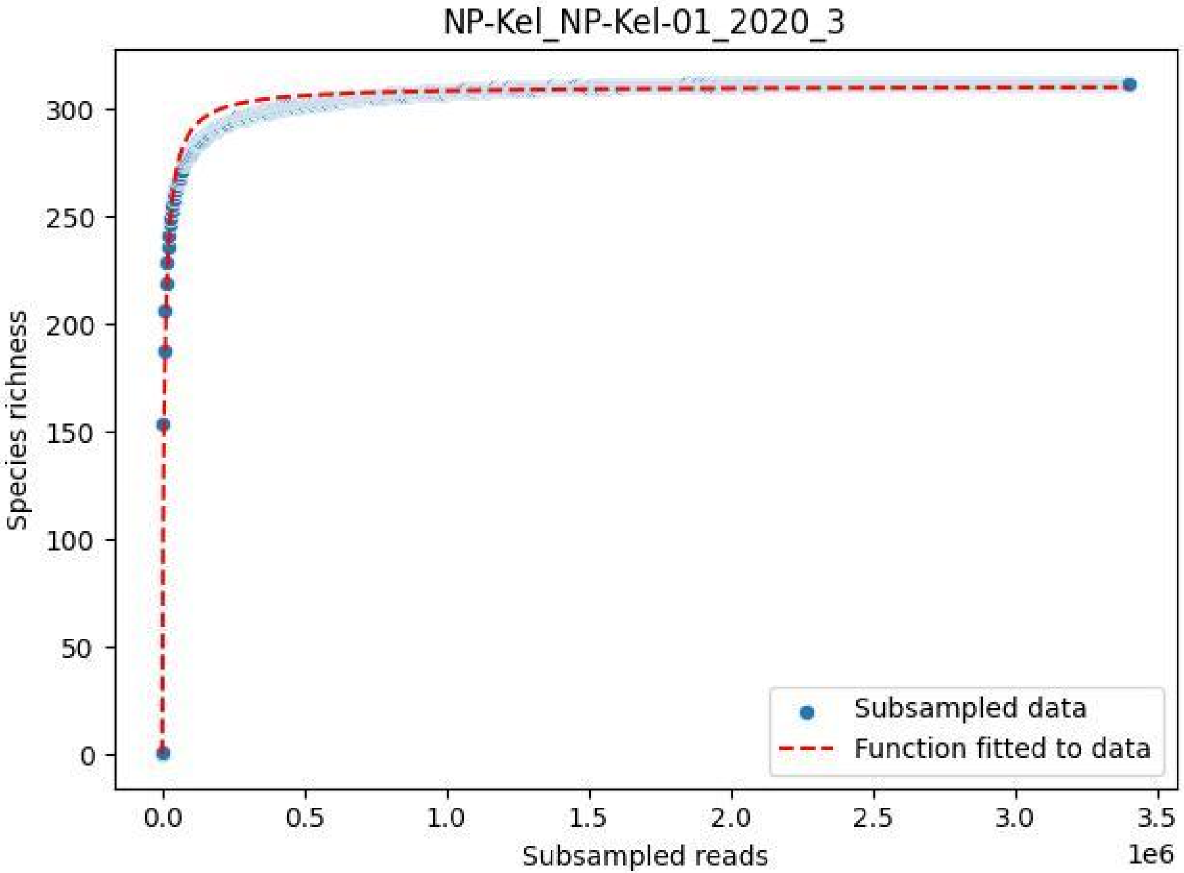

NP-Kel_NP-Kel-01_2020_3
Species richness
Subsampled reads
1e6
0
50
100
150
200
250
300
0.0
0.5
1.0
1.5
2.0
2.5
3.0
3.5
Subsampled data
Function fitted to data

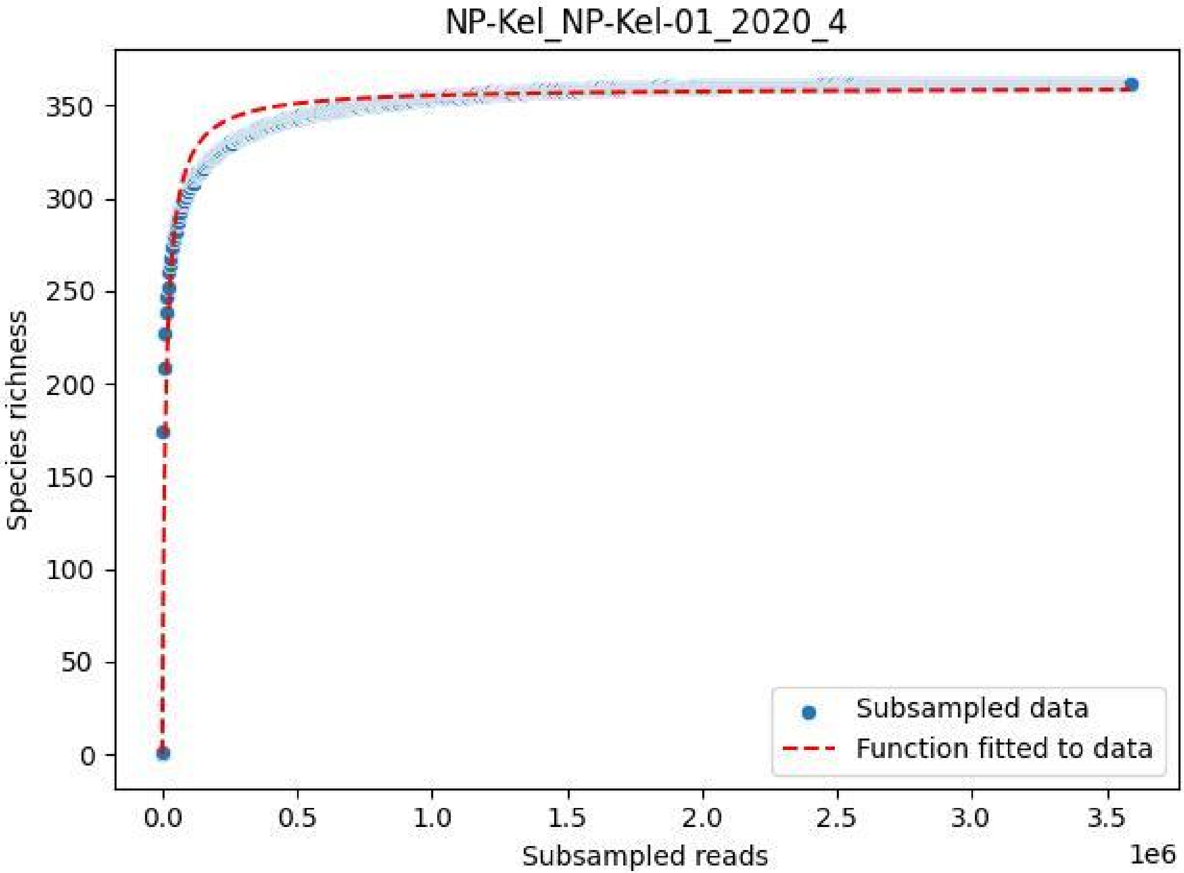
NP-Kel_NP-Kel-01_2020_4
Species richness
Subsampled reads
1e6
0
50
100
150
200
250
300
350
0.0
0.5
1.0
1.5
2.0
2.5
3.0
3.5
Subsampled data
Function fitted to data

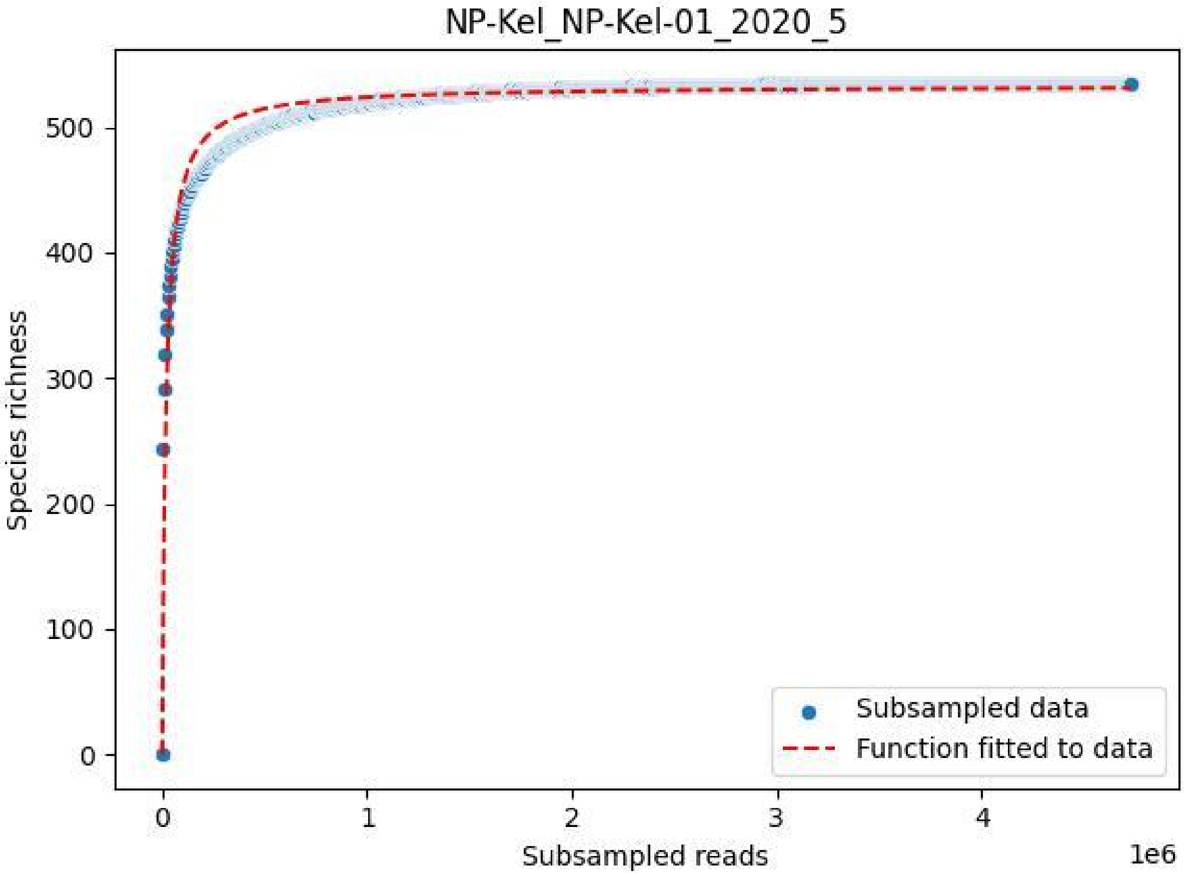

NP-Kel_NP-Kel-01_2020_5
Species richness
Subsampled reads
1e6
0
100
200
300
400
500
0
1
2
3
4
Subsampled data
Function fitted to data

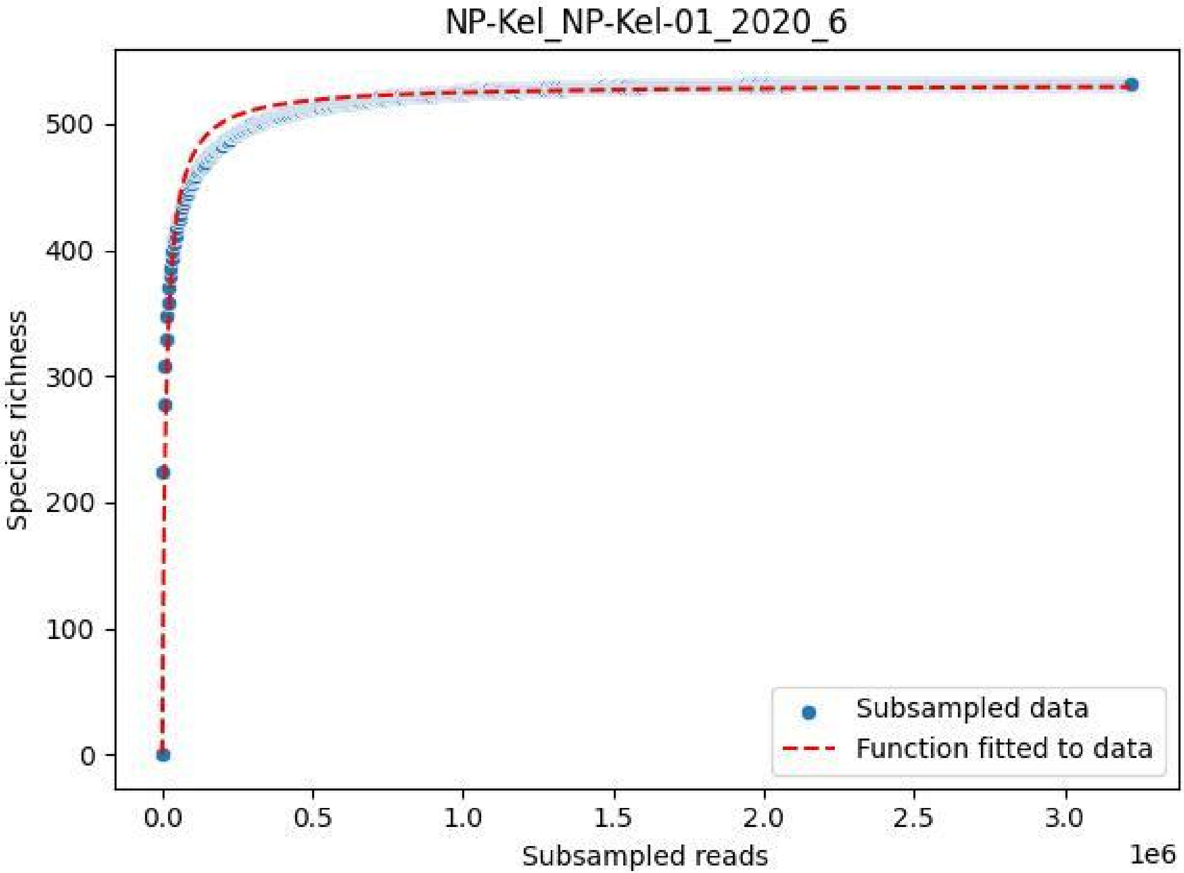
NP-Kel_NP-Kel-01_2020_6
Species richness
Subsampled reads
1e6
0
100
200
300
400
500
0.0
0.5
1.0
1.5
2.0
2.5
3.0
Subsampled data
Function fitted to data

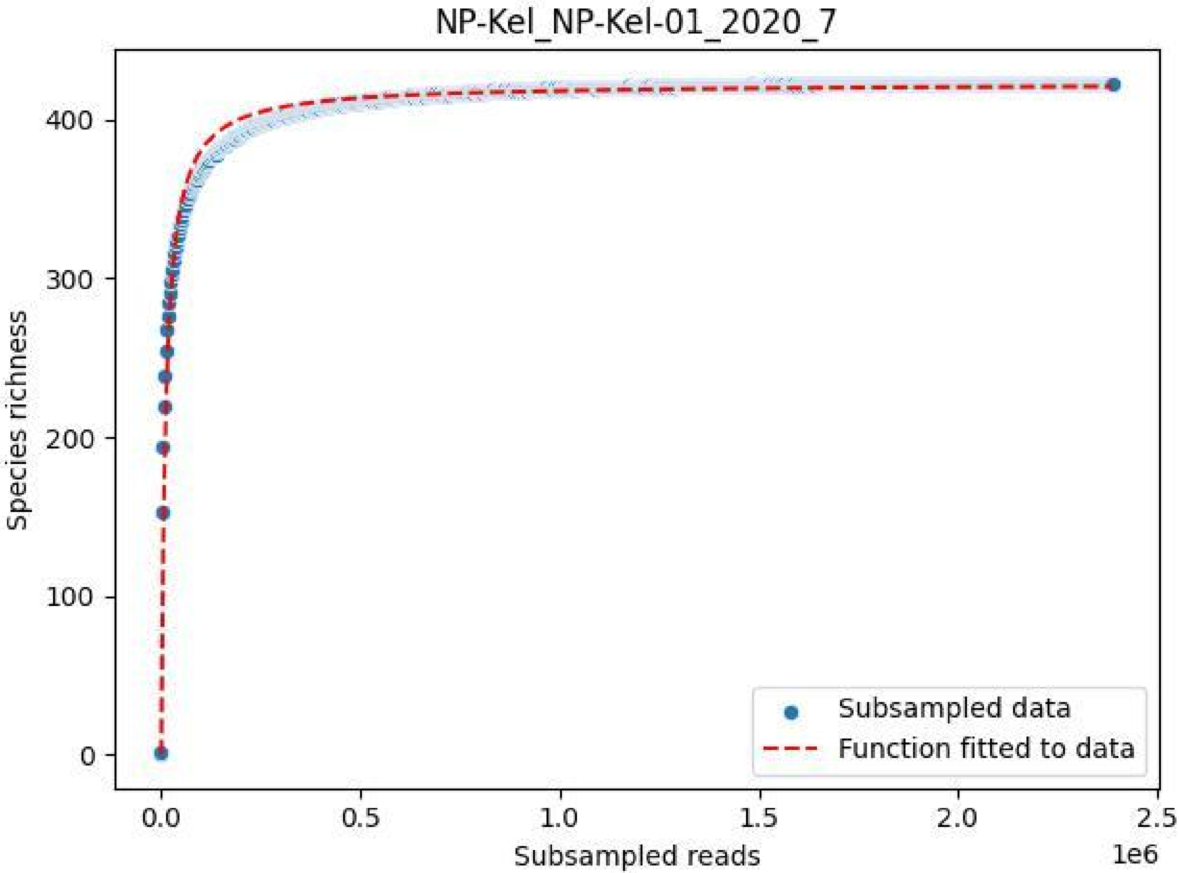

NP-Kel_NP-Kel-01_2020_7
Species richness
0
100
200
300
400
Subsampled reads
0.0
0.5
1.0
1.5
2.0
2.5
1e6
Subsampled data
Function fitted to data

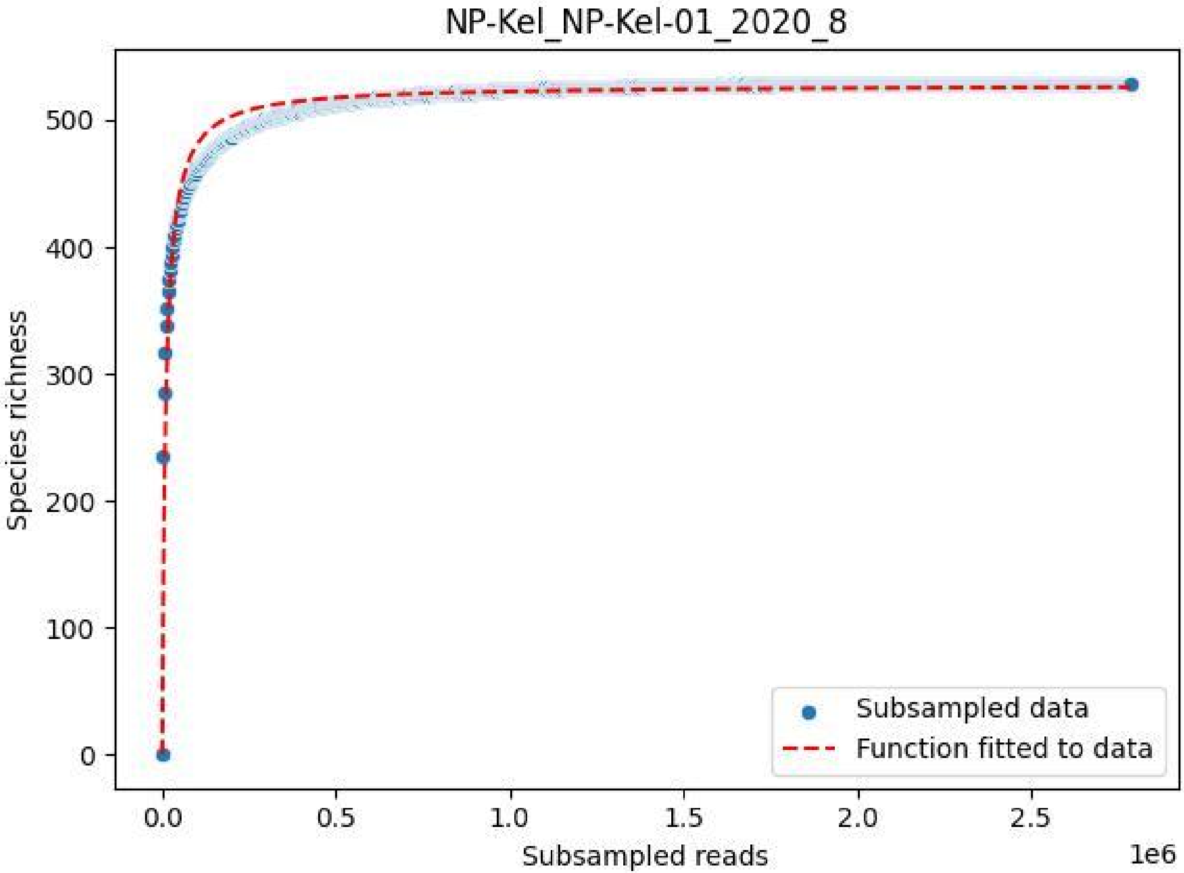

NP-Kel_NP-Kel-01_2020_8
Species richness
Subsampled reads
1e6
0
100
200
300
400
500
0.0
0.5
1.0
1.5
2.0
2.5
Subsampled data
Function fitted to data

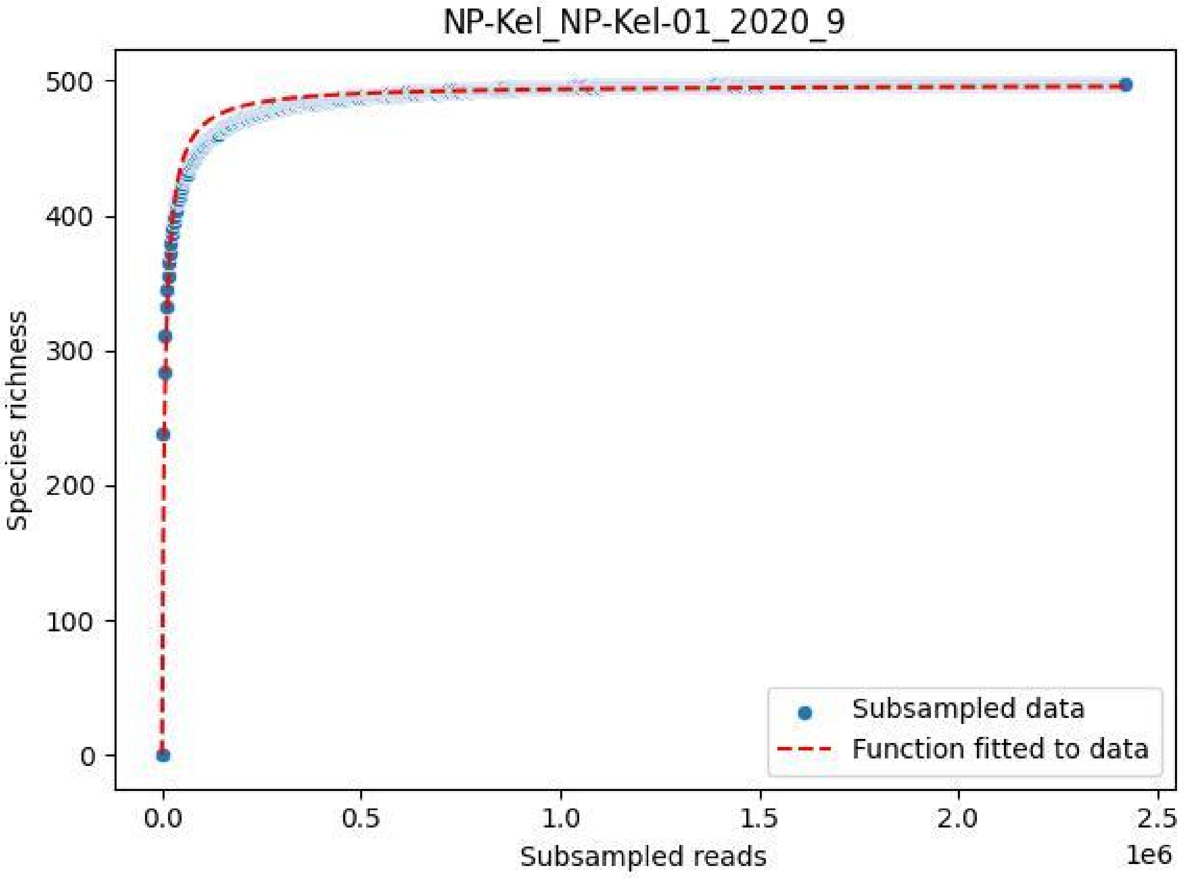
NP-Kel_NP-Kel-01_2020_9
Species richness
Subsampled reads
500
400
300
200
100
0
0.0
0.5
1.0
1.5
2.0
2.5
1e6
Subsampled data
Function fitted to data

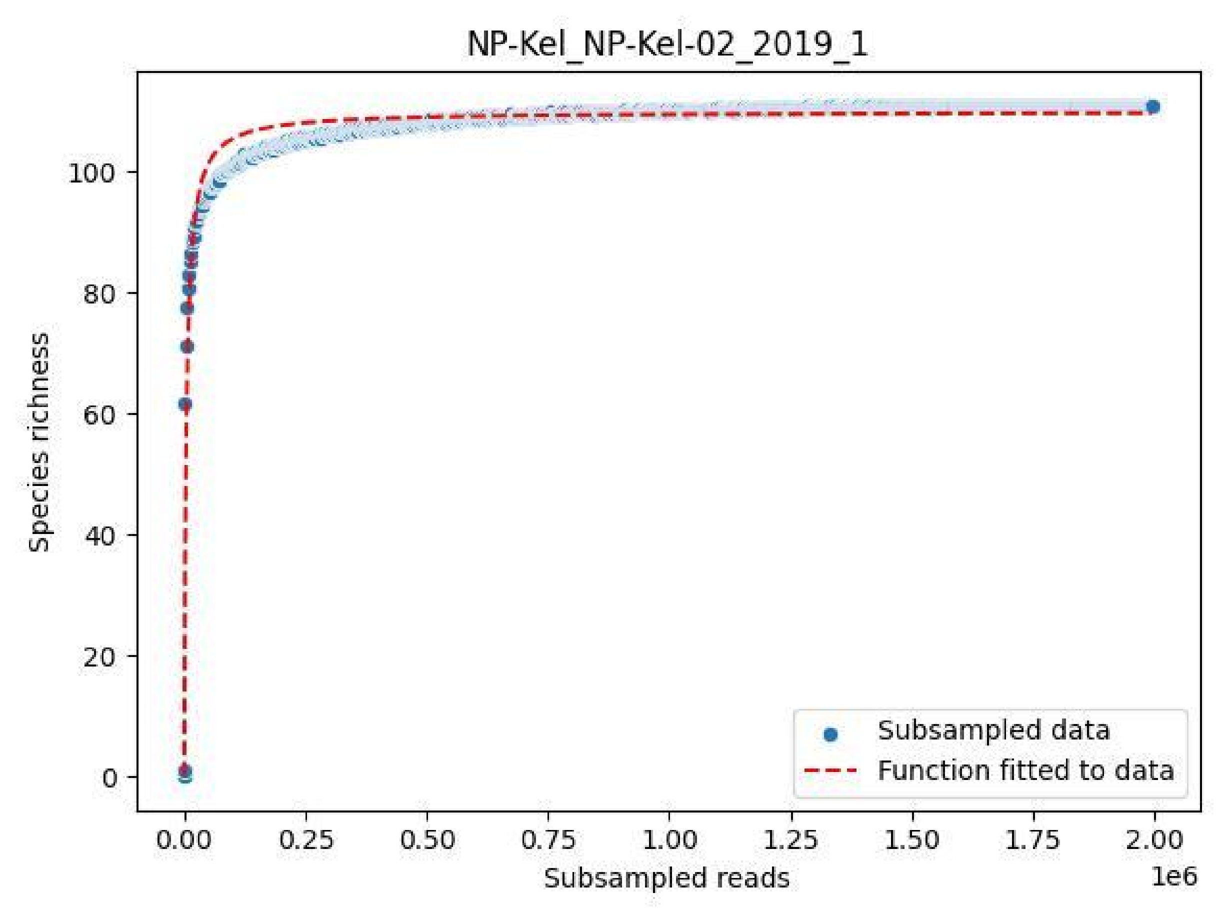

NP-Kel_NP-Kel-02_2019_1
Species richness
0
20
40
60
80
100
0.00
0.25
0.50
0.75
1.00
1.25
1.50
1.75
2.00
1e6
Subsampled reads
Subsampled data
Function fitted to data

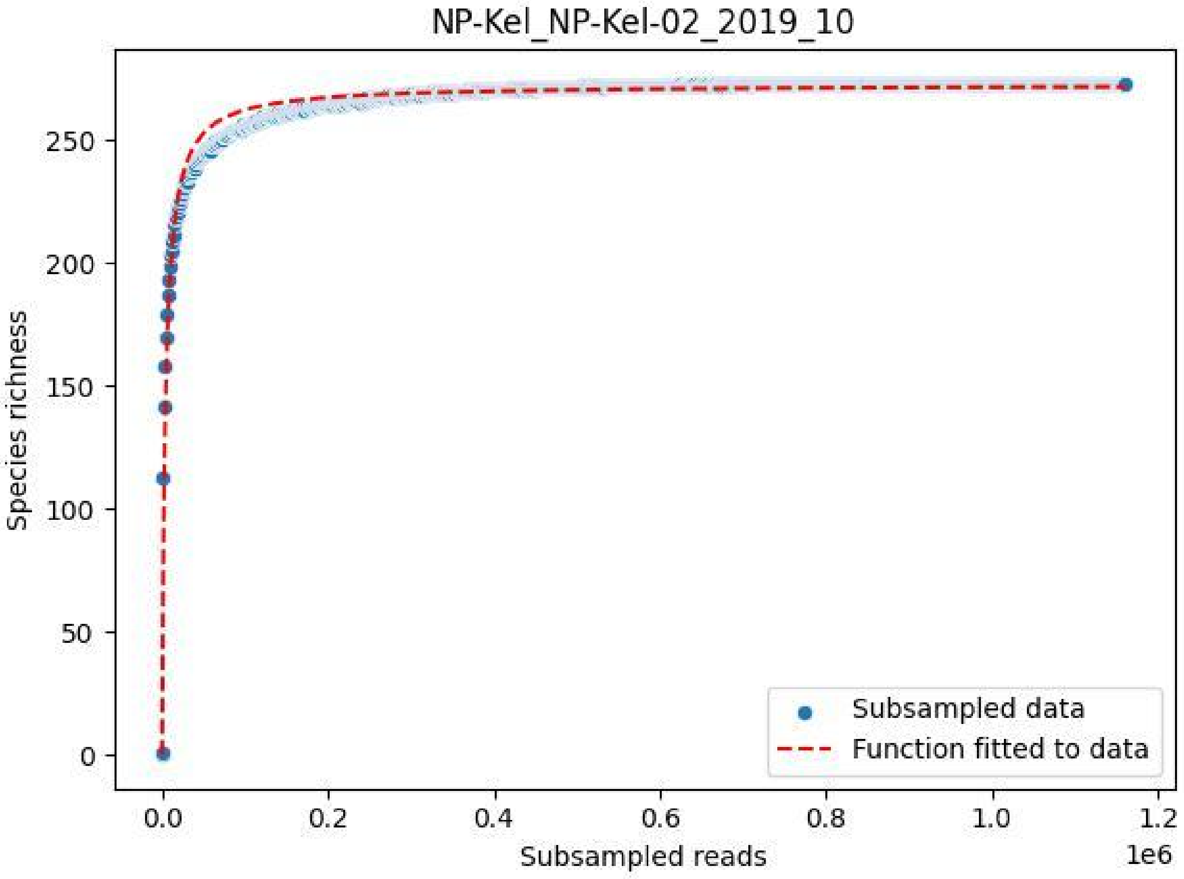
NP-Kel_NP-Kel-02_2019_10
Species richness
Subsampled reads
1e6
0
50
100
150
200
250
0.0
0.2
0.4
0.6
0.8
1.0
1.2
Subsampled data
Function fitted to data

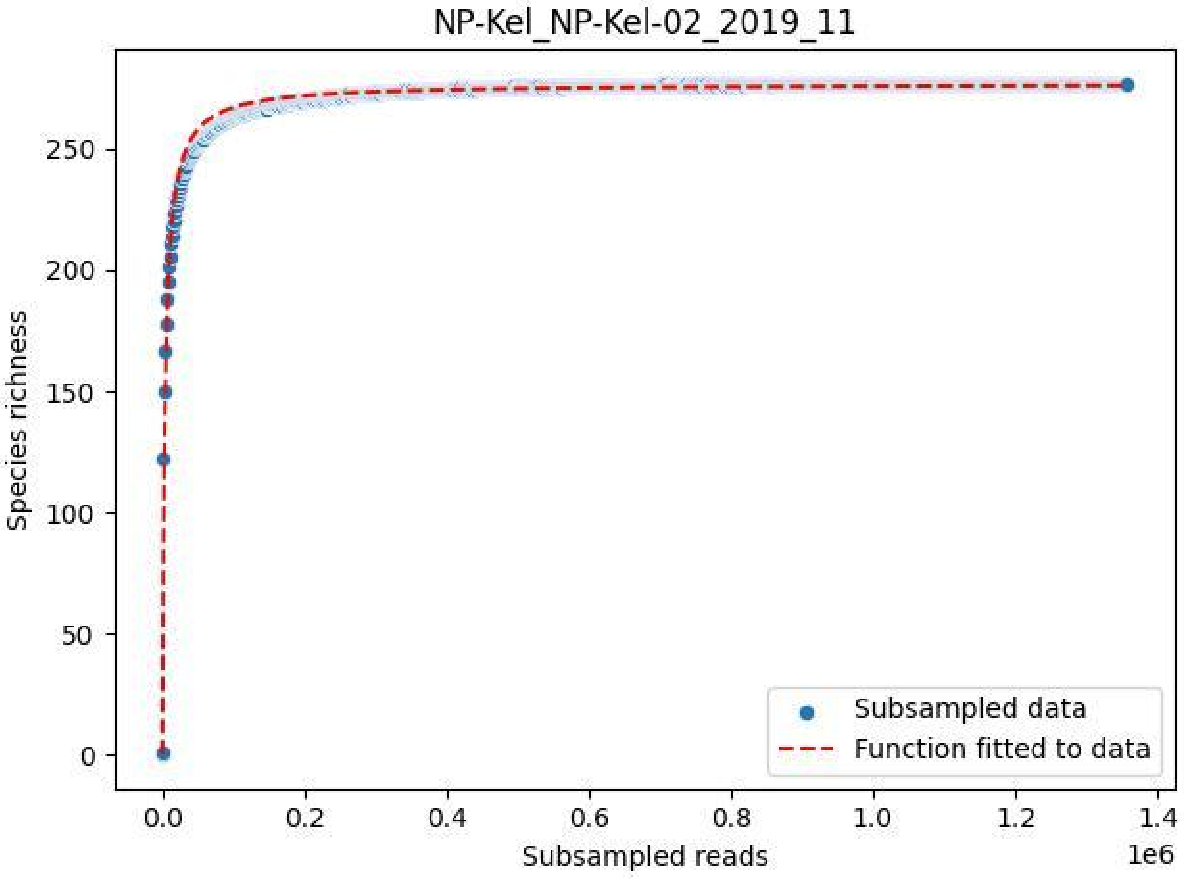

NP-Kel_NP-Kel-02_2019_11
Species richness
Subsampled reads
1e6
0
50
100
150
200
250
0.0
0.2
0.4
0.6
0.8
1.0
1.2
1.4
Subsampled data
Function fitted to data

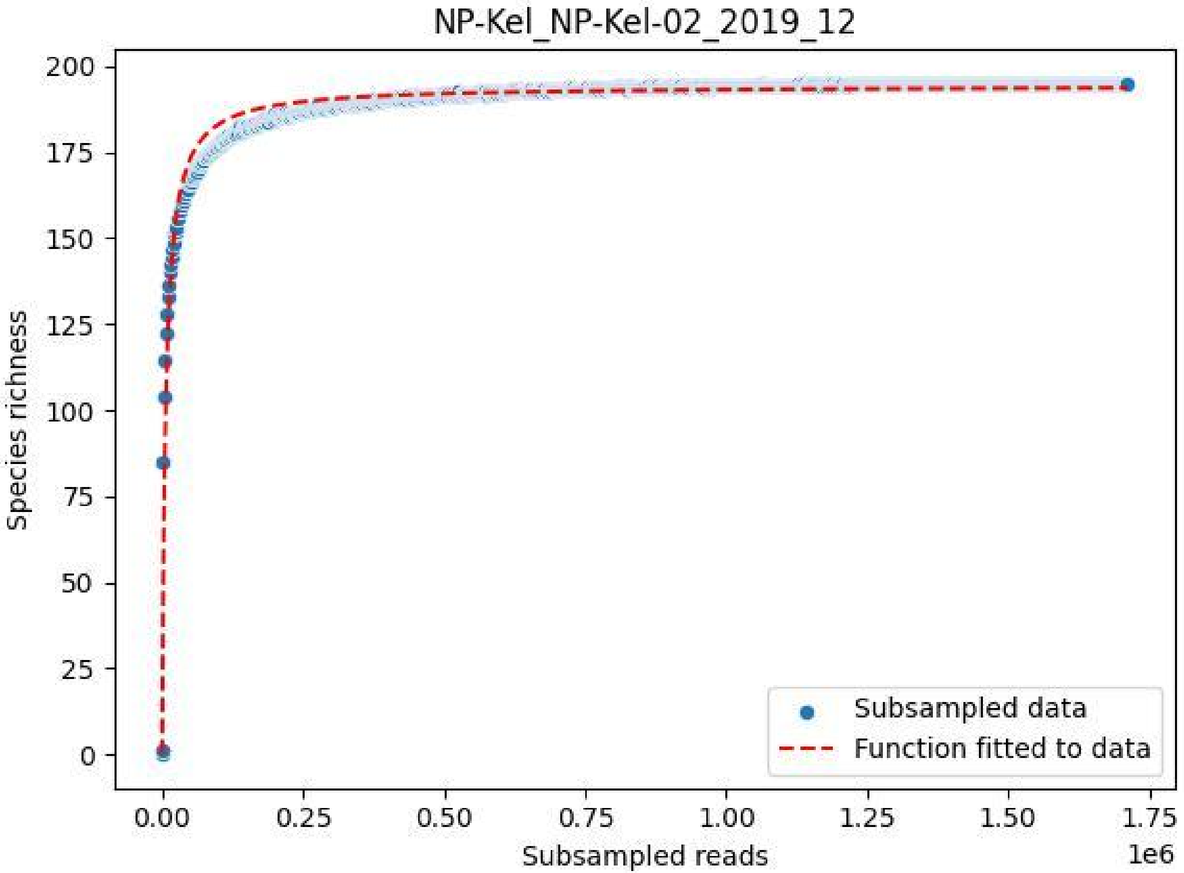

NP-Kel_NP-Kel-02_2019_12
Species richness
Subsampled reads
1e6
Subsampled data
Function fitted to data
0
25
50
75
100
125
150
175
200
0.00
0.25
0.50
0.75
1.00
1.25
1.50
1.75

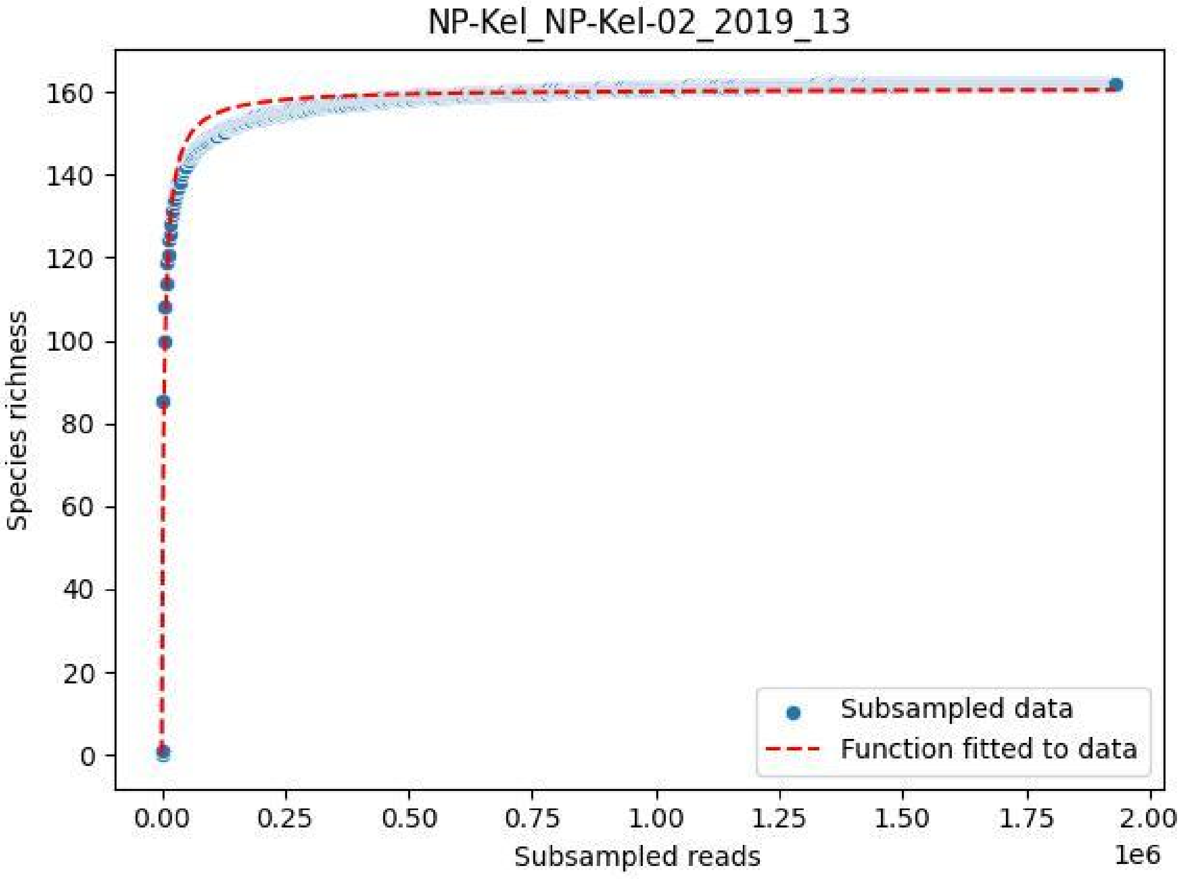
NP-Kel_NP-Kel-02_2019_13
Species richness
Subsampled reads
1e6
0.00
0.25
0.50
0.75
1.00
1.25
1.50
1.75
2.00
0
20
40
60
80
100
120
140
160
Subsampled data
Function fitted to data

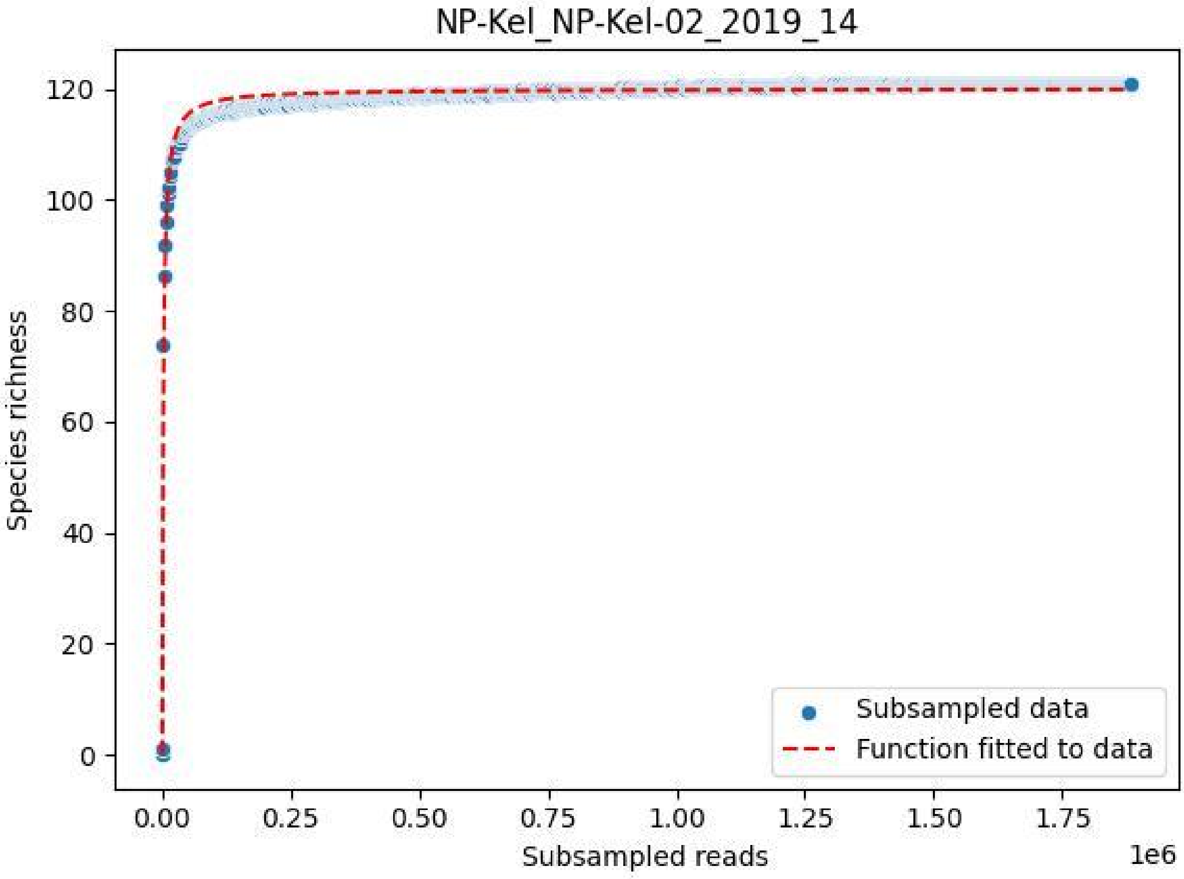

NP-Kel_NP-Kel-02_2019_14
Species richness
0
20
40
60
80
100
120
0.00
0.25
0.50
0.75
1.00
1.25
1.50
1.75
1e6
Subsampled reads
Subsampled data
Function fitted to data

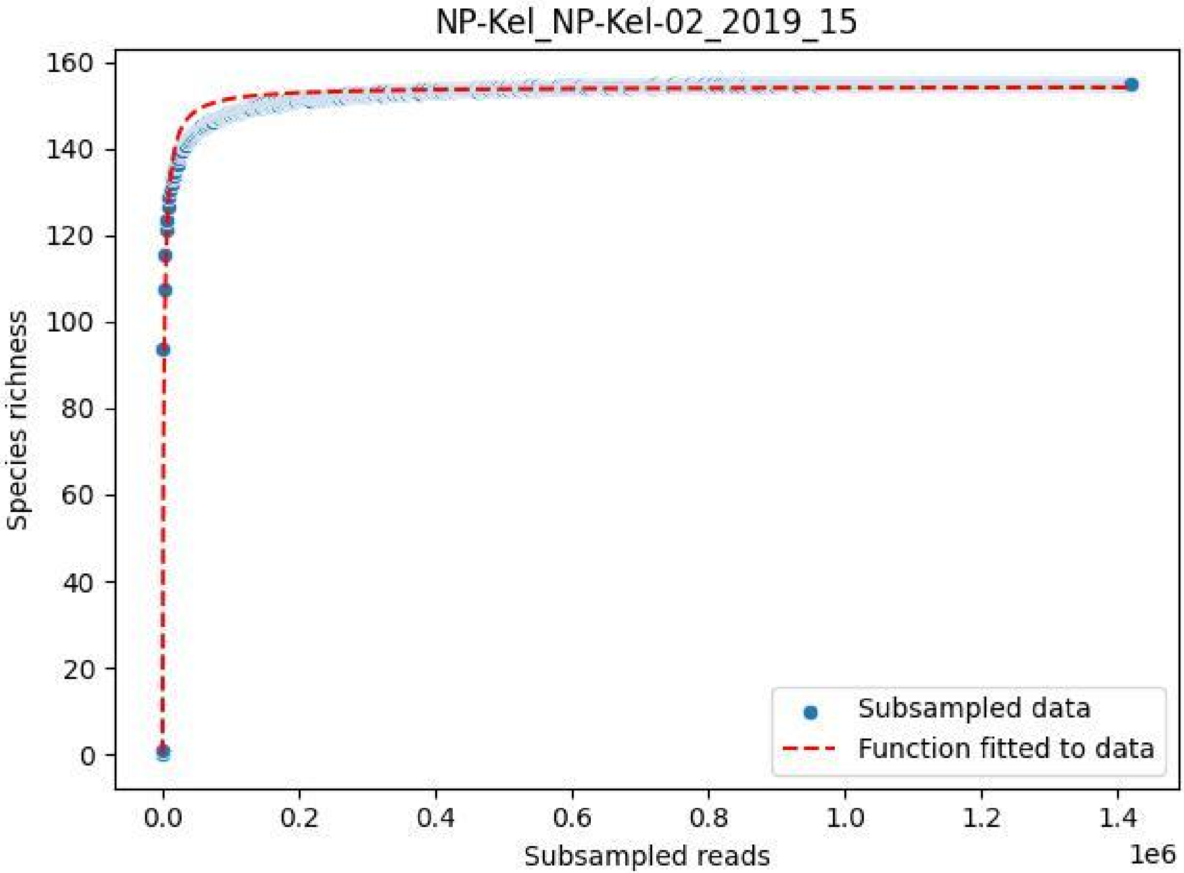

NP-Kel_NP-Kel-02_2019_15
Species richness
Subsampled reads
1e6
0
20
40
60
80
100
120
140
160
0.0
0.2
0.4
0.6
0.8
1.0
1.2
1.4
Subsampled data
Function fitted to data

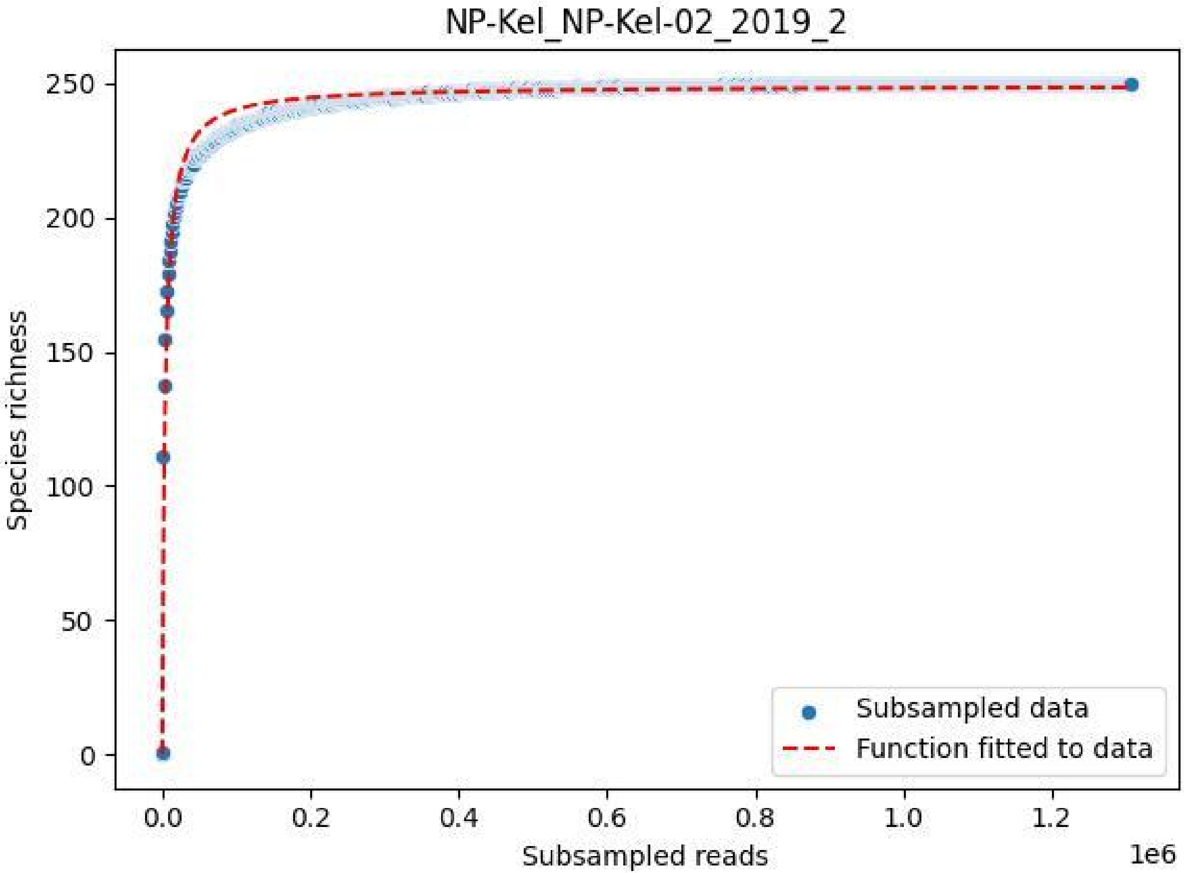
NP-Kel_NP-Kel-02_2019_2
Species richness
0
50
100
150
200
250
0.0
0.2
0.4
0.6
0.8
1.0
1.2
1e6
Subsampled reads
Subsampled data
Function fitted to data

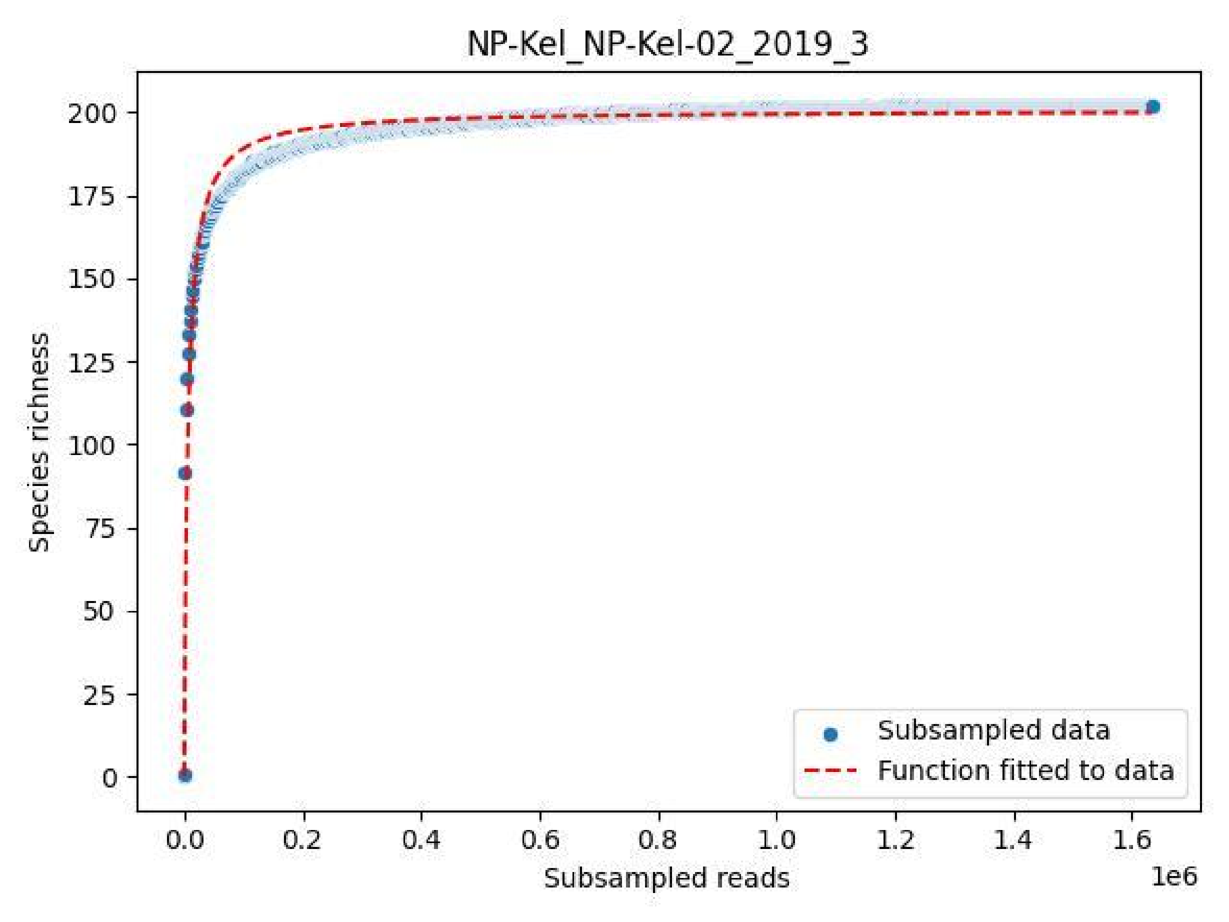
NP-Kel_NP-Kel-02_2019_3
Species richness
0
25
50
75
100
125
150
175
200
0.0
0.2
0.4
0.6
0.8
1.0
1.2
1.4
1.6
1e6
Subsampled reads
Subsampled data
Function fitted to data

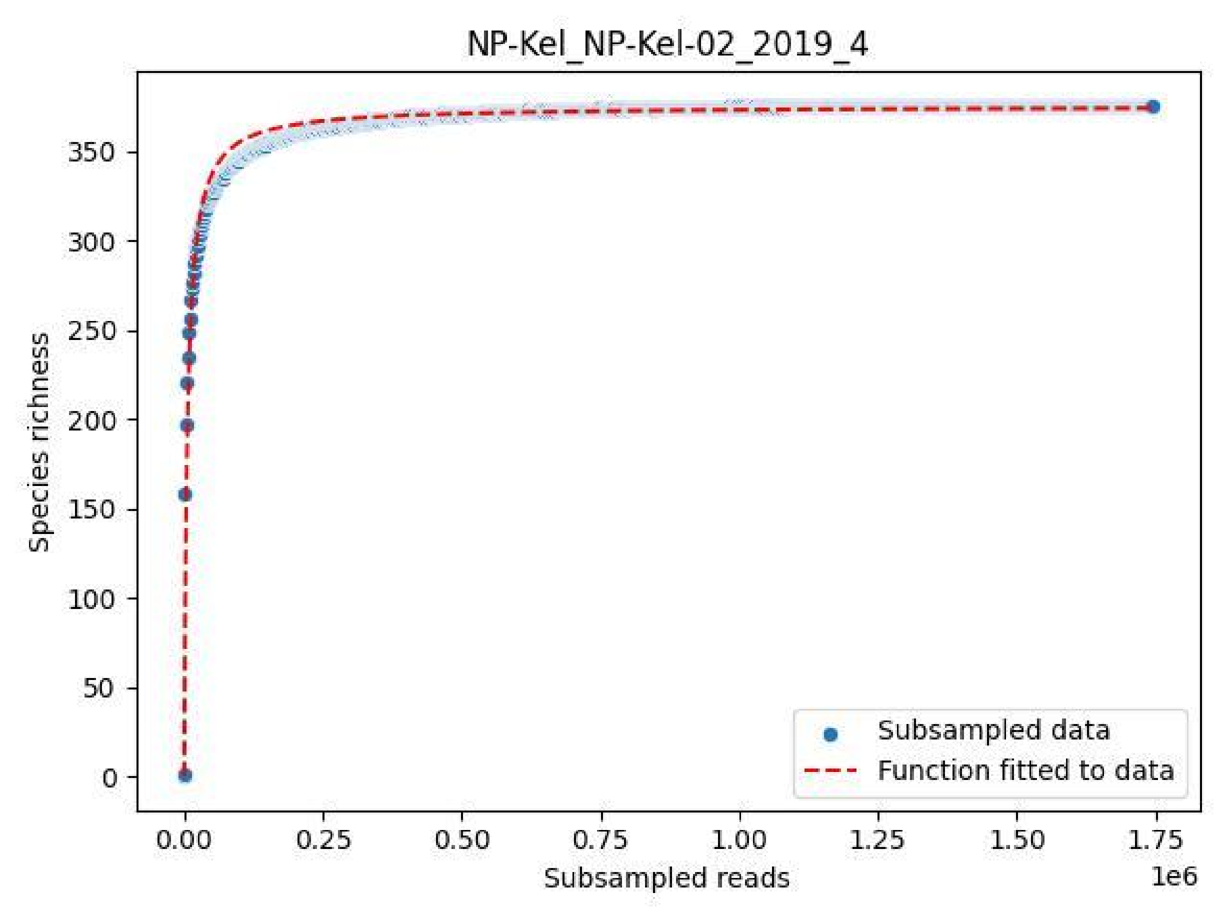
NP-Kel_NP-Kel-02_2019_4
Species richness
0
50
100
150
200
250
300
350
Subsampled reads
0.00
0.25
0.50
0.75
1.00
1.25
1.50
1.75
1e6
Subsampled data
Function fitted to data

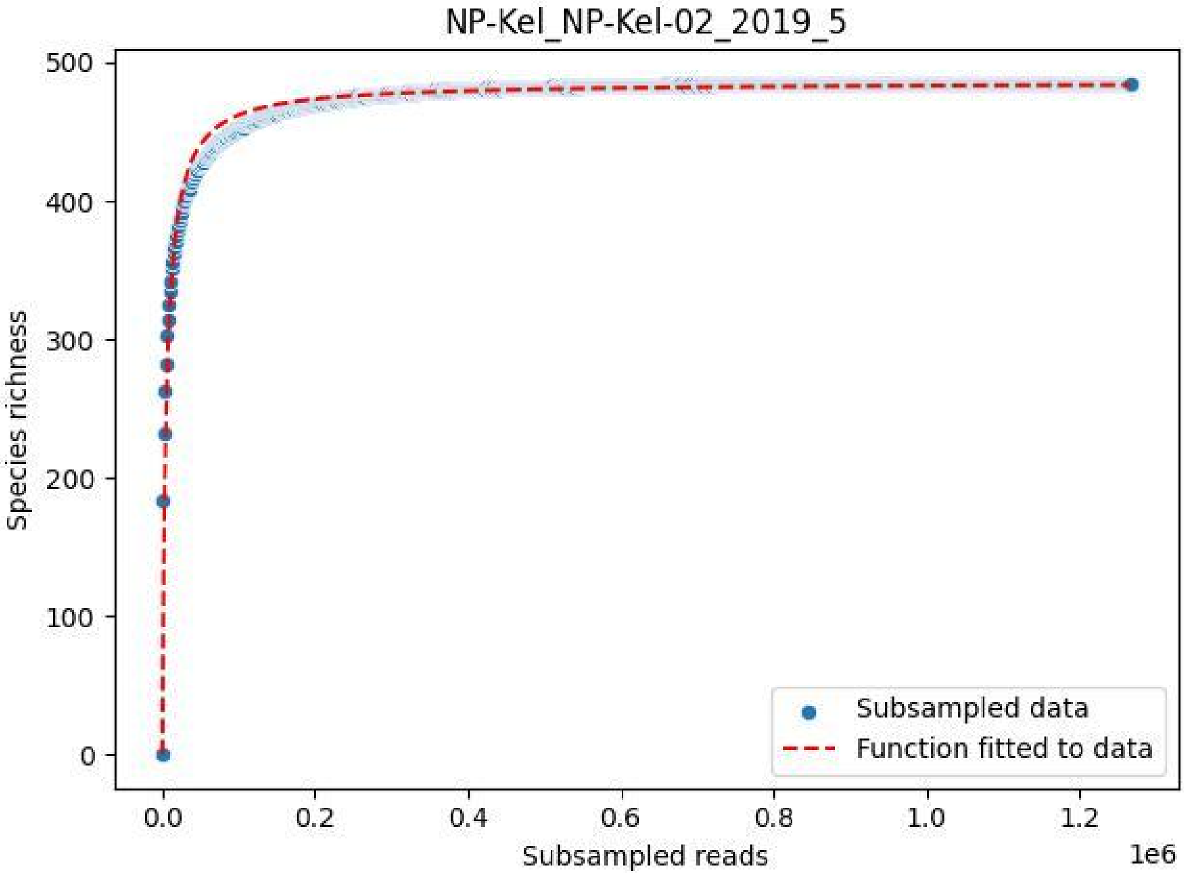
NP-Kel_NP-Kel-02_2019_5
Species richness
Subsampled reads
1e6
0
100
200
300
400
500
0.0
0.2
0.4
0.6
0.8
1.0
1.2
Subsampled data
Function fitted to data

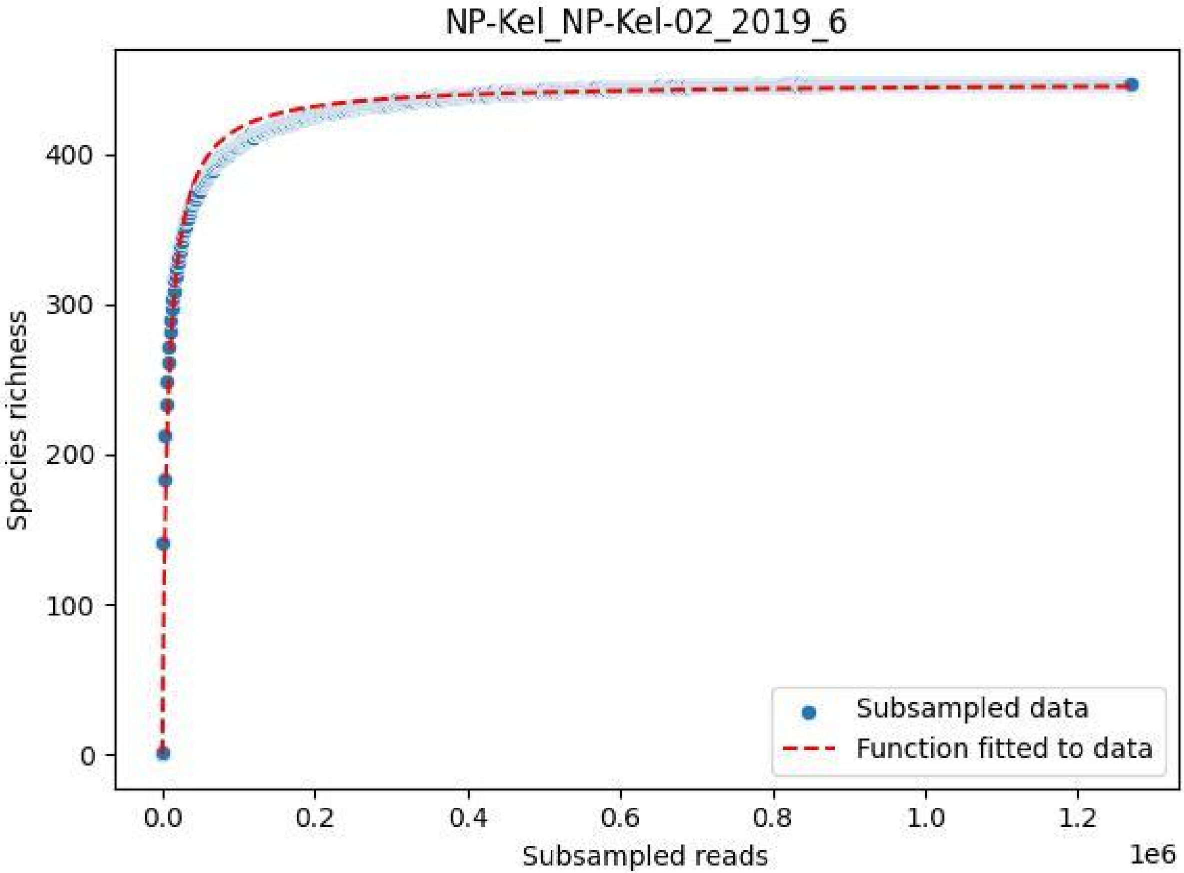

NP-Kel_NP-Kel-02_2019_6
Species richness
Subsampled reads
1e6
0
100
200
300
400
0.0
0.2
0.4
0.6
0.8
1.0
1.2
Subsampled data
Function fitted to data

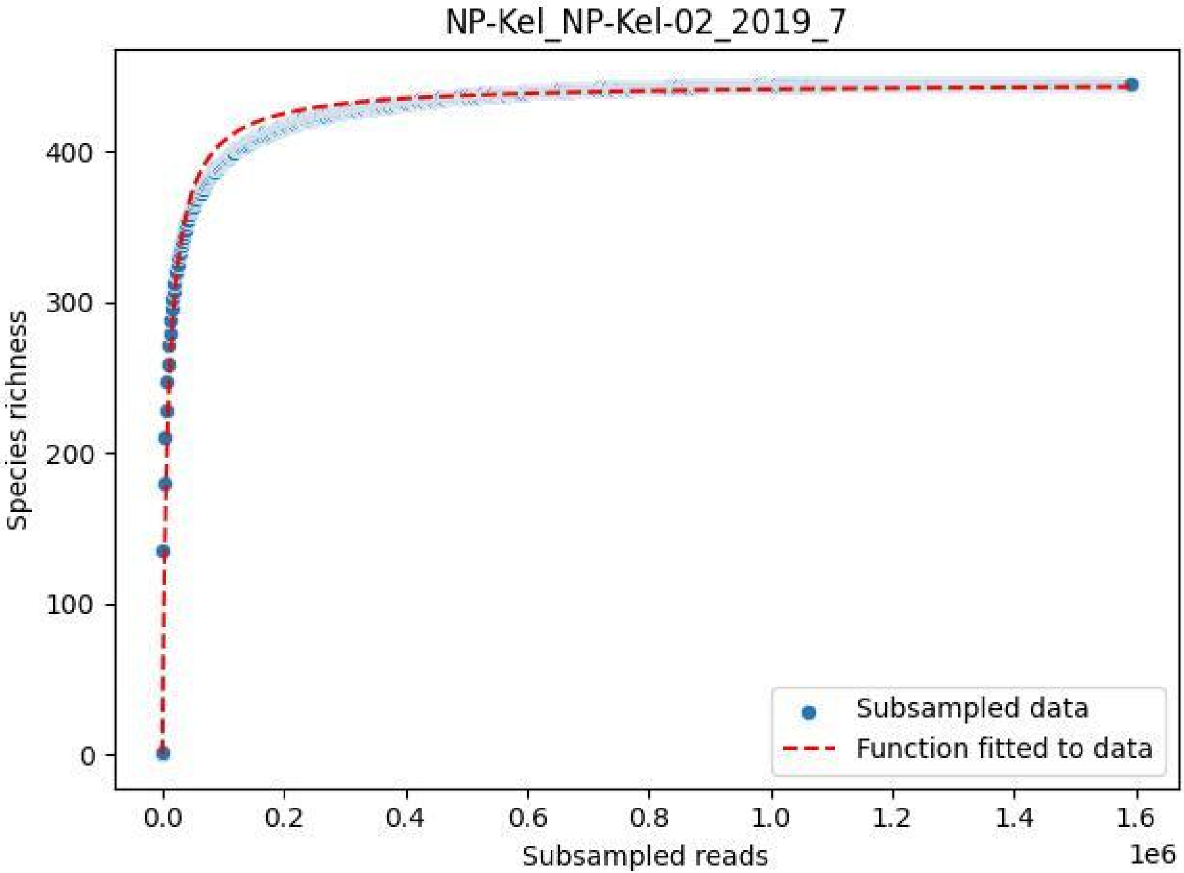

NP-Kel_NP-Kel-02_2019_7
Species richness
Subsampled reads
1e6
0
100
200
300
400
0.0
0.2
0.4
0.6
0.8
1.0
1.2
1.4
1.6
Subsampled data
Function fitted to data

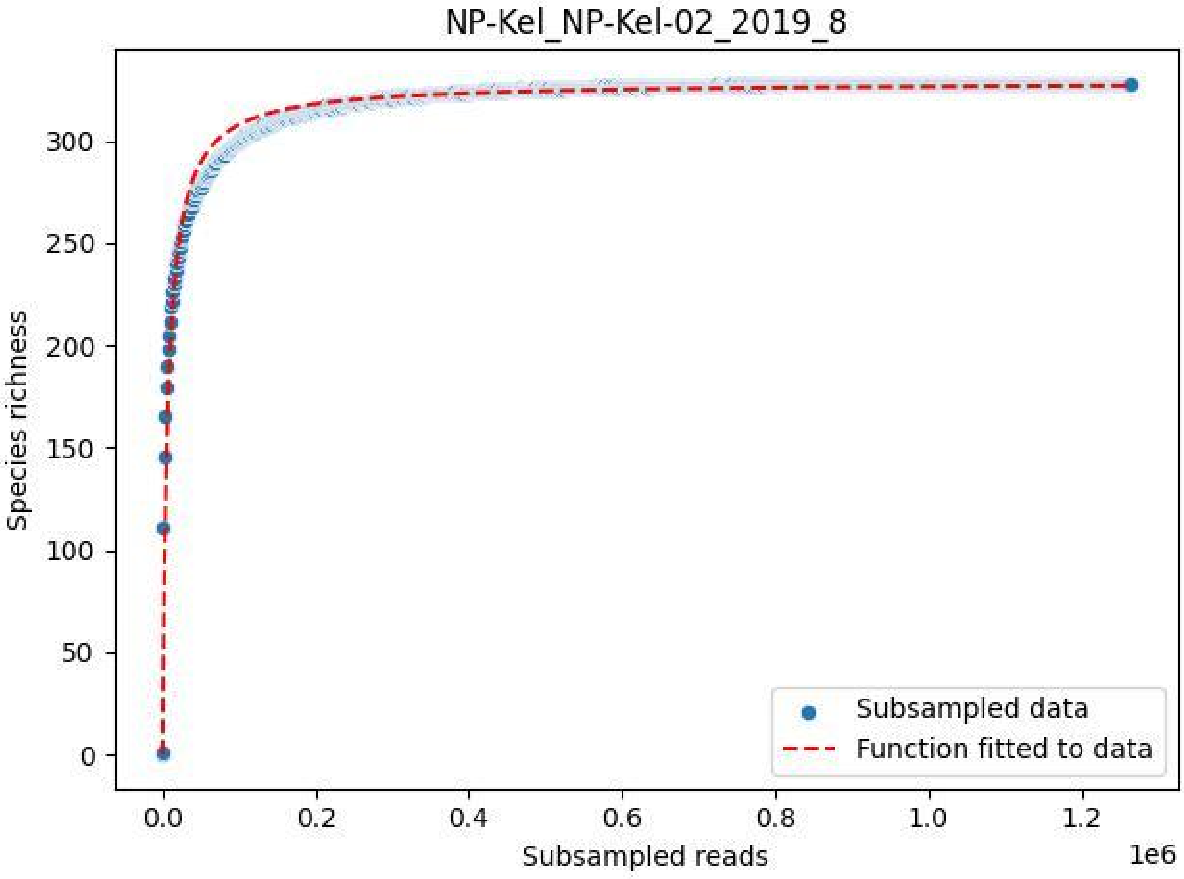

NP-Kel_NP-Kel-02_2019_8
Species richness
Subsampled reads
1e6
0
50
100
150
200
250
300
0.0
0.2
0.4
0.6
0.8
1.0
1.2
Subsampled data
Function fitted to data

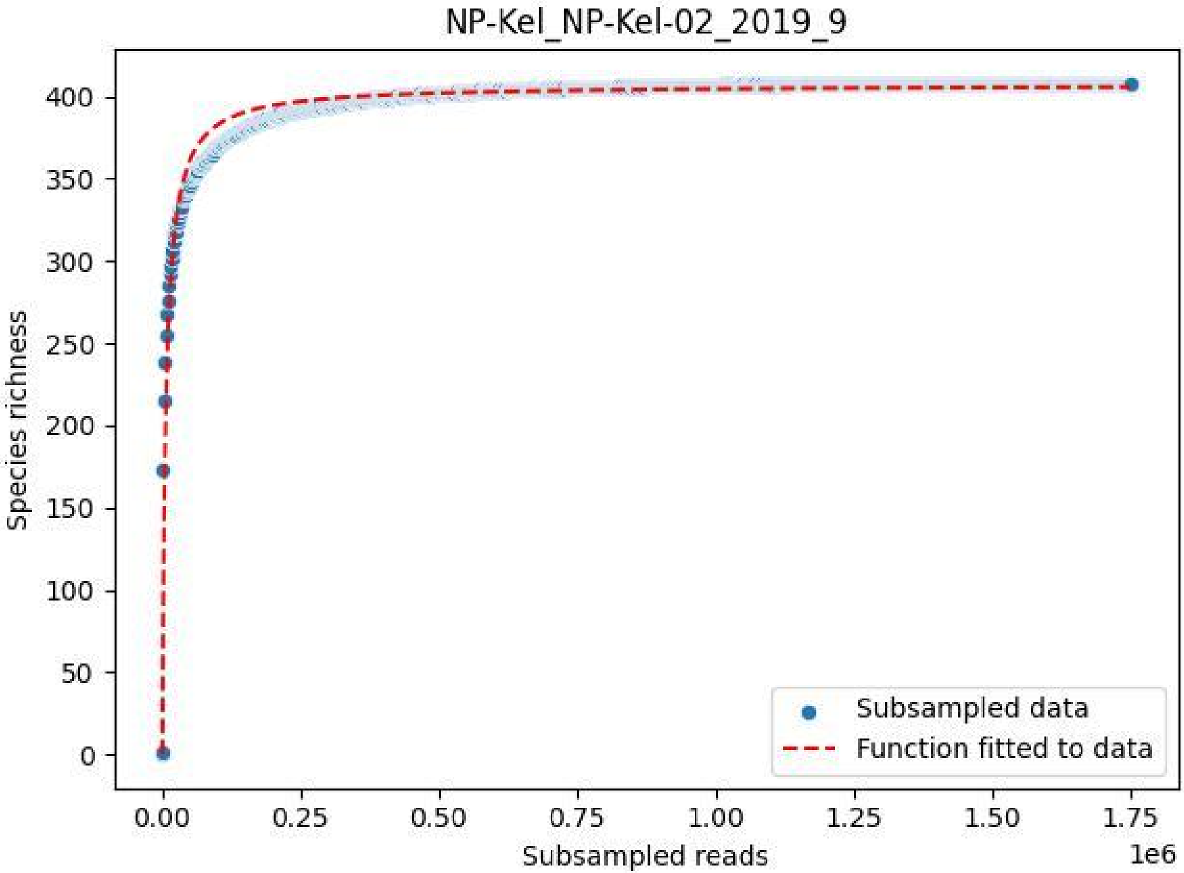

NP-Kel_NP-Kel-02_2019_9
Species richness
Subsampled reads
1e6
0
50
100
150
200
250
300
350
400
0.00
0.25
0.50
0.75
1.00
1.25
1.50
1.75
Subsampled data
Function fitted to data

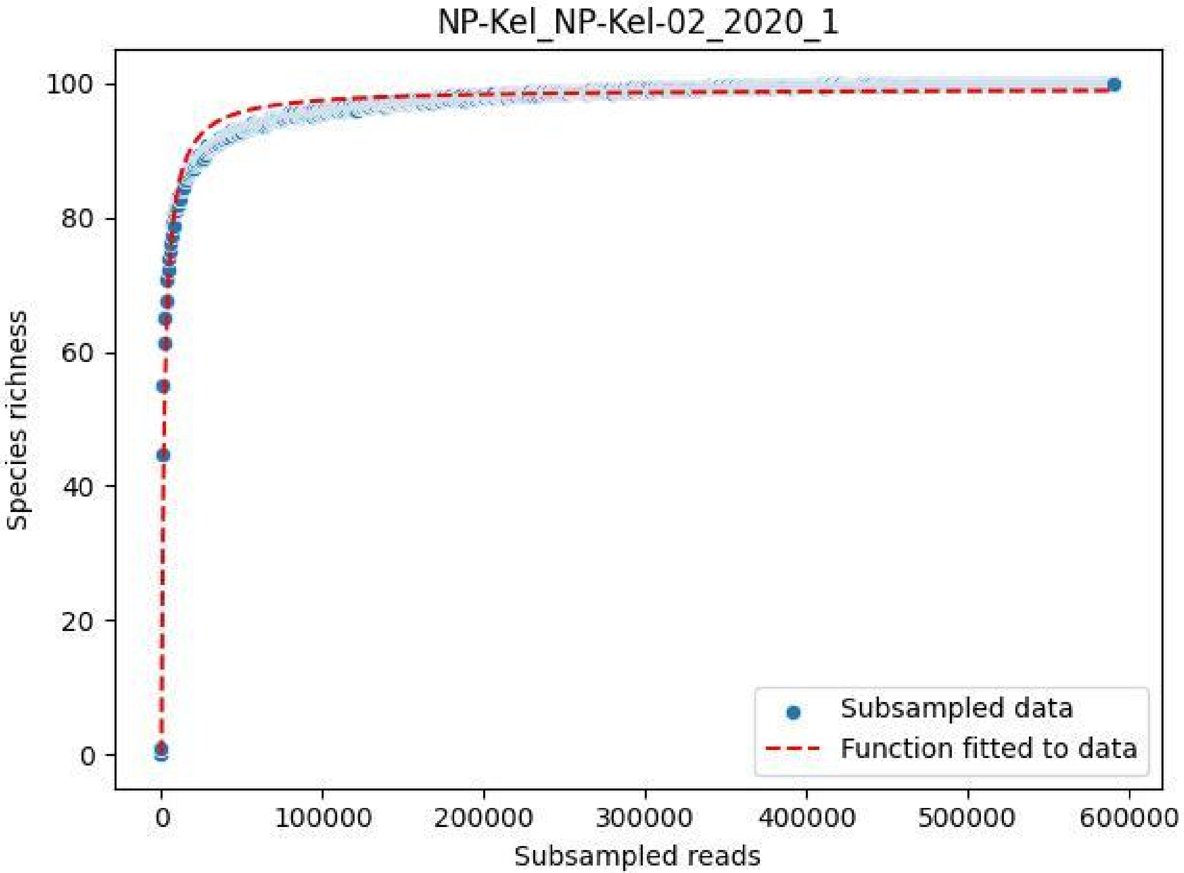

NP-Kel_NP-Kel-02_2020_1
Species richness
Subsampled reads
0
20
40
60
80
100
0
100000
200000
300000
400000
500000
600000
Subsampled data
Function fitted to data

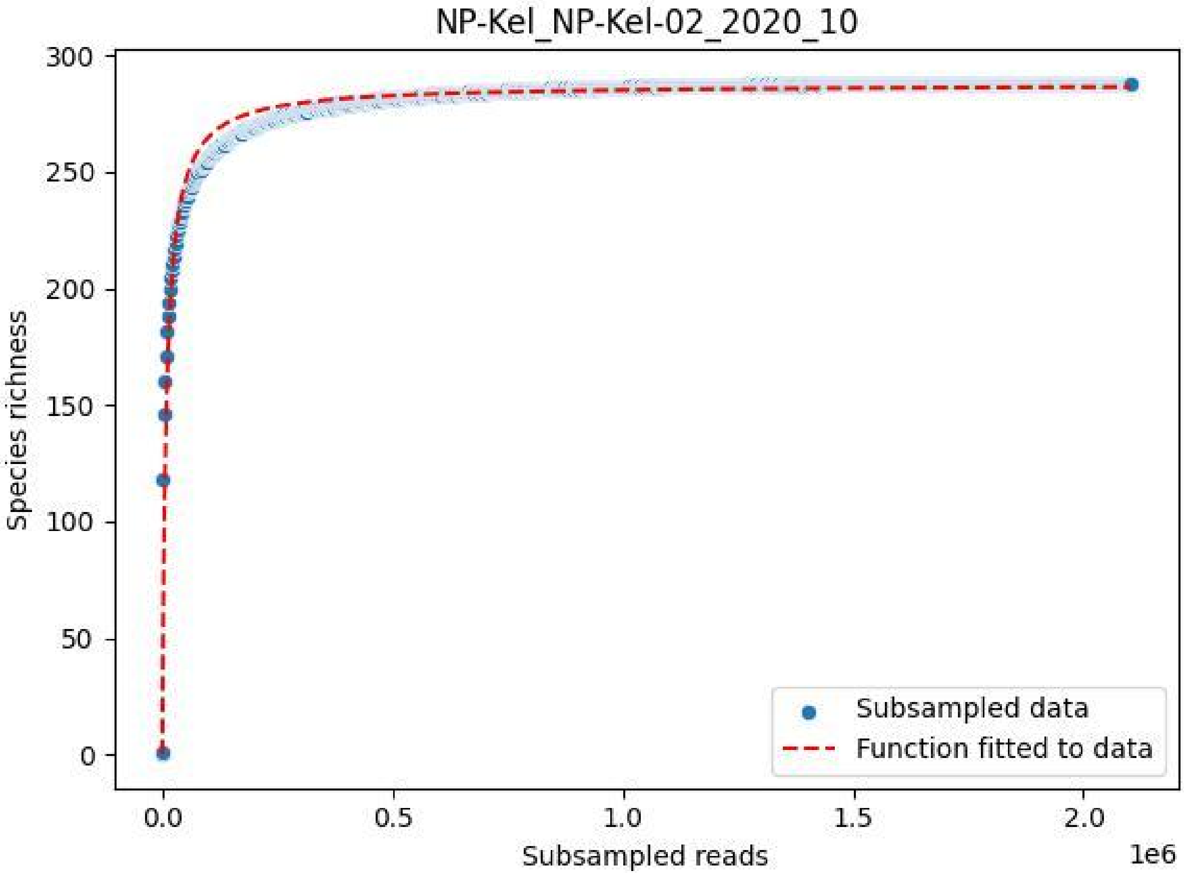
NP-Kel_NP-Kel-02_2020_10
Species richness
Subsampled reads
1e6
0
50
100
150
200
250
300
0.0
0.5
1.0
1.5
2.0
Subsampled data
Function fitted to data

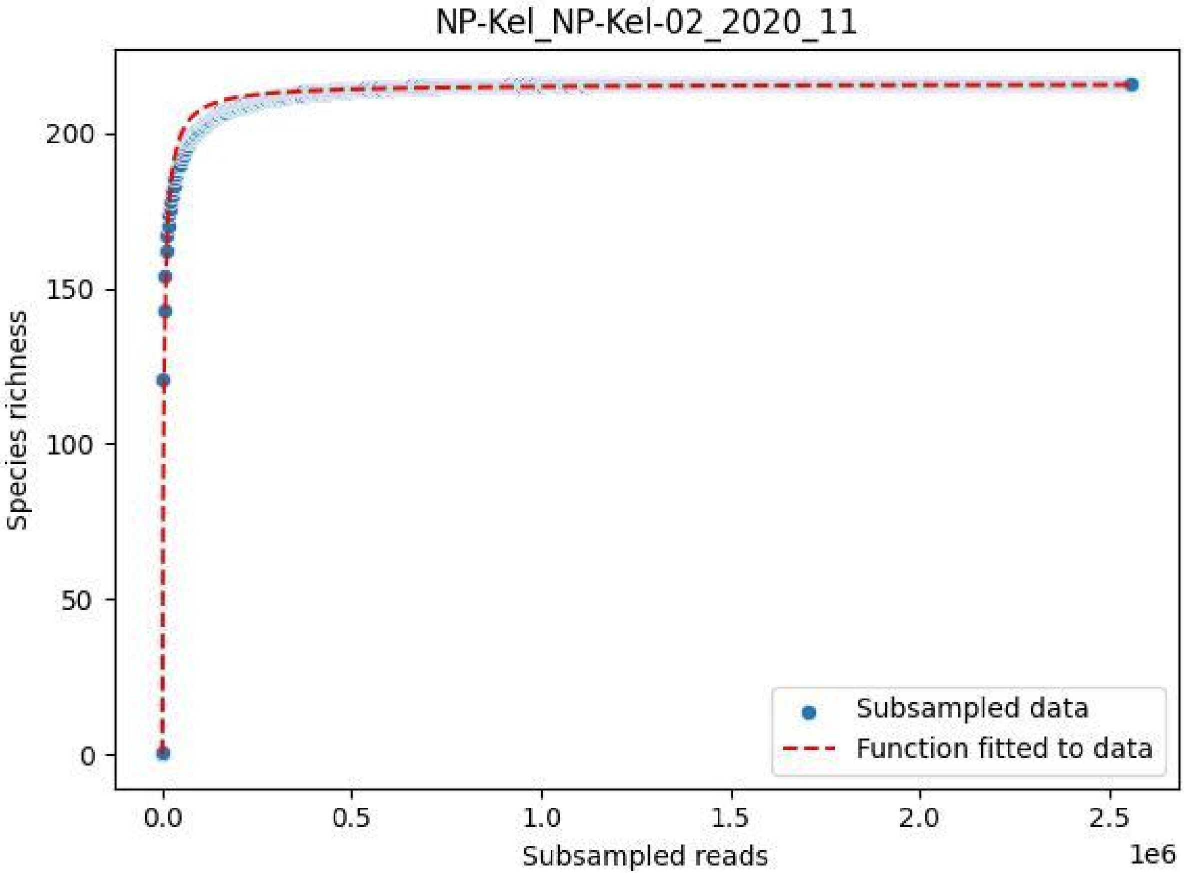

NP-Kel_NP-Kel-02_2020_11
Species richness
Subsampled reads
1e6
0
50
100
150
200
0.0
0.5
1.0
1.5
2.0
2.5
Subsampled data
Function fitted to data

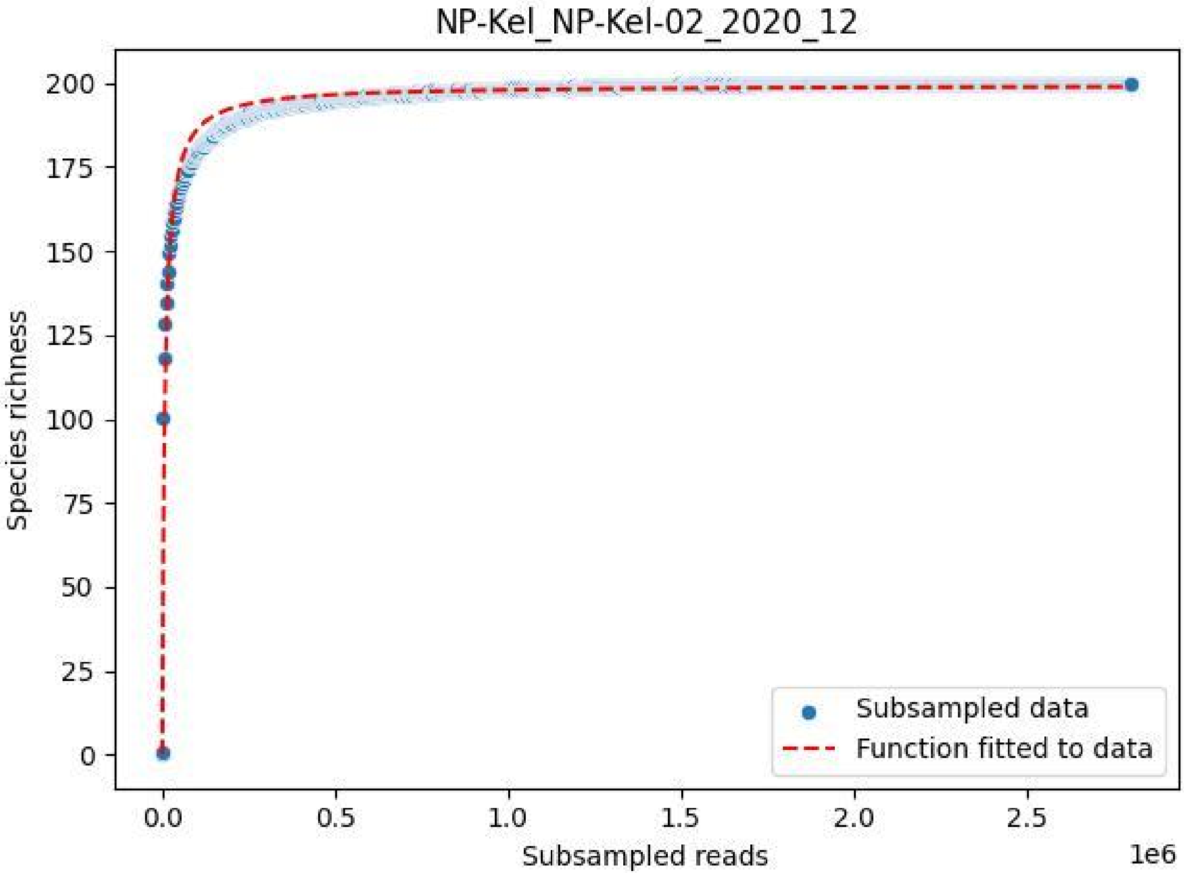
NP-Kel_NP-Kel-02_2020_12
Species richness
Subsampled reads
1e6
0
25
50
75
100
125
150
175
200
0.0
0.5
1.0
1.5
2.0
2.5
Subsampled data
Function fitted to data

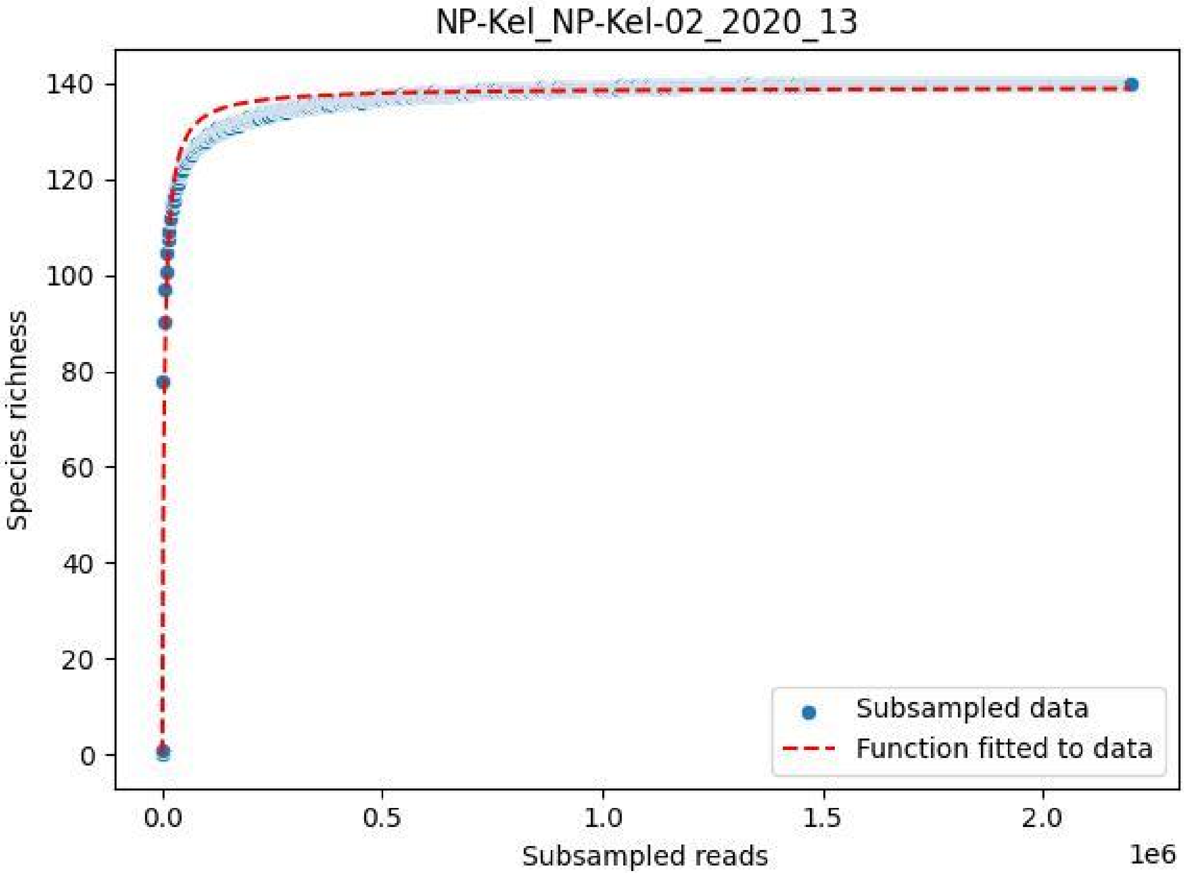

NP-Kel_NP-Kel-02_2020_13
Species richness
Subsampled reads
1e6
0
20
40
60
80
100
120
140
0.0
0.5
1.0
1.5
2.0
Subsampled data
Function fitted to data

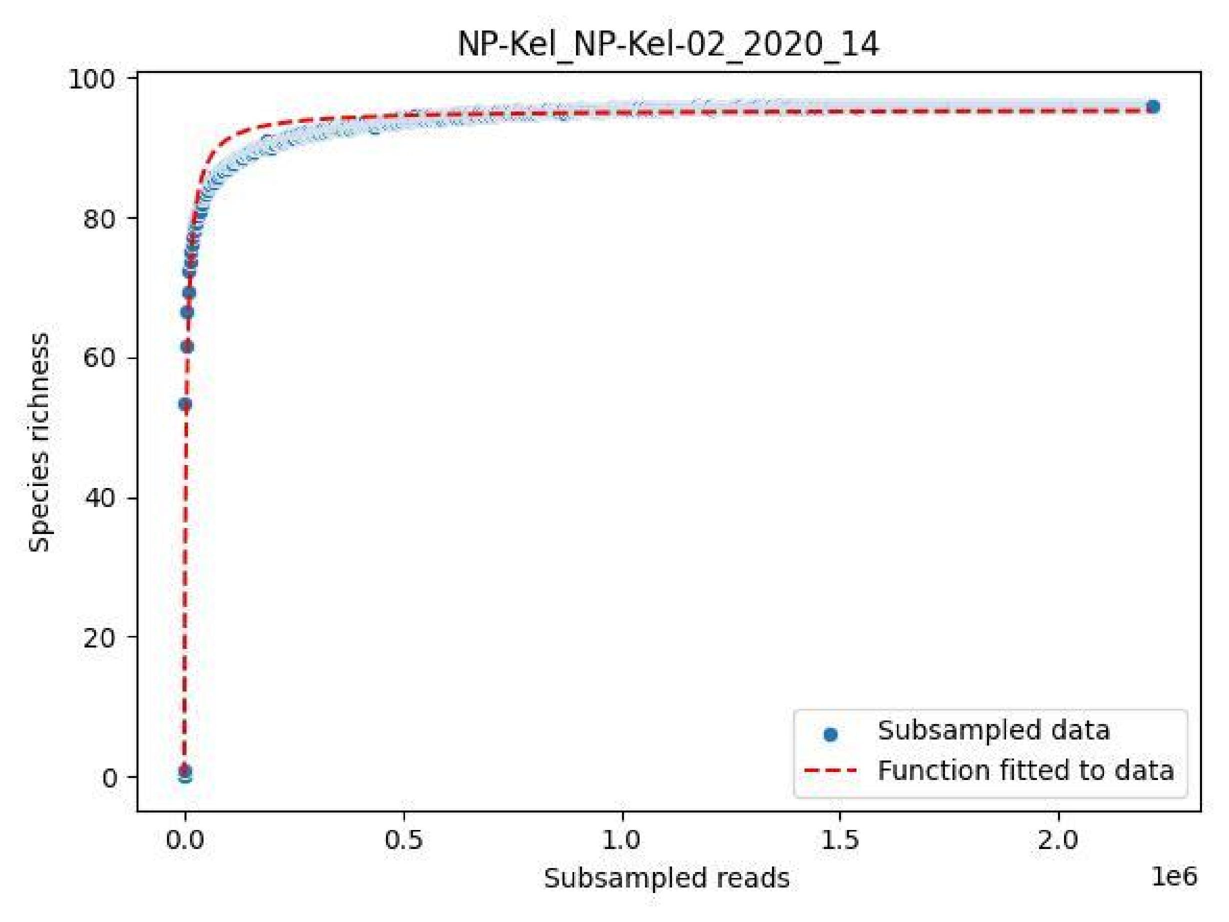

NP-Kel_NP-Kel-02_2020_14
Species richness
Subsampled reads
1e6
0
20
40
60
80
100
0.0
0.5
1.0
1.5
2.0
Subsampled data
Function fitted to data

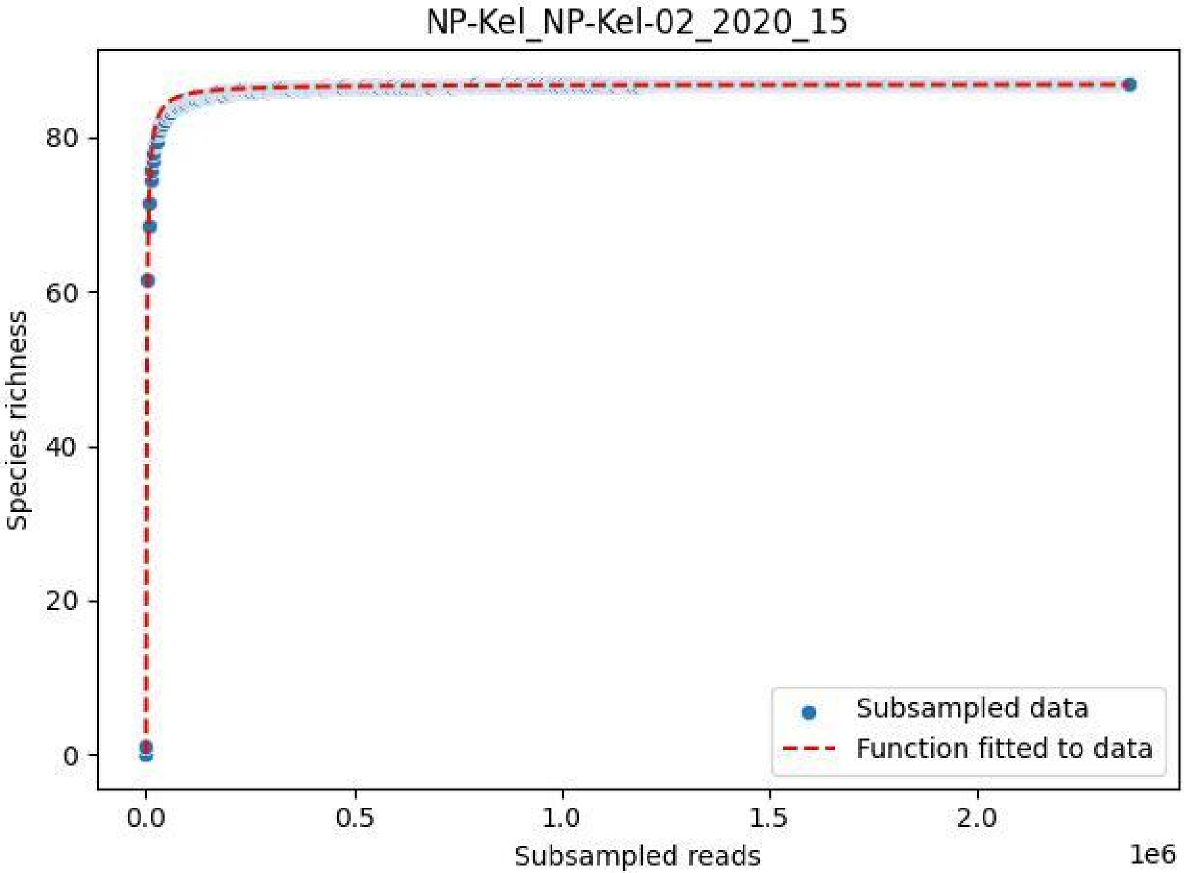
NP-Kel_NP-Kel-02_2020_15
Species richness
Subsampled reads
1e6
0
20
40
60
80
0.0
0.5
1.0
1.5
2.0
Subsampled data
Function fitted to data

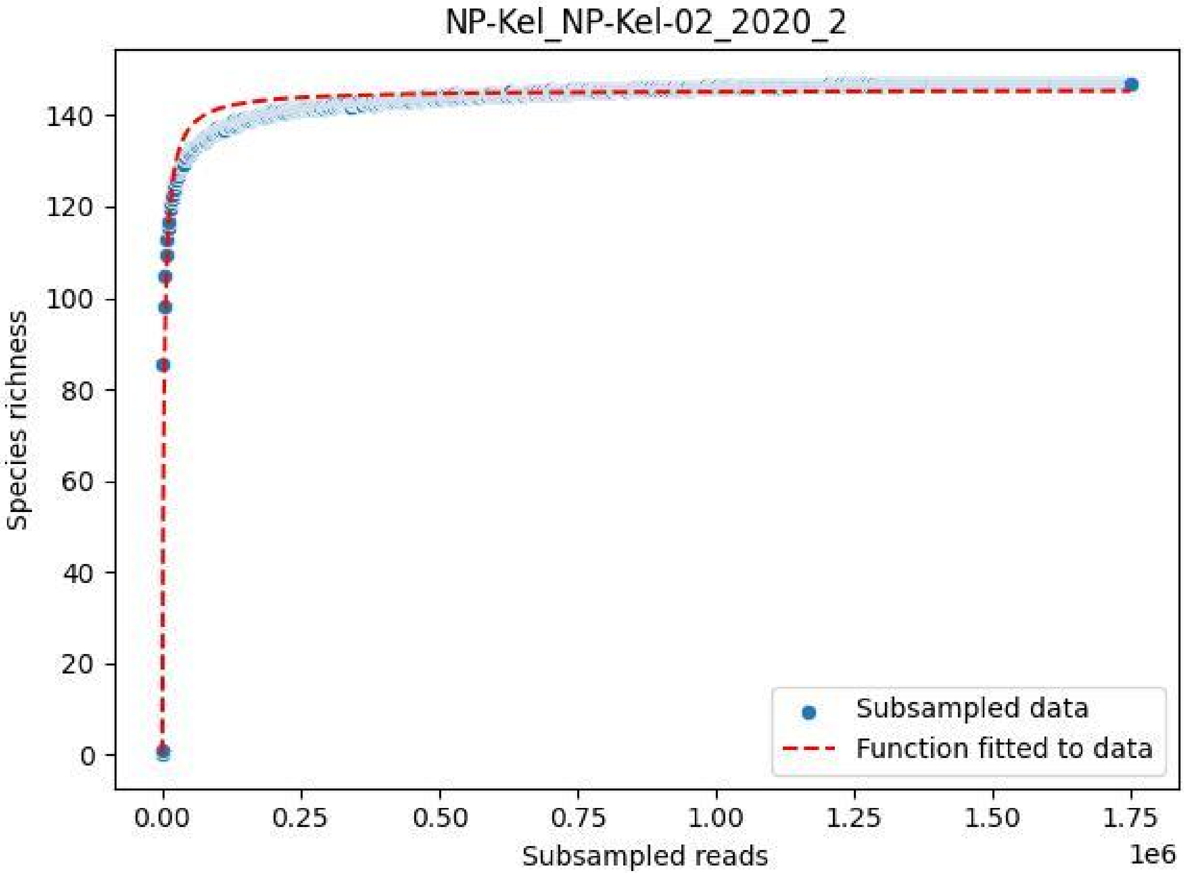
NP-Kel_NP-Kel-02_2020_2
Species richness
Subsampled reads
1e6
Subsampled data
Function fitted to data
0
20
40
60
80
100
120
140
0.00
0.25
0.50
0.75
1.00
1.25
1.50
1.75

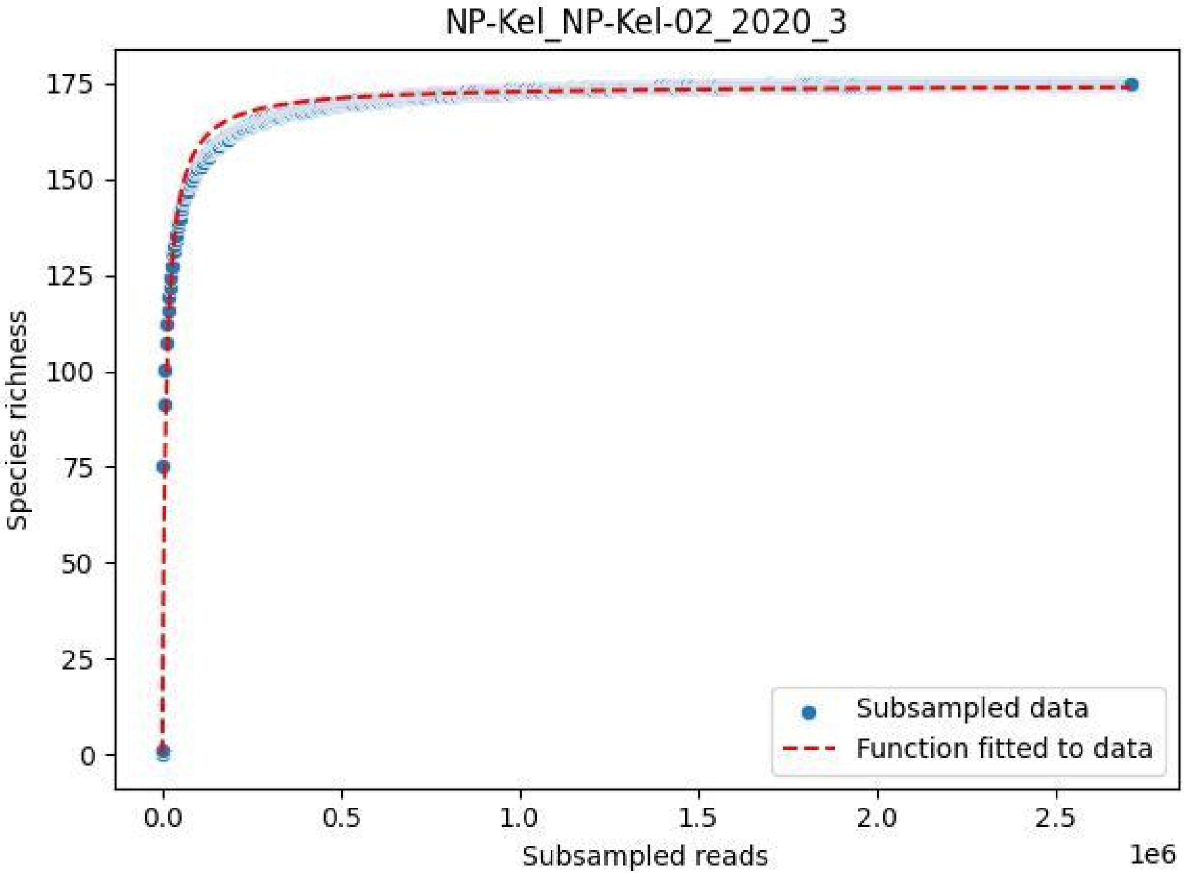

NP-Kel_NP-Kel-02_2020_3
175
150
125
100
75
50
25
0
Species richness
0.0
0.5
1.0
1.5
2.0
2.5
1e6
Subsampled reads
Subsampled data
Function fitted to data

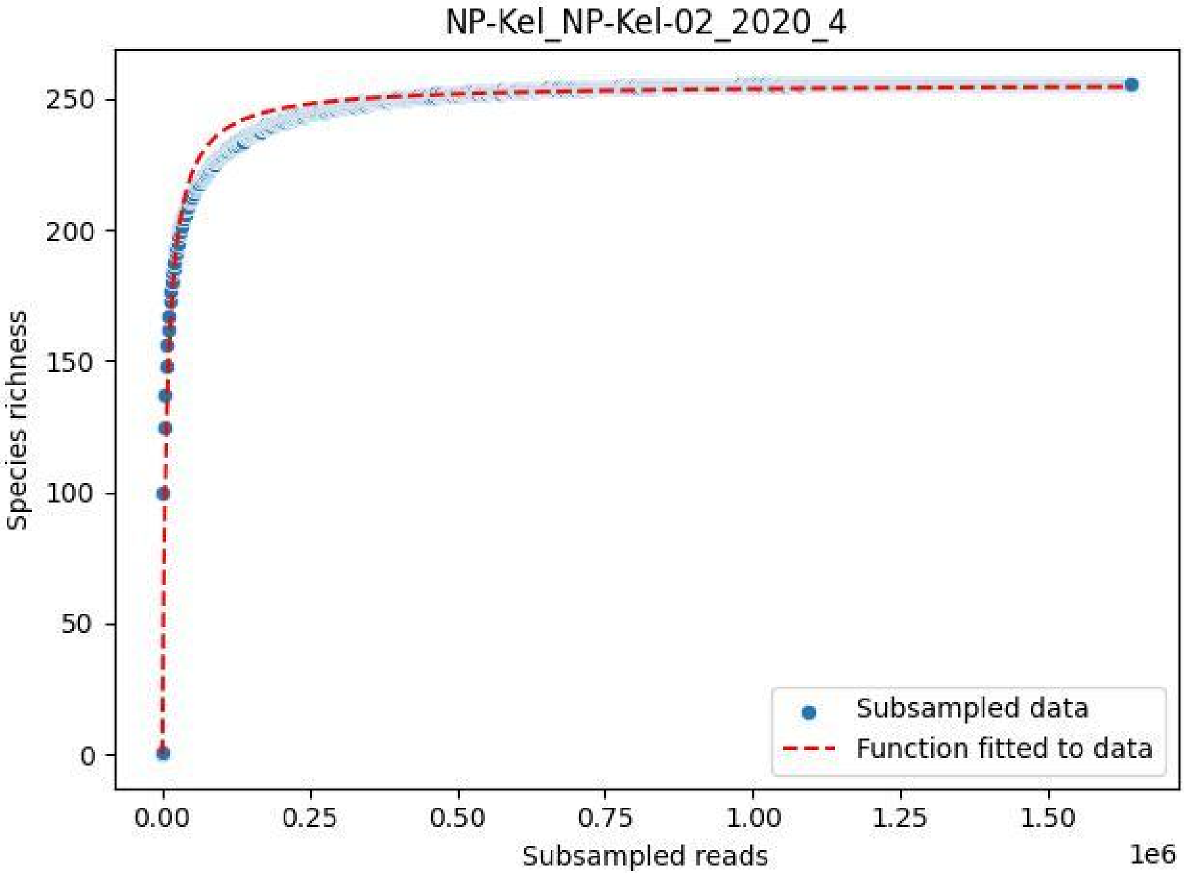

NP-Kel_NP-Kel-02_2020_4
Species richness
Subsampled reads
1e6
0
50
100
150
200
250
0.00
0.25
0.50
0.75
1.00
1.25
1.50
Subsampled data
Function fitted to data

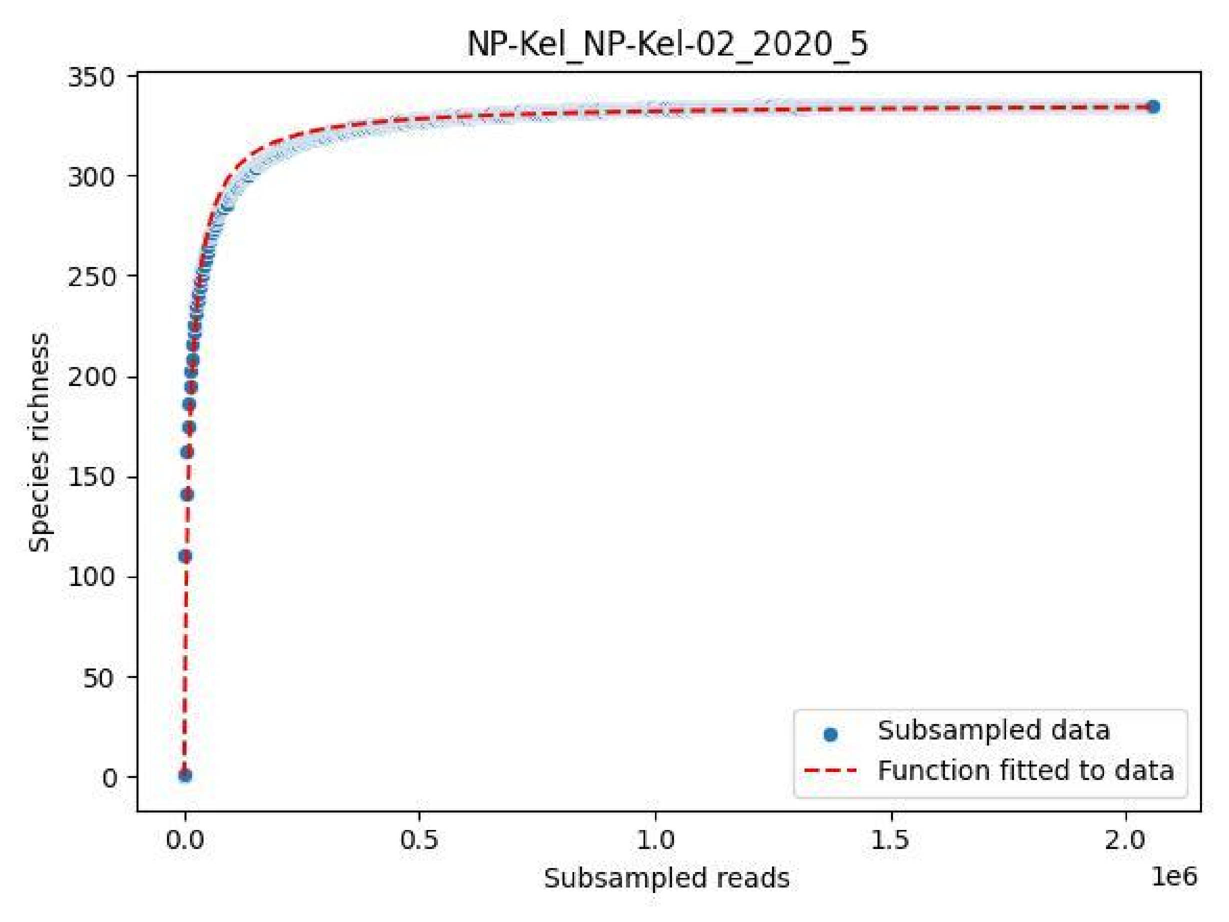
NP-Kel_NP-Kel-02_2020_5
Species richness
Subsampled reads
1e6
0
50
100
150
200
250
300
350
0.0
0.5
1.0
1.5
2.0
Subsampled data
Function fitted to data

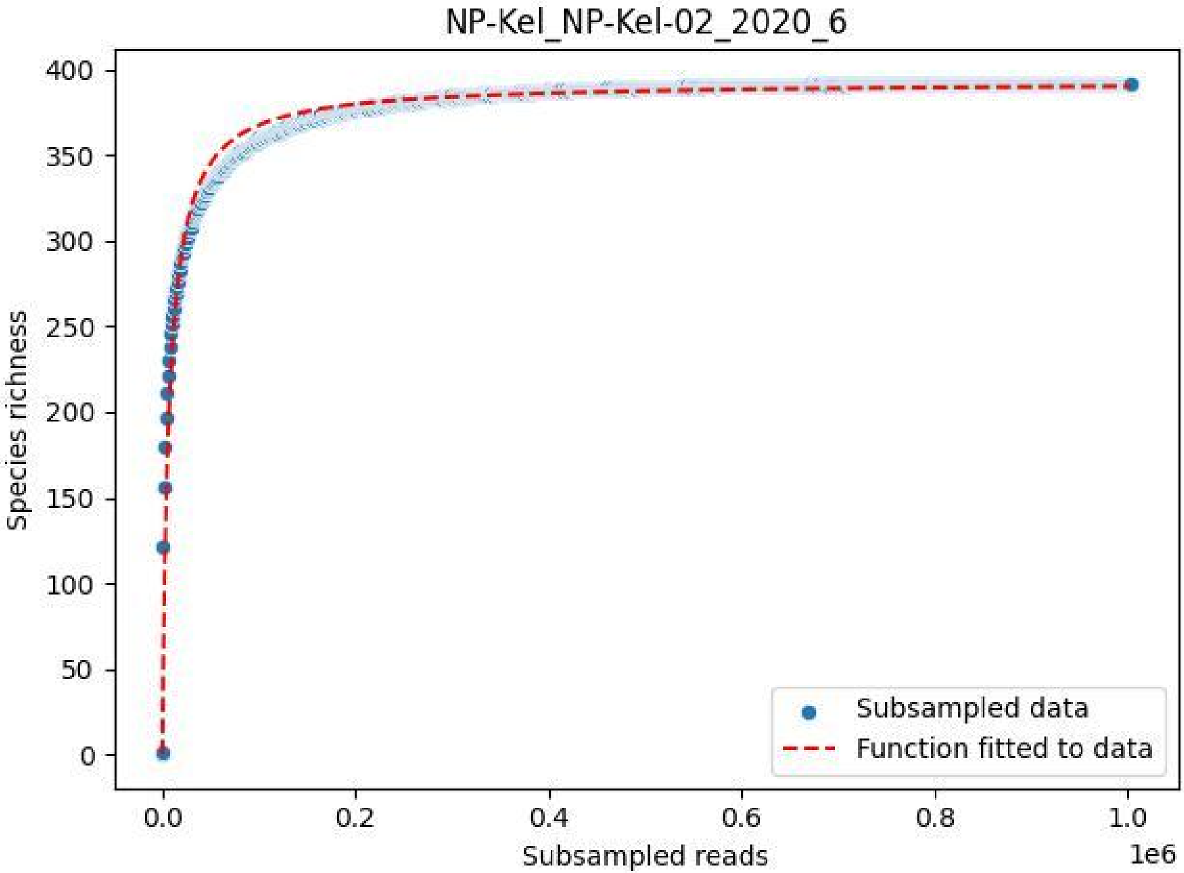

NP-Kel_NP-Kel-02_2020_6
Species richness
Subsampled reads
1e6
0
50
100
150
200
250
300
350
400
0.0
0.2
0.4
0.6
0.8
1.0
Subsampled data
Function fitted to data

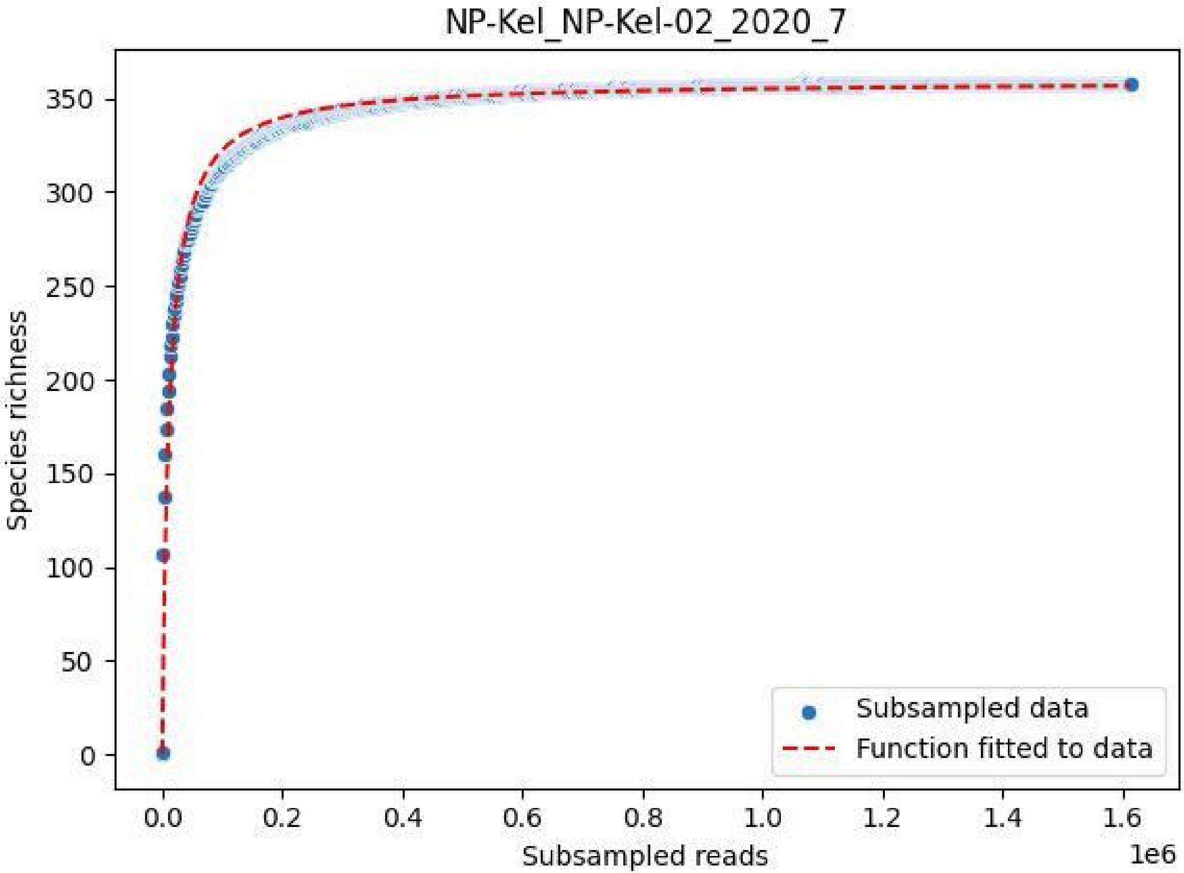
NP-Kel_NP-Kel-02_2020_7
Species richness
Subsampled reads
1e6
0
50
100
150
200
250
300
350
0.0
0.2
0.4
0.6
0.8
1.0
1.2
1.4
1.6
Subsampled data
Function fitted to data

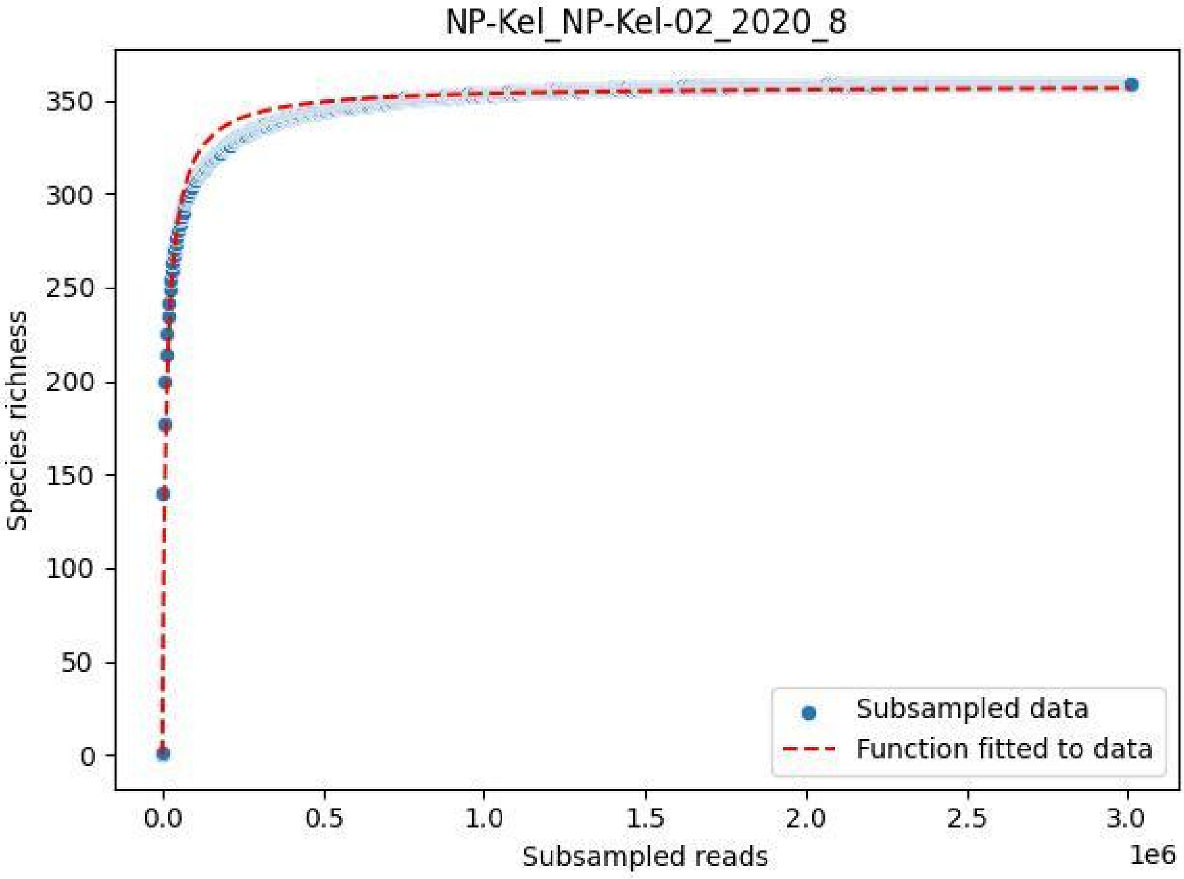

NP-Kel_NP-Kel-02_2020_8
Species richness
Subsampled reads
1e6
0
50
100
150
200
250
300
350
0.0
0.5
1.0
1.5
2.0
2.5
3.0
Subsampled data
Function fitted to data

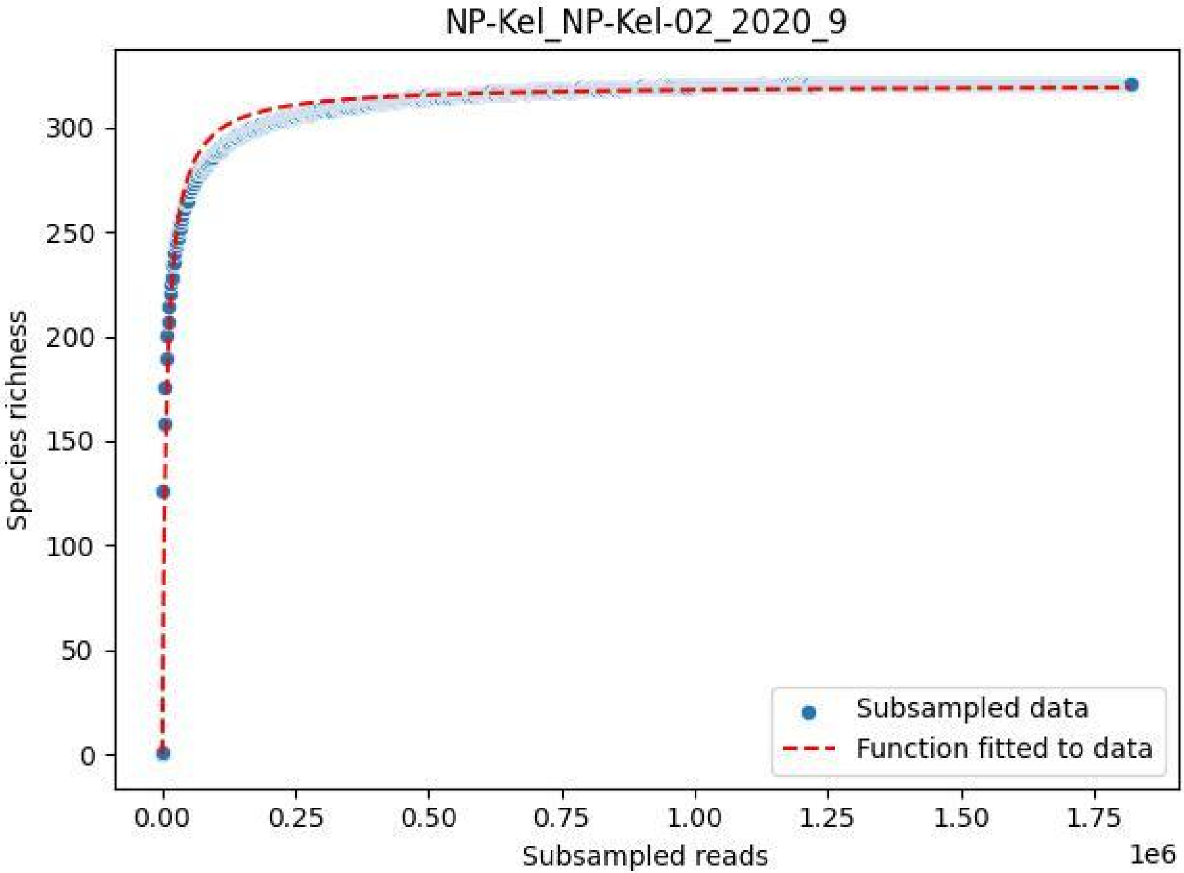

NP-Kel_NP-Kel-02_2020_9
Species richness
Subsampled reads
1e6
0
50
100
150
200
250
300
0.00
0.25
0.50
0.75
1.00
1.25
1.50
1.75
Subsampled data
Function fitted to data

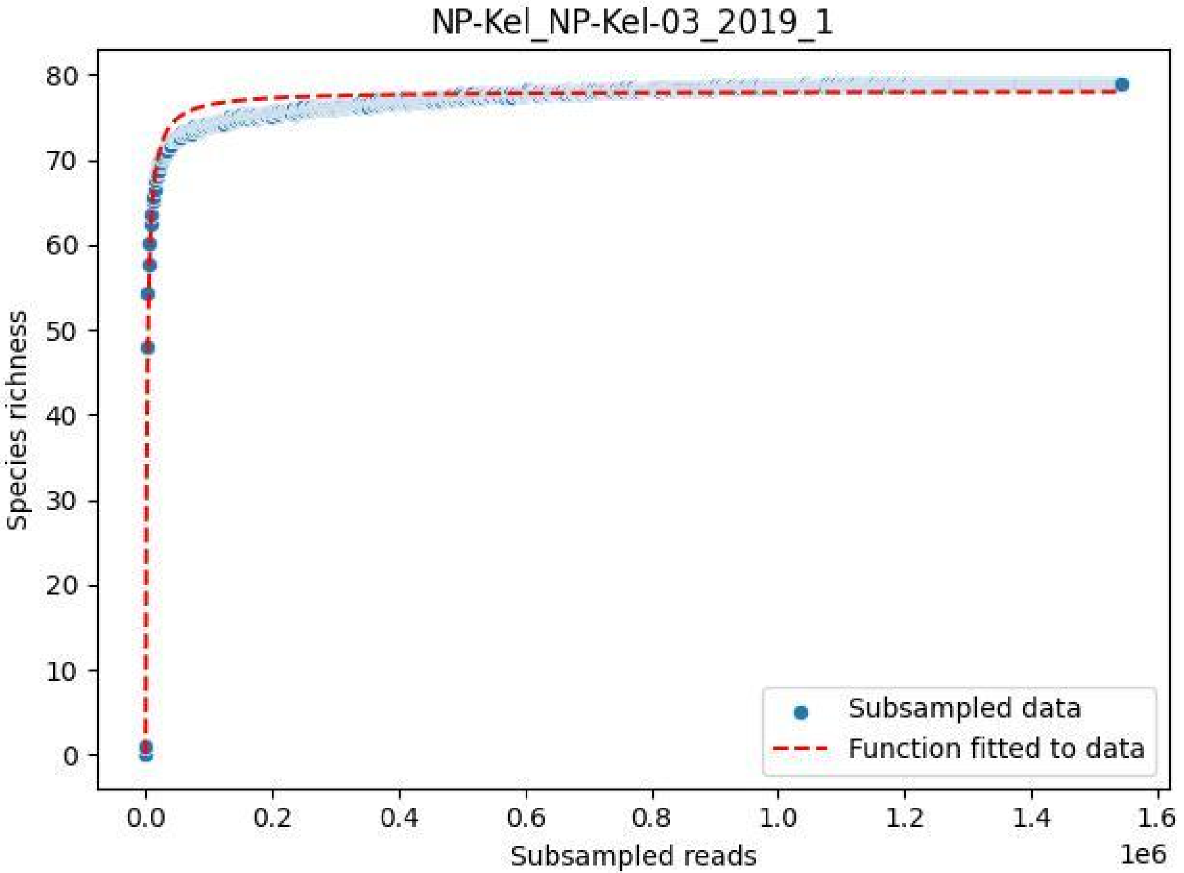

NP-Kel_NP-Kel-03_2019_1
Species richness
Subsampled reads
1e6
0
10
20
30
40
50
60
70
80
0.0
0.2
0.4
0.6
0.8
1.0
1.2
1.4
1.6
Subsampled data
Function fitted to data

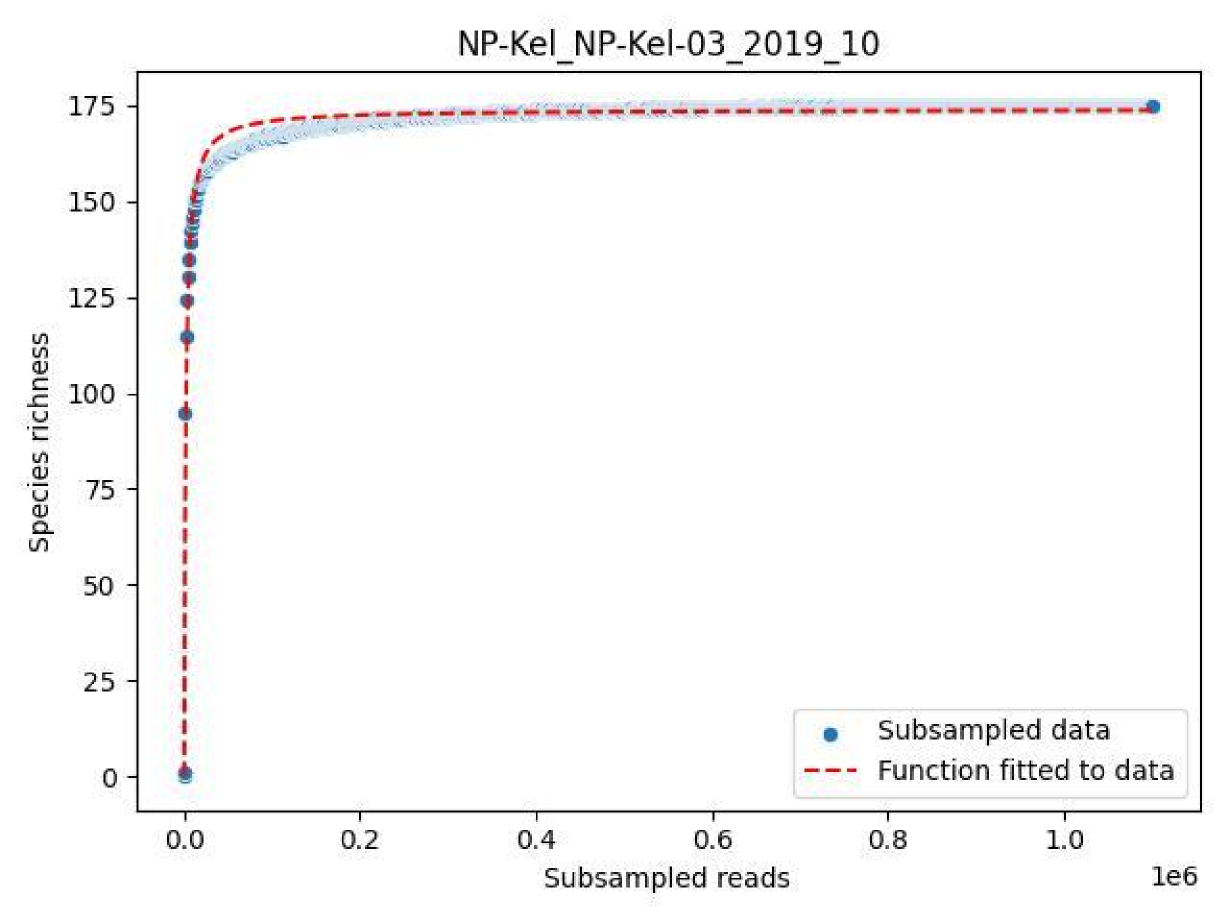
NP-Kel_NP-Kel-03_2019_10
Species richness
Subsampled reads
1e6
0
25
50
75
100
125
150
175
0.0
0.2
0.4
0.6
0.8
1.0
Subsampled data
Function fitted to data

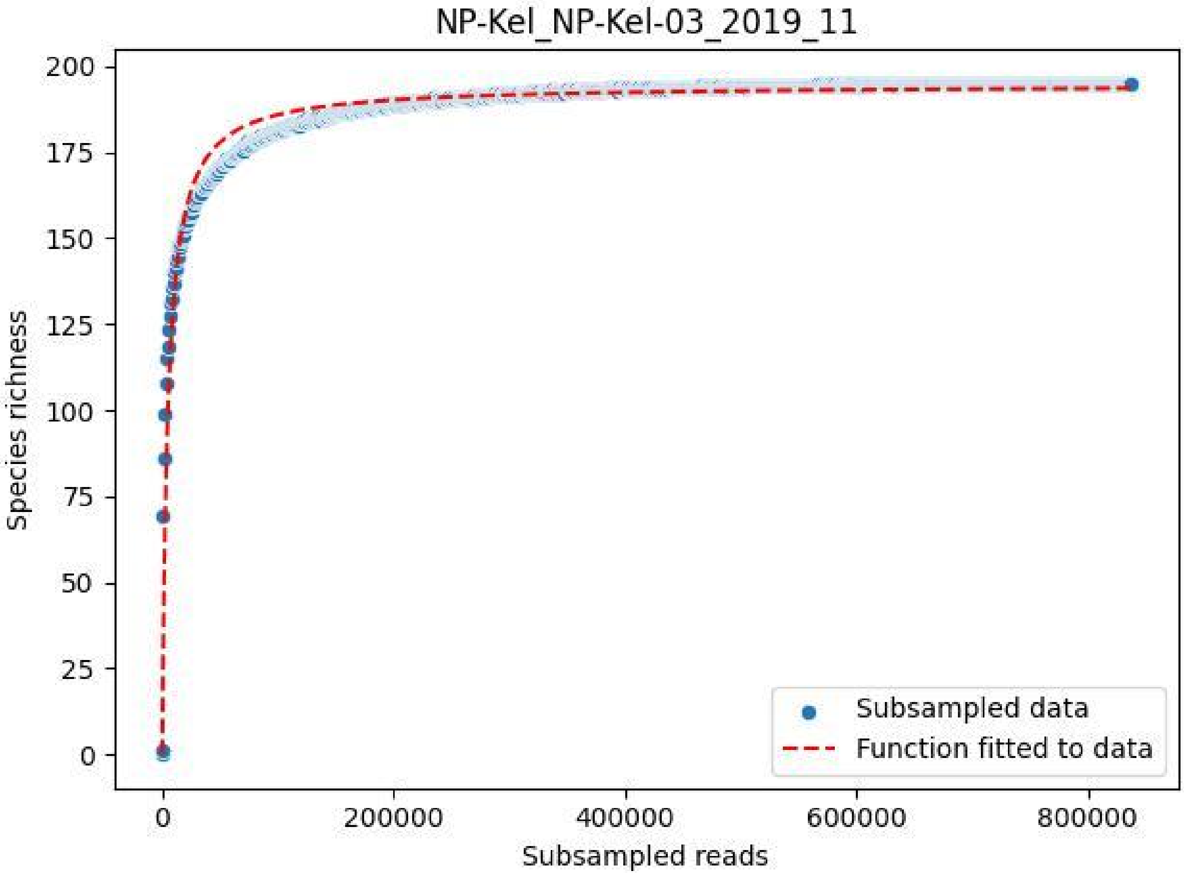

NP-Kel_NP-Kel-03_2019_11
Species richness
Subsampled reads
0
25
50
75
100
125
150
175
200
0
200000
400000
600000
800000
Subsampled data
Function fitted to data

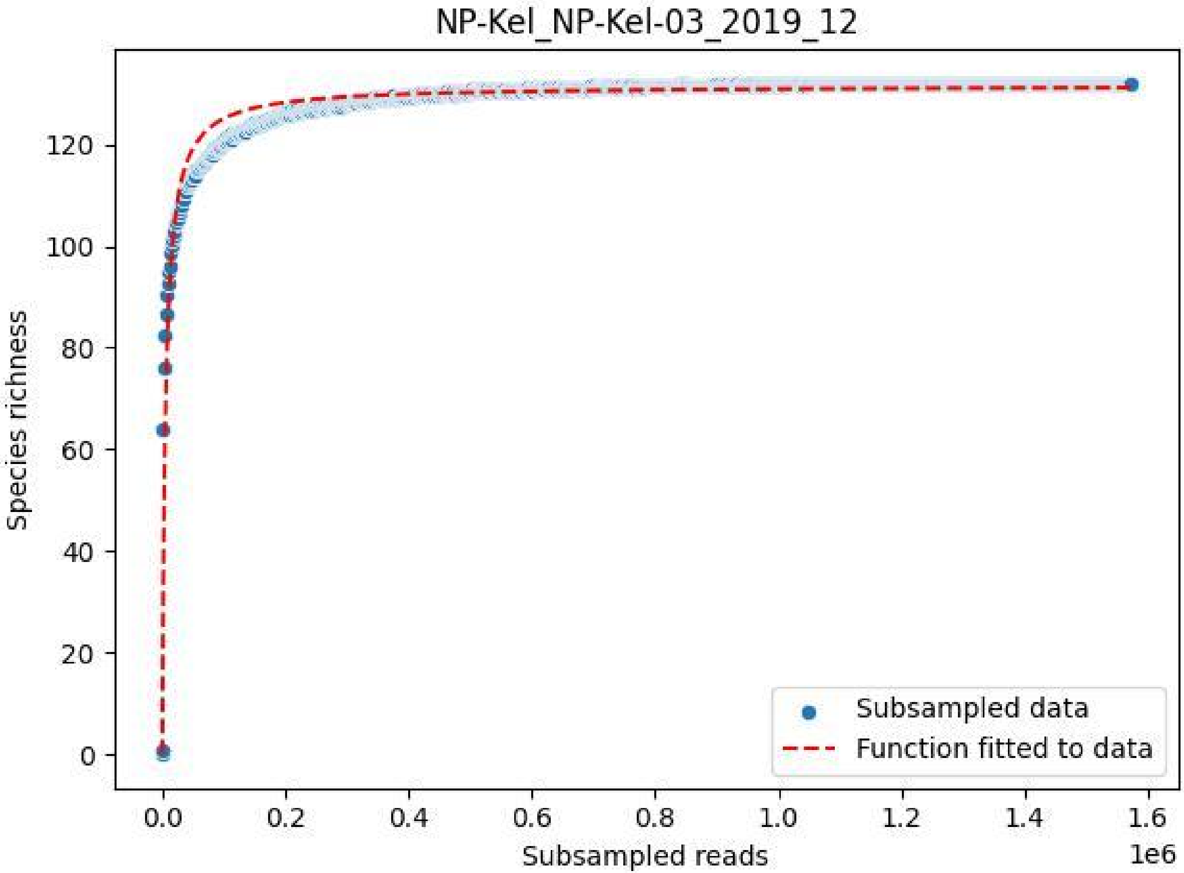
NP-Kel_NP-Kel-03_2019_12
Species richness
Subsampled reads
1e6
0
20
40
60
80
100
120
0.0
0.2
0.4
0.6
0.8
1.0
1.2
1.4
1.6
Subsampled data
Function fitted to data

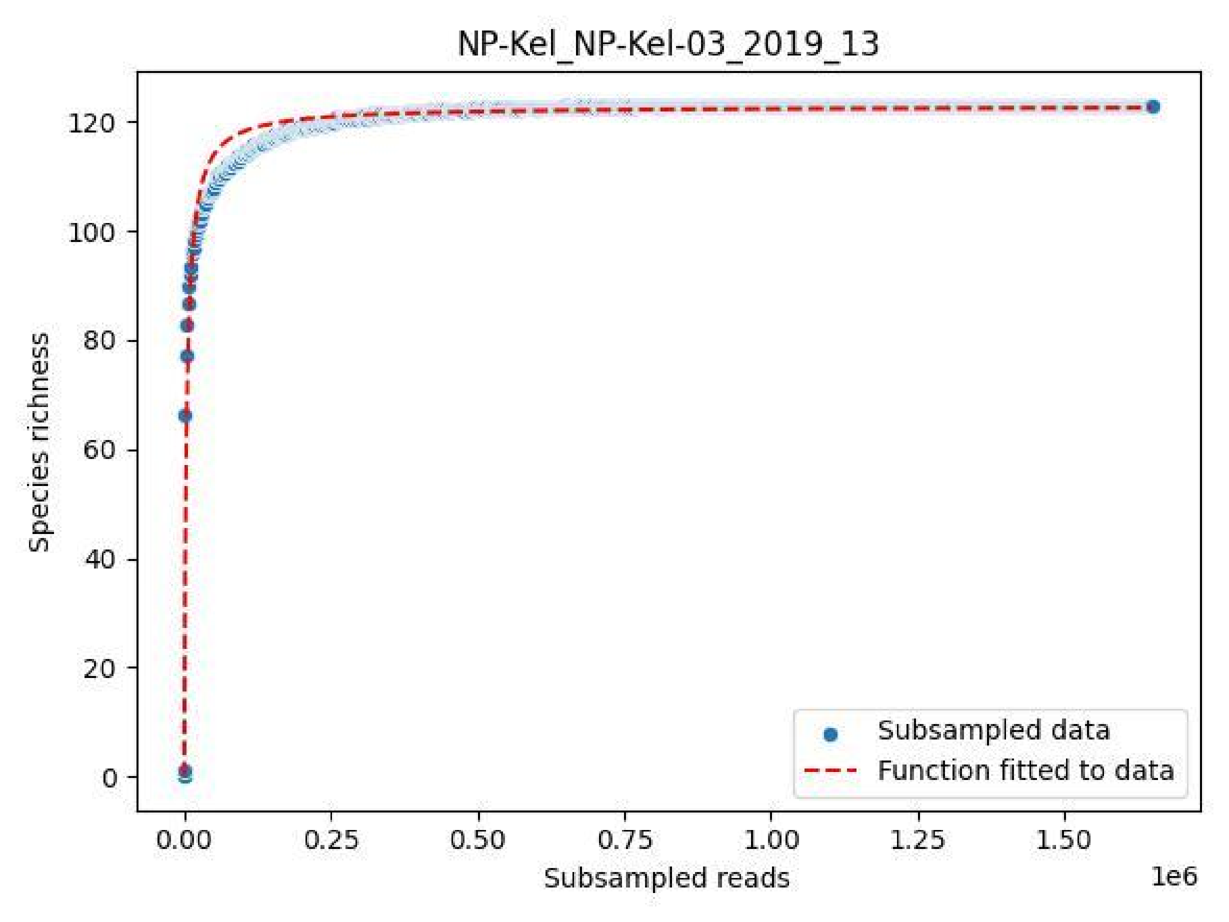
NP-Kel_NP-Kel-03_2019_13
Species richness
0
20
40
60
80
100
120
0.00
0.25
0.50
0.75
1.00
1.25
1.50
Subsampled reads
1e6
Subsampled data
Function fitted to data

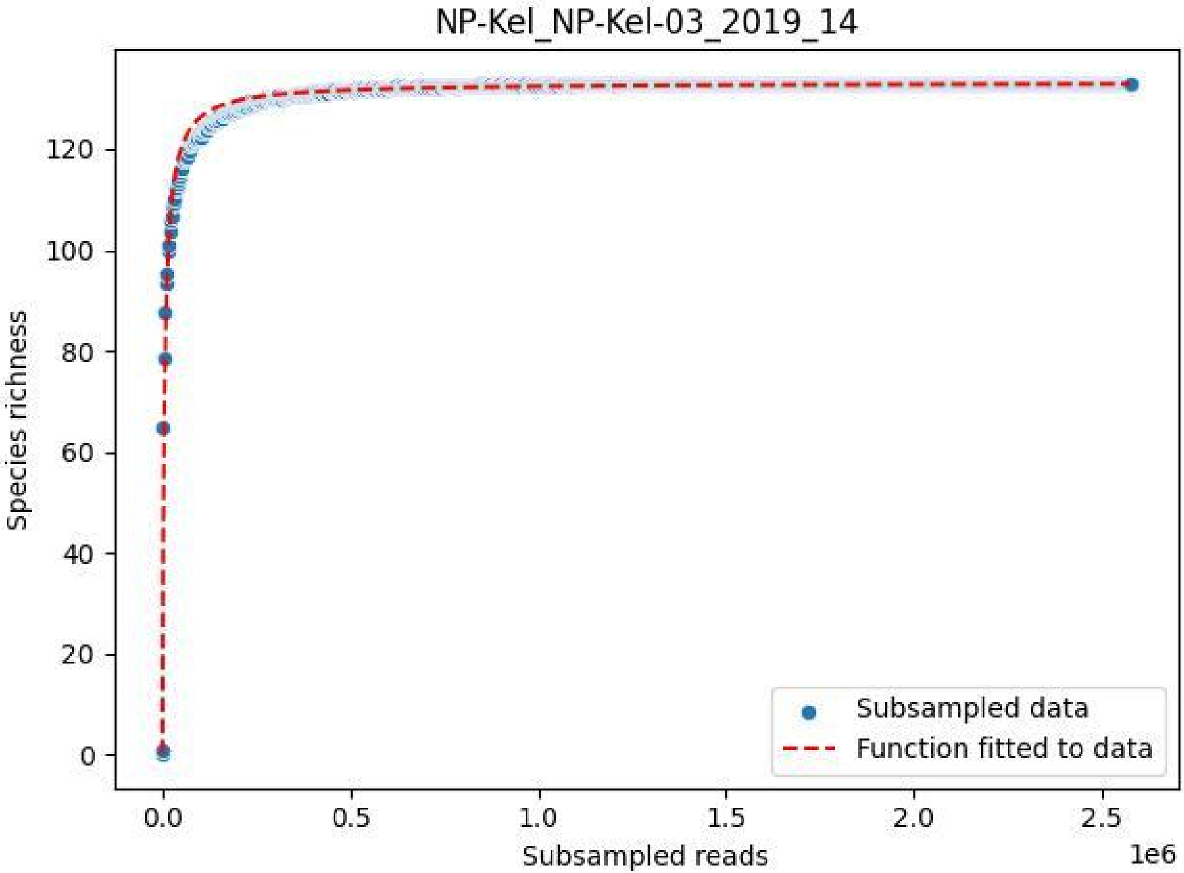

NP-Kel_NP-Kel-03_2019_14
Species richness
Subsampled reads
1e6
0
20
40
60
80
100
120
0.0
0.5
1.0
1.5
2.0
2.5
Subsampled data
Function fitted to data

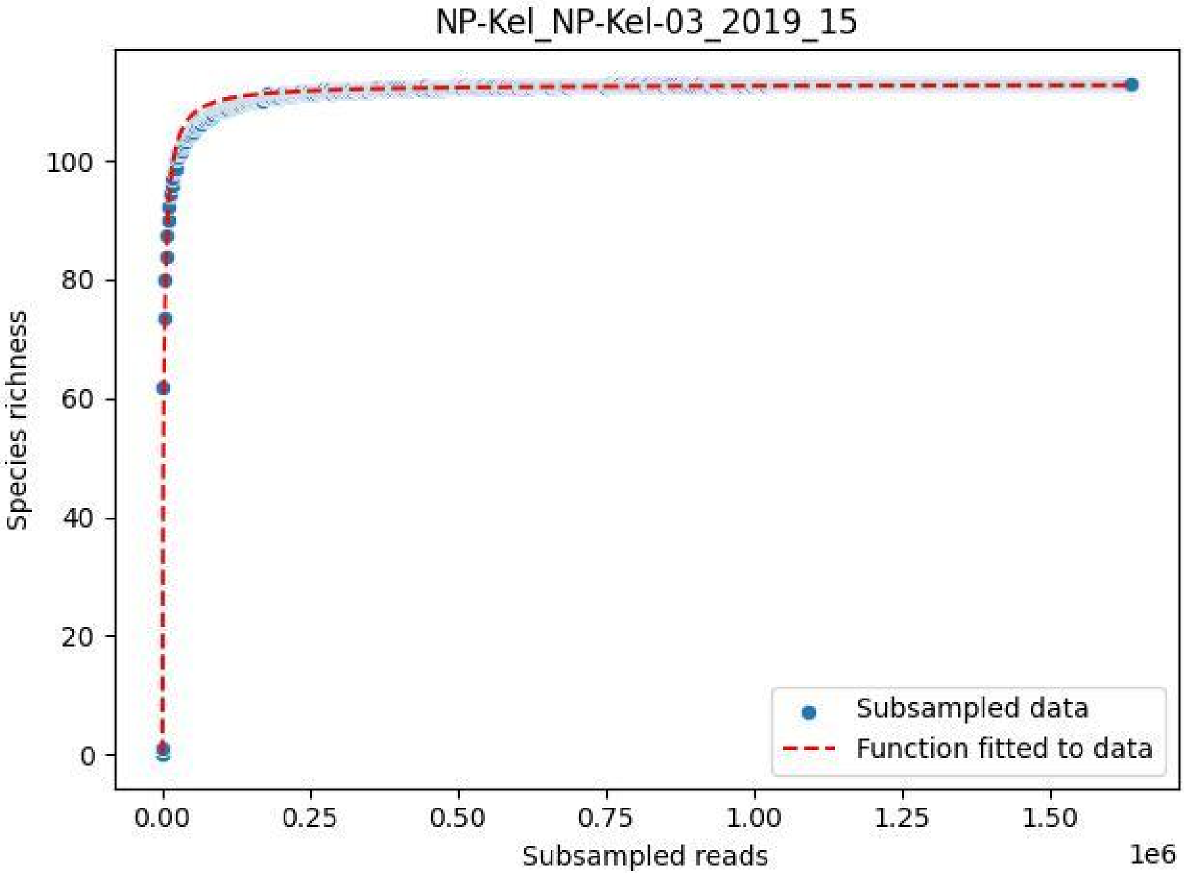
NP-Kel_NP-Kel-03_2019_15
Species richness
Subsampled reads
1e6
0
20
40
60
80
100
0.00
0.25
0.50
0.75
1.00
1.25
1.50
Subsampled data
Function fitted to data

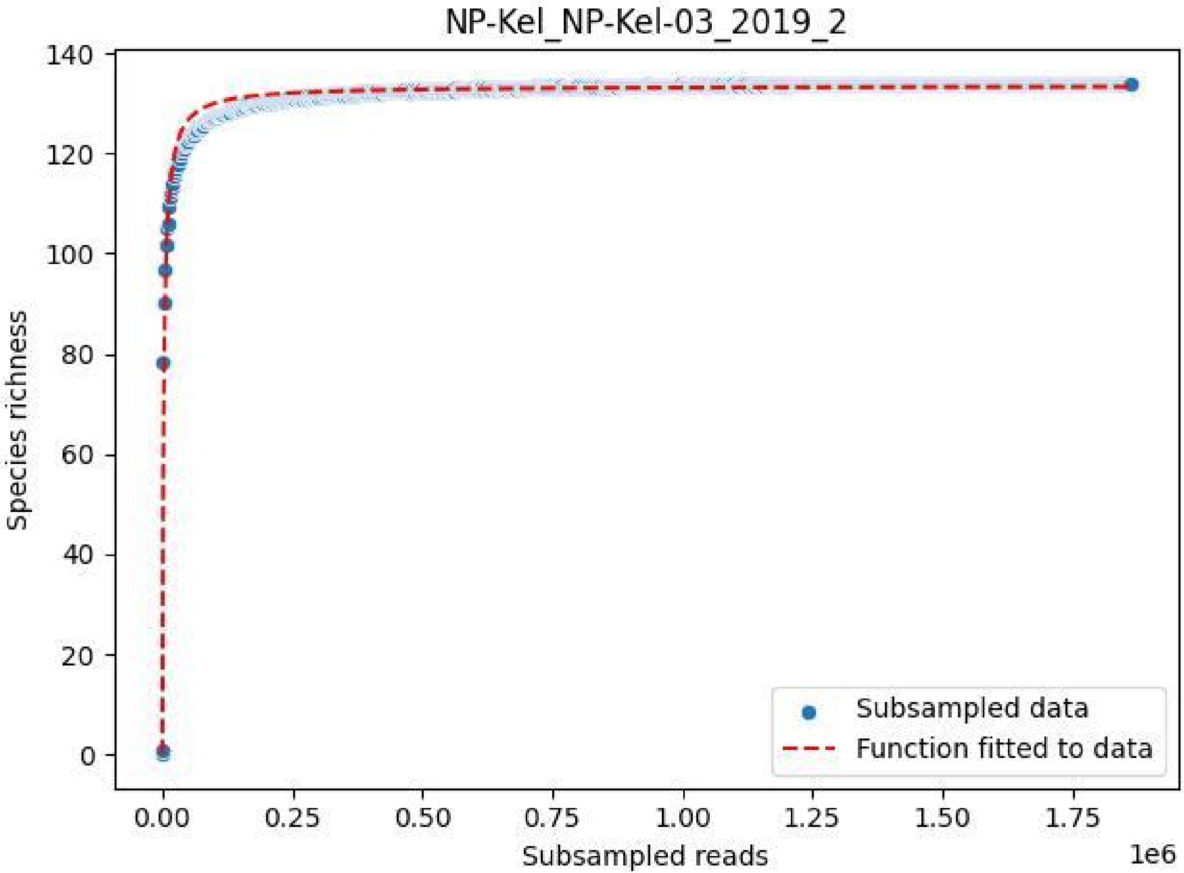
NP-Kel_NP-Kel-03_2019_2
Species richness
Subsampled reads
1e6
0
20
40
60
80
100
120
140
0.00
0.25
0.50
0.75
1.00
1.25
1.50
1.75
Subsampled data
Function fitted to data

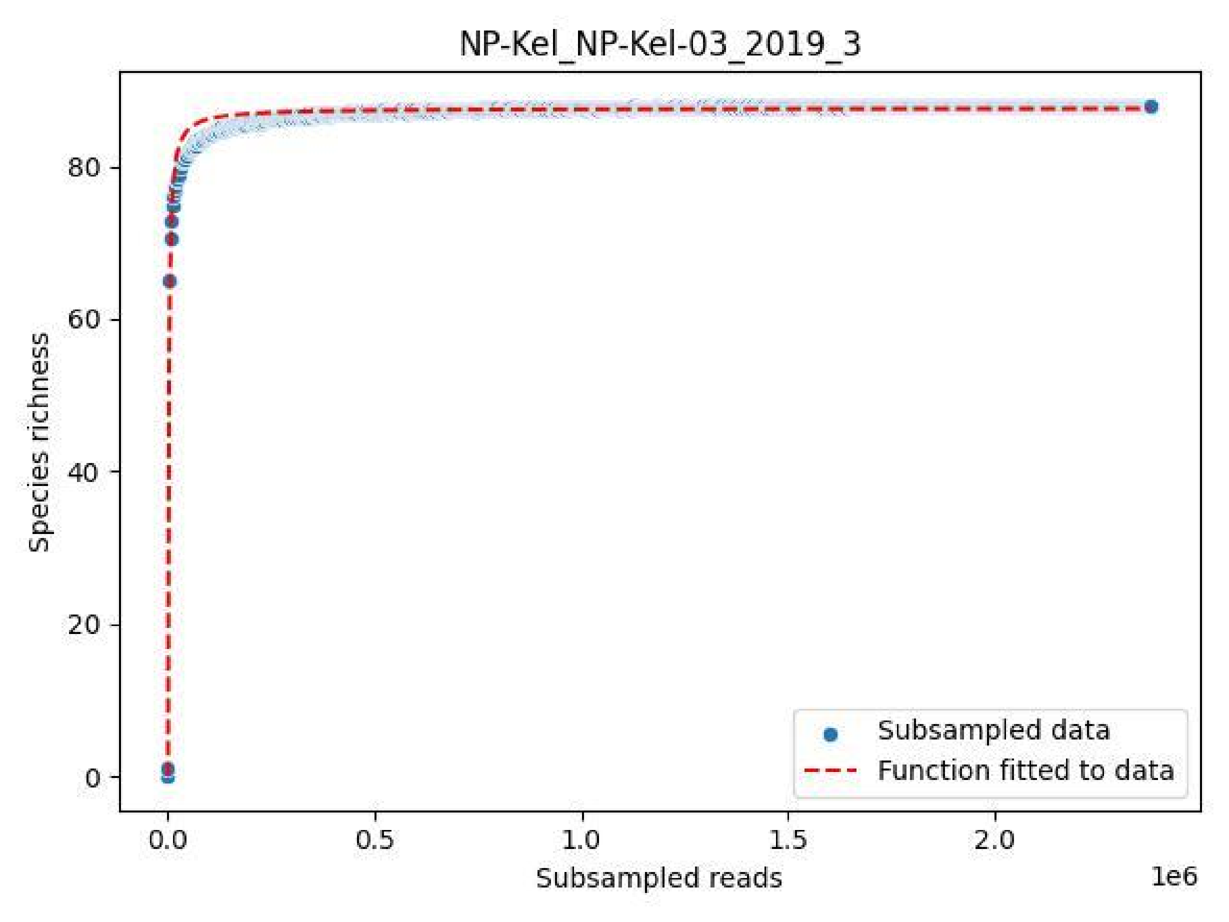
NP-Kel_NP-Kel-03_2019_3
Species richness
Subsampled reads
1e6
0
20
40
60
80
0.0
0.5
1.0
1.5
2.0
Subsampled data
Function fitted to data

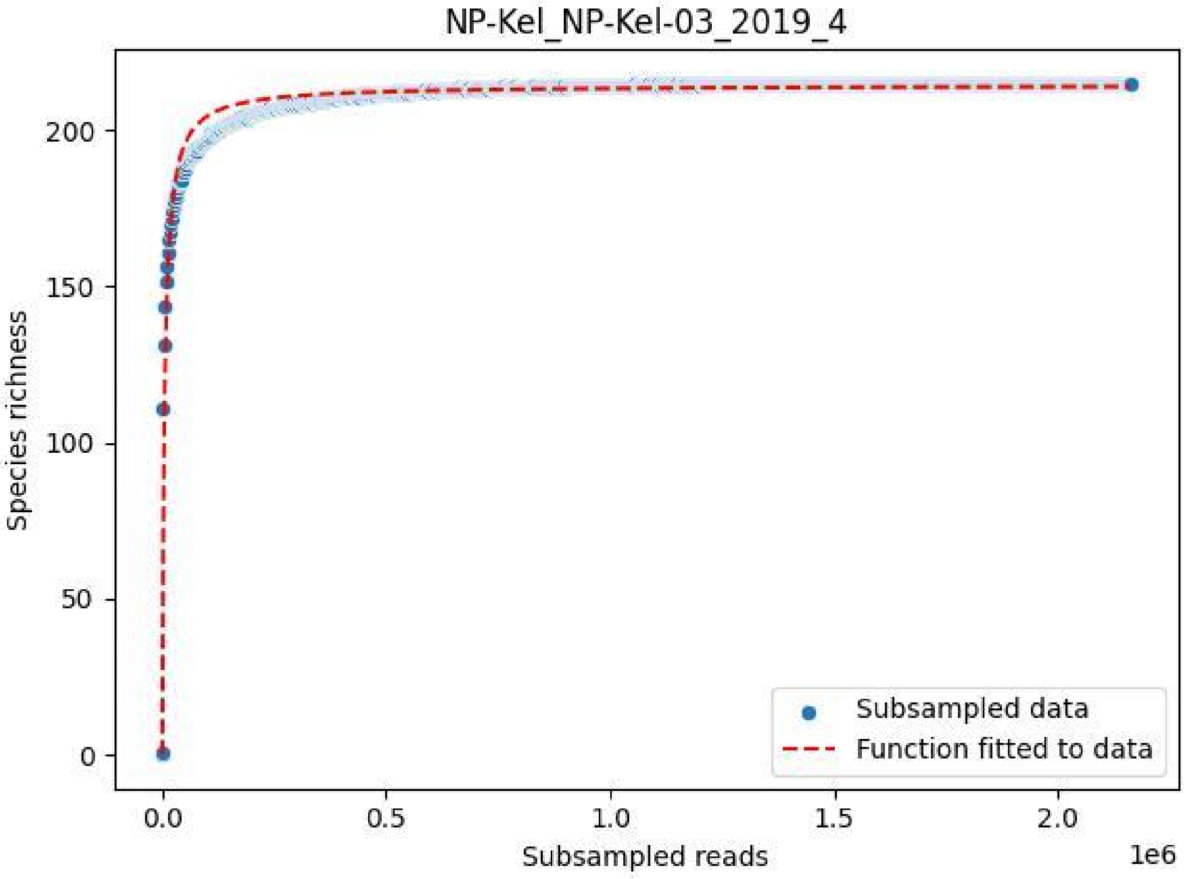

NP-Kel_NP-Kel-03_2019_4
Species richness
Subsampled reads
1e6
0
50
100
150
200
0.0
0.5
1.0
1.5
2.0
Subsampled data
Function fitted to data

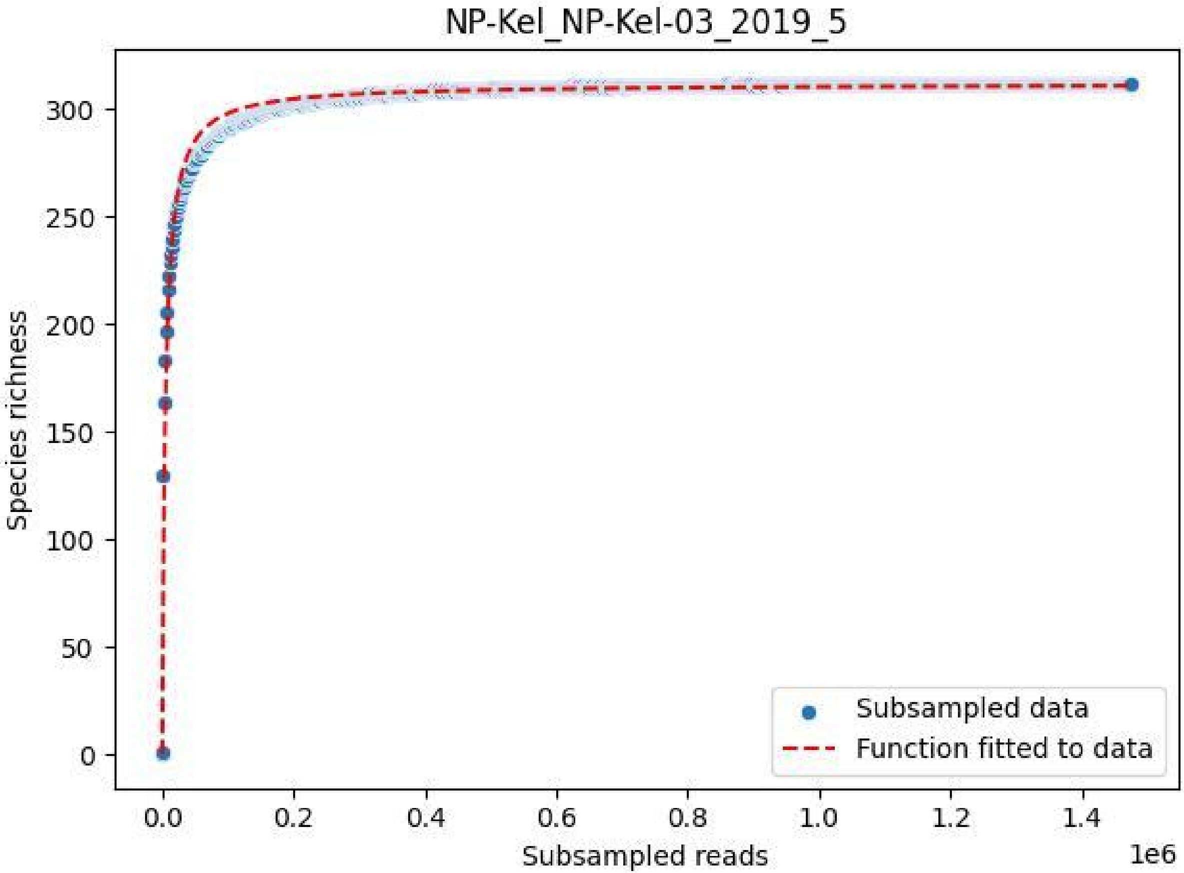

NP-Kel_NP-Kel-03_2019_5
Species richness
Subsampled reads
1e6
0
50
100
150
200
250
300
0.0
0.2
0.4
0.6
0.8
1.0
1.2
1.4
Subsampled data
Function fitted to data

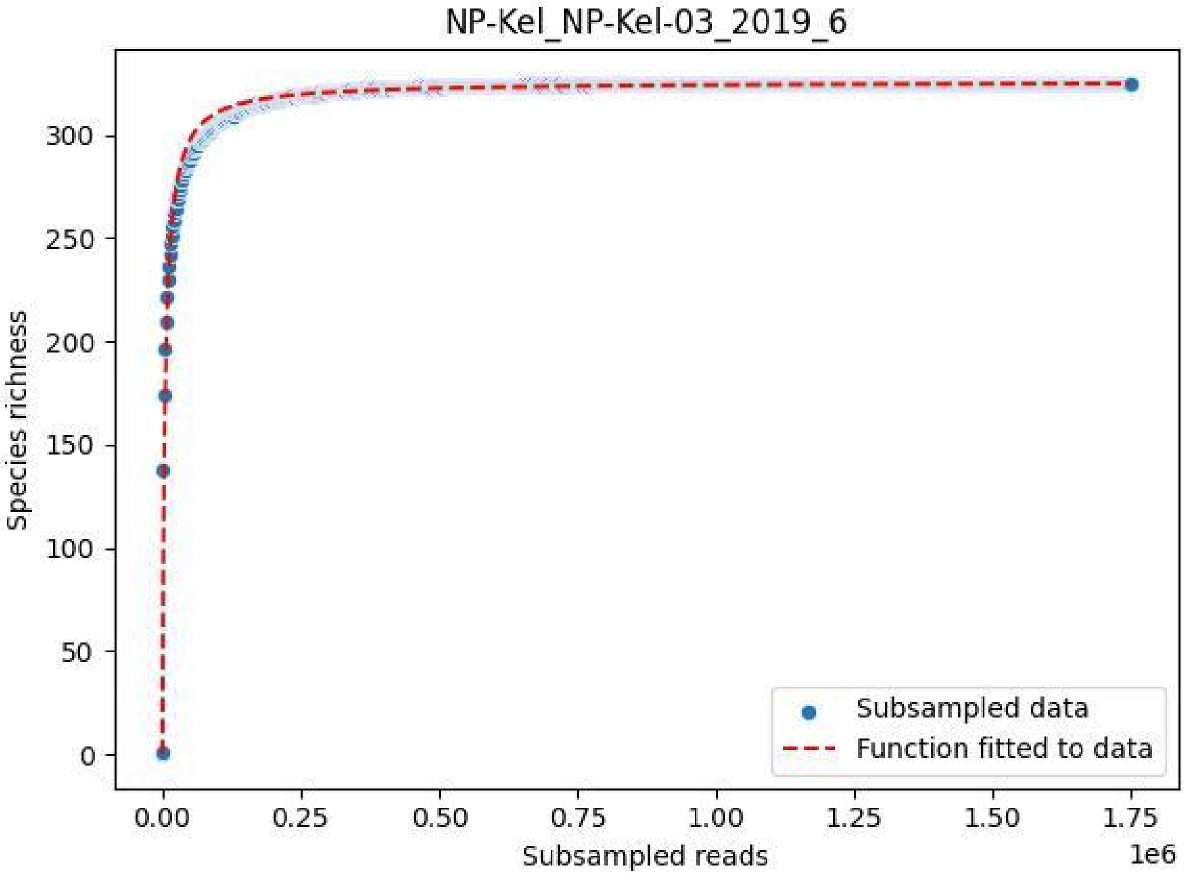
NP-Kel_NP-Kel-03_2019_6
Species richness
Subsampled reads
1e6
0
50
100
150
200
250
300
0.00
0.25
0.50
0.75
1.00
1.25
1.50
1.75
Subsampled data
Function fitted to data

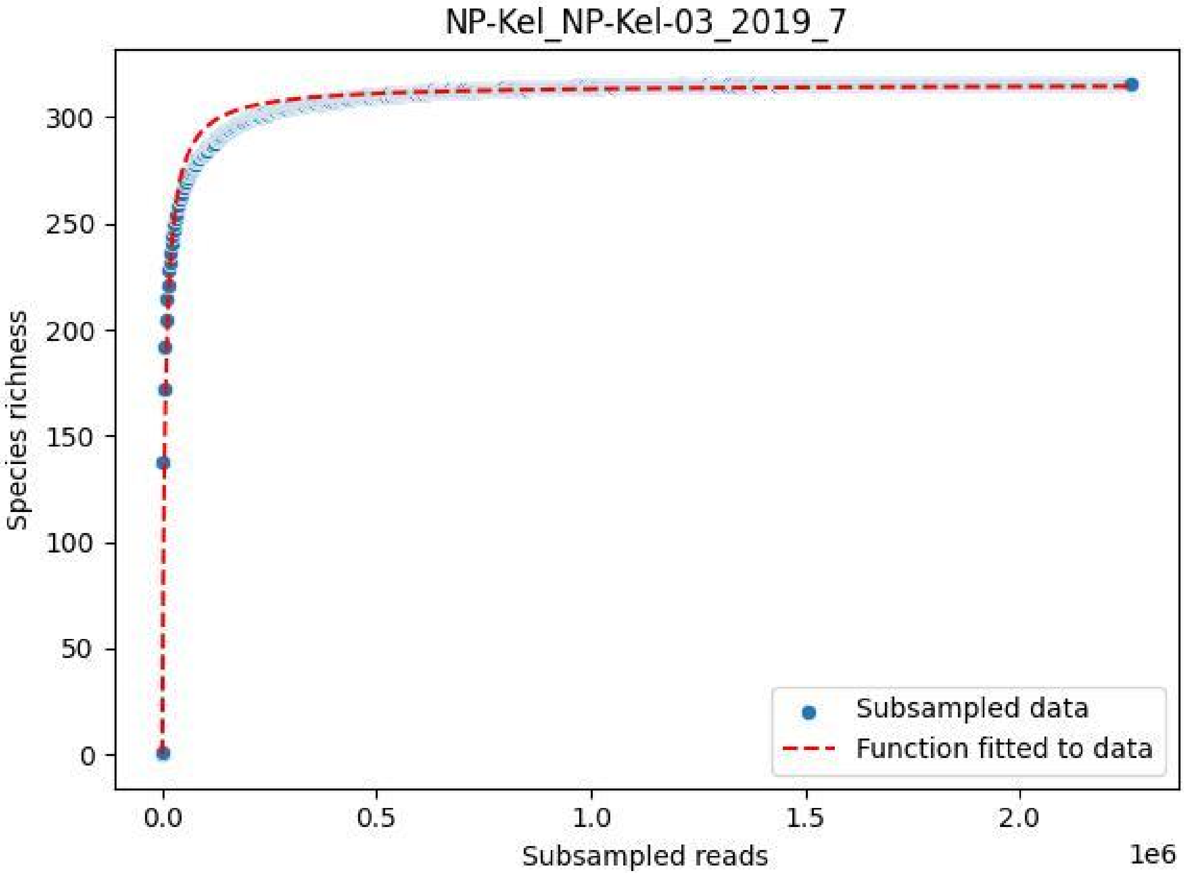

NP-Kel_NP-Kel-03_2019_7
Species richness
Subsampled reads
1e6
0
50
100
150
200
250
300
0.0
0.5
1.0
1.5
2.0
Subsampled data
Function fitted to data

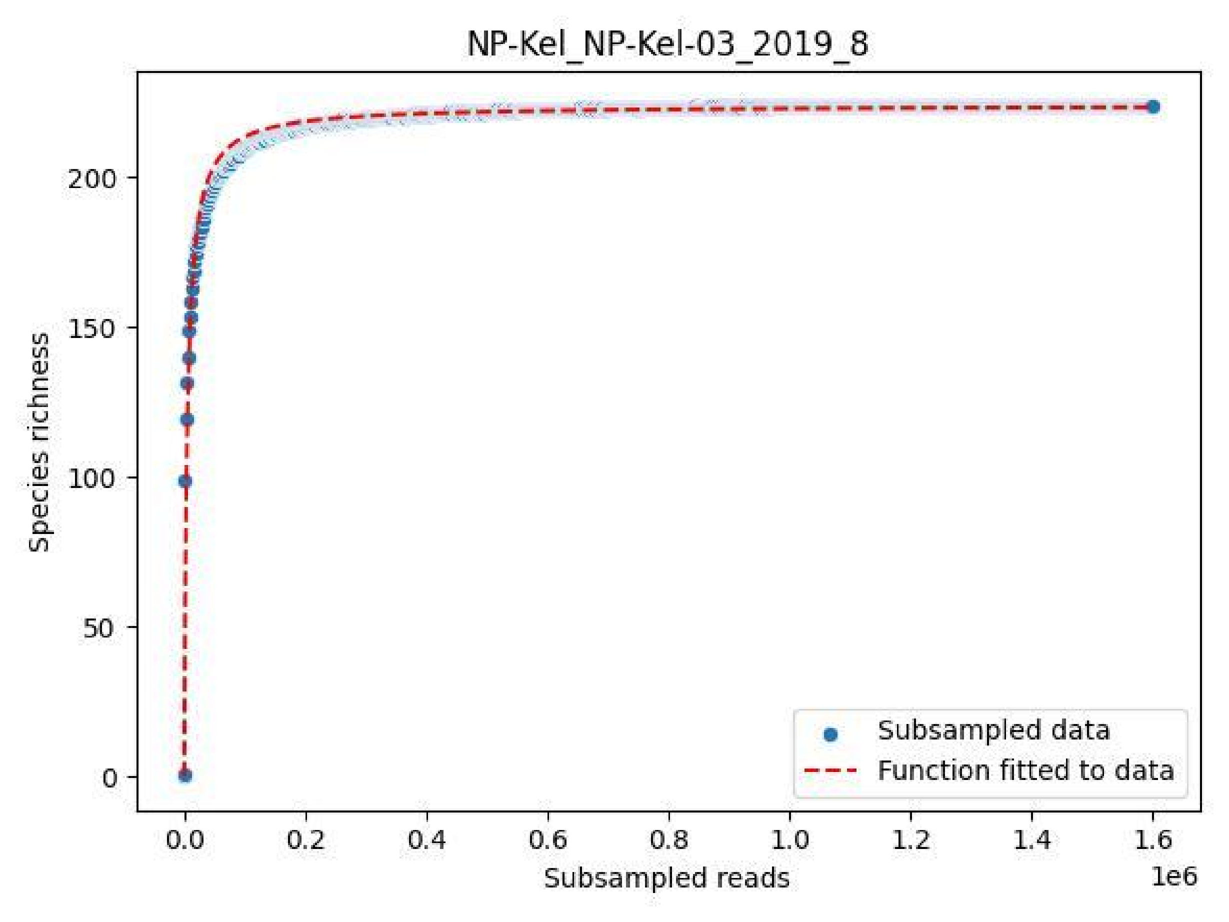

NP-Kel_NP-Kel-03_2019_8
Species richness
Subsampled reads
1e6
0
50
100
150
200
0.0
0.2
0.4
0.6
0.8
1.0
1.2
1.4
1.6
Subsampled data
Function fitted to data

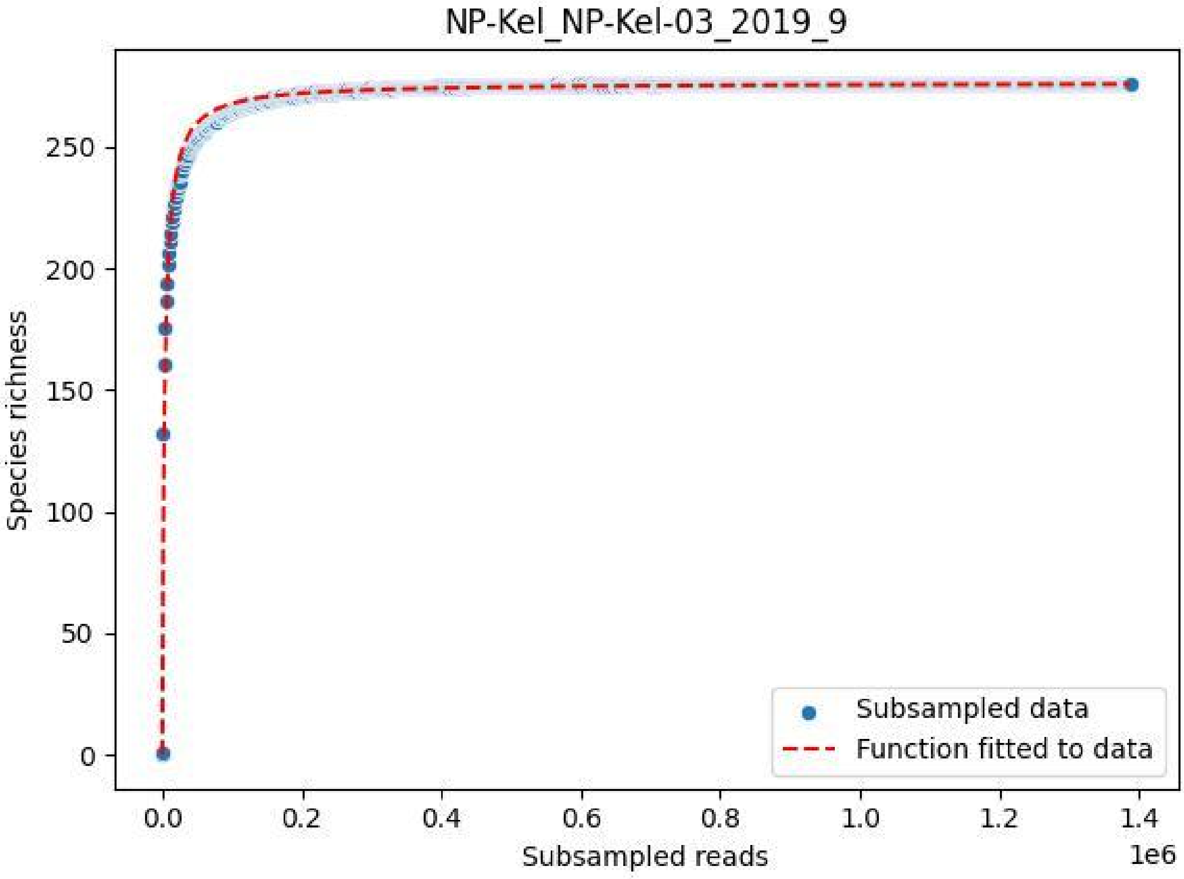
NP-Kel_NP-Kel-03_2019_9
Species richness
Subsampled reads
1e6
0
50
100
150
200
250
0.0
0.2
0.4
0.6
0.8
1.0
1.2
1.4
Subsampled data
Function fitted to data

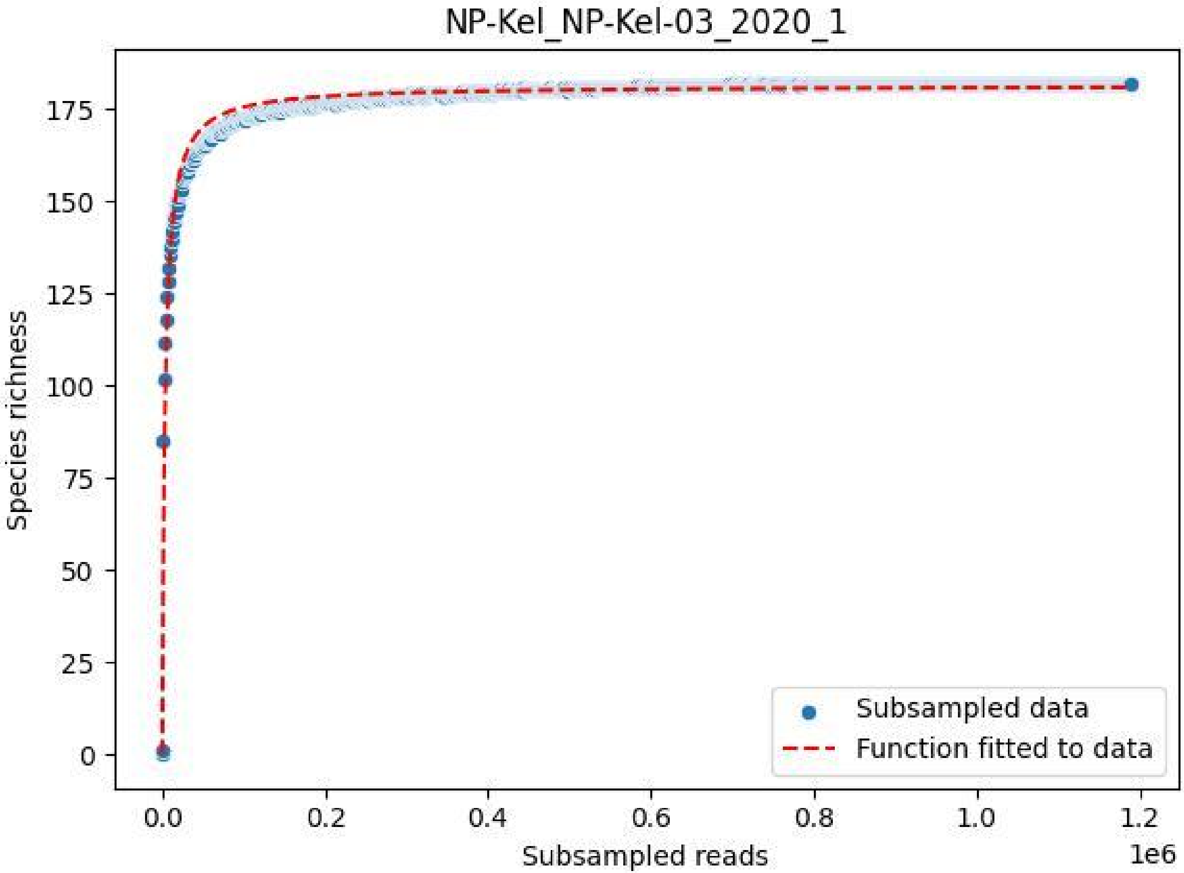

NP-Kel_NP-Kel-03_2020_1
Species richness
Subsampled reads
1e6
0
25
50
75
100
125
150
175
0.0
0.2
0.4
0.6
0.8
1.0
1.2
Subsampled data
Function fitted to data

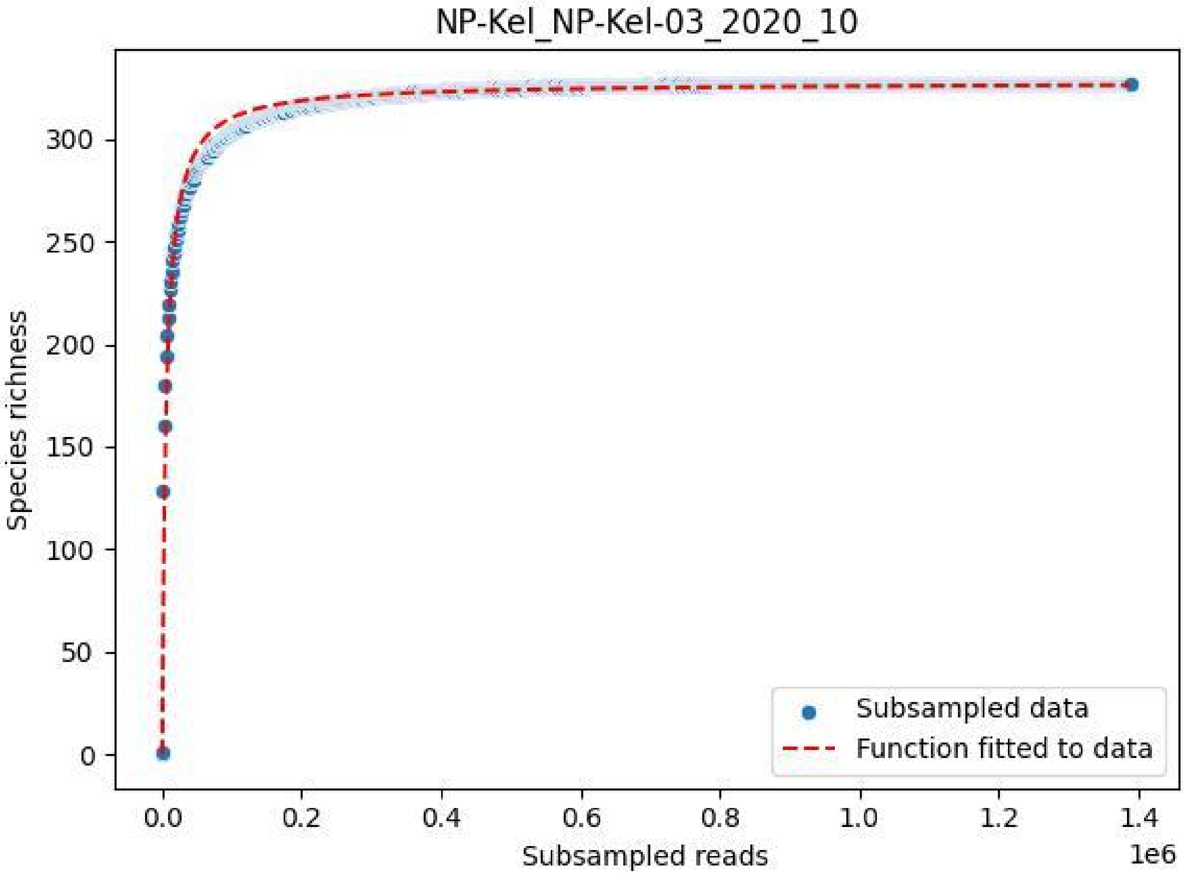

NP-Kel_NP-Kel-03_2020_10
Species richness
Subsampled reads
1e6
0
50
100
150
200
250
300
0.0
0.2
0.4
0.6
0.8
1.0
1.2
1.4
Subsampled data
Function fitted to data

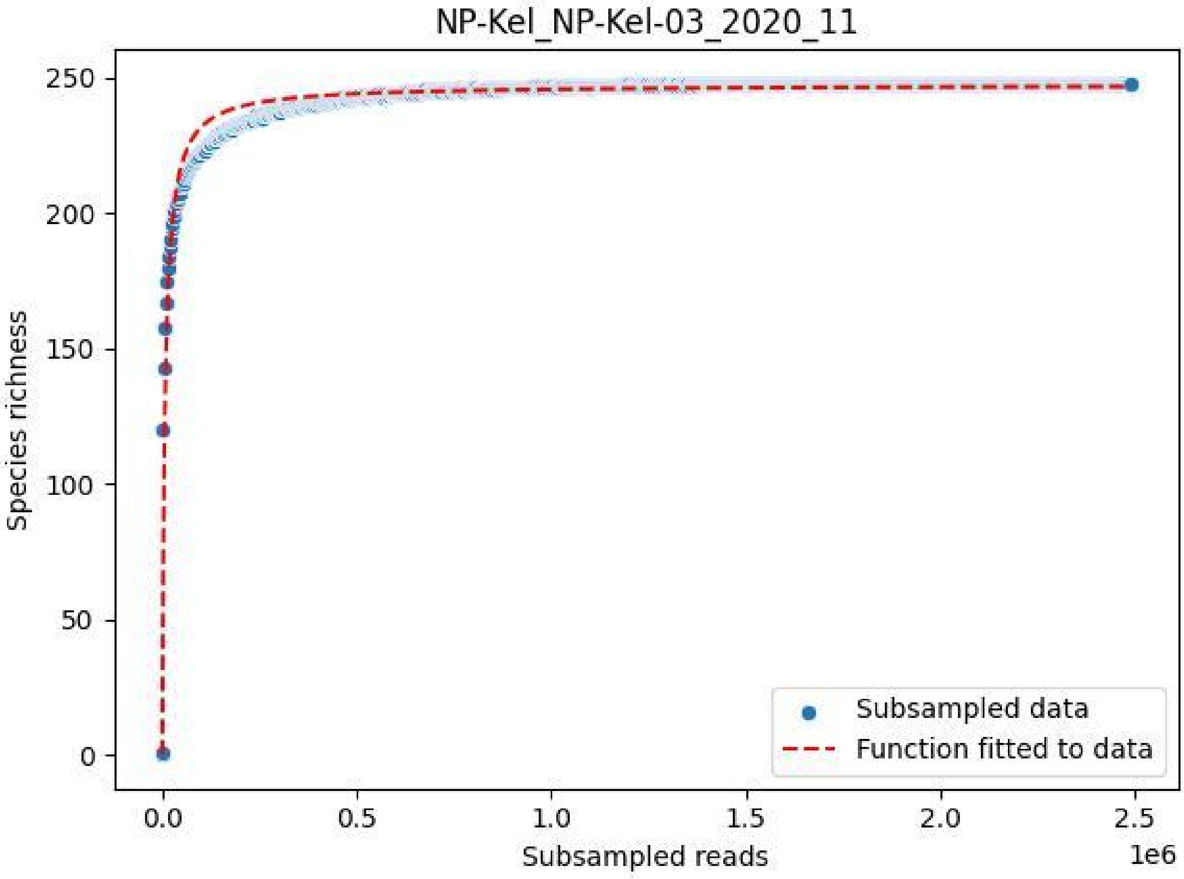
NP-Kel_NP-Kel-03_2020_11
250
200
150
100
50
0
Species richness
0.0
0.5
1.0
1.5
2.0
2.5
1e6
Subsampled reads
Subsampled data
Function fitted to data

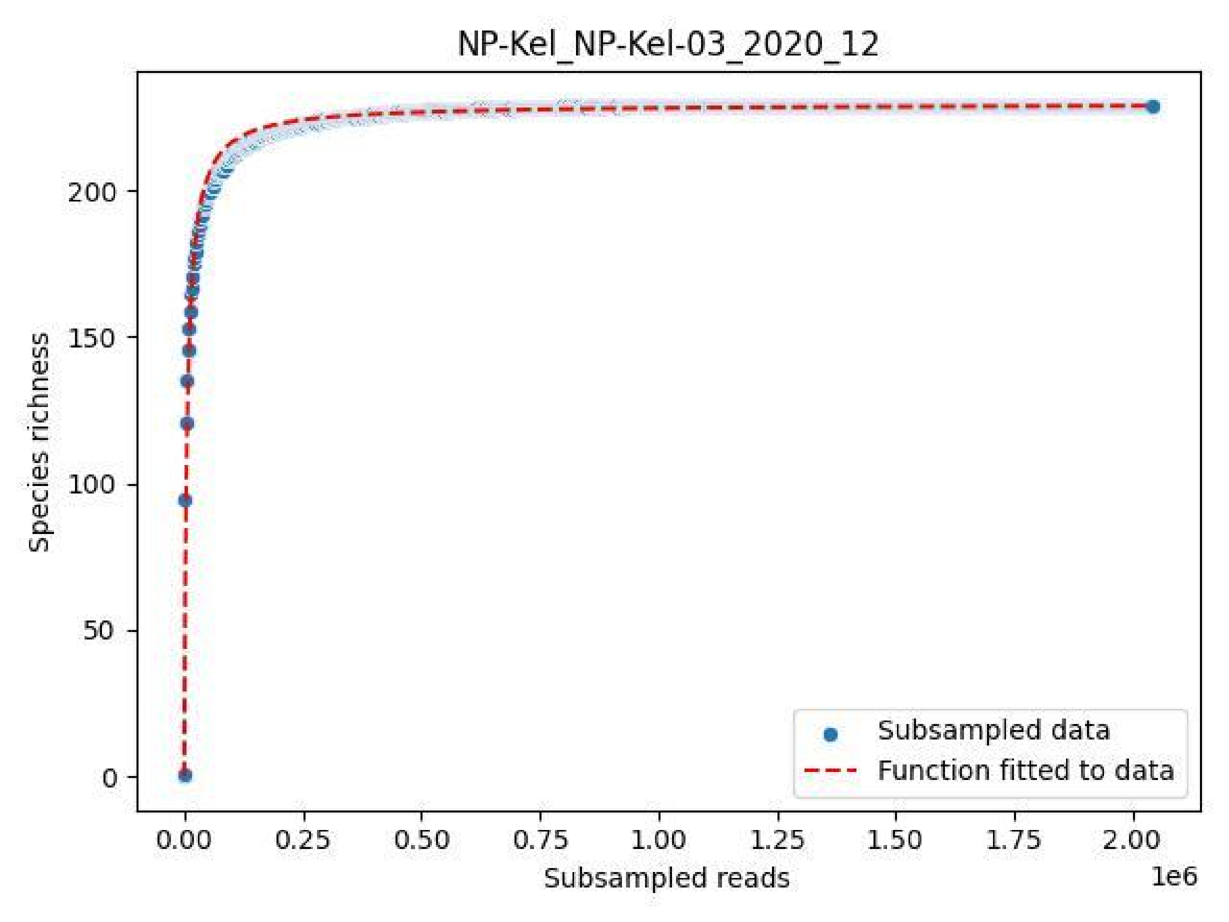

NP-Kel_NP-Kel-03_2020_12
Species richness
Subsampled reads
1e6
0
50
100
150
200
0.00
0.25
0.50
0.75
1.00
1.25
1.50
1.75
2.00
Subsampled data
Function fitted to data

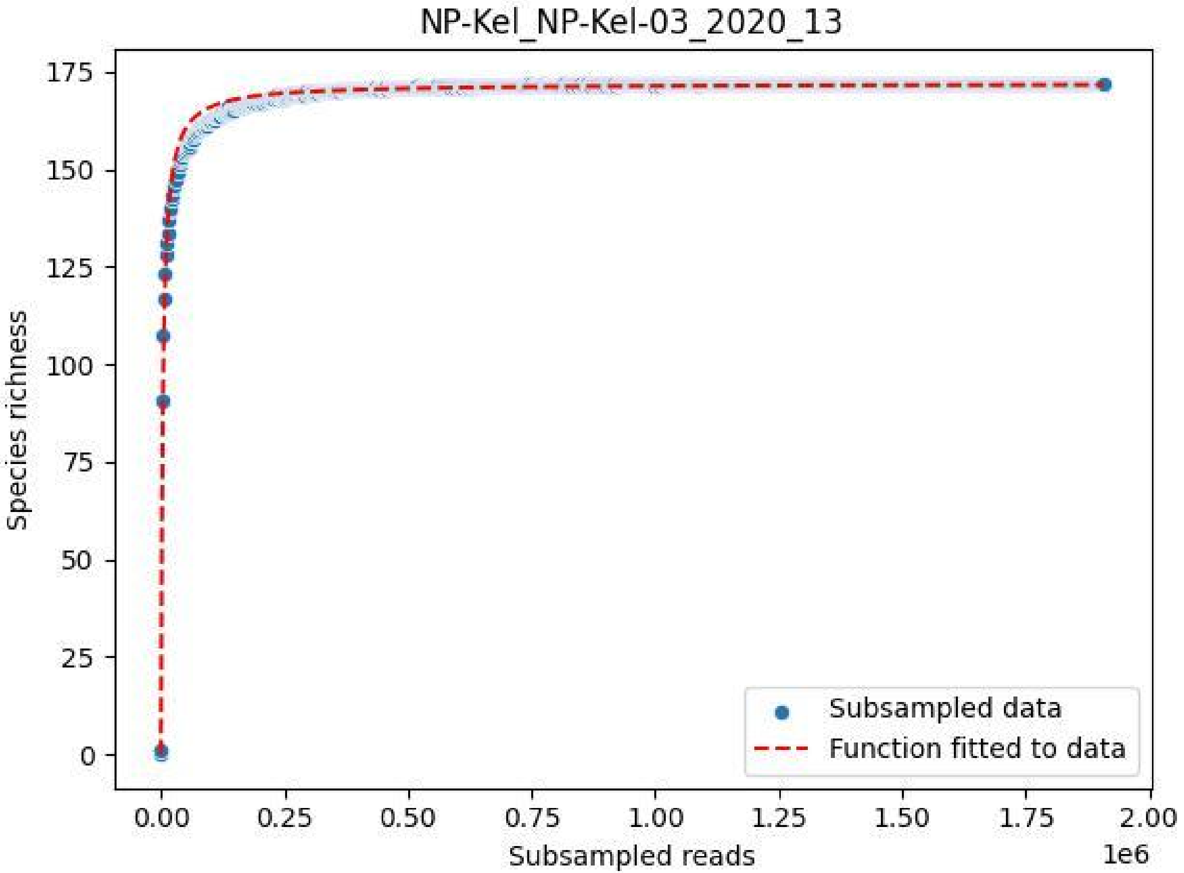

NP-Kel_NP-Kel-03_2020_13
Species richness
Subsampled reads
1e6
0
25
50
75
100
125
150
175
0.00
0.25
0.50
0.75
1.00
1.25
1.50
1.75
2.00
Subsampled data
Function fitted to data

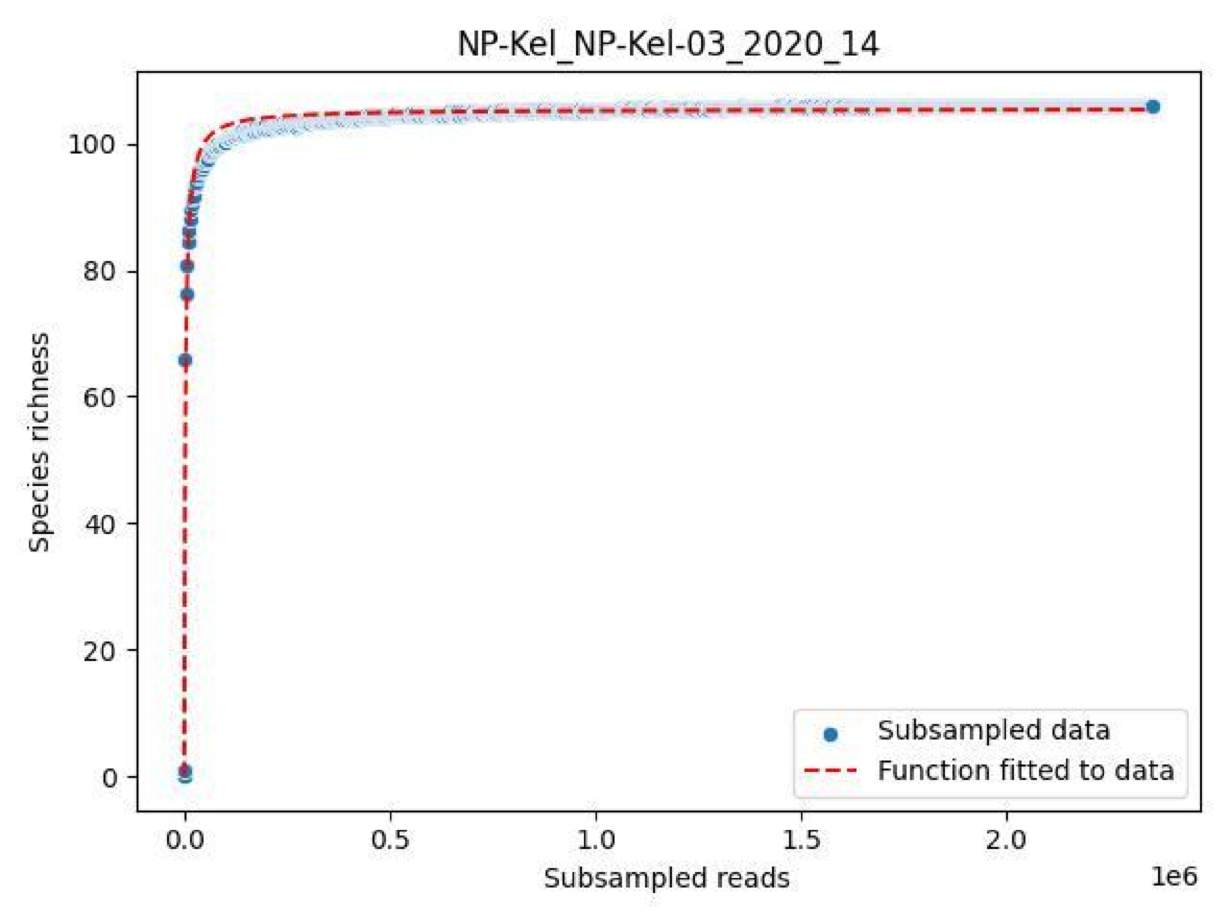

NP-Kel_NP-Kel-03_2020_14
Species richness
Subsampled reads
1e6
0
20
40
60
80
100
0.0
0.5
1.0
1.5
2.0
Subsampled data
Function fitted to data

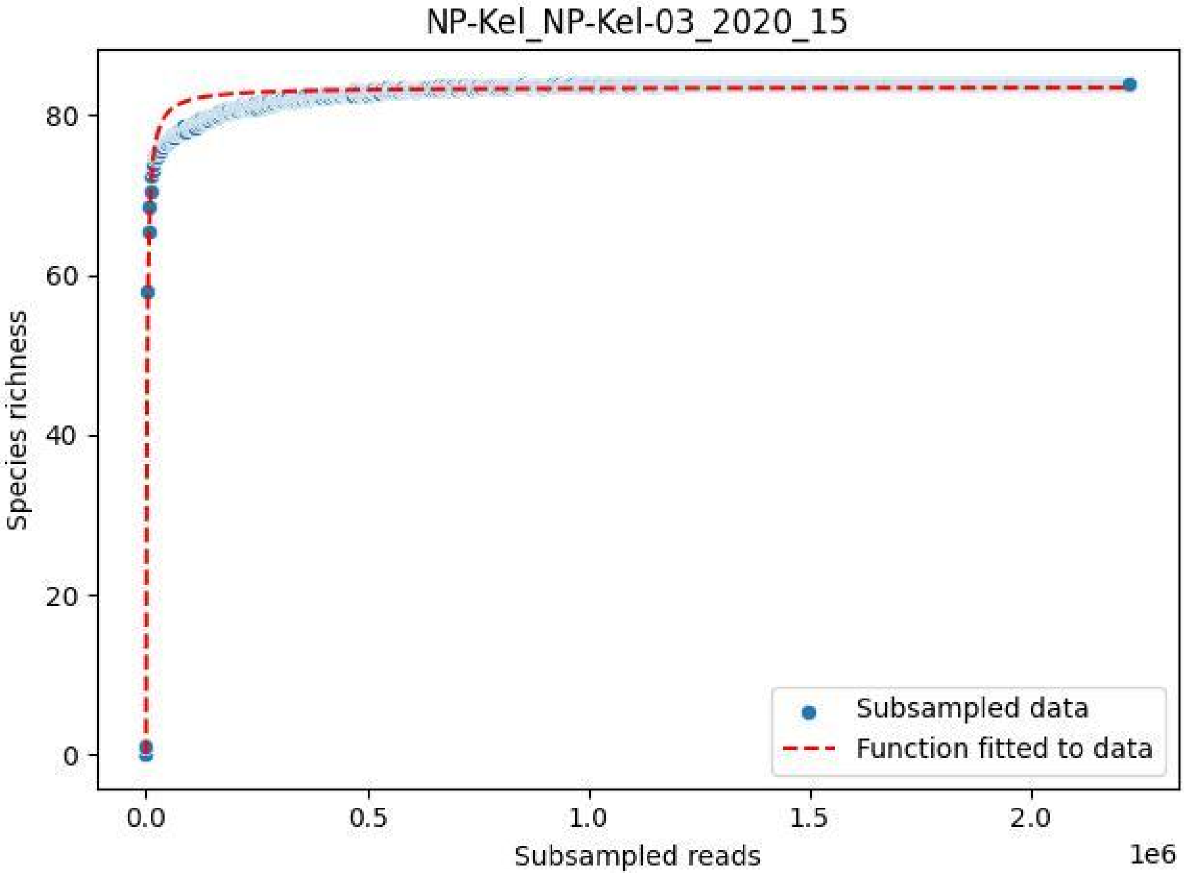
NP-Kel_NP-Kel-03_2020_15
Species richness
Subsampled reads
1e6
0
20
40
60
80
0.0
0.5
1.0
1.5
2.0
Subsampled data
Function fitted to data

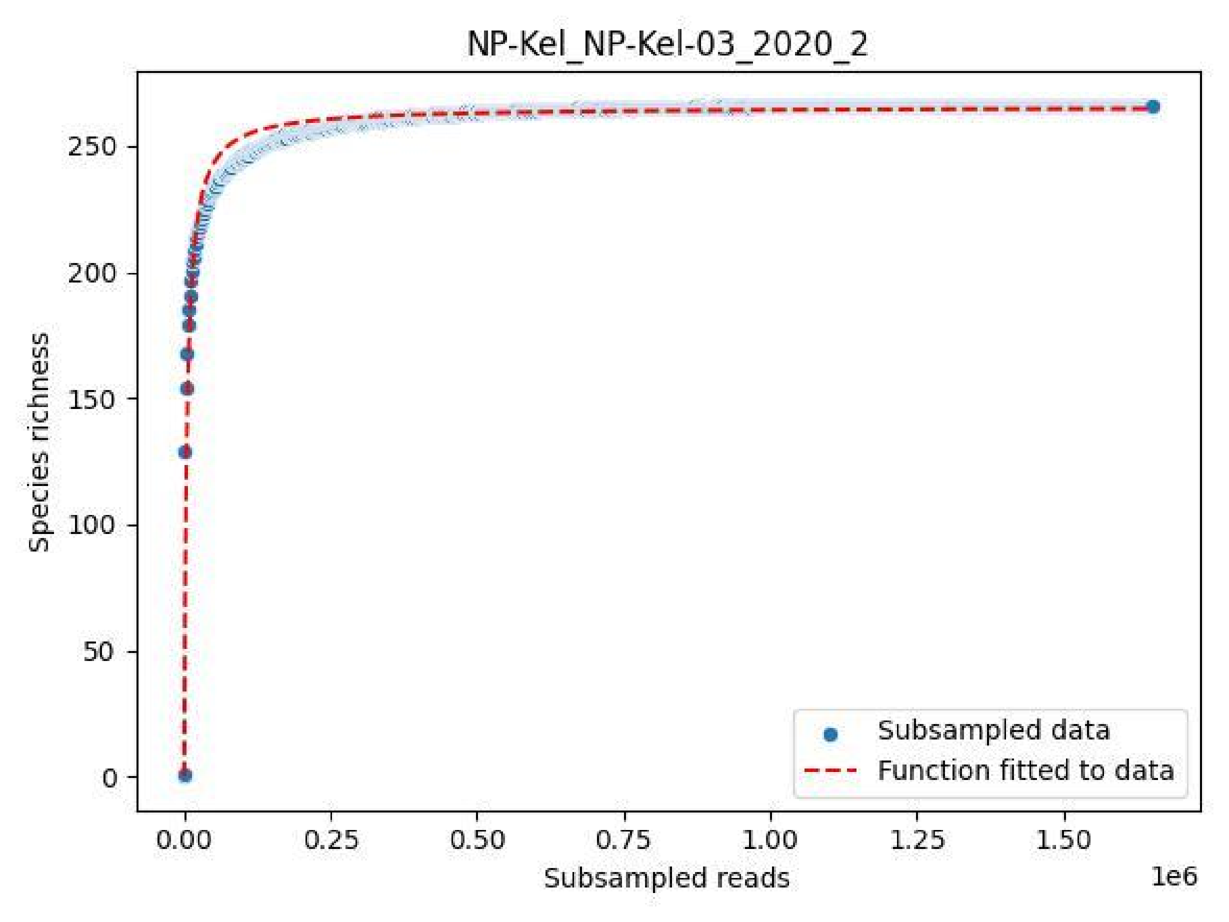
NP-Kel_NP-Kel-03_2020_2
Species richness
0
50
100
150
200
250
0.00
0.25
0.50
0.75
1.00
1.25
1.50
1e6
Subsampled reads
Subsampled data
Function fitted to data

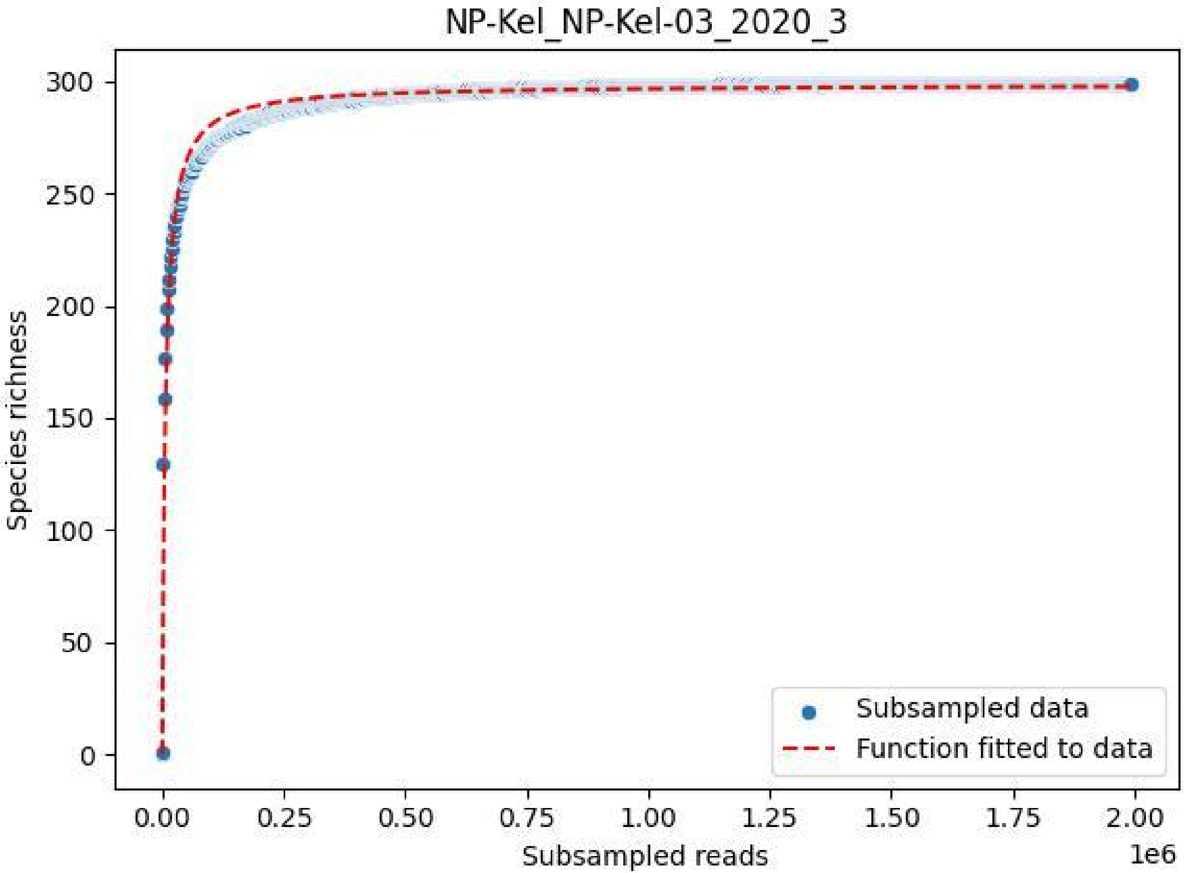

NP-Kel_NP-Kel-03_2020_3
Species richness
Subsampled reads
1e6
0
50
100
150
200
250
300
0.00
0.25
0.50
0.75
1.00
1.25
1.50
1.75
2.00
Subsampled data
Function fitted to data

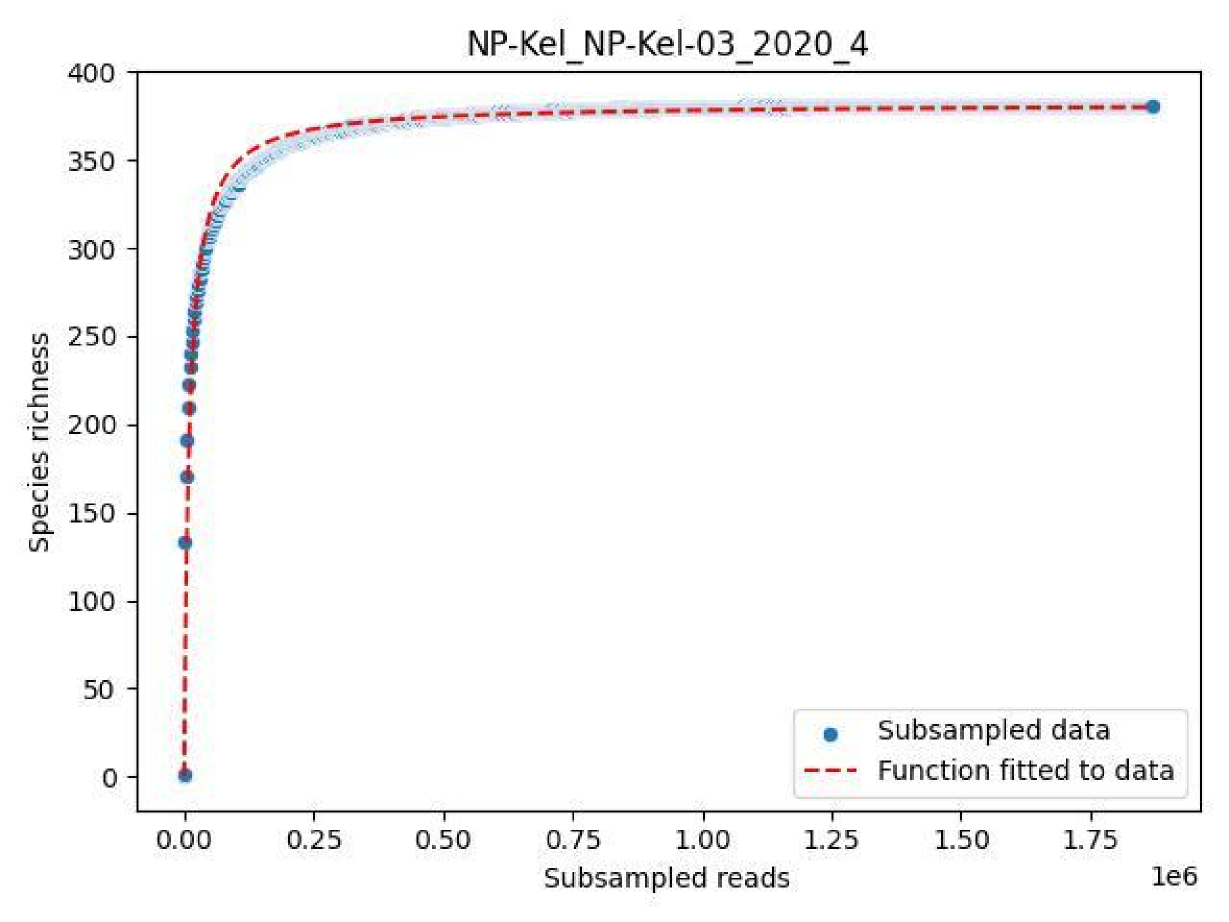

NP-Kel_NP-Kel-03_2020_4
400
350
300
250
200
150
100
50
0
Species richness
0.00
0.25
0.50
0.75
1.00
1.25
1.50
1.75
1e6
Subsampled reads
Subsampled data
Function fitted to data

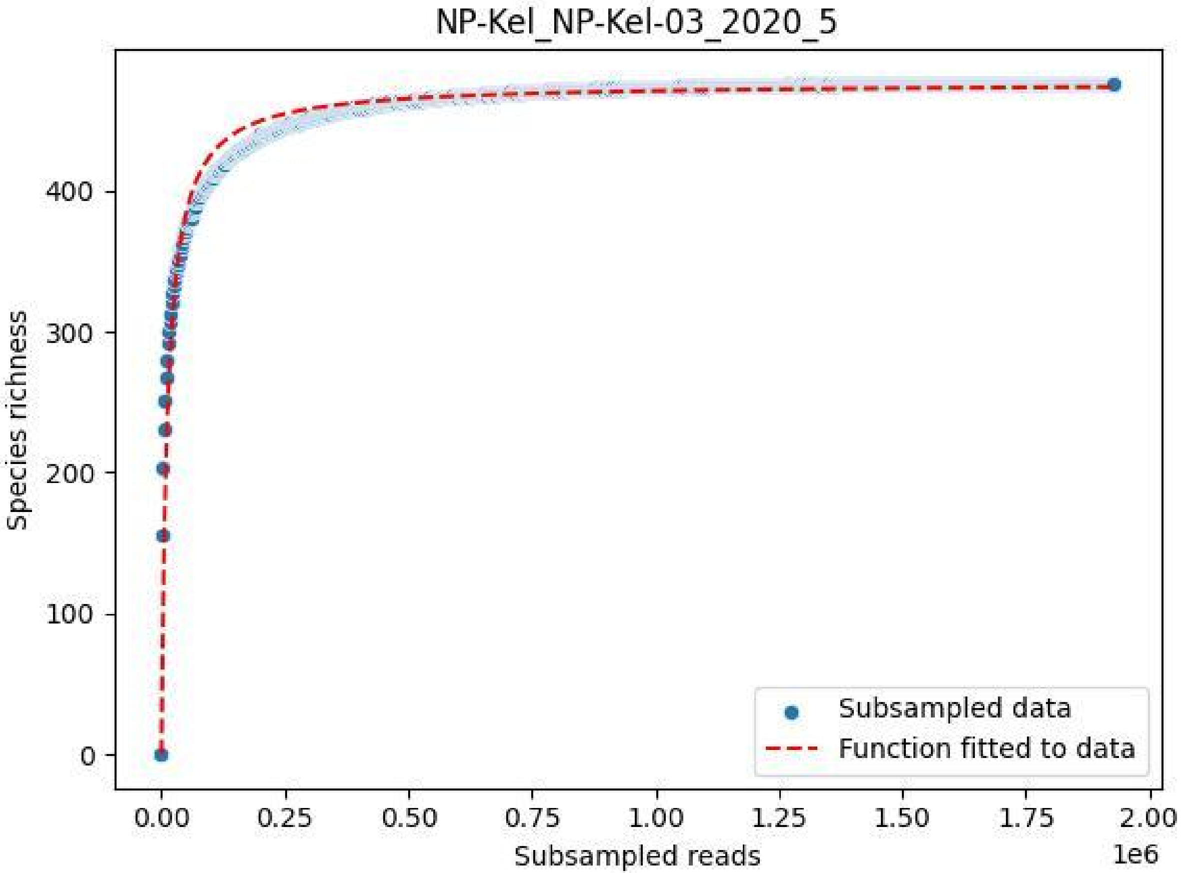

NP-Kel_NP-Kel-03_2020_5
Species richness
0
100
200
300
400
Subsampled reads
0.00
0.25
0.50
0.75
1.00
1.25
1.50
1.75
2.00
1e6
Subsampled data
Function fitted to data

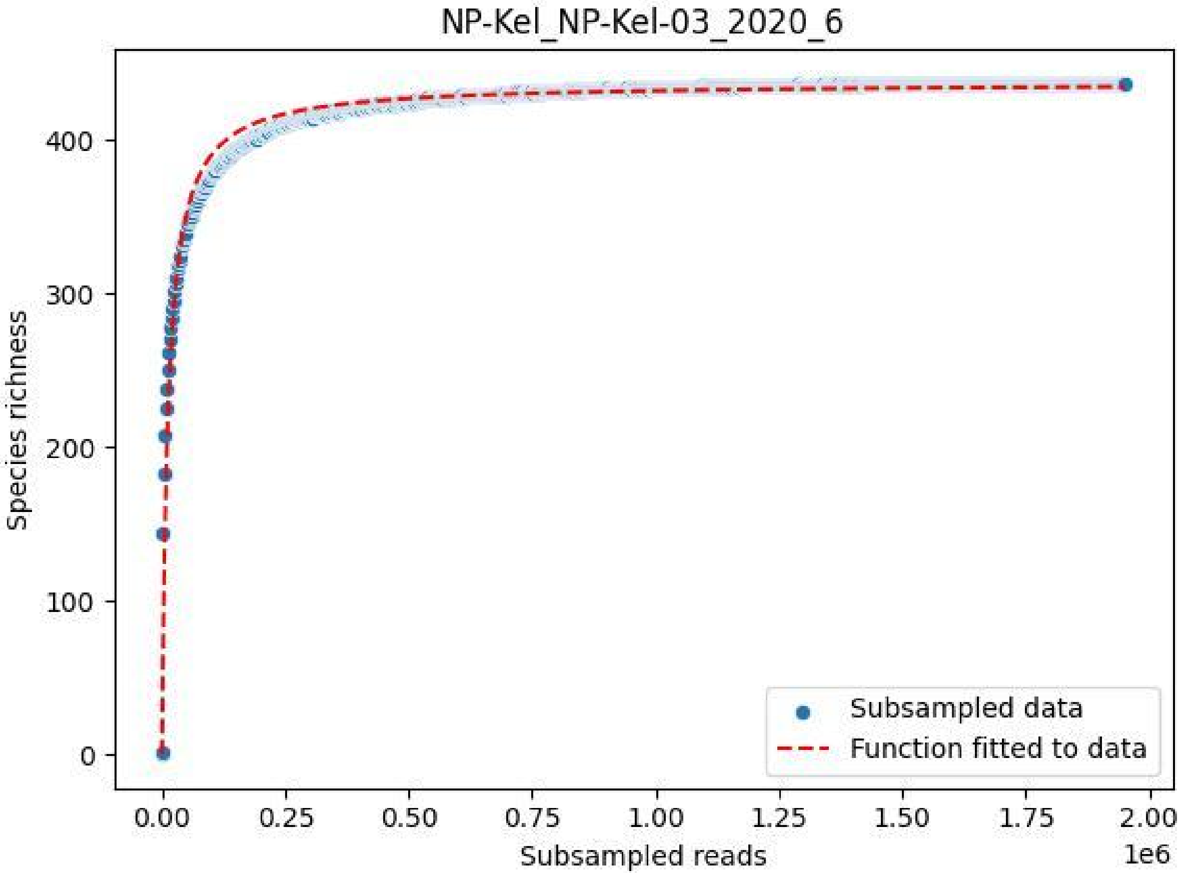
NP-Kel_NP-Kel-03_2020_6
Species richness
Subsampled reads
1e6
0
100
200
300
400
0.00
0.25
0.50
0.75
1.00
1.25
1.50
1.75
2.00
Subsampled data
Function fitted to data

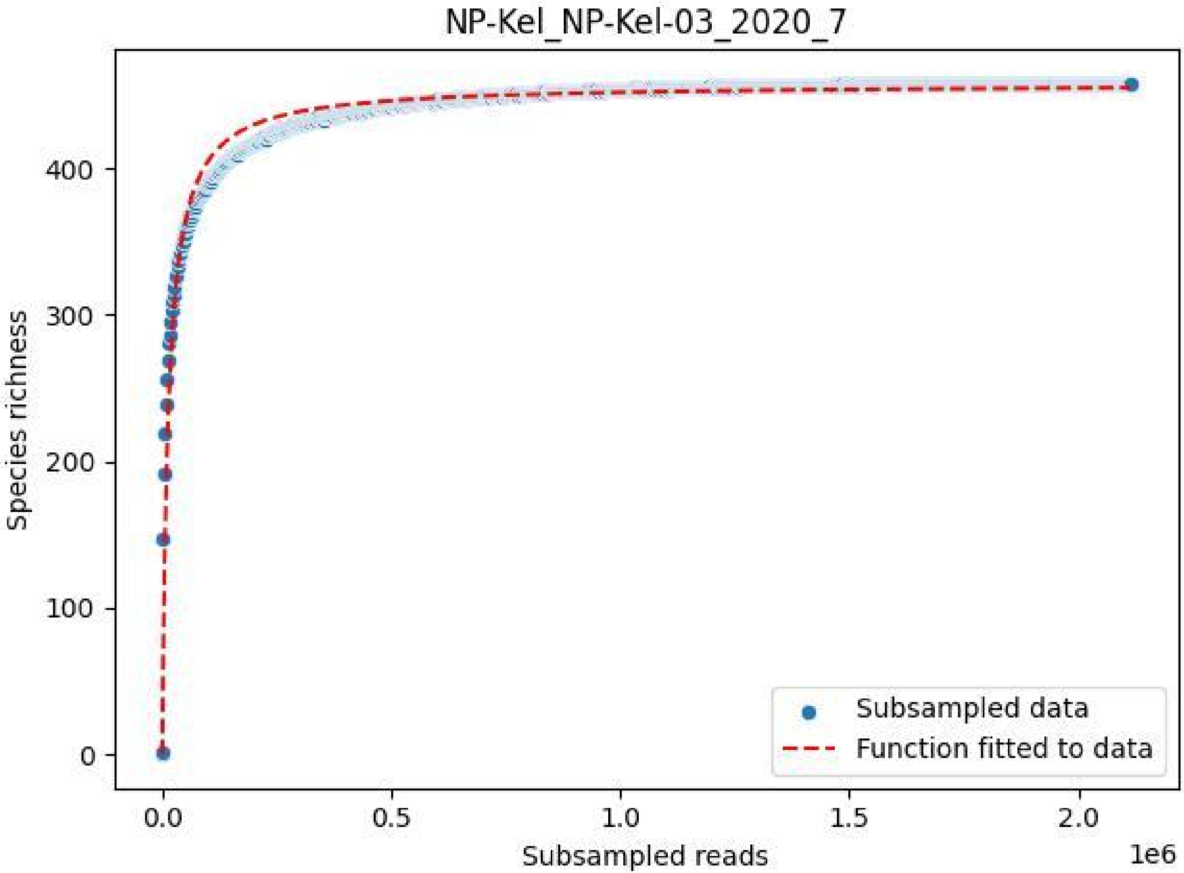

NP-Kel_NP-Kel-03_2020_7
Species richness
Subsampled reads
1e6
0
100
200
300
400
0.0
0.5
1.0
1.5
2.0
Subsampled data
Function fitted to data

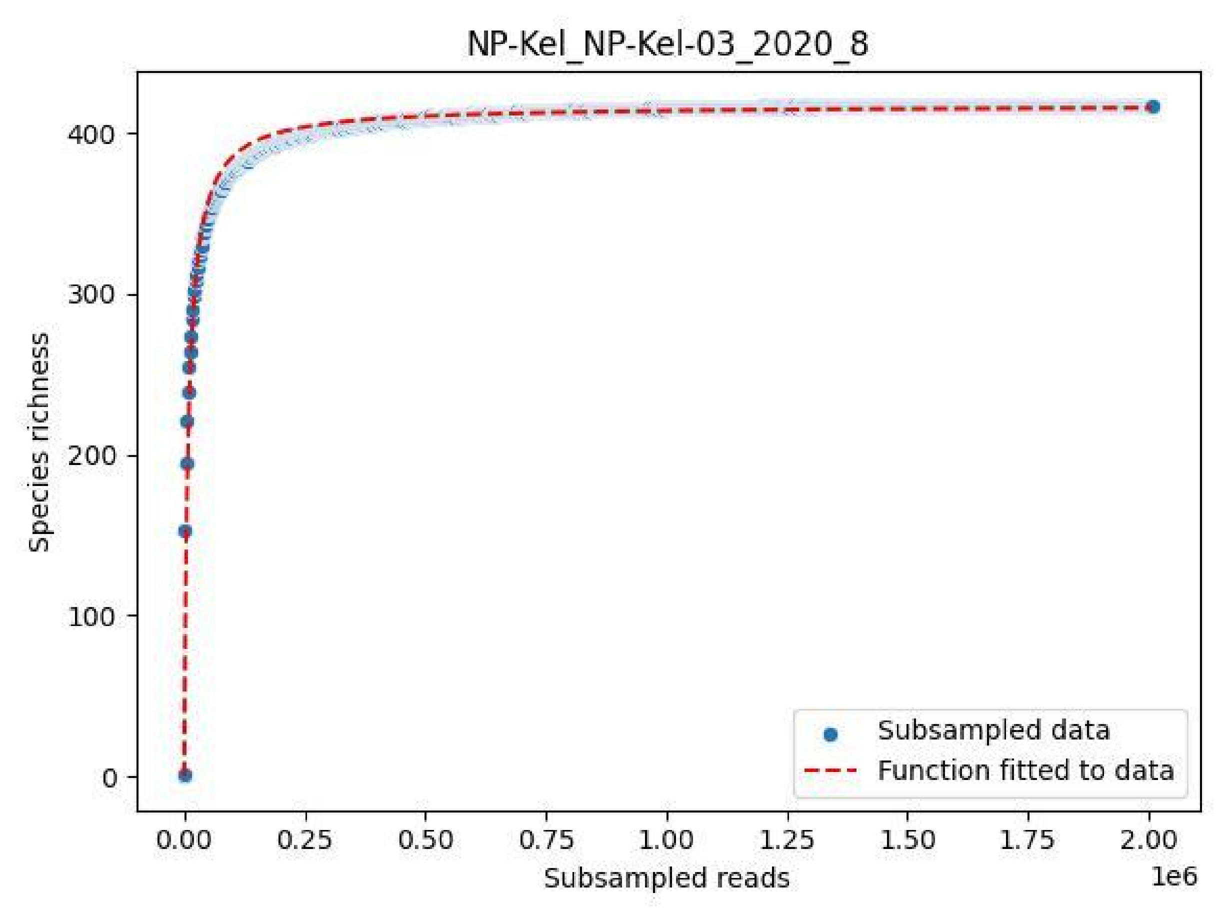

NP-Kel_NP-Kel-03_2020_8
Species richness
Subsampled reads
1e6
0
100
200
300
400
0.00
0.25
0.50
0.75
1.00
1.25
1.50
1.75
2.00
Subsampled data
Function fitted to data

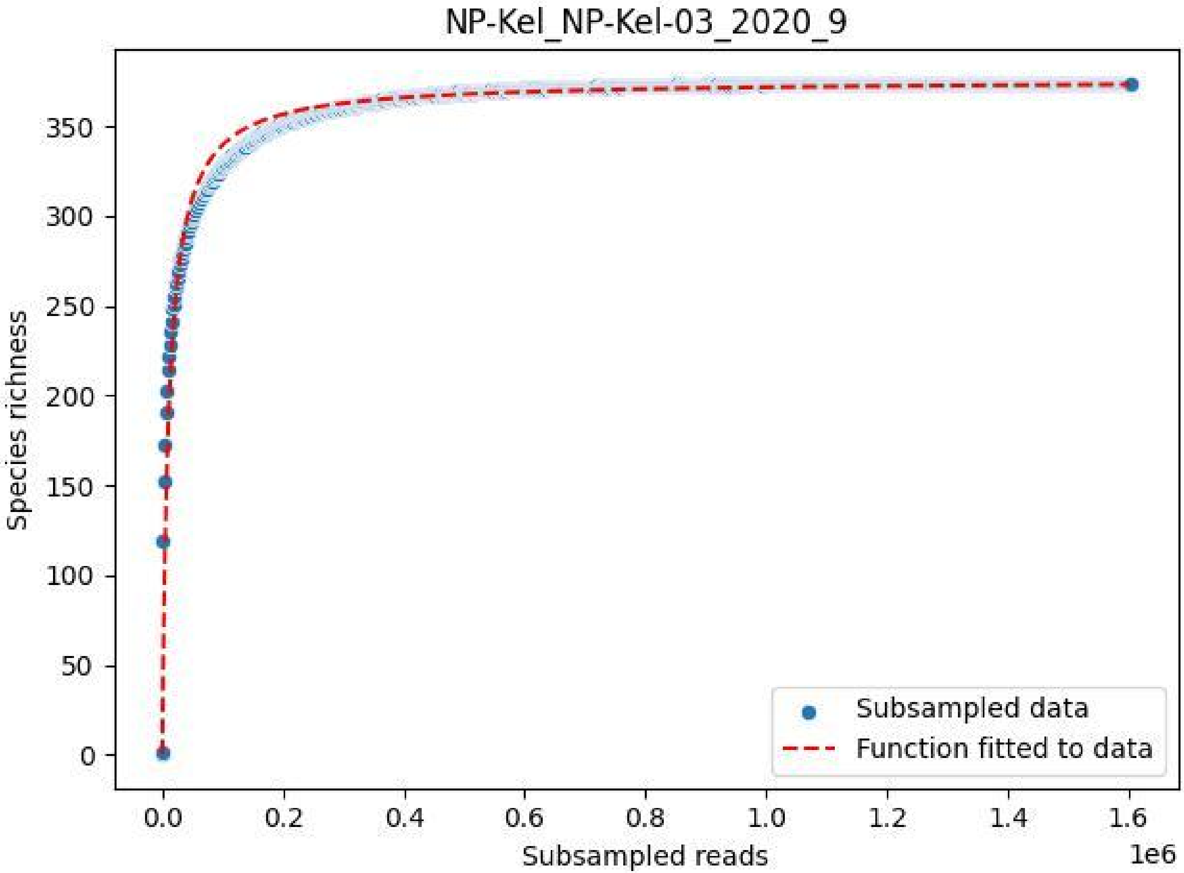

NP-Kel_NP-Kel-03_2020_9
Species richness
0
50
100
150
200
250
300
350
Subsampled reads
0.0
0.2
0.4
0.6
0.8
1.0
1.2
1.4
1.6
1e6
Subsampled data
Function fitted to data

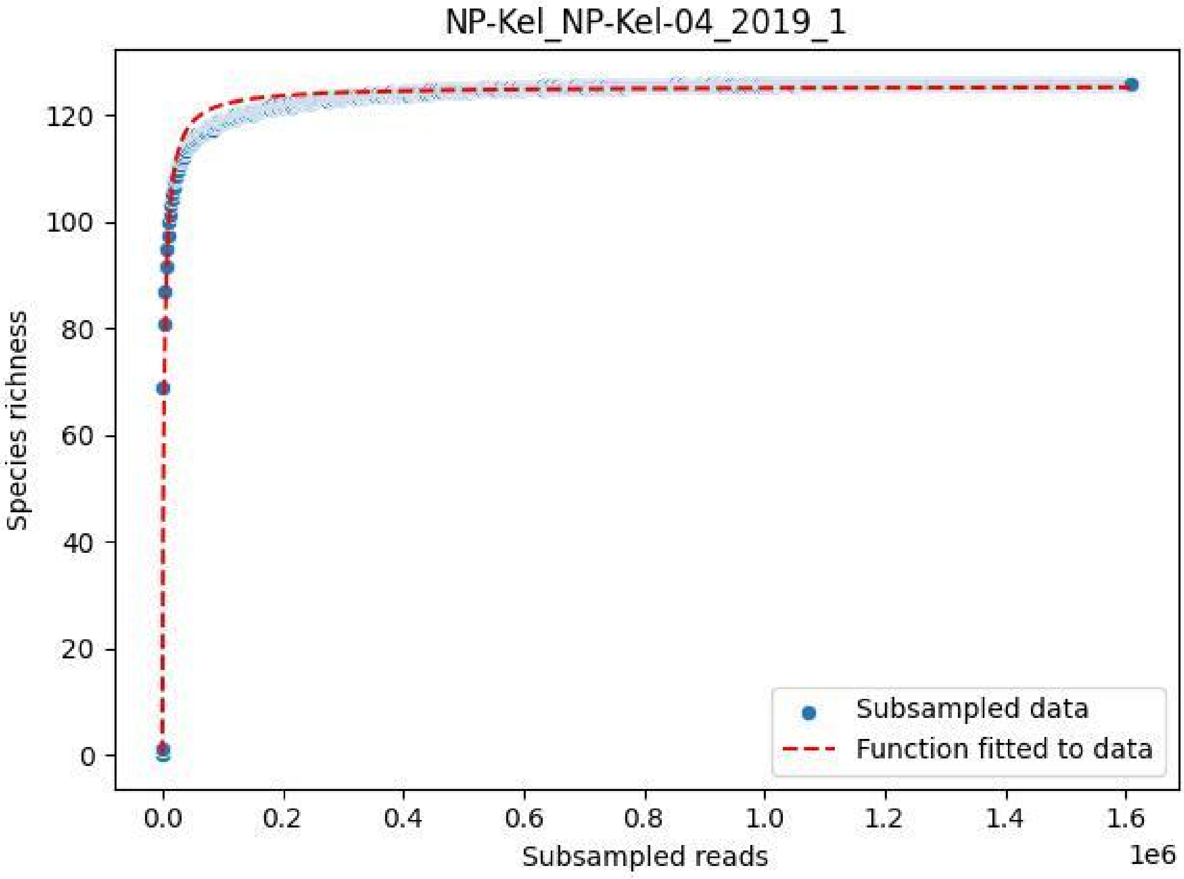

NP-Kel_NP-Kel-04_2019_1
Species richness
Subsampled reads
1e6
0
20
40
60
80
100
120
0.0
0.2
0.4
0.6
0.8
1.0
1.2
1.4
1.6
Subsampled data
Function fitted to data

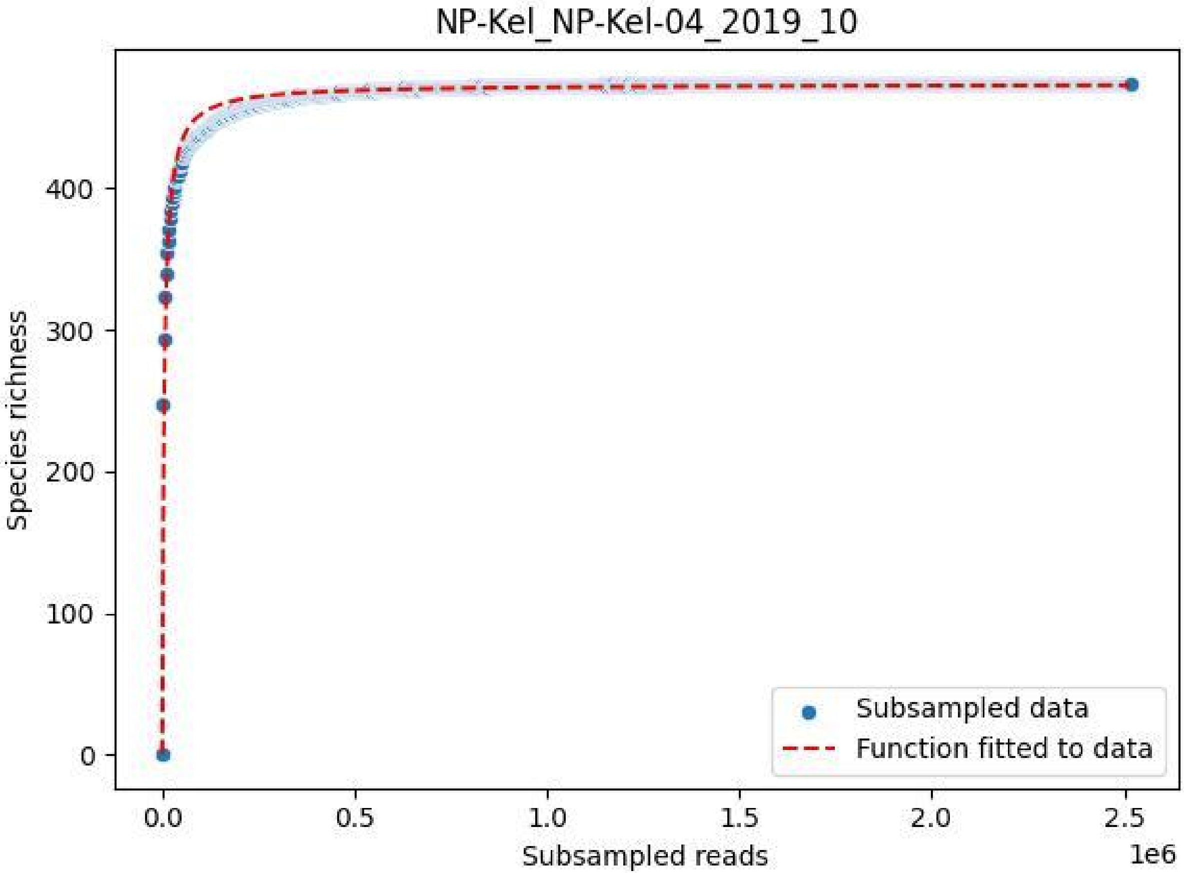
NP-Kel_NP-Kel-04_2019_10
Species richness
Subsampled reads
1e6
0
100
200
300
400
0.0
0.5
1.0
1.5
2.0
2.5
Subsampled data
Function fitted to data

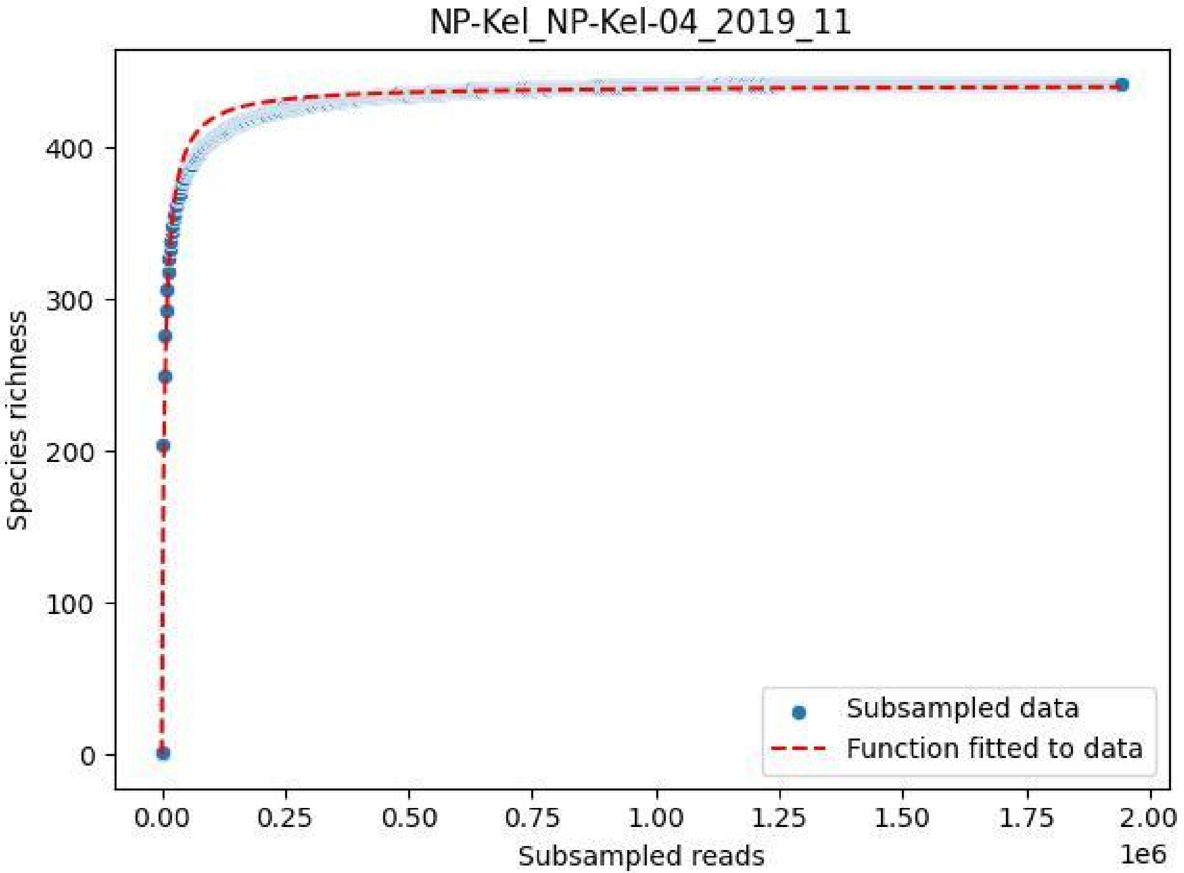

NP-Kel_NP-Kel-04_2019_11
Species richness
Subsampled reads
1e6
0
100
200
300
400
0.00
0.25
0.50
0.75
1.00
1.25
1.50
1.75
2.00
Subsampled data
Function fitted to data

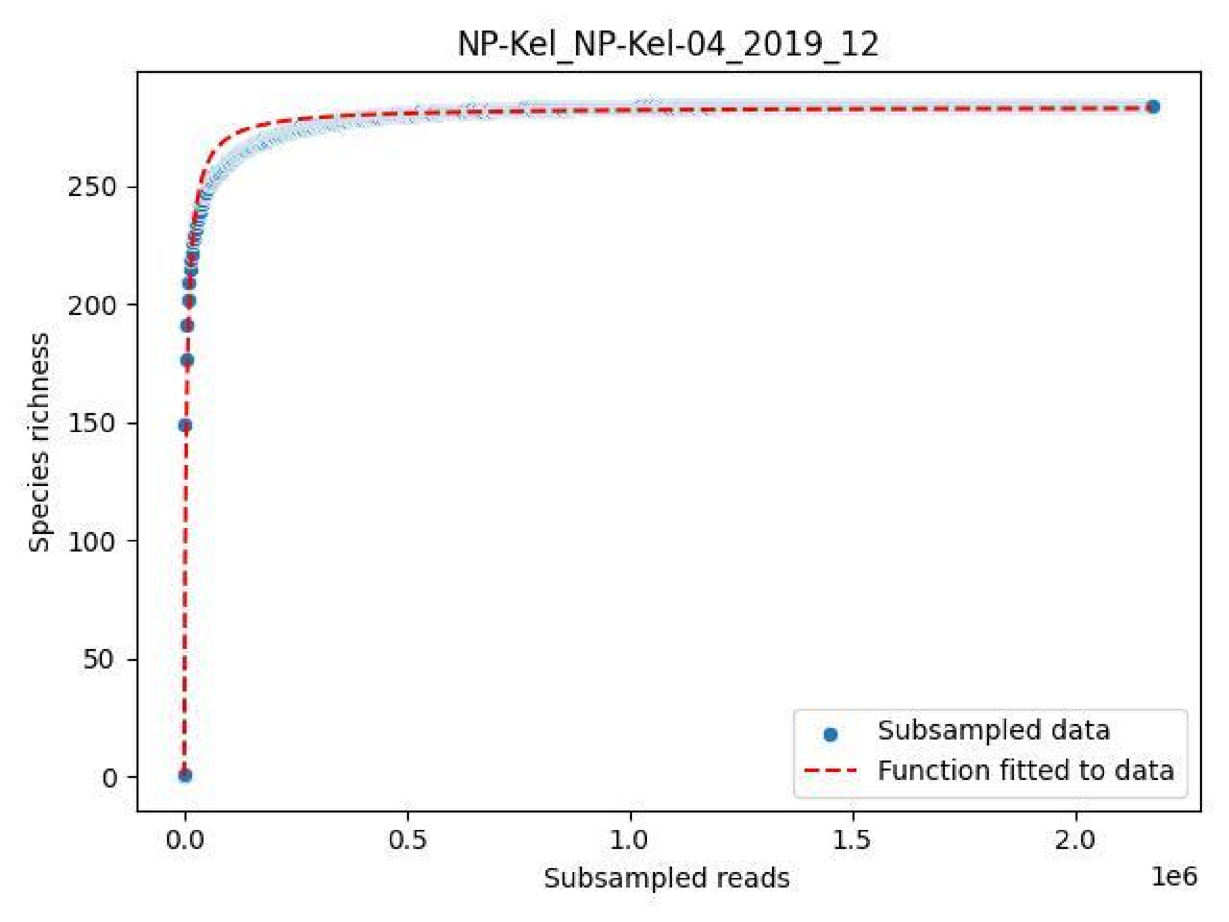

NP-Kel_NP-Kel-04_2019_12
Species richness
Subsampled reads
1e6
0
50
100
150
200
250
0.0
0.5
1.0
1.5
2.0
Subsampled data
Function fitted to data

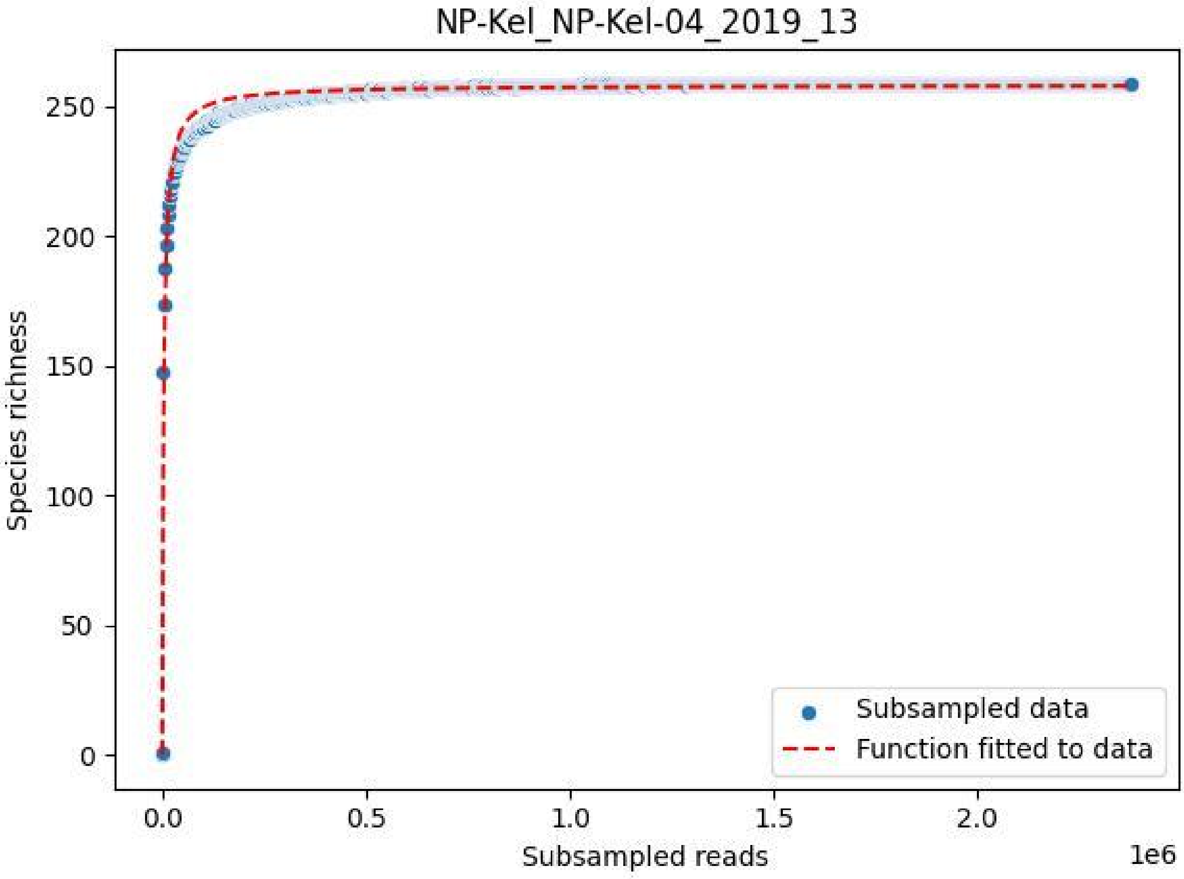
NP-Kel_NP-Kel-04_2019_13
Species richness
Subsampled reads
1e6
0.0
0.5
1.0
1.5
2.0
0
50
100
150
200
250
Subsampled data
Function fitted to data

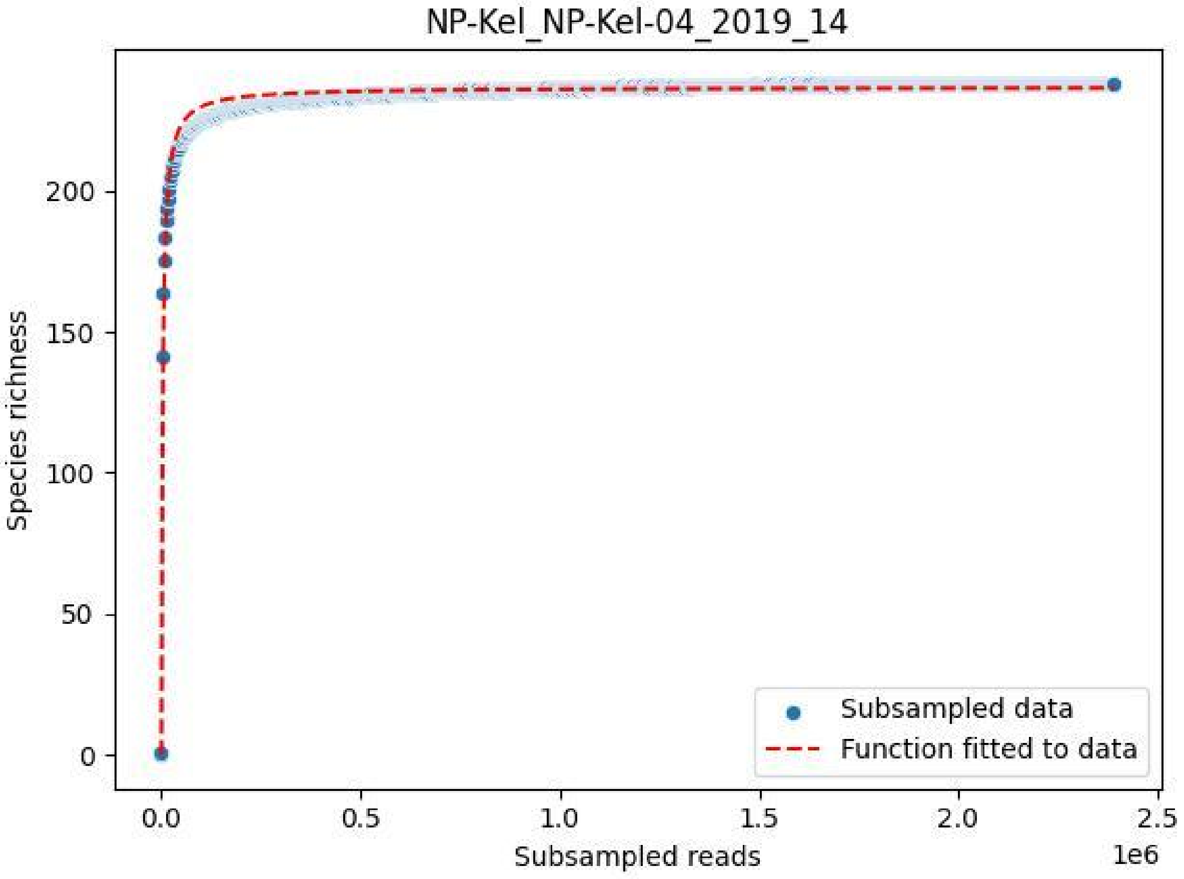
NP-Kel_NP-Kel-04_2019_14
Species richness
Subsampled reads
0
50
100
150
200
0.0
0.5
1.0
1.5
2.0
2.5
1e6
Subsampled data
Function fitted to data

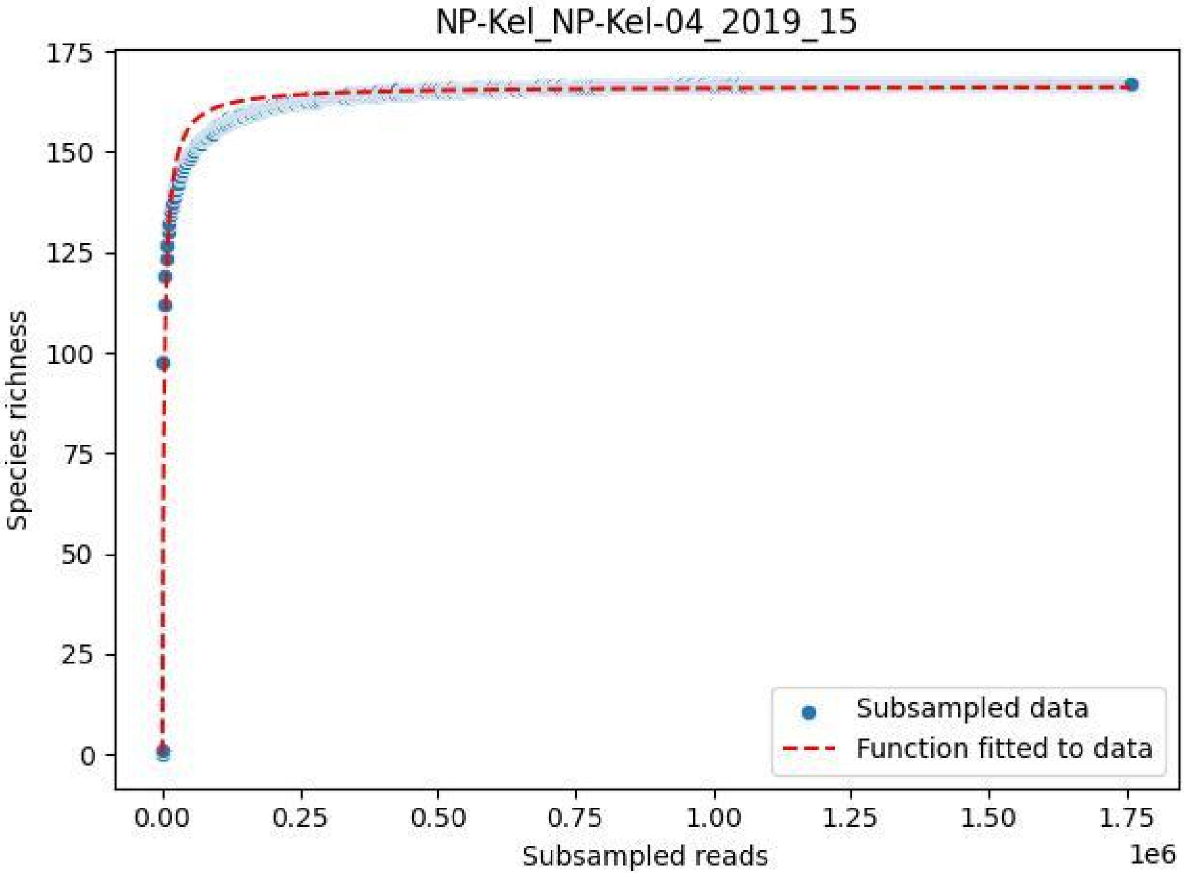
NP-Kel_NP-Kel-04_2019_15
Species richness
Subsampled reads
1e6
0
25
50
75
100
125
150
175
0.00
0.25
0.50
0.75
1.00
1.25
1.50
1.75
Subsampled data
Function fitted to data

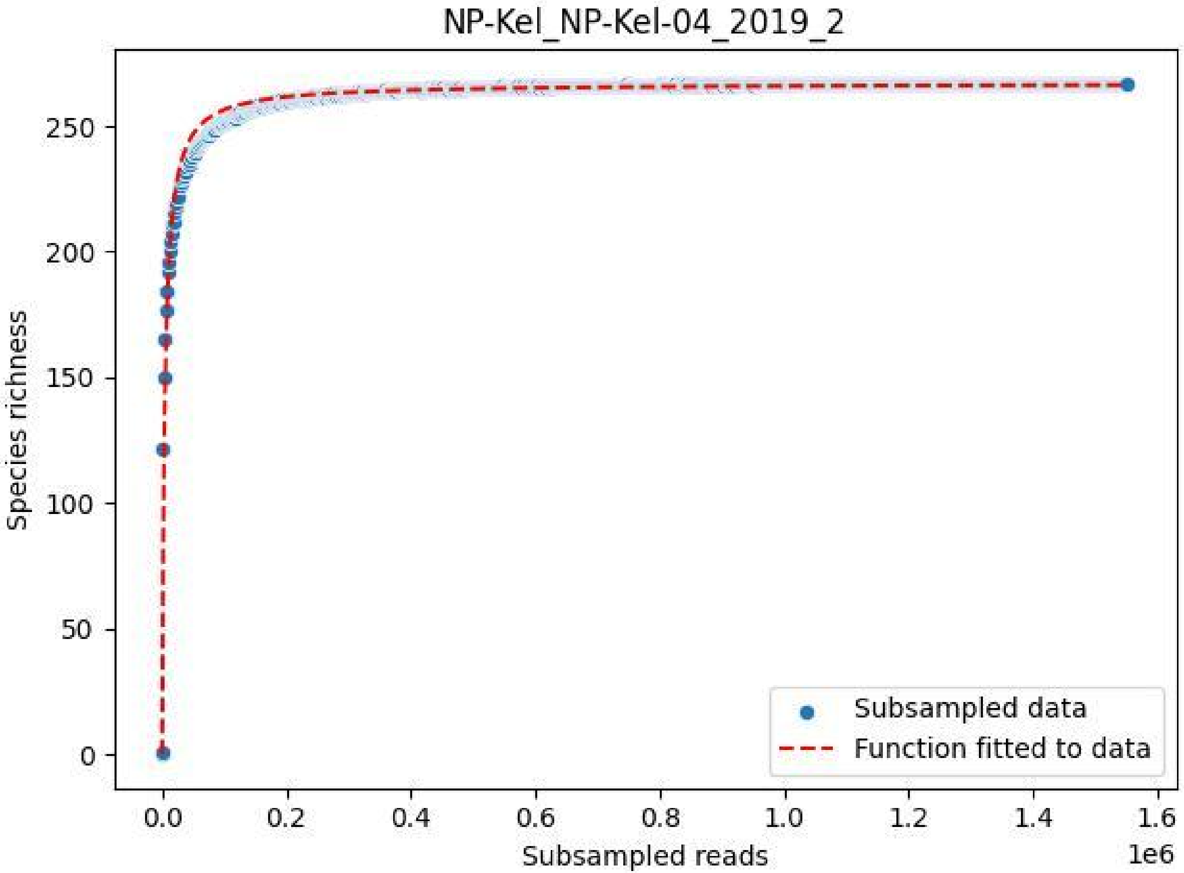
NP-Kel_NP-Kel-04_2019_2
Species richness
Subsampled reads
1e6
0
50
100
150
200
250
0.0
0.2
0.4
0.6
0.8
1.0
1.2
1.4
1.6
Subsampled data
Function fitted to data

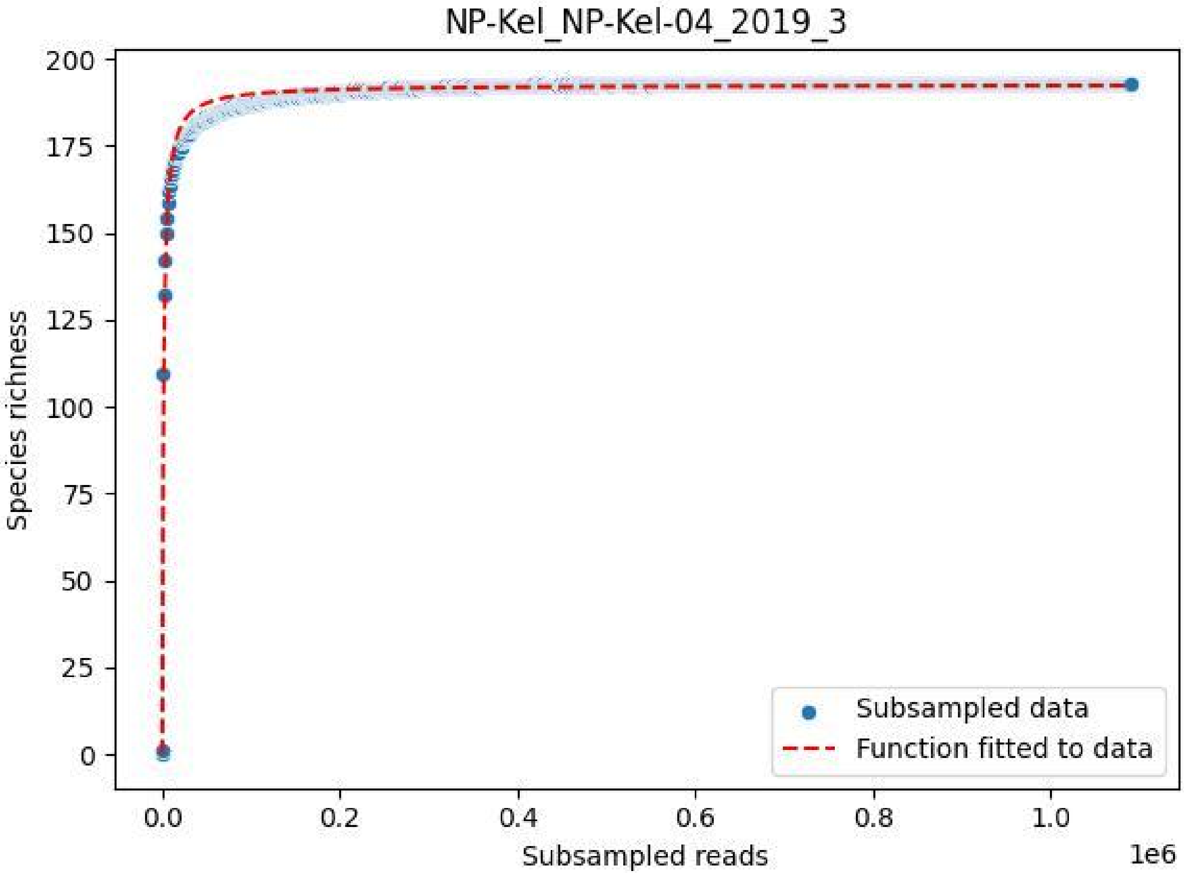

NP-Kel_NP-Kel-04_2019_3
Species richness
Subsampled reads
1e6
Subsampled data
Function fitted to data
0
25
50
75
100
125
150
175
200
0.0
0.2
0.4
0.6
0.8
1.0

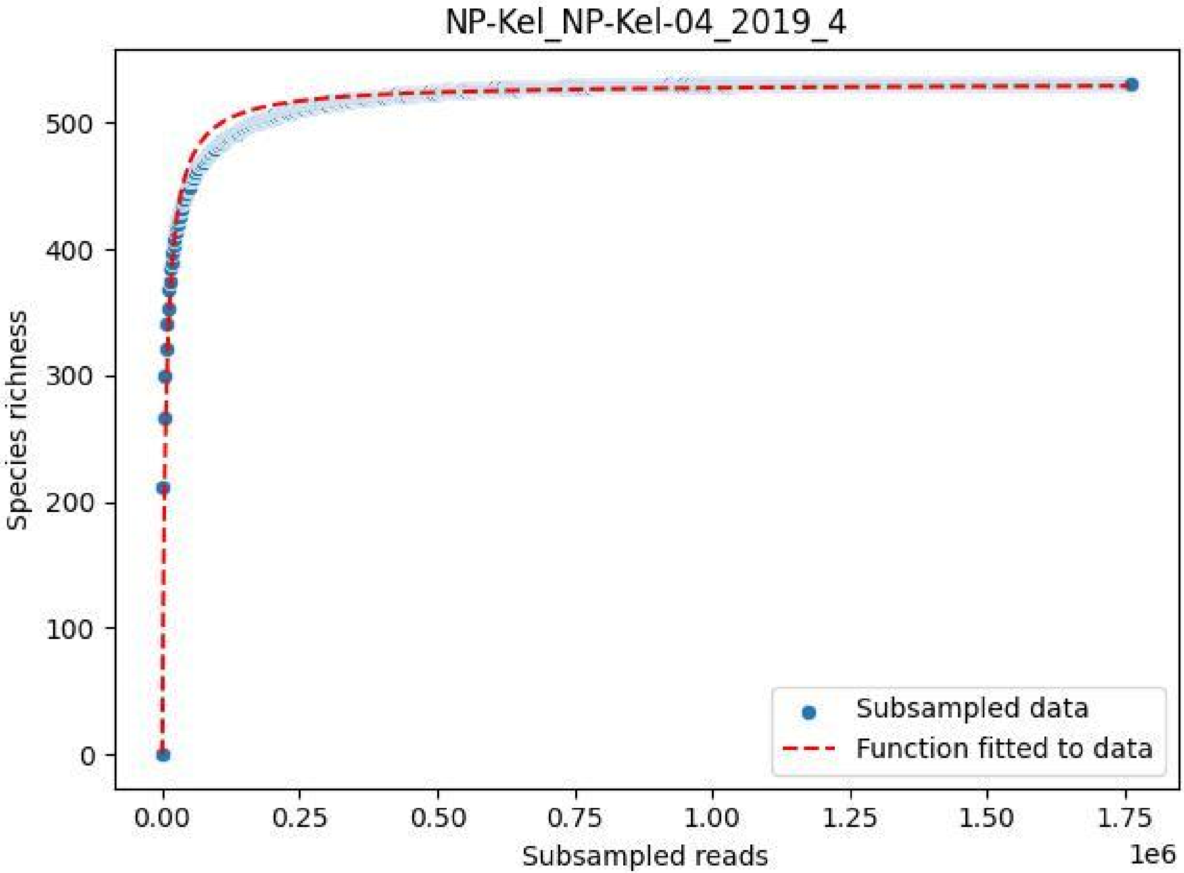
NP-Kel_NP-Kel-04_2019_4
Species richness
Subsampled reads
1e6
Subsampled data
Function fitted to data
0
100
200
300
400
500
0.00
0.25
0.50
0.75
1.00
1.25
1.50
1.75

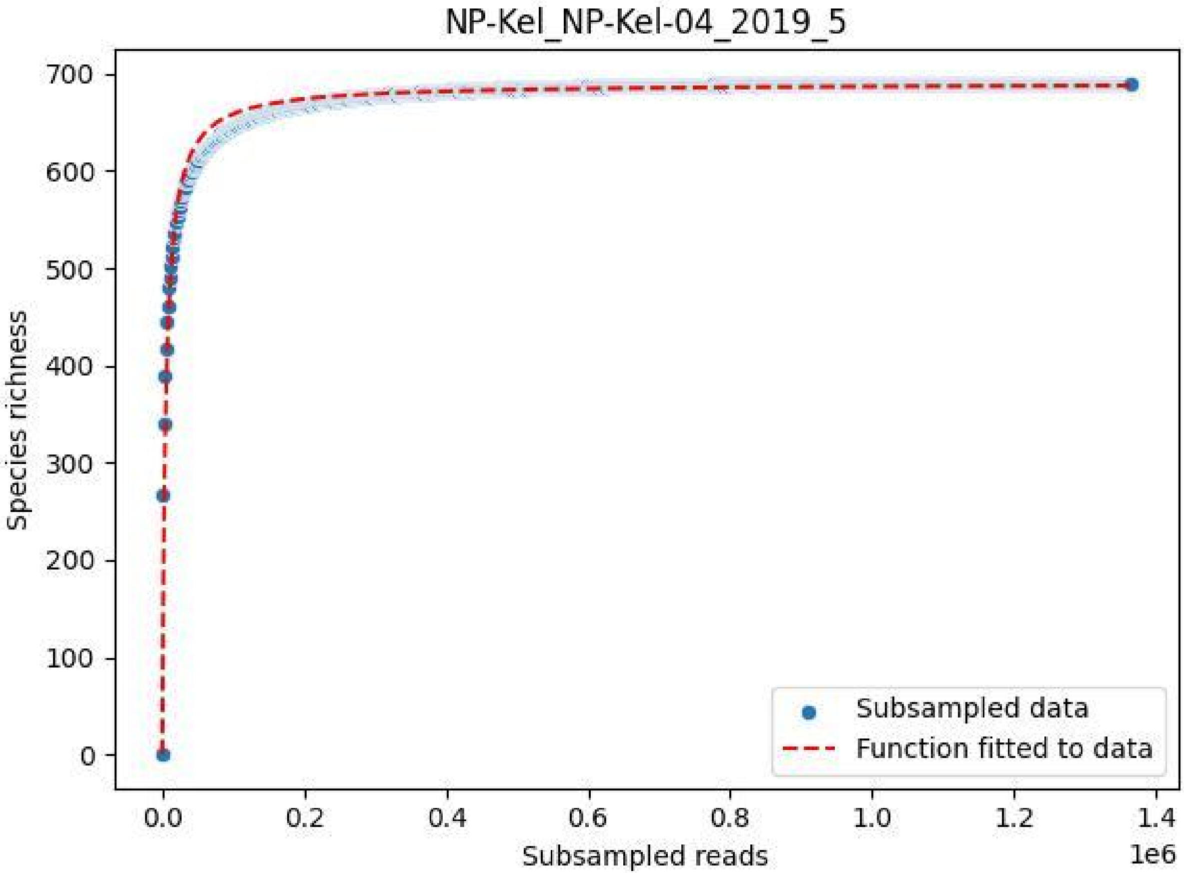
NP-Kel_NP-Kel-04_2019_5
Species richness
0
100
200
300
400
500
600
700
Subsampled reads
0.0
0.2
0.4
0.6
0.8
1.0
1.2
1.4
1e6
Subsampled data
Function fitted to data

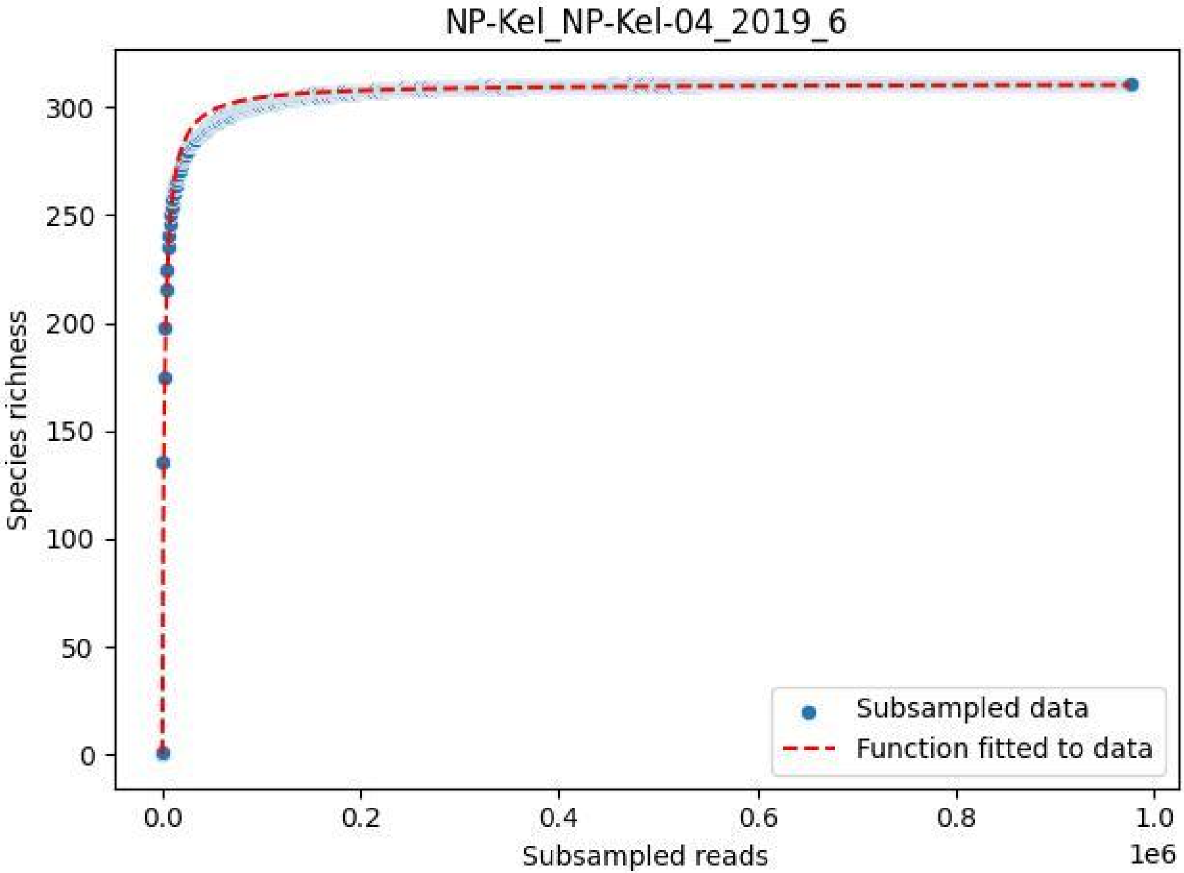
NP-Kel_NP-Kel-04_2019_6
Species richness
Subsampled reads
1e6
0
50
100
150
200
250
300
0.0
0.2
0.4
0.6
0.8
1.0
Subsampled data
Function fitted to data

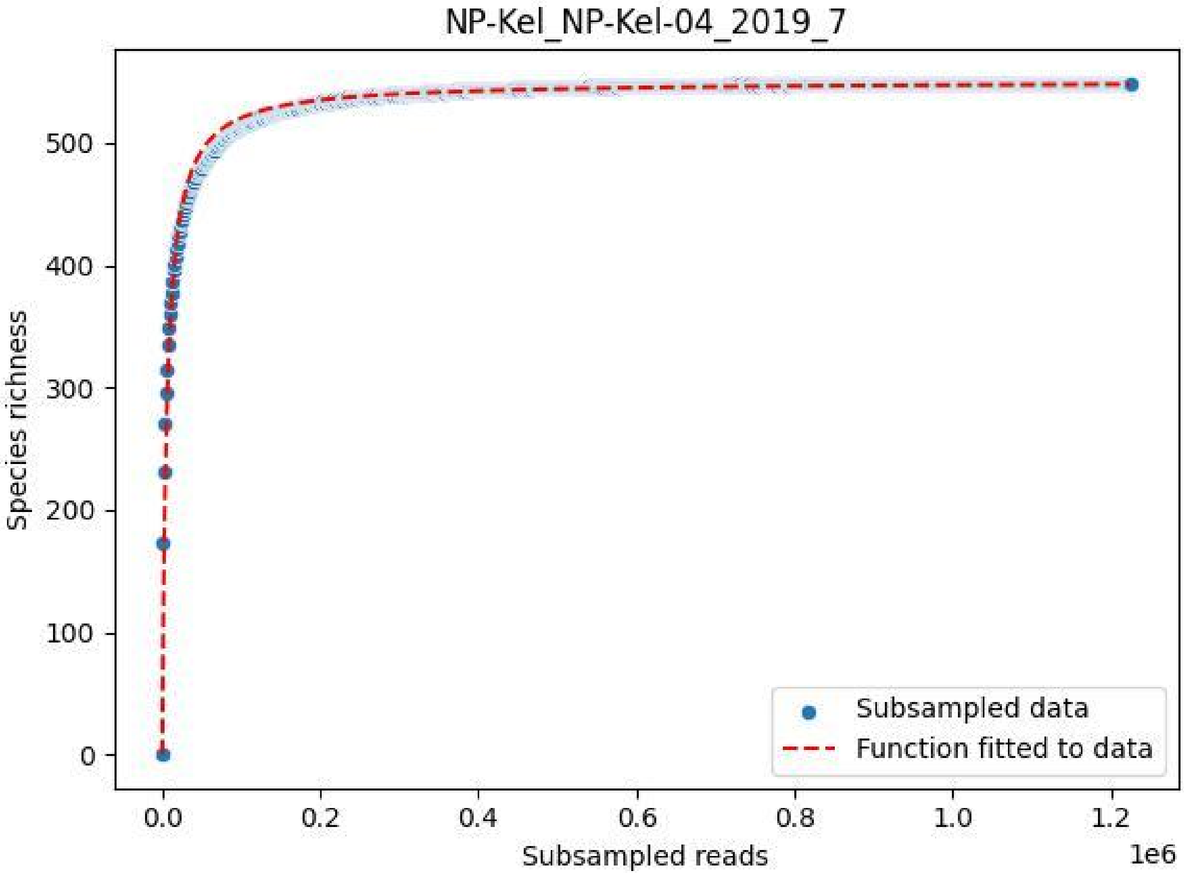

NP-Kel_NP-Kel-04_2019_7
Species richness
Subsampled reads
1e6
0
100
200
300
400
500
0.0
0.2
0.4
0.6
0.8
1.0
1.2
Subsampled data
Function fitted to data

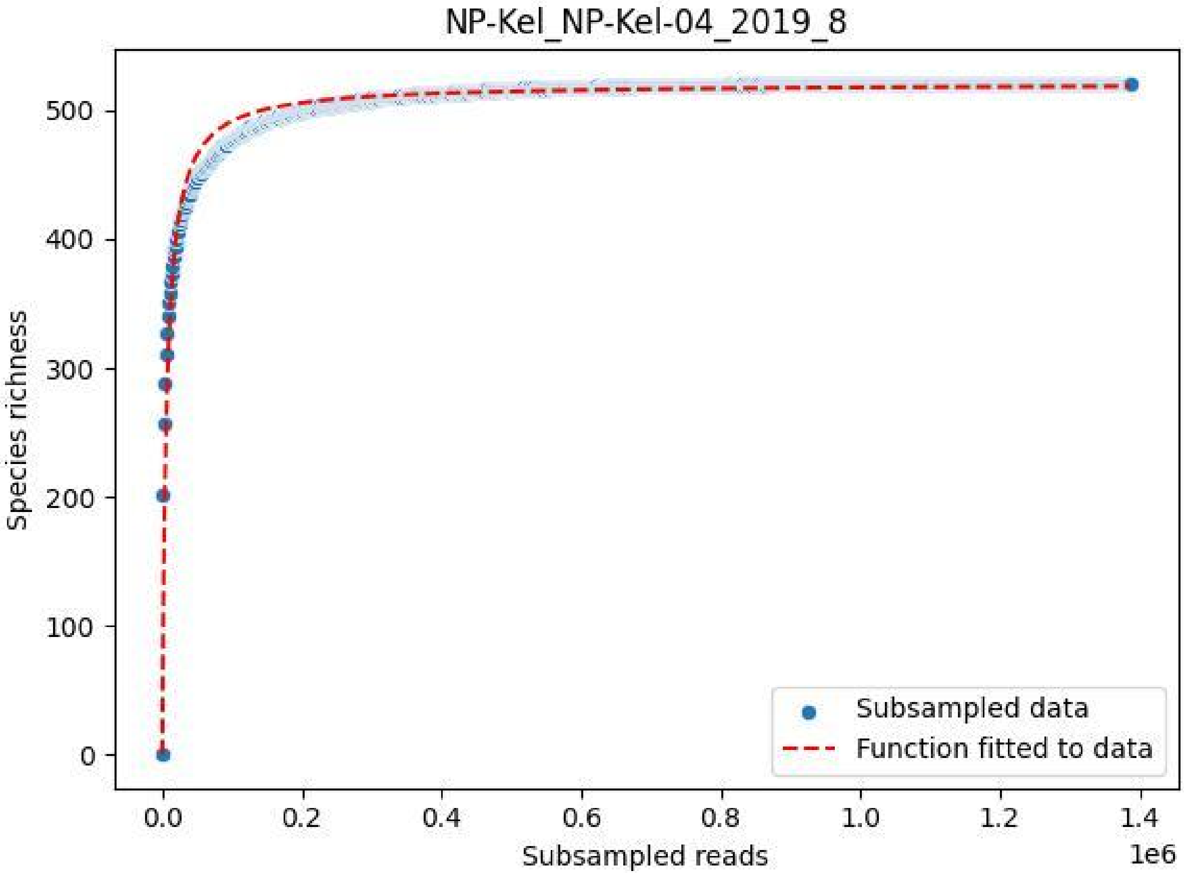

NP-Kel_NP-Kel-04_2019_8
Species richness
Subsampled reads
1e6
0
100
200
300
400
500
0.0
0.2
0.4
0.6
0.8
1.0
1.2
1.4
Subsampled data
Function fitted to data

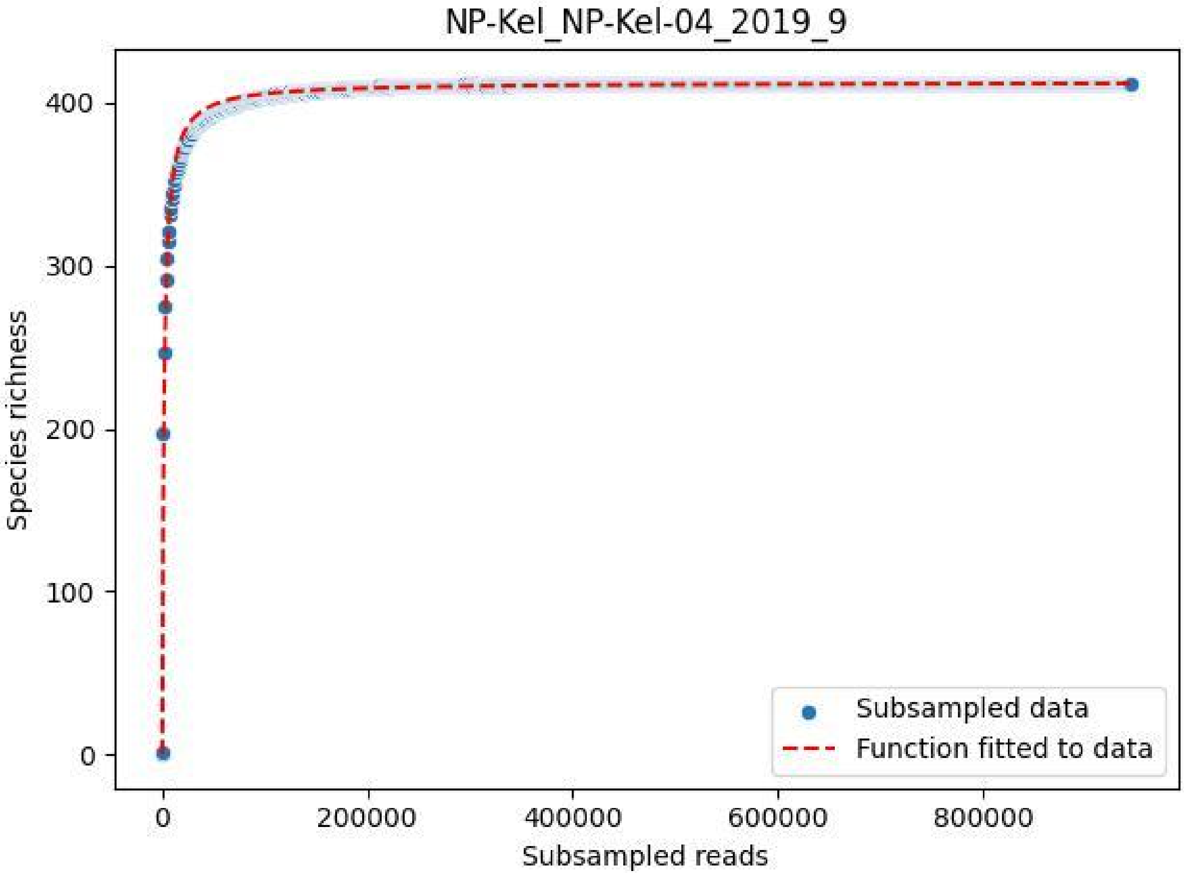
NP-Kel_NP-Kel-04_2019_9
Species richness
400
300
200
100
0
0
200000
400000
600000
800000
Subsampled reads
Subsampled data
Function fitted to data

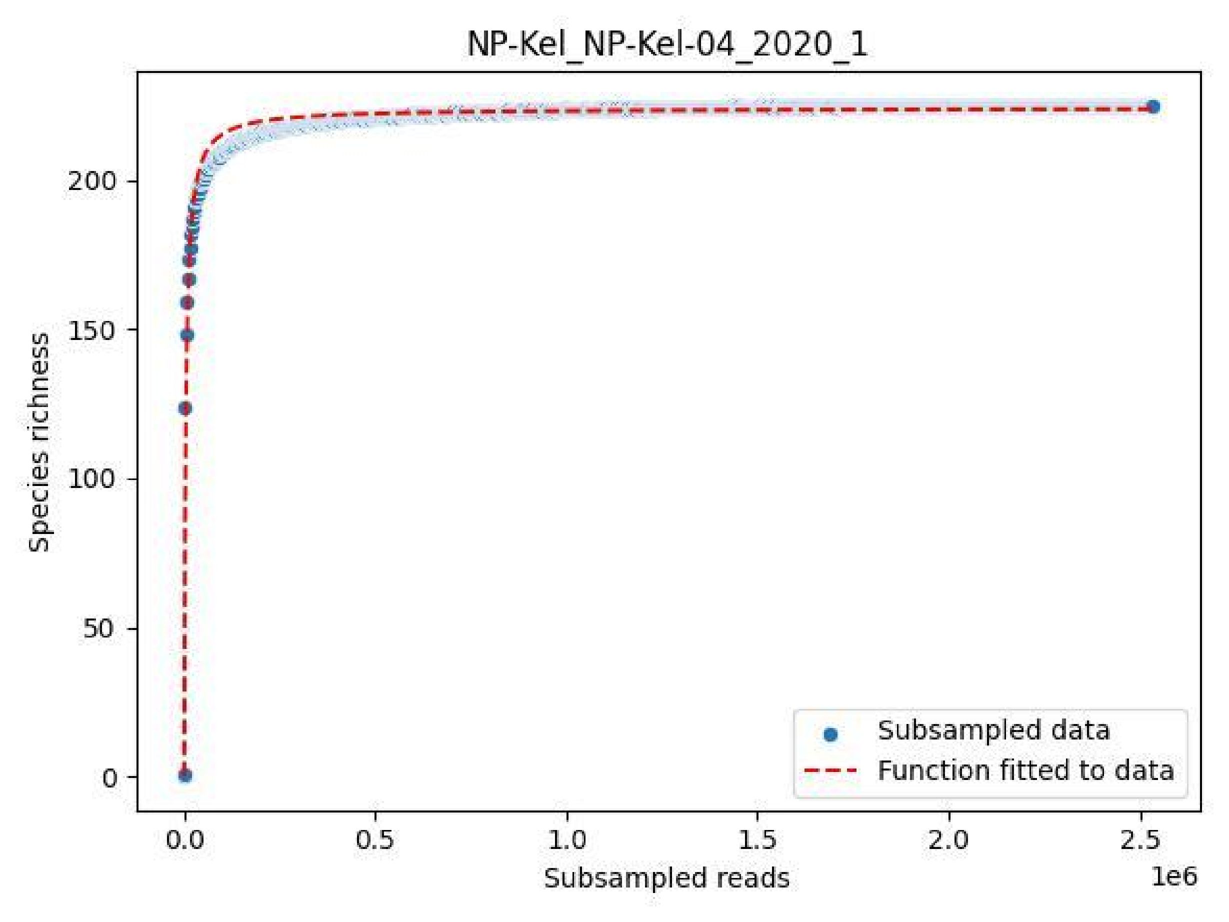
NP-Kel_NP-Kel-04_2020_1
Species richness
Subsampled reads
1e6
0
50
100
150
200
0.0
0.5
1.0
1.5
2.0
2.5
Subsampled data
Function fitted to data

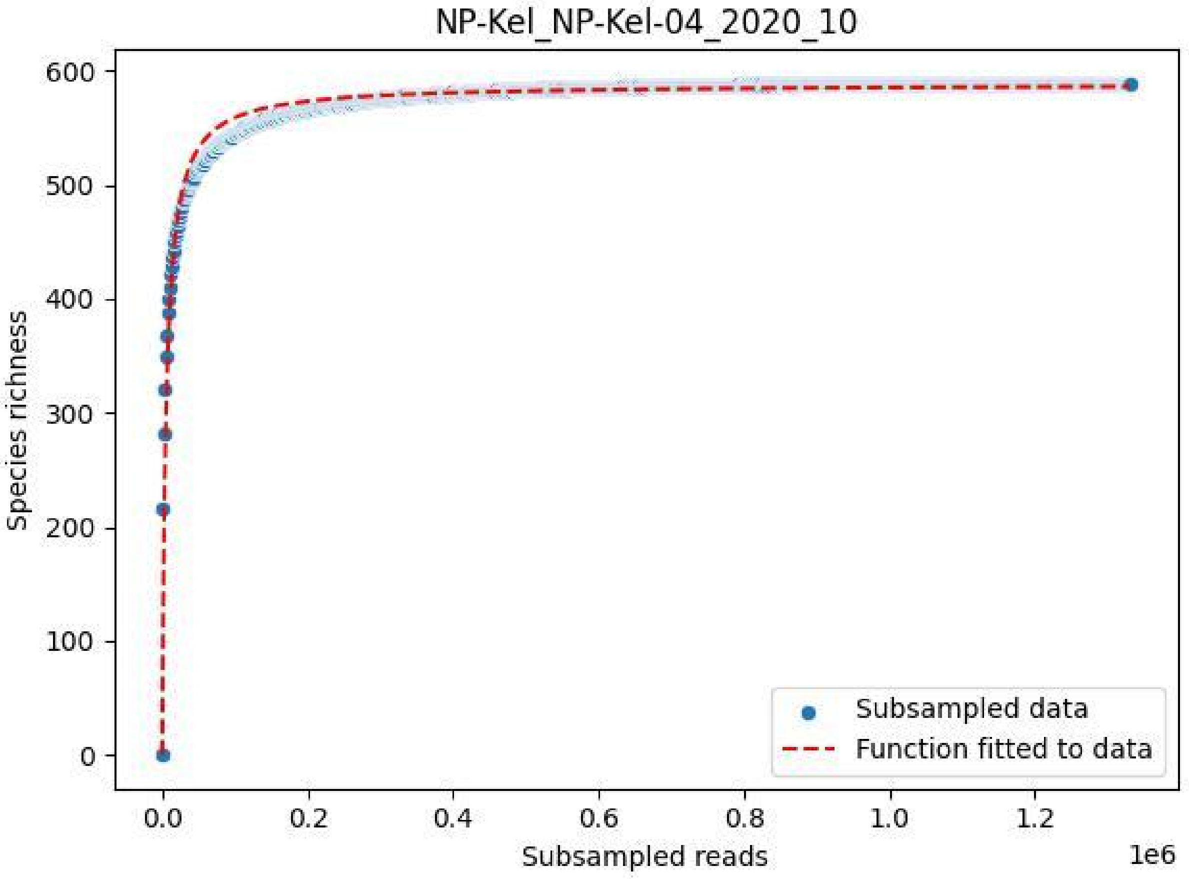

NP-Kel_NP-Kel-04_2020_10
Species richness
Subsampled reads
1e6
0
100
200
300
400
500
600
0.0
0.2
0.4
0.6
0.8
1.0
1.2
Subsampled data
Function fitted to data

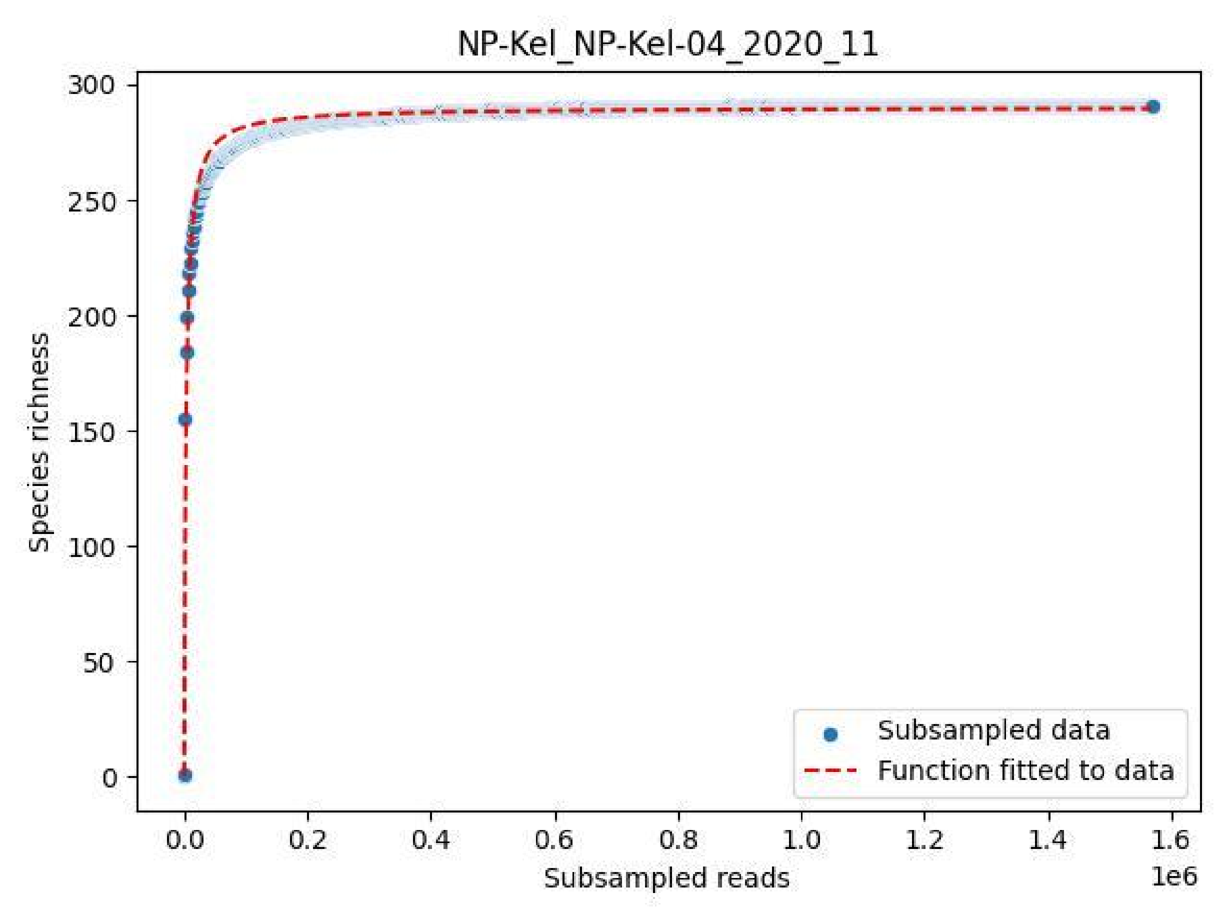

NP-Kel_NP-Kel-04_2020_11
300
250
200
150
100
50
0
Species richness
0.0
0.2
0.4
0.6
0.8
1.0
1.2
1.4
1.6
1e6
Subsampled reads
Subsampled data
Function fitted to data

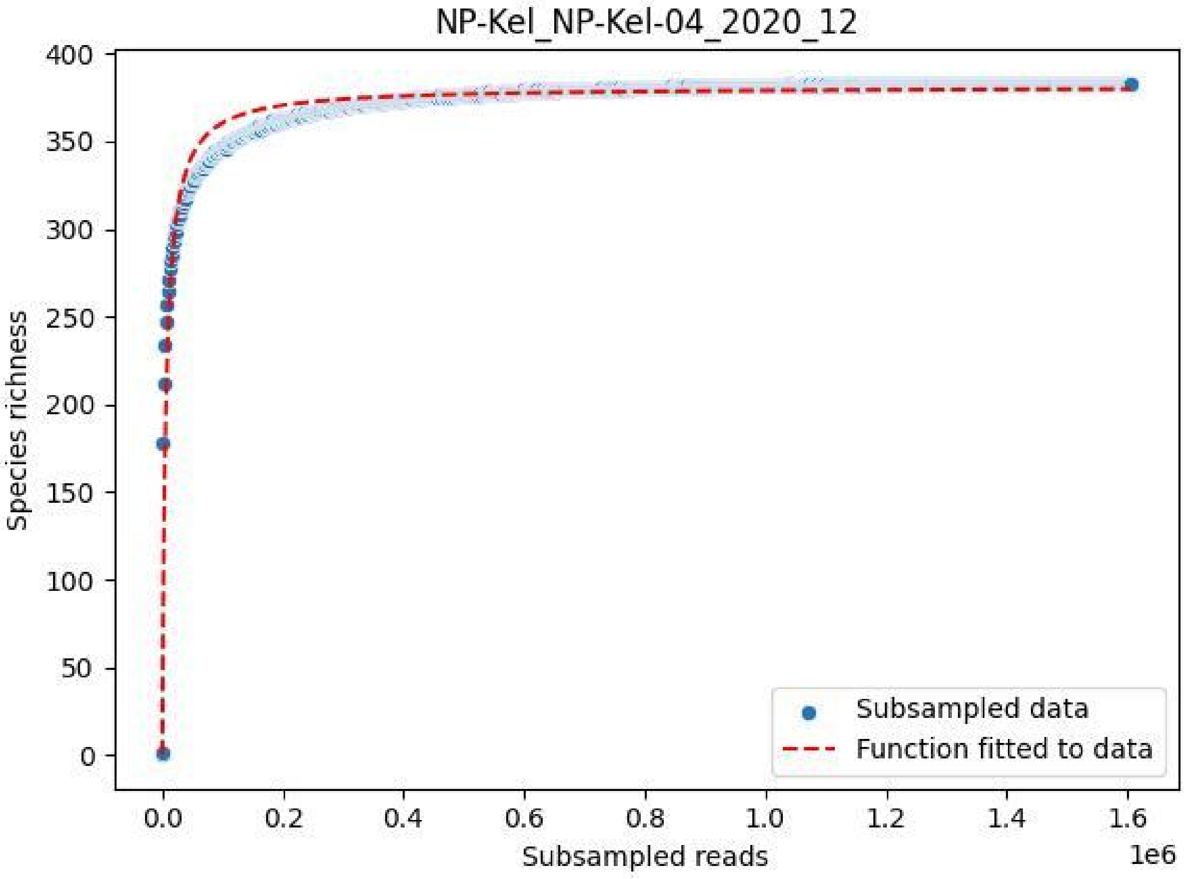

NP-Kel_NP-Kel-04_2020_12
Species richness
Subsampled reads
1e6
0
50
100
150
200
250
300
350
400
0.0
0.2
0.4
0.6
0.8
1.0
1.2
1.4
1.6
Subsampled data
Function fitted to data

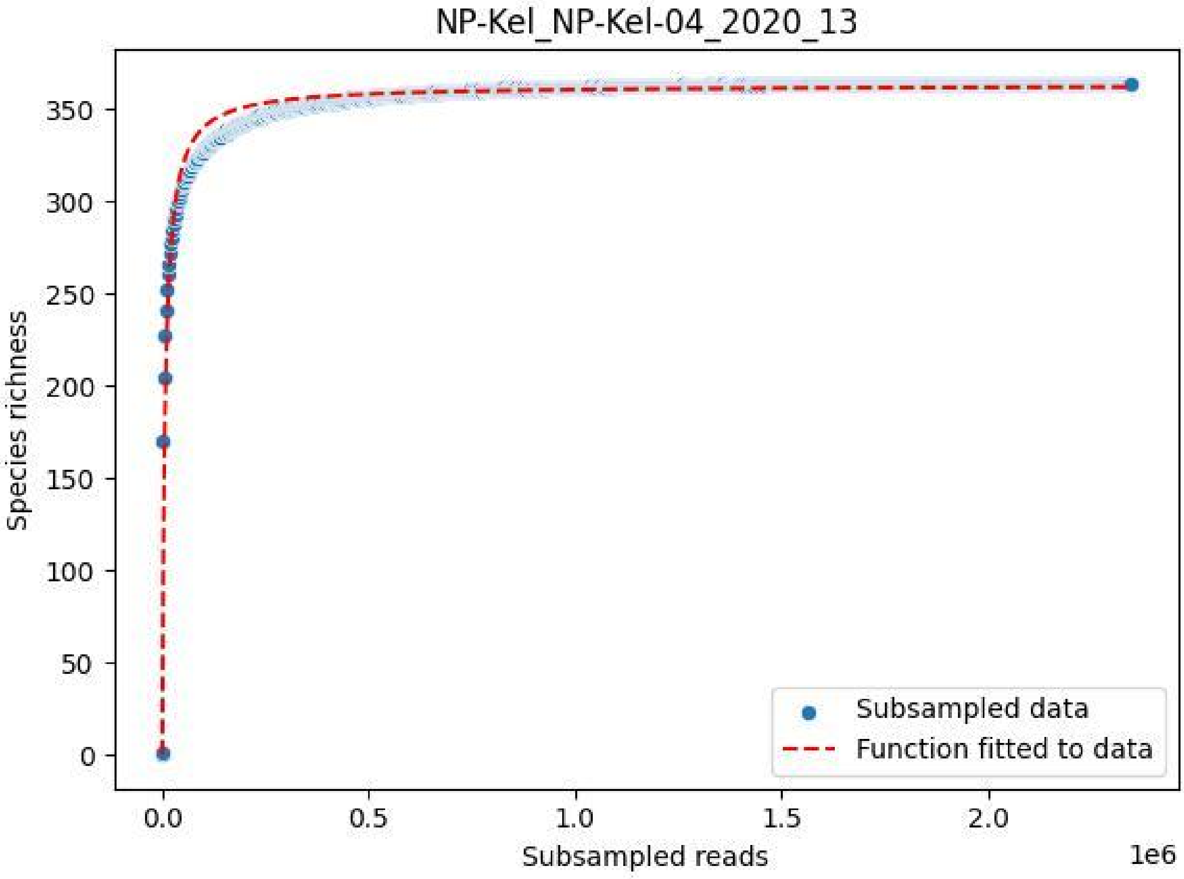

NP-Kel_NP-Kel-04_2020_13
Species richness
350
300
250
200
150
100
50
0
0.0
0.5
1.0
1.5
2.0
1e6
Subsampled reads
Subsampled data
Function fitted to data

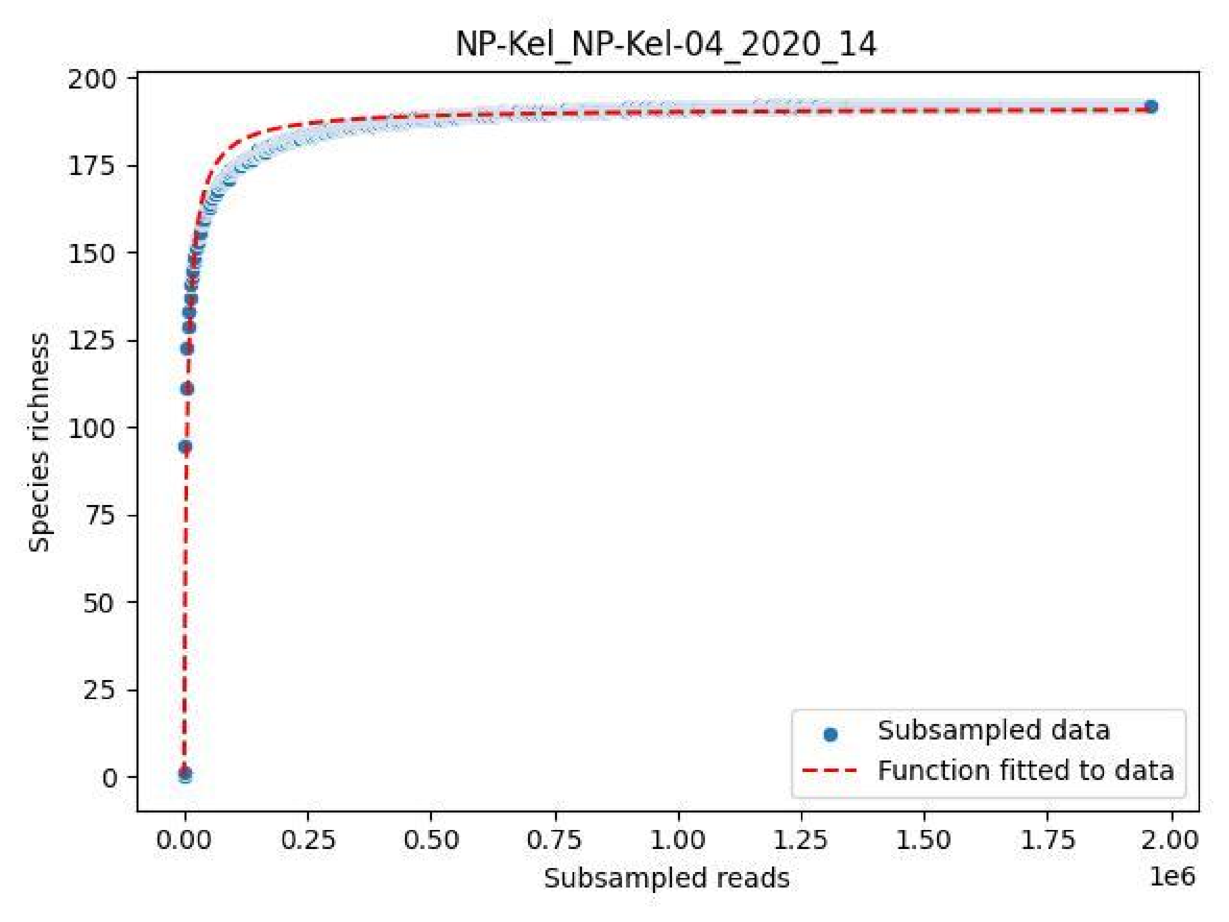

NP-Kel_NP-Kel-04_2020_14
Species richness
Subsampled reads
1e6
0
25
50
75
100
125
150
175
200
0.00
0.25
0.50
0.75
1.00
1.25
1.50
1.75
2.00
Subsampled data
Function fitted to data

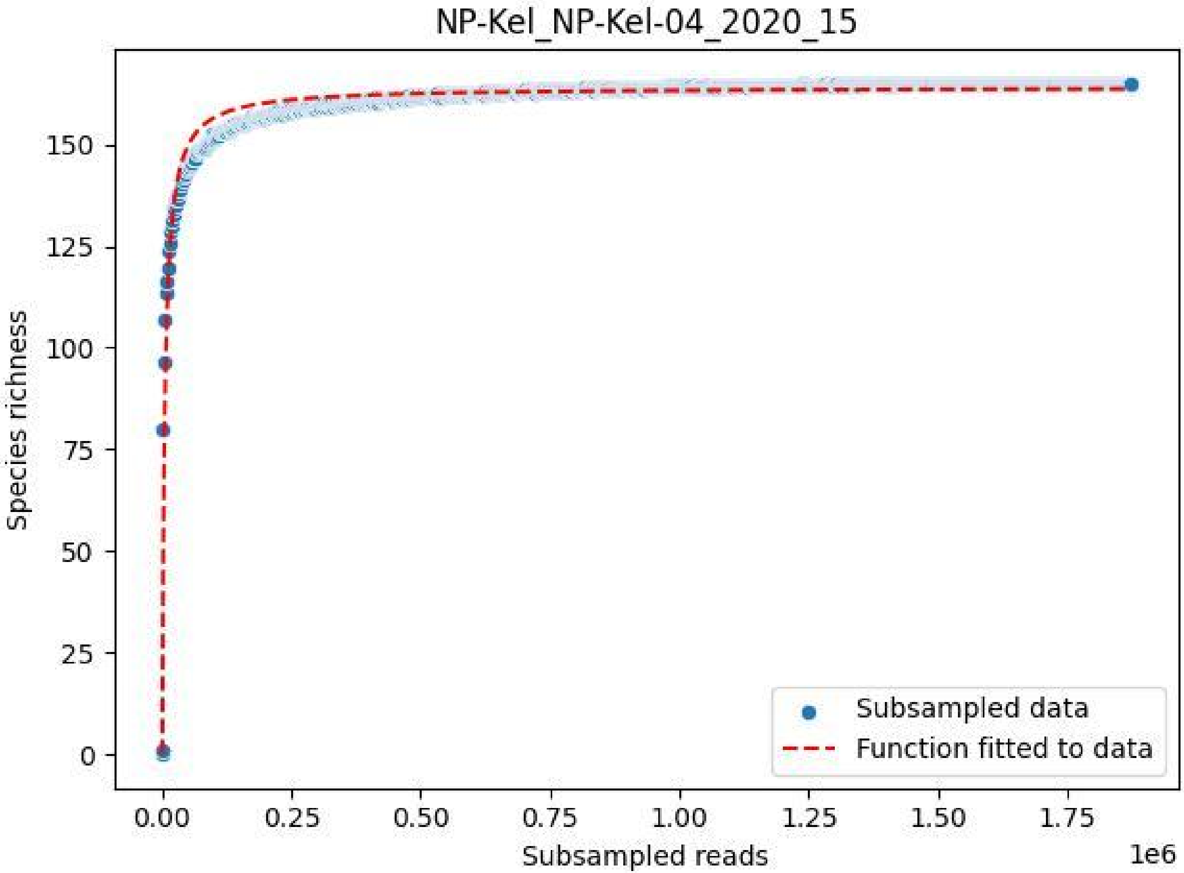
NP-Kel_NP-Kel-04_2020_15
Species richness
0
25
50
75
100
125
150
0.00
0.25
0.50
0.75
1.00
1.25
1.50
1.75
1e6
Subsampled reads
Subsampled data
Function fitted to data

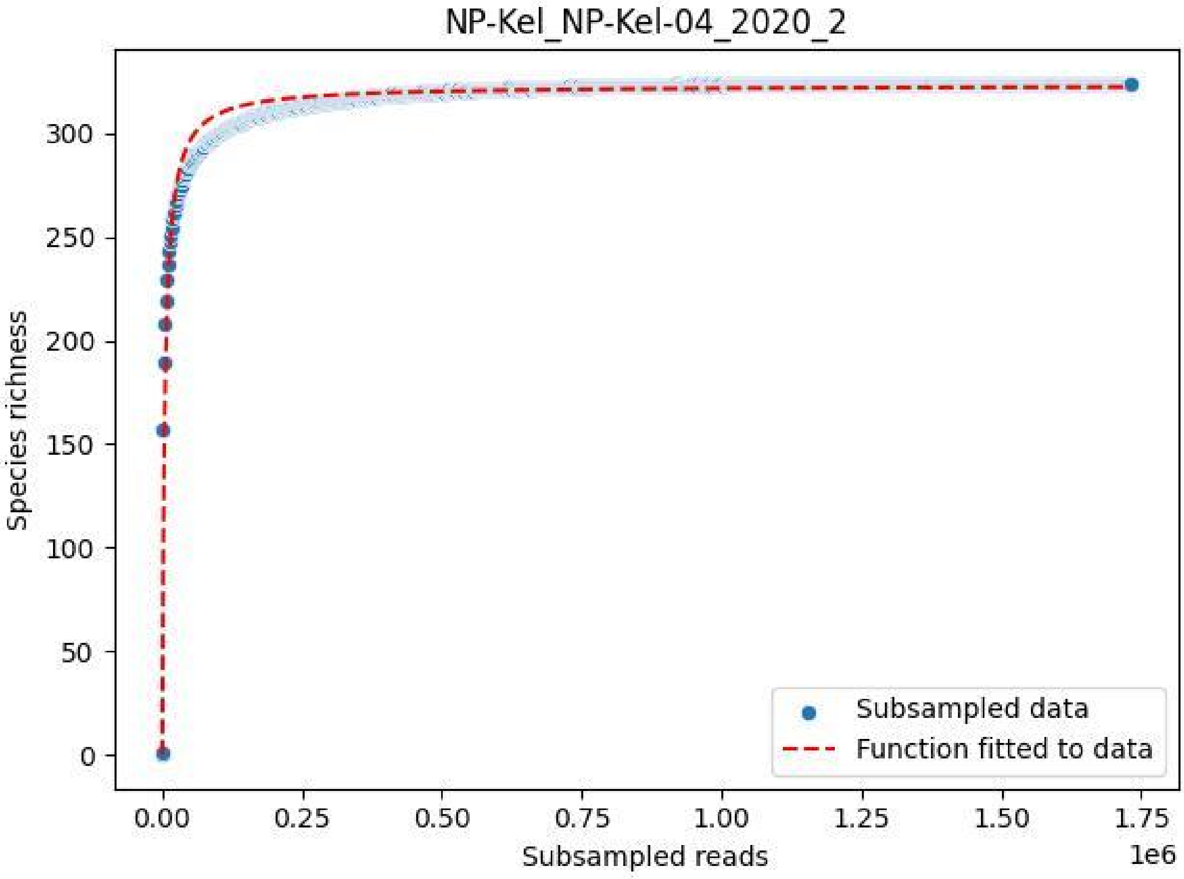
NP-Kel_NP-Kel-04_2020_2
Species richness
Subsampled reads
1e6
0
50
100
150
200
250
300
0.00
0.25
0.50
0.75
1.00
1.25
1.50
1.75
Subsampled data
Function fitted to data

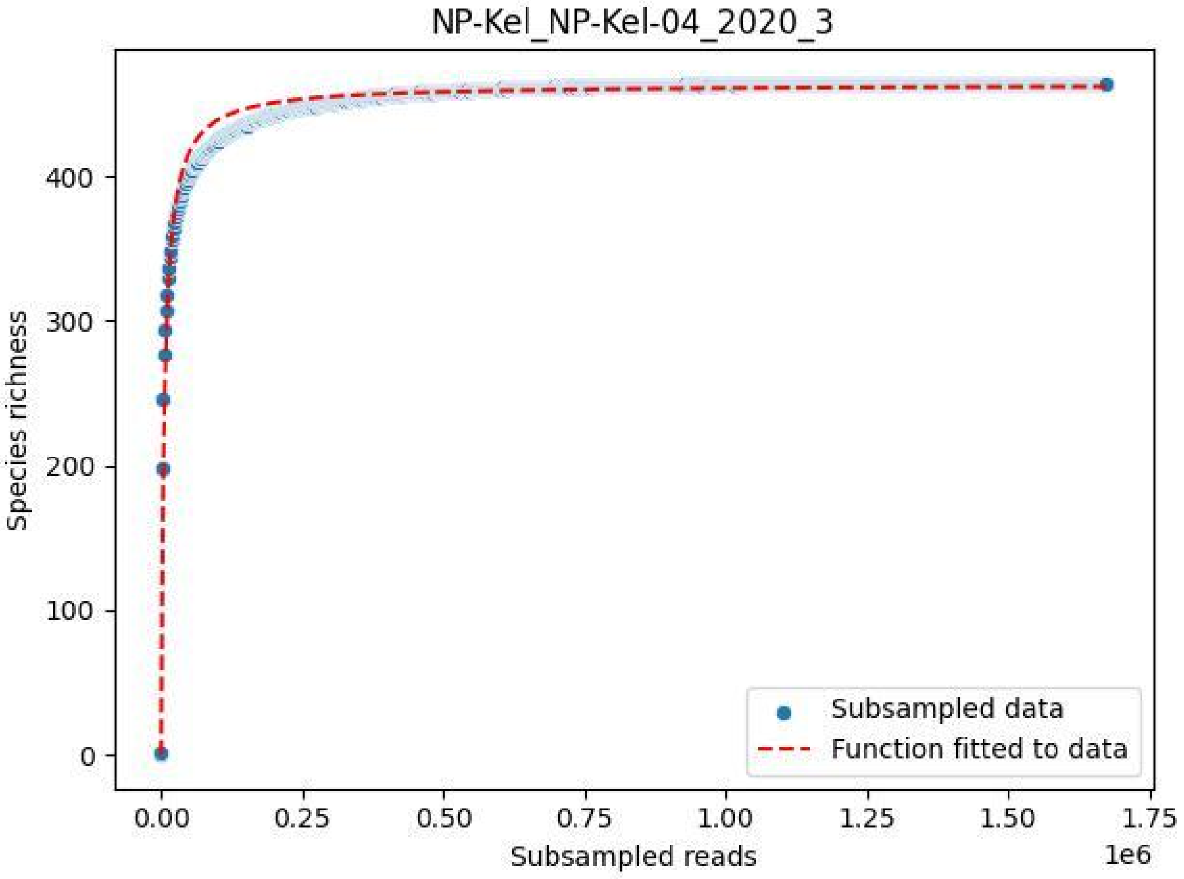

NP-Kel_NP-Kel-04_2020_3
Species richness
Subsampled reads
1e6
0
100
200
300
400
0.00
0.25
0.50
0.75
1.00
1.25
1.50
1.75
Subsampled data
Function fitted to data

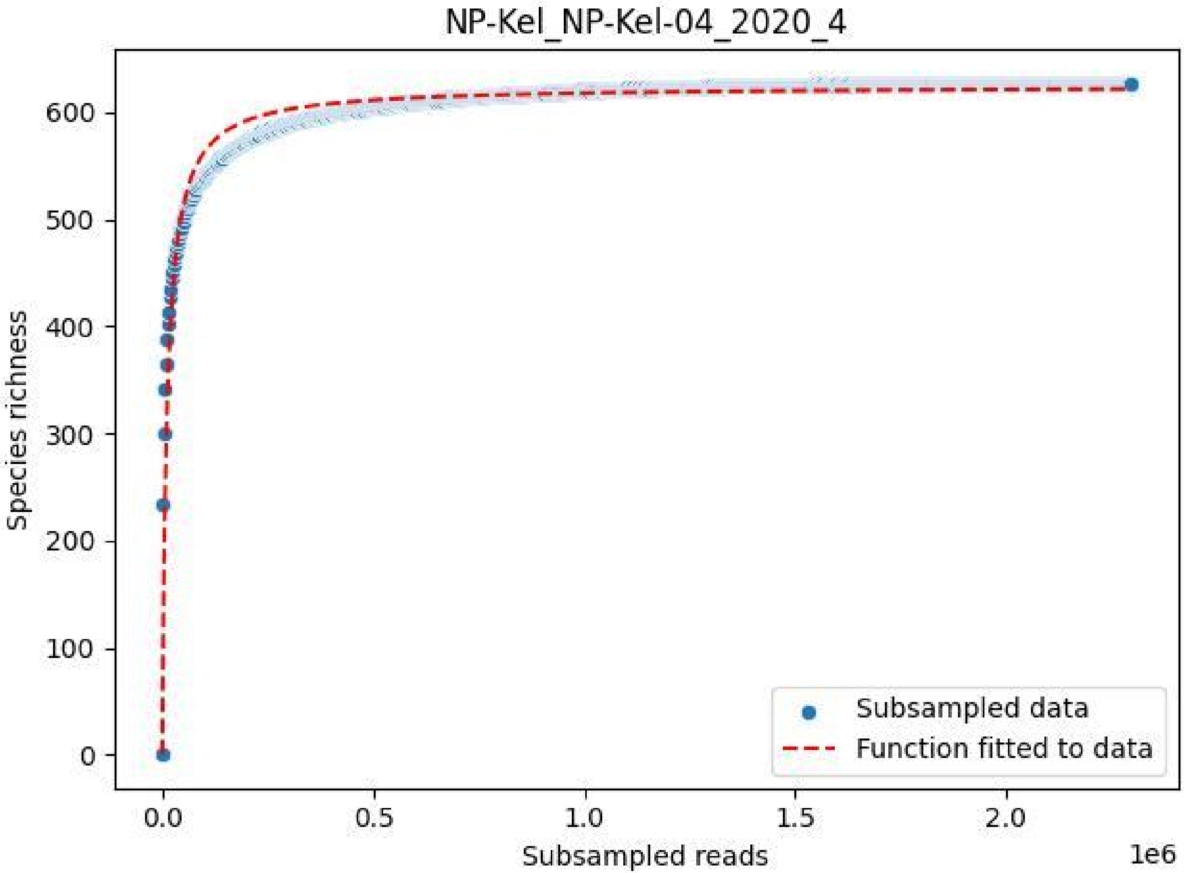

NP-Kel_NP-Kel-04_2020_4
Species richness
0
100
200
300
400
500
600
0.0
0.5
1.0
1.5
2.0
1e6
Subsampled reads
Subsampled data
Function fitted to data

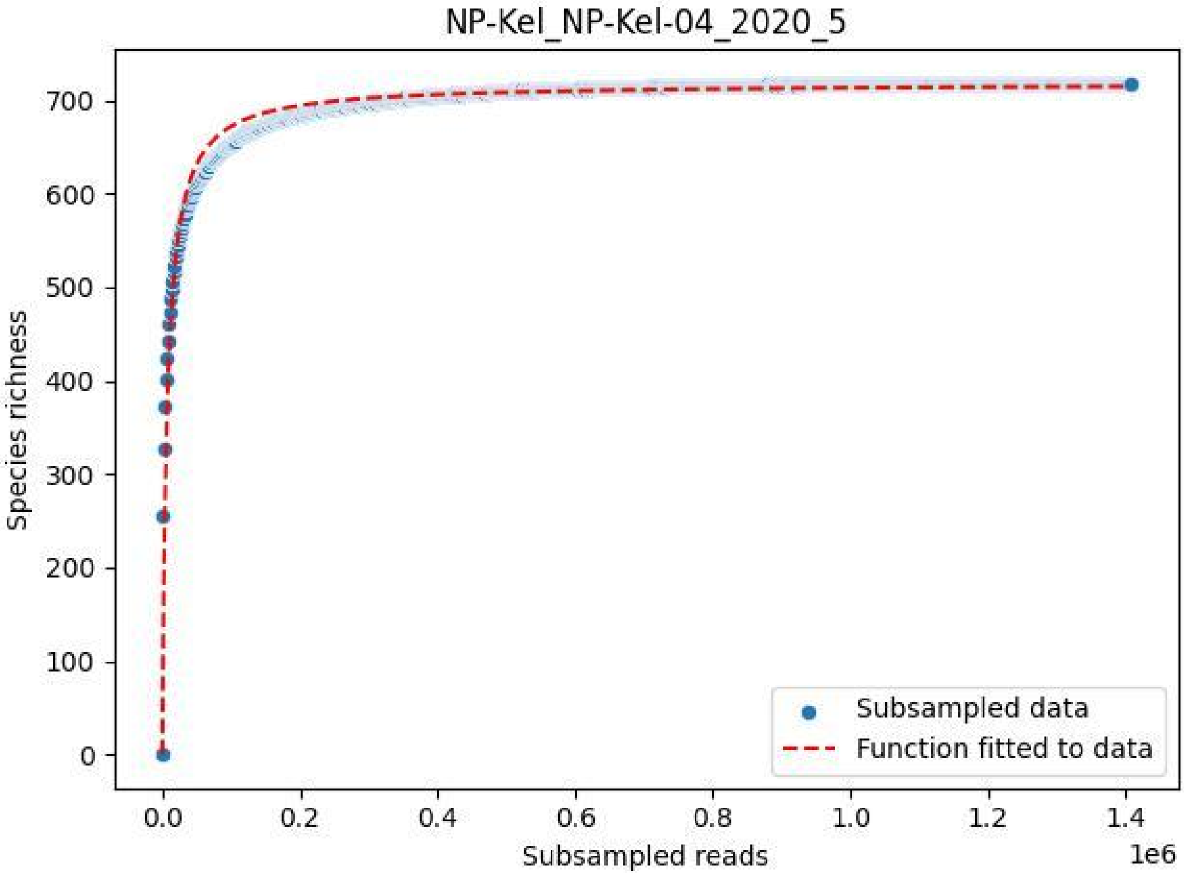
NP-Kel_NP-Kel-04_2020_5
Species richness
Subsampled reads
1e6
Subsampled data
Function fitted to data
0
100
200
300
400
500
600
700
0.0
0.2
0.4
0.6
0.8
1.0
1.2
1.4

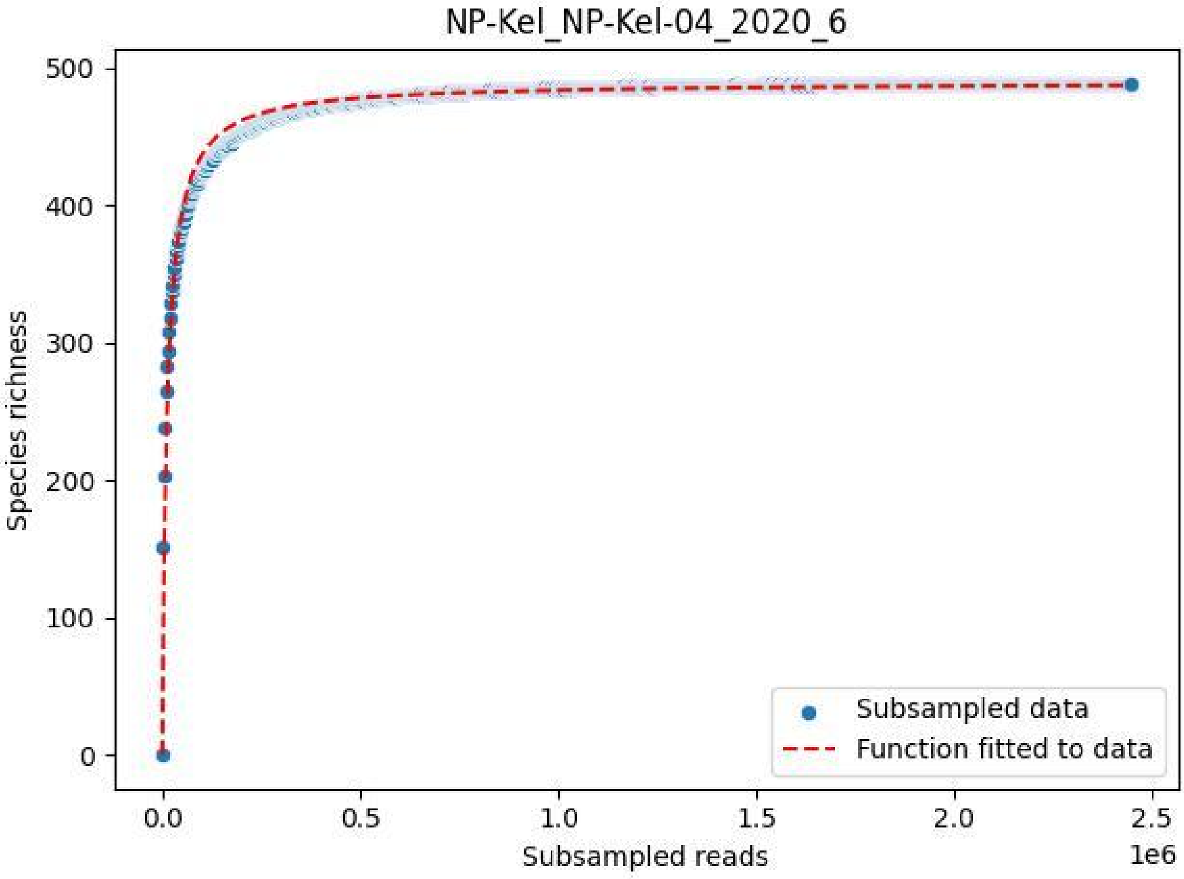

NP-Kel_NP-Kel-04_2020_6
Species richness
Subsampled reads
1e6
0
100
200
300
400
500
0.0
0.5
1.0
1.5
2.0
2.5
Subsampled data
Function fitted to data

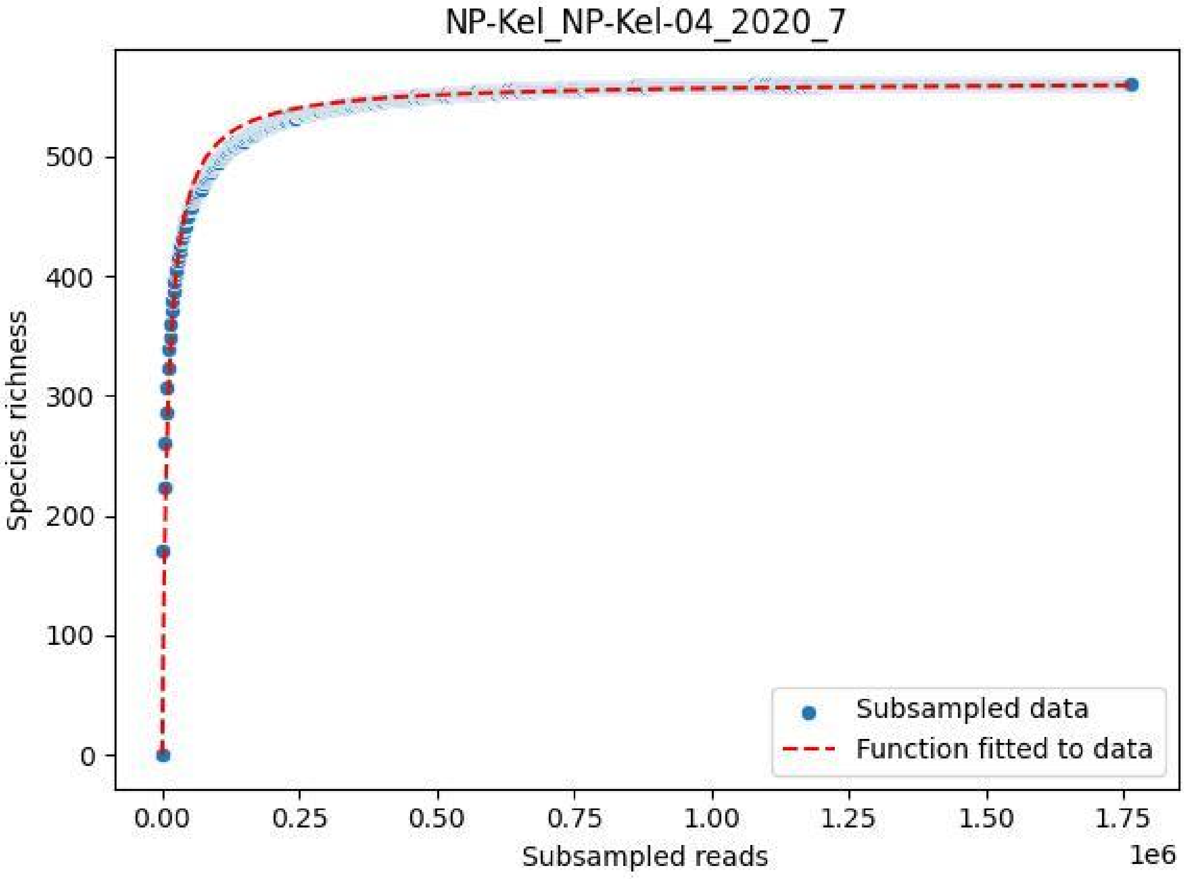
NP-Kel_NP-Kel-04_2020_7
Species richness
0
100
200
300
400
500
0.00
0.25
0.50
0.75
1.00
1.25
1.50
1.75
1e6
Subsampled reads
Subsampled data
Function fitted to data

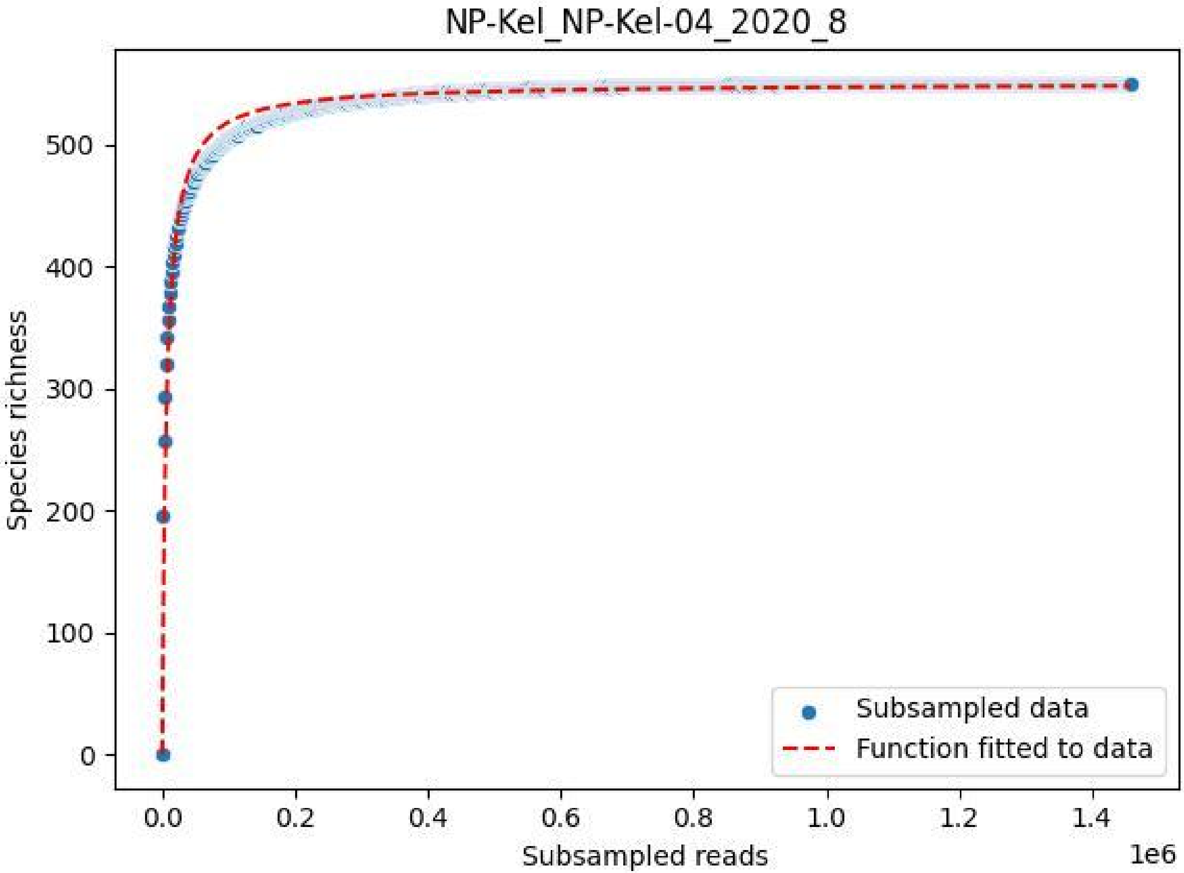
NP-Kel_NP-Kel-04_2020_8
Species richness
0
100
200
300
400
500
0.0
0.2
0.4
0.6
0.8
1.0
1.2
1.4
1e6
Subsampled reads
Subsampled data
Function fitted to data

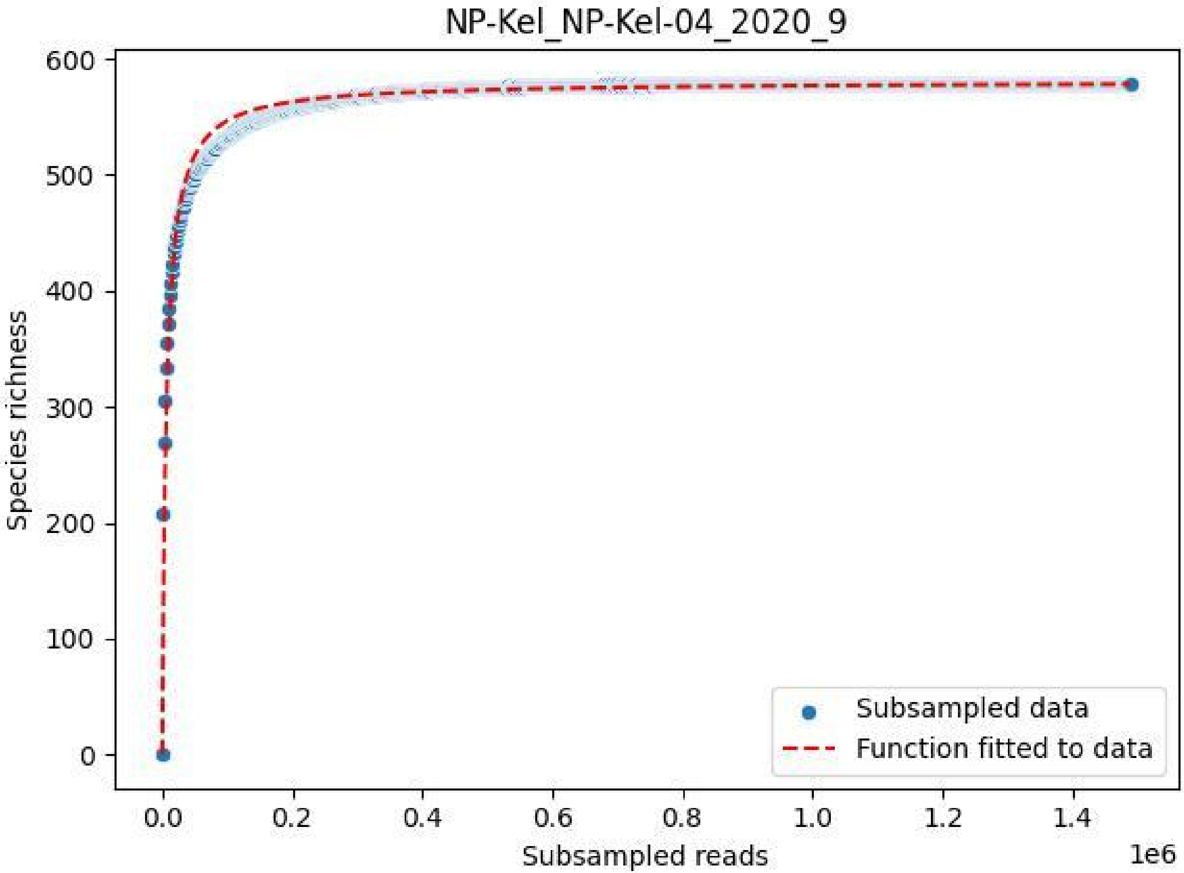

NP-Kel_NP-Kel-04_2020_9
Species richness
0
100
200
300
400
500
600
Subsampled reads
0.0
0.2
0.4
0.6
0.8
1.0
1.2
1.4
1e6
Subsampled data
Function fitted to data

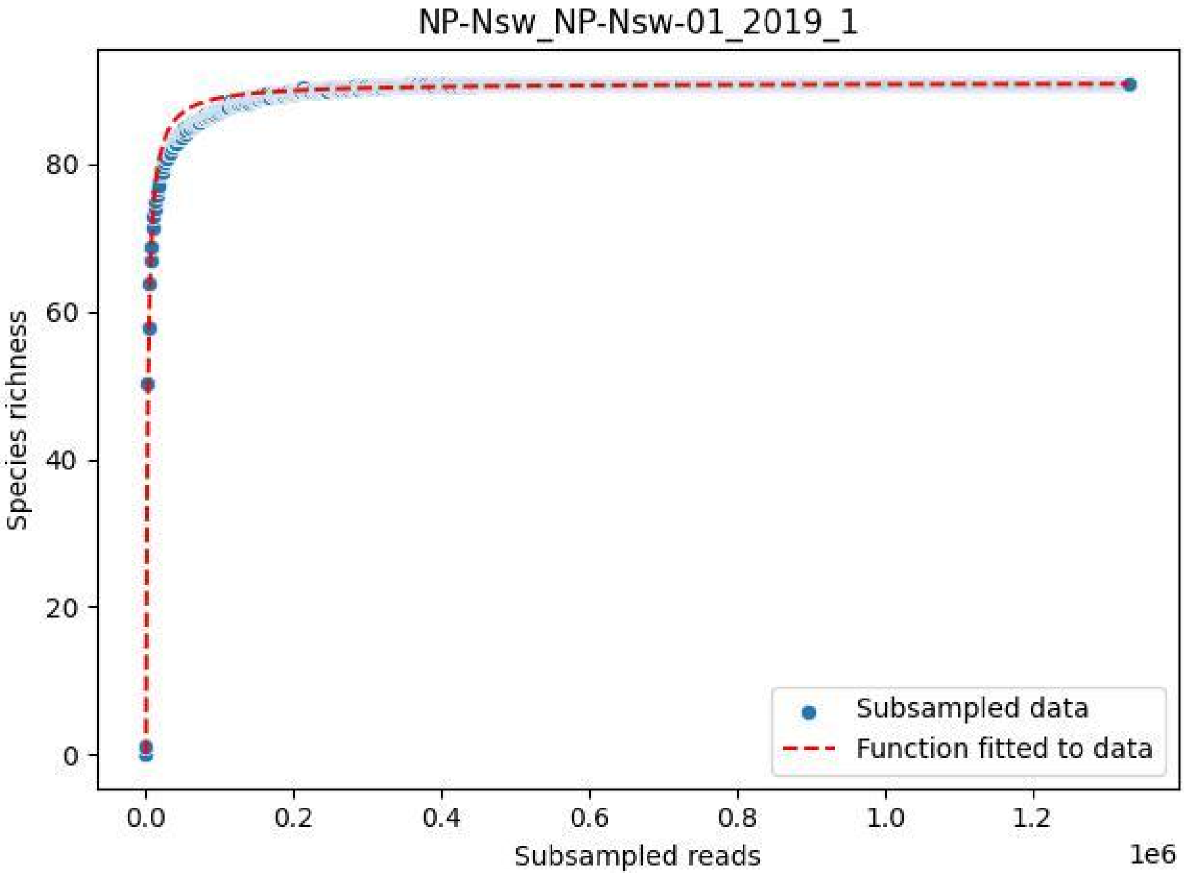

NP-Nsw_NP-Nsw-01_2019_1
Species richness
Subsampled reads
1e6
0
20
40
60
80
0.0
0.2
0.4
0.6
0.8
1.0
1.2
Subsampled data
Function fitted to data

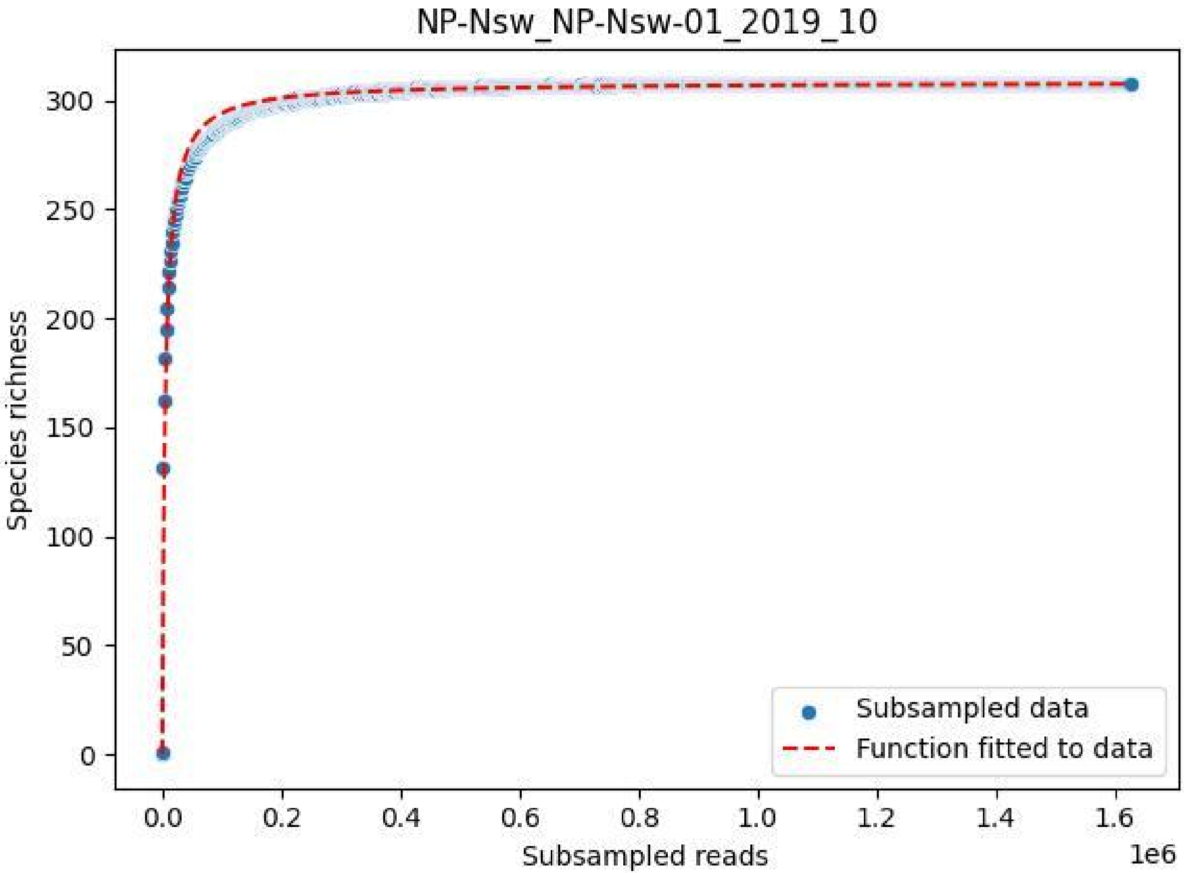
NP-Nsw_NP-Nsw-01_2019_10
Species richness
Subsampled reads
1e6
0
50
100
150
200
250
300
0.0
0.2
0.4
0.6
0.8
1.0
1.2
1.4
1.6
Subsampled data
Function fitted to data

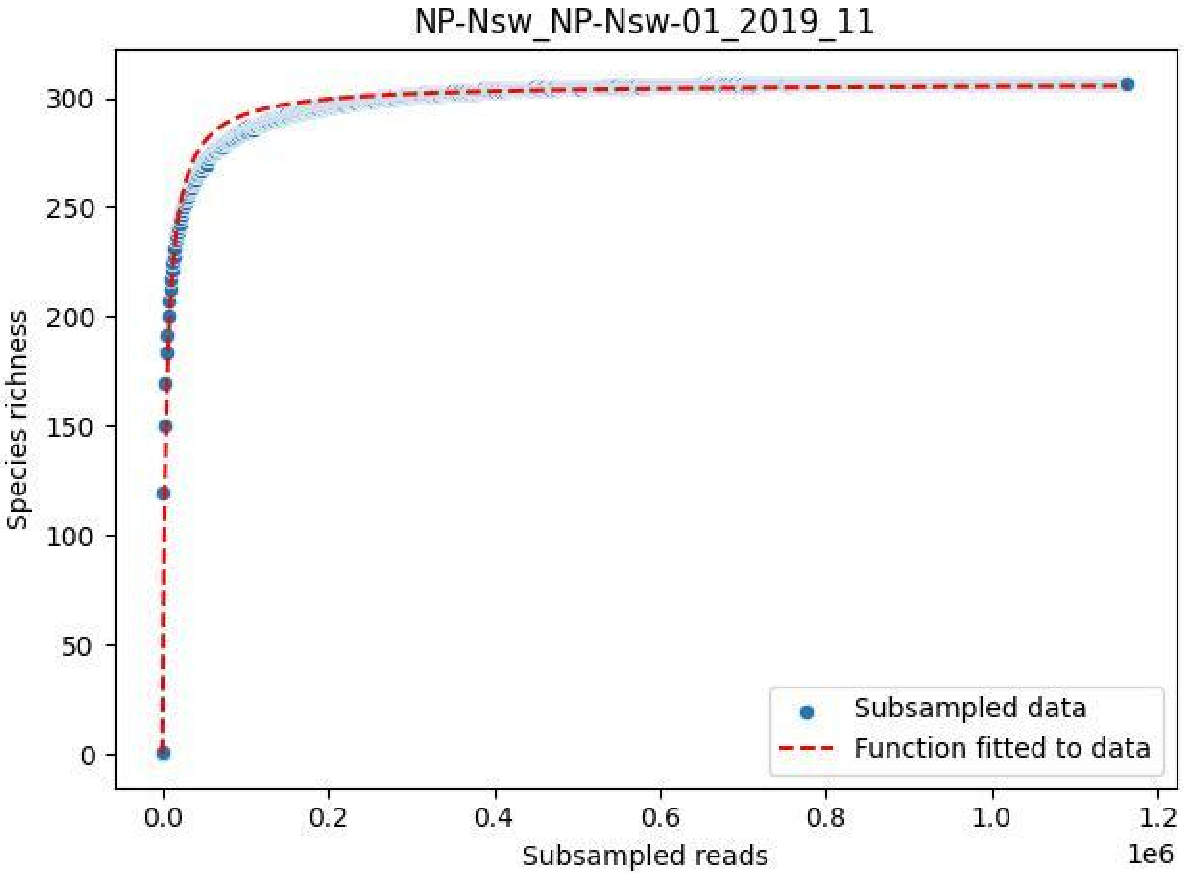

NP-Nsw_NP-Nsw-01_2019_11
Species richness
Subsampled reads
1e6
0
50
100
150
200
250
300
0.0
0.2
0.4
0.6
0.8
1.0
1.2
Subsampled data
Function fitted to data

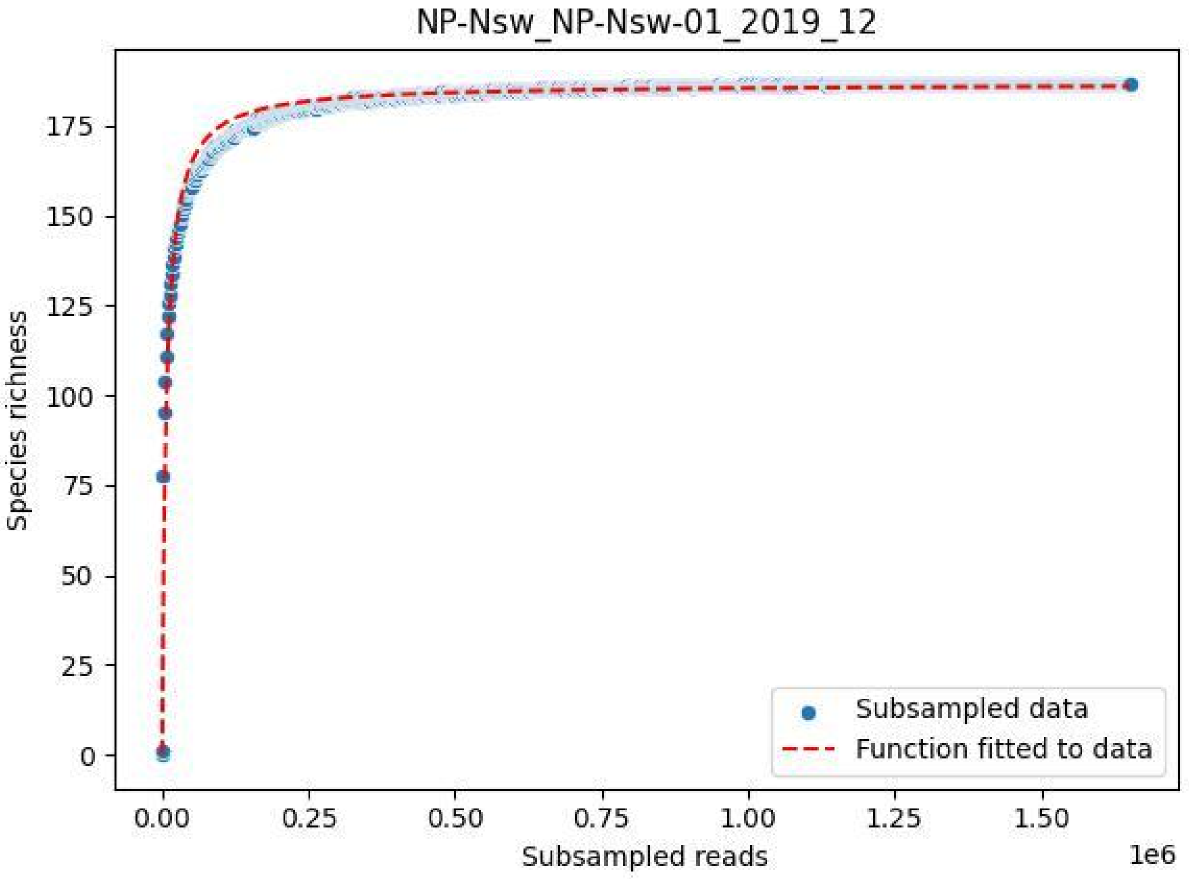
NP-Nsw_NP-Nsw-01_2019_12
Species richness
Subsampled reads
1e6
0
25
50
75
100
125
150
175
0.00
0.25
0.50
0.75
1.00
1.25
1.50
Subsampled data
Function fitted to data

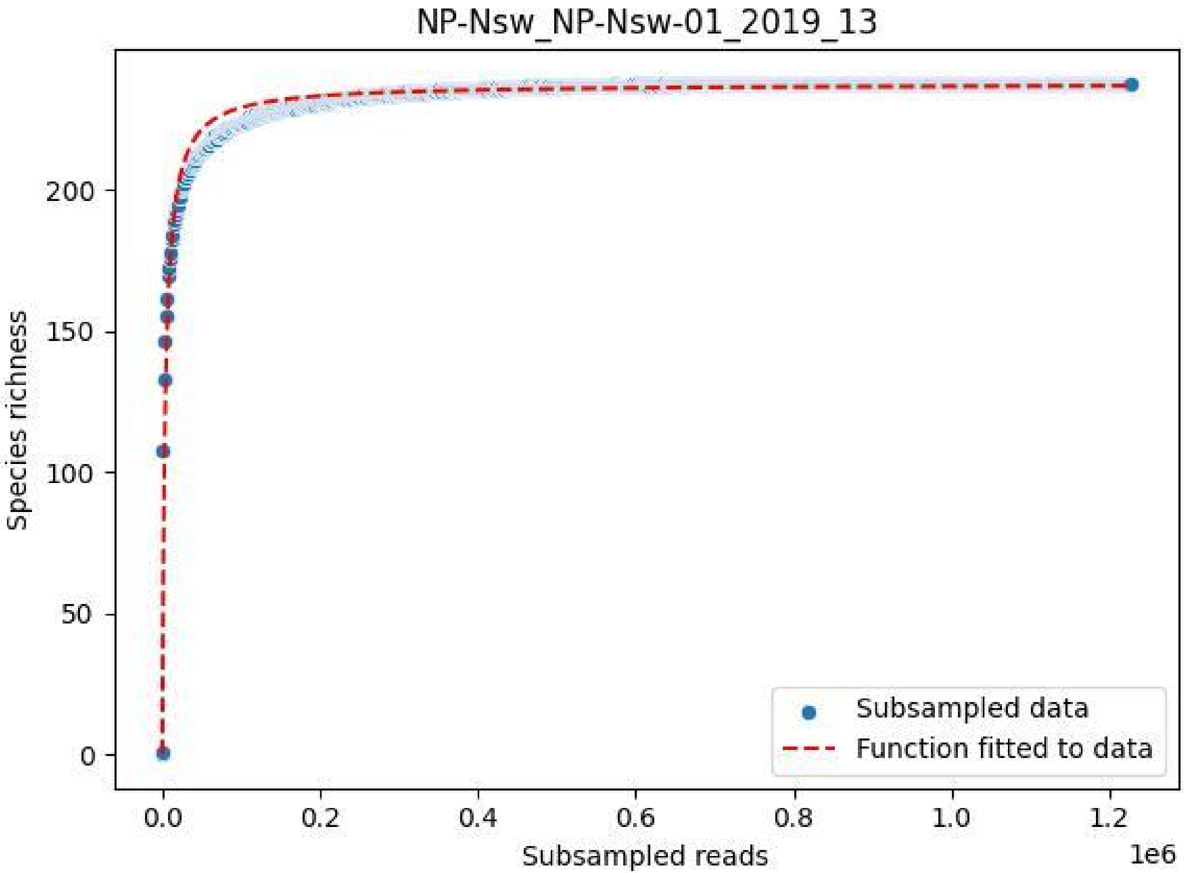
NP-Nsw_NP-Nsw-01_2019_13
Species richness
Subsampled reads
1e6
0
50
100
150
200
0.0
0.2
0.4
0.6
0.8
1.0
1.2
Subsampled data
Function fitted to data

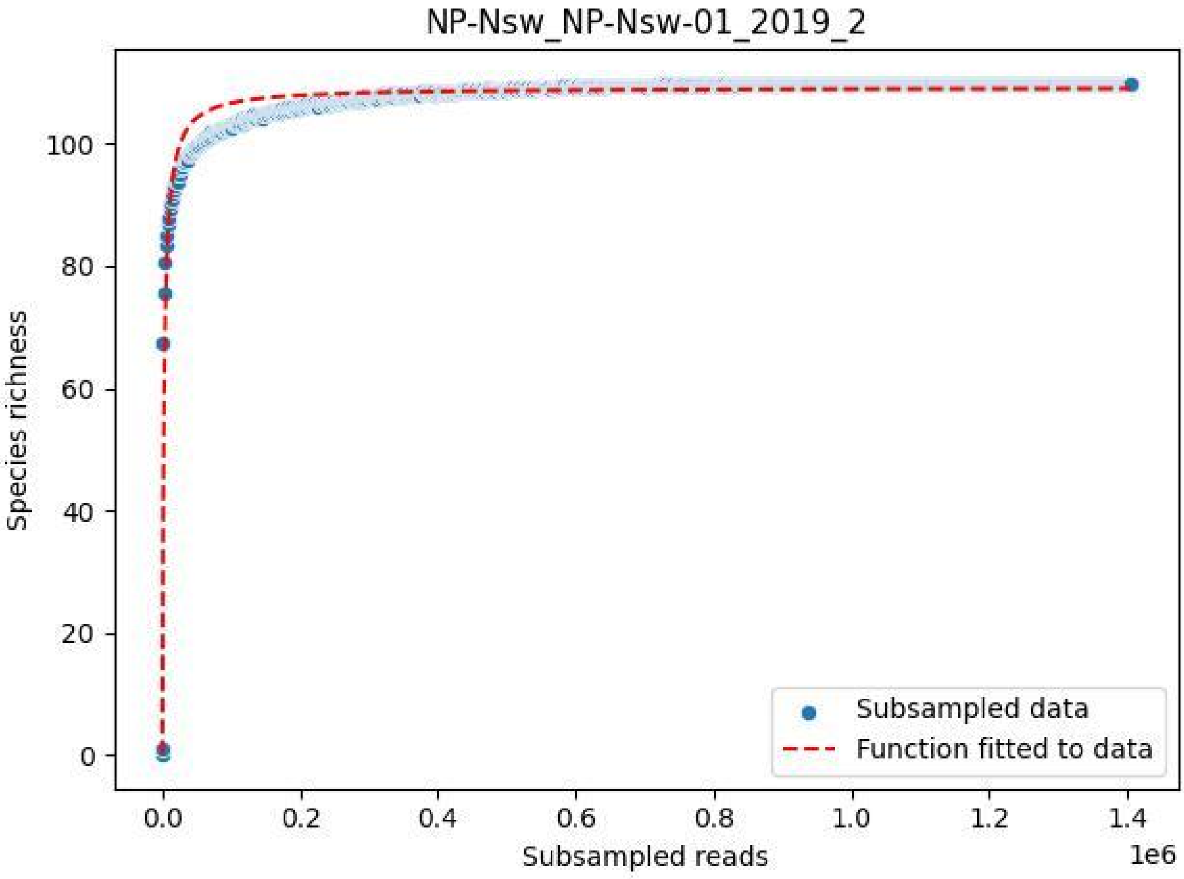
NP-Nsw_NP-Nsw-01_2019_2
Species richness
Subsampled reads
1e6
0
20
40
60
80
100
0.0
0.2
0.4
0.6
0.8
1.0
1.2
1.4
Subsampled data
Function fitted to data

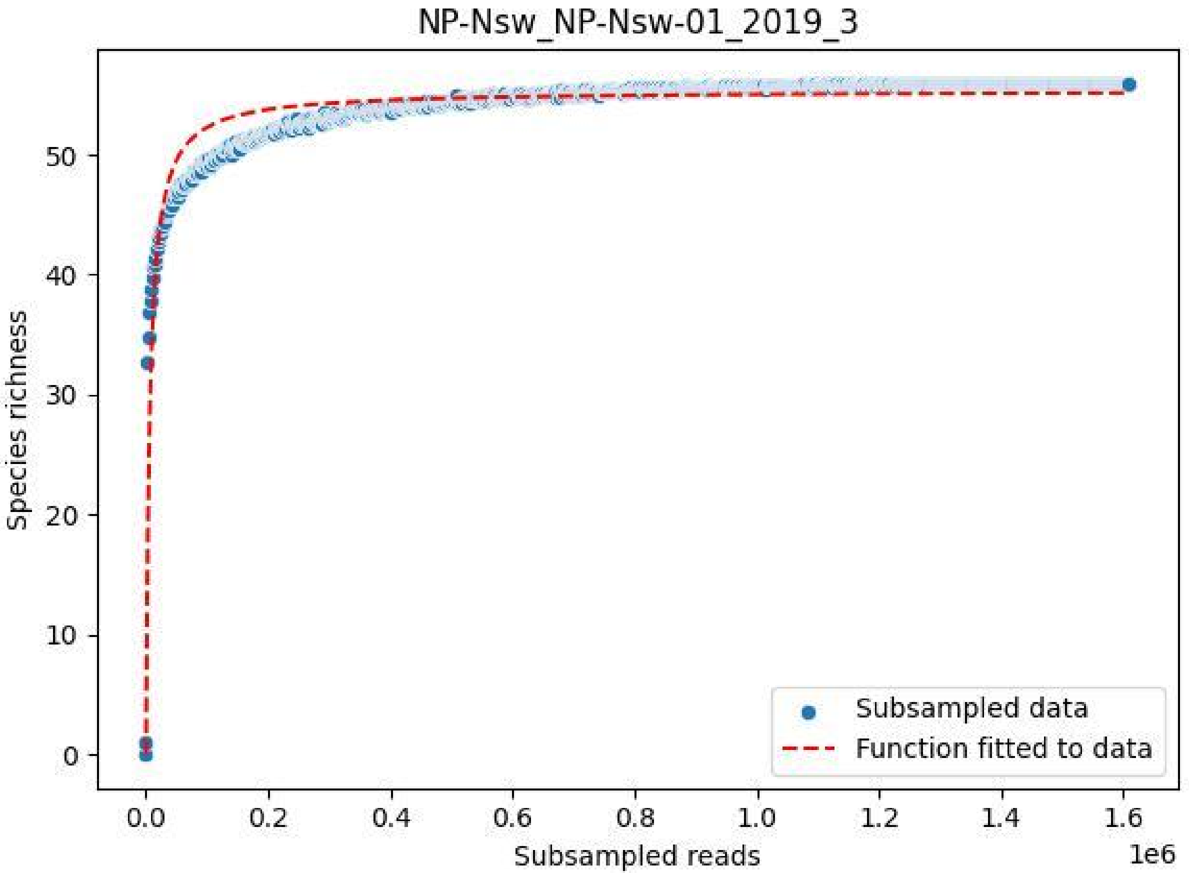
NP-Nsw_NP-Nsw-01_2019_3
Species richness
Subsampled reads
1e6
0
10
20
30
40
50
0.0
0.2
0.4
0.6
0.8
1.0
1.2
1.4
1.6
Subsampled data
Function fitted to data

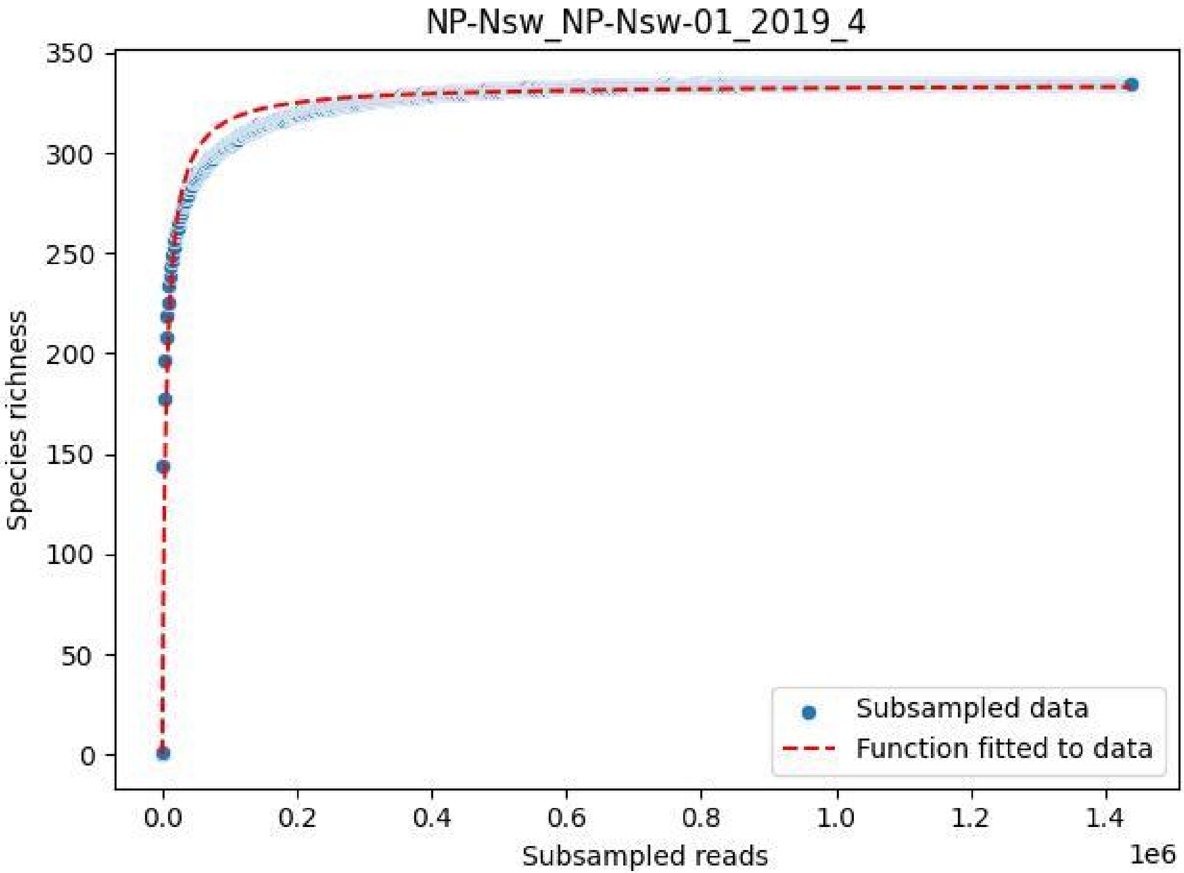

NP-Nsw_NP-Nsw-01_2019_4
Species richness
Subsampled reads
1e6
0
50
100
150
200
250
300
350
0.0
0.2
0.4
0.6
0.8
1.0
1.2
1.4
Subsampled data
Function fitted to data

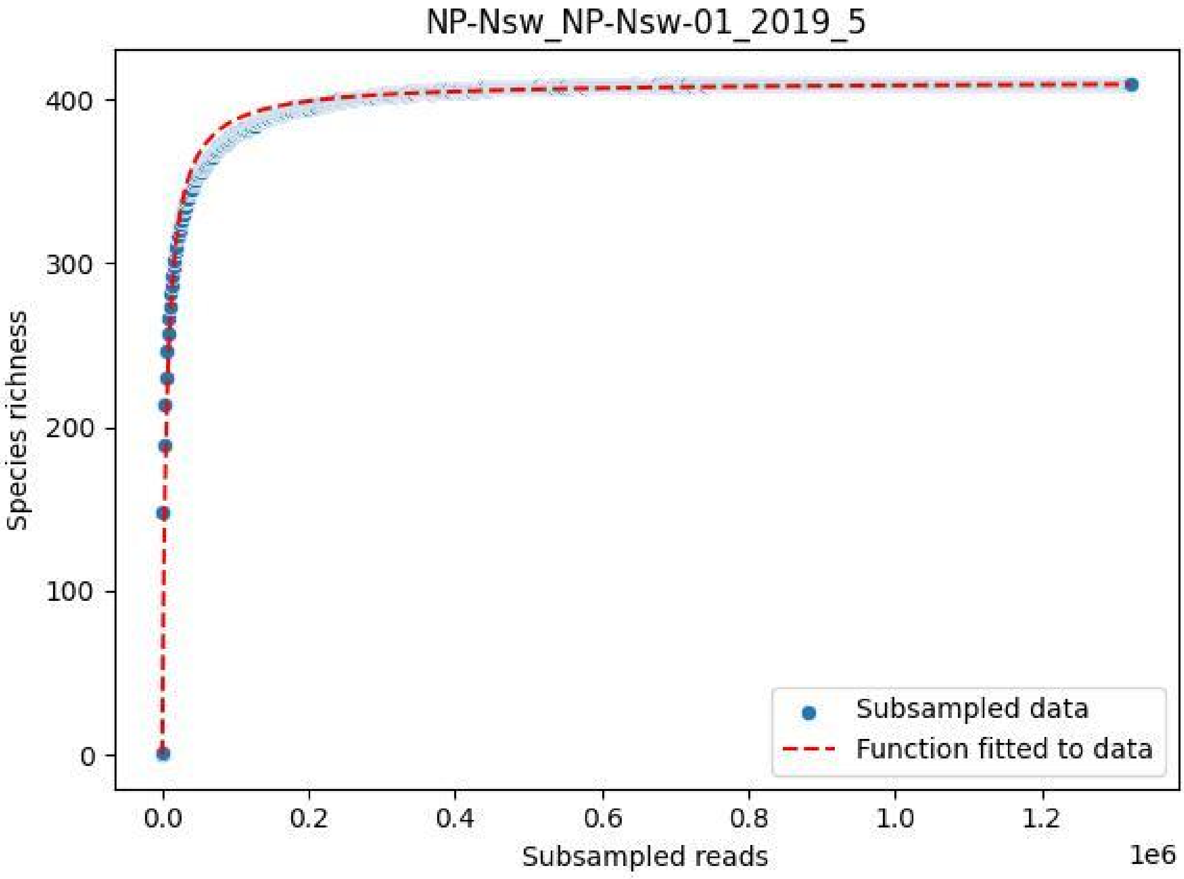

NP-Nsw_NP-Nsw-01_2019_5
Species richness
Subsampled reads
1e6
0
100
200
300
400
0.0
0.2
0.4
0.6
0.8
1.0
1.2
Subsampled data
Function fitted to data

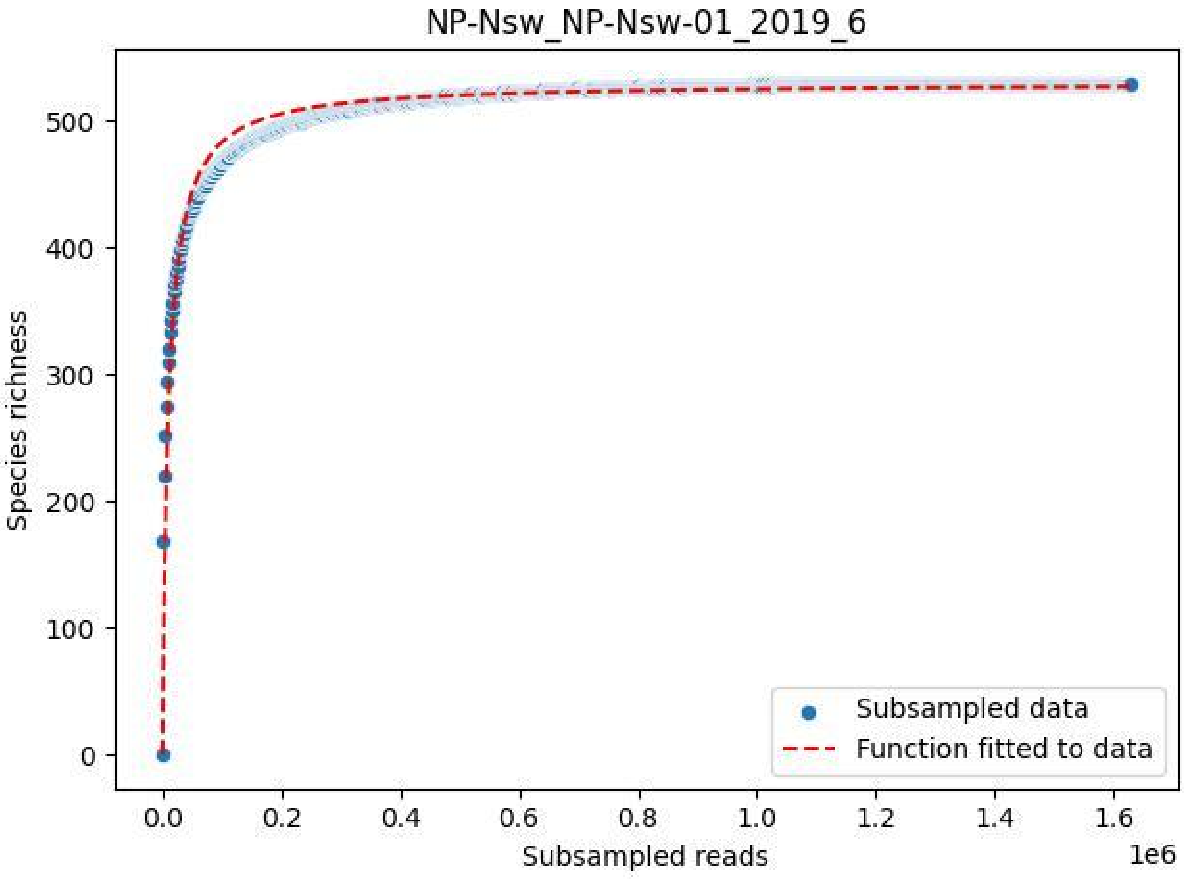
NP-Nsw_NP-Nsw-01_2019_6
Species richness
Subsampled reads
1e6
0
100
200
300
400
500
0.0
0.2
0.4
0.6
0.8
1.0
1.2
1.4
1.6
Subsampled data
Function fitted to data

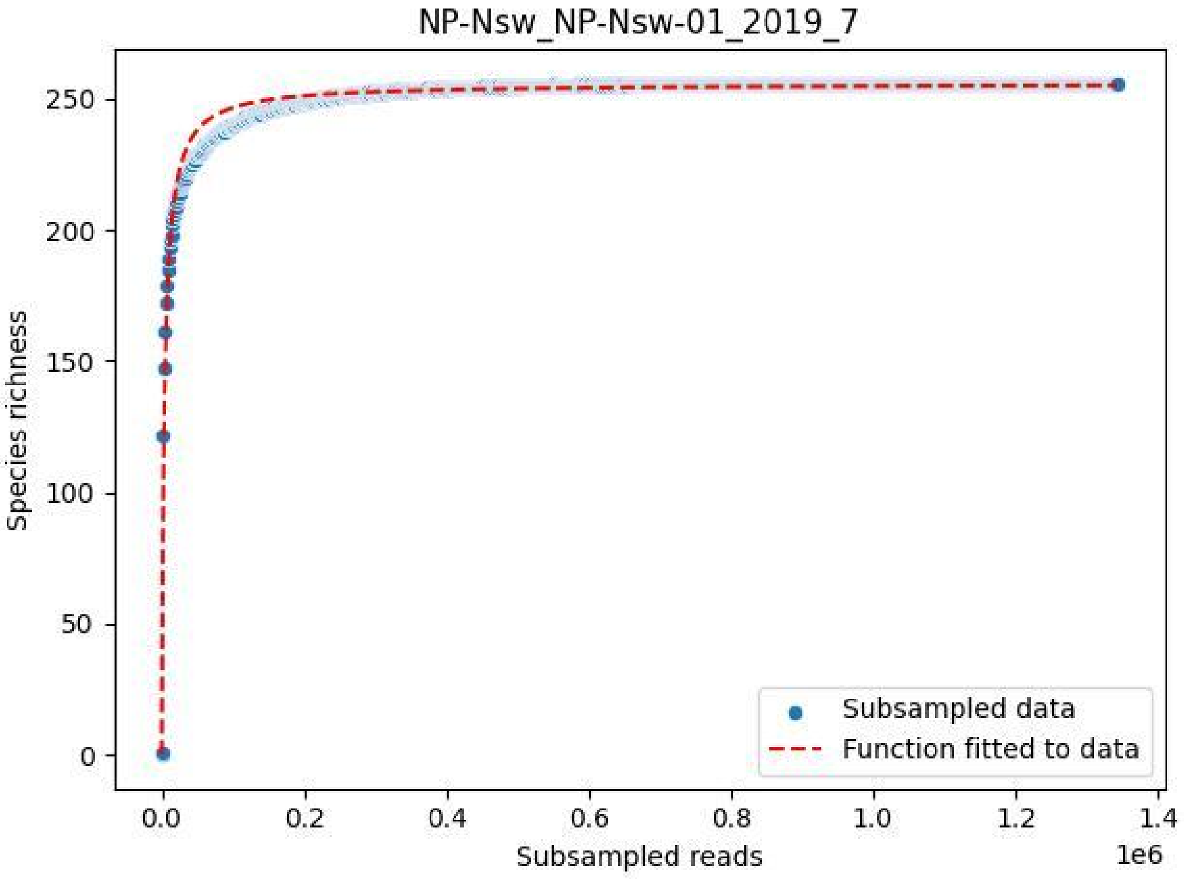

NP-Nsw_NP-Nsw-01_2019_7
Species richness
Subsampled reads
1e6
0
50
100
150
200
250
0.0
0.2
0.4
0.6
0.8
1.0
1.2
1.4
Subsampled data
Function fitted to data

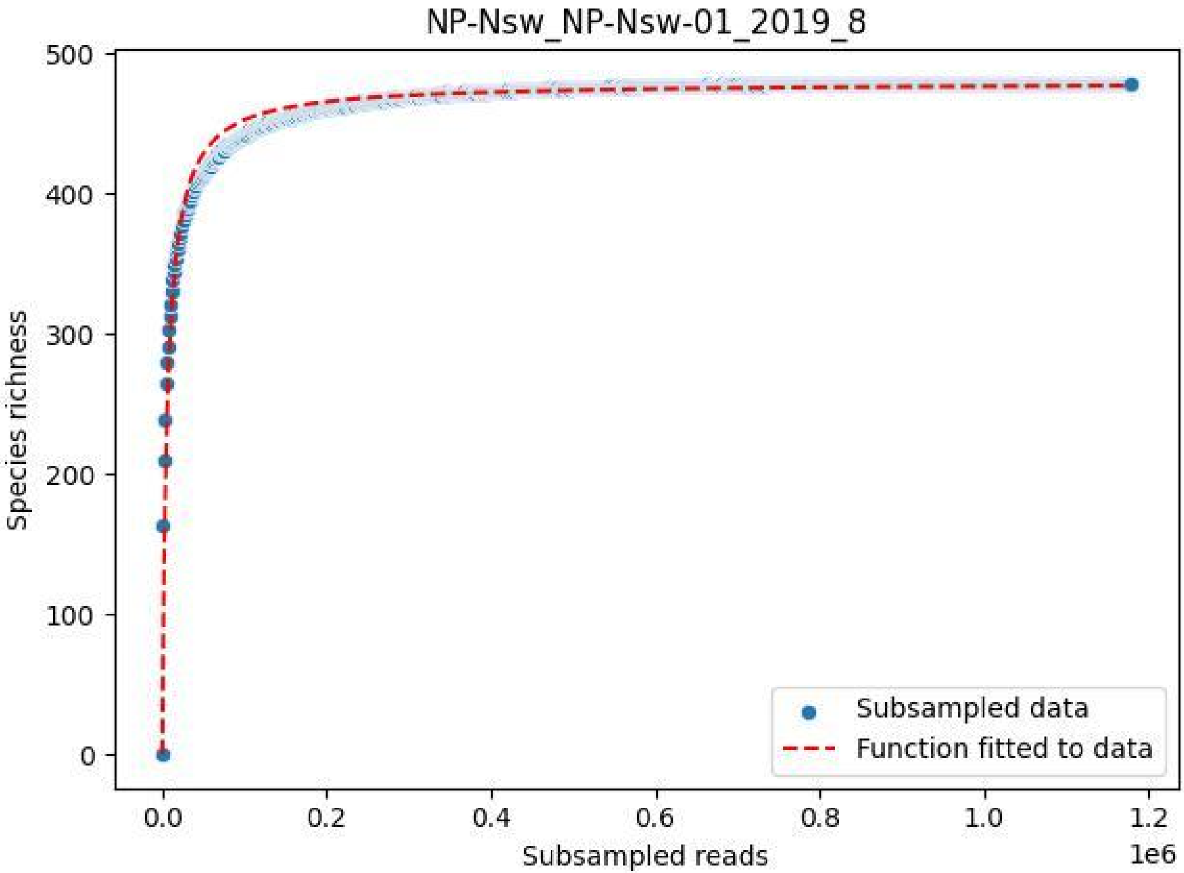

NP-Nsw_NP-Nsw-01_2019_8
Species richness
Subsampled reads
1e6
0
100
200
300
400
500
0.0
0.2
0.4
0.6
0.8
1.0
1.2
Subsampled data
Function fitted to data

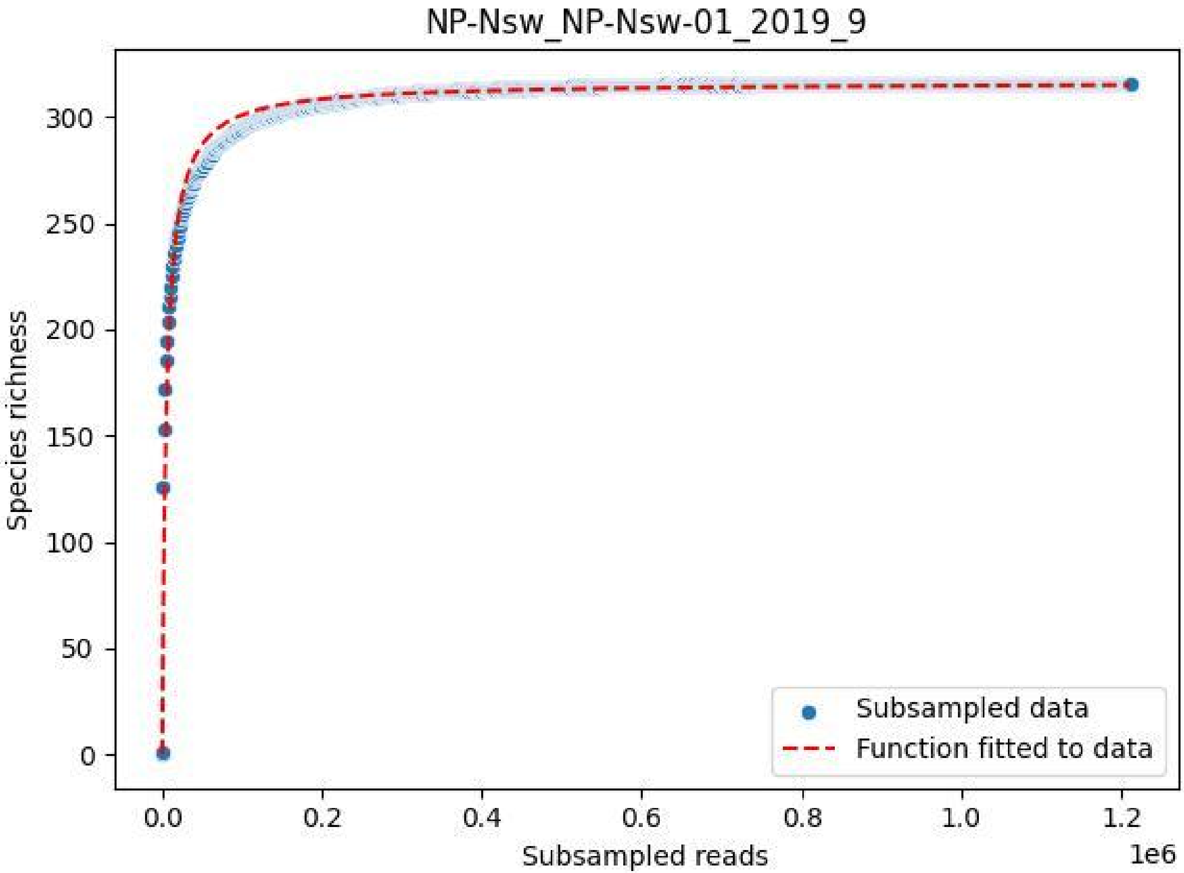

NP-Nsw_NP-Nsw-01_2019_9
Species richness
Subsampled reads
1e6
Subsampled data
Function fitted to data
0
50
100
150
200
250
300
0.0
0.2
0.4
0.6
0.8
1.0
1.2

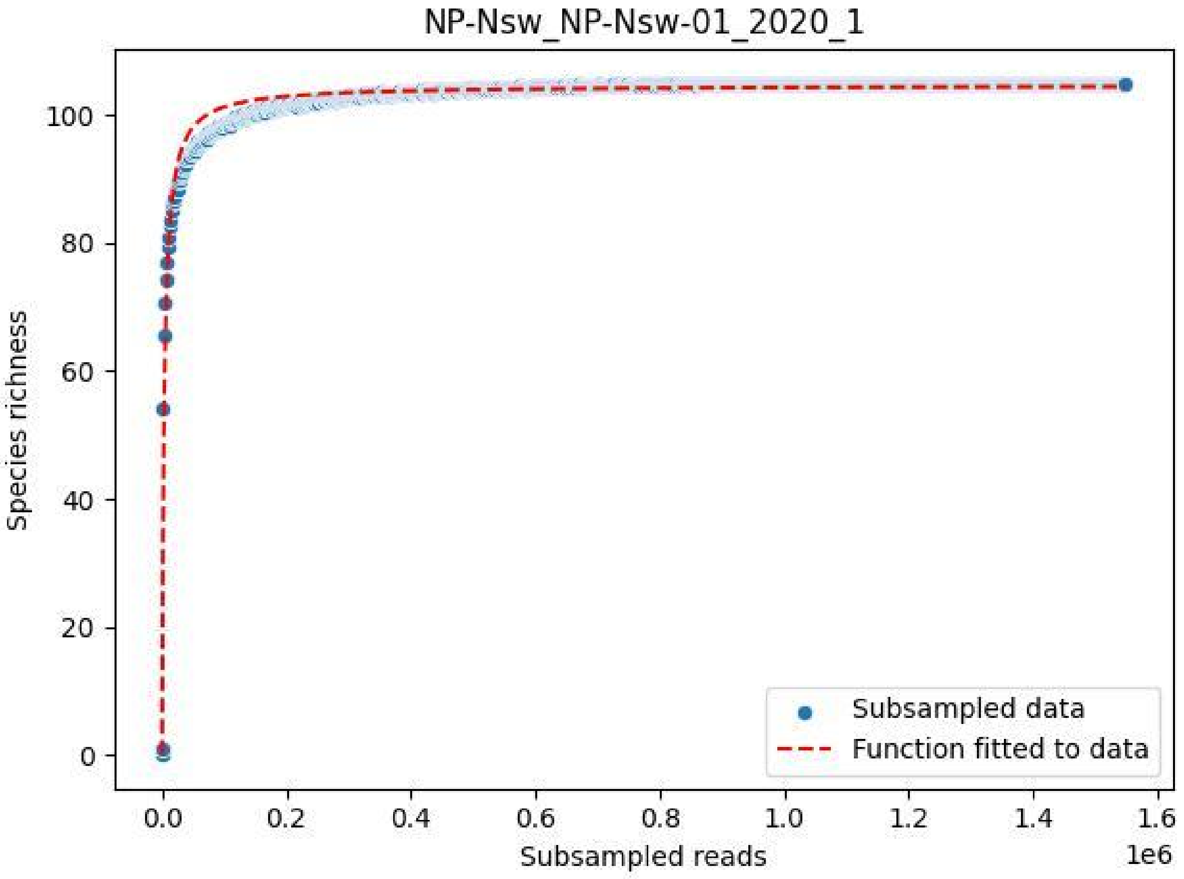
NP-Nsw_NP-Nsw-01_2020_1
Species richness
0
20
40
60
80
100
0.0
0.2
0.4
0.6
0.8
1.0
1.2
1.4
1.6
1e6
Subsampled reads
Subsampled data
Function fitted to data

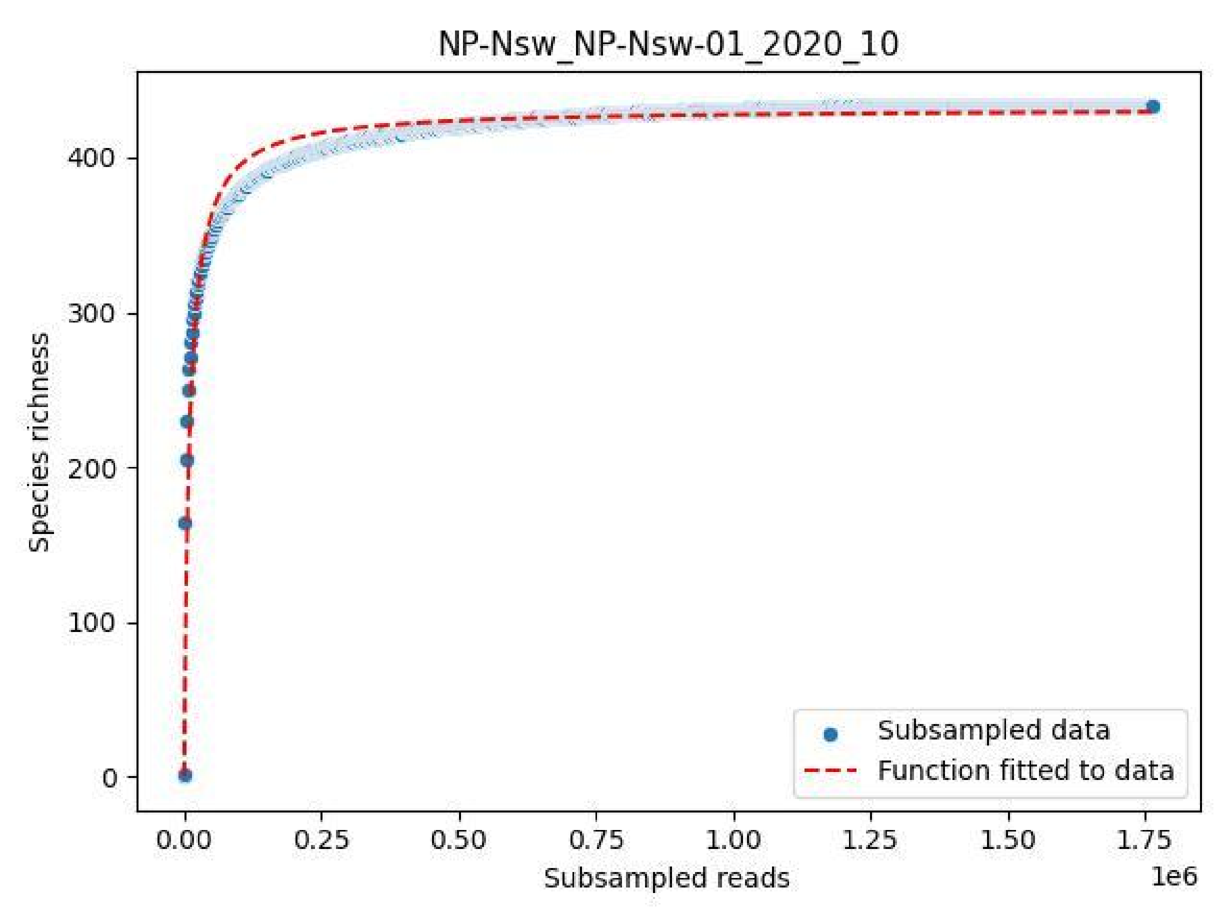
NP-Nsw_NP-Nsw-01_2020_10
Species richness
0
100
200
300
400
0.00
0.25
0.50
0.75
1.00
1.25
1.50
1.75
1e6
Subsampled reads
Subsampled data
Function fitted to data

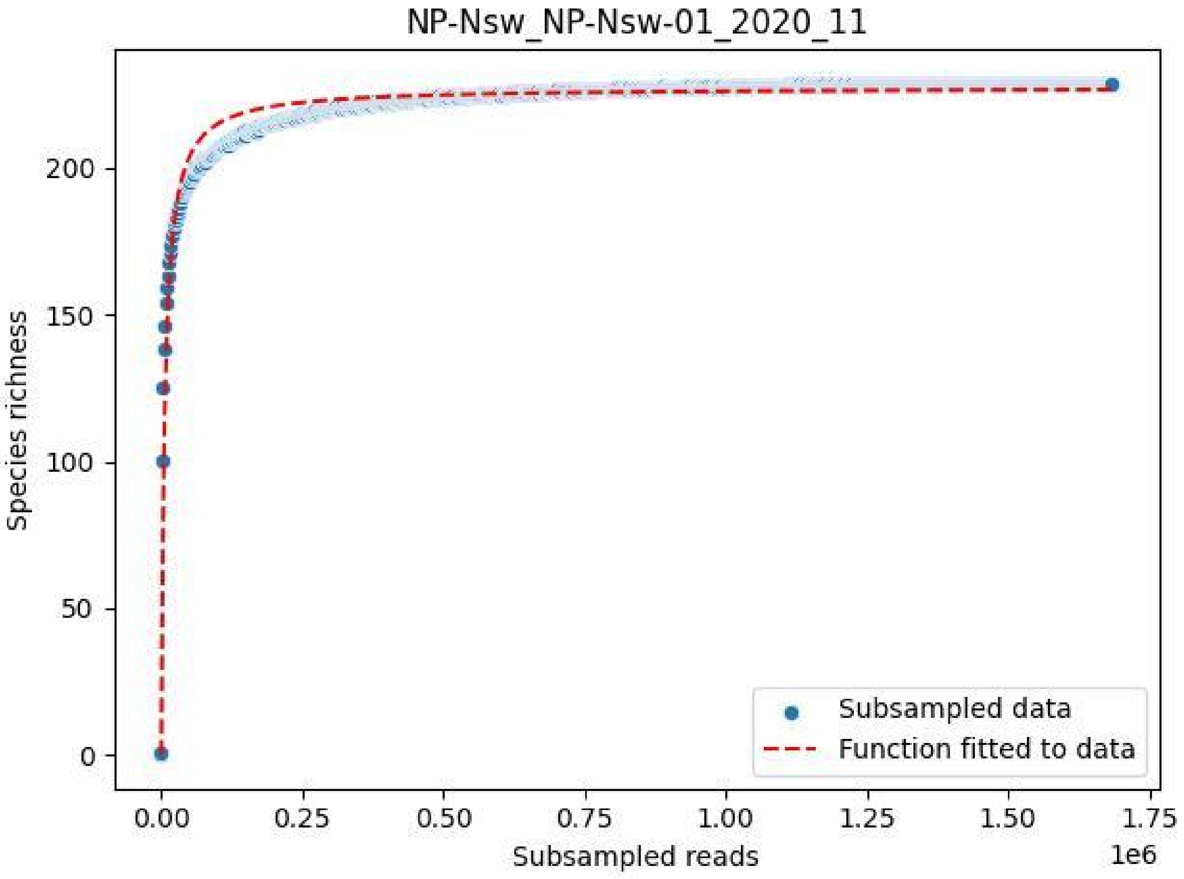
NP-Nsw_NP-Nsw-01_2020_11
Species richness
Subsampled reads
1e6
0
50
100
150
200
0.00
0.25
0.50
0.75
1.00
1.25
1.50
1.75
Subsampled data
Function fitted to data

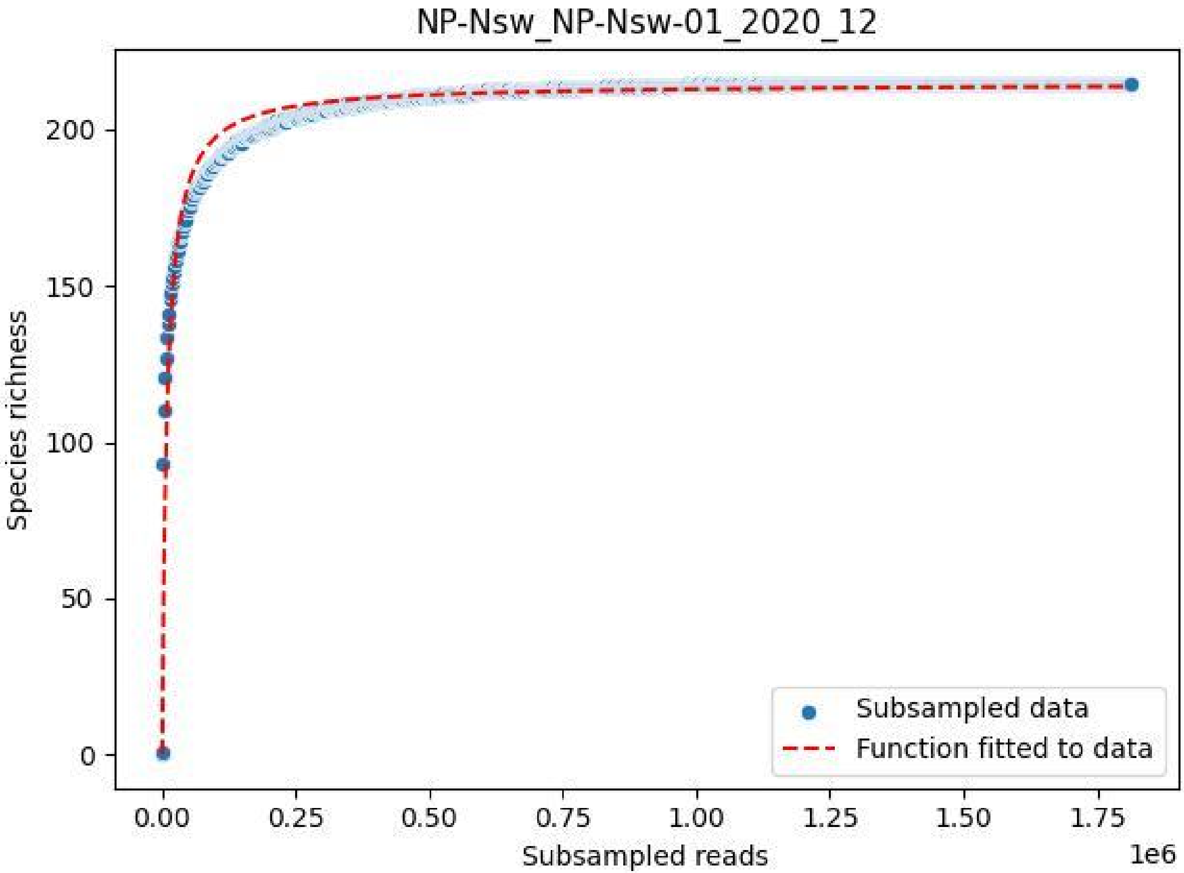
NP-Nsw_NP-Nsw-01_2020_12
Species richness
Subsampled reads
1e6
0
50
100
150
200
0.00
0.25
0.50
0.75
1.00
1.25
1.50
1.75
Subsampled data
Function fitted to data

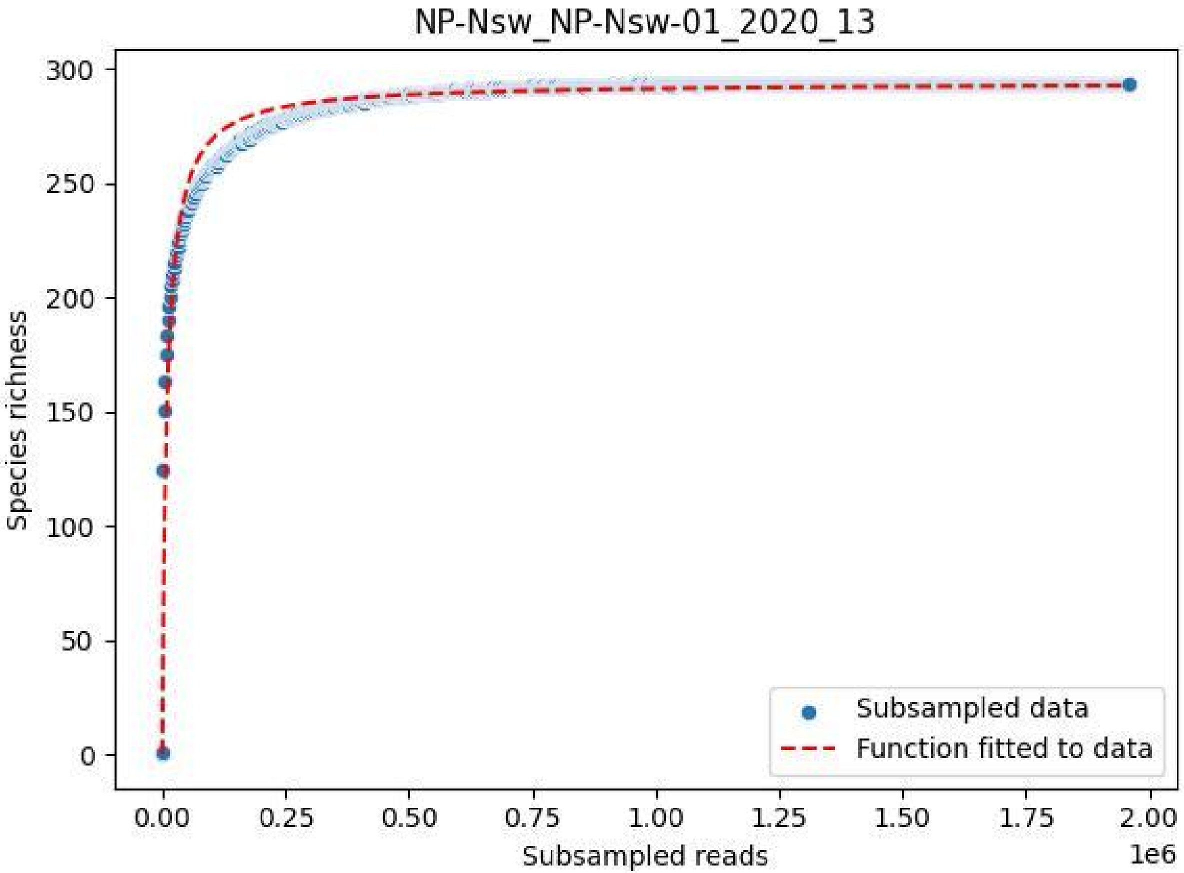

NP-Nsw_NP-Nsw-01_2020_13
Species richness
Subsampled reads
1e6
0
50
100
150
200
250
300
0.00
0.25
0.50
0.75
1.00
1.25
1.50
1.75
2.00
Subsampled data
Function fitted to data

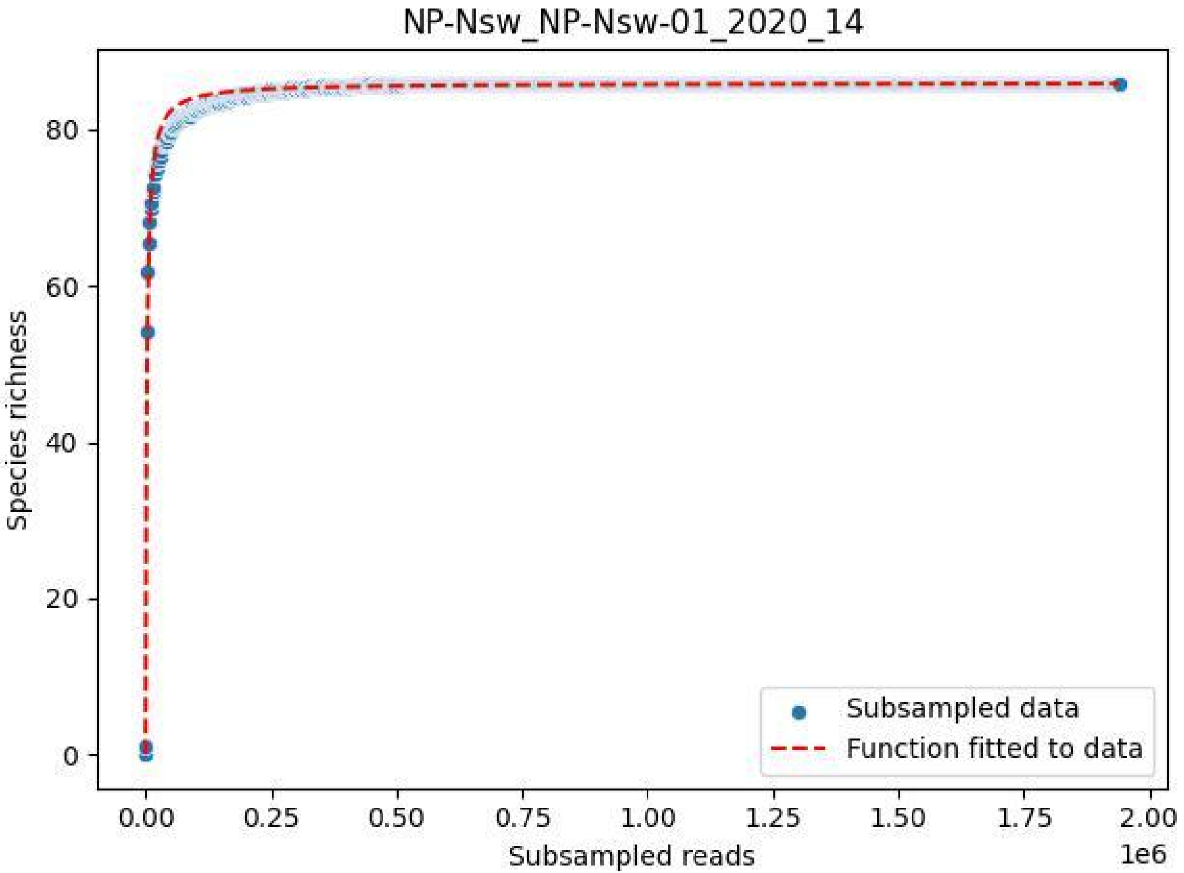
NP-Nsw_NP-Nsw-01_2020_14
Species richness
0
20
40
60
80
0.00
0.25
0.50
0.75
1.00
1.25
1.50
1.75
2.00
1e6
Subsampled reads
Subsampled data
Function fitted to data

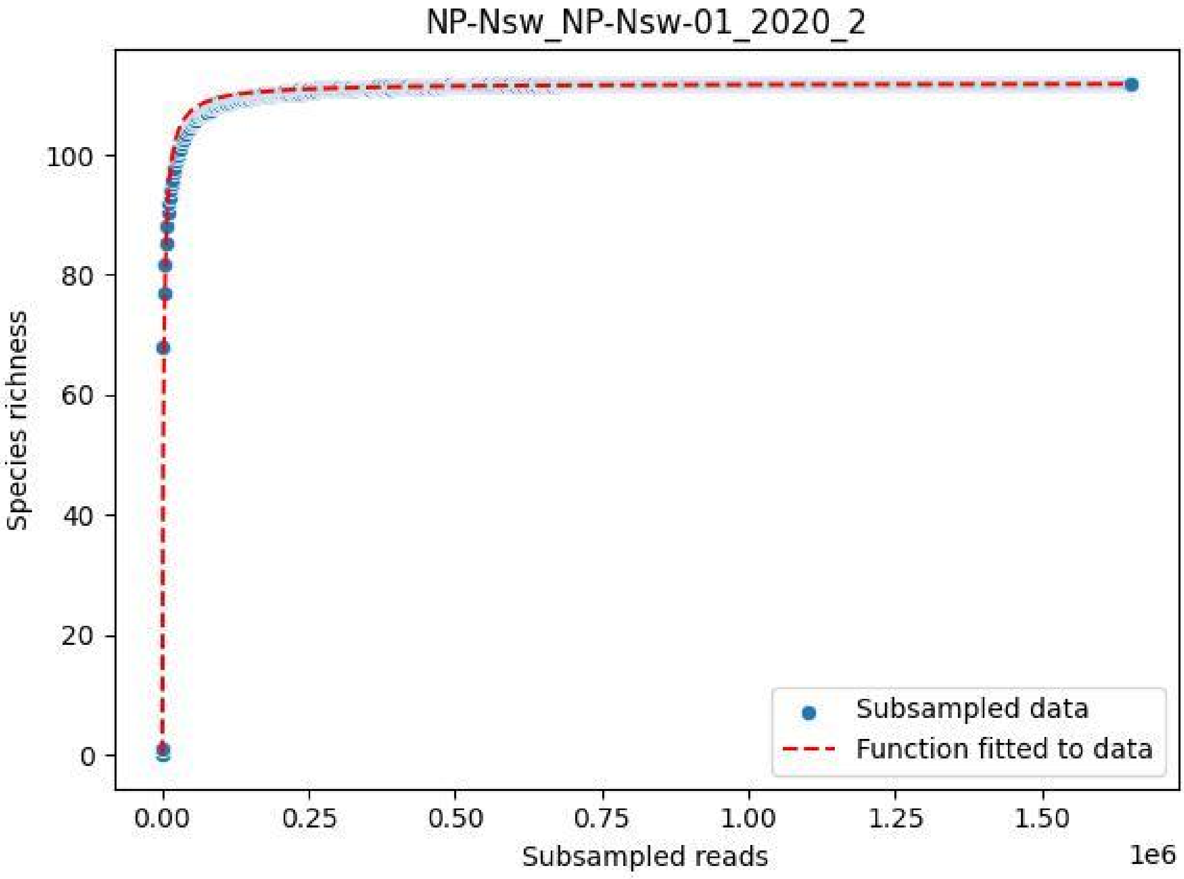
NP-Nsw_NP-Nsw-01_2020_2
Species richness
0
20
40
60
80
100
0.00
0.25
0.50
0.75
1.00
1.25
1.50
1e6
Subsampled reads
Subsampled data
Function fitted to data

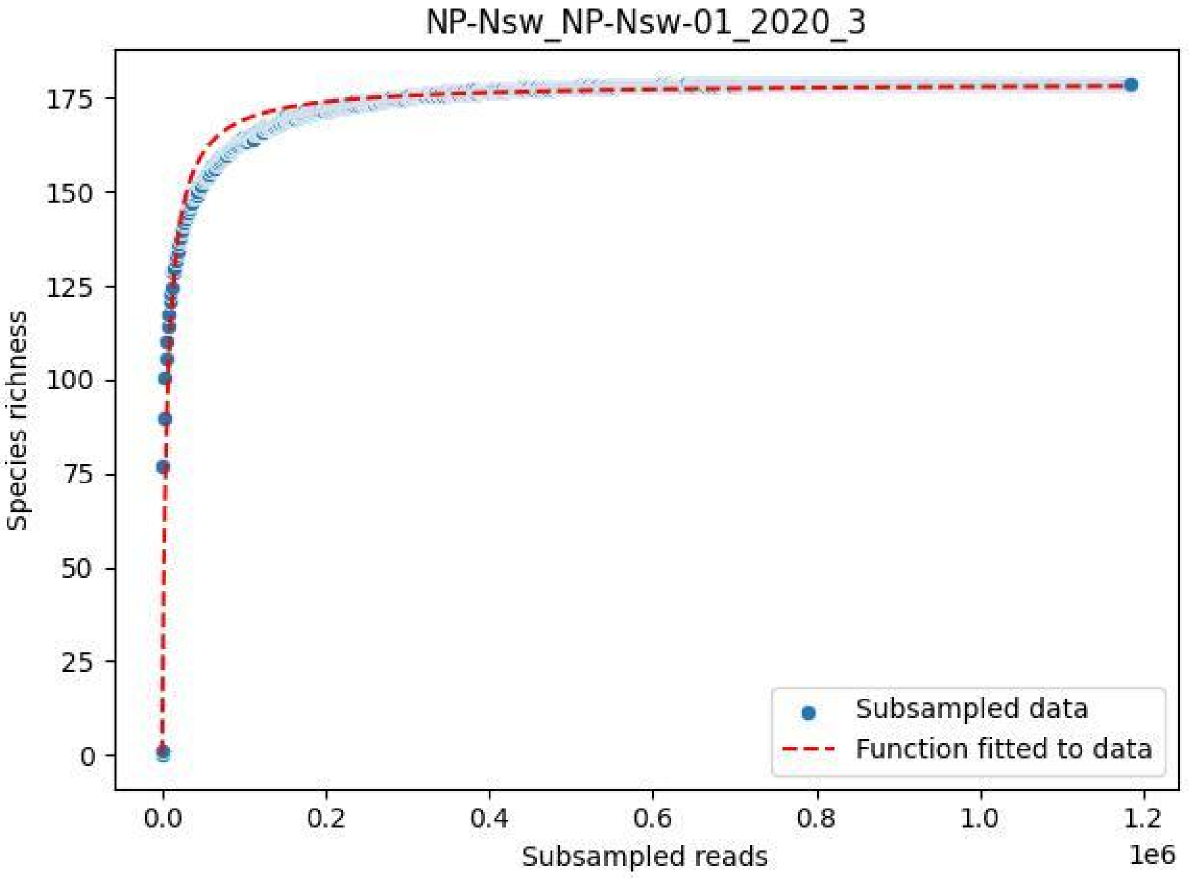

NP-Nsw_NP-Nsw-01_2020_3
Species richness
Subsampled reads
1e6
0
25
50
75
100
125
150
175
0.0
0.2
0.4
0.6
0.8
1.0
1.2
Subsampled data
Function fitted to data

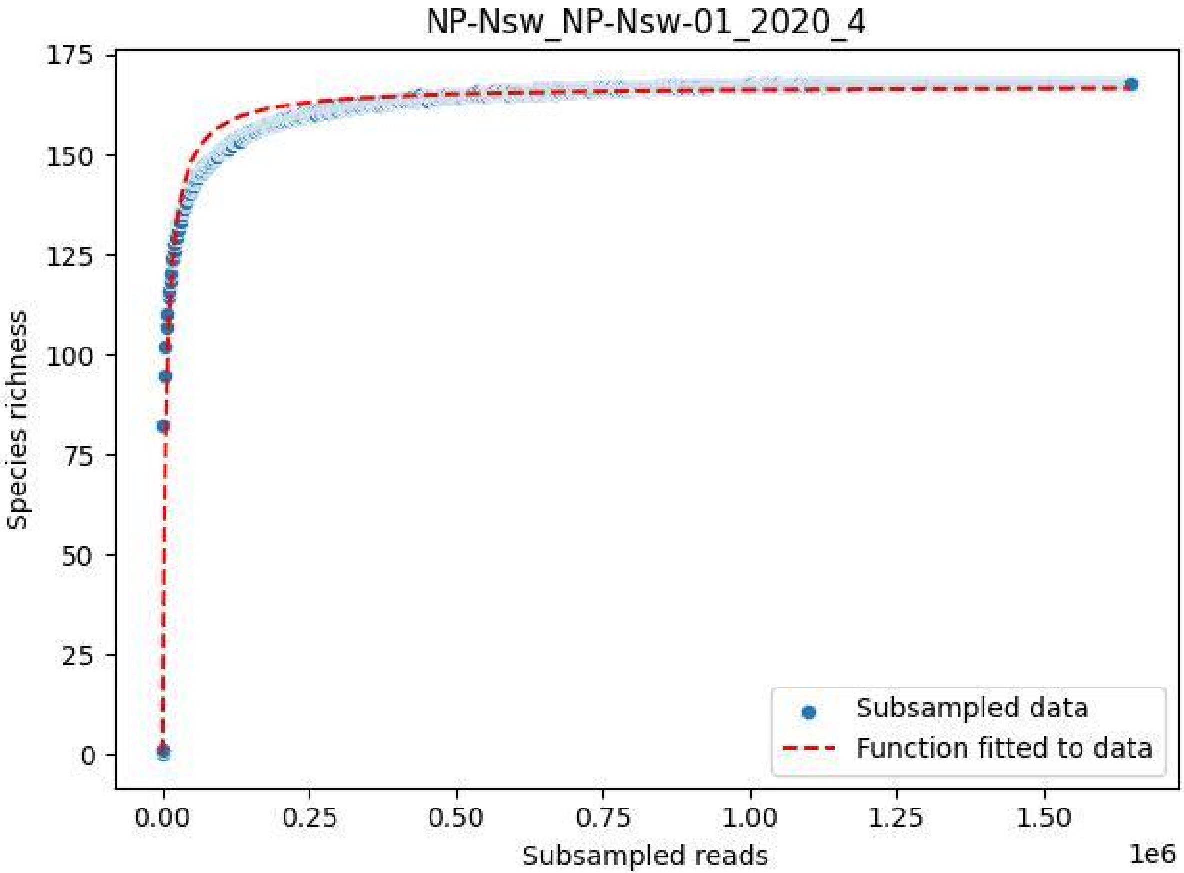

NP-Nsw_NP-Nsw-01_2020_4
Species richness
Subsampled reads
1e6
0
25
50
75
100
125
150
175
0.00
0.25
0.50
0.75
1.00
1.25
1.50
Subsampled data
Function fitted to data

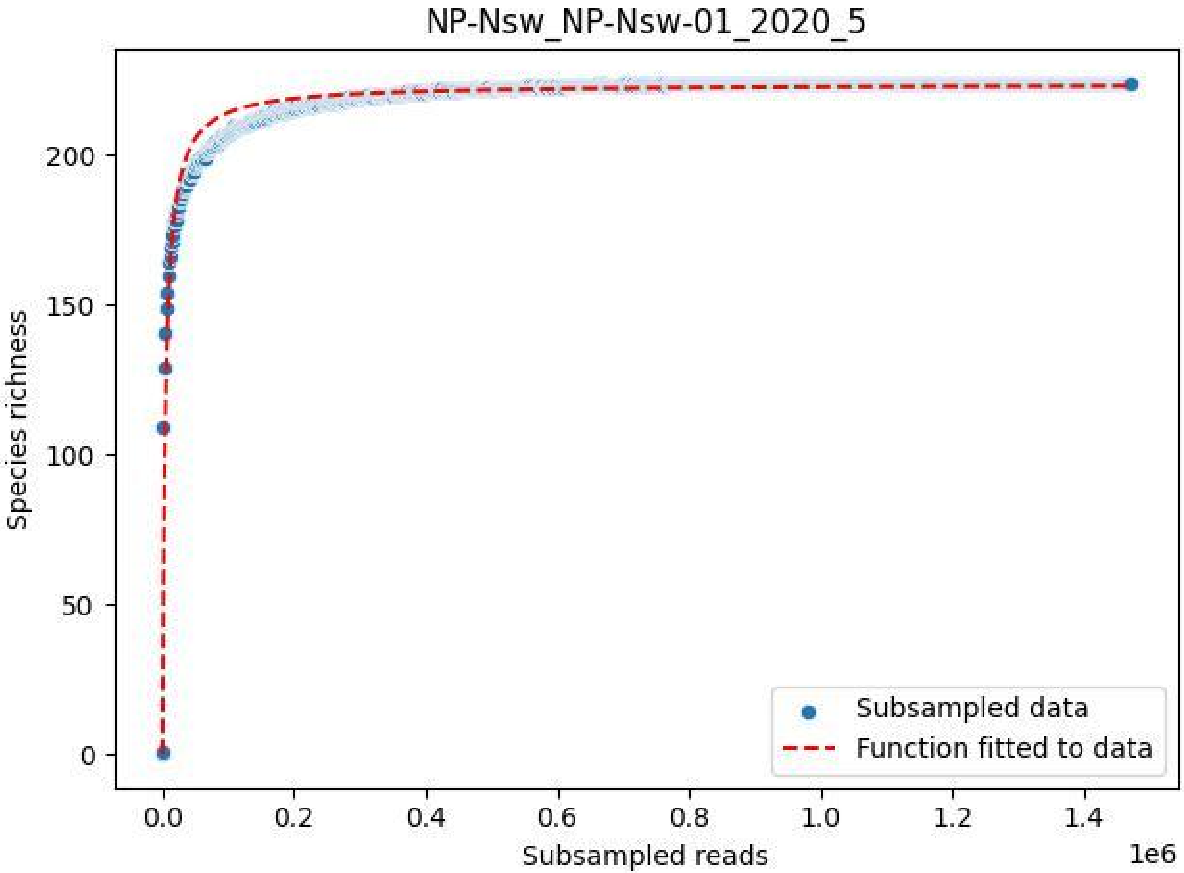

NP-Nsw_NP-Nsw-01_2020_5
Species richness
Subsampled reads
1e6
0
50
100
150
200
0.0
0.2
0.4
0.6
0.8
1.0
1.2
1.4
Subsampled data
Function fitted to data

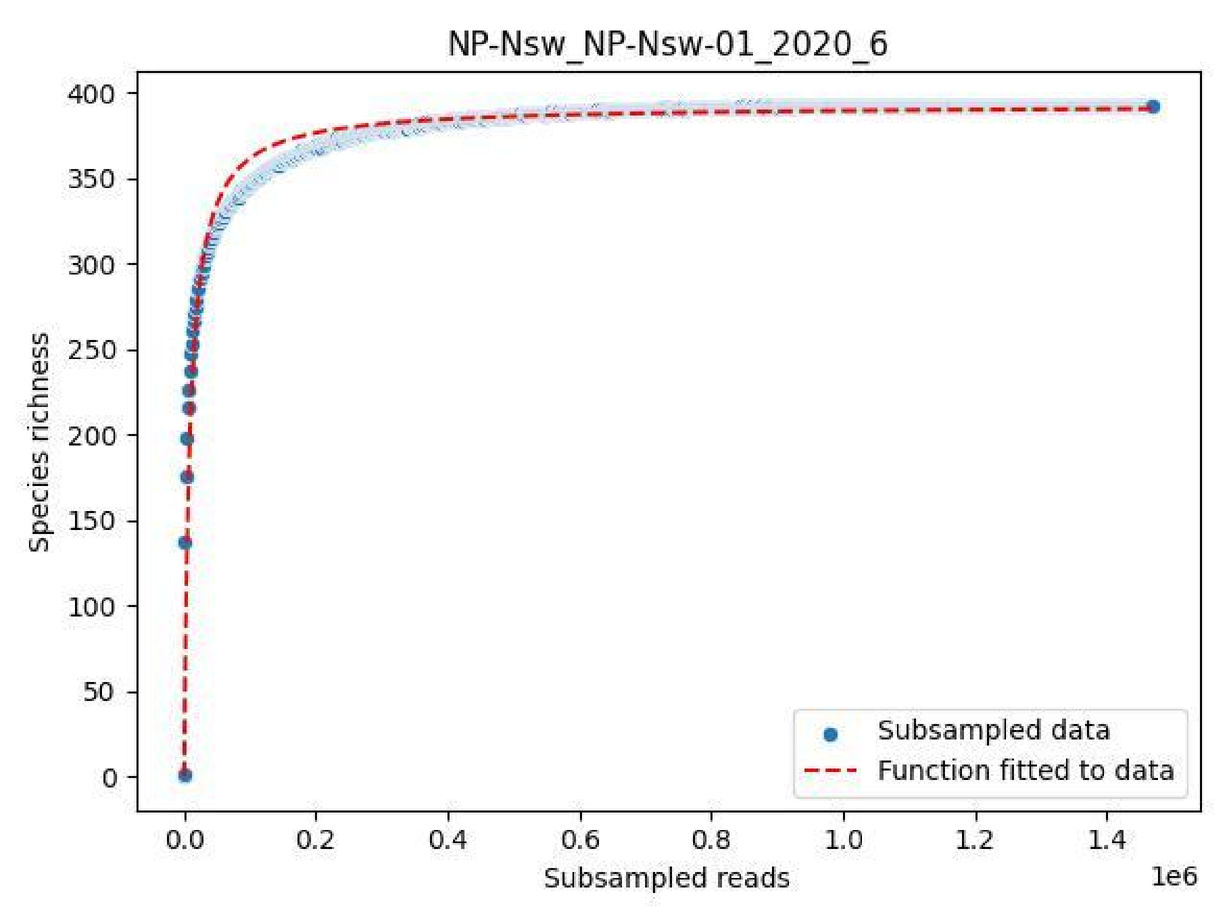

NP-Nsw_NP-Nsw-01_2020_6
Species richness
Subsampled reads
1e6
0.0
0.2
0.4
0.6
0.8
1.0
1.2
1.4
0
50
100
150
200
250
300
350
400
Subsampled data
Function fitted to data

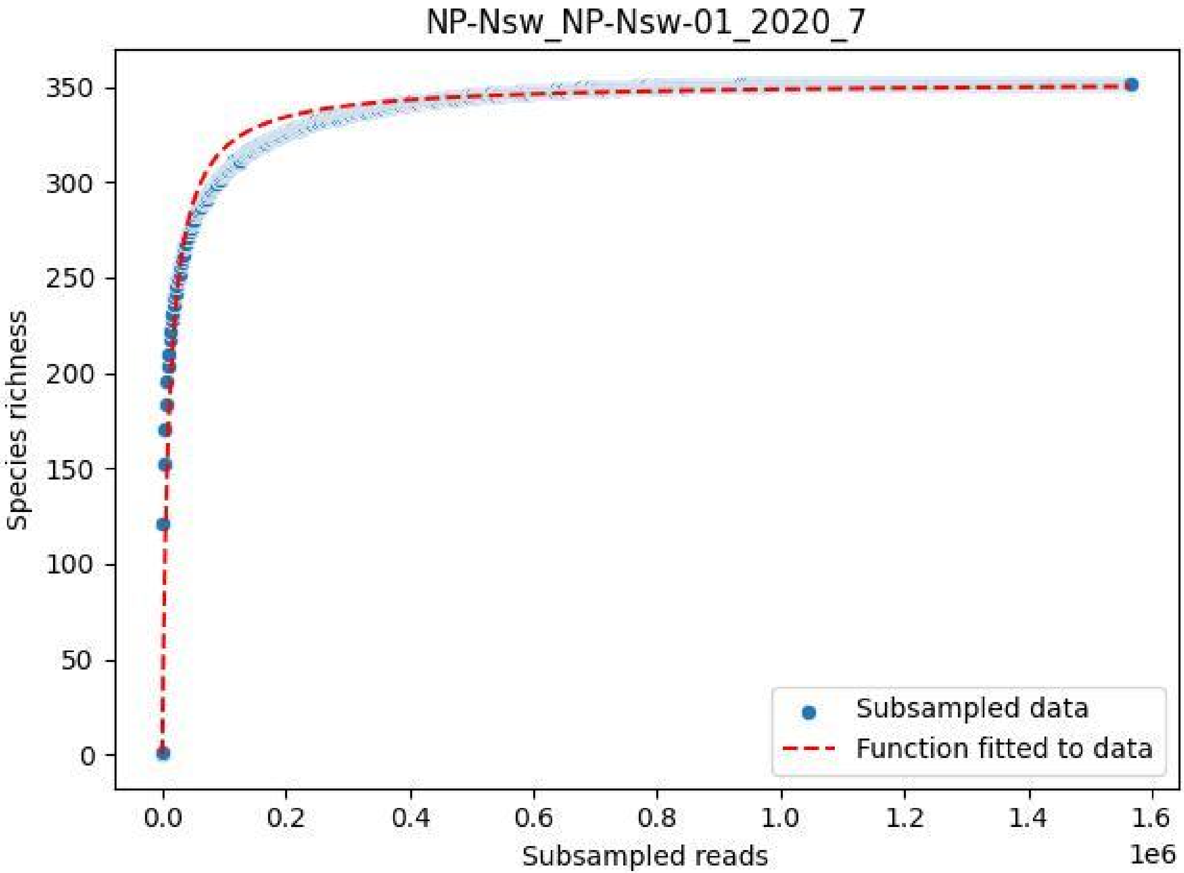

NP-Nsw_NP-Nsw-01_2020_7
Species richness
Subsampled reads
1e6
0
50
100
150
200
250
300
350
0.0
0.2
0.4
0.6
0.8
1.0
1.2
1.4
1.6
Subsampled data
Function fitted to data

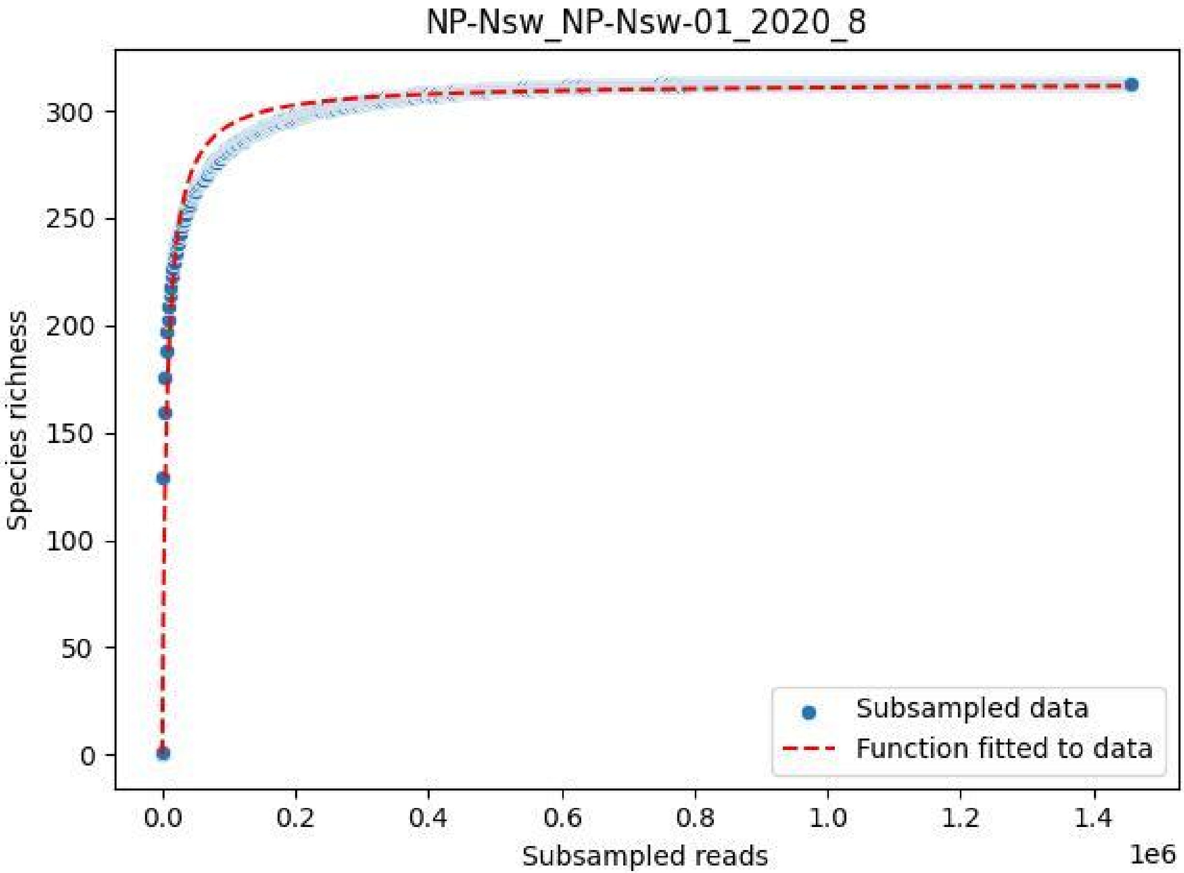

NP-Nsw_NP-Nsw-01_2020_8
Species richness
Subsampled reads
1e6
0
50
100
150
200
250
300
0.0
0.2
0.4
0.6
0.8
1.0
1.2
1.4
Subsampled data
Function fitted to data

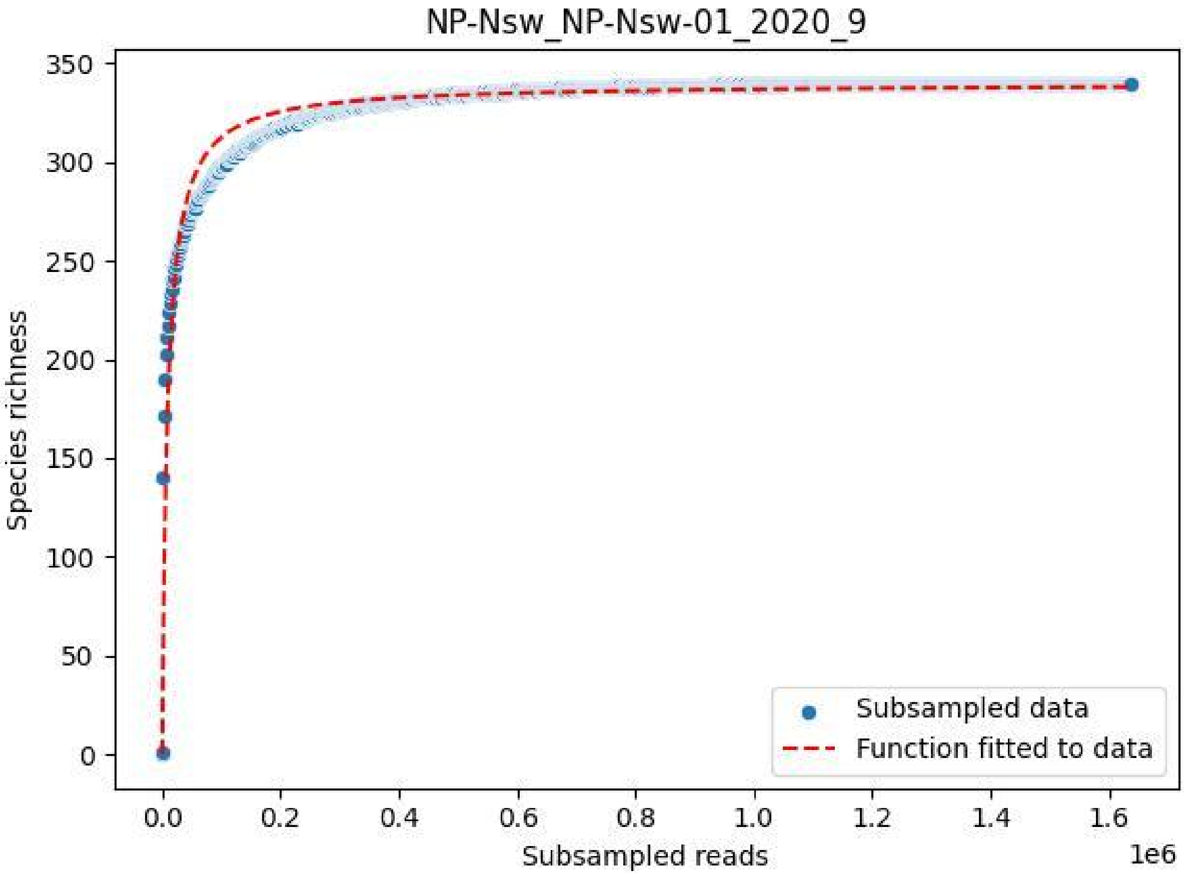
NP-Nsw_NP-Nsw-01_2020_9
Species richness
Subsampled reads
1e6
0
50
100
150
200
250
300
350
0.0
0.2
0.4
0.6
0.8
1.0
1.2
1.4
1.6
Subsampled data
Function fitted to data

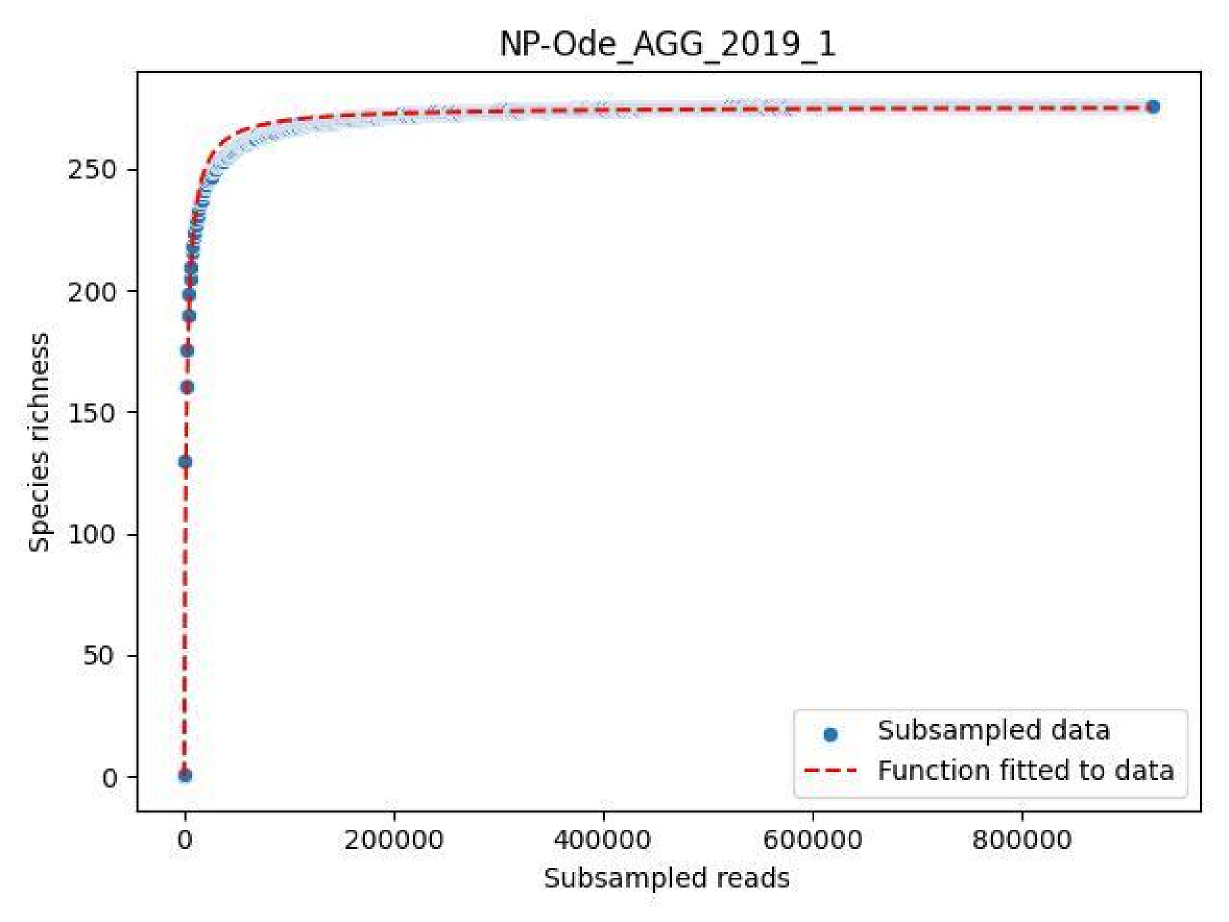

NP-Ode_AGG_2019_1
Species richness
Subsampled reads
0
50
100
150
200
250
0
200000
400000
600000
800000
Subsampled data
Function fitted to data

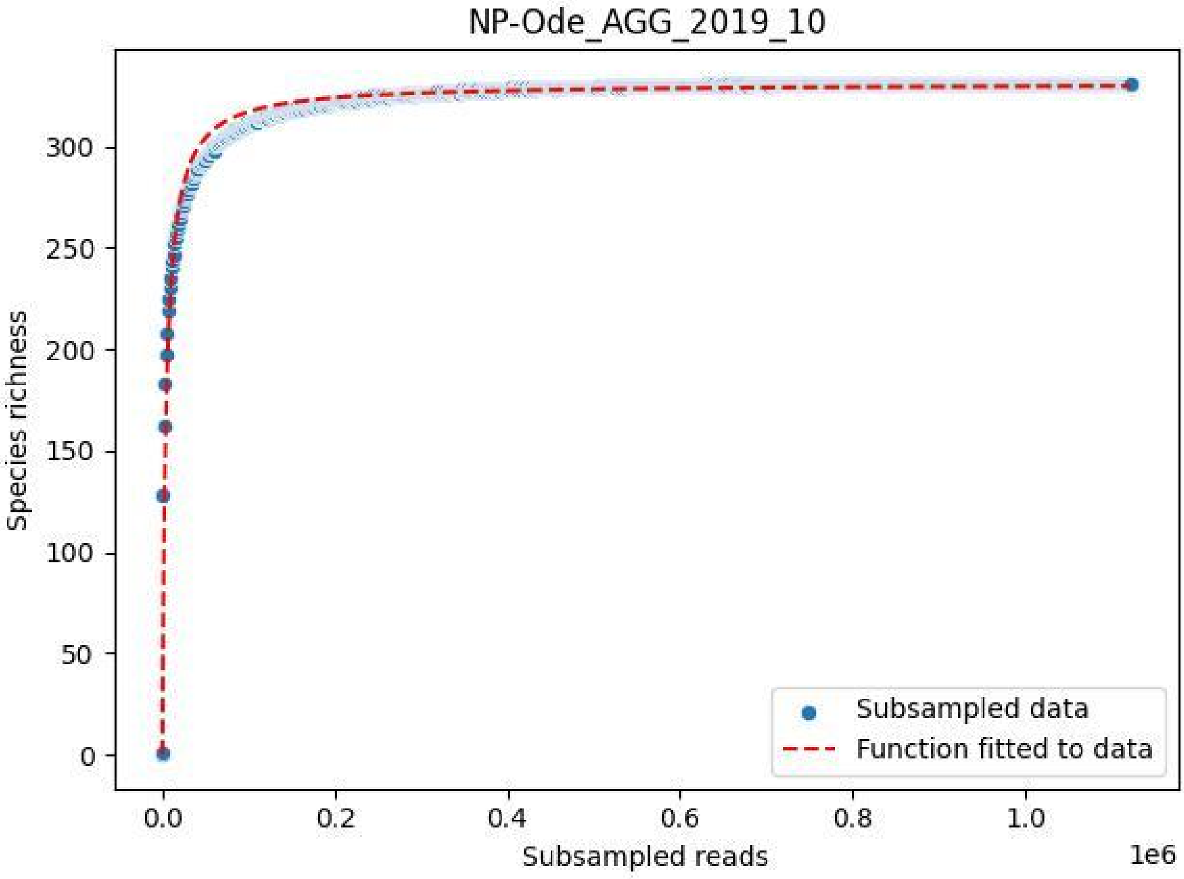
NP-Ode_AGG_2019_10
Species richness
Subsampled reads
1e6
Subsampled data
Function fitted to data
0
50
100
150
200
250
300
0.0
0.2
0.4
0.6
0.8
1.0

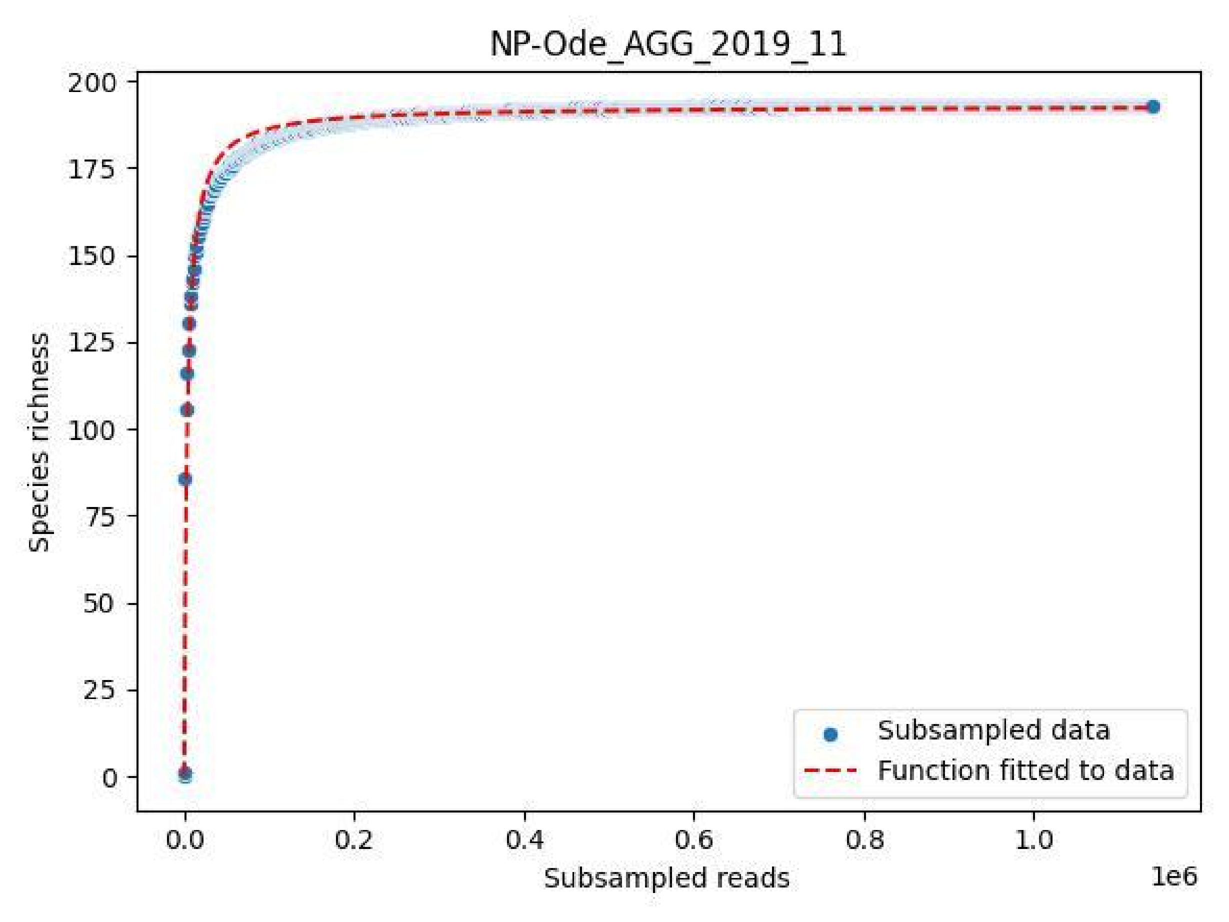

NP-Ode_AGG_2019_11
Species richness
Subsampled reads
1e6
200
175
150
125
100
75
50
25
0
0.0
0.2
0.4
0.6
0.8
1.0
Subsampled data
Function fitted to data

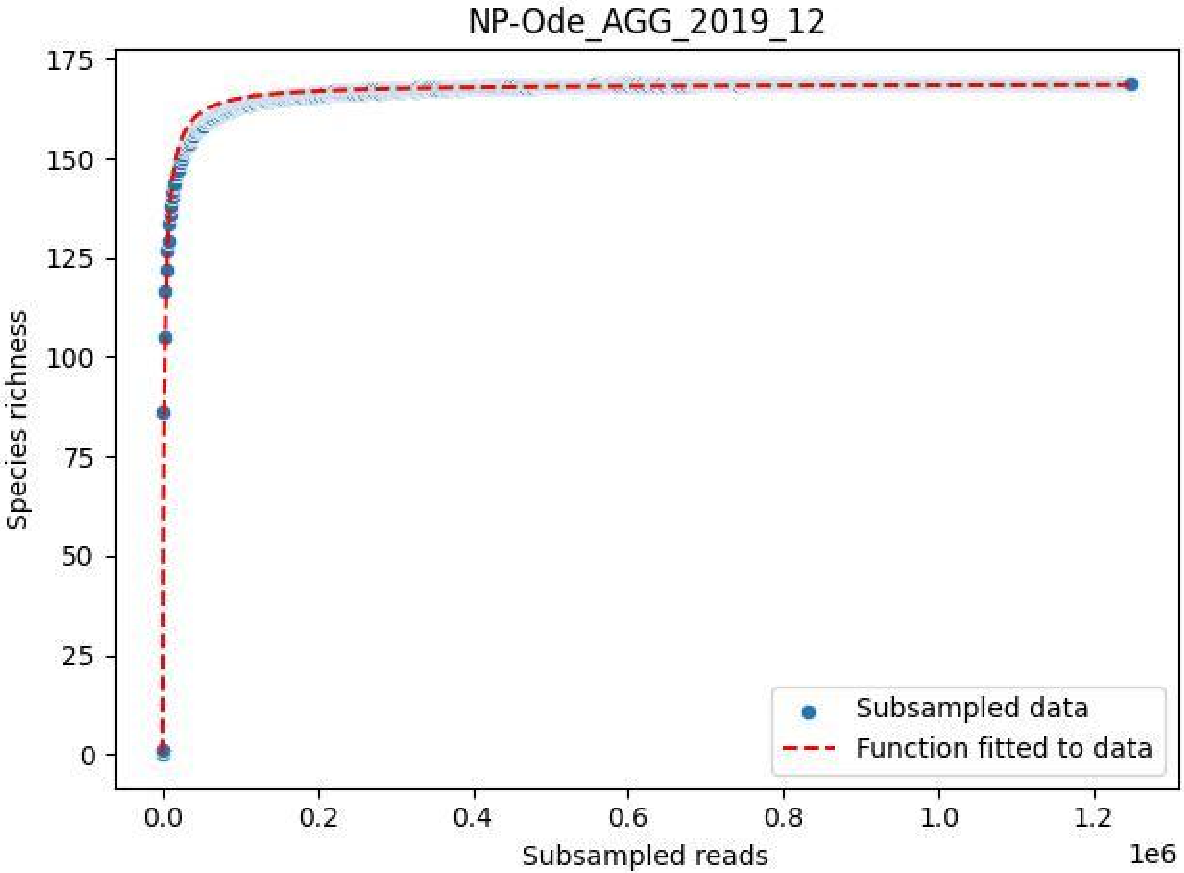

NP-Ode_AGG_2019_12
Species richness
Subsampled reads
1e6
Subsampled data
Function fitted to data
0
25
50
75
100
125
150
175
0.0
0.2
0.4
0.6
0.8
1.0
1.2

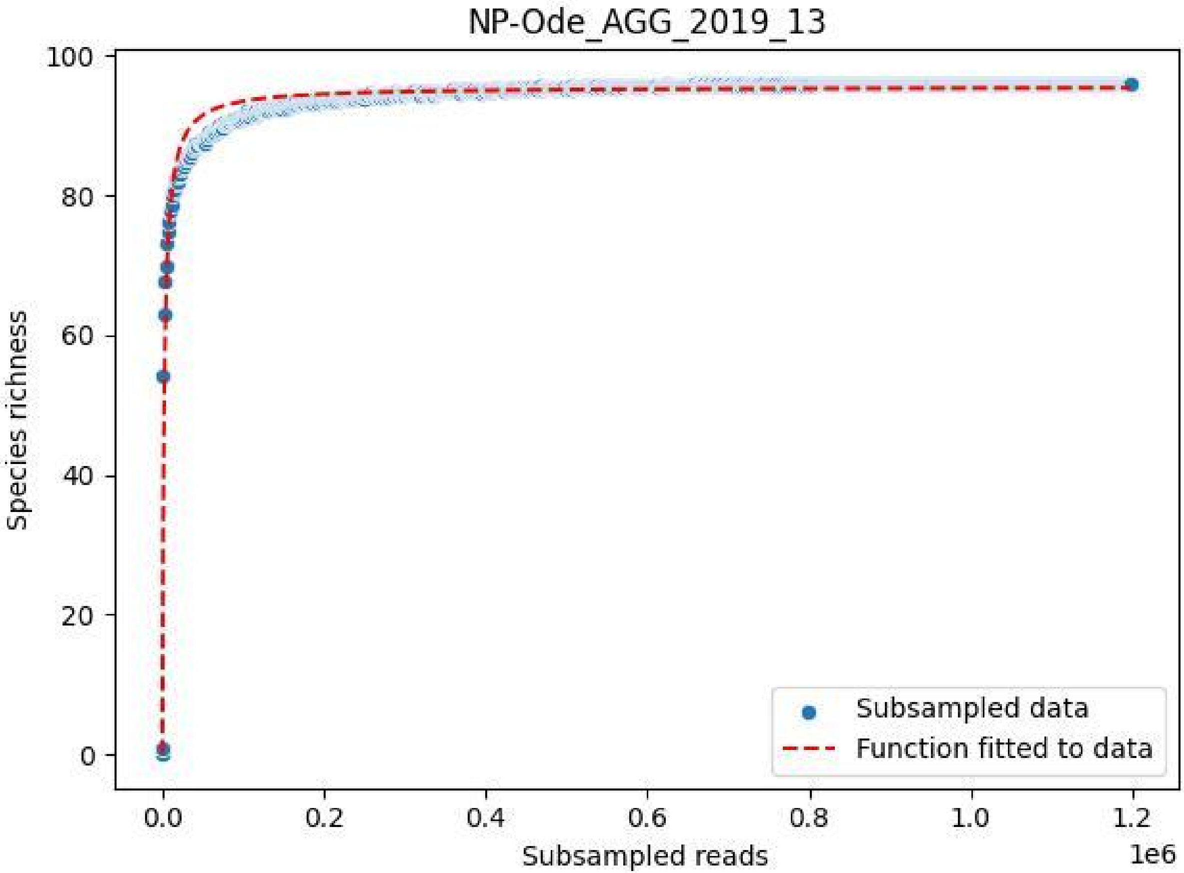
NP-Ode_AGG_2019_13
Species richness
Subsampled reads
1e6
0
20
40
60
80
100
0.0
0.2
0.4
0.6
0.8
1.0
1.2
Subsampled data
Function fitted to data

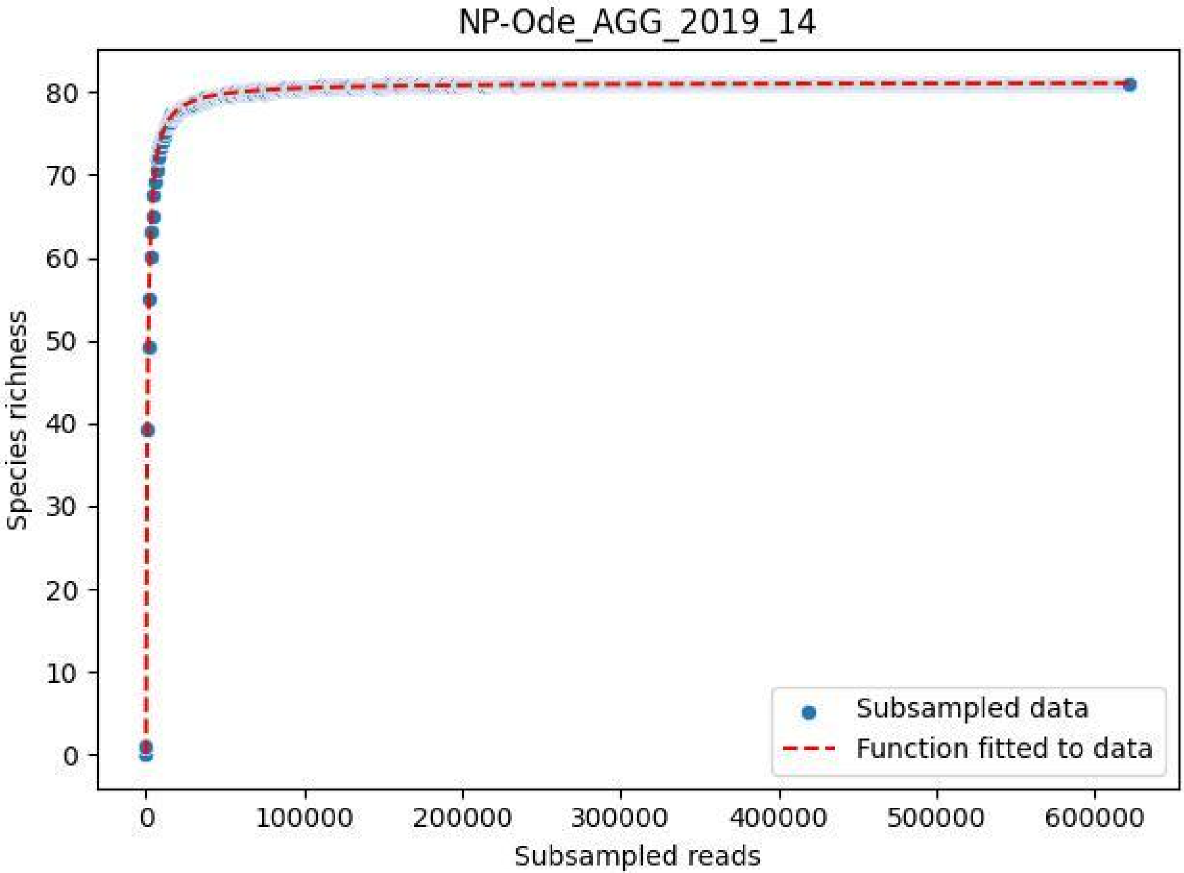

NP-Ode_AGG_2019_14
Species richness
Subsampled reads
0
10
20
30
40
50
60
70
80
0
100000
200000
300000
400000
500000
600000
Subsampled data
Function fitted to data

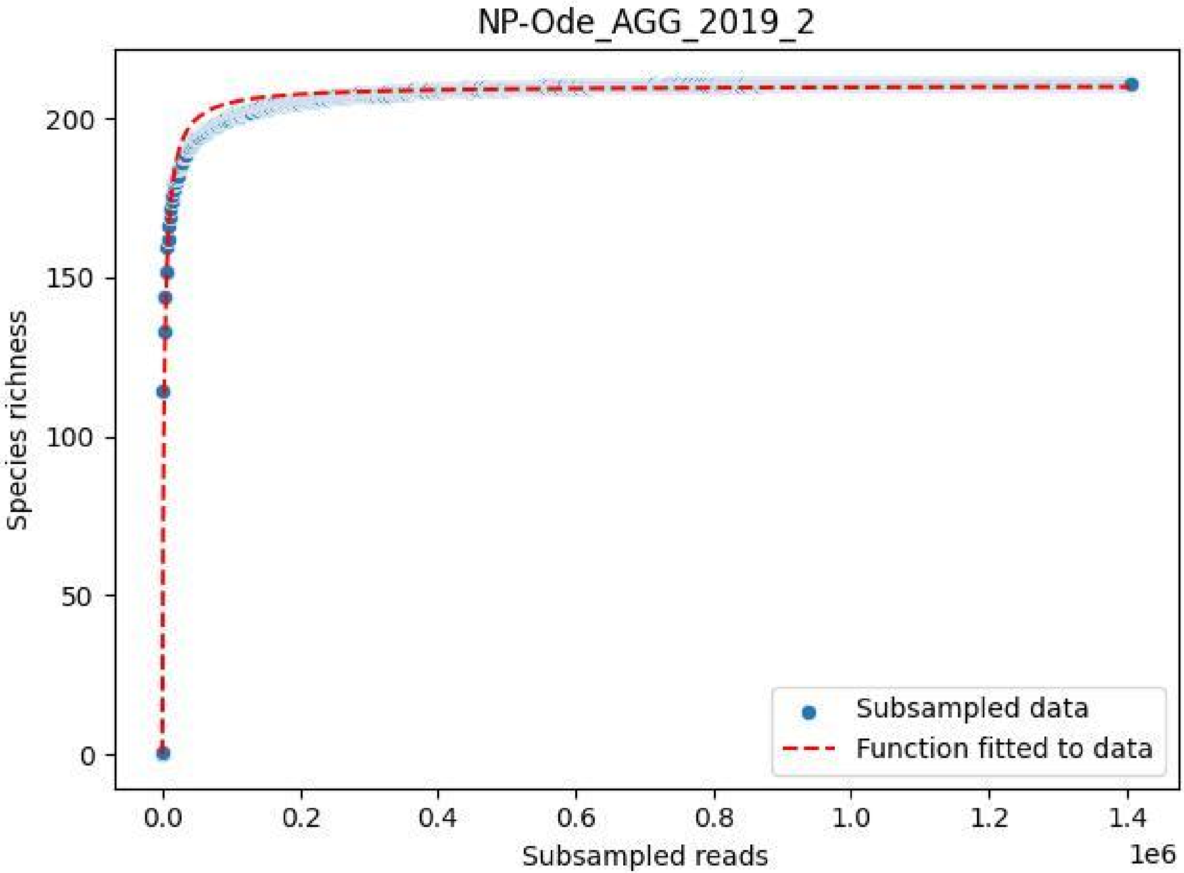

NP-Ode_AGG_2019_2
Species richness
Subsampled reads
1e6
0
50
100
150
200
0.0
0.2
0.4
0.6
0.8
1.0
1.2
1.4
Subsampled data
Function fitted to data

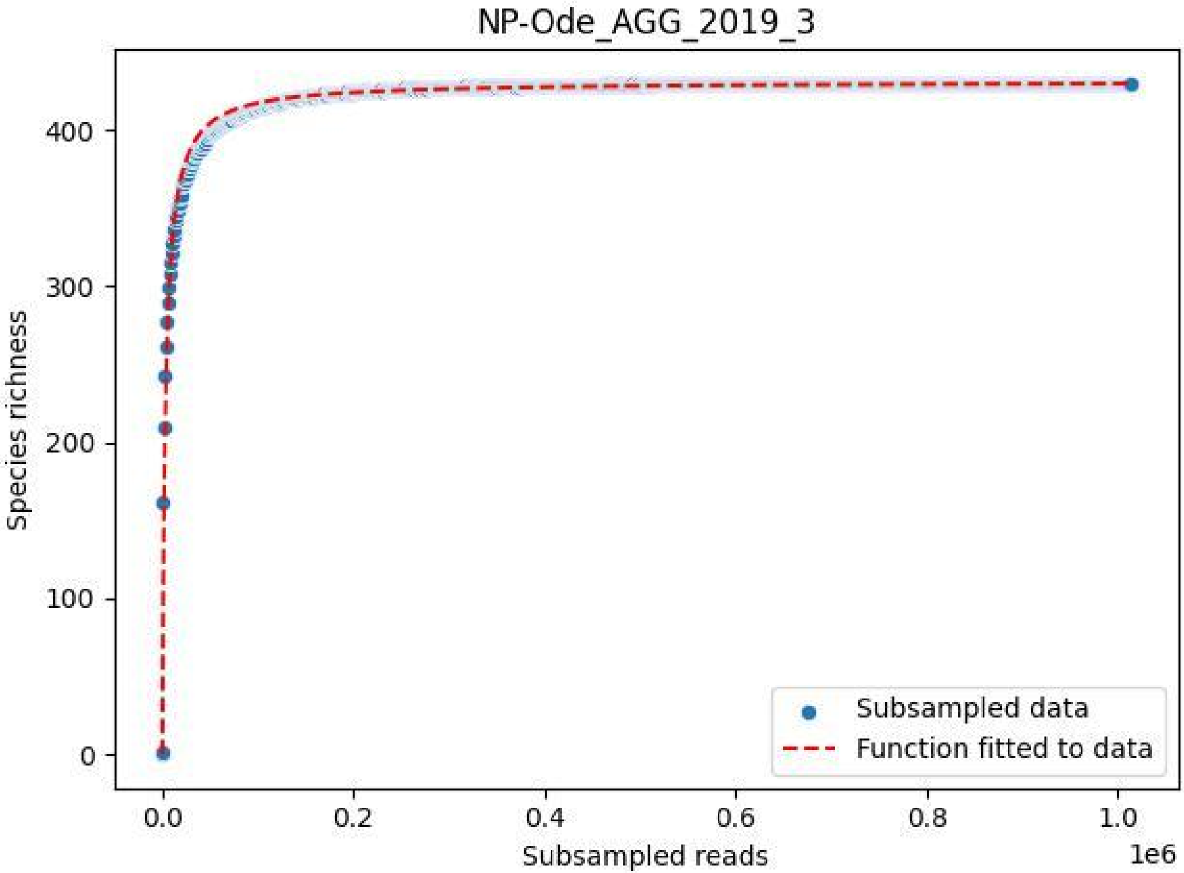

NP-Ode_AGG_2019_3
Species richness
Subsampled reads
1e6
0
100
200
300
400
0.0
0.2
0.4
0.6
0.8
1.0
Subsampled data
Function fitted to data

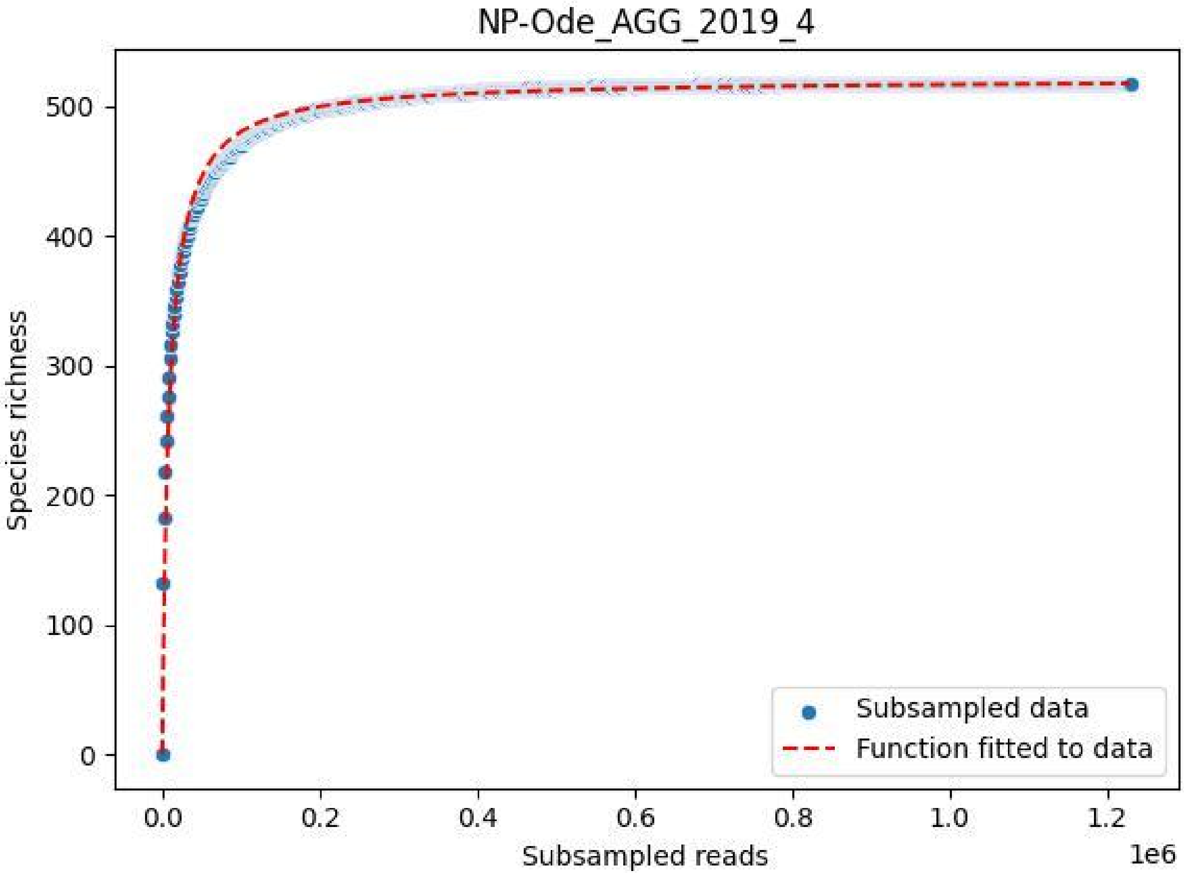

NP-Ode_AGG_2019_4
Species richness
0
100
200
300
400
500
0.0
0.2
0.4
0.6
0.8
1.0
1.2
1e6
Subsampled reads
Subsampled data
Function fitted to data

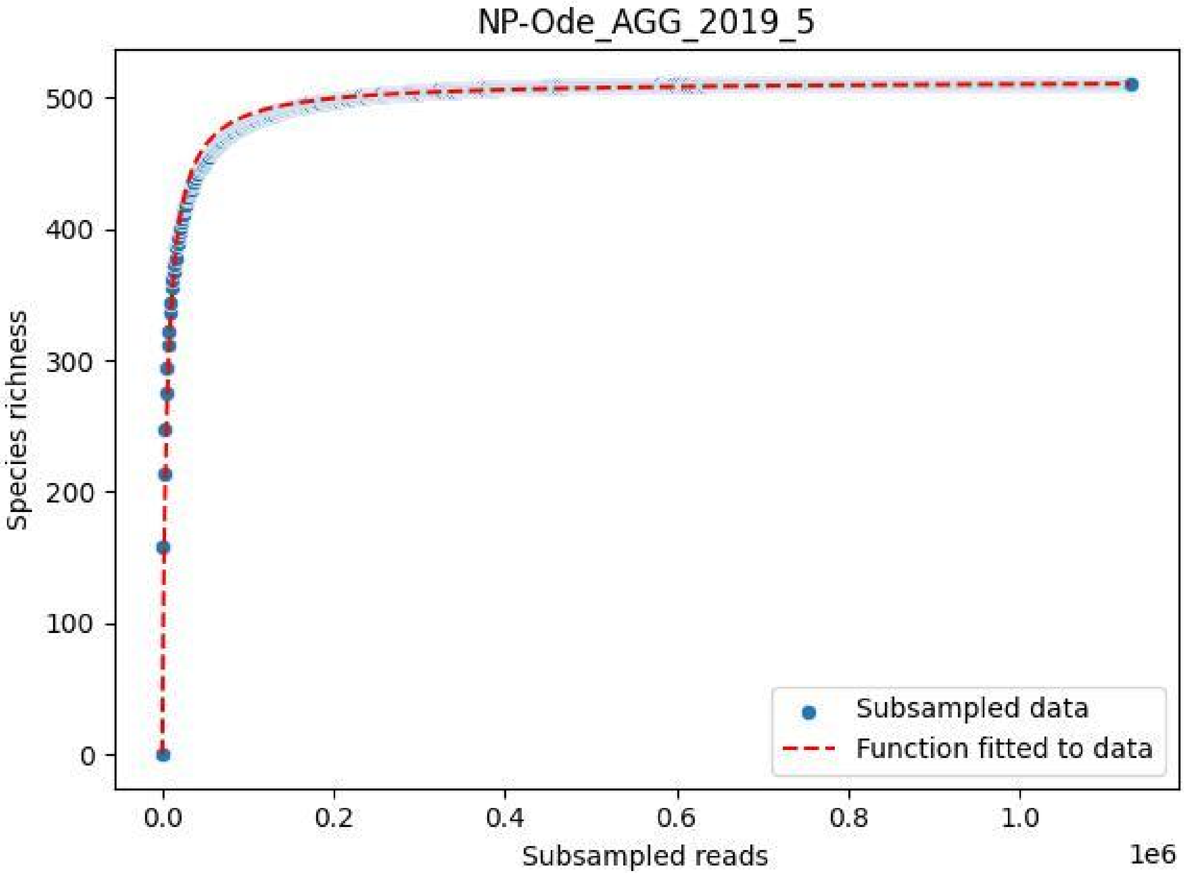

NP-Ode_AGG_2019_5
Species richness
500
400
300
200
100
0
0.0
0.2
0.4
0.6
0.8
1.0
1e6
Subsampled reads
Subsampled data
Function fitted to data

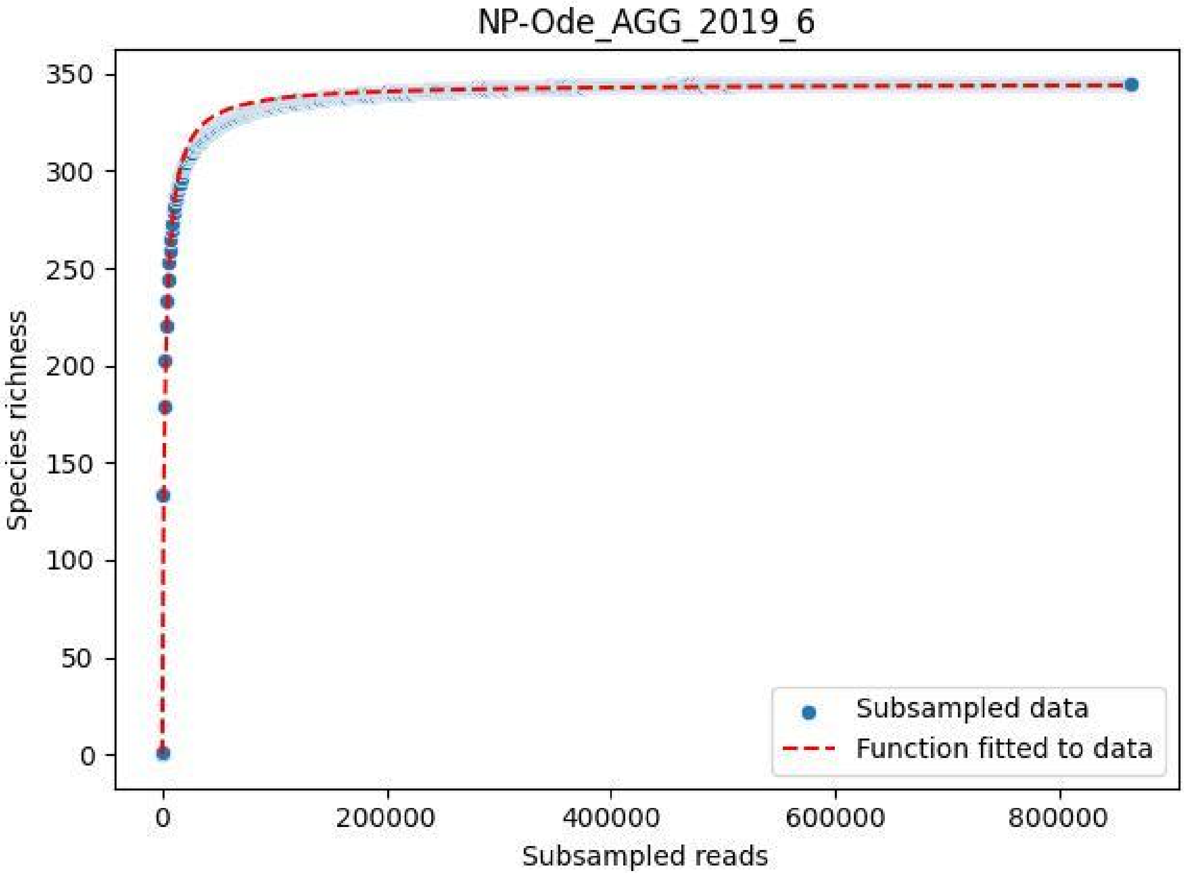
NP-Ode_AGG_2019_6
Species richness
Subsampled reads
0
50
100
150
200
250
300
350
0
200000
400000
600000
800000
Subsampled data
Function fitted to data

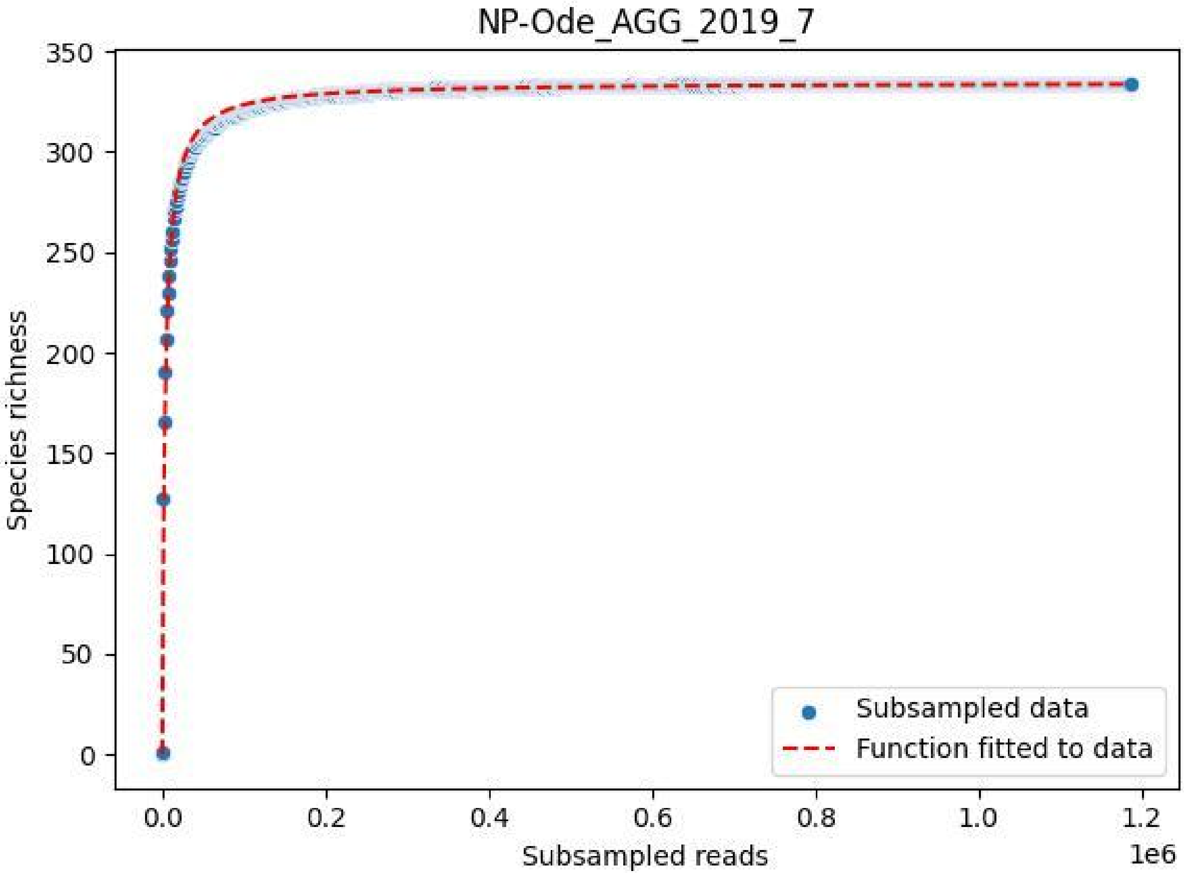

NP-Ode_AGG_2019_7
Species richness
Subsampled reads
1e6
0
50
100
150
200
250
300
350
0.0
0.2
0.4
0.6
0.8
1.0
1.2
Subsampled data
Function fitted to data

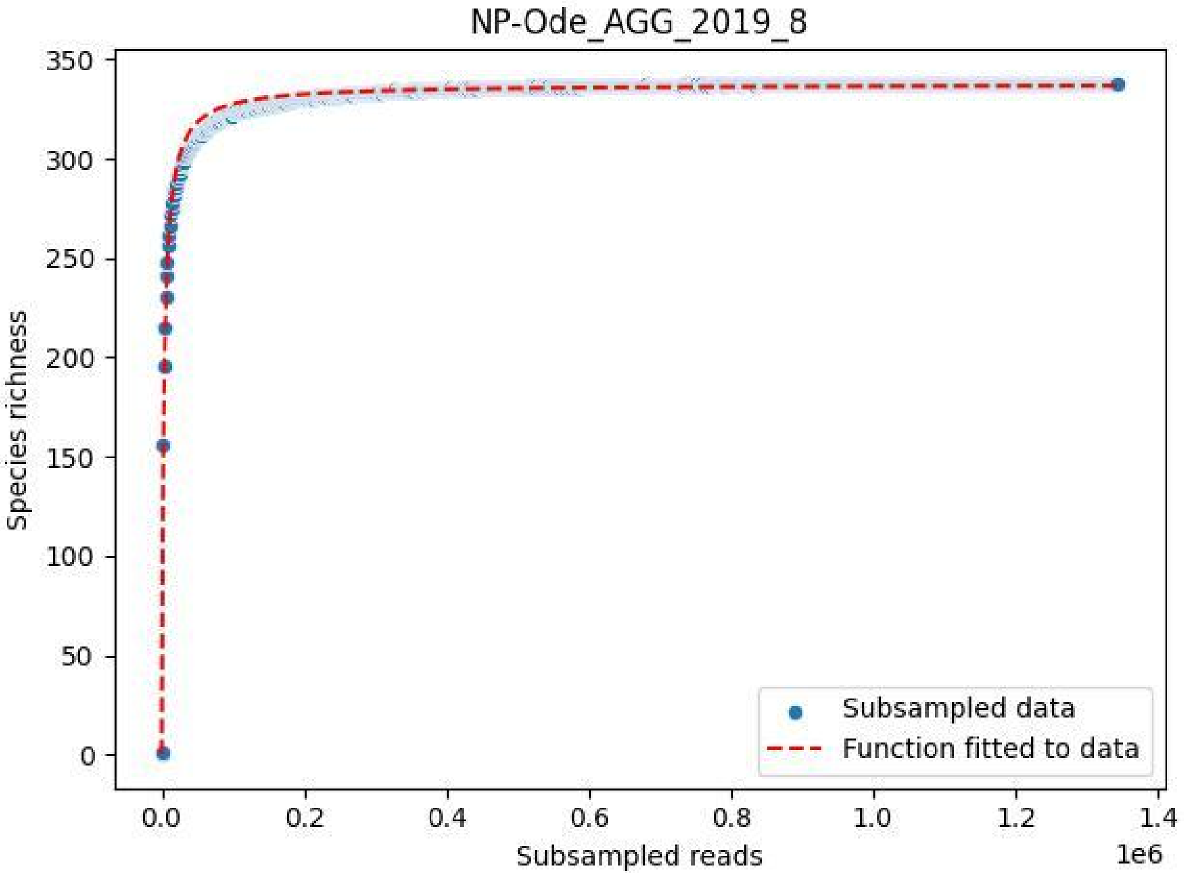
NP-Ode_AGG_2019_8
Species richness
Subsampled reads
1e6
0
50
100
150
200
250
300
350
0.0
0.2
0.4
0.6
0.8
1.0
1.2
1.4
Subsampled data
Function fitted to data

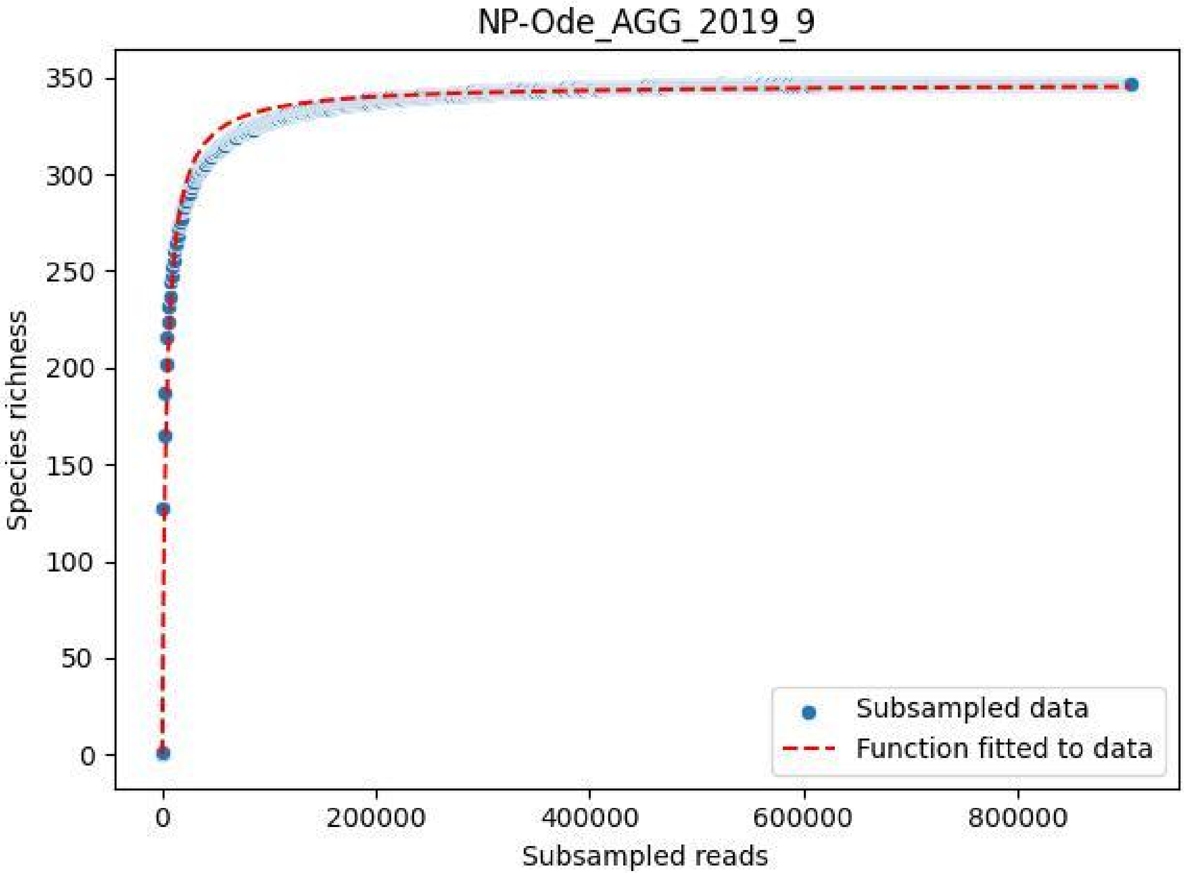
NP-Ode_AGG_2019_9
Species richness
Subsampled reads
0
50
100
150
200
250
300
350
0
200000
400000
600000
800000
Subsampled data
Function fitted to data

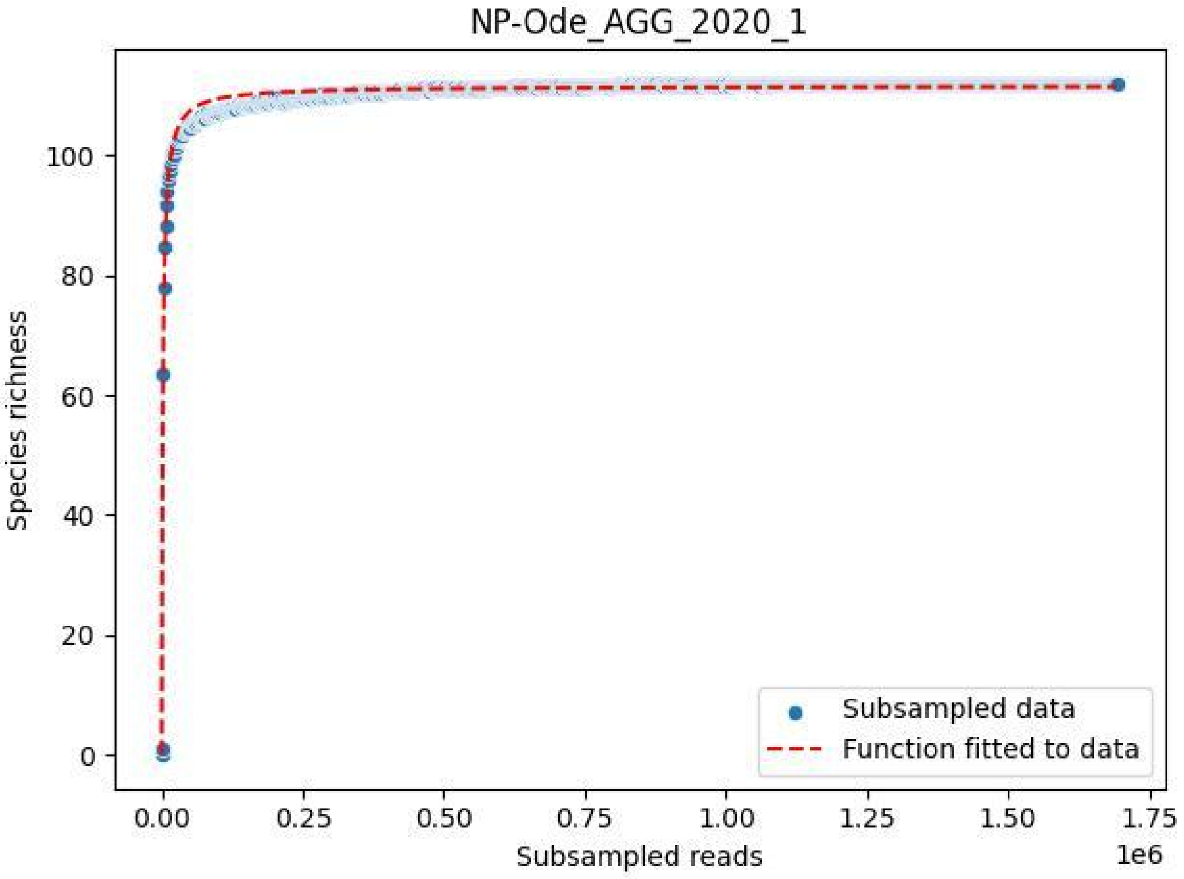

NP-Ode_AGG_2020_1
Species richness
Subsampled reads
1e6
0
20
40
60
80
100
0.00
0.25
0.50
0.75
1.00
1.25
1.50
1.75
Subsampled data
Function fitted to data

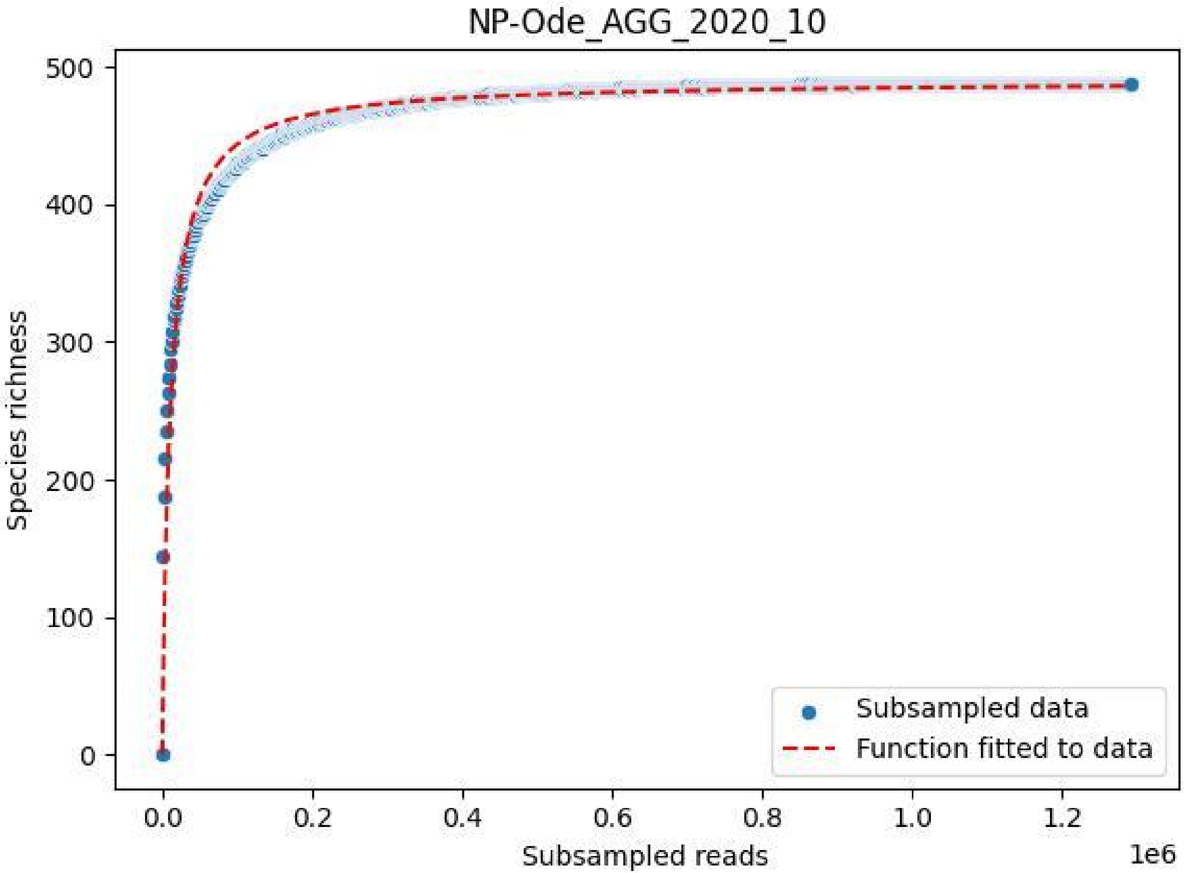

NP-Ode_AGG_2020_10
Species richness
Subsampled reads
1e6
500
400
300
200
100
0
0.0
0.2
0.4
0.6
0.8
1.0
1.2
Subsampled data
Function fitted to data

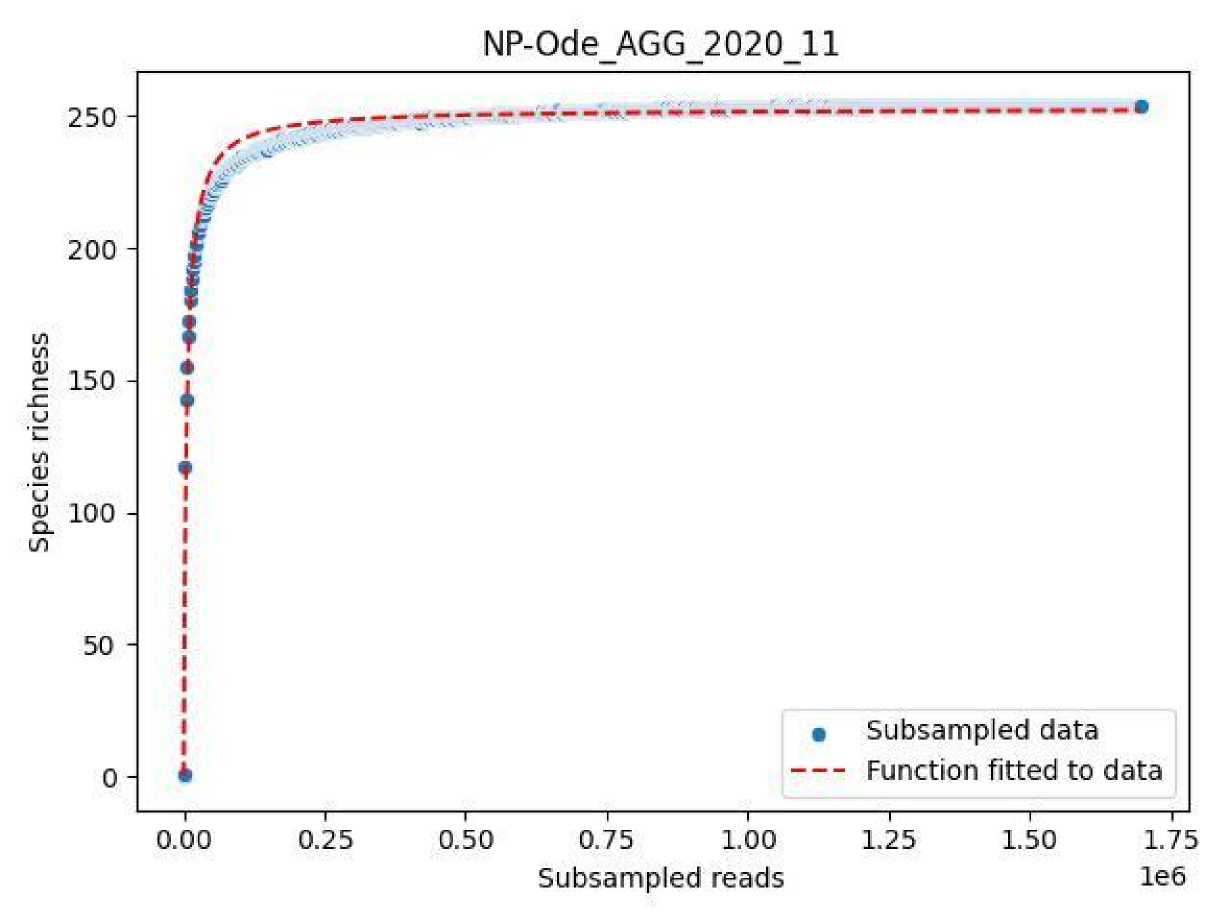

NP-Ode_AGG_2020_11
Species richness
0
50
100
150
200
250
0.00
0.25
0.50
0.75
1.00
1.25
1.50
1.75
1e6
Subsampled reads
Subsampled data
Function fitted to data

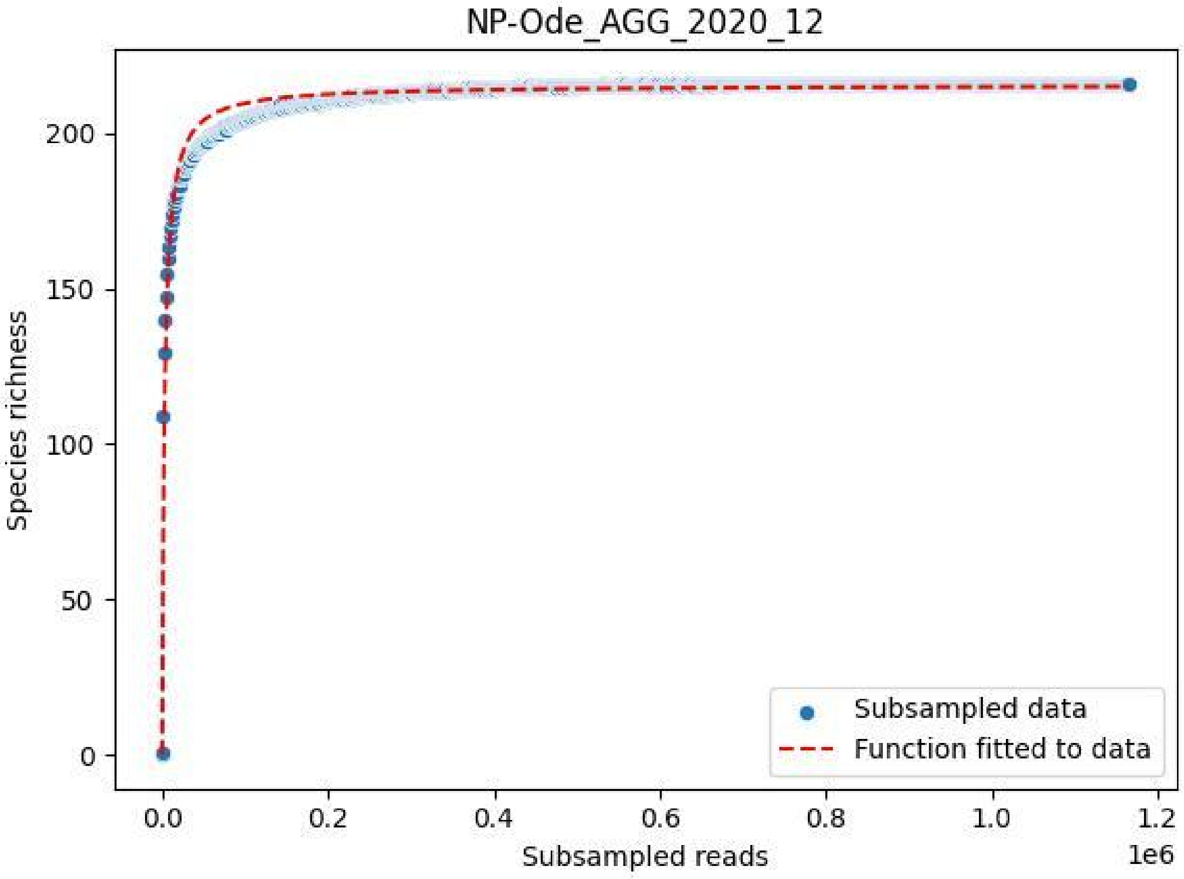

NP-Ode_AGG_2020_12
Species richness
Subsampled reads
1e6
0
50
100
150
200
0.0
0.2
0.4
0.6
0.8
1.0
1.2
Subsampled data
Function fitted to data

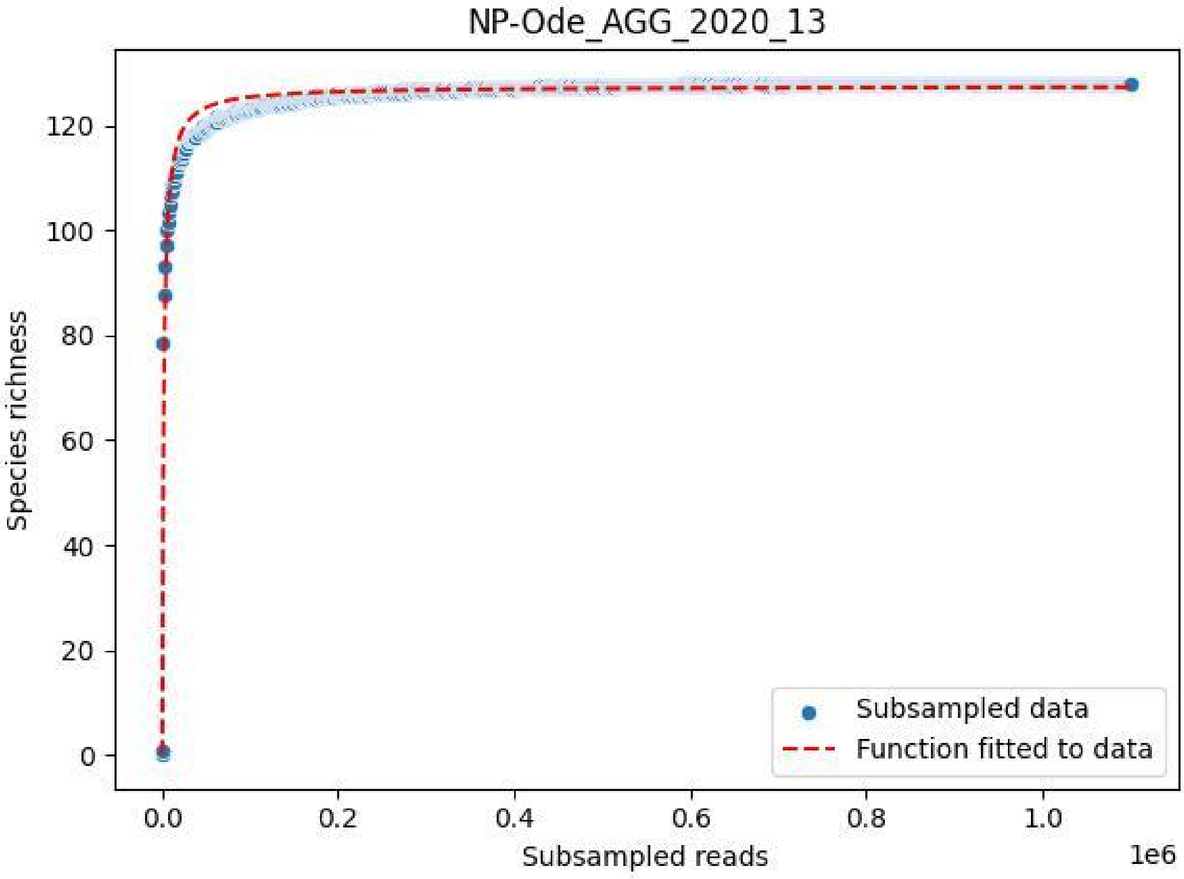
NP-Ode_AGG_2020_13
Species richness
Subsampled reads
1e6
0
20
40
60
80
100
120
0.0
0.2
0.4
0.6
0.8
1.0
Subsampled data
Function fitted to data

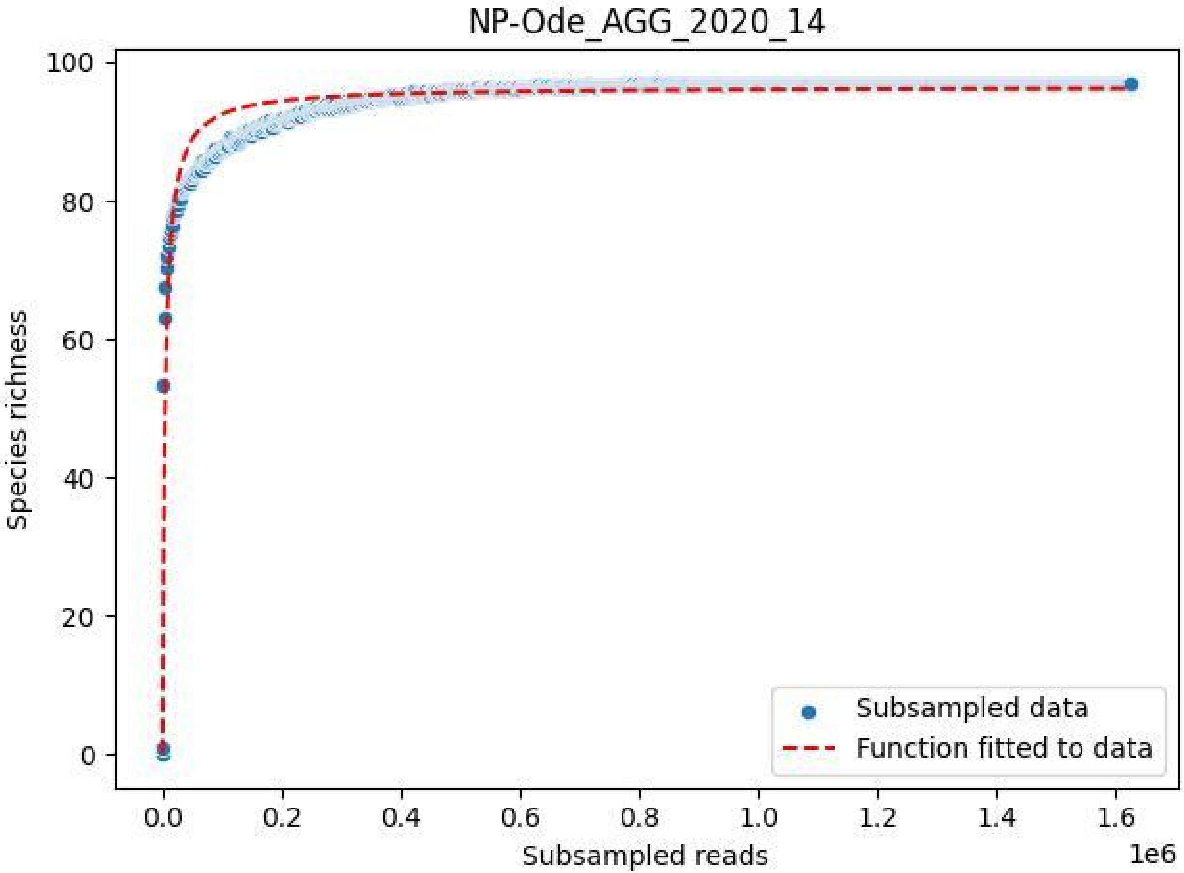
NP-Ode_AGG_2020_14
Species richness
Subsampled reads
1e6
0
20
40
60
80
100
0.0
0.2
0.4
0.6
0.8
1.0
1.2
1.4
1.6
Subsampled data
Function fitted to data

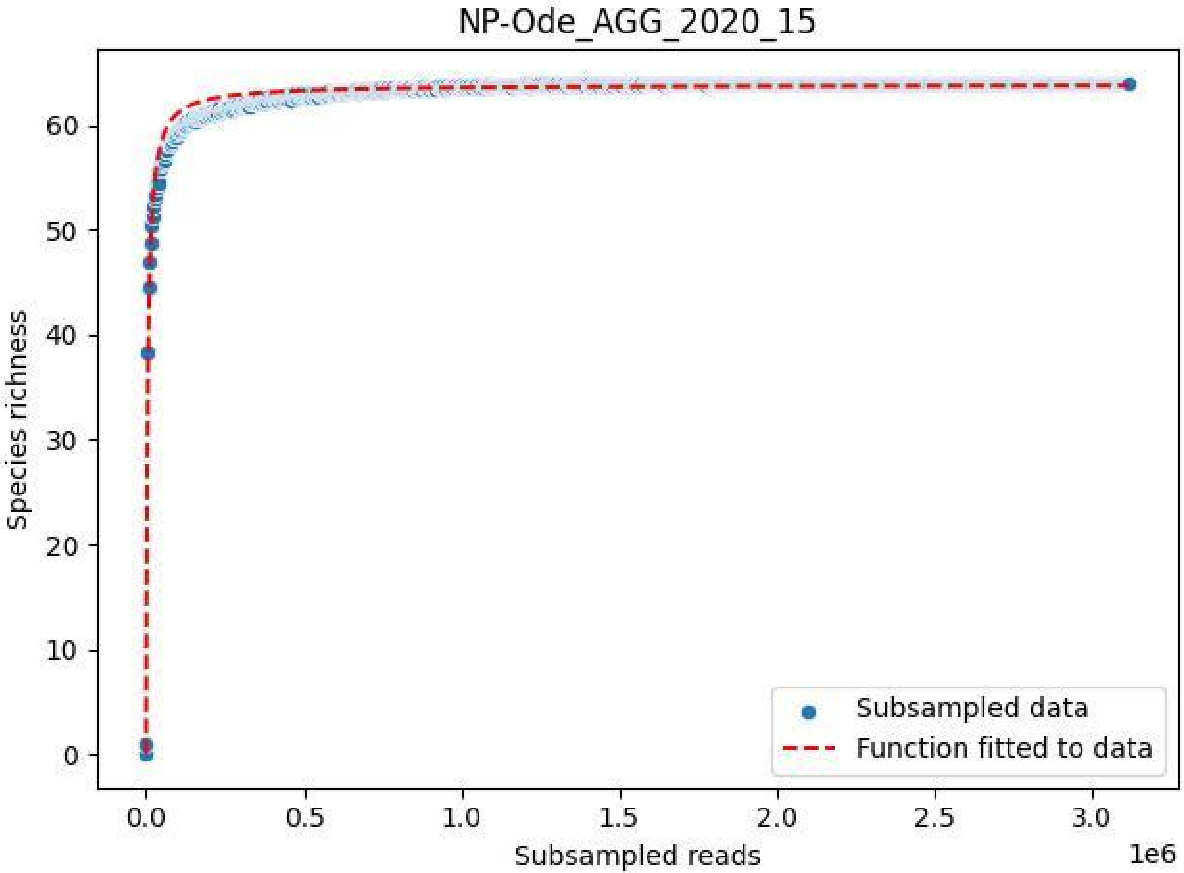

NP-Ode_AGG_2020_15
Species richness
Subsampled reads
1e6
0
10
20
30
40
50
60
0.0
0.5
1.0
1.5
2.0
2.5
3.0
Subsampled data
Function fitted to data

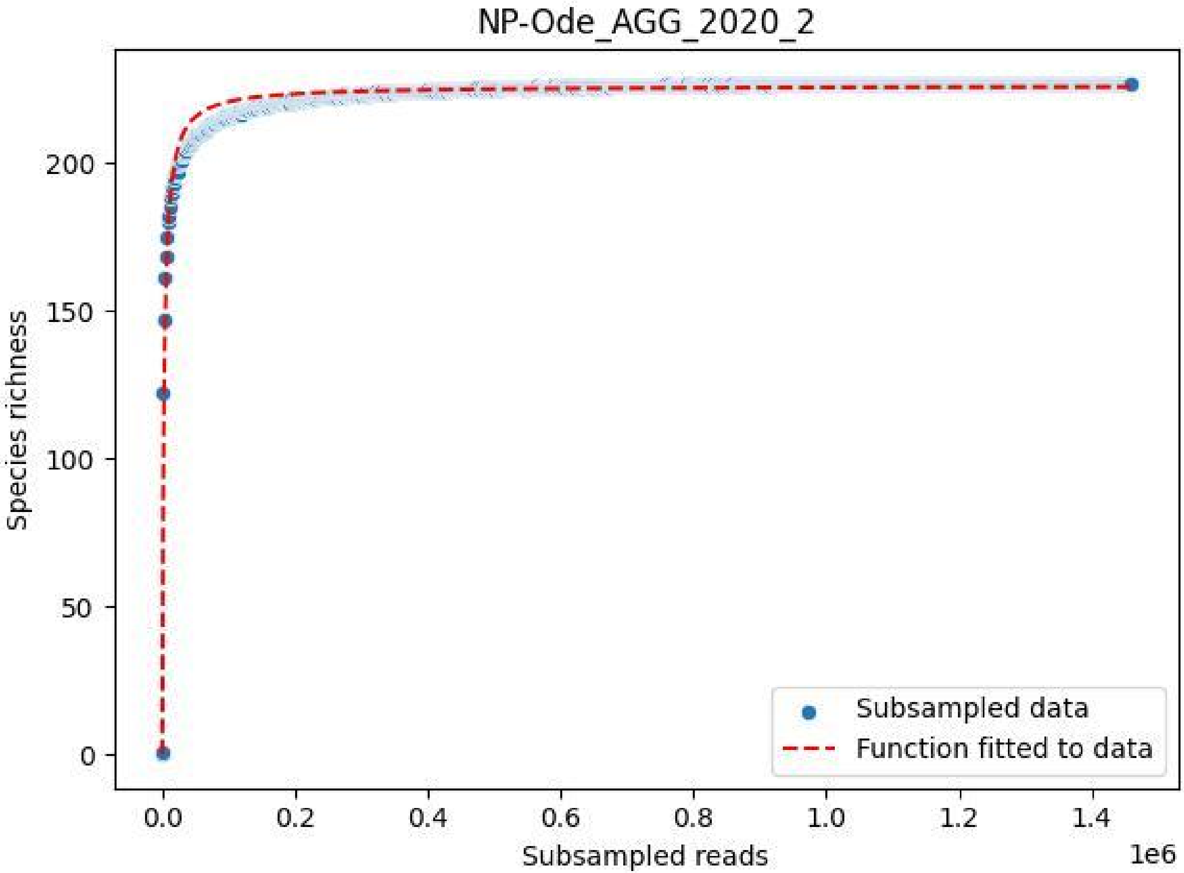
NP-Ode_AGG_2020_2
Species richness
Subsampled reads
1e6
0
50
100
150
200
0.0
0.2
0.4
0.6
0.8
1.0
1.2
1.4
Subsampled data
Function fitted to data

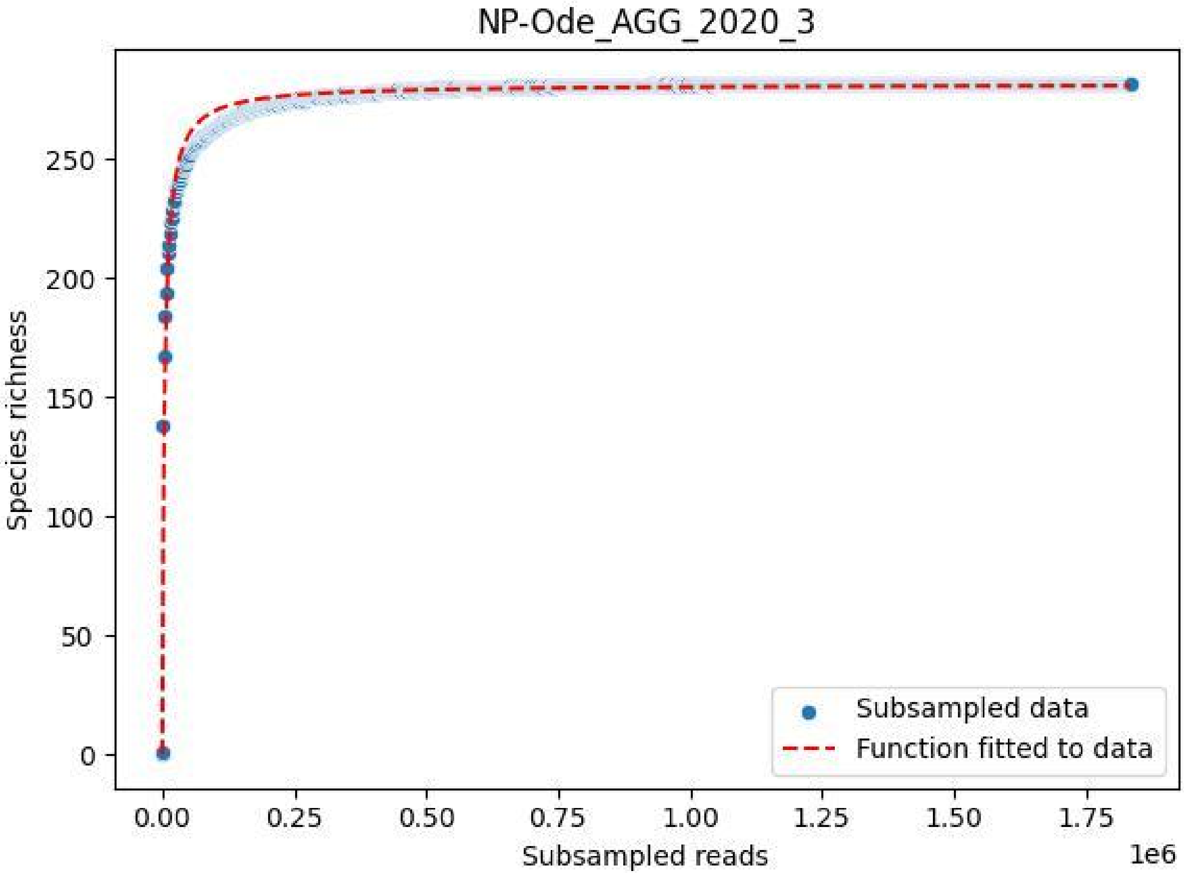
NP-Ode_AGG_2020_3
Species richness
0
50
100
150
200
250
0.00
0.25
0.50
0.75
1.00
1.25
1.50
1.75
1e6
Subsampled reads
Subsampled data
Function fitted to data

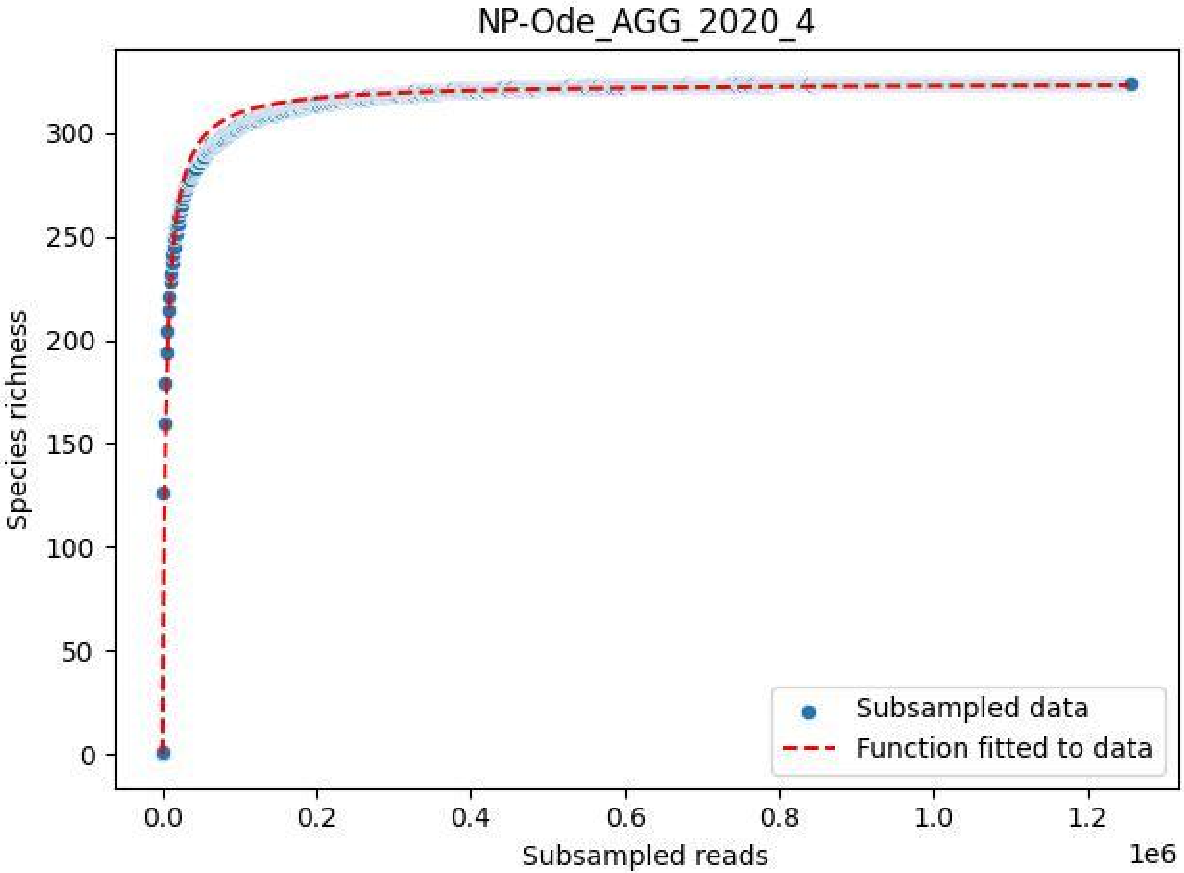

NP-Ode_AGG_2020_4
Species richness
Subsampled reads
1e6
0
50
100
150
200
250
300
0.0
0.2
0.4
0.6
0.8
1.0
1.2
Subsampled data
Function fitted to data

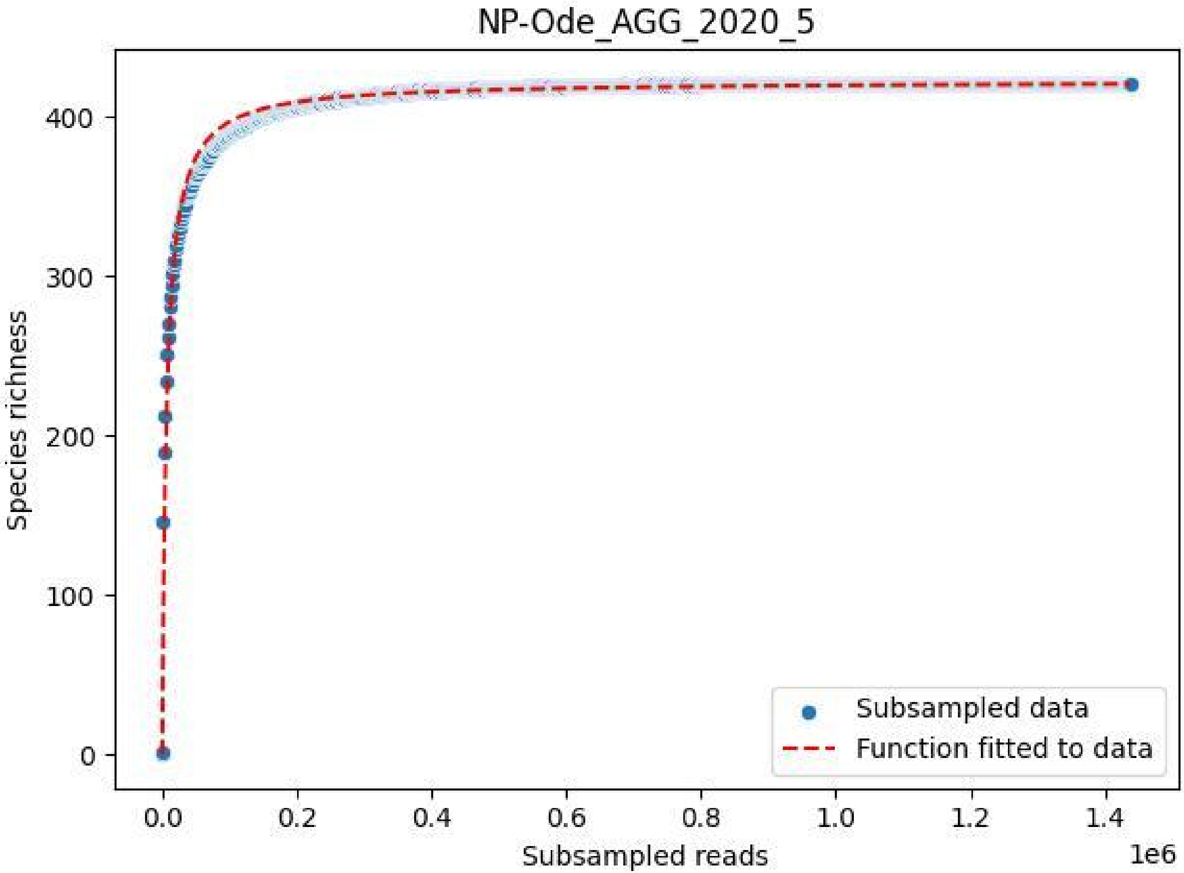
NP-Ode_AGG_2020_5
Species richness
400
300
200
100
0
0.0
0.2
0.4
0.6
0.8
1.0
1.2
1.4
Subsampled reads
1e6
Subsampled data
Function fitted to data

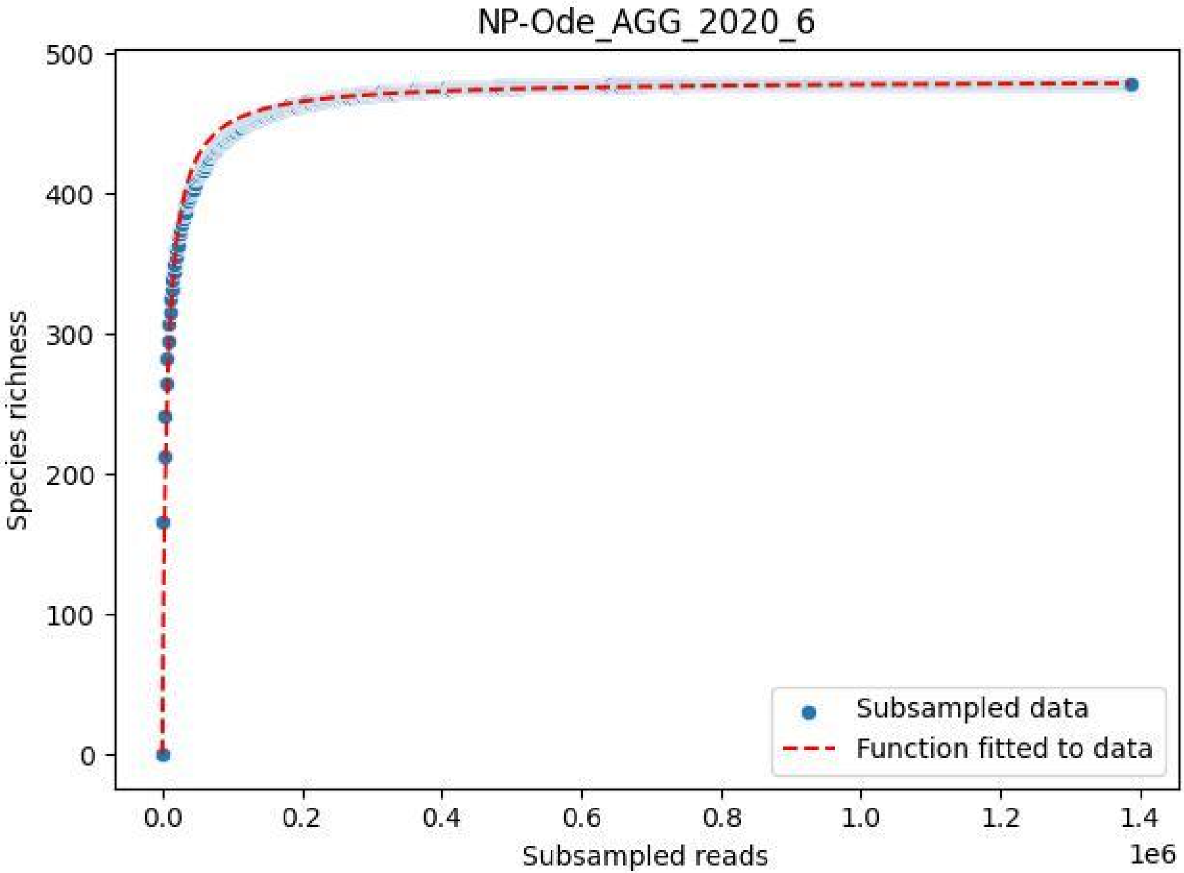

NP-Ode_AGG_2020_6
Species richness
Subsampled reads
1e6
0
100
200
300
400
500
0.0
0.2
0.4
0.6
0.8
1.0
1.2
1.4
Subsampled data
Function fitted to data

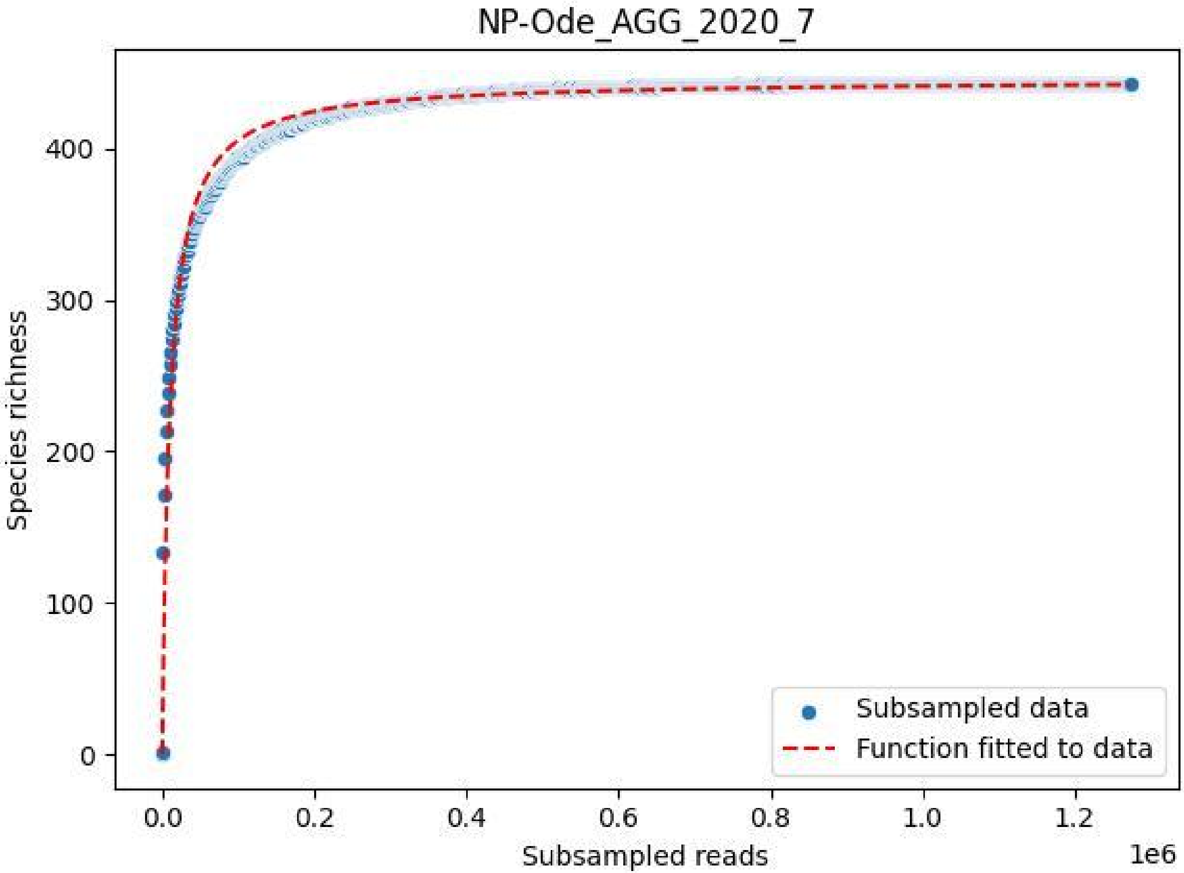
NP-Ode_AGG_2020_7
Species richness
Subsampled reads
1e6
0
100
200
300
400
0.0
0.2
0.4
0.6
0.8
1.0
1.2
Subsampled data
Function fitted to data

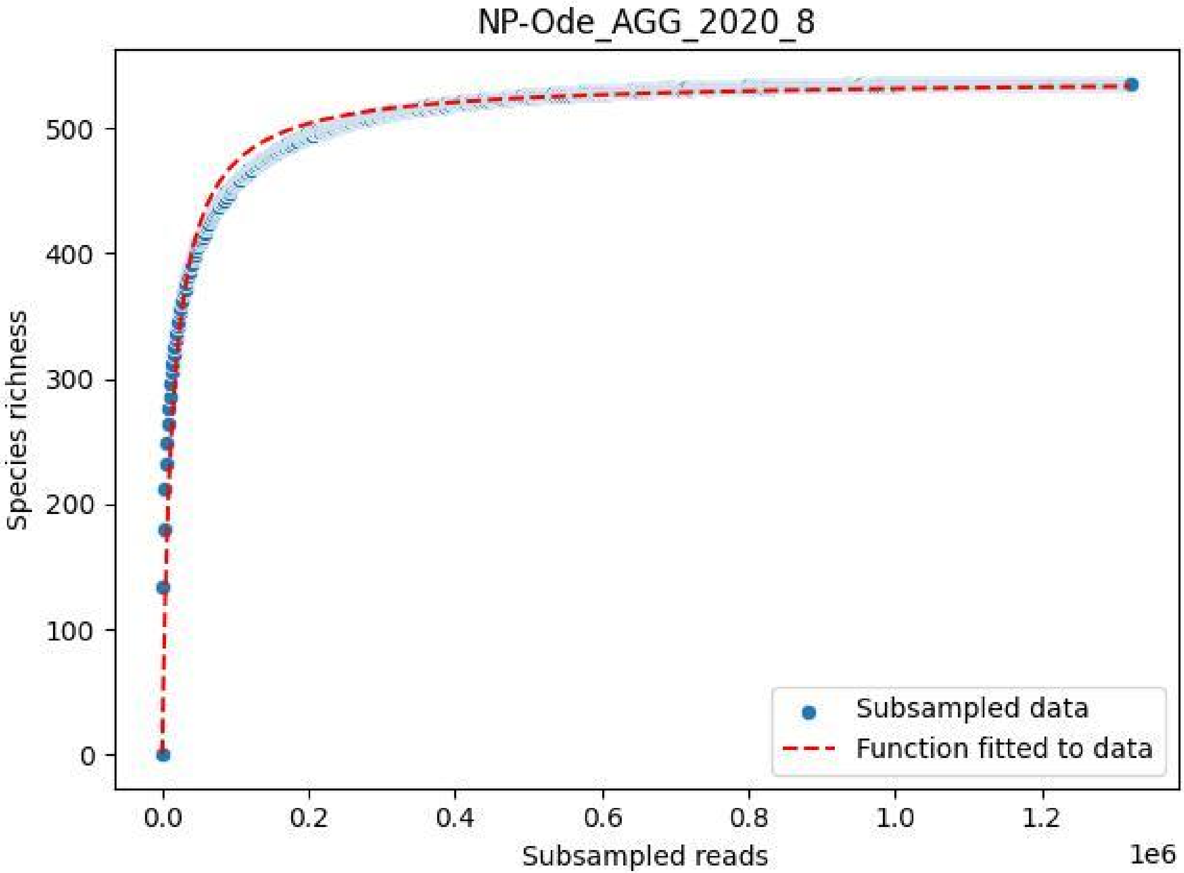

NP-Ode_AGG_2020_8
Species richness
Subsampled reads
1e6
0
100
200
300
400
500
0.0
0.2
0.4
0.6
0.8
1.0
1.2
Subsampled data
Function fitted to data

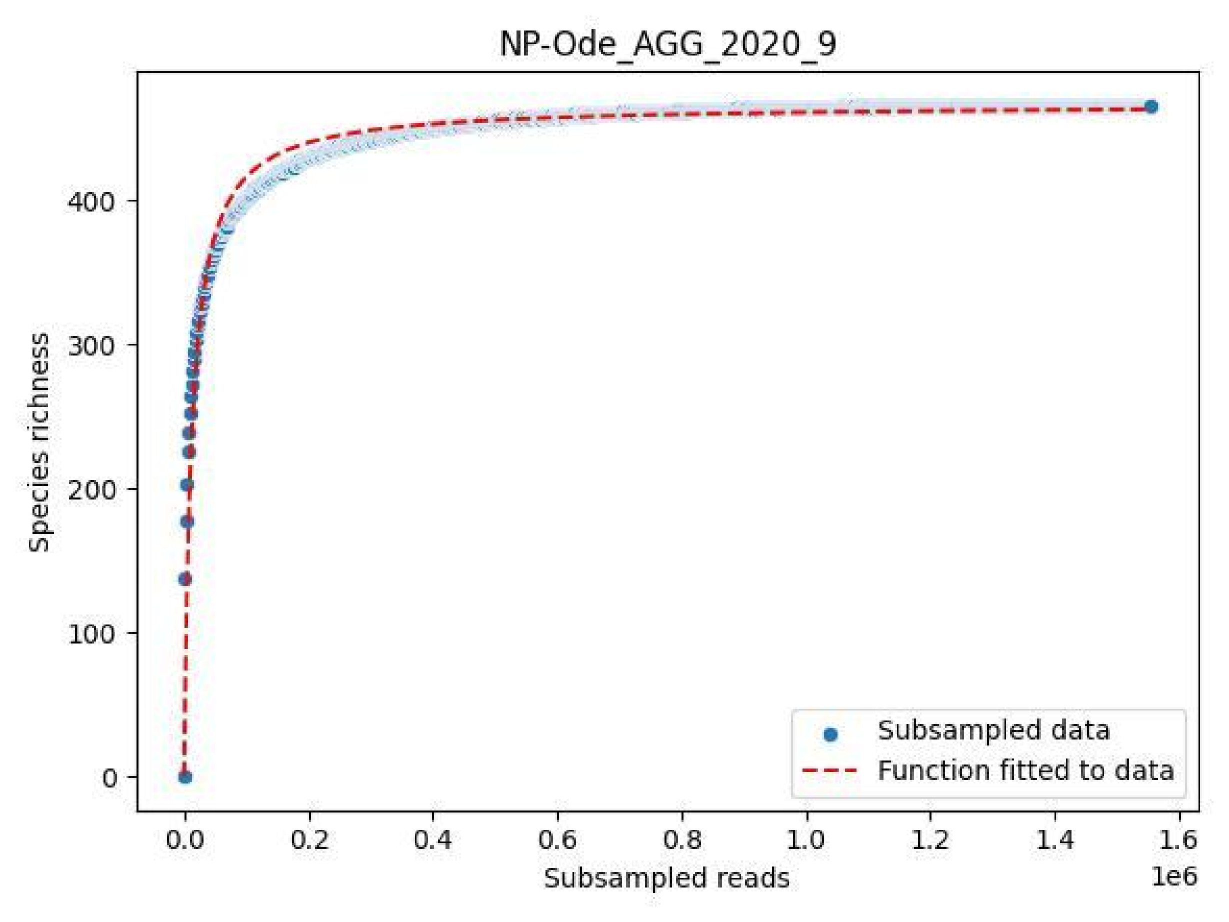

NP-Ode_AGG_2020_9
Species richness
Subsampled reads
1e6
0
100
200
300
400
0.0
0.2
0.4
0.6
0.8
1.0
1.2
1.4
1.6
Subsampled data
Function fitted to data

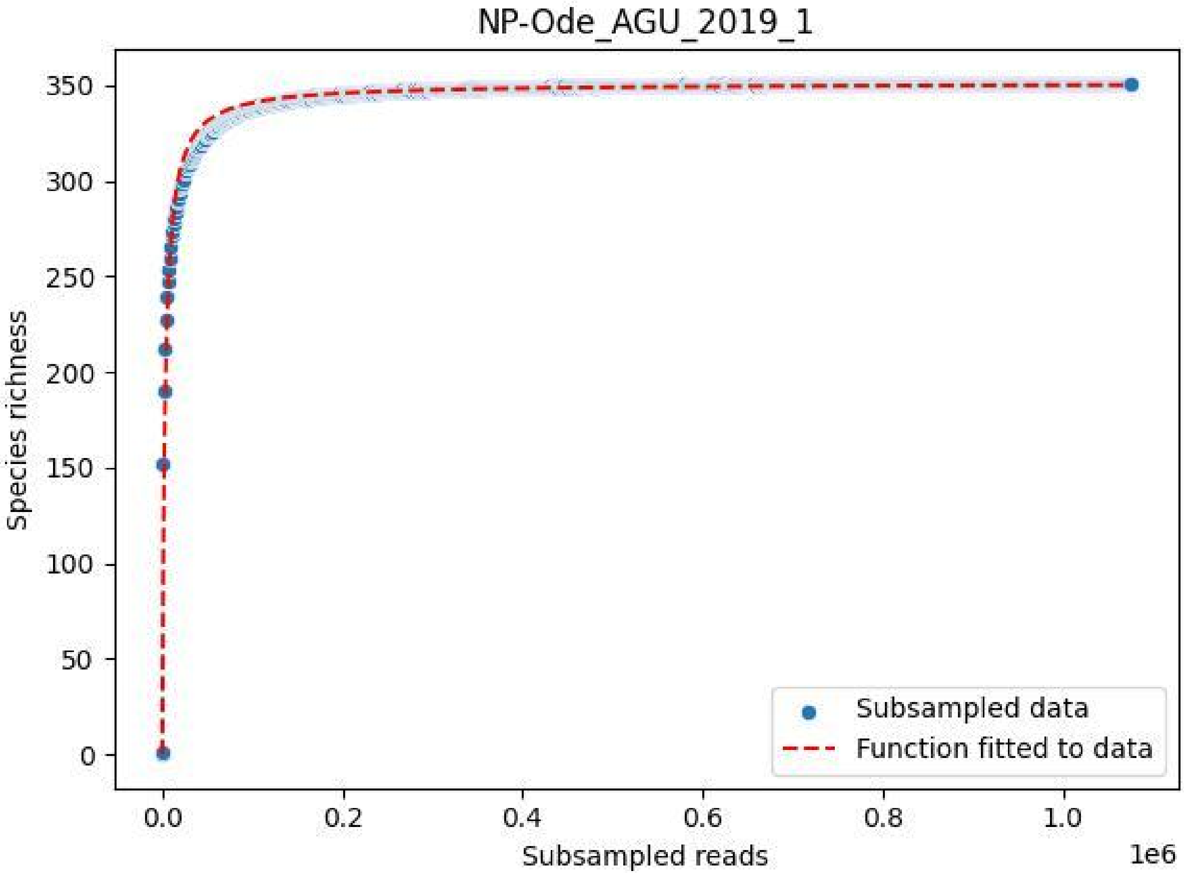

NP-Ode_AGU_2019_1
Species richness
350
300
250
200
150
100
50
0
Subsampled reads
0.0
0.2
0.4
0.6
0.8
1.0
1e6
Subsampled data
Function fitted to data

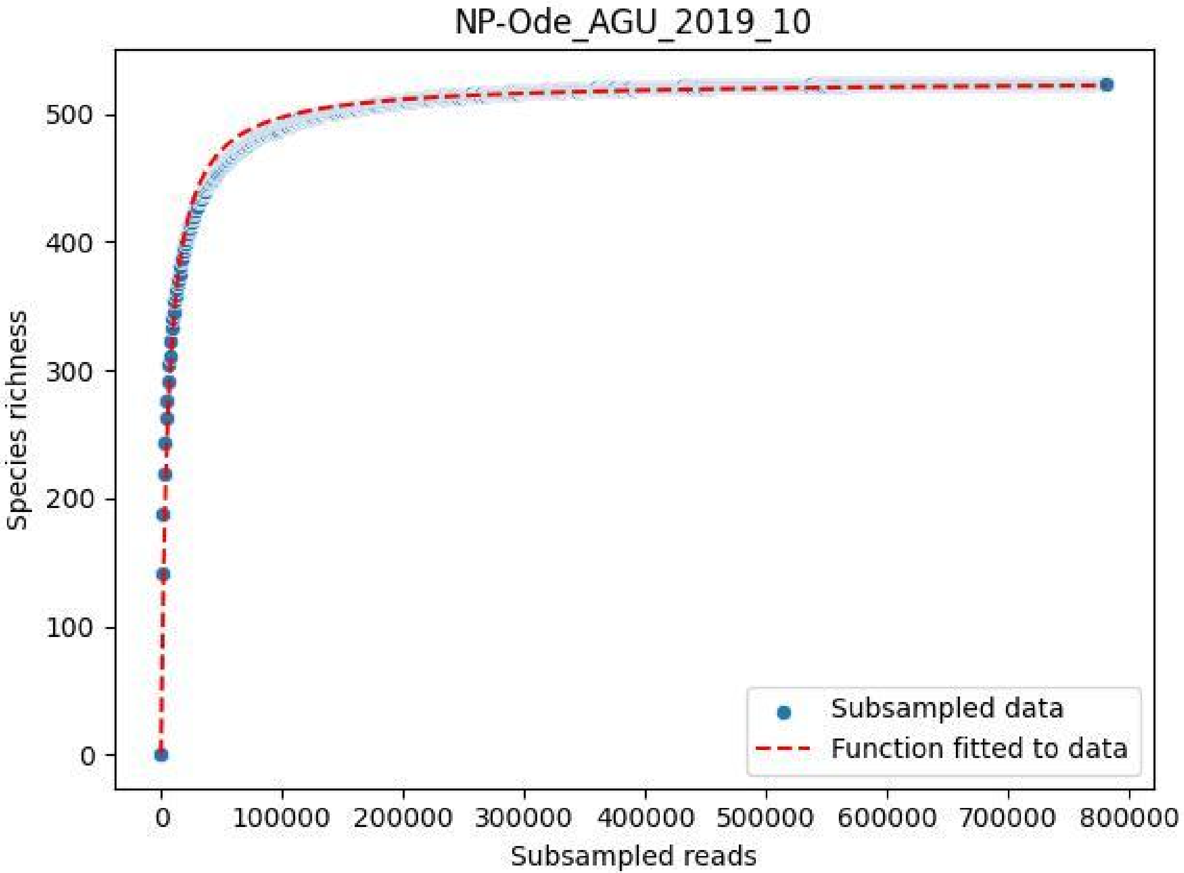

NP-Ode_AGU_2019_10
Species richness
Subsampled reads
0
100
200
300
400
500
100000
200000
300000
400000
500000
600000
700000
800000
Subsampled data
Function fitted to data

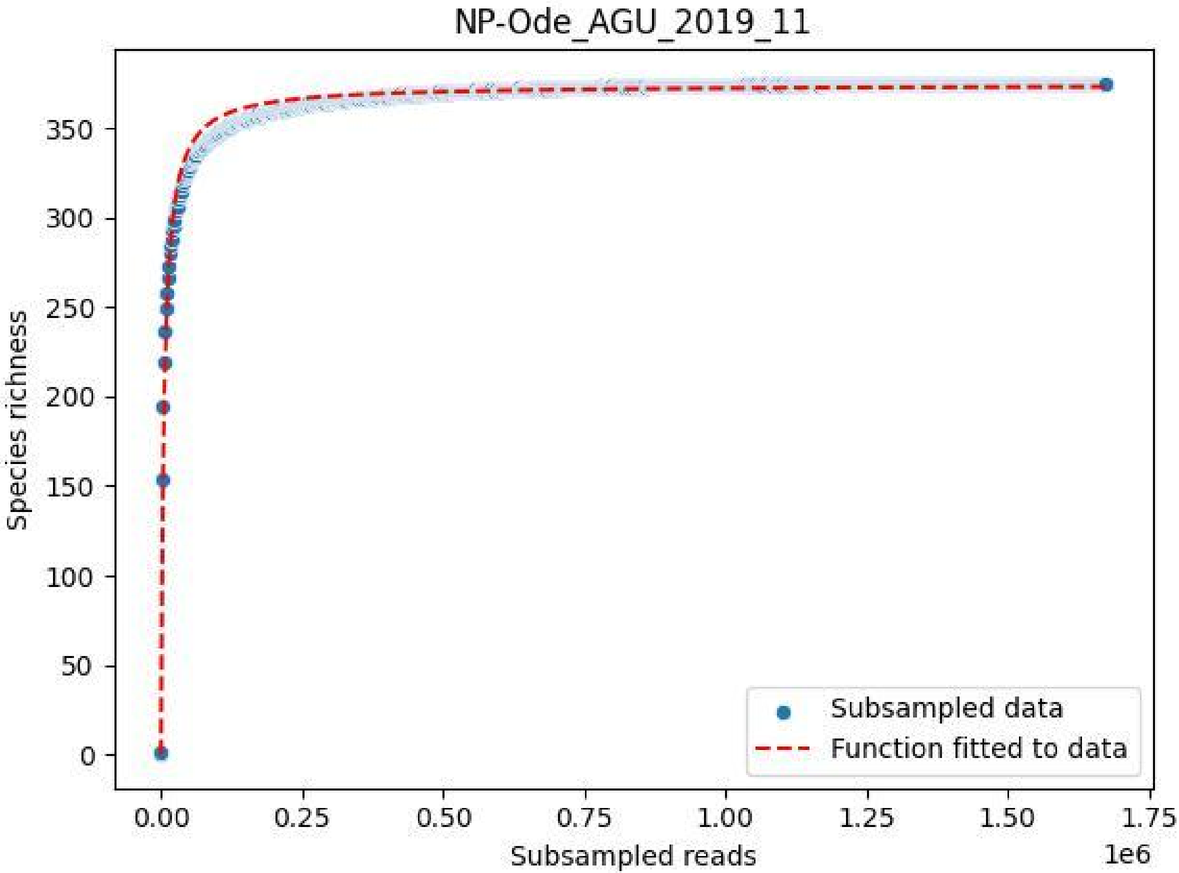

NP-Ode_AGU_2019_11
Species richness
Subsampled reads
1e6
0
50
100
150
200
250
300
350
0.00
0.25
0.50
0.75
1.00
1.25
1.50
1.75
Subsampled data
Function fitted to data

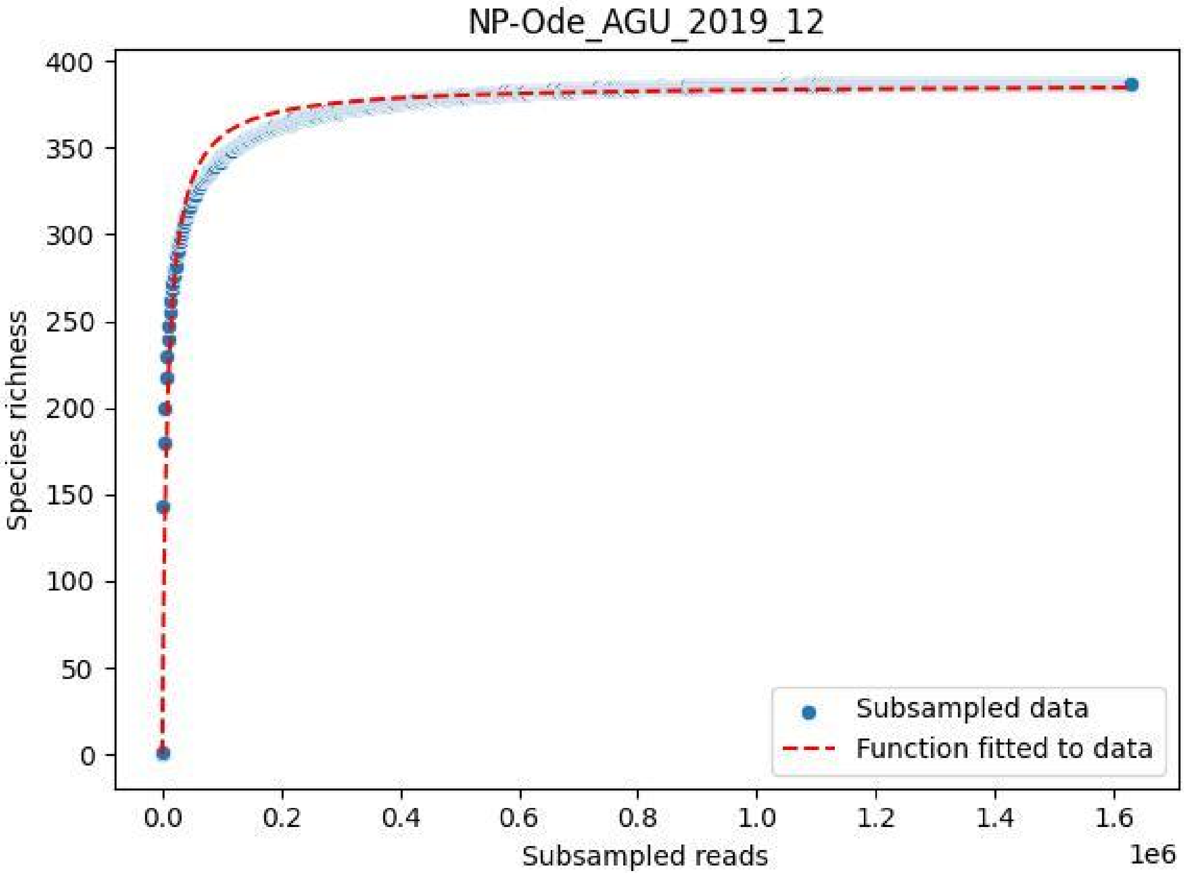
NP-Ode_AGU_2019_12
Species richness
Subsampled reads
1e6
Subsampled data
Function fitted to data
0
50
100
150
200
250
300
350
400
0.0
0.2
0.4
0.6
0.8
1.0
1.2
1.4
1.6

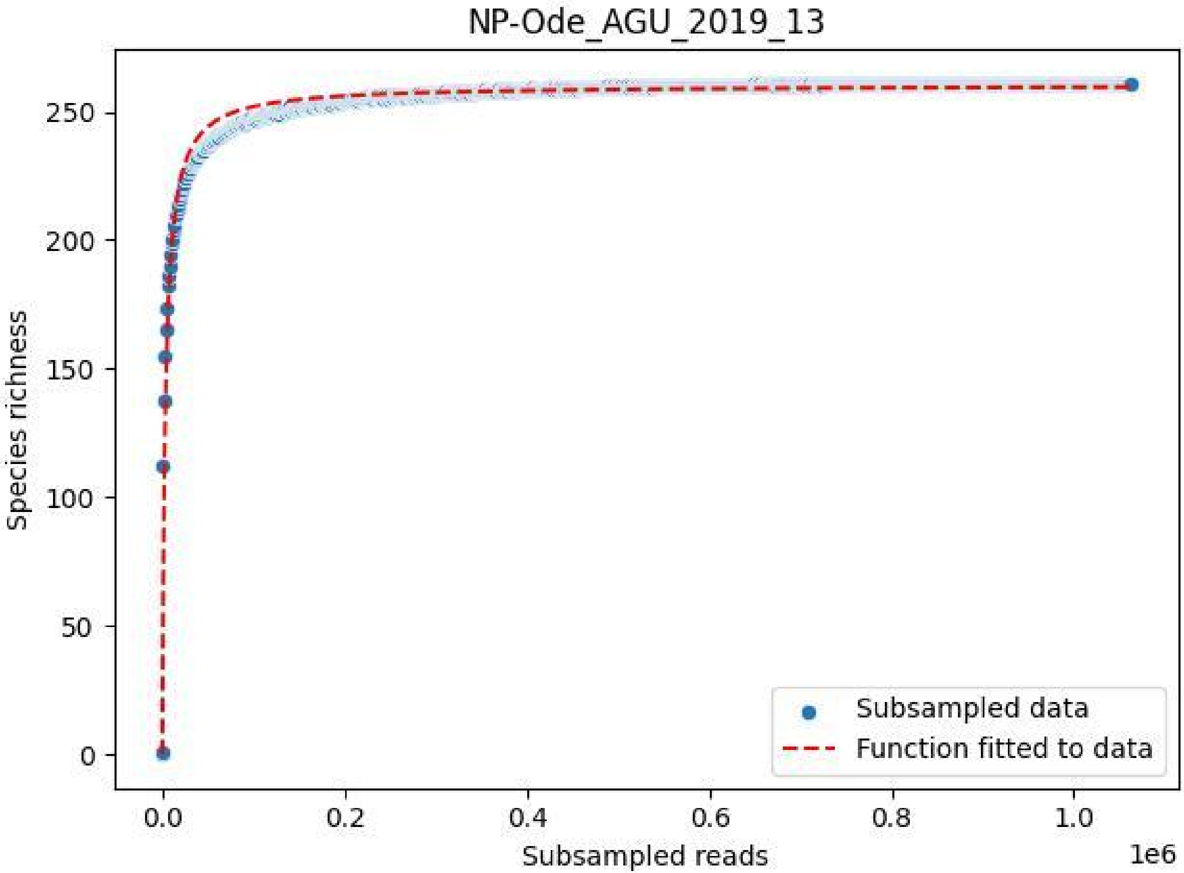
NP-Ode_AGU_2019_13
Species richness
Subsampled reads
1e6
0
50
100
150
200
250
0.0
0.2
0.4
0.6
0.8
1.0
Subsampled data
Function fitted to data

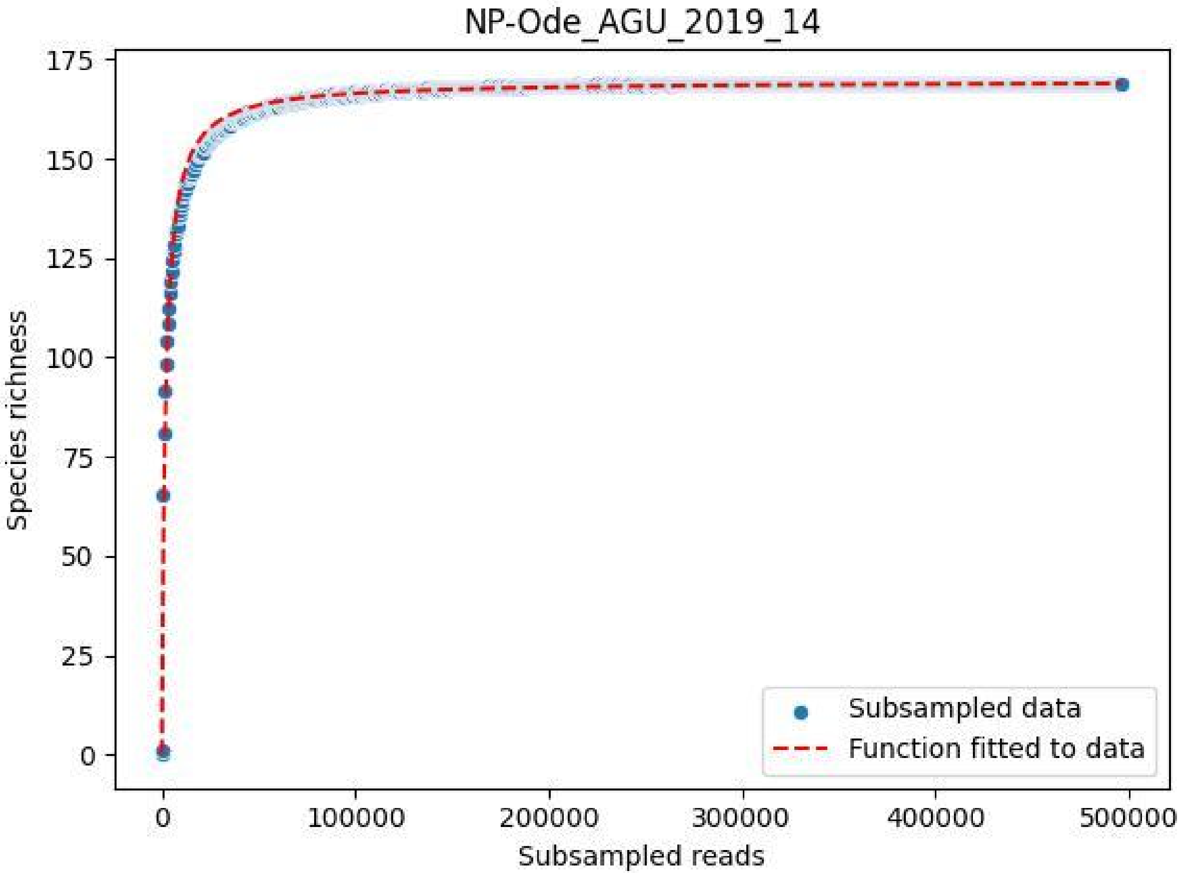
NP-Ode_AGU_2019_14
Species richness
Subsampled reads
0
25
50
75
100
125
150
175
0
100000
200000
300000
400000
500000
Subsampled data
Function fitted to data

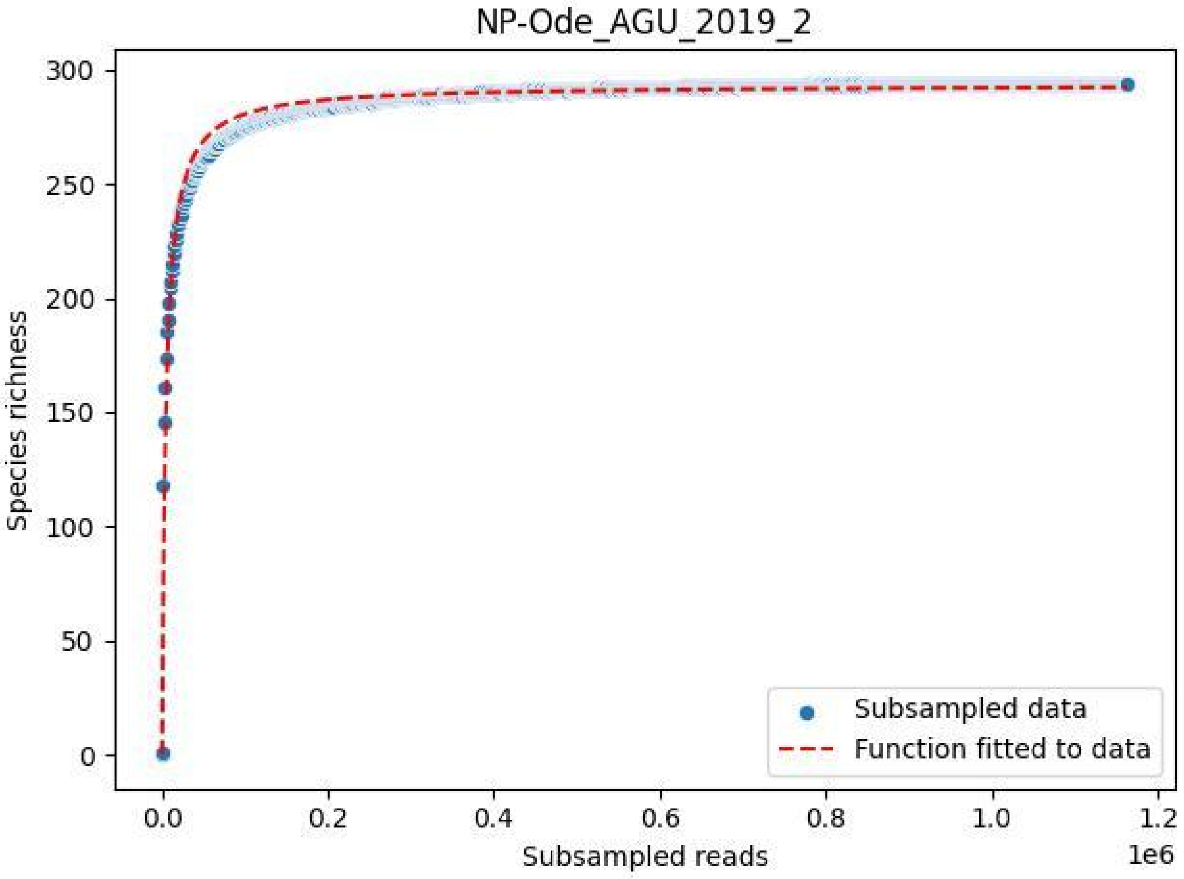
NP-Ode_AGU_2019_2
Species richness
Subsampled reads
1e6
300
250
200
150
100
50
0
0.0
0.2
0.4
0.6
0.8
1.0
1.2
Subsampled data
Function fitted to data

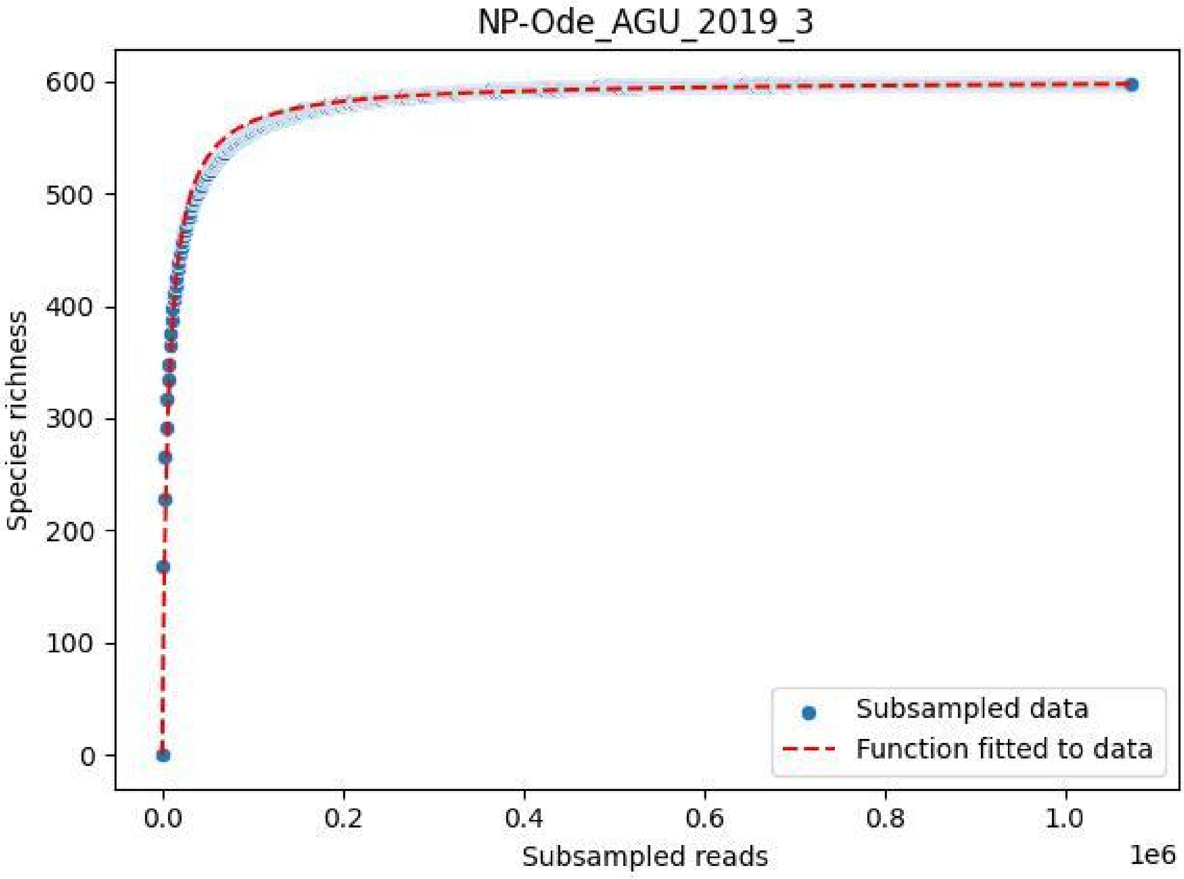

NP-Ode_AGU_2019_3
Species richness
Subsampled reads
1e6
0
100
200
300
400
500
600
0.0
0.2
0.4
0.6
0.8
1.0
Subsampled data
Function fitted to data

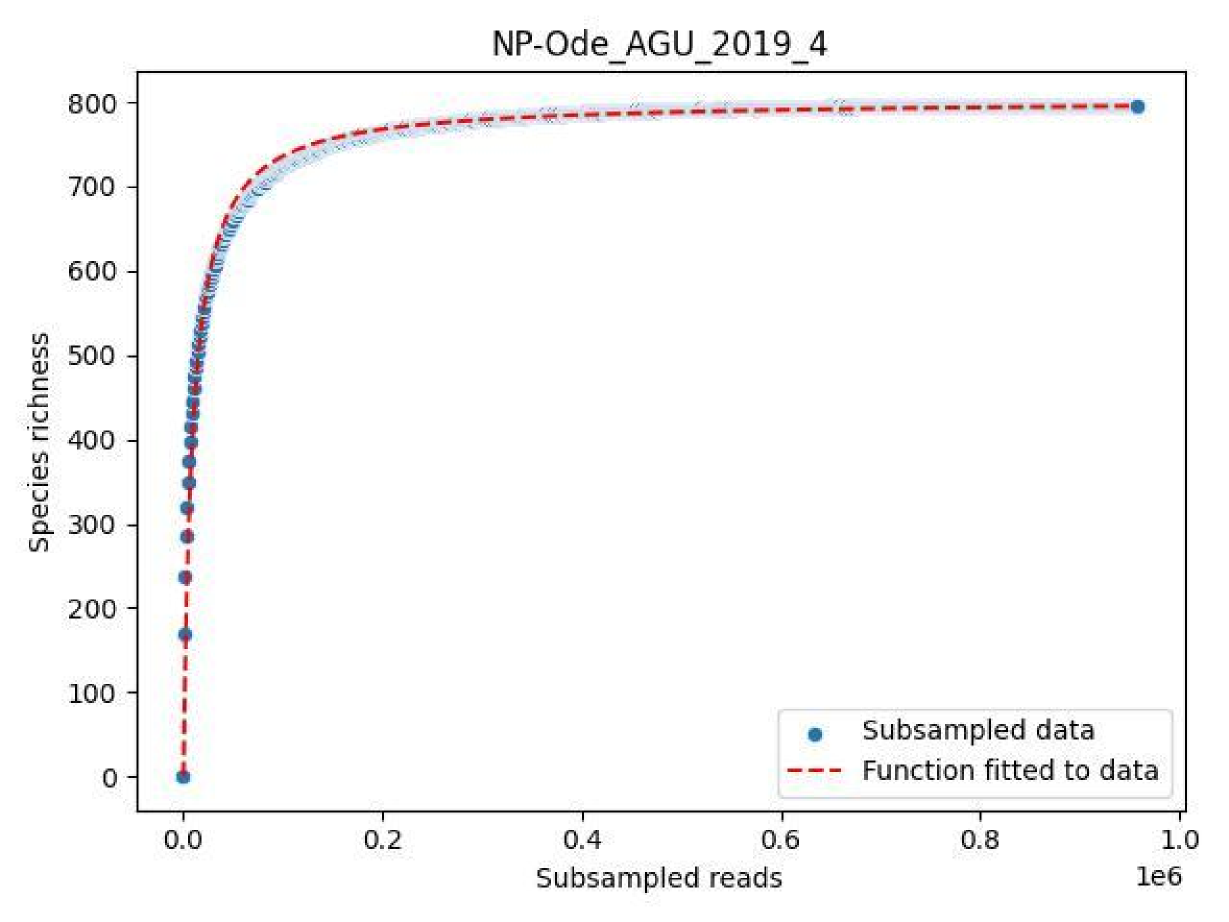
NP-Ode_AGU_2019_4
Species richness
0
100
200
300
400
500
600
700
800
0.0
0.2
0.4
0.6
0.8
1.0
1e6
Subsampled reads
Subsampled data
Function fitted to data

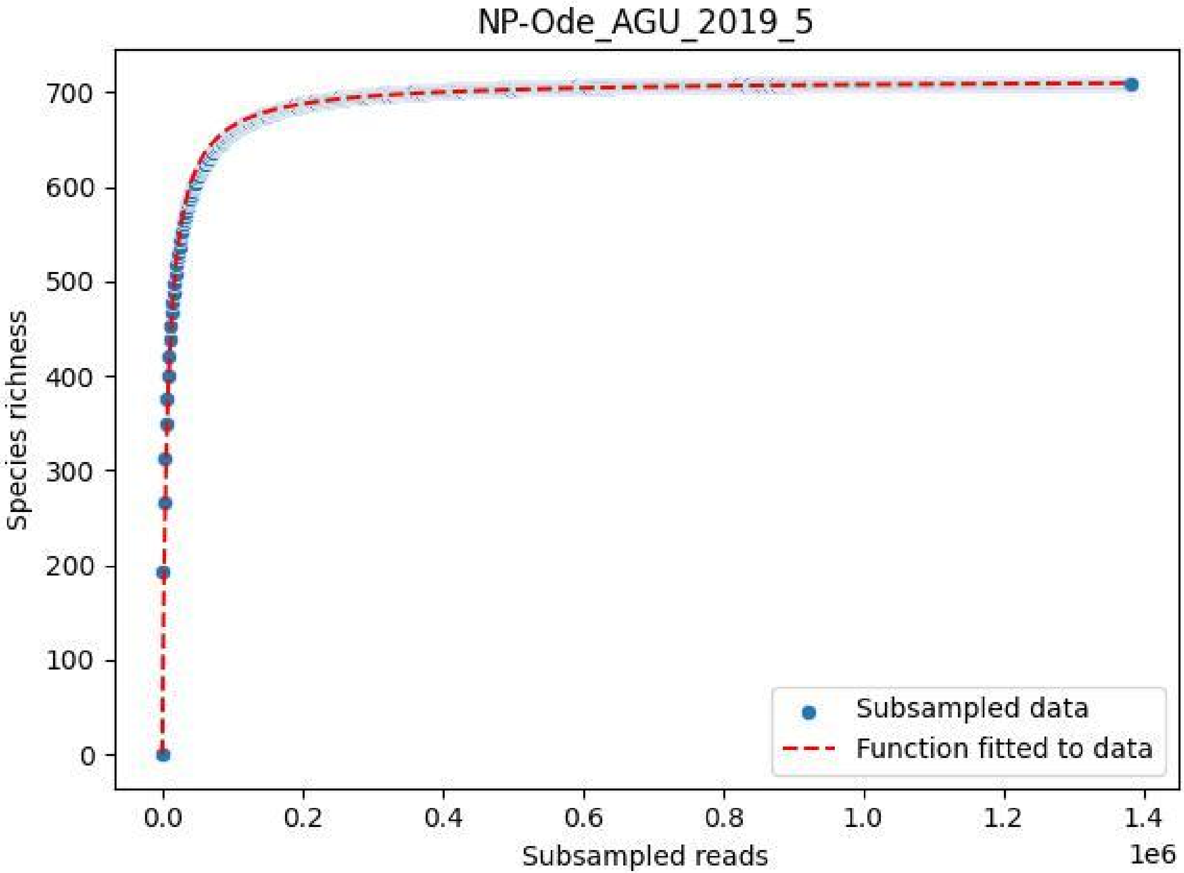
NP-Ode_AGU_2019_5
Species richness
0
100
200
300
400
500
600
700
0.0
0.2
0.4
0.6
0.8
1.0
1.2
1.4
1e6
Subsampled reads
Subsampled data
Function fitted to data

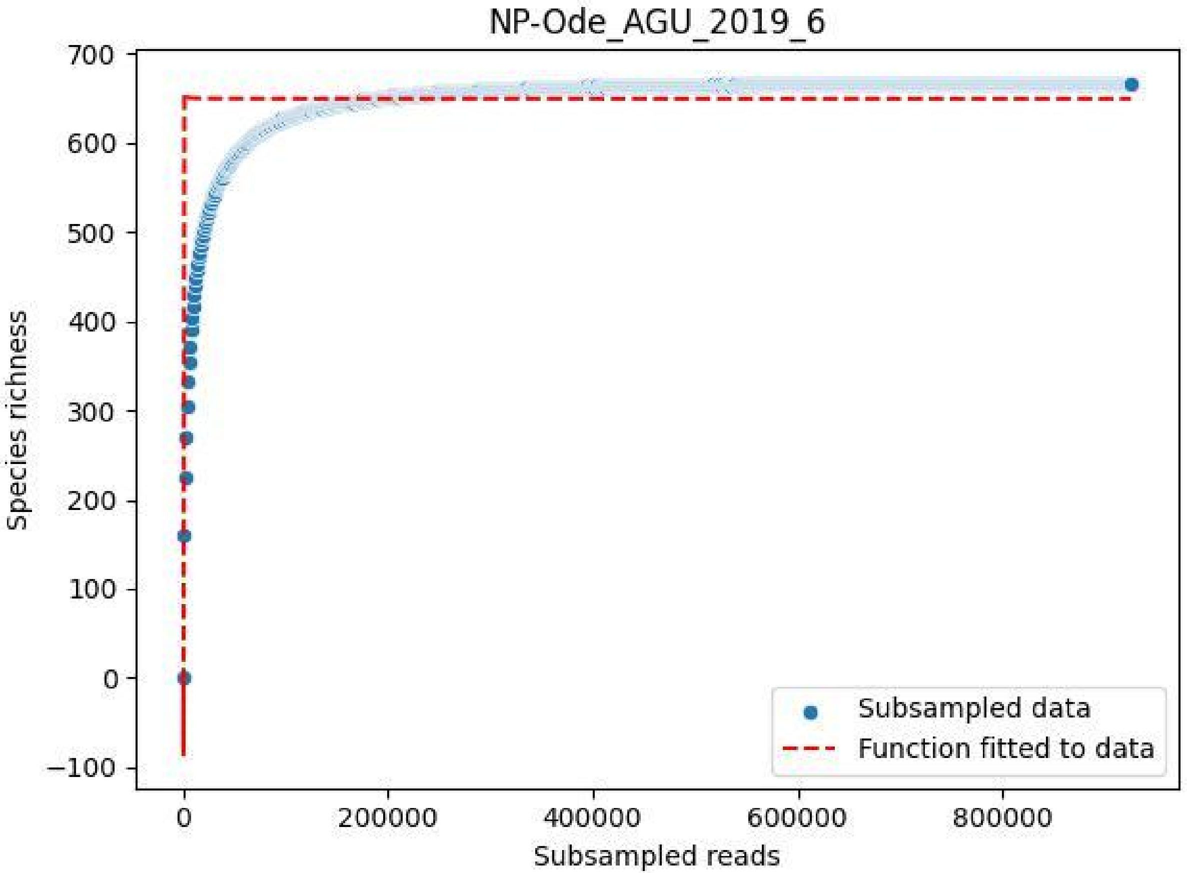

NP-Ode_AGU_2019_6
Species richness
Subsampled reads
700
600
500
400
300
200
100
0
−100
0
200000
400000
600000
800000
Subsampled data
Function fitted to data

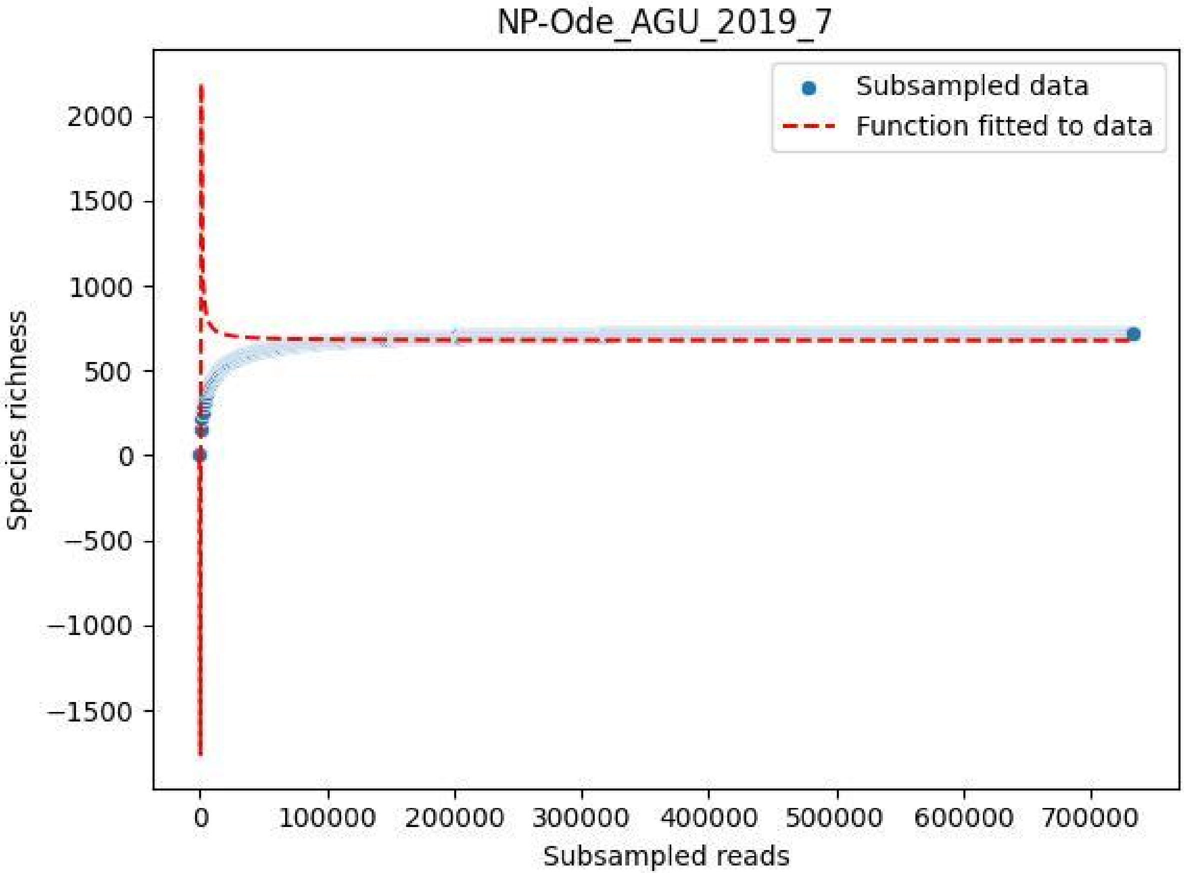

NP-Ode_AGU_2019_7
Subsampled data
Function fitted to data
Species richness
Subsampled reads
2000
1500
1000
500
0
−500
−1000
−1500
0
100000
200000
300000
400000
500000
600000
700000

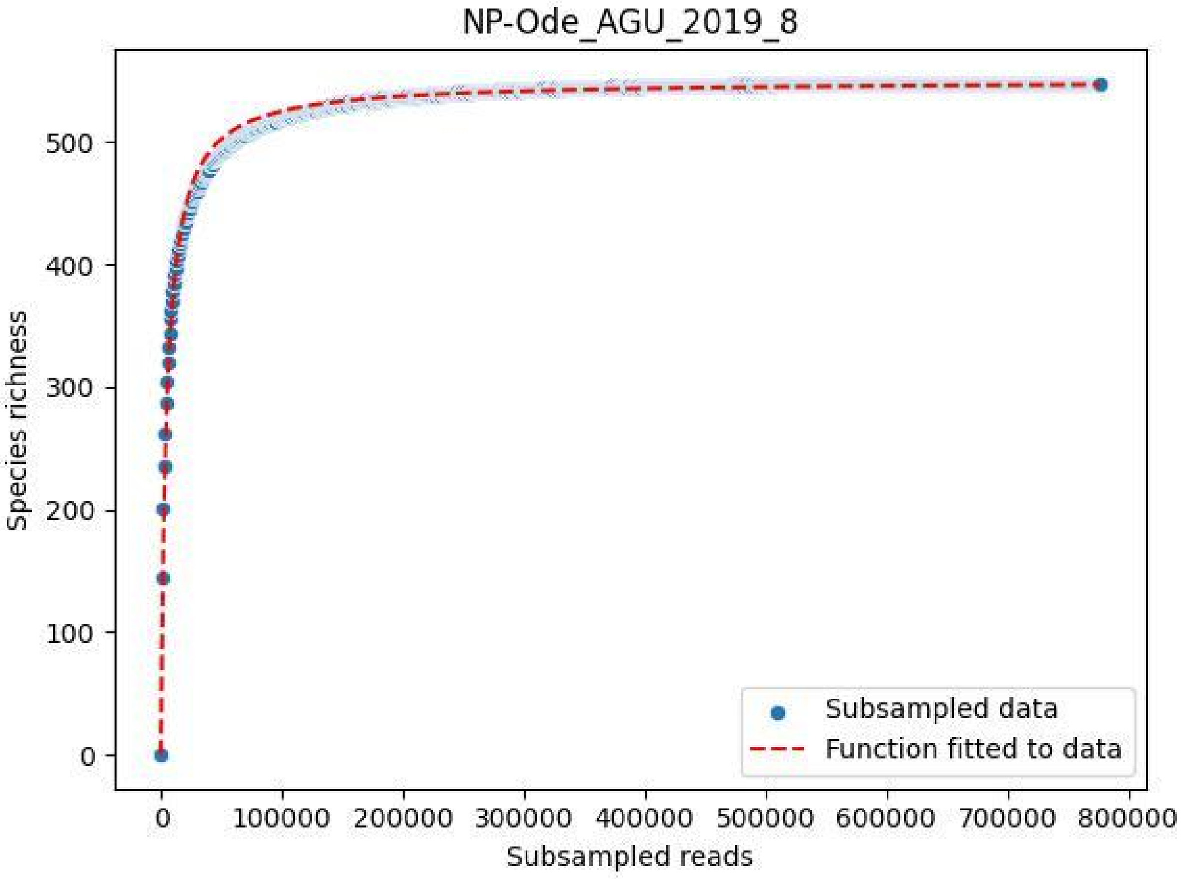
NP-Ode_AGU_2019_8
Species richness
Subsampled reads
0
100
200
300
400
500
0
100000
200000
300000
400000
500000
600000
700000
800000
Subsampled data
Function fitted to data

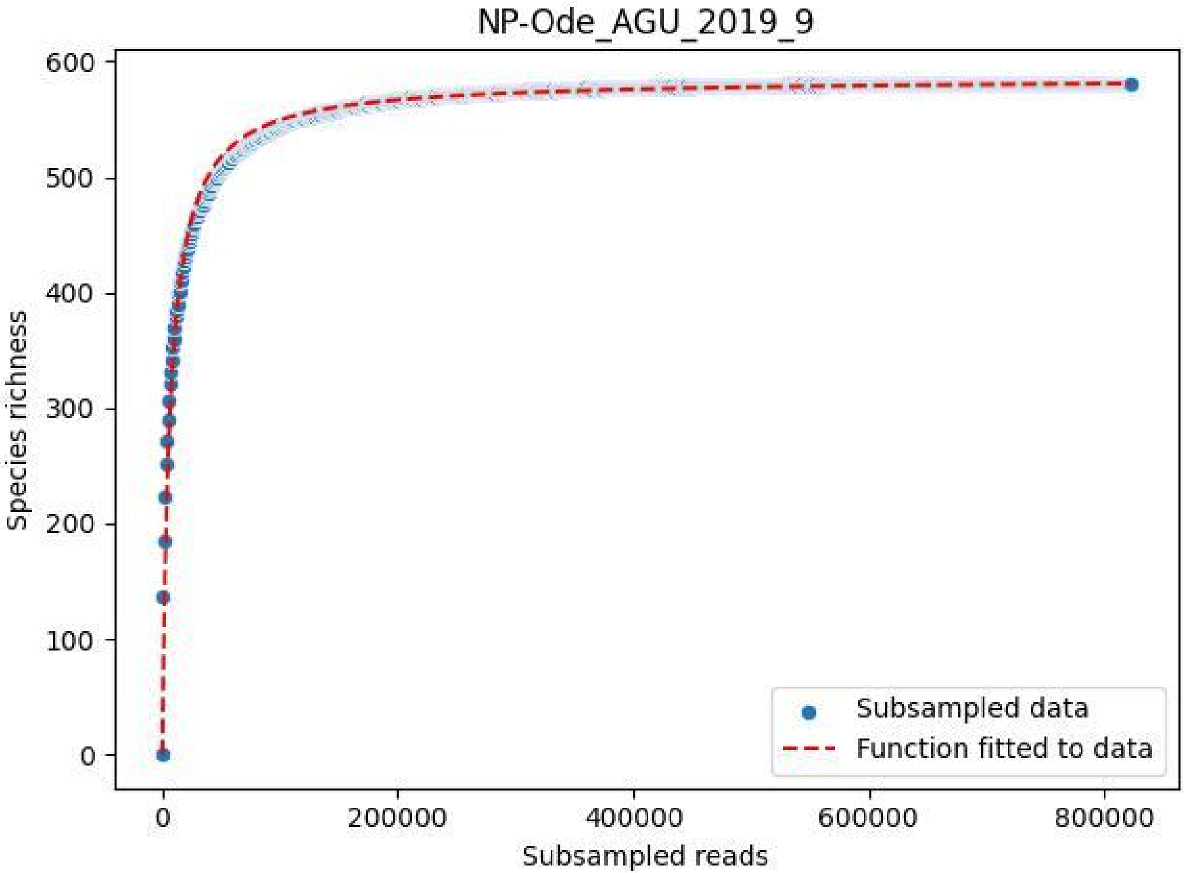
NP-Ode_AGU_2019_9
Species richness
Subsampled reads
0
100
200
300
400
500
600
0
200000
400000
600000
800000
Subsampled data
Function fitted to data

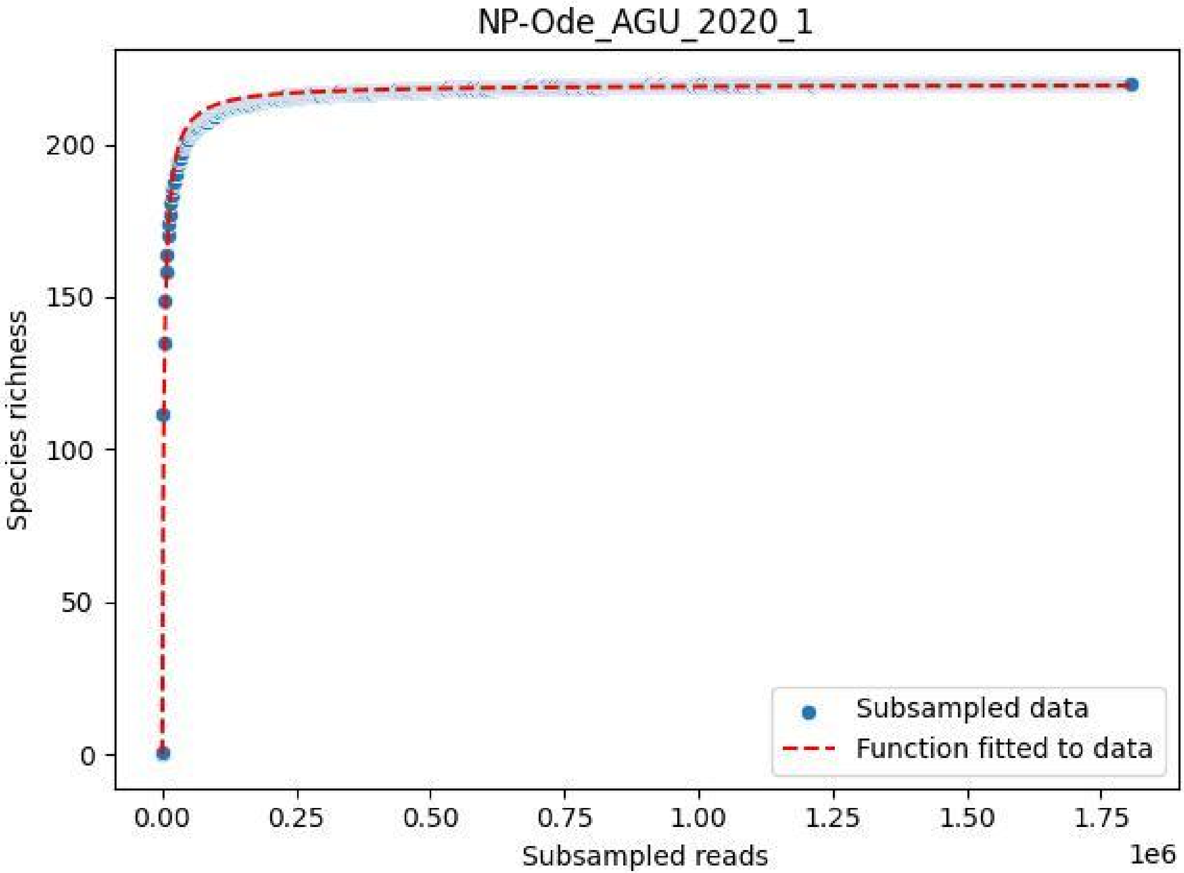

NP-Ode_AGU_2020_1
Species richness
Subsampled reads
1e6
0
50
100
150
200
0.00
0.25
0.50
0.75
1.00
1.25
1.50
1.75
Subsampled data
Function fitted to data

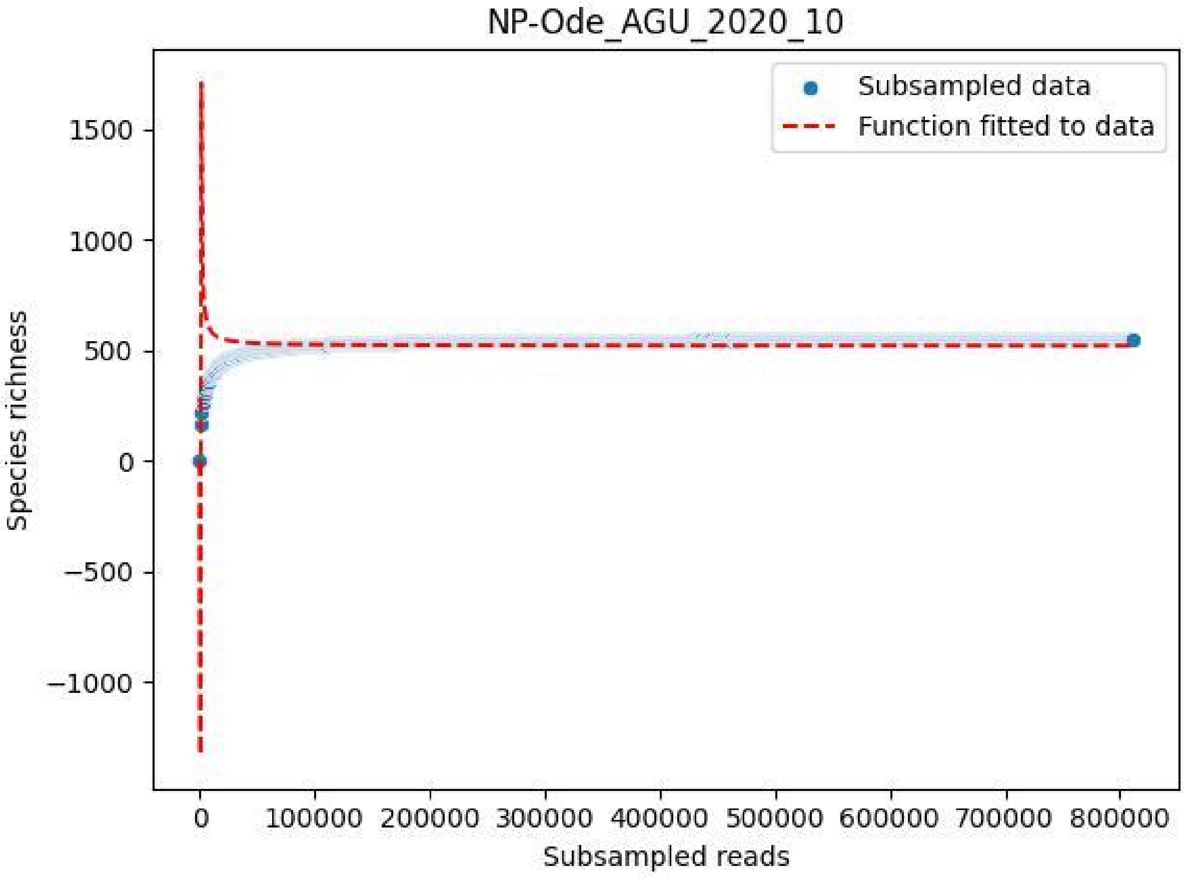

NP-Ode_AGU_2020_10
Subsampled data
Function fitted to data
Species richness
1500
1000
500
0
−500
−1000
0
100000
200000
300000
400000
500000
600000
700000
800000
Subsampled reads

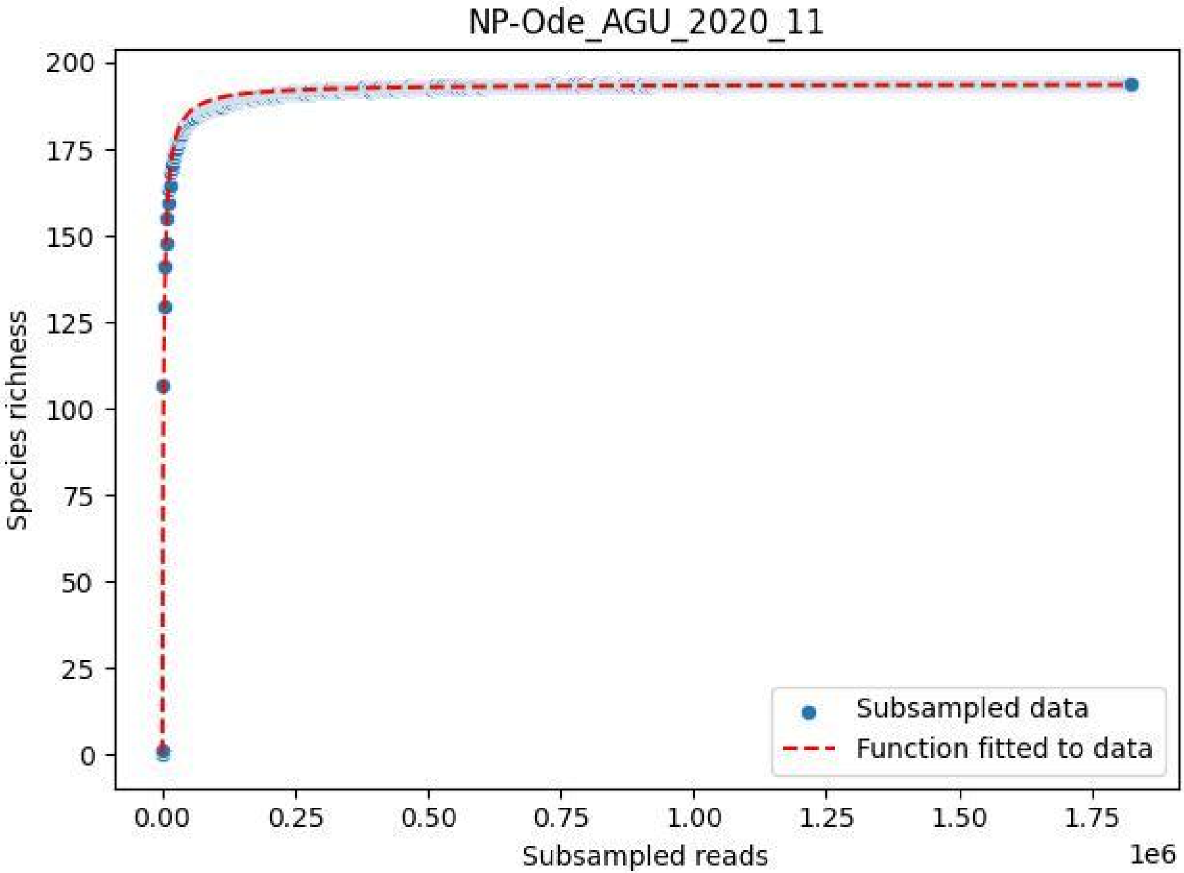
NP-Ode_AGU_2020_11
Species richness
200
175
150
125
100
75
50
25
0
0.00
0.25
0.50
0.75
1.00
1.25
1.50
1.75
1e6
Subsampled reads
Subsampled data
Function fitted to data

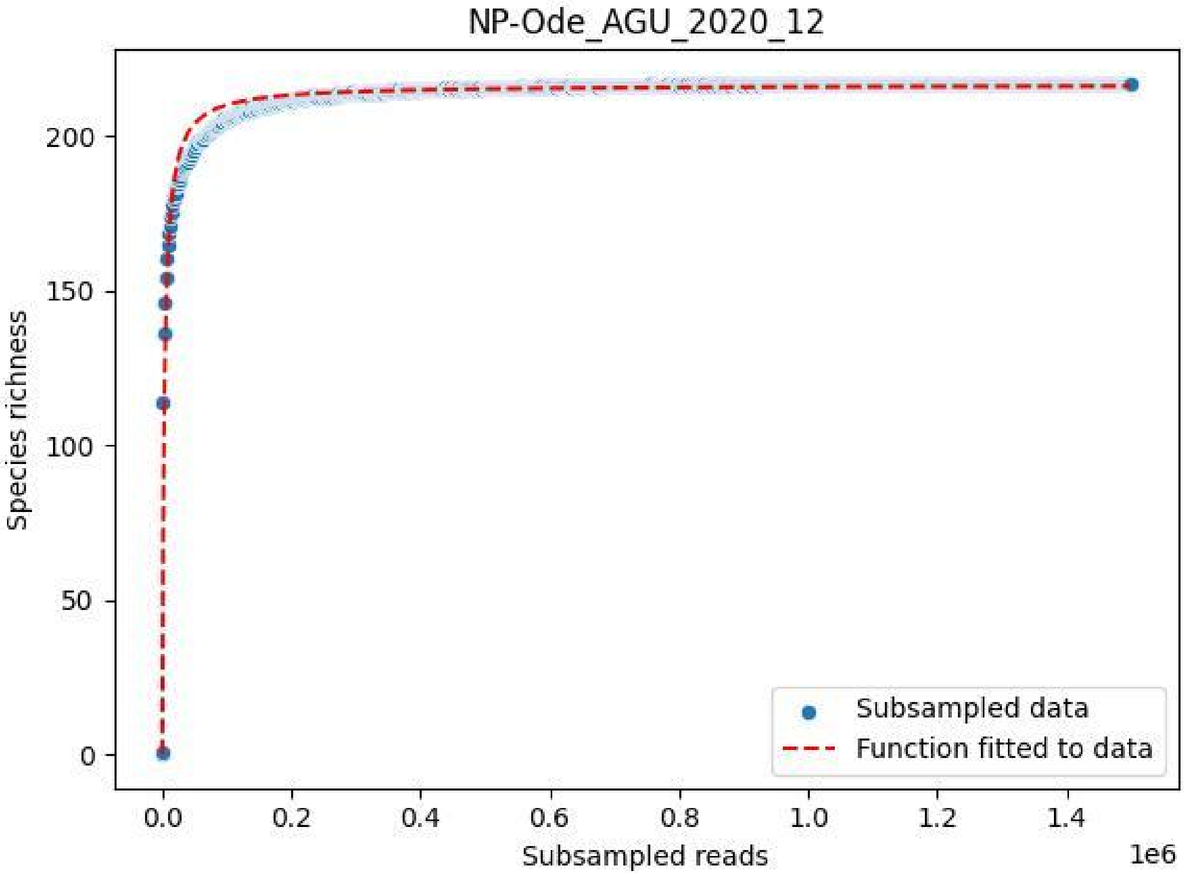
NP-Ode_AGU_2020_12
Species richness
Subsampled reads
1e6
0
50
100
150
200
0.0
0.2
0.4
0.6
0.8
1.0
1.2
1.4
Subsampled data
Function fitted to data

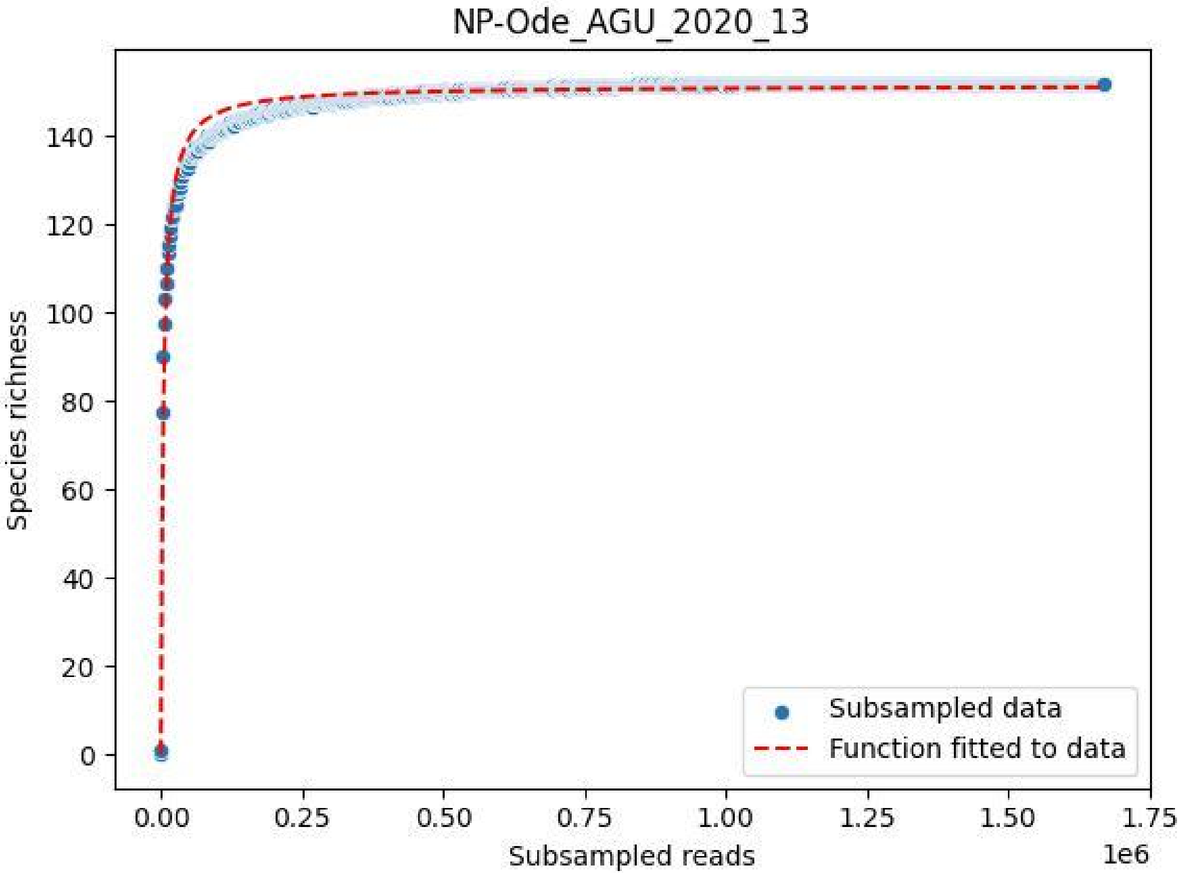

NP-Ode_AGU_2020_13
Species richness
Subsampled reads
1e6
Subsampled data
Function fitted to data
0
20
40
60
80
100
120
140
0.00
0.25
0.50
0.75
1.00
1.25
1.50
1.75

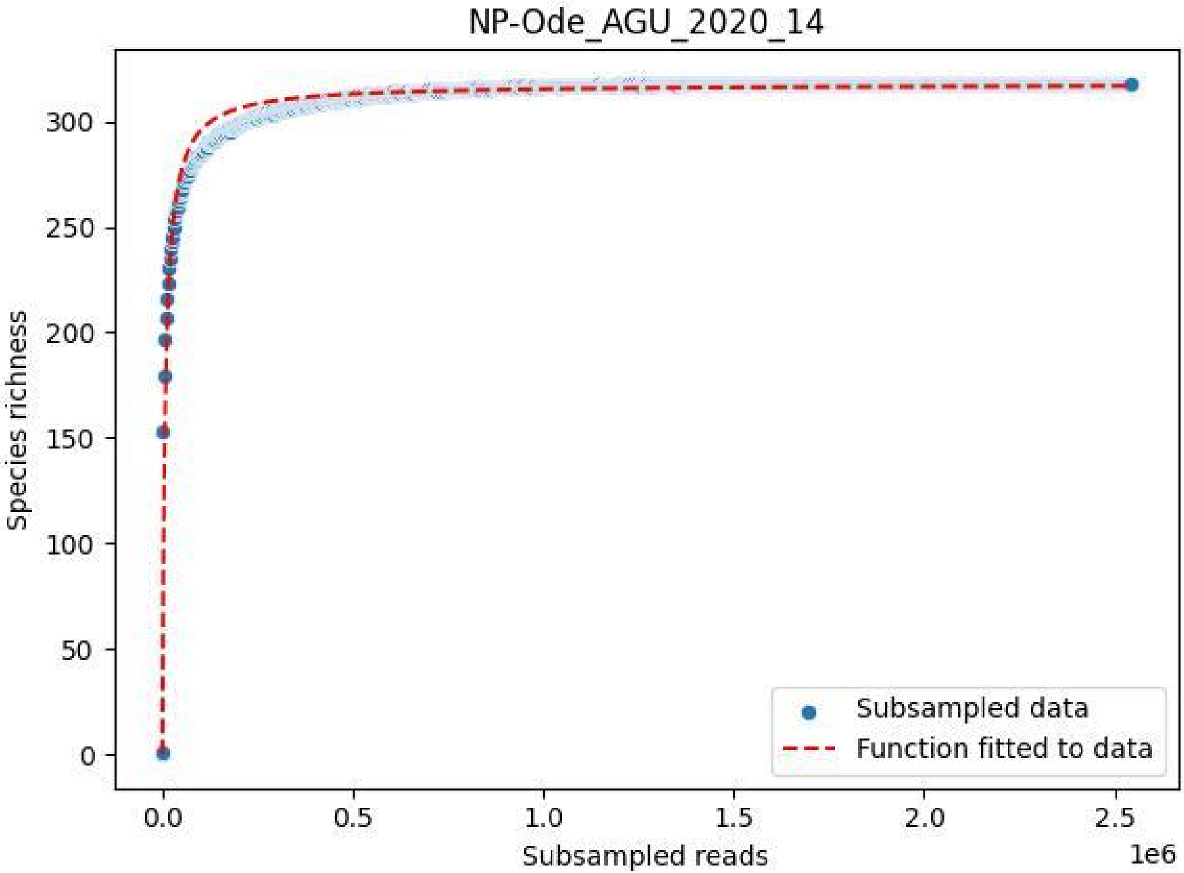
NP-Ode_AGU_2020_14
Species richness
300
250
200
150
100
50
0
0.0
0.5
1.0
1.5
2.0
2.5
1e6
Subsampled reads
Subsampled data
Function fitted to data

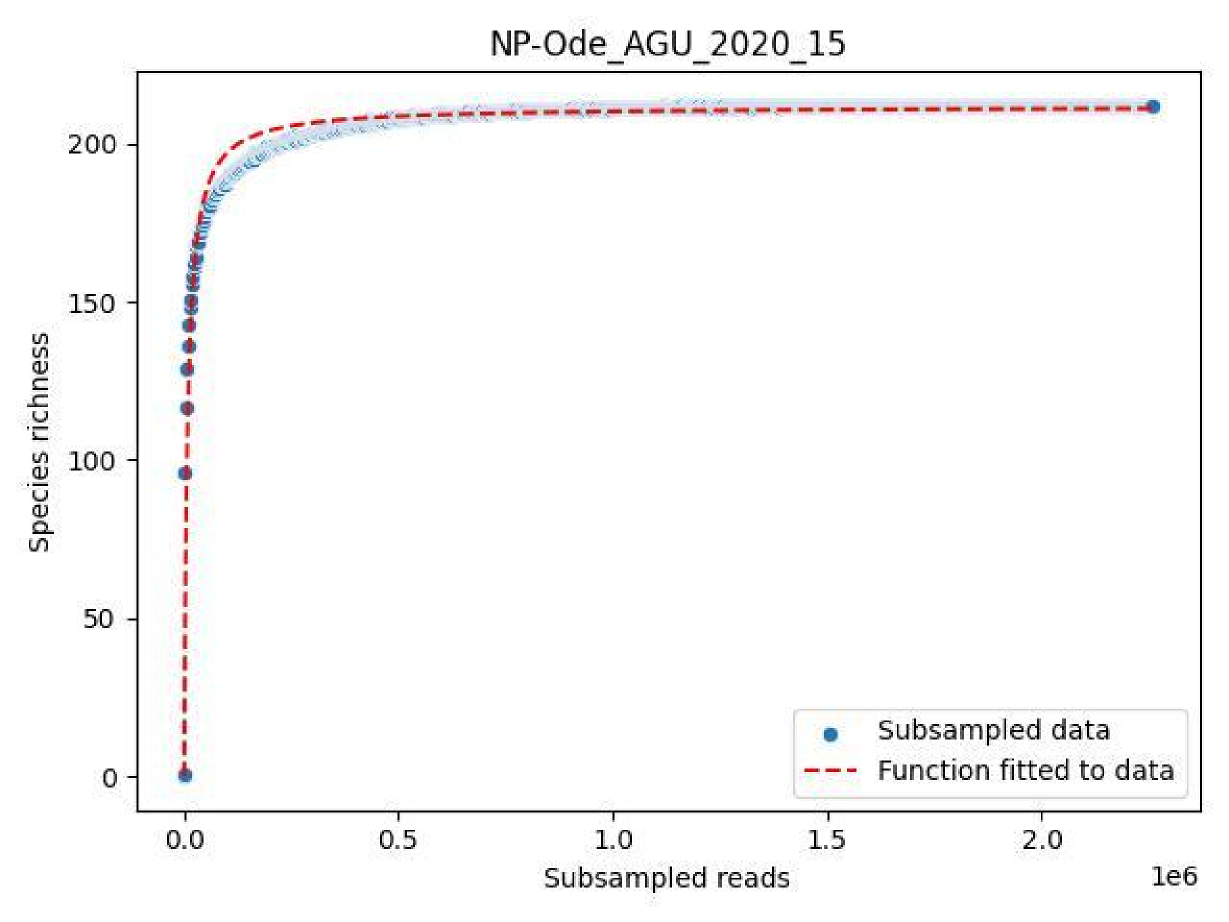
NP-Ode_AGU_2020_15
Species richness
0
50
100
150
200
0.0
0.5
1.0
1.5
2.0
1e6
Subsampled reads
Subsampled data
Function fitted to data

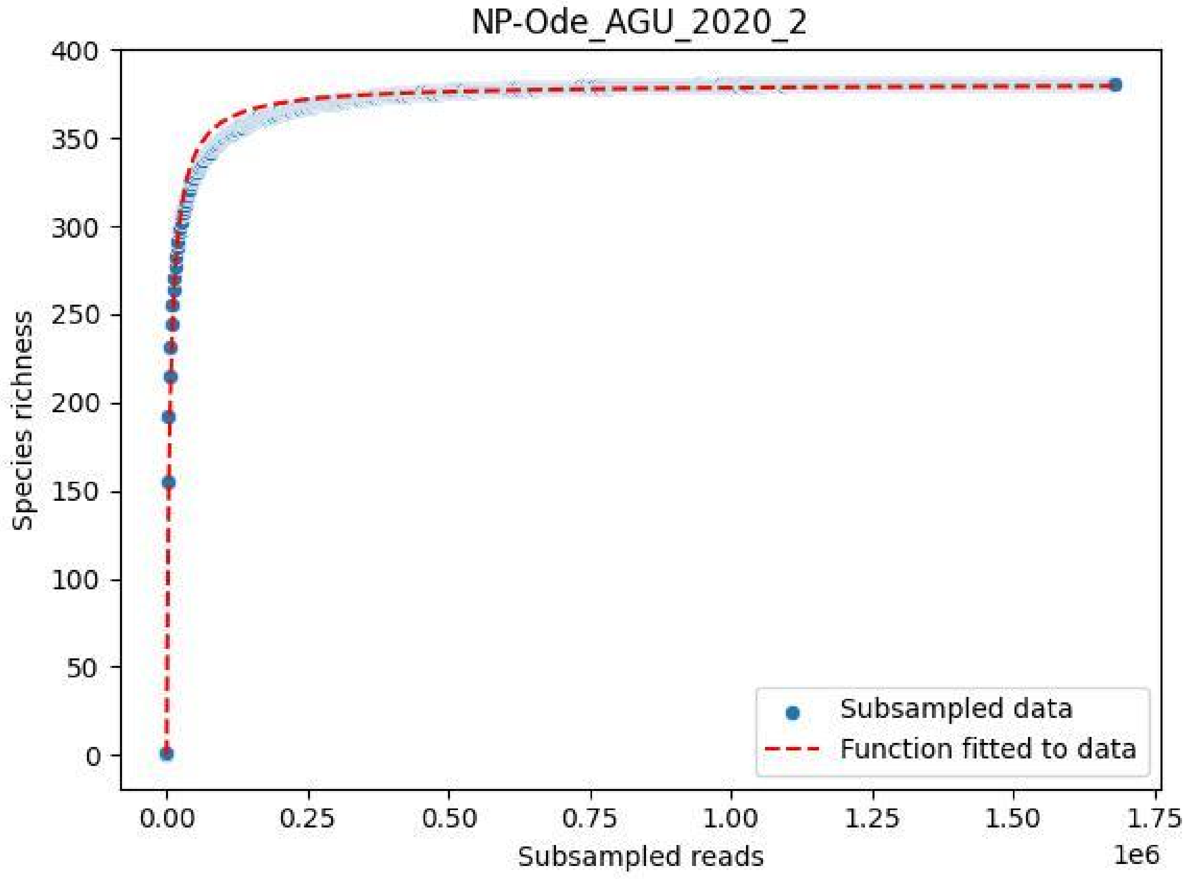

NP-Ode_AGU_2020_2
Species richness
Subsampled reads
1e6
0
50
100
150
200
250
300
350
400
0.00
0.25
0.50
0.75
1.00
1.25
1.50
1.75
Subsampled data
Function fitted to data

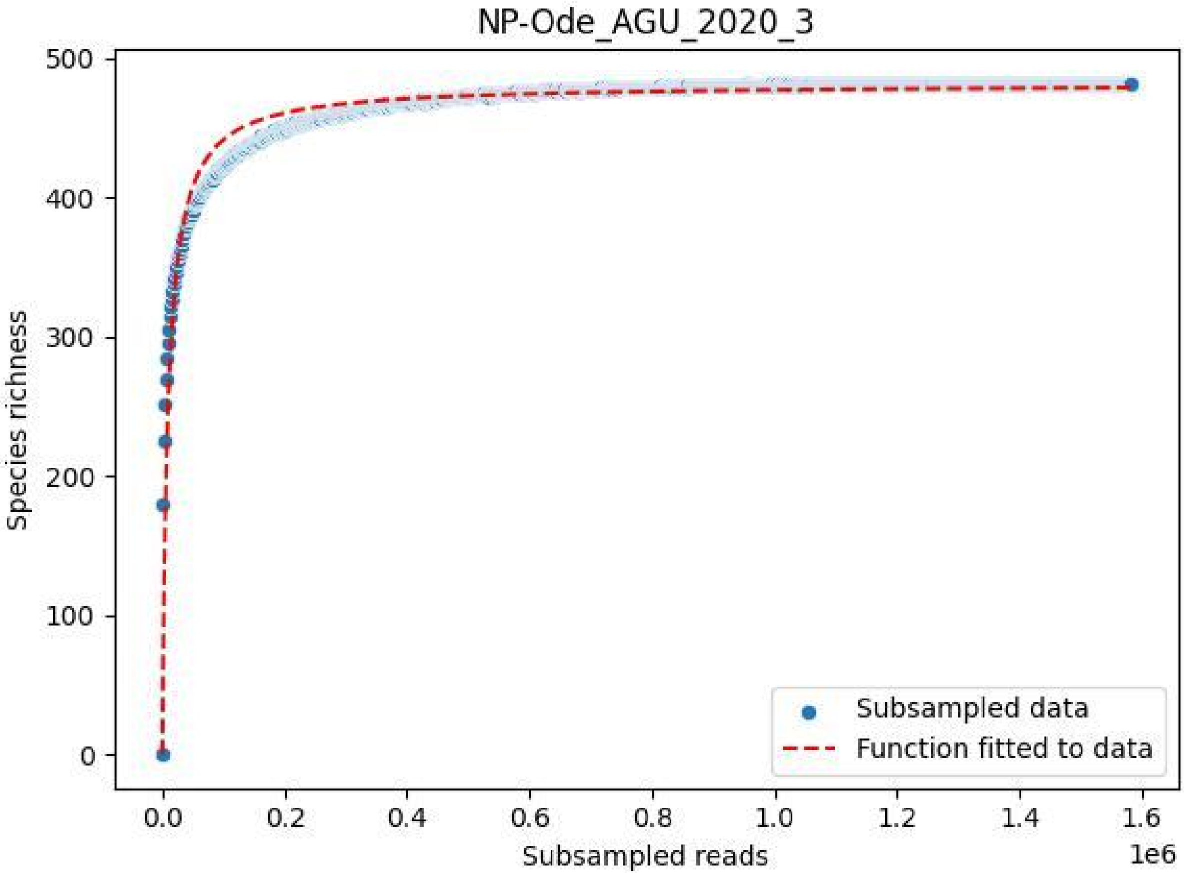
NP-Ode_AGU_2020_3
Species richness
500
400
300
200
100
0
0.0
0.2
0.4
0.6
0.8
1.0
1.2
1.4
1.6
1e6
Subsampled reads
Subsampled data
Function fitted to data

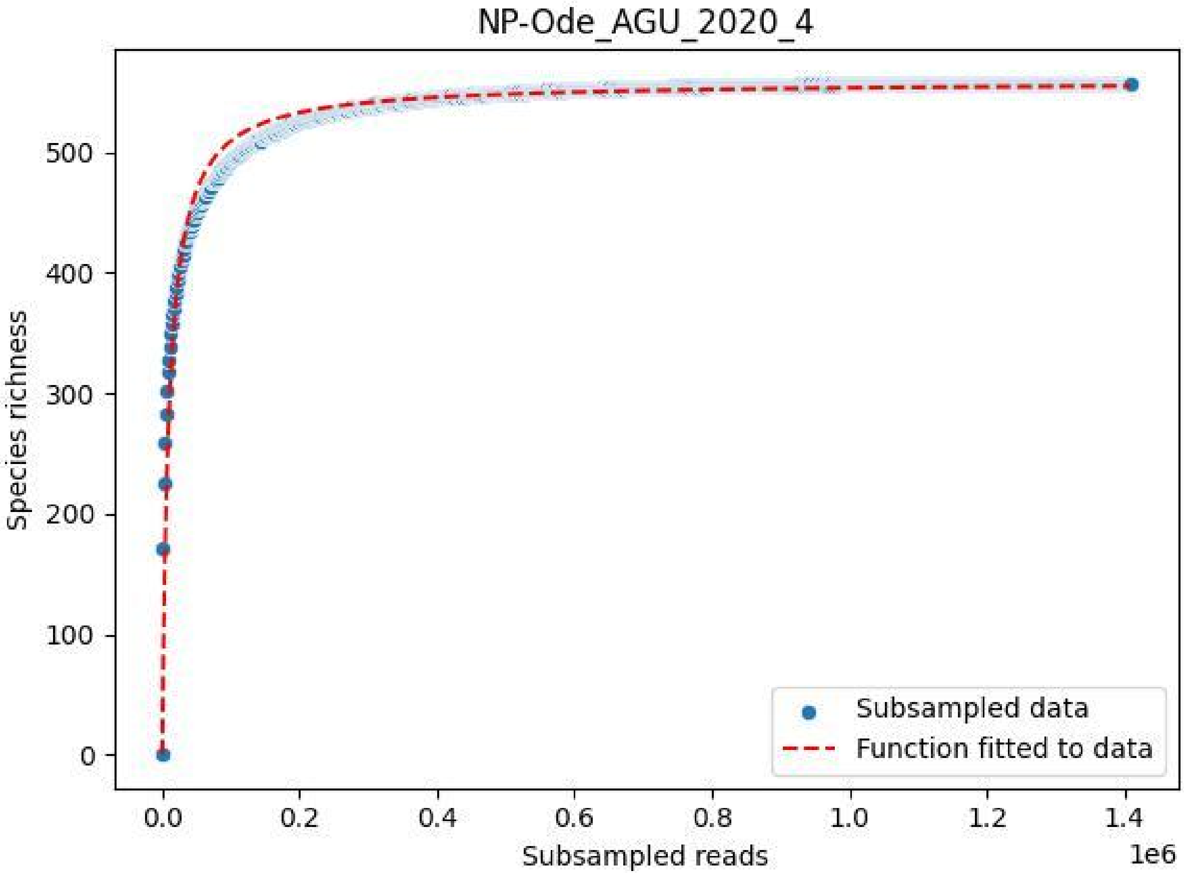

NP-Ode_AGU_2020_4
Species richness
Subsampled reads
1e6
0
100
200
300
400
500
0.0
0.2
0.4
0.6
0.8
1.0
1.2
1.4
Subsampled data
Function fitted to data

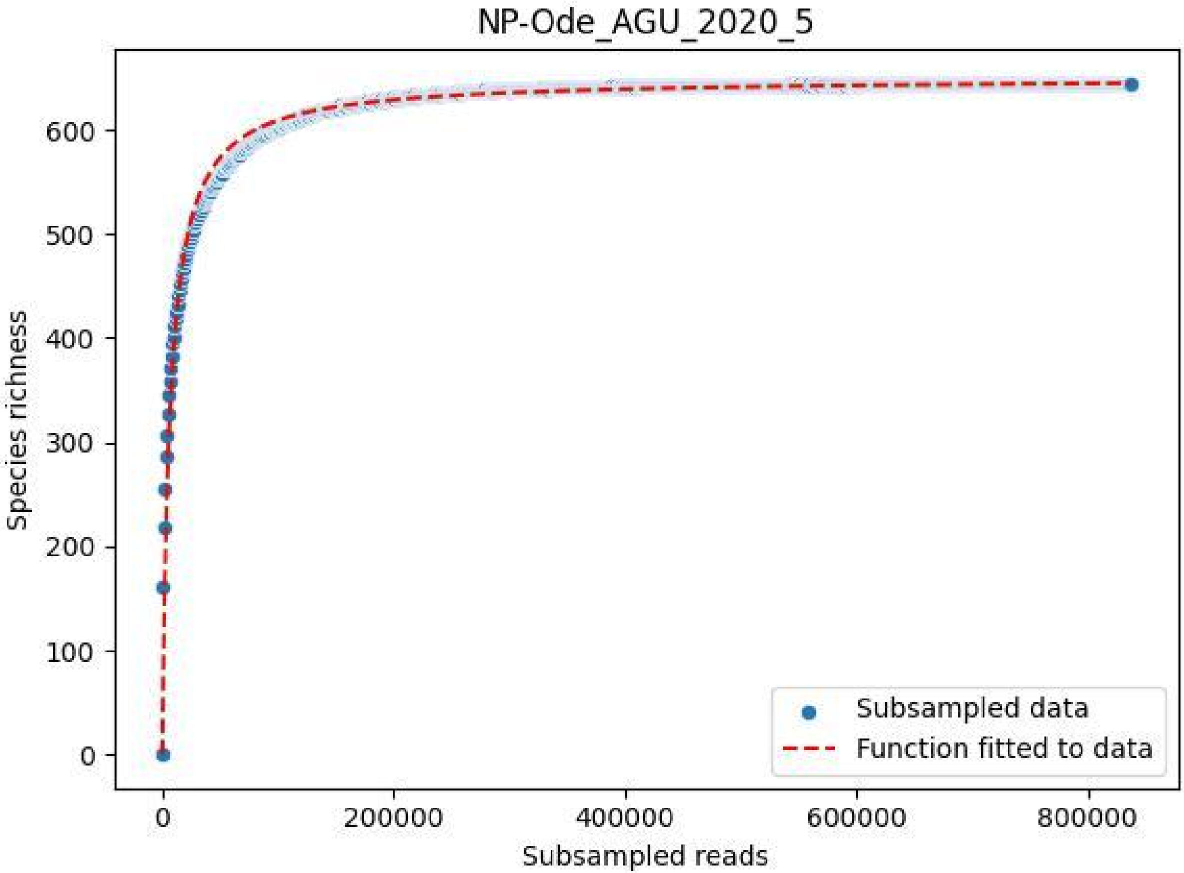
NP-Ode_AGU_2020_5
Species richness
0
100
200
300
400
500
600
0
200000
400000
600000
800000
Subsampled reads
Subsampled data
Function fitted to data

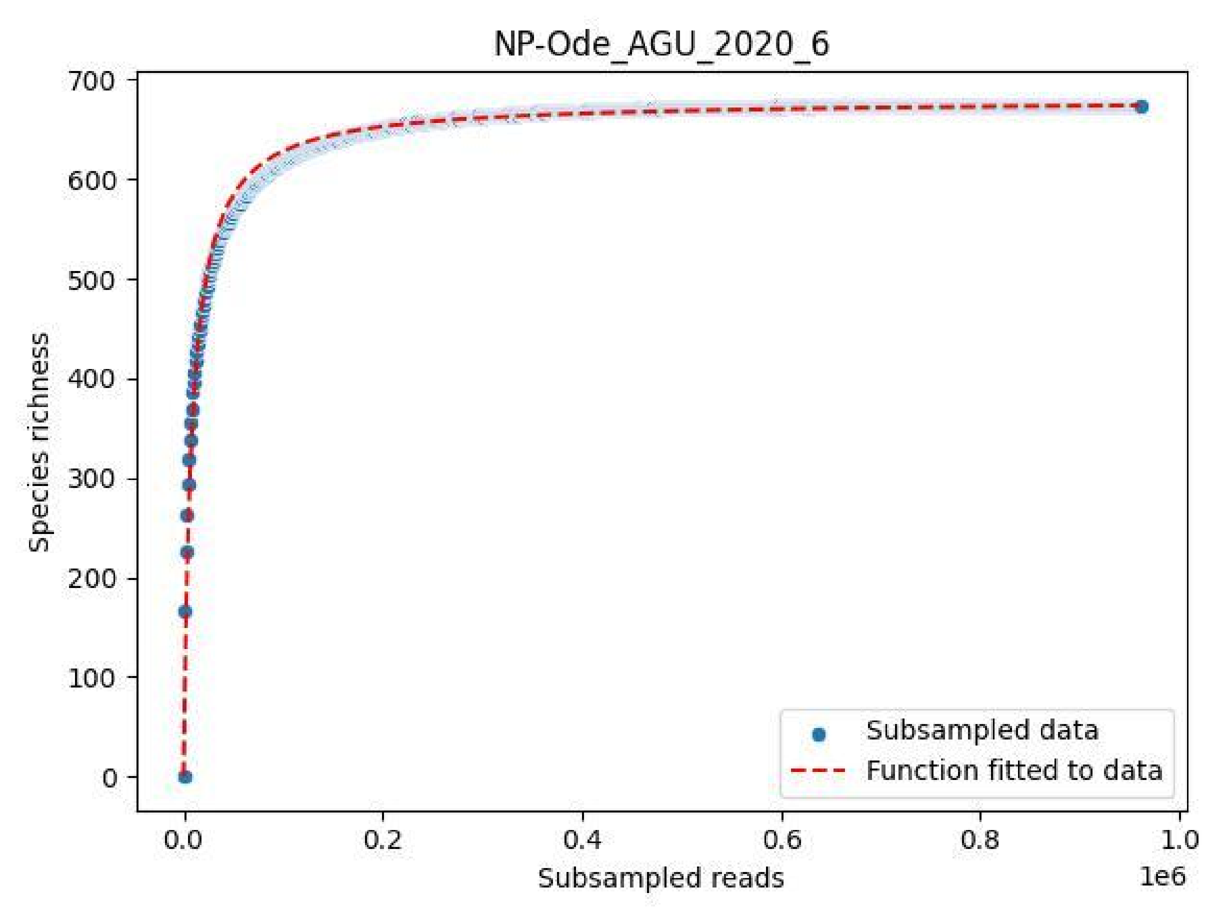

NP-Ode_AGU_2020_6
Species richness
Subsampled reads
1e6
0
100
200
300
400
500
600
700
0.0
0.2
0.4
0.6
0.8
1.0
Subsampled data
Function fitted to data

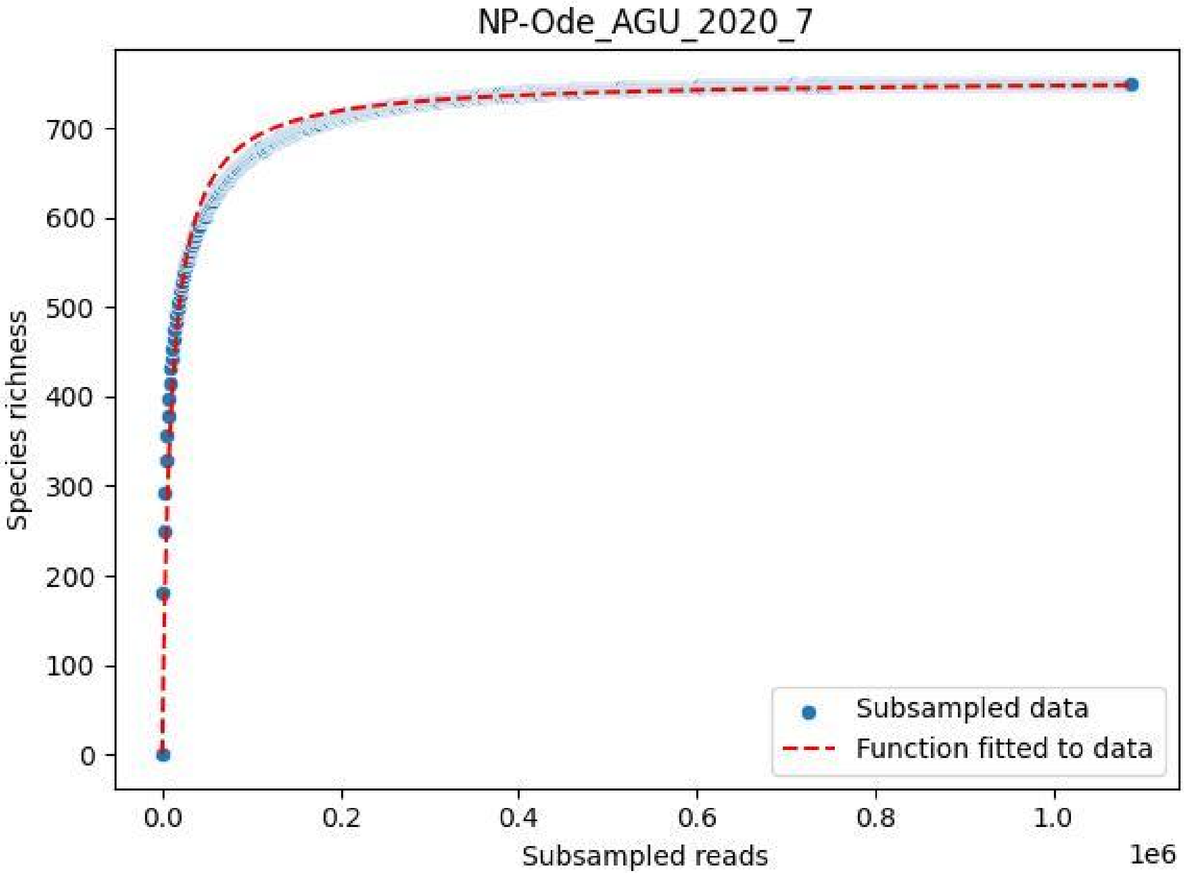

NP-Ode_AGU_2020_7
Species richness
0
100
200
300
400
500
600
700
Subsampled reads
0.0
0.2
0.4
0.6
0.8
1.0
1e6
Subsampled data
Function fitted to data

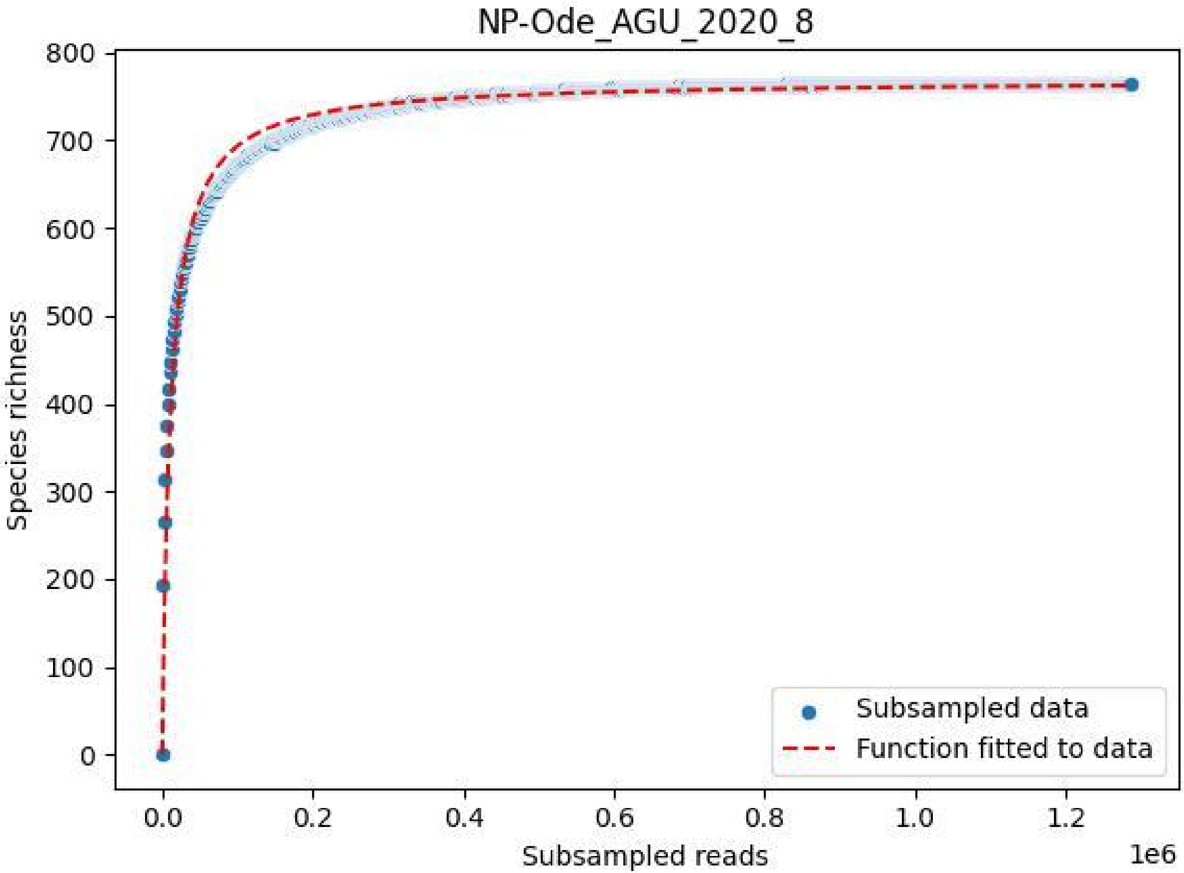

NP-Ode_AGU_2020_8
Species richness
Subsampled reads
1e6
Subsampled data
Function fitted to data
0
100
200
300
400
500
600
700
800
0.0
0.2
0.4
0.6
0.8
1.0
1.2

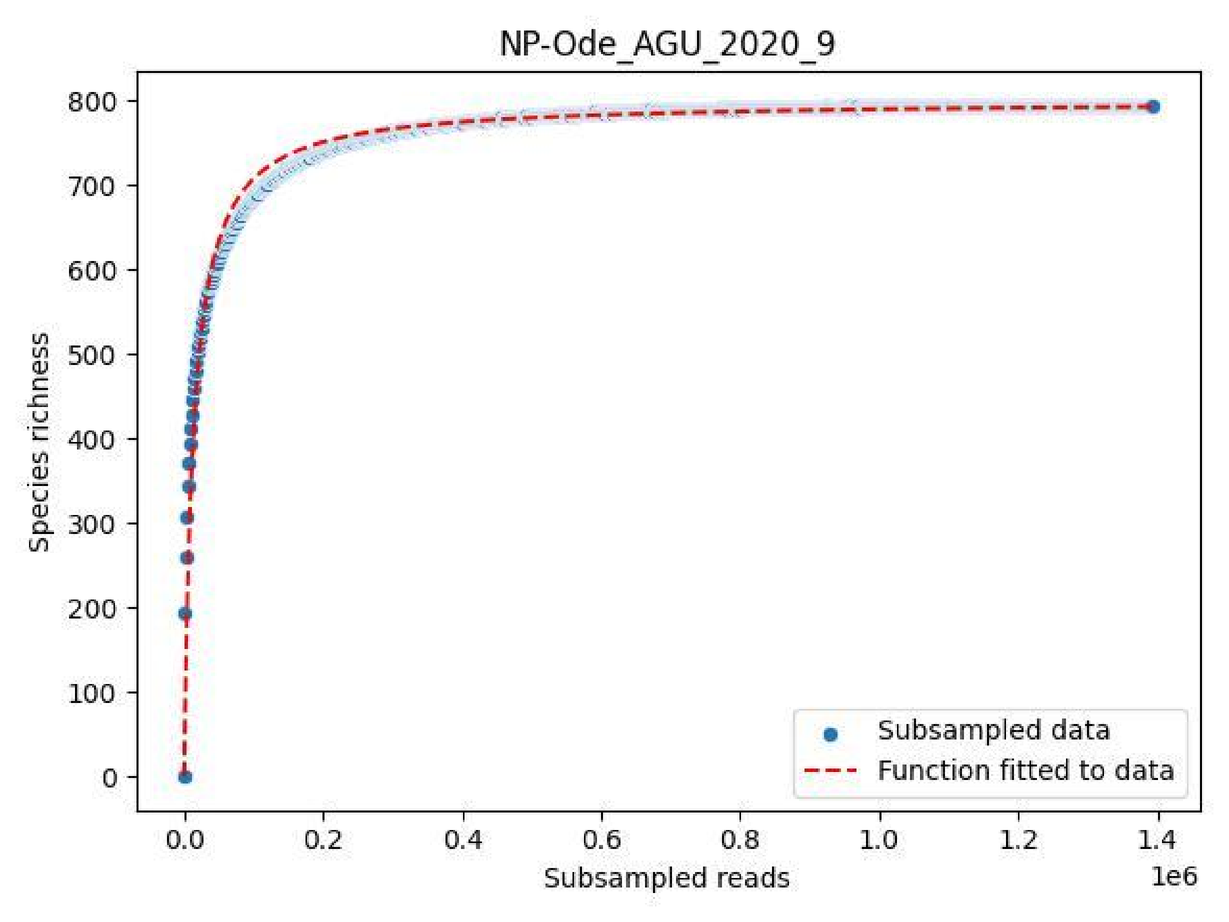

NP-Ode_AGU_2020_9
Species richness
0
100
200
300
400
500
600
700
800
0.0
0.2
0.4
0.6
0.8
1.0
1.2
1.4
1e6
Subsampled reads
Subsampled data
Function fitted to data

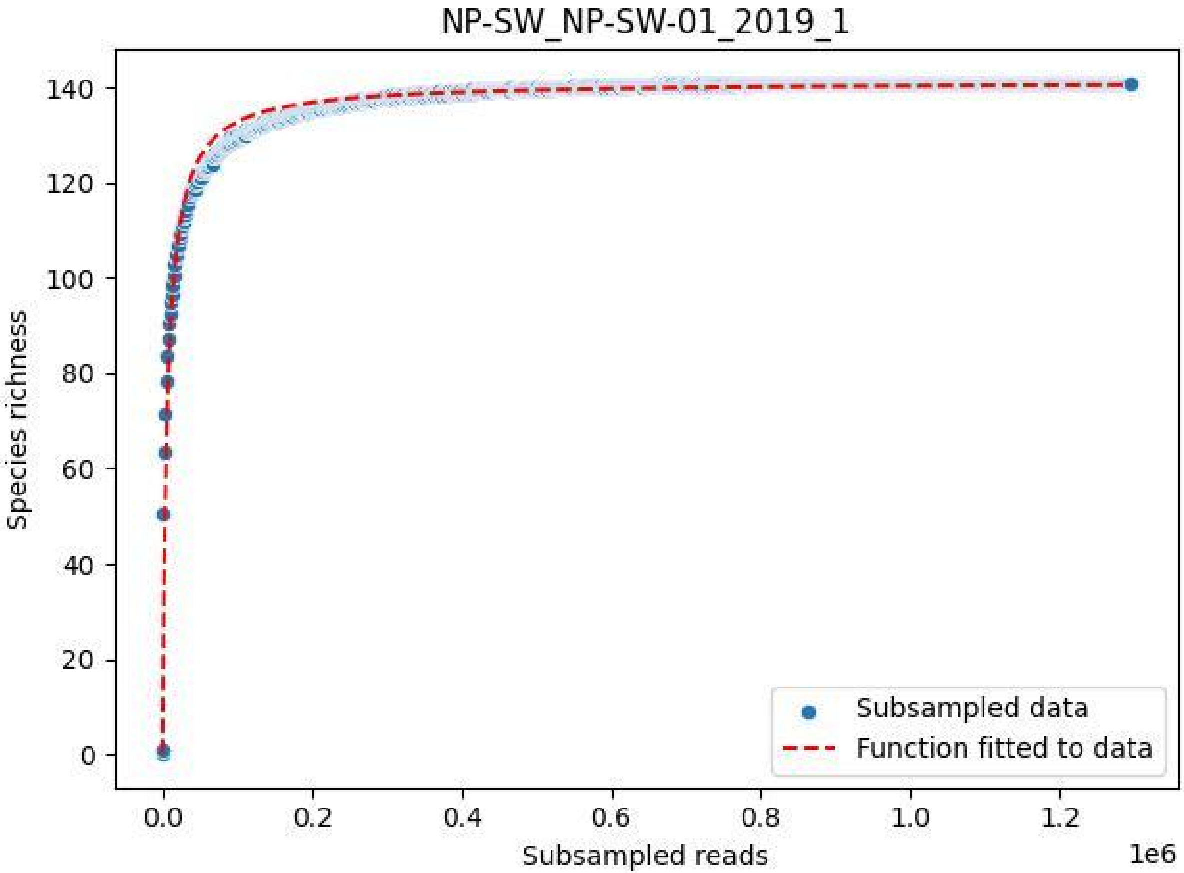

NP-SW_NP-SW-01_2019_1
Species richness
Subsampled reads
1e6
0
20
40
60
80
100
120
140
0.0
0.2
0.4
0.6
0.8
1.0
1.2
Subsampled data
Function fitted to data

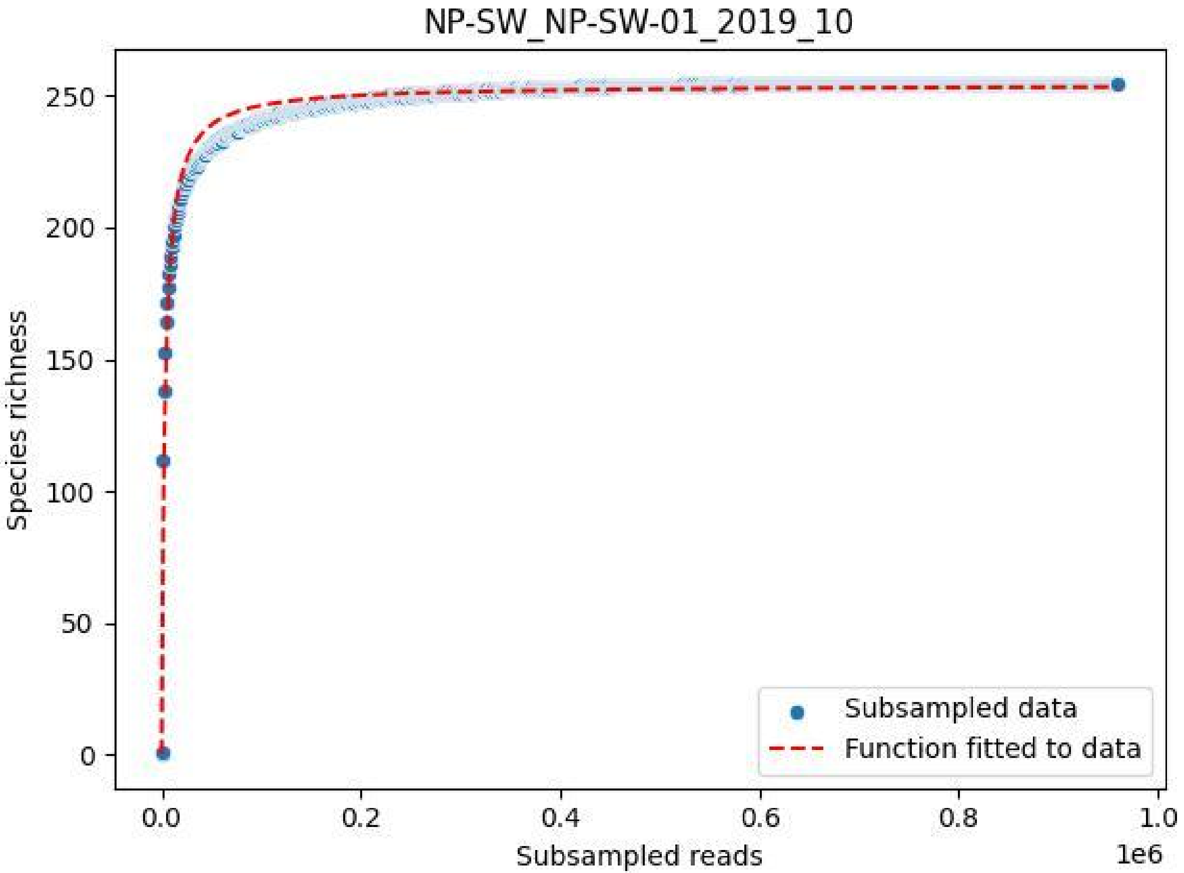
NP-SW_NP-SW-01_2019_10
Species richness
Subsampled reads
1e6
0
50
100
150
200
250
0.0
0.2
0.4
0.6
0.8
1.0
Subsampled data
Function fitted to data

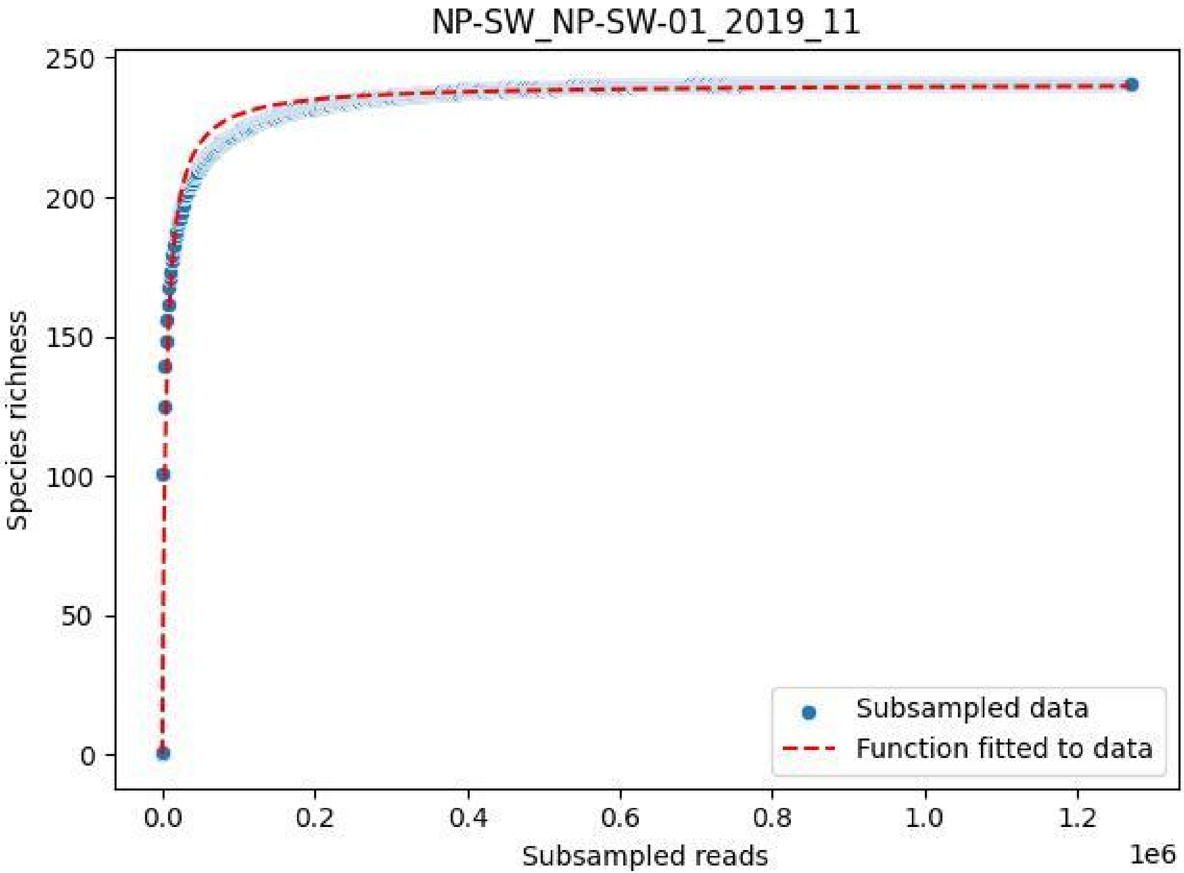

NP-SW_NP-SW-01_2019_11
Species richness
Subsampled reads
1e6
0
50
100
150
200
250
0.0
0.2
0.4
0.6
0.8
1.0
1.2
Subsampled data
Function fitted to data

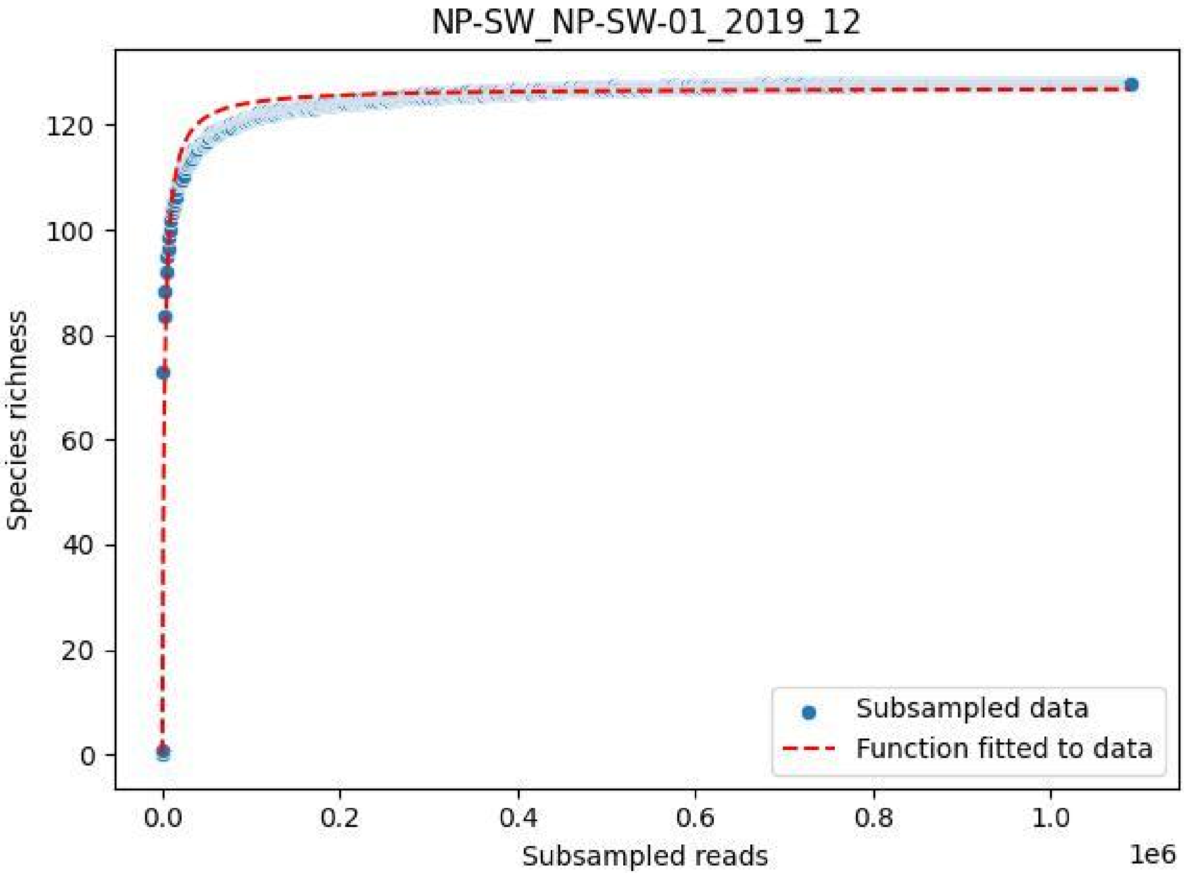

NP-SW_NP-SW-01_2019_12
Species richness
Subsampled reads
1e6
0.0
0.2
0.4
0.6
0.8
1.0
0
20
40
60
80
100
120
Subsampled data
Function fitted to data

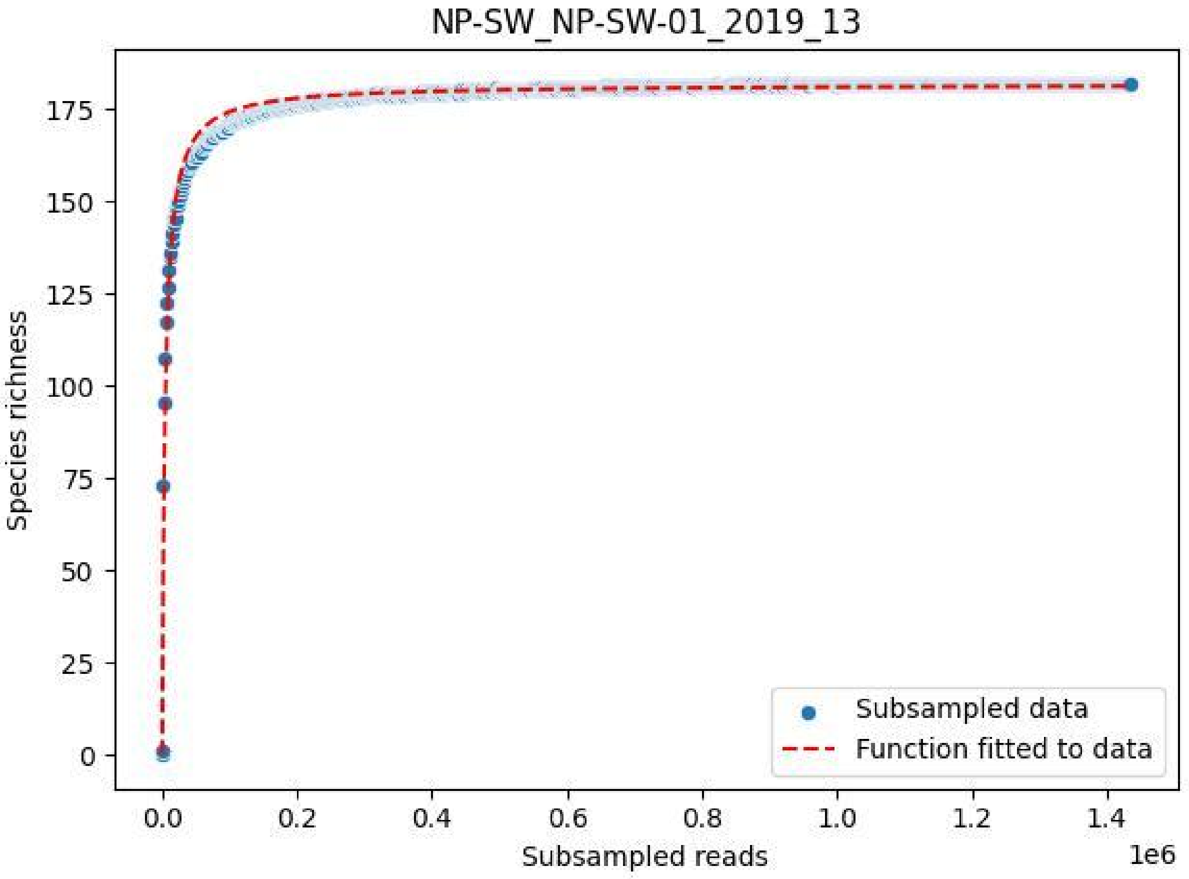
NP-SW_NP-SW-01_2019_13
Species richness
Subsampled reads
1e6
0
25
50
75
100
125
150
175
0.0
0.2
0.4
0.6
0.8
1.0
1.2
1.4
Subsampled data
Function fitted to data

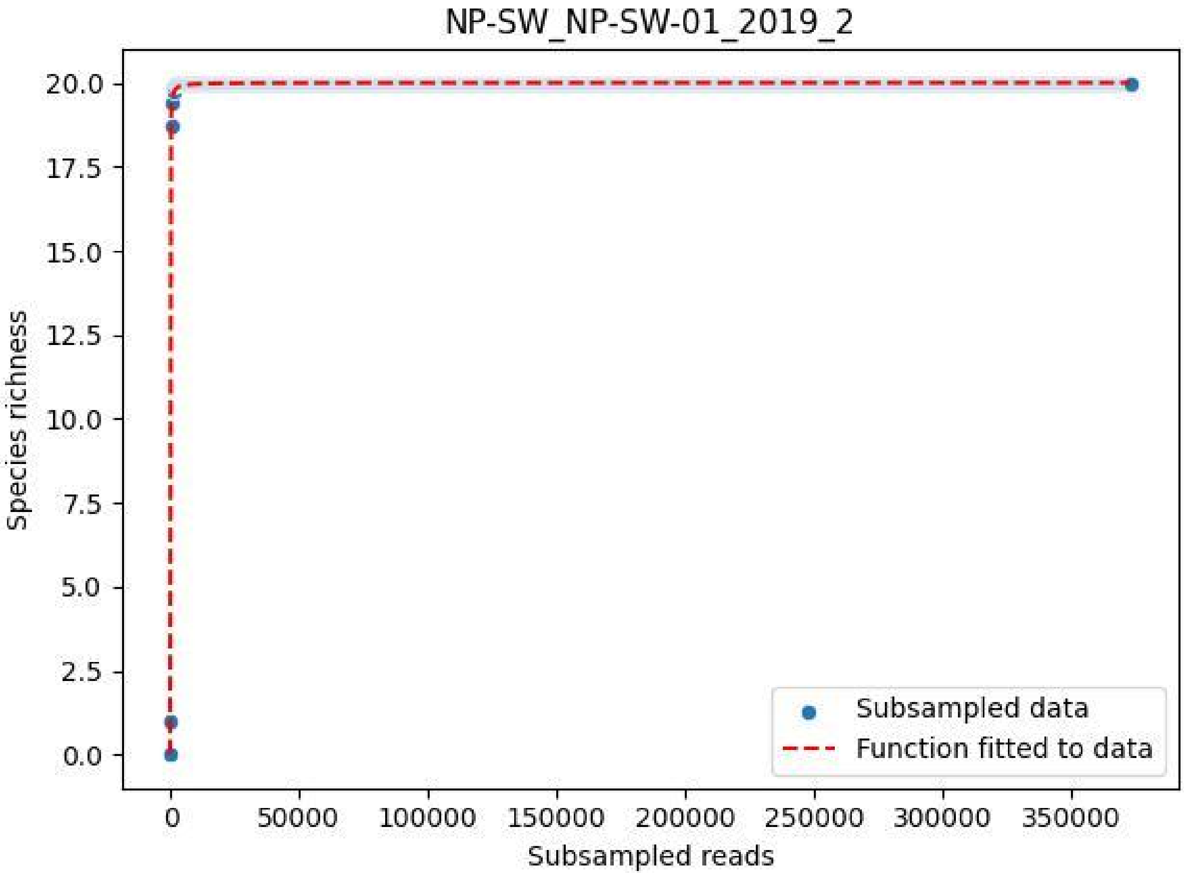

NP-SW_NP-SW-01_2019_2
Species richness
Subsampled reads
0.0
2.5
5.0
7.5
10.0
12.5
15.0
17.5
20.0
0
50000
100000
150000
200000
250000
300000
350000
Subsampled data
Function fitted to data

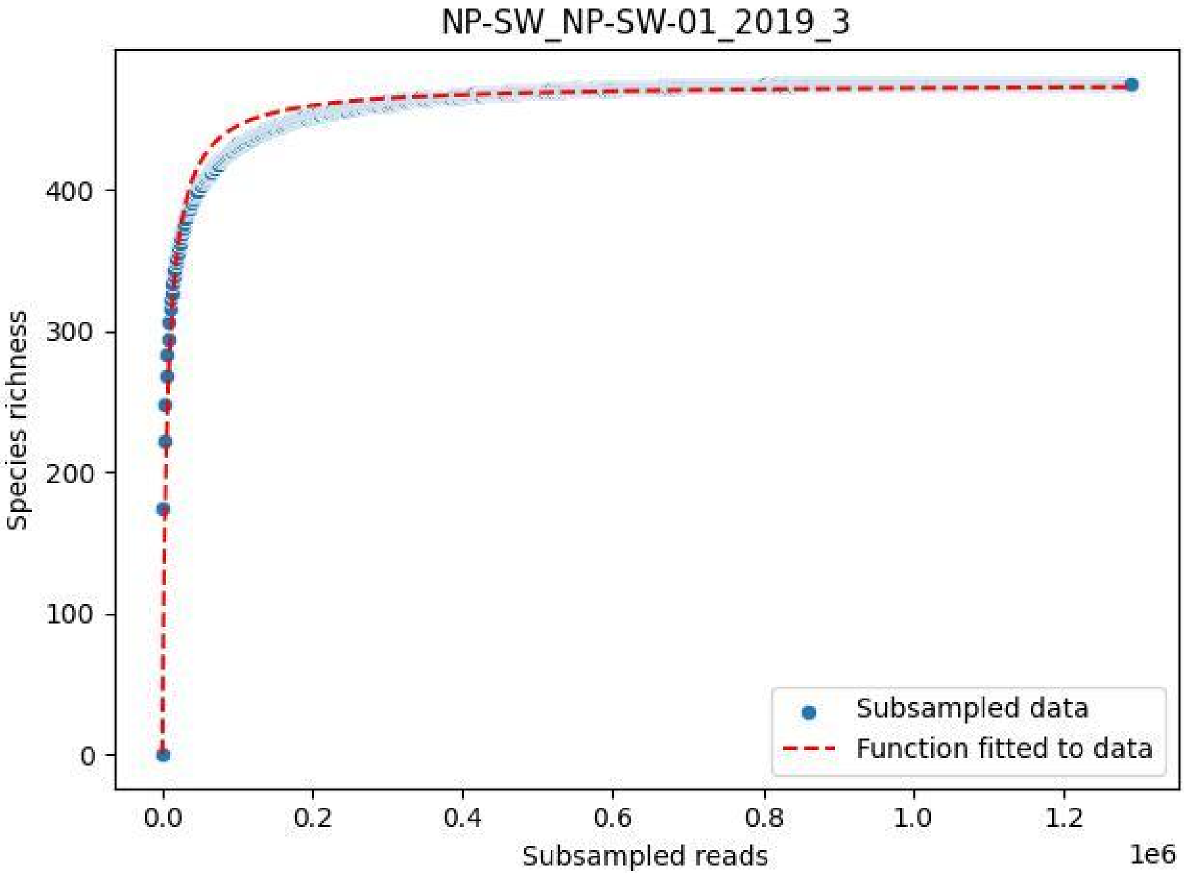
NP-SW_NP-SW-01_2019_3
Species richness
Subsampled reads
1e6
0
100
200
300
400
0.0
0.2
0.4
0.6
0.8
1.0
1.2
Subsampled data
Function fitted to data

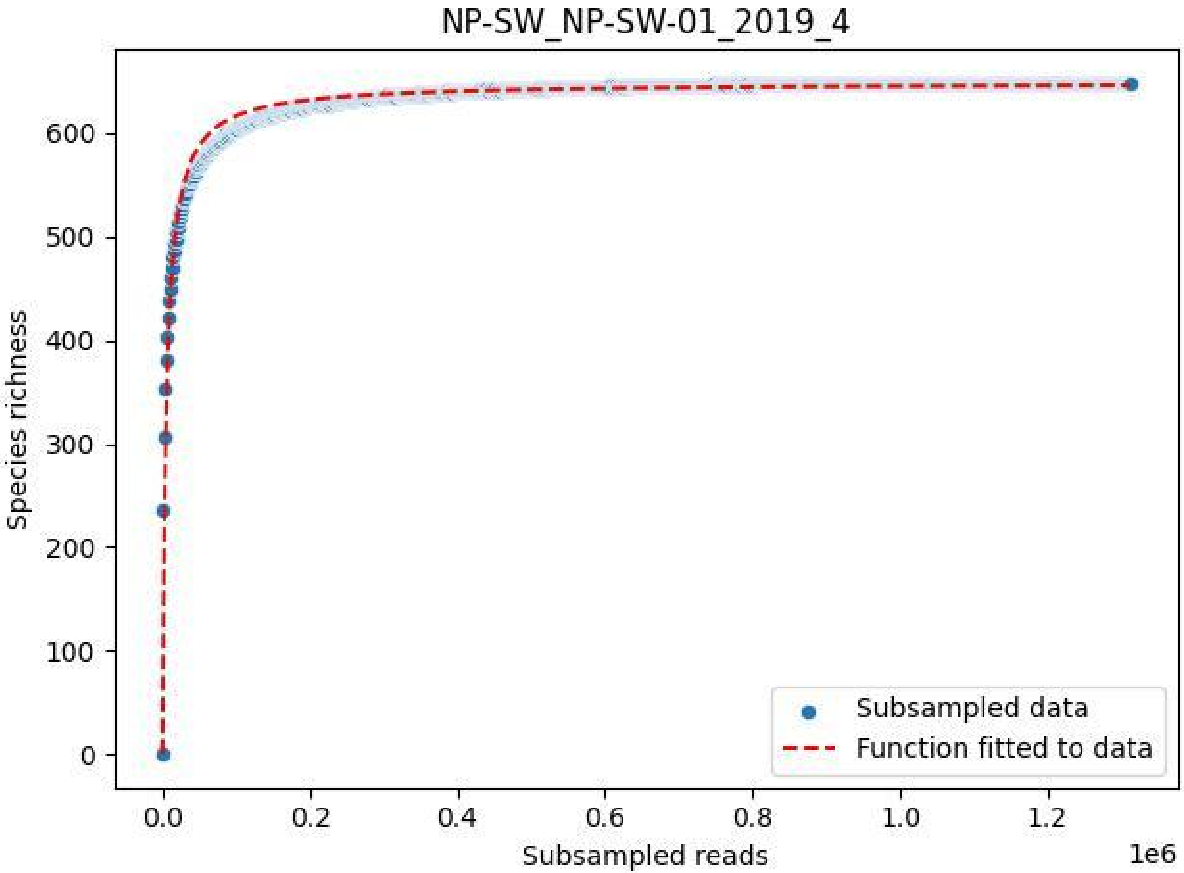

NP-SW_NP-SW-01_2019_4
Species richness
Subsampled reads
1e6
0
100
200
300
400
500
600
0.0
0.2
0.4
0.6
0.8
1.0
1.2
Subsampled data
Function fitted to data

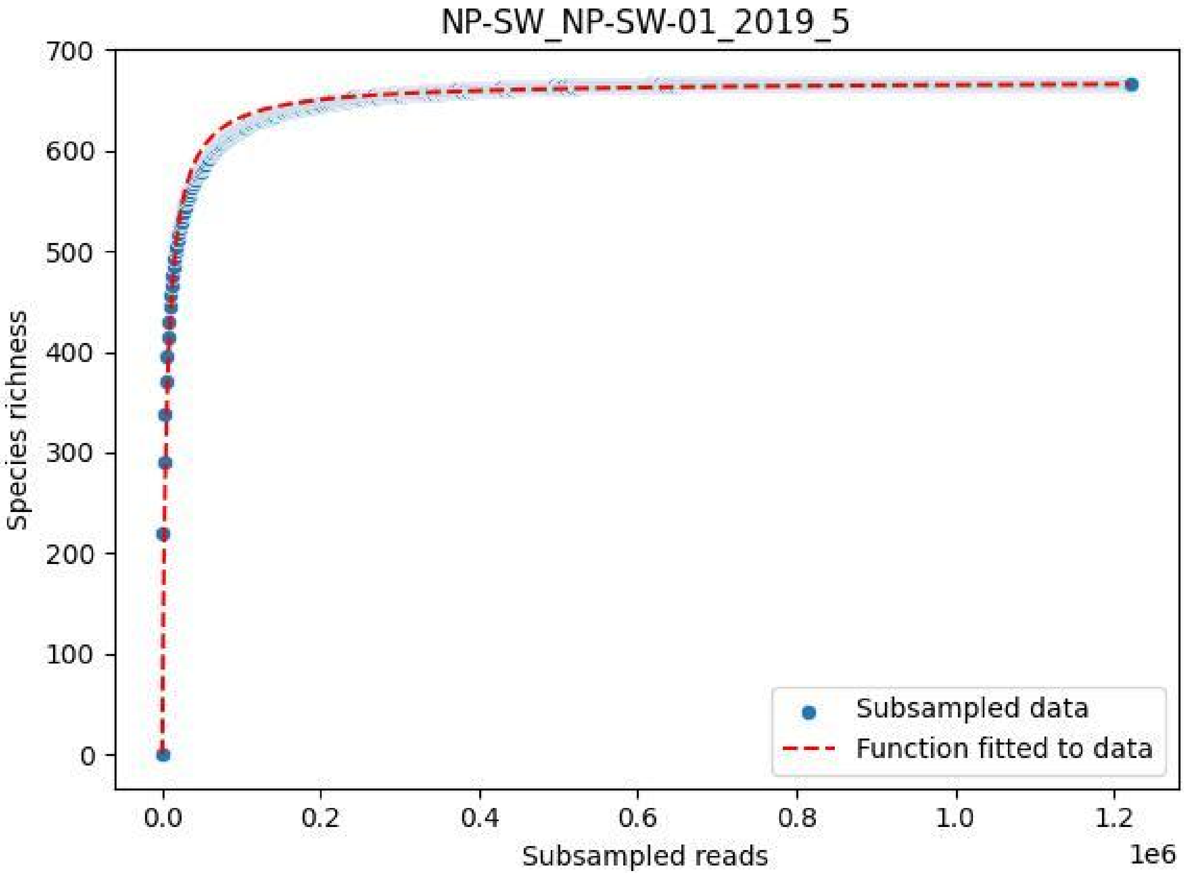
NP-SW_NP-SW-01_2019_5
Species richness
Subsampled reads
1e6
0
100
200
300
400
500
600
700
0.0
0.2
0.4
0.6
0.8
1.0
1.2
Subsampled data
Function fitted to data

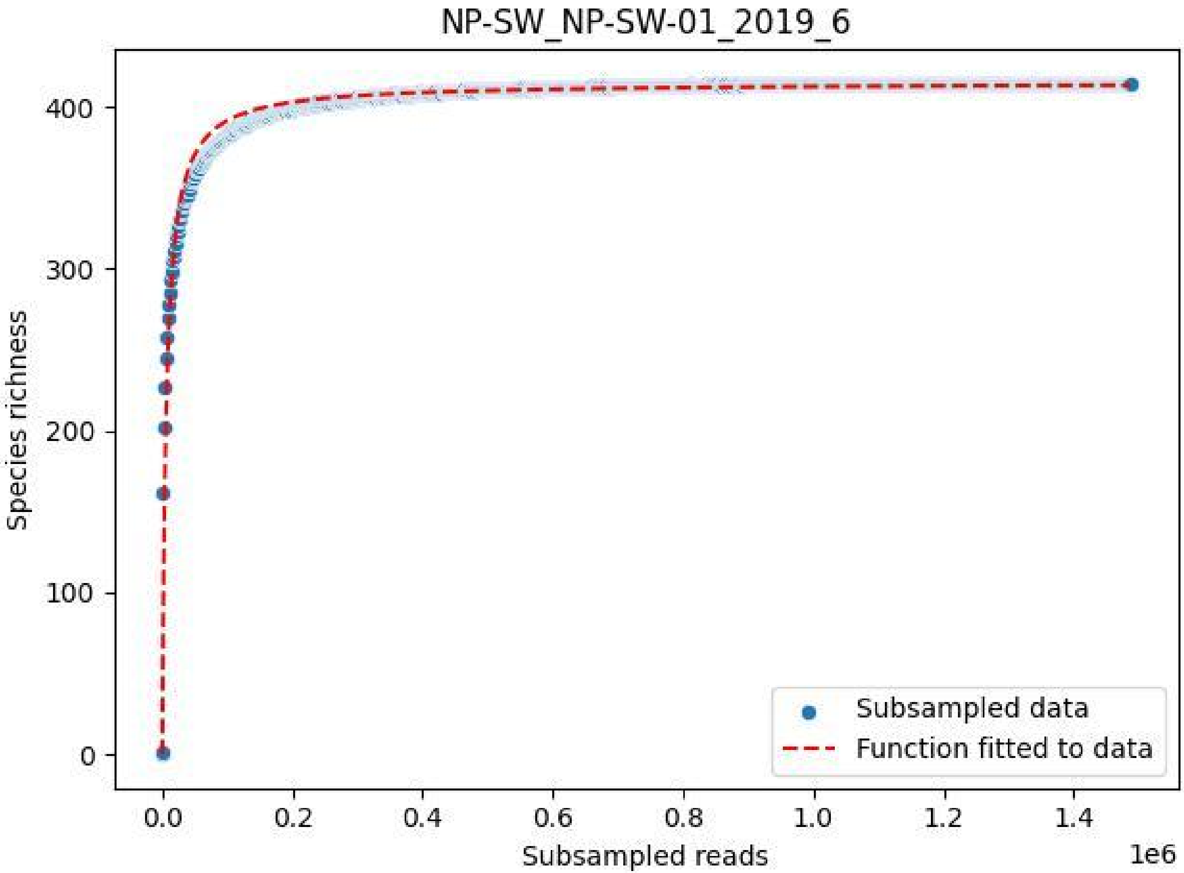
NP-SW_NP-SW-01_2019_6
Species richness
Subsampled reads
1e6
0
100
200
300
400
0.0
0.2
0.4
0.6
0.8
1.0
1.2
1.4
Subsampled data
Function fitted to data

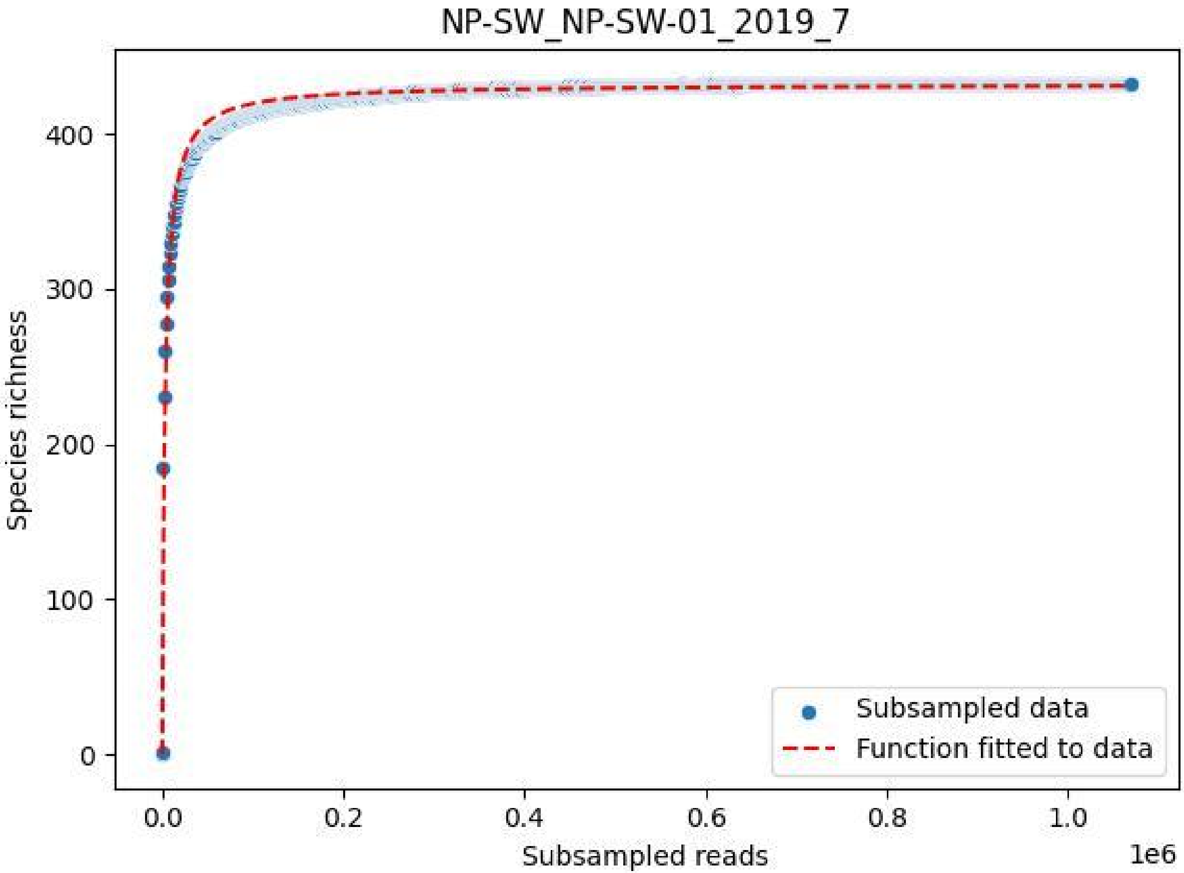
NP-SW_NP-SW-01_2019_7
Species richness
Subsampled reads
1e6
0
100
200
300
400
0.0
0.2
0.4
0.6
0.8
1.0
Subsampled data
Function fitted to data

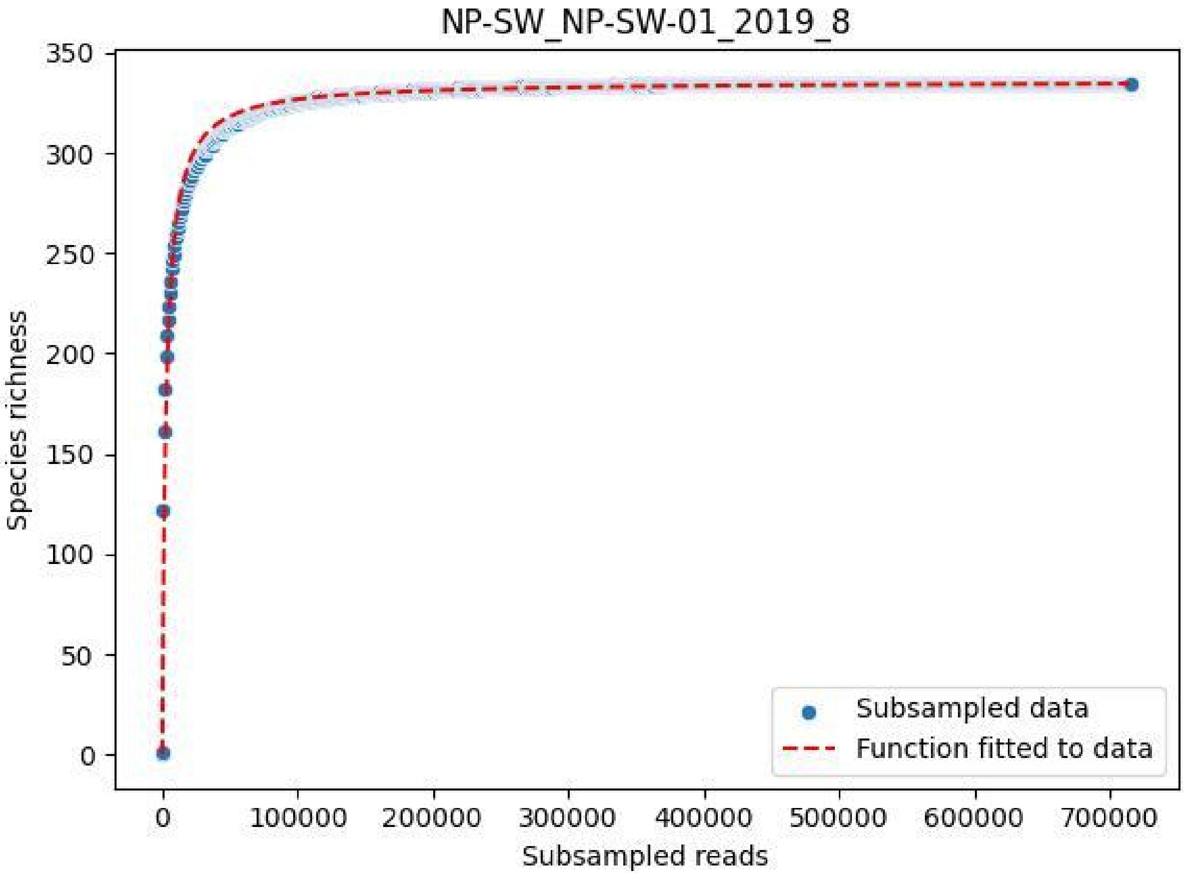
NP-SW_NP-SW-01_2019_8
Species richness
Subsampled reads
0
50
100
150
200
250
300
350
0
100000
200000
300000
400000
500000
600000
700000
Subsampled data
Function fitted to data

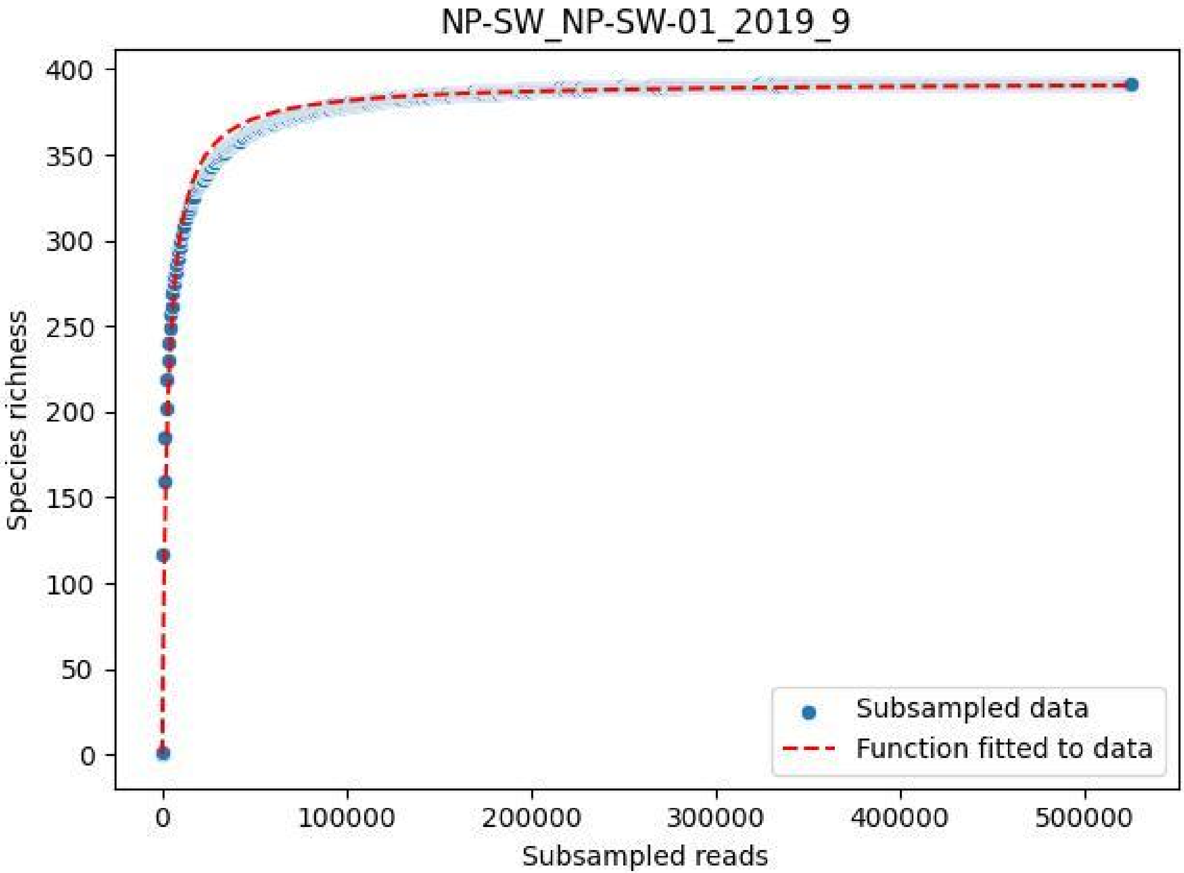
NP-SW_NP-SW-01_2019_9
Species richness
Subsampled reads
0
50
100
150
200
250
300
350
400
0
100000
200000
300000
400000
500000
Subsampled data
Function fitted to data

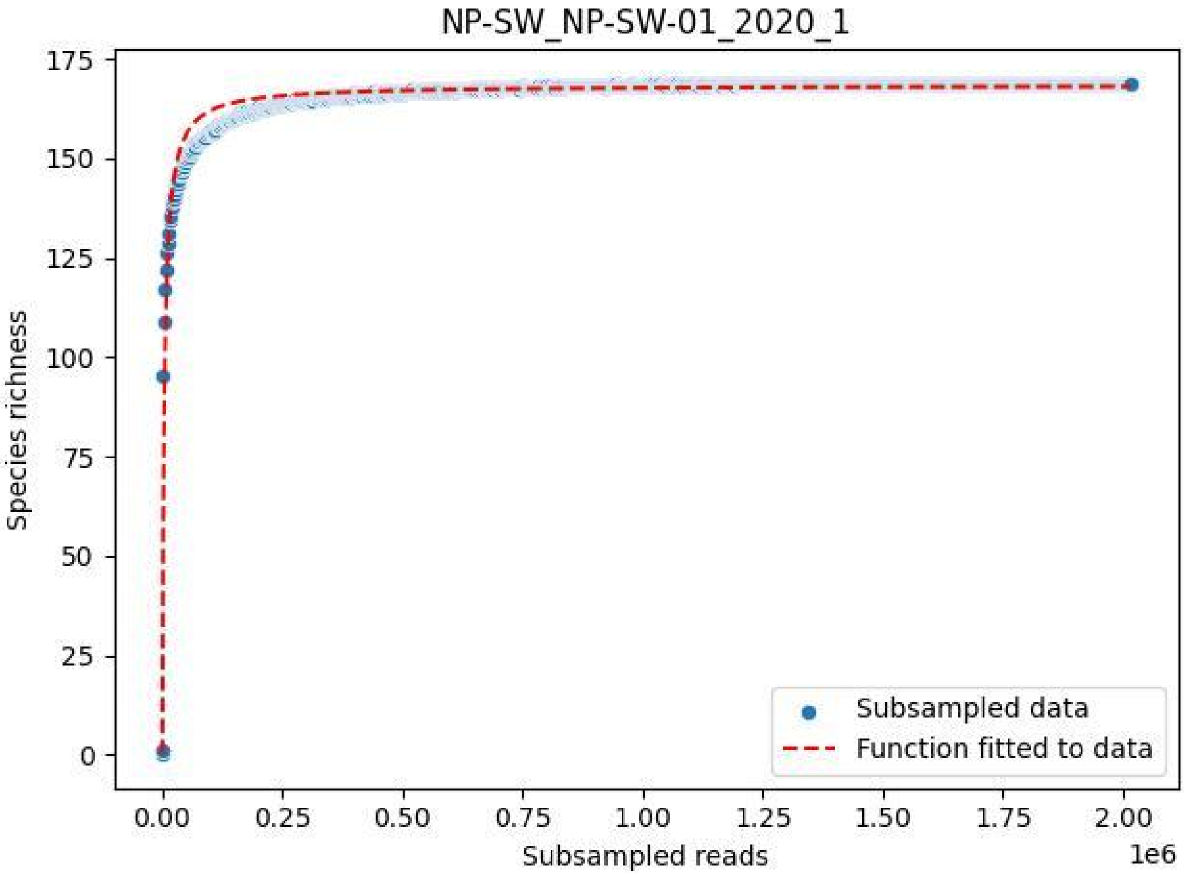
NP-SW_NP-SW-01_2020_1
Species richness
Subsampled reads
1e6
0
25
50
75
100
125
150
175
0.00
0.25
0.50
0.75
1.00
1.25
1.50
1.75
2.00
Subsampled data
Function fitted to data

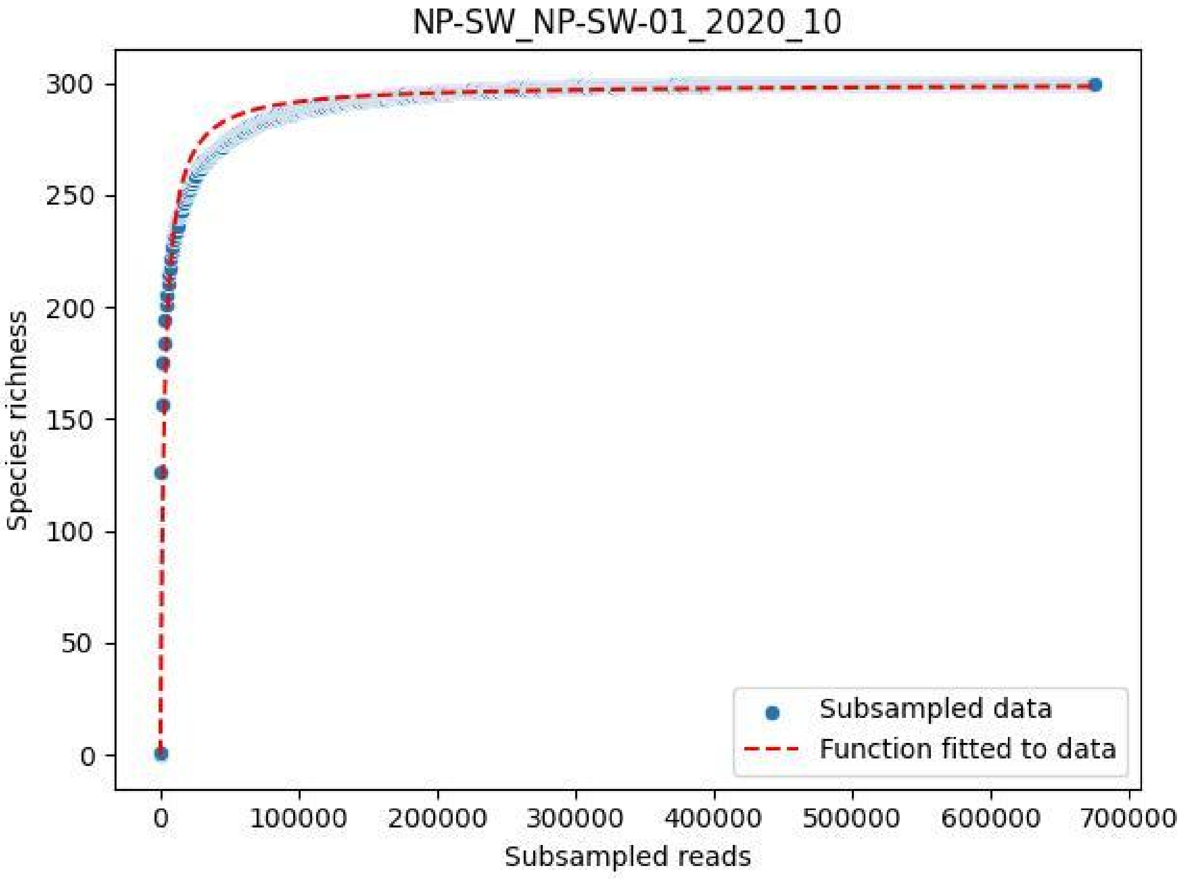

NP-SW_NP-SW-01_2020_10
Species richness
Subsampled reads
0
50
100
150
200
250
300
0
100000
200000
300000
400000
500000
600000
700000
Subsampled data
Function fitted to data

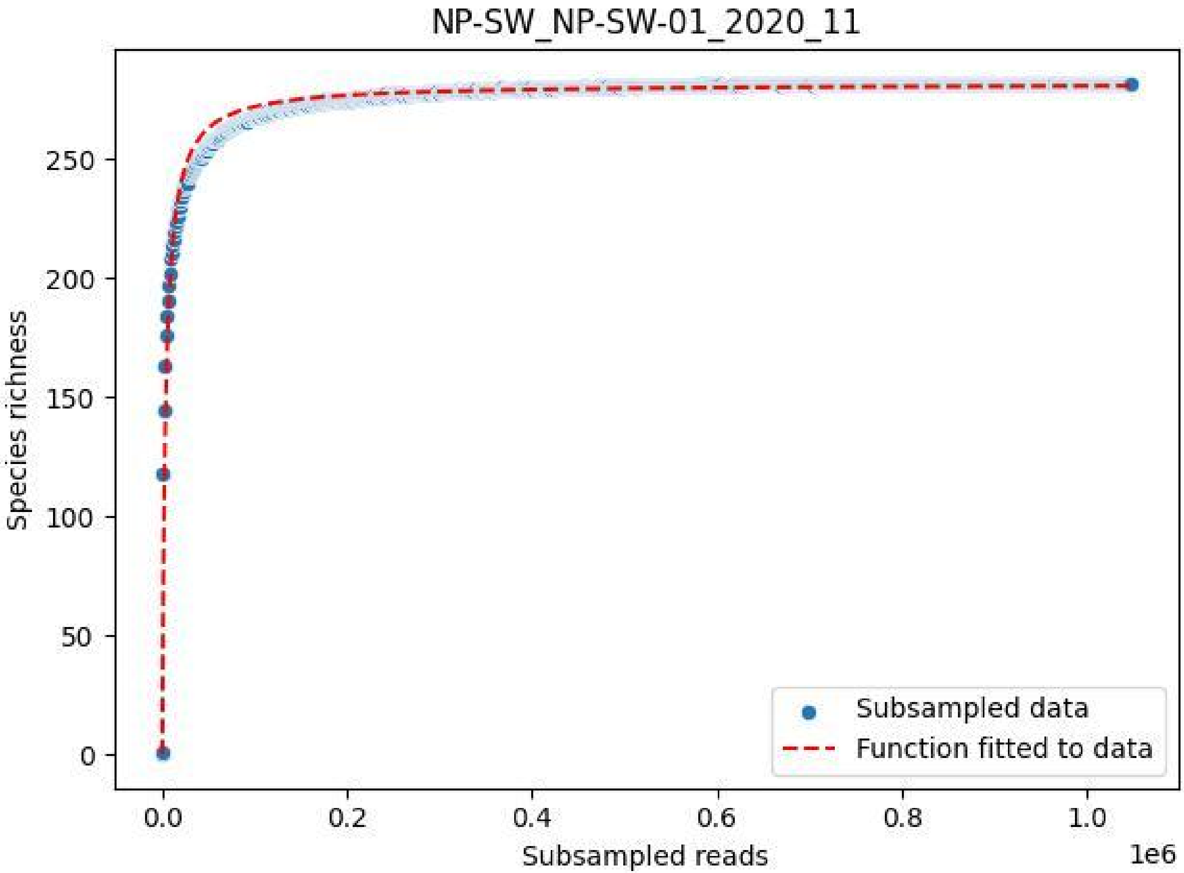

NP-SW_NP-SW-01_2020_11
Species richness
Subsampled reads
1e6
0
50
100
150
200
250
0.0
0.2
0.4
0.6
0.8
1.0
Subsampled data
Function fitted to data

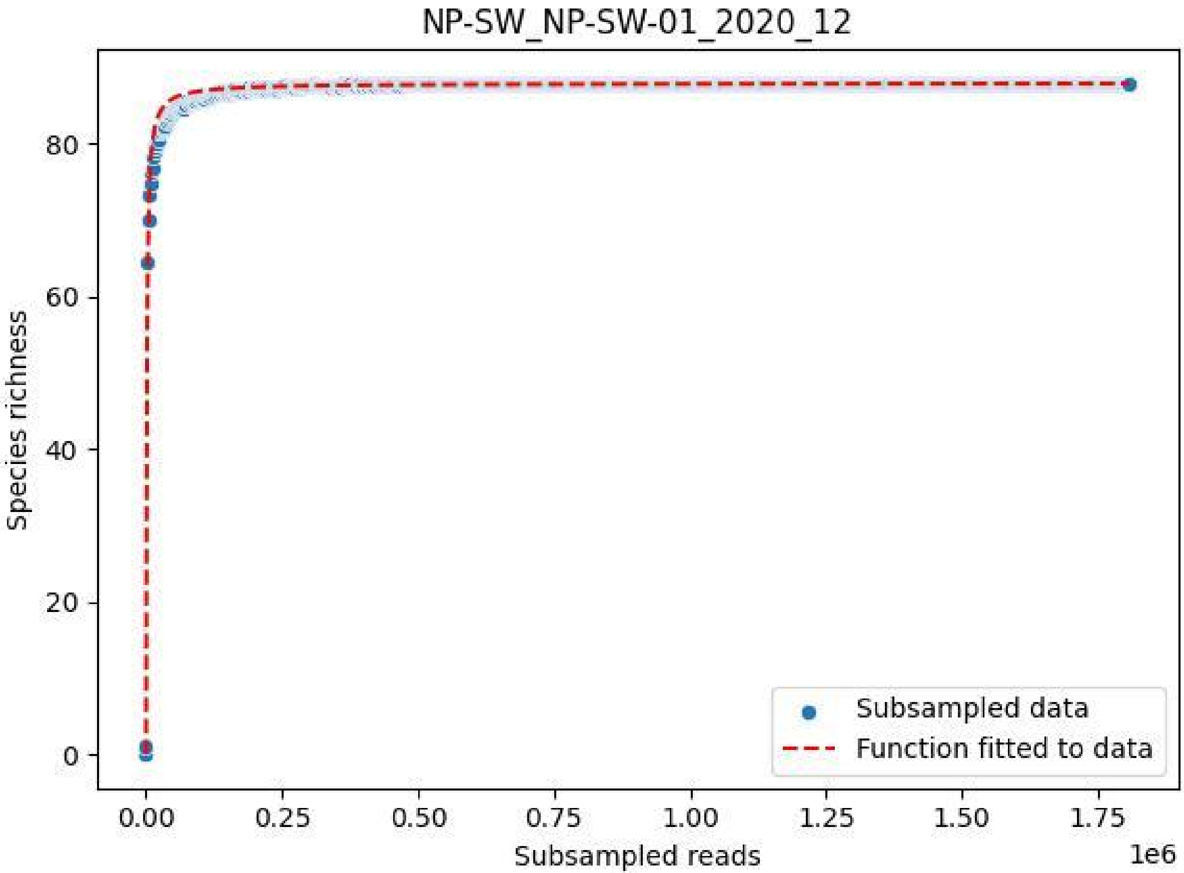
NP-SW_NP-SW-01_2020_12
Species richness
0
20
40
60
80
0.00
0.25
0.50
0.75
1.00
1.25
1.50
1.75
1e6
Subsampled reads
Subsampled data
Function fitted to data

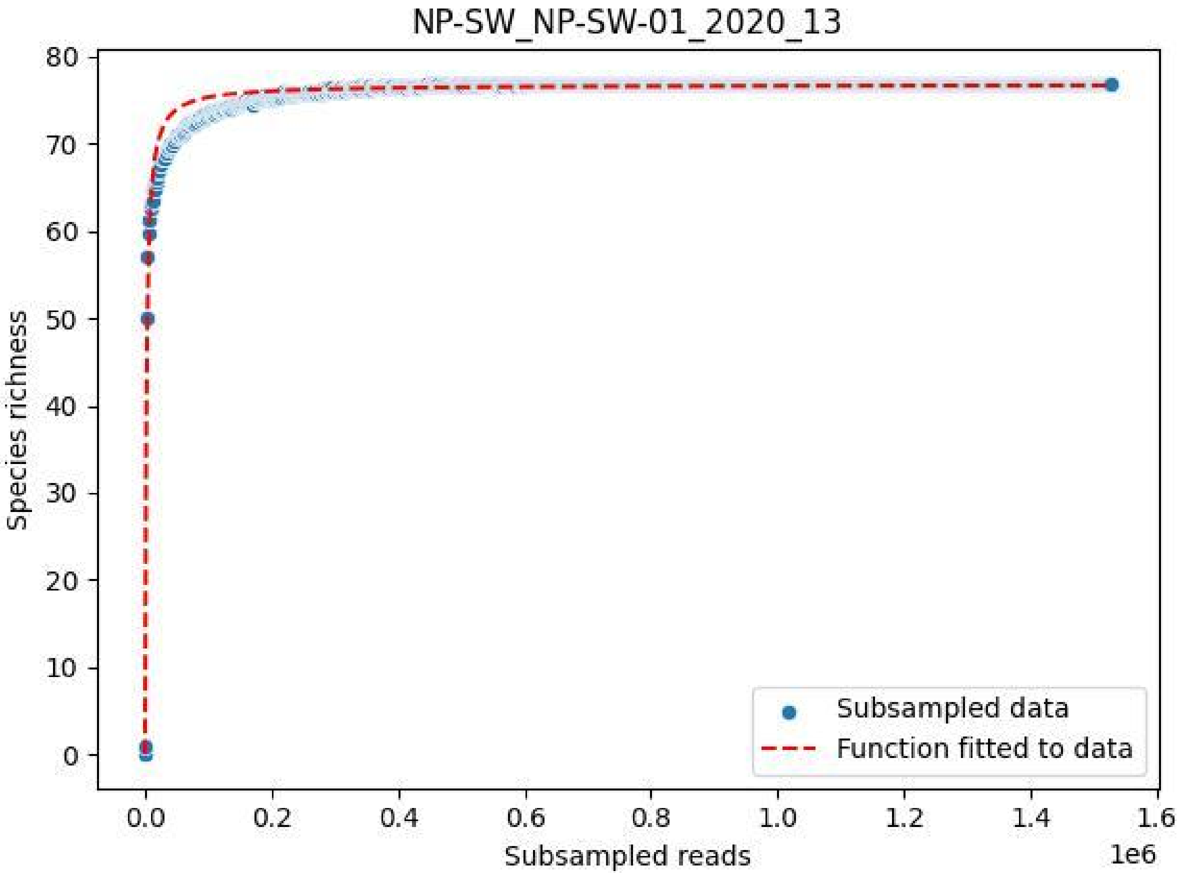

NP-SW_NP-SW-01_2020_13
Species richness
Subsampled reads
1e6
0
10
20
30
40
50
60
70
80
0.0
0.2
0.4
0.6
0.8
1.0
1.2
1.4
1.6
Subsampled data
Function fitted to data

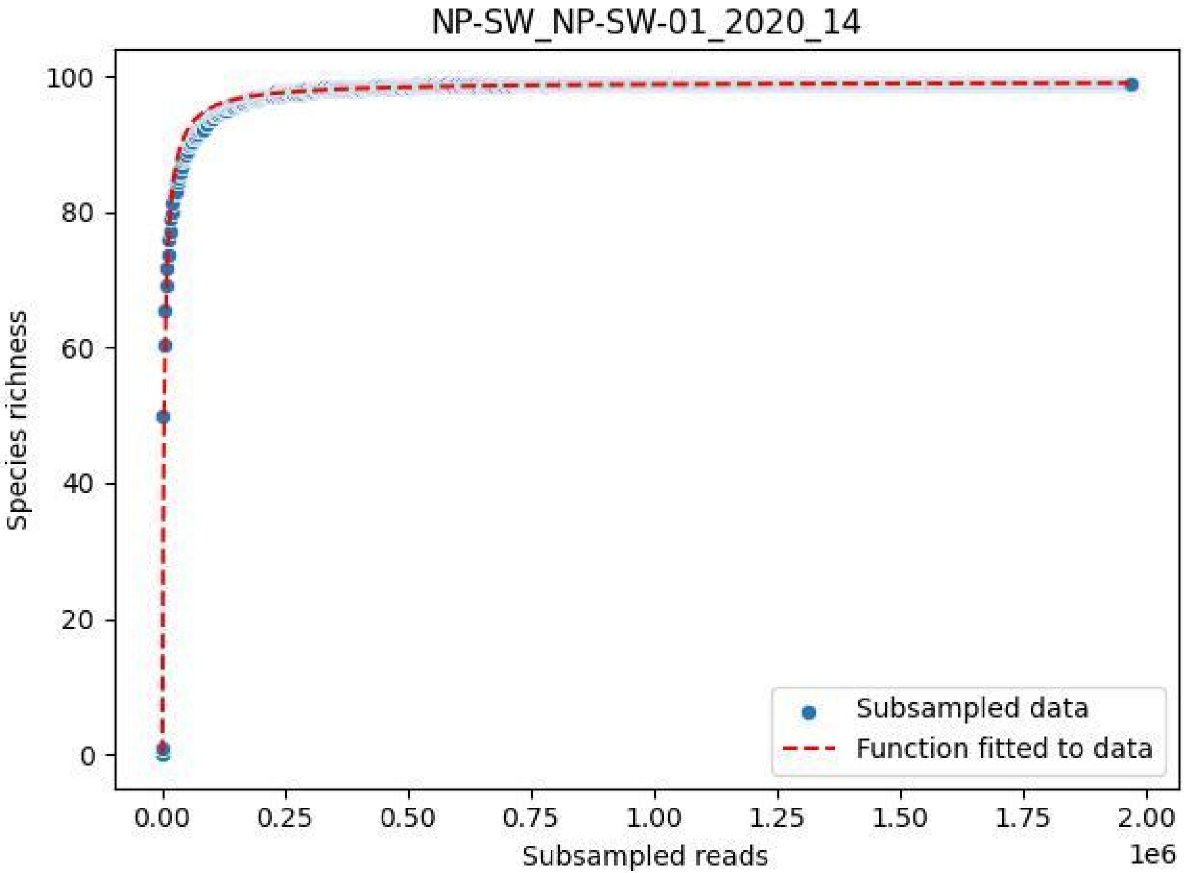

NP-SW_NP-SW-01_2020_14
100
80
60
40
20
0
Species richness
0.00
0.25
0.50
0.75
1.00
1.25
1.50
1.75
2.00
1e6
Subsampled reads
Subsampled data
Function fitted to data

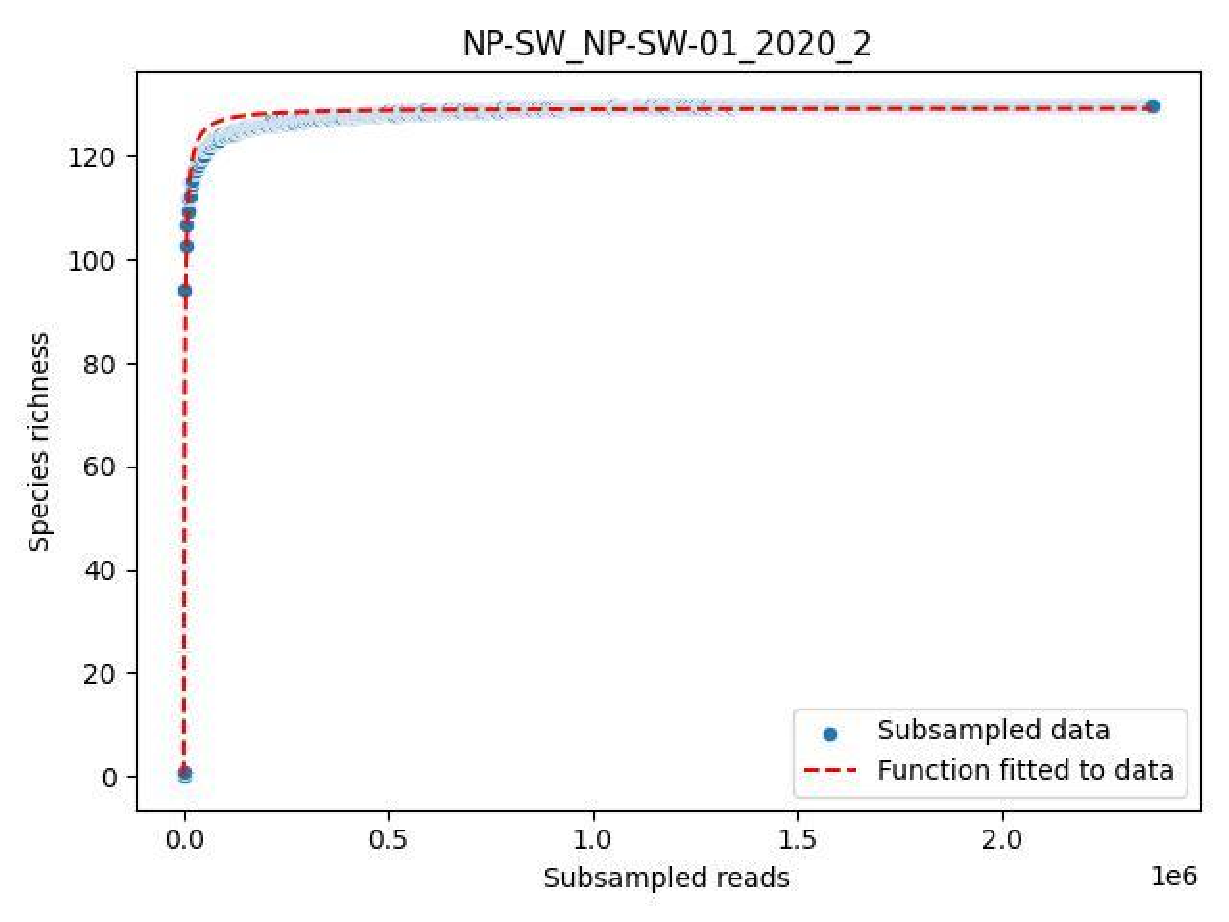
NP-SW_NP-SW-01_2020_2
Species richness
0
20
40
60
80
100
120
0.0
0.5
1.0
1.5
2.0
1e6
Subsampled reads
Subsampled data
Function fitted to data

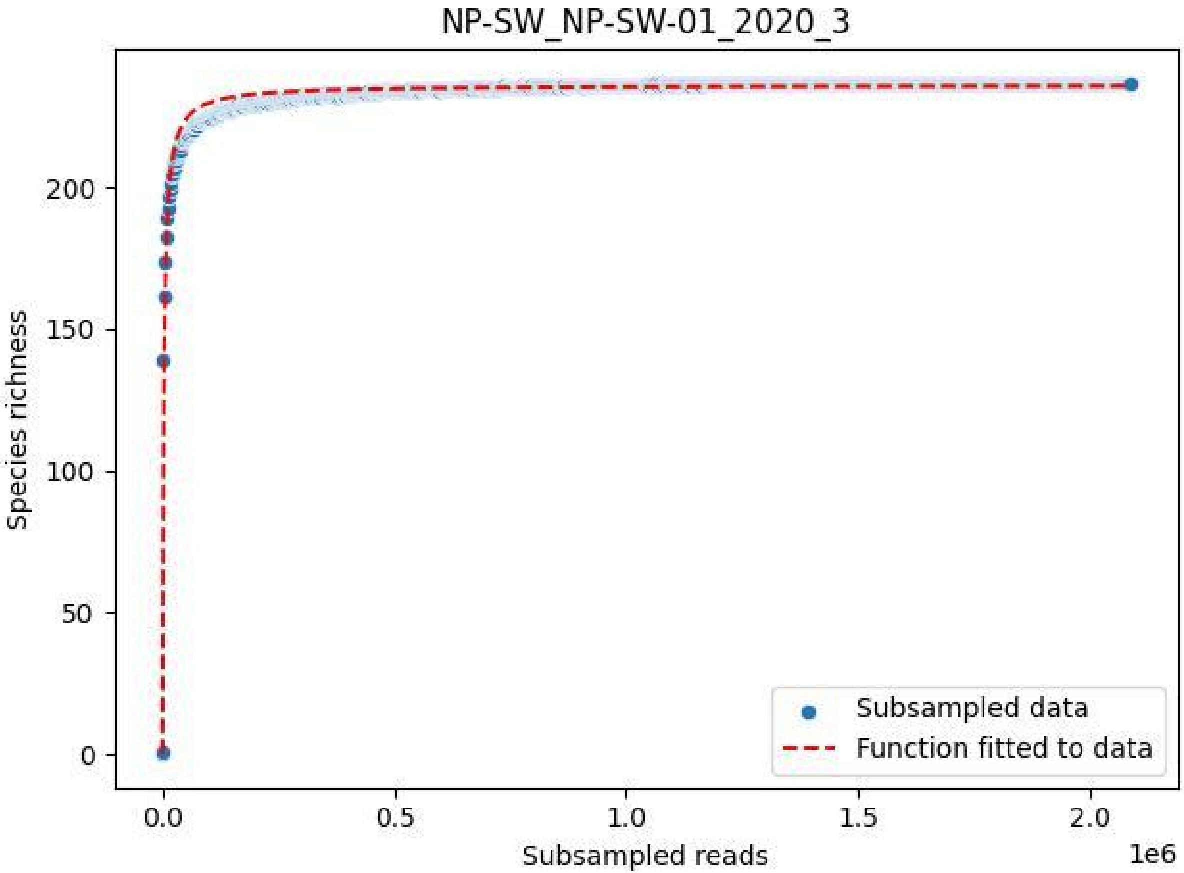

NP-SW_NP-SW-01_2020_3
Species richness
Subsampled reads
1e6
0
50
100
150
200
0.0
0.5
1.0
1.5
2.0
Subsampled data
Function fitted to data

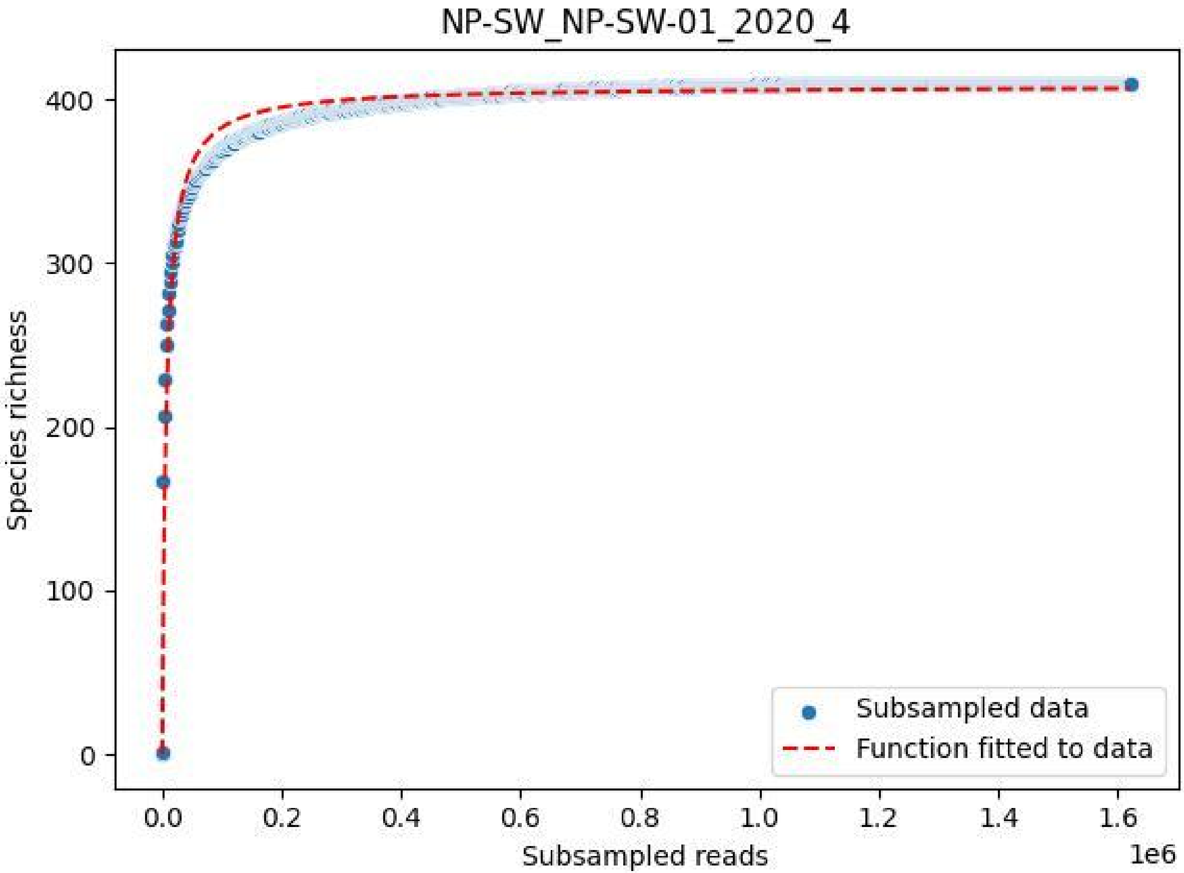
NP-SW_NP-SW-01_2020_4
Species richness
0
100
200
300
400
0.0
0.2
0.4
0.6
0.8
1.0
1.2
1.4
1.6
1e6
Subsampled reads
Subsampled data
Function fitted to data

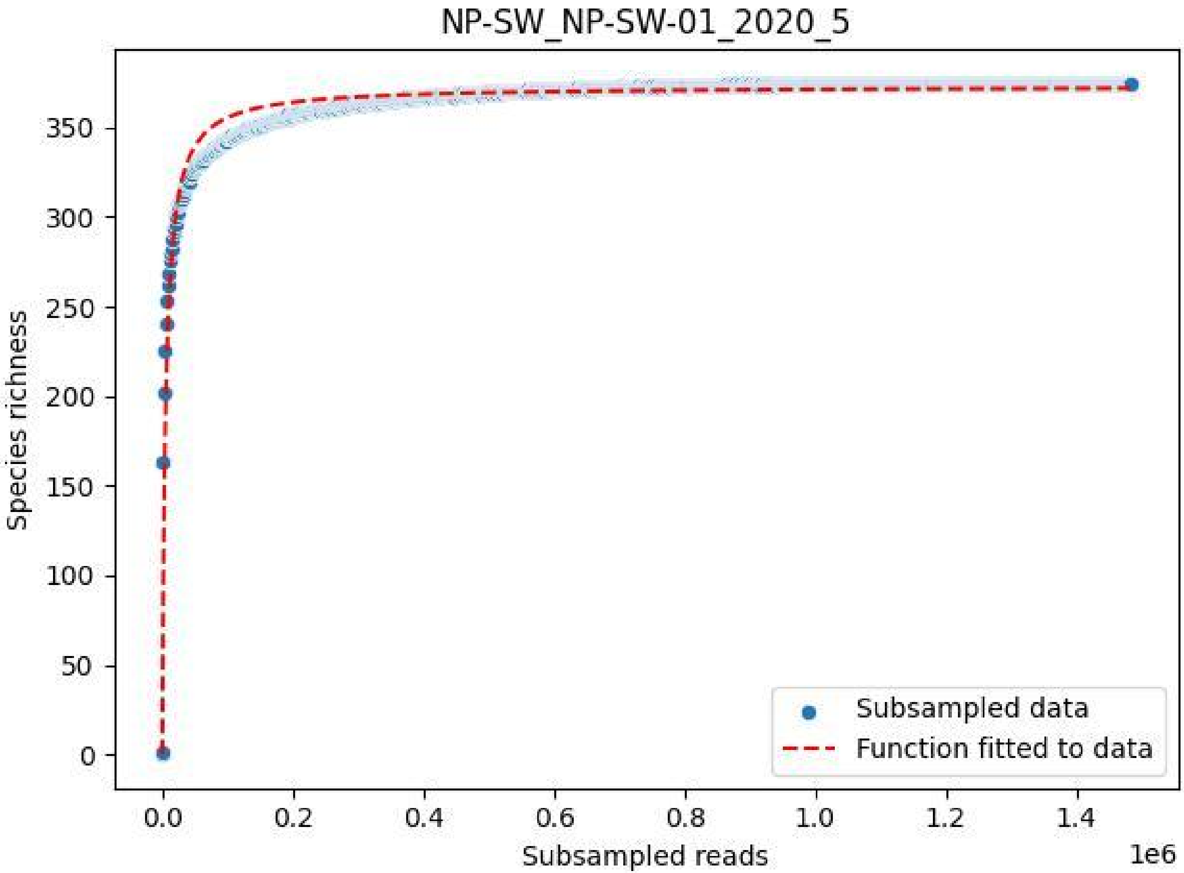

NP-SW_NP-SW-01_2020_5
Species richness
Subsampled reads
1e6
0
50
100
150
200
250
300
350
0.0
0.2
0.4
0.6
0.8
1.0
1.2
1.4
Subsampled data
Function fitted to data

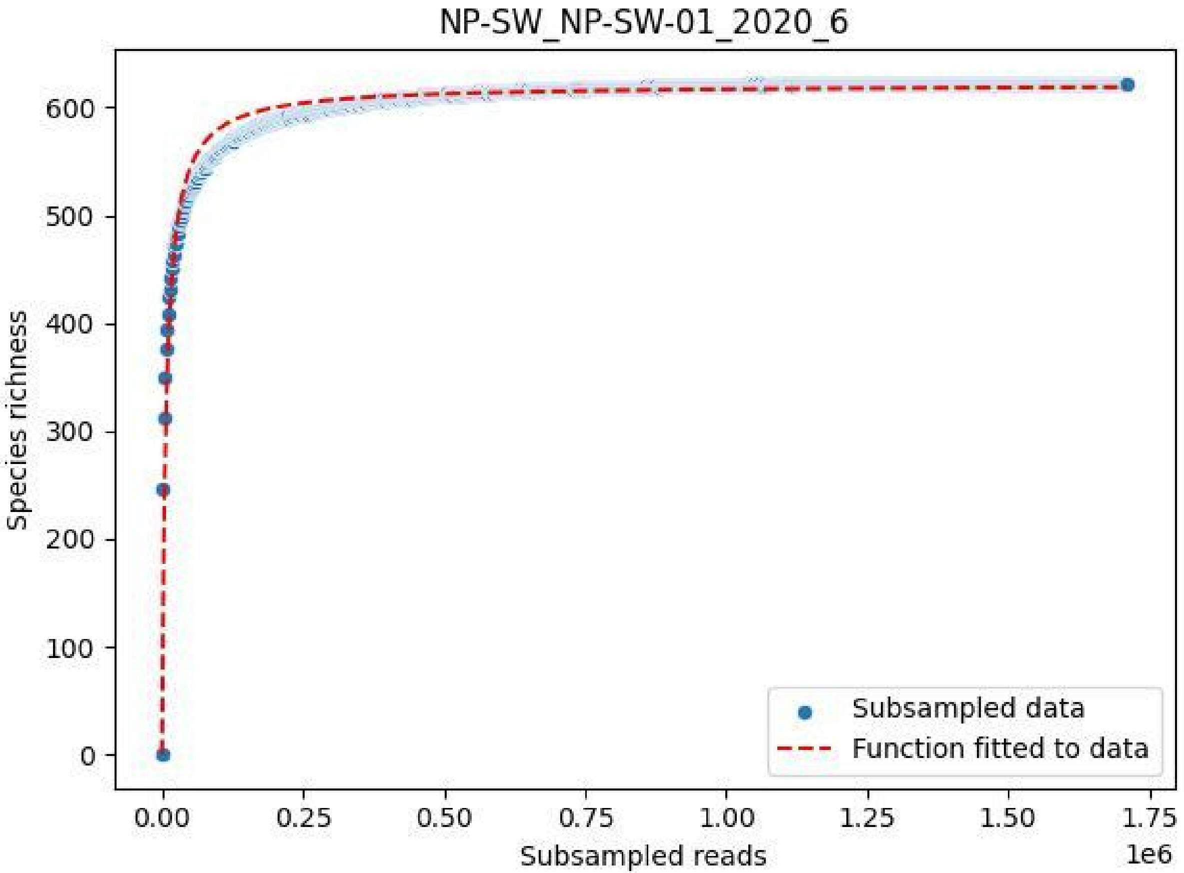

NP-SW_NP-SW-01_2020_6
Species richness
Subsampled reads
1e6
0
100
200
300
400
500
600
0.00
0.25
0.50
0.75
1.00
1.25
1.50
1.75
Subsampled data
Function fitted to data

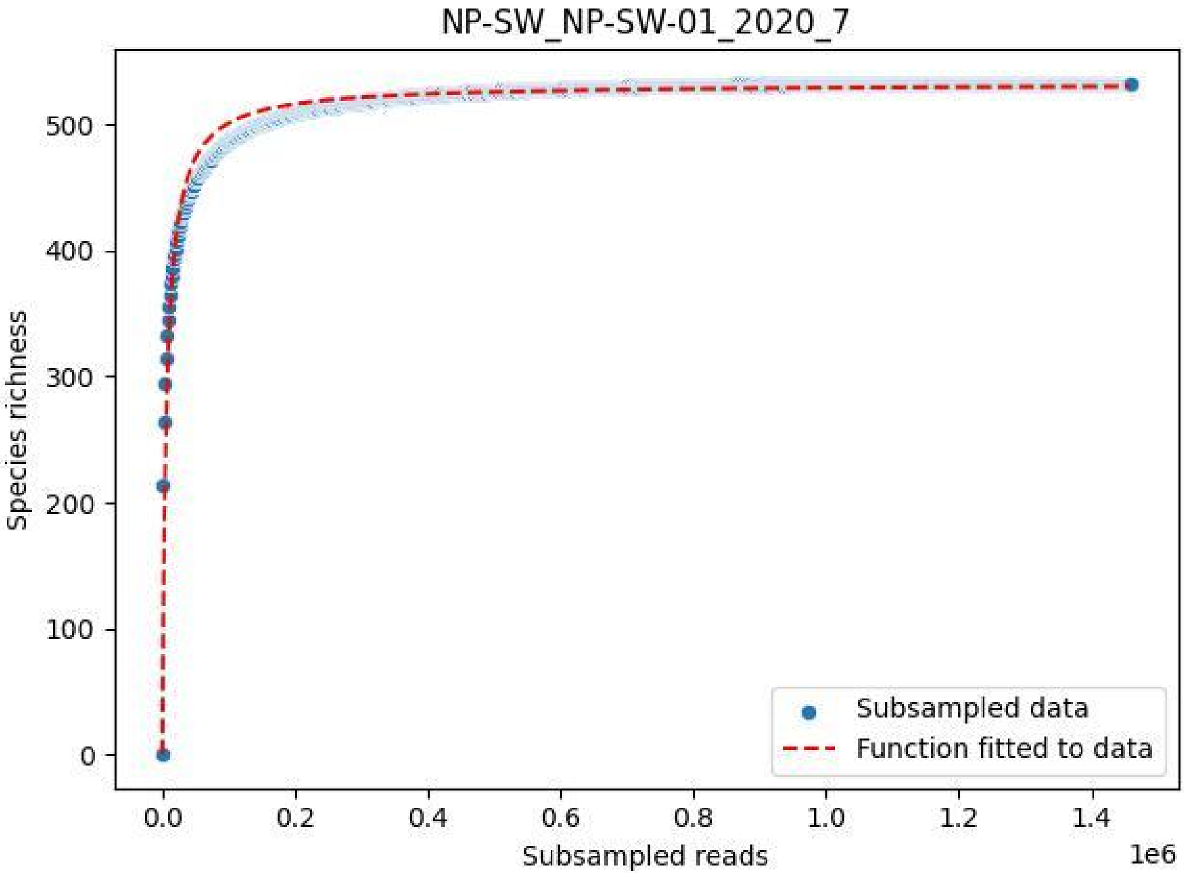
NP-SW_NP-SW-01_2020_7
Species richness
Subsampled reads
1e6
0
100
200
300
400
500
0.0
0.2
0.4
0.6
0.8
1.0
1.2
1.4
Subsampled data
Function fitted to data

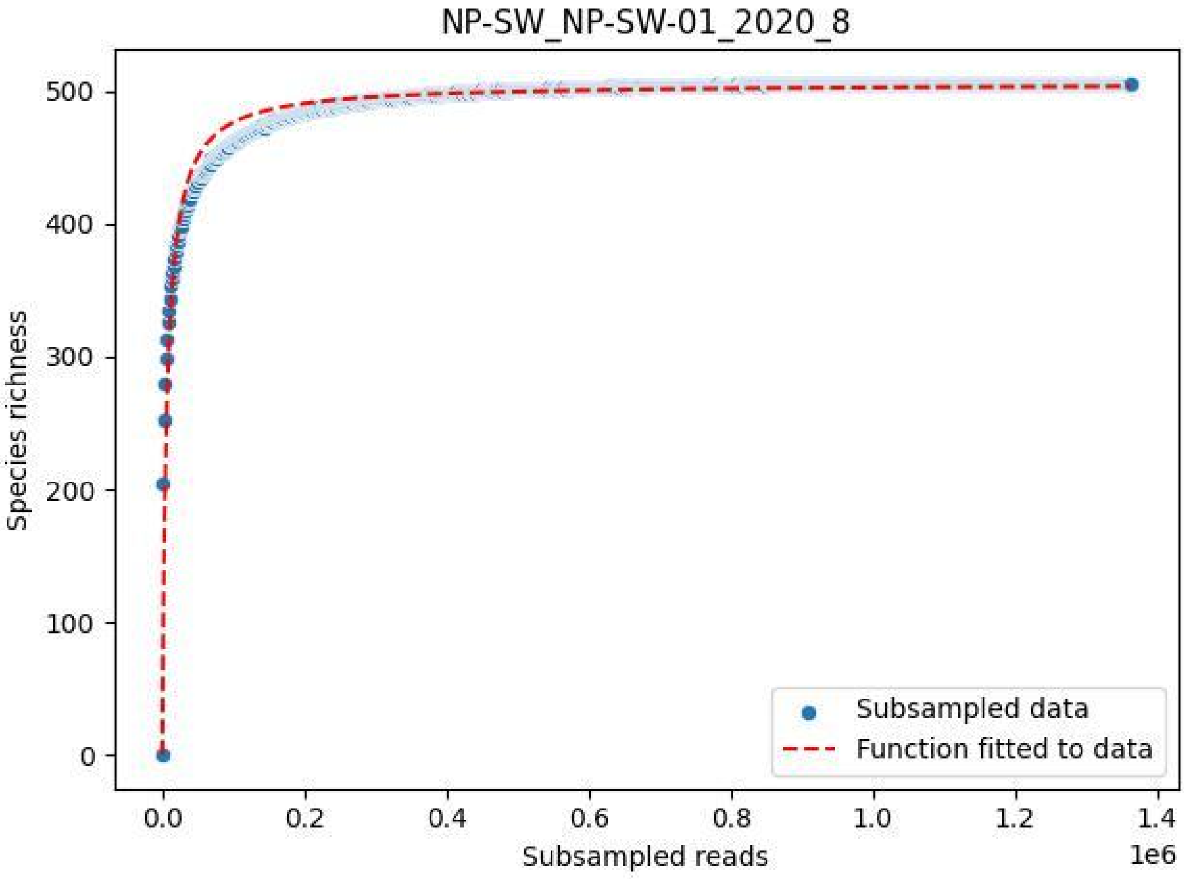
NP-SW_NP-SW-01_2020_8
Species richness
Subsampled reads
1e6
0
100
200
300
400
500
0.0
0.2
0.4
0.6
0.8
1.0
1.2
1.4
Subsampled data
Function fitted to data

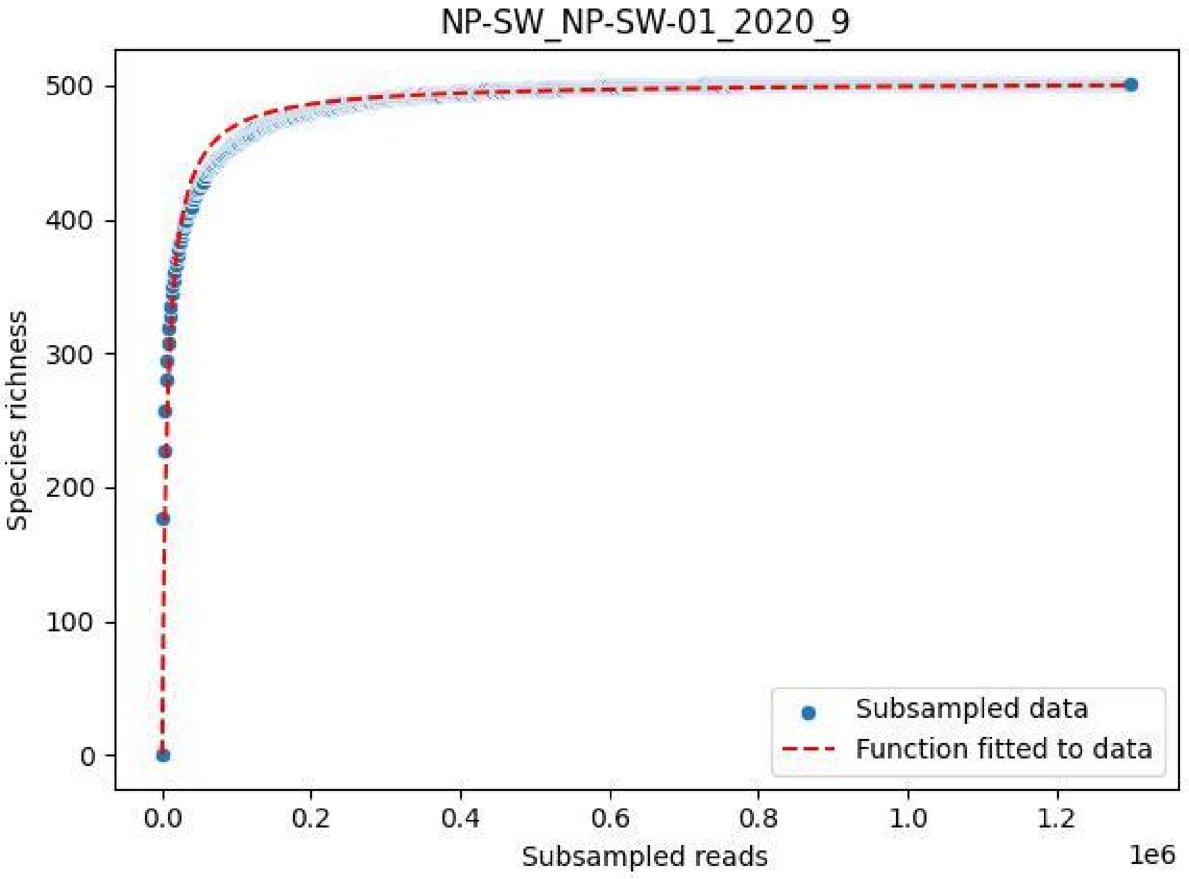

NP-SW_NP-SW-01_2020_9
Species richness
Subsampled reads
1e6
0
100
200
300
400
500
0.0
0.2
0.4
0.6
0.8
1.0
1.2
Subsampled data
Function fitted to data

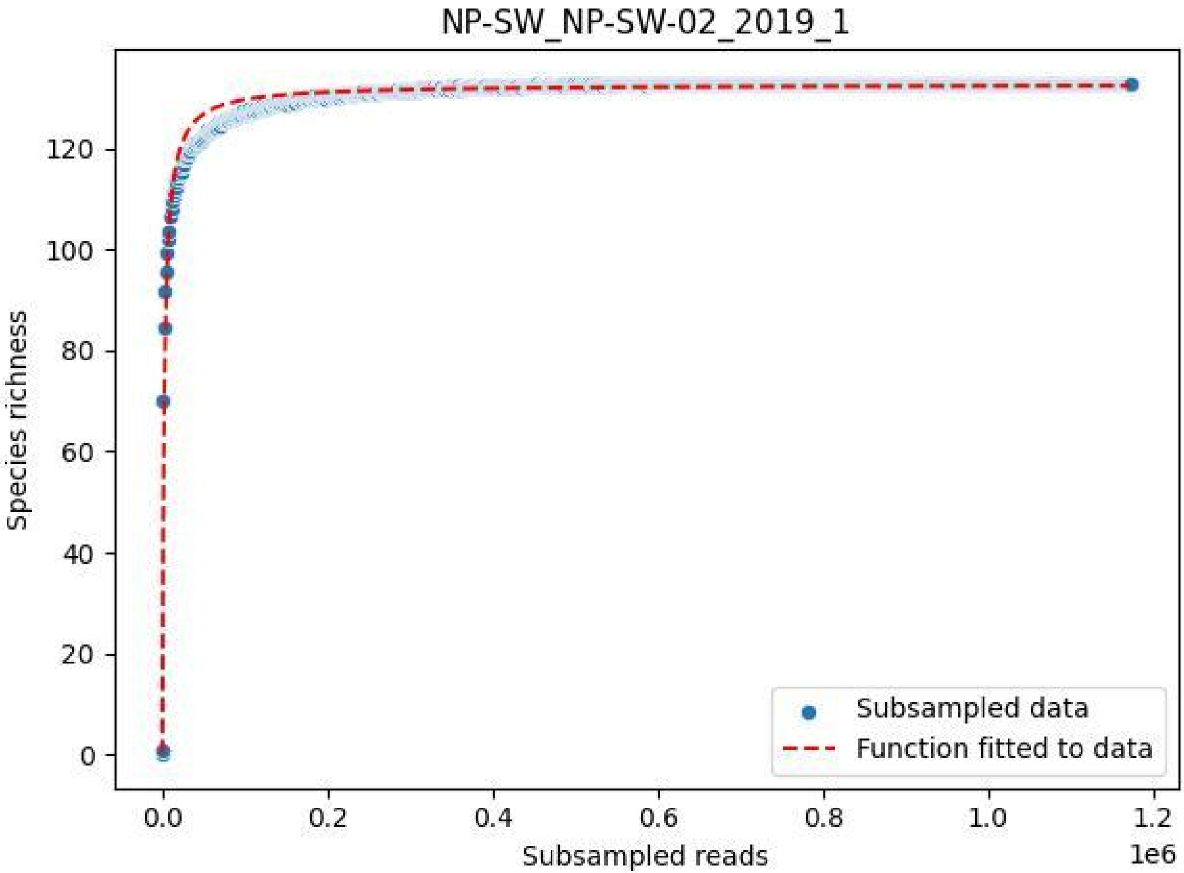
NP-SW_NP-SW-02_2019_1
Species richness
Subsampled reads
1e6
0
20
40
60
80
100
120
0.0
0.2
0.4
0.6
0.8
1.0
1.2
Subsampled data
Function fitted to data

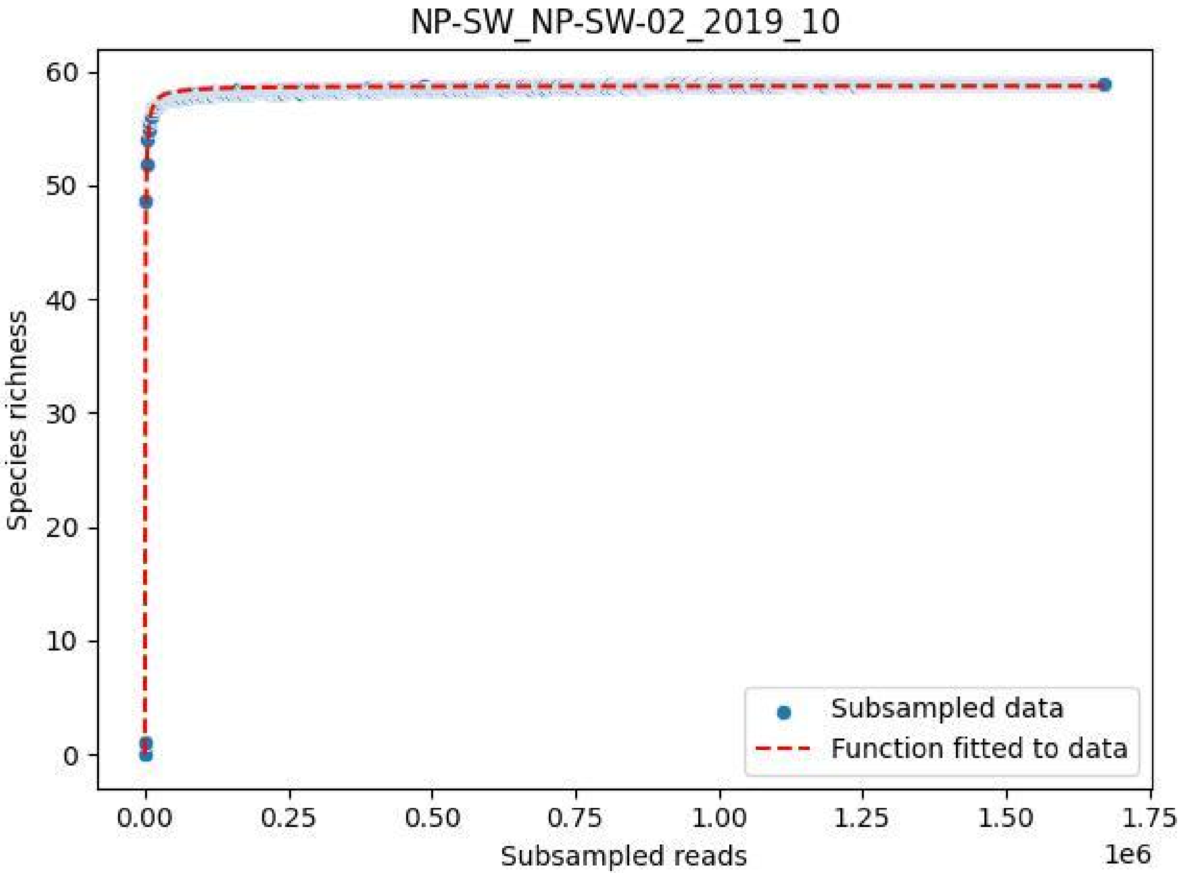

NP-SW_NP-SW-02_2019_10
Species richness
Subsampled reads
1e6
0
10
20
30
40
50
60
0.00
0.25
0.50
0.75
1.00
1.25
1.50
1.75
Subsampled data
Function fitted to data

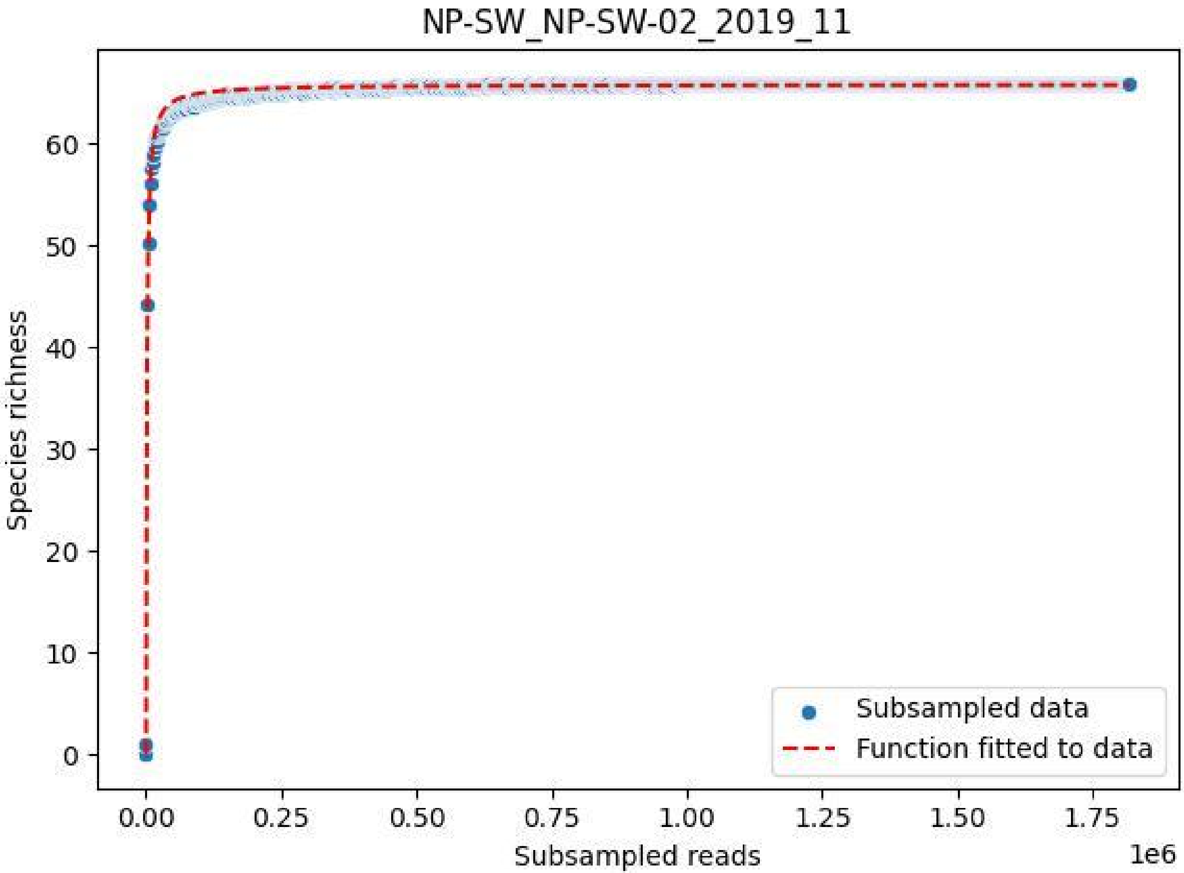
NP-SW_NP-SW-02_2019_11
Species richness
Subsampled reads
1e6
0
10
20
30
40
50
60
0.00
0.25
0.50
0.75
1.00
1.25
1.50
1.75
Subsampled data
Function fitted to data

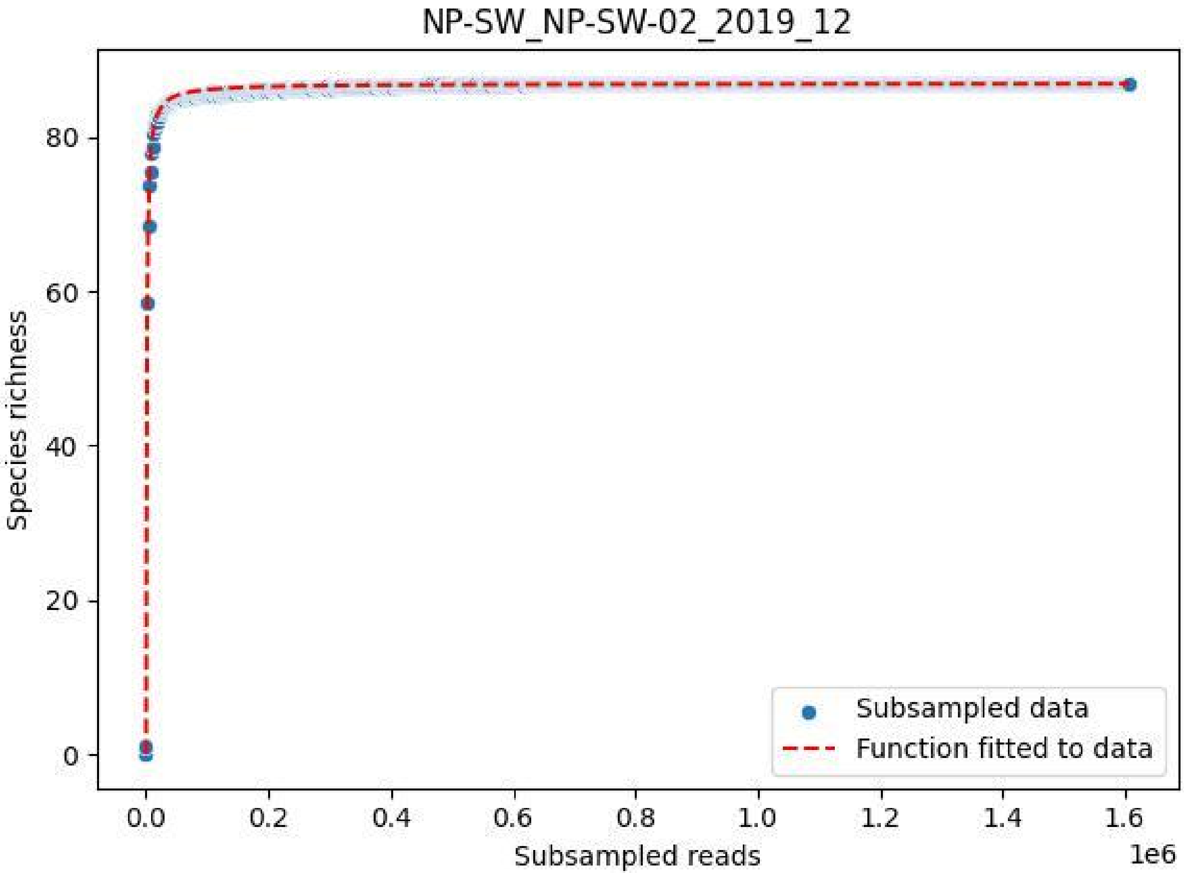

NP-SW_NP-SW-02_2019_12
Species richness
Subsampled reads
1e6
0
20
40
60
80
0.0
0.2
0.4
0.6
0.8
1.0
1.2
1.4
1.6
Subsampled data
Function fitted to data

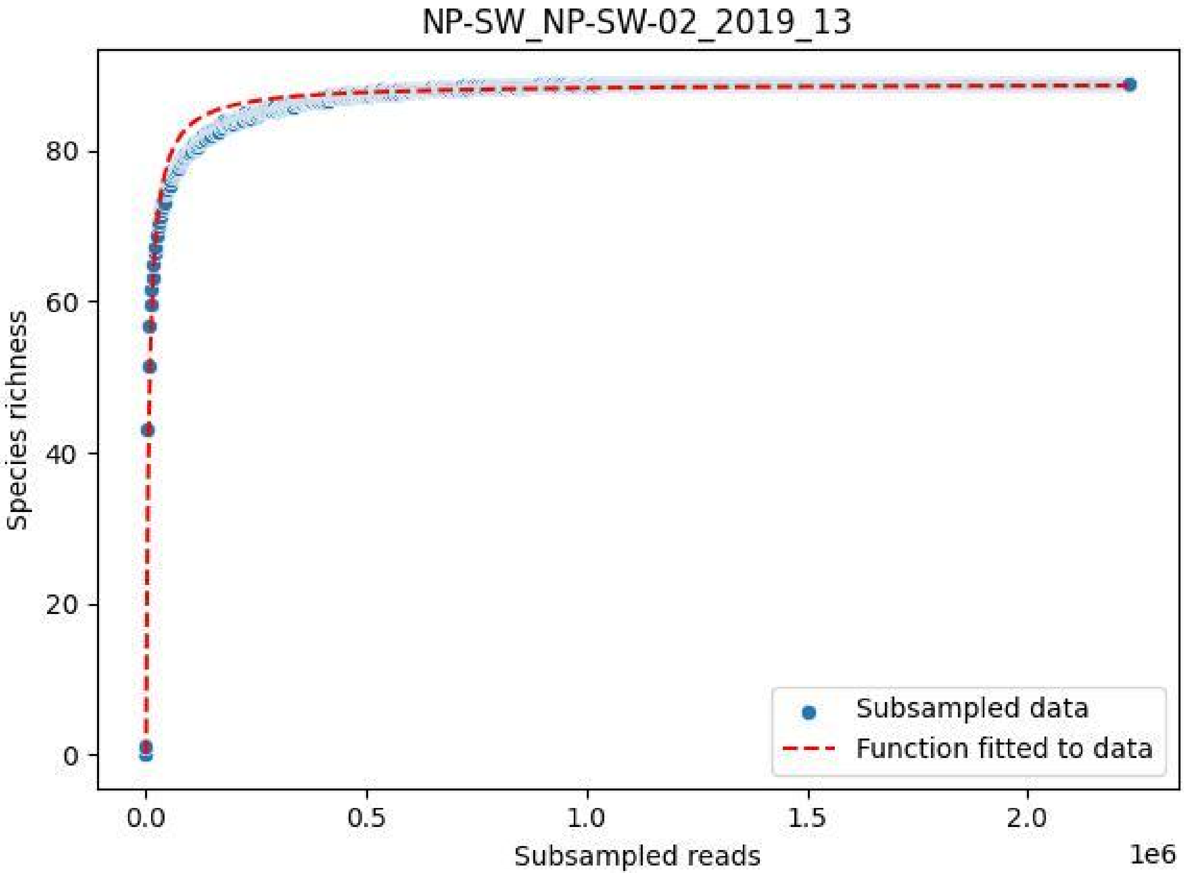
NP-SW_NP-SW-02_2019_13
Species richness
Subsampled reads
1e6
0
20
40
60
80
0.0
0.5
1.0
1.5
2.0
Subsampled data
Function fitted to data

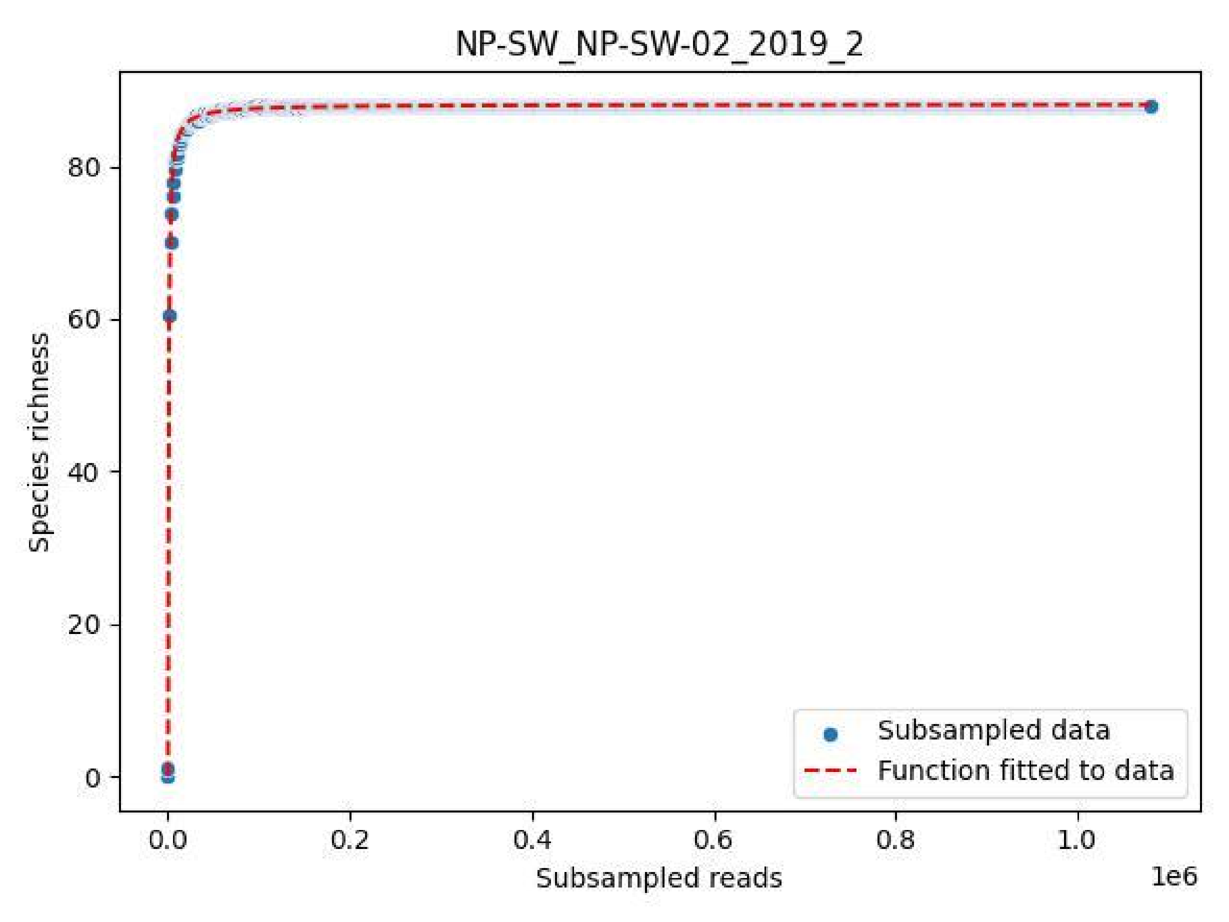

NP-SW_NP-SW-02_2019_2
Species richness
Subsampled reads
1e6
0
20
40
60
80
0.0
0.2
0.4
0.6
0.8
1.0
Subsampled data
Function fitted to data

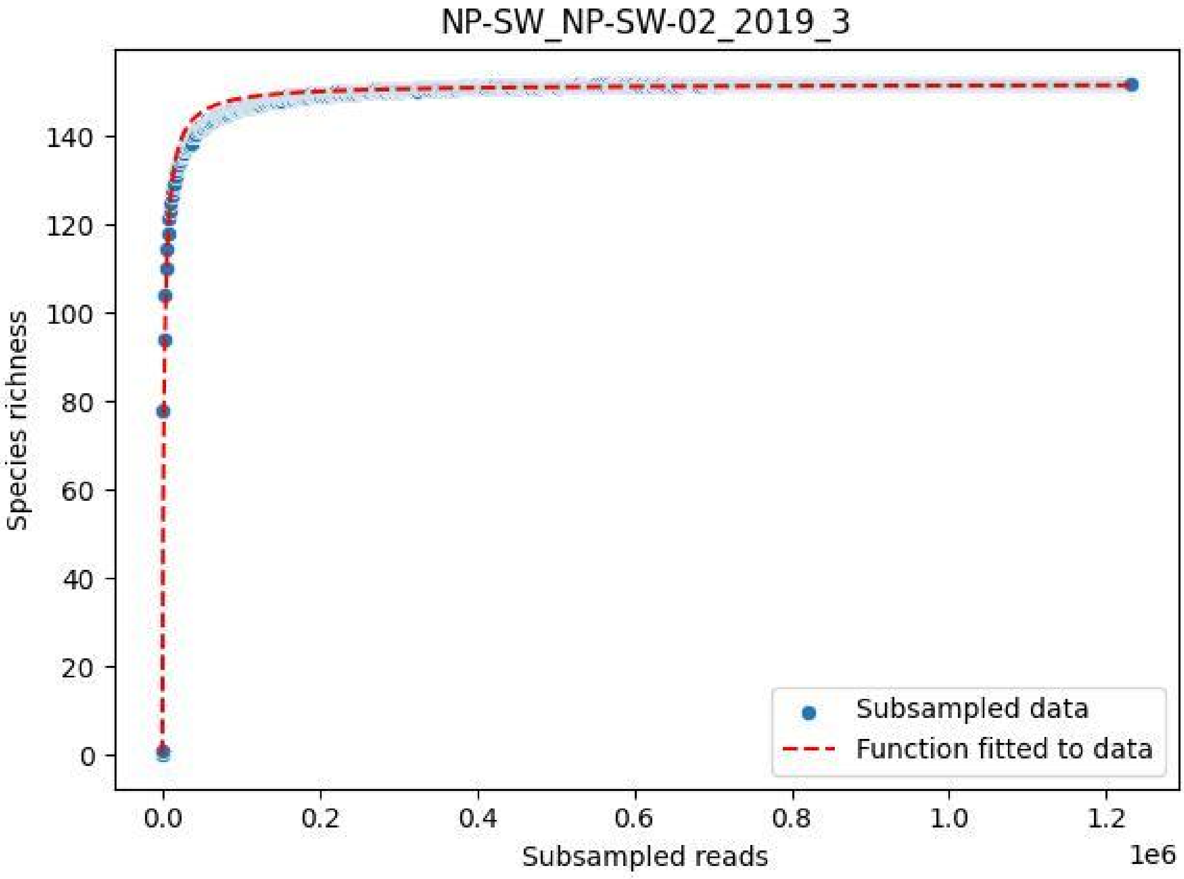

NP-SW_NP-SW-02_2019_3
Species richness
Subsampled reads
1e6
0
20
40
60
80
100
120
140
0.0
0.2
0.4
0.6
0.8
1.0
1.2
Subsampled data
Function fitted to data

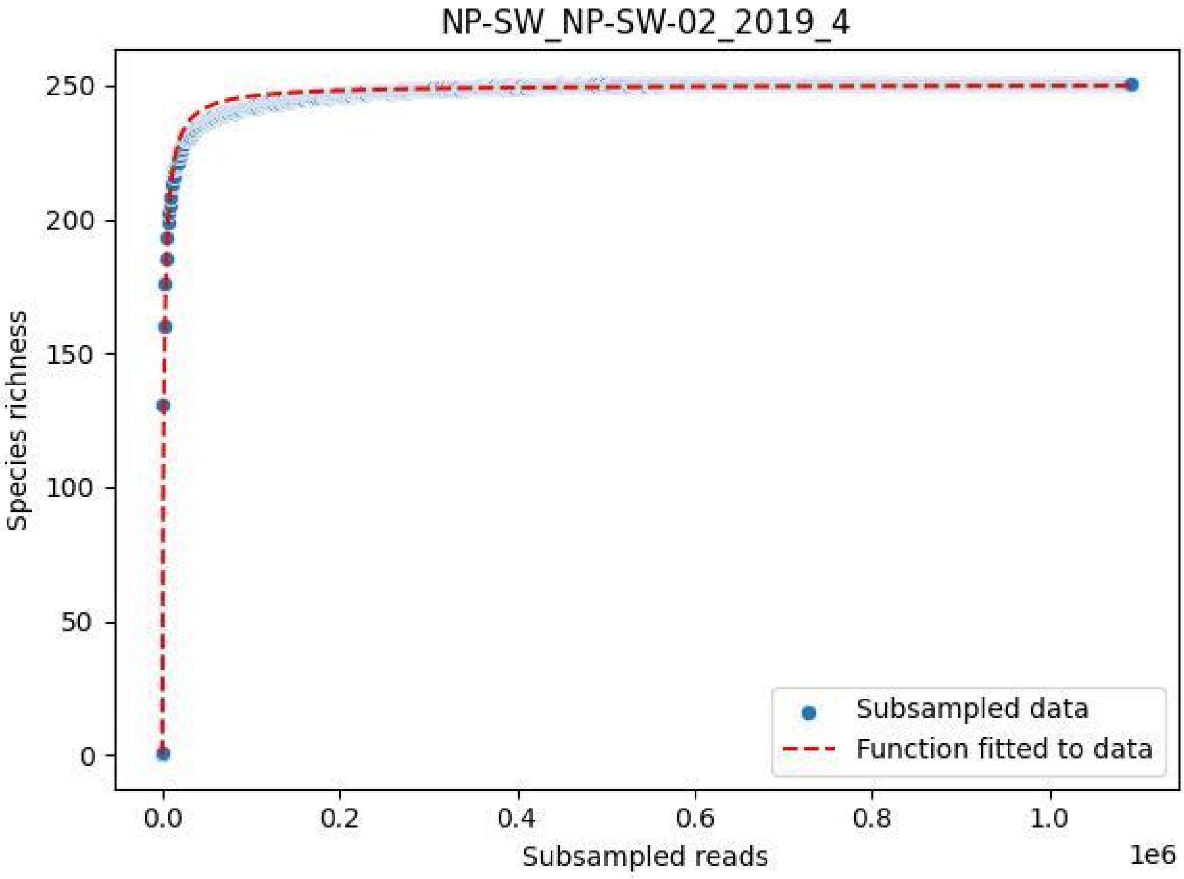
NP-SW_NP-SW-02_2019_4
Species richness
Subsampled reads
1e6
0
50
100
150
200
250
0.0
0.2
0.4
0.6
0.8
1.0
Subsampled data
Function fitted to data

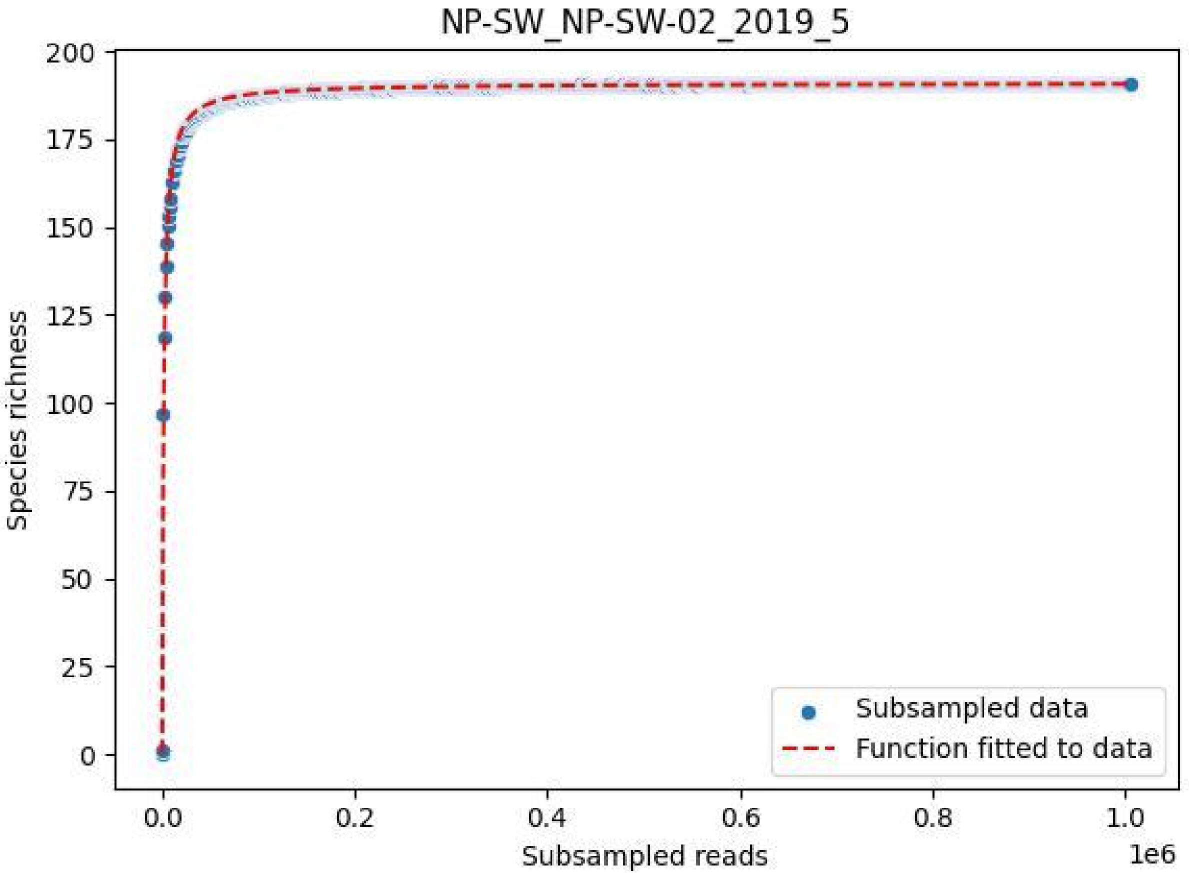

NP-SW_NP-SW-02_2019_5
Species richness
Subsampled reads
1e6
200
175
150
125
100
75
50
25
0
0.0
0.2
0.4
0.6
0.8
1.0
Subsampled data
Function fitted to data

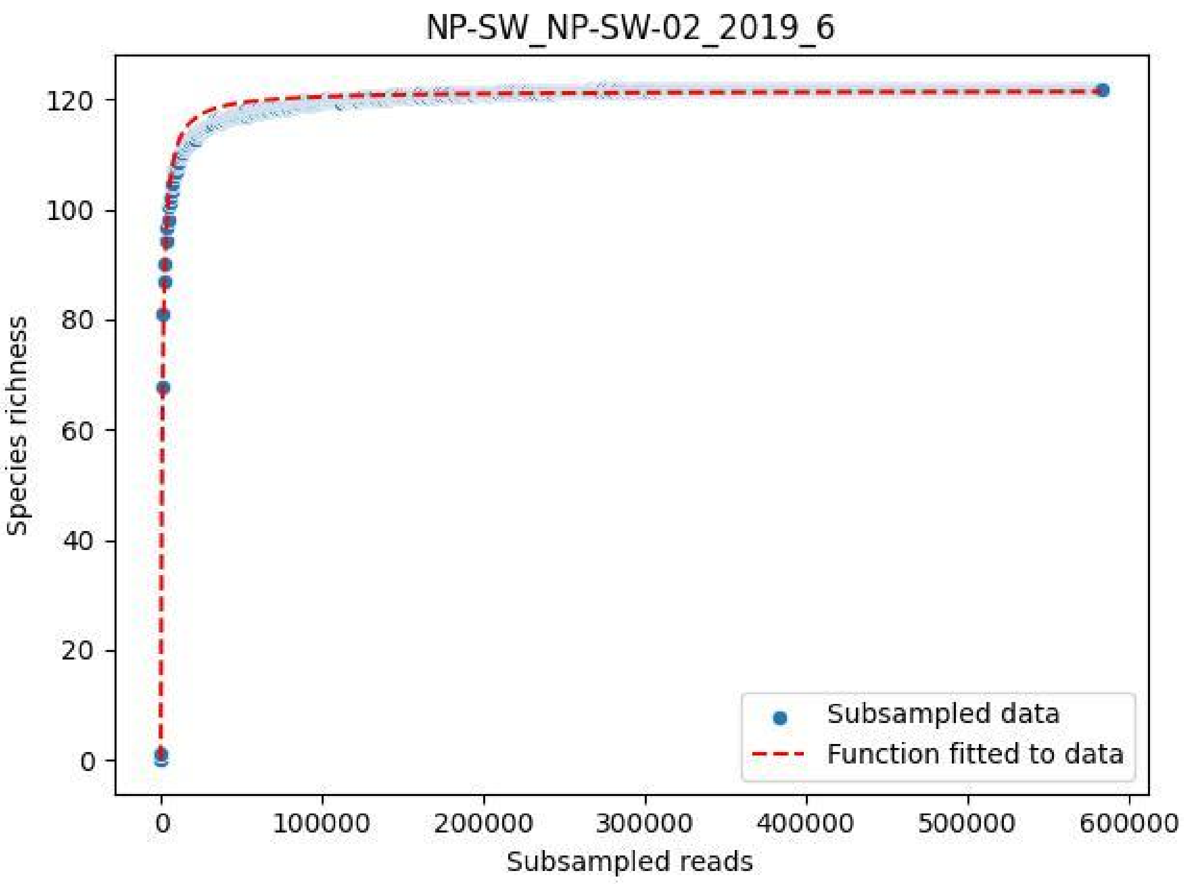

NP-SW_NP-SW-02_2019_6
Species richness
0
20
40
60
80
100
120
Subsampled reads
0
100000
200000
300000
400000
500000
600000
Subsampled data
Function fitted to data

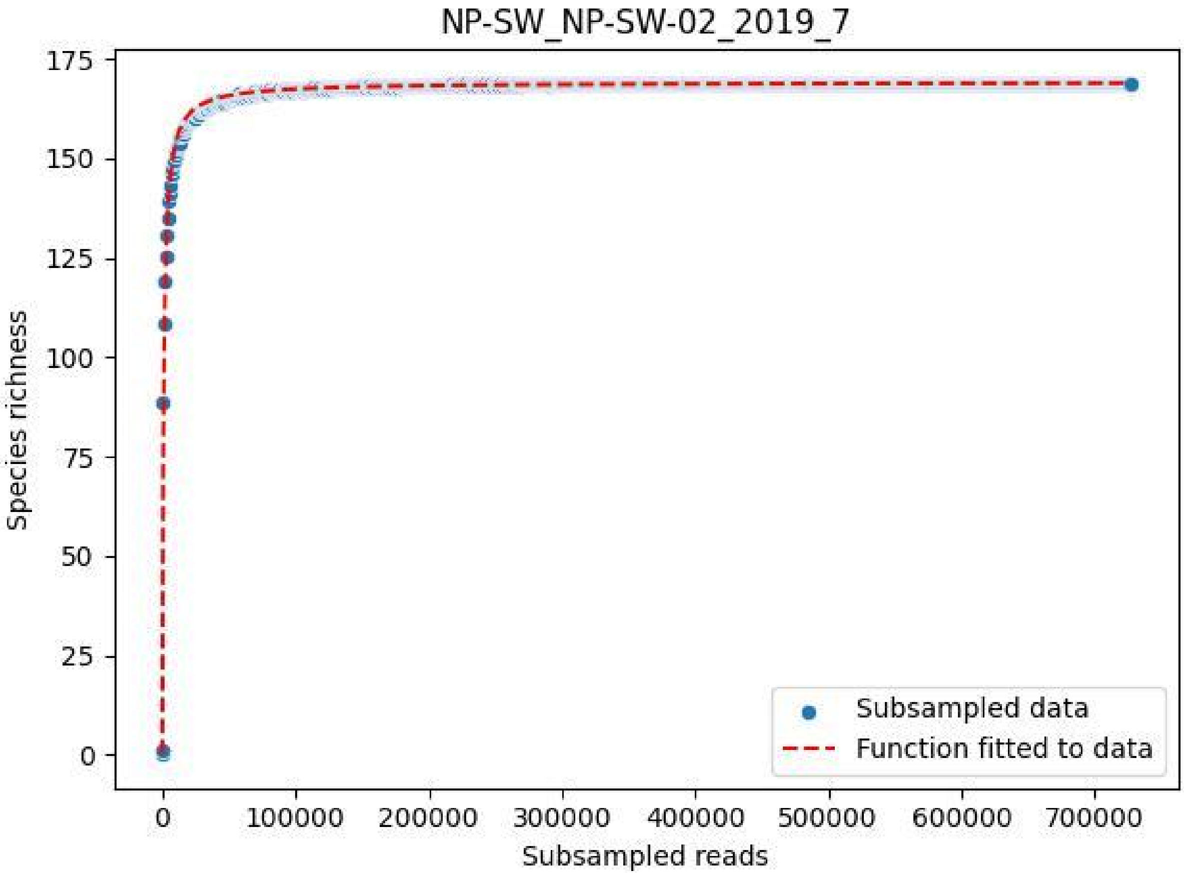
NP-SW_NP-SW-02_2019_7
Species richness
Subsampled reads
0
25
50
75
100
125
150
175
0
100000
200000
300000
400000
500000
600000
700000
Subsampled data
Function fitted to data

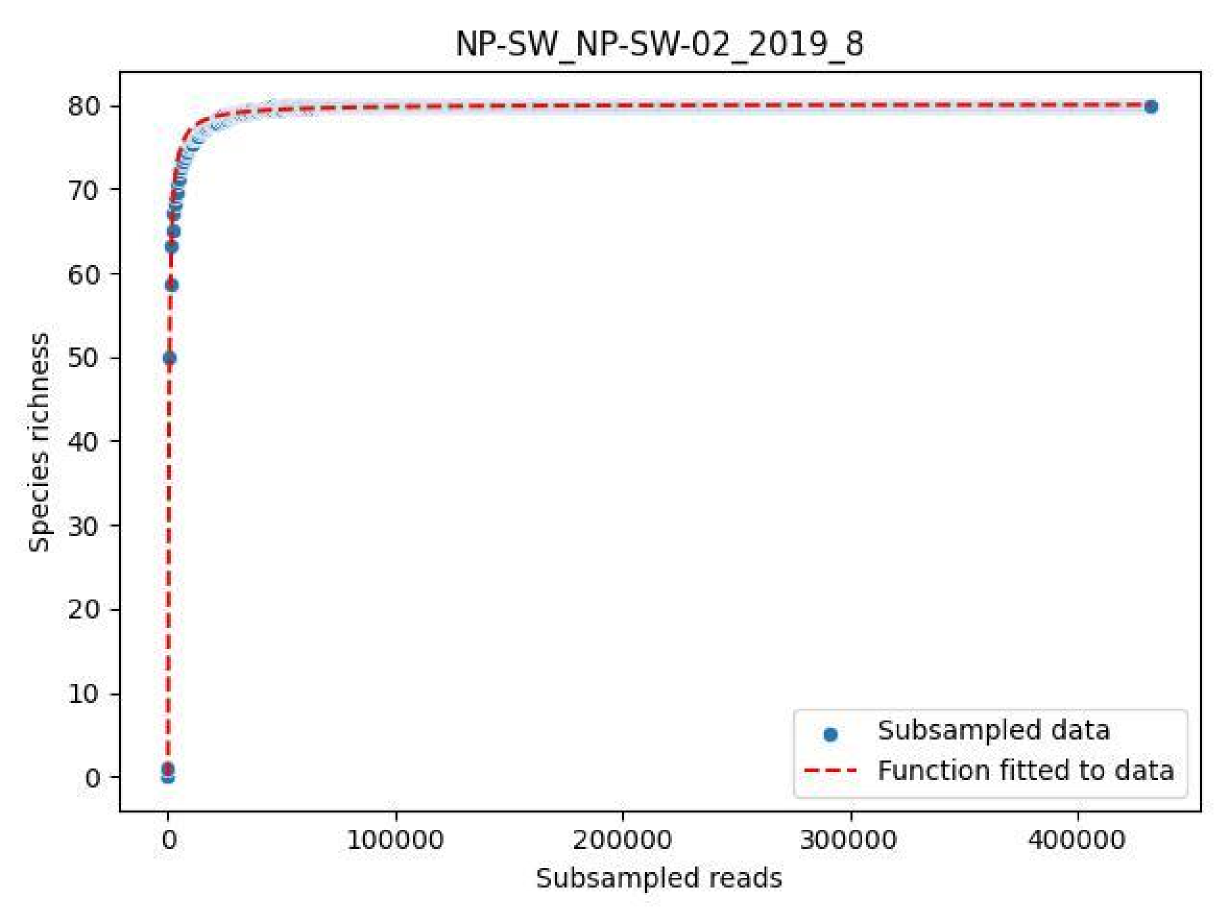

NP-SW_NP-SW-02_2019_8
Species richness
Subsampled reads
0
10
20
30
40
50
60
70
80
0
100000
200000
300000
400000
Subsampled data
Function fitted to data

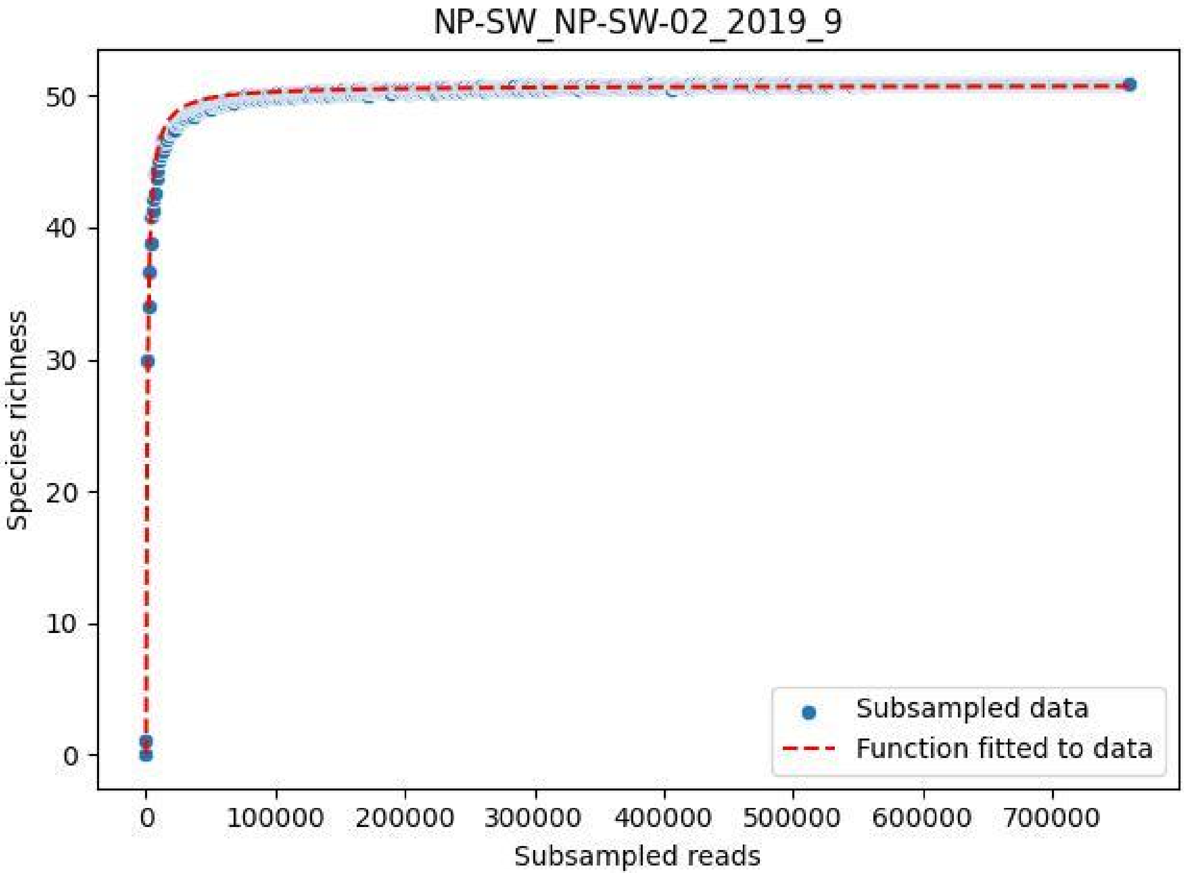
NP-SW_NP-SW-02_2019_9
Species richness
Subsampled reads
0
10
20
30
40
50
0
100000
200000
300000
400000
500000
600000
700000
Subsampled data
Function fitted to data

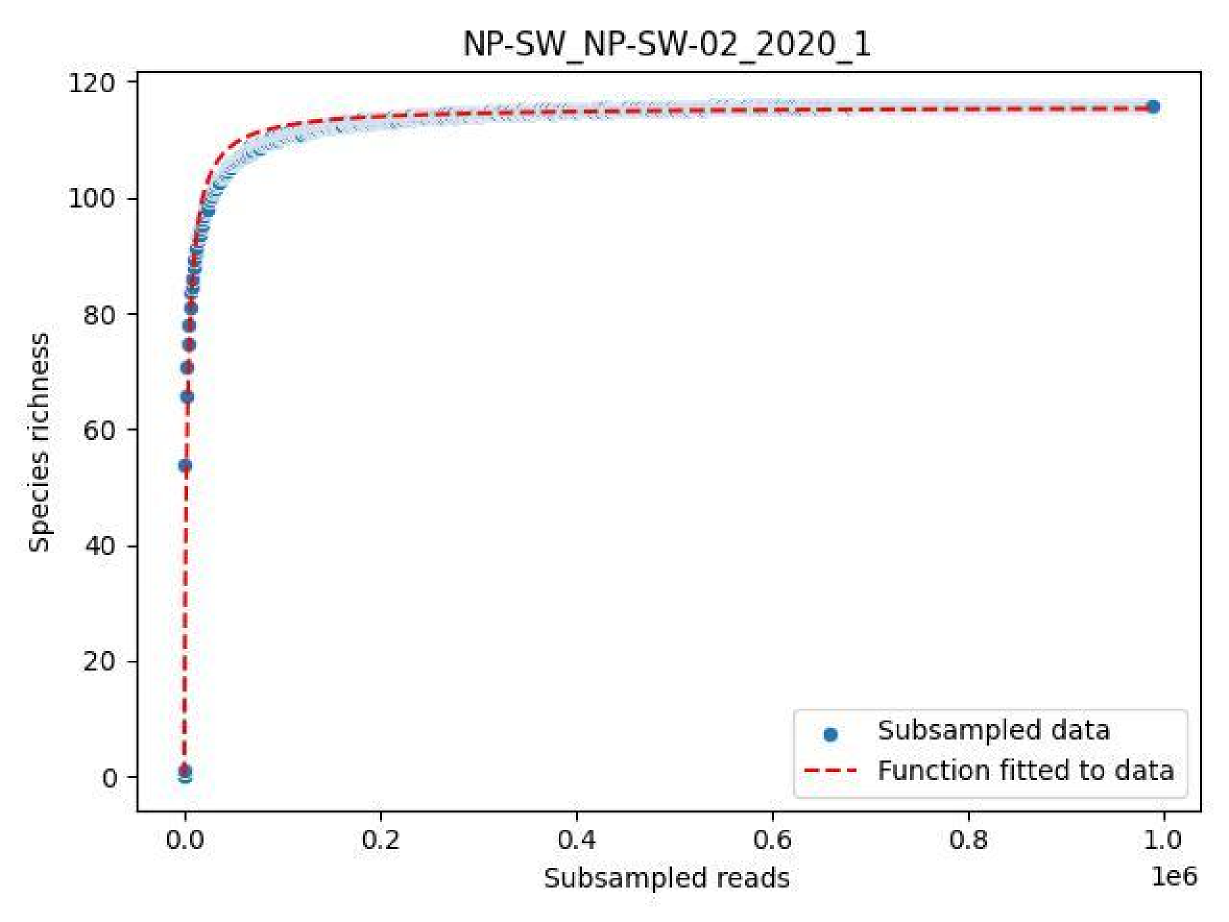
NP-SW_NP-SW-02_2020_1
Species richness
Subsampled reads
1e6
0
20
40
60
80
100
120
0.0
0.2
0.4
0.6
0.8
1.0
Subsampled data
Function fitted to data

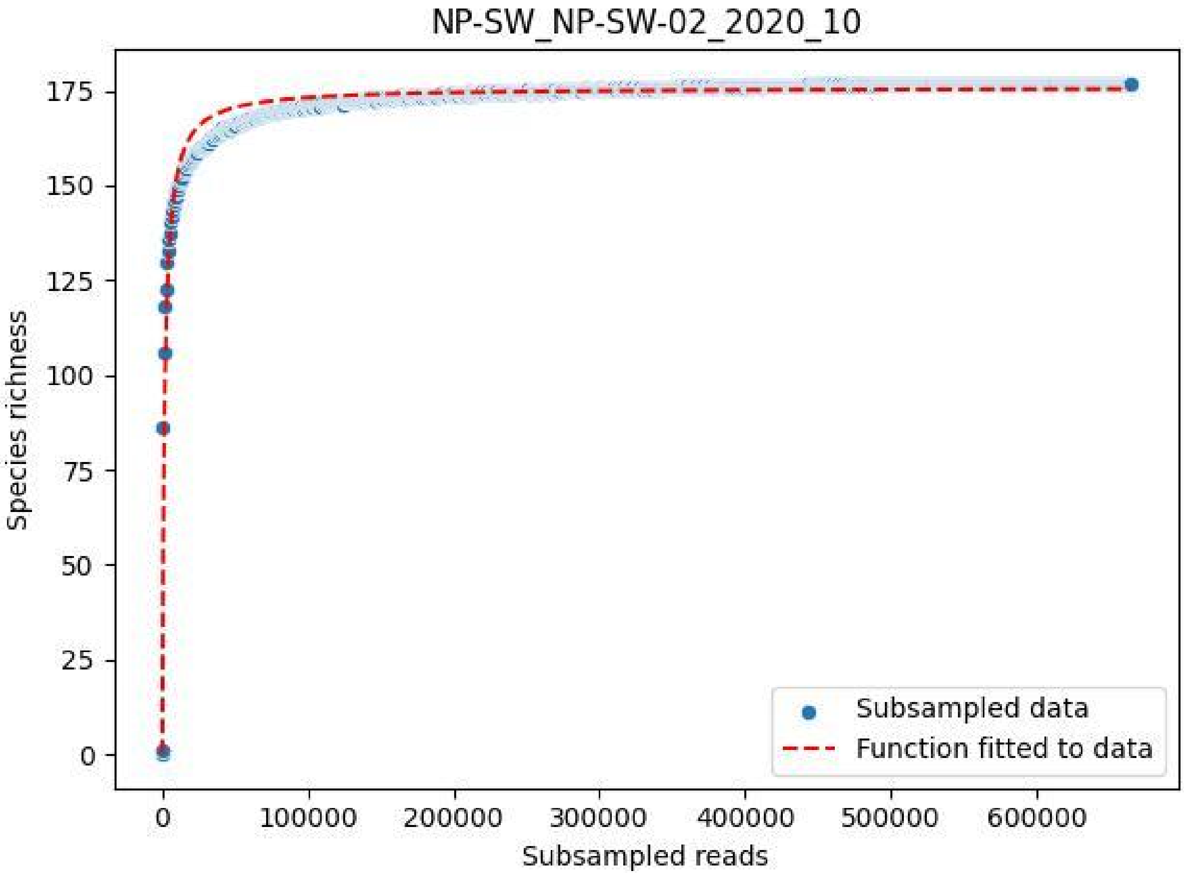

NP-SW_NP-SW-02_2020_10
Species richness
Subsampled reads
0
25
50
75
100
125
150
175
0
100000
200000
300000
400000
500000
600000
Subsampled data
Function fitted to data

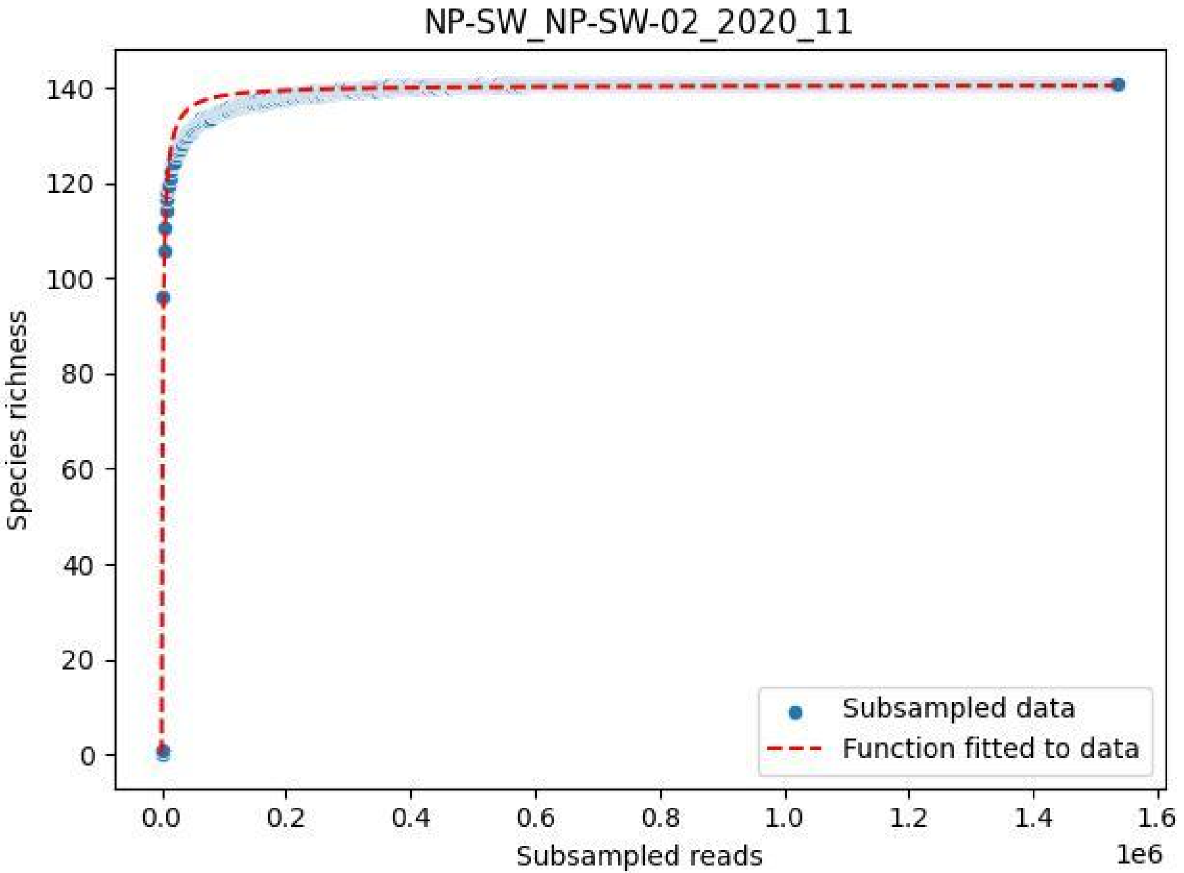

NP-SW_NP-SW-02_2020_11
Species richness
Subsampled reads
1e6
0
20
40
60
80
100
120
140
0.0
0.2
0.4
0.6
0.8
1.0
1.2
1.4
1.6
Subsampled data
Function fitted to data

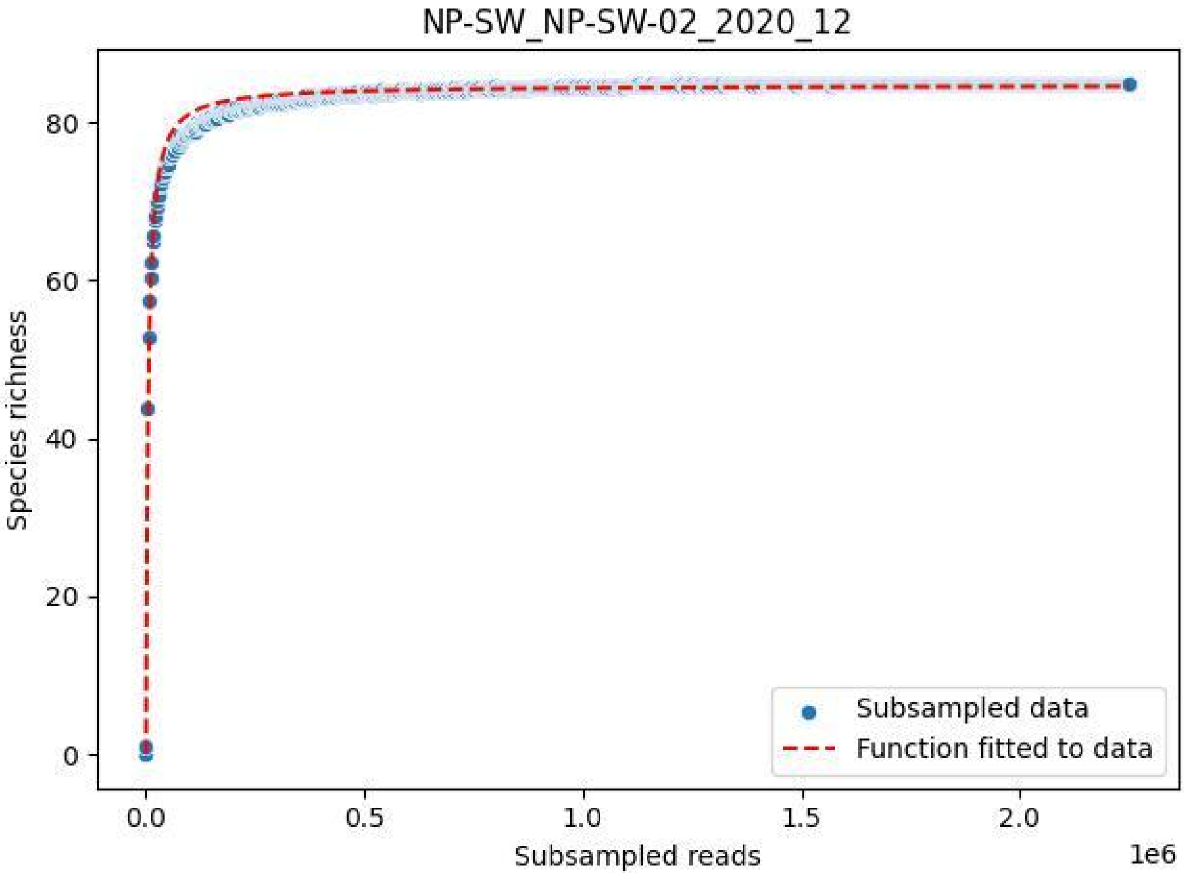

NP-SW_NP-SW-02_2020_12
Species richness
Subsampled reads
1e6
0
20
40
60
80
0.0
0.5
1.0
1.5
2.0
Subsampled data
Function fitted to data

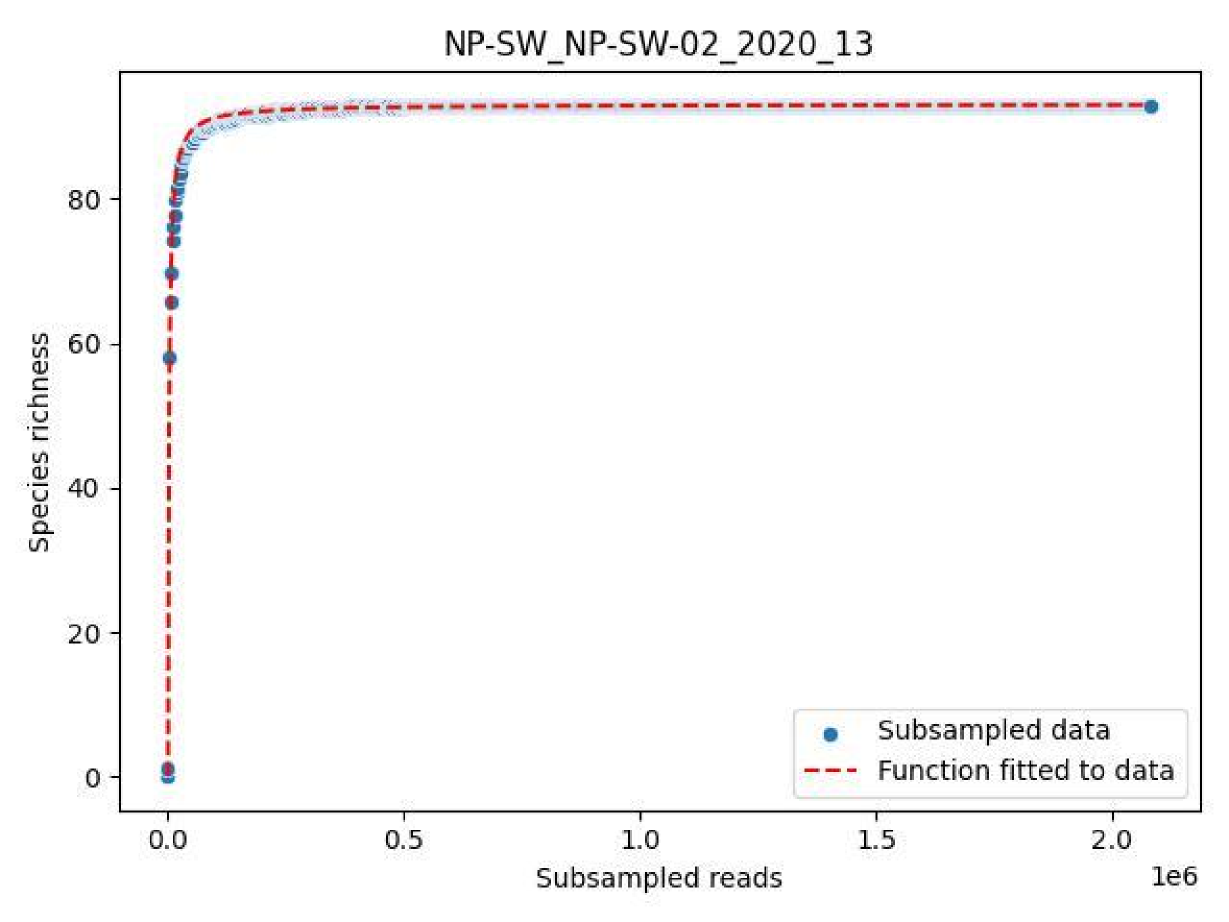

NP-SW_NP-SW-02_2020_13
Species richness
Subsampled reads
1e6
0
20
40
60
80
0.0
0.5
1.0
1.5
2.0
Subsampled data
Function fitted to data

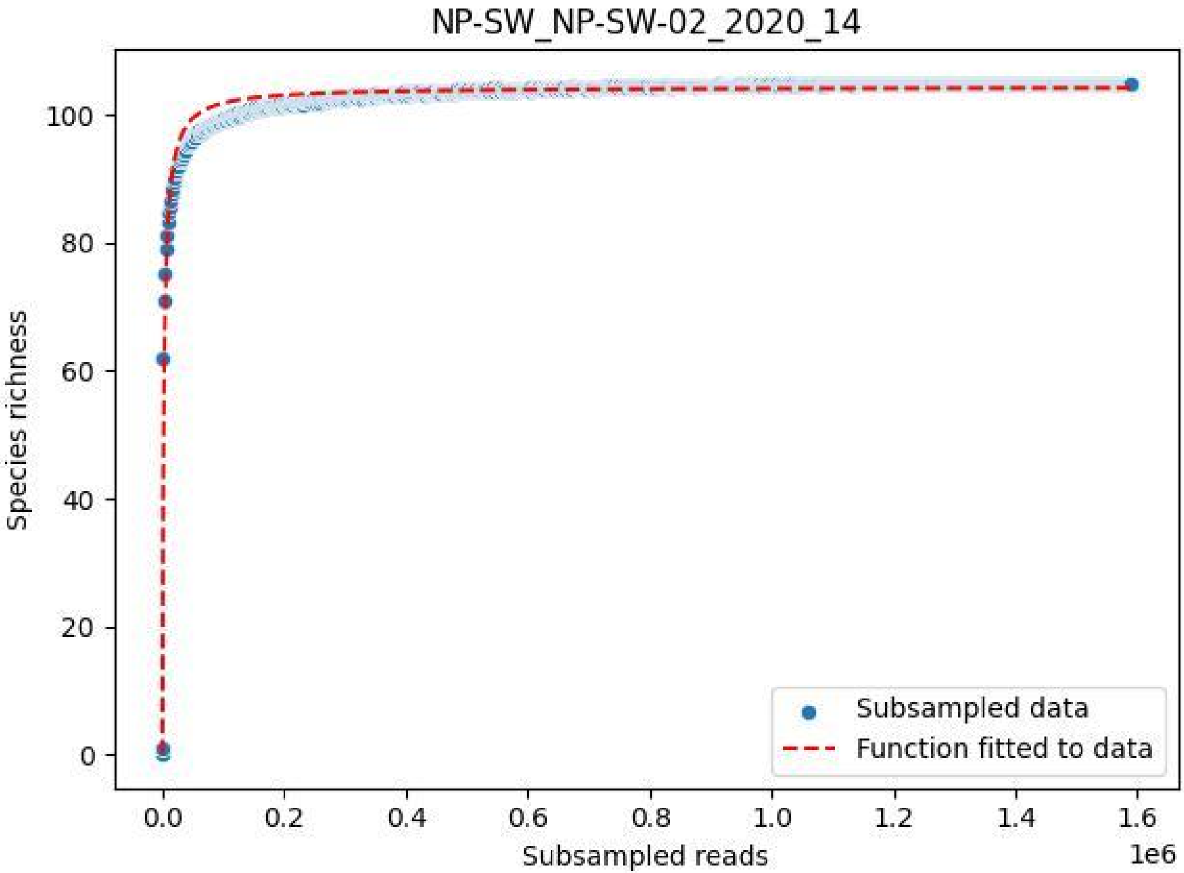

NP-SW_NP-SW-02_2020_14
Species richness
Subsampled reads
1e6
0
20
40
60
80
100
0.0
0.2
0.4
0.6
0.8
1.0
1.2
1.4
1.6
Subsampled data
Function fitted to data

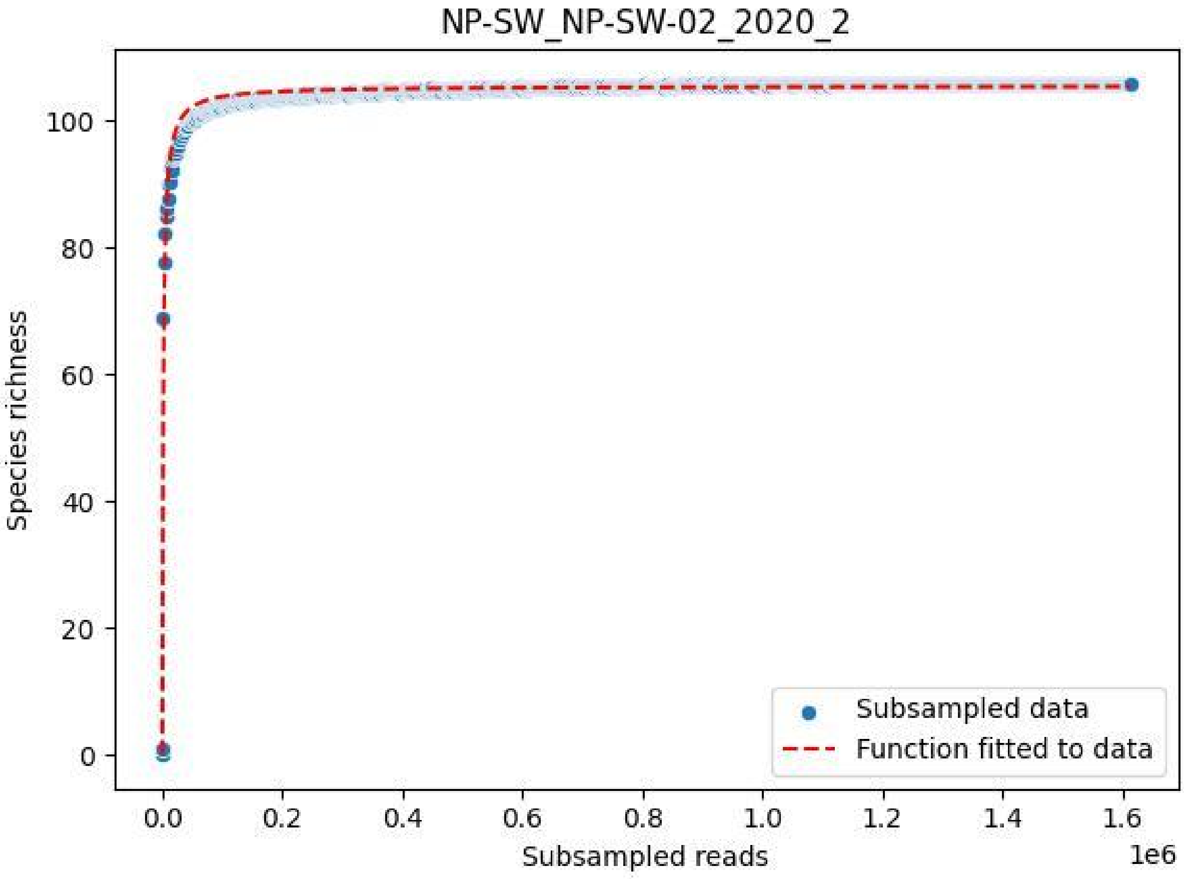

NP-SW_NP-SW-02_2020_2
Species richness
Subsampled reads
1e6
0
20
40
60
80
100
0.0
0.2
0.4
0.6
0.8
1.0
1.2
1.4
1.6
Subsampled data
Function fitted to data

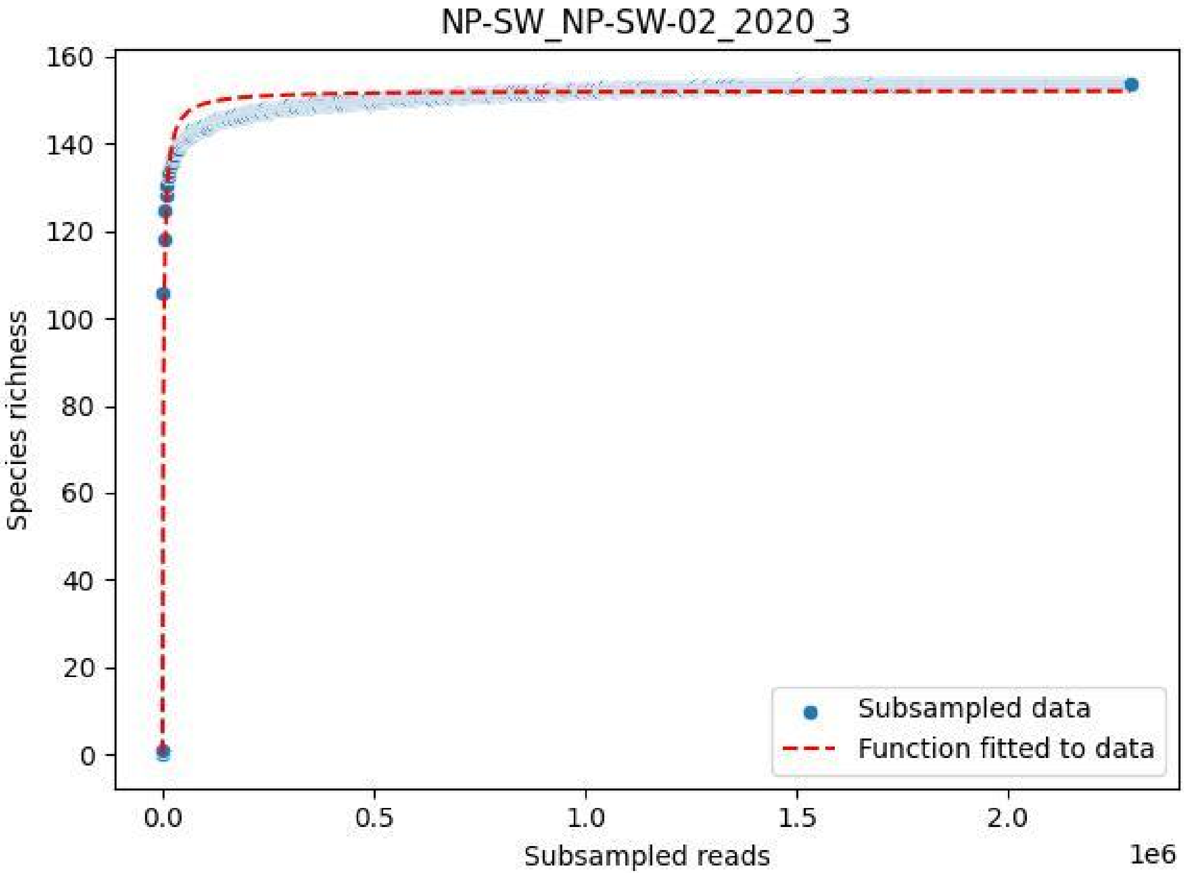
NP-SW_NP-SW-02_2020_3
Species richness
Subsampled reads
1e6
0.0
0.5
1.0
1.5
2.0
0
20
40
60
80
100
120
140
160
Subsampled data
Function fitted to data

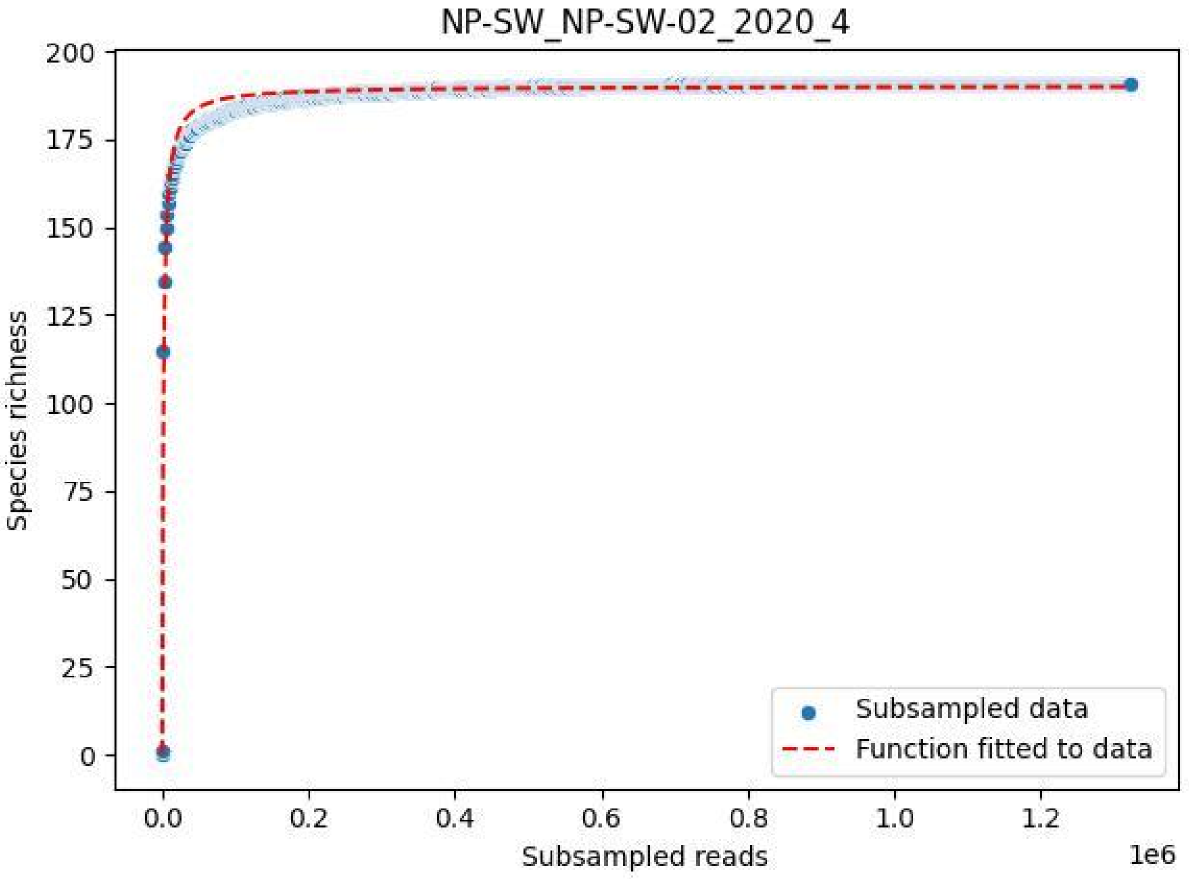
NP-SW_NP-SW-02_2020_4
Species richness
Subsampled reads
1e6
0
25
50
75
100
125
150
175
200
0.0
0.2
0.4
0.6
0.8
1.0
1.2
Subsampled data
Function fitted to data

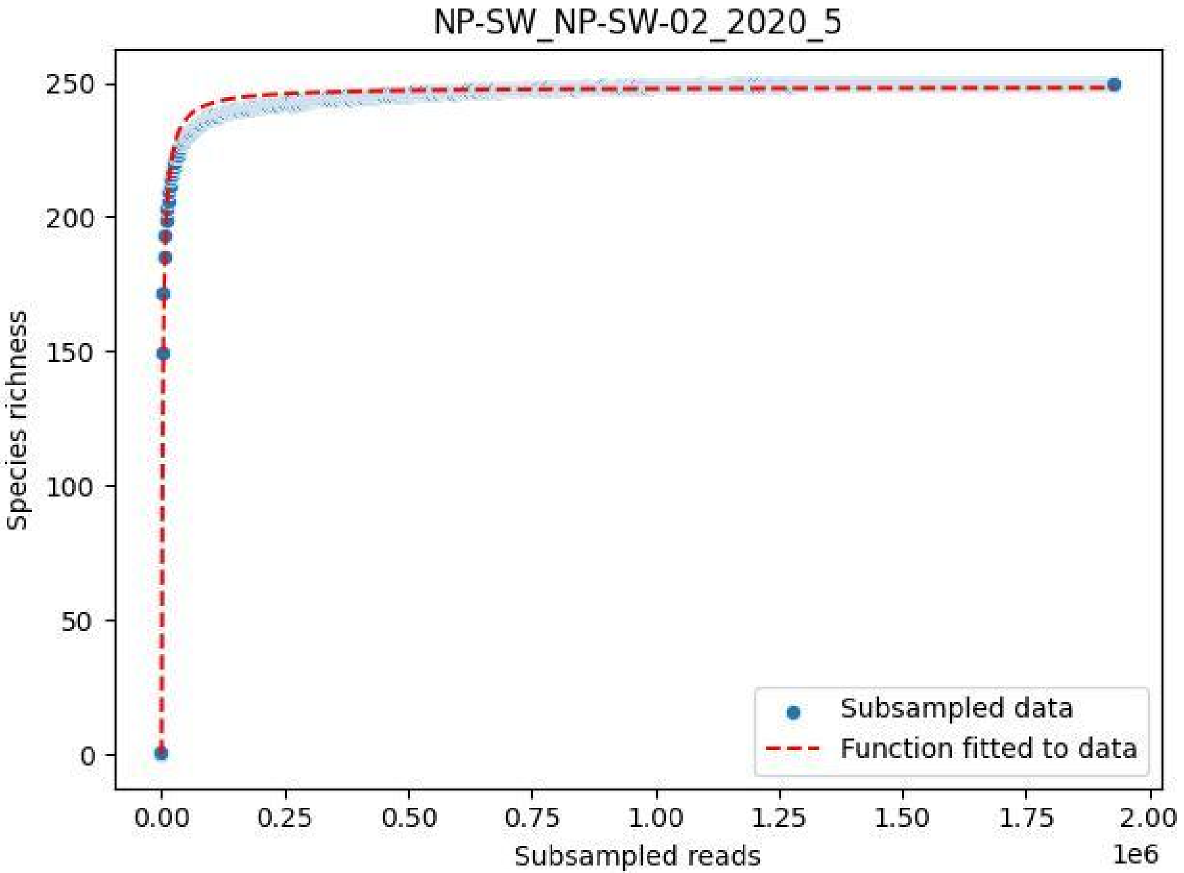
NP-SW_NP-SW-02_2020_5
Species richness
Subsampled reads
1e6
Subsampled data
Function fitted to data
0
50
100
150
200
250
0.00
0.25
0.50
0.75
1.00
1.25
1.50
1.75
2.00

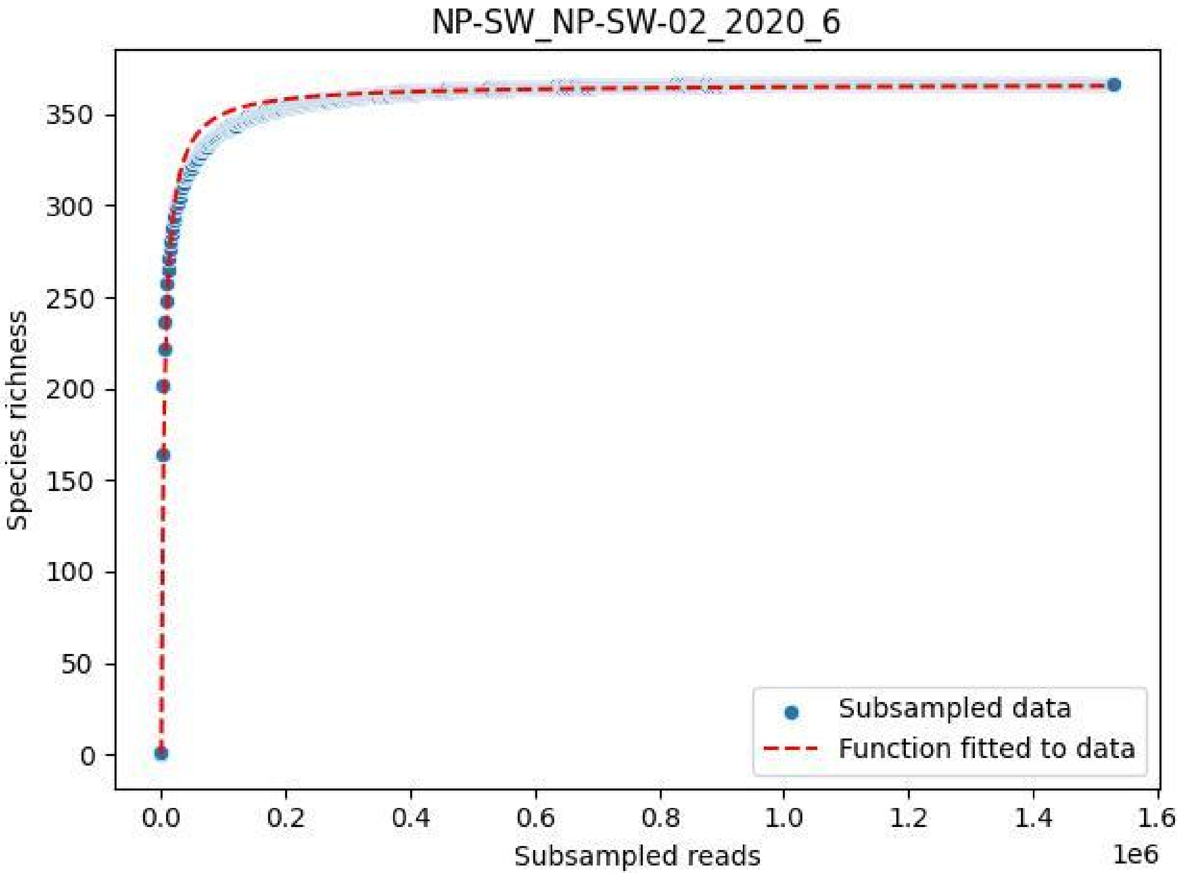

NP-SW_NP-SW-02_2020_6
Species richness
Subsampled reads
1e6
Subsampled data
Function fitted to data
0
50
100
150
200
250
300
350
0.0
0.2
0.4
0.6
0.8
1.0
1.2
1.4
1.6

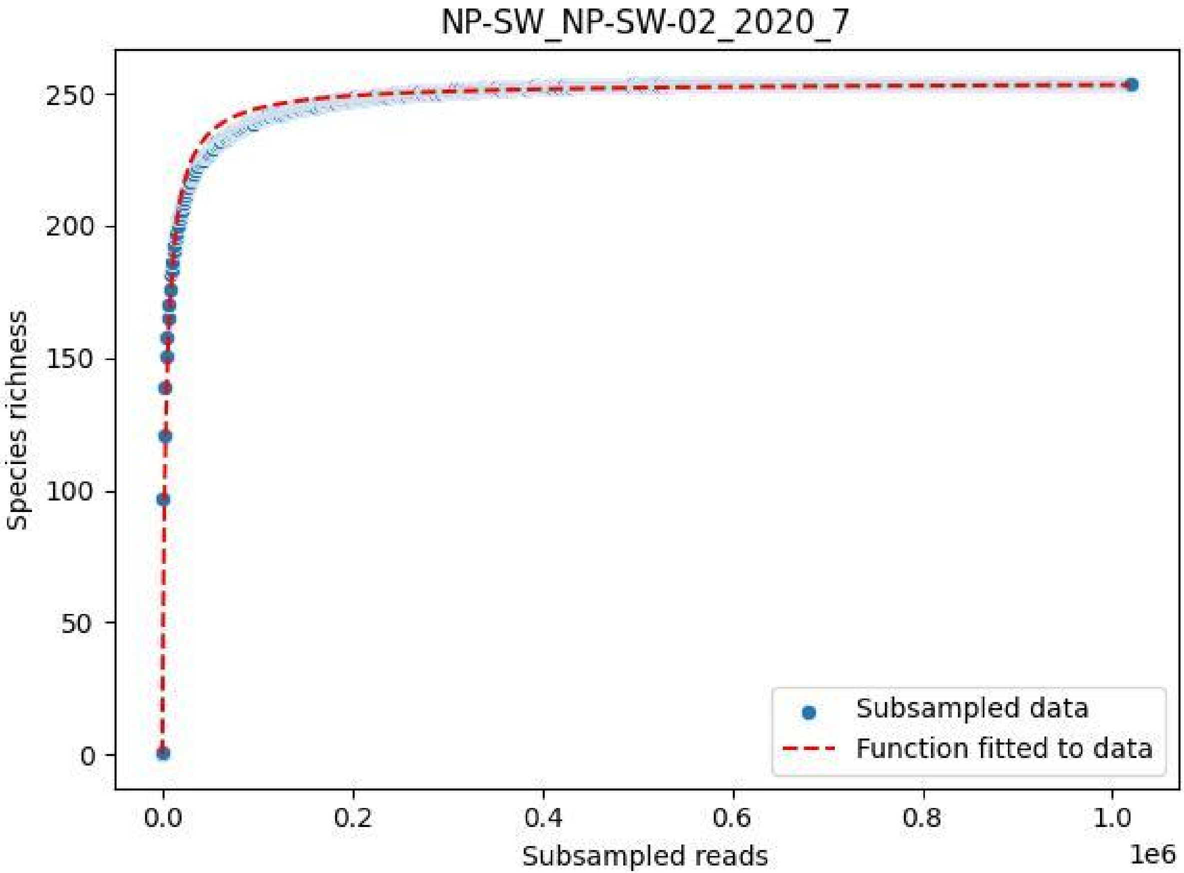

NP-SW_NP-SW-02_2020_7
Species richness
Subsampled reads
1e6
Subsampled data
Function fitted to data
0
50
100
150
200
250
0.0
0.2
0.4
0.6
0.8
1.0

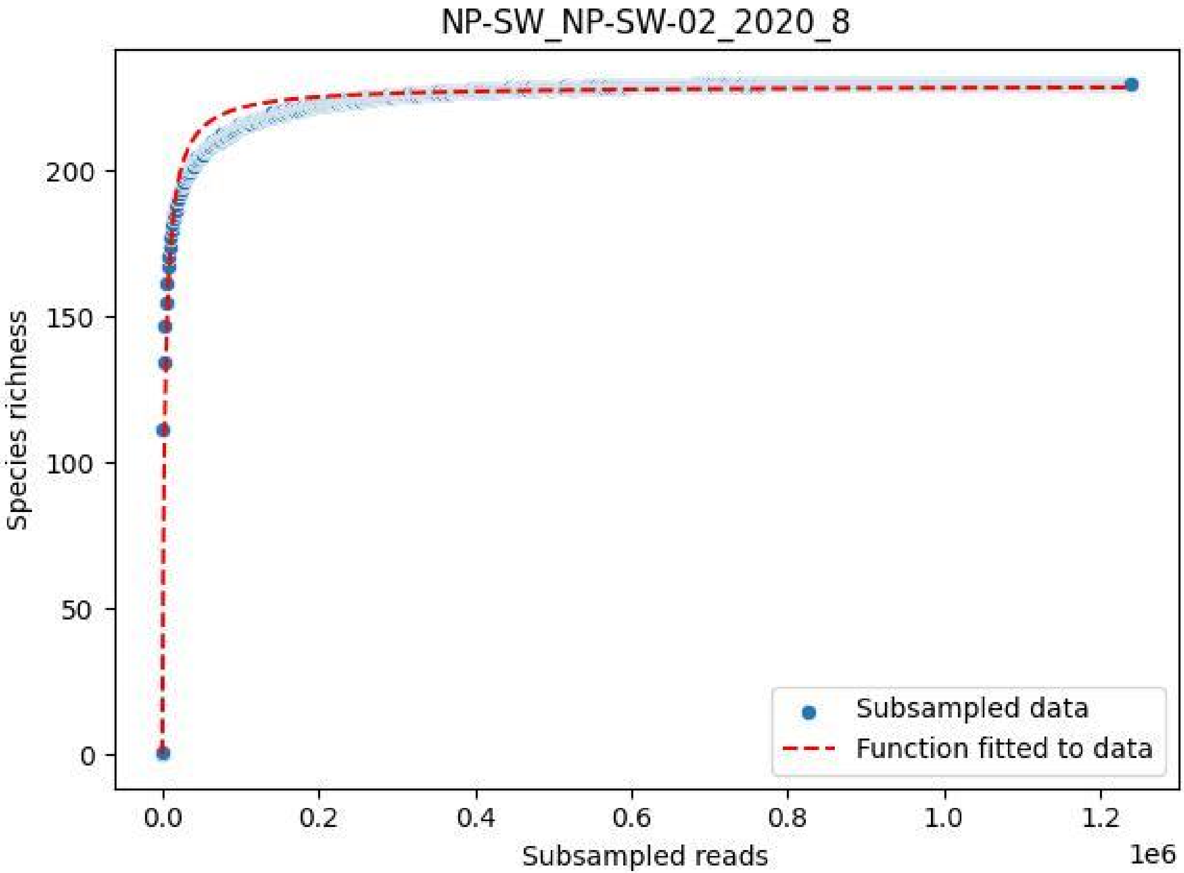

NP-SW_NP-SW-02_2020_8
Species richness
Subsampled reads
1e6
0
50
100
150
200
0.0
0.2
0.4
0.6
0.8
1.0
1.2
Subsampled data
Function fitted to data

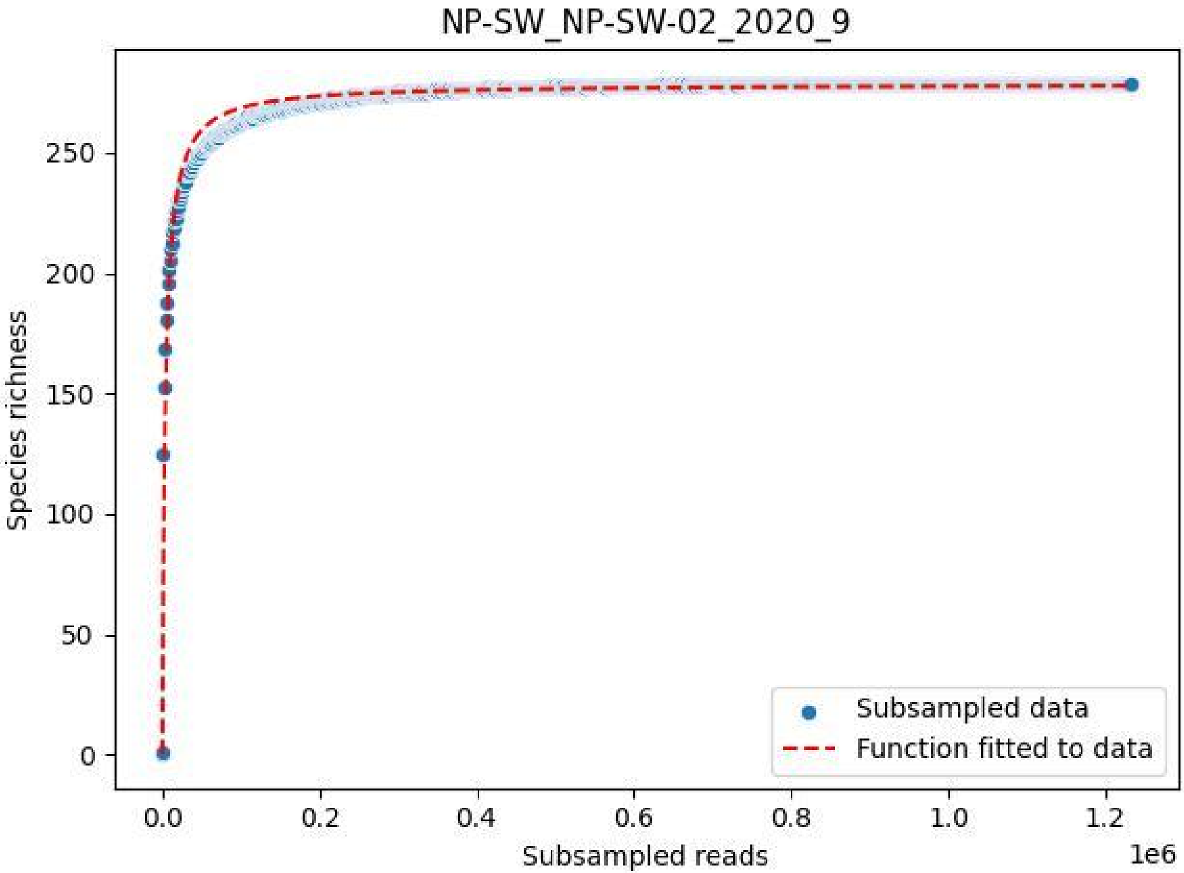

NP-SW_NP-SW-02_2020_9
Species richness
Subsampled reads
1e6
0
50
100
150
200
250
0.0
0.2
0.4
0.6
0.8
1.0
1.2
Subsampled data
Function fitted to data

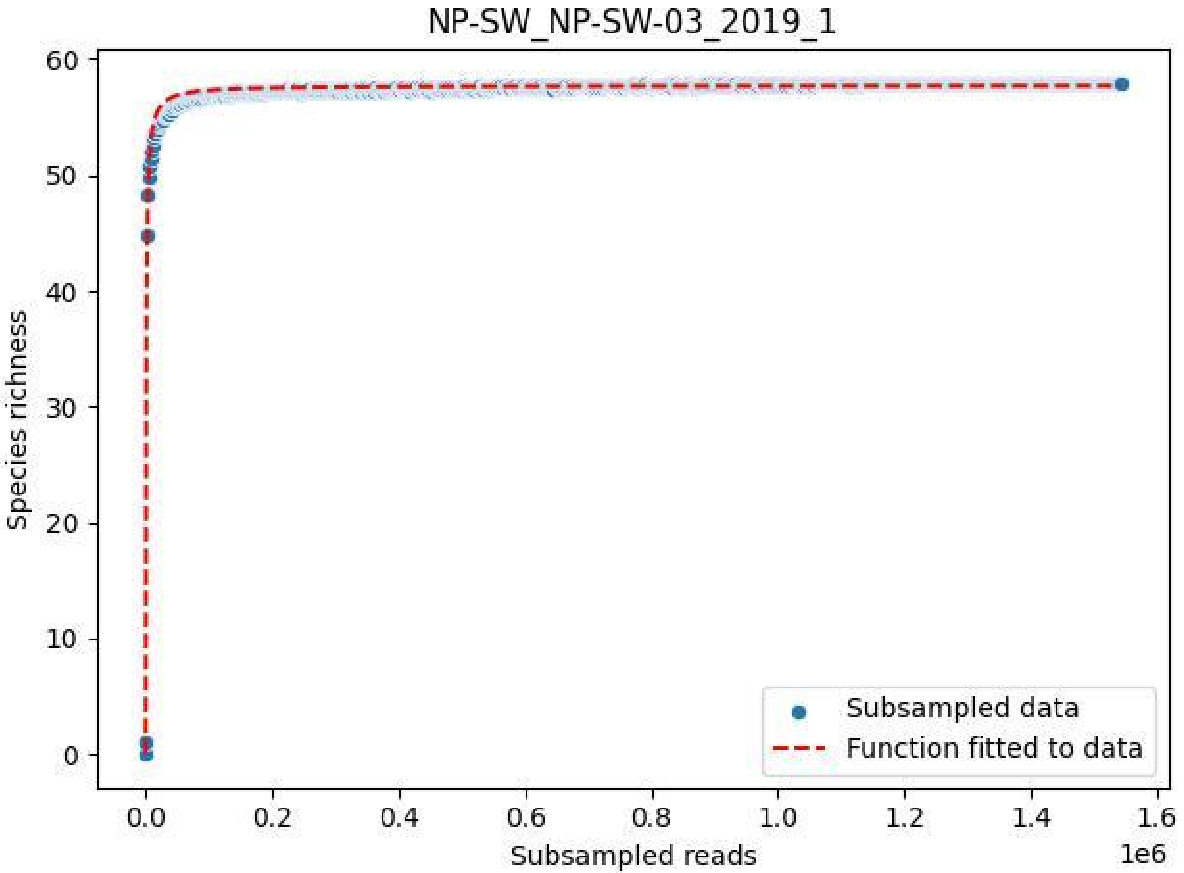
NP-SW_NP-SW-03_2019_1
Species richness
Subsampled reads
1e6
0
10
20
30
40
50
60
0.0
0.2
0.4
0.6
0.8
1.0
1.2
1.4
1.6
Subsampled data
Function fitted to data

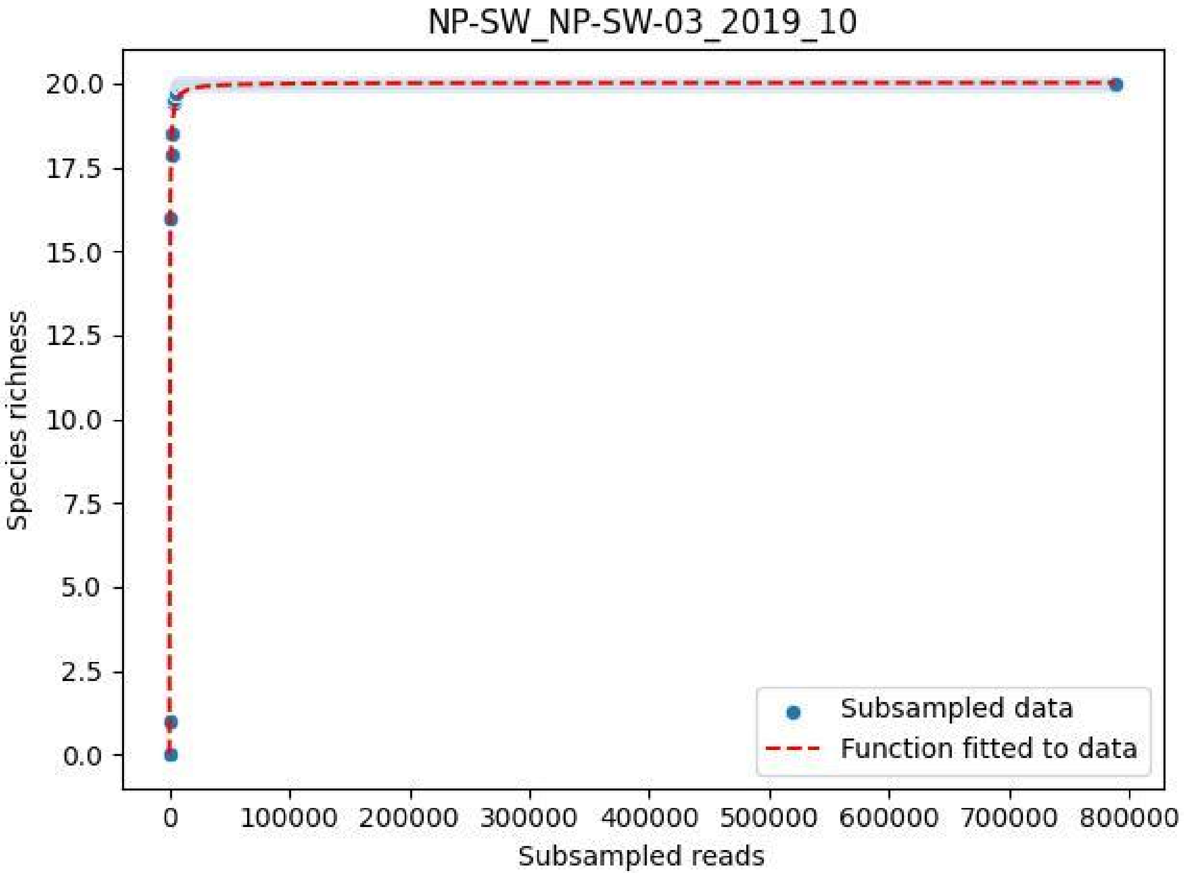

NP-SW_NP-SW-03_2019_10
Species richness
Subsampled reads
0.0
2.5
5.0
7.5
10.0
12.5
15.0
17.5
20.0
0
100000
200000
300000
400000
500000
600000
700000
800000
Subsampled data
Function fitted to data

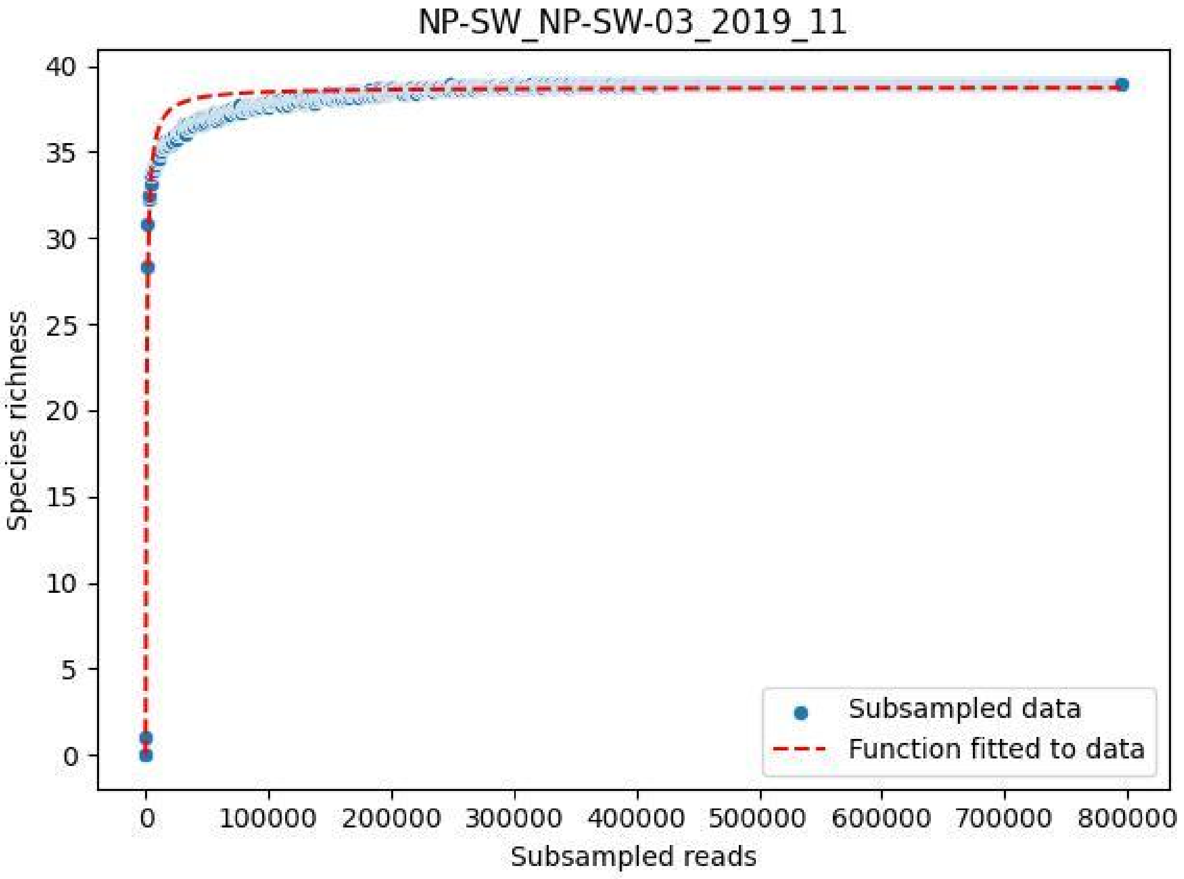
NP-SW_NP-SW-03_2019_11
Species richness
Subsampled reads
0
5
10
15
20
25
30
35
40
0
100000
200000
300000
400000
500000
600000
700000
800000
Subsampled data
Function fitted to data

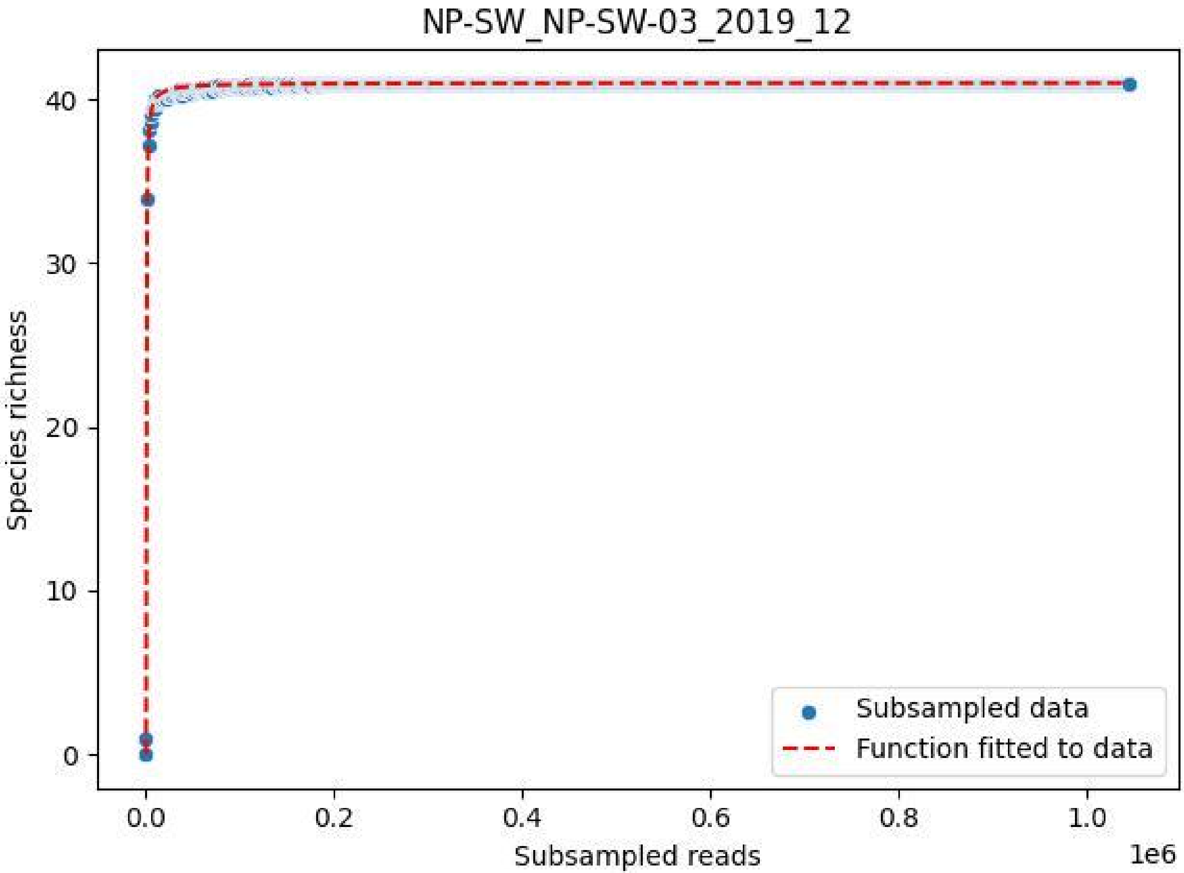

NP-SW_NP-SW-03_2019_12
Species richness
Subsampled reads
1e6
0
10
20
30
40
0.0
0.2
0.4
0.6
0.8
1.0
Subsampled data
Function fitted to data

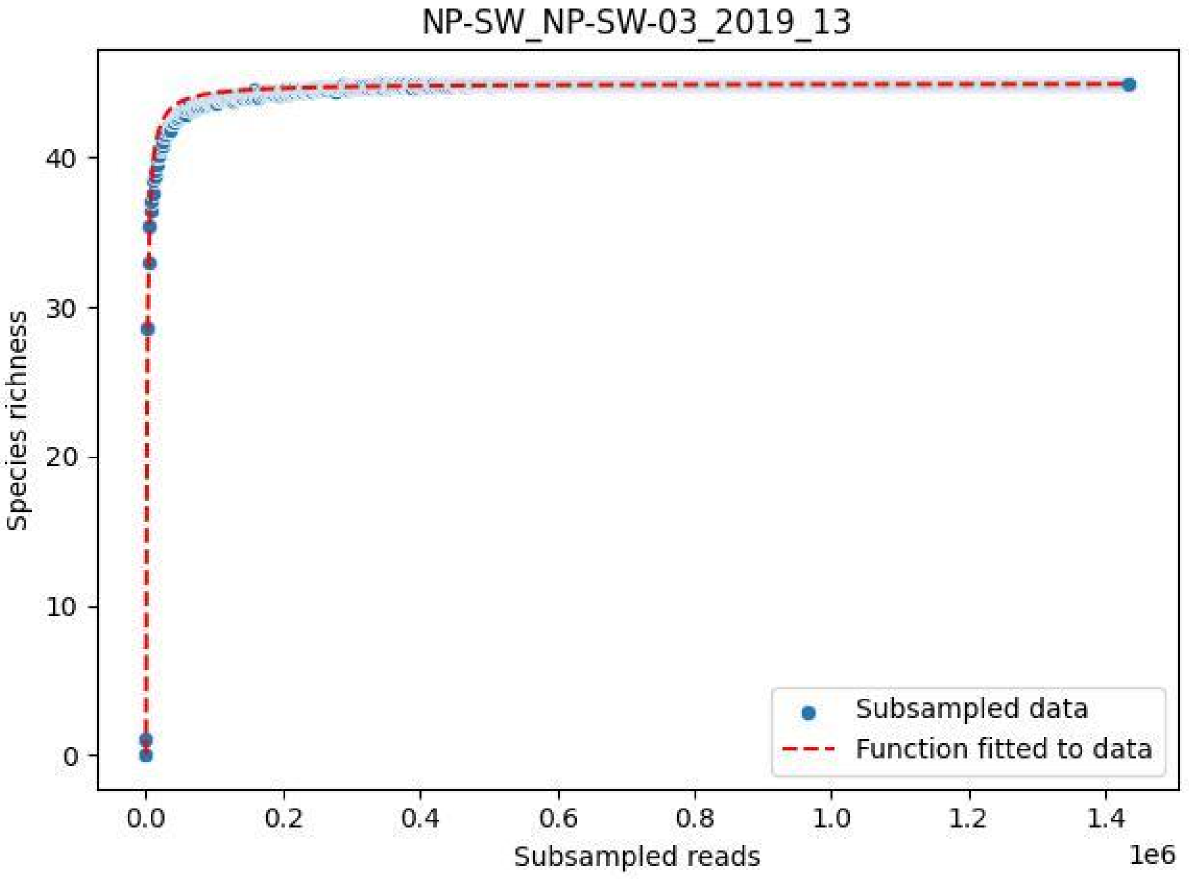

NP-SW_NP-SW-03_2019_13
Species richness
0
10
20
30
40
Subsampled reads
0.0
0.2
0.4
0.6
0.8
1.0
1.2
1.4
1e6
Subsampled data
Function fitted to data

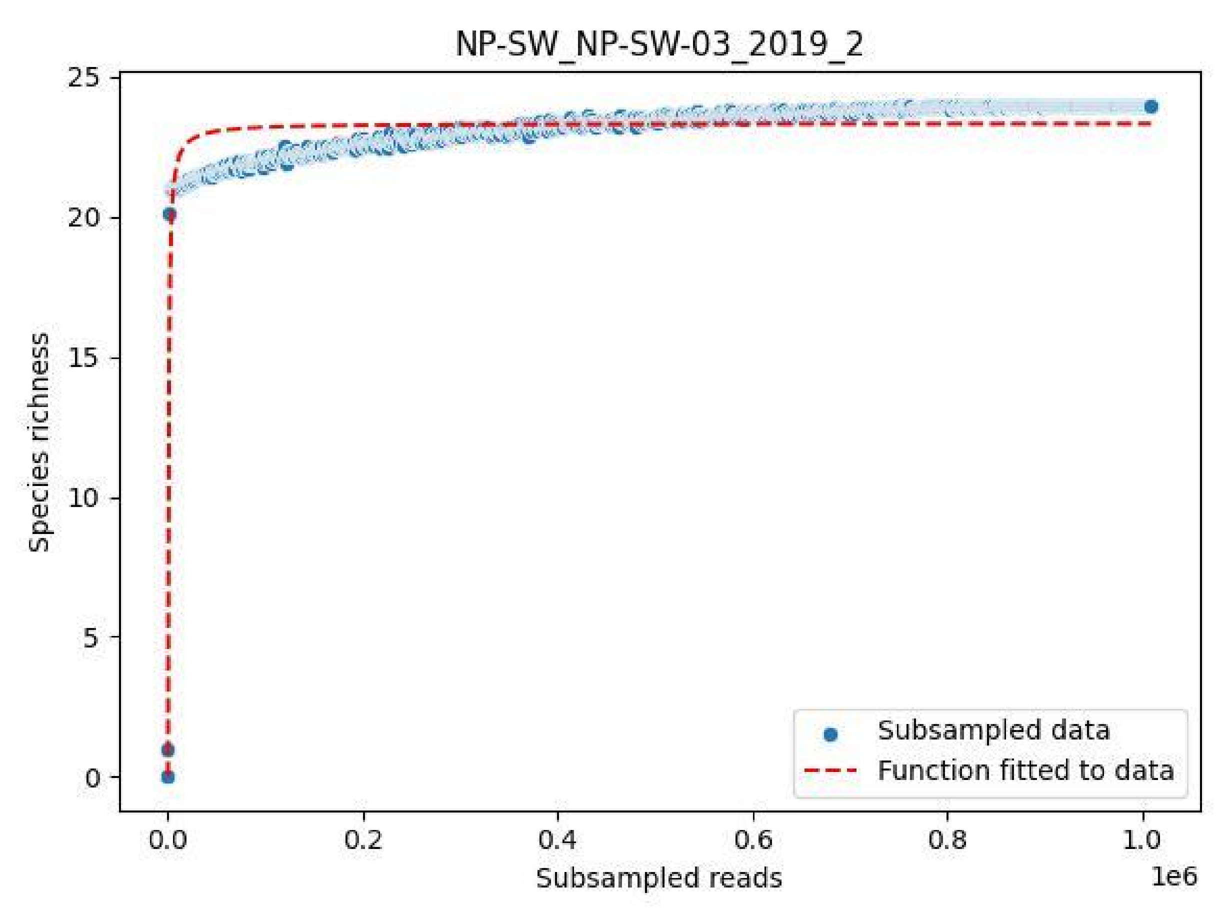
NP-SW_NP-SW-03_2019_2
Species richness
Subsampled reads
1e6
0
5
10
15
20
25
0.0
0.2
0.4
0.6
0.8
1.0
Subsampled data
Function fitted to data

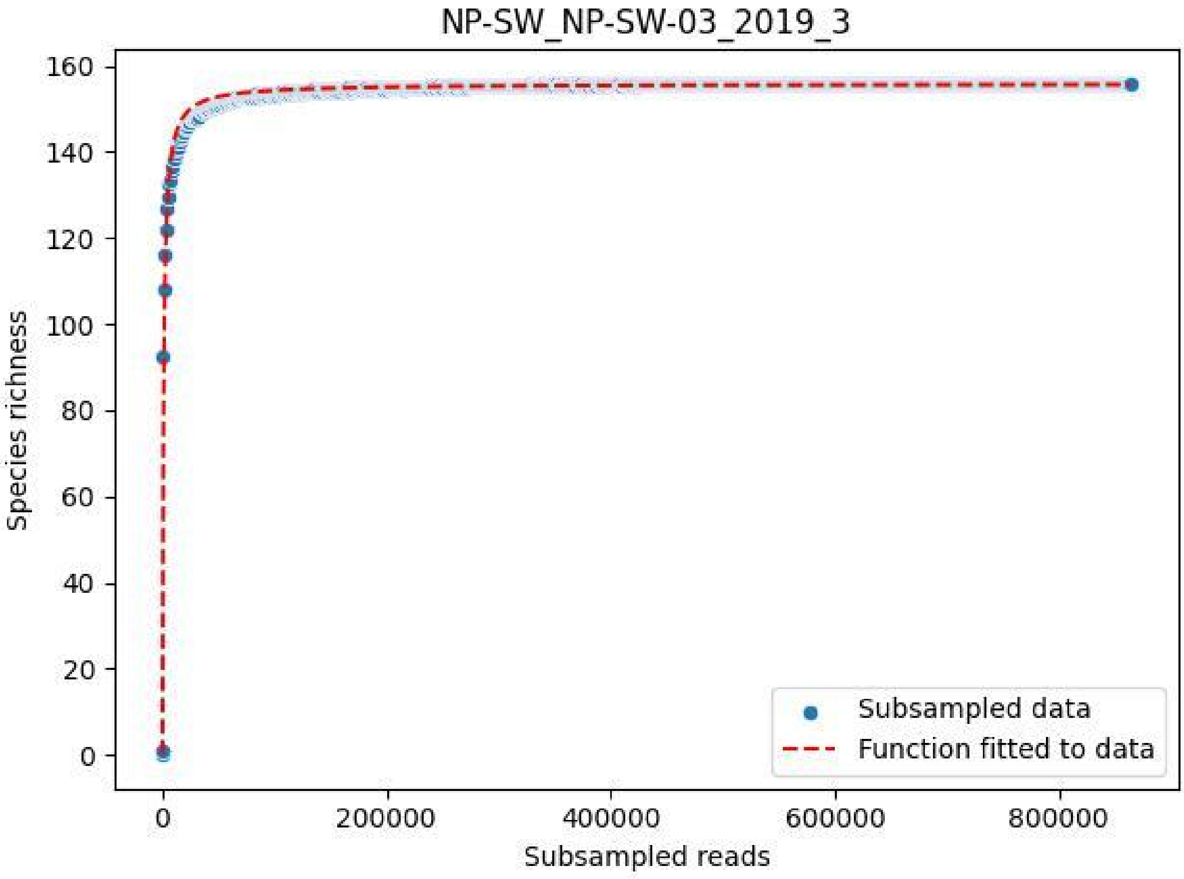

NP-SW_NP-SW-03_2019_3
Species richness
Subsampled reads
0
20
40
60
80
100
120
140
160
0
200000
400000
600000
800000
Subsampled data
Function fitted to data

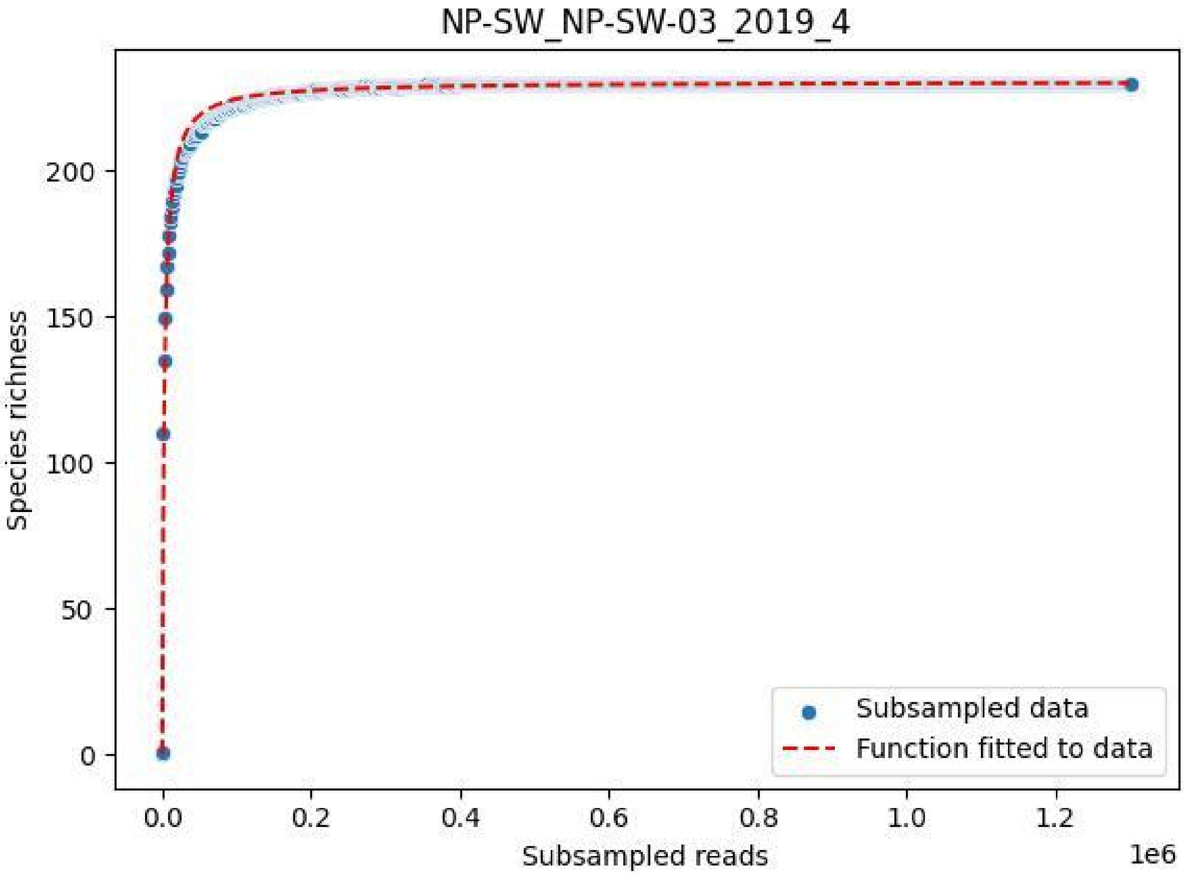
NP-SW_NP-SW-03_2019_4
Species richness
Subsampled reads
1e6
Subsampled data
Function fitted to data
0
50
100
150
200
0.0
0.2
0.4
0.6
0.8
1.0
1.2

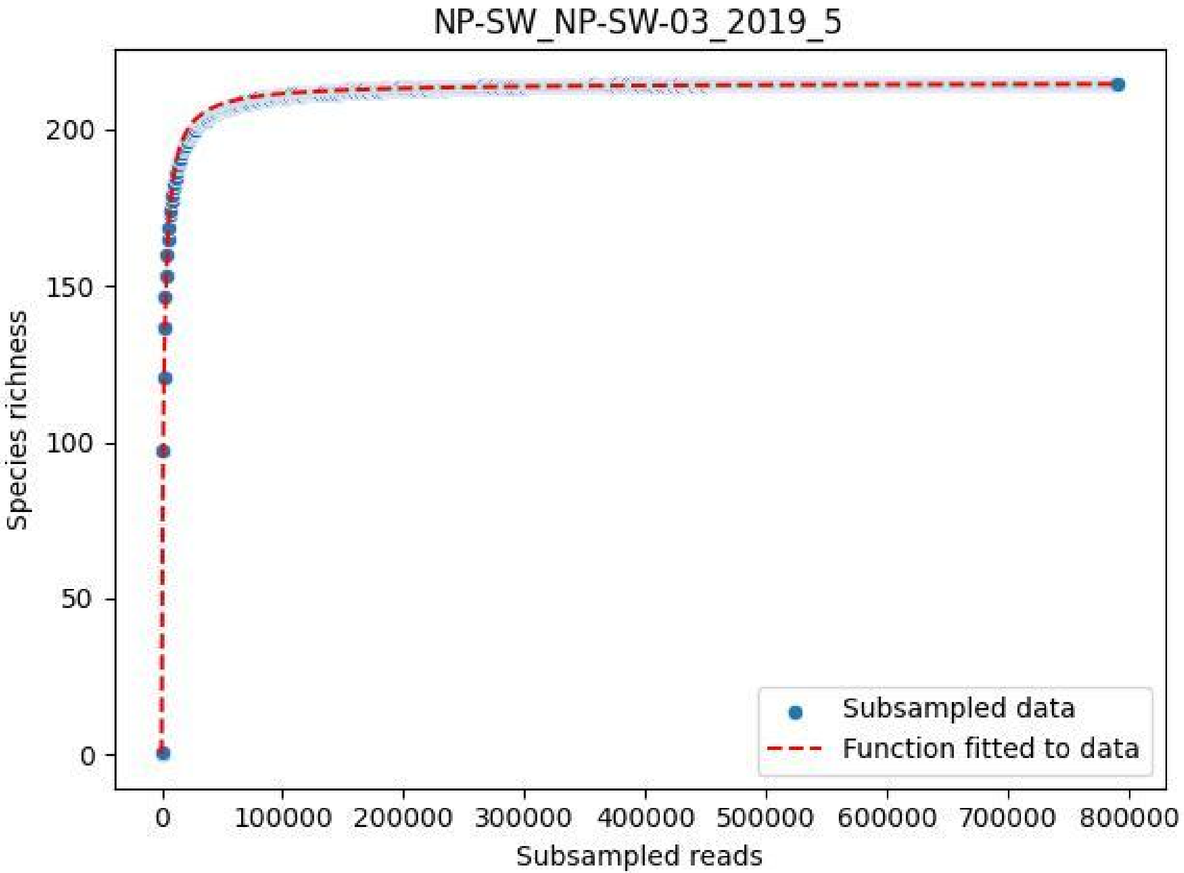

NP-SW_NP-SW-03_2019_5
Species richness
0
50
100
150
200
Subsampled reads
0
100000
200000
300000
400000
500000
600000
700000
800000
Subsampled data
Function fitted to data

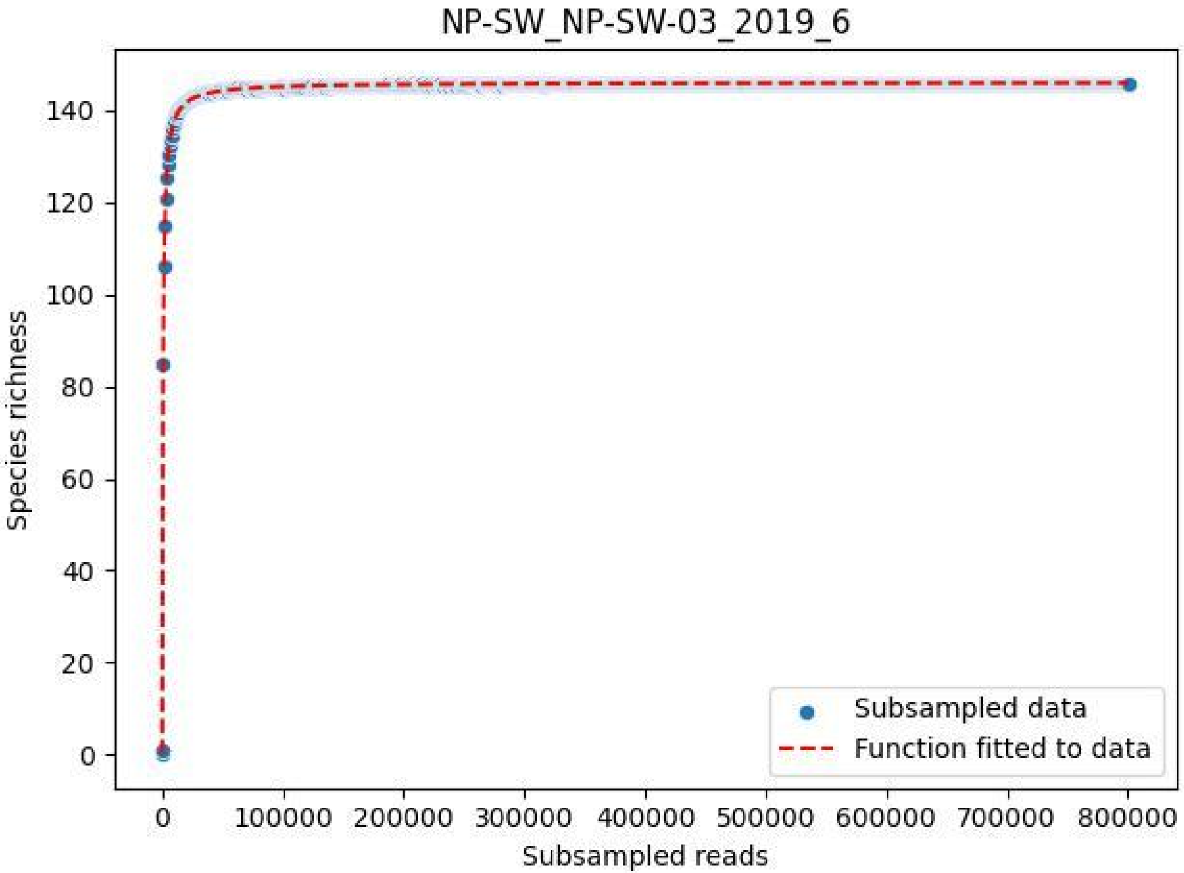
NP-SW_NP-SW-03_2019_6
Species richness
Subsampled reads
0
20
40
60
80
100
120
140
0
100000
200000
300000
400000
500000
600000
700000
800000
Subsampled data
Function fitted to data

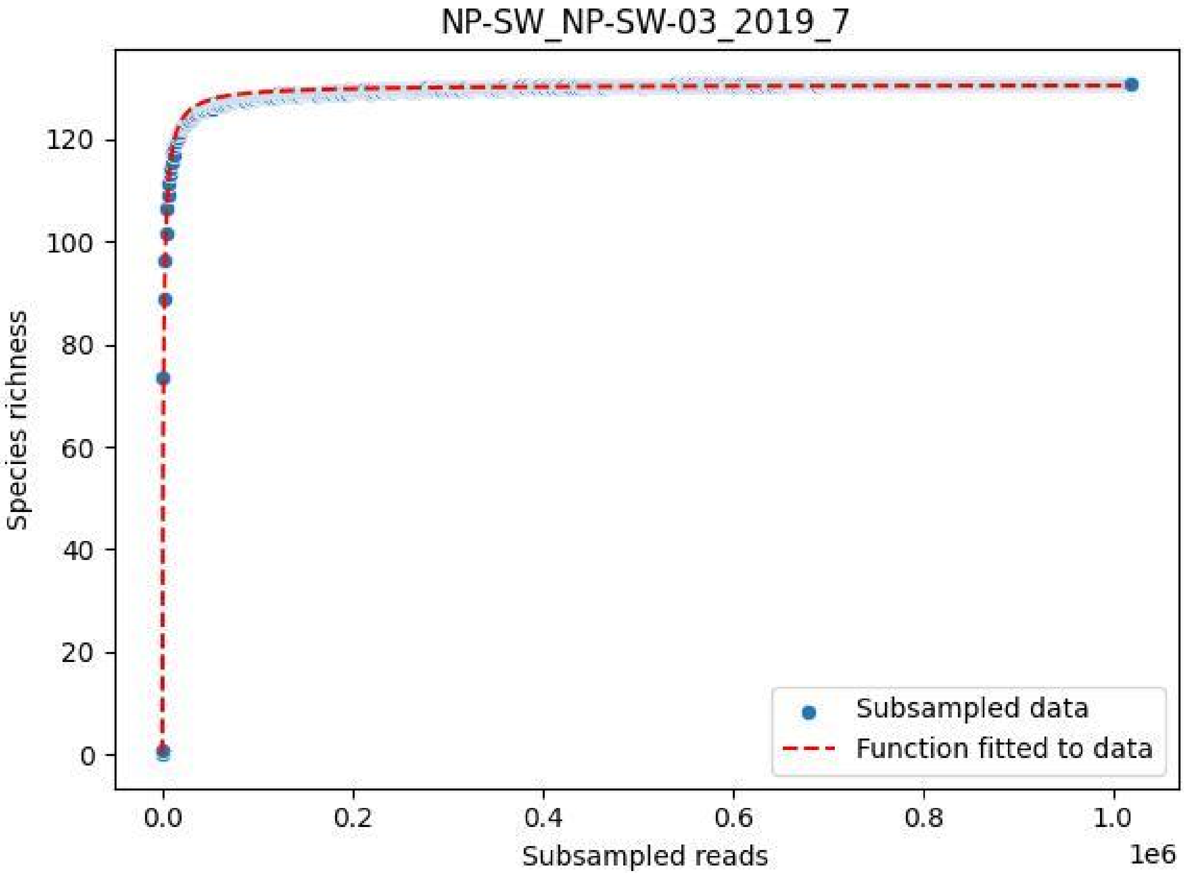
NP-SW_NP-SW-03_2019_7
Species richness
Subsampled reads
1e6
0
20
40
60
80
100
120
0.0
0.2
0.4
0.6
0.8
1.0
Subsampled data
Function fitted to data

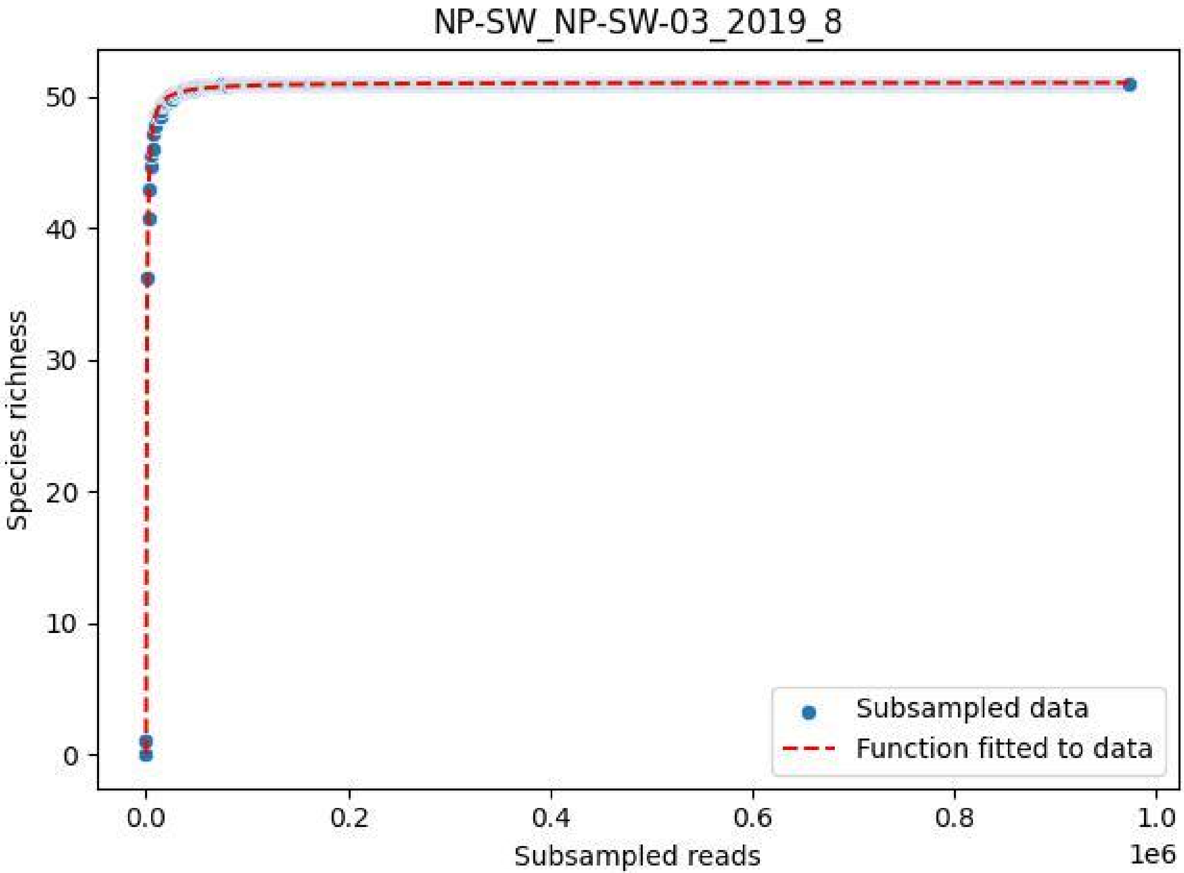
NP-SW_NP-SW-03_2019_8
Species richness
Subsampled reads
1e6
0
10
20
30
40
50
0.0
0.2
0.4
0.6
0.8
1.0
Subsampled data
Function fitted to data

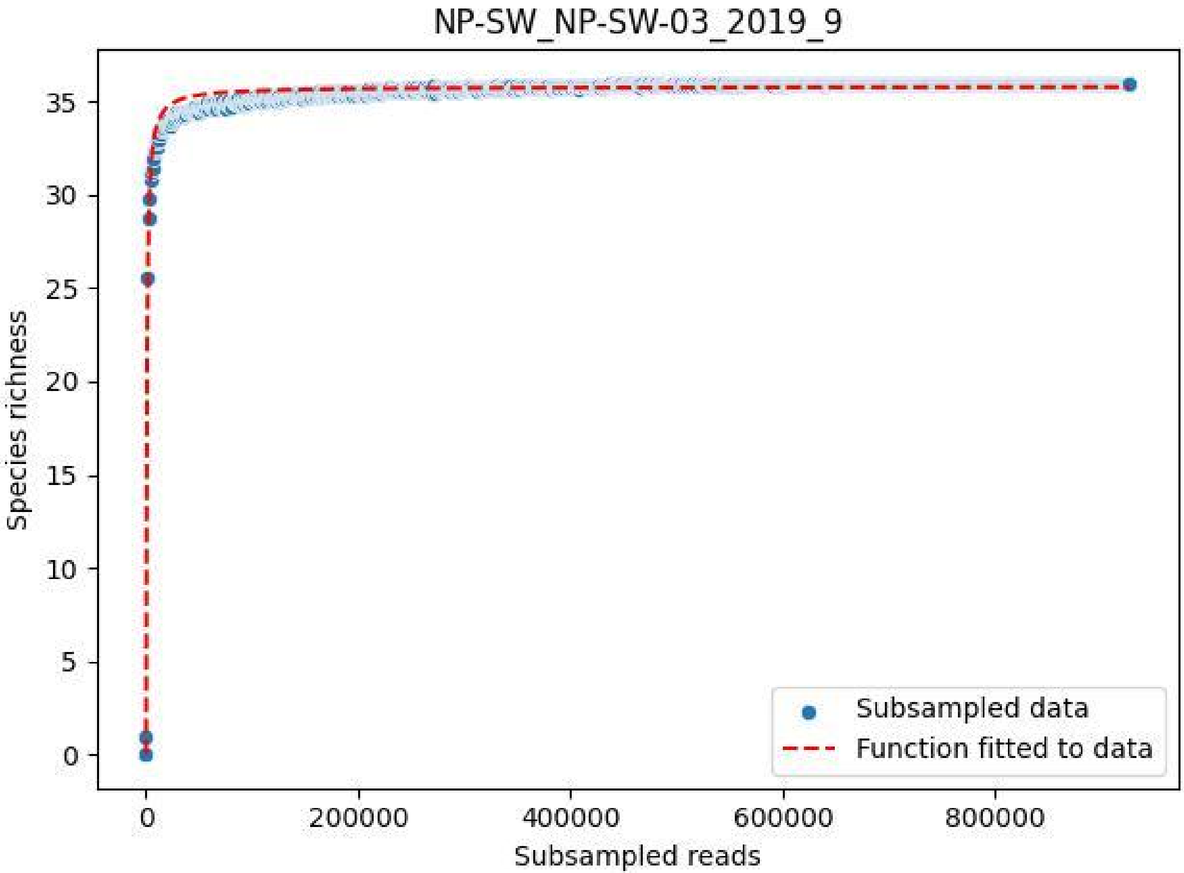

NP-SW_NP-SW-03_2019_9
Species richness
Subsampled reads
0
5
10
15
20
25
30
35
0
200000
400000
600000
800000
Subsampled data
Function fitted to data

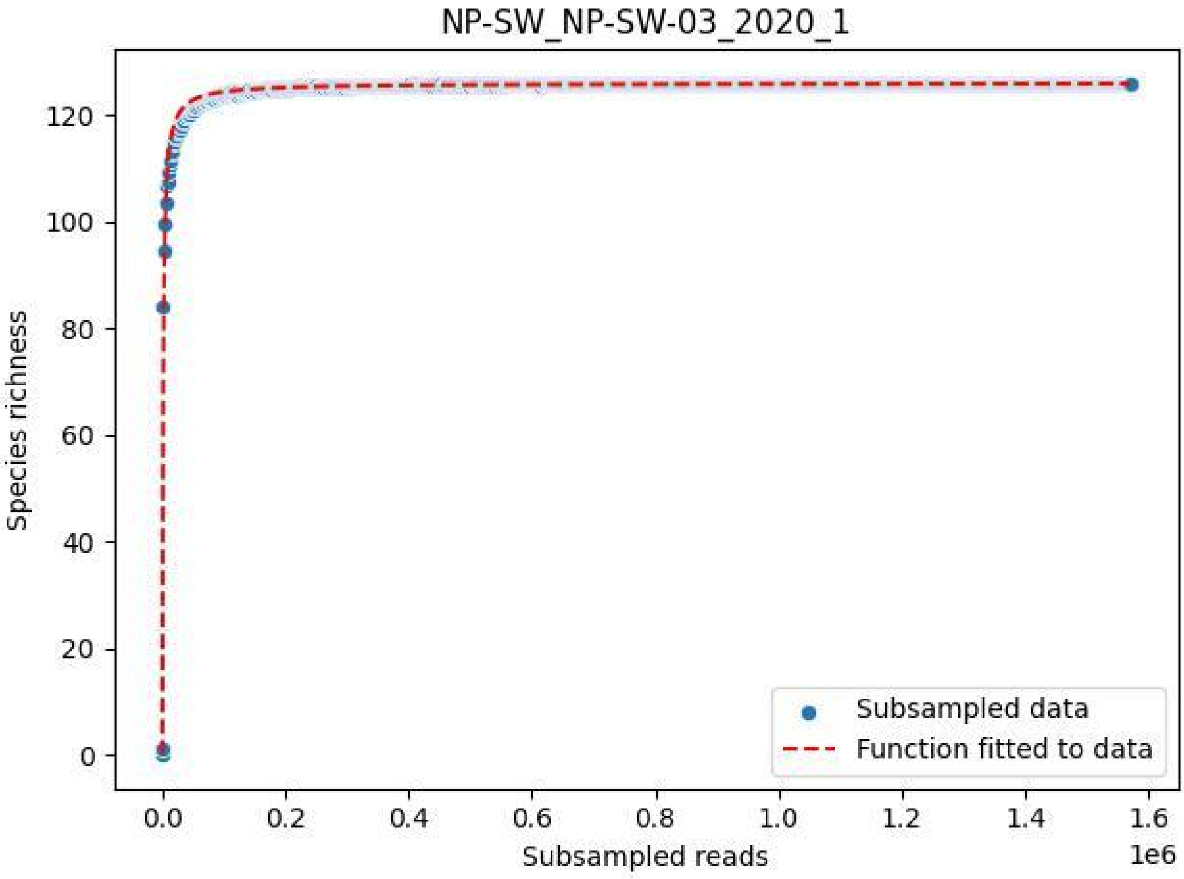

NP-SW_NP-SW-03_2020_1
Species richness
Subsampled reads
1e6
0
20
40
60
80
100
120
0.0
0.2
0.4
0.6
0.8
1.0
1.2
1.4
1.6
Subsampled data
Function fitted to data

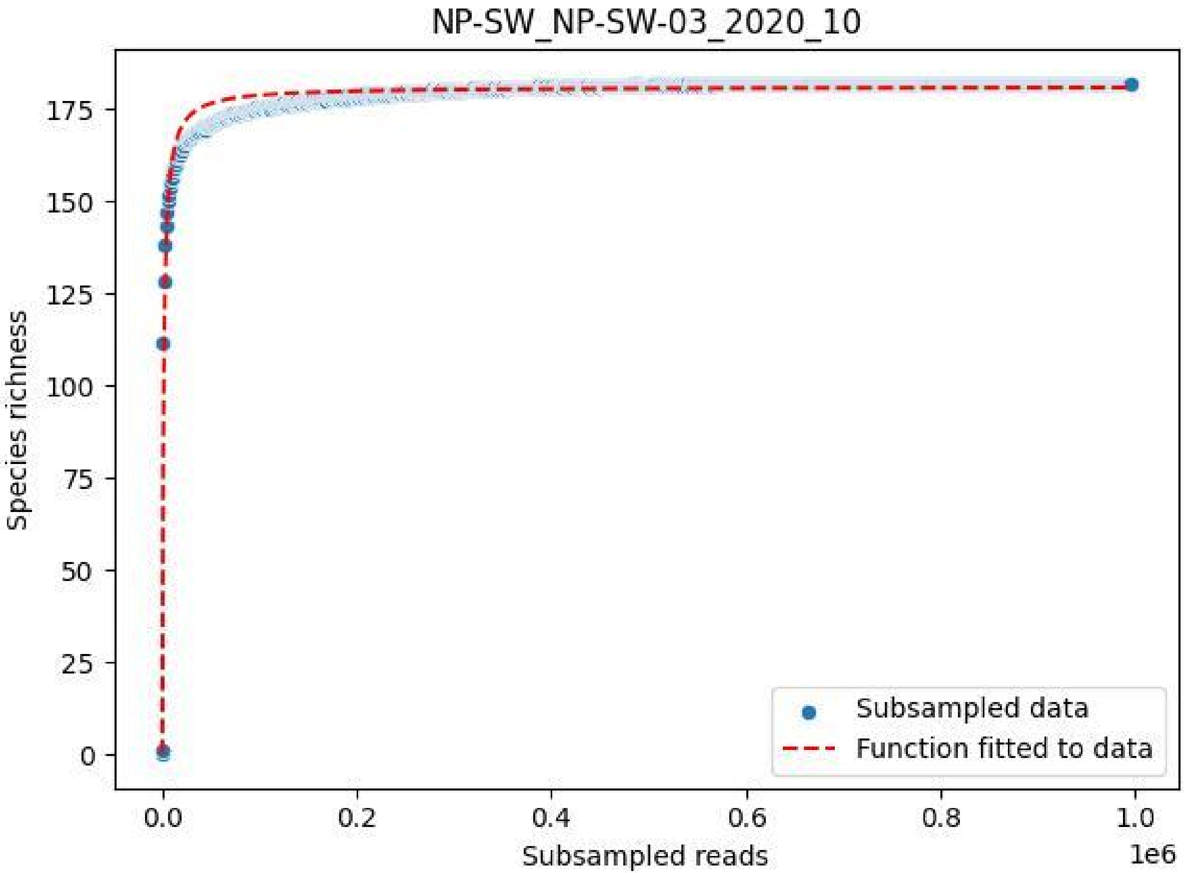

NP-SW_NP-SW-03_2020_10
Species richness
Subsampled reads
1e6
0
25
50
75
100
125
150
175
0.0
0.2
0.4
0.6
0.8
1.0
Subsampled data
Function fitted to data

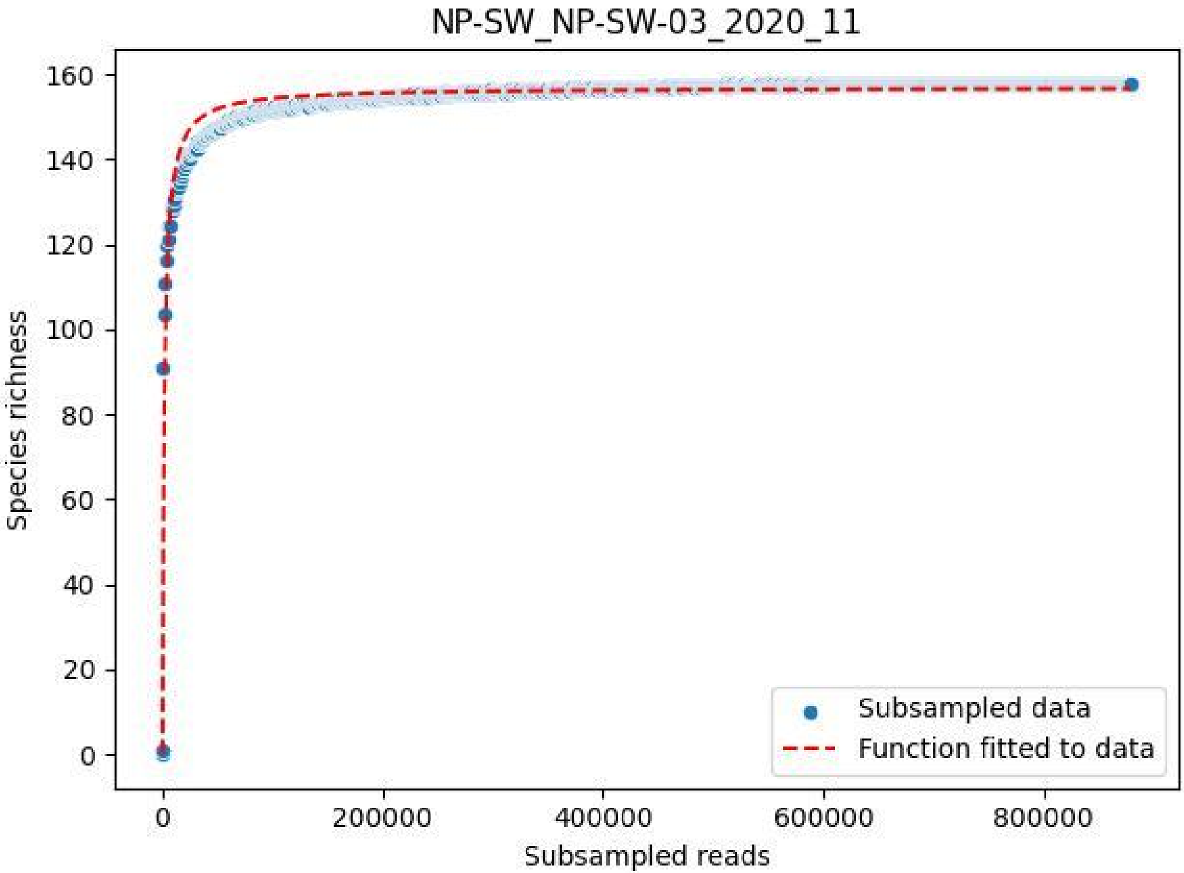
NP-SW_NP-SW-03_2020_11
Species richness
Subsampled reads
0
20
40
60
80
100
120
140
160
0
200000
400000
600000
800000
Subsampled data
Function fitted to data

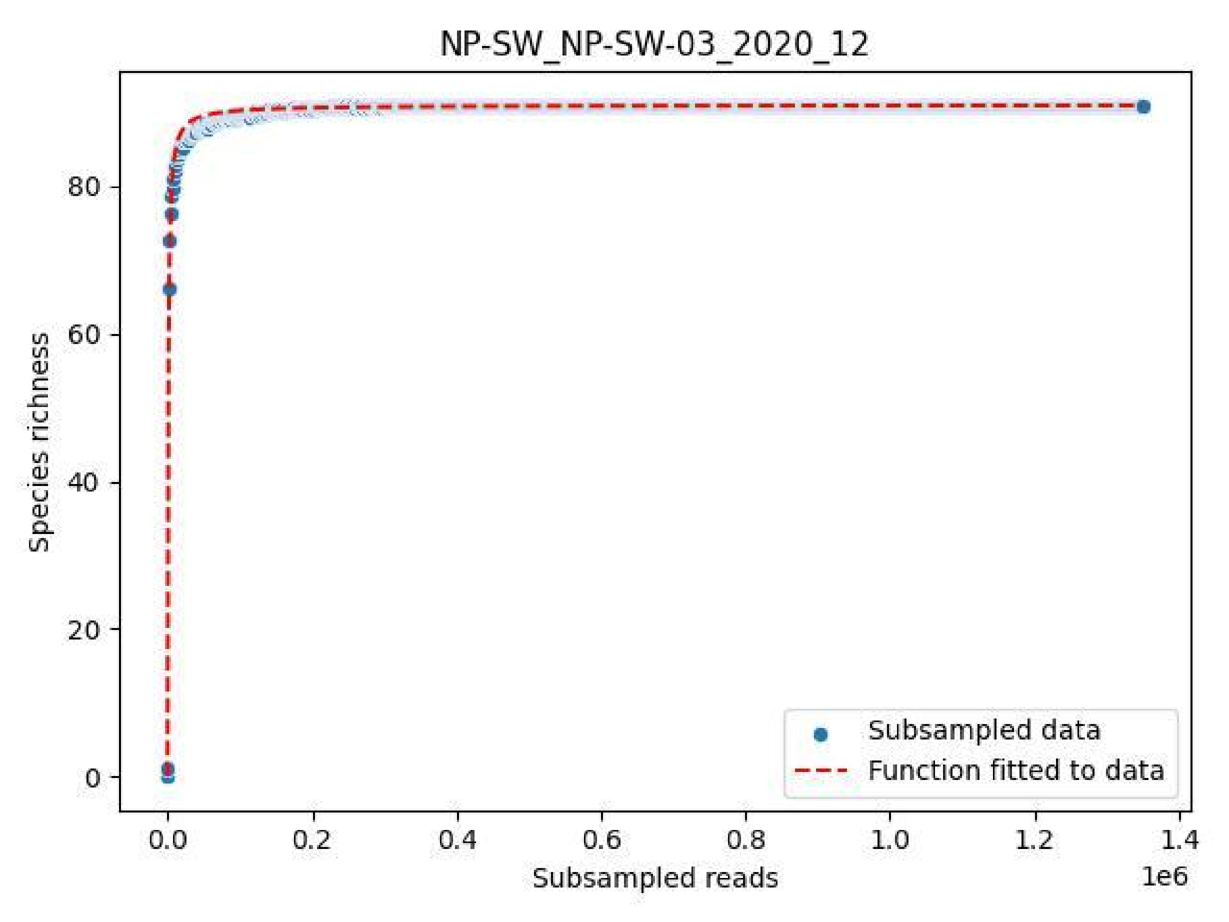

NP-SW_NP-SW-03_2020_12
Species richness
Subsampled reads
1e6
0
20
40
60
80
0.0
0.2
0.4
0.6
0.8
1.0
1.2
1.4
Subsampled data
Function fitted to data

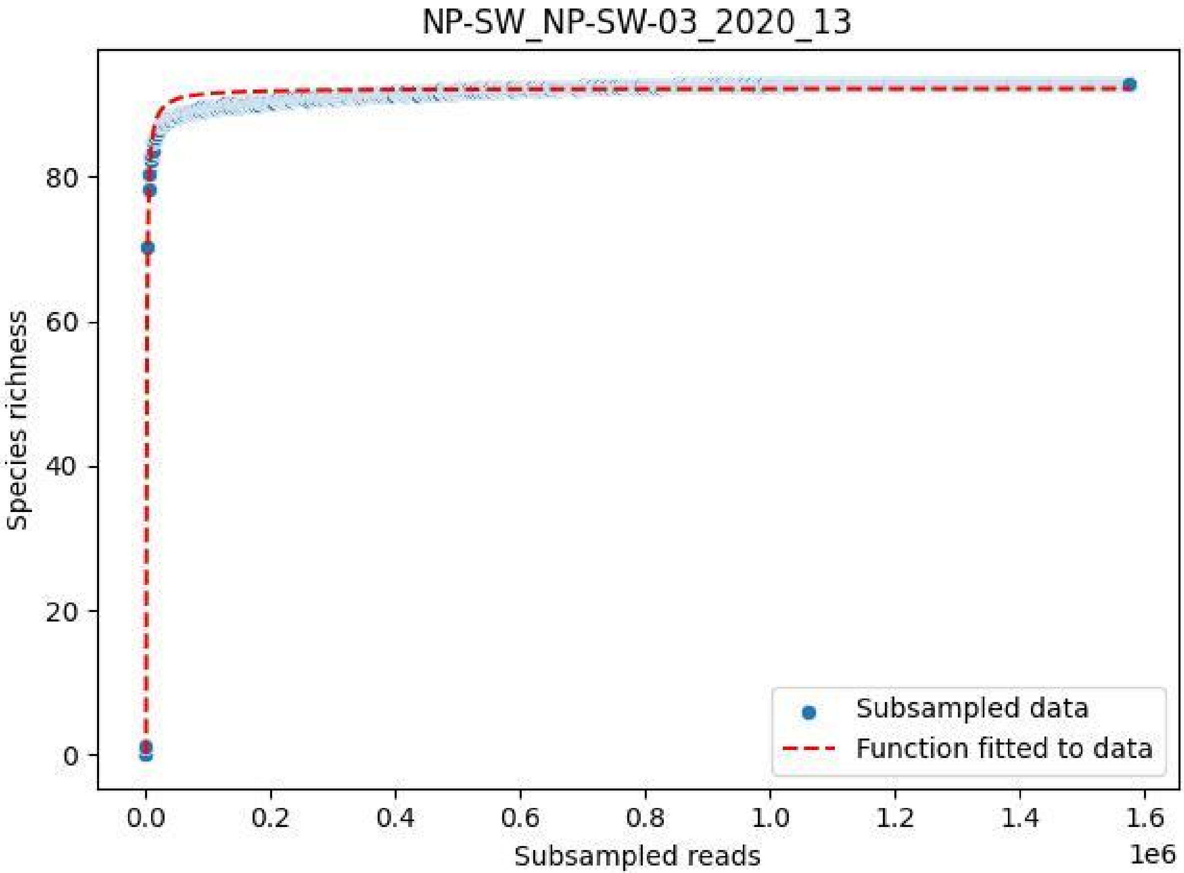

NP-SW_NP-SW-03_2020_13
Species richness
Subsampled reads
1e6
0
20
40
60
80
0.0
0.2
0.4
0.6
0.8
1.0
1.2
1.4
1.6
Subsampled data
Function fitted to data

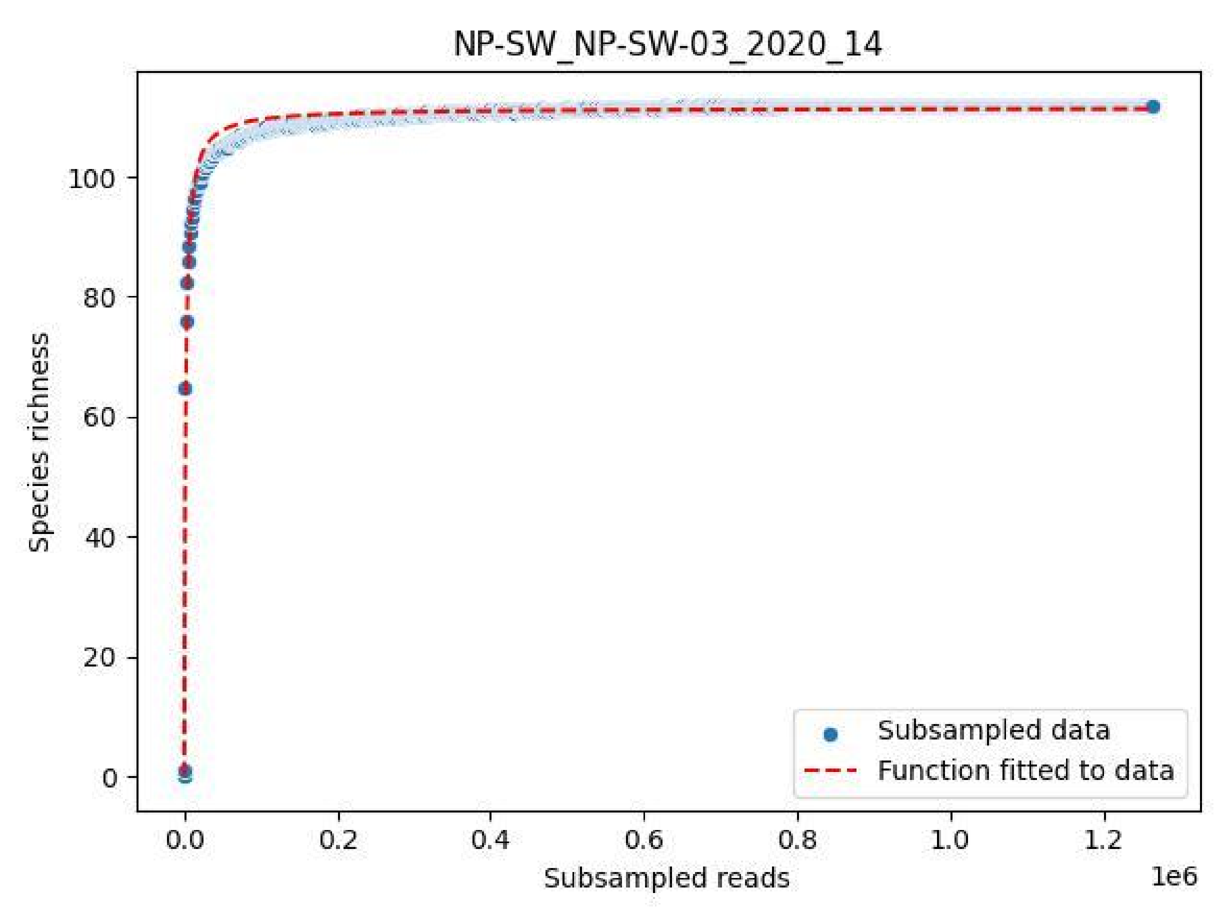

NP-SW_NP-SW-03_2020_14
Species richness
Subsampled reads
1e6
0
20
40
60
80
100
0.0
0.2
0.4
0.6
0.8
1.0
1.2
Subsampled data
Function fitted to data

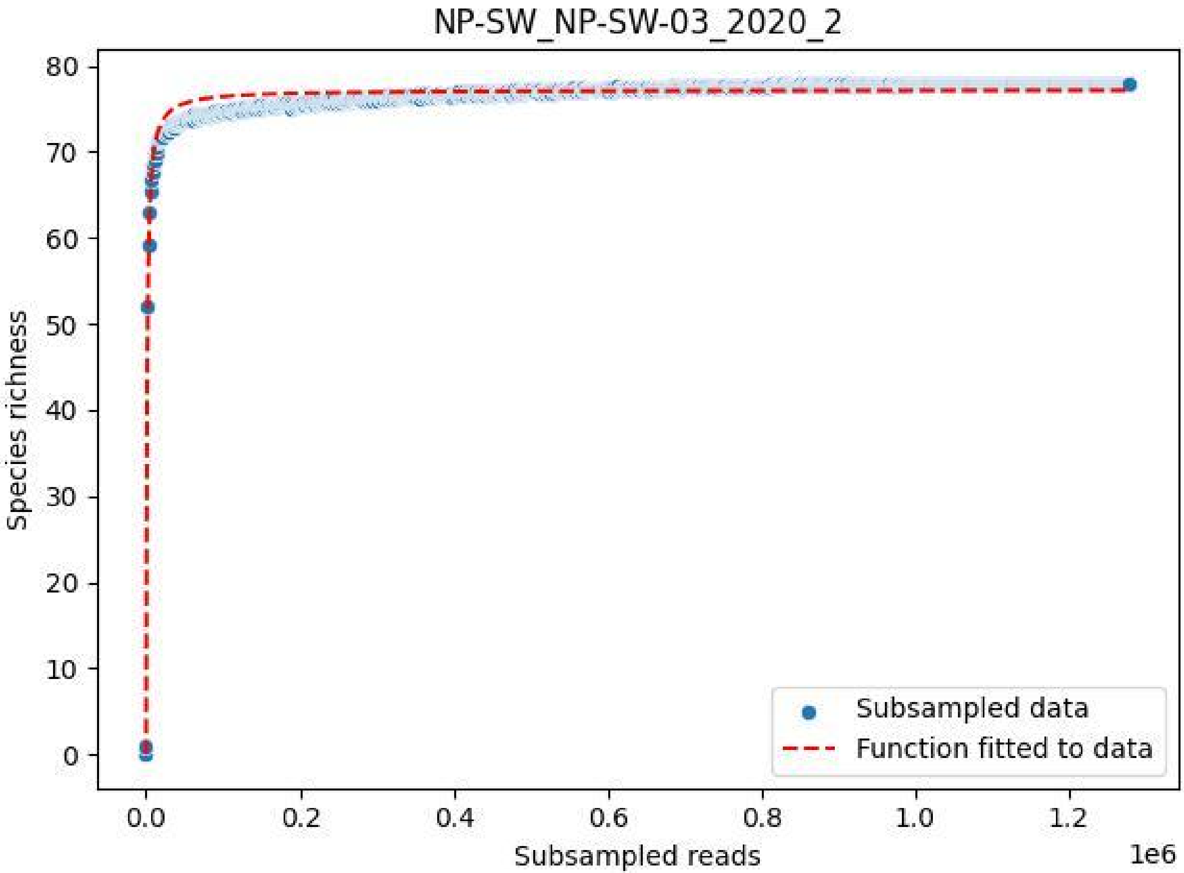

NP-SW_NP-SW-03_2020_2
Species richness
Subsampled reads
1e6
0
10
20
30
40
50
60
70
80
0.0
0.2
0.4
0.6
0.8
1.0
1.2
Subsampled data
Function fitted to data

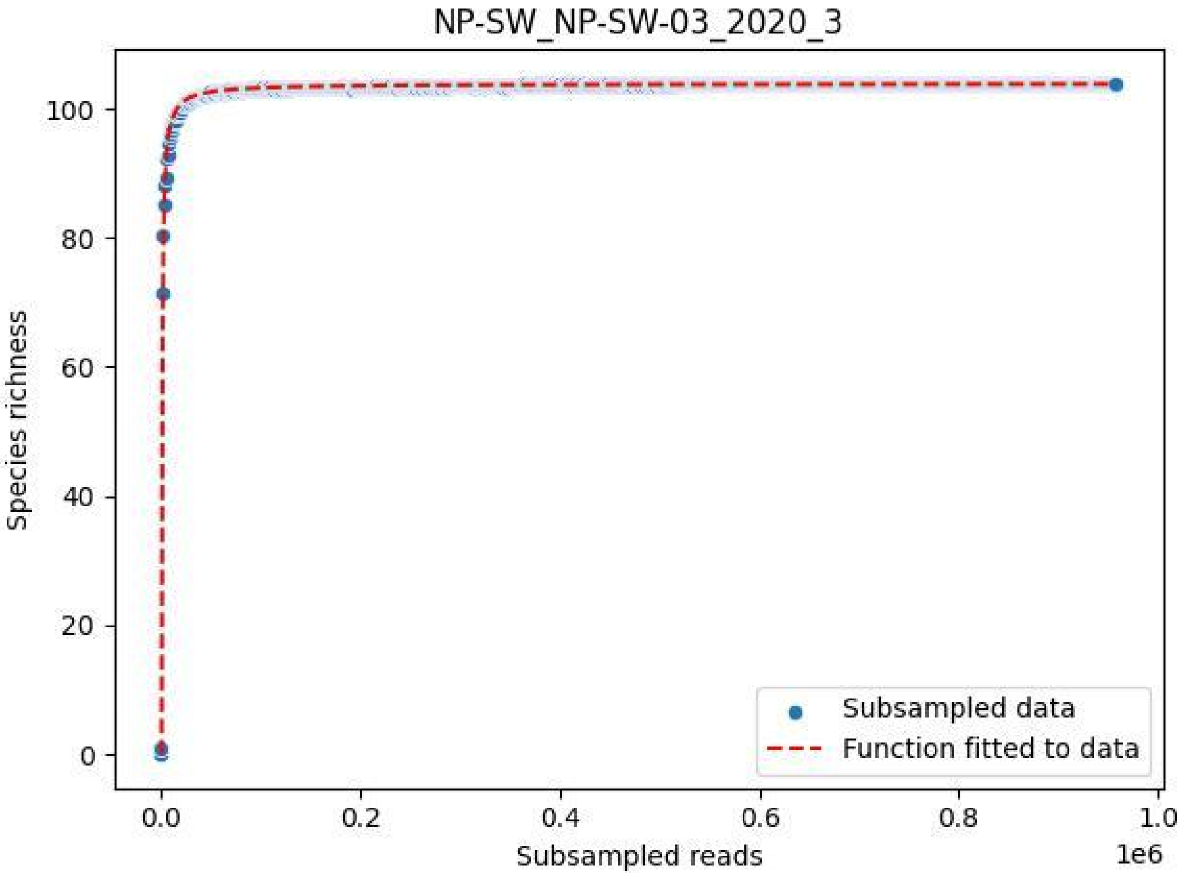
NP-SW_NP-SW-03_2020_3
Species richness
Subsampled reads
1e6
0.0
0.2
0.4
0.6
0.8
1.0
0
20
40
60
80
100
Subsampled data
Function fitted to data

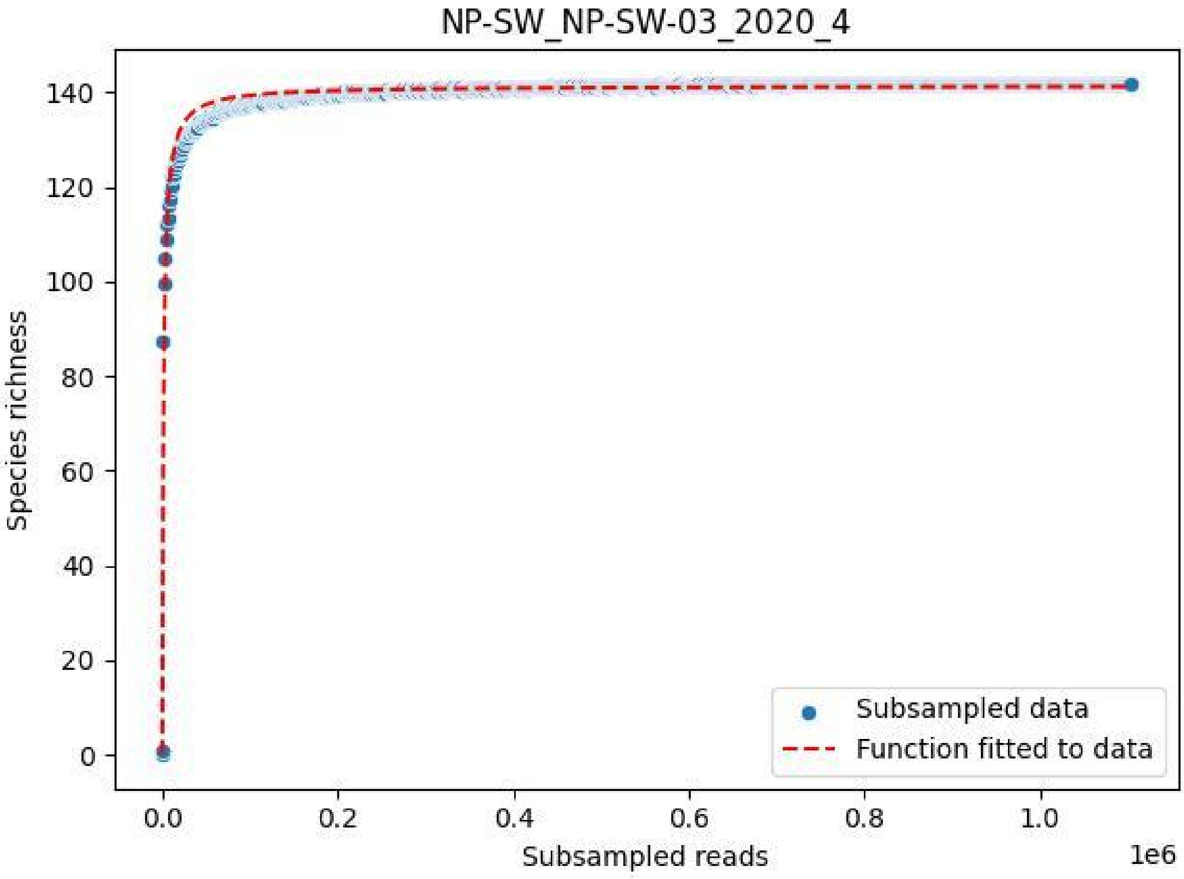

NP-SW_NP-SW-03_2020_4
Species richness
Subsampled reads
1e6
0
20
40
60
80
100
120
140
0.0
0.2
0.4
0.6
0.8
1.0
Subsampled data
Function fitted to data

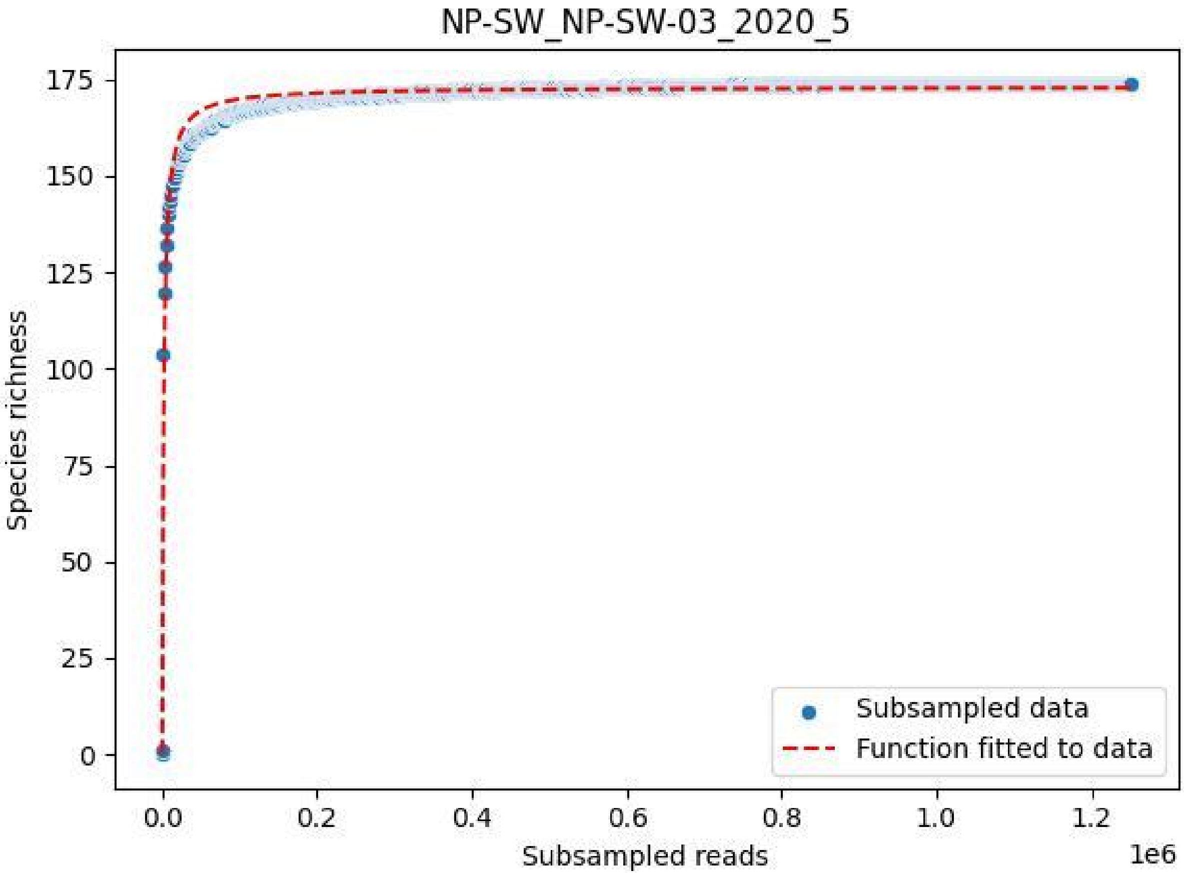

NP-SW_NP-SW-03_2020_5
175
150
125
100
75
50
25
0
Species richness
0.0
0.2
0.4
0.6
0.8
1.0
1.2
1e6
Subsampled reads
Subsampled data
Function fitted to data

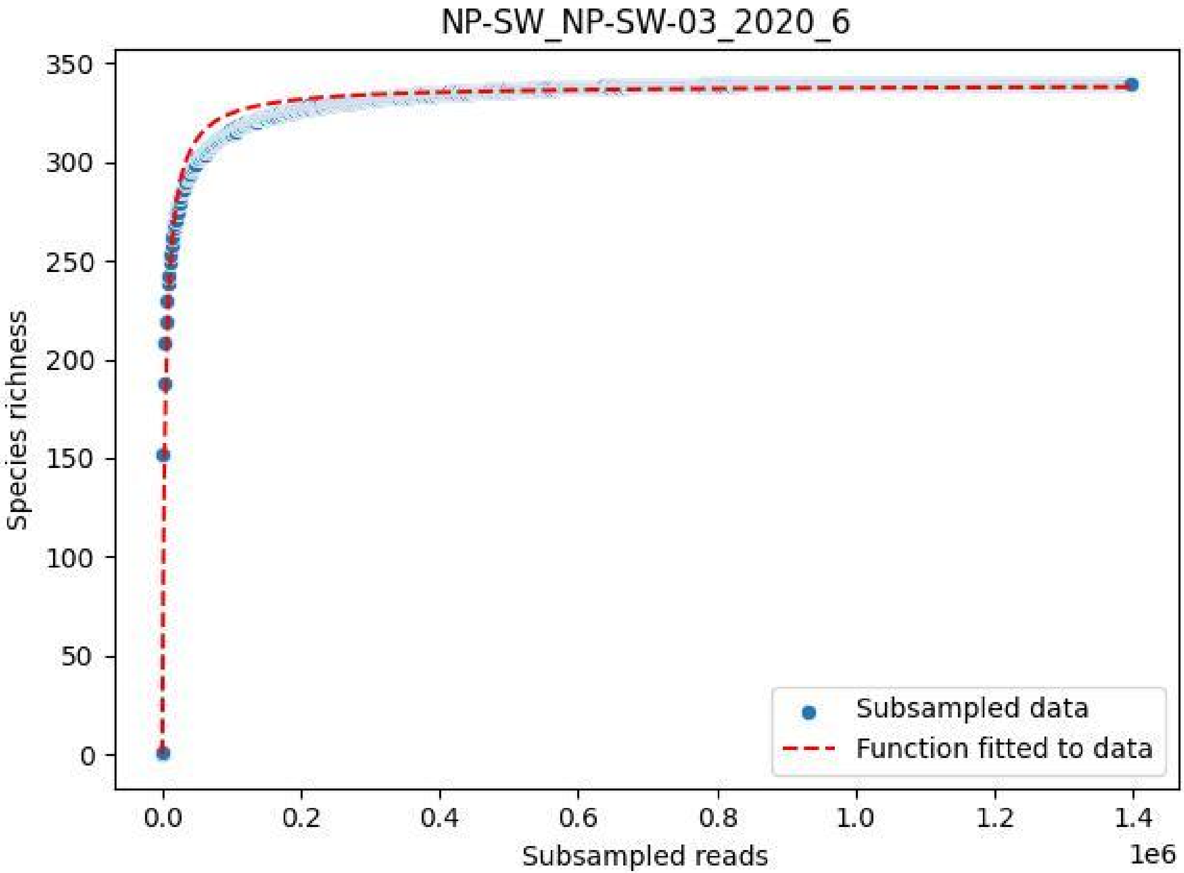

NP-SW_NP-SW-03_2020_6
Species richness
Subsampled reads
1e6
0
50
100
150
200
250
300
350
0.0
0.2
0.4
0.6
0.8
1.0
1.2
1.4
Subsampled data
Function fitted to data

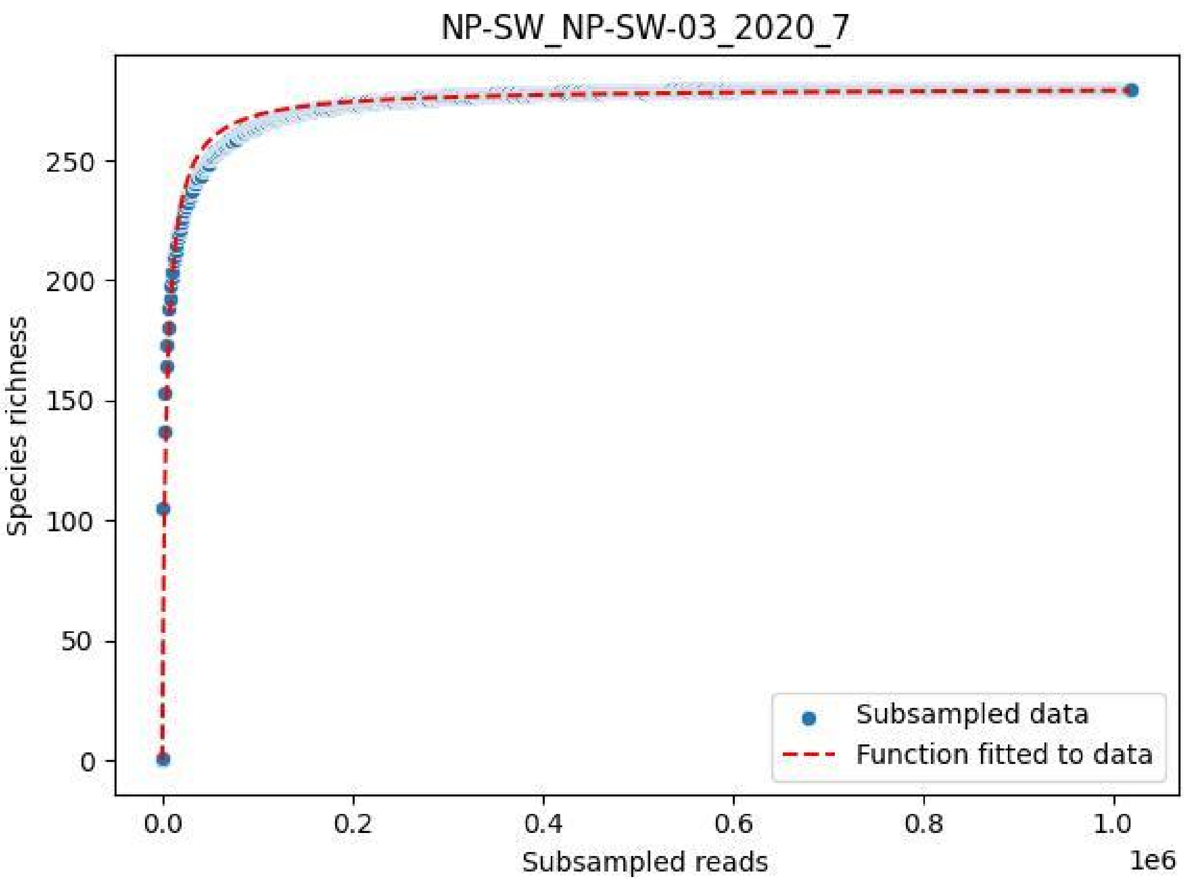
NP-SW_NP-SW-03_2020_7
Species richness
Subsampled reads
1e6
0.0
0.2
0.4
0.6
0.8
1.0
0
50
100
150
200
250
Subsampled data
Function fitted to data

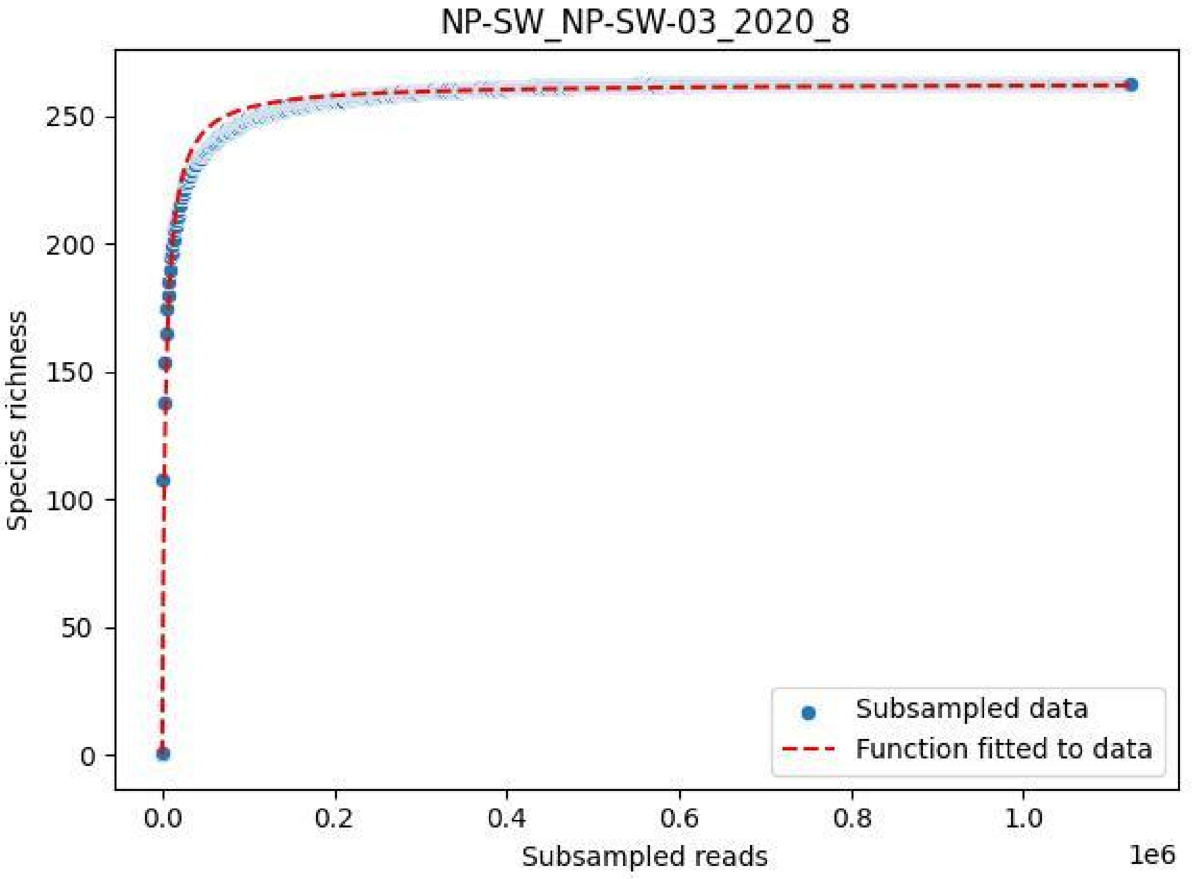

NP-SW_NP-SW-03_2020_8
Species richness
250
200
150
100
50
0
0.0
0.2
0.4
0.6
0.8
1.0
1e6
Subsampled reads
Subsampled data
Function fitted to data

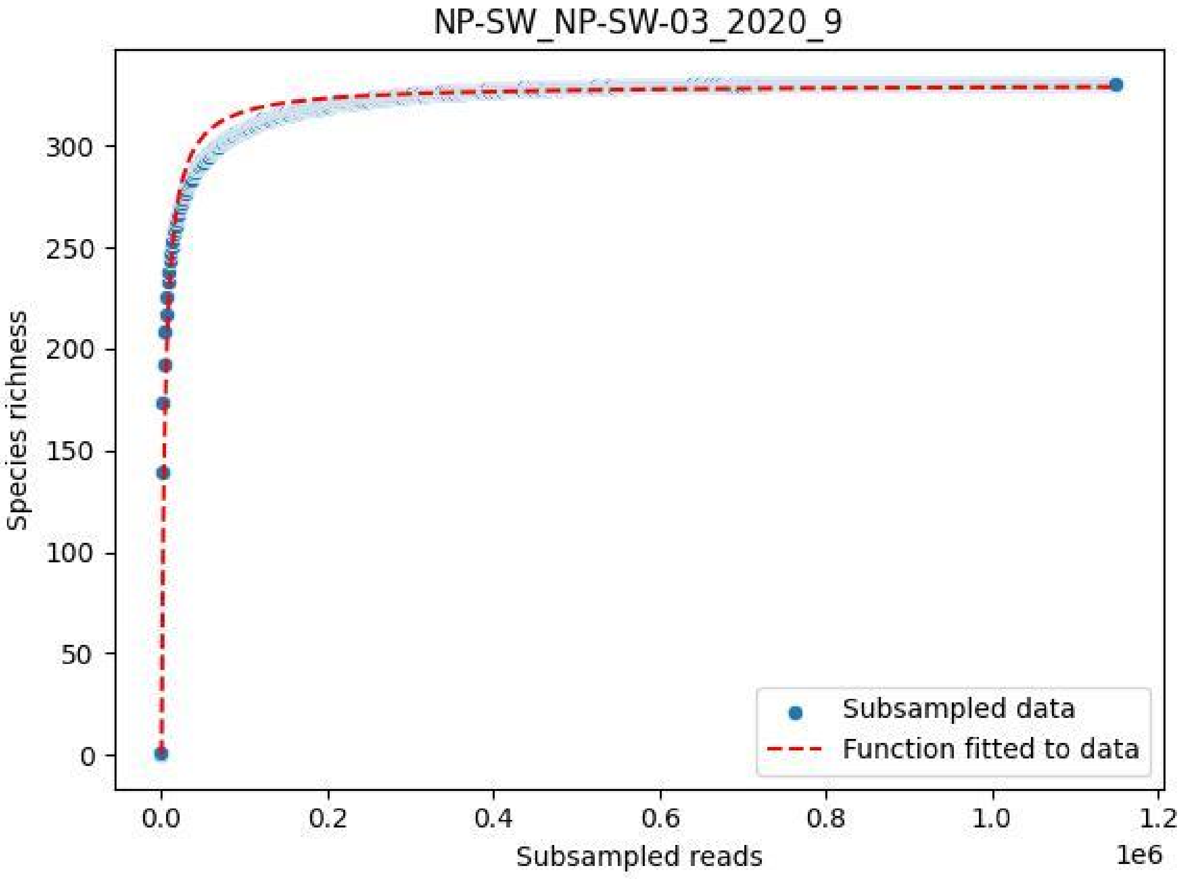
NP-SW_NP-SW-03_2020_9
Species richness
0
50
100
150
200
250
300
Subsampled reads
0.0
0.2
0.4
0.6
0.8
1.0
1.2
1e6
Subsampled data
Function fitted to data

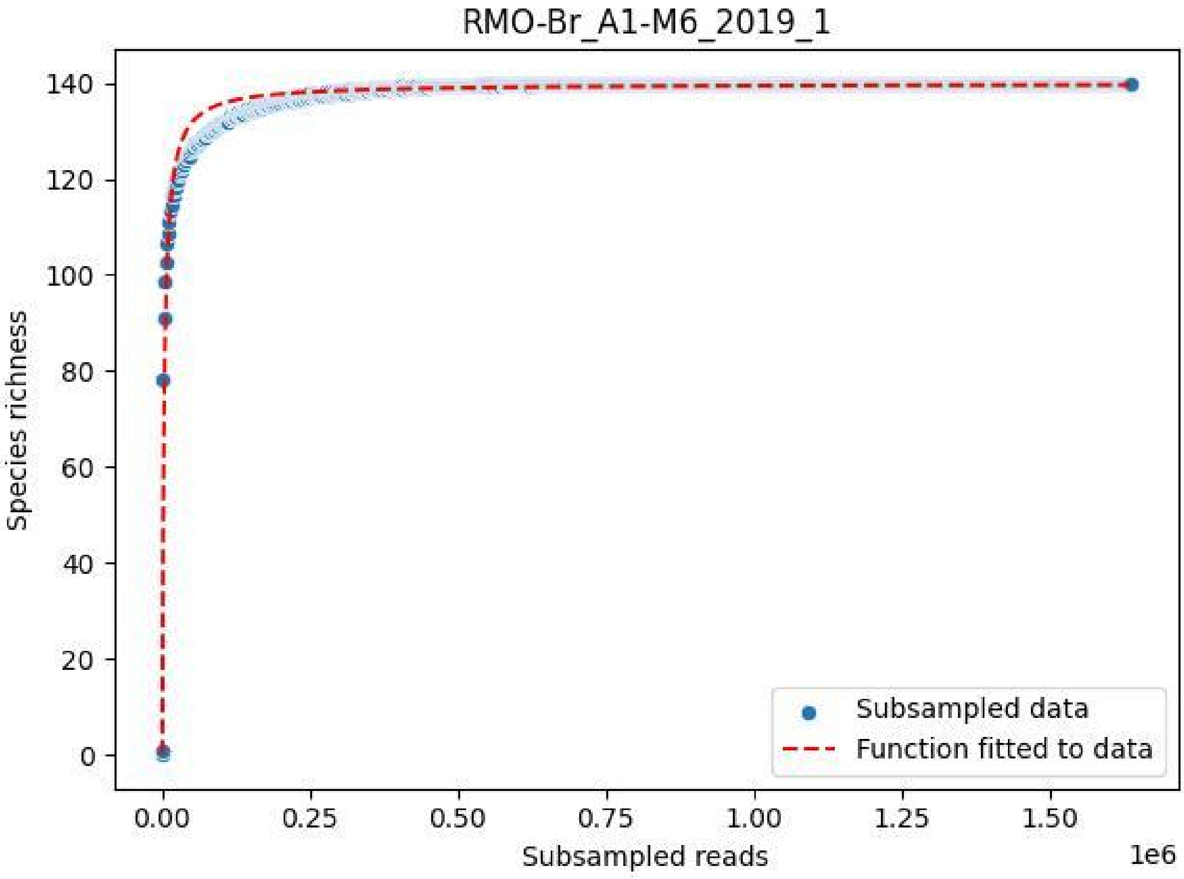
RMO-Br_A1-M6_2019_1
Species richness
Subsampled reads
1e6
0
20
40
60
80
100
120
140
0.00
0.25
0.50
0.75
1.00
1.25
1.50
Subsampled data
Function fitted to data

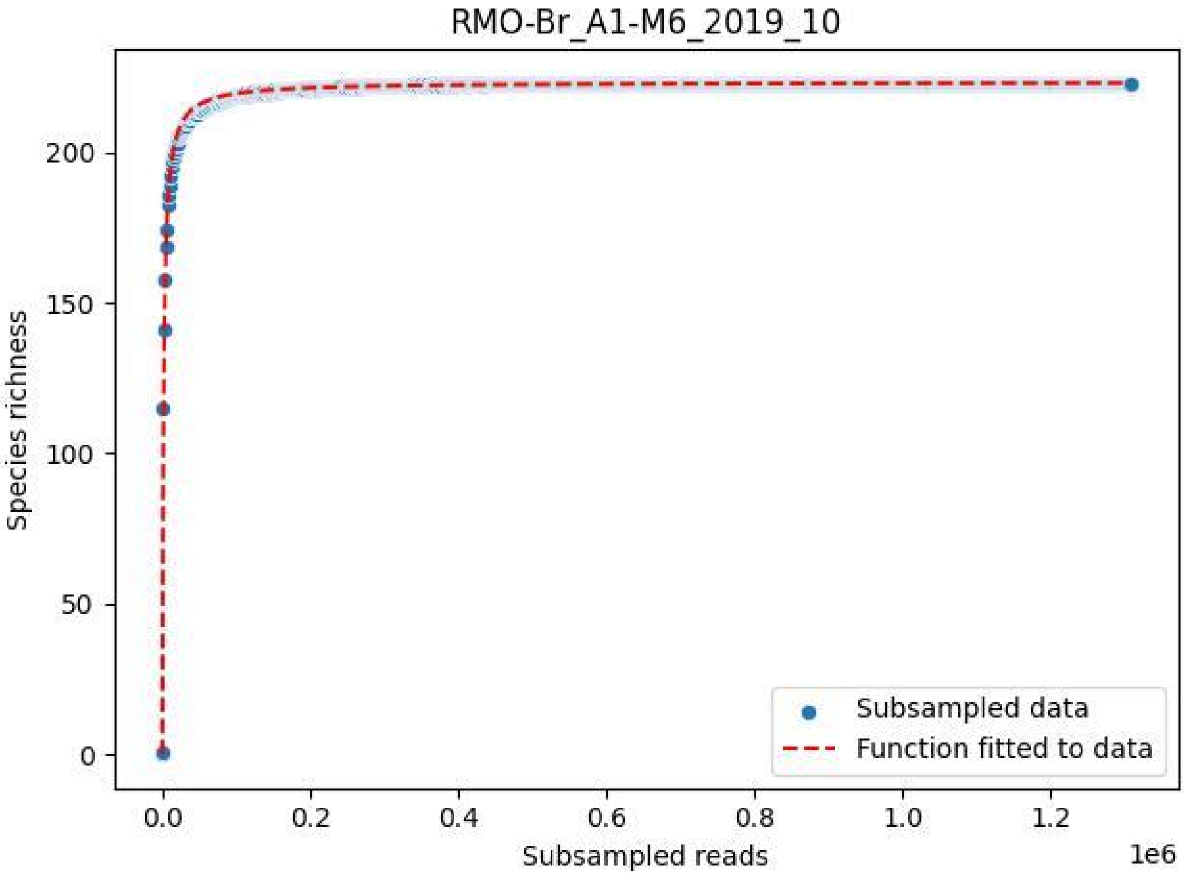

RMO-Br_A1-M6_2019_10
Species richness
Subsampled reads
1e6
0
50
100
150
200
0.0
0.2
0.4
0.6
0.8
1.0
1.2
Subsampled data
Function fitted to data

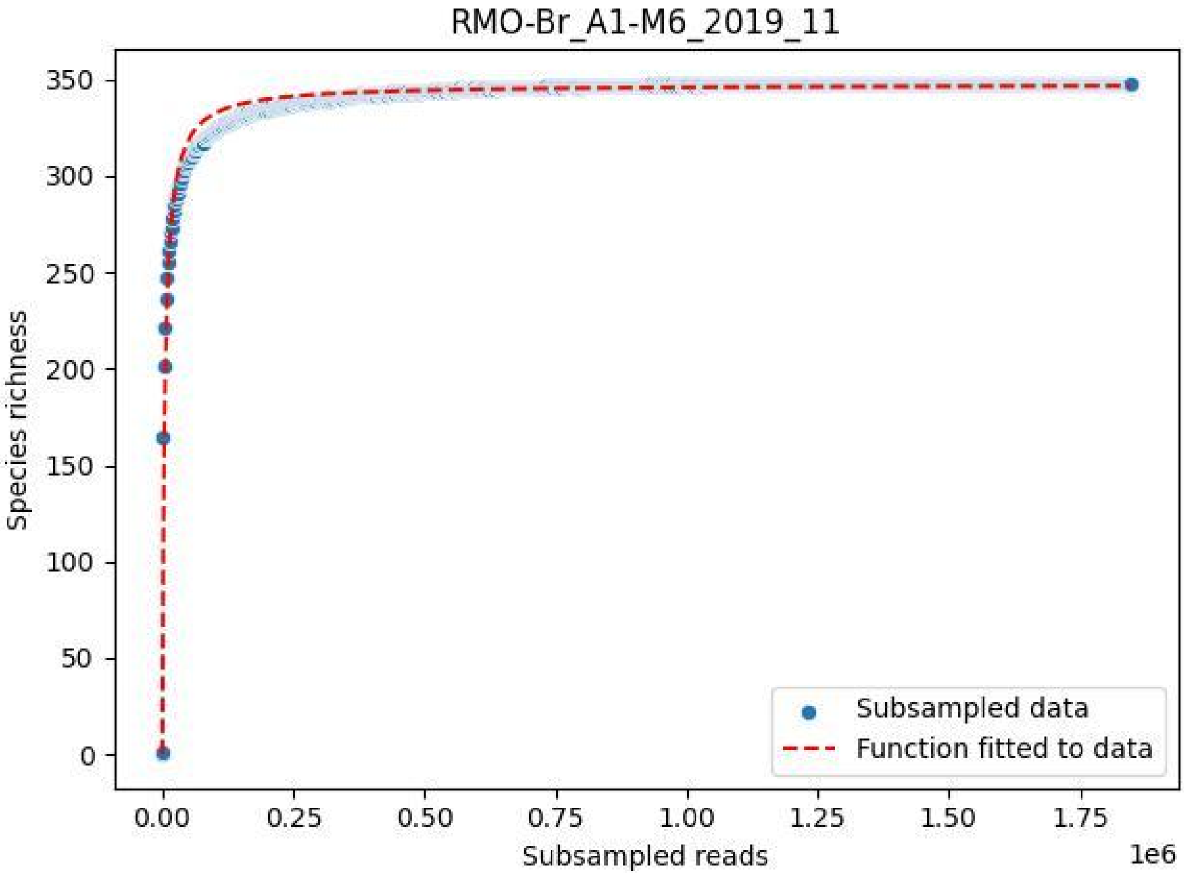
RMO-Br_A1-M6_2019_11
Species richness
Subsampled reads
1e6
0
50
100
150
200
250
300
350
0.00
0.25
0.50
0.75
1.00
1.25
1.50
1.75
Subsampled data
Function fitted to data

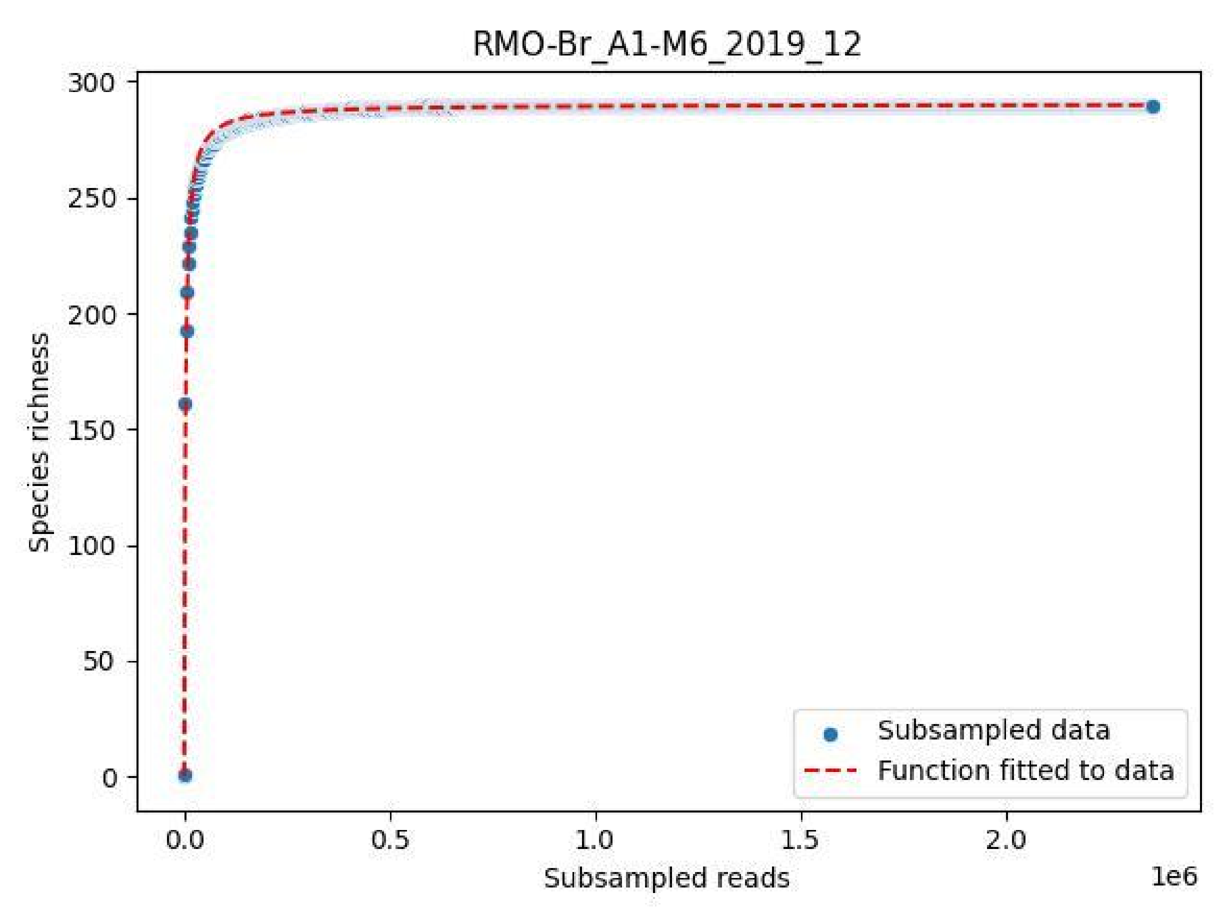
RMO-Br_A1-M6_2019_12
Species richness
Subsampled reads
1e6
0
50
100
150
200
250
300
0.0
0.5
1.0
1.5
2.0
Subsampled data
Function fitted to data

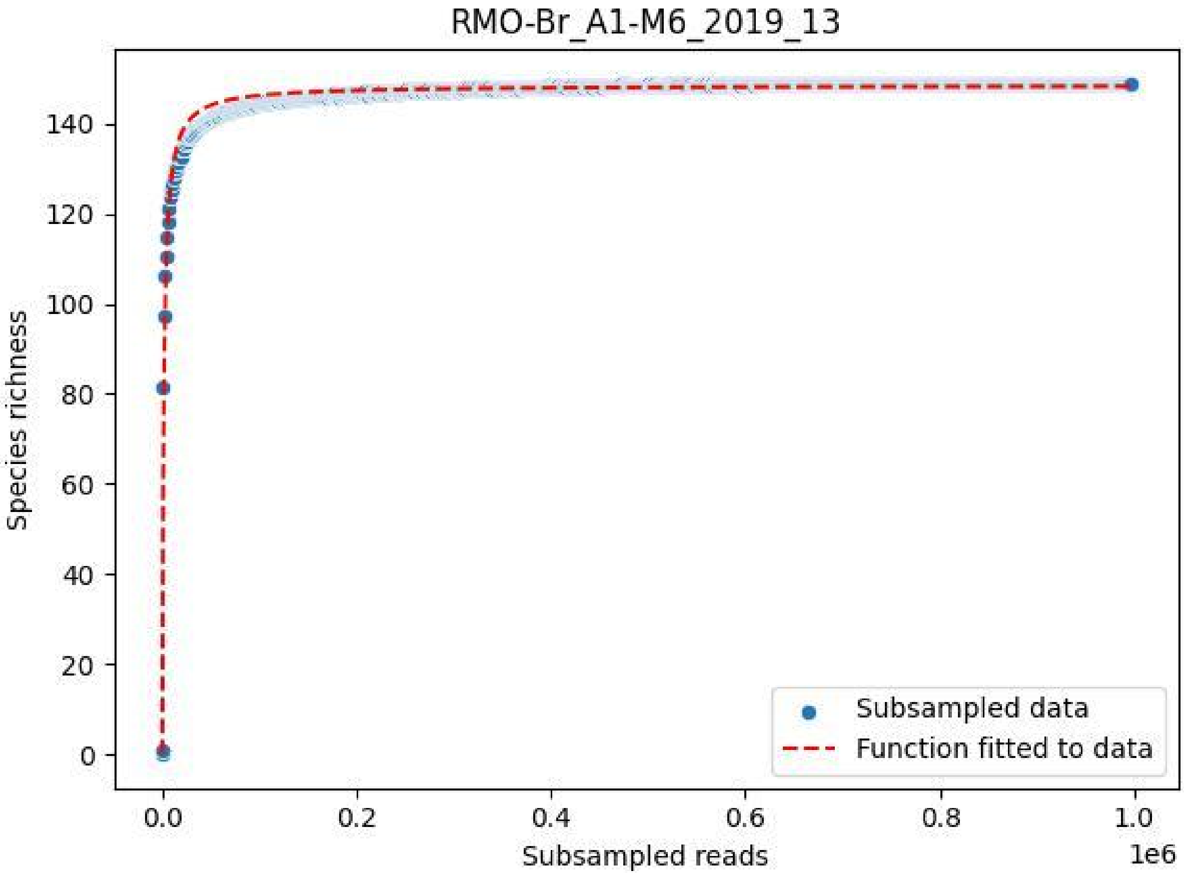

RMO-Br_A1-M6_2019_13
Species richness
Subsampled reads
1e6
0.0
0.2
0.4
0.6
0.8
1.0
0
20
40
60
80
100
120
140
Subsampled data
Function fitted to data

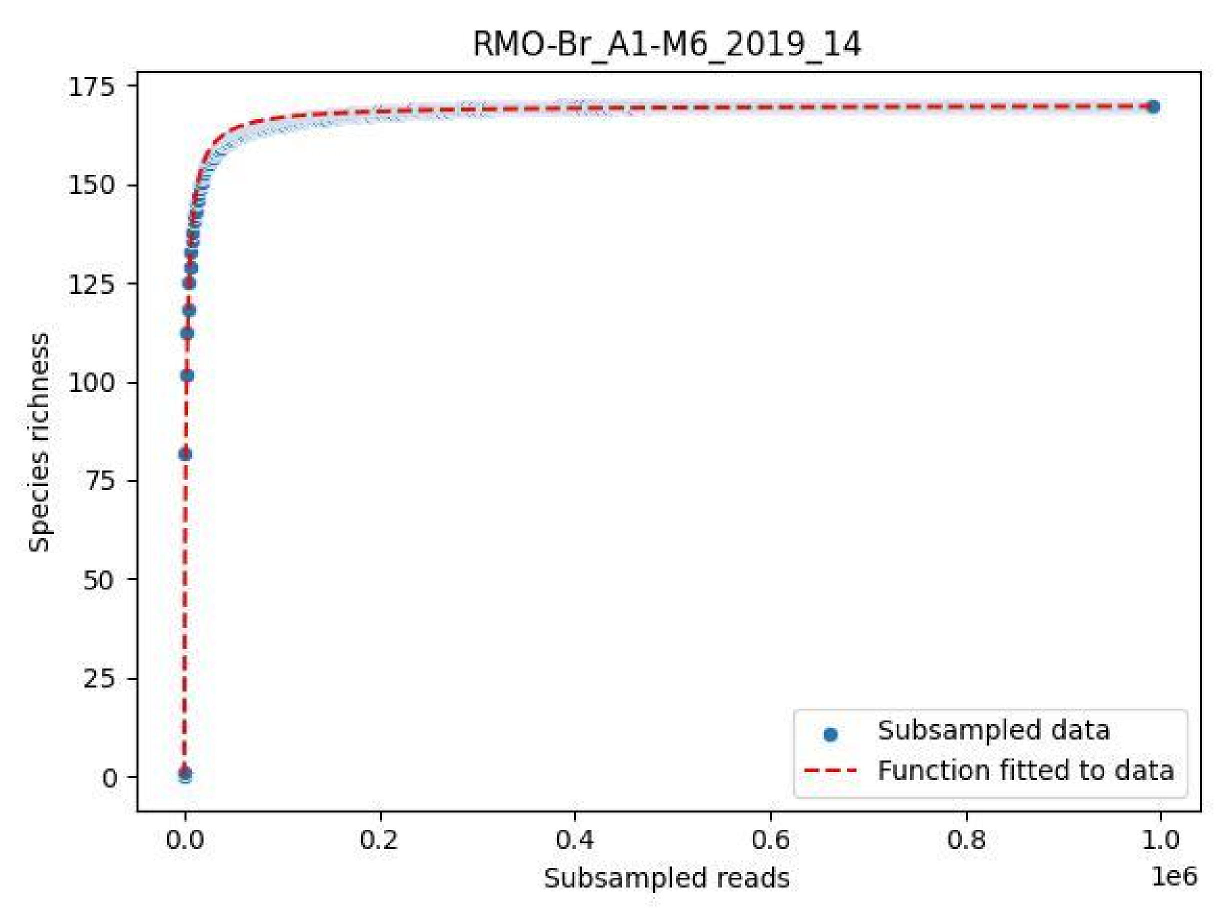
RMO-Br_A1-M6_2019_14
Species richness
Subsampled reads
1e6
0.0
0.2
0.4
0.6
0.8
1.0
0
25
50
75
100
125
150
175
Subsampled data
Function fitted to data

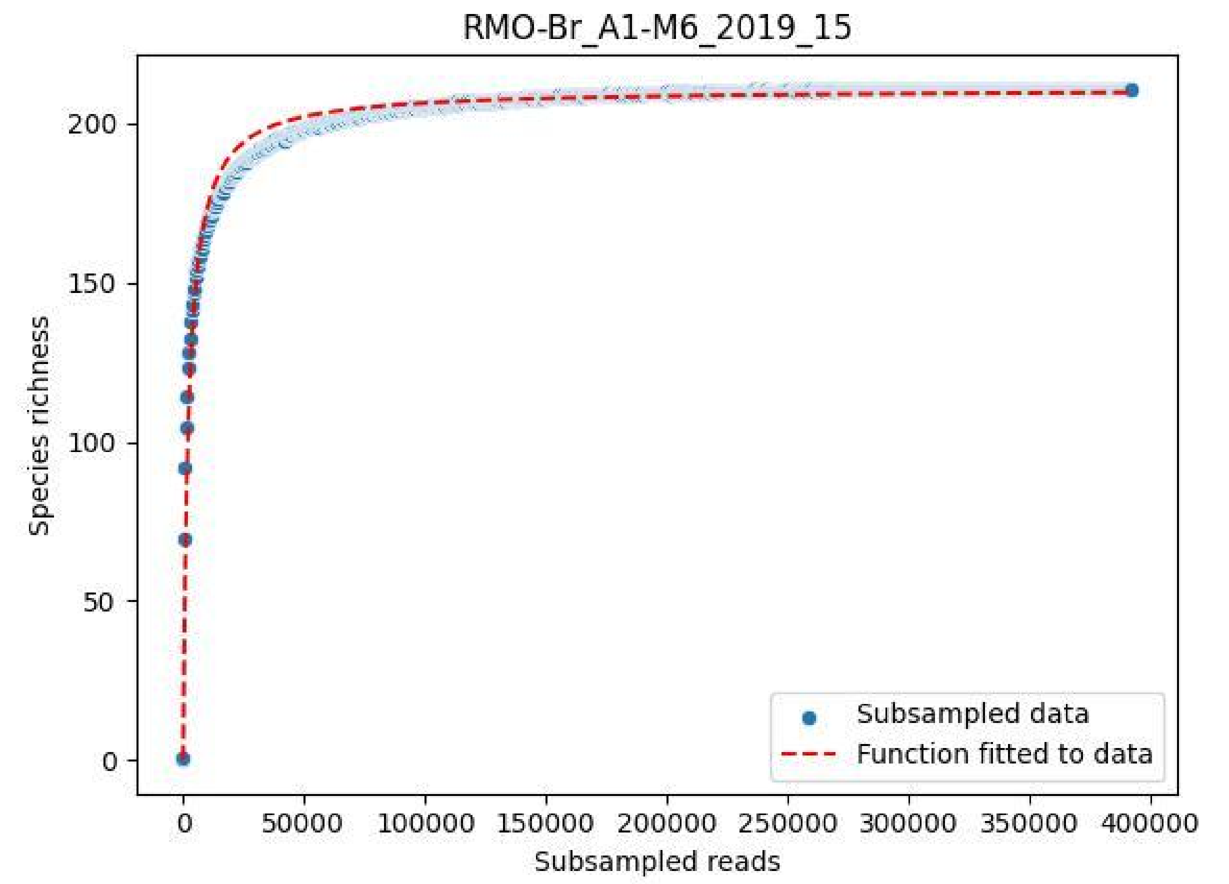
RMO-Br_A1-M6_2019_15
Species richness
Subsampled reads
0
50
100
150
200
0
50000
100000
150000
200000
250000
300000
350000
400000
Subsampled data
Function fitted to data

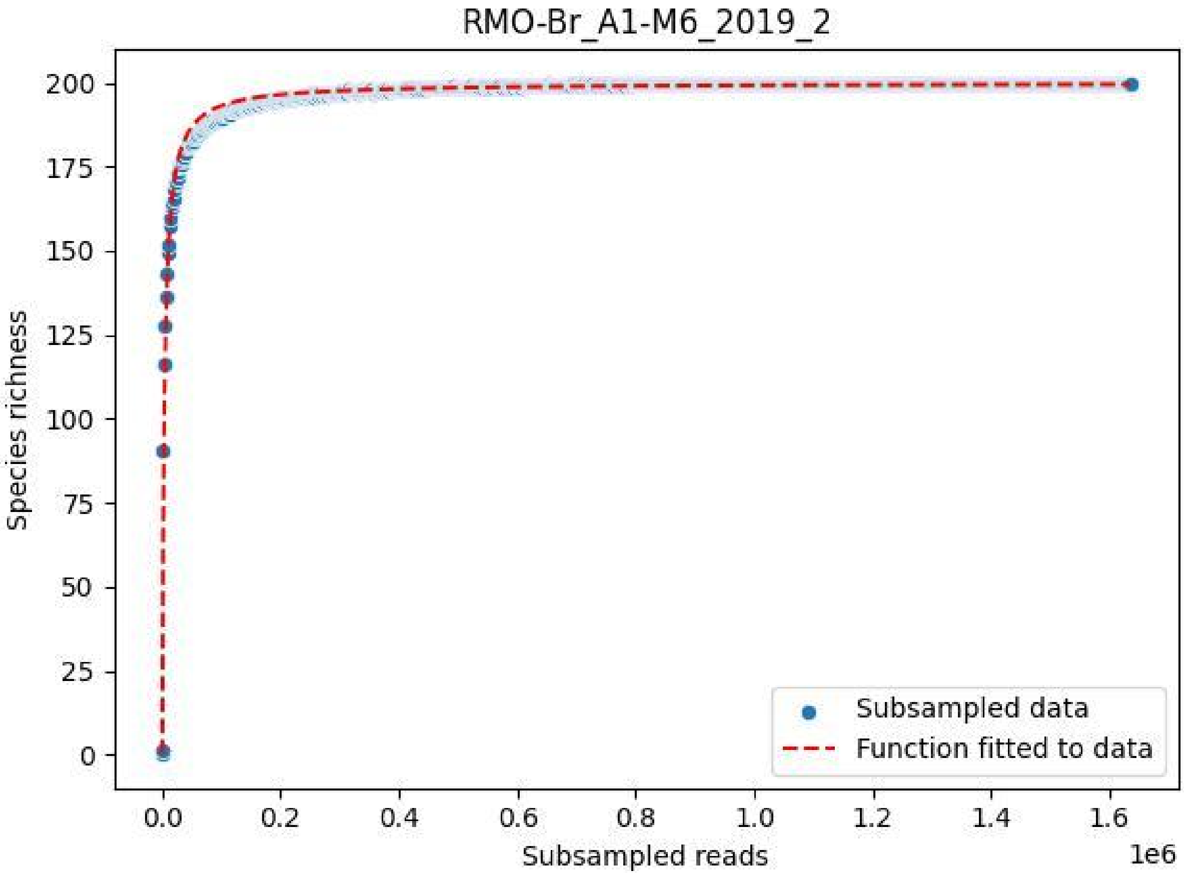

RMO-Br_A1-M6_2019_2
Species richness
0
25
50
75
100
125
150
175
200
0.0
0.2
0.4
0.6
0.8
1.0
1.2
1.4
1.6
1e6
Subsampled reads
Subsampled data
Function fitted to data

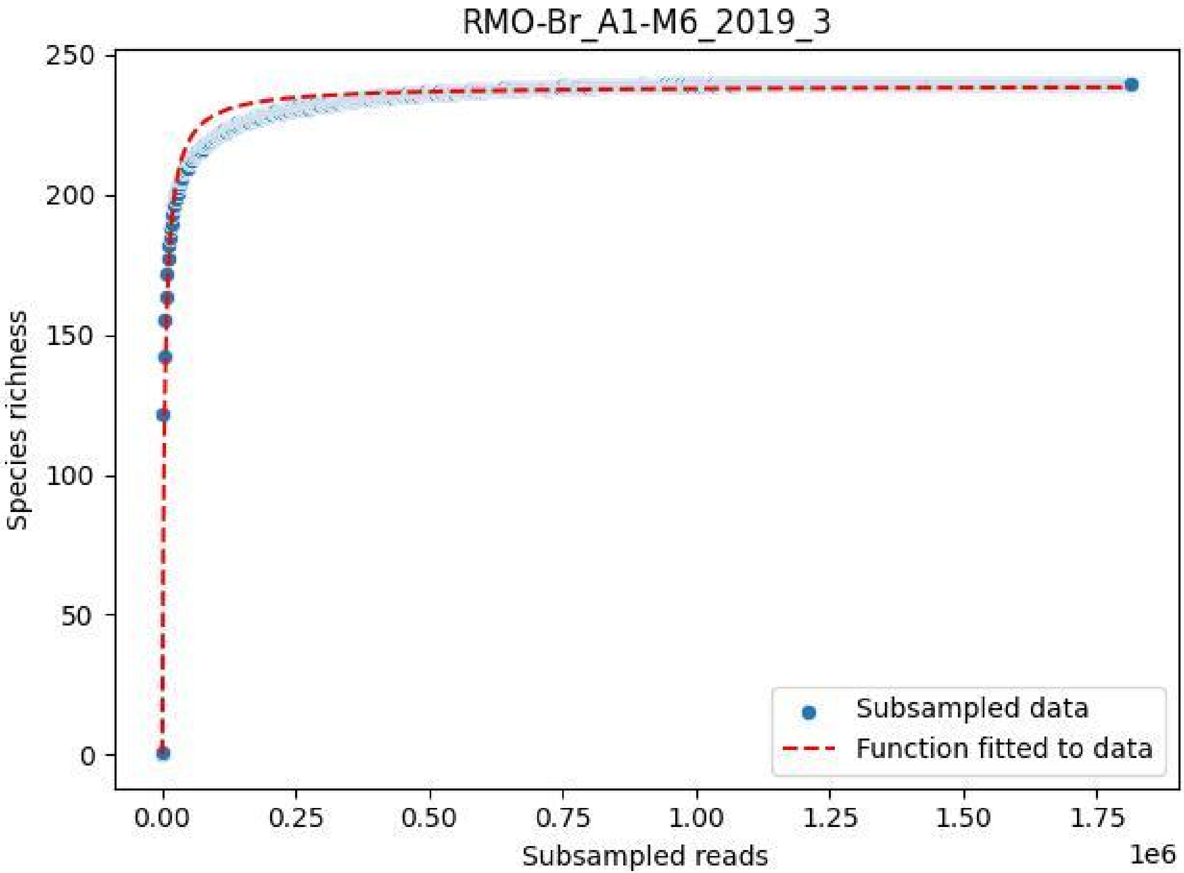

RMO-Br_A1-M6_2019_3
Species richness
Subsampled reads
1e6
250
200
150
100
50
0
0.00
0.25
0.50
0.75
1.00
1.25
1.50
1.75
Subsampled data
Function fitted to data

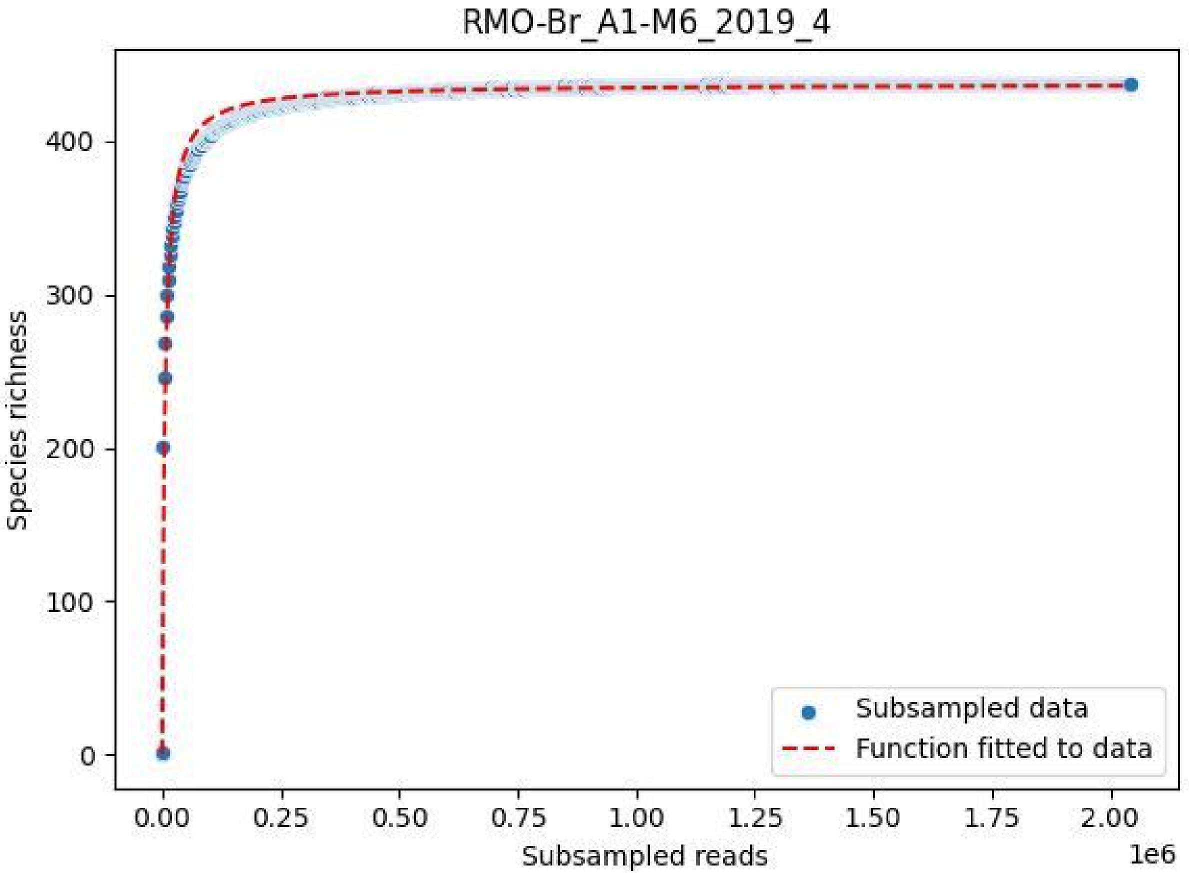

RMO-Br_A1-M6_2019_4
Species richness
0
100
200
300
400
0.00
0.25
0.50
0.75
1.00
1.25
1.50
1.75
2.00
1e6
Subsampled reads
Subsampled data
Function fitted to data

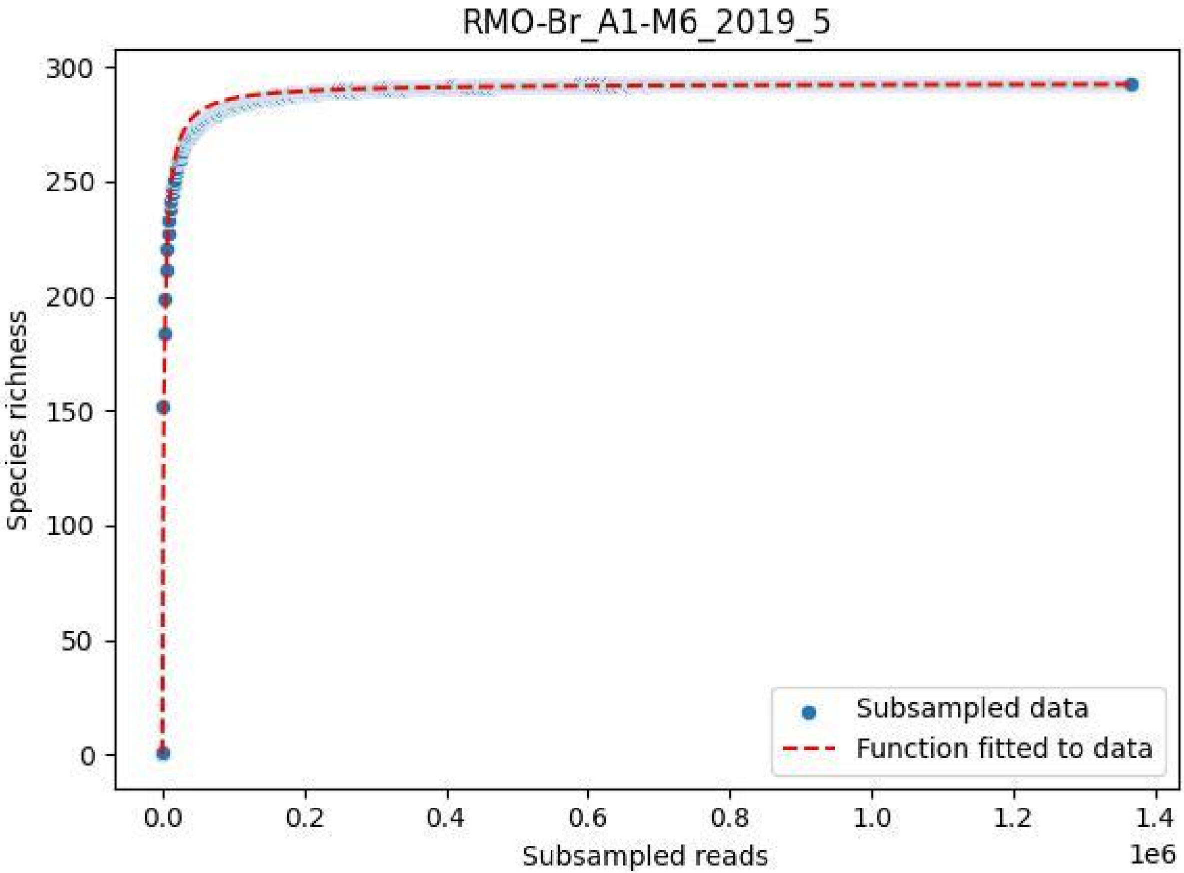

RMO-Br_A1-M6_2019_5
Species richness
300
250
200
150
100
50
0
0.0
0.2
0.4
0.6
0.8
1.0
1.2
1.4
Subsampled reads
1e6
Subsampled data
Function fitted to data

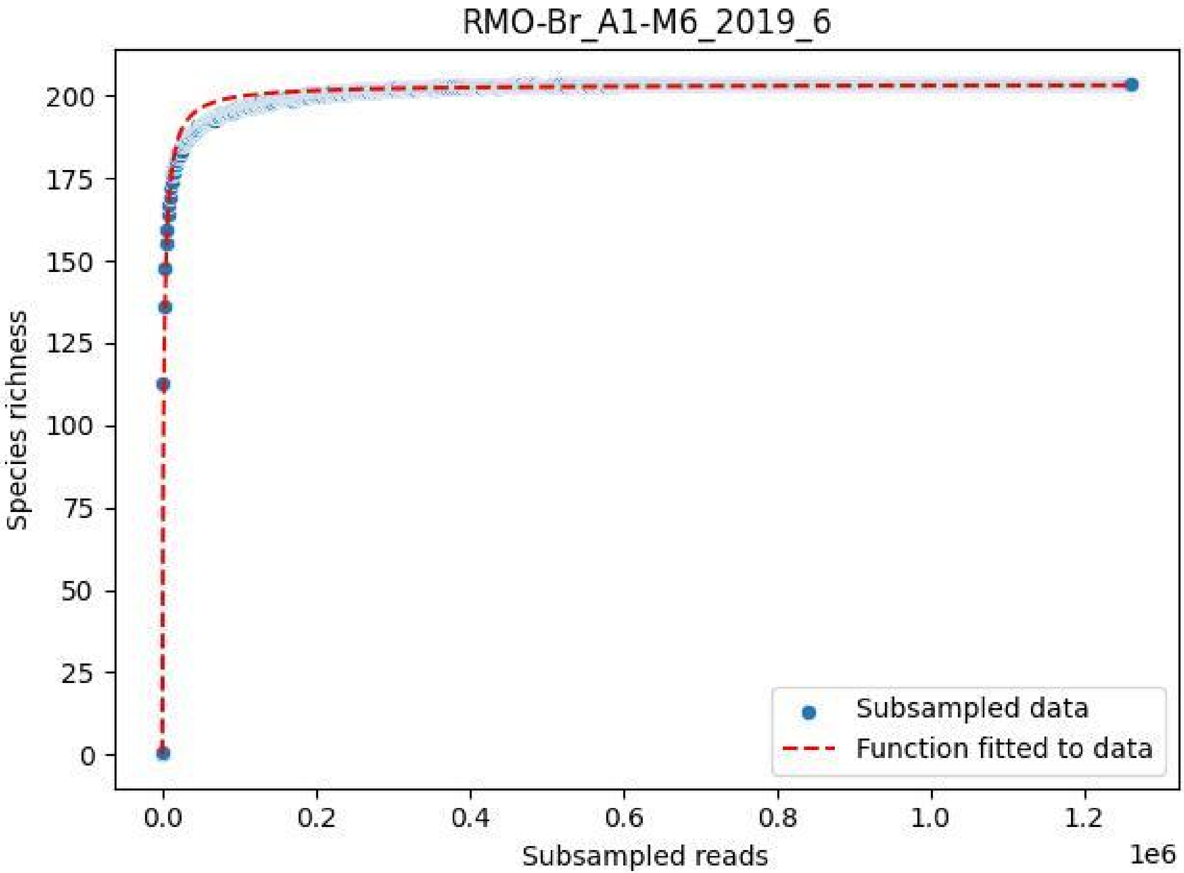
RMO-Br_A1-M6_2019_6
Species richness
Subsampled reads
1e6
0
25
50
75
100
125
150
175
200
0.0
0.2
0.4
0.6
0.8
1.0
1.2
Subsampled data
Function fitted to data

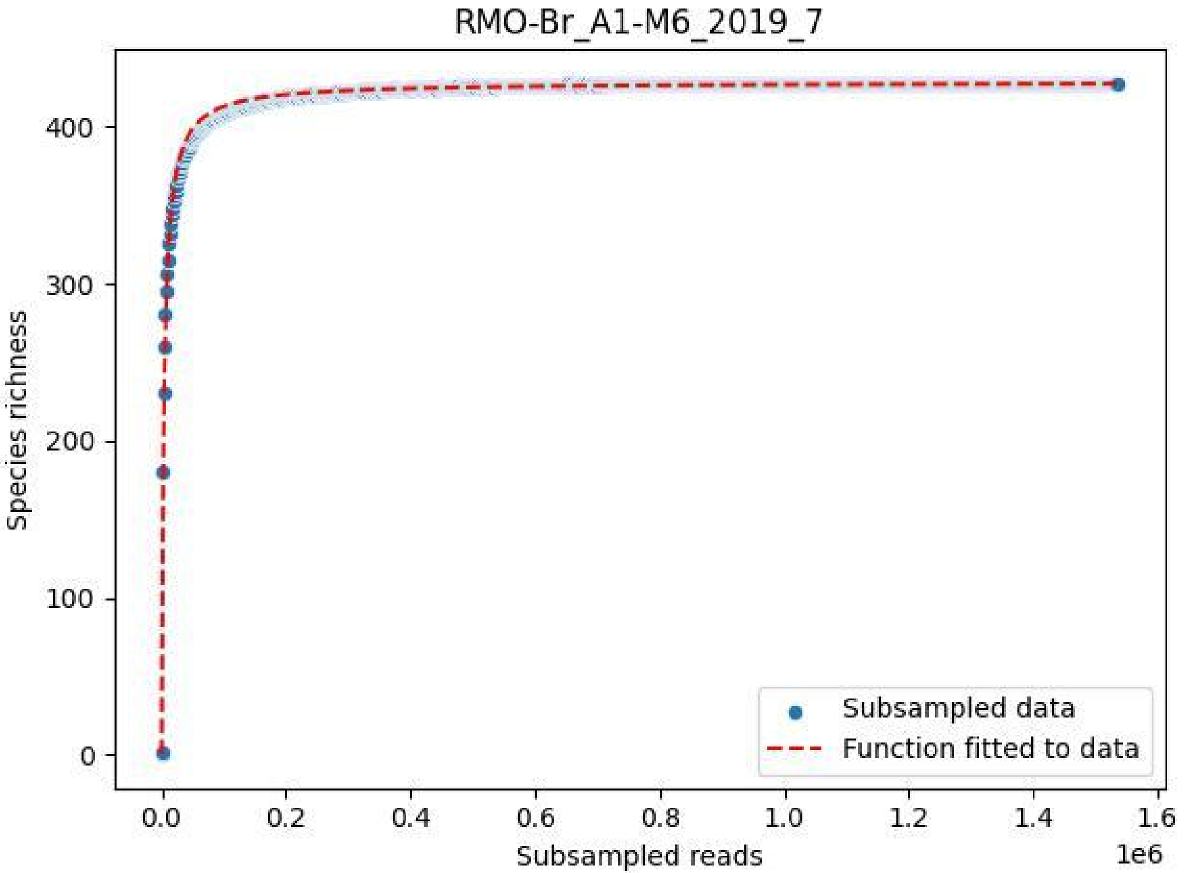

RMO-Br_A1-M6_2019_7
Species richness
Subsampled reads
1e6
0
100
200
300
400
0.0
0.2
0.4
0.6
0.8
1.0
1.2
1.4
1.6
Subsampled data
Function fitted to data

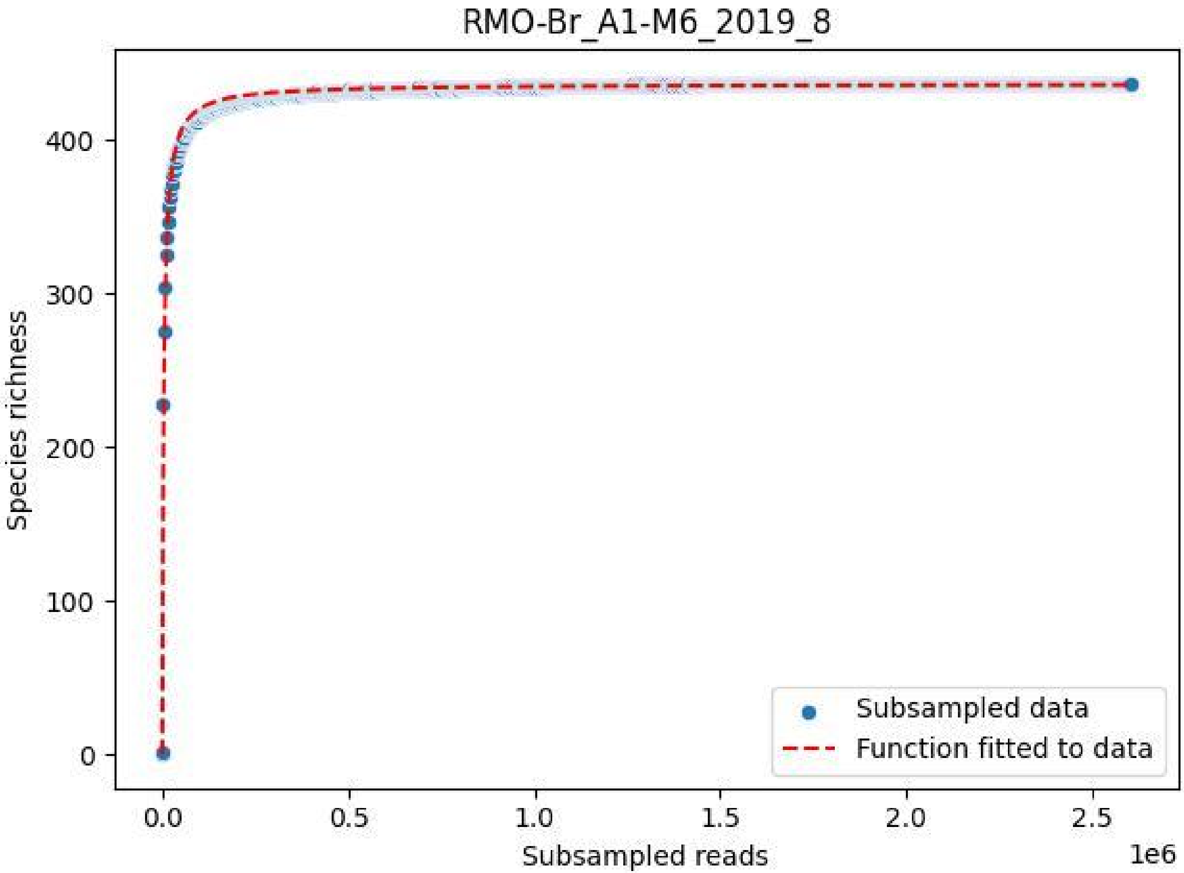

RMO-Br_A1-M6_2019_8
Species richness
Subsampled reads
1e6
0
100
200
300
400
0.0
0.5
1.0
1.5
2.0
2.5
Subsampled data
Function fitted to data

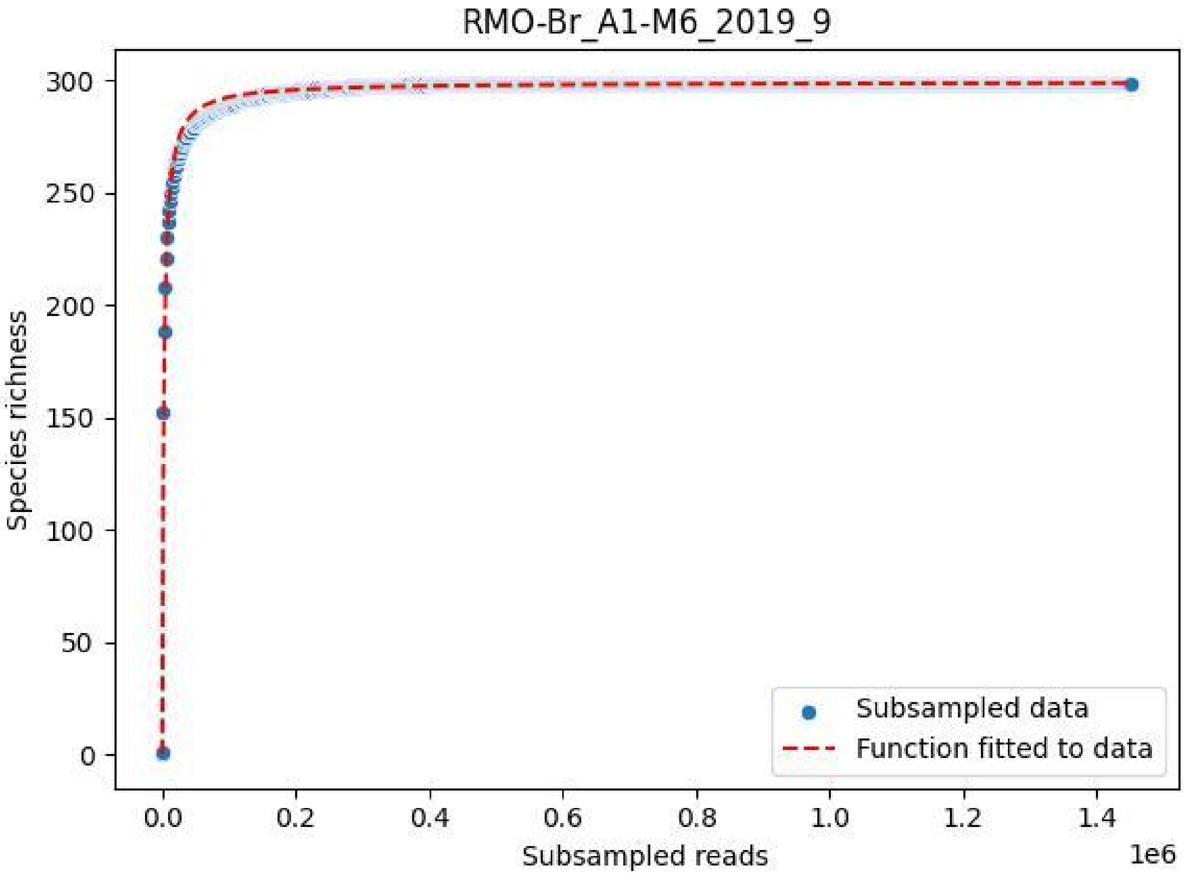

RMO-Br_A1-M6_2019_9
Species richness
Subsampled reads
1e6
0
50
100
150
200
250
300
0.0
0.2
0.4
0.6
0.8
1.0
1.2
1.4
Subsampled data
Function fitted to data

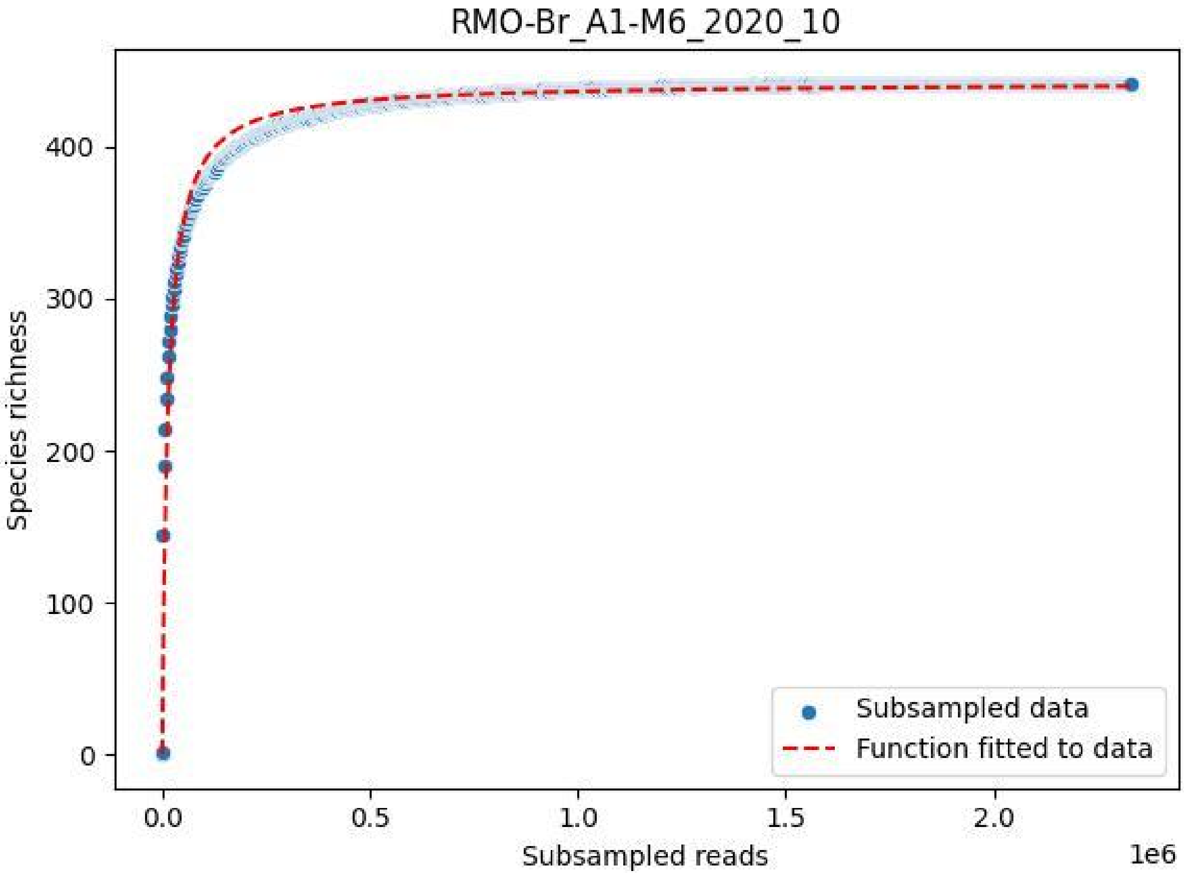

RMO-Br_A1-M6_2020_10
Species richness
0
100
200
300
400
0.0
0.5
1.0
1.5
2.0
1e6
Subsampled reads
Subsampled data
Function fitted to data

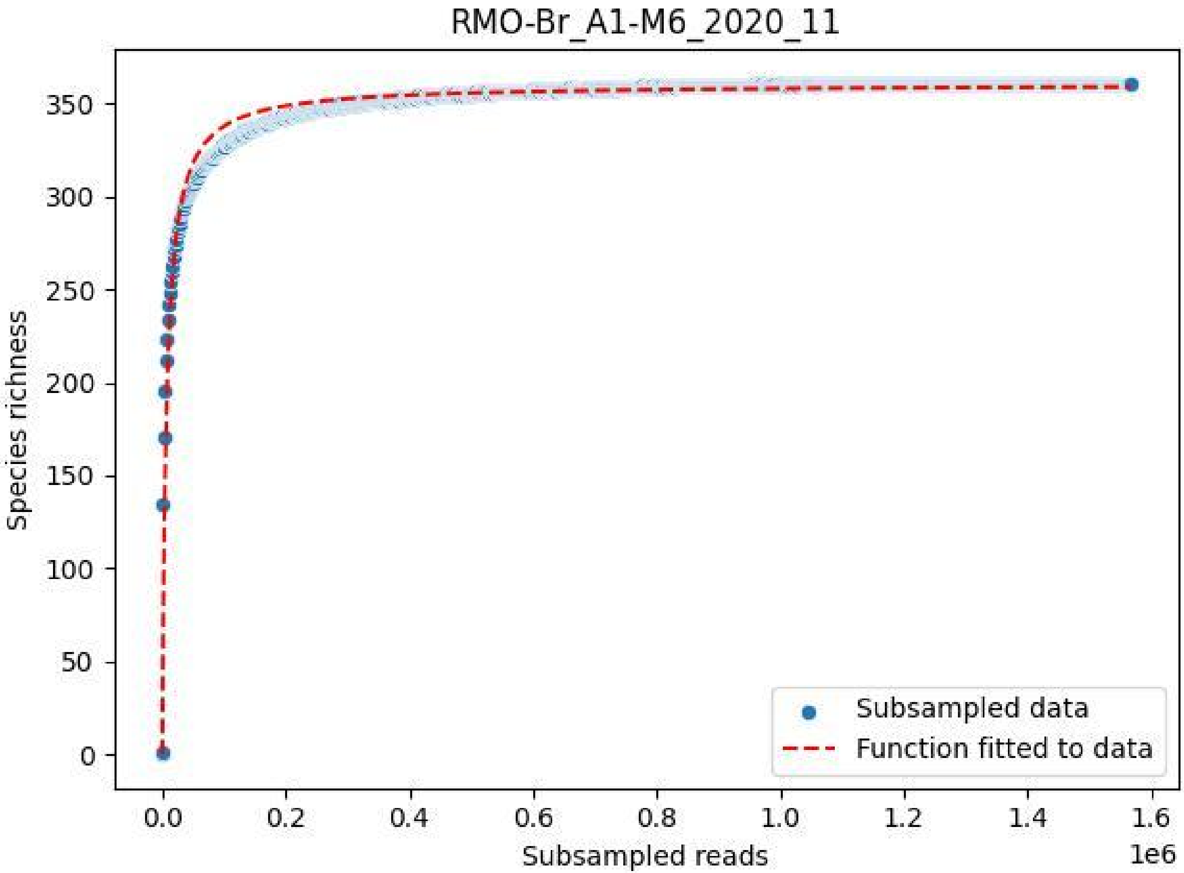

RMO-Br_A1-M6_2020_11
Species richness
Subsampled reads
1e6
0
50
100
150
200
250
300
350
0.0
0.2
0.4
0.6
0.8
1.0
1.2
1.4
1.6
Subsampled data
Function fitted to data

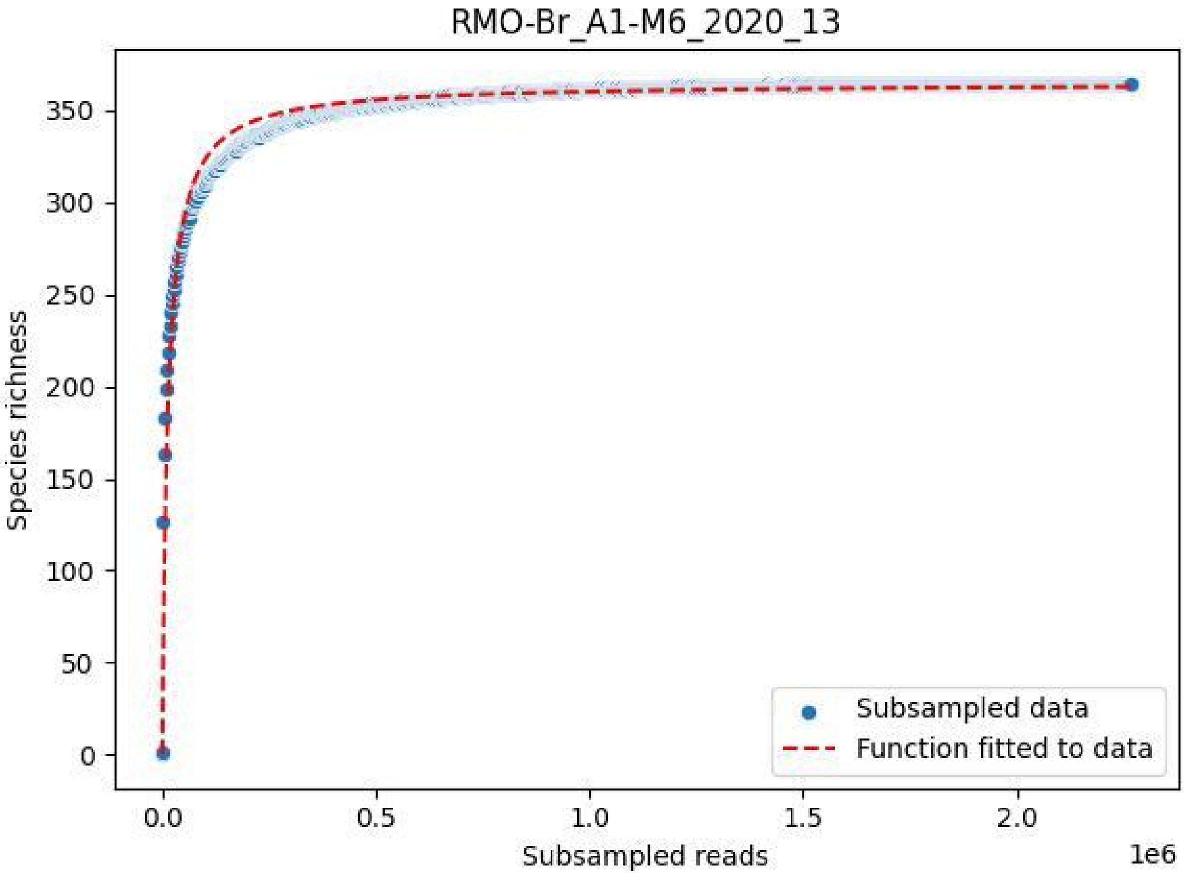

RMO-Br_A1-M6_2020_13
Species richness
Subsampled reads
1e6
0
50
100
150
200
250
300
350
0.0
0.5
1.0
1.5
2.0
Subsampled data
Function fitted to data

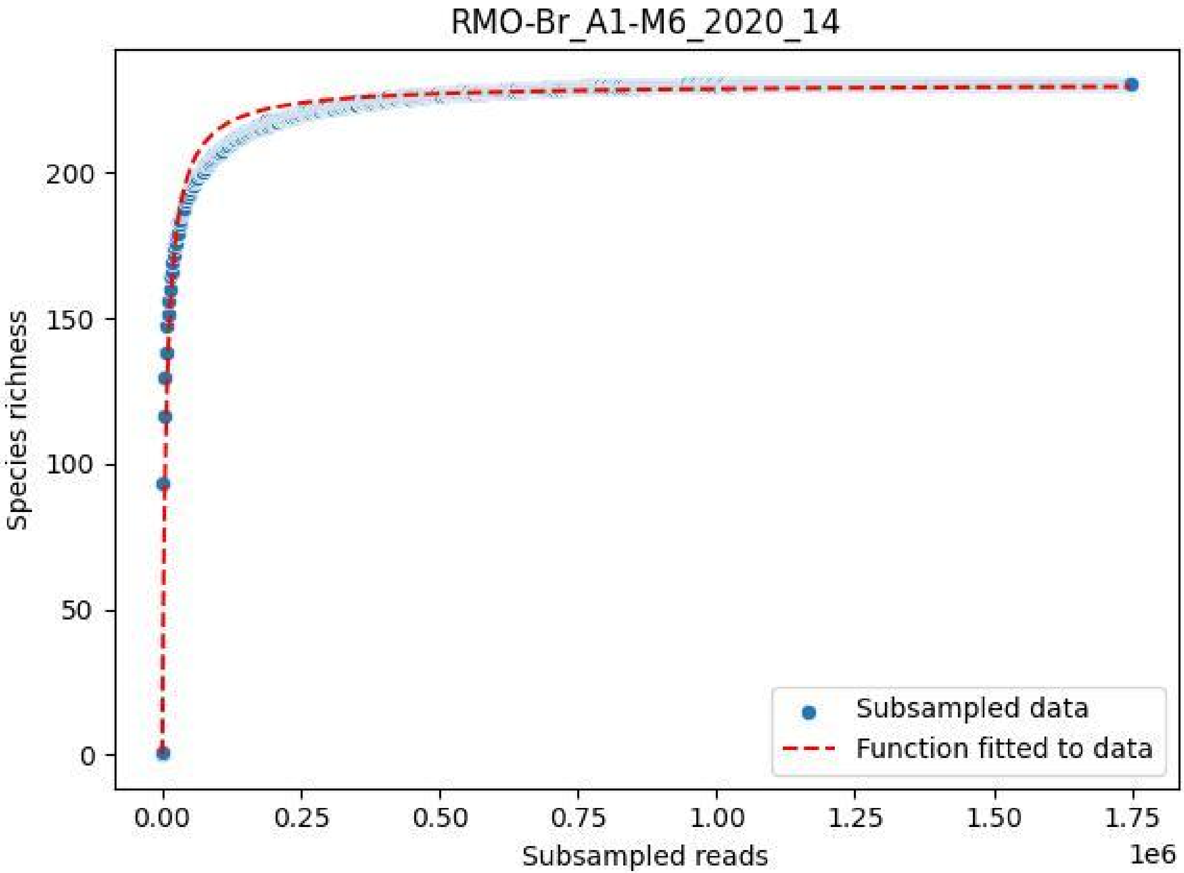

RMO-Br_A1-M6_2020_14
Species richness
Subsampled reads
1e6
0
50
100
150
200
0.00
0.25
0.50
0.75
1.00
1.25
1.50
1.75
Subsampled data
Function fitted to data

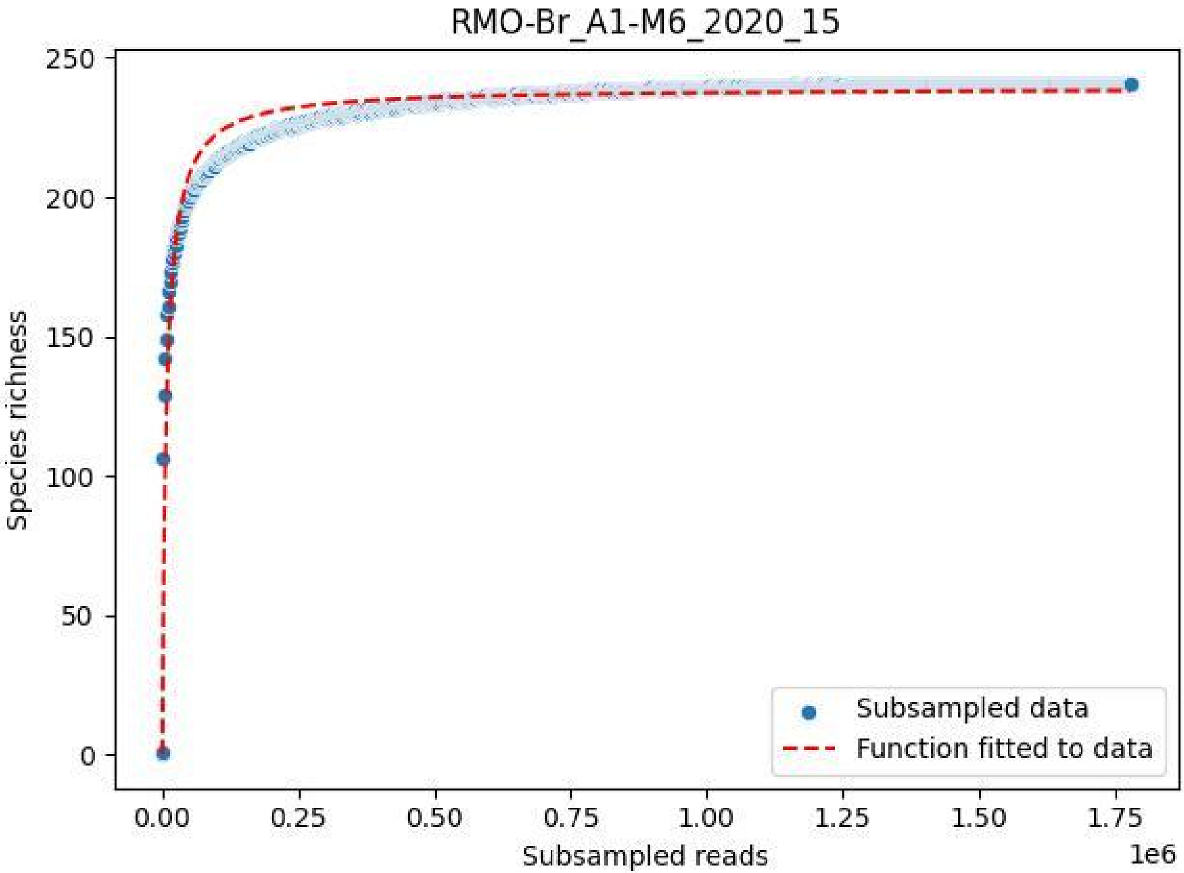
RMO-Br_A1-M6_2020_15
Species richness
Subsampled reads
1e6
250
200
150
100
50
0
0.00
0.25
0.50
0.75
1.00
1.25
1.50
1.75
Subsampled data
Function fitted to data

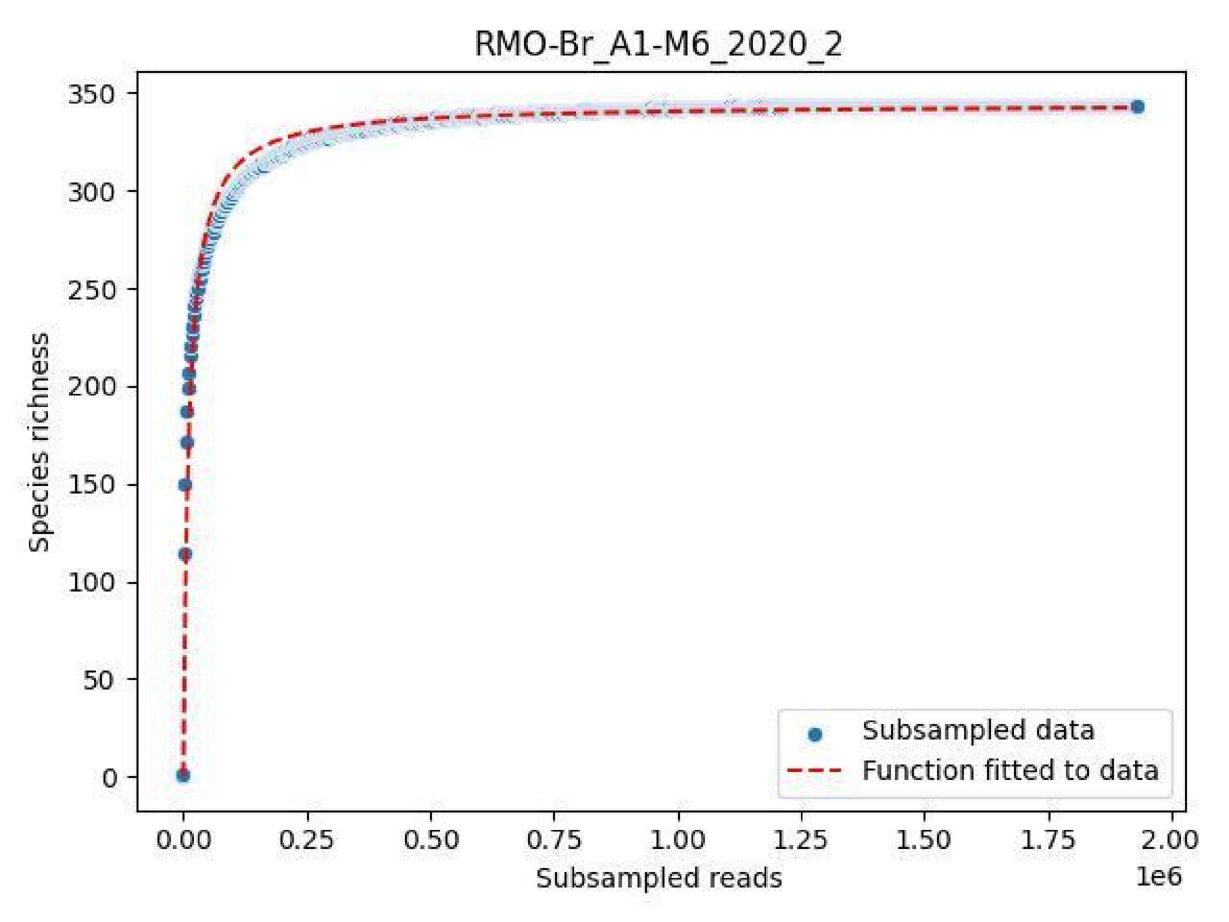
RMO-Br_A1-M6_2020_2
Species richness
Subsampled reads
1e6
0
50
100
150
200
250
300
350
0.00
0.25
0.50
0.75
1.00
1.25
1.50
1.75
2.00
Subsampled data
Function fitted to data

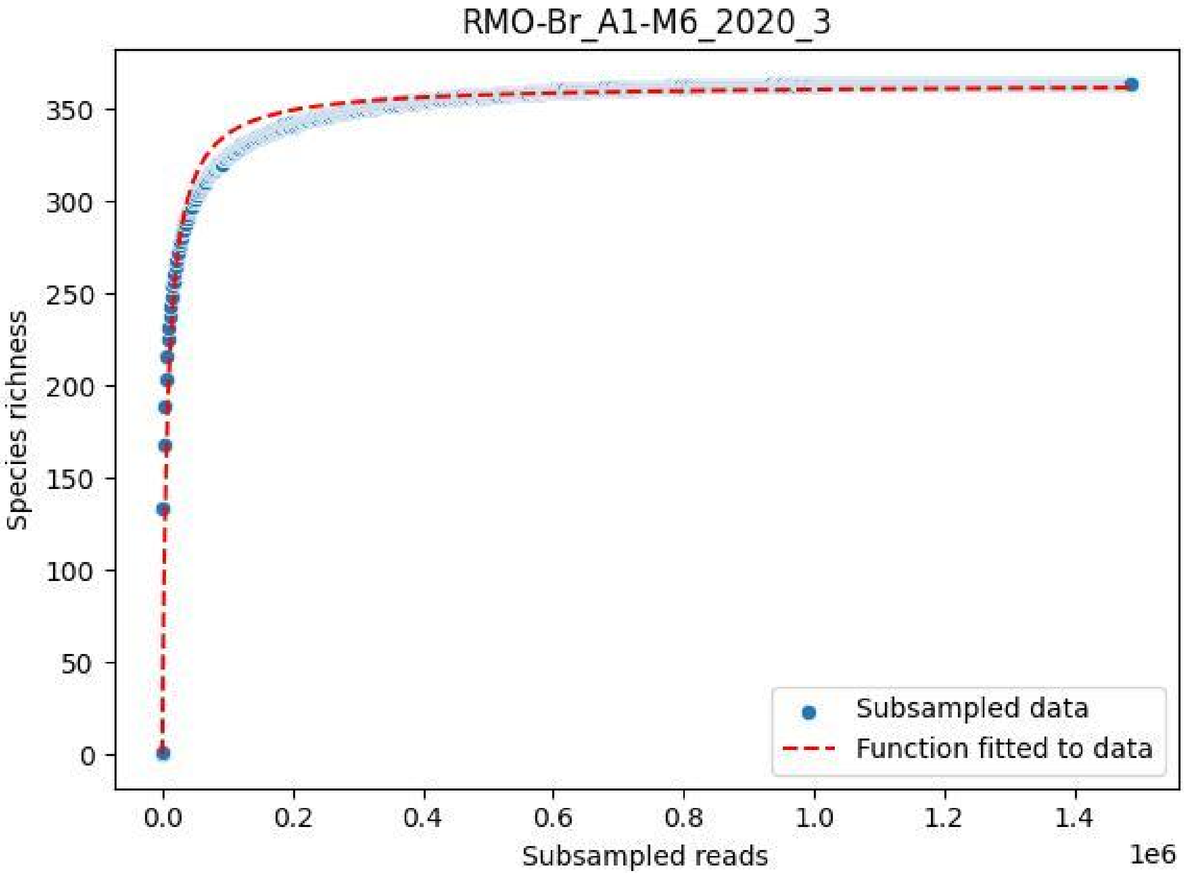

RMO-Br_A1-M6_2020_3
Species richness
Subsampled reads
1e6
0
50
100
150
200
250
300
350
0.0
0.2
0.4
0.6
0.8
1.0
1.2
1.4
Subsampled data
Function fitted to data

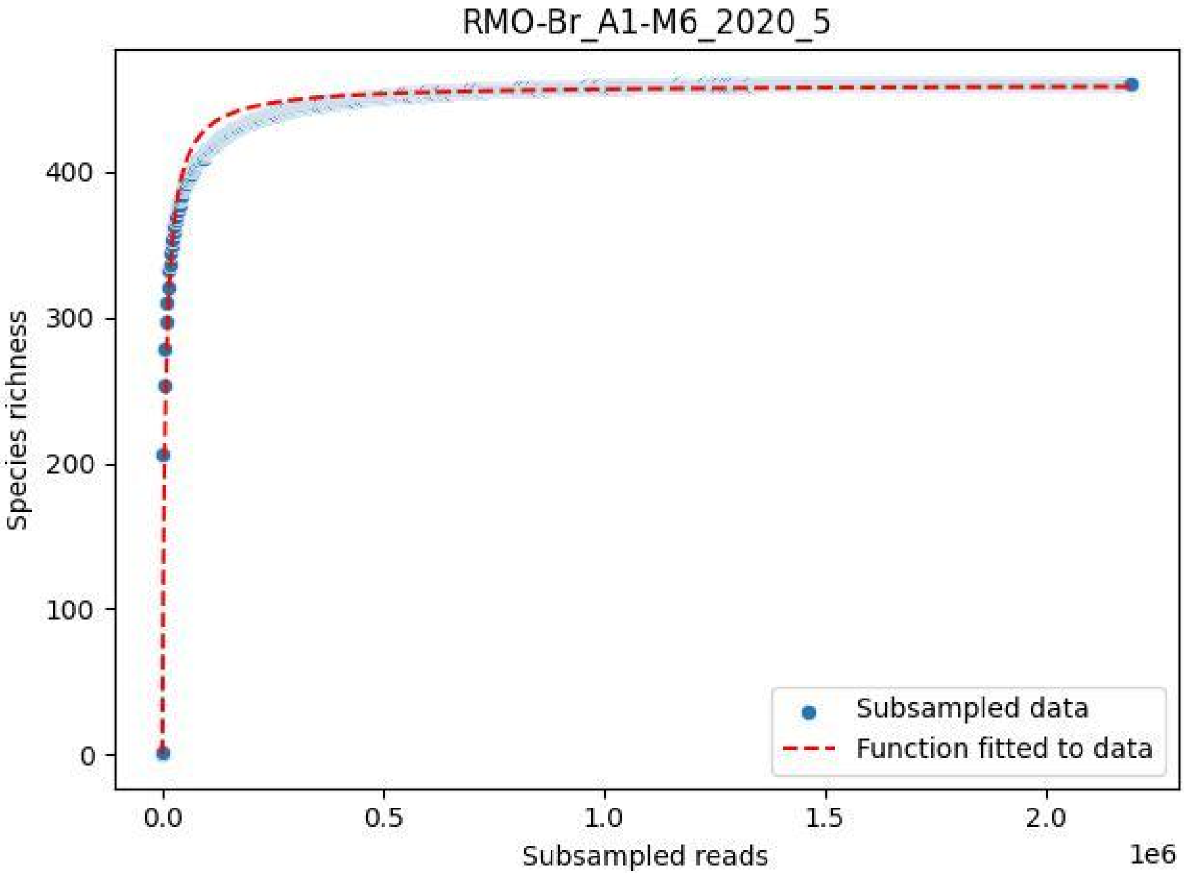
RMO-Br_A1-M6_2020_5
Species richness
Subsampled reads
1e6
0
100
200
300
400
0.0
0.5
1.0
1.5
2.0
Subsampled data
Function fitted to data

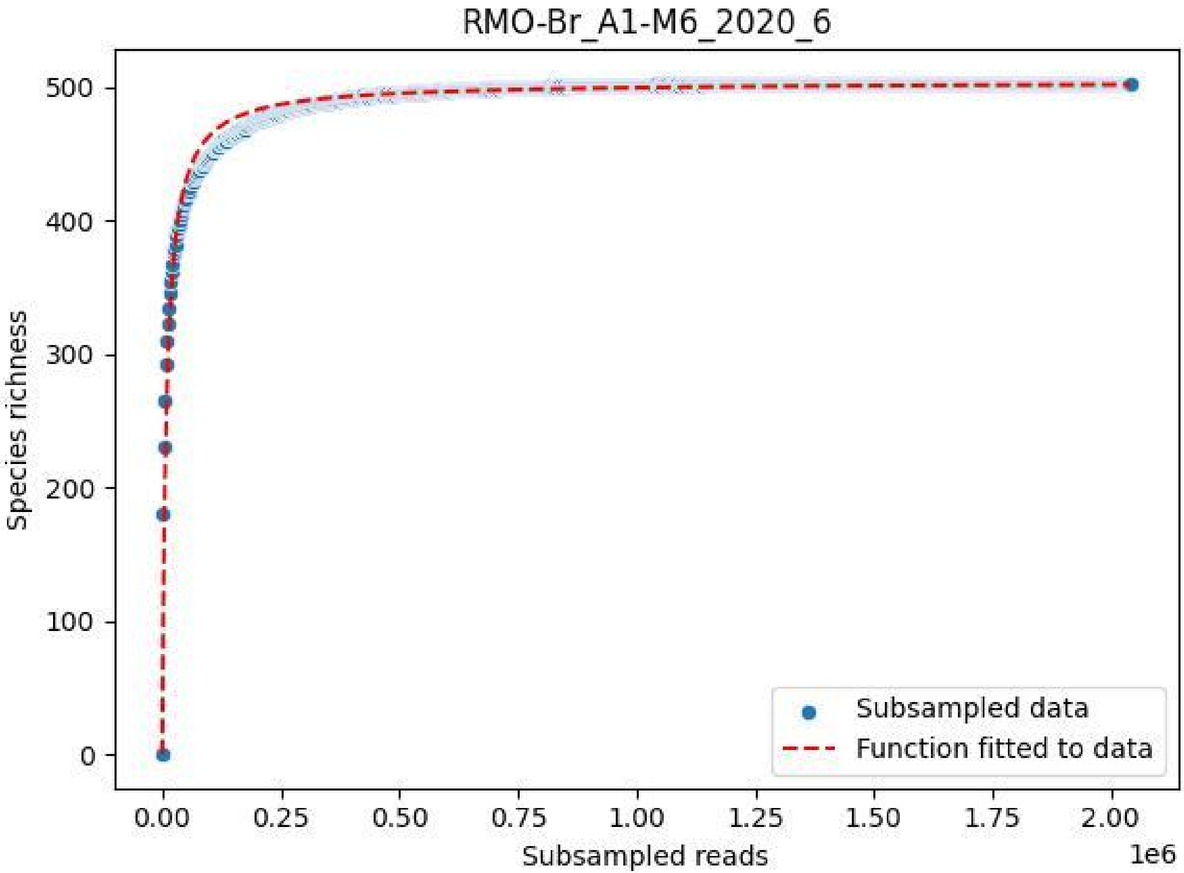
RMO-Br_A1-M6_2020_6
Species richness
Subsampled reads
1e6
0
100
200
300
400
500
0.00
0.25
0.50
0.75
1.00
1.25
1.50
1.75
2.00
Subsampled data
Function fitted to data

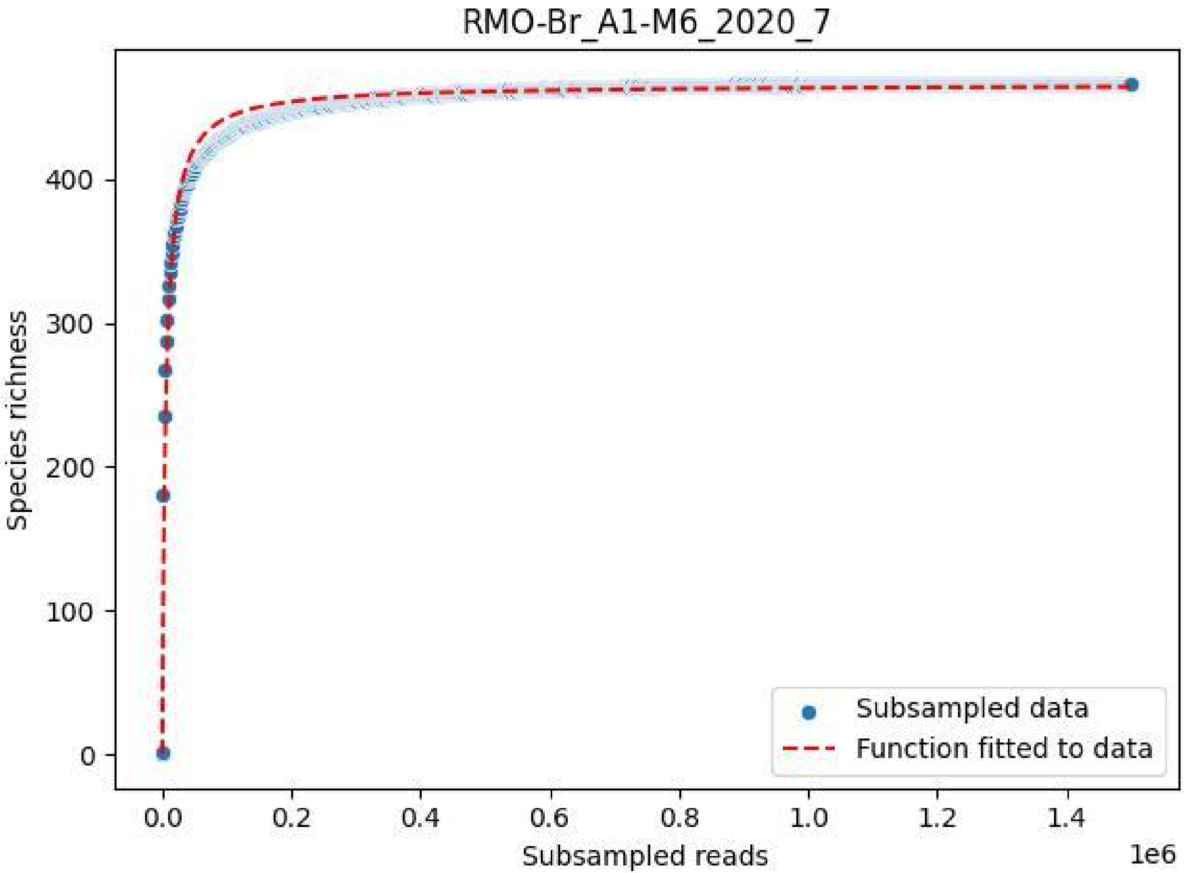

RMO-Br_A1-M6_2020_7
Species richness
400
300
200
100
0
0.0
0.2
0.4
0.6
0.8
1.0
1.2
1.4
Subsampled reads
1e6
Subsampled data
Function fitted to data

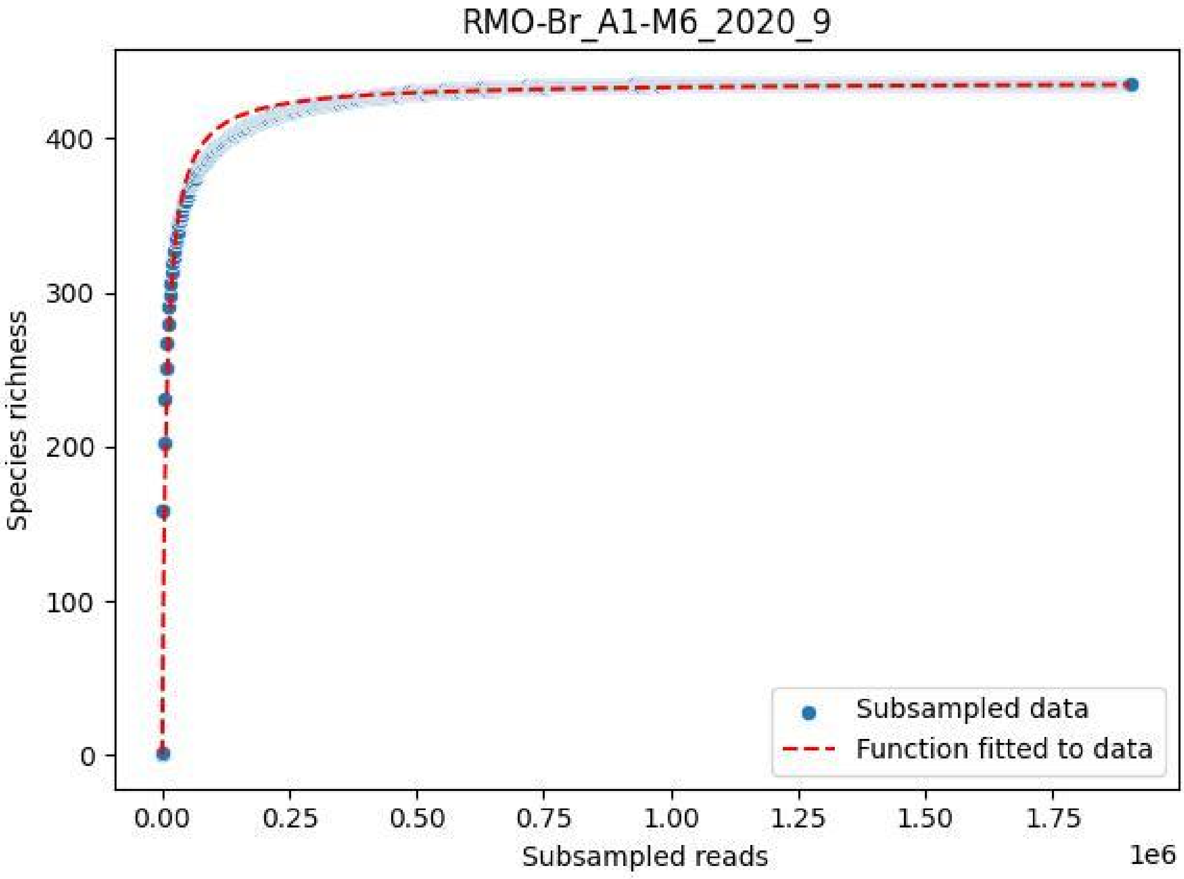
RMO-Br_A1-M6_2020_9
Species richness
Subsampled reads
1e6
0
100
200
300
400
0.00
0.25
0.50
0.75
1.00
1.25
1.50
1.75
Subsampled data
Function fitted to data

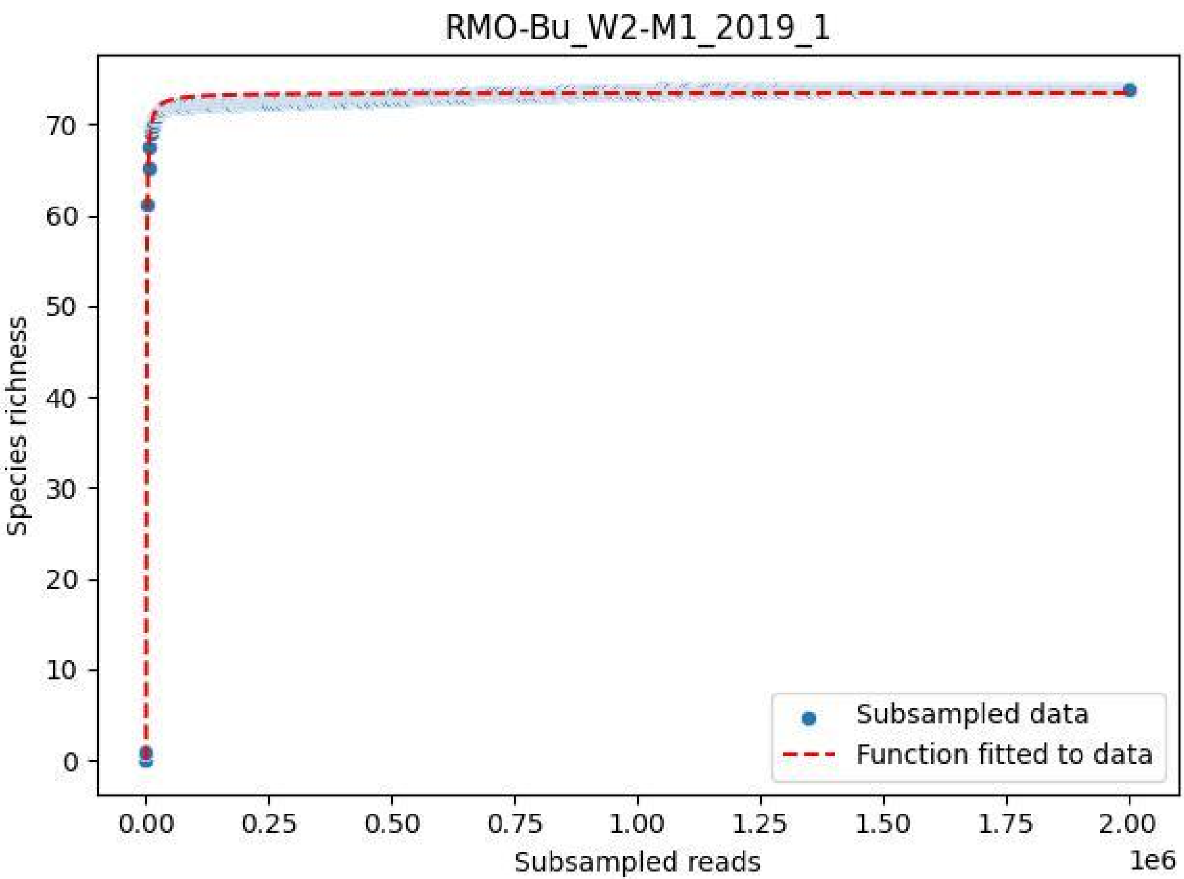
RMO-Bu_W2-M1_2019_1
Species richness
0
10
20
30
40
50
60
70
0.00
0.25
0.50
0.75
1.00
1.25
1.50
1.75
2.00
1e6
Subsampled reads
Subsampled data
Function fitted to data

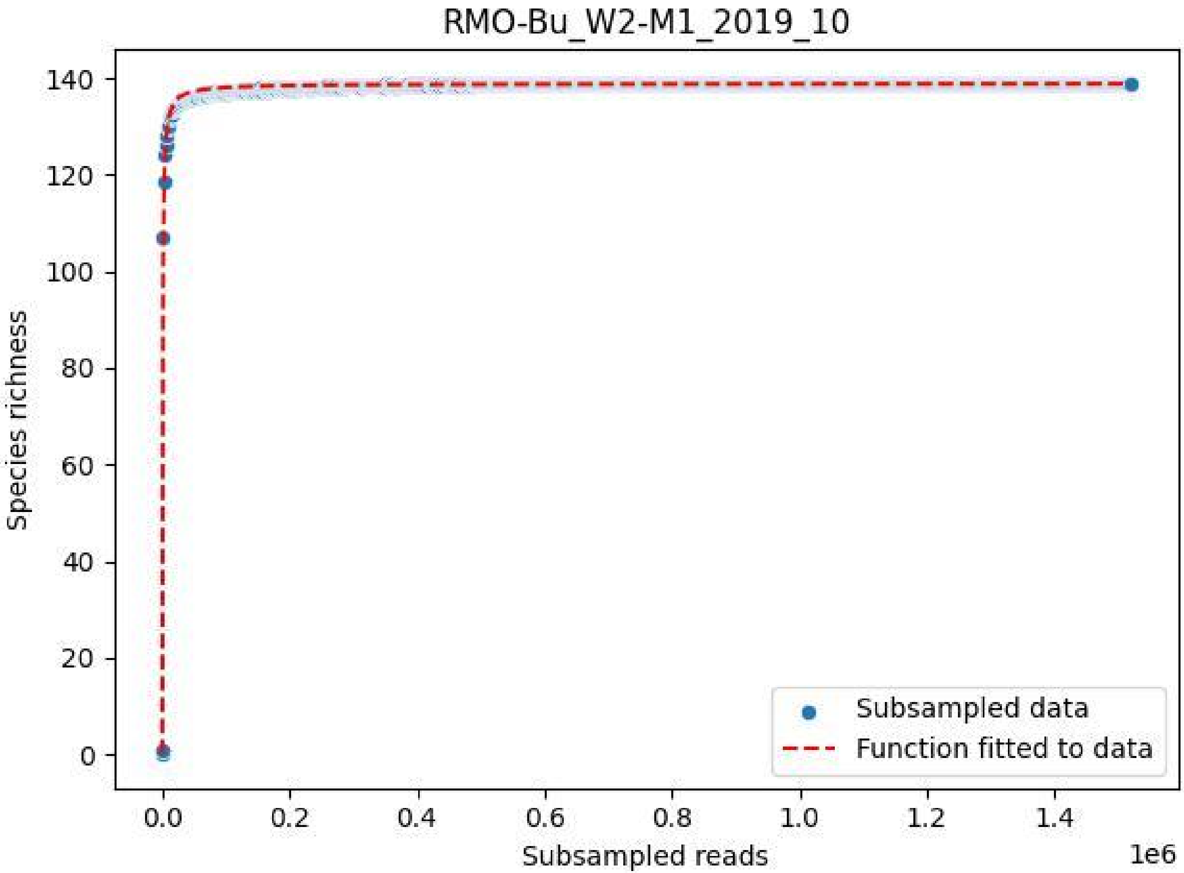

RMO-Bu_W2-M1_2019_10
Species richness
Subsampled reads
1e6
0
20
40
60
80
100
120
140
0.0
0.2
0.4
0.6
0.8
1.0
1.2
1.4
Subsampled data
Function fitted to data

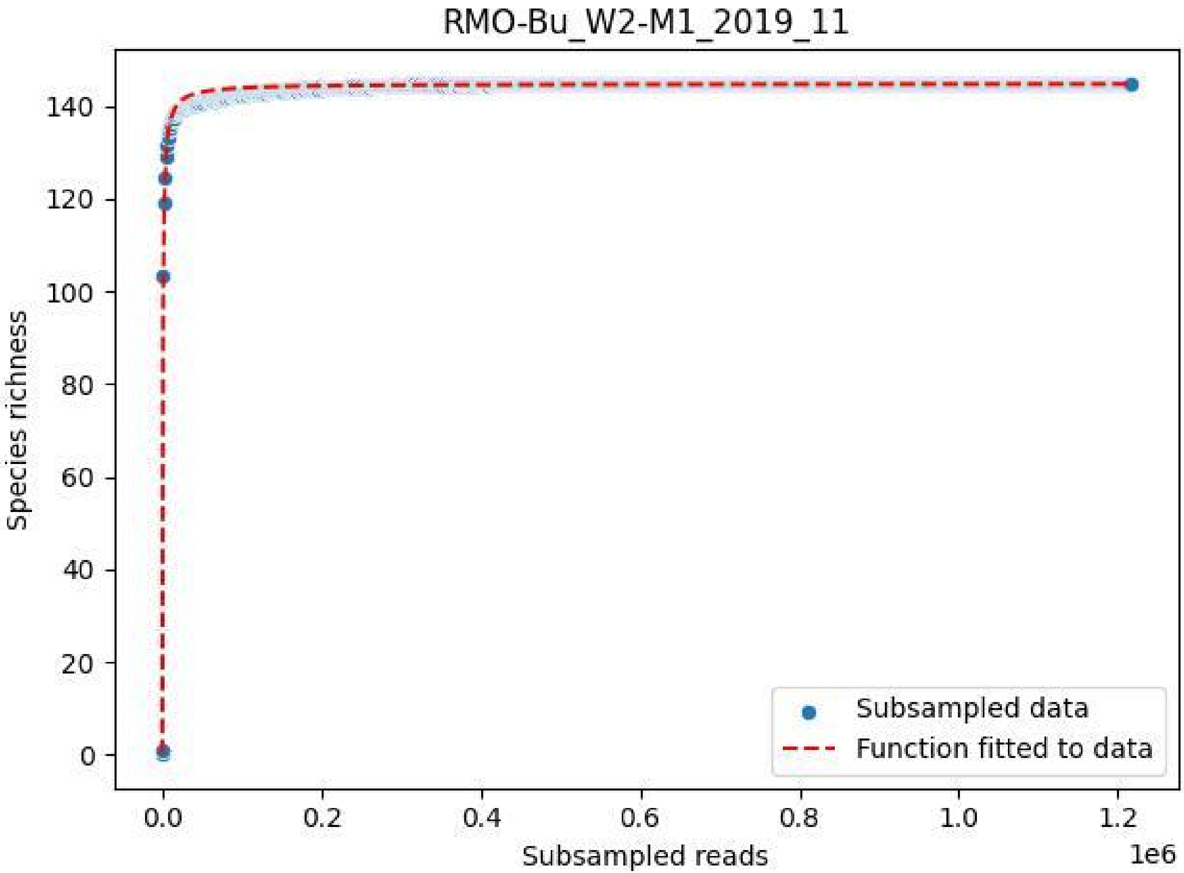
RMO-Bu_W2-M1_2019_11
Species richness
Subsampled reads
1e6
0
20
40
60
80
100
120
140
0.0
0.2
0.4
0.6
0.8
1.0
1.2
Subsampled data
Function fitted to data

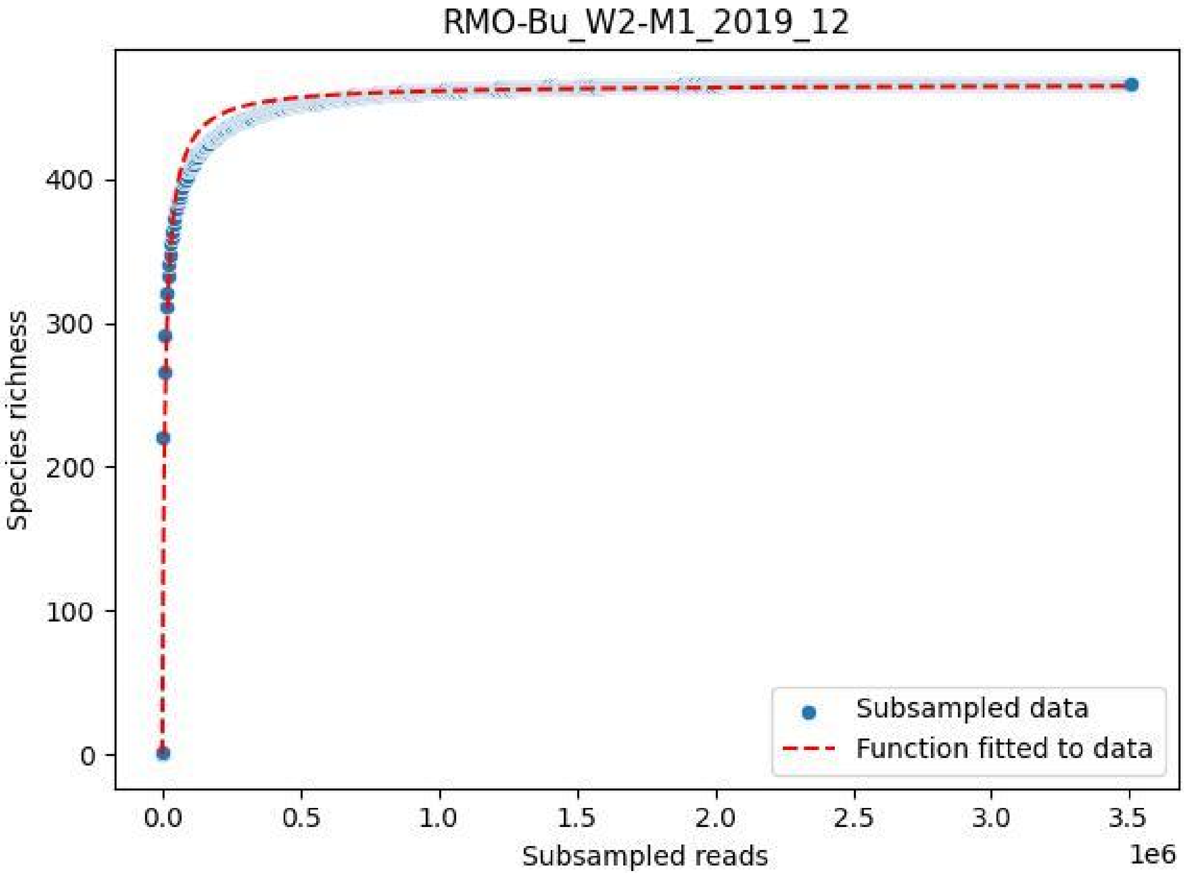

RMO-Bu_W2-M1_2019_12
Species richness
Subsampled reads
1e6
0
100
200
300
400
0.0
0.5
1.0
1.5
2.0
2.5
3.0
3.5
Subsampled data
Function fitted to data

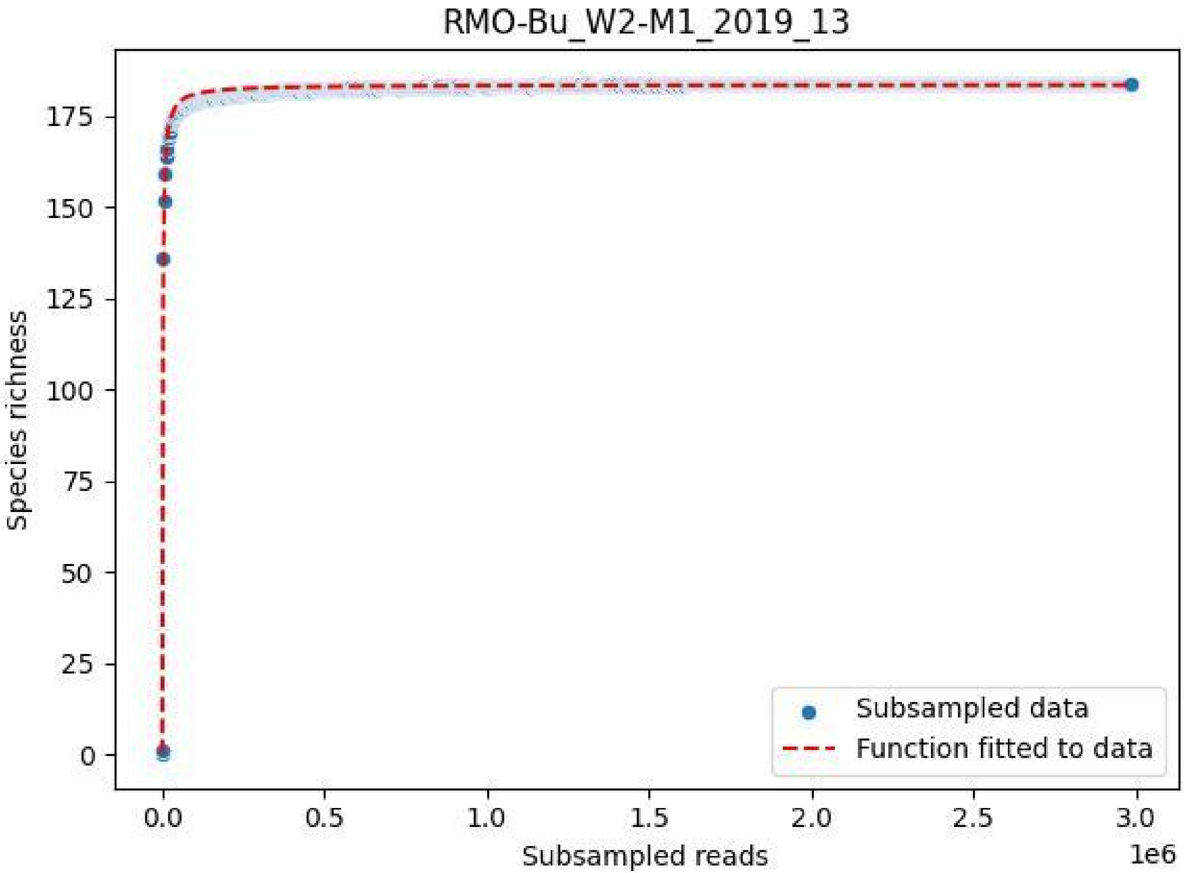
RMO-Bu_W2-M1_2019_13
Species richness
Subsampled reads
1e6
0
25
50
75
100
125
150
175
0.0
0.5
1.0
1.5
2.0
2.5
3.0
Subsampled data
Function fitted to data

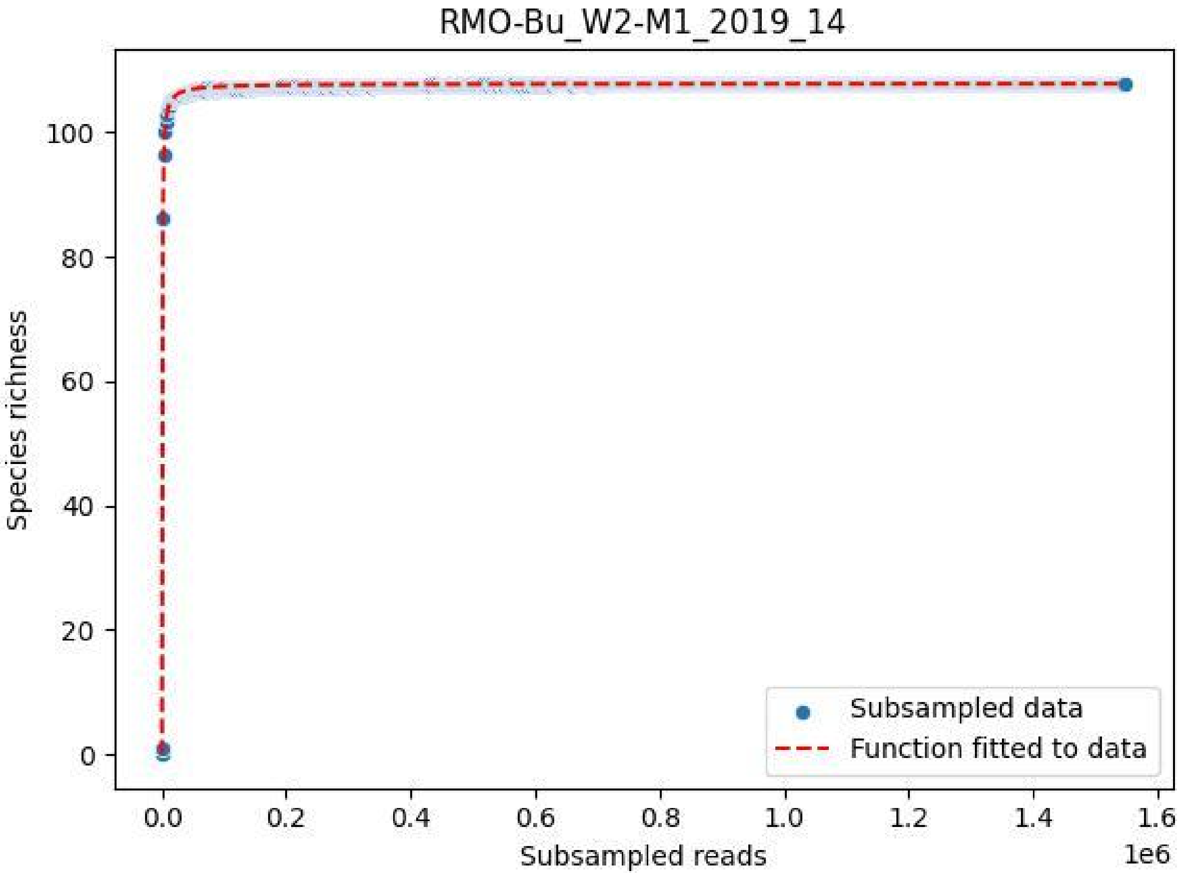
RMO-Bu_W2-M1_2019_14
Species richness
Subsampled reads
1e6
0
20
40
60
80
100
0.0
0.2
0.4
0.6
0.8
1.0
1.2
1.4
1.6
Subsampled data
Function fitted to data

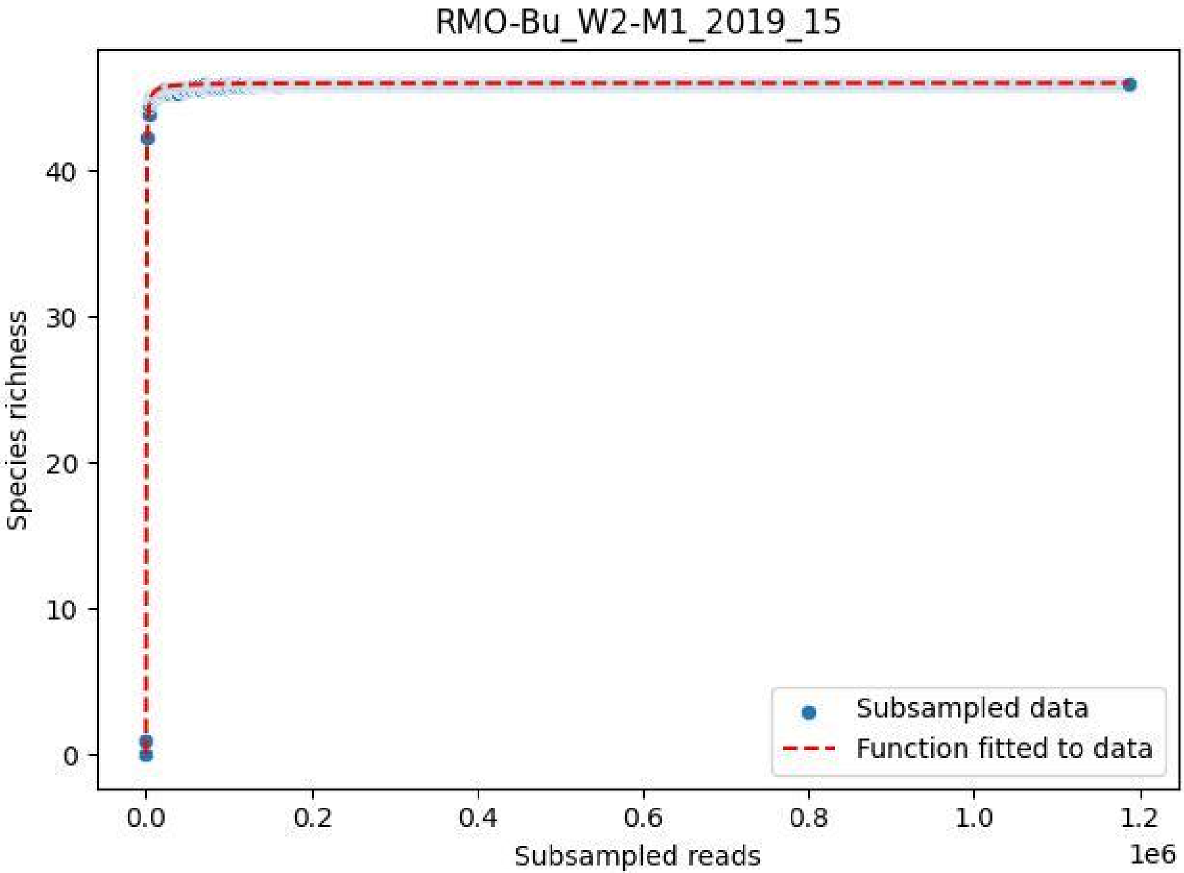

RMO-Bu_W2-M1_2019_15
Species richness
Subsampled reads
1e6
0
10
20
30
40
0.0
0.2
0.4
0.6
0.8
1.0
1.2
Subsampled data
Function fitted to data

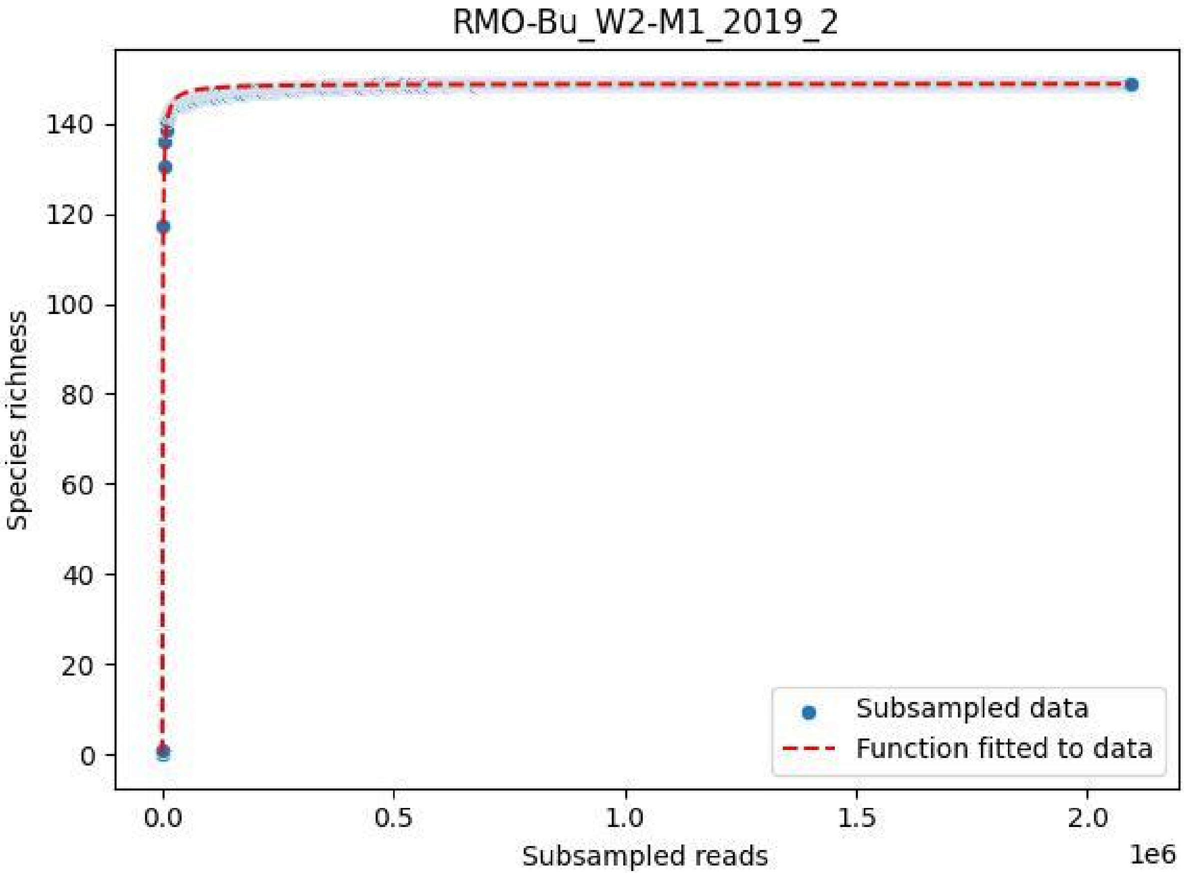
RMO-Bu_W2-M1_2019_2
Species richness
0
20
40
60
80
100
120
140
0.0
0.5
1.0
1.5
2.0
1e6
Subsampled reads
Subsampled data
Function fitted to data

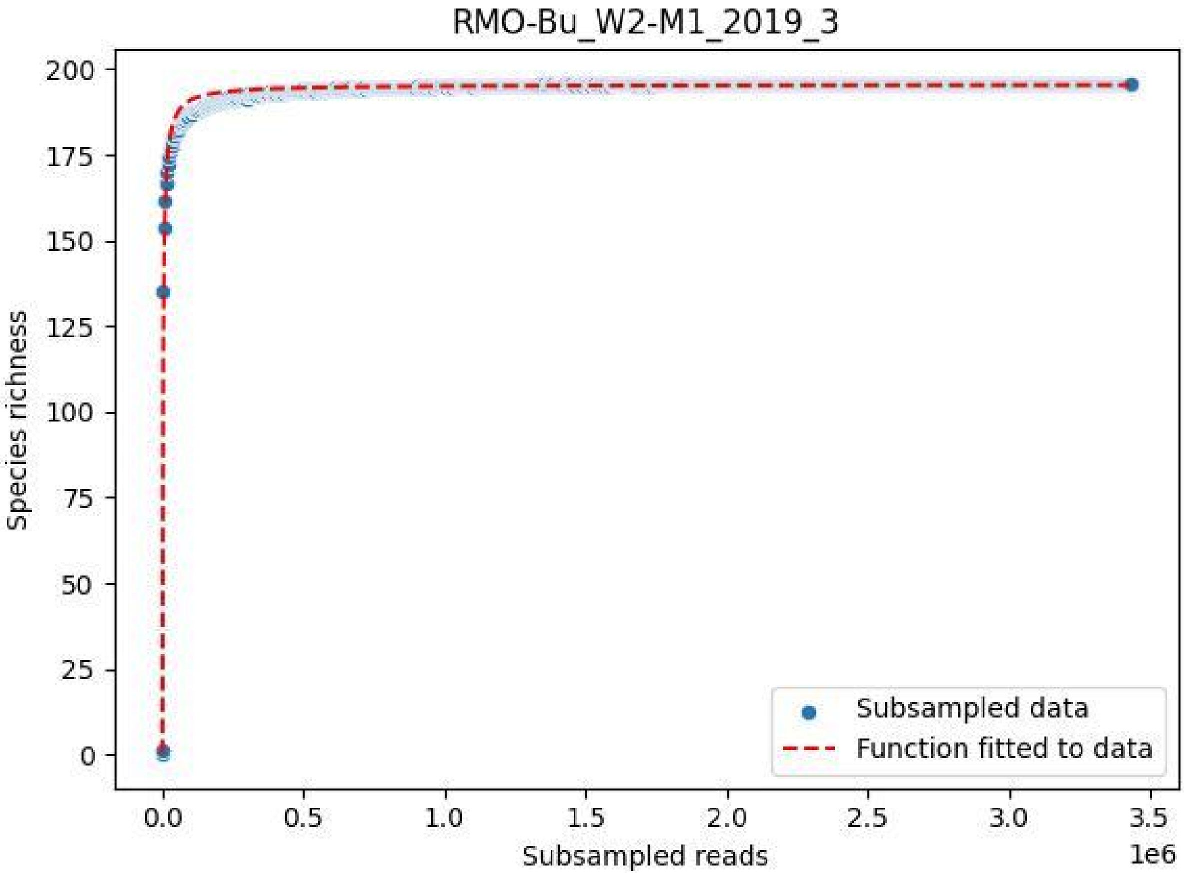

RMO-Bu_W2-M1_2019_3
Species richness
Subsampled reads
1e6
0
25
50
75
100
125
150
175
200
0.0
0.5
1.0
1.5
2.0
2.5
3.0
3.5
Subsampled data
Function fitted to data

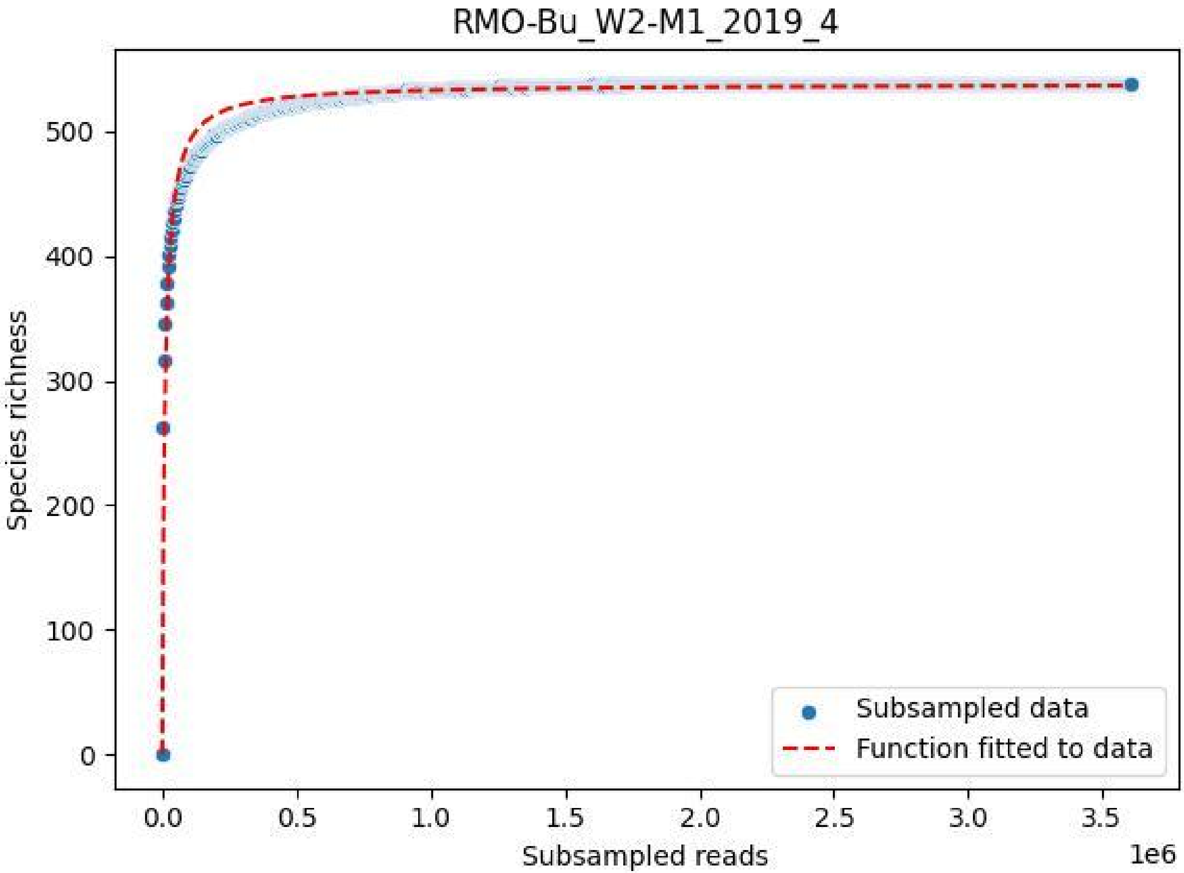

RMO-Bu_W2-M1_2019_4
Species richness
0
100
200
300
400
500
0.0
0.5
1.0
1.5
2.0
2.5
3.0
3.5
1e6
Subsampled reads
Subsampled data
Function fitted to data

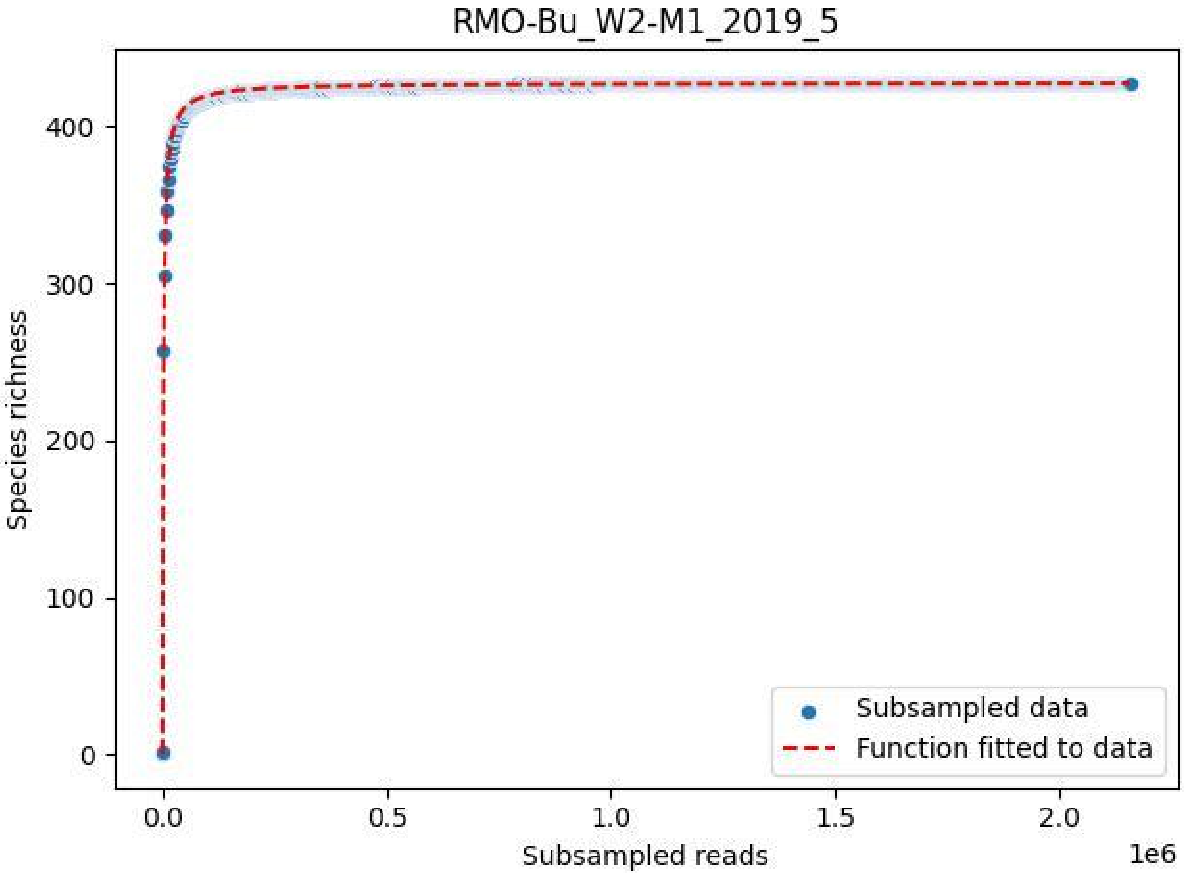

RMO-Bu_W2-M1_2019_5
Species richness
0
100
200
300
400
0.0
0.5
1.0
1.5
2.0
1e6
Subsampled reads
Subsampled data
Function fitted to data

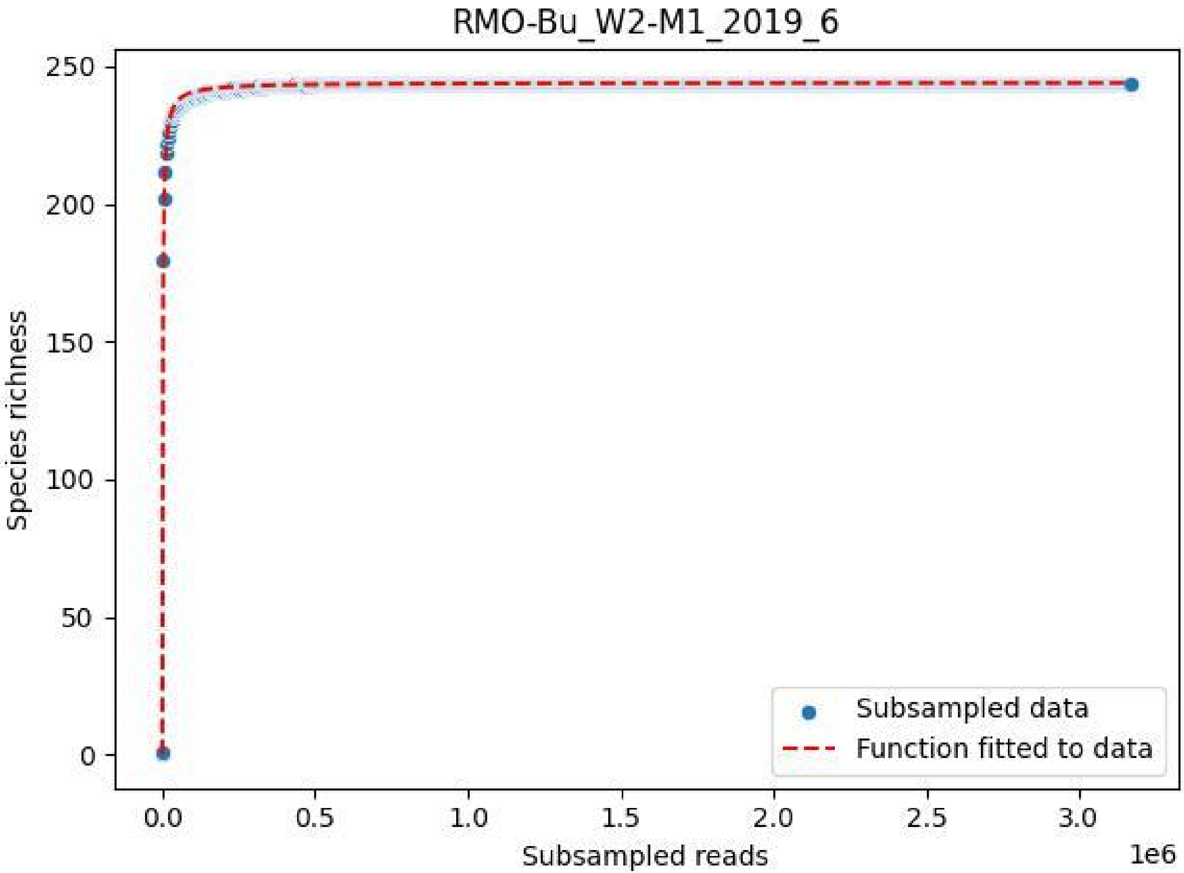

RMO-Bu_W2-M1_2019_6
Species richness
Subsampled reads
1e6
Subsampled data
Function fitted to data
0
50
100
150
200
250
0.0
0.5
1.0
1.5
2.0
2.5
3.0

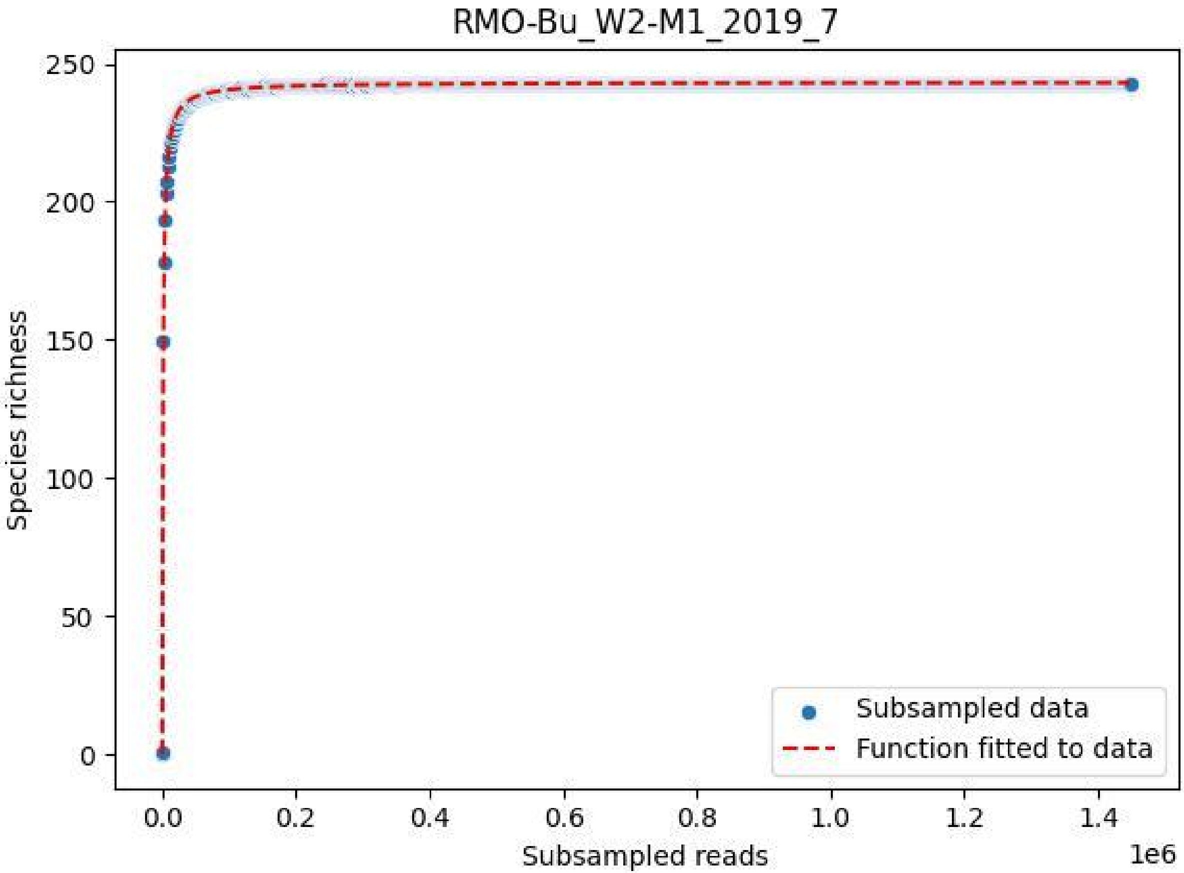
RMO-Bu_W2-M1_2019_7
Species richness
Subsampled reads
250
200
150
100
50
0
0.0
0.2
0.4
0.6
0.8
1.0
1.2
1.4
1e6
Subsampled data
Function fitted to data

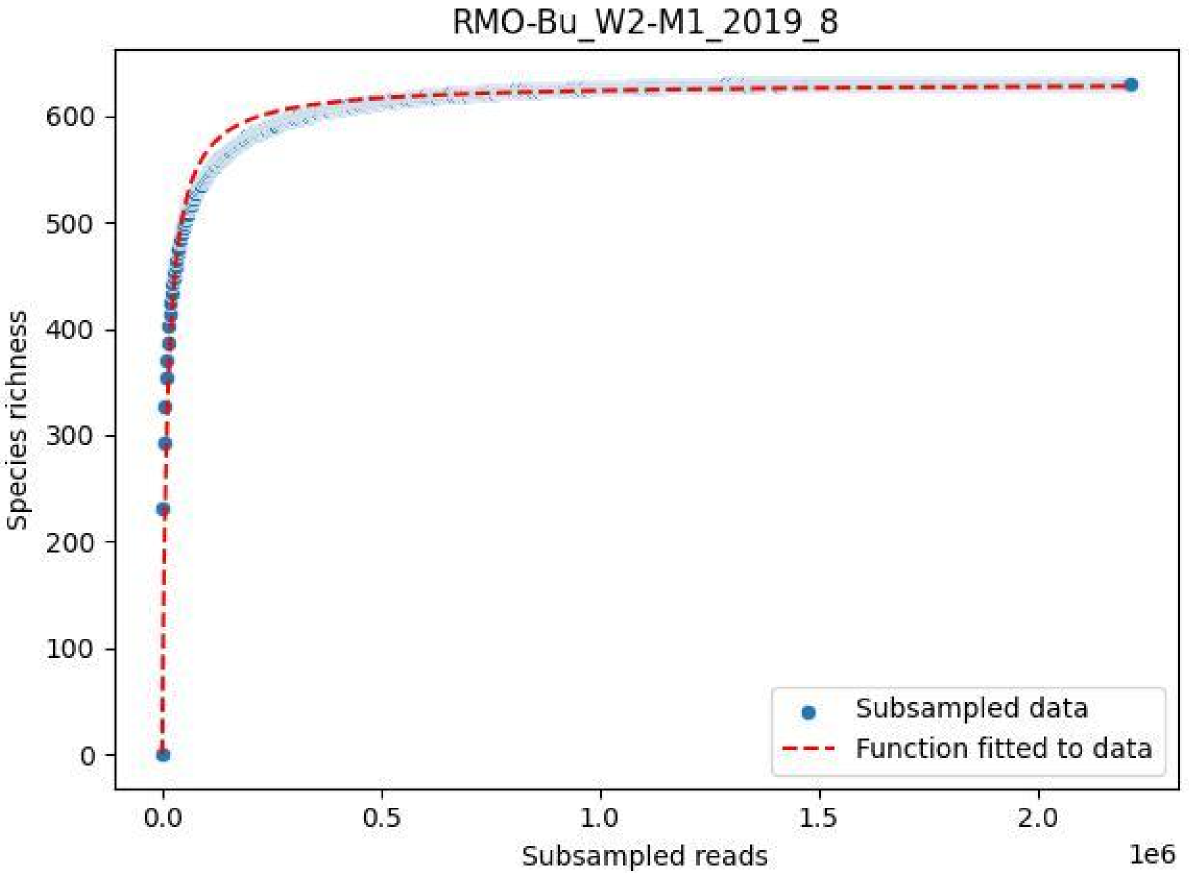
RMO-Bu_W2-M1_2019_8
Species richness
Subsampled reads
1e6
0
100
200
300
400
500
600
0.0
0.5
1.0
1.5
2.0
Subsampled data
Function fitted to data

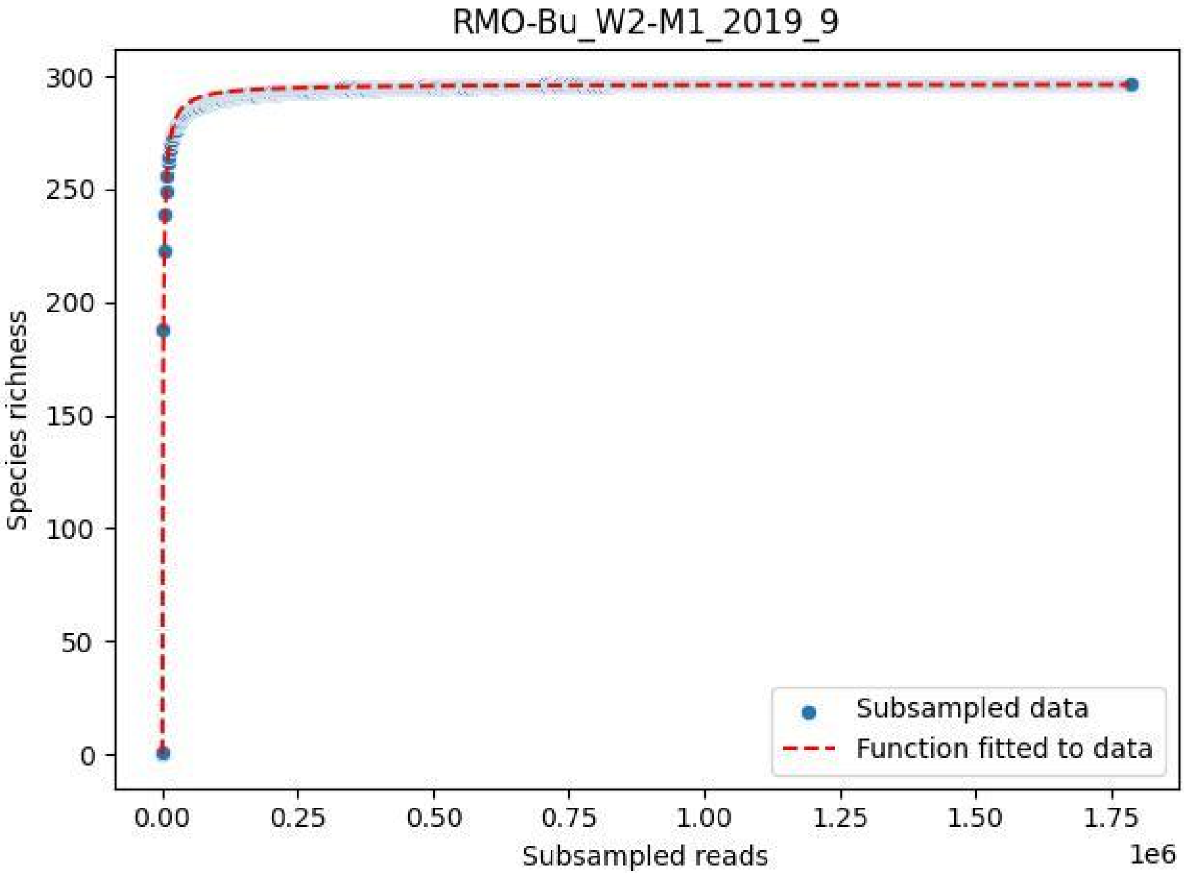
RMO-Bu_W2-M1_2019_9
Species richness
0
50
100
150
200
250
300
0.00
0.25
0.50
0.75
1.00
1.25
1.50
1.75
Subsampled reads
1e6
Subsampled data
Function fitted to data

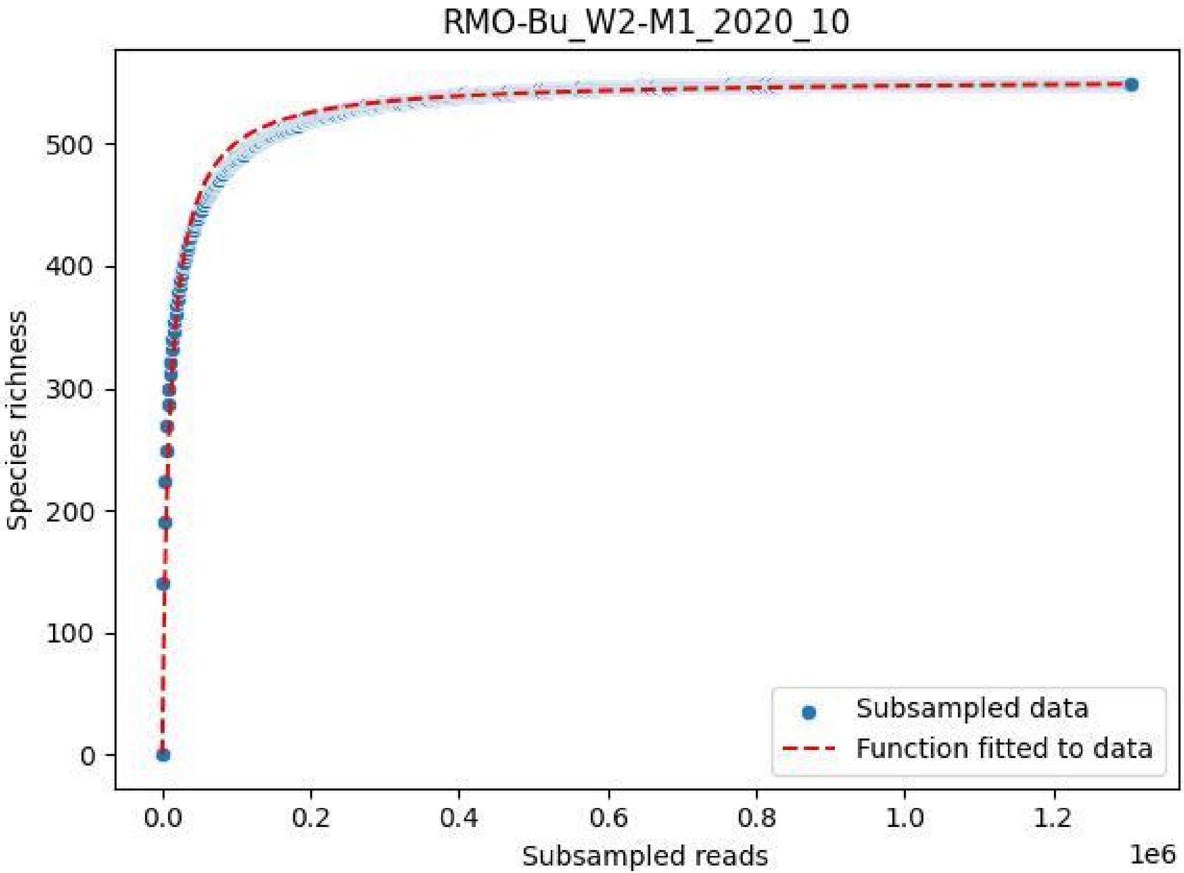
RMO-Bu_W2-M1_2020_10
Species richness
Subsampled reads
1e6
0
100
200
300
400
500
0.0
0.2
0.4
0.6
0.8
1.0
1.2
Subsampled data
Function fitted to data

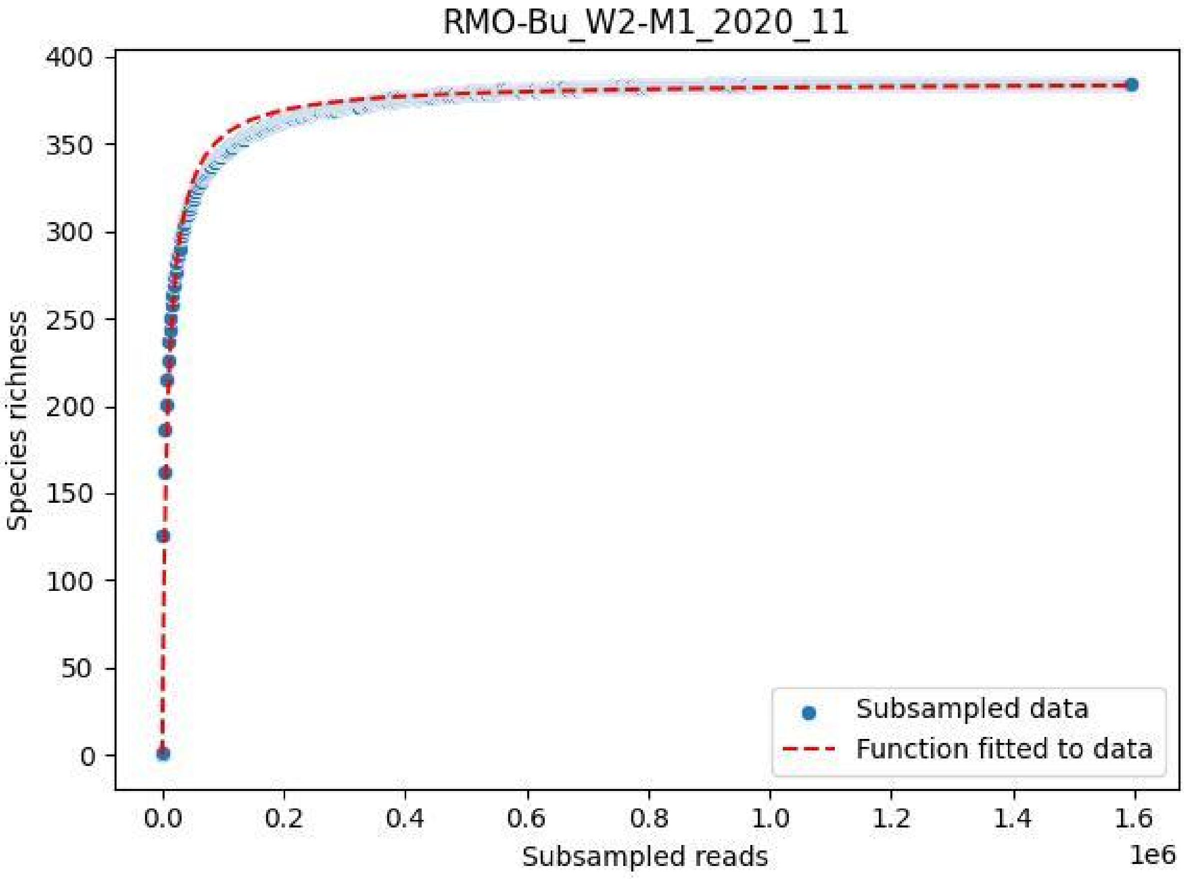

RMO-Bu_W2-M1_2020_11
Species richness
400
350
300
250
200
150
100
50
0
0.0
0.2
0.4
0.6
0.8
1.0
1.2
1.4
1.6
1e6
Subsampled reads
Subsampled data
Function fitted to data

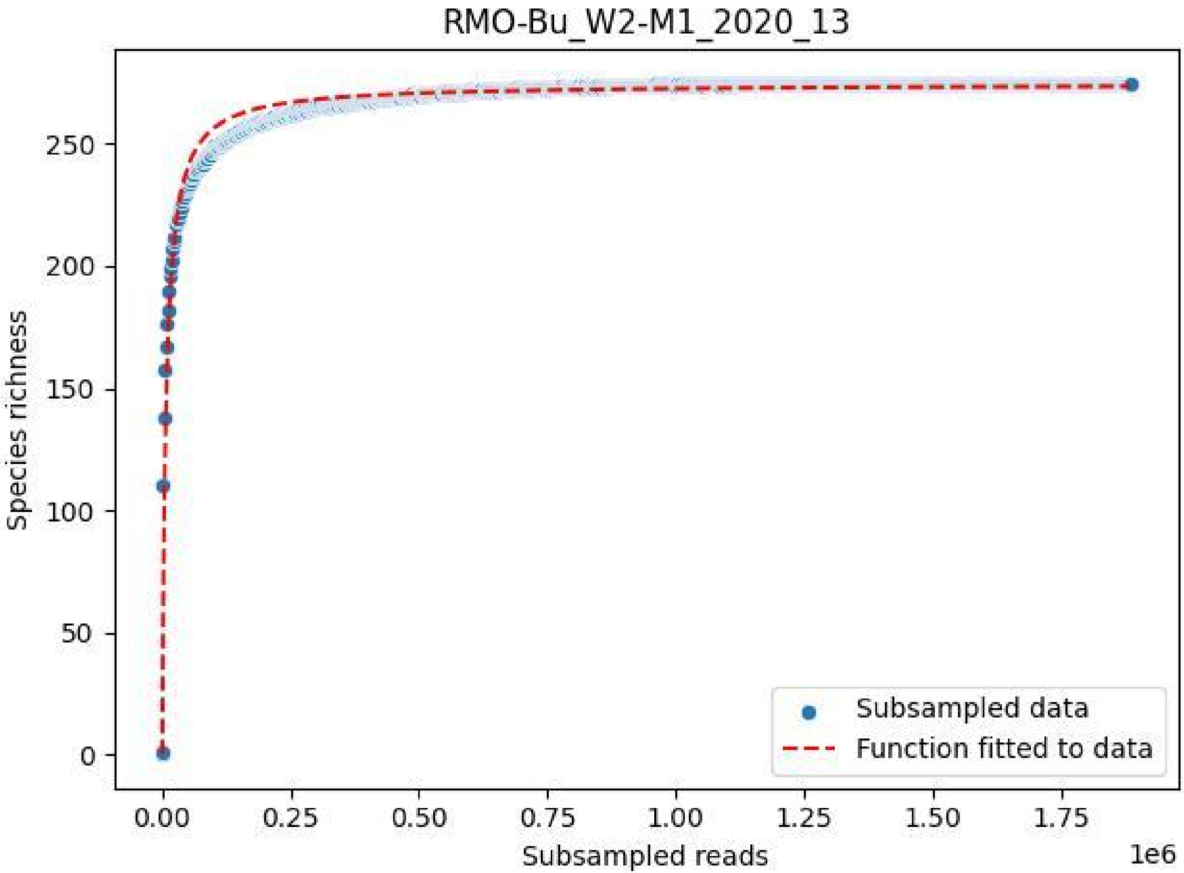

RMO-Bu_W2-M1_2020_13
Species richness
Subsampled reads
1e6
0
50
100
150
200
250
0.00
0.25
0.50
0.75
1.00
1.25
1.50
1.75
Subsampled data
Function fitted to data

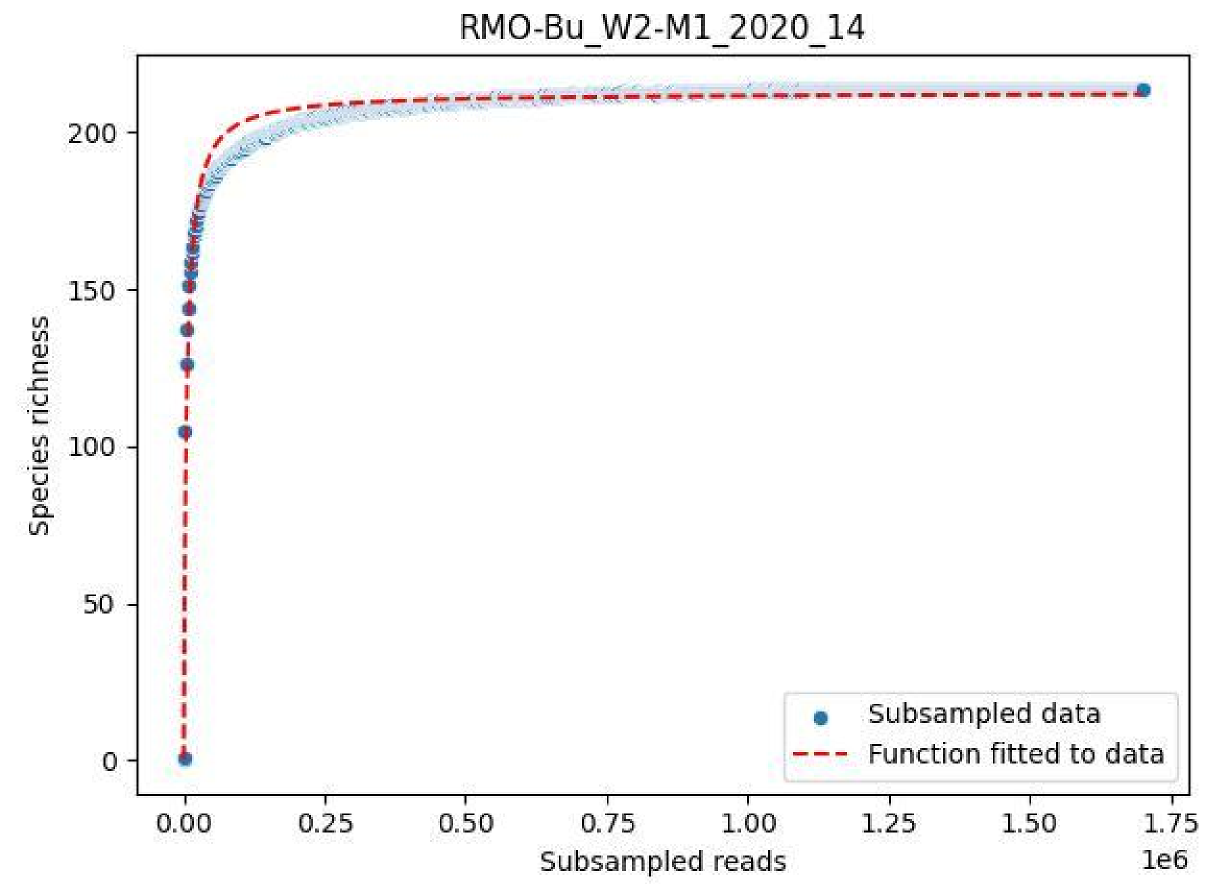
RMO-Bu_W2-M1_2020_14
Species richness
Subsampled reads
1e6
0
50
100
150
200
0.00
0.25
0.50
0.75
1.00
1.25
1.50
1.75
Subsampled data
Function fitted to data

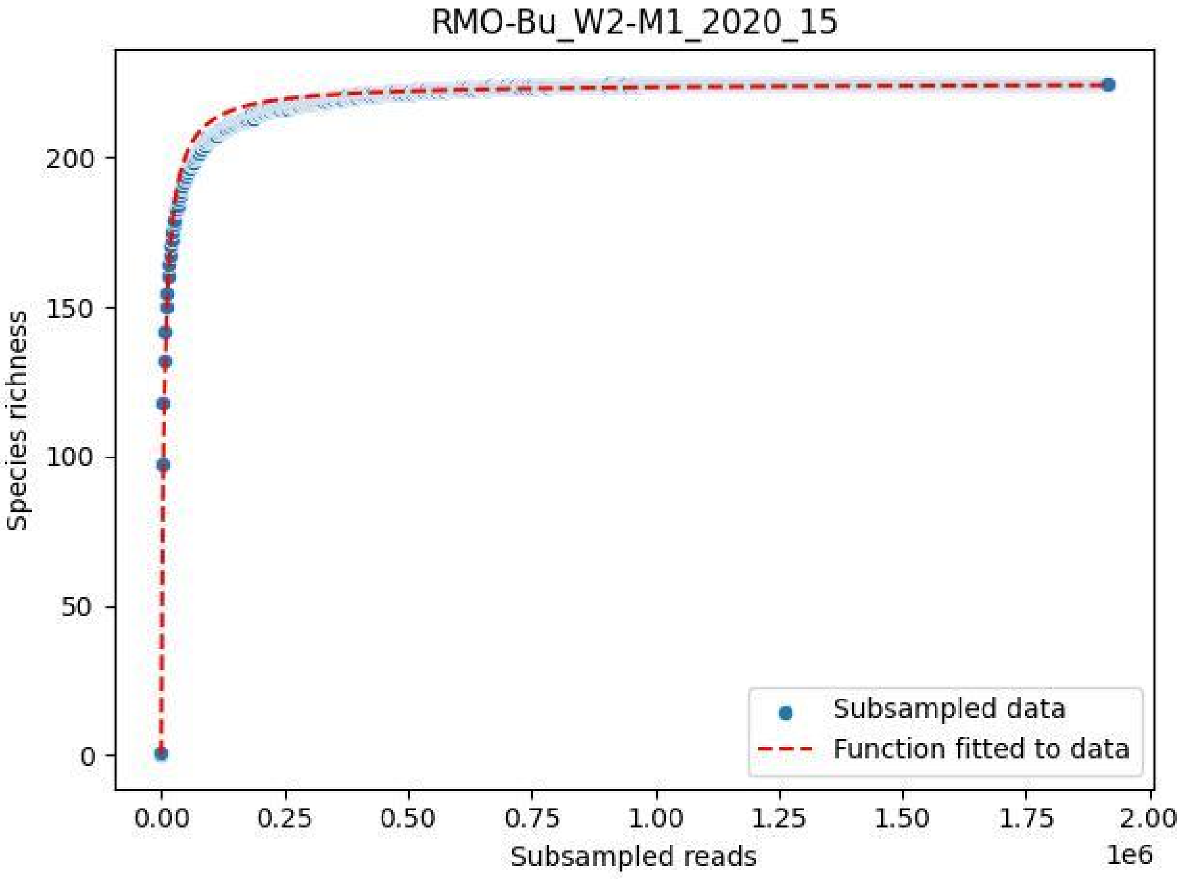

RMO-Bu_W2-M1_2020_15
Species richness
0
50
100
150
200
0.00
0.25
0.50
0.75
1.00
1.25
1.50
1.75
2.00
1e6
Subsampled reads
Subsampled data
Function fitted to data

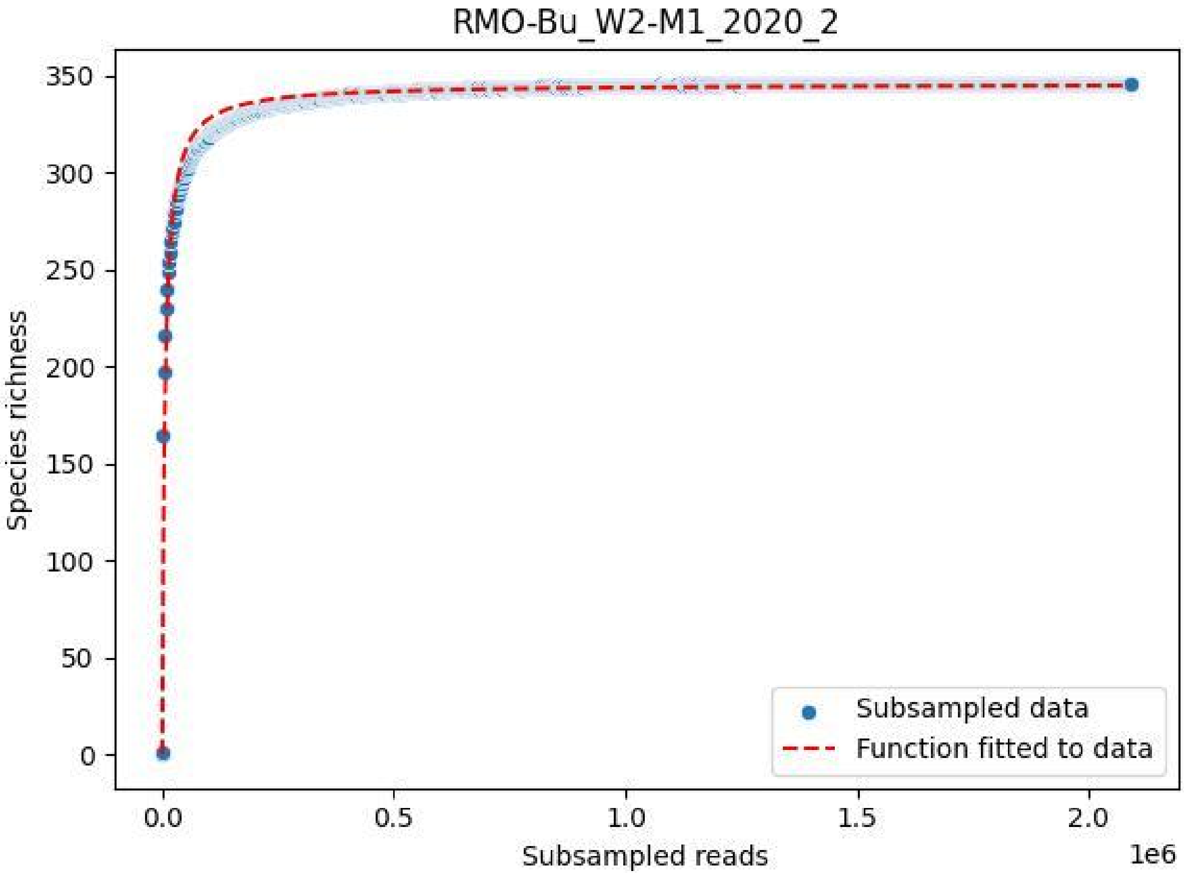
RMO-Bu_W2-M1_2020_2
350
300
250
200
150
100
50
0
Species richness
0.0
0.5
1.0
1.5
2.0
1e6
Subsampled reads
Subsampled data
Function fitted to data

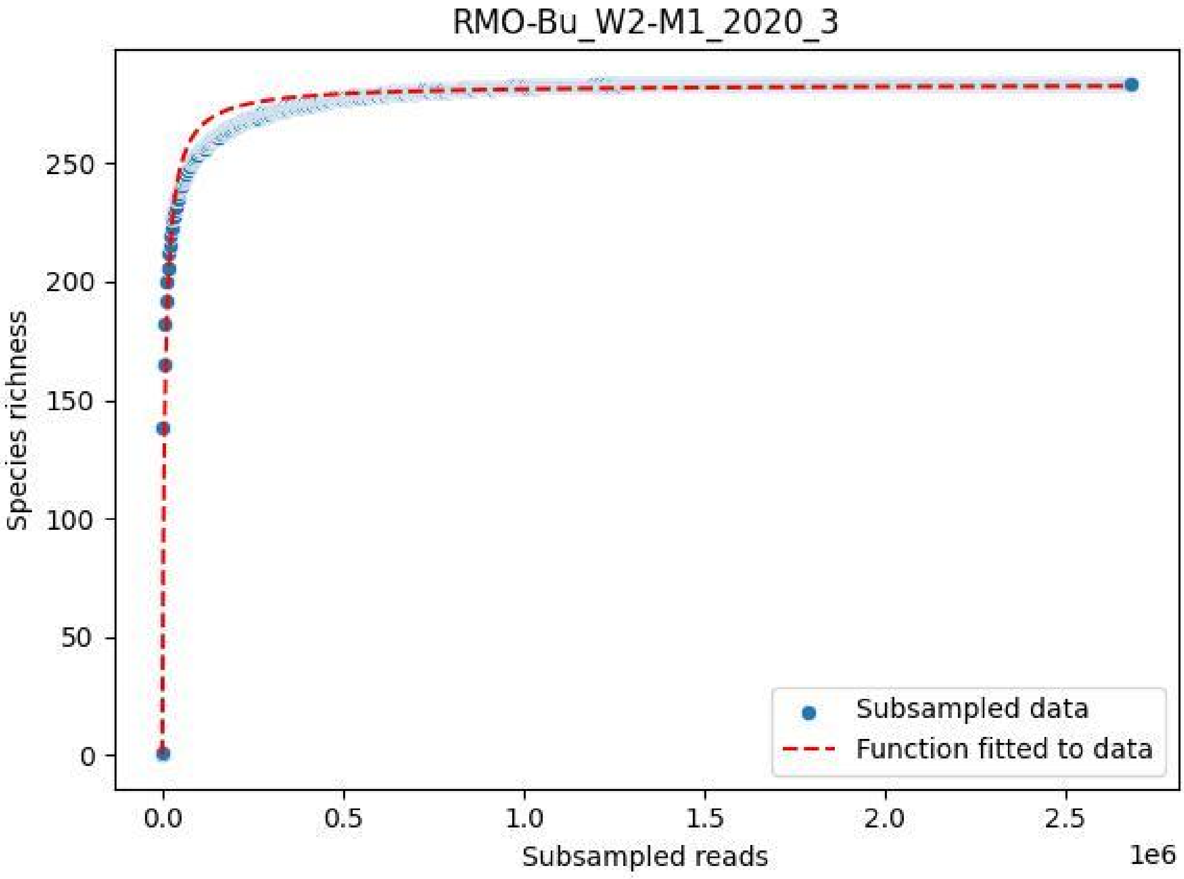
RMO-Bu_W2-M1_2020_3
Species richness
0
50
100
150
200
250
0.0
0.5
1.0
1.5
2.0
2.5
1e6
Subsampled reads
Subsampled data
Function fitted to data

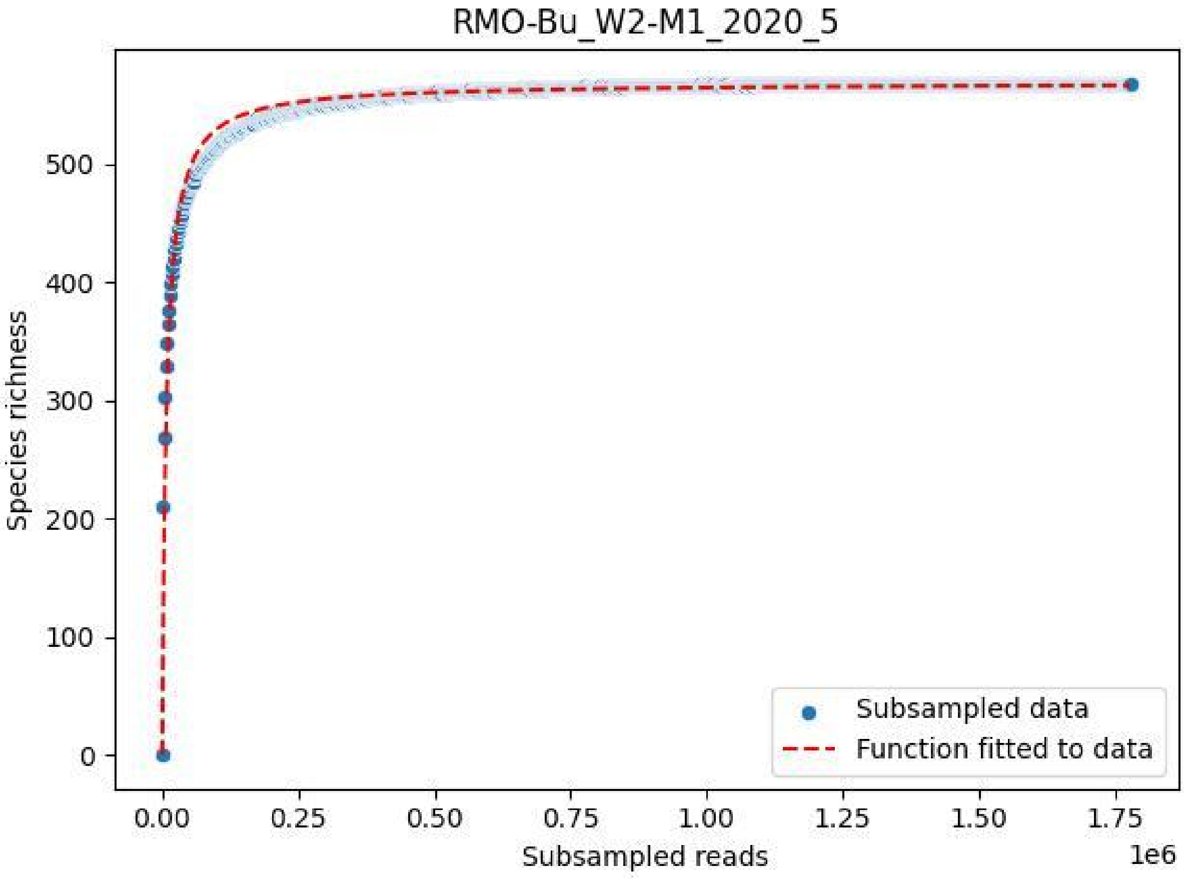

RMO-Bu_W2-M1_2020_5
Species richness
0
100
200
300
400
500
0.00
0.25
0.50
0.75
1.00
1.25
1.50
1.75
1e6
Subsampled reads
Subsampled data
Function fitted to data

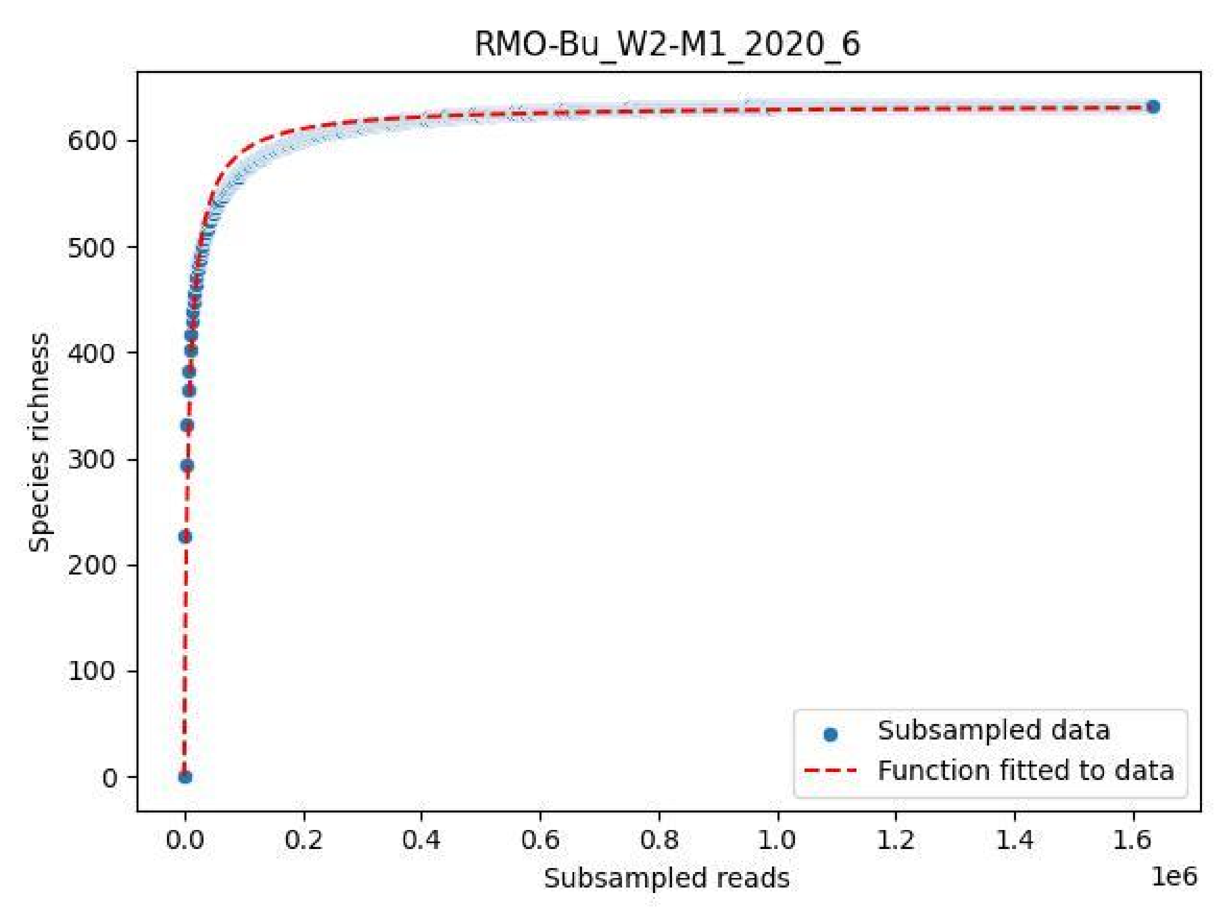

RMO-Bu_W2-M1_2020_6
Species richness
Subsampled reads
1e6
0
100
200
300
400
500
600
0.0
0.2
0.4
0.6
0.8
1.0
1.2
1.4
1.6
Subsampled data
Function fitted to data

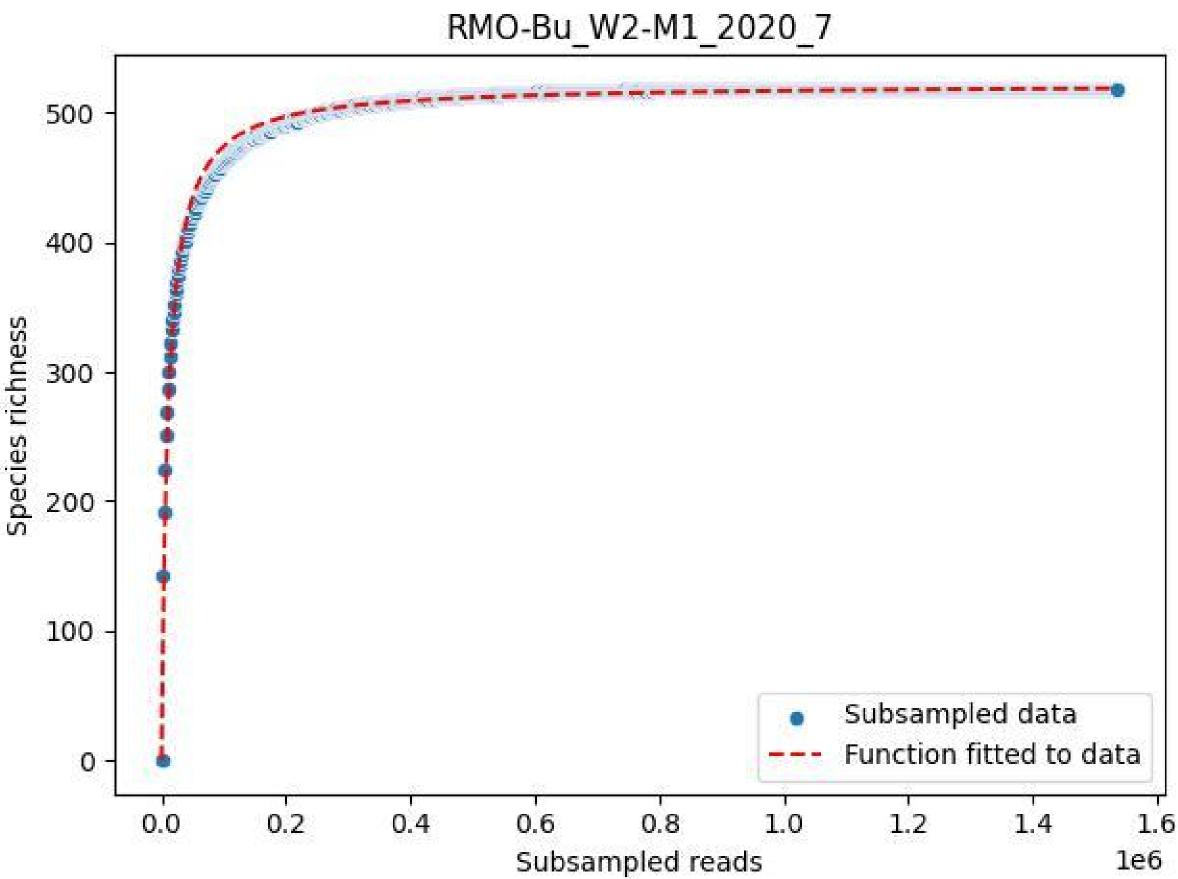
RMO-Bu_W2-M1_2020_7
Species richness
Subsampled reads
1e6
0
100
200
300
400
500
0.0
0.2
0.4
0.6
0.8
1.0
1.2
1.4
1.6
Subsampled data
Function fitted to data

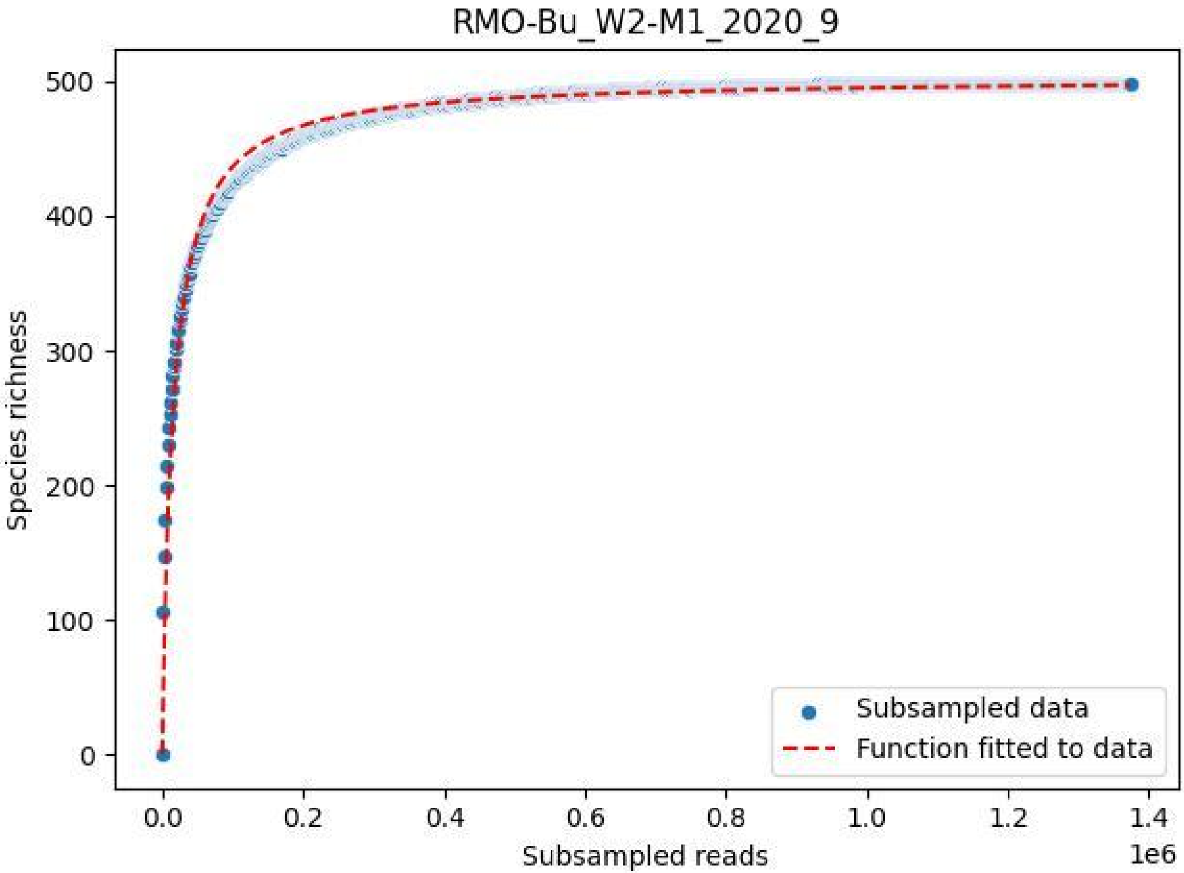

RMO-Bu_W2-M1_2020_9
Species richness
Subsampled reads
1e6
0
100
200
300
400
500
0.0
0.2
0.4
0.6
0.8
1.0
1.2
1.4
Subsampled data
Function fitted to data

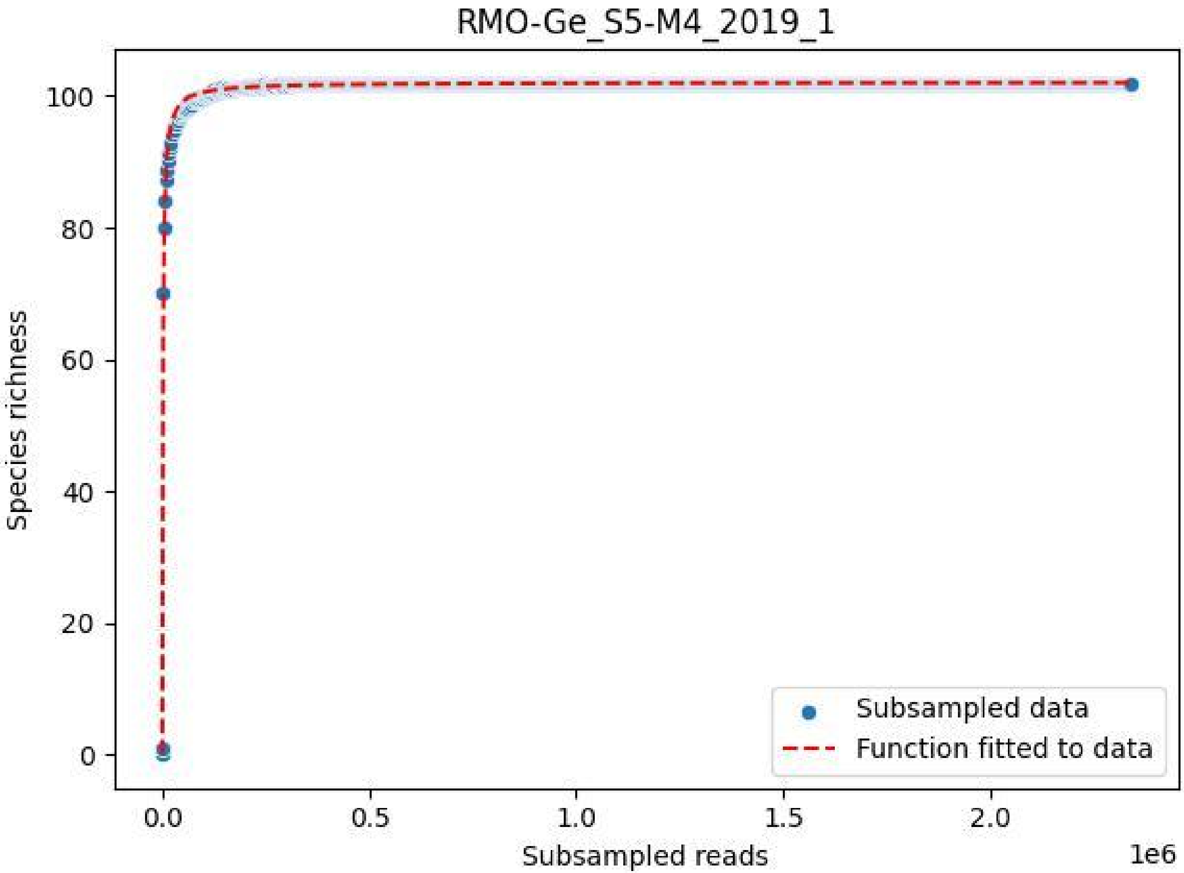

RMO-Ge_S5-M4_2019_1
Species richness
Subsampled reads
1e6
0
20
40
60
80
100
0.0
0.5
1.0
1.5
2.0
Subsampled data
Function fitted to data

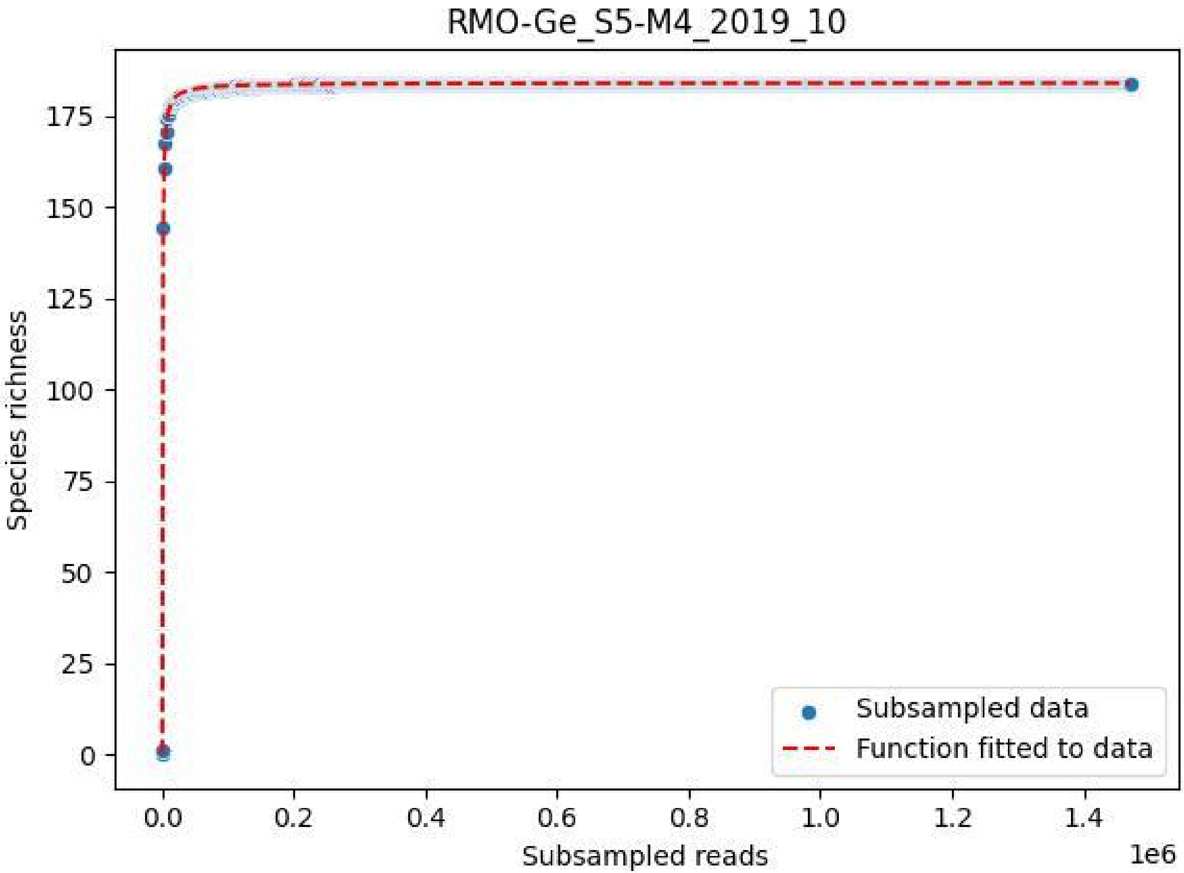

RMO-Ge_S5-M4_2019_10
Species richness
0
25
50
75
100
125
150
175
0.0
0.2
0.4
0.6
0.8
1.0
1.2
1.4
1e6
Subsampled reads
Subsampled data
Function fitted to data

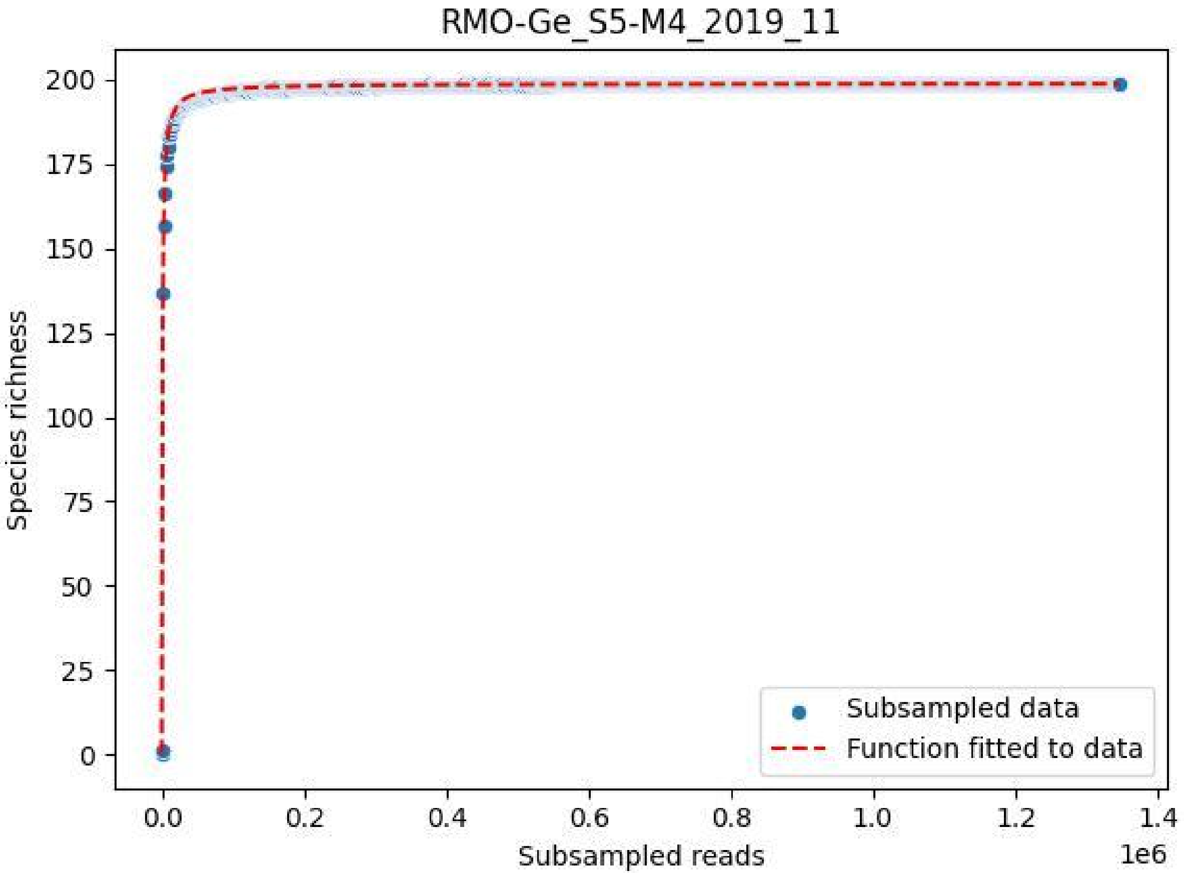
RMO-Ge_S5-M4_2019_11
Species richness
Subsampled reads
1e6
Subsampled data
Function fitted to data
0
25
50
75
100
125
150
175
200
0.0
0.2
0.4
0.6
0.8
1.0
1.2
1.4

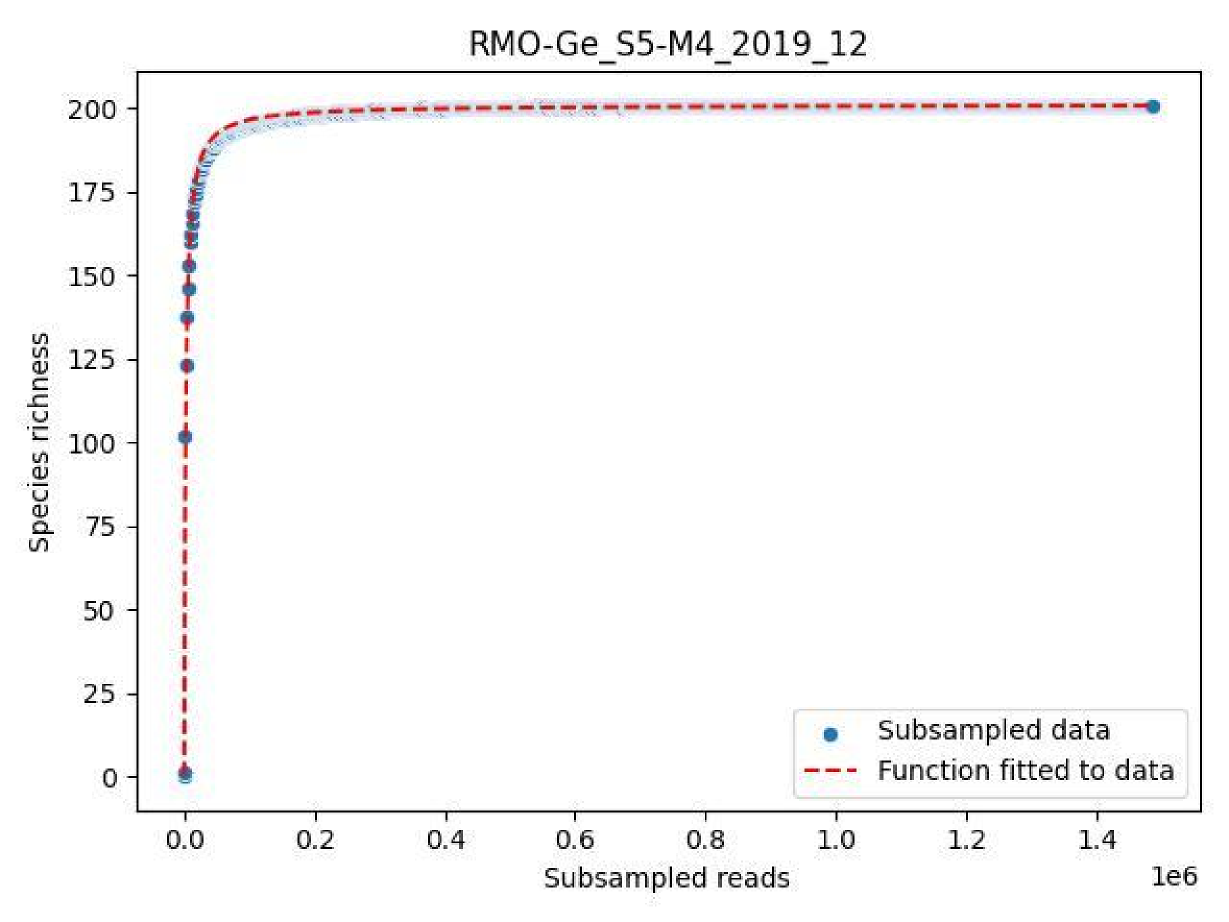

RMO-Ge_S5-M4_2019_12
Species richness
0
25
50
75
100
125
150
175
200
0.0
0.2
0.4
0.6
0.8
1.0
1.2
1.4
1e6
Subsampled reads
Subsampled data
Function fitted to data

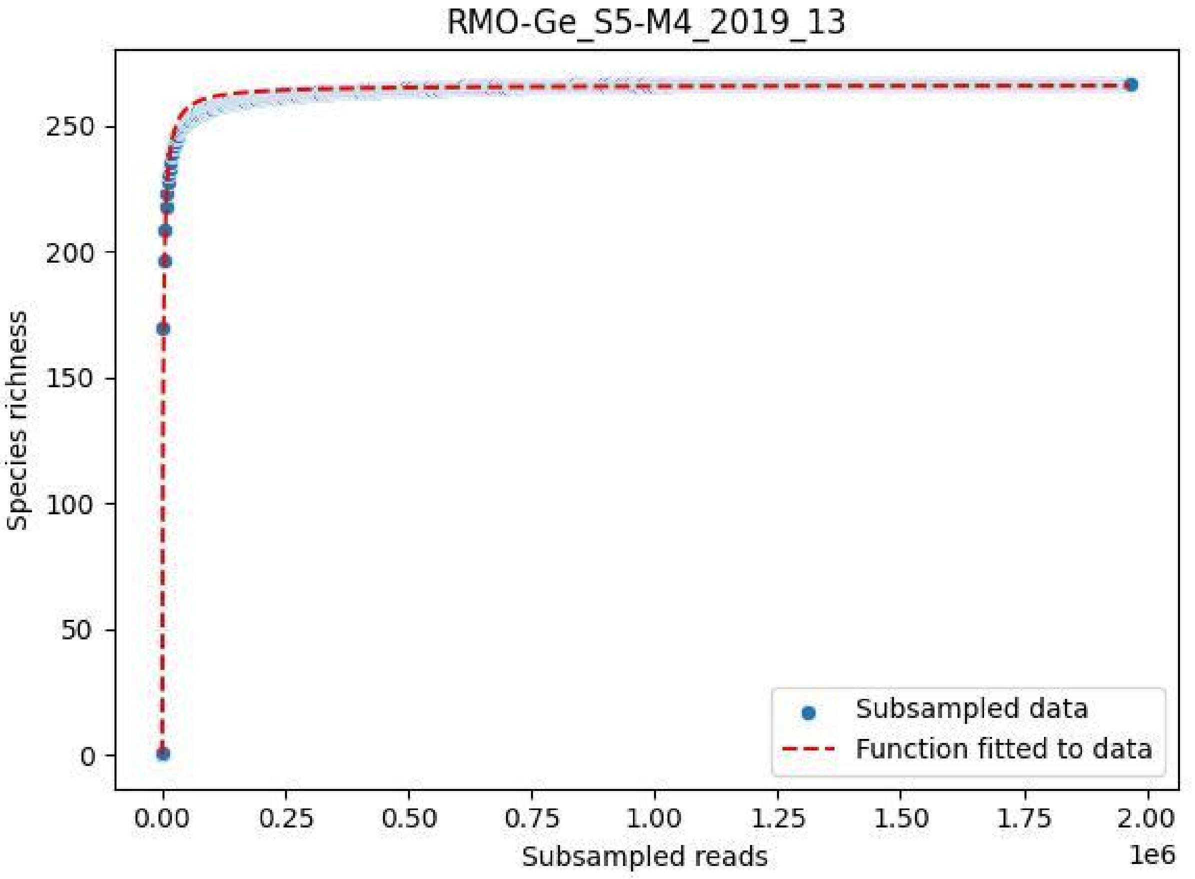

RMO-Ge_S5-M4_2019_13
Species richness
Subsampled reads
1e6
0
50
100
150
200
250
0.00
0.25
0.50
0.75
1.00
1.25
1.50
1.75
2.00
Subsampled data
Function fitted to data

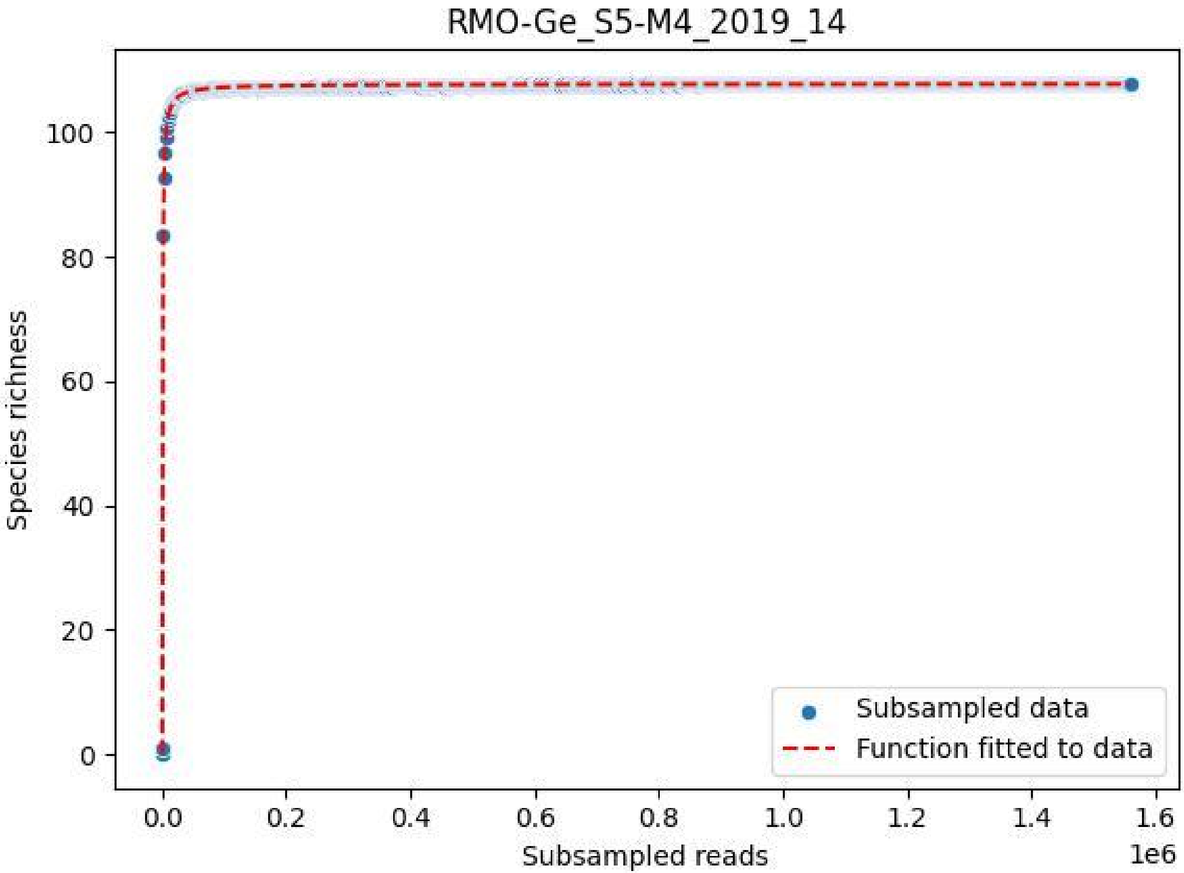
RMO-Ge_S5-M4_2019_14
Species richness
Subsampled reads
1e6
0
20
40
60
80
100
0.0
0.2
0.4
0.6
0.8
1.0
1.2
1.4
1.6
Subsampled data
Function fitted to data

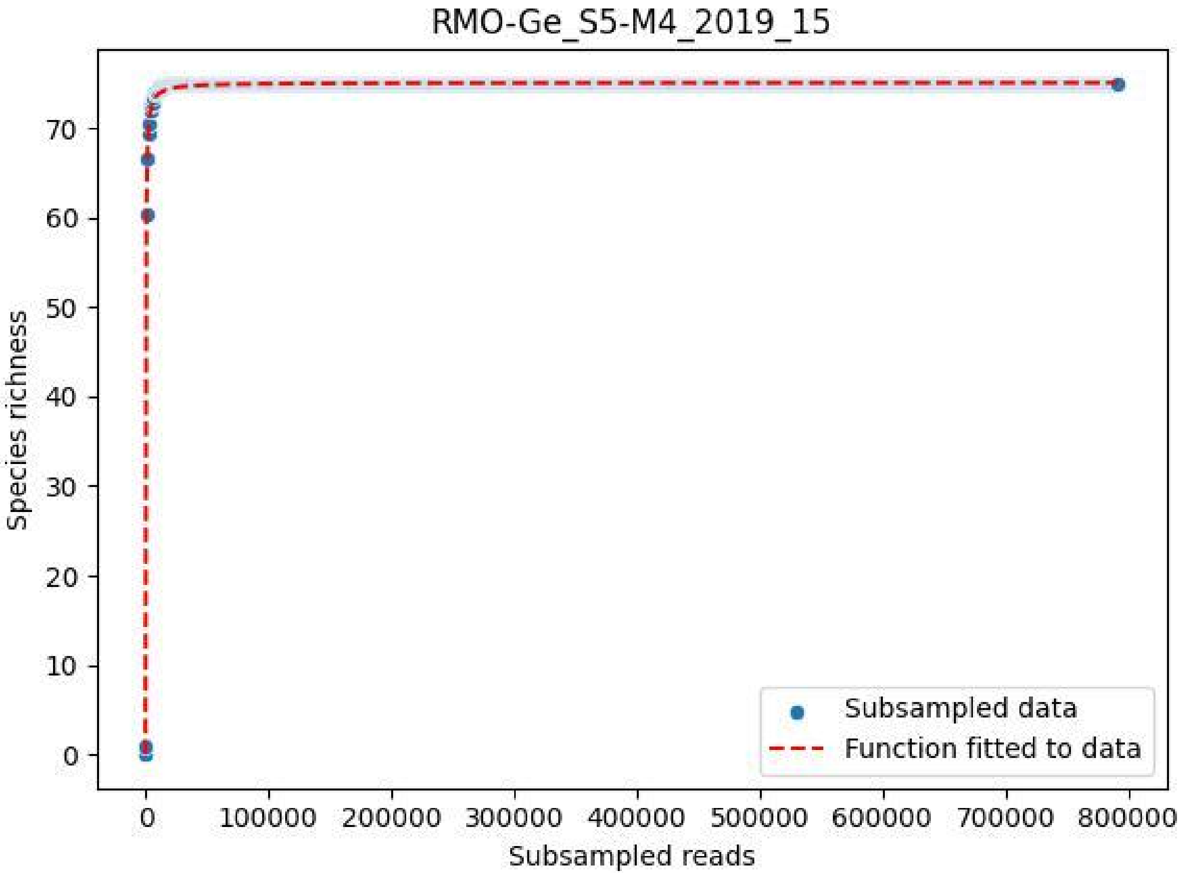

RMO-Ge_S5-M4_2019_15
Species richness
0
10
20
30
40
50
60
70
0
100000
200000
300000
400000
500000
600000
700000
800000
Subsampled reads
Subsampled data
Function fitted to data

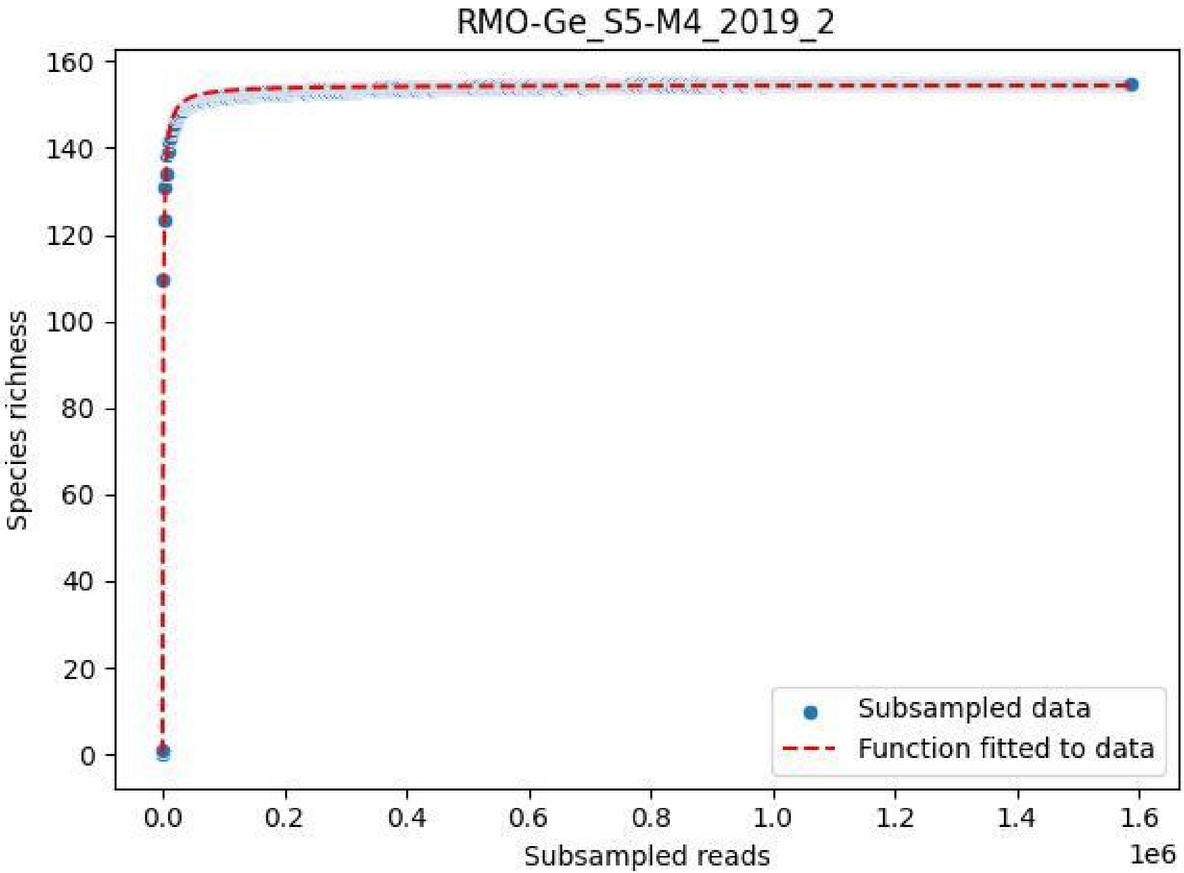

RMO-Ge_S5-M4_2019_2
Species richness
Subsampled reads
1e6
0
20
40
60
80
100
120
140
160
0.0
0.2
0.4
0.6
0.8
1.0
1.2
1.4
1.6
Subsampled data
Function fitted to data

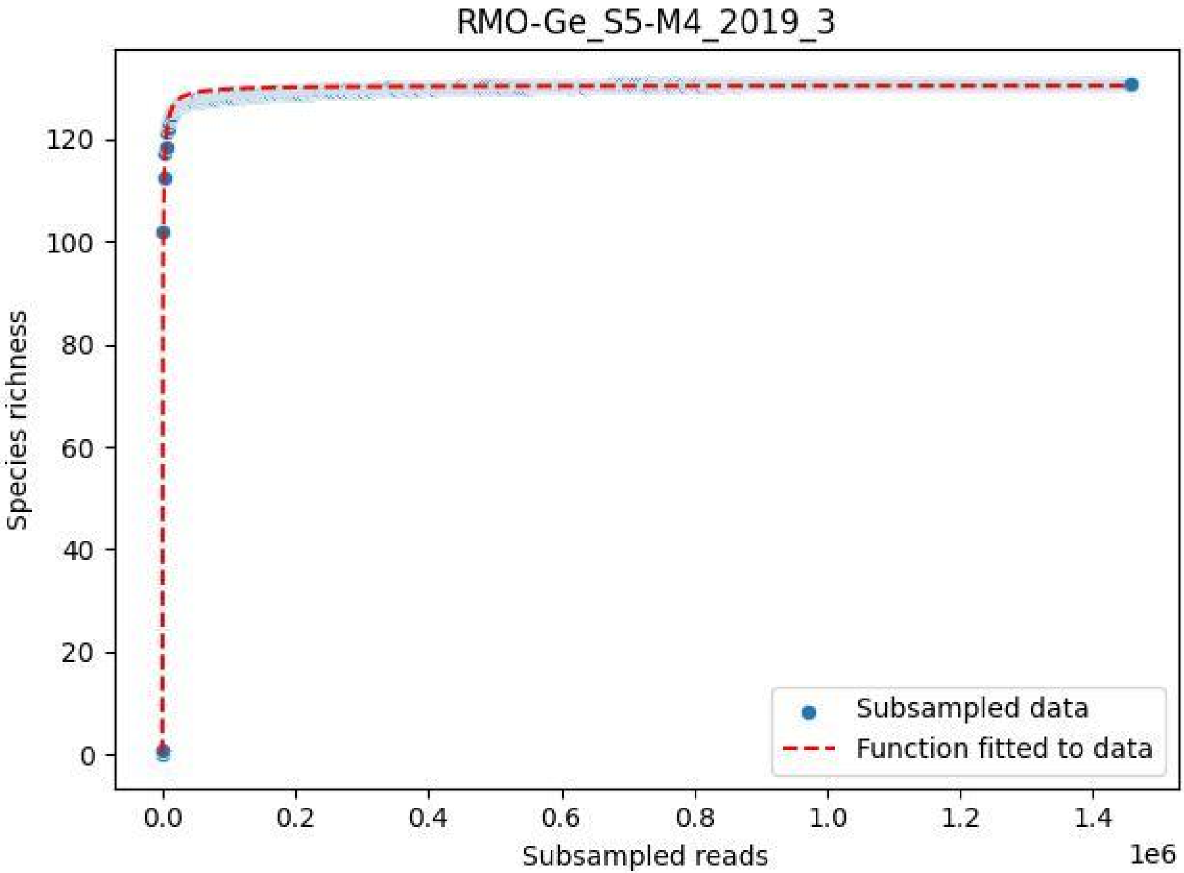
RMO-Ge_S5-M4_2019_3
Species richness
Subsampled reads
1e6
0
20
40
60
80
100
120
0.0
0.2
0.4
0.6
0.8
1.0
1.2
1.4
Subsampled data
Function fitted to data

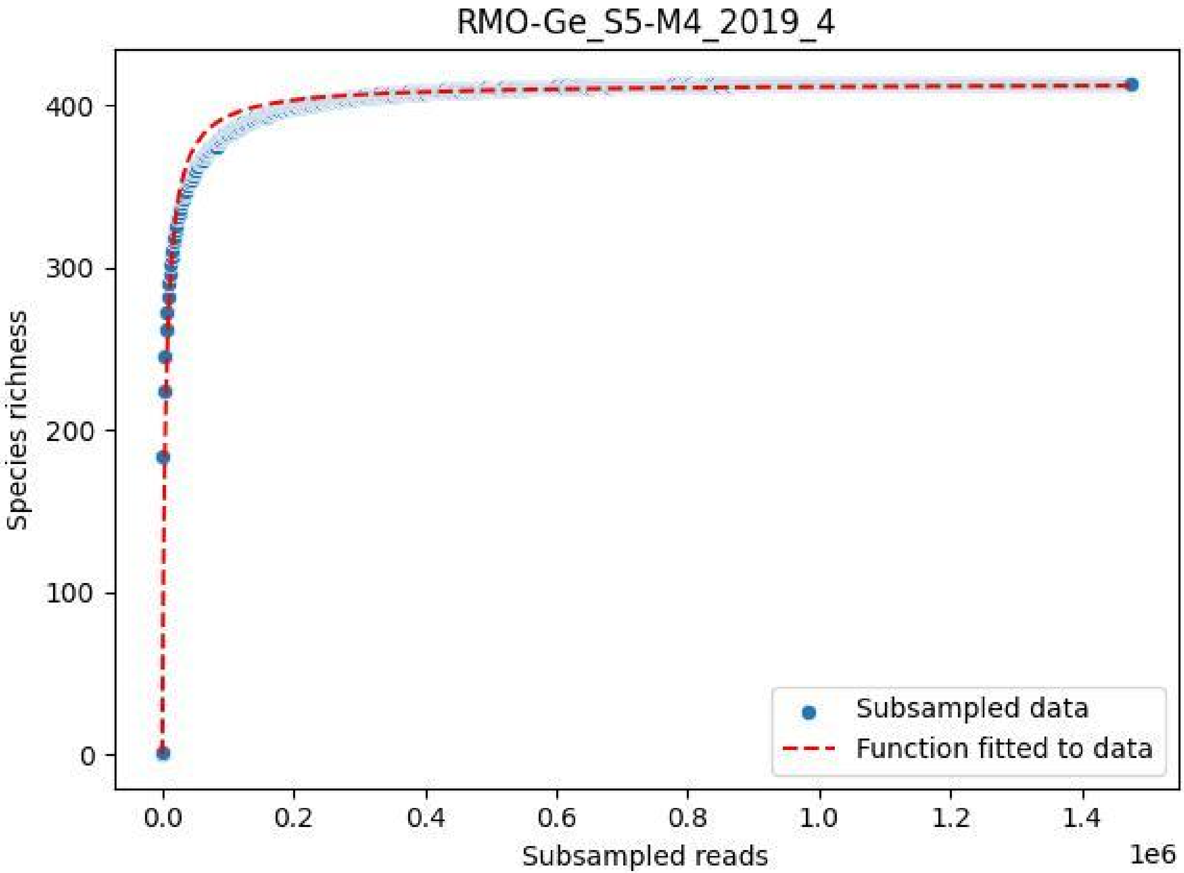

RMO-Ge_S5-M4_2019_4
Species richness
Subsampled reads
1e6
0
100
200
300
400
0.0
0.2
0.4
0.6
0.8
1.0
1.2
1.4
Subsampled data
Function fitted to data

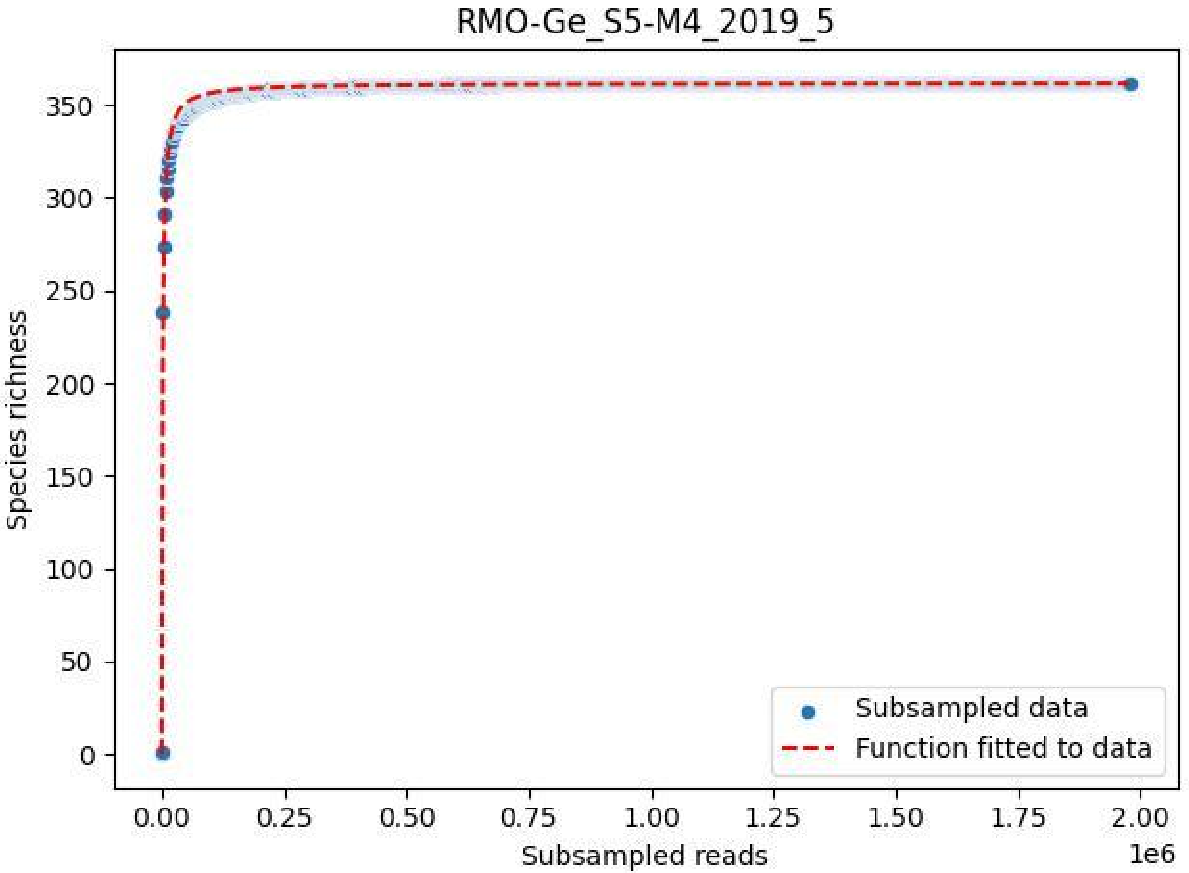

RMO-Ge_S5-M4_2019_5
Species richness
Subsampled reads
1e6
Subsampled data
Function fitted to data
0
50
100
150
200
250
300
350
0.00
0.25
0.50
0.75
1.00
1.25
1.50
1.75
2.00

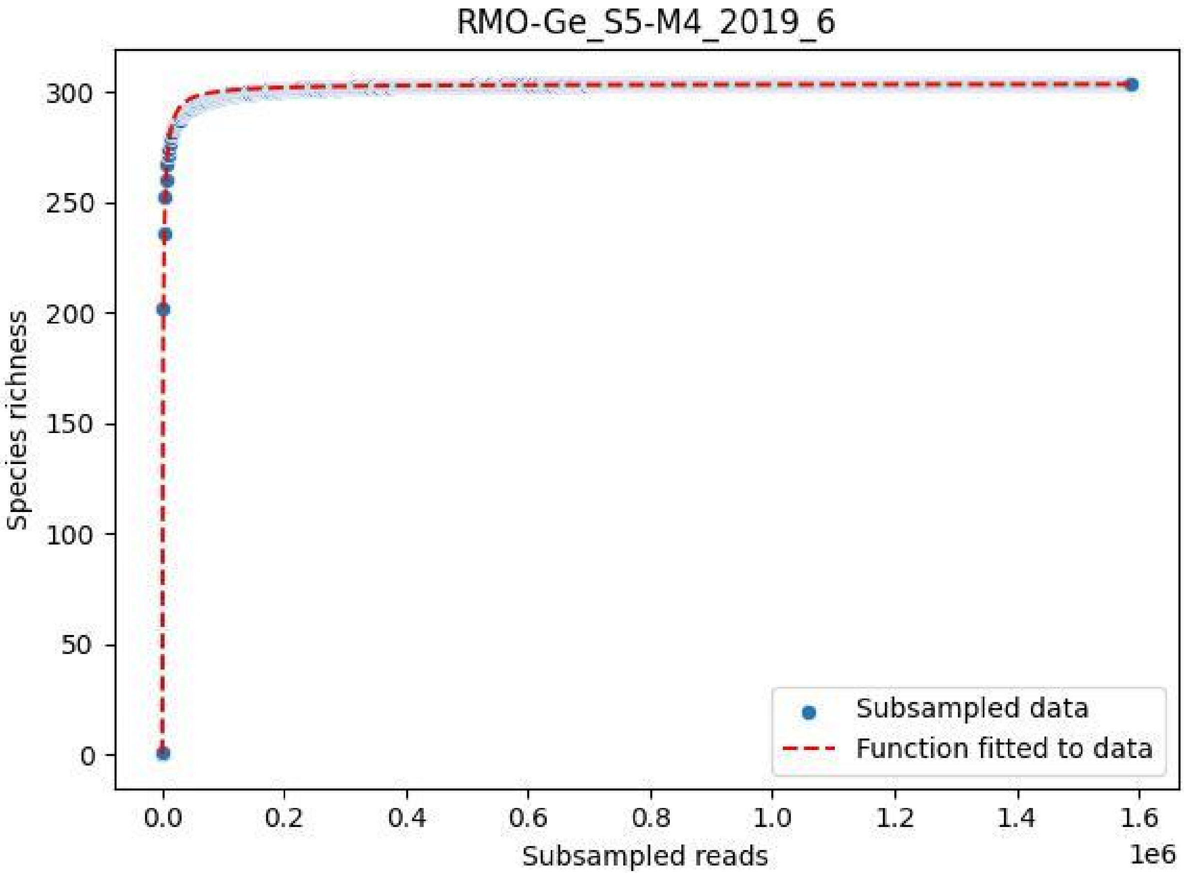
RMO-Ge_S5-M4_2019_6
Species richness
300
250
200
150
100
50
0
0.0
0.2
0.4
0.6
0.8
1.0
1.2
1.4
1.6
1e6
Subsampled reads
Subsampled data
Function fitted to data

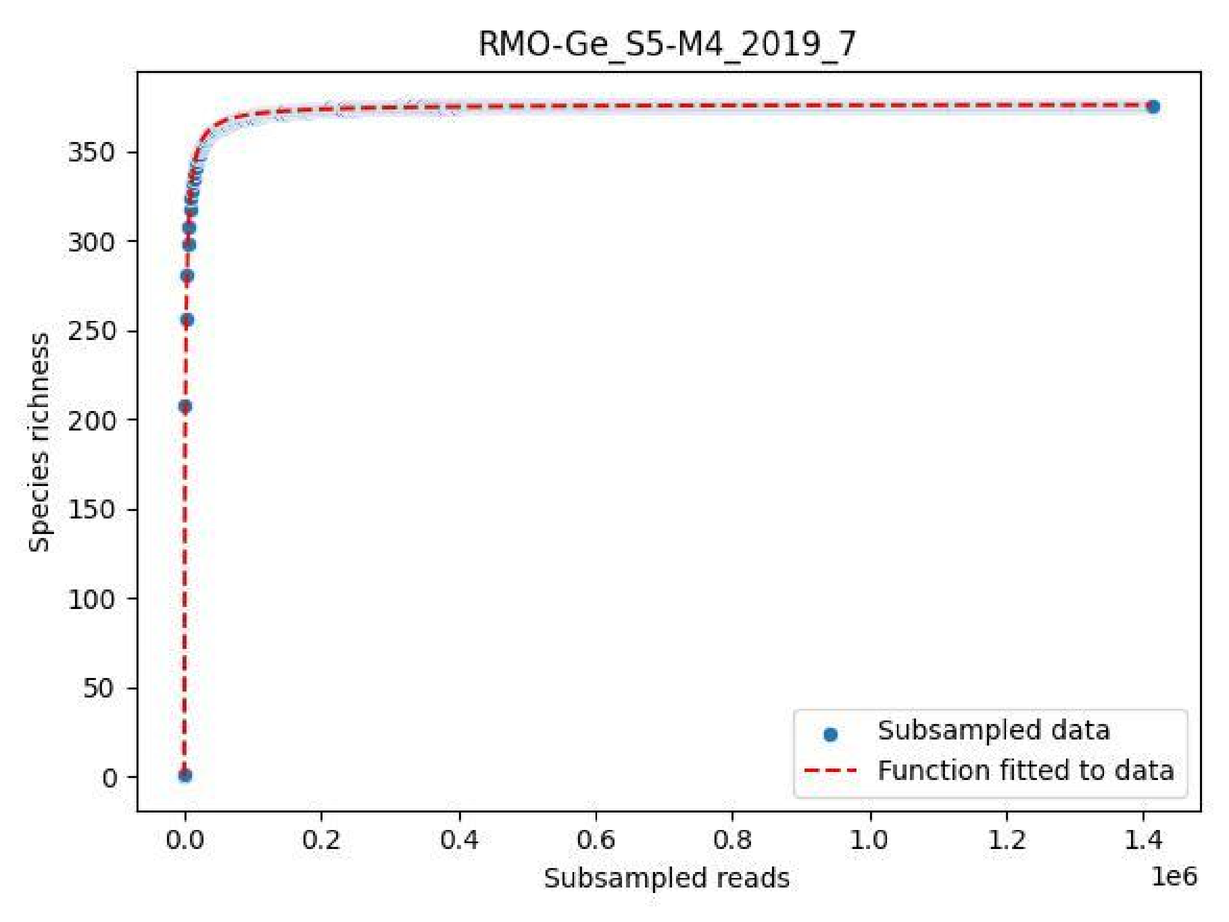

RMO-Ge_S5-M4_2019_7
Species richness
Subsampled reads
1e6
Subsampled data
Function fitted to data
0
50
100
150
200
250
300
350
0.0
0.2
0.4
0.6
0.8
1.0
1.2
1.4

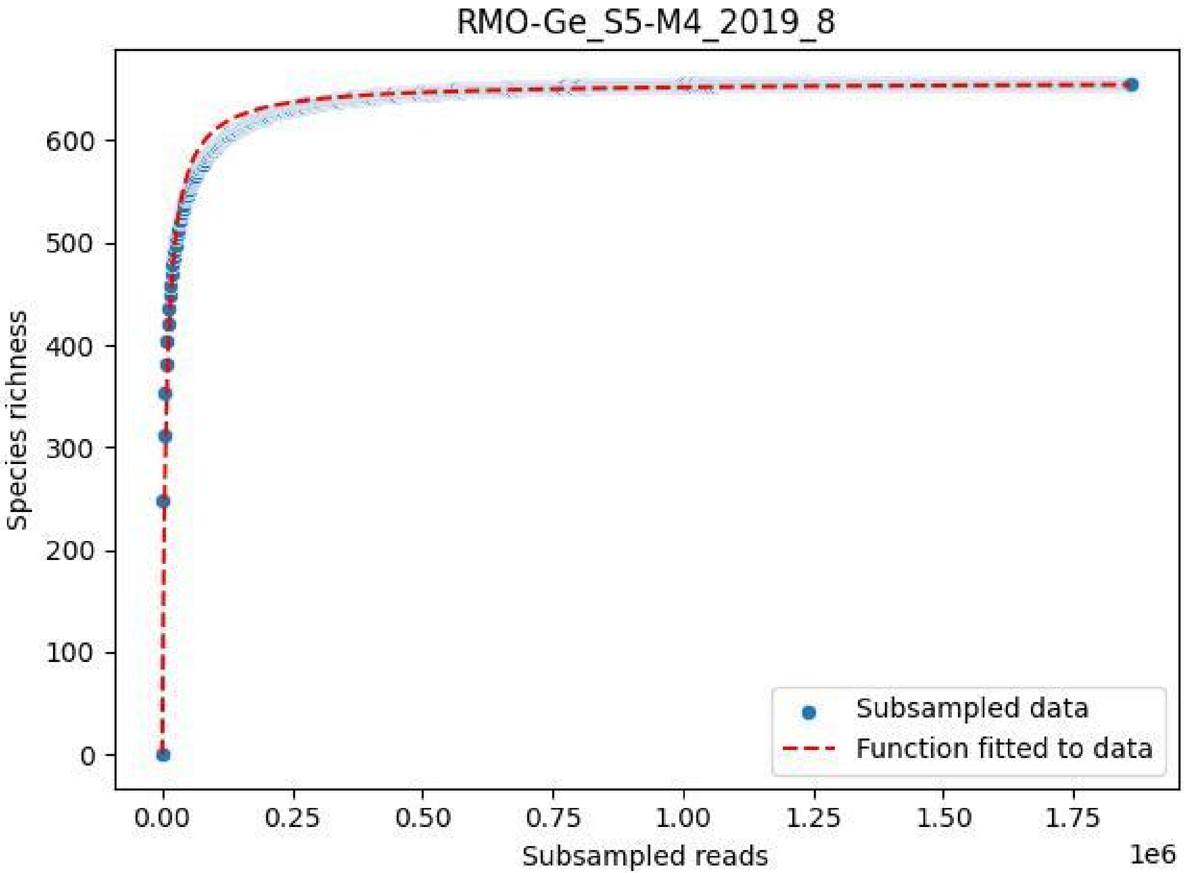

RMO-Ge_S5-M4_2019_8
Species richness
Subsampled reads
1e6
0
100
200
300
400
500
600
0.00
0.25
0.50
0.75
1.00
1.25
1.50
1.75
Subsampled data
Function fitted to data

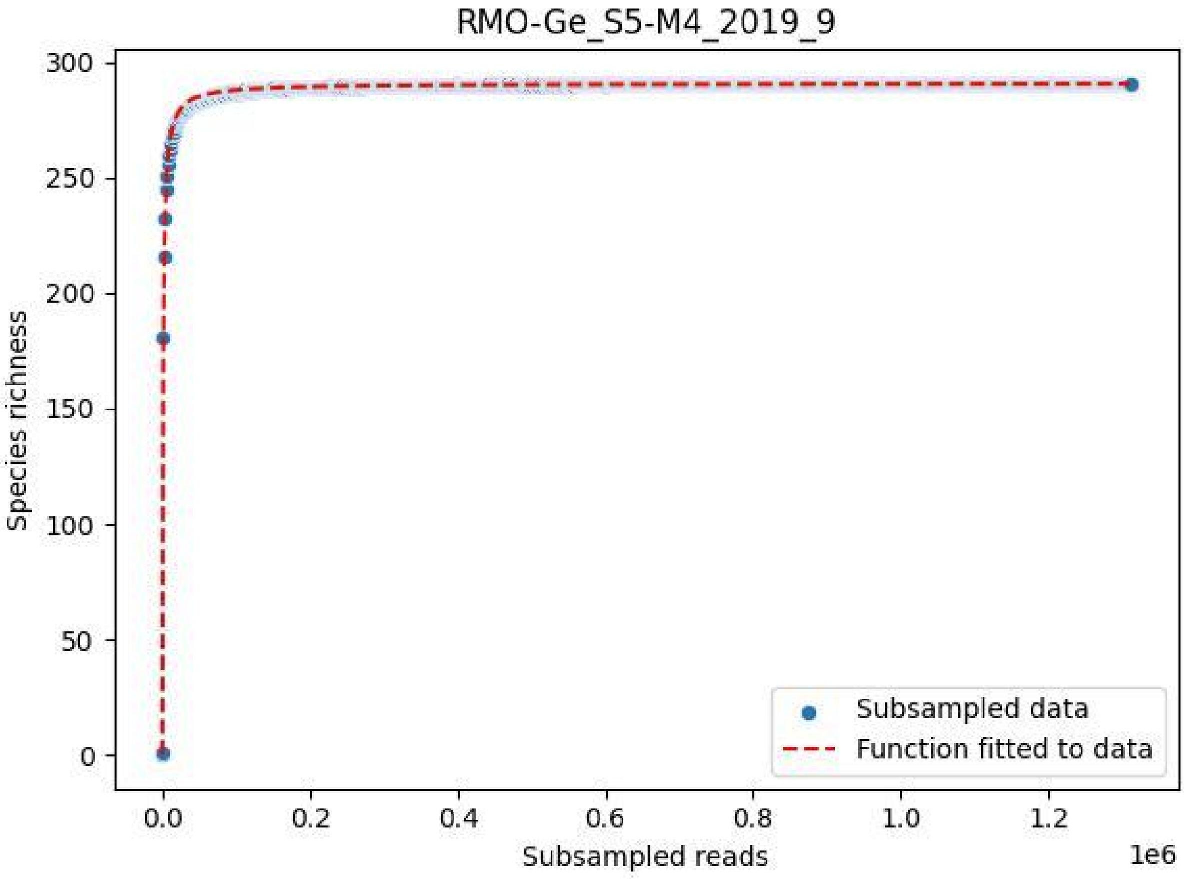
RMO-Ge_S5-M4_2019_9
Species richness
300
250
200
150
100
50
0
0.0
0.2
0.4
0.6
0.8
1.0
1.2
1e6
Subsampled reads
Subsampled data
Function fitted to data

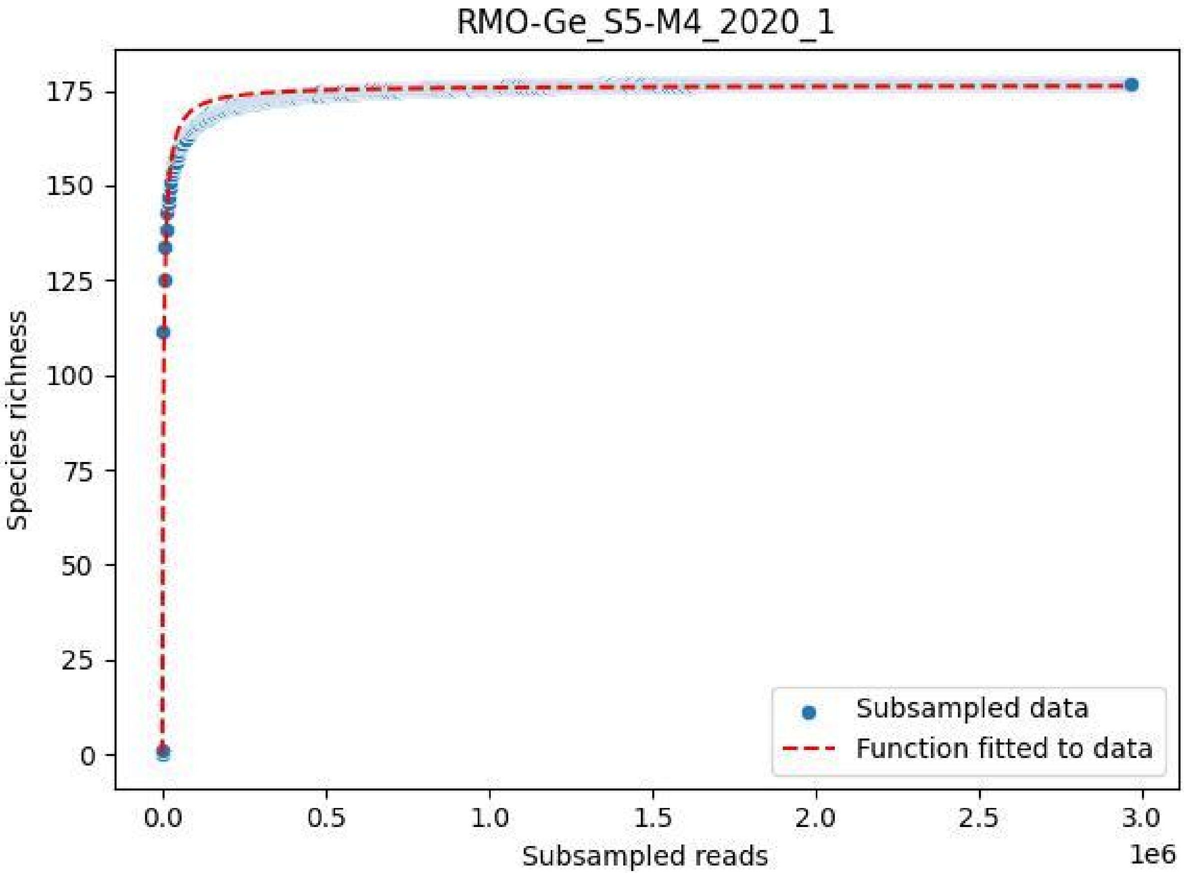

RMO-Ge_S5-M4_2020_1
Species richness
Subsampled reads
1e6
0
25
50
75
100
125
150
175
0.0
0.5
1.0
1.5
2.0
2.5
3.0
Subsampled data
Function fitted to data

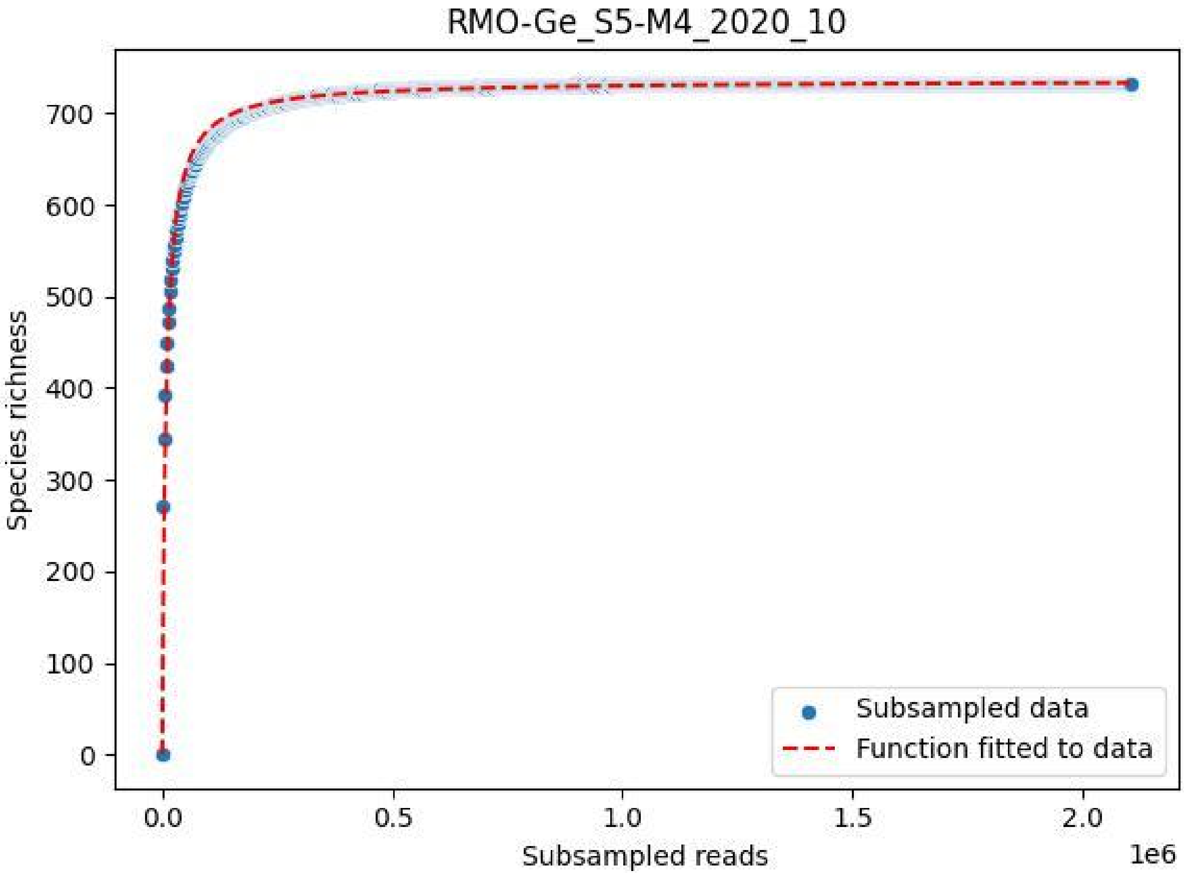

RMO-Ge_S5-M4_2020_10
Species richness
Subsampled reads
1e6
0
100
200
300
400
500
600
700
0.0
0.5
1.0
1.5
2.0
Subsampled data
Function fitted to data

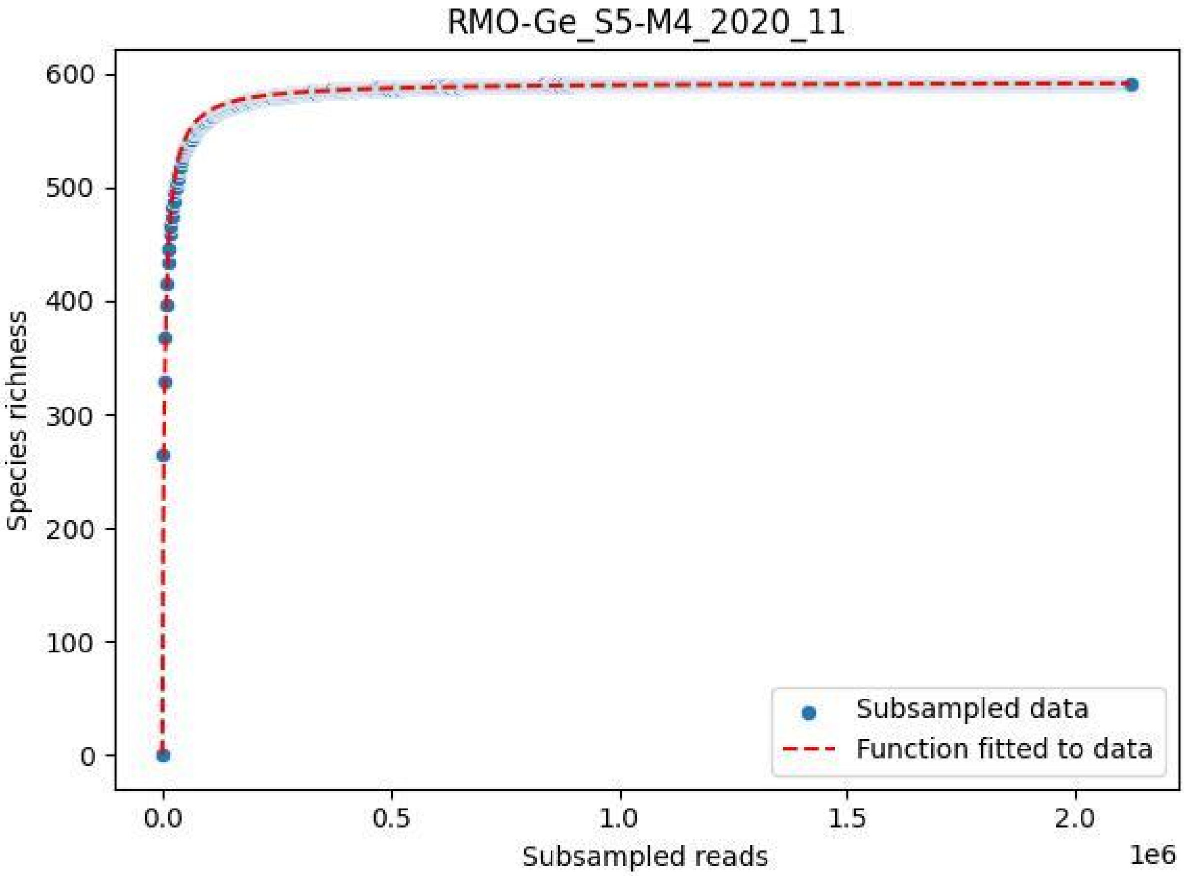
RMO-Ge_S5-M4_2020_11
Species richness
Subsampled reads
1e6
0
100
200
300
400
500
600
0.0
0.5
1.0
1.5
2.0
Subsampled data
Function fitted to data

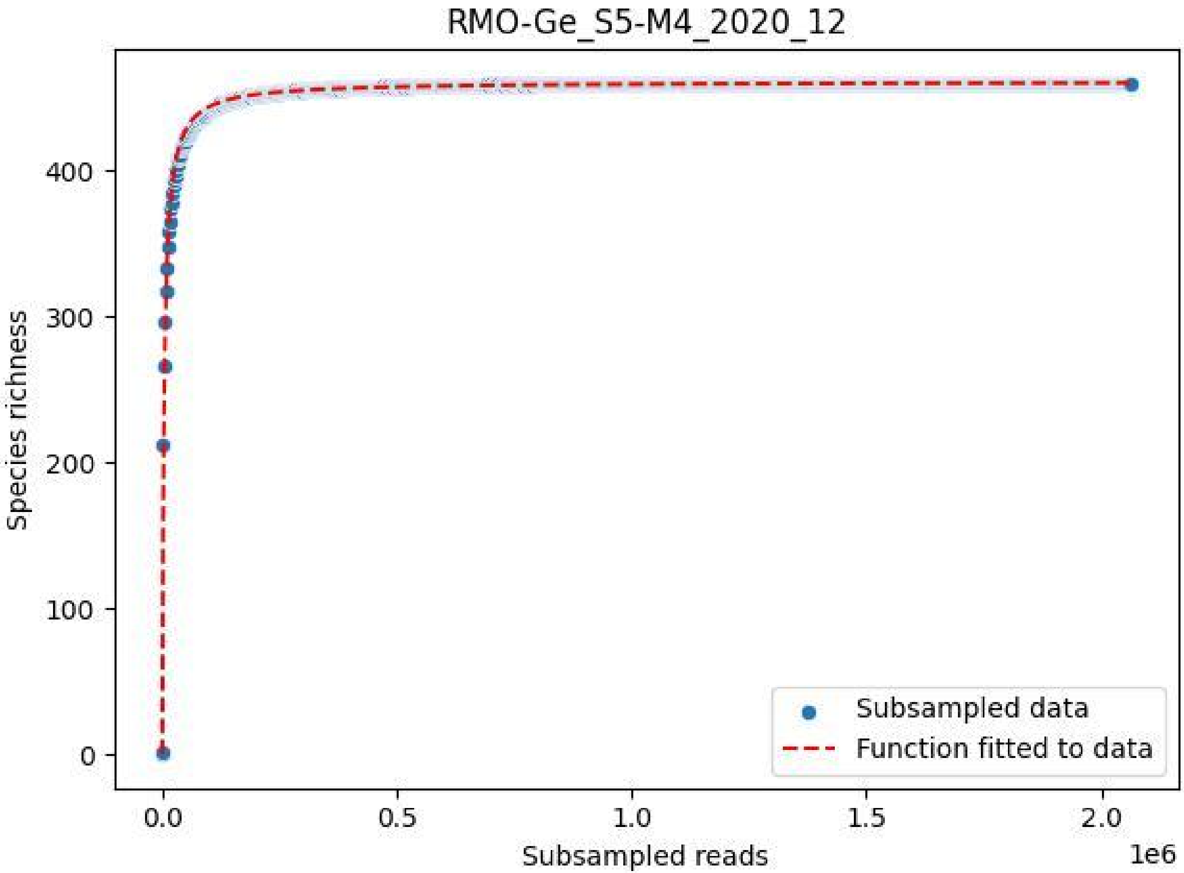

RMO-Ge_S5-M4_2020_12
Species richness
0
100
200
300
400
0.0
0.5
1.0
1.5
2.0
1e6
Subsampled reads
Subsampled data
Function fitted to data

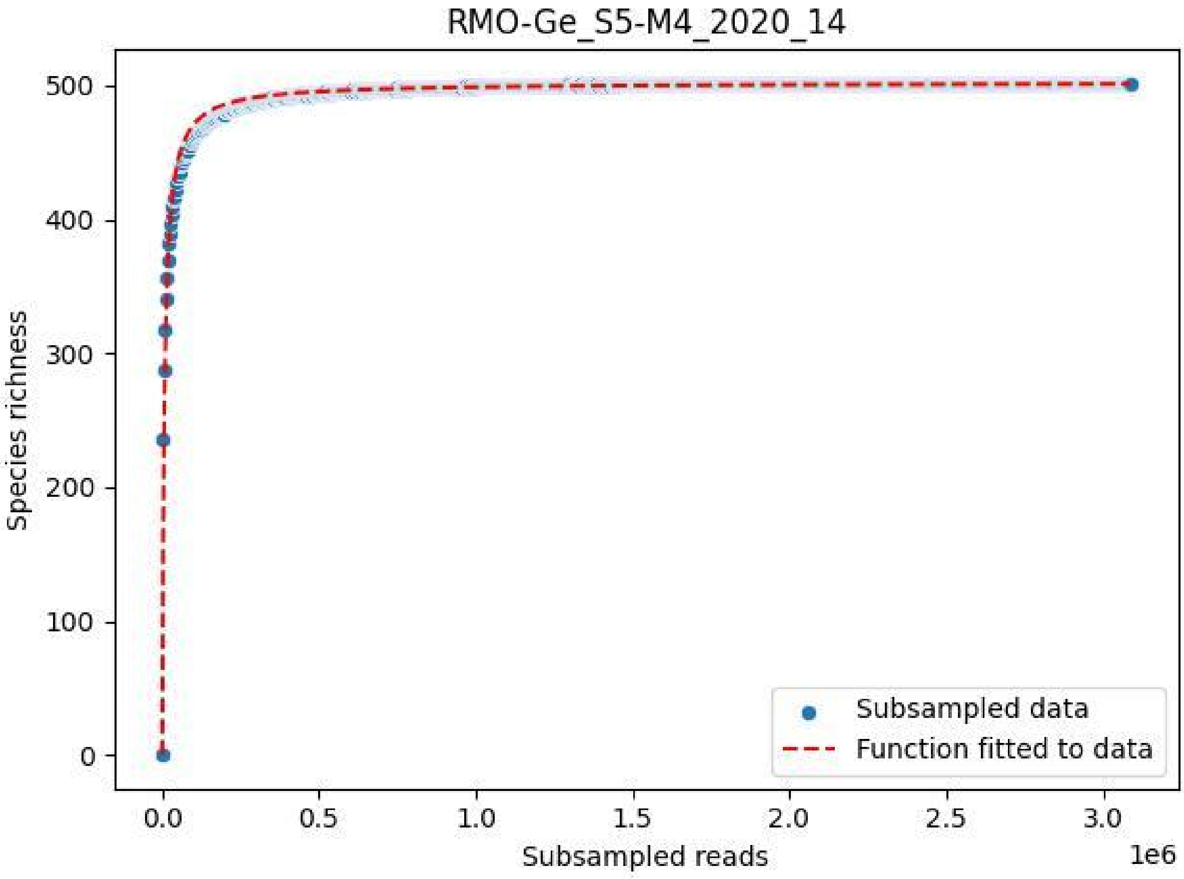

RMO-Ge_S5-M4_2020_14
Species richness
0
100
200
300
400
500
0.0
0.5
1.0
1.5
2.0
2.5
3.0
1e6
Subsampled reads
Subsampled data
Function fitted to data

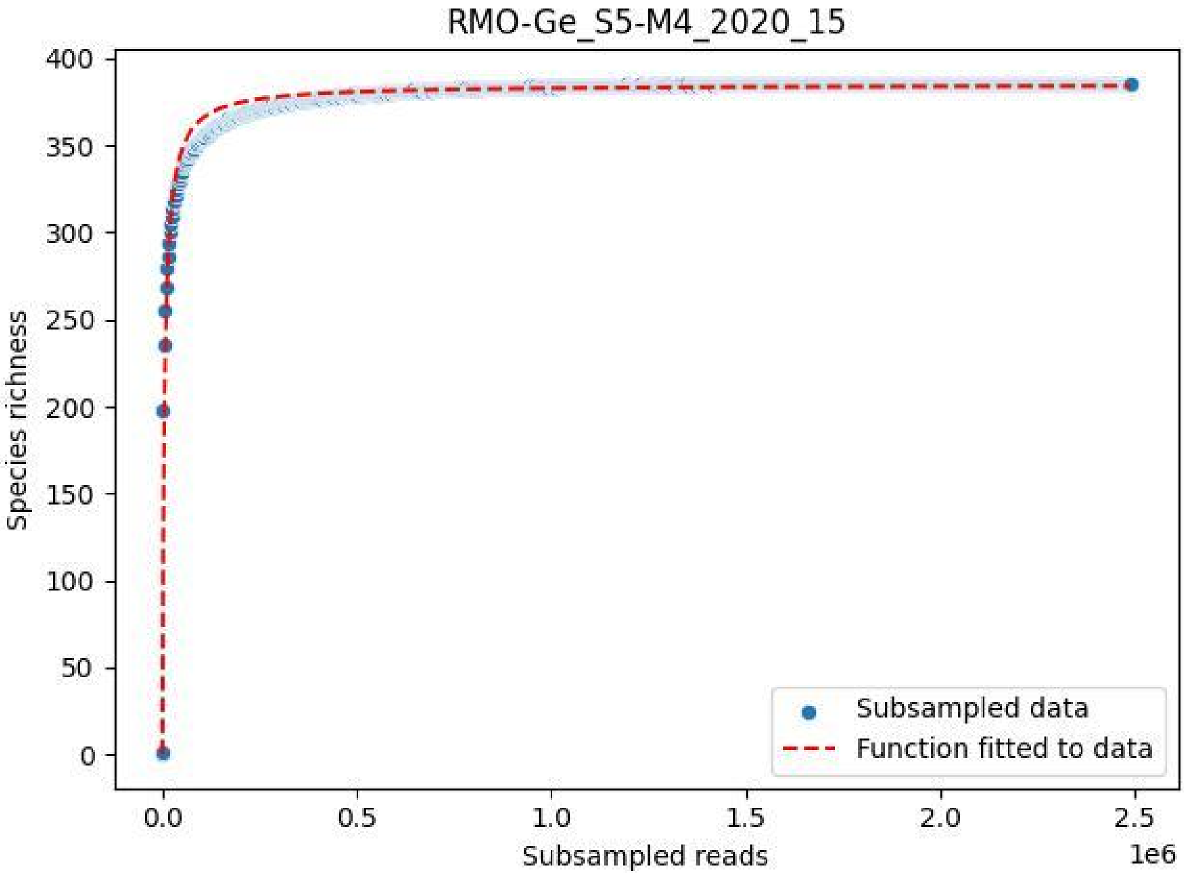

RMO-Ge_S5-M4_2020_15
Species richness
Subsampled reads
1e6
0
50
100
150
200
250
300
350
400
0.0
0.5
1.0
1.5
2.0
2.5
Subsampled data
Function fitted to data

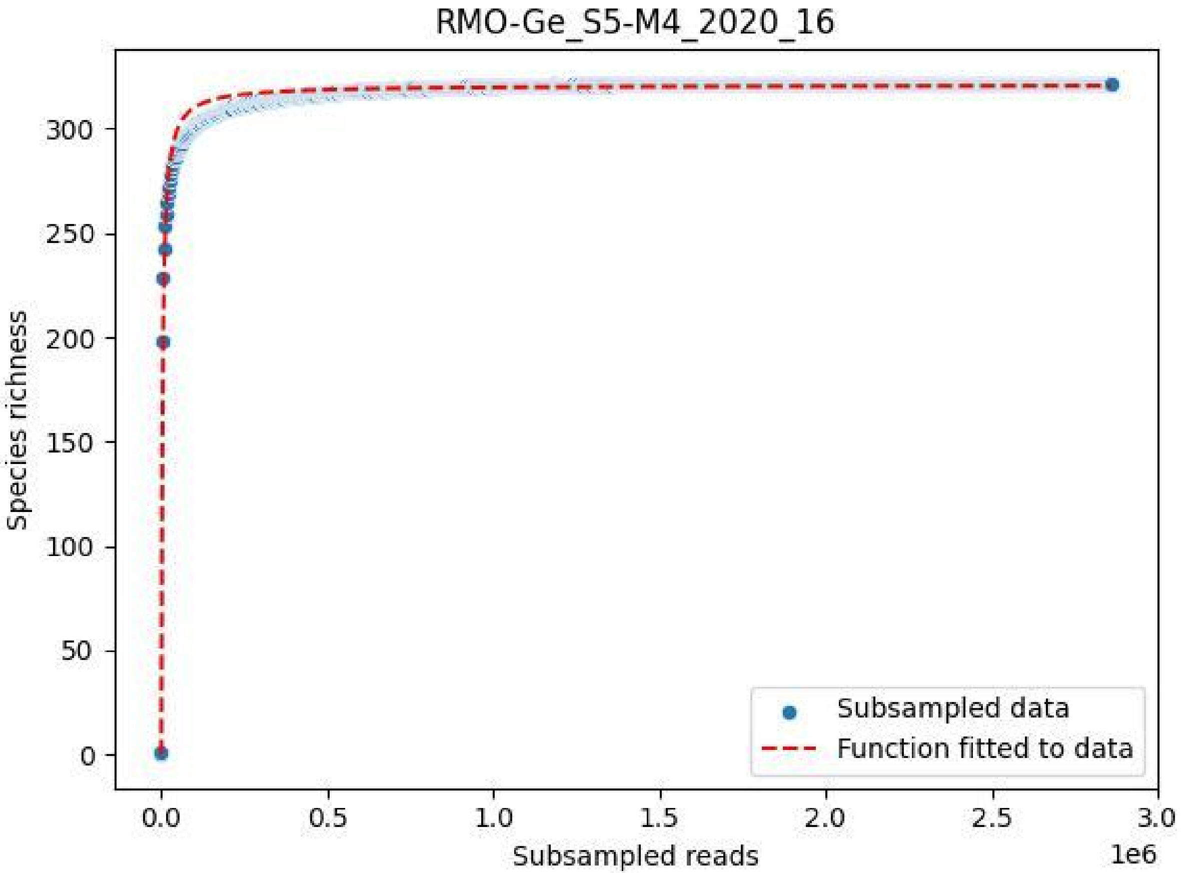
RMO-Ge_S5-M4_2020_16
Species richness
Subsampled reads
1e6
0
50
100
150
200
250
300
0.0
0.5
1.0
1.5
2.0
2.5
3.0
Subsampled data
Function fitted to data

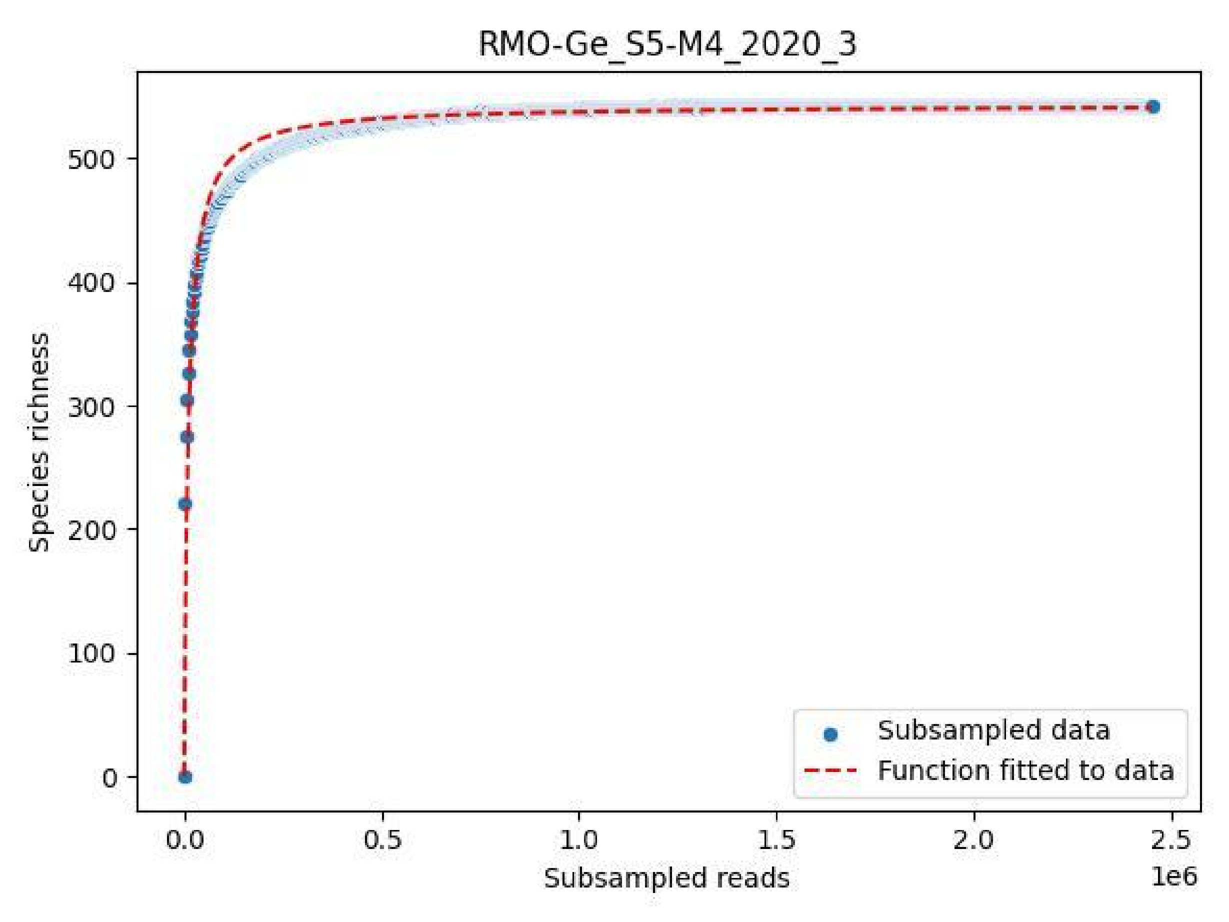

RMO-Ge_S5-M4_2020_3
Species richness
Subsampled reads
1e6
0
100
200
300
400
500
0.0
0.5
1.0
1.5
2.0
2.5
Subsampled data
Function fitted to data

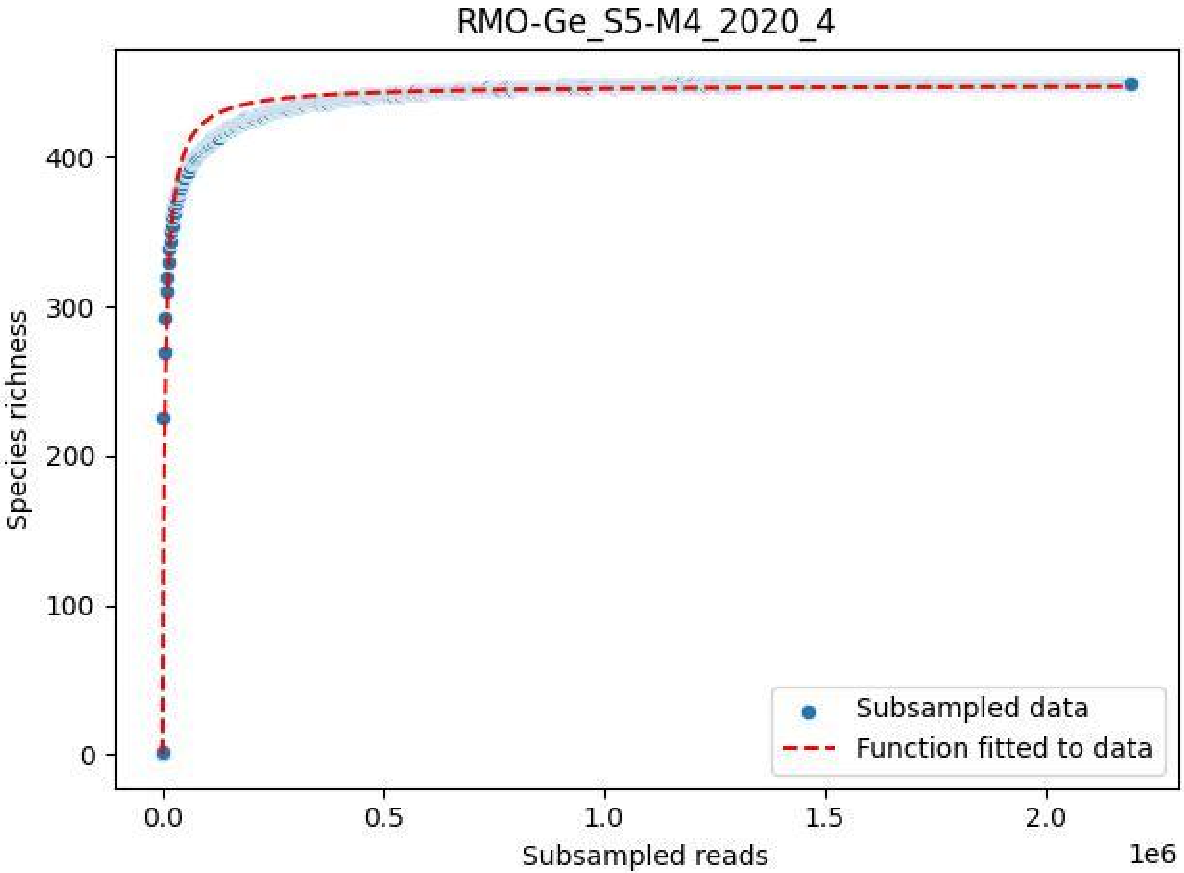

RMO-Ge_S5-M4_2020_4
Species richness
Subsampled reads
1e6
0
100
200
300
400
0.0
0.5
1.0
1.5
2.0
Subsampled data
Function fitted to data

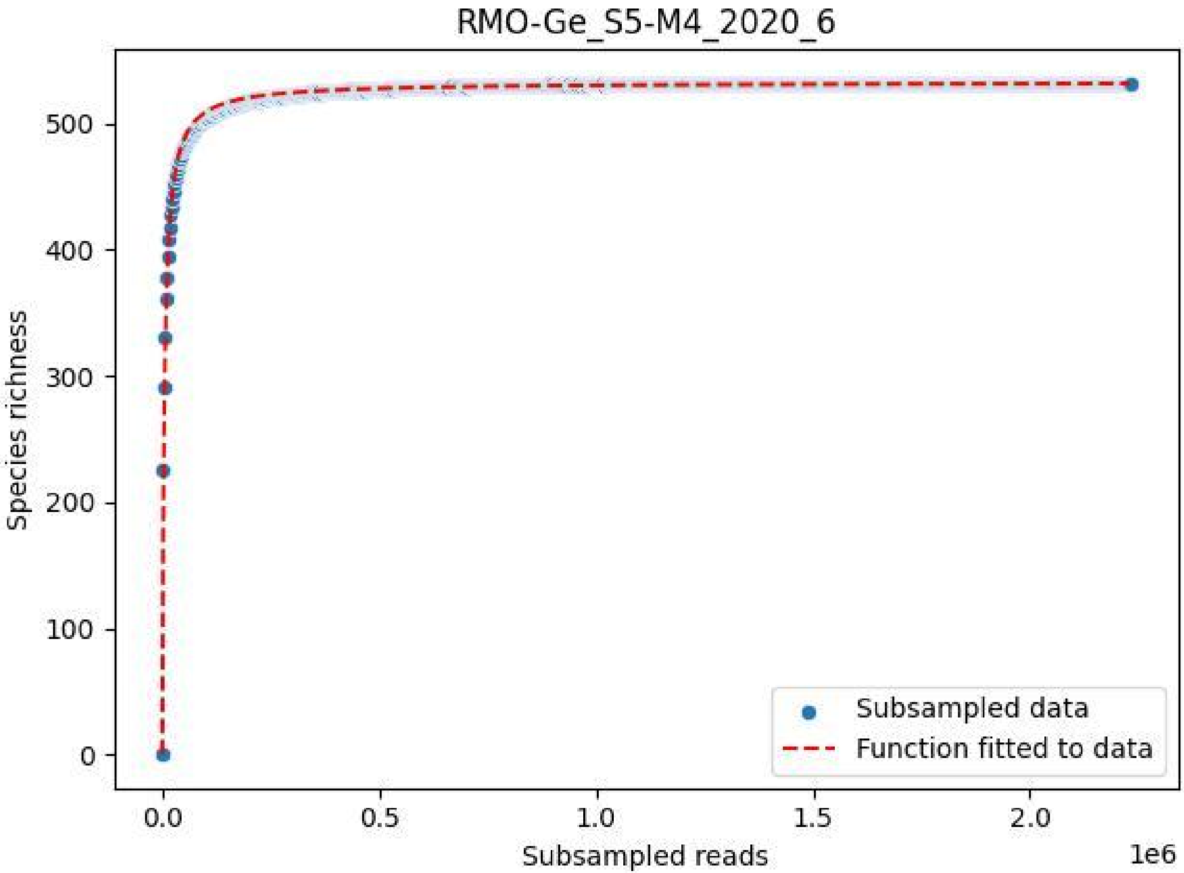
RMO-Ge_S5-M4_2020_6
Species richness
Subsampled reads
1e6
0
100
200
300
400
500
0.0
0.5
1.0
1.5
2.0
Subsampled data
Function fitted to data

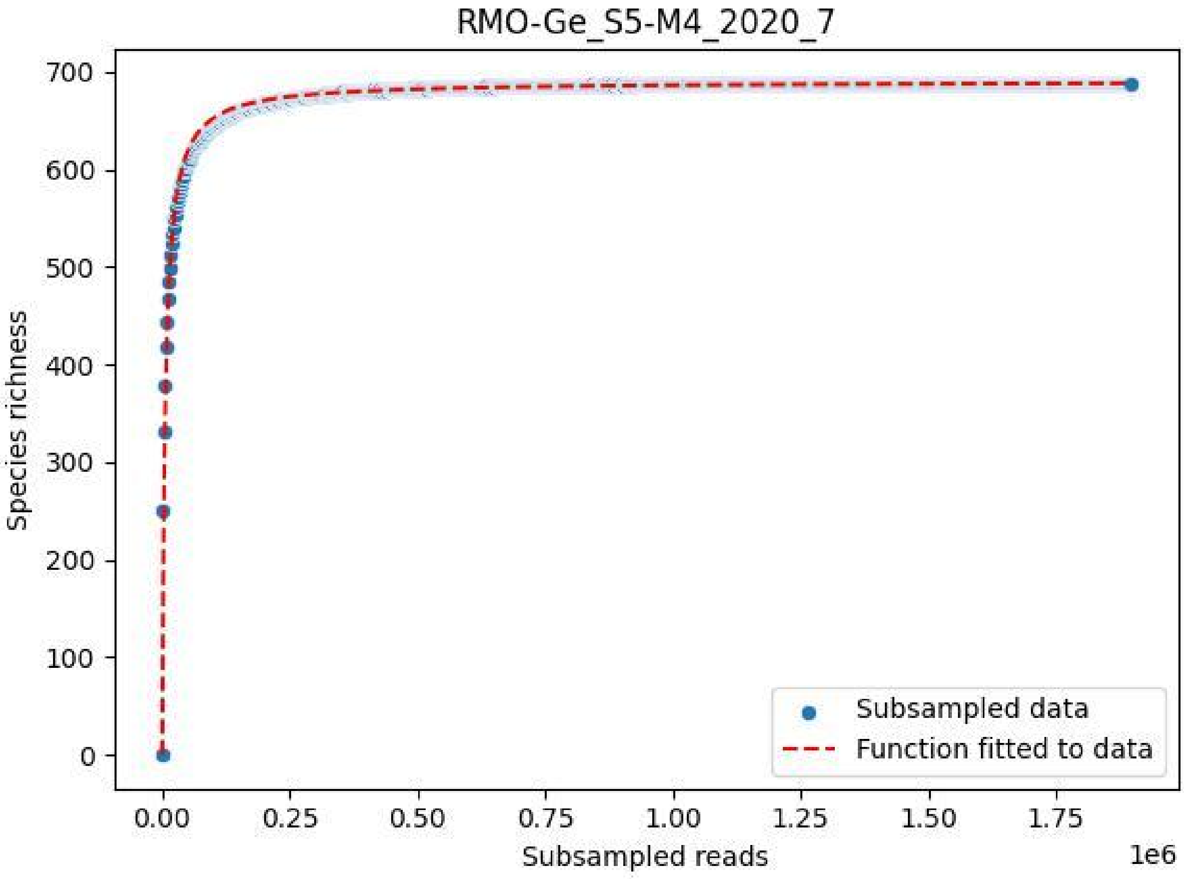
RMO-Ge_S5-M4_2020_7
Species richness
Subsampled reads
1e6
0
100
200
300
400
500
600
700
0.00
0.25
0.50
0.75
1.00
1.25
1.50
1.75
Subsampled data
Function fitted to data

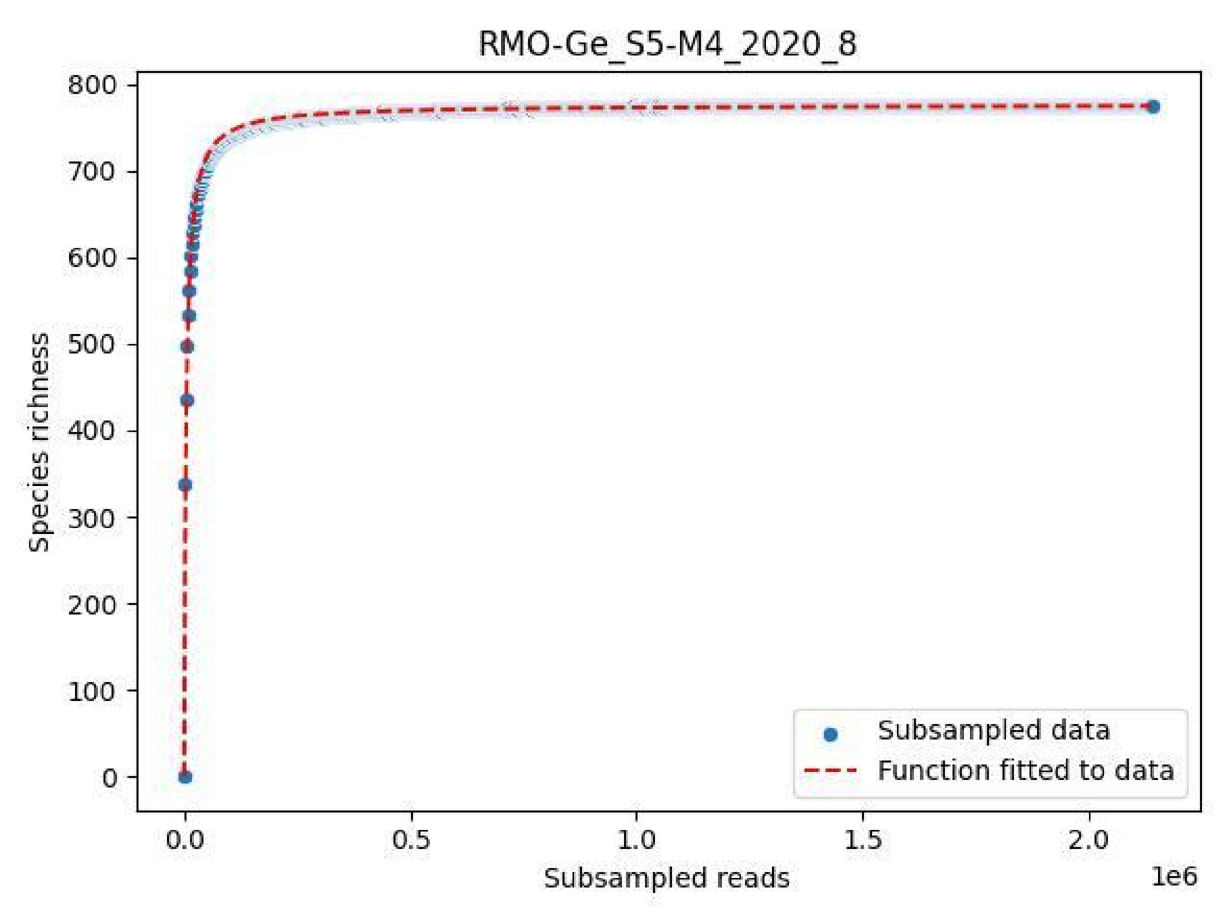

RMO-Ge_S5-M4_2020_8
800
700
600
500
400
300
200
100
0
Species richness
0.0
0.5
1.0
1.5
2.0
1e6
Subsampled reads
Subsampled data
Function fitted to data

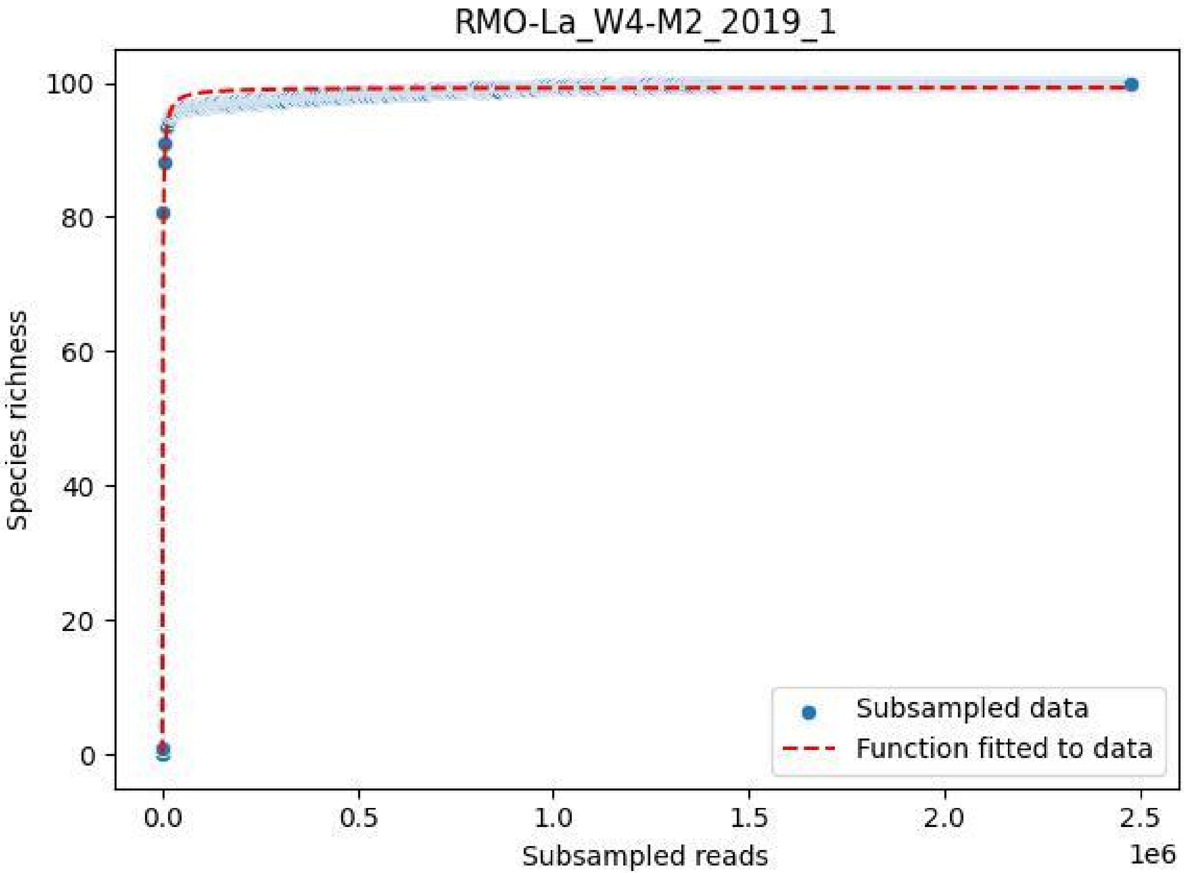
RMO-La_W4-M2_2019_1
Species richness
Subsampled reads
1e6
100
80
60
40
20
0
0.0
0.5
1.0
1.5
2.0
2.5
Subsampled data
Function fitted to data

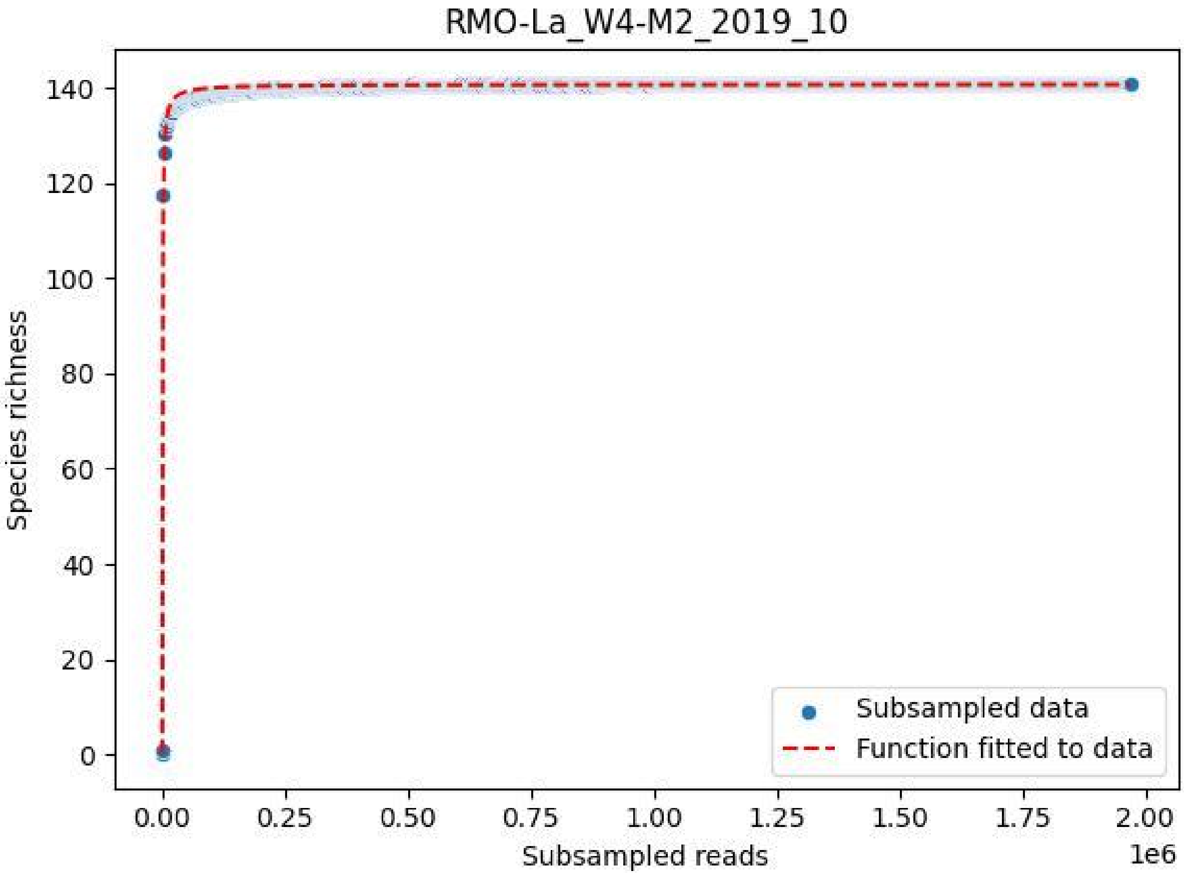
RMO-La_W4-M2_2019_10
Species richness
Subsampled reads
1e6
0
20
40
60
80
100
120
140
0.00
0.25
0.50
0.75
1.00
1.25
1.50
1.75
2.00
Subsampled data
Function fitted to data

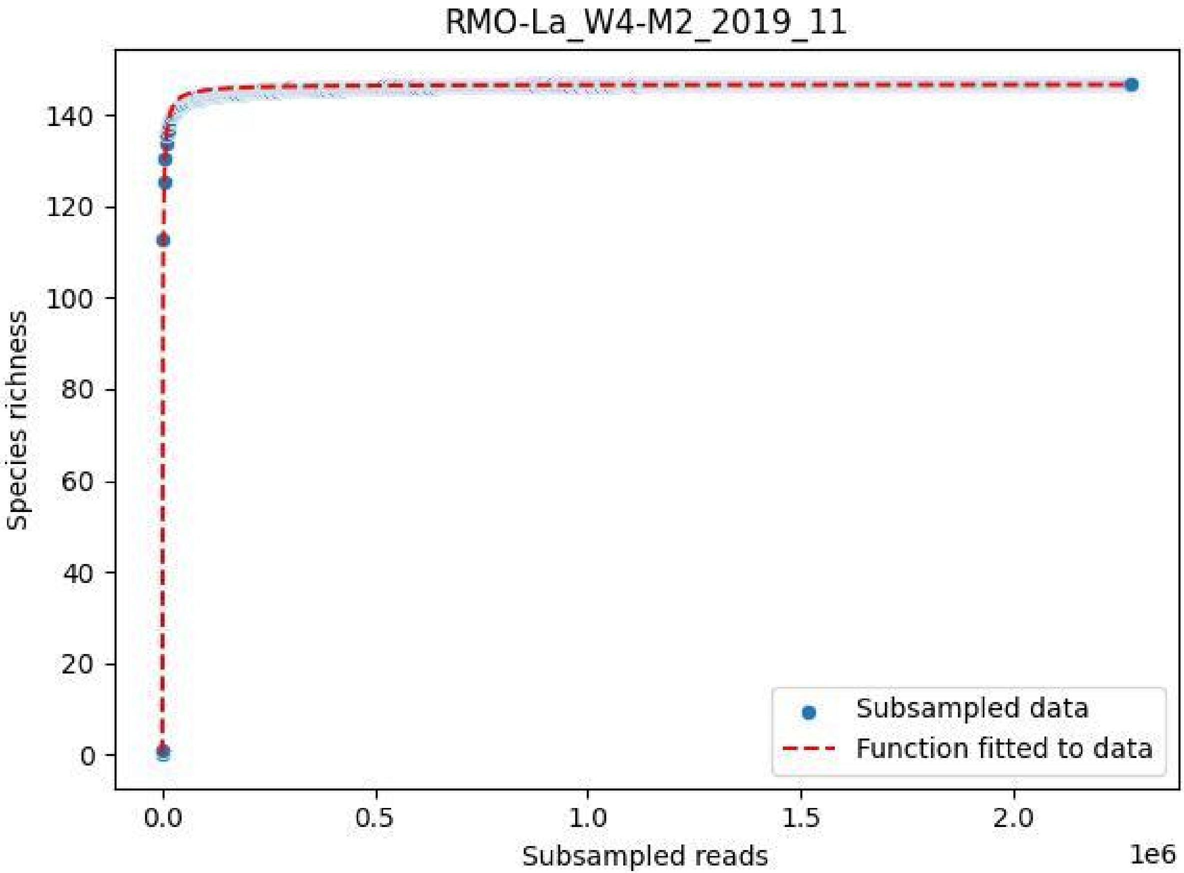

RMO-La_W4-M2_2019_11
Species richness
Subsampled reads
1e6
0
20
40
60
80
100
120
140
0.0
0.5
1.0
1.5
2.0
Subsampled data
Function fitted to data

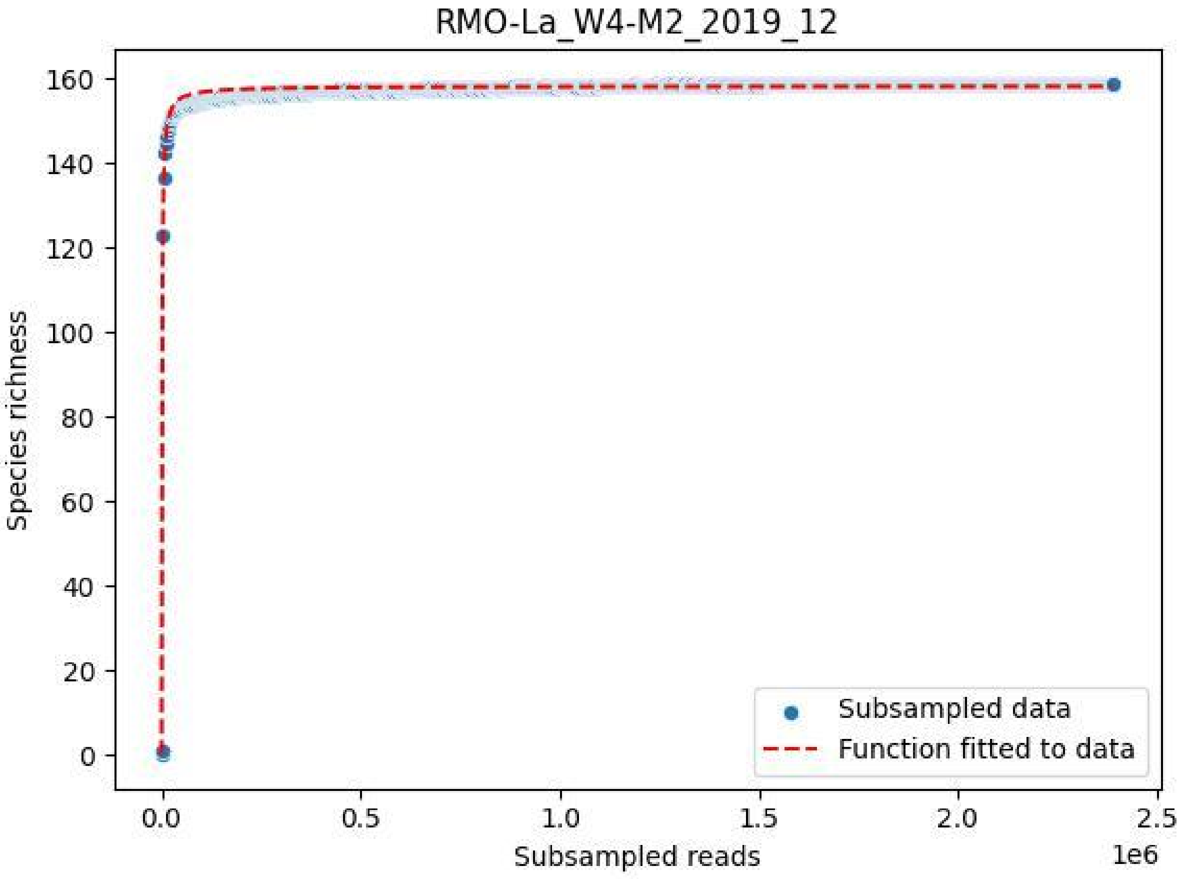

RMO-La_W4-M2_2019_12
Species richness
Subsampled reads
1e6
Subsampled data
Function fitted to data
0
20
40
60
80
100
120
140
160
0.0
0.5
1.0
1.5
2.0
2.5

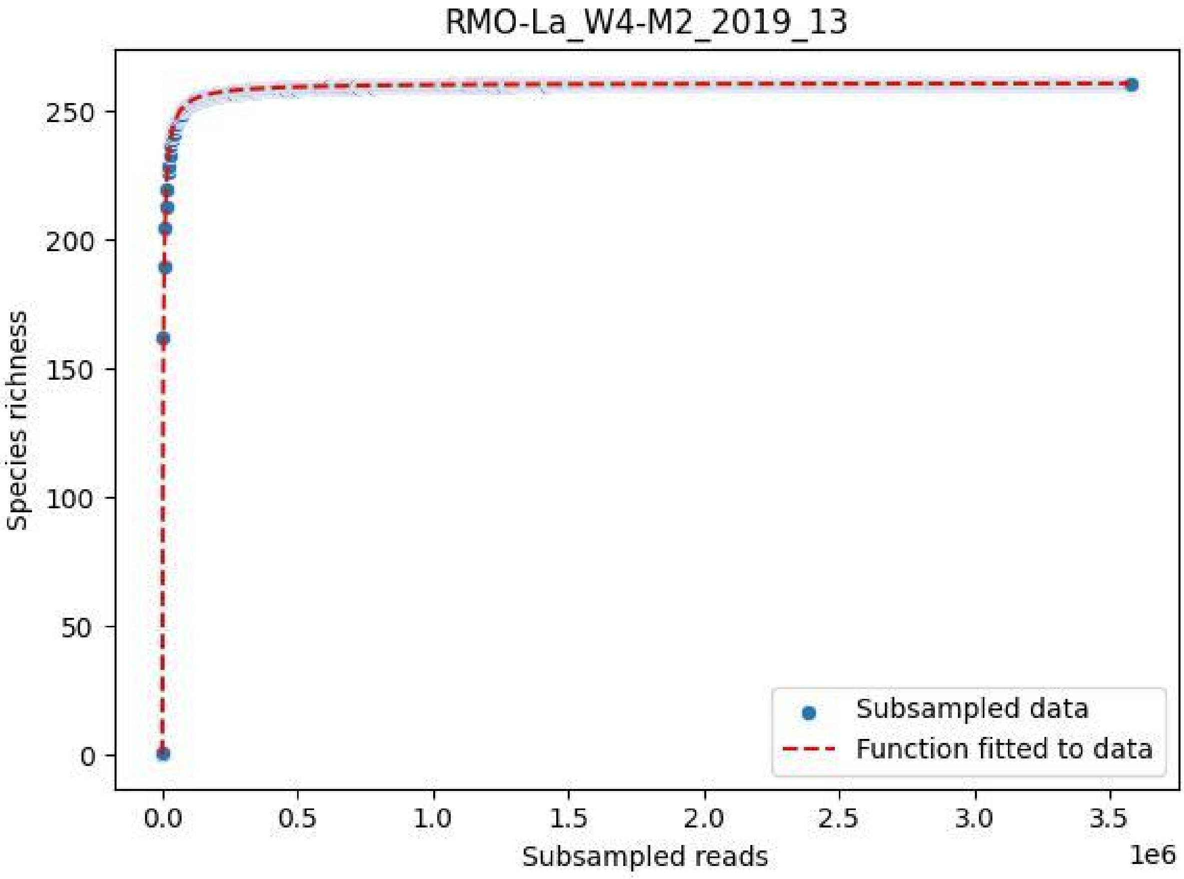

RMO-La_W4-M2_2019_13
Species richness
0
50
100
150
200
250
0.0
0.5
1.0
1.5
2.0
2.5
3.0
3.5
1e6
Subsampled reads
Subsampled data
Function fitted to data

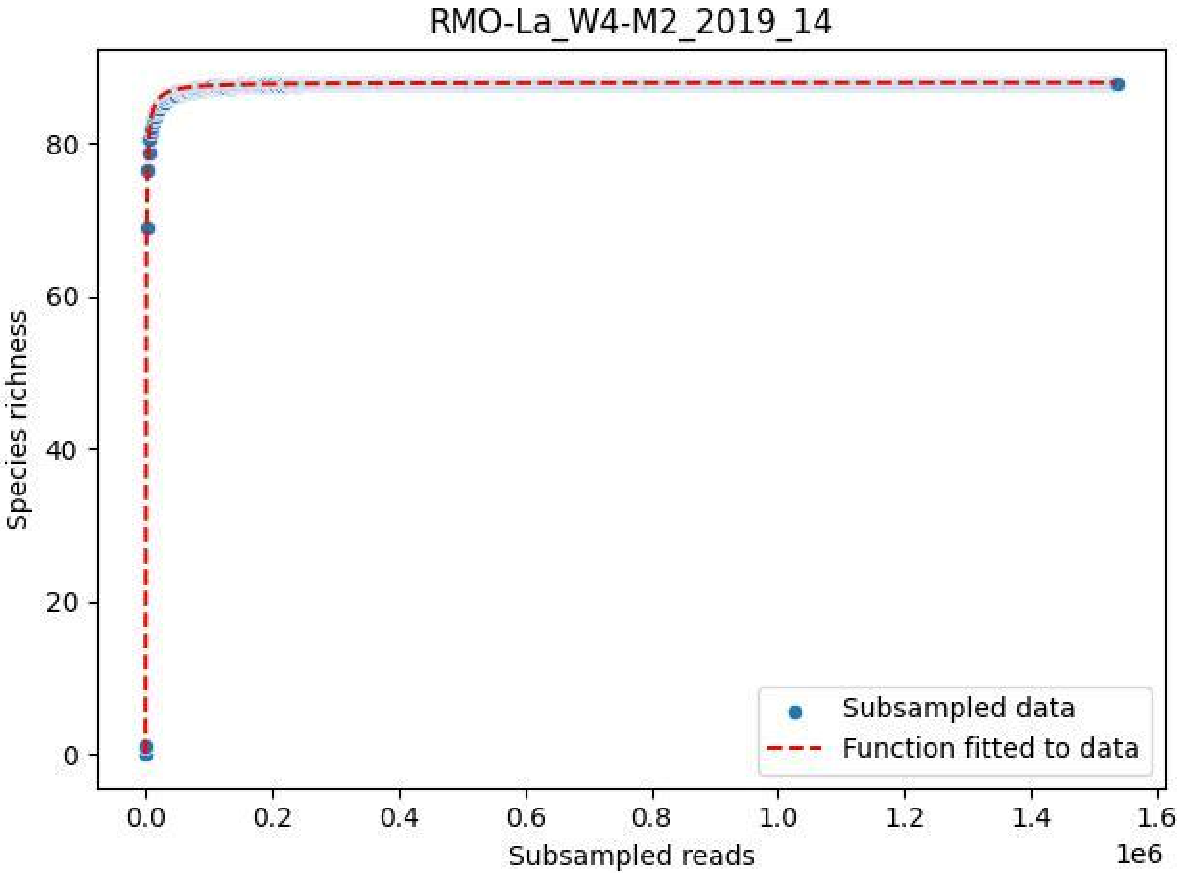
RMO-La_W4-M2_2019_14
Species richness
Subsampled reads
1e6
0
20
40
60
80
0.0
0.2
0.4
0.6
0.8
1.0
1.2
1.4
1.6
Subsampled data
Function fitted to data

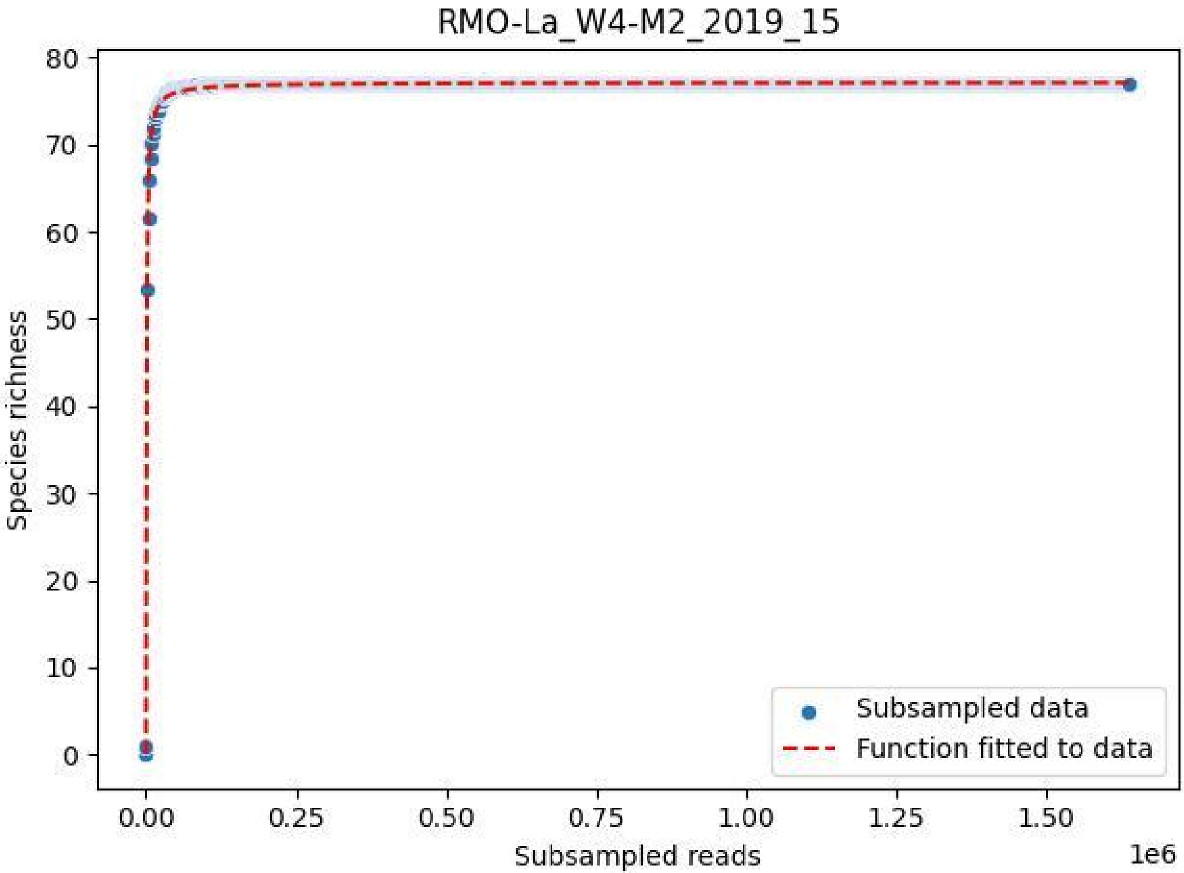
RMO-La_W4-M2_2019_15
Species richness
Subsampled reads
1e6
0
10
20
30
40
50
60
70
80
0.00
0.25
0.50
0.75
1.00
1.25
1.50
Subsampled data
Function fitted to data

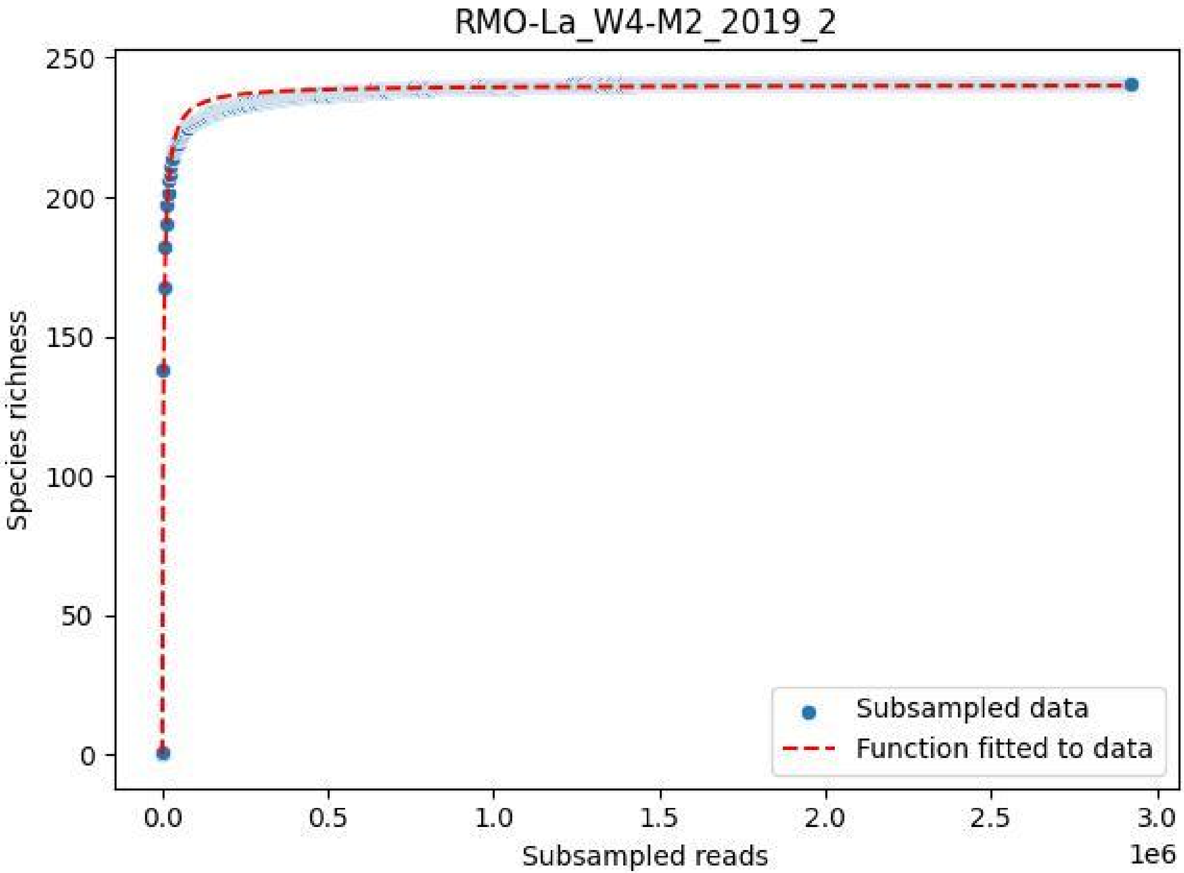

RMO-La_W4-M2_2019_2
Species richness
Subsampled reads
1e6
0
50
100
150
200
250
0.0
0.5
1.0
1.5
2.0
2.5
3.0
Subsampled data
Function fitted to data

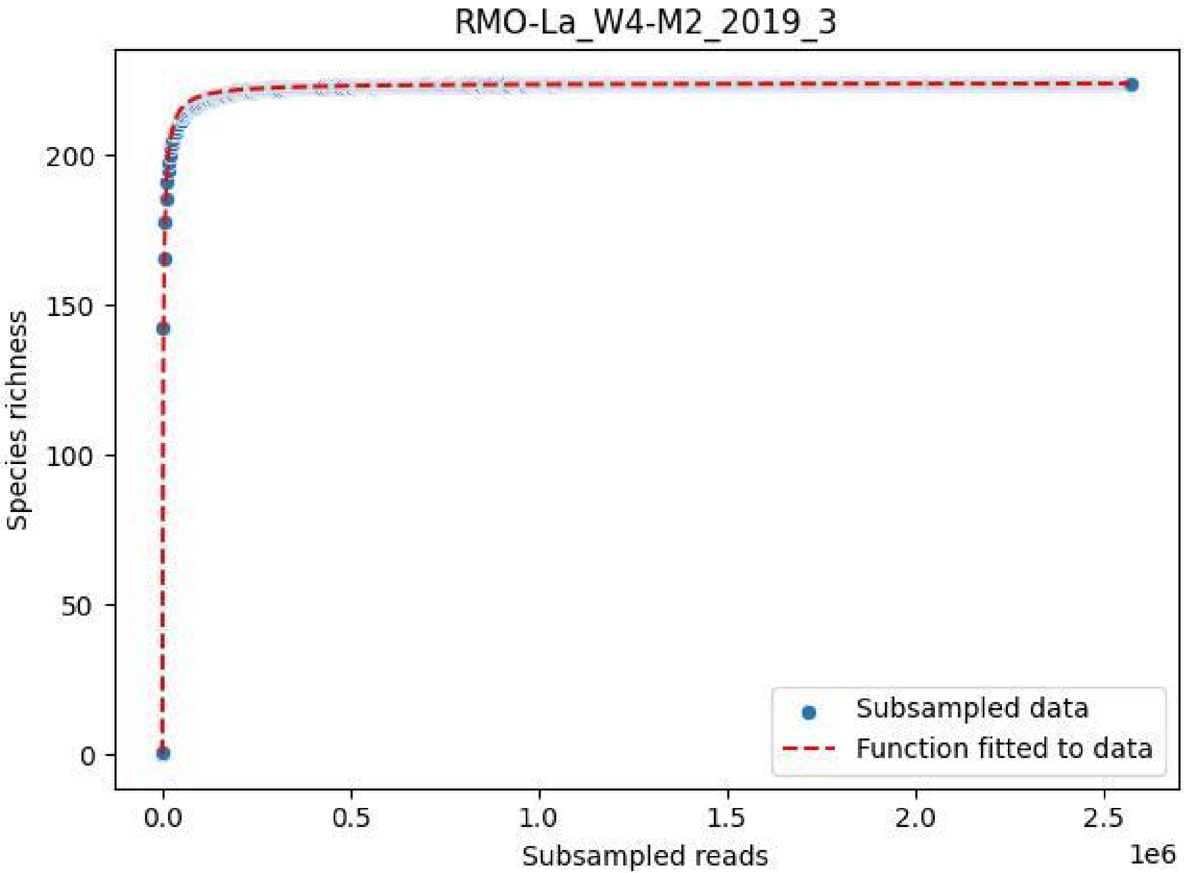

RMO-La_W4-M2_2019_3
Species richness
Subsampled reads
1e6
0
50
100
150
200
0.0
0.5
1.0
1.5
2.0
2.5
Subsampled data
Function fitted to data

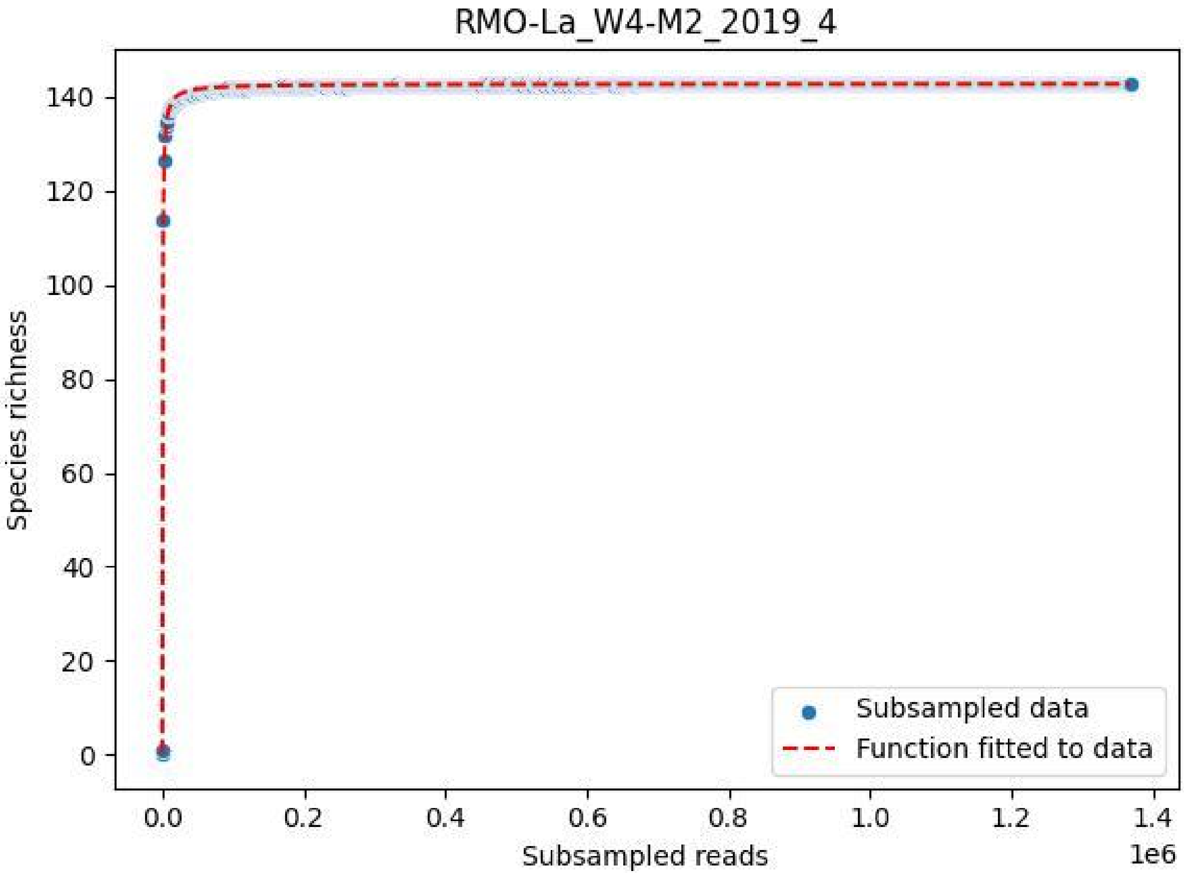

RMO-La_W4-M2_2019_4
Species richness
Subsampled reads
1e6
0
20
40
60
80
100
120
140
0.0
0.2
0.4
0.6
0.8
1.0
1.2
1.4
Subsampled data
Function fitted to data

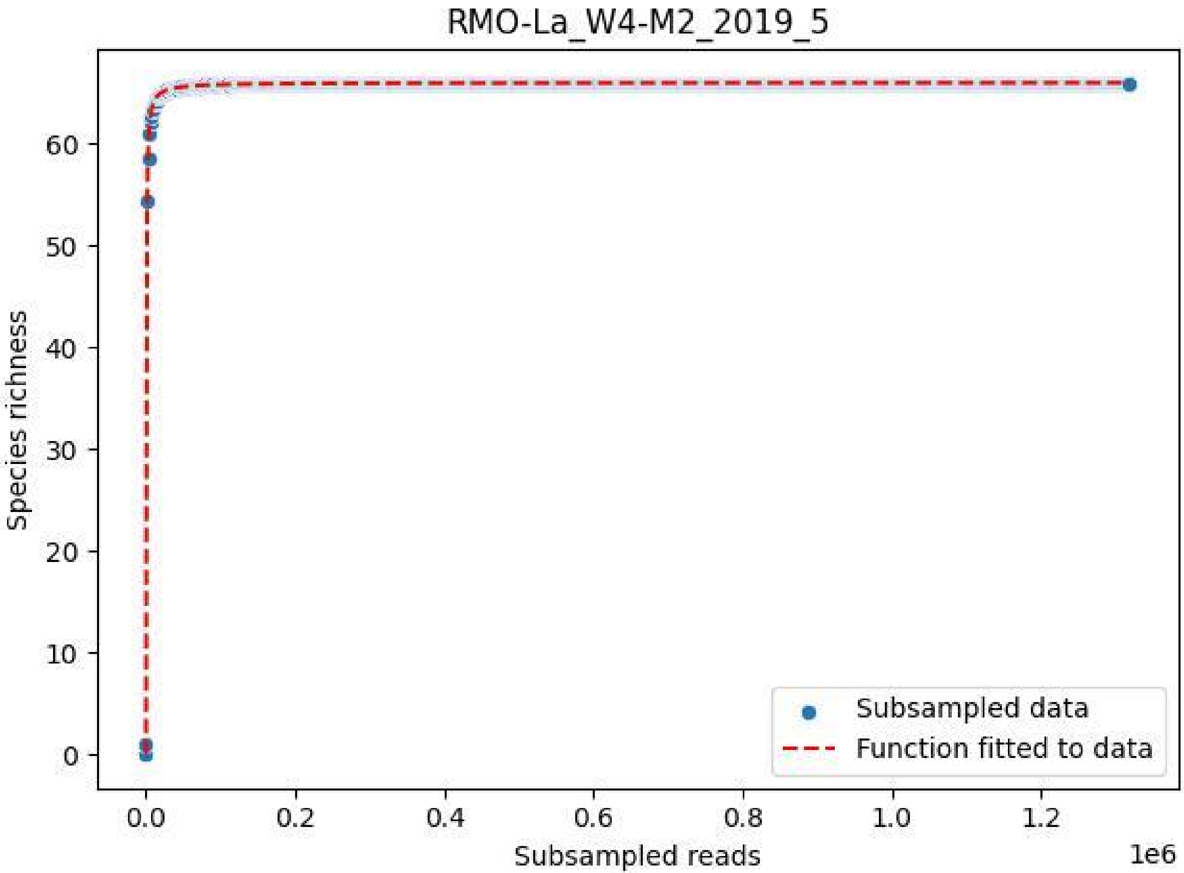

RMO-La_W4-M2_2019_5
Species richness
0
10
20
30
40
50
60
0.0
0.2
0.4
0.6
0.8
1.0
1.2
Subsampled reads
1e6
Subsampled data
Function fitted to data

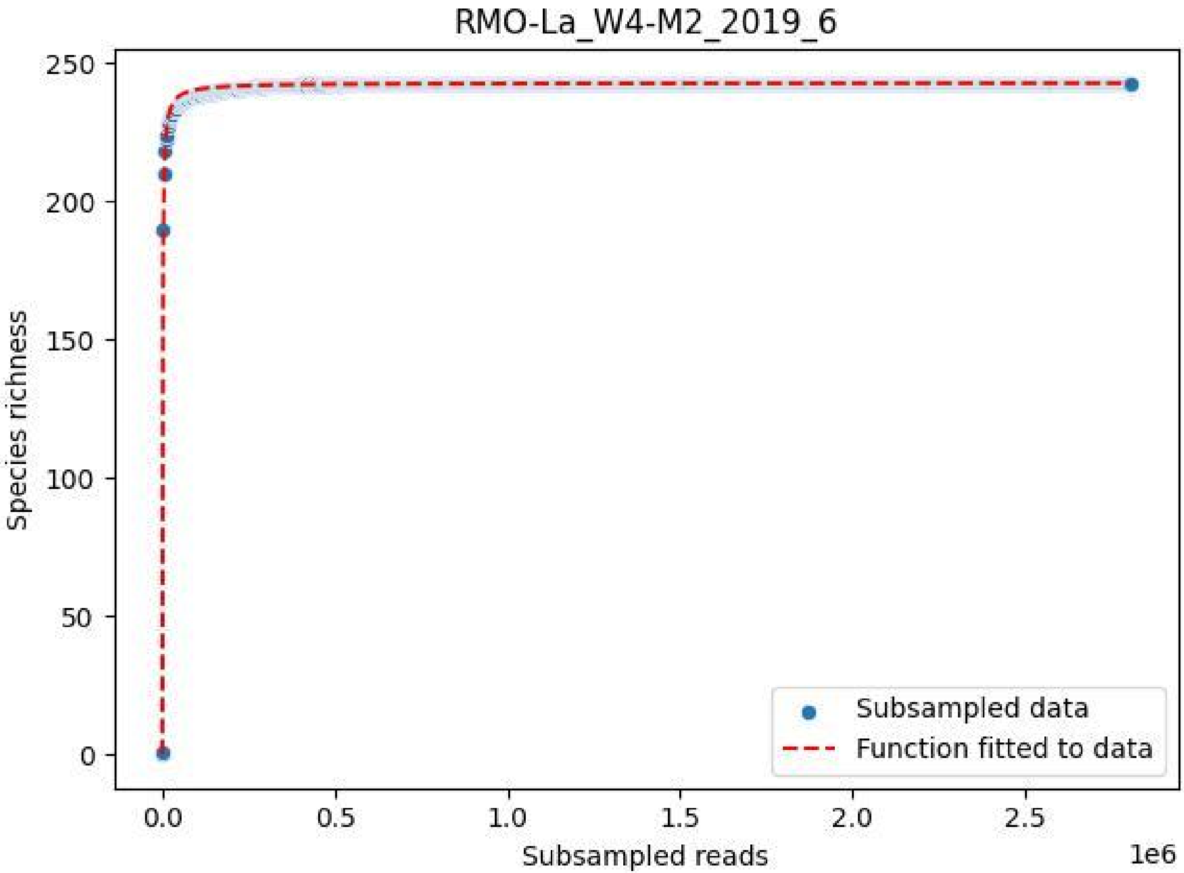

RMO-La_W4-M2_2019_6
Species richness
Subsampled reads
1e6
0
50
100
150
200
250
0.0
0.5
1.0
1.5
2.0
2.5
Subsampled data
Function fitted to data

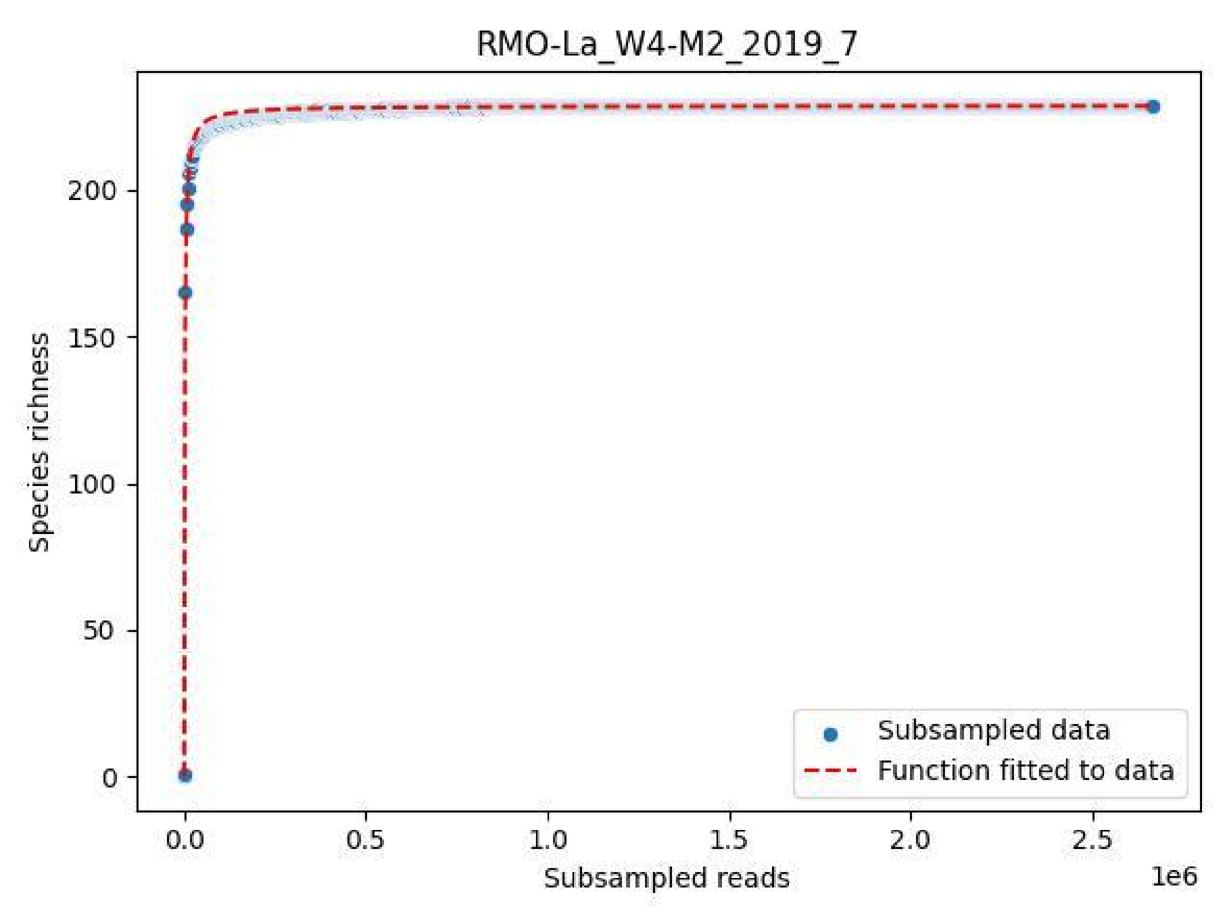

RMO-La_W4-M2_2019_7
Species richness
Subsampled reads
1e6
0
50
100
150
200
0.0
0.5
1.0
1.5
2.0
2.5
Subsampled data
Function fitted to data

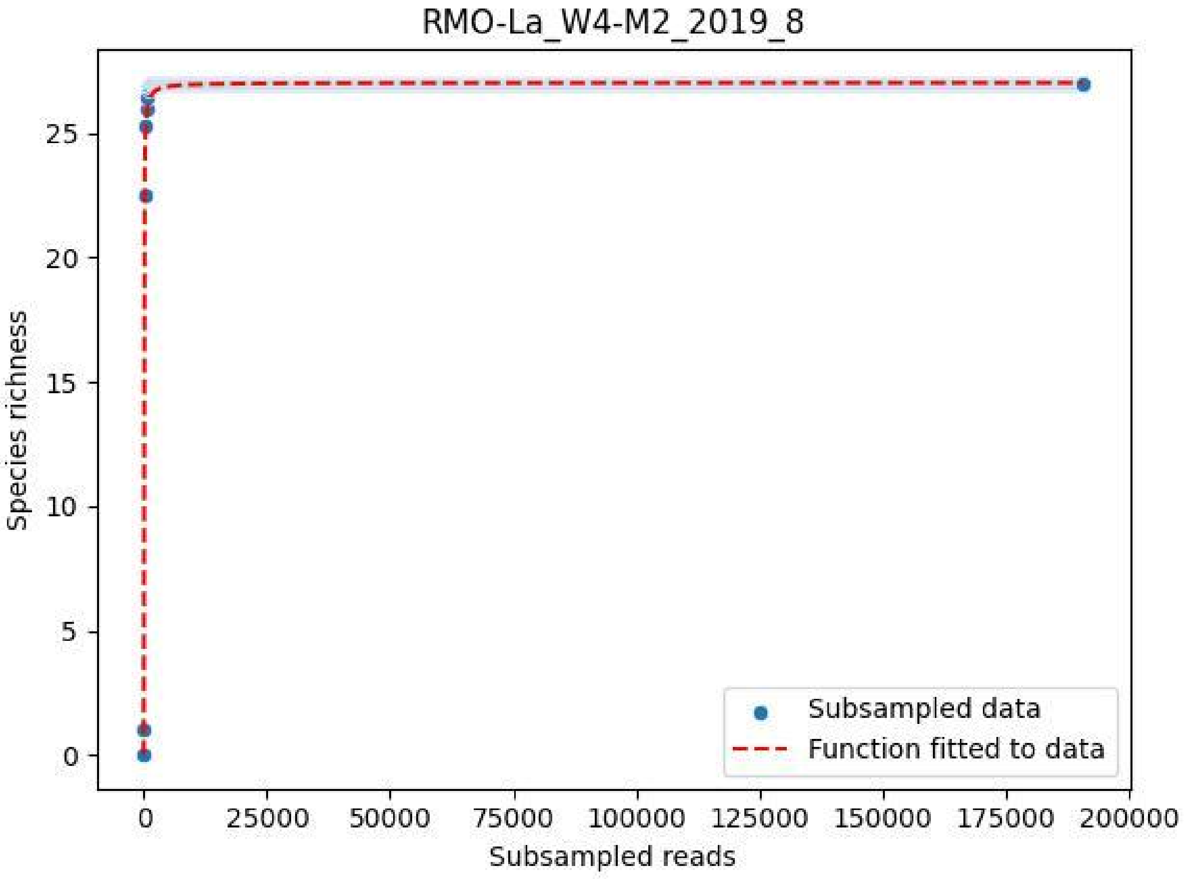

RMO-La_W4-M2_2019_8
Species richness
Subsampled reads
0
5
10
15
20
25
0
25000
50000
75000
100000
125000
150000
175000
200000
Subsampled data
Function fitted to data

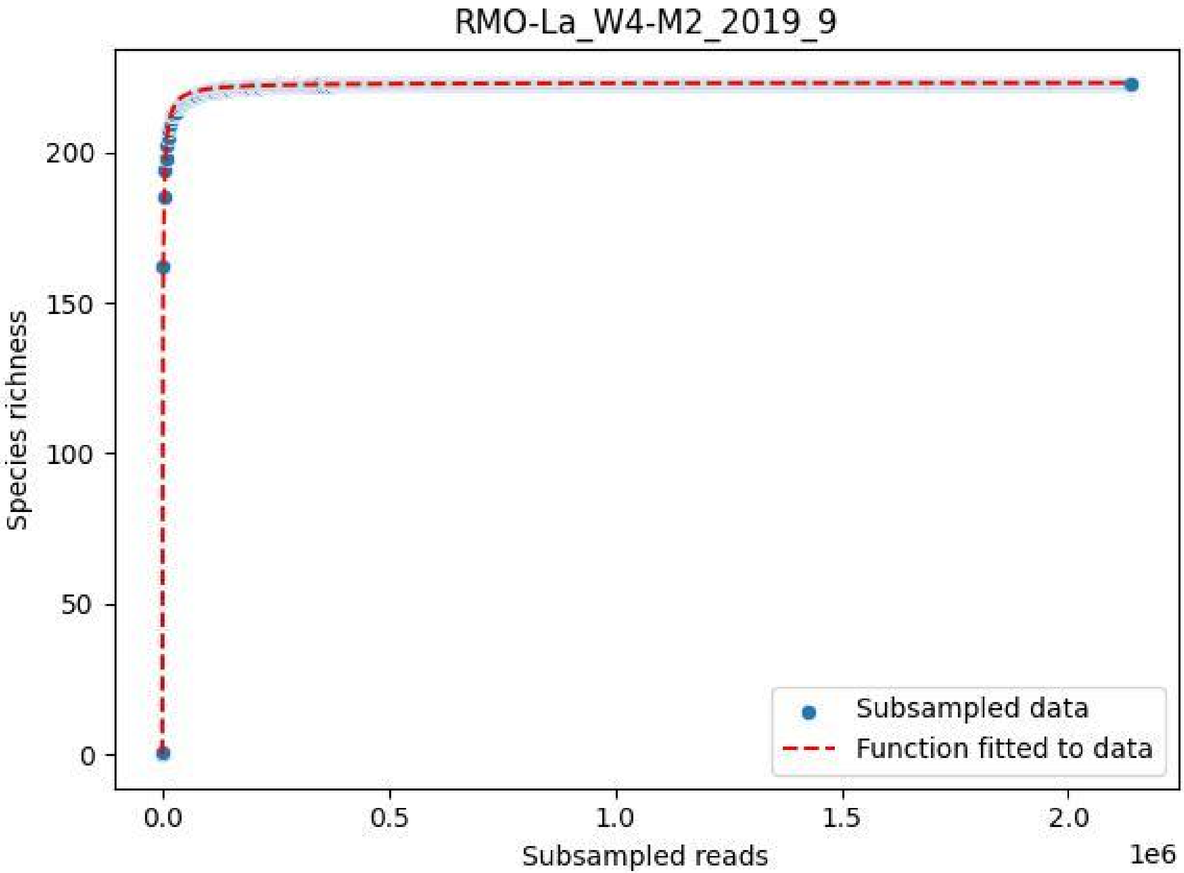
RMO-La_W4-M2_2019_9
Species richness
200
150
100
50
0
0.0
0.5
1.0
1.5
2.0
1e6
Subsampled reads
Subsampled data
Function fitted to data

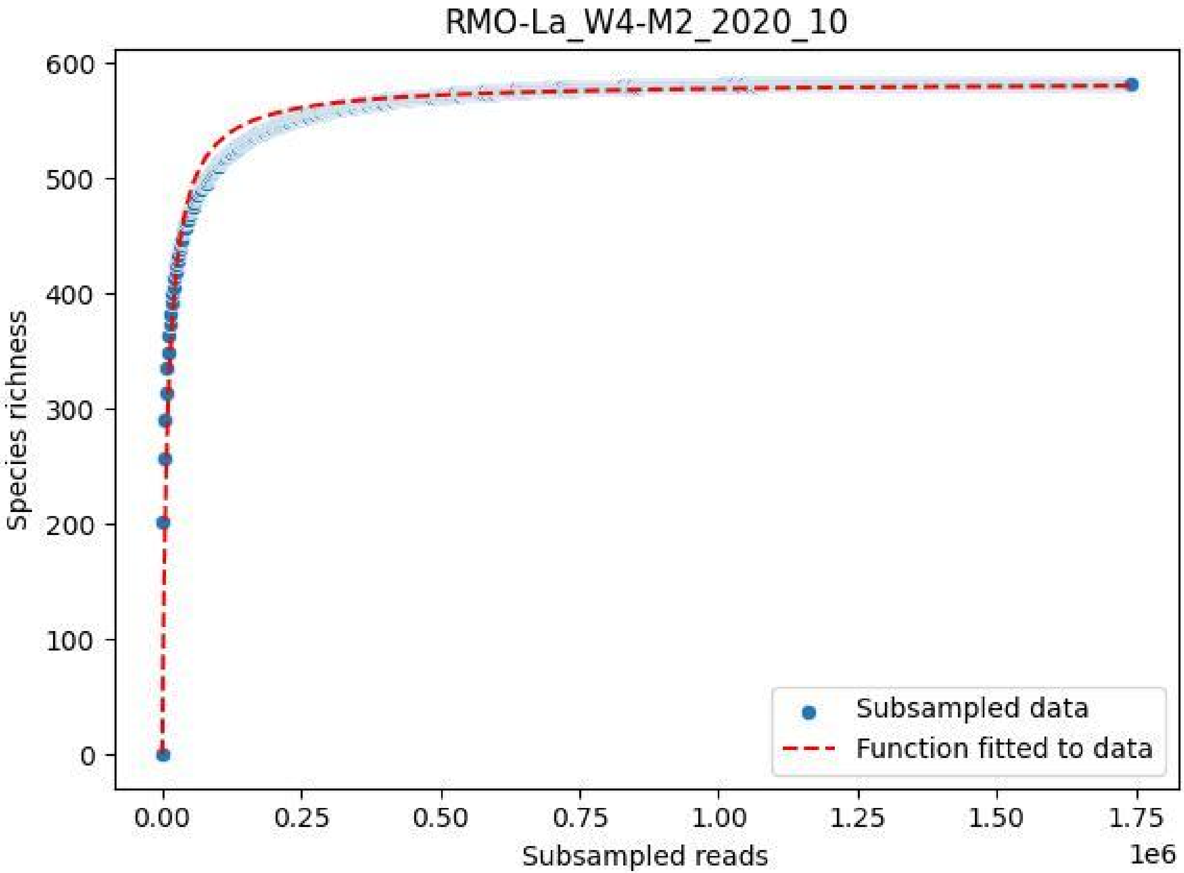
RMO-La_W4-M2_2020_10
Species richness
Subsampled reads
1e6
0
100
200
300
400
500
600
0.00
0.25
0.50
0.75
1.00
1.25
1.50
1.75
Subsampled data
Function fitted to data

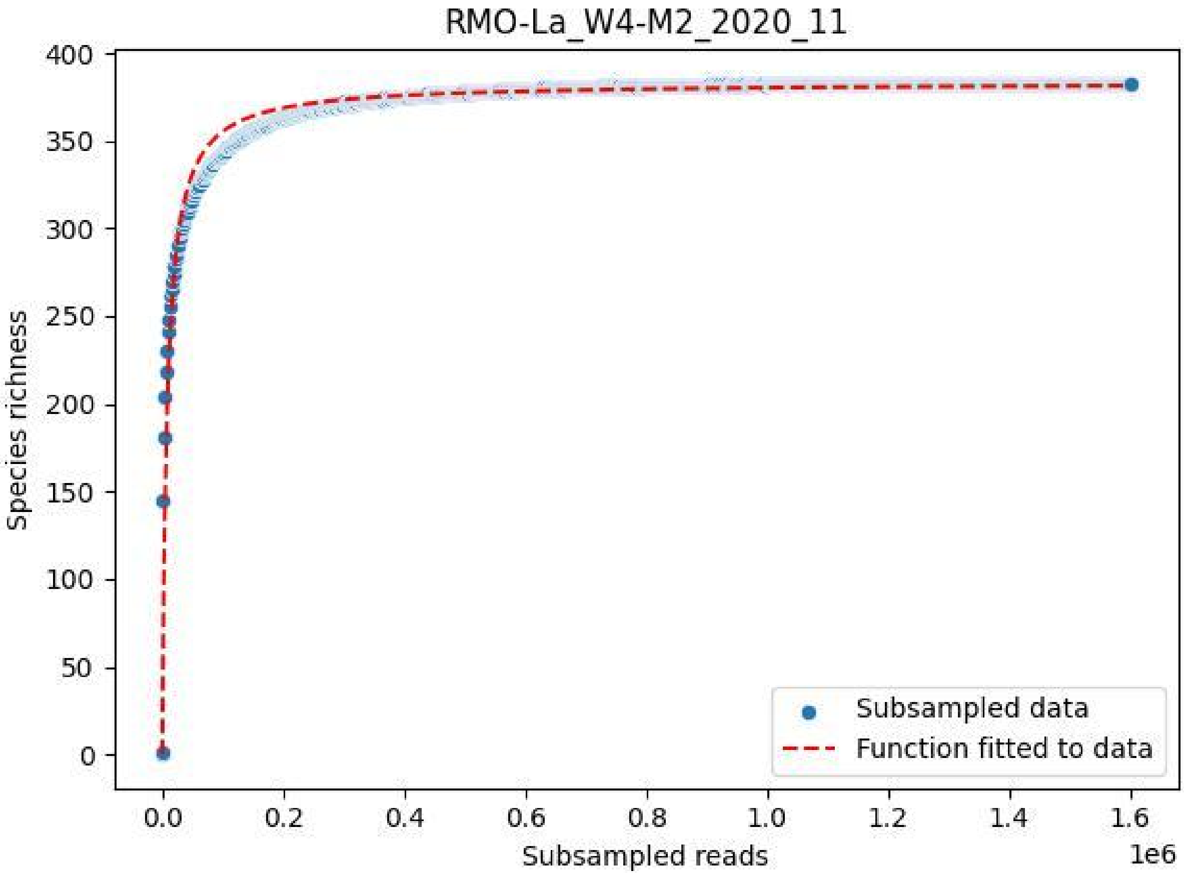
RMO-La_W4-M2_2020_11
Species richness
Subsampled reads
1e6
400
350
300
250
200
150
100
50
0
0.0
0.2
0.4
0.6
0.8
1.0
1.2
1.4
1.6
Subsampled data
Function fitted to data

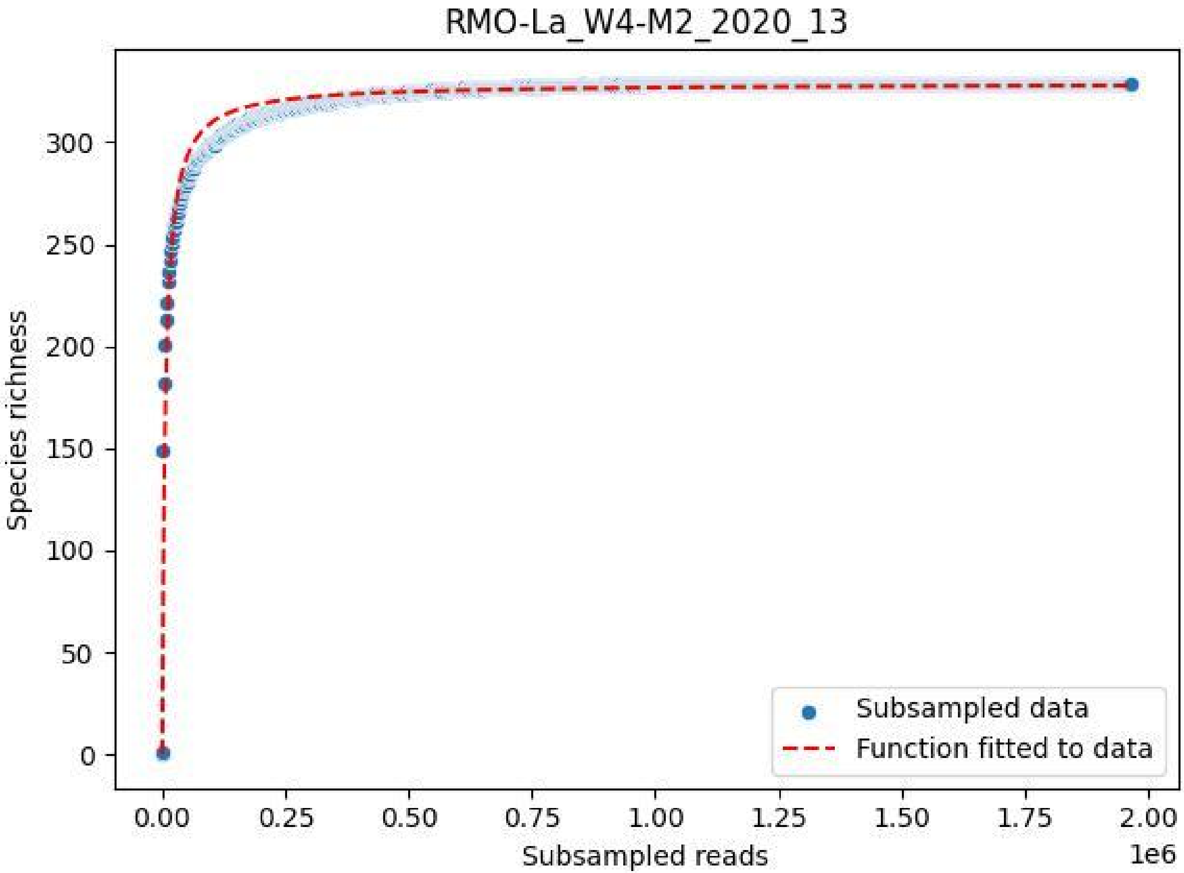

RMO-La_W4-M2_2020_13
Species richness
300
250
200
150
100
50
0
0.00
0.25
0.50
0.75
1.00
1.25
1.50
1.75
2.00
Subsampled reads
1e6
Subsampled data
Function fitted to data

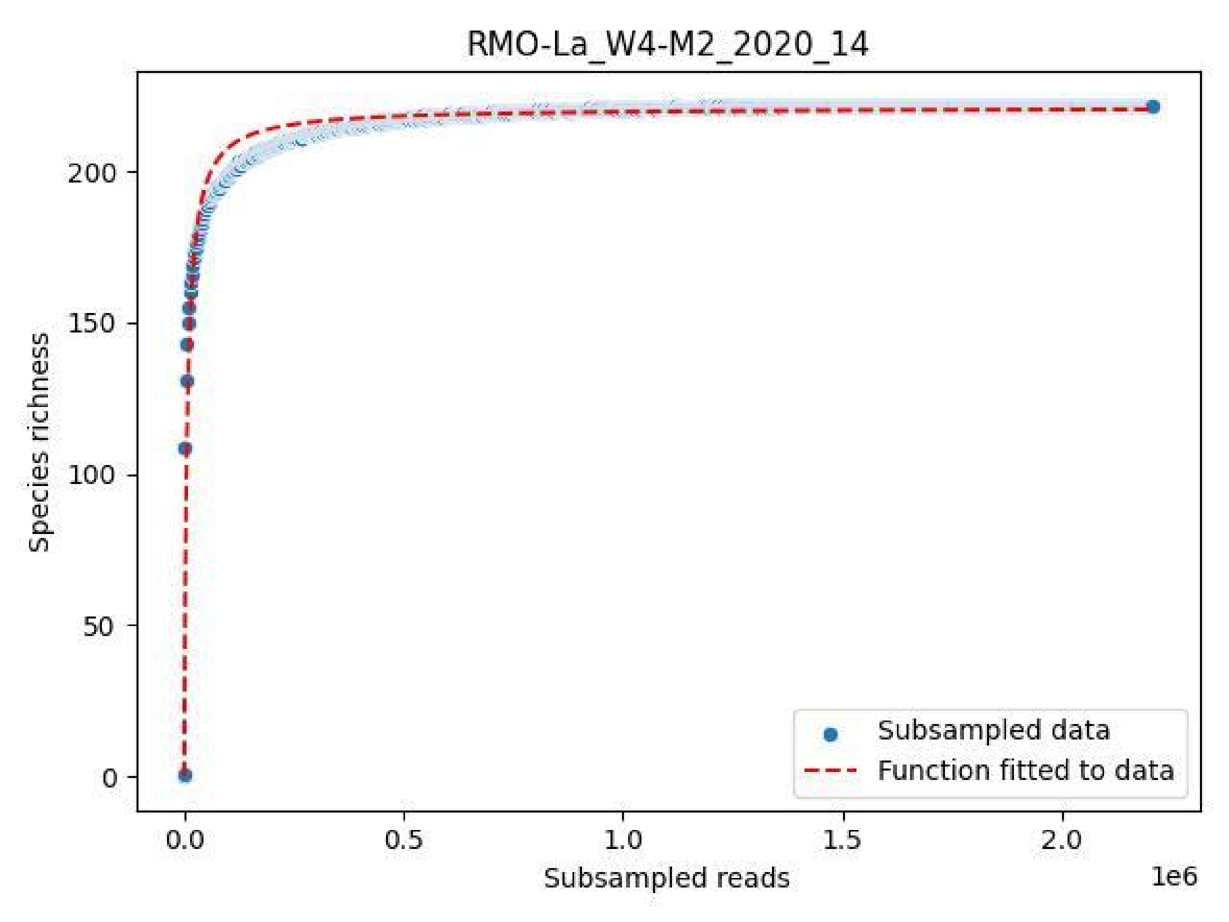

RMO-La_W4-M2_2020_14
Species richness
Subsampled reads
1e6
0
50
100
150
200
0.0
0.5
1.0
1.5
2.0
Subsampled data
Function fitted to data

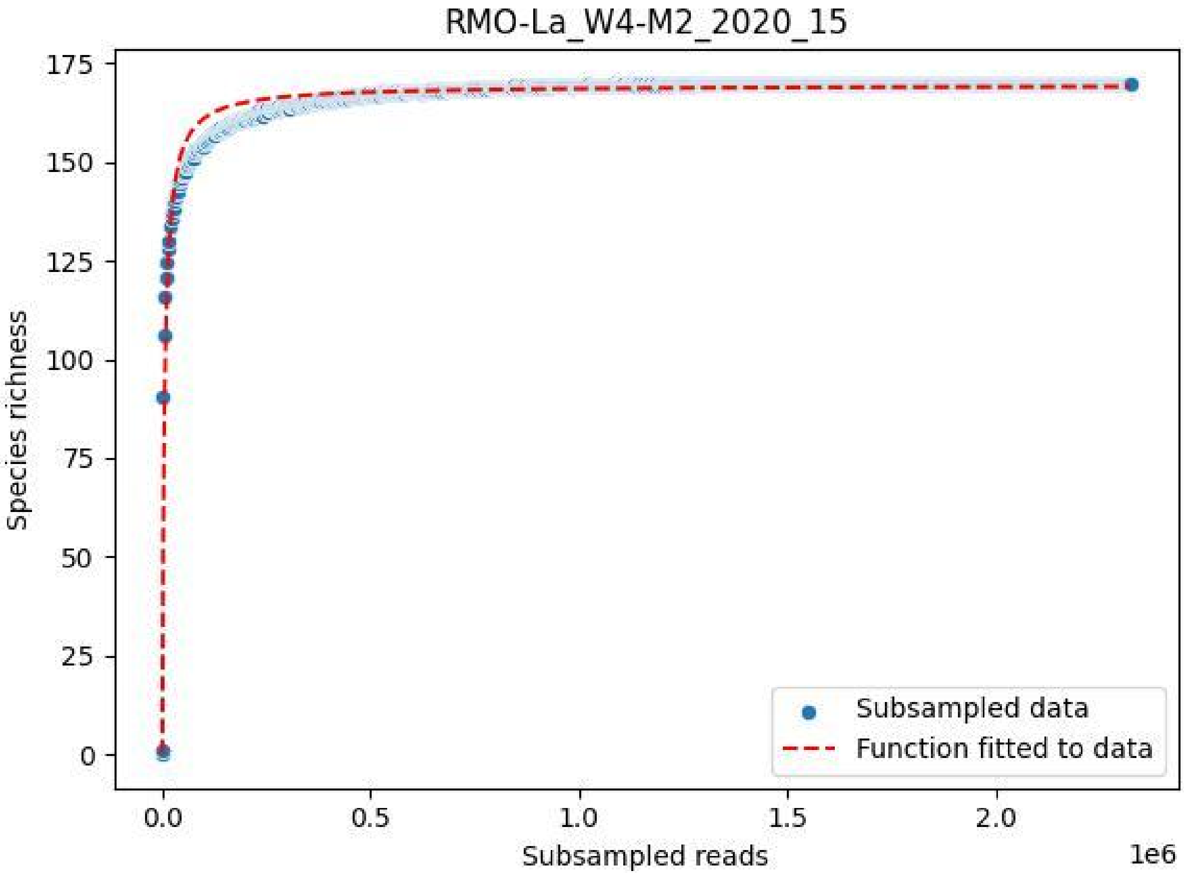

RMO-La_W4-M2_2020_15
Species richness
Subsampled reads
1e6
0
25
50
75
100
125
150
175
0.0
0.5
1.0
1.5
2.0
Subsampled data
Function fitted to data

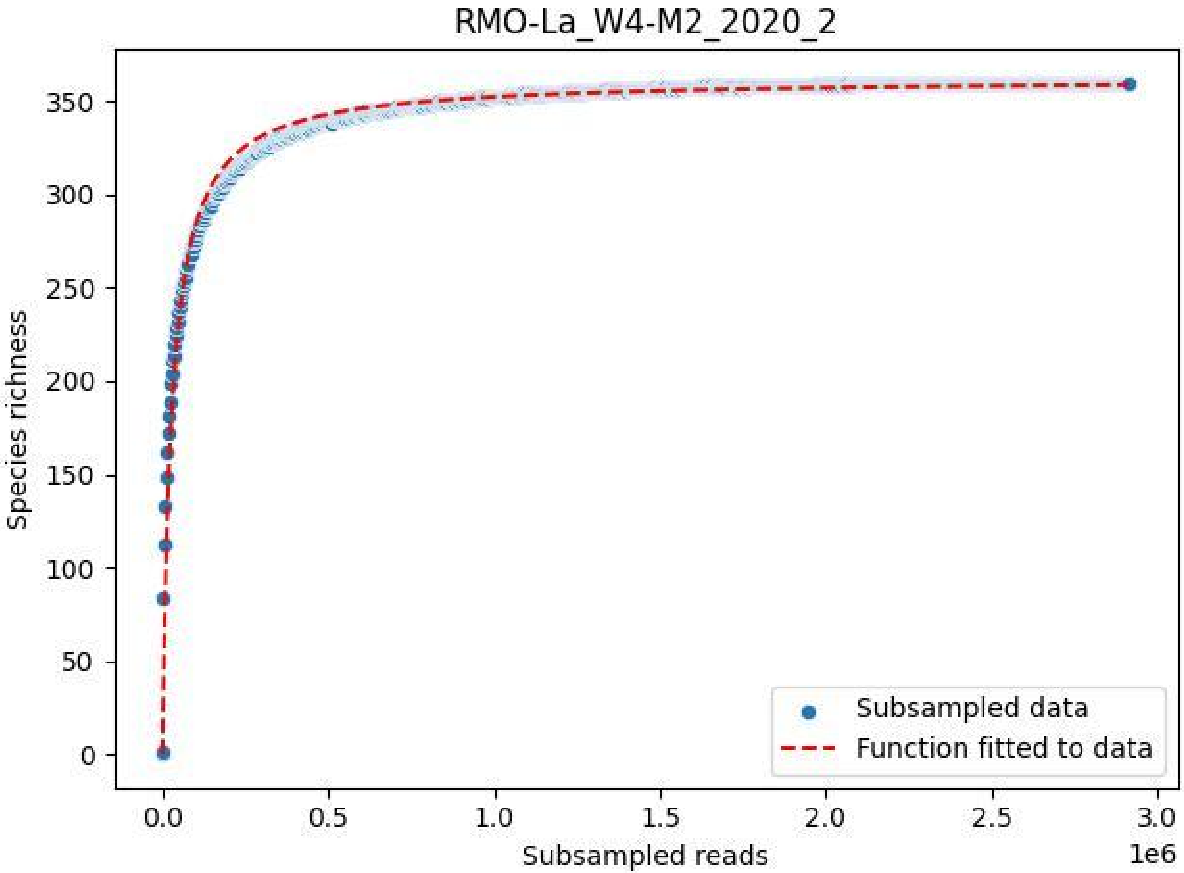

RMO-La_W4-M2_2020_2
Species richness
Subsampled reads
1e6
0
50
100
150
200
250
300
350
0.0
0.5
1.0
1.5
2.0
2.5
3.0
Subsampled data
Function fitted to data

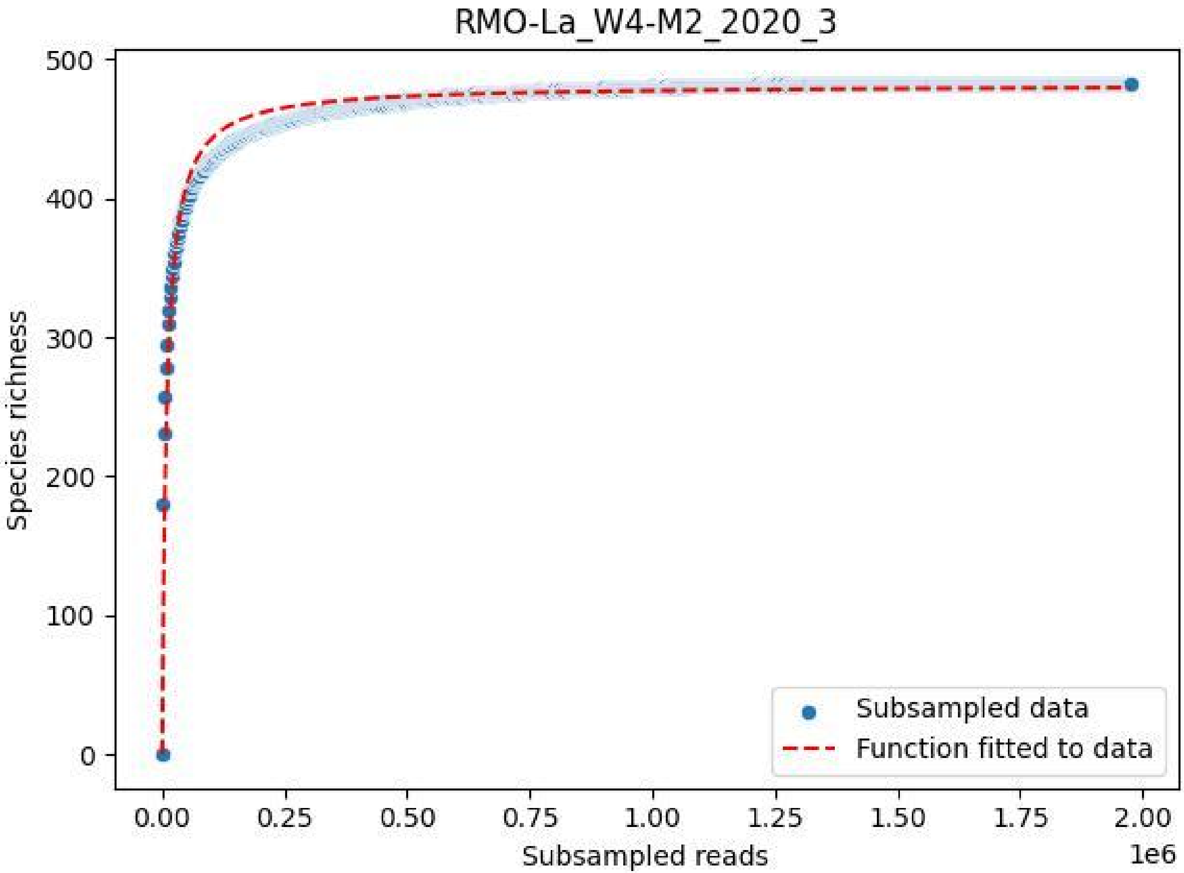

RMO-La_W4-M2_2020_3
Species richness
500
400
300
200
100
0
0.00
0.25
0.50
0.75
1.00
1.25
1.50
1.75
2.00
1e6
Subsampled reads
Subsampled data
Function fitted to data

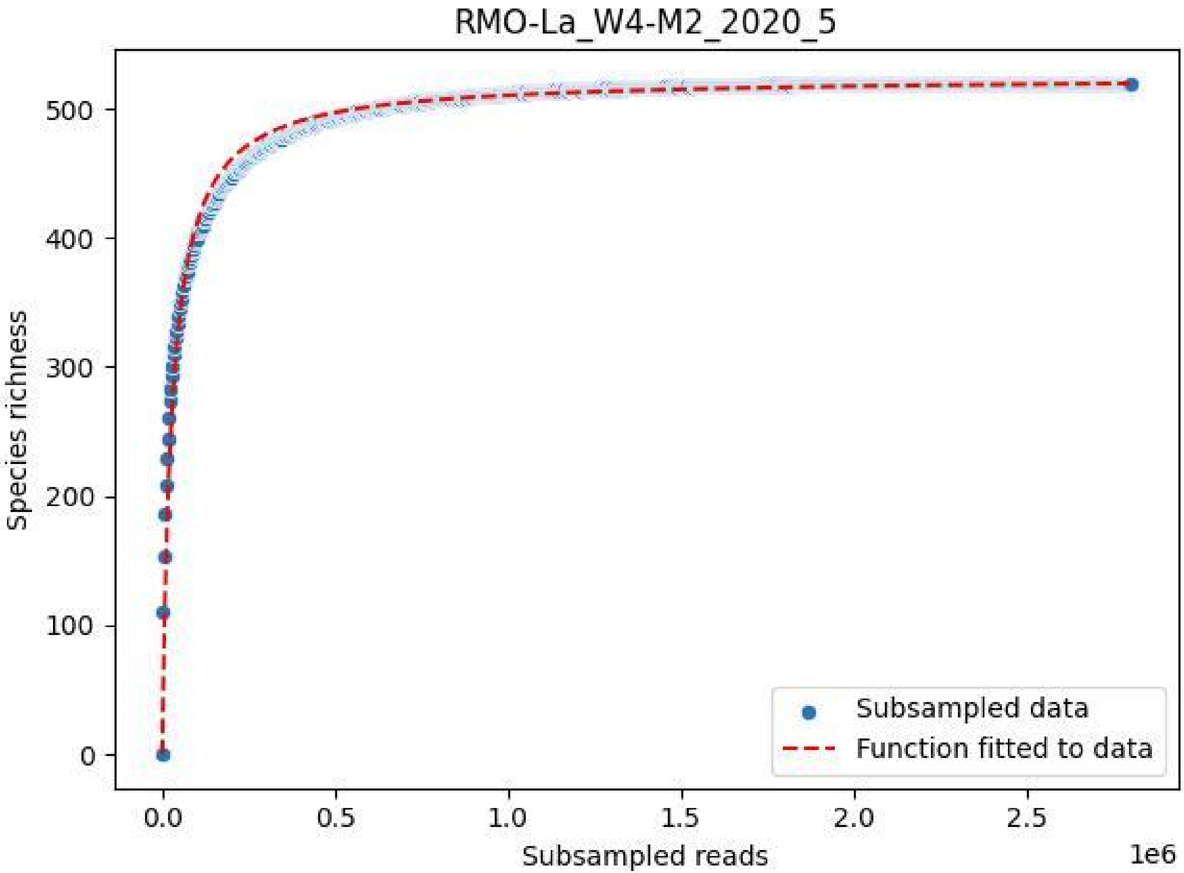
RMO-La_W4-M2_2020_5
Species richness
Subsampled reads
1e6
0
100
200
300
400
500
0.0
0.5
1.0
1.5
2.0
2.5
Subsampled data
Function fitted to data

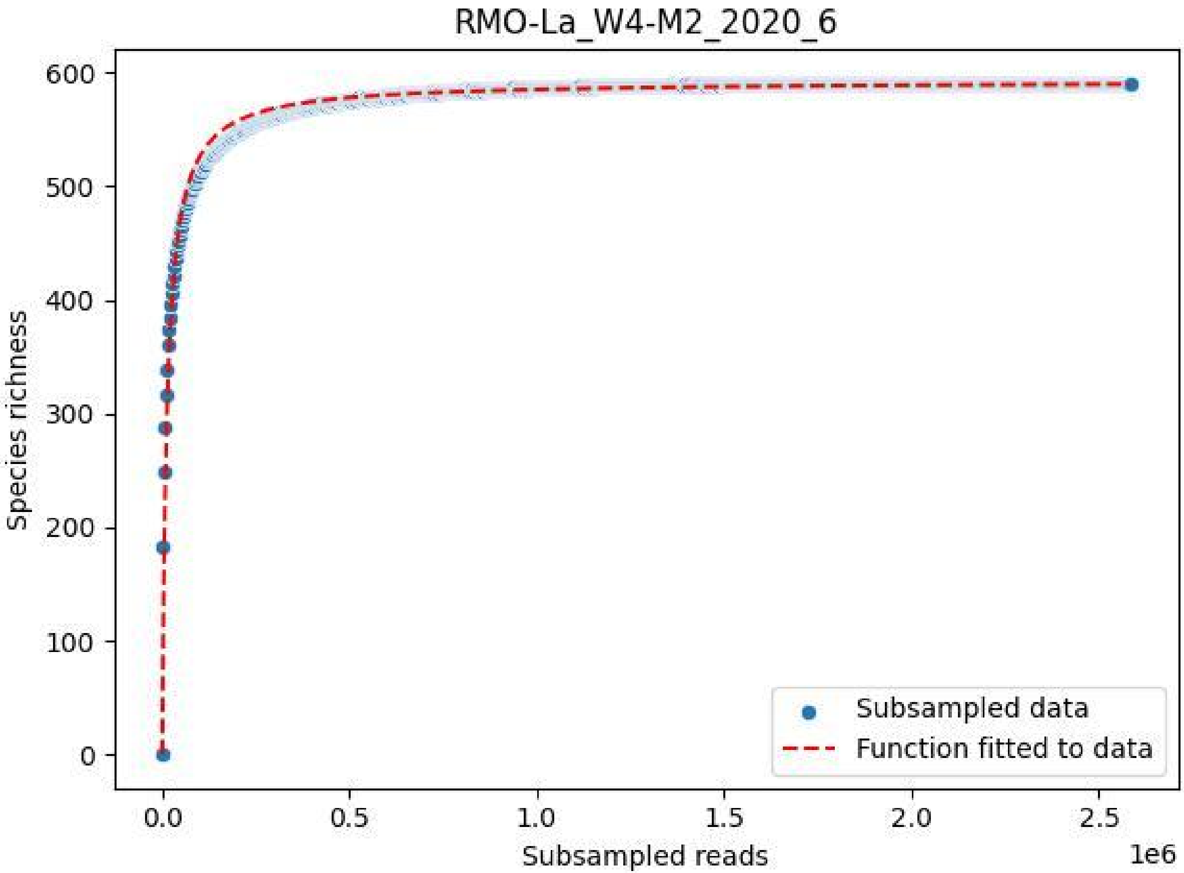
RMO-La_W4-M2_2020_6
600
500
400
300
200
100
0
Species richness
0.0
0.5
1.0
1.5
2.0
2.5
1e6
Subsampled reads
Subsampled data
Function fitted to data

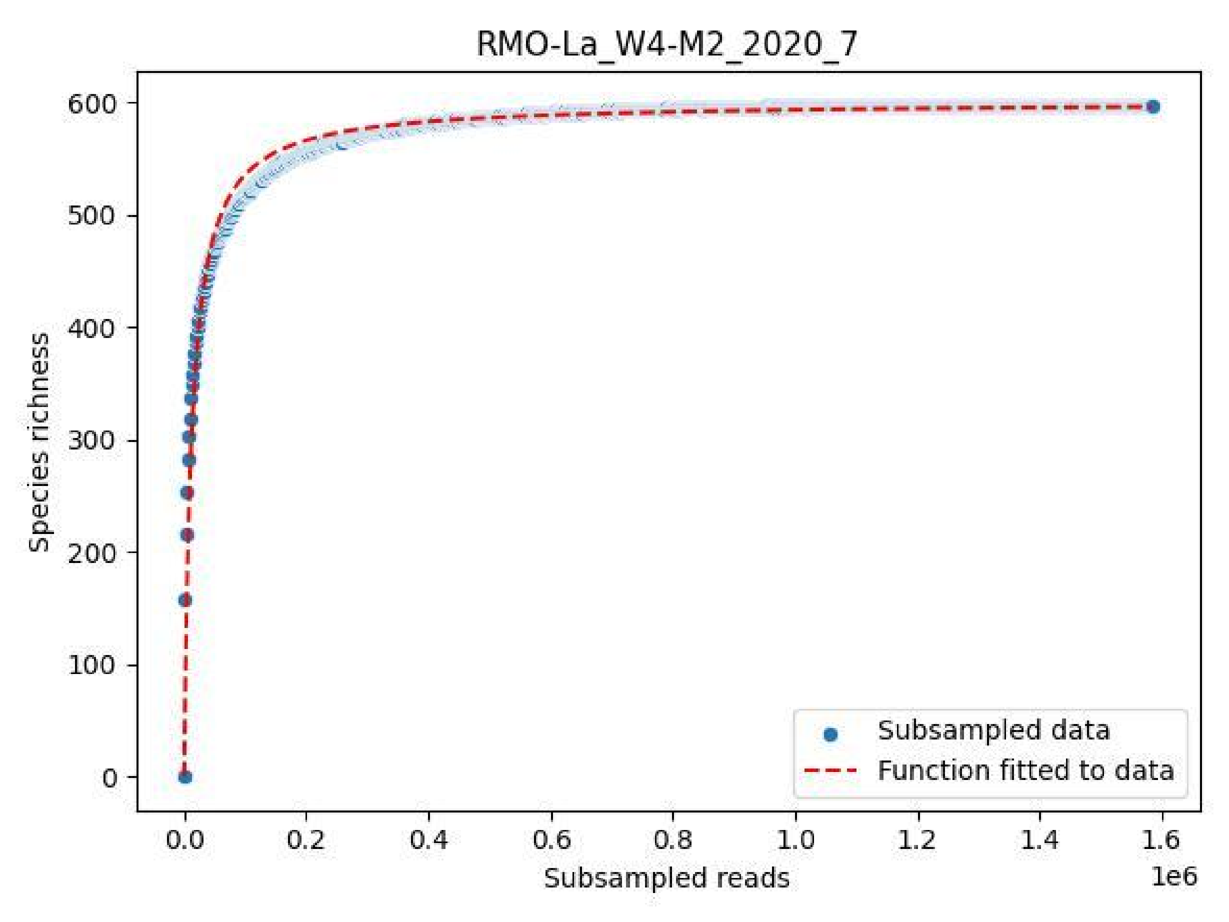
RMO-La_W4-M2_2020_7
Species richness
Subsampled reads
1e6
0
100
200
300
400
500
600
0.0
0.2
0.4
0.6
0.8
1.0
1.2
1.4
1.6
Subsampled data
Function fitted to data

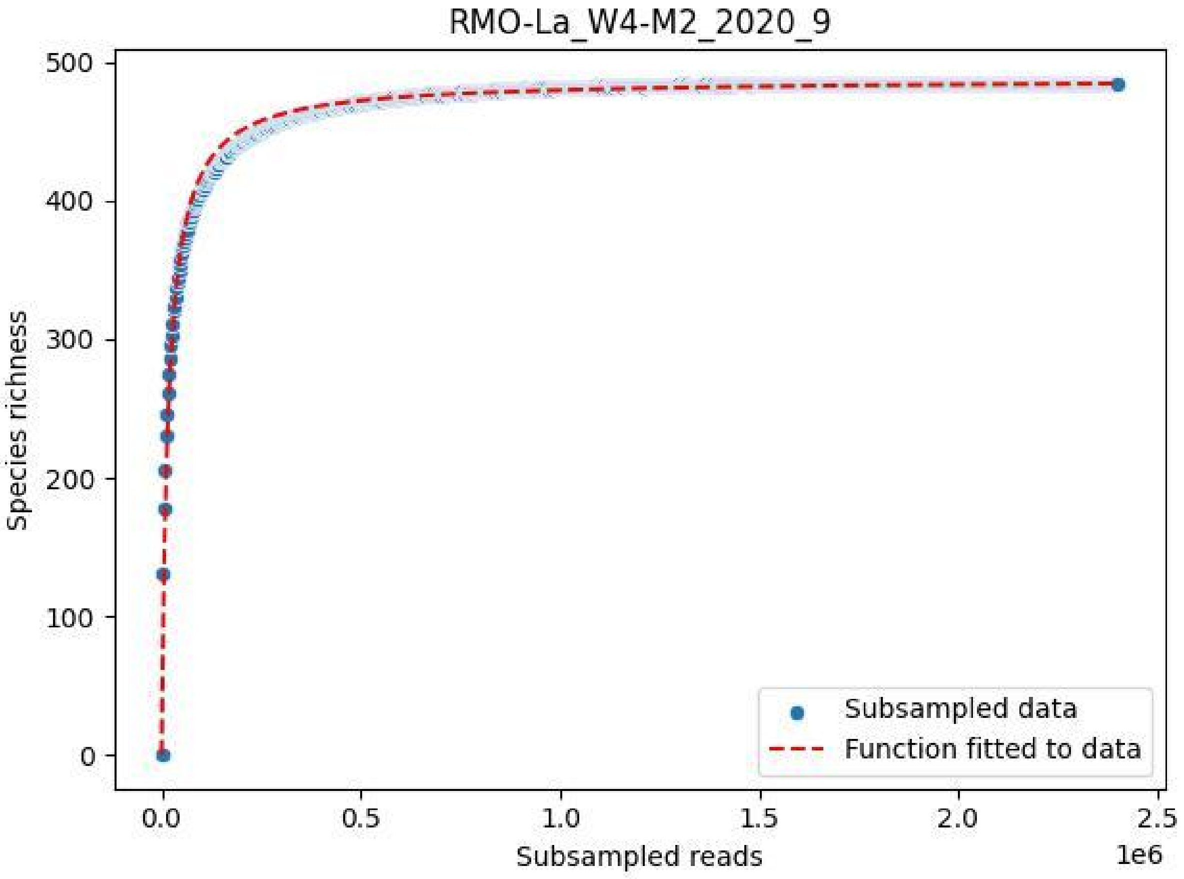
RMO-La_W4-M2_2020_9
Species richness
Subsampled reads
1e6
0
100
200
300
400
500
0.0
0.5
1.0
1.5
2.0
2.5
Subsampled data
Function fitted to data

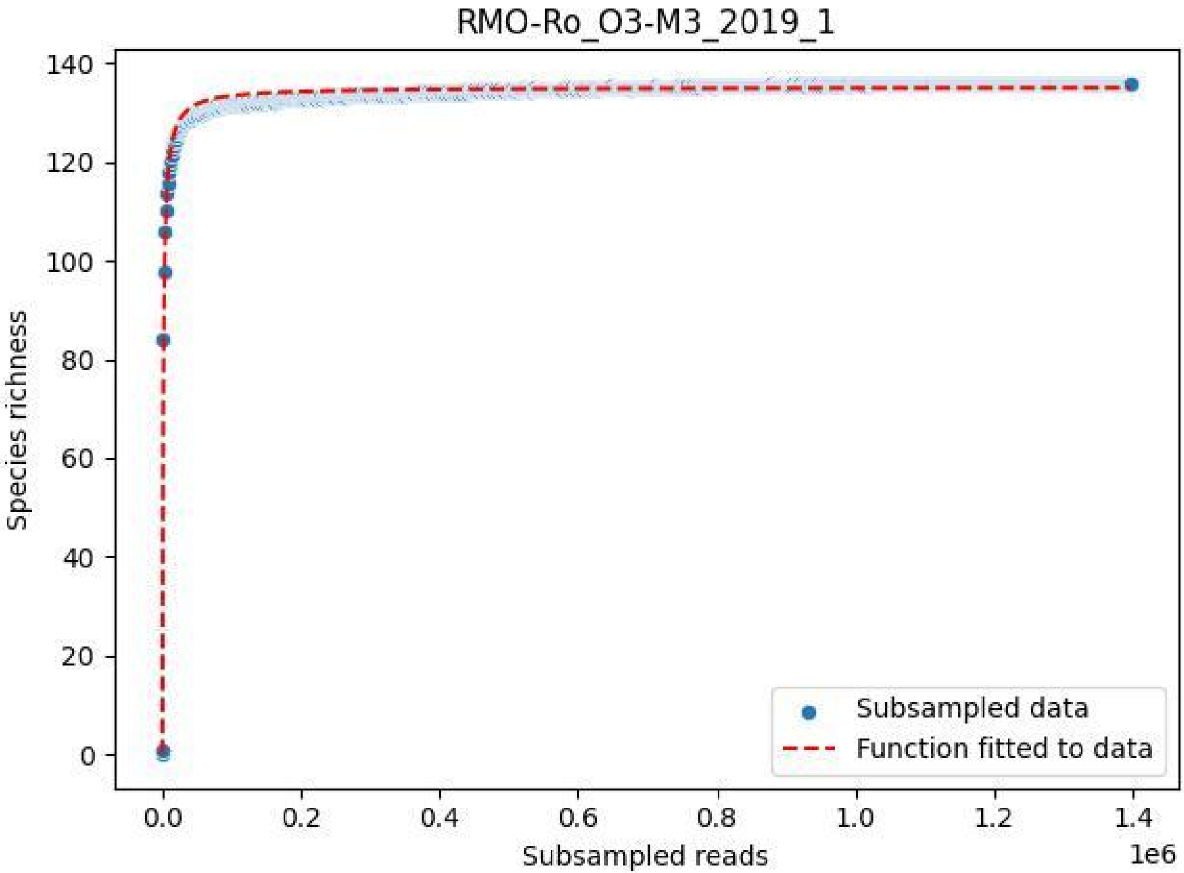

RMO-Ro_O3-M3_2019_1
Species richness
Subsampled reads
1e6
Subsampled data
Function fitted to data
0
20
40
60
80
100
120
140
0.0
0.2
0.4
0.6
0.8
1.0
1.2
1.4

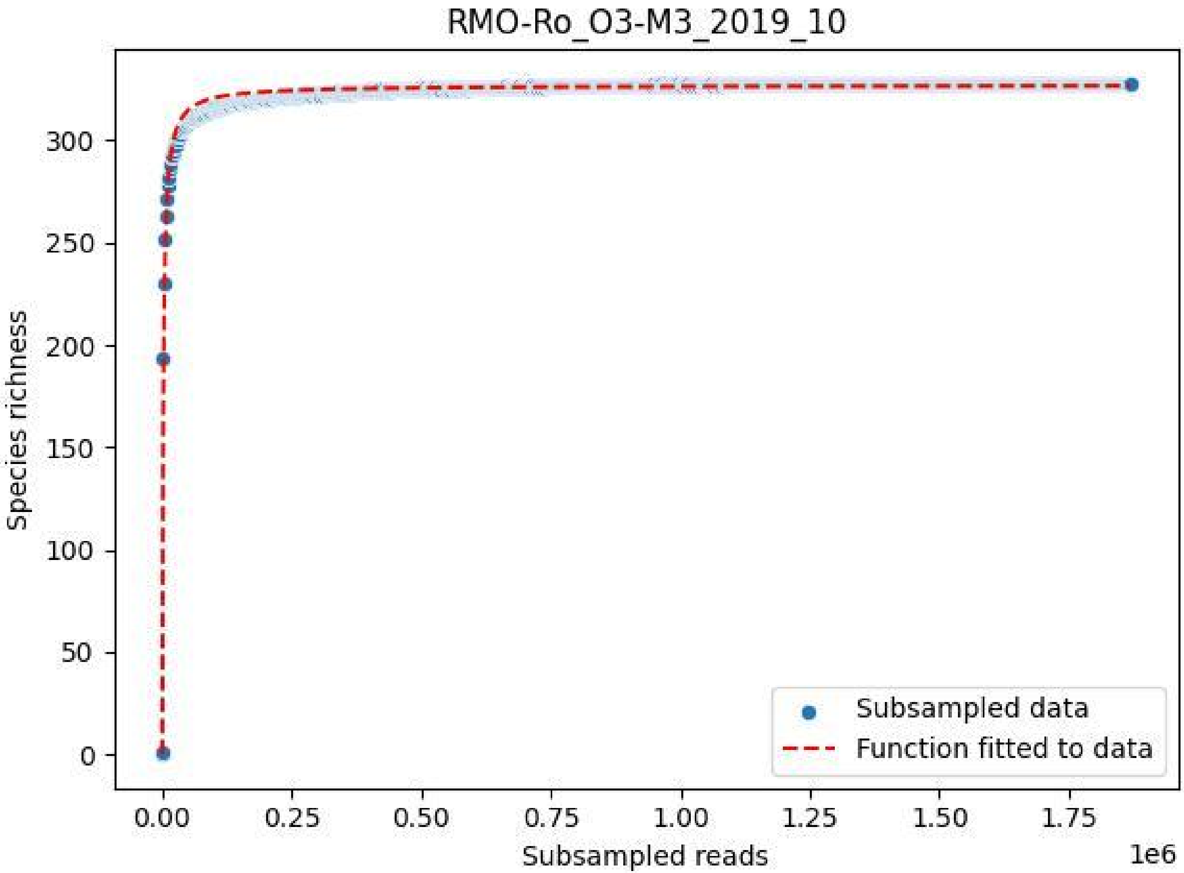
RMO-Ro_O3-M3_2019_10
Species richness
Subsampled reads
1e6
0
50
100
150
200
250
300
0.00
0.25
0.50
0.75
1.00
1.25
1.50
1.75
Subsampled data
Function fitted to data

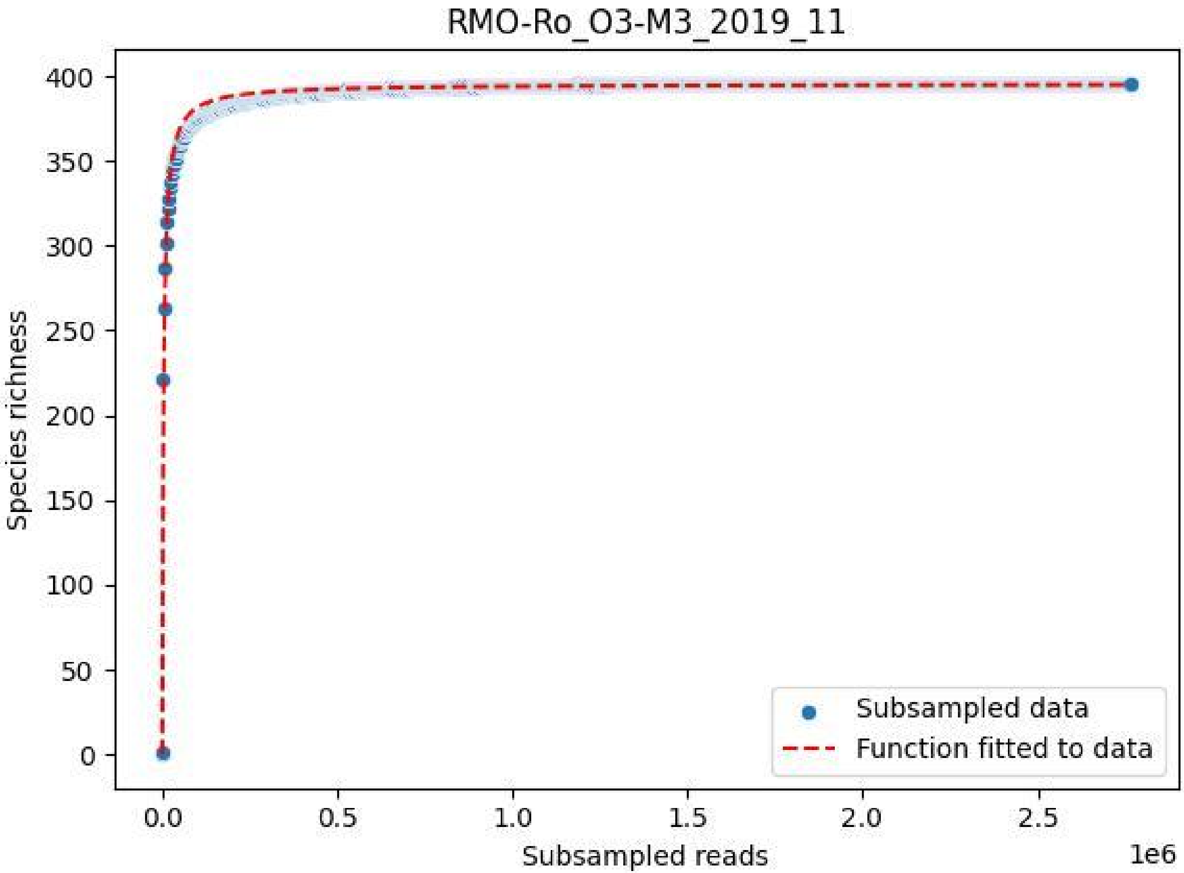

RMO-Ro_O3-M3_2019_11
Species richness
Subsampled reads
1e6
0
50
100
150
200
250
300
350
400
0.0
0.5
1.0
1.5
2.0
2.5
Subsampled data
Function fitted to data

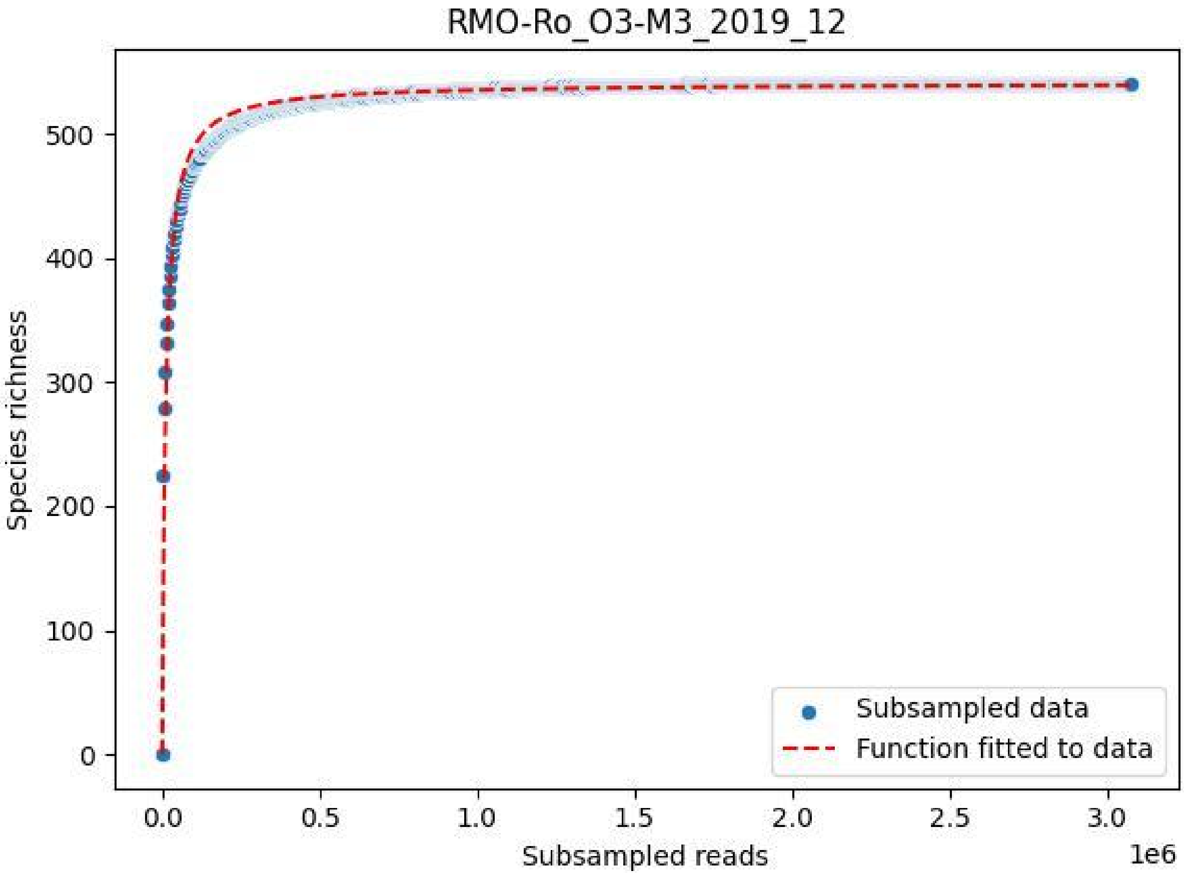

RMO-Ro_O3-M3_2019_12
Species richness
Subsampled reads
1e6
0
100
200
300
400
500
0.0
0.5
1.0
1.5
2.0
2.5
3.0
Subsampled data
Function fitted to data

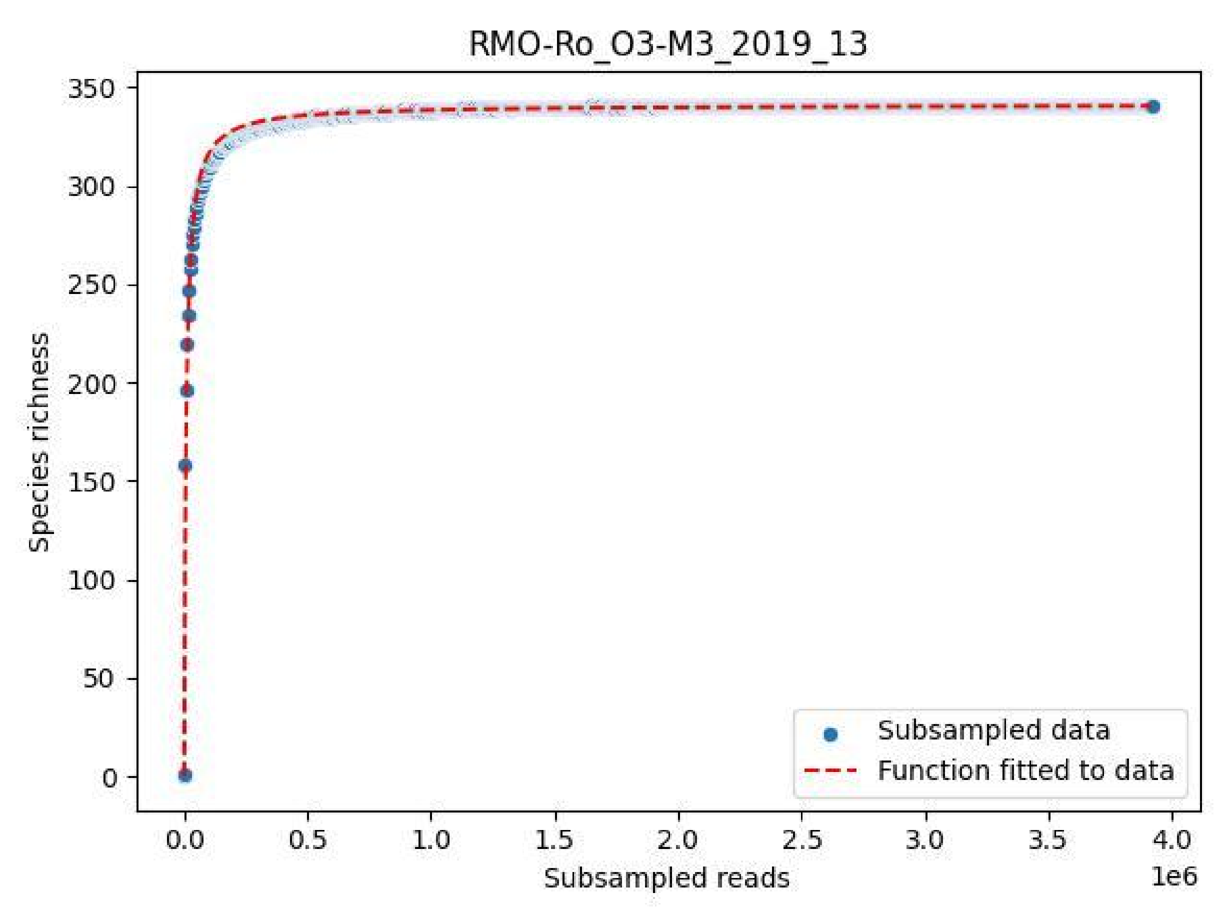
RMO-Ro_O3-M3_2019_13
Species richness
0
50
100
150
200
250
300
350
Subsampled reads
0.0
0.5
1.0
1.5
2.0
2.5
3.0
3.5
4.0
1e6
Subsampled data
Function fitted to data

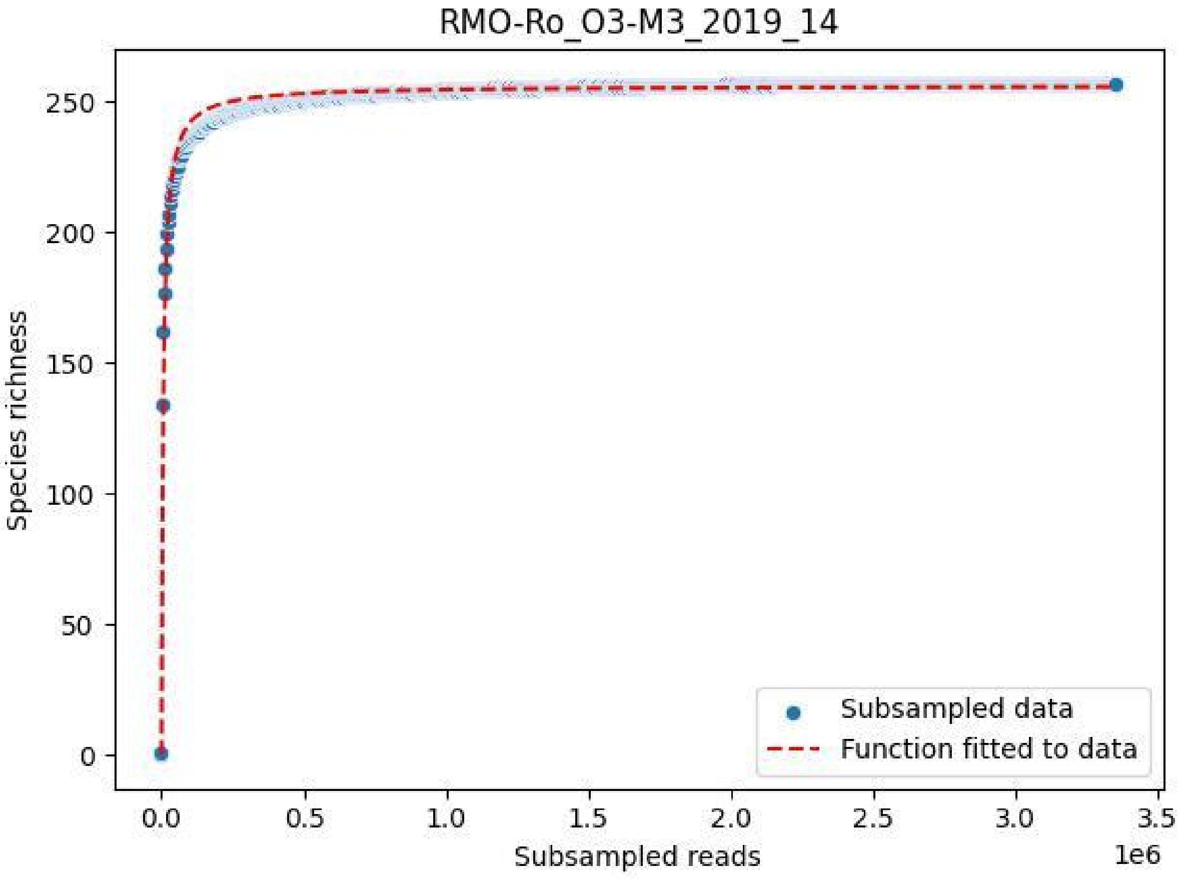
RMO-Ro_O3-M3_2019_14
Species richness
Subsampled reads
1e6
0
50
100
150
200
250
0.0
0.5
1.0
1.5
2.0
2.5
3.0
3.5
Subsampled data
Function fitted to data

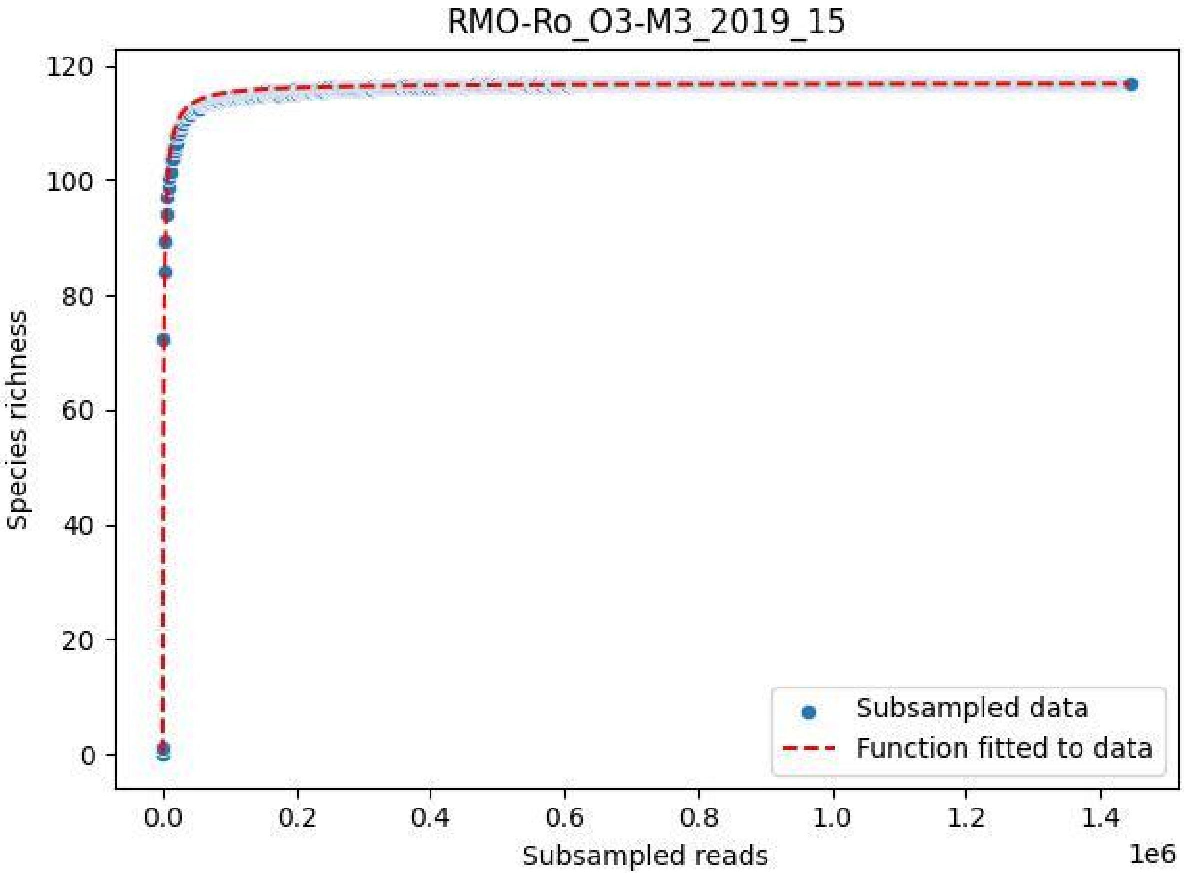

RMO-Ro_O3-M3_2019_15
Species richness
Subsampled reads
1e6
Subsampled data
Function fitted to data
0
20
40
60
80
100
120
0.0
0.2
0.4
0.6
0.8
1.0
1.2
1.4

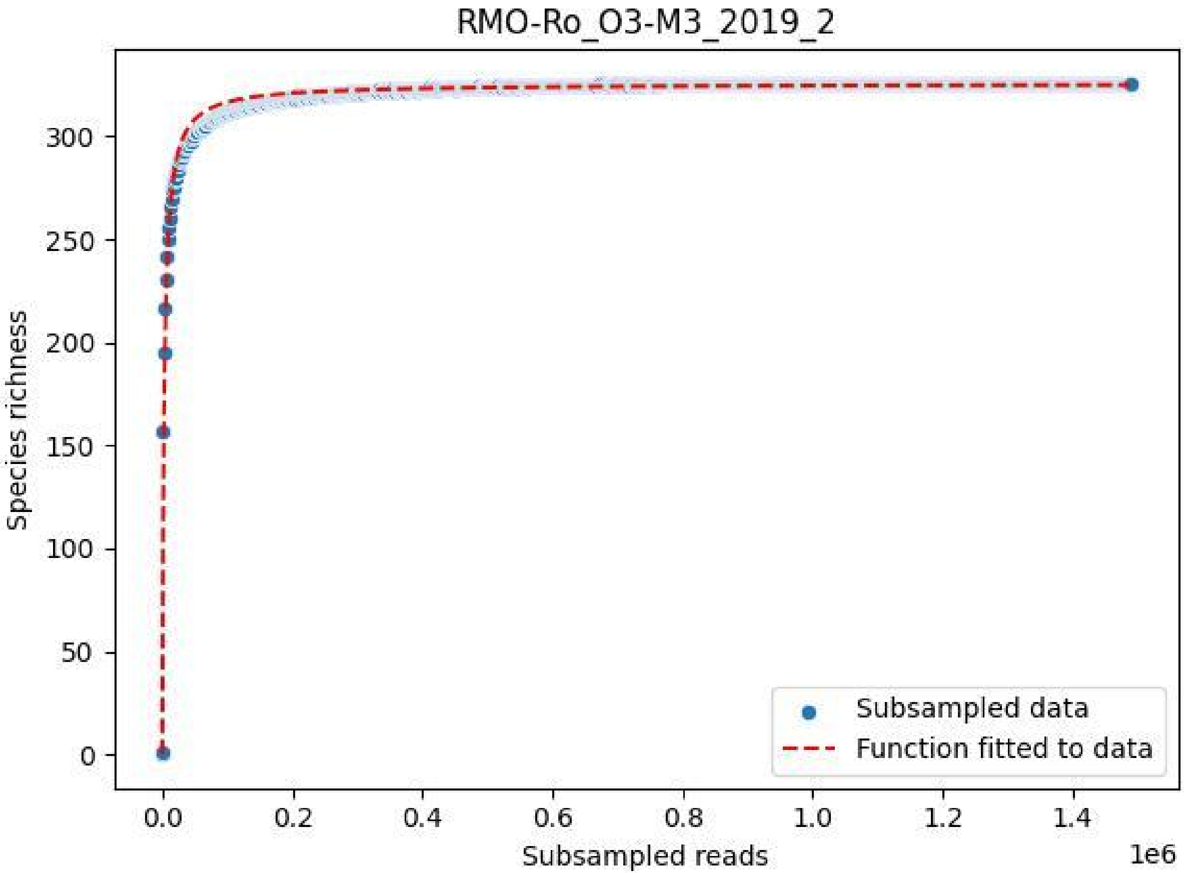
RMO-Ro_O3-M3_2019_2
Species richness
Subsampled reads
1e6
0
50
100
150
200
250
300
0.0
0.2
0.4
0.6
0.8
1.0
1.2
1.4
Subsampled data
Function fitted to data

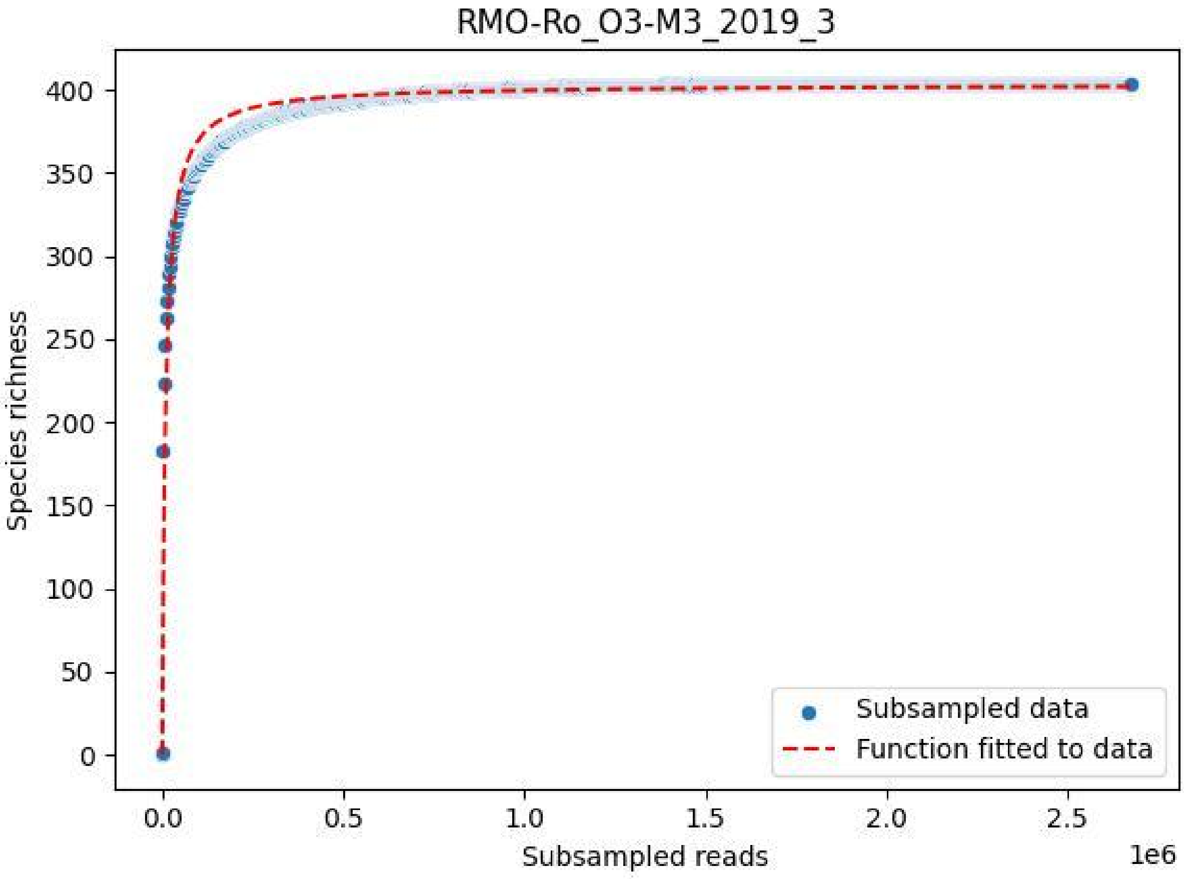

RMO-Ro_O3-M3_2019_3
Species richness
Subsampled reads
1e6
0
50
100
150
200
250
300
350
400
0.0
0.5
1.0
1.5
2.0
2.5
Subsampled data
Function fitted to data

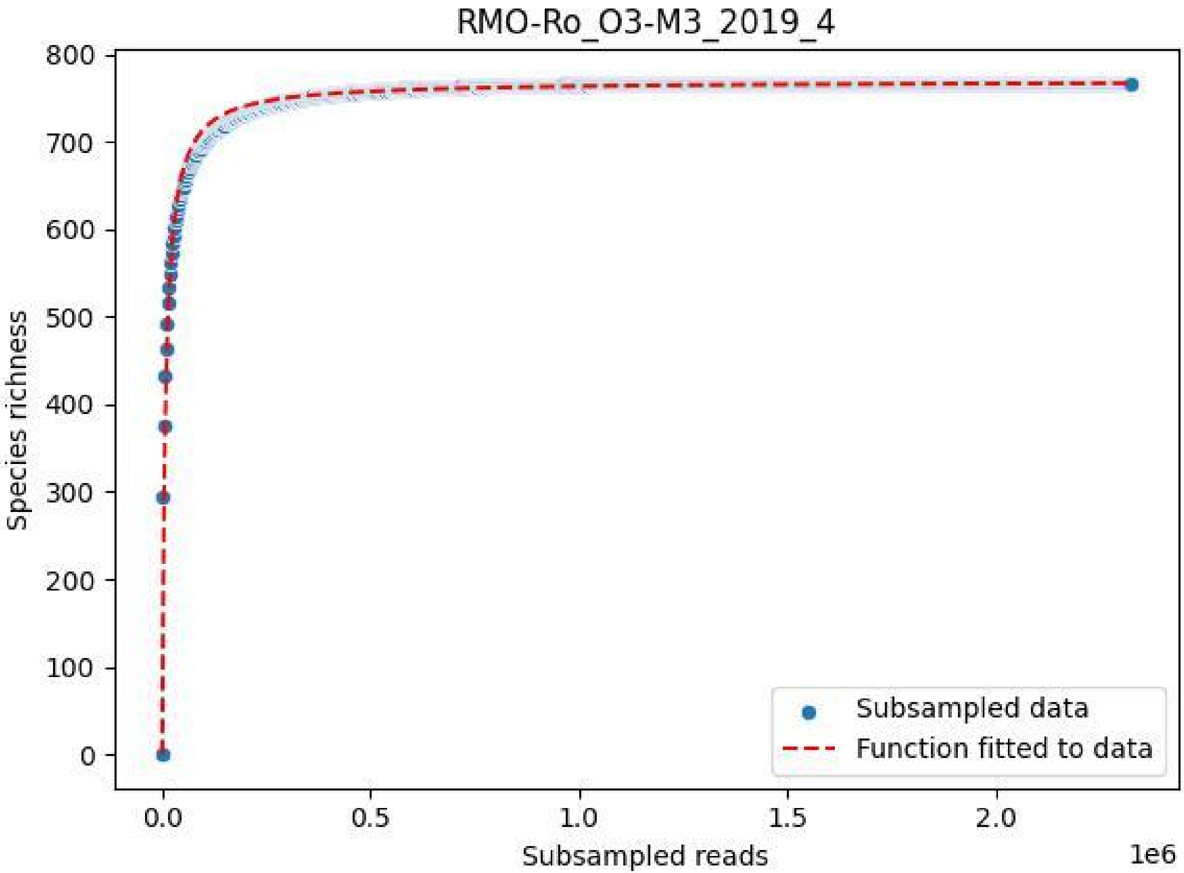

RMO-Ro_O3-M3_2019_4
800
700
600
500
400
300
200
100
0
Species richness
0.0
0.5
1.0
1.5
2.0
1e6
Subsampled reads
Subsampled data
Function fitted to data

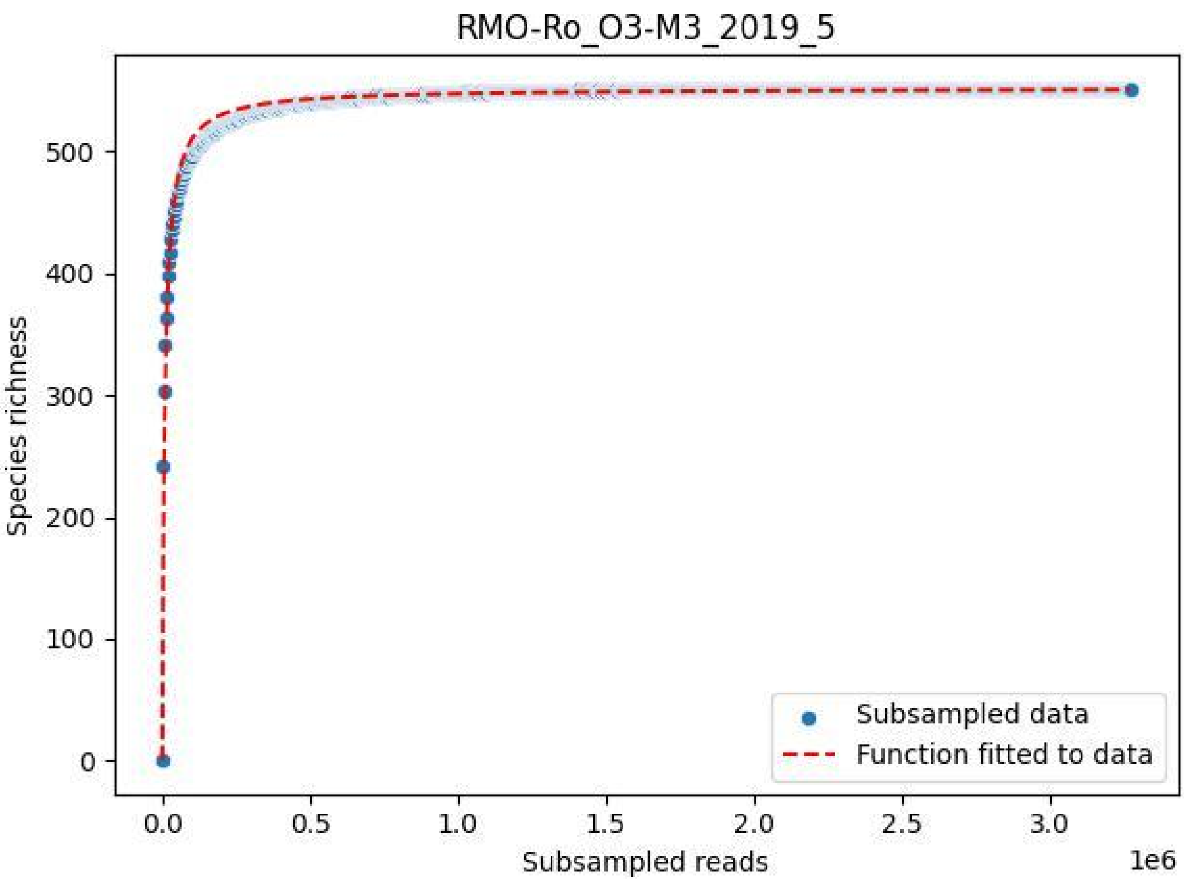
RMO-Ro_O3-M3_2019_5
Species richness
Subsampled reads
1e6
0
100
200
300
400
500
0.0
0.5
1.0
1.5
2.0
2.5
3.0
Subsampled data
Function fitted to data

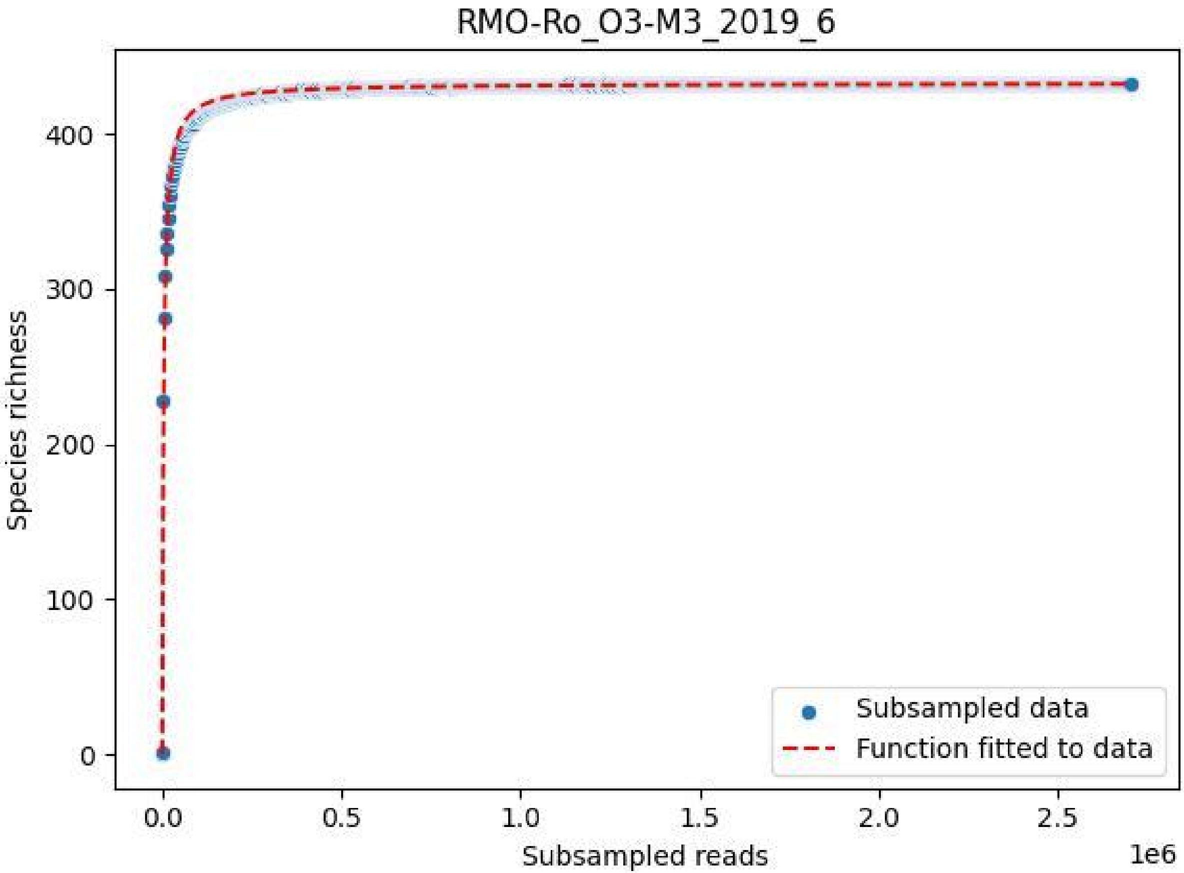
RMO-Ro_O3-M3_2019_6
Species richness
Subsampled reads
1e6
0
100
200
300
400
0.0
0.5
1.0
1.5
2.0
2.5
Subsampled data
Function fitted to data

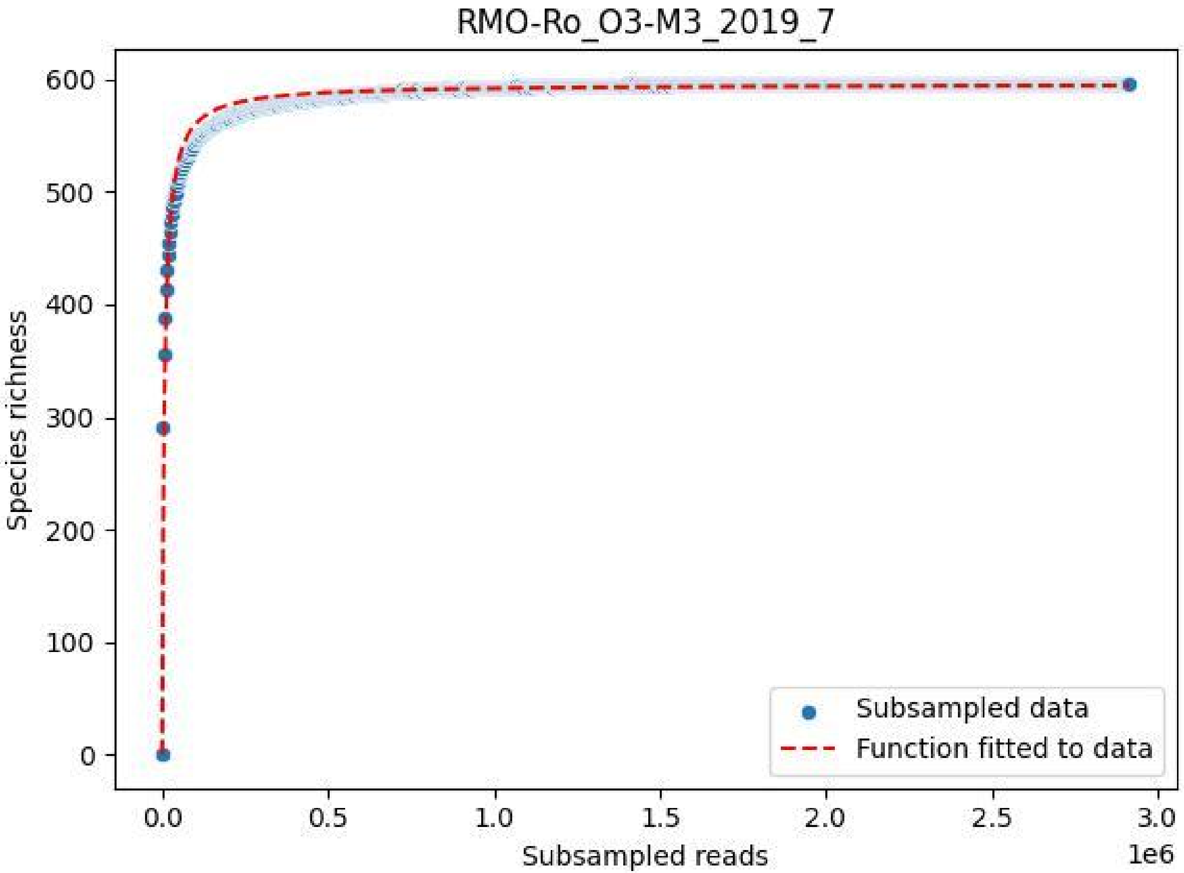
RMO-Ro_O3-M3_2019_7
Species richness
Subsampled reads
1e6
0
100
200
300
400
500
600
0.0
0.5
1.0
1.5
2.0
2.5
3.0
Subsampled data
Function fitted to data

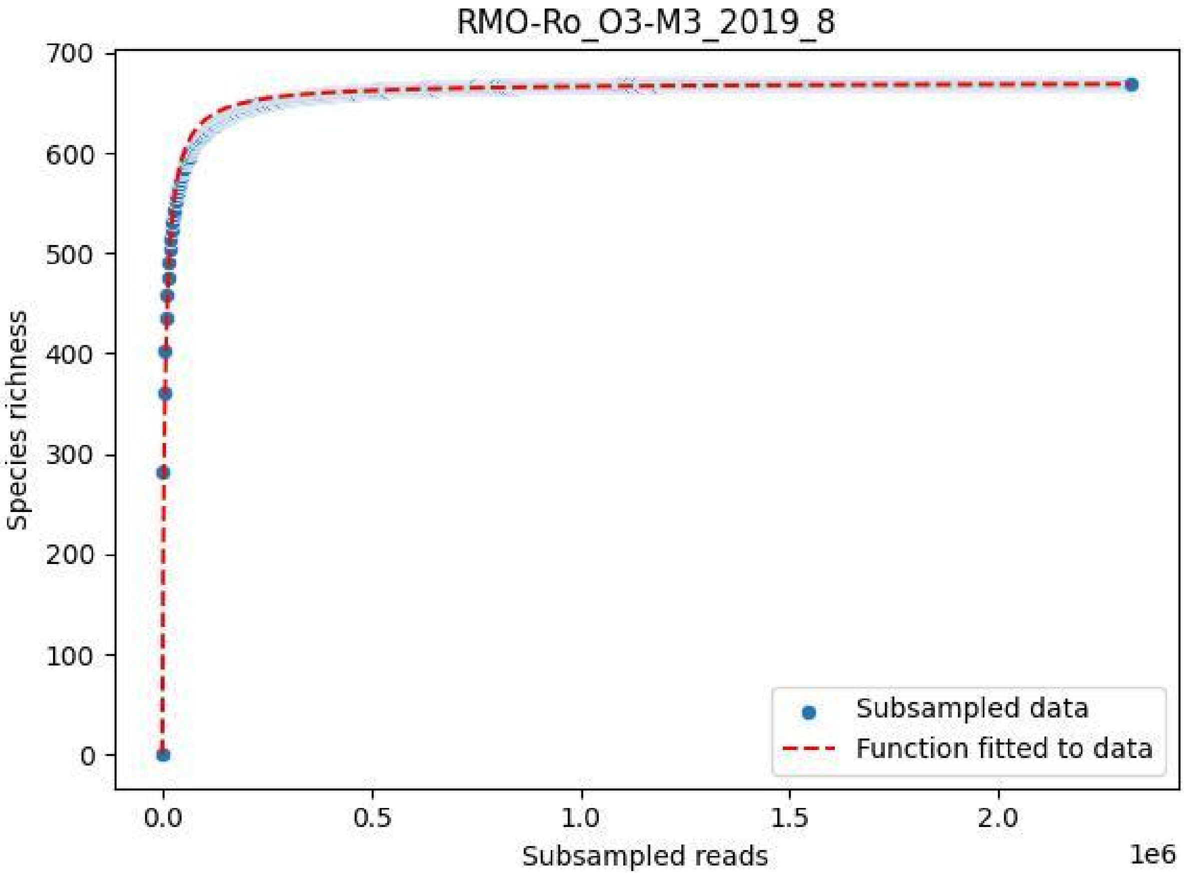
RMO-Ro_O3-M3_2019_8
700
600
500
400
300
200
100
0
Species richness
0.0
0.5
1.0
1.5
2.0
1e6
Subsampled reads
Subsampled data
Function fitted to data

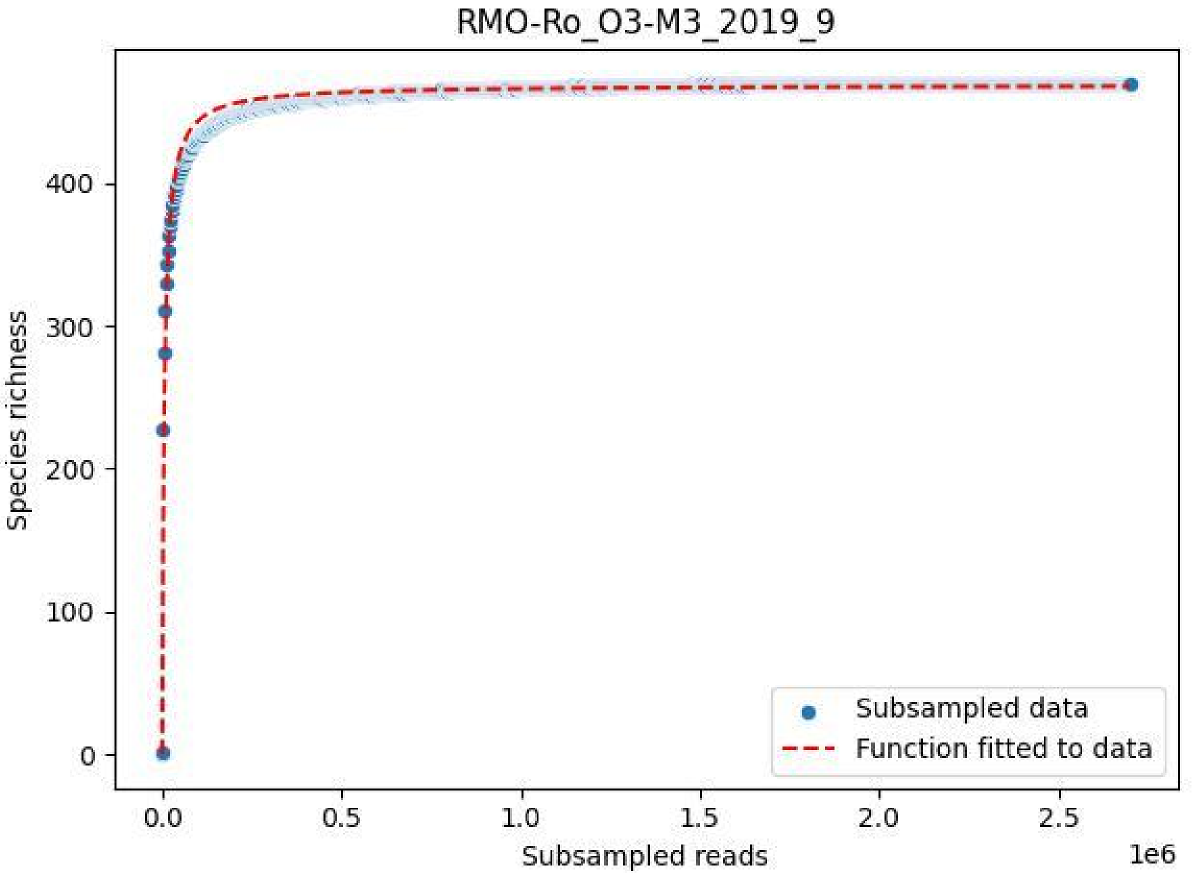

RMO-Ro_O3-M3_2019_9
Species richness
Subsampled reads
1e6
0
100
200
300
400
0.0
0.5
1.0
1.5
2.0
2.5
Subsampled data
Function fitted to data

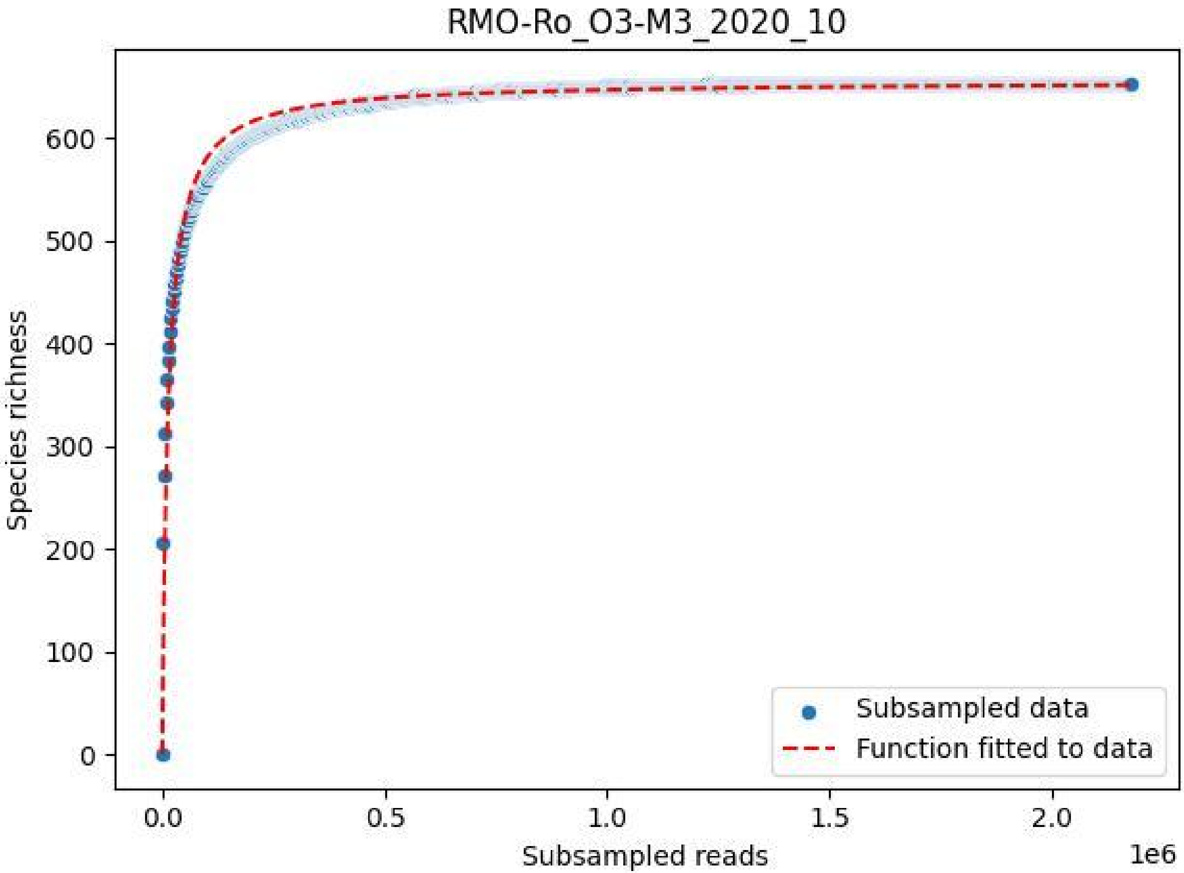
RMO-Ro_O3-M3_2020_10
Species richness
0
100
200
300
400
500
600
Subsampled reads
0.0
0.5
1.0
1.5
2.0
1e6
Subsampled data
Function fitted to data

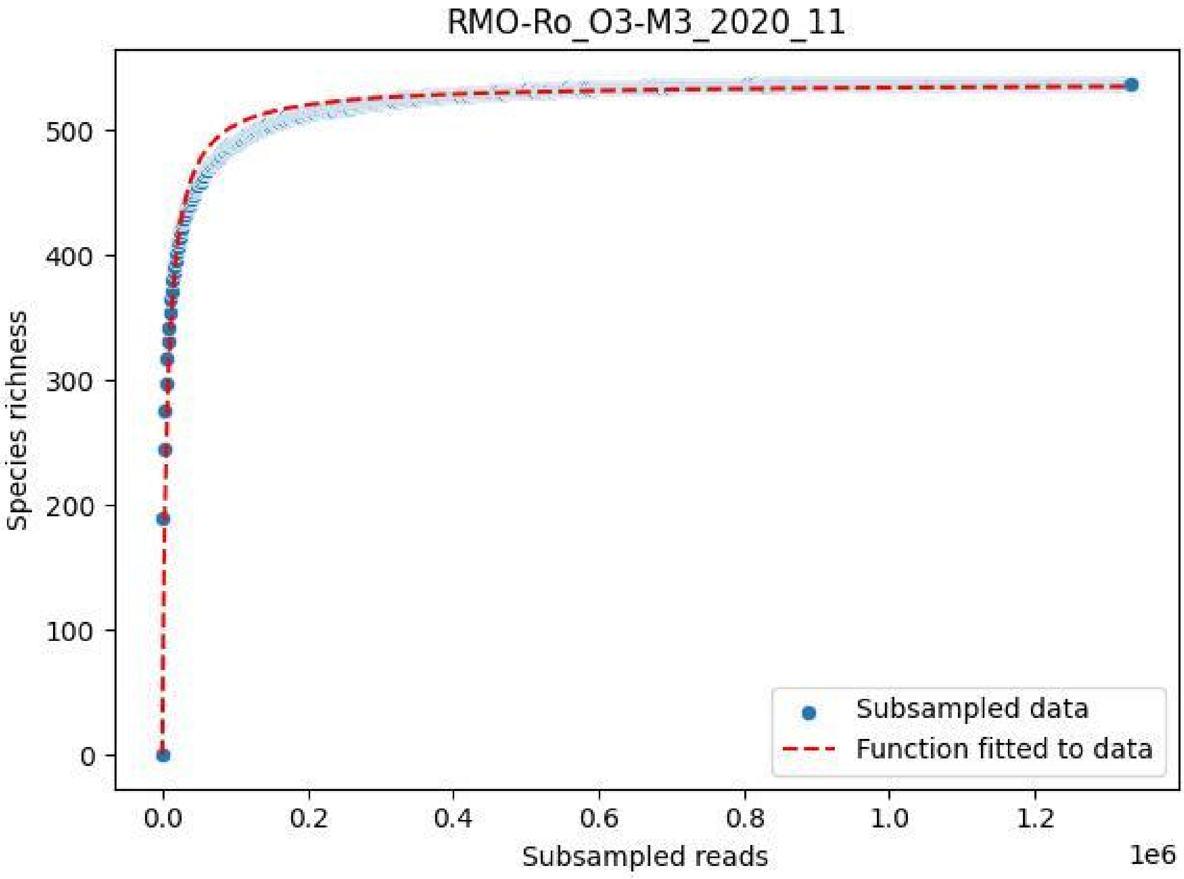

RMO-Ro_O3-M3_2020_11
Species richness
Subsampled reads
1e6
0
100
200
300
400
500
0.0
0.2
0.4
0.6
0.8
1.0
1.2
Subsampled data
Function fitted to data

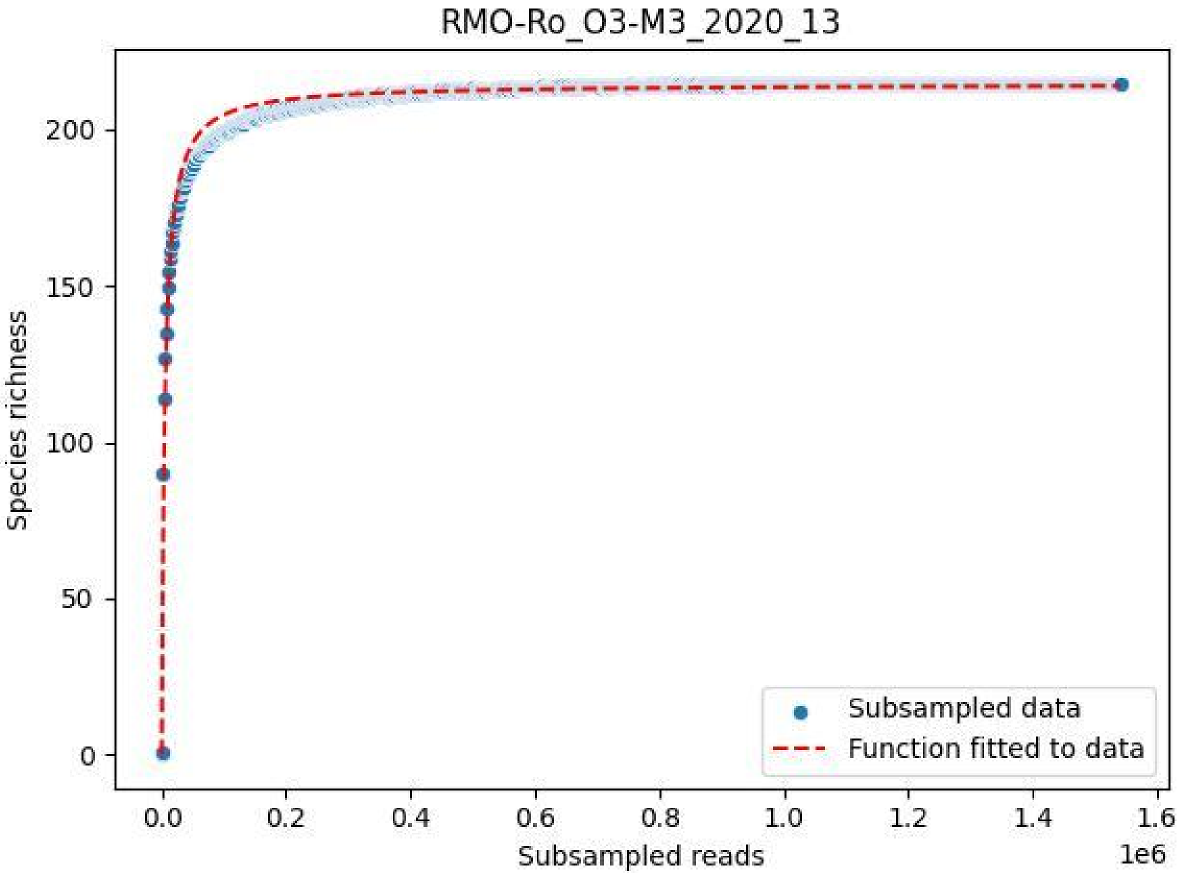

RMO-Ro_O3-M3_2020_13
Species richness
0
50
100
150
200
0.0
0.2
0.4
0.6
0.8
1.0
1.2
1.4
1.6
1e6
Subsampled reads
Subsampled data
Function fitted to data

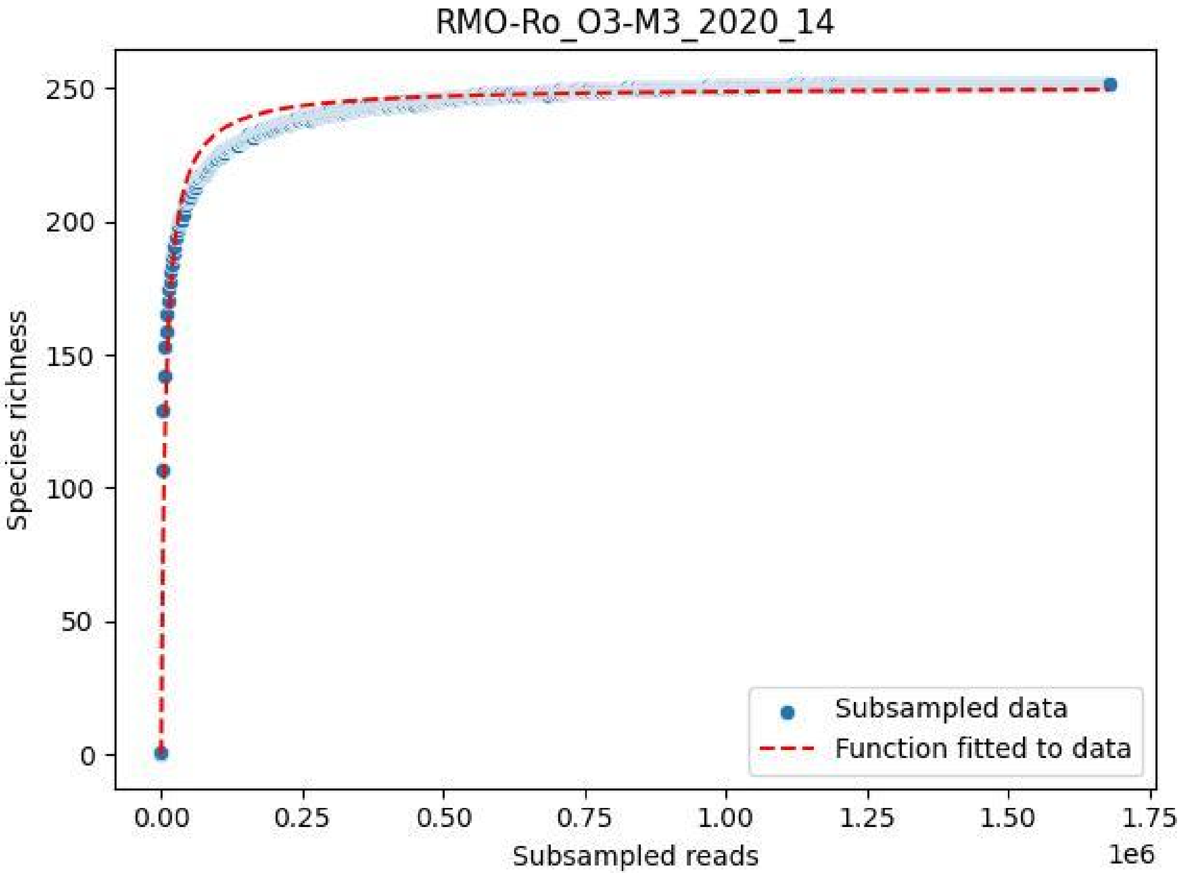
RMO-Ro_O3-M3_2020_14
Species richness
250
200
150
100
50
0
0.00
0.25
0.50
0.75
1.00
1.25
1.50
1.75
1e6
Subsampled reads
Subsampled data
Function fitted to data

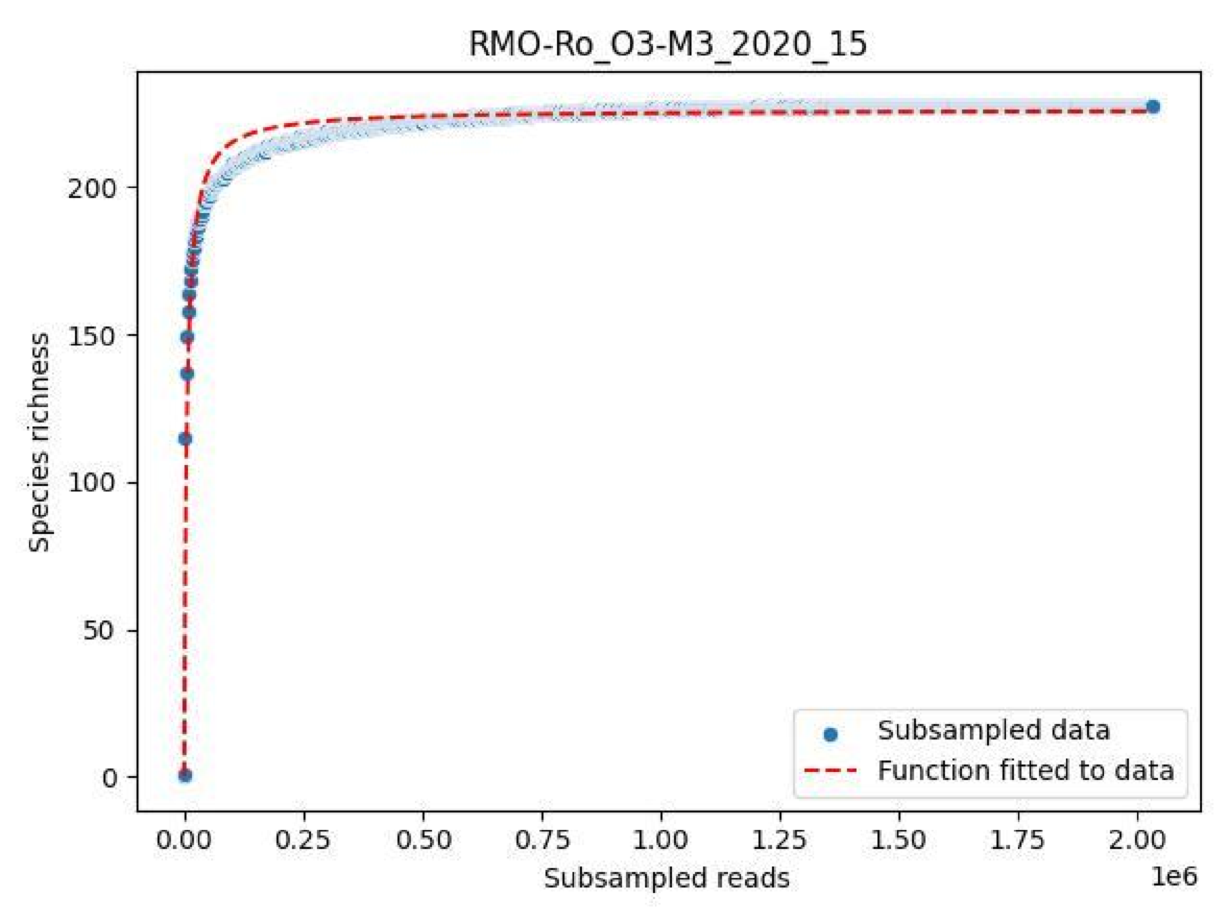

RMO-Ro_O3-M3_2020_15
Species richness
Subsampled reads
1e6
0
50
100
150
200
0.00
0.25
0.50
0.75
1.00
1.25
1.50
1.75
2.00
Subsampled data
Function fitted to data

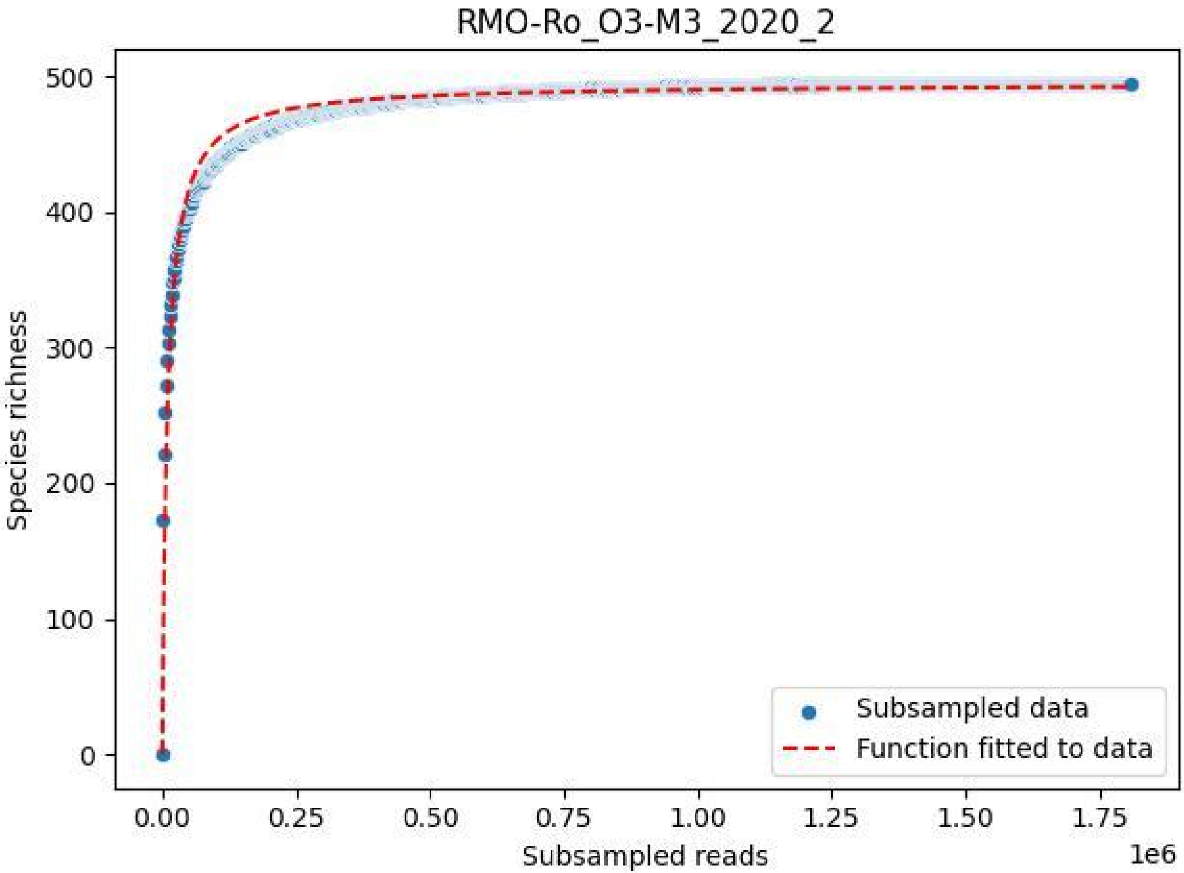

RMO-Ro_O3-M3_2020_2
Species richness
500
400
300
200
100
0
0.00
0.25
0.50
0.75
1.00
1.25
1.50
1.75
Subsampled reads
1e6
Subsampled data
Function fitted to data

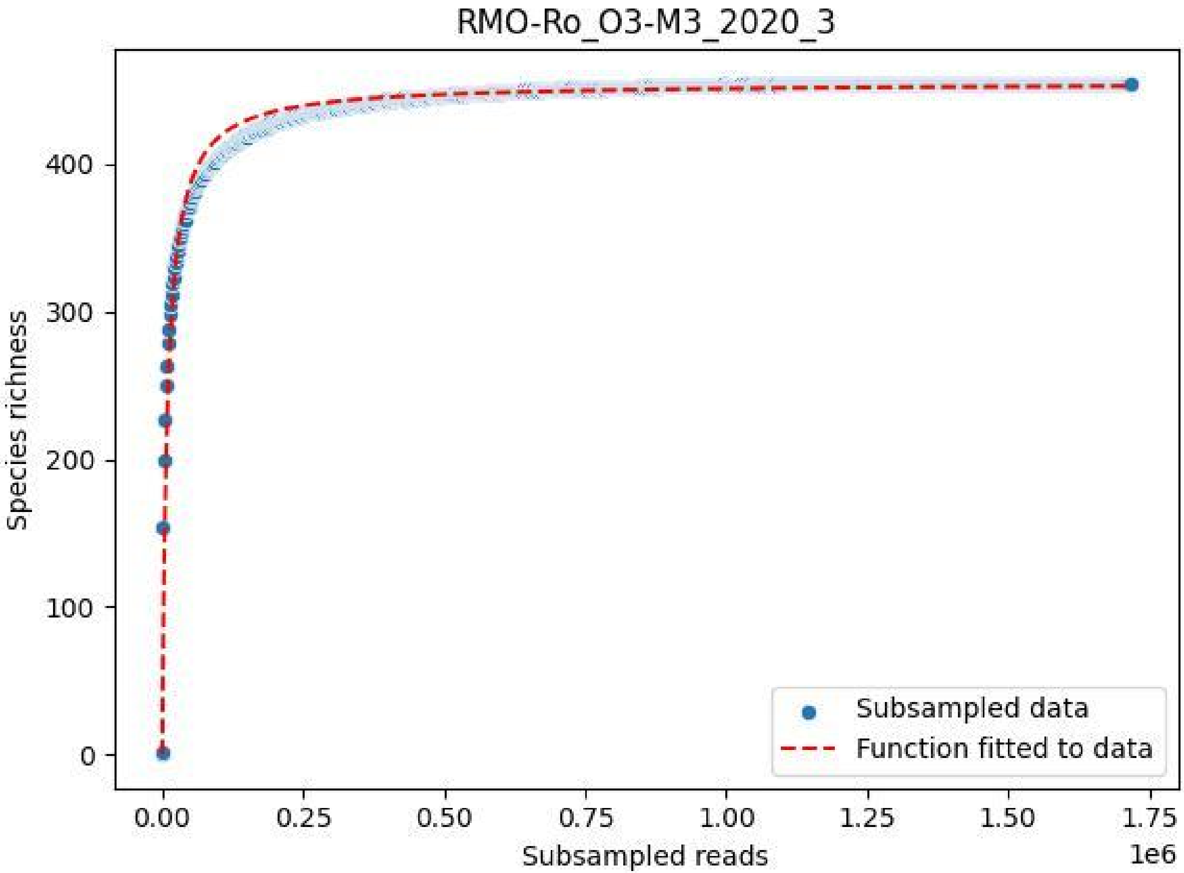

RMO-Ro_O3-M3_2020_3
Species richness
Subsampled reads
1e6
Subsampled data
Function fitted to data
0
100
200
300
400
0.00
0.25
0.50
0.75
1.00
1.25
1.50
1.75

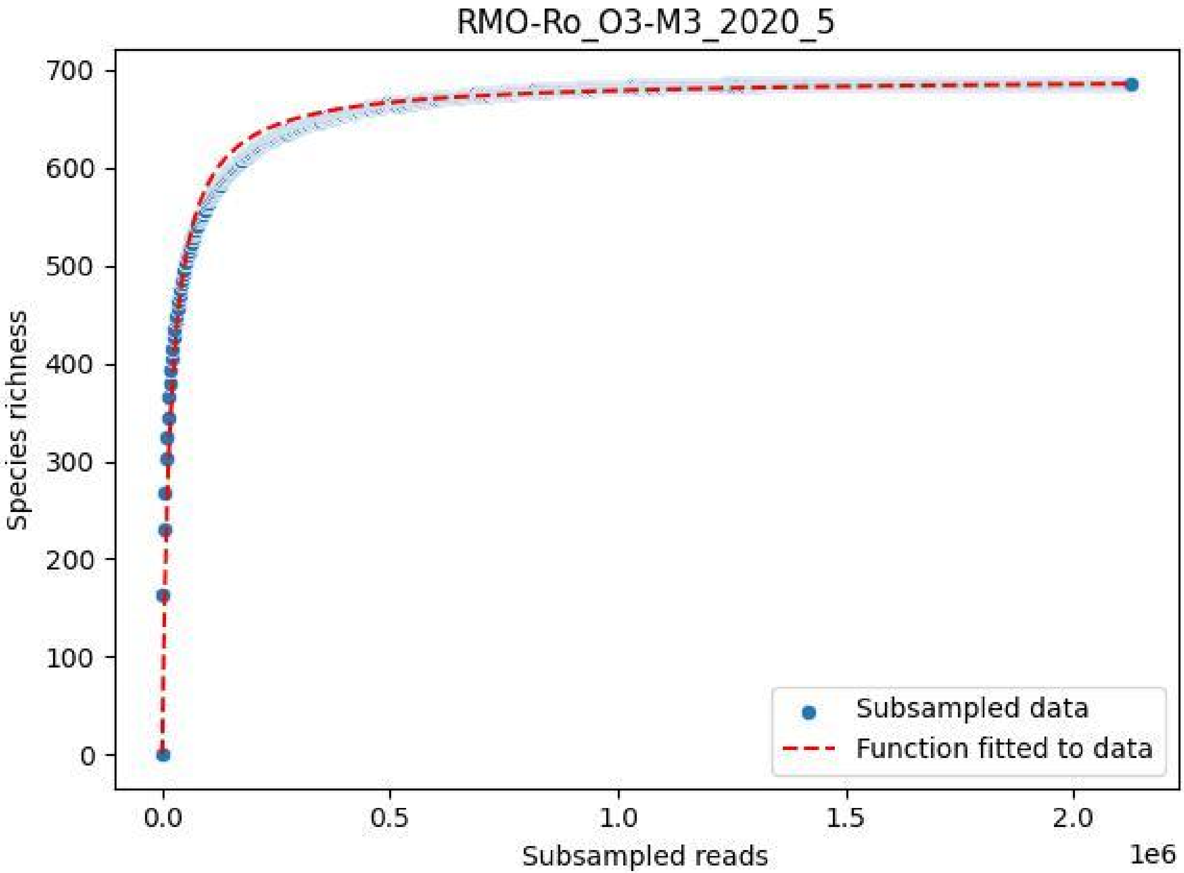

RMO-Ro_O3-M3_2020_5
Species richness
0
100
200
300
400
500
600
700
Subsampled reads
0.0
0.5
1.0
1.5
2.0
1e6
Subsampled data
Function fitted to data

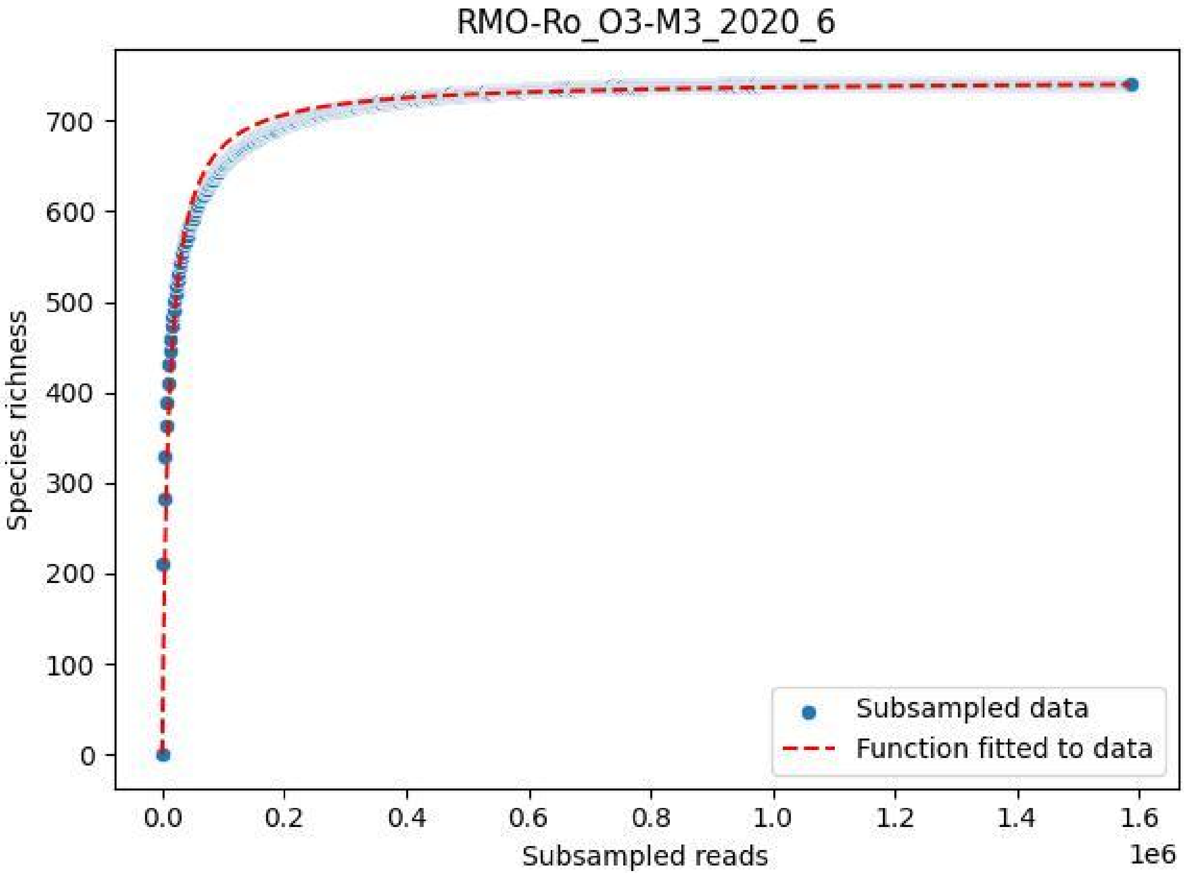

RMO-Ro_O3-M3_2020_6
Species richness
Subsampled reads
1e6
0
100
200
300
400
500
600
700
0.0
0.2
0.4
0.6
0.8
1.0
1.2
1.4
1.6
Subsampled data
Function fitted to data

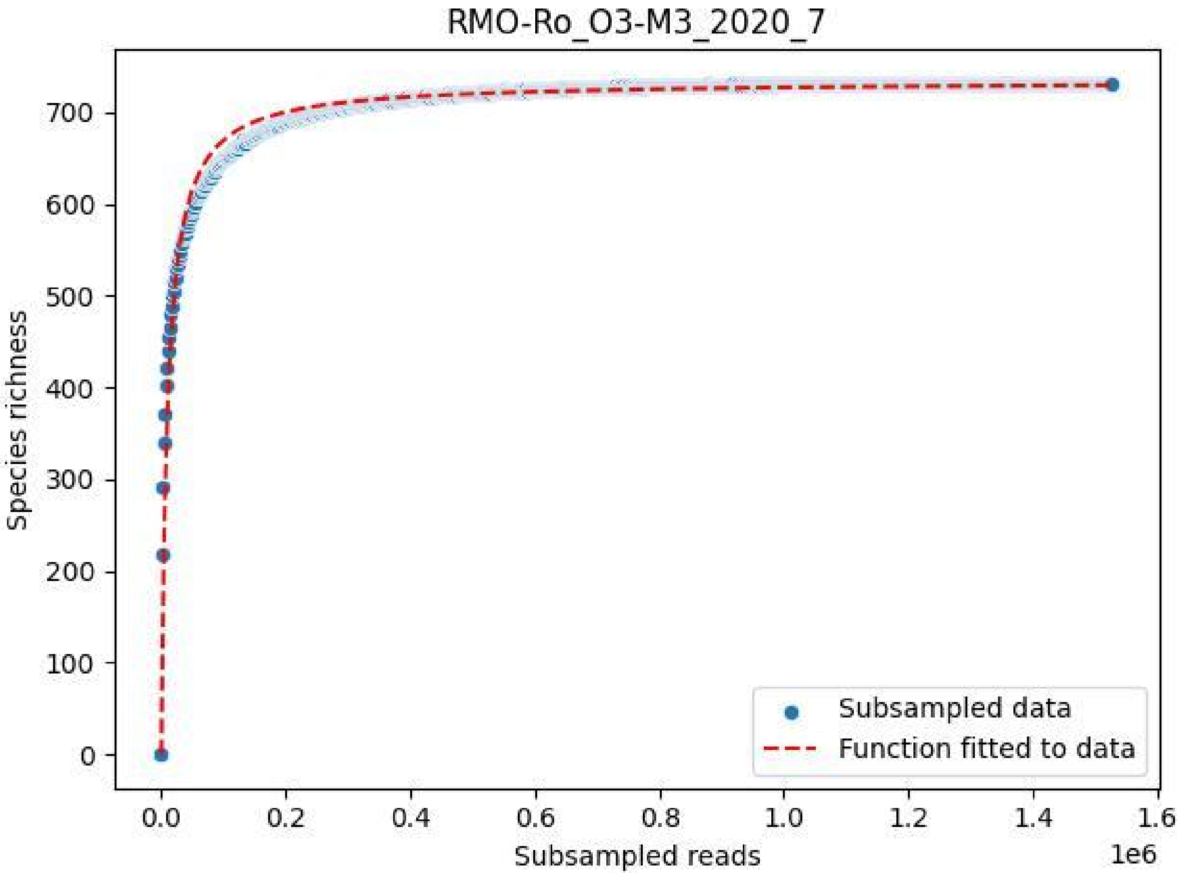
RMO-Ro_O3-M3_2020_7
Species richness
Subsampled reads
1e6
0
100
200
300
400
500
600
700
0.0
0.2
0.4
0.6
0.8
1.0
1.2
1.4
1.6
Subsampled data
Function fitted to data

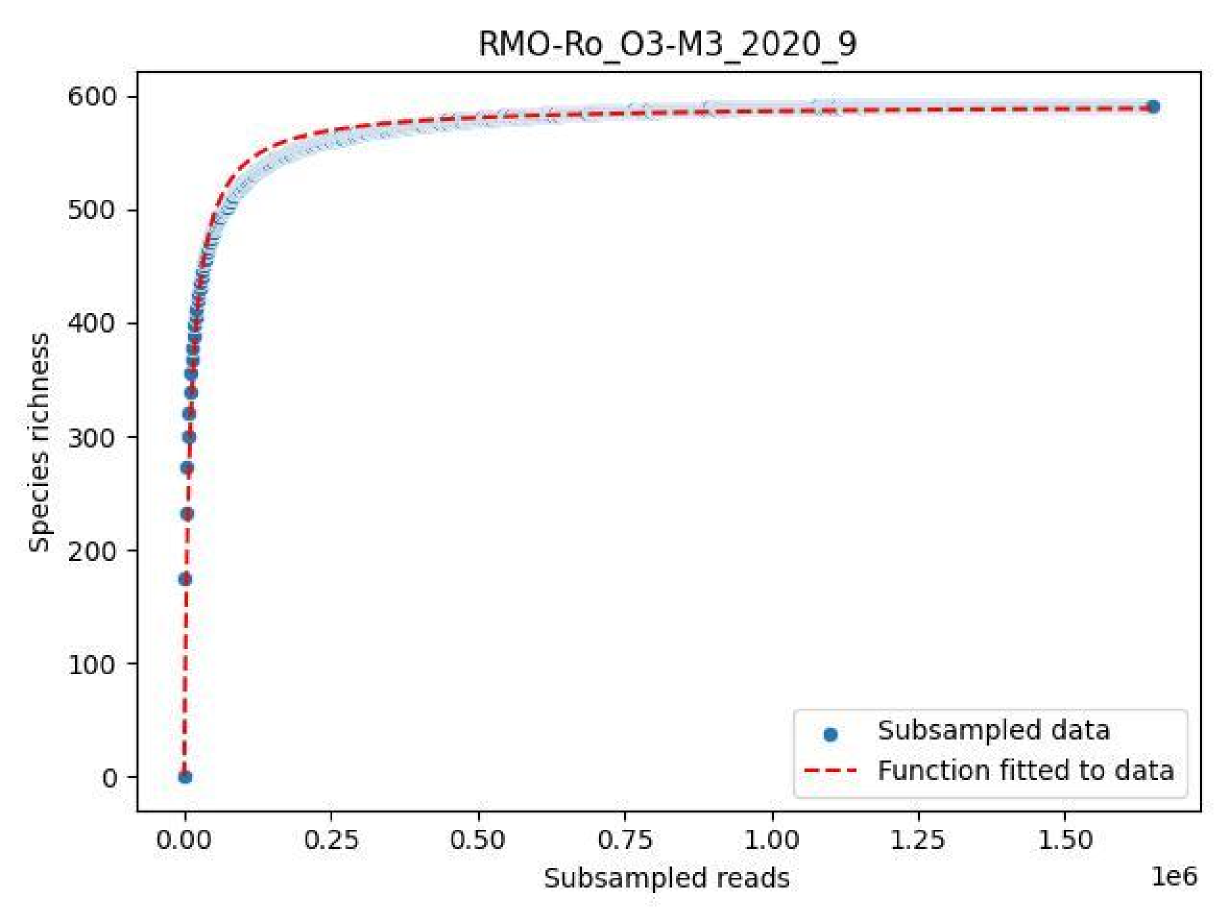
RMO-Ro_O3-M3_2020_9
600
500
400
300
200
100
0
Species richness
0.00
0.25
0.50
0.75
1.00
1.25
1.50
1e6
Subsampled reads
Subsampled data
Function fitted to data

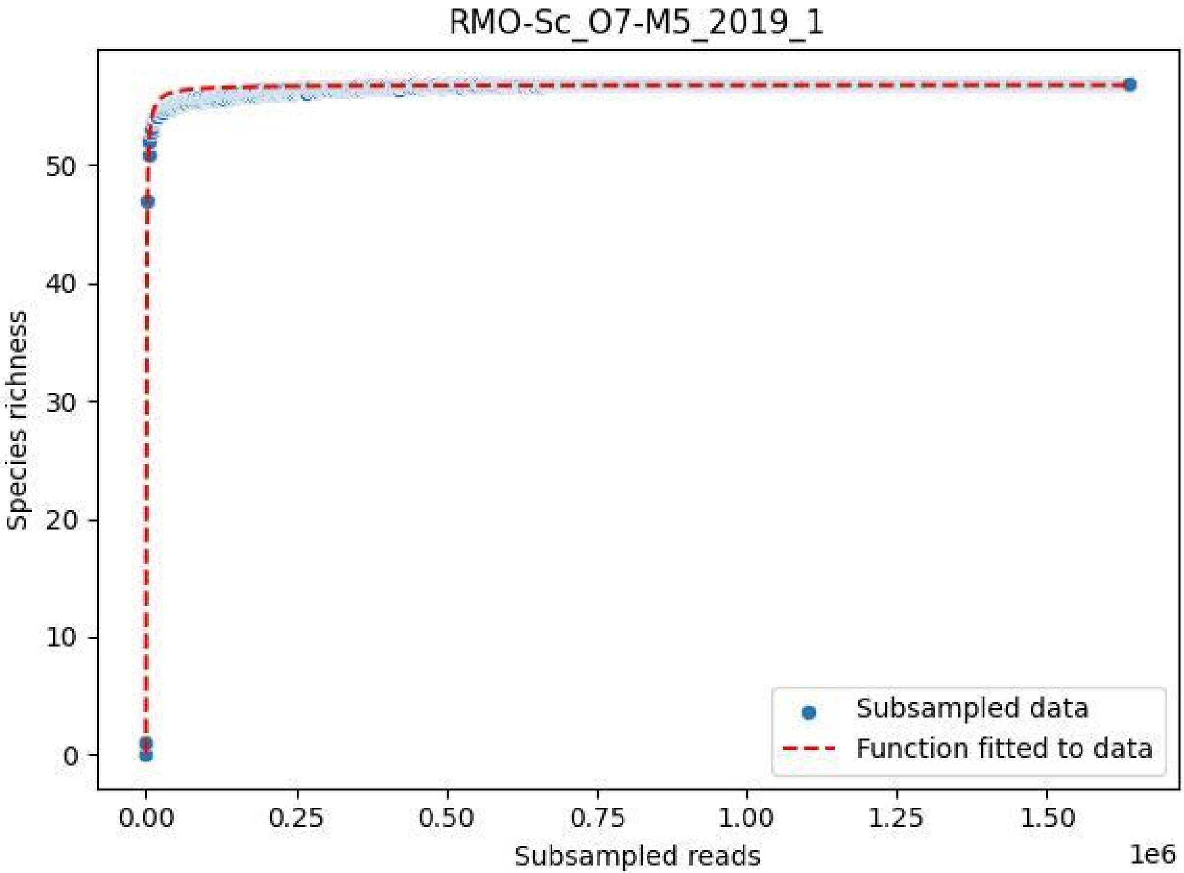

RMO-Sc_O7-M5_2019_1
Species richness
Subsampled reads
1e6
0
10
20
30
40
50
0.00
0.25
0.50
0.75
1.00
1.25
1.50
Subsampled data
Function fitted to data

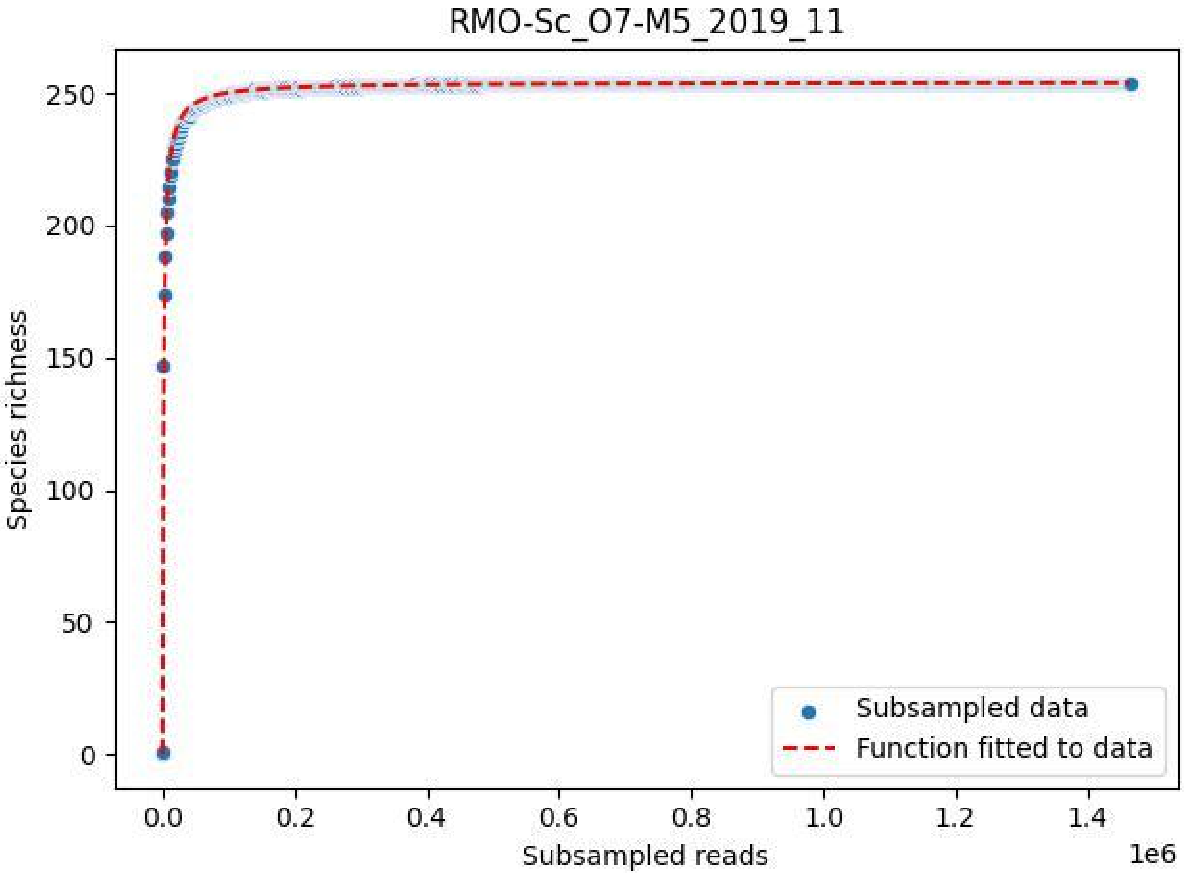

RMO-Sc_O7-M5_2019_11
Species richness
Subsampled reads
1e6
0
50
100
150
200
250
0.0
0.2
0.4
0.6
0.8
1.0
1.2
1.4
Subsampled data
Function fitted to data

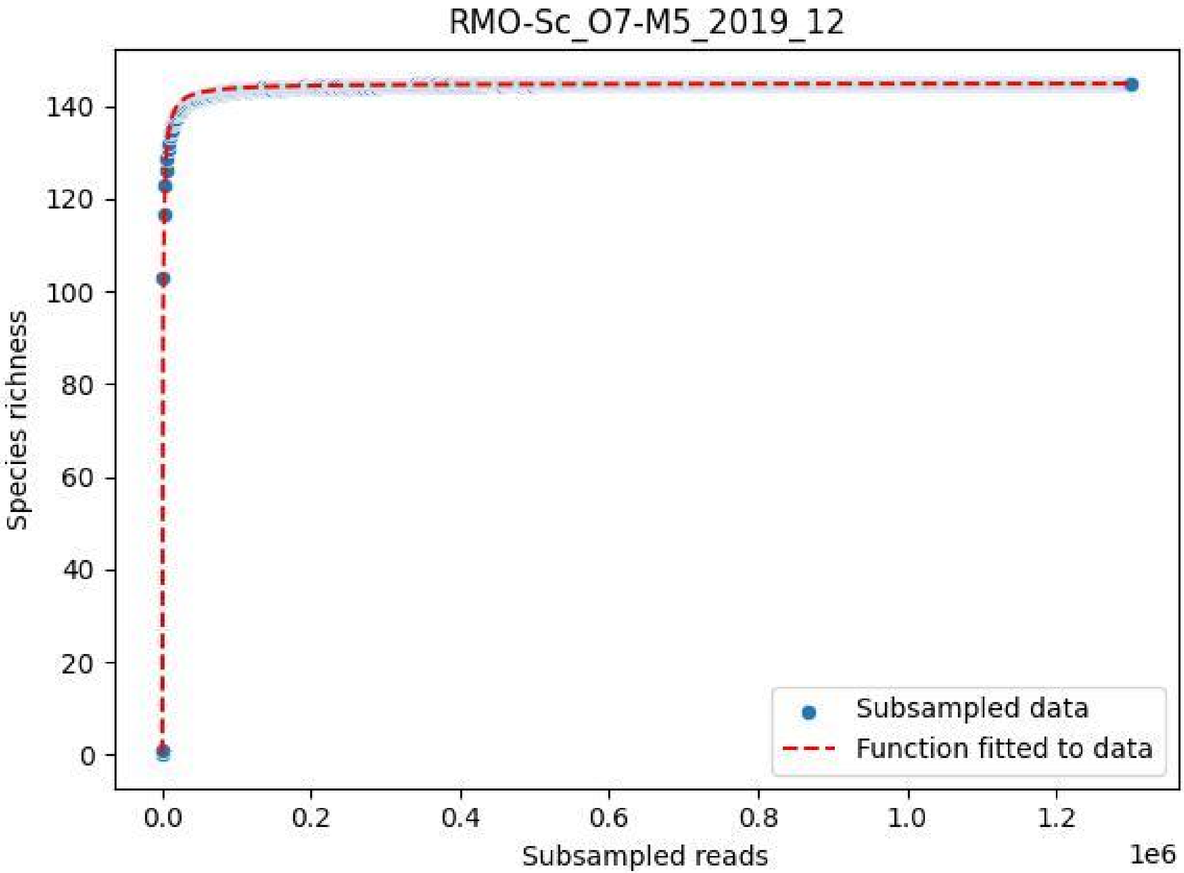

RMO-Sc_O7-M5_2019_12
Species richness
0
20
40
60
80
100
120
140
0.0
0.2
0.4
0.6
0.8
1.0
1.2
1e6
Subsampled reads
Subsampled data
Function fitted to data

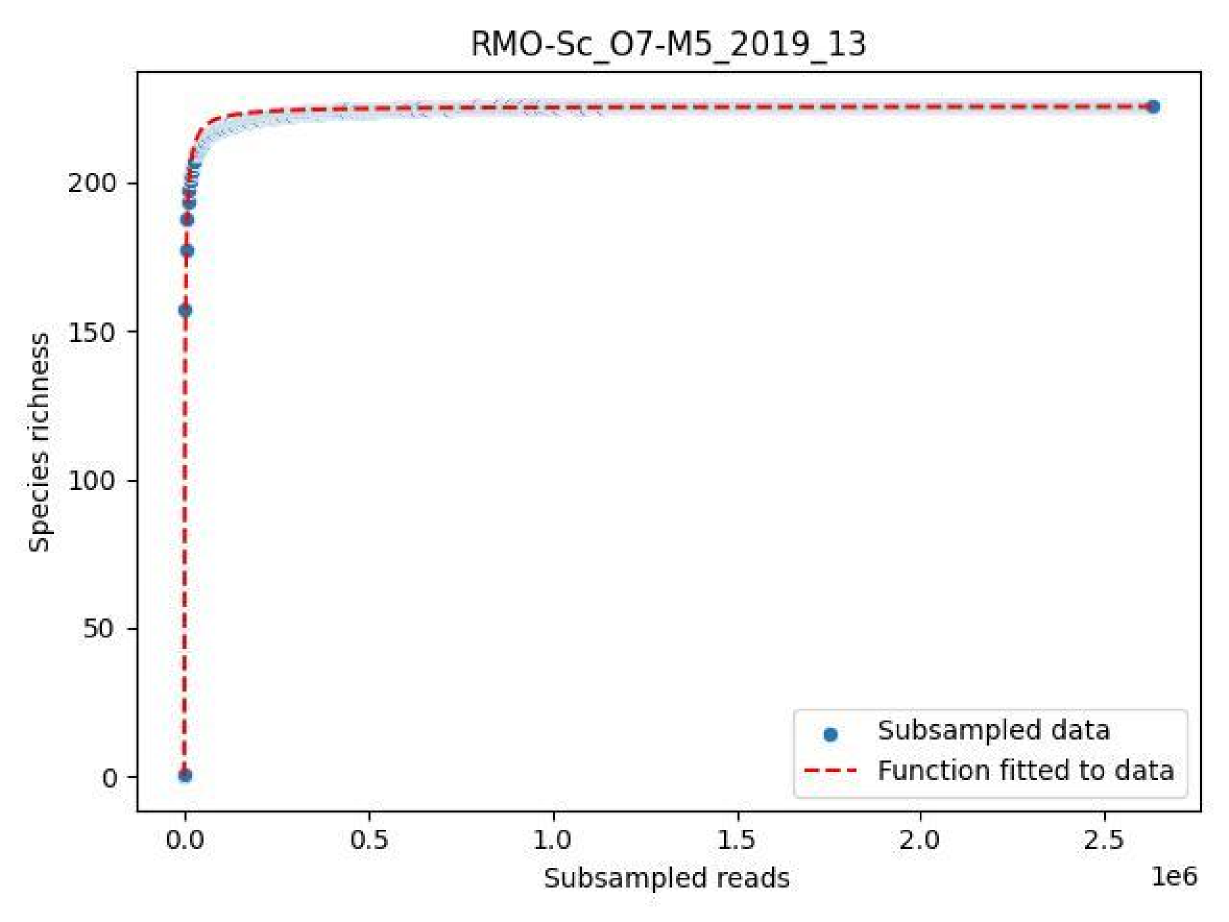
RMO-Sc_O7-M5_2019_13
Species richness
Subsampled reads
1e6
0
50
100
150
200
0.0
0.5
1.0
1.5
2.0
2.5
Subsampled data
Function fitted to data

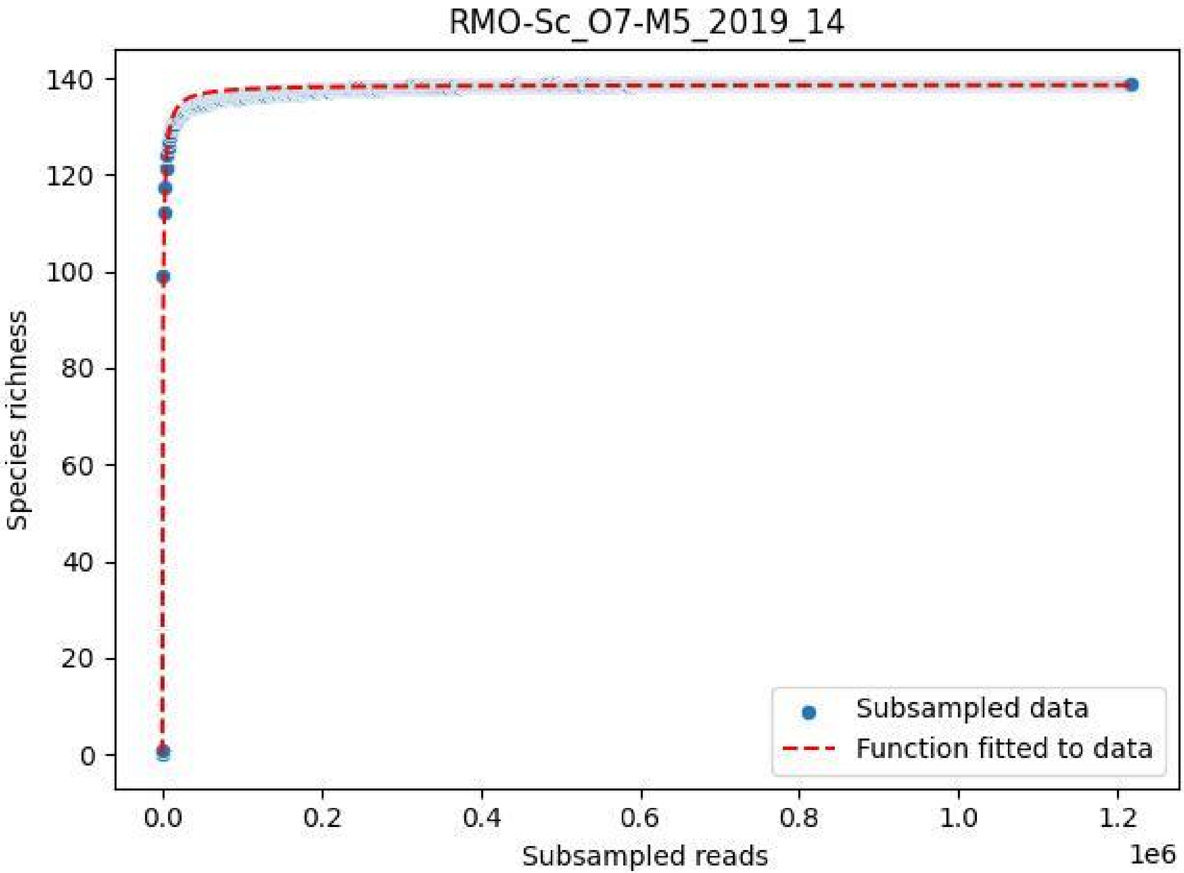
RMO-Sc_O7-M5_2019_14
Species richness
Subsampled reads
1e6
0
20
40
60
80
100
120
140
0.0
0.2
0.4
0.6
0.8
1.0
1.2
Subsampled data
Function fitted to data

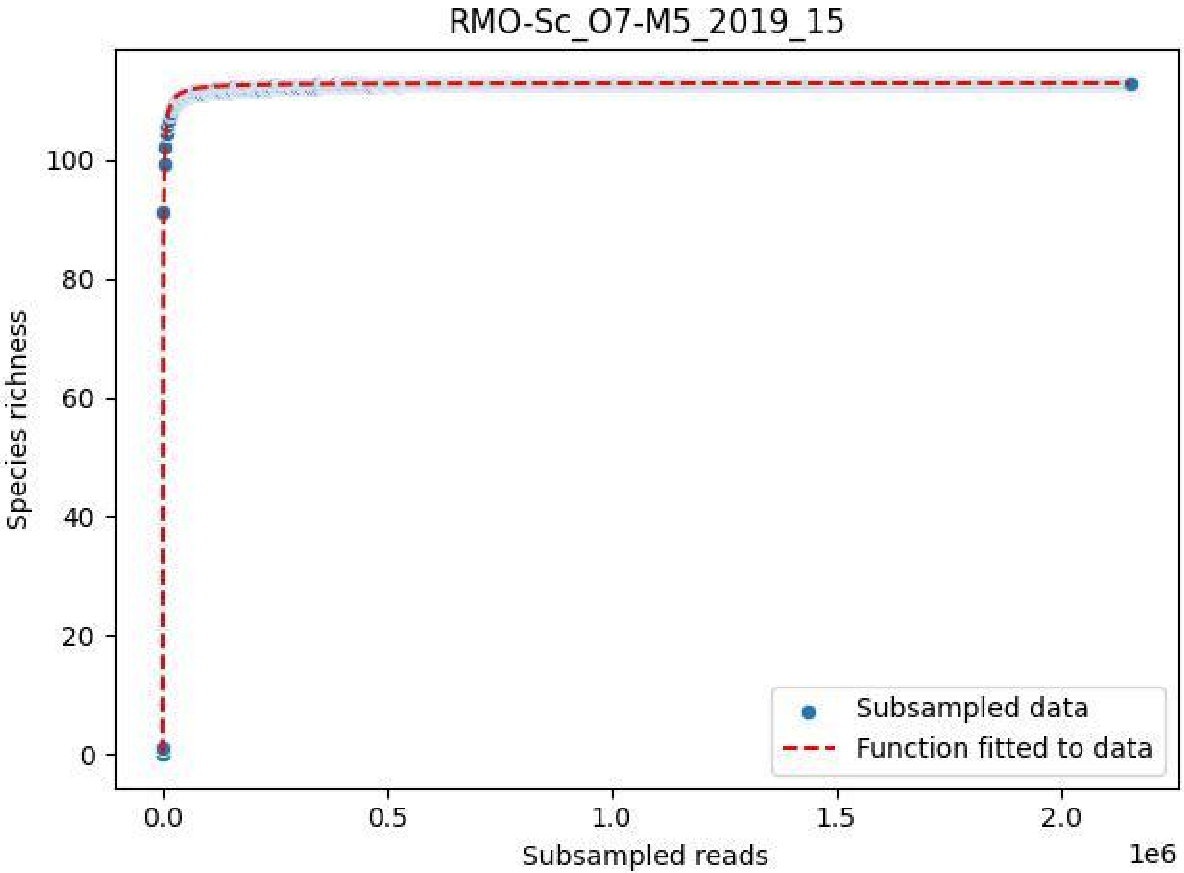

RMO-Sc_O7-M5_2019_15
Species richness
Subsampled reads
1e6
0
20
40
60
80
100
0.0
0.5
1.0
1.5
2.0
Subsampled data
Function fitted to data

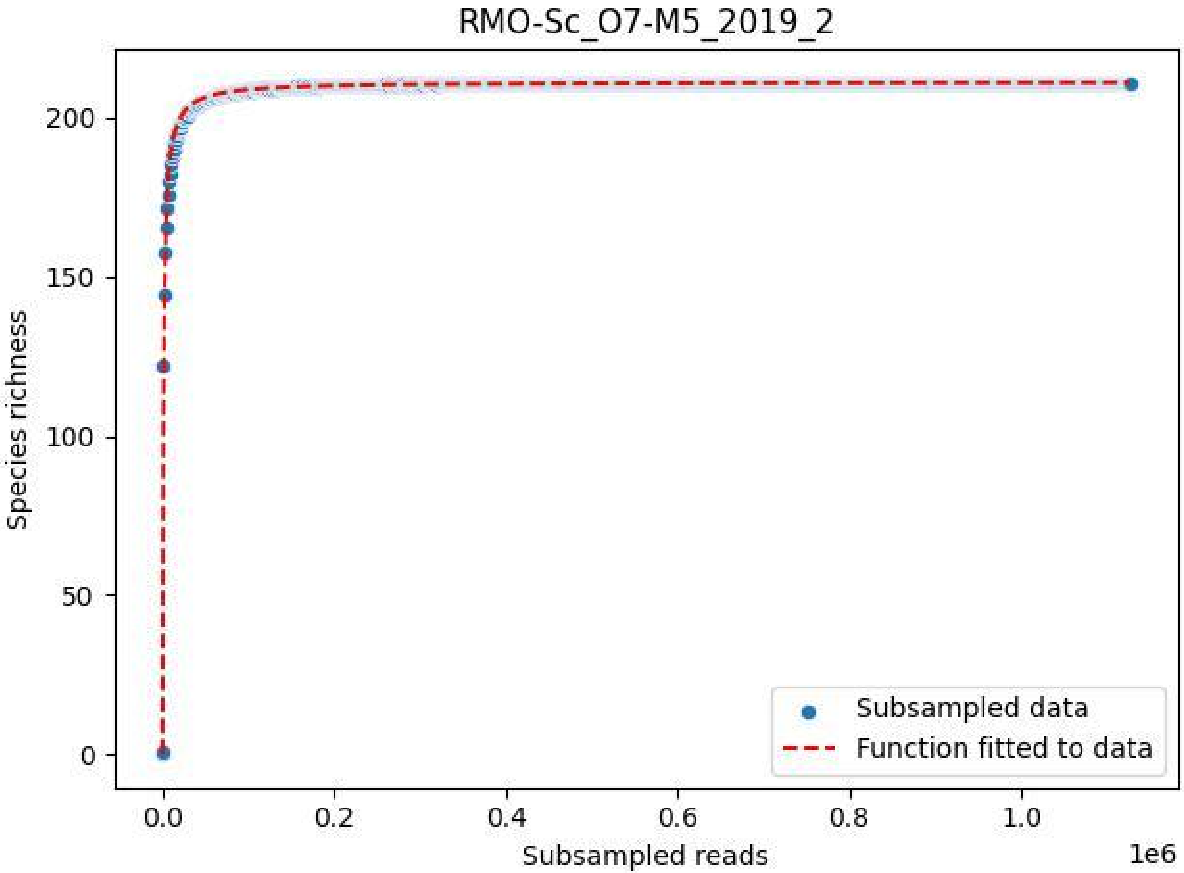
RMO-Sc_O7-M5_2019_2
Species richness
Subsampled reads
1e6
0
50
100
150
200
0.0
0.2
0.4
0.6
0.8
1.0
Subsampled data
Function fitted to data

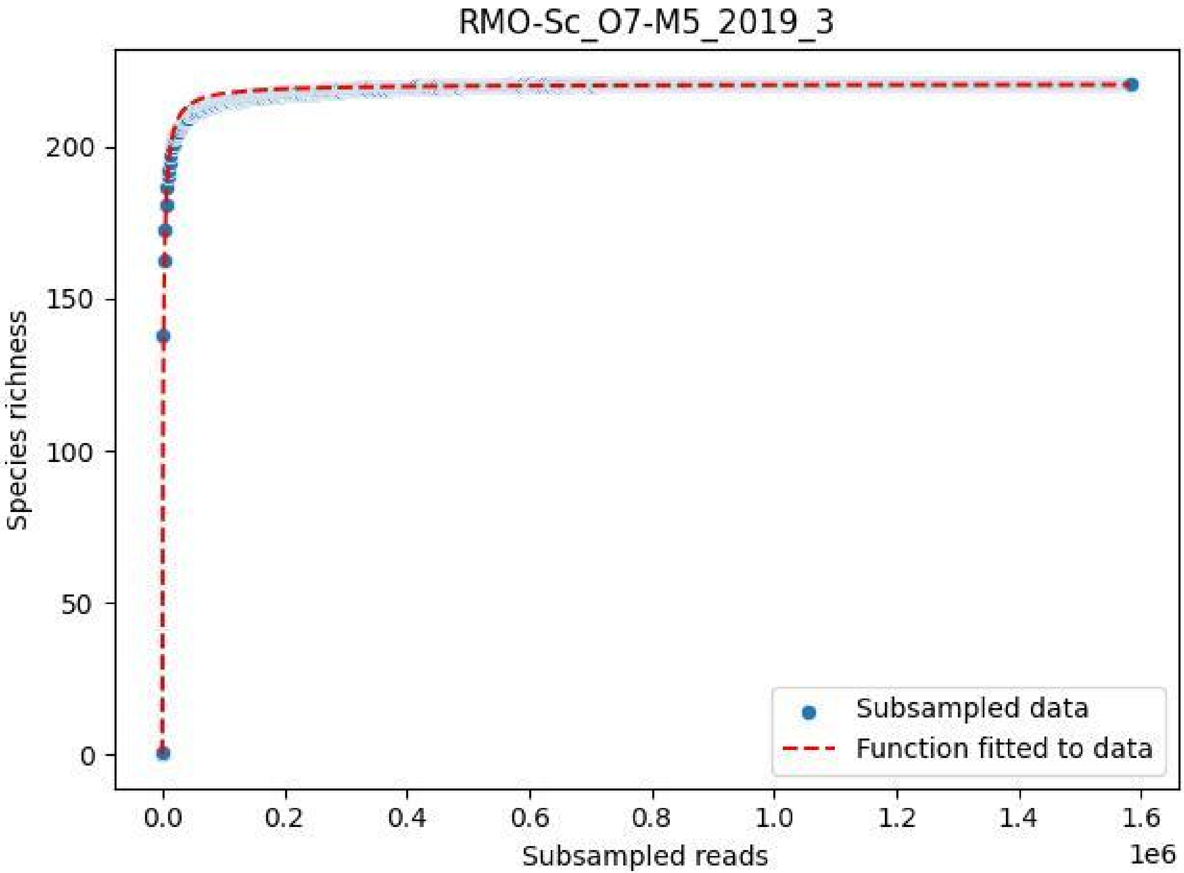
RMO-Sc_O7-M5_2019_3
Species richness
Subsampled reads
1e6
0
50
100
150
200
0.0
0.2
0.4
0.6
0.8
1.0
1.2
1.4
1.6
Subsampled data
Function fitted to data

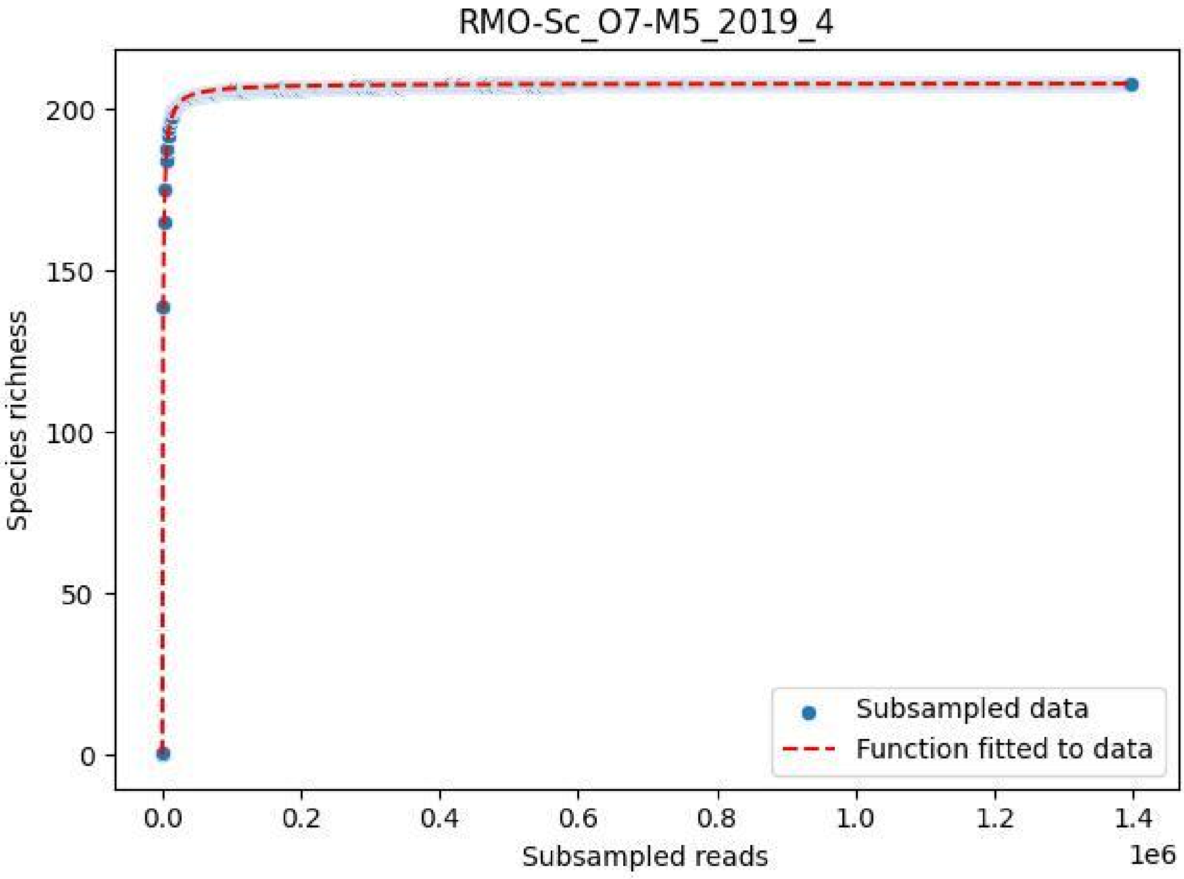

RMO-Sc_O7-M5_2019_4
Species richness
0
50
100
150
200
0.0
0.2
0.4
0.6
0.8
1.0
1.2
1.4
1e6
Subsampled reads
Subsampled data
Function fitted to data

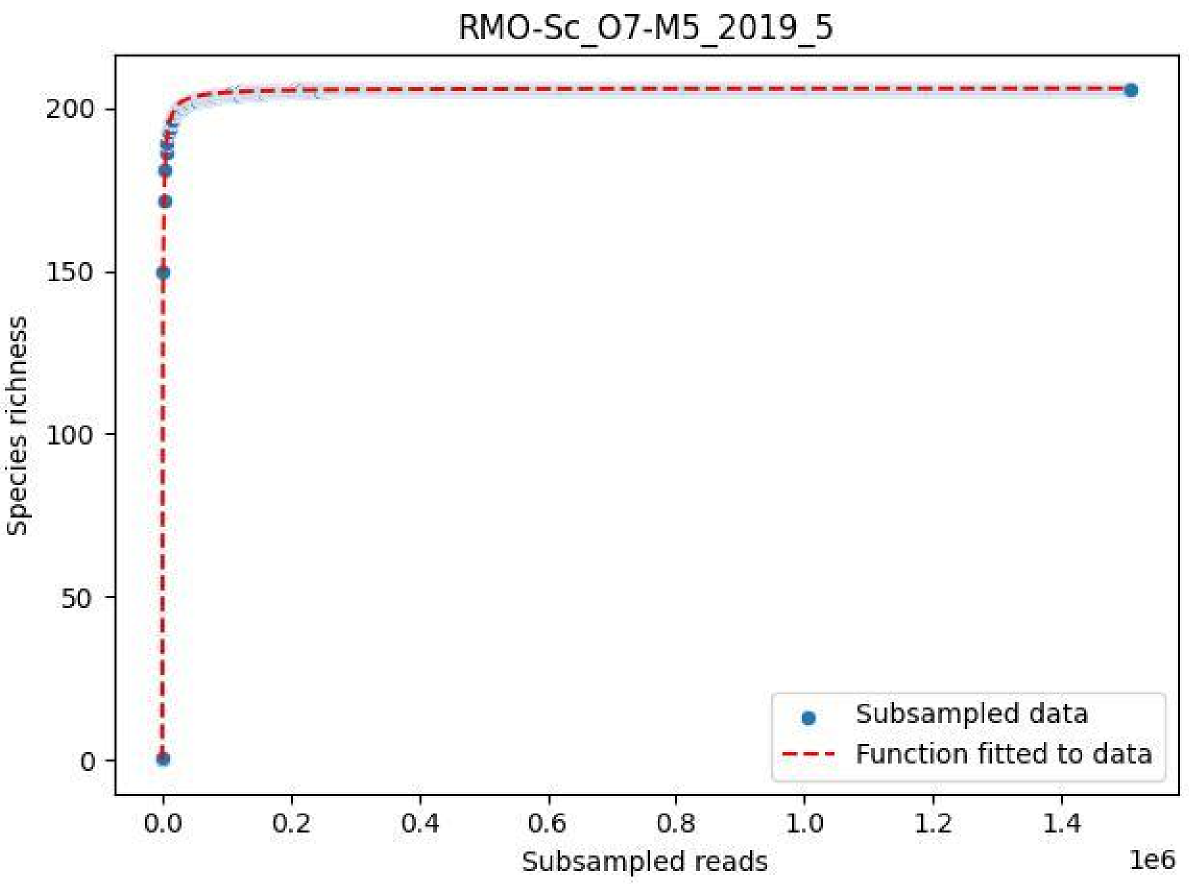

RMO-Sc_O7-M5_2019_5
Species richness
Subsampled reads
1e6
0
50
100
150
200
0.0
0.2
0.4
0.6
0.8
1.0
1.2
1.4
Subsampled data
Function fitted to data

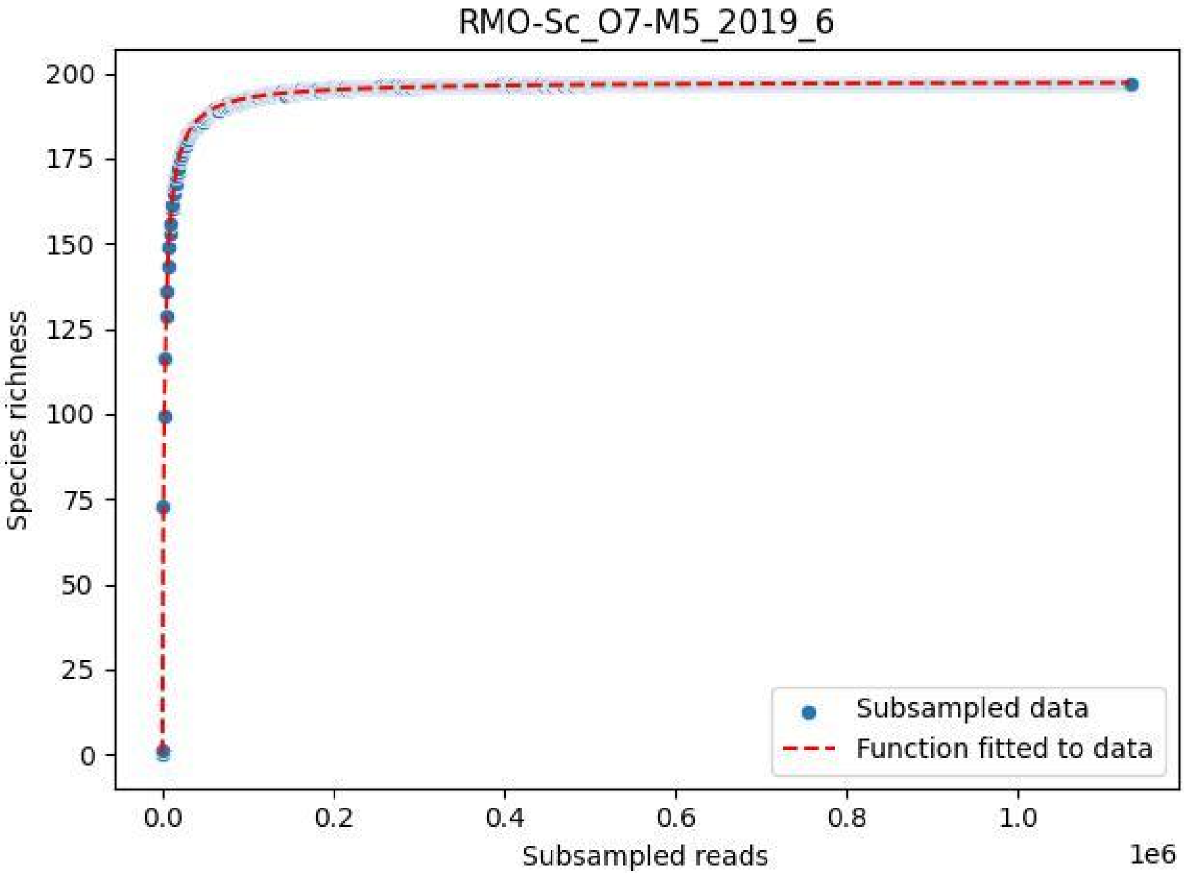

RMO-Sc_O7-M5_2019_6
Species richness
Subsampled reads
1e6
200
175
150
125
100
75
50
25
0
0.0
0.2
0.4
0.6
0.8
1.0
Subsampled data
Function fitted to data

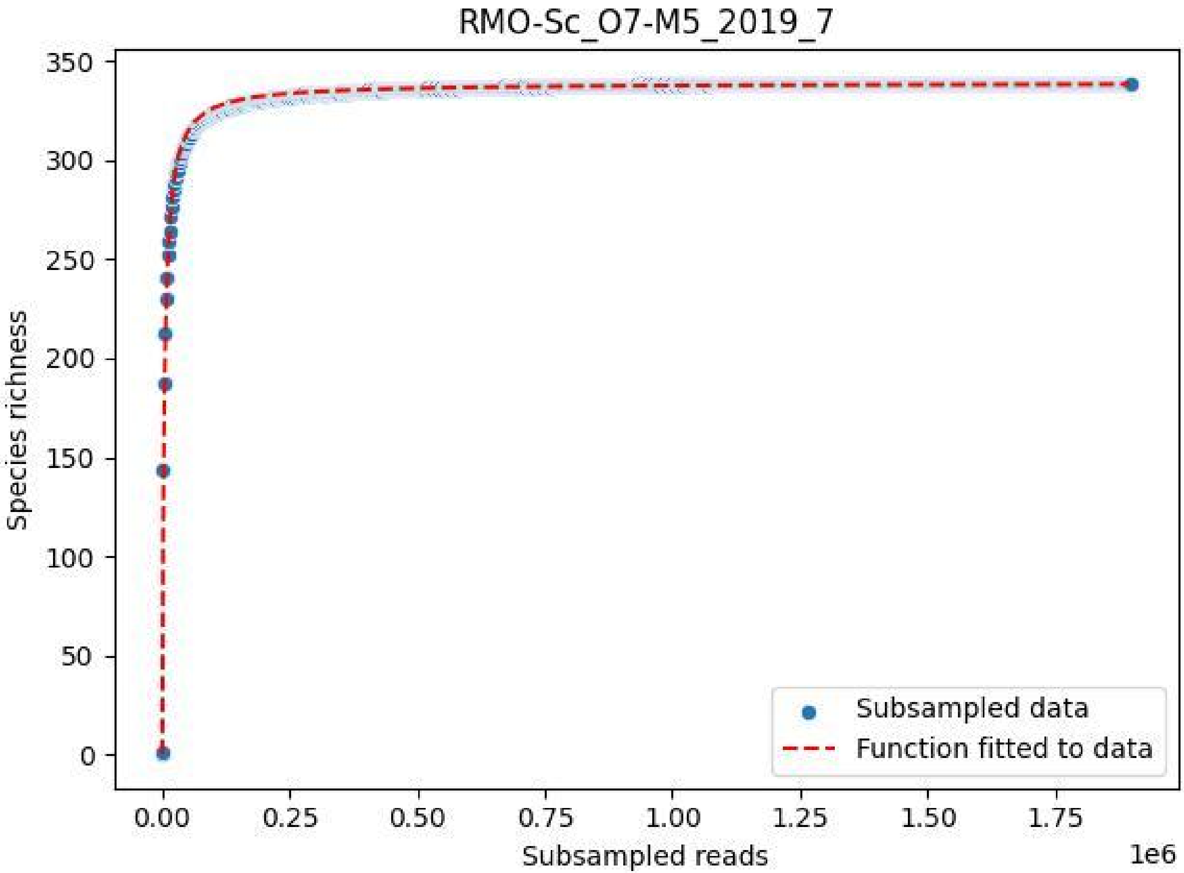
RMO-Sc_O7-M5_2019_7
Species richness
Subsampled reads
1e6
0
50
100
150
200
250
300
350
0.00
0.25
0.50
0.75
1.00
1.25
1.50
1.75
Subsampled data
Function fitted to data

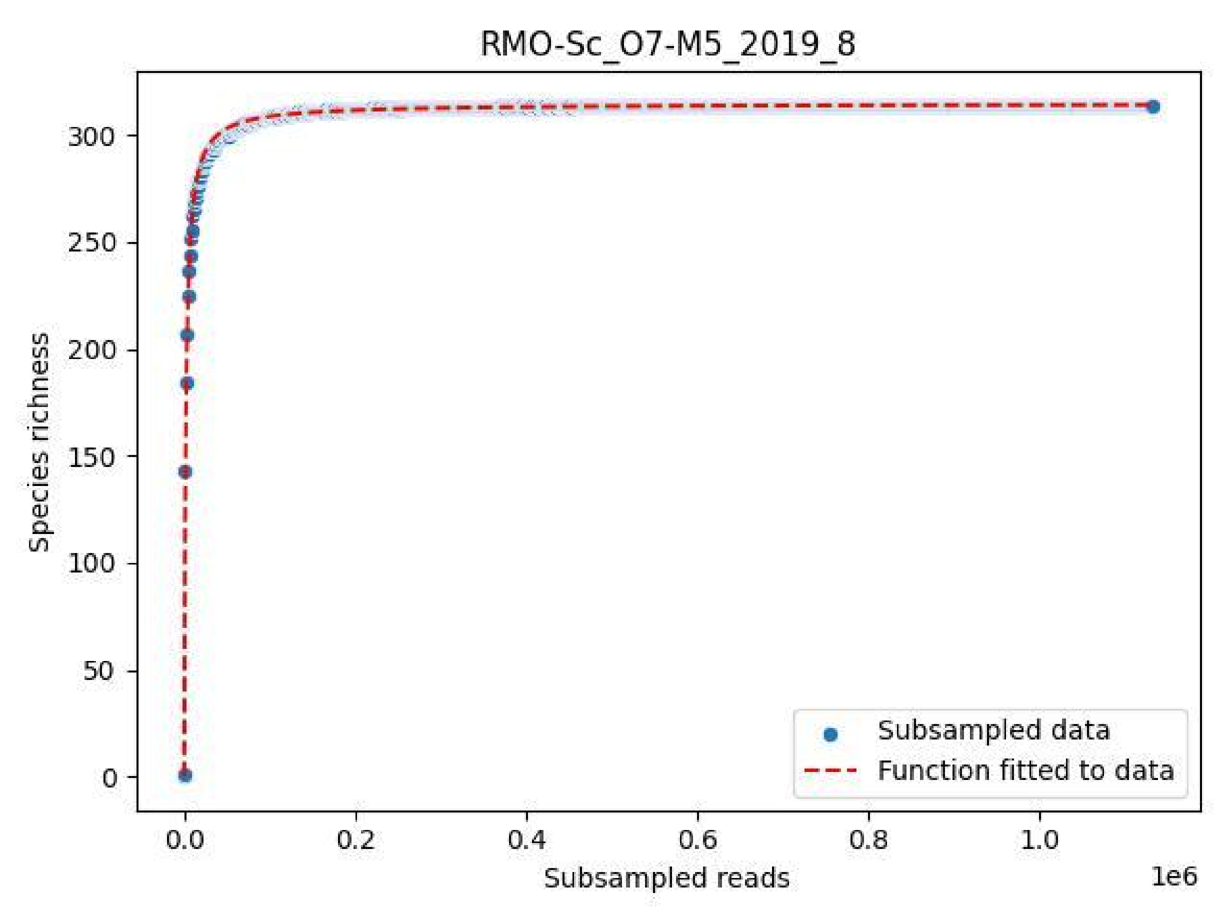

RMO-Sc_O7-M5_2019_8
Species richness
300
250
200
150
100
50
0
0.0
0.2
0.4
0.6
0.8
1.0
Subsampled reads
1e6
Subsampled data
Function fitted to data

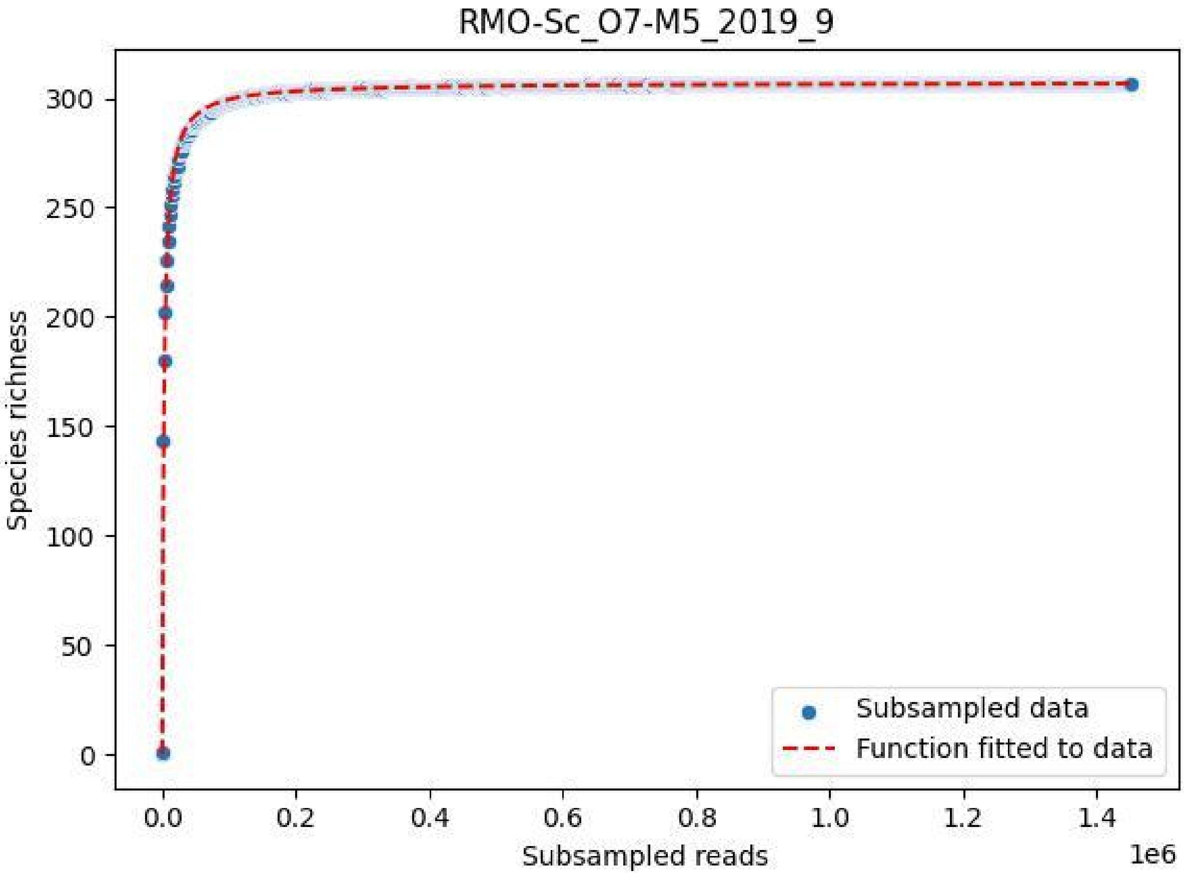

RMO-Sc_O7-M5_2019_9
Species richness
Subsampled reads
1e6
Subsampled data
Function fitted to data
0
50
100
150
200
250
300
0.0
0.2
0.4
0.6
0.8
1.0
1.2
1.4

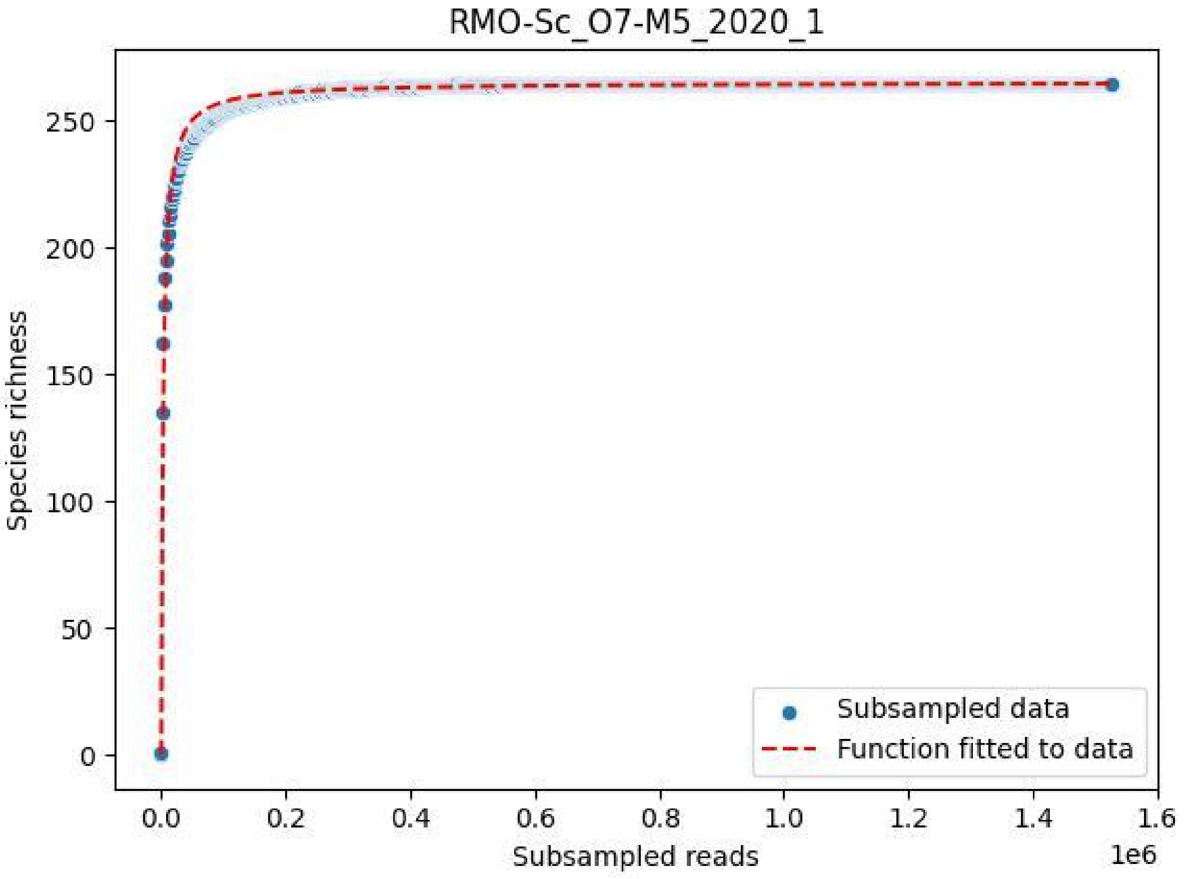

RMO-Sc_O7-M5_2020_1
Species richness
250
200
150
100
50
0
0.0
0.2
0.4
0.6
0.8
1.0
1.2
1.4
1.6
Subsampled reads
1e6
Subsampled data
Function fitted to data

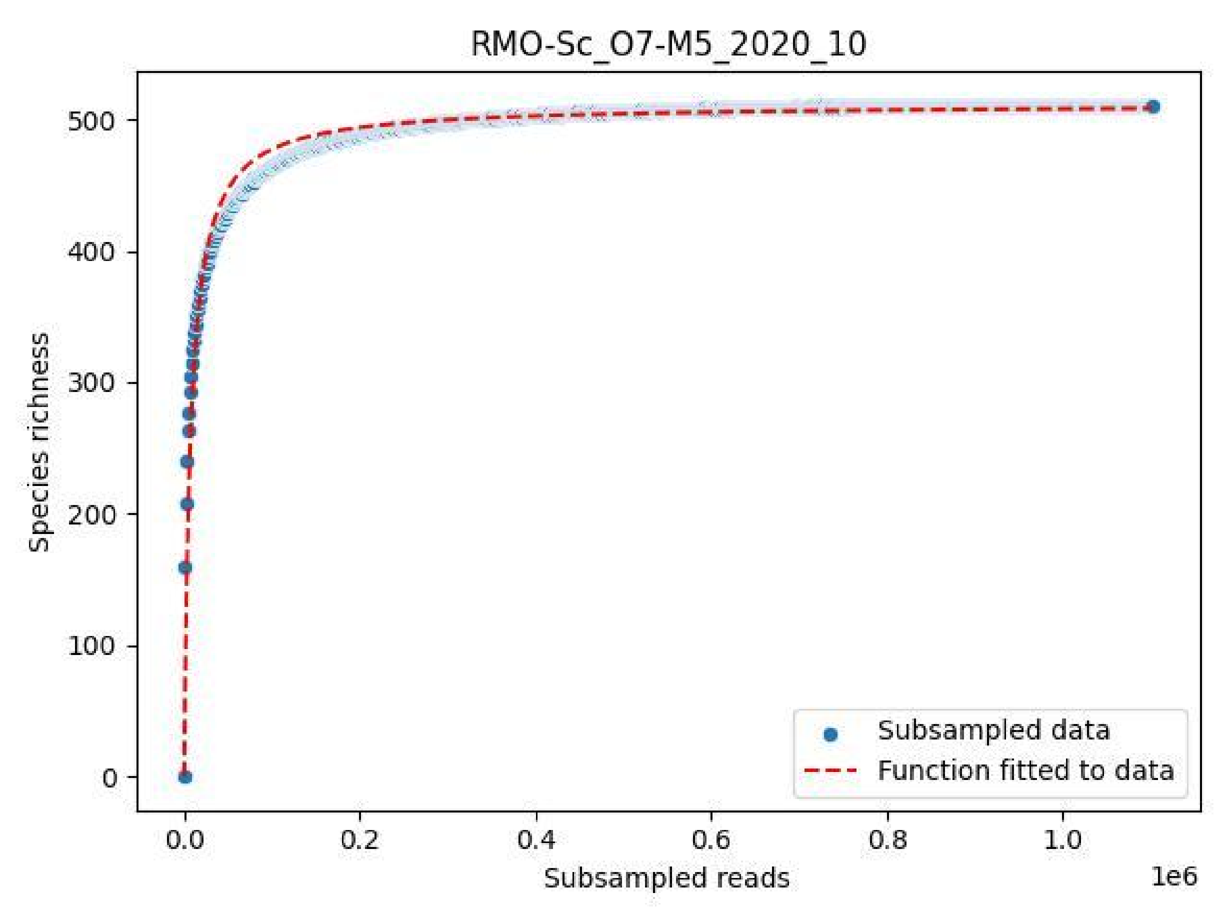
RMO-Sc_O7-M5_2020_10
Species richness
Subsampled reads
1e6
0
100
200
300
400
500
0.0
0.2
0.4
0.6
0.8
1.0
Subsampled data
Function fitted to data

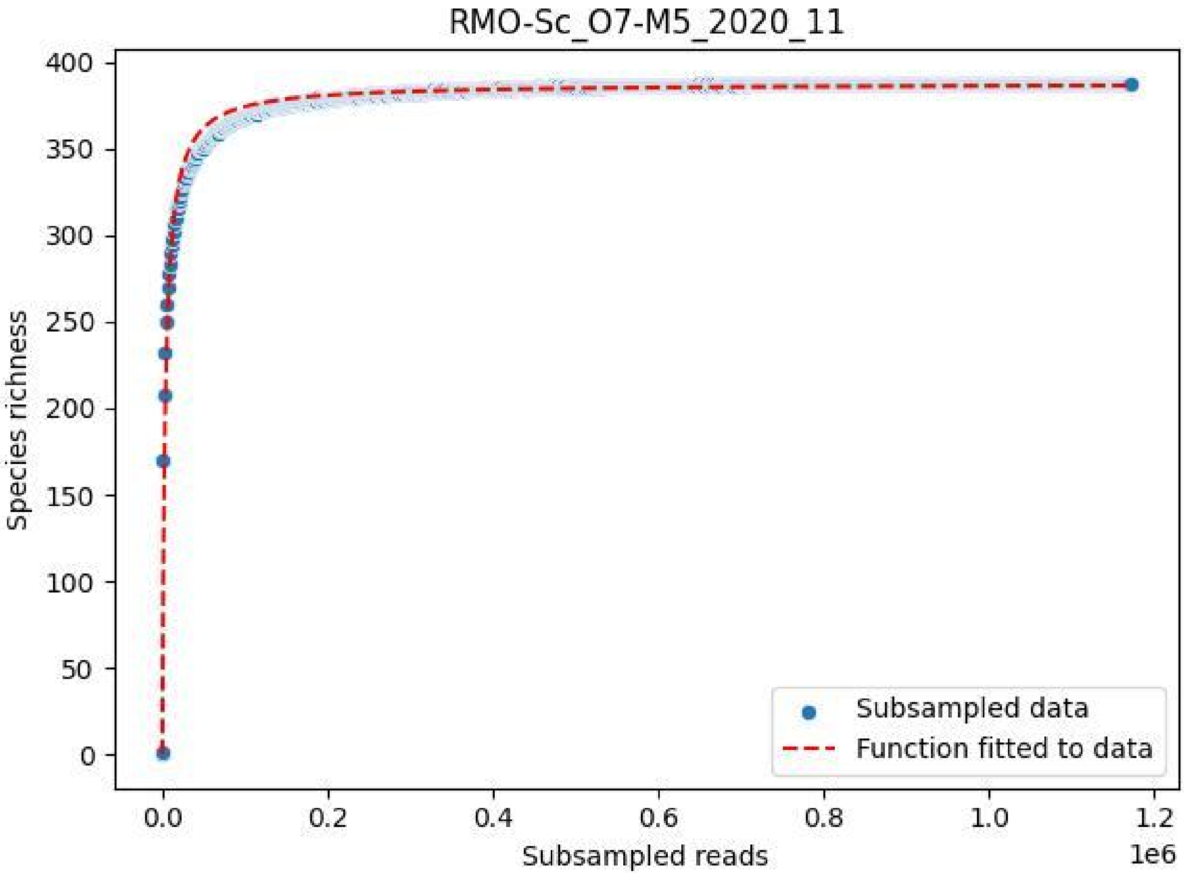

RMO-Sc_O7-M5_2020_11
Species richness
Subsampled reads
1e6
0
50
100
150
200
250
300
350
400
0.0
0.2
0.4
0.6
0.8
1.0
1.2
Subsampled data
Function fitted to data

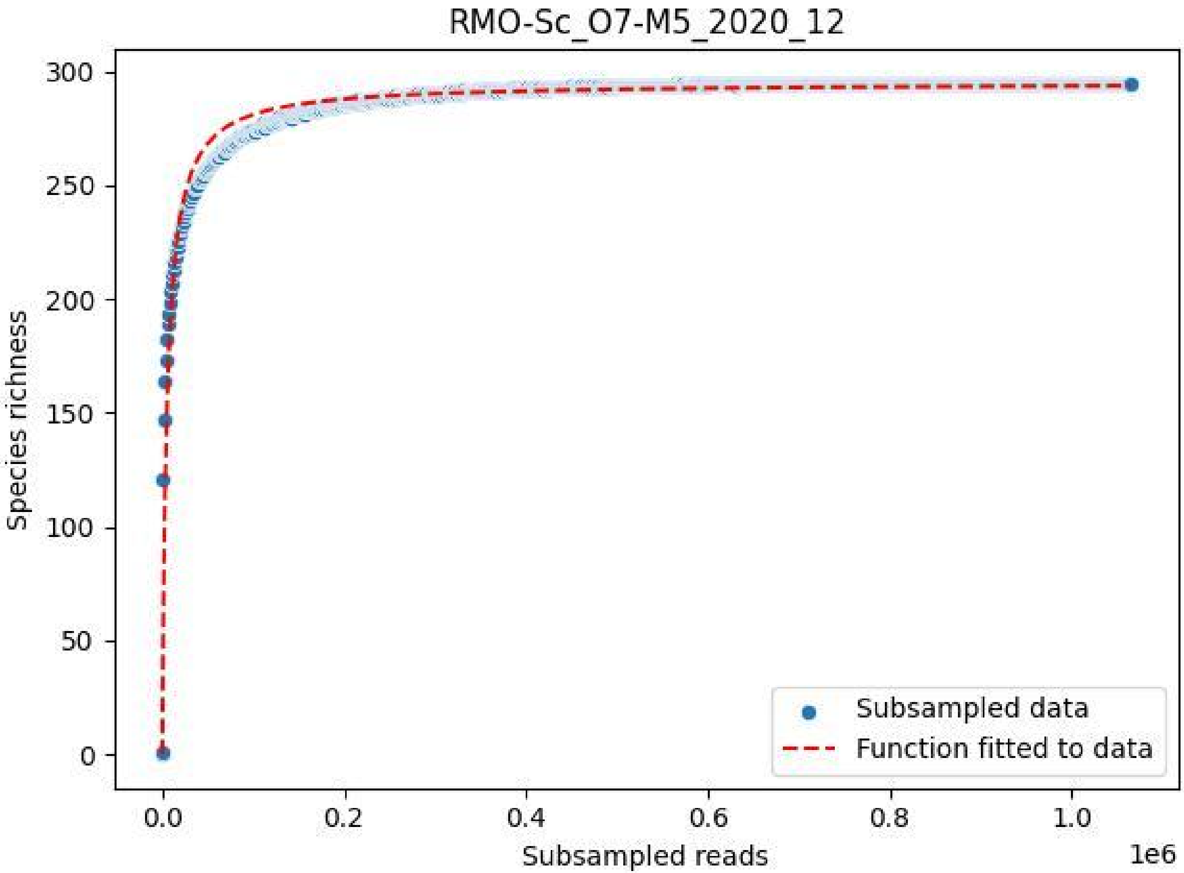

RMO-Sc_O7-M5_2020_12
Species richness
Subsampled reads
1e6
Subsampled data
Function fitted to data
0
50
100
150
200
250
300
0.0
0.2
0.4
0.6
0.8
1.0

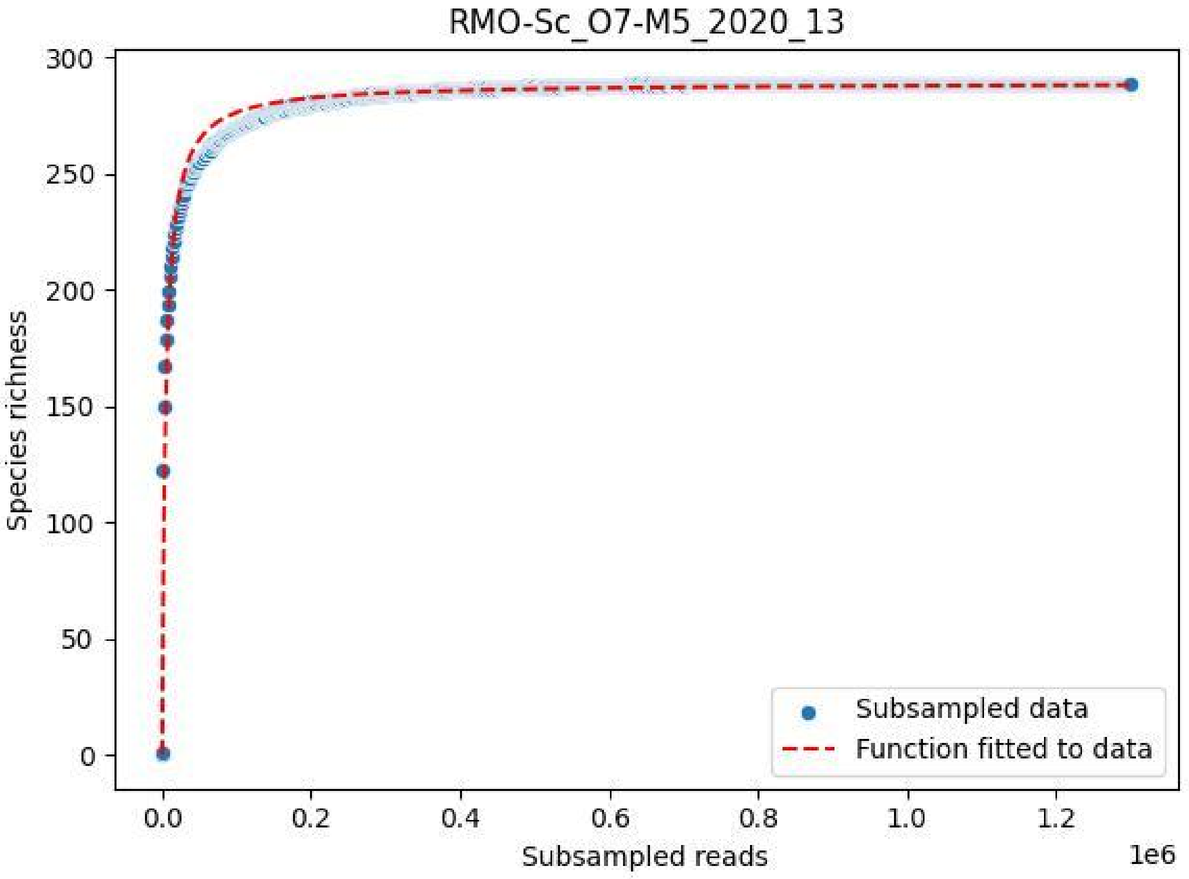

RMO-Sc_O7-M5_2020_13
Species richness
Subsampled reads
1e6
0
50
100
150
200
250
300
0.0
0.2
0.4
0.6
0.8
1.0
1.2
Subsampled data
Function fitted to data

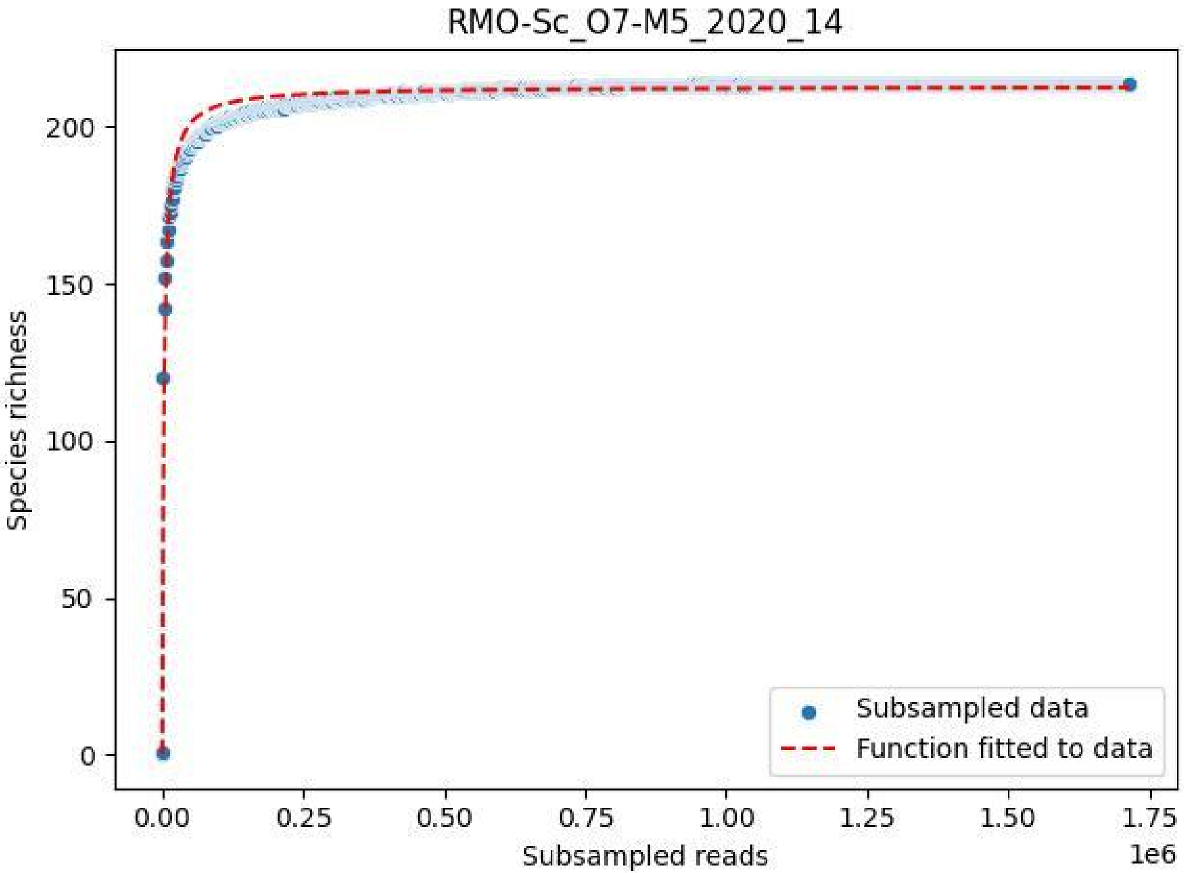
RMO-Sc_O7-M5_2020_14
Species richness
Subsampled reads
1e6
0
50
100
150
200
0.00
0.25
0.50
0.75
1.00
1.25
1.50
1.75
Subsampled data
Function fitted to data

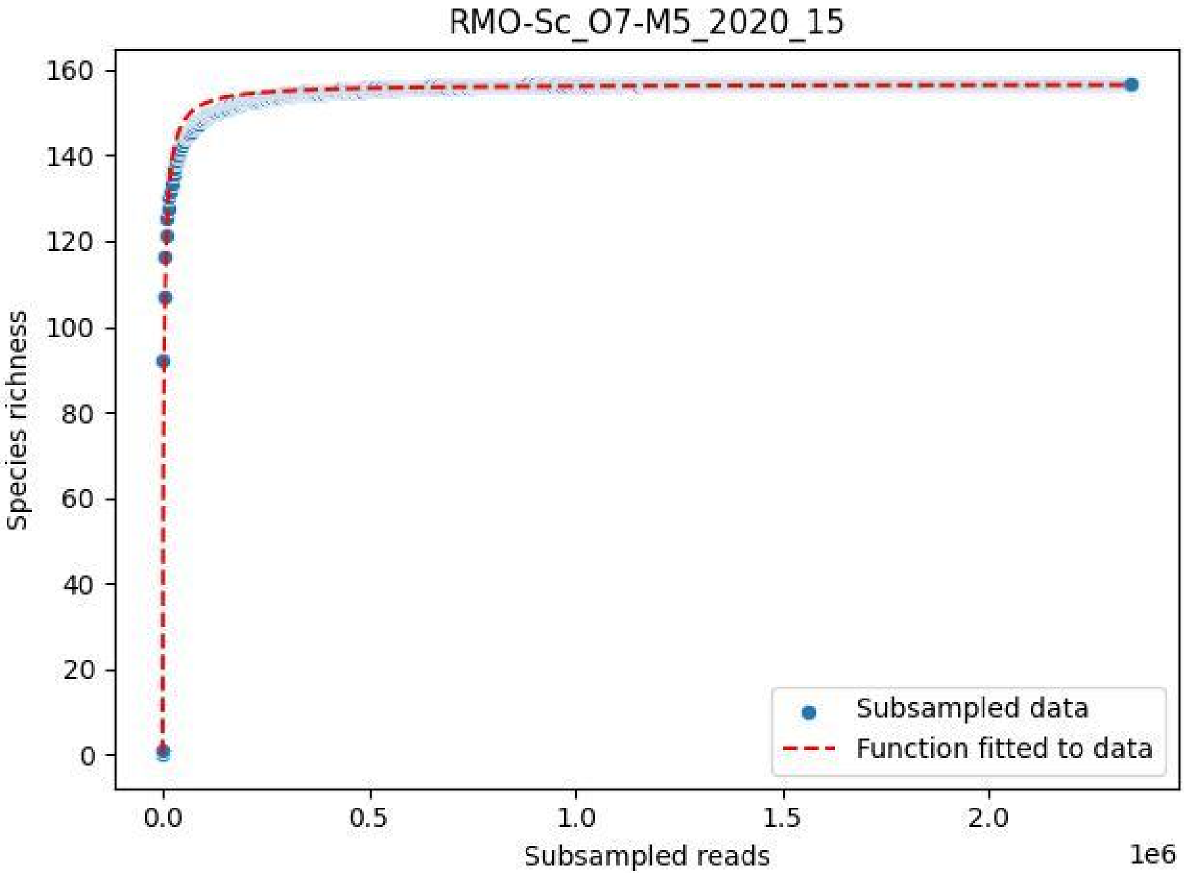

RMO-Sc_O7-M5_2020_15
Species richness
Subsampled reads
1e6
0
20
40
60
80
100
120
140
160
0.0
0.5
1.0
1.5
2.0
Subsampled data
Function fitted to data

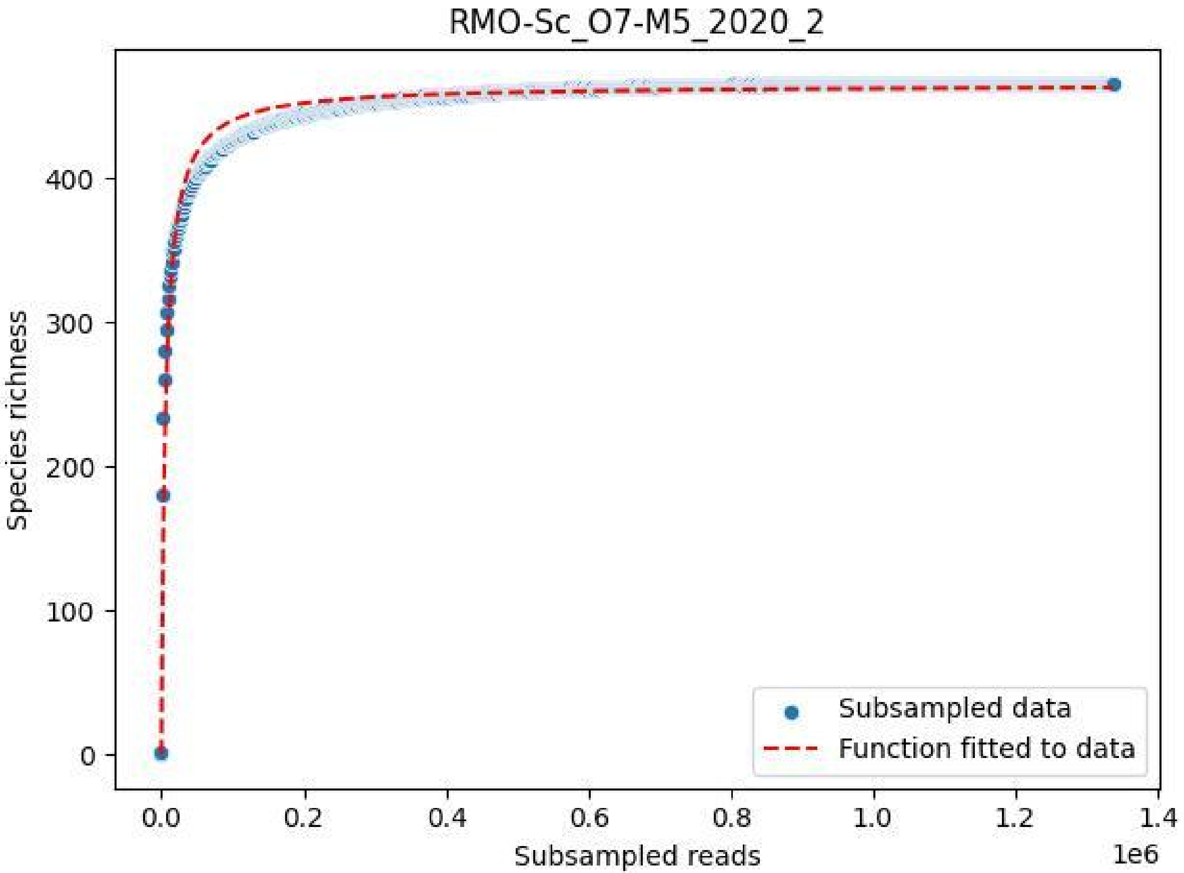

RMO-Sc_O7-M5_2020_2
Species richness
400
300
200
100
0
0.0
0.2
0.4
0.6
0.8
1.0
1.2
1.4
1e6
Subsampled reads
Subsampled data
Function fitted to data

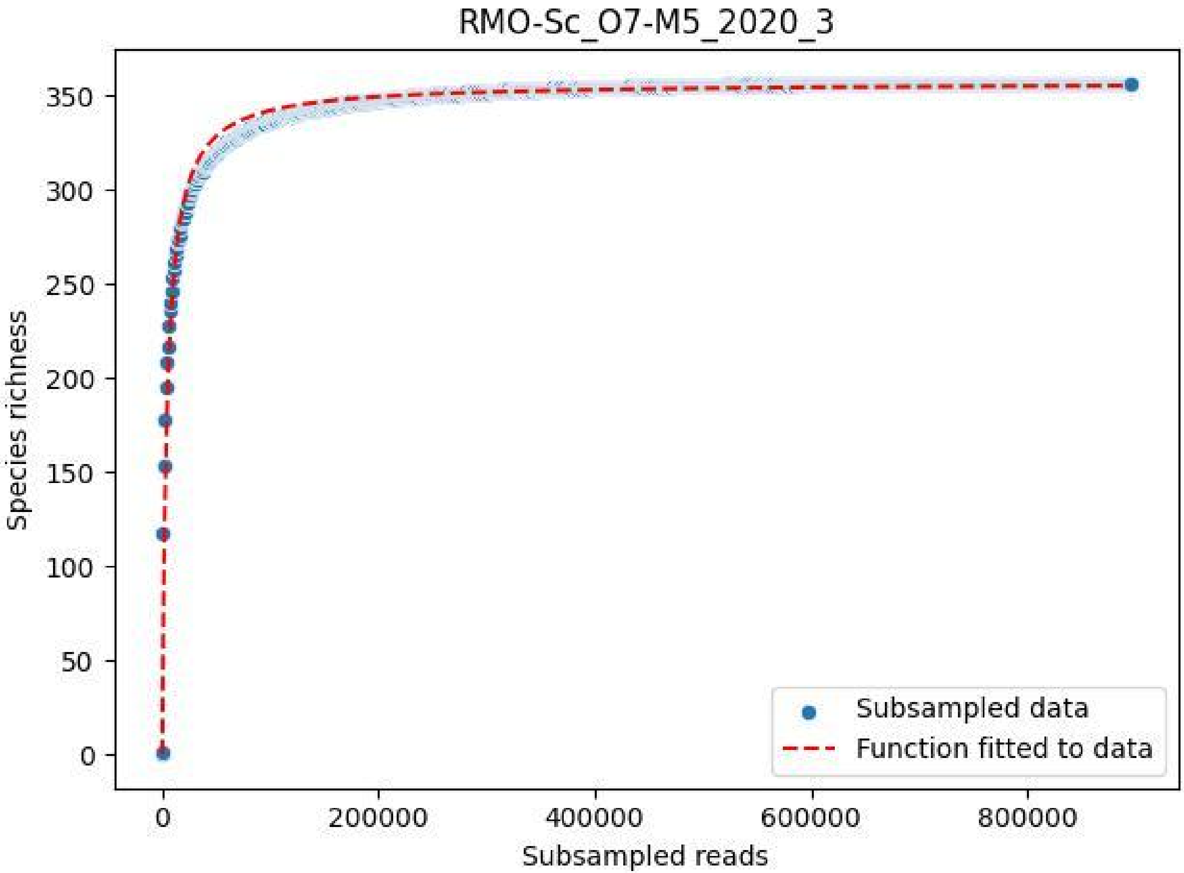

RMO-Sc_O7-M5_2020_3
Species richness
Subsampled reads
0
50
100
150
200
250
300
350
0
200000
400000
600000
800000
Subsampled data
Function fitted to data

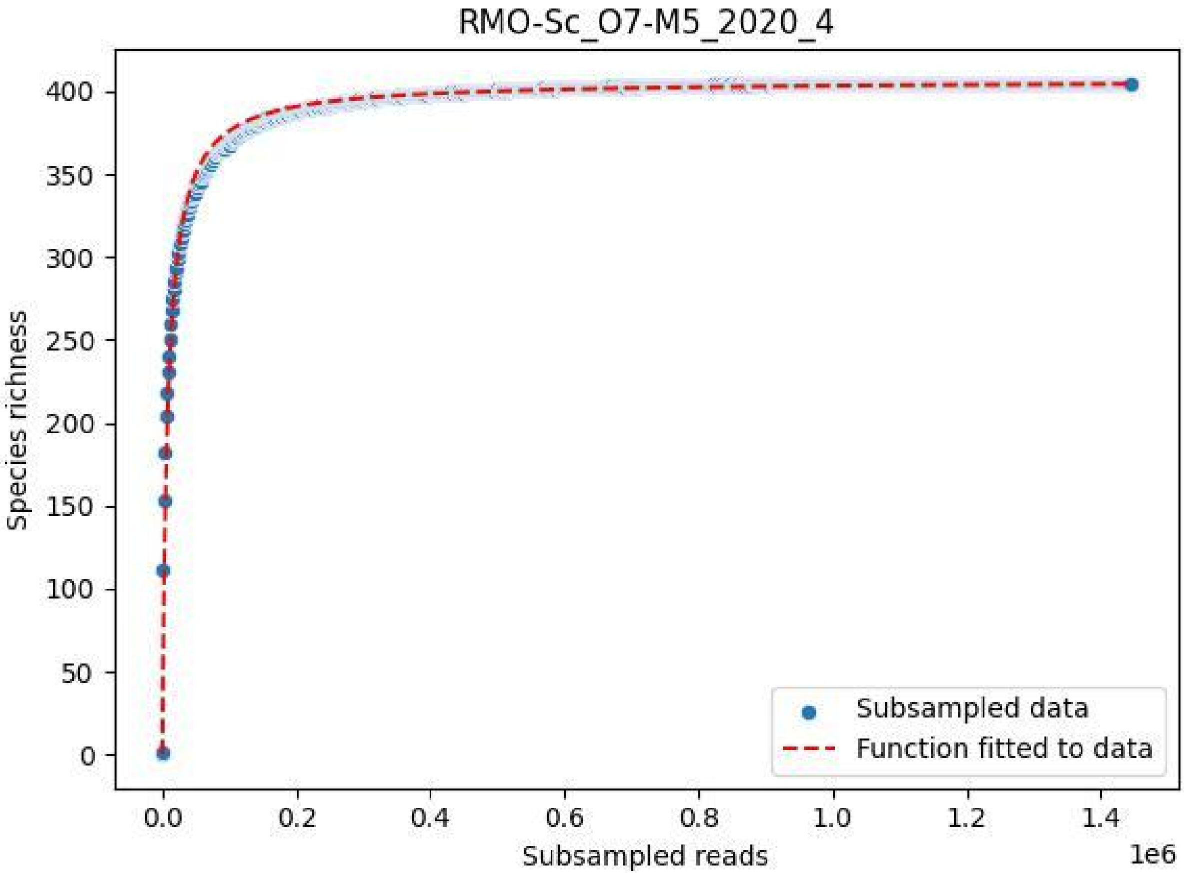
RMO-Sc_O7-M5_2020_4
Species richness
Subsampled reads
1e6
0.0
0.2
0.4
0.6
0.8
1.0
1.2
1.4
0
50
100
150
200
250
300
350
400
Subsampled data
Function fitted to data

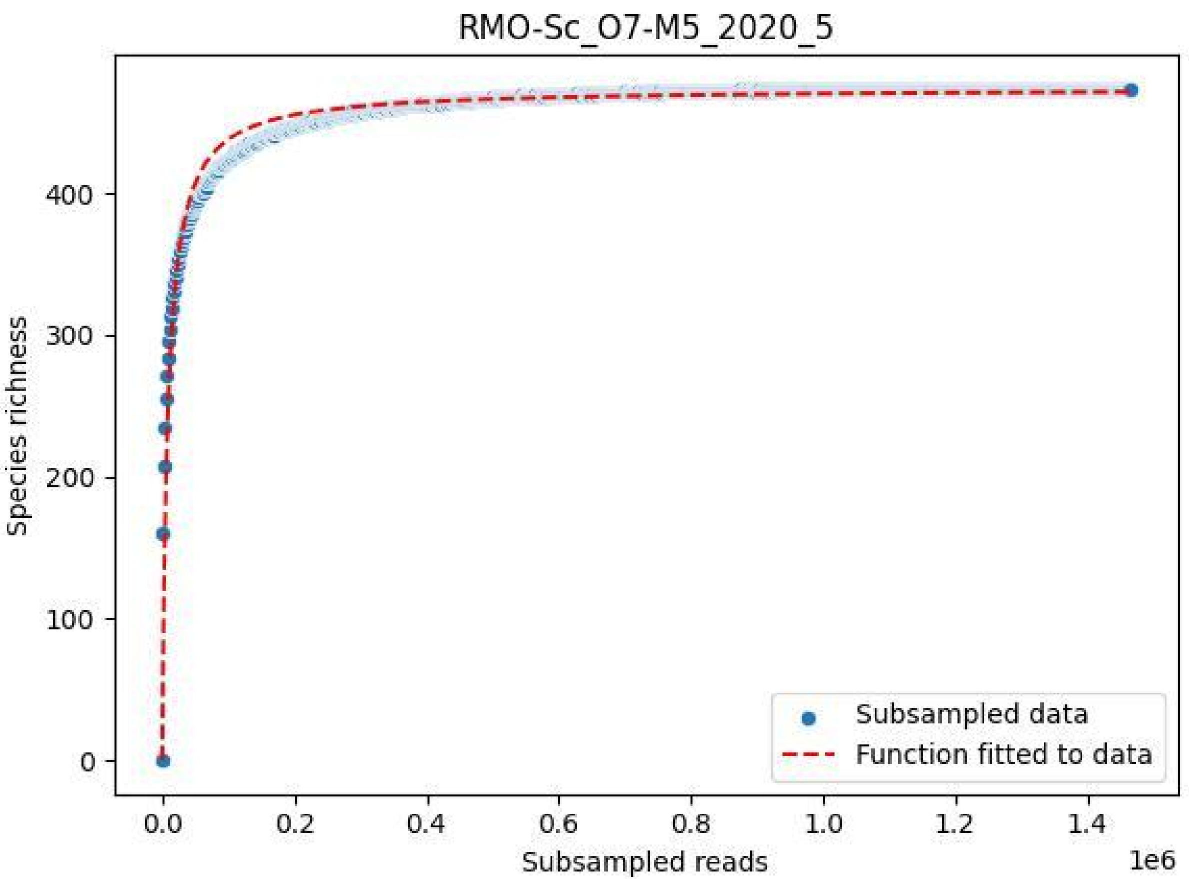
RMO-Sc_O7-M5_2020_5
Species richness
Subsampled reads
1e6
0
100
200
300
400
0.0
0.2
0.4
0.6
0.8
1.0
1.2
1.4
Subsampled data
Function fitted to data

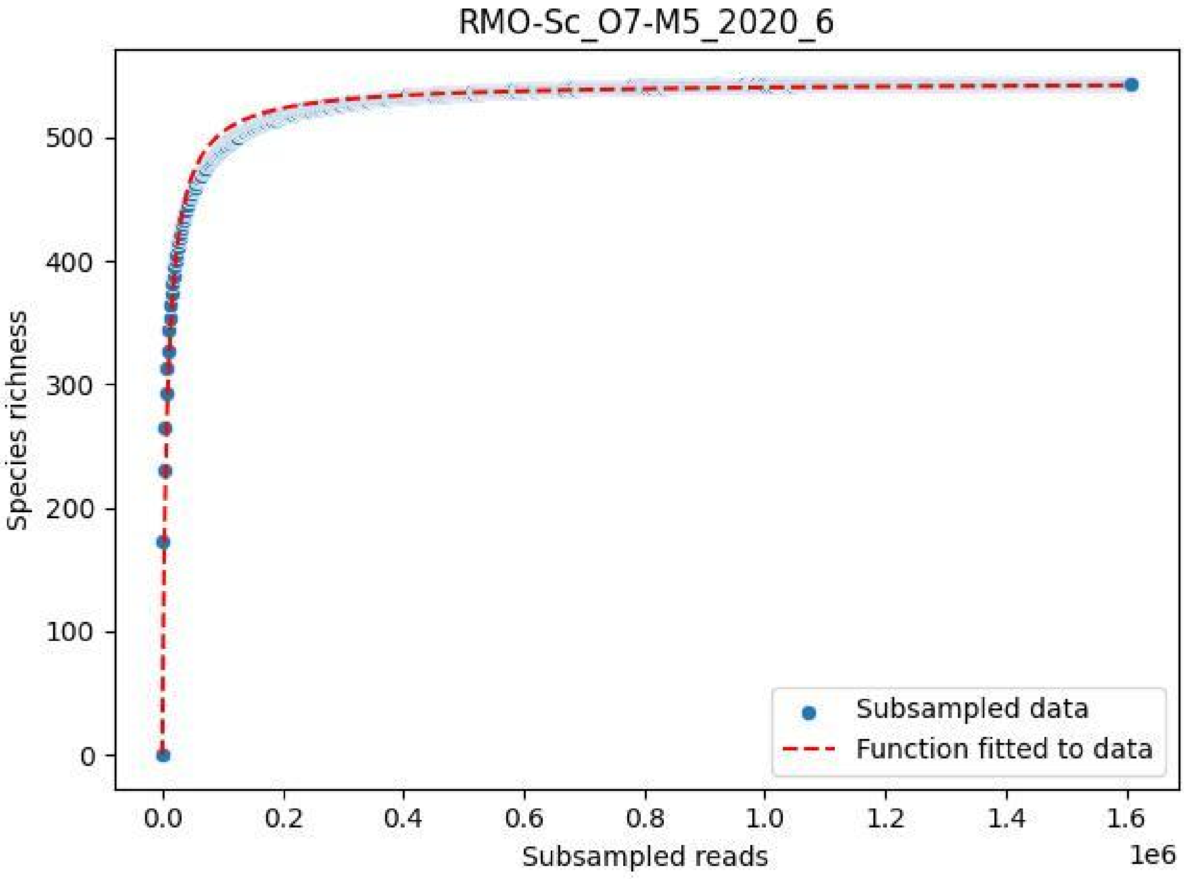
RMO-Sc_O7-M5_2020_6
Species richness
Subsampled reads
1e6
0
100
200
300
400
500
0.0
0.2
0.4
0.6
0.8
1.0
1.2
1.4
1.6
Subsampled data
Function fitted to data

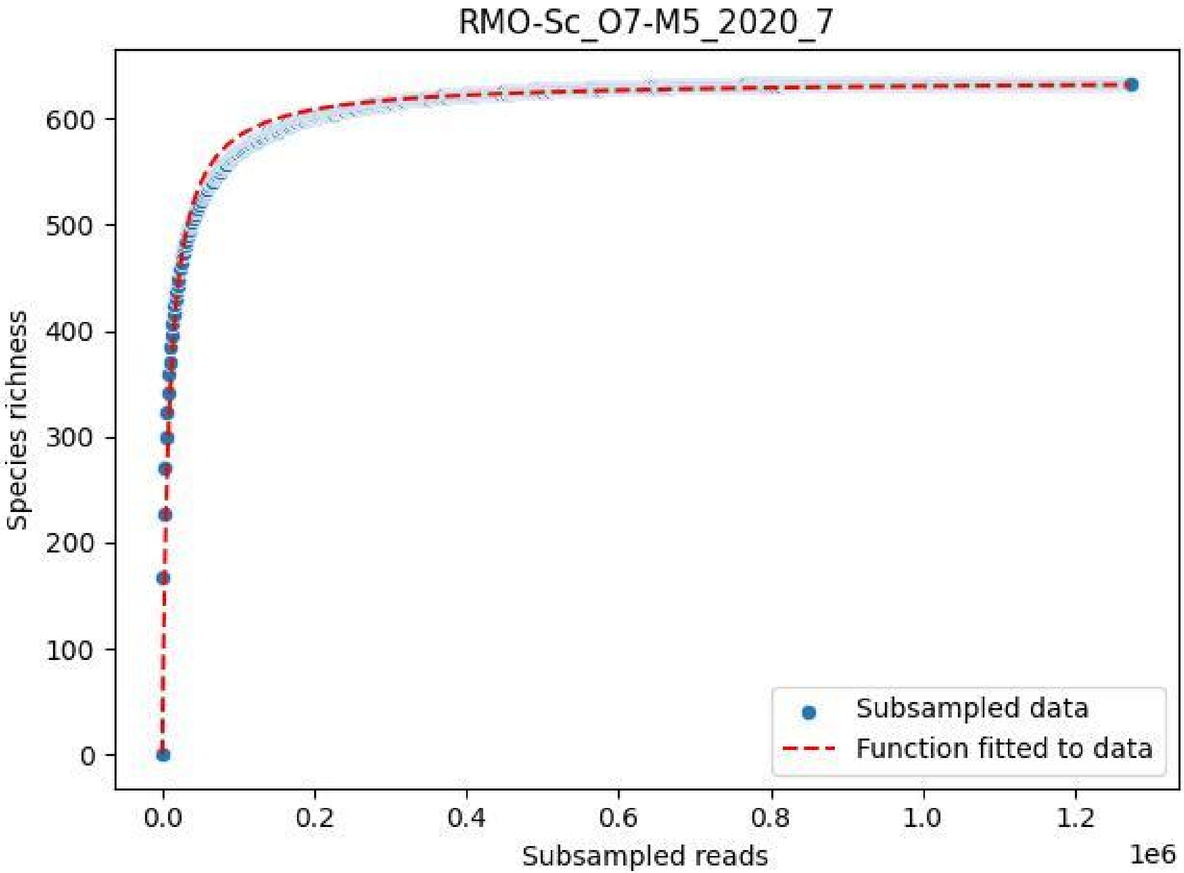

RMO-Sc_O7-M5_2020_7
Species richness
Subsampled reads
1e6
0
100
200
300
400
500
600
0.0
0.2
0.4
0.6
0.8
1.0
1.2
Subsampled data
Function fitted to data

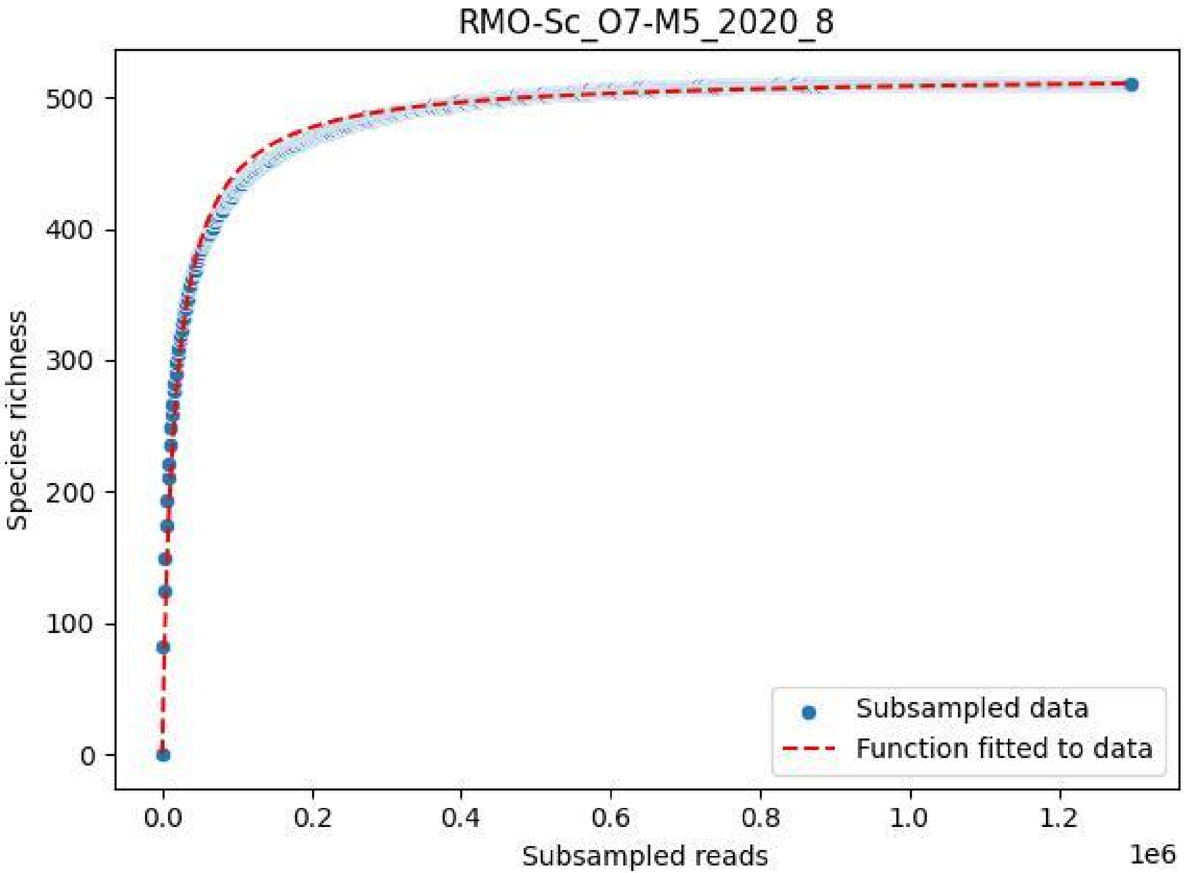

RMO-Sc_O7-M5_2020_8
Species richness
Subsampled reads
1e6
0
100
200
300
400
500
0.0
0.2
0.4
0.6
0.8
1.0
1.2
Subsampled data
Function fitted to data

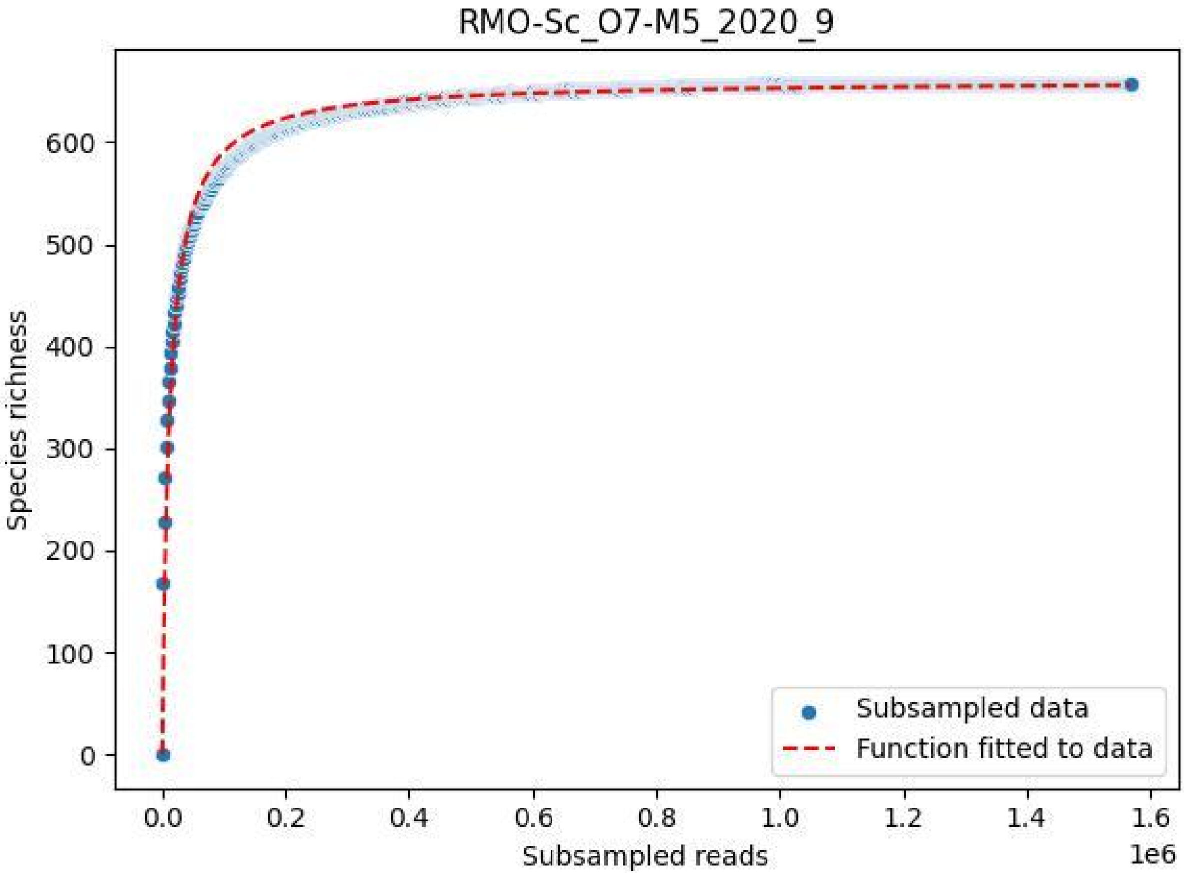
RMO-Sc_O7-M5_2020_9
Species richness
Subsampled reads
1e6
0
100
200
300
400
500
600
0.0
0.2
0.4
0.6
0.8
1.0
1.2
1.4
1.6
Subsampled data
Function fitted to data

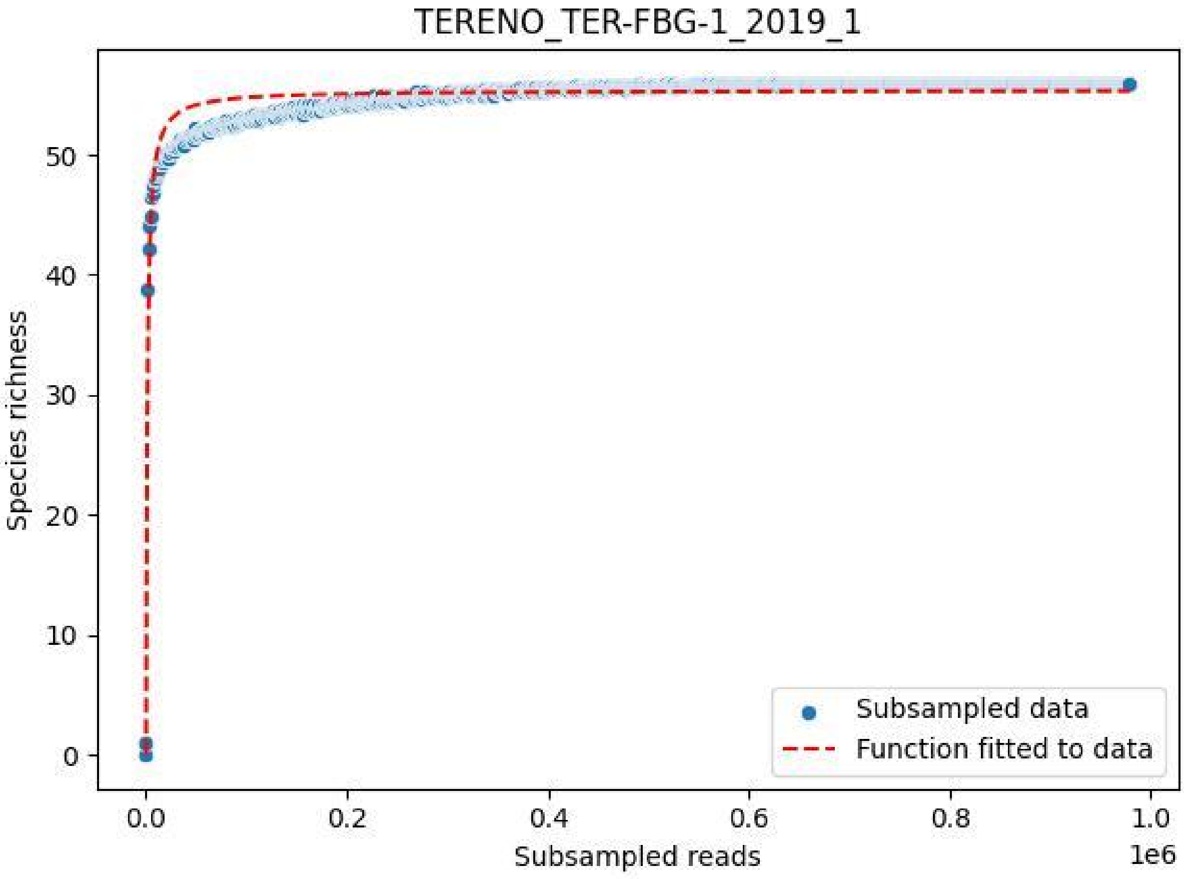

TERENO_TER-FBG-1_2019_1
Species richness
0
10
20
30
40
50
Subsampled reads
0.0
0.2
0.4
0.6
0.8
1.0
1e6
Subsampled data
Function fitted to data

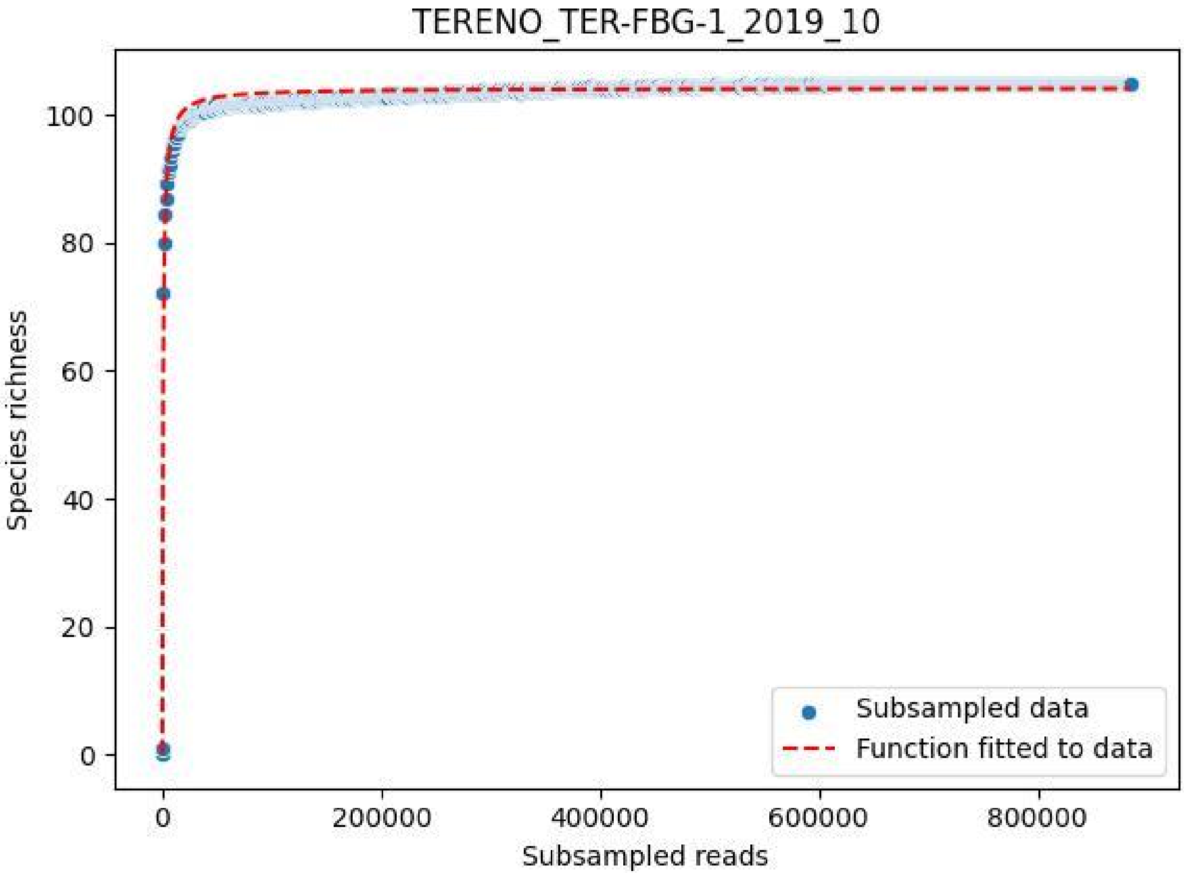

TERENO_TER-FBG-1_2019_10
Species richness
Subsampled reads
0
20
40
60
80
100
0
200000
400000
600000
800000
Subsampled data
Function fitted to data

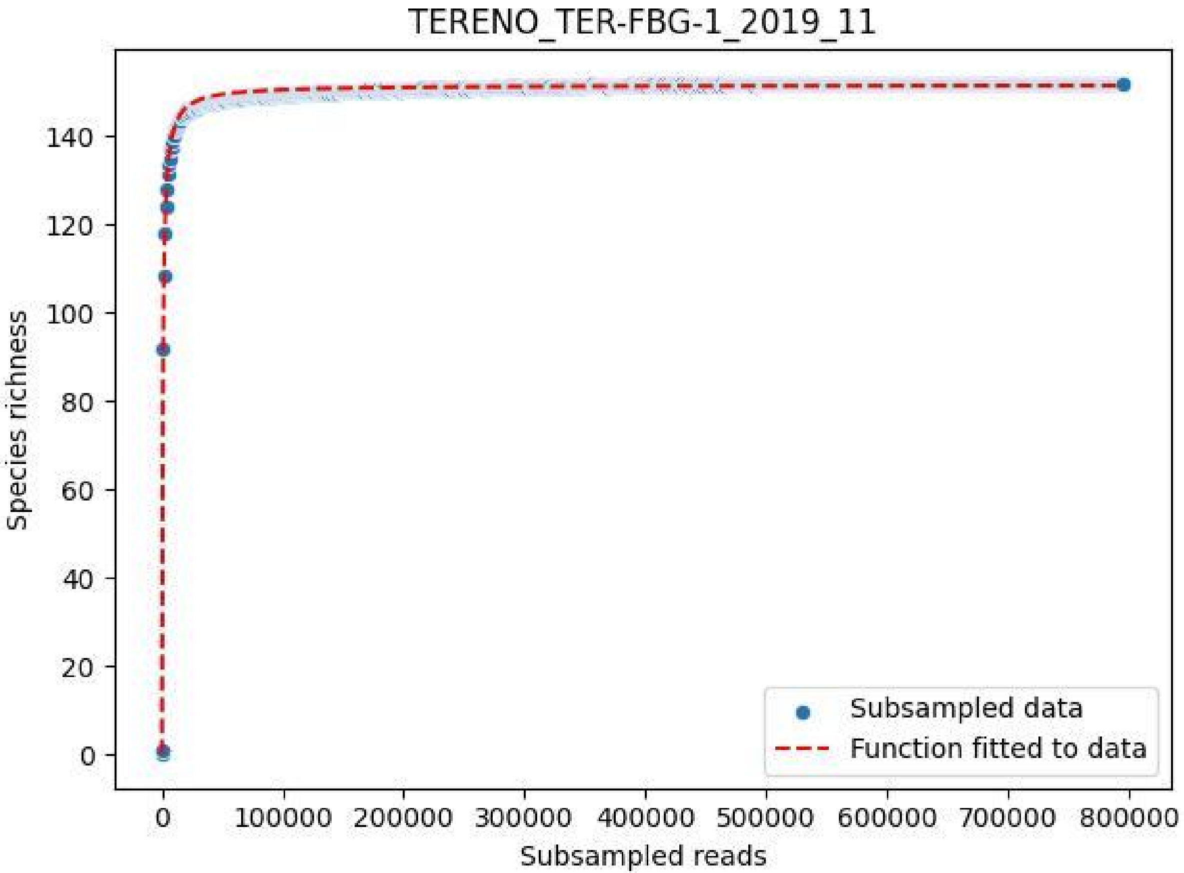
TERENO_TER-FBG-1_2019_11
Species richness
0
20
40
60
80
100
120
140
Subsampled reads
0
100000
200000
300000
400000
500000
600000
700000
800000
Subsampled data
Function fitted to data

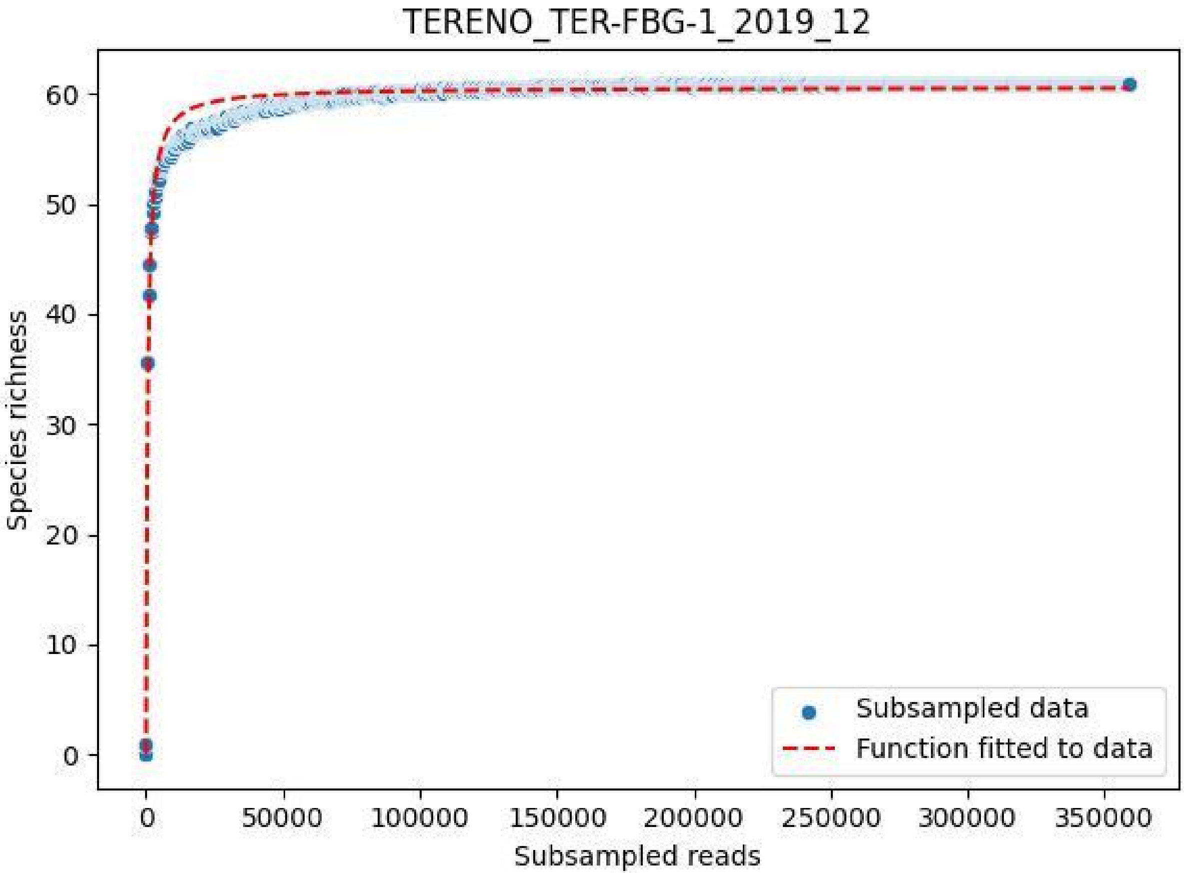
TERENO_TER-FBG-1_2019_12
Species richness
0
10
20
30
40
50
60
Subsampled reads
0
50000
100000
150000
200000
250000
300000
350000
Subsampled data
Function fitted to data

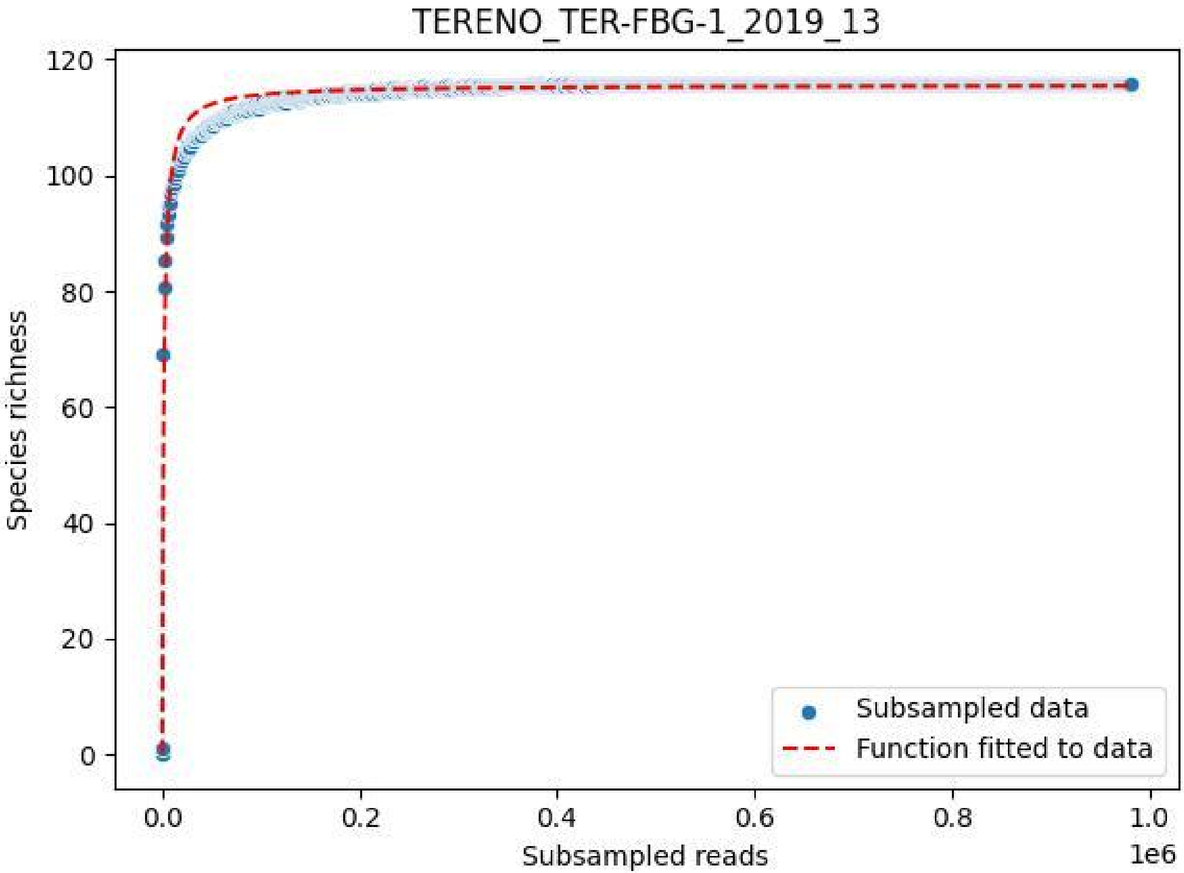
TERENO_TER-FBG-1_2019_13
Species richness
0
20
40
60
80
100
120
0.0
0.2
0.4
0.6
0.8
1.0
1e6
Subsampled reads
Subsampled data
Function fitted to data

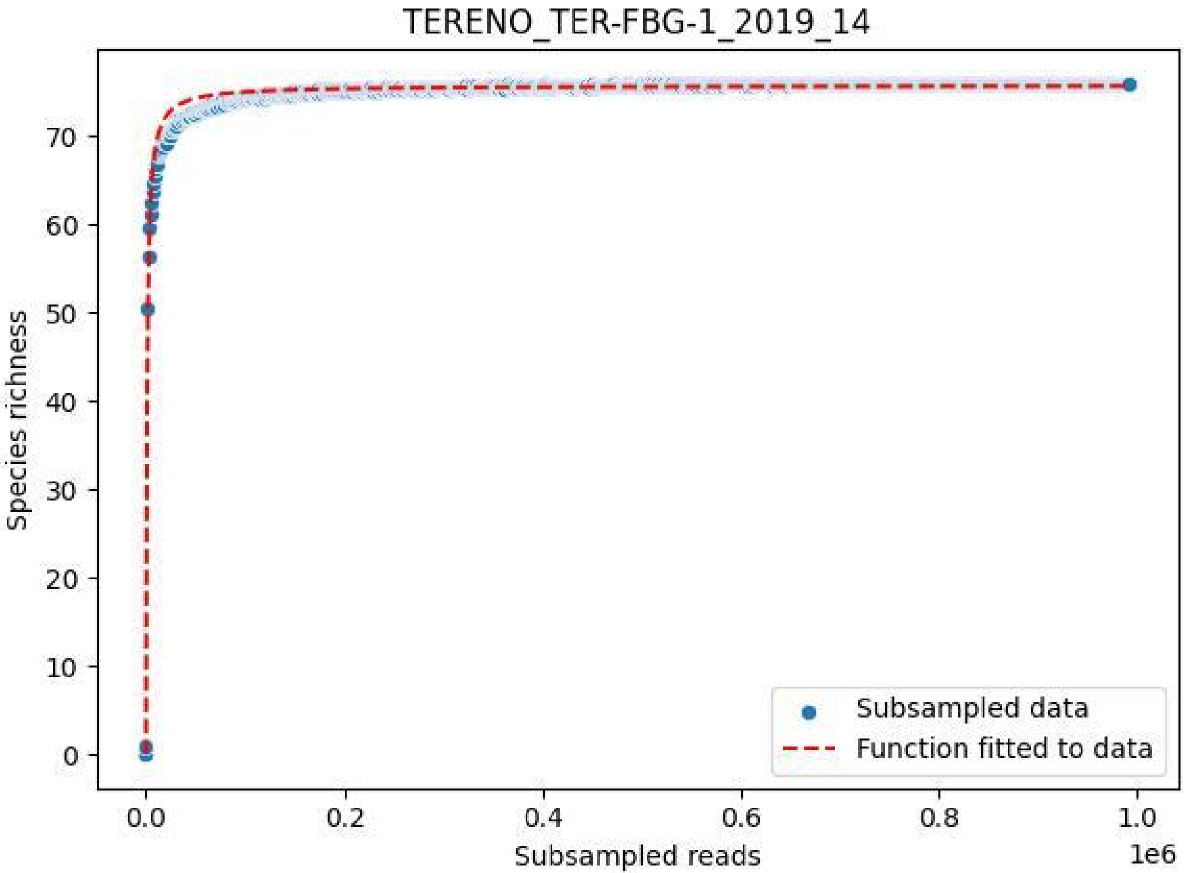

TERENO_TER-FBG-1_2019_14
Species richness
Subsampled reads
1e6
0
10
20
30
40
50
60
70
0.0
0.2
0.4
0.6
0.8
1.0
Subsampled data
Function fitted to data

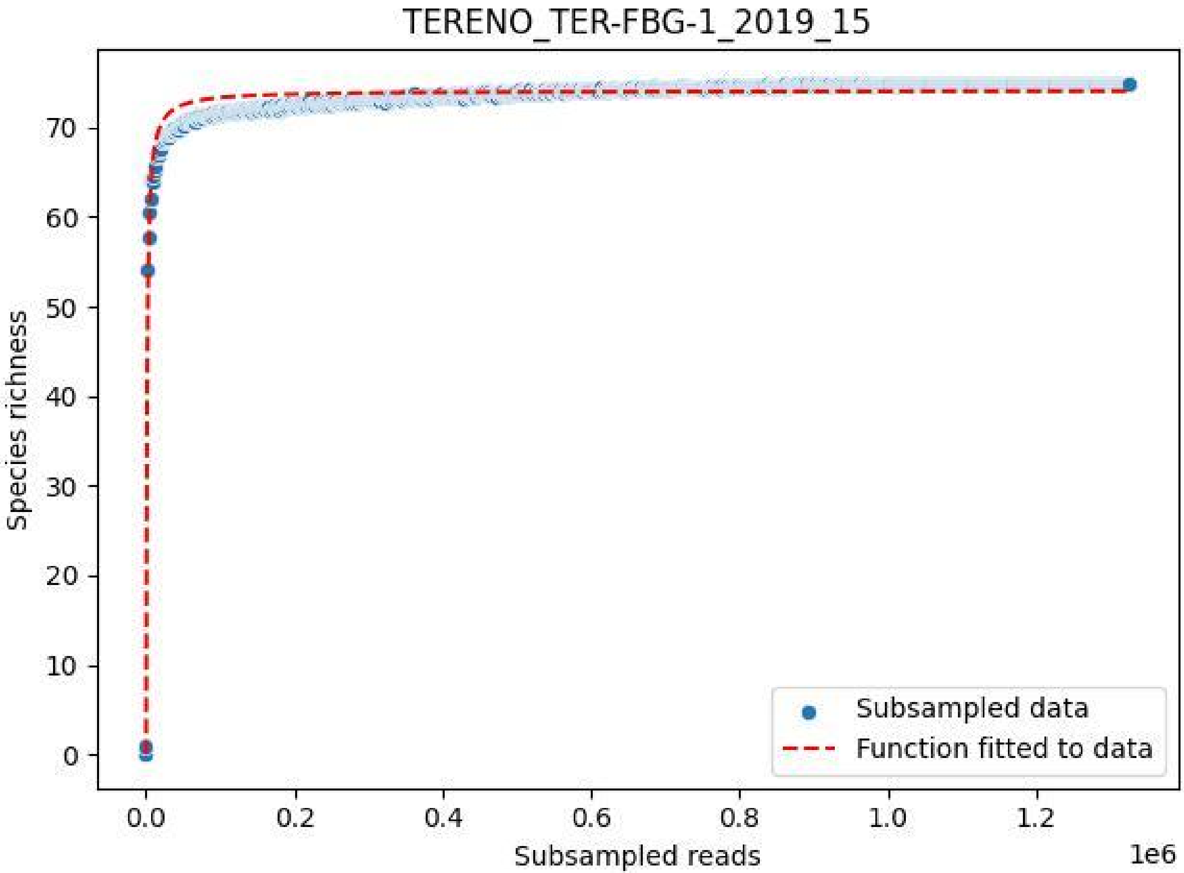

TERENO_TER-FBG-1_2019_15
Species richness
Subsampled reads
1e6
0
10
20
30
40
50
60
70
0.0
0.2
0.4
0.6
0.8
1.0
1.2
Subsampled data
Function fitted to data

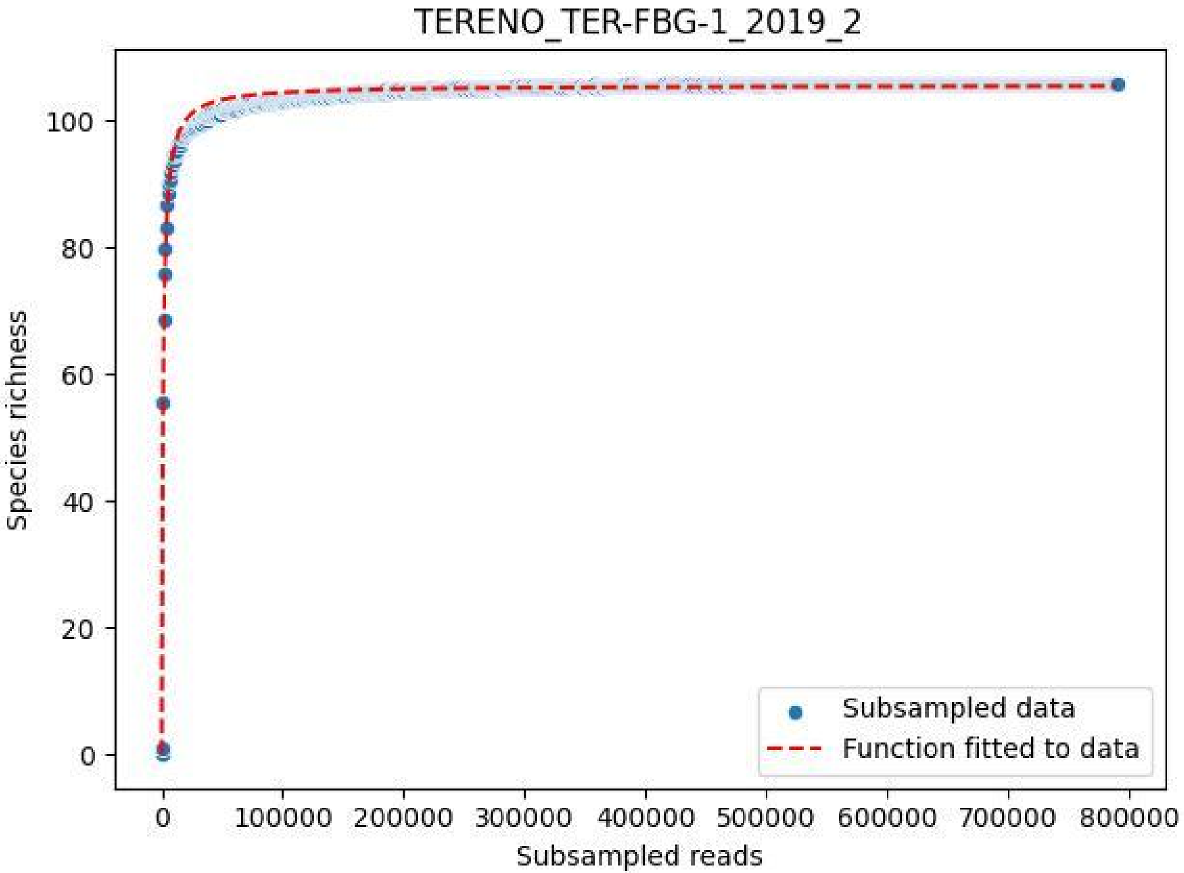

TERENO_TER-FBG-1_2019_2
Species richness
Subsampled reads
0
20
40
60
80
100
0
100000
200000
300000
400000
500000
600000
700000
800000
Subsampled data
Function fitted to data

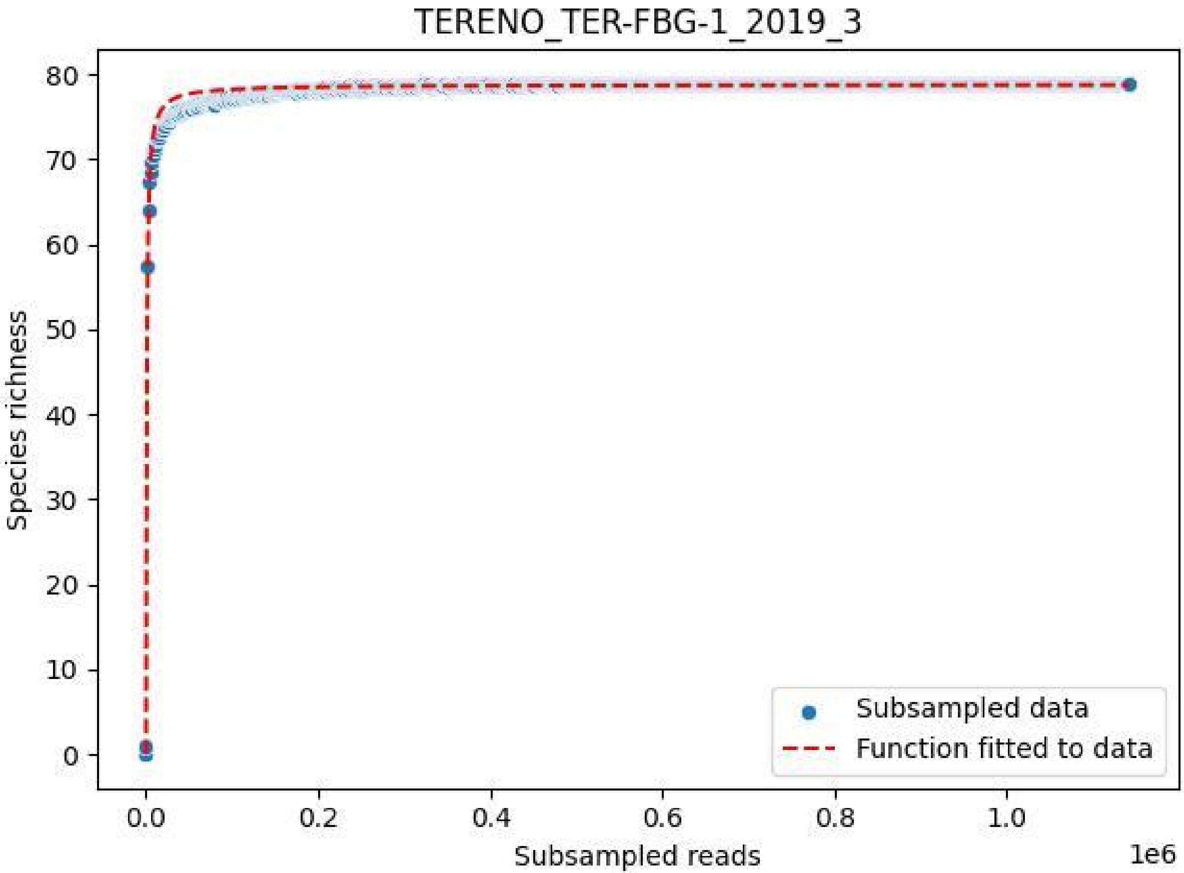

TERENO_TER-FBG-1_2019_3
Species richness
Subsampled reads
1e6
0
10
20
30
40
50
60
70
80
0.0
0.2
0.4
0.6
0.8
1.0
Subsampled data
Function fitted to data

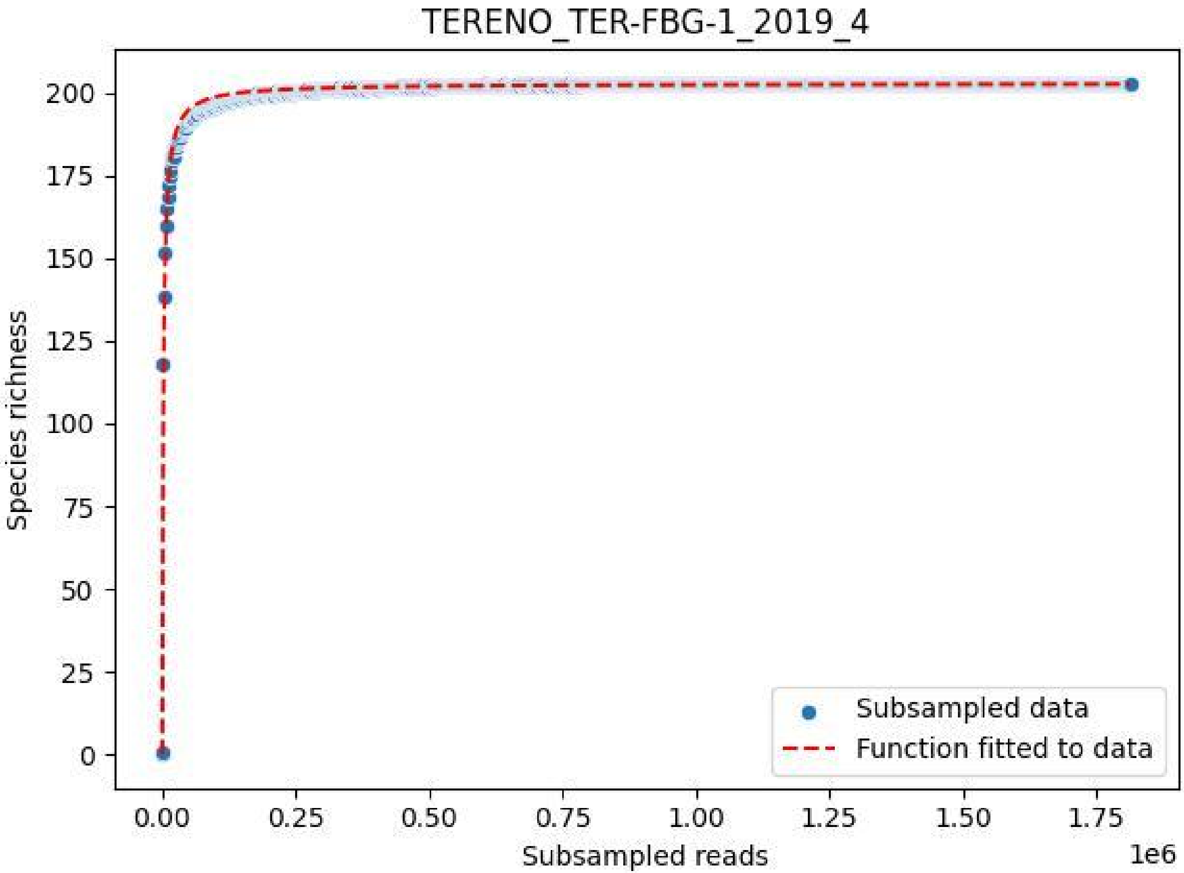
TERENO_TER-FBG-1_2019_4
Species richness
Subsampled reads
1e6
0
25
50
75
100
125
150
175
200
0.00
0.25
0.50
0.75
1.00
1.25
1.50
1.75
Subsampled data
Function fitted to data

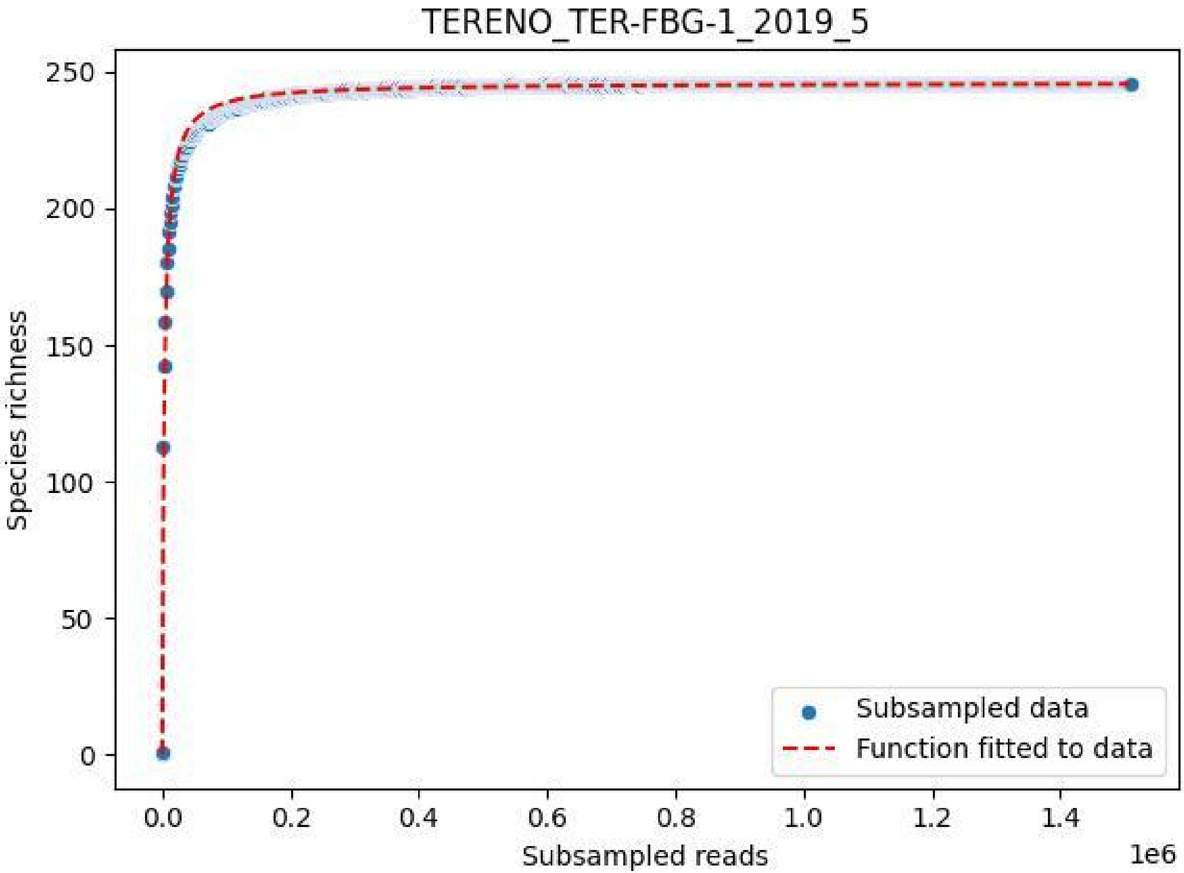
TERENO_TER-FBG-1_2019_5
Species richness
250
200
150
100
50
0
0.0
0.2
0.4
0.6
0.8
1.0
1.2
1.4
1e6
Subsampled reads
Subsampled data
Function fitted to data

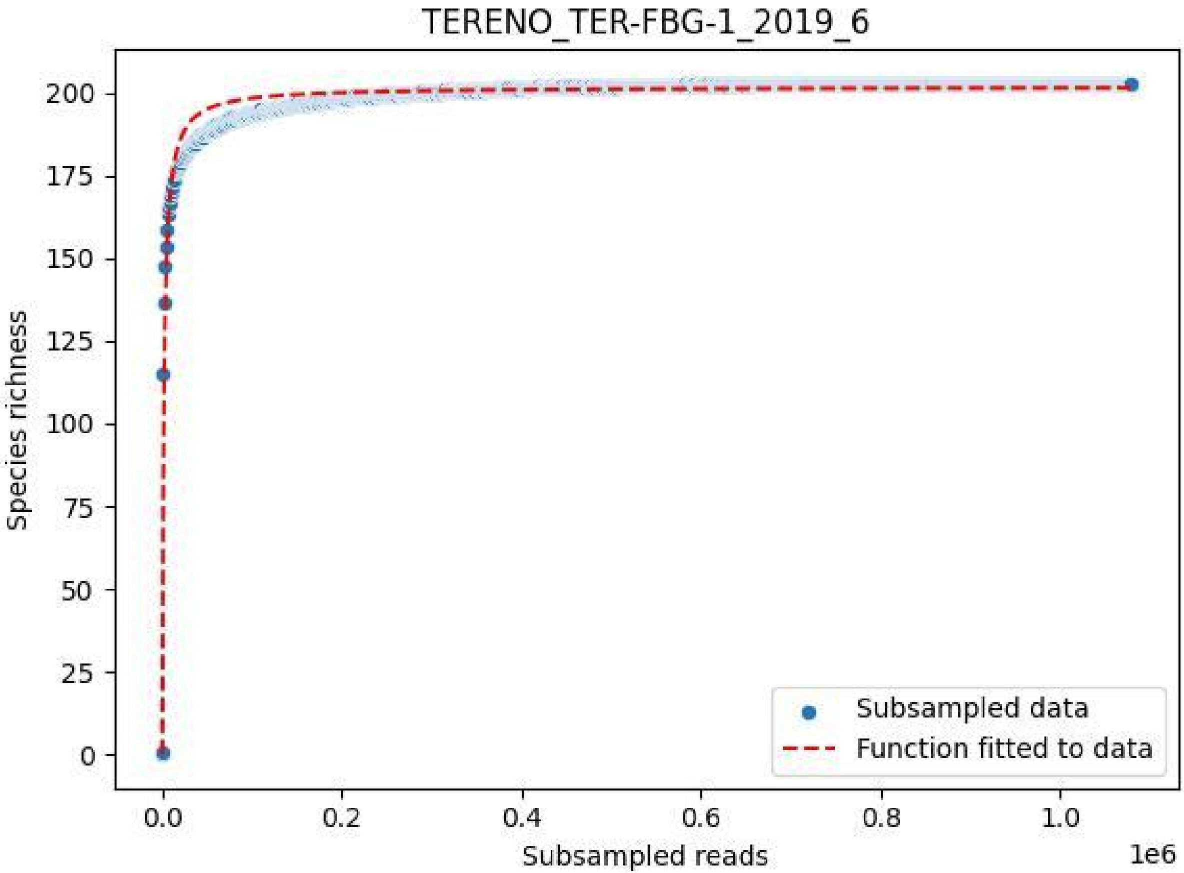
TERENO_TER-FBG-1_2019_6
Species richness
0
25
50
75
100
125
150
175
200
0.0
0.2
0.4
0.6
0.8
1.0
1e6
Subsampled reads
Subsampled data
Function fitted to data

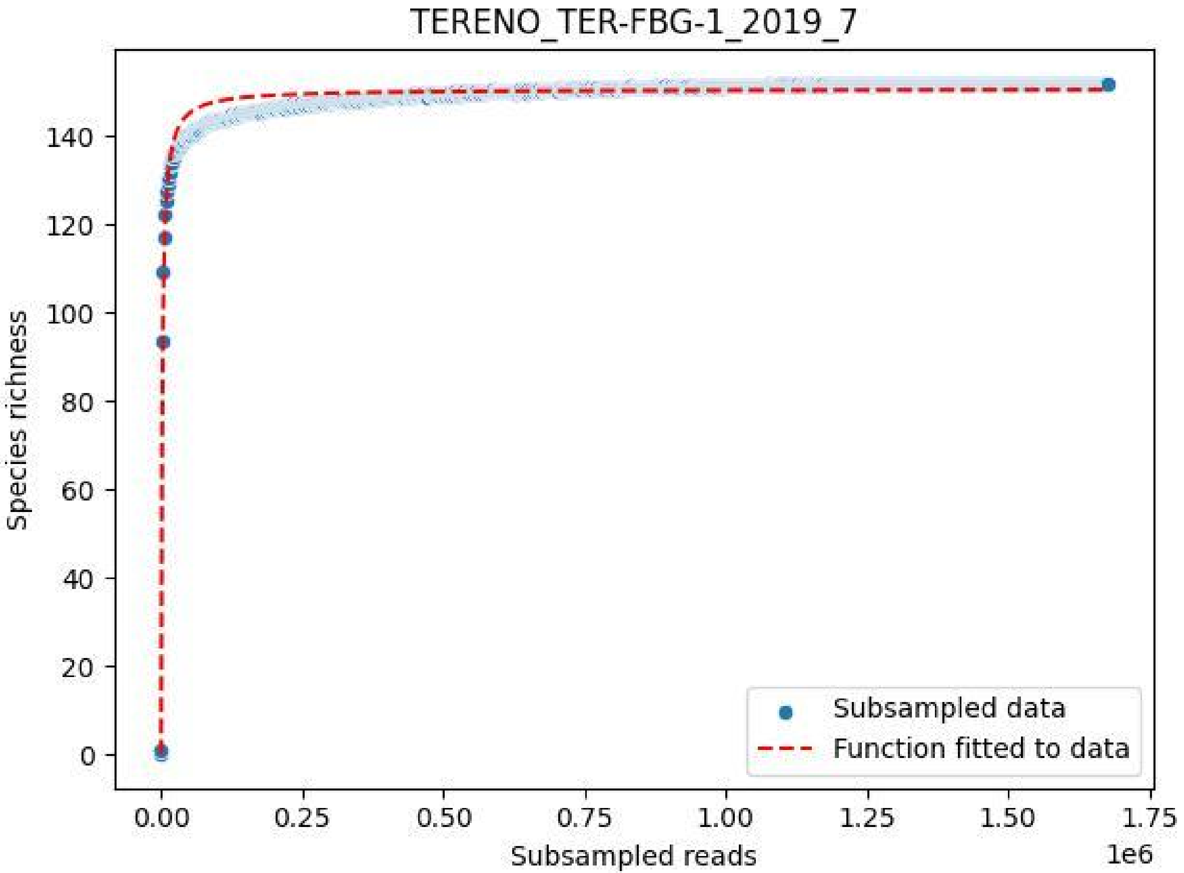
TERENO_TER-FBG-1_2019_7
Species richness
0
20
40
60
80
100
120
140
0.00
0.25
0.50
0.75
1.00
1.25
1.50
1.75
1e6
Subsampled reads
Subsampled data
Function fitted to data

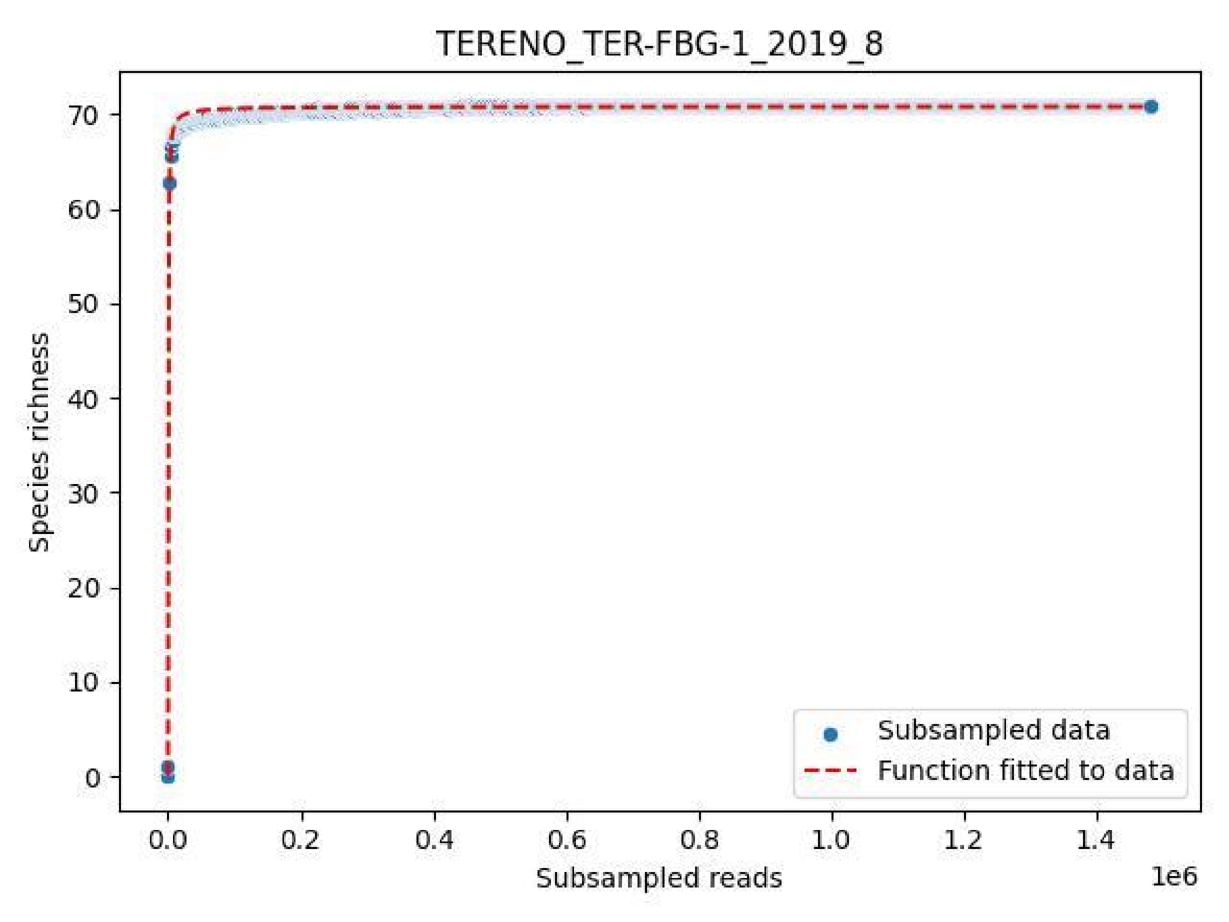

TERENO_TER-FBG-1_2019_8
Species richness
Subsampled reads
1e6
0
10
20
30
40
50
60
70
0.0
0.2
0.4
0.6
0.8
1.0
1.2
1.4
Subsampled data
Function fitted to data

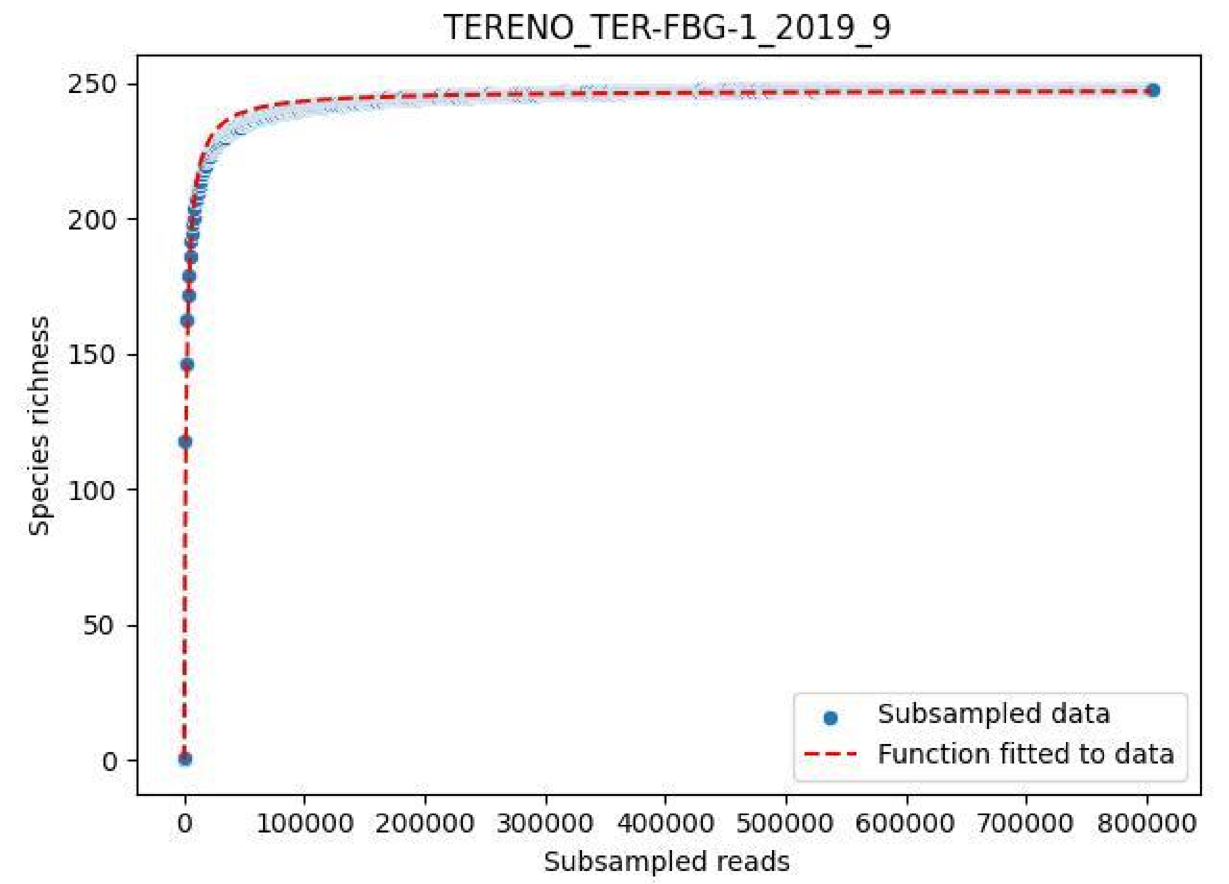
TERENO_TER-FBG-1_2019_9
Species richness
Subsampled reads
0
50
100
150
200
250
0
100000
200000
300000
400000
500000
600000
700000
800000
Subsampled data
Function fitted to data

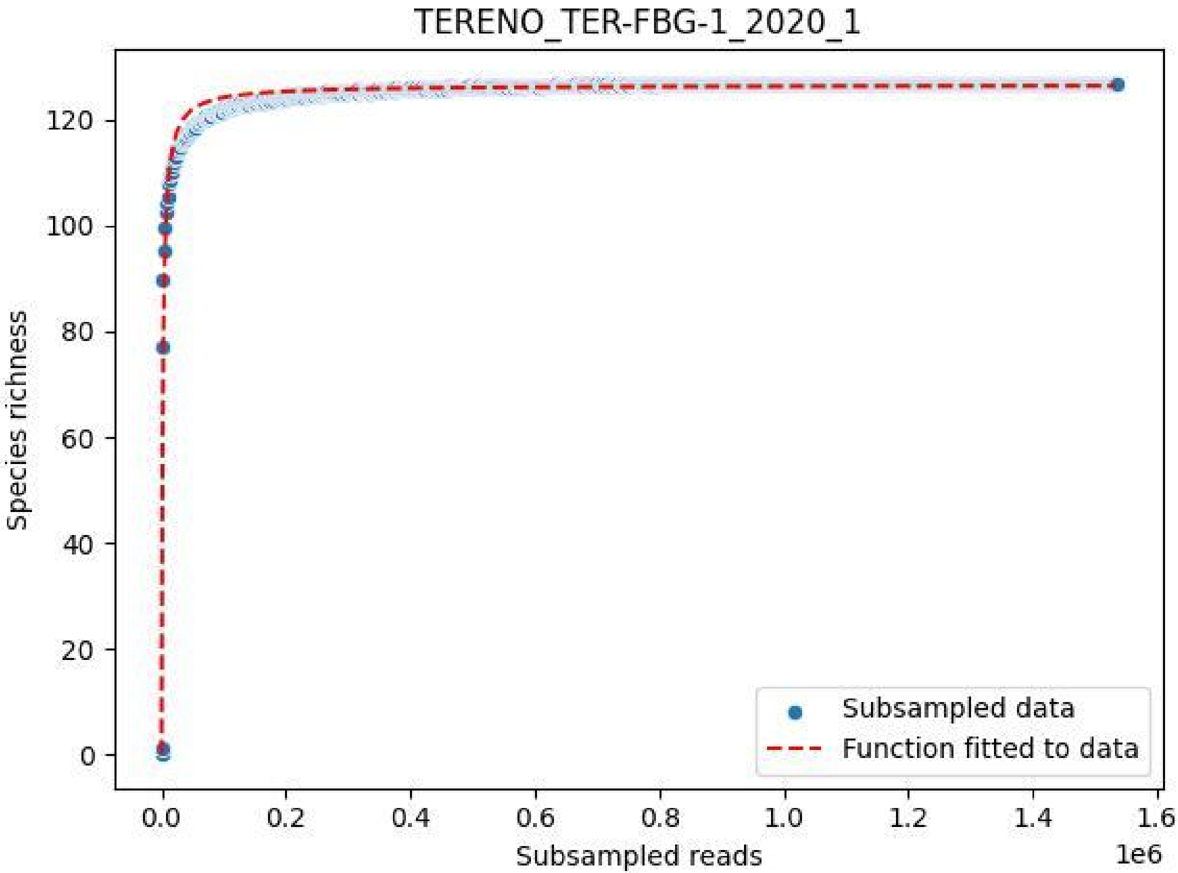

TERENO_TER-FBG-1_2020_1
Species richness
Subsampled reads
1e6
0
20
40
60
80
100
120
0.0
0.2
0.4
0.6
0.8
1.0
1.2
1.4
1.6
Subsampled data
Function fitted to data

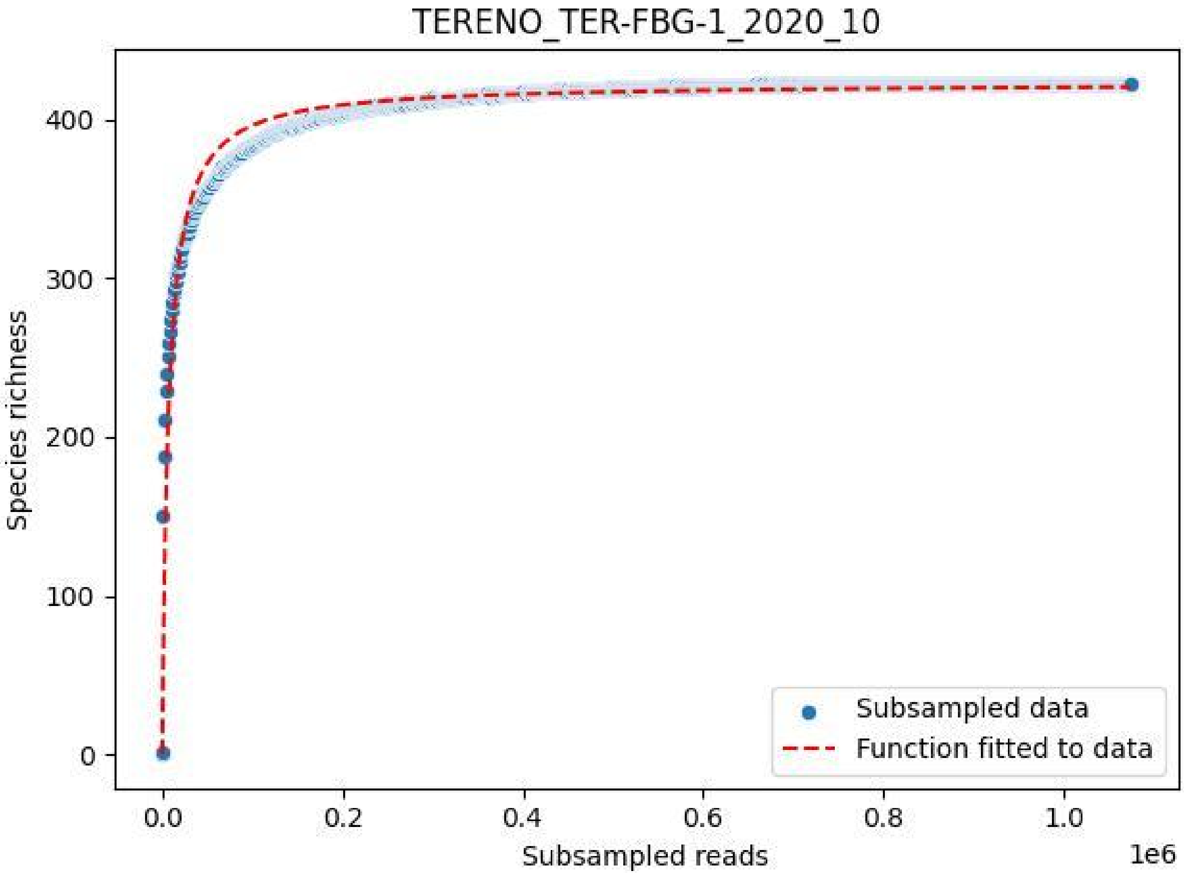

TERENO_TER-FBG-1_2020_10
Species richness
400
300
200
100
0
0.0
0.2
0.4
0.6
0.8
1.0
1e6
Subsampled reads
Subsampled data
Function fitted to data

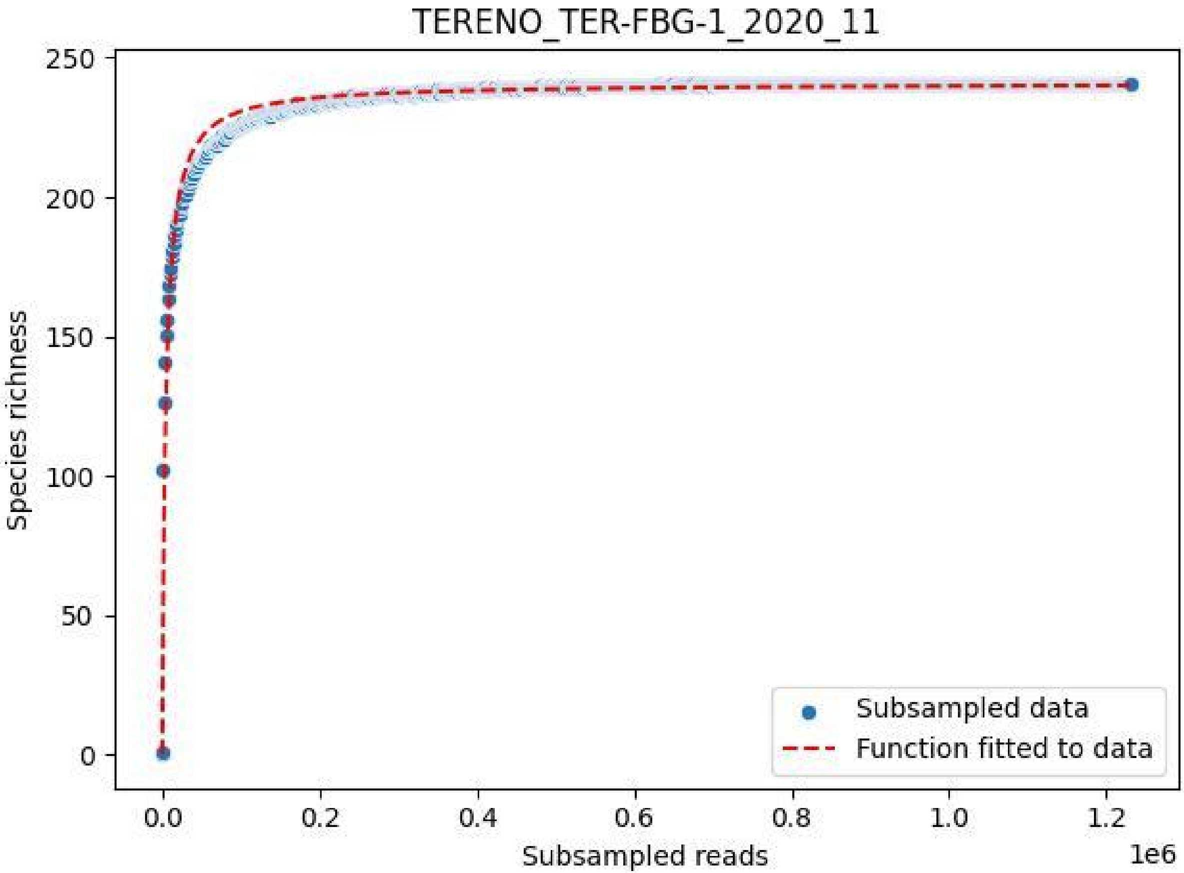

TERENO_TER-FBG-1_2020_11
250
200
150
100
50
0
Species richness
0.0
0.2
0.4
0.6
0.8
1.0
1.2
1e6
Subsampled reads
Subsampled data
Function fitted to data

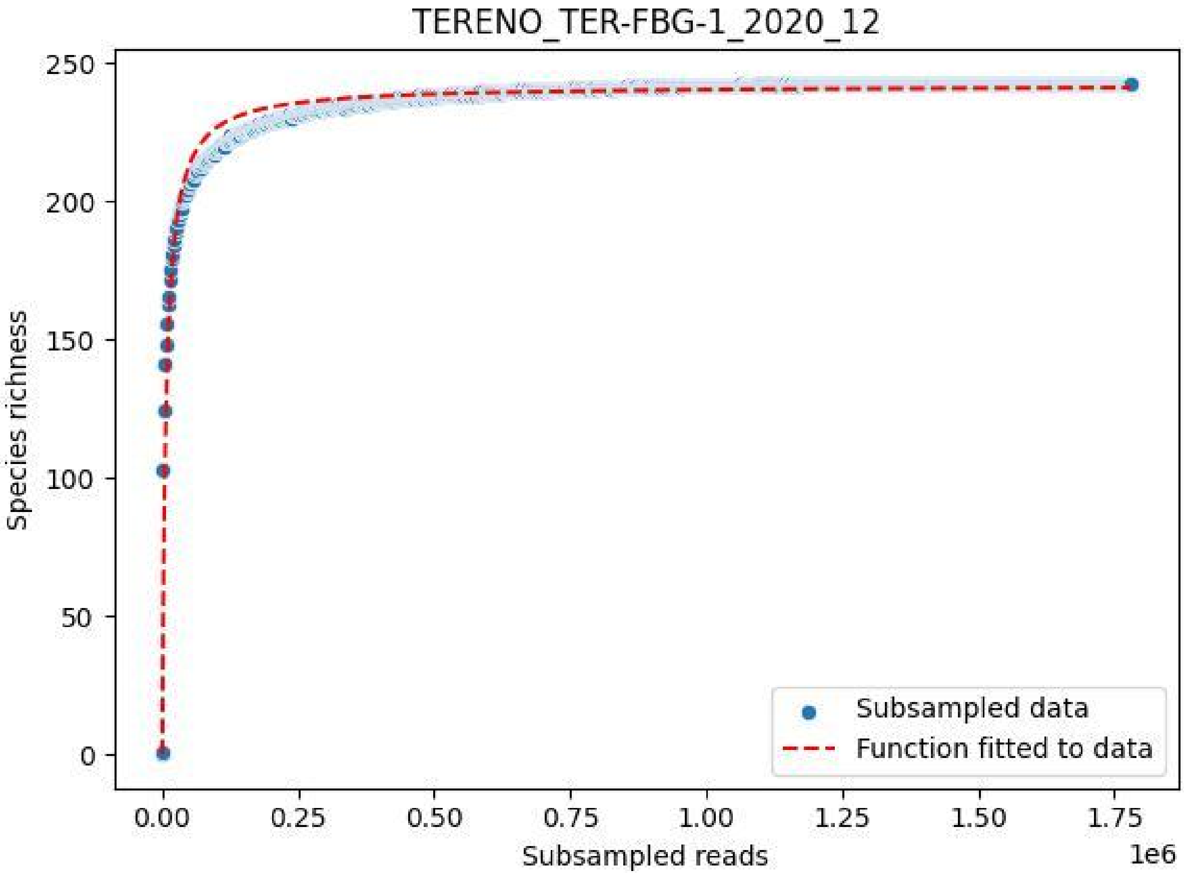
TERENO_TER-FBG-1_2020_12
Species richness
Subsampled reads
1e6
0
50
100
150
200
250
0.00
0.25
0.50
0.75
1.00
1.25
1.50
1.75
Subsampled data
Function fitted to data

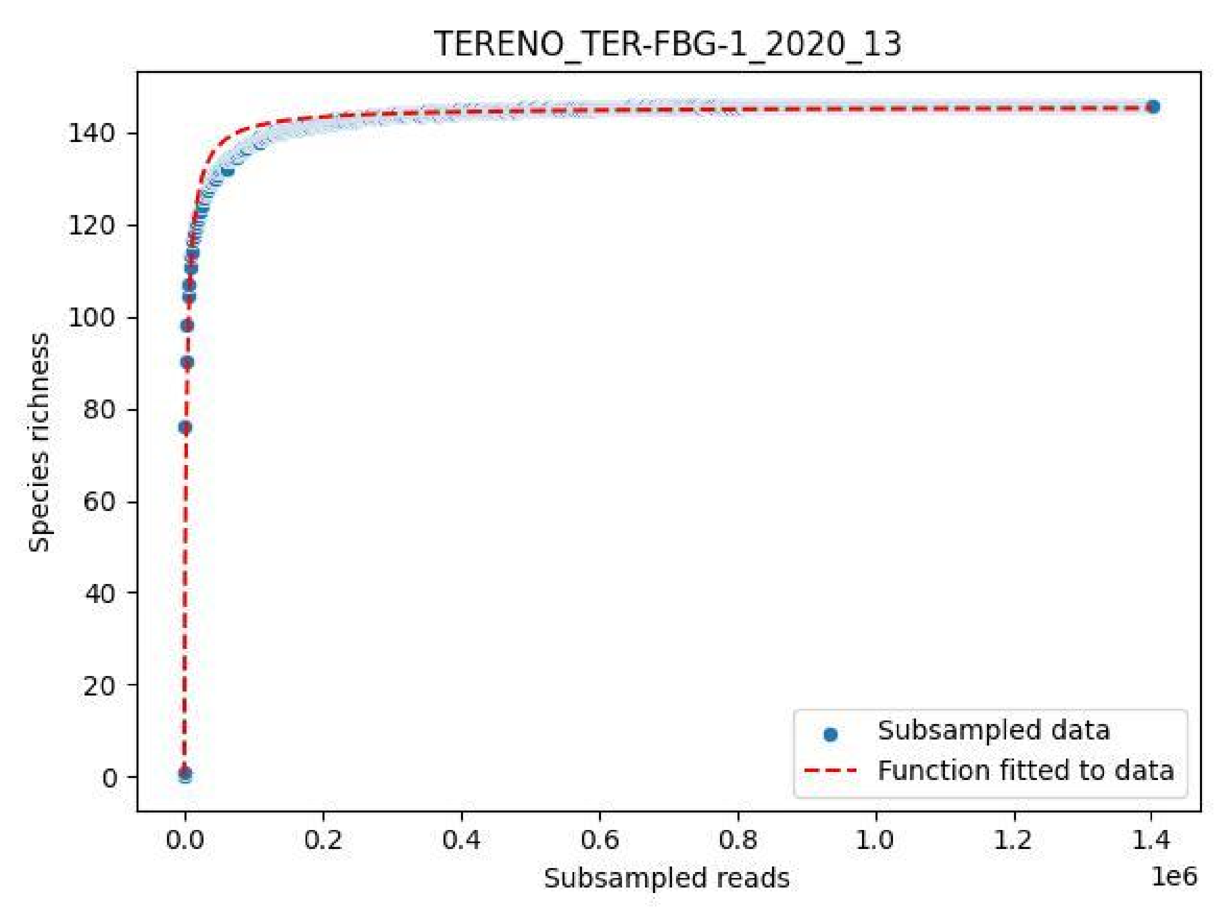
TERENO_TER-FBG-1_2020_13
Species richness
Subsampled reads
1e6
0
20
40
60
80
100
120
140
0.0
0.2
0.4
0.6
0.8
1.0
1.2
1.4
Subsampled data
Function fitted to data

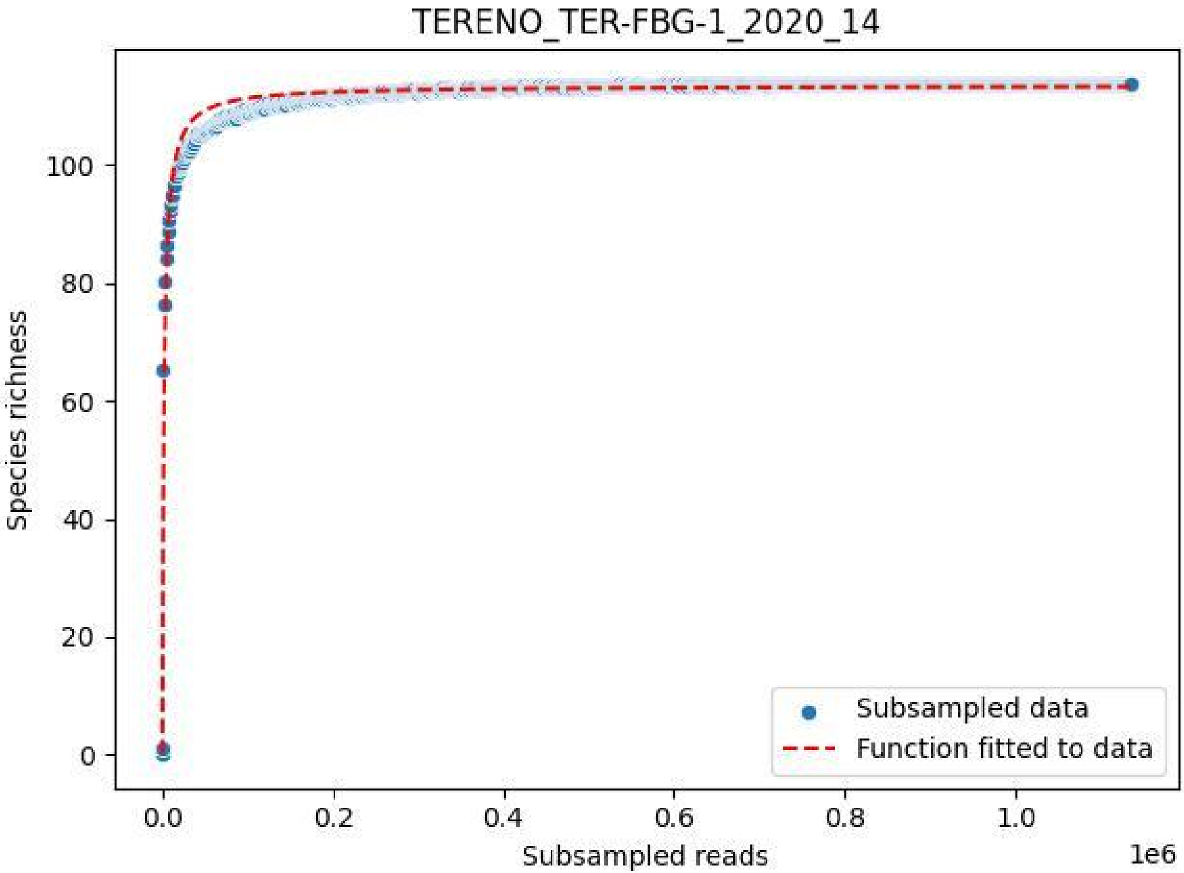

TERENO_TER-FBG-1_2020_14
Species richness
0
20
40
60
80
100
0.0
0.2
0.4
0.6
0.8
1.0
1e6
Subsampled reads
Subsampled data
Function fitted to data

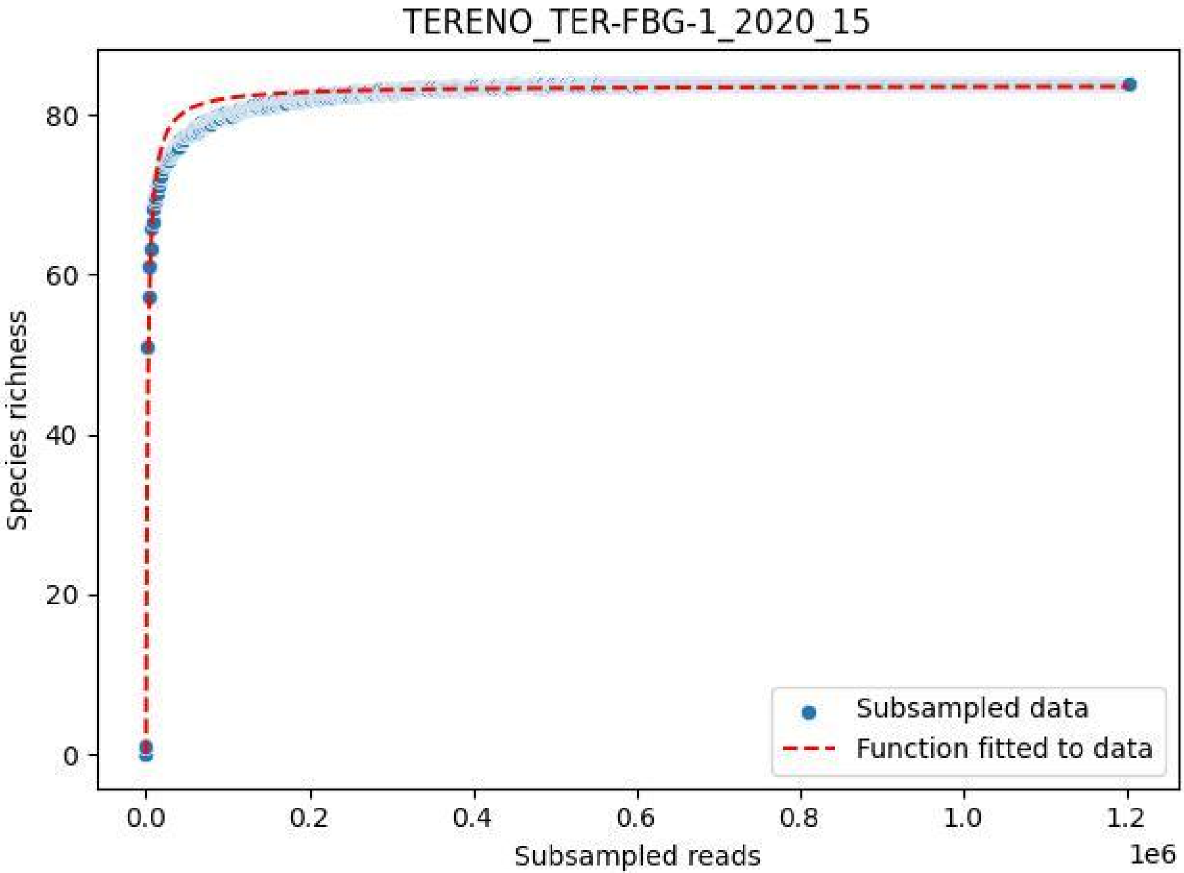

TERENO_TER-FBG-1_2020_15
Species richness
Subsampled reads
1e6
0
20
40
60
80
0.0
0.2
0.4
0.6
0.8
1.0
1.2
Subsampled data
Function fitted to data

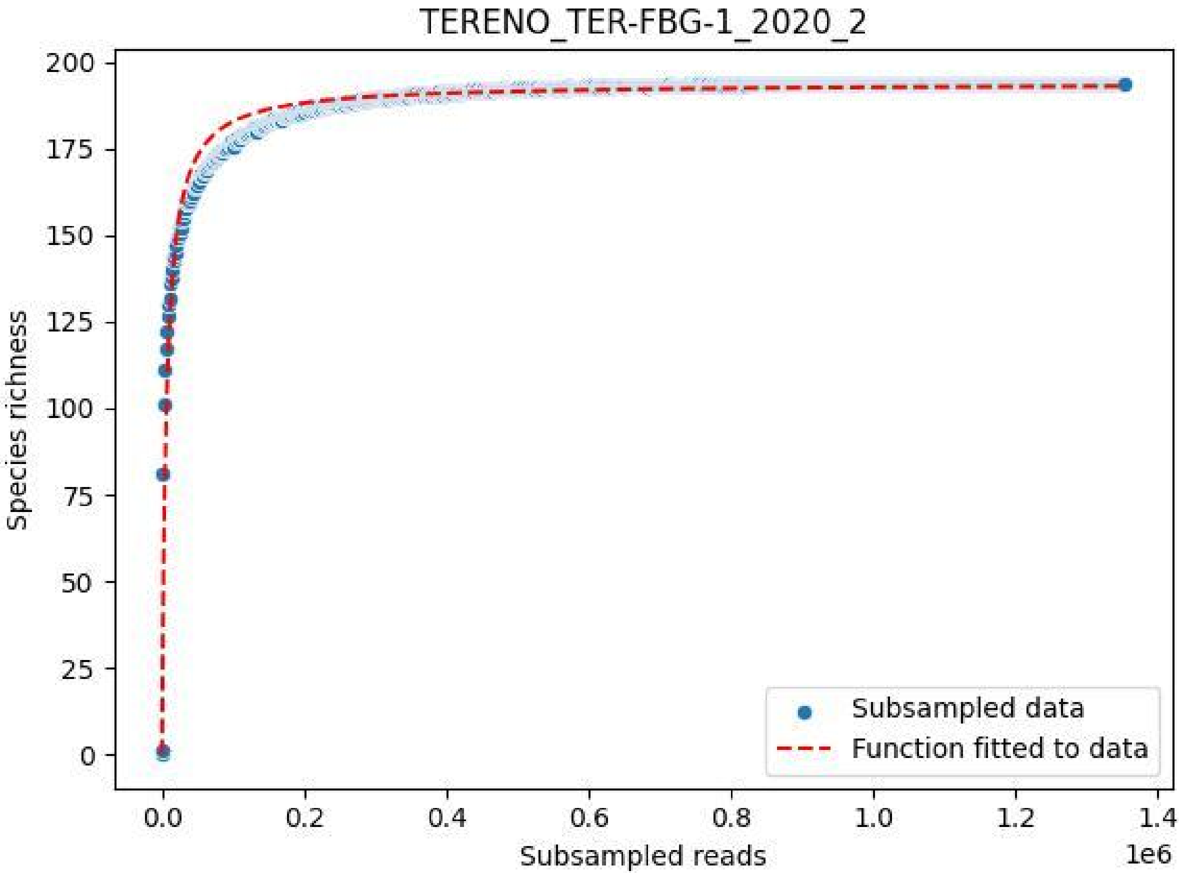
TERENO_TER-FBG-1_2020_2
Species richness
200
175
150
125
100
75
50
25
0
0.0
0.2
0.4
0.6
0.8
1.0
1.2
1.4
1e6
Subsampled reads
Subsampled data
Function fitted to data

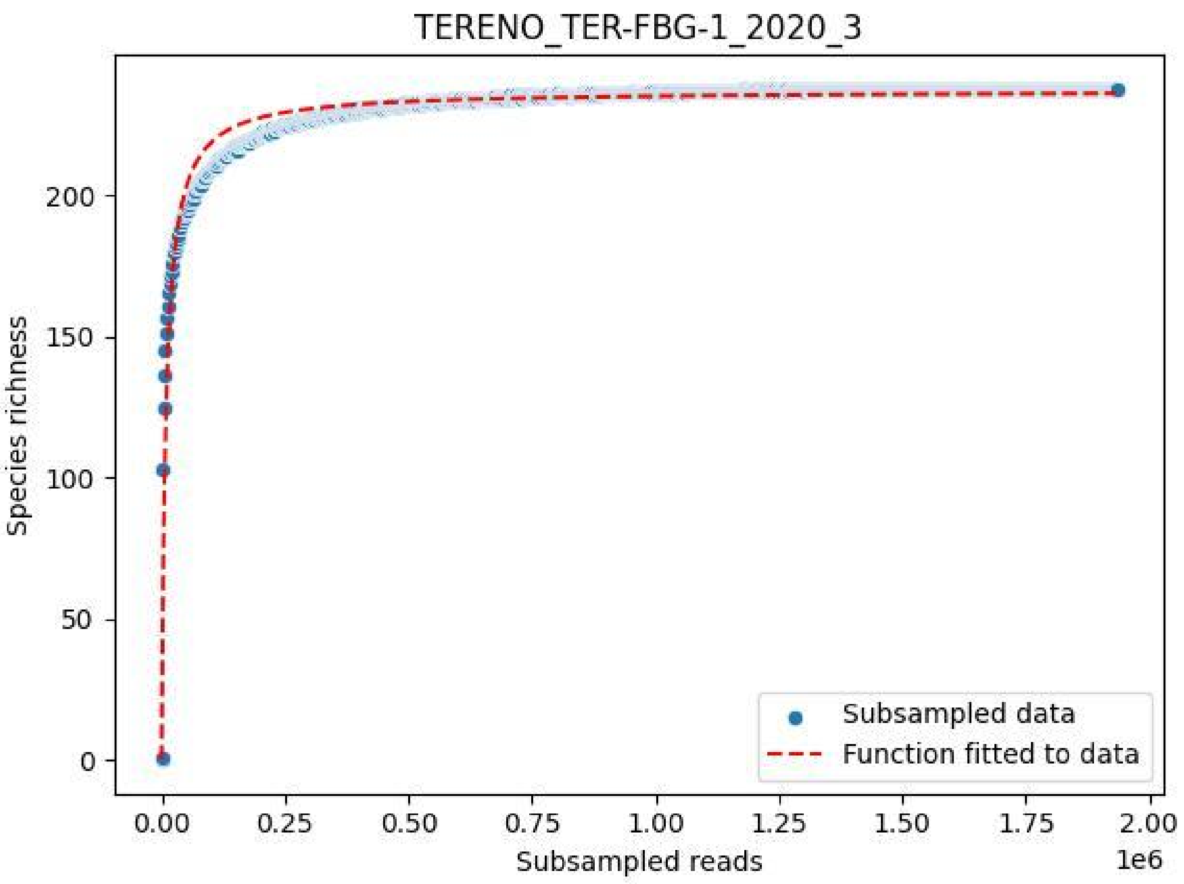

TERENO_TER-FBG-1_2020_3
Species richness
0
50
100
150
200
Subsampled reads
0.00
0.25
0.50
0.75
1.00
1.25
1.50
1.75
2.00
1e6
Subsampled data
Function fitted to data

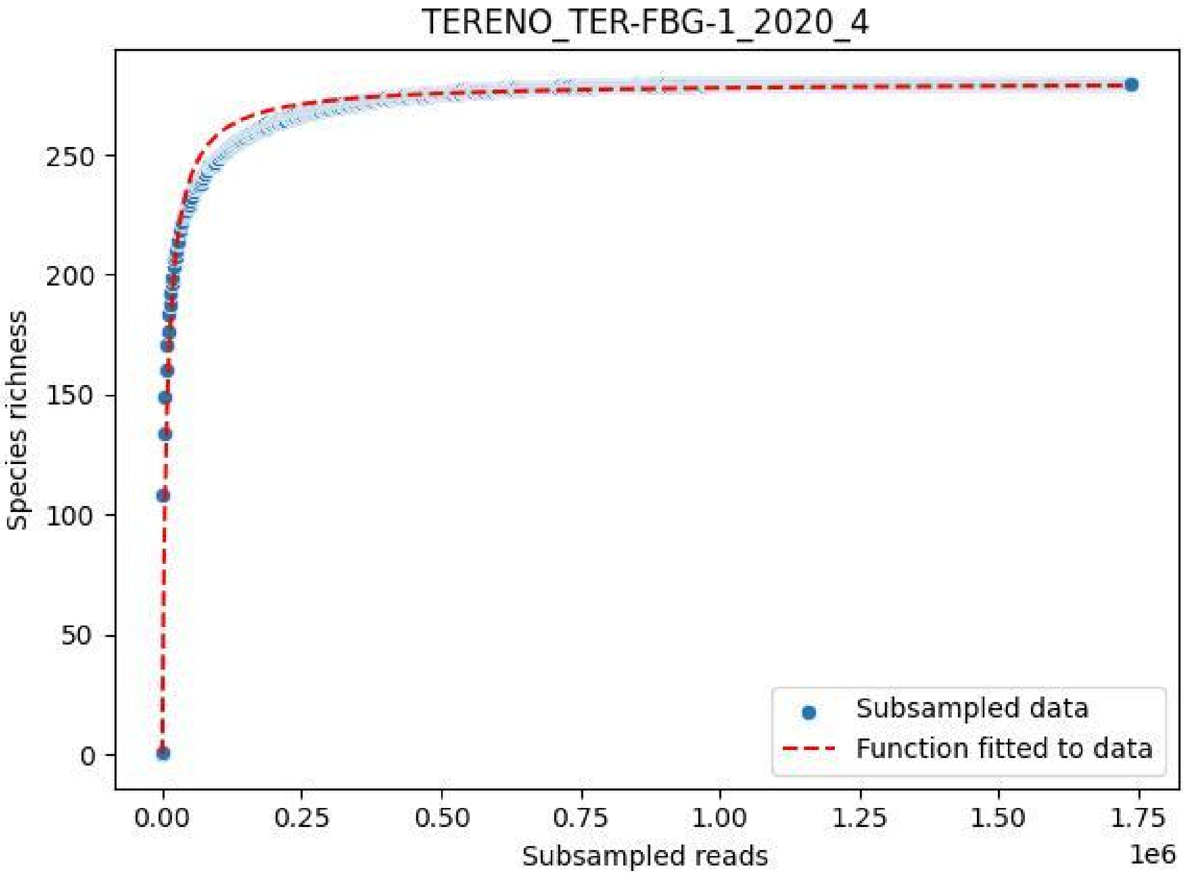

TERENO_TER-FBG-1_2020_4
Species richness
Subsampled reads
1e6
0
50
100
150
200
250
0.00
0.25
0.50
0.75
1.00
1.25
1.50
1.75
Subsampled data
Function fitted to data

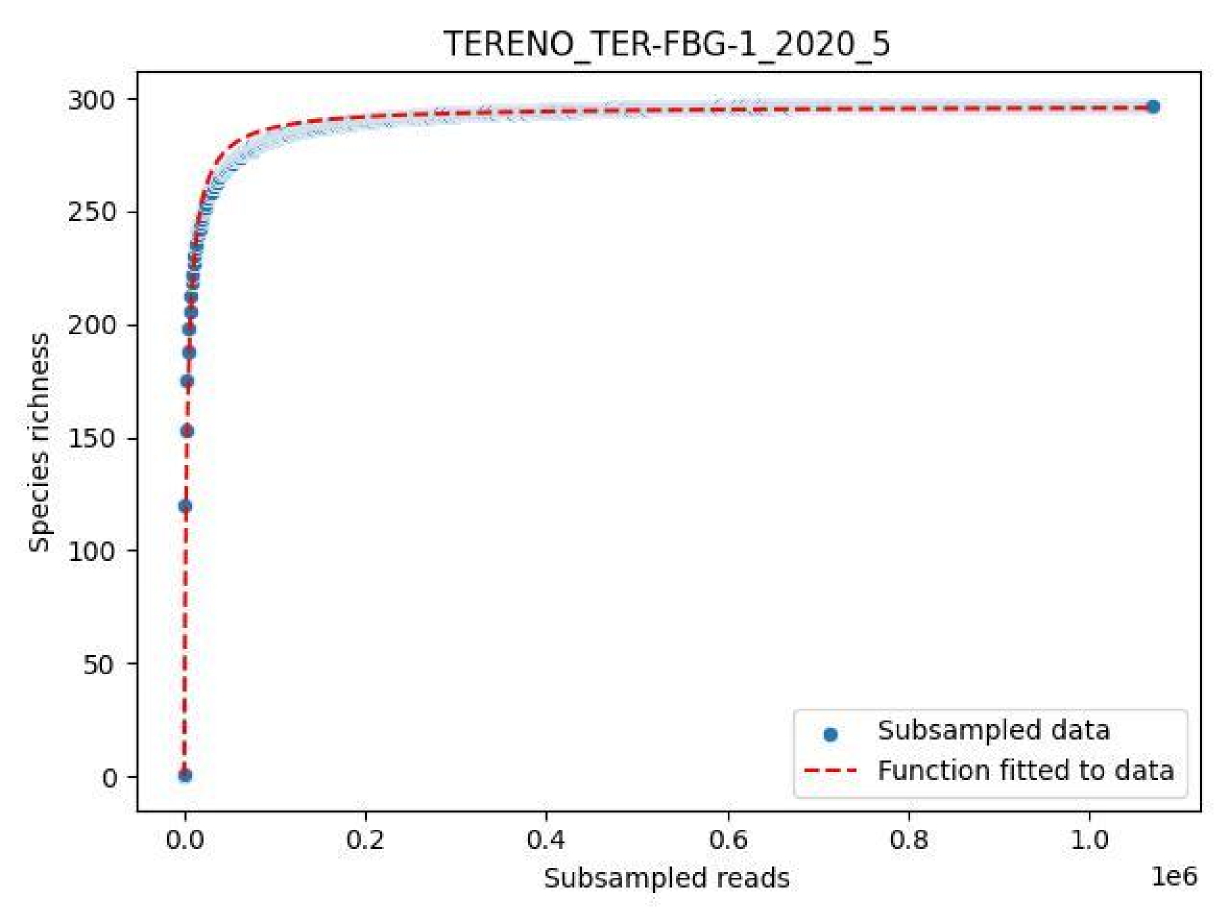

TERENO_TER-FBG-1_2020_5
Species richness
Subsampled reads
1e6
0
50
100
150
200
250
300
0.0
0.2
0.4
0.6
0.8
1.0
Subsampled data
Function fitted to data

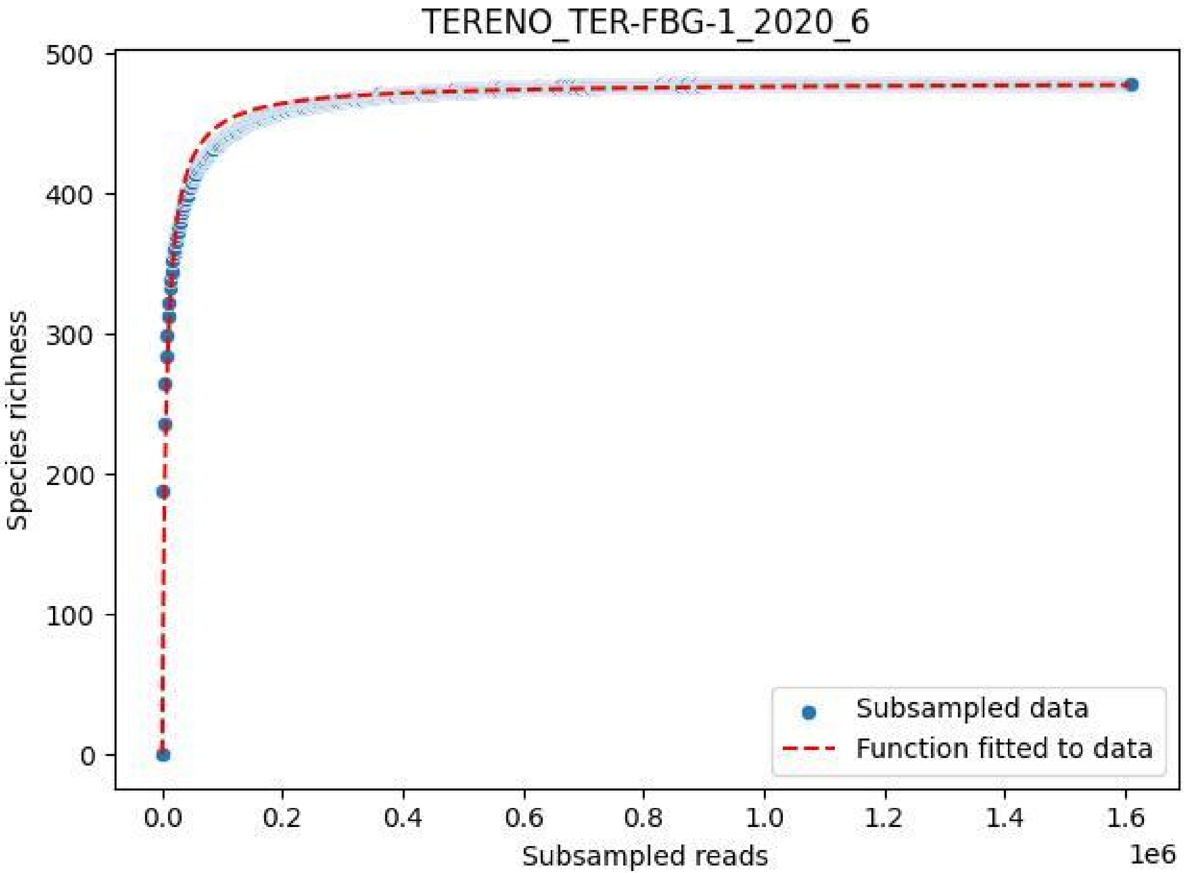

TERENO_TER-FBG-1_2020_6
Species richness
Subsampled reads
1e6
0
100
200
300
400
500
0.0
0.2
0.4
0.6
0.8
1.0
1.2
1.4
1.6
Subsampled data
Function fitted to data

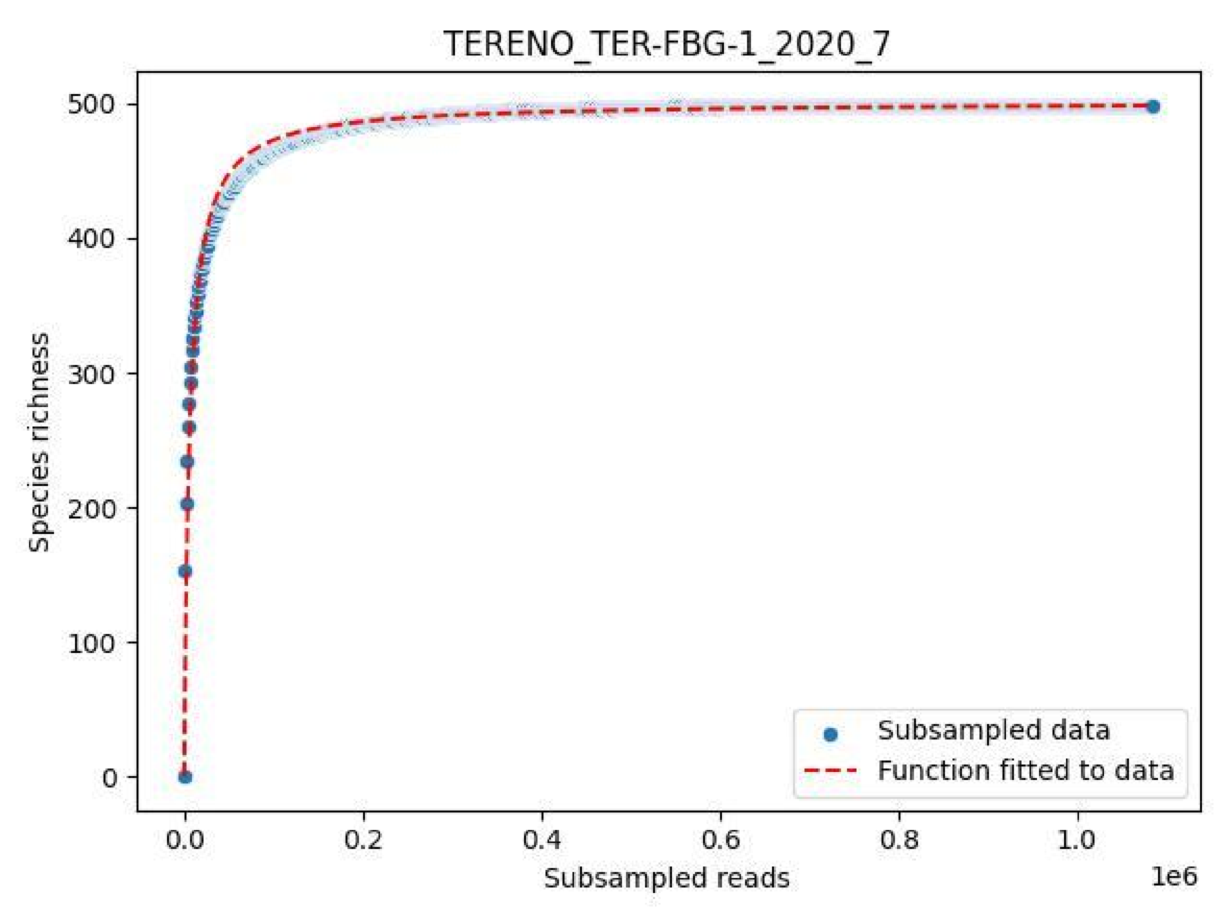

TERENO_TER-FBG-1_2020_7
Species richness
Subsampled reads
1e6
0
100
200
300
400
500
0.0
0.2
0.4
0.6
0.8
1.0
Subsampled data
Function fitted to data

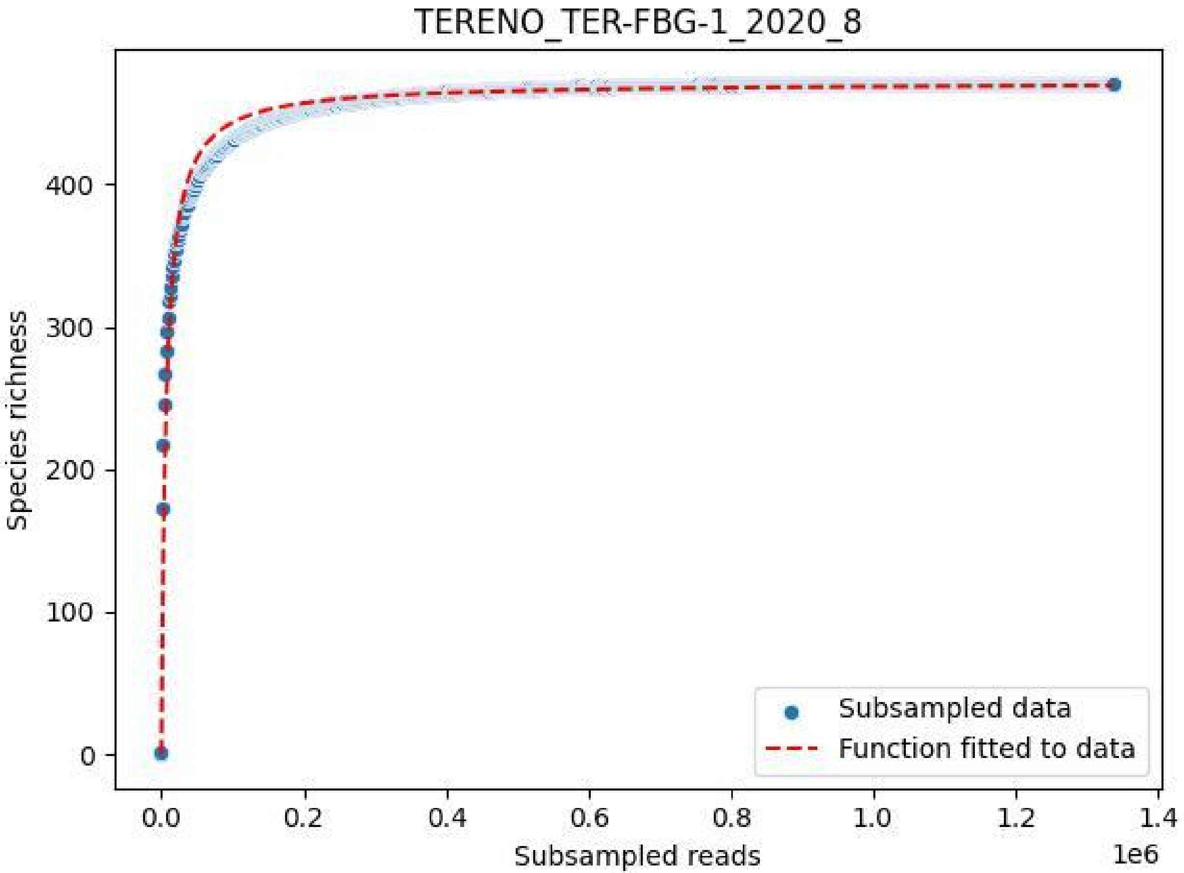

TERENO_TER-FBG-1_2020_8
Species richness
Subsampled reads
1e6
0
100
200
300
400
0.0
0.2
0.4
0.6
0.8
1.0
1.2
1.4
Subsampled data
Function fitted to data

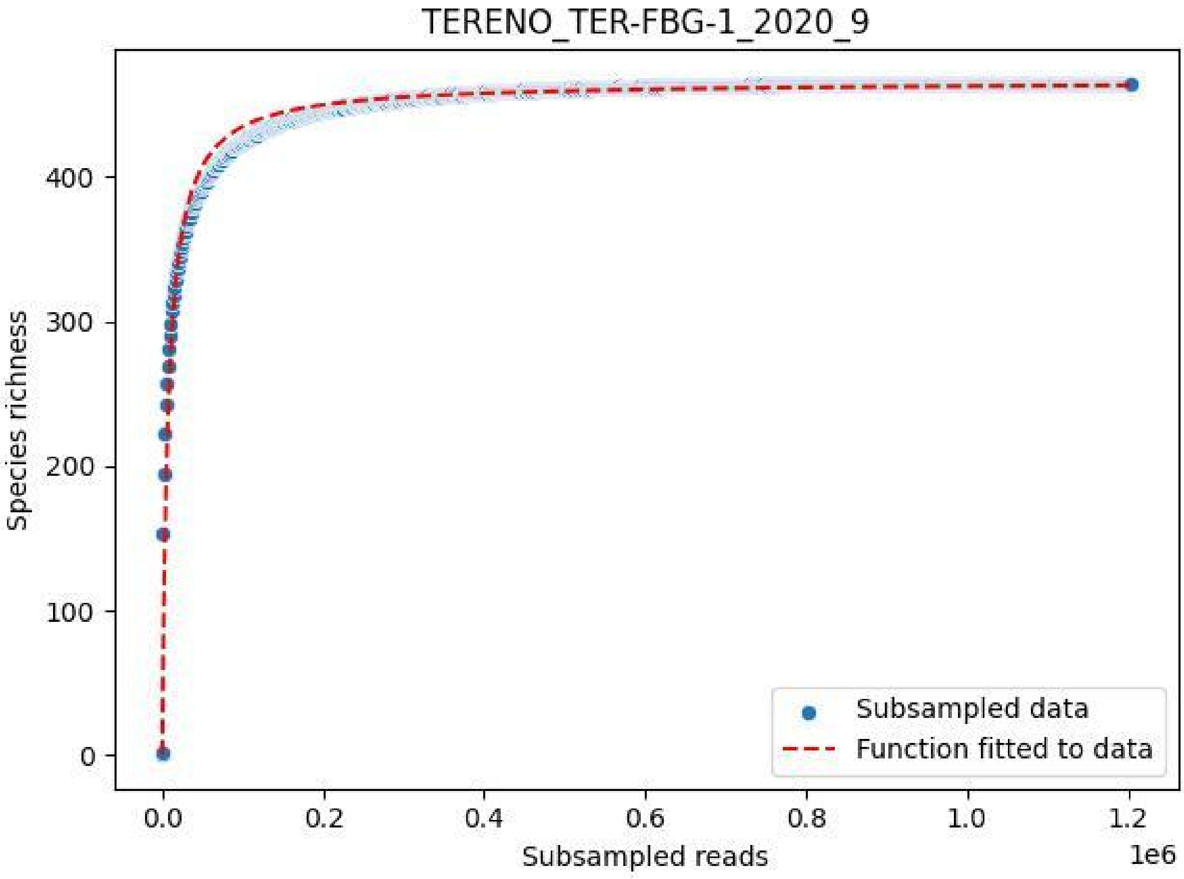
TERENO_TER-FBG-1_2020_9
Species richness
Subsampled reads
1e6
0
100
200
300
400
0.0
0.2
0.4
0.6
0.8
1.0
1.2
Subsampled data
Function fitted to data

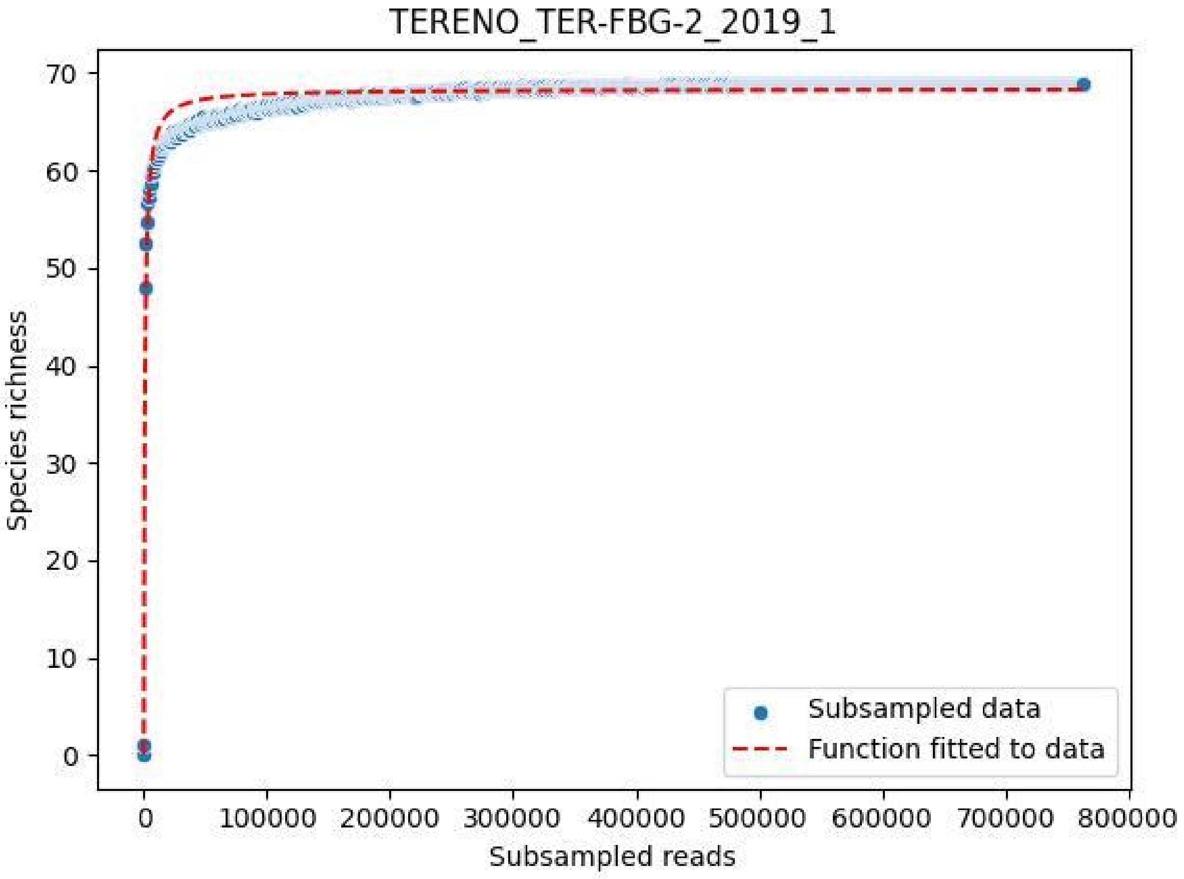
TERENO_TER-FBG-2_2019_1
Species richness
Subsampled reads
0
10
20
30
40
50
60
70
0
100000
200000
300000
400000
500000
600000
700000
800000
Subsampled data
Function fitted to data

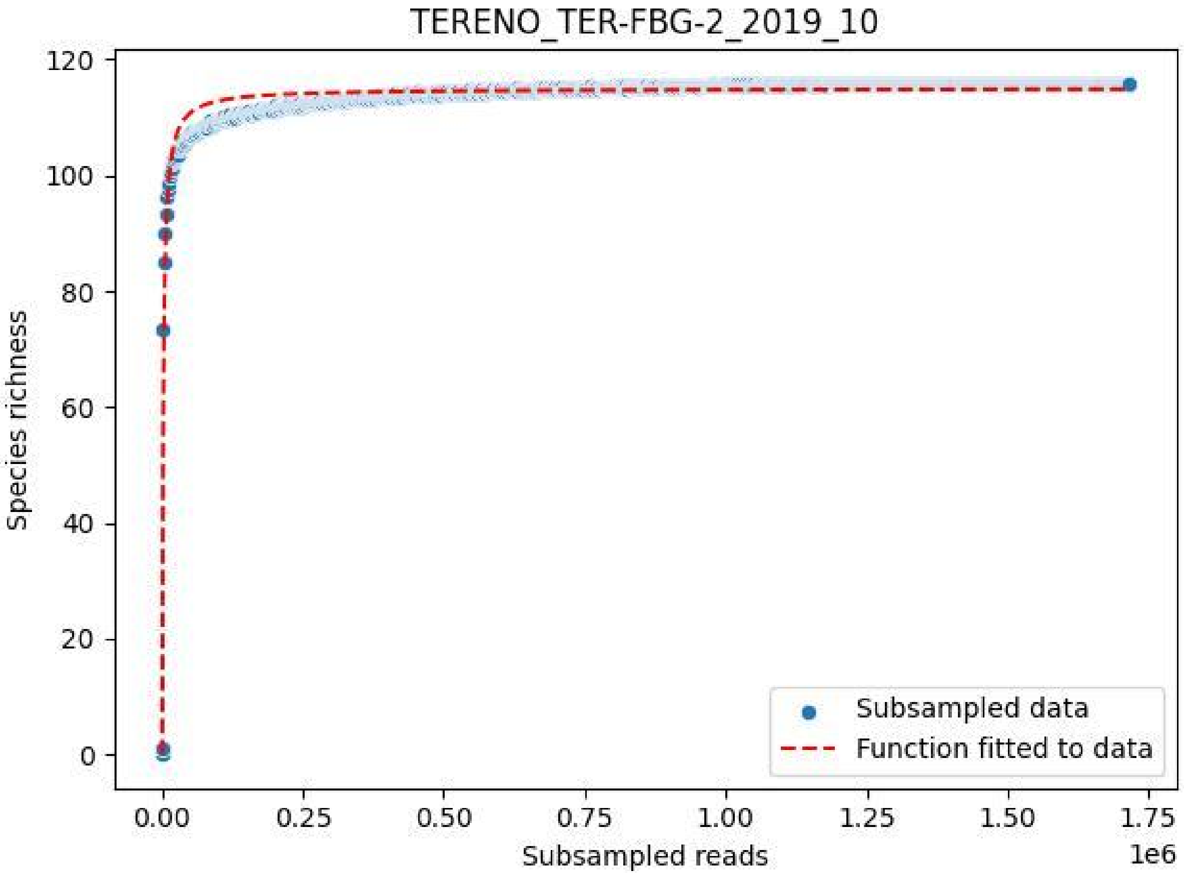
TERENO_TER-FBG-2_2019_10
Species richness
Subsampled reads
1e6
0
20
40
60
80
100
120
0.00
0.25
0.50
0.75
1.00
1.25
1.50
1.75
Subsampled data
Function fitted to data

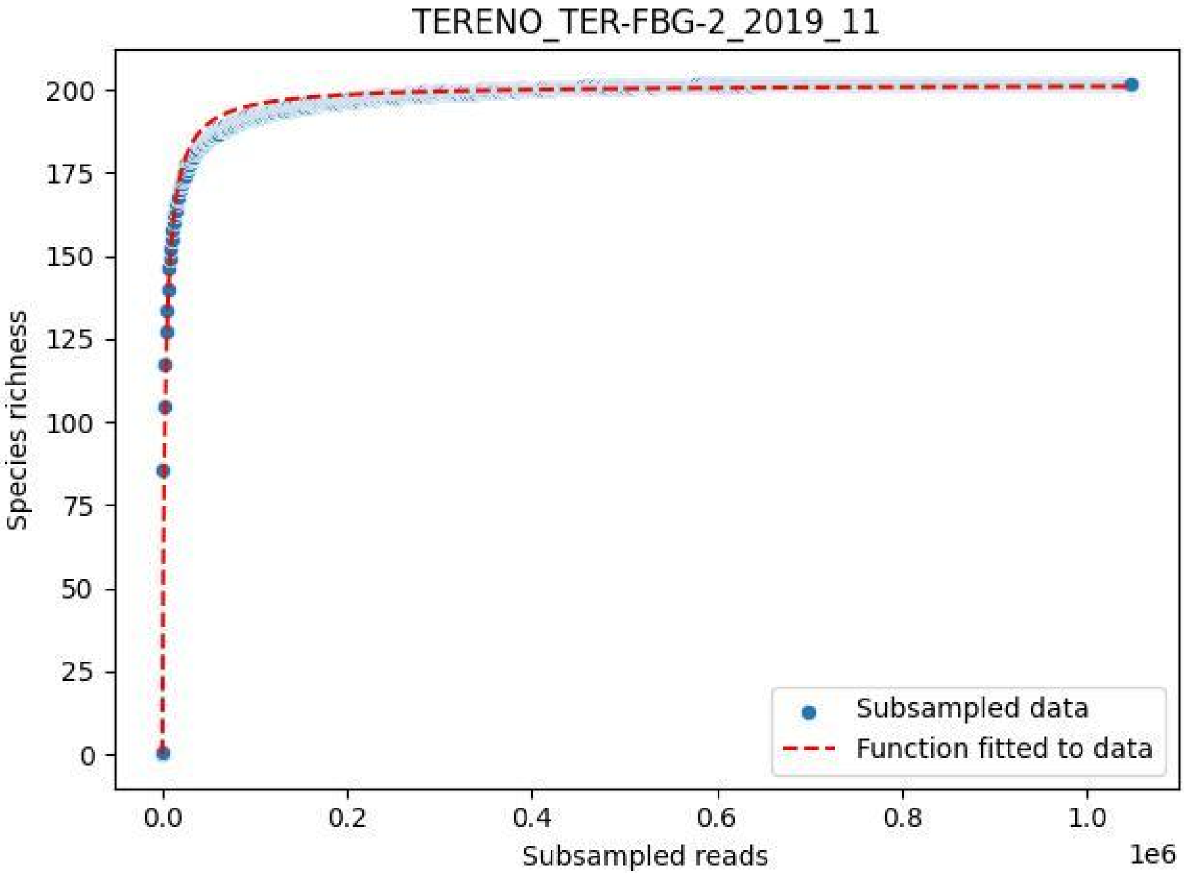
TERENO_TER-FBG-2_2019_11
Species richness
Subsampled reads
1e6
0
25
50
75
100
125
150
175
200
0.0
0.2
0.4
0.6
0.8
1.0
Subsampled data
Function fitted to data

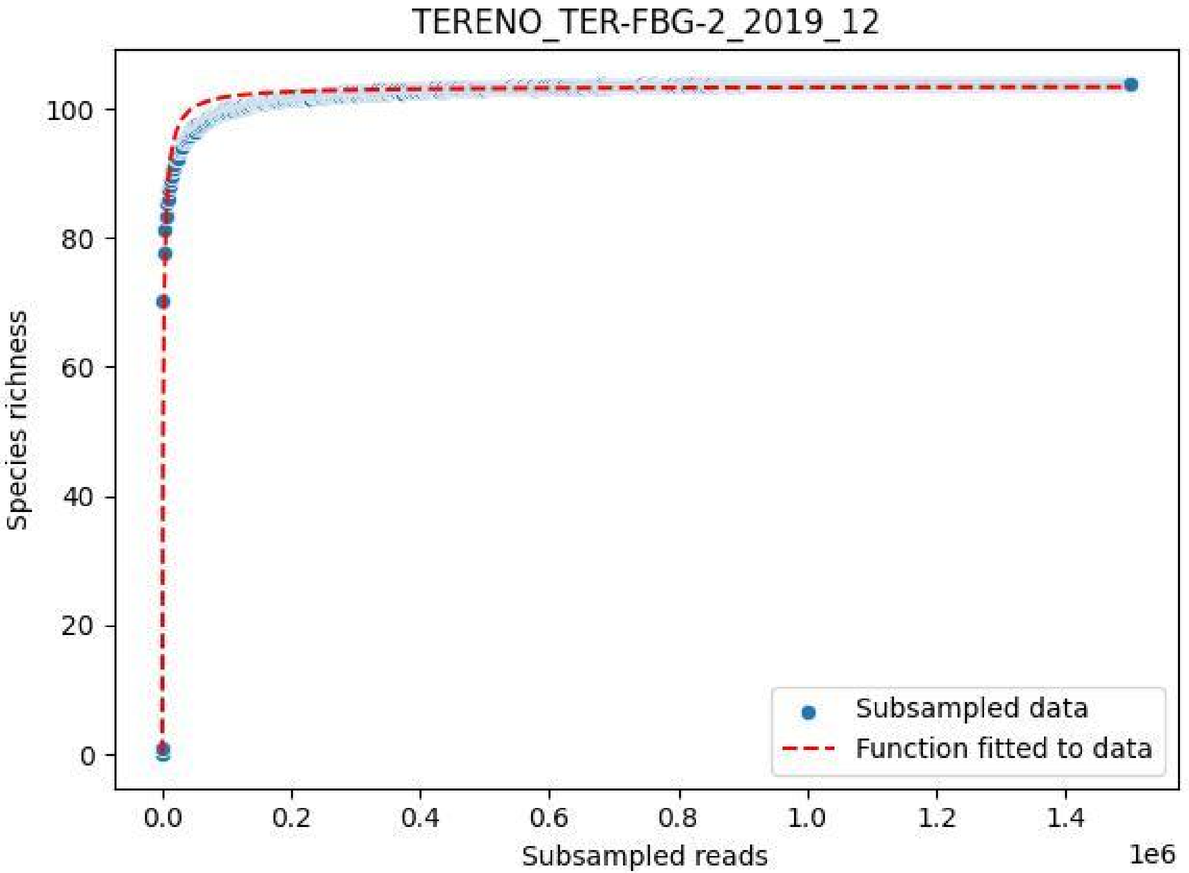

TERENO_TER-FBG-2_2019_12
Species richness
Subsampled reads
1e6
0
20
40
60
80
100
0.0
0.2
0.4
0.6
0.8
1.0
1.2
1.4
Subsampled data
Function fitted to data

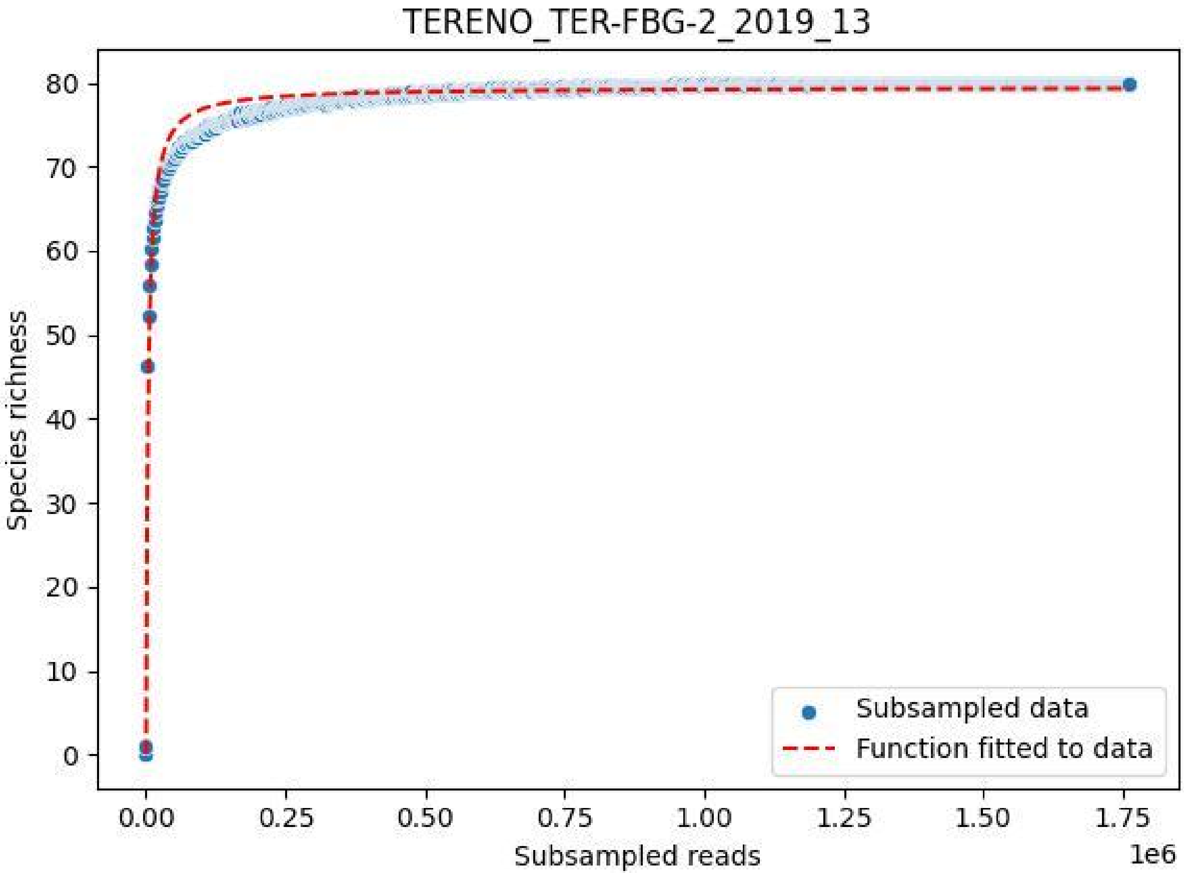

TERENO_TER-FBG-2_2019_13
Species richness
Subsampled reads
1e6
0
10
20
30
40
50
60
70
80
0.00
0.25
0.50
0.75
1.00
1.25
1.50
1.75
Subsampled data
Function fitted to data

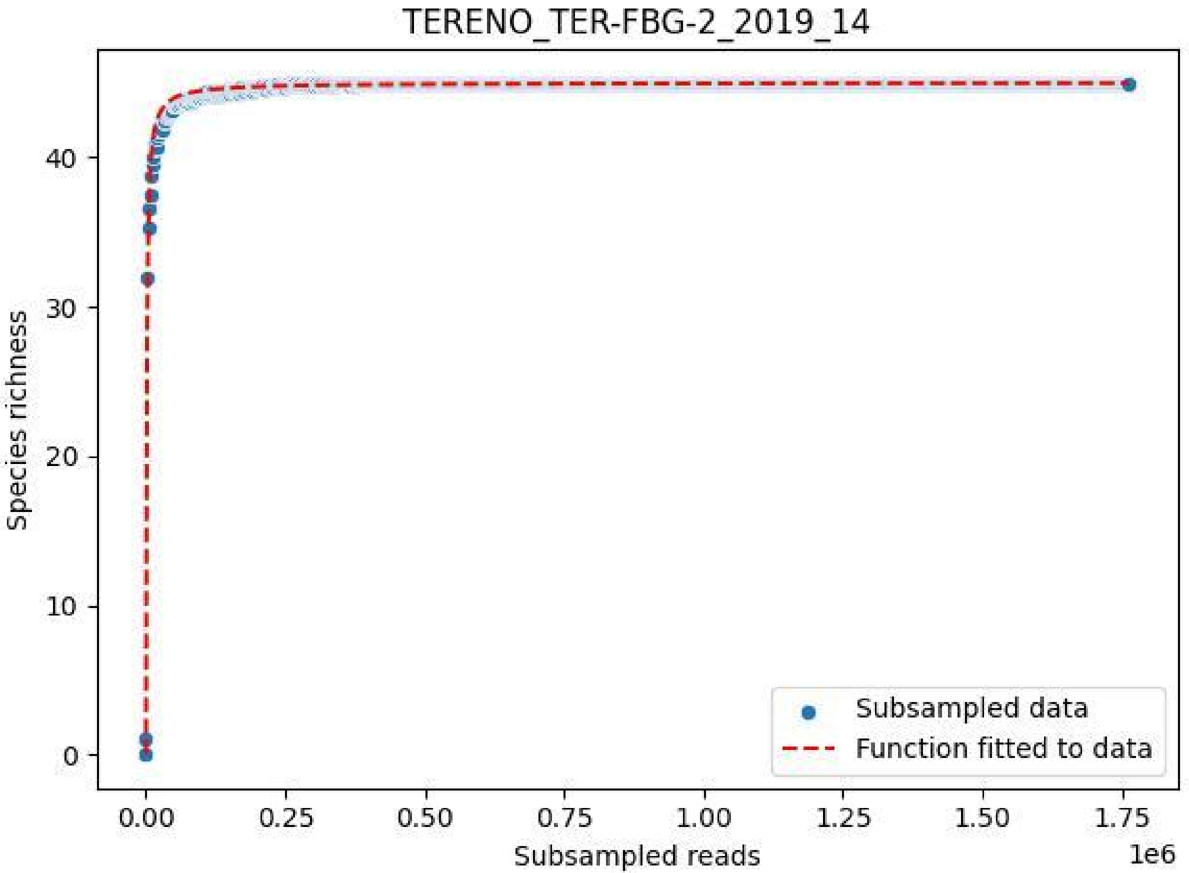

TERENO_TER-FBG-2_2019_14
Species richness
0
10
20
30
40
0.00
0.25
0.50
0.75
1.00
1.25
1.50
1.75
1e6
Subsampled reads
Subsampled data
Function fitted to data

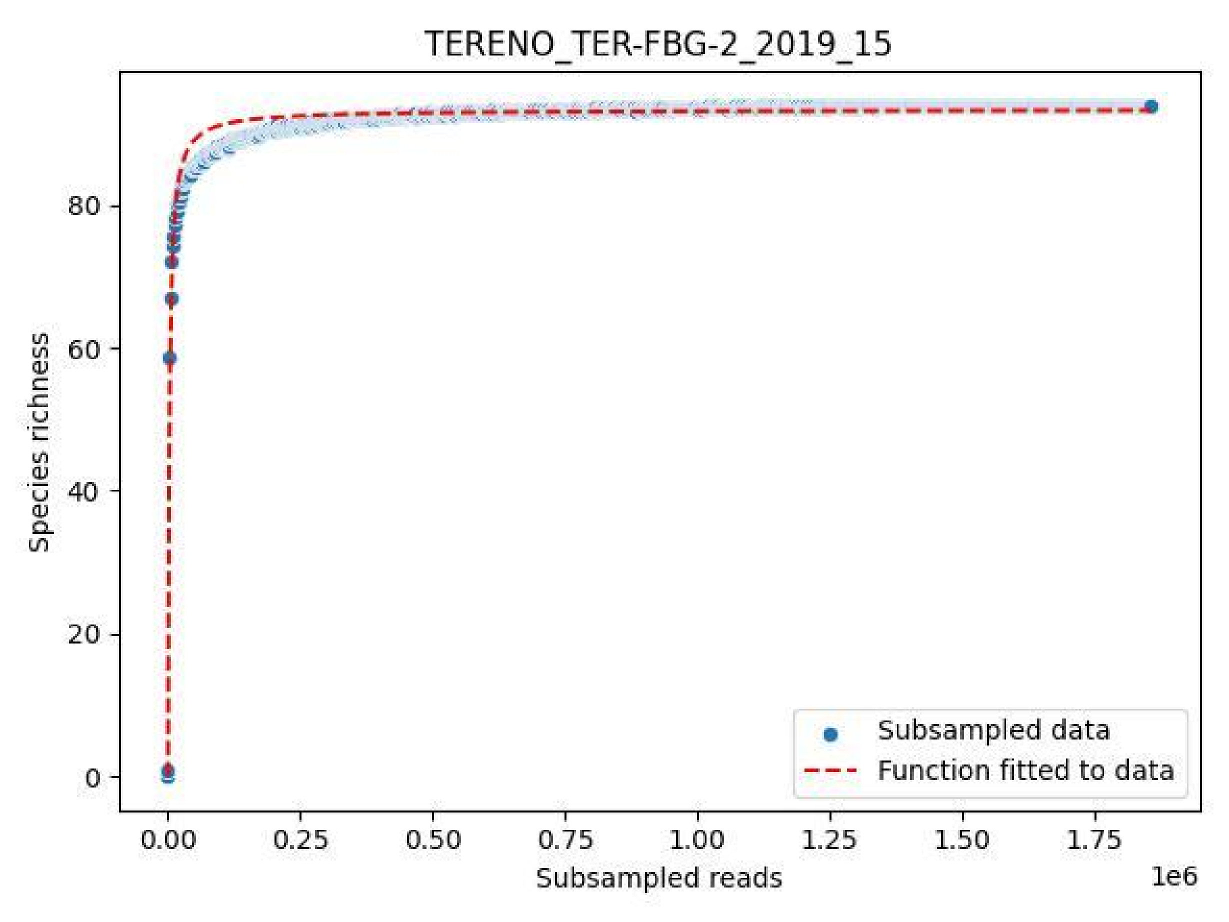

TERENO_TER-FBG-2_2019_15
Species richness
0
20
40
60
80
0.00
0.25
0.50
0.75
1.00
1.25
1.50
1.75
1e6
Subsampled reads
Subsampled data
Function fitted to data

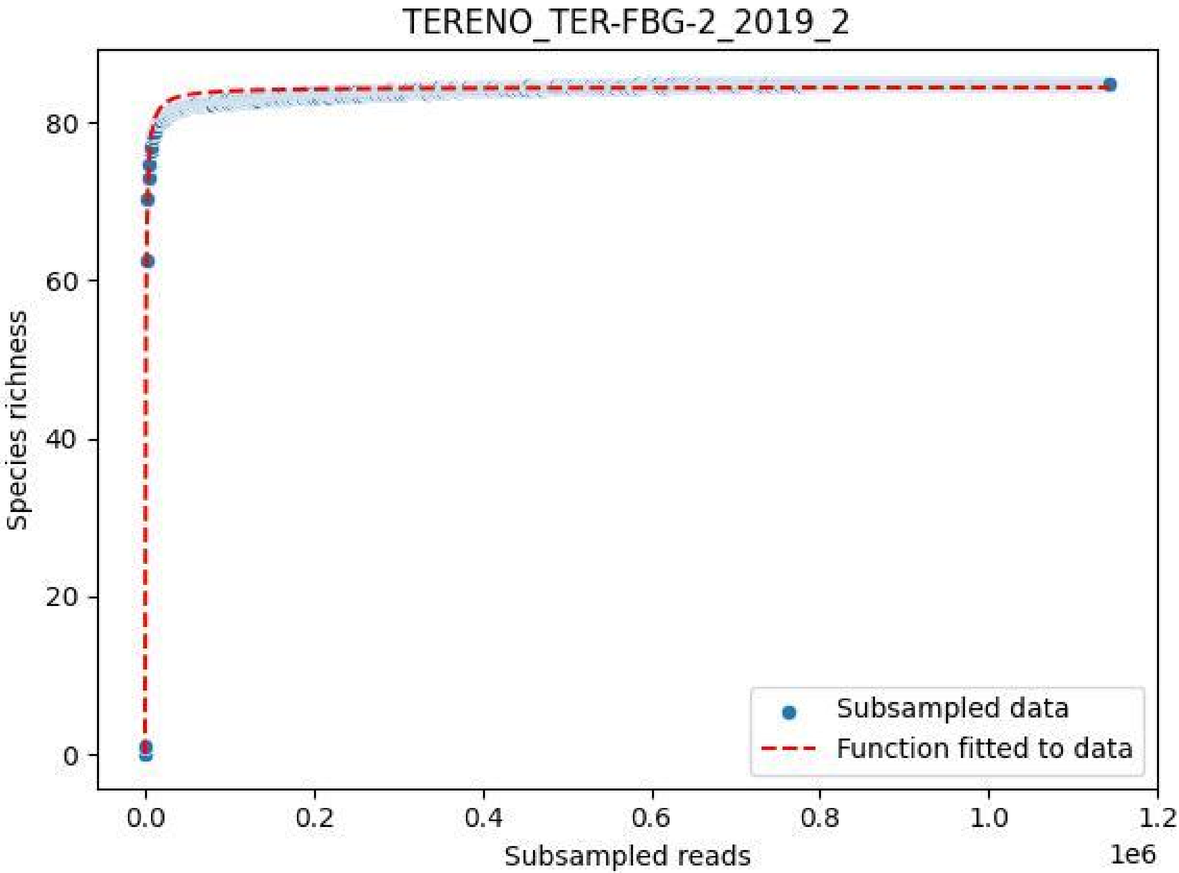

TERENO_TER-FBG-2_2019_2
Species richness
Subsampled reads
1e6
0.0
0.2
0.4
0.6
0.8
1.0
1.2
0
20
40
60
80
Subsampled data
Function fitted to data

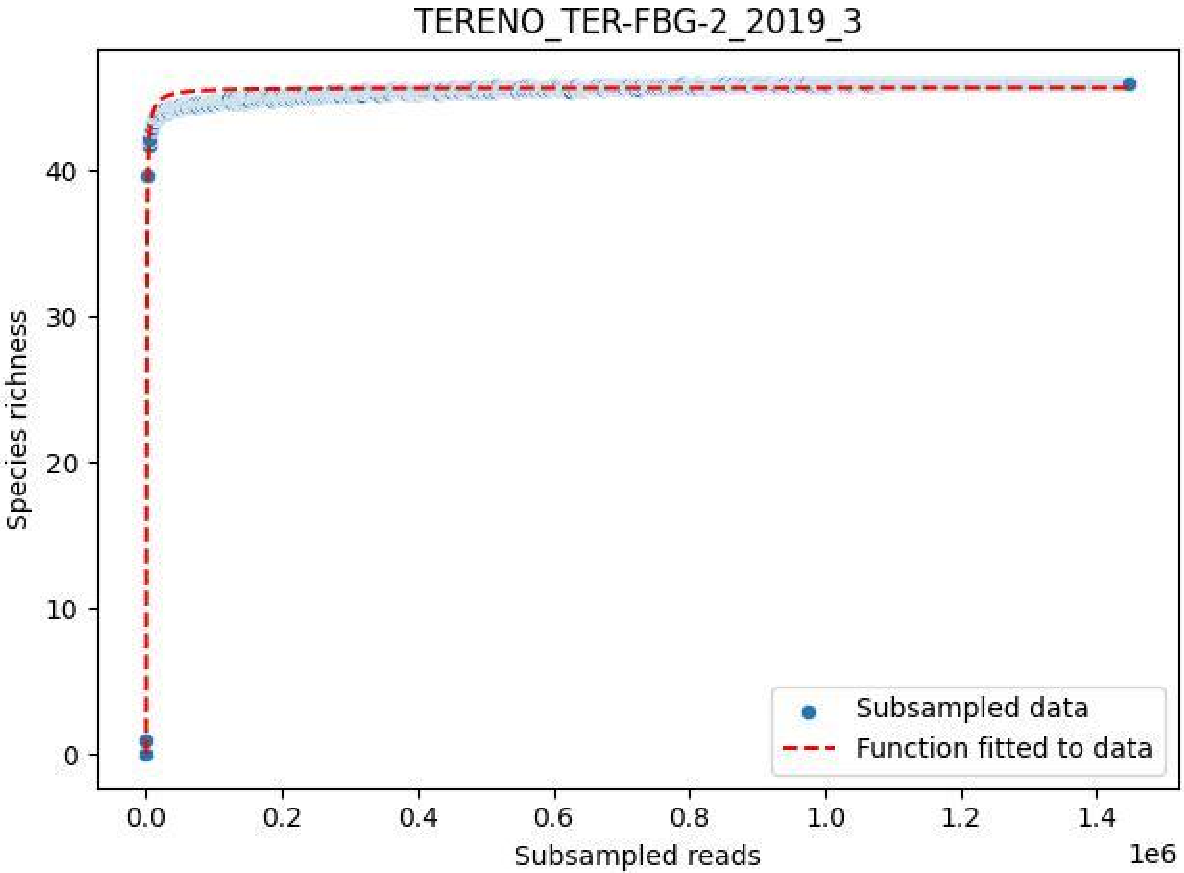

TERENO_TER-FBG-2_2019_3
Species richness
Subsampled reads
1e6
0
10
20
30
40
0.0
0.2
0.4
0.6
0.8
1.0
1.2
1.4
Subsampled data
Function fitted to data

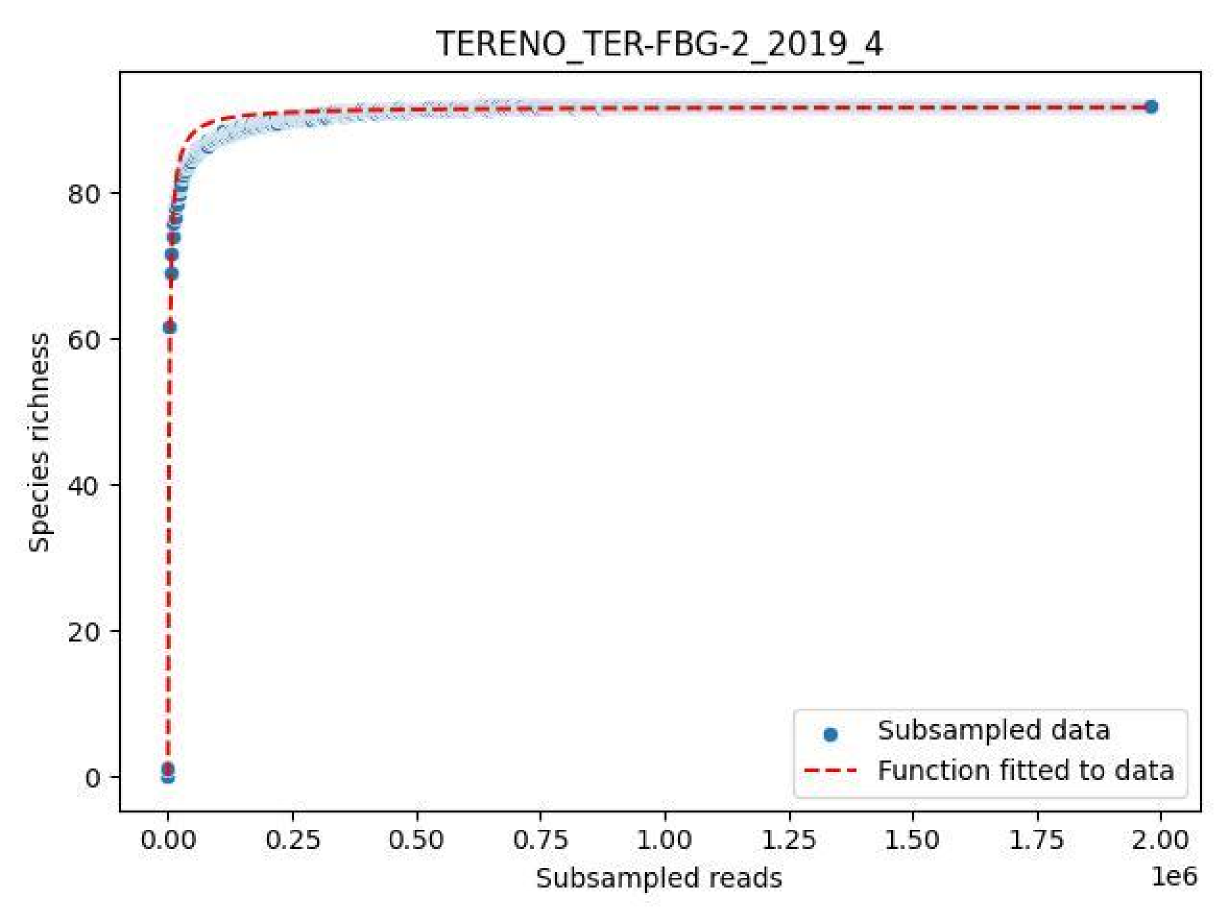

TERENO_TER-FBG-2_2019_4
Species richness
0
20
40
60
80
0.00
0.25
0.50
0.75
1.00
1.25
1.50
1.75
2.00
1e6
Subsampled reads
Subsampled data
Function fitted to data

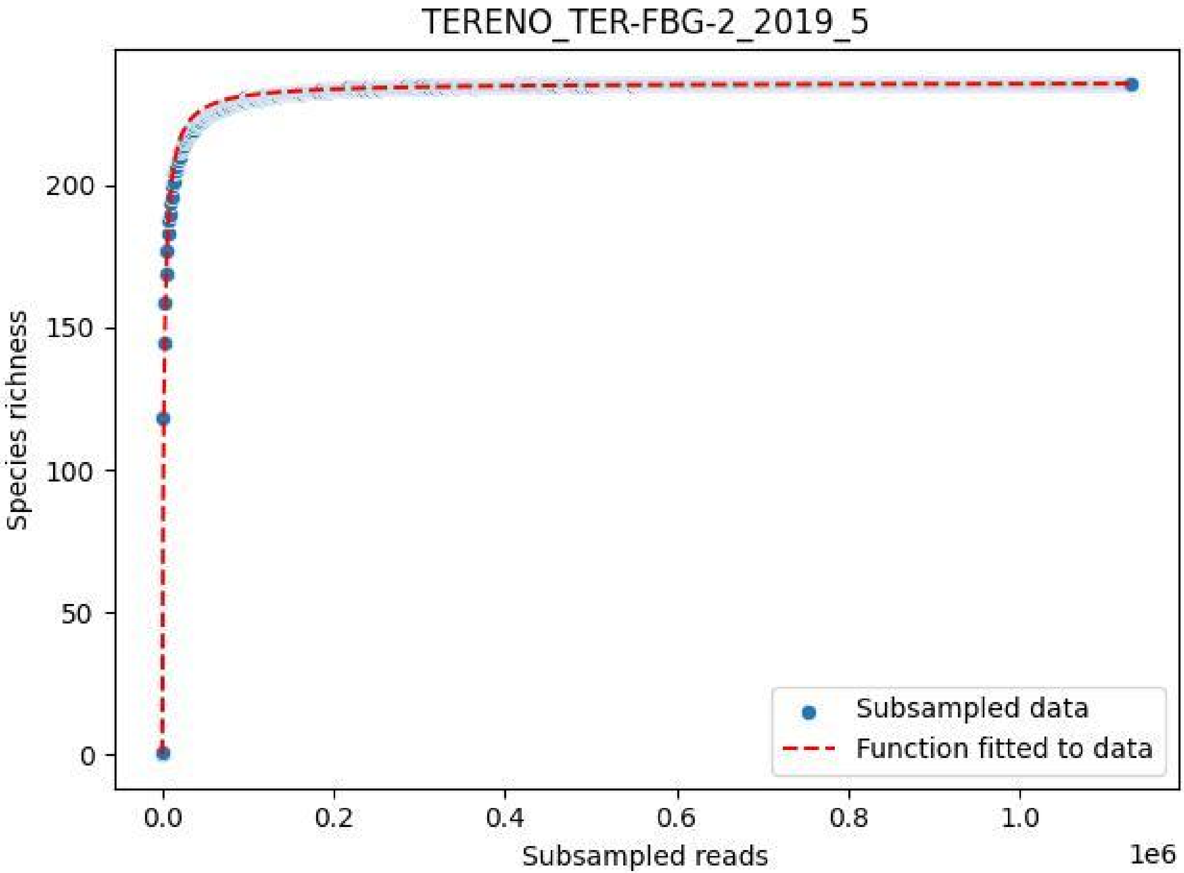

TERENO_TER-FBG-2_2019_5
Species richness
Subsampled reads
1e6
0
50
100
150
200
0.0
0.2
0.4
0.6
0.8
1.0
Subsampled data
Function fitted to data

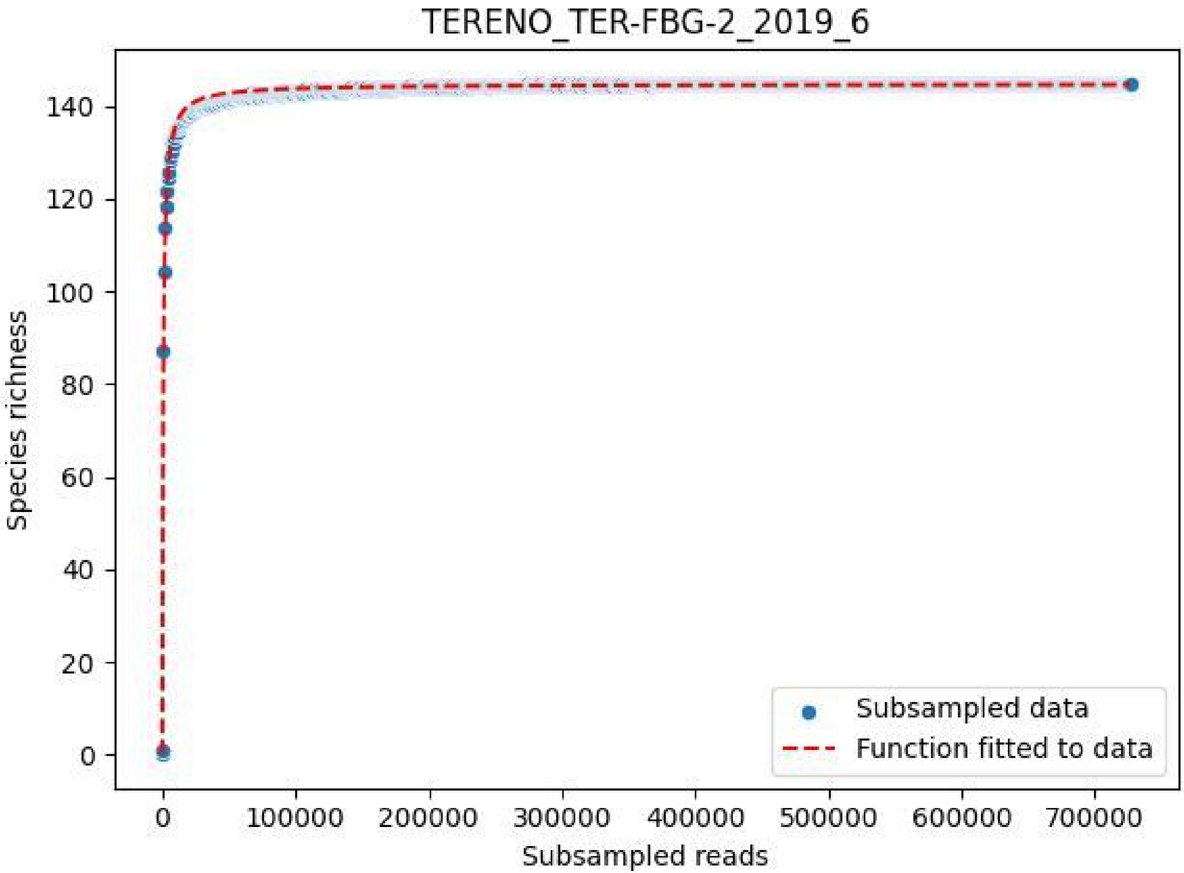

TERENO_TER-FBG-2_2019_6
Species richness
Subsampled reads
0
20
40
60
80
100
120
140
0
100000
200000
300000
400000
500000
600000
700000
Subsampled data
Function fitted to data

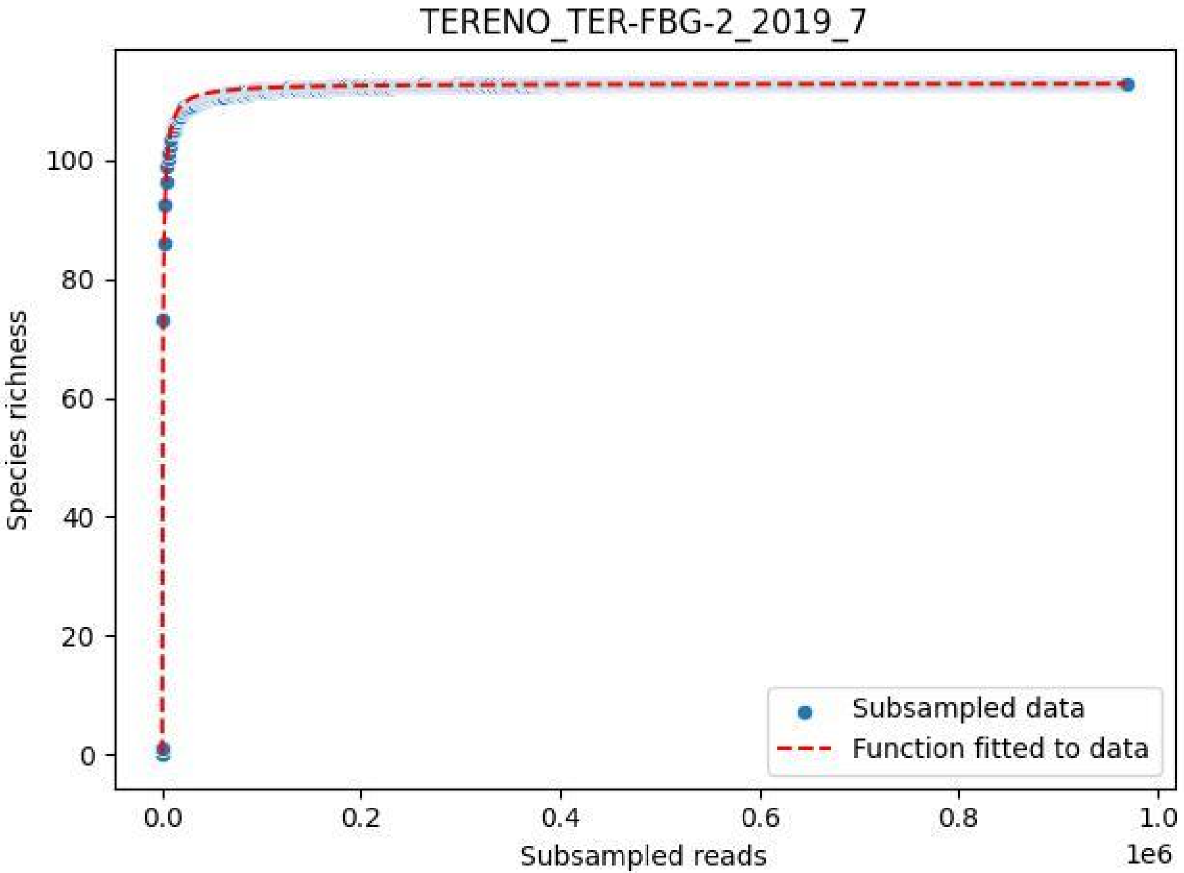

TERENO_TER-FBG-2_2019_7
Species richness
0
20
40
60
80
100
0.0
0.2
0.4
0.6
0.8
1.0
1e6
Subsampled reads
Subsampled data
Function fitted to data

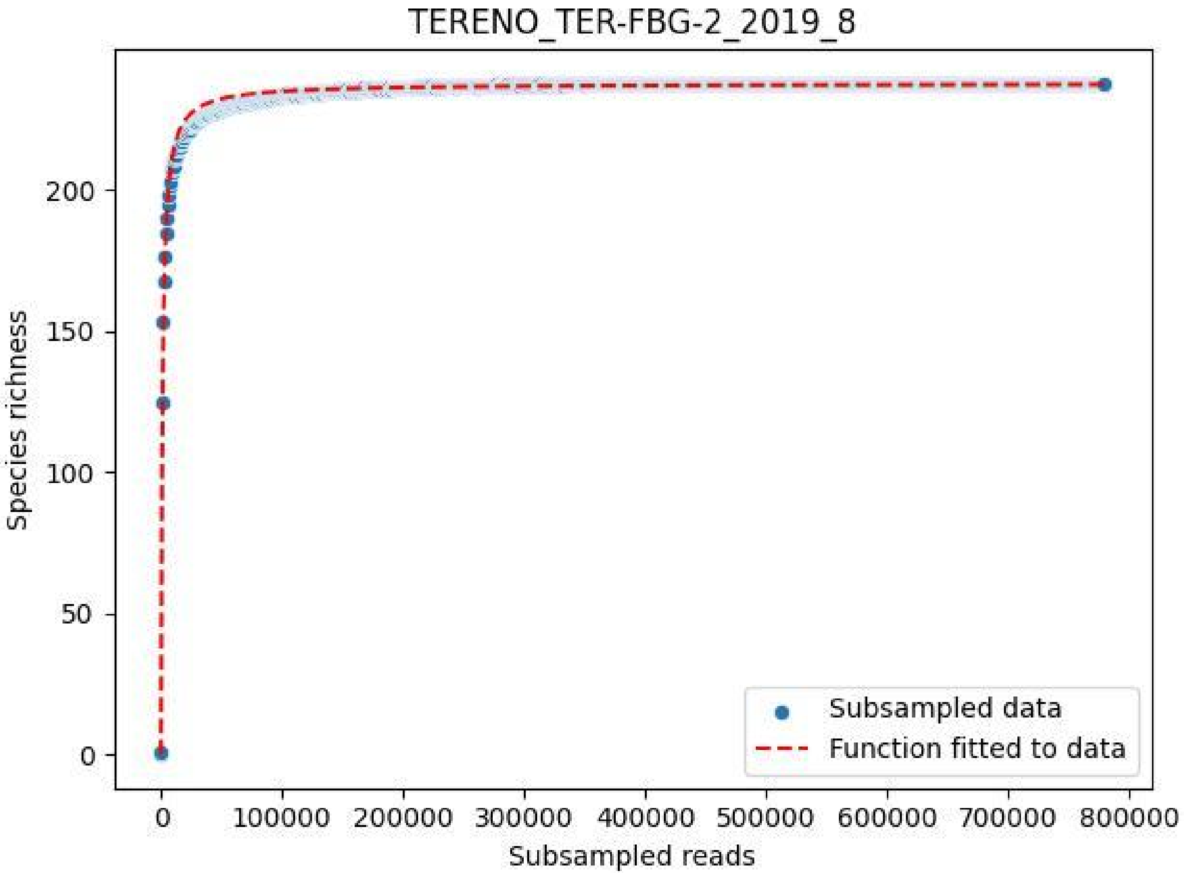

TERENO_TER-FBG-2_2019_8
Species richness
Subsampled reads
0
50
100
150
200
0
100000
200000
300000
400000
500000
600000
700000
800000
Subsampled data
Function fitted to data

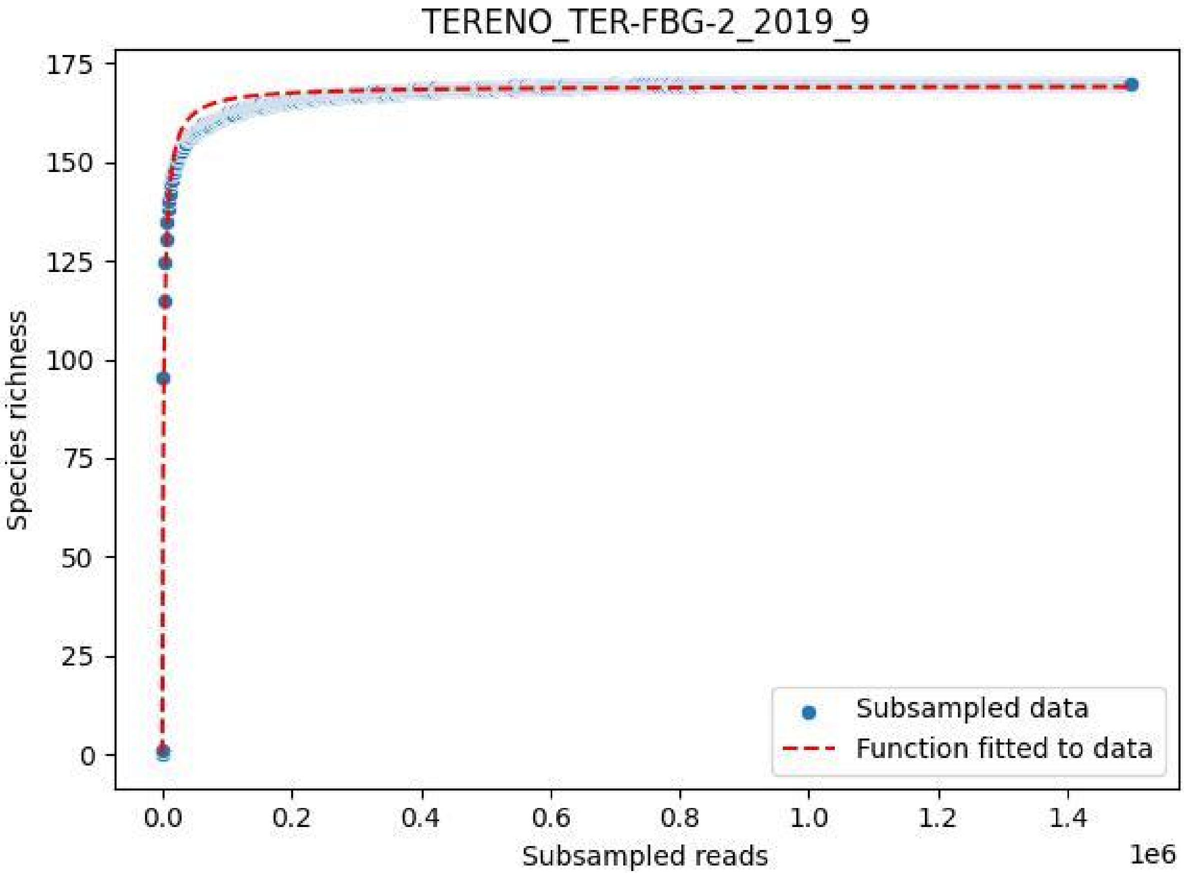

TERENO_TER-FBG-2_2019_9
Species richness
Subsampled reads
1e6
0
25
50
75
100
125
150
175
0.0
0.2
0.4
0.6
0.8
1.0
1.2
1.4
Subsampled data
Function fitted to data

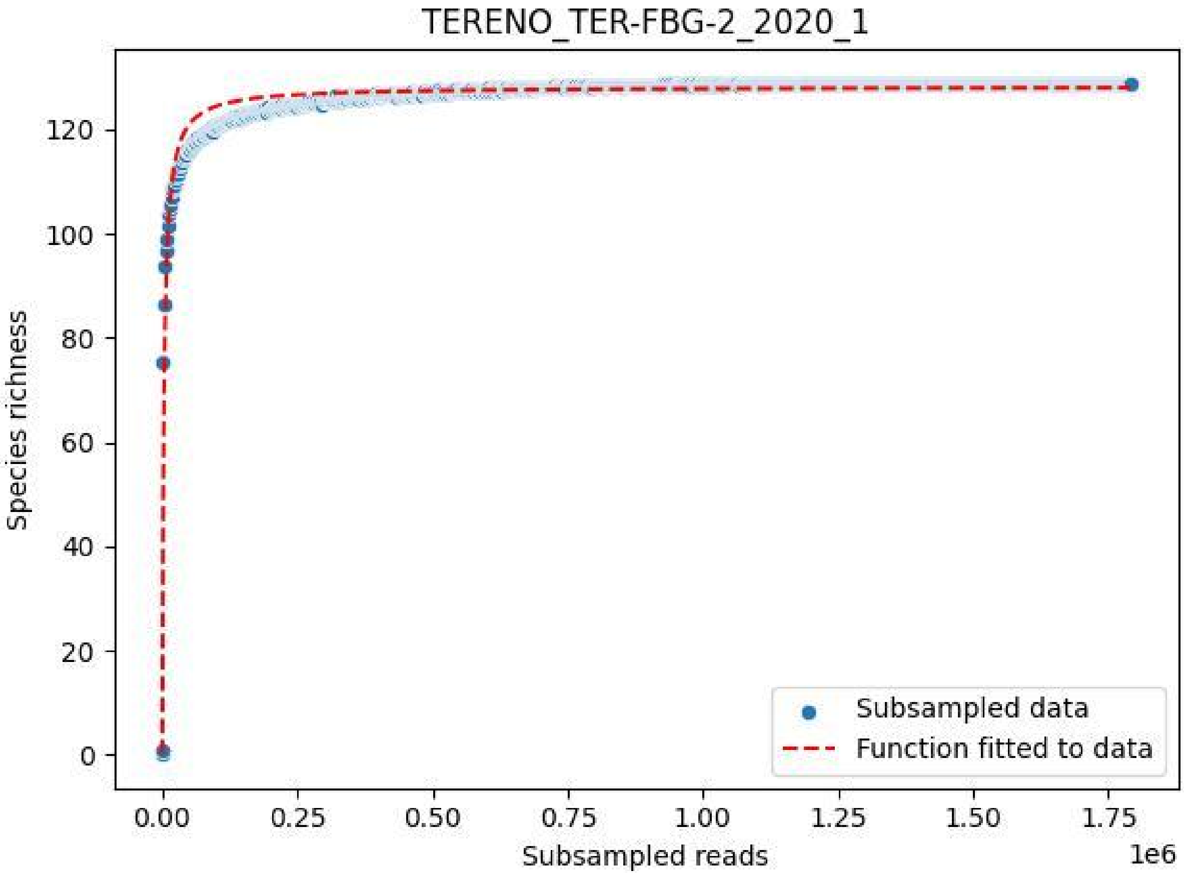
TERENO_TER-FBG-2_2020_1
Species richness
Subsampled reads
1e6
0
20
40
60
80
100
120
0.00
0.25
0.50
0.75
1.00
1.25
1.50
1.75
Subsampled data
Function fitted to data

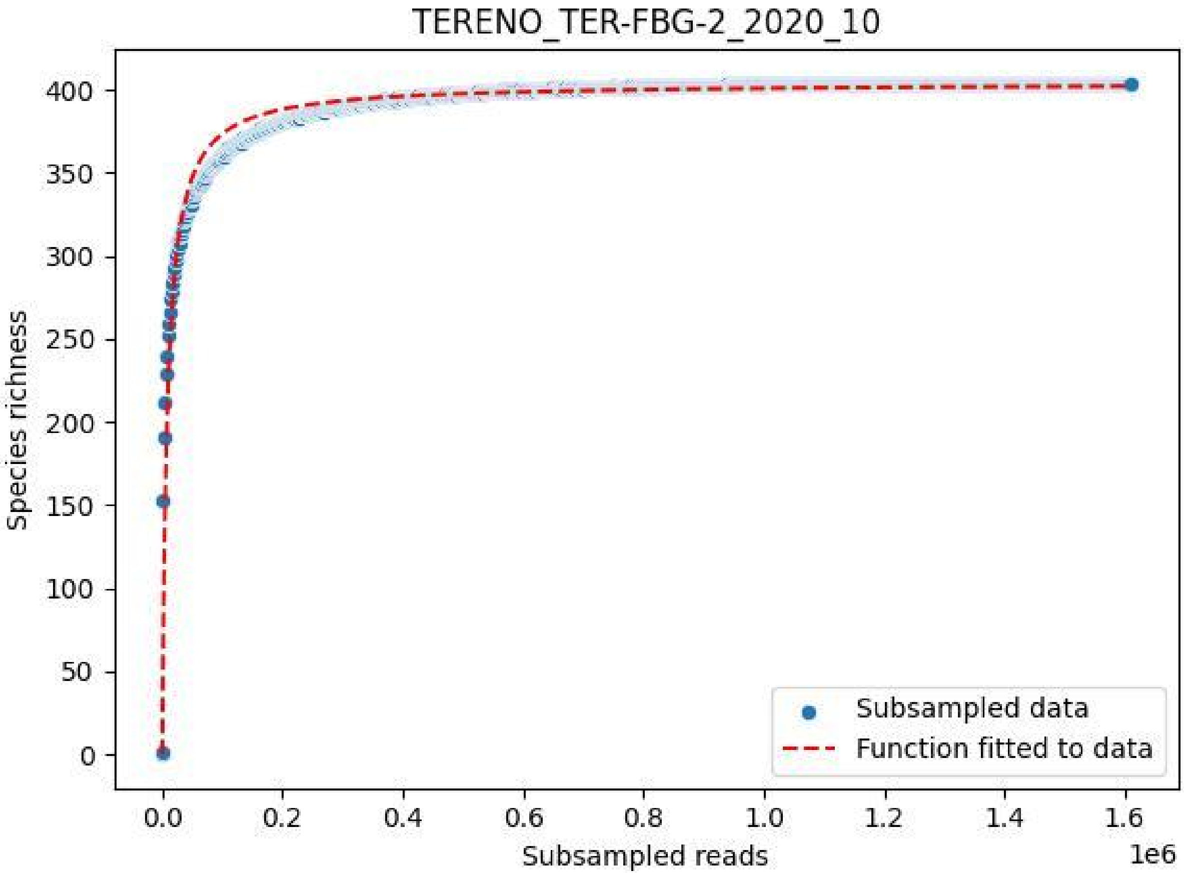

TERENO_TER-FBG-2_2020_10
Species richness
Subsampled reads
1e6
0
50
100
150
200
250
300
350
400
0.0
0.2
0.4
0.6
0.8
1.0
1.2
1.4
1.6
Subsampled data
Function fitted to data

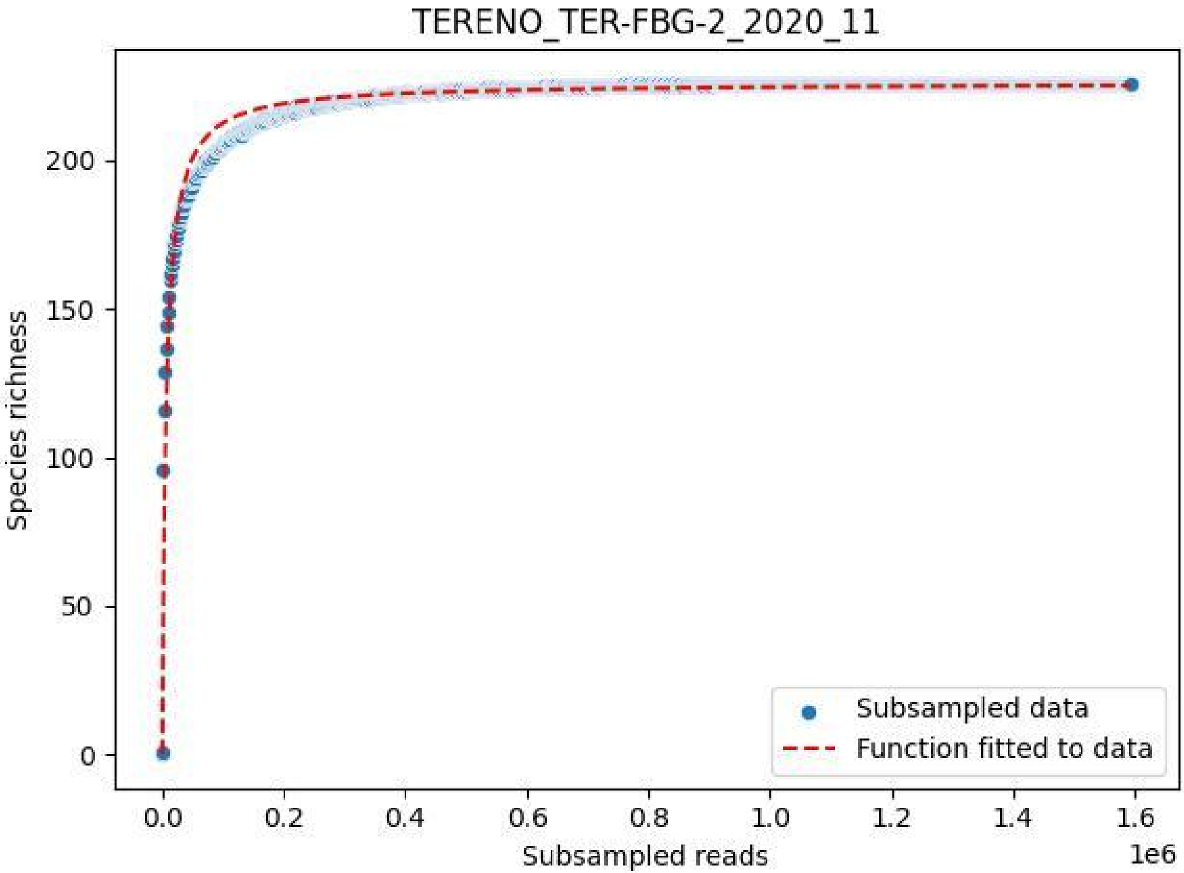

TERENO_TER-FBG-2_2020_11
Species richness
0
50
100
150
200
0.0
0.2
0.4
0.6
0.8
1.0
1.2
1.4
1.6
1e6
Subsampled reads
Subsampled data
Function fitted to data

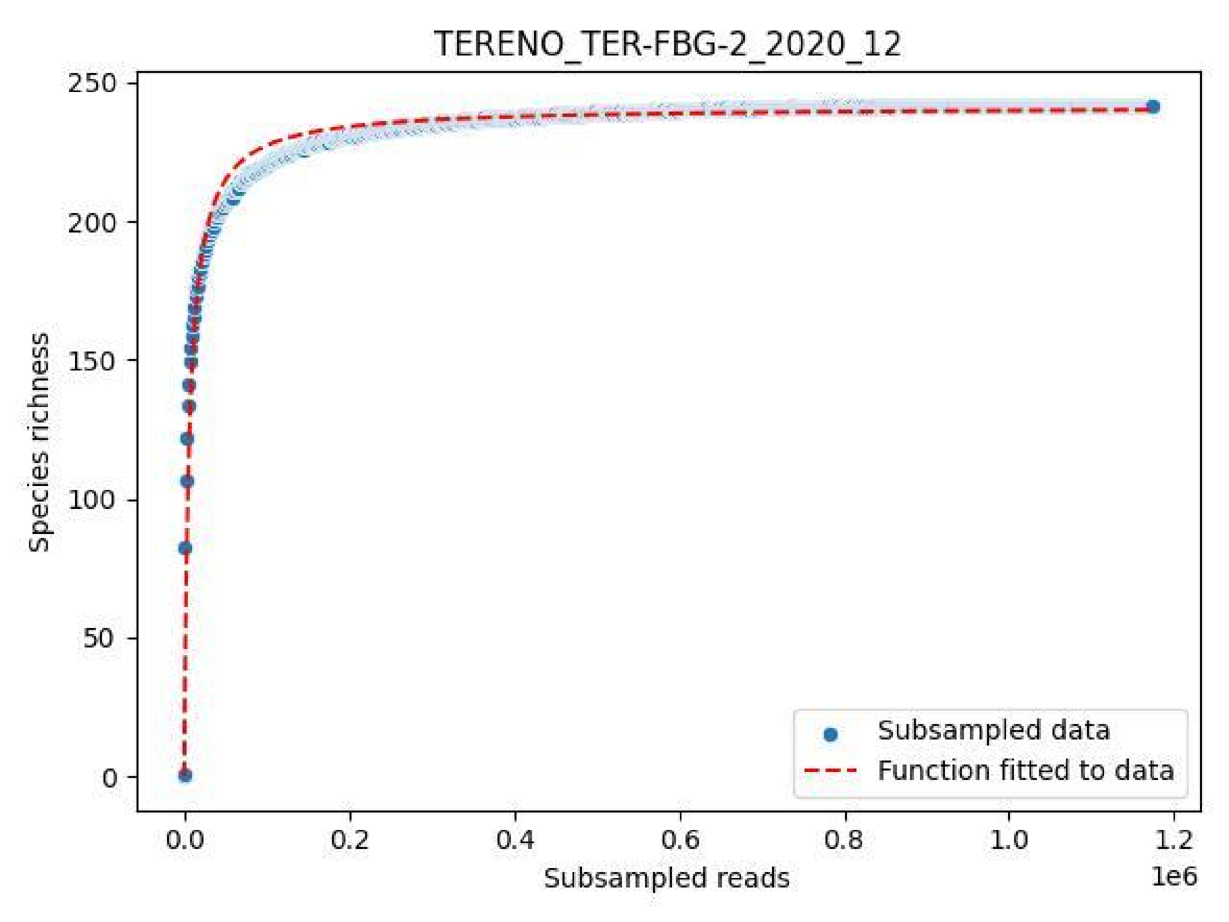

TERENO_TER-FBG-2_2020_12
250
200
150
100
50
0
Species richness
0.0
0.2
0.4
0.6
0.8
1.0
1.2
1e6
Subsampled reads
Subsampled data
Function fitted to data

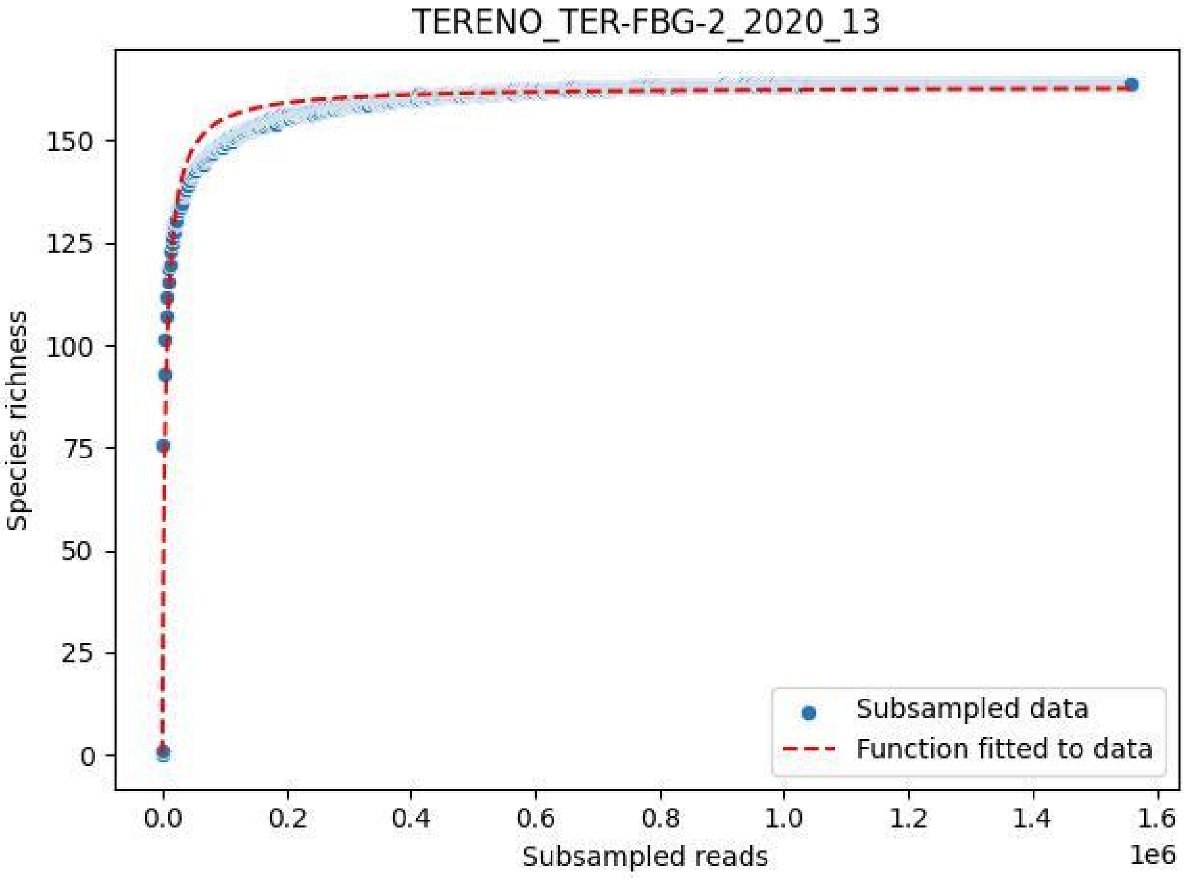
TERENO_TER-FBG-2_2020_13
Species richness
0
25
50
75
100
125
150
0.0
0.2
0.4
0.6
0.8
1.0
1.2
1.4
1.6
1e6
Subsampled reads
Subsampled data
Function fitted to data

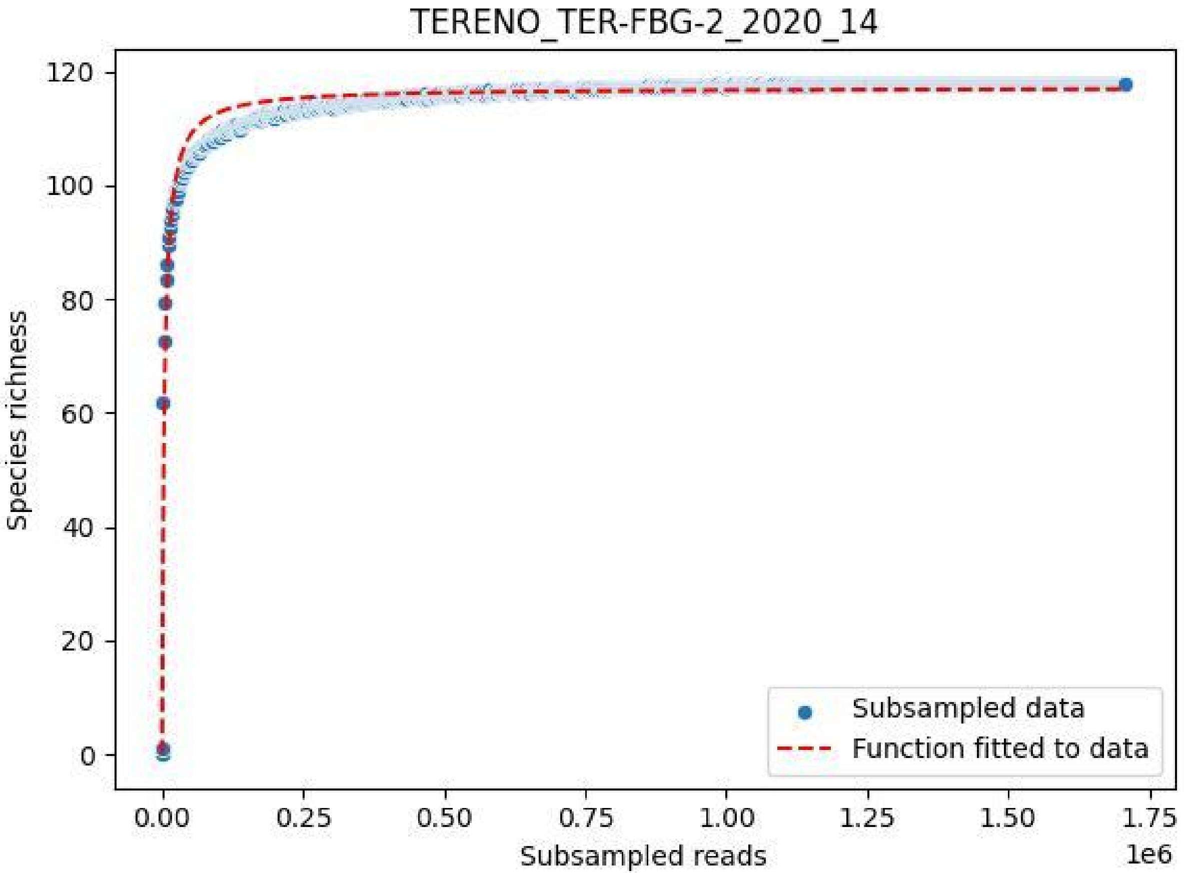

TERENO_TER-FBG-2_2020_14
Species richness
Subsampled reads
1e6
0.00
0.25
0.50
0.75
1.00
1.25
1.50
1.75
0
20
40
60
80
100
120
Subsampled data
Function fitted to data

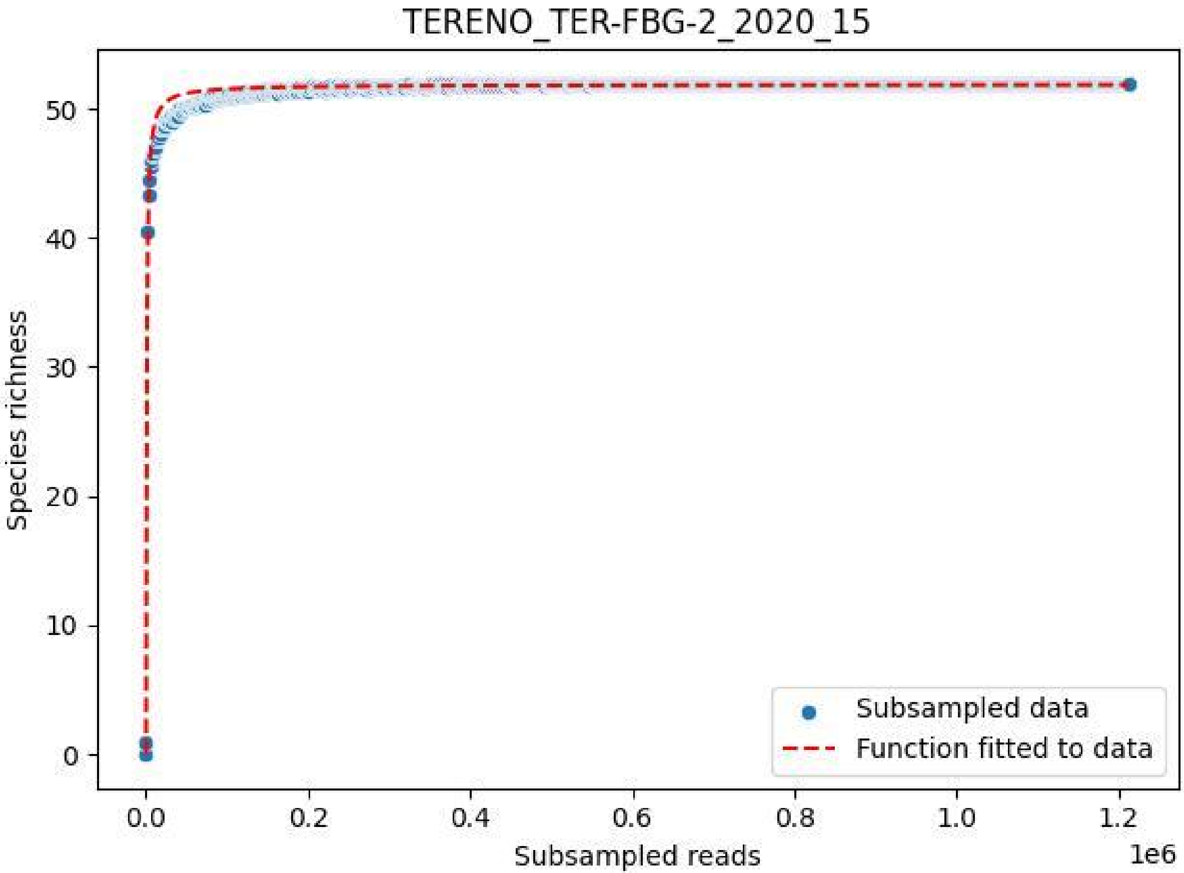
TERENO_TER-FBG-2_2020_15
Species richness
Subsampled reads
1e6
0
10
20
30
40
50
0.0
0.2
0.4
0.6
0.8
1.0
1.2
Subsampled data
Function fitted to data

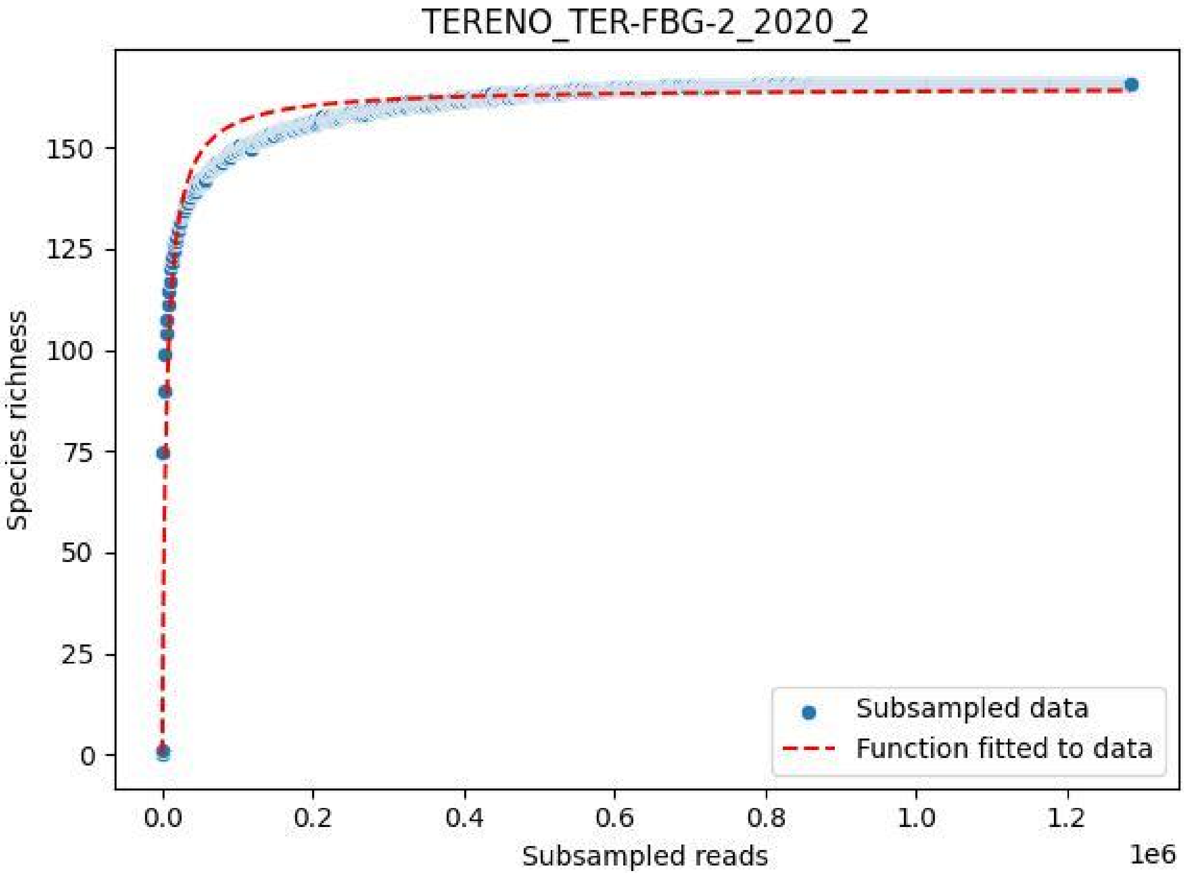

TERENO_TER-FBG-2_2020_2
Species richness
150
125
100
75
50
25
0
0.0
0.2
0.4
0.6
0.8
1.0
1.2
1e6
Subsampled reads
Subsampled data
Function fitted to data

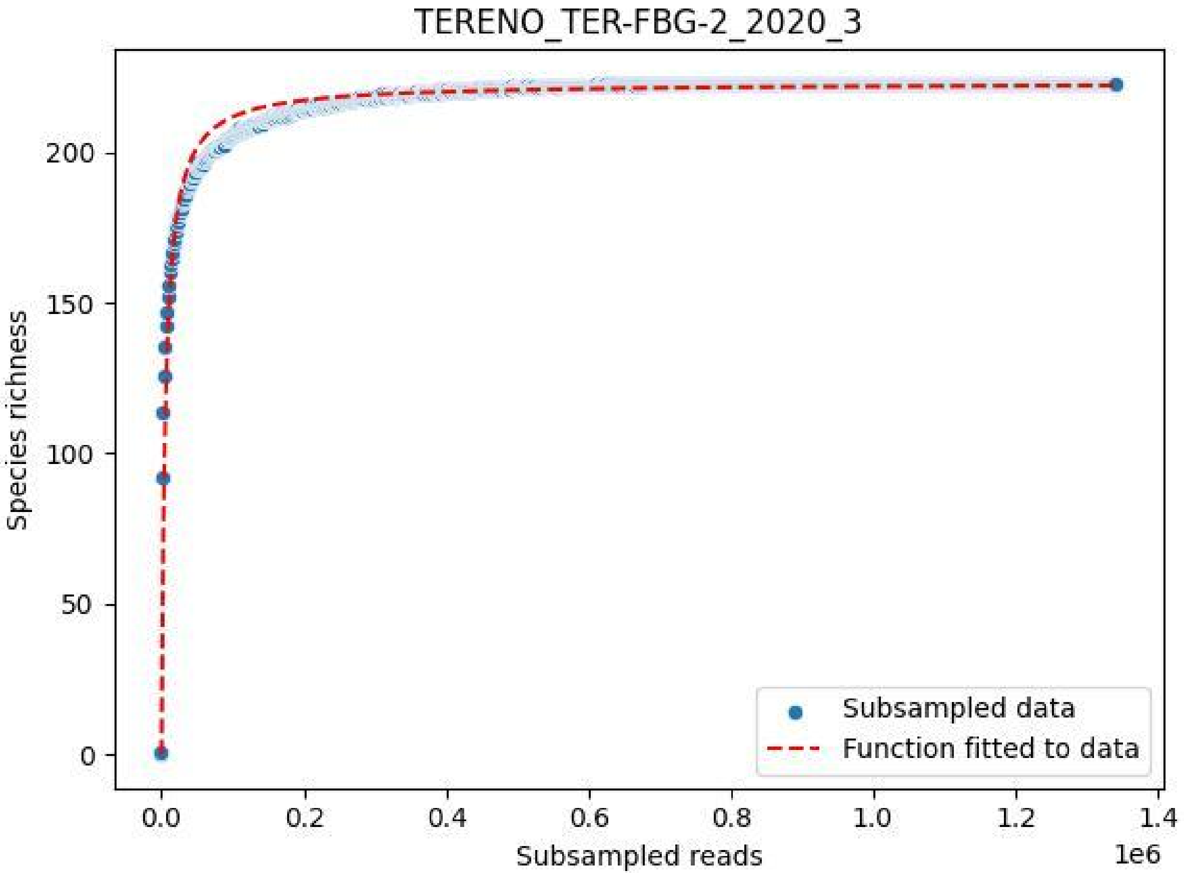

TERENO_TER-FBG-2_2020_3
Species richness
Subsampled reads
1e6
0
50
100
150
200
0.0
0.2
0.4
0.6
0.8
1.0
1.2
1.4
Subsampled data
Function fitted to data

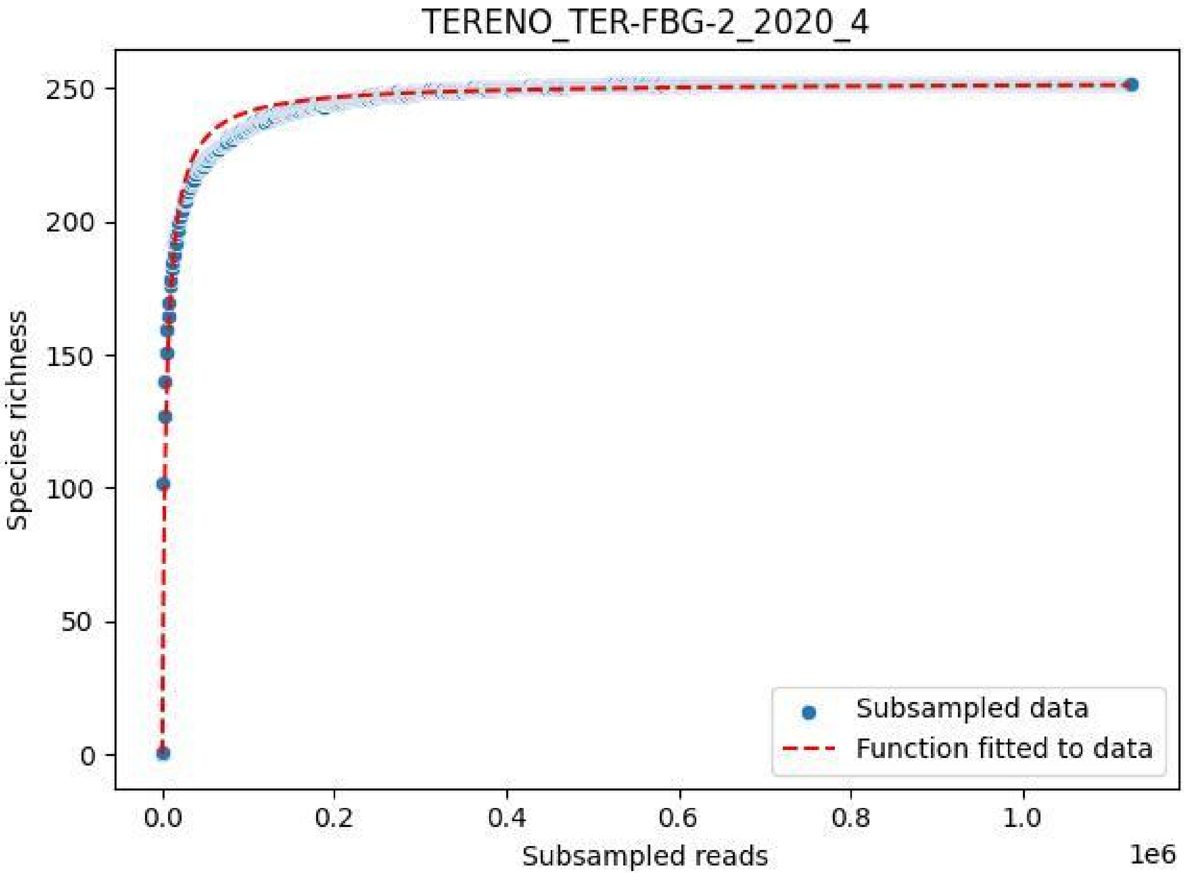

TERENO_TER-FBG-2_2020_4
Species richness
Subsampled reads
1e6
0
50
100
150
200
250
0.0
0.2
0.4
0.6
0.8
1.0
Subsampled data
Function fitted to data

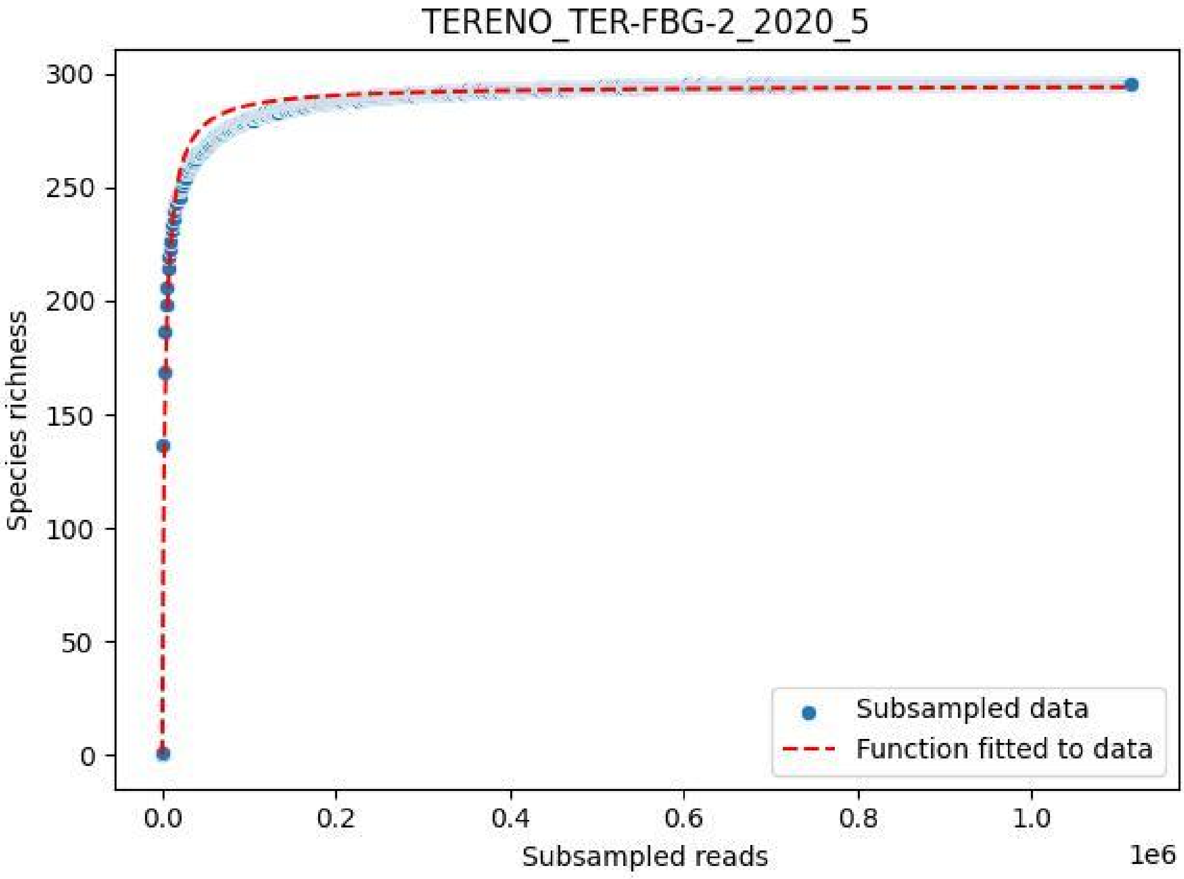
TERENO_TER-FBG-2_2020_5
Species richness
Subsampled reads
1e6
0
50
100
150
200
250
300
0.0
0.2
0.4
0.6
0.8
1.0
Subsampled data
Function fitted to data

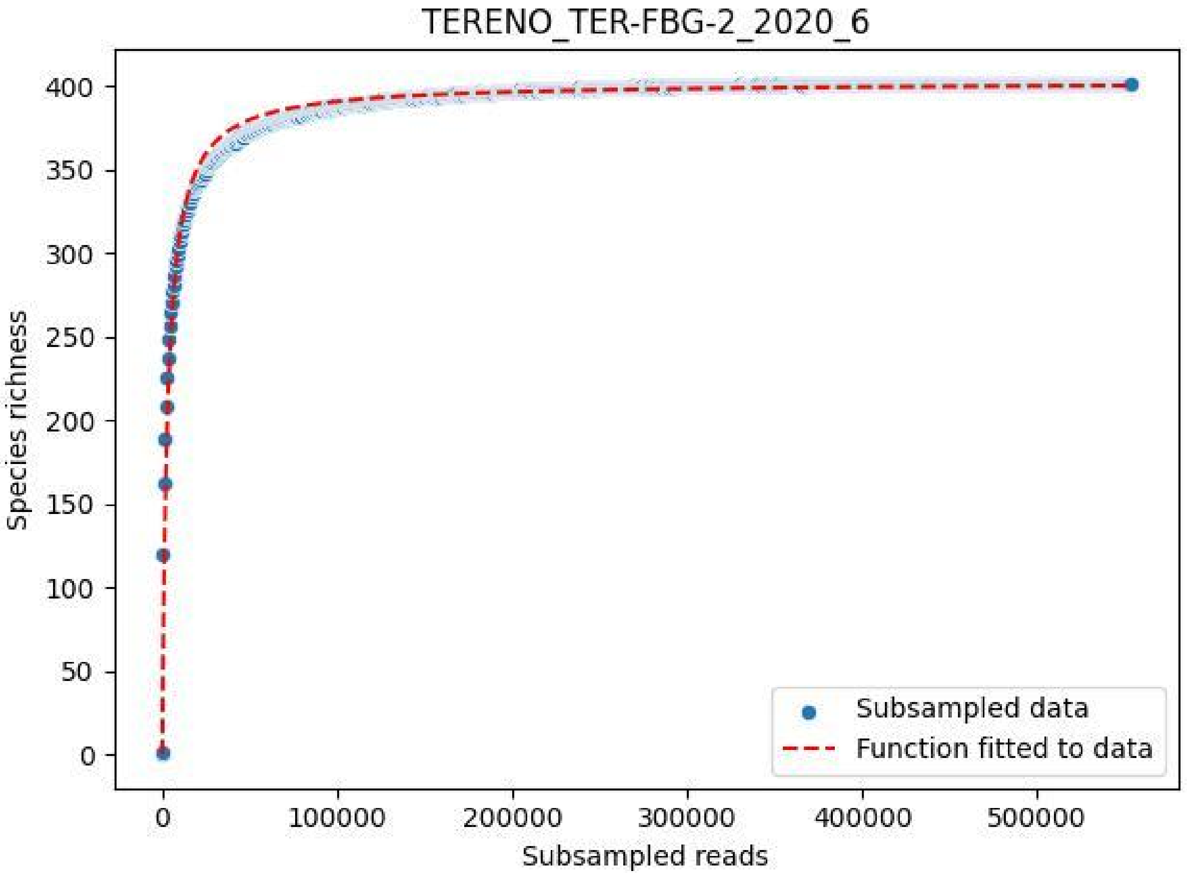

TERENO_TER-FBG-2_2020_6
Species richness
Subsampled reads
Subsampled data
Function fitted to data
0
50
100
150
200
250
300
350
400
0
100000
200000
300000
400000
500000

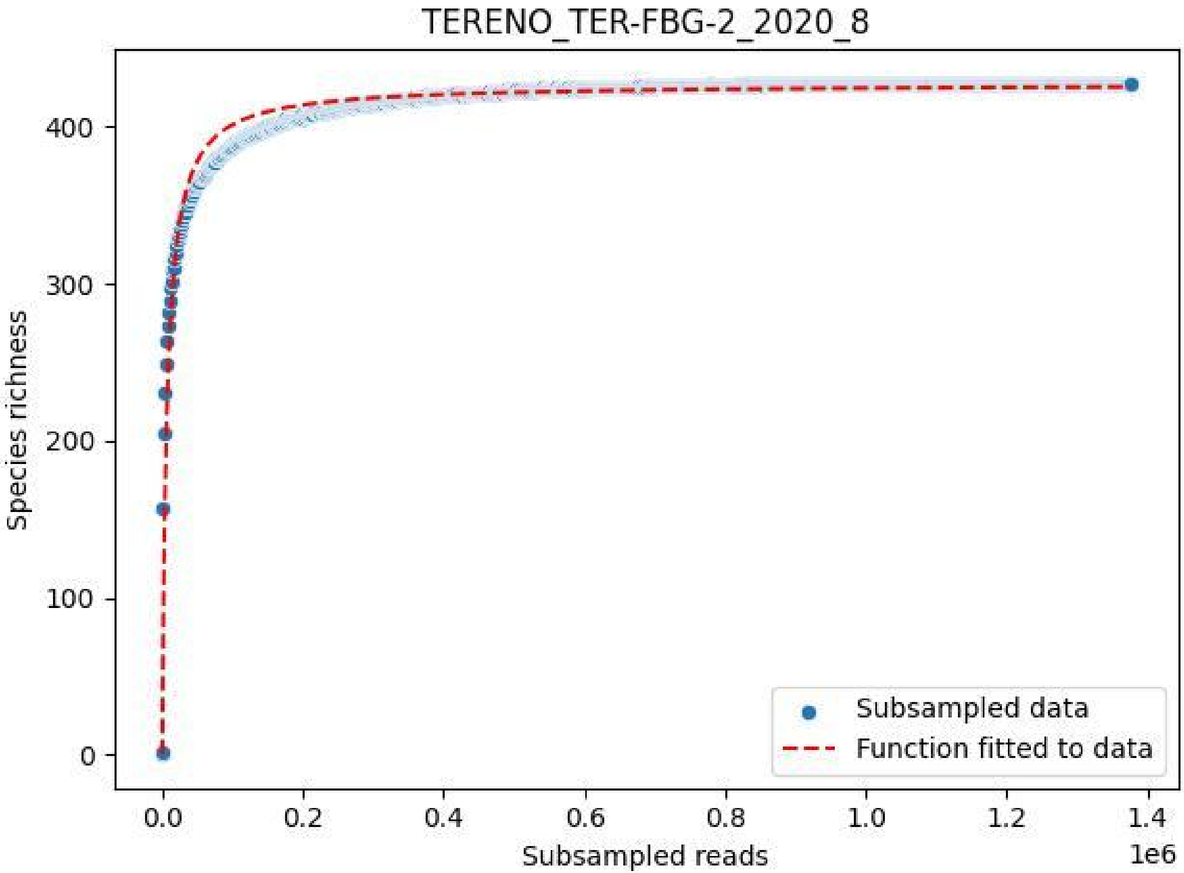
TERENO_TER-FBG-2_2020_8
Species richness
Subsampled reads
1e6
0
100
200
300
400
0.0
0.2
0.4
0.6
0.8
1.0
1.2
1.4
Subsampled data
Function fitted to data

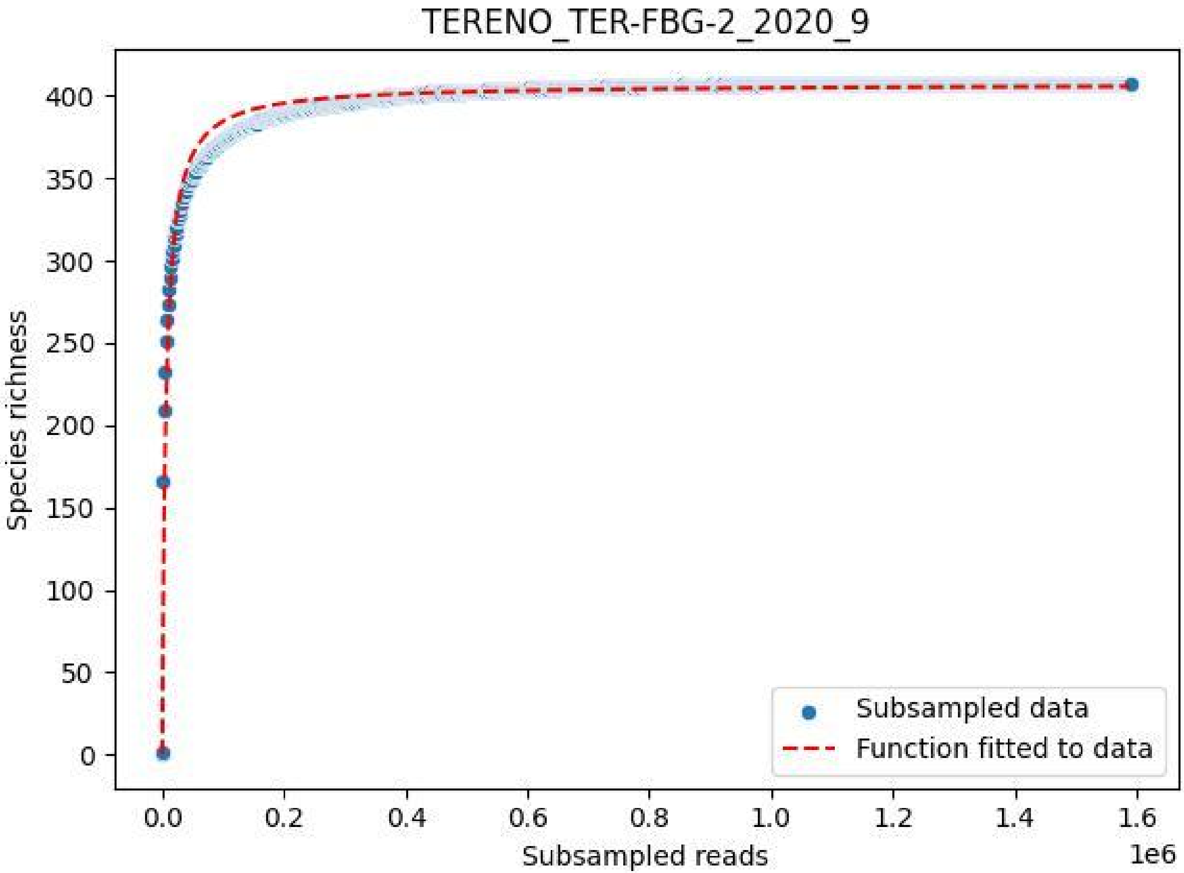

TERENO_TER-FBG-2_2020_9
Species richness
Subsampled reads
1e6
0
50
100
150
200
250
300
350
400
0.0
0.2
0.4
0.6
0.8
1.0
1.2
1.4
1.6
Subsampled data
Function fitted to data

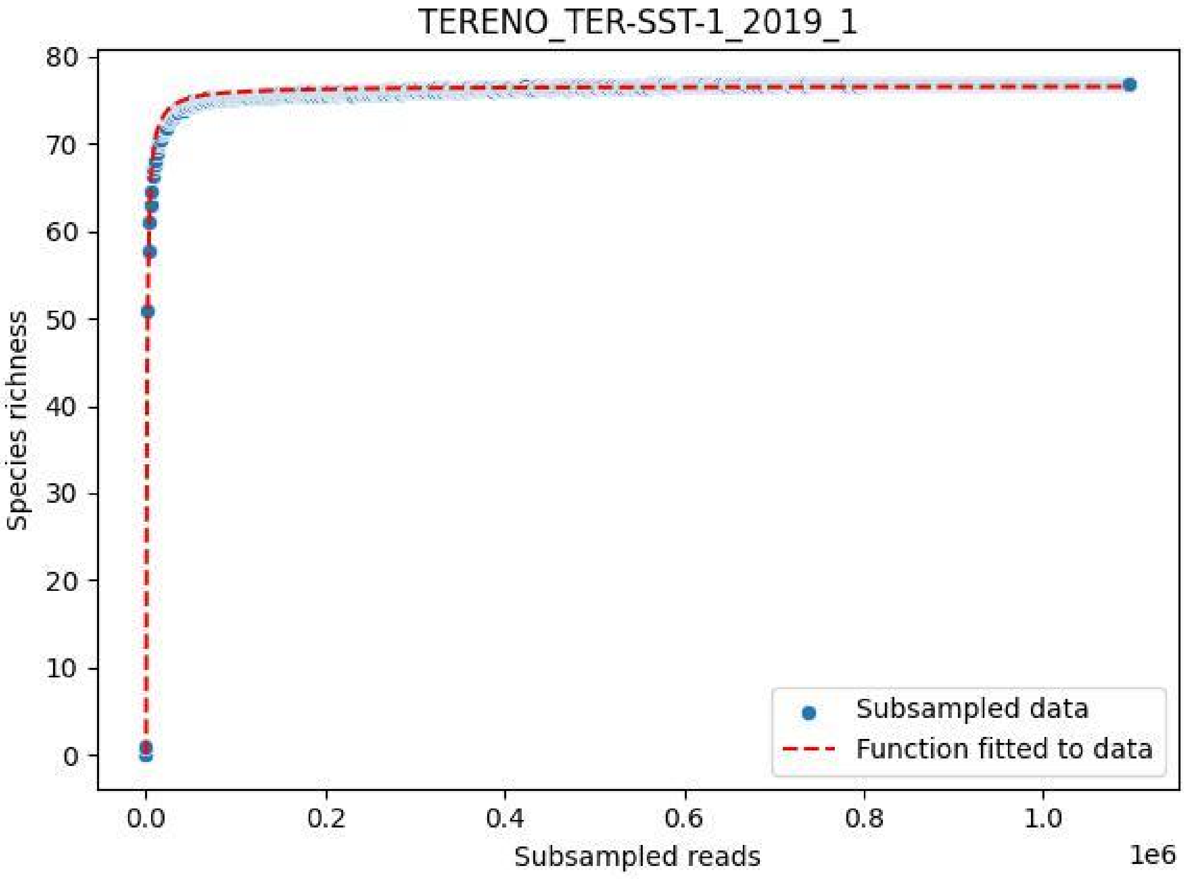

TERENO_TER-SST-1_2019_1
Species richness
0
10
20
30
40
50
60
70
80
0.0
0.2
0.4
0.6
0.8
1.0
1e6
Subsampled reads
Subsampled data
Function fitted to data

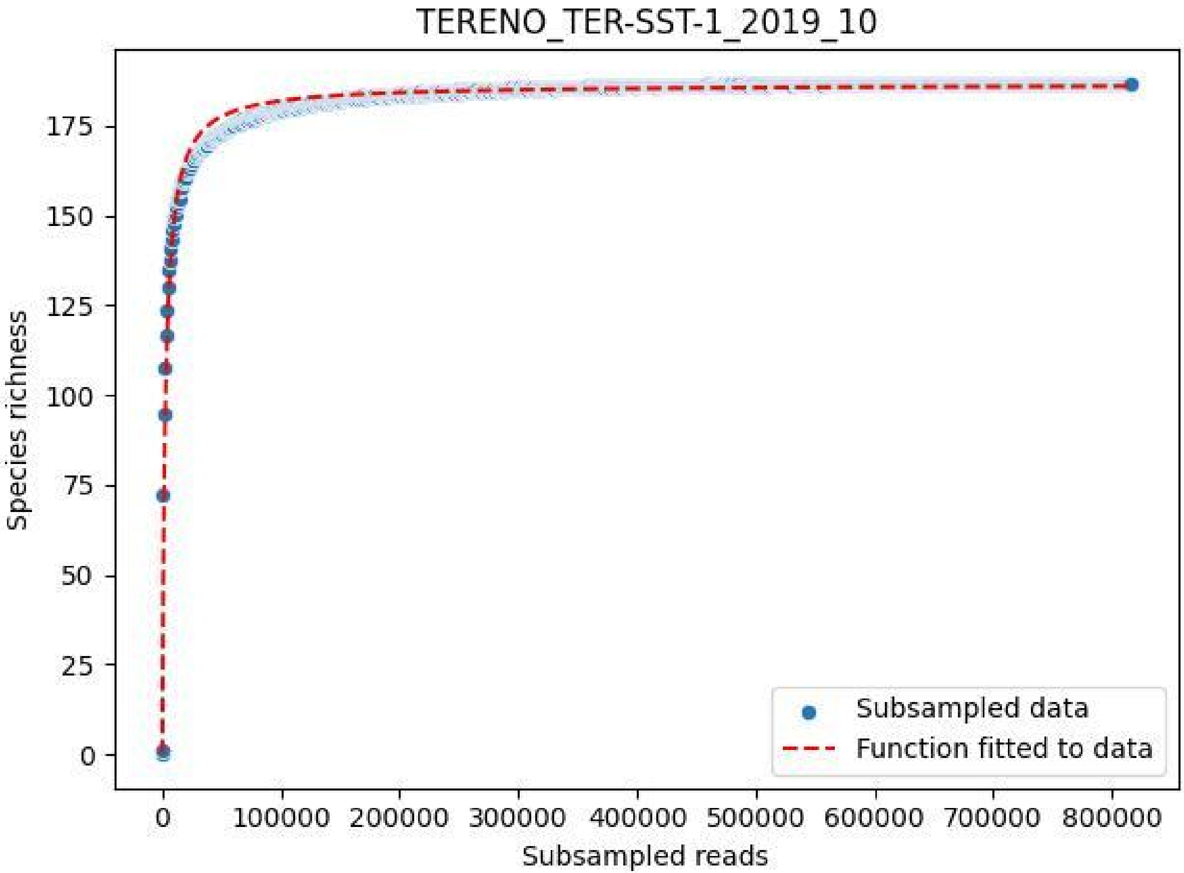
TERENO_TER-SST-1_2019_10
Species richness
Subsampled reads
0
25
50
75
100
125
150
175
0
100000
200000
300000
400000
500000
600000
700000
800000
Subsampled data
Function fitted to data

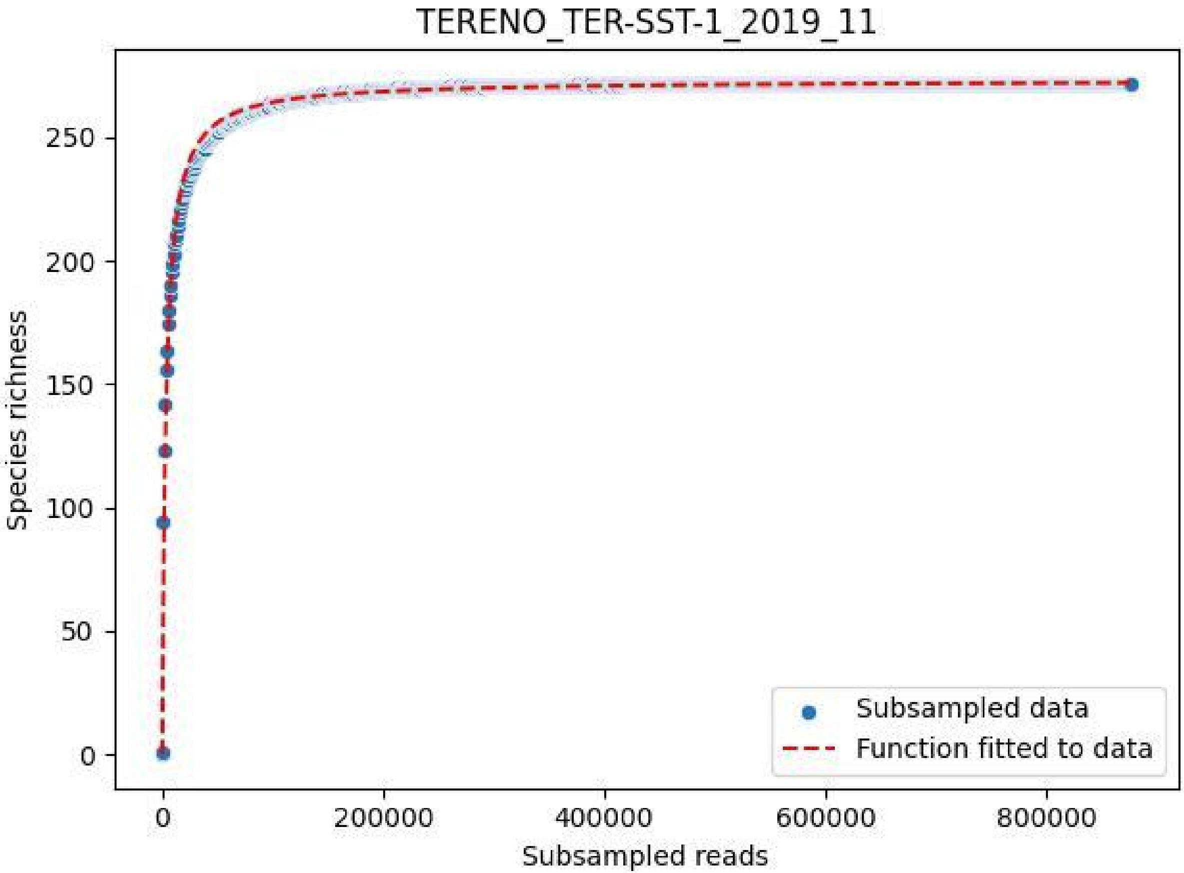
TERENO_TER-SST-1_2019_11
Species richness
0
50
100
150
200
250
0
200000
400000
600000
800000
Subsampled reads
Subsampled data
Function fitted to data

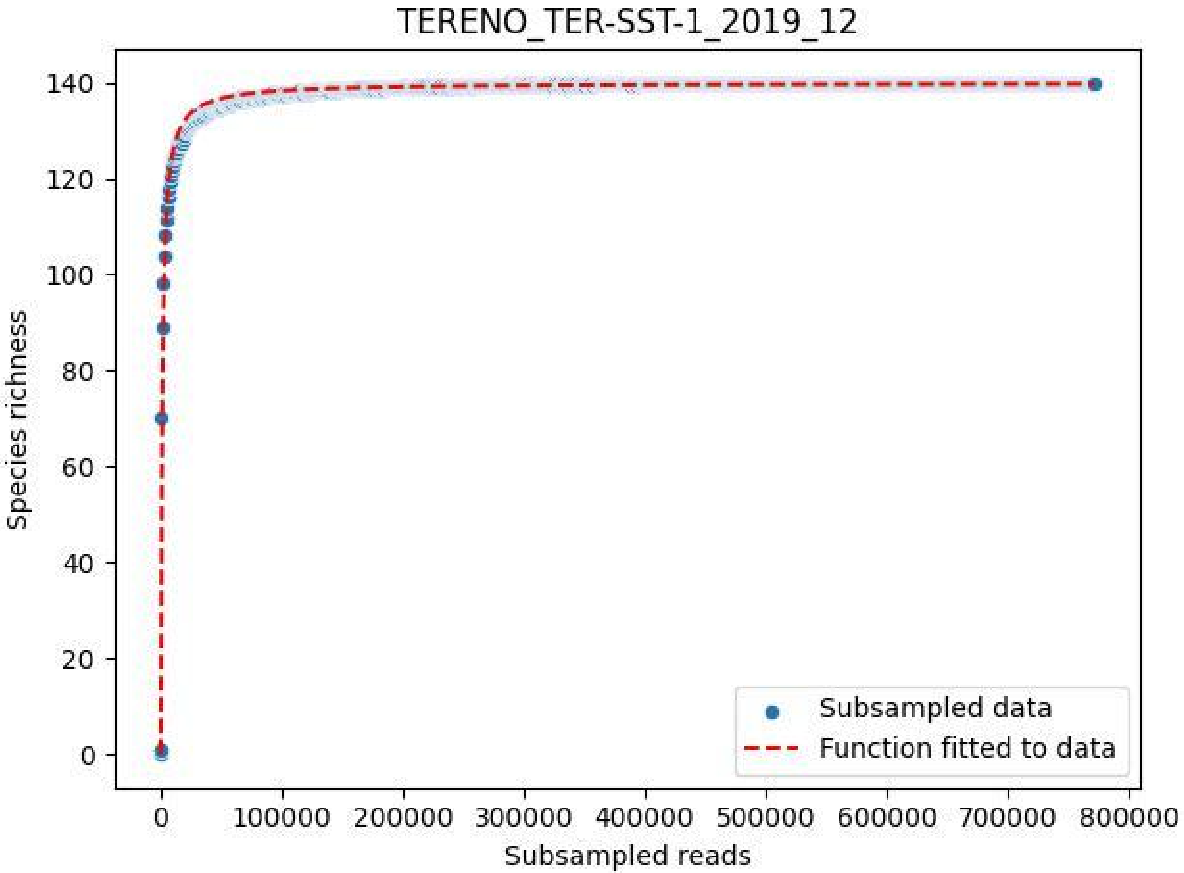
TERENO_TER-SST-1_2019_12
Species richness
Subsampled reads
0
20
40
60
80
100
120
140
0
100000
200000
300000
400000
500000
600000
700000
800000
Subsampled data
Function fitted to data

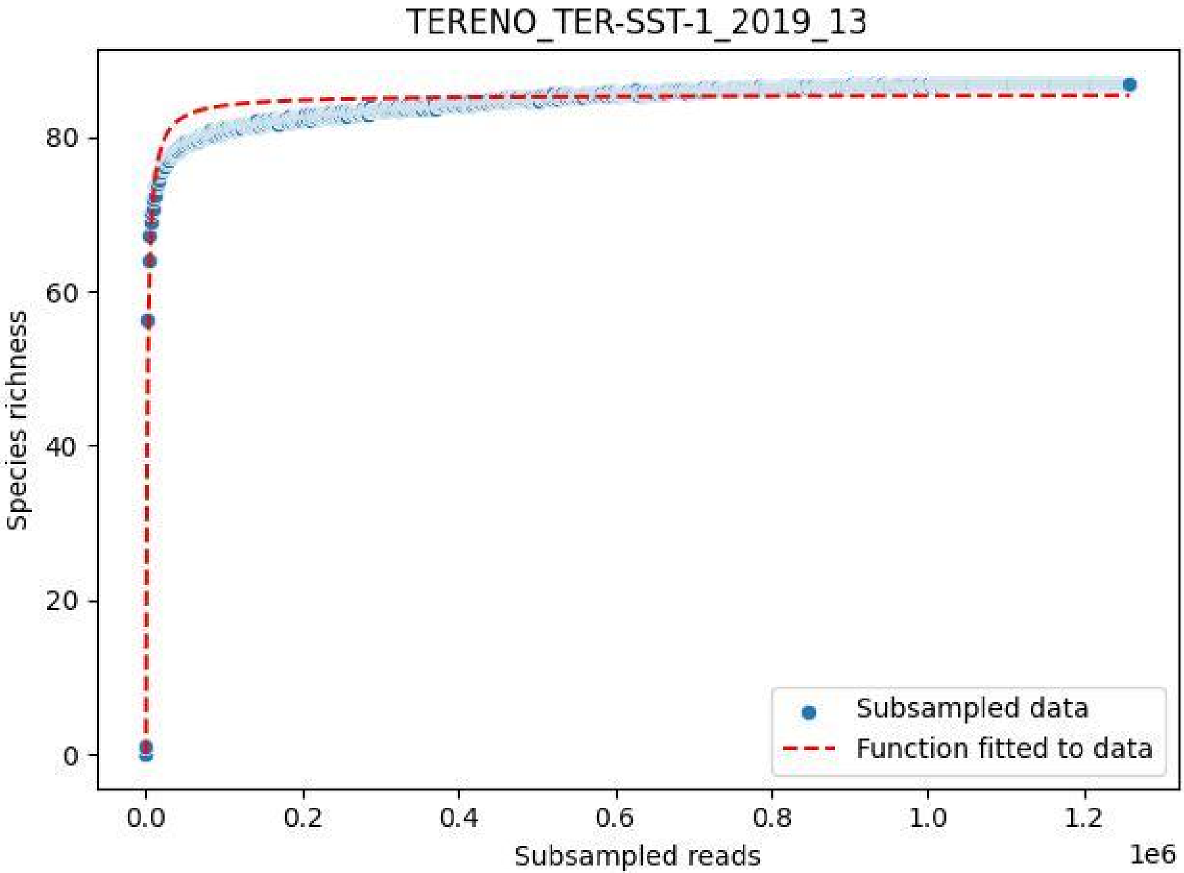
TERENO_TER-SST-1_2019_13
Species richness
Subsampled reads
1e6
Subsampled data
Function fitted to data
0
20
40
60
80
0.0
0.2
0.4
0.6
0.8
1.0
1.2

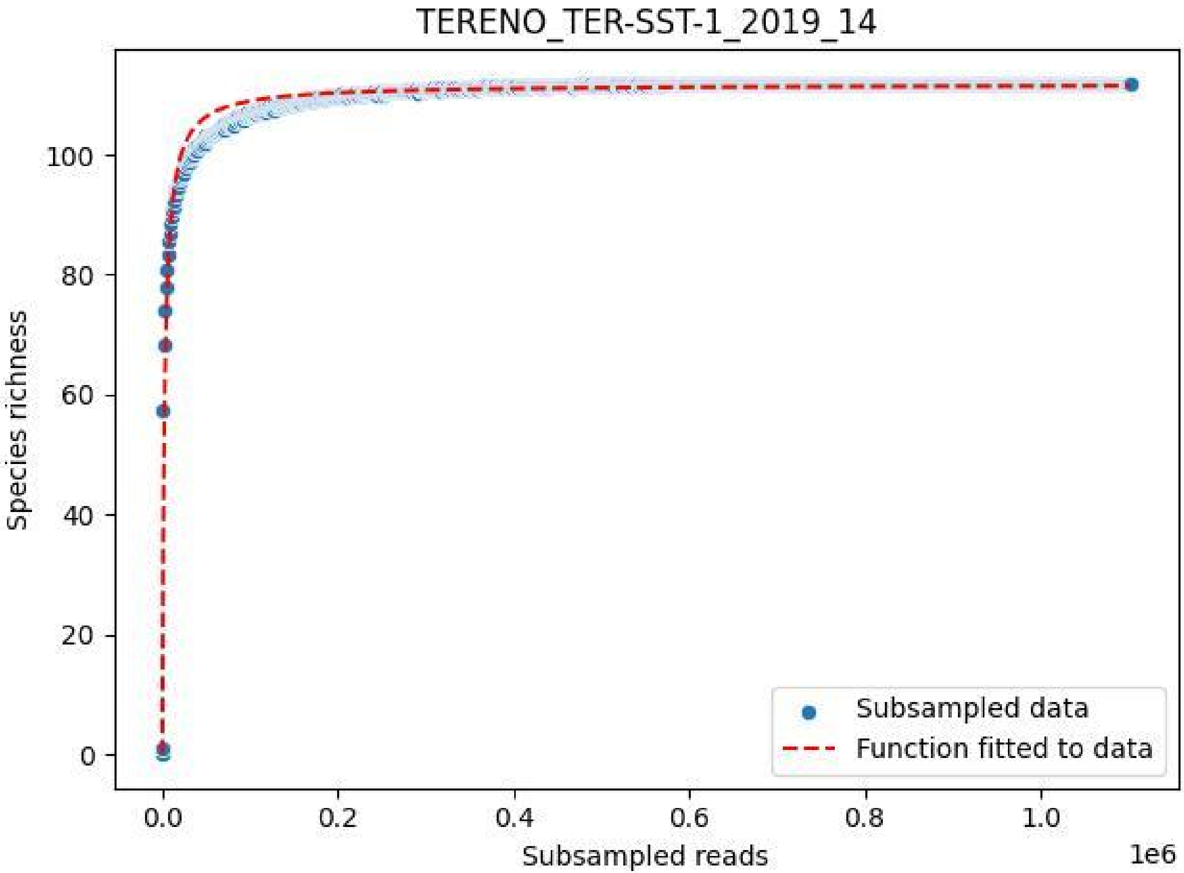

TERENO_TER-SST-1_2019_14
Species richness
Subsampled reads
1e6
0
20
40
60
80
100
0.0
0.2
0.4
0.6
0.8
1.0
Subsampled data
Function fitted to data

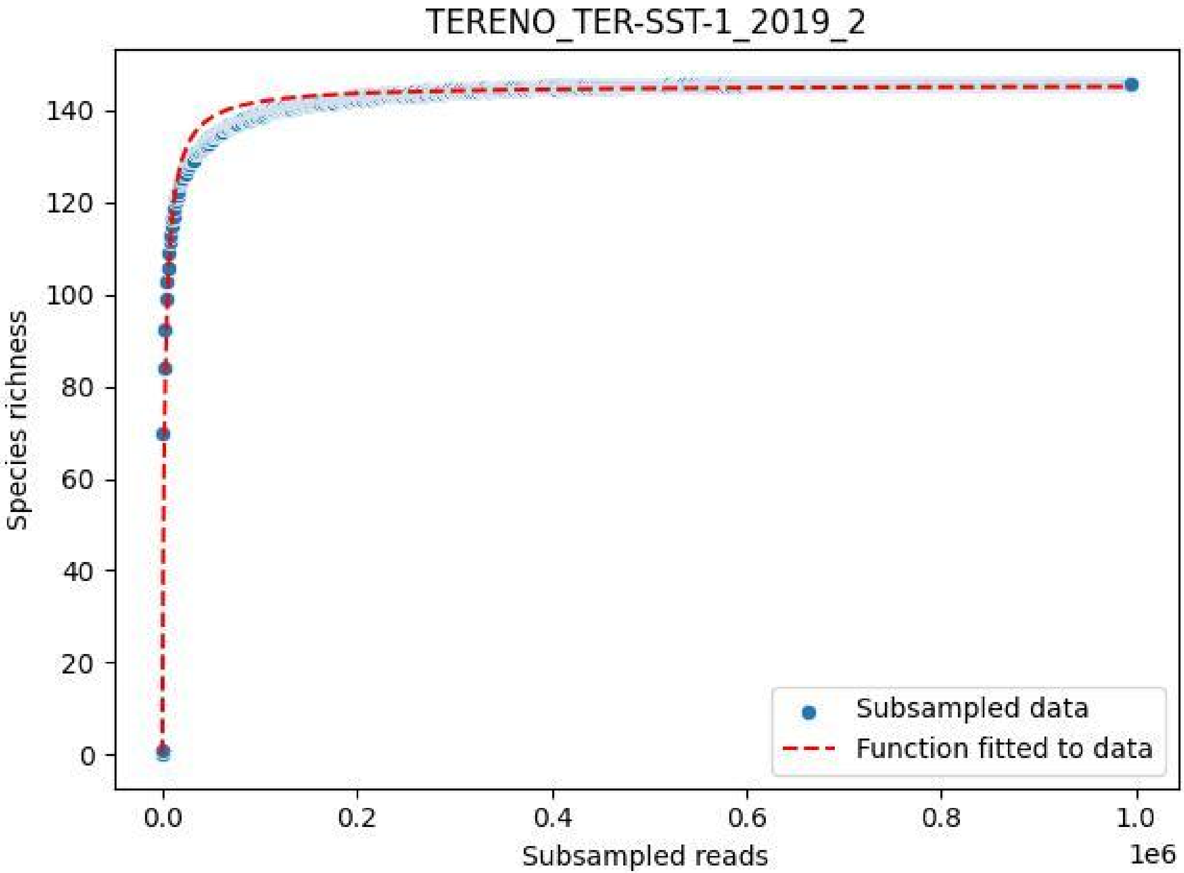

TERENO_TER-SST-1_2019_2
Species richness
Subsampled reads
1e6
0
20
40
60
80
100
120
140
0.0
0.2
0.4
0.6
0.8
1.0
Subsampled data
Function fitted to data

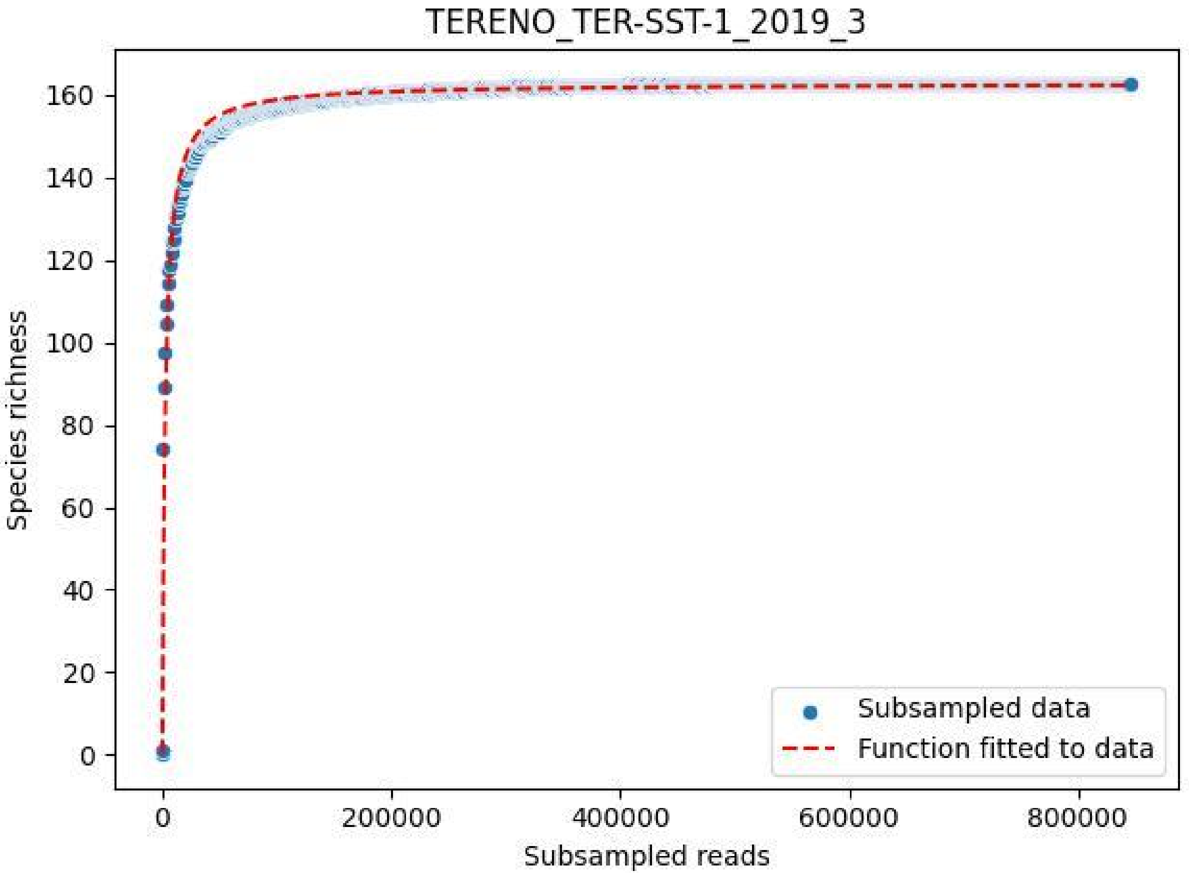
TERENO_TER-SST-1_2019_3
Species richness
Subsampled reads
0
20
40
60
80
100
120
140
160
0
200000
400000
600000
800000
Subsampled data
Function fitted to data

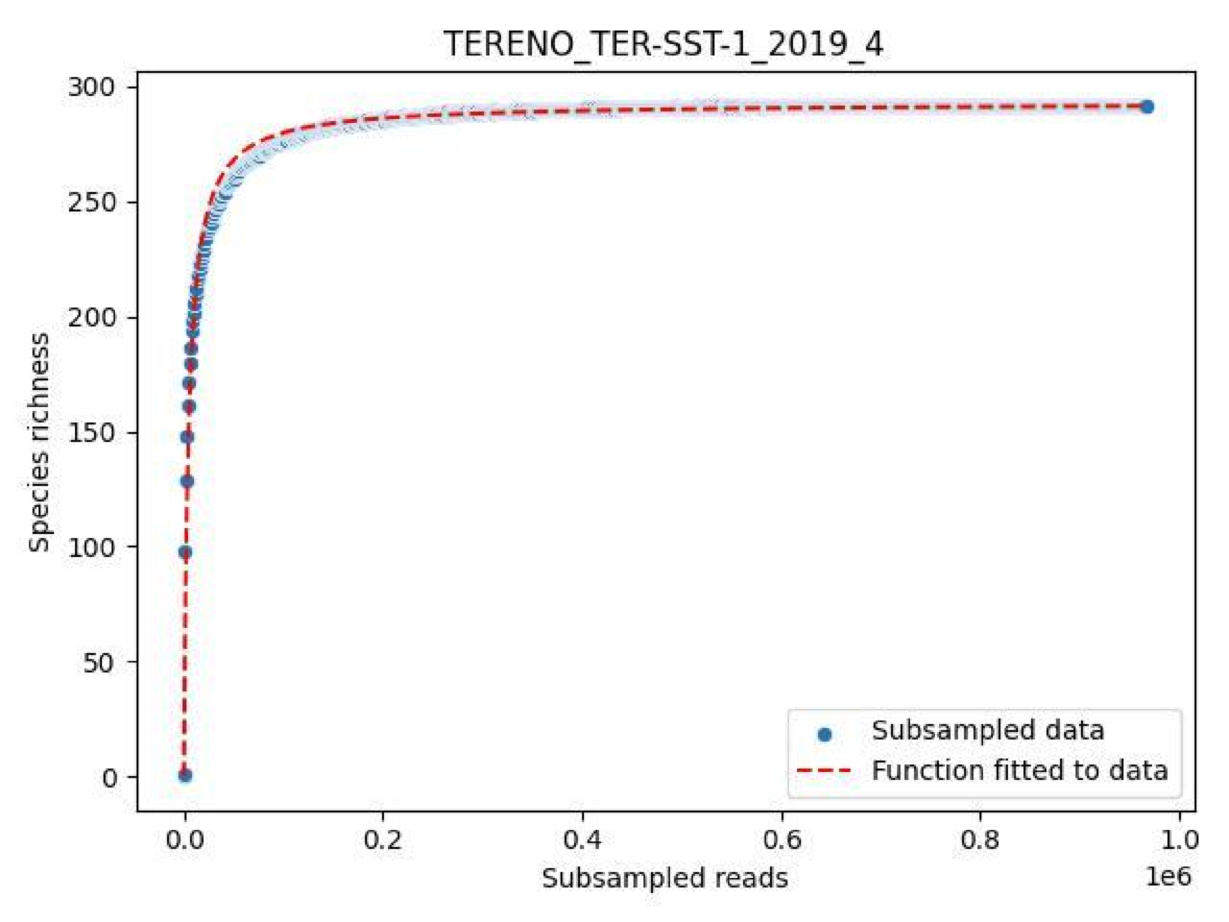

TERENO_TER-SST-1_2019_4
Species richness
Subsampled reads
1e6
0
50
100
150
200
250
300
0.0
0.2
0.4
0.6
0.8
1.0
Subsampled data
Function fitted to data

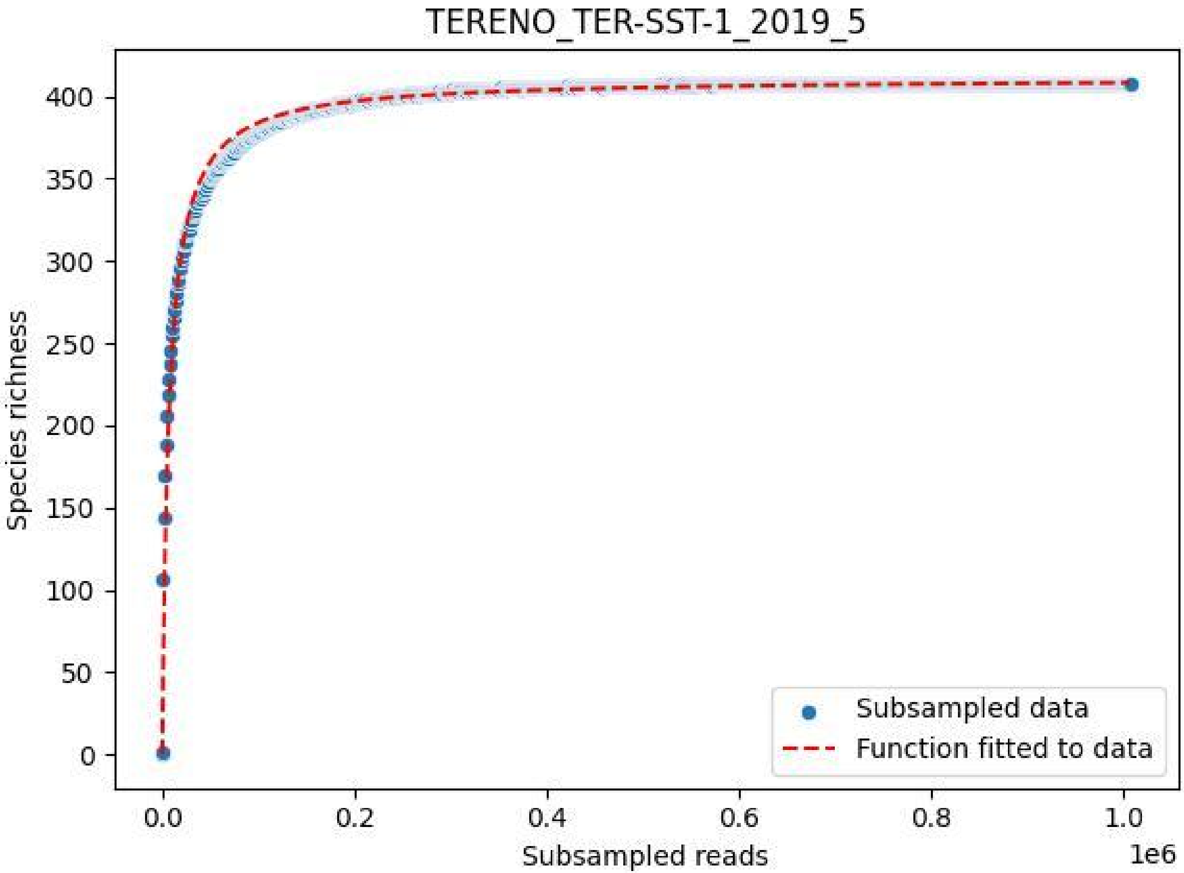

TERENO_TER-SST-1_2019_5
Species richness
0
50
100
150
200
250
300
350
400
0.0
0.2
0.4
0.6
0.8
1.0
1e6
Subsampled reads
Subsampled data
Function fitted to data

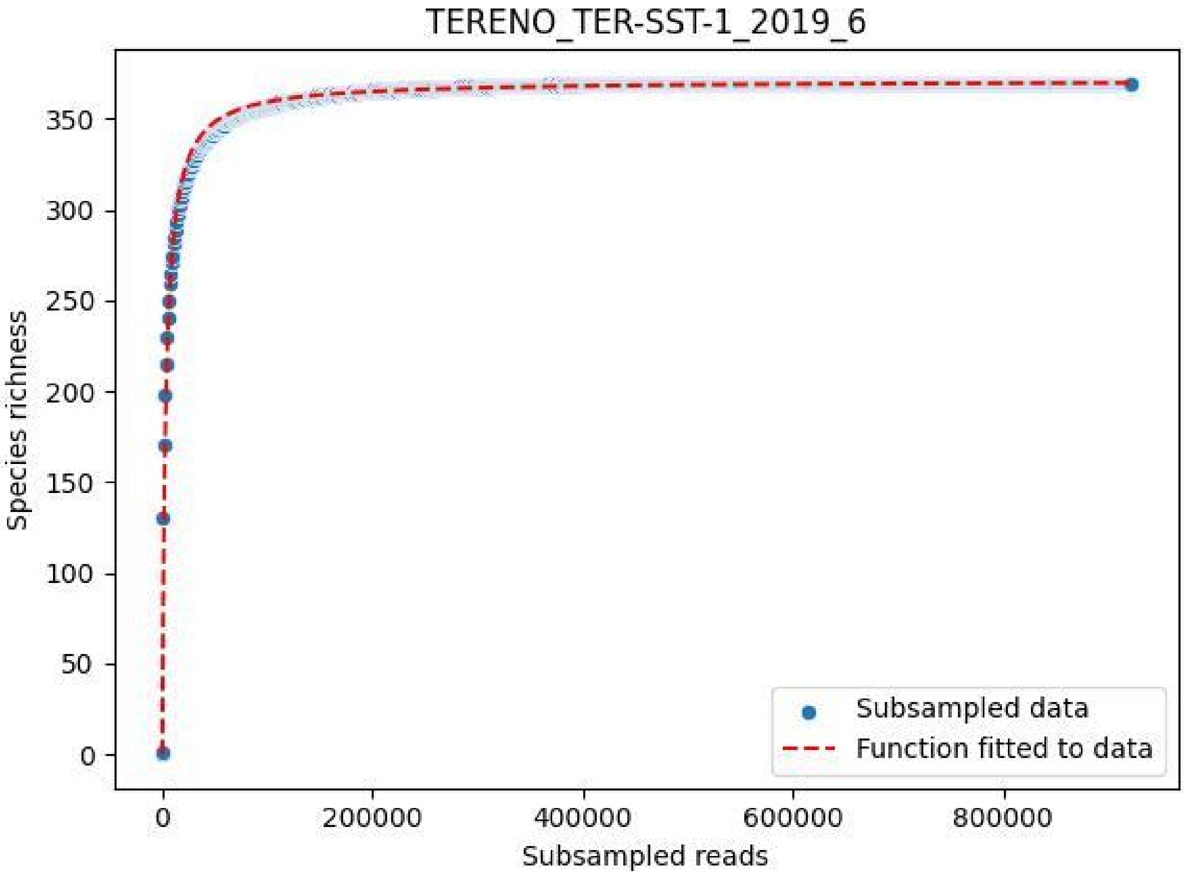

TERENO_TER-SST-1_2019_6
Species richness
Subsampled reads
0
50
100
150
200
250
300
350
0
200000
400000
600000
800000
Subsampled data
Function fitted to data

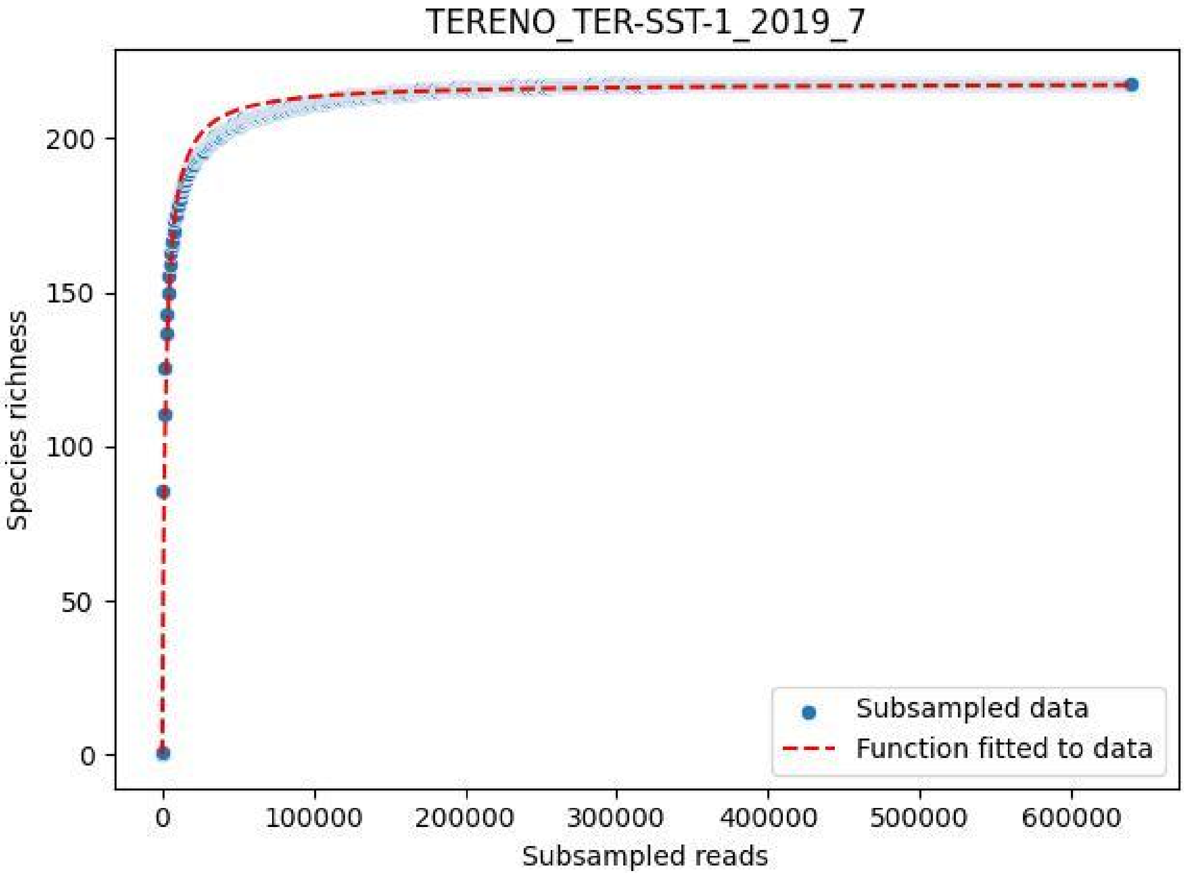

TERENO_TER-SST-1_2019_7
Species richness
Subsampled reads
0
50
100
150
200
0
100000
200000
300000
400000
500000
600000
Subsampled data
Function fitted to data

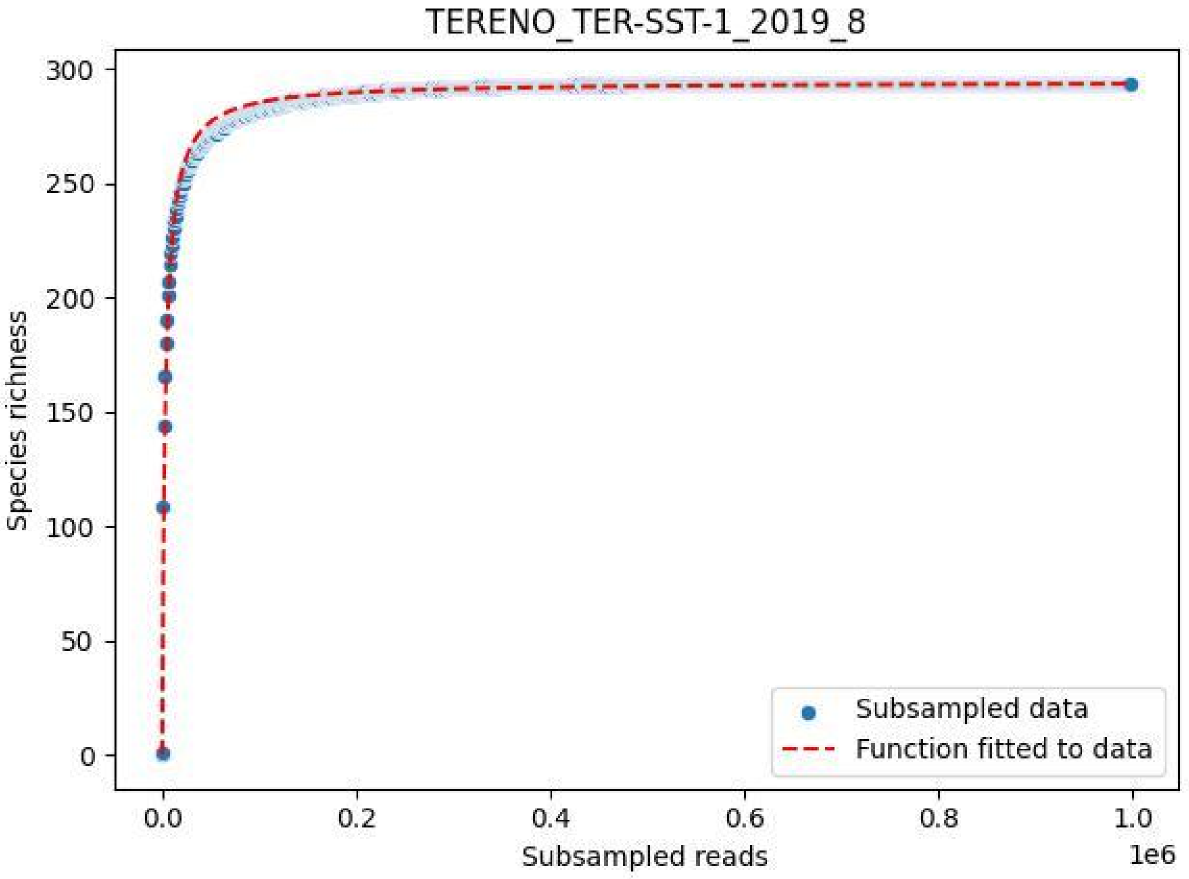

TERENO_TER-SST-1_2019_8
Species richness
Subsampled reads
1e6
0.0
0.2
0.4
0.6
0.8
1.0
0
50
100
150
200
250
300
Subsampled data
Function fitted to data

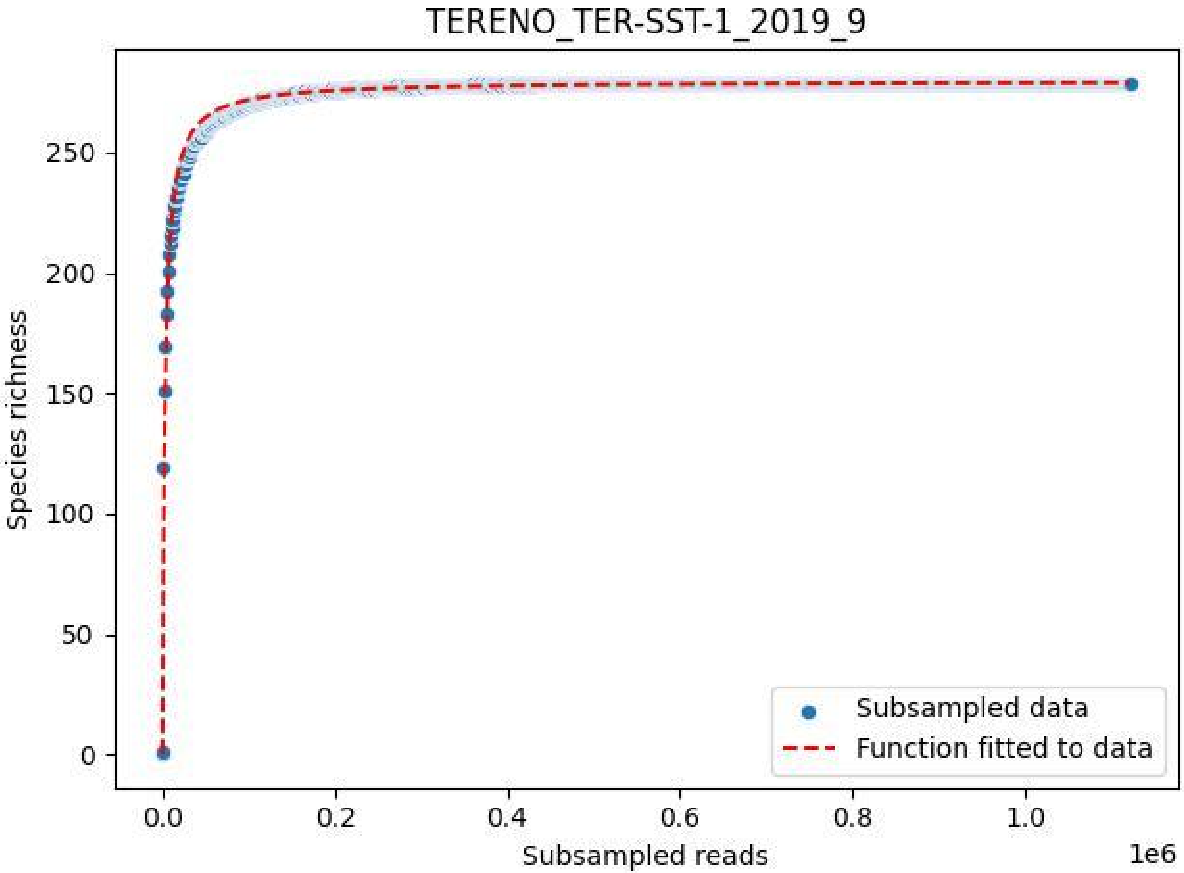

TERENO_TER-SST-1_2019_9
Species richness
0
50
100
150
200
250
0.0
0.2
0.4
0.6
0.8
1.0
1e6
Subsampled reads
Subsampled data
Function fitted to data

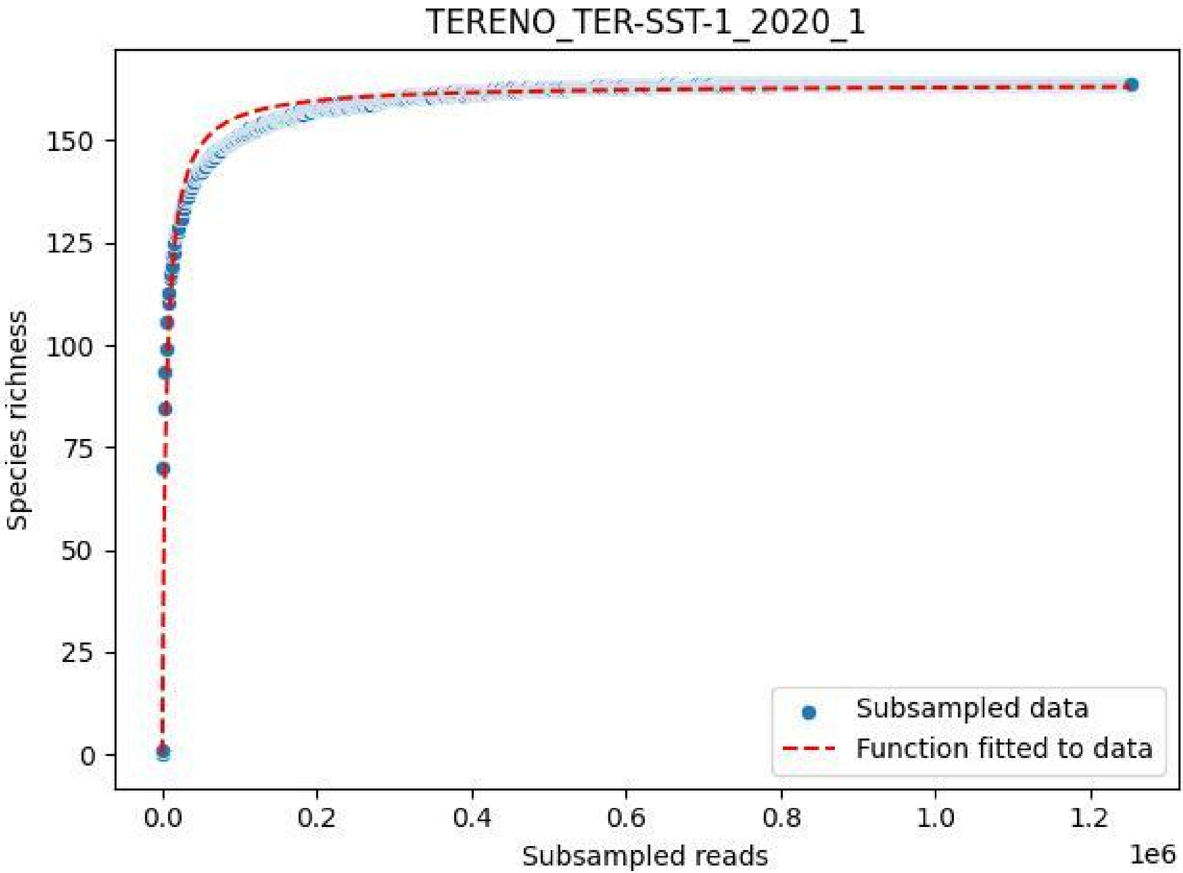

TERENO_TER-SST-1_2020_1
Species richness
Subsampled reads
1e6
0
25
50
75
100
125
150
0.0
0.2
0.4
0.6
0.8
1.0
1.2
Subsampled data
Function fitted to data

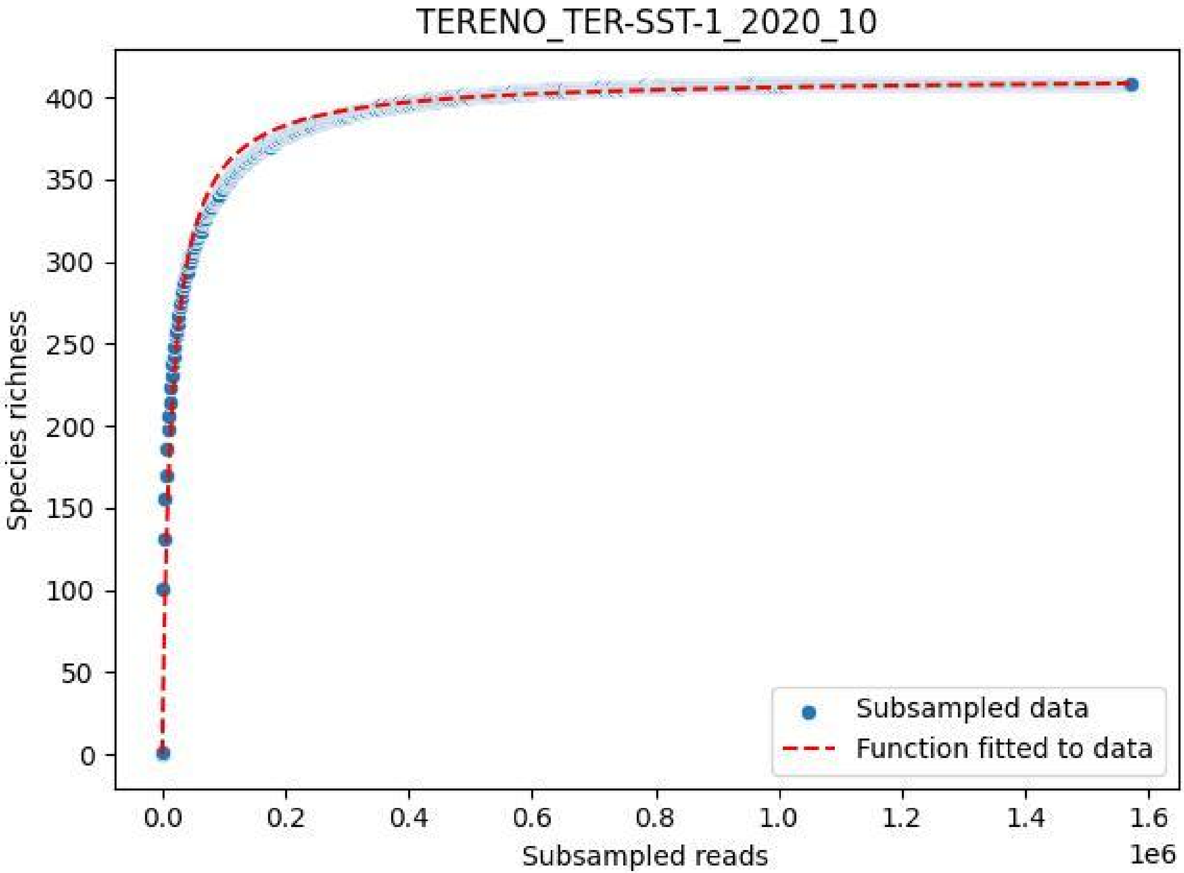

TERENO_TER-SST-1_2020_10
Species richness
Subsampled reads
1e6
Subsampled data
Function fitted to data
0
50
100
150
200
250
300
350
400
0.0
0.2
0.4
0.6
0.8
1.0
1.2
1.4
1.6

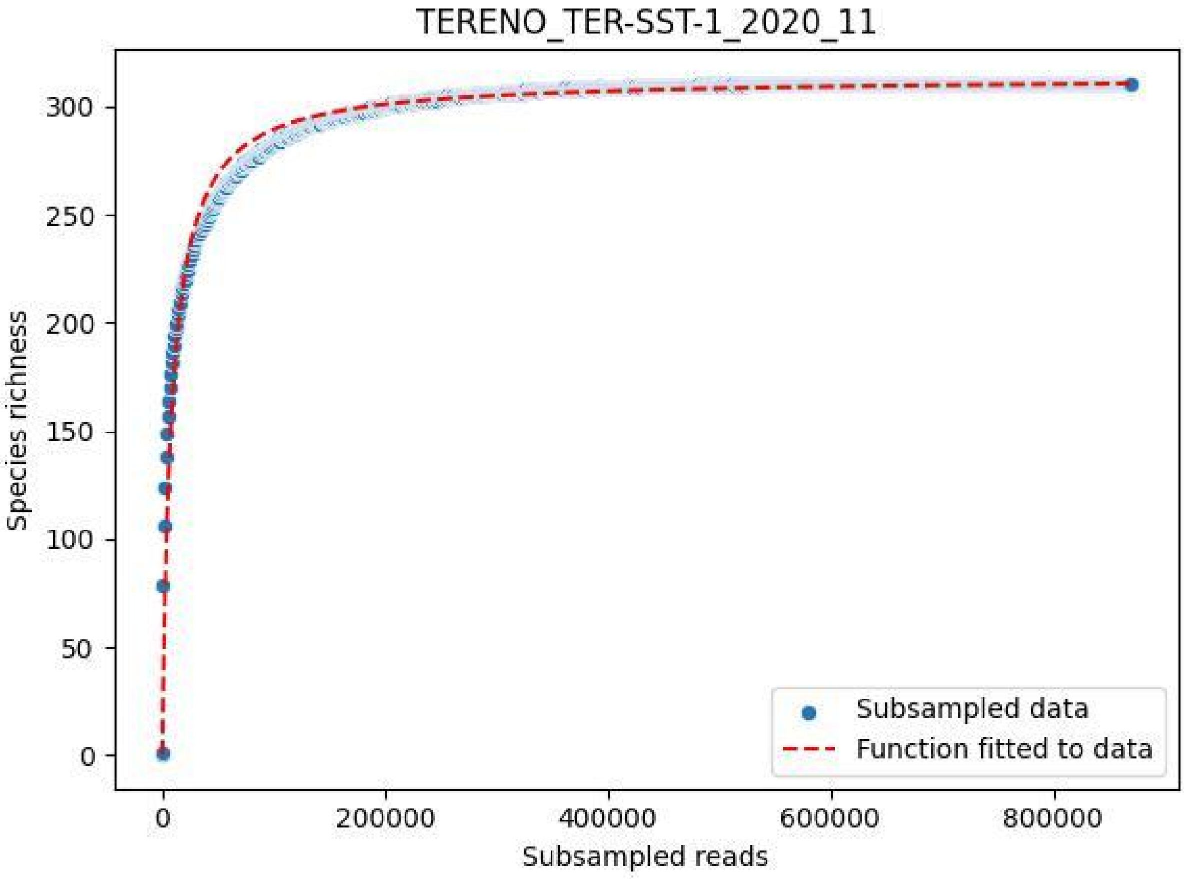

TERENO_TER-SST-1_2020_11
Species richness
Subsampled reads
0
50
100
150
200
250
300
0
200000
400000
600000
800000
Subsampled data
Function fitted to data

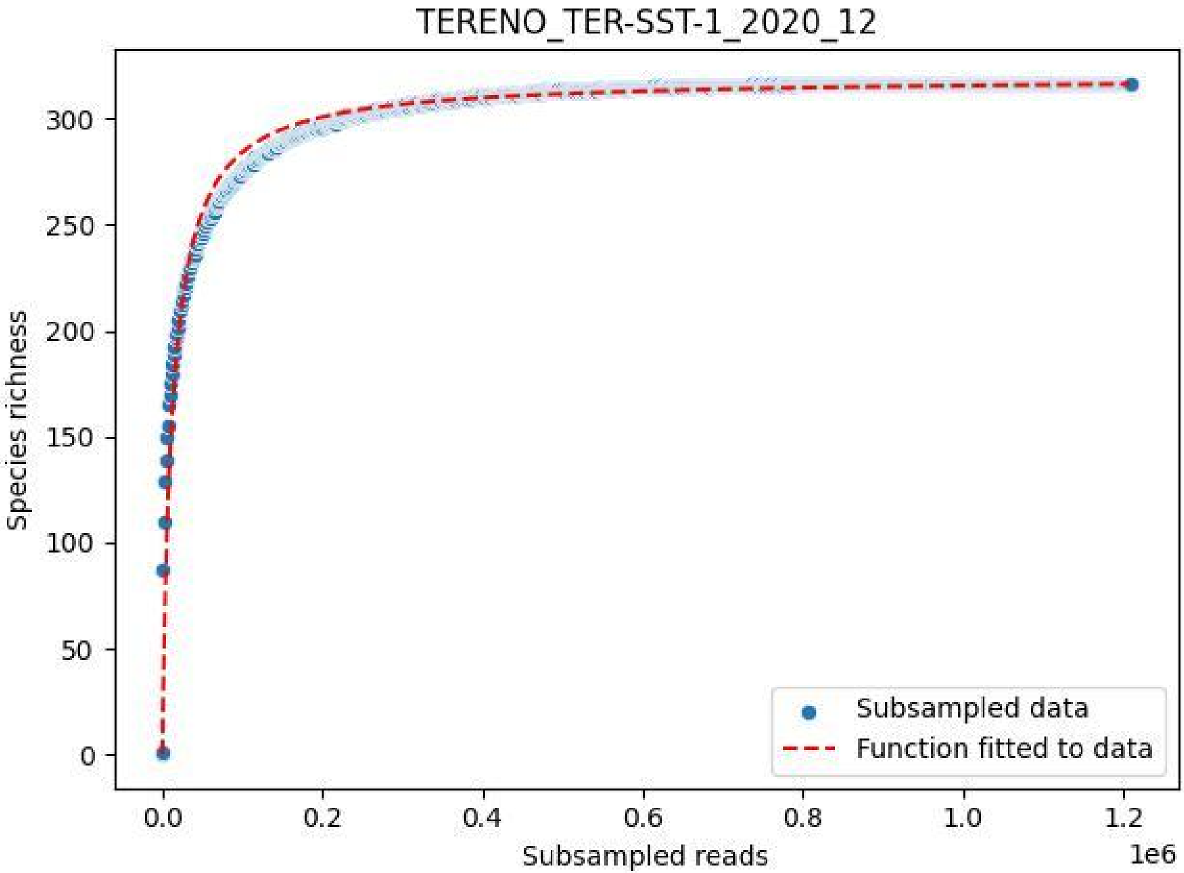
TERENO_TER-SST-1_2020_12
Species richness
Subsampled reads
1e6
0
50
100
150
200
250
300
0.0
0.2
0.4
0.6
0.8
1.0
1.2
Subsampled data
Function fitted to data

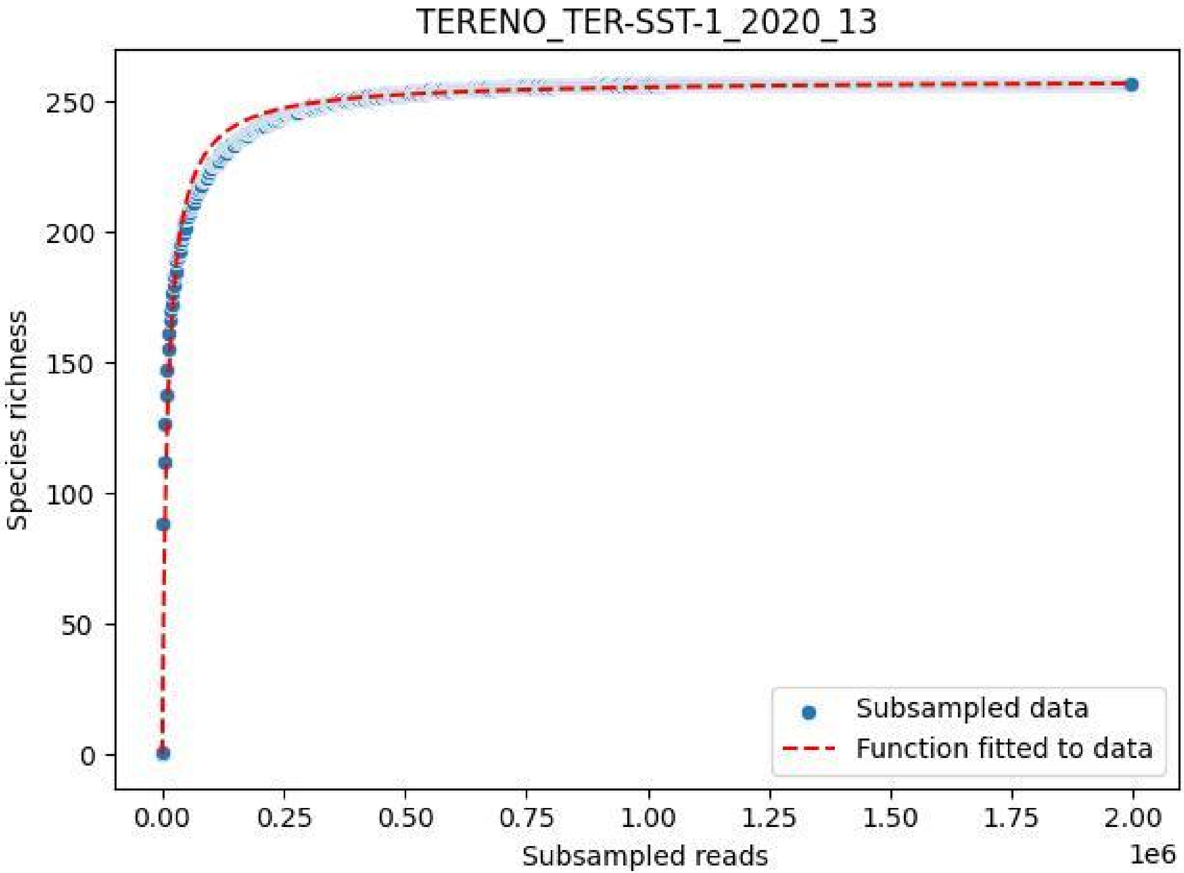

TERENO_TER-SST-1_2020_13
Species richness
Subsampled reads
1e6
0
50
100
150
200
250
0.00
0.25
0.50
0.75
1.00
1.25
1.50
1.75
2.00
Subsampled data
Function fitted to data

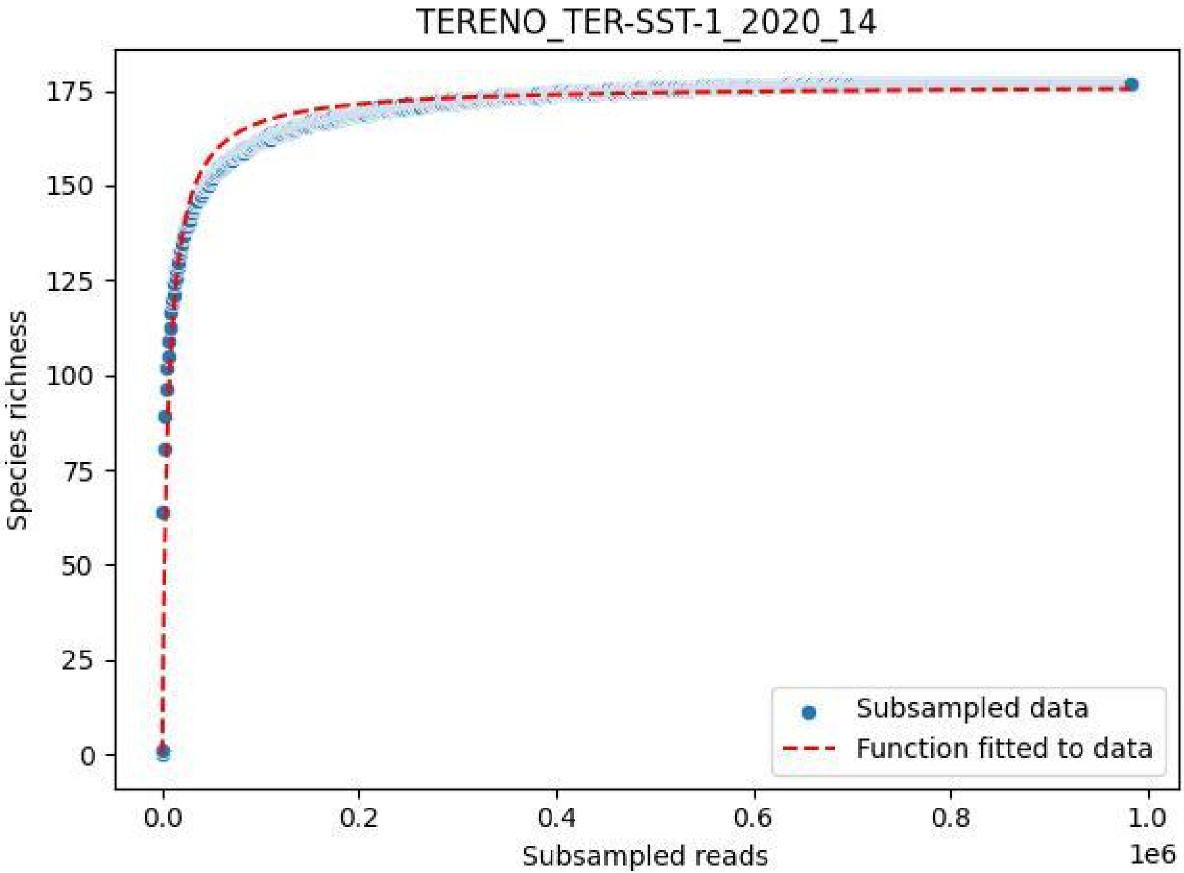

TERENO_TER-SST-1_2020_14
Species richness
0
25
50
75
100
125
150
175
Subsampled reads
0.0
0.2
0.4
0.6
0.8
1.0
1e6
Subsampled data
Function fitted to data

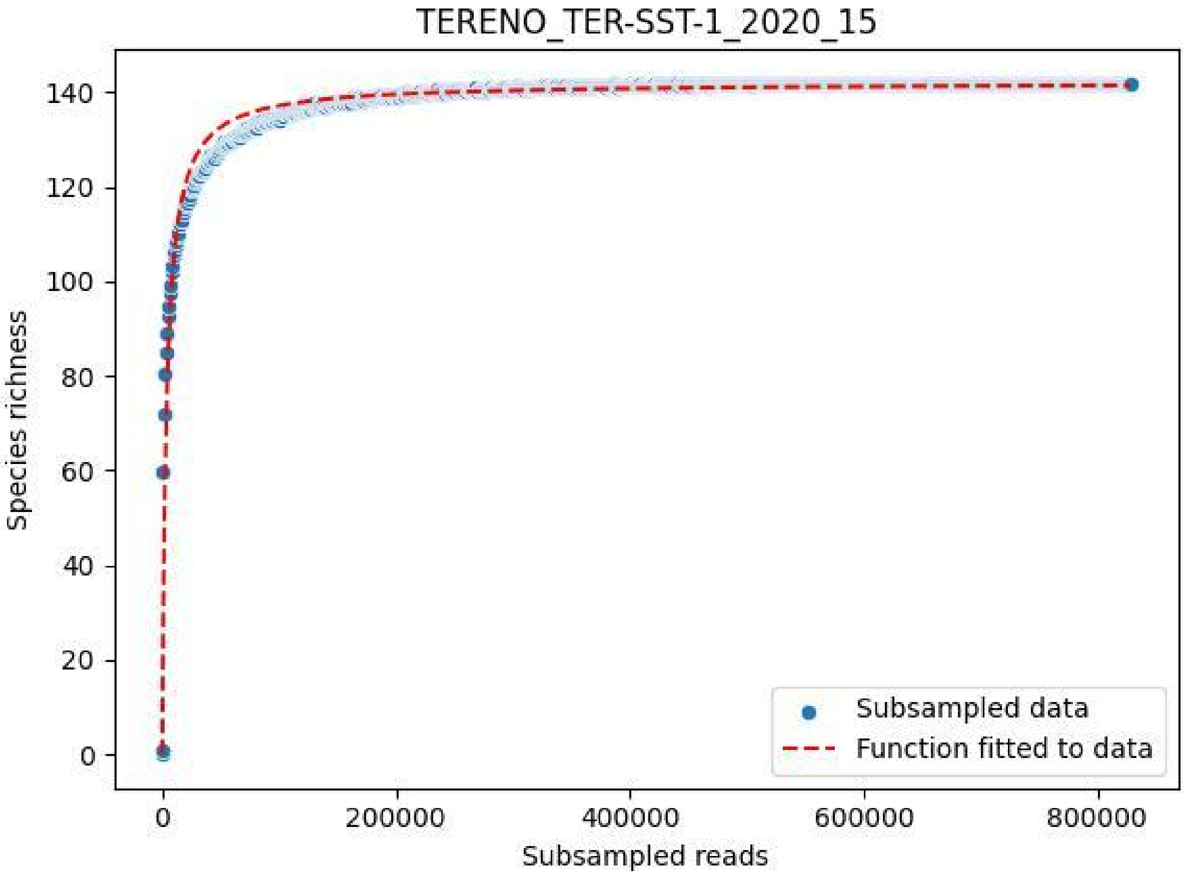
TERENO_TER-SST-1_2020_15
Species richness
Subsampled reads
0
20
40
60
80
100
120
140
0
200000
400000
600000
800000
Subsampled data
Function fitted to data

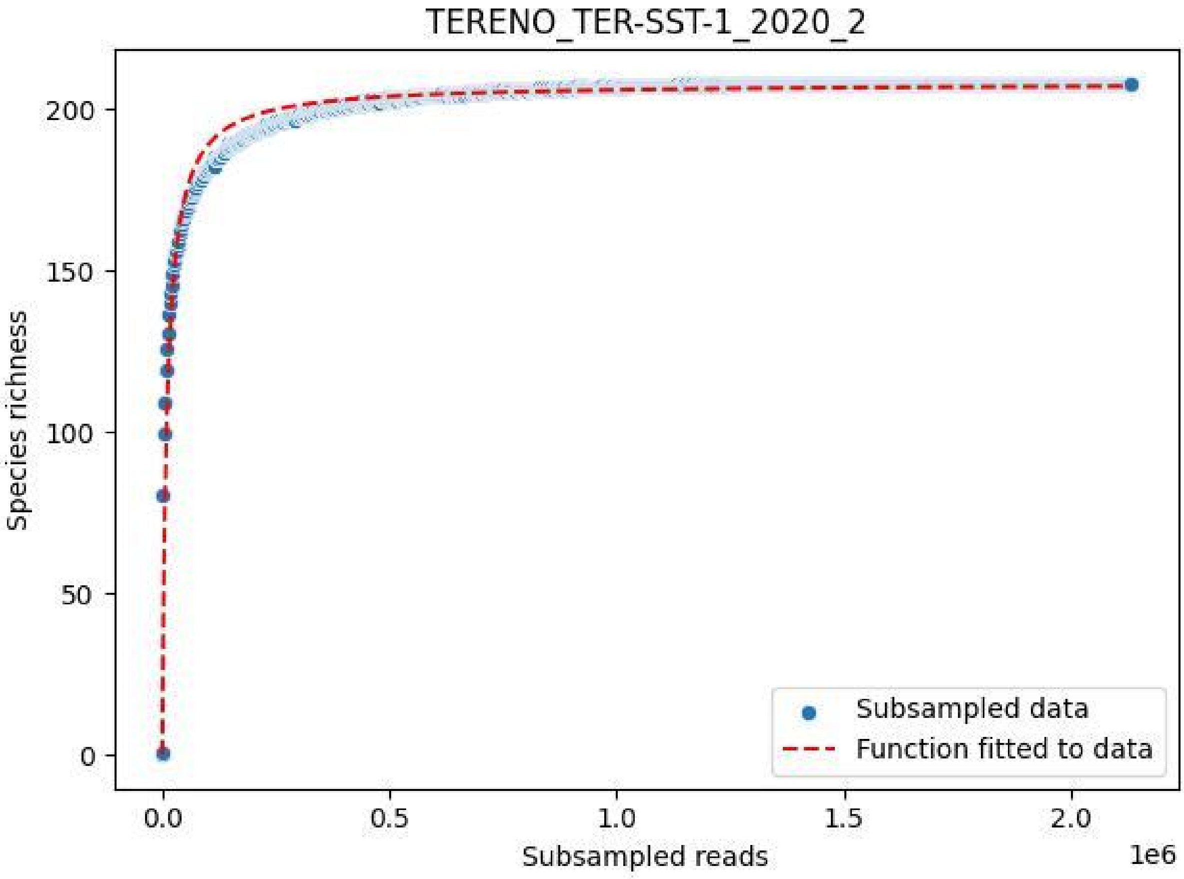
TERENO_TER-SST-1_2020_2
Species richness
Subsampled reads
1e6
0
50
100
150
200
0.0
0.5
1.0
1.5
2.0
Subsampled data
Function fitted to data

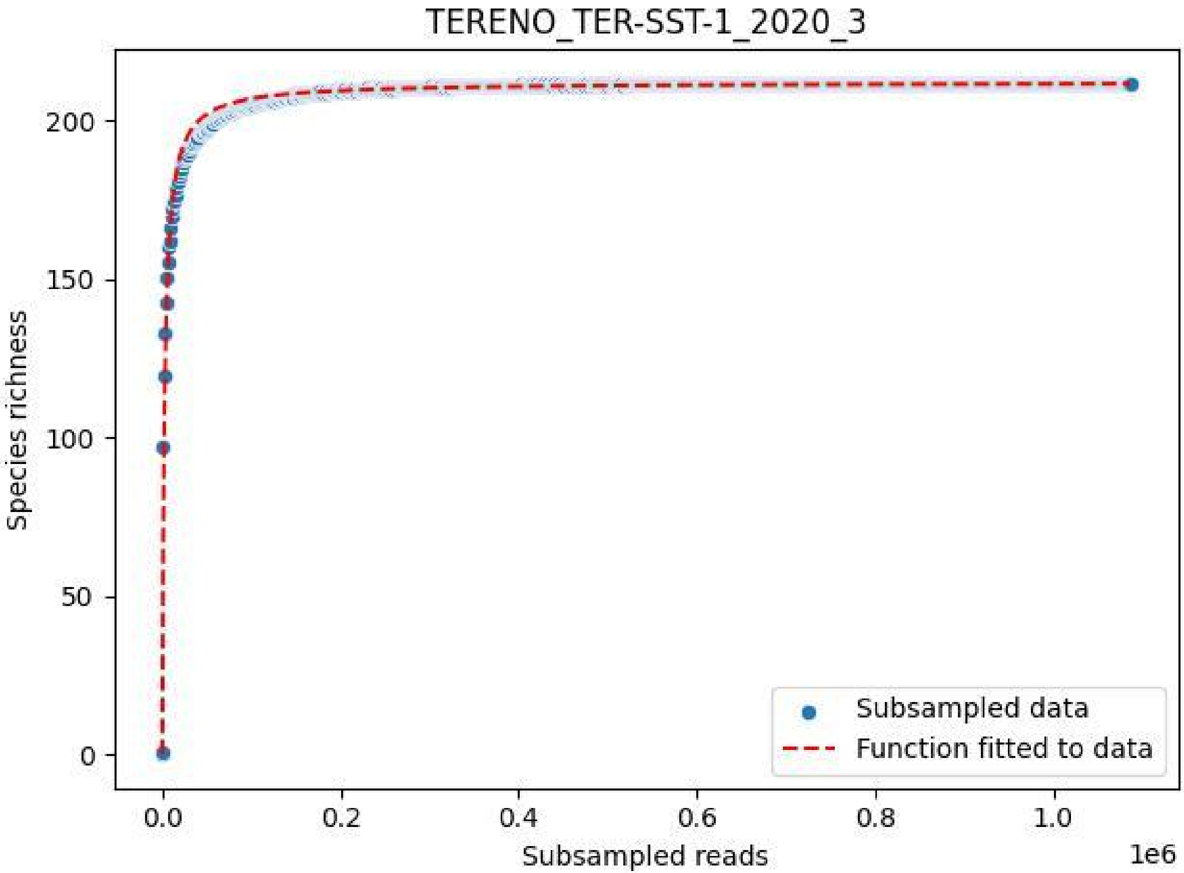

TERENO_TER-SST-1_2020_3
Species richness
Subsampled reads
1e6
0
50
100
150
200
0.0
0.2
0.4
0.6
0.8
1.0
Subsampled data
Function fitted to data

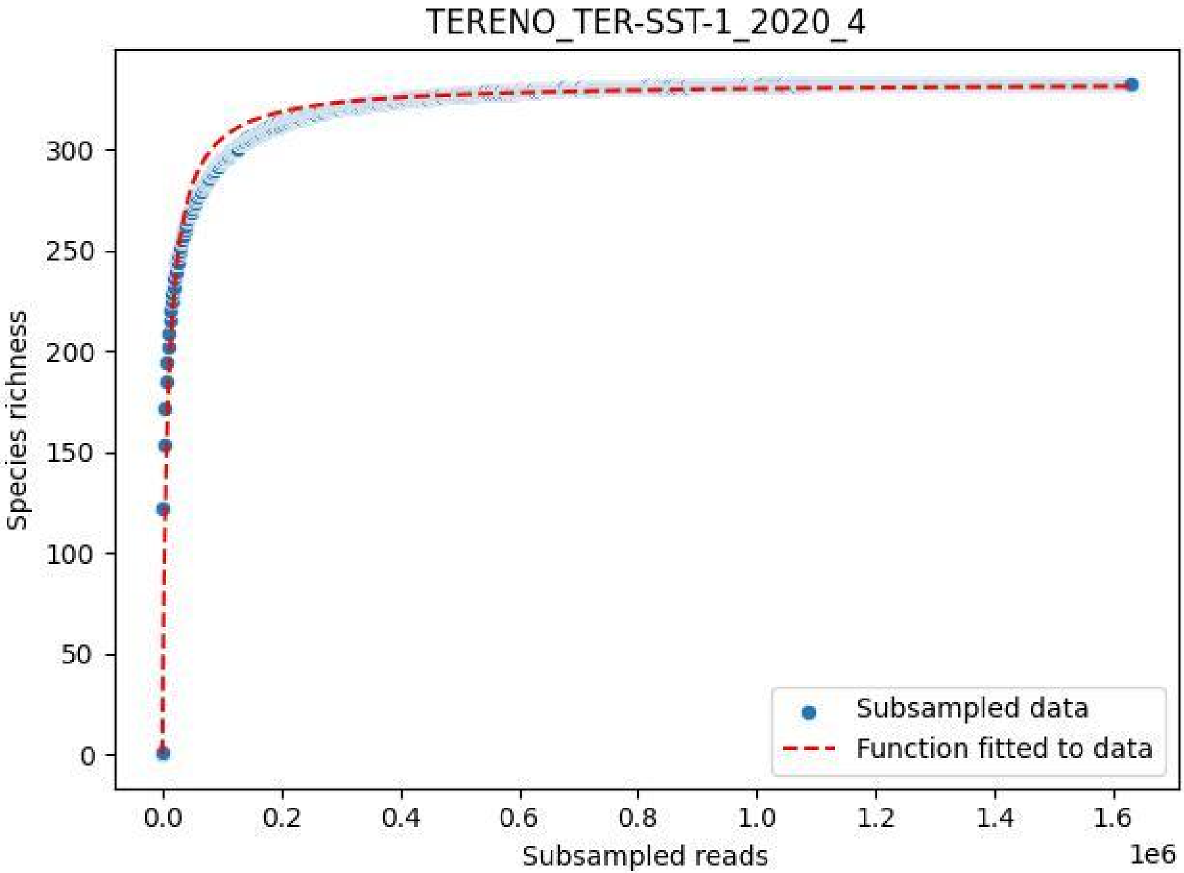
TERENO_TER-SST-1_2020_4
Species richness
Subsampled reads
1e6
0
50
100
150
200
250
300
0.0
0.2
0.4
0.6
0.8
1.0
1.2
1.4
1.6
Subsampled data
Function fitted to data

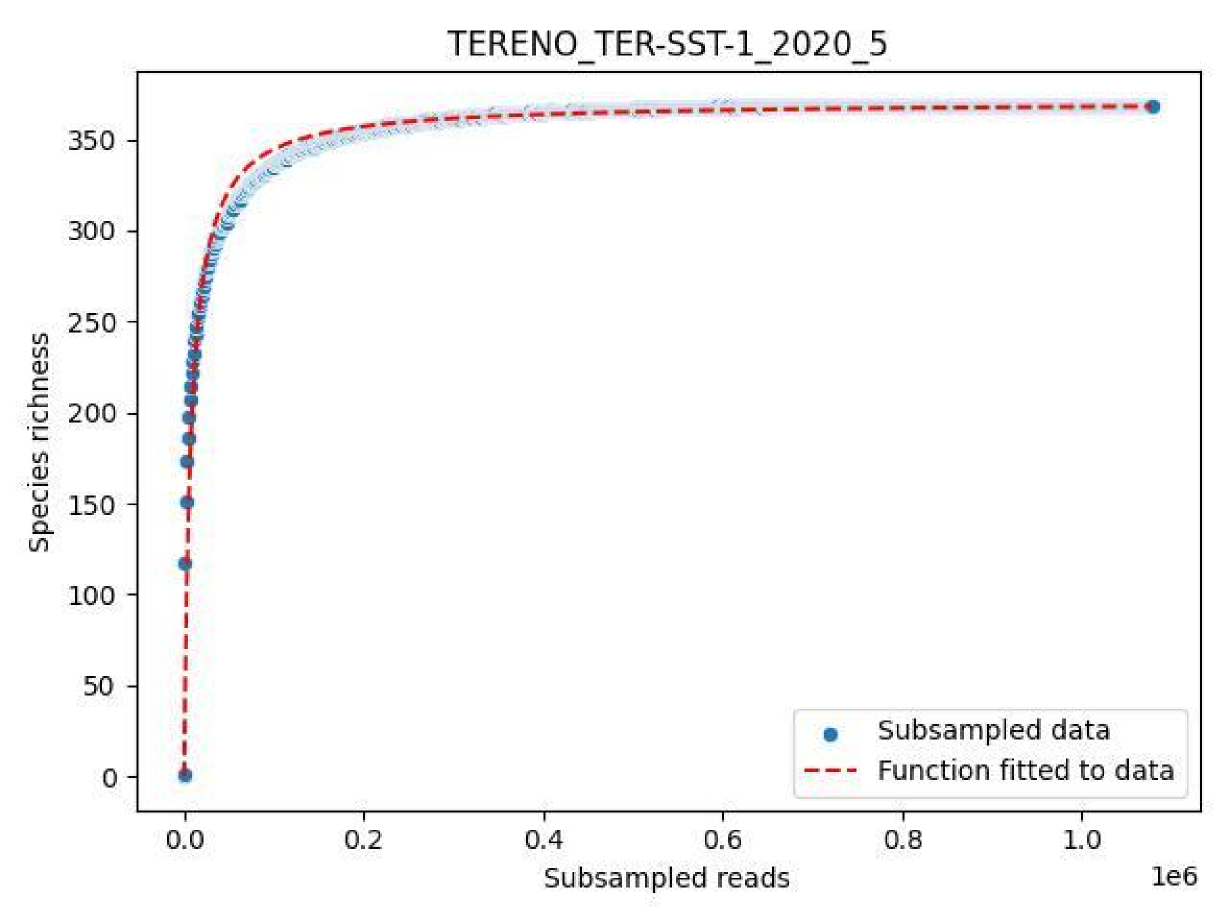
TERENO_TER-SST-1_2020_5
Species richness
Subsampled reads
1e6
0
50
100
150
200
250
300
350
0.0
0.2
0.4
0.6
0.8
1.0
Subsampled data
Function fitted to data

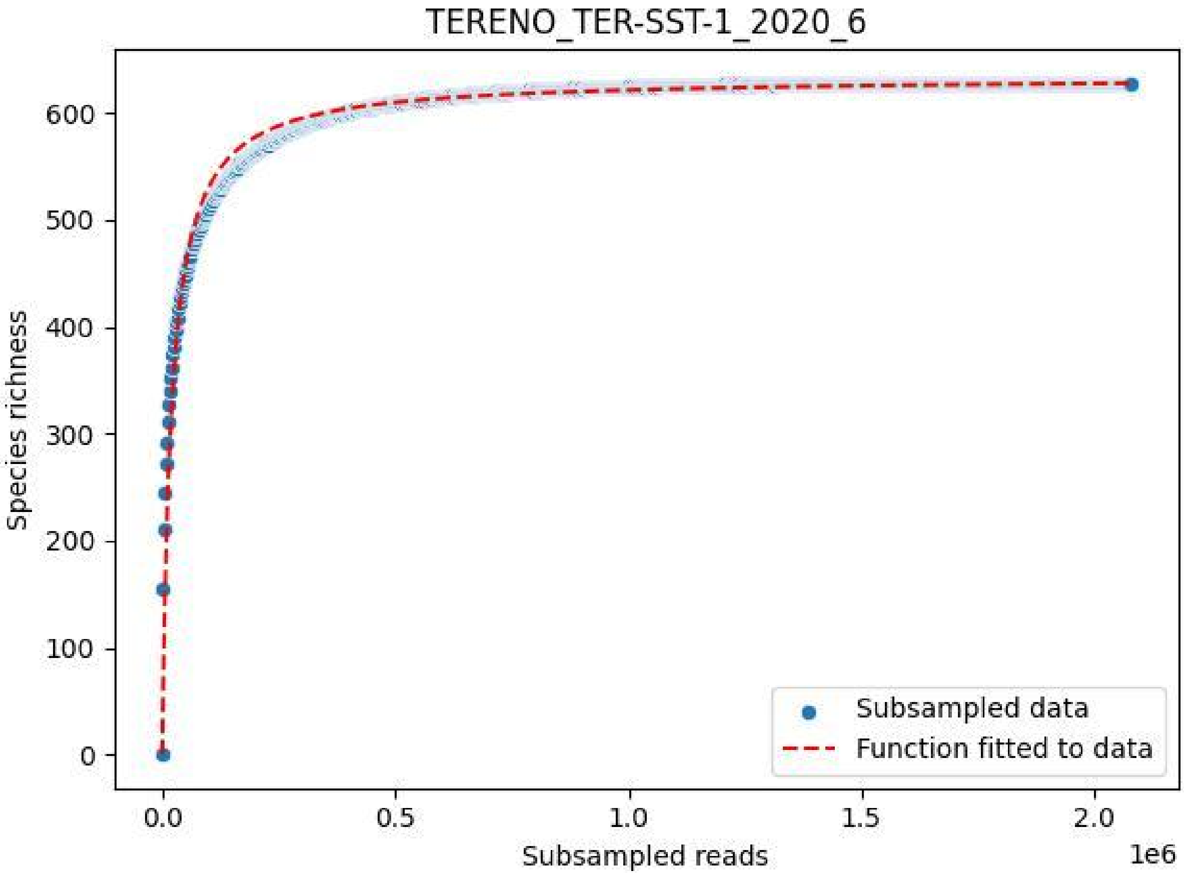

TERENO_TER-SST-1_2020_6
Species richness
Subsampled reads
1e6
Subsampled data
Function fitted to data
0
100
200
300
400
500
600
0.0
0.5
1.0
1.5
2.0

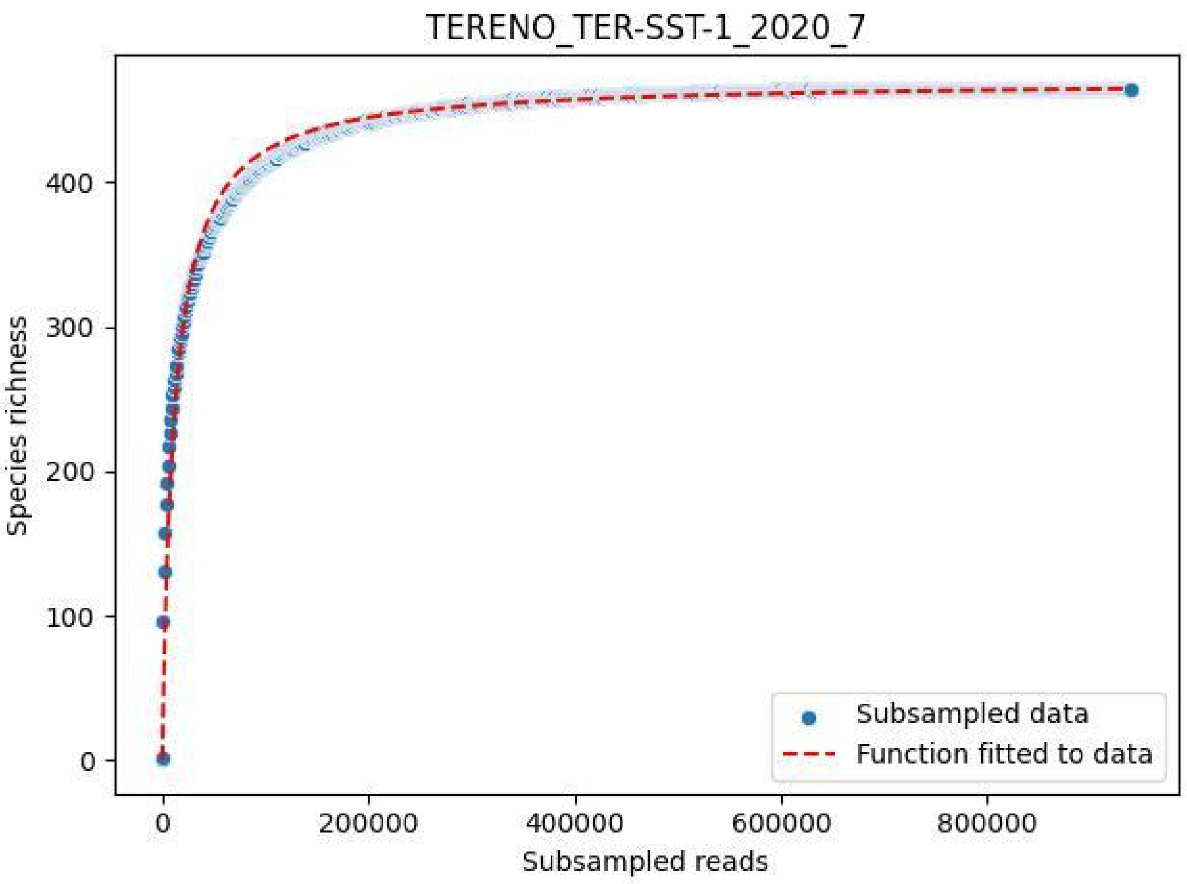

TERENO_TER-SST-1_2020_7
Species richness
Subsampled reads
0
100
200
300
400
0
200000
400000
600000
800000
Subsampled data
Function fitted to data

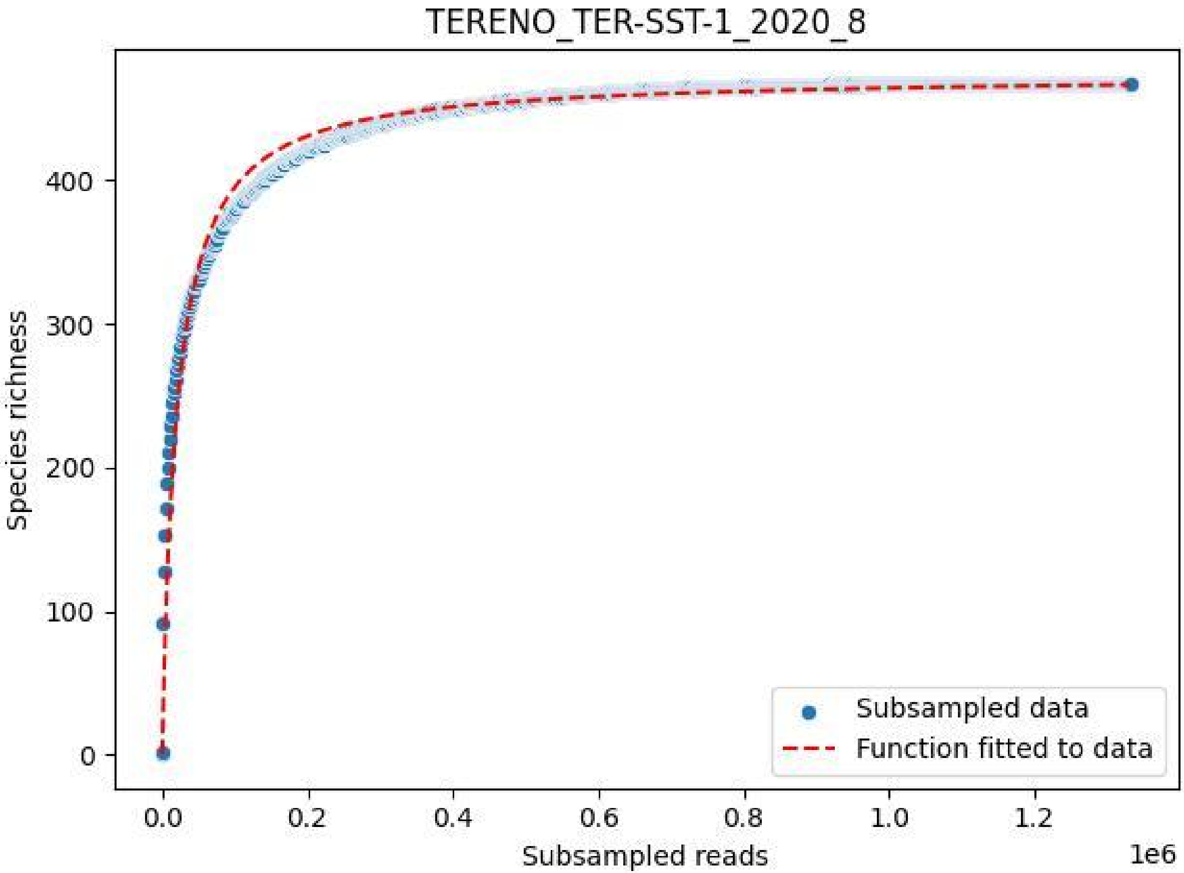
TERENO_TER-SST-1_2020_8
Species richness
Subsampled reads
1e6
0
100
200
300
400
0.0
0.2
0.4
0.6
0.8
1.0
1.2
Subsampled data
Function fitted to data

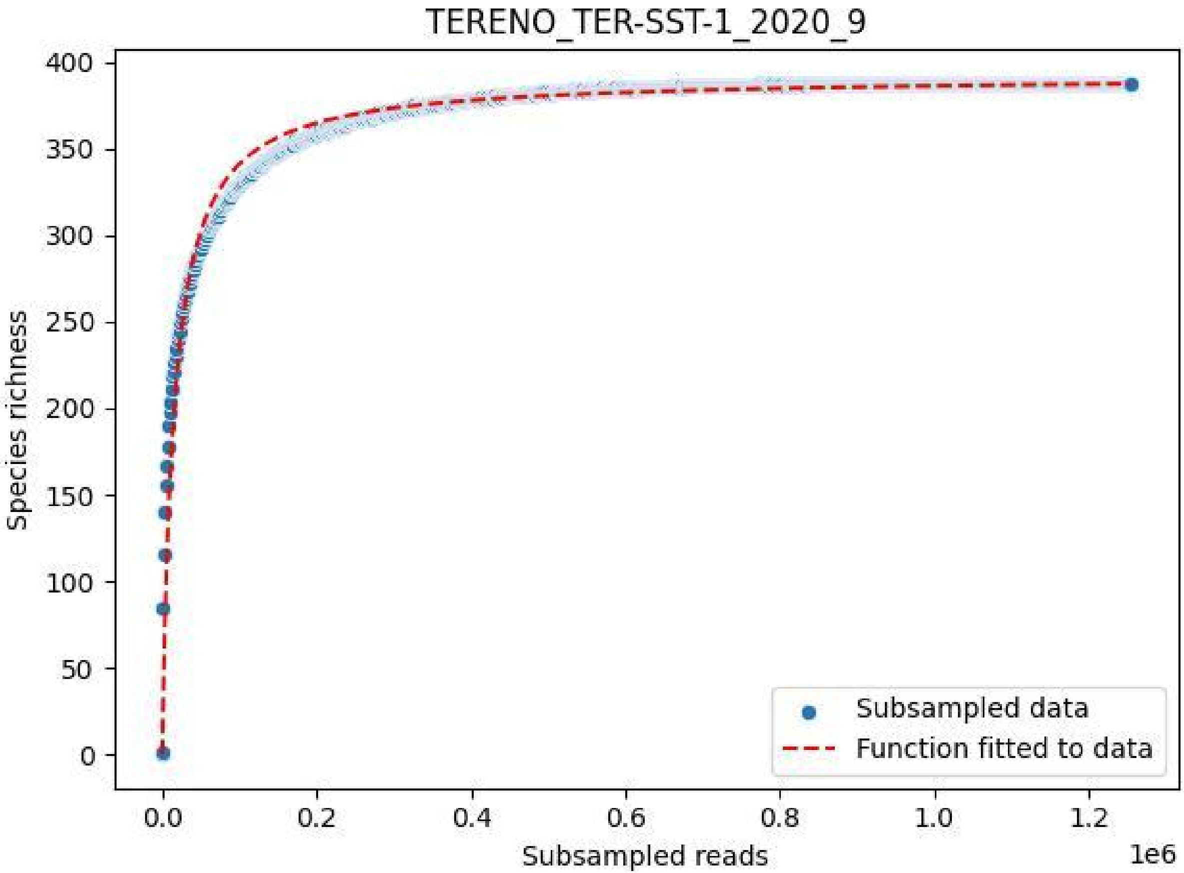

TERENO_TER-SST-1_2020_9
Species richness
Subsampled reads
1e6
0
50
100
150
200
250
300
350
400
0.0
0.2
0.4
0.6
0.8
1.0
1.2
Subsampled data
Function fitted to data

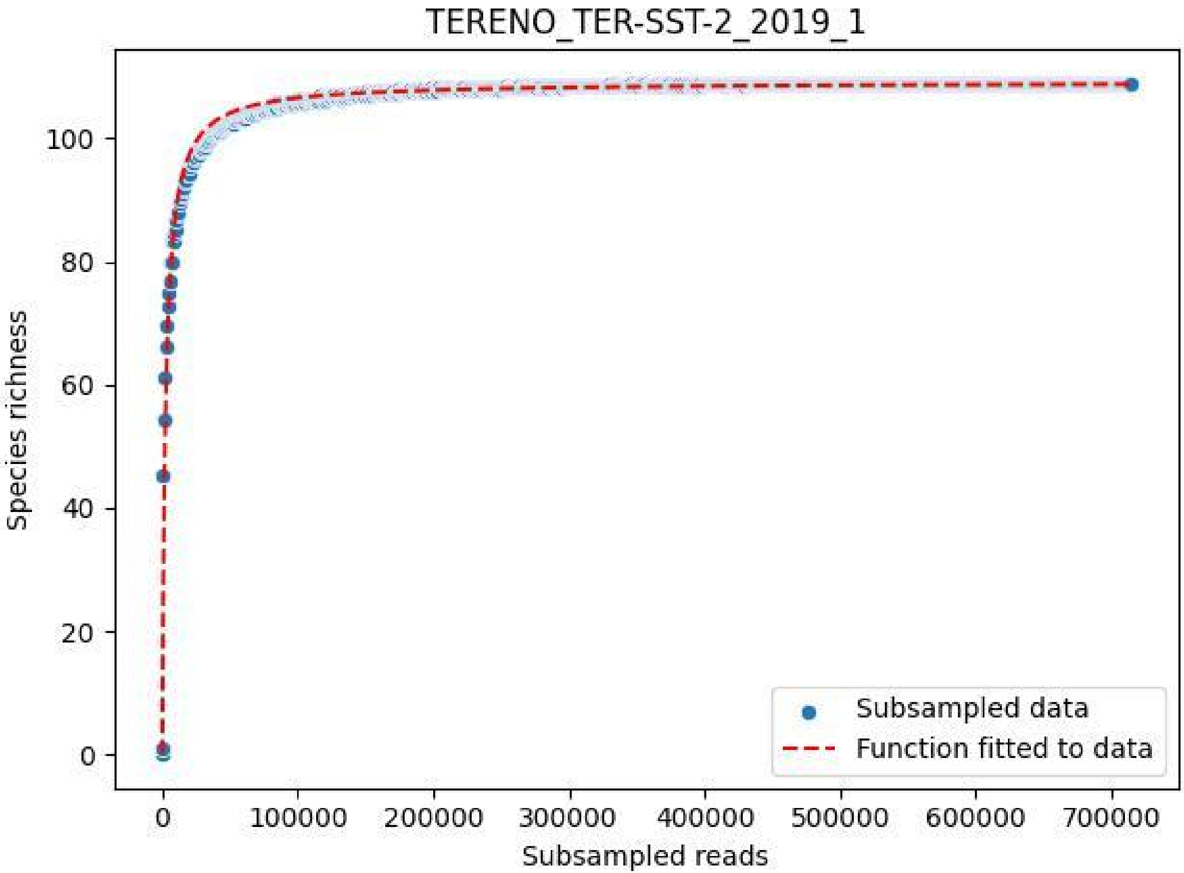
TERENO_TER-SST-2_2019_1
Species richness
Subsampled reads
0
20
40
60
80
100
0
100000
200000
300000
400000
500000
600000
700000
Subsampled data
Function fitted to data

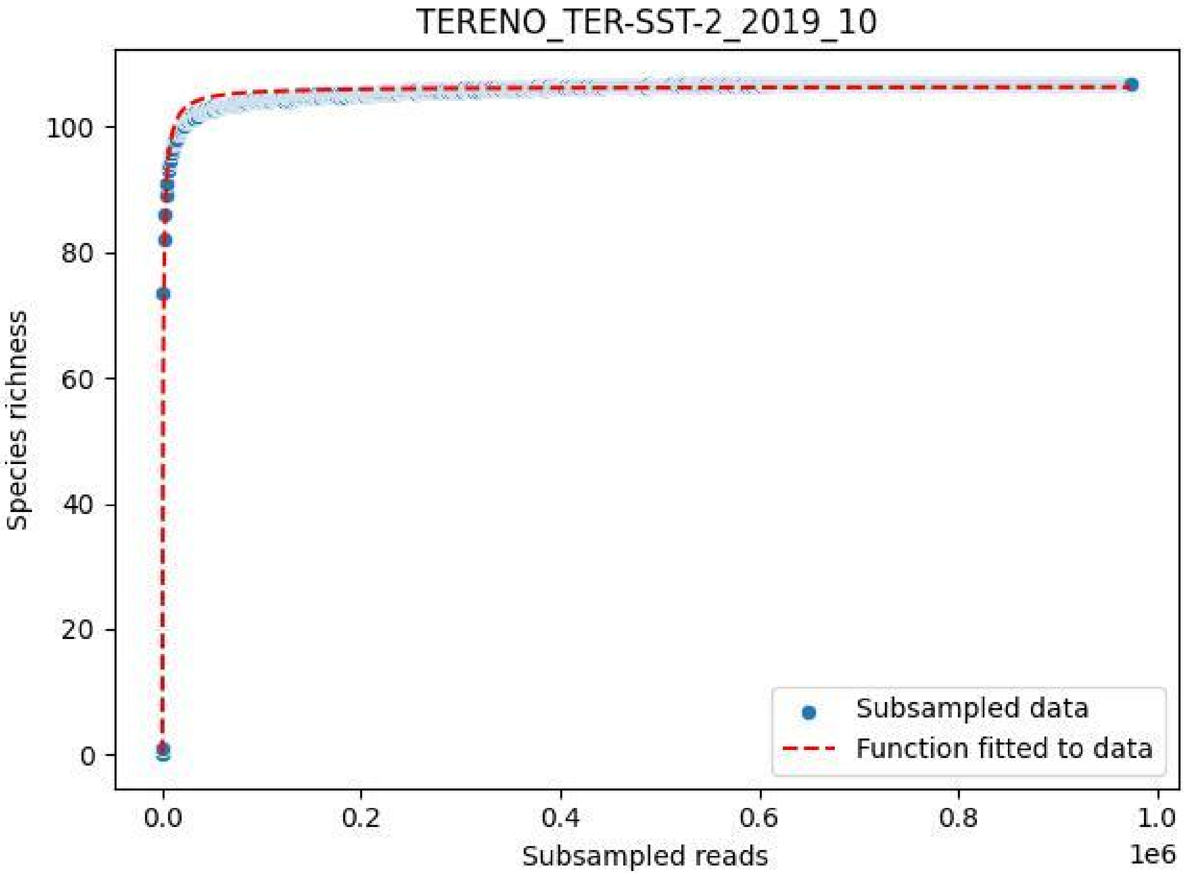
TERENO_TER-SST-2_2019_10
Species richness
Subsampled reads
1e6
0
20
40
60
80
100
0.0
0.2
0.4
0.6
0.8
1.0
Subsampled data
Function fitted to data

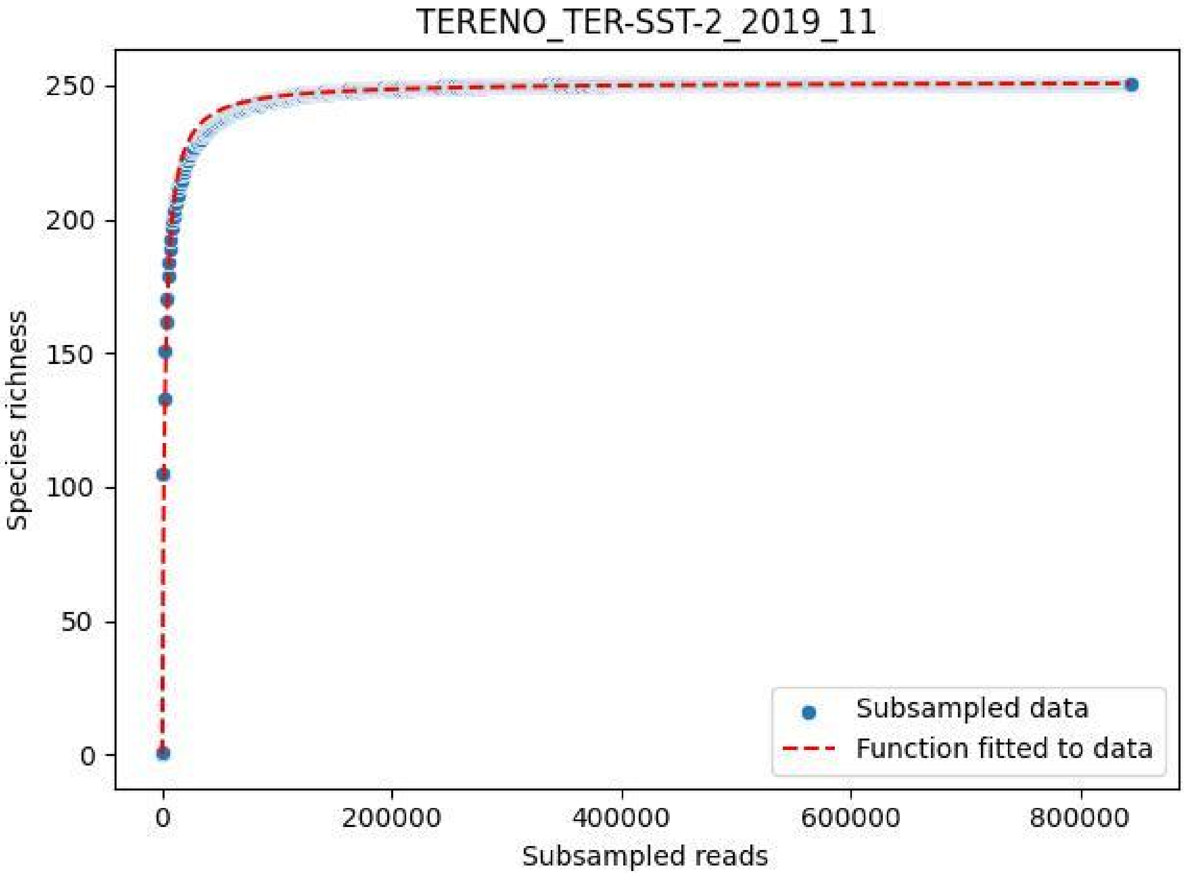
TERENO_TER-SST-2_2019_11
Species richness
Subsampled reads
250
200
150
100
50
0
0
200000
400000
600000
800000
Subsampled data
Function fitted to data

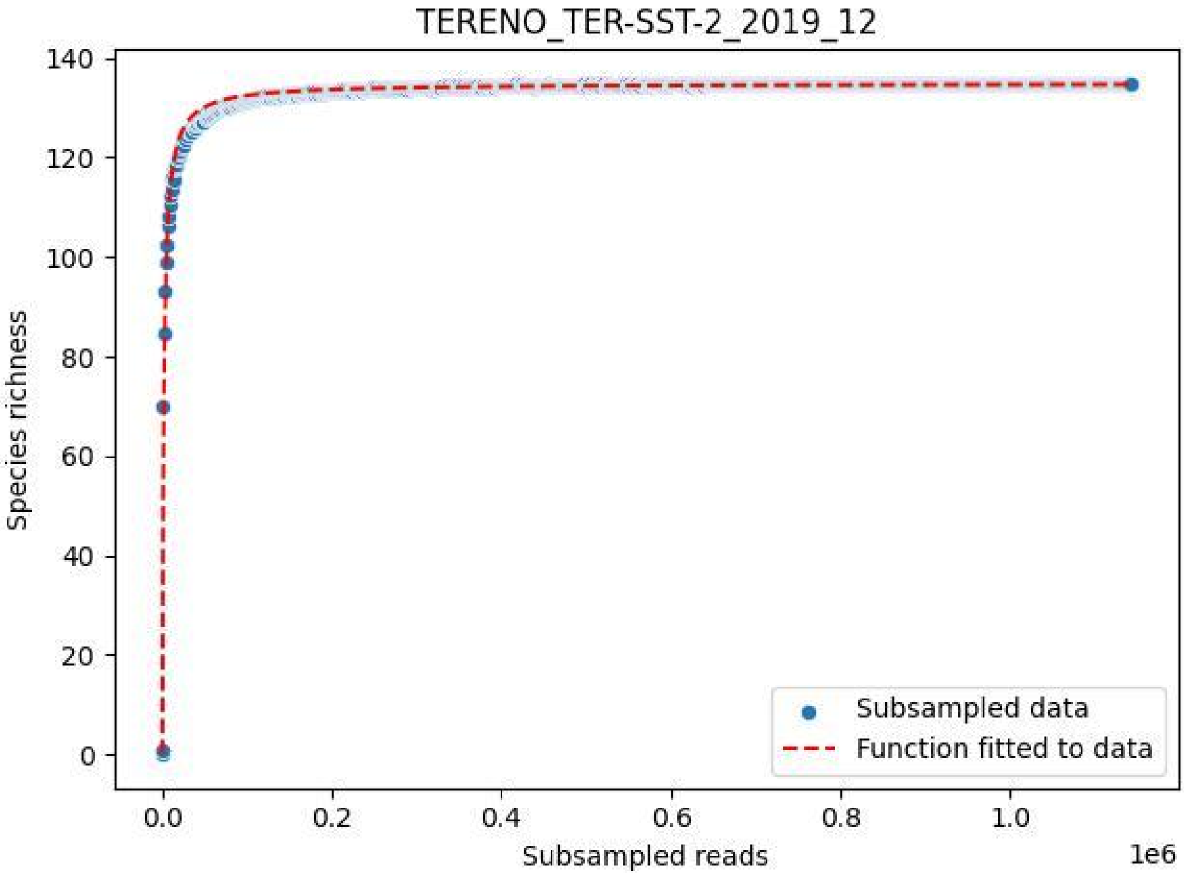
TERENO_TER-SST-2_2019_12
Species richness
140
120
100
80
60
40
20
0
0.0
0.2
0.4
0.6
0.8
1.0
1e6
Subsampled reads
Subsampled data
Function fitted to data

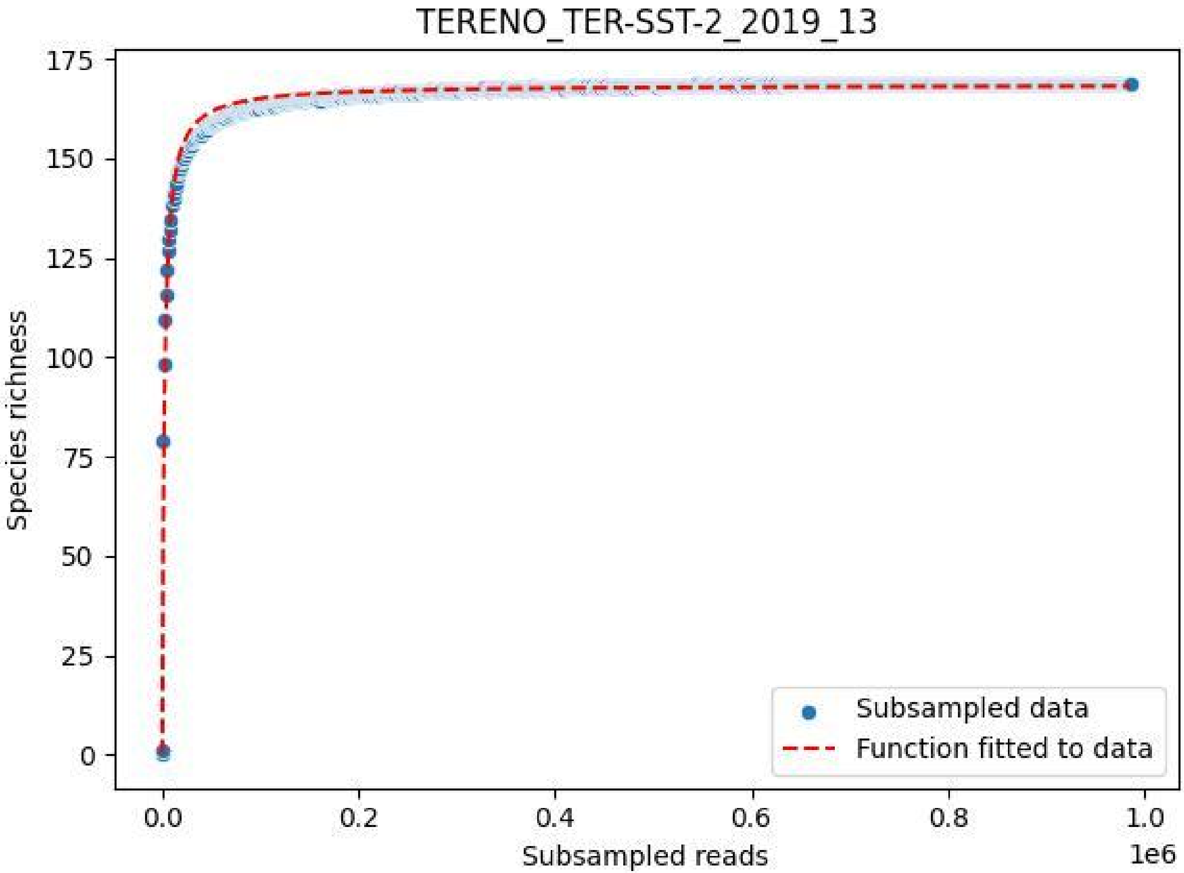
TERENO_TER-SST-2_2019_13
Species richness
Subsampled reads
1e6
0
25
50
75
100
125
150
175
0.0
0.2
0.4
0.6
0.8
1.0
Subsampled data
Function fitted to data

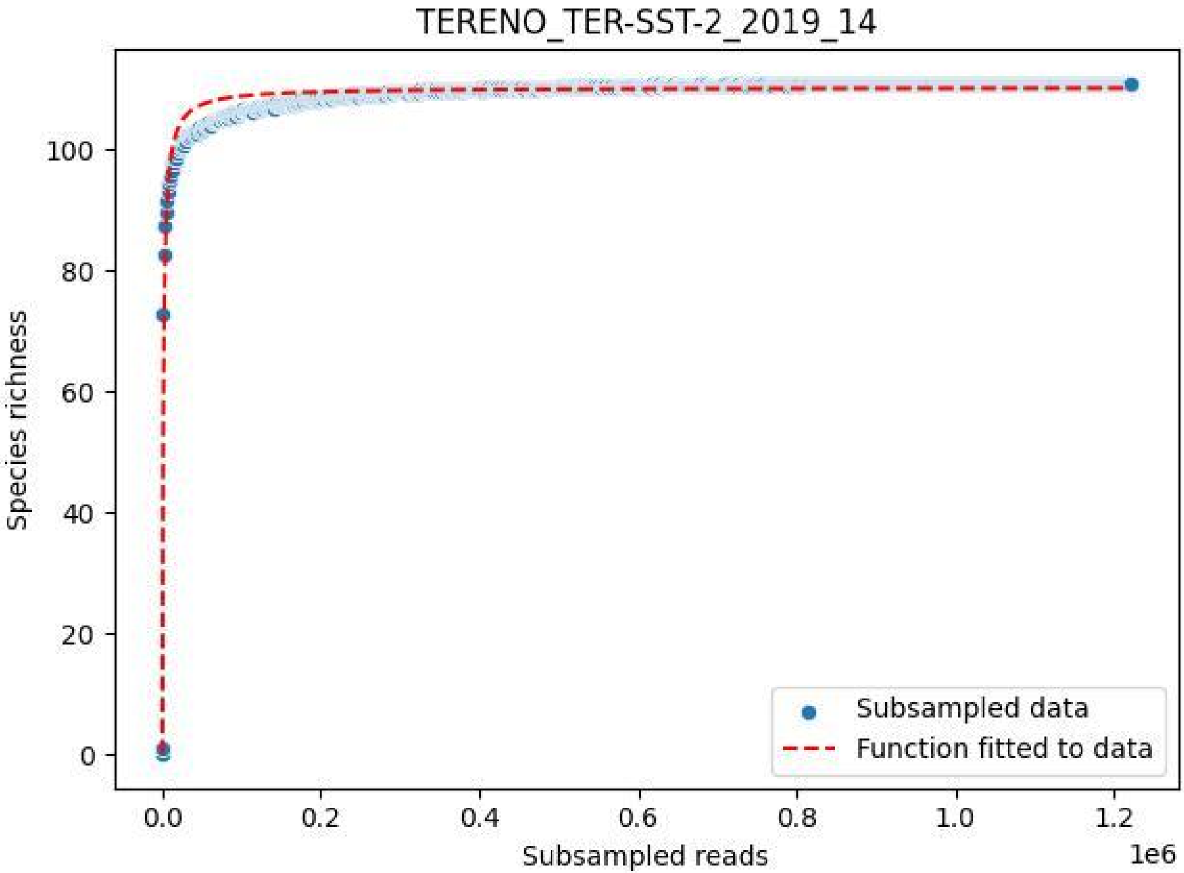
TERENO_TER-SST-2_2019_14
Species richness
Subsampled reads
1e6
Subsampled data
Function fitted to data
0
20
40
60
80
100
0.0
0.2
0.4
0.6
0.8
1.0
1.2

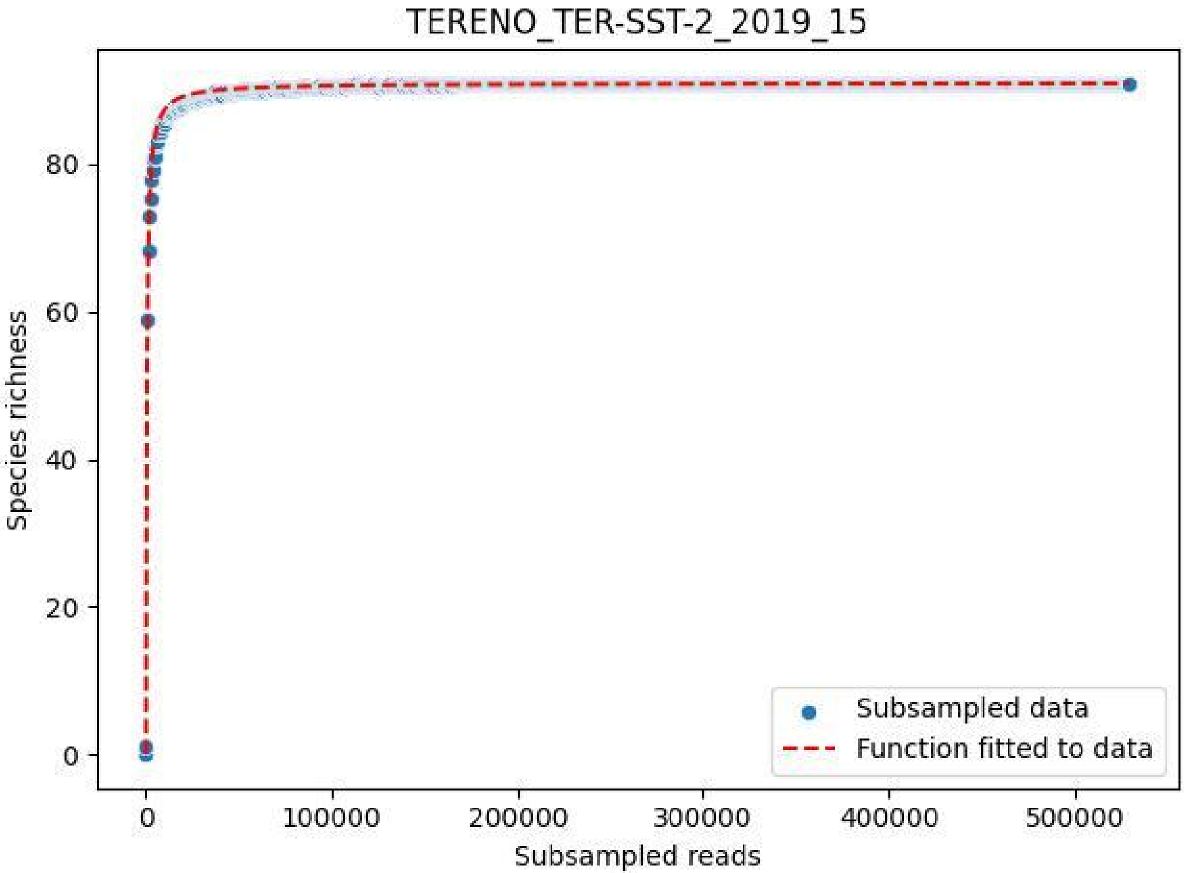

TERENO_TER-SST-2_2019_15
Species richness
Subsampled reads
0
20
40
60
80
0
100000
200000
300000
400000
500000
Subsampled data
Function fitted to data

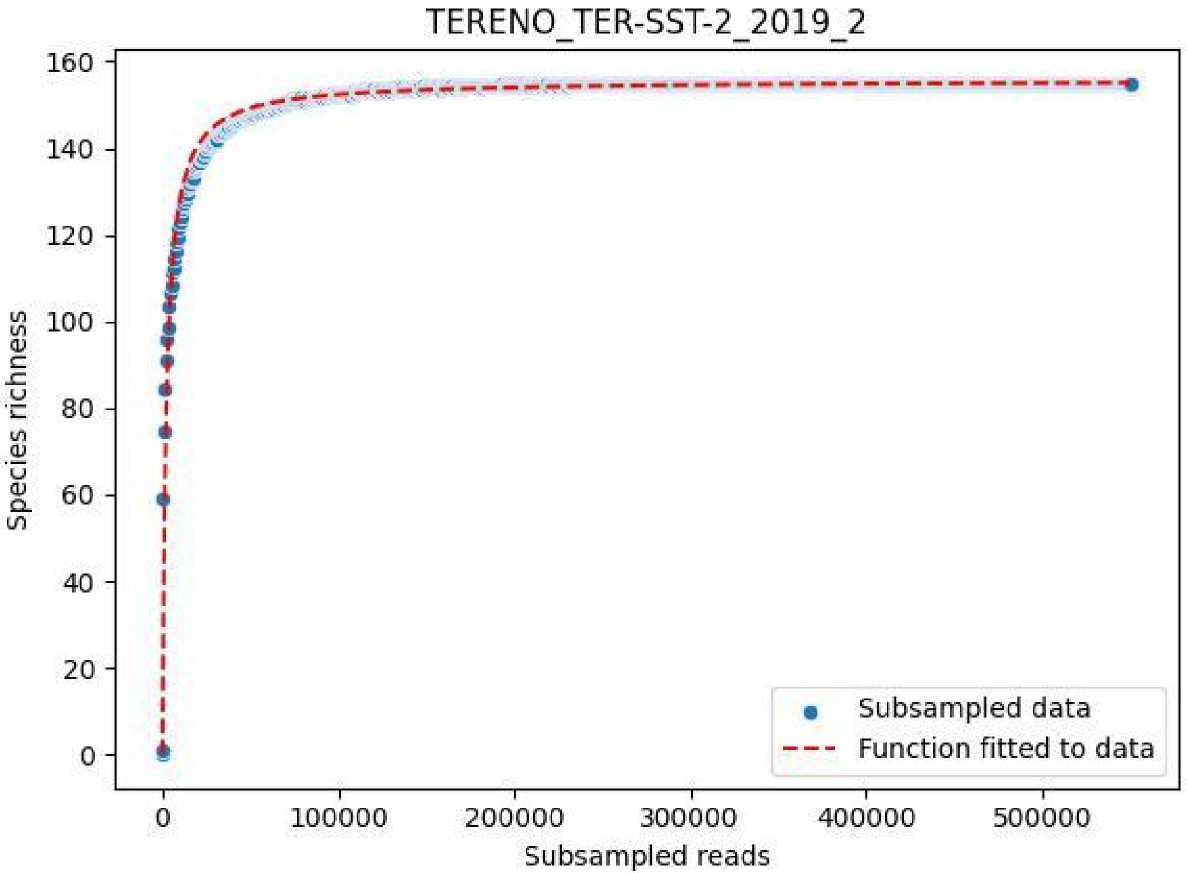
TERENO_TER-SST-2_2019_2
Species richness
Subsampled reads
0
20
40
60
80
100
120
140
160
0
100000
200000
300000
400000
500000
Subsampled data
Function fitted to data

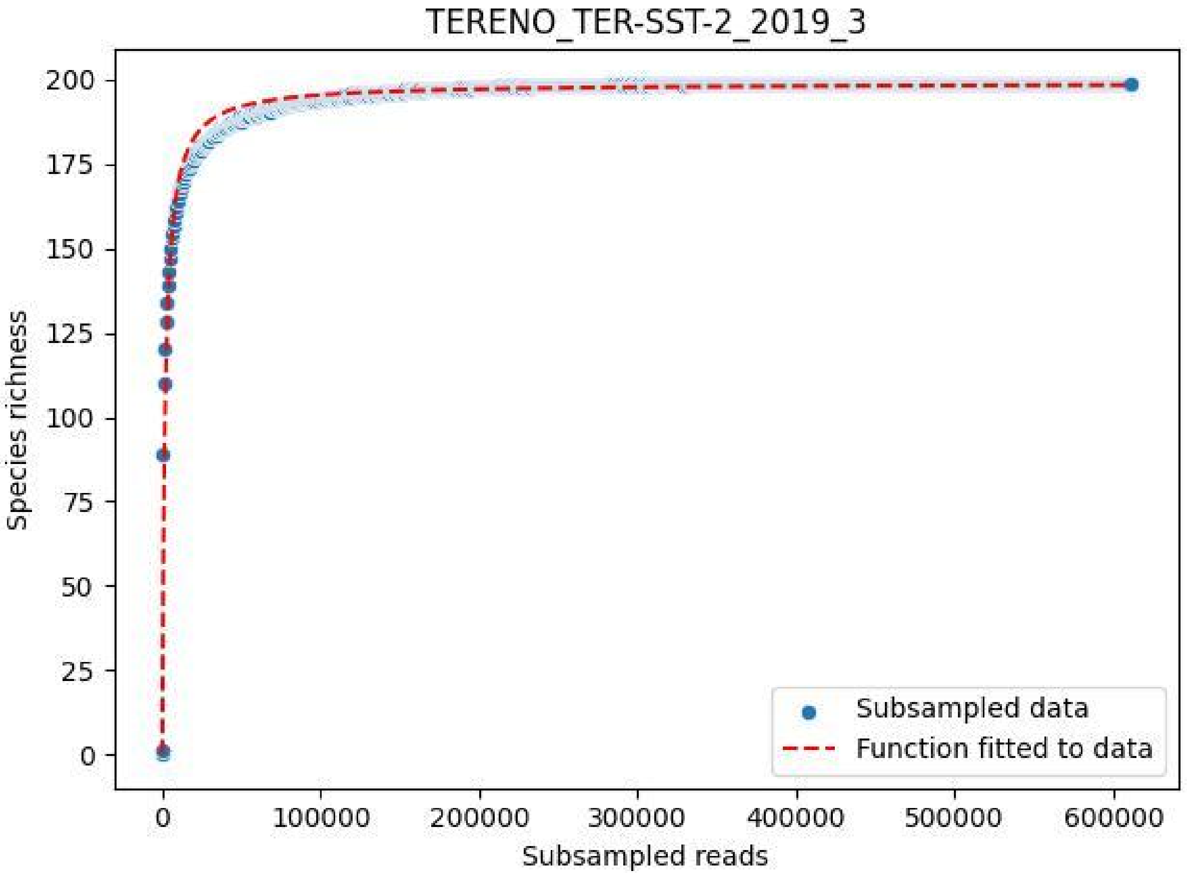
TERENO_TER-SST-2_2019_3
Species richness
Subsampled reads
0
25
50
75
100
125
150
175
200
0
100000
200000
300000
400000
500000
600000
Subsampled data
Function fitted to data

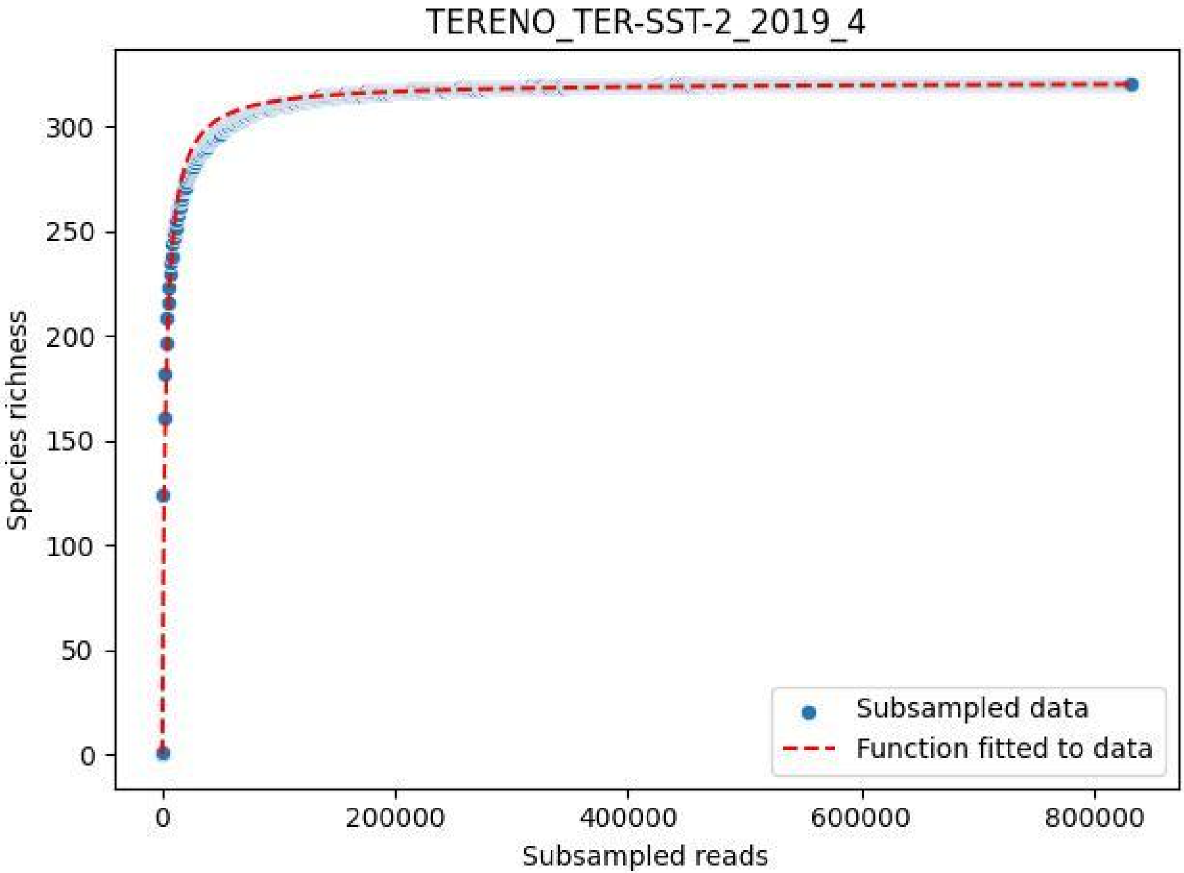

TERENO_TER-SST-2_2019_4
Species richness
Subsampled reads
0
50
100
150
200
250
300
0
200000
400000
600000
800000
Subsampled data
Function fitted to data

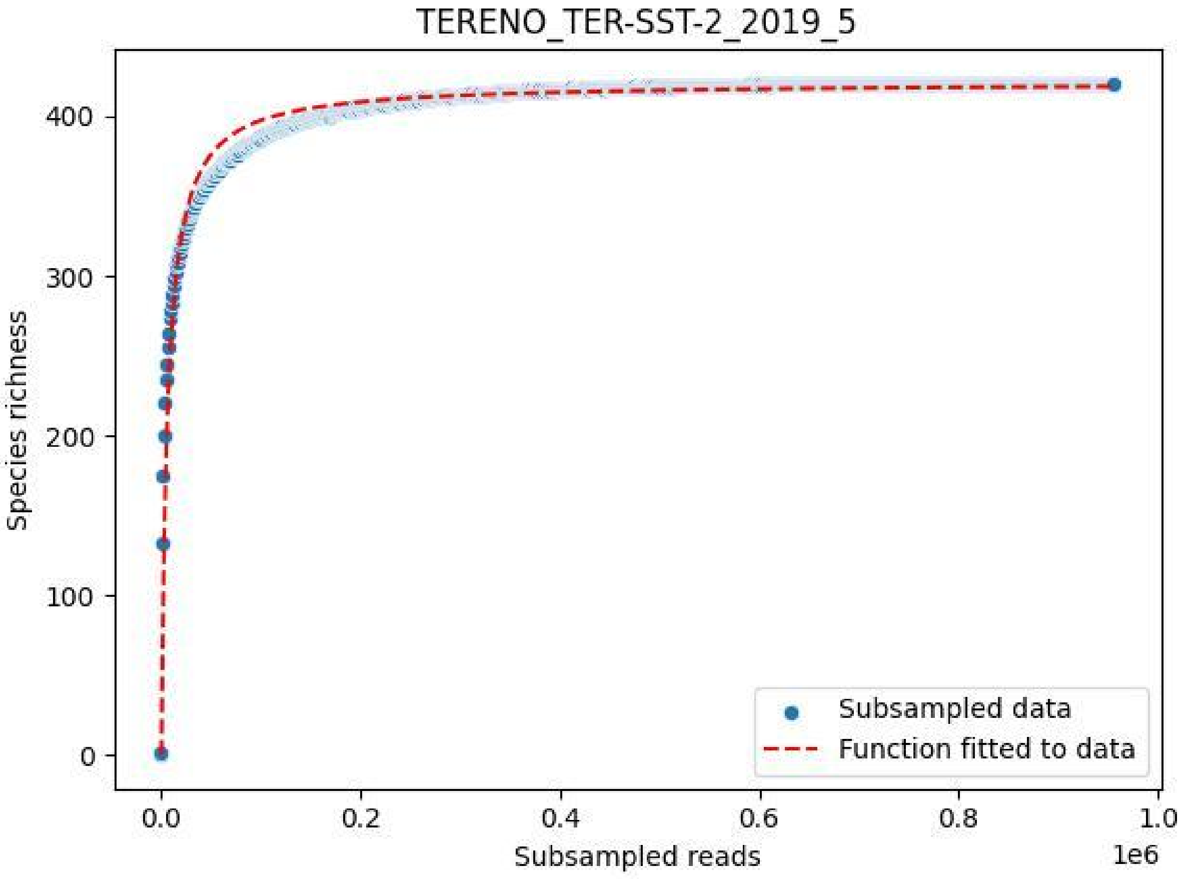
TERENO_TER-SST-2_2019_5
Species richness
Subsampled reads
1e6
0
100
200
300
400
0.0
0.2
0.4
0.6
0.8
1.0
Subsampled data
Function fitted to data

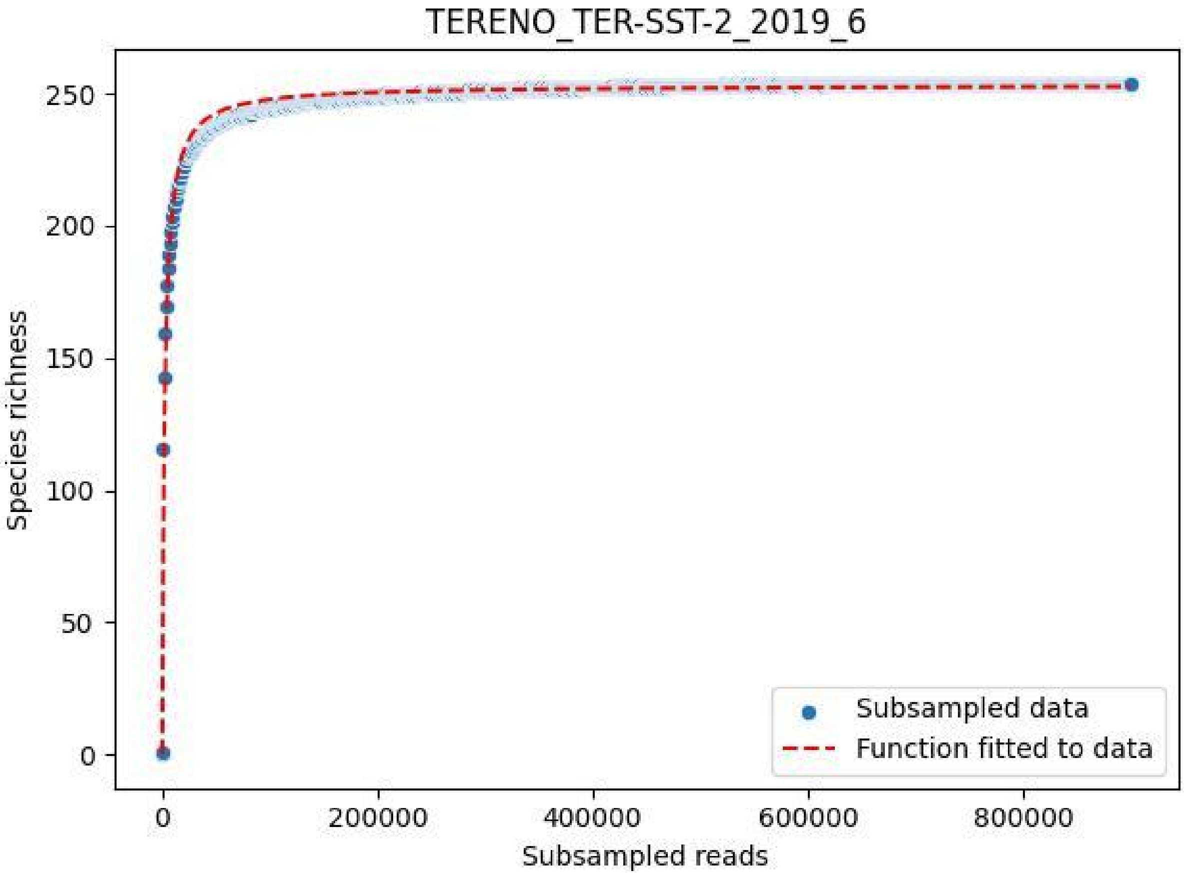

TERENO_TER-SST-2_2019_6
Species richness
Subsampled reads
0
50
100
150
200
250
0
200000
400000
600000
800000
Subsampled data
Function fitted to data

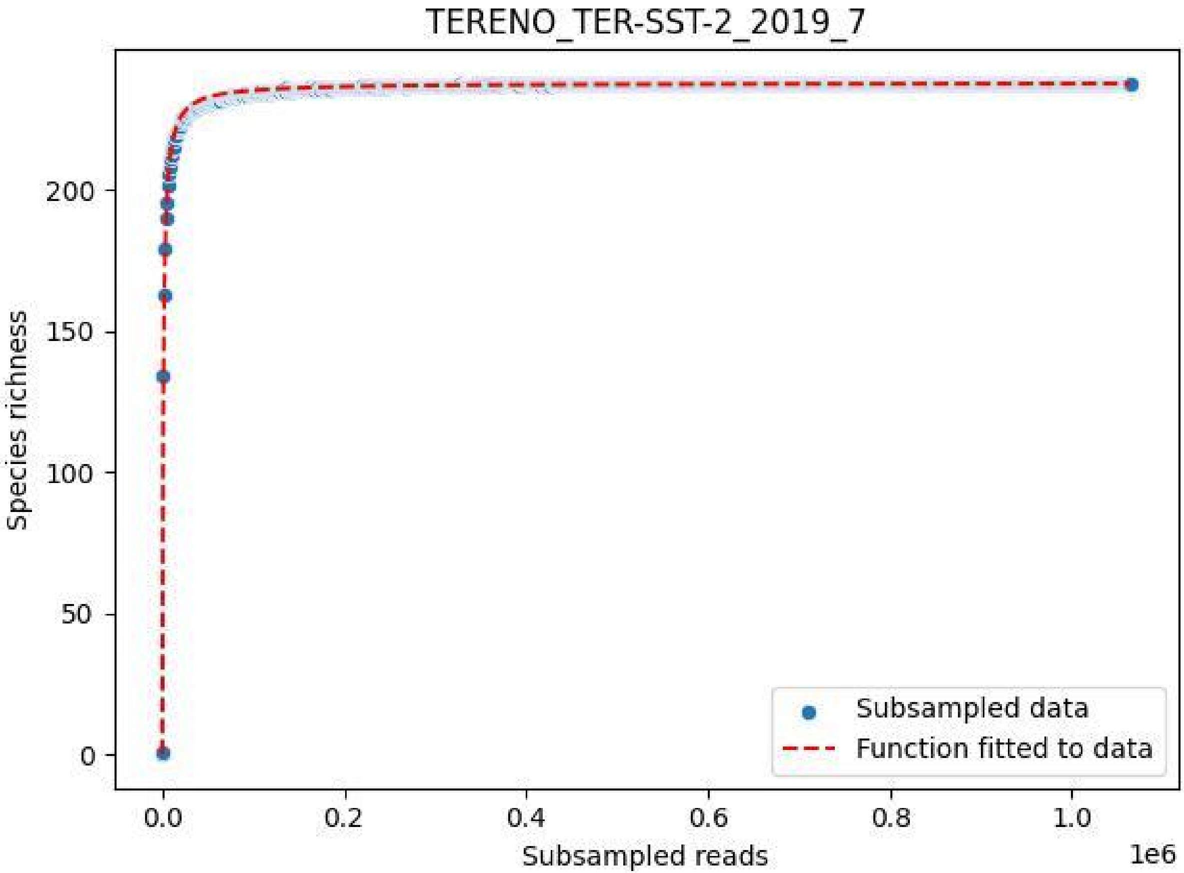

TERENO_TER-SST-2_2019_7
Species richness
Subsampled reads
1e6
0
50
100
150
200
0.0
0.2
0.4
0.6
0.8
1.0
Subsampled data
Function fitted to data

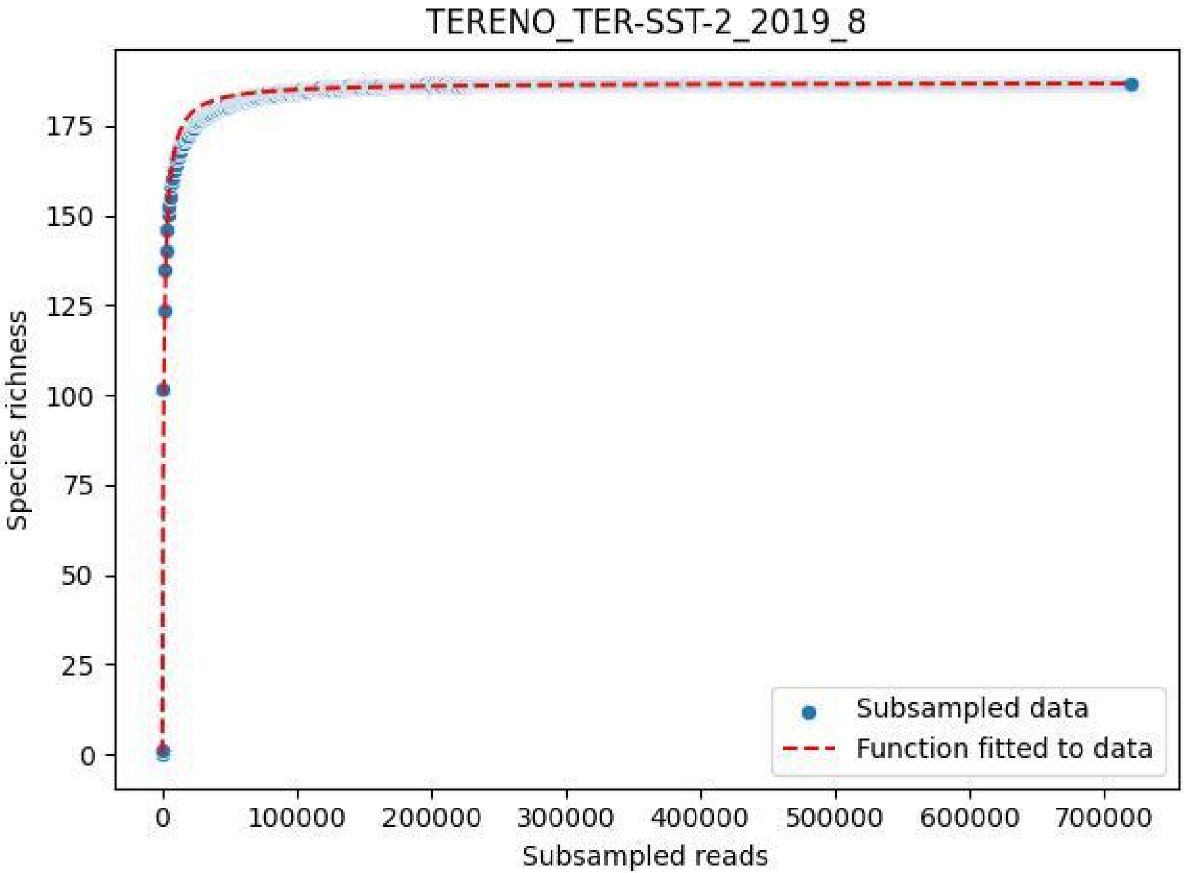

TERENO_TER-SST-2_2019_8
Species richness
Subsampled reads
175
150
125
100
75
50
25
0
0
100000
200000
300000
400000
500000
600000
700000
Subsampled data
Function fitted to data

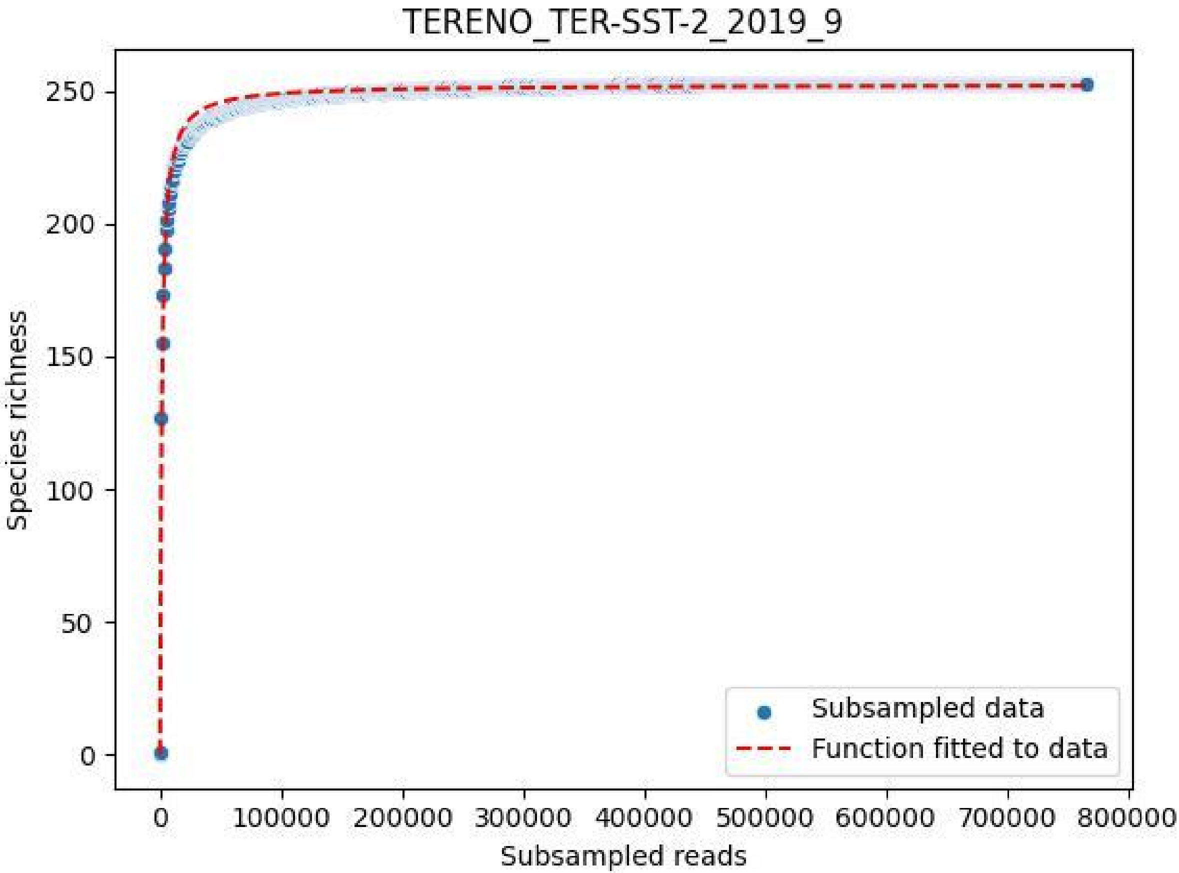

TERENO_TER-SST-2_2019_9
Species richness
Subsampled reads
0
50
100
150
200
250
0
100000
200000
300000
400000
500000
600000
700000
800000
Subsampled data
Function fitted to data

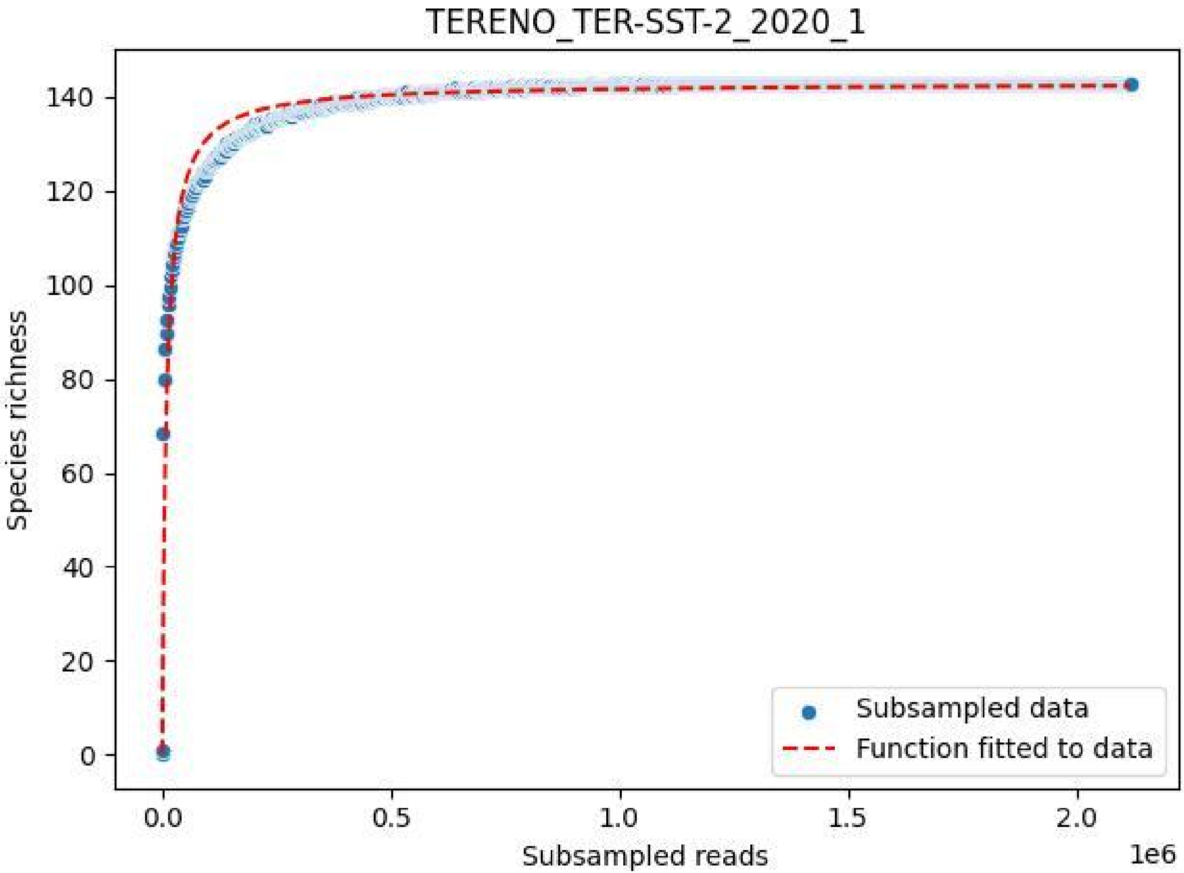

TERENO_TER-SST-2_2020_1
Species richness
0
20
40
60
80
100
120
140
Subsampled reads
0.0
0.5
1.0
1.5
2.0
1e6
Subsampled data
Function fitted to data

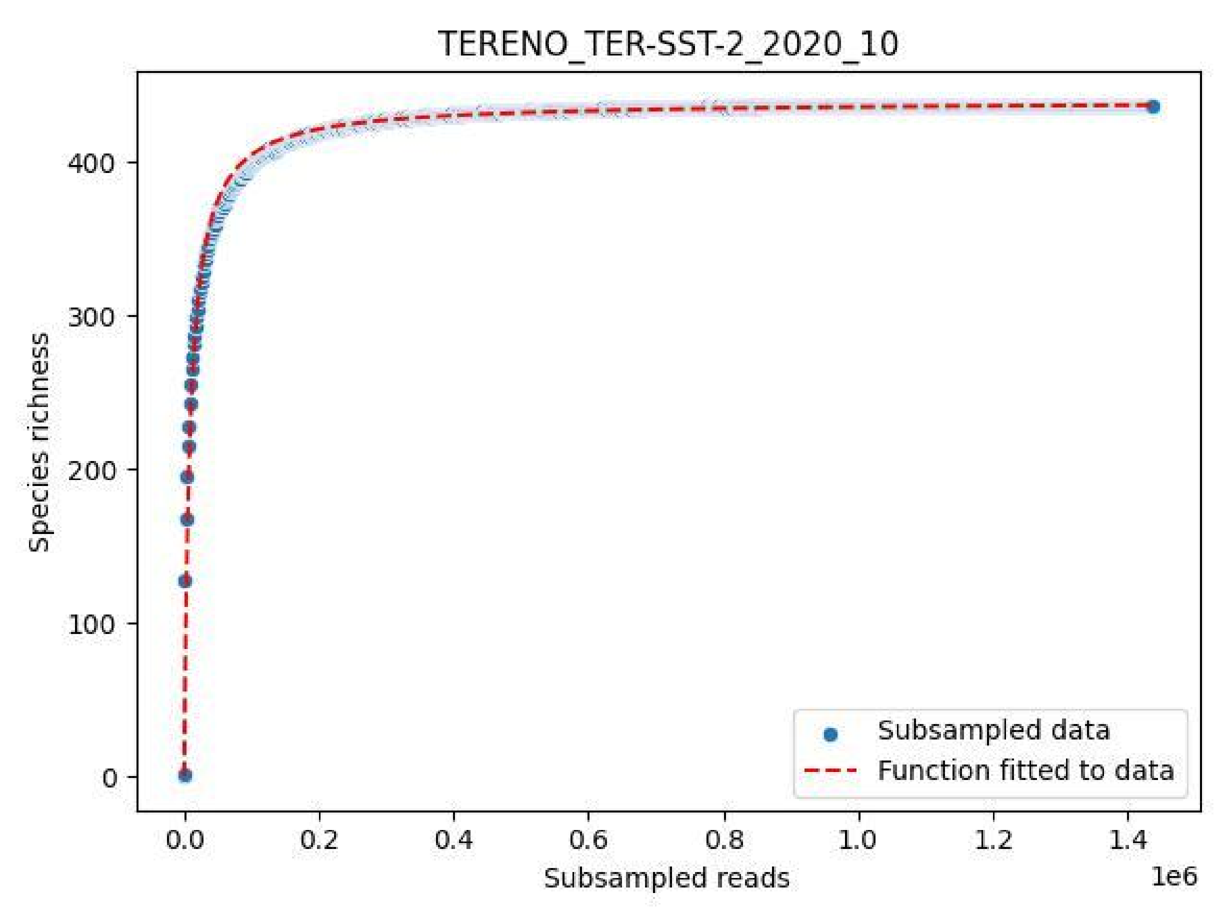

TERENO_TER-SST-2_2020_10
Species richness
Subsampled reads
1e6
0
100
200
300
400
0.0
0.2
0.4
0.6
0.8
1.0
1.2
1.4
Subsampled data
Function fitted to data

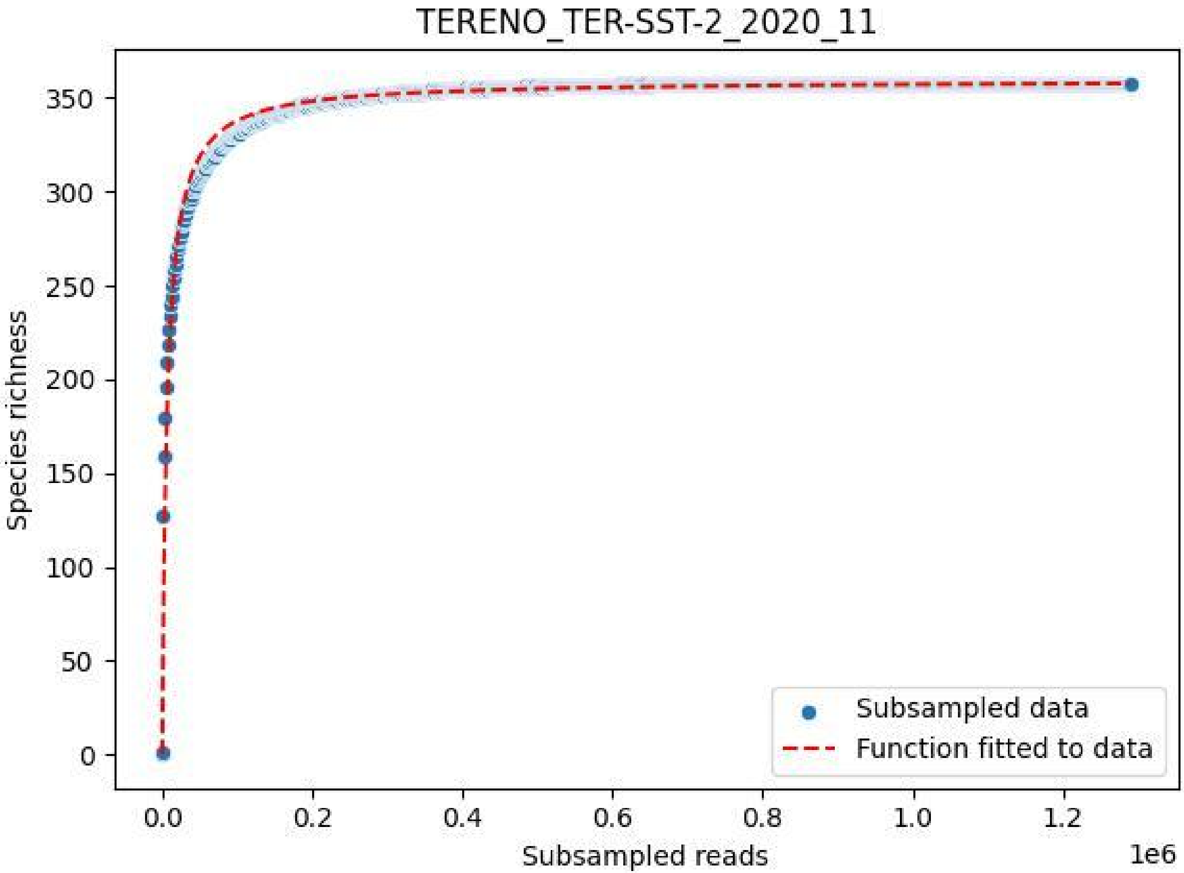
TERENO_TER-SST-2_2020_11
Species richness
0
50
100
150
200
250
300
350
0.0
0.2
0.4
0.6
0.8
1.0
1.2
1e6
Subsampled reads
Subsampled data
Function fitted to data

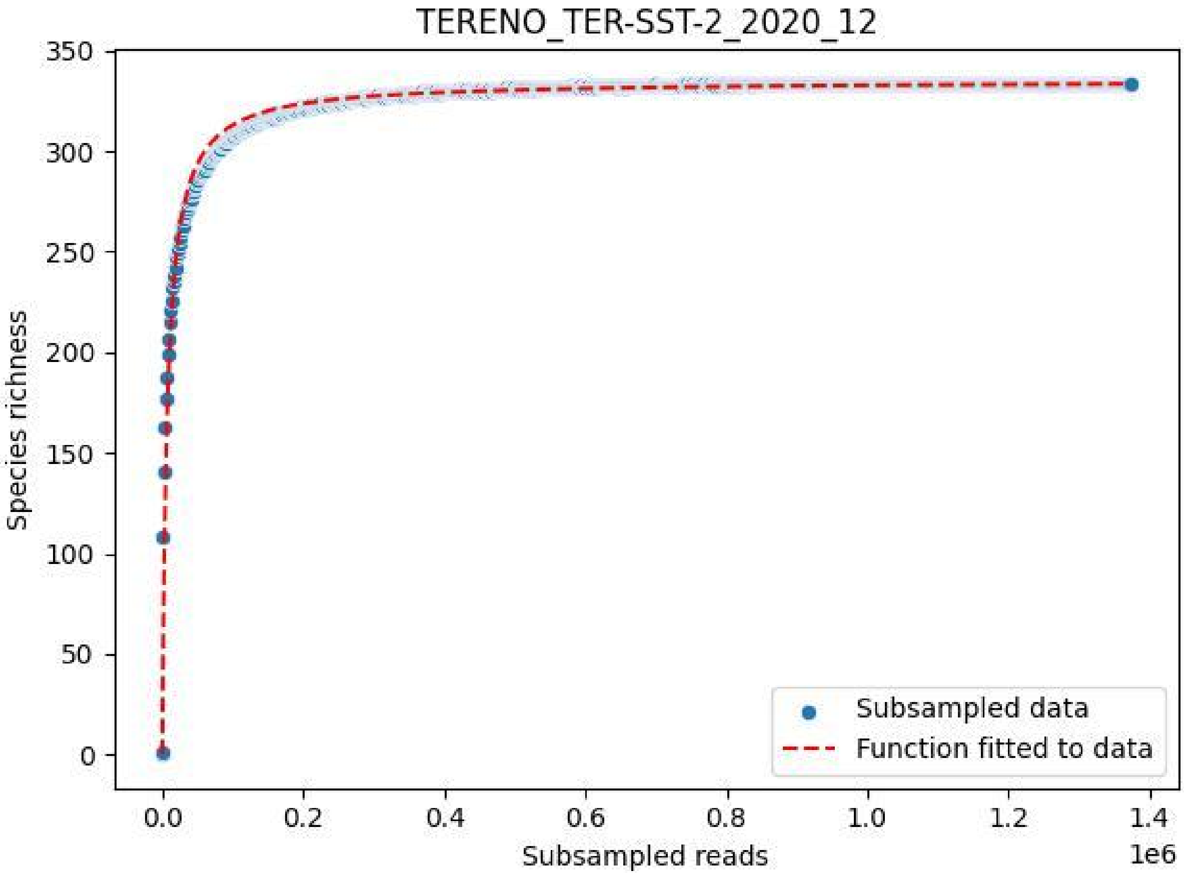

TERENO_TER-SST-2_2020_12
Species richness
Subsampled reads
1e6
Subsampled data
Function fitted to data
0
50
100
150
200
250
300
350
0.0
0.2
0.4
0.6
0.8
1.0
1.2
1.4

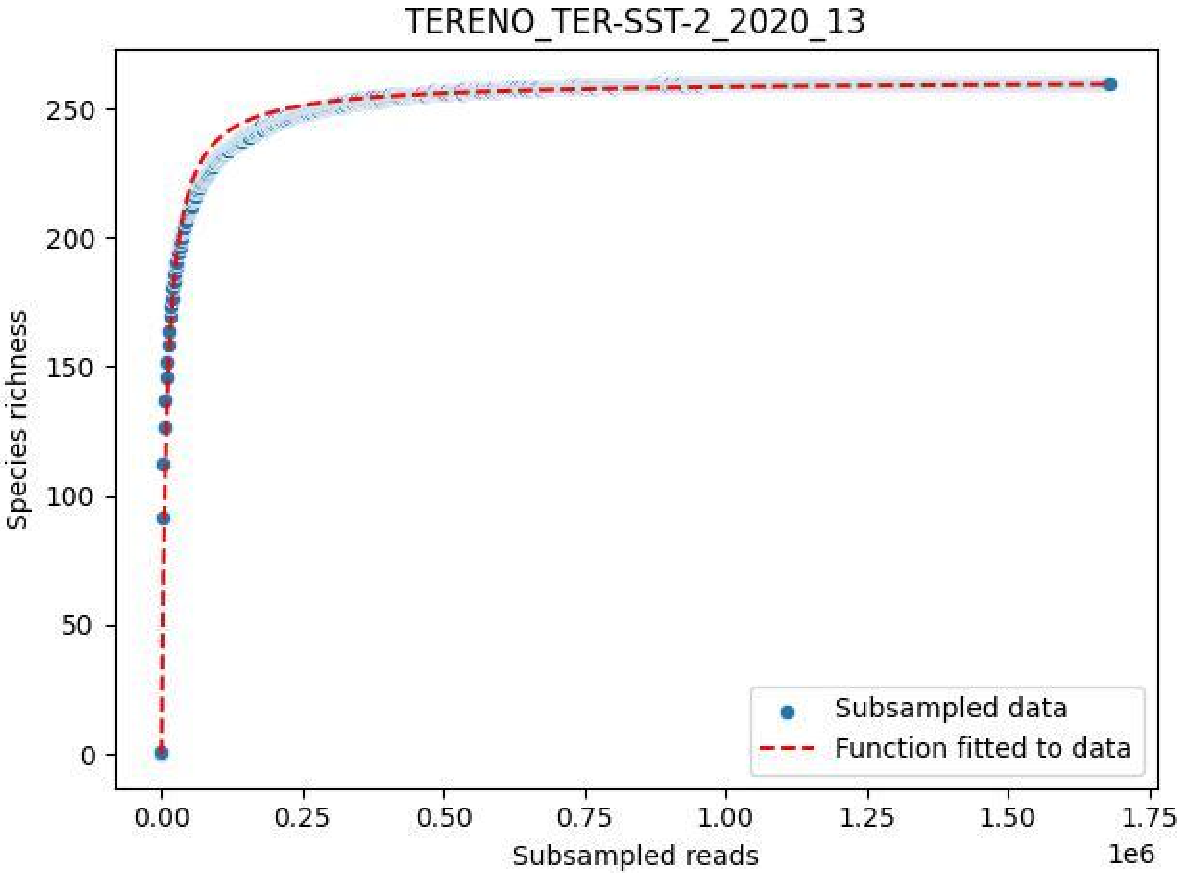

TERENO_TER-SST-2_2020_13
Species richness
Subsampled reads
1e6
0
50
100
150
200
250
0.00
0.25
0.50
0.75
1.00
1.25
1.50
1.75
Subsampled data
Function fitted to data

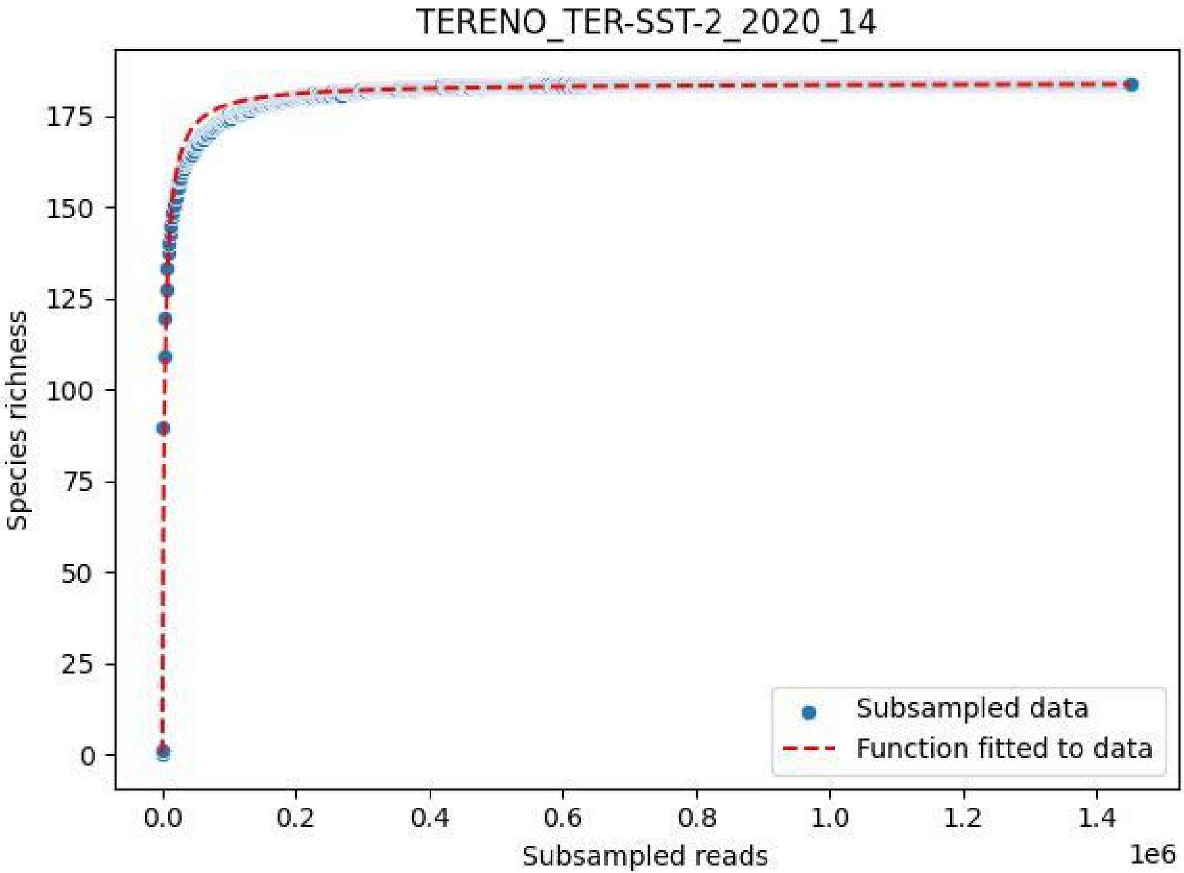

TERENO_TER-SST-2_2020_14
Species richness
Subsampled reads
1e6
0
25
50
75
100
125
150
175
0.0
0.2
0.4
0.6
0.8
1.0
1.2
1.4
Subsampled data
Function fitted to data

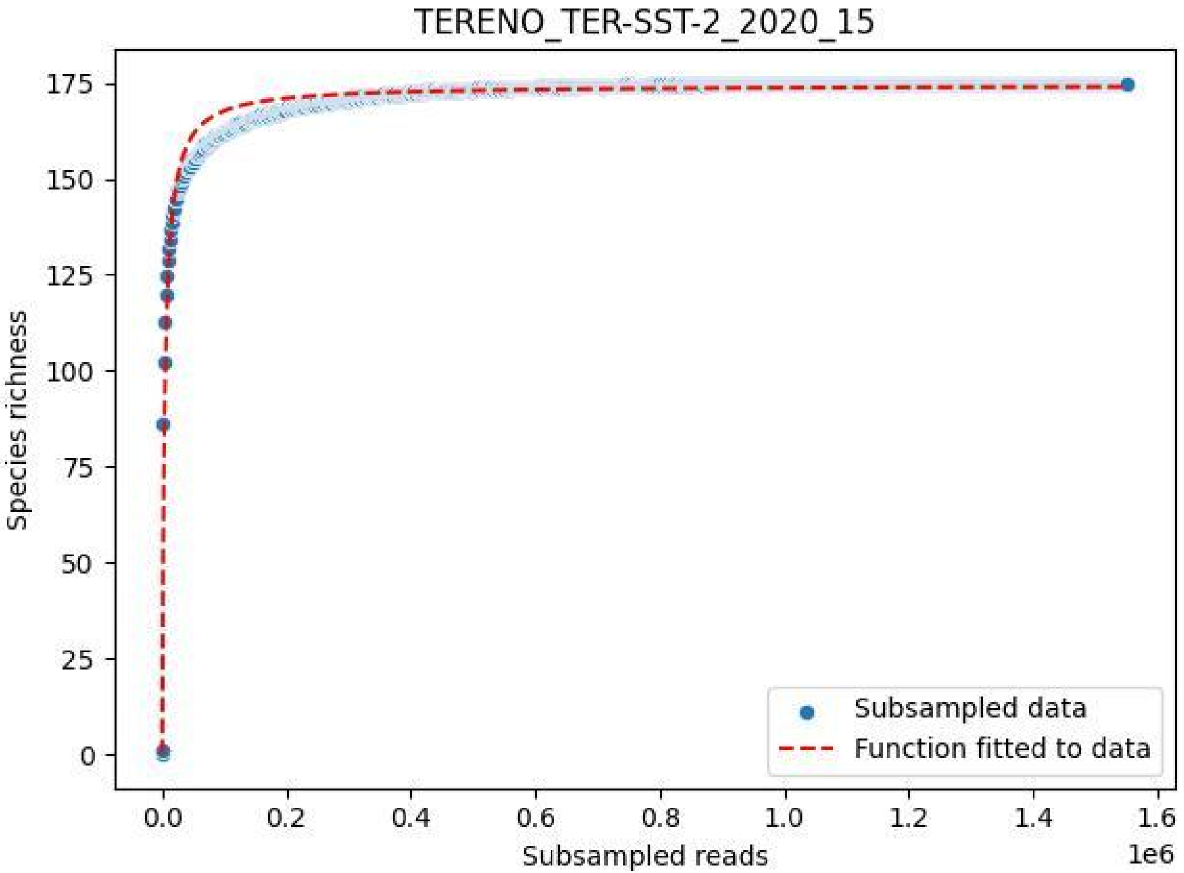
TERENO_TER-SST-2_2020_15
Species richness
Subsampled reads
1e6
0
25
50
75
100
125
150
175
0.0
0.2
0.4
0.6
0.8
1.0
1.2
1.4
1.6
Subsampled data
Function fitted to data

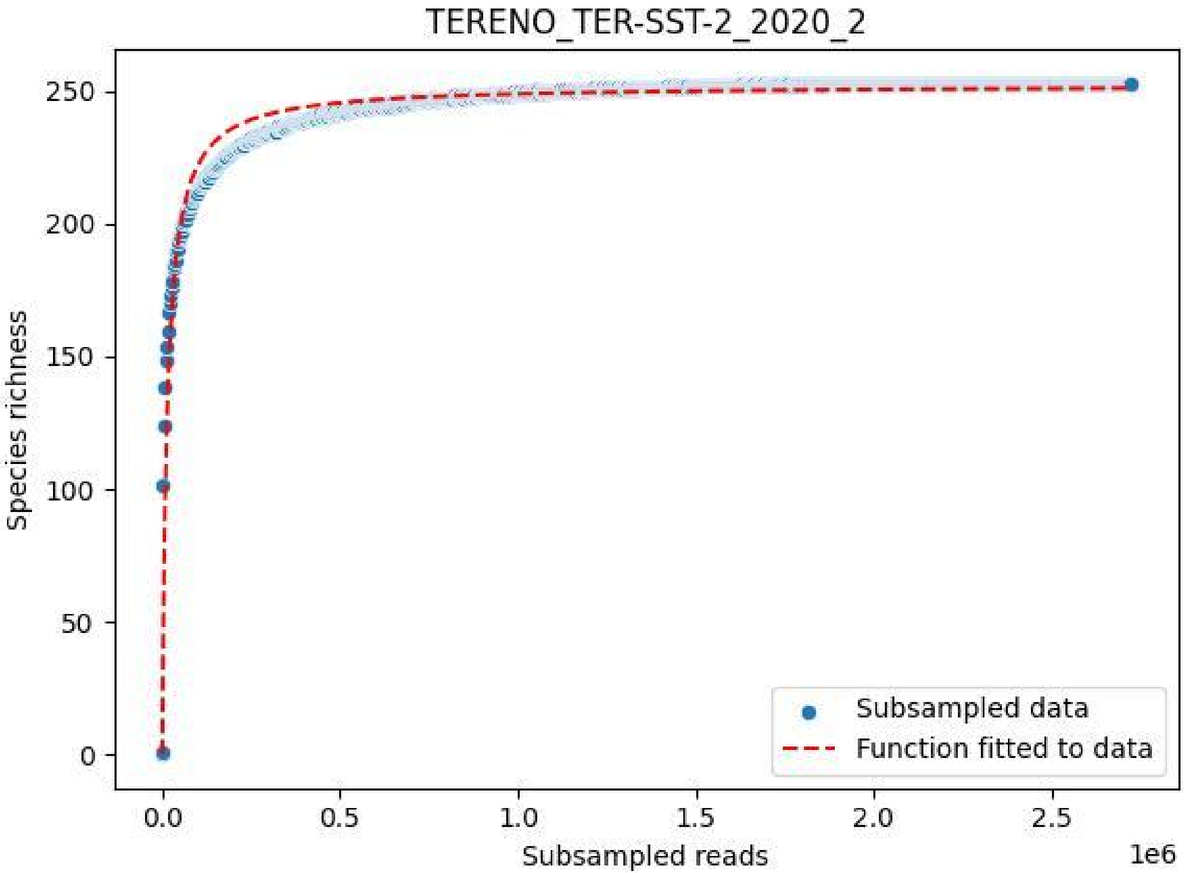

TERENO_TER-SST-2_2020_2
Species richness
Subsampled reads
1e6
0
50
100
150
200
250
0.0
0.5
1.0
1.5
2.0
2.5
Subsampled data
Function fitted to data

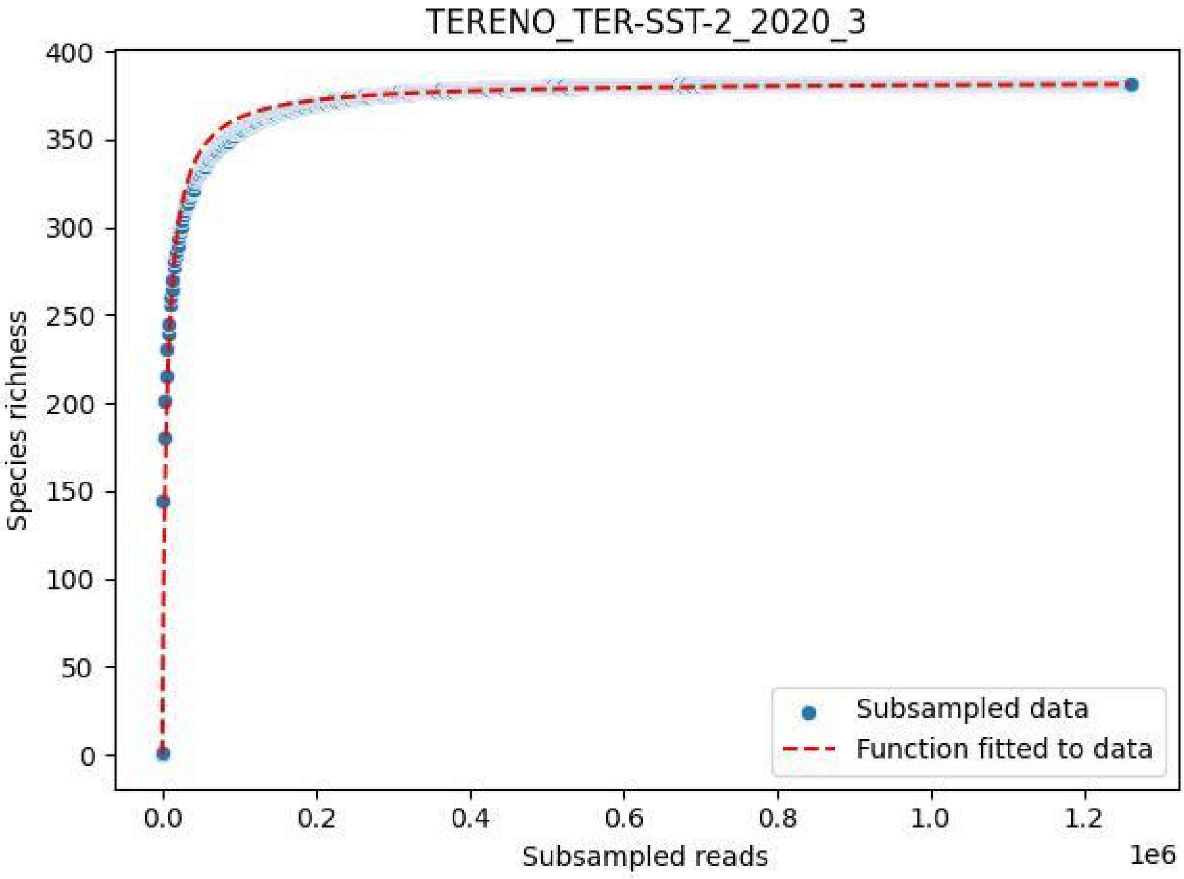

TERENO_TER-SST-2_2020_3
Species richness
Subsampled reads
1e6
0
50
100
150
200
250
300
350
400
0.0
0.2
0.4
0.6
0.8
1.0
1.2
Subsampled data
Function fitted to data

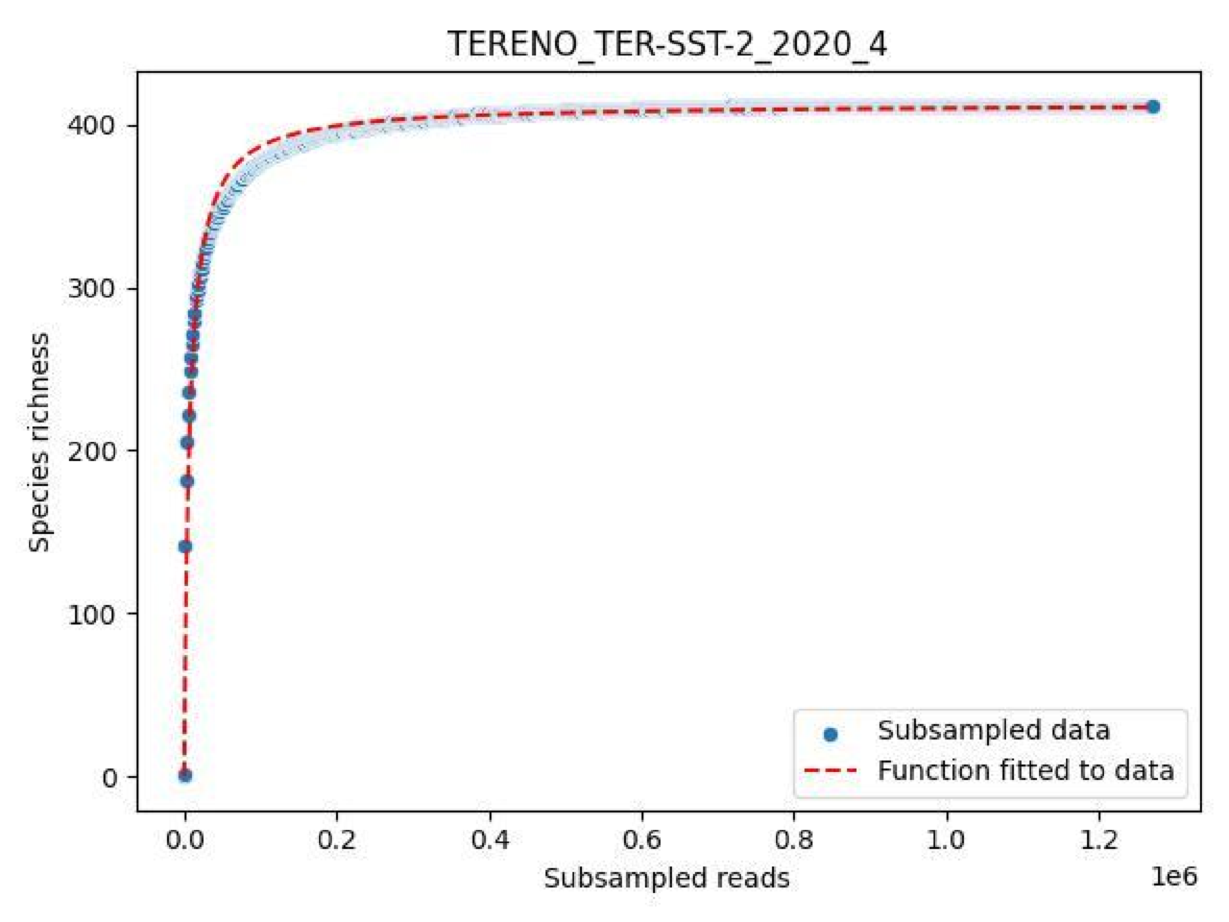

TERENO_TER-SST-2_2020_4
Species richness
Subsampled reads
1e6
0
100
200
300
400
0.0
0.2
0.4
0.6
0.8
1.0
1.2
Subsampled data
Function fitted to data

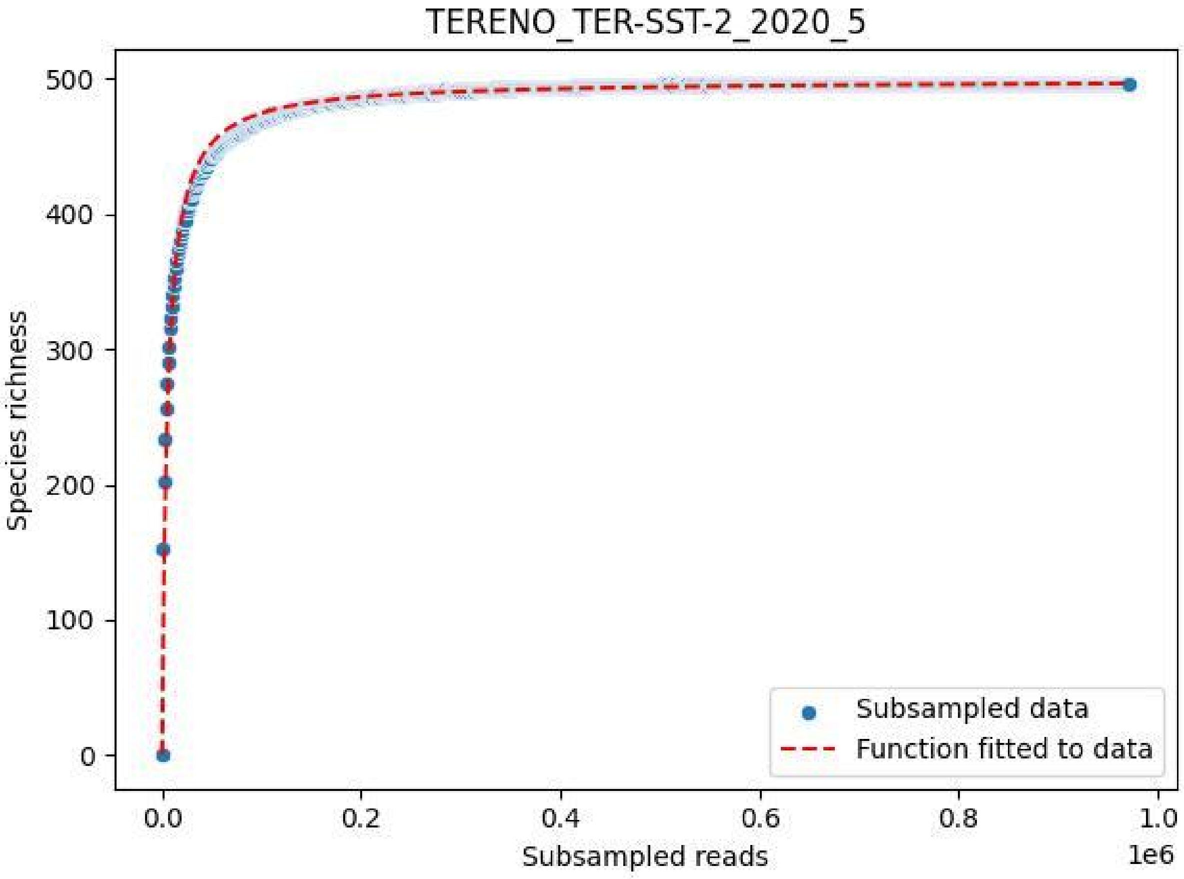
TERENO_TER-SST-2_2020_5
Species richness
Subsampled reads
1e6
0
100
200
300
400
500
0.0
0.2
0.4
0.6
0.8
1.0
Subsampled data
Function fitted to data

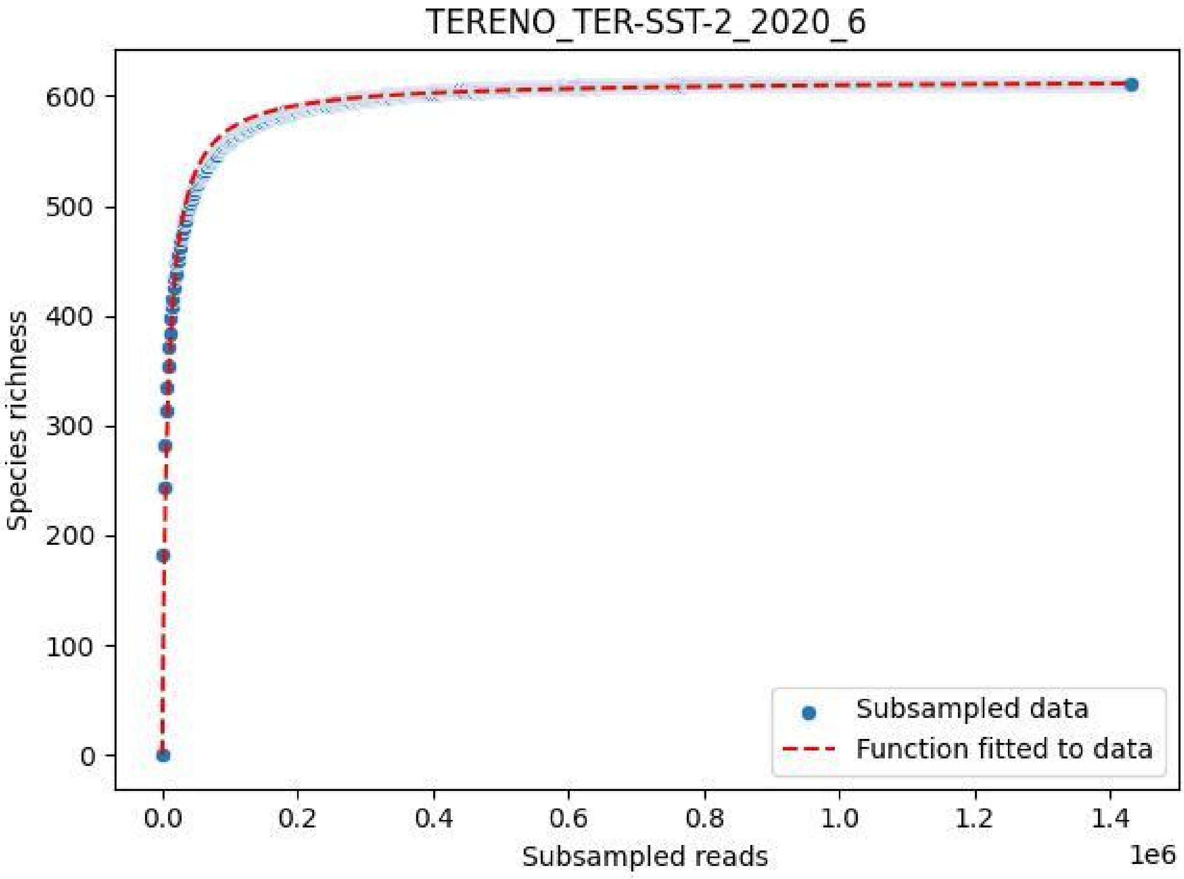

TERENO_TER-SST-2_2020_6
Species richness
0
100
200
300
400
500
600
0.0
0.2
0.4
0.6
0.8
1.0
1.2
1.4
1e6
Subsampled reads
Subsampled data
Function fitted to data

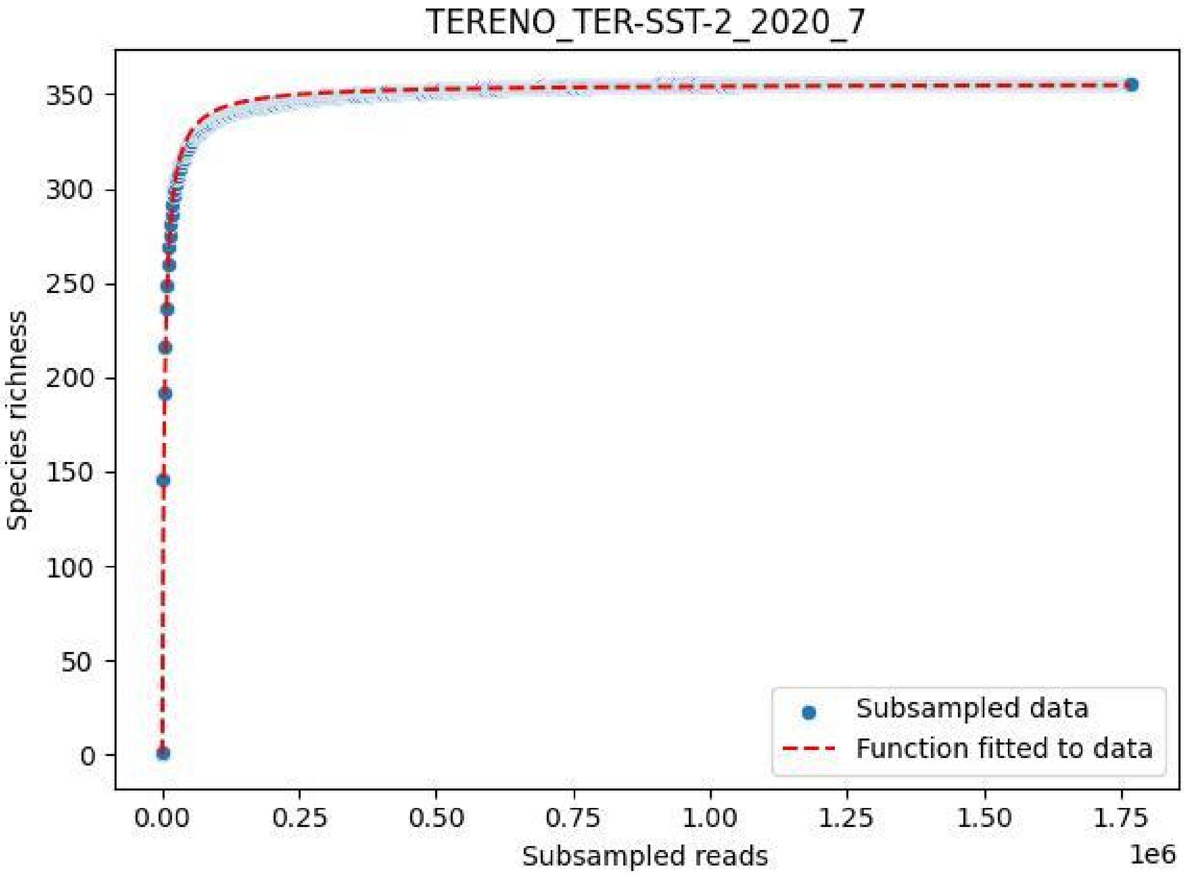

TERENO_TER-SST-2_2020_7
Species richness
0
50
100
150
200
250
300
350
0.00
0.25
0.50
0.75
1.00
1.25
1.50
1.75
1e6
Subsampled reads
Subsampled data
Function fitted to data

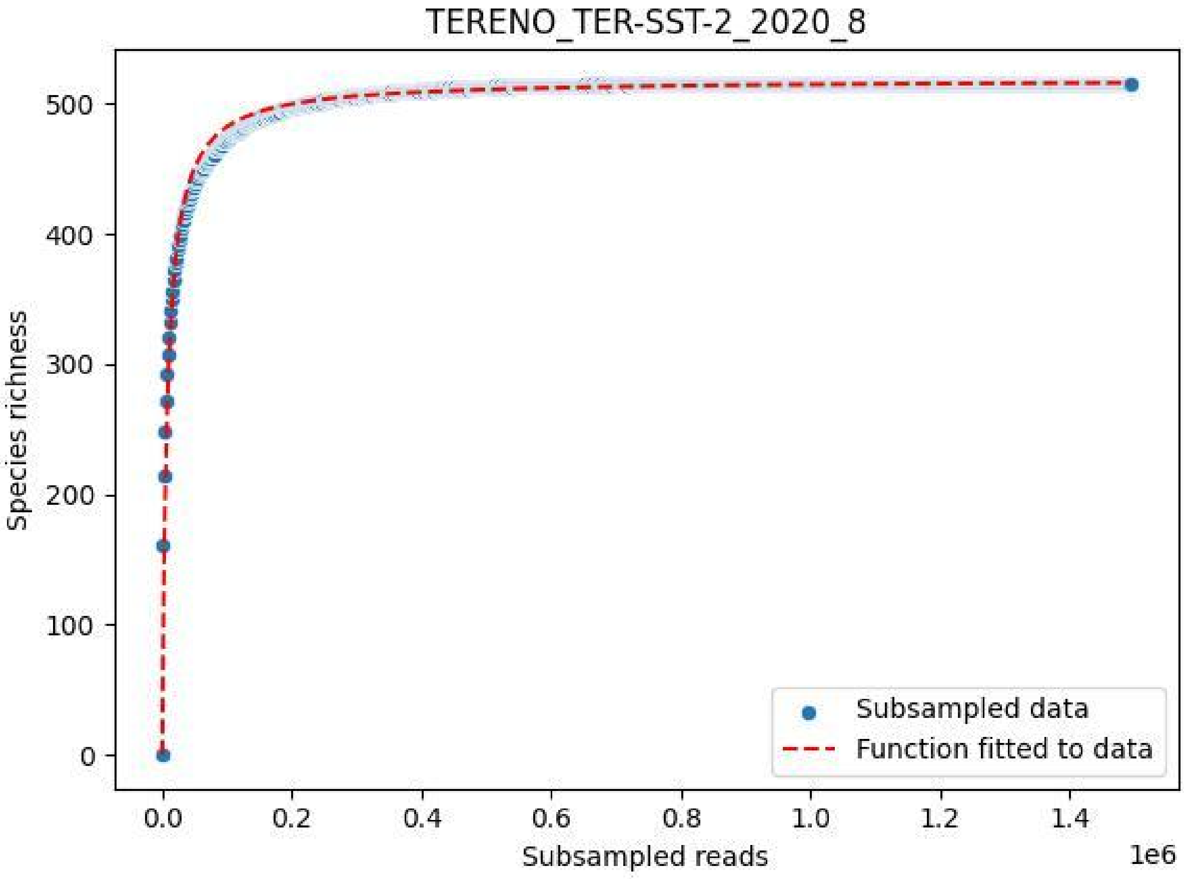
TERENO_TER-SST-2_2020_8
Species richness
0
100
200
300
400
500
0.0
0.2
0.4
0.6
0.8
1.0
1.2
1.4
1e6
Subsampled reads
Subsampled data
Function fitted to data

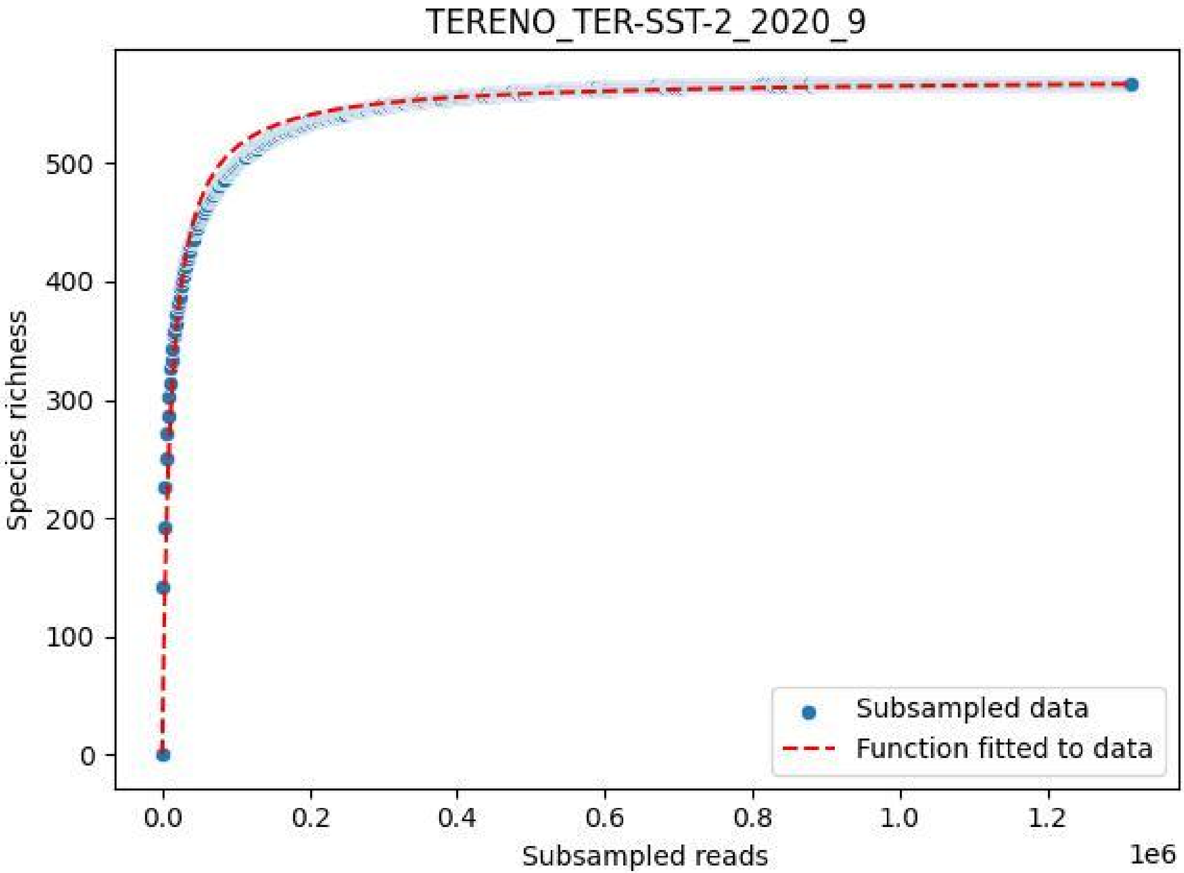

TERENO_TER-SST-2_2020_9
Species richness
Subsampled reads
1e6
0
100
200
300
400
500
0.0
0.2
0.4
0.6
0.8
1.0
1.2
Subsampled data
Function fitted to data

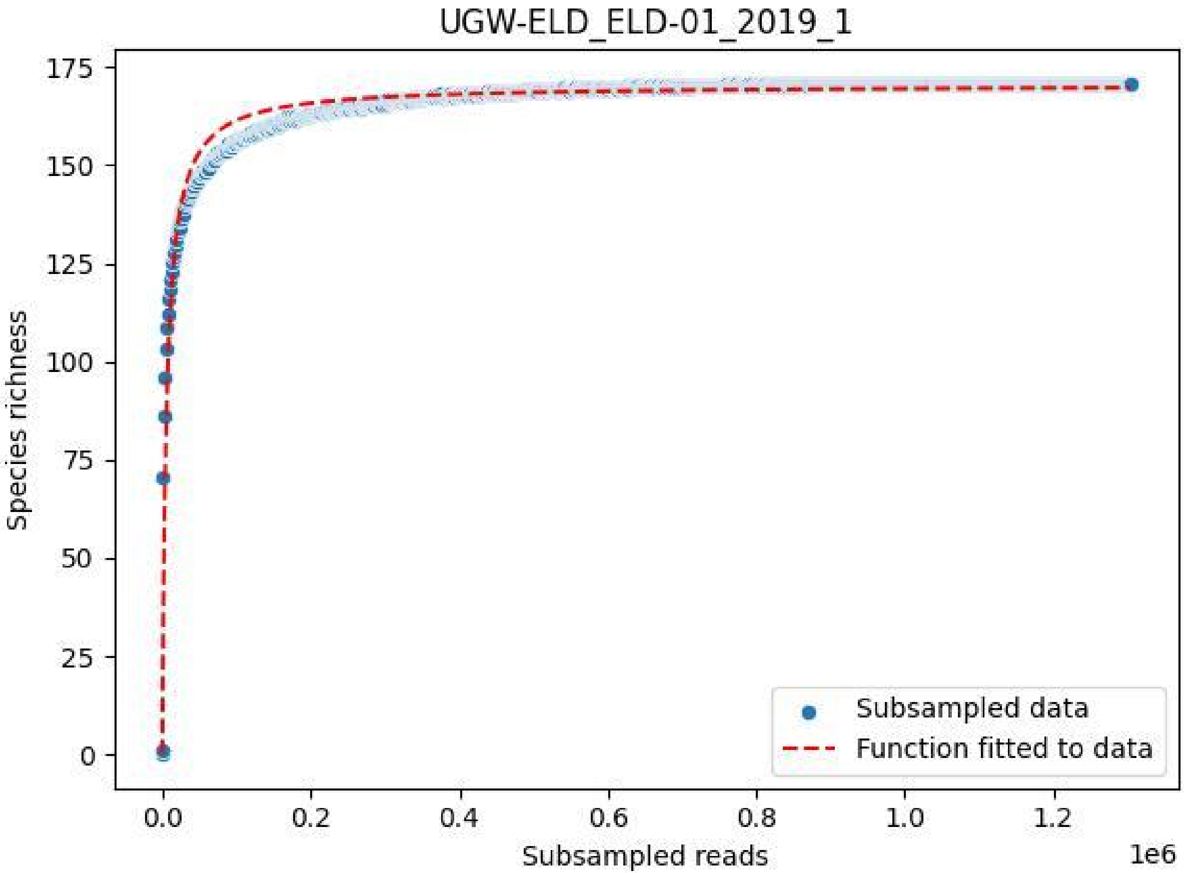

UGW-ELD_ELD-01_2019_1
Species richness
175
150
125
100
75
50
25
0
0.0
0.2
0.4
0.6
0.8
1.0
1.2
1e6
Subsampled reads
Subsampled data
Function fitted to data

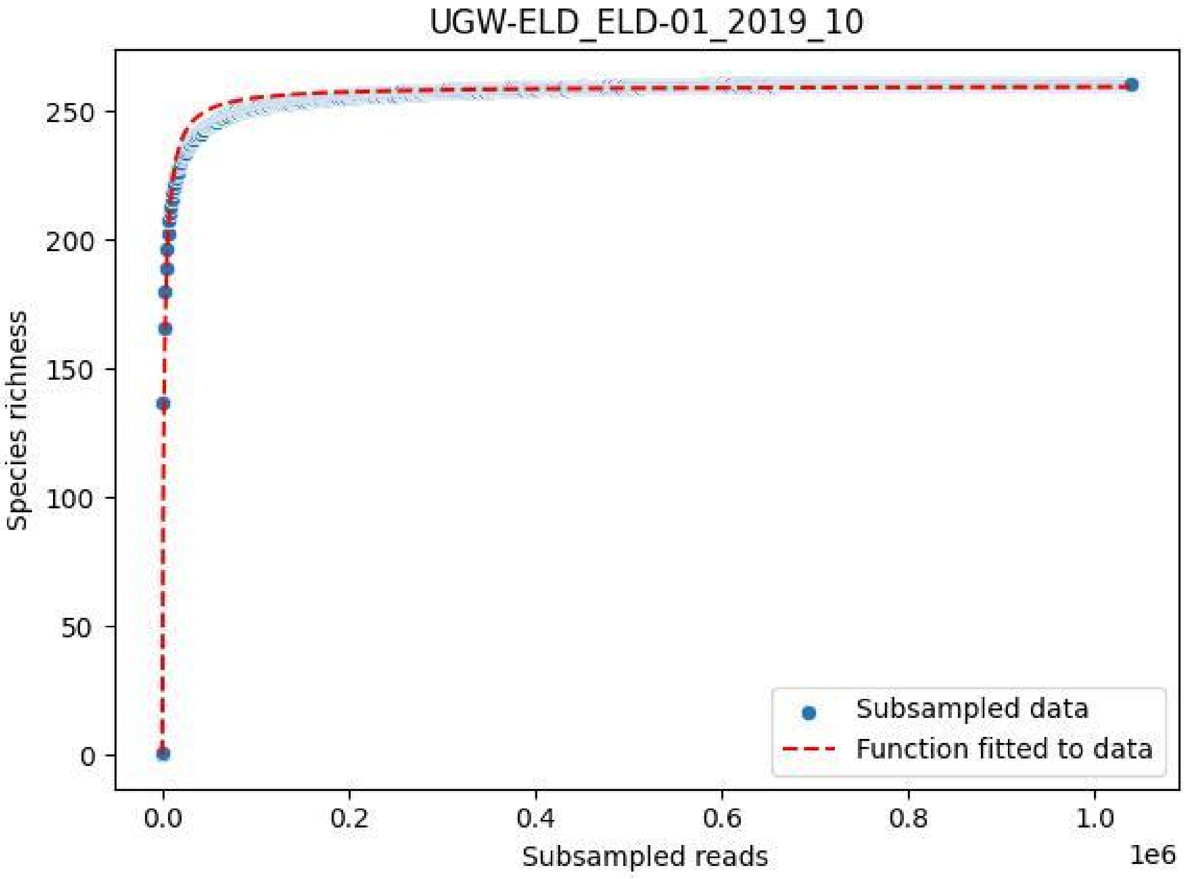
UGW-ELD_ELD-01_2019_10
Species richness
Subsampled reads
1e6
Subsampled data
Function fitted to data
0
50
100
150
200
250
0.0
0.2
0.4
0.6
0.8
1.0

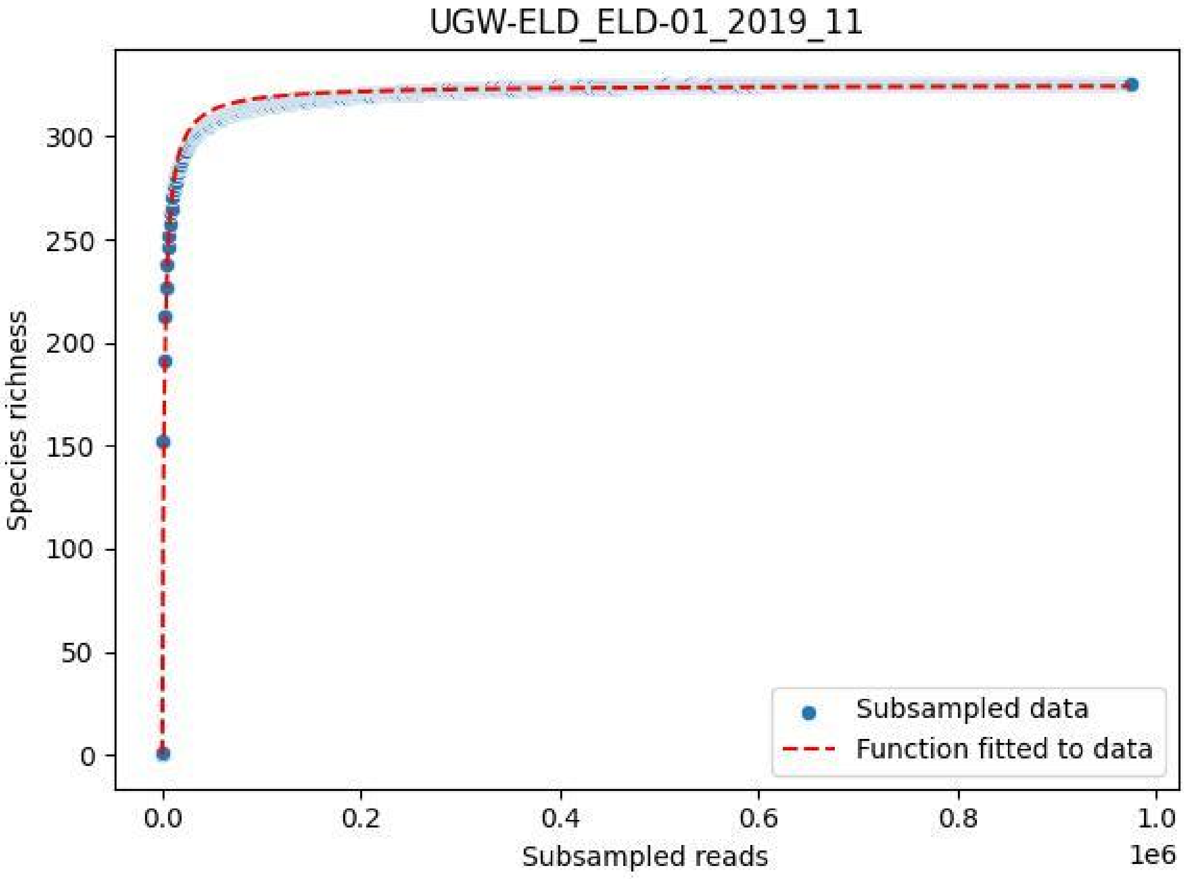

UGW-ELD_ELD-01_2019_11
300
250
200
150
100
50
0
Species richness
0.0
0.2
0.4
0.6
0.8
1.0
1e6
Subsampled reads
Subsampled data
Function fitted to data

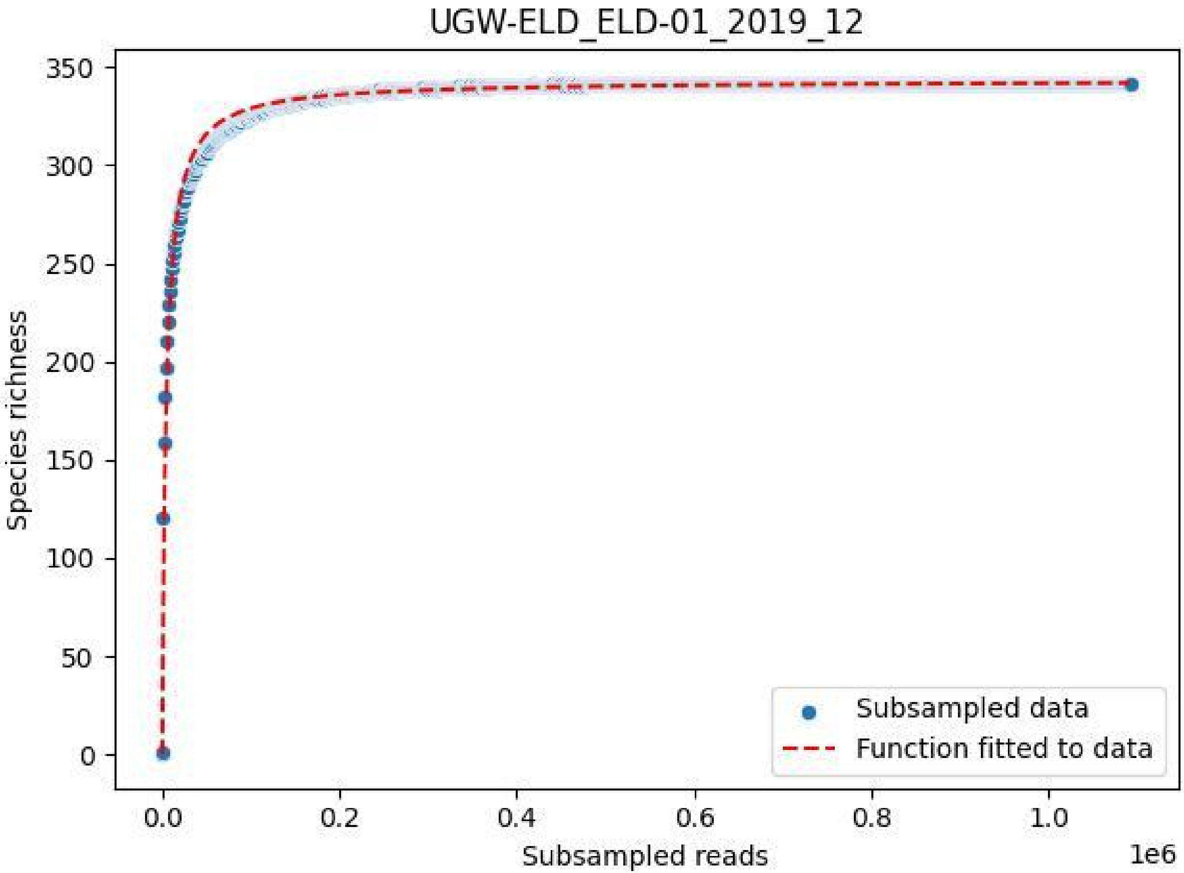

UGW-ELD_ELD-01_2019_12
Species richness
Subsampled reads
1e6
0
50
100
150
200
250
300
350
0.0
0.2
0.4
0.6
0.8
1.0
Subsampled data
Function fitted to data

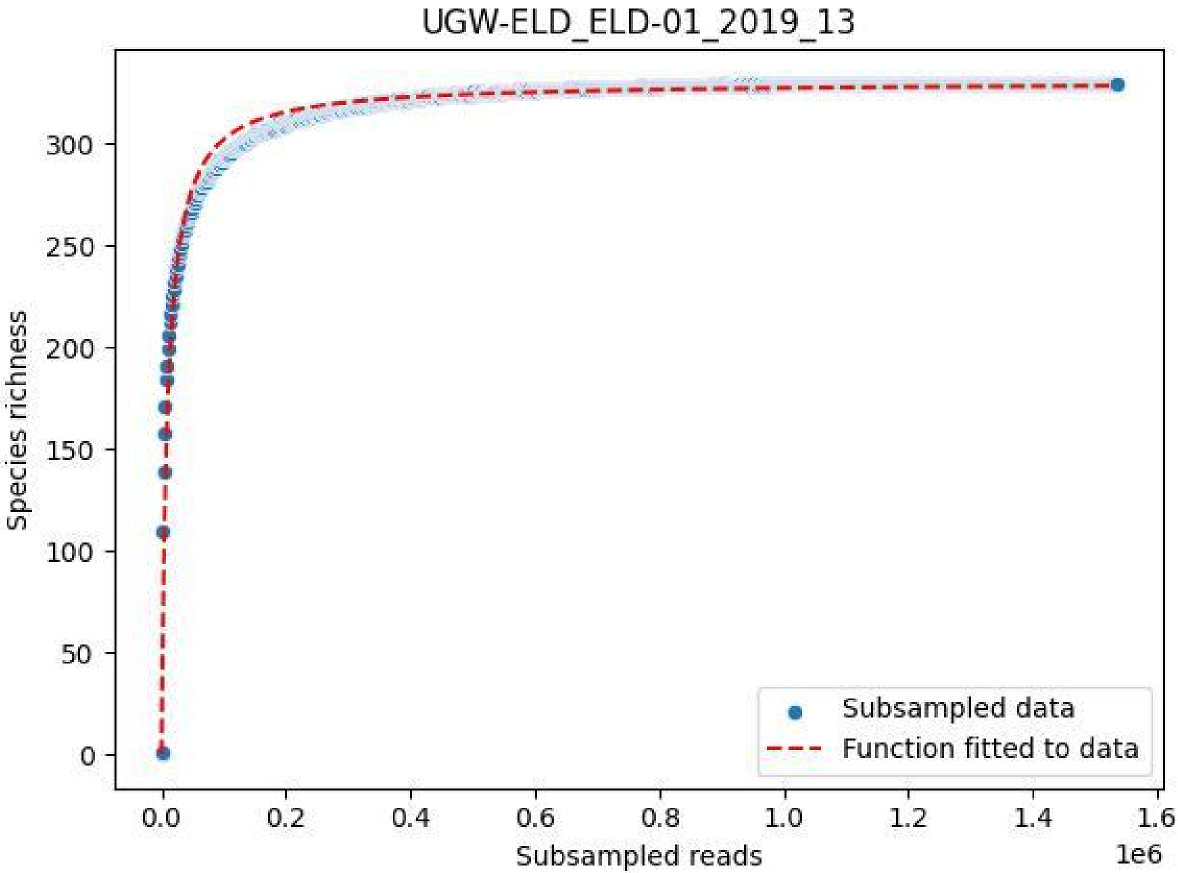
UGW-ELD_ELD-01_2019_13
Species richness
Subsampled reads
1e6
0
50
100
150
200
250
300
0.0
0.2
0.4
0.6
0.8
1.0
1.2
1.4
1.6
Subsampled data
Function fitted to data

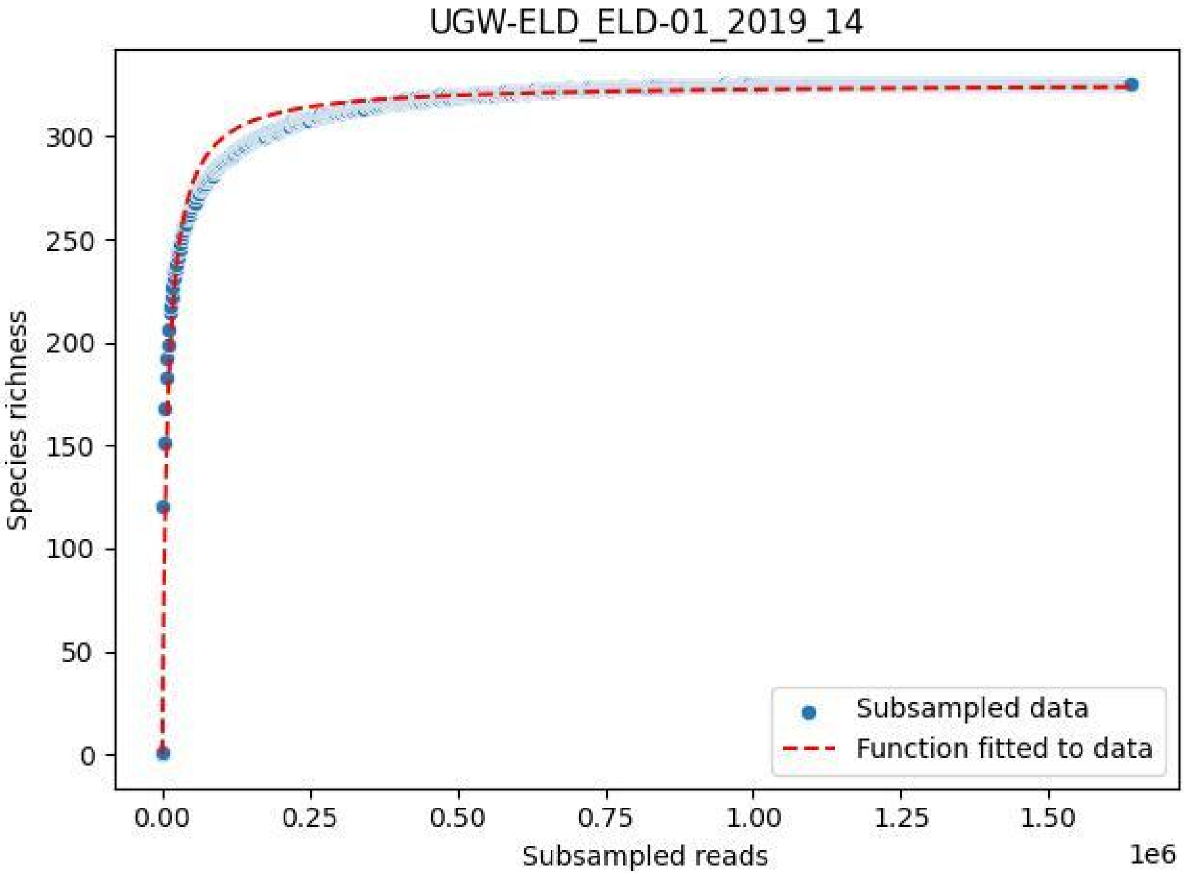
UGW-ELD_ELD-01_2019_14
Species richness
Subsampled reads
1e6
0
50
100
150
200
250
300
0.00
0.25
0.50
0.75
1.00
1.25
1.50
Subsampled data
Function fitted to data

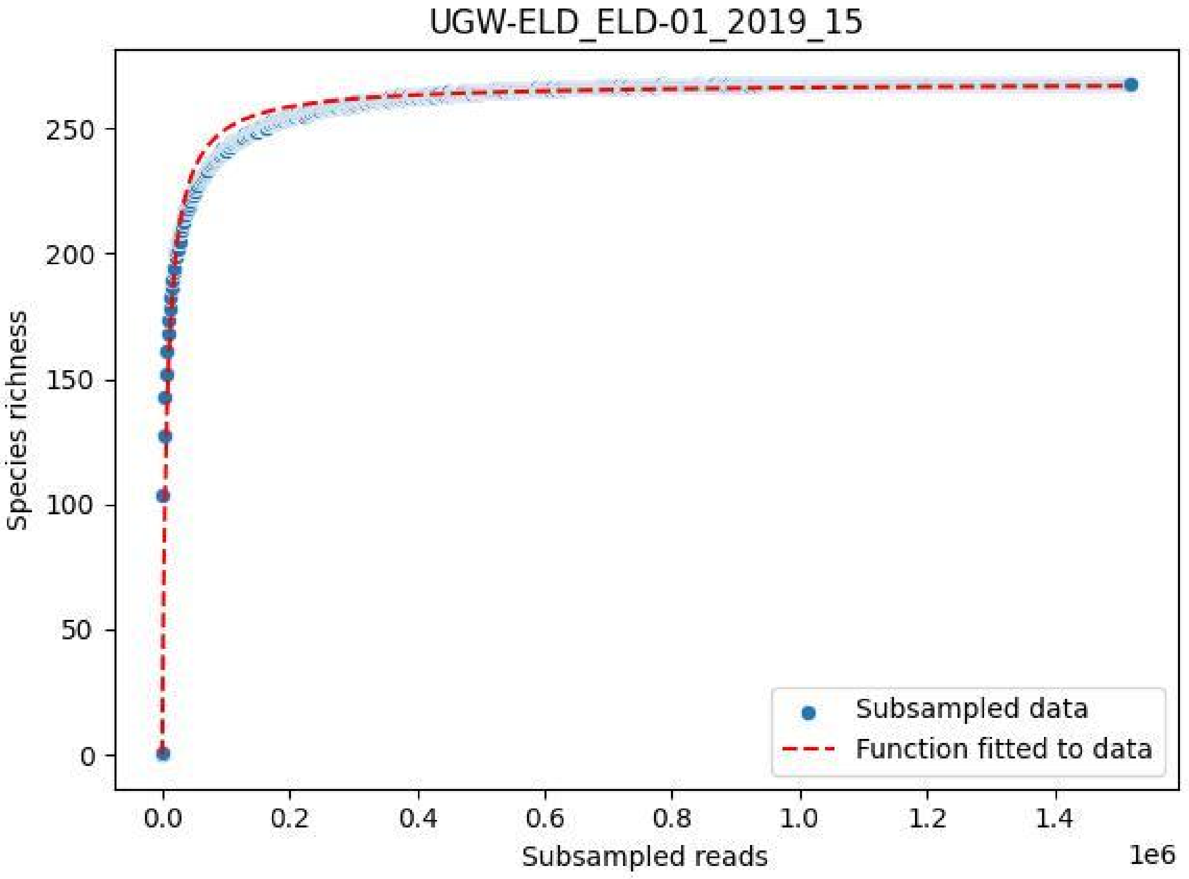

UGW-ELD_ELD-01_2019_15
Species richness
Subsampled reads
1e6
0
50
100
150
200
250
0.0
0.2
0.4
0.6
0.8
1.0
1.2
1.4
Subsampled data
Function fitted to data

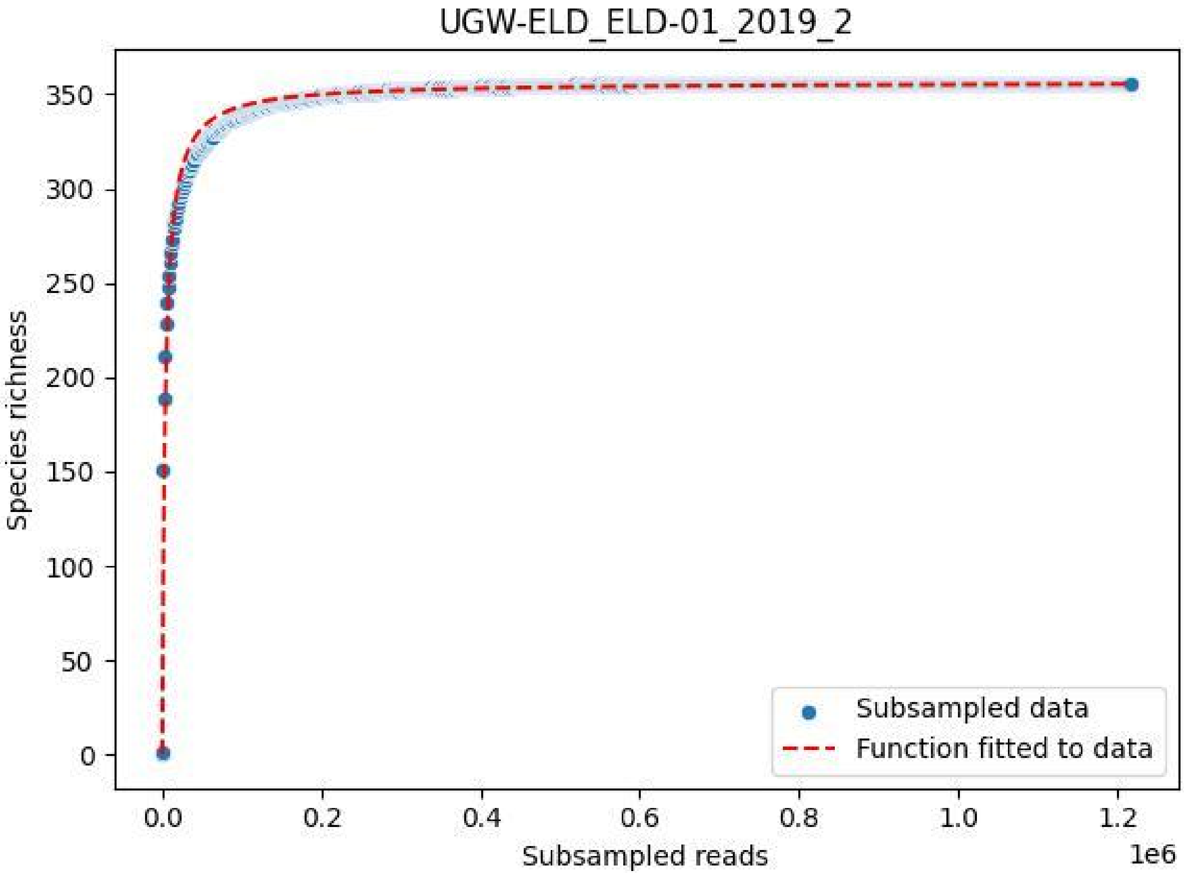

UGW-ELD_ELD-01_2019_2
Species richness
Subsampled reads
1e6
Subsampled data
Function fitted to data
0
50
100
150
200
250
300
350
0.0
0.2
0.4
0.6
0.8
1.0
1.2

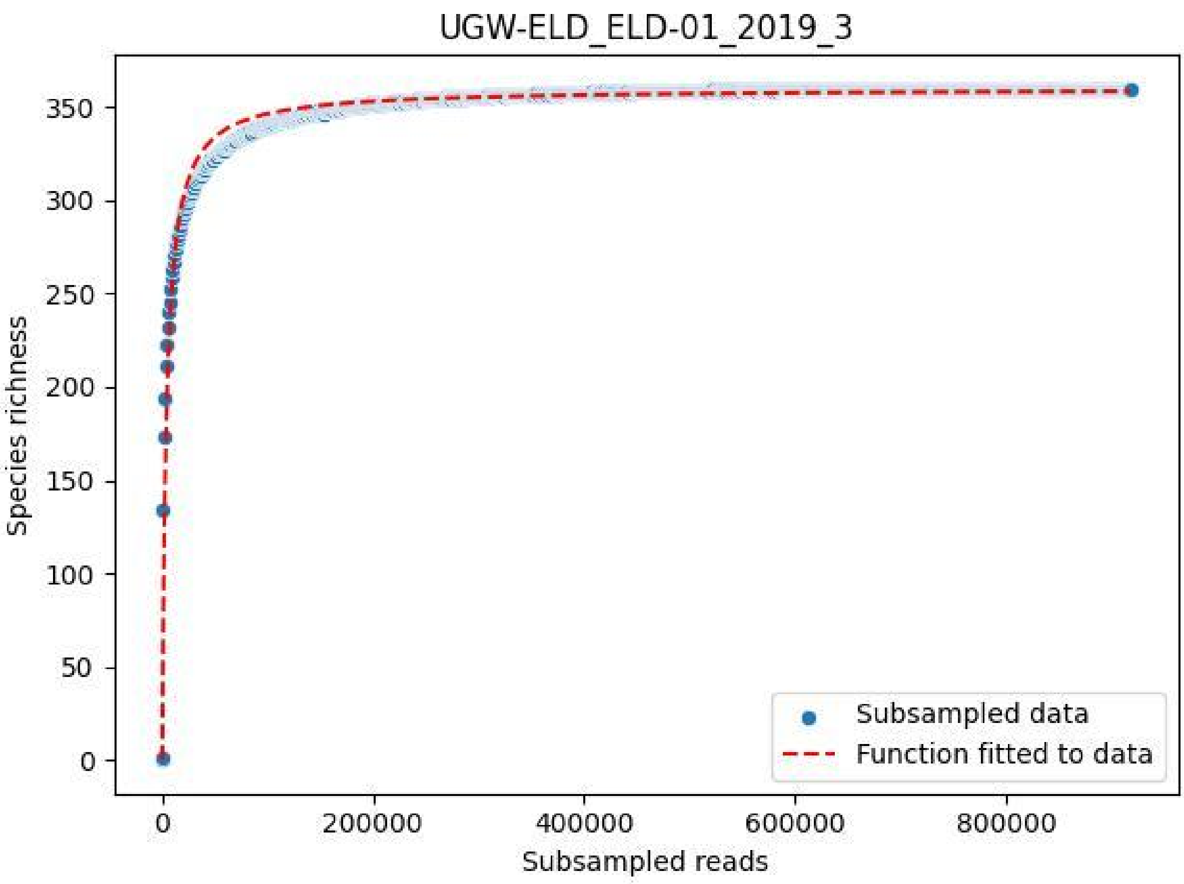

UGW-ELD_ELD-01_2019_3
Species richness
Subsampled reads
Subsampled data
Function fitted to data
0
50
100
150
200
250
300
350
0
200000
400000
600000
800000

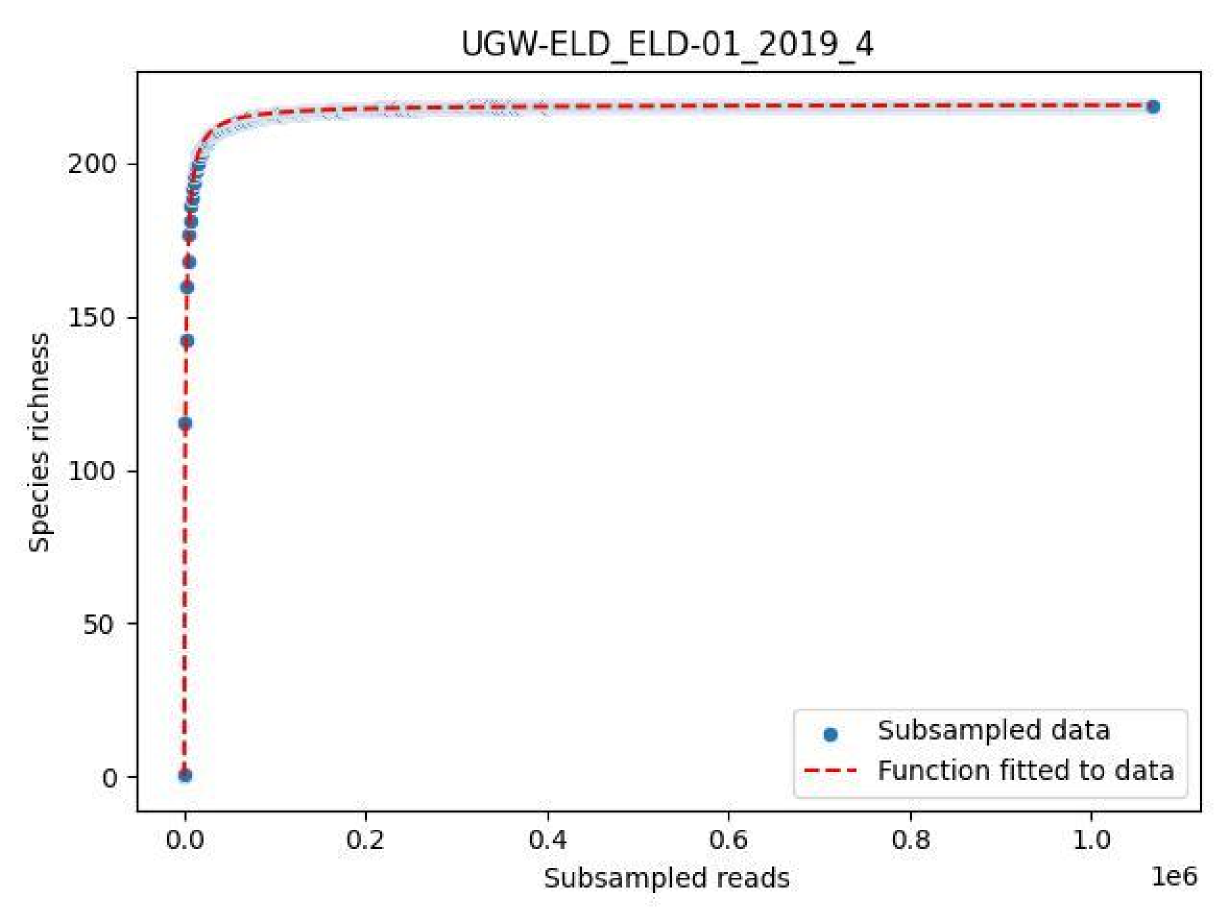

UGW-ELD_ELD-01_2019_4
Species richness
200
150
100
50
0
0.0
0.2
0.4
0.6
0.8
1.0
1e6
Subsampled reads
Subsampled data
Function fitted to data

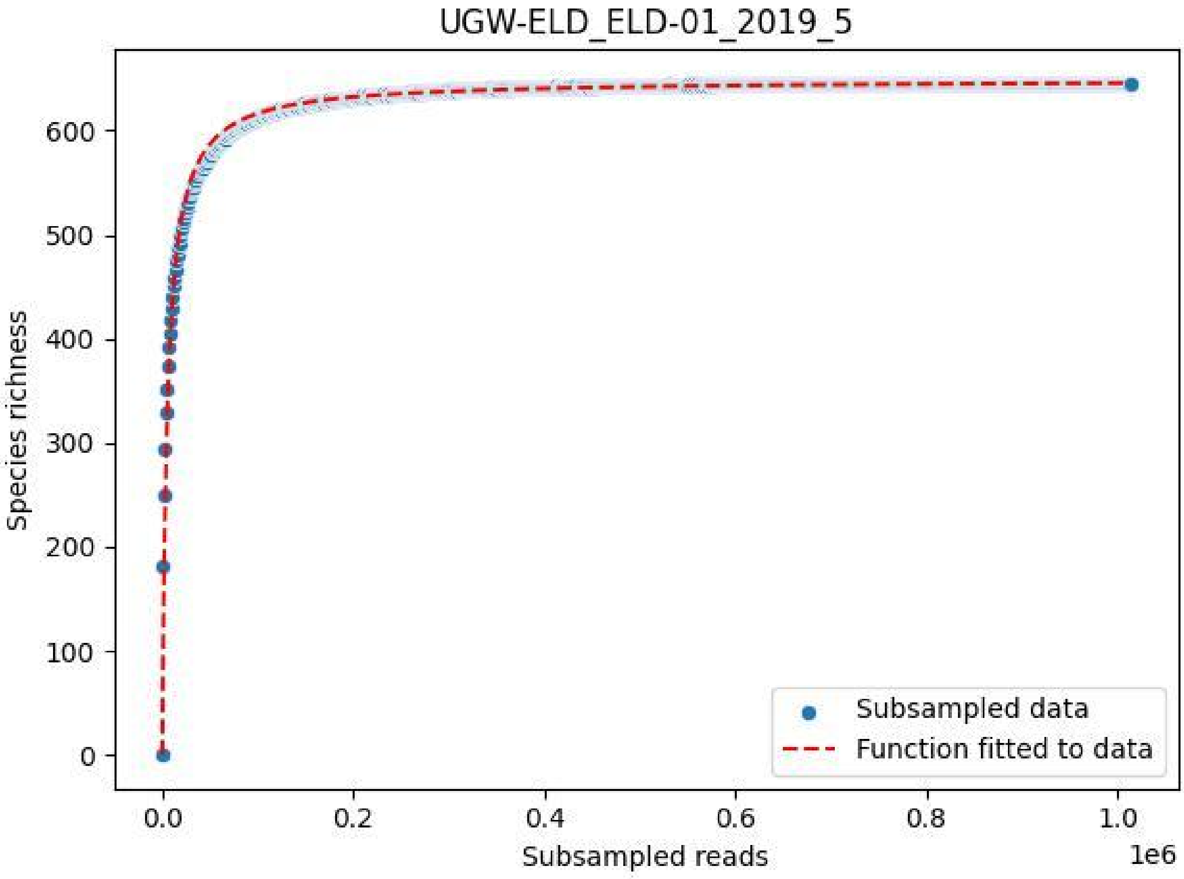

UGW-ELD_ELD-01_2019_5
Species richness
Subsampled reads
1e6
0
100
200
300
400
500
600
0.0
0.2
0.4
0.6
0.8
1.0
Subsampled data
Function fitted to data

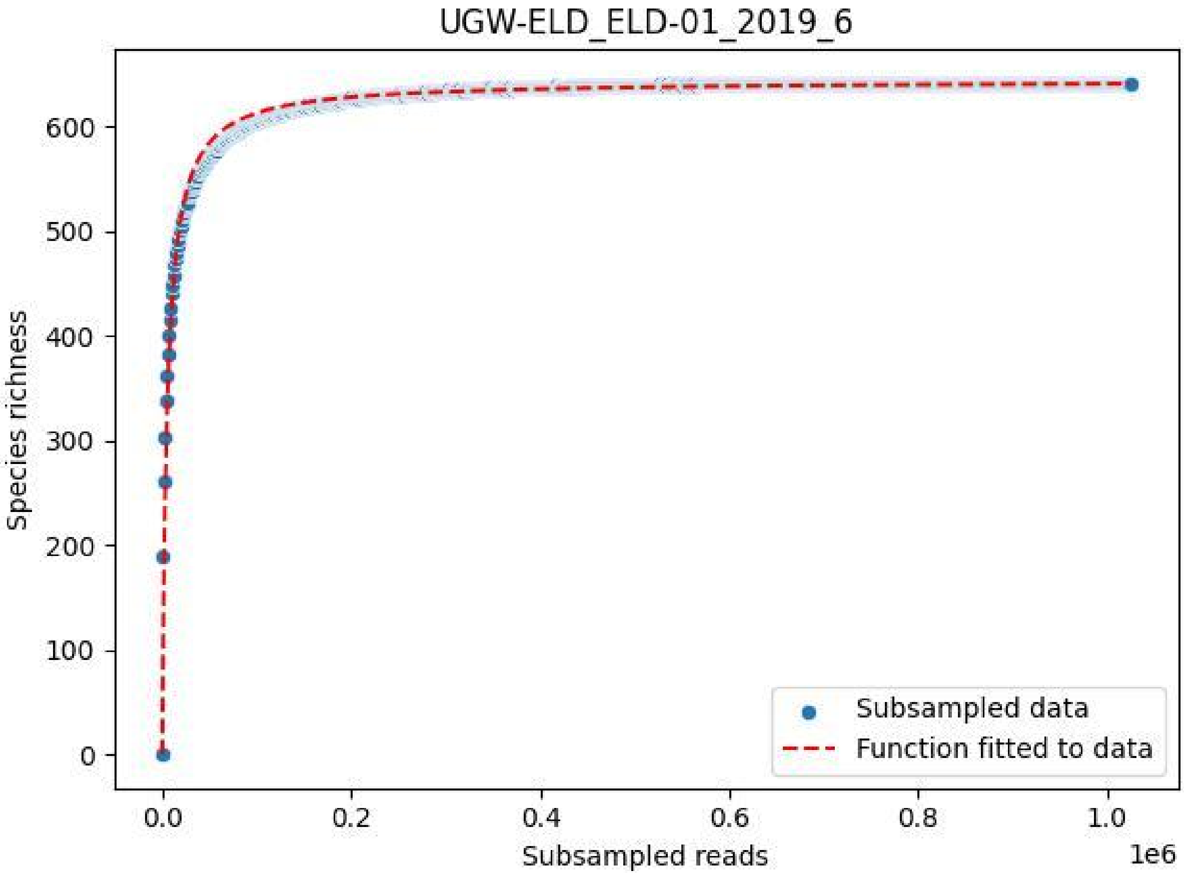
UGW-ELD_ELD-01_2019_6
Species richness
0
100
200
300
400
500
600
0.0
0.2
0.4
0.6
0.8
1.0
1e6
Subsampled reads
Subsampled data
Function fitted to data

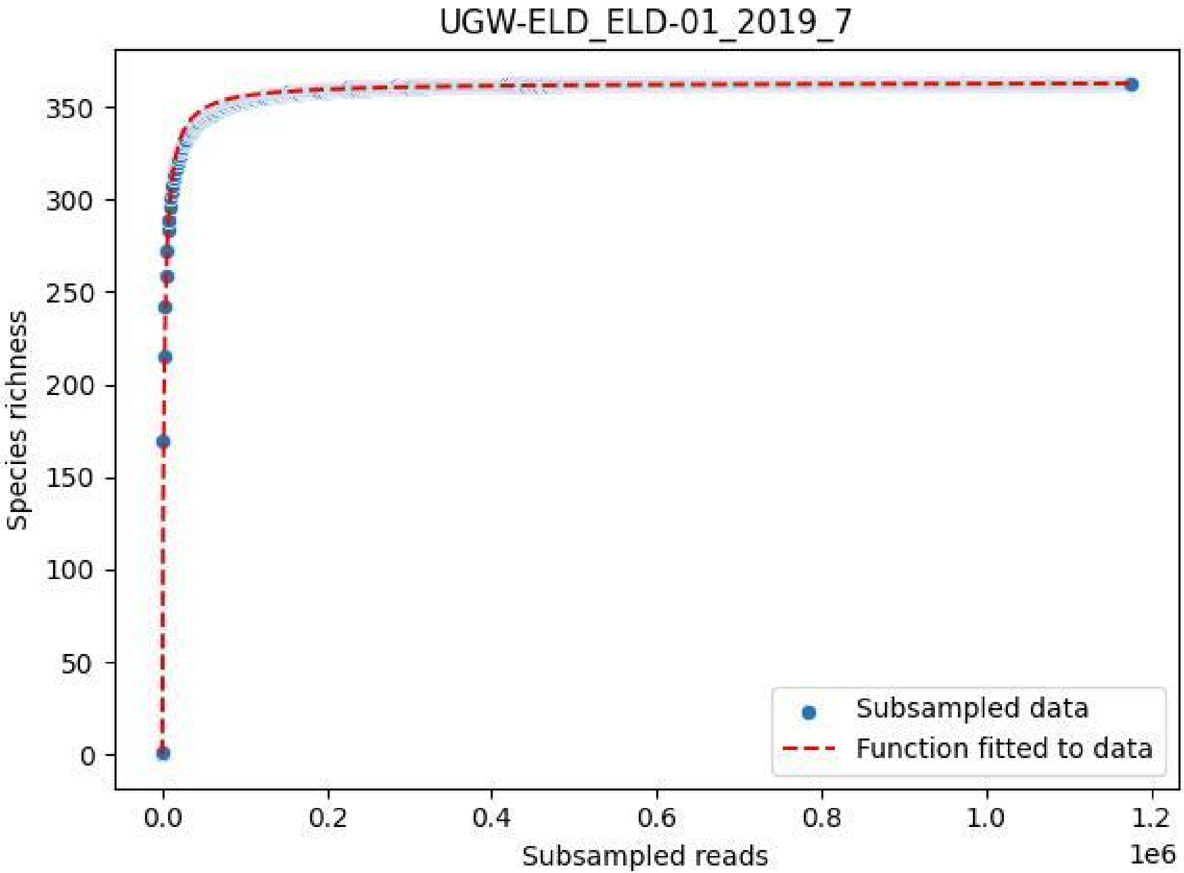

UGW-ELD_ELD-01_2019_7
Species richness
Subsampled reads
1e6
0
50
100
150
200
250
300
350
0.0
0.2
0.4
0.6
0.8
1.0
1.2
Subsampled data
Function fitted to data

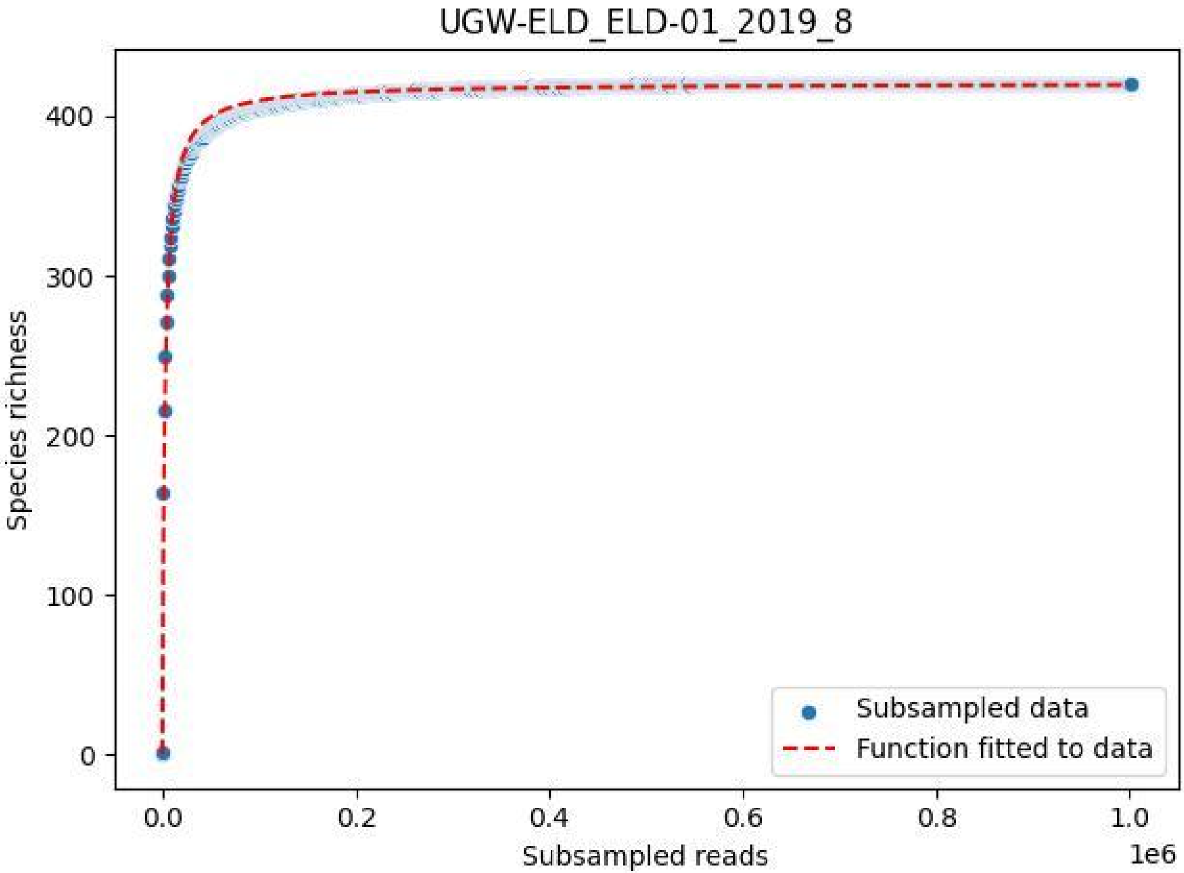

UGW-ELD_ELD-01_2019_8
Species richness
Subsampled reads
1e6
0
100
200
300
400
0.0
0.2
0.4
0.6
0.8
1.0
Subsampled data
Function fitted to data

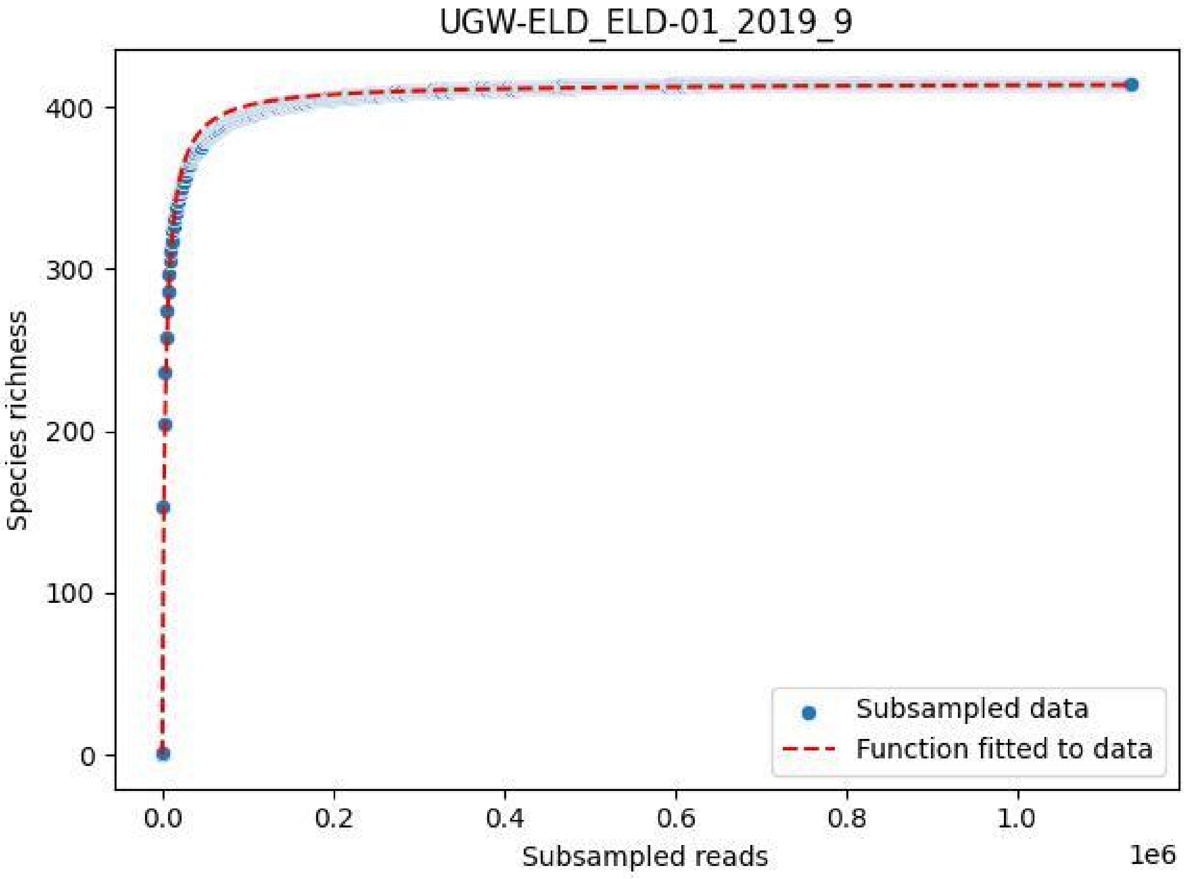
UGW-ELD_ELD-01_2019_9
Species richness
Subsampled reads
1e6
0
100
200
300
400
0.0
0.2
0.4
0.6
0.8
1.0
Subsampled data
Function fitted to data

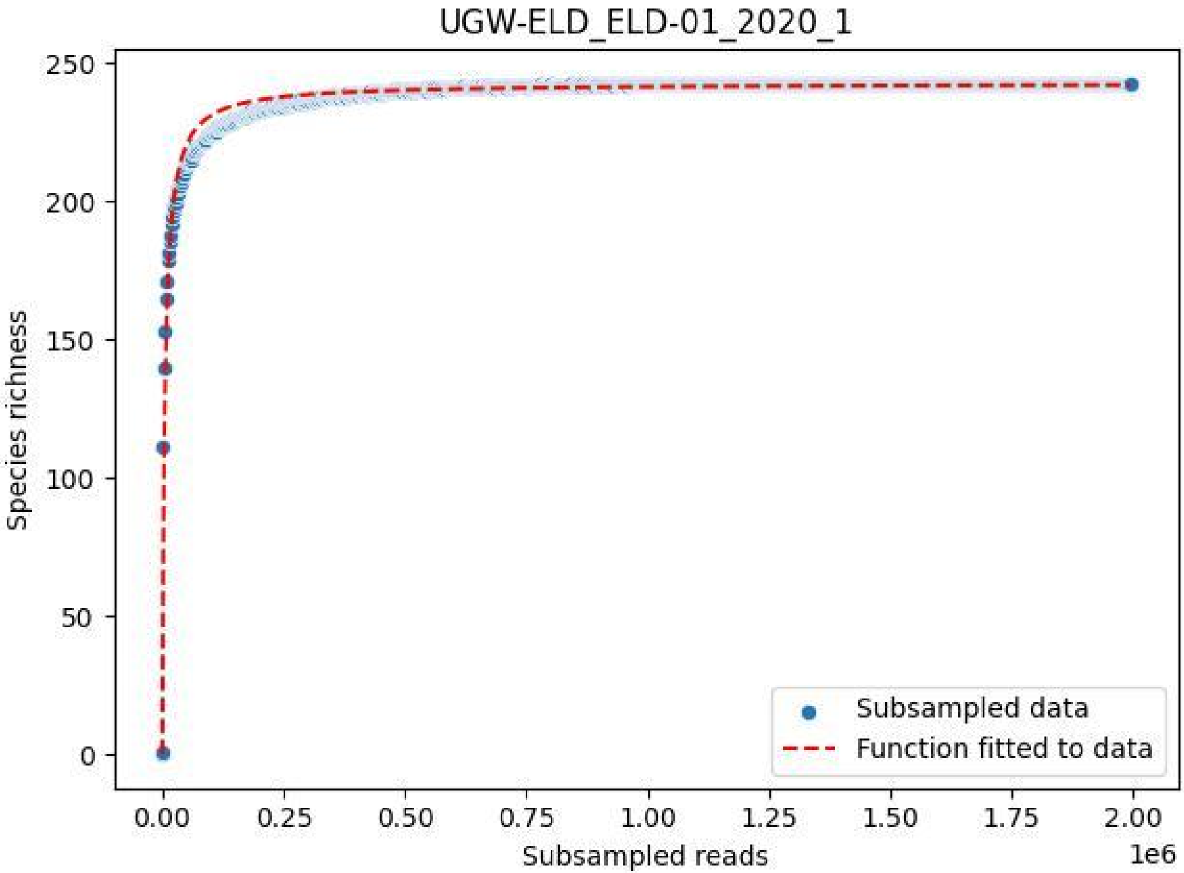
UGW-ELD_ELD-01_2020_1
Species richness
Subsampled reads
1e6
250
200
150
100
50
0
0.00
0.25
0.50
0.75
1.00
1.25
1.50
1.75
2.00
Subsampled data
Function fitted to data

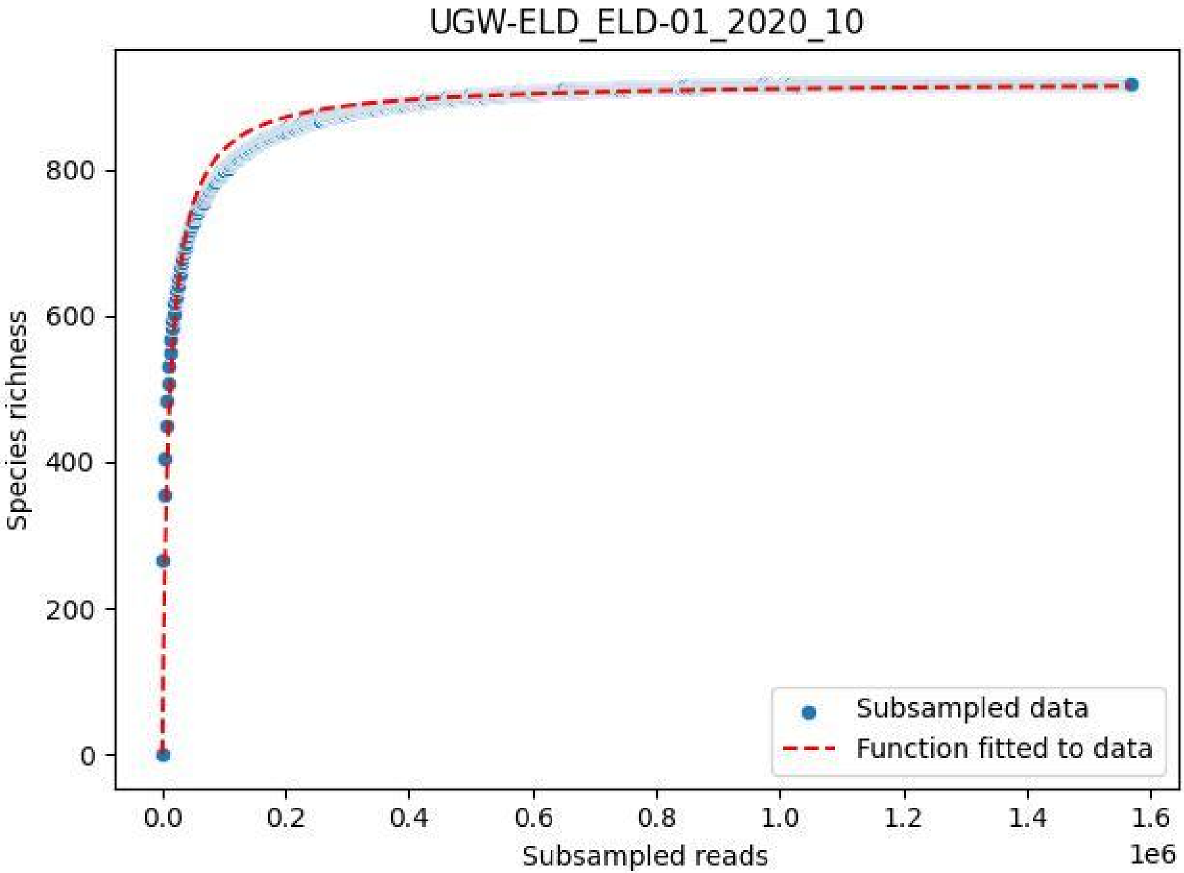
UGW-ELD_ELD-01_2020_10
Species richness
Subsampled reads
1e6
0
200
400
600
800
0.0
0.2
0.4
0.6
0.8
1.0
1.2
1.4
1.6
Subsampled data
Function fitted to data

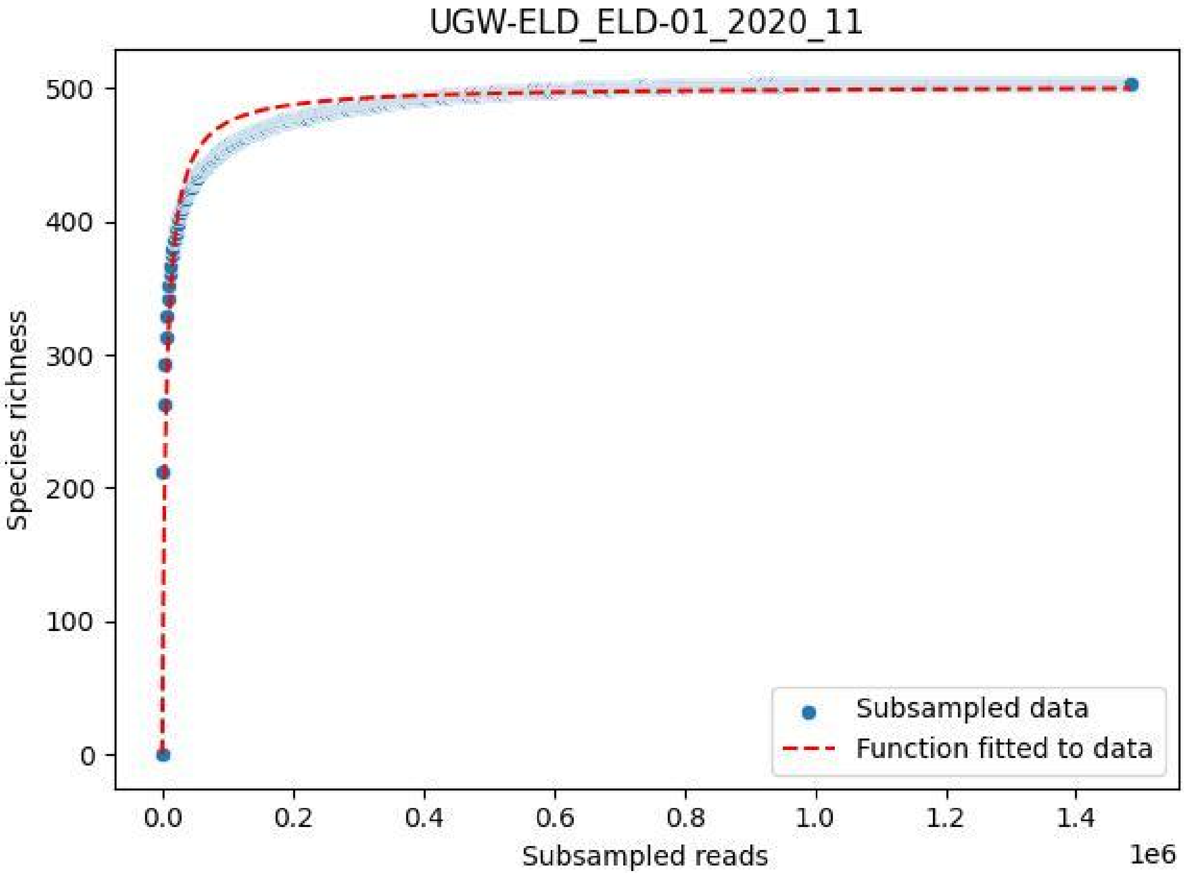

UGW-ELD_ELD-01_2020_11
Species richness
Subsampled reads
1e6
Subsampled data
Function fitted to data
0
100
200
300
400
500
0.0
0.2
0.4
0.6
0.8
1.0
1.2
1.4

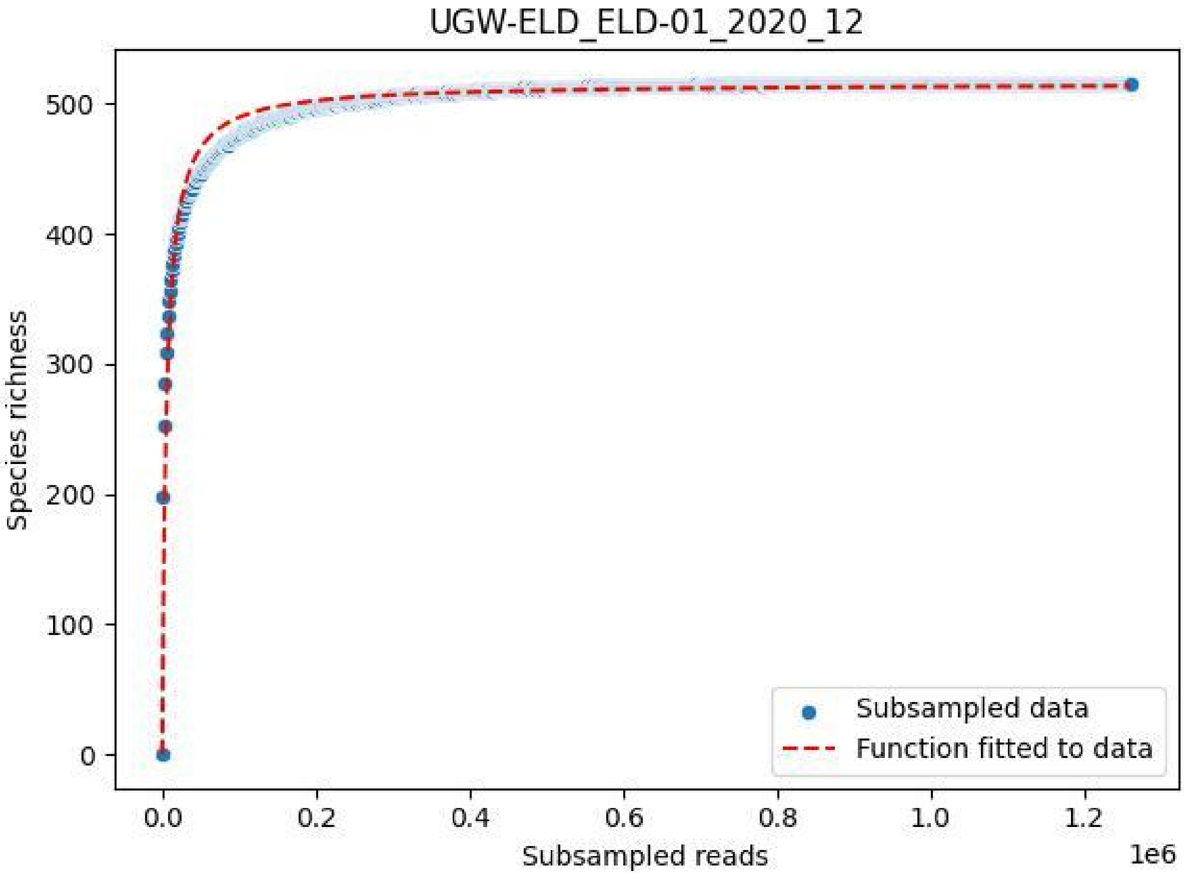
UGW-ELD_ELD-01_2020_12
Species richness
Subsampled reads
1e6
0
100
200
300
400
500
0.0
0.2
0.4
0.6
0.8
1.0
1.2
Subsampled data
Function fitted to data

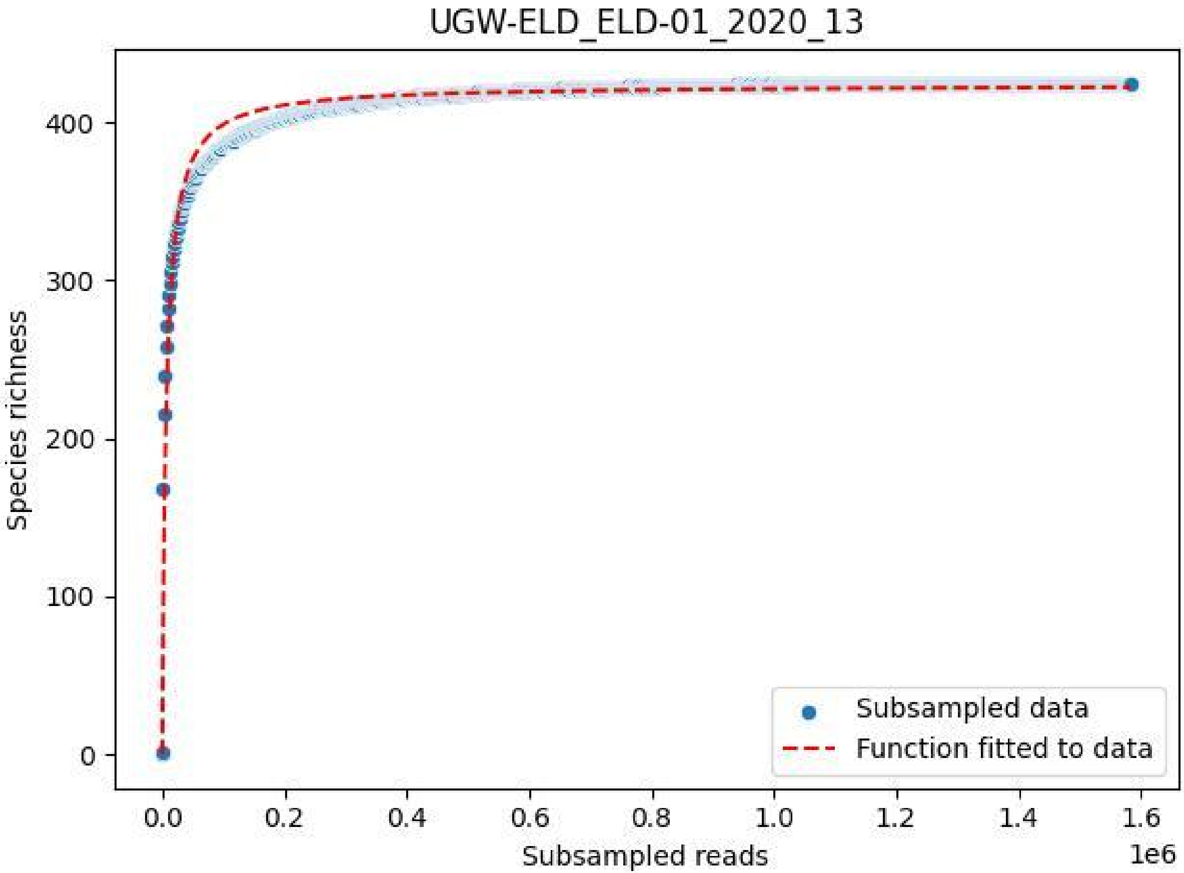
UGW-ELD_ELD-01_2020_13
Species richness
Subsampled reads
1e6
0
100
200
300
400
0.0
0.2
0.4
0.6
0.8
1.0
1.2
1.4
1.6
Subsampled data
Function fitted to data

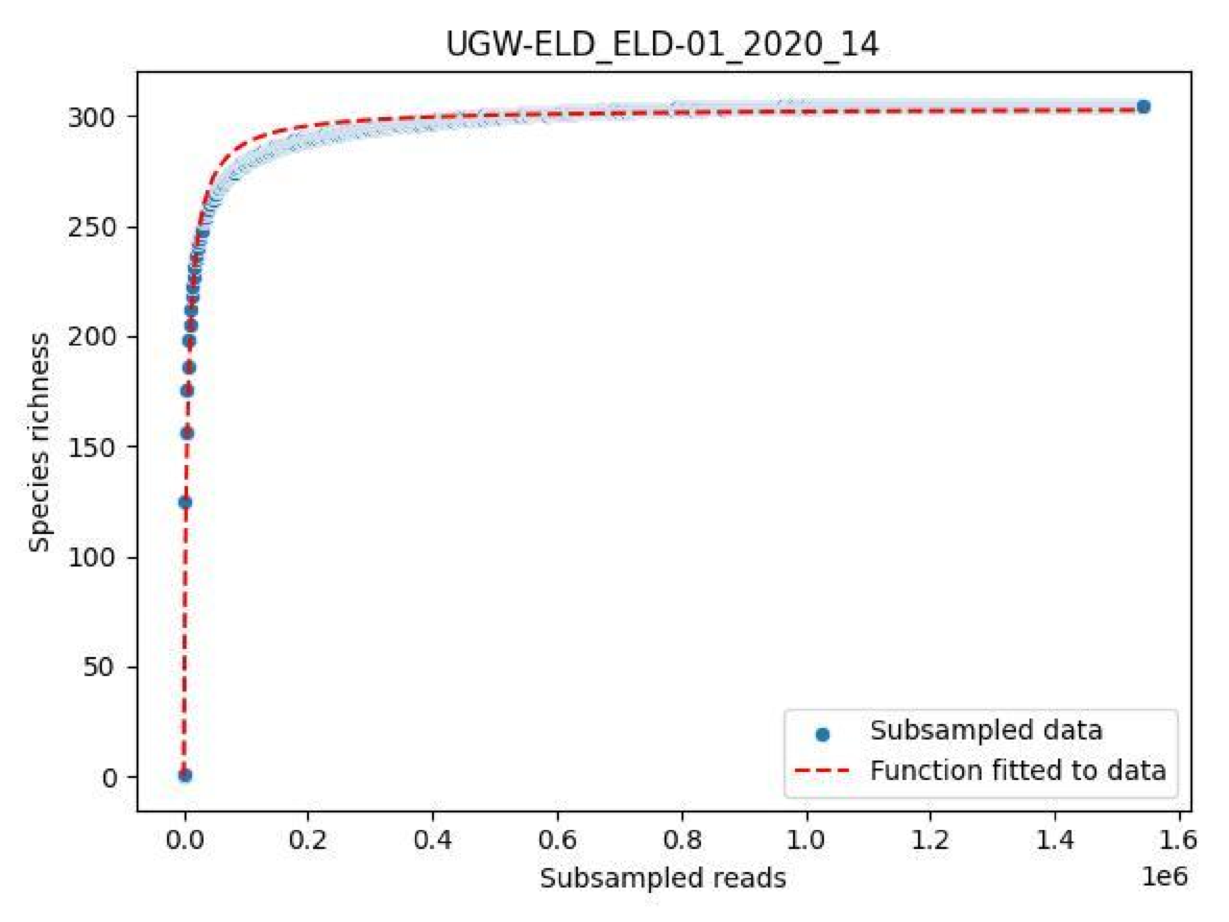
UGW-ELD_ELD-01_2020_14
Species richness
Subsampled reads
1e6
0
50
100
150
200
250
300
0.0
0.2
0.4
0.6
0.8
1.0
1.2
1.4
1.6
Subsampled data
Function fitted to data

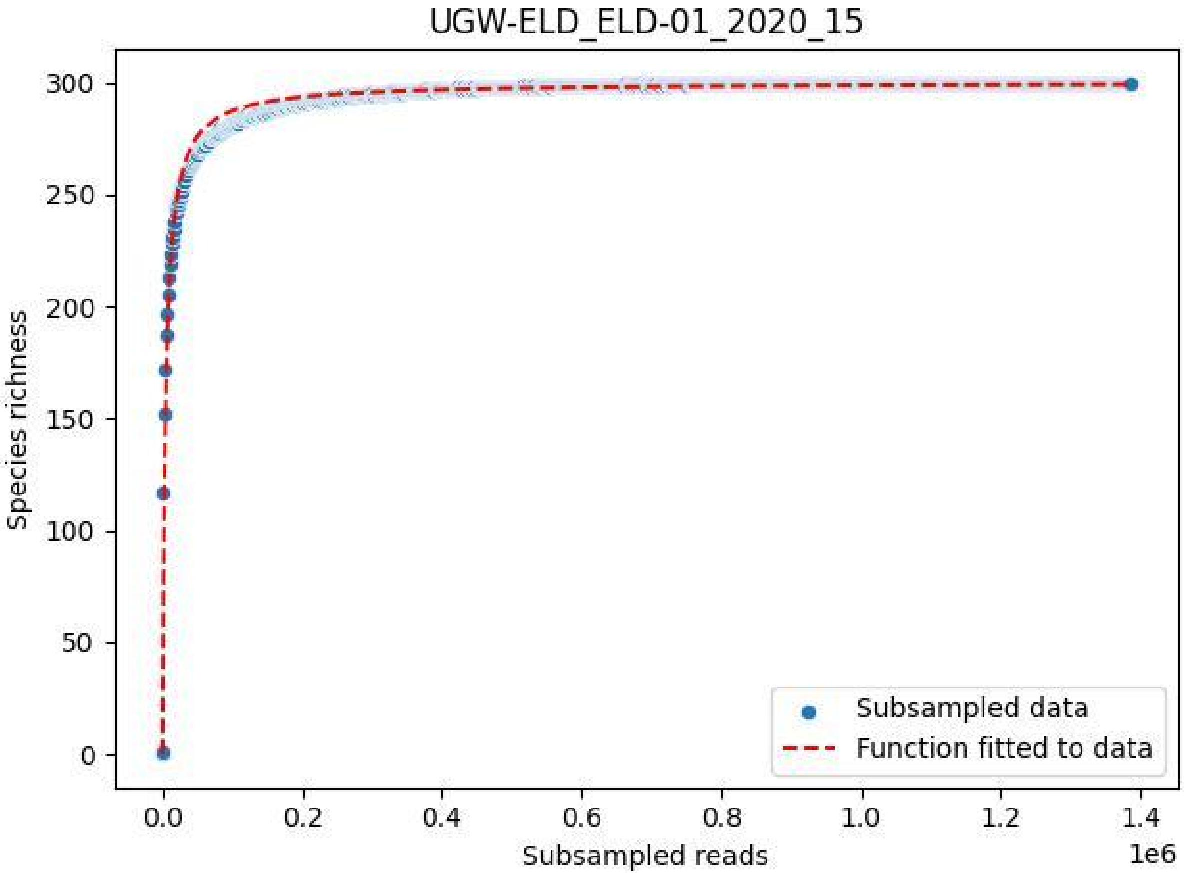

UGW-ELD_ELD-01_2020_15
Species richness
Subsampled reads
1e6
0
50
100
150
200
250
300
0.0
0.2
0.4
0.6
0.8
1.0
1.2
1.4
Subsampled data
Function fitted to data

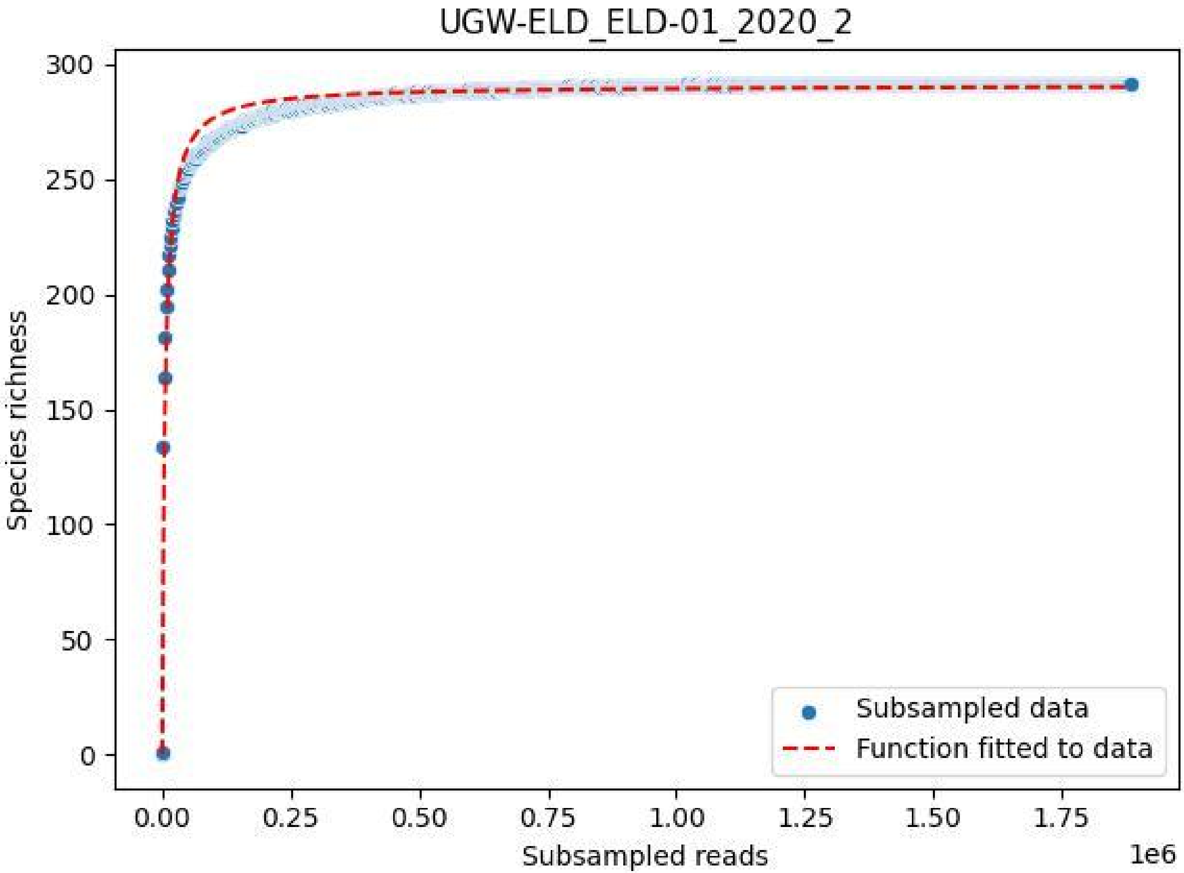
UGW-ELD_ELD-01_2020_2
Species richness
300
250
200
150
100
50
0
0.00
0.25
0.50
0.75
1.00
1.25
1.50
1.75
1e6
Subsampled reads
Subsampled data
Function fitted to data

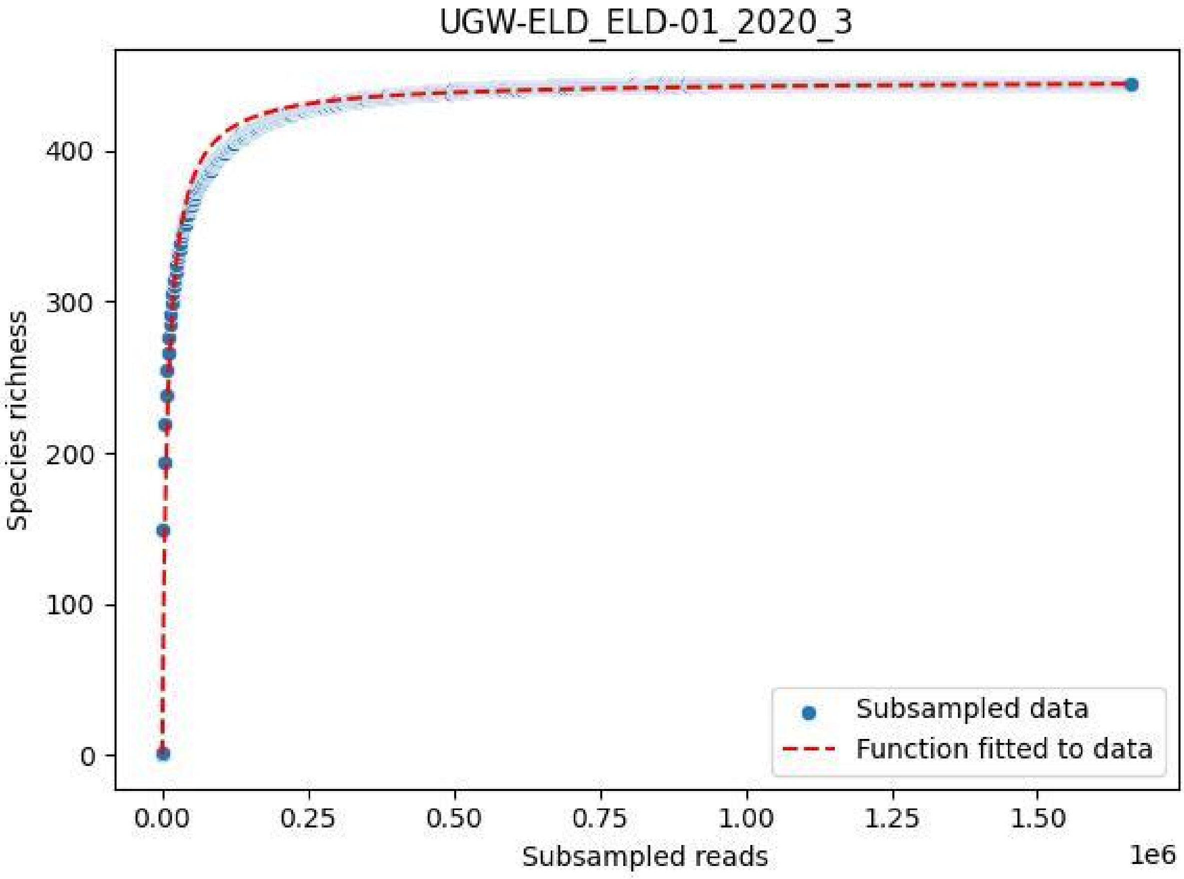

UGW-ELD_ELD-01_2020_3
Species richness
Subsampled reads
1e6
0
100
200
300
400
0.00
0.25
0.50
0.75
1.00
1.25
1.50
Subsampled data
Function fitted to data

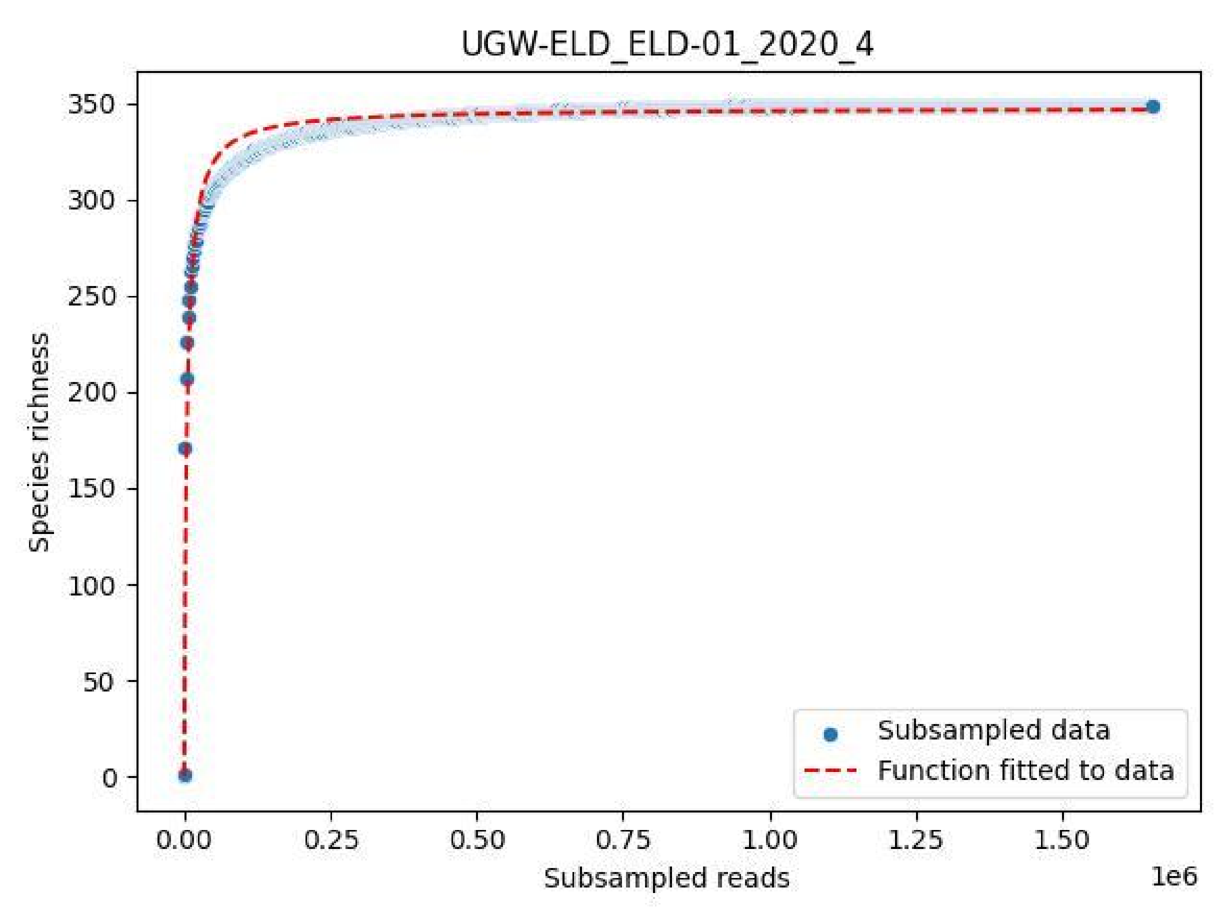
UGW-ELD_ELD-01_2020_4
Species richness
Subsampled reads
1e6
0
50
100
150
200
250
300
350
0.00
0.25
0.50
0.75
1.00
1.25
1.50
Subsampled data
Function fitted to data

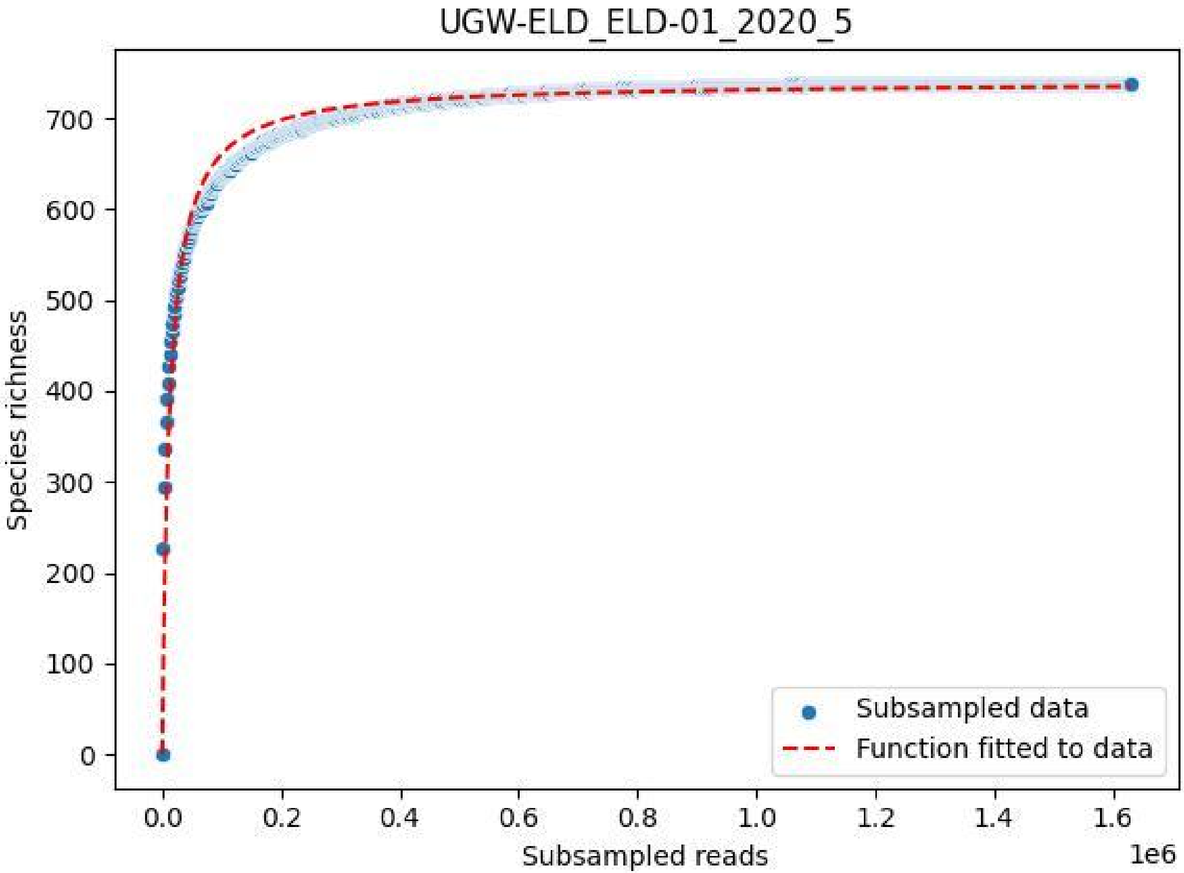
UGW-ELD_ELD-01_2020_5
Species richness
Subsampled reads
1e6
Subsampled data
Function fitted to data
0
100
200
300
400
500
600
700
0.0
0.2
0.4
0.6
0.8
1.0
1.2
1.4
1.6

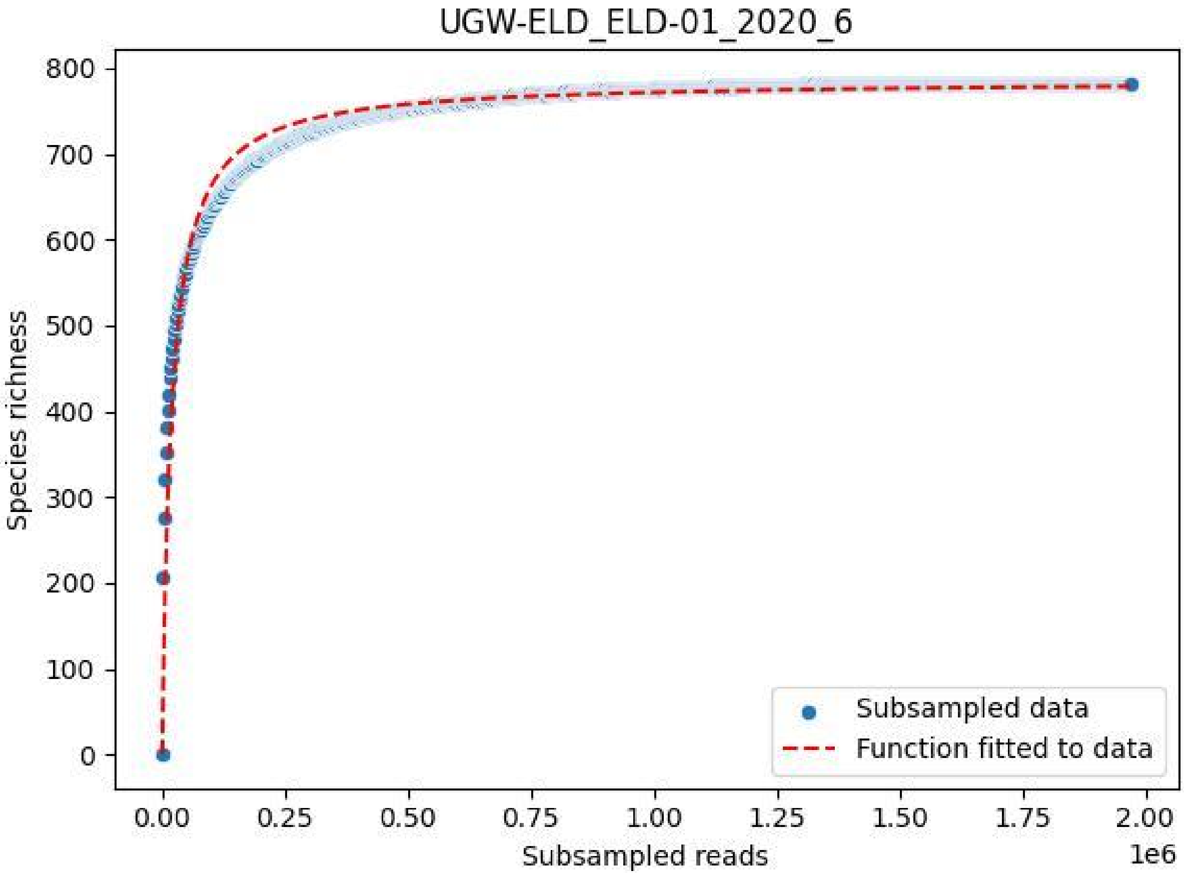
UGW-ELD_ELD-01_2020_6
Species richness
Subsampled reads
1e6
0
100
200
300
400
500
600
700
800
0.00
0.25
0.50
0.75
1.00
1.25
1.50
1.75
2.00
Subsampled data
Function fitted to data

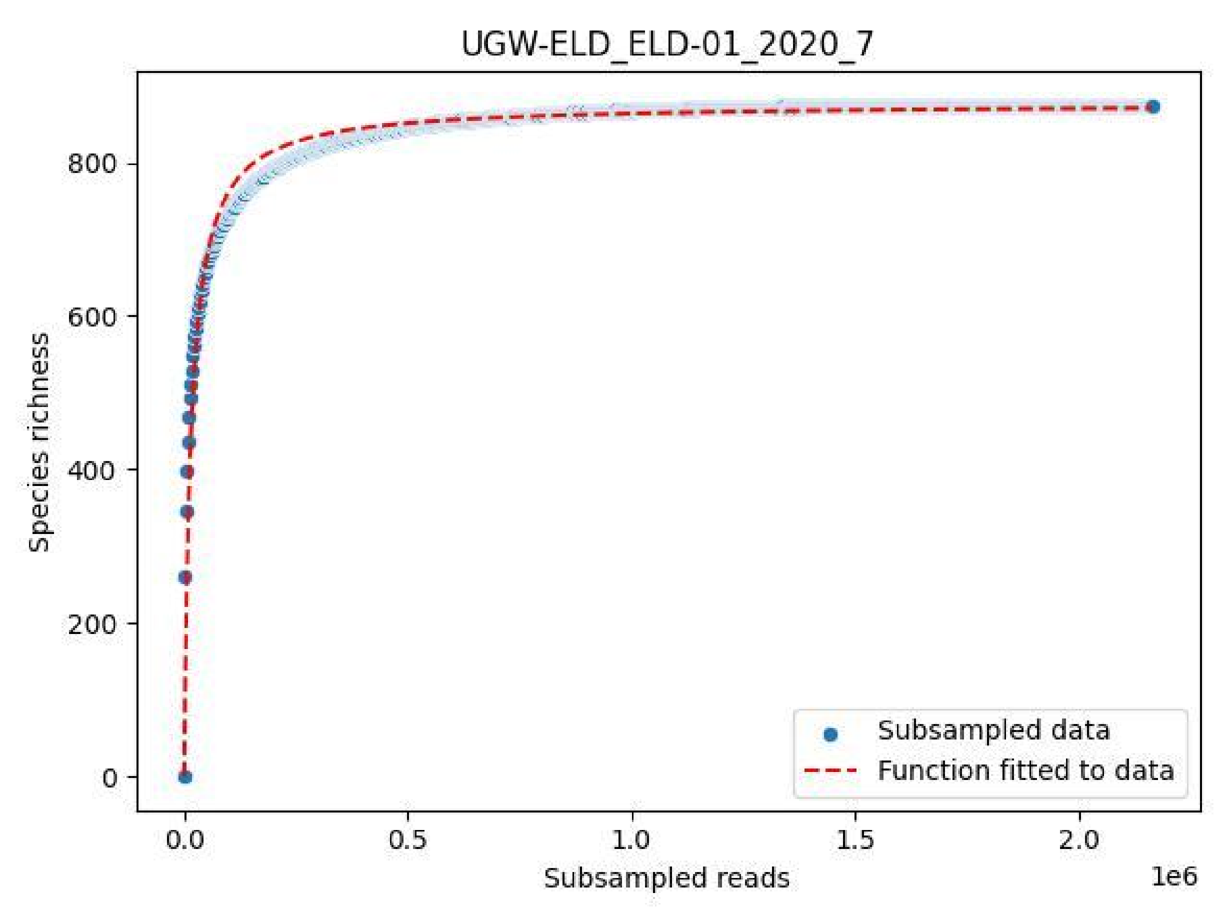
UGW-ELD_ELD-01_2020_7
Species richness
800
600
400
200
0
0.0
0.5
1.0
1.5
2.0
Subsampled reads
1e6
Subsampled data
Function fitted to data

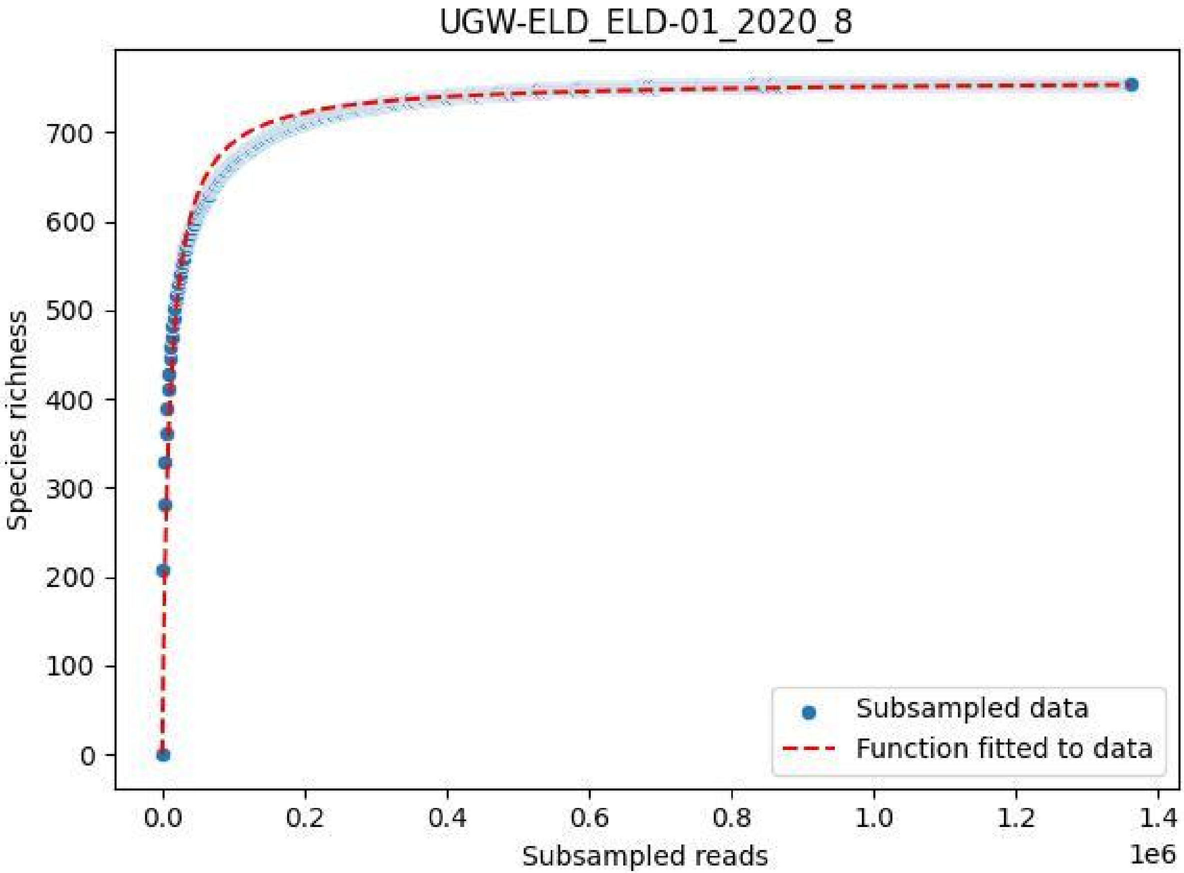

UGW-ELD_ELD-01_2020_8
Species richness
0
100
200
300
400
500
600
700
0.0
0.2
0.4
0.6
0.8
1.0
1.2
1.4
1e6
Subsampled reads
Subsampled data
Function fitted to data

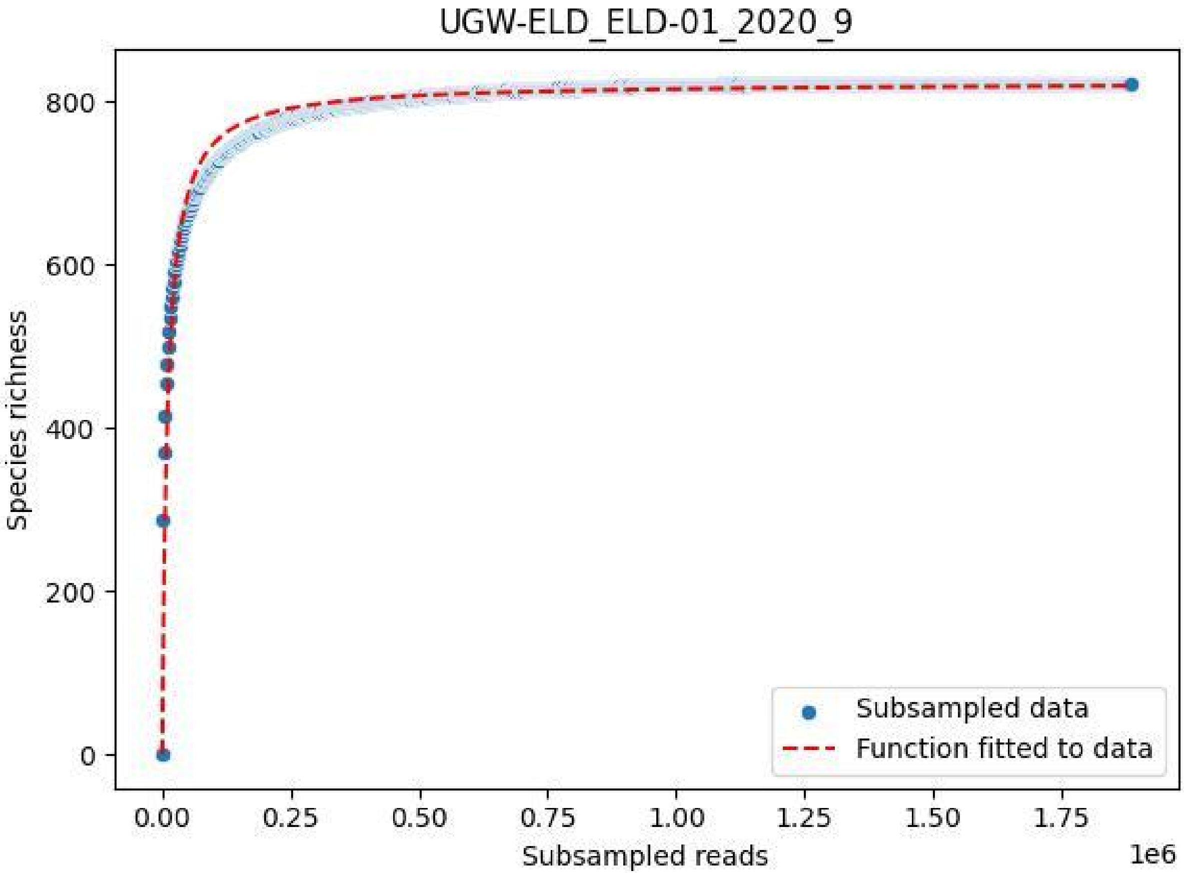

UGW-ELD_ELD-01_2020_9
Species richness
0
200
400
600
800
0.00
0.25
0.50
0.75
1.00
1.25
1.50
1.75
1e6
Subsampled reads
Subsampled data
Function fitted to data

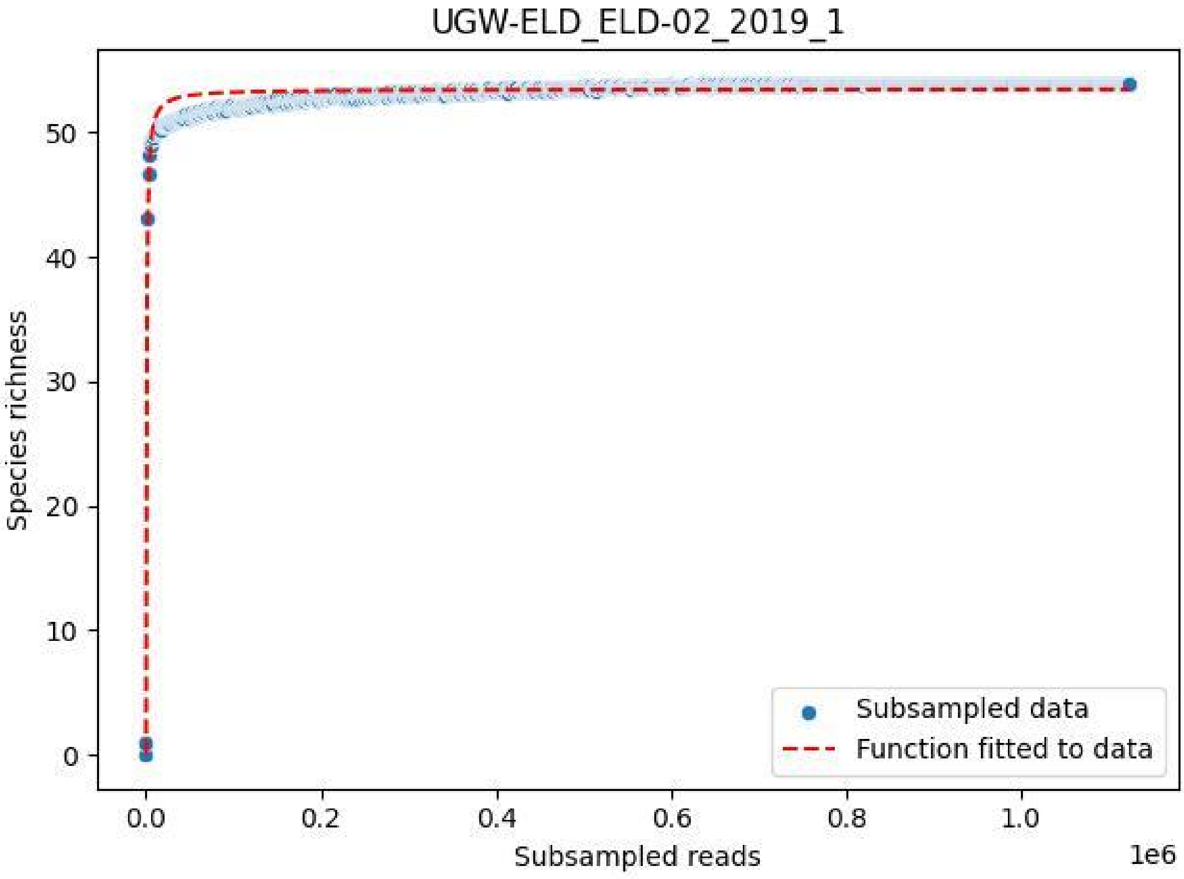
UGW-ELD_ELD-02_2019_1
Species richness
Subsampled reads
1e6
0
10
20
30
40
50
0.0
0.2
0.4
0.6
0.8
1.0
Subsampled data
Function fitted to data

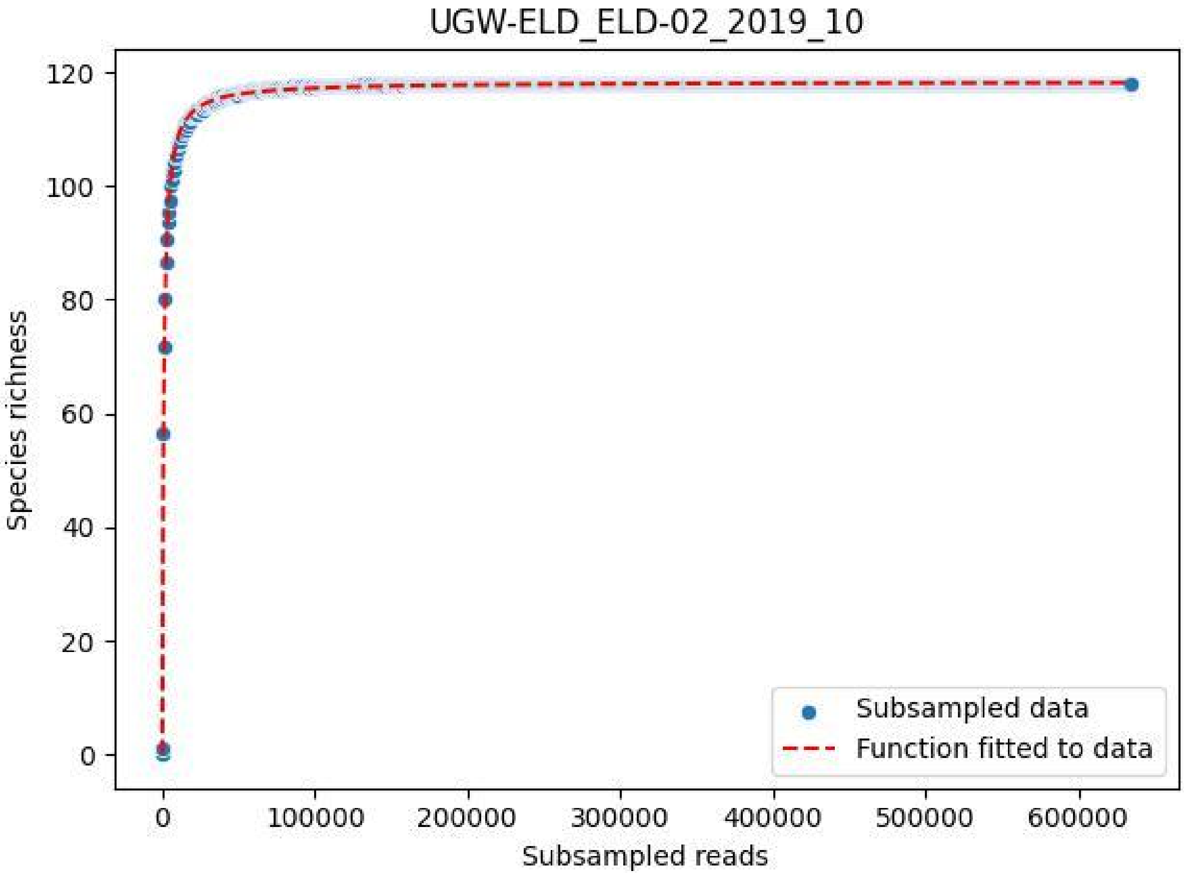

UGW-ELD_ELD-02_2019_10
Species richness
Subsampled reads
0
20
40
60
80
100
120
0
100000
200000
300000
400000
500000
600000
Subsampled data
Function fitted to data

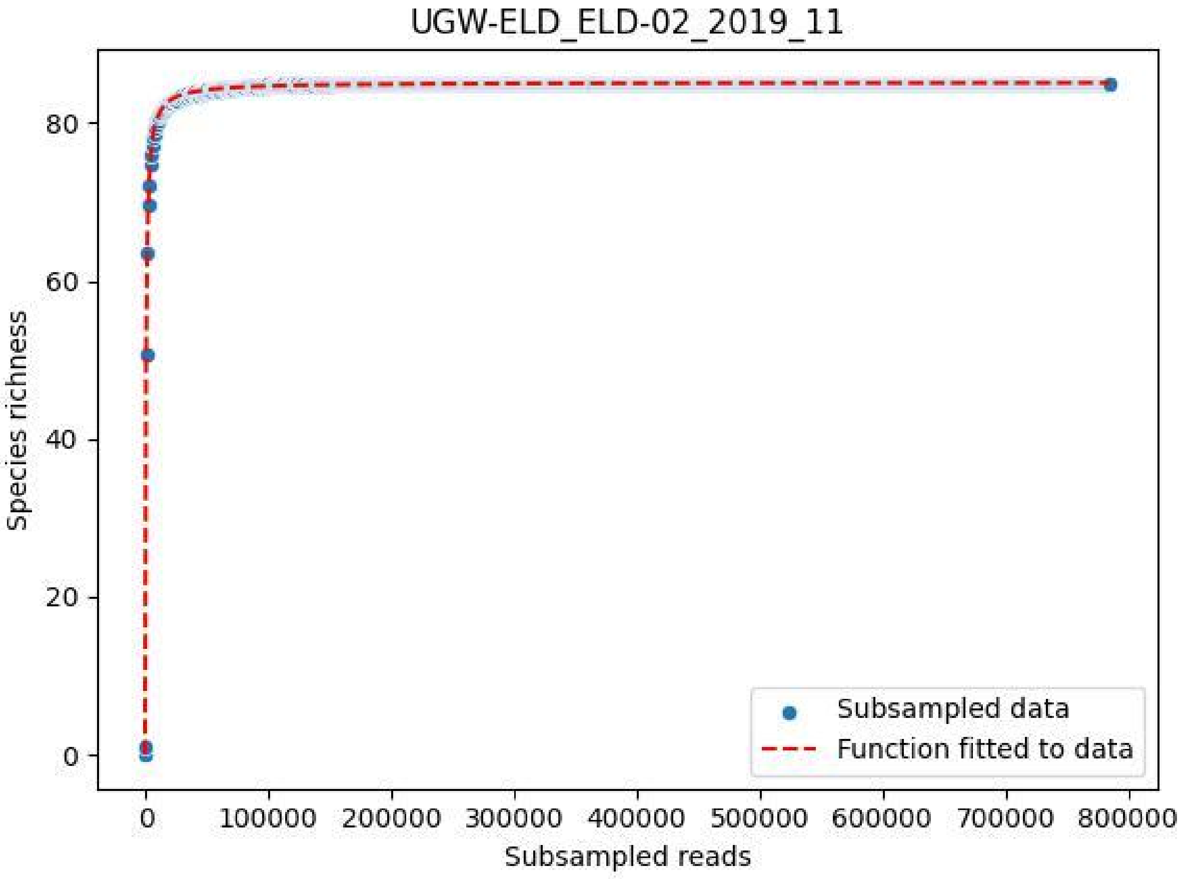
UGW-ELD_ELD-02_2019_11
Species richness
Subsampled reads
0
20
40
60
80
0
100000
200000
300000
400000
500000
600000
700000
800000
Subsampled data
Function fitted to data

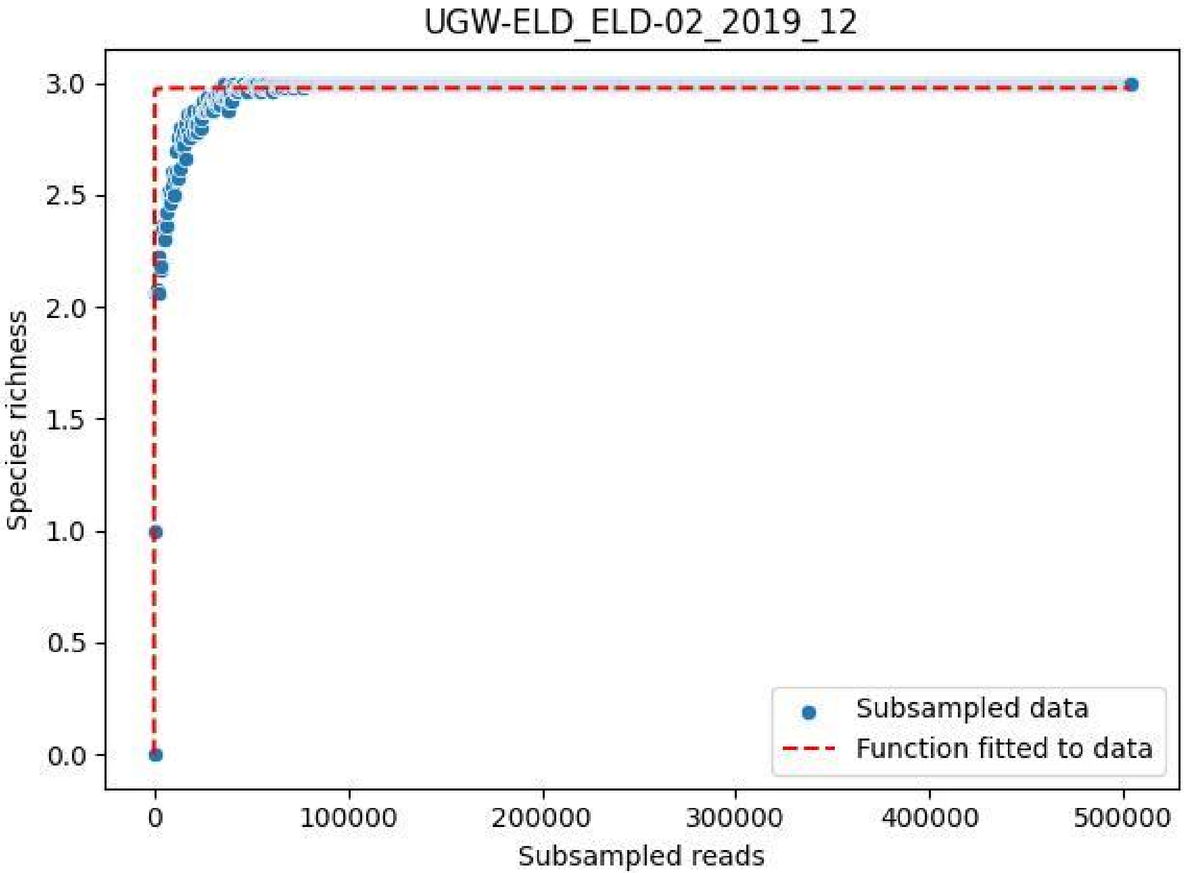
UGW-ELD_ELD-02_2019_12
Species richness
Subsampled reads
3.0
2.5
2.0
1.5
1.0
0.5
0.0
0
100000
200000
300000
400000
500000
Subsampled data
Function fitted to data

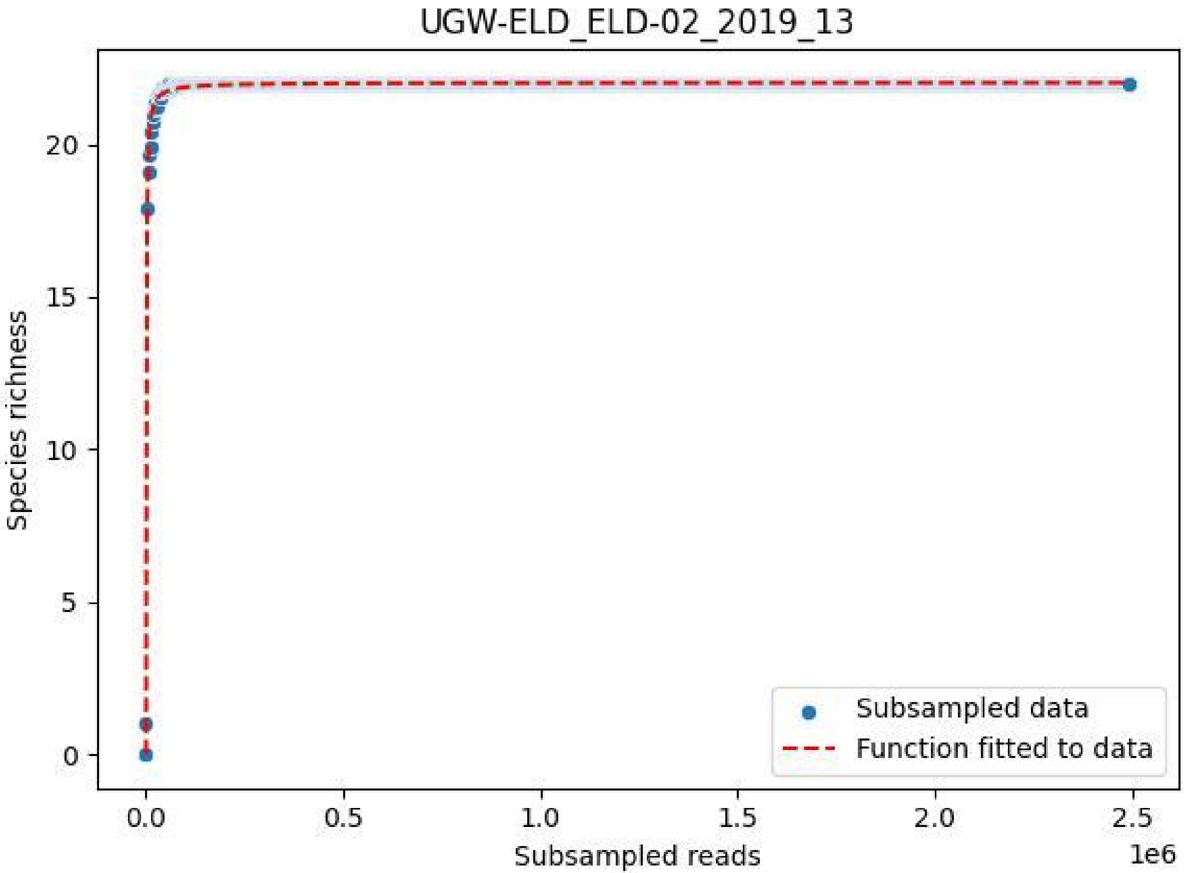

UGW-ELD_ELD-02_2019_13
Species richness
Subsampled reads
1e6
0
5
10
15
20
0.0
0.5
1.0
1.5
2.0
2.5
Subsampled data
Function fitted to data

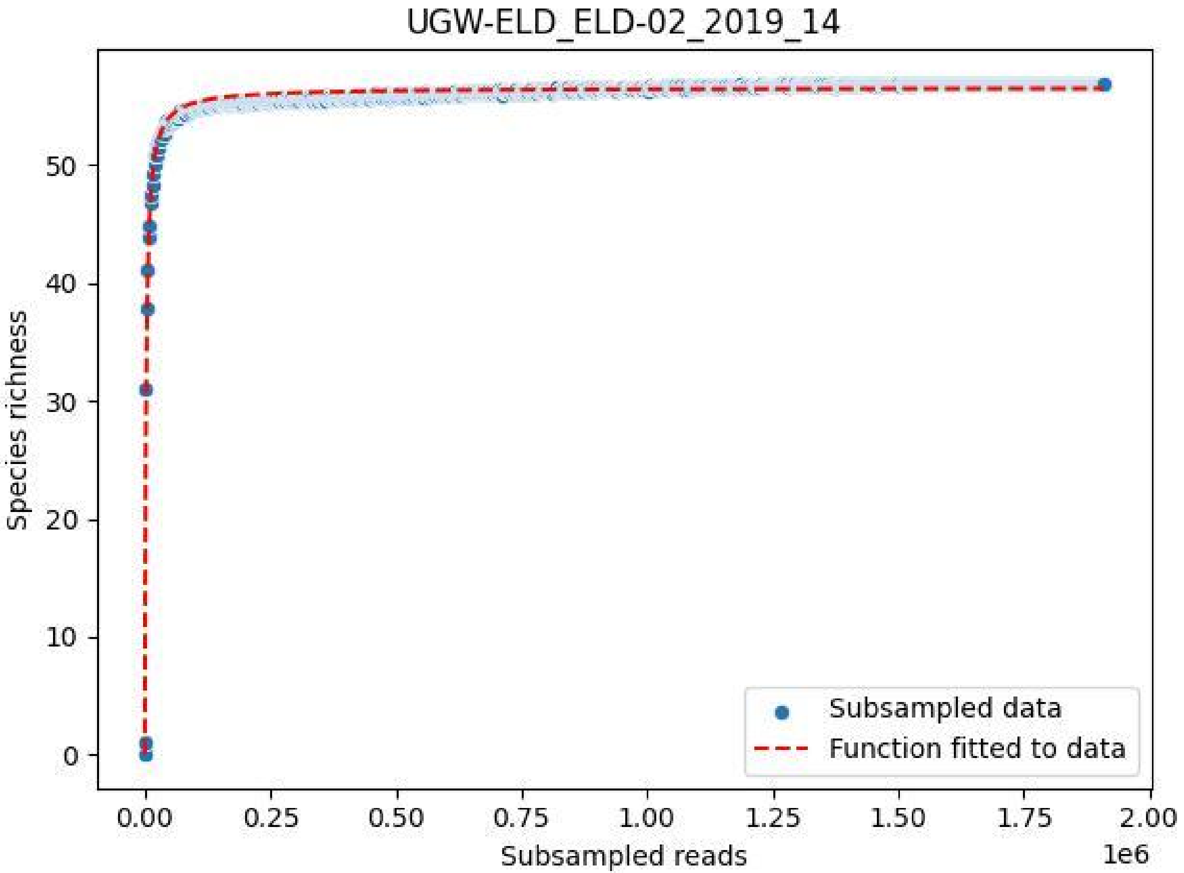
UGW-ELD_ELD-02_2019_14
Species richness
0
10
20
30
40
50
0.00
0.25
0.50
0.75
1.00
1.25
1.50
1.75
2.00
1e6
Subsampled reads
Subsampled data
Function fitted to data

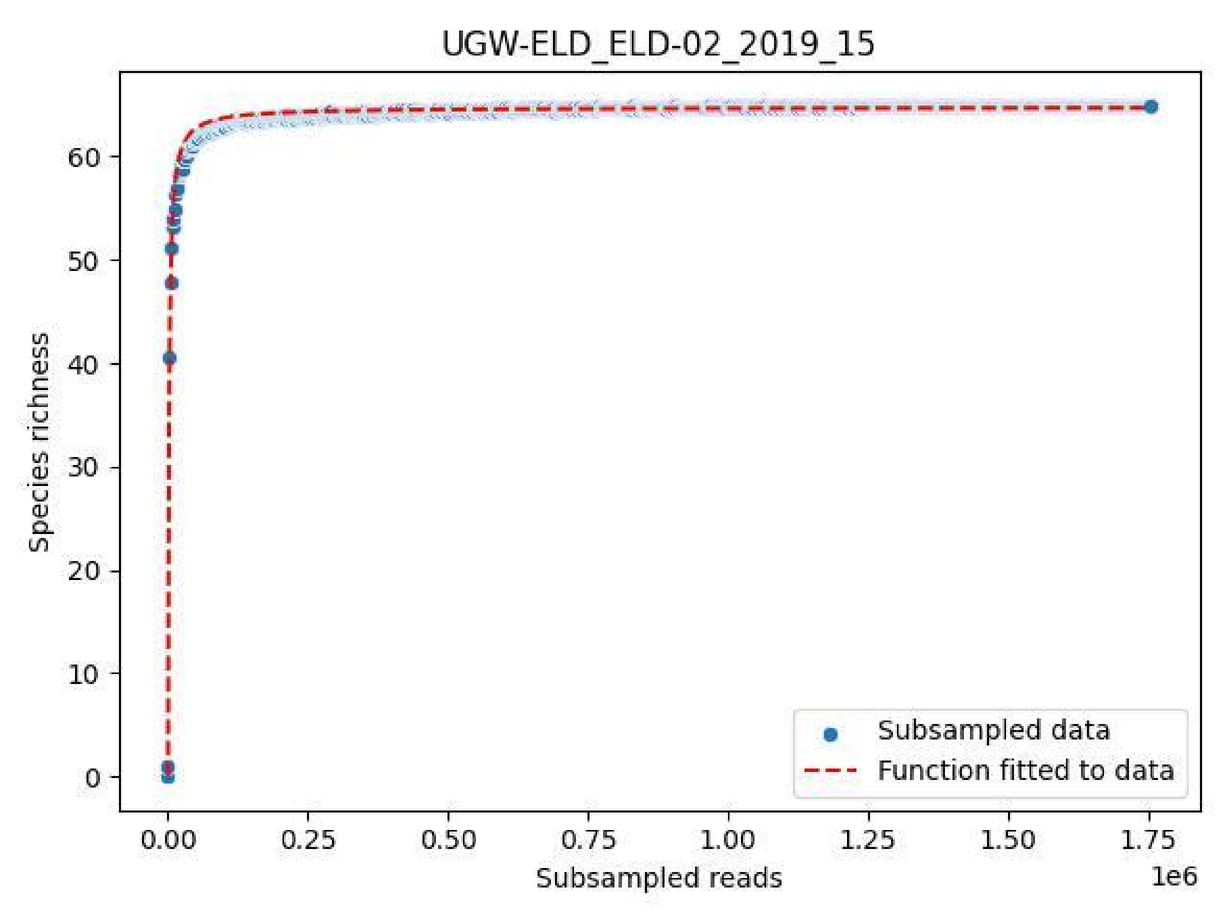

UGW-ELD_ELD-02_2019_15
Species richness
Subsampled reads
1e6
0
10
20
30
40
50
60
0.00
0.25
0.50
0.75
1.00
1.25
1.50
1.75
Subsampled data
Function fitted to data

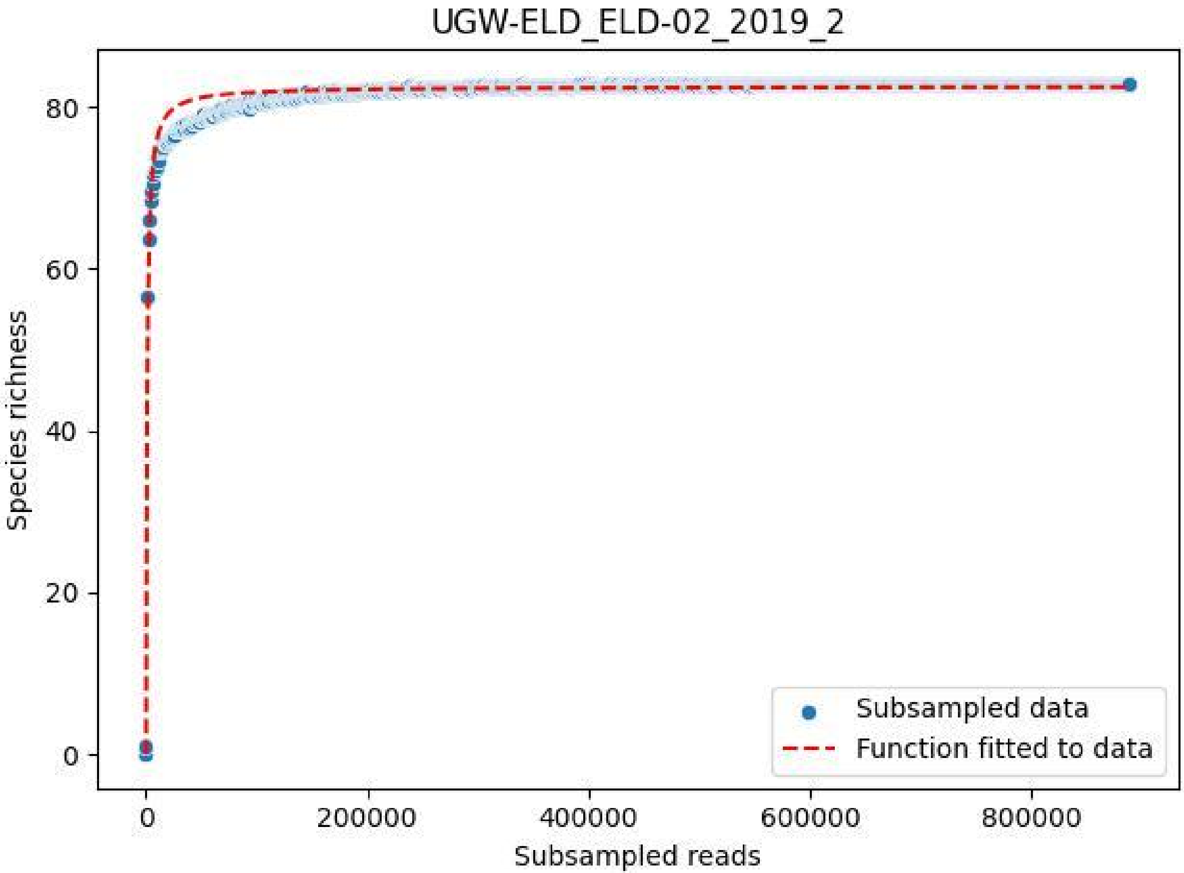

UGW-ELD_ELD-02_2019_2
Species richness
Subsampled reads
0
20
40
60
80
0
200000
400000
600000
800000
Subsampled data
Function fitted to data

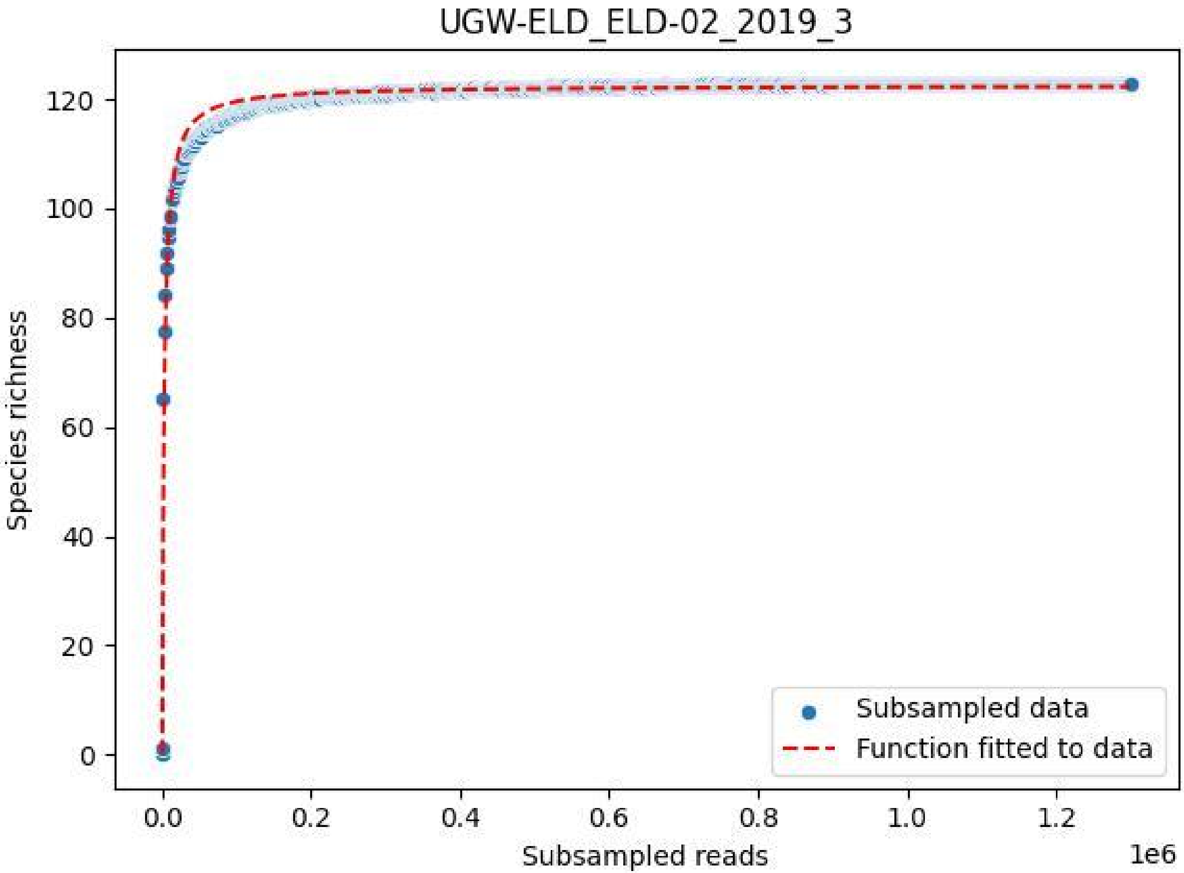
UGW-ELD_ELD-02_2019_3
Species richness
Subsampled reads
1e6
0
20
40
60
80
100
120
0.0
0.2
0.4
0.6
0.8
1.0
1.2
Subsampled data
Function fitted to data

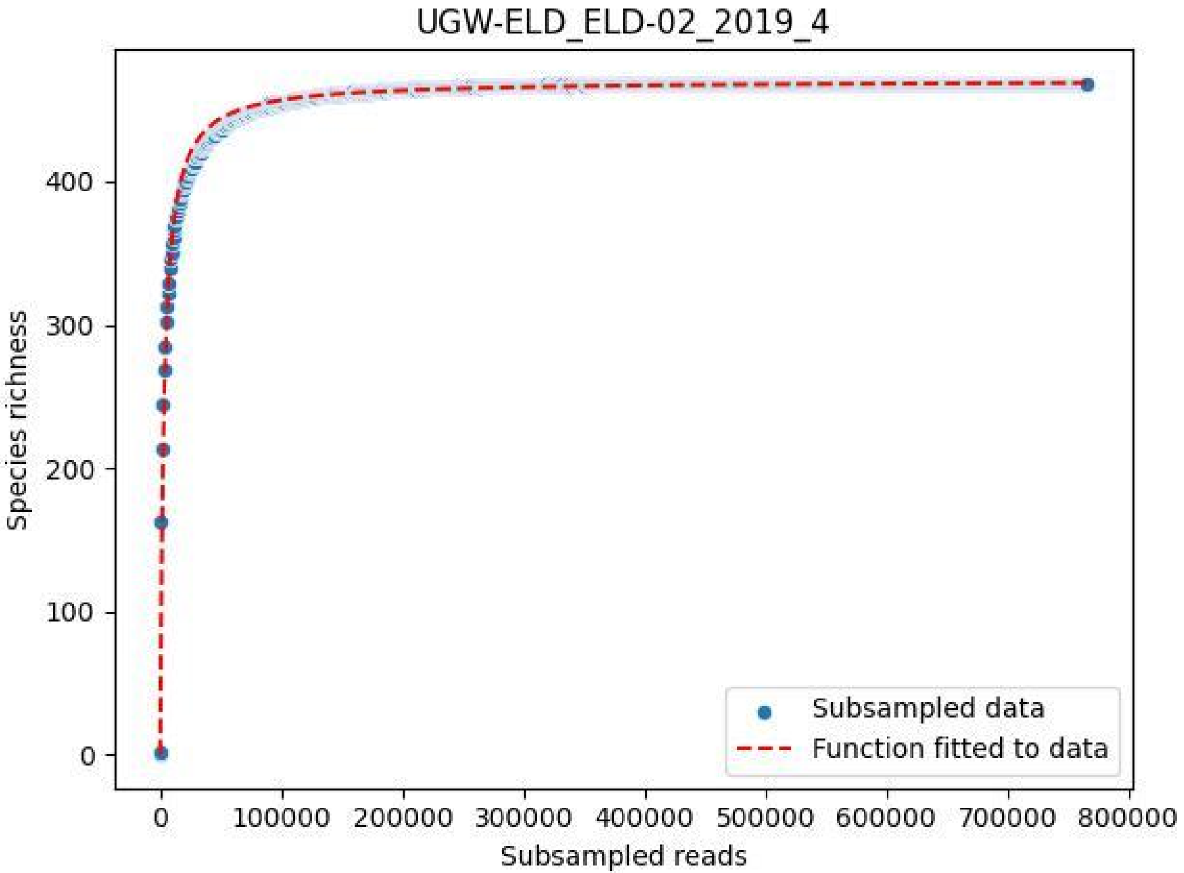
UGW-ELD_ELD-02_2019_4
Species richness
Subsampled reads
0
100
200
300
400
0
100000
200000
300000
400000
500000
600000
700000
800000
Subsampled data
Function fitted to data

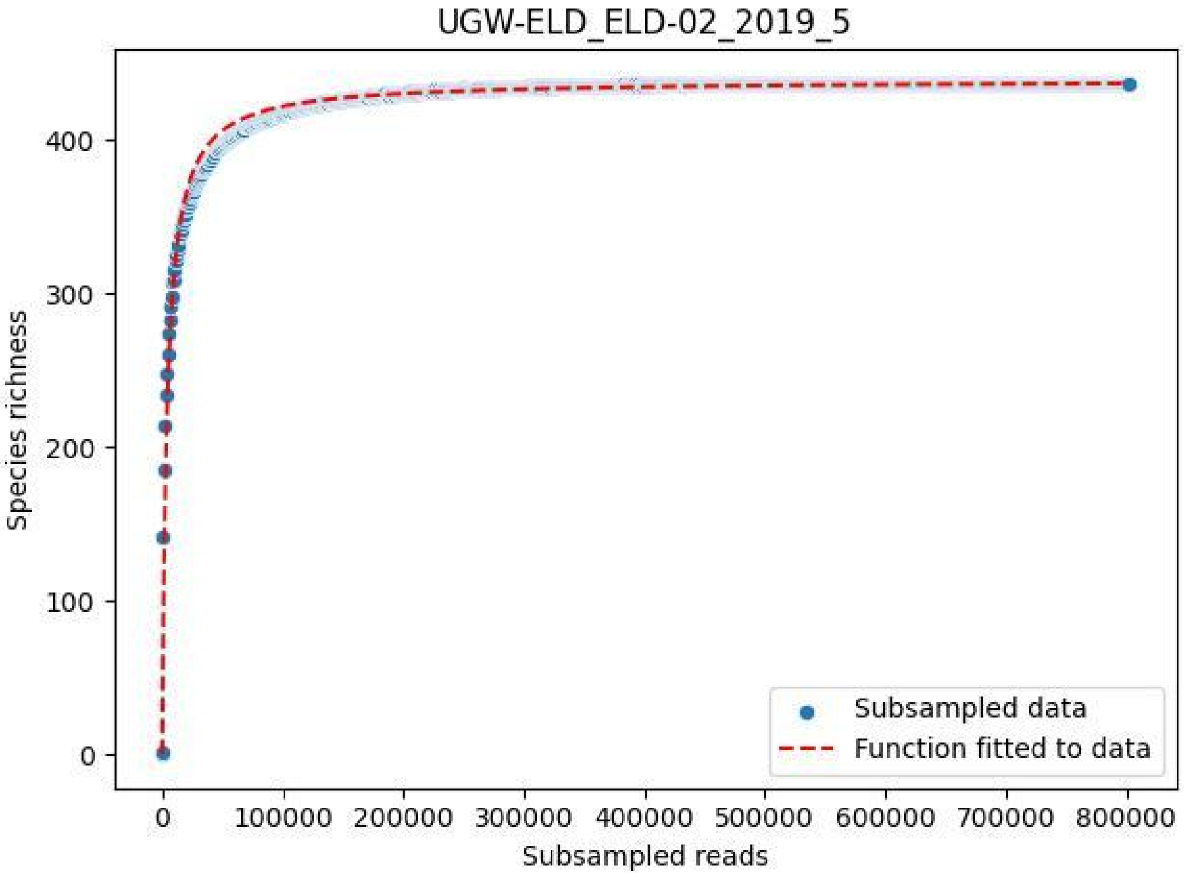
UGW-ELD_ELD-02_2019_5
Species richness
Subsampled reads
0
100
200
300
400
100000
200000
300000
400000
500000
600000
700000
800000
Subsampled data
Function fitted to data

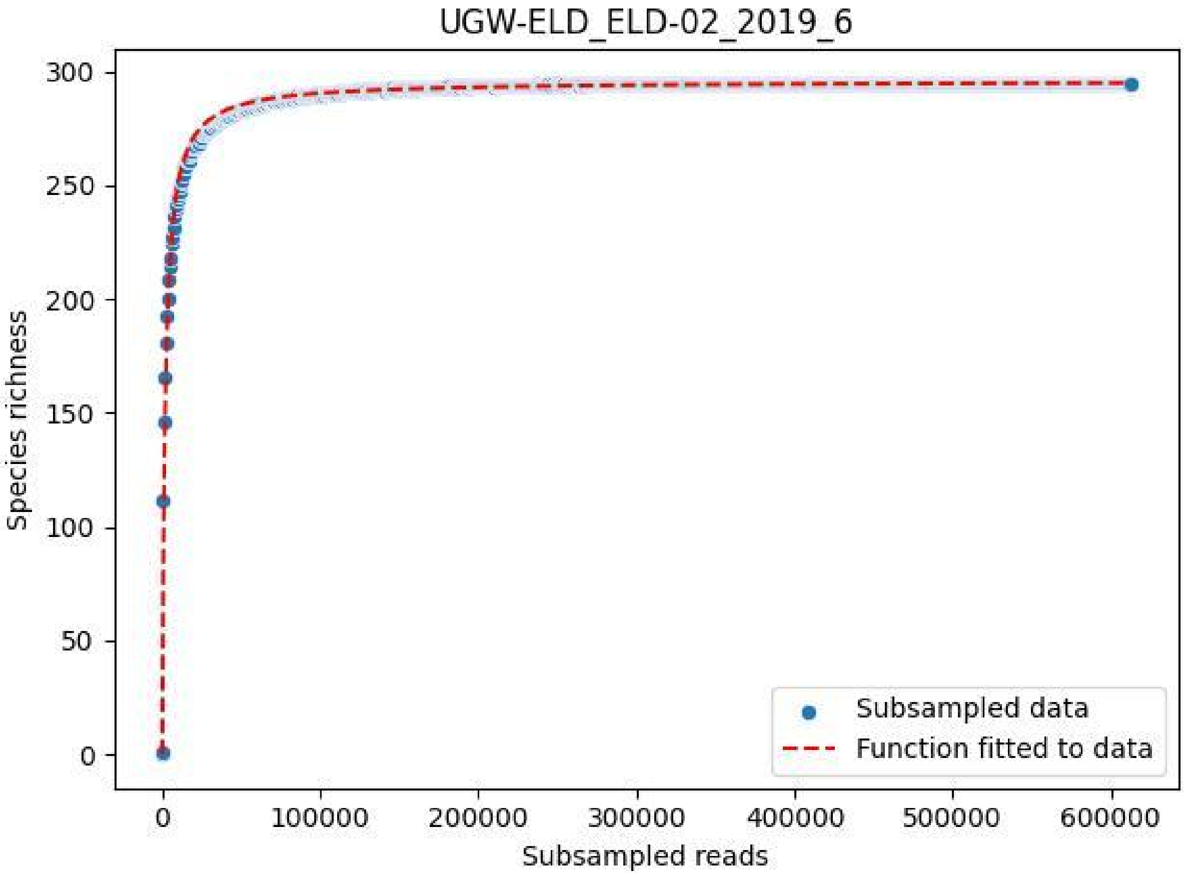

UGW-ELD_ELD-02_2019_6
Species richness
Subsampled reads
0
50
100
150
200
250
300
0
100000
200000
300000
400000
500000
600000
Subsampled data
Function fitted to data

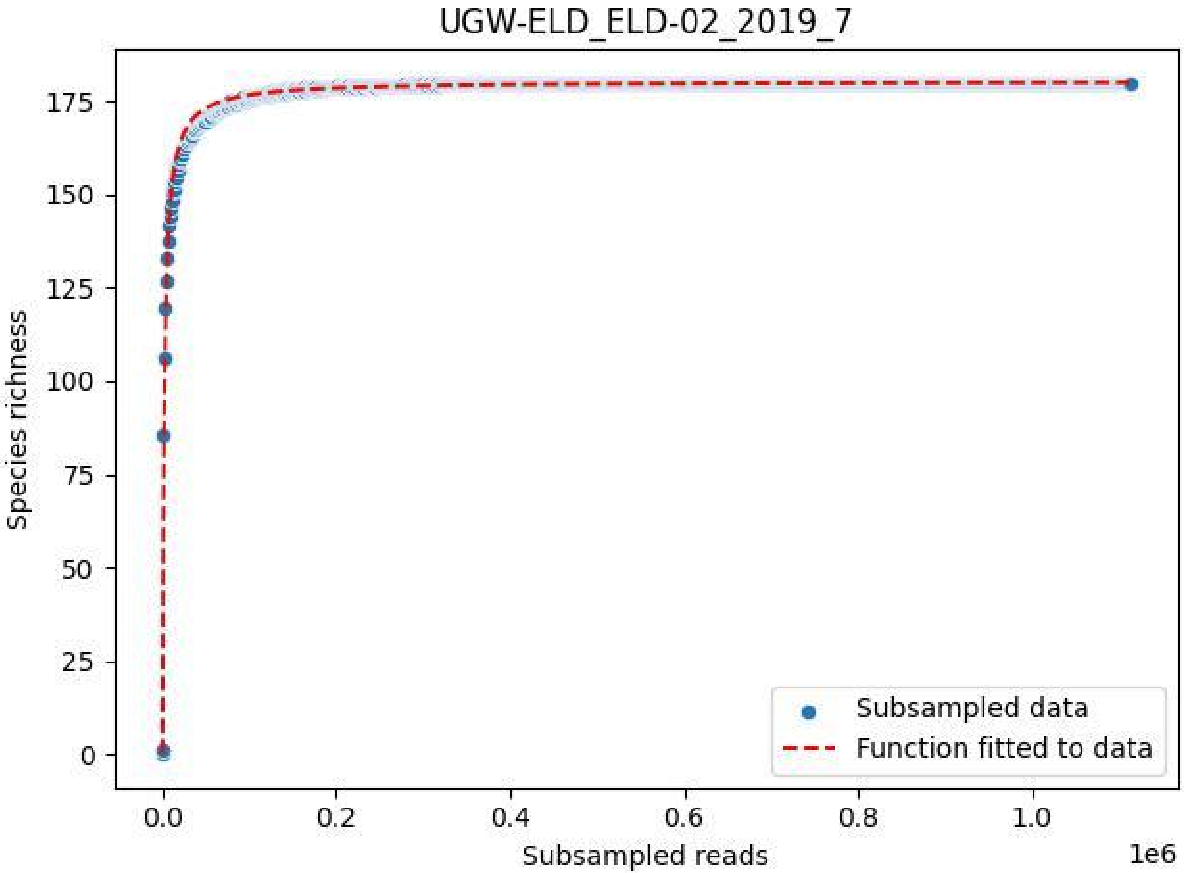

UGW-ELD_ELD-02_2019_7
Species richness
0
25
50
75
100
125
150
175
0.0
0.2
0.4
0.6
0.8
1.0
1e6
Subsampled reads
Subsampled data
Function fitted to data

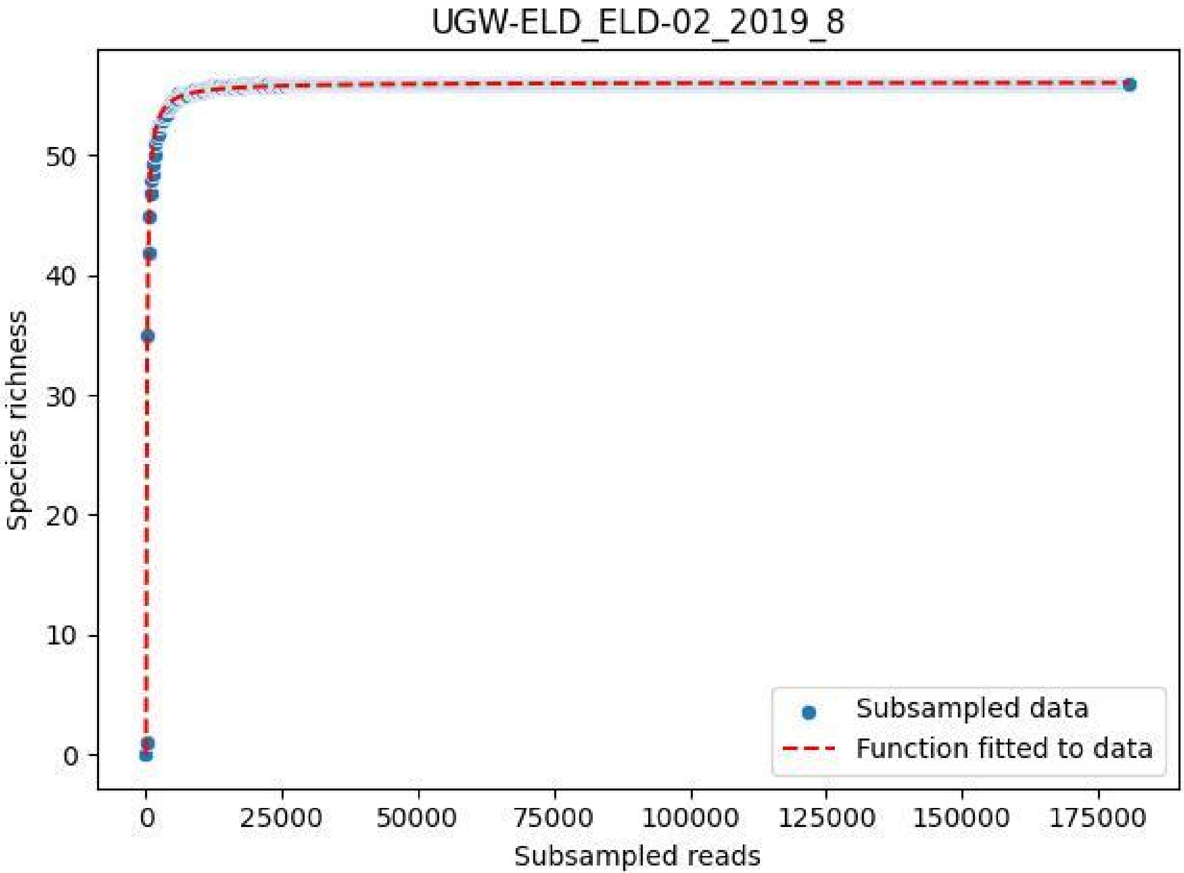

UGW-ELD_ELD-02_2019_8
Species richness
0
10
20
30
40
50
0
25000
50000
75000
100000
125000
150000
175000
Subsampled reads
Subsampled data
Function fitted to data

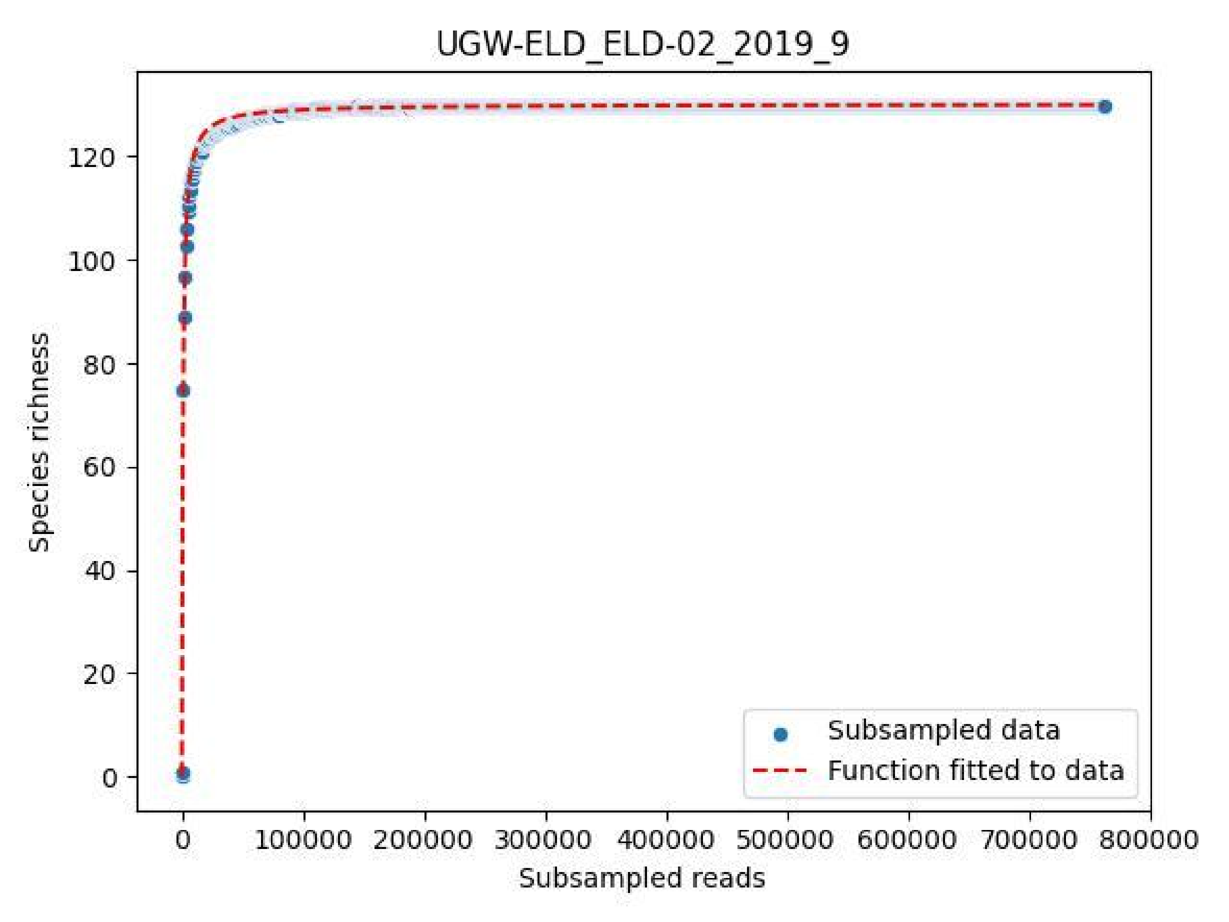
UGW-ELD_ELD-02_2019_9
Species richness
0
20
40
60
80
100
120
0
100000
200000
300000
400000
500000
600000
700000
800000
Subsampled reads
Subsampled data
Function fitted to data

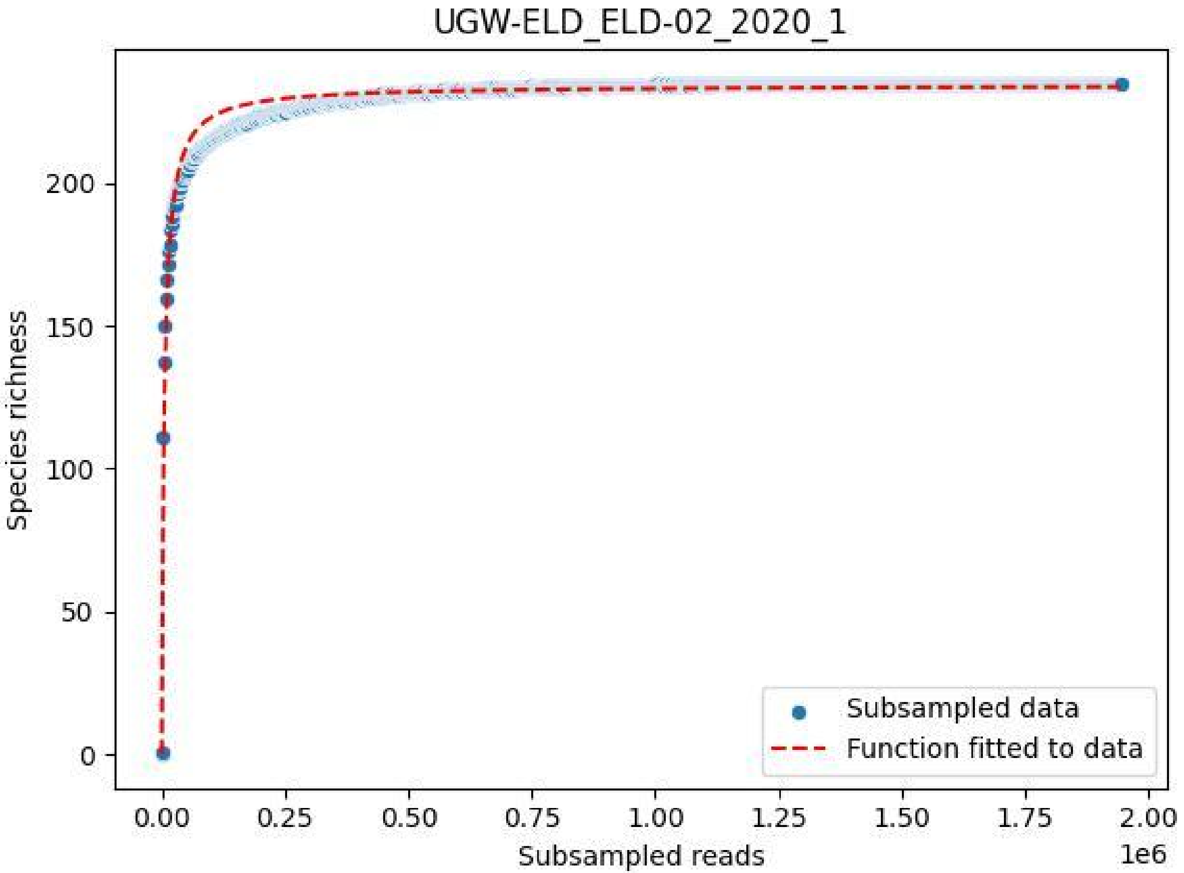

UGW-ELD_ELD-02_2020_1
Species richness
0
50
100
150
200
0.00
0.25
0.50
0.75
1.00
1.25
1.50
1.75
2.00
1e6
Subsampled reads
Subsampled data
Function fitted to data

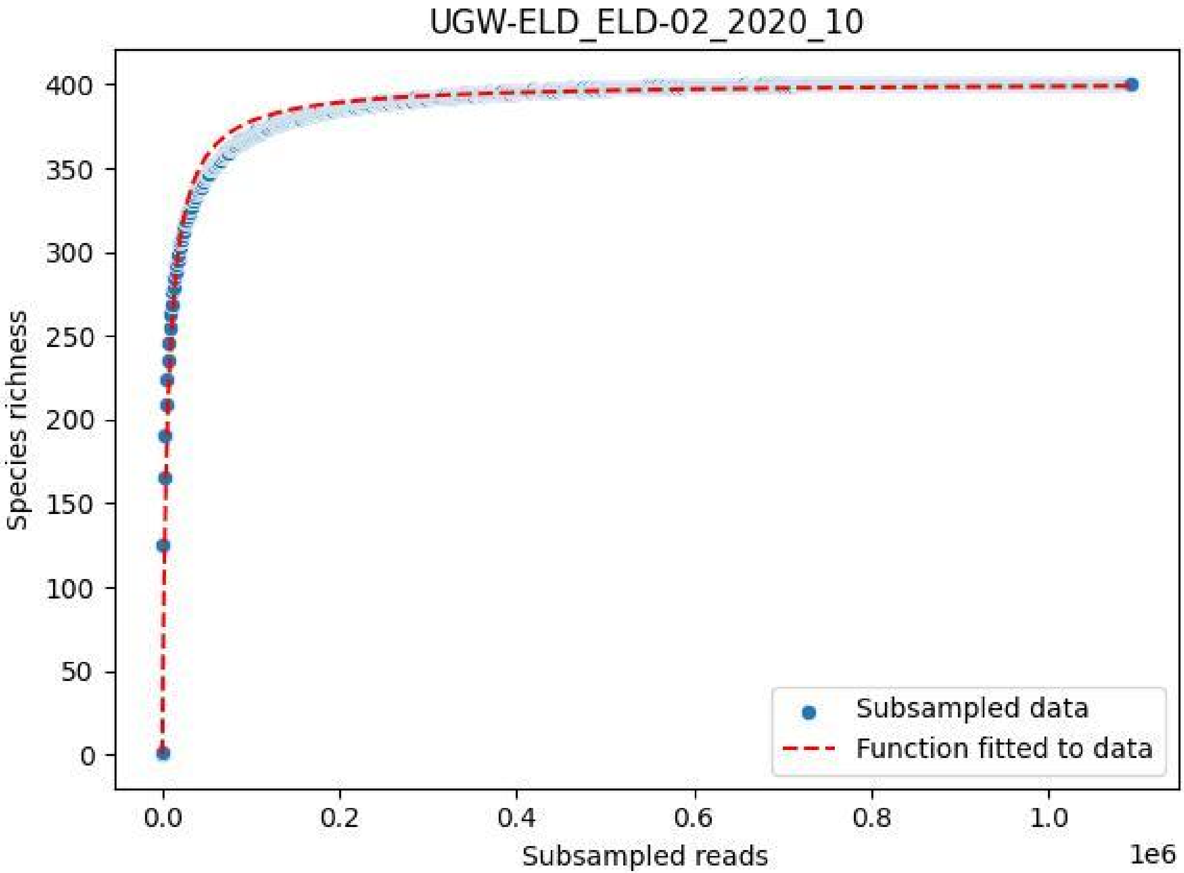
UGW-ELD_ELD-02_2020_10
Species richness
Subsampled reads
1e6
0
50
100
150
200
250
300
350
400
0.0
0.2
0.4
0.6
0.8
1.0
Subsampled data
Function fitted to data

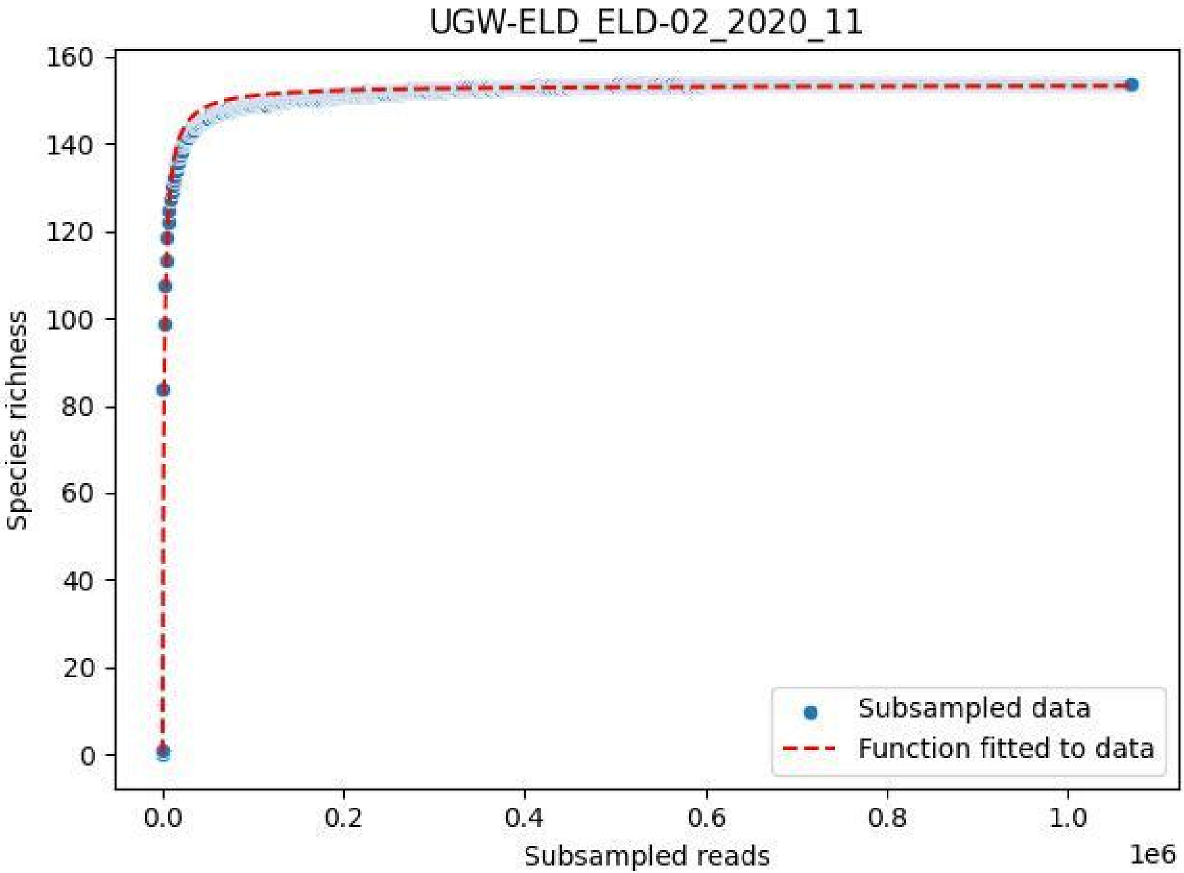

UGW-ELD_ELD-02_2020_11
Species richness
Subsampled reads
1e6
0
20
40
60
80
100
120
140
160
0.0
0.2
0.4
0.6
0.8
1.0
Subsampled data
Function fitted to data

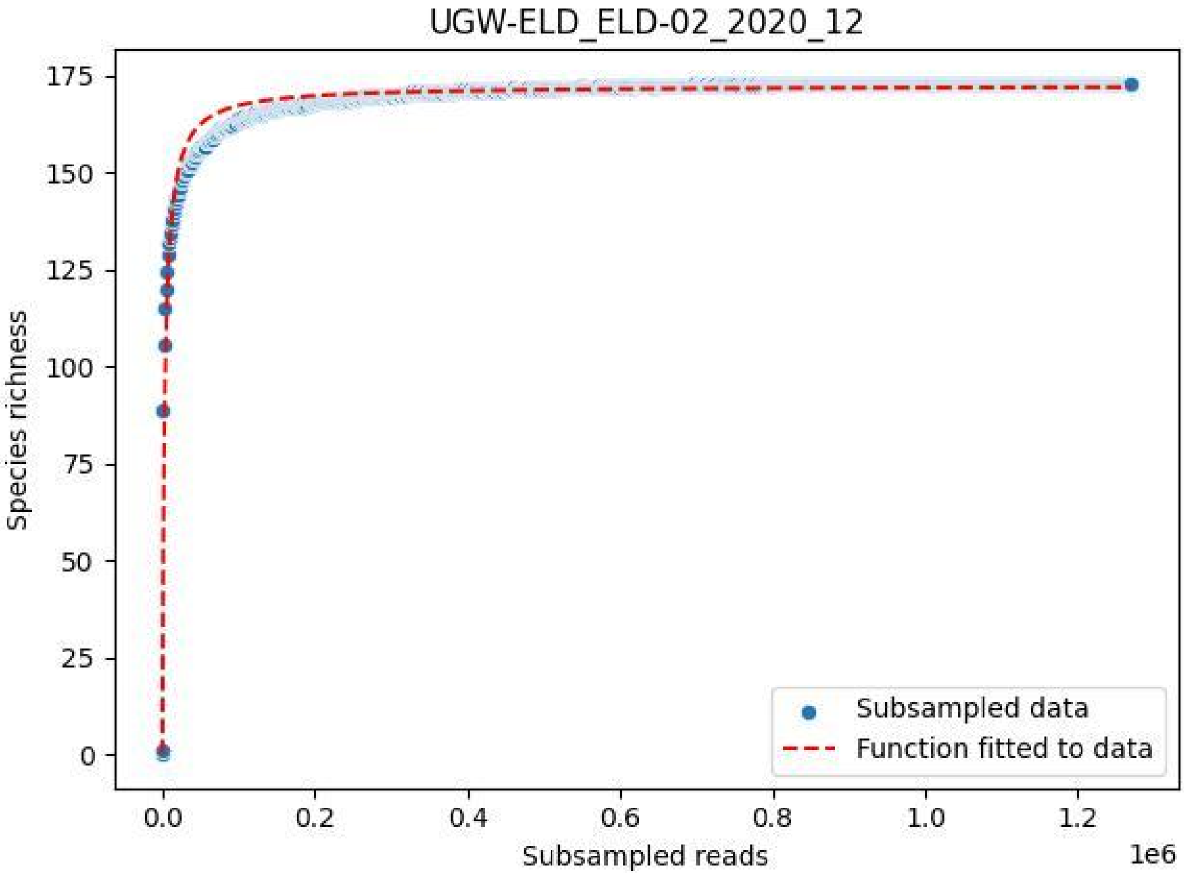
UGW-ELD_ELD-02_2020_12
Species richness
Subsampled reads
1e6
Subsampled data
Function fitted to data
0
25
50
75
100
125
150
175
0.0
0.2
0.4
0.6
0.8
1.0
1.2

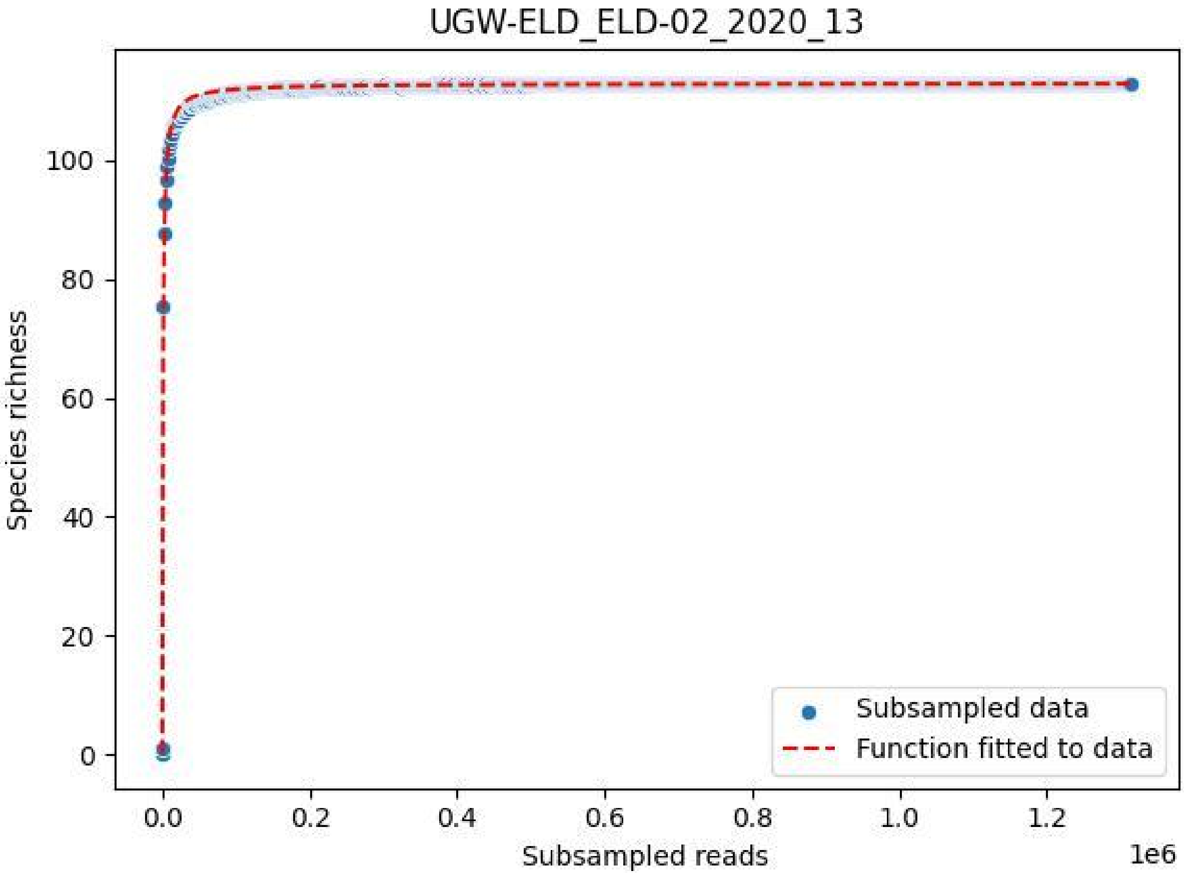

UGW-ELD_ELD-02_2020_13
Species richness
Subsampled reads
1e6
0
20
40
60
80
100
0.0
0.2
0.4
0.6
0.8
1.0
1.2
Subsampled data
Function fitted to data

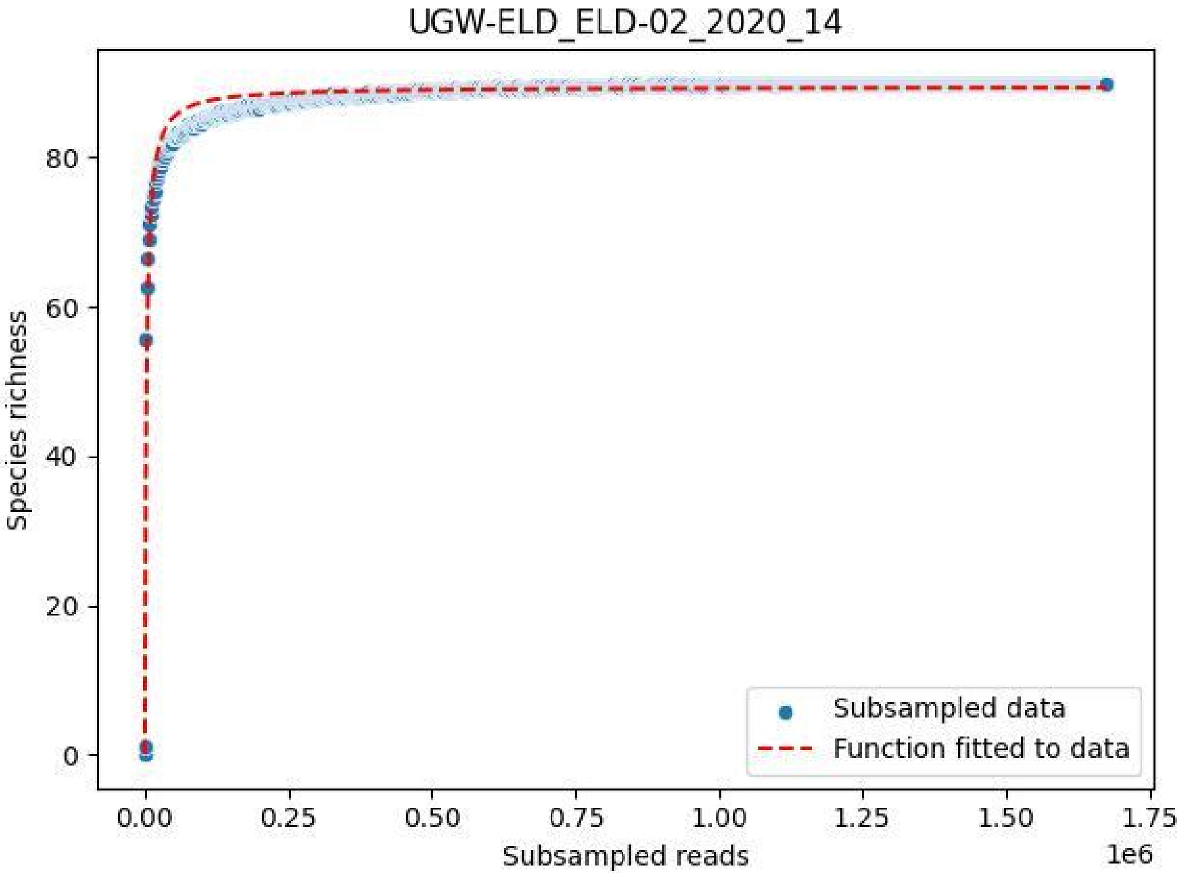

UGW-ELD_ELD-02_2020_14
Species richness
Subsampled reads
1e6
0.00
0.25
0.50
0.75
1.00
1.25
1.50
1.75
0
20
40
60
80
Subsampled data
Function fitted to data

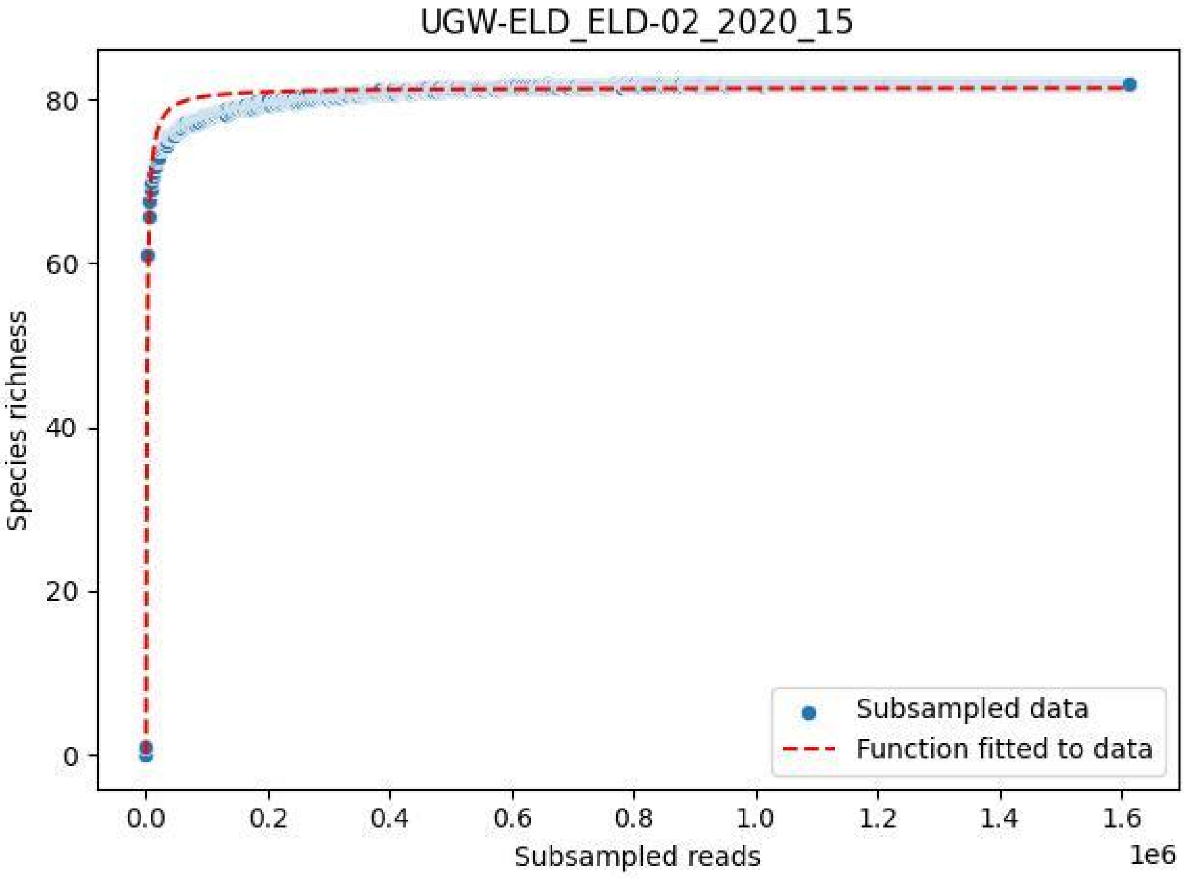
UGW-ELD_ELD-02_2020_15
Species richness
0
20
40
60
80
0.0
0.2
0.4
0.6
0.8
1.0
1.2
1.4
1.6
1e6
Subsampled reads
Subsampled data
Function fitted to data

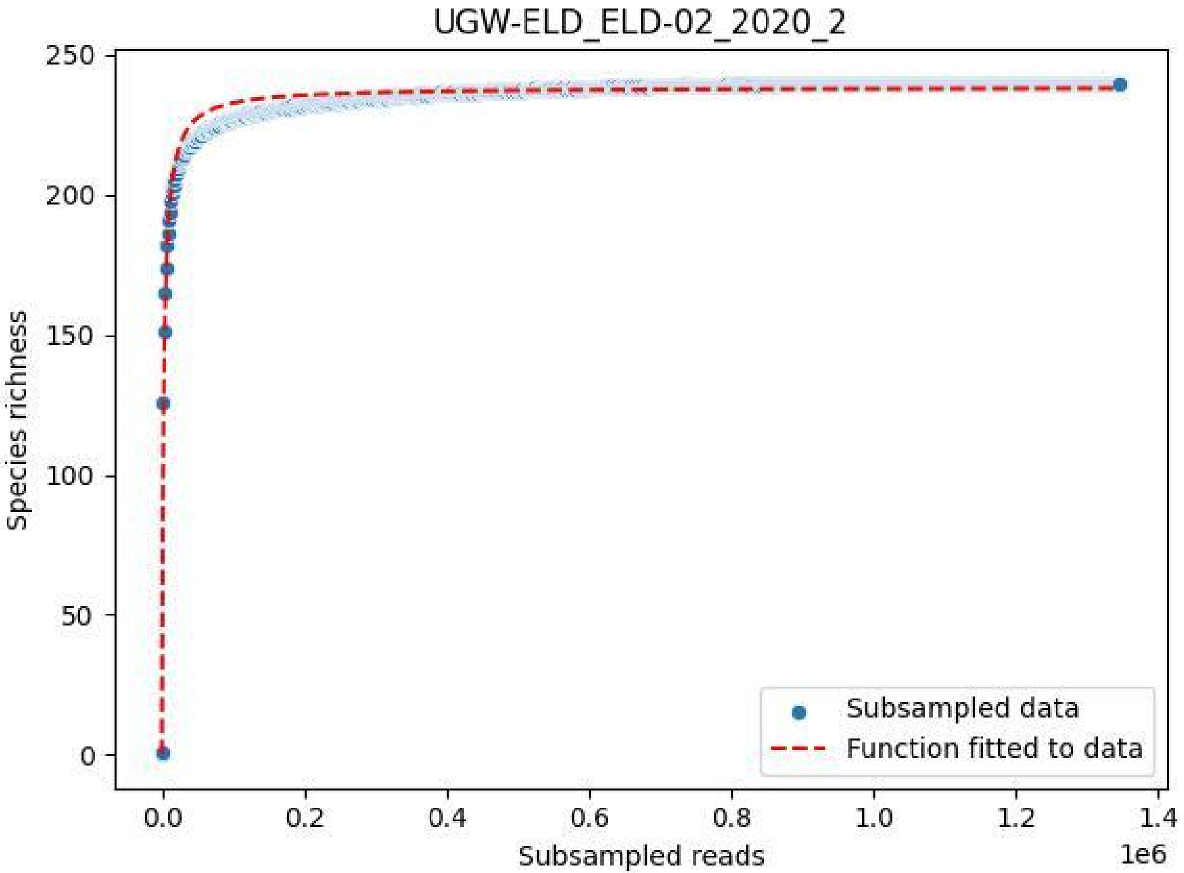
UGW-ELD_ELD-02_2020_2
Species richness
Subsampled reads
1e6
0
50
100
150
200
250
0.0
0.2
0.4
0.6
0.8
1.0
1.2
1.4
Subsampled data
Function fitted to data

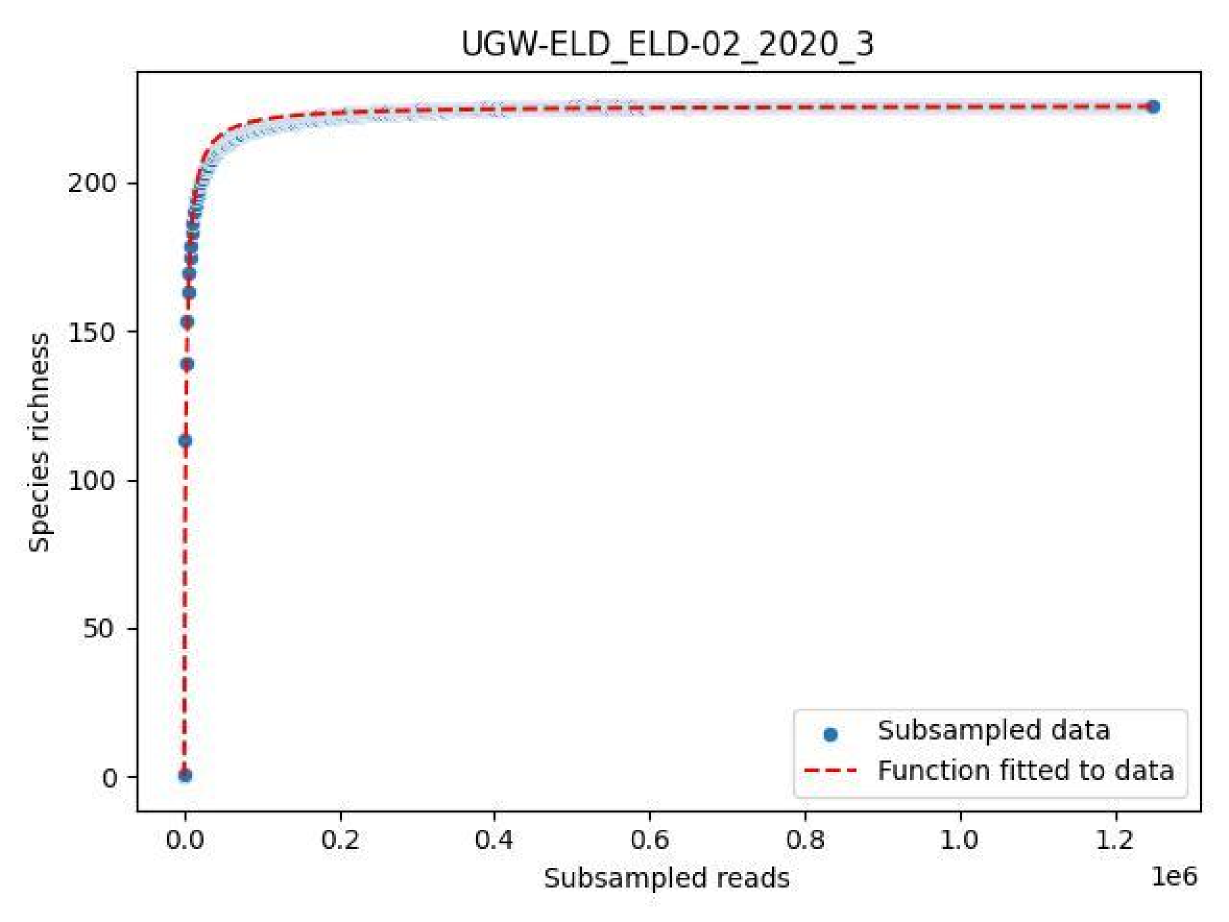
UGW-ELD_ELD-02_2020_3
Species richness
Subsampled reads
1e6
0
50
100
150
200
0.0
0.2
0.4
0.6
0.8
1.0
1.2
Subsampled data
Function fitted to data

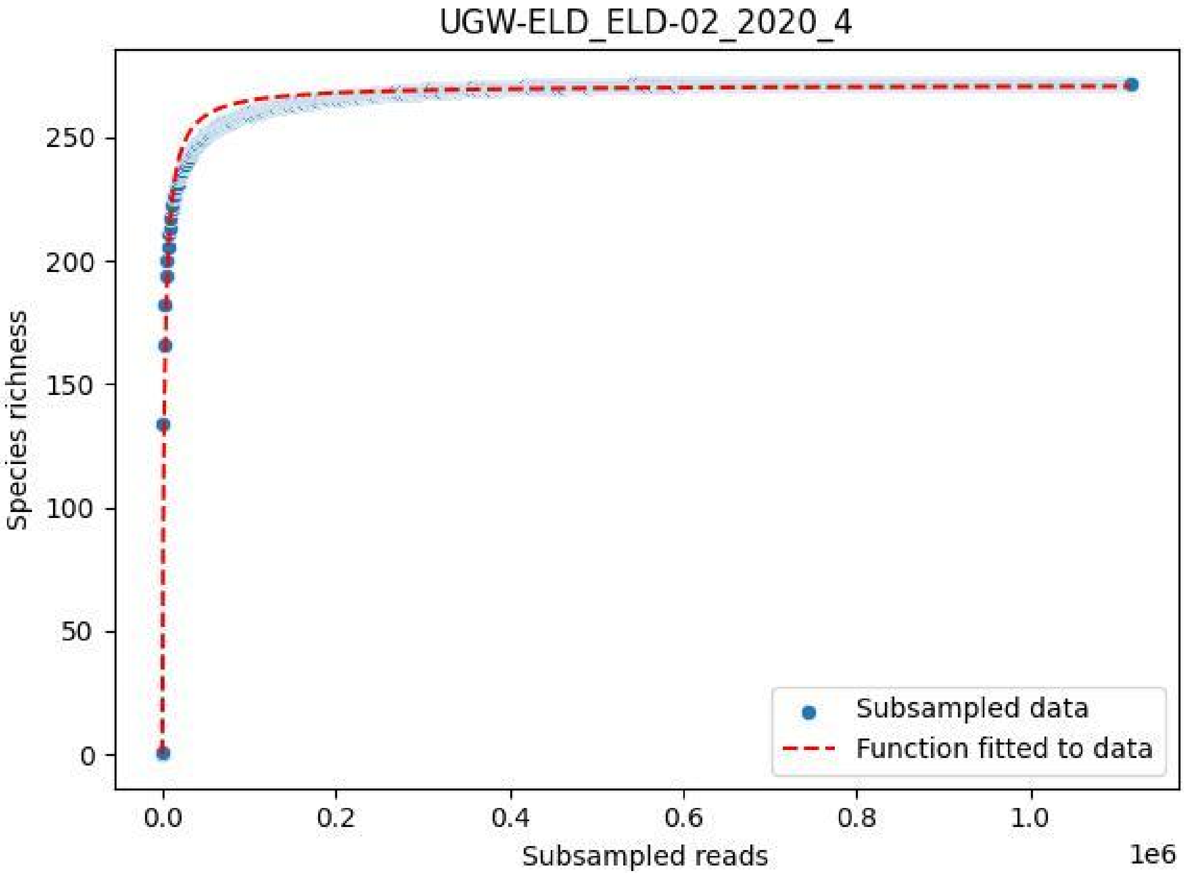

UGW-ELD_ELD-02_2020_4
Species richness
Subsampled reads
1e6
0
50
100
150
200
250
0.0
0.2
0.4
0.6
0.8
1.0
Subsampled data
Function fitted to data

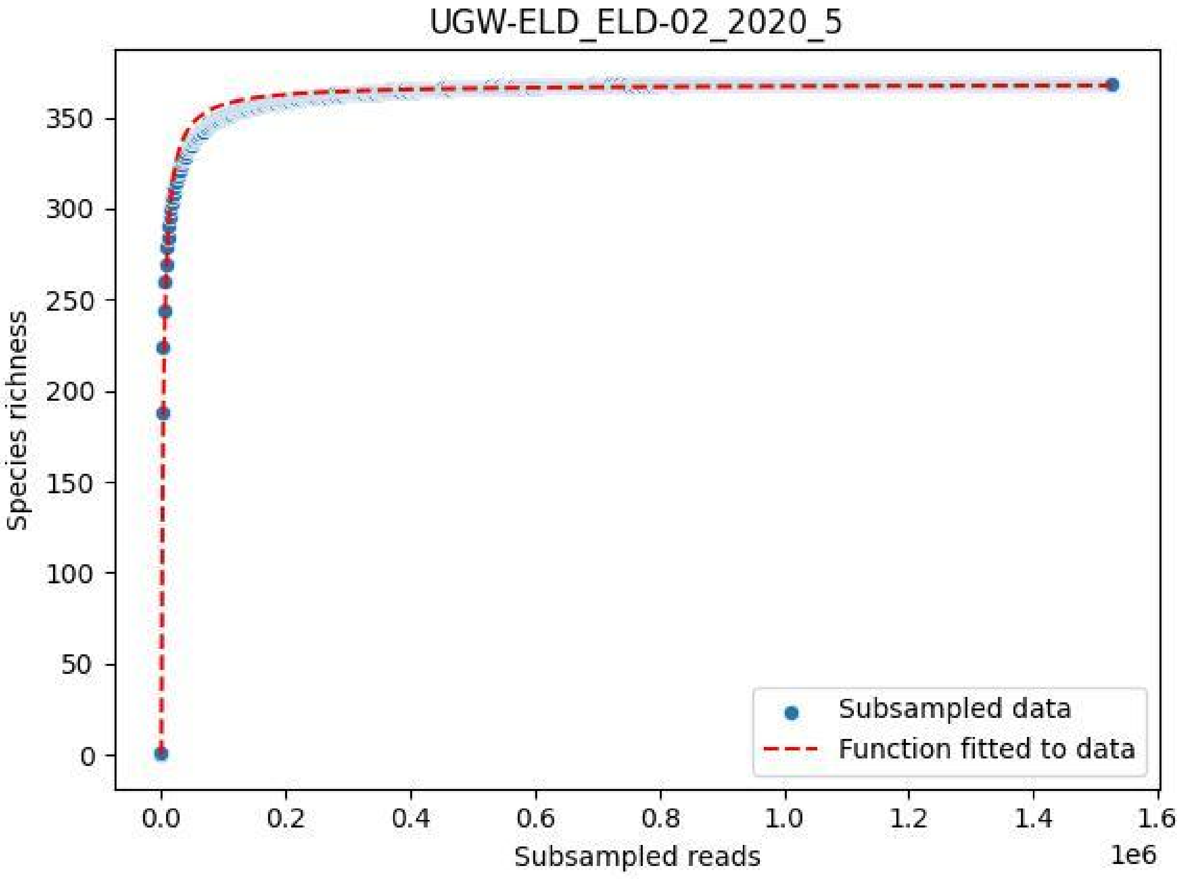
UGW-ELD_ELD-02_2020_5
Species richness
Subsampled reads
1e6
Subsampled data
Function fitted to data
0
50
100
150
200
250
300
350
0.0
0.2
0.4
0.6
0.8
1.0
1.2
1.4
1.6

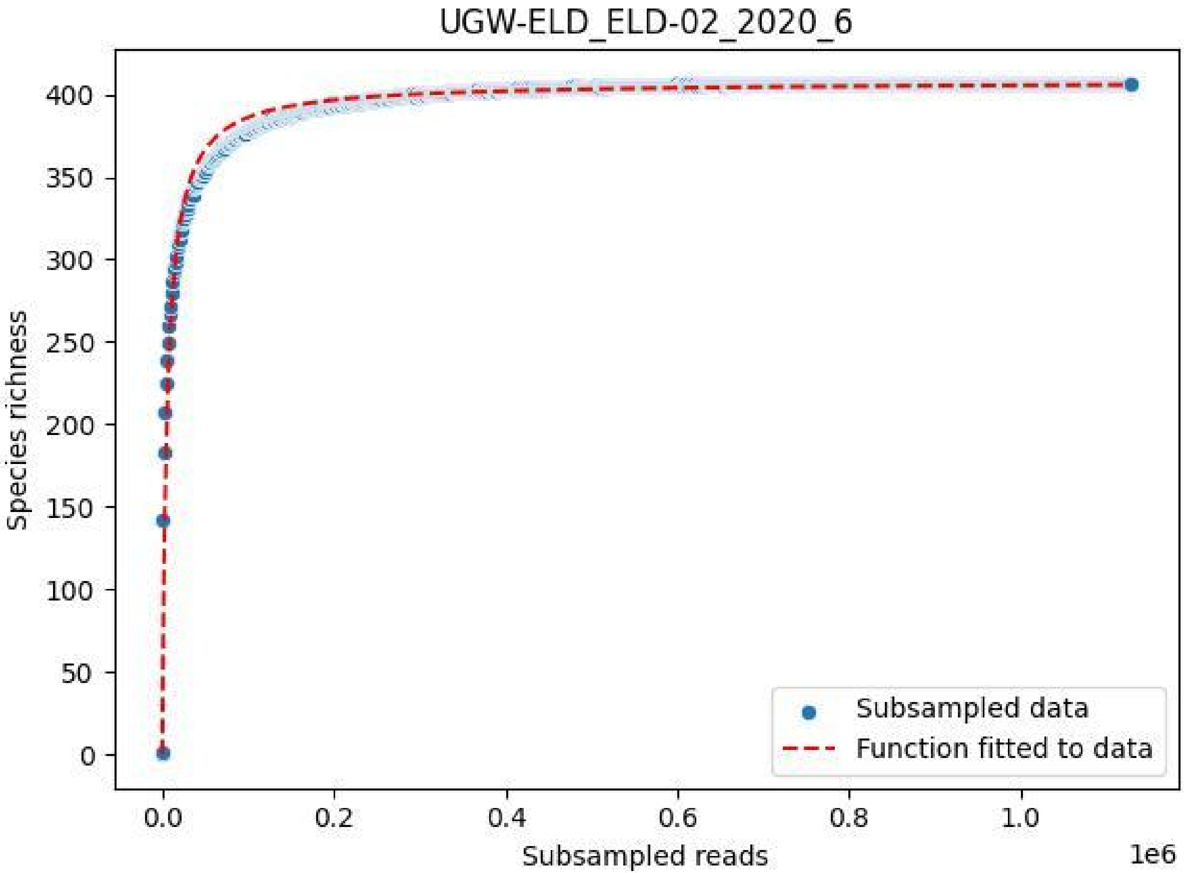

UGW-ELD_ELD-02_2020_6
Species richness
Subsampled reads
1e6
0
50
100
150
200
250
300
350
400
0.0
0.2
0.4
0.6
0.8
1.0
Subsampled data
Function fitted to data

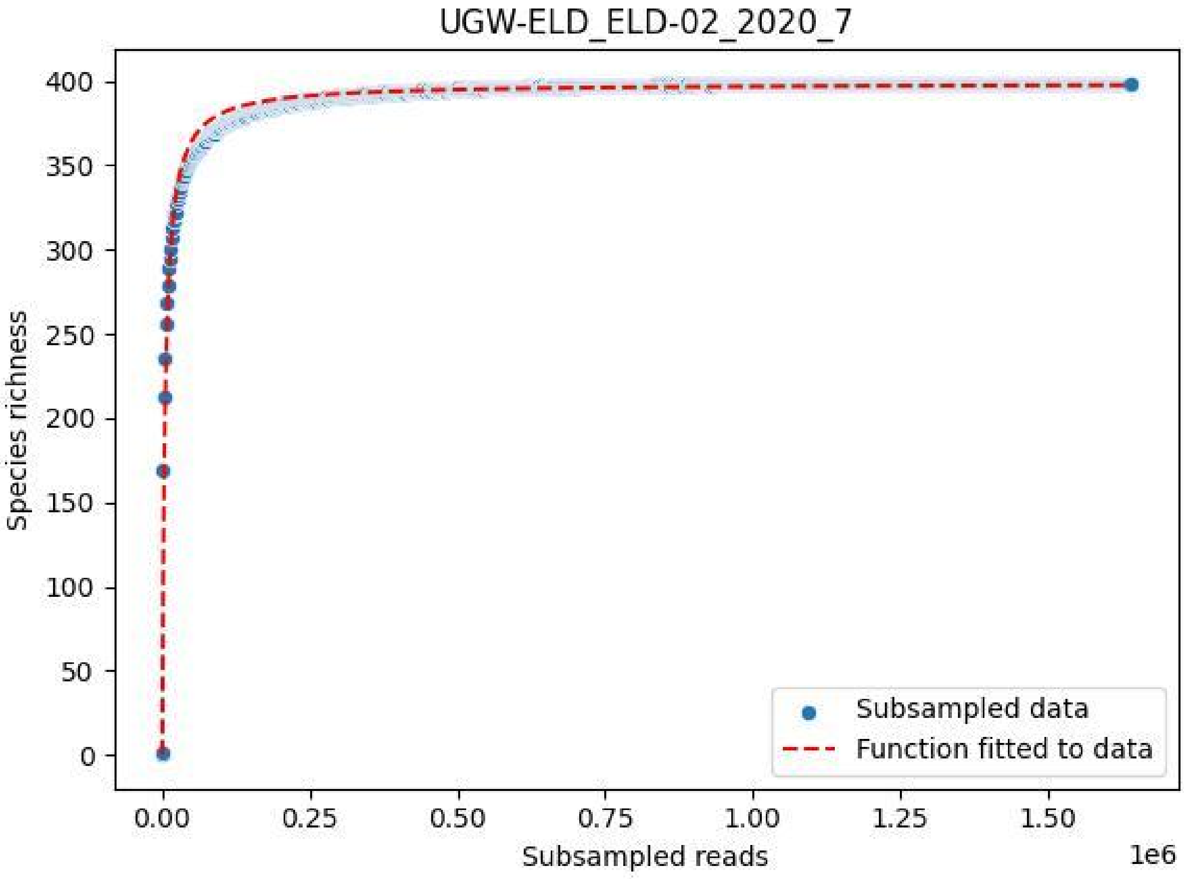
UGW-ELD_ELD-02_2020_7
Species richness
400
350
300
250
200
150
100
50
0
0.00
0.25
0.50
0.75
1.00
1.25
1.50
1e6
Subsampled reads
Subsampled data
Function fitted to data

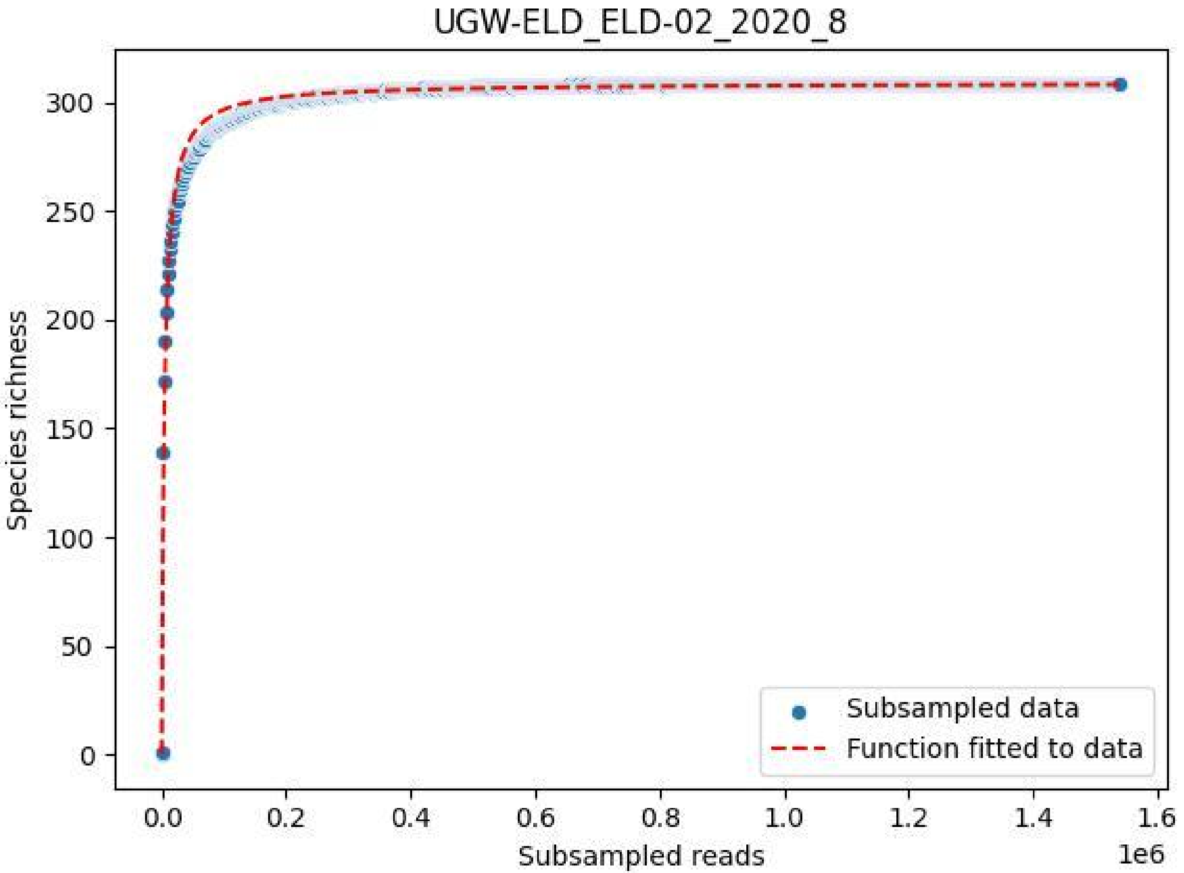
UGW-ELD_ELD-02_2020_8
Species richness
Subsampled reads
1e6
0
50
100
150
200
250
300
0.0
0.2
0.4
0.6
0.8
1.0
1.2
1.4
1.6
Subsampled data
Function fitted to data

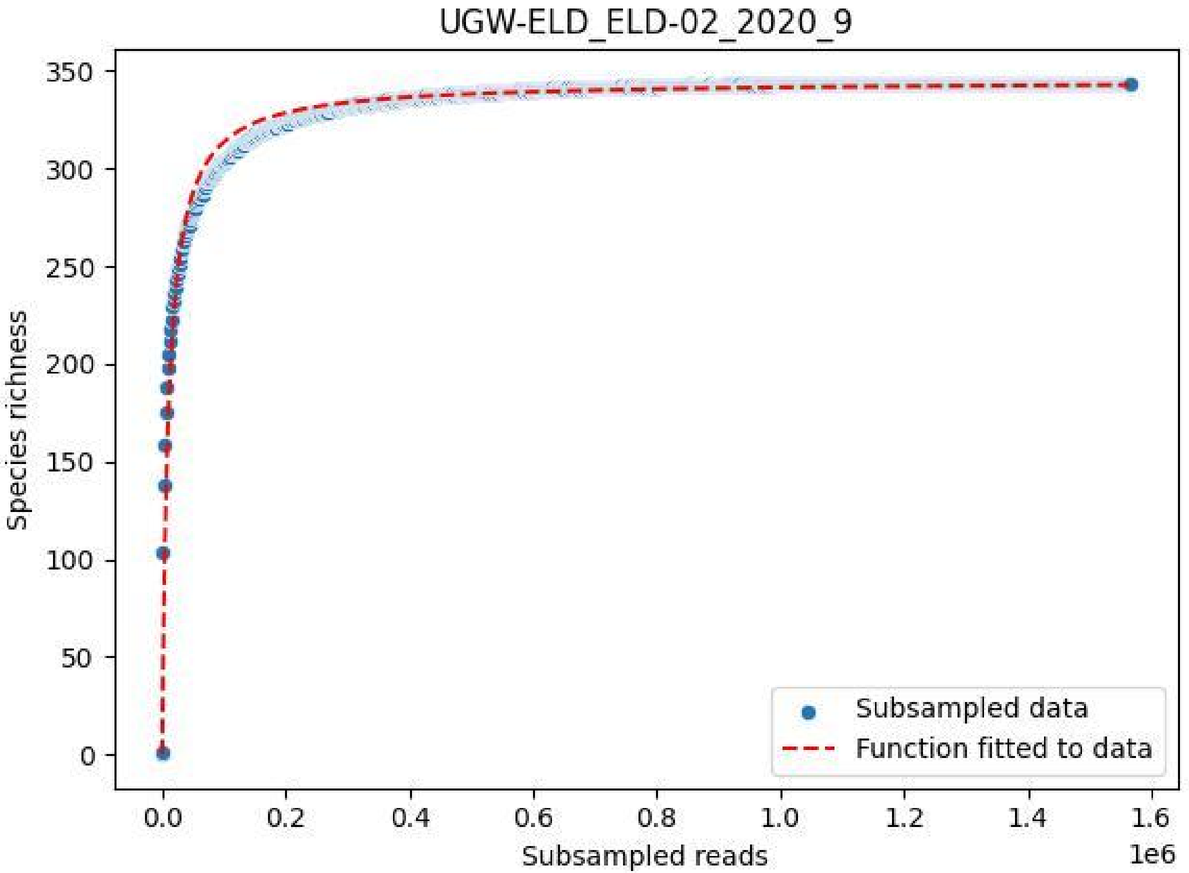
UGW-ELD_ELD-02_2020_9
Species richness
Subsampled reads
1e6
Subsampled data
Function fitted to data
0
50
100
150
200
250
300
350
0.0
0.2
0.4
0.6
0.8
1.0
1.2
1.4
1.6

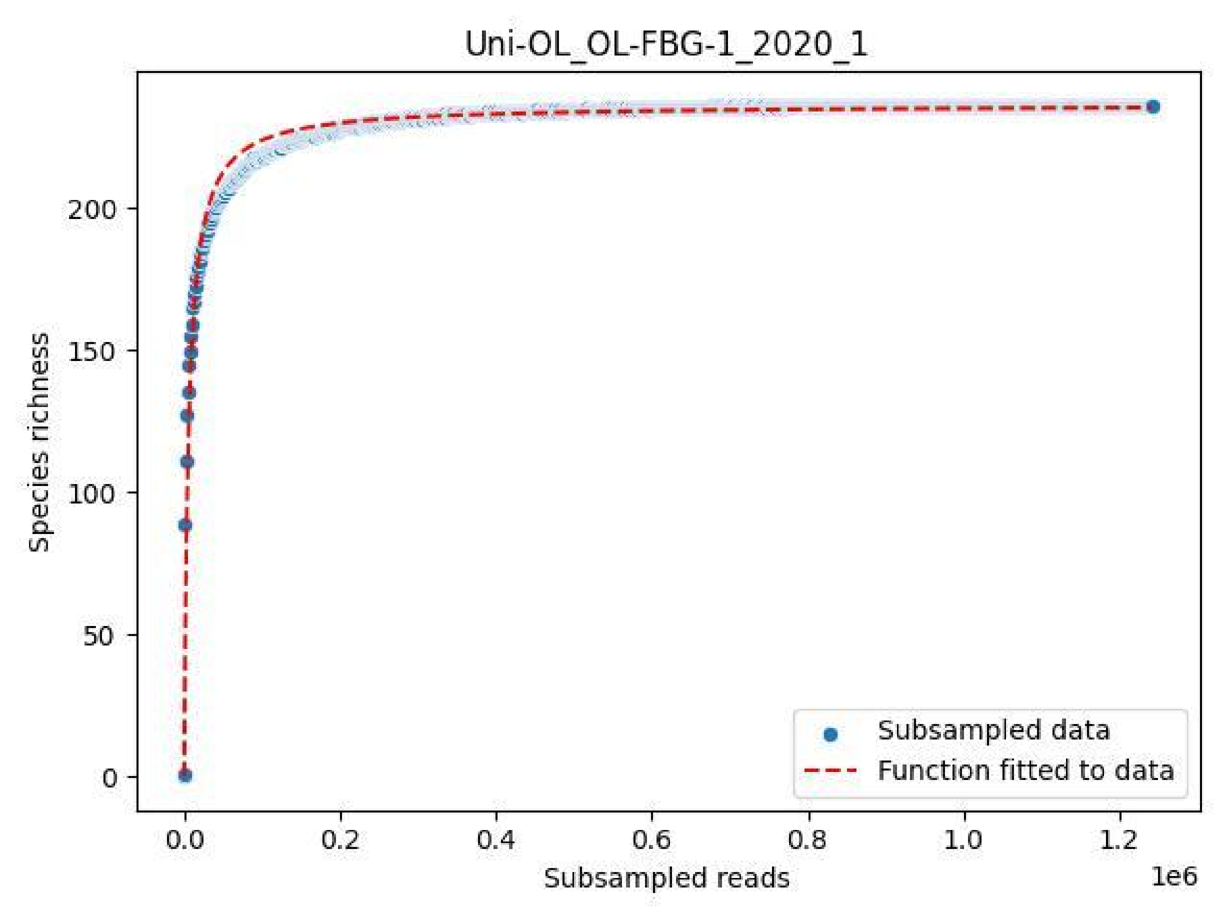

Uni-OL_OL-FBG-1_2020_1
Species richness
Subsampled reads
1e6
0
50
100
150
200
0.0
0.2
0.4
0.6
0.8
1.0
1.2
Subsampled data
Function fitted to data

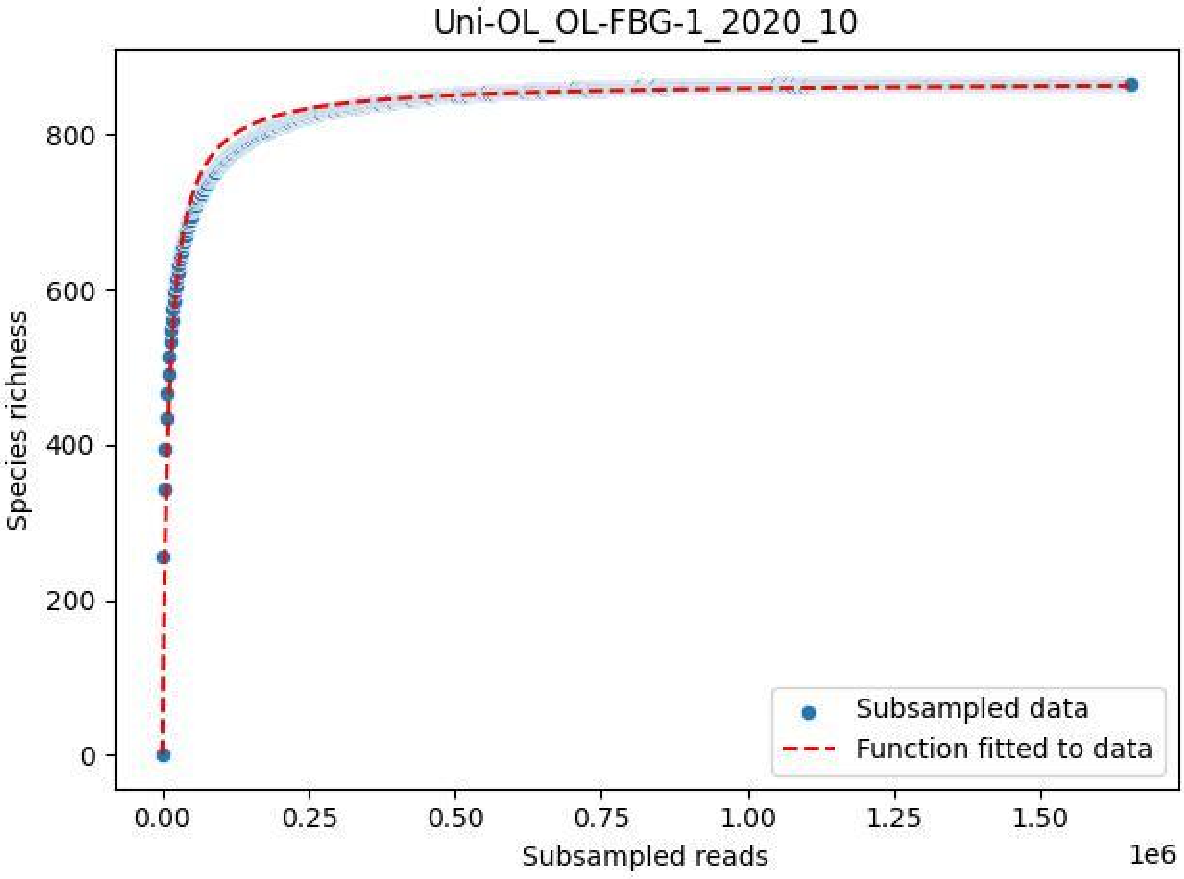
Uni-OL_OL-FBG-1_2020_10
Species richness
0
200
400
600
800
0.00
0.25
0.50
0.75
1.00
1.25
1.50
1e6
Subsampled reads
Subsampled data
Function fitted to data

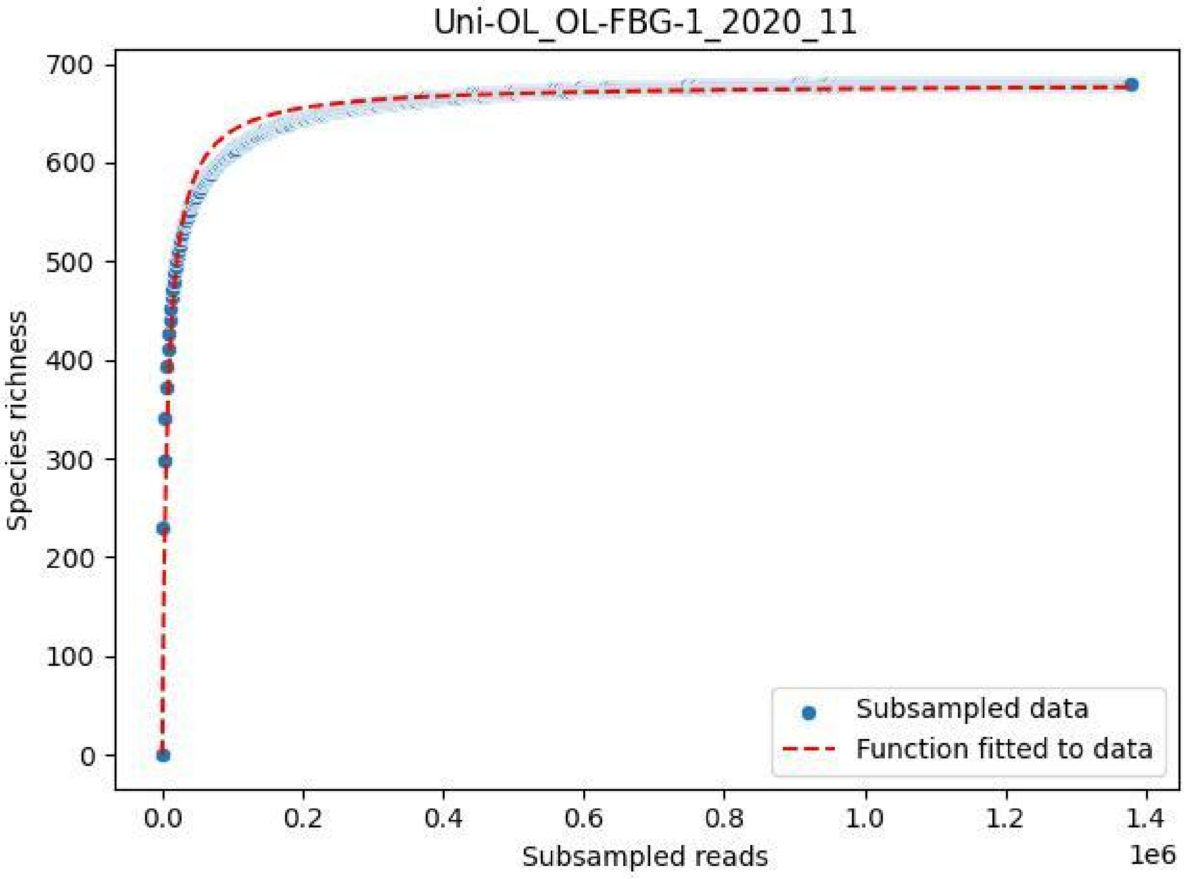

Uni-OL_OL-FBG-1_2020_11
700
600
500
400
300
200
100
0
Species richness
0.0
0.2
0.4
0.6
0.8
1.0
1.2
1.4
1e6
Subsampled reads
Subsampled data
Function fitted to data

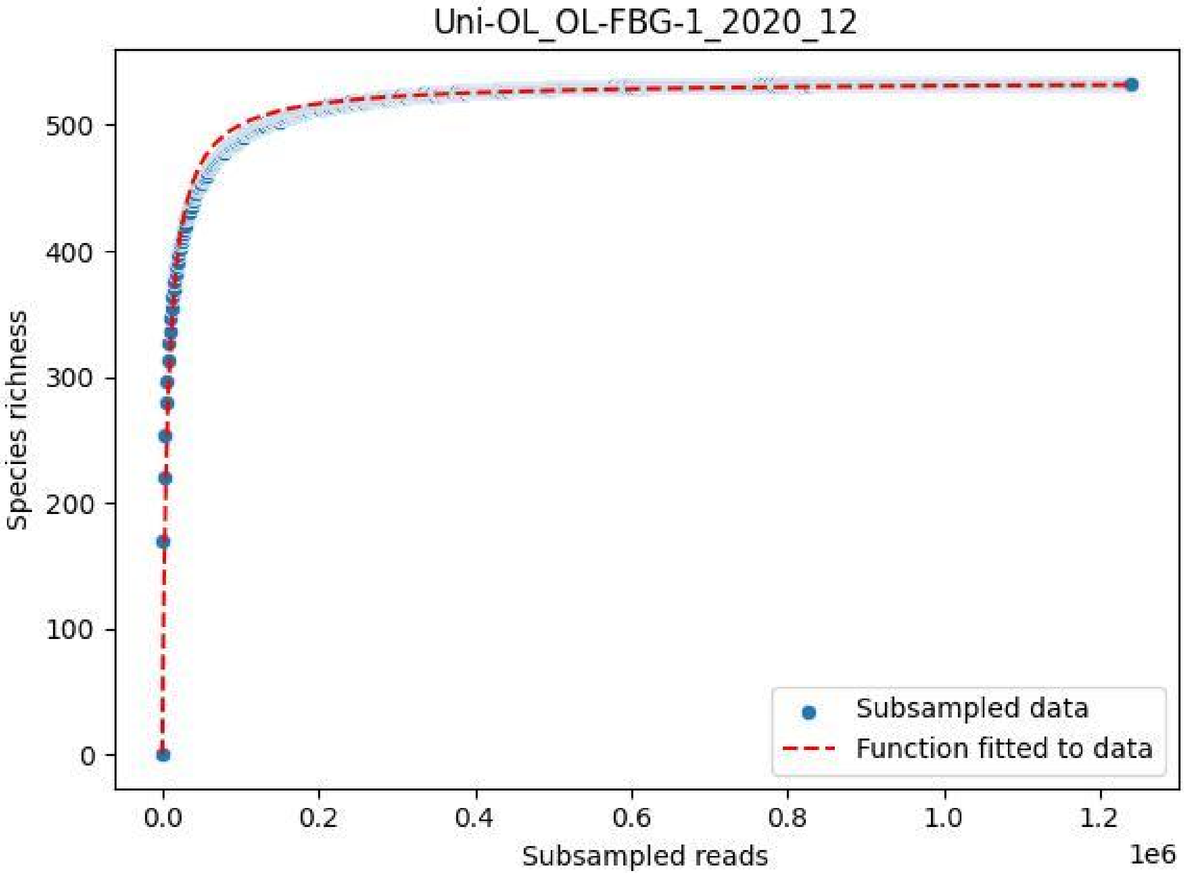

Uni-OL_OL-FBG-1_2020_12
Species richness
0
100
200
300
400
500
0.0
0.2
0.4
0.6
0.8
1.0
1.2
1e6
Subsampled reads
Subsampled data
Function fitted to data

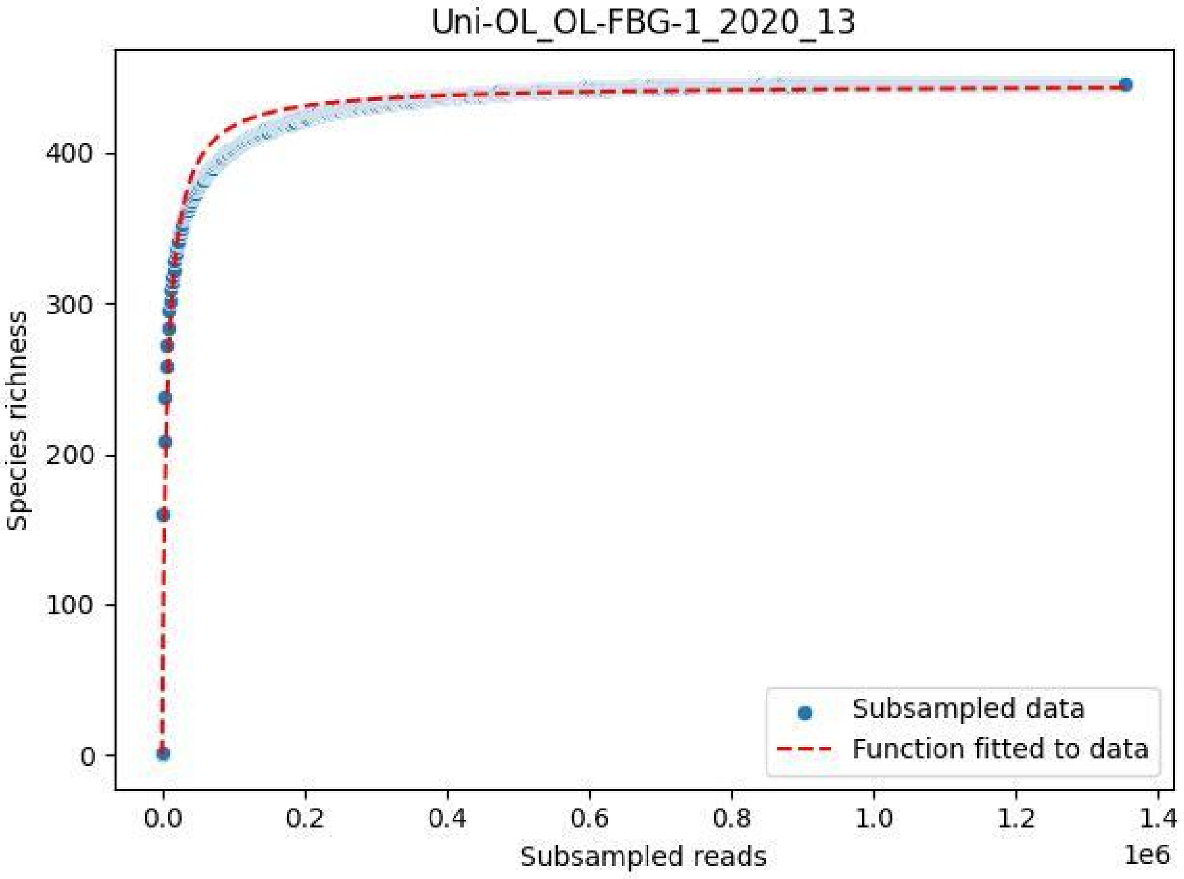
Uni-OL_OL-FBG-1_2020_13
Species richness
Subsampled reads
1e6
0
100
200
300
400
0.0
0.2
0.4
0.6
0.8
1.0
1.2
1.4
Subsampled data
Function fitted to data

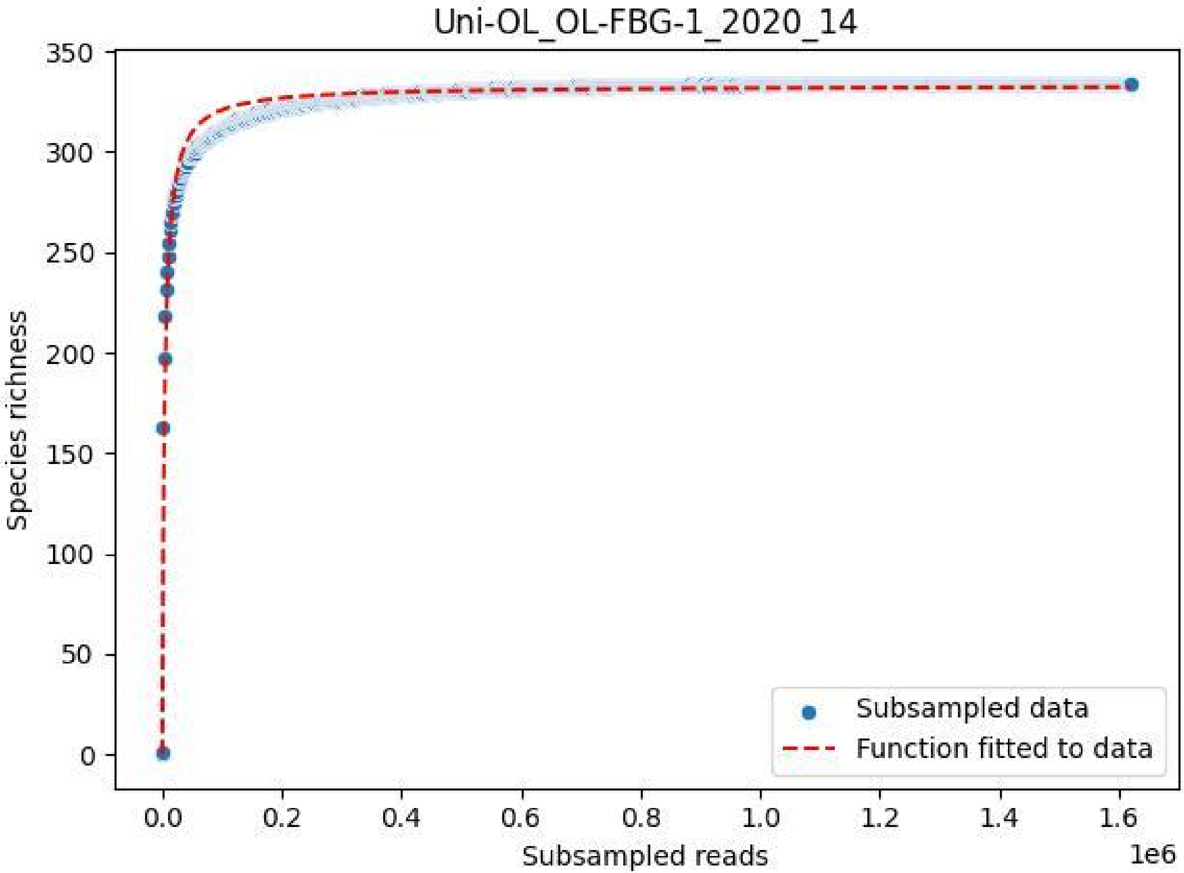

Uni-OL_OL-FBG-1_2020_14
Species richness
Subsampled reads
1e6
0
50
100
150
200
250
300
350
0.0
0.2
0.4
0.6
0.8
1.0
1.2
1.4
1.6
Subsampled data
Function fitted to data

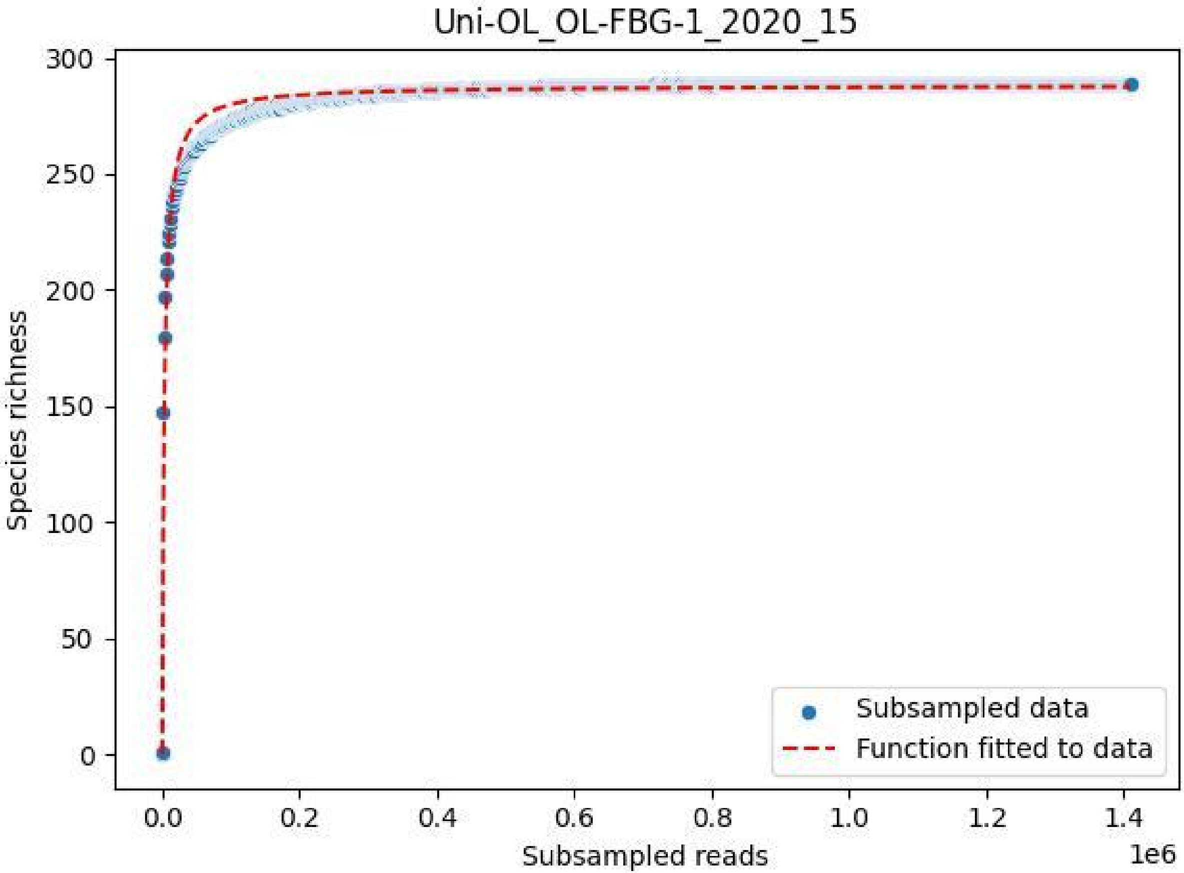
Uni-OL_OL-FBG-1_2020_15
Species richness
Subsampled reads
1e6
0
50
100
150
200
250
300
0.0
0.2
0.4
0.6
0.8
1.0
1.2
1.4
Subsampled data
Function fitted to data

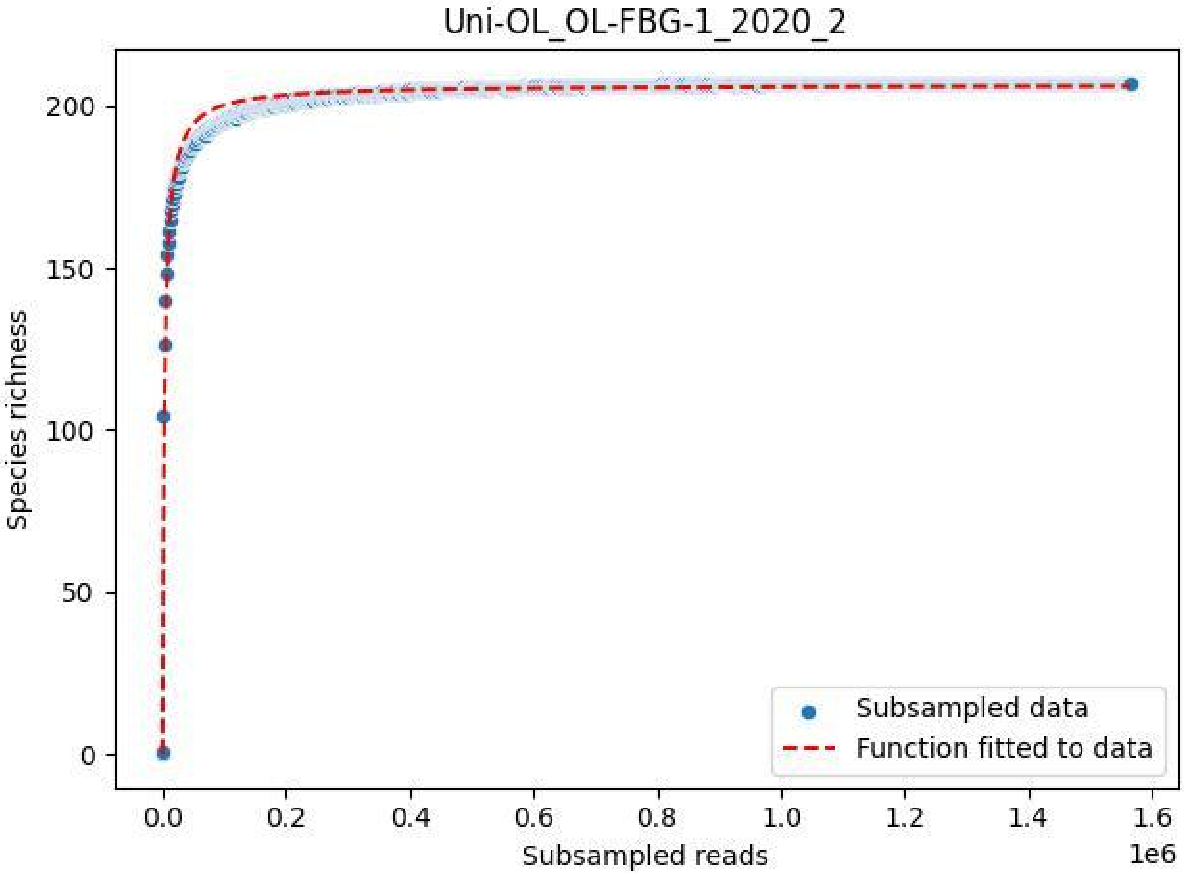

Uni-OL_OL-FBG-1_2020_2
Species richness
Subsampled reads
1e6
0
50
100
150
200
0.0
0.2
0.4
0.6
0.8
1.0
1.2
1.4
1.6
Subsampled data
Function fitted to data

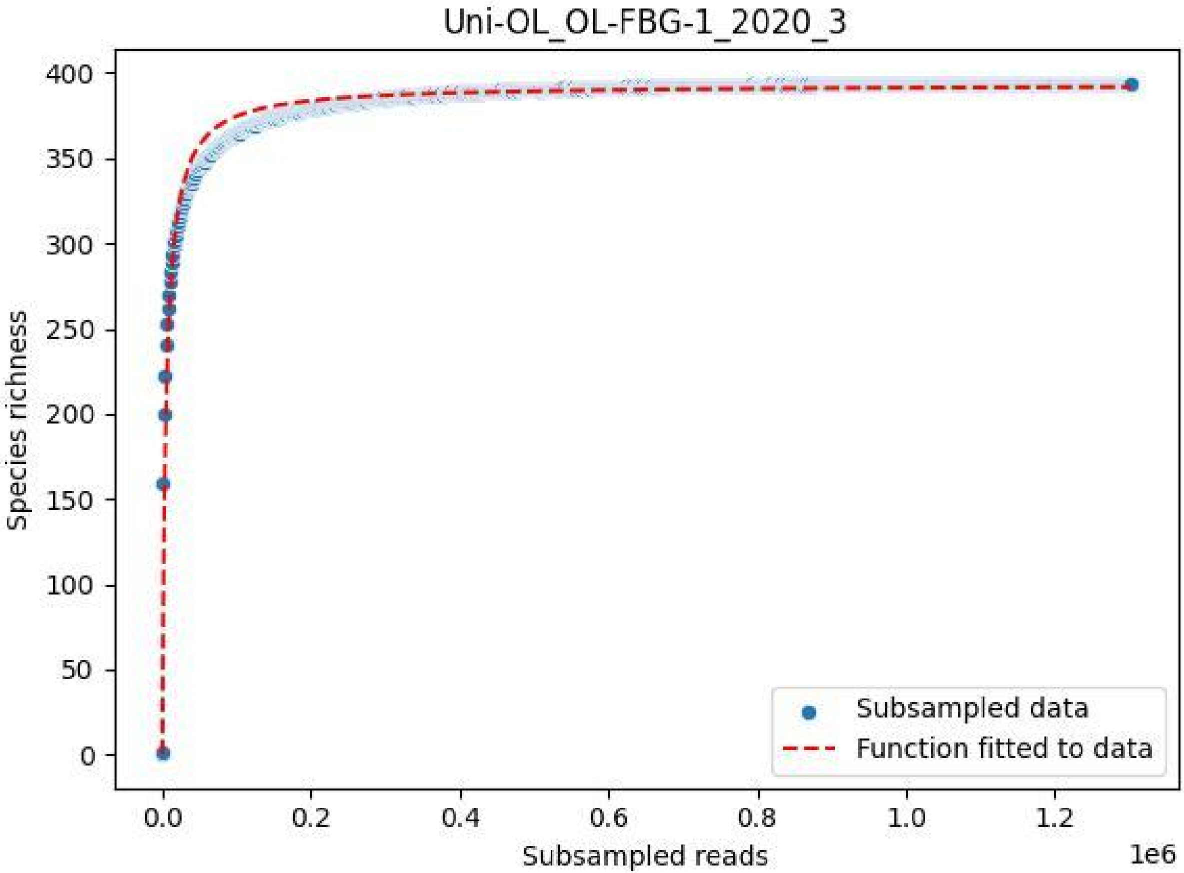

Uni-OL_OL-FBG-1_2020_3
Species richness
Subsampled reads
1e6
0
50
100
150
200
250
300
350
400
0.0
0.2
0.4
0.6
0.8
1.0
1.2
Subsampled data
Function fitted to data

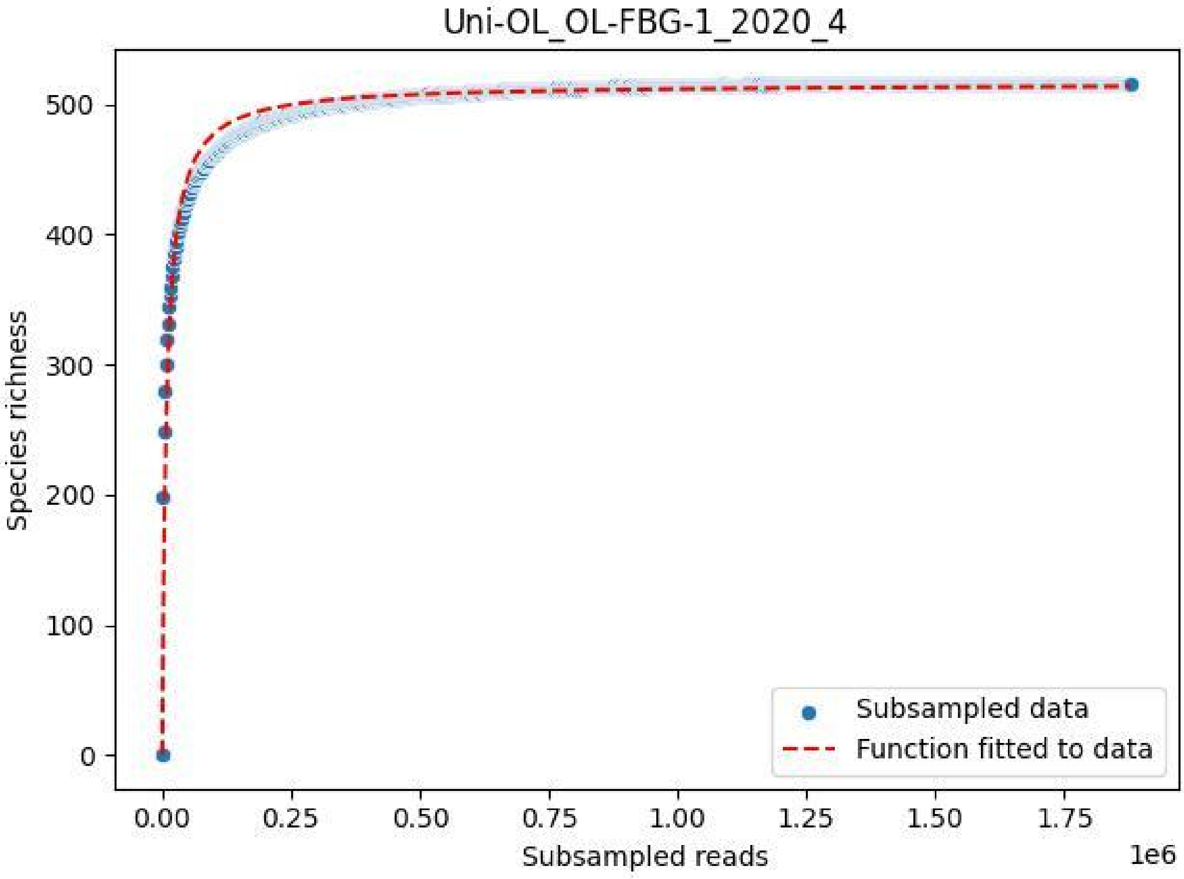

Uni-OL_OL-FBG-1_2020_4
Species richness
0
100
200
300
400
500
0.00
0.25
0.50
0.75
1.00
1.25
1.50
1.75
1e6
Subsampled reads
Subsampled data
Function fitted to data

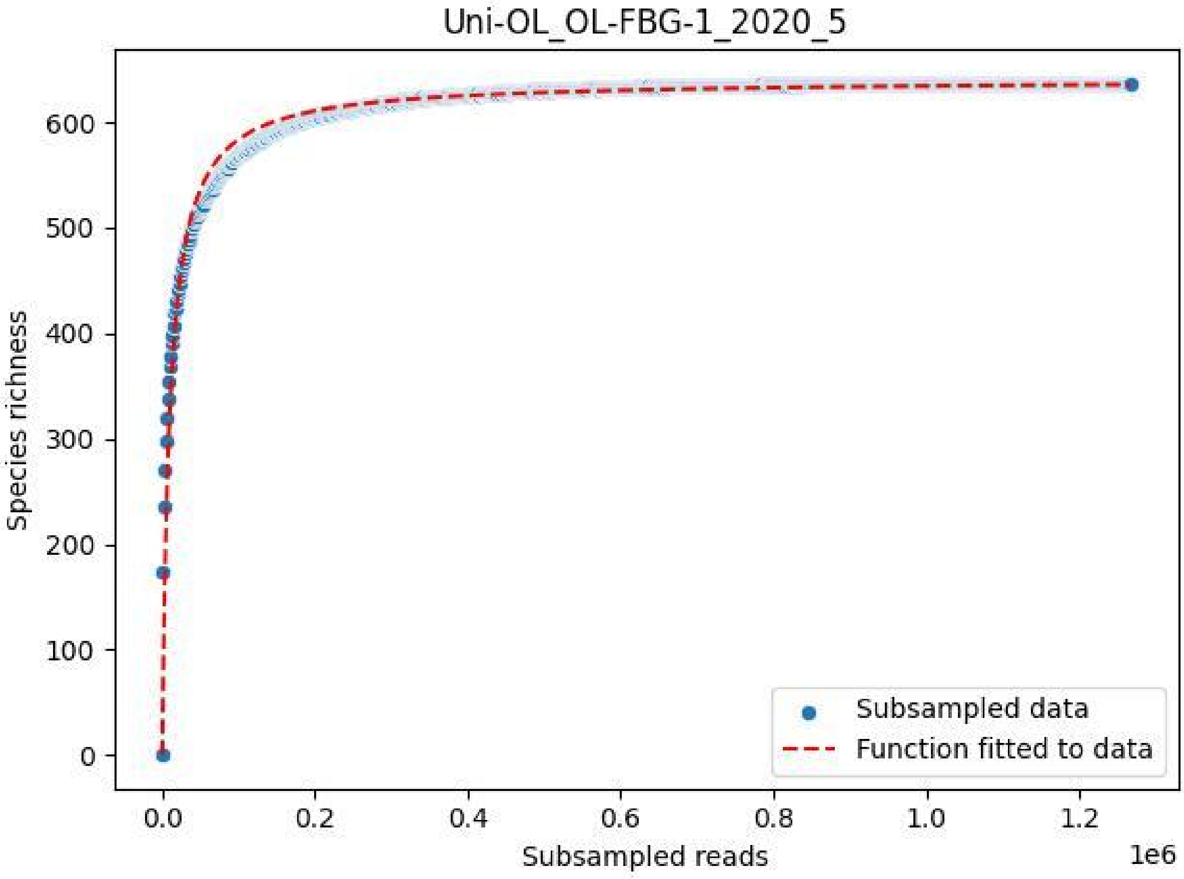
Uni-OL_OL-FBG-1_2020_5
Species richness
Subsampled reads
1e6
0.0
0.2
0.4
0.6
0.8
1.0
1.2
0
100
200
300
400
500
600
Subsampled data
Function fitted to data

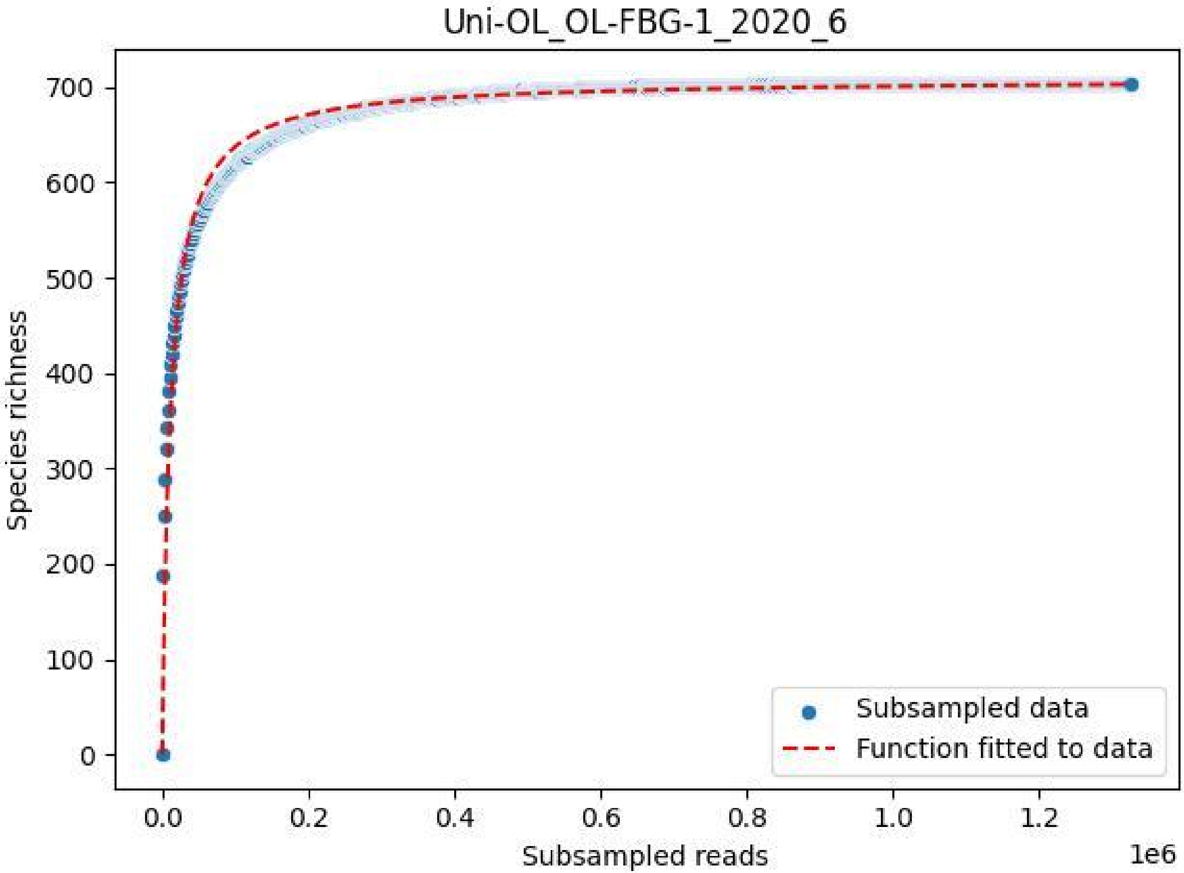
Uni-OL_OL-FBG-1_2020_6
Species richness
Subsampled reads
1e6
0
100
200
300
400
500
600
700
0.0
0.2
0.4
0.6
0.8
1.0
1.2
Subsampled data
Function fitted to data

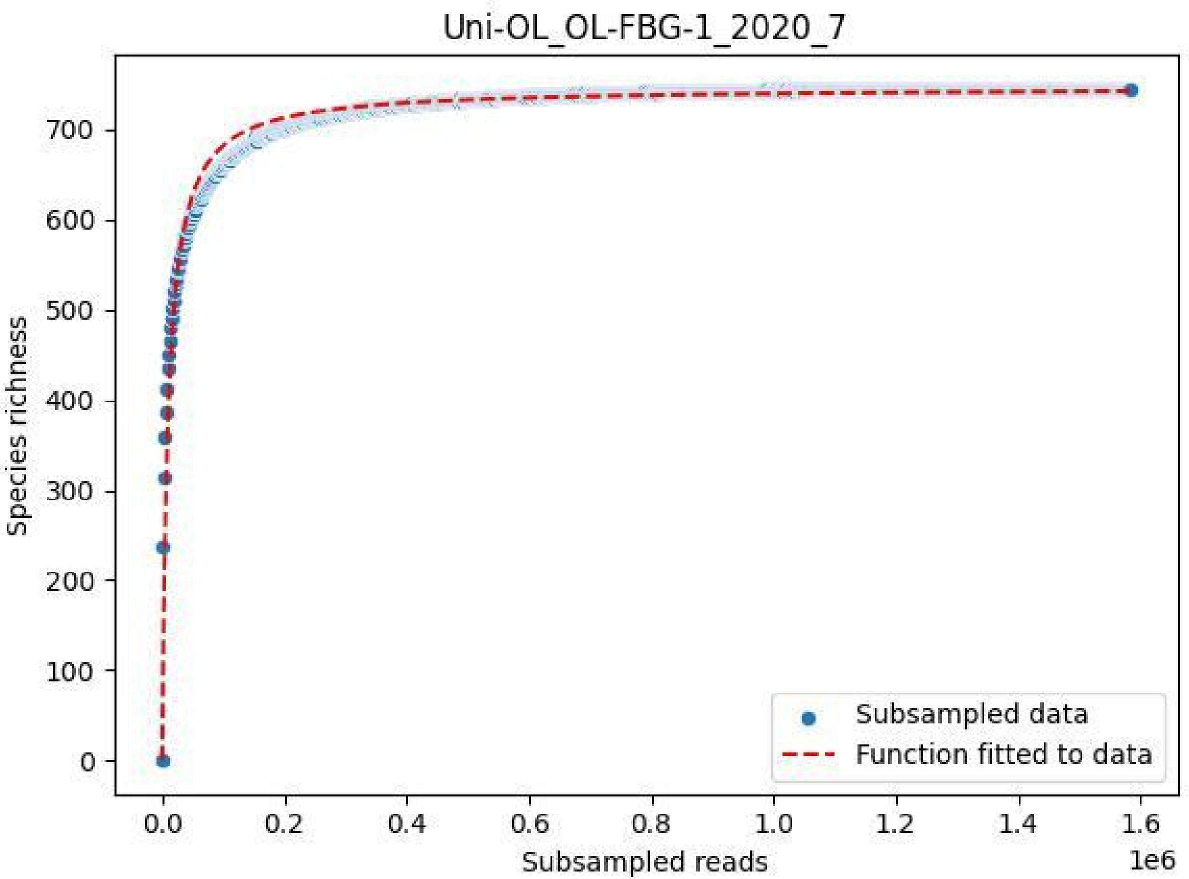
Uni-OL_OL-FBG-1_2020_7
Species richness
700
600
500
400
300
200
100
0
0.0
0.2
0.4
0.6
0.8
1.0
1.2
1.4
1.6
1e6
Subsampled reads
Subsampled data
Function fitted to data

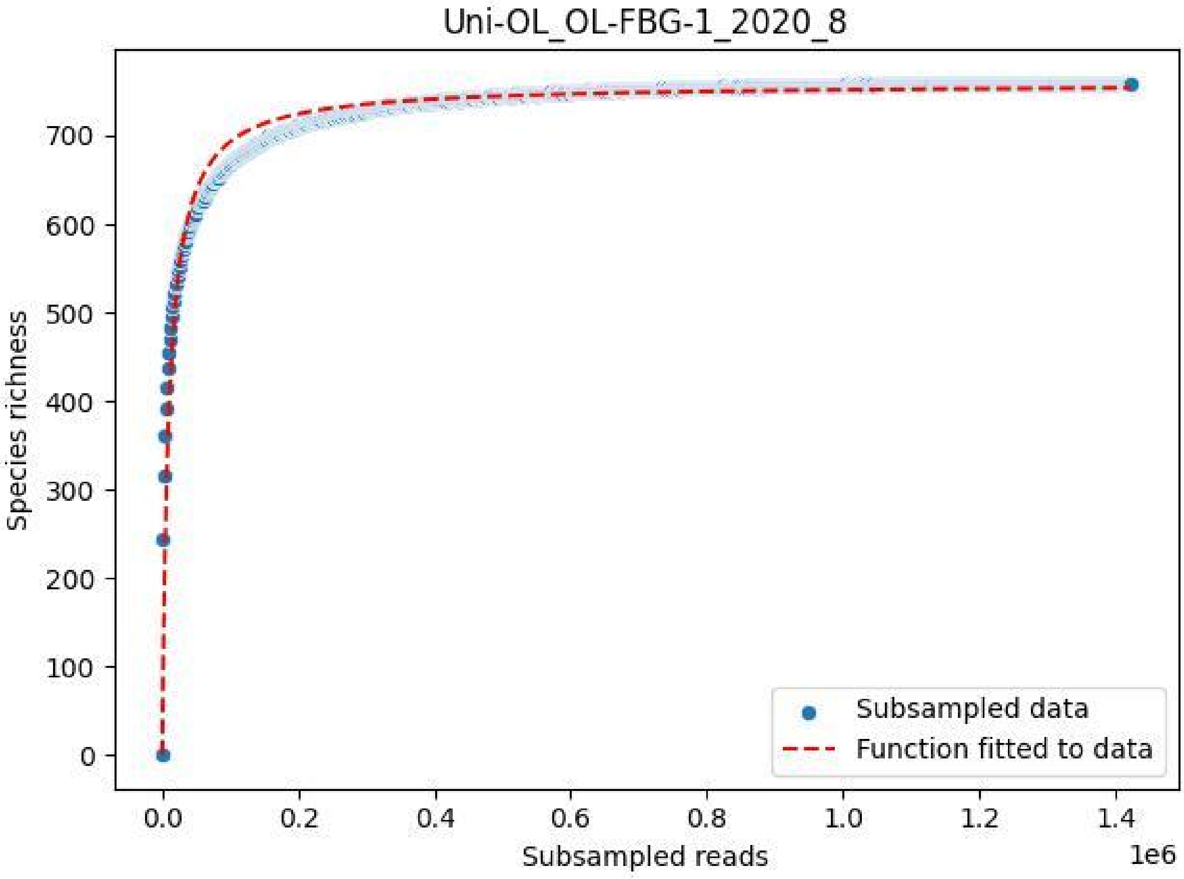

Uni-OL_OL-FBG-1_2020_8
Species richness
Subsampled reads
1e6
Subsampled data
Function fitted to data
0
100
200
300
400
500
600
700
0.0
0.2
0.4
0.6
0.8
1.0
1.2
1.4

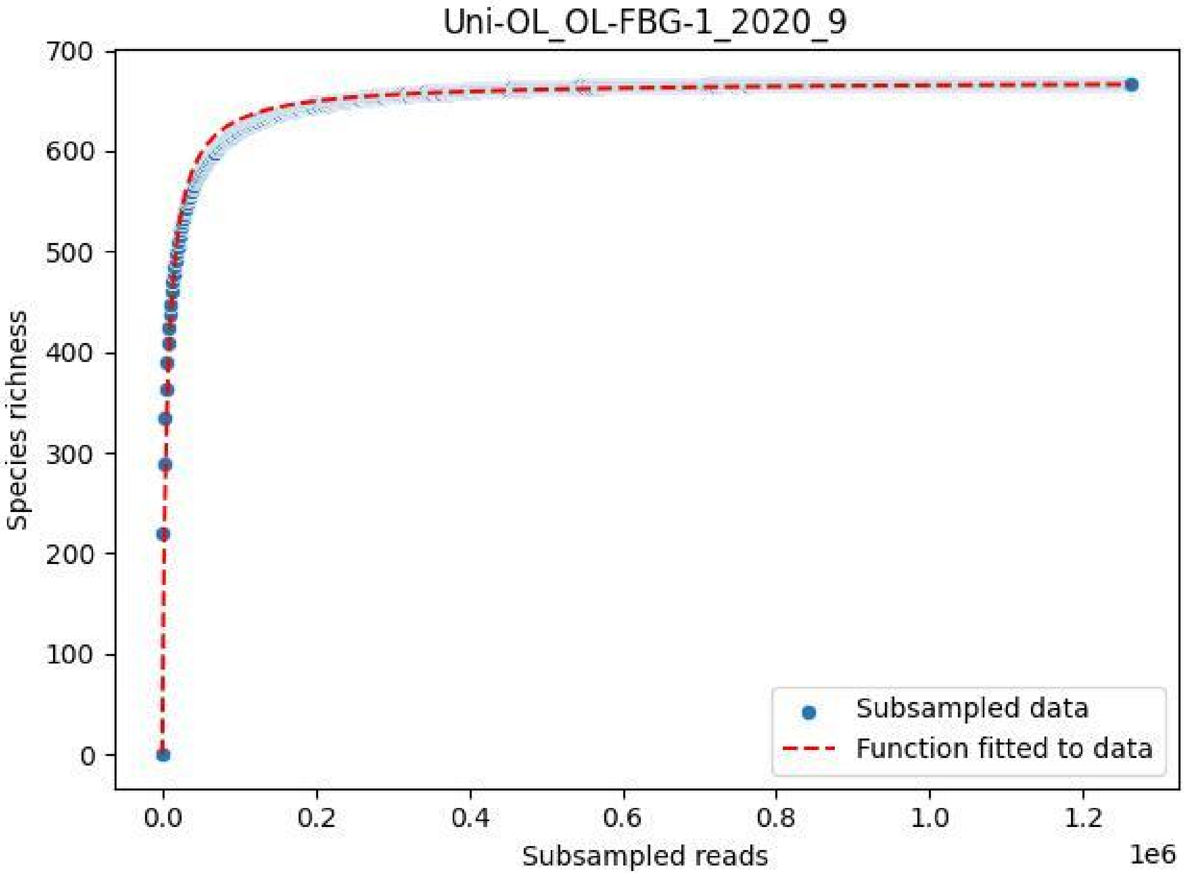

Uni-OL_OL-FBG-1_2020_9
Species richness
Subsampled reads
1e6
0
100
200
300
400
500
600
700
0.0
0.2
0.4
0.6
0.8
1.0
1.2
Subsampled data
Function fitted to data
