## Supplementary material for "Upscaling biodiversity monitoring: Metabarcoding estimates 31,846 insect species from Malaise traps across Germany": Information S1.docx

Supporting information 1:

Orthoptera was the only insect order where the number of OTUs per validated species was much higher compared to all other orders. For this reason, we excluced the order from downstream analysis. A likely reason for this observation is a high number of mitochondrial pseudogenes (“Numts”), i.e. formerly mitochondrial genes that were integrated into the nuclear genome and lost their function (Bensasson et al. 2001). Numts present a technical challenge when using our approach to estimate insect diversity. In this study for the order Orthoptera we detected about 6,000 unassigned orthopteran OTUs, while only 78 species are known from Germany. This likely occurred because Orthoptera exhibit high rates of Numts (Bensasson et al. 2000, 2001, Song et al. 2008). The often large genome sizes of orthopterans is considered one key factor leading to non-functional mitochondrial gene copies in the nuclear genome, potentially leading to inflated diversity estimates based on OTUs. If identifying insects with high Numts is critical, then metabarcoding could be combined with other approaches, including image-based approaches in combination with high-throughput single specimen barcoding as well as machine learning (Chua et al. 2023, de Waard et al. 2019, Hartop et al. 2022, Wührl et al. 2022). While metabarcoding is likely to outcompete those methods in time and costs for large-scale monitoring, single-specimen methods could be feasible if they target a particular problem (Srivathsan et al. 2023), such as specific taxa that are difficult for metabarcoding to resolve.

Bensasson D, Zhang D-X, Hewitt GM (2000) Frequent assimilation of mitochondrial DNA by grasshopper nuclear genomes. Molecular Biology and Evolution 17: 406–415. <https://doi.org/10.1093/oxfordjournals.molbev.a026320>

Bensasson D, Zhang D-X, Hartl DL, Hewitt GM (2001) Mitochondrial pseudogenes: evolution’s misplaced witnesses. Trends in Ecology & Evolution 16: 314–321. <https://doi.org/10.1016/S0169-5347(01)02151-6>

Chua PYS, Bourlat SJ, Ferguson C, Korlevic P, Zhao L, Ekrem T, Meier R, Lawniczak MKN (2023) Future of DNA-based insect monitoring. *Trends in Genetics* **0**,. doi:[10.1016/j.tig.2023.02.012](https://doi.org/10.1016/j.tig.2023.02.012).

deWaard JR, Levesque-Beaudin V, deWaard SL, Ivanova NV, McKeown JTA, Miskie R, Naik S, Perez KHJ, Ratnasingham S, Sobel CN, Sones JE, Steinke C, Telfer AC, Young AD, Young MR, Zakharov EV, Hebert PDN (2019) Expedited assessment of terrestrial arthropod diversity by coupling Malaise traps with DNA barcoding. *Genome* **62**, 85–95. doi:[10.1139/gen-2018-0093](https://doi.org/10.1139/gen-2018-0093).

Hartop E, Srivathsan A, Ronquist F, Meier R (2022) Towards large-scale integrative taxonomy (LIT): Resolving the data conundrum for dark taxa. *Systematic Biology* **71**, 1404–1422. doi:[10.1093/sysbio/syac033](https://doi.org/10.1093/sysbio/syac033).

Song H, Buhay JE, Whiting MF, Crandall KA (2008) Many species in one: DNA barcoding overestimates the number of species when nuclear mitochondrial pseudogenes are coamplified. *Proceedings of the National Academy of Sciences of the United States of America* **105**, 13486–13491. doi:[10.1073/pnas.0803076105](https://doi.org/10.1073/pnas.0803076105).

Hartop E, Srivathsan A, Ronquist F, Meier R (2022) Towards large-scale integrative taxonomy (LIT): Resolving the data conundrum for dark taxa. *Systematic Biology* **71**, 1404–1422. doi:[10.1093/sysbio/syac033](https://doi.org/10.1093/sysbio/syac033).

Wührl L, Pylatiuk C, Giersch M, Lapp F, von Rintelen T, Balke M, Schmidt S, Cerretti P, Meier R (2022) DiversityScanner: Robotic handling of small invertebrates with machine learning methods. *Molecular Ecology Resources* **22**, 1626–1638. doi:[10.1111/1755-0998.13567](https://doi.org/10.1111/1755-0998.13567).
