## Supplementary material for "Upscaling biodiversity monitoring: Metabarcoding estimates 31,846 insect species from Malaise traps across Germany": Information S2.pdf

**Title:** Land cover drives flying insect diversity, with protected areas insufficiently covering biodiversity hotspots, pollinators, and threatened species

**Authors:** James S. Sinclair[1*], Dominik Buchner[2], Mark O. Gessner[3,4], Jörg Müller[5,6], Steffen U. Pauls[7,8,9], Stefan Stoll[10,11], Ellen A. R. Welti[12], Claus Bässler[6,13], Jörn Buse[14], Frank Dziock[15], Julian Enss[16,17], Thomas Hörren[17], Robert Künast[18], Yuanheng Li[2], Andreas Marten[19], Carsten Morkel[20], Ronny Richter[21,22], Tobias Scharnweber[23], Sebastian Seibold[24,25,26], Martin Sorg[17], Sönke Twietmeyer[27], Dirk Weis[28], Wolfgang Weisser[29], Benedikt Wiggering[30], Martin Wilmking[23], Gerhard Zotz[31], Mark Frenzel[32], Florian Leese[2,33], Peter Haase[1,11]

[1]Department of River Ecology and Conservation, Senckenberg Research Institute and Natural History Museum Frankfurt, Gelnhausen 63571 Germany

[2]Aquatic Ecosystem Research, University of Duisburg-Essen, Universitätsstraße 5, Essen 45141 Germany

[3]Department of Plankton and Microbial Ecology, Leibniz Institute of Freshwater Ecology & Inland Fisheries (IGB), Zur alten Fischerhütte 2, Stechlin 16775 Germany

[4]Department of Ecology, Berlin Institute of Technology (TU Berlin), Ernst-Reuter-Platz 1, Berlin 10587 Germany

[5]Field Station Fabrikschleichach, Department of Animal Ecology and Tropical Biology, Biocenter University of Würzburg, Glashüttenstraße 5, Rauhenebrach 96181 Germany

[6]Nationalpark Bavarian Forest, Freyunger Str. 2, Grafenau 94481 Germany

[7]Senckenberg Research Institute and Nature Museum Frankfurt, Senckenberganlage 25,

Frankfurt am Main 60388 Germany
[8]Institute of Insect Biotechnology, Justus-Liebig-University Gießen, Heinrich-Buff-Ring 26-32, Gießen 35392 Germany
[9]LOEWE Centre for Translational Biodiversity Genomics, Senckenberganlage 25, Frankfurt am Main 60388 Germany
[10]University of Applied Sciences Trier, Environmental Campus Birkenfeld, Campusallee, Hoppstädten-Weiersbach 55768 Germany
[11]Faculty of Biology, University of Duisburg-Essen, Universitätsstraße 5, Essen 45141 Germany
[12]Conservation Ecology Center, Smithsonian National Zoo and Conservation Biology Institute, Front Royal, Virginia 22630 USA
[13]Ecology of Fungi, University Bayreuth, Universitätsstraße 30, Bayreuth 95440 Germany
[14]Department for Ecological Monitoring, Research and Species Protection, Black Forest National Park, Kniebisstraße 67, Freudenstadt 72250 Germany
[15]University of Applied Sciences HTW Dresden, Pillnitzer Platz 2, Dresden 01326 Germany
[16]Aquatic Ecology, Faculty of Biology, University of Duisburg-Essen, Universitätsstraße 5, Essen 45141 Germany
[17]Entomological Society Krefeld (EVK), Magdeburger Straße 38-40, Krefeld 47800 Germany
[18]Terrestrial Ecology Research Group, Department of Life Science Systems, School of Life Sciences, Technical University of Munich, Hans-Carl-von-Carlowitz-Platz 2, Freising 85354 Germany
[19]Harz National Park, Lindenallee 35, Wernigerode 38855 Germany
[20]Kellerwald-Edersee National Park, Laustrasse 8, Bad Wildungen 34537 Germany
[21]German Centre for Integrative Biodiversity Research (iDiv) Halle-Jena-Leipzig, Puschstraße 4,

Leipzig 04103 Germany
[22]Systematic Botany and Functional Biodiversity, Institute for Biology, Leipzig University, Johannisallee 21, Leipzig 04103 Germany
[23]Institute for Botany and Landscape Ecology, University Greifswald, Soldmannstraße 15, Greifswald 14487 Germany
[24]Berchtesgaden National Park, Doktorberg 6, 83471 Berchtesgaden, Germany
[25]Technical University of Munich, Ecosystem Dynamics and Forest Management, Department of Life Science Systems, School of Life Sciences, Hans-Carl-von-Carlowitz-Platz 2, Freising 85354 Germany
[26]TUD Dresden University of Technology, Forest Zoology, Pienner Str. 7, Tharandt 01737 Germany
[27]Department of Research and Documentation, Eifel National Park, Urftseestraße 34, Schleiden 53937 Germany
[28]Biosphärenreservat Oberlausitzer Heide- und Teichlandschaft, Warthaer Dorfstraße 29, Malschwitz 02694 Germany
[29]Terrestrial Ecology Research Group, Department of Life Science Systems, School of Life Sciences, Technical University of Munich, Hans-Carl-von-Carlowitz-Platz 2, Freising 85354 Germany
[30]Lower Saxon Wadden Sea National Park Authority, Virchowstraße 1, Wilhelmshaven D-26382 Germany
[31]Institute for Biology and Environmental Sciences, Carl von Ossietzky University Oldenburg, Ammerländer Heerstraße 114-118, Oldenburg 26129 Germany
[32]Helmholtz Centre for Environmental Research UFZ, Department of Community Ecology, Th.-

Lieser-Straße 4, Halle 06120 Germany
[33]Centre for Water and Environmental Research (ZWU), University of Duisburg-Essen, Universitätsstraße 3, Essen 45141 Germany

**Abstract**

Insects and their ecosystem functions are declining in many regions. Ameliorating these losses requires identifying the principal drivers across different taxa, and determining which insects are covered by protected areas. Here, we used biomass and state-of-the-art DNA metabarcoding data for 31,846 flying insect species to examine responses to land cover, weather, and climatic gradients, and differences in site protection status across Germany during 2019 and 2020. Average insect biomass was ~2.0 g day$^{-1}$, which matched reports from previous and subsequent years, indicating a potential recent plateau in biomass declines. Insect biomass, diversity, taxonomic composition, and trait composition were primarily related to land cover. Specifically, as heterogeneous low vegetation increased and forest cover decreased, insect biomass increased by 75% in 2019 and 58% in 2020, and species richness by 64% and 32%. Surveyed sites with higher protection status tended to be forested. Consequently, many non-forest-dwelling insects, including most pollinators and threatened species, occurred outside protected areas. Our results highlight the value of low vegetation and land cover heterogeneity for promoting insect biomass and diversity, with forests providing valuable contributions by supporting unique communities. Additionally, better protecting insects requires improved management of unforested areas where many biodiversity hotspots and key taxa occur.

### Introduction

Losses of key ecological functions, such as pollination and organic matter decomposition[1,2], are linked to anthropogenic declines in insect biomass and species richness[3,4,5,6,7]. However, most research has been limited to specific taxonomic or functional groups (e.g., pollinators[5,8,9]), or to metrics that summarize insect communities, such as overall biomass or richness[4,10,11,12]. This focus on taxonomic subsets or summary metrics makes it difficult to determine how entire insect communities are responding to environmental change because unstudied taxa are missed (including taxa that may be increasing[13]), and summary metrics can mask important compositional changes[14]. Additionally, studies focusing on different taxonomic subsets or summary metrics can make it difficult to identify key drivers. For example, changes in land cover, weather, and climate are often important in structuring insect communities[5,6,7], but different taxa can vary in their responses to each of these drivers[13,15]. The identity of the examined taxonomic group may therefore explain why some studies find effects of just land cover or just meteorological factors[16,17], whereas others find both to be important[18,19]. Similarly, differences in which summary metrics are examined may further exacerbate variability in insect community responses to a given environmental driver, such as biomass responding positively versus richness responding negatively to warming[20]. Studies encompassing a wide variety of insect taxa and multiple summary metrics are needed to determine how entire insect communities are responding to environmental change.

In addition to disentangling key drivers, information on a variety of insect taxa can help determine which insects are better protected from environmental change, and can better track harmful invaders. Protected areas, such as nature reserves and national parks, are typically designated to protect wildlife, but the location and design of these areas rarely considers insects[21]. Consequently, little is known about which insects are covered by current protected areas[21] and whether these areas are protecting vulnerable native species, threatened species, and key functional groups (e.g., pollinators, dung beetles). Native and threatened insects are also at risk of further decline owing to ongoing spread of invasive insects[22], which can be sources of costly ecosystem disservices, especially when they become pests or disease vectors[23]. Changes in invasive species can be easily missed if insect assessments are limited to taxonomic subsets or summary community metrics (e.g., overall biomass or richness[24]), warranting more comprehensive examinations of change across the broader insect community.

At present, obtaining a comprehensive perspective of insect communities is hindered by insufficient knowledge of species diversity and distribution. Insects are by far the most diverse animal group on Earth, comprising millions of known and as yet unknown and cryptic species[25,26]. This extremely high diversity presents challenges in terms of the needed expertise, time, and cost to identify all insects in a given community, particularly when samples are collected over large spatiotemporal scales that can encompass thousands of species within a single biogeographic region[27,28]. Genetic identification methods, such as DNA metabarcoding (hereafter 'metabarcoding'), offer a promising solution to this issue. Metabarcoding can allow for faster and more cost-effective species identification than traditional approaches based on morphological traits, enabling identification of thousands of species within a few weeks[29]. Additionally, metabarcoding can help detect unknown taxa by defining putative species (e.g., via Operational Taxonomic Units or OTUs), can resolve species that are difficult to distinguish morphologically, and can more effectively track the presence of invasive species[30]. Consequently, metabarcoding can expand the scope of insect community analyses to encompass a wider variety of taxa, thus broadening our perspective on insect responses to environmental change.

Here, we take advantage of recent advances in metabarcoding technology[29] to analyze spatial patterns and drivers of flying insect communities across Germany. Insects were collected during 2019 and 2020 from a Malaise trap program encompassing 75 sites distributed across Germany, initiated by LTER-D, a German network for long-term ecological research (Fig. 1; https://www.ufz.de/lter-d/index.php?de=46285; ref. 31). The resulting dataset provides one of the most comprehensive broad-scale assessments of insect communities, comprising 31,846 species. Of these species, 10,803 have assigned scientific species names and are referred to as ‘validated species’, whereas the additional 21,403 species were estimated from OTUs[29] and are referred to as ‘plausible species’. Our principal questions were how key characteristics of flying insect diversity differed (1) in response to spatial land cover, weather, and climatic gradients and (2) between protected versus unprotected areas. We examined these relationships for insect biomass, species richness, taxonomic composition, occurrence of threatened species, and functional composition via changes in pollinators, invasive species, and larval feeding traits. Examining changes in feeding trait composition can also help identify potential drivers, such as indicating variation in available resources[32]. We expected lower insect biomass and species richness in sites experiencing more intense anthropogenic land use, and warmer and drier conditions, given the known negative effects of these stressors on insects[2,6,7,12]. Conversely, we expected higher values in protected areas, which restrict human activity and land modification[33] and can help maintain the environmental conditions required by certain insects (e.g., lower temperatures[34]).

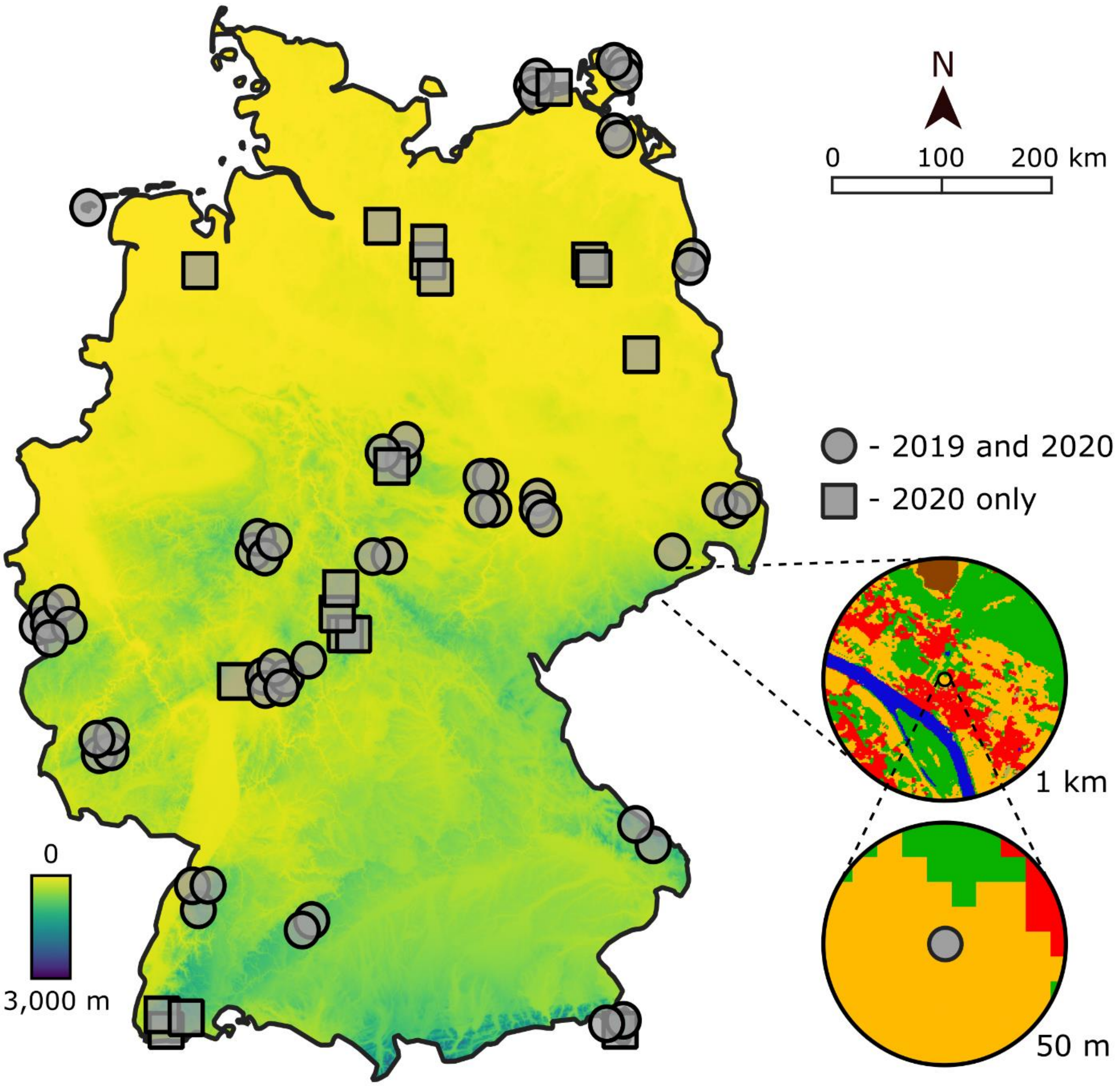


**Fig. 1: Locations of 75 Malaise trap sampling sites across Germany.** Elevation is indicated in the map by a yellow to purple color gradient (sites encompass a gradient of 1 to 1,400 m). The insets illustrate how we quantified fine-scale (50 m radius around each trap) versus broad-scale (1 km) land cover (green = forest, orange = low vegetation, red = urban, brown = agricultural, bare soil = not shown). Note that some trap positions are shifted slightly to distinguish proximate traps.

### Results

*Validated and plausible species*

We report the results for the validated species here, whereas the results for the plausible species estimated from OTUs are provided in Supplementary Information 1, given the greater uncertainty in the exact number of plausible species (further detailed in ref. 29). The principal patterns are the same in both. Of the 10,803 validated species, the majority belonged to five major orders: Diptera (36%), Hymenoptera (22%), Coleoptera (17%), Lepidoptera (16%), and Hemiptera (6%). All other orders each comprised about 1% or less of all species. The five major orders encompassed 359 different families, with 51% of species belonging to just 20 families, particularly families from the Hymenoptera (e.g., Ichneumonidae) and Diptera (e.g., Chironomidae, Mycetophilidae, and Sciaridae; Data S1).

*Seasonal dynamics of biomass, richness, and turnover*

Across all sampling periods, insect biomass averaged 2.0 ± 1.2 g $day^{-1}$ (mean ± SD) per site in 2019 and 2.3 ± 1.2 g $day^{-1}$ in 2020. We also caught an average of 1,229 ± 494 different species per site in 2019 and 1,700 ± 557 in 2020. Mid-April to September was the most consistent sampling period across sites, which encompassed the period of highest biomass and species richness, and lowest temporal turnover (i.e., temporal β-diversity), based on generalized additive mixed models (Fig. 2). Biomass peaked around 3.9 g $day^{-1}$ on Julian day 183 in 2019 (early July; $n$ = 856, edf = 3.9) and at 3.8 g $day^{-1}$ on Julian day 200 in 2020 ($n$ = 1,084, edf = 3.9; Fig. 2a). Total species richness mirrored changes in biomass, although richness peaked earlier and lower in 2019 (287 species on Julian day 178; $n$ = 775, edf = 3.8) compared to 2020 (506 species on Julian day 195; $n$ = 1,040, edf = 3.9; Fig. 2b). Temporal turnover was high throughout

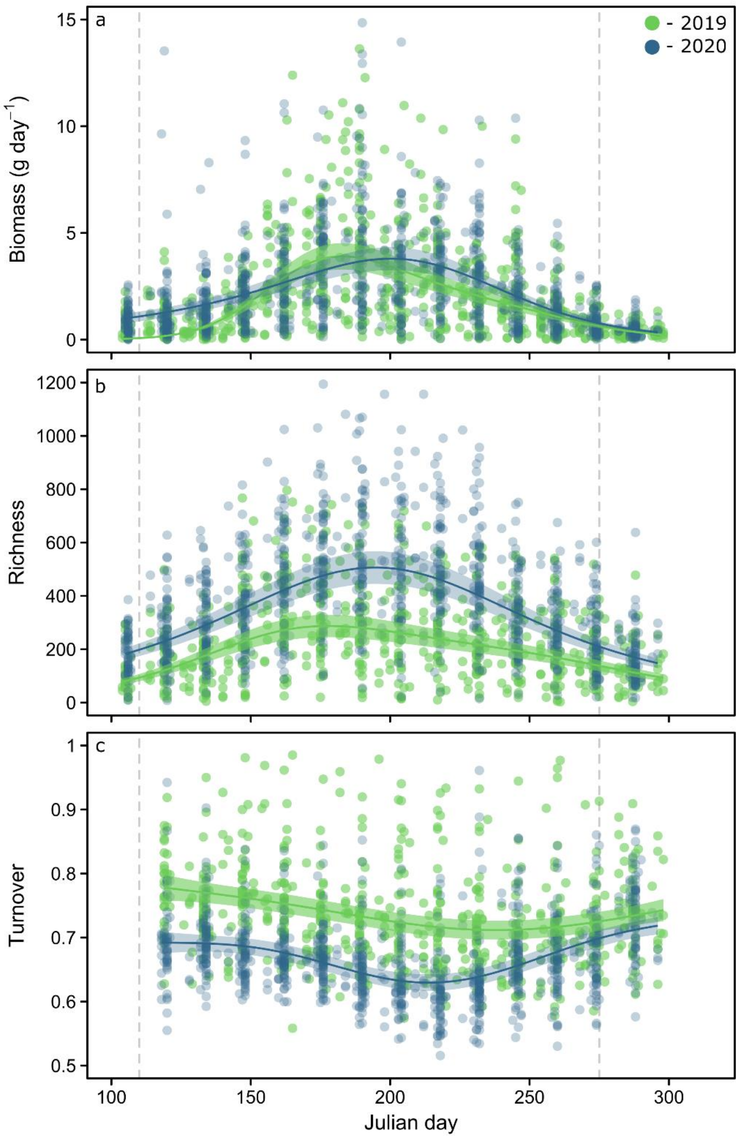


**Fig. 2: Seasonal trends in biomass, richness, and temporal turnover.** Seasonal trends in (**a**) biomass (g day$^{-1}$), (**b**) total species richness, and (**c**) temporal turnover (i.e., change in species composition between successive sampling periods) of flying insects caught in Malaise traps during the growing season in 2019 (green) and 2020 (blue). Best-fit lines (solid lines) and 95% confidence intervals (shaded areas) are based on estimates from generalized additive mixed models. Gray dashed lines indicate the period considered in all other analyses, specifically Julian days 110 (mid-April) to 273 (end of September).

the year (always > 0.6) and reached its lowest values somewhat later in the year than biomass and richness, at 0.71 on Julian day 238 in 2019 ($n$ = 716, edf = 3.0) and at 0.63 on Julian day 214 in 2020 ($n$ = 945, edf = 3.8; Fig. 2c). The species richness of the five major insect orders (Coleoptera, Diptera, Hemiptera, Hymenoptera, and Lepidoptera), pollinators, and threatened species generally followed similar seasonal patterns as total species richness, although invasive species tended to increase through the year (Supplementary Information 2).

*Drivers of site-level insect biomass, diversity, and composition*

Surveyed sites encompassed several land cover gradients, specifically agricultural (planted crops), bare soil (typically rocky areas or barren agricultural land), low vegetation (e.g., grassland, meadows, gardens, hedges, etc.), forest (trees), and urban land cover types, and gradients in land cover heterogeneity (i.e., the diversity of different land cover types around the traps; Table 1). Additionally, the survey network captured gradients in temperature (mean annual values from 5.0–11.2°C), relative humidity (72–85%), and precipitation (mean annual values from 1.4–4.9 mm day$^{-1}$), with similar gradients in temperature anomalies (ΔT; mean ± SD of 3.1 ± 1.4°C in 2019 and 0.5 ± 0.4°C in 2020), humidity anomalies (ΔH; of -3.8 ± 1.9% and -4.3 ± 2.5%), and precipitation anomalies (ΔP; of -0.34 ± 0.35 mm day$^{-1}$ and -0.43 ± 0.46 mm day$^{-1}$).

**Table 1: Gradient of land cover encompassed by the survey network.** Values show the mean ± the standard deviation of the proportion of each land cover type, and land cover heterogeneity (*H*), across sites within finer and broader spatial scales (radius of 50 m and 1 km, respectively). Additionally, the range of values within each spatial scale are shown, and we indicate the number of sites that are dominated by each land cover type (as a percentage of sites with > 50% cover out of 75 total sites).

| **Cover type** | **50 m** (mean ± SD) | **1 km** (mean ± SD) | **50 m** (range) | **1 km** (range) | **50 m** (%) | **1 km** (%) |
|---|---|---|---|---|---|---|
| Agricultural | 0.11 ± 0.25 | 0.13 ± 0.22 | 0–1 | 0–0.96 | 11 | 7 |
| Bare soil | 0.30 ± 0.09 | 0.02 ± 0.04 | 0–0.56 | 0–0.20 | 1 | 0 |
| Forest | 0.49 ± 0.41 | 0.57 ± 0.30 | 0–1 | 0–1 | 44 | 55 |
| Low vegetation | 0.34 ± 0.37 | 0.20 ± 0.16 | 0–1 | 0–0.63 | 36 | 11 |
| Urban | 0.03 ± 0.01 | 0.05 ± 0.09 | 0–0.66 | 0–0.58 | 3 | 3 |
| *H* | 0.42 ± 0.35 | 0.77 ± 0.39 | 0–1.25 | 0.06–1.4 | – | – |

Given some covariation among our land cover, weather, and climate gradients, such as forests tending to be colder and wetter (see Data S2), we used a combination of variation partitioning, Redundancy Analysis (RDA), and stepwise permutation tests to determine the degree to which insects were related to the individual effects of land cover, the individual effects of weather and climate, and the covarying effects of both. Land cover, weather, and climate explained 42% of the total site-level variability in insect biomass, species richness, and temporal turnover (hereafter collectively referred to as ‘insect biomass and diversity’; Fig. 3), and 29% of compositional differences among key insect ‘groups’, including the five major orders, and pollinator, threatened, and invasive species (Fig. 4). Additionally, these predictors explained 25% of the compositional differences among 86 insect groups comprising 125 families, and 36%

of trait composition (Supplementary Information 3). The individual effect of land cover was the primary driver in all these relationships, always accounting for between 48–72% of the total explained variation across the four variation partitioning analyses we used (48%, 59%, 72%, and 69%, respectively). The individual effect of weather and climate was of lesser importance, explaining about 24–31% of the total (26%, 31%, 24%, and 25%, respectively), with the covarying effect of land cover, weather, and climate comprising the remainder (26%, 10%, 4%, and 6%, respectively).

The only individual land cover, weather, or climate predictors that always explained a significant portion of the variation in insect biomass, diversity, and composition were low vegetation cover within a 50 m radius around the traps, bare soil within 1 km, land cover heterogeneity within 50 m, and relative humidity, which were primarily represented along RDA axis 1 in all ordinations (Figs. 3, 4, and Supplementary Information 3). Forest and urban cover within 1 km were also important in three out of the four ordinations (Table 2).

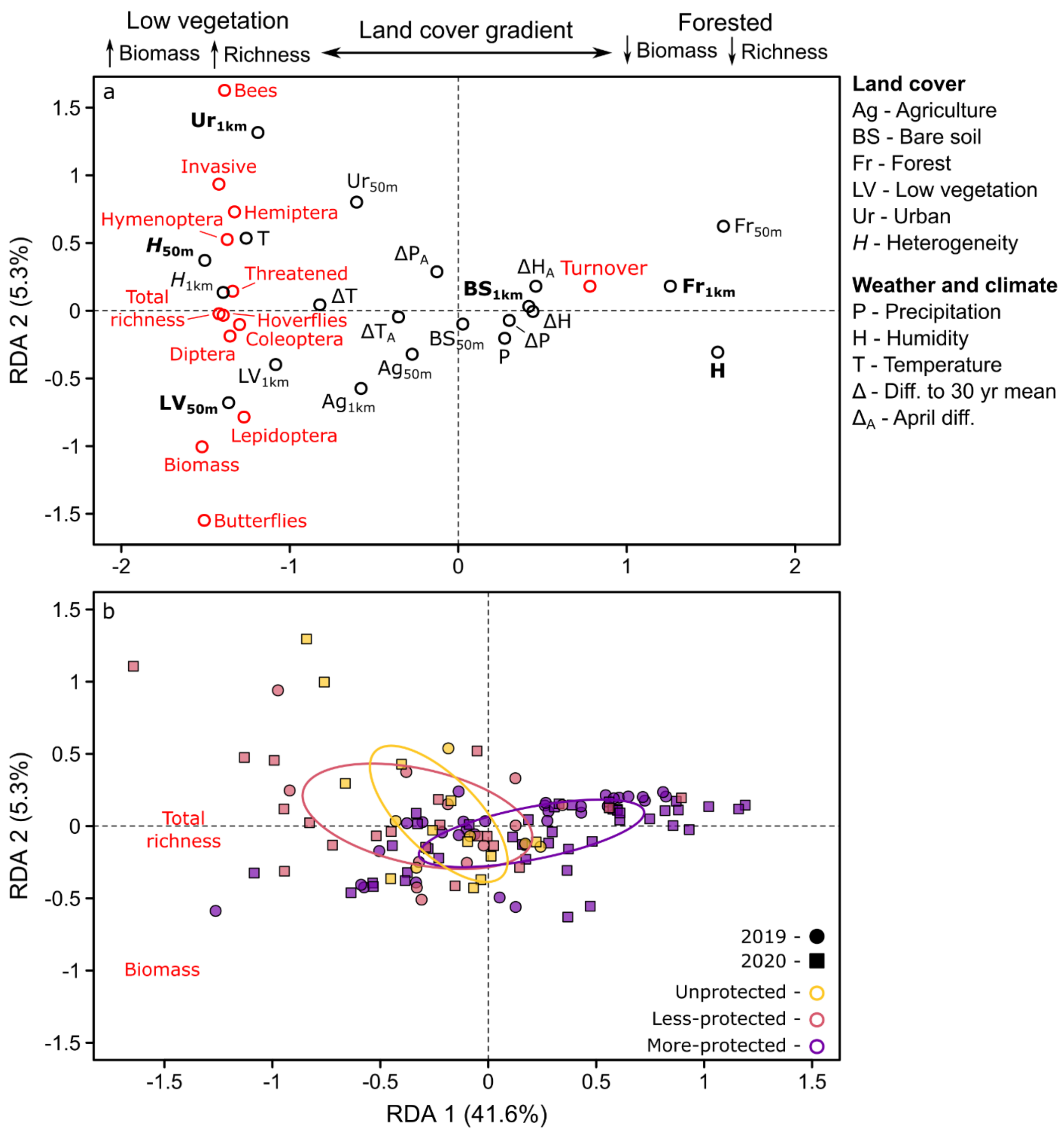


**Fig. 3: Insect biomass and diversity.** Insect biomass, richness, and temporal turnover (red text) in relation to (a) land cover, weather, and climate (black text), and (b) in more-protected (purple), less-protected (pink), and unprotected (yellow) areas, during 2019 (circles) and 2020 (squares) based on Redundancy Analysis (RDA). More proximate red and black text in (a) indicate stronger relationships, with bold black text indicating predictors that consistently

explained the most variation based on a stepwise model selection procedure (Table 1). Pollinator groups (bees, butterflies, and hoverflies) are shown separately, and their corresponding orders (Hymenoptera, Lepidoptera, and Diptera) include these groups. Land cover predictors included agricultural (Ag; i.e., planted crops), bare soil (BS), forest (Fr), low vegetation (LV), and urban (Ur) land, in addition to land cover heterogeneity (*H*), within finer (50 m) and broader (1 km) spatial scales. Weather and climate predictors included precipitation (P), relative humidity (H), and temperature (T), and differences in these values (Δ) and their values from the previous April ($\Delta_A$) compared to the previous 30 years. Differences between (b) more-protected, less-protected, and unprotected areas are shown using colored ellipses that indicate the central tendency for each group (based on standard deviations).

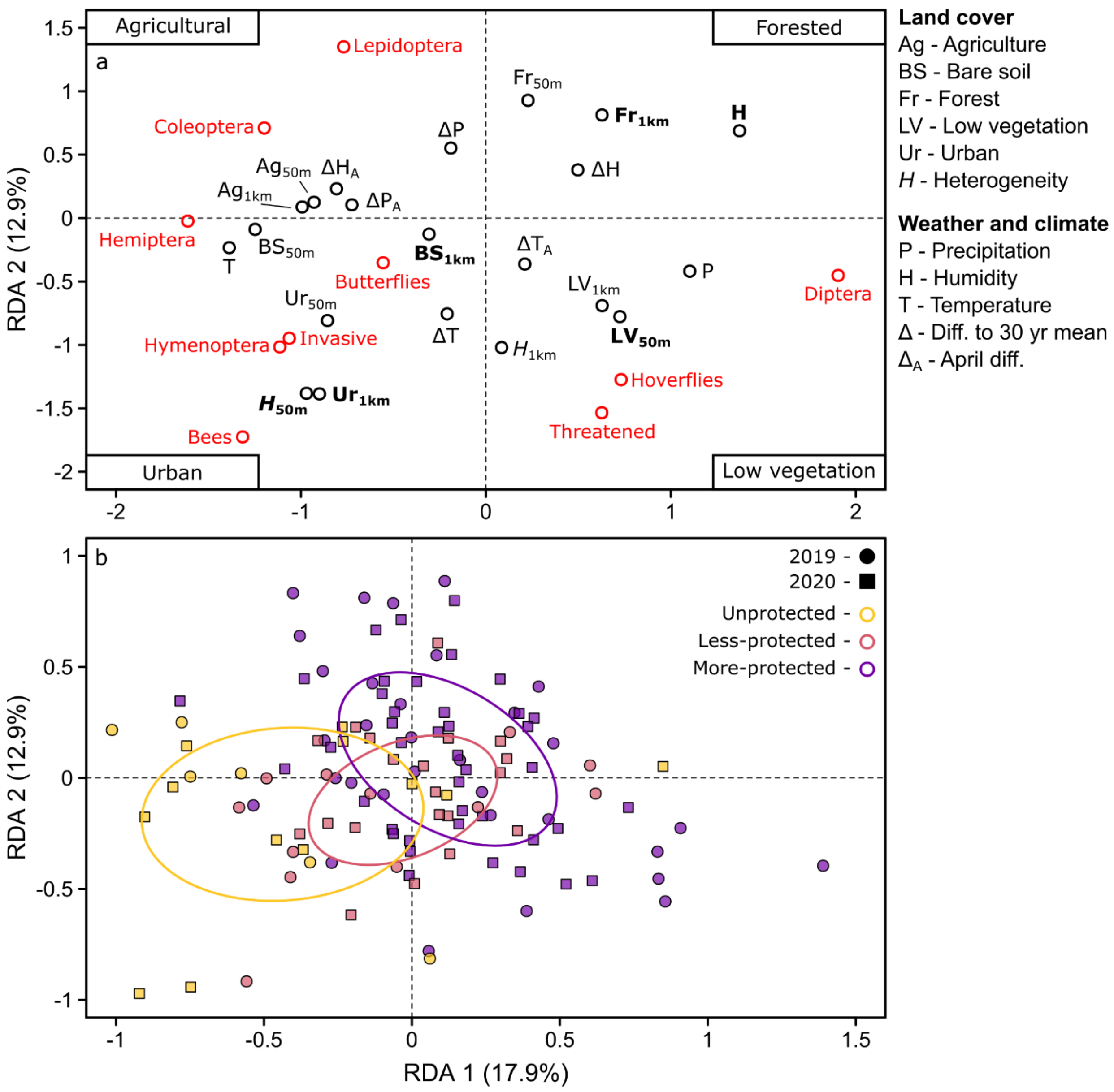


**Fig. 4: Insect group composition.** Composition of different insect groups expressed as proportional richness (red text) in relation to (a) land cover, weather, and climate (black text), and (b) in more-protected (purple), less-protected (pink), and unprotected (yellow) areas, during 2019 (circles) and 2020 (squares) based on Redundancy Analysis (RDA). More proximate red and black text in (a) indicate stronger relationships, with bold black text indicating predictors that consistently explained the most variation based on a stepwise model selection procedure

(Table 1). Pollinator groups (bees, butterflies, and hoverflies) are shown separately, and their corresponding orders (Hymenoptera, Lepidoptera, and Diptera) include these groups. Land cover predictors included agricultural (Ag; i.e., planted crops), bare soil (BS), forest (Fr), low vegetation (LV), and urban (Ur) land, in addition to land cover heterogeneity (*H*), within finer (50 m) and broader (1 km) spatial scales. Weather and climate predictors included precipitation (P), relative humidity (H), and temperature (T), and differences in these values (Δ) and their values from the previous April ($\Delta_A$) compared to the previous 30 years. Differences between (b) more-protected, less-protected, and unprotected areas are shown using colored ellipses that indicate the central tendency for each group (based on standard deviations).

**Table 2: Important land cover, weather, and climate predictors.** Six predictors consistently explained the most variation (based on stepwise permutation tests) across four Redundancy Analyses of: (i) insect biomass, richness, and turnover; (ii) group composition; (iii) family-level community composition; and (iv) feeding trait composition. Values show percentages of the total explained variation (adjusted-$R^2$) accounted for by each predictor, with land cover heterogeneity represented as '*H*'. Note that relative humidity reflects both short-term weather and longer-term climatic gradients (see *Methods*).

| Predictor | Insect biomass, richness, & turnover | Group composition | Family-level composition | Feeding trait composition |
|---|---|---|---|---|
| Bare soil (1 km) | 10% | 4% | 5% | 3% |
| Forest (1 km) | 4% | – | 34% | 46% |
| Humidity | 28% | 23% | 10% | 9% |
| Low vegetation (50 m) | 24% | 12% | 12% | 14% |
| Urban (1 km) | 11% | 15% | – | 3% |
| *H* (50 m) | 5% | 6% | 6% | 2% |

*Patterns in insect biomass, diversity, and composition*

Sites with more low vegetation exhibited higher total insect biomass and richness, and lower temporal turnover. In contrast, when low vegetation was scarce (typically in forested areas), biomass and richness were lower and turnover was higher (Fig. 3a). Humidity also tended to be higher in forested sites versus lower in low vegetation sites. Community composition at the low vegetation sites was characterized by certain Diptera families, such as hoverflies (lower right of Fig. 4a), and a mixture of multiple phytophagous guilds, such as miners (e.g., Agromyzidae) and stem feeders (e.g., Chloropidae), in addition to parasites (e.g., Tachinidae; Supplementary Information 3). Conversely, forested sites were characterized by a different set of primarily mycetophagous (i.e., fungus-feeding) Diptera, such as fungus gnats (e.g., Mycetophilidae), and specific Lepidoptera families (right and upper sides of Fig. 4a), including certain moths (e.g., Adelidae or Geometridae; Supplementary Information 3). Composition was generally similar between urban and agricultural areas because both were located on the left side of Fig. 4. However, there were some differences, such as bees, butterflies, and invasive species accounting for greater proportions of communities in urban areas where low vegetation was common (lower part of Fig. 4a), and more Coleoptera and Lepidoptera in agricultural areas with less low vegetation (upper left of Fig. 4a). Sensitivity analyses indicated these results were robust and not affected by 11 traps placed under forest canopies (Supplementary Information 4), where biomass and richness of flying insects might be underrepresented, nor by the addition of 19 new sites in 2020 (Fig. 1; Supplementary Information 5).

*Insect biomass and richness effect sizes*

Of the important predictors (Table 2), variability in biomass was best explained by forest cover within a 1 km radius around the traps ($R^2_{adj}$ = 0.23 out of a maximum of 0.41), with all other predictors providing minor contributions (see Supplementary Information 6 for model details). Biomass tended to decrease by 75%, from 3.6 to 0.9 g $day^{-1}$, during 2019 as forest cover increased from 0% to 98% across sites ($n$ = 56, edf = 2.6, $P$ = 0.014), with a similar decrease of 58%, from 3.8 to 1.6 g $day^{-1}$, during 2020 ($n$ = 75, edf = 1.0, $P < 0.001$; Fig. 5a).

Variability in total species richness was best explained by low vegetation cover within a 50 m radius around the traps ($R^2_{adj}$ = 0.34 out of a maximum of 0.44), with all other predictors providing minor contributions (Supplementary Information 6). Richness tended to increase by 64%, from 197 to 323 species, during 2019 as low vegetation cover increased from 0% to 99%

across sites ($n$ = 56, edf = 1.7, $P$ = 0.0018), with a 32% increase, from 301 to 396 species, occurring during 2020 ($n$ = 73, edf = 3.7, $P$ < 0.001; Fig. 5b). These relationships were not linear, with richness tending to plateau and even slightly decline once the proportion of low vegetation was above 50%.

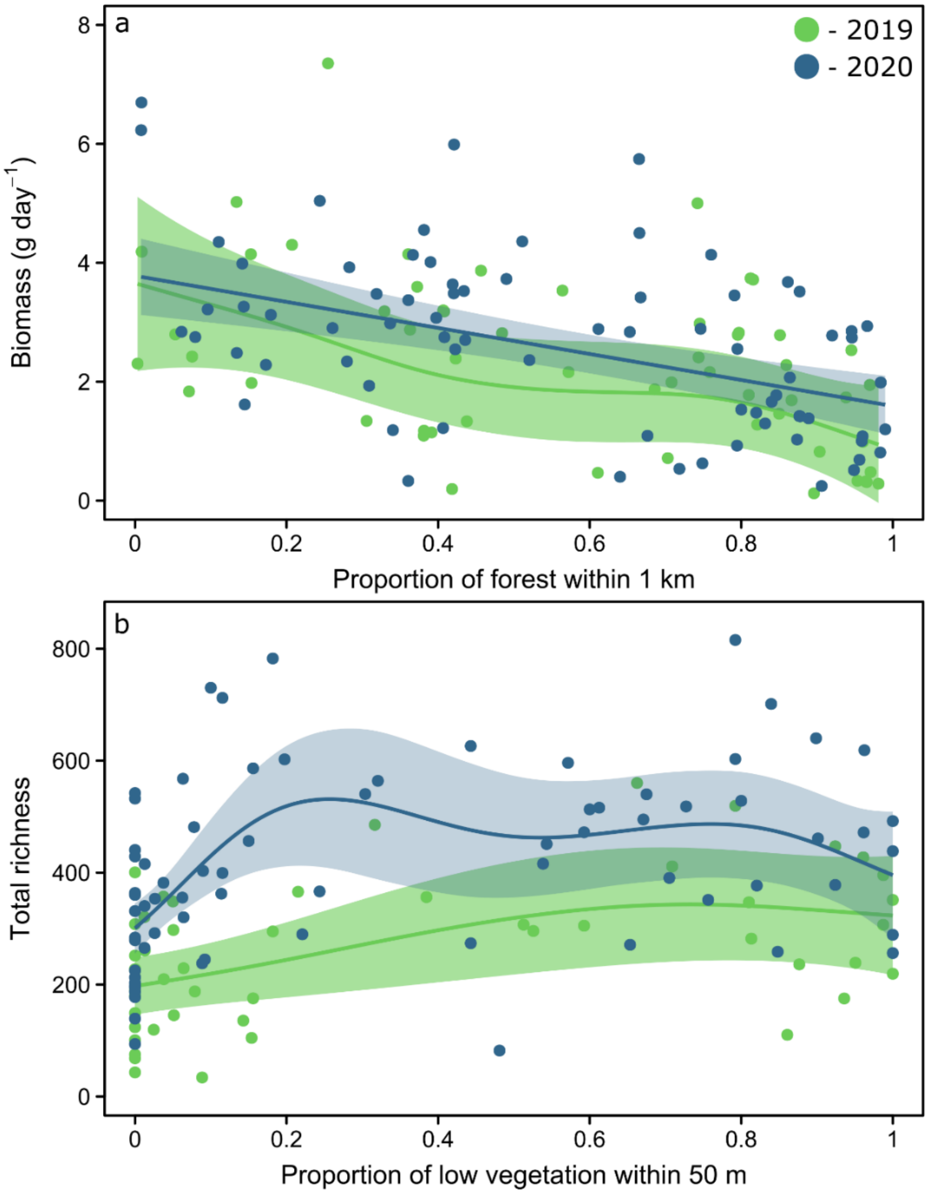


**Fig. 5: Insect biomass in relation to forest cover, and insect richness in relation to low vegetation.** Relationships between insect (a) biomass (g $day^{-1}$) and the proportion of forest cover within a 1 km radius, and (b) total species richness and the proportion of low vegetation cover

within a 50 m radius, during 2019 (green) and 2020 (blue). Best-fit lines and 95% confidence intervals were determined based on the respective generalized additive mixed models.

*Protected areas*

We found differences among the three protected area categories in insect biomass and diversity (PERMANOVA, $n$ = 123, $F_{2,120}$ = 11.1, $R^2_{adj}$ = 0.16, $P$ < 0.001; Fig. 3b), group composition (PERMANOVA, $n$ = 123, $F_{2,120}$ = 10.7, $R^2_{adj}$ = 0.15, $P$ < 0.001; Fig. 4b), family-level community composition (PERMANOVA, $n$ = 123, $F_{2,120}$ = 13.1, $R^2_{adj}$ = 0.18, $P$ < 0.001; Fig. 5a), and trait composition (PERMANOVA, $n$ = 123, $F_{2,120}$ = 7.1, $R^2_{adj}$ = 0.11, $P$ < 0.001; Fig. 6b). More-protected sites were associated with humid forests, whereas unprotected sites were characterized by more low vegetation, urban, and agricultural cover and lower humidity. Less-protected sites tended to be located between the more-protected and unprotected sites in the ordinations, except for insect biomass and diversity where less-protected and unprotected sites overlapped (Fig. 3b). Given these associations, more-protected sites were characterized by the same compositional patterns reported above for forested habitats, such as lower biomass and richness, higher turnover, and dominated by fungus-feeding Diptera. Conversely, less-protected and particularly unprotected sites were characterized by higher biomass, higher richness, lower turnover, and a mixture of insect groups and their associated feeding traits, including pollinators and threatened species.

### Discussion

Our results on the spatial distribution of 31,846 flying insect species (10,803 validated plus 21,403 plausible species) have basic implications for understanding the drivers of insect biomass, diversity, and community composition, and applied implications for improving insect protection. Overall insect biomass across Germany was 2.0 g day$^{-1}$ in 2019 and 2.3 g day$^{-1}$ in 2020, which are in the range of biomass values reported from the last decade (2007–2016) considered in a landmark study[4] that found insect biomass declined in two subregions of Germany by over 75% from 1989 to 2016. Furthermore, two recent studies from other regions of Germany reported that insect biomass had not declined further by 2020 and 2021[35], or found biomass may even have increased during 2019, 2020, and particularly 2022[12]. We acknowledge the difficulties in verifying temporal trends with only a few years of data, and difficulties in comparing studies with different sampling methods (e.g., trap designs[36]), site locations, and site protection status. However, the results from these studies, combined with our own, suggest that the decline in biomass emerging in ref. 4 has not continued in recent years. Declines may have plateaued because insect biomass has reached its lowest possible levels, or due to changes in other drivers, such as the abatement of poor weather conditions particularly during the growing period[12]. Our conclusion of a potential plateau still requires further scrutiny informed by consistent insect monitoring following the same methods and the same sites over the coming years.

At the site-level, the positive relationship we found between insect biomass/richness and low vegetation (e.g., grassland, meadows, gardens, etc.) indicated that open areas may be 'biodiversity hotspots' for a wide variety of flying insects. This inferred bolstering effect of low vegetation is supported by other research on individual insect groups, such as bees, whereby low vegetation benefits insect biomass and richness by diversifying nesting habitat and food resources[37,38]. Furthermore, given that the relationship we found applied broadly across the insect community, our results suggest that these beneficial effects can apply to diverse groups of pollinating and non-pollinating flying insects, including beetles, flies, true bugs, wasps, butterflies, and moths. Enhancing the variety of forbs in low vegetation areas could therefore improve overall insect biomass and diversity in depauperate sites.

Lower insect biomass and diversity in forests compared to urban and agricultural areas was contrary to our expectations. One possible explanation for this pattern is that Malaise traps targeting flying insects can underrepresent other groups, such as ground-dwelling, soil-dwelling, and forest canopy[39] insects. Missed canopy species may partly explain the lower diversity we found in forested areas, although we partly controlled for this issue by placing most forest traps in gaps or along forest edges, following procedures used in other studies[15,20]. An alternative (or additional) explanation is that the lower land cover heterogeneity in forests may be reducing insect diversity, with the opposite pattern in urban or agricultural sites. This interpretation is consistent with numerous studies on various insect and non-insect taxa showing that higher landscape heterogeneity can foster biodiversity, including in highly fragmented urban and agricultural landscapes, owing to higher habitat and resource diversity[40]. Our feeding-trait results lend further support to this inferred linkage between land cover heterogeneity and habitat/resource diversity. Specifically, the dominance of forest communities by fewer feeding guilds than found in other habitat types, with a prominence of fungus-feeding flies, indicated a more homogeneous vegetation structure and resource supply, which may be further exacerbated by German forest management practices that favored conifers for timber production[41]. Conversely, the more even mixture of different plant feeding guilds in urban and agricultural areas indicated a more heterogeneous vegetation structure, likely facilitated by mixtures of natural vegetation (e.g., meadows, remnant forest patches) and anthropogenic vegetation (e.g., gardens, crops).

The general pattern of homogeneous land cover in forests versus heterogeneous cover in urban and agricultural areas undoubtedly varies among studies and regions. For example, our surveyed sites captured a wide forest cover gradient, ranging from 0–100% cover at both finer and broader spatial scales, with many sites on both ends of the gradient. Consequently, sites with the highest forest cover were entirely dominated by trees, resulting in more homogenous land cover. In contrast, the urban and agricultural gradients were narrower, with only a few sites dominated (> 50%) by these land cover types, meaning that most sites with higher urban or agricultural values exhibited a mixture of different land covers. Wider forest cover gradients versus narrower urban/agricultural gradients are not atypical for insect monitoring studies (e.g., ref. 20) that rarely sample heavily urbanized or agricultural sites[42]. More extensive sampling of such sites might have altered or even reversed our results given that other studies report higher insect diversity in forests compared to heavily urbanized[43,44] or agricultural areas[15,20]. Thus, seemingly opposite conclusions may be reached about the influence of certain land cover types on insects, depending on the studied extent of land cover gradients and associated habitat heterogeneity.

Irrespective of potential caveats of the effectiveness of insect trapping in forests and the extent of land cover gradients, forested habitats clearly played an important role for regional biodiversity by supporting unique insect communities. Forests can be an underappreciated habitat for some insects (e.g., pollinators[45]) compared to low vegetation areas, but our forested sites harbored a variety of specific forest-dwelling insects, such as fungus-feeding flies, beetles, and moths. Furthermore, insects in nearby non-forested areas can also depend on forests during parts of the year or during particular life stages[46]. Consequently, our biomass and diversity results must not be misconstrued to mean that forests play a marginal role for insects compared to other land cover types. On the contrary, we found that forests contributed critically to regional diversity by supporting insects that do not occur elsewhere.

While low to moderate urbanization did not reduce insect biomass or diversity, these sites exhibited more invasive species, suggesting potential negative impacts of urban centers on insect communities. Urban areas promote biological invasions by acting as hubs for the introduction and spread of non-natives[47], and by offering unexploited niches for new species[48]. The horticultural trade is also a common vector for insect introductions[49], which could partly explain the association between urban areas and invasive insects given the greater presence of ornamental vegetation in lawns, gardens, and parks. These results highlight that, although low to moderate urbanization may increase habitat heterogeneity and thus benefit insect diversity, this effect can come with the detriment of promoting invasive insects. Such detriments could be mitigated by efforts targeted at identifying invasive species and controlling their spread[50].

Somewhat different seasonal trends in biomass and richness between years potentially reflected differences in temperature. The earlier peak for biomass and richness in 2019, and the lower richness peak, may reflect higher temperatures in this year (ΔT of +3.1°C in 2019 versus +0.5°C in 2020), particularly given that warming can shift phenology to an earlier date[51] and can reduce richness when species' temperature tolerances are exceeded[7]. Conversely, we found no evidence for strong effects of temperature or precipitation on site-level community differences, with only humidity exhibiting consistent relationships. Consistent site-level relationships to humidity likely reflect patterns in insect activity. Specifically, because Malaise traps record activity-abundance, wetter conditions that reduce insect flight activity also reduce the number of captured specimens[52,53], thus explaining why insect biomass and diversity were lower at higher humidity. The weaker effects we observed for temperature and precipitation may have occurred because, despite the wide temperature and elevational gradients in the present study, spatial differences in these environmental factors may be less important in temperate regions[20]. Responses may be stronger where organisms are closer to their environmental limits, for example in Mediterranean, tropical[54], or higher-elevation climate zones[55]. Our focus on within-year spatial patterns may also upweight the influence of land cover variability, which exhibits high spatial variation, while downweighting the influence of weather or climate, which can exert stronger changes across years or seasons. For example, temperature effects on insects may be more detectable in long-term temporal analyses across decades[12,56], or in seasonal analyses where the effects of short-term extreme events can be more evident[31]. The effects of temperature and precipitation may therefore become more evident as more temporal data is collected across regions.

Contrary to expectations, key functional insect groups (i.e., pollinators) and threatened insects tended to occur outside more-protected areas. Protected areas were primarily characterized by higher forest cover and thus lower insect biomass and diversity. This association with forests likely occurred because protected areas tend to be located in regions with little value for settlement and agriculture, such as forested, cooler, mountainous areas[52,53,54]. Additionally, protected areas are typically designated to prioritize charismatic vertebrates or plants instead of insects[21], and many charismatic species may primarily persist in more remote, forested areas. These biases mean that protected areas tend to favor forest-dwelling species[60] at the expense of other taxa, including many insects[61]. Efforts to expand protected areas to meet the Kunming-Montreal Global Biodiversity Framework target of 30% protected land by 2030[62] should therefore specifically consider insects and non-forested, warmer, low elevation areas. Protecting insects in such regions, which are often impacted by urbanization and agriculture, may require establishing and expanding protected areas that allow for some human activity. This could include IUCN category V/VI protected areas or the outer transition zones of biosphere reserves[63]. However, the benefits of any protection afforded by these types of multi-use areas are debated owing to inevitable tradeoffs across diverging ecological, socioeconomic, and cultural needs that challenge the prioritization of biodiversity[64]. An additional option could be to prioritize alternative approaches to enhance insect diversity in unprotected areas, such as via urban green spaces[65] and agro-ecological land management[66,67]. Regardless of how best to achieve protection, our results clearly show a high diversity of insects and important taxa outside more-protected areas, highlighting the need to consider insects when designating protected areas and to navigate tradeoffs between insect conservation and human needs.

The key insights from our study are how the relationships we have shown play out across the insect community at large. This includes that low vegetation and land cover heterogeneity can benefit the broader insect community regardless of whether total biomass, overall community diversity, or the diversity of multiple individual insect groups are considered. Similarly, we show the wide variety of insects left unprotected because of existing biases in the designation of protected areas. These results highlight that conservation efforts aimed at expanding protected areas and improving insect habitat in urban and agricultural sites will not only benefit particular species, but likely insect communities as a whole. Our work also informs why different studies can reveal different spatiotemporal patterns and drivers, such as missing heavily urbanized or agricultural areas potentially driving variability among studies in whether these stressors have positive or negative effects on insects. Likewise, the lack of weather and climate effects in our analyses suggest that spatially focused studies, such as ours, may upweight the influence of land cover and downweight the importance of meteorological variables. These differences may explain why spatial studies can find stronger effects of land cover[20], whereas temporal studies can find stronger effects of weather or climate[12]. In conclusion, our work highlights the value of a broader, genetics-based perspective of insect community change for identifying key drivers, informing future protection efforts, and reconciling different findings across studies. Such a broad perspective is essential to improving our understanding of, and ability to ameliorate, ongoing insect biodiversity loss.

### Methods

*Insect sampling and composition*

Townes-type Malaise traps with black roofs and an opening measuring 1.16 $m^2$ on each side were set up at 56 sites across Germany during 2019, with 19 sites added in 2020 (Fig. 1). Traps were generally exposed for 14 days, emptied, and then reset, although shorter and longer exposure periods (ranging from 7–29 days) were occasionally necessary owing to logistical constraints. Traps were primarily placed in open areas (typically agricultural fields or grasslands), adjacent to forest edges or hedgerows, or within forest clearings, but 11 traps were placed under forest canopies. Trap openings usually faced east-west and sample bottles were aligned according to the conditions on site in different cardinal directions, often with the sample bottle towards the south, southwest or southeast.

All captured insects were preserved in 80% denatured ethanol (1% methyl ethyl ketone) and transported to the lab to determine wet biomass following methods in ref. 31, and species identity via metabarcoding. Insect DNA sequences of mitochondrial cytochrome oxidase I were assigned to Operational Taxonomic Units (OTUs) based on a 97% similarity threshold (see ref. 29 for detailed metabarcoding methods). OTUs could not always be assigned to species names because of incomplete reference data or conflicting matches in the databases. Therefore, OTUs were divided into two groups. The first group included OTUs that could be unambiguously matched to a barcode with a species name. We checked the validity of these identifications based on three criteria: (i) expert assessments from taxonomists in Germany on whether a given species is present when and where it was caught; (ii) records of species detections within 200 km of the site from the Global Biodiversity Information Facility (https://www.gbif.org/); and (iii) the German Barcode of Life database. Identifications were considered valid if they met two of the three criteria and are termed ‘validated species’. The second group included OTUs that could only be resolved to genus or family level for the reasons stated above, but these taxa can still be used to estimate the likely species richness of each insect family (detailed in Supplementary Information 1). We refer to these OTUs as ‘plausible species’.

We quantified total insect biomass (g), species richness, and temporal turnover during each exposure period in 2019 and 2020. Temporal turnover was calculated as shifts in species presence/absence based on Jaccard’s index, which ranges from 0 (no turnover) to 1. We quantified insect composition as the richness of validated species within the five most important insect orders in our dataset – all Coleoptera, Diptera, Hemiptera, Hymenoptera, and Lepidoptera – and the richness of 86 insect groups comprising 125 families and > 90% of all species (see Supplementary Information 3 for the families within each group). We represented functional composition by separately quantifying the richness of bees, butterflies, and hoverflies to represent pollinators (families listed in Supplementary Information 3), and by examining patterns in larval feeding-trait composition based on traits provided by ref. 68, which were available at the family level for 89% of our identified families. Lastly, we calculated the richness of threatened and invasive species. Threatened species were assigned using a country-level Red List of species considered endangered (categories 1, 2, 3, R, and G) in Germany[69]. Invasive species were determined based on species considered non-native and invasive or potentially invasive in Germany (sources in Data S3). ‘Potentially invasive’ species are those that are invasive elsewhere in Europe, but only small populations or no populations have yet been detected in Germany.

*Land cover*

To represent differences in land cover and land cover heterogeneity among sites, we extracted data from a 10 m resolution raster map of agricultural, bare soil, forest, low vegetation, and urban land cover types in Germany derived from Sentinel-2 satellite imagery[70]. Note that satellite imagery often classifies tree-lined impervious surfaces as ‘forest’, thus urban cover values tend to be lower than their true values. We extracted the proportions of each land cover type within finer-scale (radius of 50 m) and broader-scale (radius of 1 km) areas around each site. We selected these radii to capture both fine-scale and broad-scale impacts of land cover, given that mobile taxa are expected to integrate land use-changes across larger areas, whereas less mobile insects should be more strongly influenced by land use in the immediate site surroundings. We considered including land cover within a radius of 500 m and 5 km, but these cover values were generally correlated to the 1 km values (mean $r$ of 0.91 and 0.68, respectively). In addition to the five land cover types, we included a measure of land cover heterogeneity, which can also influence insect diversity[71]. We calculated land cover heterogeneity for each site within a radius of 50 m and 1 km using the Shannon diversity index ($H$), which ranged from 0 (100% cover of one land cover type) to a potential maximum of 1.6 (equal distribution of all five land cover types).

*Weather and climate*

We obtained gridded data on daily mean temperature (°C, 5x5 km resolution), daily total precipitation (mm, 1x1 km resolution), and daily mean relative humidity (%, 5x5 km resolution) from the German Weather Service (https://opendata.dwd.de), which is interpolated from stations across Central Europe. We used this data to calculate weather or climate values at three time scales: (1) weather during each exposure period, with daily temperature and humidity averaged across days and precipitation summed across days then divided by the number of days (mm $day^{-1}$); (2) weather averaged across the 12 months prior to each exposure period to reflect conditions during hatching for different species; and (3) longer-term climate normals as mean annual values averaged across the previous 30 years. These different scales of shorter- and longer-term temperature, precipitation, and humidity variables were generally correlated to the other variables within their respective groups (*r* ranging from 0.62–0.92). For example, sites that were warmer and wetter in 2019 or 2020 also tended to be warmer and wetter during the previous year and during the previous 30 years. Consequently, we used only the exposure period values from 2019 and 2020, which thus represent both weather and climatic conditions given the positive correlations across the three time periods. Additionally, we calculated anomalies from climate normals for temperature (ΔT), precipitation (ΔP), and relative humidity (ΔH) based on the differences between these values during each exposure period and the same periods averaged across the previous 30 years. We also calculated anomalies only for April in the year before sampling compared to April normals from the previous 30 years, given that anomalies during April of the previous year can affect insect biomass in the following year[12].

*Protected areas*

We quantified differences in protection status across sites based on data from Protected Planet (https://www.protectedplanet.net) and ref. 72. We included four types of protected areas in Germany that potentially affect terrestrial insects: national parks, nature reserves, biosphere reserves, and landscape protection areas. We grouped sites within national parks, nature reserves, and the inner two zones of biosphere reserves into a single 'more-protected' category. These areas all restrict land conversion for anthropogenic purposes and thus experience reduced anthropogenic pressures[33,72]. We grouped sites within landscape protection areas and the outer transition zone of biosphere reserves into a 'less-protected' category because these protection areas allow for more extensive human activity, including the use of approved pesticides and insecticides, and even overlap with urban areas in some cases. All other sites were considered 'unprotected'.

*Statistical analyses*

To quantify and control for seasonal changes, we first examined seasonal patterns in biomass, species richness, and temporal turnover. We did so by relating these variables to the final Julian day in each exposure period for each site using generalized additive mixed models performed with the *mgcv* package in R[73]. All models included a smoothed predictor for Julian day which varied by year (using thin-plate regression splines and a basis dimension of $k = 5$ to restrict the results to unimodal relationships), a fixed categorical term for year, and a random term for each site to control for repeated measures. Additionally, biomass was converted to a daily rate (g $day^{-1}$) to account for differences in trap exposure length. We controlled the effects of exposure length on richness using an offset, which allowed us to model the raw count data using non-normal distributions. Biomass and turnover were modeled using Gaussian distributions with log-link functions, whereas richness for all taxa was modeled using Poisson distributions with log-link functions.

To analyze site-level community, land cover, weather, and climate relationships, we used a reduced dataset of biomass, richness, and temporal turnover at each site averaged across samples collected from late April (after Julian day 110) to September (Julian day 273). This period encompassed an average of 11.6 samplings per trap and included 98% of all identified species. We then conducted four Redundancy Analyses (RDAs) using the *vegan* package in R[74]. We chose this method because RDA is useful for relating numerous response variables to numerous predictors, and because the method allows for correlations among predictors. The first RDA examined patterns of average insect biomass (g $day^{-1}$), species richness (species $day^{-1}$), and temporal turnover. The second RDA examined compositional patterns in the proportion of total species richness that comprised different insect groups, which included the five most important insect orders, and pollinator, threatened, and invasive species. The third RDA examined changes in the proportional richness of the 86 insect groups representing 125 families. The final RDA examined changes in feeding-trait composition, which was quantified for each community and trait as the average of the trait values of each identified family weighted by their proportional richness. We also weighted each trait to ensure they all contributed equally to the ordinations. All response variables in these RDAs were converted to *z*-scores by subtracting each value from its respective mean and dividing by the standard deviation.

The predictors for the above RDAs included the 12 land cover variables (five cover categories and cover heterogeneity at two spatial scales) and nine weather and climate variables (average site temperature, precipitation, humidity and the six corresponding anomaly values).

We also controlled for differences between sampling years by including year as a conditioning variable, and we controlled for positive spatial autocorrelation in the compositional RDAs using Moran's Eigenvector Maps based on Gabriel graphs[75]. We checked for the need to control elevation, latitude, and longitude, but these variables generally did not improve the models because these differences were already accounted for by the meteorological variables (see correlations in Data S2).

To determine the relative contributions of land cover versus weather and climate in explaining insect diversity and composition in the RDAs, we used variation partitioning conducted via the *varpart* function in the *vegan* package. We first used RDAs to extract the residuals of all response data in relation to year and the spatial autocorrelation variables to control for these effects. We then removed highly correlated predictors based on variance inflation factors > 10 prior to performing the analyses. In addition to variation partitioning, we identified the most important individual predictors using stepwise permutation tests in both forward and backward directions via the *ordiR2step* function in the *vegan* package. Important predictors were those that contributed significantly ($P < 0.05$) to improving the adjusted-$R^2$ and that contributed > 1% of explained variation.

Given the ecological importance of total insect biomass and richness, we conducted additional analyses using generalized additive mixed models to quantify the effect sizes of relationships with the important predictors identified in the RDAs. These analyses also allowed us to test for non-linear relationships, which cannot be modeled by RDA. We related total biomass (g $day^{-1}$) and total species richness (with an offset for exposure length) to smoothed terms of the land cover and climate predictors determined to be important in the stepwise permutation tests. We then determined the relative importance of these predictors using forward selection and adjusted-$R^2$. Smoothed terms used thin-plate regression splines and a basis dimension of $k = 10$, which we confirmed based on comparisons to the effective degrees of freedom (edf) and whether relationships changed if the edf was increased. We allowed the effects of the predictors to vary by year, and included a fixed categorical term for year and a random term for each site. Biomass was modeled using a Gaussian distribution with an identity-link function and richness was modeled using a negative binomial distribution with a log-link function.

To compare protection categories, we used non-parametric permutational multivariate analyses of variance (PERMANOVAs; performed with the *adonis2* function from the *vegan* package. This method determined whether the different protection categories were in significantly ($P < 0.05$) different parts of each RDA.

**Acknowledgments**

We wish to thank Michael Sachtleben and Monika Degebrodt (IGB), Thomas Hahn and Petra Möhl (Ranger Nature Park Stechlin-Ruppiner Land), Dr. Mario Schrumpf (Nature Park Stechlin-Ruppiner Land), Günter Hoenselaar (Kellerwald-Edersee National Park), Arne Beermann, Mathias Hippke, Christoph Huber, Katrin Preuss, Stephanie Puffpaff, Torsten Raab, Robin Reiter, Gregor Scheiffarth, Rhena Schumann, and Michael Tautenhahn for their assistance. Funding for authors, data collection, and processing was provided by the EU Horizon project eLTER PLUS (grant #871128). PH and SUP also acknowledge funding from the LOEWE Centre for Translational Biodiversity Genomics.

### Author contributions

JSS and PH conceived the study. JSS performed the analyses and wrote most of the manuscript with contributions from all authors. DB processed the Malaise trap samples for metabarcoding and provided the associated data, with contributions from LY and FL. TH and MS checked the taxalist for plausibility and commented on the methods. MOG, JM, SUP, SSt, EARW, CB, JB, FD, JE, TH, RK, AM, CM, RR, TS, SSe, ST, DW, WW, BW, MW, GZ, and MF set up Malaise traps in their respective regions, maintained the traps, and collected samples.

### Competing interests

The authors declare no competing interests.

### Data availability

Demultiplexed raw read data for this study has been uploaded to the European Nucleotide Archive under the accession number PRJEB71324. The data needed to repeat our community analyses will be provided on a publicly available database upon acceptance for publication. This statement will be amended to provide the link to the dataset at that time.

### Code availability

The code required to reproduce our analyses are available upon request.
