## Supplementary material for "Upscaling biodiversity monitoring: Metabarcoding estimates 31,846 insect species from Malaise traps across Germany": Script S1.py

```
import time, sys, glob
import pandas as pd
from pathlib import Path
from selenium import webdriver
from selenium.webdriver.common.keys import Keys
from selenium.webdriver.support.ui import WebDriverWait
from selenium.webdriver.support import expected_conditions as EC
from selenium.webdriver.common.by import By

## collects all phyla from a  given taxa table in parquet format
def collect_phyla(taxa_table):
    ## collect the phylum data to search for
    data = pd.read_parquet(taxa_table) ## replace with sys argv
    data = data['Phylum'].unique()

    return data

def start_browser():
    ## start chromedriver and check the statistics germany page
    options = webdriver.ChromeOptions()
    options.add_experimental_option('excludeSwitches', ['enable-logging'])
    prefs = {'download.default_directory' : '{}'.format(Path.cwd().joinpath('temp'))}
    options.add_experimental_option('prefs', prefs)
    browser = webdriver.Chrome(options = options)
    browser.maximize_window()

    return browser

def download_existing_data(browser, phylum):

    ## get german statistics first
    browser.get('https://bolgermany.de/gbol1/ergebnisse/results#StatisticsDE')
    browser.refresh()
    time.sleep(2)
    WebDriverWait(browser, 120).until(EC.presence_of_element_located((By.XPATH, '//*[@id="choiceTaxaDE"]')))
    search_input = browser.find_element_by_xpath('//*[@id="choiceTaxaDE"]')
    search_input.send_keys(phylum)
    time.sleep(5)
    search_input.send_keys(Keys.TAB)
    time.sleep(1)

    ## click the "anzeigen" button, check if the search term is available
    show_button = browser.find_element_by_xpath('//*[@id="opDE"]')

    if show_button.is_enabled():
        show_button.click()

        ## wait for the csv download to appear, download via clicking on the csv button
        WebDriverWait(browser, 120).until(EC.element_to_be_clickable((By.XPATH, '//*[@id="datatableDE_filter"]/div/a[1]'))).click()
        time.sleep(2)
    else:
        print('{} not in database.'.format(phylum))

def download_missing_data(browser, phylum):

    ## get german statistics first
    browser.get('https://bolgermany.de/gbol1/ergebnisse/results#Missing')
    browser.refresh()
    time.sleep(2)
    WebDriverWait(browser, 120).until(EC.presence_of_element_located((By.XPATH, '//*[@id="choiceTaxaMI"]')))
    search_input = browser.find_element_by_xpath('//*[@id="choiceTaxaMI"]')
    search_input.send_keys(phylum)
    time.sleep(5)
    search_input.send_keys(Keys.TAB)
    time.sleep(1)

    ## click the "anzeigen" button, check if the search term is available
    show_button = browser.find_element_by_xpath('//*[@id="opMI"]')

    if show_button.is_enabled():
        show_button.click()

        ## wait for the csv download to appear, download via clicking on the csv button, wait for the download to finish
        WebDriverWait(browser, 120).until(EC.element_to_be_clickable((By.XPATH, '//*[@id="datatableMI_filter"]/div/a[1]'))).click()
        time.sleep(2)
    else:
        print('{} not in database.'.format(phylum))

def generate_database():
    ## collect all input files
    files = glob.glob('{}'.format(Path.cwd().joinpath('temp', '*.csv')))

    ## collect the dataframes from the input files here
    out = []

    for file in files:
        ## extract status and phylum from the file name
        status, phylum = Path(file).stem.split(' ')[-2:]

        ## collect the data
        data = pd.read_csv(file)

        if status == 'fehlenden':
            specs = data['Art']
            status = ['missing' for i in specs]
            phylum = [phylum for i in specs]
            output = pd.DataFrame(zip(status, phylum , specs), columns = ['Status', 'Phylum', 'Species'])
            out.append(output)
        elif status == 'gesammelten':
            specs = data['Art']
            status = ['collected' for i in specs]
            phylum = [phylum for i in specs]
            output = pd.DataFrame(zip(status, phylum , specs), columns = ['Status', 'Phylum', 'Species'])
            out.append(output)

    out = pd.concat(out, axis = 0)
    out.to_excel('GBOL Liste.xlsx', index = False)

def main():
    ## save the phyla
    phyla = collect_phyla(sys.argv[1])

    ## start chromium browser
    browser = start_browser()

    ## run the downloads
    for phylum in phyla:
        download_existing_data(browser, phylum)
        download_missing_data(browser, phylum)

    generate_database()

if __name__ == '__main__':
    main()
```
