## Supplementary figures and images for "Upscaling biodiversity monitoring: Metabarcoding estimates 31,846 insect species from Malaise traps across Germany"

### Figure S1.JPG

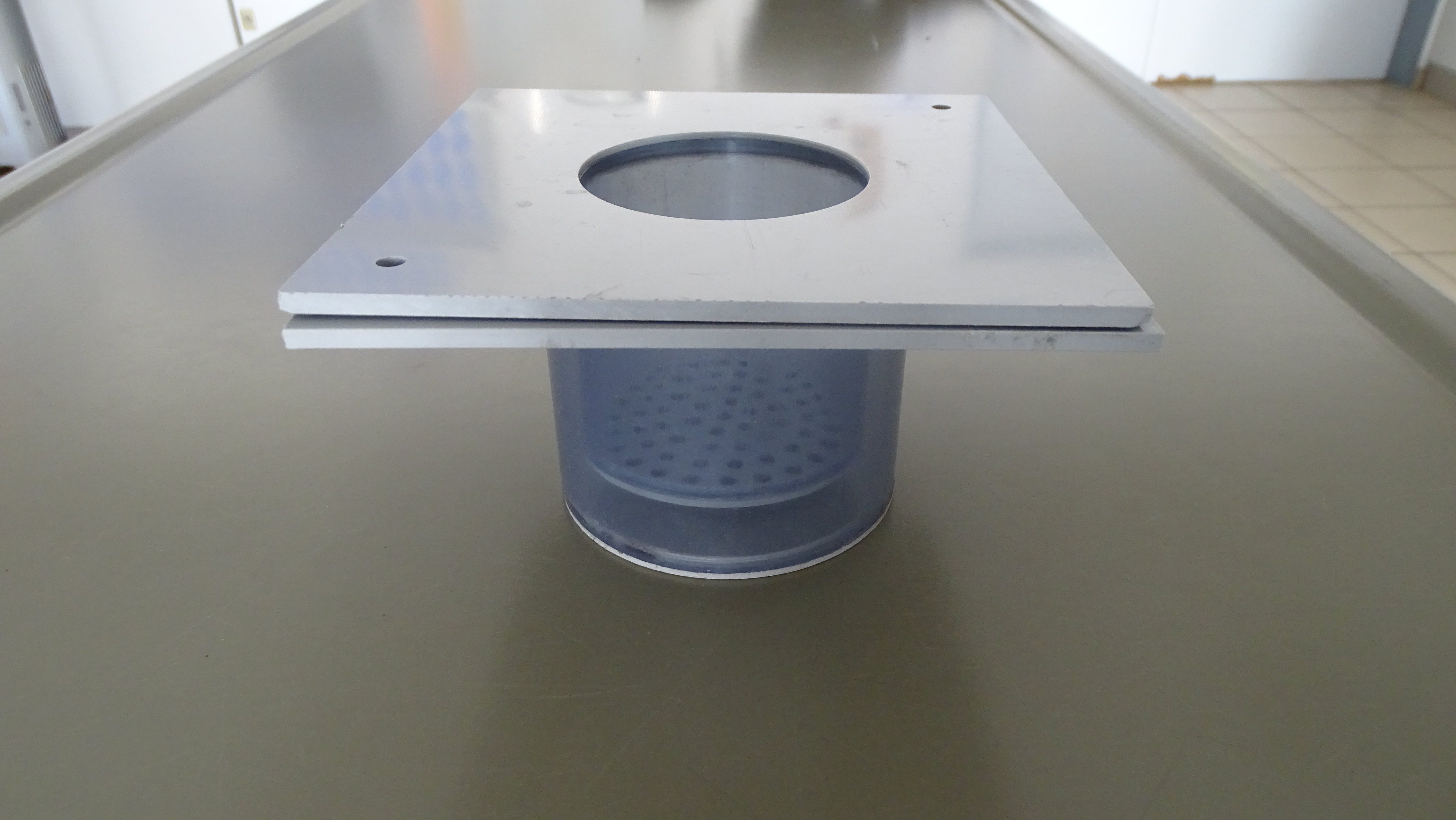

### Figure S2.png

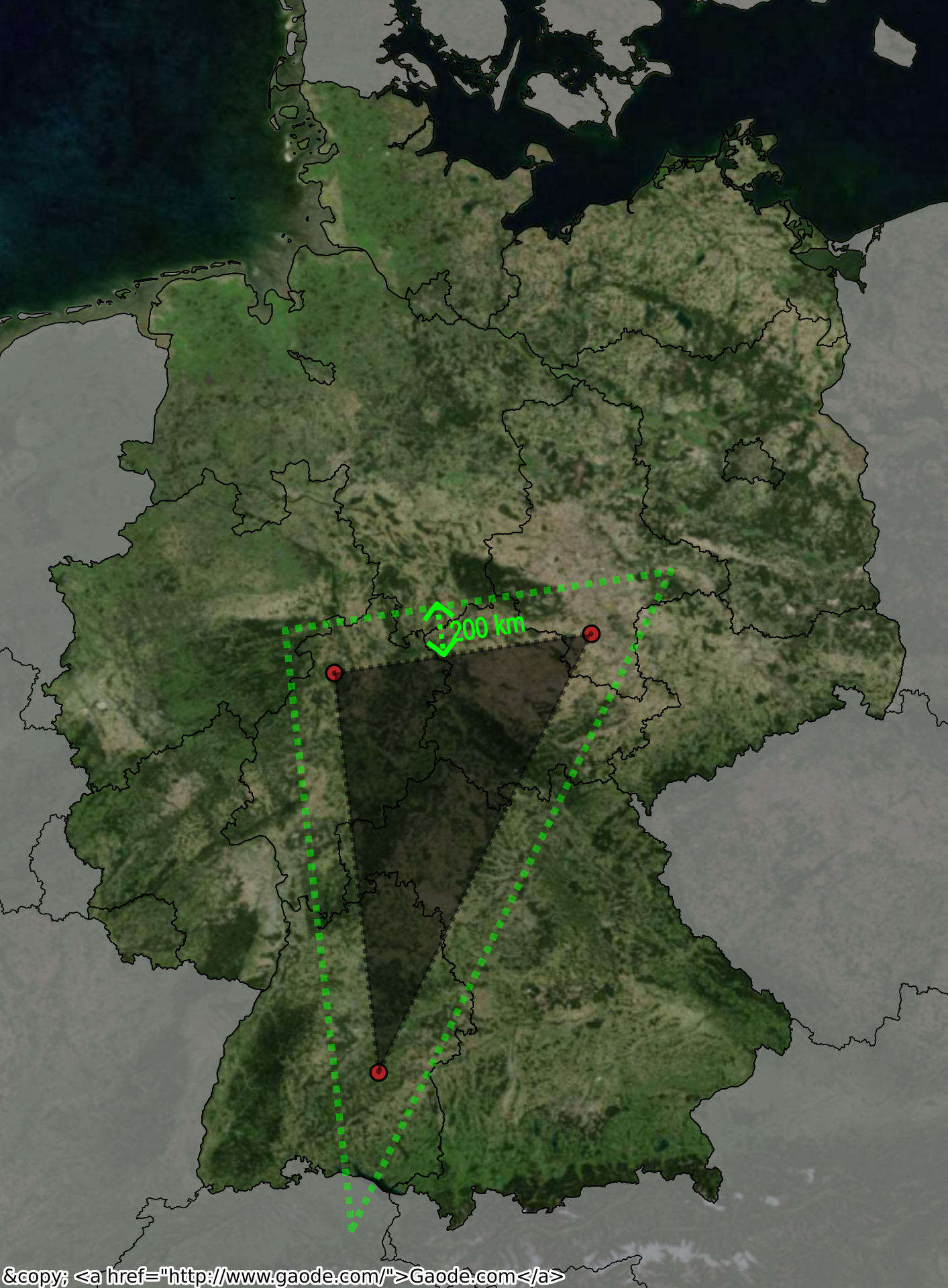
